## Supplemental Table S4 for "Transcriptomic responses of the human kidney to acute injury at single cell resolution"

**Podocytes**

*This means that positive fold changes imply higher mean value in non-COVID AKI than in COVID AKI and vice versa.

| | baseMean | log2 fold change non-COVID/COVID AKI* | lfcSE | stat | pvalue | padj |
|---|---|---|---|---|---|---|
| AL669831.5 | 12.2210027 | -0.487727741 | 0.43328269 | -1.1256571 | 0.26031069 | 0.99999872 |
| SAMD11 | 48.102533 | -0.660567997 | 0.71134411 | -0.9286195 | 0.35308632 | 0.99999872 |
| NOC2L | 12.6479149 | 0.508272397 | 0.55688334 | 0.91270893 | 0.36139565 | 0.99999872 |
| HES4 | 23.03128 | -0.47704999 | 0.84605413 | -0.5638528 | 0.57285431 | 0.99999872 |
| ISG15 | 576.585799 | -2.130823618 | 2.26227461 | -0.9418943 | 0.34624675 | 0.99999872 |
| AGRN | 111.659355 | -0.268162288 | 0.58391677 | -0.4592474 | 0.64605648 | 0.99999872 |
| C1orf159 | 12.7788718 | -0.190758877 | 0.48737104 | -0.3914038 | 0.69549878 | 0.99999872 |
| SDF4 | 51.4293417 | 0.34319402 | 0.63618507 | 0.53945626 | 0.58957207 | 0.99999872 |
| B3GALT6 | 4.97465606 | -0.201965117 | 0.85454699 | -0.2363417 | 0.8131675 | 0.99999872 |
| UBE2J2 | 22.2247154 | 0.121794267 | 0.51263636 | 0.23758413 | 0.81220366 | 0.99999872 |
| ACAP3 | 7.91076114 | -0.4486896 | 0.55172702 | -0.8132457 | 0.41607723 | 0.99999872 |
| INTS11 | 19.0675457 | -0.174858223 | 0.38760943 | -0.4511196 | 0.65190333 | 0.99999872 |
| DVL1 | 11.8569733 | -0.202269312 | 0.46558803 | -0.4344384 | 0.66397013 | 0.99999872 |
| MXRA8 | 79.7406274 | -0.238877276 | 0.4664218 | -0.5121486 | 0.60854701 | 0.99999872 |
| AURKAIP1 | 43.2543672 | -0.497629078 | 0.585493 | -0.8499317 | 0.39536304 | 0.99999872 |
| CCNL2 | 55.3029639 | -0.299908623 | 0.24072019 | -1.2458806 | 0.21280823 | 0.99999872 |
| MRPL20 | 46.4876818 | 0.279632107 | 0.31917254 | 0.87611582 | 0.38096708 | 0.99999872 |
| ANKRD65 | 16.1822483 | 1.181997496 | 0.53652802 | 2.20304895 | 0.0275913 | 0.89154528 |
| VWA1 | 70.8850489 | -0.041137946 | 0.6178723 | -0.06658 | 0.94691606 | 0.99999872 |
| ATAD3C | 5.82868676 | -0.324103131 | 0.83203067 | -0.3895327 | 0.69688215 | 0.99999872 |
| ATAD3B | 9.264103 | -0.818936379 | 0.62281162 | -1.3149022 | 0.18854277 | 0.99999872 |
| ATAD3A | 5.95827862 | -0.689155029 | 0.6196966 | -1.1120846 | 0.26610179 | 0.99999872 |
| SSU72 | 39.5444282 | -0.442217849 | 0.39711654 | -1.113572 | 0.26546285 | 0.99999872 |
| MIB2 | 9.51409055 | 0.06291301 | 0.56188067 | 0.11196863 | 0.91084828 | 0.99999872 |
| CDK11B | 9.02427123 | -0.163113718 | 0.55533421 | -0.2937217 | 0.76897056 | 0.99999872 |
| SLC35E2B | 14.235477 | -0.783705835 | 0.53282861 | -1.4708404 | 0.14133429 | 0.99999872 |
| CDK11A | 4.78124249 | 0.095552932 | 0.64982597 | 0.14704388 | 0.88309739 | 0.99999872 |
| SLC35E2A | 10.9568293 | 0.530106983 | 0.60927563 | 0.87006104 | 0.38426705 | 0.99999872 |
| NADK | 11.4037512 | -0.234101732 | 0.49108471 | -0.4767034 | 0.63357337 | 0.99999872 |
| GNB1 | 155.392165 | 0.007305721 | 0.33693624 | 0.0216828 | 0.98270099 | 0.99999872 |
| PRKCZ | 21.5365712 | -0.397321092 | 0.33545859 | -1.1844117 | 0.23625012 | 0.99999872 |
| FAAP20 | 19.4240643 | -0.152057295 | 0.46783737 | -0.3250217 | 0.74516465 | 0.99999872 |
| SKI | 39.465746 | -0.355547548 | 0.34729599 | -1.0237594 | 0.30594892 | 0.99999872 |
| MORN1 | 7.74445844 | -0.357265771 | 0.48852187 | -0.7313199 | 0.46458377 | 0.99999872 |
| RER1 | 38.6532152 | 0.21312992 | 0.36718876 | 0.58043694 | 0.56162 | 0.99999872 |
| PEX10 | 10.6140949 | -0.049873271 | 0.58296661 | -0.0855508 | 0.9318235 | 0.99999872 |
| PANK4 | 3.8199875 | 1.134545109 | 0.85342113 | 1.32940828 | 0.18371331 | NA |
| TNFRSF14 | 24.6732356 | -0.659240148 | 0.50714468 | -1.2999055 | 0.19363337 | 0.99999872 |
| FAM213B | 5.21454943 | -0.0158431 | 0.83611385 | -0.0189485 | 0.98488219 | 0.99999872 |
| ARHGEF16 | 20.5426354 | -0.806477115 | 0.41036584 | -1.9652638 | 0.04938372 | 0.97937581 |
| TPRG1L | 18.2253749 | 0.244791006 | 0.5222016 | 0.46876725 | 0.63923601 | 0.99999872 |
| WRAP73 | 6.32002732 | -0.427627452 | 0.63678316 | -0.6715433 | 0.5018745 | 0.99999872 |
| SMIM1 | 7.79233443 | 0.717997334 | 0.67992118 | 1.05600083 | 0.29096783 | 0.99999872 |
| LRRC47 | 13.8788741 | 0.248550552 | 0.58278837 | 0.42648509 | 0.66975441 | 0.99999872 |
| CEP104 | 26.122262 | 0.29431477 | 0.37685446 | 0.78097728 | 0.43481586 | 0.99999872 |
| C1orf174 | 6.00334306 | -0.950173925 | 0.56112205 | -1.6933462 | 0.09038958 | 0.99999872 |
| NPHP4 | 12.1683926 | 0.330343436 | 0.43614605 | 0.75741471 | 0.44880144 | 0.99999872 |
| RPL22 | 12.8474251 | 0.170370672 | 0.59748741 | 0.28514521 | 0.7755329 | 0.99999872 |
| RNF207 | 3.99756485 | -1.222728366 | 0.93053611 | -1.3140042 | 0.18884479 | NA |
| ICMT | 24.7736965 | -0.458948035 | 0.4850382 | -0.9462101 | 0.34404144 | 0.99999872 |
| ACOT7 | 14.1538094 | -1.12855315 | 0.5533293 | -2.0395688 | 0.0413933 | 0.95980711 |
| PLEKHG5 | 5.73621805 | 0.110824621 | 0.65560738 | 0.16904114 | 0.86576428 | 0.99999872 |
| NOL9 | 15.4076178 | -0.335821388 | 0.5810381 | -0.5779679 | 0.56328577 | 0.99999872 |
| ZBTB48 | 7.92418947 | -0.294852275 | 0.58855194 | -0.5009792 | 0.61638577 | 0.99999872 |
| KLHL21 | 12.5252922 | -0.536393867 | 0.58439398 | -0.9178634 | 0.35869036 | 0.99999872 |
| PHF13 | 9.7206617 | 0.615270004 | 0.54615451 | 1.12654934 | 0.25993307 | 0.99999872 |
| THAP3 | 7.77959471 | -0.332254121 | 0.66188586 | -0.501981 | 0.61568089 | 0.99999872 |
| DNAJC11 | 25.8794764 | -0.360407462 | 0.31224291 | -1.1542535 | 0.24839626 | 0.99999872 |
| CAMTA1 | 117.945979 | -0.132542847 | 0.28053394 | -0.4724663 | 0.63659396 | 0.99999872 |

| | | | | | | |
|---|---|---|---|---|---|---|
| VAMP3 | 28.1995901 | 0.552935265 | 0.41399589 | 1.33560568 | 0.18167819 | 0.99999872 |
| PER3 | 43.3187237 | -1.017162367 | 0.58009156 | -1.7534514 | 0.07952455 | 0.99999872 |
| PARK7 | 105.601704 | 0.703496206 | 0.42921453 | 1.63903168 | 0.10120666 | 0.99999872 |
| ERRFI1 | 63.9620585 | -0.102456609 | 0.54643904 | -0.1874987 | 0.85126964 | 0.99999872 |
| SLC45A1 | 74.3844856 | -0.155372862 | 0.54525539 | -0.2849543 | 0.77567916 | 0.99999872 |
| RERE | 604.968307 | -0.342065479 | 0.28215866 | -1.2123161 | 0.22539141 | 0.99999872 |
| ENO1 | 83.4090663 | -0.732057243 | 0.49401579 | -1.4818499 | 0.13838024 | 0.99999872 |
| GPR157 | 4.99166341 | 0.608504221 | 0.70150387 | 0.86742818 | 0.38570745 | 0.99999872 |
| H6PD | 26.1647951 | 0.249293384 | 0.38501108 | 0.64749665 | 0.51731056 | 0.99999872 |
| SPSB1 | 63.2622571 | -0.224989887 | 0.55005781 | -0.4090295 | 0.68251799 | 0.99999872 |
| SLC25A33 | 20.5517187 | -0.240559191 | 0.39547786 | -0.6082747 | 0.54300528 | 0.99999872 |
| TMEM201 | 4.60937763 | 0.468612066 | 0.73974292 | 0.63347962 | 0.52642049 | 0.99999872 |
| PIK3CD | 7.59914313 | 0.393285082 | 0.91851369 | 0.42817553 | 0.66852333 | 0.99999872 |
| CLSTN1 | 63.059107 | 0.082535549 | 0.44216706 | 0.18666146 | 0.85192608 | 0.99999872 |
| CTNNBIP1 | 89.0023943 | -0.262090463 | 0.69601866 | -0.3765567 | 0.7065031 | 0.99999872 |
| LZIC | 19.7340143 | -0.030321628 | 0.60408138 | -0.0501946 | 0.95996731 | 0.99999872 |
| NMNAT1 | 10.7964506 | -0.086227214 | 0.53116345 | -0.1623365 | 0.87104087 | 0.99999872 |
| UBE4B | 63.6233995 | -0.181484282 | 0.31516258 | -0.5758434 | 0.56472105 | 0.99999872 |
| KIF1B | 139.965804 | 0.319645678 | 0.32582041 | 0.98104866 | 0.32656875 | 0.99999872 |
| PGD | 23.3618586 | 0.34105095 | 0.46324576 | 0.73622034 | 0.46159662 | 0.99999872 |
| DFFA | 17.5786131 | 0.061604042 | 0.40864444 | 0.15075218 | 0.88017121 | 0.99999872 |
| PEX14 | 28.8394318 | -0.173300493 | 0.35748804 | -0.4847728 | 0.62783749 | 0.99999872 |
| CASZ1 | 171.502854 | -0.421457009 | 0.54207539 | -0.7774878 | 0.43687102 | 0.99999872 |
| AL139423.1 | 6.89682419 | 0.158675434 | 0.82216718 | 0.19299655 | 0.84696167 | 0.99999872 |
| SRM | 13.320911 | -0.361072003 | 0.48612544 | -0.7427548 | 0.45763015 | 0.99999872 |
| EXOSC10 | 10.8966615 | -0.922492054 | 0.45517055 | -2.0266954 | 0.04269357 | 0.95980711 |
| MTOR | 20.2158639 | -0.187898372 | 0.40511761 | -0.4638119 | 0.64278251 | 0.99999872 |
| UBIAD1 | 4.973932 | 0.016387646 | 0.59495442 | 0.02754437 | 0.97802555 | 0.99999872 |
| FBXO2 | 36.4529185 | 0.351343192 | 0.81307393 | 0.43211715 | 0.66565627 | 0.99999872 |
| FBXO44 | 7.86079415 | -0.15584108 | 0.55857178 | -0.2789992 | 0.78024544 | 0.99999872 |
| MAD2L2 | 5.19183553 | -0.329429737 | 0.77818556 | -0.4233306 | 0.67205409 | 0.99999872 |
| AGTRAP | 24.5349099 | 0.168312611 | 0.64668256 | 0.26027084 | 0.79465487 | 0.99999872 |
| MTHFR | 12.4434938 | -0.492735521 | 0.42905261 | -1.1484268 | 0.25079241 | 0.99999872 |
| CLCN6 | 7.30640911 | -0.285541323 | 0.59512065 | -0.4798041 | 0.6313667 | 0.99999872 |
| KIAA2013 | 29.1524218 | -0.232406955 | 0.37936564 | -0.6126199 | 0.54012767 | 0.99999872 |
| PLOD1 | 19.02825 | 0.091909878 | 0.53936671 | 0.17040332 | 0.86469296 | 0.99999872 |
| MFN2 | 13.1685749 | -0.16186609 | 0.53142587 | -0.3045883 | 0.76067975 | 0.99999872 |
| MIIP | 12.5764277 | 0.192133229 | 0.75523289 | 0.25440262 | 0.79918452 | 0.99999872 |
| TNFRSF8 | 8.9067448 | -0.268929932 | 0.81908843 | -0.3283283 | 0.74266343 | 0.99999872 |
| VPS13D | 91.5978601 | -0.354275116 | 0.21844299 | -1.6218196 | 0.10484198 | 0.99999872 |
| DHRS3 | 139.893866 | -0.880744663 | 0.37962518 | -2.3200375 | 0.02033885 | 0.8671024 |
| PDPN | 27.6830468 | 0.23758888 | 0.69971088 | 0.33955293 | 0.73419323 | 0.99999872 |
| PRDM2 | 54.6463598 | 0.052422893 | 0.29499933 | 0.17770513 | 0.85895456 | 0.99999872 |
| KAZN | 130.507684 | -0.626363679 | 0.38090308 | -1.6444175 | 0.10009 | 0.99999872 |
| TMEM51 | 11.141284 | -1.108503593 | 0.59394175 | -1.8663507 | 0.06199232 | 0.99999872 |
| EFHD2 | 4.42275693 | 0.577025723 | 0.7768177 | 0.74280713 | 0.45759846 | NA |
| CASP9 | 8.30982991 | -0.733583821 | 0.62794531 | -1.1682288 | 0.24271447 | 0.99999872 |
| DNAJC16 | 30.946139 | -0.177705716 | 0.34192318 | -0.5197241 | 0.60325589 | 0.99999872 |
| DDI2 | 23.2920273 | -0.030651952 | 0.43447376 | -0.0705496 | 0.94375622 | 0.99999872 |
| PLEKHM2 | 31.9278701 | -0.344536529 | 0.30608699 | -1.1256164 | 0.26032794 | 0.99999872 |
| FBLIM1 | 13.7438073 | 0.809835495 | 0.95730643 | 0.84595222 | 0.3975794 | 0.99999872 |
| AL450998.2 | 6.26872963 | -0.525201898 | 0.59881448 | -0.8770695 | 0.3804489 | 0.99999872 |
| SPEN | 63.1413115 | -0.020511561 | 0.30848406 | -0.0664915 | 0.94698654 | 0.99999872 |
| ZBTB17 | 7.38681746 | -0.857275508 | 0.52390353 | -1.6363232 | 0.10177196 | 0.99999872 |
| EPHA2 | 18.7853984 | -0.178351104 | 0.56515956 | -0.3155766 | 0.75232394 | 0.99999872 |
| FBXO42 | 36.9729752 | -0.262734364 | 0.33427322 | -0.7859869 | 0.43187518 | 0.99999872 |
| SZRD1 | 28.5561025 | -0.39236909 | 0.4282138 | -0.9162925 | 0.35951351 | 0.99999872 |
| NECAP2 | 20.7084455 | -0.474856957 | 0.39129834 | -1.213542 | 0.22492269 | 0.99999872 |
| NBPF1 | 8.49175222 | 0.664599601 | 0.63734295 | 1.04276607 | 0.29705665 | 0.99999872 |
| CROCC | 15.0714924 | -0.395394511 | 0.53737148 | -0.7357936 | 0.4618563 | 0.99999872 |
| ATP13A2 | 12.2818467 | 0.102553088 | 0.48191938 | 0.21280134 | 0.83148192 | 0.99999872 |

| | | | | | | |
|---|---|---|---|---|---|---|
| SDHB | 22.3246908 | 0.129592312 | 0.40638918 | 0.31888721 | 0.74981204 | 0.99999872 |
| RCC2 | 7.64756047 | -0.398665964 | 0.60823361 | -0.6554488 | 0.51217887 | 0.99999872 |
| ARHGEF10L | 52.0211701 | 0.051496207 | 0.54077131 | 0.09522733 | 0.92413426 | 0.99999872 |
| KLHDC7A | 5.53259563 | -0.93146056 | 1.0572719 | -0.8810038 | 0.37831576 | 0.99999872 |
| IFFO2 | 9.51712498 | 0.009824685 | 0.64724867 | 0.01517915 | 0.98788926 | 0.99999872 |
| UBR4 | 54.6860696 | -0.020071938 | 0.24022378 | -0.0835552 | 0.93341012 | 0.99999872 |
| AL035413.1 | 10.2161481 | 0.265821231 | 0.55960094 | 0.47501927 | 0.63477324 | 0.99999872 |
| EMC1 | 19.4238022 | 0.613721422 | 0.41764246 | 1.46949002 | 0.14169993 | 0.99999872 |
| MRTO4 | 7.02979189 | -0.54328277 | 0.76181558 | -0.7131421 | 0.47575783 | 0.99999872 |
| AKR7A2 | 12.208665 | 0.452880808 | 0.5362595 | 0.84451801 | 0.39838001 | 0.99999872 |
| CAPZB | 91.4814484 | -0.13757648 | 0.38663737 | -0.3558282 | 0.72196923 | 0.99999872 |
| NBL1 | 6.87673746 | 2.049774825 | 1.36179555 | 1.50520012 | 0.13227263 | 0.99999872 |
| TMCO4 | 24.4862261 | -0.574125495 | 0.47230778 | -1.2155749 | 0.22414689 | 0.99999872 |
| OTUD3 | 7.70891851 | 0.002567394 | 0.54946806 | 0.00467251 | 0.99627189 | 0.99999872 |
| CAMK2N1 | 12.1397728 | -0.027052873 | 0.63904979 | -0.042333 | 0.96623327 | 0.99999872 |
| MUL1 | 7.54281127 | 0.489887956 | 0.58753665 | 0.83379982 | 0.4043938 | 0.99999872 |
| PINK1 | 20.1388838 | -0.330051227 | 0.49741832 | -0.6635285 | 0.50699214 | 0.99999872 |
| DDOST | 21.7839367 | 0.517366467 | 0.41993221 | 1.23202378 | 0.2179402 | 0.99999872 |
| HP1BP3 | 89.2265828 | 0.007894788 | 0.24010672 | 0.03288033 | 0.97377002 | 0.99999872 |
| EIF4G3 | 176.298837 | 0.064965642 | 0.40442726 | 0.16063616 | 0.87237997 | 0.99999872 |
| ECE1 | 8.08574456 | -1.06941307 | 0.68869661 | -1.5528072 | 0.1204692 | 0.99999872 |
| USP48 | 46.8855083 | 0.141303962 | 0.33213054 | 0.42544706 | 0.67051081 | 0.99999872 |
| HSPG2 | 8.28057351 | -0.292681012 | 0.76620295 | -0.3819889 | 0.70246961 | 0.99999872 |
| CDC42 | 82.867487 | 0.155428353 | 0.34447509 | 0.45120346 | 0.65184292 | 0.99999872 |
| ZBTB40 | 10.4370573 | 0.30543079 | 0.72274635 | 0.42259749 | 0.67258896 | 0.99999872 |
| KDM1A | 25.4079449 | 0.314670451 | 0.3237538 | 0.97194366 | 0.33107858 | 0.99999872 |
| AL031428.1 | 7.34617643 | 0.260973533 | 0.72931423 | 0.35783414 | 0.72046744 | 0.99999872 |
| LUZP1 | 34.8118878 | -0.180563562 | 0.3344859 | -0.5398241 | 0.58931832 | 0.99999872 |
| LINC01355 | 8.37746869 | -0.012880514 | 0.52264575 | -0.0246448 | 0.98033826 | 0.99999872 |
| HNRNPR | 64.9481678 | -0.166828036 | 0.24009987 | -0.6948277 | 0.4871633 | 0.99999872 |
| ASAP3 | 7.13099654 | -0.754084872 | 0.80849166 | -0.9327058 | 0.35097188 | 0.99999872 |
| ID3 | 20.7891561 | -3.026780929 | 0.96046576 | -3.1513679 | 0.00162508 | 0.3314549 |
| RPL11 | 101.413158 | 0.390963122 | 0.39365562 | 0.99316028 | 0.32063185 | 0.99999872 |
| ELOA | 12.5323722 | -0.128762995 | 0.58475001 | -0.2202018 | 0.82571401 | 0.99999872 |
| ELOA-AS1 | 9.74930753 | 0.218990141 | 0.49886902 | 0.43897322 | 0.66068094 | 0.99999872 |
| PITHD1 | 20.4624707 | 0.452937265 | 0.43753754 | 1.03519635 | 0.30057722 | 0.99999872 |
| LYPLA2 | 10.1181569 | -0.025638015 | 0.52409317 | -0.0489188 | 0.960984 | 0.99999872 |
| HMGCL | 18.3757661 | -0.001232743 | 0.48630165 | -0.0025349 | 0.99797742 | 0.99999872 |
| FUCA1 | 12.4737631 | 0.127460777 | 0.42243937 | 0.30172561 | 0.76286124 | 0.99999872 |
| PNRC2 | 45.2312421 | 0.293605787 | 0.48750759 | 0.60225891 | 0.54700181 | 0.99999872 |
| SRSF10 | 66.3776333 | 0.346654905 | 0.26741009 | 1.29634192 | 0.19485771 | 0.99999872 |
| IFNLR1 | 6.4834538 | 1.173854616 | 0.73068082 | 1.60652174 | 0.10815932 | 0.99999872 |
| STPG1 | 6.60822466 | 0.285288634 | 0.58899942 | 0.48436149 | 0.62812935 | 0.99999872 |
| NIPAL3 | 28.4705669 | 0.010403557 | 0.36657289 | 0.0283806 | 0.9773586 | 0.99999872 |
| RCAN3 | 12.0121421 | 0.033781373 | 0.64105394 | 0.05269662 | 0.95797364 | 0.99999872 |
| SRRM1 | 56.70328 | 0.23234918 | 0.25434181 | 0.91353122 | 0.36096322 | 0.99999872 |
| CLIC4 | 37.0204692 | -0.48005842 | 0.58561573 | -0.8197499 | 0.41235872 | 0.99999872 |
| SYF2 | 17.6931446 | 0.103211545 | 0.39234486 | 0.26306333 | 0.79250177 | 0.99999872 |
| RSRP1 | 122.654638 | 0.472675426 | 0.35992807 | 1.31324968 | 0.18909883 | 0.99999872 |
| TMEM50A | 59.0617116 | 0.023379599 | 0.51722769 | 0.04520175 | 0.9639465 | 0.99999872 |
| MACO1 | 34.4545101 | -0.269211147 | 0.44077996 | -0.6107609 | 0.54135791 | 0.99999872 |
| LDLRAP1 | 7.40867803 | -0.571661055 | 0.54486173 | -1.0491856 | 0.29409272 | 0.99999872 |
| MAN1C1 | 10.7304857 | -0.139834637 | 0.63298099 | -0.2209144 | 0.82515906 | 0.99999872 |
| MTFR1L | 21.8660939 | 0.029570467 | 0.38974589 | 0.07587115 | 0.93952161 | 0.99999872 |
| AL033528.2 | 25.0188007 | 0.145887409 | 0.48800279 | 0.2989479 | 0.7649798 | 0.99999872 |
| PAQR7 | 49.1961145 | -0.80818957 | 0.40293311 | -2.0057661 | 0.04488121 | 0.95980711 |
| STMN1 | 38.3423636 | 0.916762566 | 0.34688584 | 2.64283656 | 0.00822147 | 0.66265058 |
| PAFAH2 | 9.6786605 | 0.169272851 | 0.44967142 | 0.37643676 | 0.70659222 | 0.99999872 |
| PDIK1L | 5.0856321 | 0.139272937 | 0.6113756 | 0.22780258 | 0.81979972 | 0.99999872 |
| ZNF593 | 11.4706942 | -0.498903544 | 0.68690181 | -0.7263098 | 0.46764885 | 0.99999872 |
| CEP85 | 3.84187604 | 0.756589312 | 0.80737468 | 0.93709814 | 0.34870808 | NA |

| | | | | | | |
|---|---|---|---|---|---|---|
| SH3BGRL3 | 41.0964851 | 0.112850471 | 0.37618181 | 0.29998918 | 0.76418541 | 0.99999872 |
| DHDDS | 13.8081258 | -0.266770768 | 0.43818962 | -0.6088021 | 0.54265561 | 0.99999872 |
| HMGN2 | 48.0033246 | 0.200972907 | 0.50553382 | 0.39754592 | 0.69096493 | 0.99999872 |
| ARID1A | 74.231916 | -0.339151944 | 0.26911253 | -1.2602607 | 0.20757533 | 0.99999872 |
| PIGV | 5.2564584 | -0.265089603 | 0.61224768 | -0.4329777 | 0.66503096 | 0.99999872 |
| ZDHHC18 | 8.03704423 | 0.721705305 | 0.59680888 | 1.20927374 | 0.2265577 | 0.99999872 |
| GPN2 | 9.4638356 | -0.743669139 | 0.60667201 | -1.2258175 | 0.22026738 | 0.99999872 |
| NUDC | 27.4711789 | -0.429264808 | 0.34971866 | -1.2274575 | 0.21965067 | 0.99999872 |
| SLC9A1 | 5.38510809 | 0.220411102 | 0.67445508 | 0.32679879 | 0.74382008 | 0.99999872 |
| WDTC1 | 27.1644267 | 0.077983939 | 0.47307096 | 0.16484618 | 0.86906506 | 0.99999872 |
| TMEM222 | 16.105494 | -0.348654159 | 0.39949963 | -0.8727271 | 0.38281184 | 0.99999872 |
| MAP3K6 | 4.0284086 | 0.28341916 | 0.8050776 | 0.35203956 | 0.7248086 | NA |
| WASF2 | 138.379961 | 0.063406169 | 0.28548477 | 0.22210001 | 0.82423603 | 0.99999872 |
| AHDC1 | 44.5368943 | -0.247951436 | 0.33016493 | -0.7509927 | 0.45265708 | 0.99999872 |
| IFI6 | 126.040823 | 1.003856005 | 1.84945686 | 0.54278422 | 0.58727838 | 0.99999872 |
| FAM76A | 18.187141 | -0.23885135 | 0.35415856 | -0.6744192 | 0.50004481 | 0.99999872 |
| STX12 | 40.1097829 | -0.022015842 | 0.26860159 | -0.0819647 | 0.9346748 | 0.99999872 |
| PPP1R8 | 14.2724052 | -0.077307274 | 0.42785425 | -0.180686 | 0.85661405 | 0.99999872 |
| THEMIS2 | 4.71514989 | 0.049754053 | 1.0123254 | 0.04914828 | 0.96080113 | 0.99999872 |
| RPA2 | 11.5584277 | -0.160584933 | 0.43503879 | -0.3691279 | 0.71203243 | 0.99999872 |
| EYA3 | 37.4168585 | 0.100004535 | 0.30912588 | 0.32350748 | 0.74631094 | 0.99999872 |
| DNAJC8 | 23.3431236 | 0.248816674 | 0.3475622 | 0.71589106 | 0.47405862 | 0.99999872 |
| ATP5IF1 | 89.5001939 | 0.661445654 | 0.45990395 | 1.43822565 | 0.15037004 | 0.99999872 |
| SESN2 | 13.3948309 | -0.225885311 | 0.5687332 | -0.3971727 | 0.6912401 | 0.99999872 |
| PHACTR4 | 342.631931 | -0.025463599 | 0.33221193 | -0.0766487 | 0.93890305 | 0.99999872 |
| RCC1 | 16.4370484 | 0.092017641 | 0.56430134 | 0.16306472 | 0.87046748 | 0.99999872 |
| TRNAU1AP | 18.6327006 | 0.302262462 | 0.46080853 | 0.65593938 | 0.51186313 | 0.99999872 |
| SNHG12 | 4.51011518 | -0.406332855 | 0.67692732 | -0.6002607 | 0.54833252 | NA |
| TAF12 | 9.30337747 | 0.046896818 | 0.56562312 | 0.08291178 | 0.93392169 | 0.99999872 |
| GMEB1 | 15.7949686 | -1.219893726 | 0.42249534 | -2.8873542 | 0.00388497 | 0.4904418 |
| YTHDF2 | 33.207824 | 0.261772101 | 0.31078368 | 0.84229681 | 0.39962184 | 0.99999872 |
| EPB41 | 43.7496362 | 0.300552742 | 0.38917933 | 0.77227314 | 0.43995267 | 0.99999872 |
| SRSF4 | 76.7943165 | -0.398815519 | 0.25240391 | -1.5800687 | 0.11409113 | 0.99999872 |
| MECR | 7.7805407 | -0.205250234 | 0.54520701 | -0.3764629 | 0.70657276 | 0.99999872 |
| SDC3 | 16.2262017 | -0.318473071 | 0.63333313 | -0.5028524 | 0.61506807 | 0.99999872 |
| PUM1 | 111.402776 | -0.338268289 | 0.24644151 | -1.3726109 | 0.16987335 | 0.99999872 |
| SNRNP40 | 14.4639463 | -0.417472053 | 0.48254663 | -0.8651434 | 0.38696006 | 0.99999872 |
| ZCCHC17 | 14.1121394 | 0.03404453 | 0.48783306 | 0.06978726 | 0.94436299 | 0.99999872 |
| SERINC2 | 8.94391186 | -1.041048241 | 0.46407339 | -2.2432836 | 0.02487853 | 0.88365618 |
| TINAGL1 | 6.60025938 | -1.340719051 | 0.76775048 | -1.7462953 | 0.08075965 | 0.99999872 |
| PEF1 | 16.58359 | 0.013174589 | 0.4621489 | 0.02850724 | 0.97725759 | 0.99999872 |
| PTP4A2 | 10.3840246 | 0.228051647 | 0.57067906 | 0.39961454 | 0.68944045 | 0.99999872 |
| KHDRBS1 | 66.8241069 | -0.119811606 | 0.34822871 | -0.3440601 | 0.73080109 | 0.99999872 |
| TMEM39B | 7.71962577 | -0.423928786 | 0.54515708 | -0.7776269 | 0.43678902 | 0.99999872 |
| KPNA6 | 72.8651021 | -0.202558225 | 0.36743976 | -0.5512692 | 0.58144914 | 0.99999872 |
| TXLNA | 26.6034543 | -0.23478133 | 0.36294664 | -0.6468756 | 0.51771245 | 0.99999872 |
| TMEM234 | 8.00140073 | 0.647892782 | 0.59553881 | 1.08791027 | 0.27663472 | 0.99999872 |
| EIF3I | 16.0110414 | 0.952165179 | 0.53928317 | 1.76561263 | 0.07746086 | 0.99999872 |
| HDAC1 | 21.2644297 | -0.180811674 | 0.50280877 | -0.3596033 | 0.71914384 | 0.99999872 |
| MARCKSL1 | 6.61558513 | -0.609160765 | 0.72868205 | -0.8359761 | 0.40316836 | 0.99999872 |
| TSSK3 | 5.37495152 | 0.080372794 | 0.85390767 | 0.09412352 | 0.92501104 | 0.99999872 |
| BSDC1 | 26.7282646 | -0.291556832 | 0.39717256 | -0.734081 | 0.46289937 | 0.99999872 |
| ZBTB8A | 8.41840522 | 0.466681087 | 0.61474309 | 0.75914816 | 0.44776393 | 0.99999872 |
| ZBTB8OS | 29.8020681 | 0.032398181 | 0.32499732 | 0.09968753 | 0.9205924 | 0.99999872 |
| RBBP4 | 38.4263843 | 0.063354345 | 0.28547273 | 0.22192783 | 0.82437006 | 0.99999872 |
| SYNC | 8.22391893 | 1.246235702 | 0.67424667 | 1.84833795 | 0.06455347 | 0.99999872 |
| KIAA1522 | 11.7233515 | 0.154570199 | 0.47705929 | 0.32400627 | 0.74593329 | 0.99999872 |
| YARS | 18.2446613 | -0.607566289 | 0.61190321 | -0.9929124 | 0.32075265 | 0.99999872 |
| S100PBP | 28.6646328 | -0.233387728 | 0.3715661 | -0.628119 | 0.52992599 | 0.99999872 |
| TMEM54 | 8.27839486 | 0.262371123 | 0.63994862 | 0.40998779 | 0.6818149 | 0.99999872 |
| RNF19B | 28.6847341 | -1.096441022 | 0.80375314 | -1.3641515 | 0.1725199 | 0.99999872 |

| | | | | | | |
|---|---|---|---|---|---|---|
| AL020995.1 | 4.01618727 | 0.460336052 | 0.80098066 | 0.57471557 | 0.56548368 | NA |
| AK2 | 25.9243049 | 0.047068668 | 0.46425269 | 0.10138588 | 0.91924414 | 0.99999872 |
| ZNF362 | 14.189581 | 0.037048621 | 0.65800037 | 0.05630486 | 0.95509894 | 0.99999872 |
| AL513327.1 | 5.87852242 | 0.358593629 | 0.8507492 | 0.42150334 | 0.67338757 | 0.99999872 |
| PHC2 | 33.5205373 | -0.520730635 | 0.51294994 | -1.0151685 | 0.3100255 | 0.99999872 |
| CSMD2 | 22.0967081 | -1.309957163 | 0.57631778 | -2.2729772 | 2.30E-02 | 0.88337526 |
| SMIM12 | 14.3375191 | 0.193981012 | 0.43721086 | 0.44367839 | 0.65727512 | 0.99999872 |
| ZMYM1 | 16.7552379 | 0.304714276 | 0.41753875 | 0.72978682 | 0.4655205 | 0.99999872 |
| SFPQ | 91.9793267 | -0.447318724 | 0.23916706 | -1.8703191 | 0.06143952 | 0.99999872 |
| ZMYM4 | 156.132932 | 0.006269102 | 0.20019902 | 0.03131435 | 0.97501885 | 0.99999872 |
| KIAA0319L | 97.9773044 | -0.405129029 | 0.27198541 | -1.4895249 | 0.13634921 | 0.99999872 |
| NCDN | 5.01188263 | 0.089733194 | 0.74454197 | 0.12052134 | 0.90407018 | 0.99999872 |
| PSMB2 | 31.2706012 | 0.894030401 | 0.40502833 | 2.20732808 | 0.02729115 | 0.89154528 |
| AGO4 | 33.729653 | -0.140042429 | 0.46431059 | -0.3016137 | 0.76294657 | 0.99999872 |
| AGO1 | 10.3481779 | 0.043171934 | 0.54709823 | 0.07891076 | 0.93710361 | 0.99999872 |
| AGO3 | 78.1713761 | 0.008571061 | 0.25972564 | 0.03300044 | 0.97367424 | 0.99999872 |
| ADPRHL2 | 19.4782074 | -0.237178171 | 0.50241772 | -0.4720737 | 0.63687422 | 0.99999872 |
| TRAPPC3 | 14.9038777 | 0.386656118 | 0.51608061 | 0.74921652 | 0.45372671 | 0.99999872 |
| MAP7D1 | 22.5469038 | 0.124101442 | 0.37590721 | 0.3301385 | 0.74129531 | 0.99999872 |
| THRAP3 | 75.0348093 | -0.215083279 | 0.25065514 | -0.8580844 | 0.39084583 | 0.99999872 |
| EVA1B | 86.9759889 | -0.218456763 | 0.37719246 | -0.5791652 | 0.56247768 | 0.99999872 |
| STK40 | 31.3949181 | -0.091825579 | 0.50993646 | -0.1800726 | 0.85709558 | 0.99999872 |
| LSM10 | 18.692129 | 1.407088252 | 0.49126013 | 2.86424272 | 0.00418008 | 0.50343486 |
| OSCP1 | 6.71445582 | -0.13832293 | 0.73711515 | -0.1876544 | 0.85114755 | 0.99999872 |
| MRPS15 | 11.5558987 | 0.27223441 | 0.47446339 | 0.57377327 | 0.56612124 | 0.99999872 |
| MEAF6 | 28.3251749 | 0.327615817 | 0.45494683 | 0.72011892 | 0.47145178 | 0.99999872 |
| SNIP1 | 10.923813 | 0.181374908 | 0.64572693 | 0.28088485 | 0.77879872 | 0.99999872 |
| DNALI1 | 4.91030896 | -0.193005223 | 0.69626011 | -0.2772028 | 0.78162442 | 0.99999872 |
| GNL2 | 8.34008605 | 0.255810584 | 0.82339933 | 0.31067621 | 0.75604678 | 0.99999872 |
| C1orf109 | 5.26250307 | 0.564013163 | 0.76519495 | 0.73708428 | 0.4610711 | 0.99999872 |
| C1orf122 | 45.8071053 | 0.483318691 | 0.41313763 | 1.16987334 | 0.24205195 | 0.99999872 |
| MTF1 | 11.021096 | -0.330268913 | 0.46462842 | -0.7108237 | 0.47719347 | 0.99999872 |
| INPP5B | 20.183457 | 0.410177487 | 0.71175275 | 0.5762921 | 0.56441776 | 0.99999872 |
| SF3A3 | 12.9087344 | 0.029631679 | 0.52777177 | 0.05614487 | 0.9552264 | 0.99999872 |
| UTP11 | 4.26442033 | 0.783853401 | 0.66841633 | 1.17270235 | 0.24091519 | NA |
| RRAGC | 19.0188211 | -0.003478329 | 0.64537836 | -0.0053896 | 0.99569975 | 0.99999872 |
| AL139260.1 | 17.3962156 | 0.415682154 | 0.7991161 | 0.52017742 | 0.60293992 | 0.99999872 |
| MYCBP | 5.50528546 | 1.081402352 | 0.8010772 | 1.34993526 | 0.17703675 | 0.99999872 |
| RHBDL2 | 4.18561746 | 0.306863718 | 1.01622096 | 0.30196555 | 0.76267833 | NA |
| AKIRIN1 | 32.8211363 | 0.257988573 | 0.29845633 | 0.86440978 | 0.38736282 | 0.99999872 |
| NDUFS5 | 69.7878377 | 0.108105855 | 0.45070133 | 0.23986141 | 0.8104377 | 0.99999872 |
| MACF1 | 659.951503 | -0.064923749 | 0.24873694 | -0.2610137 | 0.79408194 | 0.99999872 |
| PABPC4 | 11.6073331 | 0.056039154 | 0.50147094 | 0.11174956 | 0.91102199 | 0.99999872 |
| PPIE | 16.1871374 | 0.10601215 | 0.36587528 | 0.28974942 | 0.77200794 | 0.99999872 |
| BMP8B | 4.24973186 | -1.286670062 | 0.95936931 | -1.3411624 | 0.17986772 | NA |
| TRIT1 | 16.6701542 | 0.057127017 | 0.39162345 | 0.14587231 | 0.88402219 | 0.99999872 |
| CAP1 | 59.9163941 | 0.123682491 | 0.45701265 | 0.27063253 | 0.78667367 | 0.99999872 |
| PPT1 | 25.698367 | 0.770045289 | 0.36125118 | 2.13160632 | 0.03303922 | 0.92959829 |
| RLF | 39.836749 | -0.004526596 | 0.27353701 | -0.0165484 | 0.9867969 | 0.99999872 |
| ZMPSTE24 | 20.8167088 | -0.283178581 | 0.38381861 | -0.7377927 | 0.46064041 | 0.99999872 |
| SMAP2 | 12.9316183 | -0.045091846 | 0.50211779 | -0.0898033 | 0.92844351 | 0.99999872 |
| AL603839.3 | 13.7805477 | -0.730796841 | 0.46639009 | -1.5669219 | 0.11713295 | 0.99999872 |
| ZNF684 | 5.56979583 | 0.942445202 | 0.74170273 | 1.27065085 | 0.20385289 | 0.99999872 |
| NFYC | 25.2960779 | 0.287030478 | 0.37383271 | 0.76780461 | 0.44260328 | 0.99999872 |
| CTPS1 | 5.61800406 | -0.730801313 | 0.86819736 | -0.8417456 | 0.39993037 | 0.99999872 |
| SCMH1 | 105.044928 | -0.192311726 | 0.28348991 | -0.6783724 | 0.49753561 | 0.99999872 |
| HIVEP3 | 64.1803364 | -0.68591882 | 0.36962533 | -1.8557138 | 0.06349436 | 0.99999872 |
| FOXJ3 | 68.2760289 | -0.261467786 | 0.28801361 | -0.9078313 | 0.36396733 | 0.99999872 |
| PPCS | 21.4540855 | -0.026689258 | 0.35569536 | -0.075034 | 0.94018763 | 0.99999872 |
| CCDC30 | 38.3659936 | -0.126805809 | 0.35907179 | -0.3531489 | 0.72397681 | 0.99999872 |
| PPIH | 7.29667594 | -0.153673501 | 0.58527499 | -0.2625663 | 0.79288486 | 0.99999872 |

| | | | | | | |
|---|---|---|---|---|---|---|
| YBX1 | 47.4667151 | 0.570488949 | 0.40201125 | 1.41908701 | 0.15587365 | 0.99999872 |
| P3H1 | 12.8261345 | 0.264493363 | 0.47535181 | 0.55641602 | 0.57792649 | 0.99999872 |
| SVBP | 6.95450467 | -0.672629208 | 0.65037175 | -1.0342227 | 0.30103208 | 0.99999872 |
| ERMAP | 11.1694156 | 0.037382639 | 0.54876512 | 0.06812138 | 0.94568901 | 0.99999872 |
| SLC2A1 | 6.43208238 | -0.993674166 | 1.01368714 | -0.9802572 | 0.32695915 | 0.99999872 |
| EBNA1BP2 | 18.5305363 | 0.542663885 | 0.42653921 | 1.27224854 | 0.20328482 | 0.99999872 |
| ELOVL1 | 16.300821 | 0.138390477 | 0.74118091 | 0.18671619 | 0.85188316 | 0.99999872 |
| MED8 | 7.98988811 | 0.706659187 | 0.64962473 | 1.08779601 | 0.27668517 | 0.99999872 |
| SZT2 | 15.2802745 | -0.699545635 | 0.42149923 | -1.6596605 | 0.09698276 | 0.99999872 |
| HYI | 6.41059847 | -0.34058411 | 0.58696329 | -0.5802477 | 0.56174758 | 0.99999872 |
| PTPRF | 51.6050482 | 0.908904607 | 0.46924944 | 1.93693276 | 0.05275356 | 0.99999872 |
| IPO13 | 6.21566397 | 0.041728546 | 0.77896504 | 0.05356922 | 0.95727838 | 0.99999872 |
| ATP6V0B | 63.3139883 | 0.612188715 | 0.40367431 | 1.51654117 | 0.12938257 | 0.99999872 |
| B4GALT2 | 16.0790004 | -0.101479358 | 0.63843345 | -0.1589506 | 0.87370782 | 0.99999872 |
| CCDC24 | 9.28263693 | 0.398513583 | 0.51895473 | 0.76791588 | 0.44253716 | 0.99999872 |
| DMAP1 | 8.33835496 | 0.391213676 | 0.70040092 | 0.55855677 | 0.57646425 | 0.99999872 |
| ERI3 | 46.6033715 | -0.077261634 | 0.29639529 | -0.2606709 | 0.79434629 | 0.99999872 |
| RNF220 | 56.2134005 | 0.209716882 | 0.62293019 | 0.33666193 | 0.73637176 | 0.99999872 |
| TMEM53 | 8.85900713 | 0.090336724 | 0.56354475 | 0.16030089 | 0.87264406 | 0.99999872 |
| RPS8 | 81.2350447 | 0.74419231 | 0.3982169 | 1.86881148 | 0.06164905 | 0.99999872 |
| EIF2B3 | 37.2970042 | -0.004199295 | 0.3457644 | -0.012145 | 0.99030996 | 0.99999872 |
| HECTD3 | 5.69418522 | 0.218327277 | 0.58003646 | 0.37640268 | 0.70661755 | 0.99999872 |
| UROD | 8.20865931 | 0.471369124 | 0.68153708 | 0.69162653 | 0.4891719 | 0.99999872 |
| ZSWIM5 | 94.0712385 | 0.412856997 | 0.47157275 | 0.87548951 | 0.38130761 | 0.99999872 |
| TESK2 | 6.10642201 | 0.566524924 | 0.67021932 | 0.8452829 | 0.39795291 | 0.99999872 |
| PRDX1 | 148.480159 | 0.465164803 | 0.5242915 | 0.88722553 | 0.37495749 | 0.99999872 |
| AKR1A1 | 11.6324543 | 0.320019731 | 0.60855368 | 0.52586936 | 0.59897898 | 0.99999872 |
| NASP | 32.6282565 | 0.21619196 | 0.36563354 | 0.59128044 | 0.55433254 | 0.99999872 |
| GPBP1L1 | 80.5530052 | 0.071796404 | 0.23253904 | 0.3087499 | 0.75751178 | 0.99999872 |
| TMEM69 | 6.10166035 | 0.069002006 | 0.71260017 | 0.09683131 | 0.92286036 | 0.99999872 |
| IPP | 24.5339468 | -0.533528021 | 0.3928475 | -1.3581047 | 0.17443048 | 0.99999872 |
| MAST2 | 24.7761103 | 0.16617381 | 0.36144447 | 0.45974922 | 0.64569624 | 0.99999872 |
| TSPAN1 | 17.2960885 | 0.48923313 | 0.68075392 | 0.7186637 | 0.47234816 | 0.99999872 |
| POMGNT1 | 15.0042124 | 0.367686847 | 0.41299516 | 0.89029336 | 0.37330838 | 0.99999872 |
| LRRC41 | 25.1422329 | 0.047851696 | 0.39973159 | 0.11970957 | 0.90471322 | 0.99999872 |
| UQCRH | 29.7015785 | 0.341434499 | 0.50587259 | 0.67494168 | 0.49971282 | 0.99999872 |
| NSUN4 | 13.9858762 | -0.41310258 | 0.40242569 | -1.0265313 | 0.3046412 | 0.99999872 |
| FAAH | 7.8121169 | -0.807555229 | 0.63065678 | -1.2804988 | 0.20036978 | 0.99999872 |
| MKNK1 | 16.8200461 | -0.31056284 | 0.48904211 | -0.6350431 | 0.52540027 | 0.99999872 |
| ATPAF1 | 18.6854472 | -0.031670526 | 0.45859134 | -0.0690605 | 0.9449415 | 0.99999872 |
| EFCAB14-AS | 4.31407159 | 2.193511477 | 1.14354596 | 1.91816644 | 0.05508991 | NA |
| EFCAB14 | 50.8823862 | 0.011957572 | 0.3579959 | 0.03340142 | 0.97335447 | 0.99999872 |
| CYP4A11 | 7.72318477 | 1.029521804 | 2.45213514 | 0.41984709 | 0.67459716 | 0.99999872 |
| CYP4A22-AS: | 6.20460507 | 0.163815711 | 0.80827321 | 0.20267369 | 0.83939009 | 0.99999872 |
| PDZK1IP1 | 138.51237 | 0.198672225 | 0.59563106 | 0.33354913 | 0.73871981 | 0.99999872 |
| CMPK1 | 35.0365983 | 0.36987955 | 0.36772563 | 1.00585741 | 0.31448416 | 0.99999872 |
| LINC01389 | 6.76538572 | -1.392963682 | 1.05137448 | -1.3248977 | 0.18520507 | 0.99999872 |
| FOXD2 | 31.5642238 | -0.266346079 | 0.40784291 | -0.6530605 | 0.5137173 | 0.99999872 |
| TRABD2B | 31.201323 | -0.955134236 | 0.7465425 | -1.2794104 | 0.20075258 | 0.99999872 |
| SPATA6 | 76.6483502 | 0.485997075 | 0.36206952 | 1.34227557 | 0.17950666 | 0.99999872 |
| AGBL4 | 47.0902108 | 0.381405991 | 0.58287525 | 0.6543527 | 0.5128846 | 0.99999872 |
| FAF1 | 124.405064 | -0.482389949 | 0.25256119 | -1.9099924 | 0.05613419 | 0.99999872 |
| RNF11 | 69.7228021 | 0.246365328 | 0.36869485 | 0.6682093 | 0.504 | 0.99999872 |
| TTC39A | 7.7614123 | -0.830739841 | 0.76378019 | -1.0876687 | 0.27674137 | 0.99999872 |
| EPS15 | 144.06287 | 0.142811033 | 0.18585822 | 0.76838695 | 0.44225733 | 0.99999872 |
| OSBPL9 | 141.206848 | 0.068437683 | 0.26525889 | 0.25800335 | 0.79640432 | 0.99999872 |
| NRDC | 68.4321465 | 0.1171355 | 0.29104939 | 0.40245919 | 0.68734612 | 0.99999872 |
| TXNDC12 | 20.1076411 | 0.26715671 | 0.33236143 | 0.80381382 | 0.42150451 | 0.99999872 |
| BTF3L4 | 30.8017797 | 0.094388595 | 0.35957603 | 0.26249969 | 0.79293623 | 0.99999872 |
| ZFYVE9 | 127.209638 | -0.38229707 | 0.31819189 | -1.2014671 | 0.22957008 | 0.99999872 |
| CC2D1B | 15.614848 | -0.396562825 | 0.44706556 | -0.8870351 | 0.37506002 | 0.99999872 |

| | | | | | | |
|---|---|---|---|---|---|---|
| PRPF38A | 15.1726366 | -0.233300838 | 0.51495169 | -0.4530538 | 0.65050998 | 0.99999872 |
| TUT4 | 95.2941005 | 0.383759595 | 0.26802831 | 1.43178754 | 0.15220463 | 0.99999872 |
| GPX7 | 4.97333929 | 0.478893388 | 0.72545954 | 0.66012418 | 0.50917414 | 0.99999872 |
| COA7 | 4.7717975 | -0.419408502 | 0.7235437 | -0.5796588 | 0.56214471 | 0.99999872 |
| ZYG11B | 52.8937591 | 0.108444269 | 0.33914291 | 0.3197598 | 0.74915042 | 0.99999872 |
| ECHDC2 | 95.4946037 | 0.397663739 | 0.332854 | 1.1947092 | 0.23220067 | 0.99999872 |
| SCP2 | 54.3120902 | 0.504939239 | 0.26060527 | 1.9375634 | 0.05267651 | 0.99999872 |
| CPT2 | 5.88710376 | 1.143372319 | 0.61713836 | 1.85270013 | 0.06392535 | 0.99999872 |
| C1orf123 | 10.3515228 | 0.222677536 | 0.52128189 | 0.42717298 | 0.66925334 | 0.99999872 |
| MAGOH | 11.6281211 | 0.409645958 | 0.52264857 | 0.78378854 | 0.4331642 | 0.99999872 |
| NDC1 | 12.6300039 | -0.347906984 | 0.44255654 | -0.78613 | 0.43179131 | 0.99999872 |
| YIPF1 | 10.2557011 | -0.596849078 | 0.47259737 | -1.2629124 | 0.20662066 | 0.99999872 |
| HSPB11 | 26.5984869 | 0.452715056 | 0.43331633 | 1.04476805 | 0.29613019 | 0.99999872 |
| LRRC42 | 10.3862069 | 0.089992535 | 0.48489129 | 0.18559322 | 0.85276377 | 0.99999872 |
| TMEM59 | 100.403567 | 0.769443686 | 0.28062763 | 2.74186721 | 0.0061091 | 0.58725857 |
| TCEANC2 | 6.16039867 | -0.304820446 | 0.60680936 | -0.5023331 | 0.61543321 | 0.99999872 |
| MRPL37 | 9.80523259 | 0.417229697 | 0.55910055 | 0.74625163 | 0.45551543 | 0.99999872 |
| SSBP3 | 121.117686 | -0.139422609 | 0.3672956 | -0.3795924 | 0.70424802 | 0.99999872 |
| ACOT11 | 24.8846061 | -0.303214513 | 0.43262823 | -0.7008662 | 0.4833865 | 0.99999872 |
| DHCR24 | 6.31386586 | -0.341215405 | 0.55838549 | -0.611075 | 0.54114994 | 0.99999872 |
| USP24 | 76.2592174 | 0.11350561 | 0.27431357 | 0.41378051 | 0.67903486 | 0.99999872 |
| PLPP3 | 36.8245411 | -1.311465385 | 0.67638647 | -1.9389291 | 0.05250998 | 0.99999872 |
| PRKAA2 | 47.1315139 | 0.184021416 | 0.3731773 | 0.49312061 | 0.62192737 | 0.99999872 |
| DAB1 | 37.893238 | -0.087375501 | 0.80863907 | -0.1080525 | 0.91395402 | 0.99999872 |
| MYSM1 | 30.449252 | 0.068313987 | 0.40850051 | 0.1672311 | 0.86718822 | 0.99999872 |
| JUN | 23.1031873 | -0.390239529 | 0.63703075 | -0.6125914 | 0.54014658 | 0.99999872 |
| LINC01135 | 7.7588782 | 0.917193798 | 0.77551707 | 1.18268679 | 0.2369333 | 0.99999872 |
| FGGY | 51.446051 | -0.393293714 | 0.49109174 | -0.8008559 | 0.42321507 | 0.99999872 |
| HOOK1 | 14.5868172 | 0.073197067 | 0.449393 | 0.16287986 | 0.87061303 | 0.99999872 |
| NFIA | 37.1323747 | -0.085032955 | 0.62963586 | -0.135051 | 0.89257155 | 0.99999872 |
| TM2D1 | 57.2606574 | 0.047858802 | 0.2995323 | 0.15977843 | 0.87305561 | 0.99999872 |
| PATJ | 192.421797 | -0.255195614 | 0.28985559 | -0.8804233 | 0.37863004 | 0.99999872 |
| USP1 | 14.980012 | -0.022142064 | 0.42166378 | -0.0525112 | 0.95812138 | 0.99999872 |
| DOCK7 | 61.699101 | 0.259840248 | 0.32755695 | 0.7932674 | 0.42762204 | 0.99999872 |
| ATG4C | 15.0500457 | 0.289723253 | 0.40321502 | 0.7185329 | 0.47242877 | 0.99999872 |
| ALG6 | 8.72867747 | 0.174990298 | 0.61798223 | 0.28316397 | 0.77705115 | 0.99999872 |
| ITGB3BP | 28.3880445 | 0.743996256 | 0.64586586 | 1.15193618 | 0.2493473 | 0.99999872 |
| EFCAB7 | 12.7685671 | -0.147411247 | 0.59068435 | -0.2495601 | 0.80292756 | 0.99999872 |
| PGM1 | 14.1200345 | 0.091193929 | 0.40792491 | 0.22355567 | 0.82310306 | 0.99999872 |
| ROR1 | 18.1307141 | -1.122687489 | 0.80244163 | -1.3990893 | 0.16178621 | 0.99999872 |
| CACHD1 | 8.29845662 | -0.312516917 | 0.61928128 | -0.5046445 | 0.61380852 | 0.99999872 |
| RAVER2 | 9.59194105 | 0.217305615 | 0.67387586 | 0.32247128 | 0.74709569 | 0.99999872 |
| JAK1 | 53.249092 | 0.059380017 | 0.29735677 | 0.19969284 | 0.84172081 | 0.99999872 |
| AK4 | 26.0389851 | 0.901870075 | 0.64259822 | 1.40347427 | 0.16047546 | 0.99999872 |
| DNAJC6 | 37.0194011 | -0.127291251 | 0.5646045 | -0.2254521 | 0.82162761 | 0.99999872 |
| AL139294.1 | 4.59784863 | 0.884367754 | 0.94967714 | 0.9312299 | 0.35173465 | 0.99999872 |
| LEPR | 119.984916 | 1.337615494 | 0.3869427 | 3.4568826 | 0.00054646 | 0.21771468 |
| PDE4B | 50.1575875 | -0.283095417 | 0.44540432 | -0.635592 | 0.5250424 | 0.99999872 |
| SGIP1 | 304.686952 | 0.067811493 | 0.25636276 | 0.26451383 | 7.91E-01 | 1.00E+00 |
| WDR78 | 29.0346428 | 0.359091453 | 0.37894004 | 0.94762078 | 0.34332254 | 0.99999872 |
| MIER1 | 61.1041704 | -0.04260913 | 0.25668688 | -0.1659965 | 0.86815968 | 0.99999872 |
| SLC35D1 | 18.469216 | 0.082238095 | 0.4161522 | 0.19761543 | 0.84334596 | 0.99999872 |
| SERBP1 | 55.2825759 | 0.141931175 | 0.50308271 | 0.28212294 | 0.77784924 | 0.99999872 |
| GADD45A | 119.80671 | 1.121183307 | 0.71757964 | 1.56245139 | 0.11818169 | 0.99999872 |
| GNG12 | 66.9874913 | -0.019608428 | 0.32397403 | -0.0605247 | 0.95173775 | 0.99999872 |
| GNG12-AS1 | 16.0367625 | -0.45283491 | 0.73395957 | -0.6169753 | 0.53725103 | 0.99999872 |
| WLS | 13.7625718 | -0.07405559 | 0.71084744 | -0.1041793 | 0.91702706 | 0.99999872 |
| LRRC40 | 17.2982427 | -0.013123909 | 0.50327202 | -0.0260772 | 0.97919579 | 0.99999872 |
| SRSF11 | 108.804636 | 0.395646712 | 0.21847808 | 1.81092179 | 0.07015296 | 0.99999872 |
| ANKRD13C | 47.9548 | 0.047620857 | 0.36937447 | 0.12892298 | 0.89741859 | 0.99999872 |
| HHLA3 | 4.52529117 | -0.29190682 | 0.69336149 | -0.4210024 | 0.67375336 | NA |

| | | | | | | | |
|---|---|---|---|---|---|---|---|
| PTGER3 | 28.9509106 | | 0.047472186 | 0.7126482 | 0.06661377 | 0.94688918 | 0.99999872 |
| ZRANB2-AS1 | 5.20047894 | | 1.24247138 | 0.80388574 | 1.54558206 | 0.12220554 | 0.99999872 |
| ZRANB2 | 103.782052 | | 0.118056704 | 0.2484895 | 0.47509736 | 0.63471758 | 0.99999872 |
| NEGR1 | 11.2308061 | | -0.111780632 | 0.99244441 | -0.1126316 | 0.91032261 | 0.99999872 |
| CRYZ | 17.8606217 | | 0.004036132 | 0.48401104 | 0.00833893 | 0.99334658 | 0.99999872 |
| TYW3 | 13.2619833 | | -0.034880573 | 0.43162044 | -0.0808131 | 0.93559061 | 0.99999872 |
| ACADM | 31.103861 | | 0.598273727 | 0.31354849 | 1.90807401 | 0.05638165 | 0.99999872 |
| RABGGTB | 11.6464637 | | 0.197866343 | 0.61431322 | 0.32209358 | 0.7473818 | 0.99999872 |
| AC092813.2 | 336.670773 | | -0.440202565 | 0.80251022 | -0.548532 | 0.58332664 | 0.99999872 |
| ST6GALNAC3 | 2498.45019 | | -0.13099517 | 0.53311798 | -0.2457152 | 0.80590273 | 0.99999872 |
| PIGK | 35.992404 | | 0.649745264 | 0.39820033 | 1.6317045 | 0.10274175 | 0.99999872 |
| AC118549.1 | 101.189275 | | -0.005360594 | 0.27377723 | -0.0195801 | 0.98437831 | 0.99999872 |
| USP33 | 68.2693792 | | 0.125989338 | 0.27640466 | 0.45581482 | 0.64852315 | 0.99999872 |
| MIGA1 | 51.4334206 | | 0.496133793 | 0.26164883 | 1.89618198 | 0.05793599 | 0.99999872 |
| NEXN | 8.22309274 | | 1.597420645 | 0.9144275 | 1.74690792 | 0.08065331 | 0.99999872 |
| FUBP1 | 49.5871516 | | -0.006759222 | 0.32652103 | -0.0207007 | 0.98348439 | 0.99999872 |
| GIPC2 | 12.8195723 | | 0.102699021 | 0.49353893 | 0.20808697 | 0.83516106 | 0.99999872 |
| IFI44L | 51.6199489 | | -4.314767031 | 2.72677607 | -1.5823694 | 0.11356526 | 0.99999872 |
| IFI44 | 23.2152533 | | -2.185311256 | 1.6495371 | -1.3248027 | 0.18523659 | 0.99999872 |
| LINC01781 | 9.6889829 | | 1.449310297 | 1.31345492 | 1.10343361 | 0.26983891 | 0.99999872 |
| ADGRL2 | 17.0760782 | | 0.29245693 | 0.79173126 | 0.36938914 | 0.71183769 | 0.99999872 |
| LINC01362 | 6.50425612 | | 1.672459952 | 0.9128921 | 1.83204559 | 0.06694463 | 0.99999872 |
| LINC01725 | 21.1363406 | | 0.559952538 | 0.88157934 | 0.63516976 | 0.5253177 | 0.99999872 |
| TTLL7 | 49.4590234 | | 1.127151725 | 0.40855465 | 2.75887624 | 0.00580005 | 0.58410373 |
| PRKACB | 70.9612506 | | 0.093710612 | 0.34843966 | 0.26894359 | 0.78797309 | 0.99999872 |
| RPF1 | 10.7362948 | | 0.23552425 | 0.45022799 | 0.52312218 | 0.60088923 | 0.99999872 |
| GNG5 | 54.5627997 | | 0.613901942 | 0.42261046 | 1.45264258 | 0.14632302 | 0.99999872 |
| SPATA1 | 9.64414947 | | 0.658518262 | 0.6827984 | 0.96444025 | 0.33482525 | 0.99999872 |
| CTBS | 20.6605463 | | -0.007851275 | 0.49944168 | -0.0157201 | 0.98745769 | 0.99999872 |
| SSX2IP | 5.26580228 | | 0.735415538 | 0.69274628 | 1.06159436 | 0.28841988 | 0.99999872 |
| WDR63 | 4.52971091 | | 1.470356729 | 1.08514246 | 1.35498958 | 0.17542088 | NA |
| MCOLN3 | 7.65573377 | | 2.162677933 | 1.03005694 | 2.09957125 | 0.03576657 | 0.94734101 |
| SYDE2 | 23.5906676 | | 0.627332185 | 0.56957922 | 1.10139585 | 0.27072441 | 0.99999872 |
| C1orf52 | 23.9496989 | | 0.214677993 | 0.34804949 | 0.61680307 | 0.53736462 | 0.99999872 |
| BCL10 | 8.10843665 | | 1.277431235 | 0.57783282 | 2.21072806 | 0.02705467 | 0.89154528 |
| AL078459.1 | 11.725642 | | -0.673362805 | 0.81278829 | -0.8284603 | 0.40740989 | 0.99999872 |
| DDAH1 | 22.753541 | | -1.062940648 | 0.47580449 | -2.2339862 | 0.02548398 | 0.88365618 |
| CYR61 | 34.1735452 | | -2.605627134 | 0.70607234 | -3.6903119 | 0.00022398 | 0.12623956 |
| ZNHIT6 | 24.533242 | | -0.282168783 | 0.35343013 | -0.7983722 | 0.42465454 | 0.99999872 |
| AL136382.1 | 6.32707021 | | -0.225640978 | 0.68631171 | -0.3287733 | 0.74232703 | 0.99999872 |
| ODF2L | 72.589042 | | 0.197212618 | 0.46032272 | 0.42842251 | 0.66834354 | 0.99999872 |
| SH3GLB1 | 76.2347249 | | 0.02600764 | 0.26994574 | 0.09634395 | 0.9232474 | 0.99999872 |
| SELENOF | 55.3571578 | | 0.331335878 | 0.25943802 | 1.27712924 | 0.20155662 | 0.99999872 |
| HS2ST1 | 28.763886 | | -0.006290013 | 0.49045444 | -0.0128249 | 0.98976752 | 0.99999872 |
| LMO4 | 21.4819248 | | -0.124164603 | 0.44229968 | -0.2807251 | 0.77892129 | 0.99999872 |
| PKN2-AS1 | 9.96812225 | | 1.204393686 | 0.88081832 | 1.36735767 | 0.17151322 | 0.99999872 |
| PKN2 | 96.0010777 | | -0.052287476 | 0.20554184 | -0.2543885 | 0.79919545 | 0.99999872 |
| GTF2B | 20.7592 | | -0.305550762 | 0.45892795 | -0.6657924 | 0.50554377 | 0.99999872 |
| KYAT3 | 29.2127421 | | 0.277430038 | 0.42909309 | 0.64654977 | 0.51792338 | 0.99999872 |
| GBP3 | 13.3057609 | | -1.248877455 | 1.01164512 | -1.2345015 | 0.21701608 | 0.99999872 |
| GBP1 | 15.2456317 | | -1.578472035 | 1.75678244 | -0.8985017 | 0.36891813 | 0.99999872 |
| GBP2 | 6.48904618 | | 0.050838018 | 0.69382876 | 0.07327171 | 0.94158991 | 0.99999872 |
| GBP4 | 8.19847909 | | 0.703996857 | 2.94812618 | 0.23879468 | 0.81126479 | 0.99999872 |
| LRRC8B | 8.69137781 | | 0.408673263 | 0.62491482 | 0.65396635 | 0.51313348 | 0.99999872 |
| ZNF326 | 23.9537154 | | -0.331481102 | 0.33366347 | -0.9934594 | 0.32048614 | 0.99999872 |
| ZNF644 | 90.8141812 | | 0.014405884 | 0.24771141 | 0.05815592 | 0.95362444 | 0.99999872 |
| HFM1 | 71.7352793 | | 1.085368575 | 0.8471839 | 1.28114873 | 0.20014143 | 0.99999872 |
| CDC7 | 10.5152089 | | -0.801898166 | 0.66507541 | -1.2057252 | 0.22792347 | 0.99999872 |
| TGFBR3 | 1002.82591 | | 0.595865698 | 0.30797592 | 1.93478017 | 0.05301728 | 0.99999872 |
| BTBD8 | 23.640145 | | 0.516073053 | 0.50183183 | 1.02837848 | 0.30377183 | 0.99999872 |
| GLMN | 22.54868 | | 0.249149403 | 0.35718759 | 0.69753097 | 0.48547056 | 0.99999872 |

| | | | | | | |
|---|---|---|---|---|---|---|
| RPAP2 | 46.338024 | -0.314808747 | 0.28298511 | -1.1124569 | 0.26594173 | 0.99999872 |
| EVI5 | 110.725965 | -0.026973381 | 0.30346056 | -0.088886 | 0.92917255 | 0.99999872 |
| RPL5 | 47.8087098 | 0.402044411 | 0.3564382 | 1.12794984 | 0.25934111 | 0.99999872 |
| FAM69A | 5.3864156 | 0.454082835 | 0.730298 | 0.62177746 | 0.53408821 | 0.99999872 |
| MTF2 | 40.5662422 | 0.228001334 | 0.35000817 | 0.65141717 | 0.51477723 | 0.99999872 |
| TMED5 | 53.6987738 | 1.127567149 | 0.32497752 | 3.46967735 | 0.00052108 | 0.21771468 |
| CCDC18 | 11.2595629 | -0.711342227 | 0.72278405 | -0.9841698 | 0.32503203 | 0.99999872 |
| DR1 | 21.5314269 | 0.685223192 | 0.37908121 | 1.80758943 | 0.07067042 | 0.99999872 |
| FNBP1L | 251.548443 | 0.369924655 | 0.24109441 | 1.53435598 | 0.1249421 | 0.99999872 |
| BCAR3 | 206.687615 | -0.038921881 | 0.41368979 | -0.0940847 | 0.92504187 | 0.99999872 |
| DNTTIP2 | 6.90367559 | -0.035127789 | 0.65881689 | -0.0533195 | 0.95747734 | 0.99999872 |
| GCLM | 31.7696581 | -0.033726043 | 0.39395699 | -0.0856084 | 0.93177769 | 0.99999872 |
| ARHGAP29 | 412.214199 | 0.166055577 | 0.50489861 | 0.32888895 | 0.74223962 | 0.99999872 |
| ABCD3 | 46.9208013 | 0.276069771 | 0.31664228 | 0.87186643 | 0.38328125 | 0.99999872 |
| F3 | 261.443993 | -1.171366309 | 0.57526025 | -2.0362372 | 0.04172655 | 0.95980711 |
| SLC44A3 | 10.00247 | -0.34457683 | 0.52627421 | -0.6547477 | 0.51263021 | 0.99999872 |
| CNN3 | 85.320097 | 0.12106735 | 0.43495859 | 0.27834224 | 0.78074965 | 0.99999872 |
| AC105942.1 | 8.1771009 | -0.175225991 | 0.52171767 | -0.3358636 | 0.7369737 | 0.99999872 |
| ALG14 | 19.8099969 | 0.448530701 | 0.39020479 | 1.14947513 | 0.25036011 | 0.99999872 |
| PTBP2 | 81.6429558 | 0.149439921 | 0.30723591 | 0.48640121 | 0.62668273 | 0.99999872 |
| DPYD | 129.271407 | -0.17822189 | 0.2749654 | -0.6481612 | 0.51688072 | 0.99999872 |
| DPYD-AS1 | 5.34808674 | 0.364338073 | 0.79133221 | 0.46041102 | 0.64522122 | 0.99999872 |
| SNX7 | 14.8628823 | 0.295700828 | 0.46677923 | 0.63349183 | 0.52641251 | 0.99999872 |
| PALMD | 9.79899413 | -0.721871427 | 0.63337601 | -1.1397202 | 0.25440289 | 0.99999872 |
| AGL | 17.9171567 | 0.43319205 | 0.55643018 | 0.77852005 | 0.43626249 | 0.99999872 |
| TRMT13 | 17.097212 | 0.008014277 | 0.5080323 | 0.01577513 | 0.98741379 | 0.99999872 |
| DBT | 21.0792571 | -0.295481977 | 0.37924836 | -0.7791253 | 0.43590593 | 0.99999872 |
| RTCA | 16.8462928 | -0.006254457 | 0.44761429 | -0.0139729 | 0.98885162 | 0.99999872 |
| CDC14A | 740.116004 | -0.082849318 | 0.40200231 | -0.2060916 | 0.83671932 | 0.99999872 |
| AL589990.1 | 7.87104852 | 1.962242653 | 1.04138301 | 1.88426605 | 0.059529 | 0.99999872 |
| VCAM1 | 18.1254361 | 1.278219668 | 0.71412757 | 1.78990383 | 0.07346937 | 0.99999872 |
| EXTL2 | 12.9247355 | 0.251687391 | 0.44092407 | 0.57081799 | 0.56812303 | 0.99999872 |
| SLC30A7 | 34.367935 | -0.138338426 | 0.32209733 | -0.4294926 | 0.66756477 | 0.99999872 |
| DPH5 | 8.98073984 | -0.26176863 | 0.49015762 | -0.5340499 | 0.59330701 | 0.99999872 |
| AC093157.1 | 8.0896503 | 0.440377094 | 0.6149644 | 0.71610177 | 0.47392851 | 0.99999872 |
| RNPC3 | 32.6817404 | -0.194419317 | 0.33849789 | -0.574359 | 0.56572488 | 0.99999872 |
| AMY2B | 7.79299685 | -0.781240199 | 0.67594366 | -1.1557771 | 0.24777235 | 0.99999872 |
| PRMT6 | 17.295663 | -0.290030802 | 0.56834208 | -0.5103103 | 0.60983411 | 0.99999872 |
| NTNG1 | 1584.55003 | -0.473633481 | 0.31439241 | -1.5065042 | 0.1319378 | 0.99999872 |
| VAV3 | 21.4632364 | 1.019608659 | 0.88317172 | 1.15448518 | 0.24830131 | 0.99999872 |
| SLC25A24 | 34.802609 | 0.102293653 | 0.35274221 | 0.2899955 | 0.77181968 | 0.99999872 |
| PRPF38B | 30.7124601 | -0.801353376 | 0.31833803 | -2.5173033 | 0.0118257 | 0.72670507 |
| STXBP3 | 48.0161434 | 0.198761478 | 0.43553437 | 0.45636233 | 0.64812945 | 0.99999872 |
| AL449266.1 | 8.70360912 | -0.157360866 | 0.52396594 | -0.3003265 | 0.76392811 | 0.99999872 |
| GPSM2 | 9.81224327 | 0.565423896 | 0.64494791 | 0.87669699 | 0.38065124 | 0.99999872 |
| WDR47 | 10.1984925 | 0.258951472 | 0.51669446 | 0.50116944 | 0.61625188 | 0.99999872 |
| TAF13 | 6.67927044 | -0.061103832 | 0.62629236 | -0.0975644 | 0.9222782 | 0.99999872 |
| TMEM167B | 17.3276424 | 0.553533966 | 0.42866024 | 1.29131167 | 0.19659563 | 0.99999872 |
| SARS | 35.1090798 | 0.532137831 | 0.46709434 | 1.1392513 | 0.25459835 | 0.99999872 |
| CELSR2 | 13.153792 | 0.110629508 | 0.71661723 | 0.1543774 | 0.87731217 | 0.99999872 |
| SORT1 | 197.079229 | 0.288763022 | 0.36669163 | 0.78748191 | 0.43099982 | 0.99999872 |
| PSMA5 | 27.9175304 | -0.386338522 | 0.51000841 | -0.757514 | 0.44874197 | 0.99999872 |
| CYB561D1 | 8.38591682 | -0.15741107 | 0.55128879 | -0.2855329 | 0.77523593 | 0.99999872 |
| AMIGO1 | 4.23040142 | 0.566669536 | 0.76805793 | 0.73779531 | 0.46063884 | NA |
| GNAI3 | 46.6256204 | 0.59906466 | 0.28949502 | 2.0693436 | 0.03851386 | 0.95980711 |
| AMPD2 | 11.4367844 | 0.524501278 | 0.55612068 | 0.9431429 | 0.34560781 | 0.99999872 |
| GSTM4 | 5.70846727 | 1.415668919 | 0.83118376 | 1.70319607 | 0.08853138 | 0.99999872 |
| GSTM3 | 9.27673317 | 0.529641674 | 0.7474309 | 0.70861624 | 0.47856265 | 0.99999872 |
| CSF1 | 5.29180149 | -0.647188447 | 0.74938106 | -0.8636306 | 0.38779082 | 0.99999872 |
| AHCYL1 | 55.9631948 | 0.478446276 | 0.25073695 | 1.90816019 | 0.05637051 | 0.99999872 |
| STRIP1 | 12.3268135 | -0.264612511 | 0.43649153 | -0.606226 | 0.5443647 | 0.99999872 |

| | | | | | | |
|---|---|---|---|---|---|---|
| KCNC4 | 12.3873407 | -0.382059239 | 0.42839067 | -0.8918477 | 0.37247457 | 0.99999872 |
| RBM15 | 10.3395006 | 0.083662049 | 0.51598102 | 0.16214172 | 0.87119425 | 0.99999872 |
| SLC16A4 | 24.796774 | 0.06268903 | 0.42009756 | 0.14922493 | 0.88137615 | 0.99999872 |
| AL355488.1 | 5.17421206 | 0.695601746 | 0.69300147 | 1.00375219 | 0.31549807 | 0.99999872 |
| LAMTOR5 | 51.2048797 | 0.670667271 | 0.35118961 | 1.90970136 | 0.05617168 | 0.99999872 |
| LRIF1 | 13.313248 | -0.5380279 | 0.51003738 | -1.0548793 | 0.29148051 | 0.99999872 |
| DRAM2 | 17.0694197 | 0.371465203 | 0.38178642 | 0.97296598 | 0.33057021 | 0.99999872 |
| CEPT1 | 19.376248 | 0.36852939 | 0.45709453 | 0.80624327 | 0.42010259 | 0.99999872 |
| WDR77 | 6.74213161 | -0.73265073 | 0.63825015 | -1.1479053 | 0.25100766 | 0.99999872 |
| ATP5PB | 28.6090757 | 0.738009453 | 0.40805565 | 1.80859999 | 0.07051317 | 0.99999872 |
| RAP1A | 64.7974991 | -0.070037469 | 0.26468891 | -0.264603 | 0.79131534 | 0.99999872 |
| DDX20 | 5.27152541 | 0.271705668 | 0.67480016 | 0.40264612 | 0.68720857 | 0.99999872 |
| KCND3 | 56.3266575 | -0.668738448 | 0.94206168 | -0.7098669 | 0.47778665 | 0.99999872 |
| AL445426.1 | 5.45679569 | -0.22184072 | 0.77187005 | -0.2874068 | 0.77380084 | 0.99999872 |
| CTTNBP2NL | 24.5471629 | 0.356331194 | 0.48535619 | 0.73416431 | 0.4628486 | 0.99999872 |
| WNT2B | 8.77342807 | -0.405853611 | 0.58784286 | -0.6904117 | 0.4899353 | 0.99999872 |
| ST7L | 30.1996529 | 0.033327282 | 0.39685968 | 0.08397749 | 0.93307433 | 0.99999872 |
| CAPZA1 | 51.2260529 | 0.072651856 | 0.44457084 | 0.1634202 | 0.8701876 | 0.99999872 |
| MOV10 | 9.81249855 | 0.201153275 | 0.94032827 | 0.21391814 | 0.83061089 | 0.99999872 |
| RHOC | 27.0371042 | 0.102767736 | 0.48583988 | 0.21152594 | 0.83247689 | 0.99999872 |
| SLC16A1-AS1 | 6.89119735 | -0.15959044 | 0.95569088 | -0.1669896 | 0.86737823 | 0.99999872 |
| LRIG2 | 25.6502357 | -0.289879216 | 0.32293191 | -0.8976481 | 0.36937319 | 0.99999872 |
| MAGI3 | 109.893429 | 0.455363625 | 0.57771052 | 0.78822111 | 0.43056738 | 0.99999872 |
| PHTF1 | 41.4871203 | -0.462568007 | 0.36098607 | -1.2814012 | 0.20005277 | 0.99999872 |
| RSBN1 | 36.4695539 | 0.457869711 | 0.45599013 | 1.00412199 | 0.31531982 | 0.99999872 |
| HIPK1 | 33.6588193 | 0.280117783 | 0.41387427 | 0.67681854 | 0.49852109 | 0.99999872 |
| TRIM33 | 73.3939897 | -0.354861704 | 0.25856052 | -1.3724512 | 0.16992301 | 0.99999872 |
| BCAS2 | 12.8313895 | 0.139552325 | 0.40997178 | 0.34039495 | 0.73355912 | 0.99999872 |
| DENND2C | 5.27641645 | 0.641605753 | 0.67971713 | 0.94393053 | 0.34520515 | 0.99999872 |
| NRAS | 13.5391427 | -0.361863409 | 0.48958559 | -0.7391218 | 0.45983301 | 0.99999872 |
| CSDE1 | 100.77669 | 0.024504951 | 0.30892206 | 0.07932406 | 0.93677487 | 0.99999872 |
| SIKE1 | 17.7366564 | -0.062759599 | 0.45710582 | -0.1372977 | 0.89079545 | 0.99999872 |
| TSPAN2 | 119.634612 | 0.519064084 | 0.67141968 | 0.77308441 | 0.43947243 | 0.99999872 |
| NGF | 27.8425211 | 0.697830605 | 0.51871144 | 1.34531562 | 0.17852333 | 0.99999872 |
| VANGL1 | 8.09048333 | -0.12175045 | 0.51576943 | -0.236056 | 0.81338924 | 0.99999872 |
| SLC22A15 | 13.1045829 | 0.626325391 | 0.56653904 | 1.1055291 | 0.26893039 | 0.99999872 |
| ATP1A1 | 167.242481 | -0.086697336 | 0.35616904 | -0.2434163 | 0.80768295 | 0.99999872 |
| LINC01762 | 19.3912131 | 1.119635212 | 1.10837875 | 1.01015579 | 0.31242066 | 0.99999872 |
| CD58 | 18.2394372 | 0.304101994 | 0.5259653 | 0.57817881 | 0.5631434 | 0.99999872 |
| IGSF3 | 5.48639799 | -0.725117556 | 0.71763616 | -1.0104251 | 0.31229169 | 0.99999872 |
| TTF2 | 6.58080385 | 0.514040219 | 0.85544414 | 0.60090448 | 0.54790361 | 0.99999872 |
| MAN1A2 | 101.262104 | -0.068151167 | 0.34452692 | -0.1978109 | 0.84319305 | 0.99999872 |
| GDAP2 | 28.8352792 | -0.087850919 | 0.3846732 | -0.2283781 | 0.81935234 | 0.99999872 |
| WDR3 | 7.29414125 | -0.476607614 | 0.51587771 | -0.9238771 | 0.3555503 | 0.99999872 |
| SPAG17 | 19.2673787 | -0.27723743 | 0.49864147 | -0.5559855 | 0.57822077 | 0.99999872 |
| WARS2 | 16.3423294 | -0.503225337 | 0.52086115 | -0.966141 | 0.33397361 | 0.99999872 |
| AL359915.2 | 45.3647389 | -0.344901675 | 0.38617789 | -0.8931161 | 0.37179502 | 0.99999872 |
| ZNF697 | 6.26329694 | 0.4476075 | 1.06263421 | 0.42122444 | 0.6735912 | 0.99999872 |
| PHGDH | 4.83928125 | 0.429508106 | 0.74017244 | 0.58028114 | 0.56172504 | 0.99999872 |
| NOTCH2 | 151.892561 | 0.250700877 | 0.23008878 | 1.08958325 | 0.27589676 | 0.99999872 |
| SEC22B | 43.6613965 | 0.058828492 | 0.30698934 | 0.19163041 | 0.84803172 | 0.99999872 |
| NBPF26 | 14.8703082 | 0.155757482 | 0.64532414 | 0.24136317 | 0.80927365 | 0.99999872 |
| LINC00623 | 46.5422193 | -0.326080762 | 0.63072875 | -0.5169905 | 0.6051628 | 0.99999872 |
| SRGAP2C | 279.062313 | 0.137763212 | 0.31591882 | 0.43607156 | 0.66278481 | 0.99999872 |
| SRGAP2-AS1 | 16.0660439 | 0.960308487 | 0.45656948 | 2.10331292 | 0.03543842 | 0.94734101 |
| AC244021.1 | 101.12455 | 0.208888624 | 0.51501057 | 0.40560065 | 0.68503605 | 0.99999872 |
| NBPF15 | 19.9623402 | -0.009717495 | 0.45083542 | -0.0215544 | 0.98280339 | 0.99999872 |
| FP700111.1 | 24.7077163 | 0.410750879 | 0.45866605 | 0.89553365 | 0.37050188 | 0.99999872 |
| SRGAP2B | 233.649801 | 0.190980949 | 0.31234352 | 0.6114452 | 0.54090488 | 0.99999872 |
| GPR89A | 8.70474878 | -0.46292684 | 0.53819268 | -0.8601508 | 0.38970595 | 0.99999872 |
| PDZK1 | 19.4151512 | 0.440222126 | 0.97914822 | 0.44959702 | 0.65300103 | 0.99999872 |

| | | | | | | |
|---|---|---|---|---|---|---|
| RNF115 | 99.2517291 | -0.259338737 | 0.22762967 | -1.1393011 | 0.25457758 | 0.99999872 |
| POLR3C | 9.19712858 | -0.40064284 | 0.54387838 | -0.7366405 | 0.461341 | 0.99999872 |
| PIAS3 | 9.22639501 | -0.701520783 | 0.45716572 | -1.5345 | 0.1249067 | 0.99999872 |
| ITGA10 | 6.98505112 | -1.330655289 | 1.14101938 | -1.1661987 | 0.24353413 | 0.99999872 |
| PEX11B | 9.38791206 | -0.443320235 | 0.55680152 | -0.7961908 | 0.42592115 | 0.99999872 |
| RBM8A | 24.5232435 | -0.028747189 | 0.44696576 | -0.0643163 | 0.94871837 | 0.99999872 |
| LIX1L | 23.8562241 | -0.064598166 | 0.38056843 | -0.1697413 | 0.86521362 | 0.99999872 |
| POLR3GL | 6.92630603 | 0.805020528 | 0.56253107 | 1.43106856 | 0.15241057 | 0.99999872 |
| TXNIP | 413.855631 | -0.238640536 | 0.53480534 | -0.4462194 | 0.65543874 | 0.99999872 |
| NBPF12 | 20.037276 | -0.280970579 | 0.46950338 | -0.5984421 | 0.54954499 | 0.99999872 |
| PRKAB2 | 9.17563045 | 0.529668985 | 0.52103912 | 1.01656279 | 0.30936146 | 0.99999872 |
| AC242426.2 | 23.6181805 | 0.385569682 | 0.45971775 | 0.83870959 | 0.4016323 | 0.99999872 |
| FMO5 | 11.2757789 | 1.431232536 | 0.86628401 | 1.65215162 | 0.09850365 | 0.99999872 |
| CHD1L | 21.1862045 | 0.013807199 | 0.37988939 | 0.03634531 | 0.97100702 | 0.99999872 |
| BCL9 | 17.232772 | 0.630284256 | 0.37432096 | 1.68380702 | 0.09221897 | 0.99999872 |
| ACP6 | 16.6626204 | -0.083980588 | 0.48020621 | -0.1748844 | 0.86117045 | 0.99999872 |
| GPR89B | 10.4369087 | -0.364213663 | 0.56606434 | -0.6434139 | 0.51995557 | 0.99999872 |
| NBPF11 | 14.8829653 | -0.309178985 | 0.57699683 | -0.5358417 | 0.59206793 | 0.99999872 |
| AC239809.3 | 8.33318568 | -1.77245648 | 0.60134026 | -2.9475101 | 0.00320344 | 0.44319076 |
| NBPF14 | 23.6186277 | 0.192366449 | 0.34347118 | 0.56006577 | 0.57543458 | 0.99999872 |
| PDE4DIP | 352.970501 | -0.007998102 | 0.38521255 | -0.0207628 | 0.98343485 | 0.99999872 |
| NBPF9 | 8.42443448 | -0.350674986 | 0.55191806 | -0.6353751 | 0.52518381 | 0.99999872 |
| AC245297.3 | 229.815261 | 0.24894774 | 0.30680658 | 0.81141591 | 0.41712686 | 0.99999872 |
| NBPF19 | 24.418476 | 1.010393998 | 0.47501248 | 2.12708936 | 0.03341266 | 0.92959829 |
| HIST2H2AC | 4.51004014 | -0.501631285 | 0.8719658 | -0.5752878 | 0.56509667 | NA |
| BOLA1 | 5.98833222 | -0.064865811 | 0.6392193 | -0.1014766 | 0.91917212 | 0.99999872 |
| SF3B4 | 11.4275652 | -0.65987744 | 0.50180272 | -1.3150137 | 0.18850531 | 0.99999872 |
| OTUD7B | 45.2126741 | -0.639231682 | 0.34576197 | -1.8487622 | 0.06449215 | 0.99999872 |
| VPS45 | 39.2501007 | -0.200129595 | 0.39479268 | -0.5069233 | 0.61220866 | 0.99999872 |
| PLEKHO1 | 29.5627778 | 0.118731056 | 0.59595201 | 0.19922922 | 0.84208344 | 0.99999872 |
| ANP32E | 15.1393367 | 0.046590002 | 0.50389314 | 0.09246008 | 0.9263325 | 0.99999872 |
| APH1A | 25.4511937 | 0.164032364 | 0.3912237 | 0.41928023 | 0.67501134 | 0.99999872 |
| C1orf54 | 5.63174048 | 0.177421187 | 0.73616885 | 0.24100611 | 0.80955038 | 0.99999872 |
| MRPS21 | 46.3185597 | 0.331428707 | 0.2882583 | 1.14976294 | 0.25024152 | 0.99999872 |
| PRPF3 | 22.0955263 | -0.530869918 | 0.43389303 | -1.2235041 | 0.22113935 | 0.99999872 |
| RPRD2 | 90.784016 | -0.155778146 | 0.30012794 | -0.5190391 | 0.60373345 | 0.99999872 |
| TARS2 | 7.55999154 | -0.774473316 | 0.51460552 | -1.5049845 | 0.13232807 | 0.99999872 |
| ECM1 | 42.2162917 | 0.598954821 | 1.11853351 | 0.53548223 | 0.59231644 | 0.99999872 |
| ADAMTSL4 | 8.45550432 | -0.577485321 | 1.15384438 | -0.500488 | 0.61673147 | 0.99999872 |
| MCL1 | 68.8892689 | -0.276947817 | 0.43195434 | -0.6411507 | 0.52142479 | 0.99999872 |
| ENSA | 76.3557839 | -0.089508434 | 0.25418385 | -0.3521405 | 0.72473288 | 0.99999872 |
| GOLPH3L | 26.0119701 | -0.101514747 | 0.35406392 | -0.2867131 | 0.77433204 | 0.99999872 |
| ARNT | 46.9307451 | 0.138049398 | 0.4432042 | 0.31148035 | 0.75543548 | 0.99999872 |
| SETDB1 | 20.9090009 | -0.183441249 | 0.34463856 | -0.5322714 | 0.59453804 | 0.99999872 |
| CERS2 | 37.6428175 | 0.34025379 | 0.34246884 | 0.99353211 | 0.32045071 | 0.99999872 |
| MINDY1 | 6.54613876 | -0.290184434 | 0.53778808 | -0.5395888 | 0.58948063 | 0.99999872 |
| PRUNE1 | 9.25613149 | 0.228798989 | 0.54835539 | 0.41724581 | 0.67649862 | 0.99999872 |
| C1orf56 | 9.01605834 | 0.841896672 | 0.70146144 | 1.20020378 | 0.23006021 | 0.99999872 |
| CDC42SE1 | 23.8197346 | -0.536859353 | 0.38122706 | -1.4082404 | 0.15905989 | 0.99999872 |
| GABPB2 | 29.1551497 | -0.528622811 | 0.31507917 | -1.677746 | 0.0933967 | 0.99999872 |
| SCNM1 | 5.37390043 | 0.490762702 | 0.81438629 | 0.6026166 | 0.54676377 | 0.99999872 |
| VPS72 | 8.91496684 | 0.178597505 | 0.488364 | 0.36570572 | 0.71458468 | 0.99999872 |
| PIP5K1A | 94.454942 | -0.105135371 | 0.34253483 | -0.3069334 | 0.75889408 | 0.99999872 |
| PSMD4 | 24.9672184 | 0.093748382 | 0.43898335 | 0.21355794 | 0.8308918 | 0.99999872 |
| ZNF687 | 3.79280109 | -0.268139743 | 0.72447641 | -0.3701152 | 0.71129664 | NA |
| PI4KB | 24.5640358 | -0.161485945 | 0.35868306 | -0.450219 | 0.65255251 | 0.99999872 |
| PSMB4 | 18.8885299 | 0.612480831 | 0.60258131 | 1.01642851 | 0.30942537 | 0.99999872 |
| POGZ | 152.718728 | -0.21815148 | 0.25683429 | -0.8493861 | 0.39566647 | 0.99999872 |
| TUFT1 | 5.05688033 | 1.050600047 | 1.00166627 | 1.04885237 | 0.29424607 | 0.99999872 |
| SNX27 | 36.0755473 | 0.03627999 | 0.36232676 | 0.10013058 | 0.92024066 | 0.99999872 |
| MRPL9 | 7.3877579 | 0.920044786 | 0.53209493 | 1.72909896 | 0.08379139 | 0.99999872 |

| | | | | | | |
|---|---|---|---|---|---|---|
| RORC | 10.4529286 | -0.182469113 | 0.65391686 | -0.2790402 | 0.78021395 | 0.99999872 |
| THEM4 | 16.3138388 | 0.095852584 | 0.4782888 | 0.20040734 | 0.84116202 | 0.99999872 |
| S100A10 | 181.798877 | -0.002927461 | 0.4738264 | -0.0061783 | 9.95E-01 | 1.00E+00 |
| S100A11 | 175.541989 | -0.033285952 | 0.48372453 | -0.0688118 | 0.94513943 | 0.99999872 |
| FLG-AS1 | 24.9536563 | 0.002497367 | 0.48946559 | 0.00510223 | 0.99592902 | 0.99999872 |
| S100A6 | 1132.44508 | -0.112858658 | 0.73469767 | -0.1536124 | 0.87791538 | 0.99999872 |
| S100A4 | 109.838061 | -0.359156277 | 1.13789524 | -0.3156321 | 0.75228176 | 0.99999872 |
| S100A2 | 7.86529726 | -1.837022307 | 0.75945268 | -2.4188766 | 0.01556852 | 0.79608149 |
| S100A16 | 72.4638741 | -0.785652562 | 0.44525196 | -1.7645123 | 0.07764577 | 0.99999872 |
| S100A13 | 50.5262761 | -0.475293331 | 0.40794118 | -1.1651026 | 0.24397747 | 0.99999872 |
| S100A1 | 5.19285587 | -0.063426426 | 0.83176604 | -0.0762551 | 0.93921612 | 0.99999872 |
| CHTOP | 24.1659145 | -0.039072243 | 0.35545203 | -0.1099227 | 0.91247069 | 0.99999872 |
| SNAPIN | 8.38916547 | 0.621078855 | 0.63045668 | 0.98512535 | 0.3245625 | 0.99999872 |
| ILF2 | 22.6919649 | 0.539967398 | 0.5065902 | 1.065886 | 0.28647518 | 0.99999872 |
| NPR1 | 78.7882736 | 0.068102801 | 0.57320496 | 0.11881056 | 0.90542544 | 0.99999872 |
| INTS3 | 41.7984309 | 0.247650428 | 0.29386037 | 0.84274865 | 0.39936903 | 0.99999872 |
| SLC27A3 | 4.96937757 | -0.223623363 | 0.88534225 | -0.2525841 | 0.80058963 | 0.99999872 |
| GATAD2B | 72.9442497 | -0.239161752 | 0.33117676 | -0.7221574 | 0.4701977 | 0.99999872 |
| DENND4B | 10.3888881 | 0.066415132 | 0.47572106 | 0.1396094 | 0.88896861 | 0.99999872 |
| CRTC2 | 9.22663483 | -0.562337481 | 0.55740673 | -1.0088459 | 0.31304856 | 0.99999872 |
| SLC39A1 | 31.7259555 | 0.143372828 | 0.3707023 | 0.38676002 | 0.69893388 | 0.99999872 |
| JTB | 28.7772718 | 0.343479932 | 0.51559981 | 0.66617544 | 0.50529897 | 0.99999872 |
| RAB13 | 15.934395 | 0.037528759 | 0.52848938 | 0.07101138 | 0.9433887 | 0.99999872 |
| RPS27 | 23.6138305 | 0.831353654 | 0.45036776 | 1.84594396 | 0.06490034 | 0.99999872 |
| TPM3 | 60.1684172 | -0.058983819 | 0.36757936 | -0.1604655 | 0.87251437 | 0.99999872 |
| C1orf43 | 40.9304102 | 0.097141997 | 0.30389999 | 0.3196512 | 0.74923276 | 0.99999872 |
| UBAP2L | 29.5331683 | -0.123454427 | 0.34942567 | -0.3533067 | 0.72385853 | 0.99999872 |
| HAX1 | 13.8580493 | 0.375537973 | 0.51078517 | 0.73521707 | 0.46220731 | 0.99999872 |
| ATP8B2 | 6.18764045 | -0.523073748 | 0.7065478 | -0.7403232 | 0.45910389 | 0.99999872 |
| IL6R | 35.9384641 | -0.511479715 | 0.59843493 | -0.8546956 | 0.39271968 | 0.99999872 |
| UBE2Q1 | 26.6111113 | 0.260427573 | 0.50210244 | 0.51867419 | 0.60398796 | 0.99999872 |
| ADAR | 52.7637831 | -0.438133299 | 0.55120188 | -0.794869 | 0.4266897 | 0.99999872 |
| PMVK | 23.4179286 | 0.268133796 | 0.39701044 | 0.67538223 | 0.49943295 | 0.99999872 |
| PBXIP1 | 23.5951388 | 0.57935732 | 0.45616642 | 1.27005691 | 0.20406436 | 0.99999872 |
| PYGO2 | 11.1101658 | -0.547309512 | 0.46839913 | -1.1684682 | 0.24261794 | 0.99999872 |
| SHC1 | 4.22280341 | -0.437117711 | 0.96334692 | -0.453749 | 0.65000949 | NA |
| CKS1B | 5.5128193 | 0.479754034 | 0.66587093 | 0.72049104 | 0.47122272 | 0.99999872 |
| FLAD1 | 5.18713436 | -0.195989371 | 0.85089415 | -0.2303334 | 0.81783268 | 0.99999872 |
| ZBTB7B | 5.62761983 | -0.023511767 | 0.65630773 | -0.0358243 | 0.97142245 | 0.99999872 |
| ADAM15 | 12.3389337 | -0.310260616 | 0.59246581 | -0.5236768 | 0.60050333 | 0.99999872 |
| EFNA1 | 16.589111 | -0.071916356 | 0.6792648 | -0.1058738 | 0.91568247 | 0.99999872 |
| SLC50A1 | 7.97866112 | -0.14680088 | 0.59131124 | -0.2482633 | 0.80393069 | 0.99999872 |
| DPM3 | 21.4913576 | 0.145402143 | 0.46459475 | 0.31296553 | 0.75430685 | 0.99999872 |
| THBS3 | 6.01312349 | -0.326904614 | 0.61833379 | -0.5286863 | 0.59702307 | 0.99999872 |
| MTX1 | 7.08554307 | -0.289999095 | 0.64651107 | -0.4485601 | 0.65374899 | 0.99999872 |
| GBA | 14.5163407 | -0.519168081 | 0.55297256 | -0.9388677 | 0.34779867 | 0.99999872 |
| FAM189B | 6.41935713 | -0.235029458 | 0.59497716 | -0.3950227 | 0.69282617 | 0.99999872 |
| SCAMP3 | 20.6747637 | 0.037775909 | 0.42036613 | 0.0898643 | 0.92839505 | 0.99999872 |
| CLK2 | 10.374843 | -0.220114134 | 0.4819318 | -0.456733 | 0.647863 | 0.99999872 |
| FDPS | 23.3489291 | 0.284290338 | 0.42868345 | 0.66317077 | 0.50722119 | 0.99999872 |
| RUSC1 | 8.51011288 | -0.340096139 | 0.58601709 | -0.5803519 | 0.56167732 | 0.99999872 |
| ASH1L | 169.137242 | -0.279666188 | 0.29786737 | -0.938895 | 0.34778465 | 0.99999872 |
| MSTO1 | 4.48632793 | -0.95329012 | 0.63204805 | -1.5082558 | 0.13148908 | NA |
| YY1AP1 | 40.8844811 | -0.221560843 | 0.2737085 | -0.8094774 | 0.41824061 | 0.99999872 |
| DAP3 | 33.4325947 | 0.054005174 | 0.36745075 | 0.14697255 | 0.88315369 | 0.99999872 |
| GON4L | 30.3880285 | -0.537783206 | 0.38054381 | -1.4131965 | 0.15759794 | 0.99999872 |
| RIT1 | 8.46551781 | 0.626285001 | 0.59646281 | 1.04999841 | 0.29371884 | 0.99999872 |
| KHDC4 | 28.0723009 | -0.560152278 | 0.34391688 | -1.6287432 | 0.10336739 | 0.99999872 |
| ARHGEF2 | 15.0014674 | 0.072394009 | 0.40834436 | 0.17728666 | 0.85928323 | 0.99999872 |
| SSR2 | 34.8437901 | 0.88990302 | 0.50999336 | 1.74493061 | 0.08099695 | 0.99999872 |
| UBQLN4 | 7.20040921 | -0.034620974 | 0.63399367 | -0.0546078 | 0.95645096 | 0.99999872 |

| | | | | | | |
|---|---|---|---|---|---|---|
| LAMTOR2 | 11.6177217 | 0.307351108 | 0.5207982 | 0.59015393 | 0.55508745 | 0.99999872 |
| LMNA | 68.5538061 | -0.260347995 | 0.38444667 | -0.6772018 | 0.4982779 | 0.99999872 |
| SLC25A44 | 7.52264655 | -0.185156474 | 0.50974716 | -0.363232 | 0.71643159 | 0.99999872 |
| SMG5 | 25.6159276 | -0.31959252 | 0.35758401 | -0.8937551 | 0.37145295 | 0.99999872 |
| GLMP | 15.2293578 | 0.148449301 | 0.47827227 | 0.31038659 | 0.75626699 | 0.99999872 |
| CCT3 | 34.5286151 | 0.130474846 | 0.42919159 | 0.3040014 | 0.76112682 | 0.99999872 |
| MEF2D | 27.5721971 | -0.194867772 | 0.61144405 | -0.3187009 | 0.74995333 | 0.99999872 |
| NAXE | 8.9260302 | 0.493750574 | 0.61491305 | 0.80295999 | 0.42199786 | 0.99999872 |
| GPATCH4 | 6.31857654 | -0.251748927 | 0.75682173 | -0.3326397 | 0.7394063 | 0.99999872 |
| NES | 267.046714 | -0.432846075 | 0.70607456 | -0.6130317 | 0.53985539 | 0.99999872 |
| ISG20L2 | 8.80572501 | -0.723345144 | 0.57222988 | -1.2640814 | 0.20620081 | 0.99999872 |
| RRNAD1 | 7.20787905 | 1.10623713 | 0.65258534 | 1.69516086 | 0.09004491 | 0.99999872 |
| MRPL24 | 7.32589221 | 0.421852806 | 0.68534498 | 0.61553352 | 0.53820244 | 0.99999872 |
| HDGF | 33.1583782 | -0.577722351 | 0.36848709 | -1.5678225 | 0.11692256 | 0.99999872 |
| PRCC | 22.7142566 | -0.473330352 | 0.34642094 | -1.3663445 | 0.17183086 | 0.99999872 |
| ARHGEF11 | 37.4781227 | -0.411334502 | 0.29832957 | -1.3787923 | 0.16795881 | 0.99999872 |
| ETV3 | 4.46151413 | -0.278956471 | 0.70646892 | -0.3948602 | 0.69294605 | NA |
| KIRREL1 | 578.826662 | -0.014537942 | 0.39569831 | -0.03674 | 0.97069234 | 0.99999872 |
| IFI16 | 58.9014235 | 0.209499367 | 1.42862197 | 0.14664437 | 0.88341273 | 0.99999872 |
| DUSP23 | 44.4024775 | -0.224303033 | 0.51166522 | -0.4383785 | 0.66111193 | 0.99999872 |
| SLAMF8 | 21.8155206 | -0.386765573 | 0.7398833 | -0.5227386 | 0.60115616 | 0.99999872 |
| AL590560.2 | 6.36671658 | -0.047004167 | 0.63475614 | -0.0740507 | 0.94097 | 0.99999872 |
| CFAP45 | 9.49803429 | 0.298621517 | 0.87873881 | 0.33982967 | 0.73398481 | 0.99999872 |
| TAGLN2 | 59.9267045 | -0.647936446 | 0.55291605 | -1.1718532 | 0.241256 | 0.99999872 |
| IGSF8 | 18.0190587 | 0.417039993 | 0.47553452 | 0.87699205 | 0.38049096 | 0.99999872 |
| PEA15 | 104.316209 | -0.193392887 | 0.55387421 | -0.3491639 | 0.72696626 | 0.99999872 |
| DCAF8 | 28.030959 | -0.196682745 | 0.32555492 | -0.6041461 | 0.54574649 | 0.99999872 |
| AL139011.2 | 6.14628026 | 0.180133274 | 0.74010877 | 0.24338757 | 0.80770517 | 0.99999872 |
| COPA | 56.5235815 | 0.313219135 | 0.28419993 | 1.10210842 | 0.27041454 | 0.99999872 |
| NCSTN | 51.0152527 | -0.004274449 | 0.58149012 | -0.0073509 | 0.99413492 | 0.99999872 |
| F11R | 11.9652626 | -0.120250187 | 0.60587793 | -0.1984726 | 0.8426753 | 0.99999872 |
| AL591806.3 | 9.53676354 | -0.190876289 | 0.5606389 | -0.3404621 | 0.73350857 | 0.99999872 |
| USF1 | 5.97201906 | -1.113865578 | 0.72008508 | -1.5468527 | 0.12189876 | 0.99999872 |
| KLHDC9 | 4.58821104 | 0.901329312 | 0.76972559 | 1.17097485 | 0.24160889 | NA |
| PFDN2 | 26.79264 | 0.32462847 | 0.42525174 | 0.76337952 | 0.44523709 | 0.99999872 |
| NIT1 | 19.6045047 | 0.428498685 | 0.37935284 | 1.12955182 | 0.25866512 | 0.99999872 |
| DEDD | 9.06814936 | -0.482887405 | 0.52159658 | -0.9257871 | 0.35455661 | 0.99999872 |
| UFC1 | 30.5097128 | 0.879152339 | 0.37290504 | 2.35757695 | 0.01839464 | 0.85282771 |
| PPOX | 5.8929274 | -0.55465063 | 0.87820826 | -0.6315707 | 0.52766742 | 0.99999872 |
| B4GALT3 | 12.4264787 | 0.688474896 | 0.49345071 | 1.39522526 | 0.16294792 | 0.99999872 |
| ADAMTS4 | 6.10476129 | 0.388828314 | 1.52915791 | 0.2542761 | 0.79928226 | 0.99999872 |
| NDUFS2 | 21.8902886 | 0.984350017 | 0.4936909 | 1.99385893 | 0.04616747 | 0.95980711 |
| TOMM40L | 5.58488383 | 0.14982331 | 0.65994235 | 0.22702485 | 0.82040442 | 0.99999872 |
| SDHC | 38.1578859 | 0.520539359 | 0.43760566 | 1.18951697 | 0.2342363 | 0.99999872 |
| AL592295.4 | 5.27855486 | -0.294101932 | 0.56296623 | -0.5224149 | 0.6013815 | 0.99999872 |
| DUSP12 | 7.45930933 | 0.711714118 | 0.57123953 | 1.24591188 | 0.21279675 | 0.99999872 |
| ATF6 | 75.2722595 | 0.156633258 | 0.31063518 | 0.50423542 | 0.61409595 | 0.99999872 |
| NOS1AP | 169.085755 | -0.484592083 | 0.43309239 | -1.1189116 | 0.26317787 | 0.99999872 |
| C1orf226 | 10.341644 | -0.621658624 | 0.42931697 | -1.4480178 | 0.14761206 | 0.99999872 |
| UHMK1 | 51.0505285 | -0.138718177 | 0.36452577 | -0.3805442 | 0.70354147 | 0.99999872 |
| UAP1 | 10.4366355 | -0.340250237 | 0.60745199 | -0.560127 | 0.57539284 | 0.99999872 |
| HSD17B7 | 8.30294417 | 0.529532061 | 0.56132469 | 0.94336143 | 0.34549606 | 0.99999872 |
| PBX1 | 74.022809 | -0.25654733 | 0.37540577 | -0.6833867 | 0.49436249 | 0.99999872 |
| MGST3 | 69.7889092 | 0.626201848 | 0.33690694 | 1.85867896 | 0.06307266 | 0.99999872 |
| ALDH9A1 | 23.6182004 | 0.841953618 | 0.40116333 | 2.09878012 | 0.03583629 | 0.94734101 |
| TMCO1 | 47.9266687 | 0.294620695 | 0.29078425 | 1.01319345 | 0.31096777 | 0.99999872 |
| UCK2 | 15.9535492 | -0.213055619 | 0.53755806 | -0.3963397 | 0.69185442 | 0.99999872 |
| POGK | 17.0622745 | -0.252436218 | 0.3892735 | -0.6484804 | 0.51667431 | 0.99999872 |
| TADA1 | 6.28675358 | 0.431863384 | 0.53493308 | 0.80732226 | 0.41948084 | 0.99999872 |
| ILDR2 | 57.4352738 | 0.142744908 | 0.56573999 | 0.25231539 | 0.80079729 | 0.99999872 |
| MAEL | 8.71677813 | -0.368546977 | 0.90520118 | -0.4071437 | 0.68390243 | 0.99999872 |

| | | | | | | |
|---|---|---|---|---|---|---|
| POU2F1 | 68.9464951 | -0.079252329 | 0.27214771 | -0.2912107 | 0.77089017 | 0.99999872 |
| CREG1 | 15.9302871 | 0.946153548 | 0.48409649 | 1.95447306 | 0.0506453 | 0.98819634 |
| MPZL1 | 36.4234293 | -0.360566568 | 0.38642762 | -0.9330766 | 0.35078042 | 0.99999872 |
| MPC2 | 23.1823259 | 1.048654637 | 0.38027879 | 2.75759432 | 0.00582284 | 0.58410373 |
| DCAF6 | 107.344551 | 0.083979936 | 0.25542994 | 0.32877874 | 0.74232293 | 0.99999872 |
| TIPRL | 17.1094217 | 0.384992251 | 0.41629686 | 0.9248022 | 0.3550688 | 0.99999872 |
| SFT2D2 | 23.4209205 | 0.027345461 | 0.3440154 | 0.07948906 | 0.93664363 | 0.99999872 |
| ATP1B1 | 230.00172 | -0.465325819 | 0.63299325 | -0.7351197 | 0.46226659 | 0.99999872 |
| NME7 | 58.6011196 | 0.147142104 | 0.30924084 | 0.47581718 | 0.63420463 | 0.99999872 |
| BLZF1 | 22.8394963 | 0.147009046 | 0.46566613 | 0.31569624 | 0.75223308 | 0.99999872 |
| SLC19A2 | 7.37282041 | 0.087335348 | 0.58195132 | 0.15007329 | 0.88070679 | 0.99999872 |
| F5 | 66.8173247 | 0.2550792 | 0.56572512 | 0.45088894 | 0.6520696 | 0.99999872 |
| C1orf112 | 5.2189432 | -0.680008869 | 0.63771199 | -1.066326 | 0.2862763 | 0.99999872 |
| SCYL3 | 11.4713554 | 0.021316129 | 0.42564907 | 0.05007912 | 0.96005934 | 0.99999872 |
| KIFAP3 | 44.1518899 | 0.283358561 | 0.26438049 | 1.07178317 | 0.28381744 | 0.99999872 |
| FMO4 | 12.4037638 | -0.426368489 | 0.69245813 | -0.6157318 | 0.53807155 | 0.99999872 |
| PRRC2C | 122.475978 | -0.218469817 | 0.28885091 | -0.7563411 | 0.4494447 | 0.99999872 |
| VAMP4 | 15.8579865 | 0.191118453 | 0.47210289 | 0.40482373 | 0.68560708 | 0.99999872 |
| METTL13 | 5.22816228 | -0.197698517 | 0.67604072 | -0.2924358 | 0.76995343 | 0.99999872 |
| DNM3 | 31.0345472 | 0.496016115 | 0.59814708 | 0.82925442 | 0.40696045 | 0.99999872 |
| PIGC | 26.0049672 | 0.395741123 | 0.3623965 | 1.09201144 | 0.27482807 | 0.99999872 |
| SUCO | 56.7838897 | 0.254954305 | 0.35338967 | 0.72145375 | 0.47063039 | 0.99999872 |
| AL645568.1 | 15.3363371 | -0.134787678 | 0.6443315 | -0.20919 | 0.83429995 | 0.99999872 |
| PRDX6 | 115.425267 | 0.414115794 | 0.39123466 | 1.05848443 | 0.28983464 | 0.99999872 |
| KLHL20 | 23.2063509 | -0.320474871 | 0.34049829 | -0.9411938 | 0.34660556 | 0.99999872 |
| DARS2 | 7.08934149 | -0.325389904 | 0.70267959 | -0.4630701 | 0.64331413 | 0.99999872 |
| ZBTB37 | 17.7093864 | -0.294218091 | 0.37704943 | -0.780317 | 0.43520429 | 0.99999872 |
| RC3H1 | 72.6124611 | 0.116412077 | 0.30418931 | 0.38269615 | 0.70194507 | 0.99999872 |
| RABGAP1L | 680.187164 | 0.221455745 | 0.44397761 | 0.49879935 | 0.61792074 | 0.99999872 |
| CACYBP | 10.38516 | 0.459899171 | 0.71001285 | 0.64773359 | 0.51715727 | 0.99999872 |
| MRPS14 | 13.2664342 | 0.356123451 | 0.42520623 | 0.83753112 | 0.40229409 | 0.99999872 |
| KIAA0040 | 85.9532955 | -0.465652106 | 0.67018757 | -0.6948086 | 0.48717524 | 0.99999872 |
| COP1 | 113.311524 | -0.062459011 | 0.23699722 | -0.2635432 | 0.79213191 | 0.99999872 |
| RASAL2 | 119.739164 | -0.02901841 | 0.34492531 | -0.0841295 | 0.93295343 | 0.99999872 |
| RALGPS2 | 31.4581802 | 0.331947552 | 0.57978633 | 0.57253429 | 0.56696007 | 0.99999872 |
| ANGPTL1 | 9.54576408 | 0.133618088 | 0.93386625 | 0.14308054 | 0.88622657 | 0.99999872 |
| FAM20B | 62.6654127 | 0.056287915 | 0.47818927 | 0.11771053 | 0.90629702 | 0.99999872 |
| TOR3A | 12.6486099 | 0.091315779 | 0.57822045 | 0.15792554 | 0.87451547 | 0.99999872 |
| ABL2 | 54.4007473 | 0.263873068 | 0.3535434 | 0.74636683 | 0.45544585 | 0.99999872 |
| SOAT1 | 16.4211497 | -0.134168462 | 0.52178796 | -0.2571322 | 0.79707675 | 0.99999872 |
| AXDND1 | 34.7410553 | 1.791596967 | 0.56326716 | 3.18072327 | 0.00146908 | 0.32751082 |
| AL160286.3 | 11.6891068 | 1.26631398 | 1.02774001 | 1.23213455 | 0.21789883 | 0.99999872 |
| NPHS2 | 1793.17469 | 0.536765207 | 0.44016648 | 1.21945953 | 0.22266983 | 0.99999872 |
| TDRD5 | 29.0759937 | -0.4744352 | 0.59703508 | -0.7946521 | 0.42681588 | 0.99999872 |
| TOR1AIP2 | 59.5136265 | 0.43499649 | 0.25618971 | 1.69794676 | 0.08951781 | 0.99999872 |
| TOR1AIP1 | 32.4343367 | -0.269954844 | 0.3350372 | -0.8057459 | 0.42038938 | 0.99999872 |
| CEP350 | 71.3860278 | 0.176012067 | 0.26694218 | 0.65936402 | 0.50966204 | 0.99999872 |
| QSOX1 | 33.0931525 | 0.402851019 | 0.52335175 | 0.76975193 | 0.44144706 | 0.99999872 |
| ACBD6 | 52.8378346 | -0.088303357 | 0.34933982 | -0.2527721 | 0.80044434 | 0.99999872 |
| XPR1 | 51.7660315 | -0.127381227 | 0.35090889 | -0.3630037 | 0.71660212 | 0.99999872 |
| STX6 | 12.2321517 | 0.149640043 | 0.43594563 | 0.34325392 | 0.73140745 | 0.99999872 |
| MR1 | 14.9232734 | 0.06185796 | 0.49894823 | 0.12397671 | 0.90133372 | 0.99999872 |
| IER5 | 4.92858781 | 0.852187396 | 1.2977141 | 0.65668347 | 0.51138447 | 0.99999872 |
| GLUL | 105.4777 | 0.712709893 | 0.50211202 | 1.41942407 | 0.15577542 | 0.99999872 |
| RNASEL | 8.11192851 | -0.212778377 | 0.54938407 | -0.3873035 | 0.69853153 | 0.99999872 |
| DHX9 | 45.4378873 | 0.026914138 | 0.26465518 | 0.10169511 | 0.91899868 | 0.99999872 |
| LAMC1 | 39.5670976 | -0.025883136 | 0.42197621 | -0.0613379 | 0.95109011 | 0.99999872 |
| SMG7 | 49.8946896 | -0.228845141 | 0.28554847 | -0.8014231 | 0.42288674 | 0.99999872 |
| ARPC5 | 73.781554 | 0.190711809 | 0.44381391 | 0.42971121 | 0.66740573 | 0.99999872 |
| RGL1 | 60.3474846 | -0.292894348 | 0.59941023 | -0.4886376 | 0.62509832 | 0.99999872 |
| COLGALT2 | 28.8202083 | 0.096714483 | 0.53998755 | 0.17910502 | 0.85785524 | 0.99999872 |

| | | | | | | | |
|---|---|---|---|---|---|---|---|
| TSEN15 | 52.0739822 | | 0.430605845 | 0.38875171 | 1.10766289 | 0.26800743 | 0.99999872 |
| C1orf21 | 204.938925 | | -0.670677817 | 0.65766922 | -1.0197798 | 0.30783289 | 0.99999872 |
| EDEM3 | 62.6850706 | | 0.365203936 | 0.30934119 | 1.1805862 | 0.23776715 | 0.99999872 |
| RNF2 | 16.8249938 | | 0.672946869 | 0.51623807 | 1.30355916 | 0.19238393 | 0.99999872 |
| TRMT1L | 19.7712727 | | 0.399724059 | 0.43419703 | 0.92060523 | 0.35725657 | 0.99999872 |
| SWT1 | 17.7344583 | | -0.322727353 | 0.51358082 | -0.6283867 | 0.52975065 | 0.99999872 |
| IVNS1ABP | 38.4192087 | | -1.045824272 | 0.42313134 | -2.4716304 | 0.01344985 | 0.74507289 |
| TPR | 42.3354407 | | -0.20613822 | 0.27438375 | -0.7512771 | 0.45248592 | 0.99999872 |
| ODR4 | 25.582664 | | 0.144734232 | 0.36792492 | 0.3933798 | 0.69403899 | 0.99999872 |
| AL596220.1 | 5.76327842 | | 0.860384671 | 0.62462884 | 1.37743348 | 0.16837827 | 0.99999872 |
| ERVMER61-1 | 30.2913104 | | -0.902749372 | 0.46949132 | -1.9228244 | 0.05450211 | 0.99999872 |
| AL390957.1 | 7.75398779 | | -0.249061852 | 0.76571319 | -0.3252678 | 0.74497839 | 0.99999872 |
| AL136454.1 | 24.5369946 | | -0.155480526 | 0.59721497 | -0.2603426 | 0.79459948 | 0.99999872 |
| UCHL5 | 19.4749739 | | 0.031662297 | 0.38003599 | 0.08331394 | 0.93360192 | 0.99999872 |
| TROVE2 | 76.5815378 | | 0.521801604 | 0.37133724 | 1.40519601 | 0.159963 | 0.99999872 |
| GLRX2 | 10.8211022 | | 0.235790915 | 0.50429015 | 0.46756994 | 0.64009216 | 0.99999872 |
| CDC73 | 70.4549569 | | -0.149811966 | 0.28971169 | -0.5171071 | 0.60508141 | 0.99999872 |
| B3GALT2 | 16.3180084 | | -0.423690757 | 0.76734323 | -0.5521529 | 0.58084363 | 0.99999872 |
| LINC01031 | 19.7711403 | | -0.454510709 | 0.6372691 | -0.7132163 | 0.47571192 | 0.99999872 |
| AL357793.1 | 11.0200464 | | 0.605530882 | 0.60395563 | 1.00260822 | 0.31604993 | 0.99999872 |
| AL356272.1 | 6.61413263 | | 0.822480699 | 0.6487312 | 1.26782972 | 0.20485876 | 0.99999872 |
| KCNT2 | 150.738673 | | 0.361223766 | 0.72515647 | 0.49813217 | 6.18E-01 | 0.99999872 |
| CFH | 68.3230404 | | 1.025219808 | 0.6189875 | 1.65628515 | 0.09766409 | 0.99999872 |
| ZBTB41 | 23.5404092 | | 0.098331921 | 0.4125197 | 0.23836903 | 0.81159488 | 0.99999872 |
| CRB1 | 7.58867926 | | -0.116795275 | 0.89655766 | -0.1302708 | 0.89635219 | 0.99999872 |
| DENND1B | 57.3362204 | | -0.03133903 | 0.27946358 | -0.1121399 | 0.91071245 | 0.99999872 |
| NEK7 | 90.4895756 | | 0.203825774 | 0.23281657 | 0.87547797 | 0.38131389 | 0.99999872 |
| MIR181A1HG | 8.29232939 | | 0.674799281 | 1.02385047 | 0.65907992 | 0.50984445 | 0.99999872 |
| LINC00862 | 6.87084582 | | 0.666910346 | 0.86387536 | 0.77199834 | 0.44011541 | 0.99999872 |
| ZNF281 | 9.60664696 | | 0.430980349 | 0.49020911 | 0.87917654 | 0.37930556 | 0.99999872 |
| DDX59 | 10.6523352 | | -0.398959151 | 0.46174555 | -0.8640238 | 0.38757481 | 0.99999872 |
| CAMSAP2 | 135.483303 | | 0.2602094 | 0.19523453 | 1.33280422 | 0.18259606 | 0.99999872 |
| TMEM9 | 18.4163195 | | 0.300700492 | 0.49857853 | 0.60311561 | 0.54643178 | 0.99999872 |
| TNNT2 | 133.90212 | | 0.490319812 | 0.76465311 | 0.6412317 | 0.52137216 | 0.99999872 |
| TNNI1 | 58.4086274 | | 0.438296499 | 0.94433292 | 0.46413345 | 0.64255214 | 0.99999872 |
| PHLDA3 | 11.2028237 | | 0.501058533 | 0.64548661 | 0.77624931 | 0.4376018 | 0.99999872 |
| CSRP1 | 108.265857 | | -0.10002446 | 0.52110933 | -0.1919452 | 0.84778509 | 0.99999872 |
| AC096677.1 | 7.9561329 | | 0.280342525 | 0.59574502 | 0.47057469 | 0.63794449 | 0.99999872 |
| NAV1 | 15.1112977 | | -0.975458057 | 0.92669345 | -1.0526222 | 0.2925142 | 0.99999872 |
| IPO9-AS1 | 5.0829248 | | -0.413500326 | 1.21155783 | -0.3412964 | 0.73288046 | 0.99999872 |
| IPO9 | 29.285479 | | 0.084317745 | 0.30054891 | 0.28054584 | 0.77905876 | 0.99999872 |
| SHISA4 | 7.6398556 | | 0.189522466 | 0.69406077 | 0.27306322 | 0.78480462 | 0.99999872 |
| TIMM17A | 13.792263 | | -0.199209348 | 0.5598392 | -0.3558332 | 0.72196552 | 0.99999872 |
| RNPEP | 15.9832748 | | -0.02953592 | 0.39090368 | -0.0755581 | 0.93977071 | 0.99999872 |
| ELF3 | 12.8497662 | | 0.306293374 | 0.68159285 | 0.4493788 | 0.65315842 | 0.99999872 |
| ARL8A | 25.123684 | | -0.295322073 | 0.38084937 | -0.7754301 | 0.43808554 | 0.99999872 |
| PTPN7 | 33.4410623 | | -0.405672997 | 0.36300759 | -1.1175331 | 0.26376646 | 0.99999872 |
| PPP1R12B | 66.9795045 | | 0.183477272 | 0.48560225 | 0.37783448 | 0.70555356 | 0.99999872 |
| KDM5B | 49.5822865 | | 0.644644939 | 0.2903169 | 2.22048713 | 0.02638572 | 0.88440681 |
| RABIF | 6.3042286 | | -0.407571114 | 0.68895825 | -0.5915759 | 0.5541346 | 0.99999872 |
| KLHL12 | 14.0563146 | | -0.445963999 | 0.46230797 | -0.964647 | 0.33472165 | 0.99999872 |
| ADIPOR1 | 15.5281074 | | 0.11844043 | 0.42063077 | 0.28157814 | 0.778267 | 0.99999872 |
| CYB5R1 | 15.8569553 | | 0.249614275 | 0.48411891 | 0.51560529 | 0.60613011 | 0.99999872 |
| TMEM183A | 7.31313906 | | 0.844424996 | 0.64205133 | 1.31519857 | 0.18844318 | 0.99999872 |
| PPFIA4 | 60.9968032 | | -0.378362482 | 0.64020173 | -0.5910051 | 0.55451701 | 0.99999872 |
| ADORA1 | 76.307887 | | -0.037710925 | 0.68251108 | -0.0552532 | 0.95593674 | 0.99999872 |
| CHI3L1 | 356.758073 | | 0.881732495 | 0.60307673 | 1.46205689 | 0.14372562 | 0.99999872 |
| BTG2 | 48.025666 | | 0.724883918 | 0.79736033 | 0.90910457 | 0.36329493 | 0.99999872 |
| ATP2B4 | 41.0261954 | | 0.407361237 | 0.53043292 | 0.7679788 | 0.44249978 | 0.99999872 |
| SNRPE | 8.73282406 | | 1.274865511 | 0.59864655 | 2.12957965 | 0.03320633 | 0.92959829 |
| SOX13 | 16.1496922 | | -0.093212742 | 0.49809982 | -0.1871367 | 0.85155348 | 0.99999872 |

| | | | | | | |
|---|---|---|---|---|---|---|
| PLEKHA6 | 18.6741153 | 0.184251469 | 0.41293064 | 0.4462044 | 0.65544961 | 0.99999872 |
| PPP1R15B | 13.277756 | 0.316693038 | 0.39965083 | 0.79242433 | 0.4281133 | 0.99999872 |
| MDM4 | 47.4444171 | -0.168095741 | 0.28866375 | -0.5823237 | 0.56034867 | 0.99999872 |
| LRRN2 | 12.6218822 | -0.726231776 | 0.65157371 | -1.1145812 | 0.26502995 | 0.99999872 |
| NFASC | 627.851936 | 0.728497227 | 0.50666047 | 1.43784107 | 0.15047915 | 0.99999872 |
| RBBP5 | 13.7550622 | -0.690433183 | 0.4695813 | -1.4703166 | 0.14147602 | 0.99999872 |
| DSTYK | 54.5154925 | -0.189214398 | 0.3689097 | -0.5129017 | 0.60802011 | 0.99999872 |
| NUAK2 | 14.0573974 | -0.041268611 | 0.75482076 | -0.0546734 | 0.95639867 | 0.99999872 |
| CDK18 | 7.48174313 | -0.655649794 | 0.66029367 | -0.992967 | 0.32072606 | 0.99999872 |
| ELK4 | 25.3298598 | -0.085478823 | 0.39826081 | -0.2146303 | 0.83005559 | 0.99999872 |
| NUCKS1 | 139.167057 | -0.02094712 | 0.3068444 | -0.0682663 | 0.94557368 | 0.99999872 |
| RAB29 | 7.64758212 | 0.207075258 | 0.95754286 | 0.21625691 | 0.8287875 | 0.99999872 |
| SLC41A1 | 9.87758959 | 0.729355867 | 0.5229965 | 1.39457121 | 0.16314518 | 0.99999872 |
| RHEX | 31.8728281 | 0.560633217 | 1.06369067 | 0.52706415 | 0.59814904 | 0.99999872 |
| SRGAP2 | 372.392626 | -0.047085431 | 0.30800244 | -0.1528736 | 0.87849798 | 0.99999872 |
| RASSF5 | 8.54511806 | -0.643761759 | 0.69963664 | -0.9201373 | 0.35750102 | 0.99999872 |
| EIF2D | 8.89905044 | 0.408124642 | 0.53880708 | 0.75745968 | 0.44877451 | 0.99999872 |
| DYRK3 | 8.26724928 | 0.974693838 | 0.58730096 | 1.65961559 | 0.09699181 | 0.99999872 |
| MAPKAPK2 | 34.9085421 | -0.675133627 | 0.38523547 | -1.752522 | 0.07968409 | 0.99999872 |
| PIGR | 18.4896086 | 2.082620246 | 1.71800322 | 1.21223303 | 0.22542319 | 0.99999872 |
| PFKFB2 | 8.74302032 | -0.358711483 | 0.77493003 | -0.4628953 | 0.6434394 | 0.99999872 |
| CD55 | 46.5053025 | 0.118424025 | 0.52539396 | 0.22540043 | 0.82166778 | 0.99999872 |
| CR1 | 235.815543 | 0.56611795 | 0.56702828 | 0.99839456 | 0.31808807 | 0.99999872 |
| CR1L | 5.8940625 | -0.696198046 | 0.59674913 | -1.1666511 | 0.24335129 | 0.99999872 |
| CD46 | 139.39855 | 0.434591381 | 0.36752381 | 1.18248496 | 0.23701333 | 0.99999872 |
| MIR29B2CHC | 83.3911457 | 0.136490349 | 0.42918326 | 0.31802347 | 0.75046713 | 0.99999872 |
| PLXNA2 | 32.5517492 | -2.589405531 | 0.41476833 | -6.2430165 | 4.29E-10 | 4.50E-06 |
| AL031316.1 | 10.3273697 | -0.1406681 | 0.54565858 | -0.2577951 | 0.79656506 | 0.99999872 |
| G0S2 | 344.399327 | 1.031692928 | 0.94009649 | 1.09743301 | 0.27245215 | 0.99999872 |
| HSD11B1 | 5.55479525 | -0.487347969 | 0.92438906 | -0.5272109 | 0.59804715 | 0.99999872 |
| UTP25 | 4.17257153 | 0.439846801 | 0.89552967 | 0.49115827 | 0.62331451 | NA |
| HHAT | 8.70610767 | 0.583323584 | 0.94781271 | 0.61544182 | 0.53826298 | 0.99999872 |
| RCOR3 | 64.6395788 | -0.182311896 | 0.33849168 | -0.5386008 | 0.59016233 | 0.99999872 |
| TRAF5 | 39.3076865 | 0.552267622 | 0.44407058 | 1.2436483 | 0.21362903 | 0.99999872 |
| LINC00467 | 11.0254925 | -0.851304835 | 0.50324826 | -1.69162 | 0.09071844 | 0.99999872 |
| SLC30A1 | 20.887884 | 0.023648855 | 0.54789185 | 0.04316336 | 0.96557131 | 0.99999872 |
| LPGAT1 | 49.7928742 | -0.108197844 | 0.28850976 | -0.3750232 | 0.70764324 | 0.99999872 |
| INTS7 | 13.3894557 | 0.149445106 | 0.4490828 | 0.33277851 | 0.73930148 | 0.99999872 |
| PPP2R5A | 62.4306585 | -0.18125541 | 0.34080306 | -0.5318479 | 0.59483131 | 0.99999872 |
| AL360091.3 | 9.02604695 | 0.724767258 | 1.07703764 | 0.67292658 | 0.500994 | 0.99999872 |
| TMEM206 | 5.75459644 | 0.636289266 | 0.74114756 | 0.85851901 | 0.39060594 | 0.99999872 |
| NENF | 19.9530947 | 0.153060937 | 0.41537385 | 0.36848958 | 0.71250821 | 0.99999872 |
| ATF3 | 4.78611235 | 0.933955888 | 1.10400835 | 0.84596813 | 0.39757052 | 0.99999872 |
| NSL1 | 27.3773982 | -0.18563435 | 0.35218135 | -0.5270987 | 0.59812501 | 0.99999872 |
| TATDN3 | 8.16215227 | -0.578598654 | 0.68259544 | -0.8476451 | 0.39663567 | 0.99999872 |
| FLVCR1 | 14.5312661 | 0.020882051 | 0.3787937 | 0.05512777 | 0.95603667 | 0.99999872 |
| ANGEL2 | 23.7676849 | 0.33934941 | 0.38582512 | 0.87954203 | 0.37910745 | 0.99999872 |
| RPS6KC1 | 75.85705 | 0.211618992 | 0.37351575 | 0.56655976 | 0.57101332 | 0.99999872 |
| PROX1 | 4.56899895 | -0.018520129 | 0.83800204 | -0.0221003 | 0.98236792 | NA |
| SMYD2 | 21.3324926 | -0.804706369 | 0.46363752 | -1.7356369 | 0.08262806 | 0.99999872 |
| PTPN14 | 385.289457 | -0.205935672 | 0.24060953 | -0.8558916 | 0.39205776 | 0.99999872 |
| KCTD3 | 42.115296 | 0.202858395 | 0.3267379 | 0.6208597 | 0.53469194 | 0.99999872 |
| ESRRG | 179.673965 | -0.53149481 | 0.66909723 | -0.7943462 | 0.42699394 | 0.99999872 |
| GPATCH2 | 60.9183069 | -0.064025524 | 0.29240048 | -0.2189652 | 0.82667717 | 0.99999872 |
| SPATA17 | 10.531001 | -0.488940706 | 0.58251075 | -0.8393677 | 0.401263 | 0.99999872 |
| RRP15 | 15.278035 | 0.421825132 | 0.55270918 | 0.76319546 | 0.44534684 | 0.99999872 |
| TGFB2 | 8.55477524 | 0.007232869 | 1.00265876 | 0.00721369 | 0.99424436 | 0.99999872 |
| LYPLAL1 | 56.6295641 | -0.154129029 | 0.45981428 | -0.3351984 | 0.7374754 | 0.99999872 |
| EPRS | 31.7739253 | 0.065400257 | 0.32523546 | 0.20108587 | 0.84063143 | 0.99999872 |
| BPNT1 | 6.36437816 | 1.440053202 | 0.77517113 | 1.85772296 | 0.06320837 | 0.99999872 |
| IARS2 | 32.9176508 | 0.076629808 | 0.32435539 | 0.23625261 | 0.81323666 | 0.99999872 |

| | | | | | | |
|---|---|---|---|---|---|---|
| RAB3GAP2 | 108.267088 | -1.46E-05 | 0.26288941 | -5.54E-05 | 0.99995578 | 0.99999872 |
| MARK1 | 45.5351111 | -0.027521425 | 0.49124705 | -0.0560236 | 0.95532301 | 0.99999872 |
| C1orf115 | 26.3413327 | -0.126442415 | 0.59109825 | -0.213911 | 0.83061647 | 0.99999872 |
| 2-Mar | 45.3661507 | 0.539352306 | 0.30123939 | 1.79044416 | 0.07338254 | 0.99999872 |
| MIA3 | 26.0374899 | 0.108643888 | 0.34261278 | 0.31710402 | 0.75116468 | 0.99999872 |
| AIDA | 15.6335308 | -0.115837589 | 0.39654407 | -0.2921178 | 0.77019654 | 0.99999872 |
| BROX | 25.6358368 | 0.030163683 | 0.37056042 | 0.08140018 | 0.93512371 | 0.99999872 |
| FAM177B | 8.08881513 | -0.071927619 | 0.76432196 | -0.0941064 | 0.9250246 | 0.99999872 |
| AC093152.1 | 14.9810181 | -0.150406844 | 0.54303288 | -0.2769756 | 0.78179886 | 0.99999872 |
| DISP1 | 8.28577321 | 1.074223359 | 0.63205111 | 1.69958306 | 0.08920938 | 0.99999872 |
| SUSD4 | 8.26698636 | -0.973238339 | 0.82825148 | -1.1750517 | 0.23997402 | 0.99999872 |
| CAPN2 | 314.703878 | -0.125166205 | 0.3116049 | -0.4016824 | 0.68791778 | 0.99999872 |
| TP53BP2 | 24.9288963 | -0.033650412 | 0.35536878 | -0.0946915 | 0.92455985 | 0.99999872 |
| FBXO28 | 28.5897718 | -0.159282212 | 0.35382445 | -0.450173 | 0.65258571 | 0.99999872 |
| DEGS1 | 19.1247684 | 0.326581642 | 0.49906946 | 0.65438114 | 0.51286628 | 0.99999872 |
| NVL | 22.1535173 | -0.241843335 | 0.39090204 | -0.6186802 | 0.53612708 | 0.99999872 |
| CNIH4 | 25.1145747 | 0.225579535 | 0.36594301 | 0.61643352 | 0.53760844 | 0.99999872 |
| WDR26 | 107.522186 | -0.263363951 | 0.33179001 | -0.793767 | 0.4273311 | 0.99999872 |
| CNIH3 | 36.2503687 | -0.02538276 | 0.41791442 | -0.0607367 | 0.95156887 | 0.99999872 |
| DNAH14 | 61.3141807 | -0.784294631 | 0.55157726 | -1.4219126 | 0.15505164 | 0.99999872 |
| LBR | 6.81323564 | -0.489888668 | 0.67360878 | -0.7272599 | 0.46706674 | 0.99999872 |
| ENAH | 11.8419745 | -0.350724281 | 0.50680397 | -0.6920314 | 0.48891758 | 0.99999872 |
| SRP9 | 55.742322 | 0.46985059 | 0.3268045 | 1.43771149 | 0.15051593 | 0.99999872 |
| EPHX1 | 35.6939546 | -0.186434273 | 0.60036911 | -0.3105328 | 0.75615586 | 0.99999872 |
| AL591895.1 | 5.85036038 | -0.144937147 | 0.64069636 | -0.2262182 | 0.82103175 | 0.99999872 |
| TMEM63A | 9.45831068 | 0.409995232 | 0.55568054 | 0.73782543 | 0.46062054 | 0.99999872 |
| SDE2 | 7.49821373 | 0.350446533 | 0.58301035 | 0.6010983 | 0.54777451 | 0.99999872 |
| H3F3A | 9.74587383 | 0.724193044 | 0.58507737 | 1.23777312 | 0.2158002 | 0.99999872 |
| ACBD3 | 56.4977061 | 0.088667598 | 0.28751434 | 0.30839366 | 0.75778281 | 0.99999872 |
| PARP1 | 26.5381026 | 0.153617188 | 0.37185045 | 0.41311551 | 0.67952199 | 0.99999872 |
| ITPKB | 7.36672378 | -1.007170586 | 0.67243187 | -1.4978032 | 0.1341844 | 0.99999872 |
| PSEN2 | 10.6044727 | 0.635741109 | 0.52701405 | 1.20630771 | 0.22769886 | 0.99999872 |
| COQ8A | 11.6855147 | 0.205152467 | 0.74187885 | 0.27653095 | 0.78214029 | 0.99999872 |
| CDC42BPA | 202.832651 | -0.113648877 | 0.29242101 | -0.3886481 | 0.69753646 | 0.99999872 |
| ZNF678 | 22.3791879 | -0.282346096 | 0.39295738 | -0.7185158 | 0.47243929 | 0.99999872 |
| SNAP47 | 13.2791961 | -0.079497199 | 0.55951284 | -0.1420829 | 0.88701454 | 0.99999872 |
| JMJD4 | 5.22806584 | -0.129258525 | 0.60742851 | -0.2127963 | 0.83148587 | 0.99999872 |
| ARF1 | 110.745539 | 0.073089472 | 0.39979957 | 0.18281528 | 0.85494296 | 0.99999872 |
| C1orf35 | 7.1050396 | 0.141846977 | 0.53122638 | 0.26701795 | 0.78945534 | 0.99999872 |
| MRPL55 | 13.9031202 | 0.3287268 | 0.46235914 | 0.71097718 | 0.47709838 | 0.99999872 |
| GUK1 | 41.4659227 | 0.087725436 | 0.57552256 | 0.15242745 | 0.87884981 | 0.99999872 |
| OBSCN | 21.1866645 | -0.74023403 | 0.46670124 | -1.5860983 | 0.11271702 | 0.99999872 |
| TRIM11 | 6.80732707 | 0.094446781 | 0.63455566 | 0.14883924 | 0.88168049 | 0.99999872 |
| RNF187 | 27.3108296 | -0.087322794 | 0.35730344 | -0.2443939 | 0.80692574 | 0.99999872 |
| RHOU | 7.69956963 | 1.088846905 | 0.91245454 | 1.19331633 | 0.23274551 | 0.99999872 |
| RAB4A | 22.1284148 | 0.481944076 | 0.41327133 | 1.16616865 | 0.24354626 | 0.99999872 |
| CCSAP | 4.71934092 | 0.431855958 | 0.8501902 | 0.50795217 | 0.61148689 | 0.99999872 |
| NUP133 | 19.9349254 | 0.313255412 | 0.37623082 | 0.83261498 | 0.40506191 | 0.99999872 |
| ABCB10 | 12.541913 | 0.5183153 | 0.48701121 | 1.06427797 | 0.28720279 | 0.99999872 |
| TAF5L | 7.14713306 | 0.206423697 | 0.57987132 | 0.3559819 | 0.72185412 | 0.99999872 |
| GALNT2 | 35.7429025 | -0.154225577 | 0.27482906 | -0.5611691 | 0.57468227 | 0.99999872 |
| COG2 | 10.7167924 | 0.068967552 | 0.54641794 | 0.12621758 | 0.89955969 | 0.99999872 |
| C1orf198 | 15.6009875 | -0.533712557 | 0.47454823 | -1.1246751 | 0.26072676 | 0.99999872 |
| TTC13 | 17.9029493 | 0.232411739 | 0.48410929 | 0.48008114 | 0.6311697 | 0.99999872 |
| ARV1 | 11.6035627 | 0.738370955 | 0.50176826 | 1.4715378 | 0.14114573 | 0.99999872 |
| FAM89A | 158.924018 | 0.703922215 | 0.53632385 | 1.3124947 | 0.18935327 | 0.99999872 |
| C1orf131 | 5.4267533 | 0.320066376 | 0.59922272 | 0.53413591 | 0.59324751 | 0.99999872 |
| GNPAT | 22.1196429 | -0.064907245 | 0.36772517 | -0.1765102 | 0.85989313 | 0.99999872 |
| EXOC8 | 6.35505562 | 0.511265661 | 0.61322519 | 0.83373233 | 0.40443184 | 0.99999872 |
| SPRTN | 5.94459453 | 0.671437872 | 0.59899101 | 1.12094816 | 0.26230993 | 0.99999872 |
| EGLN1 | 42.9267718 | 0.219746712 | 0.29639724 | 0.74139258 | 0.45845544 | 0.99999872 |

| | | | | | | |
|---|---|---|---|---|---|---|
| DISC1 | 7.48768301 | 0.219861198 | 0.65656825 | 0.33486419 | 0.73772753 | 0.99999872 |
| SIPA1L2 | 19.6297304 | 0.01490675 | 0.41954858 | 0.03553045 | 0.97165677 | 0.99999872 |
| NTPCR | 10.7487643 | -0.555574553 | 0.4116558 | -1.3496094 | 0.1771413 | 0.99999872 |
| PCNX2 | 46.0709303 | -0.303498111 | 0.37769819 | -0.8035466 | 0.42165886 | 0.99999872 |
| MAP3K21 | 16.8515464 | 0.34195808 | 0.53762036 | 0.63605865 | 0.5247382 | 0.99999872 |
| COA6 | 7.88191583 | 0.206472023 | 0.499459 | 0.41339133 | 0.67931992 | 0.99999872 |
| TARBP1 | 17.0637084 | -0.432812789 | 0.63013263 | -0.6868598 | 0.49217107 | 0.99999872 |
| IRF2BP2 | 28.557885 | 0.33019285 | 0.51715685 | 0.63847718 | 0.5231631 | 0.99999872 |
| TOMM20 | 47.1822664 | 0.551622114 | 0.416212 | 1.32533929 | 0.18505864 | 0.99999872 |
| ARID4B | 122.060297 | -0.017706497 | 0.26314637 | -0.0672876 | 0.94635272 | 0.99999872 |
| B3GALNT2 | 12.4196935 | -0.042924111 | 0.5576501 | -0.0769732 | 0.93864487 | 0.99999872 |
| LYST | 56.9919881 | 0.736540444 | 0.4951193 | 1.48760195 | 0.1368559 | 0.99999872 |
| NID1 | 5.26226662 | 1.045910515 | 0.94099243 | 1.11149727 | 0.26635436 | 0.99999872 |
| GPR137B | 10.4754058 | -0.009844224 | 0.59203531 | -0.0166278 | 0.98673357 | 0.99999872 |
| ERO1B | 31.1490094 | 0.407238374 | 0.41013195 | 0.99294477 | 0.32073687 | 0.99999872 |
| EDARADD | 9.30325703 | -1.001617743 | 0.55980451 | -1.7892277 | 0.07357815 | 0.99999872 |
| LGALS8 | 111.027129 | -0.20860791 | 0.49884021 | -0.4181858 | 0.67581125 | 0.99999872 |
| HEATR1 | 17.4368455 | 0.037049791 | 0.37625462 | 0.09847 | 0.9215591 | 0.99999872 |
| MTR | 66.6290526 | -0.120901858 | 0.35165166 | -0.3438114 | 0.73098811 | 0.99999872 |
| MT1HL1 | 7.26721492 | -0.326061897 | 2.82451015 | -0.1154402 | 0.90809625 | 0.99999872 |
| RYR2 | 47.2601797 | -0.8879127 | 1.11488469 | -0.7964166 | 0.42578992 | 0.99999872 |
| FMN2 | 843.162557 | 0.038316808 | 0.26257791 | 0.14592548 | 0.88398022 | 0.99999872 |
| RGS7 | 15.7430319 | -1.922175362 | 0.64422176 | -2.9837169 | 0.0028477 | 0.43717075 |
| FH | 10.3790381 | 0.262718968 | 0.50356899 | 0.52171395 | 0.60186951 | 0.99999872 |
| OPN3 | 10.1774311 | 0.332996344 | 0.62910201 | 0.52932011 | 0.59658341 | 0.99999872 |
| CEP170 | 38.8971167 | 0.161359641 | 0.38825848 | 0.4155985 | 0.67770382 | 0.99999872 |
| SDCCAG8 | 77.5861423 | -0.152042282 | 0.29765174 | -0.510806 | 0.60948694 | 0.99999872 |
| AKT3 | 345.714262 | 0.012169872 | 0.32366589 | 0.03760011 | 0.97000652 | 0.99999872 |
| ZBTB18 | 9.60335886 | -0.30597435 | 0.67787196 | -0.4513748 | 0.65171941 | 0.99999872 |
| ADSS | 18.2220476 | -0.108769996 | 0.41968156 | -0.2591727 | 0.79550201 | 0.99999872 |
| CATSPERE | 17.915944 | -1.02296909 | 0.50392708 | -2.0299943 | 0.04235712 | 0.95980711 |
| DESI2 | 27.1083861 | -0.155395833 | 0.34326701 | -0.4526967 | 0.65076718 | 0.99999872 |
| COX20 | 31.7644747 | 0.019145413 | 0.41163807 | 0.04651031 | 0.96290352 | 0.99999872 |
| HNRNPU | 143.519946 | -0.147350796 | 0.26348599 | -0.5592358 | 0.57600081 | 0.99999872 |
| EFCAB2 | 55.3182127 | -0.64265512 | 0.51213705 | -1.2548499 | 0.20953326 | 0.99999872 |
| KIF26B | 13.3731484 | 0.436101425 | 0.76540551 | 0.5697652 | 0.56883696 | 0.99999872 |
| SMYD3 | 139.862297 | -0.277726593 | 0.25911495 | -1.0718277 | 0.28379741 | 0.99999872 |
| TFB2M | 5.95236905 | -0.096329353 | 0.75723944 | -0.1272112 | 0.89877322 | 0.99999872 |
| CNST | 36.4098189 | 0.041842355 | 0.32562611 | 0.12849816 | 0.89775476 | 0.99999872 |
| SCCPDH | 20.4442858 | 0.795920669 | 0.45662583 | 1.74304784 | 0.08132526 | 0.99999872 |
| AHCTF1 | 49.6459949 | -0.001447633 | 0.34716417 | -0.0041699 | 0.99667293 | 0.99999872 |
| ZNF124 | 57.8580105 | 0.181498357 | 0.36541442 | 0.49669183 | 0.61940639 | 0.99999872 |
| AL390728.6 | 13.4391981 | 0.414308804 | 0.44276146 | 0.93573819 | 0.349408 | 0.99999872 |
| ZNF496 | 38.9824289 | -0.331777698 | 0.4804843 | -0.6905068 | 0.48987551 | 0.99999872 |
| SH3BP5L | 6.57883856 | -0.611126228 | 0.53393837 | -1.1445632 | 0.25239013 | 0.99999872 |
| ZNF672 | 6.56412617 | 0.096957806 | 0.53216892 | 0.18219366 | 0.85543075 | 0.99999872 |
| ZNF692 | 7.81102502 | -0.458779837 | 0.52043464 | -0.8815321 | 0.37802989 | 0.99999872 |
| FAM110C | 7.58894502 | 0.105750585 | 0.75019272 | 0.14096456 | 0.88789794 | 0.99999872 |
| SH3YL1 | 27.7860687 | 0.157418158 | 0.40648161 | 0.38727006 | 0.69855629 | 0.99999872 |
| ACP1 | 18.8758446 | 0.535856399 | 0.39302551 | 1.36341378 | 0.17275214 | 0.99999872 |
| TMEM18 | 19.6112794 | 0.225002649 | 0.65021425 | 0.34604386 | 0.72930975 | 0.99999872 |
| EIPR1 | 19.4909314 | -0.530279347 | 0.419802 | -1.2631654 | 0.20652975 | 0.99999872 |
| TRAPPC12 | 39.3583306 | -0.07718942 | 0.33490375 | -0.2304824 | 0.81771694 | 0.99999872 |
| ADI1 | 33.6672723 | 0.427158457 | 0.43732932 | 0.97674324 | 0.32869628 | 0.99999872 |
| AC108488.2 | 6.33903239 | -0.527873258 | 0.6289356 | -0.8393121 | 0.40129419 | 0.99999872 |
| RPS7 | 62.8599933 | 0.993291749 | 0.43148303 | 2.30204131 | 0.02133284 | 0.87186485 |
| NRIR | 3.68258655 | -5.886571052 | 3.7785225 | -1.5579029 | 0.11925627 | NA |
| CMPK2 | 11.5795258 | -1.188317967 | 2.74898967 | -0.4322744 | 0.66554196 | 0.99999872 |
| RSAD2 | 27.8162473 | -2.675187148 | 1.97437669 | -1.3549528 | 0.17543261 | 0.99999872 |
| ID2 | 10.4860729 | -1.675654146 | 0.71149864 | -2.3551052 | 0.01851746 | 0.85473999 |
| KIDINS220 | 78.9812585 | 0.451594194 | 0.28885294 | 1.56340519 | 0.11795732 | 0.99999872 |

| | | | | | | | |
|---|---|---|---|---|---|---|---|
| MBOAT2 | 80.7478611 | | 0.861444762 | 0.43148766 | 1.99645283 | 0.04588466 | 0.95980711 |
| ASAP2 | 36.9282969 | | 0.133869071 | 0.46532294 | 0.28769068 | 0.77358353 | 0.99999872 |
| ITGB1BP1 | 35.2970436 | | 0.079412339 | 0.3829125 | 0.2073903 | 0.83570506 | 0.99999872 |
| CPSF3 | 8.56069709 | | -0.07310536 | 0.52237999 | -0.1399467 | 0.8887021 | 0.99999872 |
| IAH1 | 28.2967795 | | 0.144552964 | 0.4180639 | 0.34576763 | 0.72951735 | 0.99999872 |
| ADAM17 | 45.0650897 | | 0.082953599 | 0.41309247 | 0.20081121 | 0.8408462 | 0.99999872 |
| YWHAQ | 75.4199451 | | 0.421950404 | 0.34517071 | 1.22243977 | 0.22154137 | 0.99999872 |
| TAF1B | 27.2852417 | | -0.045477508 | 0.48847534 | -0.0931009 | 0.92582338 | 0.99999872 |
| GRHL1 | 13.5399375 | | -0.064456705 | 0.49648308 | -0.1298266 | 0.89670362 | 0.99999872 |
| AC104794.2 | 5.48369913 | | 0.412071439 | 0.68284099 | 0.60346617 | 0.54619861 | 0.99999872 |
| KLF11 | 11.5362055 | | 0.190724193 | 0.62708655 | 0.30414333 | 0.7610187 | 0.99999872 |
| CYS1 | 26.5799674 | | 0.182267834 | 0.5037809 | 0.36179981 | 0.71750164 | 0.99999872 |
| HPCAL1 | 15.7546984 | | -0.27382878 | 0.41622421 | -0.6578877 | 0.51061031 | 0.99999872 |
| ODC1 | 27.2968531 | | 0.484134464 | 0.45537212 | 1.06316229 | 0.28770836 | 0.99999872 |
| NOL10 | 30.5125419 | | 0.016406133 | 0.42121252 | 0.03894978 | 0.96893043 | 0.99999872 |
| PDIA6 | 72.549171 | | 1.24052215 | 0.47125849 | 2.63236035 | 0.00847939 | 0.66769402 |
| PQLC3 | 18.1346133 | | -0.07427551 | 0.60017373 | -0.1237567 | 0.90150793 | 0.99999872 |
| ROCK2 | 87.7996343 | | 0.16274909 | 0.29895626 | 0.54439097 | 0.58617246 | 0.99999872 |
| E2F6 | 11.3942675 | | 0.95351808 | 0.675862 | 1.41081771 | 0.15829837 | 0.99999872 |
| LPIN1 | 38.2008252 | | 0.027212459 | 0.29330116 | 0.09277992 | 0.9260784 | 0.99999872 |
| MIR3681HG | 20.8200155 | | -0.561798737 | 0.66393157 | -0.8461696 | 0.39745812 | 0.99999872 |
| TRIB2 | 124.920138 | | -0.64648793 | 0.45873208 | -1.4092931 | 0.15874853 | 0.99999872 |
| NBAS | 82.4378686 | | 0.041644051 | 0.22252074 | 0.18714683 | 0.85154551 | 0.99999872 |
| DDX1 | 31.4327158 | | 0.057710725 | 0.32539176 | 0.17735767 | 0.85922746 | 0.99999872 |
| SMC6 | 17.7921343 | | 0.169630468 | 0.37402847 | 0.45352287 | 0.65017228 | 0.99999872 |
| GEN1 | 6.68538265 | | -1.381991918 | 0.75095631 | -1.8403093 | 0.06572283 | 0.99999872 |
| KCNS3 | 5.7017511 | | 0.908790726 | 0.64514225 | 1.4086672 | 0.1589336 | 0.99999872 |
| RDH14 | 10.4914034 | | 0.934617942 | 0.58983055 | 1.5845533 | 0.11306786 | 0.99999872 |
| LINC01376 | 43.5540676 | | -0.215406908 | 0.39073145 | -0.5512914 | 0.5814339 | 0.99999872 |
| TTC32 | 11.1946539 | | -0.341261267 | 0.45465563 | -0.7505929 | 0.45289773 | 0.99999872 |
| WDR35 | 36.1477235 | | -0.095455224 | 0.27487876 | -0.347263 | 0.72839374 | 0.99999872 |
| MATN3 | 5.9482228 | | -0.129776723 | 0.85623829 | -0.1515661 | 0.87952915 | 0.99999872 |
| LAPTM4A | 103.153495 | | 0.470897458 | 0.37087806 | 1.26968269 | 0.20419768 | 0.99999872 |
| SDC1 | 5.27701212 | | -0.536879992 | 0.79469034 | -0.6755839 | 0.49930487 | 0.99999872 |
| PUM2 | 96.9242146 | | -0.035255107 | 0.25943817 | -0.1358902 | 0.89190808 | 0.99999872 |
| RHOB | 13.4381568 | | 0.183352661 | 0.58760491 | 0.31203392 | 0.75501475 | 0.99999872 |
| HS1BP3 | 8.86685848 | | 0.524855139 | 0.65152478 | 0.80557971 | 0.42048523 | 0.99999872 |
| LDAH | 42.472952 | | 0.090550396 | 0.35065495 | 0.25823219 | 0.79622771 | 0.99999872 |
| AC018742.1 | 152.51804 | | 0.83857324 | 0.79516065 | 1.054596 | 0.29161014 | 0.99999872 |
| KLHL29 | 693.284767 | | -0.429703273 | 0.50538138 | -0.8502554 | 0.39518308 | 0.99999872 |
| ATAD2B | 67.5476717 | | -0.053747548 | 0.26922344 | -0.1996392 | 0.84176278 | 0.99999872 |
| UBXN2A | 28.82251 | | -0.075876009 | 0.32017797 | -0.2369807 | 0.81267174 | 0.99999872 |
| FKBP1B | 7.26071812 | | -0.451202126 | 0.95695281 | -0.4714988 | 0.63728456 | 0.99999872 |
| SF3B6 | 27.043742 | | 0.903883357 | 0.37090909 | 2.43694041 | 0.01481212 | 0.78384542 |
| FAM228B | 22.8406876 | | 0.089220275 | 0.42783988 | 0.2085366 | 0.83481 | 0.99999872 |
| TP53I3 | 8.01476518 | | -0.202547825 | 0.74584549 | -0.2715681 | 0.78595415 | 0.99999872 |
| ITSN2 | 72.0076737 | | -0.185450863 | 0.32293754 | -0.5742623 | 0.5657903 | 0.99999872 |
| NCOA1 | 230.505957 | | 0.266690365 | 0.36752815 | 0.72563249 | 0.4680641 | 0.99999872 |
| PTRHD1 | 7.19217871 | | 0.886607765 | 0.57768294 | 1.53476535 | 0.12484148 | 0.99999872 |
| DNAJC27 | 9.66051715 | | 0.014126851 | 0.51227774 | 0.02757655 | 0.97799989 | 0.99999872 |
| DNAJC27-AS | 11.2227141 | | 0.139402749 | 0.73046865 | 0.19084015 | 0.84865083 | 0.99999872 |
| DNMT3A | 37.3964433 | | 0.289702846 | 0.34490755 | 0.83994347 | 0.40094008 | 0.99999872 |
| DTNB | 37.5373208 | | 0.081353727 | 0.48605724 | 0.16737479 | 0.86707516 | 0.99999872 |
| ASXL2 | 46.3125912 | | 0.507022082 | 0.27729972 | 1.82842621 | 0.06748561 | 0.99999872 |
| RAB10 | 106.811372 | | -0.163231622 | 0.22268654 | -0.7330107 | 0.46355188 | 0.99999872 |
| HADHA | 38.4514185 | | 0.602130662 | 0.31908199 | 1.88707193 | 0.05915065 | 0.99999872 |
| HADHB | 38.1465899 | | 0.934519027 | 0.38730889 | 2.412852 | 0.01582825 | 0.79608149 |
| SELENOI | 16.4808203 | | -0.3129232 | 0.46151636 | -0.6780327 | 0.49775094 | 0.99999872 |
| MAPRE3 | 12.7262067 | | 0.274356887 | 0.52561448 | 0.52197361 | 0.6016887 | 0.99999872 |
| TMEM214 | 14.2254841 | | 0.443875219 | 0.53587027 | 0.82832589 | 0.40748596 | 0.99999872 |
| AGBL5 | 14.0106643 | | 0.08335181 | 0.41592755 | 0.20039983 | 0.84116789 | 0.99999872 |

| | | | | | | |
|---|---|---|---|---|---|---|
| OST4 | 42.8624268 | 1.080387019 | 0.53410046 | 2.02281613 | 0.04309211 | 0.95980711 |
| EMILIN1 | 9.58369997 | 1.341061628 | 0.679291 | 1.97420784 | 0.04835811 | 0.97937581 |
| KHK | 4.2234055 | 0.240388875 | 0.90728473 | 0.26495417 | 0.79104477 | NA |
| PREB | 7.37630669 | 1.252597918 | 0.68107547 | 1.83914703 | 0.06589356 | 0.99999872 |
| ATRAID | 35.0273815 | 0.713736321 | 0.42476009 | 1.68032812 | 0.09289349 | 0.99999872 |
| CAD | 6.581632 | -0.312280487 | 0.54764418 | -0.5702252 | 0.56852499 | 0.99999872 |
| TRIM54 | 12.2728777 | -0.140706677 | 0.9170121 | -0.1534404 | 0.87805101 | 0.99999872 |
| MPV17 | 8.52148823 | -0.222253886 | 0.55056188 | -0.4036856 | 0.68644395 | 0.99999872 |
| GTF3C2 | 19.0771572 | -0.589537639 | 0.35070086 | -1.6810271 | 0.09275765 | 0.99999872 |
| EIF2B4 | 6.63249292 | -0.226079593 | 0.70842724 | -0.3191289 | 0.74962878 | 0.99999872 |
| SNX17 | 20.4479516 | 0.232060356 | 0.51971847 | 0.44651166 | 0.65522769 | 0.99999872 |
| ZNF513 | 4.0370515 | 0.013644203 | 0.7762695 | 0.01757663 | 0.9859766 | NA |
| PPM1G | 37.7678145 | -0.025325861 | 0.43071256 | -0.0587999 | 0.95311148 | 0.99999872 |
| NRBP1 | 20.3343459 | -0.080309338 | 0.42077099 | -0.1908623 | 0.84863344 | 0.99999872 |
| IFT172 | 11.456886 | 0.561326481 | 0.50243739 | 1.11720683 | 0.2639059 | 0.99999872 |
| ZNF512 | 22.854275 | -0.1560191 | 0.41279275 | -0.3779599 | 0.70546039 | 0.99999872 |
| GPN1 | 6.9823429 | -0.155302251 | 0.55012397 | -0.2823041 | 0.77771034 | 0.99999872 |
| SUPT7L | 12.3324119 | 0.260357371 | 0.53548692 | 0.48620678 | 0.62682056 | 0.99999872 |
| SLC4A1AP | 11.3556172 | 0.069977823 | 0.45027032 | 0.15541292 | 0.8764958 | 0.99999872 |
| MRPL33 | 43.7433035 | 0.576171477 | 0.32056316 | 1.79737273 | 0.07227647 | 0.99999872 |
| RBKS | 17.5267072 | -0.278070378 | 0.49825894 | -0.5580841 | 0.57678697 | 0.99999872 |
| BABAM2 | 117.484481 | -0.30084391 | 0.30947898 | -0.972098 | 0.33100178 | 0.99999872 |
| FOSL2 | 47.4091614 | -0.256242159 | 0.67919095 | -0.3772756 | 0.70596882 | 0.99999872 |
| PPP1CB | 24.8885585 | 0.031775499 | 0.34115388 | 0.09314125 | 0.92579135 | 0.99999872 |
| TRMT61B | 16.3543553 | 0.503408216 | 0.45223699 | 1.11315136 | 0.26564343 | 0.99999872 |
| WDR43 | 24.7717233 | -0.135637638 | 0.46732802 | -0.2902408 | 0.77163206 | 0.99999872 |
| TOGARAM2 | 16.2727135 | 0.189826758 | 0.82535653 | 0.22999365 | 0.81809671 | 0.99999872 |
| C2orf71 | 7.50967127 | 1.933264923 | 0.96843293 | 1.99628168 | 0.04590327 | 0.95980711 |
| CLIP4 | 200.734805 | 0.100018634 | 0.40094011 | 0.24946029 | 0.80300476 | 0.99999872 |
| ALK | 8.73139304 | 0.951342533 | 0.71622117 | 1.3282804 | 0.18408549 | 0.99999872 |
| YPEL5 | 35.0530984 | 0.529618922 | 0.33474774 | 1.58214338 | 0.11361684 | 0.99999872 |
| LBH | 59.3486416 | -0.143008408 | 0.67371599 | -0.2122681 | 0.83189789 | 0.99999872 |
| LCLAT1 | 47.307656 | 0.064275821 | 0.33485029 | 0.19195391 | 0.84777831 | 0.99999872 |
| GALNT14 | 24.8091689 | -0.432019869 | 0.96474465 | -0.4478075 | 0.65429215 | 0.99999872 |
| MEMO1 | 7.14976135 | 0.514376201 | 0.63505267 | 0.80997408 | 0.41795507 | 0.99999872 |
| DPY30 | 20.2743193 | 0.649266197 | 0.38699748 | 1.67770134 | 0.09340542 | 0.99999872 |
| SPAST | 20.9152424 | 0.28230102 | 0.42095792 | 0.67061578 | 0.50246533 | 0.99999872 |
| SLC30A6 | 24.1927976 | -0.163499452 | 0.34584157 | -0.4727582 | 0.63638568 | 0.99999872 |
| YIPF4 | 53.5725176 | -0.01914937 | 0.27754524 | -0.0689955 | 0.94499321 | 0.99999872 |
| BIRC6 | 198.149934 | -0.04454339 | 0.17613107 | -0.2528991 | 0.80034618 | 0.99999872 |
| TTC27 | 24.1000362 | -0.178475576 | 0.37859726 | -0.4714128 | 0.63734601 | 0.99999872 |
| LINC00486 | 38.7613961 | -0.222413637 | 0.40940448 | -0.5432614 | 0.58694986 | 0.99999872 |
| RASGRP3 | 30.1011923 | 0.175799193 | 0.5833679 | 0.30135219 | 0.76314595 | 0.99999872 |
| AC020594.1 | 10.4804508 | 0.562914226 | 0.79252598 | 0.71027857 | 0.4775314 | 0.99999872 |
| FAM98A | 43.9964414 | 0.793022449 | 0.53030687 | 1.49540293 | 0.13480932 | 0.99999872 |
| LINC01320 | 52.8845758 | -0.252551077 | 0.65311953 | -0.3866843 | 0.69898993 | 0.99999872 |
| AC012593.1 | 23.0579436 | 0.324791035 | 1.04575067 | 0.31058171 | 0.75611863 | 0.99999872 |
| AC009414.2 | 15.2351794 | -0.818283082 | 0.732472 | -1.1171527 | 0.26392903 | 0.99999872 |
| CRIM1 | 1579.77597 | 0.466329447 | 0.36630146 | 1.27307559 | 0.20299122 | 0.99999872 |
| FEZ2 | 247.322963 | 1.107646424 | 0.39179134 | 2.82713348 | 0.00469667 | 0.52915868 |
| STRN | 38.8006258 | 0.008763768 | 0.34328742 | 0.02552895 | 0.97963306 | 0.99999872 |
| HEATR5B | 32.1156547 | 0.274747451 | 0.3722784 | 0.7380161 | 0.46050467 | 0.99999872 |
| GPATCH11 | 5.94545987 | 0.557204581 | 0.6460272 | 0.86250947 | 0.38840722 | 0.99999872 |
| EIF2AK2 | 42.4764592 | -0.589815966 | 0.678575 | -0.8691979 | 0.3847389 | 0.99999872 |
| CEBPZOS | 32.7146927 | 0.403371688 | 0.3790501 | 1.06416458 | 0.28725415 | 0.99999872 |
| CEBPZ | 21.9029516 | -0.004996849 | 0.49710135 | -0.010052 | 0.99197982 | 0.99999872 |
| NDUFAF7 | 10.6822794 | 0.033259087 | 0.50578706 | 0.06575709 | 0.94757122 | 0.99999872 |
| PRKD3 | 15.2859595 | 1.370286272 | 0.52278074 | 2.62114909 | 0.00876339 | 0.67516788 |
| AC006369.1 | 52.9008443 | 0.76437642 | 0.36165468 | 2.11355323 | 0.03455344 | 0.94284111 |
| CDC42EP3 | 493.188435 | -0.541806998 | 0.31488585 | -1.7206458 | 0.08531512 | 0.99999872 |
| RMDN2 | 19.8348304 | -0.328557425 | 0.70495302 | -0.46607 | 0.64116543 | 0.99999872 |

| | | | | | | |
|---|---|---|---|---|---|---|
| CYP1B1 | 62.1750301 | -1.512266125 | 0.92087054 | -1.6422136 | 0.10054574 | 0.99999872 |
| CYP1B1-AS1 | 8.65410018 | 2.427966022 | 0.80479802 | 3.01686382 | 0.00255405 | 0.41171209 |
| ATL2 | 32.5519846 | 0.668317521 | 0.33485997 | 1.9958119 | 0.0459544 | 0.95980711 |
| AC011247.1 | 10.6143488 | -0.203812199 | 0.74084587 | -0.2751074 | 0.78323371 | 0.99999872 |
| HNRNPLL | 46.1669683 | 0.539967954 | 0.39958942 | 1.35130693 | 0.17659713 | 0.99999872 |
| GALM | 7.28025952 | 0.687138111 | 0.75231814 | 0.91336108 | 0.36105267 | 0.99999872 |
| SRSF7 | 36.4975555 | 0.842880439 | 0.3963023 | 2.12686232 | 0.03343152 | 0.92959829 |
| GEMIN6 | 5.63362694 | 0.246020018 | 0.74187008 | 0.33162143 | 0.74017513 | 0.99999872 |
| DHX57 | 18.4729234 | -0.461798794 | 0.36375354 | -1.2695376 | 0.20424939 | 0.99999872 |
| MORN2 | 7.89946158 | 0.646406306 | 0.5407937 | 1.19529185 | 0.23197302 | 0.99999872 |
| SOS1 | 117.943467 | 0.178807565 | 0.25182694 | 0.71004145 | 0.47767843 | 0.99999872 |
| MAP4K3 | 171.093224 | -0.172972929 | 0.25674594 | -0.6737124 | 0.50049416 | 0.99999872 |
| AC007388.1 | 124.573713 | -0.124418323 | 0.2582244 | -0.4818225 | 0.62993206 | 0.99999872 |
| TMEM178A | 130.848648 | 0.445556758 | 0.59268958 | 0.75175399 | 0.45219902 | 0.99999872 |
| THUMPD2 | 22.5144779 | 0.642310739 | 0.43833085 | 1.46535598 | 0.14282382 | 0.99999872 |
| SLC8A1-AS1 | 29.3908587 | 1.293464439 | 0.92586444 | 1.39703437 | 0.16240324 | 0.99999872 |
| SLC8A1 | 30.4262713 | 1.542839408 | 1.80208785 | 0.85613995 | 0.39192037 | 0.99999872 |
| PKDCC | 41.3845883 | 0.129859069 | 0.83483728 | 0.15555015 | 0.87638762 | 0.99999872 |
| EML4 | 205.759634 | 0.295486441 | 0.36521582 | 0.8090735 | 0.41847287 | 0.99999872 |
| COX7A2L | 42.2512107 | 0.619340169 | 0.34552357 | 1.79246867 | 0.07305792 | 0.99999872 |
| MTA3 | 37.3183485 | 0.021891319 | 0.31512693 | 0.06946826 | 0.9446169 | 0.99999872 |
| ZFP36L2 | 98.3187519 | -0.483402016 | 0.44615492 | -1.0834847 | 0.27859335 | 0.99999872 |
| THADA | 66.9698969 | -0.650913019 | 0.29371225 | -2.2161589 | 0.02668062 | 0.89031704 |
| PLEKHH2 | 115.700657 | 0.458980983 | 0.36474116 | 1.25837453 | 0.20825634 | 0.99999872 |
| DYNC2LI1 | 31.2589203 | 0.342253339 | 0.35305622 | 0.96940182 | 0.33234474 | 0.99999872 |
| LRPPRC | 45.530626 | -0.162284578 | 0.30834412 | -0.52631 | 0.59867287 | 0.99999872 |
| PPM1B | 52.2520311 | -0.362679886 | 0.24238184 | -1.4963162 | 0.13457126 | 0.99999872 |
| SLC3A1 | 27.1634497 | 0.751201954 | 0.85529133 | 0.87829951 | 0.3797812 | 0.99999872 |
| PREPL | 56.1112775 | 0.534925523 | 0.38192768 | 1.40059374 | 0.16133559 | 0.99999872 |
| CAMKMT | 103.512293 | -0.24874184 | 0.27110181 | -0.9175219 | 0.35886923 | 0.99999872 |
| SRBD1 | 34.5012318 | -0.248315166 | 0.28553423 | -0.8696511 | 0.38449109 | 0.99999872 |
| PRKCE | 62.5382057 | -0.069781396 | 0.43787465 | -0.1593639 | 0.87338221 | 0.99999872 |
| EPAS1 | 20.4053913 | -0.04731779 | 0.68022045 | -0.0695624 | 0.94454194 | 0.99999872 |
| RHOQ | 84.7303581 | 0.02942561 | 0.48060609 | 0.06122605 | 0.95117919 | 0.99999872 |
| AC018682.1 | 9.62071328 | 0.983921369 | 0.74355101 | 1.32327352 | 0.18574443 | 0.99999872 |
| PIGF | 18.5473592 | 0.868983593 | 0.39996554 | 2.17264613 | 0.02980696 | 0.90266962 |
| CRIPT | 11.9000135 | -0.238597583 | 0.46755928 | -0.5103044 | 0.60983819 | 0.99999872 |
| SOCS5 | 41.2121666 | -0.116339252 | 0.49262011 | -0.2361642 | 0.81330523 | 0.99999872 |
| MCFD2 | 34.9378373 | 0.355494822 | 0.46613855 | 0.76263767 | 0.44567951 | 0.99999872 |
| TTC7A | 16.2501536 | -0.566439032 | 0.37727996 | -1.5013759 | 0.13325837 | 0.99999872 |
| CALM2 | 7.91546698 | 0.851122795 | 0.61072681 | 1.39362279 | 0.16343154 | 0.99999872 |
| EPCAM | 14.9259318 | 0.166017877 | 0.68443187 | 0.24256304 | 0.80834391 | 0.99999872 |
| MSH2 | 19.4829968 | -0.163652871 | 0.54900954 | -0.2980875 | 0.76563639 | 0.99999872 |
| MSH6 | 19.7001205 | 0.207416032 | 0.50993978 | 0.40674613 | 0.68419446 | 0.99999872 |
| FBXO11 | 104.004819 | -0.147175407 | 0.2580895 | -0.5702495 | 0.56850848 | 0.99999872 |
| FOXN2 | 30.1055709 | -0.121023108 | 0.35330243 | -0.3425482 | 0.73193839 | 0.99999872 |
| PPP1R21 | 56.546179 | -0.396350149 | 0.30771912 | -1.2880257 | 0.197737 | 0.99999872 |
| STON1-GTF2 | 19.3262289 | 0.711495038 | 0.65205774 | 1.09115342 | 0.27520538 | 0.99999872 |
| AC073082.1 | 4.36214159 | 3.443402119 | 1.52016181 | 2.26515499 | 0.02350317 | NA |
| AC009975.1 | 38.8084143 | 1.773384037 | 1.47967924 | 1.1984922 | 0.23072546 | 0.99999872 |
| GPR75-ASB3 | 18.6566538 | 0.510024678 | 0.50946141 | 1.00110562 | 0.31677575 | 0.99999872 |
| ERLEC1 | 31.7049165 | -0.00780514 | 0.34853547 | -0.0223941 | 0.98213358 | 0.99999872 |
| PSME4 | 56.1049064 | -0.207038452 | 0.2722187 | -0.7605592 | 0.44692037 | 0.99999872 |
| ACYP2 | 108.728603 | -0.37865082 | 0.28686563 | -1.3199588 | 0.18684877 | 0.99999872 |
| SPTBN1 | 111.008175 | -0.283100451 | 0.28335352 | -0.9991069 | 0.31774291 | 0.99999872 |
| AC093110.1 | 4.5949472 | 1.326054031 | 1.029831 | 1.28764238 | 0.19787048 | 0.99999872 |
| RTN4 | 234.175971 | 0.476930314 | 0.30682633 | 1.55439826 | 0.12008945 | 0.99999872 |
| CLHC1 | 16.508585 | -0.137824986 | 0.6612193 | -0.2084407 | 0.83488491 | 0.99999872 |
| RPS27A | 115.081956 | 0.908662189 | 0.42960856 | 2.11509331 | 0.034422 | 0.9417067 |
| MTIF2 | 9.44440672 | -0.365584491 | 0.48293437 | -0.7570066 | 0.44904593 | 0.99999872 |
| AC012358.3 | 15.2387855 | 0.170398428 | 0.53708709 | 0.31726405 | 0.75104325 | 0.99999872 |

| | | | | | | |
|---|---|---|---|---|---|---|
| CCDC88A | 49.5105463 | 0.036025099 | 0.30574 | 0.1178292 | 0.90620299 | 0.99999872 |
| CFAP36 | 24.3761805 | 0.433981728 | 0.36942905 | 1.17473632 | 0.24010022 | 0.99999872 |
| PPP4R3B | 58.4666151 | -0.221227487 | 0.27810858 | -0.7954716 | 0.4263392 | 0.99999872 |
| PNPT1 | 33.3375077 | -0.865900272 | 0.86057234 | -1.0061912 | 0.31432363 | 0.99999872 |
| EFEMP1 | 99.4025141 | 0.684897354 | 0.4777267 | 1.43365935 | 0.1516695 | 0.99999872 |
| VRK2 | 49.0471359 | 0.257966756 | 0.37229781 | 0.69290431 | 0.48836961 | 0.99999872 |
| FANCL | 44.9440371 | 0.229622586 | 0.39359526 | 0.58339774 | 0.55962558 | 0.99999872 |
| PAPOLG | 11.9013068 | 0.037394934 | 0.43554298 | 0.08585819 | 0.93157915 | 0.99999872 |
| REL | 15.0770017 | 0.173092928 | 0.44840089 | 0.38602272 | 0.69947985 | 0.99999872 |
| PUS10 | 10.2151921 | 0.661568018 | 0.74783875 | 0.88463993 | 0.37635087 | 0.99999872 |
| PEX13 | 18.7092372 | -0.002009137 | 0.36878224 | -0.005448 | 0.99565312 | 0.99999872 |
| KIAA1841 | 22.6909347 | -0.169809318 | 0.46150609 | -0.367946 | 0.7129135 | 0.99999872 |
| C2orf74 | 12.5800204 | -1.013177124 | 0.48585937 | -2.08533 | 0.03703936 | 0.95590735 |
| USP34 | 224.770335 | -0.202588132 | 0.23633737 | -0.8571989 | 0.39133498 | 0.99999872 |
| XPO1 | 111.863006 | 0.299331927 | 0.45263644 | 0.66130761 | 0.50841506 | 0.99999872 |
| FAM161A | 8.08391923 | 0.397807287 | 0.55429485 | 0.71768173 | 0.47295355 | 0.99999872 |
| CCT4 | 34.3990981 | -0.099175293 | 0.3607855 | -0.2748871 | 0.78340295 | 0.99999872 |
| COMMD1 | 81.9421945 | -0.040445858 | 0.21425873 | -0.1887711 | 0.85027221 | 0.99999872 |
| AC018462.1 | 4.834145 | 1.387611831 | 0.86824547 | 1.59817917 | 0.11000311 | 0.99999872 |
| B3GNT2 | 5.04733574 | 0.707441777 | 0.77440195 | 0.91353306 | 0.36096226 | 0.99999872 |
| EHBP1 | 127.010232 | 0.033859122 | 0.2628595 | 0.12881072 | 0.89750742 | 0.99999872 |
| WDPCP | 91.6641763 | -0.074146198 | 0.30579392 | -0.2424711 | 0.80841512 | 0.99999872 |
| MDH1 | 30.9103125 | 0.86877064 | 0.39795645 | 2.18307969 | 0.02902994 | 0.90266962 |
| UGP2 | 38.2505094 | 0.745030709 | 0.36417384 | 2.04581063 | 0.04077501 | 0.95980711 |
| VPS54 | 69.4222272 | 0.003779856 | 0.23764792 | 0.01590528 | 0.98730996 | 0.99999872 |
| PELI1 | 9.75184076 | -0.485541207 | 0.6289656 | -0.7719678 | 0.44013348 | 0.99999872 |
| LGALSL | 4.75542139 | 1.148389955 | 0.73960394 | 1.55270935 | 0.12049259 | 0.99999872 |
| AFTPH | 35.5400326 | -0.049386157 | 0.30505524 | -0.1618925 | 0.87139051 | 0.99999872 |
| SERTAD2 | 23.1248775 | 0.171959453 | 0.42488604 | 0.404719 | 0.68568408 | 0.99999872 |
| CEP68 | 19.9758748 | 0.737225528 | 0.50265066 | 1.46667574 | 0.14246429 | 0.99999872 |
| RAB1A | 136.689881 | 0.436152644 | 0.22005646 | 1.98200337 | 0.04747886 | 0.97354899 |
| ACTR2 | 79.909406 | 0.165300633 | 0.28708392 | 0.57579203 | 0.56475576 | 0.99999872 |
| SPRED2 | 42.6853386 | -0.185873683 | 0.40527664 | -0.4586341 | 0.64649695 | 0.99999872 |
| MEIS1 | 36.3234057 | -0.92975069 | 0.45575612 | -2.040018 | 0.04134853 | 0.95980711 |
| ETAA1 | 14.8692926 | 0.04552592 | 0.43370441 | 0.10496993 | 0.91639967 | 0.99999872 |
| C1D | 23.1227506 | 0.228436885 | 0.39397202 | 0.57983023 | 0.56202911 | 0.99999872 |
| WDR92 | 7.42400985 | -0.406907653 | 0.58217667 | -0.6989419 | 0.48458836 | 0.99999872 |
| PNO1 | 8.868555 | -0.527507991 | 0.58582755 | -0.9004493 | 0.36788121 | 0.99999872 |
| APLF | 22.2096109 | 0.31437409 | 0.51511948 | 0.61029354 | 0.54166738 | 0.99999872 |
| ARHGAP25 | 8.16507312 | -0.469717457 | 0.80145729 | -0.5860792 | 0.55782228 | 0.99999872 |
| ANTXR1 | 9.79862163 | -0.062060933 | 0.67352223 | -0.0921439 | 0.92658375 | 0.99999872 |
| GFPT1 | 49.4586725 | 0.025184098 | 0.29387499 | 0.08569664 | 0.93170758 | 0.99999872 |
| NFU1 | 15.9725284 | 0.057107777 | 0.38560922 | 0.14809754 | 0.88226579 | 0.99999872 |
| AAK1 | 72.7380361 | -0.103218254 | 0.29162794 | -0.3539381 | 0.72338524 | 0.99999872 |
| ANXA4 | 67.8647807 | -0.230807799 | 0.4730787 | -0.4878846 | 0.62563161 | 0.99999872 |
| GMCL1 | 17.4520476 | 0.214042105 | 0.41059578 | 0.5212964 | 0.60216031 | 0.99999872 |
| MXD1 | 13.0475335 | -0.333324516 | 0.55821743 | -0.5971231 | 0.5504252 | 0.99999872 |
| PCBP1 | 65.8434845 | 0.326463487 | 0.42564584 | 0.76698385 | 0.44309112 | 0.99999872 |
| C2orf42 | 9.92618301 | -0.688032512 | 0.56204711 | -1.2241545 | 0.22089394 | 0.99999872 |
| PCYOX1 | 112.649602 | 1.013882092 | 0.82408258 | 1.23031614 | 0.21857874 | 0.99999872 |
| SNRPG | 25.6873478 | 0.783100118 | 0.38428422 | 2.03781492 | 0.04156845 | 0.95980711 |
| FAM136A | 8.56851267 | -0.034997087 | 0.52594781 | -0.066541 | 0.94694713 | 0.99999872 |
| NAGK | 12.8023605 | -0.132403204 | 0.42150172 | -0.3141226 | 0.75342795 | 0.99999872 |
| MCEE | 15.0065621 | -0.005284721 | 0.47530257 | -0.0111186 | 0.99112879 | 0.99999872 |
| MPHOSPH10 | 14.5085191 | -0.200481345 | 0.37814873 | -0.5301653 | 0.59599731 | 0.99999872 |
| PAIP2B | 6.10939506 | 0.07575343 | 0.81187497 | 0.09330677 | 0.92565985 | 0.99999872 |
| ZNF638 | 172.279876 | 0.096209187 | 0.22300742 | 0.43141697 | 0.66616521 | 0.99999872 |
| EXOC6B | 184.969414 | 0.121796225 | 0.29029099 | 0.41956599 | 0.67480254 | 0.99999872 |
| SPR | 4.90848671 | 0.40380324 | 0.72726777 | 0.55523324 | 0.57873514 | 0.99999872 |
| SFXN5 | 25.6252533 | -0.583178513 | 0.58555089 | -0.9959485 | 0.31927518 | 0.99999872 |
| SMYD5 | 4.29103744 | -0.264470144 | 0.92468607 | -0.2860107 | 0.7748699 | NA |

| | | | | | | |
|---|---|---|---|---|---|---|
| CCT7 | 30.3077649 | 0.59183529 | 0.5039493 | 1.17439452 | 0.24023704 | 0.99999872 |
| ALMS1 | 42.7755104 | -0.088326771 | 0.30599103 | -0.288658 | 0.77284308 | 0.99999872 |
| NAT8 | 10.10007 | 1.422476282 | 2.21533974 | 0.642103 | 0.52080631 | 0.99999872 |
| TPRKB | 6.99595035 | 0.888004517 | 0.59902828 | 1.48240835 | 0.13823167 | 0.99999872 |
| DUSP11 | 8.92179485 | 0.040173463 | 0.48627725 | 0.08261432 | 0.93415822 | 0.99999872 |
| STAMBP | 19.4633383 | -0.222196282 | 0.33425788 | -0.6647451 | 0.50621356 | 0.99999872 |
| DGUOK | 27.2907758 | -0.232467784 | 0.43615853 | -0.5329892 | 0.59404106 | 0.99999872 |
| TET3 | 59.9978258 | -0.169100674 | 0.41170271 | -0.4107349 | 0.68126693 | 0.99999872 |
| BOLA3 | 10.0975205 | 1.031192876 | 0.63270366 | 1.62981968 | 0.10313961 | 0.99999872 |
| MOB1A | 51.3783034 | 0.255161671 | 0.27530629 | 0.92682835 | 0.35401566 | 0.99999872 |
| MTHFD2 | 48.1553513 | 0.415512213 | 0.53843558 | 0.77170274 | 0.44029051 | 0.99999872 |
| AC006030.1 | 11.9904131 | 1.172314905 | 0.51340197 | 2.28342501 | 0.02240535 | 0.87985246 |
| WDR54 | 3.7389453 | 0.819115279 | 0.8307397 | 0.98600714 | 0.32412961 | NA |
| RTKN | 4.37379499 | -0.189424358 | 0.80712492 | -0.2346903 | 0.81444915 | NA |
| MOGS | 17.4327917 | 0.290605916 | 0.52696279 | 0.55147331 | 0.58130926 | 0.99999872 |
| CCDC142 | 4.31512765 | -0.668328165 | 0.64132967 | -1.0420977 | 0.29736639 | NA |
| AUP1 | 14.4476401 | 0.223270873 | 0.53894728 | 0.4142722 | 0.67867477 | 0.99999872 |
| HTRA2 | 7.04300441 | 0.372889825 | 0.63270391 | 0.58935913 | 0.55562039 | 0.99999872 |
| DOK1 | 6.72418768 | 0.380336121 | 0.55394731 | 0.6865926 | 0.49233949 | 0.99999872 |
| POLE4 | 14.6581134 | -0.109866188 | 0.54444847 | -0.2017935 | 0.84007813 | 0.99999872 |
| TACR1 | 10.1092049 | 0.36613913 | 0.72180804 | 0.50725277 | 0.61197747 | 0.99999872 |
| AC007681.1 | 4.74216236 | 0.13900267 | 0.79937519 | 0.17388915 | 0.86195258 | 0.99999872 |
| EVA1A | 7.62680058 | 0.170036003 | 1.04504743 | 0.16270649 | 0.87074953 | 0.99999872 |
| MRPL19 | 35.6585593 | 0.020787389 | 0.38954517 | 0.05336323 | 0.9574425 | 0.99999872 |
| GCFC2 | 31.4169102 | 0.408169453 | 0.33120937 | 1.23236082 | 0.21781433 | 0.99999872 |
| SUCLG1 | 32.7888169 | 0.354808496 | 0.41804983 | 0.84872297 | 0.39603546 | 0.99999872 |
| DNAH6 | 11.1465106 | -0.368508728 | 0.67440802 | -0.5464181 | 0.58477859 | 0.99999872 |
| TMSB10 | 316.08741 | -0.366371945 | 0.59315252 | -0.617669 | 0.53679353 | 0.99999872 |
| KCMF1 | 58.7492586 | -0.243176735 | 0.27315768 | -0.8902431 | 0.37333538 | 0.99999872 |
| TCF7L1 | 335.874124 | -0.527337393 | 0.33181805 | -1.5892366 | 0.11200698 | 0.99999872 |
| TGOLN2 | 119.991786 | 0.270398147 | 0.43781646 | 0.61760618 | 0.53683497 | 0.99999872 |
| RETSAT | 5.23891998 | -0.168235033 | 0.61496713 | -0.2735675 | 0.784417 | 0.99999872 |
| ELMOD3 | 19.9620449 | 0.240171985 | 0.38603733 | 0.62214705 | 0.53384518 | 0.99999872 |
| MAT2A | 35.176523 | -0.285297532 | 0.43346128 | -0.6581846 | 0.51041953 | 0.99999872 |
| GGCX | 25.8857176 | 0.594732574 | 0.48862469 | 1.21715622 | 0.22354478 | 0.99999872 |
| VAMP8 | 52.1987252 | -0.332923514 | 0.45776278 | -0.7272839 | 0.46705204 | 0.99999872 |
| VAMP5 | 207.357802 | -0.050076002 | 0.5023866 | -0.0996762 | 0.92060137 | 0.99999872 |
| RNF181 | 13.8830511 | 0.65301002 | 0.51290684 | 1.27315522 | 0.20296297 | 0.99999872 |
| TMEM150A | 8.79114564 | 0.670380711 | 0.52297247 | 1.28186615 | 0.1998896 | 0.99999872 |
| USP39 | 33.8579472 | 0.345649668 | 0.32726567 | 1.05617455 | 0.29088848 | 0.99999872 |
| C2orf68 | 13.4836511 | 0.099599237 | 0.44661243 | 0.22301045 | 0.82352738 | 0.99999872 |
| ATOH8 | 21.5857465 | -1.479163616 | 0.65641898 | -2.2533834 | 0.02423499 | 0.88337526 |
| POLR1A | 12.0614665 | -0.205263232 | 0.44888815 | -0.4572703 | 0.64747675 | 0.99999872 |
| PTCD3 | 29.846325 | -0.016064074 | 0.32469505 | -0.0494743 | 0.96054128 | 0.99999872 |
| IMMT | 38.6970342 | -0.143128005 | 0.36499134 | -0.3921408 | 0.69495416 | 0.99999872 |
| MRPL35 | 10.6311321 | 0.645754617 | 0.51275022 | 1.25939412 | 0.20788801 | 0.99999872 |
| KDM3A | 19.3837821 | -0.045241371 | 0.49454104 | -0.0914815 | 0.92710998 | 0.99999872 |
| CHMP3 | 21.0263372 | 0.260268994 | 0.39858515 | 0.65298216 | 0.51376778 | 0.99999872 |
| RMND5A | 29.3814932 | 0.038532072 | 0.50162829 | 0.07681399 | 0.93877152 | 0.99999872 |
| PLGLB1 | 10.4961815 | -0.131963807 | 0.54198914 | -0.2434805 | 0.80763316 | 0.99999872 |
| CYTOR | 27.1739804 | -0.989101987 | 0.45845057 | -2.1574889 | 0.03096759 | 0.91534437 |
| KRCC1 | 44.7998172 | 0.245018095 | 0.29442834 | 0.83218244 | 0.40530597 | 0.99999872 |
| THNSL2 | 13.3071903 | 0.476270612 | 0.81100921 | 0.58725672 | 0.5570313 | 0.99999872 |
| EIF2AK3 | 38.64134 | -0.339444001 | 0.34586013 | -0.9814488 | 0.32637148 | 0.99999872 |
| RPIA | 15.187676 | 0.275860643 | 0.48036935 | 0.57426779 | 0.5657866 | 0.99999872 |
| MAL | 13.881838 | -0.03057159 | 0.71482828 | -0.0427677 | 0.96588668 | 0.99999872 |
| MRPS5 | 27.2614867 | 0.108540424 | 0.34979774 | 0.31029481 | 0.75633678 | 0.99999872 |
| ZNF514 | 3.14147092 | 1.355004466 | 0.83150191 | 1.62958672 | 0.10318887 | NA |
| AC092835.1 | 7.83335363 | 0.27299364 | 0.52616951 | 0.51883212 | 0.60387782 | 0.99999872 |
| FAHD2A | 8.18581873 | 0.504454918 | 0.53572662 | 0.9416275 | 0.34638338 | 0.99999872 |
| ANKRD36C | 33.4140375 | 0.470717837 | 0.43659123 | 1.07816603 | 0.28095967 | 0.99999872 |

| | | | | | | |
|---|---|---|---|---|---|---|
| STARD7 | 50.049201 | 0.037251689 | 0.31071925 | 0.11988857 | 0.90457142 | 0.99999872 |
| TMEM127 | 15.4267609 | 0.349271505 | 0.50586966 | 0.69043773 | 0.48991895 | 0.99999872 |
| CIAO1 | 19.4426138 | -0.013267467 | 0.40737806 | -0.0325679 | 0.97401913 | 0.99999872 |
| SNRNP200 | 26.1127964 | -0.392863529 | 0.3840207 | -1.023027 | 0.3062951 | 0.99999872 |
| KANSL3 | 32.0493112 | -0.470516182 | 0.34301539 | -1.3717057 | 0.17015506 | 0.99999872 |
| LMAN2L | 10.5825162 | 0.643592319 | 0.52340876 | 1.22961703 | 0.21884055 | 0.99999872 |
| CNNM4 | 6.32074411 | -0.299641274 | 0.66642187 | -0.449627 | 0.65297941 | 0.99999872 |
| ANKRD23 | 9.31880458 | 0.070145821 | 0.49119942 | 0.14280518 | 0.88644405 | 0.99999872 |
| SEMA4C | 4.19086846 | 0.704003706 | 0.8385064 | 0.83959252 | 0.40113689 | NA |
| ANKRD36 | 44.5338816 | -0.097327904 | 0.40000048 | -0.2433195 | 0.80775792 | 0.99999872 |
| AC092683.1 | 34.6134059 | -0.751139843 | 0.42891265 | -1.7512653 | 0.07990022 | 0.99999872 |
| ANKRD36B | 12.8308202 | -0.010832736 | 0.45376306 | -0.0238731 | 0.98095382 | 0.99999872 |
| COX5B | 56.0842106 | 0.766445034 | 0.58870849 | 1.30190926 | 0.19294741 | 0.99999872 |
| ACTR1B | 19.3622487 | -0.131928359 | 0.65860446 | -0.200315 | 0.84123422 | 0.99999872 |
| TMEM131 | 79.3890249 | -0.018995709 | 0.29697844 | -0.0639633 | 0.94899948 | 0.99999872 |
| INPP4A | 27.4062471 | 0.104903797 | 0.35521973 | 0.29532086 | 0.76774878 | 0.99999872 |
| COA5 | 13.3937414 | 0.30864988 | 0.45222255 | 0.68251767 | 0.49491168 | 0.99999872 |
| UNC50 | 23.6118411 | 0.83180979 | 0.43606482 | 1.90753703 | 0.05645108 | 0.99999872 |
| MGAT4A | 26.176736 | 0.278478257 | 0.60468806 | 0.46053209 | 0.64513434 | 0.99999872 |
| TSGA10 | 19.4168077 | -0.079931199 | 0.63480491 | -0.1259146 | 0.89979953 | 0.99999872 |
| AC079447.1 | 21.8700453 | 0.306941529 | 0.37647495 | 0.815304 | 0.41489832 | 0.99999872 |
| MITD1 | 12.333708 | 0.761752062 | 0.51980566 | 1.4654555 | 0.14279669 | 0.99999872 |
| TXNDC9 | 14.1766917 | 0.392805356 | 0.53166723 | 0.73881807 | 0.46001748 | 0.99999872 |
| EIF5B | 47.7350887 | -0.31229821 | 0.40826221 | -0.7649452 | 0.44430419 | 0.99999872 |
| REV1 | 61.6002572 | -0.328832369 | 0.3061809 | -1.0739807 | 0.28283135 | 0.99999872 |
| LONRF2 | 6.62162184 | 0.052203713 | 0.70659151 | 0.07388104 | 0.94110505 | 0.99999872 |
| NPAS2 | 20.3672766 | 0.010030361 | 0.42162456 | 0.02378979 | 0.98102028 | 0.99999872 |
| RPL31 | 28.6637829 | 0.532906569 | 0.4258626 | 1.25135798 | 0.2108039 | 0.99999872 |
| TBC1D8 | 26.2824594 | 0.132633333 | 0.45302879 | 0.29277021 | 0.7696978 | 0.99999872 |
| CNOT11 | 14.7469046 | -0.025862883 | 0.49600723 | -0.0521421 | 0.95841543 | 0.99999872 |
| RNF149 | 56.7947757 | -0.521154638 | 0.51867417 | -1.0047823 | 0.31500168 | 0.99999872 |
| MAP4K4 | 39.1186883 | -0.562346954 | 0.41973882 | -1.3397544 | 0.18032519 | 0.99999872 |
| IL1R1 | 11.8323871 | -0.62090635 | 0.5057282 | -1.2277471 | 0.2195419 | 0.99999872 |
| MFSD9 | 4.33969921 | -0.287334707 | 0.75111788 | -0.3825428 | 0.70205882 | NA |
| TMEM182 | 6.34073796 | 0.348954665 | 0.73590782 | 0.47418258 | 0.63536972 | 0.99999872 |
| POU3F3 | 8.51747676 | -0.867191967 | 0.81654922 | -1.0620204 | 0.28822641 | 0.99999872 |
| MRPS9 | 19.9953614 | -0.24758509 | 0.408699 | -0.6057883 | 0.54465531 | 0.99999872 |
| TGFBRAP1 | 12.2302366 | 0.163145439 | 0.49167316 | 0.33181685 | 0.74002756 | 0.99999872 |
| C2orf49 | 10.5494292 | 0.892436491 | 0.46886117 | 1.90341311 | 0.05698666 | 0.99999872 |
| FHL2 | 6.17855269 | -0.003181023 | 1.00465235 | -0.0031663 | 0.99747367 | 0.99999872 |
| NCK2 | 105.98024 | 0.086619697 | 0.49829964 | 0.17383054 | 0.86199864 | 0.99999872 |
| UXS1 | 18.8982364 | 0.121373399 | 0.39556278 | 0.30683726 | 0.75896724 | 0.99999872 |
| SULT1C2 | 13.2144943 | 0.455670498 | 1.35527929 | 0.33621889 | 0.73670581 | 0.99999872 |
| GCC2 | 66.4118984 | 0.051891711 | 0.37547496 | 0.13820285 | 0.8900801 | 0.99999872 |
| GCC2-AS1 | 6.14295669 | 0.770753078 | 0.83286898 | 0.92541936 | 0.35474781 | 0.99999872 |
| LIMS1 | 84.7702097 | 0.075933355 | 0.33090008 | 0.22947518 | 0.81849961 | 0.99999872 |
| RANBP2 | 54.848854 | -0.206445547 | 0.37515029 | -0.5503009 | 0.582113 | 0.99999872 |
| CCDC138 | 5.35584036 | 0.701640256 | 0.67477482 | 1.03981393 | 0.29842636 | 0.99999872 |
| 10-Sep | 94.0071353 | -0.287279275 | 0.34358929 | -0.8361124 | 0.40309166 | 0.99999872 |
| SOWAHC | 52.6924885 | -0.313720377 | 0.4190677 | -0.748615 | 0.45408927 | 0.99999872 |
| NPHP1 | 19.6528131 | -0.185618636 | 0.51155253 | -0.3628535 | 0.71671431 | 0.99999872 |
| SMIM37 | 30.1792974 | 0.582407658 | 0.52014977 | 1.11969224 | 0.26284493 | 0.99999872 |
| MIR4435-2H | 100.49013 | -0.560658175 | 0.41827485 | -1.3404061 | 0.18011334 | 0.99999872 |
| AC017002.5 | 5.45769659 | -0.047756151 | 1.14653616 | -0.0416525 | 0.96677568 | 0.99999872 |
| ANAPC1 | 34.2564499 | 0.020301815 | 0.3231371 | 0.06282725 | 0.94990407 | 0.99999872 |
| TMEM87B | 45.8838295 | -0.08432244 | 0.36398845 | -0.2316624 | 0.81680023 | 0.99999872 |
| ZC3H8 | 14.699874 | 0.016503088 | 0.48855537 | 0.03377936 | 0.97305309 | 0.99999872 |
| ZC3H6 | 33.8541542 | 0.171702015 | 0.4315759 | 0.39784894 | 0.69074154 | 0.99999872 |
| TTL | 13.4563751 | -0.987607037 | 0.53913836 | -1.8318248 | 0.06697752 | 0.99999872 |
| POLR1B | 7.03634866 | -0.821066994 | 0.77168489 | -1.0639926 | 0.28733207 | 0.99999872 |
| CHCHD5 | 9.33891905 | 0.78697685 | 0.53586717 | 1.46860433 | 0.14194014 | 0.99999872 |

| | | | | | | |
|---|---|---|---|---|---|---|
| SLC20A1 | 20.0865413 | -0.983914149 | 0.52946527 | -1.8583167 | 0.06312406 | 0.99999872 |
| PSD4 | 6.66443238 | -0.516773805 | 0.71272858 | -0.725064 | 0.4684128 | 0.99999872 |
| PAX8 | 107.569831 | -0.731472078 | 0.60047373 | -1.2181583 | 0.22316381 | 0.99999872 |
| CBWD2 | 14.1834609 | -0.769084352 | 0.42198226 | -1.8225514 | 0.06837136 | 0.99999872 |
| RABL2A | 9.90392224 | 0.995726561 | 0.54833436 | 1.8159113 | 0.06938398 | 0.99999872 |
| SLC35F5 | 97.85791 | 0.087606249 | 0.30631686 | 0.28599878 | 0.77487906 | 0.99999872 |
| ACTR3 | 94.6666488 | 0.123571081 | 0.26960271 | 0.45834511 | 0.64670452 | 0.99999872 |
| DDX18 | 26.7675092 | -0.289108013 | 0.30996944 | -0.9326984 | 0.35097569 | 0.99999872 |
| CCDC93 | 31.7600012 | -0.133226561 | 0.42202397 | -0.3156848 | 0.75224175 | 0.99999872 |
| INSIG2 | 27.4604641 | 0.94534938 | 0.36936158 | 2.55941449 | 0.01048486 | 0.70947852 |
| STEAP3 | 24.0016054 | -0.465750122 | 0.52154629 | -0.8930178 | 3.72E-01 | 0.99999872 |
| C2orf76 | 10.9619625 | 0.713160674 | 0.58093463 | 1.22760915 | 0.21959372 | 0.99999872 |
| DBI | 46.7099685 | 0.443004402 | 0.35905975 | 1.23379021 | 0.21728109 | 0.99999872 |
| TMEM37 | 8.35027115 | 0.759353443 | 1.15952443 | 0.65488352 | 0.51254275 | 0.99999872 |
| CFAP221 | 15.9263715 | -0.713056982 | 0.69760429 | -1.0221511 | 0.3067094 | 0.99999872 |
| PTPN4 | 68.2750275 | -0.258154999 | 0.46929586 | -0.5500901 | 0.58225759 | 0.99999872 |
| EPB41L5 | 552.536765 | -0.303681707 | 0.32244011 | -0.9418236 | 0.34628296 | 0.99999872 |
| TMEM185B | 3.33323665 | 0.531809271 | 0.88206047 | 0.60291702 | 0.54656389 | NA |
| RALB | 11.1317761 | -0.443117484 | 0.46393211 | -0.9551343 | 0.33950977 | 0.99999872 |
| GLI2 | 10.0412879 | -0.822845218 | 0.76243963 | -1.0792267 | 0.28048667 | 0.99999872 |
| TFCP2L1 | 20.6395661 | 0.139254583 | 0.85943294 | 0.16203077 | 0.87128162 | 0.99999872 |
| CLASP1 | 117.793644 | 0.185985553 | 0.29576782 | 0.62882281 | 0.52946507 | 0.99999872 |
| NIFK-AS1 | 11.5432099 | -0.010138214 | 0.5973429 | -0.0169722 | 0.98645881 | 0.99999872 |
| NIFK | 8.23380787 | 0.413624813 | 0.65014974 | 0.6361993 | 0.52464653 | 0.99999872 |
| TSN | 16.253728 | 0.200309438 | 0.46053249 | 0.43495181 | 0.66359741 | 0.99999872 |
| BIN1 | 5.50009758 | 0.487463984 | 0.93792897 | 0.51972378 | 0.60325611 | 0.99999872 |
| ERCC3 | 15.4611642 | -0.382636215 | 0.38119506 | -1.0037806 | 0.31548437 | 0.99999872 |
| MAP3K2 | 122.042228 | -0.043156491 | 0.31492909 | -0.1370356 | 0.89100267 | 0.99999872 |
| IWS1 | 44.4090896 | -0.248161776 | 0.28618468 | -0.8671386 | 0.38586608 | 0.99999872 |
| AC010976.1 | 6.56530645 | 1.827441017 | 1.02777265 | 1.77805959 | 0.07539407 | 0.99999872 |
| LIMS2 | 77.7015595 | 0.317502118 | 0.52286092 | 0.6072401 | 0.54369158 | 0.99999872 |
| WDR33 | 65.405987 | -0.020486697 | 0.31364194 | -0.0653187 | 0.94792022 | 0.99999872 |
| SFT2D3 | 4.94006867 | 1.23565277 | 0.81637024 | 1.51359361 | 0.13012894 | 0.99999872 |
| POLR2D | 7.87053318 | 0.029832651 | 0.50238102 | 0.05938252 | 0.95264743 | 0.99999872 |
| AMMECR1L | 11.8781581 | 0.299154775 | 0.46642792 | 0.64137407 | 0.52127968 | 0.99999872 |
| SAP130 | 19.806989 | -0.459305944 | 0.45321551 | -1.0134383 | 0.31085086 | 0.99999872 |
| UGGT1 | 38.0954206 | -0.032702591 | 0.39232374 | -0.0833561 | 0.93356837 | 0.99999872 |
| HS6ST1 | 22.4271228 | 0.269984459 | 0.50449274 | 0.53516025 | 0.59253905 | 0.99999872 |
| SMPD4 | 16.372058 | -0.455294915 | 0.42882491 | -1.0617268 | 0.28835972 | 0.99999872 |
| MZT2B | 45.7001561 | 0.366754326 | 0.44147747 | 0.83074303 | 0.40611881 | 0.99999872 |
| CCDC115 | 15.1726393 | 0.219457532 | 0.39451671 | 0.55626929 | 0.57802677 | 0.99999872 |
| IMP4 | 9.90160977 | 0.284073402 | 0.57411002 | 0.49480656 | 0.62073668 | 0.99999872 |
| PTPN18 | 18.0426448 | 0.210072557 | 0.49978762 | 0.42032366 | 0.67424903 | 0.99999872 |
| FAM168B | 38.2633742 | -0.128215122 | 0.43928677 | -0.2918711 | 0.77038517 | 0.99999872 |
| PLEKHB2 | 37.9842524 | 0.233613972 | 0.32167728 | 0.72623709 | 0.46769344 | 0.99999872 |
| MZT2A | 16.144325 | 0.351162779 | 0.40325596 | 0.87081856 | 0.38385323 | 0.99999872 |
| NCKAP5 | 42.1444236 | 0.990077207 | 0.67600796 | 1.464594 | 0.14303172 | 0.99999872 |
| MGAT5 | 829.880599 | -0.250404278 | 0.23224736 | -1.0781792 | 0.28095377 | 0.99999872 |
| CCNT2-AS1 | 13.1357778 | -0.325087003 | 0.64123868 | -0.5069672 | 0.61217781 | 0.99999872 |
| ACMSD | 10.4703751 | 1.916007014 | 1.32420702 | 1.44690897 | 0.14792242 | 0.99999872 |
| CCNT2 | 43.5656487 | -0.042149656 | 0.29786155 | -0.1415075 | 0.887469 | 0.99999872 |
| RAB3GAP1 | 102.572759 | 0.139095643 | 0.22004203 | 0.63213215 | 0.52730053 | 0.99999872 |
| ZRANB3 | 28.7389708 | -0.373480375 | 0.32402225 | -1.152638 | 0.24905898 | 0.99999872 |
| R3HDM1 | 60.3864752 | -0.123703888 | 0.30836333 | -0.4011628 | 0.68830029 | 0.99999872 |
| UBXN4 | 83.0011742 | 0.237085943 | 0.24583025 | 0.96442949 | 0.33483064 | 0.99999872 |
| MCM6 | 46.0760143 | -0.045555995 | 0.40473367 | -0.112558 | 0.91038102 | 0.99999872 |
| DARS | 48.6909433 | -0.157467246 | 0.33352075 | -0.4721363 | 0.63682952 | 0.99999872 |
| THSD7B | 15.1037138 | 0.706465474 | 1.91443818 | 0.36901974 | 0.71211301 | 0.99999872 |
| HNMT | 13.3606076 | -0.704206459 | 0.50033449 | -1.4074714 | 0.15928766 | 0.99999872 |
| AC114763.1 | 16.0964266 | -1.038704288 | 0.46550036 | -2.2313716 | 0.02565652 | 0.88365618 |
| SPOPL | 44.6846299 | 0.021315977 | 0.41550181 | 0.05130177 | 0.95908506 | 0.99999872 |

| | | | | | | |
|---|---|---|---|---|---|---|
| LRP1B | 28.1851575 | -0.345038349 | 1.13996669 | -0.3026741 | 0.76213827 | 0.99999872 |
| GTDC1 | 123.705217 | 0.216861521 | 0.25013303 | 0.86698476 | 0.38595036 | 0.99999872 |
| ZEB2 | 96.8013062 | 0.285874971 | 0.28688369 | 0.99648389 | 0.31901509 | 0.99999872 |
| ACVR2A | 27.7096558 | -0.057515339 | 0.34236545 | -0.167994 | 0.86658801 | 0.99999872 |
| ORC4 | 53.138865 | 0.087256089 | 0.38622945 | 0.22591775 | 0.82126539 | 0.99999872 |
| MBD5 | 323.747353 | 0.08685583 | 0.2981569 | 0.29130914 | 0.77081489 | 0.99999872 |
| EPC2 | 57.2658345 | -0.233279438 | 0.31801999 | -0.733537 | 0.46323095 | 0.99999872 |
| MMADHC | 16.332693 | 0.858703015 | 0.44501086 | 1.92962261 | 0.05365362 | 0.99999872 |
| AC007364.1 | 16.7182095 | -0.27973149 | 0.65968279 | -0.4240394 | 0.67153708 | 0.99999872 |
| RND3 | 5.26118264 | -2.175624272 | 0.86673035 | -2.5101512 | 0.01206795 | 0.72670507 |
| RBM43 | 12.8366081 | -0.001344507 | 0.67168071 | -0.0020017 | 0.99840287 | 0.99999872 |
| NMI | 10.7314194 | -0.48027056 | 1.09277909 | -0.4394946 | 0.66030316 | 0.99999872 |
| RIF1 | 59.4645854 | -0.131611392 | 0.26872257 | -0.4897668 | 0.62429893 | 0.99999872 |
| ARL5A | 20.4634004 | 0.435546024 | 0.49629039 | 0.87760318 | 0.38015911 | 0.99999872 |
| CACNB4 | 7.41483488 | -0.156636888 | 0.62888289 | -0.2490716 | 0.80330537 | 0.99999872 |
| STAM2 | 39.2987698 | -0.064001392 | 0.3247649 | -0.1970699 | 0.84377282 | 0.99999872 |
| FMNL2 | 247.001612 | -0.108438369 | 0.2817174 | -0.384919 | 0.70029746 | 0.99999872 |
| PRPF40A | 45.6437249 | 0.017163733 | 0.25962714 | 0.06610917 | 0.94729091 | 0.99999872 |
| ARL6IP6 | 27.8359566 | 0.600070137 | 0.37859444 | 1.5849946 | 0.11296756 | 0.99999872 |
| KCNJ3 | 14.8162189 | -0.074604798 | 0.85844837 | -0.0869066 | 0.93074578 | 0.99999872 |
| LINC01876 | 22.2103338 | 0.308232564 | 1.00714024 | 0.30604731 | 0.75956862 | 0.99999872 |
| GPD2 | 26.8874732 | -0.055507267 | 0.40451884 | -0.137218 | 0.89085848 | 0.99999872 |
| ACVR1 | 31.1400538 | -0.456850135 | 0.522082 | -0.8750544 | 0.38154432 | 0.99999872 |
| UPP2 | 13.2365614 | 1.731132697 | 0.6999437 | 2.47324564 | 0.01338921 | 0.74507289 |
| CCDC148 | 120.919091 | 0.054189906 | 0.41303827 | 0.13119827 | 0.89561846 | 0.99999872 |
| PKP4 | 310.356462 | -0.555239867 | 0.37898982 | -1.4650522 | 0.14290668 | 0.99999872 |
| PKP4-AS1 | 8.88909606 | 0.189871218 | 0.74535644 | 0.25473882 | 0.79892483 | 0.99999872 |
| TANC1 | 397.578 | 0.28414453 | 0.2637145 | 1.07747026 | 0.28127023 | 0.99999872 |
| WDSUB1 | 50.0421164 | 0.345899908 | 0.40602773 | 0.85191203 | 0.39426292 | 0.99999872 |
| BAZ2B | 212.849926 | -0.160733742 | 0.26281441 | -0.6115865 | 0.54081138 | 0.99999872 |
| AC009961.1 | 12.456218 | 0.445551925 | 0.57054758 | 0.78091983 | 0.43484965 | 0.99999872 |
| 7-Mar | 88.7275309 | 0.107458197 | 0.20816509 | 0.51621623 | 0.6057034 | 0.99999872 |
| CD302 | 5.33113931 | 1.3028874 | 0.77596964 | 1.6790443 | 0.09314341 | 0.99999872 |
| PLA2R1 | 2663.86674 | 0.501607258 | 0.39006988 | 1.28594203 | 0.1984633 | 0.99999872 |
| ITGB6 | 42.6879592 | -1.459484598 | 0.70200409 | -2.0790258 | 0.03761498 | 0.95980711 |
| LINC02478 | 6.37238102 | 0.373752987 | 0.70373003 | 0.5311028 | 0.59534755 | 0.99999872 |
| RBMS1 | 543.609738 | 0.208548553 | 0.25470926 | 0.81877098 | 0.41291709 | 0.99999872 |
| TANK | 57.1828682 | -0.109098553 | 0.4008322 | -0.2721801 | 0.78548353 | 0.99999872 |
| PSMD14 | 31.1949888 | 0.13228382 | 0.35326535 | 0.37446022 | 0.70806195 | 0.99999872 |
| DPP4 | 32.0871052 | 1.109473046 | 1.81205742 | 0.61227257 | 0.54035744 | 0.99999872 |
| IFIH1 | 19.1055066 | -0.521484755 | 1.13914008 | -0.4577881 | 0.6471047 | 0.99999872 |
| GCA | 18.3846716 | 0.015461592 | 0.58332317 | 0.02650605 | 0.97885371 | 0.99999872 |
| FIGN | 203.170557 | -0.366941729 | 0.35935421 | -1.0211143 | 0.30720028 | 0.99999872 |
| GRB14 | 10.8573803 | -0.011189921 | 0.63004618 | -0.0177605 | 0.98582993 | 0.99999872 |
| COBLL1 | 102.813405 | 0.070607647 | 0.57280759 | 0.12326591 | 0.90189654 | 0.99999872 |
| AC019197.1 | 90.4472899 | 2.380858963 | 1.03862784 | 2.29231191 | 0.02188765 | 0.8732058 |
| SCN2A | 11.6214156 | 2.180656171 | 1.18017353 | 1.84774198 | 0.06463968 | 0.99999872 |
| CSRNP3 | 17.0877313 | 0.243768439 | 0.78027068 | 0.31241522 | 0.75472498 | 0.99999872 |
| TTC21B | 33.230025 | -0.223548409 | 0.32304817 | -0.6919971 | 0.48893913 | 0.99999872 |
| AC010127.1 | 6.80692257 | 1.18050639 | 0.6874476 | 1.7172311 | 0.08593696 | 0.99999872 |
| SCN9A | 30.9486821 | -0.119746739 | 0.46598957 | -0.256973 | 0.79719962 | 0.99999872 |
| SCN7A | 33.8138497 | 0.520692657 | 0.85086683 | 0.61195553 | 0.54056718 | 0.99999872 |
| STK39 | 33.3786644 | -0.525611895 | 0.38332588 | -1.3711881 | 0.17031631 | 0.99999872 |
| CERS6 | 422.041484 | 0.430477246 | 0.4036086 | 1.06657105 | 0.28616558 | 0.99999872 |
| LRP2 | 65.1423763 | 1.505808805 | 1.17997886 | 1.27613202 | 0.20190885 | 0.99999872 |
| FASTKD1 | 23.0781019 | 0.225964944 | 0.37911351 | 0.59603506 | 0.55115181 | 0.99999872 |
| PPIG | 63.1303441 | -0.075977529 | 0.2637471 | -0.2880696 | 0.77329344 | 0.99999872 |
| KLHL23 | 23.9718481 | -0.256280732 | 0.49187786 | -0.5210251 | 0.60234925 | 0.99999872 |
| SSB | 88.0309966 | 0.147053604 | 0.69491482 | 0.21161386 | 0.8324083 | 0.99999872 |
| METTL5 | 16.096895 | 0.438270737 | 0.41734711 | 1.05013482 | 0.29365613 | 0.99999872 |
| UBR3 | 136.241033 | 0.058672932 | 0.28561319 | 0.20542795 | 0.83723779 | 0.99999872 |

| | | | | | | |
|---|---|---|---|---|---|---|
| GORASP2 | 27.0247824 | 0.015177617 | 0.32759746 | 0.04633008 | 0.96304716 | 0.99999872 |
| TLK1 | 77.2603643 | 0.004836053 | 0.33594079 | 0.01439555 | 0.98851441 | 0.99999872 |
| METTL8 | 16.0861139 | -0.229428701 | 0.45530588 | -0.5039002 | 0.61433154 | 0.99999872 |
| DCAF17 | 12.7737961 | -0.55233658 | 0.4988935 | -1.1071232 | 0.26824065 | 0.99999872 |
| DYNC1I2 | 64.7009441 | 0.059036775 | 0.25691017 | 0.2297954 | 0.81825076 | 0.99999872 |
| SLC25A12 | 40.5725878 | -0.074360439 | 0.36871479 | -0.2016747 | 0.84017106 | 0.99999872 |
| HAT1 | 15.2231649 | -0.025100697 | 0.48686424 | -0.0515558 | 0.9588826 | 0.99999872 |
| METAP1D | 5.07912503 | -0.752086055 | 1.04938513 | -0.7166921 | 0.47356409 | 0.99999872 |
| ITGA6 | 25.9991271 | 0.953871547 | 0.57010202 | 1.67315937 | 0.09429593 | 0.99999872 |
| PDK1 | 11.7958609 | -0.091517535 | 0.54097044 | -0.1691729 | 0.86566065 | 0.99999872 |
| RAPGEF4 | 7.20472698 | 0.265728957 | 0.7154408 | 0.37141991 | 0.7103248 | 0.99999872 |
| MAP3K20 | 88.3691286 | -0.078976523 | 0.55571082 | -0.142118 | 0.88698678 | 0.99999872 |
| MAP3K20-AS | 4.54017196 | 1.265251256 | 0.82960075 | 1.52513272 | 0.127226 | NA |
| SP3 | 47.2741244 | -0.063458163 | 0.3341495 | -0.1899095 | 0.84938005 | 0.99999872 |
| OLA1 | 66.1700929 | -0.231350848 | 0.36411312 | -0.6353818 | 0.52517944 | 0.99999872 |
| CIR1 | 28.7349237 | -0.051907973 | 0.32836578 | -0.1580797 | 0.87439398 | 0.99999872 |
| SCRN3 | 10.5681858 | -0.142069937 | 0.62682867 | -0.2266488 | 0.82069688 | 0.99999872 |
| GPR155 | 19.7379729 | 0.288216313 | 0.53915715 | 0.53456828 | 0.59294842 | 0.99999872 |
| AC010894.3 | 87.2314681 | 0.693114879 | 0.52212224 | 1.32749541 | 0.18434486 | 0.99999872 |
| WIPF1 | 26.5768979 | -0.299408374 | 0.41057254 | -0.729246 | 0.46585121 | 0.99999872 |
| CHN1 | 42.1560463 | 0.643826794 | 0.43823126 | 1.46914848 | 0.14179252 | 0.99999872 |
| ATF2 | 55.2786423 | 0.368664797 | 0.26085983 | 1.41326778 | 0.157577 | 0.99999872 |
| ATP5MC3 | 55.1951033 | 1.297694554 | 0.38027318 | 3.41253239 | 0.00064362 | 0.23254757 |
| LNPK | 21.9142581 | 0.466323311 | 0.42705532 | 1.09195058 | 0.27485482 | 0.99999872 |
| HOXD-AS2 | 4.65740293 | 0.506571202 | 0.83335085 | 0.60787266 | 0.54327193 | 0.99999872 |
| HOXD8 | 7.80812008 | 0.647558525 | 0.58400076 | 1.10883164 | 0.26750282 | 0.99999872 |
| HOXD3 | 8.52588543 | 0.001335541 | 0.73499339 | 0.00181708 | 0.99855018 | 0.99999872 |
| HAGLR | 16.3890552 | -0.268455983 | 0.59025029 | -0.4548172 | 0.64924076 | 0.99999872 |
| HOXD1 | 9.59787792 | -0.186874879 | 0.71614489 | -0.2609456 | 0.79413444 | 0.99999872 |
| MTX2 | 52.051364 | 0.140712181 | 0.33260297 | 0.42306352 | 0.67224892 | 0.99999872 |
| HNRNPA3 | 99.4499727 | -0.020790985 | 0.25813692 | -0.0805425 | 0.93580582 | 0.99999872 |
| NFE2L2 | 198.240511 | 0.136134011 | 0.24560429 | 0.5542819 | 0.57938593 | 0.99999872 |
| AGPS | 48.5862535 | 0.260866166 | 0.3019217 | 0.86401927 | 0.38757731 | 0.99999872 |
| TTC30A | 4.19825116 | 0.224218739 | 0.72616857 | 0.30876955 | 0.75749684 | NA |
| RBM45 | 12.5667263 | -0.015748214 | 0.47056972 | -0.0334663 | 0.97330276 | 0.99999872 |
| AC009948.1 | 16.2693828 | -0.420147246 | 0.69083684 | -0.6081715 | 0.54307377 | 0.99999872 |
| PRKRA | 29.1848081 | 0.290019547 | 0.3141417 | 0.92321251 | 0.35589647 | 0.99999872 |
| PJVK | 9.4049757 | -0.208259005 | 0.60569981 | -0.343832 | 0.7309726 | 0.99999872 |
| FKBP7 | 4.49664614 | 0.807792615 | 0.86887477 | 0.9296997 | 0.35252659 | NA |
| PLEKHA3 | 32.1617121 | 0.528597022 | 0.29097746 | 1.81662531 | 0.0692745 | 0.99999872 |
| TTN-AS1 | 46.3343405 | 0.095901694 | 0.40937329 | 0.23426466 | 0.81477952 | 0.99999872 |
| SESTD1 | 37.2522609 | 0.410869252 | 0.37814005 | 1.08655312 | 0.27723435 | 0.99999872 |
| CWC22 | 17.2089164 | -0.071434753 | 0.37288596 | -0.1915727 | 0.84807697 | 0.99999872 |
| UBE2E3 | 79.2265591 | 0.265108749 | 0.27325739 | 0.97017961 | 0.33195697 | 0.99999872 |
| SSFA2 | 33.2786609 | -0.488267287 | 0.45208912 | -1.0800244 | 0.28013131 | 0.99999872 |
| PDE1A | 70.8795268 | -0.140295238 | 0.99929268 | -0.1403945 | 0.88834827 | 0.99999872 |
| DNAJC10 | 51.850996 | -0.0716014 | 0.31029164 | -0.2307552 | 0.81750501 | 0.99999872 |
| NCKAP1 | 136.902616 | 0.276750194 | 0.24261368 | 1.14070319 | 0.25399346 | 0.99999872 |
| NUP35 | 9.62161953 | -0.48136177 | 0.66577312 | -0.7230117 | 0.46967268 | 0.99999872 |
| ZNF804A | 367.665875 | -0.935713962 | 0.67661084 | -1.3829426 | 0.16668247 | 0.99999872 |
| FSIP2 | 11.9630366 | 0.140136153 | 0.89154958 | 0.15718268 | 0.87510088 | 0.99999872 |
| FSIP2-AS1 | 7.53107251 | 0.300451424 | 0.70756615 | 0.42462662 | 0.67110888 | 0.99999872 |
| LINC01473 | 12.847179 | 0.134717831 | 0.48127917 | 0.27991619 | 0.77954181 | 0.99999872 |
| ZC3H15 | 34.7366899 | -0.040087417 | 0.32291063 | -0.124144 | 0.90120126 | 0.99999872 |
| ITGAV | 637.920035 | 0.544028809 | 0.42243432 | 1.28784236 | 0.19780084 | 0.99999872 |
| AC017101.1 | 34.4830407 | 1.77450324 | 0.59422735 | 2.9862362 | 0.00282434 | 0.43717075 |
| FAM171B | 6.97983059 | 0.78931171 | 0.62803014 | 1.25680547 | 0.20882409 | 0.99999872 |
| AC007319.1 | 106.681482 | 1.438550244 | 0.9562141 | 1.50442276 | 0.13247255 | 0.99999872 |
| CALCRL | 9.74559128 | 0.641417865 | 0.55775633 | 1.14999656 | 0.25014529 | 0.99999872 |
| TFPI | 116.096145 | 0.020285335 | 0.3516996 | 0.05767802 | 0.9540051 | 0.99999872 |
| LINC01090 | 11.0661106 | 2.082907819 | 0.68141124 | 3.0567559 | 0.00223746 | 0.37212934 |

| | | | | | | |
|---|---|---|---|---|---|---|
| GULP1 | 329.242428 | -0.023562188 | 0.31956438 | -0.0737322 | 0.94122347 | 0.99999872 |
| COL3A1 | 27.2206474 | -0.000386297 | 0.88468248 | -0.0004367 | 0.9996516 | 0.99999872 |
| WDR75 | 15.6275494 | -0.215447139 | 0.40052462 | -0.5379123 | 0.59063756 | 0.99999872 |
| SLC40A1 | 15.1041535 | 0.888528715 | 0.69008316 | 1.2875676 | 0.19789652 | 0.99999872 |
| ASNSD1 | 12.2816795 | 0.461693865 | 0.54738897 | 0.84344751 | 0.39897821 | 0.99999872 |
| ANKAR | 54.7345311 | -0.367114847 | 0.47269075 | -0.7766491 | 0.43736582 | 0.99999872 |
| OSGEPL1 | 5.07054743 | -0.141755072 | 0.76739137 | -0.1847233 | 0.85344606 | 0.99999872 |
| ORMDL1 | 35.0736512 | 0.555639087 | 0.30668144 | 1.81177933 | 0.0700203 | 0.99999872 |
| PMS1 | 24.6467515 | -0.267576419 | 0.39822588 | -0.6719212 | 0.50163385 | 0.99999872 |
| C2orf88 | 14.0954642 | 0.994060005 | 0.48391538 | 2.05420213 | 0.03995613 | 0.95980711 |
| HIBCH | 67.6542673 | 0.221285031 | 0.35545988 | 0.62253166 | 0.53359233 | 0.99999872 |
| INPP1 | 5.27972461 | -0.246957508 | 0.66304389 | -0.3724603 | 0.70955019 | 0.99999872 |
| MFSD6 | 125.952637 | 0.21194803 | 0.50008409 | 0.42382478 | 0.6716936 | 0.99999872 |
| NEMP2 | 6.53935328 | -0.356625424 | 0.59066192 | -0.6037725 | 0.5459949 | 0.99999872 |
| NAB1 | 26.0520639 | 0.404731814 | 0.32002574 | 1.26468521 | 0.20598419 | 0.99999872 |
| GLS | 383.665036 | 0.044484205 | 0.36569631 | 0.12164248 | 0.90318217 | 0.99999872 |
| STAT1 | 122.970851 | -0.52658259 | 0.66708452 | -0.7893791 | 0.42989046 | 0.99999872 |
| MYO1B | 336.177247 | 0.415681626 | 0.46477558 | 0.89437062 | 0.37112363 | 0.99999872 |
| NABP1 | 9.06942835 | -1.004658306 | 1.06148034 | -0.9464691 | 0.34390939 | 0.99999872 |
| SLC39A10 | 36.1739002 | 0.706546822 | 0.45981962 | 1.5365739 | 0.12439769 | 0.99999872 |
| DNAH7 | 23.2258733 | -0.006733647 | 0.60094288 | -0.0112051 | 0.99105978 | 0.99999872 |
| GTF3C3 | 13.507575 | -0.098128146 | 0.4351059 | -0.225527 | 0.82156929 | 0.99999872 |
| PGAP1 | 33.1664814 | -0.446890544 | 0.60977903 | -0.7328729 | 0.46363594 | 0.99999872 |
| ANKRD44 | 197.100118 | 0.178611967 | 0.34673945 | 0.51511868 | 0.60647009 | 0.99999872 |
| SF3B1 | 62.5449236 | 0.288690777 | 0.24819171 | 1.16317654 | 0.24475788 | 0.99999872 |
| COQ10B | 17.4134243 | 0.341925883 | 0.43617675 | 0.78391589 | 0.43308947 | 0.99999872 |
| HSPD1 | 36.750388 | 0.071789884 | 0.42357812 | 0.1694844 | 0.86541564 | 0.99999872 |
| MOB4 | 15.7681701 | 0.499470233 | 0.43608003 | 1.1453637 | 0.25205853 | 0.99999872 |
| AC011997.1 | 5.78465709 | -0.122046898 | 1.05998904 | -0.1151398 | 0.90833434 | 0.99999872 |
| PLCL1 | 86.1006576 | -0.058391069 | 0.65763392 | -0.0887896 | 0.9292491 | 0.99999872 |
| FTCDNL1 | 7.44971826 | -0.466483395 | 0.68436811 | -0.6816264 | 0.4954752 | 0.99999872 |
| C2orf69 | 8.50370028 | 0.484543374 | 0.52680886 | 0.91977074 | 0.35769258 | 0.99999872 |
| TYW5 | 10.6183194 | 0.144866538 | 0.48057809 | 0.30144224 | 0.76307729 | 0.99999872 |
| MAIP1 | 7.19879714 | 0.757251197 | 0.56577501 | 1.33843167 | 0.18075576 | 0.99999872 |
| SPATS2L | 231.522711 | -0.518439229 | 0.4152737 | -1.2484278 | 0.21187443 | 0.99999872 |
| KCTD18 | 8.23277861 | -0.064256343 | 0.53825988 | -0.1193779 | 0.90497596 | 0.99999872 |
| AOX1 | 33.5391043 | -1.254667472 | 0.58778986 | -2.1345511 | 0.0327977 | 0.92959829 |
| BZW1 | 19.5646721 | -0.530308291 | 0.37625522 | -1.4094377 | 0.15870578 | 0.99999872 |
| CLK1 | 43.144963 | 0.439756824 | 0.43645583 | 1.00756318 | 0.31366421 | 0.99999872 |
| PPIL3 | 7.9828508 | -0.422458589 | 0.52745202 | -0.8009422 | 0.4231651 | 0.99999872 |
| NIF3L1 | 5.99877666 | -0.176568738 | 0.55232698 | -0.3196815 | 0.74920976 | 0.99999872 |
| ORC2 | 29.5486018 | -0.562089886 | 0.42866325 | -1.3112621 | 0.18976922 | 0.99999872 |
| AC005037.1 | 10.062001 | -0.336067566 | 0.5595526 | -0.6006005 | 0.54810611 | 0.99999872 |
| FAM126B | 56.6575345 | -0.372724185 | 0.33010253 | -1.1291164 | 0.25884873 | 0.99999872 |
| NDUFB3 | 31.8203402 | 1.104860867 | 0.39187331 | 2.81943382 | 0.00481084 | 0.5356861 |
| CFLAR | 146.236759 | -0.498988961 | 0.51310078 | -0.972497 | 0.33080336 | 0.99999872 |
| CFLAR-AS1 | 9.62682601 | 1.530089834 | 0.99907994 | 1.5314989 | 0.12564614 | 0.99999872 |
| CASP8 | 7.23580157 | 0.004062929 | 0.9659576 | 0.00420612 | 0.99664402 | 0.99999872 |
| TRAK2 | 40.5297859 | -0.175187007 | 0.42824459 | -0.4090817 | 0.68247974 | 0.99999872 |
| STRADB | 18.0883673 | 0.425372717 | 0.66463 | 0.64001432 | 0.52216329 | 0.99999872 |
| ALS2 | 28.8641398 | 0.173252462 | 0.38861409 | 0.44582136 | 0.65572629 | 0.99999872 |
| SUMO1 | 34.0586494 | 0.305783405 | 0.3166131 | 0.96579519 | 0.33414668 | 0.99999872 |
| NOP58 | 29.4203118 | -0.275055282 | 0.34541668 | -0.7962999 | 0.42585772 | 0.99999872 |
| BMPR2 | 214.898827 | -0.146680129 | 0.35381778 | -0.414564 | 0.67846107 | 0.99999872 |
| FAM117B | 11.0596996 | -0.134936689 | 0.48676585 | -0.2772107 | 0.78161834 | 0.99999872 |
| ICA1L | 28.533683 | 0.061391623 | 0.32683525 | 0.18783661 | 0.85100474 | 0.99999872 |
| WDR12 | 12.3651063 | 0.373808975 | 0.54062809 | 0.69143461 | 0.48929246 | 0.99999872 |
| CARF | 25.9393415 | -0.045689021 | 0.3968889 | -0.1151179 | 0.90835166 | 0.99999872 |
| NBEAL1 | 255.679446 | 0.19400377 | 0.18931455 | 1.02476946 | 0.30547198 | 0.99999872 |
| CYP20A1 | 50.7886426 | -0.442070524 | 0.36839286 | -1.1999975 | 0.23014029 | 0.99999872 |
| ABI2 | 93.3989663 | 0.06281988 | 0.24964065 | 0.25164123 | 0.80131839 | 0.99999872 |

| | | | | | | |
|---|---|---|---|---|---|---|
| RAPH1 | 48.3588339 | 0.088111808 | 0.31970042 | 0.27560742 | 0.78284961 | 0.99999872 |
| PARD3B | 1868.53506 | -0.267040244 | 0.30905513 | -0.8640538 | 0.38755836 | 0.99999872 |
| INO80D | 124.917304 | 0.427180746 | 0.81047296 | 0.52707588 | 0.5981409 | 0.99999872 |
| NDUFS1 | 48.1702332 | -0.037411023 | 0.29753286 | -0.1257375 | 0.89993976 | 0.99999872 |
| EEF1B2 | 23.4921814 | 0.415535947 | 0.41861402 | 0.99264698 | 0.32088202 | 0.99999872 |
| ZDBF2 | 19.2890622 | -0.020140929 | 0.37805406 | -0.0532753 | 0.95751259 | 0.99999872 |
| FASTKD2 | 14.1785568 | -0.199610621 | 0.57698877 | -0.3459524 | 0.72937852 | 0.99999872 |
| CPO | 4.50791788 | -0.328576536 | 0.78186742 | -0.4202458 | 0.67430587 | NA |
| KLF7 | 374.10306 | 0.097612597 | 0.32887385 | 0.29680863 | 0.76661262 | 0.99999872 |
| AC007879.1 | 8.0324921 | 1.816525948 | 0.76999248 | 2.35914766 | 0.01831697 | 0.85282771 |
| AC009226.1 | 5.20280058 | -0.270413649 | 1.03973961 | -0.2600782 | 0.79480342 | 0.99999872 |
| CREB1 | 51.1708704 | -0.226181991 | 0.29402144 | -0.7692704 | 0.4417328 | 0.99999872 |
| METTL21A | 28.8267675 | 0.088955895 | 0.34665454 | 0.25661252 | 0.79747791 | 0.99999872 |
| CCNYL1 | 20.2817295 | -0.635866534 | 0.39612427 | -1.6052198 | 0.10844544 | 0.99999872 |
| FZD5 | 5.87740295 | -0.505525261 | 0.70372043 | -0.7183609 | 0.47253477 | 0.99999872 |
| PLEKHM3 | 28.4773521 | 0.006196186 | 0.35024177 | 0.01769117 | 0.98588523 | 0.99999872 |
| IDH1 | 8.46936813 | 0.269037317 | 0.55735471 | 0.48270395 | 0.62930596 | 0.99999872 |
| PIKFYVE | 45.2879479 | 0.324084258 | 0.28816039 | 1.12466622 | 0.2607305 | 0.99999872 |
| PTH2R | 75.7296825 | 0.666303968 | 0.73061573 | 0.91197594 | 0.36178139 | 0.99999872 |
| MAP2 | 24.5855091 | -0.100916531 | 0.72650692 | -0.1389065 | 8.90E-01 | 0.99999872 |
| RPE | 17.8228591 | 0.696092812 | 0.45281392 | 1.5372602 | 0.1242296 | 0.99999872 |
| KANSL1L | 124.875869 | 0.215917797 | 0.30495095 | 0.70804108 | 0.47891975 | 0.99999872 |
| AC007038.1 | 19.6171587 | -0.435623381 | 0.57279252 | -0.7605256 | 0.44694047 | 0.99999872 |
| LANCL1 | 53.9263082 | 0.551942685 | 0.4024911 | 1.37131649 | 0.17027632 | 0.99999872 |
| ERBB4 | 836.620568 | 0.263651316 | 0.23754732 | 1.10988967 | 0.26704657 | 0.99999872 |
| IKZF2 | 41.8526159 | 0.356112255 | 0.55867815 | 0.63741933 | 0.52385174 | 0.99999872 |
| SPAG16 | 148.122045 | 0.647874623 | 0.39044199 | 1.65933642 | 0.09704802 | 0.99999872 |
| AC068051.1 | 7.16805838 | 2.092392589 | 0.90848433 | 2.30316861 | 0.02126936 | 0.87186485 |
| BARD1 | 9.63875616 | 0.366367251 | 0.6359812 | 0.57606617 | 0.56457046 | 0.99999872 |
| AC016708.1 | 8.7184807 | 0.462477959 | 0.65887438 | 0.7019213 | 0.48272825 | 0.99999872 |
| ATIC | 23.9652256 | -0.234480673 | 0.44511556 | -0.5267861 | 0.59834216 | 0.99999872 |
| PECR | 5.84336542 | -0.205688002 | 0.63507738 | -0.3238786 | 0.74602991 | 0.99999872 |
| XRCC5 | 81.2042349 | 0.016102443 | 0.25869986 | 0.06224373 | 0.95036874 | 0.99999872 |
| SMARCAL1 | 15.6422993 | 0.027696077 | 0.44026799 | 0.06290732 | 0.94984031 | 0.99999872 |
| RPL37A | 66.9373594 | 0.61281944 | 0.38256938 | 1.60185177 | 0.10918839 | 0.99999872 |
| IGFBP2 | 1167.78975 | 0.565825025 | 0.54525925 | 1.03771742 | 0.29940163 | 0.99999872 |
| IGFBP5 | 18.7392072 | -0.865177082 | 1.00663821 | -0.8594717 | 0.39008031 | 0.99999872 |
| TNS1 | 23.6381671 | -0.501352125 | 0.89301564 | -0.5614147 | 0.57451485 | 0.99999872 |
| ARPC2 | 61.715796 | -0.251350072 | 0.36218634 | -0.69398 | 0.48769476 | 0.99999872 |
| AAMP | 14.9327103 | 0.503282578 | 0.44909743 | 1.12065344 | 0.26243541 | 0.99999872 |
| PNKD | 43.4178646 | 0.546050133 | 0.29997489 | 1.82031946 | 0.06871037 | 0.99999872 |
| TMBIM1 | 35.1811965 | 0.429668286 | 0.53577388 | 0.80195826 | 0.4225771 | 0.99999872 |
| CTDSP1 | 28.4498787 | -0.084052891 | 0.40742786 | -0.2063013 | 0.83655558 | 0.99999872 |
| USP37 | 22.7981793 | 0.273015172 | 0.43257863 | 0.63113422 | 0.52795277 | 0.99999872 |
| CNOT9 | 15.0321737 | -0.485764364 | 0.40748927 | -1.1920912 | 0.23322549 | 0.99999872 |
| ZNF142 | 6.68505588 | 0.1612325 | 0.54703881 | 0.29473686 | 0.76819491 | 0.99999872 |
| RNF25 | 5.73462075 | -1.060407212 | 0.63632683 | -1.6664506 | 0.0956237 | 0.99999872 |
| STK36 | 7.31967983 | -1.398986785 | 0.55641699 | -2.5142776 | 0.01192765 | 0.72670507 |
| TTLL4 | 4.230667 | -0.302299856 | 0.7447787 | -0.4058922 | 0.68482182 | NA |
| CYP27A1 | 13.1730378 | 0.03587378 | 0.54004377 | 0.06642754 | 0.94703744 | 0.99999872 |
| CNPPD1 | 13.8786314 | -0.693069722 | 0.51334559 | -1.3501036 | 0.17698276 | 0.99999872 |
| RETREG2 | 22.7337811 | -0.023493418 | 0.37326607 | -0.0629401 | 0.94981417 | 0.99999872 |
| ZFAND2B | 11.9438486 | 0.065595116 | 0.43447633 | 0.15097512 | 0.87999534 | 0.99999872 |
| ANKZF1 | 9.92670352 | -0.337645538 | 0.55411744 | -0.6093393 | 0.54229956 | 0.99999872 |
| STK16 | 4.61822305 | 0.3715121 | 0.69875344 | 0.53167838 | 0.59494877 | 0.99999872 |
| TUBA4A | 5.75649311 | 0.313933623 | 0.65940247 | 0.47608803 | 0.63401166 | 0.99999872 |
| DNAJB2 | 23.3759363 | -0.495151981 | 0.48894408 | -1.0126966 | 0.31120513 | 0.99999872 |
| DNPEP | 10.7049407 | -0.411574795 | 0.61426507 | -0.670028 | 0.50283994 | 0.99999872 |
| GMPPA | 5.54053156 | -0.805808052 | 0.68702209 | -1.1728998 | 0.240836 | 0.99999872 |
| CHPF | 32.3545035 | 0.328823394 | 0.59648527 | 0.55126825 | 0.5814498 | 0.99999872 |
| OBSL1 | 15.3615789 | -0.04822591 | 0.46988273 | -0.1026339 | 0.91825352 | 0.99999872 |

| | | | | | | |
|---|---|---|---|---|---|---|
| EPHA4 | 29.349914 | -0.783388586 | 0.94601561 | -0.8280927 | 0.40761802 | 0.99999872 |
| SGPP2 | 10.4998484 | 0.284473481 | 0.8780182 | 0.32399497 | 0.74594184 | 0.99999872 |
| FARSB | 17.9887062 | 0.046499446 | 0.40774064 | 0.11404173 | 0.90920472 | 0.99999872 |
| ACSL3 | 70.6017522 | 0.3827895 | 0.41130006 | 0.93068186 | 0.35201816 | 0.99999872 |
| AP1S3 | 53.809792 | 0.135342721 | 0.30713034 | 0.44066867 | 0.65945288 | 0.99999872 |
| WDFY1 | 46.3081704 | -0.145300093 | 0.38353618 | -0.3788432 | 0.70480427 | 0.99999872 |
| MRPL44 | 7.90975681 | 0.672859378 | 0.67397491 | 0.99834484 | 0.31811217 | 0.99999872 |
| SERPINE2 | 9.17622633 | -0.569275937 | 0.83042282 | -0.6855254 | 0.49301244 | 0.99999872 |
| CUL3 | 90.1461077 | -0.106214525 | 0.21071979 | -0.5040558 | 0.61422219 | 0.99999872 |
| IRS1 | 13.1052569 | 0.534983654 | 0.56896135 | 0.94028119 | 0.34707335 | 0.99999872 |
| RHBDD1 | 58.0915933 | -0.211764705 | 0.38634392 | -0.5481249 | 0.58360618 | 0.99999872 |
| COL4A4 | 600.731885 | 0.00421372 | 0.49477725 | 0.0085164 | 0.99320498 | 0.99999872 |
| COL4A3 | 1303.90159 | -0.062187792 | 0.29195637 | -0.2130037 | 0.83132406 | 0.99999872 |
| AC097662.1 | 149.819499 | 1.466071895 | 0.47305411 | 3.09916324 | 0.00194068 | 0.36957035 |
| MFF | 27.6275975 | 0.437013679 | 0.35689215 | 1.22449788 | 0.22076447 | 0.99999872 |
| AGFG1 | 77.9189516 | -0.119199952 | 0.27006034 | -0.4413827 | 0.65893599 | 0.99999872 |
| TRIP12 | 123.699673 | -0.10265733 | 0.21770443 | -0.4715445 | 0.63725194 | 0.99999872 |
| FBXO36 | 9.5854807 | 0.360644565 | 0.4669931 | 0.77226958 | 0.43995478 | 0.99999872 |
| AC009950.1 | 3.98044971 | 0.17224013 | 1.09124146 | 0.1578387 | 0.87458391 | NA |
| SP110 | 21.9546689 | -1.218403961 | 1.18469837 | -1.0284508 | 0.30373783 | 0.99999872 |
| SP140L | 8.75674871 | -0.492291148 | 0.84664188 | -0.5814633 | 0.56092827 | 0.99999872 |
| SP100 | 81.8738936 | -0.638423154 | 0.54074038 | -1.1806463 | 0.23774325 | 0.99999872 |
| AC010149.1 | 4.03430244 | -0.470870114 | 0.75032653 | -0.6275536 | 0.53029641 | NA |
| CAB39 | 52.0727765 | -0.076645231 | 0.28498927 | -0.2689408 | 0.78797527 | 0.99999872 |
| ITM2C | 40.6257141 | -0.296881068 | 0.50746786 | -0.5850244 | 0.55853133 | 0.99999872 |
| PSMD1 | 79.2570963 | 0.013079198 | 0.21277639 | 0.06146922 | 0.95098553 | 0.99999872 |
| ARMC9 | 55.975054 | -0.005655991 | 0.25865782 | -0.0218667 | 0.9825543 | 0.99999872 |
| B3GNT7 | 10.7275675 | 0.262037938 | 0.74372306 | 0.35233268 | 0.72458878 | 0.99999872 |
| NCL | 60.6641832 | -0.228214972 | 0.38498258 | -0.592793 | 0.5533197 | 0.99999872 |
| PTMA | 316.880455 | -0.31552584 | 0.54871954 | -0.5750221 | 0.56527638 | 0.99999872 |
| PDE6D | 13.7320441 | 0.239814889 | 0.46014462 | 0.52117287 | 0.60224635 | 0.99999872 |
| COPS7B | 11.3907932 | 0.241392214 | 0.51882618 | 0.46526606 | 0.64174093 | 0.99999872 |
| DIS3L2 | 57.9804258 | -0.04212288 | 0.32679344 | -0.1288976 | 0.8974387 | 0.99999872 |
| EIF4E2 | 17.6836856 | -0.112910837 | 0.37993446 | -0.297185 | 0.76632524 | 0.99999872 |
| EFHD1 | 23.2766146 | 0.602741099 | 0.54577062 | 1.1043854 | 0.26942599 | 0.99999872 |
| GIGYF2 | 76.0999685 | 0.087527626 | 0.23791127 | 0.3679003 | 0.71294758 | 0.99999872 |
| ATG16L1 | 8.18242468 | 0.042454753 | 0.4950064 | 0.08576607 | 0.93165238 | 0.99999872 |
| DGKD | 36.3535591 | 0.384895287 | 0.45745173 | 0.84138995 | 0.40012952 | 0.99999872 |
| USP40 | 51.7037527 | -0.00018301 | 0.30577279 | -0.0005985 | 0.99952245 | 0.99999872 |
| SH3BP4 | 22.2180048 | -0.805649073 | 0.39149613 | -2.0578724 | 0.03960239 | 0.95980711 |
| AGAP1 | 117.729876 | -0.415143855 | 0.30711714 | -1.3517443 | 0.17645712 | 0.99999872 |
| IQCA1 | 10.4477375 | -0.666221765 | 0.56522106 | -1.1786924 | 0.23852069 | 0.99999872 |
| COPS8 | 23.44104 | 0.543517812 | 0.35666305 | 1.52389715 | 0.12753441 | 0.99999872 |
| RAB17 | 5.98832359 | 0.628015291 | 0.99881013 | 0.62876344 | 0.52950394 | 0.99999872 |
| LRRFIP1 | 169.35037 | -0.100632637 | 0.3120662 | -0.3224721 | 0.74709508 | 0.99999872 |
| RAMP1 | 103.627863 | 0.593992102 | 0.91610324 | 0.64838992 | 0.51673279 | 0.99999872 |
| ILKAP | 35.8524601 | -0.050445928 | 0.47362817 | -0.1065096 | 0.91517807 | 0.99999872 |
| PER2 | 7.35222441 | -1.058887042 | 0.52580431 | -2.0138425 | 0.04402608 | 0.95980711 |
| TRAF3IP1 | 21.9967632 | -0.598184876 | 0.3944943 | -1.5163334 | 0.12943508 | 0.99999872 |
| ASB1 | 9.30578255 | 0.426064402 | 0.48595653 | 0.87675414 | 0.38062019 | 0.99999872 |
| HDAC4 | 50.4192597 | -0.173032112 | 0.37685244 | -0.4591508 | 0.64612586 | 0.99999872 |
| NDUFA10 | 38.0902977 | 0.426267411 | 0.36173849 | 1.17838556 | 0.23864293 | 0.99999872 |
| COPS9 | 25.4463575 | 0.611997158 | 0.62184089 | 0.98417001 | 0.32503193 | 0.99999872 |
| AC124861.1 | 12.2914266 | -0.524719877 | 0.68643014 | -0.7644185 | 0.4446179 | 0.99999872 |
| GPC1 | 16.2345747 | -0.529627038 | 0.5627278 | -0.941178 | 0.34661364 | 0.99999872 |
| ANKMY1 | 17.4032084 | -0.50151471 | 0.42081249 | -1.1917772 | 0.23334863 | 0.99999872 |
| RNPEPL1 | 24.8018235 | -0.46254173 | 0.43896054 | -1.0537205 | 0.29201089 | 0.99999872 |
| CAPN10 | 5.12179068 | 0.723330293 | 0.65745186 | 1.10020267 | 0.27124383 | 0.99999872 |
| SNED1 | 34.8590004 | -0.044759145 | 0.59666425 | -0.0750156 | 9.40E-01 | 0.99999872 |
| AC005237.1 | 5.00128849 | 0.207696542 | 0.81622434 | 0.25446012 | 0.79914011 | 0.99999872 |
| MTERF4 | 22.475983 | -0.214995042 | 0.3829712 | -0.561387 | 0.57453377 | 0.99999872 |

| | | | | | | |
|---|---|---|---|---|---|---|
| PPP1R7 | 21.5240482 | 0.017938941 | 0.34150854 | 0.05252853 | 0.95810757 | 0.99999872 |
| HDLBP | 84.925135 | 0.074105051 | 0.30048272 | 0.24662001 | 0.80520232 | 0.99999872 |
| 2-Sep | 152.732142 | 0.167276008 | 0.32573302 | 0.51353715 | 0.60757563 | 0.99999872 |
| FARP2 | 66.4635336 | 0.162188568 | 0.3406733 | 0.47608242 | 0.63401566 | 0.99999872 |
| STK25 | 22.7888448 | -0.146582229 | 0.34529275 | -0.4245158 | 0.67118968 | 0.99999872 |
| BOK | 19.3549971 | -1.034606375 | 0.60810355 | -1.7013655 | 0.08887438 | 0.99999872 |
| THAP4 | 13.7905746 | -0.011985528 | 0.42977472 | -0.0278879 | 0.97775153 | 0.99999872 |
| ATG4B | 32.0722748 | -0.1239002 | 0.46157313 | -0.2684303 | 0.78836814 | 0.99999872 |
| DTYMK | 8.53849958 | 0.660591519 | 0.60946253 | 1.08389194 | 0.27841271 | 0.99999872 |
| ING5 | 20.6924016 | 0.174850855 | 0.41680068 | 0.41950713 | 0.67484554 | 0.99999872 |
| D2HGDH | 30.5517391 | 0.330927496 | 0.40158292 | 0.82405769 | 0.40990678 | 0.99999872 |
| LINC01266 | 8.36157035 | -0.077653473 | 0.73077185 | -0.1062623 | 0.91537427 | 0.99999872 |
| CNTN4 | 166.103712 | -0.379054015 | 1.3692072 | -0.276842 | 0.78190145 | 0.99999872 |
| TRNT1 | 14.771921 | 0.316028353 | 0.45058709 | 0.70137019 | 0.48307202 | 0.99999872 |
| CRBN | 30.6834806 | 0.482544364 | 0.40948633 | 1.17841385 | 0.23863166 | 0.99999872 |
| AC034195.1 | 39.7439024 | 0.538615829 | 0.50741956 | 1.06148022 | 0.28847172 | 0.99999872 |
| SUMF1 | 130.706004 | -0.369681468 | 0.36434781 | -1.0146389 | 0.31027798 | 0.99999872 |
| ITPR1 | 27.945813 | 0.155462256 | 0.4102179 | 0.37897482 | 0.70470656 | 0.99999872 |
| AC018816.1 | 6.79896231 | -0.91966181 | 0.68894065 | -1.3348926 | 0.18191149 | 0.99999872 |
| BHLHE40 | 17.2130101 | -1.385389147 | 0.80159056 | -1.7283002 | 0.08393441 | 0.99999872 |
| ARL8B | 52.7137506 | -0.037078216 | 0.27714128 | -0.1337881 | 0.89357011 | 0.99999872 |
| AC026202.2 | 6.59884379 | 0.361459992 | 0.74839345 | 0.48298123 | 0.62910906 | 0.99999872 |
| EDEM1 | 16.6676409 | 0.378784766 | 0.4545573 | 0.83330477 | 0.40467287 | 0.99999872 |
| LMCD1-AS1 | 41.0749818 | 0.044386214 | 0.58647594 | 0.07568292 | 0.93967136 | 0.99999872 |
| LMCD1 | 10.2964249 | -0.102573403 | 0.73668638 | -0.1392362 | 0.88926351 | 0.99999872 |
| RAD18 | 11.2267602 | -0.88136376 | 0.48564692 | -1.8148242 | 0.06955094 | 0.99999872 |
| SRGAP3 | 14.9952516 | -0.005926394 | 0.44902391 | -0.0131984 | 0.98946951 | 0.99999872 |
| THUMPD3-A! | 41.8618228 | -0.552533253 | 0.33232449 | -1.6626318 | 0.09638617 | 0.99999872 |
| THUMPD3 | 9.83455962 | 0.066640217 | 0.46141607 | 0.14442543 | 0.88516453 | 0.99999872 |
| SETD5 | 126.684034 | -0.023814497 | 0.17883873 | -0.1331619 | 0.89406538 | 0.99999872 |
| MTMR14 | 15.5187608 | -0.35072117 | 0.40738146 | -0.8609159 | 0.38928437 | 0.99999872 |
| OGG1 | 9.51483039 | 0.155842334 | 0.46139442 | 0.3377638 | 0.73554119 | 0.99999872 |
| CAMK1 | 4.91844251 | 0.94690674 | 0.7024725 | 1.34796271 | 0.17767038 | 0.99999872 |
| TADA3 | 14.9703677 | -0.180434376 | 0.41217283 | -0.4377639 | 0.66155746 | 0.99999872 |
| RPUSD3 | 7.20556262 | -0.367607429 | 0.51313816 | -0.7163908 | 0.4737501 | 0.99999872 |
| JAGN1 | 11.4977909 | 0.939231879 | 0.49959696 | 1.87997915 | 0.06011092 | 0.99999872 |
| IL17RC | 7.27013497 | -0.127512398 | 0.53351236 | -0.2390055 | 0.81110131 | 0.99999872 |
| CRELD1 | 29.9646125 | 0.925361969 | 0.5362425 | 1.72564086 | 0.08441204 | 0.99999872 |
| EMC3 | 22.6385602 | 0.051072497 | 0.37201365 | 0.13728662 | 0.89080425 | 0.99999872 |
| BRK1 | 39.0024497 | 0.691645403 | 0.33910624 | 2.03961271 | 0.04138891 | 0.95980711 |
| VHL | 11.7691067 | -0.472116388 | 0.48999332 | -0.963516 | 0.33528866 | 0.99999872 |
| SEC13 | 19.6725499 | 0.082361636 | 0.38641172 | 0.21314477 | 0.83121405 | 0.99999872 |
| ATG7 | 81.3182152 | -0.797048164 | 0.3448209 | -2.3114845 | 0.02080611 | 0.8671024 |
| VGLL4 | 179.596882 | 0.155587677 | 0.33376128 | 0.46616455 | 0.64109773 | 0.99999872 |
| TAMM41 | 20.6399039 | -0.204389818 | 0.40368594 | -0.506309 | 0.61263976 | 0.99999872 |
| PPARG | 12.2112498 | -0.181711062 | 0.70708661 | -0.2569856 | 0.7971899 | 0.99999872 |
| TSEN2 | 20.5227229 | -0.93003118 | 0.47225127 | -1.9693567 | 0.04891215 | 0.97937581 |
| MKRN2OS | 4.69464948 | 0.527482812 | 1.04591981 | 0.50432433 | 0.61403348 | 0.99999872 |
| MKRN2 | 13.3716722 | 0.850022327 | 0.55207378 | 1.53968974 | 0.123636 | 0.99999872 |
| RAF1 | 70.2550682 | -0.160899139 | 0.2344736 | -0.6862143 | 0.49257798 | 0.99999872 |
| CAND2 | 21.6047166 | 0.461642819 | 0.46056122 | 1.00234844 | 0.31617534 | 0.99999872 |
| RPL32 | 94.1279647 | 0.508998754 | 0.39138945 | 1.30049177 | 0.19343248 | 0.99999872 |
| IQSEC1 | 42.4147508 | -0.157244973 | 0.58196341 | -0.2701974 | 0.78700843 | 0.99999872 |
| NUP210 | 9.63685055 | 0.014845959 | 0.47503744 | 0.03125219 | 0.97506842 | 0.99999872 |
| HDAC11 | 5.73368238 | -0.06485625 | 0.57330275 | -0.1131274 | 0.90992955 | 0.99999872 |
| XPC | 25.5497431 | -0.299017983 | 0.31436217 | -0.9511895 | 0.3415082 | 0.99999872 |
| LSM3 | 23.8356838 | 0.795189236 | 0.48227365 | 1.6488341 | 0.09918163 | 0.99999872 |
| SLC6A6 | 138.063407 | -0.357481236 | 0.72777552 | -0.4911971 | 0.62328704 | 0.99999872 |
| CCDC174 | 14.6317804 | -0.12935698 | 0.63179744 | -0.2047444 | 0.83777184 | 0.99999872 |
| FGD5 | 6.04563543 | 0.01439219 | 0.88587309 | 0.01624633 | 0.98703787 | 0.99999872 |
| FGD5-AS1 | 49.2864844 | -0.208869835 | 0.30559862 | -0.6834777 | 0.49430505 | 0.99999872 |

| | | | | | | |
|---|---|---|---|---|---|---|
| NR2C2 | 47.5819859 | -0.164128376 | 0.36610083 | -0.4483147 | 0.65392611 | 0.99999872 |
| MRPS25 | 16.0340482 | -0.074025306 | 0.46164027 | -0.1603528 | 0.87260318 | 0.99999872 |
| RBSN | 13.0123763 | -0.021564101 | 0.42941099 | -0.0502179 | 0.95994878 | 0.99999872 |
| CAPN7 | 63.1238263 | 0.174853296 | 0.27529158 | 0.63515671 | 0.52532621 | 0.99999872 |
| SH3BP5-AS1 | 4.87745665 | -0.02843341 | 0.76057813 | -0.0373839 | 0.97017887 | 0.99999872 |
| SH3BP5 | 73.6476457 | 0.288533411 | 0.48666322 | 0.59288107 | 0.55326075 | 0.99999872 |
| METTL6 | 16.0642792 | -0.66214947 | 0.42805589 | -1.5468762 | 0.1218931 | 0.99999872 |
| EAF1 | 7.8425962 | -0.293464459 | 0.57332795 | -0.5118614 | 0.608748 | 0.99999872 |
| COLQ | 9.89218247 | -0.187708496 | 0.53960342 | -0.3478638 | 0.72794247 | 0.99999872 |
| HACL1 | 19.2243659 | 0.168194956 | 0.37352917 | 0.450286 | 0.65250423 | 0.99999872 |
| BTD | 16.1103662 | 0.406378777 | 0.3916756 | 1.03753918 | 0.29948465 | 0.99999872 |
| ANKRD28 | 103.184834 | 0.206913601 | 0.24489972 | 0.84489114 | 0.39817162 | 0.99999872 |
| DPH3 | 8.27530463 | 0.459240658 | 0.65623366 | 0.69981271 | 0.48404427 | 0.99999872 |
| OXNAD1 | 15.1624855 | -0.252388964 | 0.46525761 | -0.5424714 | 0.58749378 | 0.99999872 |
| RFTN1 | 8.66015457 | 1.216123353 | 0.98285046 | 1.23734322 | 0.21595969 | 0.99999872 |
| PLCL2 | 44.6125332 | -0.194598202 | 0.45730866 | -0.4255292 | 0.67045091 | 0.99999872 |
| TBC1D5 | 268.402748 | 0.062455145 | 0.24343175 | 0.25656122 | 0.79751752 | 0.99999872 |
| RAB5A | 49.3403308 | 0.197887709 | 0.29302944 | 0.67531683 | 0.4994745 | 0.99999872 |
| KAT2B | 54.2877477 | 0.856782704 | 0.49730184 | 1.72286252 | 0.08491339 | 0.99999872 |
| ZNF385D | 10.0156974 | -1.963511993 | 0.61725608 | -3.1810331 | 0.00146751 | 0.32751082 |
| UBE2E2 | 214.786133 | 0.355360885 | 0.34866245 | 1.01921181 | 0.30810242 | 0.99999872 |
| UBE2E1 | 77.7970632 | 0.164294037 | 0.36077365 | 0.45539367 | 0.64882605 | 0.99999872 |
| NKIRAS1 | 9.03758962 | -0.079059387 | 0.59410246 | -0.1330737 | 0.89413513 | 0.99999872 |
| RPL15 | 152.334105 | 0.706862568 | 0.46642017 | 1.51550599 | 0.12964431 | 0.99999872 |
| NR1D2 | 30.3054682 | -0.166871186 | 0.37344241 | -0.4468458 | 0.65498637 | 0.99999872 |
| THRB | 28.120401 | 0.053286024 | 0.70086613 | 0.07602882 | 0.93939617 | 0.99999872 |
| TOP2B | 55.4684864 | -0.032487978 | 0.29017094 | -0.1119615 | 0.91085392 | 0.99999872 |
| NGLY1 | 28.8812898 | -0.093363704 | 0.39749579 | -0.2348797 | 0.81430208 | 0.99999872 |
| OXSM | 3.84307895 | 0.405867012 | 0.70301771 | 0.57732117 | 0.56372251 | NA |
| NEK10 | 19.6882066 | -0.29961903 | 1.13419187 | -0.2641696 | 0.79164922 | 0.99999872 |
| SLC4A7 | 34.0936054 | -0.387813538 | 0.94919977 | -0.4085689 | 0.68285603 | 0.99999872 |
| CMC1 | 40.5768372 | 0.143469548 | 0.43225184 | 0.33191194 | 0.73995575 | 0.99999872 |
| AZI2 | 33.8231238 | 0.089616572 | 0.4567208 | 0.19621741 | 0.84444 | 0.99999872 |
| RBMS3 | 39.4438729 | -0.518935655 | 0.47663729 | -1.0887433 | 0.2762671 | 0.99999872 |
| RBMS3-AS3 | 16.7385931 | -0.122743457 | 0.62768739 | -0.1955487 | 0.84496341 | 0.99999872 |
| TGFBR2 | 38.0735182 | -0.080076434 | 0.51458392 | -0.1556139 | 0.87633733 | 0.99999872 |
| STT3B | 77.136643 | 0.363556757 | 0.32953466 | 1.10324284 | 0.26992172 | 0.99999872 |
| OSBPL10 | 106.049346 | -0.233471279 | 0.33286611 | -0.701397 | 0.48305532 | 0.99999872 |
| GPD1L | 19.0922931 | 0.827660623 | 0.715044 | 1.15749608 | 0.24706974 | 0.99999872 |
| CMTM8 | 5.77650554 | -0.893945868 | 0.63050751 | -1.4178195 | 0.15624346 | 0.99999872 |
| CMTM7 | 13.7010277 | 0.029177948 | 0.41981089 | 0.0695026 | 0.94458956 | 0.99999872 |
| CMTM6 | 37.2677378 | 0.485900689 | 0.32355034 | 1.5017777 | 0.13315453 | 0.99999872 |
| DYNC1LI1 | 22.9799532 | -0.198173152 | 0.32639402 | -0.6071593 | 0.54374523 | 0.99999872 |
| CNOT10 | 25.6317548 | -0.12737185 | 0.34471183 | -0.3695024 | 0.71175328 | 0.99999872 |
| GLB1 | 28.5768902 | 0.214742973 | 0.31589129 | 0.67980023 | 0.49663096 | 0.99999872 |
| CRTAP | 26.3650132 | 0.247790157 | 0.52035453 | 0.47619487 | 0.63393556 | 0.99999872 |
| FBXL2 | 63.2925635 | 0.574770269 | 0.28064655 | 2.04802186 | 0.04055786 | 0.95980711 |
| UBP1 | 29.5279131 | 0.397664209 | 0.32665716 | 1.21737485 | 0.22346163 | 0.99999872 |
| CLASP2 | 88.3354767 | 0.15122414 | 0.2926073 | 0.51681602 | 0.60528459 | 0.99999872 |
| PDCD6IP | 89.3454058 | 0.058390007 | 0.2143803 | 0.27236647 | 0.78534025 | 0.99999872 |
| LINC01811 | 34.6682238 | -1.018878763 | 2.10428308 | -0.4841928 | 0.62824903 | 0.99999872 |
| STAC | 21.4646641 | -0.006277003 | 0.62440487 | -0.0100528 | 0.99197918 | 0.99999872 |
| TRANK1 | 17.3621989 | -0.128538484 | 0.9447268 | -0.1360589 | 0.89177472 | 0.99999872 |
| EPM2AIP1 | 32.8968552 | 0.392237256 | 0.34820601 | 1.12645171 | 0.25997437 | 0.99999872 |
| MLH1 | 10.650378 | -0.271356004 | 0.47029641 | -0.5769893 | 0.56394667 | 0.99999872 |
| LRRFIP2 | 91.3382009 | 0.145708006 | 0.26977897 | 0.54010141 | 0.5891271 | 0.99999872 |
| GOLGA4 | 233.422779 | -0.217917566 | 0.2957821 | -0.7367503 | 0.46127419 | 0.99999872 |
| ITGA9 | 51.5950212 | 0.393364236 | 0.4472511 | 0.87951542 | 0.37912188 | 0.99999872 |
| CTDSPL | 587.781206 | -0.095296203 | 0.44767202 | -0.2128706 | 0.83142791 | 0.99999872 |
| VILL | 14.5467378 | 0.274036324 | 0.79721839 | 0.3437406 | 0.73104138 | 0.99999872 |
| PLCD1 | 11.1022304 | 0.005660086 | 0.58066963 | 0.00974751 | 0.99222273 | 0.99999872 |

| | | | | | | |
|---|---|---|---|---|---|---|
| ACAA1 | 19.1733723 | 0.552199409 | 0.48296533 | 1.14335207 | 0.25289244 | 0.99999872 |
| MYD88 | 5.00480661 | -0.301205328 | 1.29702186 | -0.2322284 | 0.81636061 | 0.99999872 |
| OXSR1 | 47.4627728 | 0.136282813 | 0.26204555 | 0.52007299 | 0.6030127 | 0.99999872 |
| XYLB | 7.44910141 | -0.259402552 | 0.83488535 | -0.3107044 | 0.75602535 | 0.99999872 |
| ACVR2B | 14.5930556 | 0.203086801 | 0.59264631 | 0.34267792 | 0.73184078 | 0.99999872 |
| EXOG | 21.1977786 | 0.198781123 | 0.45750386 | 0.43449059 | 0.66393223 | 0.99999872 |
| WDR48 | 32.3171572 | 0.257158362 | 0.4359575 | 0.58987025 | 0.55527764 | 0.99999872 |
| GORASP1 | 10.1376291 | 0.218567414 | 0.54609667 | 0.40023576 | 0.68898288 | 0.99999872 |
| TTC21A | 4.60288742 | 1.551918768 | 1.04003597 | 1.49217797 | 0.1356525 | 0.99999872 |
| CSRNP1 | 8.89259382 | -0.342377941 | 0.70143625 | -0.4881098 | 0.62547204 | 0.99999872 |
| SLC25A38 | 9.7580045 | -0.11578516 | 0.49039339 | -0.2361067 | 0.81334988 | 0.99999872 |
| RPSA | 61.3542276 | 0.402996354 | 0.47706028 | 0.84474934 | 0.39825081 | 0.99999872 |
| MYRIP | 31.7950657 | 0.480482849 | 0.7383226 | 0.6507763 | 0.5151909 | 0.99999872 |
| EIF1B | 27.770972 | 0.627065742 | 0.31596607 | 1.98459836 | 0.04718918 | 0.96950623 |
| ENTPD3-AS1 | 7.4194874 | -0.032023649 | 0.63804229 | -0.0501905 | 0.9599706 | 0.99999872 |
| RPL14 | 101.708213 | 0.514442094 | 0.5983059 | 0.85983122 | 0.38988209 | 0.99999872 |
| ZNF621 | 20.3797091 | -0.575852474 | 0.4319759 | -1.3330662 | 0.18251008 | 0.99999872 |
| CTNNB1 | 85.4996685 | 0.289535139 | 0.25917749 | 1.11713072 | 0.26393844 | 0.99999872 |
| ULK4 | 57.9557292 | -0.449379684 | 0.41639423 | -1.0792169 | 0.28049105 | 0.99999872 |
| TRAK1 | 74.260198 | 0.252138154 | 0.24803202 | 1.01655484 | 0.30936525 | 0.99999872 |
| SEC22C | 29.2082104 | 0.046163649 | 0.3897306 | 0.11845015 | 0.90571099 | 0.99999872 |
| SS18L2 | 8.12999909 | 0.695759646 | 0.53853001 | 1.29196078 | 0.19637073 | 0.99999872 |
| NKTR | 87.2761344 | -0.049426879 | 0.23323816 | -0.2119159 | 0.83217263 | 0.99999872 |
| AC006059.1 | 14.3907511 | 0.712072877 | 0.47587799 | 1.49633498 | 0.13456638 | 0.99999872 |
| AC099329.3 | 6.77822333 | 0.685614682 | 0.64625779 | 1.06089968 | 0.2887355 | 0.99999872 |
| POMGNT2 | 4.72818792 | 0.44767869 | 0.65890337 | 0.67942996 | 0.49686547 | 0.99999872 |
| SNRK | 40.6859792 | -0.102819612 | 0.32352761 | -0.3178078 | 0.75063071 | 0.99999872 |
| ANO10 | 57.1164905 | -0.123281786 | 0.36407682 | -0.3386148 | 0.73489996 | 0.99999872 |
| ABHD5 | 15.2871254 | -0.047483131 | 0.58422632 | -0.0812752 | 0.93522307 | 0.99999872 |
| TCAIM | 71.7207794 | 0.449503412 | 0.40285342 | 1.11579892 | 0.26450822 | 0.99999872 |
| ZNF445 | 14.0650075 | -0.406141793 | 0.45613184 | -0.8904044 | 0.37324879 | 0.99999872 |
| AC099669.1 | 20.1418602 | 0.034564077 | 0.46972202 | 0.07358411 | 0.94134132 | 0.99999872 |
| ZNF197 | 18.1944651 | -0.181744641 | 0.48073662 | -0.3780545 | 0.70539011 | 0.99999872 |
| KIAA1143 | 15.1621044 | 0.067892607 | 0.45058365 | 0.15067703 | 0.88023049 | 0.99999872 |
| ZDHHC3 | 22.5771882 | 0.33494706 | 0.3906674 | 0.85737141 | 0.39123966 | 0.99999872 |
| EXOSC7 | 7.55634814 | 0.319109123 | 0.67950572 | 0.46961948 | 0.6386269 | 0.99999872 |
| LARS2 | 18.7732424 | -0.428290745 | 0.41652593 | -1.0282451 | 0.30383454 | 0.99999872 |
| LIMD1 | 17.4547538 | -0.875780518 | 0.67197921 | -1.3032851 | 0.19247743 | 0.99999872 |
| SACM1L | 31.3539516 | 0.145470075 | 0.33004493 | 0.44075841 | 0.65938791 | 0.99999872 |
| LZTFL1 | 8.19027616 | 0.124295342 | 0.57001425 | 0.21805655 | 0.82738505 | 0.99999872 |
| FYCO1 | 16.1782944 | 0.054018 | 0.37435672 | 0.14429552 | 0.88526711 | 0.99999872 |
| LTF | 33.6247452 | 1.696899447 | 1.0347979 | 1.63983658 | 0.10103915 | 0.99999872 |
| LRRC2 | 239.674997 | -0.080827179 | 0.53575408 | -0.1508662 | 0.88008127 | 0.99999872 |
| ALS2CL | 549.638089 | -0.034941733 | 0.58888763 | -0.0593351 | 0.95268517 | 0.99999872 |
| TMIE | 8.43223013 | -0.354742376 | 0.59873998 | -0.5924815 | 0.55352819 | 0.99999872 |
| MYL3 | 8.9389579 | 1.332968215 | 0.76345575 | 1.74596656 | 0.08081676 | 0.99999872 |
| PTH1R | 412.133291 | 0.430021656 | 0.6374214 | 0.67462695 | 0.4999128 | 0.99999872 |
| CCDC12 | 39.4213043 | -0.435838514 | 0.32414822 | -1.3445655 | 0.1787656 | 0.99999872 |
| NBEAL2 | 4.99891797 | -1.22721533 | 0.64904965 | -1.8907881 | 0.05865264 | 0.99999872 |
| SETD2 | 106.148653 | -0.148343891 | 0.24170824 | -0.6137312 | 0.53939295 | 0.99999872 |
| KIF9-AS1 | 26.4226569 | 0.046667979 | 0.40882616 | 0.11415116 | 0.90911797 | 0.99999872 |
| KIF9 | 5.43306867 | 0.729176539 | 0.91583522 | 0.79618749 | 0.42592307 | 0.99999872 |
| KLHL18 | 14.9364997 | -0.254300586 | 0.46685035 | -0.5447154 | 0.58594925 | 0.99999872 |
| PTPN23 | 15.2848171 | -0.147550569 | 0.48315057 | -0.3053925 | 0.76006722 | 0.99999872 |
| SCAP | 34.8731611 | -0.156499583 | 0.38254681 | -0.4090992 | 0.68246688 | 0.99999872 |
| ELP6 | 11.4484242 | -0.106676393 | 0.53484086 | -0.1994545 | 0.84190727 | 0.99999872 |
| SMARCC1 | 65.6131788 | -0.270228254 | 0.28154265 | -0.9598129 | 0.33714941 | 0.99999872 |
| DHX30 | 24.9920159 | -0.110957031 | 0.34125762 | -0.3251416 | 0.74507394 | 0.99999872 |
| MAP4 | 165.381818 | -0.130514273 | 0.27432369 | -0.4757674 | 0.63424008 | 0.99999872 |
| ZNF589 | 4.31874501 | 0.676329681 | 0.74826968 | 0.9038582 | 0.3660706 | NA |
| NME6 | 6.9266103 | -0.670547612 | 0.61480902 | -1.09066 | 0.27542251 | 0.99999872 |

| | | | | | | |
|---|---|---|---|---|---|---|
| PLXNB1 | 39.6685485 | -0.270006354 | 0.39421657 | -0.6849188 | 0.49339515 | 0.99999872 |
| TMA7 | 63.6530016 | 0.356663995 | 0.45412958 | 0.78537936 | 0.43223118 | 0.99999872 |
| SHISA5 | 32.3052466 | -0.139065709 | 0.50783048 | -0.2738428 | 0.78420545 | 0.99999872 |
| PFKFB4 | 6.73994125 | -0.792256453 | 0.81790652 | -0.9686394 | 0.33272515 | 0.99999872 |
| COL7A1 | 18.3464045 | -0.92677304 | 0.6352906 | -1.4588175 | 0.14461534 | 0.99999872 |
| UQCRC1 | 21.9346605 | 0.413566166 | 0.50335945 | 0.82161201 | 0.41129775 | 0.99999872 |
| SLC26A6 | 5.9618994 | -0.562028876 | 0.62552013 | -0.8984985 | 0.36891987 | 0.99999872 |
| NCKIPSD | 13.2647845 | 0.248938826 | 0.44173135 | 0.56355255 | 0.57305869 | 0.99999872 |
| IP6K2 | 63.516508 | -0.171629064 | 0.23886364 | -0.7185232 | 0.47243476 | 0.99999872 |
| PRKAR2A | 63.8398166 | -0.330822679 | 0.3700215 | -0.8940634 | 0.37128797 | 0.99999872 |
| SLC25A20 | 10.1298732 | 0.794136855 | 0.53052173 | 1.4968979 | 0.13441983 | 0.99999872 |
| ARIH2 | 54.1114097 | -0.007511985 | 0.24721792 | -0.0303861 | 0.97575914 | 0.99999872 |
| P4HTM | 42.5327134 | 0.56085012 | 0.36812933 | 1.52351383 | 0.1276302 | 0.99999872 |
| WDR6 | 29.4864565 | -0.731321565 | 0.31611505 | -2.3134664 | 0.02069701 | 0.8671024 |
| DALRD3 | 7.8962573 | 0.488826215 | 0.51768723 | 0.9442501 | 0.34504186 | 0.99999872 |
| NDUFAF3 | 24.0046032 | 0.63224558 | 0.52350206 | 1.20772318 | 0.22715376 | 0.99999872 |
| IMPDH2 | 6.31677605 | -0.802234412 | 0.77506711 | -1.0350515 | 0.30064484 | 0.99999872 |
| QRICH1 | 34.6505547 | -0.350918469 | 0.3563547 | -0.9847449 | 0.3247494 | 0.99999872 |
| QARS | 8.07115481 | 0.687979106 | 0.58675761 | 1.17250989 | 0.2409924 | 0.99999872 |
| USP19 | 6.19140004 | -0.058968105 | 0.63935606 | -0.0922305 | 0.92651494 | 0.99999872 |
| LAMB2 | 108.835419 | 0.106699671 | 0.64887586 | 0.16443773 | 0.86938657 | 0.99999872 |
| C3orf62 | 5.14653435 | -0.33629753 | 0.70623305 | -0.4761849 | 0.63394264 | 0.99999872 |
| USP4 | 25.4333683 | -0.276870377 | 0.33701996 | -0.8215252 | 0.4113472 | 0.99999872 |
| RHOA | 125.100458 | 0.128949423 | 0.44147306 | 0.292089 | 0.77021858 | 0.99999872 |
| TCTA | 13.6931123 | 0.126216505 | 0.40060819 | 0.31506222 | 0.75271441 | 0.99999872 |
| DAG1 | 158.392221 | -0.031119677 | 0.62391884 | -0.0498778 | 0.9602198 | 0.99999872 |
| APEH | 13.438829 | -0.16951941 | 0.51928044 | -0.3264506 | 0.74408346 | 0.99999872 |
| MST1 | 5.64037638 | 0.299937849 | 0.75826106 | 0.39556014 | 0.69242955 | 0.99999872 |
| RNF123 | 9.35254691 | 0.107697821 | 0.53180171 | 0.202515 | 0.83951414 | 0.99999872 |
| IP6K1 | 47.8926987 | -0.458740375 | 0.25954186 | -1.7675005 | 0.07714444 | 0.99999872 |
| UBA7 | 7.27723386 | -1.10680088 | 0.92262391 | -1.199623 | 0.23028578 | 0.99999872 |
| RBM6 | 145.644246 | -0.558796627 | 0.27108427 | -2.0613392 | 0.03927069 | 0.95980711 |
| RBM5 | 45.1202471 | -0.148957077 | 0.26862522 | -0.5545164 | 0.57922552 | 0.99999872 |
| GNAI2 | 54.7708163 | -0.10321207 | 0.49348331 | -0.2091501 | 0.83433109 | 0.99999872 |
| SEMA3B | 180.16163 | -0.081379641 | 0.66608596 | -0.1221759 | 0.90275972 | 0.99999872 |
| IFRD2 | 6.32512375 | 0.538121062 | 0.65928432 | 0.81622002 | 0.41437431 | 0.99999872 |
| HYAL2 | 17.4866925 | 0.171365545 | 0.41474071 | 0.41318719 | 0.67946948 | 0.99999872 |
| TUSC2 | 11.2931113 | -0.070933466 | 0.58877606 | -0.1204761 | 0.90410598 | 0.99999872 |
| RASSF1 | 5.08990122 | -0.481468556 | 0.73440169 | -0.6555929 | 0.51208608 | 0.99999872 |
| TMEM115 | 15.2917317 | 0.034006954 | 0.59016077 | 0.0576232 | 0.95404877 | 0.99999872 |
| HEMK1 | 6.44701106 | -0.106507578 | 0.63342642 | -0.1681451 | 0.86646909 | 0.99999872 |
| CISH | 3.91686707 | 0.402834126 | 1.15022849 | 0.35022096 | 0.72617288 | NA |
| MAPKAPK3 | 4.57569583 | -0.205101878 | 0.59830835 | -0.342803 | 0.7317467 | NA |
| DOCK3 | 52.4873297 | 0.539675464 | 0.47596396 | 1.13385782 | 0.25685419 | 0.99999872 |
| MANF | 19.2797646 | 1.303065493 | 0.67411373 | 1.93300543 | 0.05323553 | 0.99999872 |
| RBM15B | 8.56602982 | 0.488771538 | 0.720906 | 0.67799622 | 0.49777409 | 0.99999872 |
| DCAF1 | 16.7233205 | -0.167286605 | 0.4357903 | -0.3838695 | 0.70107517 | 0.99999872 |
| RAD54L2 | 22.3392311 | -0.358385982 | 0.36586765 | -0.9795509 | 0.32730786 | 0.99999872 |
| TEX264 | 24.4514673 | -0.306955994 | 0.44754497 | -0.6858663 | 0.49279745 | 0.99999872 |
| PCBP4 | 7.60805382 | 0.496576607 | 0.68013337 | 0.73011651 | 0.46531897 | 0.99999872 |
| RPL29 | 119.914289 | 0.661501632 | 0.39015192 | 1.69549755 | 0.08998107 | 0.99999872 |
| ALAS1 | 5.1619066 | 0.901493062 | 0.70450236 | 1.27961681 | 0.20067994 | 0.99999872 |
| WDR82 | 30.0085288 | -0.157132801 | 0.32619834 | -0.4817094 | 0.63001241 | 0.99999872 |
| DNAH1 | 3.70556956 | 0.331250023 | 0.71520918 | 0.46315124 | 0.64325596 | NA |
| BAP1 | 12.8910657 | -0.064745906 | 0.50484524 | -0.128249 | 0.89795191 | 0.99999872 |
| PHF7 | 5.73299132 | 0.067652525 | 0.5955963 | 0.11358789 | 0.90956449 | 0.99999872 |
| SEMA3G | 88.7711423 | -0.078479728 | 0.76122773 | -0.1030963 | 0.91788658 | 0.99999872 |
| TNNC1 | 24.4750196 | 1.434397498 | 0.87787772 | 1.6339377 | 0.10227193 | 0.99999872 |
| NISCH | 30.915269 | 0.033023177 | 0.3670125 | 0.08997834 | 0.92830442 | 0.99999872 |
| SMIM4 | 12.9797622 | 0.614154117 | 0.44524834 | 1.37935185 | 0.1677863 | 0.99999872 |
| PBRM1 | 70.4627522 | -0.024427569 | 0.2995616 | -0.0815444 | 0.93500902 | 0.99999872 |

| | | | | | | |
|---|---|---|---|---|---|---|
| GNL3 | 14.9188387 | 0.368795968 | 0.54094374 | 0.681764 | 0.49538819 | 0.99999872 |
| GLT8D1 | 18.5810224 | 0.928065516 | 0.42716066 | 2.17263808 | 0.02980757 | 0.90266962 |
| SPCS1 | 63.4567749 | 0.794354777 | 0.30797903 | 2.57924955 | 0.00990152 | 0.70848663 |
| NEK4 | 17.2300751 | 0.027410436 | 0.46305691 | 0.05919453 | 0.95279717 | 0.99999872 |
| SFMBT1 | 15.3419876 | -0.23436753 | 0.58469672 | -0.4008361 | 0.68854083 | 0.99999872 |
| AC096887.1 | 7.24865652 | 0.04869484 | 0.77085356 | 0.06317003 | 0.94963111 | 0.99999872 |
| PRKCD | 10.6660781 | -0.026991867 | 0.50360004 | -0.0535978 | 0.95725559 | 0.99999872 |
| TKT | 21.7469154 | -1.32E-05 | 0.38607681 | -3.43E-05 | 0.99997265 | 0.99999872 |
| DCP1A | 36.1300151 | -0.198810225 | 0.30741621 | -0.6467135 | 0.51781736 | 0.99999872 |
| CACNA1D | 19.3681711 | -0.448243971 | 0.9016337 | -0.4971464 | 0.6190858 | 0.99999872 |
| CHDH | 5.4097516 | 1.956691785 | 1.35712004 | 1.44179713 | 0.14935961 | 0.99999872 |
| IL17RB | 10.1818947 | 0.972323056 | 1.31516288 | 0.73931759 | 0.45971417 | 0.99999872 |
| ACTR8 | 6.67471796 | -0.138238264 | 0.64465555 | -0.2144374 | 0.83020598 | 0.99999872 |
| SELENOK | 30.930624 | 1.01879228 | 0.48619647 | 2.0954333 | 0.03613249 | 0.94734101 |
| CACNA2D3 | 32.0652758 | -1.222359726 | 0.54338528 | -2.2495267 | 0.024479 | 0.88337526 |
| ERC2 | 61.7077563 | 0.116462184 | 0.355952 | 0.32718508 | 0.7435279 | 0.99999872 |
| CCDC66 | 39.8137872 | 0.65324297 | 0.3163979 | 2.06462484 | 0.03895851 | 0.95980711 |
| FAM208A | 67.2047408 | 0.258357967 | 0.26140835 | 0.98833098 | 0.32299058 | 0.99999872 |
| ARHGEF3 | 599.789426 | -0.321223073 | 0.3401913 | -0.9442425 | 0.34504576 | 0.99999872 |
| IL17RD | 56.8399691 | 0.449984822 | 0.65274633 | 0.68937166 | 0.49058941 | 0.99999872 |
| APPL1 | 36.0275355 | 0.194005977 | 0.3231928 | 0.60027938 | 0.54832006 | 0.99999872 |
| PDE12 | 10.4517737 | 0.148679348 | 0.45072293 | 0.32986861 | 0.74149924 | 0.99999872 |
| ARF4 | 42.2481262 | 0.405504207 | 0.28451091 | 1.42526768 | 0.15407984 | 0.99999872 |
| DENND6A | 54.8569963 | -0.230319235 | 0.30785587 | -0.7481398 | 0.45437582 | 0.99999872 |
| DENND6A-A! | 10.6660413 | 0.140090784 | 0.65272721 | 0.21462378 | 0.83006065 | 0.99999872 |
| SLMAP | 73.1979596 | -0.214955246 | 0.2591343 | -0.8295129 | 0.40681424 | 0.99999872 |
| FLNB | 15.9670953 | -0.488290256 | 0.60969578 | -0.8008752 | 0.42320387 | 0.99999872 |
| ABHD6 | 5.86492194 | 0.509219882 | 0.58669683 | 0.86794381 | 0.38542509 | 0.99999872 |
| RPP14 | 5.40209623 | 0.021816874 | 0.57538303 | 0.03791713 | 0.96975375 | 0.99999872 |
| PXK | 13.1547888 | 0.267260439 | 0.45574001 | 0.58643181 | 0.55758537 | 0.99999872 |
| PDHB | 16.5386515 | 0.728716935 | 0.51668675 | 1.41036506 | 0.15843192 | 0.99999872 |
| KCTD6 | 5.07788734 | 1.187609953 | 0.75408286 | 1.57490644 | 0.11527804 | 0.99999872 |
| ACOX2 | 5.19435106 | 0.599701549 | 0.71565507 | 0.83797569 | 0.40204436 | 0.99999872 |
| C3orf67 | 6.73706329 | -0.153479175 | 0.76301966 | -0.2011471 | 0.84058357 | 0.99999872 |
| FHIT | 247.524721 | 0.575913045 | 0.57106555 | 1.00848851 | 0.31322 | 0.99999872 |
| PTPRG | 428.384323 | 0.223780379 | 0.50205963 | 0.4457247 | 0.65579612 | 0.99999872 |
| THOC7 | 20.5291236 | 1.045640324 | 0.38836457 | 2.69241942 | 0.00709357 | 0.62327327 |
| ATXN7 | 8.16826177 | -0.396739646 | 0.50181511 | -0.7906092 | 0.42917207 | 0.99999872 |
| PSMD6-AS2 | 8.98198606 | 0.420853886 | 0.58311952 | 0.72172835 | 0.47046151 | 0.99999872 |
| PSMD6 | 13.2240198 | 0.808571039 | 0.47861498 | 1.68939769 | 0.09114324 | 0.99999872 |
| PRICKLE2 | 92.7045362 | 0.357858977 | 0.39737467 | 0.9005581 | 0.36782332 | 0.99999872 |
| ADAMTS9 | 536.04315 | 0.095894427 | 0.36873738 | 0.26006158 | 0.79481627 | 0.99999872 |
| ADAMTS9-A! | 187.44498 | 0.258590955 | 0.42029069 | 0.61526691 | 0.53837847 | 0.99999872 |
| ADAMTS9-A! | 764.269622 | 0.927783808 | 0.6395869 | 1.45059851 | 0.14689169 | 0.99999872 |
| MAGI1 | 88.9503899 | -0.615986645 | 0.57095471 | -1.0788713 | 0.28064511 | 0.99999872 |
| SLC25A26 | 32.9366433 | 0.095317556 | 0.39994805 | 0.23832484 | 0.81162916 | 0.99999872 |
| LRIG1 | 21.3105023 | 0.434052604 | 0.67410839 | 0.64389141 | 0.51964585 | 0.99999872 |
| SUCLG2 | 89.3857902 | -0.363679119 | 0.21930688 | -1.6583115 | 0.09725461 | 0.99999872 |
| SUCLG2-AS1 | 39.0884487 | -0.506487342 | 0.49331409 | -1.0267036 | 0.30456006 | 0.99999872 |
| FAM19A1 | 10.4074952 | -0.592355597 | 0.88664245 | -0.6680885 | 0.50407712 | 0.99999872 |
| EOGT | 10.1351742 | 0.064192144 | 0.56317858 | 0.11398186 | 0.90925217 | 0.99999872 |
| AC109587.1 | 3.6165596 | 1.931028423 | 0.89598801 | 2.15519449 | 0.03114662 | NA |
| TMF1 | 42.6442082 | -0.020035457 | 0.28686302 | -0.0698433 | 0.94431839 | 0.99999872 |
| UBA3 | 26.4123503 | -0.059018817 | 0.36373393 | -0.1622582 | 0.87110252 | 0.99999872 |
| ARL6IP5 | 45.259024 | 0.929202508 | 0.37789299 | 2.45890378 | 0.0139362 | 0.75772455 |
| FRMD4B | 31.4952417 | -0.421026271 | 0.48811864 | -0.8625491 | 0.38838545 | 0.99999872 |
| MITF | 35.2481439 | -0.304640481 | 0.56280126 | -0.5412932 | 0.58830553 | 0.99999872 |
| FOXP1 | 229.550409 | -0.613180332 | 0.33511671 | -1.8297516 | 0.06728709 | 0.99999872 |
| EIF4E3 | 16.7426383 | 0.143856324 | 0.55056789 | 0.26128716 | 0.79387106 | 0.99999872 |
| RYBP | 50.5297652 | -0.065355382 | 0.36180789 | -0.1806356 | 0.85665362 | 0.99999872 |
| SHQ1 | 19.2871442 | 0.644712437 | 0.4704893 | 1.37030201 | 0.17059265 | 0.99999872 |

| | | | | | | |
|---|---|---|---|---|---|---|
| GXYLT2 | 9.08023426 | 1.277351129 | 0.55327442 | 2.30871169 | 0.02095958 | 0.8671024 |
| PPP4R2 | 53.4641087 | 0.17235713 | 0.25304941 | 0.68112045 | 0.49579528 | 0.99999872 |
| PDZRN3 | 12.1747632 | 1.204080089 | 0.98783882 | 1.2189034 | 0.22288086 | 0.99999872 |
| CNTN3 | 40.2709975 | -0.188741528 | 0.78235351 | -0.2412484 | 0.80936261 | 0.99999872 |
| ZNF717 | 24.5077288 | -0.384785519 | 0.37637139 | -1.0223559 | 0.30661247 | 0.99999872 |
| ROBO2 | 288.393199 | 0.128858855 | 0.45080285 | 0.28584304 | 0.77499834 | 0.99999872 |
| ROBO1 | 155.254925 | 0.23345035 | 0.32263333 | 0.7235779 | 0.46932491 | 0.99999872 |
| GBE1 | 117.768973 | -0.210500681 | 0.43040319 | -0.4890779 | 0.62478656 | 0.99999872 |
| CHMP2B | 18.7471615 | 0.381277911 | 0.40591927 | 0.93929492 | 0.34757935 | 0.99999872 |
| CGGBP1 | 60.0566033 | 0.034164655 | 0.27710881 | 0.12328968 | 0.90187772 | 0.99999872 |
| ZNF654 | 49.9539389 | 0.199233026 | 0.31436737 | 0.6337586 | 0.52623838 | 0.99999872 |
| C3orf38 | 21.0973172 | 0.058188288 | 0.33202536 | 0.17525254 | 0.8608812 | 0.99999872 |
| PROS1 | 24.0561336 | 0.399611384 | 0.58097251 | 0.68783183 | 0.49155869 | 0.99999872 |
| ARL13B | 64.083959 | 0.396783196 | 0.26690747 | 1.48659456 | 0.13712193 | 0.99999872 |
| NSUN3 | 21.2139191 | -0.483824808 | 0.37338088 | -1.2957943 | 0.19504637 | 0.99999872 |
| MTRNR2L12 | 47.9241545 | -0.188225916 | 0.53594036 | -0.3512068 | 0.72543319 | 0.99999872 |
| ARL6 | 11.5236153 | 0.785945878 | 0.49527848 | 1.5868767 | 0.11254057 | 0.99999872 |
| CRYBG3 | 16.7869535 | 0.600050951 | 0.45939174 | 1.30618576 | 0.19148939 | 0.99999872 |
| RIOX2 | 15.3217369 | -0.506298498 | 0.45928002 | -1.1023743 | 0.27029897 | 0.99999872 |
| CLDND1 | 25.8925323 | 0.436534377 | 0.33829269 | 1.2904044 | 0.19691029 | 0.99999872 |
| CPOX | 7.38037508 | 0.205934563 | 0.66298994 | 0.31061491 | 0.75609339 | 0.99999872 |
| ST3GAL6 | 518.666056 | 0.223905533 | 0.47105811 | 0.47532466 | 0.63455559 | 0.99999872 |
| DCBLD2 | 119.044661 | 0.174164501 | 0.37363214 | 0.46613898 | 0.64111603 | 0.99999872 |
| CMSS1 | 41.0957622 | 0.132064994 | 0.47825742 | 0.27613789 | 0.78244216 | 0.99999872 |
| FILIP1L | 12.2087084 | 0.67579673 | 0.61165113 | 1.10487285 | 0.26921469 | 0.99999872 |
| TBC1D23 | 26.6769037 | 0.000876382 | 0.35428603 | 0.00247366 | 0.99802631 | 0.99999872 |
| NIT2 | 11.9217536 | -0.358357185 | 0.44340468 | -0.8081944 | 0.41897868 | 0.99999872 |
| TOMM70 | 23.116137 | 0.320609033 | 0.38871903 | 0.82478348 | 0.40949453 | 0.99999872 |
| TMEM45A | 9.9607342 | 0.596784078 | 0.55783014 | 1.06983117 | 0.28469531 | 0.99999872 |
| TFG | 50.3801921 | 0.374352577 | 0.38485561 | 0.97270915 | 0.33069788 | 0.99999872 |
| IMPG2 | 6.1035615 | -0.650581024 | 1.27164663 | -0.5116052 | 0.60892735 | 0.99999872 |
| SENP7 | 93.6424339 | 0.216236255 | 0.30500844 | 0.70895172 | 0.47835444 | 0.99999872 |
| TRMT10C | 14.0633593 | 0.155000663 | 0.43044864 | 0.36009096 | 0.71877911 | 0.99999872 |
| PCNP | 56.0285154 | 0.05939157 | 0.28967517 | 0.20502817 | 0.83755011 | 0.99999872 |
| ZBTB11 | 13.9361036 | 0.184087122 | 0.43828172 | 0.42002008 | 0.67447079 | 0.99999872 |
| RPL24 | 50.1311872 | 0.788105882 | 0.42849926 | 1.83922344 | 0.06588233 | 0.99999872 |
| CEP97 | 10.405204 | 0.104425435 | 0.47858937 | 0.21819422 | 0.82727779 | 0.99999872 |
| NXPE3 | 15.0832151 | 0.085804781 | 0.48826778 | 0.17573304 | 0.86050368 | 0.99999872 |
| NFKBIZ | 16.3719841 | -0.455243226 | 0.67769752 | -0.6717499 | 0.50174294 | 0.99999872 |
| ALCAM | 91.9224329 | 0.151022505 | 0.41359249 | 0.36514808 | 0.71500087 | 0.99999872 |
| CBLB | 855.464309 | -0.14942362 | 0.2264121 | -0.6599631 | 0.50927754 | 0.99999872 |
| LINC00882 | 16.1230131 | -0.232204809 | 0.59556923 | -0.3898872 | 0.69661997 | 0.99999872 |
| DUBR | 23.0191168 | 1.003553945 | 0.53116863 | 1.88933209 | 0.05884734 | 0.99999872 |
| BBX | 151.219418 | 0.249113745 | 0.25390496 | 0.9811299 | 0.32652869 | 0.99999872 |
| CD47 | 134.737803 | 0.324306508 | 0.36315917 | 0.89301478 | 0.37184926 | 0.99999872 |
| IFT57 | 20.0364968 | 0.577293944 | 0.37289291 | 1.54814942 | 0.12158633 | 0.99999872 |
| DZIP3 | 14.9779129 | -0.050776427 | 0.59541136 | -0.0852796 | 0.93203913 | 0.99999872 |
| NECTIN3 | 34.9635356 | 0.272099669 | 0.43368419 | 0.62741432 | 0.53038769 | 0.99999872 |
| PHLDB2 | 106.679624 | -0.0420706 | 0.42166296 | -0.0997731 | 0.9205245 | 0.99999872 |
| ABHD10 | 7.46812518 | 0.216573883 | 0.7022535 | 0.30839844 | 0.75777917 | 0.99999872 |
| CD200 | 62.0788646 | 0.463458398 | 0.53907736 | 0.85972521 | 0.38994054 | 0.99999872 |
| ATG3 | 15.0581973 | 0.331615489 | 0.41336843 | 0.80222743 | 0.42242141 | 0.99999872 |
| SLC35A5 | 13.8509725 | 0.081212144 | 0.40964347 | 0.19825079 | 0.84284885 | 0.99999872 |
| GTPBP8 | 5.96195983 | 0.083712644 | 0.59172415 | 0.14147241 | 0.88749675 | 0.99999872 |
| NEPRO | 15.7210656 | -0.137724133 | 0.52774286 | -0.2609683 | 0.79411699 | 0.99999872 |
| AC078785.1 | 5.18154396 | 0.383569609 | 0.89743579 | 0.42740619 | 0.6690835 | 0.99999872 |
| CFAP44 | 22.252767 | 0.40181207 | 0.40993022 | 0.98019627 | 0.32698925 | 0.99999872 |
| SPICE1 | 14.8216028 | -0.131467424 | 0.50028357 | -0.2627858 | 0.79271567 | 0.99999872 |
| USF3 | 25.3047164 | 0.177611745 | 0.34776491 | 0.51072359 | 0.60954462 | 0.99999872 |
| NAA50 | 28.8755772 | -0.052409462 | 0.30882958 | -0.1697035 | 0.86524332 | 0.99999872 |
| ATP6V1A | 45.041306 | 0.258785155 | 0.30164057 | 0.85792555 | 0.39093358 | 0.99999872 |

| | | | | | | |
|---|---|---|---|---|---|---|
| GRAMD1C | 9.53458955 | 0.182889503 | 0.72241171 | 0.2531652 | 0.80014056 | 0.99999872 |
| ZDHHC23 | 4.03383835 | -0.735611318 | 0.73646089 | -0.9988464 | 0.3178691 | NA |
| CCDC191 | 31.626677 | -0.536878155 | 0.4921807 | -1.0908151 | 0.27535424 | 0.99999872 |
| QTRT2 | 15.107541 | 0.2617693 | 0.4797372 | 0.54565146 | 0.58530555 | 0.99999872 |
| ZBTB20 | 1108.26057 | 0.080807527 | 0.22760895 | 0.35502789 | 0.7225687 | 0.99999872 |
| ZBTB20-AS5 | 20.1159891 | 1.033750673 | 0.62192359 | 1.66218275 | 0.09647614 | 0.99999872 |
| LSAMP | 22.7105253 | -0.590554992 | 0.68476549 | -0.8624193 | 0.38845681 | 0.99999872 |
| B4GALT4 | 11.8363566 | -0.095090683 | 0.42925791 | -0.2215234 | 0.8246849 | 0.99999872 |
| ARHGAP31 | 11.3745277 | 0.659160792 | 0.63136014 | 1.04403295 | 0.29647015 | 0.99999872 |
| TMEM39A | 18.464202 | 0.410061685 | 0.40919767 | 1.00211148 | 0.31628975 | 0.99999872 |
| POGLUT1 | 27.9219301 | 0.547368726 | 0.52729036 | 1.03807839 | 0.29923356 | 0.99999872 |
| TIMMDC1 | 28.0417782 | 0.162660394 | 0.43976522 | 0.36988008 | 0.71147185 | 0.99999872 |
| COX17 | 24.1898973 | 0.201553358 | 0.51930315 | 0.38812274 | 0.69792521 | 0.99999872 |
| MAATS1 | 13.5822103 | 0.972115843 | 0.4983357 | 1.95072485 | 0.05108979 | 0.98949866 |
| GSK3B | 109.519361 | 0.166323518 | 0.26283966 | 0.63279461 | 0.52686778 | 0.99999872 |
| LRRC58 | 34.157917 | 0.452020211 | 0.37486079 | 1.20583486 | 0.22788117 | 0.99999872 |
| FSTL1 | 144.898732 | 0.463086617 | 0.63798688 | 0.72585602 | 0.46792704 | 0.99999872 |
| NDUFB4 | 35.2247584 | 0.911220287 | 0.39910316 | 2.28316982 | 0.02242037 | 0.87985246 |
| HGD | 8.4442638 | 0.013097596 | 0.8624386 | 0.0151867 | 0.98788323 | 0.99999872 |
| RABL3 | 14.2586931 | 0.257442548 | 0.42195015 | 0.6101255 | 0.54177867 | 0.99999872 |
| GTF2E1 | 6.11168886 | 0.205008527 | 0.56049211 | 0.36576523 | 0.71454026 | 0.99999872 |
| GOLGB1 | 123.56588 | -0.240242519 | 0.3081112 | -0.7797267 | 0.43555178 | 0.99999872 |
| IQCB1 | 27.2395603 | -0.140425749 | 0.38008824 | -0.3694557 | 0.71178812 | 0.99999872 |
| CCDC58 | 4.35943194 | 0.288991256 | 0.88572828 | 0.3262753 | 0.74421608 | NA |
| FAM162A | 25.2594478 | 0.292885677 | 0.57222044 | 0.51184065 | 0.60876254 | 0.99999872 |
| AC083798.2 | 12.3178255 | 0.616986147 | 0.49201425 | 1.25400056 | 0.2098418 | 0.99999872 |
| KPNA1 | 51.1015534 | 0.230600628 | 0.26574983 | 0.86773574 | 0.38553902 | 0.99999872 |
| PARP9 | 43.8609229 | -0.715277473 | 0.81482504 | -0.8778295 | 0.38003624 | 0.99999872 |
| DTX3L | 23.9473422 | -0.51307105 | 0.78783023 | -0.6512457 | 0.51488789 | 0.99999872 |
| PARP14 | 79.6265884 | -1.154664597 | 0.91146774 | -1.2668189 | 0.20522004 | 0.99999872 |
| HSPBAP1 | 12.193086 | -0.288410531 | 0.50954004 | -0.5660213 | 0.57137928 | 0.99999872 |
| DIRC2 | 21.7317466 | -0.453781675 | 0.37361224 | -1.2145793 | 0.2245266 | 0.99999872 |
| PDIA5 | 10.906781 | -0.506913653 | 0.46663747 | -1.0863115 | 0.2773412 | 0.99999872 |
| SEC22A | 33.0414181 | -0.08035739 | 0.34175673 | -0.2351304 | 0.81410754 | 0.99999872 |
| ADCY5 | 7.01670188 | -0.684249566 | 0.63846425 | -1.0717116 | 0.28384958 | 0.99999872 |
| HACD2 | 20.1327695 | 0.868971645 | 0.45293553 | 1.91853274 | 0.05504349 | 0.99999872 |
| MYLK-AS1 | 9.00776748 | 0.295310532 | 0.67460515 | 0.43775316 | 0.66156523 | 0.99999872 |
| MYLK | 102.840163 | -0.757357699 | 0.44361246 | -1.7072507 | 0.08777546 | 0.99999872 |
| CCDC14 | 32.0438852 | 0.229683056 | 0.46470796 | 0.49425246 | 0.62112789 | 0.99999872 |
| KALRN | 25.8413177 | -0.516111462 | 1.00089103 | -0.515652 | 0.60609748 | 0.99999872 |
| UMPS | 10.1927234 | -0.634857564 | 0.67951583 | -0.9342793 | 0.35015985 | 0.99999872 |
| ITGB5 | 207.914457 | -0.184614526 | 0.30771261 | -0.5999576 | 0.54853448 | 0.99999872 |
| ITGB5-AS1 | 4.78611432 | 0.77207948 | 0.88309795 | 0.87428521 | 0.38196295 | 0.99999872 |
| HEG1 | 21.4924281 | 0.160021285 | 0.55400116 | 0.28884648 | 0.77269886 | 0.99999872 |
| ZNF148 | 151.989748 | -0.095087076 | 0.20938408 | -0.4541275 | 0.64973704 | 0.99999872 |
| SNX4 | 45.0049994 | -0.170348067 | 0.35771284 | -0.4762146 | 0.63392152 | 0.99999872 |
| OSBPL11 | 12.0052334 | -0.383212752 | 0.44852694 | -0.8543807 | 0.39289409 | 0.99999872 |
| SLC41A3 | 8.30238123 | 0.403432804 | 0.54970545 | 0.73390722 | 0.46300528 | 0.99999872 |
| ALDH1L1 | 10.3269977 | 1.095237349 | 1.2236659 | 0.89504606 | 0.37076246 | 0.99999872 |
| AC016924.1 | 5.6310806 | -0.420312923 | 1.00718278 | -0.4173154 | 0.6764477 | 0.99999872 |
| KLF15 | 9.77523333 | -0.288762804 | 0.67008059 | -0.4309374 | 0.66651387 | 0.99999872 |
| ZXDC | 39.2490486 | 0.220248334 | 0.32393038 | 0.6799249 | 0.49655201 | 0.99999872 |
| TXNRD3 | 5.30069545 | 0.212582544 | 0.63235729 | 0.33617474 | 0.7367391 | 0.99999872 |
| CHCHD6 | 19.8580482 | 0.703773041 | 0.45429773 | 1.54914498 | 0.12134687 | 0.99999872 |
| PLXNA1 | 14.9481401 | -0.587332394 | 0.8283336 | -0.709053 | 0.47829161 | 0.99999872 |
| TPRA1 | 11.3950473 | -0.221926467 | 0.62186071 | -0.3568749 | 0.72118547 | 0.99999872 |
| ABTB1 | 8.54572448 | 0.673888045 | 0.73288981 | 0.91949436 | 0.35783705 | 0.99999872 |
| MGLL | 17.1145747 | -0.419303988 | 0.42946703 | -0.9763357 | 0.32889814 | 0.99999872 |
| SEC61A1 | 51.9133064 | -0.097169416 | 0.50556962 | -0.1921979 | 0.8475872 | 0.99999872 |
| RUVBL1 | 14.9190571 | -0.027673438 | 0.47606794 | -0.0581292 | 0.95364573 | 0.99999872 |
| EEFSEC | 29.1660728 | -0.075279426 | 0.38680445 | -0.1946188 | 0.84569137 | 0.99999872 |

| | | | | | | |
|---|---|---|---|---|---|---|
| RPN1 | 62.8105908 | 0.556024578 | 0.46961935 | 1.18398993 | 0.23641705 | 0.99999872 |
| RAB7A | 88.7392628 | -0.068856334 | 0.22800473 | -0.3019952 | 0.76265572 | 0.99999872 |
| ACAD9 | 14.3355769 | -0.522870987 | 0.42021121 | -1.2443052 | 0.21338726 | 0.99999872 |
| CNBP | 85.6793027 | 0.460020273 | 0.4032129 | 1.14088679 | 0.25391704 | 0.99999872 |
| COPG1 | 19.3829701 | -0.102068022 | 0.42756481 | -0.2387194 | 0.81132316 | 0.99999872 |
| HMCES | 38.2747123 | 0.313058518 | 0.50383593 | 0.62135012 | 0.53436928 | 0.99999872 |
| H1FX | 46.300657 | -0.343090915 | 0.76598107 | -0.4479104 | 0.65421784 | 0.99999872 |
| MBD4 | 15.4747129 | 0.185895952 | 0.38409391 | 0.48398568 | 0.62839603 | 0.99999872 |
| IFT122 | 16.7288208 | -6.22E-05 | 0.45967565 | -0.0001354 | 0.99989195 | 0.99999872 |
| PLXND1 | 11.7705038 | 0.514271405 | 0.81266352 | 0.63282206 | 0.52684985 | 0.99999872 |
| TMCC1 | 113.364173 | -0.10851147 | 0.24448417 | -0.4438384 | 0.6571594 | 0.99999872 |
| PIK3R4 | 18.3140743 | 0.167990146 | 0.42587731 | 0.39445667 | 0.69324391 | 0.99999872 |
| ATP2C1 | 69.0347334 | 0.282384341 | 0.29009797 | 0.97341028 | 0.33034943 | 0.99999872 |
| NEK11 | 48.5128496 | 0.218250205 | 0.39295854 | 0.55540263 | 0.5786193 | 0.99999872 |
| NUDT16 | 8.87769921 | -0.699880069 | 0.53569028 | -1.3065013 | 0.19138213 | 0.99999872 |
| MRPL3 | 20.9195719 | -0.018273355 | 0.45515923 | -0.0401472 | 0.9679758 | 0.99999872 |
| CPNE4 | 26.4194142 | -0.426321653 | 0.65105752 | -0.6548141 | 0.51258744 | 0.99999872 |
| DNAJC13 | 44.3130674 | 0.023961394 | 0.32688303 | 0.07330266 | 0.94156528 | 0.99999872 |
| UBA5 | 17.7275283 | 0.567881394 | 0.42611585 | 1.33269251 | 0.18263273 | 0.99999872 |
| TMEM108 | 5.26214127 | 1.622639731 | 0.85334012 | 1.90151581 | 0.05723448 | 0.99999872 |
| CDV3 | 64.6217378 | -0.430484394 | 0.36212927 | -1.1887589 | 0.23453455 | 0.99999872 |
| TOPBP1 | 19.1856436 | -0.159061456 | 0.44022283 | -0.3613203 | 0.71786 | 0.99999872 |
| SRPRB | 13.0628389 | 0.825336408 | 0.64533095 | 1.27893511 | 0.20091991 | 0.99999872 |
| RAB6B | 7.13569057 | 0.274122301 | 0.78821173 | 0.3477775 | 0.72800729 | 0.99999872 |
| RYK | 59.2524042 | 0.508090766 | 0.27428546 | 1.85241596 | 0.06396612 | 0.99999872 |
| AMOTL2 | 79.4912354 | -0.703606383 | 0.48783583 | -1.4423016 | 0.14921732 | 0.99999872 |
| ANAPC13 | 11.8628305 | 0.393867662 | 0.50909012 | 0.77366982 | 0.43912607 | 0.99999872 |
| CEP63 | 23.0235883 | -0.028050268 | 0.37385962 | -0.0750289 | 0.94019174 | 0.99999872 |
| PPP2R3A | 86.5356533 | 0.074858515 | 0.35848565 | 0.20881872 | 0.83458976 | 0.99999872 |
| MSL2 | 40.486077 | 0.242840167 | 0.37157473 | 0.65354328 | 0.5134061 | 0.99999872 |
| PCCB | 17.3077096 | -0.212409926 | 0.40454978 | -0.5250526 | 0.59954661 | 0.99999872 |
| STAG1 | 165.799149 | -0.549690461 | 0.23514409 | -2.337675 | 0.01940411 | 0.8671024 |
| NCK1 | 19.6800582 | 0.026031398 | 0.49652021 | 0.05242767 | 0.95818793 | 0.99999872 |
| DBR1 | 4.57472711 | 0.155568564 | 0.68112534 | 0.22839932 | 0.81933582 | NA |
| ARMC8 | 45.1742442 | -0.351940651 | 0.33059055 | -1.0645817 | 0.28706527 | 0.99999872 |
| MRAS | 4.74271894 | 0.403993357 | 0.67598951 | 0.59763258 | 0.55008512 | 0.99999872 |
| CEP70 | 24.4812113 | 0.265049577 | 0.43782673 | 0.6053755 | 0.54492953 | 0.99999872 |
| FAIM | 7.17549827 | -0.712579291 | 0.62758929 | -1.135423 | 0.25619814 | 0.99999872 |
| PIK3CB | 58.4757969 | 0.062247814 | 0.33404407 | 0.18634612 | 0.85217335 | 0.99999872 |
| MRPS22 | 6.99999226 | 1.010783628 | 0.62809694 | 1.60927966 | 0.10755521 | 0.99999872 |
| COPB2 | 38.8942148 | 0.130748369 | 0.26490669 | 0.49356386 | 0.62161423 | 0.99999872 |
| AC097103.2 | 54.8641219 | -0.588997764 | 0.4668417 | -1.2616649 | 0.20706941 | 0.99999872 |
| NMNAT3 | 12.4973244 | 0.169493576 | 0.59114974 | 0.28671852 | 0.77432786 | 0.99999872 |
| CLSTN2 | 64.6986882 | -0.9223837 | 1.10238915 | -0.8367133 | 0.40275372 | 0.99999872 |
| SLC25A36 | 67.0231586 | 0.171904353 | 0.3034185 | 0.56655858 | 0.57101412 | 0.99999872 |
| ZBTB38 | 117.900554 | 0.172775897 | 0.299367 | 0.57713741 | 0.56384664 | 0.99999872 |
| RASA2 | 63.254258 | -0.231925884 | 0.25553919 | -0.9075942 | 0.36409267 | 0.99999872 |
| RNF7 | 32.3414237 | 0.747320027 | 0.44472588 | 1.68040598 | 0.09287835 | 0.99999872 |
| ATP1B3 | 30.3054134 | -0.195213075 | 0.4967389 | -0.3929893 | 0.69432738 | 0.99999872 |
| TFDP2 | 73.8483152 | 0.25619424 | 0.32286011 | 0.79351469 | 0.42747801 | 0.99999872 |
| GK5 | 27.1337876 | -0.216979515 | 0.3257169 | -0.6661598 | 0.50530895 | 0.99999872 |
| XRN1 | 88.8234889 | -0.295061252 | 0.37172789 | -0.793756 | 0.42733748 | 0.99999872 |
| ATR | 57.3868666 | -0.401650133 | 0.29517752 | -1.3607071 | 0.17360628 | 0.99999872 |
| PLS1 | 6.03486882 | 0.409066339 | 0.69411494 | 0.58933516 | 0.55563646 | 0.99999872 |
| TRPC1 | 54.7787233 | 0.162136036 | 0.51175846 | 0.31682141 | 0.75137912 | 0.99999872 |
| PCOLCE2 | 388.062469 | 0.213523996 | 0.64847504 | 0.32927096 | 0.7419509 | 0.99999872 |
| U2SURP | 66.0340326 | -0.022795365 | 0.28384854 | -0.0803082 | 0.93599214 | 0.99999872 |
| CHST2 | 66.8023395 | -0.92547623 | 0.85238556 | -1.0857484 | 2.78E-01 | 1.00E+00 |
| SLC9A9 | 18.6312597 | 0.189001068 | 0.66670045 | 0.28348724 | 0.77680336 | 0.99999872 |
| C3orf58 | 23.7854297 | 0.505647762 | 0.36030438 | 1.40339056 | 0.16050041 | 0.99999872 |
| AC107021.1 | 10.7564402 | 1.078709134 | 0.77816727 | 1.38621757 | 0.16568048 | 0.99999872 |

| | | | | | | |
|---|---|---|---|---|---|---|
| LNCSRLR | 5.69789923 | -0.035548721 | 0.81002113 | -0.0438862 | 0.96499514 | 0.99999872 |
| PLOD2 | 123.134985 | 0.712728702 | 0.32692006 | 2.18013147 | 0.02924772 | 0.90266962 |
| PLSCR4 | 32.9753848 | 0.574312806 | 0.38242031 | 1.50178427 | 0.13315283 | 0.99999872 |
| PLSCR1 | 86.5126067 | -0.102810991 | 0.83510931 | -0.1231108 | 0.90201934 | 0.99999872 |
| GYG1 | 22.4123056 | 0.174402068 | 0.34150815 | 0.51068201 | 0.60957374 | 0.99999872 |
| HLTF | 26.3759808 | 0.571445689 | 0.34516981 | 1.65554947 | 0.09781309 | 0.99999872 |
| HPS3 | 28.7213507 | 0.371898877 | 0.30184833 | 1.23207202 | 0.21792218 | 0.99999872 |
| CP | 9.57003718 | 2.018305785 | 1.10404844 | 1.82809531 | 0.06753525 | 0.99999872 |
| TM4SF1 | 14.2526498 | -2.350438276 | 0.62830007 | -3.7409486 | 0.00018333 | 0.12623956 |
| WWTR1 | 146.838713 | -0.236021949 | 0.2096213 | -1.1259445 | 0.26018902 | 0.99999872 |
| COMMD2 | 25.4519375 | 0.173420135 | 0.32521049 | 0.53325504 | 0.59385704 | 0.99999872 |
| ANKUB1 | 23.1403494 | 0.765979935 | 0.60007426 | 1.27647525 | 0.20178757 | 0.99999872 |
| RNF13 | 74.0259123 | -0.047879599 | 0.33199793 | -0.1442166 | 0.88532947 | 0.99999872 |
| TSC22D2 | 57.8492744 | -0.034866812 | 0.29566672 | -0.1179261 | 0.90612624 | 0.99999872 |
| SERP1 | 49.1969819 | 0.637314953 | 0.30484679 | 2.09060741 | 0.03656327 | 0.95300982 |
| EIF2A | 22.1405992 | 0.37988707 | 0.43514338 | 0.87301586 | 0.38265443 | 0.99999872 |
| SELENOT | 44.2142902 | 0.358550976 | 0.42945315 | 0.83490125 | 0.40377331 | 0.99999872 |
| SIAH2 | 7.74799868 | -0.148770589 | 0.61350129 | -0.2424943 | 0.80839714 | 0.99999872 |
| MED12L | 8.5631116 | -0.8211379 | 0.6692069 | -1.2270314 | 0.21981078 | 0.99999872 |
| MBNL1 | 187.6251 | 0.383877379 | 0.23814866 | 1.61192331 | 0.10697863 | 0.99999872 |
| RAP2B | 7.96533323 | 0.11144524 | 0.48452715 | 0.23000824 | 0.81808536 | 0.99999872 |
| C3orf79 | 9.35343355 | 1.636947976 | 0.97512612 | 1.67870386 | 0.09320977 | 0.99999872 |
| ARHGEF26 | 241.262989 | -0.113052886 | 0.31770916 | -0.3558377 | 0.72196214 | 0.99999872 |
| DHX36 | 76.5430284 | -0.340929688 | 0.3065701 | -1.1120774 | 0.26610485 | 0.99999872 |
| MME | 1512.20495 | -0.085933757 | 0.45934432 | -0.1870792 | 0.85159855 | 0.99999872 |
| MME-AS1 | 43.3984494 | 1.838399623 | 0.81004324 | 2.269508 | 0.02323745 | 0.88337526 |
| GMPS | 25.2888789 | 0.339971292 | 0.35725417 | 0.95162302 | 0.3412882 | 0.99999872 |
| SSR3 | 43.360739 | 0.463949354 | 0.28835656 | 1.60894331 | 0.10762874 | 0.99999872 |
| TIPARP | 33.3803105 | -0.657587076 | 0.42738795 | -1.5386187 | 0.12389742 | 0.99999872 |
| LINC00886 | 36.150456 | -0.29107994 | 0.5157455 | -0.5643868 | 0.57249094 | 0.99999872 |
| LEKR1 | 12.933098 | -0.579060781 | 0.57579022 | -1.0056801 | 0.31456947 | 0.99999872 |
| CCNL1 | 37.9073091 | -0.02209876 | 0.40507854 | -0.0545543 | 0.95649358 | 0.99999872 |
| VEPH1 | 5.77821337 | -0.384415652 | 0.93854928 | -0.4095849 | 0.68211045 | 0.99999872 |
| RSRC1 | 68.5044373 | -0.16908015 | 0.26153345 | -0.6464953 | 0.51795863 | 0.99999872 |
| MLF1 | 7.04968393 | 0.759051767 | 0.88756771 | 0.85520435 | 0.39243803 | 0.99999872 |
| GFM1 | 18.1506107 | 0.058835422 | 0.40525629 | 0.14518078 | 0.88456814 | 0.99999872 |
| LXN | 6.67305846 | 1.115502175 | 0.71324018 | 1.56399234 | 0.11781937 | 0.99999872 |
| RARRES1 | 5.82498225 | 1.14019486 | 0.70210617 | 1.62396359 | 0.10438358 | 0.99999872 |
| MFSD1 | 14.7720522 | 0.314376276 | 0.47673956 | 0.65942981 | 0.50961981 | 0.99999872 |
| IQCJ-SCHIP1 | 628.491331 | -0.556229773 | 0.95612931 | -0.5817516 | 0.560734 | 0.99999872 |
| SMC4 | 7.11656326 | 0.058765543 | 0.53083196 | 0.11070461 | 0.91185059 | 0.99999872 |
| KPNA4 | 59.0264867 | -0.151517023 | 0.37633199 | -0.4026153 | 0.68723124 | 0.99999872 |
| PPM1L | 60.6643913 | 0.0755146 | 0.57108383 | 0.13223032 | 0.89480212 | 0.99999872 |
| B3GALNT1 | 11.7498401 | 0.635217872 | 0.6490348 | 0.97871157 | 0.32772251 | 0.99999872 |
| NMD3 | 43.5534547 | -0.253448176 | 0.40060674 | -0.6326608 | 0.52695518 | 0.99999872 |
| LINC01322 | 7.46023593 | -0.002703644 | 0.58494889 | -0.004622 | 0.99631218 | 0.99999872 |
| SERPINI2 | 39.8229128 | -0.321995654 | 0.69764873 | -0.4615441 | 0.64440829 | 0.99999872 |
| WDR49 | 98.1205598 | -0.435600063 | 0.71582284 | -0.6085305 | 0.54283566 | 0.99999872 |
| PDCD10 | 39.1260869 | 0.63299904 | 0.29841493 | 2.12120435 | 0.03390461 | 0.93734168 |
| SERPINI1 | 9.1530639 | 0.96248077 | 0.5579121 | 1.72514769 | 0.08450086 | 0.99999872 |
| GOLIM4 | 317.272394 | 0.410952578 | 0.45669722 | 0.89983595 | 0.36820756 | 0.99999872 |
| MECOM | 189.268912 | -0.416183028 | 0.61563441 | -0.676023 | 0.49902603 | 0.99999872 |
| MYNN | 9.86337055 | 0.350456428 | 0.58976374 | 0.5942319 | 0.55235702 | 0.99999872 |
| SEC62 | 76.1072793 | 0.431895137 | 0.29060357 | 1.4862004 | 0.13722612 | 0.99999872 |
| GPR160 | 6.90922345 | 0.187375575 | 0.87656079 | 0.21376221 | 0.83073249 | 0.99999872 |
| PHC3 | 108.057802 | -0.08216474 | 0.24941585 | -0.3294287 | 0.74183167 | 0.99999872 |
| PRKCI | 162.852787 | -0.960792164 | 0.27876529 | -3.4465989 | 0.00056769 | 0.21771468 |
| SKIL | 47.9905946 | -0.420061086 | 0.31640875 | -1.32759 | 0.1843136 | 0.99999872 |
| AC026316.5 | 89.6523629 | 0.509527096 | 0.6181967 | 0.82421517 | 0.40981731 | 0.99999872 |
| SLC7A14 | 10.7797711 | 1.456476821 | 1.05264801 | 1.38363138 | 0.16647136 | 0.99999872 |
| RPL22L1 | 18.6559911 | 1.215544976 | 0.60714832 | 2.00205605 | 0.0452787 | 0.95980711 |

| | | | | | | |
|---|---|---|---|---|---|---|
| EIF5A2 | 16.2046722 | 0.338325094 | 0.46935331 | 0.72083244 | 0.47101261 | 0.99999872 |
| TNIK | 74.2879171 | -0.103435988 | 0.62210519 | -0.1662677 | 0.8679463 | 0.99999872 |
| PLD1 | 39.3904158 | 0.300729455 | 0.38392703 | 0.78329848 | 0.43345186 | 0.99999872 |
| FNDC3B | 744.26282 | -0.197859567 | 0.30913013 | -0.6400527 | 0.52213835 | 0.99999872 |
| TNFSF10 | 24.4170354 | -1.060495287 | 0.77732808 | -1.3642828 | 0.17247859 | 0.99999872 |
| NCEH1 | 15.947072 | -0.402786004 | 0.69650819 | -0.5782933 | 0.56306613 | 0.99999872 |
| ECT2 | 7.34440833 | 0.190710862 | 0.52756679 | 0.36149141 | 0.71773213 | 0.99999872 |
| NLGN1 | 55.6585809 | 0.126394884 | 0.69650854 | 0.18146925 | 0.85599926 | 0.99999872 |
| NAALADL2 | 201.136687 | -0.214225824 | 0.23779263 | -0.9008935 | 0.36764496 | 0.99999872 |
| TBL1XR1 | 141.50607 | -0.364365752 | 0.30996634 | -1.175501 | 0.23979433 | 0.99999872 |
| KCNMB2-AS1 | 6.40453817 | -0.061092346 | 0.81306037 | -0.0751388 | 0.94010431 | 0.99999872 |
| ZMAT3 | 28.3214024 | -0.026840352 | 0.57553267 | -0.0466357 | 0.9628036 | 0.99999872 |
| PIK3CA | 62.276021 | 0.35638068 | 0.40776372 | 0.87398821 | 0.38212467 | 0.99999872 |
| KCNMB3 | 9.76973976 | -0.195630918 | 0.52968655 | -0.3693334 | 0.71187926 | 0.99999872 |
| ZNF639 | 9.24005345 | 0.263552793 | 0.48542106 | 0.54293646 | 0.58717355 | 0.99999872 |
| MFN1 | 38.1016327 | 0.148058653 | 0.35576604 | 0.4161686 | 0.67728664 | 0.99999872 |
| GNB4 | 11.56706 | 0.221573945 | 0.82054682 | 0.27003206 | 0.78713559 | 0.99999872 |
| ACTL6A | 11.4329553 | 0.16433783 | 0.43017549 | 0.38202509 | 0.70244275 | 0.99999872 |
| MRPL47 | 8.18607013 | 0.086265745 | 0.64377593 | 0.13399964 | 0.89340286 | 0.99999872 |
| NDUFB5 | 23.0327811 | 0.972259874 | 0.33590563 | 2.89444353 | 0.00379831 | 0.48535019 |
| USP13 | 16.9442448 | -0.288602563 | 0.50852582 | -0.5675278 | 0.57035561 | 0.99999872 |
| PEX5L | 20.2239898 | -0.679661995 | 1.02654433 | -0.6620873 | 0.50791526 | 0.99999872 |
| TTC14 | 72.6314524 | 0.311600855 | 0.29252183 | 1.06522258 | 0.28677522 | 0.99999872 |
| FXR1 | 77.5144273 | 0.454269226 | 0.28680762 | 1.5838813 | 0.11322073 | 0.99999872 |
| DNAJC19 | 19.1950568 | 0.219094763 | 0.44164958 | 0.4960828 | 0.61983599 | 0.99999872 |
| ATP11B | 65.7361372 | -0.124756444 | 0.26654391 | -0.4680521 | 0.63974731 | 0.99999872 |
| DCUN1D1 | 40.8680388 | -0.006917103 | 0.42791353 | -0.0161647 | 0.98710298 | 0.99999872 |
| MCCC1 | 31.7806408 | -0.344393289 | 0.47454491 | -0.7257338 | 0.46800197 | 0.99999872 |
| LAMP3 | 8.19097929 | -0.196673991 | 2.71482098 | -0.0724446 | 0.94224813 | 0.99999872 |
| MCF2L2 | 15.8697168 | 1.138545092 | 0.66849955 | 1.70313516 | 0.08854278 | 0.99999872 |
| B3GNT5 | 8.75123614 | -0.087906874 | 0.56610258 | -0.1552844 | 0.87659715 | 0.99999872 |
| KLHL24 | 40.2670264 | 0.636721169 | 0.52486972 | 1.21310326 | 0.22509034 | 0.99999872 |
| YEATS2 | 40.4996202 | -0.100461703 | 0.44704374 | -0.2247245 | 0.82219357 | 0.99999872 |
| ABCC5 | 19.8581451 | -0.473250366 | 0.40199945 | -1.1772413 | 0.2390992 | 0.99999872 |
| EIF2B5 | 12.8171665 | -0.596170184 | 0.45920944 | -1.2982533 | 0.1942003 | 0.99999872 |
| DVL3 | 19.6097359 | -0.4515967 | 0.34751376 | -1.2995074 | 0.19376986 | 0.99999872 |
| AP2M1 | 52.57197 | 0.069815044 | 0.47686689 | 0.14640363 | 0.88360276 | 0.99999872 |
| ABCF3 | 6.17790305 | 0.231199087 | 0.64676663 | 0.3574691 | 0.72074065 | 0.99999872 |
| ALG3 | 5.06740179 | -0.641153286 | 0.81666251 | -0.7850897 | 0.432401 | 0.99999872 |
| PSMD2 | 26.7283783 | 0.02808431 | 0.45014226 | 0.06238985 | 0.95025237 | 0.99999872 |
| EIF4G1 | 36.1099165 | -0.321940857 | 0.35912223 | -0.896466 | 0.37000394 | 0.99999872 |
| FAM131A | 5.9777893 | 0.529154388 | 0.75836849 | 0.69775366 | 0.48533126 | 0.99999872 |
| POLR2H | 19.5777452 | 0.204918818 | 0.50813131 | 0.40327926 | 0.6867428 | 0.99999872 |
| CHRD | 3.92355122 | 0.834405661 | 0.85272078 | 0.97852155 | 0.32781644 | NA |
| MAGEF1 | 17.3584952 | 0.453505894 | 0.40473851 | 1.12049108 | 0.26250455 | 0.99999872 |
| VPS8 | 54.7232196 | -0.143277092 | 0.29933252 | -0.4786553 | 0.63218389 | 0.99999872 |
| EHHADH | 16.3370803 | -0.606843432 | 0.70222025 | -0.8641782 | 0.38749001 | 0.99999872 |
| MAP3K13 | 51.0047609 | 0.403613272 | 0.48660056 | 0.82945501 | 0.40684698 | 0.99999872 |
| TMEM41A | 8.77445651 | 0.320967378 | 0.5330315 | 0.60215462 | 0.54707122 | 0.99999872 |
| SENP2 | 25.1079998 | -0.162010951 | 0.35228509 | -0.4598859 | 0.64559811 | 0.99999872 |
| IGF2BP2 | 76.079404 | 0.034960777 | 0.26748581 | 0.13070143 | 0.8960115 | 0.99999872 |
| TRA2B | 46.9562401 | 0.194441459 | 0.31756915 | 0.61228069 | 0.54035206 | 0.99999872 |
| ETV5 | 6.35738323 | -0.617174243 | 0.53753139 | -1.1481641 | 0.25090083 | 0.99999872 |
| TBCCD1 | 6.28732781 | -0.401539163 | 0.61874174 | -0.6489608 | 0.51636369 | 0.99999872 |
| AC068631.2 | 6.52884093 | -0.299923311 | 0.58777486 | -0.510269 | 0.60986299 | 0.99999872 |
| EIF4A2 | 89.646024 | 0.776594302 | 0.37013605 | 2.09813202 | 0.03589349 | 0.94734101 |
| ST6GAL1 | 200.231654 | 0.001773758 | 0.50741911 | 0.00349565 | 9.97E-01 | 0.99999872 |
| RPL39L | 4.77326288 | 0.200707185 | 0.68847732 | 0.29152331 | 0.77065112 | 0.99999872 |
| RTP4 | 5.92962026 | -1.248672613 | 1.55252985 | -0.8042825 | 0.42123383 | 0.99999872 |
| BCL6 | 31.5699692 | 0.139462789 | 0.81485036 | 0.17115141 | 0.86410471 | 0.99999872 |
| LPP | 255.287689 | 0.062136347 | 0.32190766 | 0.19302538 | 0.84693909 | 0.99999872 |

| | | | | | | |
|---|---|---|---|---|---|---|
| TPRG1 | 10.97731 | -0.121664632 | 0.6011121 | -0.2023992 | 0.83960462 | 0.99999872 |
| P3H2 | 484.348855 | -0.152846853 | 0.37645382 | -0.4060175 | 0.68472972 | 0.99999872 |
| P3H2-AS1 | 14.3630589 | -0.537298667 | 0.64532057 | -0.8326074 | 0.4050662 | 0.99999872 |
| IL1RAP | 99.0023003 | 0.518527618 | 0.43275961 | 1.19818858 | 0.23084361 | 0.99999872 |
| CCDC50 | 65.5764902 | 0.049249329 | 0.31958284 | 0.15410505 | 0.87752691 | 0.99999872 |
| MB21D2 | 34.7140315 | 0.249746505 | 0.47922996 | 0.52114126 | 0.60226837 | 0.99999872 |
| OPA1 | 50.0978266 | 0.025526362 | 0.26426048 | 0.09659546 | 0.92304766 | 0.99999872 |
| HES1 | 19.4056511 | -0.119544408 | 0.53255502 | -0.2244733 | 0.822389 | 0.99999872 |
| ATP13A3 | 94.5702956 | 0.34971331 | 0.42089891 | 0.83087245 | 0.40604569 | 0.99999872 |
| LSG1 | 12.7384785 | -0.186601625 | 0.52301158 | -0.356783 | 0.72125429 | 0.99999872 |
| FAM43A | 14.8349647 | 0.595275483 | 0.9893211 | 0.60170098 | 0.54737319 | 0.99999872 |
| XXYLT1 | 8.20069376 | -0.419663592 | 0.53912737 | -0.7784127 | 0.43632575 | 0.99999872 |
| ACAP2 | 82.4011021 | 0.046057147 | 0.2429987 | 0.1895366 | 0.84967228 | 0.99999872 |
| PPP1R2 | 23.2580622 | 0.452394267 | 0.33311603 | 1.35806814 | 0.17444206 | 0.99999872 |
| APOD | 226.839796 | 0.611927869 | 1.46349667 | 0.41812727 | 0.67585407 | 0.99999872 |
| MUC20-OT1 | 82.7491501 | 0.008078926 | 0.48625666 | 0.01661453 | 0.98674413 | 0.99999872 |
| TNK2 | 7.90918038 | 0.237201674 | 0.60258579 | 0.39363967 | 0.69384709 | 0.99999872 |
| TFRC | 10.3795241 | -0.338383202 | 0.61636973 | -0.5489939 | 0.58300966 | 0.99999872 |
| PCYT1A | 16.7153244 | -0.291771885 | 0.40351982 | -0.723067 | 0.4696387 | 0.99999872 |
| UBXN7 | 38.9621737 | -0.049588778 | 0.33884864 | -0.1463449 | 0.88364911 | 0.99999872 |
| RNF168 | 19.3016518 | -0.271903832 | 0.35118388 | -0.7742492 | 0.43878345 | 0.99999872 |
| PIGX | 14.8980609 | 0.06417915 | 0.48847006 | 0.13138809 | 0.89546831 | 0.99999872 |
| PAK2 | 52.868892 | 0.026479603 | 0.25928749 | 0.10212449 | 0.91865786 | 0.99999872 |
| SENP5 | 52.5573891 | -0.562979158 | 0.24476843 | -2.3000481 | 0.0214455 | 0.87186485 |
| NCBP2 | 17.9774971 | -0.226241708 | 0.42809659 | -0.5284829 | 0.59716424 | 0.99999872 |
| NCBP2-AS2 | 14.4720734 | 0.237266293 | 0.44114288 | 0.53784455 | 0.59068437 | 0.99999872 |
| PIGZ | 5.98158032 | 1.225485398 | 0.6782291 | 1.80689003 | 0.07077943 | 0.99999872 |
| DLG1 | 132.387186 | 0.034526211 | 0.21946604 | 0.15731915 | 0.87499333 | 0.99999872 |
| RUBCN | 14.4426174 | -0.238836722 | 0.46907111 | -0.5091695 | 0.61063339 | 0.99999872 |
| FYTTD1 | 27.7627907 | -0.319454001 | 0.35235143 | -0.9066346 | 0.36460006 | 0.99999872 |
| LRCH3 | 85.8514991 | 0.040457563 | 0.29368515 | 0.13775829 | 0.89043145 | 0.99999872 |
| IQCG | 13.1103217 | 0.055038893 | 0.44984894 | 0.12234972 | 0.90262205 | 0.99999872 |
| RPL35A | 67.0965362 | 0.742976296 | 0.37005396 | 2.00775121 | 0.04466974 | 0.95980711 |
| LMLN | 13.1068382 | 0.511949763 | 0.49745489 | 1.02913806 | 0.3034148 | 0.99999872 |
| ZNF595 | 18.6745574 | 0.830227281 | 0.55935239 | 1.48426518 | 0.13773858 | 0.99999872 |
| ZNF718 | 50.2237832 | 0.010149954 | 0.2898165 | 0.035022 | 0.9720622 | 0.99999872 |
| ZNF141 | 24.5092621 | 0.105913665 | 0.36537587 | 0.28987592 | 0.77191117 | 0.99999872 |
| AC092574.2 | 4.9255863 | 2.027469668 | 1.31721616 | 1.53920801 | 0.12375352 | 0.99999872 |
| ZNF721 | 90.9990774 | -0.465553511 | 0.27348332 | -1.7023104 | 0.08869719 | 0.99999872 |
| PIGG | 19.7007555 | -0.308257875 | 0.47320862 | -0.6514207 | 0.51477498 | 0.99999872 |
| ATP5ME | 17.9357806 | 0.630869173 | 0.59929631 | 1.05268322 | 0.2924862 | 0.99999872 |
| PCGF3 | 24.3718591 | 0.000879936 | 0.350858 | 0.00250795 | 0.99799894 | 0.99999872 |
| AC139887.2 | 55.5530822 | -0.089783579 | 0.31263298 | -0.2871852 | 0.77397049 | 0.99999872 |
| GAK | 36.6078307 | -0.588327939 | 0.42894334 | -1.371575 | 0.17019579 | 0.99999872 |
| TMEM175 | 9.38762879 | 0.237419021 | 0.63088449 | 0.37632724 | 0.70667363 | 0.99999872 |
| IDUA | 6.32036451 | -0.977302051 | 0.57218029 | -1.7080317 | 0.08763047 | 0.99999872 |
| FGFRL1 | 8.50824694 | 0.683760929 | 0.54142365 | 1.26289446 | 0.20662711 | 0.99999872 |
| AC092535.3 | 9.33710372 | -0.340392443 | 0.60658399 | -0.5611629 | 0.57468648 | 0.99999872 |
| SPON2 | 14.7911415 | -0.555960002 | 0.66976036 | -0.8300879 | 0.40648906 | 0.99999872 |
| CTBP1-AS | 16.0882165 | 1.066880493 | 0.76487453 | 1.3948438 | 0.16306295 | 0.99999872 |
| CTBP1 | 33.0587867 | -0.189534959 | 0.45237933 | -0.4189735 | 0.67523549 | 0.99999872 |
| CTBP1-DT | 8.39669528 | -0.049060532 | 0.61828436 | -0.0793495 | 0.93675466 | 0.99999872 |
| MAEA | 20.9106229 | -0.118181761 | 0.36081873 | -0.3275378 | 0.74326118 | 0.99999872 |
| UVSSA | 23.2523122 | 0.115442949 | 0.45204724 | 0.25537806 | 0.79843111 | 0.99999872 |
| SLBP | 27.9786836 | -0.155231438 | 0.35292772 | -0.4398392 | 0.66005359 | 0.99999872 |
| TMEM129 | 12.8060071 | 0.174908236 | 0.52372166 | 0.33397174 | 0.73840088 | 0.99999872 |
| LETM1 | 13.1737891 | -0.40348788 | 0.53827286 | -0.7495973 | 0.45349728 | 0.99999872 |
| NSD2 | 54.2553675 | 0.037638869 | 0.363312 | 0.1035993 | 0.91748734 | 0.99999872 |
| NELFA | 16.8178403 | -0.031586844 | 0.49762785 | -0.0634748 | 0.9493884 | 0.99999872 |
| C4orf48 | 5.73719697 | 2.023576979 | 1.29128224 | 1.56710665 | 0.11708977 | 0.99999872 |
| POLN | 6.31791939 | -0.452875533 | 0.64756778 | -0.6993485 | 0.48433428 | 0.99999872 |

| | | | | | | |
|---|---|---|---|---|---|---|
| MXD4 | 60.2169516 | -0.302521662 | 0.42961719 | -0.7041656 | 0.48132963 | 0.99999872 |
| ZFYVE28 | 39.4922564 | -0.198638502 | 0.48864641 | -0.4065076 | 0.68436964 | 0.99999872 |
| RNF4 | 21.5742874 | 0.217141044 | 0.33734941 | 0.64366808 | 0.51979069 | 0.99999872 |
| FAM193A | 48.8718703 | -0.19844181 | 0.34802079 | -0.570201 | 0.56854136 | 0.99999872 |
| TNIP2 | 12.3107253 | -0.44162471 | 0.49368409 | -0.8945492 | 0.37102812 | 0.99999872 |
| SH3BP2 | 9.43799042 | -0.593525729 | 0.67834128 | -0.8749663 | 0.38159227 | 0.99999872 |
| ADD1 | 84.5143779 | -0.166444656 | 0.33791725 | -0.4925604 | 0.62232323 | 0.99999872 |
| MFSD10 | 22.8401063 | 0.185271176 | 0.60093277 | 0.308306 | 0.7578495 | 0.99999872 |
| NOP14-AS1 | 5.56218159 | -0.331472391 | 0.66202473 | -0.5006949 | 0.61658588 | 0.99999872 |
| NOP14 | 7.93162344 | -0.897481423 | 0.75010247 | -1.1964784 | 0.23150991 | 0.99999872 |
| GRK4 | 7.97793053 | 0.276729833 | 0.69180939 | 0.40000878 | 0.68915005 | 0.99999872 |
| HTT | 49.8975665 | -0.090676181 | 0.36597169 | -0.2477683 | 0.80431369 | 0.99999872 |
| RGS12 | 22.3778656 | -0.381215414 | 0.525617 | -0.7252722 | 0.46828503 | 0.99999872 |
| DOK7 | 9.97682321 | -1.873354784 | 0.61167833 | -3.062647 | 0.00219389 | 0.37076694 |
| LRPAP1 | 60.7877306 | 0.431464669 | 0.34600446 | 1.24699162 | 0.21240057 | 0.99999872 |
| TMEM128 | 19.3700816 | 0.449969908 | 0.49136519 | 0.91575455 | 0.35979565 | 0.99999872 |
| STX18 | 39.5503437 | 0.025060251 | 0.36251215 | 0.06912941 | 0.94488661 | 0.99999872 |
| STX18-AS1 | 24.6906166 | -0.592495588 | 0.53819583 | -1.1008922 | 0.27094358 | 0.99999872 |
| EVC2 | 15.3702105 | -0.591200481 | 0.41272275 | -1.4324398 | 0.152018 | 0.99999872 |
| EVC | 26.1039064 | -0.462102497 | 0.43632571 | -1.0590769 | 0.28956474 | 0.99999872 |
| CRMP1 | 7.12485798 | 0.687308319 | 0.85106486 | 0.8075863 | 0.41932878 | 0.99999872 |
| WFS1 | 35.7758178 | 0.489203438 | 0.56324924 | 0.86853812 | 0.38509982 | 0.99999872 |
| MAN2B2 | 24.9186623 | 0.667834823 | 0.54773608 | 1.21926389 | 0.22274405 | 0.99999872 |
| MRFAP1 | 62.7651756 | 0.442523069 | 0.30390745 | 1.45611131 | 0.14536185 | 0.99999872 |
| MRFAP1L1 | 10.5752222 | 0.095032203 | 0.54420054 | 0.17462718 | 0.86137259 | 0.99999872 |
| BLOC1S4 | 6.09889461 | 0.797791649 | 0.65866004 | 1.21123433 | 0.22580561 | 0.99999872 |
| KIAA0232 | 96.3763243 | -0.018753917 | 0.29609066 | -0.0633384 | 0.94949701 | 0.99999872 |
| TBC1D14 | 29.4097525 | -0.44946349 | 0.35040342 | -1.2827029 | 0.19959618 | 0.99999872 |
| TADA2B | 8.51174574 | -0.563736405 | 0.52784018 | -1.0680059 | 0.28551787 | 0.99999872 |
| GRPEL1 | 11.4178872 | 0.224211777 | 0.48818036 | 0.45928061 | 0.64603267 | 0.99999872 |
| SORCS2 | 8.98426513 | 0.50439985 | 0.69602319 | 0.72468828 | 0.46864328 | 0.99999872 |
| AFAP1 | 39.9229675 | -0.221727869 | 0.32588583 | -0.6803851 | 0.49626065 | 0.99999872 |
| ABLIM2 | 438.595801 | -0.523934184 | 0.36193215 | -1.4476033 | 1.48E-01 | 1.00E+00 |
| SH3TC1 | 6.16696782 | 0.140704367 | 0.8996548 | 0.15639817 | 0.87571918 | 0.99999872 |
| ACOX3 | 16.1517573 | 0.792533014 | 0.49147554 | 1.61255842 | 0.10684048 | 0.99999872 |
| TRMT44 | 10.7882801 | -0.601817731 | 0.5135322 | -1.1719182 | 0.2412299 | 0.99999872 |
| SLC2A9 | 14.7672689 | 0.3510878 | 0.75509314 | 0.4649596 | 0.64196039 | 0.99999872 |
| WDR1 | 33.8506907 | 0.108226079 | 0.46118415 | 0.23466999 | 0.81446488 | 0.99999872 |
| ZNF518B | 9.50416832 | -0.505974194 | 0.60630821 | -0.8345165 | 0.40399 | 0.99999872 |
| AC005699.1 | 8.81454656 | -1.126837914 | 0.76020207 | -1.4822874 | 0.13826385 | 0.99999872 |
| RAB28 | 52.1432027 | 0.185504339 | 0.25782175 | 0.71950617 | 0.4718291 | 0.99999872 |
| BOD1L1 | 44.1306614 | -0.15520425 | 0.2851291 | -0.5443297 | 0.58621458 | 0.99999872 |
| CPEB2-DT | 7.87769175 | 0.933740503 | 0.63956843 | 1.45995403 | 0.14430271 | 0.99999872 |
| CPEB2 | 60.1816892 | 0.24079059 | 0.42591478 | 0.56534922 | 0.57183626 | 0.99999872 |
| AC098829.1 | 155.290209 | 1.661391942 | 0.41059905 | 4.04626345 | 5.20E-05 | 0.06816157 |
| C1QTNF7 | 17.287413 | 2.965901968 | 0.80490924 | 3.68476569 | 0.00022891 | 0.12623956 |
| CC2D2A | 44.9521285 | 1.004598264 | 0.45252274 | 2.2199951 | 0.0264191 | 0.88440681 |
| FBXL5 | 106.216219 | 0.367944488 | 0.26399527 | 1.39375408 | 0.16339188 | 0.99999872 |
| FAM200B | 41.6106317 | 1.0043648 | 0.39259362 | 2.55828103 | 0.0105191 | 0.70947852 |
| BST1 | 33.3657706 | 1.379185144 | 0.53837689 | 2.56174654 | 0.01041473 | 0.70947852 |
| CD38 | 5.94698826 | 1.032005148 | 1.38758482 | 0.74374203 | 0.45703256 | 0.99999872 |
| TAPT1 | 37.0891193 | 0.115225992 | 0.34495428 | 0.33403265 | 0.73835492 | 0.99999872 |
| TAPT1-AS1 | 73.8928991 | 0.139226226 | 0.49846022 | 0.27931261 | 0.78000493 | 0.99999872 |
| LDB2 | 18.4329611 | -1.40447772 | 0.61442896 | -2.285826 | 0.02226444 | 0.87985246 |
| QDPR | 17.613492 | 0.242978698 | 0.45124935 | 0.53845773 | 0.59026108 | 0.99999872 |
| LAP3 | 33.4780797 | -0.475812946 | 1.19128881 | -0.3994102 | 0.68959095 | 0.99999872 |
| AC006160.1 | 3.71681508 | 0.864091879 | 1.67229962 | 0.51670877 | 0.60535947 | NA |
| MED28 | 19.7674818 | 0.284385935 | 0.38194161 | 0.74457962 | 0.45652589 | 0.99999872 |
| FAM184B | 25.3273357 | -0.008325893 | 0.67855246 | -0.0122701 | 0.99021014 | 0.99999872 |
| DCAF16 | 29.7481083 | -0.013397518 | 0.31235359 | -0.0428922 | 0.9657875 | 0.99999872 |
| LCORL | 67.5914647 | -0.156701146 | 0.33761553 | -0.4641408 | 0.64254687 | 0.99999872 |

| | | | | | | |
|---|---|---|---|---|---|---|
| AC024230.1 | 9.89448845 | 1.574798637 | 0.84774479 | 1.85763292 | 0.06322116 | 0.99999872 |
| SLIT2 | 13.1800314 | 0.022311293 | 0.84510878 | 0.0264005 | 0.9789379 | 0.99999872 |
| PACRGL | 26.3031618 | 0.187047643 | 0.31502699 | 0.59375117 | 0.55267855 | 0.99999872 |
| KCNIP4 | 260.397107 | 0.434202578 | 0.95692297 | 0.45374872 | 0.65000969 | 0.99999872 |
| ADGRA3 | 46.0541977 | -0.426806493 | 0.29725541 | -1.4358241 | 0.15105239 | 0.99999872 |
| PPARGC1A | 28.5630979 | -0.894073577 | 0.71129352 | -1.2569685 | 0.20876503 | 0.99999872 |
| DHX15 | 56.3876477 | -0.261558038 | 0.26938514 | -0.9709446 | 0.33157588 | 0.99999872 |
| CCDC149 | 12.6570187 | -0.537567337 | 0.45133275 | -1.1910665 | 0.23362747 | 0.99999872 |
| SEPSECS | 9.53840327 | -0.085872332 | 0.56826365 | -0.1511135 | 0.87988615 | 0.99999872 |
| AC104662.2 | 8.29930995 | 0.584373541 | 0.67456198 | 0.86630074 | 0.38632526 | 0.99999872 |
| PI4K2B | 4.25666206 | -0.517515673 | 0.83671657 | -0.6185077 | 0.5362407 | NA |
| ZCCHC4 | 9.82925419 | -0.884268581 | 0.51078629 | -1.7311909 | 0.08341772 | 0.99999872 |
| ANAPC4 | 29.8530566 | -0.215291035 | 0.35200381 | -0.6116156 | 0.54079209 | 0.99999872 |
| SLC34A2 | 5.84947263 | -0.470052448 | 0.89104778 | -0.5275278 | 0.59782714 | 0.99999872 |
| SEL1L3 | 7.28924338 | -1.249704767 | 0.71002621 | -1.7600826 | 0.0783938 | 0.99999872 |
| SMIM20 | 10.1697444 | 0.940751908 | 0.52428675 | 1.79434614 | 0.07275794 | 0.99999872 |
| RBPJ | 58.7297541 | 0.132733792 | 0.27955536 | 0.47480324 | 0.63492723 | 0.99999872 |
| TBC1D19 | 52.1123554 | -0.056335465 | 0.281131 | -0.2003887 | 0.84117663 | 0.99999872 |
| STIM2 | 44.7092316 | -0.377439726 | 0.35726597 | -1.0564671 | 0.29075488 | 0.99999872 |
| PCDH7 | 24.7734286 | -1.701444569 | 0.8508674 | -1.9996589 | 0.04553711 | 0.95980711 |
| ARAP2 | 79.288575 | 0.13867381 | 0.38870624 | 0.35675736 | 0.72127346 | 0.99999872 |
| C4orf19 | 44.4278879 | 0.81417893 | 0.48423297 | 1.68137855 | 0.09268941 | 0.99999872 |
| RELL1 | 41.4080574 | 0.580638594 | 0.34590303 | 1.67861668 | 0.09322677 | 0.99999872 |
| PGM2 | 6.60676767 | 0.234423451 | 0.77384034 | 0.30293517 | 0.76193927 | 0.99999872 |
| TBC1D1 | 216.07506 | -0.158402497 | 0.34619713 | -0.45755 | 0.64727577 | 0.99999872 |
| KLF3 | 18.3845734 | -0.418833468 | 0.38718069 | -1.0817519 | 0.27936277 | 0.99999872 |
| FAM114A1 | 50.3869036 | 0.082947796 | 0.50766065 | 0.16339221 | 0.87020964 | 0.99999872 |
| KLHL5 | 18.2214589 | -0.221713027 | 0.36181242 | -0.6127845 | 0.54001887 | 0.99999872 |
| WDR19 | 18.6521026 | -0.023675609 | 0.49514849 | -0.0478152 | 0.96186355 | 0.99999872 |
| RFC1 | 77.4016351 | 0.326221677 | 0.34679956 | 0.94066347 | 0.34687734 | 0.99999872 |
| RPL9 | 7.67032085 | 0.467467365 | 0.63022748 | 0.74174387 | 0.45824254 | 0.99999872 |
| LIAS | 7.80730851 | -0.813926874 | 0.53193572 | -1.5301226 | 0.12598638 | 0.99999872 |
| UGDH | 14.513527 | 0.709889586 | 0.53493808 | 1.32705002 | 0.18449214 | 0.99999872 |
| UGDH-AS1 | 20.7083672 | 0.171743511 | 0.39122966 | 0.43898387 | 0.66067322 | 0.99999872 |
| SMIM14 | 60.3661466 | -0.394551093 | 0.366585 | -1.0762882 | 0.28179841 | 0.99999872 |
| UBE2K | 126.960683 | -0.162829079 | 0.28470687 | -0.5719183 | 0.5673773 | 0.99999872 |
| PDS5A | 97.2412038 | -0.361720036 | 0.24102514 | -1.5007564 | 0.13341857 | 0.99999872 |
| N4BP2 | 30.9867811 | -0.372632887 | 0.37136268 | -1.0034204 | 0.31565807 | 0.99999872 |
| RBM47 | 60.6282789 | -0.85608118 | 0.54708702 | -1.5647989 | 0.11763007 | 0.99999872 |
| NSUN7 | 15.4613588 | -0.451537662 | 0.57451828 | -0.7859413 | 0.43190186 | 0.99999872 |
| APBB2 | 682.37362 | -0.387333783 | 0.36004084 | -1.0758051 | 0.28201442 | 0.99999872 |
| AC131953.1 | 5.38920832 | 1.677207772 | 0.8561807 | 1.95894134 | 0.05011966 | 0.98159583 |
| UCHL1 | 16.7024119 | 0.081029102 | 0.49877377 | 0.16245662 | 0.87094629 | 0.99999872 |
| LIMCH1 | 31.6791377 | 0.905097729 | 0.52132792 | 1.73613898 | 0.08253926 | 0.99999872 |
| TMEM33 | 27.5649332 | -0.055416586 | 0.37183292 | -0.1490363 | 0.88152502 | 0.99999872 |
| SLC30A9 | 61.4316827 | -0.427916378 | 0.27219724 | -1.572082 | 0.11593154 | 0.99999872 |
| SHISA3 | 21.4346027 | 2.12073635 | 1.25849864 | 1.68513202 | 0.0919631 | 0.99999872 |
| ATP8A1 | 32.5056856 | 0.15702393 | 0.66898687 | 0.234719 | 0.81442684 | 0.99999872 |
| KCTD8 | 20.1121928 | 1.162588263 | 0.93214047 | 1.24722432 | 0.21231526 | 0.99999872 |
| GUF1 | 10.9839679 | -0.538879839 | 0.47653198 | -1.1308367 | 0.25812384 | 0.99999872 |
| GNPDA2 | 14.6419304 | 0.123327029 | 0.55168773 | 0.22354499 | 0.82311137 | 0.99999872 |
| GABRA2 | 34.8425907 | 0.906375624 | 0.72828442 | 1.24453523 | 0.21330264 | 0.99999872 |
| COMMD8 | 16.1484474 | 0.16556571 | 0.4299576 | 0.38507451 | 0.70018221 | 0.99999872 |
| ATP10D | 87.6628303 | -0.000767066 | 0.29958047 | -0.0025605 | 0.99795704 | 0.99999872 |
| NFXL1 | 16.429478 | 0.298001505 | 0.39934771 | 0.74622065 | 0.45553414 | 0.99999872 |
| TXK | 10.4689898 | -0.407612998 | 0.53363173 | -0.763847 | 0.44495842 | 0.99999872 |
| TEC | 13.3231956 | 0.464889728 | 0.50306769 | 0.92410969 | 0.35542921 | 0.99999872 |
| SLAIN2 | 54.2218718 | -0.329614886 | 0.2886925 | -1.1417508 | 0.25355763 | 0.99999872 |
| FRYL | 94.1550207 | 0.024864761 | 0.24179714 | 0.10283315 | 0.9180954 | 0.99999872 |
| OCIAD1 | 75.8212609 | 0.285106845 | 0.29297908 | 0.97313037 | 0.33048851 | 0.99999872 |
| OCIAD2 | 12.3370762 | -0.269348964 | 0.59813164 | -0.4503172 | 0.65248174 | 0.99999872 |

| | | | | | | |
|---|---|---|---|---|---|---|
| DCUN1D4 | 54.5256335 | 0.614493955 | 0.28583296 | 2.14983586 | 0.0315682 | 0.92859542 |
| LRRC66 | 4.60630223 | 0.404847794 | 0.73509865 | 0.55073941 | 0.58181233 | 0.99999872 |
| SGCB | 78.1236185 | 1.022854693 | 0.39856553 | 2.56634007 | 0.0102778 | 0.70849208 |
| SPATA18 | 38.0452356 | 1.001990096 | 0.63801423 | 1.57048237 | 0.11630293 | 0.99999872 |
| USP46 | 117.309083 | 0.117911053 | 0.52857588 | 0.22307309 | 0.82347862 | 0.99999872 |
| ERVMER34-1 | 14.3102485 | 0.591929549 | 0.63789057 | 0.92794842 | 0.35343433 | 0.99999872 |
| RASL11B | 157.775027 | 0.723435091 | 0.65396202 | 1.1062341 | 0.26862521 | 0.99999872 |
| SCFD2 | 65.5828819 | -0.395134569 | 0.40501805 | -0.9755974 | 0.329264 | 0.99999872 |
| AC023154.1 | 110.148596 | 0.111927418 | 0.25479254 | 0.43928844 | 0.66045255 | 0.99999872 |
| AC058822.1 | 33.3830347 | 0.479369322 | 0.63989098 | 0.74914217 | 0.45377152 | 0.99999872 |
| LNX1 | 10.4698336 | -0.304393124 | 0.9243345 | -0.3293106 | 0.74192095 | 0.99999872 |
| CHIC2 | 49.1117522 | 0.003282933 | 0.39939667 | 0.00821973 | 0.99344168 | 0.99999872 |
| SRD5A3 | 6.92557073 | 0.840841161 | 0.74072111 | 1.13516565 | 0.25630592 | 0.99999872 |
| TMEM165 | 80.5449822 | -0.000488937 | 0.3551455 | -0.0013767 | 0.99890153 | 0.99999872 |
| CLOCK | 57.2427185 | 0.42237329 | 0.3260937 | 1.29525131 | 0.19523355 | 0.99999872 |
| EXOC1 | 45.4711575 | 0.168476449 | 0.38272663 | 0.44020048 | 0.65979191 | 0.99999872 |
| CEP135 | 20.8750555 | -0.685302875 | 0.38330869 | -1.7878615 | 0.07379835 | 0.99999872 |
| KIAA1211 | 117.074524 | 0.301961801 | 0.48900427 | 0.61750341 | 0.53690273 | 0.99999872 |
| AASDH | 19.8510238 | 0.287251664 | 0.45008711 | 0.63821348 | 0.52333472 | 0.99999872 |
| PAICS | 12.1845942 | -0.410304487 | 0.41695407 | -0.984052 | 0.32508995 | 0.99999872 |
| SRP72 | 29.3807517 | 0.035775177 | 0.35213855 | 0.10159404 | 0.91907891 | 0.99999872 |
| REST | 32.6596276 | -0.495649751 | 0.33292585 | -1.4887692 | 0.13654815 | 0.99999872 |
| NOA1 | 5.41451256 | 0.006557444 | 0.83173686 | 0.00788404 | 0.99370951 | 0.99999872 |
| POLR2B | 34.293763 | -0.195275897 | 0.34985133 | -0.5581682 | 0.57672952 | 0.99999872 |
| IGFBP7 | 1214.41452 | 0.409822528 | 0.24292196 | 1.68705429 | 9.16E-02 | 0.99999872 |
| AC116049.2 | 15.6941716 | -0.027753774 | 0.97529415 | -0.0284568 | 0.9772978 | 0.99999872 |
| AC104806.2 | 21.6894214 | 0.579104641 | 0.57187056 | 1.01264986 | 0.31122743 | 0.99999872 |
| CENPC | 40.9294784 | -0.015231268 | 0.42103509 | -0.0361758 | 0.97114221 | 0.99999872 |
| UBA6 | 50.9036476 | 0.065215335 | 0.45489825 | 0.14336246 | 0.88600393 | 0.99999872 |
| UBA6-AS1 | 117.069767 | -0.947723325 | 0.44559863 | -2.1268542 | 0.0334322 | 0.92959829 |
| YTHDC1 | 49.9278003 | -0.433556538 | 0.28132764 | -1.5411089 | 0.12329028 | 0.99999872 |
| UGT2A3 | 4.59756613 | -0.277695927 | 1.2791876 | -0.2170877 | 0.82813998 | 0.99999872 |
| UGT2B7 | 38.0331071 | 0.492916221 | 0.85051183 | 0.57955246 | 0.56221646 | 0.99999872 |
| UTP3 | 14.7529845 | 0.250171533 | 0.41139369 | 0.60810736 | 0.54311627 | 0.99999872 |
| RUFY3 | 113.130409 | -0.145346984 | 0.28257026 | -0.5143747 | 0.60699007 | 0.99999872 |
| GRSF1 | 21.8343565 | -0.374860363 | 0.34551087 | -1.0849452 | 0.27794592 | 0.99999872 |
| MOB1B | 41.2327469 | 0.045696067 | 0.35296322 | 0.1294641 | 0.89699043 | 0.99999872 |
| DCK | 5.26186335 | 0.527888052 | 0.75626128 | 0.69802337 | 0.48516258 | 0.99999872 |
| SLC4A4 | 30.6742779 | 0.982953912 | 1.05109669 | 0.93516983 | 0.34970079 | 0.99999872 |
| COX18 | 7.45547593 | 0.344682833 | 0.62336888 | 0.55293558 | 0.58030752 | 0.99999872 |
| ANKRD17 | 215.195733 | -0.04396878 | 0.29280588 | -0.1501636 | 0.88063556 | 0.99999872 |
| AC053527.1 | 33.071581 | 0.556292077 | 0.54964534 | 1.01209278 | 0.31149369 | 0.99999872 |
| RASSF6 | 6.94838463 | -0.322765144 | 0.63365875 | -0.5093674 | 0.61049469 | 0.99999872 |
| MTHFD2L | 31.1831391 | -0.08384406 | 0.43912156 | -0.1909359 | 0.84857583 | 0.99999872 |
| BTC | 31.7617923 | 0.294075373 | 0.49785624 | 0.59068331 | 0.55473263 | 0.99999872 |
| RCHY1 | 29.5021281 | 0.284982301 | 0.41986688 | 0.67874443 | 0.49729981 | 0.99999872 |
| THAP6 | 15.1759777 | -0.040790601 | 0.40638038 | -0.1003754 | 0.92004628 | 0.99999872 |
| CDKL2 | 7.31510412 | -0.301036221 | 0.8314631 | -0.362056 | 0.71731017 | 0.99999872 |
| G3BP2 | 47.9132589 | 0.346568854 | 0.26564924 | 1.30461076 | 0.19202542 | 0.99999872 |
| USO1 | 103.836889 | 0.314952632 | 0.24316822 | 1.29520476 | 0.1952496 | 0.99999872 |
| NAAA | 15.8825381 | 0.583455976 | 0.40488431 | 1.44104367 | 0.14957235 | 0.99999872 |
| SDAD1 | 16.0682762 | -0.506899678 | 0.43490935 | -1.1655295 | 0.24380473 | 0.99999872 |
| CXCL10 | 13.5009524 | -8.500402577 | 4.64824678 | -1.8287331 | 0.06743961 | 0.99999872 |
| NUP54 | 23.7620088 | 0.149038149 | 0.37337051 | 0.39916957 | 0.68976826 | 0.99999872 |
| SCARB2 | 142.353146 | -0.086161245 | 0.54749987 | -0.1573722 | 0.87495154 | 0.99999872 |
| SHROOM3 | 135.1734 | -0.067056623 | 0.38916689 | -0.1723081 | 0.86319528 | 0.99999872 |
| 11-Sep | 211.256147 | 0.215803455 | 0.33029313 | 0.65336948 | 0.51351811 | 0.99999872 |
| CCNI | 144.658616 | 0.279374846 | 0.36476875 | 0.76589578 | 0.44373832 | 0.99999872 |
| CCNG2 | 15.8763352 | 1.078727093 | 0.7223559 | 1.49334573 | 0.13534671 | 0.99999872 |
| CNOT6L | 47.5308057 | 0.25409898 | 0.37837651 | 0.67155062 | 0.50186982 | 0.99999872 |
| MRPL1 | 26.3227565 | 0.050529219 | 0.38971985 | 0.12965524 | 0.8968392 | 0.99999872 |

| | | | | | | |
|---|---|---|---|---|---|---|
| FRAS1 | 70.7733659 | 0.493579778 | 0.8203384 | 0.60167826 | 0.54738833 | 0.99999872 |
| BMP2K | 52.3688921 | 0.38632918 | 0.50103787 | 0.77105784 | 0.44067265 | 0.99999872 |
| PAQR3 | 16.8890943 | 0.944593564 | 0.41178602 | 2.29389419 | 0.02179658 | 0.8732058 |
| LINC01088 | 12.6871155 | -0.443659376 | 0.81249568 | -0.5460452 | 0.58503486 | 0.99999872 |
| ANTXR2 | 30.0410501 | 0.308926757 | 0.40219268 | 0.76810637 | 0.44242399 | 0.99999872 |
| CFAP299 | 7.24779561 | -0.231762963 | 0.87736976 | -0.2641565 | 0.7916593 | 0.99999872 |
| PRKG2 | 21.2947429 | -1.21639278 | 1.00935337 | -1.2051208 | 0.22815666 | 0.99999872 |
| RASGEF1B | 285.471462 | -0.228357896 | 0.80785658 | -0.2826713 | 0.77742879 | 0.99999872 |
| HNRNPD | 65.1570302 | 0.170723853 | 0.27496916 | 0.62088364 | 0.53467619 | 0.99999872 |
| HNRNPDL | 78.6803591 | 0.173077304 | 0.34090569 | 0.50769849 | 0.61166481 | 0.99999872 |
| ENOPH1 | 22.0807127 | 0.472971573 | 0.42257176 | 1.11926924 | 0.2630253 | 0.99999872 |
| TMEM150C | 254.074129 | -0.39969001 | 0.48013813 | -0.832448 | 0.40515614 | 0.99999872 |
| SCD5 | 109.195374 | -0.202281626 | 0.48875077 | -0.4138748 | 0.6789658 | 0.99999872 |
| SEC31A | 107.268836 | 0.23679693 | 0.27705259 | 0.8547003 | 0.39271709 | 0.99999872 |
| LIN54 | 17.636963 | -0.263635011 | 0.43474853 | -0.6064081 | 0.54424382 | 0.99999872 |
| COPS4 | 27.2267813 | 0.213614712 | 0.35812358 | 0.59648323 | 0.55085245 | 0.99999872 |
| COQ2 | 4.93919568 | -0.801561144 | 0.67161227 | -1.1934879 | 0.23267833 | 0.99999872 |
| HELQ | 12.3078077 | 0.141171523 | 0.44332472 | 0.3184382 | 0.75015257 | 0.99999872 |
| MRPS18C | 9.68260111 | 0.083207551 | 0.55203661 | 0.15072832 | 0.88019003 | 0.99999872 |
| ABRAXAS1 | 6.84341774 | 0.069508536 | 0.71121076 | 0.09773268 | 0.92214456 | 0.99999872 |
| GPAT3 | 25.8950422 | -0.886503888 | 1.0388327 | -0.8533654 | 0.3934567 | 0.99999872 |
| CDS1 | 43.8322644 | -0.118141239 | 0.59018699 | -0.2001759 | 0.84134298 | 0.99999872 |
| WDFY3 | 163.739178 | 0.098159045 | 0.25784814 | 0.38068549 | 0.70343664 | 0.99999872 |
| ARHGAP24 | 400.932667 | 0.515717137 | 0.37343647 | 1.38100369 | 0.16727783 | 0.99999872 |
| MAPK10 | 308.713535 | -0.221482868 | 0.3418749 | -0.6478477 | 0.51708346 | 0.99999872 |
| PTPN13 | 351.935681 | 0.129450836 | 0.46787155 | 0.27668029 | 0.78202561 | 0.99999872 |
| AC093827.5 | 5.39175912 | -0.45359012 | 0.6318852 | -0.7178363 | 0.47285824 | 0.99999872 |
| AFF1 | 115.155738 | -0.442875413 | 0.39814447 | -1.1123485 | 0.26598833 | 0.99999872 |
| KLHL8 | 16.5774021 | 0.139078008 | 0.46337019 | 0.30014449 | 0.76406695 | 0.99999872 |
| HSD17B11 | 11.4096093 | -0.847977274 | 0.54648403 | -1.5516963 | 0.1207349 | 0.99999872 |
| NUDT9 | 23.0551954 | -0.530125459 | 0.39773989 | -1.3328446 | 0.18258281 | 0.99999872 |
| SPP1 | 502.284398 | 0.178551545 | 0.64102825 | 0.27853928 | 7.81E-01 | 0.99999872 |
| PKD2 | 58.95168 | -0.103542544 | 0.39890313 | -0.2595681 | 0.79519692 | 0.99999872 |
| PPM1K | 18.0859525 | -0.763360324 | 0.78585721 | -0.9713728 | 0.33136267 | 0.99999872 |
| HERC6 | 10.0441701 | -3.206042003 | 1.34522276 | -2.3832796 | 0.01715915 | 0.82882344 |
| HERC5 | 14.4912751 | -1.314347743 | 1.15168569 | -1.1412382 | 0.2537708 | 0.99999872 |
| PYURF | 17.3265457 | 0.770232681 | 0.53071628 | 1.45130782 | 0.14669417 | 0.99999872 |
| HERC3 | 17.7932662 | 0.08432526 | 0.47176568 | 0.17874395 | 0.85813876 | 0.99999872 |
| NAP1L5 | 4.03339593 | 0.990944834 | 0.78344958 | 1.26484826 | 0.20592573 | NA |
| FAM13A-AS1 | 8.87539759 | 0.04629365 | 0.5535872 | 0.08362486 | 0.9333547 | 0.99999872 |
| FAM13A | 62.582185 | -0.425899178 | 0.32320827 | -1.3177236 | 0.18759618 | 0.99999872 |
| SNCA | 138.831833 | 0.027533937 | 0.48250245 | 0.05706486 | 0.95449352 | 0.99999872 |
| CCSER1 | 100.867109 | 0.017605075 | 0.79858005 | 0.02204547 | 0.98241168 | 0.99999872 |
| SMARCAD1 | 40.1033519 | -0.066635193 | 0.29152204 | -0.2285769 | 0.81919781 | 0.99999872 |
| PDLIM5 | 553.224582 | -0.036551208 | 0.30829352 | -0.1185598 | 0.90562414 | 0.99999872 |
| BMPR1B | 24.5772088 | 0.495845983 | 0.51242153 | 0.96765252 | 0.33321794 | 0.99999872 |
| UNC5C | 404.739313 | -0.268472778 | 0.40806219 | -0.6579212 | 0.51058875 | 0.99999872 |
| STPG2 | 83.1535284 | -0.739133533 | 0.47531081 | -1.5550531 | 0.11993343 | 0.99999872 |
| RAP1GDS1 | 73.8686424 | 0.082257595 | 0.2625388 | 0.31331595 | 0.75404063 | 0.99999872 |
| TSPAN5 | 76.3927901 | 0.345699534 | 0.51245001 | 0.67460147 | 0.499929 | 0.99999872 |
| EIF4E | 32.6105421 | -0.005309447 | 0.48014161 | -0.0110581 | 0.9911771 | 0.99999872 |
| METAP1 | 36.2821759 | 0.232268178 | 0.32136589 | 0.72275305 | 0.46983162 | 0.99999872 |
| ADH5 | 43.6439064 | 0.57103295 | 0.43890118 | 1.30105129 | 0.1932409 | 0.99999872 |
| TRMT10A | 6.01005629 | -0.013575818 | 0.7586189 | -0.0178954 | 0.98572227 | 0.99999872 |
| LAMTOR3 | 16.4921868 | 0.040479932 | 0.48286624 | 0.0838326 | 0.93318953 | 0.99999872 |
| DNAJB14 | 65.5811063 | 0.068442613 | 0.26871093 | 0.25470722 | 0.79894924 | 0.99999872 |
| H2AFZ | 25.6297784 | 1.592225787 | 0.50489927 | 3.15355135 | 0.00161297 | 0.3314549 |
| DDIT4L | 17.1750452 | -0.426530284 | 0.79471784 | -0.5367066 | 0.59147032 | 0.99999872 |
| PPP3CA | 87.0965909 | -0.443428782 | 0.30857697 | -1.4370119 | 0.15071461 | 0.99999872 |
| BANK1 | 19.8265534 | -1.078929881 | 0.70535116 | -1.5296351 | 0.12610708 | 0.99999872 |
| NFKB1 | 56.3154595 | 0.239081561 | 0.26683103 | 0.89600359 | 0.37025084 | 0.99999872 |

| | | | | | | |
|---|---|---|---|---|---|---|
| MANBA | 30.54956 | 0.550996873 | 0.42185445 | 1.3061303 | 0.19150825 | 0.99999872 |
| UBE2D3 | 127.478238 | 0.12710883 | 0.22666771 | 0.56077167 | 0.5749532 | 0.99999872 |
| AC018797.2 | 6.65429808 | 0.025130166 | 0.7075292 | 0.0355182 | 0.97166653 | 0.99999872 |
| CISD2 | 18.59099 | 0.267726321 | 0.37512071 | 0.71370713 | 0.4754083 | 0.99999872 |
| SLC9B1 | 6.41985587 | 0.771192894 | 0.78003496 | 0.98866452 | 0.32282731 | 0.99999872 |
| SLC9B2 | 9.80615598 | 0.068530877 | 0.53691913 | 0.12763724 | 0.89843606 | 0.99999872 |
| TACR3 | 17.6913554 | -0.359616077 | 0.89635366 | -0.4011989 | 0.68827371 | 0.99999872 |
| TET2 | 176.768916 | 0.085917457 | 0.49021014 | 0.17526658 | 0.86087017 | 0.99999872 |
| PPA2 | 59.6250433 | 0.443332915 | 0.42181641 | 1.05100916 | 0.29325438 | 0.99999872 |
| ARHGEF38 | 14.4572537 | 0.022609119 | 0.67277259 | 0.03360589 | 0.97319143 | 0.99999872 |
| INTS12 | 241.984336 | -0.489625553 | 0.49225216 | -0.9946641 | 0.31989966 | 0.99999872 |
| GSTCD | 50.6033371 | 0.729471928 | 0.4395604 | 1.65954878 | 0.09700526 | 0.99999872 |
| NPNT | 1143.6162 | 0.195327811 | 0.33975146 | 0.57491382 | 0.56534958 | 0.99999872 |
| AC109361.2 | 14.0268468 | 1.534946844 | 0.83856638 | 1.83044166 | 0.06718392 | 0.99999872 |
| AC109361.1 | 7.81644584 | 3.741278717 | 0.90431642 | 4.13713456 | 3.52E-05 | 0.06141329 |
| TBCK | 101.109479 | 0.445559144 | 0.29874881 | 1.49141729 | 0.13585197 | 0.99999872 |
| AIMP1 | 22.5172528 | 0.063172738 | 0.37060806 | 0.170457 | 0.86465075 | 0.99999872 |
| PAPSS1 | 87.3367871 | 0.406161969 | 0.39619115 | 1.0251667 | 0.30528454 | 0.99999872 |
| SGMS2 | 8.75073817 | -0.626726209 | 0.51740804 | -1.2112804 | 0.22578796 | 0.99999872 |
| CYP2U1 | 8.74734296 | 0.690657615 | 0.62186604 | 1.11062122 | 0.26673143 | 0.99999872 |
| HADH | 11.1025827 | 0.229233391 | 0.52919621 | 0.43317277 | 0.66488926 | 0.99999872 |
| RPL34-AS1 | 7.92710018 | 0.596786827 | 0.79095031 | 0.75451873 | 0.4505378 | 0.99999872 |
| RPL34 | 52.971398 | 0.662967342 | 0.4825739 | 1.37381516 | 0.16949907 | 0.99999872 |
| OSTC | 32.1142739 | 1.231433271 | 0.52181995 | 2.35988155 | 0.01828077 | 0.85282771 |
| SEC24B-AS1 | 9.34857539 | 0.179627181 | 0.60373279 | 0.29752762 | 0.76606372 | 0.99999872 |
| SEC24B | 76.2279817 | 0.095283707 | 0.24870357 | 0.3831216 | 0.70162961 | 0.99999872 |
| MCUB | 16.732048 | -0.455873463 | 0.60955292 | -0.7478817 | 0.45453153 | 0.99999872 |
| CASP6 | 12.8153802 | -0.18162398 | 0.44954109 | -0.4040209 | 0.68619737 | 0.99999872 |
| PLA2G12A | 10.9924831 | 0.144362263 | 0.46313176 | 0.31170884 | 0.75526181 | 0.99999872 |
| GAR1 | 10.6255932 | 0.083400474 | 0.46882759 | 0.17789157 | 0.85880813 | 0.99999872 |
| EGF | 49.5686611 | 0.078823939 | 1.50813419 | 0.05226587 | 0.95831685 | 0.99999872 |
| ELOVL6 | 23.6561021 | -0.198942456 | 0.45226511 | -0.4398802 | 0.6600239 | 0.99999872 |
| ENPEP | 899.698009 | 0.252017082 | 0.42071761 | 0.59901719 | 0.54916142 | 0.99999872 |
| FAM241A | 6.28416126 | 0.262025 | 0.68743588 | 0.38116282 | 0.70308244 | 0.99999872 |
| AP1AR | 10.3526016 | 0.723487276 | 0.54846643 | 1.31910949 | 0.18713251 | 0.99999872 |
| ALPK1 | 46.5018636 | -0.337214045 | 0.44765274 | -0.7532938 | 0.45127337 | 0.99999872 |
| LARP7 | 28.3420047 | 0.057223141 | 0.35038939 | 0.16331299 | 0.870272 | 0.99999872 |
| ANK2 | 63.5817812 | -1.206103665 | 0.58655859 | -2.0562373 | 0.03975965 | 0.95980711 |
| CAMK2D | 77.110664 | 0.332837304 | 0.247911 | 1.34256769 | 0.179412 | 0.99999872 |
| ARSJ | 14.0291663 | -0.120505795 | 0.4744482 | -0.2539915 | 0.79950215 | 0.99999872 |
| UGT8 | 20.2098765 | 0.208737868 | 0.78706015 | 0.26521209 | 0.79084608 | 0.99999872 |
| SNHG8 | 10.6660345 | 0.671677584 | 0.78678688 | 0.85369698 | 0.39327291 | 0.99999872 |
| METTL14 | 17.2094726 | 0.308261495 | 0.37230086 | 0.8279903 | 0.407676 | 0.99999872 |
| SEC24D | 31.0134832 | 0.024564278 | 0.33683847 | 0.07292599 | 0.94186501 | 0.99999872 |
| USP53 | 168.854373 | 0.006601187 | 0.36968476 | 0.01785626 | 0.98575352 | 0.99999872 |
| C4orf3 | 50.7585642 | 0.316878041 | 0.48505302 | 0.65328538 | 0.51357232 | 0.99999872 |
| PDE5A | 19.4885334 | 1.681424571 | 0.73367242 | 2.29179199 | 0.02191765 | 0.8732058 |
| AC108866.1 | 19.0595761 | -0.67426679 | 0.53022084 | -1.2716716 | 0.20348982 | 0.99999872 |
| PRDM5 | 112.791518 | 0.357092079 | 0.40160989 | 0.8891516 | 0.37392161 | 0.99999872 |
| NDNF | 408.369454 | 0.62372249 | 0.67857295 | 0.91916793 | 0.35800775 | 0.99999872 |
| ANXA5 | 84.2137895 | 0.094753578 | 0.51271036 | 0.18480918 | 0.85337871 | 0.99999872 |
| EXOSC9 | 10.8125624 | 0.149541021 | 0.49037128 | 0.30495469 | 0.76040066 | 0.99999872 |
| BBS7 | 18.5024142 | 0.525329031 | 0.46085869 | 1.13989177 | 0.25433139 | 0.99999872 |
| KIAA1109 | 115.907443 | 0.181694872 | 0.30637294 | 0.5930513 | 0.55314682 | 0.99999872 |
| SPATA5 | 43.5499007 | -0.114087786 | 0.41026376 | -0.278084 | 0.78094788 | 0.99999872 |
| SPRY1 | 12.9168856 | -2.04732402 | 0.53339915 | -3.8382589 | 0.00012391 | 0.0998713 |
| ANKRD50 | 9.00979969 | 0.334289638 | 0.6252669 | 0.53463511 | 0.5929022 | 0.99999872 |
| FAT4 | 20.4972547 | 0.295337583 | 0.66941463 | 0.44118782 | 0.65907703 | 0.99999872 |
| INTU | 32.45299 | -0.052431281 | 0.35217742 | -0.1488775 | 0.88165032 | 0.99999872 |
| HSPA4L | 13.123739 | 0.547773197 | 0.67965541 | 0.80595724 | 0.42026751 | 0.99999872 |
| MFSD8 | 27.6265508 | -0.268072338 | 0.31277282 | -0.8570832 | 0.39139889 | 0.99999872 |

| | | | | | | |
|---|---|---|---|---|---|---|
| ABHD18 | 57.2022909 | -0.358102085 | 0.34321995 | -1.0433603 | 0.29678144 | 0.99999872 |
| LARP1B | 65.3938457 | -0.22661144 | 0.28576866 | -0.7929891 | 0.42778416 | 0.99999872 |
| PGRMC2 | 41.6107212 | 0.199556312 | 0.3026457 | 0.65937269 | 0.50965647 | 0.99999872 |
| AC108062.1 | 17.4099509 | -0.179637545 | 0.46002376 | -0.3904962 | 0.69616964 | 0.99999872 |
| JADE1 | 71.6661506 | -0.020427208 | 0.33213201 | -0.0615033 | 0.9509584 | 0.99999872 |
| SCLT1 | 47.8758949 | -0.531947338 | 0.48770153 | -1.0907231 | 0.27539473 | 0.99999872 |
| C4orf33 | 7.28678486 | -0.521036169 | 0.75568162 | -0.6894916 | 0.49051393 | 0.99999872 |
| AC096711.2 | 12.4497357 | -0.296265921 | 0.50684229 | -0.5845328 | 0.55886194 | 0.99999872 |
| AC109927.1 | 26.6993012 | 1.059235676 | 0.45608644 | 2.32244499 | 0.02020899 | 0.8671024 |
| NOCT | 5.06114113 | 0.537972375 | 0.75316347 | 0.71428368 | 0.47505178 | 0.99999872 |
| ELF2 | 166.242 | -0.194426224 | 0.24412463 | -0.796422 | 0.42578679 | 0.99999872 |
| NDUFC1 | 32.5287724 | 0.993723416 | 0.4775631 | 2.0808212 | 0.03745027 | 0.95980711 |
| NAA15 | 45.069448 | 0.32899187 | 0.32441919 | 1.01409496 | 0.31053744 | 0.99999872 |
| AC097376.2 | 66.434697 | -0.242118976 | 0.38637327 | -0.6266453 | 0.5308918 | 0.99999872 |
| RAB33B | 12.9184342 | 0.84789621 | 0.48429877 | 1.75077092 | 0.07998538 | 0.99999872 |
| SETD7 | 20.967822 | -0.018831824 | 0.46153316 | -0.0408028 | 0.96745314 | 0.99999872 |
| MGST2 | 20.4201715 | -0.334046257 | 0.35690219 | -0.9359602 | 0.34929366 | 0.99999872 |
| MAML3 | 402.04993 | -0.226417377 | 0.31182925 | -0.7260941 | 0.4677811 | 0.99999872 |
| SCOC | 61.0108717 | 0.918679453 | 0.37588827 | 2.44402268 | 0.01452451 | 0.77646863 |
| SCOC-AS1 | 5.35295308 | 0.306847024 | 0.7794547 | 0.39366884 | 0.69382555 | 0.99999872 |
| ELMOD2 | 19.6714552 | -0.021183711 | 0.42012461 | -0.0504224 | 0.95978575 | 0.99999872 |
| TBC1D9 | 34.0108703 | -0.078175596 | 0.35678663 | -0.2191102 | 0.82656419 | 0.99999872 |
| RNF150 | 127.909894 | 0.377449422 | 0.44194408 | 0.85406603 | 0.39306841 | 0.99999872 |
| ZNF330 | 14.2121979 | 0.987434622 | 0.48499942 | 2.03595012 | 0.04175536 | 0.95980711 |
| IL15 | 20.1031394 | -0.198406623 | 0.36417681 | -0.5448085 | 0.58588523 | 0.99999872 |
| INPP4B | 21.3080699 | -1.095862941 | 0.95001223 | -1.1535251 | 0.24869493 | 0.99999872 |
| AC104596.1 | 12.9254088 | -1.118261728 | 0.90827774 | -1.2311892 | 0.21825212 | 0.99999872 |
| USP38 | 16.5518338 | 0.269532636 | 0.38244013 | 0.7047708 | 0.48095288 | 0.99999872 |
| GAB1 | 41.5657987 | 0.05614342 | 0.43699904 | 0.12847493 | 0.89777314 | 0.99999872 |
| SMARCA5 | 48.662237 | -0.090256042 | 0.27896974 | -0.3235335 | 0.74629128 | 0.99999872 |
| ANAPC10 | 37.4675586 | 0.391058422 | 0.30059567 | 1.30094496 | 0.1932773 | 0.99999872 |
| ABCE1 | 16.3050145 | -0.210448945 | 0.46668337 | -0.4509459 | 0.65202855 | 0.99999872 |
| OTUD4 | 41.6962951 | 0.010015254 | 0.31942972 | 0.03135354 | 0.97498759 | 0.99999872 |
| SMAD1 | 45.7195533 | -0.327762551 | 0.30267078 | -1.0829012 | 0.27885227 | 0.99999872 |
| MMAA | 6.78517457 | 0.069134849 | 0.63403974 | 0.10903867 | 0.91317182 | 0.99999872 |
| ZNF827 | 36.1140191 | 0.87520851 | 0.38504312 | 2.27301426 | 0.02302532 | 0.88337526 |
| LSM6 | 12.191101 | 0.943166967 | 0.507813 | 1.85731159 | 0.06326684 | 0.99999872 |
| SLC10A7 | 30.1865026 | 0.511451897 | 0.3708035 | 1.3793071 | 0.16780009 | 0.99999872 |
| TTC29 | 7.50639364 | 2.192919225 | 1.37944625 | 1.5897098 | 0.11190023 | 0.99999872 |
| EDNRA | 8.17980638 | 1.390412657 | 0.82118702 | 1.69317417 | 0.09042232 | 0.99999872 |
| TMEM184C | 15.5948714 | -0.208349403 | 0.43526368 | -0.478674 | 0.63217057 | 0.99999872 |
| PRMT9 | 13.6238626 | 0.437654042 | 0.48786872 | 0.89707339 | 0.36967976 | 0.99999872 |
| ARHGAP10 | 11.2560278 | -0.410629888 | 0.90335193 | -0.4545625 | 0.64942404 | 0.99999872 |
| NR3C2 | 187.179273 | 0.277120451 | 0.41736741 | 0.66397243 | 0.50670796 | 0.99999872 |
| DCLK2 | 60.8642308 | -0.031238 | 0.31636853 | -0.0987393 | 0.92134528 | 0.99999872 |
| LRBA | 208.593941 | -0.068179787 | 0.28745814 | -0.2371816 | 0.81251589 | 0.99999872 |
| RPS3A | 34.4543048 | 0.965687572 | 0.50323801 | 1.91894801 | 0.05499091 | 0.99999872 |
| SH3D19 | 91.3399211 | -0.446204197 | 0.26921205 | -1.6574451 | 0.09742951 | 0.99999872 |
| FAM160A1 | 19.7638571 | -0.33017813 | 0.73375599 | -0.4499836 | 0.6527223 | 0.99999872 |
| GATB | 6.65811706 | -0.732727288 | 0.76113345 | -0.9626791 | 0.33570858 | 0.99999872 |
| FBXW7 | 109.323407 | 0.122269474 | 0.27709211 | 0.44125932 | 0.65902528 | 0.99999872 |
| ARFIP1 | 46.7948824 | -0.089082674 | 0.26326382 | -0.338378 | 0.73507838 | 0.99999872 |
| TRIM2 | 33.4131409 | -0.134850777 | 0.56550697 | -0.23846 | 0.81152436 | 0.99999872 |
| PLRG1 | 8.95459169 | 0.298000748 | 0.52879576 | 0.56354603 | 0.57306313 | 0.99999872 |
| FGB | 8.09862854 | 2.002462586 | 2.34907467 | 0.8524474 | 0.39396582 | 0.99999872 |
| MAP9 | 24.2143647 | -0.140616781 | 0.31954875 | -0.440048 | 0.65990236 | 0.99999872 |
| GUCY1B1 | 12.7648111 | 0.892340729 | 0.72096262 | 1.23770735 | 0.21582459 | 0.99999872 |
| CTSO | 16.9967973 | -0.355008301 | 0.71669741 | -0.4953392 | 0.62036073 | 0.99999872 |
| PDGFC | 71.3435242 | 0.577188376 | 0.41405037 | 1.39400521 | 0.16331603 | 0.99999872 |
| ETFDH | 31.6803397 | 0.189748377 | 0.37726445 | 0.50295855 | 0.61499342 | 0.99999872 |
| PPID | 10.6738111 | 0.190777335 | 0.42991685 | 0.44375403 | 0.65722043 | 0.99999872 |

| | | | | | | |
|---|---|---|---|---|---|---|
| FNIP2 | 56.8279005 | -0.961570159 | 0.46819057 | -2.0538008 | 0.03999498 | 0.95980711 |
| RAPGEF2 | 144.632666 | -0.12944715 | 0.27773828 | -0.466076 | 0.6411611 | 0.99999872 |
| NAF1 | 18.5050129 | -0.016989007 | 0.4195228 | -0.040496 | 0.96769767 | 0.99999872 |
| TMA16 | 17.5298013 | 0.082690973 | 0.45649036 | 0.18114506 | 0.85625372 | 0.99999872 |
| TMEM192 | 25.9560119 | -0.295504956 | 0.49451116 | -0.5975698 | 0.55012699 | 0.99999872 |
| KLHL2 | 29.1195979 | -0.30200564 | 0.32449336 | -0.930699 | 0.3520093 | 0.99999872 |
| MSMO1 | 23.9773807 | 0.775940242 | 0.60677042 | 1.27880366 | 0.2009662 | 0.99999872 |
| DDX60 | 43.3059122 | -1.049077748 | 0.74079727 | -1.4161469 | 0.1567325 | 0.99999872 |
| DDX60L | 39.4297543 | -1.560401634 | 1.1318041 | -1.3786853 | 0.16799181 | 0.99999872 |
| PALLD | 241.349514 | -0.228634098 | 0.51937125 | -0.4402132 | 0.65978268 | 0.99999872 |
| CBR4 | 111.176847 | -0.047519038 | 0.37531532 | -0.126611 | 0.89924831 | 0.99999872 |
| SH3RF1 | 326.447446 | -0.11741589 | 0.29499501 | -0.3980267 | 0.6906105 | 0.99999872 |
| AC096741.1 | 6.693841 | 0.877656379 | 0.7238422 | 1.21249684 | 0.22532225 | 0.99999872 |
| NEK1 | 52.719379 | -0.023784054 | 0.38263182 | -0.0621591 | 0.95043612 | 0.99999872 |
| CLCN3 | 62.9467913 | 0.14625824 | 0.29562065 | 0.49474974 | 0.62077679 | 0.99999872 |
| HPF1 | 14.4096004 | 0.692790218 | 0.43835884 | 1.58041805 | 0.11401116 | 0.99999872 |
| MFAP3L | 10.2495953 | 0.005204296 | 0.53295659 | 0.00976495 | 0.99220882 | 0.99999872 |
| GALNT7 | 18.8664743 | 0.193355282 | 0.57336581 | 0.33722848 | 0.73594467 | 0.99999872 |
| HMGB2 | 9.79204624 | 0.622699904 | 0.74217126 | 0.83902455 | 0.40145553 | 0.99999872 |
| FBXO8 | 15.2396193 | 0.417603561 | 0.40580619 | 1.02907144 | 0.30344611 | 0.99999872 |
| CEP44 | 26.0817227 | 0.127573832 | 0.32779668 | 0.38918586 | 0.69713866 | 0.99999872 |
| HPGD | 180.334912 | 0.832761472 | 1.10725949 | 0.75209243 | 0.45199548 | 0.99999872 |
| WDR17 | 16.3565253 | -0.17197209 | 0.62703445 | -0.2742626 | 0.78388283 | 0.99999872 |
| SPCS3 | 64.77761 | 0.330963783 | 0.3824894 | 0.86528878 | 0.38688031 | 0.99999872 |
| AGA | 4.17500343 | 0.41596929 | 0.72553717 | 0.57332596 | 0.56642401 | NA |
| AC078881.1 | 25.1910835 | -0.531935136 | 0.47794224 | -1.1129695 | 0.26572153 | 0.99999872 |
| AC098864.1 | 21.9056728 | -0.021673903 | 0.49207198 | -0.0440462 | 9.65E-01 | 0.99999872 |
| TENM3 | 328.112325 | 0.096920591 | 0.34060711 | 0.28455246 | 0.77598704 | 0.99999872 |
| DCTD | 35.4977232 | -0.180734421 | 0.29336078 | -0.6160824 | 0.53784012 | 0.99999872 |
| WWC2 | 170.909009 | 0.01216031 | 0.28621841 | 0.04248612 | 0.96611118 | 0.99999872 |
| CDKN2AIP | 8.35152592 | 0.289597438 | 0.58896772 | 0.49170341 | 0.62292903 | 0.99999872 |
| ING2 | 6.26885483 | 0.783541628 | 0.64228409 | 1.21993 | 0.22249141 | 0.99999872 |
| RWDD4 | 11.326448 | -0.331201508 | 0.45883475 | -0.7218318 | 0.47039791 | 0.99999872 |
| TRAPPC11 | 26.9385475 | 0.099900405 | 0.39085808 | 0.25559253 | 0.79826549 | 0.99999872 |
| STOX2 | 40.4419288 | -0.574105356 | 0.45107916 | -1.2727375 | 0.20311121 | 0.99999872 |
| IRF2 | 27.8634228 | -0.119259435 | 0.44701612 | -0.26679 | 0.78963085 | 0.99999872 |
| CASP3 | 11.1567917 | -0.411001462 | 0.52347691 | -0.7851377 | 0.43237283 | 0.99999872 |
| PRIMPOL | 18.6873039 | -0.046346041 | 0.42757435 | -0.1083929 | 0.913684 | 0.99999872 |
| ACSL1 | 36.2467349 | -0.018246817 | 0.42489639 | -0.0429442 | 0.96574605 | 0.99999872 |
| SLC25A4 | 36.1613192 | 0.999252746 | 0.53292713 | 1.87502696 | 0.06078901 | 0.99999872 |
| CFAP97 | 42.5674389 | 0.374251017 | 0.31856331 | 1.17480891 | 0.24007118 | 0.99999872 |
| SNX25 | 291.735552 | -0.498682085 | 0.49454222 | -1.0083711 | 0.31327634 | 0.99999872 |
| LRP2BP | 24.7977734 | 0.474451636 | 0.63152484 | 0.75127945 | 0.45248449 | 0.99999872 |
| AC112722.1 | 47.3581187 | -0.132846755 | 0.55764997 | -0.2382261 | 0.81170577 | 0.99999872 |
| ANKRD37 | 11.8507086 | 0.393540197 | 1.12927836 | 0.34848821 | 0.72747356 | 0.99999872 |
| UFSP2 | 19.328326 | 0.560379572 | 0.38477014 | 1.45640089 | 0.14528183 | 0.99999872 |
| C4orf47 | 8.35525548 | -0.313544878 | 0.90619533 | -0.3460014 | 0.72934164 | 0.99999872 |
| SORBS2 | 26.0093951 | -0.024204788 | 0.67414046 | -0.0359047 | 0.97135838 | 0.99999872 |
| TLR3 | 12.5803142 | 0.759434735 | 0.88312367 | 0.85994155 | 0.38982127 | 0.99999872 |
| CYP4V2 | 10.4565076 | 0.025298182 | 0.53270168 | 0.04749034 | 0.96212243 | 0.99999872 |
| FAT1 | 312.718706 | 0.519145359 | 0.3874353 | 1.33995369 | 0.1802604 | 0.99999872 |
| LINC01060 | 8.80890755 | 1.42246408 | 1.22174654 | 1.16428738 | 0.24430757 | 0.99999872 |
| FRG1-DT | 10.876123 | -1.241858108 | 0.79272668 | -1.5665653 | 0.11721633 | 0.99999872 |
| FRG1 | 15.6454422 | 0.563479036 | 0.43910049 | 1.28325758 | 0.19940185 | 0.99999872 |
| CCDC127 | 9.6079715 | 0.830898217 | 0.49082007 | 1.69287744 | 0.0904788 | 0.99999872 |
| SDHA | 30.1320817 | 0.053158903 | 0.31686309 | 0.16776615 | 0.86676725 | 0.99999872 |
| PDCD6 | 15.1743429 | -0.451869861 | 0.44146862 | -1.0235605 | 0.30604289 | 0.99999872 |
| EXOC3 | 20.199845 | -0.521478552 | 0.34840784 | -1.4967475 | 0.13445898 | 0.99999872 |
| SLC9A3 | 8.82855581 | 0.263753357 | 0.72426519 | 0.36416683 | 0.71573343 | 0.99999872 |
| BRD9 | 12.560167 | -0.182002931 | 0.47948006 | -0.3795839 | 0.70425428 | 0.99999872 |
| AC116351.1 | 9.3210871 | 1.241135191 | 0.60819396 | 2.04068976 | 0.04128167 | 0.95980711 |

| | | | | | | |
|---|---|---|---|---|---|---|
| SLC12A7 | 23.0577699 | -0.242739049 | 0.42961646 | -0.5650134 | 0.57206466 | 0.99999872 |
| CLPTM1L | 24.3217395 | 0.32961329 | 0.38436807 | 0.85754597 | 0.39114322 | 0.99999872 |
| LPCAT1 | 18.4294149 | -0.959281023 | 0.50234978 | -1.9095878 | 0.0561863 | 0.99999872 |
| MRPL36 | 13.5274873 | 1.215118142 | 0.48200658 | 2.52095756 | 0.0117036 | 0.72670507 |
| NDUFS6 | 36.3435499 | 0.507027185 | 0.55410539 | 0.91503745 | 0.36017197 | 0.99999872 |
| ICE1 | 39.7177887 | 0.210803712 | 0.31611848 | 0.66685032 | 0.50486775 | 0.99999872 |
| MED10 | 19.7452782 | 0.436443012 | 0.38953483 | 1.12042102 | 0.26253439 | 0.99999872 |
| NSUN2 | 18.0860234 | 0.09869267 | 0.4418992 | 0.22333752 | 0.82327283 | 0.99999872 |
| SRD5A1 | 5.3085567 | 0.142067782 | 0.70457555 | 0.20163598 | 0.84020131 | 0.99999872 |
| TENT4A | 27.3352821 | 0.405932977 | 0.38768759 | 1.04706208 | 0.29507095 | 0.99999872 |
| MTRR | 13.2202383 | -0.579163294 | 0.59680425 | -0.970441 | 0.33182674 | 0.99999872 |
| MIR4458HG | 5.49385944 | 0.774319975 | 0.70309401 | 1.10130361 | 0.27076454 | 0.99999872 |
| AC021088.1 | 76.5651506 | 1.967738474 | 0.77292342 | 2.54583887 | 0.01090155 | 0.71391523 |
| SEMA5A | 1100.23068 | -0.006219885 | 0.29754635 | -0.0209039 | 0.9833223 | 0.99999872 |
| AC091906.1 | 9.340184 | 1.327613476 | 0.73414575 | 1.80837861 | 0.07054759 | 0.99999872 |
| SNHG18 | 9.47965764 | 0.717723071 | 0.54724995 | 1.3115087 | 0.18968596 | 0.99999872 |
| FAM173B | 8.57498569 | -0.019037625 | 0.58937903 | -0.0323012 | 0.97423189 | 0.99999872 |
| CCT5 | 29.7445213 | 0.019200806 | 0.4872946 | 0.03940287 | 0.96856919 | 0.99999872 |
| CMBL | 8.66616465 | 1.031319211 | 0.98263862 | 1.04954069 | 0.29392934 | 0.99999872 |
| 6-Mar | 125.255332 | 0.210472414 | 0.26846041 | 0.78399795 | 0.43304131 | 0.99999872 |
| ANKRD33B | 148.140922 | -0.540357746 | 0.5564605 | -0.9710622 | 0.33151731 | 0.99999872 |
| DAP | 26.0592561 | -0.191730135 | 0.38990458 | -0.491736 | 0.62290596 | 0.99999872 |
| DNAH5 | 9.03451041 | -0.205449877 | 0.90945707 | -0.2259039 | 0.82127618 | 0.99999872 |
| TRIO | 129.461008 | -0.154248037 | 0.34246262 | -0.4504084 | 0.65241599 | 0.99999872 |
| OTULIN | 17.7830716 | -0.034707892 | 0.38875902 | -0.0892787 | 0.92886044 | 0.99999872 |
| ANKH | 126.276945 | -0.662911579 | 0.68096352 | -0.9734906 | 0.33030954 | 0.99999872 |
| LINC02149 | 83.7449693 | -1.541849156 | 0.76403129 | -2.0180445 | 0.04358662 | 0.95980711 |
| FBXL7 | 333.381195 | -0.371839599 | 0.37930621 | -0.9803151 | 0.32693062 | 0.99999872 |
| ZNF622 | 6.41402183 | -0.232476517 | 0.55698033 | -0.4173873 | 0.67639514 | 0.99999872 |
| RETREG1 | 412.859695 | 0.481556989 | 0.32835267 | 1.46658467 | 0.14248907 | 0.99999872 |
| MYO10 | 30.3385586 | -0.725749141 | 0.63079668 | -1.1505278 | 0.24992654 | 0.99999872 |
| LINC02241 | 12.5033451 | 0.657752495 | 0.55672813 | 1.18146086 | 0.23741969 | 0.99999872 |
| CDH12 | 22.7161386 | -0.814090514 | 0.71146112 | -1.1442516 | 0.25251932 | 0.99999872 |
| C5orf17 | 17.2090698 | -1.763827617 | 1.0073429 | -1.7509704 | 0.07995101 | 0.99999872 |
| CDH10 | 20.2176033 | -1.072683987 | 1.1091748 | -0.9671009 | 0.33349358 | 0.99999872 |
| CDH6 | 29.8654129 | 0.647042447 | 0.91969634 | 0.70353922 | 0.48171977 | 0.99999872 |
| DROSHA | 32.4362473 | 0.088274254 | 0.30258975 | 0.29172916 | 0.77049371 | 0.99999872 |
| C5orf22 | 11.3709198 | 0.19318059 | 0.54061681 | 0.35733367 | 0.72084203 | 0.99999872 |
| PDZD2 | 56.8715457 | -0.731329746 | 0.58530697 | -1.2494807 | 0.21148932 | 0.99999872 |
| GOLPH3 | 81.6191346 | 0.091624948 | 0.28013575 | 0.32707339 | 0.74361238 | 0.99999872 |
| MTMR12 | 38.0071663 | 0.38746926 | 0.29061963 | 1.33325219 | 0.18244906 | 0.99999872 |
| ZFR | 105.138353 | -0.107377912 | 0.1972016 | -0.5445083 | 0.58609173 | 0.99999872 |
| SUB1 | 86.2805281 | 1.162096956 | 0.29978904 | 3.8763824 | 0.00010602 | 0.09257402 |
| NPR3 | 184.416851 | -0.195084704 | 0.54482758 | -0.3580669 | 0.72029327 | 0.99999872 |
| TARS | 24.9636887 | 0.465203255 | 0.43711121 | 1.0642675 | 0.28720753 | 0.99999872 |
| C1QTNF3-AN | 32.5476212 | -0.463299297 | 0.36887059 | -1.2559941 | 0.20911809 | 0.99999872 |
| RAI14 | 227.951516 | -0.193973165 | 0.39325152 | -0.4932547 | 0.62183262 | 0.99999872 |
| RAD1 | 10.8353072 | 0.604486549 | 0.56856689 | 1.06317579 | 0.28770224 | 0.99999872 |
| BRIX1 | 9.44173183 | 0.359953453 | 0.53072247 | 0.67823293 | 0.49762402 | 0.99999872 |
| DNAJC21 | 23.4631364 | 0.090321122 | 0.40261196 | 0.2243379 | 0.82249439 | 0.99999872 |
| SPEF2 | 24.9106005 | 0.004278663 | 0.58582938 | 0.0073036 | 0.99417262 | 0.99999872 |
| LMBRD2 | 22.2868814 | 0.080931056 | 0.42510114 | 0.19038071 | 0.84901081 | 0.99999872 |
| SKP2 | 4.17983132 | 0.147882788 | 0.65977361 | 0.22414171 | 0.82264704 | NA |
| NADK2 | 18.8456966 | 0.343022295 | 0.41035174 | 0.83592259 | 0.40319845 | 0.99999872 |
| RANBP3L | 12.6589547 | 0.586555644 | 1.15985329 | 0.50571538 | 0.61305647 | 0.99999872 |
| NIPBL | 182.399036 | 0.037897203 | 0.22454652 | 0.16877217 | 0.86597585 | 0.99999872 |
| CPLANE1 | 86.3852641 | -0.249650648 | 0.32656989 | -0.7644632 | 0.44459128 | 0.99999872 |
| NUP155 | 13.1562472 | -0.690192257 | 0.44519963 | -1.5502984 | 0.12106992 | 0.99999872 |
| WDR70 | 83.7075395 | -0.230201759 | 0.21476802 | -1.0718624 | 0.28378186 | 0.99999872 |
| EGFLAM | 7.79660161 | 0.482797707 | 0.94523028 | 0.51077258 | 0.60951031 | 0.99999872 |
| LIFR | 60.1058882 | -0.087614477 | 0.3862329 | -0.2268436 | 0.82054533 | 0.99999872 |

| | | | | | | |
|---|---|---|---|---|---|---|
| OSMR-AS1 | 58.6734617 | -0.146693682 | 0.61465737 | -0.2386593 | 8.11E-01 | 0.99999872 |
| OSMR | 82.430421 | -0.481933977 | 0.69103386 | -0.6974101 | 0.48554619 | 0.99999872 |
| RICTOR | 84.6579599 | -0.25058479 | 0.26638442 | -0.9406886 | 0.34686446 | 0.99999872 |
| C9 | 4.18985536 | 0.730069192 | 1.02687505 | 0.71096205 | 0.47710775 | NA |
| DAB2 | 22.0276898 | 0.599377972 | 0.7506153 | 0.79851553 | 0.42457139 | 0.99999872 |
| TTC33 | 36.3255553 | 0.528811427 | 0.36360395 | 1.45436105 | 0.14584623 | 0.99999872 |
| PTGER4 | 139.08083 | 0.269055357 | 0.46757792 | 0.57542358 | 0.56500486 | 0.99999872 |
| PRKAA1 | 38.3316431 | 0.530908494 | 0.38924181 | 1.36395547 | 0.17258158 | 0.99999872 |
| RPL37 | 40.3160364 | 0.281296985 | 0.33284951 | 0.84511762 | 0.39804517 | 0.99999872 |
| CARD6 | 5.14678065 | -0.47134455 | 0.74931079 | -0.6290374 | 0.52932454 | 0.99999872 |
| OXCT1 | 18.7360617 | -1.019326994 | 0.37088057 | -2.7483969 | 0.00598875 | 0.58725857 |
| C5orf51 | 8.58011478 | -0.186308863 | 0.46663584 | -0.3992597 | 0.68970189 | 0.99999872 |
| FBXO4 | 4.96336756 | 0.269278199 | 0.60441929 | 0.44551556 | 0.65594721 | 0.99999872 |
| GHR | 182.362022 | -1.230081741 | 0.41333745 | -2.9759745 | 0.00292059 | 0.43717075 |
| SELENOP | 88.8246449 | 0.904313641 | 0.39767643 | 2.27399354 | 0.02296637 | 0.88337526 |
| ZNF131 | 48.7498788 | -0.393094337 | 0.2967204 | -1.3247971 | 0.18523844 | 0.99999872 |
| HMGCS1 | 23.6576772 | 0.029269129 | 0.50403033 | 0.05807017 | 0.95369273 | 0.99999872 |
| CCL28 | 8.88243375 | 0.463268459 | 0.77968028 | 0.59417747 | 0.55239342 | 0.99999872 |
| TMEM267 | 25.9902478 | 0.07309448 | 0.44307353 | 0.16497144 | 0.86896646 | 0.99999872 |
| PAIP1 | 17.6484666 | 0.275127012 | 0.52003892 | 0.52905082 | 0.5967702 | 0.99999872 |
| NNT-AS1 | 19.8108401 | 0.18608541 | 0.42465251 | 0.43820631 | 0.66123674 | 0.99999872 |
| NNT | 37.9629564 | 0.241414378 | 0.33229133 | 0.72651422 | 0.4675236 | 0.99999872 |
| MRPS30-DT | 13.8650306 | 0.05787131 | 0.65872657 | 0.08785331 | 0.92999327 | 0.99999872 |
| MRPS30 | 10.4946963 | 0.641410553 | 0.53578206 | 1.19714824 | 0.23124878 | 0.99999872 |
| PARP8 | 59.1705512 | 0.227620298 | 0.48712594 | 0.46727197 | 0.64030531 | 0.99999872 |
| ITGA1 | 173.680551 | -0.445555658 | 0.34755801 | -1.2819605 | 0.19985649 | 0.99999872 |
| ITGA2 | 24.814167 | -1.933105626 | 0.43997958 | -4.3936258 | 1.11E-05 | 0.03893472 |
| MOCS2 | 18.8528543 | 0.347574321 | 0.47590156 | 0.73034919 | 0.46517676 | 0.99999872 |
| NDUFS4 | 47.6239721 | 0.391717594 | 0.38107729 | 1.02792166 | 0.30398668 | 0.99999872 |
| ARL15 | 157.463783 | 0.218711506 | 0.43760364 | 0.49979361 | 0.61722041 | 0.99999872 |
| SNX18 | 7.58910254 | 0.053166076 | 0.63312286 | 0.08397434 | 0.93307683 | 0.99999872 |
| GPX8 | 14.1602555 | 0.674836981 | 0.77610844 | 0.86951378 | 0.38456617 | 0.99999872 |
| DHX29 | 30.2415603 | 0.01245287 | 0.36699468 | 0.03393202 | 0.97293136 | 0.99999872 |
| MTREX | 49.2926826 | 0.169220928 | 0.33826166 | 0.50026636 | 0.61688754 | 0.99999872 |
| PLPP1 | 125.004542 | 0.179064268 | 0.29997142 | 0.59693776 | 0.55054893 | 0.99999872 |
| SLC38A9 | 22.18069 | 0.126259096 | 0.37831266 | 0.33374272 | 0.73857371 | 0.99999872 |
| IL6ST | 92.7888587 | 0.4222579 | 0.27975242 | 1.50939856 | 0.13119696 | 0.99999872 |
| AC008914.1 | 3.96148005 | 1.477139862 | 1.29804214 | 1.13797528 | 0.2551308 | NA |
| MAP3K1 | 27.9045525 | -0.592726972 | 0.60993195 | -0.971792 | 0.33115405 | 0.99999872 |
| MIER3 | 9.61997072 | 0.206475373 | 0.54485971 | 0.37895144 | 0.70472392 | 0.99999872 |
| GPBP1 | 88.3431839 | 0.003286882 | 0.37084695 | 0.00886318 | 0.9929283 | 0.99999872 |
| PDE4D | 517.403656 | 0.002151608 | 0.42423983 | 0.00507168 | 0.9959534 | 0.99999872 |
| ELOVL7 | 5.53497647 | -0.201901612 | 0.78074438 | -0.2586014 | 0.79594278 | 0.99999872 |
| ERCC8 | 18.7284842 | -0.103919122 | 0.38516573 | -0.2698037 | 0.78731131 | 0.99999872 |
| NDUFAF2 | 32.3820857 | 0.512283992 | 0.40468234 | 1.26589165 | 0.20555188 | 0.99999872 |
| SMIM15 | 14.3390206 | 0.546824916 | 0.42920482 | 1.27404187 | 0.20264858 | 0.99999872 |
| ZSWIM6 | 662.589592 | -0.395418542 | 0.37323625 | -1.0594323 | 0.28940295 | 0.99999872 |
| KIF2A | 37.7692147 | 0.373473158 | 0.44751191 | 0.83455468 | 0.40396849 | 0.99999872 |
| DIMT1 | 6.69127139 | 0.553838046 | 0.55181548 | 1.0036653 | 0.31553997 | 0.99999872 |
| IPO11 | 16.0153725 | -0.026846248 | 0.36971176 | -0.072614 | 0.94211329 | 0.99999872 |
| RNF180 | 43.3310492 | 0.261834531 | 0.46248717 | 0.56614443 | 0.57129561 | 0.99999872 |
| SREK1IP1 | 26.0179415 | 0.154737207 | 0.33287156 | 0.4648556 | 0.64203487 | 0.99999872 |
| CWC27 | 36.3731509 | 0.10145442 | 0.32110333 | 0.31595568 | 0.75203615 | 0.99999872 |
| PPWD1 | 23.602001 | 0.049351973 | 0.45543664 | 0.10836189 | 0.91370863 | 0.99999872 |
| TRIM23 | 20.0643731 | 0.186838995 | 0.43444671 | 0.43006194 | 0.66715059 | 0.99999872 |
| TRAPPC13 | 25.7866304 | -0.196271451 | 0.46493324 | -0.4221497 | 0.67291572 | 0.99999872 |
| SGTB | 7.91114868 | 0.58008964 | 0.61955094 | 0.93630661 | 0.34911535 | 0.99999872 |
| NLN | 29.5711825 | -0.109818339 | 0.35613317 | -0.3083631 | 0.75780604 | 0.99999872 |
| ERBIN | 240.7125 | 0.319231132 | 0.23357926 | 1.36669299 | 0.17172156 | 0.99999872 |
| SREK1 | 63.7667754 | -0.114714067 | 0.3033654 | -0.3781383 | 0.70532788 | 0.99999872 |
| MAST4 | 212.804959 | -0.103004894 | 0.31803238 | -0.3238818 | 0.74602754 | 0.99999872 |

| | | | | | | |
|---|---|---|---|---|---|---|
| PIK3R1 | 43.6639355 | -0.423077751 | 0.35369022 | -1.1961816 | 0.23162568 | 0.99999872 |
| SLC30A5 | 25.4087348 | 0.095692835 | 0.35272658 | 0.27129465 | 0.78616442 | 0.99999872 |
| MRPS36 | 20.1801702 | 0.61928542 | 0.35248155 | 1.75693002 | 0.07892974 | 0.99999872 |
| CDK7 | 13.8351144 | -0.541550464 | 0.46952682 | -1.1533962 | 0.24874778 | 0.99999872 |
| CCDC125 | 15.5597968 | -0.432324686 | 0.54399303 | -0.7947247 | 0.42677368 | 0.99999872 |
| AK6 | 17.5894523 | 0.216383339 | 0.38217564 | 0.56618821 | 0.57126584 | 0.99999872 |
| RAD17 | 21.9575456 | 0.53366618 | 0.38290381 | 1.39373431 | 0.16339785 | 0.99999872 |
| MARVELD2 | 6.71245679 | 0.88170084 | 0.72295662 | 1.21957641 | 0.22262549 | 0.99999872 |
| OCLN | 10.5289308 | -0.832789585 | 0.57580998 | -1.4462924 | 0.14809521 | 0.99999872 |
| GTF2H2C | 10.8974666 | 0.007602666 | 0.49244326 | 0.01543866 | 0.98768222 | 0.99999872 |
| NAIP | 38.5170957 | -0.018450555 | 0.42806385 | -0.0431023 | 0.96561995 | 0.99999872 |
| GTF2H2 | 12.492543 | 0.456632729 | 0.53785879 | 0.84898256 | 0.395891 | 0.99999872 |
| BDP1 | 65.3206482 | -0.134195044 | 0.27474189 | -0.4884404 | 0.62523791 | 0.99999872 |
| MCCC2 | 20.7619429 | -0.603474329 | 0.35862476 | -1.6827458 | 0.0924243 | 0.99999872 |
| MAP1B | 169.958689 | -0.70172932 | 0.45986694 | -1.5259399 | 0.12702483 | 0.99999872 |
| MRPS27 | 26.659814 | 0.162788112 | 0.47705232 | 0.34123744 | 0.73292484 | 0.99999872 |
| PTCD2 | 5.63483904 | 0.186329339 | 0.64817029 | 0.28746973 | 0.77375268 | 0.99999872 |
| TNPO1 | 124.342284 | 0.213589504 | 0.18299172 | 1.16720857 | 0.24312615 | 0.99999872 |
| FCHO2 | 42.6868243 | -0.02021331 | 0.27475698 | -0.073568 | 0.94135417 | 0.99999872 |
| FOXD1 | 15.2228363 | 0.170541655 | 0.67708106 | 0.25187775 | 0.80113556 | 0.99999872 |
| BTF3 | 53.0802845 | 1.018311648 | 0.34748311 | 2.93053568 | 0.00338378 | 0.44319076 |
| ANKRA2 | 10.4890127 | 0.723239379 | 0.50715631 | 1.42606799 | 0.15384872 | 0.99999872 |
| ARHGEF28 | 83.9912541 | -0.269878222 | 0.44584015 | -0.6053251 | 0.54496303 | 0.99999872 |
| ENC1 | 31.8964236 | 2.063817878 | 1.18178115 | 1.74636216 | 0.08074804 | 0.99999872 |
| HEXB | 24.3450179 | 0.553849086 | 0.51209353 | 1.08153892 | 0.27945746 | 0.99999872 |
| GFM2 | 10.6168493 | -0.162293117 | 0.47534564 | -0.3414213 | 0.73278646 | 0.99999872 |
| NSA2 | 12.3965379 | 0.055971619 | 0.47757216 | 0.11720034 | 0.9067013 | 0.99999872 |
| FAM169A | 18.2173729 | -0.050427003 | 0.53066761 | -0.0950256 | 0.9242945 | 0.99999872 |
| HMGCR | 16.0201742 | 0.25479865 | 0.45282375 | 0.56268836 | 0.57364711 | 0.99999872 |
| COL4A3BP | 51.3866278 | 0.039125958 | 0.28510612 | 0.13723297 | 0.89084665 | 0.99999872 |
| POLK | 75.3946776 | 0.153803138 | 0.31121888 | 0.49419604 | 0.62116774 | 0.99999872 |
| POC5 | 9.26949164 | 0.037286806 | 0.56181297 | 0.06636872 | 0.94708428 | 0.99999872 |
| IQGAP2 | 803.14777 | 0.36554514 | 0.48197044 | 0.75843892 | 0.44818827 | 0.99999872 |
| AC025188.1 | 7.04632108 | -0.346766426 | 0.69785614 | -0.4969025 | 0.61925784 | 0.99999872 |
| F2R | 212.849261 | -0.288740766 | 0.58344652 | -0.4948881 | 0.62067908 | 0.99999872 |
| AC008581.2 | 6.83117986 | 1.735818635 | 0.69479209 | 2.49832815 | 0.01247806 | 0.72670507 |
| ZBED3 | 18.1816367 | -0.263881016 | 0.44908121 | -0.587602 | 0.55679947 | 0.99999872 |
| WDR41 | 58.9006402 | 0.043537167 | 0.33095747 | 0.13154913 | 0.89534092 | 0.99999872 |
| TBCA | 93.7783599 | 0.602098119 | 0.39861921 | 1.51045938 | 0.13092625 | 0.99999872 |
| AP3B1 | 97.6724807 | -0.150772261 | 0.26336112 | -0.5724925 | 0.56698837 | 0.99999872 |
| SCAMP1-AS1 | 9.01488246 | -0.986637451 | 0.83749204 | -1.1780858 | 0.23876242 | 0.99999872 |
| SCAMP1 | 46.2208226 | 0.466188936 | 0.3205816 | 1.45419741 | 0.14589158 | 0.99999872 |
| LHFPL2 | 5.42257773 | -0.765693901 | 0.55117133 | -1.3892121 | 0.16476825 | 0.99999872 |
| ARSB | 17.7730438 | -0.826465168 | 0.45100149 | -1.8325109 | 0.06687533 | 0.99999872 |
| DMGDH | 20.1567759 | -0.214421453 | 0.81319011 | -0.2636794 | 0.792027 | 0.99999872 |
| BHMT2 | 15.1299442 | -0.069877788 | 0.91640178 | -0.0762523 | 0.93921834 | 0.99999872 |
| BHMT | 13.4727384 | 0.630998452 | 1.31780192 | 0.47882648 | 0.63206208 | 0.99999872 |
| JMY | 39.9167366 | -0.063176307 | 0.33942954 | -0.186125 | 0.85234678 | 0.99999872 |
| HOMER1 | 18.8966241 | 0.119877598 | 0.412968 | 0.29028302 | 0.77159973 | 0.99999872 |
| TENT2 | 126.127868 | 0.10949869 | 0.23401259 | 0.46791794 | 0.63984327 | 0.99999872 |
| CMYA5 | 133.615562 | 0.078579062 | 0.49516538 | 0.15869256 | 0.8739111 | 0.99999872 |
| MTX3 | 13.4160487 | 0.085649622 | 0.43366039 | 0.19750391 | 0.84343322 | 0.99999872 |
| THBS4 | 4.88433496 | 0.149598888 | 0.98656713 | 0.15163579 | 0.8794742 | 0.99999872 |
| AC012636.1 | 8.23782135 | 2.80715979 | 0.79623502 | 3.52554171 | 0.00042262 | 0.20128125 |
| SERINC5 | 340.863713 | 0.447973757 | 0.39402248 | 1.13692438 | 0.2555699 | 0.99999872 |
| AC010260.1 | 37.3119515 | 2.346275007 | 0.68206382 | 3.43996404 | 0.00058179 | 0.21771468 |
| ZFYVE16 | 50.7793396 | 0.154854987 | 0.38343532 | 0.40386208 | 0.68631414 | 0.99999872 |
| AC008771.1 | 5.25412797 | 0.0066219 | 0.67885149 | 0.00975456 | 0.99221711 | 0.99999872 |
| FAM151B | 9.73056444 | -0.313696646 | 0.53856936 | -0.5824628 | 0.56025496 | 0.99999872 |
| DHFR | 85.7485353 | -0.088993859 | 0.36996941 | -0.2405438 | 0.80990869 | 0.99999872 |
| MSH3 | 42.512104 | -0.330878918 | 0.28533703 | -1.1596074 | 0.2462087 | 0.99999872 |

| | | | | | | |
|---|---|---|---|---|---|---|
| RASGRF2 | 7.96455265 | -0.534804457 | 0.70997591 | -0.7532713 | 0.45128692 | 0.99999872 |
| CKMT2-AS1 | 23.4844058 | -0.068352914 | 0.46062962 | -0.1483902 | 0.88203485 | 0.99999872 |
| ZCCHC9 | 13.4579079 | 1.285358253 | 0.49869987 | 2.57741848 | 0.00995414 | 0.70848663 |
| SSBP2 | 218.673811 | 0.465554979 | 0.25353449 | 1.83625897 | 0.06631936 | 0.99999872 |
| ATG10 | 57.7731512 | -0.224101477 | 0.35989999 | -0.6226771 | 0.53349675 | 0.99999872 |
| RPS23 | 92.3649733 | 0.634445348 | 0.48313008 | 1.31319778 | 0.18911631 | 0.99999872 |
| ATP6AP1L | 23.5652171 | 0.096244771 | 0.54107774 | 0.17787605 | 0.85882032 | 0.99999872 |
| TMEM167A | 24.7191196 | 0.366152521 | 0.39609422 | 0.92440764 | 0.35527412 | 0.99999872 |
| XRCC4 | 20.7193356 | -0.591817182 | 0.63335271 | -0.9344196 | 0.35008749 | 0.99999872 |
| EDIL3 | 97.7129127 | 0.846263479 | 0.52702787 | 1.60572815 | 0.10833365 | 0.99999872 |
| AC113383.1 | 11.9802523 | 0.149743962 | 0.53824192 | 0.2782094 | 0.78085162 | 0.99999872 |
| COX7C | 60.1675651 | 0.98496195 | 0.39067171 | 2.52120113 | 0.0116955 | 0.72670507 |
| RASA1 | 81.445815 | 0.188393608 | 0.28204092 | 0.66796551 | 0.50415561 | 0.99999872 |
| CCNH | 97.3406558 | 0.870764704 | 0.36132186 | 2.40994194 | 0.01595506 | 0.79608149 |
| TMEM161B | 21.3187424 | 0.025042293 | 0.45038146 | 0.05560241 | 0.95565855 | 0.99999872 |
| TMEM161B-. | 57.7385199 | 0.077770815 | 0.49788196 | 0.15620332 | 0.87587276 | 0.99999872 |
| CETN3 | 11.2543055 | 0.194146768 | 0.60674573 | 0.31998044 | 0.74898316 | 0.99999872 |
| MBLAC2 | 6.19497868 | 0.216229747 | 0.64057989 | 0.33755313 | 0.73569997 | 0.99999872 |
| LYSMD3 | 16.8989522 | 0.81555462 | 0.44011069 | 1.85306704 | 0.06387275 | 0.99999872 |
| ADGRV1 | 22.0621499 | -0.301314276 | 0.63430925 | -0.4750274 | 0.63476744 | 0.99999872 |
| ARRDC3 | 29.6891797 | -0.539938411 | 0.57812466 | -0.9339481 | 0.35033068 | 0.99999872 |
| ARRDC3-AS1 | 10.9099903 | 0.151608786 | 0.5582148 | 0.27159578 | 0.78593284 | 0.99999872 |
| NR2F1-AS1 | 26.6240506 | -0.685345181 | 0.36975952 | -1.853489 | 0.06381231 | 0.99999872 |
| NR2F1 | 21.542368 | 0.575500427 | 0.59003744 | 0.97536256 | 0.32938045 | 0.99999872 |
| FAM172A | 164.585195 | -0.253923642 | 0.27393464 | -0.9269497 | 0.35395264 | 0.99999872 |
| KIAA0825 | 70.79189 | 0.096405016 | 0.26382528 | 0.36541235 | 0.71480362 | 0.99999872 |
| SLF1 | 11.8223366 | -0.696047362 | 0.50409578 | -1.3807839 | 0.1673454 | 0.99999872 |
| MCTP1 | 5.56960283 | -1.50924967 | 0.80232115 | -1.8811042 | 0.05995775 | 0.99999872 |
| TTC37 | 67.4951627 | 0.094772375 | 0.2538655 | 0.37331727 | 0.70891232 | 0.99999872 |
| ARSK | 8.56929534 | 0.45636094 | 0.62694787 | 0.72790891 | 0.46666935 | 0.99999872 |
| RHOBTB3 | 62.9787269 | 1.004561683 | 0.47659326 | 2.10779665 | 0.03504858 | 0.94734101 |
| GLRX | 18.298139 | 1.070251497 | 1.02105771 | 1.04817924 | 0.29455603 | 0.99999872 |
| ELL2 | 39.957766 | -0.025233036 | 0.32752874 | -0.0770407 | 0.93859119 | 0.99999872 |
| AC104123.1 | 35.6940283 | -0.267588831 | 0.51661584 | -0.5179648 | 0.60448282 | 0.99999872 |
| CAST | 232.980203 | 0.042189488 | 0.24575014 | 0.17167636 | 0.86369197 | 0.99999872 |
| ERAP1 | 46.0071554 | 0.074620392 | 0.31183339 | 0.23929571 | 0.81087629 | 0.99999872 |
| AC009126.1 | 8.28495266 | -0.495222535 | 0.55203777 | -0.8970809 | 0.36967576 | 0.99999872 |
| ERAP2 | 20.8148663 | 0.944500571 | 0.82138214 | 1.14989178 | 0.25018845 | 0.99999872 |
| LNPEP | 87.5946371 | 0.231192694 | 0.29143683 | 0.79328578 | 0.42761134 | 0.99999872 |
| LIX1-AS1 | 17.1988673 | -0.453669418 | 0.64572473 | -0.702574 | 0.48232126 | 0.99999872 |
| RIOK2 | 6.99705916 | 0.437947138 | 0.58698068 | 0.74610145 | 0.45560613 | 0.99999872 |
| RGMB | 4.14096014 | 0.33764297 | 0.69519999 | 0.48567747 | 0.62719586 | NA |
| CHD1 | 55.295417 | 0.082691018 | 0.29667082 | 0.27872987 | 0.78045214 | 0.99999872 |
| LINC02062 | 19.1041864 | 0.044895682 | 0.4182009 | 0.10735434 | 0.91450787 | 0.99999872 |
| FAM174A | 20.644419 | 0.749156844 | 0.39627211 | 1.89051118 | 0.05868963 | 0.99999872 |
| SLCO4C1 | 7.26108555 | 0.302481537 | 0.98750871 | 0.30630772 | 0.75937036 | 0.99999872 |
| PAM | 269.380858 | 0.099045198 | 0.2425483 | 0.40835247 | 0.68301492 | 0.99999872 |
| GIN1 | 6.41720718 | 0.109415368 | 0.59910332 | 0.18263188 | 0.85508686 | 0.99999872 |
| PPIP5K2 | 44.041249 | 0.003702765 | 0.37041889 | 0.00999616 | 0.99202435 | 0.99999872 |
| C5orf30 | 10.3142997 | 0.650140144 | 0.63950314 | 1.01663323 | 0.30932794 | 0.99999872 |
| NUDT12 | 9.26246141 | -0.218852387 | 0.47122677 | -0.4644311 | 0.64233888 | 0.99999872 |
| AC099520.1 | 20.3247921 | -0.914291035 | 1.23774522 | -0.7386747 | 0.46010457 | 0.99999872 |
| EFNA5 | 62.9610409 | -0.272937871 | 0.62773509 | -0.4347979 | 0.66370916 | 0.99999872 |
| FBXL17 | 253.518329 | 0.032439331 | 0.2530535 | 0.12819159 | 0.89799735 | 0.99999872 |
| FER | 164.888319 | 0.092201725 | 0.20544881 | 0.44878199 | 0.65358893 | 0.99999872 |
| AC008467.1 | 8.17254222 | -0.416362934 | 0.83407605 | -0.4991906 | 0.61764511 | 0.99999872 |
| PJA2 | 98.3838049 | 0.209455851 | 0.21868063 | 0.95781622 | 0.33815544 | 0.99999872 |
| MAN2A1 | 68.3050744 | 0.55424136 | 0.48610889 | 1.14015886 | 0.25422012 | 0.99999872 |
| TMEM232 | 44.924213 | -0.351393271 | 0.56802797 | -0.6186197 | 0.53616695 | 0.99999872 |
| SLC25A46 | 39.618378 | 0.273051477 | 0.33876254 | 0.80602617 | 0.42022776 | 0.99999872 |
| WDR36 | 15.2167909 | 0.073261599 | 0.52330846 | 0.13999697 | 0.88866238 | 0.99999872 |

| | | | | | | |
|---|---|---|---|---|---|---|
| STARD4 | 14.4902417 | -0.399951787 | 0.62529838 | -0.6396175 | 0.5224213 | 0.99999872 |
| NREP | 12.2384832 | -0.423129106 | 0.50072982 | -0.8450248 | 0.39809701 | 0.99999872 |
| EPB41L4A | 113.897483 | 0.450489106 | 0.45181559 | 0.99706411 | 0.31873339 | 0.99999872 |
| EPB41L4A-AS | 4.97384454 | -0.578484669 | 0.65546109 | -0.8825614 | 0.37747328 | 0.99999872 |
| APC | 43.7331227 | 0.081167608 | 0.39845592 | 0.20370537 | 0.83858375 | 0.99999872 |
| SRP19 | 21.2020666 | 0.084942673 | 0.37344223 | 0.22745867 | 0.8200671 | 0.99999872 |
| REEP5 | 85.5199447 | 0.301995482 | 0.3570863 | 0.84572128 | 0.39770825 | 0.99999872 |
| DCP2 | 35.8531859 | -0.072087495 | 0.30602908 | -0.2355577 | 0.81377593 | 0.99999872 |
| MCC | 174.275278 | 0.19953603 | 0.365546 | 0.54585751 | 0.58516389 | 0.99999872 |
| YTHDC2 | 60.7180336 | 0.049316523 | 0.262361 | 0.187972 | 0.8508986 | 0.99999872 |
| PGGT1B | 25.0636644 | -0.217900999 | 0.33377514 | -0.6528377 | 0.5138609 | 0.99999872 |
| CCDC112 | 9.06216184 | 0.001886984 | 0.54659876 | 0.00345223 | 0.99724552 | 0.99999872 |
| FEM1C | 9.14185428 | 0.719628287 | 0.56651308 | 1.27027656 | 0.20398613 | 0.99999872 |
| TMED7-TICA | 16.5896992 | 0.468195195 | 0.43136744 | 1.08537445 | 0.27775584 | 0.99999872 |
| ATG12 | 24.5521394 | 0.124077963 | 0.32790422 | 0.37839696 | 0.70513572 | 0.99999872 |
| AP3S1 | 52.2089596 | 0.2760243 | 0.2552002 | 1.08159906 | 0.27943072 | 0.99999872 |
| COMMD10 | 79.3234534 | -0.409574064 | 0.26806317 | -1.5279013 | 0.12653705 | 0.99999872 |
| SEMA6A | 9.96551437 | -1.158608781 | 0.66776401 | -1.7350572 | 0.08273067 | 0.99999872 |
| SEMA6A-AS1 | 8.83045217 | -0.612961468 | 0.47464035 | -1.291423 | 0.19655704 | 0.99999872 |
| DTWD2 | 23.9387182 | 0.221545616 | 0.41524946 | 0.53352415 | 0.5936708 | 0.99999872 |
| AC008629.1 | 11.5426357 | -0.2561964 | 0.70258407 | -0.3646487 | 0.71537362 | 0.99999872 |
| DMXL1 | 95.2821315 | 0.004514181 | 0.28112653 | 0.01605747 | 0.98718854 | 0.99999872 |
| TNFAIP8 | 16.5439862 | 0.38558198 | 0.73714821 | 0.52307253 | 0.60092378 | 0.99999872 |
| HSD17B4 | 44.8713198 | 0.618608471 | 0.42664084 | 1.44995137 | 0.14707208 | 0.99999872 |
| PRR16 | 10.6938617 | 0.438060846 | 0.85209578 | 0.51409813 | 0.60718338 | 0.99999872 |
| SRFBP1 | 45.9186349 | -0.100035204 | 0.38344512 | -0.2608853 | 0.79418094 | 0.99999872 |
| LOX | 83.87025 | 0.216424354 | 0.46485944 | 0.46556945 | 0.64152372 | 0.99999872 |
| SNX2 | 26.1154486 | 0.721866222 | 0.40834045 | 1.76780483 | 0.07709354 | 0.99999872 |
| SNX24 | 57.637923 | -0.412320278 | 0.32763092 | -1.2584901 | 0.20821455 | 0.99999872 |
| PPIC | 12.7256044 | 0.877571367 | 0.5666641 | 1.54866236 | 0.12146291 | 0.99999872 |
| CEP120 | 18.6725126 | -0.18844059 | 0.41565719 | -0.4533558 | 0.65029257 | 0.99999872 |
| CSNK1G3 | 41.7437888 | 0.464846271 | 0.30383384 | 1.52993582 | 0.12603262 | 0.99999872 |
| ZNF608 | 118.400261 | 0.26845774 | 0.28018072 | 0.9581592 | 0.33798249 | 0.99999872 |
| AC112196.1 | 3.89930628 | 1.663350189 | 0.97140351 | 1.71231643 | 0.08683837 | NA |
| GRAMD2B | 56.7899785 | -0.172836815 | 0.40804487 | -0.4235731 | 0.6718772 | 0.99999872 |
| ALDH7A1 | 55.5638109 | -0.448033572 | 0.4367055 | -1.0259398 | 0.30491995 | 0.99999872 |
| PHAX | 14.8328534 | -0.153193792 | 0.44100033 | -0.347378 | 0.72830739 | 0.99999872 |
| C5orf63 | 7.59135238 | -0.510238152 | 0.5338356 | -0.9557964 | 0.3391751 | 0.99999872 |
| PRRC1 | 23.4161606 | 0.45492585 | 0.5451612 | 0.8344795 | 0.40401083 | 0.99999872 |
| LINC01184 | 35.3046628 | -0.671029463 | 0.45702533 | -1.4682544 | 0.14203512 | 0.99999872 |
| SLC12A2 | 9.56257822 | 0.536306255 | 0.51574201 | 1.03987313 | 0.29839885 | 0.99999872 |
| SLC27A6 | 9.59750257 | 2.455212563 | 1.62952206 | 1.50670717 | 0.13188573 | 0.99999872 |
| ISOC1 | 8.26229999 | -0.002798993 | 0.61458288 | -0.0045543 | 0.99636621 | 0.99999872 |
| ADAMTS19 | 462.107097 | -0.299561705 | 0.57173852 | -0.5239488 | 0.60031416 | 0.99999872 |
| AC008591.1 | 47.4313923 | 1.351250951 | 0.81149974 | 1.66512801 | 0.09588722 | 0.99999872 |
| CHSY3 | 13.4792519 | 0.419190662 | 0.78235416 | 0.53580678 | 0.5920921 | 0.99999872 |
| HINT1 | 141.004287 | 1.103681374 | 0.43300257 | 2.54890259 | 0.01080625 | 0.71391523 |
| LYRM7 | 10.0426397 | 0.412043362 | 0.44537916 | 0.92515188 | 0.35488691 | 0.99999872 |
| CDC42SE2 | 100.793226 | -0.144786934 | 0.22205341 | -0.6520365 | 0.5143776 | 0.99999872 |
| RAPGEF6 | 4.21225876 | -0.948624823 | 1.00669675 | -0.9423144 | 0.34603171 | NA |
| P4HA2 | 18.4203279 | 0.741915764 | 0.58894151 | 1.2597444 | 0.20776158 | 0.99999872 |
| SLC22A5 | 16.3146235 | 0.946100098 | 0.71079097 | 1.33105251 | 0.18317173 | 0.99999872 |
| AC116366.3 | 88.4059081 | -0.562395004 | 0.5357851 | -1.0496653 | 0.29387204 | 0.99999872 |
| IRF1 | 23.3096857 | -0.191529469 | 0.68895558 | -0.2779997 | 0.78101256 | 0.99999872 |
| TH2LCRR | 4.32278797 | 0.740788807 | 0.85447088 | 0.86695617 | 0.38596602 | NA |
| KIF3A | 25.3813923 | 0.249667966 | 0.37791837 | 0.66063993 | 0.50884326 | 0.99999872 |
| 8-Sep | 11.6452069 | 0.370877442 | 0.6014087 | 0.61668121 | 0.53744502 | 0.99999872 |
| UQCRQ | 31.6609559 | 0.750985456 | 0.37210716 | 2.01819673 | 0.04357078 | 0.95980711 |
| AFF4 | 114.88528 | -0.131972916 | 0.18842507 | -0.7004 | 0.48367755 | 0.99999872 |
| ZCCHC10 | 12.3345908 | 0.09398992 | 0.71866871 | 0.13078337 | 0.89594668 | 0.99999872 |
| HSPA4 | 31.0449147 | -0.208910717 | 0.36697404 | -0.5692793 | 0.56916662 | 0.99999872 |

| | | | | | | |
|---|---|---|---|---|---|---|
| C5orf15 | 27.5647363 | 0.389274456 | 0.40913643 | 0.95145392 | 0.341374 | 0.99999872 |
| VDAC1 | 40.2855341 | 0.48665792 | 0.30350549 | 1.60345672 | 0.10883386 | 0.99999872 |
| SKP1 | 134.483616 | 1.055475373 | 0.37588353 | 2.80798519 | 0.00498525 | 0.5412956 |
| CDKL3 | 23.0881215 | -0.338511475 | 0.40855791 | -0.828552 | 0.40735796 | 0.99999872 |
| UBE2B | 65.8137835 | 0.204287449 | 0.24895736 | 0.82057205 | 0.41189007 | 0.99999872 |
| CDKN2AIPNL | 4.90616737 | -0.162141564 | 0.62733171 | -0.2584622 | 0.79605018 | 0.99999872 |
| JADE2 | 20.8418577 | -0.292574446 | 0.54387908 | -0.5379402 | 0.5906183 | 0.99999872 |
| SAR1B | 29.8433394 | 0.465098805 | 0.34605913 | 1.3439865 | 0.17895275 | 0.99999872 |
| SEC24A | 27.8961754 | 0.135349161 | 0.37605525 | 0.35991828 | 0.71890824 | 0.99999872 |
| CAMLG | 15.946406 | 0.968102565 | 0.40384335 | 2.397223 | 0.01651987 | 0.81253899 |
| DDX46 | 46.9378287 | -0.257699187 | 0.3131653 | -0.8228855 | 0.41057311 | 0.99999872 |
| C5orf24 | 25.7612891 | 0.062866355 | 0.34913896 | 0.18006113 | 0.85710458 | 0.99999872 |
| TXNDC15 | 36.6774438 | -0.018952155 | 0.41457816 | -0.0457143 | 0.96353796 | 0.99999872 |
| PCBD2 | 19.5407172 | -0.684179386 | 0.36044637 | -1.8981447 | 0.05767702 | 0.99999872 |
| H2AFY | 30.6750613 | -0.24364675 | 0.2857858 | -0.8525502 | 0.39390878 | 0.99999872 |
| CXCL14 | 90.6430651 | 0.162250456 | 1.63556917 | 0.09920122 | 0.92097851 | 0.99999872 |
| SMAD5 | 61.574808 | 0.297617919 | 0.36878078 | 0.80703208 | 0.419648 | 0.99999872 |
| SPOCK1 | 878.241187 | -0.046856309 | 0.44871275 | -0.1044238 | 0.916833 | 0.99999872 |
| KLHL3 | 11.6068996 | -0.176388326 | 0.70060724 | -0.2517649 | 0.80122277 | 0.99999872 |
| HNRNPA0 | 32.5525726 | 0.243485947 | 0.4816669 | 0.50550691 | 0.61320285 | 0.99999872 |
| AC106791.1 | 9.79036146 | -0.523239973 | 0.52098962 | -1.0043194 | 0.31522469 | 0.99999872 |
| PKD2L2 | 6.03424943 | 0.907652557 | 0.72278018 | 1.25577953 | 0.20919592 | 0.99999872 |
| FAM13B | 126.652667 | 0.019531667 | 0.24883362 | 0.07849288 | 0.937436 | 0.99999872 |
| NME5 | 15.8338443 | 0.240572687 | 0.63482416 | 0.37895956 | 0.70471789 | 0.99999872 |
| BRD8 | 15.6747193 | 0.182617006 | 0.38262691 | 0.47727172 | 0.63316864 | 0.99999872 |
| CDC23 | 11.9167998 | -0.144712024 | 0.42174894 | -0.3431236 | 0.73150546 | 0.99999872 |
| FAM53C | 9.30512128 | -0.144707291 | 0.51092012 | -0.2832288 | 0.77700146 | 0.99999872 |
| KDM3B | 45.6042467 | -0.044813651 | 0.3353052 | -0.1336503 | 0.89367908 | 0.99999872 |
| EGR1 | 17.8308304 | -2.002564154 | 1.39709911 | -1.433373 | 0.15175127 | 0.99999872 |
| ETF1 | 27.9134257 | -0.078622854 | 0.33730799 | -0.2330892 | 0.81569213 | 0.99999872 |
| HSPA9 | 43.3529077 | -0.259527207 | 0.40599116 | -0.6392435 | 0.52266454 | 0.99999872 |
| CTNNA1 | 270.393878 | -0.122675766 | 0.33449877 | -0.366745 | 0.7138092 | 0.99999872 |
| SIL1 | 117.160359 | 0.005618538 | 0.28172592 | 0.01994328 | 0.98408862 | 0.99999872 |
| PAIP2 | 46.3044218 | 0.539213512 | 0.30178434 | 1.78675111 | 0.07397772 | 0.99999872 |
| DNAJC18 | 8.92725243 | 0.710723335 | 0.66634298 | 1.06660287 | 0.2861512 | 0.99999872 |
| UBE2D2 | 89.8936437 | -0.523947317 | 0.30492398 | -1.7182883 | 0.08574404 | 0.99999872 |
| CXXC5 | 68.5514752 | -0.077693865 | 0.64363374 | -0.1207113 | 0.90391971 | 0.99999872 |
| PURA | 58.6194856 | -0.340829517 | 0.31223192 | -1.0915909 | 0.27501297 | 0.99999872 |
| IGIP | 9.63262359 | 0.167144951 | 0.67375544 | 0.24807956 | 0.80407285 | 0.99999872 |
| CYSTM1 | 93.1889286 | 0.33849324 | 0.28845949 | 1.17345157 | 0.24061477 | 0.99999872 |
| PFDN1 | 26.2471211 | 0.409890954 | 0.33756408 | 1.21426119 | 0.224648 | 0.99999872 |
| HBEGF | 10.5341415 | 0.050624988 | 0.72119119 | 0.07019635 | 0.94403738 | 0.99999872 |
| AC008438.1 | 4.38074865 | -0.637529864 | 0.96648822 | -0.6596354 | 0.50948782 | NA |
| SRA1 | 13.8400758 | 0.482081947 | 0.45834579 | 1.05178657 | 0.29289748 | 0.99999872 |
| SLC35A4 | 18.3819346 | 0.220602845 | 0.48802885 | 0.45202829 | 0.65124861 | 0.99999872 |
| NDUFA2 | 26.3762387 | 0.417012752 | 0.4255421 | 0.9799565 | 0.32710759 | 0.99999872 |
| IK | 10.3978038 | -0.289238233 | 0.59714867 | -0.4843655 | 0.62812647 | 0.99999872 |
| WDR55 | 5.68645783 | -0.427094516 | 0.69805362 | -0.6118363 | 0.5406461 | 0.99999872 |
| HARS | 16.5395133 | -0.421416597 | 0.37551886 | -1.1222249 | 0.26176684 | 0.99999872 |
| ZMAT2 | 13.2402896 | -0.162440505 | 0.4572253 | -0.3552745 | 0.72238393 | 0.99999872 |
| AC244517.11 | 5.76190757 | 0.939010488 | 1.00820081 | 0.93137248 | 0.35166092 | 0.99999872 |
| TAF7 | 45.3782962 | 0.367776048 | 0.4034664 | 0.91154072 | 0.36201054 | 0.99999872 |
| DIAPH1 | 30.2365884 | -0.077979435 | 0.40305695 | -0.19347 | 0.84659089 | 0.99999872 |
| HDAC3 | 11.9664856 | 0.260136974 | 0.52606349 | 0.4944973 | 0.62095502 | 0.99999872 |
| DELE1 | 18.1401466 | -0.16032966 | 0.41710426 | -0.3843875 | 0.70069127 | 0.99999872 |
| RNF14 | 23.3030942 | -0.113161677 | 0.36664752 | -0.3086389 | 0.75759625 | 0.99999872 |
| GNPDA1 | 13.3478154 | 0.033958506 | 0.64400974 | 0.0527298 | 0.95794719 | 0.99999872 |
| NDFIP1 | 90.479648 | 0.156078269 | 0.26116423 | 0.59762498 | 0.55009019 | 0.99999872 |
| SPRY4 | 7.19455403 | -1.212860858 | 0.66607002 | -1.820921 | 0.06861887 | 0.99999872 |
| SPRY4-AS1 | 67.3978595 | 1.10922178 | 0.54062277 | 2.05174816 | 0.04019414 | 0.95980711 |
| FGF1 | 1026.94526 | -0.015997001 | 0.46589536 | -0.034336 | 0.97260919 | 0.99999872 |

| | | | | | | |
|---|---|---|---|---|---|---|
| ARHGAP26 | 51.4311887 | -1.326059511 | 0.48868158 | -2.7135451 | 0.00665675 | 0.61721837 |
| NR3C1 | 80.2298828 | 0.193922764 | 0.30147578 | 0.64324493 | 0.52006519 | 0.99999872 |
| YIPF5 | 18.4546438 | 0.392410724 | 0.45791457 | 0.85695181 | 0.39147152 | 0.99999872 |
| KCTD16 | 15.847057 | -1.343890282 | 0.7153825 | -1.8785619 | 0.06030434 | 0.99999872 |
| PRELID2 | 35.1917255 | 0.367719954 | 0.6477674 | 0.56767283 | 0.57025714 | 0.99999872 |
| SH3RF2 | 5.79569956 | -0.845428009 | 0.89883396 | -0.9405831 | 0.34691856 | 0.99999872 |
| LARS | 37.4628863 | 0.093444305 | 0.32656771 | 0.28614067 | 0.77477038 | 0.99999872 |
| TCERG1 | 36.8809556 | -0.132341901 | 0.3461826 | -0.3822893 | 0.70224679 | 0.99999872 |
| PPP2R2B | 112.258015 | 0.764509999 | 0.48760411 | 1.5678908 | 0.11690662 | 0.99999872 |
| STK32A | 6.23252016 | -0.799438573 | 1.24499007 | -0.6421245 | 0.52079238 | 0.99999872 |
| DPYSL3 | 29.0196547 | 0.309383792 | 0.61768312 | 0.50087785 | 0.61645709 | 0.99999872 |
| JAKMIP2 | 43.1920296 | 0.328987878 | 0.690246 | 0.4766241 | 0.63362982 | 0.99999872 |
| AC091948.1 | 10.3348361 | -0.598872064 | 0.58896195 | -1.0168264 | 0.30923602 | 0.99999872 |
| FBXO38 | 25.5526716 | 0.155580273 | 0.38657386 | 0.40245937 | 0.68734598 | 0.99999872 |
| GRPEL2 | 5.12736986 | 0.249437762 | 0.70544726 | 0.35358811 | 0.72364759 | 0.99999872 |
| CSNK1A1 | 210.29686 | 0.131689357 | 0.22583517 | 0.58312155 | 0.55981148 | 0.99999872 |
| ARHGEF37 | 62.1290605 | -0.035837043 | 0.58367083 | -0.0613994 | 0.95104112 | 0.99999872 |
| PPARGC1B | 13.0248096 | -0.046767754 | 0.95522923 | -0.0489597 | 0.9609514 | 0.99999872 |
| SLC26A2 | 27.6227052 | 0.040850005 | 0.54855166 | 0.07446884 | 0.94063734 | 0.99999872 |
| HMGXB3 | 22.2726063 | -0.003011916 | 0.33315035 | -0.0090407 | 0.99278666 | 0.99999872 |
| PDGFRB | 140.432046 | -0.473671064 | 0.54960779 | -0.8618347 | 0.38877849 | 0.99999872 |
| CAMK2A | 12.1641558 | 0.585356399 | 1.15764417 | 0.50564449 | 0.61310624 | 0.99999872 |
| TCOF1 | 9.54902772 | -1.008805933 | 0.4945304 | -2.039927 | 0.04135759 | 0.95980711 |
| CD74 | 34.1995109 | 0.979841415 | 0.79117618 | 1.23846172 | 0.21554491 | 0.99999872 |
| RPS14 | 109.220928 | 0.410982519 | 0.57754174 | 0.71160661 | 0.47670841 | 0.99999872 |
| NDST1 | 10.4080567 | 0.053705452 | 0.49677104 | 0.10810906 | 0.91390918 | 0.99999872 |
| SYNPO | 173.332176 | -0.998170945 | 0.65686065 | -1.5196084 | 0.12860943 | 0.99999872 |
| RBM22 | 11.4757664 | -0.211448095 | 0.48273987 | -0.4380166 | 0.66137422 | 0.99999872 |
| DCTN4 | 36.4361912 | -0.385771685 | 0.29415167 | -1.311472 | 0.18969835 | 0.99999872 |
| GPX3 | 272.930273 | -0.400847988 | 0.74675776 | -0.5367845 | 0.59141648 | 0.99999872 |
| TNIP1 | 20.349157 | -0.291549252 | 0.51186422 | -0.5695832 | 0.56896044 | 0.99999872 |
| ANXA6 | 4.98331466 | 1.107080668 | 0.84823208 | 1.30516247 | 0.19183753 | 0.99999872 |
| GM2A | 4.91162124 | 0.629089022 | 0.70217804 | 0.89591099 | 0.3703003 | 0.99999872 |
| SLC36A1 | 11.8814441 | -0.715342176 | 0.59712012 | -1.1979871 | 0.23092206 | 0.99999872 |
| SPARC | 532.069853 | 0.144651705 | 0.72241707 | 0.20023295 | 0.8412984 | 0.99999872 |
| ATOX1 | 57.7815623 | 0.13748301 | 0.42292303 | 0.32507809 | 0.74512197 | 0.99999872 |
| G3BP1 | 58.2797001 | 0.071444851 | 0.30173018 | 0.23678391 | 0.81282443 | 0.99999872 |
| FAM114A2 | 22.0470813 | 0.537902564 | 0.39851582 | 1.34976466 | 0.17709149 | 0.99999872 |
| MFAP3 | 24.8184063 | 0.310978702 | 0.37125212 | 0.83764827 | 0.40222827 | 0.99999872 |
| GALNT10 | 34.3559956 | -1.240165071 | 0.43132613 | -2.8752375 | 0.00403724 | 0.49700703 |
| SAP30L-AS1 | 11.675473 | -0.349858129 | 0.45967198 | -0.7611039 | 0.44659502 | 0.99999872 |
| SAP30L | 10.6256334 | -0.064033032 | 0.47417211 | -0.1350417 | 0.89257886 | 0.99999872 |
| LARP1 | 86.3769018 | -0.325894182 | 0.34875851 | -0.9344408 | 0.35007656 | 0.99999872 |
| FAXDC2 | 31.323179 | 0.539959544 | 0.69090589 | 0.78152401 | 0.43449436 | 0.99999872 |
| CNOT8 | 18.6046359 | 0.385217276 | 0.43334098 | 0.88894726 | 0.37403143 | 0.99999872 |
| GEMIN5 | 8.55430135 | 0.219808285 | 0.50573546 | 0.43463095 | 0.66383033 | 0.99999872 |
| MRPL22 | 13.1702226 | 0.399575014 | 0.45489671 | 0.87838625 | 0.37973414 | 0.99999872 |
| HAVCR2 | 8.85432912 | 0.212330888 | 0.611606 | 0.3471694 | 0.72846406 | 0.99999872 |
| CYFIP2 | 31.7251545 | -0.098007804 | 0.65316555 | -0.1500505 | 0.88072479 | 0.99999872 |
| CLINT1 | 47.4849275 | 0.139115428 | 0.27995049 | 0.49692869 | 0.61923934 | 0.99999872 |
| EBF1 | 9.43243341 | 0.990918335 | 1.02926575 | 0.96274294 | 0.33567654 | 0.99999872 |
| RNF145 | 116.562887 | -0.282723958 | 0.39117505 | -0.7227556 | 0.46983005 | 0.99999872 |
| UBLCP1 | 20.9665166 | 0.381707412 | 0.36380726 | 1.0492023 | 0.29408502 | 0.99999872 |
| TTC1 | 32.8127483 | 0.050219279 | 0.38662103 | 0.12989278 | 0.89665126 | 0.99999872 |
| PWWP2A | 36.4735237 | 0.312534775 | 0.4039784 | 0.77364229 | 0.43914235 | 0.99999872 |
| ZBED8 | 6.14678761 | 0.46079437 | 0.73835766 | 0.62408016 | 0.53257494 | 0.99999872 |
| SLU7 | 9.60852046 | 0.524990114 | 0.56307588 | 0.93236122 | 0.35114988 | 0.99999872 |
| AC113414.1 | 12.6601534 | 0.800032012 | 0.69137025 | 1.15716869 | 0.24720345 | 0.99999872 |
| CCNG1 | 19.0256393 | 0.552247839 | 0.38126939 | 1.44844526 | 0.14749256 | 0.99999872 |
| NUDCD2 | 16.3736341 | 0.472228329 | 0.43747722 | 1.07943524 | 0.28039375 | 0.99999872 |
| MAT2B | 28.6756557 | 0.658292869 | 0.43825463 | 1.50207851 | 0.13307683 | 0.99999872 |

| | | | | | | |
|---|---|---|---|---|---|---|
| AC109466.1 | 671.751295 | -1.066291823 | 0.75497599 | -1.412352 | 0.15784635 | 0.99999872 |
| AC008415.1 | 235.893631 | 0.021995799 | 0.53282045 | 0.04128182 | 0.96707122 | 0.99999872 |
| AC114321.1 | 7.58191514 | 0.690022774 | 0.9598519 | 0.71888462 | 0.47221201 | 0.99999872 |
| TENM2 | 283.447882 | 0.137433174 | 0.48246603 | 0.28485565 | 0.77575473 | 0.99999872 |
| AC011369.1 | 5.57461716 | 1.089982419 | 1.10071542 | 0.99024907 | 0.32205239 | 0.99999872 |
| WWC1 | 40.1139661 | -1.119106337 | 0.55208974 | -2.027037 | 0.04265863 | 0.95980711 |
| RARS | 23.2425643 | -0.042966264 | 0.33660358 | -0.1276465 | 0.89842874 | 0.99999872 |
| PANK3 | 35.3824643 | 0.216274803 | 0.37082154 | 0.5832315 | 0.55973747 | 0.99999872 |
| SLIT3 | 46.4573325 | -0.211574754 | 0.70090735 | -0.3018584 | 0.76276003 | 0.99999872 |
| KCNIP1 | 15.555647 | -0.429383213 | 1.84074666 | -0.2332658 | 0.81555503 | 0.99999872 |
| RANBP17 | 38.1701626 | -0.82913197 | 0.5672237 | -1.4617372 | 0.14381325 | 0.99999872 |
| NPM1 | 28.6702781 | 0.000269127 | 0.36704227 | 0.00073323 | 0.99941497 | 0.99999872 |
| FBXW11 | 78.3867157 | -0.112773608 | 0.25164201 | -0.448151 | 0.65404426 | 0.99999872 |
| STK10 | 5.60426242 | -0.723619507 | 0.58891481 | -1.2287338 | 0.21917165 | 0.99999872 |
| UBTD2 | 15.6359842 | -0.34599486 | 0.48314258 | -0.7161341 | 0.47390857 | 0.99999872 |
| DUSP1 | 113.181218 | -0.574863343 | 0.48617173 | -1.1824286 | 0.23703569 | 0.99999872 |
| ERGIC1 | 77.6635826 | -0.432380596 | 0.31731056 | -1.3626417 | 0.17299547 | 0.99999872 |
| RPL26L1 | 6.60907524 | 0.745146364 | 0.63378153 | 1.17571486 | 0.23970884 | 0.99999872 |
| ATP6V0E1 | 93.3494091 | 0.623175292 | 0.23456797 | 2.65669384 | 0.00789111 | 0.65621445 |
| CREBRF | 66.6784691 | -0.210517966 | 0.39351502 | -0.5349681 | 0.59267194 | 0.99999872 |
| BOD1 | 9.20858956 | 0.602392734 | 0.58464572 | 1.03035516 | 0.30284332 | 0.99999872 |
| CPEB4 | 527.01446 | 0.177582496 | 0.27644635 | 0.64237599 | 0.52062909 | 0.99999872 |
| SFXN1 | 31.2843754 | 0.243547103 | 0.35514018 | 0.68577738 | 0.4928535 | 0.99999872 |
| THOC3 | 5.83992835 | -0.581292337 | 0.63248267 | -0.9190645 | 0.35806186 | 0.99999872 |
| AC139491.7 | 29.1990438 | 0.697511488 | 0.85573797 | 0.81509938 | 0.41501543 | 0.99999872 |
| SIMC1 | 32.5631119 | 0.286072496 | 0.34057336 | 0.8399732 | 0.40092341 | 0.99999872 |
| KIAA1191 | 26.0135527 | 0.088411433 | 0.45374166 | 0.19484972 | 0.8455106 | 0.99999872 |
| ARL10 | 6.94028799 | 0.895102165 | 0.57980934 | 1.54378708 | 0.12263992 | 0.99999872 |
| NOP16 | 3.08800215 | -0.297634358 | 1.0227565 | -0.2910119 | 0.77104218 | NA |
| CLTB | 35.6786038 | -0.376869081 | 0.42345138 | -0.8899938 | 0.37346924 | 0.99999872 |
| FAF2 | 31.3901268 | -0.189006819 | 0.36006245 | -0.5249279 | 0.59963333 | 0.99999872 |
| RNF44 | 11.4769974 | 0.443871442 | 0.48879699 | 0.90808956 | 0.36383091 | 0.99999872 |
| TSPAN17 | 8.06342258 | 0.173281545 | 0.68071579 | 0.25455784 | 0.79906462 | 0.99999872 |
| UIMC1 | 42.7614226 | -0.267073494 | 0.36078581 | -0.740255 | 0.45914529 | 0.99999872 |
| ZNF346 | 23.8543791 | -0.332563305 | 0.37284396 | -0.8919638 | 0.37241235 | 0.99999872 |
| FGFR4 | 34.7095111 | 0.340556104 | 0.82039468 | 0.41511252 | 0.67805953 | 0.99999872 |
| NSD1 | 119.55647 | -0.392523182 | 0.29708815 | -1.3212347 | 0.1864231 | 0.99999872 |
| PRELID1 | 24.5473065 | 0.14341887 | 0.47391962 | 0.30262277 | 0.76217736 | 0.99999872 |
| LMAN2 | 43.5123193 | 0.347508599 | 0.35403046 | 0.98157825 | 0.32630767 | 0.99999872 |
| RGS14 | 4.41229044 | 0.05936533 | 0.87831608 | 0.06758994 | 0.94611206 | NA |
| DBN1 | 16.01374 | -0.29696261 | 0.48305986 | -0.6147532 | 0.5387177 | 0.99999872 |
| PDLIM7 | 30.9573625 | -0.297335874 | 0.62490121 | -0.4758126 | 0.63420789 | 0.99999872 |
| DDX41 | 4.20408888 | 0.02317222 | 0.72665812 | 0.03188875 | 0.97456077 | NA |
| FAM193B | 35.4176951 | 0.195882617 | 0.34650544 | 0.56530892 | 0.57186367 | 0.99999872 |
| TMED9 | 81.1290293 | 0.654115139 | 0.45566291 | 1.43552421 | 0.15113777 | 0.99999872 |
| B4GALT7 | 11.2024408 | -0.087783161 | 0.55690123 | -0.1576279 | 0.87475003 | 0.99999872 |
| FAM153A | 5.19484753 | 0.144831278 | 0.99340755 | 0.14579241 | 0.88408527 | 0.99999872 |
| AC106795.3 | 4.15746337 | 0.065951977 | 0.81681992 | 0.08074237 | 0.93564684 | NA |
| RMND5B | 17.2667748 | -0.128712747 | 0.36882665 | -0.348979 | 0.72710509 | 0.99999872 |
| NHP2 | 19.1871174 | 0.211341366 | 0.43885882 | 0.48157028 | 0.63011124 | 0.99999872 |
| HNRNPAB | 23.6843488 | -0.435208505 | 0.52390542 | -0.8307005 | 0.40614283 | 0.99999872 |
| PHYKPL | 45.1819287 | -0.075570148 | 0.31528034 | -0.2396919 | 0.8105691 | 0.99999872 |
| CLK4 | 26.0336433 | 0.533066935 | 0.55720255 | 0.95668431 | 0.33872662 | 0.99999872 |
| ZNF354A | 6.50092396 | 1.003134511 | 0.69824734 | 1.43664638 | 0.1508185 | 0.99999872 |
| ZNF354B | 8.18976832 | 0.091051083 | 0.50896576 | 0.17889432 | 0.85802068 | 0.99999872 |
| ZFP2 | 6.57243849 | 0.840158362 | 0.65761589 | 1.27758222 | 0.20139677 | 0.99999872 |
| ZNF354C | 6.00149276 | 1.299066735 | 0.7871119 | 1.65042192 | 0.09885667 | 0.99999872 |
| RUFY1 | 38.3684392 | -0.263442074 | 0.34461363 | -0.7644563 | 0.44459535 | 0.99999872 |
| HNRNPH1 | 100.399445 | 0.142619106 | 0.29389294 | 0.48527572 | 0.62748078 | 0.99999872 |
| CANX | 191.663234 | 0.474799102 | 0.39461831 | 1.2031857 | 0.22890447 | 0.99999872 |
| MAML1 | 14.4214983 | -0.21714755 | 0.54829873 | -0.3960388 | 0.69207643 | 0.99999872 |

| | | | | | | |
|---|---|---|---|---|---|---|
| MGAT4B | 47.1811388 | 0.057117447 | 0.59799497 | 0.09551493 | 0.92390583 | 0.99999872 |
| SQSTM1 | 123.236809 | -0.119643704 | 0.40737218 | -0.2936963 | 0.76898998 | 0.99999872 |
| MRNIP | 15.6304277 | -0.233259245 | 0.44995432 | -0.5184065 | 0.60417468 | 0.99999872 |
| TBC1D9B | 35.1180811 | 0.158585219 | 0.3222126 | 0.49217572 | 0.62259512 | 0.99999872 |
| RNF130 | 42.9410596 | -0.118104498 | 0.40656567 | -0.290493 | 0.77143907 | 0.99999872 |
| MAPK9 | 20.273963 | -0.028028929 | 0.4507893 | -0.0621775 | 0.95042152 | 0.99999872 |
| CNOT6 | 33.4915182 | -0.50959414 | 0.4324512 | -1.1783853 | 0.23864303 | 0.99999872 |
| FLT4 | 27.8403853 | -0.285544987 | 0.635319 | -0.4494514 | 0.65310609 | 0.99999872 |
| MGAT1 | 48.6933933 | -0.29750906 | 0.4596008 | -0.6473206 | 0.51742448 | 0.99999872 |
| ZFP62 | 5.46976285 | -0.012502252 | 0.63997679 | -0.0195355 | 0.98441394 | 0.99999872 |
| TRIM41 | 7.77589654 | -0.74496972 | 0.53336981 | -1.3967227 | 0.16249697 | 0.99999872 |
| RACK1 | 58.8957939 | 0.016033293 | 0.47168587 | 0.03399146 | 0.97288396 | 0.99999872 |
| TRIM52 | 11.3535552 | -0.040627082 | 0.50575055 | -0.0803303 | 0.93597458 | 0.99999872 |
| DUSP22 | 15.46952 | 0.147386735 | 0.49344105 | 0.29869168 | 0.76517531 | 0.99999872 |
| EXOC2 | 58.8441871 | 0.043113231 | 0.24710082 | 0.17447628 | 0.86149117 | 0.99999872 |
| FOXC1 | 305.169043 | 0.117135371 | 0.35959661 | 0.32574103 | 0.7446203 | 0.99999872 |
| GMDS | 442.652513 | -0.064037157 | 0.20938835 | -0.3058296 | 0.75973439 | 0.99999872 |
| GMDS-DT | 155.640984 | -0.068127557 | 0.37244151 | -0.1829215 | 0.85485962 | 0.99999872 |
| WRNIP1 | 19.2960375 | 0.610811869 | 0.39740619 | 1.53699635 | 0.1242942 | 0.99999872 |
| SERPINB1 | 83.189791 | 0.394260264 | 0.74410667 | 0.52984374 | 0.59622027 | 0.99999872 |
| SERPINB9 | 49.157796 | 0.021153235 | 0.42904502 | 0.04930307 | 0.96067777 | 0.99999872 |
| SERPINB6 | 53.4480486 | 0.035542882 | 0.25103185 | 0.14158714 | 0.88740613 | 0.99999872 |
| NQO2 | 16.8529137 | -0.086578739 | 0.42441671 | -0.2039946 | 0.83835768 | 0.99999872 |
| RIPK1 | 40.8866507 | -0.295147516 | 0.26894762 | -1.0974164 | 0.27245942 | 0.99999872 |
| BPHL | 9.33393111 | -0.024025874 | 0.56119493 | -0.042812 | 0.9658514 | 0.99999872 |
| TUBB2A | 31.9059681 | 2.188053208 | 0.80986674 | 2.70174473 | 0.00689767 | 0.62304985 |
| PSMG4 | 17.6734449 | 0.34077995 | 0.51016017 | 0.6679862 | 0.5041424 | 0.99999872 |
| SLC22A23 | 21.1361007 | -0.196159591 | 0.71188345 | -0.2755502 | 0.7828936 | 0.99999872 |
| PXDC1 | 27.4105692 | -0.186331236 | 0.47077117 | -0.3958 | 0.69225257 | 0.99999872 |
| FAM50B | 4.99670274 | 0.364582647 | 0.78127728 | 0.4666495 | 0.64075067 | 0.99999872 |
| PRPF4B | 55.223736 | 0.208505944 | 0.25331692 | 0.82310312 | 0.41044936 | 0.99999872 |
| ECI2 | 15.8960836 | 0.385469014 | 0.41837389 | 0.92135054 | 0.35686744 | 0.99999872 |
| CDYL | 66.2093307 | -0.1833045 | 0.27165867 | -0.6747604 | 0.49982803 | 0.99999872 |
| LYRM4 | 20.0169972 | -0.37977469 | 0.4870175 | -0.7797968 | 0.43551048 | 0.99999872 |
| FARS2 | 92.3540872 | -0.034285083 | 0.25775544 | -0.133014 | 0.89418231 | 0.99999872 |
| RREB1 | 57.1686231 | -0.100436238 | 0.3568871 | -0.281423 | 0.77838598 | 0.99999872 |
| SSR1 | 57.0654244 | 0.260697114 | 0.32870273 | 0.79310906 | 0.42771429 | 0.99999872 |
| AL139095.4 | 9.13050167 | 2.187210463 | 1.01873935 | 2.14697751 | 0.03179507 | 0.92959829 |
| RIOK1 | 9.37864327 | -0.354767857 | 0.46589314 | -0.761479 | 0.44637099 | 0.99999872 |
| DSP | 9.42914477 | -0.373669991 | 0.69905823 | -0.5345334 | 0.59297253 | 0.99999872 |
| SNRNP48 | 10.2853931 | 0.397226247 | 0.55669804 | 0.71353987 | 0.47551175 | 0.99999872 |
| BLOC1S5-TXI | 5.24961957 | 0.102733282 | 0.70287447 | 0.14616164 | 0.88379379 | 0.99999872 |
| SLC35B3 | 24.4263054 | 0.090839395 | 0.37033897 | 0.24528716 | 0.80623409 | 0.99999872 |
| GCNT2 | 14.7856778 | -1.305666388 | 0.80755587 | -1.6168125 | 0.10591877 | 0.99999872 |
| PAK1IP1 | 5.00514756 | 0.079968764 | 0.73091346 | 0.10940935 | 0.91287782 | 0.99999872 |
| TMEM14C | 33.8014805 | 1.024816317 | 0.46541779 | 2.20192768 | 0.02767042 | 0.89154528 |
| AL024498.2 | 8.39428152 | 0.25494569 | 0.53990644 | 0.47220346 | 0.63678157 | 0.99999872 |
| MAK | 20.440596 | 1.457699291 | 0.59810405 | 2.43720019 | 0.01480148 | 0.78384542 |
| SYCP2L | 6.34243964 | 1.269377516 | 0.72036213 | 1.76213804 | 0.07804598 | 0.99999872 |
| SMIM13 | 12.1641446 | 0.777326663 | 0.57265229 | 1.35741474 | 0.17464947 | 0.99999872 |
| AL139807.1 | 38.770153 | 0.984506276 | 0.43017739 | 2.28860536 | 0.02210229 | 0.87722657 |
| NEDD9 | 183.523377 | -0.115413652 | 0.27273202 | -0.423176 | 0.67216684 | 0.99999872 |
| TMEM170B | 28.2710298 | 0.349413811 | 0.4566016 | 0.76524877 | 0.44412342 | 0.99999872 |
| HIVEP1 | 54.8041426 | -0.248520061 | 0.34051847 | -0.7298284 | 0.46549506 | 0.99999872 |
| PHACTR1 | 16.5484065 | -0.242325711 | 0.87222767 | -0.2778239 | 0.78114753 | 0.99999872 |
| TBC1D7 | 6.40003051 | -0.022514345 | 0.65314278 | -0.0344708 | 0.97250174 | 0.99999872 |
| GFOD1 | 22.3856613 | -0.041077386 | 0.53636959 | -0.0765841 | 0.9389544 | 0.99999872 |
| SIRT5 | 16.2296226 | -0.014471783 | 0.47625308 | -0.0303868 | 0.97575861 | 0.99999872 |
| NOL7 | 32.1978195 | 0.129802045 | 0.34019149 | 0.38155582 | 0.70279086 | 0.99999872 |
| RANBP9 | 47.8638166 | 0.238763709 | 0.32317786 | 0.7387997 | 0.46002863 | 0.99999872 |
| MCUR1 | 14.5095686 | 0.193054531 | 0.44232566 | 0.43645338 | 0.66250782 | 0.99999872 |

| | | | | | | |
|---|---|---|---|---|---|---|
| JARID2 | 106.151888 | -0.210842328 | 0.36290858 | -0.5809792 | 0.56125449 | 0.99999872 |
| DTNBP1 | 33.408309 | 0.373546172 | 0.40396092 | 0.92470868 | 0.35511746 | 0.99999872 |
| MYLIP | 75.4976267 | 0.489277229 | 0.55193574 | 0.88647499 | 0.37536162 | 0.99999872 |
| GMPR | 9.11018978 | -1.715901283 | 1.02101788 | -1.6805791 | 0.0928447 | 0.99999872 |
| ATXN1 | 134.380031 | 0.019389427 | 0.44833821 | 0.04324732 | 0.96550438 | 0.99999872 |
| CAP2 | 37.9443025 | 0.019685531 | 0.51515386 | 0.03821291 | 0.96951792 | 0.99999872 |
| FAM8A1 | 13.108516 | 0.706229279 | 0.54753177 | 1.28984164 | 0.19710565 | 0.99999872 |
| NUP153 | 40.4931818 | 0.116635917 | 0.2916182 | 0.39996103 | 0.68918522 | 0.99999872 |
| KIF13A | 158.046743 | 0.036144422 | 0.33363557 | 0.10833504 | 0.91372993 | 0.99999872 |
| TPMT | 8.19880808 | 0.041548427 | 0.5334535 | 0.07788575 | 0.93791893 | 0.99999872 |
| KDM1B | 10.1476046 | 0.071723795 | 0.58918801 | 0.12173329 | 0.90311025 | 0.99999872 |
| DEK | 42.8879161 | -0.069660215 | 0.27686707 | -0.2516017 | 0.80134898 | 0.99999872 |
| RNF144B | 17.1960207 | -0.597910154 | 0.4814123 | -1.2419918 | 0.21423957 | 0.99999872 |
| AL022068.1 | 67.1326589 | -1.037393271 | 0.68021014 | -1.525107 | 0.1272324 | 0.99999872 |
| ID4 | 20.5816974 | -2.407398373 | 0.71639666 | -3.3604266 | 0.00077822 | 0.25898594 |
| MBOAT1 | 12.7365896 | 0.769584882 | 0.61165981 | 1.25819102 | 0.20832268 | 0.99999872 |
| E2F3 | 14.9099151 | 0.723930308 | 0.69757302 | 1.03778427 | 0.2993705 | 0.99999872 |
| CDKAL1 | 138.510461 | -0.253746868 | 0.28084751 | -0.9035041 | 0.36625843 | 0.99999872 |
| SOX4 | 10.4234212 | 0.067921162 | 0.7133628 | 0.09521265 | 0.92414592 | 0.99999872 |
| CASC15 | 11.8493136 | -0.940512938 | 0.9364041 | -1.0043879 | 0.31519168 | 0.99999872 |
| DCDC2 | 78.5697799 | 0.028044502 | 0.37013837 | 0.07576762 | 0.93960398 | 0.99999872 |
| MRS2 | 26.2530677 | -0.281485509 | 0.35492587 | -0.7930825 | 0.42772974 | 0.99999872 |
| ALDH5A1 | 5.13887186 | 1.006392367 | 0.72207151 | 1.3937572 | 0.16339094 | 0.99999872 |
| TDP2 | 16.7202936 | 0.3630151 | 0.46286147 | 0.78428455 | 0.43287316 | 0.99999872 |
| ACOT13 | 17.4671725 | -0.022364597 | 0.46698536 | -0.0478914 | 0.96180277 | 0.99999872 |
| C6orf62 | 68.9566737 | -0.406940187 | 0.35151681 | -1.1576692 | 0.24699905 | 0.99999872 |
| GMNN | 6.31859491 | 0.160722274 | 0.58821332 | 0.27323807 | 0.78467021 | 0.99999872 |
| RIPOR2 | 47.3779464 | -0.296560181 | 0.454281 | -0.6528122 | 0.51387736 | 0.99999872 |
| CARMIL1 | 43.8415296 | -0.323609233 | 0.40787792 | -0.7933973 | 0.42754639 | 0.99999872 |
| SLC17A1 | 22.3211088 | 2.843843149 | 1.82525955 | 1.55804863 | 0.11922172 | 0.99999872 |
| SLC17A3 | 8.0178693 | 0.306903195 | 1.53685908 | 0.19969508 | 0.84171906 | 0.99999872 |
| TRIM38 | 37.9831622 | -0.618472324 | 0.68672389 | -0.9006128 | 0.36779423 | 0.99999872 |
| U91328.2 | 5.38334605 | 0.239612559 | 0.91592186 | 0.26160808 | 0.79362361 | 0.99999872 |
| HIST1H1C | 6.41527813 | -0.021190897 | 0.62864056 | -0.0337091 | 0.97310914 | 0.99999872 |
| HIST1H4C | 12.2988227 | 1.123760276 | 0.86370073 | 1.30109914 | 0.19322452 | 0.99999872 |
| HIST1H2AC | 22.9101893 | 0.365753653 | 0.5733049 | 0.63797406 | 0.52349057 | 0.99999872 |
| BTN3A2 | 8.95888522 | 0.257815784 | 1.11280638 | 0.23168072 | 0.81678601 | 0.99999872 |
| BTN2A2 | 4.92843621 | 0.008237581 | 0.70518833 | 0.01168139 | 0.99067981 | 0.99999872 |
| BTN3A1 | 7.71785932 | 0.655980066 | 1.33194853 | 0.49249656 | 0.62236835 | 0.99999872 |
| BTN2A1 | 11.5966988 | 0.067218355 | 0.58158154 | 0.11557856 | 0.90798656 | 0.99999872 |
| HCG11 | 4.86980371 | 0.670677831 | 0.99546016 | 0.67373649 | 0.50047886 | 0.99999872 |
| HMGN4 | 4.33441743 | 0.531635702 | 1.04696132 | 0.50778925 | 0.61160115 | NA |
| ABT1 | 7.63917699 | 0.051095094 | 0.55027371 | 0.09285396 | 0.92601958 | 0.99999872 |
| ZNF322 | 10.0976791 | 0.003172142 | 0.71367978 | 0.00444477 | 0.9964536 | 0.99999872 |
| LINC00240 | 11.3901895 | 0.417113234 | 0.69306315 | 0.60184016 | 0.54728054 | 0.99999872 |
| ZSCAN16-AS | 17.4890089 | 0.795111299 | 0.54385387 | 1.46199438 | 0.14374275 | 0.99999872 |
| ZKSCAN8 | 14.3098516 | -0.232057757 | 0.4443471 | -0.5222443 | 0.60150022 | 0.99999872 |
| ZSCAN9 | 8.01433225 | -0.349405327 | 0.66712396 | -0.5237487 | 0.60045332 | 0.99999872 |
| PGBD1 | 7.04328546 | -0.204996255 | 0.57049504 | -0.3593305 | 0.71934788 | 0.99999872 |
| ZSCAN31 | 15.2303704 | 0.416626749 | 0.55552403 | 0.74997071 | 0.45327235 | 0.99999872 |
| ZSCAN23 | 4.89673998 | 1.022576311 | 0.73664214 | 1.38815885 | 0.16508868 | 0.99999872 |
| TRIM27 | 21.9425488 | -0.111415685 | 0.5072534 | -0.219645 | 0.82614763 | 0.99999872 |
| GABBR1 | 21.8654819 | 0.128322351 | 0.51261021 | 0.25033124 | 0.8023312 | 0.99999872 |
| HLA-F | 20.8203517 | -0.120629733 | 1.21419719 | -0.0993494 | 0.92086088 | 0.99999872 |
| HLA-A | 481.869791 | -0.005551932 | 0.83112995 | -0.00668 | 0.99467019 | 0.99999872 |
| ZNRD1 | 7.29497391 | 0.0991324 | 0.64737209 | 0.15313048 | 0.87829538 | 0.99999872 |
| PPP1R11 | 12.4865102 | 0.106023391 | 0.45965309 | 0.23065958 | 0.81757927 | 0.99999872 |
| TRIM26 | 14.7461208 | -0.134390056 | 0.62504061 | -0.2150101 | 0.82975943 | 0.99999872 |
| HCG17 | 33.344494 | -0.314850127 | 0.33943068 | -0.927583 | 0.35362392 | 0.99999872 |
| HLA-E | 187.882807 | -0.159831624 | 0.81047425 | -0.1972075 | 0.84366514 | 0.99999872 |
| GNL1 | 21.4276825 | -0.179636102 | 0.41887979 | -0.4288488 | 0.66803325 | 0.99999872 |

| | | | | | | |
|---|---|---|---|---|---|---|
| PRR3 | 7.6267751 | 0.685373473 | 0.58263846 | 1.17632722 | 0.23946414 | 0.99999872 |
| ABCF1 | 19.8177975 | 0.012389479 | 0.45119038 | 0.02745954 | 0.97809321 | 0.99999872 |
| PPP1R10 | 17.9561701 | -0.300904381 | 0.41682003 | -0.7219048 | 0.47035301 | 0.99999872 |
| MRPS18B | 19.8209654 | 0.004317266 | 0.38751709 | 0.01114084 | 0.99111108 | 0.99999872 |
| ATAT1 | 5.00108063 | -0.326149289 | 0.62514783 | -0.5217155 | 0.60186846 | 0.99999872 |
| C6orf136 | 7.20037249 | -0.252292976 | 0.5949651 | -0.4240467 | 0.67153177 | 0.99999872 |
| DHX16 | 7.16073266 | -0.244542779 | 0.73355869 | -0.333365 | 0.73885879 | 0.99999872 |
| MDC1 | 8.88193844 | 0.030754921 | 0.50957852 | 0.06035364 | 0.95187398 | 0.99999872 |
| TUBB | 71.9509856 | 0.960850689 | 0.63714852 | 1.50804821 | 0.13154219 | 0.99999872 |
| FLOT1 | 41.0171306 | -0.137214021 | 0.41622573 | -0.3296625 | 0.74165498 | 0.99999872 |
| IER3 | 131.611463 | 1.241341641 | 1.00641637 | 1.23342751 | 0.21741631 | 0.99999872 |
| DDR1 | 30.9264726 | -0.197245665 | 0.35730463 | -0.5520378 | 0.58092244 | 0.99999872 |
| POU5F1 | 8.12471145 | 0.243029683 | 1.10907762 | 0.21912775 | 0.82655054 | 0.99999872 |
| HCG27 | 6.29352479 | -0.064467637 | 0.63490264 | -0.1015394 | 0.91912228 | 0.99999872 |
| HLA-C | 265.830702 | 0.680311181 | 0.95384726 | 0.71322864 | 0.47570429 | 0.99999872 |
| HLA-B | 350.180677 | 0.425708924 | 1.01817606 | 0.41810934 | 0.67586717 | 0.99999872 |
| MICA | 27.9374966 | 0.154808854 | 0.41492449 | 0.37310127 | 0.70907307 | 0.99999872 |
| NFKBIL1 | 8.76495851 | -0.368158335 | 0.55581415 | -0.6623767 | 0.50772984 | 0.99999872 |
| AIF1 | 148.510235 | 0.849081189 | 0.75656416 | 1.12228577 | 0.26174096 | 0.99999872 |
| PRRC2A | 43.5494658 | -0.338667453 | 0.39360261 | -0.8604299 | 0.38955211 | 0.99999872 |
| BAG6 | 17.0962879 | -0.414005941 | 0.42604781 | -0.9717359 | 0.33118196 | 0.99999872 |
| C6orf47 | 8.79296698 | 0.186241131 | 0.50761963 | 0.36689112 | 0.71370024 | 0.99999872 |
| GPANK1 | 4.98737696 | 0.406305614 | 0.64552566 | 0.62941822 | 0.52907529 | 0.99999872 |
| DDAH2 | 21.2500014 | -0.42088305 | 0.44372715 | -0.9485177 | 0.34286597 | 0.99999872 |
| CLIC1 | 109.066471 | 0.068146057 | 0.45120382 | 0.15103165 | 0.87995075 | 0.99999872 |
| VARS | 8.3493148 | -0.306288233 | 0.65865145 | -0.4650232 | 0.64191481 | 0.99999872 |
| LSM2 | 15.4501256 | 0.554487313 | 0.53401687 | 1.03833295 | 0.29911507 | 0.99999872 |
| HSPA1A | 69.7208083 | -0.016226458 | 0.59403947 | -0.0273155 | 0.97820813 | 0.99999872 |
| HSPA1B | 28.816143 | 0.635156722 | 0.77944092 | 0.81488758 | 0.41513667 | 0.99999872 |
| C6orf48 | 21.7664889 | 0.639097491 | 0.43267452 | 1.47708604 | 0.13965257 | 0.99999872 |
| NEU1 | 14.3244721 | 0.351595608 | 0.47927048 | 0.7336058 | 0.46318902 | 0.99999872 |
| EHMT2 | 10.049484 | 0.21931416 | 0.57960316 | 0.37838675 | 0.70514331 | 0.99999872 |
| NELFE | 9.97793396 | 0.566450014 | 0.53349661 | 1.06176872 | 0.2883407 | 0.99999872 |
| STK19 | 17.1770669 | 0.846993133 | 0.49115359 | 1.72449749 | 0.08461807 | 0.99999872 |
| ATF6B | 18.1292467 | -0.113504316 | 0.41826316 | -0.2713706 | 0.78610603 | 0.99999872 |
| AGPAT1 | 14.5749871 | 0.438266992 | 0.47510521 | 0.92246303 | 0.3562871 | 0.99999872 |
| RNF5 | 13.9461172 | 0.732266053 | 0.51599772 | 1.41912653 | 0.15586213 | 0.99999872 |
| PBX2 | 18.6377501 | -0.586700656 | 0.5885365 | -0.9968807 | 0.31882244 | 0.99999872 |
| HLA-DRA | 13.0217039 | 0.323856783 | 0.7300215 | 0.44362636 | 0.65731274 | 0.99999872 |
| HLA-DRB1 | 10.9417305 | 0.214849053 | 1.27604449 | 0.16837113 | 0.86629132 | 0.99999872 |
| PSMB8 | 32.3756236 | 0.026575864 | 0.60462981 | 0.04395394 | 0.96494112 | 0.99999872 |
| PSMB9 | 20.8128689 | -0.816125127 | 1.03215011 | -0.7907039 | 0.4291168 | 0.99999872 |
| TAP1 | 35.3997552 | 0.106757858 | 1.16084935 | 0.0919653 | 0.92672561 | 0.99999872 |
| BRD2 | 51.9145388 | -0.339055283 | 0.37180339 | -0.9119209 | 0.36181036 | 0.99999872 |
| HLA-DPA1 | 7.22312087 | 0.695312013 | 1.10150837 | 0.63123625 | 0.52788606 | 0.99999872 |
| HLA-DPB1 | 6.33323482 | 0.781164396 | 1.06622361 | 0.73264594 | 0.46377441 | 0.99999872 |
| RXRB | 8.44622411 | 0.211972187 | 0.48981831 | 0.43275676 | 0.66519149 | 0.99999872 |
| SLC39A7 | 24.9912577 | -0.12413427 | 0.47256805 | -0.2626802 | 0.79279708 | 0.99999872 |
| RING1 | 19.079539 | -0.656536009 | 0.37524307 | -1.7496286 | 0.08018242 | 0.99999872 |
| VPS52 | 11.171486 | -0.093464405 | 0.42843273 | -0.2181542 | 0.82730896 | 0.99999872 |
| RPS18 | 50.0862357 | 0.546864741 | 0.50197827 | 1.08941915 | 0.27596909 | 0.99999872 |
| B3GALT4 | 11.1585979 | 0.624693209 | 0.65923014 | 0.9476102 | 0.34332793 | 0.99999872 |
| WDR46 | 7.79173548 | 0.25182662 | 0.55183061 | 0.45634768 | 0.64813998 | 0.99999872 |
| PFDN6 | 11.5271116 | 0.328627532 | 0.60399637 | 0.54408859 | 0.58638051 | 0.99999872 |
| RGL2 | 11.0667365 | 0.044745158 | 0.53274993 | 0.08398905 | 0.93306514 | 0.99999872 |
| TAPBP | 128.932147 | -0.431401507 | 0.41144397 | -1.0485061 | 0.2944055 | 0.99999872 |
| ZBTB22 | 6.80746335 | 0.548867916 | 0.60776272 | 0.90309573 | 0.36647509 | 0.99999872 |
| DAXX | 6.405123 | 0.083473185 | 0.61365916 | 0.13602532 | 0.89180126 | 0.99999872 |
| PHF1 | 16.7744862 | -0.308954583 | 0.44299172 | -0.6974274 | 0.48553533 | 0.99999872 |
| CUTA | 32.7640077 | 0.333821093 | 0.37960482 | 0.87939109 | 0.37918926 | 0.99999872 |
| SYNGAP1 | 6.77897674 | 0.158005266 | 0.66591816 | 0.2372743 | 0.812444 | 0.99999872 |

| | | | | | | |
|---|---|---|---|---|---|---|
| ITPR3 | 8.25536212 | -1.584428161 | 0.92074042 | -1.7208196 | 0.08528356 | 0.99999872 |
| UQCC2 | 15.4752873 | 0.173865336 | 0.39551958 | 0.43958717 | 0.66023613 | 0.99999872 |
| IP6K3 | 8.98272383 | 0.805837913 | 0.90299211 | 0.89240858 | 0.37217397 | 0.99999872 |
| LEMD2 | 19.4749123 | -0.256936847 | 0.42219755 | -0.6085702 | 0.54280937 | 0.99999872 |
| SMIM29 | 11.7950533 | 1.030791857 | 0.69365743 | 1.4860244 | 0.13727267 | 0.99999872 |
| NUDT3 | 18.138421 | -0.105964229 | 0.75533561 | -0.1402876 | 0.88843275 | 0.99999872 |
| C6orf106 | 40.0622731 | -0.171114019 | 0.29019642 | -0.589649 | 0.55542602 | 0.99999872 |
| SNRPC | 20.9535312 | 0.150091399 | 0.46790649 | 0.32077221 | 0.74838302 | 0.99999872 |
| UHRF1BP1 | 30.1102723 | 0.140803651 | 0.37384055 | 0.37664093 | 0.70644047 | 0.99999872 |
| TAF11 | 15.7359139 | 0.053485485 | 0.40216275 | 0.13299463 | 0.89419763 | 0.99999872 |
| ANKS1A | 68.0999981 | -0.044706945 | 0.30948215 | -0.1444573 | 0.8851394 | 0.99999872 |
| ZNF76 | 16.0219902 | -0.100375804 | 0.42809839 | -0.234469 | 0.81462089 | 0.99999872 |
| PPARD | 12.49691 | 0.126498523 | 0.42134234 | 0.30022742 | 0.76400369 | 0.99999872 |
| RPL10A | 82.5102461 | 0.895529942 | 0.53352292 | 1.67852198 | 0.09324524 | 0.99999872 |
| TEAD3 | 36.2151987 | -0.037479123 | 0.39070484 | -0.0959269 | 0.92357859 | 0.99999872 |
| FKBP5 | 207.477875 | -0.531838045 | 0.62112321 | -0.8562521 | 0.39185836 | 0.99999872 |
| SRPK1 | 22.9912645 | -0.412123936 | 0.43055695 | -0.957188 | 0.33847238 | 0.99999872 |
| MAPK14 | 35.6131291 | -0.190839372 | 0.32026006 | -0.5958888 | 0.55124953 | 0.99999872 |
| BRPF3 | 10.0003008 | -0.976613002 | 0.62959956 | -1.5511653 | 0.12086207 | 0.99999872 |
| ETV7 | 4.193243 | 0.456440918 | 1.89034812 | 0.24145866 | 0.80919966 | NA |
| KCTD20 | 50.9967335 | 0.140710588 | 0.34483898 | 0.40804722 | 0.68323901 | 0.99999872 |
| STK38 | 63.9547706 | 0.265946754 | 0.29246527 | 0.90932763 | 0.36317721 | 0.99999872 |
| SRSF3 | 59.6787548 | -0.018178152 | 0.29659123 | -0.0612903 | 0.95112805 | 0.99999872 |
| CDKN1A | 18.4839528 | 0.454432663 | 0.6679696 | 0.68031938 | 0.49630226 | 0.99999872 |
| PPIL1 | 5.68160945 | -0.177982377 | 0.79611804 | -0.2235628 | 0.82309751 | 0.99999872 |
| C6orf89 | 35.1770798 | -0.127772961 | 0.27195684 | -0.4698281 | 0.63847785 | 0.99999872 |
| MTCH1 | 32.754109 | 0.520203916 | 0.42399211 | 1.22691887 | 0.21985309 | 0.99999872 |
| TBC1D22B | 15.5412651 | 0.142037332 | 0.42310398 | 0.33570314 | 0.73709474 | 0.99999872 |
| RNF8 | 14.6431866 | -0.006072663 | 0.41868799 | -0.014504 | 0.98842786 | 0.99999872 |
| CMTR1 | 13.0697458 | -1.074502564 | 0.6116058 | -1.7568548 | 0.07894257 | 0.99999872 |
| CCDC167 | 4.80261324 | 0.562329406 | 0.66765565 | 0.84224466 | 0.39965102 | 0.99999872 |
| ZFAND3 | 363.796641 | 0.003044871 | 0.20838334 | 0.01461187 | 0.98834183 | 0.99999872 |
| BTBD9 | 180.117282 | -0.05188952 | 0.24980999 | -0.207716 | 0.83545076 | 0.99999872 |
| GLO1 | 35.6235365 | 0.066285758 | 0.35466673 | 0.1868959 | 0.85174226 | 0.99999872 |
| SAYSD1 | 6.5514645 | 0.764236693 | 0.65675774 | 1.16365084 | 0.24456554 | 0.99999872 |
| DAAM2 | 24.2552077 | -0.392952368 | 0.51328479 | -0.765564 | 0.44393576 | 0.99999872 |
| MOCS1 | 8.3379307 | -0.872789367 | 0.59095666 | -1.4769093 | 0.13969996 | 0.99999872 |
| UNC5CL | 7.14933157 | 0.797156559 | 0.83621683 | 0.95328931 | 0.34044351 | 0.99999872 |
| OARD1 | 15.4603977 | -0.61387833 | 0.5557669 | -1.1045608 | 0.26934995 | 0.99999872 |
| NFYA | 6.8299717 | 0.069898762 | 0.61291828 | 0.11404255 | 0.90920406 | 0.99999872 |
| TFEB | 10.0193492 | 0.430589125 | 0.5225776 | 0.82397164 | 0.40995567 | 0.99999872 |
| USP49 | 14.4890484 | -0.294584957 | 0.44253589 | -0.6656747 | 0.50561903 | 0.99999872 |
| MED20 | 3.47029368 | 0.862390462 | 0.78083285 | 1.10444951 | 0.26939819 | NA |
| CCND3 | 32.12759 | -0.415175257 | 0.46821339 | -0.8867223 | 0.37522843 | 0.99999872 |
| TAF8 | 13.4234289 | -0.232254585 | 0.46780255 | -0.4964799 | 0.61955583 | 0.99999872 |
| MRPS10 | 10.26157 | -0.122850434 | 0.42930902 | -0.2861585 | 0.77475671 | 0.99999872 |
| TRERF1 | 16.397596 | 0.680428071 | 0.80504231 | 0.84520784 | 0.3979948 | 0.99999872 |
| UBR2 | 101.503772 | -0.052074002 | 0.23030485 | -0.226109 | 0.82111663 | 0.99999872 |
| TBCC | 4.46719896 | 0.25770384 | 0.63172316 | 0.40793793 | 0.68331925 | NA |
| BICRAL | 57.7268529 | -0.102405702 | 0.32131519 | -0.3187079 | 0.749948 | 0.99999872 |
| RPL7L1 | 36.6855636 | 0.065703664 | 0.45023275 | 0.14593266 | 0.88397455 | 0.99999872 |
| C6orf226 | 11.475838 | -0.040606251 | 0.44852566 | -0.0905327 | 0.92786389 | 0.99999872 |
| CNPY3 | 14.505356 | 0.046947818 | 0.54430626 | 0.08625258 | 0.93126564 | 0.99999872 |
| PEX6 | 10.3868153 | 0.411035619 | 0.48618525 | 0.84543004 | 0.39787078 | 0.99999872 |
| PPP2R5D | 8.48762337 | -0.668390718 | 0.50842976 | -1.3146176 | 0.18863845 | 0.99999872 |
| MEA1 | 11.2505307 | 0.036590326 | 0.43496694 | 0.08412208 | 0.93295937 | 0.99999872 |
| KLHDC3 | 15.4957998 | -0.124550525 | 0.59627518 | -0.2088809 | 0.83454118 | 0.99999872 |
| RRP36 | 19.8753853 | 0.004462103 | 0.47351526 | 0.00942336 | 0.99248136 | 0.99999872 |
| CUL7 | 7.4227082 | -0.044722349 | 0.54186598 | -0.082534 | 0.93422211 | 0.99999872 |
| KLC4 | 8.99326161 | 0.028800943 | 0.53696702 | 0.05363633 | 0.95722491 | 0.99999872 |
| MRPL2 | 13.6737038 | 0.388244492 | 0.52523592 | 0.73918116 | 0.459797 | 0.99999872 |

| | | | | | | |
|---|---|---|---|---|---|---|
| PTK7 | 24.2275377 | 0.034889352 | 0.5502884 | 0.06340194 | 0.94944644 | 0.99999872 |
| SRF | 11.1799599 | -0.49396628 | 0.46727043 | -1.0571315 | 0.29045159 | 0.99999872 |
| CUL9 | 9.82864752 | 0.20722874 | 0.52421889 | 0.39530957 | 0.69261444 | 0.99999872 |
| DNPH1 | 15.9303744 | 0.28636633 | 0.87642355 | 0.32674422 | 0.74386135 | 0.99999872 |
| ZNF318 | 17.4075136 | 0.378552366 | 0.46385123 | 0.81610729 | 0.41443878 | 0.99999872 |
| ABCC10 | 8.7628037 | -0.530494531 | 0.54423832 | -0.9747467 | 0.3296859 | 0.99999872 |
| TJAP1 | 11.8543731 | -0.670695764 | 0.45704684 | -1.4674552 | 0.14225226 | 0.99999872 |
| POLR1C | 16.07762 | -0.232759338 | 0.47367166 | -0.4913938 | 0.62314792 | 0.99999872 |
| YIPF3 | 28.6953961 | 0.019719706 | 0.46560794 | 0.0423526 | 0.96621762 | 0.99999872 |
| XPO5 | 43.8470399 | 0.449112827 | 0.4969193 | 0.90379429 | 0.36610449 | 0.99999872 |
| POLH | 7.63256136 | 0.669818489 | 0.55685083 | 1.20286879 | 0.2290271 | 0.99999872 |
| GTPBP2 | 11.0390002 | -0.507031653 | 0.79965989 | -0.6340591 | 0.52604223 | 0.99999872 |
| MAD2L1BP | 4.76727523 | 0.588107013 | 0.67041502 | 0.87722828 | 0.38036266 | 0.99999872 |
| MRPS18A | 15.8932469 | 0.532773168 | 0.6425031 | 0.82921493 | 0.4069828 | 0.99999872 |
| VEGFA | 1005.34766 | 0.440186799 | 0.52077227 | 0.84525775 | 0.39796695 | 0.99999872 |
| MRPL14 | 16.2273776 | 0.64761237 | 0.51145391 | 1.26621843 | 0.20543489 | 0.99999872 |
| TMEM63B | 12.1413857 | -0.510088279 | 0.55998497 | -0.9108964 | 0.36234998 | 0.99999872 |
| SLC29A1 | 16.0378829 | 0.492426209 | 0.56449273 | 0.87233401 | 0.38302619 | 0.99999872 |
| HSP90AB1 | 113.282784 | 0.140624291 | 0.45709915 | 0.30764505 | 0.75835244 | 0.99999872 |
| SLC35B2 | 12.9972692 | 0.691006644 | 0.48131946 | 1.43565076 | 0.15110174 | 0.99999872 |
| CDC5L | 19.4585402 | -0.095928552 | 0.4248602 | -0.2257885 | 0.82136591 | 0.99999872 |
| SUPT3H | 101.123793 | 0.150421243 | 0.39778262 | 0.37814936 | 0.70531964 | 0.99999872 |
| CLIC5 | 1957.78295 | 0.014636941 | 0.43002132 | 0.03403771 | 0.97284708 | 0.99999872 |
| ENPP4 | 31.9629568 | 1.141570956 | 0.51626253 | 2.21122181 | 0.02702048 | 0.89154528 |
| ENPP5 | 10.4132537 | 0.920811586 | 0.64797006 | 1.4210712 | 0.15529606 | 0.99999872 |
| RCAN2 | 6.48913539 | -1.145101829 | 0.62809332 | -1.8231396 | 0.06828225 | 0.99999872 |
| CYP39A1 | 10.2876495 | 1.259803839 | 0.76534031 | 1.64607014 | 0.09974932 | 0.99999872 |
| SLC25A27 | 23.8689803 | 0.585993694 | 0.59122505 | 0.99115167 | 0.32161152 | 0.99999872 |
| TNFRSF21 | 12.0187274 | -1.836770313 | 1.19292375 | -1.5397215 | 0.12362826 | 0.99999872 |
| CD2AP | 201.60225 | -0.085586307 | 0.28745756 | -0.2977355 | 0.76590508 | 0.99999872 |
| MUT | 15.7580804 | 0.18363821 | 0.45434613 | 0.4041813 | 0.6860794 | 0.99999872 |
| PKHD1 | 114.81957 | -0.384800735 | 0.70150481 | -0.5485361 | 0.58332383 | 0.99999872 |
| MCM3 | 8.74866348 | -0.808570694 | 0.47235047 | -1.7118025 | 0.08693308 | 0.99999872 |
| EFHC1 | 32.3074401 | 0.618451688 | 0.48873076 | 1.26542412 | 0.20571933 | 0.99999872 |
| TRAM2 | 107.022433 | 0.005175784 | 0.46016882 | 0.01124758 | 0.99102592 | 0.99999872 |
| TRAM2-AS1 | 7.14959882 | -1.363194186 | 0.58920749 | -2.3136064 | 0.02068932 | 0.8671024 |
| TMEM14A | 17.8083971 | 1.126334173 | 0.56457438 | 1.99501468 | 0.04604128 | 0.95980711 |
| GSTA4 | 25.7626071 | 0.104628498 | 0.54340265 | 0.19254322 | 0.84731671 | 0.99999872 |
| ICK | 9.86191244 | 0.24600512 | 0.45823685 | 0.53685146 | 0.59137022 | 0.99999872 |
| FBXO9 | 36.8723091 | -0.337318225 | 0.39654505 | -0.8506429 | 0.39496776 | 0.99999872 |
| ELOVL5 | 52.8281957 | 0.236136564 | 0.33987598 | 0.69477274 | 0.48719773 | 0.99999872 |
| GCLC | 20.717441 | -0.313719615 | 0.52546896 | -0.5970279 | 0.55048877 | 0.99999872 |
| LRRC1 | 63.9575892 | 0.617831408 | 0.35695962 | 1.73081597 | 0.08348459 | 0.99999872 |
| TINAG | 15.5682773 | 1.570821419 | 1.31740573 | 1.19235964 | 0.23312026 | 0.99999872 |
| HMGCLL1 | 10.7213953 | -0.720602193 | 0.64536499 | -1.1165809 | 0.26417358 | 0.99999872 |
| DST | 737.430367 | 0.055495258 | 0.29267684 | 0.18961274 | 0.8496126 | 0.99999872 |
| AL512422.1 | 6.19839618 | 1.462051069 | 0.93181636 | 1.56903348 | 0.11664014 | 0.99999872 |
| BEND6 | 8.03708124 | 0.11420594 | 0.83574017 | 0.13665245 | 0.89130552 | 0.99999872 |
| KIAA1586 | 8.67862854 | 0.174677512 | 0.48958323 | 0.35678818 | 0.72125038 | 0.99999872 |
| ZNF451 | 73.8682511 | 0.035959927 | 0.25566578 | 0.14065209 | 0.88814479 | 0.99999872 |
| ZNF451-AS1 | 10.4332478 | 1.44783989 | 0.85728181 | 1.68887276 | 0.09124382 | 0.99999872 |
| BAG2 | 6.04687519 | 0.266185 | 0.66998291 | 0.39730118 | 0.69114538 | 0.99999872 |
| RAB23 | 5.74698305 | 0.488072308 | 0.73278274 | 0.66605323 | 0.50537708 | 0.99999872 |
| PRIM2 | 41.0118932 | -0.658538553 | 0.43749116 | -1.5052614 | 0.13225689 | 0.99999872 |
| AL021368.2 | 6.15349435 | -0.273355642 | 0.65475979 | -0.41749 | 0.67632006 | 0.99999872 |
| PTP4A1 | 18.5074659 | -0.336578055 | 0.55469066 | -0.6067851 | 0.54399351 | 0.99999872 |
| PHF3 | 100.088773 | 0.276191209 | 0.29400846 | 0.93939885 | 0.347526 | 0.99999872 |
| EYS | 16.2364934 | 1.106951868 | 0.78741314 | 1.40580822 | 0.15978108 | 0.99999872 |
| LMBRD1 | 63.4652797 | 0.19889787 | 0.3334896 | 0.59641402 | 0.55089868 | 0.99999872 |
| COL19A1 | 17.9653461 | -0.239733821 | 0.5141574 | -0.4662654 | 0.64102552 | 0.99999872 |
| FAM135A | 85.9429221 | 0.095916532 | 0.28079743 | 0.34158622 | 0.73266231 | 0.99999872 |

| | | | | | | |
|---|---|---|---|---|---|---|
| SDHAF4 | 6.02229845 | 1.056007015 | 0.6054929 | 1.74404524 | 0.0811512 | 0.99999872 |
| SMAP1 | 62.9504882 | 0.027756257 | 0.25205668 | 0.11011911 | 0.91231491 | 0.99999872 |
| OGFRL1 | 67.9901794 | -0.667056489 | 0.51807139 | -1.2875764 | 0.19789346 | 0.99999872 |
| AL136164.2 | 78.9457087 | 1.178833313 | 0.38230206 | 3.08351289 | 0.00204572 | 0.36957035 |
| LINC00472 | 45.0502794 | 0.309939982 | 0.42680591 | 0.72618484 | 0.46772546 | 0.99999872 |
| AC019205.1 | 16.1166283 | -0.58169233 | 0.51452474 | -1.130543 | 0.2582475 | 0.99999872 |
| OOEP | 54.7858951 | 0.728871535 | 0.4165833 | 1.74964176 | 0.08018015 | 0.99999872 |
| MTO1 | 11.1245543 | -0.351194306 | 0.51473311 | -0.6822843 | 0.49505922 | 0.99999872 |
| EEF1A1 | 247.095267 | 0.320118844 | 0.40076832 | 0.79876284 | 0.42442794 | 0.99999872 |
| SLC17A5 | 62.8952877 | 0.439025478 | 0.52256844 | 0.84013011 | 0.40083544 | 0.99999872 |
| COX7A2 | 46.0875155 | 1.260289566 | 0.43454289 | 2.90026509 | 0.00372847 | 0.48230777 |
| TMEM30A | 92.8862835 | 0.559798553 | 0.47642441 | 1.17499972 | 0.23999483 | 0.99999872 |
| SENP6 | 180.291393 | -0.104735886 | 0.24675598 | -0.4244513 | 0.67123675 | 0.99999872 |
| MYO6 | 120.054774 | -0.146791262 | 0.34970646 | -0.4197557 | 0.67466397 | 0.99999872 |
| LINC02540 | 6.38988656 | 2.418758003 | 0.74865227 | 3.23081638 | 0.00123437 | 0.30478826 |
| MEI4 | 7.57167191 | 0.258386168 | 0.66400796 | 0.38913113 | 0.69717915 | 0.99999872 |
| IRAK1BP1 | 45.8077554 | 0.401054168 | 0.33757696 | 1.18803774 | 0.23481854 | 0.99999872 |
| PHIP | 194.691547 | -0.572267533 | 0.39093283 | -1.4638513 | 0.14323458 | 0.99999872 |
| HMGN3 | 221.554331 | -0.61808667 | 0.31418754 | -1.9672539 | 0.04915395 | 0.97937581 |
| HMGN3-AS1 | 11.4520941 | 0.43778826 | 0.63318993 | 0.69140117 | 0.48931347 | 0.99999872 |
| LCA5 | 33.4379848 | -0.115965529 | 0.44557173 | -0.2602623 | 0.79466144 | 0.99999872 |
| SH3BGRL2 | 73.0924327 | -0.195911329 | 0.46547706 | -0.4208829 | 0.6738406 | 0.99999872 |
| ELOVL4 | 16.6605501 | -0.823717331 | 0.67389907 | -1.2223156 | 0.22158831 | 0.99999872 |
| BCKDHB | 209.697504 | -0.160080315 | 0.59800006 | -0.2676928 | 7.89E-01 | 0.99999872 |
| AL359715.1 | 11.3197943 | 0.017380819 | 0.75205745 | 0.02311103 | 0.98156171 | 0.99999872 |
| TENT5A | 34.4498805 | -0.411329616 | 0.74968575 | -0.5486694 | 0.58323236 | 0.99999872 |
| IBTK | 56.450919 | -0.333332694 | 0.34252657 | -0.9731586 | 0.33047446 | 0.99999872 |
| TPBG | 22.0817711 | 0.098592312 | 0.37789439 | 0.26089911 | 0.79417031 | 0.99999872 |
| UBE3D | 28.7515589 | -0.40257974 | 0.50973326 | -0.7897851 | 0.42965327 | 0.99999872 |
| DOPEY1 | 48.305195 | 0.106192071 | 0.28602732 | 0.37126548 | 0.71043981 | 0.99999872 |
| PGM3 | 21.8234484 | 0.552197984 | 0.46057494 | 1.19893188 | 0.23055444 | 0.99999872 |
| ME1 | 35.8866322 | -0.082673976 | 0.32984173 | -0.2506474 | 0.80208672 | 0.99999872 |
| CYB5R4 | 14.8432365 | 0.304427083 | 0.47609036 | 0.63943131 | 0.52254239 | 0.99999872 |
| CEP162 | 26.3701537 | 0.476692158 | 0.44437335 | 1.07272895 | 0.28339275 | 0.99999872 |
| NT5E | 45.6737899 | 0.317241877 | 0.55085225 | 0.57591101 | 0.56467533 | 0.99999872 |
| SNX14 | 50.6900807 | 0.121707931 | 0.26225215 | 0.46408746 | 0.64258509 | 0.99999872 |
| SYNCRIP | 21.6174847 | -0.32514658 | 0.36944921 | -0.8800846 | 0.37881345 | 0.99999872 |
| ZNF292 | 130.794893 | -0.014186651 | 0.28001067 | -0.0506647 | 0.95959272 | 0.99999872 |
| SMIM8 | 19.4673835 | -0.200035624 | 0.50970427 | -0.3924543 | 0.69472258 | 0.99999872 |
| SLC35A1 | 6.68397118 | 1.213463498 | 1.06628386 | 1.13803045 | 0.25510777 | 0.99999872 |
| RARS2 | 36.837693 | -0.038517769 | 0.29172519 | -0.1320344 | 0.89495706 | 0.99999872 |
| ORC3 | 26.8497691 | -0.493896091 | 0.37564855 | -1.3147824 | 0.18858305 | 0.99999872 |
| AKIRIN2 | 12.3678721 | 0.445244718 | 0.55564639 | 0.80130947 | 0.42295251 | 0.99999872 |
| RNGTT | 54.8678816 | -0.154821523 | 0.3105724 | -0.4985038 | 0.61812899 | 0.99999872 |
| PNRC1 | 37.8226833 | 0.995050047 | 0.45351815 | 2.19406883 | 0.02823046 | 0.90182554 |
| PM20D2 | 34.5258931 | 0.630525901 | 0.46106375 | 1.36754604 | 0.17145422 | 0.99999872 |
| UBE2J1 | 23.1288881 | 0.156366627 | 0.36485936 | 0.42856686 | 0.66823847 | 0.99999872 |
| RRAGD | 11.2317126 | 0.221365294 | 0.6646732 | 0.33304381 | 0.73910122 | 0.99999872 |
| ANKRD6 | 17.4862597 | 0.092316106 | 0.54514952 | 0.16934089 | 0.86552851 | 0.99999872 |
| LYRM2 | 25.718115 | -0.113068183 | 0.40799231 | -0.2771331 | 0.78167789 | 0.99999872 |
| MDN1 | 24.0897824 | -0.476035133 | 0.40556566 | -1.173756 | 0.24049277 | 0.99999872 |
| CASP8AP2 | 21.8127349 | -0.347655925 | 0.36892089 | -0.942359 | 0.34600886 | 0.99999872 |
| BACH2 | 36.893576 | 0.644488104 | 0.55201316 | 1.16752308 | 0.24299919 | 0.99999872 |
| MAP3K7 | 36.3865903 | 0.146205224 | 0.35011578 | 0.41759107 | 0.67624613 | 0.99999872 |
| EPHA7 | 49.1353011 | 0.173616799 | 0.58121581 | 0.29871314 | 0.76515893 | 0.99999872 |
| MANEA | 40.3399179 | 0.68128599 | 0.45737393 | 1.48956016 | 0.13633992 | 0.99999872 |
| UFL1 | 39.5122699 | -0.02418612 | 0.33694344 | -0.0717809 | 0.94277623 | 0.99999872 |
| GPR63 | 8.57505946 | 1.009992674 | 0.61335377 | 1.64667231 | 0.09962542 | 0.99999872 |
| NDUFAF4 | 11.3852415 | 0.373961218 | 0.51066279 | 0.73230559 | 0.46398207 | 0.99999872 |
| KLHL32 | 7.49133474 | 0.155229252 | 0.92639406 | 0.16756288 | 0.86692718 | 0.99999872 |
| MMS22L | 15.5796857 | -0.230451252 | 0.40950053 | -0.5627618 | 0.5735971 | 0.99999872 |

| | | | | | | |
|---|---|---|---|---|---|---|
| FBXL4 | 54.524288 | 0.027219185 | 0.28931355 | 0.09408196 | 0.92504405 | 0.99999872 |
| PNISR | 135.997896 | 0.230743259 | 0.32321048 | 0.71391021 | 0.4752827 | 0.99999872 |
| AL513550.1 | 8.93068649 | 0.742612142 | 0.60819804 | 1.22100384 | 0.22208457 | 0.99999872 |
| USP45 | 33.4195926 | 0.467398327 | 0.35318769 | 1.32337093 | 0.18571205 | 0.99999872 |
| TSTD3 | 8.35673133 | -0.641876761 | 0.51430826 | -1.248039 | 0.21201678 | 0.99999872 |
| CCNC | 32.8576338 | 0.476813343 | 0.43119557 | 1.10579369 | 0.26881583 | 0.99999872 |
| SIM1 | 14.0729703 | -1.375000417 | 1.01429343 | -1.3556239 | 0.17521886 | 0.99999872 |
| ASCC3 | 93.7435872 | -0.024453912 | 0.32170873 | -0.0760126 | 0.93940909 | 0.99999872 |
| GRIK2 | 146.8447 | -0.667539853 | 0.55856923 | -1.1950888 | 0.23205233 | 0.99999872 |
| HACE1 | 32.9856132 | 0.517064905 | 0.39120054 | 1.32173873 | 0.18625517 | 0.99999872 |
| PREP | 18.0940967 | -0.47448045 | 0.37247408 | -1.2738616 | 0.20271245 | 0.99999872 |
| ATG5 | 40.0956757 | 0.156337854 | 0.31785235 | 0.49185684 | 0.62282055 | 0.99999872 |
| CRYBG1 | 12.0843005 | 0.487884143 | 0.760712 | 0.64135198 | 0.52129403 | 0.99999872 |
| RTN4IP1 | 5.60485558 | -0.136131108 | 0.72398084 | -0.1880314 | 0.85085206 | 0.99999872 |
| QRSL1 | 12.9822605 | -0.129579196 | 0.47679786 | -0.2717697 | 0.78579913 | 0.99999872 |
| LINC02532 | 39.646147 | 0.373245033 | 0.96457372 | 0.38695335 | 0.69879074 | 0.99999872 |
| CD24 | 44.4996645 | -0.190961817 | 0.54472113 | -0.350568 | 0.72591244 | 0.99999872 |
| C6orf203 | 19.1415398 | 0.363466195 | 0.41888315 | 0.86770306 | 0.38555691 | 0.99999872 |
| PDSS2 | 73.061529 | -0.196795245 | 0.29720524 | -0.6621527 | 0.50787338 | 0.99999872 |
| SOBP | 69.8075423 | 0.342197396 | 0.3134769 | 1.09161917 | 0.27500053 | 0.99999872 |
| SEC63 | 57.0826908 | -0.050749424 | 0.28764135 | -0.176433 | 0.85995378 | 0.99999872 |
| OSTM1 | 30.4733101 | 0.200235181 | 0.37154123 | 0.53893126 | 0.58993429 | 0.99999872 |
| SNX3 | 47.341811 | 0.497476239 | 0.30759316 | 1.61731892 | 0.10580946 | 0.99999872 |
| AFG1L | 21.8802967 | -0.581932836 | 0.48685672 | -1.1952856 | 0.23197546 | 0.99999872 |
| FOXO3 | 103.312765 | -0.034921845 | 0.38935338 | -0.0896919 | 0.92853205 | 0.99999872 |
| ARMC2 | 19.970858 | -0.243013958 | 0.36695637 | -0.6622421 | 0.50781608 | 0.99999872 |
| SESN1 | 22.6111693 | 0.867030922 | 0.49167791 | 1.76341238 | 0.07783096 | 0.99999872 |
| CEP57L1 | 10.8709598 | -0.133681718 | 0.567559 | -0.235538 | 0.81379118 | 0.99999872 |
| CD164 | 107.535006 | 0.35653454 | 0.34708393 | 1.02722861 | 0.30431283 | 0.99999872 |
| PPIL6 | 7.484313 | 1.159894601 | 0.61352519 | 1.89054113 | 0.05868562 | 0.99999872 |
| MICAL1 | 3.97724366 | 0.255016445 | 0.94322273 | 0.27036715 | 0.78687781 | NA |
| AK9 | 24.3990981 | -0.073138426 | 0.54802661 | -0.1334578 | 0.89383134 | 0.99999872 |
| FIG4 | 21.9706508 | 0.913055211 | 0.49495902 | 1.84470868 | 0.06507993 | 0.99999872 |
| WASF1 | 23.0420389 | 0.112856227 | 0.39302372 | 0.28714864 | 0.77399851 | 0.99999872 |
| CDC40 | 29.3460609 | 0.383782446 | 0.4040652 | 0.94980326 | 0.34221223 | 0.99999872 |
| METTL24 | 25.1365159 | -0.193114132 | 0.57376311 | -0.3365747 | 0.73643755 | 0.99999872 |
| CDK19 | 36.8594836 | 0.212875505 | 0.40748885 | 0.52240817 | 0.60138617 | 0.99999872 |
| AMD1 | 18.2322187 | 0.095921003 | 0.48967727 | 0.19588617 | 0.84469927 | 0.99999872 |
| GTF3C6 | 24.1160051 | 0.30247104 | 0.44402593 | 0.6812013 | 0.49574413 | 0.99999872 |
| RPF2 | 8.67032349 | -0.750764171 | 0.64001884 | -1.1730345 | 0.24078198 | 0.99999872 |
| SLC16A10 | 121.726798 | 0.349965484 | 0.60313484 | 0.58024419 | 0.56174996 | 0.99999872 |
| MFSD4B | 9.70865141 | -0.148295751 | 0.68405188 | -0.2167902 | 0.82837184 | 0.99999872 |
| REV3L | 94.3438236 | 0.178150968 | 0.30826787 | 0.57790963 | 0.56332514 | 0.99999872 |
| TRAF3IP2-AS | 44.2284891 | 0.170713213 | 0.40894863 | 0.41744415 | 0.67635357 | 0.99999872 |
| TRAF3IP2 | 12.6955433 | -0.111696185 | 0.55172647 | -0.2024485 | 0.83956613 | 0.99999872 |
| FYN | 390.612174 | -0.224069378 | 0.31732511 | -0.7061193 | 0.48011396 | 0.99999872 |
| WISP3 | 4.23798247 | 0.723494991 | 1.38712079 | 0.52158038 | 0.60196253 | NA |
| TUBE1 | 17.678942 | 0.717390778 | 0.58449452 | 1.22736956 | 0.21968372 | 0.99999872 |
| FAM229B | 13.251962 | -0.036357851 | 0.6239636 | -0.0582692 | 0.95353421 | 0.99999872 |
| HDAC2 | 24.4938891 | 0.077052989 | 0.43671564 | 0.17643744 | 0.85995029 | 0.99999872 |
| FRK | 10.6193145 | 0.267703572 | 0.80395493 | 0.33298331 | 0.73914688 | 0.99999872 |
| NT5DC1 | 62.0406132 | -0.216108683 | 0.35163647 | -0.6145798 | 0.53883224 | 0.99999872 |
| TSPYL4 | 11.7502473 | 0.234262684 | 0.62686348 | 0.37370607 | 0.70862301 | 0.99999872 |
| DSE | 39.3901808 | 0.122286195 | 0.45708257 | 0.26753633 | 0.78905624 | 0.99999872 |
| TSPYL1 | 52.7798883 | 0.882771793 | 0.40831167 | 2.16200481 | 0.03061781 | 0.91534437 |
| CALHM5 | 28.3113223 | -0.165189744 | 0.65203566 | -0.2533446 | 0.8000019 | 0.99999872 |
| RWDD1 | 21.3449059 | 0.17997468 | 0.43579915 | 0.41297621 | 0.67962405 | 0.99999872 |
| ZUP1 | 8.56035433 | 1.1457719 | 0.69175138 | 1.65633481 | 0.09765403 | 0.99999872 |
| KPNA5 | 43.1291209 | 0.41865375 | 0.28768063 | 1.45527264 | 0.1455938 | 0.99999872 |
| DCBLD1 | 13.5781938 | -0.058385194 | 0.58286902 | -0.1001686 | 0.92021045 | 0.99999872 |
| GOPC | 24.5145581 | 0.123903775 | 0.38383488 | 0.32280489 | 0.74684301 | 0.99999872 |

| | | | | | | |
|---|---|---|---|---|---|---|
| NUS1 | 22.9219262 | 0.577458695 | 0.42086802 | 1.372066 | 0.17004289 | 0.99999872 |
| SLC35F1 | 5.20689677 | 1.476302687 | 0.91305035 | 1.61689078 | 0.10590186 | 0.99999872 |
| CEP85L | 116.310633 | 0.113825961 | 0.39217674 | 0.29024149 | 0.7716315 | 0.99999872 |
| MCM9 | 39.9295757 | -0.282817484 | 0.36433606 | -0.7762544 | 0.43759877 | 0.99999872 |
| ASF1A | 8.77767575 | 0.654657504 | 0.59590993 | 1.09858466 | 0.27194927 | 0.99999872 |
| AL365275.1 | 11.395072 | -0.285882527 | 0.86940198 | -0.3288266 | 0.74228673 | 0.99999872 |
| FAM184A | 10.6691722 | 0.28241578 | 0.95999461 | 0.29418476 | 0.76861673 | 0.99999872 |
| MAN1A1 | 75.4346481 | 0.12775449 | 0.44997547 | 0.28391434 | 0.77647602 | 0.99999872 |
| TBC1D32 | 43.4351055 | -0.275943585 | 0.38934805 | -0.7087324 | 0.47849054 | 0.99999872 |
| GJA1 | 146.948436 | 0.731987913 | 0.67030174 | 1.09202748 | 0.27482102 | 0.99999872 |
| HSF2 | 13.9696341 | 0.633735928 | 0.47574609 | 1.33208857 | 0.18283109 | 0.99999872 |
| SERINC1 | 127.64375 | 0.206008365 | 0.42434674 | 0.48547177 | 0.62734173 | 0.99999872 |
| NKAIN2 | 19.101351 | 0.396540611 | 0.71812413 | 0.55218951 | 0.58081852 | 0.99999872 |
| RNF217-AS1 | 8.81838815 | -0.453937374 | 0.86066249 | -0.5274279 | 0.5978965 | 0.99999872 |
| RNF217 | 60.7465034 | 0.340794887 | 0.52964123 | 0.64344478 | 0.51993553 | 0.99999872 |
| HDDC2 | 13.2595611 | 0.594072402 | 0.61467429 | 0.96648325 | 0.33380243 | 0.99999872 |
| NCOA7 | 39.6533973 | -0.002520346 | 0.35613794 | -0.0070769 | 0.99435351 | 0.99999872 |
| HINT3 | 13.7032353 | -0.022327691 | 0.46981893 | -0.047524 | 0.96209558 | 0.99999872 |
| TRMT11 | 20.1523983 | -0.022634136 | 0.39786866 | -0.0568885 | 0.95463405 | 0.99999872 |
| RNF146 | 24.1569979 | 0.721193356 | 0.41961792 | 1.71869055 | 0.08567073 | 0.99999872 |
| ECHDC1 | 37.1872546 | 0.413880658 | 0.37406338 | 1.10644526 | 0.26853384 | 0.99999872 |
| PTPRK | 83.8575613 | -0.395968334 | 0.4665029 | -0.8488014 | 0.39599179 | 0.99999872 |
| LAMA2 | 7.5341648 | 1.000495104 | 0.90196383 | 1.10924083 | 0.2673263 | 0.99999872 |
| AL356124.1 | 16.5253311 | 0.845821669 | 0.7019575 | 1.20494711 | 0.22822372 | 0.99999872 |
| ARHGAP18 | 104.572025 | 0.378574944 | 0.5069447 | 0.7467776 | 0.45519782 | 0.99999872 |
| L3MBTL3 | 36.1421431 | 0.030599312 | 0.51081752 | 0.05990263 | 0.95223319 | 0.99999872 |
| EPB41L2 | 9.11455907 | -1.058164771 | 0.71914572 | -1.4714191 | 0.1411778 | 0.99999872 |
| AKAP7 | 13.0581112 | -0.406747576 | 0.94175505 | -0.4319038 | 0.66581134 | 0.99999872 |
| MED23 | 24.002878 | 0.401999337 | 0.37904858 | 1.06054832 | 0.28889522 | 0.99999872 |
| CTGF | 1932.14715 | -1.059507146 | 0.6155527 | -1.721229 | 0.08520928 | 0.99999872 |
| STX7 | 87.1321996 | 0.300468864 | 0.27984153 | 1.07371076 | 0.28295234 | 0.99999872 |
| RPS12 | 55.356627 | 0.944403964 | 0.48724984 | 1.9382335 | 0.05259474 | 0.99999872 |
| EYA4 | 25.5509922 | 1.262839213 | 0.66837726 | 1.88941081 | 0.0588368 | 0.99999872 |
| AL024497.1 | 11.526879 | -1.321738204 | 0.61735542 | -2.140968 | 0.03227662 | 0.92959829 |
| TARID | 355.99028 | -0.569838828 | 0.41007797 | -1.3895865 | 0.16465447 | 0.99999872 |
| TCF21 | 205.902318 | 0.336467968 | 0.73641429 | 0.45690038 | 0.64774265 | 0.99999872 |
| TBPL1 | 26.802317 | 0.480516654 | 0.49459842 | 0.97152889 | 0.33128497 | 0.99999872 |
| SLC2A12 | 63.1932686 | 0.839145588 | 0.37985972 | 2.20909338 | 0.02716815 | 0.89154528 |
| SGK1 | 89.6374217 | -0.087155308 | 0.83126409 | -0.1048467 | 0.91649744 | 0.99999872 |
| HBS1L | 28.9235303 | 0.225674231 | 0.34685981 | 0.65062086 | 0.51529126 | 0.99999872 |
| AHI1 | 118.447782 | -0.277078176 | 0.33574329 | -0.825268 | 0.40921948 | 0.99999872 |
| LINC00271 | 43.4337074 | -0.345561824 | 0.4837831 | -0.7142908 | 0.47504737 | 0.99999872 |
| PDE7B | 322.40724 | -0.502458928 | 0.47242895 | -1.0635651 | 0.28752578 | 0.99999872 |
| AL138828.1 | 44.739825 | -0.927187076 | 0.42504538 | -2.1813837 | 2.92E-02 | 0.90266962 |
| BCLAF1 | 77.892936 | 0.043614767 | 0.23905317 | 0.18244798 | 0.85523117 | 0.99999872 |
| MAP7 | 35.4199925 | -0.287692615 | 0.42866136 | -0.6711419 | 0.50213011 | 0.99999872 |
| MAP3K5 | 77.3282414 | 0.127447558 | 0.73635662 | 0.17307858 | 0.86258966 | 0.99999872 |
| PEX7 | 12.6335189 | 0.098164529 | 0.47671607 | 0.20591823 | 0.83685479 | 0.99999872 |
| IFNGR1 | 27.6328559 | 0.470123973 | 0.35423445 | 1.32715485 | 0.18445746 | 0.99999872 |
| TNFAIP3 | 6.21739143 | -0.70492172 | 0.70187958 | -1.0043343 | 0.31521752 | 0.99999872 |
| PERP | 44.7295887 | 0.684364304 | 0.54003627 | 1.26725619 | 0.2050637 | 0.99999872 |
| ARFGEF3 | 5.48605633 | -0.890600791 | 0.6911108 | -1.2886512 | 0.19751936 | 0.99999872 |
| HEBP2 | 31.2370961 | 0.369240355 | 0.37447161 | 0.9860303 | 0.32411824 | 0.99999872 |
| NHSL1 | 11.3724253 | -0.384355955 | 0.77276736 | -0.497376 | 0.61892391 | 0.99999872 |
| CCDC28A | 12.7988425 | 0.240436424 | 0.54576786 | 0.44054705 | 0.65954094 | 0.99999872 |
| REPS1 | 158.368778 | 0.439131031 | 0.24091481 | 1.82276477 | 0.06833903 | 0.99999872 |
| ABRACL | 39.1290858 | 0.753914843 | 0.33573471 | 2.24556716 | 0.02473175 | 0.88365618 |
| HECA | 14.2731452 | 0.219712497 | 0.43633601 | 0.50353968 | 0.61458489 | 0.99999872 |
| CITED2 | 96.6765552 | 1.245765868 | 0.45572838 | 2.73357096 | 0.00626516 | 0.59140873 |
| VTA1 | 22.8788472 | 0.061610407 | 0.36961661 | 0.16668733 | 0.86761607 | 0.99999872 |
| HIVEP2 | 192.260954 | -0.146591277 | 0.29433848 | -0.4980364 | 0.61845838 | 0.99999872 |

| | | | | | | |
|---|---|---|---|---|---|---|
| AIG1 | 82.5150249 | -0.148888615 | 0.39807021 | -0.374026 | 0.70838496 | 0.99999872 |
| ADAT2 | 4.70502429 | -0.504321654 | 0.94551154 | -0.533385 | 0.59376711 | 0.99999872 |
| PEX3 | 25.0328114 | -0.091905895 | 0.41826129 | -0.2197332 | 0.82607894 | 0.99999872 |
| FUCA2 | 17.6834188 | -0.320578501 | 0.49418555 | -0.6487007 | 0.51653187 | 0.99999872 |
| PHACTR2 | 41.7544362 | 0.327499307 | 0.44466017 | 0.73651596 | 0.46141676 | 0.99999872 |
| PLAGL1 | 20.7787158 | 0.342554309 | 0.49005858 | 0.69900685 | 0.48454775 | 0.99999872 |
| SF3B5 | 39.3068573 | 0.610044131 | 0.43244209 | 1.41069556 | 0.1583344 | 0.99999872 |
| STX11 | 18.7894347 | 0.041373837 | 0.56851905 | 0.07277476 | 0.94198536 | 0.99999872 |
| UTRN | 377.169353 | -0.07621569 | 0.29706395 | -0.2565632 | 0.79751595 | 0.99999872 |
| AL024474.2 | 8.33198033 | 1.355854484 | 0.64004287 | 2.11838072 | 0.03414284 | 0.93897283 |
| EPM2A | 17.5720404 | -0.408158987 | 0.54190771 | -0.7531891 | 0.45133628 | 0.99999872 |
| FBXO30 | 8.49054789 | 0.796749464 | 0.59134793 | 1.34734464 | 0.17786926 | 0.99999872 |
| AL356599.1 | 23.2692164 | 0.407160516 | 0.54918832 | 0.74138598 | 0.45845945 | 0.99999872 |
| SHPRH | 72.987152 | 0.13556806 | 0.34892013 | 0.38853608 | 0.69761936 | 0.99999872 |
| RAB32 | 5.10184584 | 0.440741306 | 0.7835341 | 0.56250431 | 0.57377247 | 0.99999872 |
| STXBP5 | 77.9499656 | 0.102780639 | 0.3120623 | 0.32935936 | 0.74188408 | 0.99999872 |
| SAMD5 | 28.2023994 | -1.002229502 | 0.63361937 | -1.5817533 | 0.11370591 | 0.99999872 |
| AL033504.1 | 18.4084944 | -1.190226033 | 0.69743749 | -1.7065702 | 0.08790197 | 0.99999872 |
| SASH1 | 217.33603 | 0.028560217 | 0.51992852 | 0.05493105 | 0.9561934 | 0.99999872 |
| UST | 7.52900319 | -0.476437839 | 0.86039823 | -0.5537411 | 0.57975606 | 0.99999872 |
| TAB2 | 84.448637 | 0.14410163 | 0.31552798 | 0.45670001 | 0.64788668 | 0.99999872 |
| PPIL4 | 20.3340884 | 0.371673676 | 0.38123038 | 0.97493194 | 0.32959402 | 0.99999872 |
| GINM1 | 40.2398632 | -0.029429687 | 0.30999033 | -0.0949374 | 0.92436452 | 0.99999872 |
| KATNA1 | 13.6475702 | 0.167382353 | 0.4622149 | 0.36213102 | 0.71725412 | 0.99999872 |
| LATS1 | 50.9456455 | 0.087580406 | 0.24733588 | 0.35409503 | 0.72326767 | 0.99999872 |
| NUP43 | 15.3138082 | -0.766575832 | 0.43400599 | -1.7662794 | 0.07734899 | 0.99999872 |
| PCMT1 | 68.5677958 | 0.235925681 | 0.40747687 | 0.57899159 | 0.56259485 | 0.99999872 |
| AL355312.2 | 5.93096255 | 1.045399284 | 0.84345231 | 1.23942903 | 0.21518665 | 0.99999872 |
| LRP11 | 34.7505241 | -0.550778963 | 0.59436004 | -0.9266756 | 0.35409497 | 0.99999872 |
| AL355312.3 | 5.51918871 | 1.257753657 | 0.73308297 | 1.71570438 | 0.08621616 | 0.99999872 |
| RAET1E | 21.9558963 | -0.730853799 | 0.83849945 | -0.8716211 | 0.38341512 | 0.99999872 |
| PLEKHG1 | 148.24973 | -0.177368119 | 0.3290898 | -0.5389657 | 0.58991051 | 0.99999872 |
| MTHFD1L | 36.5086624 | -0.919436241 | 0.80303429 | -1.1449527 | 0.25222877 | 0.99999872 |
| AKAP12 | 19.9039848 | -0.067681259 | 1.14016196 | -0.0593611 | 0.9526645 | 0.99999872 |
| RMND1 | 20.6919001 | -0.544175126 | 0.36492008 | -1.4912173 | 0.13590446 | 0.99999872 |
| ARMT1 | 7.97349129 | 0.722507087 | 0.5377495 | 1.34357556 | 0.17908568 | 0.99999872 |
| SYNE1 | 303.59631 | 0.290462899 | 0.40035453 | 0.72551421 | 0.46813663 | 0.99999872 |
| AL080276.2 | 5.838086 | 0.012863221 | 0.65421651 | 0.01966202 | 0.98431299 | 0.99999872 |
| MTRF1L | 9.57259638 | 0.098852535 | 0.55258714 | 0.1788904 | 0.85802376 | 0.99999872 |
| RGS17 | 3.81070239 | 1.250489126 | 1.16994741 | 1.06884216 | 0.2851408 | NA |
| IPCEF1 | 12.3384303 | -1.108699493 | 0.53114643 | -2.0873707 | 0.03685464 | 0.95584876 |
| CNKSR3 | 184.488074 | -0.144938835 | 0.50010994 | -0.2898139 | 0.77195858 | 0.99999872 |
| SCAF8 | 92.2079603 | -0.022077569 | 0.34722018 | -0.0635838 | 0.94930165 | 0.99999872 |
| TIAM2 | 70.2177596 | 0.183076188 | 0.32764722 | 0.55876008 | 0.57632547 | 0.99999872 |
| TFB1M | 23.0899527 | -0.037678481 | 0.39252749 | -0.0959894 | 0.92352899 | 0.99999872 |
| ARID1B | 337.553816 | -0.142240073 | 0.23314841 | -0.6100838 | 0.54180628 | 0.99999872 |
| TMEM242 | 11.8817927 | 0.883246929 | 0.51355771 | 1.71985916 | 0.08545805 | 0.99999872 |
| ZDHHC14 | 25.5133115 | -0.247372283 | 0.37368342 | -0.6619836 | 0.50798176 | 0.99999872 |
| SNX9 | 60.5527401 | -0.240187234 | 0.28403385 | -0.8456289 | 0.39775979 | 0.99999872 |
| SYNJ2 | 101.739148 | -0.237059119 | 0.35584936 | -0.6661783 | 0.50529715 | 0.99999872 |
| SERAC1 | 10.8725631 | 0.723194903 | 0.68506167 | 1.05566394 | 0.29112178 | 0.99999872 |
| GTF2H5 | 25.0990358 | 0.300618536 | 0.49085295 | 0.61244113 | 0.54024594 | 0.99999872 |
| TULP4 | 143.747715 | -0.263452554 | 0.37285073 | -0.7065899 | 0.47982138 | 0.99999872 |
| TMEM181 | 39.6148975 | -0.158048462 | 0.30778323 | -0.5135058 | 0.60759758 | 0.99999872 |
| DYNLT1 | 60.7422643 | -0.402250841 | 0.68000093 | -0.5915445 | 0.55415562 | 0.99999872 |
| SYTL3 | 10.3984129 | -0.28077109 | 0.84355991 | -0.3328407 | 0.73925452 | 0.99999872 |
| EZR | 91.6319351 | -0.329758584 | 0.32146846 | -1.0257883 | 0.30499138 | 0.99999872 |
| RSPH3 | 40.788684 | 0.446604981 | 0.32801612 | 1.36153366 | 0.1733451 | 0.99999872 |
| AL035530.2 | 39.4849437 | -0.376224481 | 0.47239406 | -0.7964209 | 0.42578746 | 0.99999872 |
| SOD2.1 | 13.8151105 | 0.586035138 | 0.53190805 | 1.10176024 | 0.27056592 | 0.99999872 |
| WTAP | 58.311582 | -0.334066207 | 0.31742991 | -1.0524094 | 0.29261177 | 0.99999872 |

| | | | | | | |
|---|---|---|---|---|---|---|
| ACAT2 | 5.5521821 | -0.635749714 | 0.56918906 | -1.1169394 | 0.26402022 | 0.99999872 |
| TCP1 | 20.1804545 | 0.368779235 | 0.36928609 | 0.99862749 | 0.31797518 | 0.99999872 |
| MRPL18 | 15.411542 | 1.191272804 | 0.63638468 | 1.87193823 | 0.06121515 | 0.99999872 |
| IGF2R | 95.5160284 | -0.23715514 | 0.26450236 | -0.8966088 | 0.3699277 | 0.99999872 |
| SLC22A2 | 7.83644804 | 0.169731874 | 1.09811729 | 0.15456625 | 0.87716328 | 0.99999872 |
| MAP3K4 | 33.1814266 | 0.072913321 | 0.31454016 | 0.23180926 | 0.81668617 | 0.99999872 |
| AGPAT4 | 21.0749694 | -0.085350532 | 0.47991456 | -0.1778453 | 0.8588445 | 0.99999872 |
| PRKN | 229.489985 | -0.341005044 | 0.30764103 | -1.1084511 | 0.26766703 | 0.99999872 |
| PACRG | 42.1855436 | -0.765512755 | 0.49471712 | -1.5473747 | 0.12177292 | 0.99999872 |
| QKI | 353.643633 | 0.353180244 | 0.23455862 | 1.50572275 | 0.13213836 | 0.99999872 |
| PDE10A | 327.071045 | -0.019060355 | 0.77857774 | -0.024481 | 0.98046895 | 0.99999872 |
| LINC00473 | 6.2101527 | -0.672127017 | 0.97873576 | -0.6867298 | 0.49225301 | 0.99999872 |
| SFT2D1 | 32.7953192 | 0.767770018 | 0.3393531 | 2.26245175 | 0.02366951 | 0.88337526 |
| MPC1 | 23.4480957 | 0.619427383 | 0.39028668 | 1.58710869 | 0.11248803 | 0.99999872 |
| RPS6KA2 | 62.8573305 | -1.57683998 | 0.37011607 | -4.2603932 | 2.04E-05 | 0.0534555 |
| Z94721.1 | 8.33053742 | -0.544941729 | 0.57927823 | -0.9407254 | 0.34684561 | 0.99999872 |
| FGFR1OP | 30.3598347 | 0.103759205 | 0.44880192 | 0.23119153 | 0.81716601 | 0.99999872 |
| AFDN | 78.5930134 | -0.090808336 | 0.33810958 | -0.2685766 | 0.78825549 | 0.99999872 |
| FRMD1 | 67.7400924 | -0.130897054 | 0.63763264 | -0.205286 | 0.83734868 | 0.99999872 |
| LINC02519 | 18.3626703 | 0.456106722 | 0.55919583 | 0.81564758 | 0.41470173 | 0.99999872 |
| WDR27 | 34.6314654 | -0.131916119 | 0.33400953 | -0.3949472 | 0.69288188 | 0.99999872 |
| C6orf120 | 12.6800726 | 0.049133891 | 0.475269 | 0.10338122 | 0.91766041 | 0.99999872 |
| PHF10 | 39.5246643 | 0.367689551 | 0.51847408 | 0.70917634 | 0.47821505 | 0.99999872 |
| ERMARD | 19.7876363 | -0.012666826 | 0.46145254 | -0.0274499 | 0.9781009 | 0.99999872 |
| FAM120B | 55.2511154 | -0.3330489 | 0.28561874 | -1.1660611 | 0.24358975 | 0.99999872 |
| PSMB1 | 62.7825225 | 0.715102101 | 0.30324339 | 2.35817866 | 0.01836485 | 0.85282771 |
| TBP | 11.4349313 | -0.234836748 | 0.42471379 | -0.5529294 | 0.58031175 | 0.99999872 |
| PDCD2 | 20.9901311 | 0.537113379 | 0.42875276 | 1.25273452 | 0.21030234 | 0.99999872 |
| FAM20C | 111.340524 | -0.372476882 | 0.62649783 | -0.5945382 | 0.55215221 | 0.99999872 |
| PDGFA | 27.7354146 | -0.376397528 | 0.49212813 | -0.7648364 | 0.44436895 | 0.99999872 |
| PRKAR1B | 12.8607413 | -0.475383592 | 0.52257193 | -0.9096998 | 0.36298084 | 0.99999872 |
| DNAAF5 | 15.0926659 | -0.307411768 | 0.37764808 | -0.8140165 | 0.41563549 | 0.99999872 |
| SUN1 | 39.3540624 | 0.463192566 | 0.34587009 | 1.33920964 | 0.18050243 | 0.99999872 |
| COX19 | 12.9832275 | 0.088644407 | 0.47543066 | 0.18645076 | 0.85209129 | 0.99999872 |
| C7orf50 | 33.2489341 | 0.457536685 | 0.48113119 | 0.95096035 | 0.3416245 | 0.99999872 |
| ZFAND2A | 8.17469619 | 0.746999037 | 0.50914264 | 1.46717044 | 0.1423297 | 0.99999872 |
| INTS1 | 22.979834 | -0.50586768 | 0.55856716 | -0.9056524 | 0.36511988 | 0.99999872 |
| MAFK | 30.8592579 | -1.197436955 | 0.60502294 | -1.9791596 | 0.04779804 | 0.97437325 |
| PSMG3 | 7.84559126 | -0.435924983 | 0.59018329 | -0.7386264 | 0.46013386 | 0.99999872 |
| MAD1L1 | 32.9751805 | -0.518243212 | 0.45809361 | -1.1313042 | 0.25792708 | 0.99999872 |
| MRM2 | 6.06420483 | -0.068022975 | 0.57338923 | -0.1186332 | 0.90556599 | 0.99999872 |
| SNX8 | 30.7718683 | -0.848799669 | 0.51161859 | -1.6590478 | 0.09710617 | 0.99999872 |
| EIF3B | 32.4420049 | -0.32843258 | 0.36165536 | -0.9081369 | 0.3638059 | 0.99999872 |
| CHST12 | 21.8711202 | 0.256468556 | 0.42162111 | 0.60829155 | 0.54299412 | 0.99999872 |
| BRAT1 | 9.48225725 | -0.605029458 | 0.62616914 | -0.9662397 | 0.33392427 | 0.99999872 |
| IQCE | 11.7291302 | -0.595308222 | 0.49260974 | -1.2084784 | 0.2268633 | 0.99999872 |
| TTYH3 | 8.56197365 | 0.031883267 | 0.67452959 | 0.04726741 | 0.9623001 | 0.99999872 |
| AMZ1 | 6.65385718 | 1.221544788 | 0.81977312 | 1.49010106 | 0.13619767 | 0.99999872 |
| GNA12 | 47.7611303 | -0.155396084 | 0.26207037 | -0.5929556 | 0.55321089 | 0.99999872 |
| SDK1 | 24.4652522 | 0.47079081 | 0.53438308 | 0.88099871 | 0.37831852 | 0.99999872 |
| FOXK1 | 40.9657753 | -0.65930075 | 0.34573851 | -1.9069347 | 0.05652905 | 0.99999872 |
| AP5Z1 | 8.73624782 | -0.890596607 | 0.4764458 | -1.8692506 | 0.06158795 | 0.99999872 |
| RBAK-RBAKI | 25.9075841 | -0.17280516 | 0.32530076 | -0.5312166 | 0.5952687 | 0.99999872 |
| WIPI2 | 31.1374322 | 0.381668152 | 0.48709841 | 0.7835545 | 0.43330156 | 0.99999872 |
| TNRC18 | 87.1034381 | -0.231890746 | 0.26664663 | -0.8696556 | 0.38448862 | 0.99999872 |
| FBXL18 | 9.89729756 | 0.213480874 | 0.49259262 | 0.4333822 | 0.66473713 | 0.99999872 |
| ACTB | 833.953826 | 0.07034138 | 0.65357576 | 0.10762544 | 0.91429281 | 0.99999872 |
| RNF216 | 55.2665373 | -0.087786172 | 0.29224011 | -0.3003906 | 0.76387926 | 0.99999872 |
| CCZ1 | 8.40195759 | 0.408113343 | 0.57171502 | 0.71384051 | 0.4753258 | 0.99999872 |
| PMS2 | 12.0221474 | -0.335045691 | 0.49967106 | -0.6705325 | 0.50251839 | 0.99999872 |
| AIMP2 | 12.4700804 | -0.178795064 | 0.57253161 | -0.3122885 | 0.75482125 | 0.99999872 |

| | | | | | | |
|---|---|---|---|---|---|---|
| EIF2AK1 | 36.1768651 | 0.088741015 | 0.32920805 | 0.26955907 | 0.78749949 | 0.99999872 |
| USP42 | 21.5973905 | -0.513095555 | 0.3355603 | -1.5290711 | 0.12624682 | 0.99999872 |
| CYTH3 | 24.5026982 | -0.173527356 | 0.39544011 | -0.4388208 | 0.66079137 | 0.99999872 |
| FAM220A | 4.93291572 | -0.277857859 | 0.65891725 | -0.4216885 | 0.67325237 | 0.99999872 |
| RAC1 | 96.9757458 | 0.040206045 | 0.25953849 | 0.15491362 | 0.87688942 | 0.99999872 |
| DAGLB | 5.26310942 | 0.089429717 | 0.59598572 | 0.15005346 | 0.88072244 | 0.99999872 |
| ZDHHC4 | 11.2621543 | 0.128950183 | 0.53500957 | 0.24102407 | 0.80953646 | 0.99999872 |
| C7orf26 | 6.54174169 | -0.487641836 | 0.65297848 | -0.7467962 | 0.4551866 | 0.99999872 |
| ZNF316 | 9.6304604 | -0.445402203 | 0.6073115 | -0.7333999 | 0.46331455 | 0.99999872 |
| AC073343.2 | 7.10311616 | 0.653837432 | 0.63386386 | 1.03151082 | 0.30230134 | 0.99999872 |
| ZNF12 | 15.0226097 | 0.407900991 | 0.45873791 | 0.8891809 | 0.37390586 | 0.99999872 |
| CCZ1B | 12.756549 | -0.358860072 | 0.54069259 | -0.6637044 | 0.5068795 | 0.99999872 |
| C1GALT1 | 46.63146 | 0.407436409 | 0.43417871 | 0.93840715 | 0.34803521 | 0.99999872 |
| COL28A1 | 11.0384359 | -0.700284327 | 0.57684792 | -1.2139843 | 0.22475372 | 0.99999872 |
| MIOS | 17.8889864 | -0.210443535 | 0.45487958 | -0.4626357 | 0.64362551 | 0.99999872 |
| RPA3 | 15.5133444 | 1.569416565 | 0.52669888 | 2.97972262 | 0.0028851 | 0.43717075 |
| UMAD1 | 112.803449 | 0.220679461 | 0.33318944 | 0.6623243 | 0.50776341 | 0.99999872 |
| GLCCI1 | 29.49266 | 0.514950338 | 0.36121349 | 1.42561213 | 0.15398033 | 0.99999872 |
| ICA1 | 17.3292397 | 0.527064254 | 0.73463209 | 0.71745335 | 0.47309441 | 0.99999872 |
| NDUFA4 | 86.4241406 | 1.525975383 | 0.43938296 | 3.47299627 | 0.00051468 | 0.21771468 |
| PHF14 | 66.8853129 | -0.354942113 | 0.23277564 | -1.524825 | 0.12730276 | 0.99999872 |
| THSD7A | 926.253334 | -0.509416415 | 0.55798173 | -0.9129625 | 0.36126225 | 0.99999872 |
| TMEM106B | 85.6714864 | -0.032139637 | 0.32579684 | -0.0986493 | 0.92141671 | 0.99999872 |
| SCIN | 14.432736 | -0.354814601 | 0.72870862 | -0.4869088 | 0.626323 | 0.99999872 |
| AC011287.1 | 20.2751881 | 1.193127299 | 0.98193411 | 1.21507878 | 0.22433605 | 0.99999872 |
| ISPD | 21.3594816 | -0.545901219 | 0.48704723 | -1.1208384 | 0.26235667 | 0.99999872 |
| ANKMY2 | 14.9750821 | 0.548437561 | 0.5340817 | 1.02687953 | 0.3044772 | 0.99999872 |
| BZW2 | 6.12875062 | -0.81937282 | 0.72639342 | -1.1280014 | 0.25931931 | 0.99999872 |
| AC073332.1 | 12.6829886 | -0.672274923 | 0.90166361 | -0.7455939 | 0.45591275 | 0.99999872 |
| SNX13 | 64.8795727 | 0.430138404 | 0.35036139 | 1.22769923 | 0.2195599 | 0.99999872 |
| HDAC9 | 25.9488247 | 0.670984087 | 1.08568507 | 0.61802829 | 0.53655669 | 0.99999872 |
| TWISTNB | 11.2230803 | -0.139021165 | 0.46921808 | -0.2962826 | 0.76701425 | 0.99999872 |
| MACC1 | 18.8949653 | 0.404198252 | 1.01385802 | 0.39867343 | 0.69013385 | 0.99999872 |
| MACC1-AS1 | 7.95300135 | 0.982842319 | 0.75541566 | 1.30106162 | 0.19323736 | 0.99999872 |
| ITGB8 | 862.471982 | 0.420611841 | 0.55636719 | 0.75599685 | 0.44965109 | 0.99999872 |
| SP4 | 14.5691696 | -0.281451275 | 0.44647665 | -0.630383 | 0.52844405 | 0.99999872 |
| RAPGEF5 | 9.495382 | 0.101146055 | 0.60822244 | 0.1662978 | 0.8679226 | 0.99999872 |
| STEAP1B | 43.5964912 | 0.207124608 | 0.46532979 | 0.44511358 | 0.65623768 | 0.99999872 |
| TOMM7 | 50.6495801 | 0.948721621 | 0.38243045 | 2.48076904 | 0.01310993 | 0.74263079 |
| FAM126A | 73.0162061 | 0.143510108 | 0.47670276 | 0.30104736 | 0.76337838 | 0.99999872 |
| KLHL7 | 22.2180356 | 0.221731213 | 0.39631383 | 0.55948391 | 0.57583151 | 0.99999872 |
| NUPL2 | 12.9095747 | 0.400017292 | 0.52744596 | 0.75840431 | 0.44820898 | 0.99999872 |
| GPNMB | 37.3787889 | 0.753132779 | 0.64757013 | 1.16301347 | 0.24482404 | 0.99999872 |
| MALSU1 | 36.7489245 | 0.890568653 | 0.44181187 | 2.01571918 | 0.04382935 | 0.95980711 |
| TRA2A | 105.274546 | -0.075153354 | 0.32660063 | -0.2301078 | 0.818008 | 0.99999872 |
| CCDC126 | 11.7053305 | 0.0933178 | 0.53083364 | 0.17579481 | 0.86045515 | 0.99999872 |
| FAM221A | 6.59043289 | -0.010367838 | 0.61115058 | -0.0169645 | 0.98646497 | 0.99999872 |
| MPP6 | 18.4229461 | -0.096147443 | 0.35808306 | -0.268506 | 0.78830987 | 0.99999872 |
| OSBPL3 | 23.7487913 | -1.415051464 | 0.42574728 | -3.3236888 | 0.00088835 | 0.27376937 |
| CYCS | 22.9982511 | 0.550635614 | 0.53846432 | 1.02260371 | 0.30649525 | 0.99999872 |
| C7orf31 | 5.94211819 | 0.318196082 | 0.61798554 | 0.51489244 | 0.60662819 | 0.99999872 |
| NFE2L3 | 28.148921 | -0.395071201 | 0.63575932 | -0.6214163 | 0.53432575 | 0.99999872 |
| HNRNPA2B1 | 211.147614 | -0.270464689 | 0.24987277 | -1.0824096 | 0.27907055 | 0.99999872 |
| CBX3 | 41.0616967 | -0.052733946 | 0.31239472 | -0.1688055 | 0.86594963 | 0.99999872 |
| SNX10 | 28.2444033 | -0.134883667 | 0.53777199 | -0.2508194 | 0.80195371 | 0.99999872 |
| AC004540.1 | 15.7089443 | -1.073419451 | 0.62334071 | -1.7220429 | 0.08506175 | 0.99999872 |
| SKAP2 | 81.8455259 | 0.197631086 | 0.30700521 | 0.64373855 | 0.51974499 | 0.99999872 |
| HOTAIRM1 | 29.8380994 | 0.331385464 | 0.57925454 | 0.57208954 | 0.56726131 | 0.99999872 |
| HOXA3 | 10.9683568 | 0.127741673 | 0.52732219 | 0.24224597 | 0.80858958 | 0.99999872 |
| HOXA-AS2 | 9.35357044 | 0.494349333 | 0.67094847 | 0.7367918 | 0.46124897 | 0.99999872 |
| HOXA7 | 13.0320937 | -0.284111688 | 0.73973179 | -0.3840739 | 0.70092366 | 0.99999872 |

| | | | | | | |
|---|---|---|---|---|---|---|
| HOXA9 | 27.5165296 | -0.265518476 | 0.51801863 | -0.5125655 | 0.60825529 | 0.99999872 |
| HOXA10-AS | 5.64026687 | -0.75594917 | 0.86868805 | -0.8702194 | 0.38418053 | 0.99999872 |
| HIBADH | 58.8038157 | 0.08609879 | 0.3727092 | 0.23100796 | 0.81730862 | 0.99999872 |
| TAX1BP1 | 113.845718 | 0.387269939 | 0.24374868 | 1.58880839 | 0.11210366 | 0.99999872 |
| JAZF1 | 137.242121 | 0.294161382 | 0.34016254 | 0.86476711 | 0.38716662 | 0.99999872 |
| CREB5 | 7.35812408 | -0.109892568 | 0.68704153 | -0.1599504 | 0.87292015 | 0.99999872 |
| CPVL | 8.51945671 | -0.310160171 | 0.71969151 | -0.4309627 | 0.66649552 | 0.99999872 |
| WIPF3 | 170.698941 | -0.267705517 | 0.35079723 | -0.7631346 | 0.44538311 | 0.99999872 |
| SCRN1 | 26.3693614 | -0.521829228 | 0.51203295 | -1.0191321 | 0.30814025 | 0.99999872 |
| FKBP14 | 9.60226983 | 1.041726854 | 0.64267589 | 1.62092101 | 0.10503458 | 0.99999872 |
| PLEKHA8 | 14.5159155 | 0.523675355 | 0.41787395 | 1.25318975 | 0.21013666 | 0.99999872 |
| MTURN | 12.0919602 | 0.41843162 | 0.52965474 | 0.79000826 | 0.42952294 | 0.99999872 |
| ZNRF2 | 41.3325399 | 0.223430211 | 0.46376877 | 0.48177072 | 0.62996883 | 0.99999872 |
| AC006978.2 | 45.8768616 | -0.162913822 | 0.4723938 | -0.3448687 | 0.73019311 | 0.99999872 |
| NOD1 | 38.2227535 | 0.424130616 | 0.47441339 | 0.89401063 | 0.3713162 | 0.99999872 |
| GGCT | 10.4962524 | 0.95423878 | 0.71110554 | 1.3419088 | 0.17962557 | 0.99999872 |
| AC005154.6 | 24.0793605 | -0.187019528 | 0.44458175 | -0.420664 | 0.67400047 | 0.99999872 |
| GARS | 25.0409038 | -0.521366558 | 0.62389304 | -0.8356666 | 0.4033425 | 0.99999872 |
| AQP1 | 328.842805 | 0.136881333 | 1.53878506 | 0.08895416 | 0.92911834 | 0.99999872 |
| LSM5 | 12.064148 | 0.597128127 | 0.57632981 | 1.03608752 | 0.30016131 | 0.99999872 |
| AVL9 | 59.8179046 | -0.107208562 | 0.27845799 | -0.385008 | 0.70023147 | 0.99999872 |
| KBTBD2 | 25.4672674 | 0.288655211 | 0.35810874 | 0.80605464 | 0.42021134 | 0.99999872 |
| FKBP9 | 47.0143177 | 0.086617756 | 0.48178996 | 0.17978323 | 0.85732275 | 0.99999872 |
| NT5C3A | 32.3365094 | -0.201527996 | 0.74857943 | -0.2692139 | 0.78776508 | 0.99999872 |
| RP9 | 7.10355329 | 0.115845142 | 0.59438313 | 0.19489978 | 0.84547141 | 0.99999872 |
| BBS9 | 82.2688634 | 0.300655672 | 0.31689966 | 0.94874089 | 0.34275241 | 0.99999872 |
| DPY19L1 | 23.5819315 | -0.398187517 | 0.46154812 | -0.8627216 | 0.38829057 | 0.99999872 |
| HERPUD2 | 31.5187403 | 0.163338204 | 0.32181796 | 0.50754845 | 0.61177005 | 0.99999872 |
| AC018647.2 | 4.3150129 | -0.084561492 | 0.63720442 | -0.132707 | 0.8944251 | NA |
| SEPT7-AS1 | 6.68777122 | -0.348331776 | 0.53844315 | -0.646924 | 0.51768116 | 0.99999872 |
| 7-Sep | 137.651843 | 0.178195068 | 0.32593073 | 0.54672681 | 0.58456643 | 0.99999872 |
| KIAA0895 | 19.6339292 | 0.066299718 | 0.58090485 | 0.1141318 | 0.90913331 | 0.99999872 |
| ELMO1 | 21.5884283 | 0.18206661 | 0.75757431 | 0.24032838 | 0.81007569 | 0.99999872 |
| EPDR1 | 34.9982572 | 0.404978978 | 0.50601493 | 0.8003301 | 0.42351957 | 0.99999872 |
| SFRP4 | 5.36089937 | 1.750431813 | 1.01988619 | 1.71630113 | 0.08610694 | 0.99999872 |
| STARD3NL | 44.8820204 | 0.238925945 | 0.285556 | 0.83670433 | 0.40275878 | 0.99999872 |
| VPS41 | 57.7698068 | 0.003228821 | 0.29010373 | 0.01112988 | 0.99111982 | 0.99999872 |
| POU6F2 | 11.3556968 | 0.274168116 | 0.72039902 | 0.38057814 | 0.70351631 | 0.99999872 |
| YAE1D1 | 9.66995192 | 0.478491679 | 0.48046721 | 0.99588832 | 0.31930441 | 0.99999872 |
| RALA | 52.0356279 | -0.108218778 | 0.35614841 | -0.3038587 | 0.76123557 | 0.99999872 |
| CDK13 | 119.435427 | -0.078085157 | 0.23920021 | -0.3264427 | 0.74408945 | 0.99999872 |
| MPLKIP | 22.6979575 | 0.367461035 | 0.37799431 | 0.97213378 | 0.330984 | 0.99999872 |
| SUGCT | 15.5049799 | -0.938883184 | 0.83936756 | -1.1185602 | 0.26332779 | 0.99999872 |
| GLI3 | 30.0845096 | -0.40317315 | 0.5984214 | -0.6737278 | 0.50048437 | 0.99999872 |
| MRPL32 | 18.7883346 | 0.572950382 | 0.43534507 | 1.31608331 | 0.18814608 | 0.99999872 |
| STK17A | 8.48502223 | 0.256743696 | 0.57489588 | 0.44659165 | 0.65516993 | 0.99999872 |
| COA1 | 56.6802175 | -0.266562025 | 0.34243624 | -0.7784282 | 0.43631662 | 0.99999872 |
| BLVRA | 30.6292433 | 0.836220452 | 0.41929669 | 1.99434071 | 0.04611483 | 0.95980711 |
| URGCP | 9.20790915 | 0.586167715 | 0.57458509 | 1.02015824 | 0.30765342 | 0.99999872 |
| UBE2D4 | 7.28746601 | -1.25880852 | 0.54808807 | -2.2967267 | 0.02163436 | 0.87186485 |
| DBNL | 18.5518156 | -0.04507092 | 0.47412537 | -0.0950612 | 0.92426623 | 0.99999872 |
| POLM | 8.78994353 | 0.577683266 | 0.63998814 | 0.90264683 | 0.36671336 | 0.99999872 |
| POLD2 | 10.6894497 | 0.267947636 | 0.51879063 | 0.51648511 | 0.60551564 | 0.99999872 |
| YKT6 | 9.35990621 | -0.000177234 | 0.61283611 | -0.0002892 | 0.99976925 | 0.99999872 |
| NUDCD3 | 48.4865136 | -0.350432928 | 0.29863351 | -1.1734548 | 0.24061347 | 0.99999872 |
| DDX56 | 11.1059583 | -0.137197686 | 0.46933075 | -0.2923262 | 0.77003721 | 0.99999872 |
| TMED4 | 50.2264901 | 0.451760319 | 0.427175 | 1.05755327 | 0.29025916 | 0.99999872 |
| OGDH | 46.7167704 | -0.073421624 | 0.29858647 | -0.2458974 | 0.80576169 | 0.99999872 |
| ZMIZ2 | 22.0088778 | -0.514512063 | 0.40377489 | -1.2742547 | 0.20257315 | 0.99999872 |
| PPIA | 142.811642 | 0.094120236 | 0.54005746 | 0.17427819 | 0.86164684 | 0.99999872 |
| H2AFV | 40.6082456 | -0.044165684 | 0.67688871 | -0.0652481 | 0.94797649 | 0.99999872 |

| | | | | | | |
|---|---|---|---|---|---|---|
| PURB | 27.3979323 | 0.05971795 | 0.42757319 | 0.1396672 | 0.88892294 | 0.99999872 |
| SNHG15 | 7.47397689 | 0.459127542 | 0.72445041 | 0.63375979 | 0.5262376 | 0.99999872 |
| CCM2 | 33.900526 | -0.106103372 | 0.32014462 | -0.3314233 | 0.7403248 | 0.99999872 |
| TBRG4 | 5.98019204 | 0.558256423 | 0.68635116 | 0.81336851 | 0.41600681 | 0.99999872 |
| RAMP3 | 94.7088204 | -0.95761518 | 0.61030847 | -1.5690675 | 0.11663221 | 0.99999872 |
| IGFBP3 | 7.9209592 | -2.025187808 | 1.28257061 | -1.5790069 | 0.11433448 | 0.99999872 |
| TNS3 | 315.60481 | -0.378904268 | 0.30927929 | -1.2251201 | 0.22052998 | 0.99999872 |
| HUS1 | 16.8607086 | -0.449483344 | 0.40484048 | -1.1102727 | 0.26688153 | 0.99999872 |
| UPP1 | 10.7676735 | -1.007837051 | 0.98649407 | -1.0216352 | 0.3069536 | 0.99999872 |
| GRB10 | 24.8756629 | -0.417494892 | 0.62783157 | -0.6649791 | 0.50606384 | 0.99999872 |
| COBL | 25.8345212 | -0.164324119 | 0.49805164 | -0.3299339 | 0.74144991 | 0.99999872 |
| SEC61G | 21.4939669 | 0.746075397 | 0.56791224 | 1.31371601 | 0.18894179 | 0.99999872 |
| EGFR | 92.153583 | -0.11993437 | 0.29376805 | -0.4082621 | 0.68308124 | 0.99999872 |
| LANCL2 | 20.6686169 | -0.565899043 | 0.39464959 | -1.4339279 | 0.15159285 | 0.99999872 |
| VOPP1 | 52.8208112 | 0.04828207 | 0.55863212 | 0.0864291 | 0.93112532 | 0.99999872 |
| NIPSNAP2 | 41.9032608 | 0.141886901 | 0.35529709 | 0.39934721 | 0.68963739 | 0.99999872 |
| PSPH | 5.54590584 | -0.354459501 | 0.80480157 | -0.4404309 | 0.65962503 | 0.99999872 |
| CCT6A | 27.147635 | 0.227455544 | 0.40023033 | 0.56831161 | 0.5698234 | 0.99999872 |
| SUMF2 | 31.5977174 | 0.023166929 | 0.50586487 | 0.04579667 | 0.96347231 | 0.99999872 |
| PHKG1 | 12.2843288 | 0.027708536 | 0.69655516 | 0.03977938 | 0.96826901 | 0.99999872 |
| CHCHD2 | 93.6223999 | 1.076461106 | 0.5479629 | 1.96447807 | 0.04947468 | 0.97937581 |
| ZNF727 | 10.1602265 | -0.253641669 | 0.6017167 | -0.42153 | 0.67336807 | 0.99999872 |
| ZNF736 | 43.8809564 | 0.248359936 | 0.43456898 | 0.57150867 | 0.56765489 | 0.99999872 |
| ZNF680 | 37.7777404 | 0.148052296 | 0.30641162 | 0.48318107 | 0.62896717 | 0.99999872 |
| ZNF107 | 14.2181817 | 0.254554413 | 0.56333604 | 0.45186957 | 0.65136295 | 0.99999872 |
| ZNF138 | 14.5034093 | -0.024030691 | 0.57682424 | -0.0416603 | 0.96676948 | 0.99999872 |
| ZNF117 | 18.7862111 | 0.170810048 | 0.50453288 | 0.33855087 | 0.7349481 | 0.99999872 |
| ZNF92 | 7.42917347 | -0.203931065 | 0.69138556 | -0.29496 | 0.76802446 | 0.99999872 |
| VKORC1L1 | 41.3608161 | -0.547695028 | 0.29988614 | -1.8263432 | 0.06779858 | 0.99999872 |
| GUSB | 23.7162074 | 0.69569039 | 0.46852138 | 1.4848637 | 0.13757993 | 0.99999872 |
| AC068533.4 | 5.96913218 | -1.2156336 | 0.65047222 | -1.8688478 | 0.06164399 | 0.99999872 |
| CRCP | 7.25429023 | -0.543870496 | 0.51634398 | -1.0533104 | 0.29219874 | 0.99999872 |
| AC068533.3 | 4.68074485 | 1.026533645 | 0.71630229 | 1.43310116 | 0.15182893 | 0.99999872 |
| TPST1 | 42.6107333 | -0.049498375 | 0.56236021 | -0.088019 | 0.92986159 | 0.99999872 |
| AC027644.4 | 20.2326431 | -0.187270294 | 0.35797362 | -0.5231399 | 0.60087688 | 0.99999872 |
| AC027644.3 | 11.8334617 | 0.005817279 | 0.81692486 | 0.00712095 | 0.99431835 | 0.99999872 |
| TMEM248 | 56.050897 | 0.269843681 | 0.28246751 | 0.95530875 | 0.33942158 | 0.99999872 |
| SBDS | 73.4365186 | 0.367041785 | 0.35539037 | 1.03278484 | 0.3017046 | 0.99999872 |
| TYW1 | 40.0761313 | 0.067693365 | 0.29742873 | 0.22759525 | 0.81996092 | 0.99999872 |
| AUTS2 | 201.978244 | -0.381471857 | 0.27467 | -1.388837 | 0.16488233 | 0.99999872 |
| TYW1B | 20.3916364 | 0.03970405 | 0.35084108 | 0.11316819 | 0.90989721 | 0.99999872 |
| POM121 | 39.3036379 | 0.12865762 | 0.30179683 | 0.42630541 | 0.66988531 | 0.99999872 |
| NSUN5 | 11.0495752 | -0.304896157 | 0.44088764 | -0.6915507 | 0.48921953 | 0.99999872 |
| BAZ1B | 47.4399325 | -0.254219891 | 0.31857213 | -0.7979979 | 0.42487171 | 0.99999872 |
| BCL7B | 12.5873261 | 0.055043727 | 0.51192236 | 0.10752358 | 0.91437362 | 0.99999872 |
| TBL2 | 6.51729243 | 0.230309436 | 0.56158292 | 0.41010762 | 0.681727 | 0.99999872 |
| MLXIPL | 6.1240747 | 1.525760589 | 0.89719493 | 1.70058985 | 0.08902003 | 0.99999872 |
| DNAJC30 | 6.34977811 | 0.779264012 | 0.79657076 | 0.97827343 | 0.32793911 | 0.99999872 |
| BUD23 | 15.627597 | -0.239093404 | 0.46654898 | -0.5124722 | 0.60832054 | 0.99999872 |
| CLDN3 | 5.39770497 | -0.122485352 | 0.80603886 | -0.1519596 | 0.87921879 | 0.99999872 |
| CLDN4 | 8.71883115 | -0.71300925 | 0.48791544 | -1.4613377 | 0.14392278 | 0.99999872 |
| LIMK1 | 5.38660418 | -0.207955884 | 0.65338723 | -0.3182736 | 0.75027743 | 0.99999872 |
| EIF4H | 35.1084479 | -0.002567481 | 0.29386982 | -0.0087368 | 0.99302913 | 0.99999872 |
| RFC2 | 4.50802145 | 0.2562854 | 0.67323548 | 0.3806772 | 0.70344279 | NA |
| CLIP2 | 12.9335343 | -0.107426127 | 0.51201587 | -0.2098102 | 0.83381585 | 0.99999872 |
| GTF2IRD1 | 47.1591328 | -0.659262536 | 0.45613863 | -1.4453118 | 0.14837033 | 0.99999872 |
| GTF2I | 128.688838 | 0.174343851 | 0.23337006 | 0.74707034 | 0.4550211 | 0.99999872 |
| GTF2IRD2 | 12.2975646 | -0.723799003 | 0.46859812 | -1.544605 | 0.12244184 | 0.99999872 |
| CASTOR2 | 12.1452862 | -0.518087944 | 0.71569583 | -0.723894 | 0.46913079 | 0.99999872 |
| RCC1L | 18.8231511 | -0.999725285 | 0.38423155 | -2.6018824 | 0.00927136 | 0.69734315 |
| GTF2IRD2B | 15.7763921 | -0.783830616 | 0.42471687 | -1.8455368 | 0.06495949 | 0.99999872 |

| | | | | | | |
|---|---|---|---|---|---|---|
| TRIM73 | 11.7312557 | 0.031285845 | 0.56086937 | 0.05578098 | 0.95551628 | 0.99999872 |
| POM121C | 29.7393766 | -0.042382089 | 0.31210546 | -0.1357941 | 0.89198403 | 0.99999872 |
| SPDYE5 | 7.70123698 | 0.849369877 | 0.70725587 | 1.20093718 | 0.22977557 | 0.99999872 |
| HIP1 | 98.9282647 | 0.052840009 | 0.55335081 | 0.09549098 | 0.92392486 | 0.99999872 |
| RHBDD2 | 17.2768551 | -0.174018499 | 0.57422514 | -0.3030493 | 0.76185233 | 0.99999872 |
| POR | 27.9841846 | -0.725990356 | 0.35374448 | -2.0523016 | 0.04014036 | 0.95980711 |
| TMEM120A | 15.6897305 | 0.089580599 | 0.53394786 | 0.16777031 | 0.86676398 | 0.99999872 |
| STYXL1 | 9.16847719 | -0.01723662 | 0.69916745 | -0.0246531 | 0.98033169 | 0.99999872 |
| MDH2 | 21.3190755 | 1.079621637 | 0.43709038 | 2.47001922 | 0.01351058 | 0.74507289 |
| HSPB1 | 58.6889398 | 0.57159907 | 0.50295401 | 1.13648378 | 0.25575415 | 0.99999872 |
| YWHAG | 52.2851111 | -0.1622241 | 0.36531901 | -0.4440615 | 0.65699813 | 0.99999872 |
| DTX2 | 11.6040026 | -0.722395565 | 0.59443847 | -1.2152571 | 0.22426805 | 0.99999872 |
| POMZP3 | 9.14589492 | 0.045544174 | 0.5984945 | 0.0760979 | 0.93934121 | 0.99999872 |
| CCDC146 | 25.8954724 | 0.161988322 | 0.43337216 | 0.37378571 | 0.70856375 | 0.99999872 |
| GSAP | 32.7349434 | 0.884119422 | 0.47107752 | 1.87680241 | 0.06054518 | 0.99999872 |
| PTPN12 | 65.3173998 | -0.169094256 | 0.27552556 | -0.6137153 | 0.53940346 | 0.99999872 |
| APTR | 10.2616265 | 0.445639874 | 0.54442732 | 0.81854796 | 0.41304437 | 0.99999872 |
| RSBN1L | 43.7439678 | -0.115193872 | 0.28834531 | -0.3994997 | 0.68952503 | 0.99999872 |
| TMEM60 | 6.80210185 | 0.863983188 | 0.80336765 | 1.0754518 | 0.2821725 | 0.99999872 |
| PHTF2 | 17.4919581 | 0.13815526 | 0.49220005 | 0.28068924 | 0.77894876 | 0.99999872 |
| MAGI2 | 13369.5996 | 0.162458825 | 0.33074724 | 0.49118725 | 0.62329402 | 0.99999872 |
| AC006355.2 | 67.3195273 | 1.68993455 | 0.60003842 | 2.81637724 | 0.00485686 | 0.5356861 |
| GNAI1 | 25.2039539 | -0.930340972 | 0.53692208 | -1.73273 | 0.08314367 | 0.99999872 |
| SEMA3C | 34.0278865 | -0.900373396 | 0.75652077 | -1.1901503 | 0.23398734 | 0.99999872 |
| CACNA2D1 | 78.7007689 | -0.083996348 | 0.2207949 | -0.380427 | 0.70362846 | 0.99999872 |
| PCLO | 74.709968 | -1.25971317 | 0.8545339 | -1.4741524 | 0.14044057 | 0.99999872 |
| AC003984.1 | 11.6113522 | -0.405068158 | 0.74150368 | -0.5462794 | 0.58487392 | 0.99999872 |
| KIAA1324L | 19.5874381 | -0.397297052 | 0.5811416 | -0.6836493 | 0.49419664 | 0.99999872 |
| DMTF1 | 67.8406384 | 0.504282069 | 0.45392935 | 1.11092634 | 0.26660006 | 0.99999872 |
| TMEM243 | 14.5155987 | 0.343932636 | 0.42784183 | 0.803878 | 0.42146744 | 0.99999872 |
| TP53TG1 | 6.7170326 | 1.034028709 | 0.66175845 | 1.56254705 | 0.11815917 | 0.99999872 |
| CROT | 15.7957331 | 0.214461783 | 0.51278379 | 0.41823042 | 0.67577865 | 0.99999872 |
| ABCB1 | 7.60283207 | 0.976619669 | 0.89755596 | 1.08808778 | 0.27655636 | 0.99999872 |
| SLC25A40 | 15.6280624 | 0.014314169 | 0.62388288 | 0.02294368 | 0.9816952 | 0.99999872 |
| DBF4 | 4.97930779 | 0.151949906 | 0.73382007 | 0.20706698 | 0.83595755 | 0.99999872 |
| SRI | 21.104593 | 1.034656733 | 0.47612755 | 2.17306628 | 0.02977533 | 0.90266962 |
| CFAP69 | 18.1888288 | 0.387162308 | 0.4879581 | 0.79343351 | 0.42752529 | 0.99999872 |
| GTPBP10 | 12.9805457 | -0.143992761 | 0.46897947 | -0.3070343 | 0.7588173 | 0.99999872 |
| CLDN12 | 20.2095414 | -0.03343868 | 0.42377125 | -0.0789074 | 0.93710629 | 0.99999872 |
| CDK14 | 161.369725 | -0.295925621 | 0.33452437 | -0.884616 | 0.37636376 | 0.99999872 |
| FZD1 | 9.5890847 | -1.283099 | 0.73909752 | -1.7360348 | 0.08255768 | 0.99999872 |
| AC079760.1 | 15.9401901 | 0.000142691 | 0.48388702 | 0.00029488 | 0.99976472 | 0.99999872 |
| MTERF1 | 51.7875234 | 0.248424199 | 0.35492955 | 0.69992537 | 0.48397391 | 0.99999872 |
| AKAP9 | 169.993026 | 0.173636285 | 0.21230468 | 0.81786367 | 0.41343503 | 0.99999872 |
| KRIT1 | 49.445502 | -0.066197363 | 0.2754953 | -0.2402849 | 0.81010939 | 0.99999872 |
| ANKIB1 | 88.1382181 | 0.07548005 | 0.33158221 | 0.227636 | 0.81992923 | 0.99999872 |
| GATAD1 | 20.4586708 | 0.303055083 | 0.38852741 | 0.78000954 | 0.43538526 | 0.99999872 |
| AC007566.1 | 15.7268285 | -0.172264303 | 0.55475434 | -0.3105236 | 0.75616283 | 0.99999872 |
| PEX1 | 16.0377995 | -0.033969963 | 0.52454148 | -0.0647613 | 0.94836409 | 0.99999872 |
| RBM48 | 6.31218395 | 0.626474628 | 0.57409372 | 1.09124103 | 0.27516684 | 0.99999872 |
| FAM133B | 20.273273 | 0.072567761 | 0.34757013 | 0.20878595 | 0.83461534 | 0.99999872 |
| CDK6 | 157.385195 | 0.159857185 | 0.4242865 | 0.37676708 | 0.70634671 | 0.99999872 |
| SAMD9 | 17.4353199 | -0.13751436 | 1.01990143 | -0.134831 | 0.89274547 | 0.99999872 |
| SAMD9L | 28.9419644 | -0.758872651 | 0.92201594 | -0.8230581 | 0.41047498 | 0.99999872 |
| VPS50 | 39.7435809 | -0.427882727 | 0.35809632 | -1.1948817 | 0.23213327 | 0.99999872 |
| GNGT1 | 11.3764886 | 1.642780111 | 1.05287805 | 1.56027577 | 0.11869473 | 0.99999872 |
| BET1 | 15.8043156 | 0.15681519 | 0.46591383 | 0.33657552 | 0.73643691 | 0.99999872 |
| AC002074.1 | 6.77642854 | 0.392662156 | 0.98604579 | 0.39821899 | 0.69046877 | 0.99999872 |
| CASD1 | 32.219728 | -0.154748118 | 0.41769677 | -0.3704796 | 0.7110252 | 0.99999872 |
| SGCE | 40.4802851 | 0.394268744 | 0.40820019 | 0.96587105 | 0.33410871 | 0.99999872 |
| PPP1R9A | 95.4609321 | -0.042467113 | 0.33946029 | -0.1251019 | 0.90044291 | 0.99999872 |

| | | | | | | | |
|---|---|---|---|---|---|---|---|
| AC002429.2 | 6.94275643 | | 1.317993564 | 0.88684739 | 1.486156 | 0.13723786 | 0.99999872 |
| PON2 | 87.1848329 | | 0.436655267 | 0.29387668 | 1.48584526 | 0.13732006 | 0.99999872 |
| PDK4 | 102.626026 | | 1.310680584 | 0.81461827 | 1.60895063 | 0.10762714 | 0.99999872 |
| DYNC1I1 | 127.248828 | | -0.573803178 | 0.71972259 | -0.797256 | 0.42530235 | 0.99999872 |
| SLC25A13 | 39.0499048 | | -0.072153395 | 0.38223298 | -0.1887681 | 0.85027457 | 0.99999872 |
| SEM1 | 66.7793269 | | 0.353528309 | 0.3232482 | 1.0936745 | 0.27409776 | 0.99999872 |
| SDHAF3 | 10.6621886 | | 0.32665162 | 0.49868985 | 0.65501958 | 0.51245514 | 0.99999872 |
| ASNS | 14.5171977 | | -0.666204563 | 0.81026083 | -0.82221 | 0.41095738 | 0.99999872 |
| LMTK2 | 24.4752212 | | -0.11219906 | 0.34282473 | -0.3272782 | 0.74345748 | 0.99999872 |
| TECPR1 | 11.0948479 | | -0.437706941 | 0.50857916 | -0.8606466 | 0.38943269 | 0.99999872 |
| BRI3 | 75.4540115 | | -0.173362197 | 0.29708182 | -0.5835503 | 0.55952288 | 0.99999872 |
| BAIAP2L1 | 70.4000967 | | 0.883002848 | 0.47055464 | 1.87651503 | 0.06058459 | 0.99999872 |
| TMEM130 | 9.56659448 | | 3.271806129 | 1.3199193 | 2.47879256 | 0.0131828 | 0.74263079 |
| TRRAP | 55.3443968 | | 0.102697117 | 0.29703437 | 0.34574152 | 0.72953697 | 0.99999872 |
| SMURF1 | 93.0587276 | | 0.081754344 | 0.34224255 | 0.23887837 | 0.81119989 | 0.99999872 |
| ARPC1B | 12.298811 | | 0.560634655 | 0.49857478 | 1.12447456 | 0.26081176 | 0.99999872 |
| PDAP1 | 18.5760145 | | 0.265622775 | 0.4077009 | 0.65151384 | 0.51471485 | 0.99999872 |
| BUD31 | 18.482546 | | 0.195518178 | 0.46202508 | 0.42317655 | 0.67216646 | 0.99999872 |
| CPSF4 | 14.1101185 | | 0.848697138 | 0.51364543 | 1.65230153 | 0.0984731 | 0.99999872 |
| ZNF789 | 13.758777 | | 0.170683965 | 0.49267628 | 0.34644242 | 0.72901025 | 0.99999872 |
| ZNF394 | 10.5465323 | | -0.427000096 | 0.56771179 | -0.7521424 | 0.45196544 | 0.99999872 |
| ZKSCAN5 | 8.91351041 | | -0.528983493 | 0.54876903 | -0.9639456 | 0.3350732 | 0.99999872 |
| FAM200A | 8.20804835 | | 0.773940246 | 0.64393229 | 1.20189694 | 0.22940346 | 0.99999872 |
| ZSCAN25 | 4.21411838 | | 0.350444601 | 0.68496944 | 0.51162078 | 0.60891644 | NA |
| TRIM4 | 21.9050957 | | -0.265944135 | 0.40201468 | -0.6615284 | 0.5082735 | 0.99999872 |
| ZKSCAN1 | 57.9978905 | | 0.008330552 | 0.31262007 | 0.02664753 | 0.97874087 | 0.99999872 |
| ZSCAN21 | 6.12383206 | | -0.187604179 | 0.58838839 | -0.3188441 | 0.74984472 | 0.99999872 |
| ZNF3 | 9.39225582 | | -0.330616482 | 0.60373954 | -0.5476144 | 0.58395668 | 0.99999872 |
| COPS6 | 23.5488555 | | 0.345070363 | 0.38235387 | 0.90248952 | 0.36679689 | 0.99999872 |
| MCM7 | 7.02801745 | | -1.088830067 | 0.59978552 | -1.8153657 | 0.06946773 | 0.99999872 |
| AP4M1 | 11.2152559 | | -0.467099886 | 0.46606792 | -1.0022142 | 0.31624015 | 0.99999872 |
| TAF6 | 11.4247679 | | 0.41787194 | 0.56960585 | 0.73361596 | 0.46318283 | 0.99999872 |
| CNPY4 | 5.40744608 | | 0.732327249 | 0.67719333 | 1.08141533 | 0.27951241 | 0.99999872 |
| LAMTOR4 | 64.687378 | | 0.334301723 | 0.44536851 | 0.75061822 | 0.45288245 | 0.99999872 |
| CASTOR3 | 20.0501705 | | -0.291428629 | 0.40454524 | -0.7203858 | 0.47128751 | 0.99999872 |
| SPDYE3 | 13.8244444 | | 0.420880632 | 0.50626203 | 0.8313494 | 0.40577628 | 0.99999872 |
| PILRB | 22.725233 | | -1.104732852 | 0.45164764 | -2.446006 | 0.01444486 | 0.77617042 |
| MEPCE | 11.086196 | | -0.680178978 | 0.50344332 | -1.3510537 | 0.17667822 | 0.99999872 |
| PPP1R35 | 12.9741236 | | 0.362736882 | 0.49810474 | 0.72823414 | 0.46647027 | 0.99999872 |
| TSC22D4 | 10.9716526 | | -0.375118275 | 0.49072829 | -0.7644113 | 0.44462215 | 0.99999872 |
| LRCH4 | 15.7291677 | | 0.261597464 | 0.48101266 | 0.54384736 | 0.58654652 | 0.99999872 |
| GNB2 | 42.9854881 | | 0.08417911 | 0.47289647 | 0.17800748 | 0.8587171 | 0.99999872 |
| GIGYF1 | 15.5497359 | | -0.521277546 | 0.42010724 | -1.2408202 | 0.21467218 | 0.99999872 |
| POP7 | 11.4738976 | | -0.156069078 | 0.54336623 | -0.2872263 | 0.77393904 | 0.99999872 |
| SLC12A9 | 10.63793 | | -0.593199236 | 0.44507699 | -1.3328014 | 0.18259698 | 0.99999872 |
| TRIP6 | 17.1685153 | | 0.266857629 | 0.58697527 | 0.45463181 | 0.64937415 | 0.99999872 |
| SRRT | 12.190782 | | -0.617961303 | 0.50526257 | -1.2230498 | 0.22131088 | 0.99999872 |
| TRIM56 | 37.0574552 | | -0.798050767 | 0.51459271 | -1.5508396 | 0.12094012 | 0.99999872 |
| AP1S1 | 11.7784721 | | 0.626434789 | 0.56004668 | 1.11854031 | 0.2633363 | 0.99999872 |
| PLOD3 | 25.3620699 | | 0.348481943 | 0.37917918 | 0.91904291 | 0.35807313 | 0.99999872 |
| ZNHIT1 | 29.0918857 | | -0.051892939 | 0.41117892 | -0.1262053 | 0.89956945 | 0.99999872 |
| FIS1 | 60.1560555 | | 0.711763516 | 0.32533438 | 2.18779064 | 0.02868486 | 0.90266962 |
| IFT22 | 9.02423126 | | 0.259618124 | 0.49498397 | 0.52449804 | 0.59993218 | 0.99999872 |
| CUX1 | 362.678159 | | -0.698950614 | 0.21390645 | -3.2675528 | 0.00108482 | 0.29912383 |
| PRKRIP1 | 49.3896418 | | -0.454718149 | 0.26894896 | -1.6907228 | 0.09088975 | 0.99999872 |
| ORAI2 | 12.6611224 | | -0.298337779 | 0.72816311 | -0.4097128 | 0.68201661 | 0.99999872 |
| ALKBH4 | 5.17379186 | | 0.356656528 | 0.63004759 | 0.56607871 | 0.57134027 | 0.99999872 |
| POLR2J | 17.2838032 | | 0.751976833 | 0.49283272 | 1.52582569 | 0.12705328 | 0.99999872 |
| AC105052.3 | 8.45525599 | | -0.47319284 | 0.70045743 | -0.6755483 | 0.49932747 | 0.99999872 |
| FAM185A | 28.4064366 | | 0.358043413 | 0.46944641 | 0.76269283 | 0.44564661 | 0.99999872 |
| FBXL13 | 22.3843272 | | 0.117346533 | 0.43597583 | 0.26915834 | 0.78780783 | 0.99999872 |

| | | | | | | |
|---|---|---|---|---|---|---|
| ARMC10 | 29.2985614 | 0.73472245 | 0.37500354 | 1.95924137 | 0.05008453 | 0.98159583 |
| NAPEPLD | 137.697854 | -0.054860357 | 0.49859717 | -0.1100294 | 0.91238604 | 0.99999872 |
| PMPCB | 23.0088163 | -0.066478906 | 0.37688617 | -0.1763899 | 0.85998765 | 0.99999872 |
| DNAJC2 | 14.1728666 | 0.077895324 | 0.5660731 | 0.13760647 | 0.89055144 | 0.99999872 |
| PSMC2 | 22.8839051 | 0.2804178 | 0.41132428 | 0.68174385 | 0.49540093 | 0.99999872 |
| ORC5 | 18.4174002 | 0.138937116 | 0.40464886 | 0.3433523 | 0.73133344 | 0.99999872 |
| LHFPL3 | 13.0440066 | -0.10323928 | 0.46341964 | -0.2227771 | 0.823709 | 0.99999872 |
| LINC01004 | 68.8360132 | -0.440896005 | 0.26561895 | -1.6598816 | 0.09693828 | 0.99999872 |
| KMT2E-AS1 | 5.85109496 | -0.508571928 | 0.57521884 | -0.8841364 | 0.37662257 | 0.99999872 |
| KMT2E | 145.846295 | 0.269960132 | 0.28397252 | 0.95065583 | 0.34177912 | 0.99999872 |
| SRPK2 | 110.603604 | 0.055642566 | 0.2277615 | 0.2443019 | 0.80699701 | 0.99999872 |
| PUS7 | 10.6463917 | -0.148135594 | 0.69057798 | -0.2145096 | 0.8301497 | 0.99999872 |
| RINT1 | 7.32553959 | 0.238728249 | 0.55448187 | 0.43054293 | 0.66680074 | 0.99999872 |
| EFCAB10 | 4.93893292 | 0.045136942 | 0.63080736 | 0.07155424 | 0.94295666 | 0.99999872 |
| ATXN7L1 | 57.9210674 | -0.44016956 | 0.43013777 | -1.0233223 | 0.3061555 | 0.99999872 |
| CDHR3 | 15.1655358 | 0.893288557 | 0.78074119 | 1.14415452 | 0.25255956 | 0.99999872 |
| SYPL1 | 35.2949565 | 0.517398279 | 0.40046505 | 1.2919936 | 0.19635936 | 0.99999872 |
| NAMPT | 52.8296991 | 0.559204804 | 0.7799185 | 0.71700415 | 0.47337153 | 0.99999872 |
| CCDC71L | 8.49065333 | 1.004596626 | 0.76122891 | 1.31970373 | 0.18693395 | 0.99999872 |
| PRKAR2B | 92.0682141 | 0.485954352 | 0.37446614 | 1.29772574 | 0.19438159 | 0.99999872 |
| HBP1 | 36.0464018 | 0.108166385 | 0.32695018 | 0.33083445 | 0.74076953 | 0.99999872 |
| COG5 | 96.5318296 | 0.371504367 | 0.31851759 | 1.16635432 | 0.24347122 | 0.99999872 |
| BCAP29 | 37.4763936 | 0.236247448 | 0.35170131 | 0.67172752 | 0.50175718 | 0.99999872 |
| SLC26A4 | 7.26599045 | 1.417207765 | 0.86681057 | 1.63496825 | 0.1020557 | 0.99999872 |
| CBLL1 | 9.67119673 | -0.329619655 | 0.48628662 | -0.67783 | 0.49787949 | 0.99999872 |
| SLC26A3 | 223.230793 | -0.062679067 | 0.73177147 | -0.0856539 | 0.93174156 | 0.99999872 |
| DLD | 23.7053249 | 0.302463701 | 0.37758806 | 0.80104148 | 0.42310763 | 0.99999872 |
| LAMB1 | 25.5079917 | -1.742154004 | 0.62079592 | -2.8063232 | 0.00501104 | 0.5412956 |
| NRCAM | 9.78478128 | -0.039898348 | 0.69501638 | -0.0574063 | 0.95422151 | 0.99999872 |
| PNPLA8 | 40.1798972 | 0.077652381 | 0.31671057 | 0.24518405 | 0.80631392 | 0.99999872 |
| DNAJB9 | 48.543369 | 0.995135975 | 0.62232795 | 1.59905396 | 0.10980861 | 0.99999872 |
| IMMP2L | 454.764663 | 0.341062683 | 0.33082305 | 1.03095198 | 0.30256334 | 0.99999872 |
| DOCK4 | 224.076372 | -0.303448447 | 0.2323494 | -1.3060006 | 0.19155237 | 0.99999872 |
| ZNF277 | 77.7809556 | 0.239514826 | 0.30392139 | 0.7880815 | 0.43064904 | 0.99999872 |
| AC004112.1 | 4.64653404 | 0.892930812 | 0.87595648 | 1.01937806 | 0.30802352 | 0.99999872 |
| IFRD1 | 31.8076703 | 0.431425363 | 0.32622922 | 1.32246084 | 0.18601474 | 0.99999872 |
| TMEM168 | 39.9744141 | 0.193140883 | 0.28053296 | 0.68847838 | 0.49115158 | 0.99999872 |
| BMT2 | 32.4785765 | 0.057559322 | 0.29064595 | 0.19803931 | 0.84301431 | 0.99999872 |
| SMIM30 | 13.7290215 | 0.556763441 | 0.44157131 | 1.26086869 | 0.20735616 | 0.99999872 |
| FOXP2 | 39.9758954 | -1.681041437 | 0.83624487 | -2.0102263 | 0.04440724 | 0.95980711 |
| MDFIC | 138.685437 | -0.202662702 | 0.40266271 | -0.5033064 | 0.6147489 | 0.99999872 |
| TES | 10.4894042 | 0.412333862 | 0.55587042 | 0.74178054 | 0.45822031 | 0.99999872 |
| CAV2 | 5.78111775 | 0.072880185 | 0.74038565 | 0.09843544 | 0.92158654 | 0.99999872 |
| MET | 13.4572841 | -0.980301981 | 0.75156522 | -1.3043472 | 0.19211523 | 0.99999872 |
| CAPZA2 | 78.115973 | 0.424795143 | 0.27587041 | 1.53983582 | 0.12360038 | 0.99999872 |
| ST7 | 54.7222154 | -0.418881457 | 0.40863974 | -1.0250629 | 0.30533349 | 0.99999872 |
| CFTR | 8.99404126 | 0.215308596 | 0.52632131 | 0.40908204 | 0.68247946 | 0.99999872 |
| CTTNBP2 | 182.201707 | 0.069884863 | 0.36604041 | 0.19092117 | 0.84858736 | 0.99999872 |
| LSM8 | 22.0027038 | 0.198978747 | 0.37108024 | 0.53621487 | 0.59181005 | 0.99999872 |
| TSPAN12 | 12.2023811 | 0.170792391 | 0.49839932 | 0.34268183 | 0.73183783 | 0.99999872 |
| ING3 | 9.20251401 | -0.367945033 | 0.57101866 | -0.644366 | 0.5193381 | 0.99999872 |
| FAM3C | 35.5465332 | 0.217801639 | 0.3586914 | 0.60721176 | 0.54371039 | 0.99999872 |
| AASS | 84.2748344 | -0.591629674 | 0.39082703 | -1.5137891 | 0.13007935 | 0.99999872 |
| CADPS2 | 75.7417494 | 0.426282242 | 0.73690248 | 0.5784785 | 0.56294111 | 0.99999872 |
| SLC13A1 | 12.5810099 | 3.830648911 | 2.14767032 | 1.78362986 | 0.07448383 | 0.99999872 |
| IQUB | 9.13564728 | 0.793964816 | 0.79837553 | 0.99447538 | 0.31999148 | 0.99999872 |
| NDUFA5 | 33.5516226 | 0.837199519 | 0.30117696 | 2.77975952 | 0.00543992 | 0.55881811 |
| WASL | 230.187336 | 0.213664219 | 0.26121313 | 0.8179689 | 0.41337494 | 0.99999872 |
| AC006333.2 | 5.80532888 | -0.245756031 | 0.7404879 | -0.3318839 | 0.73997691 | 0.99999872 |
| POT1 | 24.2419831 | -0.063485672 | 0.34727678 | -0.18281 | 0.85494709 | 0.99999872 |
| POT1-AS1 | 13.8360863 | -0.058825473 | 0.65015713 | -0.0904789 | 0.9279067 | 0.99999872 |

| | | | | | | |
|---|---|---|---|---|---|---|
| ZNF800 | 47.2221641 | 0.243878202 | 0.31791103 | 0.76712721 | 0.44300589 | 0.99999872 |
| GCC1 | 3.47912383 | -0.018822067 | 0.7467143 | -0.0252065 | 0.97989024 | NA |
| ARF5 | 12.9811917 | 0.211532355 | 0.592623 | 0.35694254 | 0.72113483 | 0.99999872 |
| SND1 | 90.2335489 | -0.115962622 | 0.36887649 | -0.3143671 | 0.75324226 | 0.99999872 |
| LRRC4 | 12.2106421 | -0.501974163 | 0.53513567 | -0.9380316 | 0.34822817 | 0.99999872 |
| RBM28 | 14.8145606 | -0.463351764 | 0.39543351 | -1.1717564 | 0.24129486 | 0.99999872 |
| IMPDH1 | 27.3210706 | 0.233269617 | 0.52611615 | 0.44338045 | 0.65749057 | 0.99999872 |
| HILPDA | 20.3219887 | 0.525097176 | 0.68588181 | 0.76557968 | 0.44392644 | 0.99999872 |
| METTL2B | 17.2490255 | -0.528399709 | 0.54245247 | -0.974094 | 0.33000986 | 0.99999872 |
| CALU | 59.9561208 | 0.494654224 | 0.4373661 | 1.13098438 | 0.25806167 | 0.99999872 |
| FLNC | 9.47464473 | 0.734941422 | 0.61617651 | 1.19274495 | 0.23296927 | 0.99999872 |
| KCP | 7.23986831 | -1.081821734 | 0.61846207 | -1.7492128 | 0.08025425 | 0.99999872 |
| ATP6V1F | 38.3731165 | 0.604497239 | 0.47792968 | 1.26482464 | 0.2059342 | 0.99999872 |
| TNPO3 | 83.180393 | -0.323207211 | 0.37261771 | -0.8673963 | 0.38572493 | 0.99999872 |
| TSPAN33 | 26.915007 | 0.050334795 | 0.61448305 | 0.08191405 | 0.93471506 | 0.99999872 |
| SMO | 21.0137041 | 0.295680428 | 0.76947165 | 0.38426422 | 0.70078262 | 0.99999872 |
| AHCYL2 | 51.0579391 | -0.026653053 | 0.31167527 | -0.0855155 | 0.93185161 | 0.99999872 |
| NRF1 | 59.2408548 | 0.172353016 | 0.31638312 | 0.54476047 | 0.58591827 | 0.99999872 |
| UBE2H | 106.10731 | -0.398939056 | 0.25669764 | -1.5541204 | 0.12015569 | 0.99999872 |
| ZC3HC1 | 10.2325605 | -0.544513769 | 0.52377714 | -1.0395906 | 0.29853017 | 0.99999872 |
| KLHDC10 | 58.7286989 | -0.175759742 | 0.26693101 | -0.6584463 | 0.51025137 | 0.99999872 |
| TMEM209 | 10.2555902 | -0.176267143 | 0.59304892 | -0.2972219 | 0.76629708 | 0.99999872 |
| CEP41 | 6.26120513 | 0.052327936 | 0.6388174 | 0.08191376 | 0.93471529 | 0.99999872 |
| COPG2 | 19.258629 | -0.61553591 | 0.40171719 | -1.5322618 | 0.12545784 | 0.99999872 |
| AC016831.7 | 199.032634 | -0.118753522 | 0.40385074 | -0.294053 | 0.76871741 | 0.99999872 |
| LINC00513 | 23.218458 | 0.499198706 | 0.76473079 | 0.65277704 | 0.51390003 | 0.99999872 |
| AC058791.1 | 4.02082141 | 1.101571692 | 0.90157251 | 1.22183372 | 0.22177052 | NA |
| MKLN1 | 166.758051 | -0.205243405 | 0.34379612 | -0.5969916 | 0.55051297 | 0.99999872 |
| AC008264.2 | 58.8435898 | 0.347655547 | 0.84348332 | 0.41216647 | 0.68021741 | 0.99999872 |
| PODXL | 3571.08563 | 0.101089628 | 0.49954571 | 0.20236312 | 0.83963286 | 0.99999872 |
| AC009518.1 | 7.02137843 | -2.005416297 | 0.62144236 | -3.2270351 | 0.0012508 | 0.30478826 |
| PLXNA4 | 65.528867 | -1.471339974 | 0.78045935 | -1.8852231 | 0.05939973 | 0.99999872 |
| CHCHD3 | 117.194399 | 0.142225836 | 0.23039148 | 0.61732246 | 0.53702206 | 0.99999872 |
| EXOC4 | 273.019262 | -0.036729472 | 0.29814004 | -0.1231954 | 0.90195239 | 0.99999872 |
| SLC35B4 | 13.7069849 | -0.035361524 | 0.43231775 | -0.0817952 | 0.93480956 | 0.99999872 |
| AKR1B1 | 15.9812349 | 0.264435254 | 0.43863791 | 0.60285545 | 0.54660485 | 0.99999872 |
| BPGM | 5.93885555 | 0.292320356 | 0.5985919 | 0.48834666 | 0.62530431 | 0.99999872 |
| CALD1 | 179.372478 | -0.039801546 | 0.33519844 | -0.1187403 | 0.90548115 | 0.99999872 |
| AGBL3 | 7.61913741 | 0.336292444 | 0.81544829 | 0.41240193 | 0.68004485 | 0.99999872 |
| CYREN | 11.7802471 | -0.019780353 | 0.46910714 | -0.042166 | 0.9663664 | 0.99999872 |
| TMEM140 | 13.4146967 | -0.48493154 | 0.95738651 | -0.506516 | 0.61249449 | 0.99999872 |
| WDR91 | 9.12818407 | 1.355370913 | 0.78226566 | 1.73262228 | 0.08316283 | 0.99999872 |
| CNOT4 | 79.0053874 | 0.041444092 | 0.26437807 | 0.1567607 | 0.87543345 | 0.99999872 |
| NUP205 | 16.8126717 | 0.155105513 | 0.39511955 | 0.39255338 | 0.69464938 | 0.99999872 |
| STMP1 | 44.2637508 | 0.064654649 | 0.39853784 | 0.16222964 | 0.87112502 | 0.99999872 |
| SLC13A4 | 5.79259268 | 2.548770882 | 0.93114768 | 2.73723593 | 0.00619578 | 0.5901766 |
| FAM180A | 61.5419299 | 1.470199566 | 1.09721631 | 1.33993594 | 0.18026617 | 0.99999872 |
| AC015987.1 | 3.72086887 | 1.965047599 | 1.22322915 | 1.60644275 | 0.10817666 | NA |
| MTPN | 50.6678751 | 0.352678888 | 0.32189323 | 1.09563933 | 0.27323664 | 0.99999872 |
| PTN | 4.52061187 | -0.239077491 | 1.38147964 | -0.173059 | 0.86260505 | NA |
| CREB3L2 | 92.3300611 | 0.263788876 | 0.42037027 | 0.62751554 | 0.53032135 | 0.99999872 |
| TRIM24 | 26.1878865 | 0.064950206 | 0.45487824 | 0.14278592 | 0.88645926 | 0.99999872 |
| KIAA1549 | 10.1267233 | 0.057847413 | 0.57901014 | 0.09990743 | 0.92041782 | 0.99999872 |
| ZC3HAV1 | 34.1748583 | -0.167200911 | 0.52653812 | -0.3175476 | 0.75082813 | 0.99999872 |
| TTC26 | 10.6411432 | -0.116069786 | 0.47748439 | -0.243086 | 0.80793875 | 0.99999872 |
| UBN2 | 46.709625 | 0.047342294 | 0.33365172 | 0.14189135 | 0.88716582 | 0.99999872 |
| LUC7L2 | 25.1704518 | -0.019007378 | 0.38953655 | -0.0487949 | 0.96108278 | 0.99999872 |
| HIPK2 | 351.939342 | -1.135373667 | 0.36068586 | -3.1478186 | 0.00164494 | 0.3314549 |
| TBXAS1 | 67.6975852 | -1.160480555 | 0.36261228 | -3.2003344 | 0.00137268 | 0.32688551 |
| PARP12 | 21.8315551 | -0.650172308 | 0.98013943 | -0.6633468 | 0.5071085 | 0.99999872 |
| KDM7A | 31.3322695 | 0.001715219 | 0.38702118 | 0.00443185 | 0.99646391 | 0.99999872 |

| | | | | | | |
|---|---|---|---|---|---|---|
| SLC37A3 | 21.8336167 | 0.386214492 | 0.39956905 | 0.9665776 | 0.33375524 | 0.99999872 |
| MKRN1 | 23.066804 | 0.093317896 | 0.37718554 | 0.24740582 | 0.80459417 | 0.99999872 |
| NDUFB2 | 63.1141853 | 0.540965181 | 0.41688021 | 1.29765137 | 0.19440716 | 0.99999872 |
| BRAF | 148.520649 | -0.114012879 | 0.19580835 | -0.5822677 | 0.56038638 | 0.99999872 |
| MRPS33 | 10.4110119 | -0.01519774 | 0.47348659 | -0.0320975 | 0.97439429 | 0.99999872 |
| TMEM178B | 8.51434963 | -0.646064734 | 1.08064645 | -0.5978502 | 0.54993987 | 0.99999872 |
| AGK | 19.079063 | -0.236855209 | 0.38227435 | -0.6195948 | 0.53552457 | 0.99999872 |
| KIAA1147 | 11.7476646 | 1.138178402 | 0.5938917 | 1.91647466 | 0.05530471 | 0.99999872 |
| SSBP1 | 36.877984 | 0.145983182 | 0.36042548 | 0.40503014 | 0.68545536 | 0.99999872 |
| MGAM | 8.26580967 | 0.542217897 | 0.86934891 | 0.62370573 | 0.53282086 | 0.99999872 |
| GSTK1 | 26.0425178 | 0.461850027 | 0.51734645 | 0.89272871 | 0.37200247 | 0.99999872 |
| AC073342.1 | 4.94632565 | 0.527488927 | 0.63279833 | 0.83358141 | 0.4045169 | 0.99999872 |
| TMEM139 | 8.38333275 | -0.395495988 | 0.64015749 | -0.6178105 | 0.53670029 | 0.99999872 |
| CASP2 | 10.2774384 | 0.322795171 | 0.52867627 | 0.61057245 | 0.54148266 | 0.99999872 |
| ZYX | 18.1232479 | -0.490283774 | 0.55096947 | -0.8898565 | 0.37354293 | 0.99999872 |
| TCAF2 | 19.5814538 | 0.683375082 | 0.72602763 | 0.94125218 | 0.34657564 | 0.99999872 |
| TCAF1 | 58.3788586 | -0.272549442 | 0.45388807 | -0.6004772 | 0.54818824 | 0.99999872 |
| AC004889.1 | 9.29204966 | 0.385199801 | 0.56390978 | 0.68308764 | 0.49455147 | 0.99999872 |
| AC074386.1 | 14.4333586 | -1.151196436 | 0.42790112 | -2.6903328 | 0.00713808 | 0.62327327 |
| TPK1 | 31.5820697 | -0.372657388 | 0.37590493 | -0.9913607 | 0.32150947 | 0.99999872 |
| CNTNAP2 | 21.5433427 | -1.183215256 | 0.50773337 | -2.330387 | 0.0197857 | 0.8671024 |
| CUL1 | 45.5410028 | -0.221173639 | 0.27684018 | -0.7989217 | 0.42433579 | 0.99999872 |
| PDIA4 | 26.6203901 | 0.39329617 | 0.44585267 | 0.88212137 | 0.37771118 | 0.99999872 |
| ZNF786 | 5.56107202 | -0.02268203 | 0.72343764 | -0.0313531 | 0.97498792 | 0.99999872 |
| ZNF398 | 13.8809496 | -0.236184363 | 0.39058709 | -0.6046907 | 0.54538456 | 0.99999872 |
| ZNF282 | 10.6329081 | -0.351027228 | 0.49195192 | -0.7135397 | 0.47551185 | 0.99999872 |
| ZNF212 | 3.49832703 | -0.167521134 | 0.975272 | -0.1717686 | 0.86361943 | NA |
| ZNF783 | 9.81773608 | -0.046563214 | 0.56067275 | -0.0830488 | 0.93381272 | 0.99999872 |
| ZNF777 | 5.74542408 | -0.727925046 | 0.63327525 | -1.1494608 | 0.25036604 | 0.99999872 |
| ZNF746 | 10.1377698 | 0.639886389 | 0.48769284 | 1.31206845 | 0.18949704 | 0.99999872 |
| ZNF467 | 7.81562685 | -0.265490269 | 0.59520159 | -0.446051 | 0.6555604 | 0.99999872 |
| ZNF862 | 20.3172458 | 0.509488705 | 0.41929992 | 1.21509375 | 0.22433034 | 0.99999872 |
| ATP6V0E2 | 8.19425379 | 1.082210484 | 0.68808551 | 1.57278488 | 0.11576864 | 0.99999872 |
| AC092666.1 | 6.85584439 | 0.762164226 | 0.70860274 | 1.07558746 | 0.2821118 | 0.99999872 |
| ACTR3C | 25.0375833 | -0.438213648 | 0.4478554 | -0.9784713 | 0.32784129 | 0.99999872 |
| LRRC61 | 8.08395764 | -0.042105215 | 0.66372545 | -0.0634377 | 0.94941797 | 0.99999872 |
| RARRES2 | 118.113867 | 1.078240822 | 0.53034723 | 2.03308468 | 0.04204397 | 0.95980711 |
| AC073111.5 | 4.56325671 | -0.763233441 | 0.85700553 | -0.8905817 | 0.37315362 | NA |
| TMEM176B | 17.3312286 | 0.729164238 | 0.79243257 | 0.92015935 | 0.35748949 | 0.99999872 |
| TMEM176A | 11.9258669 | 0.620518684 | 0.69006711 | 0.89921498 | 0.36853816 | 0.99999872 |
| KCNH2 | 8.29308679 | -0.455045918 | 0.6436145 | -0.7070163 | 0.47955638 | 0.99999872 |
| ABCB8 | 11.4902605 | -0.425107678 | 0.52047325 | -0.8167714 | 0.41405906 | 0.99999872 |
| SLC4A2 | 10.5676682 | -0.441962625 | 0.50346462 | -0.8778425 | 0.38002921 | 0.99999872 |
| FASTK | 8.70448087 | -0.330585825 | 0.642782 | -0.5143047 | 0.60703895 | 0.99999872 |
| TMUB1 | 16.0216266 | -0.26698351 | 0.44714213 | -0.5970887 | 0.55044816 | 0.99999872 |
| AGAP3 | 23.6510458 | -0.305300677 | 0.41749196 | -0.7312732 | 0.4646123 | 0.99999872 |
| CHPF2 | 9.66023396 | -0.23014407 | 0.69486703 | -0.3312059 | 0.74048895 | 0.99999872 |
| SMARCD3 | 11.8489527 | 0.414317159 | 0.56117495 | 0.73830301 | 0.46033034 | 0.99999872 |
| NUB1 | 32.6792805 | -0.164336818 | 0.68649047 | -0.2393869 | 0.8108056 | 0.99999872 |
| RHEB | 108.150809 | 0.649031978 | 0.26925909 | 2.41043661 | 0.01593344 | 0.79608149 |
| PRKAG2 | 27.7189137 | -0.581340772 | 0.69869258 | -0.8320409 | 0.40538588 | 0.99999872 |
| GALNT11 | 46.9077587 | 0.109107592 | 0.36376434 | 0.29994032 | 0.76422268 | 0.99999872 |
| KMT2C | 187.369228 | -0.046027694 | 0.25223189 | -0.1824817 | 0.85520475 | 0.99999872 |
| ACTR3B | 9.74544597 | -0.004576848 | 0.61282799 | -0.0074684 | 0.99404113 | 0.99999872 |
| DPP6 | 1174.66166 | -0.401069925 | 0.33672325 | -1.1910966 | 0.23361566 | 0.99999872 |
| AC006019.3 | 6.78666835 | 0.698325513 | 0.61925233 | 1.12769137 | 0.25945028 | 0.99999872 |
| PAXIP1-AS2 | 4.41460587 | 0.417004098 | 0.74779749 | 0.55764308 | 0.57708813 | NA |
| PAXIP1 | 11.6937491 | 0.739602147 | 0.47220706 | 1.56626661 | 0.11728622 | 0.99999872 |
| PAXIP1-AS1 | 6.3374154 | 0.708528807 | 0.75734568 | 0.93554215 | 0.34950897 | 0.99999872 |
| INSIG1 | 14.1040906 | -0.206523085 | 0.49467922 | -0.4174889 | 0.67632084 | 0.99999872 |
| AC009403.1 | 11.0350817 | 0.123908767 | 0.48614917 | 0.25487808 | 0.79881726 | 0.99999872 |

| | | | | | | |
|---|---|---|---|---|---|---|
| RBM33 | 78.5499148 | -0.1551514 | 0.24051294 | -0.6450855 | 0.5188718 | 0.99999872 |
| LINC01006 | 25.534305 | -0.841468202 | 0.3237755 | -2.5989249 | 0.00935162 | 0.69734315 |
| LMBR1 | 48.9021281 | 0.087341314 | 0.27222122 | 0.32084682 | 0.74832647 | 0.99999872 |
| NOM1 | 12.728447 | -0.246322958 | 0.45834243 | -0.5374212 | 0.59097668 | 0.99999872 |
| UBE3C | 68.1426972 | -0.28204402 | 0.27454922 | -1.0272986 | 0.30427989 | 0.99999872 |
| DNAJB6 | 54.612002 | 0.11319742 | 0.28690461 | 0.39454723 | 0.69317706 | 0.99999872 |
| PTPRN2 | 11.2546698 | 0.055387575 | 0.51697263 | 0.10713831 | 0.91467925 | 0.99999872 |
| ESYT2 | 101.010561 | 0.234695505 | 0.27123886 | 0.86527243 | 0.38688928 | 0.99999872 |
| WDR60 | 64.422196 | -0.222002193 | 0.37878536 | -0.5860897 | 0.55781524 | 0.99999872 |
| GTPBP6 | 16.2149442 | -0.176269364 | 0.3791847 | -0.4648641 | 0.64202876 | 0.99999872 |
| LINC00685 | 8.47309791 | 0.208915821 | 0.60752412 | 0.34388071 | 0.730936 | 0.99999872 |
| IL3RA | 6.39002898 | -0.35627343 | 0.67138146 | -0.5306572 | 0.59565633 | 0.99999872 |
| SLC25A6 | 52.5055711 | 0.479341242 | 0.42601184 | 1.12518291 | 0.26051153 | 0.99999872 |
| LINC00106 | 16.2220342 | -0.481943267 | 0.50998849 | -0.9450081 | 0.34465473 | 0.99999872 |
| ASMTL-AS1 | 9.65587039 | 0.035460797 | 0.60336676 | 0.05877155 | 0.95313407 | 0.99999872 |
| ASMTL | 24.0312592 | 0.042656707 | 0.51596482 | 0.08267367 | 0.93411102 | 0.99999872 |
| AKAP17A | 14.0944952 | 0.54599005 | 0.4136854 | 1.31981948 | 0.1868953 | 0.99999872 |
| DHRSX | 60.8737168 | -0.24607237 | 0.23419388 | -1.0507208 | 0.29338685 | 0.99999872 |
| CD99 | 35.1372255 | 0.514942729 | 0.52513319 | 0.98059452 | 0.32679274 | 0.99999872 |
| ARSD | 22.1034568 | -0.094459649 | 0.35650397 | -0.264961 | 0.7910395 | 0.99999872 |
| PRKX | 15.6658087 | -0.220089212 | 0.48397964 | -0.4547489 | 0.64928989 | 0.99999872 |
| PUDP | 20.5275982 | -0.671819918 | 0.49212215 | -1.3651487 | 0.17220632 | 0.99999872 |
| STS | 10.1609483 | -0.39405477 | 0.49220378 | -0.8005927 | 0.42336746 | 0.99999872 |
| PNPLA4 | 6.35020283 | -0.269884844 | 0.77345994 | -0.3489319 | 0.72714044 | 0.99999872 |
| TBL1X | 108.029819 | 0.043967339 | 0.31735273 | 0.13854407 | 0.88981044 | 0.99999872 |
| SHROOM2 | 26.99606 | 0.174767477 | 0.55586948 | 0.3144038 | 0.75321437 | 0.99999872 |
| WWC3 | 28.3202612 | -0.351181565 | 0.49353139 | -0.7115689 | 0.4767318 | 0.99999872 |
| MID1 | 13.7568976 | 0.445364459 | 0.73196936 | 0.60844686 | 0.54289114 | 0.99999872 |
| ARHGAP6 | 62.4124992 | 0.1979311 | 0.62599666 | 0.31618555 | 0.75186167 | 0.99999872 |
| MSL3 | 4.14438242 | 0.134311687 | 0.74503204 | 0.18027639 | 0.85693559 | NA |
| PRPS2 | 6.20508174 | 0.616045083 | 0.67441417 | 0.91345216 | 0.36100478 | 0.99999872 |
| TMSB4X | 1094.27771 | 0.366221735 | 0.57682934 | 0.6348875 | 0.52550179 | 0.99999872 |
| RAB9A | 16.2093141 | 0.116009873 | 0.42277589 | 0.2744004 | 0.78377694 | 0.99999872 |
| TRAPPC2 | 6.63962212 | -0.144604696 | 0.61947182 | -0.2334322 | 0.81542578 | 0.99999872 |
| OFD1 | 25.4581566 | -0.3793951 | 0.54442903 | -0.6968679 | 0.48588549 | 0.99999872 |
| GPM6B | 8.06420615 | 0.199439101 | 0.88603373 | 0.225092 | 0.82190771 | 0.99999872 |
| GEMIN8 | 16.4970883 | -0.265476083 | 0.4177769 | -0.6354494 | 0.52513535 | 0.99999872 |
| MOSPD2 | 24.5105573 | -0.410938674 | 0.37473658 | -1.0966068 | 0.27281332 | 0.99999872 |
| PIR | 23.3582535 | 0.547211789 | 0.44621101 | 1.22635206 | 0.22006622 | 0.99999872 |
| BMX | 5.34685546 | -1.057380373 | 0.67313809 | -1.5708224 | 0.11622392 | 0.99999872 |
| CLTRN | 11.1745266 | 1.160212866 | 1.32487492 | 0.87571502 | 0.38118498 | 0.99999872 |
| CA5B | 19.9312391 | -0.9094115 | 0.49153165 | -1.8501586 | 0.0642907 | 0.99999872 |
| ZRSR2 | 9.67695148 | -0.381730131 | 0.60039316 | -0.6358003 | 0.52490662 | 0.99999872 |
| AP1S2 | 12.990048 | 0.785095647 | 0.50874241 | 1.54320857 | 0.12278018 | 0.99999872 |
| CTPS2 | 25.2704385 | -0.291920548 | 0.4847391 | -0.602222 | 0.54702637 | 0.99999872 |
| SYAP1 | 23.9054407 | -0.034066106 | 0.35405973 | -0.0962157 | 0.92334926 | 0.99999872 |
| TXLNG | 10.9728302 | 0.085232128 | 0.52833679 | 0.16132158 | 0.87184012 | 0.99999872 |
| RBBP7 | 27.6576471 | 0.187790175 | 0.32439165 | 0.57889954 | 0.56265696 | 0.99999872 |
| REPS2 | 14.985771 | 0.966019502 | 0.58110745 | 1.66237673 | 0.09643726 | 0.99999872 |
| NHS | 59.2848734 | -0.234110948 | 0.64343788 | -0.3638439 | 0.71597458 | 0.99999872 |
| SCML1 | 15.2275938 | -0.114497767 | 0.4309737 | -0.2656723 | 0.79049161 | 0.99999872 |
| RAI2 | 5.23197585 | 0.80636544 | 0.70634202 | 1.14160763 | 0.25361715 | 0.99999872 |
| CDKL5 | 14.1785138 | 0.07927622 | 0.44480585 | 0.17822657 | 0.85854505 | 0.99999872 |
| PHKA2 | 16.5286331 | -0.34078308 | 0.41889567 | -0.8135273 | 0.41591578 | 0.99999872 |
| PDHA1 | 16.1165437 | 0.372056362 | 0.44042404 | 0.84476851 | 0.3982401 | 0.99999872 |
| SH3KBP1 | 18.091164 | -0.08451272 | 0.77432799 | -0.1091433 | 0.91308882 | 0.99999872 |
| BCLAF3 | 24.3468259 | 0.021335245 | 0.46339559 | 0.04604111 | 0.96327749 | 0.99999872 |
| EIF1AX | 54.5111513 | 0.450977144 | 0.30859025 | 1.46141085 | 0.14390272 | 0.99999872 |
| RPS6KA3 | 36.2637405 | -0.01575773 | 0.31100239 | -0.0506676 | 0.95959043 | 0.99999872 |
| MBTPS2 | 18.9931856 | -0.194943036 | 0.37428936 | -0.5208351 | 0.60248165 | 0.99999872 |
| SMS | 25.8760352 | -0.338980035 | 0.30583512 | -1.1083751 | 0.26769986 | 0.99999872 |

| | | | | | | |
|---|---|---|---|---|---|---|
| PRDX4 | 9.67420983 | 1.032047269 | 0.55664931 | 1.85403495 | 0.06373416 | 0.99999872 |
| ACOT9 | 18.9180302 | -0.587274254 | 0.39630568 | -1.4818719 | 0.13837438 | 0.99999872 |
| SAT1 | 164.425283 | -0.349029841 | 0.64005615 | -0.5453113 | 5.86E-01 | 1.00E+00 |
| APOO | 171.230023 | 0.330239607 | 0.34399376 | 0.96001627 | 0.33704703 | 0.99999872 |
| KLHL15 | 10.8602517 | 0.650491226 | 0.61936137 | 1.05026121 | 0.29359804 | 0.99999872 |
| EIF2S3 | 21.117118 | 0.302248945 | 0.4800127 | 0.62966864 | 0.52891141 | 0.99999872 |
| ZFX | 45.4147499 | -0.074900863 | 0.42116995 | -0.17784 | 0.85884863 | 0.99999872 |
| POLA1 | 24.6006939 | -0.345544745 | 0.37287955 | -0.9266927 | 0.35408612 | 0.99999872 |
| GK | 24.7667982 | 0.778159964 | 0.48392152 | 1.60802926 | 0.10782877 | 0.99999872 |
| TAB3 | 28.2345814 | 0.222766544 | 0.41275776 | 0.53970286 | 0.58940197 | 0.99999872 |
| DMD | 149.029331 | 0.270971096 | 0.47264395 | 0.57330914 | 0.5664354 | 0.99999872 |
| AL591501.1 | 14.2276869 | -0.547195513 | 0.70543599 | -0.7756841 | 4.38E-01 | 0.99999872 |
| TMEM47 | 42.4165363 | 0.147737604 | 0.3185897 | 0.46372373 | 0.64284569 | 0.99999872 |
| AF241726.2 | 9.86464091 | 0.112838622 | 0.65501764 | 0.17226807 | 0.86322679 | 0.99999872 |
| DYNLT3 | 37.4432003 | 0.175413363 | 0.48183165 | 0.36405529 | 0.71581672 | 0.99999872 |
| RPGR | 11.5451443 | -0.48897671 | 0.54228766 | -0.9016925 | 0.36722024 | 0.99999872 |
| MID1IP1 | 11.6851945 | -0.227971357 | 0.54331276 | -0.4195951 | 0.67478129 | 0.99999872 |
| BCOR | 19.6414228 | -0.221732635 | 0.49116557 | -0.4514417 | 0.65167121 | 0.99999872 |
| ATP6AP2 | 61.7312937 | 0.573828112 | 0.38911801 | 1.47468916 | 0.14029613 | 0.99999872 |
| CXorf38 | 23.1599671 | -0.283509096 | 0.42832143 | -0.6619073 | 0.50803061 | 0.99999872 |
| MED14 | 45.58845 | -0.071893681 | 0.42350051 | -0.1697606 | 0.86519845 | 0.99999872 |
| USP9X | 102.237722 | -0.173655263 | 0.31504077 | -0.5512152 | 0.58148616 | 0.99999872 |
| DDX3X | 65.2390618 | 0.182958186 | 0.23968776 | 0.76331886 | 0.44527326 | 0.99999872 |
| CASK | 55.5164549 | -0.358910896 | 0.32477912 | -1.1050923 | 0.26911961 | 0.99999872 |
| MAOA | 37.9970579 | -0.116410271 | 0.47831357 | -0.2433765 | 8.08E-01 | 0.99999872 |
| MAOB | 20.0922767 | -0.561013654 | 0.68910702 | -0.8141169 | 0.415578 | 0.99999872 |
| FUNDC1 | 10.789605 | 0.334632886 | 0.70799709 | 0.47264726 | 0.63646486 | 0.99999872 |
| KDM6A | 93.9166545 | -0.144398099 | 0.57325844 | -0.2518901 | 0.80112605 | 0.99999872 |
| KRBOX4 | 15.4107012 | 0.460683819 | 0.44432461 | 1.03681814 | 0.29982062 | 0.99999872 |
| ZNF674 | 4.67583681 | 0.09527999 | 0.87450201 | 0.10895343 | 0.91323943 | 0.99999872 |
| SLC9A7 | 6.92486675 | 0.924805363 | 0.59633337 | 1.55081939 | 0.12094497 | 0.99999872 |
| RP2 | 11.2409653 | -0.788650339 | 0.50013669 | -1.5768696 | 0.11482553 | 0.99999872 |
| JADE3 | 10.4309389 | 0.101240414 | 0.57015383 | 0.17756684 | 0.85906317 | 0.99999872 |
| RGN | 7.88270266 | 2.09429463 | 1.19527865 | 1.75213925 | 0.07974987 | 0.99999872 |
| NDUFB11 | 45.5812321 | 0.727578051 | 0.43206999 | 1.68393561 | 0.09219411 | 0.99999872 |
| RBM10 | 26.6403016 | -0.314790836 | 0.4570778 | -0.688703 | 0.49101021 | 0.99999872 |
| UBA1 | 52.8178119 | 0.050629964 | 0.45726533 | 0.11072338 | 0.9118357 | 0.99999872 |
| CDK16 | 10.9413022 | 0.270277514 | 0.52761345 | 0.51226426 | 0.60846607 | 0.99999872 |
| USP11 | 22.2185473 | 0.001377487 | 0.3975566 | 0.00346488 | 0.99723543 | 0.99999872 |
| ZNF41 | 6.18792297 | 0.089079246 | 0.61333613 | 0.14523724 | 0.88452357 | 0.99999872 |
| ARAF | 14.6519462 | 0.017828059 | 0.49135557 | 0.03628342 | 0.97105637 | 0.99999872 |
| SYN1 | 9.31501833 | -0.395625853 | 0.58154094 | -0.6803061 | 0.49631066 | 0.99999872 |
| TIMP1 | 445.439596 | -0.345684968 | 0.61210706 | -0.5647459 | 5.72E-01 | 0.99999872 |
| ELK1 | 13.6378004 | 0.786289294 | 0.56566037 | 1.39003779 | 0.1645174 | 0.99999872 |
| UXT | 7.1416785 | 0.862387405 | 0.53431192 | 1.6140149 | 0.10652419 | 0.99999872 |
| ZNF81 | 17.7592965 | -0.148493366 | 0.54345268 | -0.2732407 | 0.78466823 | 0.99999872 |
| ZNF182 | 7.99620803 | -0.628906154 | 0.57133605 | -1.100764 | 0.27099939 | 0.99999872 |
| ZNF630 | 11.6431997 | 0.949623865 | 0.64028776 | 1.48312045 | 0.13804241 | 0.99999872 |
| EBP | 9.31744279 | -0.525012445 | 0.4534 | -1.1579454 | 0.24688632 | 0.99999872 |
| TBC1D25 | 5.12516419 | 0.482552509 | 0.61960076 | 0.778812 | 0.43609047 | 0.99999872 |
| RBM3 | 37.9072781 | -0.520815001 | 0.66967547 | -0.7777125 | 0.43673851 | 0.99999872 |
| WDR13 | 27.8255647 | -0.058396946 | 0.43154298 | -0.1353213 | 0.89235786 | 0.99999872 |
| HDAC6 | 12.1289173 | -0.014867123 | 0.4591879 | -0.032377 | 0.97417141 | 0.99999872 |
| PCSK1N | 13.5611304 | 0.428002572 | 0.88611545 | 0.48300994 | 0.62908868 | 0.99999872 |
| TIMM17B | 10.7226423 | 1.078828119 | 0.48513549 | 2.22376662 | 0.02616415 | 0.88434828 |
| PQBP1 | 6.13903514 | 0.06920025 | 0.6259125 | 0.11055898 | 0.91196607 | 0.99999872 |
| SLC35A2 | 6.42211503 | 0.237337186 | 0.69429496 | 0.34183913 | 0.73247196 | 0.99999872 |
| OTUD5 | 28.924868 | -0.438203003 | 0.34930018 | -1.254517 | 0.20965416 | 0.99999872 |
| GRIPAP1 | 21.2262783 | -0.698849001 | 0.33987111 | -2.0562177 | 0.03976153 | 0.95980711 |
| TFE3 | 25.7398291 | -0.106660722 | 0.33605173 | -0.3173938 | 0.75094484 | 0.99999872 |
| PRAF2 | 5.92988086 | 0.266457194 | 0.81842895 | 0.32557156 | 0.74474853 | 0.99999872 |

| | | | | | | |
|---|---|---|---|---|---|---|
| WDR45 | 14.0769175 | -0.343340696 | 0.44612126 | -0.769613 | 0.44152952 | 0.99999872 |
| GPKOW | 3.40022281 | 0.928984134 | 0.79812702 | 1.16395524 | 0.24444215 | NA |
| PLP2 | 23.2253035 | 0.378872555 | 0.61756872 | 0.61349052 | 0.53955205 | 0.99999872 |
| CCDC22 | 5.35384286 | 0.598545566 | 0.69780711 | 0.85775217 | 0.39102933 | 0.99999872 |
| PPP1R3F | 7.98781747 | 0.931786073 | 0.60697423 | 1.53513283 | 0.1247512 | 0.99999872 |
| CLCN5 | 17.8145413 | -0.108626968 | 0.77115143 | -0.1408633 | 0.8879779 | 0.99999872 |
| SHROOM4 | 10.8927519 | -0.648792784 | 0.84482802 | -0.7679584 | 0.4425119 | 0.99999872 |
| GSPT2 | 4.38623102 | -0.415745401 | 0.69914644 | -0.5946471 | 0.55207939 | NA |
| MAGED1 | 56.0948225 | -0.215609362 | 0.2529313 | -0.8524424 | 0.3939686 | 0.99999872 |
| FAM156A | 6.86769405 | 0.783402558 | 0.6146163 | 1.27462055 | 0.20244358 | 0.99999872 |
| GPR173 | 32.1644688 | 0.700498989 | 0.47179266 | 1.48476024 | 0.13760735 | 0.99999872 |
| TSPYL2 | 9.13803891 | -0.850715154 | 0.72288935 | -1.1768262 | 0.23926487 | 0.99999872 |
| KANTR | 25.0679033 | -0.118511225 | 0.47229021 | -0.2509288 | 0.80186914 | 0.99999872 |
| KDM5C | 16.3540629 | -0.12004098 | 0.49383335 | -0.2430799 | 0.80794347 | 0.99999872 |
| IQSEC2 | 56.9839342 | -0.415751383 | 0.35207102 | -1.1808736 | 0.23765293 | 0.99999872 |
| SMC1A | 18.5935493 | -0.422165823 | 0.40393379 | -1.0451362 | 0.29596003 | 0.99999872 |
| HSD17B10 | 16.1104049 | 0.585990202 | 0.43907999 | 1.33458645 | 0.18201173 | 0.99999872 |
| HUWE1 | 76.304103 | -0.116822366 | 0.31676251 | -0.3688011 | 0.71227598 | 0.99999872 |
| PHF8 | 16.2464496 | 0.051292464 | 0.43285779 | 0.11849726 | 0.90567366 | 0.99999872 |
| FAM120C | 16.2473075 | -0.313991685 | 0.4894419 | -0.64153 | 0.52117837 | 0.99999872 |
| TSR2 | 8.73536083 | 0.394254344 | 0.53945284 | 0.73084117 | 0.46487618 | 0.99999872 |
| FGD1 | 5.93544927 | 1.327709225 | 0.82996699 | 1.59971328 | 0.1096622 | 0.99999872 |
| MAGED2 | 39.7533314 | 0.300477528 | 0.48806551 | 0.61564999 | 0.53812555 | 0.99999872 |
| APEX2 | 3.56436864 | 0.017948469 | 0.75334909 | 0.0238249 | 0.98099228 | NA |
| FAM104B | 6.44976582 | -0.24567133 | 0.59477411 | -0.4130498 | 0.67957012 | 0.99999872 |
| MAGEH1 | 8.56215118 | 2.235776392 | 0.79883542 | 2.79879476 | 0.00512937 | 0.54842417 |
| RRAGB | 22.9698564 | 0.319178774 | 0.38911985 | 0.82025827 | 0.41206889 | 0.99999872 |
| AL445472.1 | 30.1575773 | -0.550209408 | 0.32744445 | -1.6803137 | 0.09289629 | 0.99999872 |
| KLF8 | 10.3750403 | 0.468208798 | 0.48495806 | 0.96546246 | 0.33431323 | 0.99999872 |
| UBQLN2 | 16.6118652 | 0.445532303 | 0.43823167 | 1.01665931 | 0.30931553 | 0.99999872 |
| NBDY | 50.9599455 | 0.260907493 | 0.27251757 | 0.95739698 | 0.33836692 | 0.99999872 |
| SPIN3 | 9.71571017 | 0.071635664 | 0.48253079 | 0.14845822 | 0.88198115 | 0.99999872 |
| FAAH2 | 17.6162467 | -0.381298918 | 0.45994103 | -0.829017 | 0.40709481 | 0.99999872 |
| SPIN4 | 5.40381537 | 0.384608071 | 0.77952325 | 0.49338884 | 0.62173787 | 0.99999872 |
| ARHGEF9 | 17.8355846 | -0.195332018 | 0.37839542 | -0.5162114 | 0.60570679 | 0.99999872 |
| MTMR8 | 8.51127623 | -0.02271344 | 0.62142406 | -0.0365506 | 0.97084331 | 0.99999872 |
| ZC4H2 | 27.3019801 | 0.414943009 | 0.49226043 | 0.84293391 | 0.39926541 | 0.99999872 |
| ZC3H12B | 18.0769599 | 0.155879557 | 0.57212802 | 0.27245573 | 0.78527163 | 0.99999872 |
| LAS1L | 7.93767867 | -0.287295949 | 0.53985691 | -0.5321705 | 0.59460788 | 0.99999872 |
| MSN | 39.691738 | -0.133234227 | 0.42063535 | -0.3167452 | 0.75143695 | 0.99999872 |
| OPHN1 | 29.8491782 | -0.011468634 | 0.35345534 | -0.0324472 | 0.97411543 | 0.99999872 |
| YIPF6 | 35.3872596 | 0.298036156 | 0.33430658 | 0.89150552 | 0.37265803 | 0.99999872 |
| EFNB1 | 98.4660054 | -0.654677733 | 0.52289902 | -1.2520156 | 0.21056417 | 0.99999872 |
| PJA1 | 9.95771703 | 0.422187753 | 0.59094319 | 0.71443035 | 0.47496111 | 0.99999872 |
| EDA | 30.381896 | 0.672134542 | 0.38541433 | 1.74392723 | 0.08117178 | 0.99999872 |
| IGBP1 | 15.4175826 | 0.220616109 | 0.40625248 | 0.54305172 | 0.58709419 | 0.99999872 |
| PDZD11 | 7.6421985 | 0.448505382 | 0.59836766 | 0.74954817 | 0.45352688 | 0.99999872 |
| SNX12 | 11.0105511 | 0.217713957 | 0.46293793 | 0.47028758 | 0.63814957 | 0.99999872 |
| MED12 | 7.46336881 | -0.670643716 | 0.57329881 | -1.1697978 | 0.24208233 | 0.99999872 |
| ZMYM3 | 8.25377889 | 0.016487078 | 0.51437939 | 0.03205237 | 0.97443029 | 0.99999872 |
| NONO | 28.8397318 | 0.014242676 | 0.33897863 | 0.04201644 | 0.96648559 | 0.99999872 |
| TAF1 | 39.3418206 | -0.587220309 | 0.34921432 | -1.681547 | 0.09265671 | 0.99999872 |
| OGT | 59.5011632 | -0.239280595 | 0.28588007 | -0.8369964 | 0.40259459 | 0.99999872 |
| NHSL2 | 12.3047094 | -0.091220524 | 0.69272911 | -0.1316828 | 0.89523517 | 0.99999872 |
| PIN4 | 18.8059821 | -0.363047507 | 0.41133281 | -0.8826126 | 0.37744563 | 0.99999872 |
| RPS4X | 80.3458769 | 0.697080562 | 0.39077403 | 1.78384567 | 0.07444875 | 0.99999872 |
| HDAC8 | 77.9245659 | -0.215774241 | 0.287143 | -0.7514522 | 0.45238056 | 0.99999872 |
| PHKA1 | 6.41115794 | 0.571902014 | 0.75596489 | 0.75651928 | 0.44933791 | 0.99999872 |
| NAP1L2 | 6.10249021 | 0.288300576 | 0.74176628 | 0.38866768 | 0.697522 | 0.99999872 |
| CHIC1 | 40.8503269 | 0.193400855 | 0.3681933 | 0.52526989 | 0.59939558 | 0.99999872 |
| XIST | 126.618471 | 3.872161047 | 4.55732147 | 0.84965721 | 0.39551569 | 0.99999872 |

| | | | | | | |
|---|---|---|---|---|---|---|
| JPX | 126.849851 | 0.253932418 | 0.5006337 | 0.50722198 | 0.61199907 | 0.99999872 |
| FTX | 794.584752 | 0.281544741 | 0.26011344 | 1.08239214 | 0.27907832 | 0.99999872 |
| SLC16A2 | 13.1387689 | 0.516329605 | 0.52762497 | 0.97859206 | 0.32778158 | 0.99999872 |
| RLIM | 23.8719664 | 0.221307719 | 0.46234667 | 0.47866187 | 0.6321792 | 0.99999872 |
| NEXMIF | 34.9146084 | 0.421006439 | 0.78091113 | 0.53912209 | 0.58980262 | 0.99999872 |
| ABCB7 | 27.576342 | 0.006959804 | 0.32691742 | 0.02128918 | 0.98301498 | 0.99999872 |
| UPRT | 6.83005756 | -0.468904849 | 0.53958884 | -0.869004 | 0.38484496 | 0.99999872 |
| ZDHHC15 | 11.0747057 | -0.377858357 | 0.64685796 | -0.5841442 | 0.55912327 | 0.99999872 |
| PBDC1 | 6.61717445 | 0.993334632 | 0.71093953 | 1.39721396 | 0.16234924 | 0.99999872 |
| ATRX | 157.810038 | -0.099127879 | 0.27591301 | -0.3592722 | 0.71939146 | 0.99999872 |
| MAGT1 | 59.3341747 | 0.505032551 | 0.50775918 | 0.99463008 | 0.31991621 | 0.99999872 |
| COX7B | 47.0405737 | 0.373220883 | 0.42170542 | 0.88502747 | 0.37614182 | 0.99999872 |
| PGK1 | 71.97979 | 0.323805238 | 0.36029274 | 0.89872819 | 0.36879746 | 0.99999872 |
| TAF9B | 7.77817756 | 0.252491398 | 0.54296458 | 0.4650237 | 0.64191448 | 0.99999872 |
| BRWD3 | 37.1855173 | -0.100029923 | 0.33678235 | -0.2970165 | 0.7664539 | 0.99999872 |
| SH3BGRL | 53.3952309 | 0.386327076 | 0.31466924 | 1.2277243 | 0.21955048 | 0.99999872 |
| RPS6KA6 | 39.575818 | 0.294873788 | 0.46266936 | 0.63733157 | 0.52390889 | 0.99999872 |
| HDX | 13.4922361 | 0.051132794 | 0.49347894 | 0.10361697 | 0.91747332 | 0.99999872 |
| APOOL | 23.977557 | 0.091065313 | 0.33269829 | 0.27371741 | 0.7843018 | 0.99999872 |
| CHM | 41.9349474 | 0.285295052 | 0.33537893 | 0.85066482 | 0.39495557 | 0.99999872 |
| DACH2 | 279.679585 | -0.984401854 | 0.80080747 | -1.2292616 | 0.21897375 | 0.99999872 |
| DIAPH2 | 93.8590648 | 0.011942818 | 0.29039467 | 0.04112616 | 0.96719532 | 0.99999872 |
| DIAPH2-AS1 | 7.5521055 | 2.287457781 | 0.91727127 | 2.49376369 | 0.01263967 | 0.72768363 |
| TSPAN6 | 10.9076083 | 0.67285741 | 0.61767033 | 1.08934714 | 0.27600083 | 0.99999872 |
| SRPX2 | 6.26319471 | 0.448214222 | 0.83532601 | 0.536574 | 0.5915619 | 0.99999872 |
| SYTL4 | 34.1991878 | 0.617673515 | 0.37584528 | 1.64342494 | 0.10029504 | 0.99999872 |
| TRMT2B | 15.2060918 | -0.294298052 | 0.48854775 | -0.6023936 | 0.54691215 | 0.99999872 |
| HNRNPH2 | 23.6664171 | 0.286959016 | 0.33737915 | 0.8505535 | 0.39501743 | 0.99999872 |
| ARMCX4 | 63.7973117 | -0.425966479 | 0.3723471 | -1.1440038 | 0.25262208 | 0.99999872 |
| ARMCX1 | 10.0472193 | 0.41840275 | 0.54454676 | 0.76835045 | 0.44227901 | 0.99999872 |
| ARMCX6 | 10.0637302 | 0.387119359 | 0.69154321 | 0.55979056 | 0.5756223 | 0.99999872 |
| ARMCX3 | 44.3566066 | 0.001734617 | 0.45760535 | 0.00379064 | 0.99697551 | 0.99999872 |
| ARMCX2 | 13.5528751 | 0.91900869 | 0.51274803 | 1.79232028 | 0.07308168 | 0.99999872 |
| ZMAT1 | 50.6677591 | -0.389148431 | 0.41487963 | -0.9379791 | 0.34825514 | 0.99999872 |
| TCEAL2 | 29.8659894 | -0.262488224 | 0.46005172 | -0.5705624 | 0.5682963 | 0.99999872 |
| BEX5 | 5.44006254 | 0.516038887 | 0.69432368 | 0.74322524 | 0.45734532 | 0.99999872 |
| GPRASP1 | 4.86461704 | 0.705660186 | 0.69604373 | 1.01381588 | 0.31067062 | 0.99999872 |
| BHLHB9 | 6.8091511 | 0.118469416 | 0.702367 | 0.16867167 | 0.8660549 | 0.99999872 |
| LINC00630 | 19.2247364 | 0.087960602 | 0.45759425 | 0.19222401 | 0.84756674 | 0.99999872 |
| BEX4 | 66.2103548 | 1.155119652 | 0.44954053 | 2.56955618 | 0.01018289 | 0.70849208 |
| TCEAL8 | 24.0213193 | 0.742543396 | 0.40836207 | 1.81834564 | 0.06901132 | 0.99999872 |
| BEX2 | 20.9504128 | 1.131876686 | 0.75571341 | 1.49775917 | 0.13419583 | 0.99999872 |
| TCEAL9 | 57.7001126 | 0.228699196 | 0.40778145 | 0.56083766 | 0.57490821 | 0.99999872 |
| BEX3 | 131.468917 | 0.479276875 | 0.40586552 | 1.18087606 | 0.23765196 | 0.99999872 |
| TCEAL4 | 50.3184741 | 0.19364558 | 0.34763892 | 0.55703079 | 0.57750639 | 0.99999872 |
| TCEAL3 | 30.7941436 | -0.305678384 | 0.41519136 | -0.7362349 | 0.46158774 | 0.99999872 |
| TCEAL1 | 6.98958845 | 0.576535748 | 0.5516978 | 1.04502092 | 0.2960133 | 0.99999872 |
| MORF4L2 | 47.2865114 | 0.446585113 | 0.35697949 | 1.25101057 | 0.21093062 | 0.99999872 |
| SLC25A53 | 13.6911717 | 0.00446837 | 0.42159824 | 0.01059864 | 0.99154366 | 0.99999872 |
| FAM199X | 33.648286 | 0.16715409 | 0.3276478 | 0.51016393 | 0.60993662 | 0.99999872 |
| IL1RAPL2 | 7.72041034 | 1.137434166 | 0.82233254 | 1.3831803 | 0.16660959 | 0.99999872 |
| RNF128 | 7.34096327 | 1.153424095 | 0.96270645 | 1.19810571 | 0.23087587 | 0.99999872 |
| TBC1D8B | 13.2350959 | 1.195647201 | 0.49192241 | 2.43056056 | 0.01507549 | 0.78681466 |
| MORC4 | 11.373668 | 1.300911597 | 0.49161412 | 2.6462047 | 0.00814005 | 0.66265058 |
| RBM41 | 23.3566322 | 0.24475696 | 0.41846552 | 0.58489158 | 0.55862062 | 0.99999872 |
| PRPS1 | 8.8899599 | 0.916951627 | 0.58878188 | 1.55737066 | 0.11938251 | 0.99999872 |
| TSC22D3 | 170.394669 | 0.357151046 | 0.3821301 | 0.93463207 | 0.34997795 | 0.99999872 |
| MID2 | 12.546864 | 0.450472632 | 0.53035639 | 0.84937721 | 0.39567143 | 0.99999872 |
| PSMD10 | 6.39625472 | -0.003468627 | 0.65210187 | -0.0053191 | 0.99575595 | 0.99999872 |
| ATG4A | 5.39761117 | 1.412901764 | 0.77231673 | 1.82943306 | 0.06733476 | 0.99999872 |
| COL4A6 | 12.6515317 | -0.791027203 | 0.69981106 | -1.130344 | 0.25833132 | 0.99999872 |

| | | | | | | |
|---|---|---|---|---|---|---|
| COL4A5 | 195.267143 | -0.051304857 | 0.28926535 | -0.1773626 | 0.85922357 | 0.99999872 |
| ACSL4 | 100.607704 | 0.523201352 | 0.26966186 | 1.94021266 | 0.05235385 | 0.99999872 |
| TMEM164 | 31.374876 | -0.654351812 | 0.40943649 | -1.5981766 | 0.11000368 | 0.99999872 |
| AMMECR1 | 23.5269939 | 0.243964864 | 0.34670194 | 0.7036732 | 0.48163631 | 0.99999872 |
| ALG13 | 47.2154358 | 0.146714847 | 0.40644287 | 0.36097287 | 0.71811973 | 0.99999872 |
| AMOT | 19.3626877 | -0.429399524 | 0.8124308 | -0.5285367 | 0.59712686 | 0.99999872 |
| LRCH2 | 14.5104179 | 0.094061223 | 0.46635012 | 0.20169658 | 0.84015394 | 0.99999872 |
| PLS3-AS1 | 6.07981968 | 0.734048702 | 0.84999217 | 0.86359466 | 0.3878106 | 0.99999872 |
| PLS3 | 122.593121 | 0.294375627 | 0.4653874 | 0.63253888 | 0.52703481 | 0.99999872 |
| DANT2 | 99.9466237 | -0.502510788 | 0.4106813 | -1.2236028 | 0.2211021 | 0.99999872 |
| KLHL13 | 25.1572602 | -1.009819313 | 0.66706645 | -1.5138212 | 0.1300712 | 0.99999872 |
| WDR44 | 30.4422418 | -0.217062352 | 0.33832046 | -0.641588 | 0.52114072 | 0.99999872 |
| IL13RA1 | 60.0546588 | -0.059049 | 0.57380508 | -0.1029078 | 0.91803617 | 0.99999872 |
| LONRF3 | 11.2017477 | -1.275145243 | 0.73962424 | -1.7240447 | 0.08469978 | 0.99999872 |
| PGRMC1 | 54.126031 | 0.321404351 | 0.45625925 | 0.7044336 | 0.48116278 | 0.99999872 |
| SLC25A43 | 31.4851148 | -0.085875744 | 0.31237628 | -0.2749112 | 0.78338444 | 0.99999872 |
| SLC25A5 | 28.6211931 | 0.741989293 | 0.40344182 | 1.83914818 | 0.06589339 | 0.99999872 |
| CXorf56 | 6.15192948 | 0.098462689 | 0.60990497 | 0.16143939 | 0.87174734 | 0.99999872 |
| UBE2A | 18.2345659 | 0.646207746 | 0.43618445 | 1.4815011 | 0.13847309 | 0.99999872 |
| NKRF | 4.60830307 | 0.21453225 | 0.67977851 | 0.3155914 | 0.75231267 | 0.99999872 |
| 6-Sep | 8.41433991 | -0.647492412 | 0.52063225 | -1.2436656 | 0.21362266 | 0.99999872 |
| RPL39 | 5.24190269 | 0.727046312 | 0.77250751 | 0.94115112 | 0.34662742 | 0.99999872 |
| UPF3B | 15.7890136 | -0.194247125 | 0.42727382 | -0.4546198 | 0.64938281 | 0.99999872 |
| NDUFA1 | 40.9129675 | 0.374497174 | 0.40935437 | 0.91484836 | 0.36027124 | 0.99999872 |
| NKAP | 11.1640351 | 0.175931916 | 0.5581729 | 0.31519251 | 0.75261549 | 0.99999872 |
| RHOXF1-AS1 | 9.3767426 | 0.902137747 | 0.62914237 | 1.4339167 | 0.15159603 | 0.99999872 |
| ZBTB33 | 4.63696161 | 0.300642646 | 0.66753356 | 0.45037833 | 0.65243767 | 0.99999872 |
| LAMP2 | 63.4035984 | 0.471806161 | 0.31408993 | 1.50213718 | 0.13306168 | 0.99999872 |
| CUL4B | 19.9768238 | 0.132559686 | 0.34821809 | 0.38068007 | 0.70344066 | 0.99999872 |
| MCTS1 | 18.3732624 | 0.164551905 | 0.39460801 | 0.41700092 | 0.67667773 | 0.99999872 |
| C1GALT1C1 | 12.699059 | -0.113012202 | 0.670874 | -0.1684552 | 0.8662252 | 0.99999872 |
| THOC2 | 60.4177809 | 0.176342365 | 0.27382255 | 0.64400234 | 0.51957392 | 0.99999872 |
| XIAP | 68.6248055 | -0.076462559 | 0.28598376 | -0.2673668 | 0.78918677 | 0.99999872 |
| STAG2 | 62.3710165 | -0.084368299 | 0.29042976 | -0.2904947 | 0.77143782 | 0.99999872 |
| TENM1 | 8.75576872 | 0.69940833 | 0.7398769 | 0.94530365 | 0.34450388 | 0.99999872 |
| SMARCA1 | 32.5726602 | 0.565391153 | 0.33564581 | 1.68448743 | 0.09208751 | 0.99999872 |
| OCRL | 10.8354991 | 0.105744035 | 0.47028799 | 0.22484953 | 0.82209633 | 0.99999872 |
| APLN | 26.5937421 | 0.609770252 | 0.89303741 | 0.68280483 | 0.49473018 | 0.99999872 |
| ZDHHC9 | 13.3913643 | -0.644362339 | 0.43918691 | -1.4671711 | 0.14232951 | 0.99999872 |
| UTP14A | 6.1493454 | -0.865899526 | 0.67881268 | -1.2756089 | 0.20209379 | 0.99999872 |
| BCORL1 | 8.43629769 | 0.529336863 | 0.56470256 | 0.93737288 | 0.34856679 | 0.99999872 |
| ELF4 | 8.15133475 | -0.388406368 | 0.73242529 | -0.5303017 | 0.59590278 | 0.99999872 |
| AIFM1 | 13.7118714 | 0.026050991 | 0.41523992 | 0.0627372 | 0.94997577 | 0.99999872 |
| ZNF280C | 6.61686573 | -0.070860248 | 0.54676096 | -0.1296 | 0.89688286 | 0.99999872 |
| SLC25A14 | 7.27100301 | 0.275971011 | 0.78986644 | 0.34938946 | 0.72679694 | 0.99999872 |
| RBMX2 | 7.45770153 | -0.434310327 | 0.70691825 | -0.6143714 | 0.53896996 | 0.99999872 |
| ENOX2 | 24.4381064 | -0.998977197 | 0.45887985 | -2.1769908 | 0.02948126 | 0.90266962 |
| RAP2C | 16.4940189 | 0.200994758 | 0.5464589 | 0.36781313 | 0.71301258 | 0.99999872 |
| RAP2C-AS1 | 37.3432891 | -0.461876604 | 0.41830791 | -1.1041546 | 0.26952608 | 0.99999872 |
| MBNL3 | 66.3021998 | -0.136793856 | 0.36824338 | -0.3714768 | 0.71028246 | 0.99999872 |
| GPC4 | 18.4020076 | 0.064700506 | 0.38955562 | 0.16608798 | 0.86808771 | 0.99999872 |
| GPC3 | 11.7055912 | -1.365045234 | 0.60458726 | -2.2578134 | 0.02395729 | 0.88337526 |
| PHF6 | 13.6624004 | -0.298434237 | 0.40060538 | -0.7449581 | 0.45629703 | 0.99999872 |
| HPRT1 | 25.5135255 | 0.295219098 | 0.34248992 | 0.86197894 | 0.38869911 | 0.99999872 |
| MIR503HG | 39.0707995 | 0.717110022 | 0.6664462 | 1.07602088 | 0.28191792 | 0.99999872 |
| FAM122B | 11.3745601 | 0.154043399 | 0.48350036 | 0.31860038 | 0.75002957 | 0.99999872 |
| FAM122C | 5.84536813 | 0.087570643 | 0.62921423 | 0.13917461 | 0.88931217 | 0.99999872 |
| MOSPD1 | 4.94192062 | 0.105722931 | 0.74937389 | 0.14108168 | 0.88780541 | 0.99999872 |
| RTL8B | 5.80922171 | 0.724202297 | 0.68895366 | 1.05116257 | 0.29318393 | 0.99999872 |
| RTL8C | 38.156373 | 0.667082767 | 0.46626291 | 1.43070091 | 0.15251595 | 0.99999872 |
| RTL8A | 27.0599898 | 0.356750184 | 0.42150073 | 0.84638093 | 0.39734027 | 0.99999872 |

| | | | | | | | |
|---|---|---|---|---|---|---|---|
| ZNF75D | 17.8780125 | | -0.474701243 | 0.45619223 | -1.0405728 | 0.29807384 | 0.99999872 |
| ZNF449 | 5.36215137 | | 0.550203796 | 0.6074682 | 0.90573268 | 0.36507736 | 0.99999872 |
| INTS6L | 14.1047555 | | 0.435984892 | 0.54275323 | 0.80328383 | 0.4218107 | 0.99999872 |
| MMGT1 | 10.6663551 | | 0.351365939 | 0.63908338 | 0.54979671 | 0.58245881 | 0.99999872 |
| SLC9A6 | 11.4199444 | | 0.16334471 | 0.53075119 | 0.30776136 | 0.75826392 | 0.99999872 |
| FHL1 | 45.8703382 | | 0.833207585 | 0.50027256 | 1.66550728 | 0.09581159 | 0.99999872 |
| MAP7D3 | 10.1836706 | | -0.19112531 | 0.54028526 | -0.3537489 | 0.72352708 | 0.99999872 |
| HTATSF1 | 12.2283906 | | -0.509495764 | 0.46935263 | -1.0855287 | 0.27768755 | 0.99999872 |
| AL683813.1 | 16.8617337 | | 0.354440683 | 0.47675614 | 0.7434423 | 0.45721394 | 0.99999872 |
| RBMX | 55.8548047 | | 0.216678746 | 0.25734872 | 0.84196551 | 0.39980726 | 0.99999872 |
| FGF13 | 75.3884002 | | -0.462926945 | 0.98115176 | -0.4718199 | 0.63705533 | 0.99999872 |
| ATP11C | 58.1194405 | | 0.90321271 | 0.4763291 | 1.89619468 | 0.05793431 | 0.99999872 |
| LDOC1 | 4.56168965 | | 1.287529048 | 0.89068606 | 1.44554755 | 0.14830415 | NA |
| SLITRK2 | 18.9356173 | | -1.193875098 | 0.61479979 | -1.9418925 | 0.05215012 | 0.99999872 |
| FMR1 | 25.4912464 | | -0.221968155 | 0.45028014 | -0.4929557 | 0.6220439 | 0.99999872 |
| IDS | 34.6767545 | | 0.32005772 | 0.55402948 | 0.57769078 | 0.56347291 | 0.99999872 |
| TMEM185A | 14.5668611 | | -0.298197741 | 0.43929958 | -0.6788027 | 0.49726289 | 0.99999872 |
| LINC00894 | 98.4509785 | | -0.273657427 | 0.37693969 | -0.7259979 | 0.46784006 | 0.99999872 |
| MAMLD1 | 11.1961795 | | -0.49469094 | 0.49662787 | -0.9960998 | 0.31920164 | 0.99999872 |
| MTM1 | 19.1067718 | | 0.330399648 | 0.40136564 | 0.82318867 | 0.41040071 | 0.99999872 |
| MTMR1 | 22.5763514 | | 0.170827291 | 0.34449509 | 0.49587729 | 0.61998099 | 0.99999872 |
| CD99L2 | 24.0228614 | | -0.431574634 | 0.41745881 | -1.0338137 | 0.30122326 | 0.99999872 |
| VMA21 | 17.1329612 | | 0.525929928 | 0.57679983 | 0.91180666 | 0.3618705 | 0.99999872 |
| CETN2 | 14.801482 | | 0.213423539 | 0.59459125 | 0.35894161 | 0.71963877 | 0.99999872 |
| ZNF185 | 58.6171598 | | -0.182131492 | 0.48139601 | -0.3783403 | 0.70517784 | 0.99999872 |
| BGN | 183.501305 | | -0.7660675 | 0.56386791 | -1.3585939 | 0.17427531 | 0.99999872 |
| CCNQ | 7.70219857 | | 0.082851445 | 0.74403509 | 0.11135422 | 0.91133546 | 0.99999872 |
| SLC6A8 | 8.59112068 | | -0.34751419 | 0.6055239 | -0.5739066 | 0.56603098 | 0.99999872 |
| BCAP31 | 31.0468434 | | -0.208950399 | 0.44041158 | -0.4744435 | 0.6351837 | 0.99999872 |
| IDH3G | 7.97574585 | | 0.342599515 | 0.49886613 | 0.68675642 | 0.49223624 | 0.99999872 |
| SSR4 | 30.7382174 | | 1.05596458 | 0.58290052 | 1.81156911 | 0.0700528 | 0.99999872 |
| HCFC1 | 12.8626378 | | 0.674000588 | 0.4464841 | 1.50957355 | 0.13115228 | 0.99999872 |
| TMEM187 | 4.32289698 | | 0.454968425 | 0.79400841 | 0.57300202 | 0.56664333 | NA |
| IRAK1 | 9.61095944 | | -0.415584386 | 0.66737929 | -0.6227109 | 0.53347449 | 0.99999872 |
| MECP2 | 56.5417618 | | -0.14967819 | 0.30976647 | -0.4831969 | 0.62895596 | 0.99999872 |
| FLNA | 144.621285 | | 0.191573667 | 0.52143199 | 0.36739914 | 0.71332131 | 0.99999872 |
| EMD | 17.9720564 | | 0.130233012 | 0.3760994 | 0.34627286 | 0.72913767 | 0.99999872 |
| RPL10 | 152.66321 | | 0.767841954 | 0.53623296 | 1.43191861 | 0.15216712 | 0.99999872 |
| DNASE1L1 | 5.66876696 | | 0.191117867 | 0.66107055 | 0.28910358 | 0.77250211 | 0.99999872 |
| TAZ | 13.0440684 | | 0.103349486 | 0.50874138 | 0.20314739 | 0.83901983 | 0.99999872 |
| ATP6AP1 | 52.3288052 | | 0.755348372 | 0.53945659 | 1.40020232 | 0.16145274 | 0.99999872 |
| GDI1 | 31.3917929 | | 0.048005353 | 0.41073461 | 0.11687682 | 0.90695767 | 0.99999872 |
| FAM50A | 16.561732 | | -0.034944511 | 0.62844255 | -0.0556049 | 0.95565653 | 0.99999872 |
| PLXNA3 | 22.6468852 | | 0.112477307 | 0.63146239 | 0.17812194 | 0.85862721 | 0.99999872 |
| LAGE3 | 3.46932789 | | -0.141738778 | 0.73709245 | -0.1922944 | 0.84751158 | NA |
| UBL4A | 9.18646364 | | 0.181277945 | 0.59248061 | 0.30596435 | 0.75963179 | 0.99999872 |
| SLC10A3 | 4.61941506 | | -0.425685068 | 0.70552953 | -0.6033554 | 0.54627227 | 0.99999872 |
| FAM3A | 13.4449704 | | -0.27090463 | 0.52351916 | -0.5174684 | 0.60482921 | 0.99999872 |
| G6PD | 5.10847113 | | -0.433540894 | 0.5991121 | -0.723639 | 0.46928738 | 0.99999872 |
| IKBKG | 6.50277739 | | -0.034327562 | 0.59723907 | -0.0574771 | 0.95416516 | 0.99999872 |
| DKC1 | 8.4969025 | | 0.019420096 | 0.49615658 | 0.03914106 | 0.96877792 | 0.99999872 |
| MPP1 | 4.95444001 | | 0.748816752 | 0.70386185 | 1.06386893 | 0.28738808 | 0.99999872 |
| F8 | 7.50688856 | | 0.247258946 | 0.55936273 | 0.44203686 | 0.65846253 | 0.99999872 |
| F8A1 | 4.67025875 | | -0.882252006 | 0.63734746 | -1.3842559 | 0.1662801 | 0.99999872 |
| FUNDC2 | 21.5717727 | | 0.308270188 | 0.41419712 | 0.74425961 | 0.45671943 | 0.99999872 |
| BRCC3 | 11.6082893 | | -0.110835663 | 0.47578404 | -0.2329537 | 0.81579734 | 0.99999872 |
| VBP1 | 14.54422 | | 1.149756879 | 0.46024136 | 2.49816072 | 0.01248396 | 0.72670507 |
| CLIC2 | 5.43847191 | | -0.49968896 | 0.82170499 | -0.6081124 | 0.54311296 | 0.99999872 |
| TMLHE-AS1 | 9.18117492 | | 1.640779681 | 0.70348365 | 2.33236364 | 0.01968157 | 0.8671024 |
| TMLHE | 23.1410817 | | -0.00636832 | 0.37985893 | -0.016765 | 0.98662412 | 0.99999872 |
| VAMP7 | 13.6229292 | | -0.268591119 | 0.42572219 | -0.630907 | 0.52810132 | 0.99999872 |

| | | | | | | |
|---|---|---|---|---|---|---|
| ZNF596 | 9.64094305 | -0.519919119 | 0.48741185 | -1.0666936 | 0.2861102 | 0.99999872 |
| FBXO25 | 27.4190588 | 0.133449323 | 0.49741363 | 0.26828642 | 0.78847886 | 0.99999872 |
| TDRP | 18.0600916 | 0.313858205 | 0.39068994 | 0.80334345 | 0.42177625 | 0.99999872 |
| ERICH1 | 47.8848209 | -0.687747756 | 0.33786302 | -2.0355816 | 0.04179239 | 0.95980711 |
| AC100810.1 | 7.89146149 | 0.604158062 | 0.74940347 | 0.8061853 | 0.42013601 | 0.99999872 |
| KBTBD11 | 29.7823483 | -1.08037759 | 0.41680067 | -2.5920726 | 0.00953996 | 0.69901919 |
| AC246817.1 | 15.7084795 | -0.947312575 | 0.52631069 | -1.7999113 | 0.07187464 | 0.99999872 |
| CSMD1 | 29.5968763 | 1.798581592 | 1.09625299 | 1.64066288 | 0.10086742 | 0.99999872 |
| MCPH1 | 103.247469 | 0.0012212 | 0.28066222 | 0.00435114 | 0.9965283 | 0.99999872 |
| ANGPT2 | 14.4139504 | 1.062508814 | 0.65037381 | 1.63368941 | 0.10232408 | 0.99999872 |
| MCPH1-AS1 | 11.2445311 | 1.258660604 | 0.55818826 | 2.25490339 | 0.02413939 | 0.88337526 |
| AGPAT5 | 10.8757191 | -0.370160947 | 0.45572679 | -0.8122431 | 0.41665214 | 0.99999872 |
| DEFB1 | 38.7193772 | 0.168275877 | 0.95954015 | 0.17537138 | 0.86078783 | 0.99999872 |
| FAM85B | 46.0971767 | 0.111182142 | 0.60543881 | 0.18363894 | 0.85429671 | 0.99999872 |
| MFHAS1 | 42.718005 | -0.166964168 | 0.45235596 | -0.3690991 | 0.71205388 | 0.99999872 |
| ERI1 | 14.2720797 | 0.489518736 | 0.44657489 | 1.09616271 | 0.27300758 | 0.99999872 |
| PPP1R3B | 22.5003778 | 0.22810369 | 0.48973769 | 0.46576707 | 0.64138224 | 0.99999872 |
| TNKS | 129.191281 | 0.006411851 | 0.27719706 | 0.02313102 | 0.98154576 | 0.99999872 |
| MSRA | 45.8960368 | -0.242446125 | 0.52397672 | -0.462704 | 0.64357655 | 0.99999872 |
| XKR6 | 131.821225 | 0.122368309 | 0.25464718 | 0.48054061 | 0.63084304 | 0.99999872 |
| MTMR9 | 27.0209859 | 0.316269428 | 0.32966333 | 0.95937098 | 0.33737189 | 0.99999872 |
| NEIL2 | 4.21036748 | -0.392071184 | 0.67366981 | -0.5819931 | 0.56057133 | NA |
| FDFT1 | 32.6937472 | -0.159196735 | 0.32975018 | -0.4827798 | 0.62925208 | 0.99999872 |
| CTSB | 47.2092315 | -0.001562397 | 0.37008672 | -0.0042217 | 0.99663158 | 0.99999872 |
| FAM66D | 17.3360893 | -0.49691511 | 1.01869481 | -0.4877959 | 0.62569445 | 0.99999872 |
| AC145124.1 | 6.653755 | 0.578416061 | 0.81583186 | 0.7089893 | 0.47833112 | 0.99999872 |
| LONRF1 | 37.0176611 | -0.220458392 | 0.36123301 | -0.6102941 | 0.54166698 | 0.99999872 |
| TRMT9B | 11.8222227 | 0.525247312 | 0.92581084 | 0.56733762 | 0.57048483 | 0.99999872 |
| DLC1 | 124.504947 | 0.192460331 | 0.30093683 | 0.63953732 | 0.52247345 | 0.99999872 |
| TUSC3 | 175.986029 | 0.314505521 | 0.37927614 | 0.82922569 | 0.40697671 | 0.99999872 |
| MICU3 | 32.8224484 | 0.683722675 | 0.45782905 | 1.4934017 | 0.13533207 | 0.99999872 |
| ZDHHC2 | 23.6160368 | 0.350531544 | 0.40217095 | 0.87159837 | 0.38342752 | 0.99999872 |
| CNOT7 | 16.3753261 | 0.564948383 | 0.38957875 | 1.45015194 | 0.14701615 | 0.99999872 |
| VPS37A | 42.0192612 | 0.598445042 | 0.27604998 | 2.16788655 | 0.03016732 | 0.90831371 |
| MTUS1 | 20.9953956 | 0.181064709 | 0.74507706 | 0.24301474 | 0.80799398 | 0.99999872 |
| PCM1 | 143.478726 | 0.039374263 | 0.2168419 | 0.18158051 | 0.85591194 | 0.99999872 |
| ASAH1 | 62.5247263 | 0.900726838 | 0.32363783 | 2.78313213 | 0.00538369 | 0.55881811 |
| PSD3 | 135.627801 | 0.242741682 | 0.28523177 | 0.85103311 | 0.39475096 | 0.99999872 |
| SH2D4A | 18.0298867 | -0.618536983 | 0.47418465 | -1.3044222 | 0.19208967 | 0.99999872 |
| CSGALNACT1 | 28.1629119 | -2.107312176 | 0.81121018 | -2.5977388 | 0.00938398 | 0.69734315 |
| INTS10 | 37.5551163 | 0.12319811 | 0.49133841 | 0.25073983 | 0.80201526 | 0.99999872 |
| LPL | 41.772861 | 0.508296159 | 0.962301 | 0.52820912 | 0.5973542 | 0.99999872 |
| ATP6V1B2 | 16.121252 | 0.397052675 | 0.52044709 | 0.76290689 | 0.44551892 | 0.99999872 |
| XPO7 | 38.3670254 | -0.04106878 | 0.35635644 | -0.1152464 | 0.90824986 | 0.99999872 |
| DMTN | 9.8273275 | -0.354305485 | 0.51929069 | -0.6822874 | 0.49505725 | 0.99999872 |
| FAM160B2 | 13.6344585 | 0.139003669 | 0.45782975 | 0.30361432 | 0.76142174 | 0.99999872 |
| BMP1 | 53.1676363 | -0.537349049 | 0.58702557 | -0.9153759 | 0.35999434 | 0.99999872 |
| PHYHIP | 123.2924 | -0.056768918 | 0.40135902 | -0.1414417 | 0.88752098 | 0.99999872 |
| POLR3D | 4.68143952 | 0.40497055 | 0.71540919 | 0.56606842 | 0.57134727 | 0.99999872 |
| SLC39A14 | 56.4591411 | 0.537365659 | 0.52655667 | 1.02052769 | 0.30747827 | 0.99999872 |
| PPP3CC | 61.5152108 | -0.046995706 | 0.28132373 | -0.1670521 | 0.86732909 | 0.99999872 |
| SORBS3 | 23.6653691 | 0.033425909 | 0.32393322 | 0.10318765 | 0.91781404 | 0.99999872 |
| PDLIM2 | 7.02242106 | -1.242574293 | 0.56948812 | -2.1819144 | 0.02911585 | 0.90266962 |
| C8orf58 | 17.8442967 | -0.54129554 | 0.55951912 | -0.9674299 | 0.33332916 | 0.99999872 |
| CCAR2 | 25.5179844 | -0.601999156 | 0.65153618 | -0.9239689 | 0.35550252 | 0.99999872 |
| BIN3 | 14.3818778 | -0.019033496 | 0.42645603 | -0.0446318 | 0.9644008 | 0.99999872 |
| PEBP4 | 22.4539308 | -0.158548096 | 0.53290085 | -0.2975189 | 0.76607035 | 0.99999872 |
| RHOBTB2 | 46.6154676 | -0.517105586 | 0.63145853 | -0.8189066 | 0.41283968 | 0.99999872 |
| TNFRSF10B | 13.5954841 | -0.42986901 | 0.5129765 | -0.8379897 | 0.40203651 | 0.99999872 |
| CHMP7 | 9.29675539 | -0.510033953 | 0.45887328 | -1.1114919 | 0.26635666 | 0.99999872 |
| R3HCC1 | 6.57193202 | -0.683461483 | 0.61943336 | -1.1033656 | 0.26986842 | 0.99999872 |

| | | | | | | |
|---|---|---|---|---|---|---|
| ENTPD4 | 35.0587985 | -0.105651801 | 0.45895297 | -0.2302018 | 0.81793496 | 0.99999872 |
| SLC25A37 | 111.924678 | -0.011392274 | 0.40724376 | -0.0279741 | 0.97768281 | 0.99999872 |
| ADAM28 | 72.8300667 | 0.870649111 | 0.49333822 | 1.7648118 | 0.0775954 | 0.99999872 |
| AC120193.1 | 13.5758844 | 2.212203935 | 0.50003918 | 4.42406123 | 9.69E-06 | 0.03893472 |
| DOCK5 | 438.753891 | 0.225852875 | 0.33500539 | 0.67417684 | 0.50019889 | 0.99999872 |
| KCTD9 | 41.0378584 | 0.683622669 | 0.41332375 | 1.65396415 | 0.0981348 | 0.99999872 |
| PPP2R2A | 79.2064266 | 0.115389906 | 0.341835 | 0.33756024 | 0.73569461 | 0.99999872 |
| BNIP3L | 162.514298 | 0.567110341 | 0.4012734 | 1.41327667 | 0.15757439 | 0.99999872 |
| PNMA2 | 22.81442 | -0.492196434 | 0.42118306 | -1.1686045 | 0.24256301 | 0.99999872 |
| DPYSL2 | 93.1714104 | -0.244428601 | 0.36596431 | -0.6679028 | 0.50419561 | 0.99999872 |
| TRIM35 | 12.5358234 | -0.246640403 | 0.5695652 | -0.4330328 | 0.66499096 | 0.99999872 |
| PTK2B | 8.78365875 | -0.670464718 | 0.95512788 | -0.7019633 | 0.48270205 | 0.99999872 |
| EPHX2 | 16.4472597 | 0.418240094 | 0.44322526 | 0.94362875 | 0.3453594 | 0.99999872 |
| CLU | 32.6957282 | 1.534944038 | 0.77992237 | 1.96807285 | 0.04905966 | 0.97937581 |
| CCDC25 | 12.9654139 | -0.395356424 | 0.40608981 | -0.9735689 | 0.33027062 | 0.99999872 |
| ELP3 | 23.6636546 | -0.661800368 | 0.35216613 | -1.8792278 | 0.0602134 | 0.99999872 |
| ZNF395 | 10.9183628 | 0.143925354 | 0.45200941 | 0.31841229 | 0.75017222 | 0.99999872 |
| FBXO16 | 12.6329526 | -0.815889107 | 0.56608048 | -1.4412952 | 0.14950129 | 0.99999872 |
| FZD3 | 6.79841081 | 0.764819663 | 0.65795128 | 1.16242598 | 0.24506248 | 0.99999872 |
| EXTL3 | 16.9381446 | -0.138412649 | 0.58237725 | -0.2376684 | 0.81213832 | 0.99999872 |
| INTS9 | 21.0601193 | -0.331193144 | 0.35704762 | -0.9275882 | 0.35362124 | 0.99999872 |
| HMBOX1 | 71.7268794 | 0.131120426 | 0.3040937 | 0.43118429 | 0.66633437 | 0.99999872 |
| KIF13B | 38.3401024 | 0.015979717 | 0.51545184 | 0.03100138 | 0.97526844 | 0.99999872 |
| SARAF | 98.1917835 | 0.451925288 | 0.41260101 | 1.09530825 | 0.27338162 | 0.99999872 |
| LEPROTL1 | 25.3075174 | 0.360140916 | 0.36986425 | 0.97371107 | 0.33020002 | 0.99999872 |
| DCTN6 | 18.1737334 | 0.267479234 | 0.52954374 | 0.50511263 | 0.61347973 | 0.99999872 |
| RBPMS | 715.09197 | -0.616163954 | 0.23879992 | -2.5802519 | 0.00987283 | 0.70848663 |
| GTF2E2 | 32.9491655 | 0.027984856 | 0.33482367 | 0.08358088 | 0.93338967 | 0.99999872 |
| GSR | 15.510148 | -0.542797999 | 0.42622528 | -1.2735002 | 0.20284058 | 0.99999872 |
| UBXN8 | 3.35903812 | 1.925335902 | 1.00964046 | 1.90695201 | 0.0565268 | NA |
| PPP2CB | 109.061068 | 0.315026654 | 0.24969309 | 1.26165548 | 0.20707278 | 0.99999872 |
| PURG | 6.036959 | 0.77754165 | 0.9047118 | 0.85943574 | 0.39010016 | 0.99999872 |
| WRN | 63.6059265 | -0.111945185 | 0.2508347 | -0.4462907 | 0.6553873 | 0.99999872 |
| NRG1 | 14.126179 | -0.361440425 | 1.08442975 | -0.3333 | 0.73890785 | 0.99999872 |
| FUT10 | 21.6593278 | -0.250227612 | 0.50109139 | -0.4993652 | 0.61752212 | 0.99999872 |
| TTI2 | 7.38372174 | 0.248485874 | 0.51085168 | 0.48641491 | 0.62667302 | 0.99999872 |
| MAK16 | 4.06923364 | -0.775780916 | 0.71622153 | -1.0831578 | 0.2787384 | NA |
| UNC5D | 141.151901 | -1.987456694 | 1.35544267 | -1.4662787 | 0.14257238 | 0.99999872 |
| ZNF703 | 10.0012391 | -0.990001116 | 0.96667735 | -1.0241278 | 0.30577493 | 0.99999872 |
| ERLIN2 | 23.3055639 | -0.238328501 | 0.34681093 | -0.6872001 | 0.49195665 | 0.99999872 |
| PLPBP | 12.7075104 | 0.422763624 | 0.44375304 | 0.95270022 | 0.34074198 | 0.99999872 |
| RAB11FIP1 | 25.9252444 | -0.054564321 | 0.31928382 | -0.170896 | 0.86430556 | 0.99999872 |
| EIF4EBP1 | 9.51965509 | 0.541170084 | 1.09298088 | 0.49513225 | 0.62050677 | 0.99999872 |
| ASH2L | 17.2285928 | -0.317033948 | 0.57395428 | -0.5523679 | 0.58069629 | 0.99999872 |
| LSM1 | 18.9778736 | 0.279265865 | 0.4481351 | 0.62317338 | 0.53317059 | 0.99999872 |
| BAG4 | 15.9770625 | 0.032546968 | 0.473636 | 0.06871726 | 0.94521468 | 0.99999872 |
| DDHD2 | 26.9544737 | -0.580541965 | 0.3943124 | -1.4722894 | 0.14094275 | 0.99999872 |
| PLPP5 | 9.87425764 | 0.06956142 | 0.55790942 | 0.12468228 | 0.90077508 | 0.99999872 |
| NSD3 | 98.3109764 | 0.042535847 | 0.27723831 | 0.15342702 | 0.87806154 | 0.99999872 |
| FGFR1 | 75.8776997 | -0.488516661 | 0.3277849 | -1.4903574 | 0.13613028 | 0.99999872 |
| TACC1 | 50.1660248 | 0.019764281 | 0.51086192 | 0.03868811 | 0.96913905 | 0.99999872 |
| PLEKHA2 | 72.7538215 | -0.158021883 | 0.39894125 | -0.3961031 | 0.69202894 | 0.99999872 |
| AC108863.1 | 5.57205035 | 1.657282864 | 0.93190307 | 1.77838546 | 0.07534057 | 0.99999872 |
| TM2D2 | 13.2665895 | 1.126044774 | 0.65951069 | 1.70739427 | 0.08774879 | 0.99999872 |
| ADAM9 | 50.96813 | 0.558964428 | 0.34194838 | 1.63464565 | 0.10212335 | 0.99999872 |
| ADAM32 | 10.60601 | 0.649276141 | 0.56814625 | 1.14279755 | 0.25312266 | 0.99999872 |
| TCIM | 16.0442002 | -0.188549079 | 0.99227624 | -0.1900167 | 0.84929603 | 0.99999872 |
| SFRP1 | 4.96862757 | -0.72565859 | 0.73271237 | -0.9903731 | 0.32199181 | 0.99999872 |
| GOLGA7 | 21.7626776 | 0.767863489 | 0.3854775 | 1.99198002 | 0.04637325 | 0.96217606 |
| GPAT4 | 20.3502549 | -0.30609934 | 0.37861443 | -0.8084725 | 0.41881866 | 0.99999872 |
| KAT6A | 84.6395208 | 0.273411682 | 0.265996 | 1.02787892 | 0.30400678 | 0.99999872 |

| | | | | | | |
|---|---|---|---|---|---|---|
| AP3M2 | 8.76955146 | 0.481229971 | 0.51733751 | 0.93020507 | 0.35226492 | 0.99999872 |
| IKBKB | 25.0500089 | -0.184704455 | 0.39618702 | -0.4662052 | 0.64106862 | 0.99999872 |
| POLB | 5.41241529 | 0.027177505 | 0.65891561 | 0.04124581 | 0.96709994 | 0.99999872 |
| VDAC3 | 23.9005904 | 0.365562644 | 0.33761371 | 1.08278376 | 0.27890441 | 0.99999872 |
| SLC20A2 | 93.5695786 | -0.890644049 | 0.33974114 | -2.6215372 | 0.00875342 | 0.67516788 |
| SMIM19 | 21.4131402 | 0.075276378 | 0.40810413 | 0.18445385 | 0.85365743 | 0.99999872 |
| THAP1 | 5.12465516 | -0.052951941 | 0.59857771 | -0.0884629 | 0.92950874 | 0.99999872 |
| RNF170 | 29.0264099 | 0.10629014 | 0.33539247 | 0.31691272 | 0.75130983 | 0.99999872 |
| HOOK3 | 47.8343586 | 0.04058771 | 0.3184632 | 0.12744867 | 0.8985853 | 0.99999872 |
| FNTA | 23.8367418 | 0.571046653 | 0.34686614 | 1.64630266 | 0.09970146 | 0.99999872 |
| HGSNAT | 60.8876729 | -0.021667929 | 0.36729985 | -0.0589925 | 0.9529581 | 0.99999872 |
| SPIDR | 199.508834 | -0.387691572 | 0.38439204 | -1.0085838 | 0.31317429 | 0.99999872 |
| CEBPD | 191.68344 | 0.317903563 | 0.6186585 | 0.51385952 | 6.07E-01 | 1.00E+00 |
| PRKDC | 63.8139341 | 0.049651074 | 0.28253479 | 0.17573437 | 0.86050263 | 0.99999872 |
| UBE2V2 | 29.7842581 | 0.12622145 | 0.30283256 | 0.41680277 | 0.67682268 | 0.99999872 |
| PXDNL | 20.3481574 | 0.421759788 | 0.54471891 | 0.7742705 | 0.43877084 | 0.99999872 |
| PCMTD1 | 180.055469 | 0.642701987 | 0.24380775 | 2.63610152 | 0.00838646 | 0.66570732 |
| AC064807.1 | 12.0104032 | 0.282754449 | 0.54008726 | 0.52353475 | 0.60060217 | 0.99999872 |
| RB1CC1 | 76.5389521 | -0.002845491 | 0.30521478 | -0.0093229 | 0.9925615 | 0.99999872 |
| ATP6V1H | 47.7412071 | -0.14320804 | 0.41161233 | -0.3479197 | 0.72790048 | 0.99999872 |
| TCEA1 | 39.3288069 | 0.492516397 | 0.33754189 | 1.45912675 | 0.14453022 | 0.99999872 |
| LYPLA1 | 32.2184127 | 0.043548079 | 0.37648116 | 0.11567134 | 0.90791302 | 0.99999872 |
| MRPL15 | 10.0872834 | -0.408043156 | 0.49297763 | -0.8277113 | 0.40783402 | 0.99999872 |
| XKR4 | 415.746023 | -0.033231097 | 0.92190341 | -0.0360462 | 9.71E-01 | 0.99999872 |
| AC022679.2 | 5.16483001 | 0.525674155 | 1.30930741 | 0.40149025 | 0.68805922 | 0.99999872 |
| TMEM68 | 15.8013686 | 0.311644237 | 0.39086931 | 0.79731059 | 0.42527067 | 0.99999872 |
| TGS1 | 14.5013595 | -0.098751516 | 0.37645759 | -0.2623178 | 0.79307646 | 0.99999872 |
| LYN | 10.1887379 | 0.103351429 | 0.51060975 | 0.20240787 | 0.83959788 | 0.99999872 |
| RPS20 | 41.8821055 | 0.730792774 | 0.41567862 | 1.75807161 | 0.07873533 | 0.99999872 |
| PLAG1 | 14.6560419 | 1.384774265 | 0.61491361 | 2.2519818 | 0.02432342 | 0.88337526 |
| CHCHD7 | 7.42651999 | -0.105321064 | 0.56745015 | -0.1856041 | 0.85275525 | 0.99999872 |
| IMPAD1 | 64.1496562 | 0.324813723 | 0.37501593 | 0.86613314 | 0.38641715 | 0.99999872 |
| FAM110B | 12.447401 | -0.007118736 | 0.78508125 | -0.0090675 | 0.99276527 | 0.99999872 |
| UBXN2B | 27.325505 | 0.10851366 | 0.36823897 | 0.29468271 | 0.76823627 | 0.99999872 |
| SDCBP | 28.9869001 | 0.45556601 | 0.36164274 | 1.25971286 | 0.20777296 | 0.99999872 |
| NSMAF | 26.515127 | -0.702452178 | 0.3191564 | -2.2009653 | 0.02773848 | 0.89154528 |
| TOX | 19.7303667 | 0.610976848 | 1.16602427 | 0.52398296 | 0.60029039 | 0.99999872 |
| LINC01301 | 10.3740155 | 1.376863321 | 1.04536425 | 1.31711345 | 0.1878006 | 0.99999872 |
| RAB2A | 140.161583 | 0.074074269 | 0.2915813 | 0.25404328 | 0.79946213 | 0.99999872 |
| CHD7 | 138.81116 | -0.682527295 | 0.23700523 | -2.8797984 | 0.0039793 | 0.49636972 |
| CLVS1 | 5.02416745 | -0.571618195 | 0.73898241 | -0.7735207 | 0.43921428 | 0.99999872 |
| ASPH | 67.8051456 | 0.151458642 | 0.24922506 | 0.60771836 | 0.54337428 | 0.99999872 |
| YTHDF3 | 48.0461211 | 0.123889517 | 0.28427541 | 0.43580807 | 0.66297599 | 0.99999872 |
| CYP7B1 | 36.9262606 | 0.122322313 | 0.89151803 | 0.13720677 | 0.89086736 | 0.99999872 |
| ARMC1 | 19.4668673 | -0.021998838 | 0.34442373 | -0.0638714 | 0.9490726 | 0.99999872 |
| MTFR1 | 26.2668808 | 0.283636917 | 0.3435734 | 0.82554969 | 0.40905958 | 0.99999872 |
| PDE7A | 27.6807957 | 0.68074432 | 0.5418078 | 1.25643139 | 0.20895961 | 0.99999872 |
| ADHFE1 | 15.6011315 | -0.149350703 | 0.62232597 | -0.2399879 | 0.81033964 | 0.99999872 |
| VCPIP1 | 24.5708633 | 0.466975423 | 0.36788149 | 1.26936372 | 0.20431137 | 0.99999872 |
| SGK3 | 66.2266238 | 0.121912937 | 0.34689509 | 0.35144037 | 0.725258 | 0.99999872 |
| COPS5 | 5.54977174 | 0.718212165 | 0.72156349 | 0.99535547 | 0.31956341 | 0.99999872 |
| CSPP1 | 166.401581 | 0.357749673 | 0.30593785 | 1.16935408 | 0.242261 | 0.99999872 |
| ARFGEF1 | 137.856381 | 0.16966825 | 0.20122263 | 0.84318673 | 0.39912402 | 0.99999872 |
| SULF1 | 348.929933 | -0.038457249 | 0.47059065 | -0.0817212 | 0.93486839 | 0.99999872 |
| NCOA2 | 180.746845 | -0.067960233 | 0.35884938 | -0.1893837 | 0.84979208 | 0.99999872 |
| TRAM1 | 67.7248186 | 0.383211674 | 0.42061093 | 0.91108349 | 0.36225138 | 0.99999872 |
| LACTB2 | 7.59648585 | 0.21727965 | 0.75007258 | 0.28967817 | 0.77206246 | 0.99999872 |
| TERF1 | 35.2676977 | -0.016702679 | 0.33724858 | -0.0495263 | 0.96049987 | 0.99999872 |
| AC022893.1 | 12.1101408 | 0.360594055 | 0.73715569 | 0.48916947 | 0.62472172 | 0.99999872 |
| SBSPON | 158.835466 | -0.068056521 | 0.74189224 | -0.0917337 | 0.92690962 | 0.99999872 |
| RPL7 | 72.2586359 | 1.207602929 | 0.38117375 | 3.16811673 | 0.0015343 | 0.3280895 |

| | | | | | | |
|---|---|---|---|---|---|---|
| RDH10 | 14.2748252 | -0.586892096 | 0.71028447 | -0.8262775 | 0.40864668 | 0.99999872 |
| RDH10-AS1 | 4.64459498 | 0.004939338 | 0.92119872 | 0.00536186 | 0.99572188 | 0.99999872 |
| STAU2 | 69.2322568 | 0.098573049 | 0.36519676 | 0.26991764 | 0.78722361 | 0.99999872 |
| AC027018.1 | 6.24136416 | 1.779237504 | 0.76486986 | 2.32619638 | 0.02000808 | 0.8671024 |
| ELOC | 30.4587757 | 0.219584678 | 0.34001566 | 0.64580755 | 0.518404 | 0.99999872 |
| TMEM70 | 9.41278301 | 0.760532192 | 0.57749454 | 1.31695131 | 0.18785495 | 0.99999872 |
| JPH1 | 34.9104677 | -0.42828884 | 0.57360016 | -0.7466679 | 0.45526404 | 0.99999872 |
| GDAP1 | 20.8635067 | -0.504216806 | 0.49919415 | -1.0100615 | 0.31246581 | 0.99999872 |
| PEX2 | 21.7291974 | 0.395321548 | 0.35570815 | 1.11136489 | 0.26641132 | 0.99999872 |
| ZC2HC1A | 46.8286085 | 0.621672378 | 0.29084508 | 2.1374691 | 0.03255986 | 0.92959829 |
| IL7 | 14.4868541 | 1.065890068 | 0.63985629 | 1.66582728 | 0.09574782 | 0.99999872 |
| TPD52 | 57.3002844 | 0.283549932 | 0.30595048 | 0.92678374 | 0.35403883 | 0.99999872 |
| ZBTB10 | 44.8285512 | 0.569534279 | 0.42424523 | 1.34246479 | 0.17944534 | 0.99999872 |
| ZNF704 | 127.705695 | 0.25341348 | 0.36014484 | 0.70364324 | 0.48165498 | 0.99999872 |
| PAG1 | 8.13270996 | 0.673500278 | 0.8106044 | 0.83086186 | 0.40605167 | 0.99999872 |
| FABP5 | 15.9799592 | 0.382943728 | 0.92430889 | 0.41430276 | 0.67865239 | 0.99999872 |
| IMPA1 | 13.4425747 | 0.142366863 | 0.42987668 | 0.33118071 | 0.74050799 | 0.99999872 |
| ZFAND1 | 14.6817433 | 0.236086547 | 0.41964044 | 0.56259246 | 0.57371243 | 0.99999872 |
| CHMP4C | 13.8581972 | -0.284896052 | 0.42329027 | -0.6730513 | 0.50091468 | 0.99999872 |
| SNX16 | 7.95479168 | 0.94326416 | 0.64105359 | 1.47142794 | 0.14117542 | 0.99999872 |
| RALYL | 14.4943653 | 1.234294481 | 0.68995211 | 1.78895674 | 0.07362178 | 0.99999872 |
| LRRCC1 | 8.31595019 | 0.303358309 | 0.58081692 | 0.52229593 | 0.60146431 | 0.99999872 |
| E2F5 | 10.8058803 | 0.383521431 | 0.48973019 | 0.78312802 | 0.43355194 | 0.99999872 |
| C8orf59 | 18.1231594 | 0.025635165 | 0.47032062 | 0.05450572 | 0.95653225 | 0.99999872 |
| CA13 | 9.63704124 | -0.454645598 | 0.51125095 | -0.8892807 | 0.37385224 | 0.99999872 |
| CA3 | 10.7506973 | 0.091971185 | 0.46639513 | 0.19719585 | 0.84367428 | 0.99999872 |
| CA2 | 32.759413 | 0.499945164 | 0.50823937 | 0.98368051 | 0.32527263 | 0.99999872 |
| WWP1 | 191.486516 | -0.115573899 | 0.3858612 | -0.2995219 | 0.76454183 | 0.99999872 |
| RMDN1 | 33.9760891 | -0.092440759 | 0.36482412 | -0.2533844 | 0.79997115 | 0.99999872 |
| CPNE3 | 40.8648527 | 0.481342355 | 0.3566746 | 1.34952799 | 0.17716744 | 0.99999872 |
| CNGB3 | 24.8904488 | 1.183474998 | 1.09689772 | 1.07892922 | 0.28061928 | 0.99999872 |
| MMP16 | 25.2558835 | -1.381944364 | 0.95067845 | -1.4536401 | 0.1460461 | 0.99999872 |
| AC090568.2 | 9.70812379 | -1.780602983 | 0.92577006 | -1.923375 | 0.05443297 | 0.99999872 |
| AF117829.1 | 80.6695522 | 0.00962363 | 0.39343005 | 0.02446084 | 0.98048502 | 0.99999872 |
| RIPK2 | 7.20196778 | 0.21101388 | 0.57612099 | 0.36626661 | 0.71416614 | 0.99999872 |
| OSGIN2 | 12.4366523 | -0.154900849 | 0.40012535 | -0.3871308 | 0.69865937 | 0.99999872 |
| NBN | 21.0684988 | -0.333865289 | 0.54664237 | -0.6107563 | 0.5413609 | 0.99999872 |
| DECR1 | 30.4967757 | 0.339563044 | 0.4125756 | 0.82303229 | 0.41048963 | 0.99999872 |
| TMEM64 | 47.480919 | 0.325765347 | 0.37338136 | 0.87247352 | 0.38295011 | 0.99999872 |
| NECAB1 | 250.945502 | 0.332422059 | 0.44682279 | 0.74396845 | 0.45689557 | 0.99999872 |
| AC103770.1 | 4.42222165 | 2.688710879 | 1.02306762 | 2.62808716 | 0.00858665 | NA |
| PIP4P2 | 17.2175031 | 0.459912648 | 0.43593051 | 1.05501367 | 0.29141907 | 0.99999872 |
| OTUD6B-AS1 | 21.8196499 | 0.578704951 | 0.42527348 | 1.36078307 | 0.17358225 | 0.99999872 |
| OTUD6B | 11.5621921 | 0.417422994 | 0.48114219 | 0.86756682 | 0.38563152 | 0.99999872 |
| LRRC69 | 14.7074586 | 0.602094615 | 0.40805065 | 1.47553893 | 0.14006771 | 0.99999872 |
| RUNX1T1 | 32.6337754 | 0.717302563 | 0.45097148 | 1.59057191 | 0.11170595 | 0.99999872 |
| AC117834.1 | 5.31920674 | 1.294143886 | 0.6964804 | 1.85811961 | 0.06315203 | 0.99999872 |
| TRIQK | 38.121943 | 0.746784262 | 0.3185213 | 2.34453477 | 0.01905083 | 0.8671024 |
| LINC00535 | 10.1873729 | -0.333696917 | 0.90926589 | -0.366996 | 0.71362203 | 0.99999872 |
| FAM92A | 11.5749969 | 0.600607997 | 0.52039832 | 1.15413132 | 0.24844634 | 0.99999872 |
| RBM12B | 18.1350001 | 0.139224212 | 0.51760655 | 0.26897691 | 0.78794745 | 0.99999872 |
| TMEM67 | 25.2031099 | -0.114792234 | 0.35414456 | -0.3241395 | 0.74583244 | 0.99999872 |
| PDP1 | 4.62930778 | 1.214832437 | 0.78415716 | 1.54922062 | 0.12132869 | 0.99999872 |
| VIRMA | 26.8238674 | 0.020581871 | 0.30214145 | 0.06811999 | 0.94569012 | 0.99999872 |
| ESRP1 | 8.2430641 | 0.251429277 | 0.73563036 | 0.34178752 | 0.7325108 | 0.99999872 |
| DPY19L4 | 34.1272377 | -0.134493063 | 0.4441724 | -0.3027947 | 0.7620463 | 0.99999872 |
| INTS8 | 25.144642 | -0.070301042 | 0.39688505 | -0.177132 | 0.85940471 | 0.99999872 |
| NDUFAF6 | 25.9729765 | -0.074753715 | 0.47112251 | -0.1586715 | 0.87392769 | 0.99999872 |
| TP53INP1 | 20.6859303 | 0.229569376 | 0.70787549 | 0.32430757 | 0.74570519 | 0.99999872 |
| PLEKHF2 | 10.6325127 | 0.263352583 | 0.61321795 | 0.42946 | 0.66758849 | 0.99999872 |
| C8orf37-AS1 | 14.5880908 | 1.075615209 | 0.82405128 | 1.30527704 | 0.19179853 | 0.99999872 |

| | | | | | | |
|---|---|---|---|---|---|---|
| UQCRB | 36.3474702 | 0.402611882 | 0.38605364 | 1.04289104 | 0.29699876 | 0.99999872 |
| MTERF3 | 7.77944492 | 0.711901043 | 0.60517972 | 1.17634649 | 0.23945644 | 0.99999872 |
| PTDSS1 | 27.4958403 | 0.08997617 | 0.43355562 | 0.20753086 | 0.8355953 | 0.99999872 |
| SDC2 | 110.533393 | 0.041388108 | 0.44602402 | 0.09279345 | 0.92606765 | 0.99999872 |
| CPQ | 138.39815 | 0.165931949 | 0.25985961 | 0.63854459 | 0.52311924 | 0.99999872 |
| TSPYL5 | 16.1384419 | -0.213740186 | 0.46117087 | -0.4634729 | 0.64302546 | 0.99999872 |
| MTDH | 138.782679 | -0.02924004 | 0.23793093 | -0.122893 | 0.90219185 | 0.99999872 |
| LAPTM4B | 37.1838646 | 0.420294167 | 0.443963 | 0.94668737 | 0.34379811 | 0.99999872 |
| MATN2 | 8.89645458 | -0.04842879 | 1.07330995 | -0.045121 | 0.96401088 | 0.99999872 |
| RPL30 | 78.769612 | 0.616439834 | 0.34727866 | 1.77505818 | 0.07588828 | 0.99999872 |
| RIDA | 5.73879391 | 0.235778819 | 0.6641681 | 0.35499871 | 0.72259056 | 0.99999872 |
| POP1 | 4.46737677 | 0.124116015 | 0.72054787 | 0.17225228 | 0.8632392 | NA |
| NIPAL2 | 16.2396286 | -0.007399081 | 0.61663735 | -0.0119991 | 0.99042635 | 0.99999872 |
| STK3 | 122.03985 | -0.312906532 | 0.32276394 | -0.9694594 | 0.33231603 | 0.99999872 |
| AC104986.2 | 19.8791373 | 0.09631336 | 0.36773904 | 0.26190681 | 0.79339329 | 0.99999872 |
| VPS13B | 178.33244 | -0.003183249 | 0.1914562 | -0.0166265 | 0.98673457 | 0.99999872 |
| COX6C | 65.545992 | 0.677235552 | 0.38870678 | 1.74227873 | 0.08145968 | 0.99999872 |
| POLR2K | 28.6491429 | 0.87754061 | 0.47534342 | 1.84611919 | 0.0648749 | 0.99999872 |
| SPAG1 | 13.0793827 | -0.064184211 | 0.68202037 | -0.0941089 | 0.92502262 | 0.99999872 |
| RNF19A | 109.993182 | -0.256462865 | 0.31901485 | -0.8039214 | 0.42144238 | 0.99999872 |
| ANKRD46 | 16.1533659 | 0.138041565 | 0.38843893 | 0.35537521 | 0.72230852 | 0.99999872 |
| PABPC1 | 67.2367509 | -0.298467366 | 0.34755265 | -0.8587688 | 0.39046809 | 0.99999872 |
| YWHAZ | 135.146245 | 0.32213187 | 0.25399399 | 1.26826573 | 0.20470307 | 0.99999872 |
| ZNF706 | 36.1593042 | 0.575071263 | 0.28694157 | 2.00414063 | 0.045055 | 0.95980711 |
| RRM2B | 16.7544448 | 0.312182686 | 0.39122461 | 0.79796281 | 0.42489207 | 0.99999872 |
| UBR5 | 126.667634 | 0.266932535 | 0.2534834 | 1.05305726 | 0.29231475 | 0.99999872 |
| KLF10 | 36.4332789 | 0.178542978 | 0.67573731 | 0.2642195 | 0.79161078 | 0.99999872 |
| AZIN1 | 23.9135337 | -0.182013761 | 0.35719832 | -0.5095594 | 0.61036017 | 0.99999872 |
| AZIN1-AS1 | 35.7621208 | -0.713165601 | 0.37157688 | -1.9192949 | 0.05494703 | 0.99999872 |
| ATP6V1C1 | 33.3856491 | 0.289333773 | 0.34065073 | 0.84935609 | 0.39568318 | 0.99999872 |
| BAALC-AS1 | 16.0086956 | -0.284070692 | 0.50324905 | -0.5644734 | 0.57243201 | 0.99999872 |
| FZD6 | 28.3021125 | -0.170418183 | 0.32399894 | -0.5259838 | 0.59889948 | 0.99999872 |
| SLC25A32 | 18.2112379 | 0.244113734 | 0.4078796 | 0.59849458 | 0.54950997 | 0.99999872 |
| DCAF13 | 8.29853268 | -0.40918895 | 0.55559182 | -0.736492 | 0.46143131 | 0.99999872 |
| DPYS | 20.9397019 | 1.714661188 | 1.93234435 | 0.88734764 | 0.37489177 | 0.99999872 |
| LRP12 | 21.747891 | -0.136176361 | 0.49875802 | -0.2730309 | 0.78482945 | 0.99999872 |
| ZFPM2 | 159.781915 | 0.173183918 | 0.43526768 | 0.39787911 | 0.6907193 | 0.99999872 |
| OXR1 | 118.233837 | -0.257786104 | 0.43998559 | -0.5858967 | 0.55794494 | 0.99999872 |
| AC090579.1 | 13.6072143 | -0.127735092 | 0.83012078 | -0.1538753 | 0.87770805 | 0.99999872 |
| EIF3E | 43.3609267 | 0.264782332 | 0.36191499 | 0.73161471 | 0.46440377 | 0.99999872 |
| EMC2 | 42.0496261 | 0.220395318 | 0.3450722 | 0.63869335 | 0.52302244 | 0.99999872 |
| NUDCD1 | 46.6166351 | -0.024664437 | 0.3754253 | -0.0656973 | 0.9476188 | 0.99999872 |
| ENY2 | 36.3621657 | 0.547572482 | 0.38896878 | 1.40775431 | 0.15920384 | 0.99999872 |
| EBAG9 | 16.395967 | 0.385190099 | 0.43445002 | 0.88661546 | 0.37528597 | 0.99999872 |
| SYBU | 14.2076455 | -0.169945997 | 0.45232681 | -0.3757151 | 0.70712874 | 0.99999872 |
| TRPS1 | 120.496364 | 0.345044446 | 0.37429836 | 0.92184332 | 0.35661031 | 0.99999872 |
| EIF3H | 53.3793518 | 0.302469734 | 0.3539387 | 0.85458226 | 0.39278245 | 0.99999872 |
| UTP23 | 11.0269849 | 0.285273513 | 0.62120922 | 0.45922293 | 0.64607409 | 0.99999872 |
| RAD21 | 55.2777834 | 0.373641011 | 0.39083837 | 0.95599878 | 0.33907285 | 0.99999872 |
| MED30 | 10.5933948 | 0.256035763 | 0.60435754 | 0.42364949 | 0.67182146 | 0.99999872 |
| EXT1 | 106.474646 | 0.08923031 | 0.49502085 | 0.18025566 | 0.85695186 | 0.99999872 |
| SAMD12 | 71.6331466 | -0.119610524 | 0.2838834 | -0.4213368 | 0.67350916 | 0.99999872 |
| AC023590.1 | 6.14961211 | 1.881913047 | 1.1052383 | 1.70272154 | 0.08862019 | 0.99999872 |
| TNFRSF11B | 13.6527595 | -1.0246414 | 0.6976809 | -1.468639 | 0.14193072 | 0.99999872 |
| MAL2 | 164.215464 | -0.169847679 | 0.54697721 | -0.3105206 | 0.75616512 | 0.99999872 |
| MAL2-AS1 | 5.79902909 | 1.884426527 | 0.89868448 | 2.09687223 | 0.03600489 | 0.94734101 |
| TAF2 | 36.1151875 | -0.228837681 | 0.3525095 | -0.6491674 | 0.51623017 | 0.99999872 |
| DEPTOR | 69.1732524 | -0.524253211 | 0.40305134 | -1.3007107 | 0.19335748 | 0.99999872 |
| AC091563.1 | 7.4173957 | 0.121088773 | 0.68039085 | 0.17796943 | 0.85874698 | 0.99999872 |
| MRPL13 | 13.9331029 | 1.190543068 | 0.51289496 | 2.32122199 | 0.02027487 | 0.8671024 |
| MTBP | 5.96237052 | -0.85881298 | 0.67937227 | -1.2641272 | 0.20618437 | 0.99999872 |

| | | | | | | |
|---|---|---|---|---|---|---|
| SNTB1 | 13.0069209 | -0.721952685 | 0.81904279 | -0.8814591 | 0.37806941 | 0.99999872 |
| ZHX2 | 67.0274428 | -0.239044806 | 0.33848656 | -0.7062165 | 0.48005349 | 0.99999872 |
| DERL1 | 17.7229556 | 0.262483406 | 0.4388294 | 0.59814453 | 0.5497435 | 0.99999872 |
| C8orf76 | 4.64381449 | 0.128388616 | 0.71724363 | 0.1790028 | 0.8579355 | 0.99999872 |
| WDYHV1 | 8.70323988 | 0.416957568 | 0.56157051 | 0.7424848 | 0.45779366 | 0.99999872 |
| FBXO32 | 8.09795858 | -0.12936776 | 0.67835521 | -0.190708 | 0.84875438 | 0.99999872 |
| FAM91A1 | 29.3155951 | 0.133764271 | 0.30838058 | 0.4337636 | 0.66446012 | 0.99999872 |
| TMEM65 | 26.278467 | 0.084575441 | 0.47025791 | 0.17984906 | 0.85727107 | 0.99999872 |
| RNF139 | 11.5393797 | 0.619118833 | 0.49191389 | 1.25859188 | 0.20817778 | 0.99999872 |
| TATDN1 | 25.3358992 | 0.425699006 | 0.43874516 | 0.97026485 | 0.3319145 | 0.99999872 |
| NDUFB9 | 47.8458747 | 0.615393943 | 0.2842842 | 2.16471383 | 0.03040961 | 0.91298529 |
| MTSS1 | 776.138615 | -0.794127288 | 0.46569761 | -1.7052423 | 0.08814923 | 0.99999872 |
| SQLE | 22.9420382 | -0.224687015 | 0.55701741 | -0.4033752 | 0.68667222 | 0.99999872 |
| WASHC5 | 20.043303 | 0.229580344 | 0.42389221 | 0.54160076 | 0.58809357 | 0.99999872 |
| NSMCE2 | 102.154003 | -0.291299389 | 0.33708967 | -0.86416 | 0.38750003 | 0.99999872 |
| FAM84B | 5.88480343 | -1.243702392 | 0.61430606 | -2.0245648 | 0.04291208 | 0.95980711 |
| PCAT1 | 30.5380253 | -1.169126288 | 0.69835379 | -1.6741175 | 0.09410751 | 0.99999872 |
| MYC | 6.23653876 | -0.440393742 | 2.30809332 | -0.1908041 | 0.84867905 | 0.99999872 |
| PVT1 | 14.9232911 | -0.130371774 | 0.96752016 | -0.1347484 | 0.89281082 | 0.99999872 |
| FAM49B | 61.8736282 | -0.230322411 | 0.23724257 | -0.9708309 | 0.33163251 | 0.99999872 |
| ASAP1 | 63.4718902 | -0.023720964 | 0.43370296 | -0.054694 | 0.95638222 | 0.99999872 |
| EFR3A | 63.6417733 | -0.054124097 | 0.24298987 | -0.2227422 | 0.82373616 | 0.99999872 |
| LRRC6 | 19.7959016 | 0.829945357 | 0.45986538 | 1.80475719 | 0.07111269 | 0.99999872 |
| PHF20L1 | 43.7056059 | 0.188825627 | 0.31995027 | 0.59017181 | 0.55507547 | 0.99999872 |
| NDRG1 | 183.593941 | 0.068652701 | 0.34024767 | 0.20177273 | 0.8400944 | 0.99999872 |
| ST3GAL1 | 186.87957 | -0.937347424 | 0.44101857 | -2.1254149 | 0.03355201 | 0.93004754 |
| ZFAT | 19.2100001 | -0.720509492 | 0.55254394 | -1.3039859 | 0.1922384 | 0.99999872 |
| AC083843.3 | 6.4049703 | -0.421349162 | 0.64424521 | -0.6540199 | 0.51309901 | 0.99999872 |
| KHDRBS3 | 14.1892639 | -0.018291206 | 0.54206813 | -0.0337434 | 0.97308179 | 0.99999872 |
| TRAPPC9 | 127.809167 | -0.067948349 | 0.29027757 | -0.2340806 | 0.8149224 | 0.99999872 |
| CHRAC1 | 16.738475 | -0.233693454 | 0.58280331 | -0.4009817 | 0.68843361 | 0.99999872 |
| AGO2 | 48.5398193 | 0.224613389 | 0.43561097 | 0.5156284 | 0.60611397 | 0.99999872 |
| PTK2 | 241.941813 | -0.162528967 | 0.21137507 | -0.7689127 | 0.44194516 | 0.99999872 |
| DENND3 | 6.52891255 | -0.079806527 | 0.75698323 | -0.1054271 | 0.91603693 | 0.99999872 |
| SLC45A4 | 12.2764527 | -0.47617742 | 0.61908279 | -0.769166 | 0.44179478 | 0.99999872 |
| PTP4A3 | 4.81182064 | -0.510234831 | 0.9307443 | -0.5482009 | 0.58355399 | 0.99999872 |
| TSNARE1 | 10.8490942 | -0.422166763 | 0.61012912 | -0.6919302 | 0.48898117 | 0.99999872 |
| JRK | 11.3412453 | 0.43038349 | 0.59629944 | 0.72175733 | 0.47044369 | 0.99999872 |
| THEM6 | 6.9721571 | -0.648336295 | 0.70479766 | -0.91989 | 0.35763026 | 0.99999872 |
| LY6E | 160.332784 | -0.880628578 | 1.02494065 | -0.8591996 | 0.39023042 | 0.99999872 |
| GLI4 | 5.34343687 | -0.20719106 | 0.61480135 | -0.3370049 | 0.73611322 | 0.99999872 |
| RHPN1 | 14.0720535 | 0.102363249 | 0.46288522 | 0.22114175 | 0.82498206 | 0.99999872 |
| ZC3H3 | 10.487616 | -0.189547377 | 0.51370642 | -0.36898 | 0.71214266 | 0.99999872 |
| GSDMD | 18.9386944 | 0.080539497 | 0.61277533 | 0.13143397 | 0.89543201 | 0.99999872 |
| EEF1D | 49.7886491 | -0.259271679 | 0.40018021 | -0.6478873 | 0.51705784 | 0.99999872 |
| TIGD5 | 4.34247763 | 0.828960854 | 0.71456858 | 1.16008578 | 0.24601388 | NA |
| TSTA3 | 6.86698036 | 0.650640751 | 0.56335006 | 1.15494929 | 0.24811119 | 0.99999872 |
| ZNF623 | 9.47663987 | -0.081029184 | 0.57277272 | -0.1414683 | 0.8875 | 0.99999872 |
| FAM83H | 7.59776949 | -0.800083346 | 0.53565335 | -1.4936588 | 0.13526482 | 0.99999872 |
| SCRIB | 5.35549919 | -0.240915722 | 0.63277686 | -0.3807278 | 0.70340526 | 0.99999872 |
| PUF60 | 24.1566977 | 0.026469396 | 0.47341235 | 0.05591193 | 0.95541197 | 0.99999872 |
| NRBP2 | 17.1088654 | -0.022705938 | 0.45263641 | -0.0501637 | 0.9599919 | 0.99999872 |
| PLEC | 73.2606914 | -0.439365107 | 0.48640596 | -0.9032889 | 0.36637259 | 0.99999872 |
| PARP10 | 14.5845765 | -1.088041 | 0.96686234 | -1.1253319 | 0.26044843 | 0.99999872 |
| GRINA | 24.5407705 | 0.485285909 | 0.64151879 | 0.75646406 | 0.44937101 | 0.99999872 |
| EXOSC4 | 6.40521368 | 0.885949351 | 0.75724934 | 1.16995724 | 0.24201818 | 0.99999872 |
| GPAA1 | 25.1174098 | 0.791239074 | 0.67159383 | 1.1781512 | 0.23873633 | 0.99999872 |
| CYC1 | 18.4805407 | 1.607129186 | 0.6537378 | 2.45836968 | 0.01395694 | 0.75772455 |
| SHARPIN | 13.4189965 | 0.134122076 | 0.45097741 | 0.29740309 | 0.76615878 | 0.99999872 |
| MAF1 | 25.2430727 | 0.282940671 | 0.42084956 | 0.67230835 | 0.50138742 | 0.99999872 |
| HGH1 | 4.75864144 | -0.450817143 | 0.70647007 | -0.6381263 | 0.52339146 | 0.99999872 |

| | | | | | | |
|---|---|---|---|---|---|---|
| MROH1 | 46.9887967 | -0.696635145 | 0.33231375 | -2.0963175 | 0.03605403 | 0.94734101 |
| HSF1 | 21.8470607 | -0.255789062 | 0.37214595 | -0.6873353 | 0.49187143 | 0.99999872 |
| DGAT1 | 7.57198408 | 0.380608568 | 0.5617017 | 0.6775991 | 0.49802592 | 0.99999872 |
| SLC52A2 | 8.50787147 | 0.460611867 | 0.65622916 | 0.70190704 | 0.48273714 | 0.99999872 |
| CPSF1 | 5.46874191 | -0.741837148 | 0.62996489 | -1.1775849 | 0.23896213 | 0.99999872 |
| VPS28 | 47.8100354 | 0.465645958 | 0.35330136 | 1.31798517 | 0.18750861 | 0.99999872 |
| CYHR1 | 26.2820135 | -0.188777432 | 0.31050001 | -0.6079788 | 0.54320152 | 0.99999872 |
| KIFC2 | 7.66446772 | -0.024885102 | 0.57338426 | -0.0434004 | 0.96538236 | 0.99999872 |
| PPP1R16A | 18.5475951 | -0.127567547 | 0.44772327 | -0.284925 | 0.77570161 | 0.99999872 |
| LRRC14 | 6.25061813 | -0.279446112 | 0.64118856 | -0.4358252 | 0.66296358 | 0.99999872 |
| ARHGAP39 | 4.9843388 | 0.281896701 | 0.65083727 | 0.43312932 | 0.66492083 | 0.99999872 |
| ZNF251 | 23.047593 | -0.551319764 | 0.50561195 | -1.090401 | 0.27553655 | 0.99999872 |
| RPL8 | 100.434597 | 0.578270336 | 0.42119679 | 1.37292198 | 0.1697766 | 0.99999872 |
| ZNF7 | 8.36293239 | 0.45882332 | 0.70644766 | 0.64947957 | 0.51602845 | 0.99999872 |
| COMMD5 | 10.2270335 | 0.605612866 | 0.63824636 | 0.94887006 | 0.3426867 | 0.99999872 |
| ZNF250 | 16.8361382 | 1.135720269 | 0.70849181 | 1.60301115 | 0.10893219 | 0.99999872 |
| ZNF16 | 15.642208 | -0.030697405 | 0.51053947 | -0.0601274 | 0.95205418 | 0.99999872 |
| C8orf33 | 12.3823854 | -0.261313959 | 0.45818752 | -0.570321 | 0.56846001 | 0.99999872 |
| WASHC1 | 122.08578 | -0.395393806 | 0.29853709 | -1.3244378 | 0.18535768 | 0.99999872 |
| CBWD1 | 9.44952944 | 0.192784292 | 0.47717095 | 0.40401515 | 0.68620158 | 0.99999872 |
| DOCK8 | 11.4312123 | -0.454348222 | 0.80990253 | -0.5609912 | 0.57480351 | 0.99999872 |
| KANK1 | 614.52349 | 0.090004392 | 0.42589908 | 0.21132798 | 0.83263135 | 0.99999872 |
| SMARCA2 | 88.7726429 | -0.299389367 | 0.21952344 | -1.363815 | 0.17262579 | 0.99999872 |
| VLDLR-AS1 | 11.4314277 | -0.862923483 | 0.7506425 | -1.1495798 | 2.50E-01 | 0.99999872 |
| VLDLR | 8.94617284 | 0.801799249 | 0.83535047 | 0.95983576 | 0.33713788 | 0.99999872 |
| PUM3 | 49.9811358 | -0.495483184 | 0.43638434 | -1.1354284 | 0.25619586 | 0.99999872 |
| RFX3 | 95.5141491 | 0.379390913 | 0.50047462 | 0.75806224 | 0.44841372 | 0.99999872 |
| RFX3-AS1 | 62.3882339 | -0.806367251 | 0.46357648 | -1.7394481 | 0.08195597 | 0.99999872 |
| GLIS3 | 80.1347714 | -0.489482497 | 0.65968 | -0.7419999 | 0.45808739 | 0.99999872 |
| SLC1A1 | 23.3280159 | -0.52931098 | 0.9899524 | -0.5346833 | 0.5928689 | 0.99999872 |
| SPATA6L | 22.8838725 | -0.047505238 | 0.59070333 | -0.0804215 | 0.93590204 | 0.99999872 |
| CDC37L1 | 19.5442961 | 0.789617094 | 0.58851948 | 1.34170085 | 0.17969301 | 0.99999872 |
| AK3 | 64.89062 | 0.301598976 | 0.28460008 | 1.05972905 | 0.28926788 | 0.99999872 |
| RCL1 | 23.9893425 | -0.178062758 | 0.46917507 | -0.3795231 | 0.70429948 | 0.99999872 |
| JAK2 | 92.1952331 | -0.265721487 | 0.36127771 | -0.7355048 | 0.46203213 | 0.99999872 |
| PLGRKT | 14.1546059 | -0.202586598 | 0.65713935 | -0.3082856 | 0.75786502 | 0.99999872 |
| CD274 | 6.2552131 | -0.933295768 | 1.28656328 | -0.7254177 | 0.46819582 | 0.99999872 |
| RIC1 | 74.3015134 | 0.006623269 | 0.31968102 | 0.02071837 | 0.98347032 | 0.99999872 |
| ERMP1 | 38.2979371 | -0.200566878 | 0.47007129 | -0.4266733 | 0.66961729 | 0.99999872 |
| KIAA2026 | 81.7235426 | -0.034374491 | 0.30181907 | -0.1138911 | 0.90932416 | 0.99999872 |
| RANBP6 | 8.29388861 | 0.464314264 | 0.49910866 | 0.93028693 | 0.35222254 | 0.99999872 |
| UHRF2 | 44.4193211 | -0.365763845 | 0.34062025 | -1.0738171 | 0.28290467 | 0.99999872 |
| GLDC | 8.71583943 | -0.016627039 | 0.8491975 | -0.0195797 | 0.98437865 | 0.99999872 |
| AL162411.1 | 8.08468361 | 0.589615167 | 0.74082704 | 0.79588775 | 0.42609728 | 0.99999872 |
| KDM4C | 122.97996 | 0.211039837 | 0.21367944 | 0.98764692 | 0.3233256 | 0.99999872 |
| DMAC1 | 27.1971977 | 0.816958928 | 0.35560519 | 2.29737627 | 0.02159731 | 0.87186485 |
| PTPRD | 910.861167 | 0.181813167 | 0.42411777 | 0.42868556 | 0.66815207 | 0.99999872 |
| MPDZ | 88.2422259 | -0.221884455 | 0.44231068 | -0.5016484 | 0.61591485 | 0.99999872 |
| NFIB | 97.1371064 | -0.427365493 | 0.6076993 | -0.7032516 | 0.48189898 | 0.99999872 |
| ZDHHC21 | 87.2895934 | 0.246862463 | 0.28109762 | 0.87820901 | 0.3798303 | 0.99999872 |
| FREM1 | 11.3638272 | -0.44681805 | 0.91082301 | -0.4905652 | 0.62373402 | 0.99999872 |
| TTC39B | 43.364106 | -0.245689211 | 0.69103697 | -0.355537 | 0.72218733 | 0.99999872 |
| SNAPC3 | 35.4452991 | 0.312564404 | 0.31404447 | 0.99528707 | 0.31959667 | 0.99999872 |
| PSIP1 | 39.4327891 | 0.140225268 | 0.33803666 | 0.41482267 | 0.67827172 | 0.99999872 |
| CCDC171 | 30.3383348 | -0.484152207 | 0.58623542 | -0.8258665 | 0.40887982 | 0.99999872 |
| BNC2 | 35.5771569 | -1.022545059 | 0.75327316 | -1.3574691 | 0.17463219 | 0.99999872 |
| CNTLN | 54.174515 | 0.150943614 | 0.34862612 | 0.43296702 | 0.66503873 | 0.99999872 |
| SH3GL2 | 7.47082583 | -1.351038218 | 0.73589103 | -1.8359216 | 0.06636926 | 0.99999872 |
| RRAGA | 21.3443816 | 0.24493382 | 0.38374359 | 0.63827468 | 0.52329489 | 0.99999872 |
| HAUS6 | 9.13147802 | 0.027941893 | 0.49306843 | 0.0566694 | 0.95480855 | 0.99999872 |
| PLIN2 | 21.9338796 | -0.319578516 | 0.94393176 | -0.338561 | 0.73494044 | 0.99999872 |

| | | | | | | |
|---|---|---|---|---|---|---|
| DENND4C | 81.9064838 | 0.02920296 | 0.25601452 | 0.1140676 | 0.90918421 | 0.99999872 |
| RPS6 | 102.64152 | 0.693040695 | 0.46816412 | 1.48033707 | 0.13878331 | 0.99999872 |
| ACER2 | 11.0077779 | 0.123756234 | 0.97752047 | 0.12660219 | 0.89925526 | 0.99999872 |
| MLLT3 | 87.6663224 | 0.377791956 | 0.50836849 | 0.7431459 | 0.45739335 | 0.99999872 |
| FOCAD | 58.5485472 | 0.183912759 | 0.31955797 | 0.57552236 | 0.56493807 | 0.99999872 |
| HACD4 | 9.70295665 | 0.692521354 | 0.52613762 | 1.31623613 | 0.1880948 | 0.99999872 |
| KLHL9 | 30.0145257 | 0.130812352 | 0.38777164 | 0.33734378 | 0.73585776 | 0.99999872 |
| AL391117.1 | 12.350635 | 0.593103737 | 1.2147973 | 0.48823268 | 0.62538504 | 0.99999872 |
| TUSC1 | 6.7647266 | 0.377764015 | 0.60210058 | 0.62741015 | 0.53039042 | 0.99999872 |
| CAAP1 | 26.168412 | 0.075196616 | 0.34688466 | 0.216777 | 0.82838214 | 0.99999872 |
| PLAA | 21.6422501 | 0.092418618 | 0.33318811 | 0.2773767 | 0.78149087 | 0.99999872 |
| IFT74 | 30.9761443 | 0.346382256 | 0.36146829 | 0.95826458 | 0.33792936 | 0.99999872 |
| LRRC19 | 3.70249531 | 1.124374761 | 1.07167832 | 1.04917188 | 0.29409902 | NA |
| MOB3B | 16.3164606 | -0.232257027 | 0.52682791 | -0.4408594 | 0.6593148 | 0.99999872 |
| C9orf72 | 8.1320202 | 0.422400001 | 0.56094079 | 0.75302066 | 0.4514375 | 0.99999872 |
| LINGO2 | 34.2099262 | -0.754424497 | 0.61620993 | -1.2242978 | 0.22083989 | 0.99999872 |
| ACO1 | 24.5629327 | 0.73231912 | 0.48054039 | 1.52394917 | 0.12752141 | 0.99999872 |
| DDX58 | 48.3289434 | -0.736910453 | 1.04489435 | -0.7052488 | 0.48065544 | 0.99999872 |
| TOPORS | 16.4011166 | 0.47847932 | 0.39152463 | 1.22209253 | 0.22167264 | 0.99999872 |
| SMIM27 | 4.5728564 | 1.458452059 | 0.79434572 | 1.83604195 | 0.06635145 | NA |
| NDUFB6 | 21.1480194 | 0.595327851 | 0.37210127 | 1.59990816 | 0.10961896 | 0.99999872 |
| APTX | 24.6865517 | -0.027919048 | 0.3727048 | -0.0749093 | 0.94028689 | 0.99999872 |
| DNAJA1 | 58.2008659 | 0.353491742 | 0.47331503 | 0.74684242 | 0.45515869 | 0.99999872 |
| SMU1 | 19.4251017 | 0.444757391 | 0.37198291 | 1.19563932 | 0.23183734 | 0.99999872 |
| B4GALT1 | 33.3842383 | -0.17210767 | 0.35416775 | -0.4859496 | 0.62700291 | 0.99999872 |
| BAG1 | 24.6241088 | -0.208490012 | 0.33506713 | -0.6222335 | 0.53378831 | 0.99999872 |
| CHMP5 | 35.7491837 | 0.422797545 | 0.46188393 | 0.91537618 | 0.35999418 | 0.99999872 |
| NFX1 | 43.0475558 | -0.65514044 | 0.32270343 | -2.0301626 | 0.04234002 | 0.95980711 |
| AQP3 | 27.2721825 | -0.002015205 | 0.46623858 | -0.0043223 | 0.99655135 | 0.99999872 |
| UBE2R2 | 114.130643 | -0.151278766 | 0.31117368 | -0.4861554 | 0.62685699 | 0.99999872 |
| UBAP2 | 102.188289 | -0.028078704 | 0.38131231 | -0.073637 | 0.94129921 | 0.99999872 |
| DCAF12 | 12.1007906 | -0.034809605 | 0.42573716 | -0.0817631 | 0.93483507 | 0.99999872 |
| UBAP1 | 65.809084 | 0.10389219 | 0.27901279 | 0.37235637 | 0.70962753 | 0.99999872 |
| NUDT2 | 8.35352728 | 0.511427515 | 0.60403314 | 0.84668784 | 0.39716914 | 0.99999872 |
| FAM219A | 25.3896716 | 0.157033236 | 0.3692296 | 0.42529969 | 0.67061822 | 0.99999872 |
| CNTFR | 6.47319706 | -1.40179044 | 0.81718235 | -1.7153949 | 0.08627285 | 0.99999872 |
| RPP25L | 7.29641972 | -0.801513836 | 0.63177759 | -1.2686646 | 0.20456073 | 0.99999872 |
| DCTN3 | 17.7474868 | 0.855424182 | 0.50093671 | 1.70764922 | 0.08770145 | 0.99999872 |
| SIGMAR1 | 6.00672526 | -0.049646972 | 0.62267784 | -0.0797314 | 0.93645089 | 0.99999872 |
| IL11RA | 7.45528036 | 0.388266937 | 0.57194847 | 0.6788495 | 0.49723322 | 0.99999872 |
| VCP | 65.4271108 | -0.085489926 | 0.37583419 | -0.2274671 | 0.82006052 | 0.99999872 |
| PIGO | 8.34617309 | 0.27698175 | 0.62625886 | 0.44227997 | 0.65828662 | 0.99999872 |
| STOML2 | 14.5013123 | 0.445552435 | 0.56555032 | 0.78782102 | 0.4308014 | 0.99999872 |
| FAM214B | 7.48375071 | -0.096022841 | 0.64222776 | -0.1495152 | 0.88114708 | 0.99999872 |
| UNC13B | 48.9415819 | -0.122840402 | 0.27323046 | -0.4495853 | 0.65300948 | 0.99999872 |
| RUSC2 | 87.1820838 | -0.295238015 | 0.4641275 | -0.636114 | 0.5247021 | 0.99999872 |
| TESK1 | 9.89028549 | 0.055384575 | 0.48274728 | 0.11472789 | 0.90866081 | 0.99999872 |
| CCDC107 | 24.9303339 | 0.528645493 | 0.48624339 | 1.08720345 | 0.27694691 | 0.99999872 |
| TPM2 | 80.5842162 | 0.498266538 | 0.5312408 | 0.93792973 | 0.34828052 | 0.99999872 |
| TLN1 | 77.5740094 | 0.153479942 | 0.38815658 | 0.3954073 | 0.69254233 | 0.99999872 |
| CREB3 | 12.4927719 | 0.476996728 | 0.5517153 | 0.86457042 | 0.38727462 | 0.99999872 |
| GBA2 | 11.8765448 | -0.356333516 | 0.41370883 | -0.8613147 | 0.38906472 | 0.99999872 |
| RGP1 | 7.97166561 | -0.634070511 | 0.48772735 | -1.3000512 | 0.19358343 | 0.99999872 |
| HINT2 | 9.86665073 | -0.110435188 | 0.71285752 | -0.154919 | 0.87688516 | 0.99999872 |
| TMEM8B | 9.02907576 | 0.254723353 | 0.63879656 | 0.39875505 | 0.6900737 | 0.99999872 |
| RECK | 29.7374912 | 0.424072349 | 0.36049985 | 1.17634542 | 0.23945687 | 0.99999872 |
| CLTA | 87.3694748 | -0.288205445 | 0.33475171 | -0.8609529 | 0.389264 | 0.99999872 |
| GNE | 21.4128643 | -0.028677598 | 0.51000637 | -0.0562299 | 0.95515868 | 0.99999872 |
| RNF38 | 91.7038006 | -0.122564878 | 0.23752988 | -0.5159977 | 0.60585599 | 0.99999872 |
| EBLN3P | 28.344347 | 0.646975397 | 0.32517025 | 1.98965126 | 0.04662937 | 0.96425074 |
| ZCCHC7 | 136.983334 | 0.00080692 | 0.26149975 | 0.00308574 | 0.99753794 | 0.99999872 |

| | | | | | | |
|---|---|---|---|---|---|---|
| GRHPR | 29.0816346 | 1.022671957 | 0.34458854 | 2.96780608 | 0.00299933 | 0.44263424 |
| ZBTB5 | 11.2789508 | 0.0291126 | 0.53063656 | 0.05486354 | 0.95624718 | 0.99999872 |
| AL138752.2 | 18.8068731 | 0.435299527 | 0.48154553 | 0.90396338 | 0.36601482 | 0.99999872 |
| TRMT10B | 20.0458399 | -0.019250292 | 0.45551259 | -0.0422607 | 0.96629085 | 0.99999872 |
| DCAF10 | 70.1485643 | 0.334742679 | 0.25003912 | 1.33876124 | 0.18064841 | 0.99999872 |
| ALDH1B1 | 5.66515563 | -0.289330202 | 0.90161962 | -0.3209005 | 0.74828578 | 0.99999872 |
| GLIDR | 21.7433704 | -0.61555048 | 0.52956705 | -1.1623655 | 0.24508702 | 0.99999872 |
| CNTNAP3B | 24.5684898 | 0.56291197 | 0.96951412 | 0.58061245 | 0.56150168 | 0.99999872 |
| AL512625.3 | 7.70376852 | -0.210404888 | 0.60687568 | -0.3467018 | 0.72881536 | 0.99999872 |
| CBWD5 | 13.2992827 | -0.897638147 | 0.44081346 | -2.036322 | 0.04171803 | 0.95980711 |
| ZNF658 | 6.2705183 | 0.73019159 | 0.78294948 | 0.93261648 | 0.35101802 | 0.99999872 |
| PGM5 | 11.6729338 | 0.820782282 | 0.69100457 | 1.1878102 | 0.23490819 | 0.99999872 |
| AL161457.2 | 5.50923614 | -0.830389379 | 0.65414875 | -1.2694198 | 0.20429138 | 0.99999872 |
| FAM189A2 | 8.9793557 | 0.666989187 | 0.56477864 | 1.18097452 | 0.23761285 | 0.99999872 |
| APBA1 | 99.6764878 | 0.021785229 | 0.55375077 | 0.03934122 | 0.96861835 | 0.99999872 |
| PTAR1 | 45.485299 | 0.263974952 | 0.32779142 | 0.80531379 | 0.42063863 | 0.99999872 |
| MAMDC2-AS | 20.9722789 | -0.395547257 | 0.43054281 | -0.9187176 | 0.3582433 | 0.99999872 |
| MAMDC2 | 59.6043229 | 1.039478026 | 0.34610932 | 3.00332288 | 0.00267049 | 0.42396043 |
| SMC5-AS1 | 19.2725477 | -0.530162752 | 0.48661139 | -1.0894993 | 0.27593378 | 0.99999872 |
| SMC5 | 69.7030796 | -0.166652635 | 0.26049551 | -0.6397524 | 0.52233357 | 0.99999872 |
| KLF9 | 56.8342689 | 0.012887251 | 0.43894774 | 0.02935942 | 0.97657794 | 0.99999872 |
| TRPM3 | 61.4424545 | 0.463682209 | 0.6885964 | 0.67337298 | 0.50071004 | 0.99999872 |
| CEMIP2 | 134.952412 | 0.499501025 | 0.58812682 | 0.84930836 | 0.39570973 | 0.99999872 |
| ABHD17B | 14.031442 | 0.533028722 | 0.6661226 | 0.80019612 | 0.42359718 | 0.99999872 |
| C9orf85 | 22.891618 | 0.091636579 | 0.40741011 | 0.22492465 | 0.82203789 | 0.99999872 |
| GDA | 21.1064052 | 2.02798535 | 1.04014456 | 1.9497149 | 0.05121011 | 0.98999913 |
| LINC01504 | 6.57660255 | -0.827382324 | 1.39267617 | -0.5940953 | 0.5524484 | 0.99999872 |
| ZFAND5 | 107.344768 | 0.672764209 | 0.37212478 | 1.80789954 | 0.07062214 | 0.99999872 |
| TMC1 | 43.5938201 | 2.042979614 | 0.84451072 | 2.41912809 | 0.01555776 | 0.79608149 |
| ALDH1A1 | 11.0587646 | 0.853403916 | 0.98247733 | 0.86862454 | 0.38505253 | 0.99999872 |
| ANXA1 | 80.4381452 | -0.091348468 | 0.85232444 | -0.1071757 | 0.91464959 | 0.99999872 |
| CARNMT1 | 18.0030618 | 0.175388354 | 0.37764361 | 0.46442823 | 0.64234097 | 0.99999872 |
| NMRK1 | 37.2646752 | 0.482994831 | 0.39403278 | 1.22577323 | 0.22028402 | 0.99999872 |
| OSTF1 | 32.4750595 | 0.772558521 | 0.41410351 | 1.86561692 | 0.06209498 | 0.99999872 |
| PCSK5 | 18.0513284 | -0.366775799 | 0.7213744 | -0.5084403 | 0.61114462 | 0.99999872 |
| RFK | 14.2446715 | 0.511775857 | 0.4300634 | 1.19000096 | 0.23404602 | 0.99999872 |
| PRUNE2 | 38.5172137 | -0.72423839 | 0.66607047 | -1.08733 | 0.27689101 | 0.99999872 |
| PCA3 | 11.8916573 | -0.17127371 | 1.04922646 | -0.1632381 | 0.87033099 | 0.99999872 |
| VPS13A | 91.3054552 | -0.079125645 | 0.2937819 | -0.2693346 | 0.78767217 | 0.99999872 |
| GNAQ | 205.830281 | -0.003089003 | 0.27452495 | -0.0112522 | 0.99102225 | 0.99999872 |
| CEP78 | 11.5428842 | -0.489129854 | 0.53287586 | -0.9179058 | 0.35866817 | 0.99999872 |
| TLE4 | 109.888326 | 0.886558189 | 0.37103088 | 2.38944582 | 0.01687381 | 0.82234329 |
| TLE1 | 21.8846105 | -0.676513171 | 0.80076911 | -0.8448293 | 3.98E-01 | 0.99999872 |
| AL591368.1 | 4.20162757 | 0.600274108 | 1.52070525 | 0.39473403 | 0.69303919 | NA |
| FRMD3 | 553.856403 | 0.244781254 | 0.50836374 | 0.48150809 | 0.63015543 | 0.99999872 |
| UBQLN1 | 52.478604 | 0.403220714 | 0.29972669 | 1.34529466 | 0.1785301 | 0.99999872 |
| GKAP1 | 16.8187738 | 0.196161202 | 0.5658658 | 0.34665676 | 0.7288492 | 0.99999872 |
| KIF27 | 17.2750331 | -0.047905048 | 0.41712046 | -0.114847 | 0.90856637 | 0.99999872 |
| C9orf64 | 5.51598953 | -0.573355551 | 0.66706134 | -0.8595245 | 0.39005122 | 0.99999872 |
| HNRNPK | 112.812618 | 0.086269806 | 0.29165509 | 0.29579393 | 0.76738746 | 0.99999872 |
| RMI1 | 9.44188891 | -0.097932644 | 0.54619225 | -0.1793007 | 0.85770161 | 0.99999872 |
| NTRK2 | 26.8823358 | -0.924044622 | 0.48428307 | -1.9080672 | 0.05638253 | 0.99999872 |
| AGTPBP1 | 40.0285739 | 0.136549895 | 0.30274787 | 0.45103503 | 0.6519643 | 0.99999872 |
| NAA35 | 44.0407616 | -0.237595435 | 0.28824402 | -0.8242857 | 0.40977723 | 0.99999872 |
| GOLM1 | 9.48745586 | -0.857723998 | 0.52908827 | -1.6211359 | 0.10498849 | 0.99999872 |
| ISCA1 | 14.9110814 | 0.482412092 | 0.47875201 | 1.00764506 | 0.31362489 | 0.99999872 |
| TUT7 | 28.4889611 | -0.025761845 | 0.40366743 | -0.0638195 | 0.94911397 | 0.99999872 |
| AL353613.1 | 5.76682807 | 0.816840534 | 0.79584 | 1.02638788 | 0.30470879 | 0.99999872 |
| GAS1 | 101.321877 | -0.145656625 | 0.73137008 | -0.1991558 | 0.84214084 | 0.99999872 |
| AL513318.2 | 18.7463112 | 0.599456517 | 0.62765203 | 0.9550778 | 0.33953836 | 0.99999872 |
| DAPK1 | 184.532123 | -0.026212722 | 0.31071809 | -0.0843618 | 0.93276882 | 0.99999872 |

| | | | | | | |
|---|---|---|---|---|---|---|
| CTSL | 79.6963417 | -0.184088642 | 0.5717639 | -0.3219662 | 0.74747831 | 0.99999872 |
| SPIN1 | 71.5416961 | 0.081684849 | 0.29105086 | 0.28065489 | 0.77897511 | 0.99999872 |
| SECISBP2 | 43.9456552 | -0.277917173 | 0.29043915 | -0.9568861 | 0.33862476 | 0.99999872 |
| SEMA4D | 27.5750562 | -0.325219429 | 0.37096303 | -0.8766896 | 0.38065526 | 0.99999872 |
| AUH | 61.3202252 | -0.036687956 | 0.33173403 | -0.1105945 | 0.91193792 | 0.99999872 |
| NFIL3 | 8.07961353 | 0.388268562 | 0.9999012 | 0.38830693 | 0.69778891 | 0.99999872 |
| SPTLC1 | 39.0206877 | -0.399755414 | 0.37949702 | -1.0533822 | 0.29216585 | 0.99999872 |
| IARS | 75.454376 | 0.198403804 | 0.29980129 | 0.66178437 | 0.50810943 | 0.99999872 |
| NOL8 | 17.0687248 | 0.000899143 | 0.4657533 | 0.00193051 | 0.99845967 | 0.99999872 |
| CENPP | 53.0841297 | -0.35653331 | 0.38652617 | -0.9224041 | 0.35631786 | 0.99999872 |
| OMD | 4.85491194 | 2.048796025 | 1.11853253 | 1.83168211 | 0.0669988 | 0.99999872 |
| ECM2 | 12.1208191 | 0.21306594 | 0.69816417 | 0.30518028 | 0.76022885 | 0.99999872 |
| IPPK | 10.7562224 | 0.194851365 | 0.46466241 | 0.41933964 | 0.67496793 | 0.99999872 |
| BICD2 | 21.9420529 | 0.461592927 | 0.50388089 | 0.91607547 | 0.35962731 | 0.99999872 |
| ZNF484 | 8.93279181 | -0.005463646 | 0.49291478 | -0.0110844 | 0.99115614 | 0.99999872 |
| CARD19 | 5.50088252 | 0.798469964 | 0.70215238 | 1.13717476 | 0.25546524 | 0.99999872 |
| NINJ1 | 23.6382482 | 0.182226994 | 0.45020205 | 0.40476713 | 0.68564869 | 0.99999872 |
| FAM120AOS | 14.3871823 | 0.055837492 | 0.48558188 | 0.11499089 | 0.90845234 | 0.99999872 |
| FAM120A | 72.9808892 | 0.265495255 | 0.28341921 | 0.93675816 | 0.34888297 | 0.99999872 |
| PHF2 | 20.7817327 | -0.683972818 | 0.42173997 | -1.621788 | 0.10484875 | 0.99999872 |
| PTPDC1 | 22.9132927 | 0.19075562 | 0.42209014 | 0.451931 | 0.65131869 | 0.99999872 |
| AL158152.2 | 46.4585545 | -0.273770835 | 0.36572149 | -0.7485774 | 0.45411198 | 0.99999872 |
| ZNF169 | 6.96064076 | -0.176179672 | 0.74774847 | -0.2356135 | 0.81373256 | 0.99999872 |
| MFSD14B | 32.5228766 | -0.169960651 | 0.31657859 | -0.5368672 | 0.59135937 | 0.99999872 |
| FBP1 | 7.25186375 | -0.149785535 | 0.80464849 | -0.1861503 | 0.85232692 | 0.99999872 |
| C9orf3 | 334.380961 | -0.106640312 | 0.19868317 | -0.5367355 | 0.59145033 | 0.99999872 |
| FANCC | 61.2181066 | -0.111436531 | 0.29683394 | -0.3754171 | 0.70735031 | 0.99999872 |
| PTCH1 | 9.99099602 | 0.163654369 | 0.55617295 | 0.29425086 | 0.76856622 | 0.99999872 |
| LINC00476 | 28.0990179 | -0.613802784 | 0.50120265 | -1.2246599 | 0.22070339 | 0.99999872 |
| ERCC6L2 | 55.6808332 | -0.289212918 | 0.27594629 | -1.0480768 | 0.29460321 | 0.99999872 |
| SLC35D2 | 10.7825446 | -0.233232628 | 0.59990304 | -0.3887839 | 0.69743603 | 0.99999872 |
| HABP4 | 33.6727658 | -0.3175804 | 0.37096507 | -0.8560925 | 0.39194664 | 0.99999872 |
| CDC14B | 26.0632203 | 0.557773901 | 0.4614782 | 1.20866792 | 0.22679045 | 0.99999872 |
| AAED1 | 13.312079 | 0.082807607 | 0.5089309 | 0.16270894 | 0.87074761 | 0.99999872 |
| ZNF510 | 6.83249201 | 0.750763702 | 0.5916667 | 1.26889632 | 0.20447804 | 0.99999872 |
| ZNF782 | 26.938669 | -0.074979144 | 0.37176661 | -0.2016834 | 0.84016426 | 0.99999872 |
| MFSD14C | 73.8203126 | -0.259063325 | 0.24944503 | -1.0385588 | 0.29900999 | 0.99999872 |
| CTSV | 9.42865796 | -1.418695412 | 1.04253105 | -1.3608184 | 0.17357109 | 0.99999872 |
| CCDC180 | 8.61052872 | -0.485784458 | 0.69931361 | -0.694659 | 0.48726906 | 0.99999872 |
| TDRD7 | 27.5345678 | 0.257183999 | 0.4692806 | 0.54803885 | 0.58366523 | 0.99999872 |
| TMOD1 | 42.0183948 | 1.127805822 | 1.12434197 | 1.00308078 | 0.31582189 | 0.99999872 |
| TSTD2 | 19.2275082 | -0.17345629 | 0.43538526 | -0.3983972 | 0.69033739 | 0.99999872 |
| NCBP1 | 19.6823403 | -0.022550781 | 0.51734843 | -0.0435892 | 0.9652319 | 0.99999872 |
| XPA | 19.3396586 | 0.299044527 | 0.34577591 | 0.86485067 | 0.38712075 | 0.99999872 |
| TRMO | 6.74879507 | -0.132468618 | 0.74569391 | -0.1776448 | 0.85900197 | 0.99999872 |
| ANP32B | 26.6071713 | -0.393209723 | 0.35302196 | -1.1138393 | 0.26534814 | 0.99999872 |
| NANS | 6.43295208 | 0.388738316 | 0.5626847 | 0.69086349 | 0.48965133 | 0.99999872 |
| TRIM14 | 13.2292375 | -0.762858485 | 0.68946971 | -1.1064423 | 0.26853511 | 0.99999872 |
| CORO2A | 9.78825032 | 0.230007919 | 0.69725069 | 0.32987836 | 0.74149187 | 0.99999872 |
| ANKS6 | 7.93819129 | 0.228214419 | 0.53868251 | 0.42365292 | 0.67181895 | 0.99999872 |
| TGFBR1 | 22.1810536 | -0.151813559 | 0.44321396 | -0.3425288 | 0.73195295 | 0.99999872 |
| ALG2 | 13.6897106 | 0.053377024 | 0.47622825 | 0.11208286 | 0.91075771 | 0.99999872 |
| SEC61B | 56.9581561 | 0.944768648 | 0.38489366 | 2.45462252 | 0.01410326 | 0.76172142 |
| AL359710.1 | 20.2000463 | 0.157180128 | 0.6433978 | 0.24429696 | 0.80700084 | 0.99999872 |
| STX17 | 25.3607771 | 0.066524835 | 0.46850335 | 0.14199436 | 0.88708446 | 0.99999872 |
| ERP44 | 54.4362897 | 0.169951504 | 0.31007083 | 0.54810543 | 0.58361951 | 0.99999872 |
| INVS | 65.7548112 | 0.019273172 | 0.35658383 | 0.05404948 | 0.95689574 | 0.99999872 |
| TEX10 | 11.314026 | 0.806244749 | 0.51282513 | 1.5721631 | 0.11591273 | 0.99999872 |
| MRPL50 | 13.947137 | 0.62839151 | 0.47633369 | 1.31922541 | 0.18709377 | 0.99999872 |
| ZNF189 | 16.4701765 | -0.331362669 | 0.4630019 | -0.7156832 | 0.474187 | 0.99999872 |
| ALDOB | 36.1907977 | 1.353536814 | 1.84893494 | 0.73206298 | 0.46413013 | 0.99999872 |

| | | | | | | |
|---|---|---|---|---|---|---|
| TMEM246 | 14.0650651 | 0.214606869 | 0.49054061 | 0.43749052 | 0.66175565 | 0.99999872 |
| RNF20 | 9.60304422 | -0.60245681 | 0.57568298 | -1.046508 | 0.29532658 | 0.99999872 |
| SMC2 | 7.1545125 | 0.299119478 | 0.59693428 | 0.50109282 | 0.6163058 | 0.99999872 |
| NIPSNAP3A | 19.5058743 | 0.179334641 | 0.42927352 | 0.41776312 | 0.67612032 | 0.99999872 |
| ABCA1 | 20.3663767 | 1.332330502 | 1.17562272 | 1.13329768 | 0.25708927 | 0.99999872 |
| SLC44A1 | 66.4872873 | -0.347516523 | 0.32364468 | -1.0737594 | 0.28293052 | 0.99999872 |
| FSD1L | 53.0517871 | 0.07483431 | 0.31193138 | 0.23990632 | 0.81040288 | 0.99999872 |
| FKTN | 20.8962884 | 0.489046057 | 0.42751999 | 1.1439139 | 0.25265935 | 0.99999872 |
| TMEM38B | 21.3483294 | 0.753513561 | 0.58543092 | 1.28710927 | 0.19805621 | 0.99999872 |
| LINC01505 | 11.4027043 | 0.815826227 | 0.75157075 | 1.08549491 | 0.27770252 | 0.99999872 |
| ZNF462 | 109.736497 | -0.065354676 | 0.29537989 | -0.2212563 | 0.82489284 | 0.99999872 |
| AL807761.4 | 5.98347547 | 1.340903916 | 0.92944491 | 1.44269326 | 0.1491069 | 0.99999872 |
| RAD23B | 121.008685 | 0.269861112 | 0.2508021 | 1.07599225 | 0.28193072 | 0.99999872 |
| ELP1 | 18.8548365 | 0.408628082 | 0.40266081 | 1.0148196 | 0.31019183 | 0.99999872 |
| FAM206A | 15.6283285 | 0.501539569 | 0.56121907 | 0.89366096 | 0.37150332 | 0.99999872 |
| CTNNAL1 | 39.7468157 | 0.014731524 | 0.64966333 | 0.02267563 | 0.98190902 | 0.99999872 |
| TMEM245 | 188.015823 | 0.285739022 | 0.33781287 | 0.84585002 | 0.39763641 | 0.99999872 |
| PTPN3 | 40.2959809 | -0.495582554 | 0.32349995 | -1.5319401 | 0.12553721 | 0.99999872 |
| PALM2-AKAF | 7.78277543 | 0.735072796 | 0.82003415 | 0.89639291 | 0.37004295 | 0.99999872 |
| TXN | 52.949554 | 0.860698343 | 0.41929377 | 2.05273344 | 0.04009844 | 0.95980711 |
| ECPAS | 87.7161549 | 0.087064518 | 0.3117414 | 0.27928442 | 0.78002656 | 0.99999872 |
| ZNF483 | 34.0660686 | 0.175126265 | 0.45169123 | 0.38771234 | 0.69822892 | 0.99999872 |
| PTGR1 | 49.4714671 | 0.186145759 | 0.45106603 | 0.41267963 | 0.67984135 | 0.99999872 |
| C9orf84 | 14.4329923 | 1.552174207 | 0.80230358 | 1.934647 | 0.05303363 | 0.99999872 |
| UGCG | 207.530454 | -0.036403941 | 0.48079554 | -0.0757161 | 0.939645 | 0.99999872 |
| SUSD1 | 33.997859 | -0.31471729 | 0.34884182 | -0.9021776 | 0.36696251 | 0.99999872 |
| PTBP3 | 301.282896 | -0.074214729 | 0.35591244 | -0.2085196 | 0.83482326 | 0.99999872 |
| HSDL2 | 58.3594786 | 0.222029133 | 0.33805933 | 0.65677564 | 0.51132519 | 0.99999872 |
| C9orf147 | 4.52545377 | 0.988267491 | 0.79422334 | 1.24431937 | 0.21338204 | NA |
| KIAA1958 | 60.4481187 | 0.212977446 | 0.32131363 | 0.66283352 | 0.50743718 | 0.99999872 |
| INIP | 16.6393749 | 0.227852089 | 0.44588669 | 0.51100895 | 0.60934479 | 0.99999872 |
| SNX30 | 14.4259538 | -0.079251149 | 0.56179445 | -0.1410679 | 0.88781632 | 0.99999872 |
| SLC31A2 | 12.3610451 | 1.214640006 | 0.66807038 | 1.8181318 | 0.06904399 | 0.99999872 |
| FKBP15 | 16.3084391 | 0.28422539 | 0.37969041 | 0.74857142 | 0.45411556 | 0.99999872 |
| SLC31A1 | 7.67979714 | 0.162003953 | 0.69180676 | 0.23417515 | 0.814849 | 0.99999872 |
| CDC26 | 10.4465099 | 1.328845464 | 0.52818887 | 2.51585282 | 0.01187448 | 0.72670507 |
| PRPF4 | 7.09764818 | -0.245962312 | 0.56008563 | -0.4391513 | 0.66055194 | 0.99999872 |
| HDHD3 | 9.58377541 | -0.329057496 | 0.70610967 | -0.4660147 | 0.64120497 | 0.99999872 |
| ALAD | 6.3294663 | -0.391626151 | 0.60735817 | -0.6448026 | 0.51905509 | 0.99999872 |
| POLE3 | 5.29643319 | -0.195621353 | 0.71676951 | -0.2729209 | 0.78491405 | 0.99999872 |
| RGS3 | 11.4999979 | -0.401530668 | 0.75922749 | -0.5288674 | 0.59689744 | 0.99999872 |
| ZNF618 | 21.3097794 | 0.464173564 | 0.71452348 | 0.64962675 | 0.51593335 | 0.99999872 |
| KIF12 | 9.04395857 | -0.27663133 | 0.75160652 | -0.3680534 | 0.71283342 | 0.99999872 |
| COL27A1 | 12.0847225 | -0.53424161 | 0.51122873 | -1.0450148 | 0.29601611 | 0.99999872 |
| ATP6V1G1 | 92.1492788 | 1.017860828 | 0.46730667 | 2.17814315 | 0.02939538 | 0.90266962 |
| TMEM268 | 4.57828639 | -0.256322953 | 1.17349192 | -0.2184275 | 0.82709601 | NA |
| PAPPA | 97.4235485 | 0.583528507 | 0.54922933 | 1.06244964 | 2.88E-01 | 0.99999872 |
| ASTN2 | 44.9069395 | -0.340982061 | 0.57004286 | -0.5981692 | 0.54972706 | 0.99999872 |
| TRIM32 | 4.61182814 | 0.088386664 | 0.69321078 | 0.1275033 | 0.89854206 | 0.99999872 |
| CDK5RAP2 | 44.8904144 | -0.029320519 | 0.34312147 | -0.0854523 | 0.93190181 | 0.99999872 |
| MEGF9 | 138.24521 | -0.035384293 | 0.38588721 | -0.0916959 | 0.92693962 | 0.99999872 |
| FBXW2 | 23.5800182 | -0.250277049 | 0.33975685 | -0.7366358 | 0.46134388 | 0.99999872 |
| PSMD5 | 19.0718846 | 0.25942294 | 0.35803707 | 0.72457006 | 0.46871583 | 0.99999872 |
| PHF19 | 4.27861558 | -0.197825522 | 0.70011157 | -0.2825629 | 0.77751196 | NA |
| C5 | 10.0879711 | -0.34075137 | 0.7213854 | -0.4723569 | 0.63667207 | 0.99999872 |
| CNTRL | 30.5117639 | -0.158394752 | 0.35449103 | -0.446823 | 0.65500285 | 0.99999872 |
| RAB14 | 78.6508551 | 0.211826036 | 0.36474235 | 0.58075526 | 0.56140541 | 0.99999872 |
| GSN | 148.130595 | -0.145281916 | 0.42284479 | -0.3435821 | 0.73116056 | 0.99999872 |
| STOM | 13.4649955 | 0.000252036 | 0.48582052 | 0.00051878 | 0.99958607 | 0.99999872 |
| DAB2IP | 58.4621506 | -0.249635855 | 0.40024625 | -0.6237057 | 0.5328209 | 0.99999872 |
| TTLL11 | 12.8533054 | 0.275833918 | 0.58518045 | 0.47136557 | 0.6373797 | 0.99999872 |

| | | | | | | |
|---|---|---|---|---|---|---|
| NDUFA8 | 15.7759874 | 0.832090147 | 0.40396299 | 2.05981776 | 0.03941597 | 0.95980711 |
| RBM18 | 13.6234878 | -0.333314716 | 0.44191961 | -0.7542429 | 0.4507034 | 0.99999872 |
| MRRF | 18.8905183 | -0.631956006 | 0.42859861 | -1.4744705 | 0.14035495 | 0.99999872 |
| PDCL | 11.5349189 | 0.857845011 | 0.52952283 | 1.62003404 | 0.10522497 | 0.99999872 |
| RC3H2 | 51.2374504 | 0.179736437 | 0.28696449 | 0.62633686 | 0.53109402 | 0.99999872 |
| ZBTB6 | 5.02380954 | 0.551293831 | 0.77374155 | 0.71250384 | 0.47615283 | 0.99999872 |
| ZBTB26 | 8.53233011 | -0.321013297 | 0.52679672 | -0.6093684 | 0.54228025 | 0.99999872 |
| RABGAP1 | 73.1746123 | -0.090801209 | 0.26013535 | -0.3490537 | 0.72704899 | 0.99999872 |
| STRBP | 29.9899586 | -0.038516987 | 0.43388072 | -0.0887732 | 0.92926214 | 0.99999872 |
| CRB2 | 180.745447 | -0.130099484 | 0.59368765 | -0.2191379 | 0.8265426 | 0.99999872 |
| DENND1A | 123.579287 | -0.58207386 | 0.27741444 | -2.0982104 | 0.03588656 | 0.94734101 |
| PSMB7 | 30.3013237 | 0.032156379 | 0.36582839 | 0.08790017 | 0.92995602 | 0.99999872 |
| NR6A1 | 28.8837661 | 0.725423883 | 0.82572296 | 0.8785318 | 0.37965519 | 0.99999872 |
| MIR181A2H( | 10.3349824 | -0.519203089 | 0.61485891 | -0.8444264 | 0.39843118 | 0.99999872 |
| OLFML2A | 18.2597604 | -0.734202615 | 0.67530345 | -1.0872188 | 0.27694012 | 0.99999872 |
| RPL35 | 61.5414926 | 0.19769941 | 0.55906223 | 0.35362684 | 0.72361856 | 0.99999872 |
| ARPC5L | 15.7766384 | -0.391749665 | 0.44991371 | -0.8707218 | 0.38390608 | 0.99999872 |
| GOLGA1 | 16.4687192 | -0.370873105 | 0.4365239 | -0.8496055 | 0.39554445 | 0.99999872 |
| SCAI | 16.6193349 | 0.036295318 | 0.44922126 | 0.08079608 | 0.93560412 | 0.99999872 |
| PPP6C | 33.5809356 | -0.398779539 | 0.37693948 | -1.0579405 | 0.29008257 | 0.99999872 |
| RABEPK | 8.94418258 | 0.110688565 | 0.56050638 | 0.19747958 | 0.84345226 | 0.99999872 |
| HSPA5 | 145.727941 | 1.131957444 | 0.70306053 | 1.61004267 | 0.10738854 | 0.99999872 |
| GAPVD1 | 50.9557385 | 0.132767273 | 0.24480209 | 0.54234533 | 0.58758063 | 0.99999872 |
| MAPKAP1 | 77.5248121 | -0.046071523 | 0.2763273 | -0.1667281 | 0.867584 | 0.99999872 |
| PBX3 | 75.273126 | 0.307204222 | 0.44492664 | 0.69046039 | 0.48990471 | 0.99999872 |
| MVB12B | 20.6427671 | 0.008580293 | 0.39037743 | 0.02197948 | 0.98246432 | 0.99999872 |
| AL161908.1 | 10.7762055 | 0.154537364 | 0.5220671 | 0.29601054 | 0.76722203 | 0.99999872 |
| LMX1B | 129.315562 | -0.575841955 | 0.58125959 | -0.9906795 | 0.32184211 | 0.99999872 |
| ZBTB43 | 19.5074708 | -0.155759287 | 0.37779293 | -0.4122875 | 0.68012874 | 0.99999872 |
| ZBTB34 | 5.7637237 | 0.147399957 | 0.63248966 | 0.23304722 | 0.81572474 | 0.99999872 |
| RALGPS1 | 21.5481607 | -0.149964178 | 0.37080493 | -0.4044288 | 0.68589746 | 0.99999872 |
| GARNL3 | 11.1318307 | 0.407443309 | 0.57685453 | 0.70631899 | 0.47998979 | 0.99999872 |
| SLC2A8 | 4.25045589 | -0.462276671 | 0.65061872 | -0.7105186 | 0.47738263 | NA |
| RPL12 | 16.2815146 | 0.076478803 | 0.73763742 | 0.10368075 | 0.9174227 | 0.99999872 |
| LRSAM1 | 16.0938988 | -0.090872189 | 0.47634295 | -0.1907705 | 0.84870539 | 0.99999872 |
| FAM129B | 16.7876872 | -0.440499726 | 0.61291759 | -0.7186932 | 0.47232995 | 0.99999872 |
| STXBP1 | 31.0790556 | 0.023055031 | 0.32787595 | 0.07031632 | 0.94394189 | 0.99999872 |
| PTRH1 | 7.59024535 | 0.749783735 | 0.54105445 | 1.3857824 | 0.16581336 | 0.99999872 |
| CDK9 | 30.9410728 | -0.202555116 | 0.39151827 | -0.517358 | 0.60490626 | 0.99999872 |
| FPGS | 19.5044273 | -0.416808387 | 0.40641691 | -1.0255685 | 0.30509501 | 0.99999872 |
| ENG | 10.3229372 | 0.00818837 | 0.75326939 | 0.01087044 | 0.99132681 | 0.99999872 |
| AK1 | 4.7451071 | 0.752910793 | 0.79220826 | 0.95039503 | 0.34191157 | 0.99999872 |
| DPM2 | 10.7257165 | -0.477589023 | 0.63583752 | -0.751118 | 0.45258163 | 0.99999872 |
| FAM102A | 10.1475481 | -0.524078684 | 0.594892 | -0.8809644 | 0.37833708 | 0.99999872 |
| SLC25A25 | 9.77950834 | 0.471937437 | 0.50827777 | 0.92850301 | 0.35314671 | 0.99999872 |
| PTGES2 | 10.077051 | -0.519169403 | 0.47857346 | -1.084827 | 0.2779983 | 0.99999872 |
| C9orf16 | 20.418946 | 0.098403192 | 0.42491063 | 0.23158562 | 0.81685988 | 0.99999872 |
| CIZ1 | 12.9467026 | -0.714794346 | 0.41386122 | -1.7271354 | 0.08414336 | 0.99999872 |
| DNM1 | 10.3279782 | -0.189850275 | 0.65352374 | -0.2905025 | 0.77143185 | 0.99999872 |
| GOLGA2 | 36.9196341 | -0.32588145 | 0.50801241 | -0.6414832 | 0.52120876 | 0.99999872 |
| SWI5 | 16.0424565 | 0.371673185 | 0.53428612 | 0.69564447 | 0.48665151 | 0.99999872 |
| TRUB2 | 5.69696831 | -0.595602219 | 0.64022818 | -0.9302968 | 0.35221744 | 0.99999872 |
| COQ4 | 25.3291447 | -0.055545256 | 0.34664908 | -0.1602348 | 0.87269609 | 0.99999872 |
| SLC27A4 | 20.6148234 | -0.276084146 | 0.44641023 | -0.6184539 | 0.53627618 | 0.99999872 |
| URM1 | 10.4664752 | -0.013073834 | 0.60052752 | -0.0217706 | 0.98263096 | 0.99999872 |
| ODF2 | 7.57353005 | -0.703709967 | 0.53469941 | -1.3160852 | 0.18814546 | 0.99999872 |
| GLE1 | 13.626803 | -0.360959499 | 0.4313639 | -0.8367865 | 0.40271258 | 0.99999872 |
| AL356481.1 | 9.42595653 | -0.844720058 | 0.78442983 | -1.0768587 | 0.28154341 | 0.99999872 |
| SPTAN1 | 152.910038 | -0.570367643 | 0.27520458 | -2.0725224 | 0.03821675 | 0.95980711 |
| WDR34 | 8.53548504 | -0.298079003 | 0.58212338 | -0.5120547 | 0.60861274 | 0.99999872 |
| SET | 56.0171833 | -0.221476952 | 0.33467835 | -0.6617606 | 0.50812465 | 0.99999872 |

| | | | | | | |
|---|---|---|---|---|---|---|
| ZDHHC12 | 18.6026028 | -0.548471541 | 0.42283626 | -1.2971251 | 0.19458814 | 0.99999872 |
| ZER1 | 36.6194359 | -0.115180524 | 0.43519374 | -0.2646649 | 0.79126759 | 0.99999872 |
| TBC1D13 | 11.3600499 | -0.017256734 | 0.56278083 | -0.0306633 | 0.97553804 | 0.99999872 |
| ENDOG | 9.95148956 | 0.332570882 | 0.49848468 | 0.66716369 | 0.50466758 | 0.99999872 |
| SPOUT1 | 6.56326243 | -0.729352368 | 0.66007755 | -1.1049495 | 0.26918147 | 0.99999872 |
| KYAT1 | 29.2971755 | -0.04384616 | 0.61412188 | -0.0713965 | 0.94308219 | 0.99999872 |
| LRRC8A | 23.6325453 | -0.743504695 | 0.57733654 | -1.2878185 | 0.19780914 | 0.99999872 |
| DOLK | 4.06878711 | -0.67955801 | 0.69264093 | -0.9811115 | 0.32653774 | NA |
| NUP188 | 7.30923548 | 0.68764159 | 0.55139811 | 1.24708732 | 0.21236548 | 0.99999872 |
| SH3GLB2 | 21.589482 | -0.055767383 | 0.41658622 | -0.1338676 | 0.8935073 | 0.99999872 |
| MIGA2 | 6.75299102 | -0.309643535 | 0.66126633 | -0.4682584 | 0.63959979 | 0.99999872 |
| CRAT | 4.6326204 | -0.231198764 | 0.69701254 | -0.3316996 | 0.74011612 | 0.99999872 |
| PTPA | 28.7789011 | -0.627840004 | 0.41419895 | -1.5157933 | 0.12957162 | 0.99999872 |
| IER5L | 12.4529634 | -1.565619002 | 0.97464463 | -1.6063486 | 0.10819735 | 0.99999872 |
| LINC01503 | 14.4157779 | -0.653283336 | 0.84373056 | -0.7742796 | 0.43876548 | 0.99999872 |
| NTMT1 | 13.1352404 | 0.958140484 | 0.70537473 | 1.35834251 | 0.17435503 | 0.99999872 |
| ASB6 | 5.28033483 | 0.196617945 | 0.70112379 | 0.28043257 | 0.77914565 | 0.99999872 |
| TOR1B | 10.2560538 | -0.340320774 | 0.46174378 | -0.7370338 | 0.4611018 | 0.99999872 |
| TOR1A | 7.63733846 | 0.322295681 | 0.62243932 | 0.51779454 | 0.60460163 | 0.99999872 |
| C9orf78 | 16.1598482 | -0.000975297 | 0.41893981 | -0.002328 | 0.99814252 | 0.99999872 |
| USP20 | 8.53701128 | 0.267426504 | 0.63624021 | 0.42032317 | 0.67424938 | 0.99999872 |
| FNBP1 | 201.111763 | -0.191044133 | 0.40410761 | -0.4727556 | 0.63638756 | 0.99999872 |
| GPR107 | 40.7339889 | 0.061339689 | 0.31437871 | 0.19511401 | 0.8453037 | 0.99999872 |
| NCS1 | 14.8256631 | 0.345709081 | 0.54295654 | 0.63671594 | 0.52430989 | 0.99999872 |
| ASS1 | 25.5082395 | -0.335638548 | 0.55202868 | -0.6080093 | 0.54318133 | 0.99999872 |
| FUBP3 | 39.9218183 | -0.09899668 | 0.32732314 | -0.3024433 | 0.76231418 | 0.99999872 |
| EXOSC2 | 3.29218785 | -0.216291995 | 0.93714071 | -0.2307999 | 0.81747025 | NA |
| ABL1 | 74.9075882 | -0.057024483 | 0.2285435 | -0.2495126 | 0.8029643 | 0.99999872 |
| AIF1L | 108.556369 | -0.403406545 | 0.51642175 | -0.7811572 | 0.43471007 | 0.99999872 |
| NUP214 | 46.8549524 | -0.552751834 | 0.26091571 | -2.1185073 | 0.03413213 | 0.93897283 |
| AL157938.2 | 8.79817951 | 0.800253655 | 0.7442684 | 1.07522187 | 0.28227541 | 0.99999872 |
| PRRC2B | 103.327588 | -0.168696184 | 0.27212766 | -0.6199156 | 0.53531335 | 0.99999872 |
| POMT1 | 13.5869794 | -0.10118806 | 0.45380315 | -0.2229779 | 0.82355273 | 0.99999872 |
| UCK1 | 11.9437385 | -0.135538751 | 0.5642347 | -0.240217 | 0.81016206 | 0.99999872 |
| RAPGEF1 | 84.8110028 | -0.195827764 | 0.26131987 | -0.7493795 | 0.45362848 | 0.99999872 |
| MED27 | 102.583733 | -0.405777505 | 0.24879097 | -1.6309977 | 1.03E-01 | 0.99999872 |
| NTNG2 | 6.55108325 | 0.026781758 | 0.6508644 | 0.04114798 | 0.96717792 | 0.99999872 |
| SETX | 104.260891 | -0.259475881 | 0.28201048 | -0.9200931 | 0.35752413 | 0.99999872 |
| TTF1 | 12.8973318 | 0.137841409 | 0.43852753 | 0.31432784 | 0.75327206 | 0.99999872 |
| DDX31 | 11.8497314 | -0.086196866 | 0.57957405 | -0.1487245 | 0.88177102 | 0.99999872 |
| GTF3C4 | 14.1548809 | -0.051357584 | 0.40969259 | -0.1253564 | 0.90024141 | 0.99999872 |
| AK8 | 6.7702264 | -0.095243509 | 0.69043144 | -0.1379478 | 0.89028166 | 0.99999872 |
| SPACA9 | 5.0588554 | 1.512740274 | 0.77336074 | 1.95606034 | 0.05045805 | 0.9863795 |
| TSC1 | 20.5735897 | -0.252772669 | 0.44331072 | -0.570193 | 0.5685468 | 0.99999872 |
| GTF3C5 | 10.8208837 | -0.438149352 | 0.44354983 | -0.9878244 | 0.32323865 | 0.99999872 |
| SURF6 | 6.8372349 | 0.298372665 | 0.58580536 | 0.50933755 | 0.61051564 | 0.99999872 |
| MED22 | 6.44180865 | 0.627476328 | 0.60006382 | 1.04568266 | 0.29570757 | 0.99999872 |
| RPL7A | 62.6550103 | 0.345766388 | 0.43828517 | 0.78890735 | 0.43016616 | 0.99999872 |
| SURF1 | 9.74183206 | 0.273639815 | 0.6077711 | 0.45023499 | 0.65254101 | 0.99999872 |
| SURF2 | 6.1572775 | -0.391409135 | 0.55232349 | -0.7086592 | 0.47853597 | 0.99999872 |
| SURF4 | 38.2541846 | 0.269938399 | 0.44720584 | 0.60361108 | 0.54610224 | 0.99999872 |
| REXO4 | 7.07892273 | 0.69735241 | 0.56780129 | 1.22816279 | 0.21938586 | 0.99999872 |
| CACFD1 | 6.77064456 | -0.25065631 | 0.63799798 | -0.3928795 | 0.6944085 | 0.99999872 |
| VAV2 | 67.1528786 | -0.81696078 | 0.42475076 | -1.9233886 | 0.05443126 | 0.99999872 |
| BRD3OS | 7.51564277 | 0.464716042 | 0.66266646 | 0.70128197 | 0.48312706 | 0.99999872 |
| BRD3 | 20.157251 | 0.096846819 | 0.41305866 | 0.23446263 | 0.81462584 | 0.99999872 |
| WDR5 | 9.9977162 | -0.13882765 | 0.64802984 | -0.2142303 | 0.83036744 | 0.99999872 |
| RXRA | 14.0216335 | -0.733629543 | 0.42517046 | -1.7254951 | 0.08443829 | 0.99999872 |
| PPP1R26 | 7.52599211 | -0.018093624 | 0.53318999 | -0.0339347 | 0.97292925 | 0.99999872 |
| MRPS2 | 8.43280515 | -0.236344487 | 0.50150903 | -0.4712667 | 0.63745032 | 0.99999872 |
| CAMSAP1 | 18.0738934 | -0.331297975 | 0.43043711 | -0.769678 | 0.44149092 | 0.99999872 |

| | | | | | | |
|---|---|---|---|---|---|---|
| UBAC1 | 9.90253985 | 0.441057285 | 0.49838309 | 0.88497643 | 0.37616935 | 0.99999872 |
| NACC2 | 38.5428806 | -0.337310224 | 0.517894 | -0.6513113 | 0.51484554 | 0.99999872 |
| TMEM250 | 7.12784021 | 0.210294502 | 0.69690979 | 0.30175283 | 0.76284049 | 0.99999872 |
| QSOX2 | 5.96011709 | 0.353242558 | 0.67280765 | 0.52502756 | 0.59956403 | 0.99999872 |
| GPSM1 | 47.2326606 | -0.409358339 | 0.44879211 | -0.9121335 | 0.36169842 | 0.99999872 |
| SNAPC4 | 4.73992257 | 0.136425426 | 0.67535471 | 0.20200559 | 0.83991235 | 0.99999872 |
| ENTR1 | 7.83983456 | -0.842200535 | 0.60080078 | -1.4017967 | 0.16097597 | 0.99999872 |
| PMPCA | 11.3548265 | -0.317999943 | 0.47534524 | -0.6689873 | 0.50350356 | 0.99999872 |
| INPP5E | 6.41501814 | 0.027696461 | 0.60951018 | 0.04544052 | 0.96375618 | 0.99999872 |
| SEC16A | 21.4072411 | -0.409015222 | 0.42986433 | -0.9514984 | 0.34135143 | 0.99999872 |
| AGPAT2 | 11.1518632 | -0.240586 | 0.56199683 | -0.4280914 | 0.66858459 | 0.99999872 |
| SNHG7 | 17.2644157 | 0.545139029 | 0.73799292 | 0.73867786 | 0.46010263 | 0.99999872 |
| RABL6 | 34.4993276 | 0.136495308 | 0.42932943 | 0.31792674 | 0.75054051 | 0.99999872 |
| PHPT1 | 53.2486388 | 0.963007769 | 0.59203168 | 1.62661527 | 0.10381883 | 0.99999872 |
| EDF1 | 52.090126 | -0.209774506 | 0.44792613 | -0.4683239 | 0.63955299 | 0.99999872 |
| TRAF2 | 6.54272744 | -0.287796338 | 0.6205747 | -0.4637578 | 0.6428213 | 0.99999872 |
| FBXW5 | 24.6008171 | -0.588826503 | 0.43674154 | -1.3482265 | 0.17758555 | 0.99999872 |
| LCNL1 | 11.2669543 | 0.8026832 | 0.72338133 | 1.10962664 | 0.26715995 | 0.99999872 |
| PAXX | 10.0432929 | 0.18848306 | 0.51044233 | 0.36925437 | 0.71193813 | 0.99999872 |
| ABCA2 | 9.20151708 | 0.573493133 | 0.83661529 | 0.68549205 | 0.49303347 | 0.99999872 |
| MAN1B1 | 23.2485589 | 0.520656143 | 0.52036666 | 1.00055631 | 0.31704136 | 0.99999872 |
| DPP7 | 18.9205376 | 0.054358751 | 0.46317182 | 0.11736196 | 0.90657323 | 0.99999872 |
| SSNA1 | 22.1052247 | 0.121752661 | 0.38925404 | 0.31278458 | 0.75444434 | 0.99999872 |
| TPRN | 5.33096981 | -0.242277923 | 0.66152799 | -0.3662399 | 0.7141861 | 0.99999872 |
| TMEM203 | 13.7531273 | 0.440698708 | 0.51191986 | 0.8608744 | 0.38930722 | 0.99999872 |
| NDOR1 | 6.3229505 | -0.346686792 | 0.65572638 | -0.5287065 | 0.59700908 | 0.99999872 |
| BX255925.3 | 6.96370757 | -1.593695127 | 0.62725079 | -2.5407622 | 0.01106111 | 0.71986533 |
| TUBB4B | 53.1917078 | 0.536284724 | 0.60450048 | 0.88715351 | 0.37499626 | 0.99999872 |
| NELFB | 17.5471024 | -0.0143965 | 0.46933387 | -0.0306743 | 0.97552927 | 0.99999872 |
| EXD3 | 24.5431976 | 0.001615127 | 0.44295344 | 0.00364627 | 0.99709071 | 0.99999872 |
| NOXA1 | 4.91127808 | -0.279303116 | 0.65652467 | -0.4254267 | 0.67052566 | 0.99999872 |
| NSMF | 6.15217303 | -0.014287719 | 0.57128011 | -0.02501 | 0.98004698 | 0.99999872 |
| PNPLA7 | 12.2968004 | 0.384123329 | 0.79433912 | 0.48357599 | 0.62868682 | 0.99999872 |
| MRPL41 | 41.694511 | 0.269105106 | 0.34186548 | 0.78716665 | 0.43118431 | 0.99999872 |
| DPH7 | 10.9068995 | 0.095645449 | 0.45475331 | 0.21032381 | 0.83341495 | 0.99999872 |
| ARRDC1 | 11.0400172 | -0.136638302 | 0.58038459 | -0.2354272 | 0.8138772 | 0.99999872 |
| ARRDC1-AS1 | 6.5472731 | -0.064310137 | 0.56579041 | -0.1136642 | 0.90950396 | 0.99999872 |
| EHMT1 | 80.0180813 | -0.15605127 | 0.27489681 | -0.5676722 | 0.57025759 | 0.99999872 |
| AC069287.3 | 6.34914222 | 0.588766602 | 1.04994738 | 0.5607582 | 0.57496239 | 0.99999872 |
| BET1L | 8.80361654 | 0.041856875 | 0.50376798 | 0.08308761 | 0.93378188 | 0.99999872 |
| RIC8A | 12.8390321 | -0.212483253 | 0.49040596 | -0.4332803 | 0.66481113 | 0.99999872 |
| SIRT3 | 7.45456111 | -0.481974489 | 0.55973772 | -0.861072 | 0.38919838 | 0.99999872 |
| PSMD13 | 25.4930139 | -0.15180906 | 0.47338461 | -0.3206886 | 0.74844637 | 0.99999872 |
| PGGHG | 10.076328 | -0.964881758 | 0.46054915 | -2.0950679 | 0.03616496 | 0.94734101 |
| IFITM2 | 304.372133 | 0.4270526 | 0.58570023 | 0.72913169 | 4.66E-01 | 1.00E+00 |
| AC136475.1 | 4.35572969 | 0.323082595 | 0.74577756 | 0.43321576 | 0.66485803 | NA |
| IFITM3 | 693.857483 | 0.291492979 | 0.70312562 | 0.41456743 | 6.78E-01 | 0.99999872 |
| SIGIRR | 16.9579384 | -0.289137543 | 0.51213965 | -0.5645678 | 0.57236779 | 0.99999872 |
| PTDSS2 | 18.1611556 | -0.179782676 | 0.43710088 | -0.4113071 | 0.68084739 | 0.99999872 |
| RNH1 | 45.6268754 | 0.568109849 | 0.3612815 | 1.5724853 | 0.11583805 | 0.99999872 |
| HRAS | 17.0776348 | -0.602704021 | 0.40667654 | -1.4820231 | 0.13833416 | 0.99999872 |
| RASSF7 | 36.808923 | 0.334210432 | 0.5164521 | 0.64712764 | 0.51754934 | 0.99999872 |
| MIR210HG | 5.84604951 | 0.274269086 | 0.62166699 | 0.44118329 | 0.65908031 | 0.99999872 |
| PHRF1 | 15.6513299 | -0.274760534 | 0.54530884 | -0.5038622 | 0.61435818 | 0.99999872 |
| IRF7 | 9.99116107 | -1.55116037 | 1.28289721 | -1.2091073 | 0.22662163 | 0.99999872 |
| DEAF1 | 28.053499 | -0.700222135 | 0.3420351 | -2.047223 | 0.04063619 | 0.95980711 |
| EPS8L2 | 26.7610381 | -0.272506798 | 0.42854085 | -0.6358946 | 0.52484514 | 0.99999872 |
| TALDO1 | 31.3401801 | 0.787365794 | 0.5588559 | 1.40888875 | 0.15886807 | 0.99999872 |
| GATD1 | 13.3764887 | -0.368022664 | 0.64425352 | -0.5712389 | 0.56783772 | 0.99999872 |
| AP006621.3 | 12.3692387 | 0.481385226 | 0.52578802 | 0.91555001 | 0.35990296 | 0.99999872 |
| SLC25A22 | 4.34060143 | -0.326679199 | 1.02759346 | -0.317907 | 0.75055545 | NA |

| | | | | | | |
|---|---|---|---|---|---|---|
| RPLP2 | 53.3533526 | 0.294273467 | 0.48862753 | 0.60224496 | 0.5470111 | 0.99999872 |
| PNPLA2 | 16.8212078 | -0.507371806 | 0.40052449 | -1.2667685 | 0.20523808 | 0.99999872 |
| CD151 | 237.519865 | 0.252843351 | 0.53424825 | 0.4732694 | 0.63602099 | 0.99999872 |
| POLR2L | 99.7348716 | 0.154185417 | 0.49174759 | 0.31354585 | 0.75386599 | 0.99999872 |
| TSPAN4 | 53.8955587 | 0.281107953 | 0.45495023 | 0.61788726 | 0.53664966 | 0.99999872 |
| CHID1 | 11.9528211 | 0.388122916 | 0.65832849 | 0.58955814 | 0.55548692 | 0.99999872 |
| AP2A2 | 45.6014168 | -0.553794225 | 0.27531598 | -2.0114859 | 0.04427415 | 0.95980711 |
| TOLLIP | 20.5369608 | -0.27747764 | 0.35166621 | -0.789037 | 0.4300904 | 0.99999872 |
| MOB2 | 15.5641258 | -0.46242123 | 0.40599058 | -1.138995 | 0.25470525 | 0.99999872 |
| DUSP8 | 6.39221684 | -0.829764753 | 0.62936946 | -1.3184065 | 0.18736762 | 0.99999872 |
| MRPL23 | 7.05566493 | 1.618970958 | 0.69693831 | 2.32297597 | 0.02018044 | 0.8671024 |
| CD81-AS1 | 5.69493588 | -0.210168387 | 0.89411084 | -0.2350585 | 0.8141633 | 0.99999872 |
| CD81 | 624.303717 | 0.797828944 | 0.54183199 | 1.47246557 | 0.1408952 | 0.99999872 |
| TSSC4 | 15.6686875 | -0.040391805 | 0.52821622 | -0.0764683 | 0.93904652 | 0.99999872 |
| KCNQ1 | 80.6362774 | 0.579411885 | 0.39799266 | 1.45583561 | 0.14543807 | 0.99999872 |
| KCNQ1OT1 | 407.061597 | -0.016757442 | 0.43651743 | -0.0383889 | 0.96937758 | 0.99999872 |
| CDKN1C | 1144.20297 | 0.022426173 | 0.58535504 | 0.03831209 | 0.96943885 | 0.99999872 |
| SLC22A18 | 6.41048536 | -0.035701302 | 0.93751567 | -0.0380808 | 0.9696233 | 0.99999872 |
| NAP1L4 | 47.6621984 | 0.418471225 | 0.38619814 | 1.08356615 | 0.27855721 | 0.99999872 |
| CARS | 22.3786409 | -0.211936706 | 0.51098792 | -0.4147587 | 0.67831853 | 0.99999872 |
| OSBPL5 | 15.5077216 | -0.223310446 | 0.38302608 | -0.5830163 | 0.55988233 | 0.99999872 |
| ZNF195 | 14.9795729 | -0.673344171 | 0.39947794 | -1.6855603 | 0.09188052 | 0.99999872 |
| AC127526.1 | 10.451039 | -0.655779498 | 0.67915259 | -0.9655849 | 0.33425192 | 0.99999872 |
| NUP98 | 46.9264745 | -0.391104998 | 0.27264453 | -1.4344869 | 0.15143337 | 0.99999872 |
| PGAP2 | 7.20562432 | -0.605610506 | 0.56569239 | -1.0705651 | 0.28436504 | 0.99999872 |
| RHOG | 10.7008175 | 0.138081602 | 0.52927581 | 0.2608878 | 0.79417903 | 0.99999872 |
| STIM1 | 39.8766758 | -0.038822199 | 0.47191822 | -0.0822647 | 0.93443624 | 0.99999872 |
| RRM1 | 8.76545683 | 0.254092401 | 0.47478877 | 0.53516936 | 0.59253275 | 0.99999872 |
| TRIM21 | 6.88309485 | -0.862287822 | 0.93062896 | -0.9265646 | 0.35415266 | 0.99999872 |
| AC104389.4 | 23.6973537 | -0.080664853 | 0.55617188 | -0.1450358 | 0.88468258 | 0.99999872 |
| TRIM5 | 10.5953957 | 0.23073937 | 1.14650694 | 0.20125423 | 0.84049979 | 0.99999872 |
| TRIM22 | 37.6861249 | -0.655629192 | 1.02477578 | -0.6397782 | 0.52231681 | 0.99999872 |
| FAM160A2 | 10.7547048 | 0.0715852 | 0.52217808 | 0.13708963 | 0.89095995 | 0.99999872 |
| SMPD1 | 31.478768 | 0.765180887 | 0.51676612 | 1.48071024 | 0.1386838 | 0.99999872 |
| APBB1 | 10.0372016 | 0.277154415 | 0.5213723 | 0.53158638 | 0.5950125 | 0.99999872 |
| ARFIP2 | 10.2438987 | -0.406722195 | 0.47802669 | -0.8508357 | 0.39486061 | 0.99999872 |
| AC084337.2 | 8.80968654 | -0.72336158 | 0.48184516 | -1.5012324 | 0.13329546 | 0.99999872 |
| DNHD1 | 12.5821103 | 0.014422091 | 0.45979347 | 0.03136646 | 0.97497729 | 0.99999872 |
| RRP8 | 4.31501987 | 0.743410993 | 0.70653106 | 1.0521986 | 0.29270844 | NA |
| ILK | 34.7685198 | 0.218712608 | 0.38110031 | 0.57389774 | 0.566037 | 0.99999872 |
| TAF10 | 11.6226154 | 0.564338935 | 0.62047709 | 0.90952421 | 0.36307349 | 0.99999872 |
| TPP1 | 30.6698851 | 0.569186249 | 0.38840408 | 1.46544868 | 0.14279854 | 0.99999872 |
| MRPL17 | 15.1127558 | 0.396382354 | 0.69041546 | 0.57412149 | 0.56588559 | 0.99999872 |
| AC087280.2 | 8.4004642 | -0.462245976 | 0.91587884 | -0.5047021 | 0.61376808 | 0.99999872 |
| SYT9 | 35.8521333 | 0.532497292 | 0.30280356 | 1.75855688 | 0.0786528 | 0.99999872 |
| AC107884.1 | 7.47407237 | 0.29143652 | 0.72278081 | 0.40321563 | 0.6867896 | 0.99999872 |
| PPFIBP2 | 13.6709028 | 0.358981846 | 0.55602973 | 0.64561628 | 5.19E-01 | 0.99999872 |
| EIF3F | 22.1474072 | 0.204120743 | 0.4339305 | 0.47039962 | 0.63806954 | 0.99999872 |
| TUB | 14.3150289 | 0.193205033 | 0.77803571 | 0.24832412 | 0.80388364 | 0.99999872 |
| RIC3 | 82.4959506 | 0.149193368 | 0.71823971 | 0.20772086 | 0.83544693 | 0.99999872 |
| STK33 | 106.242132 | 0.066253388 | 0.35661802 | 0.1857825 | 0.85261533 | 0.99999872 |
| TRIM66 | 16.9917424 | -1.303966606 | 0.42549079 | -3.0646177 | 0.00217948 | 0.37076694 |
| RPL27A | 55.6957001 | 0.590476726 | 0.4836711 | 1.22082285 | 0.2221531 | 0.99999872 |
| ST5 | 41.3248594 | -0.663617577 | 0.43366883 | -1.5302404 | 0.12595723 | 0.99999872 |
| AKIP1 | 13.999755 | 0.486454022 | 0.45773077 | 1.06275141 | 0.2878947 | 0.99999872 |
| TMEM9B | 39.3081248 | 0.370756223 | 0.50336911 | 0.73654941 | 0.46139641 | 0.99999872 |
| TMEM9B-AS | 9.01708817 | -0.481948906 | 0.57928901 | -0.8319663 | 0.40542799 | 0.99999872 |
| DENND5A | 37.3800147 | 0.234073386 | 0.51583558 | 0.45377519 | 0.64999065 | 0.99999872 |
| TMEM41B | 35.3749517 | -0.21796019 | 0.35516789 | -0.6136821 | 0.53942539 | 0.99999872 |
| IPO7 | 36.6769066 | -0.313524856 | 0.44434743 | -0.7055849 | 0.48044629 | 0.99999872 |
| ZNF143 | 8.41709647 | -0.091330888 | 0.56073736 | -0.1628764 | 0.87061575 | 0.99999872 |

| | | | | | | |
|---|---|---|---|---|---|---|
| WEE1 | 10.5249455 | -0.483028123 | 0.51579696 | -0.9364695 | 0.34903151 | 0.99999872 |
| SWAP70 | 30.9926734 | -0.237745701 | 0.32666088 | -0.7278059 | 0.46673244 | 0.99999872 |
| SBF2-AS1 | 16.0148921 | -0.12216844 | 0.45297229 | -0.269704 | 0.78738798 | 0.99999872 |
| SBF2 | 419.773133 | 0.018751978 | 0.17711992 | 0.10587165 | 0.91568419 | 0.99999872 |
| AC011092.2 | 16.9642505 | 0.598766761 | 0.57056943 | 1.04941962 | 0.29398503 | 0.99999872 |
| AC100763.1 | 4.09204098 | 1.138928039 | 0.75251545 | 1.51349455 | 0.13015408 | NA |
| ADM | 60.5888775 | 0.368020859 | 1.08845115 | 0.33811426 | 0.73527708 | 0.99999872 |
| AMPD3 | 12.914611 | -1.199491397 | 0.73028314 | -1.6425018 | 0.10048605 | 0.99999872 |
| MTRNR2L8 | 20.9683874 | -0.49972236 | 0.62372019 | -0.8011964 | 0.42301797 | 0.99999872 |
| RNF141 | 16.3202985 | 0.271929687 | 0.38465799 | 0.70693887 | 0.47960447 | 0.99999872 |
| CTR9 | 18.6888436 | 0.297432832 | 0.38928465 | 0.76404973 | 0.44483761 | 0.99999872 |
| EIF4G2 | 77.0488015 | 0.381869783 | 0.33654449 | 1.13467844 | 0.25651007 | 0.99999872 |
| ZBED5 | 29.7612819 | 0.278125275 | 0.34352593 | 0.80961945 | 0.41815893 | 0.99999872 |
| ZBED5-AS1 | 11.2508526 | -0.198509105 | 0.68712863 | -0.2888966 | 0.77266053 | 0.99999872 |
| GALNT18 | 23.3778318 | -1.401376462 | 0.65298699 | -2.1461016 | 0.03186487 | 0.92959829 |
| USP47 | 113.06876 | -0.043929968 | 0.22985386 | -0.1911213 | 0.84843056 | 0.99999872 |
| DKK3 | 6.07305383 | 3.540755709 | 1.24783999 | 2.8375078 | 0.00454672 | 0.52915868 |
| MICAL2 | 38.3854805 | 0.48949363 | 0.65521435 | 0.74707403 | 0.45501887 | 0.99999872 |
| PARVA | 203.76296 | 0.165633629 | 0.28068877 | 0.59009711 | 0.55512555 | 0.99999872 |
| TEAD1 | 347.498595 | -0.506501445 | 0.32865614 | -1.5411288 | 0.12328544 | 0.99999872 |
| ARNTL | 10.9408806 | 0.756368621 | 0.67050055 | 1.12806562 | 0.25929221 | 0.99999872 |
| BTBD10 | 29.0774935 | 0.037476447 | 0.32948129 | 0.11374378 | 0.90944091 | 0.99999872 |
| FAR1 | 71.0497046 | 0.205915998 | 0.25448613 | 0.80914428 | 0.41843216 | 0.99999872 |
| RRAS2 | 58.0567013 | -0.588271989 | 0.27556522 | -2.1347832 | 0.03277873 | 0.92959829 |
| COPB1 | 41.3122345 | 0.258725987 | 0.31299762 | 0.82660687 | 0.40845992 | 0.99999872 |
| PSMA1 | 11.0018117 | 0.689846735 | 0.78864661 | 0.87472224 | 0.38172506 | 0.99999872 |
| PDE3B | 10.7613596 | -0.454365862 | 0.64563297 | -0.7037526 | 0.48158687 | 0.99999872 |
| CYP2R1 | 9.45070457 | -0.382596655 | 0.67709287 | -0.5650579 | 0.57203441 | 0.99999872 |
| SOX6 | 301.267826 | -0.091764512 | 0.38299639 | -0.2395963 | 0.81064325 | 0.99999872 |
| C11orf58 | 65.9364044 | -0.054911826 | 0.2444034 | -0.224677 | 0.82223055 | 0.99999872 |
| PLEKHA7 | 99.0216872 | 0.370151765 | 0.54589927 | 0.67805873 | 0.49773446 | 0.99999872 |
| RPS13 | 52.5518802 | 0.812570418 | 0.57187497 | 1.42088823 | 0.15534925 | 0.99999872 |
| PIK3C2A | 78.3792199 | 0.254587474 | 0.27116793 | 0.93885541 | 0.34780499 | 0.99999872 |
| NUCB2 | 23.7106281 | 0.882824524 | 0.41231933 | 2.14111847 | 0.03226449 | 0.92959829 |
| SERGEF | 37.3648868 | 0.471202303 | 0.3881024 | 1.21411849 | 0.22470248 | 0.99999872 |
| SAAL1 | 4.52282692 | -0.32441101 | 0.83264505 | -0.389615 | 0.69682125 | NA |
| SAA1 | 10.3185645 | 4.336489971 | 4.73571035 | 0.9157 | 0.35982426 | 0.99999872 |
| HPS5 | 56.8817525 | -0.207726292 | 0.58156853 | -0.3571828 | 0.72095494 | 0.99999872 |
| GTF2H1 | 29.0782944 | -0.234010261 | 0.37410937 | -0.625513 | 0.53163444 | 0.99999872 |
| LDHA | 90.4429268 | 0.134079499 | 0.45795063 | 0.29278156 | 7.70E-01 | 0.99999872 |
| TSG101 | 40.9611355 | -0.214851149 | 0.29668227 | -0.7241793 | 0.46895568 | 0.99999872 |
| UEVLD | 26.1374511 | -0.172869867 | 0.36295459 | -0.4762851 | 0.63387127 | 0.99999872 |
| SPTY2D1OS | 7.70497833 | 0.411266276 | 0.51234332 | 0.80271619 | 0.4221388 | 0.99999872 |
| SPTY2D1 | 11.0427268 | 0.118601453 | 0.4925471 | 0.2407921 | 0.80971625 | 0.99999872 |
| ZDHHC13 | 19.908252 | -0.030545895 | 0.40334028 | -0.0757323 | 0.93963206 | 0.99999872 |
| NAV2 | 114.933667 | -0.376027059 | 0.99084767 | -0.3795004 | 0.70431633 | 0.99999872 |
| HTATIP2 | 19.78346 | 0.537594709 | 0.39251158 | 1.36962764 | 0.17080317 | 0.99999872 |
| PRMT3 | 20.9938292 | 0.289348898 | 0.40876139 | 0.70786748 | 0.47902756 | 0.99999872 |
| NELL1 | 106.711905 | 1.503233637 | 1.56837106 | 0.95846811 | 0.33782677 | 0.99999872 |
| ANO5 | 21.6300872 | 0.24070828 | 0.54605431 | 0.44081381 | 0.6593478 | 0.99999872 |
| FANCF | 6.26059689 | 0.532440175 | 0.69263479 | 0.76871705 | 0.4420613 | 0.99999872 |
| GAS2 | 8.2912436 | 0.211427145 | 0.70676961 | 0.29914578 | 0.76482882 | 0.99999872 |
| SVIP | 31.0180611 | 0.351067147 | 0.3696035 | 0.949848 | 0.34218949 | 0.99999872 |
| AC006299.1 | 9.19880049 | 0.308086338 | 0.56866889 | 0.54176753 | 0.58797866 | 0.99999872 |
| SLC5A12 | 20.4993357 | 3.122183925 | 2.29701121 | 1.35923757 | 0.17407132 | 0.99999872 |
| BBOX1 | 13.9844246 | 2.57673966 | 1.27951485 | 2.01384114 | 0.04402621 | 0.95980711 |
| BBOX1-AS1 | 8.51728927 | 0.505526676 | 0.7939231 | 0.63674514 | 0.52429087 | 0.99999872 |
| LGR4 | 210.576488 | -0.327939672 | 0.24838271 | -1.3202999 | 0.1867349 | 0.99999872 |
| LIN7C | 25.7855187 | 0.353861426 | 0.3361508 | 1.05268654 | 0.29248468 | 0.99999872 |
| BDNF-AS | 60.2362268 | 0.326014833 | 0.33571674 | 0.9711009 | 0.33149803 | 0.99999872 |
| METTL15 | 105.800004 | -0.246153411 | 0.27524342 | -0.8943117 | 0.37115513 | 0.99999872 |

| | | | | | | |
|---|---|---|---|---|---|---|
| KCNA4 | 6.17339356 | 0.19424851 | 1.03167233 | 0.18828508 | 0.85065318 | 0.99999872 |
| ARL14EP | 19.2454184 | 0.598113112 | 0.51889241 | 1.15267269 | 0.24904475 | 0.99999872 |
| MPPED2 | 10.8728787 | 0.734334657 | 0.89640404 | 0.81920052 | 0.41267202 | 0.99999872 |
| DCDC1 | 16.80383 | 0.48748264 | 0.64317991 | 0.75792579 | 0.44849541 | 0.99999872 |
| DNAJC24 | 13.9739113 | -0.27179266 | 0.42309095 | -0.6423977 | 0.52061498 | 0.99999872 |
| IMMP1L | 29.3544573 | -0.126070412 | 0.38527622 | -0.3272208 | 0.74350086 | 0.99999872 |
| ELP4 | 54.1973693 | -0.194151041 | 0.31882821 | -0.6089519 | 0.54255633 | 0.99999872 |
| AC131571.1 | 5.96814955 | 1.476332034 | 0.86674887 | 1.70329848 | 0.08851223 | 0.99999872 |
| RCN1 | 30.1042952 | -0.089955124 | 0.64368917 | -0.1397493 | 0.88885806 | 0.99999872 |
| WT1 | 443.118313 | -0.175159943 | 0.51489915 | -0.340183 | 0.73371871 | 0.99999872 |
| EIF3M | 110.938547 | 0.05333583 | 0.26304891 | 0.20276013 | 0.83932253 | 0.99999872 |
| CCDC73 | 5.9480653 | 1.653113695 | 0.8373486 | 1.974224 | 0.04835628 | 0.97937581 |
| PRRG4 | 4.91382074 | -0.372166842 | 0.78022387 | -0.4770001 | 0.63336207 | 0.99999872 |
| QSER1 | 32.3762365 | 0.077887612 | 0.36542728 | 0.21314121 | 0.83121682 | 0.99999872 |
| TCP11L1 | 15.2334084 | -0.116908589 | 0.4671944 | -0.2502354 | 0.80240529 | 0.99999872 |
| CSTF3 | 32.0344869 | -0.11747461 | 0.3262321 | -0.3600952 | 0.71877595 | 0.99999872 |
| HIPK3 | 158.884328 | -0.132399215 | 0.4122803 | -0.3211388 | 0.74810518 | 0.99999872 |
| CD59 | 4.79271094 | 0.345965288 | 0.69195086 | 0.49998535 | 0.6170854 | 0.99999872 |
| FBXO3 | 31.4412452 | 0.253829471 | 0.37390761 | 0.67885612 | 0.49722903 | 0.99999872 |
| LMO2 | 25.6047495 | 1.929627823 | 0.82503953 | 2.33883076 | 0.01934419 | 0.8671024 |
| CAPRIN1 | 57.0744994 | -0.107224235 | 0.25870834 | -0.4144599 | 0.67853733 | 0.99999872 |
| NAT10 | 12.9586649 | -0.500535255 | 0.52857019 | -0.9469608 | 0.34365875 | 0.99999872 |
| ABTB2 | 21.7605006 | -0.601994101 | 0.97597694 | -0.6168118 | 0.53735886 | 0.99999872 |
| CAT | 23.2661612 | 0.679282424 | 0.3599223 | 1.88730296 | 0.05911959 | 0.99999872 |
| APIP | 14.9741792 | -0.263245586 | 0.54948545 | -0.4790765 | 0.63188418 | 0.99999872 |
| PDHX | 27.666731 | 0.198036851 | 0.34447923 | 0.57488764 | 0.56536729 | 0.99999872 |
| PAMR1 | 31.6845285 | 0.75908826 | 0.9104325 | 0.83376665 | 0.40441249 | 0.99999872 |
| FJX1 | 6.29527351 | -1.502556721 | 1.10297556 | -1.3622756 | 0.17311092 | 0.99999872 |
| TRIM44 | 106.865475 | -0.742835533 | 0.32995638 | -2.2513144 | 0.02436563 | 0.88337526 |
| LDLRAD3 | 6.88716754 | 0.689837293 | 0.80552062 | 0.85638688 | 0.39178382 | 0.99999872 |
| COMMD9 | 8.82834535 | -0.548527513 | 0.5824633 | -0.9417375 | 0.34632707 | 0.99999872 |
| TRAF6 | 12.1780104 | 0.437719771 | 0.47261961 | 0.92615661 | 0.3543646 | 0.99999872 |
| C11orf74 | 8.42539698 | 1.041254976 | 0.66590587 | 1.56366691 | 0.11789581 | 0.99999872 |
| API5 | 36.4691604 | -0.07886387 | 0.35003861 | -0.2253005 | 0.82174552 | 0.99999872 |
| TTC17 | 136.431629 | 0.044696945 | 0.26672604 | 0.16757623 | 0.86691667 | 0.99999872 |
| HSD17B12 | 69.5336722 | 0.187895339 | 0.22799651 | 0.82411497 | 0.40987423 | 0.99999872 |
| ALKBH3 | 12.617935 | -0.868417492 | 0.4373688 | -1.9855497 | 0.04708335 | 0.96949686 |
| ACCS | 8.38782542 | 0.011315399 | 0.92851729 | 0.01218652 | 0.9902768 | 0.99999872 |
| EXT2 | 38.7236514 | 0.126845291 | 0.38291554 | 0.3312618 | 0.74044674 | 0.99999872 |
| TP53I11 | 4.81478691 | 0.376770803 | 0.69955562 | 0.53858591 | 0.59017261 | 0.99999872 |
| PRDM11 | 10.3450921 | -0.25813005 | 0.5384854 | -0.4793631 | 0.63168032 | 0.99999872 |
| CHST1 | 21.4544723 | 0.810920706 | 0.63151338 | 1.28409108 | 0.19911009 | 0.99999872 |
| SLC35C1 | 4.22923805 | 0.362242447 | 0.76806825 | 0.47162794 | 0.63719238 | NA |
| CRY2 | 26.0700179 | -0.108862438 | 0.39924055 | -0.2726738 | 0.78510398 | 0.99999872 |
| MAPK8IP1 | 5.79911289 | -0.656518044 | 0.86338556 | -0.7603996 | 0.44701576 | 0.99999872 |
| PEX16 | 5.81956956 | -0.9501453 | 0.64434355 | -1.4745943 | 0.14032166 | 0.99999872 |
| PHF21A | 117.653897 | 0.399850626 | 0.23325198 | 1.71424326 | 0.08648406 | 0.99999872 |
| DGKZ | 7.77314203 | -0.90072868 | 0.52176723 | -1.7263037 | 0.0842928 | 0.99999872 |
| MDK | 15.9780978 | -0.998529429 | 3.26498191 | -0.30583 | 0.75973408 | 0.99999872 |
| AMBRA1 | 154.612028 | 0.449000599 | 0.3837543 | 1.17002103 | 0.24199251 | 0.99999872 |
| ATG13 | 21.1055999 | -0.187787788 | 0.38842327 | -0.4834617 | 0.62876792 | 0.99999872 |
| ARHGAP1 | 18.1528581 | -0.347537655 | 0.37452389 | -0.9279452 | 0.35343599 | 0.99999872 |
| CKAP5 | 25.3667067 | -0.097430362 | 0.39328705 | -0.2477335 | 0.80434064 | 0.99999872 |
| LRP4 | 21.5141606 | 0.294945983 | 0.58179371 | 0.50695974 | 0.61218307 | 0.99999872 |
| C11orf49 | 79.2201442 | -0.283599238 | 0.25267961 | -1.1223669 | 0.26170646 | 0.99999872 |
| ARFGAP2 | 19.5214655 | -0.624806182 | 0.35479578 | -1.7610305 | 0.07823324 | 0.99999872 |
| PACSIN3 | 11.7145155 | 0.217841511 | 0.57245 | 0.38054242 | 0.70354281 | 0.99999872 |
| DDB2 | 17.642123 | 0.003749309 | 0.46555341 | 0.00805344 | 0.99357435 | 0.99999872 |
| ACP2 | 7.5843071 | -0.08590837 | 0.6896066 | -0.1245759 | 0.9008593 | 0.99999872 |
| MADD | 17.7788046 | -0.532046727 | 0.44213775 | -1.2033506 | 0.22884067 | 0.99999872 |
| SLC39A13 | 18.5593253 | -0.669284931 | 0.50805759 | -1.3173407 | 0.18772445 | 0.99999872 |

| | | | | | | |
|---|---|---|---|---|---|---|
| PSMC3 | 10.5547048 | -0.380456647 | 0.67571741 | -0.5630411 | 0.57340692 | 0.99999872 |
| CELF1 | 102.155571 | -0.346304707 | 0.28757645 | -1.2042179 | 0.22850538 | 0.99999872 |
| NDUFS3 | 16.7998094 | 0.621649794 | 0.45406053 | 1.36909015 | 0.1709711 | 0.99999872 |
| KBTBD4 | 5.13066461 | 0.600023737 | 0.73960368 | 0.81127738 | 0.4172064 | 0.99999872 |
| MTCH2 | 22.823394 | -0.196974599 | 0.4770429 | -0.4129075 | 0.67967437 | 0.99999872 |
| FNBP4 | 74.5841472 | 0.069378144 | 0.30046303 | 0.2309041 | 0.81738931 | 0.99999872 |
| NUP160 | 31.2927905 | -0.369985713 | 0.29069969 | -1.272742 | 0.2031096 | 0.99999872 |
| PTPRJ | 50.3987187 | 0.100069955 | 0.59338029 | 0.16864388 | 0.86607676 | 0.99999872 |
| TNKS1BP1 | 24.9961584 | -0.559799176 | 0.41860306 | -1.3373031 | 0.18112371 | 0.99999872 |
| SSRP1 | 15.5911494 | -0.080082417 | 0.4019011 | -0.199259 | 0.84206014 | 0.99999872 |
| TIMM10 | 8.5108417 | 0.680255848 | 0.78650203 | 0.86491302 | 0.38708653 | 0.99999872 |
| UBE2L6 | 35.2325534 | 0.027301735 | 0.98112242 | 0.02782704 | 0.9778001 | 0.99999872 |
| SERPING1 | 73.867081 | 0.054562751 | 0.6155063 | 0.08864694 | 0.9293625 | 0.99999872 |
| CLP1 | 4.58890068 | 0.529393506 | 0.7799384 | 0.67876323 | 0.49728789 | NA |
| ZDHHC5 | 21.7139538 | -0.176476431 | 0.38053402 | -0.4637599 | 0.64281975 | 0.99999872 |
| MED19 | 7.57935237 | 0.262564921 | 0.67070531 | 0.39147583 | 0.69544555 | 0.99999872 |
| CTNND1 | 108.159263 | -0.181050945 | 0.28355917 | -0.6384944 | 0.52315189 | 0.99999872 |
| LPXN | 10.9954778 | -0.284234854 | 0.66105727 | -0.4299701 | 0.6672174 | 0.99999872 |
| GLYAT | 10.2762169 | 2.095024958 | 2.21012922 | 0.94791967 | 0.34317035 | 0.99999872 |
| FAM111A | 16.1573145 | 0.061193907 | 0.65583967 | 0.0933062 | 0.92566031 | 0.99999872 |
| DTX4 | 13.3864973 | -0.124520147 | 0.66209532 | -0.1880698 | 0.85082192 | 0.99999872 |
| OSBP | 27.9514727 | -0.344695967 | 0.32061757 | -1.0751 | 0.28232994 | 0.99999872 |
| PATL1 | 16.2062782 | 0.136892411 | 0.5595381 | 0.24465253 | 0.80672549 | 0.99999872 |
| STX3 | 17.8504249 | 0.033397989 | 0.60247977 | 0.05543421 | 0.95579254 | 0.99999872 |
| MRPL16 | 15.1416115 | 0.048076245 | 0.4196739 | 0.1145562 | 0.9087969 | 0.99999872 |
| PRPF19 | 13.4318473 | -0.00831431 | 0.52896696 | -0.015718 | 0.98745936 | 0.99999872 |
| TMEM109 | 30.1397129 | 0.87586502 | 0.55153616 | 1.5880464 | 0.11227585 | 0.99999872 |
| SLC15A3 | 8.72366902 | -1.995858149 | 1.13729193 | -1.7549216 | 0.07927272 | 0.99999872 |
| VPS37C | 18.7374941 | 0.101901487 | 0.40801419 | 0.24974986 | 0.80278079 | 0.99999872 |
| DDB1 | 32.0659081 | 0.285580021 | 0.37822466 | 0.75505394 | 0.45021662 | 0.99999872 |
| TKFC | 6.50771841 | -0.769901408 | 0.54771218 | -1.4056679 | 0.15982278 | 0.99999872 |
| CYB561A3 | 7.09943155 | -0.520343238 | 0.580878 | -0.8957875 | 0.37036628 | 0.99999872 |
| TMEM138 | 6.43282563 | -0.593356545 | 0.66032599 | -0.8985812 | 0.36887576 | 0.99999872 |
| TMEM216 | 6.63064214 | 0.382799344 | 0.62170272 | 0.61572731 | 0.53807451 | 0.99999872 |
| CPSF7 | 34.0413018 | -0.498804997 | 0.43734699 | -1.1405246 | 0.25406782 | 0.99999872 |
| AP003108.2 | 18.1874305 | -0.371371545 | 0.46260092 | -0.8027903 | 0.42209594 | 0.99999872 |
| TMEM258 | 20.1692317 | 0.875386847 | 0.56629315 | 1.54581923 | 0.12214823 | 0.99999872 |
| FADS1 | 10.1639794 | -0.832187838 | 0.81742772 | -1.0180568 | 0.30865097 | 0.99999872 |
| FADS3 | 9.169502 | -0.714576438 | 0.53909131 | -1.3255202 | 0.18499867 | 0.99999872 |
| RAB3IL1 | 9.17454605 | -0.190682716 | 0.4532399 | -0.4207104 | 0.6739666 | 0.99999872 |
| FTH1 | 430.253299 | 0.807765774 | 0.55666105 | 1.45109089 | 0.14675455 | 0.99999872 |
| ASRGL1 | 9.37377933 | -0.275126461 | 0.74639719 | -0.368606 | 0.71242145 | 0.99999872 |
| AHNAK | 177.507286 | -0.74595267 | 0.4364922 | -1.7089714 | 0.08745625 | 0.99999872 |
| EML3 | 10.3322482 | -0.98743355 | 0.63932708 | -1.5444889 | 0.12246995 | 0.99999872 |
| B3GAT3 | 23.4643293 | -0.016352445 | 0.44022336 | -0.0371458 | 0.97036876 | 0.99999872 |
| GANAB | 57.0665752 | 0.377307712 | 0.34718726 | 1.08675563 | 0.27714482 | 0.99999872 |
| UQCC3 | 10.8080285 | -0.164971151 | 0.4734224 | -0.348465 | 0.72749097 | 0.99999872 |
| UBXN1 | 17.322695 | 0.801608098 | 0.40374449 | 1.98543413 | 0.04709619 | 0.96949686 |
| TTC9C | 11.2677866 | 0.556519079 | 0.44929312 | 1.23865481 | 0.21547336 | 0.99999872 |
| POLR2G | 11.180883 | 0.333871519 | 0.64679378 | 0.5161947 | 0.60571843 | 0.99999872 |
| TAF6L | 4.71462233 | -0.370316056 | 0.69956494 | -0.5293519 | 0.59656133 | 0.99999872 |
| TMEM223 | 11.2628318 | 0.417580852 | 0.45904012 | 0.90968269 | 0.36298988 | 0.99999872 |
| TMEM179B | 23.2111209 | 0.137939166 | 0.50418324 | 0.27358935 | 0.78440022 | 0.99999872 |
| NXF1 | 7.10413739 | 0.187499339 | 0.70323551 | 0.26662382 | 0.78975881 | 0.99999872 |
| STX5 | 21.3268406 | 0.093250243 | 0.36980241 | 0.25216234 | 0.80091558 | 0.99999872 |
| WDR74 | 4.01673346 | -0.745389377 | 0.88384969 | -0.8433441 | 0.39903605 | NA |
| SLC3A2 | 67.950794 | -0.414165173 | 0.59517649 | -0.6958695 | 0.48651055 | 0.99999872 |
| SLC22A6 | 8.9226318 | 2.438446074 | 2.07189679 | 1.17691484 | 0.2392295 | 0.99999872 |
| RARRES3 | 23.1594332 | -0.05063849 | 0.99382634 | -0.0509531 | 0.95936293 | 0.99999872 |
| PLA2G16 | 47.8087218 | 0.603699044 | 0.50522802 | 1.19490413 | 0.23212449 | 0.99999872 |
| ATL3 | 39.4622729 | 0.122131646 | 0.34061135 | 0.35856599 | 0.71991979 | 0.99999872 |

| | | | | | | |
|---|---|---|---|---|---|---|
| RTN3 | 59.8335111 | 0.291095602 | 0.31178699 | 0.93363613 | 0.35049162 | 0.99999872 |
| C11orf95 | 7.88224781 | 0.268781224 | 0.68377262 | 0.39308568 | 0.6942562 | 0.99999872 |
| MARK2 | 29.7143463 | -0.080610662 | 0.38872336 | -0.2073728 | 0.8357187 | 0.99999872 |
| NAA40 | 8.62680279 | -0.352086236 | 0.5329087 | -0.6606877 | 0.5088126 | 0.99999872 |
| OTUB1 | 16.0082678 | -0.355294399 | 0.39383791 | -0.9021336 | 0.36698592 | 0.99999872 |
| STIP1 | 17.0374393 | -0.225818019 | 0.48292836 | -0.4676015 | 0.6400696 | 0.99999872 |
| TRPT1 | 5.44749635 | -0.550426813 | 0.66173876 | -0.8317887 | 0.40552823 | 0.99999872 |
| NUDT22 | 7.26602781 | 0.332795272 | 0.63239866 | 0.52624285 | 0.59871949 | 0.99999872 |
| DNAJC4 | 5.11467428 | 0.73012421 | 0.78106783 | 0.93477696 | 0.34990326 | 0.99999872 |
| VEGFB | 22.2394211 | 0.886499829 | 0.42310476 | 2.0952254 | 0.03615096 | 0.94734101 |
| FKBP2 | 36.2074854 | 0.084697046 | 0.35606262 | 0.23787121 | 0.81198098 | 0.99999872 |
| PPP1R14B | 10.4518327 | 0.165451813 | 0.6748199 | 0.24517922 | 0.80631767 | 0.99999872 |
| BAD | 16.6641278 | 0.282918471 | 0.51640364 | 0.54786304 | 0.58378595 | 0.99999872 |
| GPR137 | 12.6532968 | -0.089325118 | 0.5425844 | -0.164629 | 0.86923602 | 0.99999872 |
| ESRRA | 16.0476245 | -0.160632918 | 0.41205292 | -0.3898357 | 0.69665808 | 0.99999872 |
| TRMT112 | 43.0219263 | 0.560078724 | 0.34047532 | 1.64499065 | 0.09997174 | 0.99999872 |
| PRDX5 | 29.5978555 | 0.278224134 | 0.50290939 | 0.55322915 | 0.58010651 | 0.99999872 |
| RPS6KA4 | 4.2526954 | -0.225183911 | 0.70897499 | -0.317619 | 0.75077398 | NA |
| SLC22A12 | 5.3577222 | 1.234823922 | 1.37076535 | 0.90082809 | 0.36767973 | 0.99999872 |
| SF1 | 62.292163 | -0.32304243 | 0.25779945 | -1.2530765 | 0.21017786 | 0.99999872 |
| MAP4K2 | 5.18319944 | -0.151929198 | 0.63779012 | -0.2382119 | 0.81171675 | 0.99999872 |
| MEN1 | 4.55622638 | 0.177189999 | 0.67822959 | 0.26125372 | 0.79389686 | NA |
| EHD1 | 38.3290124 | -0.334726017 | 0.46615264 | -0.718061 | 0.4727197 | 0.99999872 |
| ATG2A | 6.99354008 | -0.207331023 | 0.56865591 | -0.3645984 | 0.71541123 | 0.99999872 |
| PPP2R5B | 6.19052 | -0.226162159 | 0.72078084 | -0.3137738 | 0.75369283 | 0.99999872 |
| SAC3D1 | 5.95137292 | 0.956358963 | 0.66018356 | 1.44862585 | 0.1474421 | 0.99999872 |
| ZFPL1 | 10.4826765 | 0.029044664 | 0.5681743 | 0.05111928 | 0.95923047 | 0.99999872 |
| VPS51 | 24.880152 | -0.094264075 | 0.41573669 | -0.2267399 | 0.82062604 | 0.99999872 |
| TM7SF2 | 8.87322623 | 0.06066668 | 0.67318196 | 0.09011929 | 0.92819242 | 0.99999872 |
| ZNHIT2 | 4.12363383 | 0.065816476 | 0.81270834 | 0.08098413 | 0.93545458 | NA |
| FAU | 87.6196485 | 0.196933286 | 0.51000348 | 0.38614106 | 0.6993922 | 0.99999872 |
| SYVN1 | 12.0651543 | -0.106978843 | 0.49107391 | -0.2178467 | 0.82754854 | 0.99999872 |
| MRPL49 | 13.4317219 | 0.444716903 | 0.50293826 | 0.88423757 | 0.37656798 | 0.99999872 |
| CAPN1 | 21.7046922 | 0.11615923 | 0.43841391 | 0.26495334 | 0.79104541 | 0.99999872 |
| POLA2 | 44.5117005 | 0.871981857 | 0.7308912 | 1.1930392 | 0.23285402 | 0.99999872 |
| CDC42EP2 | 25.2103455 | 0.747577108 | 0.52559433 | 1.42234621 | 0.15492577 | 0.99999872 |
| DPF2 | 16.9034775 | -0.319516392 | 0.40002708 | -0.7987369 | 0.42444298 | 0.99999872 |
| FRMD8 | 12.6059695 | -0.49513176 | 0.43919297 | -1.1273672 | 0.25958725 | 0.99999872 |
| NEAT1 | 3850.95771 | 1.062114623 | 0.48180078 | 2.20446845 | 0.02749142 | 0.89154528 |
| MALAT1 | 28169.7029 | 0.298432239 | 0.32532008 | 0.91734958 | 0.35895948 | 0.99999872 |
| SCYL1 | 15.5224343 | -0.418464104 | 0.5131963 | -0.8154075 | 0.4148391 | 0.99999872 |
| LTBP3 | 43.1885239 | -0.219495944 | 0.52740067 | -0.4161844 | 0.67727506 | 0.99999872 |
| SSSCA1 | 11.6770056 | 0.46967836 | 0.57803745 | 0.81253967 | 0.41648204 | 0.99999872 |
| FAM89B | 11.4939233 | 0.843777787 | 0.75490448 | 1.11772788 | 0.26368323 | 0.99999872 |
| EHBP1L1 | 6.83244587 | -1.144637474 | 0.88975047 | -1.2864702 | 0.19827901 | 0.99999872 |
| MAP3K11 | 14.9732866 | -0.128387771 | 0.4602362 | -0.2789606 | 0.78027506 | 0.99999872 |
| PCNX3 | 9.07190457 | -0.168717019 | 0.52393472 | -0.3220192 | 0.74743818 | 0.99999872 |
| SIPA1 | 6.83560392 | -0.235104189 | 0.56878267 | -0.4133463 | 0.67935294 | 0.99999872 |
| RELA | 17.3040856 | -0.345979477 | 0.52942868 | -0.6534959 | 0.51343663 | 0.99999872 |
| KAT5 | 17.032275 | -0.623005668 | 0.38686889 | -1.6103793 | 0.10731507 | 0.99999872 |
| RNASEH2C | 16.8268172 | 1.002980685 | 0.6131949 | 1.63566377 | 0.10190998 | 0.99999872 |
| CFL1 | 110.598009 | 0.10841329 | 0.46702325 | 0.23213682 | 0.81643175 | 0.99999872 |
| MUS81 | 8.2502213 | 0.946151948 | 0.58648176 | 1.61326748 | 0.10668641 | 0.99999872 |
| EFEMP2 | 35.6399697 | 0.167940955 | 0.48545708 | 0.34594398 | 0.72938481 | 0.99999872 |
| FIBP | 5.29394918 | 0.588478932 | 0.87464816 | 0.6728179 | 0.50106315 | 0.99999872 |
| CCDC85B | 16.9960451 | -0.406109817 | 0.61332098 | -0.6621489 | 0.50787579 | 0.99999872 |
| C11orf68 | 8.70449772 | -0.406985579 | 0.54465805 | -0.7472314 | 0.45492392 | 0.99999872 |
| DRAP1 | 16.7143827 | 0.070208343 | 0.49683955 | 0.14130989 | 0.88762514 | 0.99999872 |
| SART1 | 7.46024293 | -0.95224036 | 0.58576045 | -1.6256481 | 0.10402454 | 0.99999872 |
| EIF1AD | 5.44134937 | 0.181138178 | 0.58558874 | 0.30932661 | 0.75707309 | 0.99999872 |
| BANF1 | 25.2586024 | 0.684180003 | 0.53672066 | 1.27474131 | 0.20240082 | 0.99999872 |

| | | | | | | |
|---|---|---|---|---|---|---|
| SF3B2 | 32.6475994 | -0.274033412 | 0.4064596 | -0.6741959 | 0.50018675 | 0.99999872 |
| PACS1 | 78.7048856 | -0.104437465 | 0.26665166 | -0.3916625 | 0.69530757 | 0.99999872 |
| KLC2 | 7.93829014 | -0.336925756 | 0.82613938 | -0.4078316 | 0.6833973 | 0.99999872 |
| RAB1B | 26.9725105 | -0.081475918 | 0.598505 | -0.1361324 | 0.89171662 | 0.99999872 |
| YIF1A | 9.97613905 | 0.06601446 | 0.60338834 | 0.10940626 | 0.91288027 | 0.99999872 |
| BRMS1 | 6.36071497 | -0.489164918 | 0.68434752 | -0.7147902 | 0.47473867 | 0.99999872 |
| B4GAT1 | 52.476269 | 0.362142604 | 0.70977689 | 0.51022034 | 0.60989711 | 0.99999872 |
| MRPL11 | 8.82380054 | 0.594869047 | 0.53730342 | 1.10713802 | 0.26823426 | 0.99999872 |
| ZDHHC24 | 12.6979123 | -0.020507421 | 0.45039406 | -0.0455322 | 0.96368313 | 0.99999872 |
| CTSF | 20.2831217 | 0.88773384 | 0.43308575 | 2.04978769 | 0.04038515 | 0.95980711 |
| CCS | 15.5044621 | 0.122489847 | 0.4005384 | 0.30581299 | 0.75974703 | 0.99999872 |
| RBM14-RBM | 14.0721123 | -0.152476583 | 0.44400559 | -0.3434114 | 0.73128898 | 0.99999872 |
| RBM4 | 11.4245238 | -0.268182491 | 0.63836902 | -0.4201057 | 0.67440821 | 0.99999872 |
| RBM4B | 10.459693 | 1.075973897 | 0.58991004 | 1.82396269 | 0.06815771 | 0.99999872 |
| C11orf80 | 27.8299839 | 0.951435776 | 0.55672635 | 1.70898284 | 0.08745413 | 0.99999872 |
| RCE1 | 7.08604173 | 0.689691873 | 0.63128631 | 1.09251835 | 0.27460533 | 0.99999872 |
| PC | 29.3950695 | 0.343670482 | 0.50866533 | 0.67563182 | 0.49927443 | 0.99999872 |
| LRFN4 | 23.3467226 | -0.639634705 | 0.46880338 | -1.3643986 | 0.17244213 | 0.99999872 |
| KDM2A | 114.574552 | -0.501042159 | 0.26377141 | -1.8995317 | 0.0574946 | 0.99999872 |
| GRK2 | 8.68868385 | -0.2861914 | 0.52449456 | -0.5456518 | 0.58530531 | 0.99999872 |
| ANKRD13D | 15.3250269 | -0.115145944 | 0.41434901 | -0.277896 | 0.78109217 | 0.99999872 |
| SSH3 | 11.2158898 | -0.705414843 | 0.57715873 | -1.2222198 | 0.22162451 | 0.99999872 |
| RAD9A | 38.1292982 | -0.064601408 | 0.33420651 | -0.1932979 | 0.84672569 | 0.99999872 |
| POLD4 | 4.59503878 | -1.57332821 | 0.95677151 | -1.6444137 | 0.10009077 | 0.99999872 |
| PPP1CA | 27.163634 | -0.432724541 | 0.50194423 | -0.8620968 | 0.38863423 | 0.99999872 |
| RPS6KB2 | 9.4164336 | -0.489438729 | 0.47165755 | -1.0376994 | 0.29941005 | 0.99999872 |
| CORO1B | 16.6106641 | 0.457159418 | 0.49760363 | 0.91872203 | 0.35824098 | 0.99999872 |
| TMEM134 | 23.1786809 | 0.468049126 | 0.39924977 | 1.1723216 | 0.24106796 | 0.99999872 |
| AIP | 11.6741682 | -0.218509839 | 0.43926399 | -0.4974454 | 0.61887501 | 0.99999872 |
| PITPNM1 | 5.12643786 | -0.012321552 | 0.74560243 | -0.0165256 | 0.98681505 | 0.99999872 |
| CDK2AP2 | 26.4277112 | -0.035031183 | 0.54761377 | -0.0639706 | 0.94899363 | 0.99999872 |
| GSTP1 | 154.438342 | 0.745019199 | 0.61102373 | 1.21929668 | 0.22273161 | 0.99999872 |
| NDUFV1 | 19.7885614 | -0.036036993 | 0.4361885 | -0.0826179 | 0.93415534 | 0.99999872 |
| NUDT8 | 4.58359516 | 0.596611142 | 0.73017311 | 0.81708178 | 0.41388169 | NA |
| UNC93B1 | 25.5172028 | -1.298128882 | 0.85836143 | -1.5123337 | 0.130449 | 0.99999872 |
| ALDH3B1 | 18.95974 | -0.121379648 | 0.50588611 | -0.2399347 | 0.81038086 | 0.99999872 |
| NDUFS8 | 26.8655868 | -0.154400467 | 0.59429559 | -0.2598042 | 0.79501484 | 0.99999872 |
| TCIRG1 | 6.29738502 | -0.993695142 | 0.78162364 | -1.2713218 | 0.2036142 | 0.99999872 |
| CHKA | 25.4600956 | -0.45306347 | 0.50914094 | -0.8898587 | 0.37354178 | 0.99999872 |
| KMT5B | 63.5154187 | -0.170345842 | 0.29354745 | -0.5803009 | 0.56171174 | 0.99999872 |
| C11orf24 | 6.16431568 | 0.748747001 | 0.65724226 | 1.13922529 | 0.2546092 | 0.99999872 |
| LRP5 | 8.4926145 | 0.02979026 | 0.73556597 | 0.04049978 | 0.96769468 | 0.99999872 |
| PPP6R3 | 140.050498 | -0.049141418 | 0.19812731 | -0.2480295 | 0.80411158 | 0.99999872 |
| CPT1A | 51.9408489 | -0.078627337 | 0.37901957 | -0.2074493 | 0.835659 | 0.99999872 |
| MRPL21 | 8.01336688 | 0.62044292 | 0.61695399 | 1.00565509 | 0.31458151 | 0.99999872 |
| IGHMBP2 | 8.26854673 | -0.948941653 | 0.54161762 | -1.7520509 | 0.07976505 | 0.99999872 |
| MRGPRF | 123.285513 | 0.425760278 | 0.69939544 | 0.60875473 | 0.54268703 | 0.99999872 |
| TPCN2 | 20.5541145 | 1.247545754 | 0.43674756 | 2.85644586 | 0.00428413 | 0.50494083 |
| CCND1 | 9.36849027 | -0.830412266 | 0.68392518 | -1.2141858 | 0.22467677 | 0.99999872 |
| ORAOV1 | 11.6630261 | -0.544799427 | 0.42890518 | -1.2702095 | 0.20401002 | 0.99999872 |
| FADD | 8.31216925 | -0.652982371 | 0.5266949 | -1.2397735 | 0.21505919 | 0.99999872 |
| PPFIA1 | 53.3240898 | -0.524481307 | 0.32484579 | -1.6145547 | 0.10640716 | 0.99999872 |
| AP002336.2 | 18.1155588 | 0.078178352 | 0.66199435 | 0.11809519 | 0.90599223 | 0.99999872 |
| AP000487.1 | 4.84965168 | 0.409909623 | 0.66998584 | 0.61181834 | 0.54065796 | 0.99999872 |
| CTTN | 56.8771591 | -0.333236104 | 0.26207405 | -1.2715342 | 0.20353868 | 0.99999872 |
| SHANK2 | 88.4574251 | -0.340447042 | 0.55734734 | -0.6108346 | 0.54130908 | 0.99999872 |
| DHCR7 | 12.3909501 | 0.130133961 | 0.62856381 | 0.20703381 | 0.83598345 | 0.99999872 |
| NADSYN1 | 11.470857 | -0.079073513 | 0.48551879 | -0.162864 | 0.87062555 | 0.99999872 |
| AP002495.2 | 15.4429625 | -0.167619381 | 0.5721387 | -0.2929698 | 0.76954521 | 0.99999872 |
| RNF121 | 19.4768336 | 0.136479162 | 0.41603367 | 0.32804836 | 0.7428751 | 0.99999872 |
| IL18BP | 10.4619192 | -0.476290845 | 1.32479775 | -0.3595197 | 0.71920637 | 0.99999872 |

| | | | | | | |
|---|---|---|---|---|---|---|
| NUMA1 | 46.7977413 | 0.019631562 | 0.41032628 | 0.04784378 | 0.96184074 | 0.99999872 |
| LAMTOR1 | 30.181185 | 0.490316389 | 0.4070029 | 1.20469998 | 0.22831914 | 0.99999872 |
| ANAPC15 | 4.25829138 | 0.643016231 | 0.73983757 | 0.86913163 | 0.38477514 | NA |
| FOLR1 | 6.30363933 | 1.864628178 | 0.83741391 | 2.2266506 | 0.02597063 | 0.88365618 |
| INPPL1 | 28.3650899 | -0.263280874 | 0.39172085 | -0.6721135 | 0.50151144 | 0.99999872 |
| CLPB | 8.55973941 | -0.707984665 | 0.59159764 | -1.1967334 | 0.23141048 | 0.99999872 |
| ARAP1 | 16.2803091 | -0.313106856 | 0.4397839 | -0.7119562 | 0.47649192 | 0.99999872 |
| ATG16L2 | 6.9341139 | -0.006860975 | 0.67731244 | -0.0101297 | 0.9919178 | 0.99999872 |
| FCHSD2 | 175.021478 | -0.314620343 | 0.40495417 | -0.7769283 | 0.43720109 | 0.99999872 |
| ARHGEF17 | 51.2075091 | -0.166895944 | 0.39371075 | -0.423905 | 0.67163512 | 0.99999872 |
| FAM168A | 75.7064358 | 0.314316833 | 0.28378333 | 1.10759442 | 0.26803701 | 0.99999872 |
| PLEKHB1 | 5.96212992 | -0.05735253 | 0.79813985 | -0.0718577 | 0.94271512 | 0.99999872 |
| RAB6A | 75.2345003 | 0.211856854 | 0.21016604 | 1.00804512 | 0.3134328 | 0.99999872 |
| MRPL48 | 17.4675777 | -0.440987393 | 0.50644543 | -0.8707501 | 0.38389063 | 0.99999872 |
| COA4 | 24.8168521 | 0.815696481 | 0.54931637 | 1.48493023 | 0.13756231 | 0.99999872 |
| PAAF1 | 17.6690009 | -0.397552004 | 0.46389716 | -0.8569831 | 0.39145425 | 0.99999872 |
| C2CD3 | 24.7491341 | 0.024551277 | 0.33001078 | 0.07439538 | 0.94069579 | 0.99999872 |
| PPME1 | 26.4708044 | -0.050059563 | 0.35672052 | -0.1403327 | 0.88839711 | 0.99999872 |
| P4HA3 | 4.31026262 | 0.530841167 | 0.83570252 | 0.6352035 | 0.5252957 | NA |
| PGM2L1 | 15.680526 | -0.224704372 | 0.74521537 | -0.3015294 | 0.76301081 | 0.99999872 |
| POLD3 | 14.0913167 | 0.030255047 | 0.43655191 | 0.06930458 | 0.94474718 | 0.99999872 |
| RNF169 | 37.1994142 | -0.039812399 | 0.34704768 | -0.1147174 | 0.90866914 | 0.99999872 |
| XRRA1 | 37.4525181 | -0.525020046 | 0.58686502 | -0.8946181 | 0.3709913 | 0.99999872 |
| SPCS2 | 48.9062173 | 0.87751792 | 0.35174453 | 2.49475924 | 0.01260426 | 0.72768363 |
| NEU3 | 5.76453602 | -0.856463452 | 0.64787539 | -1.3219571 | 0.18618245 | 0.99999872 |
| ARRB1 | 62.8375781 | -0.130331777 | 0.46113599 | -0.282632 | 0.77745897 | 0.99999872 |
| RPS3 | 96.7667331 | 0.474066731 | 0.38952073 | 1.21705136 | 0.22358467 | 0.99999872 |
| SERPINH1 | 19.4312782 | 0.55638105 | 0.4867245 | 1.1431129 | 0.25299172 | 0.99999872 |
| MAP6 | 101.433396 | 0.194414507 | 0.69131595 | 0.28122381 | 0.77853874 | 0.99999872 |
| UVRAG | 87.009463 | 0.000352332 | 0.40928165 | 0.00086085 | 0.99931314 | 0.99999872 |
| THAP12 | 11.6458309 | -0.029492439 | 0.59408082 | -0.0496438 | 0.96040623 | 0.99999872 |
| EMSY | 36.9750902 | 0.009937997 | 0.34233115 | 0.02903036 | 0.97684038 | 0.99999872 |
| TSKU | 4.45855016 | -0.57763813 | 0.82542198 | -0.6998095 | 0.48404629 | NA |
| ACER3 | 37.3610328 | -0.135386072 | 0.30314906 | -0.446599 | 0.65516461 | 0.99999872 |
| PAK1 | 154.728861 | -0.050571864 | 0.41451279 | -0.1220031 | 0.90289653 | 0.99999872 |
| CLNS1A | 41.0174783 | 0.148471539 | 0.36109634 | 0.41116877 | 0.68094878 | 0.99999872 |
| RSF1 | 97.8211117 | 0.013431145 | 0.22976692 | 0.05845552 | 0.95338579 | 0.99999872 |
| AAMDC | 29.1312574 | -0.017586934 | 0.33909448 | -0.0518644 | 0.95863674 | 0.99999872 |
| AP002812.2 | 8.70728701 | -0.147847772 | 0.6384341 | -0.2315788 | 0.81686521 | 0.99999872 |
| INTS4 | 20.7273684 | 0.260827047 | 0.58499308 | 0.44586348 | 0.65569587 | 0.99999872 |
| ALG8 | 11.5838824 | -0.560725875 | 0.43984141 | -1.2748365 | 0.20236713 | 0.99999872 |
| KCTD21 | 6.70399652 | -0.677341736 | 0.56427337 | -1.2003787 | 0.2299923 | 0.99999872 |
| GAB2 | 76.5184074 | -0.153479228 | 0.31988906 | -0.4797889 | 0.63137748 | 0.99999872 |
| AP003086.2 | 20.2070537 | 0.479537002 | 0.67096922 | 0.71469299 | 0.47479877 | 0.99999872 |
| NARS2 | 21.5599582 | -0.360562524 | 0.46885522 | -0.7690274 | 0.44187703 | 0.99999872 |
| TENM4 | 11.6267154 | -0.579142823 | 0.85107467 | -0.6804842 | 0.49619795 | 0.99999872 |
| PRCP | 31.1338841 | 0.521399338 | 0.33144841 | 1.57309348 | 0.11569717 | 0.99999872 |
| RAB30 | 9.61987139 | 0.38049716 | 0.54050452 | 0.70396666 | 0.48145354 | 0.99999872 |
| RAB30-AS1 | 32.7593904 | -0.208537557 | 0.42016542 | -0.4963225 | 0.61966688 | 0.99999872 |
| PCF11 | 30.9014544 | -0.176776213 | 0.322929 | -0.5474151 | 0.58409358 | 0.99999872 |
| ANKRD42 | 13.8903944 | 0.182700156 | 0.45971489 | 0.39742058 | 0.69105734 | 0.99999872 |
| CCDC90B | 20.1963026 | 0.206242058 | 0.47025981 | 0.43857045 | 0.66097281 | 0.99999872 |
| AP000446.1 | 9.71563727 | -0.350357533 | 0.5977827 | -0.5860951 | 0.55781159 | 0.99999872 |
| DLG2 | 632.996219 | -0.245378588 | 0.58256059 | -0.421207 | 0.67360394 | 0.99999872 |
| AP001825.1 | 12.4987495 | 0.330738012 | 0.95325421 | 0.34695678 | 0.72862379 | 0.99999872 |
| TMEM126B | 18.0035664 | 0.368507417 | 0.50790055 | 0.72555034 | 0.46811448 | 0.99999872 |
| TMEM126A | 7.53411772 | 0.134420602 | 0.58332246 | 0.23043961 | 0.81775018 | 0.99999872 |
| CREBZF | 45.1165929 | 0.310186539 | 0.32262353 | 0.96145046 | 0.33632572 | 0.99999872 |
| SYTL2 | 11.9712237 | -0.245421545 | 0.64884206 | -0.3782454 | 0.70524828 | 0.99999872 |
| PICALM | 381.954696 | -0.085842437 | 0.23756329 | -0.3613455 | 0.71784115 | 0.99999872 |
| EED | 13.3355176 | 0.569153541 | 0.44678769 | 1.27387919 | 0.20270623 | 0.99999872 |

| | | | | | | |
|---|---|---|---|---|---|---|
| HIKESHI | 23.237991 | 0.356217174 | 0.37642559 | 0.94631498 | 0.34398796 | 0.99999872 |
| AP001831.1 | 8.151122 | 0.835408971 | 0.59817538 | 1.39659538 | 0.16253528 | 0.99999872 |
| ME3 | 20.5783445 | -0.780215776 | 0.52211851 | -1.494327 | 0.13509018 | 0.99999872 |
| PRSS23 | 20.1999733 | -0.061717626 | 0.89372157 | -0.0690569 | 0.94494434 | 0.99999872 |
| TMEM135 | 56.358056 | 0.126908528 | 0.34865753 | 0.36399193 | 0.71586404 | 0.99999872 |
| CTSC | 11.2851776 | -0.604452586 | 0.49688225 | -1.2164906 | 0.22379809 | 0.99999872 |
| NOX4 | 92.6204227 | -0.093195824 | 0.6589527 | -0.1414302 | 0.88753008 | 0.99999872 |
| CHORDC1 | 14.6653423 | 0.81874928 | 0.43828249 | 1.86808576 | 0.06175012 | 0.99999872 |
| SLC36A4 | 27.1142591 | -0.43716837 | 0.43139817 | -1.0133756 | 0.3108808 | 0.99999872 |
| SMCO4 | 5.34903743 | 1.007542484 | 0.76617892 | 1.31502245 | 0.18850236 | 0.99999872 |
| CEP295 | 17.975037 | 0.213492092 | 0.46405281 | 0.4600599 | 0.64547323 | 0.99999872 |
| TAF1D | 39.7666043 | 0.217498776 | 0.39044691 | 0.55705083 | 0.5774927 | 0.99999872 |
| PANX1 | 8.48889155 | -0.131755277 | 0.536121 | -0.2457566 | 0.80587065 | 0.99999872 |
| MRE11 | 17.7780893 | -0.246194453 | 0.41183037 | -0.5978055 | 0.54996974 | 0.99999872 |
| ANKRD49 | 6.14325177 | 0.697290412 | 0.62204318 | 1.12096786 | 0.26230154 | 0.99999872 |
| CWC15 | 17.6607877 | 0.005389754 | 0.43384834 | 0.01242313 | 0.99008803 | 0.99999872 |
| SRSF8 | 15.5069925 | 0.250599266 | 0.44487498 | 0.56330267 | 0.5732288 | 0.99999872 |
| ENDOD1 | 45.5153117 | -0.033543278 | 0.27638846 | -0.1213628 | 0.90340368 | 0.99999872 |
| AP000787.1 | 5.26240465 | 0.69253704 | 0.6960433 | 0.99496258 | 0.31975446 | 0.99999872 |
| SESN3 | 22.1401022 | -0.157995485 | 0.56562684 | -0.2793281 | 0.77999302 | 0.99999872 |
| FAM76B | 9.44398603 | 0.005810395 | 0.44526746 | 0.01304923 | 0.98958852 | 0.99999872 |
| CEP57 | 22.5531834 | 0.36225102 | 0.36156429 | 1.00189934 | 0.31639221 | 0.99999872 |
| MTMR2 | 34.9391247 | 0.363853729 | 0.3243112 | 1.12192774 | 0.26189317 | 0.99999872 |
| MAML2 | 458.816644 | -0.377685384 | 0.39637152 | -0.952857 | 0.34066253 | 0.99999872 |
| CCDC82 | 20.5073314 | 0.169659601 | 0.34167528 | 0.49655216 | 0.61950489 | 0.99999872 |
| JRKL | 6.26221832 | 0.169267682 | 0.63905684 | 0.26487109 | 0.79110877 | 0.99999872 |
| ARHGAP42 | 22.0783712 | -0.04572335 | 0.58941253 | -0.0775744 | 0.93816657 | 0.99999872 |
| PGR | 21.4680243 | 0.347628884 | 0.4267984 | 0.81450372 | 0.41535644 | 0.99999872 |
| TRPC6 | 230.745335 | -0.365295215 | 0.55260208 | -0.6610457 | 0.50858303 | 0.99999872 |
| CEP126 | 34.5336098 | -0.762316524 | 0.48056995 | -1.5862759 | 0.11267674 | 0.99999872 |
| CFAP300 | 6.46093263 | 0.128965696 | 0.54270449 | 0.23763521 | 0.81216404 | 0.99999872 |
| YAP1 | 173.818809 | -0.392379001 | 0.217518 | -1.8038921 | 0.07124822 | 0.99999872 |
| BIRC3 | 11.4582127 | 0.038298502 | 1.1120529 | 0.03443946 | 0.97252672 | 0.99999872 |
| BIRC2 | 35.6583788 | 0.062972702 | 0.39750581 | 0.15841958 | 0.87412619 | 0.99999872 |
| TMEM123 | 71.2200568 | 0.501714651 | 0.49039048 | 1.02309214 | 0.30626428 | 0.99999872 |
| MMP7 | 8.95470401 | 0.267287123 | 0.78245107 | 0.34160235 | 0.73265017 | 0.99999872 |
| DCUN1D5 | 20.0400244 | 0.540849323 | 0.46299401 | 1.16815619 | 0.24274377 | 0.99999872 |
| DYNC2H1 | 120.325526 | -0.143732826 | 0.38363873 | -0.3746567 | 0.70791579 | 0.99999872 |
| CASP4 | 22.3254127 | 0.054331283 | 0.80544585 | 0.06745492 | 0.94621955 | 0.99999872 |
| CARD16 | 3.46458777 | 0.272285683 | 1.29182609 | 0.2107758 | 0.83306222 | NA |
| MSANTD4 | 11.1919434 | -0.389546501 | 0.46010892 | -0.8466398 | 0.39719595 | 0.99999872 |
| KBTBD3 | 7.60769644 | 1.166484655 | 0.68633637 | 1.69958159 | 0.08920965 | 0.99999872 |
| AASDHPPT | 24.3968154 | 0.480834952 | 0.35899421 | 1.33939472 | 0.18044221 | 0.99999872 |
| AP000766.1 | 4.56749758 | -0.698636593 | 0.74921338 | -0.9324935 | 0.35108156 | NA |
| CWF19L2 | 35.210811 | -0.393641447 | 0.34911861 | -1.1275293 | 0.25951878 | 0.99999872 |
| ALKBH8 | 8.54654708 | -0.594999182 | 0.46563598 | -1.2778205 | 0.20131274 | 0.99999872 |
| CUL5 | 80.1674674 | -0.166656658 | 0.24234839 | -0.6876739 | 0.49165817 | 0.99999872 |
| ACAT1 | 20.2677971 | 0.274515984 | 0.4167397 | 0.6587229 | 0.51007372 | 0.99999872 |
| NPAT | 17.2269246 | 0.494021088 | 0.48327614 | 1.02223356 | 0.30667037 | 0.99999872 |
| ATM | 64.1330424 | -0.376423082 | 0.36269636 | -1.0378463 | 0.29934161 | 0.99999872 |
| C11orf65 | 12.1029369 | 0.56768642 | 1.01052509 | 0.5617737 | 0.57427021 | 0.99999872 |
| KDELC2 | 15.8093703 | -0.495593173 | 0.48073247 | -1.0309126 | 0.3025818 | 0.99999872 |
| EXPH5 | 400.858547 | -0.335036142 | 0.47563553 | -0.7043968 | 0.4811857 | 0.99999872 |
| DDX10 | 41.1762565 | -0.239444212 | 0.40558977 | -0.5903606 | 0.55494894 | 0.99999872 |
| RDX | 192.751745 | 0.28839378 | 0.34323792 | 0.84021537 | 0.40078764 | 0.99999872 |
| ZC3H12C | 30.9580599 | 0.745055935 | 0.3431567 | 2.17118281 | 0.02991736 | 0.90338345 |
| FDX1 | 24.1835265 | 0.547846332 | 0.39342164 | 1.39251704 | 0.16376589 | 0.99999872 |
| ARHGAP20 | 15.1177899 | -1.114240205 | 1.13378538 | -0.9827611 | 0.32572501 | 0.99999872 |
| SIK2 | 64.9894014 | -0.228267161 | 0.33667725 | -0.678 | 0.49777172 | 0.99999872 |
| PPP2R1B | 21.5768775 | 0.074150958 | 0.39717182 | 0.18669743 | 0.85189787 | 0.99999872 |
| C11orf1 | 6.64033721 | 0.141557958 | 0.66270442 | 0.21360648 | 0.83085395 | 0.99999872 |

| | | | | | | |
|---|---|---|---|---|---|---|
| CRYAB | 214.332159 | 0.461432825 | 0.75465624 | 0.6114477 | 0.54090323 | 0.99999872 |
| DIXDC1 | 15.0844602 | 0.458559893 | 0.5929206 | 0.77339174 | 0.43929058 | 0.99999872 |
| DLAT | 14.5923152 | 0.281773982 | 0.38621033 | 0.72958686 | 0.46564276 | 0.99999872 |
| NKAPD1 | 8.72792053 | 0.238336019 | 0.62211022 | 0.38310899 | 0.70163895 | 0.99999872 |
| TIMM8B | 16.9135575 | 0.521589036 | 0.4742416 | 1.09983823 | 0.27140261 | 0.99999872 |
| IL18 | 6.89669207 | -0.021701611 | 0.86413354 | -0.0251137 | 0.97996425 | 0.99999872 |
| PTS | 42.778836 | 0.217845239 | 0.33590954 | 0.64852353 | 5.17E-01 | 0.99999872 |
| AP002884.1 | 21.336713 | 0.356399738 | 0.5868304 | 0.60733005 | 0.5436319 | 0.99999872 |
| NCAM1 | 12.9609282 | 0.87062004 | 0.73441759 | 1.18545642 | 0.23583704 | 0.99999872 |
| TTC12 | 15.8804522 | 0.147937805 | 0.3740267 | 0.3955274 | 0.69245371 | 0.99999872 |
| ZW10 | 7.4990357 | -0.149547524 | 0.58066646 | -0.2575446 | 0.79675837 | 0.99999872 |
| USP28 | 16.3716762 | 0.411423843 | 0.39632034 | 1.03810933 | 0.29921916 | 0.99999872 |
| ZBTB16 | 54.7572777 | -0.059935951 | 0.87463518 | -0.0685268 | 0.94536629 | 0.99999872 |
| AP002518.2 | 6.15200863 | 3.269176907 | 1.6014077 | 2.04143948 | 0.04120716 | 0.95980711 |
| NNMT | 20.7688057 | 1.508408897 | 0.9516762 | 1.58500223 | 1.13E-01 | 0.99999872 |
| C11orf71 | 5.91307725 | 0.278587801 | 0.81979339 | 0.33982685 | 0.73398693 | 0.99999872 |
| REXO2 | 26.7520713 | 0.90766269 | 0.33878836 | 2.6791437 | 0.00738107 | 0.63916415 |
| CADM1 | 41.4282541 | -1.313654065 | 0.76735851 | -1.711917 | 0.08691197 | 0.99999872 |
| BUD13 | 8.04541904 | -0.078542588 | 0.57791246 | -0.1359074 | 0.89189448 | 0.99999872 |
| ZPR1 | 5.97890117 | -0.591002002 | 0.67513245 | -0.8753868 | 0.3813635 | 0.99999872 |
| SIK3 | 394.000061 | -0.075379882 | 0.33910667 | -0.2222896 | 0.82408846 | 0.99999872 |
| PAFAH1B2 | 49.2959055 | 0.202651505 | 0.32053771 | 0.6322236 | 0.52724078 | 0.99999872 |
| SIDT2 | 11.242352 | 0.272113759 | 0.54117266 | 0.50282244 | 0.61508911 | 0.99999872 |
| TAGLN | 17.8978792 | 0.089470592 | 0.82411924 | 0.1085651 | 0.91354744 | 0.99999872 |
| PCSK7 | 24.6312307 | 0.065734328 | 0.31751899 | 0.20702487 | 0.83599043 | 0.99999872 |
| RNF214 | 16.3788498 | -0.464876434 | 0.45386056 | -1.0242715 | 0.30570706 | 0.99999872 |
| BACE1 | 52.5859838 | -0.455099712 | 0.27823784 | -1.63565 | 0.10191287 | 0.99999872 |
| CEP164 | 25.88584 | -0.366045774 | 0.34948279 | -1.0473928 | 0.29491843 | 0.99999872 |
| DSCAML1 | 17.2441675 | -0.19331402 | 0.99613326 | -0.1940644 | 0.84612544 | 0.99999872 |
| FXYD2 | 15.3906762 | 0.474169031 | 0.80385122 | 0.58987163 | 0.55527671 | 0.99999872 |
| UBE4A | 32.9089956 | 0.094875328 | 0.31812632 | 0.29823163 | 0.76552638 | 0.99999872 |
| ATP5MG | 51.6276753 | 0.731494644 | 0.41818917 | 1.74919557 | 0.08025722 | 0.99999872 |
| KMT2A | 89.584226 | -0.146974026 | 0.24500357 | -0.5998852 | 0.54858272 | 0.99999872 |
| TMEM25 | 6.01988113 | 0.032015756 | 0.55620218 | 0.05756136 | 0.95409803 | 0.99999872 |
| IFT46 | 5.36448263 | 0.715803343 | 0.64087068 | 1.11692321 | 0.26402716 | 0.99999872 |
| ARCN1 | 27.3056621 | 0.308679457 | 0.46047342 | 0.67035239 | 0.50263317 | 0.99999872 |
| PHLDB1 | 33.634533 | -0.750395391 | 0.40943153 | -1.8327738 | 0.06683622 | 0.99999872 |
| AP002954.1 | 10.5940895 | 1.368765137 | 0.56993981 | 2.40159596 | 0.01632373 | 0.80846766 |
| DDX6 | 61.0591502 | 0.009805785 | 0.34806255 | 0.02817248 | 0.97752459 | 0.99999872 |
| BCL9L | 22.4487059 | -0.66564348 | 0.52697773 | -1.263134 | 0.20654103 | 0.99999872 |
| CCDC84 | 26.1066292 | -0.725618755 | 0.48358624 | -1.500495 | 0.13348622 | 0.99999872 |
| RPS25 | 75.1468201 | 0.634129458 | 0.37898189 | 1.67324475 | 0.09427912 | 0.99999872 |
| TRAPPC4 | 28.5963641 | 0.743332273 | 0.43177389 | 1.72157763 | 0.08514606 | 0.99999872 |
| SLC37A4 | 7.99918019 | 1.190300266 | 0.64241141 | 1.8528629 | 0.06390201 | 0.99999872 |
| HYOU1 | 26.0797356 | 0.413838644 | 0.67723106 | 0.61107452 | 0.54115025 | 0.99999872 |
| VPS11 | 8.56271984 | -0.409093399 | 0.47518752 | -0.8609094 | 0.38928795 | 0.99999872 |
| H2AFX | 5.43294293 | 0.136737617 | 0.6391143 | 0.21394861 | 0.83058713 | 0.99999872 |
| DPAGT1 | 6.23071595 | 0.503832239 | 0.60537212 | 0.83226866 | 0.40525732 | 0.99999872 |
| HINFP | 5.57339781 | -0.081761842 | 0.62500804 | -0.1308173 | 0.89591987 | 0.99999872 |
| NLRX1 | 6.25063949 | -1.378404062 | 0.58374165 | -2.3613255 | 0.01820974 | 0.85282771 |
| CBL | 32.3745743 | 0.078862881 | 0.39986415 | 0.19722418 | 0.84365211 | 0.99999872 |
| RNF26 | 6.59431158 | -0.105981324 | 0.55161403 | -0.1921295 | 0.84764077 | 0.99999872 |
| OAF | 4.88399698 | 0.044844953 | 0.84909982 | 0.0528147 | 0.95787955 | 0.99999872 |
| TMEM136 | 6.49069155 | 0.45046217 | 0.66067336 | 0.68182282 | 0.495351 | 0.99999872 |
| ARHGEF12 | 600.01953 | 0.489634352 | 0.35565292 | 1.37671961 | 0.16859895 | 0.99999872 |
| SC5D | 9.18522006 | -0.12795826 | 0.50085592 | -0.2554792 | 0.79835302 | 0.99999872 |
| SORL1 | 16.7908451 | -0.233317757 | 0.75981764 | -0.3070707 | 0.75878954 | 0.99999872 |
| JHY | 30.364722 | 0.028695738 | 0.39500931 | 0.07264573 | 0.94208804 | 0.99999872 |
| HSPA8 | 70.9069654 | 0.557981305 | 0.48943623 | 1.14004903 | 0.25426587 | 0.99999872 |
| VWA5A | 8.43247855 | -0.428982592 | 0.46449802 | -0.9235402 | 0.35572576 | 0.99999872 |
| TBRG1 | 25.8351857 | 0.103955176 | 0.32832552 | 0.31662229 | 0.75153022 | 0.99999872 |

| | | | | | | |
|---|---|---|---|---|---|---|
| SIAE | 12.4352177 | -0.007343467 | 0.62043465 | -0.011836 | 0.99055646 | 0.99999872 |
| MSANTD2 | 22.0633986 | -0.026381844 | 0.3800854 | -0.0694103 | 0.94466303 | 0.99999872 |
| TMEM218 | 7.04626728 | 0.083694207 | 0.59499964 | 0.14066262 | 0.88813648 | 0.99999872 |
| PKNOX2 | 16.9222937 | 0.254318492 | 0.66139075 | 0.38452079 | 0.70059249 | 0.99999872 |
| EI24 | 27.0074416 | 0.001913331 | 0.36766767 | 0.00520397 | 0.99584785 | 0.99999872 |
| STT3A | 17.2432308 | 0.513012537 | 0.43234841 | 1.18657203 | 0.23539647 | 0.99999872 |
| PUS3 | 4.50725947 | 0.253344025 | 0.73135349 | 0.34640434 | 0.72903886 | NA |
| CDON | 14.5256454 | -0.435848671 | 0.55970368 | -0.7787132 | 0.43614865 | 0.99999872 |
| RPUSD4 | 5.49747942 | -0.871389826 | 0.70428369 | -1.2372711 | 0.21598647 | 0.99999872 |
| FAM118B | 14.281737 | -0.182231777 | 0.46156255 | -0.3948149 | 0.69297949 | 0.99999872 |
| SRPRA | 27.2415468 | 0.250588021 | 0.47297986 | 0.52980696 | 0.59624578 | 0.99999872 |
| FOXRED1 | 8.35776922 | -0.218227843 | 0.61253678 | -0.356269 | 0.72163915 | 0.99999872 |
| TIRAP | 8.32953348 | -0.708717991 | 0.55979459 | -1.2660322 | 0.20550155 | 0.99999872 |
| ST3GAL4 | 12.8468643 | -0.194294514 | 0.55592658 | -0.3494967 | 0.72671644 | 0.99999872 |
| ETS1 | 13.1648269 | -0.350106918 | 0.421936 | -0.8297631 | 0.40667274 | 0.99999872 |
| KCNJ1 | 9.09476945 | 1.381027819 | 1.04004814 | 1.32784991 | 0.1842277 | 0.99999872 |
| ARHGAP32 | 39.7521041 | 0.172502777 | 0.46496634 | 0.37100057 | 0.71063711 | 0.99999872 |
| NFRKB | 13.6856674 | -0.290546536 | 0.40937884 | -0.7097253 | 0.47787448 | 0.99999872 |
| PRDM10 | 15.4292838 | -0.208996644 | 0.61703708 | -0.33871 | 0.73482819 | 0.99999872 |
| APLP2 | 192.861016 | 0.326364867 | 0.37459288 | 0.87125219 | 0.38361647 | 0.99999872 |
| ZBTB44 | 52.3753004 | -0.220768765 | 0.31013557 | -0.711846 | 0.47656015 | 0.99999872 |
| AP003486.1 | 6.21507008 | 0.015862972 | 0.70009217 | 0.02265841 | 0.98192276 | 0.99999872 |
| SNX19 | 27.3647766 | -0.244803101 | 0.38011479 | -0.6440241 | 0.51955978 | 0.99999872 |
| NTM | 10.039854 | -1.667317694 | 0.58913266 | -2.8301227 | 0.00465302 | 0.52915868 |
| OPCML | 9.75431298 | -1.659195486 | 0.66160501 | -2.5078339 | 0.01214737 | 0.72670507 |
| JAM3 | 23.371099 | 0.123473935 | 0.5065294 | 0.2437646 | 0.80741314 | 0.99999872 |
| NCAPD3 | 14.0475131 | -0.38550539 | 0.42237731 | -0.9127038 | 0.36139833 | 0.99999872 |
| VPS26B | 14.6156274 | 0.328447133 | 0.48536072 | 0.67670728 | 0.4985917 | 0.99999872 |
| THYN1 | 4.97475297 | 0.431508886 | 0.80565826 | 0.53559792 | 0.59223647 | 0.99999872 |
| ACAD8 | 6.26132845 | 0.272174283 | 0.60670099 | 0.44861355 | 0.65371046 | 0.99999872 |
| ZMYND11 | 81.4364593 | -0.24965132 | 0.27947241 | -0.893295 | 0.37169919 | 0.99999872 |
| DIP2C | 80.4187558 | -0.068410275 | 0.30789992 | -0.2221835 | 0.82417106 | 0.99999872 |
| LARP4B | 93.4716026 | -0.418622499 | 0.2375115 | -1.7625357 | 0.07797883 | 0.99999872 |
| GTPBP4 | 13.8004925 | -0.208153175 | 0.51048812 | -0.4077532 | 0.68345486 | 0.99999872 |
| IDI1 | 15.8838755 | 0.306128882 | 0.48234667 | 0.63466569 | 0.52564647 | 0.99999872 |
| WDR37 | 29.7158118 | -0.401325915 | 0.34787405 | -1.1536529 | 0.24864249 | 0.99999872 |
| PFKP | 6.74566143 | -0.377789097 | 0.88169016 | -0.4284828 | 0.66829963 | 0.99999872 |
| PITRM1 | 13.8929155 | -0.047688952 | 0.4199744 | -0.113552 | 0.9095929 | 0.99999872 |
| KLF6 | 28.938155 | -1.228344245 | 0.49503364 | -2.4813349 | 0.01308913 | 0.74263079 |
| AKR1C1 | 6.67025074 | -1.647837389 | 0.75747749 | -2.1754275 | 0.02959809 | 0.90266962 |
| AKR1C3 | 6.46675032 | 0.270423679 | 0.82187682 | 0.32903189 | 0.74213159 | 0.99999872 |
| NET1 | 4.06258492 | 0.634173266 | 0.83205151 | 0.76218029 | 0.44595241 | NA |
| ASB13 | 5.64094419 | 0.312012518 | 0.63914899 | 0.48816868 | 0.62543037 | 0.99999872 |
| FAM208B | 65.7739854 | -0.243023393 | 0.37592531 | -0.6464672 | 0.51797681 | 0.99999872 |
| GDI2 | 45.7982777 | -0.242140122 | 0.33735217 | -0.7177666 | 0.47290121 | 0.99999872 |
| ANKRD16 | 7.849937 | -0.19357067 | 0.59355257 | -0.3261222 | 0.7443319 | 0.99999872 |
| FBH1 | 32.7617327 | -0.26771351 | 0.28683406 | -0.9333393 | 0.35064479 | 0.99999872 |
| RBM17 | 27.637018 | 0.020032875 | 0.35854099 | 0.05587332 | 0.95544273 | 0.99999872 |
| PFKFB3 | 33.8666949 | -0.953501332 | 0.44206731 | -2.1569144 | 0.03101234 | 0.91534437 |
| LINP1 | 6.12904667 | -2.263184282 | 1.10696793 | -2.0444895 | 0.04090522 | 0.95980711 |
| LINC00707 | 9.11172782 | -1.56853695 | 1.00561517 | -1.5597785 | 0.11881223 | 0.99999872 |
| SFMBT2 | 410.825319 | -0.369343087 | 0.5542838 | -0.6663429 | 5.05E-01 | 0.99999872 |
| ITIH5 | 146.238797 | 0.52969286 | 0.66450506 | 0.7971239 | 0.42537908 | 0.99999872 |
| KIN | 11.7608264 | 0.113425518 | 0.50097142 | 0.22641116 | 0.82088165 | 0.99999872 |
| ATP5F1C | 38.4416905 | 0.494272243 | 0.31677529 | 1.5603245 | 0.11868322 | 0.99999872 |
| TAF3 | 86.4926816 | 0.104964239 | 0.2602673 | 0.40329399 | 0.68673196 | 0.99999872 |
| USP6NL | 31.6923894 | 0.082686645 | 0.34610688 | 0.23890494 | 0.81117929 | 0.99999872 |
| ECHDC3 | 7.08121698 | 0.544327298 | 0.68494911 | 0.79469743 | 0.42678953 | 0.99999872 |
| UPF2 | 40.1485682 | -0.235918926 | 0.31489496 | -0.7491988 | 0.45373739 | 0.99999872 |
| DHTKD1 | 21.1871731 | -1.233799732 | 0.33160192 | -3.7207255 | 0.00019865 | 0.12623956 |
| SEC61A2 | 7.94523304 | 0.945499016 | 0.69273181 | 1.36488465 | 0.17228931 | 0.99999872 |

| | | | | | | |
|---|---|---|---|---|---|---|
| NUDT5 | 21.228129 | 0.525398817 | 0.38485732 | 1.36517819 | 0.17219705 | 0.99999872 |
| CDC123 | 22.3215197 | -0.237834543 | 0.35370111 | -0.6724167 | 0.50131843 | 0.99999872 |
| CAMK1D | 9.04325694 | -0.884984092 | 0.56027513 | -1.5795527 | 0.11420934 | 0.99999872 |
| OPTN | 66.4550011 | -0.117575566 | 0.45766376 | -0.2569038 | 0.79725303 | 0.99999872 |
| PHYH | 9.31814294 | 1.021102366 | 0.53606534 | 1.90480954 | 0.05680483 | 0.99999872 |
| SEPHS1 | 18.2841587 | -0.331706302 | 0.38441239 | -0.8628918 | 0.38819695 | 0.99999872 |
| BEND7 | 26.6155309 | 0.085316288 | 0.347782 | 0.24531542 | 0.80621221 | 0.99999872 |
| PRPF18 | 4.83852919 | 0.536639358 | 0.67066374 | 0.80016158 | 0.42361718 | 0.99999872 |
| AL157392.5 | 11.5028416 | 0.080855174 | 0.50888081 | 0.15888824 | 0.87375693 | 0.99999872 |
| FRMD4A | 35.8761207 | 0.098223033 | 0.67281941 | 0.14598722 | 0.88393148 | 0.99999872 |
| FAM107B | 21.0854177 | -0.669725854 | 0.69752699 | -0.9601433 | 0.33698311 | 0.99999872 |
| DCLRE1C | 11.886213 | 0.586092784 | 0.60048264 | 0.97603619 | 0.32904653 | 0.99999872 |
| OLAH | 18.5263398 | 0.83039572 | 0.74170223 | 1.119581 | 0.26289236 | 0.99999872 |
| RPP38 | 24.1912731 | -0.041488649 | 0.46885548 | -0.0884892 | 0.92948786 | 0.99999872 |
| NMT2 | 25.7333722 | -0.56116282 | 0.42712788 | -1.3138052 | 0.18891177 | 0.99999872 |
| FAM171A1 | 15.3610575 | -0.565586787 | 0.43034367 | -1.3142677 | 0.18875613 | 0.99999872 |
| MINDY3 | 40.0499533 | 0.387522178 | 0.35922958 | 1.07875911 | 0.28069513 | 0.99999872 |
| PTER | 16.0790457 | 0.01948281 | 0.54290691 | 0.0358861 | 0.97137318 | 0.99999872 |
| RSU1 | 42.0799918 | 0.038612348 | 0.34529217 | 0.11182515 | 0.91096205 | 0.99999872 |
| CUBN | 73.5804559 | -0.001541514 | 1.01591948 | -0.0015174 | 0.99878932 | 0.99999872 |
| TRDMT1 | 12.1901632 | -0.260941143 | 0.57651911 | -0.4526149 | 0.65082606 | 0.99999872 |
| VIM | 513.986305 | 0.28567411 | 0.55298101 | 0.51660745 | 0.60543021 | 0.99999872 |
| HACD1 | 5.90042064 | 0.752823099 | 0.89752766 | 0.83877426 | 0.401596 | 0.99999872 |
| STAM | 19.5157409 | -0.082075752 | 0.40541773 | -0.2024474 | 0.839567 | 0.99999872 |
| CACNB2 | 270.032155 | -0.477974415 | 0.52617895 | -0.9083876 | 0.36367349 | 0.99999872 |
| AL450384.2 | 17.8719357 | 0.493067119 | 0.46988829 | 1.04932839 | 0.294027 | 0.99999872 |
| NSUN6 | 25.2146457 | -0.159104159 | 0.3960061 | -0.401772 | 0.68785183 | 0.99999872 |
| ARL5B | 16.4145855 | -0.015901863 | 0.45240557 | -0.0351496 | 0.97196047 | 0.99999872 |
| MALRD1 | 9.87488751 | -1.179897119 | 0.89991783 | -1.3111165 | 0.1898184 | 0.99999872 |
| PLXDC2 | 242.393239 | -0.673645643 | 0.25419563 | -2.6501071 | 0.00804663 | 0.66265058 |
| NEBL | 1621.07413 | -0.27803311 | 0.29640076 | -0.938031 | 0.34822846 | 0.99999872 |
| MLLT10 | 119.674944 | 0.210984905 | 0.20558874 | 1.02624736 | 0.304775 | 0.99999872 |
| DNAJC1 | 81.095262 | -0.071779548 | 0.4013203 | -0.1788585 | 0.85804881 | 0.99999872 |
| BMI1 | 7.74791606 | 0.017417882 | 0.64088453 | 0.02717788 | 0.97831786 | 0.99999872 |
| PIP4K2A | 28.9857608 | 0.160880935 | 0.37402093 | 0.43013885 | 0.66709464 | 0.99999872 |
| MSRB2 | 23.4407797 | 0.361121235 | 0.37359207 | 0.96661912 | 0.33373447 | 0.99999872 |
| OTUD1 | 11.8633026 | 0.331628552 | 0.65730635 | 0.50452662 | 0.61389136 | 0.99999872 |
| KIAA1217 | 78.5893791 | -0.52410887 | 0.53572193 | -0.9783226 | 0.32791479 | 0.99999872 |
| ARHGAP21 | 110.634109 | -0.15956202 | 0.27115267 | -0.5884582 | 0.55622477 | 0.99999872 |
| PDSS1 | 5.70725625 | 0.13892617 | 0.66179903 | 0.20992199 | 0.83372856 | 0.99999872 |
| ABI1 | 130.348245 | 0.469888669 | 0.35636836 | 1.31854766 | 0.18732038 | 0.99999872 |
| ANKRD26 | 48.6744736 | -0.199557523 | 0.31874737 | -0.626068 | 0.53127035 | 0.99999872 |
| YME1L1 | 52.1874454 | 0.104901764 | 0.24512965 | 0.42794401 | 0.66869189 | 0.99999872 |
| MASTL | 7.06693007 | -0.548552039 | 0.63808561 | -0.8596841 | 0.38996321 | 0.99999872 |
| ACBD5 | 18.078396 | -0.122285558 | 0.38213133 | -0.3200092 | 0.74896132 | 0.99999872 |
| RAB18 | 83.6733728 | 0.35379174 | 0.28651387 | 1.23481542 | 0.21689921 | 0.99999872 |
| MPP7 | 15.9415377 | -0.117356609 | 0.4732191 | -0.2479963 | 0.80413723 | 0.99999872 |
| WAC-AS1 | 23.1732527 | -0.711675249 | 0.35224877 | -2.0203768 | 0.04334431 | 0.95980711 |
| WAC | 149.468003 | 0.109917513 | 0.20236742 | 0.54315815 | 0.58702092 | 0.99999872 |
| BAMBI | 70.1005161 | 1.755234441 | 0.9481127 | 1.85129303 | 0.06412741 | 0.99999872 |
| SVIL | 81.4177456 | -0.347262021 | 0.46444775 | -0.747688 | 0.45464836 | 0.99999872 |
| MTPAP | 9.9612336 | 0.163017958 | 0.55436176 | 0.29406422 | 0.76870883 | 0.99999872 |
| MAP3K8 | 13.6468191 | 0.458127717 | 0.67839415 | 0.67531201 | 0.49947755 | 0.99999872 |
| ZNF438 | 35.1135765 | 0.57249748 | 0.33560568 | 1.7058635 | 0.0880335 | 0.99999872 |
| ZEB1 | 49.6843482 | -0.369086629 | 0.39399586 | -0.9367779 | 0.34887279 | 0.99999872 |
| ARHGAP12 | 47.4373181 | 0.137260871 | 0.32983065 | 0.41615559 | 0.67729615 | 0.99999872 |
| KIF5B | 80.0862191 | -0.010932661 | 0.32177256 | -0.0339764 | 0.972896 | 0.99999872 |
| EPC1 | 120.008745 | 0.154844249 | 0.28025294 | 0.55251606 | 0.58059483 | 0.99999872 |
| AL391839.1 | 4.6592533 | 1.580765233 | 0.90175371 | 1.75299 | 0.07960372 | 0.99999872 |
| CCDC7 | 57.306251 | -0.297313665 | 0.43802778 | -0.6787553 | 0.49729294 | 0.99999872 |
| ITGB1 | 230.797665 | 0.28060458 | 0.42768733 | 0.65609748 | 0.51176141 | 0.99999872 |

| | | | | | | |
|---|---|---|---|---|---|---|
| NRP1 | 47.0219275 | -0.53320521 | 0.59319248 | -0.8988738 | 0.36871986 | 0.99999872 |
| PARD3 | 959.484077 | -0.632291103 | 0.40855544 | -1.5476262 | 0.12171232 | 0.99999872 |
| CUL2 | 22.5982068 | 0.145976102 | 0.34522658 | 0.42284143 | 0.67241096 | 0.99999872 |
| CREM | 42.2029698 | -0.598984279 | 0.32331418 | -1.8526384 | 0.0639342 | 0.99999872 |
| CCNY | 116.370036 | 0.084566071 | 0.22770764 | 0.37138004 | 0.71035449 | 0.99999872 |
| AL117336.3 | 10.5932269 | 0.651569862 | 0.56500258 | 1.15321572 | 0.24882185 | 0.99999872 |
| FZD8 | 27.8624002 | 0.16463729 | 1.07638419 | 0.15295402 | 0.87843453 | 0.99999872 |
| ZNF248 | 46.8343666 | -0.247428158 | 0.4352735 | -0.568443 | 0.56973423 | 0.99999872 |
| ZNF25 | 19.3923776 | -0.054960015 | 0.37371928 | -0.1470623 | 0.88308285 | 0.99999872 |
| ZNF33A | 46.9348841 | -0.698713963 | 0.30174231 | -2.3155983 | 0.02058021 | 0.8671024 |
| AL117339.5 | 13.2436503 | 0.320117329 | 0.56274361 | 0.56885111 | 0.56945718 | 0.99999872 |
| LINC00839 | 278.608855 | 0.986569596 | 0.39365837 | 2.50615679 | 0.01220515 | 0.72670507 |
| ZNF33B | 29.0284232 | -0.934988632 | 0.36142644 | -2.5869403 | 0.00968324 | 0.70458997 |
| BMS1 | 18.5441829 | -0.023567492 | 0.37025798 | -0.0636515 | 0.94924769 | 0.99999872 |
| CSGALNACT2 | 21.8649083 | 0.573151191 | 0.38462356 | 1.49016142 | 0.1361818 | 0.99999872 |
| HNRNPF | 45.680962 | -0.047446053 | 0.3742954 | -0.126761 | 0.89912958 | 0.99999872 |
| ZNF487 | 8.78641042 | -0.129203014 | 0.52784314 | -0.2447754 | 0.80663034 | 0.99999872 |
| ZNF32 | 17.2525597 | 0.38819465 | 0.38252932 | 1.01481017 | 0.31019632 | 0.99999872 |
| CXCL12 | 6.74246043 | -1.534166782 | 1.36909297 | -1.1205717 | 0.26247023 | 0.99999872 |
| TMEM72-AS | 7.74196469 | 0.341005529 | 0.80791992 | 0.42207838 | 0.67296781 | 0.99999872 |
| RASSF4 | 21.5809021 | 0.120068152 | 0.3854406 | 0.31150883 | 0.75541383 | 0.99999872 |
| DEPP1 | 7.64858137 | 0.937385236 | 1.18251813 | 0.79270263 | 0.4279511 | 0.99999872 |
| ZNF22 | 17.4264411 | 0.387778625 | 0.42195005 | 0.91901546 | 0.35808748 | 0.99999872 |
| 8-Mar | 27.6027448 | -0.05632296 | 0.40545017 | -0.1389146 | 0.88951761 | 0.99999872 |
| ZFAND4 | 14.3455831 | -0.5939597 | 0.4111365 | -1.4446776 | 0.14854846 | 0.99999872 |
| WASHC2C | 14.0060782 | -0.432004481 | 0.46751417 | -0.9240458 | 0.3554625 | 0.99999872 |
| AGAP4 | 14.8830854 | -0.460521455 | 0.40542937 | -1.1358858 | 0.25600437 | 0.99999872 |
| TIMM23 | 15.111353 | -0.170253416 | 0.47085495 | -0.3615836 | 0.71766324 | 0.99999872 |
| NCOA4 | 31.288544 | 0.154647373 | 0.33580362 | 0.4605292 | 0.64513641 | 0.99999872 |
| AL356056.2 | 12.4315475 | -1.56549009 | 0.73101101 | -2.1415411 | 0.03223043 | 0.92959829 |
| PTPN20 | 7.63180184 | 1.193107229 | 0.70513413 | 1.69202876 | 0.09064049 | 0.99999872 |
| AGAP9 | 12.0543464 | -0.082905309 | 0.46123902 | -0.1797448 | 0.85735293 | 0.99999872 |
| MAPK8 | 69.5589308 | -0.250600064 | 0.27907291 | -0.8979735 | 0.3691997 | 0.99999872 |
| AL138760.1 | 5.35116141 | 0.42425731 | 1.17051199 | 0.36245448 | 0.71701244 | 0.99999872 |
| ERCC6 | 34.3757298 | -1.180951227 | 0.59043352 | -2.0001426 | 0.04548487 | 0.95980711 |
| PARG | 36.7849822 | 0.186595398 | 0.37944712 | 0.49175601 | 0.62289184 | 0.99999872 |
| TIMM23B | 37.407211 | -1.024774644 | 0.44065082 | -2.3255934 | 0.02004025 | 0.8671024 |
| LINC00843 | 9.61378669 | 0.106499077 | 0.62029657 | 0.17169058 | 0.86368079 | 0.99999872 |
| AGAP6 | 16.5245598 | -0.284158343 | 0.44917569 | -0.6326218 | 0.52698063 | 0.99999872 |
| WASHC2A | 17.0284747 | -0.642745101 | 0.40538837 | -1.5855045 | 0.11285175 | 0.99999872 |
| SGMS1 | 131.239196 | 0.194640326 | 0.32009378 | 0.60807281 | 0.54313919 | 0.99999872 |
| ASAH2B | 6.50692303 | 0.099613738 | 0.63968472 | 0.15572318 | 0.87625123 | 0.99999872 |
| PRKG1 | 704.232462 | -0.299231074 | 0.47576518 | -0.628947 | 0.52938377 | 0.99999872 |
| CSTF2T | 5.00666173 | -0.149027001 | 0.62252997 | -0.2393893 | 0.81080374 | 0.99999872 |
| PCDH15 | 20.2844962 | -0.267188383 | 0.72878978 | -0.3666193 | 7.14E-01 | 0.99999872 |
| IPMK | 12.6754479 | 0.282448519 | 0.72277445 | 0.39078376 | 0.69595708 | 0.99999872 |
| CISD1 | 26.2256237 | 0.414654959 | 0.3905863 | 1.06162186 | 0.28840739 | 0.99999872 |
| UBE2D1 | 14.7933227 | 0.51296314 | 0.41507765 | 1.23582452 | 0.2165238 | 0.99999872 |
| TFAM | 13.4520667 | 0.112463364 | 0.50808941 | 0.22134562 | 0.82482333 | 0.99999872 |
| BICC1 | 119.755217 | -0.985221505 | 0.71886595 | -1.3705219 | 0.17052405 | 0.99999872 |
| PHYHIPL | 10.4111741 | -0.80874494 | 0.87977087 | -0.9192677 | 0.35795557 | 0.99999872 |
| SLC16A9 | 14.3688347 | 0.298212605 | 1.7828261 | 0.1672696 | 0.86715792 | 0.99999872 |
| CCDC6 | 88.3360104 | 0.196107669 | 0.2982397 | 0.65755051 | 0.510827 | 0.99999872 |
| ANK3 | 130.432994 | -0.175246782 | 0.5194523 | -0.3373684 | 0.73583922 | 0.99999872 |
| AL592430.1 | 10.1226597 | -0.142511163 | 0.9830912 | -0.1449623 | 0.88474064 | 0.99999872 |
| RHOBTB1 | 13.2181827 | 0.033819843 | 1.04778066 | 0.0322776 | 0.97425067 | 0.99999872 |
| CABCOCO1 | 15.5845028 | -0.29145232 | 0.68958167 | -0.4226509 | 0.67254998 | 0.99999872 |
| ARID5B | 83.4506106 | 0.53259972 | 0.47798248 | 1.1142662 | 0.265165 | 0.99999872 |
| ADO | 11.8223143 | -0.106872478 | 0.47811647 | -0.2235281 | 0.82312449 | 0.99999872 |
| NRBF2 | 7.57226087 | -0.228286051 | 0.55438304 | -0.411784 | 0.68049777 | 0.99999872 |
| JMJD1C | 260.173867 | -0.092982767 | 0.28901079 | -0.3217277 | 0.74765902 | 0.99999872 |

| | | | | | | |
|---|---|---|---|---|---|---|
| REEP3 | 53.9092184 | 0.018233193 | 0.41167975 | 0.04428975 | 0.96467344 | 0.99999872 |
| LINC01515 | 37.7975258 | -0.269496476 | 0.55447808 | -0.4860363 | 0.62694143 | 0.99999872 |
| CTNNA3 | 43.740501 | -0.50150421 | 0.60012693 | -0.8356636 | 0.40334419 | 0.99999872 |
| SIRT1 | 23.7687633 | -0.043921696 | 0.35578069 | -0.1234516 | 0.9017495 | 0.99999872 |
| HERC4 | 194.43301 | 0.028823348 | 0.24370893 | 0.11826956 | 0.90585408 | 0.99999872 |
| PBLD | 15.5329244 | -0.675644681 | 0.41423582 | -1.6310629 | 0.10287705 | 0.99999872 |
| HNRNPH3 | 53.8146808 | 0.317448249 | 0.26252588 | 1.20920746 | 0.22658315 | 0.99999872 |
| RUFY2 | 43.8943375 | 0.06255388 | 0.35731158 | 0.17506816 | 0.86102608 | 0.99999872 |
| SLC25A16 | 27.8926747 | -0.448410031 | 0.33719642 | -1.3298185 | 0.18357809 | 0.99999872 |
| TET1 | 11.7178965 | -0.422707525 | 0.54807378 | -0.7712603 | 0.44055268 | 0.99999872 |
| CCAR1 | 54.7484999 | -0.201313743 | 0.36576512 | -0.5503908 | 0.58205139 | 0.99999872 |
| DDX50 | 28.5895832 | -0.405973102 | 0.30506491 | -1.3307762 | 0.18326268 | 0.99999872 |
| DDX21 | 25.9109567 | -0.470721608 | 0.59903129 | -0.7858047 | 0.43198191 | 0.99999872 |
| KIF1BP | 21.2146387 | 0.253835869 | 0.3600334 | 0.70503422 | 0.48078894 | 0.99999872 |
| VPS26A | 36.6878408 | 0.101320784 | 0.31877794 | 0.31784126 | 0.75060535 | 0.99999872 |
| SUPV3L1 | 12.8545954 | -0.273457323 | 0.48367625 | -0.5653726 | 0.57182032 | 0.99999872 |
| HKDC1 | 8.94136668 | -1.642000066 | 0.81626811 | -2.011594 | 0.04426275 | 0.95980711 |
| HK1 | 29.4644185 | -0.253752895 | 0.44816225 | -0.5662076 | 0.57125263 | 0.99999872 |
| TSPAN15 | 20.7793877 | 0.935539047 | 0.57331559 | 1.63180464 | 0.10272065 | 0.99999872 |
| H2AFY2 | 10.3841419 | 0.092235935 | 0.52830656 | 0.1745879 | 0.86140345 | 0.99999872 |
| AIFM2 | 12.0075195 | -0.19712351 | 0.48692179 | -0.4048361 | 0.685598 | 0.99999872 |
| TYSND1 | 8.84903585 | 0.017530618 | 0.62060734 | 0.02824752 | 0.97746474 | 0.99999872 |
| SAR1A | 29.9784547 | -0.103429804 | 0.33696744 | -0.306943 | 0.75888674 | 0.99999872 |
| PPA1 | 78.2397908 | -0.228231276 | 0.52310844 | -0.4362982 | 0.66262038 | 0.99999872 |
| LRRC20 | 7.49621537 | 0.334296373 | 0.54808559 | 0.60993462 | 0.54190512 | 0.99999872 |
| EIF4EBP2 | 25.3149644 | 0.143190645 | 0.34059425 | 0.42041416 | 0.67418293 | 0.99999872 |
| SGPL1 | 31.3131508 | 0.215307521 | 0.33468434 | 0.6433152 | 0.5200196 | 0.99999872 |
| PCBD1 | 31.2706746 | 0.998188846 | 0.39895799 | 2.50198985 | 0.01234975 | 0.72670507 |
| VSIR | 31.0196989 | -1.055945393 | 0.43700047 | -2.4163484 | 0.01567705 | 0.79608149 |
| PSAP | 423.952518 | 0.482295468 | 0.48515603 | 0.99410384 | 0.32017232 | 0.99999872 |
| CHST3 | 4.90903741 | -1.168295956 | 0.74593709 | -1.5662124 | 0.11729889 | 0.99999872 |
| SPOCK2 | 640.797703 | -0.473270384 | 0.57237875 | -0.8268483 | 0.40832306 | 0.99999872 |
| ASCC1 | 35.1488801 | -0.587688198 | 0.40446074 | -1.4530167 | 0.14621912 | 0.99999872 |
| ANAPC16 | 115.423617 | 0.284567472 | 0.29305909 | 0.97102421 | 0.33153622 | 0.99999872 |
| DDIT4 | 13.6791997 | 0.140698784 | 1.0050679 | 0.13998933 | 0.88866842 | 0.99999872 |
| DNAJB12 | 20.4055211 | -0.538861515 | 0.43973749 | -1.2254164 | 0.22041839 | 0.99999872 |
| MICU1 | 55.0593281 | -0.073632304 | 0.37573781 | -0.1959672 | 0.8446358 | 0.99999872 |
| MCU | 83.4289436 | -0.225165431 | 0.41134495 | -0.5473883 | 0.58411196 | 0.99999872 |
| P4HA1 | 40.5458809 | 0.522089416 | 0.43473551 | 1.20093576 | 0.22977612 | 0.99999872 |
| ECD | 15.3899634 | 0.052347186 | 0.39983072 | 0.13092337 | 0.89583593 | 0.99999872 |
| FAM149B1 | 22.1439115 | 0.134597329 | 0.42858843 | 0.31404798 | 0.7534846 | 0.99999872 |
| DNAJC9 | 19.7830094 | -0.265231825 | 0.42404776 | -0.6254763 | 0.53165849 | 0.99999872 |
| MRPS16 | 19.3910165 | 0.372710901 | 0.41034521 | 0.90828622 | 0.36372702 | 0.99999872 |
| CFAP70 | 16.1341961 | 0.057180142 | 0.59311475 | 0.09640654 | 0.9231977 | 0.99999872 |
| ANXA7 | 46.2012431 | 0.202329696 | 0.31532616 | 0.64165211 | 0.52109909 | 0.99999872 |
| PPP3CB | 48.906364 | -0.110902202 | 0.27543378 | -0.4026456 | 0.68720895 | 0.99999872 |
| USP54 | 29.41356 | -0.4061604 | 0.52401662 | -0.7750907 | 0.43828607 | 0.99999872 |
| SEC24C | 23.989127 | -0.24534383 | 0.38797447 | -0.6323711 | 0.52714443 | 0.99999872 |
| FUT11 | 31.8224467 | -0.265447665 | 0.53598012 | -0.4952565 | 0.62041904 | 0.99999872 |
| CHCHD1 | 15.350442 | 0.563493018 | 0.51037958 | 1.10406655 | 0.26956427 | 0.99999872 |
| ZSWIM8 | 20.7659861 | 0.056495352 | 0.36217933 | 0.15598723 | 0.87604308 | 0.99999872 |
| CAMK2G | 32.1969239 | 0.081740282 | 0.37879999 | 0.21578744 | 0.82915344 | 0.99999872 |
| PLAU | 9.19413081 | -0.820669761 | 0.89823764 | -0.9136444 | 0.36090375 | 0.99999872 |
| VCL | 114.679727 | 0.048173958 | 0.2428623 | 0.19835915 | 0.84276408 | 0.99999872 |
| AP3M1 | 21.073465 | -0.127437879 | 0.46276831 | -0.2753816 | 0.78302308 | 0.99999872 |
| ADK | 150.810269 | 0.197448811 | 0.33609762 | 0.58747458 | 0.55688502 | 0.99999872 |
| KAT6B | 71.6755872 | -0.260500001 | 0.36058361 | -0.7224399 | 0.47002404 | 0.99999872 |
| SAMD8 | 34.0745098 | -0.259941623 | 0.32650294 | -0.7961387 | 0.42595143 | 0.99999872 |
| VDAC2 | 46.5122442 | 1.569166774 | 0.39483436 | 3.97424068 | 7.06E-05 | 0.07163933 |
| COMTD1 | 8.74994005 | -0.566109897 | 0.78242175 | -0.7235355 | 0.46935096 | 0.99999872 |
| LRMDA | 453.25104 | -0.59349839 | 0.33883716 | -1.7515741 | 0.07984707 | 0.99999872 |

| | | | | | | |
|---|---|---|---|---|---|---|
| DLG5 | 87.6709237 | -0.682823297 | 0.65469976 | -1.0429564 | 0.2969685 | 0.99999872 |
| POLR3A | 14.4813368 | -0.655656481 | 0.39856284 | -1.6450517 | 0.09995915 | 0.99999872 |
| RPS24 | 88.7276668 | 0.42520354 | 0.38221949 | 1.11245908 | 0.26594082 | 0.99999872 |
| ZMIZ1-AS1 | 6.15865248 | -0.9832072 | 1.14766971 | -0.8566987 | 0.3916114 | 0.99999872 |
| ZMIZ1 | 65.5348508 | -0.677253245 | 0.25868577 | -2.6180538 | 0.00884329 | 0.67635009 |
| PPIF | 3.87217054 | 0.315300082 | 1.2167927 | 0.25912391 | 0.79553964 | NA |
| ZCCHC24 | 15.7630679 | 0.486420311 | 0.79007159 | 0.61566612 | 0.5381149 | 0.99999872 |
| EIF5AL1 | 4.24518242 | 0.972277249 | 0.79198489 | 1.22764621 | 0.21957981 | NA |
| NUTM2B-AS | 59.5760294 | 0.432494166 | 0.42352073 | 1.02118771 | 0.30716551 | 0.99999872 |
| TMEM254 | 9.08751506 | -0.455412753 | 0.53542762 | -0.850559 | 0.3950144 | 0.99999872 |
| ANXA11 | 51.0320733 | -0.065718388 | 0.43461944 | -0.151209 | 0.87981082 | 0.99999872 |
| FAM213A | 10.3727805 | 0.282028933 | 0.53011694 | 0.53201268 | 0.59471721 | 0.99999872 |
| TSPAN14 | 11.9571614 | -0.063296354 | 0.48169341 | -0.1314038 | 0.89545587 | 0.99999872 |
| NRG3 | 36.7151583 | 0.150736134 | 0.9902685 | 0.15221744 | 0.87901544 | 0.99999872 |
| GHITM | 35.9526141 | 0.341031052 | 0.35572693 | 0.95868775 | 0.33771607 | 0.99999872 |
| CCSER2 | 126.966607 | 0.049141047 | 0.22496948 | 0.21843428 | 0.82709076 | 0.99999872 |
| WAPL | 48.4626081 | -0.053575095 | 0.27213306 | -0.196871 | 0.84392853 | 0.99999872 |
| BMPR1A | 93.9949664 | -0.118016065 | 0.29019227 | -0.4066823 | 0.68424132 | 0.99999872 |
| ADIRF | 21.3850833 | 0.473652948 | 0.90601116 | 0.52278931 | 0.60112088 | 0.99999872 |
| GLUD1 | 42.1747329 | -0.206481778 | 0.34200848 | -0.6037329 | 0.54602122 | 0.99999872 |
| SHLD2 | 54.064357 | -0.438907074 | 0.32905996 | -1.333821 | 0.18226253 | 0.99999872 |
| NUTM2A-AS | 68.0811253 | -0.114404016 | 0.2836506 | -0.4033273 | 0.68670749 | 0.99999872 |
| MINPP1 | 11.3218472 | 0.49076106 | 0.49335069 | 0.99475093 | 0.31985742 | 0.99999872 |
| ATAD1 | 39.4989062 | 0.209201207 | 0.27935368 | 0.74887579 | 0.45393207 | 0.99999872 |
| PTEN | 77.5842686 | 0.011916875 | 0.29724328 | 0.04009132 | 0.96802032 | 0.99999872 |
| RNLS | 34.927059 | -0.637925014 | 0.43691798 | -1.4600567 | 0.1442745 | 0.99999872 |
| LIPA | 22.2639428 | 0.288077949 | 0.64195862 | 0.44874847 | 0.65361312 | 0.99999872 |
| IFIT2 | 41.7400546 | -2.82766138 | 2.05169193 | -1.3782095 | 0.1681386 | 0.99999872 |
| IFIT3 | 62.0650885 | -0.919831366 | 1.55859878 | -0.5901656 | 0.55507964 | 0.99999872 |
| IFIT1 | 97.5427548 | 0.136890909 | 1.03410179 | 0.13237663 | 0.8946864 | 0.99999872 |
| IFIT5 | 14.3755494 | 0.876968199 | 0.81401366 | 1.07733843 | 0.2813291 | 0.99999872 |
| SLC16A12 | 16.8436283 | 1.762541019 | 1.27066781 | 1.38709819 | 0.16541182 | 0.99999872 |
| PANK1 | 9.06746799 | 0.141964063 | 0.5805287 | 0.24454271 | 0.80681053 | 0.99999872 |
| RPP30 | 11.8119627 | -0.138201286 | 0.47182131 | -0.2929102 | 0.76959077 | 0.99999872 |
| PCGF5 | 70.1163114 | -0.159531874 | 0.40125876 | -0.3975785 | 0.69094088 | 0.99999872 |
| HECTD2 | 44.6220788 | 0.351886707 | 0.31582615 | 1.1141785 | 0.26520262 | 0.99999872 |
| PPP1R3C | 4.15852797 | 0.700367118 | 0.91800519 | 0.76292283 | 0.44550942 | NA |
| TNKS2 | 68.9808631 | 0.274812368 | 0.25274072 | 1.0873292 | 0.27689135 | 0.99999872 |
| BTAF1 | 54.1817526 | -0.328694761 | 0.29551396 | -1.1122817 | 0.26601706 | 0.99999872 |
| CPEB3 | 56.1845595 | -0.315753727 | 0.3764472 | -0.8387729 | 0.40159674 | 0.99999872 |
| 5-Mar | 21.506862 | 0.134565517 | 0.3757269 | 0.35814715 | 0.7202332 | 0.99999872 |
| IDE | 25.1129755 | 0.022122994 | 0.35576517 | 0.06218426 | 0.9504161 | 0.99999872 |
| EXOC6 | 48.9539435 | -0.384216802 | 0.29160706 | -1.317584 | 0.18764294 | 0.99999872 |
| MYOF | 80.5238637 | -0.265849098 | 0.37970743 | -0.700142 | 0.48383866 | 0.99999872 |
| FRA10AC1 | 22.6572483 | 0.023612566 | 0.48894663 | 0.04829273 | 0.96148295 | 0.99999872 |
| SLC35G1 | 5.07495787 | 1.524033573 | 0.84920554 | 1.79465807 | 0.0727082 | 0.99999872 |
| PLCE1 | 846.120946 | 0.131605825 | 0.43977815 | 0.29925503 | 0.76474546 | 0.99999872 |
| PLCE1-AS2 | 6.01234754 | 2.350959554 | 1.00133519 | 2.34782476 | 0.0188834 | 0.8671024 |
| NOC3L | 10.05047 | -0.324649529 | 0.57075296 | -0.5688092 | 0.56948564 | 0.99999872 |
| TBC1D12 | 13.4052984 | 0.647010158 | 0.47993182 | 1.34812931 | 0.1776168 | 0.99999872 |
| PDLIM1 | 9.24099837 | 1.079267652 | 0.74056393 | 1.4573592 | 0.14501725 | 0.99999872 |
| SORBS1 | 26.4429417 | -0.250381034 | 1.01407336 | -0.2469062 | 0.80498079 | 0.99999872 |
| ALDH18A1 | 12.0306746 | 0.219709399 | 0.58820931 | 0.37352248 | 0.70875962 | 0.99999872 |
| TCTN3 | 29.0480619 | 0.213538495 | 0.32059404 | 0.66607132 | 0.50536552 | 0.99999872 |
| ENTPD1-AS1 | 62.0155439 | 0.210082252 | 0.46247191 | 0.45425949 | 0.64964207 | 0.99999872 |
| ZNF518A | 96.3807481 | -0.063708504 | 0.31785806 | -0.2004307 | 0.84114378 | 0.99999872 |
| BLNK | 11.4636562 | 1.194998621 | 0.93772529 | 1.27435895 | 0.20253623 | 0.99999872 |
| TM9SF3 | 115.02524 | 0.418936029 | 0.26533787 | 1.57887766 | 0.11436412 | 0.99999872 |
| LCOR | 93.2959306 | -0.118785469 | 0.29584093 | -0.401518 | 0.68803876 | 0.99999872 |
| RRP12 | 12.4152461 | -0.224207585 | 0.50871872 | -0.44073 | 0.6594085 | 0.99999872 |
| PGAM1 | 67.8117402 | 0.186719019 | 0.47611217 | 0.39217443 | 0.69492933 | 0.99999872 |

| | | | | | | |
|---|---|---|---|---|---|---|
| EXOSC1 | 13.3562593 | -0.067659787 | 0.51471936 | -0.1314499 | 0.89541945 | 0.99999872 |
| ZDHHC16 | 8.60197171 | 0.046613971 | 0.51768553 | 0.09004302 | 0.92825302 | 0.99999872 |
| MMS19 | 21.4023551 | 0.163032379 | 0.34065474 | 0.47858539 | 0.63223362 | 0.99999872 |
| MORN4 | 8.89933726 | 0.128252715 | 0.51653665 | 0.24829354 | 0.8039073 | 0.99999872 |
| PI4K2A | 5.87441275 | -0.386789979 | 0.8066612 | -0.479495 | 0.63158655 | 0.99999872 |
| AVPI1 | 5.8683208 | 0.025711641 | 0.8156578 | 0.03152258 | 0.97485278 | 0.99999872 |
| MARVELD1 | 4.39687119 | -0.009298163 | 0.67659669 | -0.0137425 | 0.98903538 | NA |
| ZFYVE27 | 14.6605128 | -0.29815065 | 0.44361471 | -0.6720937 | 0.50152405 | 0.99999872 |
| R3HCC1L | 34.0623766 | -0.123820482 | 0.3846312 | -0.32192 | 0.74751329 | 0.99999872 |
| PYROXD2 | 5.1002208 | -0.005839822 | 0.63374455 | -0.0092148 | 0.99264777 | 0.99999872 |
| HPS1 | 14.0081921 | 0.058471468 | 0.48799473 | 0.11981988 | 0.90462584 | 0.99999872 |
| GOT1 | 14.5585026 | 0.143702167 | 0.58618441 | 0.24514839 | 0.80634153 | 0.99999872 |
| SLC25A28 | 20.1194998 | -0.004829851 | 0.67690151 | -0.0071352 | 0.99430695 | 0.99999872 |
| CUTC | 18.388 | 0.098329197 | 0.40663098 | 0.24181433 | 0.80892404 | 0.99999872 |
| COX15 | 11.9003379 | -0.03998662 | 0.52723854 | -0.0758416 | 0.93954511 | 0.99999872 |
| ABCC2 | 8.35988645 | -1.270600029 | 0.63435102 | -2.002992 | 0.04517815 | 0.95980711 |
| DNMBP | 13.7523817 | -0.182538638 | 0.51125039 | -0.3570435 | 0.72105923 | 0.99999872 |
| ERLIN1 | 13.0699426 | 0.140247771 | 0.49234036 | 0.28485938 | 0.77575187 | 0.99999872 |
| CHUK | 11.6410978 | 0.1130712 | 0.43339961 | 0.26089364 | 0.79417453 | 0.99999872 |
| CWF19L1 | 12.2550426 | -0.04768688 | 0.41244877 | -0.1156189 | 0.90795457 | 0.99999872 |
| BLOC1S2 | 16.8990365 | 0.109843394 | 0.4116213 | 0.26685547 | 0.78958045 | 0.99999872 |
| SCD | 12.0003389 | -0.465873928 | 0.72043673 | -0.6466549 | 0.51785533 | 0.99999872 |
| OLMALINC | 14.1050221 | -0.00964056 | 0.46824334 | -0.0205888 | 0.98357369 | 0.99999872 |
| SEC31B | 8.36093491 | -0.01615658 | 0.62860461 | -0.0257023 | 0.97949479 | 0.99999872 |
| HIF1AN | 16.0823599 | 0.004581363 | 0.46310307 | 0.00989275 | 0.99210686 | 0.99999872 |
| PAX2 | 22.1365158 | -0.556267089 | 0.76715065 | -0.725108 | 0.46838576 | 0.99999872 |
| SLF2 | 57.6951245 | -0.183900885 | 0.42207967 | -0.4357018 | 0.66305308 | 0.99999872 |
| MRPL43 | 17.2015537 | 0.376772599 | 0.39491428 | 0.95406174 | 0.34005239 | 0.99999872 |
| SEMA4G | 9.37459942 | 1.198481632 | 0.9077164 | 1.32032607 | 0.18672618 | 0.99999872 |
| LZTS2 | 46.9079491 | -0.377415245 | 0.29371159 | -1.2849859 | 0.19879722 | 0.99999872 |
| PDZD7 | 8.34110872 | 0.414768229 | 0.70918618 | 0.58485097 | 0.55864793 | 0.99999872 |
| SFXN3 | 8.20192376 | -0.058826317 | 0.67619007 | -0.0869967 | 0.93067412 | 0.99999872 |
| BTRC | 59.1493221 | -0.082952917 | 0.24403057 | -0.3399284 | 0.73391047 | 0.99999872 |
| DPCD | 8.14048177 | 0.621114359 | 0.52639308 | 1.17994401 | 0.23802249 | 0.99999872 |
| POLL | 6.94263256 | -0.771263022 | 0.51893921 | -1.48623 | 0.13721829 | 0.99999872 |
| FBXW4 | 56.8237932 | -0.050421694 | 0.25413436 | -0.1984057 | 0.84272769 | 0.99999872 |
| OGA | 94.031372 | -0.172938906 | 0.3166181 | -0.5462066 | 0.5849239 | 0.99999872 |
| ARMH3 | 56.2707138 | 0.005101109 | 0.29048058 | 0.01756093 | 0.98598913 | 0.99999872 |
| HPS6 | 3.40534855 | 0.475563584 | 0.75482038 | 0.63003543 | 0.5286714 | NA |
| LDB1 | 8.36785534 | 0.620114346 | 0.63531291 | 0.97607704 | 0.32902629 | 0.99999872 |
| PPRC1 | 7.8044019 | -0.221892547 | 1.03517233 | -0.2143532 | 0.8302716 | 0.99999872 |
| NOLC1 | 10.7650079 | 0.15714936 | 0.566563 | 0.27737314 | 0.7814936 | 0.99999872 |
| GBF1 | 73.052138 | -0.115745588 | 0.23069577 | -0.5017239 | 0.61586173 | 0.99999872 |
| NFKB2 | 6.99526776 | -0.521282571 | 0.75727101 | -0.6883699 | 0.49121987 | 0.99999872 |
| FBXL15 | 10.4202441 | 0.57108614 | 0.49041178 | 1.16450331 | 0.2442201 | 0.99999872 |
| CUEDC2 | 15.7064834 | -0.161203968 | 0.37566106 | -0.4291208 | 0.66783533 | 0.99999872 |
| ACTR1A | 15.7085427 | -0.073498678 | 0.6829474 | -0.1076198 | 0.91429727 | 0.99999872 |
| SUFU | 17.7983918 | 0.209073377 | 0.476747 | 0.43854157 | 0.66099374 | 0.99999872 |
| TRIM8 | 56.3949467 | -0.166697391 | 0.36483766 | -0.4569084 | 0.64773689 | 0.99999872 |
| ARL3 | 54.0056205 | 0.933858778 | 0.41755004 | 2.23651943 | 0.02531777 | 0.88365618 |
| SFXN2 | 20.9092025 | -0.261206398 | 0.5315408 | -0.4914136 | 0.62313392 | 0.99999872 |
| WBP1L | 89.0591718 | 0.057194543 | 0.34415731 | 0.16618721 | 0.86800962 | 0.99999872 |
| CNNM2 | 35.6133061 | 0.271383531 | 0.49945901 | 0.54335496 | 0.58688543 | 0.99999872 |
| NT5C2 | 155.010157 | -0.200564306 | 0.29565825 | -0.6783653 | 0.49754008 | 0.99999872 |
| PCGF6 | 11.0704496 | -0.085513227 | 0.42829998 | -0.1996573 | 0.84174859 | 0.99999872 |
| ATP5MD | 17.8343273 | 0.602970135 | 0.43961035 | 1.37160131 | 0.17018759 | 0.99999872 |
| PDCD11 | 7.7807637 | -0.813055351 | 0.77149243 | -1.0538734 | 0.29194088 | 0.99999872 |
| SH3PXD2A | 15.3607199 | -0.017987439 | 0.59092514 | -0.0304395 | 0.97571658 | 0.99999872 |
| STN1 | 46.0055909 | 0.616394294 | 0.60750505 | 1.01463238 | 0.3102811 | 0.99999872 |
| SLK | 204.471566 | 0.077991594 | 0.36904469 | 0.21133374 | 0.83262686 | 0.99999872 |
| CFAP43 | 8.09578958 | -0.017322337 | 0.76447811 | -0.022659 | 0.98192225 | 0.99999872 |

| | | | | | | |
|---|---|---|---|---|---|---|
| GSTO1 | 23.2131258 | 0.390802138 | 0.53647256 | 0.72846622 | 0.46632824 | 0.99999872 |
| GSTO2 | 14.4407659 | -0.687129362 | 0.59051809 | -1.1636043 | 0.24458443 | 0.99999872 |
| XPNPEP1 | 15.2586571 | -0.101321712 | 0.4204168 | -0.241003 | 0.80955279 | 0.99999872 |
| ADD3 | 196.628348 | 0.23028473 | 0.38026465 | 0.60559068 | 0.54478659 | 0.99999872 |
| MXI1 | 22.6779702 | 0.669255796 | 0.48808397 | 1.37118988 | 0.17031577 | 0.99999872 |
| SMNDC1 | 11.6946932 | 0.123517823 | 0.42795565 | 0.28862295 | 0.77286993 | 0.99999872 |
| DUSP5 | 5.01115741 | -0.589649091 | 1.10064285 | -0.5357315 | 0.5921441 | 0.99999872 |
| SMC3 | 27.6970924 | 0.135471926 | 0.32040175 | 0.42281893 | 0.67242738 | 0.99999872 |
| PDCD4 | 26.4301194 | 0.626901837 | 0.34936778 | 1.79438939 | 0.07275104 | 0.99999872 |
| BBIP1 | 19.9452852 | 0.187921774 | 0.40736062 | 0.46131552 | 0.64457225 | 0.99999872 |
| SHOC2 | 49.0727394 | 0.101988739 | 0.2897084 | 0.35203929 | 0.7248088 | 0.99999872 |
| GPAM | 8.78034779 | 0.344665675 | 0.63771974 | 0.54046574 | 0.58887588 | 0.99999872 |
| ACSL5 | 20.4345502 | 0.585468946 | 0.55223349 | 1.06018371 | 0.28906103 | 0.99999872 |
| AL157786.1 | 23.0722329 | -0.015182993 | 0.78127012 | -0.0194337 | 0.9844951 | 0.99999872 |
| ZDHHC6 | 412.650936 | 0.125712835 | 0.54121564 | 0.23227864 | 0.8163216 | 0.99999872 |
| VTI1A | 1216.2763 | -0.153085463 | 0.43840175 | -0.3491899 | 0.72694675 | 0.99999872 |
| TCF7L2 | 148.346942 | 0.07106444 | 0.2794064 | 0.25434077 | 0.7992323 | 0.99999872 |
| CASP7 | 12.0232859 | 0.259248559 | 0.65753384 | 0.39427409 | 0.69337869 | 0.99999872 |
| NHLRC2 | 54.4077026 | 0.174782401 | 0.2848685 | 0.61355468 | 0.53950964 | 0.99999872 |
| CCDC186 | 42.5875909 | 0.082385104 | 0.28995365 | 0.28413198 | 0.77630924 | 0.99999872 |
| AFAP1L2 | 67.7223061 | -0.190584381 | 0.48139871 | -0.3958972 | 0.6921809 | 0.99999872 |
| ABLIM1 | 43.9755577 | 0.112359941 | 0.53484838 | 0.21007812 | 0.8336067 | 0.99999872 |
| FAM160B1 | 28.0472514 | 0.309566299 | 0.33657815 | 0.91974567 | 0.35770568 | 0.99999872 |
| TRUB1 | 21.8741441 | 0.034456594 | 0.50359727 | 0.06842093 | 0.94545056 | 0.99999872 |
| ATRNL1 | 81.1292199 | -0.798460857 | 0.59446045 | -1.343169 | 0.17921725 | 0.99999872 |
| HSPA12A | 130.875168 | -0.192174248 | 0.40706879 | -0.4720928 | 0.63686055 | 0.99999872 |
| ENO4 | 5.44035691 | 0.41631198 | 0.676465 | 0.61542279 | 0.53827554 | 0.99999872 |
| SHTN1 | 56.9465209 | -0.213680134 | 0.26952515 | -0.7928022 | 0.42789308 | 0.99999872 |
| PDZD8 | 61.3671501 | 0.128516624 | 0.31249023 | 0.41126606 | 0.68087746 | 0.99999872 |
| EMX2OS | 13.457213 | 0.886861138 | 0.85352136 | 1.03906144 | 0.29877616 | 0.99999872 |
| EMX2 | 6.22725779 | -0.344389455 | 0.67552575 | -0.5098095 | 0.61018492 | 0.99999872 |
| RAB11FIP2 | 42.3039379 | 0.717555131 | 0.29661612 | 2.41913735 | 0.01555736 | 0.79608149 |
| CASC2 | 21.2855869 | 0.337258399 | 0.49191196 | 0.68560723 | 0.49296082 | 0.99999872 |
| FAM204A | 33.9376007 | 0.187037551 | 0.29723693 | 0.62925408 | 0.52918273 | 0.99999872 |
| CACUL1 | 39.190095 | 0.071310934 | 0.34131833 | 0.20892793 | 0.8345045 | 0.99999872 |
| EIF3A | 56.1674133 | -0.184655132 | 0.3767284 | -0.4901545 | 0.62402455 | 0.99999872 |
| FAM45A | 14.7366916 | 0.796682844 | 0.63388288 | 1.25682973 | 0.2088153 | 0.99999872 |
| SFXN4 | 10.0803389 | -0.350889372 | 0.5547272 | -0.632544 | 0.52703145 | 0.99999872 |
| PRDX3 | 33.9777403 | 0.538063131 | 0.40638632 | 1.32401881 | 0.1854968 | 0.99999872 |
| GRK5 | 196.155229 | -0.437526246 | 0.46063777 | -0.9498271 | 0.34220011 | 0.99999872 |
| RGS10 | 41.0945823 | 0.557678806 | 0.31646627 | 1.76220612 | 0.07803448 | 0.99999872 |
| TIAL1 | 55.1527407 | 0.5797562 | 0.35457061 | 1.63509379 | 0.10202939 | 0.99999872 |
| BAG3 | 15.1153146 | 0.403206394 | 0.65542574 | 0.61518243 | 0.53843425 | 0.99999872 |
| INPP5F | 20.151442 | 0.630275607 | 0.48165628 | 1.30855888 | 0.19068382 | 0.99999872 |
| MCMBP | 23.7235905 | 0.198028735 | 0.34407315 | 0.57554255 | 0.56492442 | 0.99999872 |
| SEC23IP | 24.2651114 | -0.114459427 | 0.39118726 | -0.292595 | 0.76983175 | 0.99999872 |
| WDR11 | 42.2920758 | 0.163651245 | 0.27660296 | 0.59164676 | 0.55408715 | 0.99999872 |
| FGFR2 | 22.4778694 | -0.205640025 | 0.58146346 | -0.3536594 | 0.72359414 | 0.99999872 |
| ATE1 | 60.0207853 | 0.083039983 | 0.28555334 | 0.29080376 | 0.77120141 | 0.99999872 |
| NSMCE4A | 7.31328336 | 0.067412557 | 0.52180518 | 0.12919105 | 0.89720648 | 0.99999872 |
| TACC2 | 110.109333 | 0.12067875 | 0.38817419 | 0.31088813 | 0.75588567 | 0.99999872 |
| BTBD16 | 6.68719862 | 0.503457021 | 1.03855222 | 0.48476813 | 0.62784083 | 0.99999872 |
| PLEKHA1 | 79.6572929 | 0.169520958 | 0.26501985 | 0.63965382 | 0.52239768 | 0.99999872 |
| HTRA1 | 1023.00891 | 0.612997924 | 0.57428184 | 1.06741652 | 0.28578379 | 0.99999872 |
| C10orf88 | 8.67195273 | -0.057080974 | 0.55793564 | -0.1023075 | 0.91851264 | 0.99999872 |
| IKZF5 | 11.8451441 | 0.372062494 | 0.48094338 | 0.77360976 | 0.4391616 | 0.99999872 |
| ACADSB | 19.9352321 | 0.331890238 | 0.44922598 | 0.73880464 | 0.46002563 | 0.99999872 |
| BUB3 | 19.2511458 | 0.181044533 | 0.43048253 | 0.42056186 | 0.67407505 | 0.99999872 |
| OAT | 18.3877003 | 0.588732587 | 0.49519734 | 1.1888848 | 0.234485 | 0.99999872 |
| LHPP | 12.3143232 | 0.288642935 | 0.51281686 | 0.56285773 | 0.57353176 | 0.99999872 |
| ABRAXAS2 | 13.0432733 | 0.33314444 | 0.47956171 | 0.69468523 | 0.48725259 | 0.99999872 |

| | | | | | | |
|---|---|---|---|---|---|---|
| ZRANB1 | 38.218508 | -0.111691622 | 0.49773156 | -0.2244013 | 0.82244504 | 0.99999872 |
| CTBP2 | 113.583448 | 0.026794375 | 0.31148811 | 0.08602054 | 0.93145009 | 0.99999872 |
| EDRF1 | 14.7906374 | -0.263947494 | 0.50724524 | -0.5203548 | 0.60281631 | 0.99999872 |
| UROS | 12.6293827 | 0.091656735 | 0.56629469 | 0.16185343 | 0.87142128 | 0.99999872 |
| BCCIP | 16.4691511 | 0.290616369 | 0.41799755 | 0.69525854 | 0.48689329 | 0.99999872 |
| DHX32 | 26.1861578 | -0.178247302 | 0.37112185 | -0.4802932 | 0.63101892 | 0.99999872 |
| FANK1 | 13.6879691 | -0.473404812 | 0.52102268 | -0.9086069 | 0.36355765 | 0.99999872 |
| DOCK1 | 164.816553 | -0.378025156 | 0.24019417 | -1.5738315 | 0.11552641 | 0.99999872 |
| FAM196A | 5.66838843 | 1.360165333 | 0.78081484 | 1.74198192 | 0.08151161 | 0.99999872 |
| MGMT | 57.4251334 | 0.281383377 | 0.34807316 | 0.80840297 | 0.41885864 | 0.99999872 |
| C10orf143 | 6.94865809 | -0.105704072 | 0.65416469 | -0.1615863 | 0.87163162 | 0.99999872 |
| GLRX3 | 29.7688215 | 0.110638239 | 0.32524091 | 0.34017319 | 0.73372611 | 0.99999872 |
| PPP2R2D | 34.1229003 | -0.277472852 | 0.36518549 | -0.7598135 | 0.44736609 | 0.99999872 |
| BNIP3 | 33.4206128 | 0.348368132 | 0.50562206 | 0.68898919 | 0.49083007 | 0.99999872 |
| STK32C | 7.15151386 | -0.161788148 | 0.55313888 | -0.292491 | 0.76991124 | 0.99999872 |
| LRRC27 | 7.33936705 | -0.414517903 | 0.67035543 | -0.6183554 | 0.53634109 | 0.99999872 |
| INPP5A | 17.8241891 | 0.240451514 | 0.44561435 | 0.53959554 | 0.589476 | 0.99999872 |
| TUBGCP2 | 26.6027983 | 0.292612151 | 0.3389588 | 0.8632676 | 0.38799035 | 0.99999872 |
| FUOM | 13.3434364 | 0.787927733 | 0.76265639 | 1.03313595 | 0.30154028 | 0.99999872 |
| ECHS1 | 37.8910926 | 0.365580442 | 0.53188949 | 0.68732405 | 0.49187855 | 0.99999872 |
| SCART1 | 7.80431195 | -0.019195669 | 0.82991802 | -0.0231296 | 0.9815469 | 0.99999872 |
| SLC6A12 | 8.45178441 | -0.952673883 | 0.76485123 | -1.2455676 | 0.2129232 | 0.99999872 |
| KDM5A | 69.5290594 | -0.218430594 | 0.30286454 | -0.7212155 | 0.47077695 | 0.99999872 |
| CCDC77 | 4.5345794 | -0.554232011 | 0.63788764 | -0.8688552 | 0.38492632 | NA |
| WNK1 | 120.735315 | -0.142027144 | 0.34036124 | -0.4172835 | 0.67647103 | 0.99999872 |
| RAD52 | 10.9179253 | 0.307367962 | 0.49543523 | 0.62039989 | 0.53499455 | 0.99999872 |
| ERC1 | 223.4268 | -0.045555944 | 0.22091942 | -0.2062107 | 0.83662634 | 0.99999872 |
| WNT5B | 12.8658662 | 1.008630656 | 0.90473956 | 1.11482984 | 0.26492335 | 0.99999872 |
| FBXL14 | 6.67785268 | 1.413190611 | 0.82689418 | 1.70903441 | 0.08744458 | 0.99999872 |
| ADIPOR2 | 37.2400704 | 0.15849938 | 0.39906821 | 0.39717366 | 0.69123941 | 0.99999872 |
| DCP1B | 17.1956168 | 0.025497272 | 0.44279762 | 0.05758222 | 0.95408141 | 0.99999872 |
| CACNA1C | 14.3358376 | -0.141691525 | 1.05196142 | -0.1346927 | 0.89285484 | 0.99999872 |
| FKBP4 | 13.1082651 | 0.258838482 | 0.58359907 | 0.44352107 | 0.65738888 | 0.99999872 |
| ITFG2 | 24.5239016 | -0.063876148 | 0.38439813 | -0.1661719 | 0.86802171 | 0.99999872 |
| NRIP2 | 31.7689974 | 0.149832748 | 0.83260292 | 0.17995703 | 0.8571863 | 0.99999872 |
| TULP3 | 26.4932588 | 0.145133182 | 0.3503144 | 0.41429408 | 0.67865875 | 0.99999872 |
| TEAD4 | 15.2375769 | -0.557527513 | 0.64043091 | -0.8705506 | 0.38399958 | 0.99999872 |
| TSPAN9 | 94.5108028 | -0.412656069 | 0.33996614 | -1.2138152 | 0.2248183 | 0.99999872 |
| PARP11 | 17.9439795 | 0.067720806 | 0.54328392 | 0.12465086 | 0.90079996 | 0.99999872 |
| CCND2 | 8.13172647 | 2.245930038 | 0.92870306 | 2.41835106 | 0.01559103 | 0.79608149 |
| TIGAR | 9.21761891 | -0.323136329 | 0.46092583 | -0.7010593 | 0.48326603 | 0.99999872 |
| C12orf4 | 12.4272951 | -0.226978861 | 0.49268922 | -0.4606938 | 0.64501832 | 0.99999872 |
| AC005833.2 | 5.5420565 | -0.025094824 | 0.66390455 | -0.0377988 | 0.96984807 | 0.99999872 |
| CD9 | 63.0686782 | 1.060475986 | 0.50784645 | 2.08818234 | 0.03678139 | 0.95584876 |
| TNFRSF1A | 81.6669029 | -0.324583974 | 0.35499371 | -0.9143373 | 0.36053966 | 0.99999872 |
| SCNN1A | 11.8889153 | -0.741941789 | 0.54128553 | -1.3707032 | 0.17046751 | 0.99999872 |
| LTBR | 25.6270048 | -0.146323466 | 0.45194463 | -0.3237641 | 0.7461166 | 0.99999872 |
| TAPBPL | 8.57744936 | -0.05463602 | 0.67843759 | -0.0805321 | 0.93581405 | 0.99999872 |
| MRPL51 | 25.6783539 | 0.694112762 | 0.50127583 | 1.38469225 | 0.16614659 | 0.99999872 |
| NCAPD2 | 7.44080633 | 0.396014472 | 0.61795682 | 0.64084489 | 0.52162346 | 0.99999872 |
| GAPDH | 222.181059 | 0.500890364 | 0.53040228 | 0.94435937 | 0.34498604 | 0.99999872 |
| IFFO1 | 10.1162423 | 0.207864535 | 0.59258742 | 0.35077446 | 0.72575756 | 0.99999872 |
| CHD4 | 24.5596564 | 0.006361746 | 0.56887358 | 0.01118306 | 0.9910774 | 0.99999872 |
| AC006064.2 | 5.70520959 | 0.361843102 | 0.6739388 | 0.53690796 | 0.59133119 | 0.99999872 |
| ING4 | 14.6209872 | -0.00711053 | 0.39848349 | -0.017844 | 0.98576332 | 0.99999872 |
| ZNF384 | 11.2219646 | 0.226940191 | 0.48560383 | 0.46733608 | 0.64025945 | 0.99999872 |
| COPS7A | 19.7642885 | -0.084415538 | 0.37559467 | -0.2247517 | 0.82217245 | 0.99999872 |
| MLF2 | 31.9518557 | -0.166189258 | 0.38476496 | -0.4319241 | 0.66579658 | 0.99999872 |
| PTMS | 120.990341 | -0.390255891 | 0.58815712 | -0.6635232 | 0.50699553 | 0.99999872 |
| CD4 | 5.1649992 | -0.103196594 | 0.90568695 | -0.1139429 | 0.90928306 | 0.99999872 |
| USP5 | 12.7638673 | -0.061472339 | 0.4726059 | -0.130071 | 0.89651023 | 0.99999872 |

| | | | | | | |
|---|---|---|---|---|---|---|
| TPI1 | 43.9347664 | 0.324831001 | 0.48080471 | 0.67559862 | 0.49929552 | 0.99999872 |
| LRRC23 | 16.3843475 | 0.051379771 | 0.45188285 | 0.11370153 | 0.9094744 | 0.99999872 |
| ATN1 | 39.8631478 | -0.481636562 | 0.37357134 | -1.289276 | 0.19730215 | 0.99999872 |
| C12orf57 | 24.8029398 | 0.570475191 | 0.78783354 | 0.72410625 | 0.46900051 | 0.99999872 |
| PTPN6 | 6.10142624 | -0.487097498 | 0.53690595 | -0.9072306 | 0.36428487 | 0.99999872 |
| PHB2 | 19.6898581 | 0.724565054 | 0.47606186 | 1.52199768 | 0.12800966 | 0.99999872 |
| LPCAT3 | 44.8016436 | 0.527673792 | 0.35178193 | 1.50000255 | 0.13361374 | 0.99999872 |
| C1S | 89.2570249 | 0.816789278 | 0.34380233 | 2.37575259 | 0.0175132 | 0.83791471 |
| C1R | 276.792556 | 0.611565794 | 0.43080641 | 1.41958377 | 1.56E-01 | 0.99999872 |
| C1RL | 44.6658051 | -0.220009073 | 0.63986799 | -0.3438351 | 7.31E-01 | 0.99999872 |
| RBP5 | 5.65514117 | 0.471119356 | 0.79695344 | 0.59115042 | 0.55441964 | 0.99999872 |
| CLSTN3 | 6.58210415 | 0.596465562 | 0.56418069 | 1.05722434 | 0.29040921 | 0.99999872 |
| PEX5 | 8.51514976 | 0.112039537 | 0.57581218 | 0.19457653 | 0.84572448 | 0.99999872 |
| CD163L1 | 6.98127093 | 0.232339453 | 0.58054884 | 0.40020656 | 0.68900438 | 0.99999872 |
| SLC2A3 | 16.2701968 | -0.546917324 | 0.41283278 | -1.3247914 | 0.18524034 | 0.99999872 |
| FOXJ2 | 8.21604614 | -0.397207067 | 0.51715417 | -0.7680632 | 0.44244966 | 0.99999872 |
| NECAP1 | 10.8430162 | -0.678471609 | 0.4869689 | -1.3932545 | 0.16354285 | 0.99999872 |
| FAM66C | 9.39991332 | 0.014025265 | 0.58142115 | 0.02412239 | 0.98075499 | 0.99999872 |
| LINC00937 | 7.88017537 | -0.194176158 | 0.54385878 | -0.3570342 | 0.72106624 | 0.99999872 |
| RIMKLB | 35.4374064 | -0.172977741 | 0.46295514 | -0.3736382 | 0.70867348 | 0.99999872 |
| A2ML1-AS1 | 7.96419172 | 0.394098799 | 0.71823095 | 0.54870763 | 0.58320611 | 0.99999872 |
| PHC1 | 16.85125 | -0.046389665 | 0.38539911 | -0.1203679 | 0.90419175 | 0.99999872 |
| M6PR | 29.5095681 | 0.076285004 | 0.33236458 | 0.22952207 | 0.81846317 | 0.99999872 |
| CLEC2D | 13.4651024 | -0.23463433 | 0.7049311 | -0.3328472 | 0.73924964 | 0.99999872 |
| GABARAPL1 | 35.2848546 | 0.269961895 | 0.35203186 | 0.76686778 | 0.44316013 | 0.99999872 |
| KLRD1 | 17.8390266 | 1.116885662 | 0.58164199 | 1.92022873 | 0.05482901 | 0.99999872 |
| MAGOHB | 8.68404838 | 0.155417342 | 0.52958777 | 0.29346852 | 0.76916406 | 0.99999872 |
| YBX3 | 152.073791 | 0.03585377 | 0.42652431 | 0.08406032 | 0.93300847 | 0.99999872 |
| PRH1 | 6.83743175 | 0.52971229 | 0.65087198 | 0.8138502 | 0.41573077 | 0.99999872 |
| SMIM10L1 | 9.63453098 | 0.839846178 | 0.55920359 | 1.5018612 | 0.13313296 | 0.99999872 |
| AC244131.2 | 10.4325543 | -0.545508809 | 0.82649508 | -0.6600267 | 0.5092367 | 0.99999872 |
| LINC01252 | 24.3629633 | -0.422078184 | 0.63094656 | -0.6689603 | 0.50352082 | 0.99999872 |
| ETV6 | 103.113651 | -0.261160942 | 0.34495963 | -0.7570768 | 0.44900384 | 0.99999872 |
| BCL2L14 | 20.7112806 | -0.236410352 | 0.41404036 | -0.5709838 | 0.5680106 | 0.99999872 |
| LRP6 | 50.5503783 | 0.043022497 | 0.39492523 | 0.10893834 | 0.9132514 | 0.99999872 |
| MANSC1 | 22.4151513 | 0.403721564 | 0.53464117 | 0.75512621 | 0.45017326 | 0.99999872 |
| BORCS5 | 26.4624213 | -0.14469684 | 0.39005043 | -0.3709696 | 0.71066019 | 0.99999872 |
| DUSP16 | 60.5058361 | -0.006135169 | 0.27782661 | -0.0220827 | 0.98238196 | 0.99999872 |
| CREBL2 | 22.6365388 | -0.022585148 | 0.40942151 | -0.0551636 | 0.95600816 | 0.99999872 |
| CDKN1B | 41.7598067 | 0.599714423 | 0.53957268 | 1.11146181 | 0.26636962 | 0.99999872 |
| APOLD1 | 7.36577731 | 0.398755688 | 0.85093156 | 0.46861076 | 0.63934788 | 0.99999872 |
| GPRC5A | 46.7581868 | 0.716914916 | 0.4701035 | 1.52501507 | 0.12725534 | 0.99999872 |
| GPRC5D-AS1 | 21.7956371 | 0.103160009 | 0.61694037 | 0.16721228 | 0.86720302 | 0.99999872 |
| HEBP1 | 46.3828808 | 0.400488976 | 0.47688691 | 0.83979863 | 0.4010213 | 0.99999872 |
| FAM234B | 4.8860752 | 0.278102204 | 0.79033074 | 0.35188079 | 0.72492767 | 0.99999872 |
| EMP1 | 25.4848417 | -1.29027304 | 0.81267037 | -1.5876954 | 0.11235522 | 0.99999872 |
| GRIN2B | 14.4714304 | 2.976739751 | 1.0801381 | 2.75588813 | 0.0058533 | 0.58410373 |
| ATF7IP | 77.5792794 | 0.000775186 | 0.33006842 | 0.00234856 | 0.99812612 | 0.99999872 |
| PLBD1 | 35.1698443 | 0.200055063 | 0.29447216 | 0.67936834 | 0.4969045 | 0.99999872 |
| PLBD1-AS1 | 8.54611735 | -0.879122804 | 0.67674803 | -1.2990401 | 0.19393018 | 0.99999872 |
| AC010168.1 | 10.4696151 | -0.067007482 | 0.69261754 | -0.0967453 | 0.92292868 | 0.99999872 |
| H2AFJ | 216.214879 | -0.157301098 | 0.43368054 | -0.3627119 | 0.7168201 | 0.99999872 |
| WBP11 | 25.6627484 | 0.427108198 | 0.37968086 | 1.1249137 | 0.26062561 | 0.99999872 |
| C12orf60 | 7.18023805 | 0.535744719 | 0.75236528 | 0.7120806 | 0.47641487 | 0.99999872 |
| SMCO3 | 21.6482981 | 0.170896956 | 0.84571947 | 0.20207287 | 0.83985976 | 0.99999872 |
| MGP | 322.424111 | 0.697192503 | 0.87390376 | 0.79779094 | 0.42499182 | 0.99999872 |
| RERG | 647.883893 | -0.452283415 | 0.3200223 | -1.4132872 | 0.1575713 | 0.99999872 |
| PTPRO | 4411.19941 | -0.119636905 | 0.37368313 | -0.320156 | 0.74885006 | 0.99999872 |
| EPS8 | 36.164971 | -0.608835845 | 0.82098252 | -0.7415942 | 0.45833326 | 0.99999872 |
| STRAP | 34.515296 | 0.488222433 | 0.35547659 | 1.37343063 | 0.16961851 | 0.99999872 |
| DERA | 42.5953452 | -0.398818006 | 0.35539939 | -1.1221685 | 0.2617908 | 0.99999872 |

| | | | | | | |
|---|---|---|---|---|---|---|
| MGST1 | 22.694133 | -0.379940616 | 0.94321917 | -0.4028127 | 0.68708605 | 0.99999872 |
| PIK3C2G | 10.8966854 | 3.133019744 | 1.09089731 | 2.87196578 | 0.00407927 | 0.49700703 |
| PLEKHA5 | 433.016665 | 0.158254232 | 0.34382246 | 0.46027892 | 0.64531603 | 0.99999872 |
| AEBP2 | 98.371269 | -0.250166986 | 0.3760251 | -0.6652933 | 0.5058629 | 0.99999872 |
| AC024901.1 | 36.576821 | -1.124157692 | 0.52115198 | -2.1570631 | 0.03100075 | 0.91534437 |
| LINC02398 | 44.8116277 | -0.693664316 | 0.42346087 | -1.6380836 | 0.10140425 | 0.99999872 |
| PDE3A | 155.588982 | -0.178289182 | 0.52838565 | -0.3374225 | 0.73579846 | 0.99999872 |
| PYROXD1 | 13.1884877 | 0.451172962 | 0.47894212 | 0.94201981 | 0.3461825 | 0.99999872 |
| RECQL | 12.70727 | 0.046074949 | 0.4263791 | 0.10806099 | 0.91394731 | 0.99999872 |
| GOLT1B | 16.6296381 | 0.27259882 | 0.41237055 | 0.66105307 | 0.50857828 | 0.99999872 |
| LDHB | 41.0906176 | 0.181918862 | 0.33031638 | 0.55074127 | 0.58181105 | 0.99999872 |
| CMAS | 15.6318364 | 0.598134488 | 0.42872025 | 1.39516268 | 0.16296679 | 0.99999872 |
| C2CD5 | 48.5810295 | -0.369630081 | 0.31734456 | -1.1647595 | 0.24411636 | 0.99999872 |
| ETNK1 | 63.3653826 | 0.324402776 | 0.36664719 | 0.88478184 | 0.37627431 | 0.99999872 |
| SOX5 | 20.5824853 | 0.002006499 | 1.0909995 | 0.00183914 | 0.99853258 | 0.99999872 |
| CASC1 | 8.29368412 | -0.345182553 | 0.60065539 | -0.5746765 | 0.56551009 | 0.99999872 |
| ETFRF1 | 23.5475081 | 0.53783672 | 0.44087919 | 1.2199186 | 0.22249574 | 0.99999872 |
| KRAS | 26.8502524 | 0.316948745 | 0.33261699 | 0.95289405 | 0.34064376 | 0.99999872 |
| RASSF8-AS1 | 89.0655042 | -0.665197954 | 0.50733929 | -1.3111501 | 0.18980706 | 0.99999872 |
| RASSF8 | 321.590074 | -0.106375813 | 0.43414505 | -0.2450237 | 0.80643811 | 0.99999872 |
| SSPN | 4.21881465 | 1.54107024 | 0.94041793 | 1.63870785 | 0.10127412 | NA |
| ITPR2 | 48.840171 | -0.936303172 | 0.56587781 | -1.6546031 | 0.09800504 | 0.99999872 |
| INTS13 | 7.53839707 | -0.542846416 | 0.66095509 | -0.821306 | 0.41147196 | 0.99999872 |
| FGFR1OP2 | 33.2839801 | 0.585353175 | 0.43113636 | 1.35769847 | 0.17455938 | 0.99999872 |
| TM7SF3 | 34.4739217 | 0.072797528 | 0.38548369 | 0.18884723 | 0.85021255 | 0.99999872 |
| MED21 | 37.828414 | 0.671634723 | 0.47925281 | 1.40142051 | 0.16108836 | 0.99999872 |
| STK38L | 22.6912854 | -0.136679324 | 0.55791024 | -0.2449844 | 0.80646848 | 0.99999872 |
| PPFIBP1 | 57.1490452 | -0.245854944 | 0.31301391 | -0.7854441 | 0.43219321 | 0.99999872 |
| MRPS35 | 16.296949 | 0.510768356 | 0.47670878 | 1.07144735 | 0.28396833 | 0.99999872 |
| MANSC4 | 5.40739062 | 3.453574435 | 0.95235224 | 3.62636249 | 0.00028744 | 0.15059075 |
| KLHL42 | 182.796759 | 0.582865805 | 0.27395614 | 2.12758803 | 0.03337126 | 0.92959829 |
| AC008011.2 | 38.8322475 | -0.510784558 | 0.72780586 | -0.7018143 | 0.48279498 | 0.99999872 |
| CCDC91 | 614.409161 | 0.195474327 | 0.45298315 | 0.43152671 | 0.66608544 | 0.99999872 |
| AC022364.1 | 7.72226207 | 0.520071975 | 0.65866401 | 0.78958615 | 0.4297695 | 0.99999872 |
| FAR2 | 5.51428303 | 0.465705167 | 0.7501411 | 0.62082343 | 0.53471581 | 0.99999872 |
| AC009318.1 | 6.80846671 | -1.148811054 | 0.57073029 | -2.0128791 | 0.04412735 | 0.95980711 |
| ERGIC2 | 29.4398289 | 0.099187881 | 0.31182449 | 0.31808881 | 0.75041757 | 0.99999872 |
| TMTC1 | 36.8426292 | -2.323467887 | 0.87213333 | -2.66412 | 0.007719 | 0.65139548 |
| IPO8 | 23.1641688 | 0.29772521 | 0.38159283 | 0.78021698 | 0.43526317 | 0.99999872 |
| CAPRIN2 | 20.8085039 | 0.484456575 | 0.55459072 | 0.87353892 | 0.3823694 | 0.99999872 |
| SINHCAF | 39.6482897 | 0.129600308 | 0.47631037 | 0.27209214 | 0.78555117 | 0.99999872 |
| DENND5B | 84.932964 | 0.251034799 | 0.38462199 | 0.65267927 | 0.51396307 | 0.99999872 |
| ETFBKMT | 6.79341165 | -1.268440284 | 0.62041912 | -2.0444894 | 0.04090523 | 0.95980711 |
| AMN1 | 7.88861012 | -0.557310054 | 0.54697868 | -1.0188881 | 0.3082561 | 0.99999872 |
| KIAA1551 | 19.0547602 | 0.402121736 | 0.37176666 | 1.08165088 | 0.27940769 | 0.99999872 |
| BICD1 | 166.072433 | -0.868947023 | 0.26025013 | -3.3388917 | 0.00084113 | 0.26707259 |
| FGD4 | 70.6834529 | -0.589819513 | 0.4316744 | -1.3663528 | 0.17182826 | 0.99999872 |
| DNM1L | 49.7948983 | 0.214961469 | 0.27883484 | 0.77092759 | 0.44074986 | 0.99999872 |
| YARS2 | 16.3499385 | -0.060473869 | 0.42319297 | -0.142899 | 0.88636992 | 0.99999872 |
| PKP2 | 6.642025 | 0.191978957 | 0.63747405 | 0.30115572 | 0.76329575 | 0.99999872 |
| ALG10B | 10.3261518 | 0.332480873 | 0.57181193 | 0.58145145 | 0.56093623 | 0.99999872 |
| CPNE8 | 225.449935 | -0.296932931 | 0.2730592 | -1.0874306 | 0.27684655 | 0.99999872 |
| KIF21A | 60.9782862 | 0.456002172 | 0.47191158 | 0.96628731 | 0.33390044 | 0.99999872 |
| C12orf40 | 6.78941304 | 0.876481706 | 1.05037884 | 0.83444341 | 0.40403116 | 0.99999872 |
| SLC2A13 | 135.889203 | 0.554557473 | 0.40790261 | 1.35953402 | 0.17397743 | 0.99999872 |
| LRRK2 | 34.1646178 | -0.010290506 | 0.44629983 | -0.0230574 | 0.9816045 | 0.99999872 |
| CNTN1 | 86.8144006 | 0.542909054 | 0.57640253 | 0.94189221 | 0.34624783 | 0.99999872 |
| GXYLT1 | 43.038566 | 0.052153 | 0.40931721 | 0.12741463 | 0.89861224 | 0.99999872 |
| YAF2 | 56.8950772 | -0.273301507 | 0.33551387 | -0.8145759 | 0.41531513 | 0.99999872 |
| PPHLN1 | 99.0274962 | 0.014548758 | 0.30872985 | 0.04712456 | 0.96241395 | 0.99999872 |
| ZCRB1 | 19.6889461 | 0.209858195 | 0.56087766 | 0.37416037 | 0.708285 | 0.99999872 |

| | | | | | | |
|---|---|---|---|---|---|---|
| PRICKLE1 | 56.4116767 | -0.158377382 | 0.35059463 | -0.4517393 | 0.65145679 | 0.99999872 |
| PUS7L | 13.8805662 | 0.560671745 | 0.42934143 | 1.30588782 | 0.19159071 | 0.99999872 |
| IRAK4 | 16.6511123 | 0.40286576 | 0.38459521 | 1.04750593 | 0.2948663 | 0.99999872 |
| TWF1 | 10.6363457 | 0.116537265 | 0.45813269 | 0.25437448 | 0.79920626 | 0.99999872 |
| TMEM117 | 63.189617 | -0.219383743 | 0.4349037 | -0.5044421 | 0.61395073 | 0.99999872 |
| ANO6 | 49.6755007 | 0.277560153 | 0.35300076 | 0.78628769 | 0.43169896 | 0.99999872 |
| AC008124.1 | 10.7386685 | 0.285348892 | 0.57265897 | 0.49828765 | 0.61828131 | 0.99999872 |
| ARID2 | 115.446031 | -0.003370827 | 0.26976795 | -0.0124953 | 0.99003046 | 0.99999872 |
| SCAF11 | 120.929707 | -0.142959313 | 0.25928046 | -0.5513694 | 0.58138047 | 0.99999872 |
| SLC38A1 | 11.7924837 | -0.221850968 | 0.74056885 | -0.2995683 | 0.76450645 | 0.99999872 |
| SLC38A2 | 110.362695 | -0.081202502 | 0.39318779 | -0.2065235 | 0.83638204 | 0.99999872 |
| AC008014.1 | 160.476956 | -0.426132452 | 0.61273934 | -0.6954547 | 0.48677039 | 0.99999872 |
| SLC38A4 | 5.8492193 | 2.753943958 | 1.04375427 | 2.63849838 | 0.00832741 | 0.66570732 |
| AMIGO2 | 71.0492542 | 0.105707455 | 0.68459 | 0.15440987 | 0.87728657 | 0.99999872 |
| PCED1B | 216.726221 | -0.12488436 | 0.41131045 | -0.3036255 | 0.76141319 | 0.99999872 |
| RPAP3 | 13.2820361 | 0.581336748 | 0.49139228 | 1.18304005 | 0.23679328 | 0.99999872 |
| RAPGEF3 | 85.612267 | -1.233200859 | 0.41829657 | -2.9481496 | 0.00319682 | 0.44319076 |
| SLC48A1 | 87.0595011 | -0.45129851 | 0.37828171 | -1.1930223 | 0.23286064 | 0.99999872 |
| HDAC7 | 28.400159 | -1.105709225 | 0.35799584 | -3.0886092 | 0.00201096 | 0.36957035 |
| TMEM106C | 15.4803262 | 0.602758316 | 0.59367704 | 1.01529665 | 0.30996444 | 0.99999872 |
| SENP1 | 16.1050083 | 0.043099376 | 0.39479929 | 0.10916781 | 0.91306939 | 0.99999872 |
| PFKM | 7.59614243 | 0.438751128 | 0.56132263 | 0.78163805 | 0.43442732 | 0.99999872 |
| ASB8 | 12.1431792 | 0.761573466 | 0.47417845 | 1.60609041 | 0.10825405 | 0.99999872 |
| ZNF641 | 7.41866873 | -1.212051723 | 0.55622383 | -2.1790719 | 0.02932633 | 0.90266962 |
| AC090115.1 | 6.44896543 | -0.415764036 | 0.60609408 | -0.6859728 | 0.49273028 | 0.99999872 |
| KANSL2 | 11.8955426 | -0.072390254 | 0.42844251 | -0.1689614 | 0.86582699 | 0.99999872 |
| CCNT1 | 16.4096157 | 0.345173027 | 0.36808383 | 0.93775656 | 0.34836953 | 0.99999872 |
| ADCY6 | 11.3361501 | 0.144068633 | 0.48521546 | 0.29691682 | 0.76653001 | 0.99999872 |
| DDX23 | 14.6463974 | -0.406541067 | 0.4573489 | -0.8889079 | 0.37405256 | 0.99999872 |
| CCDC65 | 12.863731 | 0.090310989 | 0.60527327 | 0.14920697 | 0.88139032 | 0.99999872 |
| DDN | 132.693251 | 3.134125763 | 1.12115387 | 2.7954466 | 0.00518281 | 0.54853984 |
| DDN-AS1 | 9.797614 | 1.332428603 | 0.84410718 | 1.57850641 | 0.11444932 | 0.99999872 |
| PRKAG1 | 24.5456348 | 0.135306816 | 0.35523564 | 0.38089314 | 0.70328255 | 0.99999872 |
| KMT2D | 20.8416781 | -0.456749378 | 0.38740426 | -1.1789994 | 0.23839842 | 0.99999872 |
| LMBR1L | 16.0070879 | -0.444031703 | 0.43517456 | -1.0203531 | 0.30756104 | 0.99999872 |
| TUBA1B | 81.1956413 | 0.987439341 | 0.69963544 | 1.41136267 | 0.1581377 | 0.99999872 |
| TUBA1A | 30.1966697 | 1.260674492 | 0.89548858 | 1.40780634 | 0.15918842 | 0.99999872 |
| TUBA1C | 13.7776991 | 0.355221332 | 0.42955212 | 0.82695746 | 0.40826117 | 0.99999872 |
| SPATS2 | 23.4591663 | 0.704901858 | 0.44521803 | 1.58327339 | 0.11335916 | 0.99999872 |
| KCNH3 | 9.57694379 | -0.465404773 | 0.81224467 | -0.5729859 | 0.56665422 | 0.99999872 |
| MCRS1 | 12.9755663 | -0.312382235 | 0.41408723 | -0.7543875 | 0.45061657 | 0.99999872 |
| PRPF40B | 22.5453892 | -0.14961117 | 0.34921064 | -0.4284267 | 0.66834047 | 0.99999872 |
| FMNL3 | 9.05501025 | -0.696627313 | 0.77730998 | -0.8962027 | 0.37014451 | 0.99999872 |
| TMBIM6 | 165.275694 | -0.050285653 | 0.37715105 | -0.1333303 | 0.89393219 | 0.99999872 |
| NCKAP5L | 11.0949239 | 0.265556176 | 0.53065622 | 0.50042978 | 0.61677249 | 0.99999872 |
| AQP2 | 37.7863599 | 0.549416274 | 0.70823693 | 0.77575208 | 0.43789538 | 0.99999872 |
| SMARCD1 | 18.2507761 | -0.245120607 | 0.38403648 | -0.6382743 | 0.52329516 | 0.99999872 |
| GPD1 | 4.92221772 | 0.13297134 | 0.73482981 | 0.18095529 | 0.85640267 | 0.99999872 |
| AC074032.1 | 38.9290063 | 0.005210832 | 0.36271303 | 0.01436626 | 0.98853777 | 0.99999872 |
| CERS5 | 31.6507821 | 0.242764418 | 0.34024067 | 0.71350793 | 0.47553151 | 0.99999872 |
| LIMA1 | 35.9141993 | 0.659697381 | 0.40066281 | 1.64651514 | 0.09965775 | 0.99999872 |
| LARP4 | 54.5197141 | -0.459275036 | 0.2614553 | -1.7566102 | 0.07898428 | 0.99999872 |
| DIP2B | 36.7059971 | -0.402101829 | 0.48531848 | -0.8285319 | 0.40736935 | 0.99999872 |
| ATF1 | 20.5275058 | 0.430538494 | 0.39277976 | 1.09613208 | 0.27302098 | 0.99999872 |
| METTL7A | 8.8996854 | 0.265154351 | 0.96963469 | 0.27345799 | 0.78450118 | 0.99999872 |
| SLC11A2 | 30.6663348 | 0.288675257 | 0.29788363 | 0.96908735 | 0.33250161 | 0.99999872 |
| LETMD1 | 18.3053805 | -0.160698276 | 0.45448537 | -0.3535829 | 0.72365146 | 0.99999872 |
| CSRNP2 | 15.645057 | 0.465378739 | 0.42137129 | 1.10443866 | 0.2694029 | 0.99999872 |
| TFCP2 | 59.2910399 | 0.030359478 | 0.30357327 | 0.10000709 | 0.9203387 | 0.99999872 |
| POU6F1 | 9.51866222 | -0.77947671 | 0.51108887 | -1.5251295 | 0.1272268 | 0.99999872 |
| DAZAP2 | 69.5147414 | 0.030032422 | 0.49825419 | 0.0602753 | 0.95193637 | 0.99999872 |

| | | | | | | |
|---|---|---|---|---|---|---|
| ACVR1B | 25.4435443 | -0.493900582 | 0.32223641 | -1.5327274 | 0.12534303 | 0.99999872 |
| ATG101 | 9.38224397 | 0.621627815 | 0.61305998 | 1.01397553 | 0.31059443 | 0.99999872 |
| KRT7 | 6.17887053 | -0.490693461 | 0.75705982 | -0.6481568 | 0.51688355 | 0.99999872 |
| KRT8 | 14.1610271 | -0.854108629 | 0.55304719 | -1.5443684 | 0.1224991 | 0.99999872 |
| KRT18 | 18.2853026 | -0.505696417 | 0.65562109 | -0.7713242 | 0.44051478 | 0.99999872 |
| EIF4B | 42.078912 | -0.111809892 | 0.34633417 | -0.3228382 | 0.7468178 | 0.99999872 |
| AC068888.1 | 24.4416841 | -0.549947215 | 0.49179831 | -1.1182373 | 0.26346565 | 0.99999872 |
| TNS2 | 79.8424073 | -0.817336173 | 0.44864935 | -1.8217706 | 0.06848981 | 0.99999872 |
| SPRYD3 | 8.86117334 | 0.012610526 | 0.55090937 | 0.02289038 | 0.98173771 | 0.99999872 |
| CSAD | 21.4922199 | -0.322397473 | 0.42103549 | -0.7657252 | 0.44383984 | 0.99999872 |
| ZNF740 | 6.04096781 | 0.189515338 | 0.62992603 | 0.30085332 | 0.76352635 | 0.99999872 |
| RARG | 5.80634795 | -0.373242047 | 0.842752 | -0.4428848 | 0.65784907 | 0.99999872 |
| MFSD5 | 9.37877307 | 0.384455054 | 0.53060978 | 0.72455328 | 0.46872613 | 0.99999872 |
| PFDN5 | 50.5702226 | 0.737421677 | 0.4664461 | 1.58093652 | 0.11389255 | 0.99999872 |
| C12orf10 | 10.447098 | -0.025366 | 0.50765461 | -0.049967 | 0.96014865 | 0.99999872 |
| AAAS | 5.07800523 | 0.246100431 | 0.63022151 | 0.39049831 | 0.69616811 | 0.99999872 |
| SP1 | 43.0072859 | 0.031903874 | 0.32065534 | 0.09949585 | 0.92074459 | 0.99999872 |
| PRR13 | 32.4385178 | 0.213363239 | 0.49215999 | 0.43352414 | 0.66463403 | 0.99999872 |
| PCBP2 | 80.6715886 | -0.30326427 | 0.22584317 | -1.3428091 | 0.17933379 | 0.99999872 |
| MAP3K12 | 10.930159 | 1.026269447 | 0.45630778 | 2.24907284 | 0.02450786 | 0.88337526 |
| ATP5MC2 | 75.3065749 | -0.049762967 | 0.35943191 | -0.1384489 | 0.88988562 | 0.99999872 |
| CALCOCO1 | 37.8574352 | 0.393238316 | 0.38779503 | 1.0140365 | 0.31056534 | 0.99999872 |
| HOXC4 | 8.54302121 | -0.056478935 | 0.66249858 | -0.0852514 | 0.93206152 | 0.99999872 |
| SMUG1 | 10.359507 | 0.225396638 | 0.51742467 | 0.43561247 | 0.66311792 | 0.99999872 |
| LINC02381 | 4.39147542 | -0.110193704 | 0.76312036 | -0.1443989 | 0.88518552 | NA |
| CBX5 | 41.5422275 | 0.166581451 | 0.38171812 | 0.43639912 | 0.66254718 | 0.99999872 |
| HNRNPA1 | 31.2420759 | -0.173660976 | 0.3875 | -0.4481574 | 0.65403964 | 0.99999872 |
| COPZ1 | 29.2529944 | 0.141114933 | 0.36983389 | 0.38156301 | 0.70278552 | 0.99999872 |
| ZNF385A | 4.76974457 | 0.307249928 | 1.04559699 | 0.2938512 | 0.76887161 | 0.99999872 |
| PPP1R1A | 5.34649761 | -0.142147645 | 0.95884552 | -0.1482487 | 0.88214646 | 0.99999872 |
| CD63 | 126.129055 | 0.427005172 | 0.5318355 | 0.80288956 | 0.42203857 | 0.99999872 |
| ORMDL2 | 9.81288118 | 0.75689679 | 0.54630019 | 1.38549611 | 0.16590082 | 0.99999872 |
| PYM1 | 6.30698422 | 0.109126244 | 0.57203162 | 0.1907696 | 0.84870611 | 0.99999872 |
| CDK2 | 6.06926928 | 1.171270904 | 0.7621746 | 1.53674881 | 0.12435483 | 0.99999872 |
| RAB5B | 32.5270951 | -0.39573233 | 0.40369941 | -0.9802648 | 0.32695542 | 0.99999872 |
| SUOX | 5.9388183 | -0.371690765 | 0.66483144 | -0.5590752 | 0.5761104 | 0.99999872 |
| IKZF4 | 11.0378489 | -1.034285143 | 0.57791287 | -1.7896905 | 0.07350369 | 0.99999872 |
| RPS26 | 14.5680772 | -0.425643822 | 0.44548297 | -0.955466 | 0.33934211 | 0.99999872 |
| ERBB3 | 8.66552781 | -1.038870528 | 0.63029982 | -1.6482165 | 0.09930826 | 0.99999872 |
| PA2G4 | 13.1382913 | 0.299509684 | 0.45087205 | 0.66428975 | 0.50650488 | 0.99999872 |
| RPL41 | 177.416993 | 1.056497298 | 0.37366846 | 2.8273655 | 0.00469327 | 0.52915868 |
| ESYT1 | 34.8917459 | -0.030625071 | 0.35170334 | -0.0870764 | 0.93061076 | 0.99999872 |
| MYL6 | 200.099441 | 0.117665429 | 0.63256641 | 0.18601277 | 0.85243475 | 0.99999872 |
| SMARCC2 | 35.6874295 | -0.271226655 | 0.38142846 | -0.7110813 | 0.47703385 | 0.99999872 |
| RNF41 | 14.1166233 | 0.378649417 | 0.43520985 | 0.87003871 | 0.38427925 | 0.99999872 |
| NABP2 | 6.58975623 | -0.394404431 | 0.62787089 | -0.6281617 | 0.52989803 | 0.99999872 |
| ANKRD52 | 10.3655658 | 0.498575093 | 0.54814022 | 0.90957582 | 0.36304625 | 0.99999872 |
| CS | 14.7730118 | 0.31165926 | 0.40668073 | 0.76634873 | 0.44346883 | 0.99999872 |
| PAN2 | 13.4280234 | 0.00063503 | 0.46481459 | 0.0013662 | 0.99890993 | 0.99999872 |
| STAT2 | 26.4518766 | -0.650063639 | 0.6764318 | -0.9610187 | 0.33654274 | 0.99999872 |
| SPRYD4 | 5.4268739 | -0.208144663 | 0.92332562 | -0.2254293 | 0.82164531 | 0.99999872 |
| RBMS2 | 62.7889542 | -0.729163692 | 0.31743766 | -2.2970296 | 0.02161708 | 0.87186485 |
| BAZ2A | 44.5260002 | -0.416664184 | 0.34998274 | -1.1905278 | 0.23383901 | 0.99999872 |
| ATP5F1B | 58.0732206 | 0.514791567 | 0.44280924 | 1.16255833 | 0.24500875 | 0.99999872 |
| PTGES3 | 78.2447127 | 0.227996018 | 0.2626368 | 0.86810386 | 0.38533748 | 0.99999872 |
| NACA | 47.4245596 | 0.998650478 | 0.34061945 | 2.93186571 | 0.00336932 | 0.44319076 |
| NEMP1 | 19.1157386 | -0.431736147 | 0.48492944 | -0.8903072 | 0.37330098 | 0.99999872 |
| NAB2 | 12.0499216 | -1.190277508 | 0.51034435 | -2.3323027 | 0.01968477 | 0.8671024 |
| STAT6 | 31.3709566 | -0.496553885 | 0.4208447 | -1.1798982 | 0.23804072 | 0.99999872 |
| LRP1 | 15.3919123 | 0.129882803 | 0.52664855 | 0.2466214 | 0.80520124 | 0.99999872 |
| SHMT2 | 5.36385572 | 0.094629292 | 0.78614586 | 0.12037116 | 0.90418913 | 0.99999872 |

| | | | | | | | |
|---|---|---|---|---|---|---|---|
| R3HDM2 | 77.420085 | | -0.517303805 | 0.37947179 | -1.3632207 | 0.17281296 | 0.99999872 |
| MARS | 15.1903151 | | -0.393031512 | 0.65867725 | -0.5966982 | 0.55070891 | 0.99999872 |
| DDIT3 | 24.9254614 | | 0.943129962 | 1.57448199 | 0.59900969 | 0.54916643 | 0.99999872 |
| MBD6 | 12.2776472 | | 0.088601348 | 0.42788845 | 0.20706646 | 0.83595795 | 0.99999872 |
| DCTN2 | 26.2660772 | | 0.535682106 | 0.39348617 | 1.36137469 | 0.17339531 | 0.99999872 |
| PIP4K2C | 12.7889058 | | 0.494815094 | 0.70750533 | 0.69938002 | 0.48431457 | 0.99999872 |
| DTX3 | 10.8730566 | | 0.505483229 | 0.58874071 | 0.85858378 | 0.39057019 | 0.99999872 |
| OS9 | 58.2469814 | | 0.260609521 | 0.37498716 | 0.69498252 | 0.48706626 | 0.99999872 |
| TSPAN31 | 13.3710137 | | 1.306942305 | 0.55285595 | 2.36398343 | 0.01807962 | 0.85282771 |
| CDK4 | 10.7950068 | | 0.066336858 | 0.50326067 | 0.13181411 | 0.89513132 | 0.99999872 |
| 9-Mar | 7.42195377 | | 0.184545953 | 0.55732792 | 0.33112634 | 0.74054906 | 0.99999872 |
| TSFM | 7.87138882 | | 0.450764002 | 0.60676411 | 0.74289826 | 0.45754328 | 0.99999872 |
| CTDSP2 | 32.276242 | | -0.039352132 | 0.30838404 | -0.1276076 | 0.89845956 | 0.99999872 |
| AC084033.3 | 5.07600636 | | 0.060657245 | 0.60018286 | 0.10106461 | 0.91949917 | 0.99999872 |
| LRIG3 | 8.81734509 | | 0.650632016 | 0.49700225 | 1.30911282 | 0.19049614 | 0.99999872 |
| SLC16A7 | 22.0108941 | | 0.208498071 | 0.53623737 | 0.38881675 | 0.69741171 | 0.99999872 |
| FAM19A2 | 22.0866225 | | 0.158288846 | 0.78895905 | 0.20062999 | 0.84098791 | 0.99999872 |
| USP15 | 74.8796164 | | 0.010246602 | 0.36517727 | 0.02805925 | 0.97761489 | 0.99999872 |
| MON2 | 115.064627 | | 0.123993176 | 0.25345588 | 0.48921009 | 0.62469296 | 0.99999872 |
| AC048341.1 | 14.8149004 | | -0.74333505 | 0.58954199 | -1.2608687 | 0.20735615 | 0.99999872 |
| PPM1H | 54.186168 | | -0.027713117 | 0.40227117 | -0.0688916 | 0.94507588 | 0.99999872 |
| DPY19L2 | 13.4854276 | | 0.403707348 | 1.07446052 | 0.37573028 | 0.70711743 | 0.99999872 |
| RXYLT1 | 13.8239133 | | 0.072249063 | 0.50080709 | 0.14426526 | 0.88529101 | 0.99999872 |
| SRGAP1 | 531.536417 | | 0.026029516 | 0.26516026 | 0.09816522 | 0.9218011 | 0.99999872 |
| AC020611.2 | 21.7503917 | | 1.654469032 | 0.64961975 | 2.54682687 | 0.01087074 | 0.71391523 |
| C12orf56 | 16.5852597 | | 0.665085967 | 0.49018283 | 1.35681203 | 0.17484094 | 0.99999872 |
| XPOT | 36.150954 | | -0.052739003 | 0.44245603 | -0.119196 | 0.90512005 | 0.99999872 |
| TBK1 | 39.9385004 | | 0.053972722 | 0.29470717 | 0.18314017 | 0.85468804 | 0.99999872 |
| RASSF3 | 6.80535412 | | -0.693337729 | 0.80847354 | -0.8575886 | 0.39111965 | 0.99999872 |
| GNS | 51.3858469 | | 0.494752304 | 0.30021786 | 1.64797758 | 0.09935727 | 0.99999872 |
| LEMD3 | 36.3919969 | | 0.323078481 | 0.35259577 | 0.91628577 | 0.35951703 | 0.99999872 |
| MSRB3 | 151.942665 | | -0.047724834 | 0.40718622 | -0.1172064 | 0.90669649 | 0.99999872 |
| LLPH | 12.5353004 | | 0.016603112 | 0.51080805 | 0.03250362 | 0.97407043 | 0.99999872 |
| IRAK3 | 36.8584619 | | 0.24332275 | 0.61337373 | 0.39669575 | 0.69159184 | 0.99999872 |
| HELB | 15.1528531 | | -0.376732067 | 0.64537274 | -0.5837434 | 0.55939297 | 0.99999872 |
| GRIP1 | 121.959478 | | 0.289680166 | 0.3179385 | 0.91112014 | 3.62E-01 | 0.99999872 |
| CAND1 | 96.0086663 | | 0.431647823 | 0.24739578 | 1.74476631 | 0.08102555 | 0.99999872 |
| MDM1 | 18.0199165 | | -0.019451455 | 0.41995635 | -0.0463178 | 0.96305695 | 0.99999872 |
| RAP1B | 97.0161536 | | 0.243124512 | 0.22096324 | 1.10029392 | 0.27120408 | 0.99999872 |
| NUP107 | 18.5828008 | | -1.071462231 | 0.36429735 | -2.9411749 | 0.0032697 | 0.44319076 |
| SLC35E3 | 16.2469684 | | -0.993850219 | 0.42504981 | -2.3381971 | 0.01937703 | 0.8671024 |
| MDM2 | 35.687227 | | -0.327028991 | 0.55858213 | -0.5854627 | 0.55823665 | 0.99999872 |
| CPM | 16.7322625 | | -1.123075967 | 0.7819761 | -1.4362024 | 0.15094475 | 0.99999872 |
| CPSF6 | 76.89309 | | -0.009246808 | 0.29350504 | -0.0315048 | 0.97486699 | 0.99999872 |
| YEATS4 | 6.65291834 | | -0.586256521 | 0.59679 | -0.9823498 | 0.32592756 | 0.99999872 |
| FRS2 | 78.5895843 | | 0.177667673 | 0.24347453 | 0.72971769 | 0.46556277 | 0.99999872 |
| CCT2 | 18.7090451 | | 0.376649777 | 0.4378558 | 0.86021419 | 0.38967098 | 0.99999872 |
| RAB3IP | 12.8546106 | | 0.237861079 | 0.58419232 | 0.40716228 | 0.6838888 | 0.99999872 |
| AC025159.1 | 37.8070119 | | -1.085274846 | 0.51736694 | -2.0976888 | 0.03593264 | 0.94734101 |
| CNOT2 | 75.609938 | | -0.032395113 | 0.23064249 | -0.140456 | 0.88829974 | 0.99999872 |
| TSPAN8 | 5.95225338 | | 0.318857739 | 0.61899188 | 0.51512427 | 0.60646619 | 0.99999872 |
| ZFC3H1 | 69.3654879 | | -0.12067897 | 0.36910564 | -0.3269497 | 0.74370595 | 0.99999872 |
| TMEM19 | 16.957774 | | 0.229538411 | 0.56271619 | 0.40791151 | 0.68333864 | 0.99999872 |
| RAB21 | 64.534825 | | 0.244409303 | 0.23330839 | 1.04758041 | 0.29483197 | 0.99999872 |
| TBC1D15 | 46.411227 | | -0.164385508 | 0.32272812 | -0.5093622 | 0.61049837 | 0.99999872 |
| TRHDE | 17.5787643 | | -0.712013394 | 0.80955773 | -0.8795091 | 0.3791253 | 0.99999872 |
| ATXN7L3B | 35.0067155 | | 0.224444339 | 0.32876039 | 0.68269885 | 0.49479716 | 0.99999872 |
| CAPS2 | 42.9925772 | | 0.707700952 | 0.47680496 | 1.4842567 | 0.13774083 | 0.99999872 |
| GLIPR1L2 | 8.75005105 | | 2.008386253 | 1.10155366 | 1.82323051 | 0.06826849 | 0.99999872 |
| KRR1 | 18.3022359 | | -0.245824627 | 0.43493793 | -0.5651947 | 0.57194132 | 0.99999872 |
| PHLDA1 | 31.6799897 | | 1.044452913 | 1.14063845 | 0.91567395 | 0.35983793 | 0.99999872 |

| | | | | | | |
|---|---|---|---|---|---|---|
| NAP1L1 | 75.5614591 | 0.263708758 | 0.34501187 | 0.76434691 | 0.44466054 | 0.99999872 |
| BBS10 | 3.77891713 | 0.432181079 | 0.72139938 | 0.59908712 | 0.54911479 | NA |
| OSBPL8 | 66.1639823 | -0.015119282 | 0.25339128 | -0.0596677 | 0.95242027 | 0.99999872 |
| ZDHHC17 | 52.1635661 | -0.025995357 | 0.30162319 | -0.0861849 | 0.93131945 | 0.99999872 |
| NAV3 | 9.59377122 | -0.028591073 | 0.73026304 | -0.0391517 | 0.96876941 | 0.99999872 |
| SYT1 | 32.8014441 | 1.438666846 | 1.12977565 | 1.27340933 | 0.20287283 | 0.99999872 |
| AC027288.3 | 74.678327 | -0.237878414 | 0.66432366 | -0.3580761 | 0.72028638 | 0.99999872 |
| PAWR | 382.490697 | -0.523728698 | 0.41026155 | -1.2765727 | 0.20175314 | 0.99999872 |
| AC073569.2 | 26.9712508 | -0.620463335 | 0.76991253 | -0.8058881 | 0.42030738 | 0.99999872 |
| PPP1R12A | 107.141704 | -0.089980439 | 0.27019501 | -0.3330204 | 0.73911892 | 0.99999872 |
| PTPRQ | 2479.13882 | -0.81441559 | 0.4151567 | -1.9617065 | 0.04979667 | 0.97937581 |
| AC074031.1 | 20.8715147 | 0.775042643 | 0.58986074 | 1.31394174 | 0.18886581 | 0.99999872 |
| ACSS3 | 14.8747414 | 0.371517549 | 0.92660576 | 0.40094457 | 0.68846093 | 0.99999872 |
| CCDC59 | 21.0383638 | 0.589401237 | 0.55253964 | 1.06671303 | 0.28610144 | 0.99999872 |
| METTL25 | 24.6166735 | -0.098315026 | 0.38331004 | -0.2564896 | 0.79757284 | 0.99999872 |
| TMTC2 | 26.3826848 | 0.73788909 | 0.67061174 | 1.10032235 | 0.2711917 | 0.99999872 |
| LRRIQ1 | 13.0998651 | 0.350600536 | 0.57053335 | 0.61451366 | 0.53887595 | 0.99999872 |
| C12orf29 | 9.87157551 | 0.394445551 | 0.49674317 | 0.79406336 | 0.42715855 | 0.99999872 |
| CEP290 | 55.4994286 | 0.287295496 | 0.27606882 | 1.04066621 | 0.29803049 | 0.99999872 |
| TMTC3 | 28.9295951 | 0.056188587 | 0.31164126 | 0.18029894 | 0.85691789 | 0.99999872 |
| KITLG | 11.6210952 | -0.498974086 | 1.02441041 | -0.4870842 | 0.62619869 | 0.99999872 |
| DUSP6 | 31.8772404 | -1.613206958 | 0.9109904 | -1.7708276 | 0.07658937 | 0.99999872 |
| POC1B | 20.3409157 | 0.223208669 | 0.39221967 | 0.56909096 | 0.56929442 | 0.99999872 |
| POC1B-AS1 | 10.5039204 | -0.55395714 | 0.47477858 | -1.1667694 | 0.24330349 | 0.99999872 |
| ATP2B1 | 57.4266485 | 0.239239701 | 0.40372809 | 0.59257631 | 0.55346474 | 0.99999872 |
| DCN | 450.288791 | 0.119582258 | 0.6312339 | 0.18944207 | 0.84974635 | 0.99999872 |
| BTG1 | 21.3110149 | -0.019725786 | 0.41896396 | -0.0470823 | 0.96244764 | 0.99999872 |
| EEA1 | 129.151385 | 0.081594985 | 0.39707271 | 0.20549129 | 0.8371883 | 0.99999872 |
| AC138123.1 | 12.1463401 | -0.798541246 | 0.77607126 | -1.0289535 | 0.30350152 | 0.99999872 |
| NUDT4 | 16.4200025 | -0.171023401 | 0.49143041 | -0.3480114 | 0.7278316 | 0.99999872 |
| UBE2N | 36.6696426 | 0.380307633 | 0.36169395 | 1.05146252 | 0.29304621 | 0.99999872 |
| MRPL42 | 28.7593608 | -0.070886522 | 0.32913701 | -0.2153709 | 0.82947818 | 0.99999872 |
| SOCS2 | 13.2061748 | 1.143041046 | 0.73356476 | 1.5582006 | 0.1191857 | 0.99999872 |
| CRADD | 113.897766 | -0.410960522 | 0.3224574 | -1.2744645 | 0.20249884 | 0.99999872 |
| AC012085.2 | 7.84613638 | 0.624573267 | 0.76659876 | 0.81473295 | 0.41522519 | 0.99999872 |
| PLXNC1 | 15.4802926 | -0.366865911 | 0.46054183 | -0.7965963 | 0.42568553 | 0.99999872 |
| CEP83 | 43.9847818 | 0.132182551 | 0.30475826 | 0.43372918 | 0.66448511 | 0.99999872 |
| TMCC3 | 150.975271 | 0.301740985 | 0.42981358 | 0.70202757 | 0.48266197 | 0.99999872 |
| NDUFA12 | 22.6551415 | 0.588284468 | 0.44661852 | 1.31719675 | 0.18777268 | 0.99999872 |
| NR2C1 | 27.8047245 | 0.142954795 | 0.39308404 | 0.36367489 | 0.7161008 | 0.99999872 |
| FGD6 | 19.9031495 | -0.762607151 | 0.41848248 | -1.8223156 | 0.06840711 | 0.99999872 |
| VEZT | 110.87252 | -0.28917427 | 0.22237155 | -1.3004104 | 0.19346036 | 0.99999872 |
| METAP2 | 44.3096622 | 0.348181302 | 0.25299955 | 1.37621311 | 0.16875567 | 0.99999872 |
| NTN4 | 33.8203612 | -0.044039436 | 0.60778106 | -0.0724594 | 0.94223633 | 0.99999872 |
| SNRPF | 11.9194524 | 0.103925851 | 0.46748901 | 0.22230651 | 0.82407528 | 0.99999872 |
| LTA4H | 12.1913552 | 0.410367572 | 0.49036122 | 0.83686792 | 0.40266681 | 0.99999872 |
| ELK3 | 5.18470608 | -1.288912167 | 0.99172853 | -1.2996623 | 0.19371674 | 0.99999872 |
| CDK17 | 61.1508156 | 0.116031229 | 0.26692381 | 0.43469793 | 0.6637817 | 0.99999872 |
| CFAP54 | 10.2347701 | 0.473032355 | 0.65833869 | 0.71852431 | 0.47243406 | 0.99999872 |
| NEDD1 | 18.7754383 | -0.108578228 | 0.38870759 | -0.2793314 | 0.77999052 | 0.99999872 |
| TMPO | 20.0072524 | 0.373059365 | 0.36217125 | 1.03006346 | 0.30298022 | 0.99999872 |
| SLC25A3 | 70.9313906 | 0.600288423 | 0.64272709 | 0.93397094 | 0.35031888 | 0.99999872 |
| IKBIP | 12.1916903 | 0.111948094 | 0.60650948 | 0.18457765 | 0.85356032 | 0.99999872 |
| APAF1 | 17.5101005 | 0.1822699 | 0.44991923 | 0.40511694 | 0.68539156 | 0.99999872 |
| ANKS1B | 117.997965 | 0.640364044 | 0.73499442 | 0.87125021 | 0.38361755 | 0.99999872 |
| UHRF1BP1L | 32.115998 | 0.108839091 | 0.42416627 | 0.25659535 | 0.79749117 | 0.99999872 |
| ACTR6 | 10.2731574 | 0.305932577 | 0.53148336 | 0.57562024 | 0.5648719 | 0.99999872 |
| SCYL2 | 38.0851844 | -0.242494881 | 0.30161171 | -0.8039969 | 0.42139876 | 0.99999872 |
| NR1H4 | 17.2176247 | 0.184030149 | 0.76773043 | 0.23970673 | 0.81055762 | 0.99999872 |
| UTP20 | 19.8713737 | -0.320827389 | 0.41901678 | -0.7656672 | 0.44387437 | 0.99999872 |
| ARL1 | 36.0499763 | 0.460684921 | 0.38058284 | 1.21047215 | 0.22609777 | 0.99999872 |

| | | | | | | |
|---|---|---|---|---|---|---|
| CHPT1 | 77.5374171 | 0.681373264 | 0.39460246 | 1.72673345 | 0.08421554 | 0.99999872 |
| SYCP3 | 4.16736036 | 0.379867919 | 0.82681326 | 0.45943617 | 0.64592098 | NA |
| GNPTAB | 70.9913965 | -0.437570689 | 0.42847184 | -1.0212356 | 0.30714284 | 0.99999872 |
| DRAM1 | 20.833539 | -0.543415682 | 0.39251567 | -1.3844433 | 0.16622276 | 0.99999872 |
| WASHC3 | 47.6810135 | 0.102521452 | 0.42797237 | 0.23955157 | 0.81067791 | 0.99999872 |
| NUP37 | 9.5587138 | 0.569004698 | 0.5036685 | 1.12972064 | 0.25859396 | 0.99999872 |
| LINC02456 | 7.86766442 | 1.349531597 | 1.38596765 | 0.97371075 | 0.33020018 | 0.99999872 |
| IGF1 | 49.7673517 | 0.062561115 | 1.52539512 | 0.04101306 | 0.96728549 | 0.99999872 |
| HSP90B1 | 238.375487 | 0.996196803 | 0.57184416 | 1.74207741 | 0.0814949 | 0.99999872 |
| C12orf73 | 8.54383773 | 0.128528983 | 0.5184619 | 0.24790439 | 0.80420838 | 0.99999872 |
| TDG | 15.3728135 | 0.027263489 | 0.41268039 | 0.06606442 | 0.94732654 | 0.99999872 |
| HCFC2 | 19.4926768 | 0.45252384 | 0.40475437 | 1.1180209 | 0.26355807 | 0.99999872 |
| NFYB | 10.9908604 | -0.129353004 | 0.44807681 | -0.2886849 | 0.77282253 | 0.99999872 |
| TXNRD1 | 201.380756 | 0.527262597 | 0.30386124 | 1.73520847 | 0.08270389 | 0.99999872 |
| CHST11 | 7.55647107 | -0.042380844 | 0.5338154 | -0.0793923 | 0.93672057 | 0.99999872 |
| SLC41A2 | 20.8734811 | 0.278128244 | 0.48488814 | 0.57359259 | 0.56624353 | 0.99999872 |
| C12orf45 | 6.84037287 | -0.097216518 | 0.6265477 | -0.1551622 | 0.87669345 | 0.99999872 |
| ALDH1L2 | 3.95411351 | 1.339355809 | 1.39021115 | 0.96341898 | 0.33533731 | NA |
| WASHC4 | 62.0926011 | 0.000518897 | 0.38951973 | 0.00133214 | 0.9989371 | 0.99999872 |
| APPL2 | 31.4340693 | 0.218386194 | 0.33986281 | 0.64257161 | 0.52050211 | 0.99999872 |
| C12orf75 | 9.77955327 | 0.740124215 | 0.61471266 | 1.20401655 | 0.22858318 | 0.99999872 |
| NUAK1 | 7.08997769 | -0.567417985 | 0.71320633 | -0.7955874 | 0.42627188 | 0.99999872 |
| CKAP4 | 39.5471043 | 0.80284055 | 0.53119138 | 1.51139603 | 0.13068758 | 0.99999872 |
| TCP11L2 | 9.24840466 | 0.385753798 | 0.67327628 | 0.57295023 | 0.5666784 | 0.99999872 |
| POLR3B | 9.92151436 | -0.215313248 | 0.52939944 | -0.4067123 | 0.68421933 | 0.99999872 |
| RIC8B | 38.464652 | 0.354333891 | 0.41795639 | 0.84777717 | 0.39656208 | 0.99999872 |
| TMEM263 | 24.9103917 | 0.743562588 | 0.32986262 | 2.25415839 | 0.02418621 | 0.88337526 |
| MTERF2 | 7.76595458 | 0.027209772 | 0.53829853 | 0.05054774 | 0.95968591 | 0.99999872 |
| CRY1 | 19.7177128 | -0.373455087 | 0.47911762 | -0.7794643 | 0.43570626 | 0.99999872 |
| PWP1 | 13.8406766 | -0.144844651 | 0.40038807 | -0.3617607 | 0.71753089 | 0.99999872 |
| PRDM4 | 20.6821656 | -0.160561484 | 0.54937641 | -0.2922613 | 0.77008682 | 0.99999872 |
| FICD | 9.8386953 | 1.274440746 | 0.72742852 | 1.75198072 | 0.07977712 | 0.99999872 |
| SART3 | 14.2307045 | -0.198539495 | 0.44601649 | -0.4451394 | 0.65621905 | 0.99999872 |
| ISCU | 38.9732514 | 1.191751979 | 0.46998916 | 2.53570097 | 0.01122225 | 0.72584414 |
| CORO1C | 18.7665176 | 0.058501731 | 0.44993076 | 0.13002385 | 0.89654756 | 0.99999872 |
| SSH1 | 23.9303046 | -0.23789382 | 0.36024693 | -0.6603632 | 0.50902078 | 0.99999872 |
| USP30 | 8.76423079 | -0.172523369 | 0.47360557 | -0.3642765 | 0.71565156 | 0.99999872 |
| UNG | 20.2115002 | 0.726188217 | 0.50290408 | 1.4439895 | 0.14874193 | 0.99999872 |
| ACACB | 49.2881382 | -0.12408465 | 0.57369484 | -0.2162903 | 0.82876144 | 0.99999872 |
| KCTD10 | 13.1812122 | -0.508027576 | 0.42928457 | -1.1834285 | 0.23663938 | 0.99999872 |
| UBE3B | 16.8834645 | 0.373156283 | 0.4114549 | 0.90691905 | 0.36444961 | 0.99999872 |
| MMAB | 12.8835334 | 0.492009111 | 0.47117985 | 1.04420661 | 0.29638981 | 0.99999872 |
| MVK | 5.4611499 | 0.106737504 | 0.64824689 | 0.16465564 | 0.86921504 | 0.99999872 |
| TRPV4 | 9.68727901 | 1.124476631 | 0.71480781 | 1.57311745 | 0.11569163 | 0.99999872 |
| GLTP | 14.3319934 | 0.262551486 | 0.45742162 | 0.57398137 | 0.56598041 | 0.99999872 |
| TCHP | 15.627864 | -0.005416717 | 0.42033801 | -0.0128866 | 0.98971829 | 0.99999872 |
| GIT2 | 32.410232 | 0.024782089 | 0.37159121 | 0.06669181 | 0.94682706 | 0.99999872 |
| ANKRD13A | 20.2184357 | -0.143737165 | 0.44713629 | -0.3214616 | 0.74786058 | 0.99999872 |
| C12orf76 | 12.1429093 | 0.298766374 | 0.43527565 | 0.68638431 | 0.4924708 | 0.99999872 |
| IFT81 | 17.2309094 | 0.02611267 | 0.40957449 | 0.06375561 | 0.94916483 | 0.99999872 |
| ATP2A2 | 93.1835487 | -0.12796605 | 0.38825447 | -0.3295932 | 0.74170733 | 0.99999872 |
| ANAPC7 | 20.7509795 | -0.530385254 | 0.36952045 | -1.435334 | 0.15119195 | 0.99999872 |
| ARPC3 | 76.3267313 | 0.749453411 | 0.28896404 | 2.59358711 | 0.00949805 | 0.69901919 |
| GPN3 | 8.43017333 | 0.927676891 | 0.60578096 | 1.53137346 | 0.12567712 | 0.99999872 |
| VPS29 | 39.757057 | 0.747223798 | 0.29866125 | 2.50191074 | 0.01235251 | 0.72670507 |
| PPTC7 | 30.888287 | 0.340444396 | 0.33831826 | 1.00628443 | 0.31427877 | 0.99999872 |
| TCTN1 | 18.2584303 | 0.461061749 | 0.39680329 | 1.16194034 | 0.2452597 | 0.99999872 |
| PPP1CC | 24.8136108 | 0.097826125 | 0.3157746 | 0.30979732 | 0.75671509 | 0.99999872 |
| SH2B3 | 9.66610224 | -0.696776531 | 0.53132344 | -1.3113981 | 0.18972331 | 0.99999872 |
| ATXN2 | 207.179214 | -0.328269769 | 0.23614155 | -1.3901398 | 0.16448643 | 0.99999872 |
| BRAP | 17.9732363 | 0.105228686 | 0.42712396 | 0.24636568 | 0.80539917 | 0.99999872 |

| | | | | | | |
|---|---|---|---|---|---|---|
| ACAD10 | 13.1990042 | -0.68236526 | 0.51836469 | -1.3163807 | 0.1880463 | 0.99999872 |
| ALDH2 | 26.8109403 | 0.361763741 | 0.42884235 | 0.84358212 | 0.39890296 | 0.99999872 |
| MAPKAPK5-A | 5.90816334 | 0.504687486 | 0.58960026 | 0.85598246 | 0.39200748 | 0.99999872 |
| MAPKAPK5 | 17.1110896 | -0.071443211 | 0.3909167 | -0.1827581 | 0.85498779 | 0.99999872 |
| TMEM116 | 12.58084 | -0.901796521 | 0.5871666 | -1.5358444 | 0.12457655 | 0.99999872 |
| ERP29 | 42.6069071 | 0.574788217 | 0.36868568 | 1.55901963 | 0.11899174 | 0.99999872 |
| NAA25 | 40.3251652 | -0.447056035 | 0.42540295 | -1.0509002 | 0.29330444 | 0.99999872 |
| TRAFD1 | 24.1329168 | 0.197494762 | 0.6312087 | 0.31288346 | 0.7543692 | 0.99999872 |
| HECTD4 | 116.786322 | 0.110143249 | 0.47051917 | 0.23408876 | 0.81491607 | 0.99999872 |
| RPL6 | 65.3967512 | 0.73322788 | 0.43222176 | 1.69641593 | 0.08980714 | 0.99999872 |
| PTPN11 | 38.3436586 | -0.281814636 | 0.41221333 | -0.6836621 | 0.49418856 | 0.99999872 |
| OAS1 | 49.2362114 | -3.467047486 | 1.48766914 | -2.3305232 | 0.01977851 | 0.8671024 |
| AC004551.1 | 5.08839831 | -1.934612453 | 1.20138892 | -1.6103132 | 0.1073295 | 0.99999872 |
| OAS3 | 12.3106195 | -3.060834614 | 2.76844754 | -1.1056141 | 0.26889359 | 0.99999872 |
| OAS2 | 12.5324577 | -4.969945723 | 2.79182446 | -1.7801784 | 0.07504676 | 0.99999872 |
| DDX54 | 13.0318472 | -0.451746966 | 0.40132193 | -1.1256473 | 0.26031482 | 0.99999872 |
| TPCN1 | 65.9023033 | 0.132585202 | 0.37909958 | 0.34973714 | 0.72653598 | 0.99999872 |
| SLC8B1 | 10.3189166 | -0.455903038 | 0.51317896 | -0.88839 | 0.37433102 | 0.99999872 |
| PLBD2 | 13.8656158 | 0.306935764 | 0.46028483 | 0.66683876 | 0.50487513 | 0.99999872 |
| SDSL | 9.72316145 | 0.14061513 | 0.81033872 | 0.17352636 | 0.86223771 | 0.99999872 |
| RBM19 | 12.4704006 | -0.915062163 | 0.49624336 | -1.8439786 | 0.06518625 | 0.99999872 |
| MED13L | 295.870916 | 0.139167894 | 0.36024687 | 0.38631257 | 0.69926519 | 0.99999872 |
| MAP1LC3B2 | 6.03097835 | 0.825948307 | 0.57602529 | 1.43387507 | 0.15160792 | 0.99999872 |
| C12orf49 | 11.8963739 | -0.39033459 | 0.39395225 | -0.990817 | 0.32177494 | 0.99999872 |
| FBXW8 | 27.3933666 | -0.391252084 | 0.32005265 | -1.2224616 | 0.22153311 | 0.99999872 |
| FBXO21 | 153.139651 | 0.257479218 | 0.45131752 | 0.57050571 | 0.56833475 | 0.99999872 |
| WSB2 | 19.7507388 | 0.139328832 | 0.41057556 | 0.33935004 | 0.73434605 | 0.99999872 |
| VSIG10 | 8.51011519 | -0.516106035 | 0.61056045 | -0.8452988 | 0.39794401 | 0.99999872 |
| PEBP1 | 187.134968 | 0.384702343 | 0.39314117 | 0.97853488 | 0.32780985 | 0.99999872 |
| TAOK3 | 66.6103352 | -0.462584819 | 0.30611125 | -1.5111657 | 0.13074624 | 0.99999872 |
| SUDS3 | 17.5678819 | 0.048005201 | 0.46926845 | 0.10229795 | 0.91852018 | 0.99999872 |
| HSPB8 | 58.0436789 | 0.635933357 | 0.49545768 | 1.28352709 | 0.19930748 | 0.99999872 |
| PRKAB1 | 10.8653364 | 0.5088998 | 0.53323201 | 0.95436843 | 0.33989718 | 0.99999872 |
| CIT | 7.30238677 | -0.262838019 | 0.59292091 | -0.4432936 | 0.65755342 | 0.99999872 |
| BICDL1 | 32.2070309 | -0.639249764 | 0.56704754 | -1.1273301 | 0.25960294 | 0.99999872 |
| RAB35 | 10.7487094 | -0.014367501 | 0.66569338 | -0.0215828 | 0.98278078 | 0.99999872 |
| GCN1 | 17.4243337 | 0.052131446 | 0.45939344 | 0.11347886 | 0.90965092 | 0.99999872 |
| RPLP0 | 92.6378619 | 0.48418879 | 0.63100794 | 0.76732599 | 0.44288772 | 0.99999872 |
| PXN-AS1 | 4.81842777 | -0.896867205 | 0.6133301 | -1.4622912 | 0.14366143 | 0.99999872 |
| PXN | 13.9842682 | -0.856836382 | 0.44295672 | -1.9343569 | 0.05306926 | 0.99999872 |
| TRIAP1 | 3.76083376 | 0.499204364 | 0.76014446 | 0.65672302 | 0.51135903 | NA |
| GATC | 9.76832889 | 0.046120027 | 0.62256361 | 0.07408083 | 0.94094607 | 0.99999872 |
| SRSF9 | 42.799664 | 0.42667375 | 0.30568134 | 1.3958122 | 0.16277105 | 0.99999872 |
| DYNLL1 | 200.757614 | 1.033685682 | 0.60179517 | 1.7176703 | 0.08585677 | 0.99999872 |
| COQ5 | 6.24826803 | -0.123056328 | 0.6649298 | -0.1850666 | 0.85317676 | 0.99999872 |
| RNF10 | 45.1861248 | 0.113806591 | 0.31036785 | 0.36668292 | 0.71385555 | 0.99999872 |
| POP5 | 16.8900193 | -0.450090783 | 0.39836151 | -1.1298551 | 0.25853729 | 0.99999872 |
| MLEC | 60.5474748 | 0.410553768 | 0.4250036 | 0.96600068 | 0.33404384 | 0.99999872 |
| UNC119B | 10.716828 | -0.297145292 | 0.50268504 | -0.5911162 | 0.55444253 | 0.99999872 |
| ACADS | 9.1039015 | -0.230879932 | 0.48025135 | -0.4807481 | 0.63069553 | 0.99999872 |
| SPPL3 | 116.72379 | -0.202869576 | 0.21008715 | -0.9656449 | 0.33422192 | 0.99999872 |
| C12orf43 | 5.44114992 | -0.706854187 | 0.80055998 | -0.8829497 | 0.37726345 | 0.99999872 |
| OASL | 8.06792561 | -1.656754785 | 4.78621197 | -0.3461516 | 0.72922881 | 0.99999872 |
| P2RX4 | 12.0012503 | -0.640263392 | 0.49926598 | -1.2824094 | 0.19969906 | 0.99999872 |
| CAMKK2 | 18.4730459 | 0.097579847 | 0.49849209 | 0.19575004 | 0.84480582 | 0.99999872 |
| ANAPC5 | 42.0926369 | -0.032019722 | 0.30808872 | -0.1039302 | 0.91722474 | 0.99999872 |
| RNF34 | 15.2051555 | -0.296827655 | 0.43567742 | -0.6813014 | 0.49568077 | 0.99999872 |
| KDM2B | 19.2939814 | 0.26675961 | 0.46723221 | 0.57093583 | 0.56804314 | 0.99999872 |
| ORAI1 | 3.90071374 | 0.312380961 | 0.68747575 | 0.45438834 | 0.64954935 | NA |
| TMEM120B | 13.6276229 | -0.629565211 | 0.52039731 | -1.209778 | 0.2263641 | 0.99999872 |
| SETD1B | 10.6673874 | 0.051228855 | 0.49252698 | 0.10401228 | 0.9171596 | 0.99999872 |

| | | | | | | |
|---|---|---|---|---|---|---|
| WDR66 | 25.3386453 | -1.221079655 | 0.55003685 | -2.2199961 | 0.02641903 | 0.88440681 |
| BCL7A | 6.07206024 | -0.199004741 | 0.9492408 | -0.2096462 | 0.83394381 | 0.99999872 |
| MLXIP | 72.9736763 | -0.036426552 | 0.43838373 | -0.0830928 | 0.93377772 | 0.99999872 |
| CLIP1 | 90.1523717 | -0.543479255 | 0.29959643 | -1.8140378 | 0.0696719 | 0.99999872 |
| ZCCHC8 | 19.8297142 | -0.063046443 | 0.46953421 | -0.1342744 | 0.89318556 | 0.99999872 |
| RSRC2 | 44.3005346 | 0.211047715 | 0.28474263 | 0.74118762 | 0.45857969 | 0.99999872 |
| DENR | 21.9732283 | 0.087169059 | 0.46877857 | 0.18594933 | 0.8524845 | 0.99999872 |
| HIP1R | 18.212889 | -1.15443516 | 0.60301911 | -1.9144255 | 0.05556581 | 0.99999872 |
| VPS37B | 16.4893615 | -0.537975258 | 0.42905051 | -1.253874 | 0.20988782 | 0.99999872 |
| ARL6IP4 | 53.2246355 | 0.015942395 | 0.34666627 | 0.04598773 | 0.96332003 | 0.99999872 |
| PITPNM2 | 10.5049187 | -0.413883405 | 0.48970076 | -0.8451762 | 0.39801249 | 0.99999872 |
| MPHOSPH9 | 20.1335793 | 0.182731459 | 0.41656041 | 0.43866736 | 0.66090258 | 0.99999872 |
| C12orf65 | 16.6165554 | -0.019083041 | 0.37856483 | -0.0504089 | 0.95979654 | 0.99999872 |
| SBNO1 | 29.8107612 | -0.075190517 | 0.32556138 | -0.2309565 | 0.81734859 | 0.99999872 |
| KMT5A | 14.8535448 | -0.251174609 | 0.52237683 | -0.4808303 | 0.63063712 | 0.99999872 |
| RILPL2 | 13.9422911 | -0.057719624 | 0.52949042 | -0.1090098 | 0.91319475 | 0.99999872 |
| SNRNP35 | 23.9768463 | -0.07174251 | 0.34665194 | -0.2069583 | 0.83604239 | 0.99999872 |
| RILPL1 | 43.5862868 | -0.104295103 | 0.35899179 | -0.2905223 | 0.77141673 | 0.99999872 |
| TMED2 | 30.4500669 | 0.733958728 | 0.48578561 | 1.51086964 | 0.13082167 | 0.99999872 |
| DDX55 | 11.0259528 | -0.689305784 | 0.64409723 | -1.070189 | 0.28453424 | 0.99999872 |
| EIF2B1 | 12.8965503 | 0.052187809 | 0.52167265 | 0.10003938 | 0.92031306 | 0.99999872 |
| GTF2H3 | 8.70563351 | 0.204037062 | 0.53811361 | 0.37917097 | 0.7045609 | 0.99999872 |
| TCTN2 | 13.4259682 | 0.470423242 | 0.60590821 | 0.77639357 | 0.43751664 | 0.99999872 |
| ATP6V0A2 | 11.120605 | -0.042338259 | 0.42362235 | -0.0999434 | 0.92038926 | 0.99999872 |
| CCDC92 | 34.4493201 | 0.14151808 | 0.3090169 | 0.45796226 | 0.64697955 | 0.99999872 |
| RFLNA | 8.01069654 | 1.100993025 | 0.73942407 | 1.48898727 | 0.13649072 | 0.99999872 |
| NCOR2 | 82.4187771 | -0.637483699 | 0.36403893 | -1.7511415 | 0.07992153 | 0.99999872 |
| SCARB1 | 4.5468393 | 1.204157013 | 0.91682739 | 1.31339555 | 0.1890497 | NA |
| UBC | 263.202337 | 0.348867628 | 0.37573788 | 0.92848671 | 0.35315516 | 0.99999872 |
| DHX37 | 3.83564102 | -1.118646225 | 0.75645158 | -1.4788074 | 0.13919179 | NA |
| BRI3BP | 10.76409 | 0.776947272 | 0.57600216 | 1.34886174 | 0.17738138 | 0.99999872 |
| AACS | 11.9325849 | -0.436490012 | 0.53462237 | -0.8164455 | 0.4142454 | 0.99999872 |
| SLC15A4 | 30.707321 | 0.603067183 | 0.3379775 | 1.78434124 | 0.07436823 | 0.99999872 |
| STX2 | 35.2525872 | 0.373788182 | 0.31324678 | 1.19327063 | 0.2327634 | 0.99999872 |
| RAN | 39.1729672 | 0.877523065 | 0.48843711 | 1.79659375 | 0.07240014 | 0.99999872 |
| SFSWAP | 39.4180476 | -0.248475292 | 0.33016488 | -0.7525794 | 0.45170269 | 0.99999872 |
| ULK1 | 27.9451514 | -0.766424152 | 0.39059482 | -1.9621974 | 0.0497395 | 0.97937581 |
| PUS1 | 4.49976537 | -0.645398332 | 0.6516512 | -0.9904046 | 0.32197641 | NA |
| EP400 | 31.9263891 | -0.263903541 | 0.30773143 | -0.8575775 | 0.39112583 | 0.99999872 |
| DDX51 | 6.5733848 | -1.125175343 | 0.55126354 | -2.0410843 | 0.04124245 | 0.95980711 |
| FBRSL1 | 18.4305333 | 0.091114028 | 0.45116425 | 0.20195312 | 0.83995337 | 0.99999872 |
| PGAM5 | 5.00742427 | -1.437565123 | 0.64533869 | -2.2276134 | 0.0259063 | 0.88365618 |
| ANKLE2 | 30.8051018 | -0.231860773 | 0.30827161 | -0.7521315 | 0.45197201 | 0.99999872 |
| GOLGA3 | 38.4108521 | -0.612666098 | 0.48254783 | -1.2696484 | 0.20420989 | 0.99999872 |
| CHFR | 35.6831537 | -0.335165182 | 0.28253853 | -1.1862636 | 0.2355182 | 0.99999872 |
| ZNF26 | 14.6976896 | -0.970136124 | 0.37975891 | -2.5546105 | 0.01063067 | 0.70947852 |
| ZNF84 | 29.0858297 | 0.061113938 | 0.34853109 | 0.17534716 | 0.86080685 | 0.99999872 |
| ZNF140 | 11.3125829 | 0.298462389 | 0.54515774 | 0.54747896 | 0.58404972 | 0.99999872 |
| ZNF891 | 10.760402 | 0.450199996 | 0.51339544 | 0.87690689 | 0.38053721 | 0.99999872 |
| ZNF268 | 8.42007955 | -0.464350333 | 0.49699387 | -0.934318 | 0.35013987 | 0.99999872 |
| MPHOSPH8 | 74.2674897 | -0.058385944 | 0.34096418 | -0.1712378 | 0.86403681 | 0.99999872 |
| PSPC1 | 62.8384381 | -0.146429945 | 0.34878699 | -0.4198263 | 0.67461236 | 0.99999872 |
| ZMYM5 | 27.4622353 | 0.165989832 | 0.4559604 | 0.36404441 | 0.71582485 | 0.99999872 |
| AL355001.2 | 4.36008058 | 0.414124091 | 0.70985793 | 0.58339011 | 0.55963072 | NA |
| ZMYM2 | 173.871598 | -0.077922419 | 0.30174518 | -0.2582392 | 0.79622234 | 0.99999872 |
| GJA3 | 16.8650131 | 0.398592004 | 0.73903808 | 0.53933893 | 0.58965301 | 0.99999872 |
| CRYL1 | 32.7878335 | 0.429669902 | 0.49510641 | 0.86783344 | 0.38548552 | 0.99999872 |
| IFT88 | 47.1314447 | -0.00370876 | 0.28896926 | -0.0128344 | 0.98975987 | 0.99999872 |
| EEF1AKMT1 | 10.9970311 | 0.733173078 | 0.62408235 | 1.17480182 | 0.24007401 | 0.99999872 |
| XPO4 | 35.5234004 | -0.006351441 | 0.32209868 | -0.0197189 | 0.98426759 | 0.99999872 |
| LATS2 | 75.039513 | 0.156816075 | 0.2883202 | 0.54389555 | 0.58651335 | 0.99999872 |

| | | | | | | |
|---|---|---|---|---|---|---|
| SAP18 | 53.4512255 | 0.655089416 | 0.33631805 | 1.94782713 | 0.05143565 | 0.9907036 |
| MRPL57 | 34.9582647 | 0.508812767 | 0.35704878 | 1.42505114 | 0.15414242 | 0.99999872 |
| ZDHHC20 | 36.4810652 | -0.56909557 | 0.29478386 | -1.930552 | 0.05353848 | 0.99999872 |
| MICU2 | 72.240708 | -0.094878578 | 0.22602395 | -0.4197722 | 0.67465185 | 0.99999872 |
| FGF9 | 27.0317541 | 0.685930561 | 0.69553749 | 0.98618776 | 0.32404098 | 0.99999872 |
| SACS | 5.86329095 | -0.358527679 | 0.57775228 | -0.6205561 | 0.53489176 | 0.99999872 |
| TNFRSF19 | 20.5956486 | 0.864132024 | 0.80716337 | 1.07057885 | 0.28435884 | 0.99999872 |
| MIPEP | 29.4547299 | 0.017249077 | 0.49454589 | 0.03487862 | 0.97217653 | 0.99999872 |
| PARP4 | 39.0352322 | -0.147436538 | 0.46702099 | -0.3156957 | 0.75223346 | 0.99999872 |
| MTMR6 | 40.0314548 | 0.482710593 | 0.41710884 | 1.1572773 | 0.24715908 | 0.99999872 |
| NUP58 | 62.0881986 | 0.601414165 | 0.28917861 | 2.0797325 | 0.03755007 | 0.95980711 |
| ATP8A2 | 48.8768553 | 1.338931153 | 0.82386738 | 1.62517802 | 0.10412463 | 0.99999872 |
| SHISA2 | 181.537893 | 0.869243272 | 0.79101297 | 1.09889889 | 0.27181217 | 0.99999872 |
| RNF6 | 14.3290475 | 1.09920374 | 0.50563776 | 2.17389569 | 0.02971297 | 0.90266962 |
| CDK8 | 41.6047674 | 0.592266803 | 0.31679273 | 1.86957195 | 0.06154328 | 0.99999872 |
| WASF3 | 16.8768937 | 0.079831069 | 0.36862582 | 0.21656397 | 0.82854817 | 0.99999872 |
| USP12 | 41.9800505 | 0.128015926 | 0.3894693 | 0.32869324 | 0.74238756 | 0.99999872 |
| RPL21 | 39.9272112 | 1.029582767 | 0.5441357 | 1.8921434 | 0.05847188 | 0.99999872 |
| GTF3A | 12.4956926 | 0.004724806 | 0.48929733 | 0.00965631 | 0.9922955 | 0.99999872 |
| MTIF3 | 24.9989528 | 0.14896523 | 0.32437891 | 0.45923216 | 0.64606746 | 0.99999872 |
| LNX2 | 77.2826491 | -0.054805005 | 0.26282022 | -0.2085266 | 0.83481782 | 0.99999872 |
| POLR1D | 25.5247326 | 0.279615276 | 0.45686592 | 0.61202918 | 0.54051845 | 0.99999872 |
| PAN3-AS1 | 4.84379858 | 0.702706696 | 0.86504599 | 0.81233449 | 0.41659973 | 0.99999872 |
| PAN3 | 303.563718 | -0.257912472 | 0.29176593 | -0.8839705 | 0.37671214 | 0.99999872 |
| FLT1 | 53.1752746 | -1.036243057 | 0.77139699 | -1.343333 | 1.79E-01 | 0.99999872 |
| POMP | 39.7496297 | 0.356694058 | 0.40540814 | 0.87983941 | 0.37894631 | 0.99999872 |
| SLC46A3 | 9.4650308 | 0.88539799 | 0.58085449 | 1.52430255 | 0.12743315 | 0.99999872 |
| SLC7A1 | 19.8447762 | -0.064966782 | 0.61115044 | -0.1063024 | 0.9153424 | 0.99999872 |
| UBL3 | 61.6161364 | 0.298768032 | 0.36090037 | 0.82784074 | 0.4077607 | 0.99999872 |
| KATNAL1 | 43.2816061 | 0.411013857 | 0.34893961 | 1.17789396 | 0.23883888 | 0.99999872 |
| HMGB1 | 139.297921 | -0.147763359 | 0.32450962 | -0.4553435 | 0.64886211 | 0.99999872 |
| USPL1 | 17.7028471 | 0.145895877 | 0.48816162 | 0.29886798 | 0.76504078 | 0.99999872 |
| HSPH1 | 23.0526165 | 0.045673062 | 0.37798734 | 0.12083225 | 0.9038239 | 0.99999872 |
| B3GLCT | 10.8887644 | -0.645732627 | 0.51411874 | -1.255999 | 0.20911633 | 0.99999872 |
| FRY | 747.230957 | -0.108121123 | 0.40188151 | -0.2690373 | 0.78790097 | 0.99999872 |
| FRY-AS1 | 16.9247019 | 0.33949633 | 0.73707871 | 0.46059712 | 0.64508768 | 0.99999872 |
| N4BP2L1 | 29.2021581 | -0.140632505 | 0.48115445 | -0.2922814 | 0.77007147 | 0.99999872 |
| N4BP2L2 | 280.863089 | -0.05896982 | 0.26390618 | -0.2234499 | 0.82318534 | 0.99999872 |
| PDS5B | 72.8901779 | 0.03598929 | 0.26991072 | 0.13333776 | 0.89392627 | 0.99999872 |
| STARD13 | 82.6780674 | -0.530622414 | 0.39523369 | -1.3425536 | 0.17941656 | 0.99999872 |
| AL139383.1 | 82.9577778 | -0.305837374 | 0.44942263 | -0.6805117 | 0.4961805 | 0.99999872 |
| RFC3 | 11.3197947 | -1.611047993 | 0.43623783 | -3.6930497 | 0.00022158 | 0.12623956 |
| NBEA | 94.9596666 | 0.257022127 | 0.37399672 | 0.68723097 | 0.49193719 | 0.99999872 |
| SPART | 48.39889 | 0.693062723 | 0.50200444 | 1.38059082 | 0.16740481 | 0.99999872 |
| RFXAP | 9.71049052 | 0.537398911 | 0.52509051 | 1.02344053 | 0.30609961 | 0.99999872 |
| SMAD9 | 4.97521996 | -1.285689354 | 1.05336238 | -1.2205575 | 0.22225361 | 0.99999872 |
| ALG5 | 18.2830973 | 0.414468945 | 0.38531733 | 1.07565612 | 0.28208108 | 0.99999872 |
| EXOSC8 | 7.49271717 | -0.223666316 | 0.53634272 | -0.4170213 | 0.67666286 | 0.99999872 |
| SUPT20H | 40.4445341 | 0.377669763 | 0.41771239 | 0.90413828 | 0.36592208 | 0.99999872 |
| LINC00571 | 9.22050881 | -0.619628966 | 0.62963005 | -0.9841159 | 0.32505851 | 0.99999872 |
| UFM1 | 29.8343919 | 0.273170552 | 0.31865251 | 0.85726784 | 0.39129688 | 0.99999872 |
| FREM2 | 10.9478689 | 0.577942836 | 0.78426835 | 0.73691974 | 0.46117116 | 0.99999872 |
| PROSER1 | 13.2202456 | 0.306120275 | 0.41392749 | 0.73955049 | 0.4595728 | 0.99999872 |
| NHLRC3 | 31.0298005 | 0.196719266 | 0.47013391 | 0.41843241 | 0.67563099 | 0.99999872 |
| LHFPL6 | 123.226248 | 0.484811489 | 0.42196466 | 1.14893862 | 0.25058129 | 0.99999872 |
| COG6 | 32.7316428 | -0.002184674 | 0.32318716 | -0.0067598 | 0.99460652 | 0.99999872 |
| FOXO1 | 57.1093896 | -0.321509136 | 0.2984567 | -1.0772388 | 0.2813736 | 0.99999872 |
| MRPS31 | 22.9261572 | 0.371254919 | 0.38229433 | 0.97112326 | 0.3314869 | 0.99999872 |
| ELF1 | 183.890211 | -0.049122792 | 0.27241413 | -0.1803239 | 0.85689826 | 0.99999872 |
| WBP4 | 23.418658 | 0.071525897 | 0.41798995 | 0.1711187 | 0.86413043 | 0.99999872 |
| KBTBD6 | 7.67823624 | 0.535158183 | 0.61329881 | 0.87258963 | 0.3828868 | 0.99999872 |

| | | | | | | |
|---|---|---|---|---|---|---|
| AL354696.2 | 6.66809659 | -1.045316549 | 0.73678847 | -1.4187472 | 0.15597275 | 0.99999872 |
| KBTBD7 | 7.27426473 | 0.540471547 | 0.59476138 | 0.90871997 | 0.36349796 | 0.99999872 |
| MTRF1 | 5.00781182 | -0.224473306 | 0.64774661 | -0.3465449 | 0.72893322 | 0.99999872 |
| NAA16 | 28.6888547 | 0.400314902 | 0.39019972 | 1.02592309 | 0.30492784 | 0.99999872 |
| VWA8 | 87.7175674 | -0.212017153 | 0.33932142 | -0.6248269 | 0.53208467 | 0.99999872 |
| DGKH | 58.7846104 | -0.288678895 | 0.42972581 | -0.6717746 | 0.50172719 | 0.99999872 |
| AKAP11 | 67.4252498 | 0.282704028 | 0.29237645 | 0.96691792 | 0.33358507 | 0.99999872 |
| EPSTI1 | 40.4103284 | -0.915529212 | 1.60726205 | -0.5696204 | 0.56893521 | 0.99999872 |
| DNAJC15 | 24.8701628 | -0.08472561 | 0.45767459 | -0.1851219 | 0.85313339 | 0.99999872 |
| ENOX1 | 17.8414363 | 2.162728692 | 0.66414643 | 3.25640339 | 0.00112833 | 0.30314552 |
| CCDC122 | 31.9750704 | 0.992494266 | 0.40018045 | 2.48011684 | 0.01313393 | 0.74263079 |
| SMIM2-AS1 | 15.4159587 | 1.155311098 | 1.10134132 | 1.04900368 | 0.29417642 | 0.99999872 |
| TSC22D1 | 236.852861 | 0.251708892 | 0.36095794 | 0.6973358 | 0.48559267 | 0.99999872 |
| NUFIP1 | 6.46020852 | -0.784765761 | 0.62233899 | -1.260994 | 0.207311 | 0.99999872 |
| AL359706.1 | 4.71376208 | 1.709148156 | 0.8656504 | 1.97440924 | 0.04833523 | 0.97937581 |
| GPALPP1 | 19.6866673 | -0.076365119 | 0.38461405 | -0.19855 | 0.84261477 | 0.99999872 |
| GTF2F2 | 27.5874418 | -0.354570923 | 0.33298587 | -1.0648227 | 0.28695615 | 0.99999872 |
| TPT1 | 184.053076 | 0.692092868 | 0.42469732 | 1.62961441 | 0.10318302 | 0.99999872 |
| SLC25A30 | 8.50285075 | -0.409629317 | 0.60449903 | -0.6776344 | 0.49800355 | 0.99999872 |
| COG3 | 27.5767574 | 0.221675194 | 0.37866379 | 0.58541428 | 0.55826919 | 0.99999872 |
| ZC3H13 | 51.2890725 | -0.336409117 | 0.2660607 | -1.2644074 | 0.20608383 | 0.99999872 |
| AL138686.1 | 25.3236524 | 0.383133071 | 0.68325624 | 0.56074581 | 0.57497083 | 0.99999872 |
| LRCH1 | 43.9432332 | 0.199976951 | 0.411025 | 0.48653233 | 0.62658979 | 0.99999872 |
| ESD | 32.9628144 | 0.599444713 | 0.40002959 | 1.49850091 | 0.13400316 | 0.99999872 |
| SUCLA2 | 38.3688625 | -0.42508719 | 0.2927816 | -1.4518918 | 0.1465317 | 0.99999872 |
| NUDT15 | 10.3547564 | 0.179856481 | 0.56139161 | 0.32037615 | 0.74868321 | 0.99999872 |
| MED4 | 70.9031363 | 0.378061147 | 0.34159442 | 1.10675447 | 0.2684001 | 0.99999872 |
| ITM2B | 1478.05734 | 0.470291915 | 0.51382563 | 0.91527531 | 0.36004712 | 0.99999872 |
| RB1 | 88.7257433 | 0.360249675 | 0.26309298 | 1.36928653 | 0.17090973 | 0.99999872 |
| LPAR6 | 5.19071344 | -0.280232934 | 0.88049508 | -0.3182675 | 0.75028206 | 0.99999872 |
| RCBTB2 | 8.7063706 | -0.78583552 | 0.51502417 | -1.5258226 | 0.12705405 | 0.99999872 |
| FNDC3A | 112.917288 | 0.014760937 | 0.29673593 | 0.04974435 | 0.96032611 | 0.99999872 |
| CDADC1 | 11.6630567 | -0.310572229 | 0.59723301 | -0.5200185 | 0.60305066 | 0.99999872 |
| CAB39L | 22.155519 | 0.065070991 | 0.69245517 | 0.09397141 | 0.92513186 | 0.99999872 |
| SETDB2 | 31.0466981 | -0.351549626 | 0.36335295 | -0.9675155 | 0.33328638 | 0.99999872 |
| PHF11 | 23.7037979 | -0.513974595 | 0.53252263 | -0.9651695 | 0.33445992 | 0.99999872 |
| RCBTB1 | 9.74851214 | -1.070813544 | 0.51034582 | -2.0982116 | 0.03588645 | 0.94734101 |
| EBPL | 10.2885367 | -0.293538318 | 0.48906563 | -0.6002023 | 0.54837142 | 0.99999872 |
| KPNA3 | 48.9408973 | 0.136914017 | 0.3099006 | 0.44179979 | 0.65863409 | 0.99999872 |
| SPRYD7 | 12.7820804 | 0.525788886 | 0.44869865 | 1.17180849 | 0.24127395 | 0.99999872 |
| DLEU2 | 21.7782745 | 0.70311567 | 0.61618846 | 1.14107245 | 0.25383978 | 0.99999872 |
| TRIM13 | 14.6752483 | -0.249285214 | 0.38693731 | -0.6442522 | 0.51941191 | 0.99999872 |
| RNASEH2B | 25.1358865 | -0.069133303 | 0.45164075 | -0.1530714 | 0.87834193 | 0.99999872 |
| INTS6 | 55.1452346 | -0.129218344 | 0.4291925 | -0.3010732 | 0.76335871 | 0.99999872 |
| INTS6-AS1 | 13.2697119 | 0.069343087 | 0.5402358 | 0.12835707 | 0.8978664 | 0.99999872 |
| WDFY2 | 25.3168344 | -0.289372231 | 0.48209839 | -0.6002348 | 0.54834976 | 0.99999872 |
| ALG11 | 5.14880053 | 0.736804137 | 0.66874302 | 1.1017747 | 0.27055963 | 0.99999872 |
| UTP14C | 10.6213657 | -0.374491769 | 0.72420748 | -0.5171056 | 0.60508242 | 0.99999872 |
| NEK3 | 12.320664 | -0.47179881 | 0.47392191 | -0.9955201 | 0.31948335 | 0.99999872 |
| VPS36 | 40.0054824 | 0.145852974 | 0.27142147 | 0.53736713 | 0.59101404 | 0.99999872 |
| CKAP2 | 5.55267838 | 0.557417935 | 0.67678088 | 0.82363133 | 0.41014907 | 0.99999872 |
| SUGT1 | 32.5077705 | -0.213178986 | 0.42385017 | -0.5029584 | 0.61499355 | 0.99999872 |
| TDRD3 | 70.1133755 | 0.575817338 | 0.2801873 | 2.05511574 | 0.03986783 | 0.95980711 |
| PCDH9 | 67.6198849 | 1.131081654 | 0.72757148 | 1.55459866 | 0.12004168 | 0.99999872 |
| DACH1 | 2206.25696 | -0.119885728 | 0.33084811 | -0.3623588 | 0.71708392 | 0.99999872 |
| MZT1 | 11.4549752 | 0.903361776 | 0.51740528 | 1.74594618 | 0.0808203 | 0.99999872 |
| DIS3 | 21.0483091 | 0.097832265 | 0.41681317 | 0.23471491 | 0.81443002 | 0.99999872 |
| PIBF1 | 83.1256012 | -0.193406187 | 0.24064079 | -0.8037132 | 0.42156261 | 0.99999872 |
| KLF12 | 71.3544127 | 0.291536089 | 0.47851346 | 0.60925368 | 0.5423563 | 0.99999872 |
| TBC1D4 | 23.7348982 | -0.414129589 | 0.70867286 | -0.5843734 | 0.5589691 | 0.99999872 |
| COMMD6 | 26.6727477 | 0.941786463 | 0.47912813 | 1.96562547 | 0.04934189 | 0.97937581 |

| | | | | | | |
|---|---|---|---|---|---|---|
| UCHL3 | 12.9959252 | -0.098035711 | 0.5271214 | -0.1859832 | 0.85245796 | 0.99999872 |
| LMO7 | 56.8487177 | -0.329516557 | 0.3747513 | -0.879294 | 3.79E-01 | 0.99999872 |
| KCTD12 | 39.0753753 | 0.20892914 | 0.81542956 | 0.25621973 | 0.79778118 | 0.99999872 |
| AC001226.2 | 10.2270173 | 0.656283207 | 0.64569723 | 1.01639464 | 0.3094415 | 0.99999872 |
| FBXL3 | 50.7376718 | 0.294583884 | 0.31440926 | 0.93694406 | 0.34878734 | 0.99999872 |
| MYCBP2 | 125.908443 | -0.067041817 | 0.24183094 | -0.277226 | 0.7816066 | 0.99999872 |
| SLAIN1 | 11.0884952 | 1.204244722 | 0.55207035 | 2.18132476 | 0.0291594 | 0.90266962 |
| RNF219 | 10.1428937 | -0.016006757 | 0.58983262 | -0.0271378 | 0.97834983 | 0.99999872 |
| RBM26 | 110.671582 | 0.262344961 | 0.25147931 | 1.04320693 | 0.29685247 | 0.99999872 |
| NDFIP2 | 42.5186118 | -0.264203446 | 0.33278372 | -0.7939194 | 0.42724236 | 0.99999872 |
| SPRY2 | 10.2252243 | -1.208878029 | 0.52184001 | -2.3165683 | 0.02052726 | 0.8671024 |
| GPC5 | 36.152832 | 1.327332043 | 1.14953963 | 1.15466402 | 0.24822804 | 0.99999872 |
| GPC6 | 834.533514 | -0.067275016 | 0.23315411 | -0.2885431 | 0.77293102 | 0.99999872 |
| TGDS | 10.0424426 | -0.082995305 | 0.54735228 | -0.1516305 | 0.87947837 | 0.99999872 |
| GPR180 | 19.5414084 | 0.151892094 | 0.58282624 | 0.260613 | 0.79439097 | 0.99999872 |
| ABCC4 | 46.7394181 | -0.148606584 | 0.63699476 | -0.2332933 | 0.81553369 | 0.99999872 |
| CLDN10 | 30.615728 | 0.844715899 | 0.75002696 | 1.12624738 | 0.26006082 | 0.99999872 |
| CLDN10-AS1 | 20.2782124 | -2.500227218 | 1.05990879 | -2.3589079 | 0.0183288 | 0.85282771 |
| DZIP1 | 7.11082846 | 0.059208153 | 0.77149616 | 0.07674459 | 0.93882673 | 0.99999872 |
| DNAJC3-DT | 5.3794202 | 0.485136184 | 0.83257819 | 0.58269144 | 0.56010104 | 0.99999872 |
| DNAJC3 | 83.857796 | 0.310189169 | 0.37861153 | 0.81928083 | 0.41262621 | 0.99999872 |
| UGGT2 | 131.658546 | -0.158181835 | 0.31298277 | -0.5054011 | 0.61327715 | 0.99999872 |
| MBNL2 | 122.60263 | 0.225574869 | 0.27894307 | 0.80867709 | 0.41870091 | 0.99999872 |
| RAP2A | 13.6980618 | 0.99320603 | 0.63623266 | 1.56107364 | 0.11850638 | 0.99999872 |
| IPO5 | 26.5480868 | 0.060668284 | 0.37321839 | 0.16255438 | 0.87086931 | 0.99999872 |
| FARP1 | 134.345494 | -0.323156634 | 0.26521905 | -1.2184518 | 0.22305233 | 0.99999872 |
| STK24 | 55.8287056 | -0.209486998 | 0.39005009 | -0.5370772 | 0.59121431 | 0.99999872 |
| DOCK9 | 71.1071229 | -0.145325748 | 0.27971003 | -0.5195586 | 0.60337127 | 0.99999872 |
| UBAC2 | 56.0595492 | -0.456873151 | 0.32387879 | -1.4106301 | 0.15835373 | 0.99999872 |
| TM9SF2 | 77.3988568 | 0.324851511 | 0.36597611 | 0.88763038 | 0.37473961 | 0.99999872 |
| CLYBL | 50.3317132 | -0.612877788 | 0.35709636 | -1.7162812 | 0.08611058 | 0.99999872 |
| PCCA | 138.349822 | -0.344336423 | 0.29326685 | -1.1741403 | 0.24033884 | 0.99999872 |
| GGACT | 20.6185449 | 0.089977708 | 0.44925185 | 0.20028344 | 0.84125891 | 0.99999872 |
| TMTC4 | 24.0921534 | 0.590519509 | 0.44045307 | 1.34070927 | 0.18001486 | 0.99999872 |
| NALCN | 40.3943583 | -0.262203141 | 0.34549732 | -0.7589151 | 0.44790333 | 0.99999872 |
| ITGBL1 | 11.0415473 | -0.245370956 | 1.07816658 | -0.2275817 | 0.81997147 | 0.99999872 |
| FGF14 | 17.8384589 | 1.309949301 | 0.92903532 | 1.41001022 | 0.15853667 | 0.99999872 |
| FGF14-AS2 | 4.81707772 | -0.576384709 | 0.64691087 | -0.8909801 | 0.37293985 | 0.99999872 |
| TPP2 | 26.6151964 | 0.212062269 | 0.34698195 | 0.61116224 | 0.54109218 | 0.99999872 |
| EFNB2 | 59.6306499 | 0.512542287 | 0.6253701 | 0.81958234 | 0.41245424 | 0.99999872 |
| ARGLU1 | 140.825152 | 0.023032688 | 0.32997901 | 0.06980046 | 0.94435248 | 0.99999872 |
| FAM155A | 22.9045511 | -1.387038615 | 0.55428305 | -2.5024013 | 0.0123354 | 0.72670507 |
| LIG4 | 11.5738262 | 0.666070414 | 0.54893557 | 1.21338542 | 0.2249825 | 0.99999872 |
| ABHD13 | 18.6535413 | 1.134110115 | 0.44375631 | 2.55570474 | 0.0105973 | 0.70947852 |
| TNFSF13B | 23.4664866 | 0.263555533 | 0.65265616 | 0.40381988 | 0.68634518 | 0.99999872 |
| IRS2 | 21.2529632 | -0.324033762 | 0.72463002 | -0.4471713 | 0.65475137 | 0.99999872 |
| COL4A1 | 33.2399628 | -0.451478367 | 0.84565637 | -0.5338792 | 0.5934251 | 0.99999872 |
| COL4A2 | 40.8266684 | 0.373265073 | 0.72848456 | 0.51238571 | 0.60838109 | 0.99999872 |
| RAB20 | 9.15964649 | 0.184154029 | 0.60914646 | 0.30231486 | 0.76241205 | 0.99999872 |
| NAXD | 13.8759588 | 0.480118781 | 0.49206794 | 0.97571644 | 0.329205 | 0.99999872 |
| CARS2 | 29.0697382 | 0.097779645 | 0.34701056 | 0.28177715 | 0.77811439 | 0.99999872 |
| ING1 | 11.7169477 | 1.244252333 | 0.70411485 | 1.76711559 | 0.07720887 | 0.99999872 |
| ANKRD10 | 96.2519008 | 0.149072169 | 0.25524806 | 0.58402861 | 0.55920106 | 0.99999872 |
| ARHGEF7 | 44.3931472 | -0.079573005 | 0.34372592 | -0.2315013 | 0.81692536 | 0.99999872 |
| TUBGCP3 | 34.3975582 | -0.318231231 | 0.28126861 | -1.1314139 | 0.25788091 | 0.99999872 |
| ATP11A | 67.6240094 | -0.367371074 | 0.31454384 | -1.1679487 | 0.24282745 | 0.99999872 |
| MCF2L | 21.8519668 | -0.442810443 | 0.7278667 | -0.6083675 | 0.54294377 | 0.99999872 |
| PCID2 | 16.5721555 | 0.473812683 | 0.37939695 | 1.24885738 | 0.21171724 | 0.99999872 |
| CUL4A | 42.5615207 | -0.280392459 | 0.28911453 | -0.9698318 | 0.33213034 | 0.99999872 |
| LAMP1 | 136.85482 | 0.351602917 | 0.3989556 | 0.8813084 | 0.37815092 | 0.99999872 |
| DCUN1D2 | 38.4646925 | 0.062621694 | 0.35396685 | 0.17691401 | 0.85957594 | 0.99999872 |

| | | | | | | |
|---|---|---|---|---|---|---|
| TMCO3 | 91.7424368 | 0.139568029 | 0.47701554 | 0.29258592 | 0.76983868 | 0.99999872 |
| TFDP1 | 18.4626281 | -1.053856119 | 0.5951652 | -1.7706951 | 0.07661141 | 0.99999872 |
| GAS6 | 47.4646382 | -0.076475029 | 0.60987071 | -0.1253955 | 0.90021047 | 0.99999872 |
| RASA3 | 5.97179548 | 0.373297735 | 0.90181036 | 0.41394261 | 0.67891614 | 0.99999872 |
| CDC16 | 34.1660378 | 0.161568842 | 0.3070508 | 0.5261958 | 0.59875217 | 0.99999872 |
| UPF3A | 42.9086777 | -0.372241473 | 0.38830776 | -0.9586249 | 0.33774777 | 0.99999872 |
| CHAMP1 | 11.2184597 | 0.413487961 | 0.49163049 | 0.84105435 | 0.40031749 | 0.99999872 |
| TTC5 | 10.4899022 | -0.192848853 | 0.49302429 | -0.3911549 | 0.69568276 | 0.99999872 |
| CCNB1IP1 | 12.5365 | 0.000232183 | 0.44412081 | 0.00052279 | 0.99958287 | 0.99999872 |
| TEP1 | 9.8753449 | -1.136853737 | 0.50977276 | -2.2301186 | 0.02573957 | 0.88365618 |
| OSGEP | 8.23210032 | -0.705565363 | 0.51428323 | -1.3719393 | 0.17008235 | 0.99999872 |
| APEX1 | 14.3043137 | -0.247792874 | 0.42536927 | -0.5825359 | 0.56020576 | 0.99999872 |
| PIP4P1 | 9.20531392 | 0.610987566 | 0.60331977 | 1.01270933 | 0.31119902 | 0.99999872 |
| PNP | 6.40235098 | 0.567277038 | 0.77752759 | 0.72959088 | 0.4656403 | 0.99999872 |
| METTL17 | 6.13399381 | 0.637383162 | 0.72965729 | 0.87353771 | 0.38237006 | 0.99999872 |
| NDRG2 | 70.8371941 | -0.200055789 | 0.57800753 | -0.3461128 | 0.72925797 | 0.99999872 |
| ARHGEF40 | 32.0659617 | -0.241974444 | 0.49349877 | -0.4903243 | 0.62390443 | 0.99999872 |
| ZNF219 | 13.1050054 | -0.097673834 | 0.56975158 | -0.1714323 | 0.86388385 | 0.99999872 |
| HNRNPC | 132.976857 | -0.001109929 | 0.24455286 | -0.0045386 | 0.99637873 | 0.99999872 |
| SUPT16H | 34.3353441 | 0.184821634 | 0.33563595 | 0.55066102 | 0.58186607 | 0.99999872 |
| CHD8 | 41.6442063 | -0.001542913 | 0.46376377 | -0.0033269 | 0.99734549 | 0.99999872 |
| RAB2B | 17.2146964 | -0.434853067 | 0.39320234 | -1.105927 | 0.26875814 | 0.99999872 |
| TOX4 | 29.9001986 | 0.065089001 | 0.34265202 | 0.18995656 | 0.84934317 | 0.99999872 |
| METTL3 | 11.1184536 | 0.554996286 | 0.52265951 | 1.06186968 | 0.28829485 | 0.99999872 |
| DAD1 | 81.1041708 | 1.04010777 | 0.4772599 | 2.17933199 | 0.02930701 | 0.90266962 |
| ABHD4 | 7.57483557 | 0.797831785 | 0.59614598 | 1.33831614 | 0.1807934 | 0.99999872 |
| OXA1L | 8.71751867 | -0.319032447 | 0.6269063 | -0.5088997 | 0.61082251 | 0.99999872 |
| MRPL52 | 13.7982829 | 0.740649359 | 0.58636991 | 1.26310943 | 0.20654985 | 0.99999872 |
| MMP14 | 5.90044451 | 0.016346126 | 0.6280351 | 0.02602741 | 0.97923548 | 0.99999872 |
| LRP10 | 72.6134773 | -0.191607407 | 0.4735444 | -0.404624 | 0.68575395 | 0.99999872 |
| RBM23 | 28.8370838 | -0.474856513 | 0.31332548 | -1.5155375 | 0.12963634 | 0.99999872 |
| PRMT5 | 9.34617896 | 0.360854216 | 0.58802422 | 0.61367237 | 0.53943185 | 0.99999872 |
| AL132780.3 | 7.90835642 | -0.917992817 | 0.83265682 | -1.1024864 | 0.27025027 | 0.99999872 |
| PSMB5 | 39.7297593 | 0.418277822 | 0.43078803 | 0.97095971 | 0.33156834 | 0.99999872 |
| ACIN1 | 40.2443927 | -0.492590339 | 0.30714636 | -1.6037642 | 0.10876603 | 0.99999872 |
| C14orf119 | 15.4418371 | 0.172347427 | 0.44435034 | 0.38786383 | 0.6981168 | 0.99999872 |
| SLC7A8 | 135.813605 | 0.314426998 | 0.57814275 | 0.54385703 | 0.58653986 | 0.99999872 |
| HOMEZ | 4.4228851 | -0.645774937 | 0.67153741 | -0.9616366 | 0.33623219 | NA |
| PPP1R3E | 6.37042435 | -0.15439999 | 0.63055366 | -0.2448642 | 0.80656161 | 0.99999872 |
| BCL2L2-PABI | 15.3958386 | -0.165391416 | 0.41455311 | -0.3989631 | 0.68992036 | 0.99999872 |
| PABPN1 | 69.3615777 | -0.070211979 | 0.40351601 | -0.1740005 | 8.62E-01 | 0.99999872 |
| SLC22A17 | 4.93233815 | 0.674271035 | 0.7143452 | 0.94390084 | 0.34522032 | 0.99999872 |
| NGDN | 8.06173157 | -2.47E-01 | 0.52084191 | -0.4751291 | 0.63469496 | 0.99999872 |
| ZFHX2 | 27.8420469 | 0.160691306 | 0.39225364 | 0.40966174 | 0.6820541 | 0.99999872 |
| THTPA | 5.46821883 | -0.124100587 | 0.78457663 | -0.1581752 | 0.87431872 | 0.99999872 |
| AP1G2 | 9.02149792 | -0.879061805 | 0.51277248 | -1.7143311 | 0.08646794 | 0.99999872 |
| DHRS4-AS1 | 9.56065078 | 0.402921417 | 0.7742609 | 0.52039489 | 0.60278837 | 0.99999872 |
| DHRS4 | 4.97530946 | 0.640864442 | 0.68346857 | 0.93766484 | 0.34841668 | 0.99999872 |
| DHRS4L2 | 14.860957 | -0.003127699 | 0.44057396 | -0.0070991 | 0.99433575 | 0.99999872 |
| PCK2 | 5.98085178 | -0.280194259 | 0.95186722 | -0.2943628 | 0.76848072 | 0.99999872 |
| PSME1 | 36.0451448 | -0.062090749 | 0.6033355 | -0.1029125 | 0.91803244 | 0.99999872 |
| EMC9 | 6.47159653 | 0.048070325 | 0.62002589 | 0.07752954 | 0.93820229 | 0.99999872 |
| PSME2 | 5.41965713 | -1.532174819 | 0.78618121 | -1.9488825 | 0.05130946 | 0.99009295 |
| AL136295.1 | 9.54165687 | 0.229915033 | 0.46344949 | 0.49609513 | 0.6198273 | 0.99999872 |
| GMPR2 | 9.00358432 | 1.025668154 | 0.60471906 | 1.69610688 | 0.08986564 | 0.99999872 |
| TINF2 | 7.84086475 | 0.044425573 | 0.63756901 | 0.06967963 | 0.94444865 | 0.99999872 |
| DHRS1 | 6.51012311 | 1.481842356 | 0.69270865 | 2.13920001 | 0.03241947 | 0.92959829 |
| KHNYN | 8.95277989 | -0.409185678 | 0.48475201 | -0.8441134 | 0.39860602 | 0.99999872 |
| SDR39U1 | 12.0443134 | -0.553808332 | 0.39545553 | -1.4004314 | 0.16138418 | 0.99999872 |
| NOVA1 | 39.5078894 | 0.833434229 | 0.56327215 | 1.47962974 | 0.13897208 | 0.99999872 |
| PRKD1 | 337.623964 | 0.040740029 | 0.35165189 | 0.11585329 | 0.90776881 | 0.99999872 |

| | | | | | | |
|---|---|---|---|---|---|---|
| G2E3 | 20.3087145 | 0.365723951 | 0.38186187 | 0.95773885 | 0.33819446 | 0.99999872 |
| SCFD1 | 73.8945235 | 0.158689876 | 0.27630379 | 0.57433116 | 0.56574373 | 0.99999872 |
| COCH | 8.32556416 | 0.766719337 | 0.6435968 | 1.19130384 | 0.23353433 | 0.99999872 |
| STRN3 | 112.670373 | -0.019675861 | 0.3258835 | -0.060377 | 0.95185539 | 0.99999872 |
| AP4S1 | 16.8965354 | -0.134135252 | 0.40560149 | -0.330707 | 0.74086582 | 0.99999872 |
| HECTD1 | 95.7437106 | -0.043174793 | 0.28933765 | -0.1492194 | 0.88138051 | 0.99999872 |
| HEATR5A | 18.5289098 | 0.81102485 | 0.43037174 | 1.88447515 | 0.05950073 | 0.99999872 |
| AL139353.1 | 8.0249102 | 0.310330713 | 0.63285816 | 0.49036377 | 0.62387651 | 0.99999872 |
| NUBPL | 53.5507401 | 0.013861393 | 0.37883648 | 0.03658938 | 0.97081241 | 0.99999872 |
| ARHGAP5 | 155.994514 | 0.187332763 | 0.2424923 | 0.77253078 | 0.43980013 | 0.99999872 |
| AKAP6 | 11.7492335 | 0.461308577 | 0.8539012 | 0.54023648 | 0.58903396 | 0.99999872 |
| NPAS3 | 1457.9805 | -0.069890856 | 0.28324077 | -0.2467542 | 0.80509844 | 0.99999872 |
| EAPP | 26.2004107 | 0.22614985 | 0.34670818 | 0.65227723 | 0.51422235 | 0.99999872 |
| SNX6 | 52.4530939 | -0.143571499 | 0.33673194 | -0.4263673 | 0.6698402 | 0.99999872 |
| CFL2 | 15.0604998 | 0.661754196 | 0.50947148 | 1.29890332 | 0.19397711 | 0.99999872 |
| BAZ1A | 27.2402041 | -0.079692145 | 0.44639553 | -0.1785236 | 0.85831177 | 0.99999872 |
| SRP54 | 41.6170135 | -0.206950059 | 0.31479087 | -0.6574208 | 0.5109104 | 0.99999872 |
| FAM177A1 | 38.7049374 | -0.076130934 | 0.34666564 | -0.2196091 | 0.82617559 | 0.99999872 |
| PPP2R3C | 19.423271 | -0.034870735 | 0.44214538 | -0.0788671 | 0.93713831 | 0.99999872 |
| NFKBIA | 57.9034354 | -0.54222521 | 0.32838934 | -1.6511657 | 0.09870475 | 0.99999872 |
| RALGAPA1 | 130.998446 | -0.16089249 | 0.2305193 | -0.6979567 | 0.48520427 | 0.99999872 |
| BRMS1L | 8.70431141 | 0.593965149 | 0.58764987 | 1.01074667 | 0.31213769 | 0.99999872 |
| MBIP | 15.4389698 | 0.893895011 | 0.48854233 | 1.82971864 | 0.06729202 | 0.99999872 |
| SLC25A21 | 6.10492629 | -0.698257795 | 0.81465019 | -0.8571259 | 0.3913753 | 0.99999872 |
| MIPOL1 | 51.5718203 | -0.115326288 | 0.34534181 | -0.3339482 | 0.73841862 | 0.99999872 |
| SEC23A | 57.2121088 | 0.148881861 | 0.27600557 | 0.53941615 | 0.58959974 | 0.99999872 |
| GEMIN2 | 4.63628728 | -0.578905405 | 0.73622754 | -0.7863132 | 0.43168405 | 0.99999872 |
| TRAPPC6B | 22.7614001 | -0.289955679 | 0.3504776 | -0.8273159 | 0.40805805 | 0.99999872 |
| PNN | 47.3833962 | -0.174675092 | 0.29831109 | -0.5855468 | 0.55818014 | 0.99999872 |
| MIA2 | 46.3677628 | -0.3204306 | 0.32047868 | -0.99985 | 0.31738311 | 0.99999872 |
| FBXO33 | 36.0842465 | 0.20806197 | 0.42339004 | 0.49141915 | 0.62313003 | 0.99999872 |
| AL049875.1 | 9.57973805 | -0.452922427 | 0.56743283 | -0.7981957 | 0.42475695 | 0.99999872 |
| KLHL28 | 27.5338687 | 0.433208622 | 0.44165728 | 0.98087056 | 0.32665658 | 0.99999872 |
| TOGARAM1 | 59.131678 | 0.140691604 | 0.29728842 | 0.47324952 | 0.63603517 | 0.99999872 |
| PRPF39 | 16.2514173 | 0.229245747 | 0.40866997 | 0.56095569 | 0.57482774 | 0.99999872 |
| FKBP3 | 17.3809041 | 0.576666622 | 0.40040475 | 1.44020926 | 0.1498082 | 0.99999872 |
| FANCM | 6.51635206 | -0.832622943 | 0.64053189 | -1.299893 | 0.19363764 | 0.99999872 |
| MIS18BP1 | 15.7677885 | 0.721612904 | 0.40567966 | 1.77877515 | 0.07527663 | 0.99999872 |
| LINC00871 | 15.4354327 | 1.169011033 | 1.04464017 | 1.11905617 | 0.26311618 | 0.99999872 |
| MDGA2 | 12.176942 | 0.144236593 | 1.21572791 | 0.11864217 | 0.90555886 | 0.99999872 |
| RPS29 | 40.3433001 | 0.732211811 | 0.43884525 | 1.66849663 | 0.09521718 | 0.99999872 |
| LRR1 | 5.73333244 | 0.149070928 | 0.6556859 | 0.22735113 | 0.82015072 | 0.99999872 |
| RPL36AL | 86.3037817 | 0.383295268 | 0.37355147 | 1.02608422 | 0.30485189 | 0.99999872 |
| MGAT2 | 11.1417854 | -0.074329509 | 0.5928227 | -0.1253824 | 0.90022085 | 0.99999872 |
| DNAAF2 | 7.29006895 | 0.057452645 | 0.53598748 | 0.10719027 | 0.91463803 | 0.99999872 |
| POLE2 | 7.84828 | 1.388636293 | 1.1914174 | 1.165533 | 0.24380331 | 0.99999872 |
| KLHDC1 | 27.45682 | 0.191620276 | 0.43021112 | 0.44540986 | 0.65602359 | 0.99999872 |
| KLHDC2 | 27.8745155 | 0.294949845 | 0.32893055 | 0.89669337 | 0.36988256 | 0.99999872 |
| NEMF | 75.5532273 | -0.324748636 | 0.24864607 | -1.3060678 | 0.19152949 | 0.99999872 |
| ARF6 | 80.2322054 | -0.029813437 | 0.40969391 | -0.07277 | 0.94198912 | 0.99999872 |
| LINC01588 | 12.6880986 | 0.082206218 | 0.5444831 | 0.15098029 | 0.87999126 | 0.99999872 |
| VCPKMT | 10.177316 | 0.433870541 | 0.48251233 | 0.89919058 | 0.36855115 | 0.99999872 |
| SOS2 | 104.197037 | -0.105759325 | 0.25635096 | -0.4125568 | 0.67993138 | 0.99999872 |
| L2HGDH | 15.8550335 | -0.62291412 | 0.45722725 | -1.3623731 | 0.17308016 | 0.99999872 |
| ATP5S | 20.1294509 | -0.532414835 | 0.38214633 | -1.3932224 | 0.16355254 | 0.99999872 |
| CDKL1 | 23.5582975 | -0.524644855 | 0.53889729 | -0.9735526 | 0.33027873 | 0.99999872 |
| MAP4K5 | 63.6147417 | 0.012128996 | 0.28827873 | 0.04207385 | 0.96643983 | 0.99999872 |
| ATL1 | 20.300507 | -0.057307083 | 0.44380564 | -0.1291265 | 0.89725753 | 0.99999872 |
| SAV1 | 39.1291009 | 0.154764879 | 0.28998044 | 0.533708 | 0.59354358 | 0.99999872 |
| NIN | 41.5009429 | 0.261480928 | 0.43465146 | 0.60158761 | 0.54744868 | 0.99999872 |
| PYGL | 5.95779311 | 1.153394018 | 0.66156264 | 1.74343886 | 0.08125698 | 0.99999872 |

| | | | | | | |
|---|---|---|---|---|---|---|
| TMX1 | 19.2437657 | 0.1429678 | 0.44632515 | 0.32032208 | 0.74872419 | 0.99999872 |
| FRMD6 | 6.17169322 | -0.429386492 | 0.53742792 | -0.7989657 | 0.42431028 | 0.99999872 |
| RTRAF | 26.4788088 | 0.499486261 | 0.3725777 | 1.34062307 | 0.18004286 | 0.99999872 |
| TXNDC16 | 10.0211464 | 0.14556768 | 0.53231089 | 0.27346365 | 0.78449683 | 0.99999872 |
| ERO1A | 33.2681129 | 0.166795734 | 0.46130448 | 0.36157406 | 0.71767035 | 0.99999872 |
| AL133453.1 | 8.75907394 | 1.315717628 | 0.74818859 | 1.75853741 | 0.07865611 | 0.99999872 |
| PSMC6 | 33.4094935 | -0.030704786 | 0.2878367 | -0.1066743 | 0.91504735 | 0.99999872 |
| STYX | 11.708272 | 0.08314278 | 0.52610986 | 0.15803312 | 0.8744307 | 0.99999872 |
| GNPNAT1 | 13.2410546 | 0.504557123 | 0.44137902 | 1.14313797 | 0.25298131 | 0.99999872 |
| FERMT2 | 263.805302 | -0.197907845 | 0.34530852 | -0.5731334 | 0.56655437 | 0.99999872 |
| DDHD1 | 29.2469682 | 0.629348113 | 0.4107401 | 1.53222955 | 0.1254658 | 0.99999872 |
| CNIH1 | 21.4797325 | 0.759165186 | 0.38556279 | 1.96897939 | 0.04895546 | 0.97937581 |
| GMFB | 18.9806308 | 0.182935196 | 0.41782426 | 0.43782808 | 0.66151091 | 0.99999872 |
| CGRRF1 | 9.01376335 | 0.770005879 | 0.52806538 | 1.4581639 | 0.14479537 | 0.99999872 |
| SAMD4A | 75.0420831 | -0.238829061 | 0.42419976 | -0.5630108 | 0.57342751 | 0.99999872 |
| SOCS4 | 16.1850318 | 0.591791752 | 0.41197271 | 1.43648291 | 0.15086498 | 0.99999872 |
| MAPK1IP1L | 46.9758203 | -0.278238829 | 0.39064862 | -0.7122483 | 0.47631102 | 0.99999872 |
| LGALS3 | 9.37339336 | -0.485225519 | 0.54825046 | -0.8850435 | 0.37613316 | 0.99999872 |
| FBXO34 | 36.2433519 | 0.308742946 | 0.29770262 | 1.03708507 | 0.29969621 | 0.99999872 |
| ATG14 | 14.5820994 | -0.032343091 | 0.52671129 | -0.0614057 | 0.95103609 | 0.99999872 |
| KTN1 | 116.27312 | 0.126025014 | 0.31609169 | 0.39869765 | 0.690116 | 0.99999872 |
| PELI2 | 86.728133 | 0.095872563 | 0.21784058 | 0.44010424 | 0.65986161 | 0.99999872 |
| TMEM260 | 26.1257089 | 0.401727304 | 0.39983866 | 1.00472351 | 0.31503 | 0.99999872 |
| EXOC5 | 50.2745862 | 0.482708691 | 0.30520715 | 1.58157728 | 0.1137461 | 0.99999872 |
| AP5M1 | 31.0739705 | 0.245767623 | 0.31461218 | 0.78117644 | 0.43469873 | 0.99999872 |
| NAA30 | 15.96963 | -0.016265851 | 0.4275872 | -0.038041 | 0.96965498 | 0.99999872 |
| CCDC198 | 15.5017364 | 0.269081147 | 0.74249625 | 0.36240068 | 0.71705263 | 0.99999872 |
| ARMH4 | 745.972154 | -0.017525453 | 0.47835838 | -0.0366367 | 0.97077471 | 0.99999872 |
| ACTR10 | 36.9966294 | 0.265172514 | 0.32879785 | 0.80649103 | 0.41995978 | 0.99999872 |
| PSMA3 | 30.9026149 | 0.14558589 | 0.54229109 | 0.26846447 | 0.78834182 | 0.99999872 |
| ARID4A | 35.2502005 | 0.28824596 | 0.39033446 | 0.73845892 | 0.46023562 | 0.99999872 |
| AL139021.1 | 4.83727233 | 0.474156491 | 0.79249316 | 0.59830988 | 0.54963318 | 0.99999872 |
| TIMM9 | 11.4639133 | 0.861660884 | 0.56905198 | 1.51420418 | 0.12997406 | 0.99999872 |
| KIAA0586 | 26.5677875 | -0.2081448 | 0.46496402 | -0.4476579 | 0.65440014 | 0.99999872 |
| DAAM1 | 18.2586151 | 0.070918575 | 0.46684093 | 0.15191165 | 0.87925662 | 0.99999872 |
| GPR135 | 11.8497613 | 1.851798469 | 0.59875452 | 3.09275074 | 0.00198311 | 0.36957035 |
| L3HYPDH | 12.5087303 | 0.991173053 | 0.59554256 | 1.66431943 | 0.09604861 | 0.99999872 |
| JKAMP | 24.1786323 | 0.771588784 | 0.37143683 | 2.07730824 | 0.03777312 | 0.95980711 |
| PCNX4 | 141.072772 | -0.364208568 | 0.35434972 | -1.0278224 | 0.30403339 | 0.99999872 |
| DHRS7 | 25.4963407 | 0.128693006 | 0.38289368 | 0.33610638 | 0.73679065 | 0.99999872 |
| PPM1A | 40.219486 | 0.001943449 | 0.3908735 | 0.00497207 | 0.99603288 | 0.99999872 |
| MNAT1 | 93.4434265 | -0.020928256 | 0.28881335 | -0.0724629 | 0.94223352 | 0.99999872 |
| TRMT5 | 5.76236528 | -0.265635175 | 0.65490716 | -0.4056074 | 0.68503111 | 0.99999872 |
| SLC38A6 | 20.0634704 | 0.094607574 | 0.39771034 | 0.2378806 | 0.8119737 | 0.99999872 |
| HIF1A | 157.939817 | 0.012868475 | 0.41547759 | 0.03097273 | 0.97529129 | 0.99999872 |
| HIF1A-AS2 | 51.3527849 | -0.542005493 | 0.66453903 | -0.8156112 | 0.41472253 | 0.99999872 |
| SNAPC1 | 16.509375 | -0.448494934 | 0.43745026 | -1.0252478 | 0.30524626 | 0.99999872 |
| PPP2R5E | 90.239061 | -0.199991846 | 0.25418642 | -0.786792 | 0.43140363 | 0.99999872 |
| WDR89 | 4.93084625 | 0.136809942 | 0.65670492 | 0.20832788 | 0.83497297 | 0.99999872 |
| SGPP1 | 29.6457838 | 0.052772351 | 0.35043462 | 0.15059115 | 0.88029825 | 0.99999872 |
| SYNE2 | 453.11967 | -0.068394129 | 0.30034269 | -0.2277203 | 0.81986368 | 0.99999872 |
| ESR2 | 29.2667458 | 1.008256413 | 0.62301426 | 1.61835208 | 0.10558674 | 0.99999872 |
| MTHFD1 | 9.59869152 | -0.273723298 | 0.48430544 | -0.5651873 | 0.57194635 | 0.99999872 |
| AL122035.1 | 9.66252966 | -0.313973571 | 0.59651445 | -0.526347 | 0.59864716 | 0.99999872 |
| ZBTB25 | 21.433337 | -0.143441312 | 0.49594785 | -0.2892266 | 0.77240798 | 0.99999872 |
| ZBTB1 | 84.8312834 | 0.439776207 | 0.32814268 | 1.34019812 | 0.18018094 | 0.99999872 |
| AL049869.3 | 9.4783033 | 0.530001154 | 0.59135188 | 0.89625343 | 0.37011742 | 0.99999872 |
| HSPA2 | 41.1901515 | 0.023331189 | 0.85955036 | 0.02714348 | 0.97834529 | 0.99999872 |
| PLEKHG3 | 55.923763 | -0.568716398 | 0.38980281 | -1.4589849 | 0.14456927 | 0.99999872 |
| SPTB | 36.7869177 | 0.108350756 | 0.646258 | 0.16765867 | 0.86685181 | 0.99999872 |
| CHURC1-FNT | 9.8326453 | -0.551744087 | 0.49221355 | -1.1209445 | 0.26231148 | 0.99999872 |

| | | | | | | |
|---|---|---|---|---|---|---|
| FNTB | 4.78396828 | -0.106426758 | 0.87519809 | -0.1216031 | 0.9032134 | 0.99999872 |
| MAX | 32.3129228 | 0.05084064 | 0.43585082 | 0.11664688 | 0.90713988 | 0.99999872 |
| FUT8 | 42.9135738 | -0.283200418 | 0.29976355 | -0.944746 | 0.34478857 | 0.99999872 |
| AL391261.2 | 7.77804711 | -0.825804533 | 0.75904228 | -1.0879559 | 0.27661457 | 0.99999872 |
| AL359232.1 | 74.1638791 | -1.426347345 | 0.48440017 | -2.9445641 | 0.0032341 | 0.44319076 |
| CCDC196 | 6.61403124 | -1.412278833 | 1.02404559 | -1.3791171 | 0.16785864 | 0.99999872 |
| GPHN | 525.740576 | -0.862958035 | 0.21164687 | -4.0773485 | 4.56E-05 | 0.06816157 |
| MPP5 | 282.654123 | 0.538793012 | 0.4111384 | 1.3104906 | 0.19002992 | 0.99999872 |
| ATP6V1D | 43.0733497 | 0.310275709 | 0.34589471 | 0.89702357 | 0.36970634 | 0.99999872 |
| EIF2S1 | 23.9406777 | 0.083284603 | 0.38136638 | 0.21838476 | 0.82712934 | 0.99999872 |
| TMEM229B | 14.4455491 | -0.462315202 | 0.49603585 | -0.9320197 | 0.35132633 | 0.99999872 |
| PLEKHH1 | 6.90909729 | -0.912495444 | 0.63425407 | -1.4386907 | 0.15023817 | 0.99999872 |
| PIGH | 8.4646471 | 0.03810315 | 0.55394944 | 0.06878453 | 0.94516113 | 0.99999872 |
| ARG2 | 8.07100275 | -0.052273834 | 0.7735519 | -0.0675764 | 0.94612286 | 0.99999872 |
| VTI1B | 19.1680096 | 0.497128946 | 0.49587202 | 1.00253477 | 0.31608538 | 0.99999872 |
| ZFYVE26 | 13.4966428 | -0.003195651 | 0.5354129 | -0.0059686 | 0.9952378 | 0.99999872 |
| RAD51B | 106.828532 | -0.266576843 | 0.59871822 | -0.4452459 | 0.65614204 | 0.99999872 |
| ZFP36L1 | 157.380002 | -0.350896736 | 0.54603925 | -0.6426218 | 0.52046952 | 0.99999872 |
| ACTN1 | 10.0511249 | -0.900127333 | 0.52385733 | -1.7182681 | 0.08574773 | 0.99999872 |
| DCAF5 | 56.9652602 | 0.331760494 | 0.38397041 | 0.8640262 | 0.3875735 | 0.99999872 |
| EXD2 | 19.0203766 | -0.07550733 | 0.43895198 | -0.1720173 | 0.86342394 | 0.99999872 |
| ERH | 38.7484574 | 0.246999612 | 0.30671835 | 0.80529781 | 0.42064785 | 0.99999872 |
| SLC39A9 | 31.461022 | -0.108004603 | 0.2859073 | -0.3777609 | 0.70560821 | 0.99999872 |
| SUSD6 | 107.962696 | 0.17972905 | 0.47070603 | 0.38182865 | 0.70258846 | 0.99999872 |
| SRSF5 | 120.250096 | 0.232057349 | 0.324661 | 0.71476818 | 0.4747523 | 0.99999872 |
| AL357153.2 | 26.8685941 | 0.330022593 | 0.48450342 | 0.68115638 | 0.49577255 | 0.99999872 |
| MED6 | 13.8146507 | 0.384491463 | 0.46238881 | 0.8315328 | 0.40567271 | 0.99999872 |
| MAP3K9 | 4.96054 | 0.584725041 | 0.7355692 | 0.79492867 | 0.426655 | 0.99999872 |
| PCNX1 | 123.816004 | 0.254781112 | 0.33338572 | 0.76422324 | 0.44473422 | 0.99999872 |
| SIPA1L1 | 115.360037 | -0.441593677 | 0.40479668 | -1.0909024 | 0.27531583 | 0.99999872 |
| DPF3 | 7.66296769 | 0.672483438 | 0.94732648 | 0.70987505 | 0.47778162 | 0.99999872 |
| ZFYVE1 | 15.4530274 | 0.06650877 | 0.44371834 | 0.14988961 | 0.88085171 | 0.99999872 |
| RBM25 | 66.9748152 | -0.435767116 | 0.2582213 | -1.6875723 | 0.09149335 | 0.99999872 |
| PSEN1 | 37.885621 | -0.32369435 | 0.28981667 | -1.1168935 | 0.26403987 | 0.99999872 |
| PAPLN | 4.1168293 | 1.075737798 | 0.92843693 | 1.15865469 | 0.24659697 | NA |
| NUMB | 95.9713231 | -0.084665928 | 0.20220811 | -0.4187069 | 0.67543036 | 0.99999872 |
| HEATR4 | 5.44800637 | 1.472924789 | 0.85555553 | 1.72160045 | 0.08514192 | 0.99999872 |
| DNAL1 | 21.7175224 | -0.068671298 | 0.45347051 | -0.151435 | 0.8796326 | 0.99999872 |
| PNMA1 | 24.8790416 | 0.35982285 | 0.52610646 | 0.68393543 | 0.49401594 | 0.99999872 |
| ELMSAN1 | 18.5046143 | -0.715115707 | 0.38233731 | -1.8703791 | 0.06143119 | 0.99999872 |
| AC005480.1 | 10.21345 | 0.120654307 | 0.75115957 | 0.16062407 | 0.8723895 | 0.99999872 |
| COQ6 | 6.89491571 | 0.324212249 | 0.71795665 | 0.45157636 | 0.65157421 | 0.99999872 |
| ENTPD5 | 14.0586412 | -0.08965405 | 0.74678039 | -0.1200541 | 0.90444029 | 0.99999872 |
| BBOF1 | 18.8883271 | 0.332655852 | 0.47974572 | 0.69340036 | 0.48805834 | 0.99999872 |
| ALDH6A1 | 20.2971778 | 0.442209817 | 0.40716783 | 1.08606276 | 0.27745123 | 0.99999872 |
| LIN52 | 28.1833832 | -0.573229842 | 0.36446844 | -1.5727832 | 0.11576902 | 0.99999872 |
| ABCD4 | 26.7253042 | -0.236173457 | 0.33502339 | -0.7049462 | 0.48084374 | 0.99999872 |
| NPC2 | 76.5387557 | 0.768263439 | 0.41215786 | 1.86400288 | 0.0623213 | 0.99999872 |
| ISCA2 | 13.6892759 | 0.492761703 | 0.62256004 | 0.79150872 | 0.42864719 | 0.99999872 |
| LTBP2 | 10.891944 | 0.50851243 | 0.83810291 | 0.60674223 | 0.54402199 | 0.99999872 |
| AREL1 | 25.5503658 | -0.127547933 | 0.32389761 | -0.3937909 | 0.69373542 | 0.99999872 |
| FCF1 | 11.150063 | -0.566004969 | 0.41628866 | -1.3596454 | 0.17394216 | 0.99999872 |
| YLPM1 | 56.907615 | 0.028731672 | 0.35252759 | 0.08150191 | 0.93504281 | 0.99999872 |
| DLST | 16.4880777 | 0.0137852 | 0.39867234 | 0.03457777 | 0.97241643 | 0.99999872 |
| EIF2B2 | 5.56886564 | 0.743776771 | 0.64119979 | 1.15997663 | 0.24605832 | 0.99999872 |
| MLH3 | 26.3165032 | -0.137001735 | 0.4421534 | -0.3098511 | 0.75667416 | 0.99999872 |
| ACYP1 | 13.4048642 | 0.385320227 | 0.46470516 | 0.82917139 | 0.40700743 | 0.99999872 |
| NEK9 | 28.3772543 | -0.119091088 | 0.31139502 | -0.3824438 | 0.70213222 | 0.99999872 |
| AL049780.2 | 11.5610516 | 2.061923173 | 0.70259341 | 2.93473173 | 0.00333836 | 0.44319076 |
| TMED10 | 85.5993498 | 0.745621264 | 0.28949662 | 2.5755785 | 0.01000726 | 0.70848663 |
| FOS | 14.8558802 | -0.536168825 | 0.93539303 | -0.5732016 | 0.56650817 | 0.99999872 |

| | | | | | | |
|---|---|---|---|---|---|---|
| JDP2 | 14.1468227 | -0.42835419 | 0.50490209 | -0.8483906 | 0.39622047 | 0.99999872 |
| TTLL5 | 49.3921706 | -0.232819016 | 0.26349808 | -0.8835701 | 0.37692835 | 0.99999872 |
| ERG28 | 12.4222879 | 0.146690385 | 0.47672421 | 0.30770492 | 0.75830688 | 0.99999872 |
| IFT43 | 36.2313161 | 0.239341633 | 0.29238798 | 0.81857548 | 0.41302866 | 0.99999872 |
| GPATCH2L | 57.535339 | -0.143497428 | 0.25669211 | -0.5590255 | 0.57614434 | 0.99999872 |
| ANGEL1 | 6.41430469 | 1.073143312 | 0.61912868 | 1.73331223 | 0.08304019 | 0.99999872 |
| IRF2BPL | 46.7127134 | 0.247577207 | 0.51124962 | 0.48425896 | 0.6282021 | 0.99999872 |
| POMT2 | 13.7323683 | -0.038282957 | 0.57834594 | -0.0661939 | 0.94722348 | 0.99999872 |
| TMED8 | 23.3641984 | -0.097242344 | 0.36447564 | -0.2668007 | 0.78962264 | 0.99999872 |
| VIPAS39 | 10.5117276 | 0.069442606 | 0.44168612 | 0.15722162 | 0.87507019 | 0.99999872 |
| AHSA1 | 14.4496708 | 0.276303283 | 0.52343848 | 0.527862 | 0.59759512 | 0.99999872 |
| SPTLC2 | 53.1642475 | 0.200824721 | 0.40101233 | 0.50079438 | 0.61651584 | 0.99999872 |
| ALKBH1 | 12.4466283 | -0.386349102 | 0.50089589 | -0.7713162 | 0.44051955 | 0.99999872 |
| SLIRP | 21.019415 | 0.649347477 | 0.37372694 | 1.73749176 | 0.08230041 | 0.99999872 |
| SNW1 | 21.8318992 | -0.375249722 | 0.37666876 | -0.9962327 | 0.31913711 | 0.99999872 |
| ADCK1 | 8.60622243 | -0.582482699 | 0.5619664 | -1.036508 | 0.29996519 | 0.99999872 |
| NRXN3 | 48.5678293 | -1.66916331 | 1.1263739 | -1.481891 | 0.13836931 | 0.99999872 |
| CEP128 | 24.3978128 | -0.328008894 | 0.56538419 | -0.5801522 | 0.56181197 | 0.99999872 |
| GTF2A1 | 27.1446977 | -0.080988426 | 0.32669462 | -0.2479025 | 0.80420981 | 0.99999872 |
| STON2 | 60.8186961 | -0.118155044 | 0.44568328 | -0.2651099 | 0.79092481 | 0.99999872 |
| SEL1L | 142.890542 | 0.591169794 | 0.35120549 | 1.68325899 | 0.09232497 | 0.99999872 |
| FLRT2 | 88.3321229 | 0.236781842 | 0.57508887 | 0.41173087 | 0.6805367 | 0.99999872 |
| GALC | 43.7662213 | 0.195258577 | 0.5075642 | 0.3846973 | 0.7004617 | 0.99999872 |
| SPATA7 | 35.4698059 | -0.004532701 | 0.39266453 | -0.0115434 | 0.99078987 | 0.99999872 |
| PTPN21 | 67.7740436 | 0.226407152 | 0.29866639 | 0.75806037 | 0.44841484 | 0.99999872 |
| ZC3H14 | 53.4447987 | 0.250338318 | 0.32558245 | 0.76889377 | 0.44195638 | 0.99999872 |
| EML5 | 8.74437336 | -0.106901509 | 0.56239345 | -0.1900831 | 0.84924399 | 0.99999872 |
| TTC8 | 20.8579884 | 0.437960503 | 0.43540734 | 1.00586386 | 0.31448107 | 0.99999872 |
| FOXN3 | 47.6181947 | 0.601368335 | 0.43928075 | 1.36898404 | 0.17100426 | 0.99999872 |
| EFCAB11 | 15.98089 | -0.55584025 | 0.46590799 | -1.1930258 | 0.23285927 | 0.99999872 |
| TDP1 | 5.26997652 | -0.102047209 | 0.64077521 | -0.1592559 | 0.87346729 | 0.99999872 |
| PSMC1 | 29.0081894 | -0.327526447 | 0.35031005 | -0.9349616 | 0.34980809 | 0.99999872 |
| NRDE2 | 18.5078803 | -0.565047091 | 0.47470585 | -1.1903099 | 0.2339246 | 0.99999872 |
| CALM1 | 86.2681104 | -0.067142023 | 0.30284222 | -0.2217063 | 0.82454254 | 0.99999872 |
| TTC7B | 23.0725988 | -0.186610093 | 0.48342305 | -0.3860182 | 0.6994832 | 0.99999872 |
| RPS6KA5 | 15.2453055 | 0.667188348 | 0.48102568 | 1.3870119 | 0.16543813 | 0.99999872 |
| DGLUCY | 36.4807177 | -0.312445653 | 0.57703843 | -0.5414642 | 0.58818766 | 0.99999872 |
| CCDC88C | 6.07503116 | -1.269205845 | 0.76760155 | -1.6534696 | 0.09823533 | 0.99999872 |
| PPP4R3A | 50.936865 | 0.032915863 | 0.26616052 | 0.12366922 | 0.90157719 | 0.99999872 |
| TC2N | 7.07739142 | -0.100543728 | 0.95794956 | -0.1049572 | 0.91640975 | 0.99999872 |
| TRIP11 | 63.2988724 | -0.051229721 | 0.26504058 | -0.1932901 | 0.84673178 | 0.99999872 |
| ATXN3 | 48.146496 | -0.435294242 | 0.35839575 | -1.2145631 | 0.22453278 | 0.99999872 |
| NDUFB1 | 22.1342395 | 0.593641559 | 0.43190009 | 1.37448817 | 0.16929018 | 0.99999872 |
| CPSF2 | 25.3878727 | -0.067000055 | 0.31623782 | -0.211866 | 0.83221155 | 0.99999872 |
| RIN3 | 18.9429785 | -0.287134832 | 0.45262957 | -0.6343705 | 0.52583908 | 0.99999872 |
| LGMN | 22.0171194 | -0.306874516 | 0.78950004 | -0.3886947 | 0.69750197 | 0.99999872 |
| GOLGA5 | 20.2728479 | 0.515422649 | 0.3681921 | 1.39987426 | 0.16155097 | 0.99999872 |
| ITPK1 | 13.2603832 | -0.526832878 | 0.39540133 | -1.3324004 | 0.18272866 | 0.99999872 |
| MOAP1 | 11.1728642 | 0.118138181 | 0.45892144 | 0.25742572 | 0.79685015 | 0.99999872 |
| GON7 | 4.95987679 | -0.222612124 | 0.64688595 | -0.3441289 | 0.73074938 | 0.99999872 |
| UBR7 | 7.26452756 | 0.032312343 | 0.59512927 | 0.05429466 | 0.9567004 | 0.99999872 |
| BTBD7 | 78.3192304 | -0.081358031 | 0.25190929 | -0.3229656 | 0.7467213 | 0.99999872 |
| UNC79 | 7.40871759 | 0.300194046 | 0.66585912 | 0.45083718 | 0.6521069 | 0.99999872 |
| DDX24 | 77.8128222 | 0.050415263 | 0.29947523 | 0.16834535 | 0.8663116 | 0.99999872 |
| IFI27L1 | 6.15691764 | 0.307682977 | 0.84804164 | 0.36281589 | 0.71674242 | 0.99999872 |
| IFI27 | 55.7616178 | 0.681073992 | 2.11001264 | 0.32278195 | 0.74686039 | 0.99999872 |
| IFI27L2 | 16.2603334 | 0.254779564 | 0.55171084 | 0.46179909 | 0.64422541 | 0.99999872 |
| SERPINA1 | 27.7611827 | 0.163577721 | 0.73927015 | 0.22126921 | 0.82488283 | 0.99999872 |
| DICER1 | 63.9046486 | 0.09978894 | 0.25731779 | 0.38780428 | 0.69816088 | 0.99999872 |
| DICER1-AS1 | 10.2196213 | -0.845086419 | 0.67432036 | -1.2532417 | 0.21011774 | 0.99999872 |
| CLMN | 21.8641183 | 0.29253671 | 0.75537624 | 0.38727285 | 0.69855422 | 0.99999872 |

| | | | | | | |
|---|---|---|---|---|---|---|
| SYNE3 | 5.6024087 | -0.140774705 | 0.70535127 | -0.199581 | 0.8418083 | 0.99999872 |
| GLRX5 | 22.8102586 | 0.313783157 | 0.43953461 | 0.71389863 | 0.47528987 | 0.99999872 |
| C14orf132 | 10.8332468 | -0.119103719 | 1.03553804 | -0.1150163 | 0.90843223 | 0.99999872 |
| ATG2B | 44.8135559 | 0.083912565 | 0.3871267 | 0.21675737 | 0.82839744 | 0.99999872 |
| GSKIP | 6.56663077 | 0.828017584 | 0.58201827 | 1.42266596 | 0.15483301 | 0.99999872 |
| PAPOLA | 92.9392111 | 0.209612356 | 0.28767409 | 0.72864524 | 0.4662187 | 0.99999872 |
| VRK1 | 10.401638 | -0.480172341 | 0.63057206 | -0.7614869 | 0.44636632 | 0.99999872 |
| SETD3 | 31.3274456 | 0.285134781 | 0.30646928 | 0.93038618 | 0.35217117 | 0.99999872 |
| CCNK | 19.0074678 | 0.372353125 | 0.37035323 | 1.00539997 | 0.3147043 | 0.99999872 |
| CCDC85C | 101.756829 | -0.689065322 | 0.57541992 | -1.1974999 | 0.23111175 | 0.99999872 |
| EML1 | 12.584613 | -0.432635611 | 0.59122988 | -0.7317553 | 0.46431792 | 0.99999872 |
| EVL | 134.837974 | 0.465292912 | 0.58891969 | 0.79007871 | 0.4294818 | 0.99999872 |
| YY1 | 81.8214751 | 0.268602338 | 0.31157849 | 0.86206959 | 0.38864922 | 0.99999872 |
| SLC25A29 | 18.3201188 | -0.023101044 | 0.6096043 | -0.0378951 | 0.96977128 | 0.99999872 |
| WARS | 35.1880988 | -1.305302619 | 0.88610658 | -1.4730763 | 0.14073046 | 0.99999872 |
| WDR25 | 13.9546079 | -0.704898091 | 0.48965755 | -1.4395736 | 0.14998807 | 0.99999872 |
| PPP2R5C | 55.8684291 | -0.137303962 | 0.25815599 | -0.5318643 | 0.59481996 | 0.99999872 |
| DYNC1H1 | 70.7466094 | -0.0770271 | 0.29921734 | -0.2574286 | 0.79684793 | 0.99999872 |
| HSP90AA1 | 162.340724 | 0.092184205 | 0.35415441 | 0.26029382 | 0.79463714 | 0.99999872 |
| WDR20 | 49.2694657 | -0.382624573 | 0.30062661 | -1.2727568 | 0.20310435 | 0.99999872 |
| MOK | 18.2462064 | 0.121025433 | 0.47151083 | 0.25667583 | 0.79742903 | 0.99999872 |
| ZNF839 | 7.48703451 | -0.522260755 | 0.51013803 | -1.0237636 | 0.30594694 | 0.99999872 |
| CINP | 7.48628623 | 0.216221274 | 0.618121 | 0.34980412 | 0.72648571 | 0.99999872 |
| TECPR2 | 26.8842725 | -0.722504426 | 0.40408116 | -1.7880181 | 0.07377309 | 0.99999872 |
| ANKRD9 | 9.7574791 | -0.300953984 | 0.50727856 | -0.5932716 | 0.55299937 | 0.99999872 |
| RCOR1 | 21.6244716 | 0.087086636 | 0.41329018 | 0.21071547 | 0.8331093 | 0.99999872 |
| TRAF3 | 44.3704807 | -0.302317912 | 0.34098797 | -0.8865941 | 0.37529749 | 0.99999872 |
| CDC42BPB | 63.4798192 | -0.523667716 | 0.25324531 | -2.0678279 | 0.03865621 | 0.95980711 |
| TNFAIP2 | 23.4427358 | -1.493610246 | 0.41702019 | -3.5816257 | 0.00034146 | 0.17037366 |
| EIF5 | 53.0678212 | 0.497868478 | 0.36825884 | 1.35195256 | 0.17639049 | 0.99999872 |
| MARK3 | 136.786883 | -0.023208991 | 0.26534095 | -0.0874686 | 0.93029907 | 0.99999872 |
| CKB | 57.4593055 | 0.689774886 | 0.60087351 | 1.14795356 | 0.25098773 | 0.99999872 |
| BAG5 | 20.3938116 | 0.361754498 | 0.39522371 | 0.91531578 | 0.36002588 | 0.99999872 |
| KLC1 | 53.5242702 | -0.296712732 | 0.43914394 | -0.6756617 | 0.49925547 | 0.99999872 |
| ZFYVE21 | 26.8111428 | 0.425802023 | 0.46009942 | 0.92545656 | 0.35472847 | 0.99999872 |
| PPP1R13B | 25.6484079 | -0.618734666 | 0.3580207 | -1.7282092 | 0.08395073 | 0.99999872 |
| ATP5MPL | 29.7040568 | 0.979602293 | 0.37048477 | 2.6441095 | 0.00819062 | 0.66265058 |
| INF2 | 112.85301 | -1.137750915 | 0.38456337 | -2.9585525 | 0.00309088 | 0.44319076 |
| SIVA1 | 17.3531904 | 0.439490305 | 0.44329036 | 0.99142761 | 0.32147683 | 0.99999872 |
| AKT1 | 18.6697107 | -0.130164803 | 0.5055806 | -0.2574561 | 0.79682671 | 0.99999872 |
| CEP170B | 11.1583241 | -1.114916704 | 0.41848093 | -2.6641995 | 0.00771718 | 0.65139548 |
| NUDT14 | 4.06937886 | 0.151550324 | 0.96391191 | 0.15722425 | 0.87506812 | NA |
| BRF1 | 12.948573 | 0.042180733 | 0.40742294 | 0.10353058 | 0.91754188 | 0.99999872 |
| BTBD6 | 12.1703403 | 0.042998981 | 0.48307235 | 0.08901147 | 0.92907279 | 0.99999872 |
| PACS2 | 25.4387257 | -0.111667643 | 0.38652856 | -0.2888988 | 0.77265881 | 0.99999872 |
| MTA1 | 17.0197728 | -0.849288815 | 0.42118114 | -2.0164455 | 0.04375341 | 0.95980711 |
| CRIP2 | 58.1490432 | 0.378730595 | 0.68173607 | 0.55553844 | 0.57852643 | 0.99999872 |
| NIPA1 | 18.3175011 | -0.095854588 | 0.43335081 | -0.221194 | 0.8249414 | 0.99999872 |
| NIPA2 | 20.8370146 | -0.067237142 | 0.4518901 | -0.1487909 | 0.88171862 | 0.99999872 |
| CYFIP1 | 42.8190748 | -0.244901057 | 0.33525368 | -0.7304948 | 0.46508781 | 0.99999872 |
| TUBGCP5 | 12.3827635 | 0.083545206 | 0.43559914 | 0.19179378 | 0.84790374 | 0.99999872 |
| NDN | 41.3365963 | 0.734712876 | 0.76415366 | 0.96147269 | 0.33631455 | 0.99999872 |
| AC124312.1 | 90.2394424 | 0.146871147 | 0.33757363 | 0.43507885 | 0.66350519 | 0.99999872 |
| PWAR6 | 22.5908457 | 0.656161195 | 0.45803873 | 1.43254522 | 0.15198785 | 0.99999872 |
| UBE3A | 82.250035 | -0.05992739 | 0.25762531 | -0.2326145 | 0.81606074 | 0.99999872 |
| ATP10A | 592.81694 | 0.104128539 | 0.51988703 | 0.2002907 | 0.84125323 | 0.99999872 |
| HERC2 | 77.1097261 | -0.289337184 | 0.23516312 | -1.230368 | 0.21855932 | 0.99999872 |
| TJP1 | 619.489146 | 0.135403064 | 0.32679605 | 0.41433507 | 0.67862873 | 0.99999872 |
| FAN1 | 15.3514451 | 0.538939019 | 0.3892009 | 1.38473218 | 0.16613437 | 0.99999872 |
| MTMR10 | 138.484577 | -0.825705937 | 0.50150455 | -1.6464575 | 0.0996696 | 0.99999872 |
| KLF13 | 34.9055455 | -0.085887558 | 0.44866406 | -0.1914295 | 0.84818907 | 0.99999872 |

| | | | | | | |
|---|---|---|---|---|---|---|
| OTUD7A | 21.8831877 | 0.175634623 | 0.56084272 | 0.31316199 | 0.75415759 | 0.99999872 |
| LINC02256 | 49.4865152 | -0.704508688 | 0.39488541 | -1.7840839 | 0.07441004 | 0.99999872 |
| AC123768.3 | 9.31538006 | -0.041762385 | 0.59112277 | -0.0706493 | 0.94367691 | 0.99999872 |
| GOLGA8N | 5.37136043 | 0.125752853 | 0.63337132 | 0.19854523 | 0.8426185 | 0.99999872 |
| FMN1 | 10.2587473 | -1.175931117 | 0.57078328 | -2.060206 | 0.03937885 | 0.95980711 |
| RYR3 | 393.80975 | -0.195194265 | 0.62487106 | -0.3123753 | 0.75475534 | 0.99999872 |
| AVEN | 28.5261963 | -0.380719813 | 0.48398566 | -0.7866345 | 0.43149587 | 0.99999872 |
| EMC7 | 33.250589 | 0.717102729 | 0.4942151 | 1.45099314 | 0.14678177 | 0.99999872 |
| KATNBL1 | 58.9743563 | 0.167229878 | 0.28255842 | 0.59184179 | 0.55395654 | 0.99999872 |
| EMC4 | 14.6115064 | 0.231523541 | 0.44096131 | 0.52504276 | 0.59955347 | 0.99999872 |
| SLC12A6 | 23.8246964 | -0.094104522 | 0.44452478 | -0.2116969 | 0.8323435 | 0.99999872 |
| NOP10 | 28.9304883 | 0.613220288 | 0.35921915 | 1.70709243 | 0.08780487 | 0.99999872 |
| LPCAT4 | 6.00385275 | -0.268154207 | 0.72576295 | -0.3694791 | 0.71177068 | 0.99999872 |
| GOLGA8A | 27.1850693 | -1.06172948 | 0.47215893 | -2.2486697 | 0.02453352 | 0.88337526 |
| GOLGA8B | 43.7040357 | 0.76875704 | 0.41951763 | 1.83247851 | 0.06688016 | 0.99999872 |
| AQR | 36.1108239 | -0.293094805 | 0.31249529 | -0.9379175 | 0.3482868 | 0.99999872 |
| ZNF770 | 20.7027769 | 0.783758647 | 0.42151841 | 1.85936991 | 0.06297472 | 0.99999872 |
| AC021231.1 | 33.6404128 | -0.331027905 | 0.48165782 | -0.6872678 | 0.49191399 | 0.99999872 |
| DPH6 | 90.2799733 | -0.663393543 | 0.3502595 | -1.8940058 | 0.05822423 | 0.99999872 |
| DPH6-DT | 40.1763004 | 0.235294998 | 0.77382071 | 0.30406914 | 0.76107522 | 0.99999872 |
| C15orf41 | 23.9322013 | -0.470392155 | 0.50787396 | -0.9261986 | 0.35434278 | 0.99999872 |
| MEIS2 | 32.5971075 | 0.549959097 | 0.44901664 | 1.22480783 | 0.22064763 | 0.99999872 |
| SPRED1 | 54.6344313 | -0.612884702 | 0.33195116 | -1.8463099 | 0.06484722 | 0.99999872 |
| FAM98B | 12.9114356 | 0.76782061 | 0.41607711 | 1.84538054 | 0.0649822 | 0.99999872 |
| RASGRP1 | 5.95396753 | 0.460002884 | 0.83433383 | 0.55134152 | 0.58139958 | 0.99999872 |
| C15orf53 | 5.36430456 | 0.445720353 | 0.79594773 | 0.55998696 | 0.57548833 | 0.99999872 |
| AC013652.1 | 54.3731286 | -0.566132665 | 0.66686247 | -0.8489497 | 0.39590931 | 0.99999872 |
| THBS1 | 87.6116443 | 0.071758706 | 0.44065739 | 0.16284467 | 0.87064074 | 0.99999872 |
| FSIP1 | 14.8093132 | 1.118499851 | 0.5016277 | 2.22974099 | 0.02576464 | 0.88365618 |
| EIF2AK4 | 28.1207142 | -0.268195893 | 0.35587971 | -0.7536139 | 0.45108111 | 0.99999872 |
| SRP14 | 161.091043 | 0.691709082 | 0.50713732 | 1.36394829 | 0.17258384 | 0.99999872 |
| CCDC9B | 5.08498931 | -0.496713382 | 0.73275903 | -0.6778673 | 0.49785584 | 0.99999872 |
| IVD | 26.2801805 | -0.124280185 | 0.37413101 | -0.3321836 | 0.73975063 | 0.99999872 |
| BAHD1 | 8.71439007 | -0.075615143 | 0.4890863 | -0.1546049 | 0.8771328 | 0.99999872 |
| CHST14 | 5.90376408 | 0.617034162 | 0.73522031 | 0.83925071 | 0.40132864 | 0.99999872 |
| CCDC32 | 20.4547695 | -0.549329258 | 0.45255098 | -1.2138506 | 0.22480479 | 0.99999872 |
| RMDN3 | 14.7983819 | -0.290919742 | 0.42101763 | -0.6909918 | 0.48957067 | 0.99999872 |
| GCHFR | 7.13746036 | 1.102425309 | 0.81120196 | 1.35900227 | 0.17414587 | 0.99999872 |
| DNAJC17 | 14.8208671 | -0.528449245 | 0.38242472 | -1.3818386 | 0.16702126 | 0.99999872 |
| ZFYVE19 | 6.4471133 | -0.806927521 | 0.57042528 | -1.4146069 | 0.15718379 | 0.99999872 |
| SPINT1 | 12.8378954 | -0.379874842 | 0.6093301 | -0.6234303 | 0.5330018 | 0.99999872 |
| VPS18 | 9.07976024 | -0.647687772 | 0.47768667 | -1.3558841 | 0.17513604 | 0.99999872 |
| INO80 | 39.5986768 | -0.701090193 | 0.3730256 | -1.8794694 | 0.06018043 | 0.99999872 |
| CHP1 | 26.5916546 | -0.019943844 | 0.36030592 | -0.0553525 | 0.95585762 | 0.99999872 |
| NDUFAF1 | 6.29533127 | -0.250926751 | 0.72667404 | -0.3453085 | 0.72986243 | 0.99999872 |
| RTF1 | 39.6546569 | -0.17232335 | 0.33550534 | -0.5136233 | 0.60751541 | 0.99999872 |
| RPAP1 | 6.22619286 | -0.040839971 | 0.56843611 | -0.0718462 | 0.94272431 | 0.99999872 |
| TYRO3 | 326.865688 | -0.313500098 | 0.37238624 | -0.8418681 | 0.39986179 | 0.99999872 |
| MGA | 42.6985976 | -0.347972555 | 0.35064089 | -0.9923901 | 0.32100726 | 0.99999872 |
| MAPKBP1 | 16.2001932 | -0.16326042 | 0.43454154 | -0.3757073 | 0.70713453 | 0.99999872 |
| EHD4 | 60.4463243 | 0.043817006 | 0.39377713 | 0.11127362 | 0.91139938 | 0.99999872 |
| VPS39 | 24.7369025 | -0.268738867 | 0.34483271 | -0.7793311 | 0.43578468 | 0.99999872 |
| TMEM87A | 45.9800289 | -0.214329449 | 0.31553413 | -0.6792592 | 0.49697367 | 0.99999872 |
| GANC | 16.5454016 | 0.193674878 | 0.60702242 | 0.31905721 | 0.74968313 | 0.99999872 |
| ZNF106 | 51.8298699 | -0.192122949 | 0.48611146 | -0.3952241 | 0.69267753 | 0.99999872 |
| SNAP23 | 33.0882495 | 0.738330783 | 0.4028449 | 1.83279169 | 0.06683356 | 0.99999872 |
| LRRC57 | 14.8431045 | 0.070684319 | 0.45359975 | 0.15582971 | 0.87616725 | 0.99999872 |
| HAUS2 | 10.3925948 | 0.915763122 | 0.54156192 | 1.69096662 | 0.09084318 | 0.99999872 |
| STARD9 | 10.8450577 | 0.430492519 | 0.54318753 | 0.79253019 | 0.42805159 | 0.99999872 |
| TTBK2 | 49.4190254 | 0.149913451 | 0.32655046 | 0.45908203 | 0.64617526 | 0.99999872 |
| UBR1 | 51.4410506 | -0.114030088 | 0.27922937 | -0.4083743 | 0.68299893 | 0.99999872 |

| | | | | | | |
|---|---|---|---|---|---|---|
| AC068724.3 | 12.3950729 | 0.839637304 | 0.50357488 | 1.66735342 | 0.09544415 | 0.99999872 |
| TMEM62 | 20.6318535 | -0.989343058 | 0.57685506 | -1.7150635 | 0.08633359 | 0.99999872 |
| CCNDBP1 | 19.4234174 | 0.787084019 | 0.45365357 | 1.73498914 | 0.08274273 | 0.99999872 |
| ADAL | 9.15586441 | -0.558390394 | 0.56844702 | -0.9823086 | 0.32594784 | 0.99999872 |
| ZSCAN29 | 4.25519368 | 0.732889377 | 0.70166404 | 1.04450183 | 0.29625327 | NA |
| TUBGCP4 | 10.1791253 | 0.125767054 | 0.46622409 | 0.26975666 | 0.78734747 | 0.99999872 |
| TP53BP1 | 48.126039 | 0.239371879 | 0.26223508 | 0.91281411 | 0.36134031 | 0.99999872 |
| MAP1A | 8.89413159 | 0.270286462 | 0.81714725 | 0.33076837 | 0.74081946 | 0.99999872 |
| PPIP5K1 | 12.3580351 | 0.539643797 | 0.55673106 | 0.96930787 | 0.3323916 | 0.99999872 |
| CATSPER2 | 15.1273703 | -0.197456285 | 0.59222097 | -0.3334166 | 0.73881986 | 0.99999872 |
| PDIA3 | 128.123653 | 0.597742719 | 0.48396643 | 1.23509129 | 0.21679653 | 0.99999872 |
| SERF2 | 218.409035 | 0.647505137 | 0.43921007 | 1.47424931 | 0.14041448 | 0.99999872 |
| MFAP1 | 13.152698 | -0.125174335 | 0.46571762 | -0.2687773 | 0.78810105 | 0.99999872 |
| FRMD5 | 6.9791657 | -0.103806531 | 0.52550131 | -0.1975381 | 0.84340647 | 0.99999872 |
| CASC4 | 108.636578 | -0.206133812 | 0.31931777 | -0.6455444 | 0.51857443 | 0.99999872 |
| AC025043.1 | 4.58760349 | 0.950598797 | 0.67859612 | 1.40083147 | 0.16126448 | NA |
| CTDSPL2 | 45.9333941 | -0.355490695 | 0.27948096 | -1.2719675 | 0.20338467 | 0.99999872 |
| EIF3J-DT | 16.3361168 | 0.287837112 | 0.50586972 | 0.56899454 | 0.56935984 | 0.99999872 |
| EIF3J | 43.9898084 | 0.362243553 | 0.29606121 | 1.22354277 | 0.22112476 | 0.99999872 |
| SPG11 | 57.0025059 | -0.295138801 | 0.26885777 | -1.0977507 | 0.27231337 | 0.99999872 |
| B2M | 930.090397 | 0.868096487 | 0.72575296 | 1.1961322 | 0.23164497 | 0.99999872 |
| TRIM69 | 6.80299668 | -0.378919246 | 0.96449604 | -0.3928676 | 0.69441727 | 0.99999872 |
| SORD | 7.59299148 | 0.476981813 | 0.97048131 | 0.49148995 | 0.62307996 | 0.99999872 |
| DUOX1 | 9.08821154 | 0.222870554 | 0.73980992 | 0.3012538 | 0.76322096 | 0.99999872 |
| GATM | 9.16108085 | 0.59680507 | 0.73168397 | 0.81565962 | 0.41469484 | 0.99999872 |
| SPATA5L1 | 4.76646556 | 0.691352905 | 0.74117023 | 0.93278558 | 0.35093069 | 0.99999872 |
| SLC30A4 | 11.980175 | 0.405895832 | 0.45960531 | 0.88313999 | 0.37716064 | 0.99999872 |
| SQOR | 67.2113865 | 0.29043475 | 0.45788007 | 0.63430311 | 0.52588303 | 0.99999872 |
| SEMA6D | 23.7222331 | -0.133313504 | 0.44363818 | -0.3005005 | 0.7637954 | 0.99999872 |
| MYEF2 | 55.9858325 | 0.160998251 | 0.32558011 | 0.49449658 | 0.62095552 | 0.99999872 |
| SLC12A1 | 67.2814925 | 3.248907996 | 1.60457283 | 2.02478064 | 0.04288989 | 0.95980711 |
| DUT | 28.3335532 | 0.665463017 | 0.38874471 | 1.71182525 | 0.08692888 | 0.99999872 |
| CEP152 | 7.89150605 | 0.276525262 | 0.76523852 | 0.36135826 | 0.71783165 | 0.99999872 |
| EID1 | 123.757742 | 0.183671781 | 0.38472076 | 0.47741583 | 0.63306604 | 0.99999872 |
| SECISBP2L | 75.5401921 | -0.182563096 | 0.35987446 | -0.5072966 | 0.61194671 | 0.99999872 |
| COPS2 | 23.2106525 | 0.131257565 | 0.42320268 | 0.31015296 | 0.75644464 | 0.99999872 |
| GALK2 | 44.634349 | -0.044000384 | 0.37238791 | -0.1181574 | 0.90594295 | 0.99999872 |
| FAM227B | 16.1905955 | 0.281056953 | 0.61712387 | 0.45543037 | 0.64879965 | 0.99999872 |
| DTWD1 | 29.1396581 | 0.14971959 | 0.45959735 | 0.32576252 | 0.74460404 | 0.99999872 |
| SLC27A2 | 9.32091653 | 0.799132199 | 1.29626189 | 0.61648978 | 0.53757132 | 0.99999872 |
| GABPB1 | 31.1161565 | -0.505206573 | 0.34622364 | -1.4591915 | 0.14451242 | 0.99999872 |
| GABPB1-AS1 | 37.1980299 | -0.186041028 | 0.75929165 | -0.2450192 | 0.80644157 | 0.99999872 |
| USP8 | 43.7229657 | 0.027292832 | 0.28867371 | 0.09454561 | 0.92467575 | 0.99999872 |
| TRPM7 | 79.0797714 | -0.224486632 | 0.33260269 | -0.6749393 | 0.49971432 | 0.99999872 |
| SPPL2A | 76.5790129 | -0.378017844 | 0.23684427 | -1.5960608 | 0.11047523 | 0.99999872 |
| AC021752.1 | 6.41699276 | 0.512292542 | 0.91767781 | 0.55824881 | 0.5766745 | 0.99999872 |
| AP4E1 | 25.0773672 | -0.124540347 | 0.39608484 | -0.3144285 | 0.75319564 | 0.99999872 |
| DMXL2 | 41.9883592 | 0.306502034 | 0.51336667 | 0.59704312 | 0.5504786 | 0.99999872 |
| LYSMD2 | 8.66412838 | 0.717686375 | 0.809865 | 0.88618026 | 0.3755204 | 0.99999872 |
| TMOD2 | 31.8899933 | 0.617532988 | 0.55384468 | 1.11499308 | 0.26485339 | 0.99999872 |
| TMOD3 | 148.853171 | -0.107434181 | 0.2567932 | -0.4183685 | 0.67567772 | 0.99999872 |
| AC090971.2 | 6.62744475 | 0.727778819 | 0.644517 | 1.12918484 | 0.25881987 | 0.99999872 |
| LEO1 | 6.46548453 | -0.31709219 | 0.62346408 | -0.5085974 | 0.61103448 | 0.99999872 |
| MAPK6 | 28.1479421 | -0.168712288 | 0.38417301 | -0.4391571 | 0.66054774 | 0.99999872 |
| GNB5 | 24.4020285 | 0.308004705 | 0.35546906 | 0.86647402 | 0.38623027 | 0.99999872 |
| CERNA1 | 4.29611103 | 1.38086898 | 0.82994439 | 1.66380903 | 0.0961506 | NA |
| MYO5C | 57.2033839 | -0.043747677 | 0.30241893 | -0.1446592 | 0.88497996 | 0.99999872 |
| MYO5A | 40.2355703 | -0.070836875 | 0.27983844 | -0.2531349 | 0.80016397 | 0.99999872 |
| ARPP19 | 64.9615776 | 0.049363519 | 0.33351164 | 0.14801138 | 0.88233379 | 0.99999872 |
| FAM214A | 58.1673952 | 0.060594392 | 0.33687736 | 0.17987078 | 0.85725402 | 0.99999872 |
| WDR72 | 45.3674407 | 1.89233139 | 1.22627041 | 1.54315995 | 0.12279197 | 0.99999872 |

| | | | | | | |
|---|---|---|---|---|---|---|
| RSL24D1 | 20.5991876 | 0.354956563 | 0.36460311 | 0.97354233 | 0.33028383 | 0.99999872 |
| RAB27A | 21.640566 | 0.052146958 | 0.63676213 | 0.08189394 | 0.93473105 | 0.99999872 |
| PIGB | 17.8019035 | -0.428255895 | 0.54415663 | -0.7870085 | 0.43127689 | 0.99999872 |
| CCPG1 | 93.114846 | 0.251493363 | 0.20073115 | 1.2528866 | 0.21024698 | 0.99999872 |
| C15orf65 | 4.28622242 | 1.291983567 | 0.9279078 | 1.39236201 | 0.1638128 | NA |
| DNAAF4 | 18.4708493 | 0.374069057 | 0.40240278 | 0.92958864 | 0.35258411 | 0.99999872 |
| PYGO1 | 16.7263833 | 0.192737316 | 0.58545946 | 0.32920694 | 0.74199928 | 0.99999872 |
| PRTG | 7.18956489 | -0.229282271 | 0.95132718 | -0.2410131 | 0.809545 | 0.99999872 |
| NEDD4 | 24.7689599 | 0.026582194 | 0.39002665 | 0.06815481 | 0.9456624 | 0.99999872 |
| RFX7 | 57.3264735 | 0.164623655 | 0.27834284 | 0.59144204 | 0.55422428 | 0.99999872 |
| TEX9 | 28.4064272 | -0.025554079 | 0.33572753 | -0.0761155 | 0.93932719 | 0.99999872 |
| ZNF280D | 38.9678243 | -0.082231519 | 0.32356726 | -0.2541404 | 0.79938708 | 0.99999872 |
| AC090517.4 | 32.8140884 | -0.502229543 | 0.48304408 | -1.0397178 | 0.29847102 | 0.99999872 |
| TCF12 | 188.15933 | -0.044230207 | 0.24438831 | -0.1809833 | 0.85638067 | 0.99999872 |
| CGNL1 | 243.589387 | -0.057951947 | 0.31094519 | -0.1863735 | 0.85215186 | 0.99999872 |
| POLR2M | 10.3323235 | -0.679619013 | 0.51634835 | -1.3162025 | 0.18810607 | 0.99999872 |
| ALDH1A2 | 32.379633 | -1.834071744 | 0.46324564 | -3.9591776 | 7.52E-05 | 0.07163933 |
| ADAM10 | 67.3619363 | -0.175776371 | 0.35601855 | -0.4937281 | 0.6214982 | 0.99999872 |
| MINDY2 | 62.6733273 | -0.180895058 | 0.32966614 | -0.548722 | 0.58319626 | 0.99999872 |
| RNF111 | 58.9872813 | 0.077867155 | 0.30273093 | 0.25721572 | 0.79701225 | 0.99999872 |
| SLTM | 76.6962617 | -0.248898597 | 0.29437241 | -0.8455228 | 0.39781898 | 0.99999872 |
| MYO1E | 862.285106 | 0.604849186 | 0.3516416 | 1.72007291 | 0.08541919 | 0.99999872 |
| FAM81A | 68.0736669 | -0.498689143 | 0.57312685 | -0.87012 | 0.38423482 | 0.99999872 |
| GTF2A2 | 79.1971002 | 1.047217099 | 0.34094199 | 3.07154037 | 0.00212957 | 0.37076694 |
| BNIP2 | 46.7598688 | 0.060598228 | 0.30960605 | 0.19572688 | 0.84482394 | 0.99999872 |
| ANXA2 | 421.64187 | 0.328261048 | 0.50574941 | 0.64905868 | 0.51630045 | 0.99999872 |
| ICE2 | 47.2861508 | -0.213771371 | 0.31464337 | -0.6794085 | 0.49687908 | 0.99999872 |
| RORA-AS1 | 16.1812896 | 0.702245415 | 0.7427558 | 0.94545934 | 0.34442442 | 0.99999872 |
| RORA | 236.689875 | 0.392666805 | 0.60953525 | 0.64420689 | 0.51944129 | 0.99999872 |
| AC012404.1 | 5.22880755 | 0.788460454 | 0.93028858 | 0.84754394 | 0.396692 | 0.99999872 |
| AC009554.1 | 24.1615682 | 1.408128639 | 0.68031669 | 2.06981346 | 0.03846982 | 0.95980711 |
| VPS13C | 160.688422 | 0.061519506 | 0.26877174 | 0.22889128 | 0.81895342 | 0.99999872 |
| TLN2 | 22.1933138 | -0.483232353 | 0.68170983 | -0.7088534 | 0.47841544 | 0.99999872 |
| TPM1 | 60.9061329 | 0.446033108 | 0.8148439 | 0.54738473 | 0.58411444 | 0.99999872 |
| LACTB | 10.7867956 | 0.352241568 | 0.46504853 | 0.75742969 | 0.44879247 | 0.99999872 |
| RPS27L | 35.1921767 | 0.630576608 | 0.49683005 | 1.26919982 | 0.2043698 | 0.99999872 |
| RAB8B | 29.6729843 | 0.144897317 | 0.42825791 | 0.33834125 | 0.73510604 | 0.99999872 |
| APH1B | 18.608975 | 0.584482672 | 0.46397102 | 1.25973961 | 0.20776331 | 0.99999872 |
| CA12 | 32.0702629 | -0.172433864 | 0.61534779 | -0.2802218 | 0.77930734 | 0.99999872 |
| USP3 | 70.2457779 | 0.17590803 | 0.29968509 | 0.58697625 | 0.55721966 | 0.99999872 |
| USP3-AS1 | 10.6261586 | 1.621304403 | 0.67200004 | 2.41265523 | 0.01583679 | 0.79608149 |
| HERC1 | 67.3318437 | 0.140157658 | 0.27524664 | 0.50920751 | 0.61060678 | 0.99999872 |
| FAM96A | 9.58990017 | 0.578041646 | 0.57979998 | 0.99696734 | 0.31878036 | 0.99999872 |
| SNX1 | 48.2301158 | -0.371671665 | 0.27715428 | -1.341028 | 0.17991135 | 0.99999872 |
| PPIB | 89.6979012 | 0.837785565 | 0.57365692 | 1.46042963 | 0.14417203 | 0.99999872 |
| CSNK1G1 | 12.5399897 | 0.639676866 | 0.45489772 | 1.40619932 | 0.15966495 | 0.99999872 |
| TRIP4 | 24.5269593 | 0.237321126 | 0.3880892 | 0.6115118 | 0.54086081 | 0.99999872 |
| ZNF609 | 226.265635 | -0.120516971 | 0.26408757 | -0.4563523 | 0.64813666 | 0.99999872 |
| OAZ2 | 33.0366991 | 0.141087138 | 0.50375683 | 0.28006992 | 0.77942386 | 0.99999872 |
| SPG21 | 21.4066934 | 0.271093403 | 0.37346117 | 0.72589448 | 0.46790346 | 0.99999872 |
| MTFMT | 9.43409728 | 0.324853292 | 0.48819409 | 0.66541833 | 0.50578297 | 0.99999872 |
| UBAP1L | 8.85575002 | 0.643613673 | 0.67014562 | 0.96040868 | 0.33684957 | 0.99999872 |
| PDCD7 | 15.9449851 | 0.202700903 | 0.49970369 | 0.4056422 | 0.68500552 | 0.99999872 |
| CLPX | 25.1710457 | -0.368335448 | 0.32966488 | -1.1173027 | 0.26386494 | 0.99999872 |
| PARP16 | 7.78279665 | -0.096050165 | 0.6268563 | -0.1532252 | 0.87822071 | 0.99999872 |
| DPP8 | 46.2024829 | -0.185826162 | 0.409348 | -0.4539564 | 0.64986018 | 0.99999872 |
| HACD3 | 34.075018 | 1.005742915 | 0.3242018 | 3.10221265 | 0.0019208 | 0.36957035 |
| INTS14 | 9.16774213 | 0.054350717 | 0.51738074 | 0.10504975 | 0.91633633 | 0.99999872 |
| SLC24A1 | 4.78148819 | -0.494202302 | 0.86702992 | -0.5699945 | 0.56868142 | 0.99999872 |
| DENND4A | 90.690337 | 0.081678647 | 0.29108943 | 0.2805964 | 0.77901998 | 0.99999872 |
| RAB11A | 64.6302762 | 0.43778478 | 0.30050429 | 1.45683369 | 0.14516229 | 0.99999872 |

| | | | | | | |
|---|---|---|---|---|---|---|
| MEGF11 | 6.88478965 | 0.064555865 | 1.17514237 | 0.05493451 | 0.95619064 | 0.99999872 |
| DIS3L | 21.2827209 | -0.133877477 | 0.37274154 | -0.3591697 | 0.71946812 | 0.99999872 |
| TIPIN | 7.70073912 | -0.069642539 | 0.794392 | -0.0876677 | 0.93014077 | 0.99999872 |
| MAP2K1 | 20.0850915 | 0.37037861 | 0.41202102 | 0.89893134 | 0.36868923 | 0.99999872 |
| SNAPC5 | 11.2956331 | 0.435976643 | 0.46672635 | 0.93411619 | 0.35024396 | 0.99999872 |
| RPL4 | 53.7931453 | 0.001693227 | 0.49289438 | 0.00343527 | 0.99725905 | 0.99999872 |
| SMAD6 | 91.6478452 | -0.84246607 | 0.49940092 | -1.6869534 | 0.09161232 | 0.99999872 |
| AC093334.1 | 7.03262775 | -0.281172534 | 0.839524 | -0.334919 | 0.73768619 | 0.99999872 |
| AC087482.1 | 9.79262762 | 0.99613456 | 0.93141906 | 1.06948054 | 0.28485319 | 0.99999872 |
| SMAD3 | 54.969002 | -1.105450967 | 0.35935845 | -3.0761791 | 0.00209672 | 0.37076694 |
| AAGAB | 40.3301711 | -0.02577477 | 0.3036755 | -0.084876 | 0.93235995 | 0.99999872 |
| IQCH | 32.0065142 | -0.375201424 | 0.55000412 | -0.6821793 | 0.49512558 | 0.99999872 |
| IQCH-AS1 | 13.2136771 | 0.354374423 | 0.61504993 | 0.57617179 | 0.56449907 | 0.99999872 |
| C15orf61 | 9.7401047 | 0.924004098 | 0.54624564 | 1.69155418 | 0.09073101 | 0.99999872 |
| MAP2K5 | 59.9999856 | 0.011390001 | 0.29536191 | 0.03856287 | 0.96923891 | 0.99999872 |
| PIAS1 | 124.090439 | -0.120627221 | 0.2668419 | -0.452055 | 0.65122935 | 0.99999872 |
| FEM1B | 32.7141066 | 0.440915496 | 0.32382348 | 1.36159211 | 0.17332665 | 0.99999872 |
| CORO2B | 178.180252 | -0.357366121 | 0.49607707 | -0.7203843 | 0.47128843 | 0.99999872 |
| ANP32A | 26.5207617 | -0.205989525 | 0.28674278 | -0.7183774 | 0.47252464 | 0.99999872 |
| SPESP1 | 3.54398286 | 1.062552434 | 0.78851069 | 1.34754347 | 0.17780526 | NA |
| GLCE | 28.2621205 | -0.347862236 | 0.52348015 | -0.6645185 | 0.50635852 | 0.99999872 |
| PAQR5 | 15.9937561 | -0.579450311 | 0.57598673 | -1.0060133 | 0.31440917 | 0.99999872 |
| RPLP1 | 138.779677 | 0.475857312 | 0.52883869 | 0.89981562 | 0.36821838 | 0.99999872 |
| DRAIC | 6.74632922 | -0.282706218 | 0.70322437 | -0.4020143 | 0.68767354 | 0.99999872 |
| TLE3 | 6.39218323 | -1.81239299 | 0.53371266 | -3.3958216 | 0.00068423 | 0.23897852 |
| UACA | 329.479932 | -0.376442024 | 0.44889365 | -0.8385996 | 0.40169404 | 0.99999872 |
| LARP6 | 4.65441088 | 0.413142161 | 0.63878446 | 0.64676301 | 0.51778534 | 0.99999872 |
| LRRC49 | 9.80504047 | -0.121079822 | 0.55791519 | -0.2170219 | 0.82819127 | 0.99999872 |
| THSD4 | 14.3529657 | -0.356682229 | 0.81388166 | -0.4382483 | 0.66120632 | 0.99999872 |
| THSD4-AS1 | 49.8256558 | 0.022360371 | 0.3784613 | 0.05908232 | 9.53E-01 | 0.99999872 |
| MYO9A | 159.568699 | -0.092879183 | 0.28986343 | -0.3204239 | 0.74864698 | 0.99999872 |
| SENP8 | 6.2875388 | -0.058093585 | 0.78670426 | -0.0738442 | 0.94113432 | 0.99999872 |
| PKM | 82.3182831 | -0.45770774 | 0.41787213 | -1.0953297 | 0.27337224 | 0.99999872 |
| PARP6 | 17.2675363 | -0.487316989 | 0.39093556 | -1.2465405 | 0.21256604 | 0.99999872 |
| AC009690.2 | 5.17506907 | 1.619085521 | 0.8744082 | 1.85163579 | 0.06407814 | 0.99999872 |
| AC009690.1 | 23.8030589 | 0.056806966 | 0.50183709 | 0.11319802 | 0.90987356 | 0.99999872 |
| ARIH1 | 147.123791 | 0.315125483 | 0.27089801 | 1.16326243 | 0.24472304 | 0.99999872 |
| BBS4 | 24.8414588 | 0.570568737 | 0.38357823 | 1.4874899 | 0.13688547 | 0.99999872 |
| ADPGK | 19.2586843 | -0.471750285 | 0.38877848 | -1.2134167 | 0.22497056 | 0.99999872 |
| NEO1 | 64.8166103 | 0.019964148 | 0.30299338 | 0.06588971 | 0.94746563 | 0.99999872 |
| NPTN | 113.464332 | 0.418200769 | 0.35387001 | 1.18179205 | 0.23728822 | 0.99999872 |
| CD276 | 7.47241462 | 0.023922872 | 0.60527795 | 0.03952378 | 0.9684728 | 0.99999872 |
| C15orf59 | 15.8487699 | 0.613836369 | 0.99418699 | 0.61742547 | 0.53695413 | 0.99999872 |
| LOXL1-AS1 | 6.74783678 | 0.754295541 | 0.7951641 | 0.94860361 | 0.34282225 | 0.99999872 |
| LOXL1 | 35.9157472 | 0.647120106 | 0.76475264 | 0.84618224 | 0.39745109 | 0.99999872 |
| PML | 11.6404257 | -0.54197231 | 0.86072294 | -0.629671 | 0.52890983 | 0.99999872 |
| UBL7 | 10.2134492 | -0.205859828 | 0.5995298 | -0.3433688 | 0.73132103 | 0.99999872 |
| ARID3B | 13.2874024 | -0.165516066 | 0.65347612 | -0.2532856 | 0.80004755 | 0.99999872 |
| CLK3 | 9.7110852 | 0.32072694 | 0.52104514 | 0.6155454 | 0.53819459 | 0.99999872 |
| EDC3 | 13.9340248 | -0.558217236 | 0.42947277 | -1.2997733 | 0.19367868 | 0.99999872 |
| CSK | 4.50490437 | 0.326003862 | 0.70439417 | 0.46281454 | 0.64349731 | NA |
| SCAMP2 | 21.4141853 | 0.102043871 | 0.45719478 | 0.22319562 | 0.82338326 | 0.99999872 |
| MPI | 6.3255146 | 0.185741797 | 0.69203431 | 0.2683997 | 0.78839167 | 0.99999872 |
| FAM219B | 9.48332462 | -0.021639732 | 0.5646855 | -0.0383217 | 0.96943116 | 0.99999872 |
| COX5A | 22.9158256 | 0.455786519 | 0.38288477 | 1.19040128 | 0.23388871 | 0.99999872 |
| PPCDC | 27.2352112 | -0.273825555 | 0.34674932 | -0.7896931 | 0.42970701 | 0.99999872 |
| COMMD4 | 15.6115091 | -0.137240728 | 0.55096362 | -0.2490922 | 0.80328948 | 0.99999872 |
| MAN2C1 | 20.3744203 | -0.649413439 | 0.38155919 | -1.7019992 | 0.08875552 | 0.99999872 |
| SIN3A | 31.5168449 | -0.170723515 | 0.30904935 | -0.5524151 | 0.58066401 | 0.99999872 |
| PTPN9 | 48.4619436 | -0.111573167 | 0.26457861 | -0.4217014 | 0.67324299 | 0.99999872 |
| SNUPN | 6.69484252 | 0.149928066 | 0.61605665 | 0.24336734 | 0.80772084 | 0.99999872 |

| | | | | | | |
|---|---|---|---|---|---|---|
| IMP3 | 15.1716278 | 0.419166669 | 0.41816317 | 1.00239978 | 0.31615055 | 0.99999872 |
| SNX33 | 8.69596052 | 0.215779766 | 0.5090072 | 0.42392282 | 0.6716221 | 0.99999872 |
| DNM1P35 | 4.98032706 | 1.489101539 | 0.78387385 | 1.89966988 | 0.05747646 | 0.99999872 |
| UBE2Q2 | 40.1525315 | 0.581282567 | 0.40635031 | 1.43049619 | 0.15257466 | 0.99999872 |
| FBXO22 | 29.9033183 | 0.340515344 | 0.3338951 | 1.01982733 | 0.30781036 | 0.99999872 |
| NRG4 | 13.553091 | -0.396791061 | 0.82215432 | -0.4826236 | 0.62936303 | 0.99999872 |
| TMEM266 | 116.129964 | -0.652430897 | 0.51475527 | -1.2674584 | 0.20499142 | 0.99999872 |
| ETFA | 39.8566621 | -0.03480525 | 0.29232571 | -0.1190633 | 0.90522524 | 0.99999872 |
| SCAPER | 107.584788 | 0.249115234 | 0.25919503 | 0.96111114 | 0.33649629 | 0.99999872 |
| RCN2 | 75.8341126 | 0.802190058 | 0.36016376 | 2.22729256 | 0.02592773 | 0.88365618 |
| TSPAN3 | 67.1219484 | -0.014903756 | 0.41526055 | -0.0358901 | 0.97136996 | 0.99999872 |
| PEAK1 | 40.7738046 | 0.879623978 | 0.32373136 | 2.71714173 | 0.00658484 | 0.61603533 |
| HMG20A | 28.5947104 | 0.324568019 | 0.38092375 | 0.85205509 | 0.39418352 | 0.99999872 |
| LINGO1 | 407.35979 | -0.310128644 | 0.61824103 | -0.5016306 | 0.61592736 | 0.99999872 |
| TBC1D2B | 52.6579874 | -0.381380897 | 0.36442076 | -1.04654 | 0.2953118 | 0.99999872 |
| IDH3A | 15.1912621 | -0.145328216 | 0.44650116 | -0.3254823 | 0.74481609 | 0.99999872 |
| DNAJA4 | 5.49529851 | -0.11165161 | 0.62462533 | -0.1787497 | 0.85813422 | 0.99999872 |
| WDR61 | 17.058582 | 0.009622525 | 0.5222297 | 0.01842585 | 0.98529913 | 0.99999872 |
| IREB2 | 43.9626536 | 0.023039519 | 0.29694877 | 0.07758752 | 0.93815617 | 0.99999872 |
| PSMA4 | 32.7498919 | 0.068128311 | 0.49265696 | 0.13828752 | 0.89001319 | 0.99999872 |
| MORF4L1 | 115.447968 | 0.478548101 | 0.3458524 | 1.38367726 | 0.1664573 | 0.99999872 |
| CTSH | 11.9187659 | -0.176751515 | 0.62278889 | -0.2838065 | 0.7765587 | 0.99999872 |
| TMED3 | 38.1986679 | 1.065156623 | 0.62191921 | 1.71269291 | 0.08676905 | 0.99999872 |
| MTHFS | 9.08771142 | -0.12355821 | 0.54158523 | -0.2281418 | 0.81953603 | 0.99999872 |
| ZFAND6 | 116.614091 | 0.210078217 | 0.25326396 | 0.82948328 | 0.40683099 | 0.99999872 |
| FAH | 9.1348979 | 0.044705297 | 0.47407895 | 0.09429927 | 0.92487143 | 0.99999872 |
| ARNT2 | 22.2670516 | -0.151373463 | 0.55090215 | -0.2747738 | 0.78349005 | 0.99999872 |
| ABHD17C | 11.6823883 | -0.441048136 | 0.48870177 | -0.9024893 | 0.36679698 | 0.99999872 |
| CEMIP | 7.02969039 | 1.913955243 | 0.87482371 | 2.1878182 | 0.02868285 | 0.90266962 |
| AC027808.1 | 9.42519579 | -0.010639883 | 0.50904872 | -0.0209015 | 0.98332423 | 0.99999872 |
| MESD | 38.0575099 | 0.303105244 | 0.41232441 | 0.7351135 | 0.46227037 | 0.99999872 |
| TLNRD1 | 9.07241966 | -0.112109962 | 0.5730961 | -0.1956216 | 0.84490637 | 0.99999872 |
| AC104041.1 | 17.6381964 | -0.111520653 | 1.01585854 | -0.1097797 | 0.91258409 | 0.99999872 |
| EFL1 | 19.7469821 | -0.48596773 | 0.39131588 | -1.2418809 | 0.21428049 | 0.99999872 |
| RPS17 | 6.4333834 | 0.68270533 | 0.88786625 | 0.76892812 | 0.44193598 | 0.99999872 |
| CPEB1 | 17.0967307 | 0.620577941 | 0.82717776 | 0.75023528 | 0.45311301 | 0.99999872 |
| AP3B2 | 4.49170098 | -0.502709369 | 0.89607757 | -0.561011 | 0.57479004 | NA |
| WHAMM | 20.1828384 | -0.384360152 | 0.5352263 | -0.7181264 | 0.47267934 | 0.99999872 |
| HOMER2 | 6.20511157 | -0.651770637 | 0.64553057 | -1.0096666 | 0.31265506 | 0.99999872 |
| C15orf40 | 13.3834589 | 0.528144935 | 0.43401973 | 1.21686849 | 0.22365426 | 0.99999872 |
| BTBD1 | 55.7839795 | -0.261340373 | 0.34232578 | -0.7634259 | 0.44520943 | 0.99999872 |
| HDGFL3 | 33.1171916 | -0.004847191 | 0.37356154 | -0.0129756 | 0.98964725 | 0.99999872 |
| WDR73 | 23.3584559 | -0.364512665 | 0.35558536 | -1.0251059 | 0.3053132 | 0.99999872 |
| SEC11A | 54.6654402 | 0.26751064 | 0.3074634 | 0.87005687 | 0.38426933 | 0.99999872 |
| ZNF592 | 22.6722103 | -0.323509126 | 0.32098981 | -1.0078486 | 0.31352716 | 0.99999872 |
| ALPK3 | 13.8945714 | -0.023453814 | 0.41865014 | -0.0560225 | 0.95532391 | 0.99999872 |
| SLC28A1 | 10.8024608 | 1.676290106 | 1.2237254 | 1.36982538 | 0.17074142 | 0.99999872 |
| PDE8A | 107.064582 | -0.239710977 | 0.35738513 | -0.6707357 | 0.5023889 | 0.99999872 |
| AKAP13 | 142.629619 | 0.219256487 | 0.29441087 | 0.74472958 | 0.45643521 | 0.99999872 |
| AC087286.2 | 5.43184719 | 0.895957444 | 0.77123366 | 1.16171984 | 0.24534928 | 0.99999872 |
| KLHL25 | 4.53406263 | 1.162305227 | 0.74633425 | 1.55735213 | 0.11938691 | NA |
| MRPL46 | 10.1582845 | 0.1684534 | 0.56468253 | 0.29831523 | 0.76546258 | 0.99999872 |
| MRPS11 | 14.5473658 | -0.253365704 | 0.42168783 | -0.6008371 | 0.54794847 | 0.99999872 |
| DET1 | 11.8055128 | 0.138174762 | 0.56703599 | 0.243679 | 0.80747944 | 0.99999872 |
| ISG20 | 16.417041 | -1.553959863 | 3.15252555 | -0.4929254 | 6.22E-01 | 0.99999872 |
| MFGE8 | 6.80733427 | -1.316370188 | 0.82408411 | -1.5973736 | 0.11018246 | 0.99999872 |
| ABHD2 | 20.5192384 | -0.287822185 | 0.40025651 | -0.7190943 | 0.4720828 | 0.99999872 |
| POLG | 10.0806392 | 0.433112336 | 0.48407698 | 0.89471789 | 0.37093792 | 0.99999872 |
| PEX11A | 4.13775665 | 1.037010611 | 0.81409188 | 1.27382503 | 0.20272543 | NA |
| ANPEP | 16.4346283 | 0.051803995 | 0.62907553 | 0.0823494 | 0.93436887 | 0.99999872 |
| AP3S2 | 8.79760742 | -0.310539392 | 0.60323017 | -0.5147942 | 0.60669684 | 0.99999872 |

| | | | | | | |
|---|---|---|---|---|---|---|
| ZNF710 | 9.4269413 | 0.012420045 | 0.64186088 | 0.01935006 | 0.98456185 | 0.99999872 |
| IDH2 | 11.3467712 | 1.327138046 | 0.86242793 | 1.53883937 | 0.12384352 | 0.99999872 |
| SEMA4B | 26.8421318 | -0.946264422 | 0.48019244 | -1.9705942 | 0.04877031 | 0.97937581 |
| CIB1 | 34.6393118 | 0.446390065 | 0.39702398 | 1.12434031 | 0.26086869 | 0.99999872 |
| AC091167.6 | 6.50470313 | -0.114059429 | 0.62166331 | -0.1834746 | 0.85442564 | 0.99999872 |
| IQGAP1 | 110.542937 | -0.198205689 | 0.28519971 | -0.6949716 | 0.48707312 | 0.99999872 |
| CRTC3 | 47.4270087 | 0.476113494 | 0.30151926 | 1.57904838 | 0.11432496 | 0.99999872 |
| FURIN | 13.9323153 | -0.23659379 | 0.5912086 | -0.4001867 | 0.68901905 | 0.99999872 |
| MAN2A2 | 13.5376245 | 1.164031134 | 0.57980895 | 2.00761152 | 0.04468459 | 0.95980711 |
| HDDC3 | 6.13965915 | 0.10970637 | 0.62369933 | 0.17589624 | 0.86037546 | 0.99999872 |
| UNC45A | 19.3851133 | 0.411564433 | 0.43307589 | 0.95032867 | 0.34194528 | 0.99999872 |
| SLCO3A1 | 12.7206375 | -0.019121385 | 0.538575 | -0.0355037 | 0.97167813 | 0.99999872 |
| FAM174B | 18.0569599 | -1.170271403 | 0.38518366 | -3.0382167 | 0.00237983 | 0.38962243 |
| AC013394.1 | 69.0923689 | -0.06299734 | 0.27750917 | -0.2270099 | 0.82041601 | 0.99999872 |
| CHD2 | 60.8170078 | -0.269583542 | 0.36634164 | -0.7358801 | 0.46180364 | 0.99999872 |
| RGMA | 8.06004922 | 0.796663882 | 0.93899066 | 0.84842577 | 0.39620089 | 0.99999872 |
| MCTP2 | 95.8423031 | 0.603424374 | 0.46696572 | 1.29222414 | 0.19627953 | 0.99999872 |
| LINC01197 | 17.1772956 | -0.005974712 | 0.58679519 | -0.0101819 | 0.99187613 | 0.99999872 |
| AC012409.2 | 21.9213394 | 0.893349976 | 0.84733912 | 1.0543004 | 0.2917454 | 0.99999872 |
| NR2F2-AS1 | 172.104491 | 0.005637369 | 0.48213799 | 0.01169244 | 0.990671 | 0.99999872 |
| NR2F2 | 82.9529235 | -0.129444384 | 0.4726292 | -0.2738815 | 0.7841757 | 0.99999872 |
| AC087477.2 | 14.330243 | -1.121035517 | 0.55705817 | -2.0124209 | 0.04417558 | 0.95980711 |
| ARRDC4 | 174.942218 | -0.070880937 | 0.68620831 | -0.1032936 | 0.91772994 | 0.99999872 |
| IGF1R | 481.36896 | -0.389461453 | 0.29562664 | -1.3174099 | 0.18770127 | 0.99999872 |
| AC036108.1 | 9.4087403 | 1.901448159 | 0.79373915 | 2.39555798 | 0.01659509 | 0.81253899 |
| TTC23 | 20.5212894 | -0.134467828 | 0.35766613 | -0.3759591 | 0.70694732 | 0.99999872 |
| LRRC28 | 37.7004681 | 0.026821561 | 0.31595423 | 0.08489065 | 0.93234832 | 0.99999872 |
| MEF2A | 101.869565 | -0.186213691 | 0.3151016 | -0.590964 | 0.55454457 | 0.99999872 |
| LYSMD4 | 4.75402509 | 1.107637314 | 0.69749442 | 1.58802319 | 0.11228109 | 0.99999872 |
| AC090825.1 | 34.215159 | 0.182297723 | 0.65496814 | 0.27833067 | 0.78075853 | 0.99999872 |
| LINS1 | 8.03454096 | 0.0017607 | 0.59108122 | 0.00297878 | 0.99762328 | 0.99999872 |
| ASB7 | 11.4777186 | 0.204046279 | 0.44834985 | 0.45510504 | 0.64903367 | 0.99999872 |
| AC087762.1 | 10.6320774 | 1.833751342 | 1.11587088 | 1.6433365 | 0.10031333 | 0.99999872 |
| CHSY1 | 28.9311843 | -0.144595778 | 0.32024146 | -0.4515211 | 0.65161402 | 0.99999872 |
| SELENOS | 61.4020278 | 0.333964437 | 0.42460479 | 0.78653008 | 0.43155701 | 0.99999872 |
| SNRPA1 | 16.1448507 | 0.082186753 | 0.41622571 | 0.19745717 | 0.84346979 | 0.99999872 |
| PCSK6 | 15.0482464 | 0.252805093 | 1.28309188 | 0.19702805 | 0.84380559 | 0.99999872 |
| TM2D3 | 31.8735746 | 0.335080924 | 0.31078761 | 1.07816692 | 0.28095927 | 0.99999872 |
| TARSL2 | 50.0965747 | -0.761612832 | 0.33773859 | -2.2550364 | 0.02413104 | 0.88337526 |
| POLR3K | 6.16457172 | 0.160652857 | 0.68361375 | 0.2350053 | 0.81420462 | 0.99999872 |
| SNRNP25 | 5.12349894 | 0.433784189 | 0.65693481 | 0.66031543 | 0.50905143 | 0.99999872 |
| RHBDF1 | 4.80337207 | -0.317847752 | 0.75145699 | -0.4229753 | 0.67231328 | 0.99999872 |
| MPG | 17.7297239 | 0.319714879 | 0.43137455 | 0.74115378 | 0.45860021 | 0.99999872 |
| NPRL3 | 19.893921 | -0.023440647 | 0.5039399 | -0.0465148 | 0.96289996 | 0.99999872 |
| LUC7L | 48.42624 | -0.34320879 | 0.39383977 | -0.8714427 | 0.38351249 | 0.99999872 |
| RGS11 | 7.35464661 | 2.661256497 | 0.9563849 | 2.78262078 | 0.00539218 | 0.55881811 |
| AXIN1 | 12.6494395 | -0.500265759 | 0.42927264 | -1.16538 | 0.2438652 | 0.99999872 |
| MRPL28 | 15.0817785 | 0.096596863 | 0.62080824 | 0.15559855 | 0.87634947 | 0.99999872 |
| TMEM8A | 11.8498235 | 0.463552632 | 0.45197802 | 1.0256088 | 0.30507602 | 0.99999872 |
| NME4 | 7.80048124 | 1.197571583 | 0.59160289 | 2.02428283 | 0.04294106 | 0.95980711 |
| RAB11FIP3 | 52.7844119 | -0.704810529 | 0.62057484 | -1.1357382 | 0.25606617 | 0.99999872 |
| CAPN15 | 7.58400149 | -0.774908586 | 0.51397182 | -1.5076869 | 0.13163468 | 0.99999872 |
| PIGQ | 16.5890079 | -0.575083315 | 0.49464599 | -1.1626159 | 0.24498536 | 0.99999872 |
| RAB40C | 21.7681026 | -0.67130232 | 0.39308796 | -1.7077662 | 0.08767974 | 0.99999872 |
| METTL26 | 12.1042463 | 0.78214106 | 0.62876267 | 1.24393686 | 0.2135228 | 0.99999872 |
| MCRIP2 | 10.8329069 | 0.311329494 | 0.50004081 | 0.62260818 | 0.53354204 | 0.99999872 |
| RHOT2 | 14.1696356 | 0.05784352 | 0.44302891 | 0.13056376 | 0.89612041 | 0.99999872 |
| STUB1 | 37.6764305 | 0.276923465 | 0.44226936 | 0.6261421 | 0.53122174 | 0.99999872 |
| JMJD8 | 26.3324435 | -0.297788514 | 0.60064476 | -0.4957814 | 0.62004863 | 0.99999872 |
| METRN | 11.6346351 | 0.293521481 | 0.53522934 | 0.54840319 | 0.58341508 | 0.99999872 |
| FAM173A | 13.1478015 | 0.214377157 | 0.54440861 | 0.39377988 | 0.69374356 | 0.99999872 |

| | | | | | | |
|---|---|---|---|---|---|---|
| NARFL | 5.35154468 | 0.081880068 | 0.79641289 | 0.10281108 | 0.91811291 | 0.99999872 |
| MSLN | 11.8922984 | 1.598264956 | 1.07242364 | 1.49032984 | 0.13613753 | 0.99999872 |
| LMF1 | 17.809234 | -0.456576139 | 0.46278191 | -0.9865903 | 0.32384353 | 0.99999872 |
| UBE2I | 31.2136266 | 0.180618202 | 0.36169557 | 0.49936526 | 0.61752209 | 0.99999872 |
| TSR3 | 16.4660502 | 0.31130676 | 0.38992624 | 0.79837345 | 0.42465381 | 0.99999872 |
| GNPTG | 12.4531256 | 0.071505895 | 0.4235542 | 0.16882348 | 0.86593549 | 0.99999872 |
| UNKL | 11.424964 | -0.080603855 | 0.77597672 | -0.1038741 | 0.91726929 | 0.99999872 |
| C16orf91 | 6.35780573 | 0.213844286 | 0.62825713 | 0.34037702 | 0.73357263 | 0.99999872 |
| CLCN7 | 17.7438015 | -0.673313038 | 0.44531092 | -1.5120066 | 0.13053219 | 0.99999872 |
| TELO2 | 7.36009621 | -1.212269209 | 0.54379854 | -2.2292616 | 0.0257965 | 0.88365618 |
| IFT140 | 18.300926 | -0.218421307 | 0.41506016 | -0.5262401 | 0.59872138 | 0.99999872 |
| JPT2 | 11.8759468 | -0.704423672 | 0.54931378 | -1.2823703 | 0.19971278 | 0.99999872 |
| MAPK8IP3 | 23.9194532 | -0.154854347 | 0.37650606 | -0.4112931 | 0.68085762 | 0.99999872 |
| NME3 | 48.6617323 | 0.552731224 | 0.39245658 | 1.40838822 | 0.15901615 | 0.99999872 |
| MRPS34 | 23.0985985 | 0.365296109 | 0.54568753 | 0.66942359 | 0.50322531 | 0.99999872 |
| SPSB3 | 20.0367039 | -0.009275975 | 0.37495225 | -0.0247391 | 0.98026308 | 0.99999872 |
| NUBP2 | 11.1652148 | 0.475045746 | 0.49772826 | 0.95442792 | 0.33986708 | 0.99999872 |
| HAGH | 30.8206285 | -0.131869683 | 0.32396558 | -0.4070484 | 0.68397242 | 0.99999872 |
| FAHD1 | 7.88797268 | -0.644462092 | 0.6597256 | -0.9768638 | 0.32863656 | 0.99999872 |
| MSRB1 | 7.34950106 | -0.870077447 | 0.54473532 | -1.5972481 | 0.11021042 | 0.99999872 |
| NDUFB10 | 30.9099383 | 0.152153043 | 0.3636533 | 0.41840138 | 0.67565368 | 0.99999872 |
| RPS2 | 61.8419026 | 0.78871551 | 0.52660138 | 1.49774676 | 0.13419906 | 0.99999872 |
| SNHG9 | 6.64917447 | 0.061302308 | 0.62494431 | 0.09809243 | 0.9218589 | 0.99999872 |
| GFER | 4.71420777 | -0.186381817 | 0.89923908 | -0.2072661 | 0.83580201 | 0.99999872 |
| ZNF598 | 6.42565351 | -0.799592415 | 0.67815483 | -1.1790706 | 0.23837007 | 0.99999872 |
| SLC9A3R2 | 68.9201796 | -1.008912908 | 0.58089194 | -1.7368341 | 0.08241646 | 0.99999872 |
| NTHL1 | 9.92639073 | -0.048898769 | 0.58776895 | -0.0831939 | 0.93369739 | 0.99999872 |
| TSC2 | 22.901423 | 0.138886264 | 0.38725763 | 0.35864048 | 0.71986406 | 0.99999872 |
| PKD1 | 33.4362876 | 0.487603966 | 0.47708539 | 1.02204757 | 0.30675839 | 0.99999872 |
| SNHG19 | 6.55487062 | 0.332754661 | 0.77604522 | 0.42878257 | 0.66808147 | 0.99999872 |
| TRAF7 | 8.21321017 | 0.132499547 | 0.52622085 | 0.25179456 | 0.80119986 | 0.99999872 |
| MLST8 | 4.6668601 | 0.005981212 | 0.74548738 | 0.00802322 | 0.99359846 | 0.99999872 |
| PGP | 17.7872172 | 0.776807155 | 0.38693996 | 2.00756507 | 0.04468953 | 0.95980711 |
| E4F1 | 6.1714115 | 0.145634576 | 0.63761814 | 0.22840407 | 0.81933213 | 0.99999872 |
| ECI1 | 23.686066 | -0.093297423 | 0.37668443 | -0.2476806 | 0.80438155 | 0.99999872 |
| RNPS1 | 32.5126565 | 0.227141344 | 0.41240835 | 0.55076806 | 0.58179268 | 0.99999872 |
| ABCA3 | 20.8597073 | 0.344007958 | 0.56697912 | 0.60673832 | 0.54402459 | 0.99999872 |
| PDPK1 | 50.4826284 | -0.073872399 | 0.26157915 | -0.2824093 | 0.77762965 | 0.99999872 |
| AC093525.4 | 4.91749462 | -1.127322544 | 0.73864356 | -1.5262064 | 0.12695847 | 0.99999872 |
| KCTD5 | 6.75573773 | -0.707662226 | 0.58665593 | -1.2062645 | 0.22771552 | 0.99999872 |
| SRRM2 | 146.392668 | -0.019580487 | 0.23502532 | -0.0833122 | 0.93360326 | 0.99999872 |
| ELOB | 88.7404803 | 0.392596479 | 0.44108913 | 0.89006155 | 0.37343284 | 0.99999872 |
| FLYWCH2 | 22.8264083 | -0.252644093 | 0.54348181 | -0.4648621 | 0.64203021 | 0.99999872 |
| FLYWCH1 | 14.3355193 | -0.036587241 | 0.44523189 | -0.0821757 | 0.934507 | 0.99999872 |
| TNFRSF12A | 30.7100536 | -1.891296108 | 0.92499027 | -2.0446659 | 0.0408878 | 0.95980711 |
| HCFC1R1 | 45.32421 | 0.064345614 | 0.46584775 | 0.13812585 | 0.89014096 | 0.99999872 |
| THOC6 | 4.87325276 | -0.580074025 | 0.64962788 | -0.8929328 | 0.37189317 | 0.99999872 |
| IL32 | 13.8918995 | -0.049697881 | 0.67905209 | -0.0731871 | 0.9416572 | 0.99999872 |
| ZNF213-AS1 | 12.1632763 | -0.539726078 | 0.43004331 | -1.2550505 | 0.20946042 | 0.99999872 |
| ZNF205 | 5.99145481 | 0.244057259 | 0.61641618 | 0.39592935 | 0.69215715 | 0.99999872 |
| ZNF213 | 17.883171 | 0.03610305 | 0.45418869 | 0.0794891 | 0.9366436 | 0.99999872 |
| ZNF263 | 11.4361628 | -0.216819008 | 0.4194225 | -0.5169465 | 0.60519349 | 0.99999872 |
| TIGD7 | 6.4884761 | 0.13409403 | 0.59443795 | 0.2255812 | 0.82152716 | 0.99999872 |
| ZNF75A | 19.8873989 | -0.199470361 | 0.44743382 | -0.4458098 | 0.65573467 | 0.99999872 |
| AC025283.3 | 6.98570474 | -0.26710789 | 0.56723965 | -0.4708907 | 0.63771877 | 0.99999872 |
| ZSCAN32 | 6.18026361 | 0.04553106 | 0.57818243 | 0.07874861 | 0.93723258 | 0.99999872 |
| NAA60 | 16.515068 | 0.153135536 | 0.42076335 | 0.36394695 | 0.71589762 | 0.99999872 |
| CLUAP1 | 24.6249283 | -0.135725012 | 0.32776858 | -0.4140879 | 0.67880971 | 0.99999872 |
| DNASE1 | 23.7780517 | 0.320177068 | 0.42196281 | 0.7587803 | 0.44798399 | 0.99999872 |
| TRAP1 | 20.5075515 | -0.048548796 | 0.36333911 | -0.1336184 | 0.89370432 | 0.99999872 |
| CREBBP | 78.5033121 | -0.067017671 | 0.33905639 | -0.1976594 | 0.84331158 | 0.99999872 |

| | | | | | | |
|---|---|---|---|---|---|---|
| ADCY9 | 41.1800518 | -0.003848489 | 0.38296266 | -0.0100493 | 0.99198199 | 0.99999872 |
| GLIS2 | 10.2080852 | -0.664848229 | 0.46939285 | -1.4164004 | 0.1566583 | 0.99999872 |
| VASN | 145.464219 | 0.221505642 | 0.50438392 | 0.43916079 | 0.66054503 | 0.99999872 |
| DNAJA3 | 14.5011854 | -0.015783916 | 0.43401894 | -0.0363669 | 0.97098982 | 0.99999872 |
| NMRAL1 | 9.70492068 | -0.999387442 | 0.53318014 | -1.8743898 | 0.06087671 | 0.99999872 |
| HMOX2 | 10.3466638 | 0.189916367 | 0.50889651 | 0.37319251 | 0.70900517 | 0.99999872 |
| CDIP1 | 8.17322512 | -0.026832273 | 0.53614062 | -0.0500471 | 0.96008487 | 0.99999872 |
| UBALD1 | 6.97672348 | -0.646939732 | 0.62131832 | -1.0412372 | 0.29776548 | 0.99999872 |
| MGRN1 | 24.0529413 | -0.649051375 | 0.32174879 | -2.0172613 | 0.04366825 | 0.95980711 |
| NUDT16L1 | 9.80343992 | 0.467367364 | 0.57338152 | 0.81510712 | 0.41501099 | 0.99999872 |
| ANKS3 | 18.0711513 | -0.461001327 | 0.37863624 | -1.2175309 | 0.22340229 | 0.99999872 |
| ZNF500 | 8.81241135 | -0.950380767 | 0.65913533 | -1.4418599 | 0.14934191 | 0.99999872 |
| ROGDI | 5.73592472 | -0.519870775 | 0.65248045 | -0.7967607 | 0.42559003 | 0.99999872 |
| GLYR1 | 35.238014 | -0.488181929 | 0.35843197 | -1.3619933 | 0.17320001 | 0.99999872 |
| UBN1 | 13.4847237 | -0.192443992 | 0.48718152 | -0.395015 | 0.69283184 | 0.99999872 |
| ALG1 | 4.96494001 | -0.367756424 | 0.63811782 | -0.5763143 | 0.56440276 | 0.99999872 |
| RBFOX1 | 562.989914 | -0.799136327 | 0.75492433 | -1.0585648 | 0.28979802 | 0.99999872 |
| METTL22 | 7.03613938 | -0.992318571 | 0.55205696 | -1.7974931 | 0.07225737 | 0.99999872 |
| ABAT | 12.6840185 | -0.231087239 | 0.56435528 | -0.4094712 | 0.68219389 | 0.99999872 |
| PMM2 | 20.1686633 | 0.142802473 | 0.35872337 | 0.39808523 | 0.69056736 | 0.99999872 |
| CARHSP1 | 75.2519598 | 0.083757767 | 0.42085846 | 0.19901647 | 0.84224986 | 0.99999872 |
| USP7 | 34.823268 | -0.224104209 | 0.32793186 | -0.6833865 | 0.49436265 | 0.99999872 |
| C16orf72 | 42.3163963 | -0.295151563 | 0.32442865 | -0.909758 | 0.36295014 | 0.99999872 |
| AC087190.1 | 5.11276078 | 0.865931468 | 0.6890746 | 1.25665853 | 0.20887731 | 0.99999872 |
| ATF7IP2 | 30.1570975 | 0.111189665 | 0.3644511 | 0.30508803 | 0.76029911 | 0.99999872 |
| EMP2 | 12.1736429 | 0.176275722 | 0.61869378 | 0.28491594 | 0.77570854 | 0.99999872 |
| NUBP1 | 6.69172644 | -0.231373249 | 0.59531997 | -0.3886536 | 0.69753241 | 0.99999872 |
| DEXI | 17.3125786 | 0.816127024 | 0.51523055 | 1.58400357 | 0.11319291 | 0.99999872 |
| CLEC16A | 181.457293 | 0.056098302 | 0.47719336 | 0.11755885 | 0.90641721 | 0.99999872 |
| AC007014.1 | 8.55123031 | 1.595224984 | 0.78958716 | 2.02032793 | 0.04334938 | 0.95980711 |
| AC007220.1 | 7.83582292 | -0.591103529 | 0.72933326 | -0.8104711 | 0.41766947 | 0.99999872 |
| AC099489.1 | 6.06752588 | 0.545906995 | 1.2657938 | 0.4312764 | 0.66626741 | 0.99999872 |
| LITAF | 91.6081313 | 0.867609709 | 0.69117874 | 1.25526098 | 0.20938403 | 0.99999872 |
| SNN | 3.97050702 | 0.635361708 | 0.91370519 | 0.69536839 | 0.48682447 | NA |
| TXNDC11 | 31.0369206 | 0.056277453 | 0.30829455 | 0.18254443 | 0.85515549 | 0.99999872 |
| ZC3H7A | 44.8433294 | 0.157799851 | 0.28358082 | 0.5564546 | 0.57790012 | 0.99999872 |
| RSL1D1 | 14.9575731 | -0.300355795 | 0.53973589 | -0.5564866 | 0.57787825 | 0.99999872 |
| AC007216.4 | 15.1564483 | -0.146358567 | 0.54614214 | -0.2679862 | 0.78870993 | 0.99999872 |
| GSPT1 | 46.6721069 | 0.408494029 | 0.42319168 | 0.96526952 | 0.33440983 | 0.99999872 |
| SNX29 | 95.3849964 | -0.364232438 | 0.52947407 | -0.6879136 | 0.49150716 | 0.99999872 |
| CPPED1 | 14.5411661 | -0.790107627 | 0.43241433 | -1.8272004 | 0.06766965 | 0.99999872 |
| ERCC4 | 9.30083673 | 0.057788908 | 0.45822708 | 0.12611413 | 0.89964159 | 0.99999872 |
| MKL2 | 35.8835337 | 0.100887819 | 0.50831251 | 0.19847597 | 0.84267268 | 0.99999872 |
| PARN | 39.7581317 | -0.423075466 | 0.33474823 | -1.2638617 | 0.20627967 | 0.99999872 |
| BFAR | 14.0769564 | 0.280094891 | 0.49333329 | 0.56775996 | 0.57019797 | 0.99999872 |
| NOMO1 | 9.65528664 | 0.443636413 | 0.71956357 | 0.6165354 | 0.53754122 | 0.99999872 |
| PDXDC1 | 89.1509276 | 0.151741432 | 0.28851729 | 0.52593532 | 0.59893315 | 0.99999872 |
| NTAN1 | 10.5160963 | 0.403991036 | 0.70581483 | 0.57237539 | 0.56706769 | 0.99999872 |
| RRN3 | 14.3625793 | -0.380499005 | 0.4478306 | -0.8496494 | 0.39552004 | 0.99999872 |
| C16orf45 | 36.5793426 | -0.354439031 | 0.31891923 | -1.1113756 | 0.26640673 | 0.99999872 |
| MARF1 | 27.5117941 | -0.734889995 | 0.36819463 | -1.9959281 | 0.04594175 | 0.95980711 |
| NDE1 | 8.90422207 | -0.530355796 | 0.4989243 | -1.0629985 | 0.28778262 | 0.99999872 |
| FOPNL | 13.2568537 | -0.307312775 | 0.45497769 | -0.6754458 | 0.49939257 | 0.99999872 |
| ABCC1 | 53.544351 | -0.650527542 | 0.34831389 | -1.8676475 | 0.06181123 | 0.99999872 |
| XYLT1 | 52.0480861 | 0.537684041 | 0.87025465 | 0.61784679 | 0.53667634 | 0.99999872 |
| RPS15A | 18.2854201 | 1.25643668 | 0.55780389 | 2.25247025 | 0.02429257 | 0.88337526 |
| SMG1 | 89.7016363 | -0.374139186 | 0.28426704 | -1.3161539 | 0.18812238 | 0.99999872 |
| COQ7 | 8.71458121 | -0.455466409 | 0.55354916 | -0.8228111 | 0.4106154 | 0.99999872 |
| GDE1 | 23.5503776 | 0.274078018 | 0.42135742 | 0.65046444 | 0.51539226 | 0.99999872 |
| CCP110 | 12.6094688 | 0.307460528 | 0.48814891 | 0.62984988 | 0.52879281 | 0.99999872 |
| VPS35L | 38.304917 | -0.593485853 | 0.28452192 | -2.0859056 | 0.03698718 | 0.95590735 |

| | | | | | | |
|---|---|---|---|---|---|---|
| KNOP1 | 7.25651616 | -0.338157655 | 0.59739313 | -0.5660555 | 0.57135607 | 0.99999872 |
| IQCK | 25.2423479 | -0.341444167 | 0.35780471 | -0.9542752 | 0.33994435 | 0.99999872 |
| UMOD | 52.7840892 | -0.558794347 | 0.72927076 | -0.7662371 | 0.44353521 | 0.99999872 |
| ACSM2A | 30.5962524 | 3.755122917 | 2.25879108 | 1.66244809 | 0.09642296 | 0.99999872 |
| ACSM2B | 31.7112123 | 2.581772579 | 1.91268727 | 1.34981427 | 0.17707557 | 0.99999872 |
| ACSM3 | 11.2519729 | 0.657209763 | 0.92612767 | 0.70963193 | 0.47793241 | 0.99999872 |
| THUMPD1 | 22.4126202 | 0.05793865 | 0.34322692 | 0.16880567 | 0.8659495 | 0.99999872 |
| ERI2 | 10.0212765 | 0.351800949 | 0.6344538 | 0.5544942 | 0.57924068 | 0.99999872 |
| LYRM1 | 10.7442486 | 0.70807255 | 0.66479852 | 1.06509345 | 0.28683365 | 0.99999872 |
| TMEM159 | 6.28273449 | 0.018019366 | 0.56030593 | 0.03215987 | 0.97434456 | 0.99999872 |
| METTL9 | 27.1180745 | 0.047520342 | 0.37454853 | 0.12687366 | 0.8990404 | 0.99999872 |
| NPIPB4 | 8.56055463 | -1.104373468 | 0.59142482 | -1.86731 | 0.06185831 | 0.99999872 |
| UQCRC2 | 28.7623811 | 0.321487598 | 0.33779138 | 0.95173418 | 0.34123181 | 0.99999872 |
| MOSMO | 70.8180544 | 0.00961033 | 0.58107512 | 0.01653888 | 0.98680449 | 0.99999872 |
| EEF2K | 13.2322096 | -0.355629143 | 0.55544745 | -0.640257 | 0.5220055 | 0.99999872 |
| POLR3E | 12.8753177 | -0.382935268 | 0.47020817 | -0.8143952 | 0.4154186 | 0.99999872 |
| CDR2 | 64.8738628 | -0.25415528 | 0.32366701 | -0.7852369 | 0.43231469 | 0.99999872 |
| NPIPB5 | 23.9985224 | 0.143010418 | 0.33823678 | 0.42281156 | 0.67243275 | 0.99999872 |
| USP31 | 35.8241867 | -0.447969909 | 0.37275032 | -1.2017962 | 0.2294425 | 0.99999872 |
| SCNN1G | 9.31517193 | -1.343992106 | 1.46293001 | -0.9186988 | 0.35825312 | 0.99999872 |
| SCNN1B | 9.20224446 | -0.148853189 | 0.97400964 | -0.1528252 | 0.87853615 | 0.99999872 |
| COG7 | 43.0340866 | -0.50916241 | 0.43848131 | -1.1611952 | 0.24556251 | 0.99999872 |
| AC008915.2 | 4.21075254 | -0.680860033 | 0.81323062 | -0.8372287 | 0.40246402 | NA |
| GGA2 | 28.6874693 | -0.366856449 | 0.38624627 | -0.9497993 | 0.34221423 | 0.99999872 |
| EARS2 | 6.12603289 | -0.025645757 | 0.57488037 | -0.0446106 | 0.96441769 | 0.99999872 |
| UBFD1 | 14.918404 | 0.147587862 | 0.49492833 | 0.29820047 | 0.76555016 | 0.99999872 |
| NDUFAB1 | 19.9503039 | 0.928561072 | 0.43871422 | 2.1165511 | 0.03429796 | 0.9407698 |
| PALB2 | 3.99585773 | 0.232023221 | 0.72284613 | 0.32098563 | 0.74822128 | NA |
| DCTN5 | 22.6374085 | -0.210958633 | 0.42789008 | -0.4930206 | 0.62199802 | 0.99999872 |
| RBBP6 | 49.7045119 | -0.081392316 | 0.34710442 | -0.2344894 | 0.81460504 | 0.99999872 |
| TNRC6A | 153.101569 | -0.216761286 | 0.25626382 | -0.8458521 | 0.39763526 | 0.99999872 |
| AC008731.1 | 4.99735557 | 0.688503813 | 0.87414243 | 0.78763345 | 0.43091114 | 0.99999872 |
| ARHGAP17 | 31.3439028 | 0.306559442 | 0.31820455 | 0.96340371 | 0.33534497 | 0.99999872 |
| LCMT1 | 22.3622877 | -0.019566997 | 0.34358986 | -0.0569487 | 0.95458606 | 0.99999872 |
| NSMCE1 | 14.179266 | 0.584398284 | 0.54840566 | 1.06563139 | 0.2865903 | 0.99999872 |
| IL4R | 33.0088809 | 0.533371783 | 0.85330836 | 0.62506335 | 0.53192948 | 0.99999872 |
| GTF3C1 | 47.3875731 | -0.369614393 | 0.33819454 | -1.0929047 | 0.27443566 | 0.99999872 |
| KIAA0556 | 47.5076 | -0.216809784 | 0.30055039 | -0.7213758 | 0.47067832 | 0.99999872 |
| XPO6 | 31.5356812 | -0.319391883 | 0.47358799 | -0.6744087 | 0.50005149 | 0.99999872 |
| NUPR1 | 583.510951 | -0.151713955 | 0.83638894 | -0.1813916 | 0.85606019 | 0.99999872 |
| SGF29 | 10.3623922 | -0.384713837 | 0.52100091 | -0.738413 | 0.46026353 | 0.99999872 |
| SULT1A2 | 8.00810219 | -0.762918739 | 0.554445 | -1.3760044 | 0.16882028 | 0.99999872 |
| SULT1A1 | 29.1907647 | -0.513101222 | 0.44233433 | -1.1599851 | 0.24605488 | 0.99999872 |
| ATXN2L | 19.7319562 | -0.194248762 | 0.43682188 | -0.4446864 | 0.65654638 | 0.99999872 |
| TUFM | 31.0517296 | 0.880697653 | 0.39903448 | 2.20707154 | 0.02730906 | 0.89154528 |
| SH2B1 | 20.487604 | -0.389697821 | 0.36687448 | -1.0622102 | 0.28814026 | 0.99999872 |
| RABEP2 | 9.12810502 | -0.612354274 | 0.53389426 | -1.146958 | 0.251399 | 0.99999872 |
| NFATC2IP | 38.7123675 | -0.316819363 | 0.31260668 | -1.013476 | 0.31083286 | 0.99999872 |
| SPNS1 | 8.11510001 | -0.102900882 | 0.57546922 | -0.1788121 | 0.85808521 | 0.99999872 |
| QPRT | 43.0242924 | 0.341373816 | 0.43356532 | 0.78736422 | 0.43106869 | 0.99999872 |
| KIF22 | 7.65847287 | -0.607579919 | 0.59315228 | -1.0243237 | 0.30568242 | 0.99999872 |
| MAZ | 16.6436627 | -0.448116746 | 0.41039357 | -1.0919195 | 0.27486848 | 0.99999872 |
| AC009133.1 | 6.79726891 | -0.826255701 | 0.55225511 | -1.4961486 | 0.13461494 | 0.99999872 |
| MVP | 35.7731564 | -0.051710291 | 0.46968517 | -0.1100956 | 0.91233352 | 0.99999872 |
| CDIPT | 23.0127108 | 0.58695265 | 0.51318792 | 1.14373825 | 0.2527322 | 0.99999872 |
| SEZ6L2 | 18.1072983 | 0.090580973 | 0.59767783 | 0.15155485 | 0.87953805 | 0.99999872 |
| KCTD13 | 12.6515388 | -0.293883602 | 0.45218138 | -0.6499242 | 0.51574122 | 0.99999872 |
| TMEM219 | 45.5915703 | 0.70431293 | 0.55019423 | 1.2801169 | 0.20050403 | 0.99999872 |
| TAOK2 | 8.2668477 | -0.431263265 | 0.60107281 | -0.7174892 | 0.47307228 | 0.99999872 |
| HIRIP3 | 5.02687385 | 0.096183516 | 0.65107925 | 0.14772935 | 0.88255637 | 0.99999872 |
| INO80E | 17.0798196 | 0.110760376 | 0.50339118 | 0.22002844 | 0.82584901 | 0.99999872 |

| | | | | | | |
|---|---|---|---|---|---|---|
| PPP4C | 17.3553793 | -0.064008231 | 0.41565384 | -0.1539941 | 0.8776144 | 0.99999872 |
| YPEL3 | 24.1369749 | 0.105133415 | 0.42928807 | 0.24490178 | 0.80653248 | 0.99999872 |
| MAPK3 | 14.0298905 | -0.065706663 | 0.5199772 | -0.1263645 | 0.8994434 | 0.99999872 |
| NPIPB13 | 6.94794304 | -0.936254938 | 0.58283652 | -1.6063766 | 0.1081912 | 0.99999872 |
| CD2BP2 | 14.4218774 | -0.54730903 | 0.42168425 | -1.2979119 | 0.19431759 | 0.99999872 |
| TBC1D10B | 8.48214096 | -0.715820183 | 0.56549692 | -1.2658251 | 0.20557572 | 0.99999872 |
| 1-Sep | 5.7476062 | 0.978900829 | 0.81440641 | 1.20198075 | 0.22937099 | 0.99999872 |
| DCTPP1 | 10.0170745 | 0.49037714 | 0.61840247 | 0.79297409 | 0.42779292 | 0.99999872 |
| SEPHS2 | 15.8435534 | 0.570343737 | 0.46482254 | 1.22701394 | 0.21981735 | 0.99999872 |
| ZNF768 | 7.15936362 | -0.678717123 | 0.55013436 | -1.2337297 | 0.21730363 | 0.99999872 |
| ZNF688 | 9.2335002 | -1.375649734 | 0.51039522 | -2.6952637 | 0.00703329 | 0.62327327 |
| ZNF785 | 10.1588574 | -0.518230928 | 0.45558471 | -1.1375073 | 0.25532628 | 0.99999872 |
| ZNF689 | 3.91840676 | 0.089485821 | 0.68743816 | 0.1301729 | 0.89642963 | NA |
| PRR14 | 11.8995235 | -0.394727233 | 0.44358139 | -0.8898643 | 0.37353877 | 0.99999872 |
| FBRS | 15.2906997 | -0.517803188 | 0.3897163 | -1.328667 | 0.18395785 | 0.99999872 |
| PHKG2 | 7.25624014 | 0.856552186 | 0.60111716 | 1.42493384 | 0.15417633 | 0.99999872 |
| RNF40 | 13.7721639 | -0.502986873 | 0.46150815 | -1.0898765 | 0.27576757 | 0.99999872 |
| ZNF629 | 6.67012051 | -0.753466337 | 0.58571562 | -1.286403 | 0.19830245 | 0.99999872 |
| BCL7C | 28.9860248 | -0.323725132 | 0.41495469 | -0.7801457 | 0.4353051 | 0.99999872 |
| MIR762HG | 6.24334219 | 0.002863263 | 0.55970413 | 0.00511567 | 0.9959183 | 0.99999872 |
| FBXL19 | 9.42877985 | -0.587137288 | 0.69763279 | -0.8416137 | 0.40000424 | 0.99999872 |
| ORAI3 | 8.14719267 | 0.079005762 | 0.67591063 | 0.11688788 | 0.9069489 | 0.99999872 |
| SETD1A | 7.52069393 | -0.330567961 | 0.50348803 | -0.6565557 | 0.51146661 | 0.99999872 |
| HSD3B7 | 4.00746187 | -1.017220907 | 0.79447212 | -1.2803733 | 0.20041387 | NA |
| STX4 | 10.2165934 | 0.722682383 | 0.50078225 | 1.44310701 | 0.14899033 | 0.99999872 |
| BCKDK | 10.6103281 | 1.209391377 | 0.56799409 | 2.12923234 | 0.03323504 | 0.92959829 |
| KAT8 | 15.7127875 | -0.101397676 | 0.52904009 | -0.1916635 | 0.8480058 | 0.99999872 |
| FUS | 59.3876112 | -0.608226629 | 0.33189272 | -1.8326001 | 0.06686206 | 0.99999872 |
| AC026471.4 | 9.08822843 | -1.109649708 | 0.64094759 | -1.7312644 | 0.08340463 | 0.99999872 |
| ARMC5 | 4.95083173 | -0.846605781 | 0.75709132 | -1.1182347 | 0.26346677 | 0.99999872 |
| TGFB1I1 | 21.0428693 | -0.256909041 | 0.53203988 | -0.4828755 | 0.62918412 | 0.99999872 |
| SLC5A2 | 4.86374473 | -0.237498597 | 0.91712072 | -0.2589611 | 0.79566525 | 0.99999872 |
| C16orf58 | 12.4013572 | 0.70154629 | 0.48563274 | 1.44460254 | 0.14856956 | 0.99999872 |
| ZNF720 | 19.0219555 | 0.275541131 | 0.40934909 | 0.67312018 | 0.50087084 | 0.99999872 |
| ZNF267 | 8.55062077 | 0.022866011 | 0.733035 | 0.03119361 | 0.97511513 | 0.99999872 |
| VPS35 | 36.045195 | 0.228071007 | 0.34989694 | 0.65182338 | 0.51451512 | 0.99999872 |
| MYLK3 | 51.9609535 | 0.825025591 | 0.81946963 | 1.00677995 | 0.31404053 | 0.99999872 |
| C16orf87 | 9.09814352 | 0.866546406 | 0.58333234 | 1.48551064 | 0.13740861 | 0.99999872 |
| GPT2 | 5.05304396 | -0.112407241 | 1.2033754 | -0.09341 | 0.92557788 | 0.99999872 |
| DNAJA2 | 18.3062869 | 0.260679445 | 0.41271857 | 0.63161549 | 0.52763816 | 0.99999872 |
| ITFG1 | 135.800232 | -0.178820785 | 0.21125944 | -0.8464511 | 0.39730114 | 0.99999872 |
| PHKB | 102.932994 | -0.116023857 | 0.28435839 | -0.4080198 | 0.68325913 | 0.99999872 |
| LONP2 | 93.9771091 | 0.12941046 | 0.22136667 | 0.58459776 | 0.55881821 | 0.99999872 |
| SIAH1 | 23.8819105 | 0.367894103 | 0.39988879 | 0.91999104 | 0.35757744 | 0.99999872 |
| N4BP1 | 59.1503701 | -0.280697583 | 0.40690451 | -0.6898365 | 0.49029701 | 0.99999872 |
| ZNF423 | 20.2549142 | -0.145295842 | 0.76696272 | -0.1894432 | 0.8497455 | 0.99999872 |
| CNEP1R1 | 9.35014401 | 0.727580307 | 0.64300276 | 1.13153528 | 0.25782986 | 0.99999872 |
| HEATR3 | 8.91563044 | -0.222079933 | 0.48459324 | -0.4582811 | 0.64675049 | 0.99999872 |
| TENT4B | 24.9426153 | 0.093160825 | 0.40638053 | 0.2292453 | 0.81867827 | 0.99999872 |
| ADCY7 | 6.22830172 | -0.780345307 | 0.79817072 | -0.9776672 | 0.32823897 | 0.99999872 |
| BRD7 | 27.289362 | -0.500979783 | 0.3233916 | -1.5491428 | 0.12134739 | 0.99999872 |
| NKD1 | 107.349806 | 0.356555844 | 0.49626959 | 0.71847207 | 0.47246626 | 0.99999872 |
| CYLD | 27.8149907 | 0.331104446 | 0.38556992 | 0.85874034 | 0.39048379 | 0.99999872 |
| SALL1 | 17.5325188 | 0.080315949 | 0.56488059 | 0.14218217 | 0.88693611 | 0.99999872 |
| AC087564.1 | 75.5369968 | -0.336218635 | 0.48291462 | -0.6962279 | 0.48628612 | 0.99999872 |
| HNRNPA1P4 | 5.46075718 | -0.918507764 | 0.61332363 | -1.4975907 | 0.13423963 | 0.99999872 |
| TOX3 | 10.3594631 | -0.381344298 | 0.66869781 | -0.570279 | 0.5684885 | 0.99999872 |
| CHD9 | 229.179895 | 0.081408827 | 0.27730816 | 0.29356809 | 0.76908796 | 0.99999872 |
| RBL2 | 48.0269298 | 0.215680691 | 0.3463041 | 0.62280721 | 0.53341122 | 0.99999872 |
| AKTIP | 12.3803025 | 0.340833057 | 0.48944382 | 0.69636809 | 0.48619834 | 0.99999872 |
| RPGRIP1L | 10.0995017 | -0.427659731 | 0.57601364 | -0.7424472 | 0.45781642 | 0.99999872 |

| | | | | | | |
|---|---|---|---|---|---|---|
| FTO | 137.144517 | 0.065662672 | 0.23636554 | 0.27780137 | 0.78116484 | 0.99999872 |
| LPCAT2 | 84.5224081 | 0.070913304 | 0.45909148 | 0.15446443 | 0.87724355 | 0.99999872 |
| AC007495.1 | 9.64286072 | -0.329218559 | 0.53717855 | -0.6128662 | 0.53996483 | 0.99999872 |
| GNAO1 | 18.6297398 | -1.300287052 | 0.86918292 | -1.4959878 | 0.13465683 | 0.99999872 |
| AMFR | 35.2103644 | 0.015314508 | 0.34999102 | 0.04375686 | 0.96509822 | 0.99999872 |
| NUDT21 | 27.2924805 | 0.004885675 | 0.33056807 | 0.01477963 | 0.98820799 | 0.99999872 |
| OGFOD1 | 12.0060573 | -0.236794838 | 0.48197045 | -0.4913057 | 0.62321024 | 0.99999872 |
| BBS2 | 24.9918089 | 0.072204654 | 0.39916544 | 0.18088904 | 0.85645467 | 0.99999872 |
| MT2A | 304.479635 | -1.81664347 | 1.03407882 | -1.7567747 | 0.07895622 | 0.99999872 |
| MT1E | 156.070017 | -2.352511929 | 0.96785707 | -2.4306398 | 0.01507219 | 0.78681466 |
| MT1M | 17.8514696 | -3.16032864 | 1.18726272 | -2.6618613 | 0.00777099 | 0.65139548 |
| MT1F | 18.562421 | -1.72313356 | 0.85021215 | -2.0267101 | 0.04269207 | 0.95980711 |
| MT1G | 155.413967 | -2.212283257 | 1.47492988 | -1.4999244 | 0.133634 | 0.99999872 |
| MT1H | 47.4033832 | -0.987837925 | 1.8455584 | -0.5352515 | 0.59247595 | 0.99999872 |
| MT1X | 69.6828609 | -1.760748054 | 0.95618894 | -1.8414227 | 0.06555963 | 0.99999872 |
| NUP93 | 9.71161373 | -0.047924616 | 0.52498807 | -0.0912871 | 0.9272645 | 0.99999872 |
| SLC12A3 | 5.65615126 | 0.435719059 | 1.19602387 | 0.36430632 | 0.71562928 | 0.99999872 |
| HERPUD1 | 70.9361847 | 0.789478212 | 0.52670671 | 1.49889531 | 0.13390079 | 0.99999872 |
| NLRC5 | 12.5015609 | -1.590418425 | 0.8229871 | -1.932495 | 0.05329844 | 0.99999872 |
| FAM192A | 25.0756612 | -0.472979612 | 0.33418308 | -1.4153308 | 0.15697152 | 0.99999872 |
| RSPRY1 | 21.8073744 | -0.268229769 | 0.41224713 | -0.6506529 | 0.5152706 | 0.99999872 |
| ARL2BP | 36.2284778 | 0.346567019 | 0.37720838 | 0.91876807 | 0.3582169 | 0.99999872 |
| CIAPIN1 | 12.2275461 | -0.131849562 | 0.46032083 | -0.2864297 | 0.77454902 | 0.99999872 |
| COQ9 | 12.1513417 | -0.354304287 | 0.47602766 | -0.7442935 | 0.45669894 | 0.99999872 |
| POLR2C | 11.0754224 | 0.854452138 | 0.62583165 | 1.36530669 | 0.17215668 | 0.99999872 |
| DOK4 | 8.04198678 | -0.009282332 | 0.67223495 | -0.0138082 | 0.98898303 | 0.99999872 |
| ADGRG1 | 11.0797241 | -0.363943262 | 0.49176245 | -0.7400794 | 0.45925182 | 0.99999872 |
| KATNB1 | 4.78168207 | 0.422371543 | 0.74064067 | 0.57027862 | 0.56848874 | 0.99999872 |
| KIFC3 | 28.3902007 | -1.238491643 | 0.61901965 | -2.0007307 | 0.04542141 | 0.95980711 |
| USB1 | 6.59183938 | 0.007188913 | 0.55974797 | 0.01284312 | 0.98975295 | 0.99999872 |
| MMP15 | 6.40413262 | 0.377227711 | 0.82697304 | 0.45615478 | 0.64827868 | 0.99999872 |
| CFAP20 | 9.80253019 | 0.505595465 | 0.48921292 | 1.03348757 | 0.30137579 | 0.99999872 |
| CSNK2A2 | 26.0144234 | 0.157593216 | 0.33971525 | 0.46389798 | 0.64272084 | 0.99999872 |
| SETD6 | 3.84149604 | 0.838995245 | 0.72409902 | 1.15867474 | 0.24658879 | NA |
| CNOT1 | 53.8231366 | 0.102158018 | 0.29567486 | 0.34550796 | 0.72971253 | 0.99999872 |
| SLC38A7 | 6.2842481 | -0.307491088 | 0.70342065 | -0.4371369 | 0.6620121 | 0.99999872 |
| GOT2 | 17.1245772 | 0.086740071 | 0.43101458 | 0.20124626 | 0.84050602 | 0.99999872 |
| AC138305.1 | 15.9577178 | -0.835642461 | 0.7891824 | -1.0588711 | 0.28965847 | 0.99999872 |
| CDH11 | 17.5327025 | -1.345702672 | 0.73080204 | -1.8414052 | 0.0655622 | 0.99999872 |
| BEAN1 | 5.82702397 | 0.443487057 | 0.5775771 | 0.76784044 | 0.44258198 | 0.99999872 |
| AC010542.3 | 12.5386385 | -0.491787076 | 0.63932925 | -0.7692235 | 0.44176065 | 0.99999872 |
| TK2 | 7.21015844 | 0.015629685 | 0.63901685 | 0.02445896 | 0.98048652 | 0.99999872 |
| CMTM3 | 18.5042628 | -0.615699291 | 0.53115245 | -1.1591762 | 0.24638437 | 0.99999872 |
| CMTM4 | 46.4649524 | -0.563237534 | 0.47465337 | -1.1866292 | 0.23537393 | 0.99999872 |
| DYNC1LI2 | 43.6407181 | -0.058061687 | 0.36223585 | -0.160287 | 0.87265502 | 0.99999872 |
| NAE1 | 21.8002712 | -0.266953157 | 0.36534178 | -0.7306943 | 0.46496589 | 0.99999872 |
| PDP2 | 8.29896432 | -0.219418981 | 0.68933058 | -0.3183073 | 0.75025182 | 0.99999872 |
| CDH16 | 19.2412972 | 0.034298958 | 0.45051843 | 0.0761322 | 0.93931393 | 0.99999872 |
| RRAD | 6.36334213 | 0.255284415 | 1.03721166 | 0.24612567 | 0.80558495 | 0.99999872 |
| FAM96B | 29.4171498 | 0.352666761 | 0.38423361 | 0.91784465 | 0.3587002 | 0.99999872 |
| CES2 | 11.4903088 | -0.432256634 | 0.45941584 | -0.9408832 | 0.34676473 | 0.99999872 |
| CBFB | 35.9484725 | -0.188789293 | 0.39653936 | -0.4760922 | 0.6340087 | 0.99999872 |
| C16orf70 | 7.82345473 | -0.212526473 | 0.60046296 | -0.3539377 | 0.72338558 | 0.99999872 |
| TRADD | 11.6045742 | -0.158466701 | 0.5633364 | -0.2813003 | 0.77848008 | 0.99999872 |
| NOL3 | 23.2842837 | 0.098667937 | 0.41770243 | 0.23621586 | 0.81326518 | 0.99999872 |
| KIAA0895L | 4.93960156 | -0.533269162 | 0.66332989 | -0.8039275 | 0.42143883 | 0.99999872 |
| E2F4 | 9.11905774 | -0.633870411 | 0.49279344 | -1.2862801 | 0.19834531 | 0.99999872 |
| LRRC29 | 5.74799304 | -0.650668102 | 0.60726682 | -1.0714699 | 0.28395821 | 0.99999872 |
| TMEM208 | 18.3405204 | 0.560724779 | 0.43608356 | 1.28581958 | 0.19850604 | 0.99999872 |
| TPPP3 | 133.047938 | 1.050886485 | 0.90243423 | 1.16450202 | 0.24422062 | 0.99999872 |
| ZDHHC1 | 10.6337776 | 0.376757315 | 0.47342621 | 0.79581 | 0.42614248 | 0.99999872 |

| | | | | | | |
|---|---|---|---|---|---|---|
| HSD11B2 | 11.5837119 | 0.770586844 | 1.19169078 | 0.64663322 | 0.51786935 | 0.99999872 |
| ATP6V0D1 | 23.4038058 | 0.387400817 | 0.43243352 | 0.89586215 | 0.37032639 | 0.99999872 |
| AC009061.2 | 8.41184008 | -0.674004639 | 0.6672285 | -1.0101556 | 0.31242072 | 0.99999872 |
| AC027682.6 | 38.1519344 | 0.001220919 | 0.61128658 | 0.00199729 | 0.99840639 | 0.99999872 |
| RIPOR1 | 399.082513 | -0.478587285 | 0.57262502 | -0.8357778 | 0.40327991 | 0.99999872 |
| AC027682.2 | 8.56018629 | 0.418253043 | 0.72366192 | 0.57796746 | 0.56328609 | 0.99999872 |
| CTCF | 27.8543566 | -0.334008907 | 0.37272632 | -0.8961238 | 0.37018667 | 0.99999872 |
| ENKD1 | 8.74781652 | 0.481392543 | 0.60128364 | 0.80060808 | 0.42335857 | 0.99999872 |
| GFOD2 | 11.8995937 | -0.170456481 | 0.48344346 | -0.3525882 | 0.72439715 | 0.99999872 |
| RANBP10 | 20.7029977 | -0.345059411 | 0.39318153 | -0.8776084 | 0.38015627 | 0.99999872 |
| THAP11 | 7.01139103 | 1.478061918 | 0.64782842 | 2.28156388 | 0.0225151 | 0.88027325 |
| NUTF2 | 8.99698595 | -0.777606988 | 0.5301724 | -1.4667059 | 0.14245608 | 0.99999872 |
| PSKH1 | 9.89952076 | -0.013123598 | 0.54222428 | -0.0242033 | 0.98069047 | 0.99999872 |
| SLC12A4 | 9.74941878 | 0.701018784 | 0.71867413 | 0.97543345 | 0.3293453 | 0.99999872 |
| DUS2 | 5.30724157 | 0.699643601 | 0.87115271 | 0.80312395 | 0.4219031 | 0.99999872 |
| NFATC3 | 134.442245 | -0.115780962 | 0.2771151 | -0.4178082 | 0.67608735 | 0.99999872 |
| PLA2G15 | 22.8961851 | 0.475188525 | 0.37603045 | 1.26369692 | 0.20633883 | 0.99999872 |
| SLC7A6 | 12.1191603 | -0.144161573 | 0.4644654 | -0.3103817 | 0.75627069 | 0.99999872 |
| SLC7A6OS | 13.8432469 | 0.152393623 | 0.4734254 | 0.32189575 | 0.74753167 | 0.99999872 |
| PRMT7 | 9.79177125 | -0.279887554 | 0.50289871 | -0.5565486 | 0.57783591 | 0.99999872 |
| ZFP90 | 17.5533251 | 0.10672479 | 0.36856943 | 0.28956496 | 0.77214908 | 0.99999872 |
| CDH1 | 7.85903603 | -0.399556805 | 0.94737073 | -0.4217534 | 0.67320504 | 0.99999872 |
| TANGO6 | 35.0937607 | -0.408095299 | 0.38923344 | -1.048459 | 0.29442719 | 0.99999872 |
| CHTF8 | 20.8835252 | -0.26544942 | 0.42234668 | -0.6285107 | 0.52966942 | 0.99999872 |
| UTP4 | 6.57393875 | -0.225823869 | 0.81680159 | -0.2764733 | 0.78218453 | 0.99999872 |
| SNTB2 | 9.71206465 | 0.0660532 | 0.54548395 | 0.12109101 | 0.90361895 | 0.99999872 |
| VPS4A | 16.9113828 | -0.206972737 | 0.43603718 | -0.4746676 | 0.63502389 | 0.99999872 |
| COG8 | 10.818386 | -0.283442144 | 0.60795712 | -0.4662206 | 0.64105759 | 0.99999872 |
| NIP7 | 4.49997335 | 1.207570317 | 1.08033481 | 1.11777414 | 0.26366347 | NA |
| TERF2 | 14.1536839 | -0.337435854 | 0.42232475 | -0.7989962 | 0.42429264 | 0.99999872 |
| CYB5B | 78.3802579 | -0.423002645 | 0.30460484 | -1.3886931 | 0.16492609 | 0.99999872 |
| NFAT5 | 431.073521 | -0.017978604 | 0.23462003 | -0.0766286 | 0.93891901 | 0.99999872 |
| NQO1 | 113.551878 | 0.230329954 | 0.51946634 | 0.44339726 | 0.65747842 | 0.99999872 |
| NOB1 | 11.2831368 | -0.112242514 | 0.48864984 | -0.2296993 | 0.81832546 | 0.99999872 |
| WWP2 | 50.3281424 | -0.890026638 | 0.3239476 | -2.7474401 | 0.00600625 | 0.58725857 |
| PDPR | 35.4721156 | -0.441051165 | 0.44933184 | -0.9815711 | 0.32631118 | 0.99999872 |
| AC009060.1 | 7.66135357 | 0.663506314 | 0.96094506 | 0.69047268 | 0.48989698 | 0.99999872 |
| EXOSC6 | 21.4300312 | -0.233378124 | 0.52662443 | -0.4431586 | 0.65765105 | 0.99999872 |
| AARS | 25.8804753 | -0.597825469 | 0.44103276 | -1.3555126 | 0.17525429 | 0.99999872 |
| DDX19A | 5.35912978 | 0.126055954 | 0.71070932 | 0.1773664 | 0.8592206 | 0.99999872 |
| ST3GAL2 | 18.6140149 | -0.295951896 | 0.39463564 | -0.7499371 | 0.4532926 | 0.99999872 |
| FUK | 5.41321416 | 0.161431709 | 0.61900204 | 0.2607935 | 0.79425176 | 0.99999872 |
| COG4 | 20.6021988 | -0.382449022 | 0.39724852 | -0.962745 | 0.33567551 | 0.99999872 |
| SF3B3 | 28.8547811 | 0.087945779 | 0.33361243 | 0.26361661 | 0.79207535 | 0.99999872 |
| MTSS1L | 38.606302 | -0.050965648 | 0.4404678 | -0.115708 | 0.90788397 | 0.99999872 |
| VAC14 | 17.740349 | -0.472203291 | 0.38813185 | -1.2166054 | 0.22375441 | 0.99999872 |
| HYDIN | 20.3679934 | -0.751086021 | 0.50878169 | -1.4762442 | 0.13987834 | 0.99999872 |
| CMTR2 | 13.1191729 | -0.232657027 | 0.59228715 | -0.3928112 | 0.69445893 | 0.99999872 |
| ZNF19 | 7.55367441 | -0.405336704 | 0.63797665 | -0.6353472 | 0.52520197 | 0.99999872 |
| PHLPP2 | 23.4872645 | -0.085362998 | 0.40095792 | -0.2128976 | 0.8314068 | 0.99999872 |
| AP1G1 | 62.204028 | 0.397459728 | 0.29998285 | 1.3249415 | 0.18519055 | 0.99999872 |
| IST1 | 47.5150975 | -0.021264359 | 0.2561979 | -0.0829997 | 0.93385175 | 0.99999872 |
| ZNF821 | 6.62481561 | 0.365201401 | 0.68152549 | 0.53585875 | 0.59205617 | 0.99999872 |
| DHODH | 5.23387616 | 0.045641235 | 0.67380255 | 0.06773681 | 0.94599515 | 0.99999872 |
| TXNL4B | 10.5493858 | -0.832205791 | 0.65573773 | -1.2691138 | 0.20440048 | 0.99999872 |
| DHX38 | 10.5563515 | -0.388386351 | 0.47127043 | -0.8241263 | 0.4098678 | 0.99999872 |
| LINC01572 | 21.3259307 | -0.360866826 | 0.6785229 | -0.5318418 | 0.59483558 | 0.99999872 |
| AC004943.2 | 13.3701826 | -0.095407077 | 0.47373871 | -0.2013918 | 0.84039225 | 0.99999872 |
| ZFHX3 | 100.512763 | -0.180141738 | 0.25613689 | -0.7033026 | 0.4818672 | 0.99999872 |
| AC138627.1 | 7.36868544 | -0.808353961 | 0.72876426 | -1.1092119 | 0.2673388 | 0.99999872 |
| PSMD7 | 26.1662552 | 0.20482196 | 0.32156723 | 0.6369491 | 0.524158 | 0.99999872 |

| | | | | | | |
|---|---|---|---|---|---|---|
| NPIPB15 | 4.91577861 | -0.418311577 | 1.21143459 | -0.3453027 | 0.72986685 | 0.99999872 |
| GLG1 | 122.75866 | -0.062244052 | 0.28708254 | -0.2168159 | 0.82835184 | 0.99999872 |
| RFWD3 | 13.5583845 | -0.362219497 | 0.54886851 | -0.6599386 | 0.50929325 | 0.99999872 |
| WDR59 | 36.8621995 | -0.307802806 | 0.37479796 | -0.82125 | 0.4115039 | 0.99999872 |
| ZNRF1 | 9.65589463 | -0.459678868 | 0.63753929 | -0.7210205 | 0.47089693 | 0.99999872 |
| ZFP1 | 4.19236295 | 1.312971877 | 0.90383994 | 1.45265973 | 0.14631825 | NA |
| BCAR1 | 31.8921407 | 0.2121685 | 0.51025107 | 0.41581196 | 0.6775476 | 0.99999872 |
| CFDP1 | 60.2403283 | -0.391447909 | 0.3122885 | -1.2534817 | 0.21003047 | 0.99999872 |
| GABARAPL2 | 74.5206414 | 1.388546957 | 0.5122575 | 2.71064252 | 0.0067153 | 0.61721837 |
| ADAT1 | 10.4719828 | -0.460590473 | 0.47190947 | -0.9760145 | 0.32905729 | 0.99999872 |
| KARS | 19.2795841 | -0.238190135 | 0.51211239 | -0.465113 | 0.64185053 | 0.99999872 |
| TERF2IP | 95.7564773 | 0.171267617 | 0.36463356 | 0.4696979 | 0.63857087 | 0.99999872 |
| AC010528.1 | 3.67587954 | 2.502591616 | 1.213747 | 2.06187255 | 0.03921987 | NA |
| NUDT7 | 7.13010609 | 0.750603537 | 0.70380831 | 1.0664886 | 0.28620283 | 0.99999872 |
| WWOX | 977.000411 | 0.18198626 | 0.40754098 | 0.44654714 | 0.65520207 | 0.99999872 |
| AC046158.1 | 5.37139846 | 1.311620911 | 1.03554882 | 1.26659495 | 0.20530016 | 0.99999872 |
| MAF | 27.9427096 | 0.054645069 | 0.55568876 | 0.09833755 | 0.92166426 | 0.99999872 |
| CDYL2 | 14.3622894 | 0.139853179 | 0.57160659 | 0.24466684 | 0.8067144 | 0.99999872 |
| CMC2 | 23.2296667 | 0.545226894 | 0.35015919 | 1.55708294 | 0.11945079 | 0.99999872 |
| ATMIN | 15.0343788 | -0.100948742 | 0.39915411 | -0.2529067 | 0.80034032 | 0.99999872 |
| GAN | 12.9831125 | 0.592228239 | 0.41448843 | 1.4288173 | 0.15305675 | 0.99999872 |
| CMIP | 82.0796816 | -0.129919531 | 0.36264076 | -0.3582596 | 0.72014906 | 0.99999872 |
| PLCG2 | 217.288109 | -1.362738037 | 0.34099188 | -3.9963944 | 6.43E-05 | 0.07163933 |
| SDR42E1 | 14.9640602 | 0.094982689 | 0.48893119 | 0.19426596 | 0.84596764 | 0.99999872 |
| MPHOSPH6 | 15.432266 | 0.335788515 | 0.46119243 | 0.72808765 | 0.46655993 | 0.99999872 |
| HSBP1 | 50.6163217 | 0.397526338 | 0.42197281 | 0.94206625 | 0.34615872 | 0.99999872 |
| OSGIN1 | 9.27511493 | -1.830655515 | 0.85188457 | -2.1489478 | 0.03163853 | 0.92859542 |
| MBTPS1 | 46.7611865 | 0.618085871 | 0.40989259 | 1.50792155 | 0.13157461 | 0.99999872 |
| HSDL1 | 10.0777266 | -0.034574071 | 0.59653693 | -0.057958 | 0.9537821 | 0.99999872 |
| TAF1C | 7.19183136 | -0.200989138 | 0.52991184 | -0.3792879 | 0.7044741 | 0.99999872 |
| TLDC1 | 14.7497196 | -0.033748086 | 0.52964861 | -0.0637179 | 0.94919487 | 0.99999872 |
| COTL1 | 28.8815133 | -0.993529774 | 0.70625047 | -1.4067669 | 0.15949652 | 0.99999872 |
| KLHL36 | 19.8523146 | 0.060113764 | 0.54667087 | 0.10996336 | 0.91243844 | 0.99999872 |
| USP10 | 44.653784 | -0.369000616 | 0.26731913 | -1.3803749 | 0.16747125 | 0.99999872 |
| ZDHHC7 | 21.0318478 | -0.219563572 | 0.33653316 | -0.6524278 | 0.51412526 | 0.99999872 |
| KIAA0513 | 5.84898804 | -1.073672995 | 0.63513428 | -1.6904661 | 0.09093881 | 0.99999872 |
| GSE1 | 30.1502818 | -0.657832423 | 0.4322253 | -1.5219665 | 0.12801748 | 0.99999872 |
| EMC8 | 12.67694 | -0.072822834 | 0.44415143 | -0.1639595 | 0.86976305 | 0.99999872 |
| COX4I1 | 91.8007256 | 0.402678784 | 0.38396122 | 1.04874857 | 0.29429385 | 0.99999872 |
| MTHFSD | 5.76147431 | -0.167943881 | 0.62145005 | -0.2702452 | 0.78697164 | 0.99999872 |
| FOXC2 | 24.9640907 | -1.456840466 | 0.41697189 | -3.4938577 | 0.00047609 | 0.21689216 |
| AC009154.1 | 6.28831089 | -0.7441677 | 0.9113741 | -0.8165337 | 0.41419493 | 0.99999872 |
| FBXO31 | 8.85870237 | -0.229891644 | 0.53704786 | -0.4280655 | 0.66860346 | 0.99999872 |
| MAP1LC3B | 44.2740192 | 0.601776213 | 0.31948612 | 1.88357546 | 0.05962242 | 0.99999872 |
| ZCCHC14 | 49.6305483 | -0.341074624 | 0.38260315 | -0.891458 | 0.37268354 | 0.99999872 |
| KLHDC4 | 19.0159853 | -0.287201023 | 0.42787012 | -0.6712341 | 0.5020714 | 0.99999872 |
| SLC7A5 | 3.8289226 | -1.354290768 | 1.22396402 | -1.1064792 | 0.26851915 | NA |
| BANP | 20.7773619 | -0.322991915 | 0.44586214 | -0.724421 | 0.46880728 | 0.99999872 |
| ZFPM1 | 15.3671556 | 0.053731343 | 0.37445757 | 0.14349114 | 0.88590231 | 0.99999872 |
| ZC3H18 | 19.3115837 | -0.645166068 | 0.4278696 | -1.5078568 | 0.13159119 | 0.99999872 |
| CYBA | 48.0986039 | 0.773842427 | 0.42619347 | 1.81570689 | 0.06941535 | 0.99999872 |
| MVD | 10.1332062 | -0.224489097 | 0.68427814 | -0.328067 | 0.74286098 | 0.99999872 |
| RNF166 | 8.49549175 | 1.057843116 | 0.71061825 | 1.48862363 | 0.13658651 | 0.99999872 |
| PIEZO1 | 79.8293497 | -0.152362674 | 0.42390852 | -0.3594235 | 0.71927831 | 0.99999872 |
| APRT | 27.7691019 | 0.514309395 | 0.44975373 | 1.14353559 | 0.25281629 | 0.99999872 |
| GALNS | 11.8346953 | -0.006845041 | 0.48918398 | -0.0139928 | 0.98883575 | 0.99999872 |
| TRAPPC2L | 7.35750782 | 0.581544286 | 0.71385026 | 0.81465865 | 0.41526773 | 0.99999872 |
| ACSF3 | 10.0341204 | -0.455211631 | 0.45156398 | -1.0080778 | 0.3134171 | 0.99999872 |
| ZNF778 | 5.69149218 | -0.999499423 | 0.64293177 | -1.5545964 | 0.12004222 | 0.99999872 |
| ANKRD11 | 156.431756 | -0.235012286 | 0.20134043 | -1.1672384 | 0.24311409 | 0.99999872 |
| SPG7 | 41.2785534 | -0.708305952 | 0.32096635 | -2.2067919 | 0.0273286 | 0.89154528 |

| | | | | | | |
|---|---|---|---|---|---|---|
| RPL13 | 157.597358 | 0.245909706 | 0.47480881 | 0.5179131 | 0.6045189 | 0.99999872 |
| DPEP1 | 6.24938955 | -1.821607026 | 0.96448244 | -1.8886886 | 0.05893356 | 0.99999872 |
| CHMP1A | 21.2521445 | -0.380392862 | 0.49645412 | -0.7662196 | 0.44354567 | 0.99999872 |
| SPATA33 | 4.70671817 | 0.400355955 | 0.64565904 | 0.62007333 | 0.53520951 | 0.99999872 |
| CDK10 | 13.0879897 | -0.378561844 | 0.42700329 | -0.8865549 | 0.3753186 | 0.99999872 |
| VPS9D1 | 8.25187496 | -0.125442958 | 0.53590835 | -0.2340754 | 0.81492645 | 0.99999872 |
| ZNF276 | 6.2077268 | -0.408114097 | 0.56778092 | -0.718788 | 0.47227157 | 0.99999872 |
| TCF25 | 92.8093783 | -0.229079717 | 0.31122962 | -0.7360473 | 0.46170193 | 0.99999872 |
| DEF8 | 9.96046226 | -0.209662734 | 0.54663736 | -0.38355 | 0.70131204 | 0.99999872 |
| GAS8 | 9.9166664 | -0.319349132 | 0.53657228 | -0.5951652 | 0.55173307 | 0.99999872 |
| FAM157C | 23.5477573 | -0.10467286 | 0.49687792 | -0.2106611 | 0.83315171 | 0.99999872 |
| RPH3AL | 27.6181265 | -0.818694425 | 0.69001919 | -1.1864807 | 0.23543254 | 0.99999872 |
| RFLNB | 17.4845461 | -1.968354228 | 1.21328513 | -1.6223344 | 0.10473176 | 0.99999872 |
| VPS53 | 51.7594938 | -0.282178911 | 0.34857415 | -0.8095233 | 0.41821418 | 0.99999872 |
| GLOD4 | 15.5061008 | 0.669319672 | 0.46580989 | 1.43689449 | 0.15074797 | 0.99999872 |
| NXN | 89.0570056 | -0.411474622 | 0.42409033 | -0.9702523 | 0.33192075 | 0.99999872 |
| TIMM22 | 8.42136591 | 0.35154728 | 0.55507533 | 0.63333256 | 0.5265165 | 0.99999872 |
| ABR | 34.8803385 | -0.510054119 | 0.49829602 | -1.0235966 | 0.30602585 | 0.99999872 |
| YWHAE | 144.178646 | 0.218786828 | 0.25154354 | 0.86977718 | 0.38442218 | 0.99999872 |
| CRK | 41.2140752 | -0.132097248 | 0.37073049 | -0.3563161 | 0.72160384 | 0.99999872 |
| MYO1C | 103.13862 | -0.477355743 | 0.5353108 | -0.8917357 | 0.37253462 | 0.99999872 |
| INPP5K | 7.82535817 | -1.002216614 | 0.59928184 | -1.6723627 | 0.09445282 | 0.99999872 |
| PITPNA | 39.4944436 | -0.321524154 | 0.30681297 | -1.0479484 | 0.29466239 | 0.99999872 |
| SLC43A2 | 14.254269 | 0.540886304 | 0.98866354 | 0.54708835 | 0.58431804 | 0.99999872 |
| PRPF8 | 46.5978858 | -0.138871029 | 0.33494042 | -0.4146141 | 0.6784244 | 0.99999872 |
| TLCD2 | 6.67981031 | -0.211606093 | 0.81576299 | -0.2593965 | 0.7953293 | 0.99999872 |
| MIR22HG | 16.1000542 | -0.583789427 | 0.48707798 | -1.1985543 | 0.23070128 | 0.99999872 |
| WDR81 | 10.1913803 | -0.201951399 | 0.47493092 | -0.4252227 | 0.67067435 | 0.99999872 |
| SMYD4 | 10.1039241 | -0.072645761 | 0.72662579 | -0.0999769 | 0.9203627 | 0.99999872 |
| RPA1 | 16.7046828 | 0.166830607 | 0.42121791 | 0.39606722 | 0.69205544 | 0.99999872 |
| SMG6 | 68.3179977 | -0.619900785 | 0.23570533 | -2.629982 | 0.00853894 | 0.66769402 |
| SRR | 6.61076748 | 0.083321023 | 0.62421939 | 0.13348035 | 0.8938135 | 0.99999872 |
| TSR1 | 9.11469733 | -1.037440707 | 0.64225369 | -1.615313 | 0.10624294 | 0.99999872 |
| SGSM2 | 9.94920475 | 0.423169986 | 0.60205746 | 0.70287308 | 0.48213485 | 0.99999872 |
| MNT | 9.31447391 | -0.198529361 | 0.65245844 | -0.3042789 | 0.76091539 | 0.99999872 |
| METTL16 | 23.8971646 | -0.536407874 | 0.30876995 | -1.7372412 | 0.0823446 | 0.99999872 |
| PAFAH1B1 | 153.755395 | -0.013636616 | 0.24505464 | -0.0556472 | 0.95562283 | 0.99999872 |
| CLUH | 12.1321025 | -0.759872554 | 0.554821 | -1.3695815 | 0.17081759 | 0.99999872 |
| CTNS | 6.45112767 | 0.156624177 | 0.56238263 | 0.27850109 | 0.78062773 | 0.99999872 |
| TAX1BP3 | 15.3813352 | -0.701387756 | 0.46480186 | -1.5090038 | 0.13129782 | 0.99999872 |
| EMC6 | 19.203876 | 1.278558897 | 0.56600738 | 2.25890851 | 0.02388908 | 0.88337526 |
| ITGAE | 10.4906776 | 0.488663066 | 0.69105146 | 0.70712978 | 0.47948583 | 0.99999872 |
| NCBP3 | 29.1668875 | 0.100669195 | 0.39865925 | 0.2525194 | 0.80063962 | 0.99999872 |
| CAMKK1 | 4.50700248 | -0.932187101 | 0.98699345 | -0.9444714 | 0.3449288 | NA |
| ZZEF1 | 36.8380581 | 0.007110425 | 0.38773912 | 0.01833817 | 0.98536908 | 0.99999872 |
| CYB5D2 | 22.5990445 | 0.23030203 | 0.44075546 | 0.52251657 | 0.60131071 | 0.99999872 |
| ANKFY1 | 36.1353604 | -0.226037056 | 0.41926831 | -0.5391227 | 0.5898022 | 0.99999872 |
| UBE2G1 | 36.972598 | -0.04665424 | 0.33254052 | -0.1402964 | 0.8884258 | 0.99999872 |
| MYBBP1A | 6.54866305 | -0.597118431 | 0.70520655 | -0.8467284 | 0.39714651 | 0.99999872 |
| PELP1 | 10.9675385 | -0.731681214 | 0.44527281 | -1.6432201 | 0.10033741 | 0.99999872 |
| MED11 | 9.75314929 | 0.117136531 | 0.68968343 | 0.16984101 | 0.86513517 | 0.99999872 |
| CXCL16 | 8.75125841 | -0.508415894 | 0.91158667 | -0.5577263 | 0.57703127 | 0.99999872 |
| PSMB6 | 28.2971628 | 1.141222691 | 0.61696199 | 1.84974553 | 0.06435023 | 0.99999872 |
| MINK1 | 17.432106 | -0.190958234 | 0.37678901 | -0.5068041 | 0.61229225 | 0.99999872 |
| SLC25A11 | 13.8412286 | 1.436354208 | 0.6036463 | 2.37946327 | 0.01733787 | 0.83333123 |
| RNF167 | 16.3746915 | 0.532363107 | 0.54199909 | 0.9822214 | 0.32599079 | 0.99999872 |
| PFN1 | 93.0388257 | -0.452784212 | 0.51878271 | -0.872782 | 0.38278191 | 0.99999872 |
| SPAG7 | 31.4491953 | -0.374209211 | 0.46136593 | -0.8110898 | 0.41731409 | 0.99999872 |
| CAMTA2 | 12.9245865 | -0.292113352 | 0.47486481 | -0.6151506 | 0.5384553 | 0.99999872 |
| KIF1C | 14.6498633 | -0.765588508 | 0.63121392 | -1.2128828 | 0.22517463 | 0.99999872 |
| ZNF594 | 6.72441542 | -0.494763656 | 0.63883299 | -0.7744804 | 0.43864673 | 0.99999872 |

| | | | | | | |
|---|---|---|---|---|---|---|
| AC087500.1 | 14.209397 | 0.04659195 | 0.47327284 | 0.09844628 | 0.92157793 | 0.99999872 |
| RABEP1 | 65.2339801 | -0.185157811 | 0.26655901 | -0.6946222 | 0.48729208 | 0.99999872 |
| NUP88 | 26.2644853 | -0.338320049 | 0.37981709 | -0.8907447 | 0.37306616 | 0.99999872 |
| RPAIN | 23.5308117 | 0.302919298 | 0.38230116 | 0.79235778 | 0.42815209 | 0.99999872 |
| C1QBP | 20.5825477 | 0.541690471 | 0.4221707 | 1.28310769 | 0.19945435 | 0.99999872 |
| DERL2 | 22.7224671 | 0.650467023 | 0.4184265 | 1.55455505 | 0.12005208 | 0.99999872 |
| MIS12 | 10.0032588 | 0.420689787 | 0.48653996 | 0.86465619 | 0.38722752 | 0.99999872 |
| NLRP1 | 40.343826 | 0.429513497 | 0.73194738 | 0.58680925 | 0.55733182 | 0.99999872 |
| PITPNM3 | 19.1787329 | -0.538251383 | 0.550984 | -0.9768911 | 0.32862306 | 0.99999872 |
| KIAA0753 | 17.157255 | -0.051593555 | 0.41226721 | -0.1251459 | 0.90040804 | 0.99999872 |
| TXNDC17 | 11.04633 | 0.039534689 | 0.61198466 | 0.06460078 | 0.94849186 | 0.99999872 |
| MED31 | 8.64866405 | 0.343254067 | 0.52279781 | 0.65657135 | 0.51145658 | 0.99999872 |
| XAF1 | 36.2153798 | -0.307956913 | 1.78782102 | -0.1722527 | 0.86323891 | 0.99999872 |
| ALOX12-AS1 | 7.86133765 | 2.871788694 | 1.30716907 | 2.19695275 | 0.02802382 | 0.89796207 |
| ACADVL | 50.7986659 | 0.204198449 | 0.31343754 | 0.65148051 | 0.51473636 | 0.99999872 |
| DVL2 | 10.3714521 | 0.998922513 | 0.50081299 | 1.99460186 | 0.04608632 | 0.95980711 |
| PHF23 | 7.71138615 | 0.77656839 | 0.55920774 | 1.38869391 | 0.16492585 | 0.99999872 |
| ELP5 | 9.05228804 | 0.204696881 | 0.4691656 | 0.43629985 | 0.66261919 | 0.99999872 |
| EIF5A | 27.8530261 | -0.70888623 | 0.65411823 | -1.083728 | 0.27848543 | 0.99999872 |
| GPS2 | 21.576886 | 0.170039893 | 0.57320503 | 0.29664759 | 0.76673557 | 0.99999872 |
| KCTD11 | 5.09308531 | -0.459442394 | 0.71411318 | -0.6433748 | 0.51998096 | 0.99999872 |
| NLGN2 | 6.21893694 | 0.4042146 | 0.64753168 | 0.62423911 | 0.53247057 | 0.99999872 |
| CHRNB1 | 8.67462102 | -0.545927624 | 0.5860693 | -0.9315069 | 0.35159139 | 0.99999872 |
| ZBTB4 | 65.1249429 | -0.106823779 | 0.35507277 | -0.3008504 | 0.76352859 | 0.99999872 |
| POLR2A | 55.9588609 | 0.056574618 | 0.45010375 | 0.12569239 | 0.89997542 | 0.99999872 |
| EIF4A1 | 28.913177 | 0.160850672 | 0.84078833 | 0.19130935 | 0.84828323 | 0.99999872 |
| CD68 | 9.44244756 | -2.705300598 | 1.52546907 | -1.7734221 | 0.07615879 | 0.99999872 |
| MPDU1 | 11.4796194 | -0.175605024 | 0.60184823 | -0.2917763 | 0.7704577 | 0.99999872 |
| FXR2 | 20.0865936 | 0.033576267 | 0.52345638 | 0.06414339 | 0.94885605 | 0.99999872 |
| SAT2 | 23.5380321 | 0.82552092 | 0.54249686 | 1.52170637 | 0.12808267 | 0.99999872 |
| TP53 | 9.56725076 | -0.380383469 | 0.62613867 | -0.6075068 | 0.54351466 | 0.99999872 |
| KDM6B | 10.5902494 | -0.720372607 | 0.58655015 | -1.2281518 | 0.21939 | 0.99999872 |
| NAA38 | 45.920674 | 0.302783742 | 0.38993864 | 0.77649074 | 0.43745929 | 0.99999872 |
| CHD3 | 91.239615 | -0.60085649 | 0.53105291 | -1.1314437 | 0.25786837 | 0.99999872 |
| TRAPPC1 | 15.91345 | 0.380494579 | 0.45943663 | 0.82817641 | 0.4075706 | 0.99999872 |
| PER1 | 44.0452804 | -0.779649258 | 0.46970355 | -1.6598752 | 0.09693957 | 0.99999872 |
| TMEM107 | 16.5912357 | 0.48165129 | 0.58503735 | 0.82328298 | 0.41034709 | 0.99999872 |
| BORCS6 | 4.4782755 | 0.938294302 | 0.89262934 | 1.05115781 | 0.29318611 | NA |
| CTC1 | 16.3445891 | -0.202993719 | 0.4632369 | -0.4382071 | 0.66123613 | 0.99999872 |
| SLC25A35 | 3.75940003 | -0.741674229 | 0.89474765 | -0.82892 | 0.40714967 | NA |
| RANGRF | 7.89081793 | 0.717768957 | 0.60715161 | 1.18219066 | 0.23713006 | 0.99999872 |
| NDEL1 | 25.4205986 | 0.147102241 | 0.43609382 | 0.33731788 | 0.73587728 | 0.99999872 |
| MYH10 | 34.2611716 | 0.618657103 | 0.39281072 | 1.57494966 | 0.11526806 | 0.99999872 |
| STX8 | 75.1563447 | 0.007810508 | 0.23242966 | 0.03360375 | 0.97319313 | 0.99999872 |
| USP43 | 5.99293125 | 0.837381067 | 0.7561456 | 1.10743363 | 0.26810649 | 0.99999872 |
| GAS7 | 30.0232469 | -1.207178469 | 0.80827496 | -1.4935245 | 0.13529994 | 0.99999872 |
| SCO1 | 14.9675317 | 0.196324547 | 0.55853481 | 0.35149921 | 0.72521386 | 0.99999872 |
| TMEM220 | 13.8985836 | -0.358204328 | 0.71746221 | -0.4992658 | 0.61759216 | 0.99999872 |
| SHISA6 | 73.9307948 | 1.007244816 | 0.8431618 | 1.19460443 | 0.23224162 | 0.99999872 |
| DNAH9 | 29.6810584 | 1.2052152 | 0.71008579 | 1.6972811 | 0.08964353 | 0.99999872 |
| MAP2K4 | 65.1572259 | -0.069162937 | 0.22609448 | -0.3059028 | 0.75967864 | 0.99999872 |
| ARHGAP44 | 20.1773494 | -0.392570106 | 0.46400622 | -0.8460449 | 0.39752769 | 0.99999872 |
| ELAC2 | 12.5060467 | 0.06101792 | 0.46876539 | 0.13016729 | 0.89643407 | 0.99999872 |
| HS3ST3A1 | 78.6036352 | -0.763297145 | 0.407081 | -1.8750498 | 0.06078587 | 0.99999872 |
| COX10 | 18.6421998 | 0.067502209 | 0.5190683 | 0.13004495 | 0.89653087 | 0.99999872 |
| HS3ST3B1 | 117.903947 | -0.59743169 | 0.49531507 | -1.206165 | 0.22775389 | 0.99999872 |
| TRIM16 | 5.27058784 | -0.141541075 | 0.64174177 | -0.2205577 | 0.82543686 | 0.99999872 |
| AC015922.4 | 4.50967966 | 0.253767884 | 0.78804091 | 0.32202374 | 0.74743471 | NA |
| ZSWIM7 | 20.5241791 | -0.051963065 | 0.38756428 | -0.134076 | 0.89334248 | 0.99999872 |
| TTC19 | 44.4785163 | 0.534383456 | 0.30578164 | 1.74759824 | 0.08053362 | 0.99999872 |
| NCOR1 | 114.448191 | -0.257600583 | 0.1883514 | -1.3676595 | 0.17141867 | 0.99999872 |

| | | | | | | |
|---|---|---|---|---|---|---|
| PIGL | 34.910534 | -0.692036504 | 0.30586177 | -2.2625793 | 0.02366163 | 0.88337526 |
| UBB | 178.627722 | 0.601853404 | 0.36180692 | 1.66346571 | 0.09621925 | 0.99999872 |
| LRRC75A | 5.47192555 | 0.412854714 | 0.83288493 | 0.49569238 | 0.62011146 | 0.99999872 |
| ZNF624 | 5.89527078 | 0.279927554 | 0.70314709 | 0.39810668 | 0.69055155 | 0.99999872 |
| AC098850.3 | 19.1577804 | 0.627199199 | 0.79058627 | 0.79333429 | 0.42758308 | 0.99999872 |
| MPRIP | 75.0652756 | 0.042423096 | 0.29498546 | 0.1438142 | 0.88564719 | 0.99999872 |
| COPS3 | 10.7744519 | -0.03456622 | 0.43949388 | -0.0786501 | 0.93731097 | 0.99999872 |
| MED9 | 6.66738873 | 0.148247336 | 0.63966773 | 0.23175678 | 0.81672692 | 0.99999872 |
| RASD1 | 12.782378 | -0.013195677 | 0.82495756 | -0.0159956 | 0.98723792 | 0.99999872 |
| PEMT | 7.53019226 | -0.216727483 | 0.64482749 | -0.3361015 | 0.73679433 | 0.99999872 |
| RAI1 | 48.02974 | -0.060667863 | 0.39366735 | -0.1541095 | 0.87752343 | 0.99999872 |
| SMCR5 | 4.3912513 | 1.204906344 | 0.92597208 | 1.30123399 | 0.19317837 | NA |
| SREBF1 | 20.9111632 | -0.039788206 | 0.72226261 | -0.0550883 | 0.95606813 | 0.99999872 |
| TOM1L2 | 267.079719 | 0.005561761 | 0.43318274 | 0.01283929 | 0.98975601 | 0.99999872 |
| ATPAF2 | 5.02691033 | 0.913083817 | 0.6921037 | 1.31928758 | 0.18707299 | 0.99999872 |
| GID4 | 5.14826175 | 0.074621603 | 0.70071257 | 0.10649388 | 0.91519051 | 0.99999872 |
| DRG2 | 8.85271272 | -0.509297934 | 0.54030565 | -0.9426108 | 0.34588002 | 0.99999872 |
| ALKBH5 | 35.4287119 | -0.003562317 | 0.3554017 | -0.0100234 | 0.99200265 | 0.99999872 |
| FLII | 20.9353911 | -0.260766661 | 0.41718422 | -0.6250636 | 0.53192934 | 0.99999872 |
| TOP3A | 8.3617687 | -0.594703204 | 0.55237849 | -1.0766227 | 0.28164887 | 0.99999872 |
| SMCR8 | 5.59940743 | 0.233411382 | 0.61982694 | 0.37657509 | 0.70648941 | 0.99999872 |
| SHMT1 | 8.67942425 | 1.169049755 | 0.90716463 | 1.28868534 | 0.1975075 | 0.99999872 |
| TVP23B | 10.0196093 | -0.054543077 | 0.51391762 | -0.1061319 | 0.91547767 | 0.99999872 |
| PRPSAP2 | 27.6463741 | -0.143647729 | 0.40673248 | -0.353175 | 0.72395728 | 0.99999872 |
| SLC5A10 | 5.28219581 | 1.44358577 | 1.00545029 | 1.43576045 | 0.15107052 | 0.99999872 |
| FAM83G | 8.5701348 | -1.332984062 | 0.5951525 | -2.2397353 | 0.02510811 | 0.88365618 |
| EPN2 | 97.086787 | 0.034063283 | 0.24914011 | 0.1367234 | 0.89124943 | 0.99999872 |
| B9D1 | 25.0801876 | 0.076859927 | 0.37543598 | 0.20472179 | 0.83778949 | 0.99999872 |
| MAPK7 | 11.3188576 | -0.329220647 | 0.55132158 | -0.5971481 | 0.55040849 | 0.99999872 |
| AC004448.2 | 5.64663572 | 0.968499961 | 0.64537973 | 1.50066684 | 0.13344175 | 0.99999872 |
| SLC47A1 | 12.7226386 | 1.608233592 | 1.06106981 | 1.5156718 | 0.12960236 | 0.99999872 |
| ALDH3A2 | 18.6885972 | -0.468667341 | 0.38658317 | -1.2123325 | 0.22538513 | 0.99999872 |
| SLC47A2 | 9.82922144 | 0.148278472 | 1.20803637 | 0.12274338 | 0.90231031 | 0.99999872 |
| ULK2 | 23.6602274 | 0.16357542 | 0.39680056 | 0.41223586 | 0.68016655 | 0.99999872 |
| AKAP10 | 51.3353818 | -0.454808173 | 0.28661557 | -1.586823 | 0.11255275 | 0.99999872 |
| SPECC1 | 9.39658987 | 0.519961082 | 0.71735665 | 0.72482925 | 0.46855679 | 0.99999872 |
| CCDC144NL-/ | 18.7771525 | 1.417050212 | 0.76802012 | 1.84506913 | 0.06502748 | 0.99999872 |
| USP22 | 56.2191788 | -0.088098689 | 0.38625596 | -0.2280837 | 0.81958117 | 0.99999872 |
| DHRS7B | 30.6091033 | -0.242026028 | 0.40404168 | -0.5990125 | 0.54916453 | 0.99999872 |
| TMEM11 | 17.3393867 | 0.527614575 | 0.52498464 | 1.00500954 | 0.31489226 | 0.99999872 |
| AC087294.1 | 8.52668055 | 1.15155316 | 0.64044468 | 1.7980525 | 0.07216869 | 0.99999872 |
| NATD1 | 17.1709673 | 0.415506779 | 0.56620106 | 0.73385023 | 0.46304002 | 0.99999872 |
| MAP2K3 | 27.1875669 | -0.286941561 | 0.45205137 | -0.6347543 | 0.52558866 | 0.99999872 |
| C17orf51 | 5.16830732 | 0.037648553 | 0.63933389 | 0.05888715 | 0.95304199 | 0.99999872 |
| WSB1 | 208.327895 | 0.132370755 | 0.37901389 | 0.3492504 | 0.72690133 | 0.99999872 |
| KSR1 | 29.1333765 | 0.021874693 | 0.50403436 | 0.04339921 | 0.96538331 | 0.99999872 |
| LGALS9 | 4.12216035 | -1.191454397 | 1.58783334 | -0.7503649 | 0.45303497 | NA |
| NLK | 443.58649 | -0.107435735 | 0.43524671 | -0.2468387 | 0.80503306 | 0.99999872 |
| IFT20 | 14.2002344 | 0.143587512 | 0.52205677 | 0.27504195 | 0.78328401 | 0.99999872 |
| TNFAIP1 | 9.89783994 | 0.231855617 | 0.54366981 | 0.42646402 | 0.66976976 | 0.99999872 |
| POLDIP2 | 13.2512599 | -0.349919949 | 0.5069636 | -0.690227 | 0.49005146 | 0.99999872 |
| TMEM199 | 7.71865756 | -0.753364256 | 0.61639478 | -1.2222106 | 0.22162798 | 0.99999872 |
| SARM1 | 13.230395 | -0.209787351 | 0.43123579 | -0.4864794 | 0.62662727 | 0.99999872 |
| AC005726.1 | 6.05711005 | -0.360260968 | 0.78069924 | -0.4614594 | 0.64446908 | 0.99999872 |
| KIAA0100 | 21.3020447 | -0.11321632 | 0.55700397 | -0.2032594 | 0.83893225 | 0.99999872 |
| SDF2 | 20.2038734 | 0.640279689 | 0.43316995 | 1.47812583 | 0.1393741 | 0.99999872 |
| SUPT6H | 29.3643157 | -0.581436822 | 0.31995794 | -1.8172289 | 0.06918207 | 0.99999872 |
| RAB34 | 23.7889194 | 0.238666504 | 0.39613019 | 0.60249511 | 0.54684462 | 0.99999872 |
| RPL23A | 64.606387 | 0.728586844 | 0.41416249 | 1.75918114 | 0.07854675 | 0.99999872 |
| TRAF4 | 14.5350463 | -0.324628669 | 0.47842643 | -0.6785341 | 0.49743308 | 0.99999872 |
| FAM222B | 72.004984 | -0.288418034 | 0.30427176 | -0.9478962 | 0.34318232 | 0.99999872 |

| | | | | | | |
|---|---|---|---|---|---|---|
| ERAL1 | 4.71462512 | 0.041699532 | 0.71145334 | 0.05861176 | 0.95326134 | 0.99999872 |
| FLOT2 | 23.0671674 | -0.528698059 | 0.41970791 | -1.259681 | 0.20778448 | 0.99999872 |
| PHF12 | 23.481429 | -0.041180921 | 0.47526264 | -0.0866488 | 0.9309507 | 0.99999872 |
| MYO18A | 8.40001379 | -0.137516081 | 0.50860823 | -0.2703772 | 0.78687006 | 0.99999872 |
| NUFIP2 | 93.1298435 | -0.009706575 | 0.27087816 | -0.0358337 | 0.97141494 | 0.99999872 |
| TAOK1 | 112.058035 | -0.18219053 | 0.3220331 | -0.5657509 | 0.5715631 | 0.99999872 |
| ABHD15-AS1 | 7.21461645 | 0.319515167 | 0.65966796 | 0.48435757 | 0.62813213 | 0.99999872 |
| GIT1 | 11.3518336 | -0.25037821 | 0.49481879 | -0.5059998 | 0.61285679 | 0.99999872 |
| SSH2 | 75.2387632 | -0.162169482 | 0.35516282 | -0.456606 | 0.64795425 | 0.99999872 |
| NSRP1 | 58.358354 | -0.088213963 | 0.27477471 | -0.3210411 | 0.74817928 | 0.99999872 |
| BLMH | 17.4472057 | 0.024764577 | 0.43464301 | 0.05697682 | 0.95456366 | 0.99999872 |
| CPD | 83.5876898 | 0.273841281 | 0.3608257 | 0.75892954 | 0.44789471 | 0.99999872 |
| GOSR1 | 37.3549669 | -0.285148655 | 0.28447325 | -1.0023742 | 0.31616289 | 0.99999872 |
| CRLF3 | 26.7070966 | 0.70372507 | 0.4324166 | 1.6274238 | 0.10364712 | 0.99999872 |
| ATAD5 | 4.70249567 | 0.831555215 | 0.81298166 | 1.02284621 | 0.30638056 | 0.99999872 |
| TEFM | 13.544311 | 0.474668964 | 0.42878019 | 1.10702168 | 0.26828455 | 0.99999872 |
| RNF135 | 12.4096505 | -0.118398315 | 0.45077487 | -0.2626551 | 0.79281643 | 0.99999872 |
| NF1 | 173.135766 | -0.100238485 | 0.26178373 | -0.3829057 | 0.70178967 | 0.99999872 |
| COPRS | 13.7827429 | -0.053686826 | 0.50506095 | -0.1062977 | 0.91534614 | 0.99999872 |
| UTP6 | 11.6888565 | -0.058408633 | 0.52612032 | -0.1110176 | 0.91160236 | 0.99999872 |
| SUZ12 | 57.062352 | -0.101211637 | 0.27068824 | -0.3739048 | 0.70847513 | 0.99999872 |
| LRRC37B | 16.8390543 | 0.490962326 | 0.4520637 | 1.08604678 | 0.2774583 | 0.99999872 |
| RHOT1 | 33.3409067 | 0.077135651 | 0.30152655 | 0.25581711 | 0.79809206 | 0.99999872 |
| RHBDL3 | 13.279518 | -0.424206023 | 0.80884489 | -0.524459 | 0.5999593 | 0.99999872 |
| C17orf75 | 8.11443131 | -0.209167431 | 0.52942377 | -0.3950851 | 0.6927801 | 0.99999872 |
| ZNF207 | 84.3744951 | 4.05E-07 | 0.25314877 | 1.60E-06 | 0.99999872 | 0.99999872 |
| PSMD11 | 36.1708832 | -0.117753558 | 0.32303906 | -0.364518 | 0.71547122 | 0.99999872 |
| MYO1D | 431.01897 | 0.135378994 | 0.25608172 | 0.52865545 | 0.59704449 | 0.99999872 |
| AC079336.4 | 8.32394193 | 0.750447584 | 0.67628057 | 1.109669 | 0.26714169 | 0.99999872 |
| AC025211.1 | 5.75364718 | 0.556752619 | 0.72434729 | 0.76862663 | 0.44211499 | 0.99999872 |
| TMEM98 | 60.5628466 | 1.071637561 | 0.45960245 | 2.33166197 | 0.01971848 | 0.8671024 |
| CCL2 | 12.2488209 | -0.360304398 | 1.12619503 | -0.3199307 | 0.74902084 | 0.99999872 |
| ZNF830 | 6.32543266 | 0.170892119 | 0.55383778 | 0.30855988 | 0.75765634 | 0.99999872 |
| LIG3 | 8.94332184 | 0.070006233 | 0.53494832 | 0.13086541 | 0.89588178 | 0.99999872 |
| RFFL | 5.14192702 | 0.324602131 | 0.89518034 | 0.36261088 | 0.71689558 | 0.99999872 |
| SLFN5 | 9.69870328 | -1.477538911 | 1.19024473 | -1.241374 | 0.2144676 | 0.99999872 |
| SLFN12 | 4.63834035 | -0.249081698 | 1.08751523 | -0.2290374 | 0.81883982 | 0.99999872 |
| LINC02001 | 8.19136601 | -0.15528101 | 0.51187222 | -0.3033589 | 0.76161634 | 0.99999872 |
| AP2B1 | 41.7713791 | 0.201015661 | 0.31914599 | 0.62985488 | 0.52878953 | 0.99999872 |
| TAF15 | 122.194791 | 0.010535419 | 0.30835919 | 0.03416606 | 0.97274473 | 0.99999872 |
| MMP28 | 19.5376737 | 0.705117144 | 0.71183561 | 0.99056177 | 0.32189961 | 0.99999872 |
| AC015849.3 | 8.53701256 | 1.209887288 | 0.86748996 | 1.39469889 | 0.16310666 | 0.99999872 |
| ZNHIT3 | 9.24063187 | 1.51201516 | 0.56123172 | 2.6941014 | 0.00705787 | 0.62327327 |
| MYO19 | 17.0237309 | -0.593786431 | 0.37674189 | -1.5761094 | 0.11500059 | 0.99999872 |
| GGNBP2 | 46.5085104 | -0.332273072 | 0.27603017 | -1.2037564 | 0.22868376 | 0.99999872 |
| AATF | 22.6103414 | -0.291631792 | 0.52785545 | -0.5524842 | 0.58061666 | 0.99999872 |
| ACACA | 64.2858503 | -0.139476123 | 0.3224957 | -0.4324899 | 0.66538541 | 0.99999872 |
| TADA2A | 16.7380015 | -0.612986126 | 0.35420798 | -1.7305825 | 0.08352626 | 0.99999872 |
| DUSP14 | 12.3835011 | 0.683482377 | 0.66243055 | 1.03177968 | 0.30217534 | 0.99999872 |
| SYNRG | 48.1436343 | -0.002335327 | 0.30932046 | -0.0075499 | 0.99397614 | 0.99999872 |
| DDX52 | 19.2720772 | 0.147085951 | 0.37950381 | 0.38757437 | 0.69833104 | 0.99999872 |
| HNF1B | 12.4250147 | -0.465695231 | 0.5342509 | -0.8716789 | 0.38338358 | 0.99999872 |
| MRPL45 | 11.3069257 | -0.481496552 | 0.47920196 | -1.0047884 | 0.31499877 | 0.99999872 |
| SOCS7 | 14.2455002 | -0.198248264 | 0.53422893 | -0.3710923 | 0.71056876 | 0.99999872 |
| ARHGAP23 | 87.6899486 | -0.73320766 | 0.38260273 | -1.9163681 | 0.05531826 | 0.99999872 |
| MLLT6 | 50.7225127 | -0.6248367 | 0.33073826 | -1.8892181 | 0.05886261 | 0.99999872 |
| CISD3 | 15.4771373 | 0.226912469 | 0.42974124 | 0.52802116 | 0.59748465 | 0.99999872 |
| PCGF2 | 12.7848325 | -0.508328627 | 0.47463225 | -1.0709947 | 0.28417181 | 0.99999872 |
| PSMB3 | 24.4235327 | 0.584809828 | 0.48836201 | 1.19749247 | 0.23111466 | 0.99999872 |
| PIP4K2B | 20.2685783 | -0.240778674 | 0.34923025 | -0.6894554 | 0.49053674 | 0.99999872 |
| CWC25 | 12.1445574 | -0.488979788 | 0.46053315 | -1.0617689 | 0.2883406 | 0.99999872 |

| | | | | | | |
|---|---|---|---|---|---|---|
| RPL23 | 49.2068338 | 0.564661178 | 0.47057064 | 1.19994986 | 0.23015882 | 0.99999872 |
| LASP1 | 26.6166502 | 0.276391241 | 0.41314308 | 0.66899641 | 0.50349777 | 0.99999872 |
| RPL19 | 103.595341 | 0.466223964 | 0.37885335 | 1.23061854 | 0.21846557 | 0.99999872 |
| FBXL20 | 68.8623593 | -0.444416193 | 0.33447295 | -1.3287059 | 0.183945 | 0.99999872 |
| MED1 | 20.0400811 | 0.011503495 | 0.36287462 | 0.03170102 | 0.97471048 | 0.99999872 |
| CDK12 | 39.423547 | -0.336535822 | 0.27770662 | -1.2118394 | 0.22557387 | 0.99999872 |
| STARD3 | 7.06127387 | 0.154944791 | 0.58186167 | 0.26629146 | 0.79001475 | 0.99999872 |
| PGAP3 | 4.82294523 | 0.235631732 | 0.69216168 | 0.34042875 | 0.73353367 | 0.99999872 |
| ERBB2 | 14.3046349 | 0.12673769 | 0.46584235 | 0.27206133 | 0.78557486 | 0.99999872 |
| MIEN1 | 13.0848659 | 0.034968866 | 0.41166749 | 0.08494444 | 0.93230556 | 0.99999872 |
| GRB7 | 6.3446405 | 0.419131344 | 0.76127938 | 0.5505618 | 0.5819341 | 0.99999872 |
| ORMDL3 | 15.7000843 | -0.037576016 | 0.53632008 | -0.0700627 | 0.94414378 | 0.99999872 |
| PSMD3 | 27.8997462 | -0.027982747 | 0.42550428 | -0.0657637 | 0.94756594 | 0.99999872 |
| MED24 | 15.884886 | 0.261789433 | 0.45009704 | 0.58162887 | 0.56081669 | 0.99999872 |
| THRA | 18.9958781 | -0.173951606 | 0.61344495 | -0.2835651 | 0.77674365 | 0.99999872 |
| MSL1 | 37.821552 | 0.405021133 | 0.47449651 | 0.85358085 | 0.39333727 | 0.99999872 |
| CASC3 | 30.2549307 | -0.748573765 | 0.37383452 | -2.0024201 | 0.04523957 | 0.95980711 |
| WIPF2 | 40.477389 | -0.220753907 | 0.30263241 | -0.7294457 | 0.46572908 | 0.99999872 |
| RARA | 11.7791384 | 0.027053801 | 0.49957993 | 0.0541531 | 0.95681319 | 0.99999872 |
| IGFBP4 | 58.5979406 | -0.723561817 | 0.50195947 | -1.4414746 | 1.49E-01 | 0.99999872 |
| KRT10 | 30.4980402 | 0.532401007 | 0.44179251 | 1.20509287 | 0.22816745 | 0.99999872 |
| KRT19 | 17.1567472 | 0.418141205 | 0.74291031 | 0.56284211 | 0.57354241 | 0.99999872 |
| EIF1 | 177.768525 | 0.526421004 | 0.51088593 | 1.03040812 | 0.30281847 | 0.99999872 |
| JUP | 63.897671 | -0.169155613 | 0.45109727 | -0.374987 | 0.70767013 | 0.99999872 |
| P3H4 | 7.89764362 | 0.709462415 | 0.65801971 | 1.07817806 | 0.2809543 | 0.99999872 |
| FKBP10 | 5.56774594 | 0.601697231 | 0.76493003 | 0.78660428 | 0.43151355 | 0.99999872 |
| NT5C3B | 12.3760833 | 0.211482332 | 0.48928982 | 0.43222304 | 0.66557932 | 0.99999872 |
| ACLY | 22.8843006 | 0.010436334 | 0.37481398 | 0.02784404 | 0.97778654 | 0.99999872 |
| CNP | 23.1559202 | -1.287888153 | 0.82431403 | -1.5623756 | 0.11819952 | 0.99999872 |
| DNAJC7 | 37.0427093 | -0.360332959 | 0.35888582 | -1.0040323 | 0.31536304 | 0.99999872 |
| NKIRAS2 | 13.2795552 | -0.698624158 | 0.50031583 | -1.3963663 | 0.16260422 | 0.99999872 |
| DHX58 | 7.2441387 | -1.013887211 | 1.10626224 | -0.9164981 | 0.35940572 | 0.99999872 |
| KAT2A | 5.29618789 | 0.701784569 | 0.6986761 | 1.00444908 | 0.3151622 | 0.99999872 |
| GHDC | 7.82416481 | 0.225329667 | 0.70842841 | 0.31806978 | 0.75043201 | 0.99999872 |
| STAT5B | 39.4668475 | 0.080479989 | 0.34175652 | 0.23548925 | 0.81382902 | 0.99999872 |
| STAT5A | 6.51734707 | -0.670309094 | 0.58456749 | -1.1466753 | 0.25151585 | 0.99999872 |
| STAT3 | 156.561803 | 0.026047461 | 0.36662838 | 0.07104595 | 9.43E-01 | 0.99999872 |
| CAVIN1 | 32.6434686 | -0.531865497 | 0.54811911 | -0.9703466 | 0.33187377 | 0.99999872 |
| ATP6V0A1 | 37.1890983 | -0.418488088 | 0.39460453 | -1.0605253 | 0.28890568 | 0.99999872 |
| NAGLU | 29.4785372 | 0.271796586 | 0.42935403 | 0.63303607 | 0.52671009 | 0.99999872 |
| COASY | 8.13799615 | 0.264262457 | 0.5261129 | 0.5022923 | 0.61546193 | 0.99999872 |
| MLX | 8.3069792 | 0.343266007 | 0.51271079 | 0.66951196 | 0.50316895 | 0.99999872 |
| RETREG3 | 30.1630828 | -0.478142547 | 0.31196757 | -1.5326675 | 0.12535781 | 0.99999872 |
| TUBG1 | 4.188687 | 1.023916955 | 0.77984555 | 1.31297403 | 0.1891917 | NA |
| TUBG2 | 10.4275978 | 0.329087974 | 0.58611295 | 0.56147535 | 0.57447353 | 0.99999872 |
| PLEKHH3 | 6.35250001 | 0.918941453 | 0.65998621 | 1.39236462 | 0.16381201 | 0.99999872 |
| EZH1 | 23.6454077 | -0.361722567 | 0.41596935 | -0.8695895 | 0.38452479 | 0.99999872 |
| VPS25 | 10.1897248 | -0.007987563 | 0.56154707 | -0.0142242 | 0.98865111 | 0.99999872 |
| COA3 | 20.990805 | 0.293778833 | 0.41324688 | 0.71090394 | 0.47714376 | 0.99999872 |
| BECN1 | 12.6439085 | -0.392548077 | 0.44664034 | -0.8788908 | 0.3794605 | 0.99999872 |
| PSME3 | 10.8818537 | -0.555656451 | 0.45891387 | -1.2108077 | 0.22596911 | 0.99999872 |
| RUNDC1 | 11.277904 | -0.522778024 | 0.46606123 | -1.1216939 | 0.26199262 | 0.99999872 |
| RPL27 | 46.9947287 | 0.515790011 | 0.43380361 | 1.18899429 | 0.23444192 | 0.99999872 |
| IFI35 | 24.5775991 | -1.42942671 | 1.55974546 | -0.9164487 | 0.3594316 | 0.99999872 |
| VAT1 | 9.8592226 | 0.072447826 | 0.6914922 | 0.10477027 | 0.9165581 | 0.99999872 |
| BRCA1 | 9.47558228 | 0.652010589 | 0.83108266 | 0.78453158 | 0.43272826 | 0.99999872 |
| NBR1 | 36.3178546 | 0.099168452 | 0.27709861 | 0.35788144 | 0.72043204 | 0.99999872 |
| TMEM106A | 7.19409439 | 0.11245412 | 0.72524399 | 0.15505695 | 0.87677642 | 0.99999872 |
| LINC00854 | 34.5817103 | -1.687845653 | 0.61546107 | -2.7424085 | 0.00609904 | 0.58725857 |
| ARL4D | 6.64808584 | 0.77167567 | 0.86402958 | 0.89311256 | 0.3717969 | 0.99999872 |
| DHX8 | 13.3889018 | -0.186588173 | 0.43317813 | -0.4307424 | 0.66665571 | 0.99999872 |

| | | | | | | |
|---|---|---|---|---|---|---|
| DUSP3 | 30.1231328 | 0.253988895 | 0.45778506 | 0.55482128 | 0.57901691 | 0.99999872 |
| MPP3 | 12.3158281 | -0.369876319 | 0.72072326 | -0.5132016 | 0.60781032 | 0.99999872 |
| TMEM101 | 12.1400227 | 0.838053995 | 0.66313517 | 1.26377552 | 0.20631061 | 0.99999872 |
| LSM12 | 14.2621904 | -0.507897426 | 0.45759828 | -1.1099199 | 0.26703355 | 0.99999872 |
| G6PC3 | 14.4647146 | -0.344447251 | 0.61366003 | -0.5612998 | 0.57459318 | 0.99999872 |
| HDAC5 | 15.7501508 | -0.074793321 | 0.48709364 | -0.1535502 | 0.87796441 | 0.99999872 |
| TMUB2 | 12.9467027 | -0.305220743 | 0.49851207 | -0.6122635 | 0.54036344 | 0.99999872 |
| ATXN7L3 | 9.41805623 | -0.496793865 | 0.49160389 | -1.0105572 | 0.3122284 | 0.99999872 |
| UBTF | 22.9181386 | -0.7574378 | 0.45648054 | -1.6592992 | 0.09705552 | 0.99999872 |
| AC003102.1 | 5.67101716 | 0.077779671 | 0.90533654 | 0.08591244 | 0.93153602 | 0.99999872 |
| SLC25A39 | 12.4179003 | 0.766154487 | 0.55037002 | 1.39207163 | 0.16390071 | 0.99999872 |
| GRN | 80.2655993 | 0.557165674 | 0.57165287 | 0.97465736 | 0.32973025 | 0.99999872 |
| GPATCH8 | 138.585511 | -0.006789966 | 0.18238874 | -0.037228 | 0.97030322 | 0.99999872 |
| FZD2 | 24.2927693 | 0.498492408 | 0.65394601 | 0.76228374 | 0.44589068 | 0.99999872 |
| CCDC43 | 6.0256336 | -0.677794901 | 0.56456434 | -1.2005627 | 0.22992087 | 0.99999872 |
| DBF4B | 7.61002962 | -1.213701826 | 0.63210171 | -1.9201053 | 0.0548446 | 0.99999872 |
| EFTUD2 | 17.91287 | -0.268506927 | 0.42723377 | -0.6284778 | 0.52969101 | 0.99999872 |
| C1QL1 | 149.535308 | 1.039690579 | 0.55097239 | 1.88701031 | 0.05915894 | 0.99999872 |
| DCAKD | 18.4819704 | -0.380467484 | 0.4392114 | -0.8662514 | 0.38635231 | 0.99999872 |
| NMT1 | 39.7322194 | -0.061683914 | 0.30111978 | -0.2048484 | 0.83769055 | 0.99999872 |
| PLCD3 | 38.3871255 | -0.08913922 | 0.40563389 | -0.2197529 | 0.82606361 | 0.99999872 |
| ACBD4 | 9.51180861 | -0.25531785 | 0.50528198 | -0.5052978 | 0.61334972 | 0.99999872 |
| HEXIM1 | 31.457426 | -0.839436091 | 0.45445934 | -1.8471093 | 0.0647313 | 0.99999872 |
| AC138150.1 | 19.9102028 | -0.010648326 | 0.55097867 | -0.0193262 | 0.98458088 | 0.99999872 |
| MAP3K14 | 13.0452211 | -0.624103864 | 0.4875772 | -1.2800103 | 0.2005415 | 0.99999872 |
| PLEKHM1 | 15.1139797 | -0.661134526 | 0.38280474 | -1.7270803 | 0.08415324 | 0.99999872 |
| LINC02210-C | 15.3937664 | 0.721922054 | 0.48309444 | 1.49437046 | 0.13507882 | 0.99999872 |
| MAPT | 41.7880095 | 0.545047336 | 0.74810757 | 0.72856813 | 0.46626588 | 0.99999872 |
| KANSL1 | 197.616816 | 0.211840158 | 0.34522152 | 0.61363543 | 0.53945626 | 0.99999872 |
| ARL17B | 51.7611641 | 1.754374283 | 0.99346469 | 1.76591508 | 0.0774101 | 0.99999872 |
| LRRC37A2 | 10.1443146 | 0.516301898 | 0.69111582 | 0.74705554 | 0.45503004 | 0.99999872 |
| ARL17A | 11.3410575 | -0.469357629 | 0.4799789 | -0.9778714 | 0.32813795 | 0.99999872 |
| NSF | 46.4053706 | 0.578670845 | 0.52791081 | 1.09615267 | 0.27301197 | 0.99999872 |
| WNT3 | 15.0776667 | 0.189994132 | 1.1305026 | 0.16806165 | 0.86653478 | 0.99999872 |
| AC005670.2 | 7.66679021 | 0.554883191 | 0.64136942 | 0.8651538 | 0.38695438 | 0.99999872 |
| CDC27 | 69.6365203 | -0.036277941 | 0.32045027 | -0.1132093 | 0.90986465 | 0.99999872 |
| AC068234.1 | 178.724815 | -0.535619713 | 0.4706373 | -1.1380732 | 0.25508991 | 0.99999872 |
| AC068234.2 | 24.0877577 | 0.756973348 | 0.39973457 | 1.89368997 | 0.05826617 | 0.99999872 |
| EFCAB13 | 86.2712781 | -0.17915266 | 0.47446695 | -0.3775872 | 0.70573725 | 0.99999872 |
| AC040934.1 | 5.6709363 | 0.306420609 | 0.62498306 | 0.49028626 | 0.62393135 | 0.99999872 |
| NPEPPS | 106.66594 | -0.104444667 | 0.19256115 | -0.5423974 | 0.58754476 | 0.99999872 |
| AC025682.1 | 14.2097228 | 0.517145342 | 0.71198496 | 0.72634307 | 0.46762848 | 0.99999872 |
| KPNB1 | 76.914669 | -0.624799138 | 0.39410288 | -1.5853706 | 0.11288215 | 0.99999872 |
| MRPL10 | 9.80641162 | -0.160276121 | 0.64492604 | -0.2485186 | 0.80373317 | 0.99999872 |
| SCRN2 | 10.8781252 | 0.308481535 | 0.48972058 | 0.62991336 | 0.52875127 | 0.99999872 |
| SP2 | 11.099968 | -0.745851698 | 0.52197866 | -1.4288931 | 0.15303497 | 0.99999872 |
| PNPO | 6.56928221 | -0.218655035 | 0.68195339 | -0.3206305 | 0.74849044 | 0.99999872 |
| CDK5RAP3 | 13.767391 | -0.236063495 | 0.50443722 | -0.467974 | 0.63980319 | 0.99999872 |
| COPZ2 | 5.53644915 | 0.055738707 | 0.58964918 | 0.09452859 | 0.92468927 | 0.99999872 |
| NFE2L1 | 105.594068 | -0.303345546 | 0.28202342 | -1.0756041 | 0.28210436 | 0.99999872 |
| CBX1 | 17.3931319 | 0.058097434 | 0.50521852 | 0.11499466 | 0.90844935 | 0.99999872 |
| SNX11 | 9.49232596 | -0.03785966 | 0.5150571 | -0.0735058 | 0.94140367 | 0.99999872 |
| SKAP1 | 49.0530799 | -0.907905675 | 0.63024579 | -1.4405581 | 0.14970957 | 0.99999872 |
| HOXB2 | 7.0447773 | 0.43964283 | 0.66998673 | 0.65619633 | 0.51169781 | 0.99999872 |
| HOXB3 | 17.1266679 | -0.653904569 | 0.46534558 | -1.4052021 | 0.1599612 | 0.99999872 |
| HOXB7 | 10.8554435 | 0.144475164 | 0.4504608 | 0.32072749 | 0.74841691 | 0.99999872 |
| CALCOCO2 | 24.7796603 | -0.121593459 | 0.47396759 | -0.2565438 | 0.79753094 | 0.99999872 |
| ATP5MC1 | 27.8626189 | 0.312841286 | 0.37241735 | 0.84002876 | 0.40089226 | 0.99999872 |
| UBE2Z | 40.6160864 | -0.228959964 | 0.37808238 | -0.6055822 | 0.54479222 | 0.99999872 |
| SNF8 | 18.1632469 | 0.197011412 | 0.36572673 | 0.53868475 | 0.5901044 | 0.99999872 |
| ZNF652 | 37.9860397 | 0.069919979 | 0.41420262 | 0.16880622 | 0.86594907 | 0.99999872 |

| | | | | | | |
|---|---|---|---|---|---|---|
| AC091180.5 | 4.80676547 | 2.218053076 | 0.87940212 | 2.52222848 | 0.01166139 | 0.72670507 |
| PHB | 26.2484237 | 0.54299138 | 0.36693597 | 1.47979872 | 0.13892697 | 0.99999872 |
| SPOP | 40.6858411 | -0.321312286 | 0.26904905 | -1.1942517 | 0.23237952 | 0.99999872 |
| SLC35B1 | 7.77167031 | 1.37016995 | 0.66203666 | 2.06962852 | 0.03848715 | 0.95980711 |
| FAM117A | 9.22633008 | -0.334418212 | 0.54396723 | -0.6147764 | 0.5387024 | 0.99999872 |
| KAT7 | 51.996851 | 0.33936727 | 0.4647355 | 0.73023745 | 0.46524505 | 0.99999872 |
| TAC4 | 5.20149177 | 0.547694518 | 0.64749703 | 0.84586414 | 0.39762854 | 0.99999872 |
| ITGA3 | 394.312611 | -0.704680999 | 0.45967452 | -1.5329999 | 0.12527588 | 0.99999872 |
| PDK2 | 9.73011286 | 1.060816745 | 0.53571137 | 1.98020205 | 0.04768083 | 0.97387863 |
| SAMD14 | 7.47625561 | 1.41796588 | 0.63264496 | 2.24132961 | 0.02500473 | 0.88365618 |
| PPP1R9B | 6.89571042 | -1.001783029 | 0.72847068 | -1.3751865 | 0.16907364 | 0.99999872 |
| XYLT2 | 10.266619 | -0.163116524 | 0.47893343 | -0.3405829 | 0.73341762 | 0.99999872 |
| MRPL27 | 16.155681 | 0.214454063 | 0.51841568 | 0.41367203 | 0.67911432 | 0.99999872 |
| LRRC59 | 11.472976 | 0.102850668 | 0.57466071 | 0.17897633 | 0.85795628 | 0.99999872 |
| ACSF2 | 13.8441562 | 1.114214997 | 0.91852446 | 1.21304881 | 2.25E-01 | 0.99999872 |
| RSAD1 | 7.33239939 | -0.182931626 | 0.55570484 | -0.3291885 | 0.74201324 | 0.99999872 |
| SPATA20 | 9.51567043 | -0.442475813 | 0.48980114 | -0.9033785 | 0.36632505 | 0.99999872 |
| ABCC3 | 11.7380391 | -0.423452718 | 0.67957952 | -0.6231099 | 0.53321231 | 0.99999872 |
| ANKRD40 | 20.9033347 | -0.377462301 | 0.68399232 | -0.5518517 | 0.58104999 | 0.99999872 |
| AC005921.2 | 9.32731574 | 0.454667033 | 0.56228902 | 0.80860024 | 0.41874513 | 0.99999872 |
| LUC7L3 | 129.180563 | -0.058203037 | 0.21310657 | -0.273117 | 0.78476324 | 0.99999872 |
| TOB1 | 31.3969772 | -0.114966321 | 0.5262701 | -0.218455 | 0.82707464 | 0.99999872 |
| SPAG9 | 91.9091665 | -0.128764437 | 0.26237182 | -0.4907708 | 0.62358854 | 0.99999872 |
| AC005920.2 | 7.30336248 | 0.54116253 | 0.72923765 | 0.74209351 | 0.45803068 | 0.99999872 |
| MBTD1 | 35.0548036 | -0.260138803 | 0.31789516 | -0.8183163 | 0.41317658 | 0.99999872 |
| UTP18 | 10.1266323 | -0.226078045 | 0.54339232 | -0.4160494 | 0.67737386 | 0.99999872 |
| CA10 | 282.176391 | 0.322163965 | 0.4292786 | 0.75047758 | 0.45296712 | 0.99999872 |
| TOM1L1 | 43.8443887 | 0.575183946 | 0.33579212 | 1.71291676 | 0.08672785 | 0.99999872 |
| COX11 | 16.2036612 | 0.530861276 | 0.45278691 | 1.17243069 | 0.24102418 | 0.99999872 |
| STXBP4 | 38.4813174 | 0.337524186 | 0.40214978 | 0.83929968 | 0.40130116 | 0.99999872 |
| HLF | 6.42459527 | -1.630139921 | 0.9100119 | -1.7913391 | 0.07323889 | 0.99999872 |
| PCTP | 7.62271567 | -0.037502236 | 0.58466915 | -0.0641427 | 0.94885663 | 0.99999872 |
| DGKE | 7.36256928 | 0.164130802 | 0.72808636 | 0.22542765 | 0.8216466 | 0.99999872 |
| TRIM25 | 6.82348696 | 0.09731883 | 0.88293454 | 0.11022202 | 0.9122333 | 0.99999872 |
| COIL | 7.08992403 | -0.076940959 | 0.54413799 | -0.1413997 | 0.88755418 | 0.99999872 |
| SCPEP1 | 16.8989126 | 1.090471838 | 0.57807112 | 1.88639735 | 0.05924143 | 0.99999872 |
| AKAP1 | 13.7656186 | -0.119833946 | 0.45588287 | -0.2628613 | 0.79265752 | 0.99999872 |
| MSI2 | 444.350888 | 0.039137918 | 0.38899221 | 0.10061363 | 0.91985718 | 0.99999872 |
| MRPS23 | 17.4848472 | 0.204599851 | 0.42684612 | 0.47932929 | 0.63170439 | 0.99999872 |
| CUEDC1 | 25.2208982 | -0.241214092 | 0.38886183 | -0.620308 | 0.53505505 | 0.99999872 |
| VEZF1 | 35.2317724 | -0.039088558 | 0.41373698 | -0.0944768 | 0.92473039 | 0.99999872 |
| AC015813.2 | 12.0960151 | 0.364027087 | 0.53918752 | 0.67514004 | 0.4995868 | 0.99999872 |
| DYNLL2 | 19.4928335 | -0.774576144 | 0.3968352 | -1.9518837 | 0.05095202 | 0.98949866 |
| MKS1 | 5.37943862 | -1.01568409 | 0.67076589 | -1.5142155 | 0.1299712 | 0.99999872 |
| TSPOAP1 | 6.93221834 | 0.839519764 | 0.64902919 | 1.29350078 | 0.19583791 | 0.99999872 |
| SUPT4H1 | 11.9720818 | 0.232534846 | 0.77234835 | 0.30107509 | 0.76335724 | 0.99999872 |
| MTMR4 | 9.49430435 | 0.173403692 | 0.61926568 | 0.28001502 | 0.77946598 | 0.99999872 |
| RAD51C | 9.80027611 | 0.379218598 | 0.53003564 | 0.71545868 | 0.47432566 | 0.99999872 |
| TRIM37 | 20.4341271 | -0.497074729 | 0.34810609 | -1.4279404 | 0.15330903 | 0.99999872 |
| SKA2 | 19.9471384 | 0.097366378 | 0.41166533 | 0.23651828 | 0.81303052 | 0.99999872 |
| YPEL2 | 21.3599691 | 0.506097954 | 0.56256809 | 0.8996208 | 0.36832208 | 0.99999872 |
| DHX40 | 18.4697964 | 0.39282588 | 0.3951453 | 0.9941302 | 0.32015948 | 0.99999872 |
| CLTC | 134.44675 | 0.34629642 | 0.3284467 | 1.05434587 | 0.2917246 | 0.99999872 |
| PTRH2 | 16.6512043 | -0.314415724 | 0.49955096 | -0.6293967 | 0.52908938 | 0.99999872 |
| VMP1 | 92.8308685 | -0.746681694 | 0.56716373 | -1.3165188 | 0.18799999 | 0.99999872 |
| TUBD1 | 13.4474239 | 0.079184242 | 0.49788768 | 0.15904037 | 0.87363707 | 0.99999872 |
| RPS6KB1 | 18.0227593 | -0.043586469 | 0.50572344 | -0.0861864 | 0.93131826 | 0.99999872 |
| HEATR6 | 6.2385946 | 0.601014605 | 0.59278746 | 1.01387875 | 0.31064061 | 0.99999872 |
| USP32 | 57.2603698 | 0.060380014 | 0.32104473 | 0.18807353 | 0.85081901 | 0.99999872 |
| APPBP2 | 99.2039599 | 0.109475054 | 0.31856885 | 0.34364644 | 0.73111219 | 0.99999872 |
| PPM1D | 23.2803013 | -0.143279055 | 0.43125545 | -0.3322371 | 0.73971024 | 0.99999872 |

| | | | | | | |
|---|---|---|---|---|---|---|
| BCAS3 | 176.675264 | -0.069805012 | 0.25349593 | -0.2753694 | 0.78303248 | 0.99999872 |
| NACA2 | 7.84812077 | 0.322839737 | 0.52977924 | 0.60938541 | 0.54226901 | 0.99999872 |
| INTS2 | 12.814366 | -0.395475033 | 0.45937855 | -0.8608914 | 0.38929787 | 0.99999872 |
| MED13 | 110.496265 | -0.175145916 | 0.24901893 | -0.7033438 | 0.48184154 | 0.99999872 |
| METTL2A | 8.88581012 | -0.071262811 | 0.494598 | -0.1440823 | 0.88543549 | 0.99999872 |
| TLK2 | 57.5181546 | -0.144244198 | 0.31460722 | -0.4584898 | 0.6466006 | 0.99999872 |
| MRC2 | 58.560262 | 0.813776823 | 0.86746056 | 0.93811392 | 0.34818587 | 0.99999872 |
| TANC2 | 58.480972 | -0.146499499 | 0.38113747 | -0.3843744 | 0.70070095 | 0.99999872 |
| CYB561 | 14.6171867 | -0.143508037 | 0.52244448 | -0.2746857 | 0.78355771 | 0.99999872 |
| DCAF7 | 39.4738028 | -0.237209802 | 0.30266675 | -0.7837326 | 0.43319702 | 0.99999872 |
| TACO1 | 6.940879 | -0.855403314 | 0.59542714 | -1.4366213 | 0.15082563 | 0.99999872 |
| MAP3K3 | 63.4225966 | 0.289148105 | 0.46533415 | 0.62137736 | 0.53435136 | 0.99999872 |
| CCDC47 | 8.78492023 | -0.162722709 | 0.4652802 | -0.3497306 | 0.72654091 | 0.99999872 |
| DDX42 | 40.7352034 | -0.601950455 | 0.3169338 | -1.899294 | 0.05752583 | 0.99999872 |
| FTSJ3 | 8.18603316 | -0.274203767 | 0.51324615 | -0.5342539 | 0.59316587 | 0.99999872 |
| PSMC5 | 14.9421495 | -0.368590986 | 0.45413211 | -0.8116382 | 0.41699924 | 0.99999872 |
| SMARCD2 | 9.60024457 | -0.456559789 | 0.51290113 | -0.8901517 | 0.37338446 | 0.99999872 |
| ERN1 | 11.9213332 | 0.073569577 | 0.49950595 | 0.14728469 | 0.88290732 | 0.99999872 |
| SNHG25 | 6.25219217 | -0.329103251 | 0.61666394 | -0.5336833 | 0.59356066 | 0.99999872 |
| TEX2 | 11.2886003 | 0.720696313 | 0.55221512 | 1.30510065 | 0.19185858 | 0.99999872 |
| POLG2 | 9.11472447 | -0.422687673 | 0.57944604 | -0.7294686 | 0.46571509 | 0.99999872 |
| DDX5 | 220.753364 | 0.261294101 | 0.29306964 | 0.89157684 | 0.37261979 | 0.99999872 |
| CEP95 | 23.9335312 | 0.472909342 | 0.4230987 | 1.11772818 | 0.2636831 | 0.99999872 |
| SMURF2 | 47.1971462 | 0.326624185 | 0.28800785 | 1.13408083 | 0.25676065 | 0.99999872 |
| LRRC37A3 | 44.8878786 | 0.322986523 | 0.36288596 | 0.89004964 | 0.37343923 | 0.99999872 |
| GNA13 | 29.003652 | 0.043932782 | 0.39713269 | 0.11062495 | 0.91191377 | 0.99999872 |
| CEP112 | 110.762615 | 0.404750779 | 0.29431873 | 1.37521243 | 0.1690656 | 0.99999872 |
| PRKCA | 99.1075875 | -0.230151433 | 0.33912906 | -0.6786544 | 0.49735686 | 0.99999872 |
| HELZ | 79.9239633 | 0.127181881 | 0.29800648 | 0.42677555 | 0.66954282 | 0.99999872 |
| PSMD12 | 22.7247811 | -0.055517876 | 0.50155906 | -0.1106906 | 0.9118617 | 0.99999872 |
| PITPNC1 | 72.0277495 | -1.371274405 | 0.72577303 | -1.8893984 | 0.05883846 | 0.99999872 |
| NOL11 | 13.3966683 | -0.030210757 | 0.56163907 | -0.0537903 | 0.9571022 | 0.99999872 |
| BPTF | 138.426044 | -0.124874746 | 0.34159071 | -0.3655683 | 0.71468721 | 0.99999872 |
| KPNA2 | 8.89306641 | 0.242820187 | 0.75864511 | 0.32007085 | 0.74891462 | 0.99999872 |
| AC005332.7 | 8.51956262 | -0.521053371 | 0.54941236 | -0.948383 | 0.34293449 | 0.99999872 |
| AMZ2 | 8.11102908 | -0.314326238 | 0.47509385 | -0.6616087 | 0.50822201 | 0.99999872 |
| ARSG | 9.08086522 | -0.181779964 | 0.77638011 | -0.2341378 | 0.81487797 | 0.99999872 |
| WIPI1 | 21.2275799 | 0.177263247 | 0.39232533 | 0.45182717 | 0.65139349 | 0.99999872 |
| PRKAR1A | 94.3688139 | 0.231196262 | 0.32770599 | 0.70549905 | 0.48049972 | 0.99999872 |
| FAM20A | 11.4491707 | 0.271802779 | 0.65582262 | 0.41444557 | 0.67854782 | 0.99999872 |
| LINC01482 | 9.62384949 | 1.324606876 | 1.22300009 | 1.08307995 | 0.27877294 | 0.99999872 |
| ABCA10 | 15.7030788 | -0.671079612 | 0.73920224 | -0.907843 | 0.36396116 | 0.99999872 |
| ABCA5 | 47.870951 | 0.102531617 | 0.33291019 | 0.30798582 | 0.75809312 | 0.99999872 |
| AC005208.1 | 11.436578 | 2.47423417 | 1.23745029 | 1.99946146 | 0.04555845 | 0.95980711 |
| KCNJ16 | 34.1392288 | -0.092427604 | 0.75239844 | -0.122844 | 0.90223066 | 0.99999872 |
| SLC39A11 | 59.5292862 | -0.047623495 | 0.41288004 | -0.1153446 | 0.90817197 | 0.99999872 |
| COG1 | 10.3259974 | 0.242486688 | 0.47450715 | 0.51102852 | 0.60933108 | 0.99999872 |
| FAM104A | 22.790882 | -0.029787239 | 0.33464429 | -0.0890116 | 0.92907265 | 0.99999872 |
| C17orf80 | 9.34609525 | 0.401296091 | 0.51561183 | 0.77829109 | 0.43639743 | 0.99999872 |
| CDC42EP4 | 30.9801447 | -0.417699946 | 0.45227523 | -0.9235526 | 0.35571932 | 0.99999872 |
| SDK2 | 7.78683559 | 0.956782317 | 1.04865362 | 0.91239118 | 0.36156283 | 0.99999872 |
| RPL38 | 32.1320638 | 1.051208354 | 0.45364175 | 2.31726547 | 0.02048927 | 0.8671024 |
| GPRC5C | 13.3410494 | -0.231295612 | 0.61941054 | -0.3734125 | 0.70884149 | 0.99999872 |
| SLC9A3R1 | 54.1119864 | 0.069755154 | 0.43479917 | 0.16043074 | 0.87254177 | 0.99999872 |
| NAT9 | 17.5945186 | -0.382552411 | 0.41514687 | -0.9214869 | 0.35679626 | 0.99999872 |
| TMEM104 | 8.40449587 | -0.34458696 | 0.5143494 | -0.6699472 | 0.50289142 | 0.99999872 |
| MRPL58 | 7.22403215 | -0.403234362 | 0.61203389 | -0.6588432 | 0.50999647 | 0.99999872 |
| KCTD2 | 24.2265921 | -0.069985279 | 0.54558377 | -0.128276 | 0.89793059 | 0.99999872 |
| ATP5PD | 17.1735775 | 1.217949904 | 0.47416998 | 2.56859347 | 0.01021122 | 0.70849208 |
| SLC16A5 | 11.5235741 | -0.452450483 | 0.53684689 | -0.8427924 | 0.39934455 | 0.99999872 |
| ARMC7 | 3.9390146 | 0.616674116 | 0.80994604 | 0.76137679 | 0.44643205 | NA |

| | | | | | | | |
|---|---|---|---|---|---|---|---|
| NT5C | 29.5028115 | | -0.099927699 | 0.40112104 | -0.2491211 | 0.80326714 | 0.99999872 |
| JPT1 | 11.8530077 | | -0.562248173 | 0.5129118 | -1.0961888 | 0.27299615 | 0.99999872 |
| SUMO2 | 15.0864769 | | 0.769014593 | 0.41522109 | 1.85206054 | 0.06401713 | 0.99999872 |
| NUP85 | 7.47134265 | | 0.269585965 | 0.64848911 | 0.41571395 | 0.67761934 | 0.99999872 |
| GGA3 | 11.8669424 | | -0.322295024 | 0.49876012 | -0.6461924 | 0.51815473 | 0.99999872 |
| MRPS7 | 16.622638 | | 0.916608124 | 0.42493148 | 2.15707275 | 0.031 | 0.91534437 |
| MIF4GD | 5.96795492 | | -0.299424813 | 0.58645587 | -0.5105667 | 0.60965453 | 0.99999872 |
| GRB2 | 44.2157988 | | -0.215724137 | 0.31190616 | -0.6916315 | 0.48916875 | 0.99999872 |
| AC011933.4 | 5.75200536 | | 0.573000352 | 0.88831116 | 0.64504464 | 0.51889826 | 0.99999872 |
| TMEM94 | 12.7660611 | | 0.05833627 | 0.5290125 | 0.1102739 | 0.91219216 | 0.99999872 |
| CASKIN2 | 6.44997787 | | -0.596900272 | 0.61874072 | -0.9647018 | 0.33469421 | 0.99999872 |
| TSEN54 | 3.75668138 | | 1.533197767 | 0.87091914 | 1.76043641 | 0.07833384 | NA |
| LLGL2 | 23.2722194 | | 0.047785983 | 0.41516086 | 0.11510233 | 0.90836401 | 0.99999872 |
| MYO15B | 11.4065374 | | -1.191744995 | 0.64373084 | -1.8513095 | 0.06412505 | 0.99999872 |
| RECQL5 | 7.7114343 | | -0.681924287 | 0.58909648 | -1.1575766 | 0.24703687 | 0.99999872 |
| SAP30BP | 20.9111626 | | -0.24932356 | 0.38005613 | -0.6560177 | 0.51181271 | 0.99999872 |
| GALK1 | 6.81450502 | | 0.55373318 | 0.68374347 | 0.80985517 | 0.41802342 | 0.99999872 |
| H3F3B | 230.825398 | | 0.781655867 | 0.47427835 | 1.64809518 | 0.09933314 | 0.99999872 |
| UNK | 24.5246147 | | -0.122367547 | 0.35980337 | -0.3400956 | 0.73378453 | 0.99999872 |
| WBP2 | 29.9699155 | | -0.076301265 | 0.31863005 | -0.2394666 | 0.81074377 | 0.99999872 |
| TRIM47 | 4.13182041 | | 2.012167683 | 1.00293509 | 2.00627907 | 0.04482648 | NA |
| TRIM65 | 6.59887571 | | 0.173887856 | 0.52699515 | 0.32996101 | 0.74142942 | 0.99999872 |
| ACOX1 | 19.0905691 | | 0.262177515 | 0.41166685 | 0.63686817 | 0.52421072 | 0.99999872 |
| TEN1 | 6.46807029 | | -0.16521796 | 0.60108155 | -0.2748678 | 0.78341781 | 0.99999872 |
| SRP68 | 18.9359702 | | -0.095207805 | 0.41549812 | -0.2291414 | 0.81875905 | 0.99999872 |
| EXOC7 | 22.6334177 | | -0.168769907 | 0.34414334 | -0.4904058 | 0.62384674 | 0.99999872 |
| UBALD2 | 12.9334135 | | -0.217046639 | 0.65490698 | -0.331416 | 0.7403303 | 0.99999872 |
| PRPSAP1 | 15.6101639 | | 1.015767238 | 0.50891576 | 1.99594377 | 0.04594004 | 0.95980711 |
| UBE2O | 19.0402072 | | -0.411879358 | 0.46484207 | -0.886063 | 0.37558358 | 0.99999872 |
| MXRA7 | 46.6887201 | | 0.394478939 | 0.51077149 | 0.77231981 | 0.43992503 | 0.99999872 |
| JMJD6 | 6.78252104 | | -0.047668488 | 0.54834658 | -0.0869313 | 0.93072611 | 0.99999872 |
| SRSF2 | 47.0734452 | | 0.15769006 | 0.33006869 | 0.47774922 | 0.6328287 | 0.99999872 |
| MFSD11 | 29.8333336 | | 0.506455765 | 0.49201334 | 1.02935373 | 0.30331348 | 0.99999872 |
| SEC14L1 | 83.2713271 | | 0.078680701 | 0.27723937 | 0.2838006 | 0.77656319 | 0.99999872 |
| 9-Sep | 51.8191087 | | -0.118776172 | 0.52354235 | -0.2268702 | 0.82052465 | 0.99999872 |
| TNRC6C | 71.2749312 | | -0.325515477 | 0.43787879 | -0.7433917 | 0.45724454 | 0.99999872 |
| SYNGR2 | 31.8305559 | | 0.197051728 | 0.53666929 | 0.36717534 | 0.71348823 | 0.99999872 |
| AFMID | 8.9012158 | | 0.072431541 | 0.52342034 | 0.13838121 | 0.88993914 | 0.99999872 |
| SOCS3 | 42.9594446 | | 0.274354903 | 0.96492635 | 0.2843273 | 7.76E-01 | 0.99999872 |
| AC061992.1 | 5.83251776 | | 0.045984861 | 1.22631248 | 0.03749849 | 0.97008755 | 0.99999872 |
| PGS1 | 25.1591622 | | -0.754204568 | 0.39175923 | -1.9251737 | 0.05420763 | 0.99999872 |
| CYTH1 | 40.2277775 | | -0.143100357 | 0.29868556 | -0.4791004 | 0.63186723 | 0.99999872 |
| USP36 | 20.9347942 | | -0.76761799 | 0.33700596 | -2.2777579 | 0.02274101 | 0.88337526 |
| TIMP2 | 193.580207 | | -0.035311819 | 0.51322133 | -0.0688043 | 0.94514542 | 0.99999872 |
| LGALS3BP | 113.254209 | | -0.07823669 | 0.79865996 | -0.09796 | 0.9219641 | 0.99999872 |
| CANT1 | 12.4560935 | | 0.172080097 | 0.67433284 | 0.2551857 | 0.79857968 | 0.99999872 |
| C1QTNF1 | 13.5067796 | | -0.557266841 | 0.84004404 | -0.6633781 | 0.50708841 | 0.99999872 |
| CBX4 | 4.1511021 | | 0.183773651 | 0.79416206 | 0.23140573 | 0.81699961 | NA |
| TBC1D16 | 19.7436264 | | 0.113566815 | 0.74208422 | 0.15303764 | 0.8783686 | 0.99999872 |
| CCDC40 | 6.08807519 | | -0.651356118 | 0.67880629 | -0.9595611 | 0.33727615 | 0.99999872 |
| GAA | 18.8936283 | | 0.293051029 | 0.51679084 | 0.56705925 | 0.57067392 | 0.99999872 |
| EIF4A3 | 11.7451964 | | 0.265330903 | 0.54038852 | 0.49100026 | 0.62342626 | 0.99999872 |
| SGSH | 13.3314484 | | 0.051006776 | 0.54771169 | 0.09312705 | 0.92580263 | 0.99999872 |
| SLC26A11 | 9.96483554 | | 0.07944539 | 0.52365844 | 0.15171223 | 0.87941391 | 0.99999872 |
| RNF213 | 123.97136 | | -0.668896743 | 0.60600468 | -1.1037815 | 0.26968794 | 0.99999872 |
| AC124319.2 | 31.8695213 | | -0.287639844 | 0.47439689 | -0.6063274 | 0.54429735 | 0.99999872 |
| ENDOV | 13.5357292 | | 0.145896368 | 0.41825324 | 0.34882305 | 0.72722216 | 0.99999872 |
| RPTOR | 48.1247409 | | -0.419007645 | 0.27095294 | -1.5464222 | 0.12200263 | 0.99999872 |
| CHMP6 | 6.6739919 | | -0.313193663 | 0.62359698 | -0.5022373 | 0.61550061 | 0.99999872 |
| BAIAP2 | 43.0477416 | | -0.512171638 | 0.42518037 | -1.2045985 | 0.22835833 | 0.99999872 |
| NDUFAF8 | 14.4817674 | | 0.360940623 | 0.52931967 | 0.68189535 | 0.49530513 | 0.99999872 |

| | | | | | | |
|---|---|---|---|---|---|---|
| SLC38A10 | 52.6635539 | 0.344510268 | 0.30725496 | 1.12125209 | 0.26218057 | 0.99999872 |
| AC110285.2 | 7.00599405 | -0.583032976 | 0.6522644 | -0.8938599 | 0.37139687 | 0.99999872 |
| BAHCC1 | 4.10765332 | -0.979303674 | 0.80480752 | -1.2168173 | 0.22367375 | NA |
| ACTG1 | 594.228316 | 0.057855699 | 0.58473132 | 0.09894407 | 0.92118267 | 0.99999872 |
| FAAP100 | 5.73963289 | -1.121537407 | 0.56375719 | -1.989398 | 0.04665729 | 0.96425074 |
| NPLOC4 | 51.2890599 | -0.372374435 | 0.31034116 | -1.1998873 | 0.2301831 | 0.99999872 |
| OXLD1 | 12.5712362 | 0.606840944 | 0.48013029 | 1.26390888 | 0.20626273 | 0.99999872 |
| CCDC137 | 9.52797897 | -0.32133119 | 0.48870867 | -0.6575107 | 0.51085258 | 0.99999872 |
| ARL16 | 14.2052415 | 0.411387908 | 0.41149158 | 0.99974807 | 0.31743244 | 0.99999872 |
| HGS | 17.6283597 | -0.422272145 | 0.40889015 | -1.0327276 | 0.30173139 | 0.99999872 |
| MCRIP1 | 24.8885744 | -0.147579483 | 0.46686947 | -0.3161044 | 0.75192328 | 0.99999872 |
| P4HB | 147.605826 | 0.526903276 | 0.51486524 | 1.02338095 | 0.30612776 | 0.99999872 |
| ARHGDIA | 68.4173018 | -0.351260125 | 0.45741291 | -0.7679279 | 0.44253004 | 0.99999872 |
| ALYREF | 3.93832764 | 0.595714083 | 0.86775248 | 0.68650231 | 0.49239641 | NA |
| ANAPC11 | 37.1994453 | 0.515160708 | 0.40209783 | 1.28118252 | 0.20012957 | 0.99999872 |
| PCYT2 | 8.50857975 | -0.367731375 | 0.71663895 | -0.5131334 | 0.60785802 | 0.99999872 |
| SIRT7 | 9.60074531 | -0.34042081 | 0.47379628 | -0.7184962 | 0.47245141 | 0.99999872 |
| MAFG | 8.13287633 | 0.093027436 | 0.48929429 | 0.19012573 | 0.84921061 | 0.99999872 |
| ASPSCR1 | 12.6354739 | -0.985090759 | 0.43968159 | -2.240464 | 0.02506082 | 0.88365618 |
| CENPX | 10.0015899 | 0.344746383 | 0.59858937 | 0.57593135 | 0.56466159 | 0.99999872 |
| DCXR | 17.0275688 | -0.745182843 | 0.53610359 | -1.3899979 | 0.16452953 | 0.99999872 |
| RFNG | 12.1049304 | -0.6566675 | 0.50298386 | -1.3055439 | 0.19170772 | 0.99999872 |
| GPS1 | 15.2645608 | -0.124295478 | 0.57694162 | -0.2154386 | 0.8294254 | 0.99999872 |
| DUS1L | 10.2625399 | -0.558041924 | 0.54589629 | -1.022249 | 0.30666308 | 0.99999872 |
| FASN | 5.54818242 | -0.926890115 | 0.63644421 | -1.4563572 | 0.1452939 | 0.99999872 |
| CCDC57 | 32.5948567 | -0.272088735 | 0.31276997 | -0.8699324 | 0.38433735 | 0.99999872 |
| SLC16A3 | 7.85154939 | 0.675554357 | 0.58887786 | 1.14718926 | 0.25130341 | 0.99999872 |
| CSNK1D | 32.4503351 | 0.074481248 | 0.36865749 | 0.20203373 | 0.83989036 | 0.99999872 |
| SECTM1 | 9.13847049 | -0.610410015 | 2.69065459 | -0.226863 | 0.82053026 | 0.99999872 |
| OGFOD3 | 17.0633234 | -0.466697071 | 0.46153953 | -1.0111746 | 0.31193285 | 0.99999872 |
| HEXDC | 12.7306879 | -0.589768528 | 0.41793003 | -1.4111657 | 0.15819576 | 0.99999872 |
| CYBC1 | 11.8079375 | -0.082457464 | 0.61410571 | -0.1342724 | 0.89318716 | 0.99999872 |
| NARF | 10.8641963 | -0.042539138 | 0.62412562 | -0.068158 | 0.94565988 | 0.99999872 |
| FOXK2 | 51.7613939 | -0.251250469 | 0.2815772 | -0.8922969 | 0.37223381 | 0.99999872 |
| WDR45B | 37.504721 | -0.391270554 | 0.29972794 | -1.305419 | 0.19175021 | 0.99999872 |
| RAB40B | 8.06520729 | -0.383525269 | 0.62323524 | -0.615378 | 0.5383051 | 0.99999872 |
| FN3KRP | 8.73391779 | -0.050821945 | 0.53980551 | -0.0941486 | 0.9249911 | 0.99999872 |
| FN3K | 6.18579284 | 0.731360165 | 0.88825013 | 0.82337187 | 0.41029656 | 0.99999872 |
| TBCD | 29.9699004 | -0.107715209 | 0.35315764 | -0.305006 | 0.76036156 | 0.99999872 |
| METRNL | 4.65402609 | 0.225861361 | 0.71609419 | 0.31540733 | 0.7524524 | 0.99999872 |
| USP14 | 49.6042638 | 0.080753905 | 0.43289063 | 0.18654574 | 0.85201681 | 0.99999872 |
| THOC1 | 27.8033231 | -0.033963482 | 0.36467215 | -0.0931343 | 0.92579688 | 0.99999872 |
| ENOSF1 | 70.5745258 | -0.445257877 | 0.38157164 | -1.166905 | 0.24324872 | 0.99999872 |
| YES1 | 43.6237481 | -0.118756209 | 0.27568247 | -0.4307717 | 0.66663437 | 0.99999872 |
| AP005328.1 | 9.36799503 | 0.963762634 | 0.94858246 | 1.01600301 | 0.30962795 | 0.99999872 |
| METTL4 | 10.4454704 | 0.666654415 | 0.75475204 | 0.88327607 | 0.37708713 | 0.99999872 |
| SMCHD1 | 79.0697297 | -0.098369357 | 0.36514276 | -0.2693997 | 0.7876221 | 0.99999872 |
| AP001011.1 | 11.070641 | 1.383614472 | 0.75140807 | 1.84136227 | 0.06556849 | 0.99999872 |
| LPIN2 | 39.3125321 | 0.346111741 | 0.35394755 | 0.97786166 | 0.32814275 | 0.99999872 |
| MYOM1 | 7.52915671 | -0.639208073 | 0.79715651 | -0.8018602 | 0.42263384 | 0.99999872 |
| MYL12A | 184.761782 | 0.318589252 | 0.60780912 | 0.52416004 | 0.60016723 | 0.99999872 |
| MYL12B | 235.885876 | 0.295922671 | 0.66397964 | 0.44568034 | 0.65582817 | 0.99999872 |
| TGIF1 | 11.1974912 | -0.598653217 | 0.65070259 | -0.9200105 | 0.35756727 | 0.99999872 |
| DLGAP1 | 19.2948084 | 0.399315938 | 0.51716606 | 0.77212325 | 0.44004143 | 0.99999872 |
| DLGAP1-AS1 | 11.8558372 | -0.348097229 | 0.41558748 | -0.8376028 | 0.40225383 | 0.99999872 |
| LINC00667 | 15.5417612 | -0.195805147 | 0.44864226 | -0.4364394 | 0.66251797 | 0.99999872 |
| ZBTB14 | 5.64364816 | -0.710419457 | 0.74652403 | -0.9516364 | 0.3412814 | 0.99999872 |
| EPB41L3 | 25.7714244 | 0.269477054 | 0.92834185 | 0.29027783 | 0.7716037 | 0.99999872 |
| TMEM200C | 44.3398403 | 0.86010788 | 0.74349284 | 1.15684756 | 0.24733465 | 0.99999872 |
| AP005433.1 | 45.998016 | 0.903516181 | 0.59400061 | 1.52106945 | 0.12824241 | 0.99999872 |
| L3MBTL4 | 46.7472111 | -0.697650341 | 0.66680112 | -1.0462645 | 0.29543893 | 0.99999872 |

| | | | | | | |
|---|---|---|---|---|---|---|
| AP001021.2 | 10.0897241 | 1.085018399 | 0.76745063 | 1.4137957 | 0.1574219 | 0.99999872 |
| ARHGAP28 | 574.190415 | 0.654662171 | 0.36268412 | 1.80504781 | 0.0710672 | 0.99999872 |
| LAMA1 | 11.0274406 | -1.933210477 | 0.59730711 | -3.2365436 | 0.00120987 | 0.30478826 |
| PTPRM | 116.525026 | -0.197586291 | 0.57471929 | -0.3437962 | 0.73099958 | 0.99999872 |
| RAB12 | 23.0831131 | 0.043163232 | 0.31290224 | 0.13794479 | 0.89028405 | 0.99999872 |
| NDUFV2 | 33.0596216 | 0.175904019 | 0.38770209 | 0.45370924 | 0.65003812 | 0.99999872 |
| NDUFV2-AS1 | 4.69524893 | 1.049086077 | 0.70330483 | 1.49165203 | 0.13579039 | 0.99999872 |
| ANKRD12 | 217.665839 | 0.100975089 | 0.21736296 | 0.46454597 | 0.64225663 | 0.99999872 |
| TWSG1 | 44.6780418 | 0.626012592 | 0.45390984 | 1.37915625 | 0.16784658 | 0.99999872 |
| RALBP1 | 46.6598754 | 0.157524364 | 0.43232109 | 0.36436891 | 0.71558255 | 0.99999872 |
| PPP4R1 | 50.8195558 | -0.025404486 | 0.43060152 | -0.0589977 | 0.95295396 | 0.99999872 |
| RAB31 | 39.8314936 | -0.113679159 | 0.57365721 | -0.1981657 | 0.84291545 | 0.99999872 |
| VAPA | 111.109106 | 0.430101932 | 0.28882922 | 1.48912193 | 0.13645526 | 0.99999872 |
| NAPG | 25.4826139 | -0.182879148 | 0.36941167 | -0.4950551 | 0.6205612 | 0.99999872 |
| GNAL | 19.2214953 | -0.065797885 | 0.54613509 | -0.1204791 | 0.90410361 | 0.99999872 |
| MPPE1 | 24.7798849 | 0.211275588 | 0.38598832 | 0.54736265 | 0.58412961 | 0.99999872 |
| IMPA2 | 13.6497151 | 0.178878986 | 0.55913584 | 0.31992044 | 0.74902864 | 0.99999872 |
| TUBB6 | 29.898099 | -0.332903367 | 0.43874301 | -0.7587662 | 0.44799243 | 0.99999872 |
| AFG3L2 | 30.6801559 | -0.440810406 | 0.34226168 | -1.2879339 | 0.19776899 | 0.99999872 |
| PRELID3A | 5.14277858 | 0.586300415 | 0.64080317 | 0.91494619 | 0.36021988 | 0.99999872 |
| SPIRE1 | 115.427268 | -0.229433576 | 0.38241959 | -0.5999525 | 0.54853792 | 0.99999872 |
| PSMG2 | 27.9266601 | 0.172999407 | 0.43105197 | 0.40134234 | 0.6881681 | 0.99999872 |
| CEP76 | 6.18350217 | 0.34576473 | 0.7000036 | 0.49394707 | 0.62134356 | 0.99999872 |
| PTPN2 | 38.035419 | -0.225462737 | 0.30759625 | -0.7329827 | 0.46356896 | 0.99999872 |
| SEH1L | 19.7955344 | 0.248190883 | 0.53420588 | 0.46459781 | 0.6422195 | 0.99999872 |
| AP002449.1 | 3.71432087 | 3.535365929 | 1.20753 | 2.92776653 | 0.00341406 | NA |
| CEP192 | 43.0875579 | 0.223669644 | 0.33486855 | 0.66793268 | 0.50417657 | 0.99999872 |
| LDLRAD4 | 20.3996704 | -0.118673943 | 0.87468632 | -0.135676 | 0.89207742 | 0.99999872 |
| FAM210A | 14.8010859 | 0.033293983 | 0.47538481 | 0.07003586 | 0.94416512 | 0.99999872 |
| RNMT | 25.8040178 | -0.140211207 | 0.32663389 | -0.429261 | 0.66773326 | 0.99999872 |
| ZNF519 | 9.25366009 | 0.018974823 | 0.58609834 | 0.03237481 | 0.97417315 | 0.99999872 |
| ANKRD30B | 7.37290775 | 0.450720715 | 0.92201242 | 0.48884452 | 0.62495178 | 0.99999872 |
| ROCK1 | 145.940771 | -0.157906306 | 0.23391151 | -0.6750686 | 0.49963221 | 0.99999872 |
| ESCO1 | 29.3189865 | 0.153305443 | 0.4220454 | 0.36324396 | 0.71642265 | 0.99999872 |
| SNRPD1 | 18.6491885 | -0.415603672 | 0.4765483 | -0.8721124 | 0.38314708 | 0.99999872 |
| ABHD3 | 49.8859498 | 0.084122611 | 0.33699385 | 0.24962655 | 0.80287616 | 0.99999872 |
| MIB1 | 298.613101 | 0.273136198 | 0.37583978 | 0.72673574 | 0.46738786 | 0.99999872 |
| RBBP8 | 24.0022839 | -0.209408981 | 0.67259594 | -0.3113444 | 0.75553882 | 0.99999872 |
| CABLES1 | 42.1361625 | -0.306941342 | 0.55460226 | -0.5534441 | 0.57995936 | 0.99999872 |
| TMEM241 | 60.5388562 | 0.37835571 | 0.44408601 | 0.85198745 | 0.39422106 | 0.99999872 |
| RIOK3 | 61.0551302 | -0.030570632 | 0.32461178 | -0.094176 | 0.92496937 | 0.99999872 |
| RMC1 | 13.975276 | -0.400073827 | 0.42529226 | -0.9407033 | 0.34685693 | 0.99999872 |
| NPC1 | 32.6734263 | -0.737342296 | 0.37379077 | -1.972607 | 0.04854035 | 0.97937581 |
| LAMA3 | 28.349367 | -0.756914863 | 1.12295044 | -0.6740412 | 0.50028512 | 0.99999872 |
| TTC39C | 6.64386044 | -0.626092086 | 0.64904401 | -0.9646373 | 0.33472649 | 0.99999872 |
| OSBPL1A | 26.1500428 | 0.606571858 | 0.41736057 | 1.45335209 | 0.14612602 | 0.99999872 |
| IMPACT | 13.1364475 | -0.036470886 | 0.49482754 | -0.0737042 | 0.94124573 | 0.99999872 |
| ZNF521 | 69.1964223 | 0.111804874 | 0.40641855 | 0.27509786 | 0.78324105 | 0.99999872 |
| SS18 | 64.3387731 | -0.037255932 | 0.28968637 | -0.1286078 | 0.89766798 | 0.99999872 |
| TAF4B | 10.1706911 | 0.213900652 | 0.49905653 | 0.42861006 | 0.66820702 | 0.99999872 |
| KCTD1 | 13.7301144 | 0.23552437 | 0.50043282 | 0.47064133 | 0.63789689 | 0.99999872 |
| CDH2 | 12.5532742 | 0.153211277 | 0.70427795 | 0.21754377 | 0.8277846 | 0.99999872 |
| DSC2 | 24.2224748 | -0.288988429 | 0.4229073 | -0.6833375 | 0.49439359 | 0.99999872 |
| DSG2 | 30.5719529 | 0.417292665 | 0.39567572 | 1.05463298 | 0.29159321 | 0.99999872 |
| TRAPPC8 | 65.0731311 | 0.133347365 | 0.26822509 | 0.49714724 | 0.61908523 | 0.99999872 |
| RNF138 | 20.1842681 | 0.278313741 | 0.40161225 | 0.69299117 | 0.4883151 | 0.99999872 |
| GAREM1 | 148.106151 | -0.412585676 | 0.38673345 | -1.0668476 | 0.28604064 | 0.99999872 |
| DTNA | 49.5906631 | -0.024260338 | 0.51695228 | -0.0469295 | 0.96256938 | 0.99999872 |
| MAPRE2 | 38.9063893 | -0.187824034 | 0.60795438 | -0.3089443 | 0.75736391 | 0.99999872 |
| ZNF397 | 16.979088 | -0.225320098 | 0.55113862 | -0.4088265 | 0.68266696 | 0.99999872 |
| ZSCAN30 | 13.1930608 | -0.043519128 | 0.43621416 | -0.0997655 | 0.92053049 | 0.99999872 |

| | | | | | | |
|---|---|---|---|---|---|---|
| ZNF24 | 29.3377347 | -0.278223168 | 0.33421189 | -0.8324754 | 0.40514067 | 0.99999872 |
| GALNT1 | 31.3262516 | 0.039392106 | 0.36674405 | 0.10741035 | 0.91446344 | 0.99999872 |
| RPRD1A | 62.5897348 | -0.150410932 | 0.24877883 | -0.604597 | 0.54544682 | 0.99999872 |
| SLC39A6 | 24.0696959 | 0.441960728 | 0.44189759 | 1.00014288 | 0.31724137 | 0.99999872 |
| ELP2 | 33.4611323 | 0.159169299 | 0.32097052 | 0.49590006 | 0.61996493 | 0.99999872 |
| MOCOS | 4.32890253 | -0.316798508 | 0.82899439 | -0.382148 | 0.70235163 | NA |
| FHOD3 | 42.2743678 | -0.46364516 | 0.51029095 | -0.9085898 | 3.64E-01 | 0.99999872 |
| TPGS2 | 35.5337083 | 0.633133436 | 0.47507651 | 1.33269784 | 0.18263098 | 0.99999872 |
| KIAA1328 | 78.3818734 | 0.131802897 | 0.30133208 | 0.43740081 | 0.66182069 | 0.99999872 |
| AC016205.1 | 13.3055133 | 0.222977858 | 0.60917286 | 0.3660338 | 0.71433985 | 0.99999872 |
| PIK3C3 | 63.6425604 | -0.074428693 | 0.27801682 | -0.2677129 | 0.78892031 | 0.99999872 |
| LINC00907 | 5.88148457 | 0.218848251 | 0.62864297 | 0.34812805 | 0.72774402 | 0.99999872 |
| SETBP1 | 62.221254 | -1.08629805 | 0.54280253 | -2.0012767 | 0.04536258 | 0.95980711 |
| EPG5 | 26.5783112 | -0.150159461 | 0.37587258 | -0.3994957 | 0.68952803 | 0.99999872 |
| ATP5F1A | 41.8535846 | 0.53239401 | 0.52521647 | 1.01366587 | 0.31074222 | 0.99999872 |
| HAUS1 | 7.94300184 | -0.486540576 | 0.5076084 | -0.9584959 | 0.33781276 | 0.99999872 |
| C18orf25 | 29.4743275 | 0.4981062 | 0.34866976 | 1.42859018 | 0.15312206 | 0.99999872 |
| PIAS2 | 44.8228437 | -0.496399541 | 0.39640937 | -1.2522397 | 0.21048254 | 0.99999872 |
| KATNAL2 | 13.0776664 | -0.021967715 | 0.5305544 | -0.0414052 | 0.96697286 | 0.99999872 |
| HDHD2 | 7.53412416 | 0.347952675 | 0.61435813 | 0.56636782 | 0.57114377 | 0.99999872 |
| SMAD2 | 123.811621 | -0.131702242 | 0.27471584 | -0.4794126 | 0.63164512 | 0.99999872 |
| ZBTB7C | 219.53696 | 0.258522593 | 0.4897897 | 0.52782367 | 0.59762173 | 0.99999872 |
| CTIF | 33.0057765 | -0.655254125 | 0.41649668 | -1.5732517 | 0.11566054 | 0.99999872 |
| DYM | 113.018915 | 0.159063671 | 0.22435709 | 0.70897546 | 0.4783397 | 0.99999872 |
| ACAA2 | 14.2653397 | 0.2319354 | 0.44896956 | 0.51659493 | 0.60543895 | 0.99999872 |
| SNHG22 | 8.56045497 | 0.993131963 | 0.64802261 | 1.53255759 | 0.1253849 | 0.99999872 |
| MYO5B | 14.6207684 | -0.497016526 | 0.82630321 | -0.6014941 | 0.54751093 | 0.99999872 |
| MBD1 | 12.6818868 | 0.01247915 | 0.43850585 | 0.02845834 | 0.97729659 | 0.99999872 |
| CXXC1 | 8.20078583 | 0.027804554 | 0.54733503 | 0.05079988 | 0.95948499 | 0.99999872 |
| ME2 | 20.4165746 | 0.044235727 | 0.38245836 | 0.11566155 | 0.90792078 | 0.99999872 |
| AC091551.1 | 6.75003047 | -0.148711871 | 0.59477672 | -0.2500297 | 0.80256435 | 0.99999872 |
| SMAD4 | 44.7678881 | 0.0777915 | 0.32285576 | 0.24094815 | 0.8095953 | 0.99999872 |
| MEX3C | 12.946993 | -0.373603626 | 0.42435425 | -0.8804051 | 0.37863991 | 0.99999872 |
| MBD2 | 81.3926594 | -0.165327359 | 0.30434861 | -0.5432171 | 0.58698036 | 0.99999872 |
| POLI | 33.1431129 | 0.584342932 | 0.39007314 | 1.49803428 | 0.13412435 | 0.99999872 |
| CCDC68 | 11.6008541 | -0.93390256 | 0.54907405 | -1.7008682 | 0.08896775 | 0.99999872 |
| TCF4 | 17.7497239 | -1.041511773 | 0.60493237 | -1.7216995 | 0.08512396 | 0.99999872 |
| AC009271.1 | 25.4444298 | -0.300344125 | 0.58069304 | -0.5172167 | 0.60500491 | 0.99999872 |
| TXNL1 | 31.446794 | 0.077121322 | 0.29695408 | 0.2597079 | 0.7950891 | 0.99999872 |
| WDR7 | 61.9101814 | 0.177608285 | 0.25401349 | 0.69920808 | 0.484422 | 0.99999872 |
| FECH | 12.2667486 | 0.580441765 | 0.46549984 | 1.24692151 | 0.21242628 | 0.99999872 |
| NARS | 28.3786083 | 0.344991885 | 0.46414064 | 0.7432917 | 0.4573051 | 0.99999872 |
| AC027097.2 | 33.122601 | 0.300942913 | 0.56278854 | 0.53473533 | 0.59283289 | 0.99999872 |
| ATP8B1 | 234.892471 | 0.019545654 | 0.35112481 | 0.05566583 | 0.95560802 | 0.99999872 |
| NEDD4L | 150.346276 | -0.0131089 | 0.40147443 | -0.0326519 | 0.97395219 | 0.99999872 |
| MALT1 | 33.2333394 | -0.265746026 | 0.32186196 | -0.8256522 | 0.40900144 | 0.99999872 |
| AC104365.1 | 4.29501416 | 1.050315728 | 0.98009163 | 1.07165054 | 0.28387702 | NA |
| ZNF532 | 49.5014852 | -0.245719885 | 0.3074149 | -0.7993103 | 0.42411053 | 0.99999872 |
| SEC11C | 14.5791326 | 0.675792494 | 0.49672296 | 1.36050183 | 0.17367117 | 0.99999872 |
| LMAN1 | 68.2174518 | 0.742633655 | 0.34405359 | 2.15848249 | 0.03089034 | 0.91534437 |
| CCBE1 | 370.198249 | -0.200375196 | 0.3670882 | -0.5458503 | 0.58516886 | 0.99999872 |
| RNF152 | 19.5720942 | 0.800788333 | 1.02576112 | 0.78067721 | 0.43499237 | 0.99999872 |
| PIGN | 39.8642709 | -0.021508619 | 0.31651326 | -0.0679549 | 0.94582156 | 0.99999872 |
| KIAA1468 | 50.5645522 | 0.161239457 | 0.33698685 | 0.47847404 | 0.63231285 | 0.99999872 |
| ZCCHC2 | 40.3624201 | -0.69871108 | 0.31380387 | -2.2265853 | 0.02597501 | 0.88365618 |
| PHLPP1 | 45.723125 | 0.041748208 | 0.35609421 | 0.11723922 | 0.90667049 | 0.99999872 |
| BCL2 | 70.3957028 | -0.189381577 | 0.47680737 | -0.3971868 | 0.69122974 | 0.99999872 |
| KDSR | 38.2352523 | -0.143907671 | 0.35226217 | -0.4085243 | 0.68288877 | 0.99999872 |
| VPS4B | 43.6209181 | 0.337028124 | 0.28062986 | 1.20097031 | 0.22976272 | 0.99999872 |
| SERPINB8 | 3.98264693 | -1.109113252 | 0.76230204 | -1.4549525 | 0.14568241 | NA |
| DSEL | 3.74631559 | 1.944693548 | 1.00348535 | 1.93793915 | 0.05263065 | NA |

| | | | | | | |
|---|---|---|---|---|---|---|
| TMX3 | 43.3523591 | 0.279016276 | 0.45296247 | 0.61598101 | 0.53790706 | 0.99999872 |
| CCDC102B | 8.69510545 | -1.272298547 | 0.56300084 | -2.2598519 | 0.02383044 | 0.88337526 |
| DOK6 | 9.54354802 | 0.224055401 | 0.85952305 | 0.26067411 | 0.79434384 | 0.99999872 |
| RTTN | 26.320388 | -0.03233781 | 0.50196374 | -0.0644226 | 0.94863373 | 0.99999872 |
| SOCS6 | 13.1858716 | 0.490686488 | 0.48760336 | 1.00632303 | 0.31426021 | 0.99999872 |
| TIMM21 | 5.39770998 | 0.565952529 | 0.60350785 | 0.93777162 | 0.34836179 | 0.99999872 |
| CYB5A | 117.951105 | 0.669097723 | 0.40554603 | 1.64986877 | 0.09896978 | 0.99999872 |
| CNDP2 | 45.2070145 | -0.155489835 | 0.42029937 | -0.3699502 | 0.7114196 | 0.99999872 |
| LINC00909 | 9.87491493 | -0.013716896 | 0.55236974 | -0.0248328 | 0.98018832 | 0.99999872 |
| ZNF407 | 85.9880643 | -0.194521589 | 0.23218871 | -0.8377737 | 0.40215782 | 0.99999872 |
| ZADH2 | 41.900564 | 0.41514425 | 0.39962779 | 1.03882728 | 0.29888507 | 0.99999872 |
| TSHZ1 | 76.1175273 | 0.393822005 | 0.45742466 | 0.8609549 | 0.38926288 | 0.99999872 |
| ZNF516 | 64.6274759 | -0.087608504 | 0.30374742 | -0.2884255 | 0.77302105 | 0.99999872 |
| ZNF236-DT | 7.37701962 | -0.289663001 | 0.54922814 | -0.5274001 | 0.59791574 | 0.99999872 |
| ZNF236 | 32.3504892 | -0.531787722 | 0.34756943 | -1.5300187 | 0.1260121 | 0.99999872 |
| MBP | 44.885321 | -0.369883974 | 0.35416264 | -1.0443901 | 0.29630492 | 0.99999872 |
| ATP9B | 74.4042925 | -0.208376091 | 0.24413656 | -0.8535227 | 0.39336951 | 0.99999872 |
| NFATC1 | 11.5337618 | -1.037063722 | 0.52666971 | -1.969097 | 0.04894196 | 0.97937581 |
| CTDP1 | 10.6862375 | -0.448620338 | 0.50059116 | -0.8961811 | 0.37015605 | 0.99999872 |
| PQLC1 | 34.663753 | 0.35671905 | 0.49510568 | 0.72049072 | 0.47122291 | 0.99999872 |
| HSBP1L1 | 34.7696449 | 0.053788675 | 0.4010106 | 0.1341328 | 0.89329756 | 0.99999872 |
| TXNL4A | 32.3986229 | 0.717776786 | 0.36593042 | 1.96151167 | 0.04981937 | 0.97937581 |
| ADNP2 | 16.3725405 | 0.197802297 | 0.37652479 | 0.52533672 | 0.59934914 | 0.99999872 |
| PARD6G-AS1 | 7.9662146 | 1.202074029 | 0.7442617 | 1.61512278 | 0.10628411 | 0.99999872 |
| PARD6G | 167.089041 | 0.405849991 | 0.49504542 | 0.81982375 | 0.41231659 | 0.99999872 |
| AC139100.1 | 8.05618363 | 0.78868373 | 0.73623259 | 1.07124262 | 0.28406035 | 0.99999872 |
| ZCCHC3 | 11.374444 | 0.21449587 | 0.50451924 | 0.42514904 | 0.67072803 | 0.99999872 |
| NRSN2-AS1 | 14.1508093 | 0.116774866 | 0.45321608 | 0.25765826 | 0.79667066 | 0.99999872 |
| SOX12 | 7.43751123 | 0.14383186 | 0.54342546 | 0.26467634 | 0.79125881 | 0.99999872 |
| NRSN2 | 17.5556433 | -0.336668815 | 0.43553279 | -0.7730045 | 0.43951971 | 0.99999872 |
| RBCK1 | 37.0450347 | -0.491969497 | 0.72374643 | -0.679754 | 0.49666027 | 0.99999872 |
| TBC1D20 | 14.5790089 | -0.240104326 | 0.41186124 | -0.5829738 | 0.55991092 | 0.99999872 |
| CSNK2A1 | 63.8255982 | 0.092429981 | 0.2296355 | 0.40250737 | 0.68731067 | 0.99999872 |
| PSMF1 | 41.8467476 | -0.269209388 | 0.48064625 | -0.5600988 | 0.57541205 | 0.99999872 |
| SNPH | 5.5833822 | 1.477749415 | 0.82662412 | 1.78769211 | 0.0738257 | 0.99999872 |
| AL136531.2 | 14.0231882 | 0.71566332 | 0.50757908 | 1.40995433 | 0.15855317 | 0.99999872 |
| SDCBP2-AS1 | 24.2893088 | -0.187207504 | 0.43137154 | -0.4339821 | 0.66430147 | 0.99999872 |
| NSFL1C | 23.1142168 | -0.471907144 | 0.35861081 | -1.3159312 | 0.18819714 | 0.99999872 |
| AL117335.1 | 6.46691183 | 0.233158341 | 0.70583387 | 0.33033034 | 0.74115037 | 0.99999872 |
| SIRPA | 115.899753 | 0.499761926 | 0.43861493 | 1.13940929 | 0.25453248 | 0.99999872 |
| PDYN-AS1 | 23.8294665 | -0.033623801 | 0.62945888 | -0.053417 | 0.95739967 | 0.99999872 |
| STK35 | 13.6503486 | -0.221415941 | 0.54008452 | -0.4099654 | 0.68183136 | 0.99999872 |
| SNRPB | 34.0175242 | -0.166582664 | 0.40132308 | -0.4150837 | 0.67808064 | 0.99999872 |
| NOP56 | 14.0500362 | -0.38342642 | 0.53244436 | -0.7201249 | 0.47144812 | 0.99999872 |
| IDH3B | 12.1717857 | 0.120453912 | 0.52763166 | 0.22829167 | 0.8194195 | 0.99999872 |
| EBF4 | 6.47356243 | 1.357817394 | 0.67244622 | 2.01922079 | 0.04346428 | 0.95980711 |
| CPXM1 | 11.9717152 | 1.510090533 | 1.51139527 | 0.99913673 | 0.31772846 | 0.99999872 |
| PCED1A | 12.3577653 | -0.045601423 | 0.53857121 | -0.0846711 | 0.93252286 | 0.99999872 |
| PTPRA | 94.3609182 | -0.101051681 | 0.25884542 | -0.3903939 | 0.69624528 | 0.99999872 |
| MRPS26 | 21.9333989 | 0.884215943 | 0.46416579 | 1.90495715 | 0.05678564 | 0.99999872 |
| UBOX5-AS1 | 7.0384059 | 0.04695541 | 0.64311751 | 0.07301218 | 0.94179643 | 0.99999872 |
| UBOX5 | 8.89469842 | -0.393476407 | 0.5062421 | -0.7772495 | 0.43701159 | 0.99999872 |
| DDRGK1 | 18.2060063 | 0.318364977 | 0.44710716 | 0.71205519 | 0.4764306 | 0.99999872 |
| ITPA | 19.9341539 | -0.348954182 | 0.3854543 | -0.9053062 | 0.36530318 | 0.99999872 |
| C20orf194 | 185.127169 | 0.255575698 | 0.28783929 | 0.8879111 | 0.37458858 | 0.99999872 |
| ATRN | 59.7196296 | 0.124271286 | 0.28231472 | 0.44018706 | 0.65980163 | 0.99999872 |
| CENPB | 9.09155474 | -0.297676992 | 0.52708612 | -0.5647597 | 0.57223723 | 0.99999872 |
| CDC25B | 23.5661994 | 0.599805794 | 0.38686975 | 1.55040757 | 0.12104372 | 0.99999872 |
| AP5S1 | 4.23753989 | -0.539109542 | 0.67373255 | -0.8001833 | 0.4236046 | NA |
| MAVS | 28.8423122 | -0.827553821 | 0.36769235 | -2.2506691 | 0.0244065 | 0.88337526 |
| PANK2 | 33.6696659 | -0.060649222 | 0.32883298 | -0.1844378 | 0.85367004 | 0.99999872 |

| | | | | | | |
|---|---|---|---|---|---|---|
| RNF24 | 43.0509351 | 0.668586211 | 0.60780016 | 1.10000992 | 0.2713278 | 0.99999872 |
| PRNP | 50.3935618 | 0.067794726 | 0.46008855 | 0.14735147 | 0.8828546 | 0.99999872 |
| SLC23A2 | 17.3580801 | -0.093272443 | 0.47508933 | -0.1963261 | 0.84435492 | 0.99999872 |
| TMEM230 | 35.2623503 | 0.297313838 | 0.3317131 | 0.89629816 | 0.37009354 | 0.99999872 |
| PCNA | 9.75375753 | 0.325268213 | 0.49973514 | 0.65088121 | 0.51512317 | 0.99999872 |
| CDS2 | 47.9213818 | -0.059572263 | 0.33336579 | -0.1786994 | 0.85817375 | 0.99999872 |
| GPCPD1 | 57.5148139 | -0.227027894 | 0.40483236 | -0.5607948 | 0.57493742 | 0.99999872 |
| SHLD1 | 20.3532708 | -1.091803327 | 0.43224934 | -2.5258646 | 0.01154139 | 0.72670507 |
| TRMT6 | 6.00436885 | -0.29151704 | 0.58930469 | -0.4946797 | 0.62082626 | 0.99999872 |
| MCM8 | 7.38033408 | -0.719887005 | 0.75778857 | -0.949984 | 0.34212039 | 0.99999872 |
| CRLS1 | 26.5149901 | 0.139244886 | 0.37869775 | 0.36769399 | 0.71310142 | 0.99999872 |
| BMP2 | 46.0092105 | 0.203304135 | 0.83367511 | 0.24386495 | 0.80733542 | 0.99999872 |
| TMX4 | 41.8708312 | -0.101395632 | 0.51107892 | -0.1983953 | 0.84273583 | 0.99999872 |
| PLCB1 | 88.0039411 | -1.910588416 | 0.94148943 | -2.0293254 | 0.04242516 | 0.95980711 |
| PLCB4 | 57.1799371 | 0.903676308 | 0.50182712 | 1.80077215 | 0.0717388 | 0.99999872 |
| ANKEF1 | 5.24637734 | 0.377078561 | 0.66340422 | 0.5683994 | 0.5697638 | 0.99999872 |
| MKKS | 9.74508583 | 0.867519891 | 0.54664732 | 1.58698279 | 0.11251654 | 0.99999872 |
| SLX4IP | 27.0606978 | -0.229846716 | 0.49246547 | -0.4667266 | 0.64069552 | 0.99999872 |
| BTBD3 | 20.9510207 | 0.207767233 | 0.37006649 | 0.56143217 | 0.57450296 | 0.99999872 |
| SPTLC3 | 14.4316841 | -0.276443663 | 0.51948954 | -0.5321448 | 0.5946257 | 0.99999872 |
| TASP1 | 57.7787479 | -0.146201746 | 0.30266049 | -0.4830553 | 0.62905649 | 0.99999872 |
| ESF1 | 18.7170807 | 0.305639837 | 0.41418553 | 0.73792979 | 0.46055712 | 0.99999872 |
| NDUFAF5 | 12.0153173 | -0.345016279 | 0.48979806 | -0.7044052 | 0.48118049 | 0.99999872 |
| MACROD2 | 413.470322 | -0.131032514 | 0.5639415 | -0.2323513 | 0.81626521 | 0.99999872 |
| FLRT3 | 256.452741 | 0.396904268 | 0.61630259 | 0.64400876 | 0.51956975 | 0.99999872 |
| KIF16B | 48.1163115 | -0.095403457 | 0.38178747 | -0.2498863 | 0.80267528 | 0.99999872 |
| SNRPB2 | 19.9066244 | 0.019660329 | 0.44646454 | 0.04403559 | 0.96487603 | 0.99999872 |
| DSTN | 277.613836 | 0.395681856 | 0.37550825 | 1.05372347 | 0.29200954 | 0.99999872 |
| RRBP1 | 57.5368772 | -0.522846371 | 0.54676559 | -0.9562533 | 0.3389443 | 0.99999872 |
| SNX5 | 21.6720799 | 0.544994014 | 0.46797025 | 1.16459117 | 0.24418452 | 0.99999872 |
| OVOL2 | 7.1609487 | 0.280507813 | 0.84338849 | 0.33259621 | 0.73943911 | 0.99999872 |
| MGME1 | 13.8472556 | 0.467693049 | 0.49068856 | 0.95313624 | 0.34052105 | 0.99999872 |
| KAT14 | 8.84461029 | -0.416956988 | 0.54834098 | -0.7603973 | 0.44701715 | 0.99999872 |
| ZNF133 | 14.3767319 | 0.087349681 | 0.44912171 | 0.19449 | 0.84579222 | 0.99999872 |
| DZANK1 | 9.64697911 | -0.616992637 | 0.58080702 | -1.0623023 | 0.28809847 | 0.99999872 |
| POLR3F | 6.41113134 | 1.102163886 | 0.78296395 | 1.40768153 | 0.15922539 | 0.99999872 |
| RBBP9 | 5.8799908 | 0.770148307 | 0.64406312 | 1.19576526 | 0.23178818 | 0.99999872 |
| SEC23B | 21.2524446 | -0.079827587 | 0.37564943 | -0.2125055 | 0.83171265 | 0.99999872 |
| DTD1 | 13.6640422 | 0.55213775 | 0.46169473 | 1.19589356 | 0.2317381 | 0.99999872 |
| SLC24A3 | 14.591447 | 1.743680257 | 0.72628014 | 2.40083702 | 0.01635762 | 0.80846766 |
| AL121761.1 | 21.2697317 | 0.385254308 | 0.54397625 | 0.70821899 | 0.47880927 | 0.99999872 |
| RIN2 | 34.1799167 | -0.246506119 | 0.53440865 | -0.461269 | 0.64460566 | 0.99999872 |
| NAA20 | 25.6419194 | 0.530204083 | 0.4358872 | 1.21637912 | 0.22384054 | 0.99999872 |
| CRNKL1 | 8.25256501 | -0.523926543 | 0.5267469 | -0.9946457 | 0.31990861 | 0.99999872 |
| RALGAPA2 | 59.9752632 | 0.091275126 | 0.33863845 | 0.26953562 | 0.78751753 | 0.99999872 |
| KIZ | 36.5652123 | 0.566338754 | 0.51728158 | 1.0948365 | 0.27358827 | 0.99999872 |
| KIZ-AS1 | 9.39805989 | 0.936295817 | 0.65668679 | 1.4257875 | 0.15392969 | 0.99999872 |
| XRN2 | 43.1652316 | -0.064454715 | 0.36288409 | -0.1776179 | 0.85902306 | 0.99999872 |
| AL118508.1 | 22.5091764 | 1.900094398 | 1.09057913 | 1.74228018 | 0.08145943 | 0.99999872 |
| NXT1 | 7.31123126 | 0.451843029 | 0.76400708 | 0.5914121 | 0.55424434 | 0.99999872 |
| GZF1 | 5.7248854 | -0.382298821 | 0.6657372 | -0.5742488 | 0.56579942 | 0.99999872 |
| NAPB | 30.9530837 | 0.447377893 | 0.36110022 | 1.23892998 | 0.21537143 | 0.99999872 |
| CST3 | 145.141238 | 0.477531313 | 0.40157065 | 1.18915891 | 0.23437714 | 0.99999872 |
| APMAP | 26.4038875 | 0.493730606 | 0.52388785 | 0.94243569 | 0.34596962 | 0.99999872 |
| ACSS1 | 17.0872764 | -0.073234315 | 0.51662942 | -0.1417541 | 0.88727428 | 0.99999872 |
| ENTPD6 | 13.0641497 | -0.838953551 | 0.52393748 | -1.6012475 | 0.10932212 | 0.99999872 |
| PYGB | 19.1198406 | -0.476833499 | 0.41988056 | -1.1356408 | 0.25610692 | 0.99999872 |
| ABHD12 | 20.4506643 | -0.276509234 | 0.42866129 | -0.645053 | 0.51889286 | 0.99999872 |
| NINL | 31.1155327 | -0.859272674 | 0.44746342 | -1.9203194 | 0.05481757 | 0.99999872 |
| ZNF337-AS1 | 9.22867049 | -0.254417661 | 0.61916411 | -0.4109051 | 0.68114216 | 0.99999872 |
| ZNF337 | 9.49734682 | 0.173353596 | 0.47612869 | 0.36408979 | 0.71579096 | 0.99999872 |

| | | | | | | |
|---|---|---|---|---|---|---|
| FAM182B | 16.7572179 | 0.526641364 | 0.4337626 | 1.2141235 | 0.22470057 | 0.99999872 |
| HM13 | 47.0642505 | -0.138025975 | 0.4365824 | -0.316151 | 0.75188788 | 0.99999872 |
| ID1 | 31.6865865 | -3.484128318 | 0.83349265 | -4.1801548 | 2.91E-05 | 0.06104707 |
| BCL2L1 | 56.3524268 | -0.85999778 | 0.31789747 | -2.7052677 | 0.00682494 | 0.62184066 |
| AL117381.1 | 9.36280694 | -0.338042686 | 0.75047534 | -0.4504381 | 0.65239458 | 0.99999872 |
| DUSP15 | 6.17274451 | -0.972982886 | 1.35523321 | -0.717945 | 0.47279121 | 0.99999872 |
| PDRG1 | 7.16669608 | 0.48957073 | 0.6793698 | 0.7206248 | 0.47114039 | 0.99999872 |
| TM9SF4 | 35.7513997 | -0.026220745 | 0.29036644 | -0.0903023 | 0.92804703 | 0.99999872 |
| PLAGL2 | 12.1646729 | -0.74746482 | 0.48114512 | -1.5535122 | 0.12030082 | 0.99999872 |
| POFUT1 | 18.9310037 | -0.224548548 | 0.59308258 | -0.3786126 | 0.70497555 | 0.99999872 |
| KIF3B | 26.1190356 | -0.33658444 | 0.39491297 | -0.8523003 | 0.39404744 | 0.99999872 |
| ASXL1 | 85.2554988 | -0.11020041 | 0.32290874 | -0.3412742 | 0.7328972 | 0.99999872 |
| NOL4L | 24.6001865 | -0.34890124 | 0.404563 | -0.8624151 | 0.38845913 | 0.99999872 |
| FO393400.1 | 7.78782335 | -0.211399518 | 0.62907315 | -0.3360492 | 0.73683378 | 0.99999872 |
| MAPRE1 | 21.687606 | 0.176476698 | 0.35597112 | 0.49576128 | 0.62006284 | 0.99999872 |
| AL035071.2 | 8.83300761 | 1.493956017 | 0.71298326 | 2.09535919 | 0.03613907 | 0.94734101 |
| AL035071.1 | 8.82844284 | -0.118641716 | 0.77287274 | -0.1535074 | 0.87799812 | 0.99999872 |
| CDK5RAP1 | 18.7694738 | -0.589243842 | 0.38403765 | -1.5343387 | 0.12494634 | 0.99999872 |
| SNTA1 | 24.187386 | 0.522532399 | 0.57296209 | 0.91198424 | 0.36177701 | 0.99999872 |
| CBFA2T2 | 66.7261445 | 0.026775544 | 0.31704512 | 0.08445342 | 0.93269594 | 0.99999872 |
| PXMP4 | 7.91290799 | -0.565855872 | 0.68668831 | -0.824036 | 0.40991912 | 0.99999872 |
| CHMP4B | 30.7903922 | 0.060279996 | 0.38399404 | 0.15698159 | 0.87525935 | 0.99999872 |
| RALY | 37.1137304 | -0.453923503 | 0.34583259 | -1.3125527 | 0.18933371 | 0.99999872 |
| EIF2S2 | 46.812292 | 0.219788147 | 0.36939906 | 0.59498838 | 0.55185124 | 0.99999872 |
| ASIP | 5.45577511 | 0.66875918 | 1.19997608 | 0.55731043 | 0.57731535 | 0.99999872 |
| AHCY | 11.3461672 | 0.002120364 | 0.445246 | 0.00476223 | 0.9962003 | 0.99999872 |
| ITCH | 102.960411 | -0.017514111 | 0.28279671 | -0.0619318 | 0.95061714 | 0.99999872 |
| DYNLRB1 | 104.72848 | 0.452885259 | 0.50139832 | 0.90324448 | 0.36639616 | 0.99999872 |
| MAP1LC3A | 16.2120902 | 0.80107329 | 0.59456524 | 1.34732614 | 0.17787522 | 0.99999872 |
| PIGU | 15.9784765 | -0.551976725 | 0.44113771 | -1.2512572 | 0.21084067 | 0.99999872 |
| NCOA6 | 63.1799713 | -0.017297599 | 0.26424329 | -0.0654609 | 0.94780705 | 0.99999872 |
| TP53INP2 | 6.16186823 | -0.376693998 | 0.73316475 | -0.5137917 | 0.6073976 | 0.99999872 |
| GGT7 | 12.628804 | 0.158106173 | 0.51031029 | 0.3098236 | 0.7566951 | 0.99999872 |
| ACSS2 | 27.2948505 | -0.039622855 | 0.36264696 | -0.1092601 | 0.91299617 | 0.99999872 |
| GSS | 15.0882301 | -0.192780159 | 0.4051737 | -0.4757963 | 0.63421949 | 0.99999872 |
| TRPC4AP | 49.2624227 | -0.018349087 | 0.29097595 | -0.0630605 | 0.94971833 | 0.99999872 |
| EDEM2 | 13.8127986 | 0.190074794 | 0.56632655 | 0.33562755 | 0.73715174 | 0.99999872 |
| PROCR | 8.35545979 | 0.359466778 | 0.69474131 | 0.51741097 | 0.6048693 | 0.99999872 |
| MMP24OS | 23.8286885 | 0.704538249 | 0.42478181 | 1.65858856 | 0.09719873 | 0.99999872 |
| MMP24 | 12.4755015 | 0.952070607 | 0.76898684 | 1.23808439 | 0.21568477 | 0.99999872 |
| UQCC1 | 33.0976717 | -0.322314252 | 0.31530275 | -1.0222374 | 0.30666856 | 0.99999872 |
| CEP250 | 6.59476556 | 0.176712549 | 0.60324698 | 0.29293566 | 0.76957133 | 0.99999872 |
| ERGIC3 | 20.1364711 | 0.195582273 | 0.47323165 | 0.41329077 | 0.67939359 | 0.99999872 |
| CPNE1 | 22.8789727 | 0.693743282 | 0.46312836 | 1.49795034 | 0.13414615 | 0.99999872 |
| ROMO1 | 20.588319 | 0.256641079 | 0.56003684 | 0.45825749 | 0.64676746 | 0.99999872 |
| RBM39 | 131.308766 | -0.027287681 | 0.24428531 | -0.1117041 | 0.91105799 | 0.99999872 |
| PHF20 | 77.7746908 | -0.293537869 | 0.336783 | -0.8715935 | 0.38343019 | 0.99999872 |
| SCAND1 | 47.1129108 | -0.475188246 | 0.51413759 | -0.9242434 | 0.35535963 | 0.99999872 |
| NORAD | 104.295634 | 0.045419354 | 0.44138448 | 0.10290202 | 0.91804073 | 0.99999872 |
| EPB41L1 | 20.9457231 | -0.668635987 | 0.54514822 | -1.2265214 | 0.22000251 | 0.99999872 |
| AAR2 | 9.68862433 | -0.300728697 | 0.48716129 | -0.6173083 | 0.53703141 | 0.99999872 |
| DLGAP4 | 60.4905423 | -0.33289524 | 0.41971101 | -0.7931535 | 0.42768842 | 0.99999872 |
| DLGAP4-AS1 | 11.8292948 | 0.288070106 | 0.44615772 | 0.64566876 | 0.51849389 | 0.99999872 |
| MYL9 | 450.562133 | 0.496059232 | 0.55775665 | 0.8893829 | 0.37379733 | 0.99999872 |
| RAB5IF | 10.370756 | 0.109177337 | 0.60978603 | 0.17904204 | 0.85790469 | 0.99999872 |
| NDRG3 | 12.7228529 | 0.562814064 | 0.59429782 | 0.94702361 | 0.34362675 | 0.99999872 |
| SOGA1 | 21.2803575 | 0.115809953 | 0.56864761 | 0.20365856 | 0.83862033 | 0.99999872 |
| SAMHD1 | 7.76736131 | -0.541851574 | 1.1323995 | -0.4784986 | 0.63229537 | 0.99999872 |
| RBL1 | 9.50455515 | -0.11794912 | 0.63596232 | -0.1854656 | 0.85286388 | 0.99999872 |
| MROH8 | 6.83034891 | -0.945060443 | 0.54347246 | -1.7389298 | 0.08204711 | 0.99999872 |
| RPN2 | 67.6238974 | 0.73887094 | 0.6632108 | 1.11408158 | 0.26524419 | 0.99999872 |

| | | | | | | |
|---|---|---|---|---|---|---|
| MANBAL | 15.0834036 | 0.07998196 | 0.4680064 | 0.17089929 | 0.86430296 | 0.99999872 |
| SRC | 5.70511251 | 0.490873551 | 0.70018343 | 0.70106422 | 0.48326294 | 0.99999872 |
| BLCAP | 33.5280522 | -0.365899832 | 0.33598295 | -1.0890428 | 0.27613499 | 0.99999872 |
| CTNNBL1 | 23.4201502 | -0.736719238 | 0.42828895 | -1.7201453 | 0.08540603 | 0.99999872 |
| TTI1 | 8.15367365 | 0.074731991 | 0.56261379 | 0.13283 | 0.89432783 | 0.99999872 |
| RPRD1B | 34.5561026 | 0.22055016 | 0.33607871 | 0.65624556 | 0.51166614 | 0.99999872 |
| TGM2 | 9.03983144 | -0.442805229 | 0.76939492 | -0.575524 | 0.56493698 | 0.99999872 |
| RALGAPB | 52.4883626 | -0.242482933 | 0.29880949 | -0.8114968 | 0.41708045 | 0.99999872 |
| DHX35 | 7.89656136 | -0.168784969 | 0.60946587 | -0.2769392 | 0.78182682 | 0.99999872 |
| MAFB | 244.608928 | 0.310409703 | 0.4541079 | 0.68355935 | 0.49425346 | 0.99999872 |
| TOP1 | 62.2717556 | -0.23992809 | 0.37633996 | -0.6375302 | 0.52377954 | 0.99999872 |
| PLCG1 | 6.76480761 | -0.140780625 | 0.5644758 | -0.2494006 | 0.80305089 | 0.99999872 |
| ZHX3 | 27.0009656 | 0.015351053 | 0.49933318 | 0.03074311 | 0.97547441 | 0.99999872 |
| LPIN3 | 8.19395699 | 1.130796471 | 0.76891067 | 1.47064738 | 0.1413865 | 0.99999872 |
| CHD6 | 86.1547215 | 0.098124202 | 0.32304717 | 0.30374574 | 0.76132161 | 0.99999872 |
| SRSF6 | 25.8686531 | 0.147302764 | 0.4036283 | 0.36494657 | 0.71515129 | 0.99999872 |
| L3MBTL1 | 7.86070815 | -0.544216873 | 0.57222692 | -0.9510508 | 0.34157859 | 0.99999872 |
| SGK2 | 5.51094208 | -0.557885836 | 0.67060838 | -0.83191 | 0.40545973 | 0.99999872 |
| IFT52 | 14.257213 | -0.18156102 | 0.41057155 | -0.4422153 | 0.65833342 | 0.99999872 |
| TOX2 | 7.57116199 | 1.257425403 | 0.97488939 | 1.2898134 | 0.19711545 | 0.99999872 |
| OSER1 | 17.419108 | -0.27994716 | 0.40462385 | -0.6918701 | 0.48901888 | 0.99999872 |
| FITM2 | 5.72443501 | 0.694283793 | 0.72491285 | 0.95774795 | 0.33818987 | 0.99999872 |
| TTPAL | 9.20875092 | -0.11059513 | 0.53570073 | -0.2064495 | 0.83643983 | 0.99999872 |
| SERINC3 | 68.7440147 | 0.083703823 | 0.38736613 | 0.21608452 | 0.82892187 | 0.99999872 |
| PKIG | 46.0630003 | 0.304876333 | 0.34253568 | 0.89005715 | 0.3734352 | 0.99999872 |
| YWHAB | 91.9248071 | 0.116497676 | 0.48416378 | 0.24061625 | 0.80985255 | 0.99999872 |
| TOMM34 | 7.05221774 | -0.423911062 | 0.67040387 | -0.6323219 | 0.52717654 | 0.99999872 |
| STK4 | 20.9327724 | -0.21582183 | 0.39793381 | -0.5423561 | 0.58757322 | 0.99999872 |
| SLPI | 53.1342668 | 2.396790162 | 1.07538553 | 2.22877292 | 0.02582902 | 0.88365618 |
| SDC4 | 164.019475 | -0.169510021 | 0.57717242 | -0.2936904 | 0.76899446 | 0.99999872 |
| PIGT | 34.8982055 | 1.019494462 | 0.6029538 | 1.69083346 | 0.09086861 | 0.99999872 |
| WFDC2 | 95.4349701 | 1.814339224 | 0.55117919 | 3.29174114 | 9.96E-04 | 0.29808178 |
| DNTTIP1 | 8.91552596 | 0.563102065 | 0.5126905 | 1.09832748 | 0.27206152 | 0.99999872 |
| SNX21 | 15.0108692 | -0.140635018 | 0.43183794 | -0.3256662 | 0.74467692 | 0.99999872 |
| ACOT8 | 13.2216292 | -0.19547939 | 0.46507289 | -0.4203199 | 0.67425177 | 0.99999872 |
| CTSA | 41.2335399 | 0.785861131 | 0.45096052 | 1.74263844 | 0.08139679 | 0.99999872 |
| PLTP | 45.8507599 | -0.387898434 | 0.56941232 | -0.6812259 | 0.49572855 | 0.99999872 |
| PCIF1 | 5.14835679 | -0.069524986 | 0.67548284 | -0.1029264 | 0.91802142 | 0.99999872 |
| ZNF335 | 9.61241177 | -0.967921526 | 0.60259517 | -1.606255 | 0.10821788 | 0.99999872 |
| NCOA5 | 24.0284097 | -0.630045399 | 0.43681411 | -1.442365 | 0.14919942 | 0.99999872 |
| CD40 | 4.75083224 | 0.331145251 | 1.0928249 | 0.30301767 | 0.7618764 | 0.99999872 |
| SLC35C2 | 16.3721842 | 0.418480695 | 0.46605721 | 0.897917 | 0.3692298 | 0.99999872 |
| ELMO2 | 14.7755955 | -0.330017292 | 0.59863327 | -0.5512846 | 0.58143861 | 0.99999872 |
| SLC13A3 | 36.9694365 | 0.675907829 | 1.10586782 | 0.61120128 | 0.54106634 | 0.99999872 |
| TP53RK | 5.28643226 | 0.500732571 | 0.79272158 | 0.6316626 | 0.52760737 | 0.99999872 |
| EYA2 | 40.4751271 | -0.159179787 | 0.74927824 | -0.2124442 | 0.83176054 | 0.99999872 |
| ZMYND8 | 86.913169 | -0.930709834 | 0.284422 | -3.272285 | 0.00106682 | 0.29912383 |
| NCOA3 | 49.8614001 | -0.251753709 | 0.28322525 | -0.8888816 | 0.37406672 | 0.99999872 |
| SULF2 | 177.838408 | 0.132568444 | 0.49917096 | 0.26557724 | 0.79056482 | 0.99999872 |
| ARFGEF2 | 53.323493 | -0.208961246 | 0.33617529 | -0.6215842 | 0.53421531 | 0.99999872 |
| CSE1L | 21.7661008 | 0.023135705 | 0.40985761 | 0.05644815 | 0.9549848 | 0.99999872 |
| STAU1 | 48.201717 | -0.091835581 | 0.37151995 | -0.2471888 | 0.80476209 | 0.99999872 |
| DDX27 | 12.1937076 | 0.492088904 | 0.45248467 | 1.08752614 | 0.27680435 | 0.99999872 |
| ZNFX1 | 29.3982129 | -0.7316704 | 0.72891447 | -1.0037809 | 0.31548425 | 0.99999872 |
| ZFAS1 | 21.0883184 | -0.098702863 | 0.5822837 | -0.1695099 | 0.86539557 | 0.99999872 |
| B4GALT5 | 47.7228794 | -0.083965989 | 0.44938035 | -0.1868484 | 0.85177951 | 0.99999872 |
| SLC9A8 | 17.9072135 | -0.269482819 | 0.42687315 | -0.6312948 | 0.52784776 | 0.99999872 |
| RNF114 | 36.502194 | 0.057595176 | 0.55416414 | 0.10393162 | 0.91722362 | 0.99999872 |
| CEBPB | 31.0015621 | 0.466855771 | 0.7345511 | 0.63556609 | 5.25E-01 | 0.99999872 |
| PTPN1 | 65.6589021 | -0.204824147 | 0.53640012 | -0.3818496 | 0.70257296 | 0.99999872 |
| PARD6B | 97.7311905 | 0.227896941 | 0.34798377 | 0.65490681 | 0.51252775 | 0.99999872 |

| | | | | | | |
|---|---|---|---|---|---|---|
| BCAS4 | 8.91963618 | 1.066713963 | 0.65852484 | 1.61985381 | 0.10526368 | 0.99999872 |
| ADNP | 89.043994 | 0.067659155 | 0.24841319 | 0.27236539 | 0.78534108 | 0.99999872 |
| DPM1 | 22.4776361 | -0.033373154 | 0.47807734 | -0.069807 | 0.94434726 | 0.99999872 |
| ATP9A | 60.4846139 | 0.26582458 | 0.32768907 | 0.81120979 | 0.4172452 | 0.99999872 |
| ZFP64 | 15.4091579 | -0.380353201 | 0.43523084 | -0.8739114 | 0.3821665 | 0.99999872 |
| TSHZ2 | 33.5886999 | -0.513848529 | 1.00229688 | -0.512671 | 0.60818149 | 0.99999872 |
| AL109930.1 | 8.95957291 | -1.395713102 | 1.02078157 | -1.3672985 | 0.17153176 | 0.99999872 |
| ZNF217 | 16.5967192 | -0.035838615 | 0.4493601 | -0.0797548 | 0.93643229 | 0.99999872 |
| CYP24A1 | 22.3998981 | -1.566863406 | 1.18347886 | -1.3239471 | 0.18552061 | 0.99999872 |
| PFDN4 | 10.802539 | 0.606025618 | 0.49160798 | 1.23274162 | 0.21767218 | 0.99999872 |
| FAM210B | 16.7165472 | 0.060965434 | 0.43329327 | 0.14070247 | 0.88810499 | 0.99999872 |
| CSTF1 | 7.09739453 | 0.400423751 | 0.55009906 | 0.72791208 | 0.46666741 | 0.99999872 |
| RTF2 | 14.1824285 | -0.103249138 | 0.46047887 | -0.2242212 | 0.82258516 | 0.99999872 |
| BMP7 | 72.8714864 | 0.199198058 | 0.73227851 | 0.27202499 | 0.7856028 | 0.99999872 |
| RAE1 | 11.9913561 | 0.010934991 | 0.43017448 | 0.0254199 | 0.97972004 | 0.99999872 |
| RBM38 | 10.7734332 | -0.770915984 | 0.52505545 | -1.4682563 | 0.14203461 | 0.99999872 |
| PCK1 | 50.3917669 | 1.509265868 | 1.01521461 | 1.48664712 | 0.13710804 | 0.99999872 |
| PMEPA1 | 97.8027463 | 0.165621142 | 0.69292018 | 0.23901908 | 0.81109079 | 0.99999872 |
| RAB22A | 24.7134257 | 0.012413323 | 0.44113979 | 0.0281392 | 0.97755113 | 0.99999872 |
| VAPB | 38.9461597 | -0.1082381 | 0.26699207 | -0.4053982 | 0.68518484 | 0.99999872 |
| NPEPL1 | 12.27013 | -0.34580482 | 0.44043135 | -0.7851503 | 0.43236543 | 0.99999872 |
| GNAS | 199.69877 | 0.450459021 | 0.39098443 | 1.15211499 | 0.24927382 | 0.99999872 |
| NELFCD | 17.8304396 | -0.155037667 | 0.38137421 | -0.4065237 | 0.68435782 | 0.99999872 |
| CTSZ | 37.2782176 | 0.450614617 | 0.3400322 | 1.32521161 | 0.18510097 | 0.99999872 |
| ATP5F1E | 110.497621 | 0.628505907 | 0.53853905 | 1.16705727 | 0.24318724 | 0.99999872 |
| PRELID3B | 9.40904428 | 0.570629735 | 0.56062995 | 1.01783669 | 0.30875557 | 0.99999872 |
| PHACTR3 | 14.9129823 | 1.838645638 | 0.8757096 | 2.09960658 | 0.03576346 | 0.94734101 |
| SYCP2 | 5.57833409 | -0.281763036 | 0.60140615 | -0.4685071 | 0.63942201 | 0.99999872 |
| FAM217B | 14.3773008 | 0.004548406 | 0.51007733 | 0.00891709 | 0.99288529 | 0.99999872 |
| PPP1R3D | 8.6534944 | 0.805251276 | 0.55372668 | 1.45423961 | 0.14587989 | 0.99999872 |
| MIR646HG | 16.1595167 | 0.367151565 | 1.16907719 | 0.31405246 | 0.7534812 | 0.99999872 |
| TAF4 | 18.2907793 | 0.244788604 | 0.447683 | 0.54679003 | 0.58452299 | 0.99999872 |
| LSM14B | 6.81636544 | 0.255857657 | 0.53271191 | 0.48029273 | 0.63101926 | 0.99999872 |
| PSMA7 | 62.7856945 | 0.280966326 | 0.34946936 | 0.80397985 | 0.42140861 | 0.99999872 |
| SS18L1 | 16.9036134 | 0.150357278 | 0.38588411 | 0.38964361 | 0.6968001 | 0.99999872 |
| MTG2 | 19.1998868 | 0.018830615 | 0.376062 | 0.05007317 | 0.96006408 | 0.99999872 |
| OSBPL2 | 14.8572336 | -0.132004107 | 0.48190319 | -0.2739225 | 0.78414421 | 0.99999872 |
| ADRM1 | 11.8974769 | -0.581303762 | 0.65035608 | -0.8938238 | 0.37141616 | 0.99999872 |
| AL354836.1 | 11.8708388 | -0.439862584 | 0.78637435 | -0.5593552 | 0.57591933 | 0.99999872 |
| LAMA5 | 93.1440878 | -0.300090298 | 0.43712781 | -0.6865047 | 0.4923949 | 0.99999872 |
| RPS21 | 6.91999058 | -0.149201247 | 0.83268901 | -0.17918 | 0.85779634 | 0.99999872 |
| SLCO4A1 | 6.73711968 | -0.941187286 | 0.79675629 | -1.1812737 | 0.237494 | 0.99999872 |
| MRGBP | 5.1544585 | -0.272587061 | 0.63201052 | -0.4313015 | 0.66624919 | 0.99999872 |
| OGFR | 17.0983148 | -0.645731971 | 0.64686405 | -0.9982499 | 0.31815819 | 0.99999872 |
| TCFL5 | 4.18347396 | -0.082829083 | 0.75275851 | -0.1100341 | 0.91238236 | NA |
| DIDO1 | 45.6429132 | 0.233369669 | 0.3097307 | 0.75345993 | 0.45117358 | 0.99999872 |
| GID8 | 18.3703617 | -0.134850236 | 0.40505377 | -0.3329193 | 0.73919516 | 0.99999872 |
| YTHDF1 | 17.8124561 | 0.204452221 | 0.41798369 | 0.48913924 | 0.62474312 | 0.99999872 |
| ARFGAP1 | 13.1137614 | -0.700750953 | 0.45285839 | -1.5473953 | 0.12176795 | 0.99999872 |
| PPDPF | 97.754822 | 0.442609056 | 0.64628599 | 0.68485015 | 0.49343848 | 0.99999872 |
| HELZ2 | 10.4132988 | -1.041024692 | 1.18386991 | -0.8793404 | 0.37921671 | 0.99999872 |
| GMEB2 | 8.09734937 | -0.870393565 | 0.53402051 | -1.6298879 | 0.10312519 | 0.99999872 |
| MHENCR | 8.60836633 | -0.158757439 | 0.68256064 | -0.232591 | 0.81607904 | 0.99999872 |
| ARFRP1 | 13.333845 | 0.203001517 | 0.47705512 | 0.42553053 | 0.67044997 | 0.99999872 |
| SLC2A4RG | 18.7649182 | -0.357510446 | 0.68444529 | -0.522336 | 0.60143638 | 0.99999872 |
| ZBTB46 | 94.6969389 | -0.60458646 | 0.56664532 | -1.0669575 | 0.28599104 | 0.99999872 |
| TPD52L2 | 21.9728213 | -0.208857837 | 0.41316909 | -0.5055021 | 0.61320623 | 0.99999872 |
| DNAJC5 | 34.3150096 | -0.093689124 | 0.29695989 | -0.3154942 | 0.75238645 | 0.99999872 |
| UCKL1 | 8.9894298 | -0.086086698 | 0.51314176 | -0.167764 | 0.86676897 | 0.99999872 |
| ZNF512B | 5.79122706 | -0.447053784 | 0.62737472 | -0.7125786 | 0.47610658 | 0.99999872 |
| PRPF6 | 32.8343203 | -0.220574845 | 0.31748785 | -0.6947505 | 0.48721167 | 0.99999872 |

| | | | | | | |
|---|---|---|---|---|---|---|
| TCEA2 | 14.4547826 | 0.197718338 | 0.44755447 | 0.44177492 | 0.65865209 | 0.99999872 |
| PCMTD2 | 97.0048958 | 0.417294135 | 0.46888495 | 0.88997126 | 0.37348132 | 0.99999872 |
| MIER2 | 7.12763029 | 0.225613214 | 0.55684809 | 0.40516115 | 0.68535906 | 0.99999872 |
| TPGS1 | 10.9492981 | -0.405010969 | 0.5146019 | -0.7870374 | 0.43125994 | 0.99999872 |
| CDC34 | 8.51502019 | -0.401090866 | 0.63895175 | -0.6277326 | 0.5301791 | 0.99999872 |
| BSG | 74.3648542 | 0.161323768 | 0.42824639 | 0.37670783 | 0.70639074 | 0.99999872 |
| RNF126 | 11.0531658 | 0.242157363 | 0.49200304 | 0.49218672 | 0.62258735 | 0.99999872 |
| PALM | 8.34788195 | 1.728182287 | 0.66461406 | 2.60027946 | 0.00931479 | 0.69734315 |
| AC006273.1 | 12.0934942 | 0.353024268 | 0.66868762 | 0.527936 | 0.59754376 | 0.99999872 |
| LINC01836 | 9.37586859 | -0.737024982 | 0.72807452 | -1.0122933 | 0.31139783 | 0.99999872 |
| PTBP1 | 56.1775258 | -0.20266813 | 0.35133902 | -0.5768449 | 0.56404422 | 0.99999872 |
| MED16 | 15.701781 | -0.058943945 | 0.38295845 | -0.1539173 | 0.87767491 | 0.99999872 |
| R3HDM4 | 12.1816485 | -0.469124268 | 0.51240785 | -0.915529 | 0.35991397 | 0.99999872 |
| ARID3A | 29.1351793 | -0.606571374 | 0.54075995 | -1.1217017 | 0.26198928 | 0.99999872 |
| WDR18 | 6.35155031 | -0.18782771 | 0.6689518 | -0.2807791 | 0.77887981 | 0.99999872 |
| TMEM259 | 54.8100014 | -0.225764866 | 0.46821402 | -0.4821831 | 0.62967591 | 0.99999872 |
| CNN2 | 29.2729273 | -0.413035081 | 0.54196272 | -0.7621098 | 0.44599449 | 0.99999872 |
| POLR2E | 15.4640536 | 0.157867565 | 0.42765057 | 0.36915084 | 0.7120153 | 0.99999872 |
| GPX4 | 45.1634129 | 0.079013523 | 0.36008636 | 0.21942937 | 0.8263156 | 0.99999872 |
| SBNO2 | 28.3698154 | -0.816326645 | 0.64719444 | -1.2613314 | 0.20718949 | 0.99999872 |
| STK11 | 17.2240192 | -0.152378531 | 0.41400116 | -0.3680631 | 0.71282622 | 0.99999872 |
| ATP5F1D | 48.3384322 | 0.084442034 | 0.47291002 | 0.17855835 | 0.8582845 | 0.99999872 |
| MIDN | 47.3242899 | -0.768408076 | 0.4516353 | -1.7013906 | 0.08886966 | 0.99999872 |
| CIRBP | 97.2806907 | -0.091915435 | 0.32729882 | -0.2808303 | 0.77884054 | 0.99999872 |
| C19orf24 | 17.0748569 | -0.056683206 | 0.53159579 | -0.1066284 | 0.91508379 | 0.99999872 |
| MUM1 | 49.5252871 | -0.088709192 | 0.37660044 | -0.2355525 | 0.8137799 | 0.99999872 |
| NDUFS7 | 16.8717353 | 0.227057718 | 0.50282497 | 0.45156412 | 0.65158302 | 0.99999872 |
| GAMT | 4.19264603 | 1.104978621 | 0.93841918 | 1.17748938 | 0.23900023 | NA |
| DAZAP1 | 23.2733388 | -0.405909652 | 0.37717718 | -1.0761776 | 0.28184782 | 0.99999872 |
| RPS15 | 132.134397 | 0.295270927 | 0.41823888 | 0.70598631 | 0.48019665 | 0.99999872 |
| C19orf25 | 14.0903773 | -0.095240037 | 0.4904255 | -0.1941988 | 0.84602023 | 0.99999872 |
| MBD3 | 17.6889437 | -1.074198152 | 0.46491178 | -2.3105419 | 0.02085817 | 0.8671024 |
| TCF3 | 9.32443693 | -0.270153981 | 0.51911915 | -0.5204084 | 0.60277894 | 0.99999872 |
| REXO1 | 10.5390716 | -0.397615996 | 0.49729233 | -0.7995619 | 0.42396468 | 0.99999872 |
| KLF16 | 9.88571803 | -0.075107988 | 0.54792881 | -0.1370762 | 0.89097058 | 0.99999872 |
| ABHD17A | 22.5989006 | 0.284600116 | 0.56423166 | 0.50440296 | 0.61397824 | 0.99999872 |
| SCAMP4 | 20.8619223 | -0.045848907 | 0.37741353 | -0.1214819 | 0.90330936 | 0.99999872 |
| CSNK1G2 | 20.4171513 | -0.751147668 | 0.56634726 | -1.3263023 | 0.18473959 | 0.99999872 |
| BTBD2 | 16.5689469 | -0.922555116 | 0.37448837 | -2.463508 | 0.01375848 | 0.75477153 |
| MKNK2 | 15.3486251 | -0.299639574 | 0.50676971 | -0.5912736 | 0.55433709 | 0.99999872 |
| MOB3A | 12.8918282 | -0.600464702 | 0.47280155 | -1.2700142 | 0.20407956 | 0.99999872 |
| AP3D1 | 31.7727951 | -0.663068797 | 0.31178659 | -2.1266751 | 0.03344708 | 0.92959829 |
| DOT1L | 14.901923 | -1.040199424 | 0.44957419 | -2.3137436 | 0.02068179 | 0.8671024 |
| PLEKHJ1 | 19.4678102 | -0.28243676 | 0.39939808 | -0.707156 | 0.47946952 | 0.99999872 |
| SF3A2 | 6.8930678 | -0.876287162 | 0.5801863 | -1.5103548 | 0.13095291 | 0.99999872 |
| LSM7 | 23.9553252 | 0.407299858 | 0.40853933 | 0.99696608 | 0.31878097 | 0.99999872 |
| SPPL2B | 12.8992766 | -0.457596699 | 0.42184412 | -1.0847531 | 0.27803104 | 0.99999872 |
| TIMM13 | 21.6134555 | 0.698539699 | 0.4905742 | 1.42392261 | 0.15446887 | 0.99999872 |
| GADD45B | 107.757883 | 0.075200394 | 1.05486595 | 0.07128905 | 0.94316771 | 0.99999872 |
| GNG7 | 13.9323938 | -0.294238069 | 0.41982071 | -0.700866 | 0.48338664 | 0.99999872 |
| SGTA | 16.8103599 | -0.179282529 | 0.44373017 | -0.404035 | 0.68618696 | 0.99999872 |
| THOP1 | 7.83324326 | -0.676436205 | 0.66003981 | -1.0248415 | 0.30543797 | 0.99999872 |
| ZNF555 | 4.26332234 | 0.19903184 | 0.72573336 | 0.27424926 | 0.78389307 | NA |
| AES | 68.7439123 | -0.270020701 | 0.34758292 | -0.7768526 | 0.43724573 | 0.99999872 |
| GNA11 | 24.4261603 | -0.441246061 | 0.44470607 | -0.9922196 | 0.32109044 | 0.99999872 |
| NCLN | 13.1106092 | -0.12597224 | 0.62993361 | -0.199977 | 0.84149856 | 0.99999872 |
| NFIC | 165.135714 | -0.64293286 | 0.36784967 | -1.7478142 | 0.08049621 | 0.99999872 |
| DOHH | 3.45947726 | -0.030326765 | 0.72614866 | -0.0417639 | 0.96668695 | NA |
| FZR1 | 10.0543381 | -0.373014547 | 0.47183637 | -0.7905591 | 0.42920131 | 0.99999872 |
| MFSD12 | 21.3148838 | -0.969425955 | 0.60264038 | -1.6086309 | 0.10769706 | 0.99999872 |
| HMG20B | 16.9632589 | -0.164643881 | 0.43537304 | -0.3781674 | 0.70530626 | 0.99999872 |

| | | | | | | |
|---|---|---|---|---|---|---|
| TBXA2R | 13.9815602 | -0.759115489 | 0.91383592 | -0.8306912 | 0.40614808 | 0.99999872 |
| CACTIN | 5.53829225 | -0.989731956 | 0.66450377 | -1.4894302 | 0.13637413 | 0.99999872 |
| PIP5K1C | 14.8015818 | -0.756969073 | 0.49920559 | -1.5163474 | 0.12943154 | 0.99999872 |
| MRPL54 | 21.1809802 | 0.042048697 | 0.46033156 | 0.09134437 | 0.92721896 | 0.99999872 |
| DAPK3 | 14.5239763 | -0.574335617 | 0.48619097 | -1.1812964 | 0.23748501 | 0.99999872 |
| EEF2 | 88.1516498 | 0.186922072 | 0.42532147 | 0.43948422 | 0.66031071 | 0.99999872 |
| PIAS4 | 7.85191794 | -1.010108825 | 0.54654719 | -1.848164 | 0.06457863 | 0.99999872 |
| ZBTB7A | 43.5253356 | -0.65308432 | 0.37813416 | -1.7271233 | 0.08414552 | 0.99999872 |
| MAP2K2 | 31.8051174 | -0.260298233 | 0.28561805 | -0.9113508 | 0.36211057 | 0.99999872 |
| SIRT6 | 5.14930996 | -1.258331195 | 0.76341127 | -1.6483005 | 0.09929102 | 0.99999872 |
| YJU2 | 6.79206404 | -1.150816569 | 0.68639094 | -1.6766197 | 0.09361687 | 0.99999872 |
| STAP2 | 3.7324849 | -0.519433112 | 1.01590114 | -0.5113028 | 0.60913904 | NA |
| SH3GL1 | 12.142607 | -0.742341098 | 0.4711211 | -1.5756906 | 0.11509712 | 0.99999872 |
| UBXN6 | 25.9127199 | -0.436547405 | 0.45964962 | -0.9497395 | 0.34224463 | 0.99999872 |
| HDGFL2 | 12.8927897 | -0.652466908 | 0.43067773 | -1.5149771 | 0.1297782 | 0.99999872 |
| PLIN5 | 7.90460617 | 0.123087685 | 0.7058782 | 0.17437525 | 0.86157057 | 0.99999872 |
| MYDGF | 43.3560499 | 1.131157902 | 0.57603859 | 1.96368426 | 0.04956672 | 0.97937581 |
| DPP9 | 14.2364361 | -0.385630657 | 0.46452982 | -0.8301526 | 0.40645249 | 0.99999872 |
| TICAM1 | 6.17337837 | -0.685504555 | 0.59882482 | -1.1447497 | 0.25231284 | 0.99999872 |
| PLIN3 | 13.2939624 | -0.080765157 | 0.47278531 | -0.1708284 | 0.8643587 | 0.99999872 |
| KDM4B | 35.6478839 | -0.46576901 | 0.48698322 | -0.9564375 | 0.33885126 | 0.99999872 |
| PTPRS | 18.1733978 | 0.507504853 | 0.43267868 | 1.17293705 | 0.24082105 | 0.99999872 |
| TINCR | 5.84796743 | 0.511977721 | 0.74042965 | 0.69146032 | 0.48927631 | 0.99999872 |
| SAFB2 | 23.3108549 | -0.574994238 | 0.32795593 | -1.7532668 | 0.07955623 | 0.99999872 |
| SAFB | 36.0383396 | -0.018866793 | 0.35557804 | -0.0530595 | 0.9576845 | 0.99999872 |
| RPL36 | 71.7169546 | 0.186157325 | 0.51353864 | 0.36249916 | 0.71697905 | 0.99999872 |
| C19orf70 | 29.8344579 | 0.025998198 | 0.47675727 | 0.05453131 | 0.95651187 | 0.99999872 |
| LONP1 | 17.1700071 | -0.561374759 | 0.44325594 | -1.2664799 | 0.20534131 | 0.99999872 |
| CAPS | 4.8526533 | 0.107080607 | 0.70789646 | 0.15126592 | 0.87976595 | 0.99999872 |
| RANBP3 | 14.9932423 | -0.24511139 | 0.38225546 | -0.641224 | 0.52137714 | 0.99999872 |
| RFX2 | 6.89769662 | 0.348840025 | 0.91997287 | 0.37918512 | 0.7045504 | 0.99999872 |
| MLLT1 | 17.471753 | -0.117195752 | 0.47159583 | -0.2485089 | 0.8037407 | 0.99999872 |
| CLPP | 15.6708098 | 0.648378537 | 0.42462655 | 1.52693829 | 0.12677636 | 0.99999872 |
| ALKBH7 | 21.825682 | 1.126399911 | 0.80752546 | 1.39487852 | 0.16305248 | 0.99999872 |
| GTF2F1 | 14.5188002 | 0.0757449 | 0.43439105 | 0.1743703 | 0.86157445 | 0.99999872 |
| KHSRP | 17.2908786 | -0.422761775 | 0.54689195 | -0.7730261 | 0.4395069 | 0.99999872 |
| SLC25A23 | 11.2041261 | 0.191941085 | 0.48565492 | 0.39522113 | 0.6926797 | 0.99999872 |
| C3 | 8.851895 | 2.574325288 | 1.23941441 | 2.07704966 | 0.03779698 | 0.95980711 |
| TRIP10 | 26.9782267 | -0.037542277 | 0.42310062 | -0.0887313 | 0.92929544 | 0.99999872 |
| ZNF557 | 4.93038571 | 0.9907097 | 0.73309179 | 1.3514129 | 0.17656321 | 0.99999872 |
| INSR | 145.833354 | 0.035945152 | 0.27693027 | 0.12979857 | 0.8967258 | 0.99999872 |
| ZNF358 | 12.7737685 | -0.447197392 | 0.6667398 | -0.6707225 | 0.50239732 | 0.99999872 |
| PNPLA6 | 18.5198643 | -0.207461013 | 0.58838779 | -0.3525923 | 0.7243941 | 0.99999872 |
| XAB2 | 5.92574056 | -0.025431889 | 0.65274546 | -0.0389614 | 0.96892115 | 0.99999872 |
| PET100 | 5.15129388 | -0.322205813 | 0.70957859 | -0.4540805 | 0.64977087 | 0.99999872 |
| STXBP2 | 8.07788302 | -0.557093288 | 0.63785475 | -0.8733858 | 0.38245282 | 0.99999872 |
| EVI5L | 6.44600472 | 0.239622779 | 0.622667 | 0.38483295 | 0.70036118 | 0.99999872 |
| MAP2K7 | 16.8606596 | -1.032771453 | 0.52622632 | -1.9625994 | 0.04969274 | 0.97937581 |
| SNAPC2 | 3.56699447 | 0.516368199 | 0.87289649 | 0.5915572 | 0.55414714 | NA |
| TIMM44 | 6.10702164 | -0.192435554 | 0.69495636 | -0.2769031 | 0.78185453 | 0.99999872 |
| ELAVL1 | 37.4255578 | -0.198454285 | 0.33553672 | -0.5914533 | 0.55421676 | 0.99999872 |
| CERS4 | 22.1607231 | -0.382032712 | 0.51016162 | -0.7488464 | 0.45394977 | 0.99999872 |
| CD320 | 14.8266346 | 0.714480492 | 0.61576267 | 1.16031797 | 0.24591937 | 0.99999872 |
| RPS28 | 27.0689765 | 0.52668247 | 0.561276 | 0.93836628 | 0.34805621 | 0.99999872 |
| KANK3 | 11.280914 | -0.117785805 | 0.52395931 | -0.2247995 | 0.82213523 | 0.99999872 |
| ANGPTL4 | 35.988963 | 1.369426493 | 0.70121848 | 1.95292414 | 0.05082858 | 0.98949866 |
| RAB11B-AS1 | 4.49119258 | -0.447966842 | 0.95586733 | -0.4686496 | 0.63932012 | NA |
| RAB11B | 33.5890285 | -0.276391414 | 0.48719593 | -0.5673106 | 0.57050318 | 0.99999872 |
| 2-Mar | 14.4563582 | -0.673503057 | 0.60870095 | -1.1064597 | 0.2685276 | 0.99999872 |
| HNRNPM | 67.4892566 | -0.229375886 | 0.23383706 | -0.9809219 | 0.32663127 | 0.99999872 |
| ZNF414 | 4.39617279 | -0.625787086 | 0.7167656 | -0.8730708 | 0.38262451 | NA |

| | | | | | | |
|---|---|---|---|---|---|---|
| ADAMTS10 | 4.96888869 | 0.377140441 | 0.67053325 | 0.56244853 | 0.57381046 | 0.99999872 |
| ZNF558 | 6.01285994 | 0.490517854 | 0.60344711 | 0.81285972 | 0.41629849 | 0.99999872 |
| ZNF317 | 7.99944769 | -0.25774615 | 0.59062178 | -0.436398 | 0.662548 | 0.99999872 |
| ZNF699 | 4.00650495 | 0.414867495 | 0.71113837 | 0.58338505 | 0.55963412 | NA |
| ZNF559-ZNF | 7.6611352 | 0.460502503 | 0.55458325 | 0.83035775 | 0.40633655 | 0.99999872 |
| ZNF266 | 12.3563283 | 0.001765106 | 0.42644179 | 0.00413915 | 0.99669745 | 0.99999872 |
| ZNF426 | 10.9834038 | -0.525011986 | 0.44585321 | -1.1775445 | 0.23897825 | 0.99999872 |
| ZNF121 | 14.8565922 | -0.313452973 | 0.44242806 | -0.7084835 | 0.47864507 | 0.99999872 |
| ZNF561 | 8.59955477 | -0.489541616 | 0.67007408 | -0.7305784 | 0.46503674 | 0.99999872 |
| ZNF562 | 18.2868035 | -0.086956896 | 0.39018825 | -0.2228588 | 0.82364538 | 0.99999872 |
| ZNF846 | 40.4883134 | -0.357333689 | 0.39008603 | -0.9160381 | 0.35964689 | 0.99999872 |
| FBXL12 | 8.69944855 | 0.770388085 | 0.51765609 | 1.48822373 | 0.1366919 | 0.99999872 |
| UBL5 | 31.7955759 | 0.826864814 | 0.46364268 | 1.78340963 | 0.07451965 | 0.99999872 |
| PIN1 | 27.8396275 | 0.261842225 | 0.39530407 | 0.66238181 | 0.50772656 | 0.99999872 |
| C19orf66 | 22.2804696 | -0.765144596 | 0.82878333 | -0.9232143 | 0.35589555 | 0.99999872 |
| EIF3G | 17.8240066 | 0.068205624 | 0.45694236 | 0.14926527 | 0.88134432 | 0.99999872 |
| DNMT1 | 7.40813912 | -0.170486484 | 0.59978488 | -0.2842461 | 0.77622182 | 0.99999872 |
| MRPL4 | 13.4033555 | 0.58101945 | 0.5501326 | 1.05614438 | 0.29090226 | 0.99999872 |
| ICAM1 | 66.7599182 | 0.252289228 | 1.45217725 | 0.1737317 | 0.86207632 | 0.99999872 |
| TYK2 | 11.5678463 | -0.499462365 | 0.45217648 | -1.104574 | 0.26934423 | 0.99999872 |
| CDC37 | 31.4660752 | -0.316355685 | 0.30483918 | -1.037779 | 0.29937298 | 0.99999872 |
| PDE4A | 17.3041979 | 0.074685985 | 0.46958197 | 0.15904781 | 0.87363121 | 0.99999872 |
| KEAP1 | 14.8162432 | 0.091616544 | 0.49057364 | 0.18675391 | 0.85185359 | 0.99999872 |
| ATG4D | 6.14358766 | -0.070284387 | 0.57971786 | -0.121239 | 0.90350177 | 0.99999872 |
| KRI1 | 6.2144285 | 0.083001841 | 0.53301886 | 0.15572027 | 0.87625352 | 0.99999872 |
| CDKN2D | 10.3915281 | 0.518171099 | 0.85052651 | 0.60923569 | 0.54236823 | 0.99999872 |
| AP1M2 | 8.61480941 | -0.27754335 | 0.60237098 | -0.4607515 | 0.64497688 | 0.99999872 |
| SLC44A2 | 87.6727736 | 0.269035165 | 0.36985533 | 0.72740649 | 0.46697698 | 0.99999872 |
| ILF3-DT | 20.2486717 | -0.510903761 | 0.3395154 | -1.5048029 | 0.13237475 | 0.99999872 |
| ILF3 | 45.5806985 | -0.043056564 | 0.27972741 | -0.1539233 | 0.87767022 | 0.99999872 |
| QTRT1 | 13.028791 | -1.490037001 | 0.52170555 | -2.8560881 | 0.00428896 | 0.50494083 |
| DNM2 | 45.8841638 | -0.786108126 | 0.35330287 | -2.2250261 | 0.02607948 | 0.88433927 |
| TMED1 | 14.106342 | 0.238987086 | 0.60540428 | 0.39475619 | 0.69302283 | 0.99999872 |
| CARM1 | 9.19531367 | -0.209162316 | 0.4914193 | -0.425629 | 0.67037819 | 0.99999872 |
| YIPF2 | 11.2860189 | 0.201410532 | 0.48900435 | 0.41187881 | 0.68042825 | 0.99999872 |
| TIMM29 | 8.50492129 | -0.879534972 | 0.57620598 | -1.5264246 | 0.12690416 | 0.99999872 |
| SMARCA4 | 39.0915233 | -0.372311907 | 0.2851378 | -1.3057263 | 0.19164566 | 0.99999872 |
| LDLR | 11.783151 | -0.603082504 | 0.80810025 | -0.7462966 | 0.45548824 | 0.99999872 |
| KANK2 | 57.62903 | -0.456272032 | 0.52728518 | -0.8653231 | 0.38686149 | 0.99999872 |
| DOCK6 | 15.7756797 | 0.174895853 | 0.56456697 | 0.30978761 | 0.75672247 | 0.99999872 |
| RAB3D | 6.31775946 | -0.349228519 | 0.62492911 | -0.558829 | 0.57627842 | 0.99999872 |
| RGL3 | 13.7249461 | 0.314312391 | 0.66422318 | 0.47320298 | 0.63606837 | 0.99999872 |
| PRKCSH | 27.3770339 | 0.133320996 | 0.60273568 | 0.22119314 | 0.82494205 | 0.99999872 |
| ECSIT | 8.99430466 | -0.587576952 | 0.51289732 | -1.1456035 | 0.25195926 | 0.99999872 |
| ELOF1 | 6.10541754 | 0.698706415 | 0.75211239 | 0.92899202 | 0.35289322 | 0.99999872 |
| ZNF627 | 7.99046409 | -0.572906417 | 0.64949398 | -0.8820812 | 0.37773291 | 0.99999872 |
| ZNF823 | 5.76560118 | 0.057228824 | 0.64775543 | 0.08834943 | 0.92959895 | 0.99999872 |
| ZNF441 | 5.71746382 | -0.549570846 | 0.87040817 | -0.6313944 | 0.52778268 | 0.99999872 |
| ZNF440 | 6.02664385 | -0.496342294 | 0.63866841 | -0.7771518 | 0.43706923 | 0.99999872 |
| ZNF439 | 20.6624852 | -0.440652719 | 0.56218653 | -0.7838194 | 0.43314608 | 0.99999872 |
| ZNF69 | 9.60988548 | -0.159353268 | 0.5561746 | -0.2865166 | 0.77448247 | 0.99999872 |
| ZNF844 | 9.56284536 | 0.016913362 | 0.77829068 | 0.02173142 | 0.9826622 | 0.99999872 |
| ZNF788P | 6.86901186 | -0.52474111 | 0.61962 | -0.8468757 | 0.39706442 | 0.99999872 |
| ZNF136 | 9.30439577 | 0.061278016 | 0.49370144 | 0.12411958 | 0.90122059 | 0.99999872 |
| ZNF44 | 73.0399994 | -0.427581339 | 0.3804735 | -1.1238137 | 0.26109206 | 0.99999872 |
| ZNF563 | 7.92888884 | -0.804791636 | 0.56285427 | -1.4298402 | 0.1527629 | 0.99999872 |
| ZNF442 | 7.82993889 | -1.293350614 | 0.64686058 | -1.9994272 | 0.04556216 | 0.95980711 |
| ZNF490 | 9.18242799 | 0.153743554 | 0.52810344 | 0.29112394 | 0.77095653 | 0.99999872 |
| ZNF791 | 19.139552 | -0.055630189 | 0.43337202 | -0.1283659 | 0.89785942 | 0.99999872 |
| WDR83 | 4.13087142 | -0.488713434 | 0.6893808 | -0.7089165 | 0.47837628 | NA |
| DHPS | 8.8181006 | 0.054635716 | 0.53474331 | 0.10217186 | 0.91862027 | 0.99999872 |

| | | | | | | | |
|---|---|---|---|---|---|---|---|
| TNPO2 | 24.7201039 | | 0.161401522 | 0.35180288 | 0.45878397 | 0.64638931 | 0.99999872 |
| TRIR | 42.7164472 | | 0.259505356 | 0.31874139 | 0.81415643 | 0.41555534 | 0.99999872 |
| ASNA1 | 9.71990445 | | 0.564892468 | 0.57378831 | 0.98449631 | 0.32487155 | 0.99999872 |
| HOOK2 | 13.526773 | | -0.32429511 | 0.60059089 | -0.5399601 | 0.58922456 | 0.99999872 |
| JUNB | 29.7980667 | | -0.506685407 | 0.67868467 | -0.7465697 | 0.45532335 | 0.99999872 |
| DNASE2 | 18.207335 | | 0.873440532 | 0.57467182 | 1.51989448 | 0.1285375 | 0.99999872 |
| GCDH | 6.90050841 | | -0.418450277 | 0.56892739 | -0.7355074 | 0.46203056 | 0.99999872 |
| FARSA | 10.5730392 | | -0.751564598 | 0.52789103 | -1.4237116 | 0.15452997 | 0.99999872 |
| CALR | 134.997097 | | 1.372464761 | 0.57590252 | 2.38315463 | 0.01716498 | 0.82882344 |
| AC092069.1 | 23.5568221 | | 2.013515281 | 0.6229063 | 3.23245289 | 0.00122732 | 0.30478826 |
| RAD23A | 29.31158 | | 0.283155668 | 0.32630954 | 0.86775174 | 0.38553026 | 0.99999872 |
| GADD45GIP1 | 55.6802256 | | -0.113849699 | 0.33250543 | -0.3423995 | 0.73205025 | 0.99999872 |
| NFIX | 9.79686325 | | -0.669242887 | 0.65766712 | -1.0176013 | 0.30886749 | 0.99999872 |
| TRMT1 | 3.92863606 | | -0.519282847 | 0.72621125 | -0.7150576 | 0.47457348 | NA |
| NACC1 | 8.3970173 | | 0.495804574 | 0.62400103 | 0.7945573 | 0.42687106 | 0.99999872 |
| STX10 | 5.41668397 | | 0.505334087 | 0.69146871 | 0.73081266 | 0.46489359 | 0.99999872 |
| IER2 | 26.5484096 | | -1.000097749 | 0.60773479 | -1.6456154 | 0.09984296 | 0.99999872 |
| CCDC130 | 14.9729217 | | -0.328393498 | 0.54942142 | -0.5977079 | 0.55003488 | 0.99999872 |
| MRI1 | 10.1867537 | | 0.437811417 | 0.5097894 | 0.85880839 | 0.39044624 | 0.99999872 |
| C19orf53 | 34.9861866 | | 0.872247776 | 0.46099613 | 1.89209347 | 0.05847853 | 0.99999872 |
| ZSWIM4 | 7.59562897 | | -0.360776564 | 0.5375662 | -0.6711296 | 0.502138 | 0.99999872 |
| CC2D1A | 7.57098197 | | -0.322814519 | 0.60601947 | -0.5326801 | 0.59425503 | 0.99999872 |
| RFX1 | 13.7980058 | | -0.547570305 | 0.51687626 | -1.0593837 | 0.28942505 | 0.99999872 |
| PRKACA | 11.0428415 | | 0.040583939 | 0.44309943 | 0.09159104 | 0.92702297 | 0.99999872 |
| AC022098.1 | 4.57682387 | | 0.598329687 | 0.80467122 | 0.74357038 | 0.45713643 | NA |
| ADGRL1 | 9.52796084 | | 0.564843574 | 0.71743381 | 0.78731106 | 0.4310998 | 0.99999872 |
| DDX39A | 4.68376879 | | -0.798647652 | 0.66786696 | -1.1958185 | 0.23176741 | 0.99999872 |
| PKN1 | 30.929506 | | -0.516959454 | 0.44625571 | -1.1584377 | 0.24668545 | 0.99999872 |
| GIPC1 | 34.348674 | | -0.995566278 | 0.30382973 | -3.2767244 | 0.00105019 | 0.29912383 |
| DNAJB1 | 11.821591 | | 0.033007633 | 0.55559741 | 0.05940926 | 0.95262614 | 0.99999872 |
| TECR | 13.9157335 | | 0.339710896 | 0.48463352 | 0.7009645 | 0.48332517 | 0.99999872 |
| NDUFB7 | 51.3146233 | | 0.32102927 | 0.57160316 | 0.56162963 | 0.57436839 | 0.99999872 |
| ADGRE3 | 11.2185487 | | 0.816792014 | 0.70863126 | 1.15263334 | 0.24906091 | 0.99999872 |
| ZNF333 | 9.37204914 | | -0.882111651 | 0.45210298 | -1.9511299 | 0.05104159 | 0.98949866 |
| SYDE1 | 18.7809376 | | -0.696439857 | 0.4426203 | -1.5734476 | 0.11561521 | 0.99999872 |
| ILVBL | 10.4884048 | | 0.083861568 | 0.48857847 | 0.17164401 | 0.86371741 | 0.99999872 |
| BRD4 | 71.9322318 | | 0.00130154 | 0.32086676 | 0.00405633 | 0.99676353 | 0.99999872 |
| AKAP8 | 17.9679317 | | -0.32806567 | 0.37855435 | -0.8666277 | 0.38614606 | 0.99999872 |
| AKAP8L | 45.0364624 | | -0.054149354 | 0.36381844 | -0.1488362 | 0.88168289 | 0.99999872 |
| WIZ | 10.1769684 | | -0.08704222 | 0.50790649 | -0.1713745 | 0.86392931 | 0.99999872 |
| TPM4 | 79.5768389 | | 0.152454471 | 0.31849374 | 0.47867337 | 0.63217101 | 0.99999872 |
| RAB8A | 13.2634977 | | -0.178154609 | 0.53836843 | -0.3309158 | 0.7407081 | 0.99999872 |
| FAM32A | 14.080874 | | -0.053283271 | 0.52283128 | -0.1019129 | 0.91882578 | 0.99999872 |
| AP1M1 | 16.6685953 | | -0.381458375 | 0.46632848 | -0.8180036 | 0.41335514 | 0.99999872 |
| EPS15L1 | 22.4705242 | | 0.155942763 | 0.356775 | 0.43708994 | 0.66204612 | 0.99999872 |
| C19orf44 | 6.17737978 | | -0.409597992 | 0.63935928 | -0.6406382 | 0.52175776 | 0.99999872 |
| SIN3B | 18.4440101 | | -0.076337857 | 0.41502645 | -0.1839349 | 0.85406451 | 0.99999872 |
| MYO9B | 26.5731672 | | -0.189910833 | 0.34728975 | -0.5468369 | 0.58449082 | 0.99999872 |
| USE1 | 7.25195729 | | -0.556438511 | 0.56420158 | -0.9862406 | 0.32401506 | 0.99999872 |
| OCEL1 | 10.0363815 | | -0.097751107 | 0.50789985 | -0.1924614 | 0.84738081 | 0.99999872 |
| NR2F6 | 19.8881303 | | -0.084096492 | 0.38476657 | -0.218565 | 0.82698895 | 0.99999872 |
| ABHD8 | 6.06443109 | | -0.06963813 | 0.59658551 | -0.1167278 | 0.90707574 | 0.99999872 |
| MRPL34 | 17.4866995 | | 0.575523578 | 0.43405265 | 1.32593035 | 0.18486277 | 0.99999872 |
| DDA1 | 11.1115569 | | -0.012437702 | 0.54172727 | -0.0229593 | 0.9816827 | 0.99999872 |
| ANO8 | 5.20589557 | | -0.366430684 | 0.6881197 | -0.5325101 | 0.59437275 | 0.99999872 |
| GTPBP3 | 6.88067957 | | -0.23810444 | 0.65264049 | -0.3648325 | 0.71523647 | 0.99999872 |
| BST2 | 340.028855 | | -0.043202711 | 0.760398 | -0.0568159 | 0.95469183 | 0.99999872 |
| MVB12A | 17.8339309 | | -0.099114667 | 0.53756817 | -0.184376 | 0.8537185 | 0.99999872 |
| SLC27A1 | 24.1742835 | | -0.080561472 | 0.46562741 | -0.173017 | 0.86263804 | 0.99999872 |
| AC010618.3 | 13.4455262 | | -0.107755497 | 0.46088094 | -0.2338033 | 0.81513767 | 0.99999872 |
| PGLS | 28.2945206 | | -0.067518435 | 0.34570671 | -0.1953055 | 0.84515376 | 0.99999872 |

| | | | | | | |
|---|---|---|---|---|---|---|
| COLGALT1 | 14.3803999 | 0.644000277 | 0.54034201 | 1.19183825 | 0.23332467 | 0.99999872 |
| MAP1S | 8.16538525 | -0.4941241 | 0.6101095 | -0.8098941 | 0.41800103 | 0.99999872 |
| RPL18A | 6.46197575 | -0.041961113 | 0.9191331 | -0.0456529 | 0.96358689 | 0.99999872 |
| CCDC124 | 10.8511341 | 0.093529368 | 0.5242307 | 0.17841261 | 0.85839895 | 0.99999872 |
| ARRDC2 | 9.96012352 | -1.20666823 | 0.70204116 | -1.7187998 | 0.08565083 | 0.99999872 |
| JUND | 91.6097292 | -0.195671017 | 0.33491265 | -0.5842449 | 0.55905554 | 0.99999872 |
| LSM4 | 23.8598681 | 0.320596817 | 0.45154973 | 0.70999227 | 0.47770893 | 0.99999872 |
| PGPEP1 | 11.071319 | 0.080533473 | 0.46024122 | 0.17498101 | 0.86109456 | 0.99999872 |
| GDF15 | 21.5866494 | -1.236429789 | 0.95377813 | -1.2963495 | 0.19485511 | 0.99999872 |
| SSBP4 | 25.6431668 | 0.0634076 | 0.33945666 | 0.18679145 | 0.85182415 | 0.99999872 |
| ELL | 16.9197862 | -0.688150249 | 0.38490093 | -1.7878633 | 0.07379806 | 0.99999872 |
| FKBP8 | 23.6383441 | 0.373677149 | 0.49055438 | 0.76174459 | 0.44621245 | 0.99999872 |
| KXD1 | 10.6042942 | 0.214757788 | 0.50326567 | 0.42672846 | 0.66957712 | 0.99999872 |
| UBA52 | 48.8075575 | 0.756276161 | 0.60157117 | 1.25716822 | 0.20869273 | 0.99999872 |
| REX1BD | 19.4044392 | 0.202585374 | 0.40842962 | 0.49601048 | 0.61988702 | 0.99999872 |
| CRTC1 | 19.9336326 | -0.490278937 | 0.39391483 | -1.2446318 | 0.21326711 | 0.99999872 |
| UPF1 | 22.0479998 | -0.310028892 | 0.4559736 | -0.6799273 | 0.4965505 | 0.99999872 |
| COPE | 19.968297 | -0.028380577 | 0.4888865 | -0.0580515 | 0.95370764 | 0.99999872 |
| DDX49 | 7.35012577 | 0.120138884 | 0.6401267 | 0.18767985 | 0.85112763 | 0.99999872 |
| SUGP2 | 37.0962628 | -0.102447915 | 0.37794869 | -0.271063 | 0.78634256 | 0.99999872 |
| ARMC6 | 5.26234695 | 0.018063535 | 0.6255654 | 0.02887553 | 0.97696386 | 0.99999872 |
| SLC25A42 | 25.9446282 | -0.17601546 | 0.64615264 | -0.2724054 | 0.78531033 | 0.99999872 |
| TMEM161A | 8.3812872 | -0.192589548 | 0.52178042 | -0.3691008 | 0.71205263 | 0.99999872 |
| RFXANK | 7.56444186 | 0.124699155 | 0.61746938 | 0.20195196 | 0.83995428 | 0.99999872 |
| SUGP1 | 10.8132893 | -0.302596042 | 0.53203738 | -0.5687496 | 0.56952609 | 0.99999872 |
| MAU2 | 25.9833115 | -0.243457264 | 0.33845729 | -0.7193146 | 0.47194711 | 0.99999872 |
| GATAD2A | 42.9608697 | -0.277842774 | 0.26472877 | -1.0495375 | 0.29393081 | 0.99999872 |
| PBX4 | 6.43998637 | -0.224951624 | 0.75965204 | -0.2961246 | 0.76713496 | 0.99999872 |
| ATP13A1 | 7.92813146 | -0.234436333 | 0.55293255 | -0.4239872 | 0.67157513 | 0.99999872 |
| ZNF14 | 7.13375142 | 0.274354634 | 0.58665534 | 0.46765897 | 0.64002848 | 0.99999872 |
| AC011477.4 | 6.63275127 | 0.003564195 | 0.73545876 | 0.00484622 | 0.99613329 | 0.99999872 |
| ZNF253 | 19.3351845 | 0.098648636 | 0.4698502 | 0.20995763 | 0.83370074 | 0.99999872 |
| ZNF682 | 10.2127109 | -1.08588316 | 0.67871073 | -1.5999204 | 0.10961624 | 0.99999872 |
| ZNF90 | 47.8925719 | 0.345391164 | 0.50721363 | 0.68095797 | 0.49589809 | 0.99999872 |
| AC011447.3 | 14.6413671 | -1.093598743 | 0.42605252 | -2.5668167 | 0.01026368 | 0.70849208 |
| ZNF737 | 16.1940543 | -0.02824628 | 0.58545081 | -0.0482471 | 0.96151935 | 0.99999872 |
| ZNF430 | 17.3882702 | 0.088501497 | 0.43117149 | 0.20525823 | 0.83737038 | 0.99999872 |
| ZNF714 | 8.53724082 | -0.866900514 | 0.60019601 | -1.4443623 | 0.14863708 | 0.99999872 |
| ZNF431 | 27.2676072 | -0.315127959 | 0.31901169 | -0.9878257 | 0.323238 | 0.99999872 |
| ZNF708 | 30.9926339 | -0.19637417 | 0.35658093 | -0.5507142 | 0.58182964 | 0.99999872 |
| ZNF493 | 9.02814244 | 0.244132533 | 0.55699248 | 0.4383049 | 0.66116527 | 0.99999872 |
| AC010615.4 | 36.243231 | 0.317407145 | 0.4064238 | 0.78097578 | 0.43481674 | 0.99999872 |
| ZNF429 | 23.9153281 | 0.008224773 | 0.35885098 | 0.02291974 | 0.98171429 | 0.99999872 |
| AC123912.1 | 5.32743304 | -0.302762048 | 0.79417144 | -0.3812301 | 0.70303253 | 0.99999872 |
| ZNF100 | 12.3495092 | 0.529914593 | 0.71937224 | 0.73663475 | 0.4613445 | 0.99999872 |
| ZNF43 | 22.406956 | -0.716734194 | 0.33605214 | -2.1328065 | 0.0329406 | 0.92959829 |
| ZNF208 | 32.1922336 | -0.162878124 | 0.56462456 | -0.2884716 | 0.7729858 | 0.99999872 |
| ZNF257 | 6.76746936 | -1.097358596 | 0.59292472 | -1.8507553 | 0.06420476 | 0.99999872 |
| ZNF676 | 3.82240996 | 0.840769305 | 0.79729731 | 1.0545242 | 0.29164299 | NA |
| ZNF98 | 15.584419 | -1.097130987 | 0.65604075 | -1.6723519 | 0.09445495 | 0.99999872 |
| ZNF91 | 94.6825293 | -0.15668365 | 0.34277755 | -0.4571001 | 0.64759911 | 0.99999872 |
| ZNF675 | 9.88092467 | 0.105439023 | 0.51480834 | 0.20481219 | 0.83771886 | 0.99999872 |
| ZNF254 | 56.5172867 | -0.305457865 | 0.30571111 | -0.9991716 | 0.31771156 | 0.99999872 |
| LINC00662 | 35.3339831 | -0.390272442 | 0.35665796 | -1.0942485 | 0.27384601 | 0.99999872 |
| AC006504.5 | 40.1238093 | 0.65995502 | 0.30806126 | 2.14228504 | 0.03217055 | 0.92959829 |
| AC020905.1 | 6.76328436 | -0.671038883 | 0.84221599 | -0.7967539 | 0.42559397 | 0.99999872 |
| AC005394.1 | 7.14803986 | 1.627724992 | 1.44615333 | 1.12555491 | 0.26035396 | 0.99999872 |
| UQCRFS1 | 30.1359199 | 0.8996406 | 0.4276121 | 2.1038708 | 0.03538972 | 0.94734101 |
| POP4 | 5.22658275 | 1.402444067 | 0.99557181 | 1.40868199 | 0.15892922 | 0.99999872 |
| C19orf12 | 20.7464372 | -0.365272176 | 0.48376554 | -0.7550603 | 0.45021278 | 0.99999872 |
| URI1 | 86.9538265 | 0.030489224 | 0.25172637 | 0.1211205 | 0.90359559 | 0.99999872 |

| | | | | | | |
|---|---|---|---|---|---|---|
| TSHZ3 | 54.1247414 | -0.069551392 | 0.38584801 | -0.1802559 | 0.85695165 | 0.99999872 |
| AC025809.1 | 21.3132235 | 0.06431296 | 0.49094642 | 0.13099792 | 0.89577695 | 0.99999872 |
| ZNF507 | 19.9823358 | 0.02620573 | 0.38067774 | 0.06883967 | 0.94511724 | 0.99999872 |
| DPY19L3 | 60.3254993 | 0.028891785 | 0.47887761 | 0.0603323 | 0.95189098 | 0.99999872 |
| PDCD5 | 13.1199662 | -0.195021543 | 0.47263407 | -0.4126269 | 0.67987997 | 0.99999872 |
| ANKRD27 | 20.0193535 | -0.305367932 | 0.38432958 | -0.7945471 | 0.42687703 | 0.99999872 |
| NUDT19 | 7.93753968 | 0.063709515 | 0.53638799 | 0.11877506 | 0.90545357 | 0.99999872 |
| CEP89 | 19.7592844 | -0.082894724 | 0.34311259 | -0.2415963 | 0.80909301 | 0.99999872 |
| RHPN2 | 11.107548 | -0.408038281 | 0.44636853 | -0.9141287 | 0.36064923 | 0.99999872 |
| GPATCH1 | 6.01737124 | 0.204339099 | 0.72049385 | 0.28360977 | 0.77670945 | 0.99999872 |
| CEBPG | 7.27908511 | 0.123131697 | 0.59837215 | 0.20577779 | 0.83696449 | 0.99999872 |
| PEPD | 25.8924655 | -0.042684114 | 0.42627036 | -0.1001339 | 0.92023802 | 0.99999872 |
| LSM14A | 91.2442182 | -0.325265759 | 0.22473683 | -1.4473184 | 0.14780776 | 0.99999872 |
| KIAA0355 | 41.6050112 | -0.142858233 | 0.37471212 | -0.381248 | 0.70301926 | 0.99999872 |
| GPI | 16.0992849 | -0.852848934 | 0.6051017 | -1.4094307 | 0.15870784 | 0.99999872 |
| UBA2 | 40.8457194 | 0.090949996 | 0.29051149 | 0.3130685 | 0.75422862 | 0.99999872 |
| WTIP | 20.4210781 | -0.666237848 | 0.45782703 | -1.4552174 | 0.14560909 | 0.99999872 |
| SCGB2B2 | 23.3960078 | -0.309818527 | 0.46075716 | -0.6724117 | 0.50132161 | 0.99999872 |
| ZNF302 | 29.3331746 | 0.285604952 | 0.39274994 | 0.72719286 | 0.46710782 | 0.99999872 |
| ZNF181 | 7.17271926 | 0.096223073 | 0.51832633 | 0.18564188 | 0.85272561 | 0.99999872 |
| GRAMD1A | 23.3098496 | -0.274109797 | 0.54298994 | -0.5048156 | 0.61368835 | 0.99999872 |
| HPN | 14.4959309 | 0.044646576 | 1.0331609 | 0.04321357 | 0.96553128 | 0.99999872 |
| LSR | 19.3303039 | -0.486887411 | 0.52283289 | -0.9312486 | 0.35172497 | 0.99999872 |
| USF2 | 22.7321373 | -0.040613442 | 0.46611466 | -0.0871319 | 0.93056669 | 0.99999872 |
| TMEM147 | 17.7570706 | 0.494361198 | 0.42512233 | 1.1628681 | 0.24488302 | 0.99999872 |
| RBM42 | 12.8463086 | 0.081329215 | 0.78683507 | 0.10336247 | 0.9176753 | 0.99999872 |
| COX6B1 | 50.631468 | 0.367474634 | 0.42461053 | 0.8654393 | 0.38679772 | 0.99999872 |
| KMT2B | 11.013387 | -0.185677237 | 0.48323966 | -0.3842343 | 0.70080482 | 0.99999872 |
| PROSER3 | 4.48636652 | -0.605395857 | 0.71895102 | -0.8420544 | 0.39975751 | NA |
| PRODH2 | 20.8330446 | -0.167992736 | 0.80106487 | -0.2097118 | 0.83389264 | 0.99999872 |
| NPHS1 | 1427.54048 | 0.147601282 | 0.7241365 | 0.20383074 | 0.83848577 | 0.99999872 |
| KIRREL2 | 9.33244298 | 1.285659284 | 0.56795269 | 2.26367321 | 0.02359422 | 0.88337526 |
| APLP1 | 17.6797975 | -0.122937497 | 0.64057336 | -0.1919179 | 0.84780651 | 0.99999872 |
| SDHAF1 | 7.82696019 | 0.35733074 | 0.52863118 | 0.67595472 | 0.4990694 | 0.99999872 |
| AC002116.2 | 4.66954103 | 0.701238738 | 0.81517082 | 0.86023533 | 0.38965933 | 0.99999872 |
| CLIP3 | 34.1581207 | -0.195838987 | 0.38550245 | -0.5080097 | 0.61144653 | 0.99999872 |
| THAP8 | 5.34857929 | -0.117725552 | 0.80723372 | -0.1458382 | 0.88404908 | 0.99999872 |
| POLR2I | 5.51640642 | -0.155134914 | 0.65311948 | -0.2375291 | 0.81224631 | 0.99999872 |
| TBCB | 28.8194307 | -0.158495216 | 0.48040819 | -0.3299178 | 0.74146207 | 0.99999872 |
| CAPNS1 | 78.7830493 | -0.16743919 | 0.54902313 | -0.3049766 | 0.760384 | 0.99999872 |
| COX7A1 | 9.57718459 | -0.168375133 | 0.75985448 | -0.2215887 | 0.82463411 | 0.99999872 |
| ZNF565 | 24.9786641 | 0.095390914 | 0.37007999 | 0.25775755 | 0.79659402 | 0.99999872 |
| ZNF146 | 34.4878453 | -0.28018592 | 0.32148979 | -0.8715236 | 0.38346835 | 0.99999872 |
| LINC00665 | 11.3240815 | -0.361514351 | 0.68013863 | -0.5315304 | 0.59505129 | 0.99999872 |
| ZFP14 | 21.5181556 | 0.183367707 | 0.36565605 | 0.50147593 | 0.61603621 | 0.99999872 |
| ZFP82 | 11.0854837 | -0.034081065 | 0.43461801 | -0.0784161 | 0.93749704 | 0.99999872 |
| ZNF566 | 14.2634894 | 0.314546078 | 0.40880084 | 0.76943599 | 0.44163453 | 0.99999872 |
| ZNF260 | 10.7464961 | -0.004460832 | 0.44603864 | -0.010001 | 0.99202049 | 0.99999872 |
| ZNF529 | 19.146294 | -0.207553981 | 0.5301949 | -0.3914673 | 0.69545184 | 0.99999872 |
| ZNF529-AS1 | 16.1052019 | 0.088517351 | 0.47688396 | 0.18561612 | 0.85274581 | 0.99999872 |
| ZNF382 | 6.91070167 | 0.227906237 | 0.54548967 | 0.4178012 | 0.67609247 | 0.99999872 |
| ZNF461 | 7.32387559 | 0.220073004 | 0.55053276 | 0.39974552 | 0.68934396 | 0.99999872 |
| ZNF567 | 8.54621007 | 0.648632511 | 0.51542295 | 1.25844709 | 0.20823011 | 0.99999872 |
| AC020928.1 | 21.0021232 | -0.23334164 | 0.39929611 | -0.5843825 | 0.55896303 | 0.99999872 |
| ZNF790 | 14.3866863 | 0.262088807 | 0.57896125 | 0.45268799 | 0.65077343 | 0.99999872 |
| ZNF345 | 26.190883 | -0.364598028 | 0.41636027 | -0.8756792 | 0.38120446 | 0.99999872 |
| ZNF568 | 13.8032322 | 0.017400318 | 0.44807251 | 0.03883371 | 0.96902297 | 0.99999872 |
| ZNF420 | 27.2707851 | 0.199787565 | 0.31228431 | 0.63976178 | 0.52232748 | 0.99999872 |
| ZNF383 | 8.99524002 | -0.237850505 | 0.63980117 | -0.3717569 | 0.71007385 | 0.99999872 |
| HKR1 | 22.0055803 | -0.160911907 | 0.367444 | -0.4379223 | 0.66144264 | 0.99999872 |
| ZNF569 | 8.40926924 | 0.144435743 | 0.54495703 | 0.26504061 | 0.79097818 | 0.99999872 |

| | | | | | | | |
|---|---|---|---|---|---|---|---|
| ZNF793 | 11.2872337 | | -0.171209161 | 0.49925836 | -0.342927 | 0.7316534 | 0.99999872 |
| ZNF540 | 7.39779934 | | 0.591222824 | 0.71402443 | 0.82801484 | 0.40766209 | 0.99999872 |
| ZNF571 | 7.16701337 | | 0.333438744 | 0.52752864 | 0.63207704 | 0.52733654 | 0.99999872 |
| ZFP30 | 16.7217173 | | 0.110488075 | 0.51144064 | 0.21603304 | 0.828962 | 0.99999872 |
| ZNF573 | 12.1118594 | | 0.363062662 | 0.53932271 | 0.67318259 | 0.50083113 | 0.99999872 |
| SIPA1L3 | 43.4522429 | | -0.132961749 | 0.39016133 | -0.3407866 | 0.73326424 | 0.99999872 |
| SPINT2 | 7.8171016 | | 0.492812165 | 0.57992577 | 0.8497849 | 0.39544469 | 0.99999872 |
| YIF1B | 6.55623111 | | 0.40759157 | 0.74467452 | 0.5473419 | 0.58414386 | 0.99999872 |
| CATSPERG | 6.16249537 | | -1.031792704 | 0.5835787 | -1.7680438 | 0.07705358 | 0.99999872 |
| PSMD8 | 36.1362566 | | 0.883797664 | 0.4430565 | 1.99477416 | 0.04606752 | 0.95980711 |
| EIF3K | 21.5499526 | | 0.718573437 | 0.48165673 | 1.49187875 | 0.13573093 | 0.99999872 |
| ACTN4 | 880.217561 | | -0.014916023 | 0.44497816 | -0.0335208 | 0.97325928 | 0.99999872 |
| AC008649.2 | 4.98735851 | | 1.563264814 | 0.95060701 | 1.64449114 | 0.10007479 | 0.99999872 |
| CAPN12 | 10.5549585 | | 0.763310645 | 0.62280787 | 1.2255957 | 0.22035086 | 0.99999872 |
| ECH1 | 32.5271619 | | 1.241959812 | 0.39626717 | 3.1341476 | 0.00172354 | 0.34074087 |
| HNRNPL | 23.9657245 | | -0.202820867 | 0.40915664 | -0.4957047 | 0.62010277 | 0.99999872 |
| SIRT2 | 13.6866768 | | 0.658749408 | 0.52798738 | 1.24766128 | 0.21215513 | 0.99999872 |
| NFKBIB | 4.45985475 | | -0.250713742 | 0.80362199 | -0.3119797 | 0.75505596 | NA |
| MRPS12 | 10.5792177 | | 0.809342807 | 0.62989176 | 1.28489189 | 0.19883007 | 0.99999872 |
| FBXO17 | 34.0711037 | | 0.00672154 | 0.33889504 | 0.01983369 | 0.98417604 | 0.99999872 |
| FBXO27 | 5.32799354 | | 0.872681738 | 0.81301734 | 1.07338637 | 0.2830978 | 0.99999872 |
| PAK4 | 11.1196468 | | 0.089635388 | 0.45312562 | 0.19781576 | 0.84318922 | 0.99999872 |
| SAMD4B | 39.99198 | | -0.421713535 | 0.35706298 | -1.181062 | 0.23757808 | 0.99999872 |
| PAF1 | 10.0758845 | | -0.105704695 | 0.51116762 | -0.2067907 | 0.83617334 | 0.99999872 |
| MED29 | 14.4162448 | | -0.437012791 | 0.59207085 | -0.7381089 | 0.46044825 | 0.99999872 |
| ZFP36 | 17.6363272 | | 0.029699372 | 0.92505143 | 0.03210564 | 0.9743878 | 0.99999872 |
| PLEKHG2 | 8.24797282 | | -0.097280772 | 0.61256334 | -0.1588093 | 0.8738191 | 0.99999872 |
| RPS16 | 54.848759 | | 0.664675729 | 0.47272543 | 1.4060503 | 0.15970919 | 0.99999872 |
| SUPT5H | 22.9373769 | | -0.833978524 | 0.36140027 | -2.3076311 | 0.02101966 | 0.8671024 |
| TIMM50 | 8.2768954 | | -0.675548978 | 0.53878349 | -1.2538413 | 0.2098997 | 0.99999872 |
| EID2B | 4.91121122 | | -0.407883463 | 0.66874389 | -0.6099248 | 0.54191165 | 0.99999872 |
| EID2 | 7.48384092 | | 0.528546105 | 0.54998783 | 0.96101417 | 0.33654504 | 0.99999872 |
| DYRK1B | 6.00671962 | | -0.343572227 | 0.61523769 | -0.5584382 | 0.57654519 | 0.99999872 |
| FBL | 6.95681484 | | 0.420434788 | 0.61423542 | 0.68448477 | 0.4936691 | 0.99999872 |
| PSMC4 | 13.5634141 | | -0.213770299 | 0.57964607 | -0.3687945 | 0.71228088 | 0.99999872 |
| ZNF546 | 7.69112769 | | 0.744775523 | 0.60796186 | 1.22503659 | 0.22056144 | 0.99999872 |
| ZNF780B | 34.1703304 | | -0.303836956 | 0.4688338 | -0.6480697 | 0.5169399 | 0.99999872 |
| ZNF780A | 8.03491725 | | -0.492326294 | 0.59092584 | -0.833144 | 0.40476355 | 0.99999872 |
| MAP3K10 | 7.47329637 | | 0.060442324 | 0.57400887 | 0.10529859 | 0.91613888 | 0.99999872 |
| AKT2 | 71.9348254 | | 0.111487718 | 0.34950124 | 0.31899091 | 0.74973341 | 0.99999872 |
| PLD3 | 128.362248 | | 0.342230391 | 0.42679106 | 0.8018687 | 0.42262891 | 0.99999872 |
| PRX | 10.2164273 | | 0.161456504 | 0.55268369 | 0.29213184 | 0.77018582 | 0.99999872 |
| SERTAD1 | 16.7028182 | | 0.069625054 | 0.57266675 | 0.1215804 | 0.90323133 | 0.99999872 |
| SERTAD3 | 9.32920356 | | -0.01176879 | 0.60607044 | -0.0194182 | 0.9845075 | 0.99999872 |
| BLVRB | 40.3395225 | | 0.551220883 | 0.45734092 | 1.20527348 | 0.22809774 | 0.99999872 |
| SPTBN4 | 5.50710432 | | 0.227314643 | 0.71455172 | 0.31812203 | 0.75039237 | 0.99999872 |
| SHKBP1 | 9.87980893 | | -0.622401787 | 0.49867199 | -1.2481186 | 0.21198763 | 0.99999872 |
| LTBP4 | 60.6870151 | | 0.037510281 | 0.66039832 | 0.05679948 | 0.95470493 | 0.99999872 |
| NUMBL | 8.59768272 | | 0.882075848 | 0.60843515 | 1.44974504 | 0.14712963 | 0.99999872 |
| COQ8B | 9.65771784 | | 0.218262904 | 0.61922338 | 0.35247846 | 0.72447947 | 0.99999872 |
| ITPKC | 7.85879294 | | 0.308393246 | 0.7480277 | 0.41227517 | 0.68013775 | 0.99999872 |
| SNRPA | 11.3168412 | | -0.332320195 | 0.46898361 | -0.7085966 | 0.47857484 | 0.99999872 |
| AC008537.1 | 15.9662511 | | 0.446518726 | 0.64891597 | 0.68809946 | 0.49139015 | 0.99999872 |
| AXL | 51.1411037 | | -1.106891633 | 0.50590547 | -2.1879416 | 0.02867386 | 0.90266962 |
| HNRNPUL1 | 23.5624986 | | -0.010885679 | 0.3485155 | -0.0312344 | 0.97508259 | 0.99999872 |
| TGFB1 | 20.2017641 | | -0.491745553 | 0.51321718 | -0.9581627 | 0.33798073 | 0.99999872 |
| CCDC97 | 6.83456045 | | 0.094517327 | 0.54796144 | 0.17248901 | 0.8630531 | 0.99999872 |
| DMAC2 | 5.86448284 | | 0.164220512 | 0.58661816 | 0.27994447 | 0.77952011 | 0.99999872 |
| RPS19 | 111.924251 | | 0.444973766 | 0.43307939 | 1.02746465 | 0.30420172 | 0.99999872 |
| ARHGEF1 | 8.55654614 | | -0.224827346 | 0.53299722 | -0.4218171 | 0.67315852 | 0.99999872 |
| RABAC1 | 79.3846797 | | 0.536271888 | 0.53648903 | 0.99959526 | 0.31750642 | 0.99999872 |

| | | | | | | |
|---|---|---|---|---|---|---|
| GRIK5 | 16.020224 | -0.4695769 | 0.5190212 | -0.9047355 | 0.36560553 | 0.99999872 |
| DEDD2 | 6.13497415 | 0.30240368 | 0.58341054 | 0.51833771 | 0.60422267 | 0.99999872 |
| GSK3A | 13.6098994 | -0.375089881 | 0.47507292 | -0.7895417 | 0.42979547 | 0.99999872 |
| CIC | 11.4884177 | -0.80436303 | 0.48117784 | -1.6716544 | 0.0945925 | 0.99999872 |
| PAFAH1B3 | 5.44215842 | -0.638668583 | 0.59386377 | -1.0754463 | 0.28217497 | 0.99999872 |
| MEGF8 | 15.8811126 | -0.096029798 | 0.48855538 | -0.1965587 | 0.84417291 | 0.99999872 |
| LIPE-AS1 | 19.3732747 | 0.077817192 | 0.46224062 | 0.1683478 | 0.86630967 | 0.99999872 |
| PHLDB3 | 7.58817775 | -0.104219309 | 0.50346453 | -0.2070043 | 0.83600652 | 0.99999872 |
| ETHE1 | 25.851453 | 0.227926414 | 0.51587076 | 0.44182852 | 0.6586133 | 0.99999872 |
| XRCC1 | 6.46806943 | 0.249820755 | 0.73177019 | 0.34139236 | 0.73280822 | 0.99999872 |
| ZNF428 | 30.9171867 | 0.297140041 | 0.49088304 | 0.60531739 | 0.54496813 | 0.99999872 |
| CADM4 | 8.04821414 | -0.08386336 | 0.55857686 | -0.1501375 | 0.8806561 | 0.99999872 |
| PLAUR | 7.2336195 | 0.052601291 | 0.50574847 | 0.10400682 | 0.91716393 | 0.99999872 |
| SMG9 | 10.2839549 | -0.579593073 | 0.46916079 | -1.2353826 | 0.21668815 | 0.99999872 |
| ZNF283 | 7.66798389 | 0.147579469 | 0.56031342 | 0.26338735 | 0.79225204 | 0.99999872 |
| ZNF45 | 4.83364036 | 0.597414268 | 0.6907168 | 0.86491927 | 0.3870831 | 0.99999872 |
| ZNF155 | 10.272234 | 0.225106597 | 0.48526751 | 0.46388145 | 0.64273268 | 0.99999872 |
| ZNF224 | 11.9068891 | 0.357565673 | 0.59855688 | 0.59737961 | 0.55025396 | 0.99999872 |
| ZNF234 | 8.20286987 | -0.247601986 | 0.5077423 | -0.4876529 | 0.62579574 | 0.99999872 |
| ZNF226 | 13.6103253 | -0.186562279 | 0.43181105 | -0.4320461 | 0.66570791 | 0.99999872 |
| ZNF227 | 8.57537619 | 0.060064399 | 0.56062879 | 0.10713756 | 0.91467985 | 0.99999872 |
| ZNF235 | 35.1535421 | 0.297630843 | 0.42454529 | 0.70105793 | 0.48326686 | 0.99999872 |
| ZNF229 | 5.93252652 | 0.185912927 | 0.71840829 | 0.25878449 | 0.79580152 | 0.99999872 |
| ZNF180 | 7.03433532 | -0.993388989 | 0.57990004 | -1.7130349 | 0.08670613 | 0.99999872 |
| AC243964.2 | 31.6360169 | 0.81017918 | 0.49352957 | 1.64160211 | 0.10067249 | 0.99999872 |
| PVR | 15.2822411 | 0.6166589 | 0.5044419 | 1.22245773 | 0.22153458 | 0.99999872 |
| CEACAM19 | 51.2212746 | -0.709811367 | 0.35467258 | -2.0013145 | 0.04535851 | 0.95980711 |
| BCL3 | 20.7650406 | -0.019543375 | 1.05568402 | -0.0185125 | 0.98522999 | 0.99999872 |
| BCAM | 482.880143 | -0.420479341 | 0.59362044 | -0.7083303 | 0.47874017 | 0.99999872 |
| NECTIN2 | 227.484453 | -0.284826458 | 0.45084667 | -0.631759 | 0.52754434 | 0.99999872 |
| TOMM40 | 9.89217392 | -0.026969313 | 0.46603405 | -0.0578698 | 0.95385231 | 0.99999872 |
| APOE | 46.2622135 | -1.004892803 | 1.10564635 | -0.9088736 | 0.36341684 | 0.99999872 |
| CLPTM1 | 28.1397676 | 0.05549185 | 0.61943741 | 0.08958427 | 0.92861758 | 0.99999872 |
| RELB | 17.0065169 | -0.872769621 | 0.54205022 | -1.6101269 | 0.10737016 | 0.99999872 |
| CLASRP | 20.8777731 | -0.540668768 | 0.35657848 | -1.5162686 | 0.12945144 | 0.99999872 |
| GEMIN7-AS1 | 5.77470759 | -1.145125987 | 0.58164048 | -1.9687866 | 0.0489776 | 0.97937581 |
| MARK4 | 28.4995597 | 0.039471079 | 0.31713813 | 0.12446021 | 0.9009509 | 0.99999872 |
| TRAPPC6A | 14.0776665 | 0.384173211 | 0.57957503 | 0.66285328 | 0.50742453 | 0.99999872 |
| PPP1R13L | 4.74570395 | -0.405991948 | 0.71581434 | -0.5671749 | 0.57059534 | 0.99999872 |
| ERCC1 | 11.1315859 | -0.029703697 | 0.48232108 | -0.0615849 | 0.9508934 | 0.99999872 |
| VASP | 12.7223956 | -0.309007988 | 0.56867819 | -0.5433794 | 0.58686864 | 0.99999872 |
| OPA3 | 14.4715421 | -0.674407932 | 0.58838145 | -1.1462087 | 0.25170881 | 0.99999872 |
| EML2 | 14.2207722 | 0.301951909 | 0.41970535 | 0.71943783 | 0.47187119 | 0.99999872 |
| SNRPD2 | 34.4768296 | 0.335409899 | 0.39020437 | 0.85957494 | 0.39002339 | 0.99999872 |
| FBXO46 | 8.15356431 | 0.358303429 | 0.56661067 | 0.63236266 | 0.52714993 | 0.99999872 |
| SIX5 | 7.11963164 | 0.08642061 | 0.60940184 | 0.14181219 | 0.88722835 | 0.99999872 |
| DM1-AS | 6.71492405 | -0.549886042 | 0.55541429 | -0.9900466 | 0.32215133 | 0.99999872 |
| DMPK | 9.71554558 | -0.051362371 | 0.56466703 | -0.0909605 | 0.92752401 | 0.99999872 |
| SYMPK | 28.9620007 | -0.296080632 | 0.30629948 | -0.9666377 | 0.33372516 | 0.99999872 |
| MYPOP | 5.44411045 | -0.240857087 | 0.58313195 | -0.4130405 | 0.67957697 | 0.99999872 |
| PPP5C | 20.137698 | -0.302824306 | 0.39960072 | -0.7578172 | 0.44856041 | 0.99999872 |
| PNMA8A | 4.62111669 | 2.0312006 | 0.83584111 | 2.43012767 | 0.01509351 | 0.78681466 |
| PPP5D1 | 16.611786 | -0.931293751 | 0.78033985 | -1.1934464 | 0.23269461 | 0.99999872 |
| CALM3 | 22.402401 | 0.120324414 | 0.45901215 | 0.26213775 | 0.79321524 | 0.99999872 |
| PRKD2 | 17.3274904 | -0.747909561 | 0.44661832 | -1.6746056 | 0.09401163 | 0.99999872 |
| STRN4 | 8.93609732 | -0.812059394 | 0.52302164 | -1.5526306 | 0.12051142 | 0.99999872 |
| FKRP | 11.6836597 | 0.308847379 | 0.49004684 | 0.63024053 | 0.52853722 | 0.99999872 |
| SLC1A5 | 29.8395446 | 0.120676455 | 0.41483536 | 0.29090205 | 0.77112624 | 0.99999872 |
| AP2S1 | 22.467083 | 0.31770699 | 0.48711518 | 0.6522215 | 0.51425829 | 0.99999872 |
| ARHGAP35 | 41.6410416 | -0.182048064 | 0.31739176 | -0.5735753 | 0.56625525 | 0.99999872 |
| TMEM160 | 20.8544002 | 0.422543678 | 0.43689311 | 0.96715574 | 0.33346618 | 0.99999872 |

| | | | | | | |
|---|---|---|---|---|---|---|
| ZC3H4 | 17.6540363 | -1.002278172 | 0.45788181 | -2.1889452 | 0.02860083 | 0.90266962 |
| SAE1 | 26.1162517 | -0.484760255 | 0.4229892 | -1.1460346 | 0.25178084 | 0.99999872 |
| INAFM1 | 5.97451807 | -0.297660936 | 0.58332789 | -0.5102807 | 0.60985486 | 0.99999872 |
| DHX34 | 8.80092202 | -1.183593634 | 0.53715549 | -2.2034469 | 0.02756326 | 0.89154528 |
| NAPA | 38.5568219 | -0.501498561 | 0.40031317 | -1.2527656 | 0.21029103 | 0.99999872 |
| EHD2 | 40.5031949 | -0.24772773 | 0.60249043 | -0.4111729 | 0.68094577 | 0.99999872 |
| NOP53 | 6.85764877 | 0.083268403 | 0.59934327 | 0.13893274 | 0.8895033 | 0.99999872 |
| SELENOW | 59.2830305 | 0.810533156 | 0.40926047 | 1.98048239 | 0.04764935 | 0.97387863 |
| PLA2G4C | 15.6377458 | -1.145270963 | 0.68129078 | -1.6810311 | 0.09275688 | 0.99999872 |
| ZSWIM9 | 6.66022991 | -0.739595384 | 0.59045736 | -1.2525805 | 0.21035841 | 0.99999872 |
| CARD8 | 11.1464845 | -0.072421347 | 0.4785626 | -0.151331 | 0.87971463 | 0.99999872 |
| EMP3 | 37.2127264 | 0.295524847 | 0.69595326 | 0.42463318 | 0.6711041 | 0.99999872 |
| KDELR1 | 31.9628441 | 0.037459289 | 0.36192248 | 0.10350086 | 0.91756546 | 0.99999872 |
| GRWD1 | 6.34451144 | -0.011924105 | 0.88641826 | -0.013452 | 0.98926717 | 0.99999872 |
| CYTH2 | 12.4397667 | -0.455099578 | 0.44127724 | -1.0313235 | 0.30238915 | 0.99999872 |
| RPL18 | 65.8269476 | 0.160753085 | 0.52214691 | 0.30786946 | 0.75818167 | 0.99999872 |
| SPHK2 | 7.62085464 | -0.161034796 | 0.52796937 | -0.3050078 | 0.76036018 | 0.99999872 |
| DBP | 10.5840753 | 0.03998979 | 0.72521964 | 0.05514163 | 0.95602563 | 0.99999872 |
| CA11 | 6.80895385 | 0.295536294 | 0.93566696 | 0.31585629 | 0.75211159 | 0.99999872 |
| BCAT2 | 15.7470075 | -0.52286587 | 0.37812065 | -1.3828017 | 0.16672569 | 0.99999872 |
| HSD17B14 | 14.0264582 | -0.278349519 | 0.66779508 | -0.4168188 | 0.67681098 | 0.99999872 |
| PLEKHA4 | 10.7965364 | -0.538384949 | 0.53264279 | -1.0107805 | 0.31212149 | 0.99999872 |
| PPP1R15A | 15.5154551 | -0.242478533 | 0.53501722 | -0.4532163 | 0.65039299 | 0.99999872 |
| NUCB1 | 133.855602 | -0.007914616 | 0.47504656 | -0.0166607 | 0.98670729 | 0.99999872 |
| BAX | 16.4377775 | 0.108156503 | 0.52367367 | 0.20653416 | 0.83637368 | 0.99999872 |
| FTL | 476.231343 | 1.039142068 | 0.70268107 | 1.47882462 | 0.13918719 | 0.99999872 |
| GYS1 | 10.4203853 | -0.691849303 | 0.47209337 | -1.4654925 | 0.14278659 | 0.99999872 |
| RUVBL2 | 7.72050824 | -0.275522129 | 0.53048843 | -0.5193744 | 0.60349966 | 0.99999872 |
| SNRNP70 | 76.128102 | -0.381980133 | 0.22844587 | -1.6720816 | 0.09450823 | 0.99999872 |
| TRPM4 | 11.9576597 | -0.008647693 | 0.66842813 | -0.0129374 | 0.98967777 | 0.99999872 |
| SLC6A16 | 18.3854892 | -0.437228812 | 0.46375719 | -0.9427968 | 0.34578483 | 0.99999872 |
| TEAD2 | 10.5676057 | -0.029241524 | 0.64815459 | -0.045115 | 0.96401561 | 0.99999872 |
| PIH1D1 | 10.4647033 | 0.474835041 | 0.46886974 | 1.01272271 | 0.31119263 | 0.99999872 |
| RPL13A | 83.8082517 | 0.327925803 | 0.50175751 | 0.65355435 | 0.51339897 | 0.99999872 |
| RPS11 | 71.3426291 | 0.494256056 | 0.45340601 | 1.09009596 | 0.27567088 | 0.99999872 |
| FCGRT | 36.9095188 | 0.641780825 | 0.31064387 | 2.06596968 | 0.03883134 | 0.95980711 |
| NOSIP | 9.41900399 | 0.394043218 | 0.49383137 | 0.79793071 | 0.4249107 | 0.99999872 |
| PRR12 | 10.739488 | -0.629234179 | 0.50007651 | -1.2582758 | 0.20829203 | 0.99999872 |
| RRAS | 56.3905873 | 0.727582939 | 0.63570194 | 1.14453472 | 0.25240195 | 0.99999872 |
| SCAF1 | 14.0156976 | -0.118850827 | 0.41069022 | -0.2893929 | 0.77228074 | 0.99999872 |
| IRF3 | 19.9868252 | -0.545172606 | 0.36785812 | -1.4820187 | 0.13833532 | 0.99999872 |
| PRMT1 | 24.3952972 | 0.149525462 | 0.44364097 | 0.3370416 | 0.73608554 | 0.99999872 |
| AP2A1 | 23.0666213 | 0.202092251 | 0.35558556 | 0.5683365 | 0.5698065 | 0.99999872 |
| MED25 | 21.7735349 | -0.393454677 | 0.47629583 | -0.8260721 | 0.40876322 | 0.99999872 |
| PTOV1 | 19.3420075 | -0.007554711 | 0.44829861 | -0.016852 | 0.98655472 | 0.99999872 |
| PNKP | 9.58408406 | -0.76257019 | 0.65466549 | -1.1648242 | 0.24409017 | 0.99999872 |
| TBC1D17 | 11.5786815 | -0.362833924 | 0.4733829 | -0.7664703 | 0.44339653 | 0.99999872 |
| ATF5 | 6.03634092 | -0.615709248 | 0.88853584 | -0.6929481 | 0.4883421 | 0.99999872 |
| VRK3 | 14.0334293 | 0.086502535 | 0.52858453 | 0.16364939 | 0.87000716 | 0.99999872 |
| MYH14 | 6.85917893 | -0.680823597 | 0.6841551 | -0.9951305 | 0.3196728 | 0.99999872 |
| KCNC3 | 7.17038748 | -0.434231861 | 0.67697554 | -0.6414292 | 0.52124388 | 0.99999872 |
| NR1H2 | 23.3908698 | -0.428671063 | 0.38626313 | -1.1097903 | 0.26708941 | 0.99999872 |
| NAPSA | 8.76236046 | -0.479433534 | 1.04763666 | -0.4576334 | 0.64721584 | 0.99999872 |
| EMC10 | 48.5503225 | 0.175206073 | 0.65817501 | 0.26619983 | 0.79008532 | 0.99999872 |
| JOSD2 | 12.8481768 | 0.29144185 | 0.72817258 | 0.40023733 | 0.68898172 | 0.99999872 |
| CLEC11A | 5.73163201 | 0.023845534 | 0.58189391 | 0.04097918 | 0.9673125 | 0.99999872 |
| C19orf48 | 6.33921174 | -1.206198176 | 0.84213837 | -1.432304 | 0.15205683 | 0.99999872 |
| KLK6 | 64.9318961 | 0.383678453 | 0.63506353 | 0.60415759 | 0.54573887 | 0.99999872 |
| AC011483.1 | 6.59880313 | 2.464895166 | 1.32565144 | 1.85938406 | 0.06297272 | 0.99999872 |
| KLK7 | 64.2776788 | 1.01457437 | 0.72646118 | 1.39659819 | 0.16253444 | 0.99999872 |
| CTU1 | 7.73910961 | -0.247810416 | 0.61590246 | -0.4023533 | 0.687424 | 0.99999872 |

| | | | | | | |
|---|---|---|---|---|---|---|
| ETFB | 32.7857366 | 0.481142041 | 0.35562918 | 1.35293184 | 0.17607741 | 0.99999872 |
| ZNF175 | 9.27140102 | 0.0087051 | 0.6310139 | 0.01379542 | 0.9889932 | 0.99999872 |
| SPACA6 | 15.6550028 | -0.810803835 | 0.59792109 | -1.3560382 | 0.17508702 | 0.99999872 |
| ZNF577 | 13.7322287 | 0.347955922 | 0.53490169 | 0.65050444 | 0.51536644 | 0.99999872 |
| ZNF350-AS1 | 9.20841679 | -0.698672637 | 0.7146315 | -0.9776684 | 0.32823836 | 0.99999872 |
| ZNF350 | 5.62367091 | 0.608887277 | 0.69183662 | 0.88010269 | 0.37880368 | 0.99999872 |
| ZNF615 | 4.84769584 | 1.315691944 | 0.91830642 | 1.43273739 | 0.1519329 | 0.99999872 |
| ZNF614 | 4.96389528 | 0.458943427 | 0.73086031 | 0.62794958 | 0.53003697 | 0.99999872 |
| ZNF432 | 17.3029138 | -0.422670277 | 0.39584243 | -1.067774 | 0.28562245 | 0.99999872 |
| AC011468.3 | 5.41072921 | 1.539175796 | 0.92076922 | 1.67161952 | 0.09459937 | 0.99999872 |
| PPP2R1A | 51.5271374 | -0.010471423 | 0.40743669 | -0.0257007 | 0.97949604 | 0.99999872 |
| ZNF766 | 12.7946903 | -0.067050044 | 0.55846452 | -0.1200614 | 0.90443449 | 0.99999872 |
| ZNF480 | 8.15451364 | 0.095437275 | 0.65358488 | 0.14602124 | 0.88390462 | 0.99999872 |
| ZNF880 | 7.70429932 | 0.144696927 | 0.73922058 | 0.19574256 | 0.84481167 | 0.99999872 |
| ZNF528 | 7.67859442 | -0.58682617 | 0.52047632 | -1.1274791 | 0.25953997 | 0.99999872 |
| ZNF578 | 4.89586117 | -0.666494052 | 0.63585391 | -1.0481874 | 0.29455228 | 0.99999872 |
| ZNF701 | 8.97297687 | -0.196486834 | 0.59378509 | -0.3309056 | 0.74071576 | 0.99999872 |
| ZNF83 | 54.3087912 | -0.229112891 | 0.26807558 | -0.8546578 | 0.39274061 | 0.99999872 |
| ZNF611 | 22.0217944 | -1.018397407 | 0.40249658 | -2.5302014 | 0.01139971 | 0.72670507 |
| ZNF600 | 4.32967138 | -0.115131943 | 0.64448739 | -0.1786411 | 0.85821951 | NA |
| ZNF28 | 4.72466615 | -0.419874613 | 0.65554676 | -0.6404953 | 0.52185065 | 0.99999872 |
| ZNF160 | 31.1157144 | -0.314874194 | 0.42749662 | -0.7365536 | 0.46139384 | 0.99999872 |
| ZNF415 | 4.94947534 | 0.007709465 | 0.67977265 | 0.01134124 | 0.99095119 | 0.99999872 |
| ZNF347 | 9.14242951 | 0.214669685 | 0.49839813 | 0.43071928 | 0.6666725 | 0.99999872 |
| ZNF677 | 10.9165189 | 0.480060271 | 0.48195287 | 0.99607307 | 0.31921465 | 0.99999872 |
| ZNF525 | 6.06547343 | -0.115534939 | 0.6257207 | -0.184643 | 0.85350907 | 0.99999872 |
| ZNF765 | 12.5421019 | 0.0118341 | 0.4622125 | 0.02560316 | 0.97957387 | 0.99999872 |
| ZNF761 | 9.13486076 | -0.676794363 | 0.51852383 | -1.3052329 | 0.19181356 | 0.99999872 |
| ZNF331 | 10.7707879 | 0.319128982 | 0.65456 | 0.48754733 | 0.62587051 | 0.99999872 |
| MYADM | 31.0071448 | -0.794898693 | 0.48710307 | -1.6318901 | 0.10270263 | 0.99999872 |
| AC008440.2 | 8.33844484 | 0.420580116 | 0.71698449 | 0.58659584 | 0.55747518 | 0.99999872 |
| NDUFA3 | 12.1733591 | 1.001687049 | 0.59698874 | 1.6778994 | 0.09336674 | 0.99999872 |
| PRPF31 | 16.2307729 | 0.033265923 | 0.36551108 | 0.09101208 | 0.92748299 | 0.99999872 |
| CNOT3 | 10.5098991 | 0.088549919 | 0.51475188 | 0.17202447 | 0.86341829 | 0.99999872 |
| LENG1 | 8.85524324 | 0.023676445 | 0.55871071 | 0.04237693 | 0.96619822 | 0.99999872 |
| TMC4 | 5.26940985 | -0.120026644 | 0.64687543 | -0.1855483 | 0.85279899 | 0.99999872 |
| MBOAT7 | 22.1138394 | -0.083377171 | 0.47086837 | -0.1770711 | 0.85945256 | 0.99999872 |
| TSEN34 | 14.665017 | 0.65584226 | 0.46066432 | 1.42368798 | 0.15453681 | 0.99999872 |
| RPS9 | 125.608315 | 0.510151465 | 0.42902215 | 1.18910285 | 0.2343992 | 0.99999872 |
| LENG8-AS1 | 6.47816893 | -0.393425161 | 0.62970959 | -0.6247724 | 0.53212046 | 0.99999872 |
| LENG8 | 74.9665441 | -0.333119837 | 0.2768309 | -1.2033333 | 0.22884737 | 0.99999872 |
| AC011476.3 | 4.41414518 | 1.273900247 | 1.04995156 | 1.2132943 | 0.22501732 | NA |
| RDH13 | 10.4924201 | -0.369179598 | 0.50142938 | -0.7362544 | 0.46157588 | 0.99999872 |
| PPP1R12C | 18.8490847 | -0.237149485 | 0.43684681 | -0.5428665 | 0.58722175 | 0.99999872 |
| PPP6R1 | 12.7434162 | -0.659836614 | 0.40925287 | -1.6122956 | 0.10689762 | 0.99999872 |
| HSPBP1 | 8.10053301 | -0.269095236 | 0.59902716 | -0.4492204 | 0.65327265 | 0.99999872 |
| KMT5C | 5.24374906 | 0.20494945 | 0.69299678 | 0.29574373 | 0.7674258 | 0.99999872 |
| RPL28 | 89.8411367 | 0.100771774 | 0.52658348 | 0.19136904 | 0.84823648 | 0.99999872 |
| UBE2S | 11.4798557 | 0.764147405 | 0.63849067 | 1.19680278 | 0.23138343 | 0.99999872 |
| ISOC2 | 12.5843278 | 0.427629757 | 0.65505325 | 0.65281679 | 0.5138744 | 0.99999872 |
| NAT14 | 9.44305426 | -0.211235596 | 0.63215503 | -0.3341516 | 0.73826519 | 0.99999872 |
| SSC5D | 9.30184496 | -0.001822171 | 0.63415469 | -0.0028734 | 0.99770737 | 0.99999872 |
| ZNF524 | 6.23339736 | -0.370152917 | 0.63278124 | -0.5849619 | 0.55857333 | 0.99999872 |
| CCDC106 | 9.31893144 | -0.718214317 | 0.5355249 | -1.3411409 | 0.17987472 | 0.99999872 |
| U2AF2 | 12.6352286 | -0.177394513 | 0.56241975 | -0.315413 | 0.75244808 | 0.99999872 |
| EPN1 | 33.0845987 | -0.484169708 | 0.33977707 | -1.4249629 | 0.15416794 | 0.99999872 |
| ZNF787 | 10.0726184 | -0.822605659 | 0.4644101 | -1.7712915 | 0.07651224 | 0.99999872 |
| ZNF444 | 13.3496584 | -0.244418817 | 0.4334686 | -0.5638674 | 0.57284437 | 0.99999872 |
| ZSCAN5A | 27.9568644 | -0.179874297 | 0.43424756 | -0.4142206 | 0.67871254 | 0.99999872 |
| ZNF582 | 5.68645337 | 0.584023529 | 0.62051025 | 0.94119885 | 0.34660297 | 0.99999872 |
| ZNF583 | 7.10728428 | 0.42323613 | 0.58227944 | 0.72686085 | 0.4673112 | 0.99999872 |

| | | | | | | |
|---|---|---|---|---|---|---|
| ZNF667 | 8.3072958 | -0.631320373 | 0.67591474 | -0.9340237 | 0.35029168 | 0.99999872 |
| ZNF667-AS1 | 16.2692114 | 0.336005585 | 0.49102934 | 0.68428821 | 0.49379319 | 0.99999872 |
| ZNF471 | 9.42342716 | 0.19387871 | 0.4866615 | 0.39838515 | 0.69034631 | 0.99999872 |
| ZFP28 | 5.12632451 | 0.146961159 | 0.68245661 | 0.2153414 | 0.82950116 | 0.99999872 |
| ZNF470 | 5.26547137 | -0.920067011 | 0.7252885 | -1.2685532 | 0.20460048 | 0.99999872 |
| ZNF264 | 18.7834956 | -0.054943822 | 0.41899165 | -0.1311335 | 0.89566974 | 0.99999872 |
| ZNF460-AS1 | 5.48286575 | 0.185387301 | 0.6746424 | 0.27479343 | 0.78347494 | 0.99999872 |
| ZNF460 | 4.03725768 | -0.042225472 | 0.81990769 | -0.0515003 | 0.95892688 | NA |
| AC005261.1 | 23.1257312 | 0.298545363 | 0.37157647 | 0.80345604 | 0.42171119 | 0.99999872 |
| AC004076.1 | 15.234104 | -0.735784223 | 0.48377593 | -1.5209194 | 0.12828005 | 0.99999872 |
| ZNF549 | 9.1338541 | 0.045480715 | 0.52212989 | 0.08710613 | 0.93058715 | 0.99999872 |
| ZNF134 | 8.17219146 | 0.749033463 | 0.59761434 | 1.25337264 | 0.21007012 | 0.99999872 |
| ZNF211 | 4.61795523 | 0.25031997 | 0.75241593 | 0.33268829 | 0.73936959 | 0.99999872 |
| AC003006.1 | 5.78550657 | -0.050704512 | 0.6621074 | -0.0765805 | 0.93895728 | 0.99999872 |
| ZNF776 | 5.93080197 | -0.453380682 | 0.65229108 | -0.6950588 | 0.48701842 | 0.99999872 |
| ZNF586 | 44.1055593 | -0.075883609 | 0.29911899 | -0.2536904 | 0.79973477 | 0.99999872 |
| ZNF552 | 7.65747733 | 0.397235283 | 0.54609734 | 0.72740747 | 0.46697638 | 0.99999872 |
| ZNF814 | 29.0319385 | -0.188744062 | 0.458742 | -0.4114384 | 0.68075112 | 0.99999872 |
| ZNF587 | 12.7271394 | -0.324463097 | 0.43101175 | -0.7527941 | 0.45157366 | 0.99999872 |
| ZNF418 | 6.05746704 | -0.116286882 | 0.63924973 | -0.1819115 | 0.85565217 | 0.99999872 |
| C19orf18 | 8.3059559 | -1.045293989 | 0.64197165 | -1.6282557 | 0.10347068 | 0.99999872 |
| ZNF606 | 7.36884577 | -0.451337617 | 0.57111916 | -0.7902687 | 0.42937084 | 0.99999872 |
| ZNF135 | 11.4044328 | 0.16993101 | 0.51687491 | 0.32876622 | 0.7423324 | 0.99999872 |
| ZSCAN18 | 20.0486375 | 0.384028035 | 0.52456635 | 0.73208667 | 0.46411567 | 0.99999872 |
| ZNF329 | 9.8530556 | 0.403430883 | 0.48102151 | 0.83869613 | 0.40163985 | 0.99999872 |
| ZNF274 | 27.5686041 | -0.323001314 | 0.4168738 | -0.774818 | 0.43844722 | 0.99999872 |
| AC020915.3 | 5.26896258 | 0.243714215 | 0.6373922 | 0.38236146 | 0.70219327 | 0.99999872 |
| ERVK3-1 | 4.9527386 | -0.199115967 | 0.80171822 | -0.2483615 | 0.80385469 | 0.99999872 |
| AC010642.2 | 5.01829076 | 0.616546532 | 0.77071311 | 0.79996891 | 0.42372881 | 0.99999872 |
| A1BG | 11.1022431 | 0.59130025 | 0.60437943 | 0.97835933 | 0.32789664 | 0.99999872 |
| A1BG-AS1 | 8.08132485 | 0.028227445 | 0.53343779 | 0.0529161 | 0.95779876 | 0.99999872 |
| RPS5 | 42.5242407 | 0.764066645 | 0.54666709 | 1.39768181 | 0.16220864 | 0.99999872 |
| ZNF584 | 5.13183964 | 2.541854799 | 1.09391142 | 2.32363859 | 0.02014487 | 0.8671024 |
| ZNF324 | 4.97209416 | 0.305791317 | 0.82865182 | 0.36902268 | 0.71211082 | 0.99999872 |
| ZNF446 | 4.99882076 | 0.015979546 | 0.7667072 | 0.02084178 | 0.98337187 | 0.99999872 |
| SLC27A5 | 9.5986648 | 0.620286703 | 0.4813894 | 1.2885342 | 0.19756007 | 0.99999872 |
| ZBTB45 | 3.78896068 | 0.417107945 | 0.80027136 | 0.52120814 | 0.60222178 | NA |
| TRIM28 | 25.7807079 | 0.046950656 | 0.47453726 | 0.09893987 | 0.92118601 | 0.99999872 |
| CHMP2A | 22.0405065 | 0.329728114 | 0.39745783 | 0.82959271 | 0.4067691 | 0.99999872 |
| UBE2M | 21.1470675 | 0.721879859 | 0.44463516 | 1.62353302 | 0.10447551 | 0.99999872 |
| MZF1-AS1 | 5.17922139 | 0.234139606 | 0.69562281 | 0.33658989 | 0.73642608 | 0.99999872 |
| MZF1 | 25.0195963 | -0.281653573 | 0.37056351 | -0.7600683 | 0.44721377 | 0.99999872 |
| RPS4Y1 | 25.4585301 | 0.508024553 | 2.37580006 | 0.21383304 | 0.83067726 | 0.99999872 |
| ZFY | 17.5252232 | 0.148257854 | 2.22768543 | 0.06655242 | 0.94693803 | 0.99999872 |
| LINC00278 | 34.2611364 | 0.307445285 | 2.40306488 | 0.12793882 | 0.89819739 | 0.99999872 |
| TTTY15 | 11.7148586 | 0.754138013 | 2.39951284 | 0.31428797 | 0.75330234 | 0.99999872 |
| USP9Y | 64.6421979 | 0.64479174 | 2.95456282 | 0.21823592 | 0.8272453 | 0.99999872 |
| DDX3Y | 22.5477855 | 0.702149966 | 2.32652012 | 0.30180266 | 0.7628025 | 0.99999872 |
| UTY | 140.560476 | 0.216713235 | 3.13643477 | 0.06909541 | 0.94491368 | 0.99999872 |
| NLGN4Y | 36.8407533 | -0.838392488 | 2.48833613 | -0.336929 | 0.73617046 | 0.99999872 |
| TTTY14 | 101.803258 | 0.637789656 | 1.97654415 | 0.32267919 | 0.74693822 | 0.99999872 |
| KDM5D | 14.8091436 | 0.35874263 | 2.52939751 | 0.14182928 | 0.88721486 | 0.99999872 |
| TTTY10 | 14.148505 | -0.299975509 | 2.4378262 | -0.1230504 | 0.90206718 | 0.99999872 |
| EIF1AY | 20.3763898 | 1.034883731 | 2.61456215 | 0.39581531 | 0.69224128 | 0.99999872 |
| AP000547.3 | 5.61963888 | -0.21296043 | 0.61237407 | -0.347762 | 0.72801893 | 0.99999872 |
| IL17RA | 11.8497723 | 0.539685672 | 0.50131336 | 1.07654357 | 0.28168422 | 0.99999872 |
| HDHD5 | 9.78018339 | 0.075397277 | 0.46850829 | 0.16093051 | 0.87214814 | 0.99999872 |
| ADA2 | 7.51784162 | 0.193078334 | 0.83769299 | 0.23048818 | 0.81771245 | 0.99999872 |
| CECR2 | 89.842917 | -0.267452154 | 0.46251549 | -0.5782556 | 0.56309159 | 0.99999872 |
| ATP6V1E1 | 54.3261882 | 0.38969437 | 0.2875702 | 1.35512778 | 0.17537685 | 0.99999872 |
| BCL2L13 | 51.5845569 | -0.142868583 | 0.30616668 | -0.4666366 | 0.64075989 | 0.99999872 |

| | | | | | | | |
|---|---|---|---|---|---|---|---|
| MICAL3 | 55.603565 | | 0.049709853 | 0.32456813 | 0.15315691 | 0.87827453 | 0.99999872 |
| PEX26 | 16.5406047 | | -0.362169924 | 0.37819648 | -0.9576237 | 0.33825253 | 0.99999872 |
| USP18 | 9.24255119 | | -1.208121807 | 1.54433727 | -0.7822914 | 0.43404332 | 0.99999872 |
| DGCR2 | 37.9832343 | | -0.620794899 | 0.33811632 | -1.8360395 | 0.06635182 | 0.99999872 |
| SLC25A1 | 30.2642145 | | 0.112642808 | 0.49084038 | 0.22948969 | 0.81848833 | 0.99999872 |
| CLTCL1 | 7.74303608 | | -0.486092051 | 0.50336586 | -0.9656834 | 0.33420264 | 0.99999872 |
| HIRA | 10.5298297 | | 0.132985582 | 0.60735178 | 0.21895973 | 0.82668142 | 0.99999872 |
| MRPL40 | 23.3726609 | | 0.08884159 | 0.4560664 | 0.19479968 | 0.84554977 | 0.99999872 |
| UFD1 | 11.2505983 | | -0.524020598 | 0.64616951 | -0.8109646 | 0.417386 | 0.99999872 |
| CLDN5 | 103.442978 | | 0.666630799 | 0.94134236 | 0.7081704 | 0.47883944 | 0.99999872 |
| GNB1L | 4.90027135 | | 1.081142366 | 0.80873851 | 1.33682562 | 0.18127956 | 0.99999872 |
| TXNRD2 | 52.4125161 | | -1.319044079 | 0.71601635 | -1.8421983 | 0.06544615 | 0.99999872 |
| COMT | 108.445311 | | 0.198391908 | 0.30112578 | 0.65883402 | 0.51000236 | 0.99999872 |
| TANGO2 | 16.1650679 | | -0.248691945 | 0.54952444 | -0.4525585 | 0.65086671 | 0.99999872 |
| DGCR8 | 17.0264913 | | -0.162786104 | 0.48541487 | -0.3353546 | 0.73735762 | 0.99999872 |
| TRMT2A | 6.48470364 | | 1.40503125 | 0.81990858 | 1.71364379 | 0.08659417 | 0.99999872 |
| RANBP1 | 15.4837724 | | 0.913405996 | 0.47095978 | 1.9394565 | 0.05244578 | 0.99999872 |
| ZDHHC8 | 12.9943157 | | 0.286911484 | 0.46583468 | 0.61590838 | 0.53795499 | 0.99999872 |
| DGCR6L | 7.52457198 | | 0.150088854 | 0.65583166 | 0.22885271 | 0.8189834 | 0.99999872 |
| KLHL22 | 22.8957253 | | -0.585643778 | 0.46915125 | -1.2483048 | 0.21191945 | 0.99999872 |
| MED15 | 50.3387929 | | -0.398886366 | 0.31810415 | -1.253949 | 0.20986055 | 0.99999872 |
| PI4KA | 59.1918282 | | -0.031314582 | 0.35926684 | -0.0871625 | 0.93054238 | 0.99999872 |
| SNAP29 | 16.8569852 | | -0.312463162 | 0.45853478 | -0.6814383 | 0.49559419 | 0.99999872 |
| CRKL | 13.7971653 | | 0.261082166 | 0.44036863 | 0.59287186 | 0.55326691 | 0.99999872 |
| LZTR1 | 8.4222508 | | 0.380309649 | 0.58056586 | 0.65506719 | 0.5124245 | 0.99999872 |
| THAP7 | 10.7749174 | | 0.07876342 | 0.43314811 | 0.18183946 | 0.85570871 | 0.99999872 |
| GGT2 | 12.5708099 | | -0.248892872 | 1.08931689 | -0.2284853 | 0.819269 | 0.99999872 |
| HIC2 | 13.4348898 | | 0.460267812 | 0.5133564 | 0.89658532 | 0.36994024 | 0.99999872 |
| UBE2L3 | 52.3075948 | | 0.157994588 | 0.30174288 | 0.52360668 | 0.60055213 | 0.99999872 |
| YDJC | 5.68589293 | | 1.202393467 | 0.80154491 | 1.50009494 | 0.13358981 | 0.99999872 |
| SDF2L1 | 15.776307 | | 0.458376045 | 0.70996534 | 0.64563158 | 0.51851798 | 0.99999872 |
| PPIL2 | 15.0046796 | | -0.805039512 | 0.39668106 | -2.0294377 | 0.04241373 | 0.95980711 |
| YPEL1 | 5.79562176 | | -0.284562126 | 0.82721122 | -0.3440018 | 0.73084495 | 0.99999872 |
| MAPK1 | 91.603579 | | 0.231775303 | 0.25664634 | 0.90309219 | 0.36647697 | 0.99999872 |
| PPM1F | 9.8743429 | | 0.466390945 | 0.56211928 | 0.82970103 | 0.40670784 | 0.99999872 |
| TOP3B | 5.41251542 | | -0.283330446 | 0.54967716 | -0.5154488 | 0.60623942 | 0.99999872 |
| AC245060.5 | 10.8146312 | | -0.41484723 | 0.84601854 | -0.4903524 | 0.62388455 | 0.99999872 |
| BCR | 21.6800201 | | -0.898377532 | 0.35491629 | -2.5312378 | 0.01136608 | 0.72670507 |
| CHCHD10 | 32.4704728 | | 0.23128062 | 0.56092555 | 0.41231964 | 0.68010516 | 0.99999872 |
| SMARCB1 | 15.355804 | | -0.052830867 | 0.37984505 | -0.1390853 | 0.88938273 | 0.99999872 |
| DDT | 40.8468717 | | 0.939549495 | 0.6591736 | 1.42534455 | 0.15405763 | 0.99999872 |
| CABIN1 | 41.9256814 | | -0.179287742 | 0.32620574 | -0.5496155 | 0.5825831 | 0.99999872 |
| GUCD1 | 12.9204115 | | -0.550826486 | 0.45154449 | -1.219872 | 0.2225134 | 0.99999872 |
| SNRPD3 | 33.224821 | | -0.034750075 | 0.34480515 | -0.1007818 | 0.91972369 | 0.99999872 |
| KIAA1671 | 70.2652382 | | -0.320898234 | 0.3639971 | -0.8815956 | 0.37799554 | 0.99999872 |
| AL022323.4 | 5.72380729 | | 1.219459949 | 0.8916886 | 1.367585 | 0.17144201 | 0.99999872 |
| LRP5L | 11.6325704 | | -0.090287738 | 0.93544529 | -0.0965185 | 0.92310881 | 0.99999872 |
| GRK3 | 90.7495899 | | 0.023652411 | 0.69983713 | 0.03379702 | 0.97303901 | 0.99999872 |
| HPS4 | 12.4630712 | | 0.290726559 | 0.55751236 | 0.52147105 | 0.60203866 | 0.99999872 |
| SRRD | 6.02700867 | | 0.699578177 | 0.71095235 | 0.9840015 | 0.32511478 | 0.99999872 |
| TFIP11 | 8.60346217 | | -0.107261689 | 0.65909818 | -0.1627401 | 0.87072309 | 0.99999872 |
| TPST2 | 4.9779191 | | 0.412275204 | 0.84037364 | 0.4905856 | 0.62371957 | 0.99999872 |
| MIATNB | 16.5281661 | | -0.803900174 | 0.47668163 | -1.6864509 | 0.09170898 | 0.99999872 |
| PITPNB | 46.4551915 | | 0.102452082 | 0.33267835 | 0.30796138 | 0.75811172 | 0.99999872 |
| TTC28 | 214.104853 | | -0.148349467 | 0.23600201 | -0.6285941 | 0.52961483 | 0.99999872 |
| HSCB | 13.0440995 | | -0.495184727 | 0.51463402 | -0.9622075 | 0.33594537 | 0.99999872 |
| CCDC117 | 10.484378 | | 0.080634612 | 0.47277681 | 0.17055534 | 0.86457341 | 0.99999872 |
| XBP1 | 31.7826282 | | 0.889815988 | 0.5616697 | 1.58423357 | 0.11314058 | 0.99999872 |
| Z93930.2 | 37.4710025 | | 0.094302861 | 0.45275474 | 0.20828686 | 0.83500499 | 0.99999872 |
| ZNRF3 | 542.654128 | | -0.317996415 | 0.33757416 | -0.9420046 | 0.34619029 | 0.99999872 |
| ZNRF3-AS1 | 6.15780148 | | 1.17713668 | 0.75467529 | 1.55979226 | 0.11880898 | 0.99999872 |

| | | | | | | |
|---|---|---|---|---|---|---|
| KREMEN1 | 25.8089653 | 0.262773299 | 0.66378446 | 0.39587142 | 0.69219988 | 0.99999872 |
| RHBDD3 | 9.7405655 | -0.749437221 | 0.45896722 | -1.6328775 | 0.10249477 | 0.99999872 |
| EWSR1 | 60.1728822 | -0.468824046 | 0.28404953 | -1.650501 | 0.0988405 | 0.99999872 |
| GAS2L1 | 7.84088779 | -0.154490145 | 0.52465337 | -0.2944614 | 0.76840539 | 0.99999872 |
| AP1B1 | 15.6503578 | -0.334754647 | 0.4017627 | -0.8332149 | 0.40472357 | 0.99999872 |
| THOC5 | 5.6960137 | -0.250624097 | 0.63700739 | -0.3934399 | 0.69399464 | 0.99999872 |
| NIPSNAP1 | 7.32855736 | 0.364121834 | 0.56545305 | 0.64394707 | 0.51960976 | 0.99999872 |
| NF2 | 16.8631049 | 0.104356876 | 0.47819808 | 0.21822939 | 0.82725038 | 0.99999872 |
| ZMAT5 | 4.89087585 | 0.109807313 | 0.60536153 | 0.1813913 | 0.85606045 | 0.99999872 |
| UQCR10 | 82.8191846 | 0.891892353 | 0.47646541 | 1.87189319 | 0.06122138 | 0.99999872 |
| ASCC2 | 23.0966697 | -0.184640812 | 0.43157709 | -0.4278281 | 0.66877627 | 0.99999872 |
| MTMR3 | 99.8211225 | 0.290498109 | 0.25170367 | 1.15412745 | 0.24844792 | 0.99999872 |
| SF3A1 | 13.4284447 | 0.112389517 | 0.59902051 | 0.18762215 | 0.85117286 | 0.99999872 |
| RNF215 | 10.3889969 | -0.678326661 | 0.64662574 | -1.0490251 | 0.29416654 | 0.99999872 |
| PES1 | 7.25978006 | -0.50325891 | 0.56971247 | -0.883356 | 0.37704397 | 0.99999872 |
| TCN2 | 8.23203329 | -0.043079453 | 1.26893616 | -0.0339493 | 0.97291761 | 0.99999872 |
| DUSP18 | 9.28016948 | -0.146569619 | 0.64506173 | -0.227218 | 0.82025425 | 0.99999872 |
| OSBP2 | 17.3654894 | 0.902448345 | 0.54191852 | 1.6652842 | 0.09585607 | 0.99999872 |
| MORC2 | 10.0494133 | -0.238728752 | 0.48189436 | -0.4953964 | 0.62032031 | 0.99999872 |
| SELENOM | 38.1111761 | 1.443402013 | 0.45471525 | 3.17429867 | 0.00150199 | 0.32787201 |
| RNF185 | 22.4827962 | 0.10114712 | 0.48139015 | 0.21011465 | 0.8335782 | 0.99999872 |
| LIMK2 | 57.2003708 | 0.079505759 | 0.41511211 | 0.1915284 | 0.84811163 | 0.99999872 |
| PIK3IP1 | 24.7966995 | 0.512677223 | 0.72578689 | 0.70637432 | 0.47995539 | 0.99999872 |
| PIK3IP1-AS1 | 4.91527224 | 0.236730386 | 0.70496028 | 0.3358067 | 0.73701664 | 0.99999872 |
| PATZ1 | 11.5643688 | -0.234399165 | 0.47757988 | -0.4908062 | 0.62356353 | 0.99999872 |
| DRG1 | 22.8475797 | -0.069250105 | 0.34796261 | -0.1990159 | 0.84225028 | 0.99999872 |
| EIF4ENIF1 | 17.3773568 | -0.113526964 | 0.38675129 | -0.29354 | 0.76910945 | 0.99999872 |
| SFI1 | 23.6793239 | -0.898634633 | 0.40794343 | -2.2028413 | 0.02760594 | 0.89154528 |
| PISD | 34.1457274 | -0.52255656 | 0.40295072 | -1.296825 | 0.19469142 | 0.99999872 |
| PRR14L | 41.6223896 | -0.392307801 | 0.32037921 | -1.2245108 | 0.2207596 | 0.99999872 |
| YWHAH | 16.2584683 | 0.848178436 | 0.60325985 | 1.40599186 | 0.15972654 | 0.99999872 |
| RTCB | 25.7462465 | 0.483872872 | 0.31308052 | 1.54552211 | 0.12222003 | 0.99999872 |
| FBXO7 | 45.7164942 | -0.005887694 | 0.31942413 | -0.0184322 | 0.98529405 | 0.99999872 |
| SYN3 | 21.7133589 | 1.68807539 | 0.50301069 | 3.35594334 | 0.00079095 | 0.25898594 |
| TIMP3 | 268.892417 | 0.382563606 | 0.6173265 | 0.61971032 | 0.53544852 | 0.99999872 |
| LARGE1 | 664.824312 | -0.729013245 | 0.27385534 | -2.6620377 | 0.00776692 | 0.65139548 |
| HMGXB4 | 14.7111263 | 0.4987577 | 0.49945143 | 0.99861101 | 0.31798317 | 0.99999872 |
| TOM1 | 13.0291472 | -0.218920207 | 0.57127177 | -0.3832155 | 0.70155997 | 0.99999872 |
| HMOX1 | 8.11136458 | -0.322402608 | 1.00789206 | -0.3198781 | 0.74906073 | 0.99999872 |
| APOL6 | 39.5123495 | -0.510228431 | 0.75261906 | -0.6779372 | 0.49781154 | 0.99999872 |
| RBFOX2 | 124.302449 | 0.115398664 | 0.26576828 | 0.43420782 | 0.66413753 | 0.99999872 |
| APOL2 | 46.8777902 | -0.613437875 | 0.81797792 | -0.7499443 | 0.45328825 | 0.99999872 |
| APOL1 | 103.915922 | -1.011913701 | 0.63263021 | -1.5995343 | 0.10970194 | 0.99999872 |
| MYH9 | 461.955052 | 0.004245172 | 0.4637134 | 0.00915473 | 0.99269568 | 0.99999872 |
| TXN2 | 17.412019 | -0.056468123 | 0.42595404 | -0.1325686 | 0.89453458 | 0.99999872 |
| EIF3D | 23.1463628 | 0.060879335 | 0.41494833 | 0.14671546 | 0.88335661 | 0.99999872 |
| IFT27 | 9.92618278 | 0.248880261 | 0.49656953 | 0.50119922 | 0.61623092 | 0.99999872 |
| TST | 18.9761632 | -0.328407764 | 0.60846155 | -0.5397346 | 0.58938006 | 0.99999872 |
| MPST | 18.5824919 | -0.297596565 | 0.42387432 | -0.7020868 | 0.48262503 | 0.99999872 |
| C1QTNF6 | 5.21453832 | -0.405855755 | 0.64084552 | -0.6333129 | 0.52652931 | 0.99999872 |
| CARD10 | 10.4935472 | -0.868081835 | 0.5443456 | -1.5947255 | 0.11077362 | 0.99999872 |
| CDC42EP1 | 11.8595999 | -1.160512663 | 0.56508282 | -2.0537037 | 0.04000438 | 0.95980711 |
| GGA1 | 20.1325481 | -0.177296159 | 0.42108781 | -0.4210432 | 0.67372354 | 0.99999872 |
| LGALS1 | 203.944424 | -0.03937843 | 0.636972 | -0.0618213 | 0.95070515 | 0.99999872 |
| TRIOBP | 32.8003686 | -0.492064402 | 0.3538819 | -1.3904763 | 0.16438428 | 0.99999872 |
| H1F0 | 29.853755 | -1.535252191 | 0.91030348 | -1.6865279 | 0.09169417 | 0.99999872 |
| ANKRD54 | 4.22309058 | 0.232568586 | 0.78311488 | 0.29697889 | 0.76648263 | NA |
| EIF3L | 22.0829544 | 0.156683006 | 0.43023145 | 0.36418306 | 0.71572132 | 0.99999872 |
| POLR2F | 12.9893695 | -0.019010379 | 0.49856008 | -0.0381306 | 0.96958358 | 0.99999872 |
| PICK1 | 8.05395342 | -0.085859498 | 0.55795692 | -0.1538819 | 0.87770282 | 0.99999872 |
| PLA2G6 | 6.08845139 | 0.152414626 | 0.79145956 | 0.19257412 | 0.84729252 | 0.99999872 |

| | | | | | | |
|---|---|---|---|---|---|---|
| MAFF | 4.68143158 | -1.304302541 | 0.91857124 | -1.4199253 | 0.15562943 | 0.99999872 |
| TMEM184B | 16.7132048 | 0.292506396 | 0.44863344 | 0.65199419 | 0.51440492 | 0.99999872 |
| CSNK1E | 12.0915372 | -0.570771948 | 0.54087578 | -1.0552736 | 0.2913002 | 0.99999872 |
| TPTEP2-CSNI | 41.0295063 | 0.058751563 | 0.34357685 | 0.17099977 | 0.86422395 | 0.99999872 |
| KDELR3 | 16.3810688 | 0.221938024 | 0.82871111 | 0.26781109 | 0.78884474 | 0.99999872 |
| DDX17 | 375.140183 | 0.168201974 | 0.22417862 | 0.75030336 | 0.45307202 | 0.99999872 |
| FAM227A | 6.19289832 | -0.163780143 | 0.80871196 | -0.2025197 | 0.83951042 | 0.99999872 |
| CBY1 | 15.2493547 | 0.150722714 | 0.3828065 | 0.39373082 | 0.69377979 | 0.99999872 |
| TOMM22 | 11.168606 | 0.22433989 | 0.60284807 | 0.37213338 | 0.70979355 | 0.99999872 |
| JOSD1 | 28.0210208 | 0.298283862 | 0.34877369 | 0.85523613 | 0.39242044 | 0.99999872 |
| GTPBP1 | 24.8430675 | -0.191524056 | 0.40599797 | -0.4717365 | 0.63711489 | 0.99999872 |
| SUN2 | 95.5409852 | 0.079929489 | 0.58089353 | 0.13759748 | 0.89055854 | 0.99999872 |
| AL021707.1 | 10.474976 | -0.567490397 | 0.53855856 | -1.0537209 | 0.29201072 | 0.99999872 |
| CBX6 | 46.4209063 | -0.160986491 | 0.56168948 | -0.2866112 | 0.77441004 | 0.99999872 |
| APOBEC3G | 4.69632547 | -1.384081979 | 1.52188537 | -0.9094522 | 0.36311149 | 0.99999872 |
| CBX7 | 31.2591631 | 0.363522649 | 0.61146628 | 0.59450972 | 0.55217124 | 0.99999872 |
| AL031846.2 | 4.80290582 | 1.139513538 | 0.81899604 | 1.39135414 | 0.16411806 | 0.99999872 |
| RPL3 | 62.1961521 | 0.201890616 | 0.50505363 | 0.39974095 | 0.68934733 | 0.99999872 |
| TAB1 | 11.889695 | 0.089418312 | 0.45880552 | 0.19489372 | 0.84547615 | 0.99999872 |
| MIEF1 | 10.1384422 | 0.163741951 | 0.53356208 | 0.30688453 | 0.75893126 | 0.99999872 |
| ATF4 | 58.7478701 | -0.182495304 | 0.43001375 | -0.4243941 | 0.67127841 | 0.99999872 |
| RPS19BP1 | 32.6531439 | 0.463825836 | 0.32816557 | 1.41338971 | 0.15754117 | 0.99999872 |
| TNRC6B | 204.784699 | -0.130694436 | 0.26535157 | -0.4925331 | 0.62234252 | 0.99999872 |
| MKL1 | 122.484654 | 0.123514967 | 0.352844 | 0.35005545 | 0.72629708 | 0.99999872 |
| SLC25A17 | 18.6831449 | 0.118863257 | 0.45908455 | 0.25891365 | 0.79570186 | 0.99999872 |
| ST13 | 60.2518298 | 0.478651649 | 0.25906684 | 1.84759906 | 0.06466037 | 0.99999872 |
| XPNPEP3 | 26.6223955 | -0.100867042 | 0.3779867 | -0.2668534 | 0.78958203 | 0.99999872 |
| RBX1 | 45.219307 | 0.015401875 | 0.38318329 | 0.04019454 | 0.96793803 | 0.99999872 |
| EP300 | 77.6532747 | -0.298735752 | 0.31391821 | -0.9516356 | 0.34128181 | 0.99999872 |
| L3MBTL2 | 9.01057055 | 0.103027881 | 0.59655318 | 0.17270528 | 0.8628831 | 0.99999872 |
| RANGAP1 | 44.0947364 | -0.58890198 | 0.77292611 | -0.7619124 | 0.4461123 | 0.99999872 |
| ZC3H7B | 65.0219742 | -0.435785144 | 0.49414019 | -0.8819059 | 0.3778277 | 0.99999872 |
| TEF | 40.6750334 | -0.429935449 | 0.53269487 | -0.8070952 | 0.41961167 | 0.99999872 |
| TOB2 | 20.7119494 | 0.314353922 | 0.4212057 | 0.74631924 | 0.45547459 | 0.99999872 |
| PHF5A | 7.47063594 | 0.248730073 | 0.50515718 | 0.49238155 | 0.62244964 | 0.99999872 |
| ACO2 | 26.9690537 | 0.14090597 | 0.33916947 | 0.41544414 | 0.6778168 | 0.99999872 |
| POLR3H | 6.62129909 | -0.095810559 | 0.59326054 | -0.1614983 | 0.87170096 | 0.99999872 |
| CSDC2 | 36.3550568 | 0.003377507 | 0.41862986 | 0.008068 | 0.99356273 | 0.99999872 |
| PMM1 | 11.1255036 | 0.353494784 | 0.5499174 | 0.64281433 | 0.52034458 | 0.99999872 |
| DESI1 | 14.0097874 | 0.3099542 | 0.51825148 | 0.59807683 | 0.54978867 | 0.99999872 |
| XRCC6 | 57.5986518 | 0.038062971 | 0.41619763 | 0.09145408 | 0.92713179 | 0.99999872 |
| SNU13 | 49.5683104 | 0.29491481 | 0.34782925 | 0.84787237 | 0.39650905 | 0.99999872 |
| SREBF2 | 74.9983105 | -0.393333623 | 0.2897285 | -1.3575938 | 0.1745926 | 0.99999872 |
| WBP2NL | 17.6170214 | -0.130966118 | 0.50965065 | -0.2569723 | 0.79720013 | 0.99999872 |
| NAGA | 7.54906316 | 0.153165755 | 0.64041358 | 0.23916694 | 0.81097614 | 0.99999872 |
| SMDT1 | 25.3101826 | 0.45071552 | 0.504775 | 0.89290381 | 0.37190869 | 0.99999872 |
| NDUFA6 | 22.672313 | 0.750300467 | 0.47962072 | 1.56436207 | 0.11773256 | 0.99999872 |
| TCF20 | 33.640141 | -0.254549518 | 0.29531841 | -0.8619494 | 0.38871538 | 0.99999872 |
| RRP7A | 12.6844063 | -0.403197931 | 0.71064874 | -0.567366 | 0.57046555 | 0.99999872 |
| POLDIP3 | 15.3158206 | -0.082425301 | 0.44346449 | -0.1858667 | 0.85254926 | 0.99999872 |
| CYB5R3 | 85.6392545 | 0.309365786 | 0.42117239 | 0.73453482 | 0.46262285 | 0.99999872 |
| A4GALT | 20.1238163 | 0.0339059 | 0.5543572 | 0.06116255 | 0.95122975 | 0.99999872 |
| ARFGAP3 | 34.1823203 | 0.119723794 | 0.33935904 | 0.35279388 | 0.72424297 | 0.99999872 |
| PACSIN2 | 36.5546876 | -0.3762878 | 0.28735806 | -1.3094736 | 0.19037397 | 0.99999872 |
| TTLL1 | 4.67972434 | -0.302459787 | 0.71638604 | -0.4222022 | 0.67287741 | 0.99999872 |
| MCAT | 5.11532037 | -0.142363544 | 0.57221531 | -0.2487937 | 0.80352039 | 0.99999872 |
| TSPO | 48.6828918 | 0.525243514 | 0.57305447 | 0.91656821 | 0.35936895 | 0.99999872 |
| TTLL12 | 6.85301112 | 0.232531832 | 0.65296448 | 0.35611712 | 0.72175286 | 0.99999872 |
| SCUBE1 | 4.86118496 | -0.184803647 | 0.86934782 | -0.2125773 | 0.83165665 | 0.99999872 |
| EFCAB6 | 22.197317 | -0.650607583 | 0.45361966 | -1.4342579 | 0.15149868 | 0.99999872 |
| SAMM50 | 5.35561266 | -0.034861599 | 0.58426241 | -0.0596677 | 0.95242029 | 0.99999872 |

| | | | | | | |
|---|---|---|---|---|---|---|
| RTL6 | 6.44064519 | -0.245787273 | 0.58888899 | -0.4173745 | 0.67640447 | 0.99999872 |
| PHF21B | 74.733749 | 0.246292145 | 0.51145747 | 0.48154961 | 0.63012593 | 0.99999872 |
| NUP50-DT | 24.8596019 | 0.186023545 | 0.49229072 | 0.37787335 | 0.70552468 | 0.99999872 |
| NUP50 | 16.4599184 | 0.368488173 | 0.39318726 | 0.93718237 | 0.34866476 | 0.99999872 |
| KIAA0930 | 13.8775529 | -0.300641855 | 0.77038484 | -0.3902489 | 0.69635249 | 0.99999872 |
| FAM118A | 15.536712 | -0.163064774 | 0.71837834 | -0.2269901 | 0.82043144 | 0.99999872 |
| ATXN10 | 132.129149 | 0.254784649 | 0.22319403 | 1.1415388 | 0.25364577 | 0.99999872 |
| MIRLET7BHG | 30.6621639 | -0.131492898 | 0.3601512 | -0.3651047 | 0.71503325 | 0.99999872 |
| PPARA | 41.8467183 | 0.032732594 | 0.43245113 | 0.07569085 | 0.93966505 | 0.99999872 |
| TTC38 | 13.1995941 | -0.135565012 | 0.54368329 | -0.2493456 | 0.8030935 | 0.99999872 |
| TRMU | 11.6812885 | 0.052870148 | 0.43795938 | 0.1207193 | 0.90391337 | 0.99999872 |
| CELSR1 | 9.43971299 | -0.943343529 | 0.92334646 | -1.0216572 | 0.30694319 | 0.99999872 |
| GRAMD4 | 9.48989511 | 0.238671558 | 0.626067 | 0.38122367 | 0.70303729 | 0.99999872 |
| CERK | 35.9116343 | 0.066780336 | 0.55151053 | 0.12108624 | 0.90362273 | 0.99999872 |
| AL118516.1 | 4.08767517 | -0.398937034 | 0.67114338 | -0.594414 | 0.55223523 | NA |
| TBC1D22A | 110.141032 | -0.546833217 | 0.22101006 | -2.4742458 | 0.01335178 | 0.74507289 |
| BRD1 | 21.8884689 | -0.211576987 | 0.39433149 | -0.536546 | 0.59158125 | 0.99999872 |
| ZBED4 | 11.0817968 | 0.513822441 | 0.56148382 | 0.91511531 | 0.3601311 | 0.99999872 |
| ALG12 | 7.30801423 | 0.09133222 | 0.54871909 | 0.16644622 | 0.8678058 | 0.99999872 |
| CRELD2 | 11.7992063 | 0.985950971 | 0.73832346 | 1.33539164 | 0.1817482 | 0.99999872 |
| PIM3 | 35.017974 | -0.197313009 | 0.41556945 | -0.4748015 | 0.63492844 | 0.99999872 |
| TRABD | 10.4630064 | -0.413666801 | 0.55080278 | -0.7510253 | 0.45263745 | 0.99999872 |
| SELENOO | 16.5182235 | -0.396192173 | 0.37771682 | -1.0489132 | 0.29421806 | 0.99999872 |
| TUBGCP6 | 4.47237504 | -0.071252273 | 0.74655398 | -0.0954416 | 0.92396411 | NA |
| PLXNB2 | 134.394687 | -0.565806563 | 0.46820914 | -1.2084483 | 0.22687485 | 0.99999872 |
| DENND6B | 14.5718016 | -0.963945675 | 0.56611472 | -1.7027391 | 0.0886169 | 0.99999872 |
| PPP6R2 | 59.7835853 | -0.475562221 | 0.25168878 | -1.8894852 | 0.05882685 | 0.99999872 |
| SBF1 | 5.53104172 | -0.180078917 | 0.72739071 | -0.2475683 | 0.8044684 | 0.99999872 |
| MIOX | 13.2178821 | -1.578405204 | 1.21510678 | -1.2989848 | 0.19394915 | 0.99999872 |
| LMF2 | 11.4875377 | 0.067334308 | 0.67300616 | 0.10005006 | 0.92030458 | 0.99999872 |
| NCAPH2 | 6.24684833 | -0.626925668 | 0.72807412 | -0.861074 | 0.38919731 | 0.99999872 |
| ARSA | 14.1618211 | -0.552531591 | 0.49994254 | -1.1051902 | 0.26907719 | 0.99999872 |
| RABL2B | 9.77993668 | -0.379424128 | 0.58036219 | -0.6537713 | 0.51325918 | 0.99999872 |
| FP671120.1 | 331.338542 | -1.45960699 | 0.38374577 | -3.8035781 | 0.00014262 | 0.10674163 |
| FP236383.1 | 203.727624 | -1.450408936 | 0.45585191 | -3.1817546 | 0.00146386 | 0.32751082 |
| CR381653.1 | 13.6718074 | -1.543564604 | 0.6140976 | -2.5135493 | 0.01195231 | 0.72670507 |
| AP001347.1 | 12.6990209 | -0.538578216 | 0.56073535 | -0.9604856 | 0.33681089 | 0.99999872 |
| RBM11 | 6.91261862 | 0.30843112 | 0.63421982 | 0.4863158 | 0.62674328 | 0.99999872 |
| HSPA13 | 38.0650337 | 0.506074951 | 0.54886229 | 0.92204357 | 0.35650585 | 0.99999872 |
| AF127577.4 | 7.10795278 | 1.201629486 | 0.66463123 | 1.80796422 | 0.07061207 | 0.99999872 |
| NRIP1 | 77.3845131 | 0.114573009 | 0.30193063 | 0.37946799 | 0.70434037 | 0.99999872 |
| AJ009632.2 | 13.7268562 | -0.34854275 | 0.99734351 | -0.3494711 | 0.72673565 | 0.99999872 |
| USP25 | 115.138305 | -0.135287692 | 0.33098219 | -0.4087461 | 0.68272597 | 0.99999872 |
| MIR99AHG | 111.595359 | 0.607029945 | 0.36930508 | 1.64370862 | 1.00E-01 | 0.99999872 |
| CXADR | 84.4746334 | 0.726329568 | 0.54080292 | 1.34305778 | 0.17925327 | 0.99999872 |
| BTG3 | 52.6025173 | 0.859691814 | 0.49832171 | 1.72517431 | 0.08449606 | 0.99999872 |
| C21orf91 | 7.30249175 | 0.196959534 | 0.69702905 | 0.28257005 | 0.77750644 | 0.99999872 |
| MRPL39 | 7.7350104 | 0.331643645 | 0.57897818 | 0.57280854 | 0.56677434 | 0.99999872 |
| ATP5PF | 41.7558131 | 0.775352264 | 0.37029834 | 2.09385832 | 0.0362726 | 0.94779136 |
| GABPA | 17.7787685 | 1.288816222 | 0.64437396 | 2.000106 | 0.04548882 | 0.95980711 |
| APP | 193.944556 | 0.126635579 | 0.231124 | 0.54791184 | 0.58375244 | 0.99999872 |
| ADAMTS1 | 53.2236289 | 0.202891567 | 0.99284214 | 0.20435431 | 0.83807663 | 0.99999872 |
| ADAMTS5 | 29.8121279 | -1.639611049 | 2.41508216 | -0.6789049 | 0.49719814 | 0.99999872 |
| AF165147.1 | 16.9076085 | 0.255348982 | 0.89458838 | 0.2854374 | 0.77530906 | 0.99999872 |
| N6AMT1 | 6.80691687 | 0.883697335 | 0.64689323 | 1.36606366 | 0.17191898 | 0.99999872 |
| LTN1 | 28.8367741 | 0.593767622 | 0.38520085 | 1.54144943 | 0.12320744 | 0.99999872 |
| USP16 | 32.1039302 | -0.488917087 | 0.34283576 | -1.4260971 | 0.15384031 | 0.99999872 |
| CCT8 | 20.5094837 | 0.329053786 | 0.3404749 | 0.96645533 | 0.33381639 | 0.99999872 |
| MAP3K7CL | 9.44713573 | -0.428278212 | 0.66239642 | -0.6465588 | 0.51791755 | 0.99999872 |
| BACH1 | 321.633547 | 0.372745284 | 0.32579526 | 1.14410899 | 0.25257844 | 0.99999872 |
| GRIK1 | 49.3126004 | 1.41900101 | 0.47918946 | 2.96125256 | 0.00306391 | 0.44319076 |

| | | | | | | |
|---|---|---|---|---|---|---|
| GRIK1-AS1 | 18.3735111 | 0.626012004 | 0.59587987 | 1.05056747 | 0.29345729 | 0.99999872 |
| SOD1 | 78.6985395 | 0.557469294 | 0.2687318 | 2.0744448 | 0.03803802 | 0.95980711 |
| SCAF4 | 29.4641029 | -0.092421462 | 0.34783054 | -0.2657083 | 0.79046387 | 0.99999872 |
| URB1 | 10.723348 | -0.700776204 | 0.52388145 | -1.3376618 | 0.18100671 | 0.99999872 |
| EVA1C | 36.723099 | -0.564502915 | 0.40729255 | -1.3859888 | 0.16575033 | 0.99999872 |
| SYNJ1 | 18.2442192 | -0.15451946 | 0.39613639 | -0.3900663 | 0.69648752 | 0.99999872 |
| PAXBP1 | 33.2471071 | -0.056698056 | 0.37595843 | -0.1508094 | 0.88012609 | 0.99999872 |
| C21orf62-AS | 8.12639799 | -0.638987944 | 0.58571348 | -1.0909565 | 0.27529201 | 0.99999872 |
| IL10RB | 3.73287846 | 0.200345577 | 0.75703253 | 0.26464593 | 0.79128224 | NA |
| IFNAR1 | 54.0509124 | 0.000855091 | 0.40589482 | 0.00210668 | 0.99831911 | 0.99999872 |
| IFNGR2 | 25.7540105 | 0.578129974 | 0.33540656 | 1.72366928 | 0.08476756 | 0.99999872 |
| TMEM50B | 43.3185744 | 0.130832406 | 0.3241143 | 0.40366132 | 0.68646179 | 0.99999872 |
| GART | 18.2252001 | 0.073671163 | 0.58920043 | 0.12503583 | 0.90049519 | 0.99999872 |
| SON | 124.415419 | 0.08204816 | 0.29377959 | 0.27928476 | 0.7800263 | 0.99999872 |
| DONSON | 6.38292238 | 0.182879987 | 0.64925165 | 0.28167812 | 0.77819033 | 0.99999872 |
| AP000311.1 | 15.3532245 | 1.291619141 | 0.5795068 | 2.22882482 | 0.02582556 | 0.88365618 |
| ITSN1 | 59.2484516 | -0.086403792 | 0.39747882 | -0.2173796 | 0.82791252 | 0.99999872 |
| LINC00649 | 16.582221 | 0.758006198 | 0.41615556 | 1.82144916 | 0.06853861 | 0.99999872 |
| MRPS6 | 62.7813645 | 0.352806491 | 0.39219194 | 0.89957609 | 0.36834589 | 0.99999872 |
| LINC00310 | 7.861009 | 0.303696823 | 0.85573861 | 0.35489438 | 0.72266872 | 0.99999872 |
| AP000317.1 | 9.11973703 | 1.018502196 | 0.641321 | 1.58813168 | 0.11225657 | 0.99999872 |
| RCAN1 | 99.2656441 | -0.389952104 | 0.48513005 | -0.8038094 | 0.42150704 | 0.99999872 |
| RUNX1 | 47.8946594 | -1.550879578 | 0.72472162 | -2.1399659 | 0.03235752 | 0.92959829 |
| SETD4 | 11.0622533 | 0.094936724 | 0.49151432 | 0.19315149 | 0.84684033 | 0.99999872 |
| CBR1 | 19.0142953 | 0.565466193 | 0.52919808 | 1.0685341 | 0.28527966 | 0.99999872 |
| DOPEY2 | 12.2965516 | -0.60643468 | 0.53123141 | -1.1415641 | 0.25363527 | 0.99999872 |
| MORC3 | 34.4773049 | 0.008901506 | 0.30009402 | 0.02966239 | 0.97633631 | 0.99999872 |
| HLCS | 57.7363439 | -0.284573304 | 0.37059525 | -0.7678817 | 0.44255748 | 0.99999872 |
| PIGP | 10.1659266 | 1.036141348 | 0.63190325 | 1.63971516 | 0.1010644 | 0.99999872 |
| TTC3 | 100.329453 | -0.123661162 | 0.3423184 | -0.361246 | 0.71791554 | 0.99999872 |
| VPS26C | 26.0395532 | 0.303875075 | 0.31114633 | 0.97663074 | 0.32875199 | 0.99999872 |
| DYRK1A | 147.410074 | -0.187102586 | 0.24904202 | -0.7512892 | 0.4524786 | 0.99999872 |
| KCNJ15 | 36.0011126 | 0.04031454 | 1.00791023 | 0.03999815 | 0.9680946 | 0.99999872 |
| ETS2 | 95.0858529 | 0.050032329 | 0.54238891 | 0.09224438 | 0.92650388 | 0.99999872 |
| PSMG1 | 7.84090114 | -0.092470601 | 0.48588786 | -0.1903126 | 0.84906415 | 0.99999872 |
| BRWD1 | 95.7747796 | 0.021956596 | 0.28472891 | 0.07711404 | 0.93853282 | 0.99999872 |
| HMGN1 | 97.3388654 | 0.155237132 | 0.33548179 | 0.46272894 | 0.64355867 | 0.99999872 |
| WRB | 33.0519402 | 0.518412179 | 0.49906549 | 1.03876582 | 0.29891366 | 0.99999872 |
| LCA5L | 6.0828246 | 0.368829135 | 0.92034565 | 0.40075067 | 0.6886037 | 0.99999872 |
| SH3BGR | 5.86834692 | 1.664896923 | 1.22342301 | 1.3608514 | 0.17356065 | 0.99999872 |
| BACE2 | 21.9766368 | -1.290533958 | 0.84105585 | -1.5344212 | 0.12492606 | 0.99999872 |
| MX1 | 83.7706671 | -1.377284131 | 2.18440176 | -0.6305086 | 0.52836187 | 0.99999872 |
| C2CD2 | 9.46774422 | -0.446764322 | 0.49065477 | -0.9105472 | 0.36253401 | 0.99999872 |
| ZBTB21 | 13.4225266 | 0.332738843 | 0.47263467 | 0.70400854 | 0.48142745 | 0.99999872 |
| SLC37A1 | 8.16712435 | -0.631457419 | 0.57889034 | -1.0908066 | 0.27535798 | 0.99999872 |
| PDE9A | 11.4202007 | -1.14223354 | 0.71469177 | -1.5982184 | 0.10999438 | 0.99999872 |
| WDR4 | 4.47780585 | -1.260376098 | 0.94023892 | -1.3404849 | 0.18008774 | NA |
| NDUFV3 | 26.9745576 | -0.418181665 | 0.36082006 | -1.1589756 | 0.24646611 | 0.99999872 |
| PKNOX1 | 31.3028094 | 0.013204652 | 0.30886406 | 0.04275231 | 0.96589898 | 0.99999872 |
| RRP1B | 10.634374 | -0.223809677 | 0.41597914 | -0.538031 | 0.59055564 | 0.99999872 |
| PDXK | 46.5241013 | -0.600009034 | 0.26676485 | -2.2492057 | 0.02449941 | 0.88337526 |
| CSTB | 75.7762611 | 0.601600416 | 0.43362556 | 1.38737304 | 0.16532804 | 0.99999872 |
| RRP1 | 4.35888297 | -0.442943205 | 0.68969538 | -0.6422302 | 0.52072372 | NA |
| AGPAT3 | 47.5495847 | -0.1759845 | 0.28483583 | -0.6178454 | 0.53667727 | 0.99999872 |
| TRAPPC10 | 74.0170967 | 0.16091309 | 0.31483587 | 0.51110151 | 0.60927998 | 0.99999872 |
| PFKL | 38.6653692 | 0.179893092 | 0.43593801 | 0.41265751 | 0.67985756 | 0.99999872 |
| C21orf2 | 16.8050717 | 0.619930465 | 0.62584343 | 0.99055201 | 0.32190438 | 0.99999872 |
| TSPEAR | 16.1711942 | -0.295898344 | 0.98109775 | -0.3015992 | 0.76295758 | 0.99999872 |
| UBE2G2 | 43.324324 | 0.100284675 | 0.30037074 | 0.33386965 | 0.73847792 | 0.99999872 |
| SUMO3 | 27.1014269 | -0.140999026 | 0.44542176 | -0.3165517 | 0.75158377 | 0.99999872 |
| PTTG1IP | 81.579178 | -0.116564509 | 0.28994901 | -0.4020173 | 0.68767131 | 0.99999872 |

| | | | | | | |
|---|---|---|---|---|---|---|
| FAM207A | 6.79338396 | -0.305239129 | 0.76216657 | -0.4004887 | 0.68879658 | 0.99999872 |
| ADARB1 | 10.1826029 | 0.270633328 | 0.74091073 | 0.36527117 | 0.714909 | 0.99999872 |
| POFUT2 | 17.4732012 | -0.196540036 | 0.5090109 | -0.3861215 | 0.69940671 | 0.99999872 |
| COL18A1 | 84.2909433 | -0.222964696 | 0.41117876 | -0.5422573 | 0.58764125 | 0.99999872 |
| COL6A1 | 11.3290321 | 0.27634613 | 0.47457198 | 0.58230604 | 0.56036056 | 0.99999872 |
| FTCD | 6.38623728 | 0.371969193 | 1.02537759 | 0.36276314 | 0.71678183 | 0.99999872 |
| LSS | 11.3311923 | -0.066641603 | 0.54145469 | -0.1230788 | 0.90204468 | 0.99999872 |
| MCM3AP | 28.1831875 | -0.203158121 | 0.3310361 | -0.6137038 | 0.53941106 | 0.99999872 |
| YBEY | 4.85013818 | -0.827717718 | 0.68293626 | -1.2119985 | 0.22551296 | 0.99999872 |
| PCNT | 16.2699026 | -0.581579246 | 0.36653274 | -1.5867047 | 0.11257953 | 0.99999872 |
| DIP2A | 31.1277216 | 0.167415781 | 0.31135347 | 0.53770328 | 0.59078192 | 0.99999872 |
| PRMT2 | 89.3690477 | 0.236106132 | 0.31624861 | 0.74658394 | 0.45531474 | 0.99999872 |
| AL592183.1 | 46.6001026 | -0.257225732 | 0.59190258 | -0.4345744 | 0.66387135 | 0.99999872 |

**Proximal tubules**

*This means that positive fold changes imply higher mean value in non-COVID AKI than in COVID AKI and vice versa.

| | baseMean | log2 fold change | lfcSE | stat | pvalue | padj |
|---|---|---|---|---|---|---|
| AL627309.1 | 46.0856473 | -0.337164275 | 0.42343382 | -0.796262 | 0.42587975 | 0.78894887 |
| AL669831.5 | 349.587562 | -0.209452394 | 0.35172544 | -0.5954997 | 0.5515095 | 0.84691246 |
| NOC2L | 274.350881 | -0.17566525 | 0.37949677 | -0.46289 | 0.64344324 | 0.8848654 |
| KLHL17 | 55.733763 | -0.066455655 | 0.46659167 | -0.1424279 | 0.88674205 | 0.97042502 |
| PLEKHN1 | 47.4948433 | -0.65953477 | 0.60802334 | -1.0847195 | 2.78E-01 | 7.04E-01 |
| HES4 | 116.388637 | -2.057516722 | 0.84398583 | -2.437857 | 0.01477462 | 0.30832605 |
| ISG15 | 1169.63657 | -5.659945281 | 1.51418758 | -3.7379419 | 0.00018553 | 0.0505134 |
| AGRN | 499.353221 | -0.306456979 | 0.4274962 | -0.7168648 | 0.47345752 | 0.81478861 |
| C1orf159 | 304.992355 | -0.119893421 | 0.43704064 | -0.2743301 | 0.78383092 | 0.93864304 |
| LINC01342 | 94.1721291 | -0.046625682 | 0.44210064 | -0.105464 | 0.91600767 | 0.97801193 |
| AL390719.2 | 109.671187 | 0.035237446 | 0.46451074 | 0.07585927 | 0.93953106 | 0.98435387 |
| SDF4 | 473.687245 | -0.096033254 | 0.32521045 | -0.2952957 | 0.76776799 | 0.93180413 |
| B3GALT6 | 61.0958485 | -0.569170918 | 0.51342954 | -1.1085668 | 0.26761713 | 0.69874881 |
| C1QTNF12 | 39.0858734 | -0.141473022 | 0.4548934 | -0.3110026 | 7.56E-01 | 0.92799282 |
| UBE2J2 | 715.129567 | -0.516686558 | 0.36073496 | -1.4323163 | 0.15205333 | 0.61001662 |
| LINC01786 | 36.2560674 | 0.308768236 | 0.61284155 | 0.50383045 | 0.61438053 | 0.87225883 |
| SCNN1D | 51.6699071 | -0.542280344 | 0.49818877 | -1.0885038 | 2.76E-01 | 7.03E-01 |
| ACAP3 | 339.851362 | -0.56170737 | 0.37832818 | -1.4847093 | 0.13762085 | 0.59474076 |
| PUSL1 | 42.683084 | -0.674765111 | 0.54484686 | -1.2384491 | 0.21554958 | 0.6596384 |
| INTS11 | 351.174265 | -0.27753178 | 0.26194183 | -1.0595169 | 0.28936445 | 0.71218341 |
| CPTP | 78.8397927 | -0.769044621 | 0.4002719 | -1.9213055 | 0.0546932 | 0.46772575 |
| DVL1 | 157.554134 | -0.476390555 | 0.31259239 | -1.5239992 | 0.12750891 | 0.58793678 |
| AURKAIP1 | 459.348943 | -0.577711037 | 0.49331389 | -1.171082 | 0.2415658 | 0.68015474 |
| CCNL2 | 1393.10989 | -0.589658552 | 0.24309177 | -2.4256623 | 0.01528048 | 0.3105046 |
| MRPL20 | 450.607117 | -0.346102392 | 0.33413229 | -1.0358244 | 0.30028405 | 0.71879123 |
| VWA1 | 45.0710487 | 0.362538894 | 0.63455429 | 0.57132841 | 0.56777705 | 0.85263558 |
| ATAD3B | 296.151961 | -0.697504519 | 0.42693812 | -1.6337368 | 0.10231413 | 0.5491072 |
| ATAD3A | 172.913781 | -0.674996439 | 0.54678215 | -1.2344888 | 0.21702082 | 0.66032006 |
| SSU72 | 531.394181 | -0.667835483 | 0.3897831 | -1.7133516 | 0.08664788 | 0.52518597 |
| MIB2 | 236.600328 | -0.686147119 | 0.29169884 | -2.352245 | 0.01866048 | 0.33968023 |
| CDK11B | 316.680707 | -0.853554518 | 0.46272634 | -1.8446205 | 0.06509276 | 0.48790819 |
| SLC35E2B | 341.622686 | -0.190324645 | 0.2935173 | -0.6484274 | 0.51670858 | 0.83258835 |
| CDK11A | 189.110388 | -0.96005401 | 0.45656625 | -2.1027705 | 0.03548584 | 0.41997369 |
| SLC35E2A | 270.882248 | -0.086025366 | 0.21312406 | -0.4036399 | 6.86E-01 | 0.90196028 |
| NADK | 495.685555 | -0.584293113 | 0.36472475 | -1.6020111 | 0.10915315 | 0.55946616 |
| GNB1 | 3046.74651 | -0.548483089 | 0.36110835 | -1.5188878 | 0.12879073 | 0.58793678 |
| PRKCZ | 1302.21601 | -0.838557078 | 0.59155434 | -1.4175487 | 0.15632257 | 0.61255389 |
| FAAP20 | 393.375962 | -0.099933165 | 0.29825879 | -0.3350552 | 0.73758343 | 0.92051862 |
| AL590822.1 | 117.713718 | 1.520092891 | 1.00641294 | 1.51040675 | 1.31E-01 | 5.89E-01 |
| SKI | 814.23026 | 0.411808408 | 0.51541635 | 0.79898205 | 0.42430082 | 0.78894887 |
| MORN1 | 383.651251 | -0.131203676 | 0.27042564 | -0.4851747 | 0.62755243 | 0.87944599 |
| RER1 | 542.356114 | -0.261562506 | 0.26017232 | -1.0053433 | 0.31473156 | 0.72719146 |
| PEX10 | 137.57292 | -0.185672388 | 0.40724887 | -0.4559187 | 0.64844842 | 0.88787189 |
| PLCH2 | 328.615729 | 2.275987922 | 0.85697571 | 2.65583714 | 0.00791118 | 0.24533611 |
| PANK4 | 64.1561829 | 0.216568438 | 0.25906611 | 0.8359582 | 0.40317842 | 0.78277148 |
| AL139246.5 | 38.6613966 | -0.655367311 | 0.67089908 | -0.9768493 | 0.32864375 | 0.73829083 |
| TNFRSF14-A | 90.456375 | -0.201501036 | 0.45779973 | -0.4401511 | 0.6598277 | 0.89094706 |
| TNFRSF14 | 1183.56461 | -0.543304114 | 0.48626486 | -1.1173008 | 0.26386574 | 0.69567438 |
| FAM213B | 60.594743 | 0.949056312 | 0.46351796 | 2.04750711 | 0.04060832 | 0.43086475 |
| ARHGEF16 | 269.400766 | 0.240071598 | 0.53185768 | 0.45138315 | 0.65171342 | 0.88935488 |
| MEGF6 | 41.4678717 | 0.33310592 | 0.73938277 | 0.45051891 | 0.65233632 | 0.88935488 |
| TPRG1L | 224.454408 | 0.043683212 | 0.32801882 | 0.13317288 | 0.89405666 | 0.97272276 |

| | | | | | | |
|---|---|---|---|---|---|---|
| WRAP73 | 121.287539 | -0.361267011 | 0.26305081 | -1.3733735 | 0.16963627 | 0.62326131 |
| SMIM1 | 39.9886605 | 0.425266525 | 0.71567155 | 0.59422024 | 0.55236481 | 0.84721285 |
| LRRC47 | 201.055471 | -0.706484259 | 0.25251047 | -2.7978415 | 0.00514454 | 0.21811586 |
| CEP104 | 625.841361 | -0.556310696 | 0.29650281 | -1.8762409 | 0.06062221 | 0.48026727 |
| DFFB | 141.907623 | -0.190838723 | 0.53776964 | -0.3548708 | 0.72268641 | 0.91592159 |
| C1orf174 | 137.493628 | -0.576121379 | 0.38211285 | -1.5077257 | 0.13162473 | 0.59017999 |
| LINC01134 | 83.5001576 | -0.494283305 | 0.54647189 | -0.9044991 | 0.36573084 | 0.76409651 |
| AJAP1 | 264.36015 | 0.218951423 | 0.45888636 | 0.47713648 | 0.63326494 | 0.88177189 |
| NPHP4 | 261.881381 | 0.275614923 | 0.32939785 | 0.83672351 | 0.402748 | 0.78260941 |
| KCNAB2 | 596.067903 | 1.66649415 | 0.6138622 | 2.71476912 | 0.0066322 | 0.22966139 |
| RPL22 | 333.392476 | -0.180556602 | 0.39612419 | -0.4558081 | 0.64852801 | 0.88787189 |
| RNF207 | 310.621842 | 0.576357558 | 0.4310846 | 1.33699408 | 0.18122457 | 0.6318513 |
| ICMT | 260.501815 | -0.044203007 | 0.2832045 | -0.1560816 | 0.87596871 | 0.96617876 |
| GPR153 | 122.554968 | 1.612801394 | 0.62855552 | 2.56588533 | 0.01029128 | 0.26767103 |
| ACOT7 | 324.982747 | -0.321270308 | 0.57931505 | -0.5545692 | 0.57918933 | 0.85805186 |
| ESPN | 617.738708 | 0.173415768 | 0.39833864 | 0.43534759 | 6.63E-01 | 8.93E-01 |
| TNFRSF25 | 35.3972948 | 0.540023128 | 0.63568843 | 0.84950914 | 0.39559805 | 0.77931337 |
| PLEKHG5 | 62.1323316 | 0.624961022 | 0.40604508 | 1.53914195 | 0.12376965 | 0.58354574 |
| NOL9 | 421.535283 | -0.352089561 | 0.33911843 | -1.0382495 | 0.29915389 | 0.71852421 |
| ZBTB48 | 300.695668 | 0.362634765 | 0.30952228 | 1.17159503 | 0.24135968 | 0.68015474 |
| KLHL21 | 268.513745 | -1.346478571 | 0.49058363 | -2.7446463 | 0.00605761 | 0.22637955 |
| PHF13 | 96.6520075 | -0.157846733 | 0.36861988 | -0.42821 | 0.66849821 | 0.8955341 |
| THAP3 | 103.520056 | -0.246123573 | 0.2831433 | -0.8692544 | 0.38470797 | 0.77317162 |
| DNAJC11 | 1142.52983 | 0.00703571 | 0.27617345 | 0.02547569 | 0.97967554 | 0.99484461 |
| CAMTA1 | 2616.82794 | 0.025695856 | 0.23742652 | 0.10822657 | 0.91381597 | 0.97742298 |
| VAMP3 | 217.67277 | -0.194150533 | 0.23031835 | -0.842966 | 0.39924746 | 0.78161291 |
| PER3 | 871.629428 | -0.321479922 | 0.74052029 | -0.4341271 | 0.66419615 | 0.89315615 |
| PARK7 | 605.352895 | -0.10924815 | 0.26827347 | -0.4072268 | 0.68384141 | 0.90059706 |
| ERRFI1 | 3637.64952 | -0.643654463 | 1.00728526 | -0.6389992 | 0.52282346 | 0.83504325 |
| AL034417.2 | 88.3456581 | 0.521544804 | 0.81193193 | 0.64235041 | 0.52064569 | 0.833897 |
| RERE | 8120.43922 | 0.051555164 | 0.25718576 | 0.20045886 | 0.84112173 | 0.95677175 |
| ENO1 | 3980.72896 | -0.283194033 | 0.43814658 | -0.6463454 | 5.18E-01 | 8.33E-01 |
| SLC2A5 | 239.041087 | 0.995263732 | 0.64961989 | 1.53207091 | 0.12550494 | 0.58564339 |
| GPR157 | 200.588286 | -0.378728969 | 0.37777391 | -1.0025281 | 0.31608859 | 0.72798933 |
| MIR34AHG | 301.231291 | 0.263974827 | 0.74218751 | 0.35567134 | 0.72208671 | 0.9157845 |
| H6PD | 593.437879 | 0.155685676 | 0.34006158 | 0.45781613 | 0.64708455 | 0.88774616 |
| SPSB1 | 2310.62964 | -1.549037973 | 0.79946469 | -1.937594 | 5.27E-02 | 4.66E-01 |
| AL928921.2 | 56.3461281 | -0.038897297 | 1.04464993 | -0.0372348 | 0.97029782 | 0.99205112 |
| SLC25A33 | 460.341772 | -0.248866698 | 0.56572508 | -0.4399075 | 0.66000411 | 0.89094706 |
| TMEM201 | 35.8283037 | -0.733744887 | 0.62360799 | -1.1766124 | 0.23935025 | 0.6772861 |
| PIK3CD | 301.339243 | -1.274899536 | 0.5044107 | -2.527503 | 0.01148768 | 0.28627263 |
| PIK3CD-AS2 | 45.5075888 | -0.026496662 | 0.39855479 | -0.0664819 | 0.9469942 | 0.98646931 |
| CLSTN1 | 943.278427 | -0.136457052 | 0.26158916 | -0.5216464 | 0.60191653 | 0.86765503 |
| CTNNBIP1 | 276.870366 | 0.297998832 | 0.29346727 | 1.01544146 | 0.30989544 | 0.72498238 |
| LZIC | 254.812571 | -0.440910513 | 0.24961681 | -1.7663495 | 0.07733724 | 0.5110562 |
| NMNAT1 | 513.518751 | 0.599427427 | 0.22172094 | 2.703522 | 0.00686089 | 0.23083391 |
| UBE4B | 1638.28681 | -0.009906842 | 0.18150037 | -0.054583 | 0.95647065 | 0.98861313 |
| KIF1B | 3502.52922 | 0.00176995 | 0.18954259 | 0.00933801 | 0.99254946 | 0.9976934 |
| AL139424.2 | 33.0584275 | -0.257676839 | 0.5547169 | -0.4645195 | 0.64227556 | 0.88441609 |
| PGD | 253.506275 | -0.896061692 | 0.47279999 | -1.8952236 | 0.05806279 | 0.4774245 |
| DFFA | 439.742699 | -0.178932035 | 0.33391786 | -0.5358564 | 0.59205778 | 0.86382106 |
| PEX14 | 1386.3095 | -0.309064129 | 0.20642137 | -1.4972487 | 0.13432855 | 0.59240843 |
| CASZ1 | 316.97194 | 0.545510515 | 0.49399239 | 1.10428931 | 0.26946766 | 0.69961625 |
| SRM | 217.695984 | -1.31176485 | 0.63030795 | -2.0811492 | 0.03742026 | 0.42215481 |

| | | | | | | |
|---|---|---|---|---|---|---|
| EXOSC10 | 366.709439 | -0.726730596 | 0.26480516 | -2.7443974 | 0.00606221 | 0.22637955 |
| AL109811.1 | 47.9816591 | -0.048713913 | 0.43622345 | -0.1116719 | 0.91108354 | 0.97697493 |
| MTOR | 718.730112 | -0.390581893 | 0.3089025 | -1.264418 | 0.20608003 | 0.65206346 |
| UBIAD1 | 89.7307323 | -0.608541095 | 0.48452799 | -1.2559462 | 0.20913548 | 0.65403148 |
| FBXO2 | 50.3068433 | -0.436921496 | 0.51982949 | -0.8405093 | 0.40062292 | 0.78232083 |
| FBXO44 | 129.380299 | 0.134801373 | 0.57687241 | 0.23367623 | 0.81523634 | 0.94906482 |
| FBXO6 | 75.9438994 | 0.046945487 | 0.45846831 | 0.10239636 | 0.91844207 | 0.97860783 |
| MAD2L2 | 38.6442171 | 0.337490892 | 0.36773069 | 0.91776644 | 0.35874115 | 0.75936492 |
| AGTRAP | 310.777093 | 0.090279826 | 0.42264751 | 0.21360548 | 0.83085472 | 0.95325673 |
| MTHFR | 846.971535 | 0.312024954 | 0.27049825 | 1.15351932 | 0.24869729 | 0.68534254 |
| CLCN6 | 497.942366 | -0.551471123 | 0.34333763 | -1.6062065 | 0.10822855 | 0.55770254 |
| NPPA-AS1 | 55.7831464 | -1.643908768 | 0.77295605 | -2.1267817 | 0.03343823 | 0.41254527 |
| KIAA2013 | 278.512354 | -0.14790116 | 0.23426782 | -0.6313336 | 0.52782239 | 0.83761342 |
| PLOD1 | 420.995552 | 0.095766612 | 0.44617254 | 0.21464031 | 0.83004776 | 0.95325673 |
| MFN2 | 407.107461 | 0.268183555 | 0.30841993 | 0.86954029 | 0.38455168 | 0.77317162 |
| MIIP | 76.1204639 | 0.017203799 | 0.37110192 | 0.04635869 | 0.96302436 | 0.99012655 |
| VPS13D | 2654.74606 | 0.269533652 | 0.21106936 | 1.27699089 | 0.20160546 | 0.64879726 |
| DHRS3 | 1091.55138 | -0.728070539 | 0.42634195 | -1.707715 | 0.08768924 | 0.5273546 |
| PRDM2 | 768.799305 | -0.228121441 | 0.30300296 | -0.7528687 | 0.45152882 | 0.80384625 |
| KAZN | 5134.10966 | 0.365767633 | 0.45545601 | 0.80308005 | 4.22E-01 | 7.89E-01 |
| TMEM51-AS | 174.734639 | 0.055734414 | 0.6267881 | 0.08892066 | 0.92914496 | 0.98272895 |
| TMEM51 | 1999.34149 | -0.337258565 | 0.41862514 | -0.8056338 | 0.42045403 | 0.7882866 |
| C1orf195 | 73.2613896 | 0.087898491 | 0.4867832 | 0.1805701 | 0.85670503 | 0.96245027 |
| FHAD1 | 90.9601447 | 1.108762556 | 0.65701404 | 1.68757816 | 0.09149223 | 0.53537574 |
| EFHD2 | 101.403181 | -0.516155756 | 0.46689679 | -1.1055029 | 0.26894174 | 0.69961625 |
| CASP9 | 158.210077 | -0.132962123 | 0.26720476 | -0.4976039 | 0.61876328 | 0.87502492 |
| DNAJC16 | 1042.31721 | 0.087783655 | 0.25124227 | 0.34939842 | 0.72679022 | 0.91639348 |
| AGMAT | 578.402442 | 0.49214159 | 0.4758275 | 1.03428571 | 3.01E-01 | 0.71879123 |
| DDI2 | 1302.31918 | -0.532786367 | 0.29065546 | -1.8330513 | 0.06679494 | 0.48884194 |
| PLEKHM2 | 631.313156 | -0.89396488 | 0.40638858 | -2.1997785 | 0.02782261 | 0.38968579 |
| SLC25A34 | 58.7090072 | 0.040054786 | 0.87851096 | 0.04559395 | 0.96363389 | 0.99012655 |
| TMEM82 | 184.48111 | 0.985155458 | 0.56548625 | 1.74213866 | 0.08148418 | 0.51593841 |
| FBLIM1 | 288.236929 | -1.346256866 | 0.57576021 | -2.3382249 | 0.01937558 | 0.34312806 |
| UQCRHL | 69.2741663 | 0.367050127 | 0.44957045 | 0.81644629 | 0.41424493 | 0.78632759 |
| AL450998.2 | 104.074751 | -0.110504591 | 0.43653968 | -0.2531376 | 0.80016191 | 0.94403774 |
| SPEN | 1895.39243 | -1.120703986 | 0.51131121 | -2.1918236 | 0.02839225 | 0.39280225 |
| ZBTB17 | 176.705671 | -0.428268379 | 0.30406348 | -1.4084835 | 0.15898796 | 0.61423843 |
| CLCNKA | 56.9832499 | 1.046080667 | 1.02225766 | 1.02330431 | 0.30616399 | 0.72169506 |
| CLCNKB | 46.3752207 | 0.310511568 | 0.75851324 | 0.40936869 | 0.68226912 | 0.89962501 |
| EPHA2 | 154.669311 | -0.577778108 | 0.5528355 | -1.0451176 | 0.29596863 | 0.71727018 |
| FBXO42 | 1333.44248 | -0.697960567 | 0.26888915 | -2.5957186 | 0.00943933 | 0.25998727 |
| SZRD1 | 443.824992 | -0.506336315 | 0.25875422 | -1.9568234 | 0.05036823 | 0.45843118 |
| SPATA21 | 41.9417663 | 0.288996266 | 0.48122367 | 0.60054458 | 0.54814336 | 0.84486687 |
| NECAP2 | 220.689704 | -0.303151685 | 0.26438667 | -1.1466224 | 0.2515377 | 0.68791444 |
| NBPF1 | 504.833923 | 0.293688483 | 0.32900117 | 0.89266699 | 0.37203553 | 0.76845856 |
| CROCC | 539.634712 | 0.189777759 | 0.4330383 | 0.43824706 | 0.6612072 | 0.89184036 |
| BX284668.5 | 40.8387058 | -0.064588676 | 0.91611931 | -0.0705025 | 0.94379373 | 0.98522232 |
| ATP13A2 | 142.219901 | -0.28460147 | 0.46253317 | -0.6153104 | 0.53834975 | 0.84049498 |
| SDHB | 718.560576 | -0.078089385 | 0.23456745 | -0.332908 | 0.73920372 | 0.92108726 |
| RCC2 | 334.686431 | -0.106424073 | 0.37531027 | -0.2835629 | 0.77674536 | 0.93549325 |
| ARHGEF10L | 3361.68556 | 0.622199508 | 0.48954683 | 1.27097035 | 0.2037392 | 0.6491523 |
| KLHDC7A | 147.462765 | -0.978225304 | 0.63924644 | -1.5302788 | 0.12594772 | 0.58564339 |
| AL080251.1 | 36.0694289 | 1.364050704 | 1.21793332 | 1.11997158 | 0.26272587 | 0.69467275 |
| IFFO2 | 246.86062 | 0.62652838 | 0.59342073 | 1.0557912 | 0.29106362 | 0.71302224 |

| | | | | | | |
|---|---|---|---|---|---|---|
| UBR4 | 2309.15304 | -0.148341504 | 0.16979694 | -0.8736406 | 0.382314 | 0.77279408 |
| AL035413.1 | 178.15109 | -0.315267611 | 0.37315372 | -0.8448733 | 0.39818159 | 0.78121522 |
| EMC1 | 193.384681 | -0.297537947 | 0.27681174 | -1.0748748 | 0.2824308 | 0.70682473 |
| MRTO4 | 198.078483 | -0.907533718 | 0.44343729 | -2.0465886 | 0.04069849 | 0.43086475 |
| AKR7A3 | 209.630387 | 0.489621896 | 0.3481797 | 1.40623332 | 1.60E-01 | 6.15E-01 |
| AKR7A2 | 318.958717 | 0.412629926 | 0.41150687 | 1.00272913 | 0.31599157 | 0.72798933 |
| PQLC2 | 114.906145 | -0.928438815 | 0.54372472 | -1.7075531 | 0.0877193 | 0.5273546 |
| CAPZB | 1608.7724 | -0.295972425 | 0.26326656 | -1.124231 | 0.26091504 | 0.69289016 |
| NBL1 | 162.154849 | 0.314264081 | 0.53059092 | 0.59229072 | 0.55365593 | 0.84727776 |
| TMCO4 | 593.614145 | -0.05021142 | 0.42436847 | -0.1183203 | 0.90581384 | 0.97633014 |
| RNF186 | 161.281597 | 0.460982234 | 1.03557709 | 0.44514526 | 0.65621478 | 0.89053559 |
| OTUD3 | 373.467373 | -0.410504061 | 0.29705873 | -1.3818953 | 0.16700386 | 0.62238224 |
| CAMK2N1 | 869.899735 | 0.897738694 | 0.32755484 | 2.74072791 | 0.00613032 | 0.22637955 |
| MUL1 | 111.372229 | -0.372282605 | 0.3425153 | -1.086908 | 0.27707749 | 0.7036882 |
| PINK1 | 330.165637 | 0.290741868 | 0.29960196 | 0.97042711 | 0.33183364 | 0.73939532 |
| PINK1-AS | 36.0856343 | -0.097381118 | 0.35674387 | -0.2729721 | 0.78487468 | 0.93884957 |
| DDOST | 226.376949 | 0.109566783 | 0.34299575 | 0.31944065 | 0.74939239 | 0.92565124 |
| KIF17 | 51.1210025 | -0.05118221 | 0.48098392 | -0.1064115 | 0.91525589 | 0.97782888 |
| HP1BP3 | 1726.17342 | -0.153234544 | 0.1778387 | -0.861649 | 0.38888068 | 0.7748665 |
| EIF4G3 | 4244.25516 | -0.245493443 | 0.35218279 | -0.6970626 | 0.48576363 | 0.81839266 |
| ECE1 | 620.350887 | 0.508932497 | 0.37582993 | 1.35415638 | 0.17568649 | 0.62699479 |
| NBPF3 | 163.138918 | 0.680075689 | 0.35969973 | 1.89067611 | 0.05866759 | 0.47778423 |
| ALPL | 426.839268 | -0.024214449 | 0.59900636 | -0.0404244 | 0.96775481 | 0.99177684 |
| RAP1GAP | 448.573902 | 0.839699089 | 0.56634944 | 1.48265192 | 0.13816691 | 0.59474076 |
| USP48 | 1225.38746 | -0.254478935 | 0.17779009 | -1.4313449 | 0.15233139 | 0.61001662 |
| HSPG2 | 192.410573 | 0.810111942 | 0.54727835 | 1.48025578 | 0.138805 | 0.59474076 |
| CDC42 | 1300.77304 | -0.388961484 | 0.23939304 | -1.6247819 | 0.10420903 | 0.5505924 |
| ZBTB40 | 523.306575 | -0.213961883 | 0.41336114 | -0.5176149 | 0.60472699 | 0.86865188 |
| EPHB2 | 84.6432939 | -1.991184391 | 0.94991806 | -2.0961644 | 0.03606761 | 0.42040764 |
| KDM1A | 638.691869 | -0.003038902 | 0.26010276 | -0.0116835 | 0.99067815 | 0.99737132 |
| AL031428.1 | 154.446868 | 0.5847623 | 0.54599441 | 1.07100419 | 0.28416755 | 0.70897061 |
| LUZP1 | 1239.86512 | 0.186730152 | 0.2890041 | 0.64611594 | 0.51820427 | 0.83325693 |
| LINC01355 | 145.31611 | 0.246729478 | 0.26400891 | 0.93454983 | 0.35002034 | 0.75455562 |
| AL109936.6 | 45.9498545 | 0.687037589 | 0.54286674 | 1.26557318 | 0.20566593 | 0.651931 |
| HNRNPR | 1000.89849 | -0.338834525 | 0.30923402 | -1.095722 | 0.27320044 | 0.70126562 |
| ZNF436 | 51.8518305 | -0.277552123 | 0.65853746 | -0.4214675 | 0.67341376 | 0.89790684 |
| ZNF436-AS1 | 37.5212939 | 0.296480764 | 0.66731164 | 0.44429131 | 0.65683198 | 0.89053559 |
| TCEA3 | 655.221299 | 0.729861601 | 0.55164438 | 1.32306542 | 0.18581362 | 0.63661432 |
| ASAP3 | 686.7801 | 1.205440135 | 0.45587933 | 2.64420882 | 0.00818821 | 0.24877162 |
| RPL11 | 1513.20199 | -0.029983067 | 0.47255992 | -0.0634482 | 0.94940962 | 0.9871942 |
| ELOA | 240.925874 | -0.364158503 | 0.35357859 | -1.0299224 | 0.30304644 | 0.72043311 |
| ELOA-AS1 | 282.493417 | 0.705185756 | 0.50731763 | 1.3900281 | 0.16452034 | 0.62065658 |
| PITHD1 | 231.377265 | -0.360957874 | 0.26115451 | -1.3821621 | 0.16692193 | 0.62238224 |
| LYPLA2 | 257.732961 | -0.305953743 | 0.40355637 | -0.7581438 | 0.44836493 | 0.80252521 |
| GALE | 463.588389 | 0.182529469 | 0.36127501 | 0.50523691 | 0.61339245 | 0.87222135 |
| HMGCL | 555.467191 | 0.161215689 | 0.31881329 | 0.5056743 | 0.61308531 | 0.87222135 |
| FUCA1 | 167.520144 | 0.219117089 | 0.24442342 | 0.89646521 | 0.37000435 | 0.76669785 |
| PNRC2 | 277.486022 | -0.456226314 | 0.23700897 | -1.9249327 | 0.05423778 | 0.46711025 |
| SRSF10 | 818.991114 | -0.280875375 | 0.21027576 | -1.3357478 | 0.18163173 | 0.6318513 |
| MYOM3 | 1052.78627 | 0.669139493 | 0.39931742 | 1.67570824 | 0.09379535 | 0.53723919 |
| IL22RA1 | 124.329842 | 1.371447723 | 0.90114763 | 1.52189018 | 0.1280366 | 0.58793678 |
| IFNLR1 | 286.49483 | 0.854760814 | 0.4068818 | 2.10075951 | 0.03566208 | 0.42040764 |
| STPG1 | 173.669537 | -0.043725319 | 0.22990969 | -0.1901848 | 0.84916436 | 0.95959925 |
| NIPAL3 | 381.982472 | -0.392317017 | 0.38895689 | -1.0086388 | 0.31314788 | 0.72717675 |

| | | | | | | |
|---|---|---|---|---|---|---|
| RCAN3 | 68.5124397 | 0.430130365 | 0.63406053 | 0.6783743 | 0.4975344 | 0.82313869 |
| NCMAP | 132.848202 | 1.512091576 | 0.75397894 | 2.00548252 | 0.04491149 | 0.44360789 |
| SRRM1 | 850.80195 | -0.161117947 | 0.20087848 | -0.8020667 | 0.42251436 | 0.78894887 |
| CLIC4 | 5837.21726 | 0.140458408 | 0.39116343 | 0.35907858 | 0.71953631 | 0.91465123 |
| SYF2 | 259.657139 | -0.235178076 | 0.27742131 | -0.8477289 | 0.39658894 | 0.77992032 |
| RSRP1 | 2485.01335 | 0.492810565 | 0.35397856 | 1.39220455 | 0.16386046 | 0.62065658 |
| RHD | 66.9984504 | 0.899496698 | 1.17911472 | 0.76285766 | 0.44554829 | 0.80125038 |
| TMEM50A | 398.93877 | -0.160621334 | 0.26687158 | -0.6018675 | 5.47E-01 | 0.84483892 |
| MACO1 | 807.712681 | -0.411267011 | 0.24335209 | -1.6900081 | 0.0910264 | 0.53537574 |
| LDLRAP1 | 87.4331411 | 0.213106616 | 0.35866761 | 0.59416186 | 0.55240386 | 0.84721285 |
| MAN1C1 | 1375.13772 | 1.667105362 | 0.85743837 | 1.94428593 | 0.05186098 | 0.46400683 |
| AL031280.1 | 129.268233 | 1.479969711 | 1.08770542 | 1.36063467 | 0.17362917 | 0.62663931 |
| SELENON | 28.7656928 | 0.83544799 | 0.56655963 | 1.47459853 | 0.14032051 | 0.59500405 |
| AL020996.1 | 33.6388432 | 0.537311842 | 0.6763517 | 0.79442669 | 0.42694707 | 0.78933193 |
| MTFR1L | 278.145694 | -0.109015825 | 0.32890457 | -0.3314512 | 7.40E-01 | 0.92173153 |
| AL033528.2 | 72.8422884 | 0.26331883 | 0.65639533 | 0.4011589 | 0.68830314 | 0.90307854 |
| PAQR7 | 229.256774 | 0.662152296 | 0.61089754 | 1.08390075 | 0.27840881 | 0.70442645 |
| STMN1 | 100.502861 | 0.410086953 | 0.63065026 | 0.6502605 | 0.51552396 | 0.83185952 |
| PAFAH2 | 256.551946 | 0.365117302 | 0.27974338 | 1.30518656 | 0.19182933 | 0.64497936 |
| SLC30A2 | 264.150669 | -0.022580686 | 0.55814479 | -0.0404567 | 0.96772904 | 0.99177684 |
| TRIM63 | 48.0191351 | 2.761884075 | 0.88778619 | 3.11097887 | 0.00186468 | 0.13738894 |
| PDIK1L | 101.595484 | 0.409345035 | 0.36253217 | 1.12912747 | 0.25884407 | 0.69241621 |
| ZNF593 | 156.993159 | -0.590129765 | 0.48188079 | -1.2246385 | 0.22071146 | 0.66263062 |
| CEP85 | 185.160252 | -0.150380549 | 0.37121702 | -0.4051014 | 0.68540294 | 0.90113631 |
| SH3BGRL3 | 367.167034 | -0.588353291 | 0.43610171 | -1.3491194 | 1.77E-01 | 0.62731853 |
| UBXN11 | 201.218861 | -0.078986123 | 0.32979661 | -0.2394995 | 0.81071829 | 0.94717173 |
| CRYBG2 | 47.567375 | 1.27494609 | 0.61015096 | 2.08955845 | 0.03665748 | 0.42192353 |
| DHDDS | 538.158692 | -1.092626284 | 0.4132874 | -2.6437445 | 0.00819945 | 0.24877162 |
| HMGN2 | 366.447533 | 0.082147589 | 0.38673048 | 0.21241561 | 0.83178281 | 0.95325673 |
| RPS6KA1 | 497.089429 | 0.025994226 | 0.29566505 | 0.08791782 | 0.92994199 | 0.98272895 |
| ARID1A | 1381.47674 | -0.006489436 | 0.24212673 | -0.0268018 | 0.97861781 | 0.99482978 |
| PIGV | 60.9156049 | 0.086087625 | 0.32207411 | 0.26729135 | 0.78924485 | 0.94005579 |
| ZDHHC18 | 218.301788 | -0.606726166 | 0.48401985 | -1.2535151 | 0.21001832 | 0.65441616 |
| SFN | 98.5810041 | -2.900832573 | 1.29711062 | -2.2363803 | 2.53E-02 | 0.38148951 |
| GPN2 | 226.688273 | -0.745280762 | 0.36011471 | -2.0695649 | 0.03849311 | 0.42447711 |
| GPATCH3 | 103.695509 | -0.622849519 | 0.3520722 | -1.769096 | 0.07687786 | 0.50988423 |
| NUDC | 565.870192 | -0.747711003 | 0.35117239 | -2.129185 | 0.03323895 | 0.41254527 |
| KDF1 | 66.4227045 | 0.2252536 | 0.35563273 | 0.63338827 | 0.52648012 | 0.83698249 |
| SLC9A1 | 124.508597 | -0.65288101 | 0.4162087 | -1.5686386 | 0.11673218 | 0.57350419 |
| WDTC1 | 518.825515 | 0.10588715 | 0.23022428 | 0.45993043 | 0.64556616 | 0.88687652 |
| TMEM222 | 183.885887 | -0.509425298 | 0.38391464 | -1.3269234 | 0.18453404 | 0.63377801 |
| MAP3K6 | 50.3463611 | 0.495830023 | 0.60731895 | 0.81642442 | 0.41425743 | 0.78632759 |
| WASF2 | 1294.42091 | -0.131834596 | 0.21797748 | -0.6048083 | 0.54530635 | 0.84449215 |
| AHDC1 | 356.090259 | 0.879184782 | 0.433211 | 2.02946087 | 0.04241137 | 0.43742111 |
| IFI6 | 687.590344 | -5.083800156 | 1.55849034 | -3.262003 | 0.00110628 | 0.11093704 |
| FAM76A | 256.318372 | -0.306655627 | 0.2269831 | -1.3510064 | 0.17669338 | 0.62709281 |
| STX12 | 893.680117 | -0.029059495 | 0.17094462 | -0.1699936 | 0.86501515 | 0.96428252 |
| PPP1R8 | 201.184393 | -0.471293937 | 0.29914475 | -1.5754712 | 0.11514773 | 0.57182344 |
| RPA2 | 189.876139 | -0.133094864 | 0.34660651 | -0.3839941 | 0.7009828 | 0.90782041 |
| SMPDL3B | 92.4768266 | 0.069065838 | 0.33385409 | 0.20687432 | 0.83610801 | 0.95468028 |
| XKR8 | 40.7080981 | -0.007588888 | 0.47298579 | -0.0160446 | 0.98719878 | 0.9963052 |
| EYA3 | 1771.07906 | -0.205635758 | 0.23698219 | -0.8677266 | 0.385544 | 0.77317162 |
| DNAJC8 | 431.024046 | -0.555274524 | 0.27573448 | -2.0138015 | 0.04403038 | 0.44360789 |
| ATP5IF1 | 840.422798 | -0.009660762 | 0.43188459 | -0.0223689 | 0.98215373 | 0.99522272 |

| | | | | | | |
|---|---|---|---|---|---|---|
| AL353622.1 | 106.818788 | 0.212370762 | 0.63733252 | 0.33321815 | 7.39E-01 | 9.21E-01 |
| SESN2 | 503.232762 | -0.214122178 | 0.38739695 | -0.5527203 | 0.58045492 | 0.85837691 |
| MED18 | 41.3647327 | -0.538551366 | 0.36905008 | -1.4592907 | 0.1444851 | 0.60101297 |
| PHACTR4 | 2747.35211 | -0.277209292 | 0.21143645 | -1.3110762 | 0.18983203 | 0.64203405 |
| RCC1 | 513.770689 | -0.448634001 | 0.47130429 | -0.9518988 | 0.3411483 | 0.74770062 |
| TRNAU1AP | 556.251472 | -0.240521337 | 0.22590677 | -1.0646929 | 0.28701491 | 0.71118494 |
| SNHG12 | 148.178538 | -0.278554203 | 0.47559843 | -0.585692 | 0.5580825 | 0.8489192 |
| TAF12 | 283.036143 | -0.366976712 | 0.34370192 | -1.067718 | 0.28564775 | 0.71059415 |
| GMEB1 | 313.096509 | -1.126654345 | 0.45799442 | -2.459974 | 0.01389471 | 0.30363654 |
| YTHDF2 | 862.383174 | -0.209921513 | 0.28725254 | -0.7307908 | 0.46490694 | 0.81002663 |
| EPB41 | 3554.63023 | -0.008295902 | 0.23832046 | -0.0348099 | 0.97223136 | 0.9924192 |
| TMEM200B | 55.2342624 | 0.404533166 | 0.46659504 | 0.86698986 | 0.38594756 | 0.77317162 |
| SRSF4 | 2948.40096 | -0.276015961 | 0.20719792 | -1.3321367 | 0.18281527 | 0.63276491 |
| MECR | 257.224284 | 0.020259191 | 0.26732273 | 0.07578551 | 0.93958974 | 0.98435387 |
| PTPRU | 60.5064737 | 0.591240372 | 0.47541688 | 1.2436251 | 2.14E-01 | 0.65834762 |
| PUM1 | 3640.17902 | -0.062533115 | 0.20987251 | -0.2979576 | 0.7657355 | 0.93106862 |
| SNRNP40 | 270.438114 | -0.282801068 | 0.22694939 | -1.2460975 | 0.2127286 | 0.65745584 |
| ZCCHC17 | 461.701984 | -0.216968408 | 0.23630773 | -0.9181604 | 0.35853489 | 0.75936492 |
| AL451070.1 | 57.2207139 | 0.525401426 | 0.44872096 | 1.17088676 | 0.2416443 | 0.68015474 |
| FABP3 | 123.647917 | -0.591761973 | 0.47317313 | -1.2506246 | 0.21107146 | 0.65561118 |
| SERINC2 | 995.322328 | -0.666734244 | 0.48241089 | -1.3820879 | 0.16694472 | 0.62238224 |
| TINAGL1 | 688.564732 | -1.054728206 | 0.36326613 | -2.9034587 | 0.00369066 | 0.19760872 |
| PEF1 | 198.328778 | -0.636094502 | 0.22290743 | -2.8536263 | 0.00432233 | 0.20749797 |
| PTP4A2 | 349.926813 | -0.295993457 | 0.31996194 | -0.9250896 | 0.35491931 | 0.75757147 |
| KHDRBS1 | 1170.57012 | -0.274391353 | 0.26516768 | -1.0347843 | 0.30076966 | 0.71879123 |
| AL445248.1 | 53.6472295 | 0.148375587 | 0.41041203 | 0.36152836 | 0.71770451 | 0.91359359 |
| TMEM39B | 138.535484 | -0.169150633 | 0.29665607 | -0.570191 | 0.56854813 | 0.85263558 |
| KPNA6 | 1177.93928 | -0.189559974 | 0.26155469 | -0.7247432 | 0.46860962 | 0.81217042 |
| TXLNA | 362.678943 | -0.419246558 | 0.39352719 | -1.065356 | 0.28671485 | 0.71118494 |
| DCDC2B | 49.6079832 | 0.070342405 | 0.55549293 | 0.1266306 | 0.89923277 | 0.97458352 |
| TMEM234 | 146.946688 | -0.241383611 | 0.28498241 | -0.8470123 | 0.39698826 | 0.78014596 |
| EIF3I | 254.366824 | -0.366906904 | 0.44334777 | -0.8275826 | 0.40790692 | 0.78481094 |
| HDAC1 | 365.505325 | 0.069684122 | 0.27269417 | 0.25553946 | 0.79830647 | 0.94320409 |
| MARCKSL1 | 61.995523 | -0.506226694 | 0.87539246 | -0.5782854 | 0.56307144 | 0.85057939 |
| TSSK3 | 61.5652107 | 0.54595095 | 0.47365619 | 1.1526313 | 0.24906175 | 0.68557761 |
| FAM229A | 78.2710494 | -0.224512548 | 0.47398154 | -0.4736736 | 0.63573267 | 0.88257807 |
| BSDC1 | 760.316375 | -0.385841321 | 0.30567531 | -1.2622587 | 0.2068557 | 0.65251041 |
| ZBTB8A | 64.8460079 | 0.28317985 | 0.6752251 | 0.41938584 | 0.67493417 | 0.89801183 |
| ZBTB8OS | 1097.15205 | -0.063151331 | 0.25426871 | -0.2483645 | 0.80385236 | 0.94543168 |
| RBBP4 | 784.010266 | -0.330268234 | 0.22450225 | -1.4711132 | 0.1412605 | 0.59581113 |
| SYNC | 131.264955 | 0.387292677 | 0.41673401 | 0.92935222 | 0.35270658 | 0.75673243 |
| KIAA1522 | 781.191298 | 0.062772427 | 0.32232365 | 0.19474968 | 0.84558892 | 0.95764192 |
| YARS | 534.422362 | -0.553895615 | 0.42897879 | -1.2911958 | 0.19663579 | 0.64558745 |
| S100PBP | 917.183343 | 0.200681284 | 0.18486681 | 1.08554522 | 0.27768025 | 0.7036882 |
| TMEM54 | 277.908649 | 0.149095146 | 0.60390447 | 0.24688531 | 0.80499698 | 0.94585363 |
| RNF19B | 1171.02839 | -1.421238718 | 0.33290351 | -4.2692212 | 1.96E-05 | 0.01647703 |
| AL020995.1 | 299.860296 | 0.032653207 | 0.77167468 | 0.04231473 | 0.9662478 | 0.99112162 |
| AK2 | 1367.78505 | -0.316457168 | 0.38363032 | -0.8249013 | 0.4094276 | 0.78542936 |
| AZIN2 | 263.024779 | -0.259077777 | 0.53177324 | -0.487196 | 0.62611946 | 0.87877458 |
| TRIM62 | 89.0110872 | 0.169226896 | 0.45063236 | 0.37553206 | 0.70726481 | 0.91002281 |
| ZNF362 | 285.220174 | 0.303818308 | 0.43592162 | 0.69695628 | 4.86E-01 | 0.81839266 |
| AL513327.1 | 358.943949 | 0.675207892 | 0.46652326 | 1.44731881 | 1.48E-01 | 0.60605027 |
| PHC2 | 1843.97616 | -0.023363091 | 0.33052377 | -0.0706851 | 0.94364842 | 0.98522232 |
| AL513327.2 | 54.0141759 | 0.386855365 | 0.58556988 | 0.66064764 | 0.50883831 | 0.82874455 |

| | | | | | | |
|---|---|---|---|---|---|---|
| ZSCAN20 | 75.3906492 | -0.875910461 | 0.46002026 | -1.9040693 | 0.05690116 | 0.4746515 |
| SMIM12 | 316.911348 | 0.203745593 | 0.32706189 | 0.62295731 | 0.53331258 | 0.8389895 |
| AC114490.3 | 52.945995 | 1.369639062 | 0.39183224 | 3.49547307 | 0.00047322 | 0.07721972 |
| ZMYM1 | 420.499334 | 0.378690365 | 0.26409232 | 1.43393174 | 0.15159174 | 0.60934546 |
| SFPQ | 1752.1317 | -0.111735993 | 0.27018211 | -0.4135581 | 0.67919778 | 0.89932032 |
| ZMYM4 | 2897.55101 | 0.064515271 | 0.13699523 | 0.47093079 | 0.63769016 | 0.8827026 |
| ZMYM4-AS1 | 57.1202676 | 0.118158327 | 0.81885094 | 0.14429772 | 0.88526537 | 0.96957206 |
| KIAA0319L | 1917.82555 | -0.125086089 | 0.22721613 | -0.5505159 | 0.58196558 | 0.85938792 |
| NCDN | 47.2022463 | -0.345238951 | 0.40812926 | -0.8459059 | 0.39760524 | 0.78063007 |
| PSMB2 | 383.649436 | -0.089922274 | 0.19996624 | -0.4496873 | 0.65293594 | 0.88949911 |
| AGO4 | 640.567217 | -0.073108385 | 0.24091697 | -0.3034588 | 7.62E-01 | 0.93000994 |
| AGO1 | 307.945891 | 0.140459167 | 0.23608913 | 0.59494128 | 0.55188272 | 0.84701114 |
| AGO3 | 2751.00442 | -0.157378075 | 0.2094239 | -0.751481 | 0.45236325 | 0.80432522 |
| AL138787.2 | 197.335439 | -1.221312282 | 0.79788553 | -1.5306861 | 0.12584699 | 0.58564339 |
| ADPRHL2 | 350.841389 | -0.446498657 | 0.39117478 | -1.1414301 | 0.253691 | 0.68891958 |
| TRAPPC3 | 275.585396 | -0.398981822 | 0.31100163 | -1.282893 | 0.19952956 | 0.64671356 |
| MAP7D1 | 164.201877 | -0.589965575 | 0.56278898 | -1.0482891 | 0.29450541 | 0.71625012 |
| THRAP3 | 1953.20779 | -0.685783363 | 0.27391287 | -2.5036551 | 0.01229178 | 0.29629837 |
| SH3D21 | 71.1340547 | 0.167573041 | 0.58250261 | 0.28767775 | 0.77359342 | 0.93415955 |
| EVA1B | 132.075588 | -0.326993202 | 0.33500479 | -0.9760852 | 3.29E-01 | 0.73829083 |
| STK40 | 901.842831 | -0.340669597 | 0.55536012 | -0.6134211 | 0.53959796 | 0.84113285 |
| LSM10 | 89.7143435 | 0.037523445 | 0.40095075 | 0.09358617 | 0.9254379 | 0.98089833 |
| OSCP1 | 145.704027 | 0.436045942 | 0.32052614 | 1.36040679 | 0.17370123 | 0.62672355 |
| MRPS15 | 211.324248 | -0.127422501 | 0.27655963 | -0.4607415 | 0.64498407 | 0.88656204 |
| LINC01137 | 121.138083 | 0.787310371 | 0.39711748 | 1.98256286 | 0.04741628 | 0.44991578 |
| ZC3H12A | 84.0790897 | 0.286343525 | 0.3865932 | 0.74068432 | 4.59E-01 | 8.07E-01 |
| MEAF6 | 282.996253 | -0.541388124 | 0.21352004 | -2.5355377 | 0.01122748 | 0.28237637 |
| SNIP1 | 163.480897 | -0.600999547 | 0.27153349 | -2.2133533 | 0.0268733 | 0.38702446 |
| DNALI1 | 96.5919636 | 0.049684235 | 0.44003991 | 0.11290848 | 0.91010312 | 0.97692341 |
| GNL2 | 230.520265 | -1.042668577 | 0.44457117 | -2.3453355 | 0.01900996 | 0.34084276 |
| C1orf109 | 113.393968 | -0.508713424 | 0.52144202 | -0.9755896 | 0.32926787 | 0.73829083 |
| YRDC | 55.8047356 | -0.617593498 | 0.46253642 | -1.3352321 | 0.18180038 | 0.6318513 |
| C1orf122 | 329.785595 | 0.080542703 | 0.39279719 | 0.20504908 | 0.83753378 | 0.95496955 |
| MTF1 | 366.462077 | -0.099216325 | 0.2907163 | -0.3412823 | 0.73289108 | 0.91952673 |
| AL929472.2 | 108.6881 | 0.845134644 | 0.56926837 | 1.48459794 | 0.13765036 | 0.59474076 |
| INPP5B | 1060.36055 | 0.405464317 | 0.31472188 | 1.28832581 | 0.19763257 | 0.64558745 |
| SF3A3 | 268.348648 | -0.316510838 | 0.27022654 | -1.1712796 | 0.2414864 | 0.68015474 |
| UTP11 | 82.9612717 | -0.327546604 | 0.34781476 | -0.9417272 | 0.34633235 | 0.75166079 |
| AL354702.2 | 96.6777858 | -0.05599059 | 0.89826832 | -0.0623317 | 0.95029869 | 0.98724754 |
| RRAGC | 430.63201 | -0.814678096 | 0.35384701 | -2.3023456 | 0.02131569 | 0.35688511 |
| AL139260.1 | 1138.55903 | -0.600204367 | 0.59944649 | -1.0012643 | 0.31669905 | 0.7285009 |
| RHBDL2 | 163.558811 | -0.056683516 | 0.51754718 | -0.1095234 | 0.91278738 | 0.977179 |
| AKIRIN1 | 544.076659 | -0.150461341 | 0.2680653 | -0.5612862 | 0.57460248 | 0.8547017 |
| NDUFS5 | 766.2847 | -0.122664907 | 0.41607552 | -0.294814 | 0.76813593 | 0.93198626 |
| MACF1 | 10348.8519 | -0.555592628 | 0.34091393 | -1.6297153 | 0.10316169 | 0.55022042 |
| PABPC4 | 306.131541 | -0.631670319 | 0.34897671 | -1.8100644 | 0.0702858 | 0.49583505 |
| PPIE | 196.696517 | -0.202797115 | 0.36457781 | -0.5562519 | 0.57803864 | 0.85714683 |
| TRIT1 | 575.776641 | -0.453289105 | 0.17868635 | -2.5367864 | 0.01118752 | 0.28188844 |
| MFSD2A | 420.076887 | 0.907648365 | 0.84234736 | 1.07752265 | 0.28124684 | 0.70625976 |
| CAP1 | 623.606548 | -0.289165348 | 0.26829922 | -1.0777718 | 0.28113559 | 0.70625976 |
| PPT1 | 124.777995 | -0.654468141 | 0.36248951 | -1.8054816 | 0.07099936 | 0.49789693 |
| RLF | 1293.03874 | -0.453497245 | 0.15798497 | -2.8705089 | 0.00409812 | 0.20352497 |
| AL050341.2 | 31.2656271 | -0.287996254 | 0.68104947 | -0.4228713 | 0.67238919 | 0.89727632 |
| ZMPSTE24 | 432.087279 | -0.561463133 | 0.36835494 | -1.5242449 | 0.12744755 | 0.58793678 |

| | | | | | | |
|---|---|---|---|---|---|---|
| SMAP2 | 257.282554 | 0.004749192 | 0.23599406 | 0.0201242 | 0.98394429 | 0.99523767 |
| AL603839.3 | 231.111999 | -0.01395898 | 0.28076714 | -0.0497173 | 0.96034768 | 0.98936382 |
| EXO5 | 69.500029 | -0.063133738 | 0.29302631 | -0.2154542 | 0.82941325 | 0.95325673 |
| ZNF684 | 290.353802 | 1.149513624 | 0.4656891 | 2.4684143 | 0.01357131 | 0.3005202 |
| NFYC | 955.677426 | -0.063435404 | 0.2312916 | -0.2742659 | 7.84E-01 | 9.39E-01 |
| CITED4 | 254.858091 | 0.082699297 | 0.58915713 | 0.14036883 | 8.88E-01 | 0.97134248 |
| CTPS1 | 429.231973 | -0.726786101 | 0.41560984 | -1.7487221 | 0.08033907 | 0.51297858 |
| SLFNL1-AS1 | 303.988986 | -0.124124103 | 0.31468685 | -0.3944369 | 0.69325851 | 0.90551691 |
| SLFNL1 | 127.924208 | 0.15981232 | 0.38566906 | 0.41437683 | 0.67859816 | 0.8992258 |
| SCMH1 | 5099.34893 | 0.532616967 | 0.3400145 | 1.56645369 | 0.11724244 | 0.57350419 |
| AC093151.3 | 410.10916 | -0.883168848 | 0.85427057 | -1.033828 | 0.30121657 | 0.71879123 |
| HIVEP3 | 832.922315 | 0.737956134 | 0.36308573 | 2.03245701 | 0.04210741 | 0.43742111 |
| FOXJ3 | 2338.03373 | -0.258506561 | 0.23917662 | -1.0808187 | 0.27977777 | 0.70446562 |
| RIMKLA | 144.428696 | -0.329112794 | 0.6043353 | -0.5445864 | 5.86E-01 | 8.61E-01 |
| ZMYND12 | 87.3992737 | 0.122627026 | 0.6936891 | 0.1767752 | 0.85968497 | 0.96283855 |
| PPCS | 267.847518 | 0.056608018 | 0.31865631 | 0.177646 | 0.859001 | 0.96274135 |
| CCDC30 | 2334.27542 | 0.540905599 | 0.41658311 | 1.29843381 | 0.19413831 | 0.64548758 |
| PPIH | 142.055834 | -1.045510114 | 0.34364017 | -3.0424561 | 0.00234656 | 0.15766817 |
| YBX1 | 745.490956 | -0.582221388 | 0.39938072 | -1.4578105 | 0.14489279 | 0.60109715 |
| P3H1 | 174.368104 | -0.421406497 | 0.35842377 | -1.1757214 | 0.23970622 | 0.67801344 |
| SVBP | 77.3892187 | -0.66532476 | 0.41229272 | -1.6137194 | 0.10658831 | 0.55467196 |
| ERMAP | 244.030599 | 0.401169638 | 0.34867741 | 1.15054669 | 0.24991877 | 0.68617881 |
| AL512353.1 | 46.1850395 | -1.120253098 | 0.65630718 | -1.7069036 | 0.08783997 | 0.52761721 |
| ZNF691 | 80.2453234 | 0.11014525 | 0.29364272 | 0.37509954 | 0.70758644 | 0.91012042 |
| SLC2A1 | 185.625844 | -1.650067688 | 0.38871589 | -4.2449195 | 2.19E-05 | 1.65E-02 |
| SLC2A1-AS1 | 55.3860421 | 1.019476675 | 0.72894814 | 1.39855859 | 0.16194539 | 0.61845523 |
| EBNA1BP2 | 1744.11873 | 0.232338162 | 0.58265337 | 0.3987588 | 6.90E-01 | 9.04E-01 |
| CFAP57 | 290.971782 | 0.567455136 | 0.70523778 | 0.80462952 | 0.4210335 | 0.78872573 |
| TMEM125 | 66.0592611 | 0.156767474 | 0.60820917 | 0.25775256 | 0.79659788 | 0.94276732 |
| C1orf210 | 66.6951003 | 0.447220604 | 0.60998542 | 0.73316606 | 0.46345716 | 0.8082072 |
| AL139289.2 | 30.6179952 | -0.026548832 | 0.68303012 | -0.0388692 | 0.96899468 | 0.99187457 |
| ELOVL1 | 152.976063 | -0.132780546 | 0.32387077 | -0.40998 | 0.6818206 | 0.89962501 |
| MED8 | 142.016401 | -0.408194271 | 0.38927027 | -1.048614 | 0.29435579 | 0.71601327 |
| SZT2 | 573.959959 | -0.234670512 | 0.26493463 | -0.8857676 | 0.37574277 | 0.7697436 |
| HYI | 134.540691 | -0.145885537 | 0.36015886 | -0.4050589 | 0.68543424 | 0.90113631 |
| PTPRF | 1074.77546 | 0.118683162 | 0.38159609 | 0.31101776 | 0.75578712 | 0.92799282 |
| KDM4A-AS1 | 41.7390119 | 0.038484104 | 0.53554416 | 0.07185982 | 0.94271347 | 0.98492559 |
| AL357079.1 | 48.731595 | -0.006780473 | 0.59556916 | -0.0113849 | 0.99091639 | 0.99737132 |
| IPO13 | 177.570386 | -0.444074309 | 0.44705648 | -0.9933293 | 0.3205495 | 0.73210121 |
| DPH2 | 47.1464484 | -0.511243635 | 0.53552289 | -0.9546625 | 0.33974839 | 0.74648557 |
| ATP6V0B | 827.244083 | -0.474864323 | 0.33363015 | -1.4233256 | 0.1546418 | 0.61119904 |
| B4GALT2 | 161.517813 | -0.460949974 | 0.43449893 | -1.0608771 | 0.28874575 | 0.71218341 |
| CCDC24 | 220.381697 | 0.505711434 | 0.54698616 | 0.92454155 | 0.35520443 | 0.75757147 |
| SLC6A9 | 38.1674706 | -0.63847353 | 0.42776816 | -1.4925691 | 0.13555002 | 0.59381224 |
| DMAP1 | 165.145021 | -0.657309562 | 0.29870652 | -2.2005196 | 0.02777005 | 0.38960498 |
| ERI3 | 1695.53504 | 0.493726922 | 0.43658919 | 1.13087299 | 0.25810855 | 0.69210282 |
| RNF220 | 232.476936 | 0.136469323 | 0.40415288 | 0.33766758 | 0.73561371 | 0.92019178 |
| TMEM53 | 196.937998 | 0.725244263 | 0.38903995 | 1.86418967 | 0.06229507 | 0.4840117 |
| ARMH1 | 148.605693 | 0.744777506 | 0.64407332 | 1.15635516 | 0.24753591 | 0.68482118 |
| RPS8 | 1185.06381 | 0.316319551 | 0.48467207 | 0.65264655 | 0.51398417 | 0.83126535 |
| PLK3 | 75.9240628 | -0.32676813 | 0.49409886 | -0.6613416 | 5.08E-01 | 8.29E-01 |
| BTBD19 | 91.7864314 | 0.156740098 | 0.42141752 | 0.37193541 | 0.70994094 | 0.91118132 |
| EIF2B3 | 614.82586 | -0.352342914 | 0.31965948 | -1.1022445 | 0.27035537 | 0.69961625 |
| HECTD3 | 178.9977 | -0.374571815 | 0.29643727 | -1.2635787 | 0.20638128 | 0.65243721 |

| | | | | | | |
|---|---|---|---|---|---|---|
| UROD | 139.028246 | -0.355370676 | 0.40443065 | -0.8786937 | 0.37956736 | 0.77095585 |
| ZSWIM5 | 1943.91963 | -0.248475738 | 0.37939177 | -0.6549318 | 0.51251166 | 0.83055441 |
| MUTYH | 135.516618 | 0.122571878 | 0.49889526 | 0.2456866 | 0.80592485 | 0.94585363 |
| TOE1 | 57.6161578 | -0.03763671 | 0.34033075 | -0.1105886 | 0.91194257 | 0.977179 |
| TESK2 | 375.213983 | 0.067952439 | 0.44192873 | 0.15376334 | 0.87779634 | 0.96658493 |
| MMACHC | 52.036798 | 0.275303719 | 0.45129202 | 0.61003453 | 0.54183893 | 0.84244399 |
| PRDX1 | 2167.03618 | 0.108335179 | 0.54926462 | 0.19723677 | 0.84364226 | 0.95731603 |
| AKR1A1 | 1390.80443 | 0.47056986 | 0.28120678 | 1.67339444 | 0.09424967 | 0.53783536 |
| NASP | 360.818776 | -0.058559437 | 0.2481987 | -0.2359377 | 0.81348099 | 0.9486459 |
| GPBP1L1 | 2105.34011 | -0.073642278 | 0.15147527 | -0.486167 | 0.62684875 | 0.87890915 |
| TMEM69 | 140.212227 | -0.268540643 | 0.28985047 | -0.9264799 | 0.35419662 | 0.75728796 |
| IPP | 368.735712 | 0.205882849 | 0.30373907 | 0.67782801 | 0.49788075 | 0.82326489 |
| MAST2 | 2172.63834 | -0.366127834 | 0.3368178 | -1.0870204 | 0.27702778 | 0.7036882 |
| TSPAN1 | 1256.13735 | 0.605609608 | 0.39182552 | 1.54561044 | 0.12219868 | 0.58058139 |
| POMGNT1 | 359.757908 | 0.262687552 | 0.16324468 | 1.60916454 | 0.10758037 | 0.55624448 |
| LURAP1 | 103.366583 | 1.503005144 | 0.50818241 | 2.95760956 | 0.00310035 | 0.18216645 |
| LRRC41 | 730.554073 | -0.645162639 | 0.42357965 | -1.52312 | 0.12772869 | 0.58793678 |
| UQCRH | 1082.58089 | -0.107672363 | 0.52074461 | -0.2067662 | 0.83619249 | 0.95468028 |
| NSUN4 | 411.804569 | 0.158676973 | 0.22613888 | 0.70167932 | 0.48287917 | 0.8177376 |
| FAAH | 462.574896 | -0.289172945 | 0.50672578 | -0.5706695 | 5.68E-01 | 0.85263558 |
| MKNK1 | 386.202289 | -0.007787147 | 0.25272326 | -0.0308129 | 0.97541872 | 0.99420467 |
| ATPAF1 | 301.210184 | 0.380056601 | 0.24265233 | 1.56625986 | 0.11728779 | 0.57350419 |
| EFCAB14 | 478.86889 | -0.173795556 | 0.25774869 | -0.674283 | 0.50013142 | 0.8251446 |
| CYP4A11 | 1795.97109 | 0.11798075 | 1.03010366 | 0.11453289 | 0.90881537 | 0.97680747 |
| CYP4X1 | 75.1851216 | 1.368884425 | 1.02862779 | 1.33078693 | 0.18325913 | 0.63276491 |
| CYP4Z1 | 113.942855 | 0.550242846 | 1.02765664 | 0.53543453 | 0.59234942 | 0.86382106 |
| CYP4A22-AS1 | 1185.76139 | 0.332461233 | 0.80336122 | 0.41383779 | 0.67899291 | 0.89932032 |
| CYP4A22 | 149.635119 | 0.227232087 | 1.18419906 | 0.19188673 | 0.84783093 | 0.95908381 |
| LINC00853 | 108.085522 | 0.074076184 | 0.796155 | 0.09304242 | 0.92586986 | 0.98098729 |
| PDZK1IP1 | 4254.56156 | -0.073960876 | 0.43207646 | -0.1711754 | 0.86408582 | 0.96393376 |
| STIL | 97.3170623 | 0.087014267 | 0.34429369 | 0.25273268 | 0.80047479 | 0.94403774 |
| CMPK1 | 977.429976 | -0.051606091 | 0.25211433 | -0.2046932 | 0.83781182 | 0.95496955 |
| LINC01389 | 32.974129 | 0.379983442 | 0.53756209 | 0.70686429 | 0.47965082 | 0.81661579 |
| TRABD2B | 8126.78479 | 1.092981167 | 0.57785429 | 1.89144771 | 0.0585646 | 0.47778423 |
| AL356289.1 | 123.887979 | 1.206332959 | 0.66000269 | 1.82776977 | 0.06758411 | 0.48963361 |
| AL109659.1 | 105.47376 | 1.204834524 | 0.76918895 | 1.56637004 | 0.11726201 | 0.57350419 |
| SLC5A9 | 286.427444 | 0.869017058 | 0.81834051 | 1.06192599 | 0.28826928 | 0.71194722 |
| SPATA6 | 1153.10411 | 1.037487104 | 0.42775601 | 2.42541795 | 0.01529077 | 0.3105046 |
| AGBL4 | 1699.72597 | 0.752033832 | 0.92822467 | 0.81018514 | 0.41783378 | 0.78679526 |
| AC099788.1 | 37.1609144 | -0.090767022 | 1.13384245 | -0.0800526 | 0.93619543 | 0.98415216 |
| FAF1 | 5176.19954 | -0.069525208 | 0.16488598 | -0.4216563 | 0.67327593 | 0.89782405 |
| RNF11 | 923.632826 | 0.069928919 | 0.21317244 | 0.32803922 | 0.74288201 | 0.92318522 |
| TTC39A | 951.605148 | 0.403491198 | 0.48781415 | 0.82714124 | 4.08E-01 | 7.85E-01 |
| TTC39A-AS1 | 52.6960732 | 0.325994026 | 0.34191178 | 0.95344485 | 3.40E-01 | 7.47E-01 |
| EPS15 | 2364.60503 | -0.056531971 | 0.19507293 | -0.2897992 | 0.77196989 | 0.93340577 |
| AC104170.1 | 714.411067 | 1.920572558 | 0.7561003 | 2.5401029 | 0.01108199 | 0.27974361 |
| OSBPL9 | 3812.06768 | -0.226265912 | 0.36921799 | -0.6128247 | 0.53999225 | 0.8414758 |
| NRDC | 1843.8654 | 0.092882489 | 0.14587675 | 0.63671894 | 0.52430794 | 0.83589803 |
| TXNDC12 | 369.280464 | 0.094792952 | 0.19257889 | 0.4922292 | 0.62255732 | 0.87714058 |
| BTF3L4 | 509.586242 | -0.178829425 | 0.19504265 | -0.9168734 | 0.35920896 | 0.75936492 |
| ZFYVE9 | 1185.0238 | -0.470210236 | 0.14213161 | -3.3082736 | 0.00093873 | 0.10293743 |
| CC2D1B | 341.290906 | -0.474362149 | 0.33949554 | -1.3972559 | 0.16233663 | 0.61910325 |
| PRPF38A | 214.559281 | -0.232545042 | 0.2286218 | -1.0171604 | 0.30907713 | 0.72449199 |
| TUT4 | 2499.18232 | -0.019335786 | 0.28768902 | -0.0672107 | 0.94641395 | 0.98620161 |

| | | | | | | |
|---|---|---|---|---|---|---|
| COA7 | 102.281322 | -0.735448182 | 0.50685332 | -1.4510079 | 0.14677765 | 0.60383734 |
| ZYG11B | 1207.36208 | 0.029438047 | 0.23393383 | 0.1258392 | 9.00E-01 | 0.97465308 |
| ZYG11A | 895.927294 | -0.435232023 | 0.53157843 | -0.8187541 | 4.13E-01 | 7.86E-01 |
| ECHDC2 | 2399.52254 | 0.554203807 | 0.35644689 | 1.55480051 | 0.11999359 | 0.57730856 |
| SCP2 | 1408.91928 | 0.521397933 | 0.30261953 | 1.72294872 | 0.0848978 | 0.52183702 |
| SLC1A7 | 22.0615483 | 4.277115179 | 1.6158698 | 2.64694296 | 0.00812231 | 0.24877162 |
| CPT2 | 206.561274 | 0.947697334 | 0.36388105 | 2.60441517 | 0.00920312 | 0.25849829 |
| C1orf123 | 98.7111118 | -0.008266938 | 0.29753057 | -0.0277852 | 0.97783349 | 0.99481657 |
| MAGOH | 210.725937 | -0.461967915 | 0.21655195 | -2.1332891 | 0.03290102 | 0.41180153 |
| LRP8 | 168.784943 | -0.846357928 | 1.0918097 | -0.7751881 | 4.38E-01 | 7.97E-01 |
| GLIS1 | 2461.5801 | 0.422690799 | 0.50152206 | 0.84281596 | 0.39933138 | 0.78161291 |
| NDC1 | 267.549962 | -0.551817621 | 0.39173521 | -1.4086495 | 0.15893882 | 0.61423843 |
| YIPF1 | 307.16964 | 0.435922678 | 0.2814478 | 1.54885801 | 0.12141586 | 0.57985101 |
| DIO1 | 911.637691 | 3.040299218 | 0.71260132 | 4.26647991 | 1.99E-05 | 0.01647703 |
| HSPB11 | 362.259224 | 0.009102528 | 0.31927958 | 0.02850958 | 0.97725572 | 0.99473325 |
| LRRC42 | 176.749687 | 0.074349297 | 0.20760512 | 0.35812844 | 0.72024719 | 0.91487613 |
| TMEM59 | 891.093304 | 0.266143164 | 0.30288046 | 0.87870694 | 0.37956019 | 0.77095585 |
| TCEANC2 | 187.79753 | 0.413937093 | 0.35018241 | 1.18206137 | 0.23718135 | 0.67463059 |
| MRPL37 | 168.121 | -0.101152844 | 0.23335036 | -0.4334806 | 0.66466569 | 0.89315615 |
| SSBP3 | 1019.75945 | -0.45738157 | 0.47190998 | -0.9692136 | 0.33243863 | 0.73988623 |
| ACOT11 | 876.387694 | 0.6829163 | 0.64883705 | 1.05252359 | 0.29255939 | 0.71399546 |
| FAM151A | 253.249605 | 0.431389124 | 0.67147182 | 0.64245306 | 5.21E-01 | 8.34E-01 |
| MROH7-TTC4 | 44.610276 | -0.338720458 | 0.66874714 | -0.5065001 | 0.61250565 | 0.87222135 |
| PARS2 | 37.358971 | 0.049413687 | 0.42885653 | 0.11522195 | 0.9082692 | 0.97677491 |
| TTC22 | 261.02799 | 0.041770914 | 0.43795829 | 0.09537647 | 0.92401581 | 0.98018878 |
| DHCR24 | 277.938214 | 0.102008743 | 0.41044868 | 0.24852983 | 0.80372449 | 0.94543168 |
| USP24 | 1877.03681 | -0.069841626 | 0.164629 | -0.4242365 | 0.67139337 | 0.89704376 |
| AC119674.2 | 126.32144 | -0.059042288 | 0.58770709 | -0.1004621 | 0.91997747 | 0.97939204 |
| PLPP3 | 2402.41493 | -0.033056117 | 0.29543261 | -0.1118905 | 0.91091019 | 0.97697493 |
| PRKAA2 | 2397.23175 | -0.276948115 | 0.18241453 | -1.518235 | 0.12895517 | 0.58793678 |
| FYB2 | 583.417273 | 1.861776051 | 0.75316843 | 2.47192525 | 0.01343876 | 0.29875272 |
| DAB1 | 572.441636 | 0.451351338 | 0.72377233 | 0.62360955 | 0.53288403 | 0.83870036 |
| TACSTD2 | 185.114448 | 0.392400187 | 0.63445922 | 0.61847977 | 0.53625913 | 0.83957783 |
| MYSM1 | 1032.58902 | -0.012439445 | 0.23028714 | -0.0540171 | 0.95692154 | 0.98861313 |
| JUN | 1395.99781 | -0.539758226 | 0.44196463 | -1.2212702 | 2.22E-01 | 6.63E-01 |
| LINC01135 | 326.76049 | 0.582131256 | 0.70252277 | 0.82862973 | 0.40731396 | 0.78481094 |
| LINC01358 | 70.375865 | 0.7678804 | 0.62990856 | 1.21903472 | 0.22283102 | 0.66398799 |
| FGGY | 7132.42434 | 0.895086869 | 0.46087735 | 1.94213682 | 0.05212054 | 0.46437412 |
| HOOK1 | 1087.44777 | 0.558483818 | 0.4383937 | 1.27393214 | 0.20268747 | 0.64881821 |
| CYP2J2 | 94.5627704 | -0.592555486 | 1.26062844 | -0.4700477 | 0.63832095 | 0.8827026 |
| LINC01748 | 87.6297963 | 2.592352573 | 1.2178579 | 2.12861663 | 0.03328599 | 0.41254527 |
| AC099792.1 | 122.445537 | 2.496191175 | 1.1064598 | 2.25601615 | 0.02406962 | 0.37387451 |
| NFIA | 7802.80343 | 0.705995427 | 0.42920429 | 1.64489368 | 0.09999174 | 0.54542315 |
| TM2D1 | 975.195387 | -0.259351918 | 0.18914349 | -1.3711914 | 0.17031531 | 0.62420107 |
| AC099791.2 | 39.8192991 | -0.960933724 | 0.59166489 | -1.6241182 | 0.10435058 | 0.55066578 |
| PATJ | 19961.4356 | 0.022465313 | 0.27069345 | 0.08299171 | 0.93385813 | 0.98355048 |
| USP1 | 144.053879 | 0.022542075 | 0.34430907 | 0.06547046 | 0.94779942 | 0.98659528 |
| DOCK7 | 1669.92594 | -0.160459865 | 0.31477213 | -0.5097652 | 0.61021599 | 0.87109254 |
| ANGPTL3 | 108.729228 | 0.776423459 | 0.83035692 | 0.93504786 | 0.34976363 | 0.75455562 |
| AC103923.1 | 93.7689019 | -0.119889061 | 0.81897556 | -0.1463891 | 0.88361426 | 0.96854864 |
| ATG4C | 401.506817 | 0.313823099 | 0.31792132 | 0.98710932 | 0.32358905 | 0.7354443 |
| ALG6 | 189.819982 | 0.22976915 | 0.30968934 | 0.74193433 | 0.45812712 | 0.80600032 |
| ITGB3BP | 595.524387 | 0.414070065 | 0.51222216 | 0.80837984 | 0.41887195 | 0.78766781 |
| EFCAB7 | 317.474511 | 0.243357598 | 0.44634068 | 0.54522836 | 0.58559647 | 0.86077718 |

| | | | | | | |
|---|---|---|---|---|---|---|
| PGM1 | 992.156339 | 0.172136691 | 0.31551988 | 0.54556528 | 0.5853648 | 0.86077718 |
| ROR1 | 2800.80739 | 0.168922885 | 0.22294722 | 0.75768107 | 0.44864194 | 0.80267462 |
| ROR1-AS1 | 51.060686 | -1.007672688 | 0.80197392 | -1.2564906 | 0.20893816 | 0.65401126 |
| UBE2U | 55.6752667 | -2.479015748 | 1.22077218 | -2.0306948 | 0.04228596 | 0.43742111 |
| CACHD1 | 1459.76866 | -0.515812591 | 0.46367412 | -1.1124464 | 0.26594628 | 0.69766997 |
| RAVER2 | 419.260516 | 0.40134993 | 0.3905993 | 1.02752343 | 0.30417406 | 0.72082426 |
| JAK1 | 2687.56057 | -0.354694053 | 0.2145694 | -1.6530505 | 0.09832059 | 0.54193402 |
| AK4 | 4875.65303 | -0.139005478 | 0.46911129 | -0.2963166 | 0.76698828 | 0.93151767 |
| DNAJC6 | 1750.7989 | 0.007360916 | 0.44584134 | 0.01651017 | 0.98682739 | 0.9962029 |
| AL139294.1 | 164.864986 | 1.01175496 | 0.79358143 | 1.27492268 | 0.20233661 | 0.64881821 |
| LEPR | 923.338322 | 1.012175217 | 0.25338899 | 3.99455085 | 6.48E-05 | 0.02490203 |
| PDE4B | 99.0275882 | 0.560819798 | 0.72650686 | 0.77194013 | 0.44014989 | 0.79826598 |
| SGIP1 | 140.60563 | -0.384868277 | 0.86855629 | -0.4431126 | 0.65768426 | 0.89053559 |
| WDR78 | 880.844749 | 0.622209118 | 0.40658721 | 1.53032141 | 0.12593719 | 0.58564339 |
| MIER1 | 1204.88914 | -0.26113764 | 0.16191539 | -1.6128031 | 0.10678729 | 0.55488138 |
| SLC35D1 | 694.878746 | 0.670357716 | 0.31764267 | 2.11041455 | 0.03482266 | 0.41686835 |
| IL12RB2 | 135.600976 | 1.058140671 | 0.48544371 | 2.17973919 | 0.0292768 | 0.39693085 |
| SERBP1 | 1042.6223 | -0.160808289 | 0.22314198 | -0.7206546 | 0.47112206 | 0.81387913 |
| LINC01702 | 91.1026936 | 1.090442412 | 0.84893003 | 1.28449032 | 1.99E-01 | 6.46E-01 |
| GADD45A | 1288.66294 | -0.399525519 | 0.39655163 | -1.0074994 | 0.31369485 | 0.72717675 |
| GNG12 | 3364.5216 | -0.01510931 | 0.22219754 | -0.0679994 | 0.94578608 | 0.98584713 |
| GNG12-AS1 | 1170.94321 | 0.695729994 | 0.36315451 | 1.91579611 | 0.05539105 | 0.46947044 |
| WLS | 1208.58879 | 0.29006179 | 0.29872193 | 0.97100935 | 0.33154362 | 0.73939532 |
| LRRC7 | 745.476636 | 0.09008408 | 0.6560514 | 0.13731253 | 0.89078376 | 0.97230741 |
| LRRC40 | 484.620986 | 0.185158362 | 0.24767315 | 0.74759157 | 0.45470655 | 0.80465616 |
| SRSF11 | 2693.23227 | -0.324386023 | 0.17843784 | -1.8179217 | 0.0690761 | 0.49273866 |
| ANKRD13C | 1191.68163 | -0.175384149 | 0.1296285 | -1.3529753 | 0.17606353 | 0.62699479 |
| HHLA3 | 79.2400854 | 0.494121532 | 0.46875177 | 1.05412195 | 0.29182709 | 0.71340715 |
| CTH | 173.178671 | 0.207434827 | 0.52027022 | 0.39870594 | 0.69010989 | 0.90416137 |
| LINC01788 | 943.075653 | 0.346845217 | 0.81861871 | 0.42369569 | 0.67178776 | 0.89704376 |
| PTGER3 | 963.6654 | -0.775578221 | 0.91876719 | -0.844151 | 0.39858505 | 0.78122921 |
| AL031429.1 | 103.902371 | 0.783721355 | 0.60594351 | 1.29339013 | 1.96E-01 | 0.64558745 |
| ZRANB2-AS1 | 81.055784 | 0.182423938 | 0.71368672 | 0.25560787 | 0.79825364 | 0.94320409 |
| ZRANB2 | 1616.67585 | -0.401106935 | 0.14737998 | -2.7215835 | 0.006497 | 0.22952148 |
| NEGR1 | 1235.27281 | 0.862430494 | 0.76364106 | 1.12936632 | 0.25874333 | 0.69241621 |
| LRRIQ3 | 80.3074574 | 1.129341671 | 0.65183453 | 1.73255883 | 0.08317411 | 0.51935659 |
| CRYZ | 1345.9038 | 0.036119993 | 0.3636371 | 0.09932978 | 0.92087644 | 0.97939204 |
| TYW3 | 321.765798 | -0.29441527 | 0.21864139 | -1.346567 | 0.17811974 | 0.62849772 |
| SLC44A5 | 201.474099 | 1.493577156 | 0.61185391 | 2.44106826 | 0.01464389 | 0.30832605 |
| ACADM | 1402.23263 | 0.552463579 | 0.4853886 | 1.1381882 | 0.25504191 | 0.68976367 |
| RABGGTB | 321.520186 | -0.520219529 | 0.39859862 | -1.3051212 | 0.19185157 | 0.64497936 |
| MSH4 | 41.481184 | 0.625537442 | 0.78000435 | 0.80196661 | 0.42257227 | 0.78894887 |
| ASB17 | 50.6950692 | 2.349133047 | 0.79831733 | 2.94260559 | 0.00325463 | 0.18622085 |
| ST6GALNAC3 | 115.840016 | 0.043406627 | 0.36757199 | 0.11809014 | 0.90599624 | 0.97637317 |
| PIGK | 447.911053 | 0.02297279 | 0.24175904 | 0.0950235 | 0.92429616 | 0.9803537 |
| AK5 | 237.262085 | -0.119036462 | 0.81212884 | -0.1465734 | 8.83E-01 | 0.96846661 |
| AC118549.1 | 3013.56603 | -0.252439067 | 0.1817045 | -1.3892835 | 0.16474656 | 0.62065658 |
| USP33 | 2070.32103 | -0.075073935 | 0.25554347 | -0.2937815 | 0.7689249 | 0.93209094 |
| MIGA1 | 1215.57319 | -0.019311727 | 0.11464314 | -0.1684508 | 0.86622865 | 0.96437968 |
| NEXN | 128.908583 | 0.642381228 | 0.43015448 | 1.49337333 | 0.13533949 | 0.59381224 |
| FUBP1 | 1213.79069 | -0.10291684 | 0.14162681 | -0.7266763 | 0.4674243 | 0.81166836 |
| GIPC2 | 2477.85834 | 0.409577527 | 0.5284194 | 0.77509933 | 0.43828097 | 0.79696434 |
| PTGFR | 302.329535 | 0.398032038 | 0.61306047 | 0.64925412 | 0.51617413 | 0.83219275 |
| IFI44L | 722.513462 | -4.985160167 | 1.7705012 | -2.8156774 | 0.00486745 | 0.21582309 |

| | | | | | | |
|---|---|---|---|---|---|---|
| IFI44 | 266.228353 | -3.993200959 | 1.60170987 | -2.4930863 | 0.01266381 | 0.29875272 |
| AL136234.1 | 545.53719 | 0.881393121 | 0.80252626 | 1.09827324 | 0.27208519 | 0.70056282 |
| LINC01781 | 4724.20893 | 0.969704801 | 0.77387712 | 1.25304751 | 0.21018842 | 0.65441616 |
| ADGRL2 | 1709.41547 | -0.135245719 | 0.32242009 | -0.4194705 | 0.6748723 | 0.89801183 |
| LINC01362 | 119.554388 | 0.326855611 | 0.80713487 | 0.40495786 | 0.68550849 | 0.90114748 |
| LINC01725 | 1728.99278 | 1.194615044 | 0.59976948 | 1.99179033 | 0.04639407 | 0.44508419 |
| TTLL7 | 1002.13997 | -0.132379022 | 0.54161804 | -0.244414 | 8.07E-01 | 0.94585363 |
| AL359504.2 | 41.6333499 | 0.401433116 | 0.77125553 | 0.52049301 | 0.60272 | 0.8677246 |
| PRKACB | 1175.34483 | 0.195346295 | 0.14227357 | 1.37303294 | 0.16974211 | 0.62326131 |
| RPF1 | 162.725268 | -0.231774014 | 0.35398859 | -0.65475 | 0.51262876 | 0.83055441 |
| GNG5 | 480.653525 | -0.249367665 | 0.30692629 | -0.8124676 | 0.41652337 | 0.78632759 |
| SPATA1 | 206.352196 | 0.847827713 | 0.65704381 | 1.29036709 | 0.19692323 | 0.64558745 |
| CTBS | 354.216155 | 0.719519351 | 0.34990562 | 2.05632409 | 0.03975129 | 0.42869465 |
| SSX2IP | 707.231447 | 0.754755574 | 0.2929002 | 2.57683527 | 0.00997095 | 0.26511053 |
| MCOLN2 | 185.834977 | 1.570216059 | 0.79744238 | 1.96906522 | 0.0489456 | 0.45368151 |
| WDR63 | 277.295925 | 0.196247752 | 0.57113606 | 0.34360946 | 0.73114001 | 0.9190831 |
| MCOLN3 | 408.900763 | -0.332537528 | 0.33265858 | -0.9996361 | 0.31748664 | 0.72939256 |
| SYDE2 | 395.669997 | 0.172962793 | 0.34292307 | 0.50437783 | 6.14E-01 | 8.72E-01 |
| C1orf52 | 311.450139 | -0.043291054 | 0.22307654 | -0.1940637 | 0.84612602 | 0.95789828 |
| BCL10 | 165.553821 | -0.105243877 | 0.24776231 | -0.4247776 | 0.67099882 | 0.89694443 |
| AL078459.1 | 483.24116 | 0.613186396 | 0.51371232 | 1.1936377 | 0.23261972 | 0.66978724 |
| DDAH1 | 3951.83172 | 0.430605161 | 0.30704305 | 1.402426 | 0.16078807 | 0.61630932 |
| CYR61 | 210.805818 | -1.980730738 | 0.55018705 | -3.6001042 | 0.00031809 | 0.06474783 |
| ZNHIT6 | 566.26334 | -0.481488199 | 0.2778794 | -1.7327236 | 0.08314481 | 0.51935659 |
| AL136382.1 | 57.5042045 | 0.795899623 | 0.88286212 | 0.90149934 | 0.36732288 | 0.76576346 |
| ODF2L | 900.711621 | 0.180507424 | 0.27960005 | 0.64559153 | 0.51854392 | 0.83325693 |
| AL356270.1 | 40.389895 | 0.358756061 | 0.67373462 | 0.53248869 | 0.59438758 | 0.86488065 |
| SH3GLB1 | 964.464868 | -0.060016615 | 0.17206429 | -0.3488034 | 7.27E-01 | 9.16E-01 |
| SELENOF | 977.121878 | 0.141688968 | 0.17371966 | 0.81561851 | 0.41471836 | 0.78632759 |
| HS2ST1 | 1773.27427 | -0.16203061 | 0.23787364 | -0.6811625 | 0.49576865 | 0.82243887 |
| LMO4 | 250.589451 | 0.487455693 | 0.3825169 | 1.27433767 | 0.20254377 | 0.64881821 |
| PKN2-AS1 | 196.620022 | 0.989490234 | 0.59330389 | 1.66776293 | 0.0953628 | 0.53824794 |
| PKN2 | 2910.11401 | 0.027958153 | 0.15725711 | 0.17778626 | 0.85889084 | 0.9626966 |
| GTF2B | 379.395074 | -0.071233526 | 0.2076342 | -0.3430722 | 0.73154413 | 0.91917457 |
| KYAT3 | 642.199707 | 0.358550692 | 0.38310254 | 0.93591312 | 0.34931792 | 0.7542101 |
| GBP3 | 228.760019 | 0.758376699 | 0.48855224 | 1.55229397 | 0.1205919 | 0.57856254 |
| GBP1 | 244.27275 | -1.03724857 | 0.84745227 | -1.2239611 | 0.22096692 | 0.66263062 |
| GBP2 | 675.022963 | -0.063912355 | 0.61009502 | -0.104758 | 9.17E-01 | 9.78E-01 |
| GBP4 | 45.9770625 | -0.191844563 | 0.90198757 | -0.2126909 | 0.83156805 | 0.95325673 |
| LRRC8B | 432.181723 | 0.415384709 | 0.39300916 | 1.0569339 | 0.29054175 | 0.71269438 |
| LRRC8D | 153.757946 | 0.453600654 | 0.36877492 | 1.23002035 | 0.21868948 | 0.66098715 |
| ZNF326 | 790.538931 | 0.045535358 | 0.19466814 | 0.23391273 | 0.81505273 | 0.94895023 |
| ZNF644 | 2626.67807 | -0.182647691 | 0.17325425 | -1.0542176 | 0.29178328 | 0.71340715 |
| HFM1 | 73.2204755 | -0.783394149 | 0.68514455 | -1.1433998 | 0.25287263 | 0.68887358 |
| CDC7 | 86.6195587 | -1.182102541 | 0.83852078 | -1.4097475 | 0.15861427 | 0.61423843 |
| TGFBR3 | 2442.85205 | -0.00958872 | 0.34771562 | -0.0275763 | 0.97800006 | 0.99482978 |
| BTBD8 | 341.701873 | 0.44346317 | 0.65195507 | 0.68020512 | 0.49637459 | 0.82243887 |
| GLMN | 416.374671 | -0.129444767 | 0.29284404 | -0.4420263 | 0.65847017 | 0.89058983 |
| RPAP2 | 1551.17269 | 0.046864397 | 0.25898059 | 0.18095718 | 0.85640118 | 0.96242445 |
| EVI5 | 2331.44276 | -0.321590238 | 0.24793502 | -1.2970747 | 0.19460548 | 0.64548758 |
| RPL5 | 975.551514 | 0.103283435 | 0.40463569 | 0.25525043 | 0.79852968 | 0.9432477 |
| FAM69A | 902.342407 | -0.238805911 | 0.37310473 | -0.6400506 | 0.52213969 | 0.83463192 |
| MTF2 | 1262.26053 | 0.232487734 | 0.27106217 | 0.85769155 | 0.39106281 | 0.77606489 |
| TMED5 | 985.719819 | -0.115935355 | 0.2784808 | -0.4163136 | 0.67718051 | 0.89864474 |

| | | | | | | |
|---|---|---|---|---|---|---|
| CCDC18 | 521.655333 | -0.130746599 | 0.31512097 | -0.4149092 | 0.67820836 | 0.89913305 |
| DR1 | 323.077177 | 0.170229153 | 0.25860409 | 0.65826165 | 0.51037002 | 0.82930674 |
| FNBP1L | 3286.16623 | 0.243241139 | 0.25867201 | 0.94034579 | 0.34704022 | 0.75207626 |
| BCAR3 | 1511.26177 | -1.303950649 | 0.60955168 | -2.1391962 | 0.03241978 | 0.4107005 |
| AL109613.1 | 67.0832788 | 0.842230005 | 0.4827338 | 1.744709 | 0.08103554 | 0.51567042 |
| DNTTIP2 | 218.540464 | -1.460789724 | 0.61687319 | -2.3680551 | 0.01788187 | 0.33530341 |
| GCLM | 1712.16936 | -0.527826497 | 0.62875461 | -0.8394793 | 4.01E-01 | 7.82E-01 |
| ARHGAP29 | 4503.79132 | -0.197821704 | 0.41953128 | -0.4715303 | 0.6372621 | 0.88267497 |
| AC097059.1 | 199.761129 | 0.903170665 | 0.58853974 | 1.53459588 | 0.12488313 | 0.58564339 |
| ABCD3 | 2978.57461 | 0.072232892 | 0.38189772 | 0.18914198 | 0.84998154 | 0.95969497 |
| SLC44A3 | 1166.96354 | 0.00950022 | 0.37187198 | 0.02554702 | 0.97961865 | 0.99484461 |
| CNN3 | 2217.91844 | -0.615690819 | 0.35881017 | -1.7159235 | 0.08617604 | 0.52402549 |
| AC105942.1 | 428.947737 | 0.229737702 | 0.37981859 | 0.60486166 | 0.54527092 | 0.84449215 |
| ALG14 | 611.092879 | -0.133689136 | 0.2790164 | -0.4791444 | 0.63183593 | 0.88123095 |
| TMEM56-RW | 49.3374301 | -0.254921468 | 0.46469508 | -0.5485779 | 0.58329513 | 0.85996253 |
| AC092802.1 | 118.497886 | 0.367674439 | 0.52659217 | 0.69821479 | 0.48504288 | 0.81839266 |
| PTBP2 | 2364.79408 | -0.108295316 | 0.21151394 | -0.5120009 | 0.60865041 | 0.86985416 |
| DPYD | 1868.90412 | 0.7110834 | 0.37985554 | 1.87198376 | 0.06120885 | 0.4813972 |
| DPYD-AS1 | 69.4745506 | 0.84473357 | 0.69400812 | 1.2171811 | 0.22353532 | 0.66449764 |
| SNX7 | 714.264119 | 0.080741487 | 0.21505639 | 0.37544333 | 0.70733078 | 0.91002281 |
| LINC01708 | 36.2060125 | -1.280270267 | 0.73127664 | -1.7507332 | 8.00E-02 | 0.51264427 |
| FRRS1 | 190.831351 | 0.475661487 | 0.55833701 | 0.85192541 | 0.39425549 | 0.77823445 |
| AGL | 677.717641 | 0.265991261 | 0.20600364 | 1.29119692 | 0.1966354 | 0.64558745 |
| AC093019.2 | 42.8512851 | 0.473691733 | 0.62721401 | 0.75523143 | 0.45011013 | 0.80344571 |
| SASS6 | 109.855945 | -0.682204409 | 0.22783489 | -2.994293 | 0.00275082 | 0.16989777 |
| TRMT13 | 443.99103 | -0.506471397 | 0.14950132 | -3.3877386 | 0.00070471 | 0.08943993 |
| LRRC39 | 64.5220094 | -0.058921307 | 0.3870475 | -0.1522328 | 0.87900336 | 0.9674841 |
| DBT | 500.595339 | 0.294622027 | 0.32231167 | 0.91409049 | 0.3606693 | 0.76009788 |
| RTCA | 524.77668 | 0.082908448 | 0.30998733 | 0.26745754 | 0.78911691 | 0.94005579 |
| CDC14A | 677.494108 | 0.246966444 | 0.37049144 | 0.6665915 | 0.5050331 | 0.82715271 |
| LINC01349 | 254.199806 | 1.813146811 | 0.62996657 | 2.87816354 | 4.00E-03 | 0.20256898 |
| VCAM1 | 2641.79014 | 2.355639329 | 0.86460234 | 2.724535 | 0.00643921 | 0.22806785 |
| EXTL2 | 327.328774 | -0.48997245 | 0.29088485 | -1.6844207 | 0.0921004 | 0.53619162 |
| SLC30A7 | 873.795934 | -0.16925245 | 0.27050146 | -0.6256988 | 0.5315125 | 0.83789294 |
| DPH5 | 252.886336 | 0.240934885 | 0.29116245 | 0.82749299 | 0.40795769 | 0.78481094 |
| AC093157.1 | 280.945921 | 0.441201711 | 0.32502401 | 1.35744346 | 0.17464035 | 0.62699479 |
| RNPC3 | 962.114711 | 0.109238999 | 0.22731711 | 0.48055776 | 0.63083084 | 0.88115748 |
| AMY2B | 209.531064 | 0.801929606 | 0.45174131 | 1.77519653 | 0.07586544 | 0.50730849 |
| VAV3 | 2961.23021 | 0.587579621 | 0.52811918 | 1.11258905 | 0.26588497 | 0.69764265 |
| SLC25A24 | 578.761588 | -0.401191101 | 0.24593303 | -1.6313022 | 0.10282656 | 0.55022042 |
| FAM102B | 117.447085 | 0.629532169 | 0.35535642 | 1.77155139 | 0.07646905 | 0.50833864 |
| PRPF38B | 794.933367 | -0.38108895 | 0.42253916 | -0.9019021 | 0.36710889 | 0.76546934 |
| STXBP3 | 694.24714 | 0.050212745 | 0.22037026 | 0.22785627 | 0.81975798 | 0.94990455 |
| AL449266.1 | 172.467499 | -0.029781493 | 0.29966739 | -0.0993818 | 0.92083511 | 0.97939204 |
| GPSM2 | 155.315495 | 0.689193012 | 0.50526663 | 1.36401845 | 0.17256176 | 0.62593809 |
| WDR47 | 302.599032 | -0.155158462 | 0.2844888 | -0.5453939 | 0.58548263 | 0.86077718 |
| TAF13 | 242.418573 | -0.865962495 | 0.46522056 | -1.861402 | 0.06268742 | 0.48421747 |
| AL356488.3 | 33.7633969 | -0.470273622 | 0.62646127 | -0.7506827 | 0.45284365 | 0.80433814 |
| TMEM167B | 165.050576 | -0.266603881 | 0.23133491 | -1.1524585 | 0.24913271 | 0.68557761 |
| KIAA1324 | 43.1885848 | 0.608527977 | 0.48943109 | 1.24333739 | 2.14E-01 | 0.65849767 |
| SARS | 419.852218 | -0.360645763 | 0.24432721 | -1.4760769 | 0.13992324 | 0.59500405 |
| CELSR2 | 60.4937478 | 1.02547436 | 0.5756137 | 1.78153224 | 0.07482554 | 0.5071704 |
| SORT1 | 730.781208 | 0.33973196 | 0.39744883 | 0.85478164 | 0.39267204 | 0.77703588 |
| PSMA5 | 949.01968 | -0.885308452 | 0.45719805 | -1.9363785 | 0.05282137 | 0.46564998 |

| | | | | | | |
|---|---|---|---|---|---|---|
| SYPL2 | 486.189949 | -0.619166969 | 0.51054976 | -1.2127456 | 0.22522711 | 0.66649135 |
| ATXN7L2 | 73.6542635 | -0.365509805 | 0.5241942 | -0.6972794 | 0.48562797 | 0.81839266 |
| CYB561D1 | 104.109539 | -0.070900407 | 0.44176842 | -0.1604922 | 0.87249333 | 0.96578281 |
| GNAI3 | 906.139675 | -0.207807965 | 0.33751726 | -0.6156958 | 0.5380953 | 0.84049498 |
| AMPD2 | 159.450547 | -0.384942701 | 0.47839176 | -0.80466 | 0.42101592 | 0.78872573 |
| GSTM4 | 124.917348 | 0.984557252 | 0.50712247 | 1.94145854 | 0.05220268 | 0.46463778 |
| GSTM3 | 92.0635042 | -0.452070509 | 0.55047647 | -0.8212349 | 0.41151246 | 0.78632759 |
| CSF1 | 136.049431 | -0.352265765 | 0.60062166 | -0.5865019 | 5.58E-01 | 8.49E-01 |
| AHCYL1 | 2884.39594 | 0.513583133 | 0.42737464 | 1.20171644 | 0.22947341 | 0.66705629 |
| STRIP1 | 293.485339 | -0.074884202 | 0.31610296 | -0.2368981 | 0.81273582 | 0.94801888 |
| KCNC4 | 44.5943306 | 0.713282179 | 0.51685012 | 1.38005615 | 0.16756936 | 0.62296534 |
| RBM15-AS1 | 54.8738691 | 0.483463791 | 0.63887246 | 0.7567454 | 0.44920241 | 0.8030804 |
| RBM15 | 337.544251 | -0.306014694 | 0.29136161 | -1.0502917 | 0.293584 | 0.71502415 |
| SLC16A4 | 2064.43858 | 0.284730684 | 0.38728122 | 0.73520394 | 0.4622153 | 0.8082072 |
| AL355488.1 | 78.7915547 | -0.242402477 | 0.30440229 | -0.7963228 | 0.42584445 | 0.78894887 |
| LAMTOR5 | 932.771473 | -0.266419126 | 0.2894004 | -0.92059 | 0.35726451 | 0.75903693 |
| LRIF1 | 205.144773 | 0.027225759 | 0.2798079 | 0.09730161 | 0.92248688 | 0.97974025 |
| DRAM2 | 527.3859 | 0.414309956 | 0.28119611 | 1.47338439 | 0.14064742 | 0.59524859 |
| CEPT1 | 1157.43388 | 0.298825111 | 0.29136134 | 1.02561688 | 0.30507221 | 0.72103276 |
| DENND2D | 468.891559 | 1.047766024 | 0.53529312 | 1.95736873 | 0.05030413 | 0.45824658 |
| CHI3L2 | 100.22535 | 1.425126903 | 0.7974765 | 1.78704563 | 7.39E-02 | 5.06E-01 |
| OVGP1 | 101.45782 | 1.150446176 | 0.53192185 | 2.16281054 | 0.03055575 | 0.40446361 |
| WDR77 | 218.162366 | -0.032836842 | 0.32891942 | -0.0998325 | 0.92047732 | 0.97939204 |
| ATP5PB | 598.037895 | 0.463218141 | 0.2690163 | 1.72189617 | 0.08508833 | 0.52183702 |
| RAP1A | 1430.73862 | -0.078901772 | 0.20145741 | -0.3916548 | 0.69531326 | 0.90589033 |
| INKA2 | 81.4325611 | 0.355206234 | 0.54299426 | 0.65416205 | 0.51300741 | 0.83073024 |
| DDX20 | 117.215377 | -0.271018452 | 0.38631217 | -0.701553 | 0.48295799 | 0.8177376 |
| KCND3 | 143.070086 | 1.593753907 | 0.5615092 | 2.83833982 | 0.00453489 | 0.20749797 |
| AL445426.1 | 37.8423947 | 0.187641457 | 0.72426551 | 0.25907827 | 0.79557485 | 0.94252227 |
| CTTNBP2NL | 1351.07195 | 0.447755359 | 0.31098294 | 1.4398068 | 0.14992207 | 0.60758225 |
| WNT2B | 591.769593 | 0.821467641 | 0.50674701 | 1.62106067 | 0.10500463 | 0.55219437 |
| ST7L | 722.666041 | 0.160405324 | 0.19811974 | 0.80963826 | 0.41814811 | 0.78697737 |
| CAPZA1 | 1188.35584 | -0.132945209 | 0.18703852 | -0.7107905 | 0.47721405 | 0.81570932 |
| MOV10 | 501.896749 | -0.197761572 | 0.40947973 | -0.4829582 | 0.62912545 | 0.88057005 |
| AL603832.1 | 95.1174404 | -0.104303471 | 0.62445395 | -0.1670315 | 0.86734527 | 0.96486219 |
| RHOC | 748.07572 | -0.201971365 | 0.50142421 | -0.4027954 | 0.68709875 | 0.90237257 |
| SLC16A1 | 298.506254 | -0.885484591 | 0.61039266 | -1.4506803 | 0.14686891 | 0.60383734 |
| SLC16A1-AS1 | 499.947993 | 0.308257386 | 0.40347854 | 0.76399945 | 0.44486757 | 0.80041177 |
| AL357055.3 | 92.1075152 | -0.040659922 | 0.64187942 | -0.0633451 | 0.9494917 | 0.9871942 |
| LRIG2 | 734.238755 | -0.302885437 | 0.37219952 | -0.8137717 | 0.41577574 | 0.78632759 |
| MAGI3 | 3240.82421 | -0.047257398 | 0.33654311 | -0.14042 | 0.88832812 | 0.97134248 |
| PHTF1 | 405.042207 | 0.011732127 | 0.35658321 | 0.03290151 | 0.97375313 | 0.99361528 |
| RSBN1 | 444.405478 | -0.02692766 | 0.15604066 | -0.1725682 | 0.86299083 | 0.96393376 |
| AP4B1 | 68.1652171 | -0.646276009 | 0.40352949 | -1.6015583 | 0.10925332 | 0.55946616 |
| HIPK1 | 555.166653 | 0.287063545 | 0.31896627 | 0.89998089 | 0.36813042 | 0.76615653 |
| SYT6 | 185.49192 | 0.501357349 | 0.67054394 | 0.74768754 | 0.45464865 | 0.80465616 |
| AL121999.1 | 622.382957 | -0.733817769 | 0.77372208 | -0.9484255 | 3.43E-01 | 0.7505827 |
| TRIM33 | 2202.90892 | -0.565466042 | 0.24971185 | -2.2644742 | 0.02354496 | 0.37138185 |
| BCAS2 | 208.99617 | -0.479528582 | 0.22614387 | -2.120458 | 0.03396744 | 0.41378378 |
| DENND2C | 116.441083 | -1.461055139 | 0.44670443 | -3.2707424 | 0.00107266 | 0.11082848 |
| NRAS | 228.75009 | -0.77451079 | 0.38088728 | -2.0334383 | 0.04200826 | 0.4368795 |
| CSDE1 | 1271.97525 | -0.011824783 | 0.17381679 | -0.0680302 | 0.94576163 | 0.98584713 |
| SIKE1 | 309.811587 | 0.015212684 | 0.32957764 | 0.04615812 | 0.96318422 | 0.99012655 |
| VANGL1 | 227.027048 | 0.174747425 | 0.3054594 | 0.57208069 | 0.56726731 | 0.85239988 |

| | | | | | | |
|---|---|---|---|---|---|---|
| SLC22A15 | 121.943929 | 1.259451963 | 0.71012046 | 1.77357508 | 0.07613346 | 0.50767193 |
| ATP1A1 | 5195.73519 | 0.46489113 | 0.22594624 | 2.05752982 | 3.96E-02 | 0.42845498 |
| LINC01762 | 791.859252 | 0.506664574 | 0.75312254 | 0.67275184 | 0.50110518 | 0.82605517 |
| AL390066.1 | 103.283869 | 0.371132294 | 0.36791152 | 1.00875419 | 0.31309253 | 0.72717675 |
| CD58 | 1148.18871 | 0.077391276 | 0.2500771 | 0.30946967 | 0.75696428 | 0.92799282 |
| IGSF3 | 835.928619 | 0.559576477 | 0.57245934 | 0.97749559 | 0.32832386 | 0.73826734 |
| PTGFRN | 122.111994 | 1.136274092 | 0.54126902 | 2.09927791 | 0.03579241 | 0.42040764 |
| TTF2 | 259.373537 | 0.23416083 | 0.22371655 | 1.04668533 | 0.29524473 | 0.71646902 |
| MAN1A2 | 1990.80207 | -0.359959145 | 0.25540862 | -1.4093461 | 0.15873284 | 0.61423843 |
| TENT5C | 121.092029 | 0.317702185 | 0.58071185 | 0.54709092 | 0.58431627 | 0.86025731 |
| GDAP2 | 768.776011 | -0.08511426 | 0.19096838 | -0.4456982 | 0.65581527 | 0.89053559 |
| WDR3 | 163.740946 | -0.406883149 | 0.37444253 | -1.0866371 | 0.27719722 | 0.7036882 |
| SPAG17 | 408.004982 | 0.647455571 | 0.69132009 | 0.93654962 | 0.34899028 | 0.75415932 |
| WARS2 | 438.502332 | -0.314267981 | 0.26878914 | -1.169199 | 0.24232348 | 0.68080966 |
| AL359915.2 | 1070.55326 | 0.185952763 | 0.41796564 | 0.44489964 | 0.65639228 | 0.89053559 |
| AL359915.1 | 345.969257 | 0.523806798 | 0.98831231 | 0.53000129 | 0.59611104 | 0.86539512 |
| LINC01780 | 80.5779907 | 0.637912696 | 1.02990306 | 0.61939101 | 0.5356588 | 0.83954482 |
| HAO2 | 960.243653 | 1.181145565 | 1.02300895 | 1.15457989 | 0.24826251 | 0.6848328 |
| ZNF697 | 729.067575 | -0.105091276 | 0.56619044 | -0.1856112 | 0.85274969 | 0.96073035 |
| PHGDH | 110.715221 | 0.208466522 | 0.56154343 | 0.37123846 | 0.71045993 | 0.91123308 |
| HMGCS2 | 66.6389811 | -0.143391801 | 1.31798077 | -0.1087966 | 0.91336384 | 0.97720308 |
| NOTCH2 | 688.290654 | 0.125447743 | 0.2761151 | 0.45433133 | 6.50E-01 | 8.89E-01 |
| SEC22B | 749.720667 | -0.494680382 | 0.2439533 | -2.0277667 | 0.04258407 | 0.43849941 |
| NBPF26 | 119.181524 | 0.01610167 | 0.68856164 | 0.0233845 | 0.98134357 | 0.9951377 |
| LINC00623 | 623.090176 | 0.636474662 | 0.50132635 | 1.26958152 | 0.20423374 | 0.64982169 |
| SRGAP2C | 64.4388036 | 0.988471207 | 0.4068859 | 2.42935724 | 0.01512562 | 0.31036958 |
| AC239800.3 | 53.9808278 | 0.828467362 | 0.69401527 | 1.19373073 | 0.23258331 | 0.66978724 |
| AC245595.1 | 43.8202746 | -0.676968292 | 0.49969367 | -1.3547666 | 0.17549193 | 0.62699479 |
| NBPF15 | 306.913239 | 0.563492641 | 0.51906556 | 1.08559051 | 0.2776602 | 0.7036882 |
| FP700111.1 | 1506.25798 | -0.088017108 | 0.28252138 | -0.3115414 | 0.75538907 | 0.92799282 |
| SRGAP2B | 119.914048 | 0.943317693 | 0.49367979 | 1.91078857 | 0.05603176 | 0.47114668 |
| NBPF20 | 43.2916966 | -0.337407463 | 0.47882975 | -0.7046502 | 0.48102797 | 0.81720798 |
| GPR89A | 416.040493 | -0.10674817 | 0.28616483 | -0.3730304 | 0.70912585 | 0.91086765 |
| PDZK1 | 5995.49376 | 0.403749157 | 0.61943087 | 0.65180664 | 0.51452592 | 0.83169608 |
| RNF115 | 1709.02961 | -0.728588603 | 0.27516259 | -2.6478476 | 0.0081006 | 0.24877162 |
| POLR3C | 277.990393 | -0.406555054 | 0.3081969 | -1.3191406 | 0.1871221 | 0.63818925 |
| PIAS3 | 64.2642165 | -0.093082253 | 0.34033365 | -0.2735029 | 0.78446663 | 0.9387711 |
| PEX11B | 101.339472 | 0.613884473 | 0.49146206 | 1.24909839 | 0.21162909 | 0.65584609 |
| RBM8A | 235.273096 | -0.459764314 | 0.23476818 | -1.9583757 | 0.05018594 | 0.45824658 |
| LIX1L | 105.307901 | 0.218410246 | 0.36737879 | 0.59450967 | 0.55217127 | 0.84721285 |
| TXNIP | 2748.02646 | 0.200027499 | 0.86724039 | 0.23064827 | 0.81758806 | 0.94954998 |
| NBPF12 | 597.531059 | 0.445525993 | 0.26058183 | 1.70973545 | 0.0873148 | 0.52706515 |
| PRKAB2 | 275.180153 | 0.114821022 | 0.2794412 | 0.41089511 | 0.68114945 | 0.89947163 |
| AC242426.2 | 494.519757 | 0.158829022 | 0.38066269 | 0.41724347 | 0.67650033 | 0.89843878 |
| FMO5 | 1203.28857 | 1.701850509 | 0.63230431 | 2.69150549 | 0.00711303 | 0.23330842 |
| CHD1L | 651.100285 | 0.62177917 | 0.23219928 | 2.67778249 | 0.00741113 | 0.23590813 |
| LINC00624 | 99.3902225 | 0.787351249 | 0.50459363 | 1.56036699 | 0.11867318 | 0.57589001 |
| BCL9 | 615.874431 | 0.125974601 | 0.2668827 | 0.47202235 | 0.63691083 | 0.88266168 |
| ACP6 | 439.433686 | 0.741164715 | 0.39435588 | 1.87943115 | 0.06018565 | 0.48004006 |
| GPR89B | 928.081787 | 0.403965243 | 0.36396588 | 1.10989867 | 2.67E-01 | 6.98E-01 |
| NBPF11 | 476.770658 | 0.294046598 | 0.28486517 | 1.03223079 | 0.30196402 | 0.71925676 |
| AC239809.3 | 164.098346 | 0.213223004 | 0.52675863 | 0.40478313 | 0.68563693 | 0.9012299 |
| NBPF14 | 214.871435 | 0.330564526 | 0.29141172 | 1.13435564 | 0.2566454 | 0.69102923 |
| PDE4DIP | 1016.5124 | 0.410945416 | 0.43417418 | 0.94649898 | 0.34389414 | 0.7507483 |

| | | | | | | |
|---|---|---|---|---|---|---|
| NBPF9 | 311.533944 | -0.220749091 | 0.2998036 | -0.7363123 | 0.46154064 | 0.80796137 |
| AC245297.3 | 1379.73988 | 0.225154161 | 0.38561941 | 0.58387663 | 0.55930332 | 0.84936251 |
| NBPF19 | 286.408268 | -0.255628026 | 0.7443242 | -0.3434364 | 0.73127018 | 0.9190831 |
| HIST2H2BE | 43.8846224 | -1.809570555 | 0.57308755 | -3.1575813 | 0.00159084 | 0.13207839 |
| BOLA1 | 56.0409246 | 0.239522922 | 0.47290238 | 0.50649549 | 0.61250886 | 0.87222135 |
| SF3B4 | 187.349801 | -0.429617786 | 0.38530197 | -1.1150158 | 0.26484364 | 0.69614236 |
| MTMR11 | 214.95422 | 0.545180747 | 0.50684365 | 1.0756389 | 0.28208878 | 0.70674299 |
| OTUD7B | 1619.86152 | -0.787090258 | 0.37391865 | -2.1049773 | 0.03529329 | 0.4198859 |
| VPS45 | 715.678598 | 0.078228454 | 0.14387997 | 0.54370636 | 0.58664356 | 0.86102615 |
| ANP32E | 225.601349 | 0.451475102 | 0.35904999 | 1.25741571 | 0.20860314 | 0.65401126 |
| APH1A | 268.233626 | -0.059466186 | 0.34386166 | -0.1729364 | 0.86270145 | 0.96391903 |
| C1orf54 | 128.941881 | 0.131155585 | 0.52090405 | 0.25178454 | 0.80120761 | 0.9444777 |
| CIART | 46.9645589 | -2.010067204 | 0.55101393 | -3.6479426 | 0.00026435 | 0.05940045 |
| MRPS21 | 646.494704 | -0.42874328 | 0.23446754 | -1.8285827 | 0.06746215 | 0.48963361 |
| PRPF3 | 526.281931 | -0.412636747 | 0.27333903 | -1.5096152 | 0.13114165 | 0.58930656 |
| RPRD2 | 3003.85019 | 0.04213453 | 0.24536196 | 0.17172397 | 0.86365454 | 0.96393376 |
| TARS2 | 151.289156 | 0.258601985 | 0.46755696 | 0.55309194 | 0.58020046 | 0.85831909 |
| ADAMTSL4 | 50.6253622 | 0.301295233 | 0.52425489 | 0.57471134 | 0.56548654 | 0.85194709 |
| MCL1 | 686.936169 | -0.328427972 | 0.31858774 | -1.030887 | 0.30259379 | 0.7202037 |
| ENSA | 1319.00386 | 0.233380787 | 0.28738768 | 0.81207652 | 0.41674773 | 0.78632759 |
| GOLPH3L | 562.753375 | 0.168640737 | 0.25321062 | 0.66600973 | 0.50540488 | 0.8272145 |
| CTSS | 48.7421593 | 0.159056609 | 0.63845924 | 0.24912571 | 0.80326355 | 0.94543168 |
| CTSK | 39.583229 | 0.555699941 | 0.42648317 | 1.30298211 | 0.19258087 | 0.64507333 |
| ARNT | 786.514977 | -0.011633155 | 0.1966815 | -0.0591472 | 0.95283488 | 0.98786081 |
| SETDB1 | 686.814838 | -0.033659678 | 0.17762191 | -0.1895018 | 0.84969952 | 0.95959925 |
| CERS2 | 721.545356 | -0.197141407 | 0.25425138 | -0.7753799 | 0.43811521 | 0.79696434 |
| ANXA9 | 72.2969556 | 0.454840802 | 0.49304936 | 0.92250562 | 0.3562649 | 0.75841231 |
| MINDY1 | 198.411152 | 0.207104232 | 0.49368449 | 0.41950727 | 0.67484544 | 0.89801183 |
| PRUNE1 | 289.845158 | -0.077589631 | 0.39931146 | -0.1943086 | 0.84593429 | 0.95784082 |
| BNIPL | 40.0978333 | 0.25806859 | 0.52721413 | 0.48949483 | 0.62449141 | 0.87829918 |
| C1orf56 | 87.1330891 | -0.045200786 | 0.52207604 | -0.0865789 | 0.93100621 | 0.98272895 |
| CDC42SE1 | 245.624475 | -0.062996039 | 0.23178909 | -0.2717817 | 0.78578986 | 0.938961 |
| GABPB2 | 415.535516 | 0.537206801 | 0.35704761 | 1.50458032 | 0.13243201 | 0.5912647 |
| SEMA6C | 170.644003 | 0.876842882 | 0.55136179 | 1.59032219 | 1.12E-01 | 0.56445264 |
| SCNM1 | 139.792694 | -0.414411 | 0.25024166 | -1.6560432 | 0.09771308 | 0.54093423 |
| LYSMD1 | 55.3838912 | -0.410326089 | 0.32597414 | -1.2587688 | 0.20811384 | 0.65352038 |
| VPS72 | 132.823935 | -0.19420217 | 0.25150911 | -0.7721477 | 0.44002697 | 0.79826598 |
| PIP5K1A | 1949.32706 | -0.298401485 | 0.24554328 | -1.2152704 | 0.22426296 | 0.66551422 |
| PSMD4 | 496.86117 | -0.212166446 | 0.17518103 | -1.2111268 | 0.22584681 | 0.66688879 |
| ZNF687 | 63.093206 | 0.293052923 | 0.38714054 | 0.7569678 | 0.44906916 | 0.80300352 |
| PI4KB | 560.97418 | -0.075148919 | 0.24757095 | -0.303545 | 0.76147458 | 0.93000994 |
| RFX5 | 51.6026537 | 0.516262418 | 0.60929571 | 0.84731011 | 0.39682229 | 0.78014596 |
| SELENBP1 | 194.168681 | 1.010241558 | 0.68942851 | 1.46533185 | 0.1428304 | 0.59824924 |
| PSMB4 | 253.183039 | -0.57371399 | 0.46943516 | -1.2221368 | 0.2216559 | 0.66293637 |
| POGZ | 2108.83886 | 0.059257789 | 0.30868901 | 0.19196599 | 0.84776885 | 0.95908381 |
| CGN | 550.852361 | -0.022747023 | 0.61585703 | -0.0369356 | 0.97053639 | 0.99205112 |
| TUFT1 | 583.671164 | -0.285952383 | 0.41442148 | -0.6900038 | 0.49019182 | 0.81989696 |
| SNX27 | 1153.76653 | -0.173535122 | 0.37966022 | -0.4570801 | 0.64761349 | 0.88774616 |
| MRPL9 | 128.831969 | -0.087659805 | 0.34430965 | -0.2545958 | 0.79903527 | 0.94343841 |
| TDRKH | 55.3094908 | 0.412082792 | 0.46825344 | 0.8800422 | 0.37883645 | 0.77061593 |
| RORC | 405.008342 | 0.439780953 | 0.71271487 | 0.61705034 | 0.53720152 | 0.84028543 |
| AL450992.2 | 43.991019 | -0.159771773 | 0.30290126 | -0.5274715 | 0.59786622 | 0.86636561 |
| THEM4 | 489.88728 | 0.080923955 | 0.26844686 | 0.30145242 | 0.76306952 | 0.93023365 |
| S100A10 | 1391.94551 | -0.613827284 | 0.59593616 | -1.0300219 | 3.03E-01 | 7.20E-01 |

| | | | | | | |
|---|---|---|---|---|---|---|
| S100A11 | 2343.0759 | -0.570685316 | 0.52319997 | -1.0907595 | 2.75E-01 | 7.02E-01 |
| FLG-AS1 | 601.86541 | 0.653774466 | 0.51335672 | 1.2735286 | 0.20283053 | 0.64881821 |
| S100A6 | 1772.16522 | -0.270235361 | 0.83106978 | -0.3251657 | 0.74505569 | 0.92389439 |
| S100A4 | 50.4720323 | -0.769379266 | 0.86100415 | -0.8935837 | 0.37154468 | 0.76820041 |
| S100A2 | 152.454395 | -1.687552645 | 0.98553783 | -1.7123165 | 0.08683837 | 0.52575431 |
| S100A16 | 448.204917 | -0.692459033 | 0.58553701 | -1.1826051 | 2.37E-01 | 0.67429706 |
| S100A14 | 76.2942827 | -0.788203914 | 0.7191228 | -1.096063 | 0.2730512 | 0.70123486 |
| S100A13 | 451.981492 | 0.213397629 | 0.32146443 | 0.66382968 | 0.50679933 | 0.82806179 |
| S100A1 | 503.896455 | 0.218485189 | 0.6610904 | 0.33049215 | 0.74102812 | 0.92213095 |
| AL162258.2 | 40.5904501 | -0.691993284 | 0.41682066 | -1.6601703 | 0.0968802 | 0.54003127 |
| CHTOP | 449.332052 | -0.483342501 | 0.23805764 | -2.0303591 | 0.04232005 | 0.43742111 |
| SNAPIN | 61.293505 | -0.090018517 | 0.46020016 | -0.1956073 | 0.84491753 | 0.95760132 |
| ILF2 | 413.854297 | 0.038332443 | 0.34991426 | 0.1095481 | 0.91276777 | 0.977179 |
| NPR1 | 274.663503 | 0.505092418 | 0.57240542 | 0.88240327 | 3.78E-01 | 0.77055888 |
| INTS3 | 834.880989 | -0.11621527 | 0.243245 | -0.4777704 | 0.63281361 | 0.88151688 |
| SLC27A3 | 98.5622501 | -0.071357647 | 0.437356 | -0.1631569 | 0.87039491 | 0.96564168 |
| GATAD2B | 2338.63479 | -0.345171958 | 0.28713863 | -1.2021091 | 0.22932126 | 0.66705629 |
| DENND4B | 282.310738 | -0.013302643 | 0.25530386 | -0.0521051 | 0.95844492 | 0.98878187 |
| CRTC2 | 166.537569 | -0.696822607 | 0.31307873 | -2.2257105 | 0.02603358 | 0.38392873 |
| SLC39A1 | 341.353767 | -0.906779221 | 0.55344988 | -1.6384125 | 0.10133566 | 0.54663831 |
| CREB3L4 | 38.3119058 | 0.166674831 | 0.45890359 | 0.36320229 | 0.71645378 | 0.91298968 |
| JTB | 322.555366 | 0.039467165 | 0.38010939 | 0.10383107 | 0.91730341 | 0.97828877 |
| RAB13 | 405.460785 | 0.030710944 | 0.3588871 | 0.08557272 | 9.32E-01 | 9.83E-01 |
| RPS27 | 385.258839 | -0.238852369 | 0.45667041 | -0.5230301 | 0.60095331 | 0.86755264 |
| NUP210L | 61.4518182 | 0.097047345 | 0.54266398 | 0.17883506 | 0.85806722 | 0.9626966 |
| TPM3 | 711.519953 | -0.440557683 | 0.37781768 | -1.1660589 | 0.24359062 | 0.68164834 |
| C1orf43 | 626.265702 | -0.141476689 | 0.30210651 | -0.4683007 | 0.63956957 | 0.88321992 |
| UBAP2L | 802.798748 | -0.367593225 | 0.24018561 | -1.5304548 | 0.1259042 | 0.58564339 |
| HAX1 | 130.008442 | 0.205991656 | 0.32974128 | 0.62470691 | 0.53216344 | 0.83804736 |
| IL6R | 1965.51257 | -1.091900455 | 0.45822043 | -2.3829153 | 0.01717615 | 0.32964126 |
| UBE2Q1 | 191.46344 | -0.635798624 | 0.3393903 | -1.8733553 | 0.06101934 | 0.48123823 |
| ADAR | 1050.77668 | -0.491759176 | 0.35469047 | -1.386446 | 0.16561077 | 0.62065658 |
| PMVK | 247.885023 | 0.361910544 | 0.3404002 | 1.06319132 | 0.2876952 | 0.7113446 |
| PBXIP1 | 114.550998 | 0.546374597 | 0.44666291 | 1.22323701 | 0.22124019 | 0.66263062 |
| PYGO2 | 87.2108963 | -0.13910851 | 0.29472469 | -0.4719948 | 0.63693052 | 0.88266168 |
| SHC1 | 52.2514846 | -0.241897584 | 0.35695465 | -0.6776704 | 0.49798072 | 0.82326489 |
| CKS1B | 50.4785552 | 0.390602076 | 0.52106057 | 0.74962893 | 0.45347822 | 0.80433814 |
| FLAD1 | 125.650885 | -0.575979884 | 0.37775339 | -1.5247511 | 0.1273212 | 0.58793678 |
| ZBTB7B | 189.419589 | 0.374484966 | 0.29029961 | 1.28999472 | 0.19705249 | 0.64558745 |
| ADAM15 | 84.4568997 | 0.217243312 | 0.49369244 | 0.44003776 | 0.65990976 | 0.89094706 |
| EFNA1 | 1188.62686 | -0.669352939 | 0.55713465 | -1.2014204 | 0.22958816 | 0.66705629 |
| SLC50A1 | 140.52751 | -0.175498129 | 0.32053835 | -0.5475106 | 5.84E-01 | 0.86025731 |
| DPM3 | 211.667458 | -0.405202269 | 0.3822633 | -1.0600083 | 0.28914082 | 0.71218341 |
| THBS3 | 206.882359 | -0.184295487 | 0.47755018 | -0.3859186 | 0.69955697 | 0.90721166 |
| MTX1 | 120.542661 | -0.121755556 | 0.52029406 | -0.234013 | 0.81497491 | 0.94895023 |
| GBA | 224.963813 | -0.351063321 | 0.26993594 | -1.3005431 | 0.19341489 | 0.64548758 |
| FAM189B | 74.0287283 | 0.517061759 | 0.3726394 | 1.38756601 | 0.16526924 | 0.62065658 |
| SCAMP3 | 298.822643 | -0.40359454 | 0.39297803 | -1.0270155 | 0.30441315 | 0.72082426 |
| CLK2 | 331.952819 | -0.079206871 | 0.24118408 | -0.3284084 | 0.74260292 | 0.92308499 |
| HCN3 | 118.069119 | 0.592308674 | 0.3552066 | 1.6675047 | 0.09541409 | 0.53824794 |
| PKLR | 196.527223 | 2.647030473 | 0.72094532 | 3.6716106 | 0.00024103 | 0.05632519 |
| FDPS | 915.954016 | 1.30073686 | 0.27636938 | 4.70651584 | 2.52E-06 | 0.00863494 |
| RUSC1-AS1 | 43.6532536 | -0.022114268 | 0.49689312 | -0.0445051 | 0.9645018 | 0.99031076 |
| RUSC1 | 142.79788 | 0.028193733 | 0.36060431 | 0.07818468 | 0.93768115 | 0.98435387 |

| | | | | | | |
|---|---|---|---|---|---|---|
| ASH1L | 5566.71047 | -0.01950996 | 0.17069476 | -0.1142974 | 0.90900208 | 0.97680747 |
| MSTO1 | 119.905576 | -1.112314274 | 0.44713358 | -2.4876554 | 0.01285883 | 0.29875272 |
| YY1AP1 | 811.394966 | 0.04741878 | 0.20945694 | 0.22638916 | 0.82089876 | 0.95072145 |
| DAP3 | 919.391008 | -0.218125804 | 0.31893379 | -0.6839219 | 0.49402449 | 0.82179535 |
| GON4L | 1863.82906 | -0.082453904 | 0.23628125 | -0.3489651 | 0.72711553 | 0.91645085 |
| RIT1 | 584.859779 | -0.387921539 | 0.30615425 | -1.2670787 | 0.20512714 | 0.65140961 |
| KHDC4 | 995.724033 | -0.226844863 | 0.25600588 | -0.8860924 | 0.37556775 | 0.7697436 |
| ARHGEF2 | 168.322503 | 0.043419871 | 0.49973088 | 0.08688651 | 0.93076172 | 0.98272895 |
| SSR2 | 423.863603 | -0.206404771 | 0.33320125 | -0.6194598 | 0.53561352 | 0.83954482 |
| UBQLN4 | 252.322803 | 0.003360737 | 0.24418604 | 0.01376302 | 0.98901905 | 0.99721497 |
| LAMTOR2 | 161.248564 | -0.421877009 | 0.3725166 | -1.1325053 | 0.25742208 | 0.691812 |
| RAB25 | 61.3915291 | -0.013753392 | 0.73990612 | -0.018588 | 0.98516976 | 0.99570284 |
| MEX3A | 60.5664447 | 0.571784063 | 0.52271468 | 1.09387412 | 0.27401019 | 0.7014484 |
| LMNA | 815.543738 | -0.391687127 | 0.34970735 | -1.1200426 | 0.26269561 | 0.69467275 |
| SLC25A44 | 433.815819 | 0.070667534 | 0.34826071 | 0.20291561 | 8.39E-01 | 9.56E-01 |
| PAQR6 | 59.3688213 | 0.055826706 | 0.75663331 | 0.07378304 | 0.94118302 | 0.98489601 |
| SMG5 | 856.437205 | 0.041751382 | 0.1722347 | 0.24240982 | 8.08E-01 | 0.94625542 |
| TMEM79 | 58.7168895 | 0.138915262 | 0.51771563 | 0.26832348 | 0.78845033 | 0.93976423 |
| GLMP | 224.351725 | -0.203590994 | 0.31343571 | -0.6495463 | 0.51598535 | 0.83210529 |
| CCT3 | 703.3649 | -0.084407892 | 0.32643775 | -0.2585727 | 0.79596494 | 0.94252227 |
| RHBG | 33.526269 | 2.211423249 | 0.84656495 | 2.61223104 | 0.00899534 | 0.2563089 |
| MEF2D | 261.045869 | -0.025853814 | 0.32832855 | -0.0787437 | 0.93723646 | 0.98435387 |
| NAXE | 158.485441 | -0.020037549 | 0.4373018 | -0.0458209 | 0.96345302 | 0.99012655 |
| GPATCH4 | 129.175567 | -0.917560225 | 0.540711 | -1.6969513 | 0.08970587 | 0.5318332 |
| ISG20L2 | 147.601819 | -0.814155945 | 0.51882364 | -1.5692345 | 0.11659331 | 0.57350419 |
| RRNAD1 | 200.079193 | -0.043888947 | 0.35484809 | -0.1236838 | 0.90156567 | 0.97491785 |
| MRPL24 | 128.941295 | -0.007464384 | 0.28655374 | -0.0260488 | 0.97921841 | 0.99482978 |
| HDGF | 737.203458 | -0.268801967 | 0.35860554 | -0.7495756 | 0.45351034 | 0.80433814 |
| PRCC | 398.228393 | -0.371924367 | 0.31808185 | -1.1692725 | 0.24229385 | 0.68080966 |
| ARHGEF11 | 939.211336 | -0.020027721 | 0.25522319 | -0.0784714 | 0.93745308 | 0.98435387 |
| ETV3 | 112.87517 | -0.061722234 | 0.32579505 | -0.1894511 | 0.84973927 | 0.95959925 |
| KIRREL1 | 211.860597 | -0.121427263 | 0.61599992 | -0.1971222 | 0.84373191 | 0.95731603 |
| IFI16 | 201.719627 | 0.849529585 | 0.76082792 | 1.11658572 | 2.64E-01 | 0.69567438 |
| DUSP23 | 1398.98015 | -0.279023876 | 0.71138251 | -0.3922276 | 0.69489003 | 0.90589033 |
| TAGLN2 | 446.403978 | -0.502076327 | 0.61484283 | -0.816593 | 0.41416108 | 0.78632759 |
| IGSF8 | 122.273493 | 0.047452662 | 0.39426497 | 0.12035729 | 0.90420013 | 0.9756941 |
| PEA15 | 157.048185 | -1.493105143 | 0.75951436 | -1.9658682 | 0.04931383 | 0.45419182 |
| DCAF8 | 292.395268 | -0.142627526 | 0.31421184 | -0.4539215 | 0.64988529 | 0.88857633 |
| AL139011.2 | 99.9780342 | 0.824978122 | 0.43984322 | 1.8756186 | 0.06070767 | 0.48026727 |
| COPA | 1407.88292 | -0.492235683 | 0.22652907 | -2.172947 | 0.0297843 | 0.39981555 |
| NCSTN | 229.98766 | -0.080052303 | 0.25579135 | -0.3129594 | 0.75431152 | 0.92799282 |
| F11R | 378.89746 | -0.135538272 | 0.36748528 | -0.3688264 | 0.71225713 | 0.91165921 |
| AL591806.3 | 163.052141 | 0.772657611 | 0.35636565 | 2.16815962 | 0.03014654 | 0.40235509 |
| USF1 | 126.984504 | -0.912738306 | 0.34822385 | -2.6211252 | 0.00876401 | 0.25397092 |
| KLHDC9 | 36.632511 | 0.553756203 | 0.68668188 | 0.80642321 | 0.41999887 | 0.78798479 |
| PFDN2 | 450.26642 | -0.66932807 | 0.35379276 | -1.8918648 | 0.05850899 | 0.47778423 |
| NIT1 | 434.64129 | 0.533228917 | 0.28155559 | 1.89386726 | 0.05824263 | 0.47775688 |
| DEDD | 188.118704 | -0.122454044 | 0.21938308 | -0.5581745 | 0.57672522 | 0.85615431 |
| UFC1 | 194.719062 | -0.0768419 | 0.31266812 | -0.2457619 | 0.80586658 | 0.94585363 |
| USP21 | 54.3006501 | 0.322088939 | 0.38047227 | 0.84655036 | 0.39724579 | 0.78031644 |
| PPOX | 90.9121221 | 0.412443073 | 0.34566147 | 1.19319946 | 0.23279127 | 0.66978724 |
| B4GALT3 | 115.126391 | 0.001045866 | 0.33901291 | 0.00308503 | 0.9975385 | 0.99972657 |
| NDUFS2 | 504.866348 | 0.094022321 | 0.3143863 | 0.29906622 | 0.76488952 | 0.93083208 |
| TOMM40L | 59.6259414 | 0.097733622 | 0.4049881 | 0.24132468 | 0.80930349 | 0.94673968 |

| | | | | | | |
|---|---|---|---|---|---|---|
| NR1I3 | 163.492312 | 1.015077422 | 0.98782622 | 1.02758704 | 0.30414412 | 0.72082426 |
| SDHC | 1104.37419 | 0.432458828 | 0.27858725 | 1.55232811 | 0.12058373 | 0.57856254 |
| AL592295.4 | 141.953867 | 1.093651873 | 0.42772682 | 2.55689336 | 0.01056116 | 0.27159813 |
| AL592295.3 | 40.5227463 | 1.924447348 | 0.66732371 | 2.88382884 | 0.00392872 | 0.20256898 |
| DUSP12 | 153.675603 | -0.325304222 | 0.28184754 | -1.1541851 | 0.24842431 | 0.68514124 |
| ATF6 | 1437.13338 | -0.102512857 | 0.15837736 | -0.6472697 | 0.51745743 | 0.83297581 |
| NOS1AP | 629.099 | -0.405560844 | 0.53625381 | -0.7562852 | 0.44947819 | 0.80311564 |
| UHMK1 | 581.00285 | -0.108213537 | 0.18469514 | -0.5859035 | 0.55794034 | 0.84889425 |
| UAP1 | 213.255338 | -0.168201579 | 0.33432329 | -0.5031106 | 0.61488654 | 0.87260075 |
| DDR2 | 227.114087 | -0.612525078 | 0.76800094 | -0.7975577 | 0.4251272 | 0.78894887 |
| HSD17B7 | 250.574579 | 1.122109361 | 0.36134564 | 3.10536293 | 0.00190046 | 0.13753904 |
| RGS5 | 98.1928975 | 0.365715783 | 0.52965698 | 0.69047666 | 0.48989448 | 0.81970016 |
| NUF2 | 56.8657807 | 0.573017703 | 0.70409371 | 0.81383727 | 0.41573818 | 0.78632759 |
| PBX1 | 5440.92206 | 0.088329132 | 0.29097945 | 0.30355798 | 7.61E-01 | 9.30E-01 |
| MGST3 | 2357.54235 | -0.265013262 | 0.26876485 | -0.9860414 | 0.32411281 | 0.73603979 |
| ALDH9A1 | 417.807707 | 0.272893504 | 0.30897332 | 0.88322677 | 0.37711376 | 0.77047225 |
| TMCO1 | 587.135396 | -0.147199962 | 0.18431264 | -0.7986428 | 0.42449755 | 0.78894887 |
| AL451074.2 | 49.4120077 | 0.577744071 | 0.57885557 | 0.99807984 | 0.31824064 | 0.73030714 |
| UCK2 | 364.376775 | -0.515768285 | 0.49989469 | -1.0317539 | 0.30218743 | 0.71961138 |
| FAM78B | 197.59303 | 0.110571004 | 0.43986668 | 0.2513739 | 0.80152504 | 0.94475052 |
| POGK | 216.580987 | -0.293526598 | 0.39490556 | -0.743283 | 0.45731035 | 0.80559735 |
| TADA1 | 219.008456 | 0.213738079 | 0.2853159 | 0.74912783 | 0.45378016 | 0.80433814 |
| ILDR2 | 321.149742 | 0.082144675 | 0.33325042 | 0.24649534 | 0.80529881 | 0.94585363 |
| MAEL | 76.8991702 | 0.527257797 | 0.91575449 | 0.57576327 | 0.5647752 | 0.85167372 |
| POU2F1 | 2335.83814 | -0.554795902 | 0.28149852 | -1.9708661 | 0.04873919 | 0.4536808 |
| CREG1 | 289.854536 | -0.41891288 | 0.42347213 | -0.9892337 | 0.32254884 | 0.73441476 |
| MPZL1 | 908.429204 | 0.124563312 | 0.31409385 | 0.39657991 | 6.92E-01 | 9.05E-01 |
| ADCY10 | 132.129861 | 1.474216739 | 0.53902947 | 2.73494644 | 0.00623904 | 0.22644975 |
| MPC2 | 543.785329 | 0.299611869 | 0.37600541 | 0.79682862 | 0.42555057 | 0.78894887 |
| DCAF6 | 2773.23775 | 0.141604735 | 0.28080763 | 0.50427667 | 0.61406697 | 0.87225883 |
| TIPRL | 346.294664 | 0.056467444 | 0.21935635 | 0.25742333 | 0.79685199 | 0.94280281 |
| SFT2D2 | 1256.79019 | -0.21438379 | 0.22620938 | -0.9477228 | 3.43E-01 | 7.51E-01 |
| TBX19 | 262.908021 | 0.710547023 | 0.33759918 | 2.104706 | 0.03531691 | 0.4198859 |
| AL023755.1 | 103.087748 | 1.707798242 | 0.71517985 | 2.38792837 | 0.01694364 | 0.32896109 |
| LINC00970 | 384.774825 | 0.577656798 | 0.75227051 | 0.76788442 | 0.44255586 | 0.79933184 |
| ATP1B1 | 4335.04536 | 0.197903394 | 0.35369779 | 0.5595268 | 0.57580225 | 0.85537243 |
| NME7 | 1670.52835 | -0.582246219 | 0.31286028 | -1.8610423 | 0.0627382 | 0.48421747 |
| BLZF1 | 494.503306 | -0.539454911 | 0.30720768 | -1.7559942 | 0.07908939 | 0.51192014 |
| CCDC181 | 79.9407568 | 0.997093614 | 0.34722504 | 2.8716063 | 0.00408391 | 0.20352497 |
| SLC19A2 | 570.028884 | -0.35628901 | 0.45762338 | -0.7785638 | 0.43623669 | 0.79555046 |
| C1orf112 | 100.983915 | -0.257462566 | 0.32326492 | -0.7964445 | 0.42577372 | 0.78894887 |
| SCYL3 | 338.424521 | 0.393324695 | 0.2768692 | 1.42061558 | 0.15542855 | 0.6118547 |
| KIFAP3 | 901.388971 | 0.21165922 | 0.26845537 | 0.78843356 | 0.43044315 | 0.79055252 |
| GORAB | 130.743819 | -0.399614644 | 0.33018397 | -1.2102788 | 0.22617195 | 0.66688879 |
| MROH9 | 68.6228414 | 0.857562507 | 0.71689376 | 1.1962198 | 0.2316108 | 0.66878058 |
| FMO2 | 109.79018 | 3.297834512 | 0.9775754 | 3.37348354 | 0.00074223 | 0.09083761 |
| FMO1 | 1218.26929 | 0.640929624 | 0.73950565 | 0.8667001 | 0.38610635 | 0.77317162 |
| FMO4 | 1260.35729 | 0.976392486 | 0.77610916 | 1.25806077 | 0.20836978 | 0.65368257 |
| PRRC2C | 2856.65936 | -0.195011832 | 0.26757829 | -0.7288029 | 0.46612225 | 0.81070139 |
| MYOCOS | 687.541224 | 0.033855282 | 0.96044256 | 0.03524967 | 0.97188066 | 0.99240251 |
| VAMP4 | 215.083097 | 0.459917103 | 0.23877376 | 1.92616265 | 0.05408407 | 0.46711025 |
| METTL13 | 141.45454 | -0.278774147 | 0.31391455 | -0.8880574 | 0.37450986 | 0.7697436 |
| DNM3 | 448.233332 | -0.357597164 | 0.30231848 | -1.1828492 | 0.23686893 | 0.67416162 |
| PIGC | 222.907507 | 0.712822907 | 0.35616456 | 2.00138641 | 0.04535076 | 0.44360789 |

| | | | | | | |
|---|---|---|---|---|---|---|
| SUCO | 1384.42593 | -0.386838352 | 0.22085338 | -1.7515618 | 7.98E-02 | 5.13E-01 |
| AL031599.1 | 191.054736 | 0.399683698 | 0.93388726 | 0.42797853 | 0.66866675 | 0.89566976 |
| AL645568.1 | 272.36452 | 0.344085438 | 0.28792229 | 1.19506357 | 0.23206219 | 0.66937637 |
| PRDX6 | 842.272322 | 0.227256592 | 0.24784671 | 0.91692398 | 0.35918248 | 0.75936492 |
| KLHL20 | 1039.08808 | -0.196921012 | 0.25116504 | -0.7840304 | 0.4330223 | 0.79350758 |
| CENPL | 49.2371814 | -0.503560919 | 0.4422442 | -1.138649 | 0.25484958 | 0.68976367 |
| DARS2 | 135.32936 | 0.15937901 | 0.26889243 | 0.59272405 | 0.55336585 | 0.84727776 |
| ZBTB37 | 301.67256 | -0.029188951 | 0.36303588 | -0.0804024 | 0.93591723 | 0.98406248 |
| RC3H1 | 2058.84163 | -0.459278998 | 0.28007846 | -1.6398226 | 0.10104205 | 0.54608983 |
| AL121983.2 | 61.8937479 | 0.365863902 | 0.58319319 | 0.62734598 | 0.53043247 | 0.83789294 |
| RABGAP1L | 10702.2175 | 0.28903937 | 0.28498374 | 1.01423108 | 0.3104725 | 0.72498238 |
| Z99127.1 | 34.699176 | 0.300432012 | 0.54990675 | 0.54633265 | 0.58483729 | 0.86062622 |
| CACYBP | 235.603114 | -0.884334135 | 0.47190181 | -1.8739791 | 0.0609333 | 0.48123823 |
| MRPS14 | 159.457793 | -0.081296748 | 0.3460189 | -0.2349489 | 0.81424842 | 0.94881072 |
| KIAA0040 | 114.264838 | 0.11600876 | 0.47098829 | 0.24630923 | 0.80544287 | 0.94585363 |
| AL513329.1 | 43.5780979 | -1.24286851 | 0.76826677 | -1.6177564 | 0.1057151 | 0.55370151 |
| COP1 | 3127.98395 | -0.038498698 | 0.22244189 | -0.1730731 | 0.862594 | 0.96391903 |
| AL359265.3 | 59.9739484 | 0.178665053 | 0.7359663 | 0.24276255 | 0.80818935 | 0.94617795 |
| PAPPA2 | 124.744655 | -0.389198702 | 0.55188333 | -0.7052192 | 0.48067386 | 0.81714875 |
| AL122019.1 | 52.9297298 | 1.061910163 | 0.88057327 | 1.2059305 | 0.22784429 | 0.66705629 |
| SEC16B | 772.996215 | 0.077554325 | 0.78027829 | 0.09939316 | 0.92082611 | 0.97939204 |
| RASAL2 | 4594.60371 | -0.876806766 | 0.3742152 | -2.3430549 | 0.01912657 | 0.34084276 |
| RALGPS2 | 1797.59609 | 0.455113574 | 0.30917991 | 1.47200241 | 0.14102023 | 0.59555904 |
| ANGPTL1 | 97.3746932 | 0.783342558 | 0.71118762 | 1.10145697 | 0.27069782 | 0.69961625 |
| FAM20B | 337.156195 | -0.368192296 | 0.31653264 | -1.1632048 | 0.2447464 | 0.68250191 |
| TOR3A | 114.431879 | -0.417710761 | 0.38217955 | -1.0929699 | 0.274407 | 0.7014484 |
| ABL2 | 1951.55655 | -0.451660083 | 0.31674098 | -1.4259604 | 0.15387978 | 0.6105654 |
| SOAT1 | 770.645397 | 0.242481384 | 0.28379534 | 0.85442341 | 3.93E-01 | 7.77E-01 |
| AXDND1 | 80.5712967 | 0.095920689 | 0.5203283 | 0.18434648 | 0.85374166 | 0.96109041 |
| NPHS2 | 37.0079044 | 0.204416083 | 0.81868853 | 0.24968724 | 0.80282922 | 0.94543168 |
| TOR1AIP2 | 985.396659 | -0.422641205 | 0.32323498 | -1.3075355 | 0.19103091 | 0.6431984 |
| TOR1AIP1 | 348.358271 | -0.007899532 | 0.21401296 | -0.0369115 | 0.9705556 | 0.99205112 |
| CEP350 | 1967.31417 | -0.028157907 | 0.12032986 | -0.234006 | 0.81498033 | 0.94895023 |
| QSOX1 | 151.556643 | 0.533113281 | 0.46645777 | 1.14289722 | 0.25308127 | 0.68887358 |
| LHX4 | 50.1668779 | 0.84271559 | 0.64932252 | 1.29783822 | 0.19434293 | 0.64548758 |
| ACBD6 | 2262.67364 | -0.441259338 | 0.24066221 | -1.8335215 | 0.06672506 | 0.48884194 |
| XPR1 | 4137.18341 | -0.356587756 | 0.22220873 | -1.6047423 | 0.10855053 | 0.55858026 |
| KIAA1614 | 132.554039 | 0.92540428 | 0.49996548 | 1.85093635 | 0.06417871 | 0.48593331 |
| AL162431.2 | 36.0914709 | -0.033780079 | 0.38482351 | -0.0877807 | 0.93005097 | 0.98272895 |
| STX6 | 375.332345 | 0.142650557 | 0.22787838 | 0.62599425 | 0.53131871 | 0.83789294 |
| MR1 | 106.029111 | 0.930676892 | 0.42746907 | 2.17717948 | 0.02946718 | 0.39817664 |
| CACNA1E | 2536.22493 | -1.460656967 | 0.83003679 | -1.7597497 | 0.07845026 | 0.51192014 |
| GLUL | 222.943555 | -0.077408939 | 0.27197776 | -0.284615 | 0.77593915 | 0.9352344 |
| RNASEL | 101.422751 | 1.114724892 | 0.54980422 | 2.02749424 | 0.04261189 | 0.43849941 |
| NPL | 2031.18879 | 0.731510891 | 0.49325364 | 1.48303191 | 0.13806593 | 0.59474076 |
| DHX9 | 1366.71548 | -0.231889731 | 0.20528621 | -1.1295923 | 0.25864805 | 0.69241621 |
| LAMC1 | 2298.14788 | -0.135799753 | 0.42566094 | -0.3190327 | 7.50E-01 | 9.26E-01 |
| LAMC2 | 100.375988 | -0.566758991 | 0.94641308 | -0.5988495 | 0.54927325 | 0.84594253 |
| SMG7 | 1560.81692 | -0.130108874 | 0.23331226 | -0.5576598 | 0.5770767 | 0.85637657 |
| ARPC5 | 468.809993 | -0.521321313 | 0.39189121 | -1.3302705 | 0.18342916 | 0.63276491 |
| RGL1 | 2905.79457 | 0.219314964 | 0.46156801 | 0.475152 | 6.35E-01 | 0.88238823 |
| TSEN15 | 438.745151 | 0.487041583 | 0.29249384 | 1.66513448 | 0.09588593 | 0.53870645 |
| C1orf21 | 105.062347 | -0.213968003 | 0.39908744 | -0.5361432 | 0.59185961 | 0.86382106 |
| EDEM3 | 903.982097 | 0.652463176 | 0.22643973 | 2.88139886 | 0.00395914 | 0.20256898 |

| | | | | | | |
|---|---|---|---|---|---|---|
| FAM129A | 67.8672094 | 0.361655223 | 0.62088977 | 0.58247895 | 5.60E-01 | 0.84944908 |
| RNF2 | 310.345176 | -0.234953867 | 0.30967819 | -0.7587033 | 0.44803006 | 0.80223612 |
| TRMT1L | 389.999121 | 0.576234948 | 0.35734927 | 1.61252587 | 0.10684756 | 0.55488138 |
| SWT1 | 624.025605 | -0.458121514 | 0.18441224 | -2.4842251 | 0.01298337 | 0.29875272 |
| IVNS1ABP | 1060.41574 | -0.598774198 | 0.35576243 | -1.6830732 | 0.09236093 | 0.53621484 |
| HMCN1 | 101.398684 | 0.798318621 | 0.55338275 | 1.44261565 | 0.14912877 | 0.60622442 |
| TPR | 1153.71686 | -0.256659468 | 0.24205362 | -1.0603414 | 0.28898931 | 0.71218341 |
| ODR4 | 726.393837 | -0.061721831 | 0.17376086 | -0.3552114 | 0.72243126 | 0.91592159 |
| AL596220.1 | 235.48205 | 0.553484681 | 0.41584292 | 1.33099461 | 0.18319078 | 0.63276491 |
| ERVMER61-1 | 96.571292 | 0.563442906 | 0.42251271 | 1.33355255 | 0.18235054 | 0.63216521 |
| BRINP3 | 128.444378 | 0.697626249 | 0.73683568 | 0.94678674 | 0.34374746 | 0.7507483 |
| AL390957.1 | 136.679617 | 0.928195612 | 0.48503575 | 1.91366433 | 0.05566306 | 0.46970412 |
| AL136454.1 | 364.439638 | -0.073495633 | 0.47120911 | -0.1559724 | 0.87605475 | 0.96617876 |
| UCHL5 | 465.219963 | -0.066331615 | 0.18833419 | -0.3522017 | 0.72468703 | 0.91600702 |
| TROVE2 | 637.946853 | 0.315290275 | 0.15036691 | 2.09680625 | 0.03601073 | 0.42040764 |
| GLRX2 | 144.95907 | -0.499482084 | 0.42954883 | -1.1628063 | 0.24490811 | 0.68250191 |
| CDC73 | 1473.4658 | -0.368336144 | 0.24766294 | -1.4872477 | 0.1369494 | 0.59474076 |
| LINC01031 | 33.7152003 | -0.546931828 | 0.76049989 | -0.7191741 | 0.47203365 | 0.81395965 |
| KCNT2 | 1213.61521 | 1.487308413 | 0.96511284 | 1.54107205 | 0.12329925 | 0.5825794 |
| ZBTB41 | 306.108459 | 0.036176012 | 0.16818485 | 0.21509673 | 0.8296919 | 0.95325673 |
| CRB1 | 81.9279552 | 1.747897286 | 0.67762176 | 2.57945861 | 0.00989553 | 0.26491807 |
| DENND1B | 1737.2304 | 0.229514169 | 0.28576813 | 0.80314824 | 0.42188906 | 0.78892549 |
| NEK7 | 1840.20583 | -0.145890678 | 0.2084221 | -0.699977 | 0.48394167 | 0.81839266 |
| MIR181A1HG | 510.034584 | 0.095882761 | 0.48460282 | 0.19785845 | 0.84315582 | 0.95716002 |
| LINC00862 | 745.072642 | -0.026960556 | 0.51331306 | -0.0525226 | 0.95811226 | 0.98878187 |
| AC104461.1 | 214.442312 | 0.651243624 | 0.71926405 | 0.90543053 | 0.36523735 | 0.76362239 |
| ZNF281 | 289.582692 | -0.074992877 | 0.26789095 | -0.2799381 | 0.77952502 | 0.93661899 |
| DDX59 | 157.167748 | -0.683750467 | 0.30773162 | -2.2219051 | 0.02628972 | 0.38417182 |
| CAMSAP2 | 1176.41888 | -0.036875858 | 0.27172813 | -0.1357087 | 0.89205161 | 0.97237481 |
| INAVA | 586.723919 | 0.26729999 | 0.39609704 | 0.67483461 | 0.49978085 | 0.82476476 |
| TMEM9 | 213.660831 | 0.588454119 | 0.41189449 | 1.42865256 | 0.15310412 | 0.61008674 |
| LAD1 | 263.227693 | -0.263233612 | 0.54979086 | -0.4787886 | 0.63208901 | 0.88123095 |
| CSRP1 | 128.095302 | -1.198846904 | 0.47515677 | -2.5230555 | 0.011634 | 0.28824618 |
| AC096677.1 | 122.254747 | -1.010896478 | 0.58946179 | -1.7149483 | 0.08635471 | 0.52421699 |
| NAV1 | 217.587978 | 0.483696734 | 0.62455328 | 0.77446832 | 0.43865389 | 0.79718009 |
| IPO9-AS1 | 70.492363 | 0.883975833 | 0.68368175 | 1.29296391 | 0.19602354 | 0.64558745 |
| IPO9 | 711.710087 | -0.177680034 | 0.27656924 | -0.6424432 | 0.52058544 | 0.833897 |
| TIMM17A | 302.326373 | -0.168274848 | 0.24359966 | -0.6907844 | 0.48970103 | 0.81970016 |
| RNPEP | 412.061182 | -0.20560567 | 0.27290686 | -0.7533914 | 0.45121477 | 0.80384625 |
| ELF3-AS1 | 34.306494 | -0.697125721 | 0.61944642 | -1.1254012 | 0.26041908 | 0.69280433 |
| ELF3 | 1644.10815 | 0.801218123 | 0.53237179 | 1.50499732 | 0.13232477 | 0.5912647 |
| ARL8A | 335.974362 | -0.454225506 | 0.2971392 | -1.5286623 | 0.12634817 | 0.58647288 |
| PPP1R12B | 2259.1508 | -0.063028916 | 0.24215252 | -0.260286 | 0.79464316 | 0.94252227 |
| KDM5B | 1273.3609 | -0.134471311 | 0.26609929 | -0.5053426 | 0.61331822 | 0.87222135 |
| RABIF | 126.007046 | -0.77349537 | 0.40884727 | -1.8918932 | 0.05850521 | 0.47778423 |
| KLHL12 | 298.189645 | -0.122328475 | 0.2927938 | -0.4177973 | 0.67609529 | 0.89835672 |
| ADIPOR1 | 177.692528 | -0.053976809 | 0.26445315 | -0.2041073 | 0.83826968 | 0.95512573 |
| CYB5R1 | 155.625627 | 0.16082661 | 0.29713874 | 0.54125091 | 0.58833465 | 0.86140264 |
| TMEM183A | 122.77472 | -0.008566435 | 0.2508438 | -0.0341505 | 0.97275716 | 0.9926729 |
| PPFIA4 | 95.6206128 | -1.370672013 | 0.98860021 | -1.3864776 | 1.66E-01 | 0.62065658 |
| BTG2 | 134.338863 | 1.12931198 | 0.61265666 | 1.8433032 | 0.06528475 | 0.48790819 |
| ATP2B4 | 549.361241 | 1.285208247 | 0.46967217 | 2.73639429 | 0.00621165 | 0.22644975 |
| SNRPE | 135.068962 | -0.245711585 | 0.27225394 | -0.9025088 | 0.36678663 | 0.76522744 |
| SOX13 | 405.587082 | 1.152880695 | 0.37006579 | 3.11533981 | 0.00183733 | 0.13687118 |

| | | | | | | |
|---|---|---|---|---|---|---|
| ETNK2 | 398.186604 | 1.006540806 | 0.71600831 | 1.40576694 | 0.15979334 | 0.61542146 |
| REN | 56.6626062 | 0.910082025 | 0.87271334 | 1.04281897 | 0.29703214 | 0.71825247 |
| GOLT1A | 100.511383 | -0.766904989 | 0.56596952 | -1.3550288 | 0.17540839 | 0.62699479 |
| PLEKHA6 | 3150.89834 | -0.397214726 | 0.35548211 | -1.1173972 | 0.26382452 | 0.69567438 |
| AL606489.1 | 326.205766 | -0.090447393 | 0.48590449 | -0.1861423 | 0.85233315 | 0.96061646 |
| PPP1R15B | 436.998586 | -0.333846609 | 0.4011867 | -0.8321478 | 0.40532555 | 0.78429302 |
| PIK3C2B | 327.064917 | 0.616033868 | 0.41979767 | 1.46745423 | 0.14225253 | 0.59738219 |
| MDM4 | 1657.41039 | -0.424614922 | 0.20832281 | -2.0382546 | 0.04152448 | 0.43367572 |
| AL512306.3 | 61.0041681 | -0.671087467 | 0.56099407 | -1.196247 | 0.23160019 | 0.66878058 |
| LRRN2 | 410.965905 | 0.908726298 | 0.96305507 | 0.94358706 | 0.34538071 | 0.75166079 |
| NFASC | 156.229282 | -0.25242376 | 0.50115535 | -0.5036837 | 6.14E-01 | 8.72E-01 |
| RBBP5 | 438.165047 | -0.094841978 | 0.25884297 | -0.3664074 | 0.71406111 | 0.91214838 |
| DSTYK | 618.399588 | 0.008656448 | 0.23868865 | 0.03626669 | 0.97106971 | 0.99220667 |
| NUAK2 | 312.619961 | -1.464481952 | 0.4910058 | -2.9826164 | 0.00285796 | 0.17333652 |
| LEMD1 | 145.762004 | 1.736347155 | 0.46757751 | 3.71349585 | 2.04E-04 | 0.05350954 |
| CDK18 | 805.380196 | -0.373616098 | 0.40641162 | -0.9193047 | 0.35793624 | 0.75936492 |
| MFSD4A | 106.895908 | 0.557989536 | 0.56928052 | 0.98016622 | 0.32700408 | 0.73745391 |
| ELK4 | 928.461729 | 0.054979412 | 0.34175192 | 0.16087521 | 0.87219169 | 0.9657206 |
| SLC45A3 | 75.9992819 | 0.551690885 | 0.59614993 | 0.92542305 | 0.35474589 | 0.75755608 |
| NUCKS1 | 1364.74334 | 0.835653017 | 0.3220499 | 2.59479361 | 0.00946477 | 0.25998727 |
| RAB29 | 787.800975 | 0.271959428 | 0.34010855 | 0.79962537 | 0.42392788 | 0.78894887 |
| SLC41A1 | 393.746606 | 0.043500668 | 0.32920268 | 0.13213947 | 8.95E-01 | 9.73E-01 |
| PM20D1 | 86.6191097 | -0.304920214 | 0.73834235 | -0.4129794 | 0.67962167 | 0.89941358 |
| SLC26A9 | 100.978984 | -0.137404569 | 0.65156338 | -0.2108844 | 8.33E-01 | 9.54E-01 |
| RHEX | 9090.91604 | -0.71838246 | 0.81796504 | -0.8782557 | 0.37980497 | 0.77095585 |
| AC244035.1 | 63.279735 | -0.378516683 | 0.78229971 | -0.4838512 | 0.62849145 | 0.88009297 |
| SRGAP2 | 230.973984 | 0.410471388 | 0.4244545 | 0.96705627 | 3.34E-01 | 0.74022334 |
| IKBKE | 67.1404867 | 0.775913207 | 0.64319287 | 1.20634609 | 0.22768407 | 0.66705629 |
| EIF2D | 187.547512 | 0.022478436 | 0.29322253 | 0.07665999 | 0.93889403 | 0.98435387 |
| DYRK3 | 140.93217 | -0.864875272 | 0.42200422 | -2.049447 | 0.04041843 | 0.43086475 |
| MAPKAPK2 | 1233.31567 | -0.52082983 | 0.3498422 | -1.4887565 | 0.13655151 | 0.5942049 |
| PIGR | 2391.42353 | 1.803839356 | 0.6551299 | 2.75340715 | 0.00589785 | 0.22637955 |
| FCAMR | 219.142892 | 2.550062487 | 0.8691261 | 2.93405352 | 0.00334567 | 0.18902009 |
| PFKFB2 | 1411.76052 | 0.275933192 | 0.59258435 | 0.46564374 | 0.64147053 | 0.88403746 |
| YOD1 | 366.392856 | -0.37488273 | 0.53379496 | -0.7022972 | 0.48249381 | 0.8177376 |
| CD55 | 92.7705793 | -0.412438817 | 0.37674476 | -1.0947433 | 0.2736291 | 0.70140695 |
| CR1L | 245.236458 | -0.353096529 | 0.84751996 | -0.4166233 | 0.67695399 | 0.89843878 |
| CD46 | 1344.07314 | 0.213362393 | 0.24390621 | 0.8747723 | 0.38169781 | 0.77237191 |
| MIR29B2CHG | 1793.75202 | 0.36531287 | 0.43279213 | 0.8440839 | 0.39862252 | 0.78122921 |
| PLXNA2 | 480.34528 | -0.257826263 | 0.21276888 | -1.2117668 | 0.22560167 | 0.66688879 |
| LAMB3 | 110.409746 | -0.109159527 | 0.90198902 | -0.1210209 | 0.90367448 | 0.97555848 |
| AL031316.1 | 159.218094 | 0.904304986 | 0.7859498 | 1.15058874 | 0.24990147 | 0.68617881 |
| G0S2 | 1076.44596 | 1.491434893 | 0.855985 | 1.74236102 | 0.08144529 | 0.51593841 |
| HSD11B1 | 57.7202443 | 1.134626667 | 1.1658411 | 0.97322583 | 0.33044108 | 0.73849818 |
| IRF6 | 129.610152 | 0.591493701 | 0.50146408 | 1.17953354 | 0.23818579 | 0.67642542 |
| UTP25 | 201.677427 | -0.80833987 | 0.4153367 | -1.9462279 | 0.05162737 | 0.4629611 |
| SYT14 | 177.647455 | 0.653746689 | 0.77693124 | 0.84144729 | 0.4000974 | 0.7818841 |
| HHAT | 219.590451 | 0.729389257 | 0.63267543 | 1.15286483 | 0.24896587 | 0.68557761 |
| RCOR3 | 1877.81223 | 0.276454864 | 0.32539852 | 0.84958857 | 0.39555387 | 0.77931337 |
| TRAF5 | 136.284676 | -0.222930576 | 0.325227 | -0.6854615 | 4.93E-01 | 8.21E-01 |
| LINC00467 | 338.586222 | 0.349746472 | 0.53682929 | 0.65150407 | 0.51472115 | 0.83169608 |
| SLC30A1 | 369.267625 | -0.846876375 | 0.40264782 | -2.1032682 | 0.03544233 | 0.4198859 |
| LPGAT1 | 4247.55879 | 0.155455179 | 0.41030469 | 0.37887742 | 7.05E-01 | 0.909554 |
| INTS7 | 346.271067 | -0.014175811 | 0.30618793 | -0.0462977 | 0.96307294 | 0.99012655 |

| | | | | | | |
|---|---|---|---|---|---|---|
| PPP2R5A | 1080.15574 | 0.253732415 | 0.23627503 | 1.07388587 | 0.28287384 | 0.70756556 |
| AL360091.3 | 172.615564 | 0.688525253 | 0.51618282 | 1.33387867 | 0.18224362 | 0.63216521 |
| TMEM206 | 518.462699 | 0.044561405 | 0.40544597 | 0.10990713 | 0.91248302 | 0.977179 |
| NENF | 286.262117 | -0.054277333 | 0.29193463 | -0.1859229 | 0.85250522 | 0.96064527 |
| ATF3 | 338.879041 | -1.402622736 | 0.59016665 | -2.3766554 | 0.0174704 | 0.33064634 |
| NSL1 | 481.282099 | 0.411916759 | 0.22807499 | 1.80605844 | 0.07090921 | 0.49767156 |
| TATDN3 | 206.6395 | 0.091743411 | 0.43892738 | 0.20901729 | 0.83443474 | 0.95393659 |
| FLVCR1-DT | 67.165561 | 0.656642694 | 0.46898328 | 1.40014095 | 0.16147111 | 0.61754591 |
| FLVCR1 | 412.565636 | 0.139943965 | 0.39457508 | 0.35467006 | 0.72283679 | 0.91592159 |
| ANGEL2 | 397.677608 | -0.120866736 | 0.23166043 | -0.521741 | 0.6018507 | 0.86765503 |
| RPS6KC1 | 1803.43175 | -0.399584317 | 0.31350309 | -1.2745785 | 0.20245845 | 0.64881821 |
| AL592402.1 | 45.2564375 | -2.721809465 | 1.20112489 | -2.2660503 | 0.0234483 | 0.37088088 |
| AC096639.1 | 53.4935163 | -3.874295602 | 1.38696435 | -2.7933635 | 0.0052163 | 0.21865406 |
| SMYD2 | 493.974466 | -0.266148249 | 0.23957632 | -1.1109122 | 2.67E-01 | 6.98E-01 |
| PTPN14 | 3632.43134 | 0.026556367 | 0.42760029 | 0.06210559 | 0.95047875 | 0.98731502 |
| KCTD3 | 1034.71496 | 0.698125714 | 0.37834601 | 1.84520437 | 0.06500782 | 0.48790819 |
| USH2A | 107.270029 | 0.059782936 | 0.55482357 | 0.10775126 | 9.14E-01 | 9.77E-01 |
| ESRRG | 11121.6418 | 0.078009755 | 0.49654116 | 0.15710632 | 0.87516106 | 0.96615698 |
| GPATCH2 | 2102.32566 | 0.12178376 | 0.25047703 | 0.48620729 | 0.6268202 | 0.87890915 |
| SPATA17 | 581.989039 | 0.569461911 | 0.43981189 | 1.29478518 | 0.19539435 | 0.64558745 |
| RRP15 | 255.216928 | -0.472402315 | 0.31060638 | -1.5209034 | 0.12828407 | 0.58793678 |
| TGFB2 | 71.7901765 | -0.093283798 | 0.58633409 | -0.1590967 | 0.87359272 | 0.96592775 |
| LYPLAL1 | 2332.34739 | 0.176279826 | 0.4261112 | 0.41369443 | 0.67909791 | 0.89932032 |
| AC096642.1 | 82.7509529 | 0.688781893 | 0.60529765 | 1.13792262 | 0.25515279 | 0.68976367 |
| AL513283.1 | 662.484853 | 0.790026807 | 1.06595271 | 0.74114621 | 0.4586048 | 0.80636987 |
| ZC3H11B | 103.713305 | -1.067090946 | 0.43667851 | -2.4436535 | 0.01453938 | 0.30802371 |
| SLC30A10 | 46.4497281 | -0.613997477 | 1.06021819 | -0.5791237 | 0.56250571 | 0.85054239 |
| EPRS | 475.512452 | -0.500297647 | 0.29847441 | -1.6761827 | 0.0937024 | 0.53694769 |
| BPNT1 | 311.445789 | 0.465165208 | 0.31425311 | 1.48022466 | 0.1388133 | 0.59474076 |
| IARS2 | 568.191906 | -0.095347033 | 0.19546417 | -0.487798 | 0.62569291 | 0.87854882 |
| RAB3GAP2 | 1963.56718 | -0.288799459 | 0.18664455 | -1.5473233 | 0.12178531 | 0.5804281 |
| MARK1 | 101.012229 | 0.207534825 | 0.897599 | 0.23121107 | 0.81715083 | 0.94954998 |
| C1orf115 | 240.584331 | -0.201338423 | 0.32689707 | -0.6159077 | 0.53795543 | 0.84040975 |
| 2-Mar | 2100.67857 | 0.950041373 | 0.56770502 | 1.67347714 | 0.0942334 | 0.53783536 |
| 1-Mar | 303.600179 | -0.032522052 | 0.51733134 | -0.062865 | 9.50E-01 | 9.87E-01 |
| DUSP10 | 53.2875211 | 0.467172743 | 0.68746361 | 0.67955996 | 0.49678313 | 0.82259077 |
| AL513314.2 | 178.897658 | -0.067983527 | 0.30898989 | -0.2200186 | 0.82585666 | 0.95192757 |
| TAF1A | 79.3147752 | -0.382253488 | 0.32921188 | -1.1611169 | 0.24559435 | 0.68310013 |
| MIA3 | 625.983373 | -0.336795956 | 0.31366235 | -1.0737532 | 0.28293331 | 0.70756556 |
| AIDA | 269.125471 | 0.17808155 | 0.20010076 | 0.88995939 | 3.73E-01 | 7.70E-01 |
| BROX | 386.175129 | 0.051180348 | 0.25826007 | 0.19817368 | 0.84290918 | 0.95716002 |
| FAM177B | 562.708535 | -0.352033258 | 0.66191078 | -0.531844 | 5.95E-01 | 0.86499581 |
| AC093152.1 | 1085.0348 | 0.242796742 | 0.16619413 | 1.46092253 | 0.1440367 | 0.60082504 |
| DISP1 | 141.272429 | 0.18444619 | 0.44965446 | 0.4101954 | 0.68166262 | 0.89962501 |
| CAPN2 | 721.0313 | -0.296657817 | 0.52834478 | -0.5614853 | 5.74E-01 | 8.55E-01 |
| TP53BP2 | 748.443339 | -0.355033366 | 0.33529796 | -1.0588593 | 2.90E-01 | 0.71218341 |
| FBXO28 | 1546.76126 | -0.018243058 | 0.23932336 | -0.0762277 | 0.93923798 | 0.98435387 |
| DEGS1 | 452.170494 | -0.113661061 | 0.4051649 | -0.2805304 | 0.77907062 | 0.93646866 |
| NVL | 787.121942 | -0.357520847 | 0.26908233 | -1.3286671 | 0.18395782 | 0.63338605 |
| CNIH4 | 523.938249 | -0.41740842 | 0.25198205 | -1.6565006 | 0.09762048 | 0.54093104 |
| WDR26 | 1563.88043 | -0.078959318 | 0.21680148 | -0.364201 | 0.71570791 | 0.91298152 |
| CNIH3 | 497.454815 | 0.75570628 | 0.37923331 | 1.99272124 | 0.04629198 | 0.44508419 |
| DNAH14 | 2326.41985 | 0.151743877 | 0.3573729 | 0.42460935 | 0.67112147 | 0.89694443 |
| LBR | 762.273881 | 0.073641779 | 0.37372442 | 0.19704835 | 0.8437897 | 0.95731603 |

| | | | | | | |
|---|---|---|---|---|---|---|
| ENAH | 994.886056 | -0.375760967 | 0.46955049 | -0.8002568 | 0.42356203 | 0.78894887 |
| SRP9 | 460.043575 | -0.133274617 | 0.32754465 | -0.4068899 | 0.68408884 | 0.9007343 |
| EPHX1 | 1081.55951 | -0.628196942 | 0.53207033 | -1.1806652 | 0.23773573 | 0.67592693 |
| AL591895.1 | 385.934399 | 0.987858291 | 0.48138671 | 2.05210962 | 0.04015901 | 0.4304996 |
| TMEM63A | 1440.95175 | 0.836881725 | 0.51332382 | 1.63031927 | 0.10303404 | 0.55022042 |
| AL117348.2 | 36.0128857 | -0.398430934 | 0.77470002 | -0.5143035 | 0.60703981 | 0.86985416 |
| SDE2 | 118.398148 | -0.269192566 | 0.26417979 | -1.0189749 | 0.3082149 | 0.72367286 |
| H3F3A | 150.773092 | 0.355894533 | 0.34993934 | 1.01701779 | 0.30914497 | 0.72449199 |
| ACBD3 | 1054.75625 | -0.118043105 | 0.28650227 | -0.4120146 | 0.68032876 | 0.89941358 |
| LIN9 | 115.629195 | 0.197603768 | 0.2527049 | 0.78195462 | 0.43424124 | 0.79442004 |
| PARP1 | 234.270592 | 0.223574842 | 0.31811385 | 0.70281391 | 0.48217173 | 0.8177376 |
| STUM | 229.406585 | -0.143477741 | 0.83848191 | -0.1711161 | 0.86413249 | 0.96393376 |
| ITPKB | 106.000192 | 0.88206052 | 0.53458778 | 1.64998257 | 0.0989465 | 0.5425667 |
| PSEN2 | 69.0849272 | -0.884623268 | 0.37375574 | -2.3668486 | 0.01794027 | 0.33548057 |
| COQ8A | 516.386223 | -0.471944094 | 0.36186612 | -1.3041953 | 0.192167 | 0.64497936 |
| CDC42BPA | 5067.6545 | 0.155897168 | 0.27205595 | 0.57303347 | 0.56662203 | 0.85226469 |
| ZNF678 | 267.408368 | 0.241664503 | 0.3333623 | 0.72493051 | 0.46849466 | 0.81217042 |
| SNAP47 | 190.153188 | 0.096145304 | 0.24536283 | 0.39184951 | 0.69516941 | 0.90589033 |
| JMJD4 | 76.7112612 | -0.666589183 | 0.50526838 | -1.3192774 | 0.18707638 | 0.63818925 |
| ARF1 | 886.298841 | -0.502980189 | 0.36950193 | -1.3612383 | 0.17343839 | 0.62663931 |
| C1orf35 | 100.373998 | -0.210370532 | 0.27441622 | -0.7666111 | 0.44331276 | 0.7993765 |
| MRPL55 | 157.705977 | -0.419172903 | 0.39393384 | -1.0640693 | 0.2872973 | 0.71118494 |
| GUK1 | 677.736614 | -0.061845255 | 0.38724424 | -0.1597061 | 0.87311262 | 0.96592775 |
| IBA57 | 60.3837331 | -0.825216631 | 0.52556489 | -1.5701518 | 0.11637981 | 0.57350419 |
| OBSCN | 433.919316 | 0.720095572 | 0.42263834 | 1.70381033 | 0.08841653 | 0.52922503 |
| TRIM11 | 108.719592 | -0.463050713 | 0.4674026 | -0.9906892 | 0.32183736 | 0.73375526 |
| RNF187 | 233.198494 | -0.694738323 | 0.45261154 | -1.5349549 | 0.1247949 | 0.58564339 |
| RHOU | 89.1242293 | 0.929368877 | 0.67005013 | 1.38701395 | 0.16543751 | 0.62065658 |
| RAB4A | 299.769234 | 0.212864153 | 0.19312051 | 1.1022348 | 0.27035961 | 0.69961625 |
| NUP133 | 489.048258 | -0.067249553 | 0.17796725 | -0.377876 | 0.70552271 | 0.90974598 |
| ABCB10 | 309.370066 | 0.205085907 | 0.24817887 | 0.82636328 | 0.40859805 | 0.78506496 |
| TAF5L | 167.039653 | -0.487759874 | 0.36561887 | -1.3340665 | 0.18218207 | 0.63216521 |
| URB2 | 94.864164 | -0.719316663 | 0.60108169 | -1.1967037 | 0.23142208 | 0.66878058 |
| GALNT2 | 3329.88693 | -0.429487527 | 0.32736038 | -1.3119716 | 0.18952971 | 0.6416112 |
| PGBD5 | 164.217971 | 0.278069853 | 0.37127142 | 0.7489665 | 0.4538774 | 0.80433814 |
| COG2 | 364.82481 | -0.154132637 | 0.23656427 | -0.6515466 | 0.51469372 | 0.83169608 |
| AGT | 602.929024 | -1.271957864 | 0.611757 | -2.0791881 | 0.03760007 | 0.42215481 |
| C1orf198 | 100.834385 | -0.832069207 | 0.41891051 | -1.9862696 | 0.0470034 | 0.44732709 |
| TTC13 | 217.690911 | 0.010829294 | 0.25262368 | 0.04286729 | 0.96580732 | 0.99081812 |
| ARV1 | 143.278041 | 0.112939409 | 0.28476988 | 0.39659885 | 0.6916633 | 0.90491747 |
| FAM89A | 87.2001458 | 0.276292055 | 0.47556581 | 0.58097543 | 5.61E-01 | 8.50E-01 |
| C1orf131 | 147.180644 | 0.372010747 | 0.33531581 | 1.10943395 | 0.26724302 | 0.69853168 |
| GNPAT | 527.105803 | 0.065934657 | 0.19355655 | 0.34064803 | 0.73336856 | 0.91961238 |
| EXOC8 | 109.435429 | -0.413183926 | 0.28757232 | -1.4368001 | 0.15077481 | 0.60873943 |
| SPRTN | 164.109125 | -0.143383202 | 0.34240046 | -0.4187588 | 0.67539243 | 0.89801183 |
| EGLN1 | 1659.76543 | -0.097667173 | 0.33410274 | -0.2923268 | 7.70E-01 | 0.93266075 |
| TSNAX-DISC1 | 78.2260334 | 0.229073109 | 0.57373354 | 0.39926742 | 0.68969617 | 0.90396495 |
| DISC1 | 229.973558 | 0.876706548 | 0.34162294 | 2.56629885 | 1.03E-02 | 0.26767103 |
| SIPA1L2 | 151.80491 | 1.200990436 | 0.67606027 | 1.77645468 | 0.07565801 | 0.50730849 |
| NTPCR | 244.895185 | 0.410931453 | 0.24718069 | 1.66247394 | 0.09641778 | 0.53969191 |
| PCNX2 | 1173.07447 | 0.381862138 | 0.39425621 | 0.96856341 | 0.33276307 | 0.73988623 |
| MAP3K21 | 1581.2042 | 0.602534314 | 0.33764093 | 1.78454167 | 0.07433569 | 0.5071704 |
| KCNK1 | 571.38169 | 0.285768704 | 0.31112503 | 0.91850117 | 0.35835655 | 0.75936492 |
| SLC35F3 | 349.182891 | -1.325227005 | 0.92315961 | -1.435534 | 0.15113499 | 0.60905477 |

| | | | | | | |
|---|---|---|---|---|---|---|
| AL355472.1 | 76.3447158 | 0.305701554 | 0.73825122 | 0.41408879 | 0.67880908 | 0.89932032 |
| COA6 | 132.348583 | -0.225150969 | 0.39209831 | -0.5742207 | 0.56581846 | 0.85199095 |
| TARBP1 | 497.443829 | -0.050475751 | 0.31576944 | -0.15985 | 0.87299922 | 0.96592775 |
| IRF2BP2 | 676.052778 | -0.308378765 | 0.43212003 | -0.7136415 | 0.47544892 | 0.81564184 |
| TOMM20 | 484.633852 | -0.144962557 | 0.34698599 | -0.4177764 | 0.67611061 | 0.89835672 |
| ARID4B | 2969.59164 | -0.024036543 | 0.20460246 | -0.1174793 | 0.90648029 | 0.97651095 |
| TBCE | 42.7308475 | 1.42682164 | 0.46987477 | 3.03659981 | 0.00239263 | 0.15843364 |
| B3GALNT2 | 611.051576 | 0.039408439 | 0.26243985 | 0.15016179 | 0.88063697 | 0.96773407 |
| GNG4 | 136.364975 | -0.12931127 | 0.79683794 | -0.1622805 | 0.87108496 | 0.9657206 |
| LYST | 916.452352 | 0.015788257 | 0.24657404 | 0.06403049 | 0.94894595 | 0.9871942 |
| NID1 | 932.175656 | 0.850868385 | 0.41071013 | 2.07170052 | 0.03829338 | 0.42447711 |
| GPR137B | 1824.42309 | -0.675052495 | 0.46734446 | -1.4444431 | 0.14861437 | 0.60622442 |
| ERO1B | 479.408336 | 0.074352104 | 0.33113371 | 0.22453801 | 0.82233869 | 0.95112619 |
| EDARADD | 71.8766589 | -0.135061127 | 0.43638833 | -0.3094976 | 0.75694306 | 0.92799282 |
| LGALS8 | 455.962752 | -0.384301381 | 0.24387716 | -1.575799 | 1.15E-01 | 0.57182344 |
| AL359921.1 | 74.4681684 | -0.575349071 | 0.58668706 | -0.9806745 | 0.32675326 | 0.73745391 |
| HEATR1 | 513.139441 | -0.715603471 | 0.34321628 | -2.0849928 | 0.03706995 | 0.42215481 |
| MTR | 2838.62144 | 0.150356342 | 0.38740226 | 0.38811426 | 0.69793148 | 0.90696517 |
| MT1HL1 | 564.572199 | -3.04104131 | 0.81426873 | -3.73469 | 0.00018795 | 0.0505134 |
| RYR2 | 107.223147 | -0.214896331 | 0.54466202 | -0.3945499 | 6.93E-01 | 9.06E-01 |
| LINC01139 | 245.879254 | -0.375216961 | 0.90032735 | -0.4167562 | 0.67685678 | 0.89843878 |
| CHRM3 | 649.730079 | -0.361575033 | 0.35292052 | -1.0245225 | 0.30558853 | 0.72122 |
| CHRM3-AS1 | 122.259647 | 2.745465844 | 1.01732252 | 2.69871727 | 0.00696073 | 0.23157938 |
| AL356361.2 | 97.9174348 | 2.790566053 | 0.94089338 | 2.96586852 | 0.00301829 | 0.17987725 |
| FMN2 | 40.4837887 | -0.164940084 | 0.57490977 | -0.2868973 | 0.77419093 | 0.93453439 |
| RGS7 | 48.3072591 | 0.089802133 | 0.48700312 | 0.18439745 | 0.85370167 | 0.96109041 |
| FH | 166.644627 | 0.453441791 | 0.22003651 | 2.06075705 | 0.03932622 | 0.42704465 |
| KMO | 1024.25548 | -0.302986935 | 0.82260689 | -0.3683253 | 0.71263069 | 0.91183303 |
| OPN3 | 201.383309 | 0.362982851 | 0.28769327 | 1.26170087 | 0.20705644 | 0.65251041 |
| PLD5 | 40.3595193 | -0.486243195 | 0.4761762 | -1.0211413 | 0.30718749 | 0.72250292 |
| CEP170 | 948.61047 | -0.109331785 | 0.27997274 | -0.3905087 | 0.69616044 | 0.90612803 |
| SDCCAG8 | 2386.63355 | 0.41592287 | 0.28589552 | 1.45480722 | 0.14572264 | 0.60203043 |
| AKT3 | 2182.21679 | 0.552920823 | 0.45410801 | 1.21759759 | 0.22337693 | 0.66426142 |
| ZBTB18 | 344.326841 | 0.139925818 | 0.34482192 | 0.40579154 | 0.68489577 | 0.90098343 |
| AL645465.1 | 141.388951 | 0.389101924 | 0.54946072 | 0.7081524 | 0.47885062 | 0.81596289 |
| ADSS | 542.840847 | 0.047905749 | 0.29778393 | 0.16087419 | 0.87219249 | 0.9657206 |
| CATSPERE | 633.818418 | -0.240448976 | 0.44920605 | -0.5352755 | 0.59245939 | 0.86382106 |
| DESI2 | 546.855744 | -0.12471805 | 0.21394085 | -0.5829558 | 0.55992309 | 0.84941263 |
| COX20 | 459.766575 | -0.184828076 | 0.25155368 | -0.7347461 | 0.46249416 | 0.8082072 |
| HNRNPU | 2545.27847 | -0.008529304 | 0.15929314 | -0.0535447 | 0.95729791 | 0.98861313 |
| EFCAB2 | 1441.63741 | 0.083185511 | 0.34474932 | 0.24129275 | 0.80932823 | 0.94673968 |
| KIF26B | 880.083438 | 1.596051421 | 0.5939523 | 2.68717103 | 0.007206 | 0.2351731 |
| SMYD3 | 5813.31607 | -0.354489442 | 0.19941366 | -1.7776588 | 0.07545991 | 0.50728736 |
| TFB2M | 180.085818 | -0.730839493 | 0.2954685 | -2.4734938 | 0.01337991 | 0.29875272 |
| CNST | 542.386109 | -0.335486313 | 0.30394736 | -1.1037645 | 0.2696953 | 0.69961625 |
| SCCPDH | 301.015904 | 0.187339392 | 0.2486047 | 0.75356334 | 0.45111146 | 0.80384625 |
| AHCTF1 | 1263.24536 | -0.383813595 | 0.27935103 | -1.3739473 | 0.16945805 | 0.62326131 |
| ZNF669 | 40.1253836 | -0.318079077 | 0.43049316 | -0.7388714 | 0.4599851 | 0.80696859 |
| ZNF124 | 575.977504 | 0.163739061 | 0.25032123 | 0.65411575 | 0.51303723 | 0.83073024 |
| AL390728.6 | 233.703152 | -0.167663299 | 0.42841498 | -0.3913572 | 0.6955332 | 0.90589033 |
| ZNF496 | 263.616097 | -0.221655383 | 0.35978598 | -0.6160757 | 0.53784459 | 0.84040975 |
| OR2T2 | 72.9789911 | -0.408679516 | 0.87119606 | -0.4691017 | 6.39E-01 | 8.83E-01 |
| OR2G6 | 210.020064 | 0.450917891 | 1.06174253 | 0.42469608 | 0.67105824 | 0.89694443 |
| AC098483.1 | 45.3515049 | -0.173648786 | 0.98175436 | -0.176876 | 8.60E-01 | 0.96283855 |

| | | | | | | |
|---|---|---|---|---|---|---|
| OR2T10 | 179.39909 | 0.264590683 | 0.82892233 | 0.3191984 | 7.50E-01 | 9.26E-01 |
| OR2T11 | 365.17913 | 0.404247352 | 0.78457383 | 0.5152445 | 6.06E-01 | 8.70E-01 |
| LYPD8 | 75.99108 | 0.323965778 | 0.52457551 | 0.61757702 | 0.5368542 | 0.83993385 |
| SH3BP5L | 200.239732 | -0.751049894 | 0.31822151 | -2.3601481 | 0.01826764 | 0.33797156 |
| ZNF672 | 112.132548 | -0.066133754 | 0.31611758 | -0.2092062 | 0.83428728 | 0.95393659 |
| ZNF692 | 224.69694 | -0.151055356 | 0.27110353 | -0.557187 | 0.57739969 | 0.85659658 |
| PGBD2 | 74.2329414 | -0.485363854 | 0.39081096 | -1.2419402 | 0.21425862 | 0.65906957 |
| FAM110C | 540.853312 | -0.21386644 | 0.56508629 | -0.3784669 | 0.7050838 | 0.909554 |
| SH3YL1 | 1688.13033 | 0.193546785 | 0.48260114 | 0.40104917 | 0.68838393 | 0.90307854 |
| ACP1 | 319.995608 | -0.265978507 | 0.29281535 | -0.9083489 | 0.36369391 | 0.76272056 |
| ALKAL2 | 221.118262 | 0.054106607 | 0.88199593 | 0.06134564 | 0.95108394 | 0.98731502 |
| TMEM18 | 90.224853 | -0.199901988 | 0.35631004 | -0.5610338 | 0.57477447 | 0.85474196 |
| LINC01115 | 78.5497134 | 1.759036602 | 1.09982566 | 1.59937767 | 0.10973671 | 0.56020898 |
| SNTG2 | 336.191243 | 1.428080859 | 0.81399407 | 1.75441187 | 0.07935996 | 0.51192014 |
| PXDN | 265.178886 | 0.86626486 | 0.74115536 | 1.16880334 | 0.24248288 | 0.68080966 |
| MYT1L | 44.1864045 | 0.928040852 | 0.97038041 | 0.95636808 | 0.3388863 | 0.74632304 |
| LINC01250 | 49.800428 | 0.44714611 | 0.43063997 | 1.03832932 | 0.29911676 | 0.71852421 |
| EIPR1 | 884.948346 | -0.196595747 | 0.22023294 | -0.8926719 | 0.37203292 | 0.76845856 |
| TRAPPC12 | 1060.14603 | -0.090880077 | 0.2284176 | -0.3978681 | 0.69072741 | 0.90453814 |
| ADI1 | 650.444844 | -0.059773773 | 0.40697999 | -0.1468715 | 0.88323343 | 0.96846661 |
| AC108488.2 | 118.895011 | -0.580565213 | 0.46972182 | -1.2359767 | 0.21646723 | 0.66032006 |
| RNASEH1 | 51.5374478 | -0.57929747 | 0.47291357 | -1.2249542 | 0.22059247 | 0.66263062 |
| RNASEH1-AS | 37.0663151 | 0.636206949 | 0.34000795 | 1.87115314 | 0.06132386 | 0.4813972 |
| RPS7 | 1238.25132 | 0.017973941 | 0.48497167 | 0.03706184 | 0.9704357 | 0.99205112 |
| COLEC11 | 190.107519 | -0.390778399 | 0.56774881 | -0.6882945 | 0.49126733 | 0.82035445 |
| ALLC | 220.033231 | -1.111685135 | 0.63916026 | -1.7392901 | 0.08198374 | 0.51595552 |
| CMPK2 | 42.4547763 | -6.659306925 | 1.43005023 | -4.6566944 | 3.21E-06 | 0.00875212 |
| RSAD2 | 68.9912924 | -5.744725955 | 1.15890432 | -4.9570321 | 7.16E-07 | 0.00591552 |
| RNF144A | 59.3926982 | -0.191498135 | 0.49413137 | -0.387545 | 0.69835279 | 0.90696517 |
| LINC00299 | 54.4684492 | 1.031309811 | 0.87460063 | 1.17917798 | 0.23832731 | 0.67642542 |
| ID2 | 444.944092 | -0.627126122 | 0.44188749 | -1.4191986 | 0.15584112 | 0.6125316 |
| KIDINS220 | 1467.59125 | -0.121781286 | 0.22100331 | -0.5510383 | 0.58160742 | 0.85915432 |
| MBOAT2 | 684.060819 | 1.374031412 | 0.52488024 | 2.61779986 | 0.00884987 | 0.25484277 |
| ASAP2 | 2225.0095 | 0.172697672 | 0.35815124 | 0.48219202 | 0.62966955 | 0.88083664 |
| ITGB1BP1 | 492.290995 | -0.20282749 | 0.3552892 | -0.57088 | 0.568081 | 0.85263558 |
| CPSF3 | 262.431662 | 0.005598285 | 0.28192995 | 0.01985701 | 0.98415744 | 0.99523767 |
| IAH1 | 308.181454 | 0.155788617 | 0.25054802 | 0.62179145 | 0.53407901 | 0.83916403 |
| ADAM17 | 1003.16784 | -0.536445267 | 0.31275093 | -1.7152475 | 0.08629985 | 0.52421699 |
| AC080162.1 | 40.0951476 | 0.079821726 | 0.4846921 | 0.16468543 | 0.86919159 | 0.96556934 |
| YWHAQ | 959.82674 | -0.269917017 | 0.34763238 | -0.7764438 | 0.43748698 | 0.79647151 |
| TAF1B | 848.454837 | -0.01650431 | 0.29478531 | -0.0559876 | 0.95535172 | 0.98840182 |
| AC010969.2 | 42.3178782 | -0.201682358 | 0.59433572 | -0.3393408 | 0.73435301 | 0.91975299 |
| GRHL1 | 314.001089 | 0.040760072 | 0.33437993 | 0.12189748 | 0.90298021 | 0.97526986 |
| KLF11 | 196.067716 | -0.602139376 | 0.30579466 | -1.9690971 | 0.04894194 | 0.45368151 |
| CYS1 | 774.774431 | 0.872134998 | 0.54367821 | 1.60413821 | 0.10868359 | 0.55858026 |
| C2orf48 | 42.016326 | -0.163115012 | 0.63509416 | -0.256836 | 0.79730541 | 0.9429392 |
| HPCAL1 | 2182.43184 | -0.11252648 | 0.43882363 | -0.2564276 | 0.7976207 | 0.94314932 |
| ODC1 | 428.595053 | 0.18036698 | 0.36591409 | 0.49292165 | 0.62206795 | 0.87713418 |
| NOL10 | 962.155409 | -0.392194472 | 0.28907298 | -1.3567317 | 0.17486649 | 0.62699479 |
| ATP6V1C2 | 36.2810835 | 0.23118432 | 0.48287519 | 0.4787662 | 0.63210496 | 0.88123095 |
| PDIA6 | 771.262651 | 0.13720085 | 0.45790331 | 0.29962843 | 0.7644606 | 0.93083208 |
| PQLC3 | 94.4962421 | 0.117742455 | 0.42203768 | 0.27898564 | 0.78025584 | 0.93716849 |
| ROCK2 | 2495.02451 | -0.069816055 | 0.21810404 | -0.3201044 | 0.74888922 | 0.92554142 |
| E2F6 | 267.892048 | -0.310834607 | 0.35793656 | -0.868407 | 0.38517158 | 0.77317162 |

| | | | | | | |
|---|---|---|---|---|---|---|
| GREB1 | 302.527001 | 0.127266702 | 0.46821188 | 0.27181434 | 7.86E-01 | 9.39E-01 |
| LPIN1 | 645.671889 | -0.524869862 | 0.74700816 | -0.7026294 | 0.48228676 | 0.8177376 |
| MIR3681HG | 63.5155984 | -0.481775731 | 0.43896323 | -1.097531 | 0.27240936 | 0.70073986 |
| TRIB2 | 45.1207268 | -0.313288397 | 0.41100075 | -0.7622575 | 0.44590635 | 0.8013686 |
| LINC00276 | 49.8062312 | 0.036984326 | 0.42804741 | 0.08640241 | 0.93114653 | 0.98272895 |
| FAM84A | 50.2078631 | 0.504679612 | 0.67588824 | 0.74669092 | 0.45525015 | 0.80465616 |
| NBAS | 2409.30176 | -0.266656348 | 0.15918318 | -1.6751541 | 0.093904 | 0.53742888 |
| DDX1 | 646.959883 | -0.019673206 | 0.19961744 | -0.0985545 | 0.92149196 | 0.97939204 |
| MYCN | 102.405751 | -0.309960494 | 0.74016063 | -0.4187746 | 6.75E-01 | 8.98E-01 |
| VSNL1 | 104.533773 | 0.534441056 | 0.68232668 | 0.78326272 | 0.43347285 | 0.79380258 |
| SMC6 | 738.485355 | 0.354270651 | 0.18775349 | 1.8868925 | 0.05917479 | 0.47855094 |
| GEN1 | 337.253914 | -0.5430894 | 0.45574527 | -1.1916512 | 0.23339804 | 0.6699868 |
| KCNS3 | 179.479463 | 0.40159057 | 0.70800684 | 0.56721284 | 0.57056959 | 0.85361246 |
| RDH14 | 138.654353 | -0.0633383 | 0.38814766 | -0.1631809 | 0.87037597 | 0.96564168 |
| LINC01376 | 1112.00827 | 0.389095328 | 0.56570629 | 0.68780449 | 0.49157591 | 0.82061027 |
| AC010096.1 | 31.820599 | -1.128313595 | 0.64231937 | -1.756624 | 0.07898193 | 0.51192014 |
| LINC00954 | 59.4316434 | -1.025027247 | 0.59549172 | -1.7213123 | 0.08519416 | 0.5221244 |
| TTC32 | 145.259248 | -0.323417714 | 0.25147068 | -1.2861051 | 0.1984064 | 0.64582202 |
| WDR35 | 696.633495 | -0.056178515 | 0.31757393 | -0.176899 | 0.85958772 | 0.96283855 |
| LAPTM4A | 2069.25579 | 0.242390894 | 0.41338795 | 0.5863521 | 0.55763893 | 0.84862404 |
| AC098828.3 | 110.770067 | 1.785843384 | 1.11740399 | 1.59820746 | 0.10999681 | 0.56111885 |
| SDC1 | 669.153478 | -0.377241629 | 0.46398949 | -0.8130392 | 0.4161956 | 0.78632759 |
| PUM2 | 3527.24479 | -0.459884285 | 0.19923672 | -2.3082306 | 0.02098631 | 0.35422078 |
| RHOB | 934.979229 | -0.288396559 | 0.46931567 | -0.6145044 | 0.53888205 | 0.84070752 |
| HS1BP3 | 367.951001 | 0.646829202 | 0.38927169 | 1.66163945 | 0.09658509 | 0.53969191 |
| GDF7 | 164.521338 | 0.696653163 | 0.55197918 | 1.26210044 | 0.20691265 | 0.65251041 |
| LDAH | 1152.39332 | 0.622100172 | 0.43056962 | 1.44483063 | 0.14850547 | 0.60622442 |
| AC018742.1 | 91.5813583 | 0.826456089 | 0.80681693 | 1.02434153 | 0.30567399 | 0.72122 |
| AC018467.1 | 1482.91744 | -0.764865063 | 0.55641737 | -1.3746247 | 0.16924782 | 0.62326131 |
| KLHL29 | 5457.37321 | -2.19750687 | 0.82566112 | -2.6615119 | 7.78E-03 | 0.24233539 |
| AC009242.1 | 42.5934294 | 1.218636685 | 0.58543755 | 2.08158273 | 0.0373806 | 0.42215481 |
| ATAD2B | 1587.76391 | -0.082348512 | 0.17429357 | -0.4724702 | 0.63659124 | 0.88266168 |
| UBXN2A | 731.186933 | 0.09738032 | 0.2381764 | 0.40885798 | 0.68264389 | 0.89962501 |
| WDCP | 37.7182106 | 0.291801071 | 0.39809992 | 0.73298449 | 0.46356789 | 0.8082072 |
| SF3B6 | 248.028581 | -0.15277077 | 0.30875089 | -0.4948027 | 0.62073941 | 0.87609696 |
| FAM228B | 335.072959 | 0.269283483 | 0.35411461 | 0.76044161 | 0.44699066 | 0.80223612 |
| TP53I3 | 230.921993 | 0.422366362 | 0.55105928 | 0.76646266 | 0.44340106 | 0.7993765 |
| ITSN2 | 2387.57745 | 0.21861263 | 0.19290207 | 1.13328297 | 0.25709544 | 0.69138851 |
| NCOA1 | 3662.45466 | 0.006900048 | 0.1465204 | 0.04709275 | 0.96243931 | 0.99012655 |
| PTRHD1 | 61.8720702 | 0.066595188 | 0.54597174 | 0.12197552 | 0.9029184 | 0.97526986 |
| CENPO | 57.9619364 | -0.645310554 | 0.41468846 | -1.5561334 | 0.11967638 | 0.57673415 |
| ADCY3 | 51.7091135 | -0.285007149 | 0.52597141 | -0.5418681 | 0.58790935 | 0.8613161 |
| DNAJC27 | 224.856206 | 0.186018246 | 0.40717146 | 0.45685482 | 0.6477754 | 0.88774616 |
| DNAJC27-AS | 177.741901 | 0.93851596 | 0.66626568 | 1.40862119 | 0.15894721 | 0.61423843 |
| DNMT3A | 784.5073 | 0.239568805 | 0.28601421 | 0.83761156 | 0.4022489 | 0.78260179 |
| DTNB | 1196.16442 | 0.02096325 | 0.26032828 | 0.08052621 | 0.93581875 | 0.98406248 |
| ASXL2 | 1681.73013 | -0.19314946 | 0.18544735 | -1.0415326 | 0.29762843 | 0.71839551 |
| KIF3C | 50.0212314 | 0.368699548 | 0.42306561 | 0.87149496 | 0.38348396 | 0.77317162 |
| RAB10 | 4166.21434 | -0.887064112 | 0.43102826 | -2.0580184 | 0.03958836 | 0.42828548 |
| HADHA | 1369.63058 | 0.058088959 | 0.22112336 | 0.26269933 | 0.79278234 | 0.9415707 |
| HADHB | 933.943766 | 0.248818963 | 0.26599519 | 0.93542655 | 0.34956852 | 0.75434303 |
| ADGRF3 | 49.2149641 | -0.289177303 | 0.62596819 | -0.4619681 | 0.64410423 | 0.88546907 |
| SELENOI | 582.258149 | -0.243085468 | 0.3409879 | -0.7128859 | 0.47591636 | 0.81570932 |
| KCNK3 | 80.6018307 | 0.476585823 | 0.54911008 | 0.86792402 | 3.85E-01 | 7.73E-01 |

| | | | | | | |
|---|---|---|---|---|---|---|
| CENPA | 151.493388 | -0.440718152 | 0.82404026 | -0.534826 | 0.59277019 | 0.86382106 |
| AC013472.2 | 39.6579224 | 0.178541926 | 0.77107568 | 0.23154916 | 0.8168882 | 0.94946888 |
| MAPRE3 | 220.00195 | 0.035406964 | 0.26177039 | 0.13525962 | 0.89240661 | 0.97237481 |
| TMEM214 | 206.6134 | -0.485156611 | 0.42388397 | -1.1445505 | 0.2523954 | 0.68874851 |
| AGBL5 | 211.866079 | 0.23512648 | 0.3088769 | 0.76123039 | 0.44651947 | 0.80205685 |
| OST4 | 341.208755 | -0.509318971 | 0.52616776 | -0.9679783 | 0.33305521 | 0.74000014 |
| KHK | 922.773511 | 0.822871377 | 0.54301608 | 1.51537202 | 1.30E-01 | 0.58838111 |
| CGREF1 | 76.6554471 | 0.548552192 | 0.46937832 | 1.16867816 | 0.24253333 | 0.68081186 |
| PREB | 118.709504 | 0.121383879 | 0.36944415 | 0.32855813 | 0.7424897 | 0.92302796 |
| SLC5A6 | 73.5380355 | -0.407193974 | 0.40910162 | -0.995337 | 0.3195724 | 0.73132207 |
| ATRAID | 199.321741 | 0.265960198 | 0.36190388 | 0.73489181 | 0.46240539 | 0.8082072 |
| CAD | 133.855842 | -0.426844198 | 0.43677067 | -0.977273 | 0.328434 | 0.73826734 |
| MPV17 | 302.396809 | -0.012194063 | 0.43162042 | -0.0282518 | 0.97746131 | 0.99473325 |
| GTF3C2 | 646.308258 | -0.224102463 | 0.261494 | -0.857008 | 0.39144044 | 0.77606943 |
| AC074117.1 | 105.916768 | -0.217943959 | 0.3454141 | -0.6309643 | 0.52806388 | 0.83761342 |
| EIF2B4 | 180.370253 | -0.387352298 | 0.28544207 | -1.3570259 | 0.17477297 | 0.62699479 |
| SNX17 | 155.171027 | 0.176519841 | 0.40333794 | 0.4376475 | 0.66164183 | 0.89184036 |
| ZNF513 | 117.217677 | -0.058449381 | 0.36501492 | -0.1601287 | 0.87277966 | 0.96592775 |
| PPM1G | 889.571633 | -0.229413528 | 0.51181071 | -0.448239 | 0.65398071 | 0.89009171 |
| NRBP1 | 559.585165 | -0.747353796 | 0.4952034 | -1.5091855 | 0.13125138 | 0.58930656 |
| KRTCAP3 | 74.7331043 | -0.270711433 | 0.40949871 | -0.6610801 | 0.50856097 | 0.82874455 |
| IFT172 | 463.017736 | 0.439349308 | 0.33896293 | 1.2961574 | 0.19492126 | 0.64558745 |
| ZNF512 | 420.485613 | -0.227583879 | 0.24815373 | -0.9171084 | 0.35908583 | 0.75936492 |
| GPN1 | 143.426053 | -1.108657432 | 0.46508041 | -2.3837973 | 0.01713504 | 0.32964126 |
| SUPT7L | 212.239889 | -0.239562204 | 0.23960626 | -0.9998161 | 0.3173995 | 0.72935371 |
| SLC4A1AP | 266.36717 | -0.226690851 | 0.21269627 | -1.0657961 | 0.28651582 | 0.71118494 |
| MRPL33 | 657.745471 | -0.214484306 | 0.28746881 | -0.7461133 | 0.45559898 | 0.80465616 |
| RBKS | 1134.44462 | 1.18158492 | 0.58698487 | 2.01297339 | 0.04411743 | 0.44360789 |
| BABAM2 | 4535.21319 | 0.626723612 | 0.36753117 | 1.70522576 | 0.08815233 | 0.52841003 |
| FOSL2 | 1007.52666 | -0.376384134 | 0.46100512 | -0.8164424 | 4.14E-01 | 0.78632759 |
| PPP1CB | 761.015531 | -0.411874451 | 0.28978672 | -1.421302 | 1.55E-01 | 0.61159057 |
| SPDYA | 86.8915893 | -0.156803357 | 0.49224532 | -0.3185472 | 0.75006992 | 0.92565124 |
| TRMT61B | 337.357517 | -0.231055338 | 0.22678092 | -1.0188482 | 0.30827503 | 0.72367286 |
| WDR43 | 926.148368 | -0.852972215 | 0.41326532 | -2.0639821 | 0.03901941 | 0.42684685 |
| CLIP4 | 1160.82901 | -0.179980145 | 0.20041661 | -0.8980301 | 3.69E-01 | 0.76620563 |
| ALK | 120.2557 | 0.463576174 | 0.47681403 | 0.97223685 | 0.33093273 | 0.73887944 |
| AC016907.2 | 49.2942262 | 0.290168488 | 0.82897742 | 0.35003184 | 0.7263148 | 0.91635631 |
| YPEL5 | 475.159439 | -0.377179124 | 0.2943219 | -1.281519 | 0.20001142 | 0.64705133 |
| LBH | 137.429014 | -0.807502756 | 0.54673816 | -1.476946 | 0.1396901 | 0.59500405 |
| LCLAT1 | 1872.2951 | 0.106986279 | 0.28078158 | 0.38103027 | 7.03E-01 | 0.90887942 |
| CAPN13 | 285.071162 | 1.169986193 | 0.56596115 | 2.06725529 | 0.0387101 | 0.42523931 |
| GALNT14 | 5171.12102 | 0.500871181 | 0.27165792 | 1.84375698 | 0.06521856 | 0.48790819 |
| MEMO1 | 109.62442 | 0.000512459 | 0.36996475 | 0.00138516 | 0.99889481 | 0.99977652 |
| DPY30 | 338.293645 | -0.239372619 | 0.23932977 | -1.0001791 | 0.31722387 | 0.72919462 |
| SPAST | 612.492062 | -0.13086178 | 0.19010719 | -0.6883579 | 0.49122744 | 0.82035445 |
| SLC30A6 | 638.21832 | -0.021399969 | 0.22287751 | -0.0960167 | 0.92350729 | 0.98012108 |
| YIPF4 | 953.325426 | -0.232189372 | 0.23149242 | -1.0030107 | 0.3158557 | 0.72798933 |
| AL133245.1 | 61.0290134 | -1.05588305 | 0.65916456 | -1.6018505 | 0.10918868 | 0.55946616 |
| BIRC6 | 5960.49356 | -0.245039577 | 0.15838791 | -1.5470851 | 0.12184273 | 0.58049991 |
| TTC27 | 692.949985 | -0.554297624 | 0.22403719 | -2.4741321 | 0.01335603 | 0.29875272 |
| LINC00486 | 1083.97624 | -0.227609746 | 0.21020905 | -1.0827781 | 0.27890695 | 0.70446562 |
| LTBP1 | 110.865735 | -0.089266376 | 0.65400569 | -0.1364917 | 0.89143255 | 0.97234681 |
| RASGRP3 | 471.97576 | -0.010765844 | 0.47941729 | -0.0224561 | 0.98208413 | 0.99522272 |
| AC020594.1 | 61.3187637 | 0.049183844 | 0.36463015 | 0.13488693 | 0.89270127 | 0.9724379 |

| | | | | | | |
|---|---|---|---|---|---|---|
| FAM98A | 293.455207 | -0.236258636 | 0.27251477 | -0.8669572 | 0.38596547 | 0.77317162 |
| LINC01317 | 111.031696 | 0.232347337 | 0.64492361 | 0.3602711 | 0.71864441 | 0.91436544 |
| AC068599.1 | 445.524677 | 1.341810662 | 0.58066151 | 2.31083106 | 0.02084219 | 0.35313211 |
| LINC01320 | 12172.2991 | 0.74370373 | 0.58367441 | 1.27417566 | 0.20260117 | 0.64881821 |
| AC073218.2 | 100.317178 | 1.159332987 | 0.73700441 | 1.57303399 | 0.11571095 | 0.57350419 |
| AC073218.1 | 63.3448147 | 1.11452402 | 1.1075297 | 1.00631524 | 0.31426395 | 0.72719146 |
| AC012593.1 | 5898.81359 | 3.611620887 | 1.07407657 | 3.36253577 | 0.0007723 | 0.09205159 |
| CRIM1 | 2579.6943 | -0.42360155 | 0.27293894 | -1.5520012 | 0.12066194 | 0.57869603 |
| FEZ2 | 731.638192 | 0.222855296 | 0.27016212 | 0.82489469 | 0.40943138 | 0.78542936 |
| STRN | 1176.14002 | -0.207329918 | 0.20526121 | -1.0100784 | 0.31245773 | 0.7265535 |
| HEATR5B | 886.243031 | -0.345944995 | 0.19658284 | -1.7597924 | 0.07844301 | 0.51192014 |
| GPATCH11 | 90.1175265 | -0.578546968 | 0.37575816 | -1.539679 | 0.12363861 | 0.58354574 |
| EIF2AK2 | 813.603837 | -0.94628488 | 0.54827407 | -1.725934 | 0.08435928 | 0.52183702 |
| CEBPZOS | 593.822675 | 0.040407802 | 0.19309596 | 0.2092628 | 0.83424309 | 0.95393659 |
| CEBPZ | 576.07915 | -0.121285859 | 0.26011241 | -0.4662825 | 0.64101332 | 0.88396933 |
| NDUFAF7 | 388.347486 | -0.091807008 | 0.14950385 | -0.6140779 | 0.53916387 | 0.84091105 |
| PRKD3 | 781.254461 | 0.281302705 | 0.22776725 | 1.23504455 | 0.21681392 | 0.66032006 |
| QPCT | 61.5204117 | 0.461749286 | 0.54953432 | 0.8402556 | 0.40076509 | 0.78232083 |
| CDC42EP3 | 119.809576 | -0.213778023 | 0.39591751 | -0.539956 | 0.5892274 | 0.8624175 |
| RMDN2 | 451.675249 | 0.17186963 | 0.37002105 | 0.46448608 | 6.42E-01 | 8.84E-01 |
| RMDN2-AS1 | 50.1234804 | 0.121140425 | 0.70133362 | 0.17272867 | 0.86286471 | 0.96391903 |
| CYP1B1 | 70.5183605 | 0.905116503 | 0.57813271 | 1.56558605 | 0.11744556 | 0.57350419 |
| CYP1B1-AS1 | 105.727962 | 0.9593408 | 0.59392273 | 1.61526199 | 0.10625397 | 0.55440878 |
| ATL2 | 1289.49207 | -0.01063355 | 0.19112551 | -0.0556365 | 9.56E-01 | 9.88E-01 |
| AC011247.1 | 94.9889006 | 1.009533403 | 0.84833968 | 1.19001083 | 0.23404214 | 0.67060722 |
| HNRNPLL | 621.336964 | -0.150621904 | 0.21450455 | -0.7021851 | 0.48256373 | 0.8177376 |
| GALM | 1885.04255 | 0.405590998 | 0.48312118 | 0.83952228 | 4.01E-01 | 7.82E-01 |
| SRSF7 | 913.144009 | -0.156865334 | 0.28675251 | -0.5470408 | 0.58435068 | 0.86025731 |
| GEMIN6 | 74.2831944 | -0.201371555 | 0.31439969 | -0.6404954 | 0.52185058 | 0.83445062 |
| DHX57 | 748.341995 | -0.14381528 | 0.25661801 | -0.5604255 | 0.57518924 | 0.85492506 |
| MORN2 | 141.759404 | 0.047243 | 0.38457752 | 0.12284389 | 0.90223072 | 0.9750023 |
| ARHGEF33 | 62.59857 | 0.758421729 | 0.60105639 | 1.26181459 | 0.20701551 | 0.65251041 |
| SOS1 | 3013.98115 | 0.042088876 | 0.1969387 | 0.21371563 | 0.83076883 | 0.95325673 |
| CDKL4 | 34.5275756 | -0.074633105 | 0.51149054 | -0.145913 | 0.88399009 | 0.96880564 |
| MAP4K3 | 2711.86459 | -0.333283311 | 0.26952183 | -1.2365726 | 0.2162458 | 0.66015172 |
| AC007388.1 | 2171.0566 | 0.273419559 | 0.41238734 | 0.66301637 | 0.50732007 | 0.8283307 |
| TMEM178A | 96.7039891 | -0.15639496 | 0.5785379 | -0.2703279 | 0.78690797 | 0.93939623 |
| THUMPD2 | 523.399007 | -0.206268411 | 0.2121786 | -0.9721452 | 0.33097831 | 0.73887944 |
| SLC8A1 | 187.324948 | 0.486920871 | 0.70488924 | 0.69077642 | 0.48970605 | 0.81970016 |
| PKDCC | 216.783222 | 1.733569194 | 0.66844115 | 2.59345075 | 0.00950181 | 0.26008591 |
| EML4 | 3006.86371 | 0.019803873 | 0.25453157 | 0.07780518 | 0.93798303 | 0.98435387 |
| COX7A2L | 717.062874 | -0.34755198 | 0.33825082 | -1.0274978 | 0.30418611 | 0.72082426 |
| MTA3 | 1349.91314 | 0.117107659 | 0.30719136 | 0.38122055 | 0.7030396 | 0.90885257 |
| OXER1 | 76.1858302 | 0.691177567 | 0.62765765 | 1.10120153 | 2.71E-01 | 7.00E-01 |
| HAAO | 513.905959 | 0.74482709 | 0.49647134 | 1.50024187 | 1.34E-01 | 5.92E-01 |
| ZFP36L2 | 1012.50355 | 0.493061089 | 0.38074181 | 1.29500118 | 0.19531982 | 0.64558745 |
| THADA | 1844.61886 | -0.358478865 | 0.21228287 | -1.6886848 | 0.09127985 | 0.53537574 |
| PLEKHH2 | 475.911505 | 0.36356472 | 0.57101 | 0.63670464 | 0.52431725 | 0.83589803 |
| DYNC2LI1 | 743.973535 | 0.605654107 | 0.48800191 | 1.24108962 | 0.21457264 | 0.65925411 |
| LRPPRC | 1826.49566 | -0.190873054 | 0.2289538 | -0.833675 | 0.40446416 | 0.78408331 |
| PPM1B | 1609.87095 | -0.118713688 | 0.22238648 | -0.533817 | 0.59346813 | 0.86444505 |
| SLC3A1 | 5402.6267 | 0.459013278 | 0.52948493 | 0.86690527 | 0.38599392 | 0.77317162 |
| PREPL | 1250.71224 | 0.071561069 | 0.37065866 | 0.19306461 | 0.84690837 | 0.95835656 |
| CAMKMT | 5225.66081 | 0.117942729 | 0.13308334 | 0.88623208 | 0.37549248 | 0.7697436 |

| | | | | | | |
|---|---|---|---|---|---|---|
| LINC01833 | 51.63066 | 0.466030917 | 0.81185052 | 0.57403538 | 0.56594386 | 0.85208617 |
| SRBD1 | 1079.98362 | 0.125521098 | 0.21189047 | 0.59238671 | 0.55359166 | 0.84727776 |
| PRKCE | 3188.28711 | -0.262149394 | 0.37287775 | -0.7030438 | 0.48202845 | 0.8177376 |
| EPAS1 | 1133.34113 | -0.067352475 | 0.41589399 | -0.1619463 | 0.87134818 | 0.9657206 |
| ATP6V1E2 | 89.724896 | -0.693536521 | 0.4107178 | -1.6885962 | 0.09129683 | 0.53537574 |
| RHOQ | 1198.00013 | -0.357169875 | 0.49945079 | -0.7151253 | 4.75E-01 | 0.81531007 |
| AC018682.1 | 229.151392 | -0.026517051 | 0.51025657 | -0.0519681 | 9.59E-01 | 9.89E-01 |
| PIGF | 364.718772 | 0.008916898 | 0.29808087 | 0.02991436 | 0.97613535 | 0.99429832 |
| CRIPT | 136.826558 | -0.2532089 | 0.32825991 | -0.7713671 | 0.44048936 | 0.79826598 |
| SOCS5 | 608.67223 | -0.386280054 | 0.21166372 | -1.8249705 | 0.06800547 | 0.4900219 |
| MCFD2 | 483.131673 | -0.38301542 | 0.35818903 | -1.0693108 | 0.28492962 | 0.70996734 |
| TTC7A | 860.553726 | 0.139584677 | 0.44185759 | 0.31590422 | 0.75207521 | 0.92679087 |
| AC106869.1 | 460.598096 | 0.385818005 | 0.66952898 | 0.57625288 | 0.56444427 | 0.85163263 |
| EPCAM | 1002.78525 | -0.359119378 | 0.36748566 | -0.9772337 | 0.32845346 | 0.73826734 |
| MSH2 | 284.126832 | 0.15454632 | 0.27483 | 0.56233424 | 0.57388831 | 0.85447394 |
| MSH6 | 215.848422 | 0.127951619 | 0.21080009 | 0.60698087 | 0.54386361 | 0.84396451 |
| FBXO11 | 2374.71339 | -0.052040625 | 0.17866445 | -0.2912758 | 0.77084042 | 0.93302012 |
| FOXN2 | 396.353604 | -0.364314373 | 0.36141885 | -1.0080115 | 0.31344892 | 0.72717675 |
| PPP1R21 | 3534.97643 | 0.253818197 | 0.39421283 | 0.64386082 | 0.51966569 | 0.8335052 |
| NRXN1 | 67.9078289 | 0.041700693 | 0.50613533 | 0.0823904 | 0.93433627 | 0.98359701 |
| AC007402.1 | 101.984745 | 0.35429153 | 0.44671275 | 0.79310817 | 0.4277148 | 0.78990661 |
| ASB3 | 64.3513578 | 0.425045623 | 0.36759287 | 1.15629452 | 0.24756071 | 0.68482118 |
| GPR75-ASB3 | 351.449309 | -0.428051792 | 0.38717669 | -1.1055722 | 0.26891171 | 0.69961625 |
| ERLEC1 | 268.955005 | 0.107572914 | 0.27994279 | 0.38426749 | 0.7007802 | 0.90772956 |
| PSME4 | 1711.50206 | -0.484592569 | 0.3549648 | -1.3651848 | 0.17219496 | 0.62592399 |
| ACYP2 | 2413.9961 | 0.079288101 | 0.25008789 | 0.31704094 | 0.75121254 | 0.92622742 |
| C2orf73 | 46.4341221 | 1.646728699 | 0.84048639 | 1.95925683 | 0.05008272 | 0.45824658 |
| SPTBN1 | 5278.30272 | -0.498277936 | 0.26795091 | -1.8595867 | 0.06294403 | 0.48524961 |
| AC092839.1 | 121.148893 | -0.141138474 | 0.60918036 | -0.2316859 | 0.81678202 | 0.94946888 |
| AC093110.1 | 127.028884 | 0.296235366 | 0.61995412 | 0.47783434 | 0.63276812 | 0.88151688 |
| EML6 | 310.3743 | 0.427873325 | 0.52206996 | 0.81957085 | 0.4124608 | 0.78632759 |
| RTN4 | 3161.12072 | -0.605387935 | 0.42640071 | -1.419763 | 0.1556767 | 0.61229857 |
| CLHC1 | 295.444493 | 0.249955628 | 0.35580385 | 0.70250963 | 0.4823614 | 0.8177376 |
| RPS27A | 1810.17792 | 0.004395064 | 0.55384333 | 0.00793557 | 0.9936684 | 0.99781778 |
| MTIF2 | 318.466172 | 0.061459002 | 0.32690039 | 0.18800529 | 0.8508725 | 0.96006828 |
| AC012358.3 | 273.004837 | 1.007263452 | 0.49963653 | 2.01599243 | 0.04380077 | 0.44242971 |
| CCDC88A | 518.902938 | 0.706303692 | 0.20189926 | 3.49829764 | 0.00046824 | 0.07721972 |
| CFAP36 | 598.79062 | 0.216287799 | 0.20397701 | 1.06035383 | 0.28898366 | 0.71218341 |
| PPP4R3B | 1652.62561 | 0.094851903 | 0.13652655 | 0.69475059 | 0.48721162 | 0.81893983 |
| PNPT1 | 549.302373 | -1.127081721 | 0.41669097 | -2.7048384 | 0.00683376 | 0.23083391 |
| EFEMP1 | 84.6929649 | 0.230595778 | 0.57218974 | 0.40300579 | 0.68694397 | 0.90225574 |
| AC011306.1 | 35.9825756 | 0.338156351 | 0.99092581 | 0.34125295 | 0.73291317 | 0.91952673 |
| CCDC85A | 70.9877584 | 1.304345601 | 0.66346637 | 1.96595586 | 0.04930371 | 0.45419182 |
| VRK2 | 812.706949 | 0.004384417 | 0.23069836 | 0.01900498 | 0.98483714 | 0.99553301 |
| FANCL | 369.884493 | -0.341603589 | 0.3319675 | -1.0290272 | 0.3034669 | 0.72082426 |
| BCL11A | 63.2462878 | 2.200360296 | 0.67379799 | 3.26560828 | 0.00109229 | 0.11093704 |
| PAPOLG | 307.302464 | -0.015290472 | 0.25817 | -0.0592264 | 0.95277181 | 0.98786081 |
| REL | 440.20918 | 0.321509375 | 0.28528594 | 1.12697238 | 0.25975416 | 0.69242517 |
| PUS10 | 645.393222 | 0.730944755 | 0.49485733 | 1.47708181 | 0.13965371 | 0.59500405 |
| PEX13 | 687.862325 | 0.031884522 | 0.20454012 | 0.15588395 | 0.8761245 | 0.96617876 |
| KIAA1841 | 172.552904 | 0.004395703 | 0.33120687 | 0.01327178 | 9.89E-01 | 0.99725454 |
| C2orf74 | 213.85587 | 0.377937665 | 0.58499567 | 0.64605207 | 0.51824563 | 0.83325693 |
| USP34 | 5032.14203 | -0.240955145 | 0.17161738 | -1.4040253 | 0.1603113 | 0.61590405 |
| AC016727.1 | 162.804002 | -0.594468035 | 0.3229347 | -1.8408305 | 0.06564641 | 0.48790819 |

| | | | | | | |
|---|---|---|---|---|---|---|
| XPO1 | 2356.08852 | 0.014452305 | 0.36920139 | 0.03914477 | 0.96877496 | 0.99187457 |
| FAM161A | 131.100471 | 0.14081835 | 0.31740427 | 0.44365613 | 0.65729121 | 0.89053559 |
| CCT4 | 510.042312 | -0.158434478 | 0.25693915 | -0.6166226 | 0.53748372 | 0.84034325 |
| COMMD1 | 2458.77787 | -0.199116602 | 0.19680466 | -1.0117474 | 3.12E-01 | 0.72600794 |
| AC018462.1 | 177.325386 | 0.158238131 | 0.63384442 | 0.24964822 | 0.8028594 | 0.94543168 |
| B3GNT2 | 396.825788 | -0.314626241 | 0.3127489 | -1.0060027 | 0.31441427 | 0.72719146 |
| EHBP1 | 4777.43276 | 0.135536343 | 0.26010065 | 0.5210919 | 0.60230275 | 0.8677246 |
| AC092567.1 | 69.1202258 | 0.579436067 | 0.69905623 | 0.82888335 | 0.40717042 | 0.78481094 |
| WDPCP | 3518.16714 | 0.568846789 | 0.28149482 | 2.02080729 | 0.04329972 | 0.44144764 |
| MDH1 | 911.704885 | 0.63702066 | 0.5046984 | 1.26218087 | 0.20688371 | 0.65251041 |
| UGP2 | 1558.1501 | 0.385964009 | 0.38512512 | 1.00217822 | 0.31625753 | 0.72798933 |
| VPS54 | 1639.53155 | 0.044026902 | 0.20595935 | 0.213765 | 0.83073032 | 0.95325673 |
| PELI1 | 852.341333 | -0.059638164 | 0.29157998 | -0.2045345 | 0.83793584 | 0.95496955 |
| AC012368.1 | 73.4566646 | 0.157946404 | 0.74487686 | 0.21204364 | 0.83207299 | 0.95325673 |
| AC008074.1 | 302.140959 | 0.340174398 | 0.39273422 | 0.86616948 | 0.38639723 | 0.77334938 |
| LGALSL | 69.2211484 | -0.185200707 | 0.4245288 | -0.43625 | 0.66265532 | 0.89232896 |
| AFTPH | 2266.00215 | -0.060751262 | 0.18037657 | -0.3368024 | 0.73626586 | 0.92019178 |
| SERTAD2 | 838.961026 | -0.288280809 | 0.37595636 | -0.7667933 | 0.44320444 | 0.7993765 |
| SLC1A4 | 69.3653463 | -1.986126115 | 0.66012655 | -3.0087051 | 0.00262364 | 0.16823326 |
| CEP68 | 241.29251 | 0.403993697 | 0.30724698 | 1.31488254 | 0.18854938 | 0.64066593 |
| RAB1A | 2979.46585 | -0.239056633 | 0.22001972 | -1.0865237 | 0.27724738 | 0.7036882 |
| ACTR2 | 1313.73396 | -0.073029861 | 0.28193895 | -0.2590272 | 0.79561424 | 0.94252227 |
| SPRED2 | 1668.55661 | -0.788067536 | 0.54008188 | -1.4591631 | 0.14452023 | 0.60101297 |
| MEIS1 | 479.573617 | 0.452258929 | 0.51145934 | 0.88425196 | 0.37656021 | 0.77035529 |
| LINC01798 | 61.3386057 | 0.253044527 | 0.75061412 | 0.33711666 | 0.73602896 | 0.92019178 |
| LINC01829 | 69.3765364 | 0.767251726 | 0.69104197 | 1.11028238 | 0.26687736 | 0.69812269 |
| ETAA1 | 206.40848 | 0.205477922 | 0.23945516 | 0.85810607 | 0.3908339 | 0.77606489 |
| C1D | 279.673891 | 0.1019353 | 0.20563998 | 0.49569787 | 0.62010759 | 0.87586466 |
| WDR92 | 77.0441851 | 0.100007647 | 0.33393627 | 0.29948123 | 0.76457289 | 0.93083208 |
| AC017083.3 | 38.9047602 | -0.379601925 | 0.45782365 | -0.8291444 | 0.40702269 | 0.78481094 |
| PNO1 | 144.514235 | -0.992590312 | 0.52510926 | -1.8902548 | 0.05872389 | 0.47778423 |
| APLF | 475.349305 | 0.681642825 | 0.32049506 | 2.12684347 | 0.03343309 | 0.41254527 |
| ARHGAP25 | 41.9458708 | 0.719331395 | 0.5942557 | 1.21047454 | 0.22609686 | 0.66688879 |
| ANTXR1 | 95.6202755 | 1.418881505 | 0.66396706 | 2.13697573 | 0.03259997 | 0.41149854 |
| GFPT1 | 1190.90941 | -0.191418093 | 0.40258703 | -0.4754701 | 0.63445194 | 0.88238823 |
| NFU1 | 351.282029 | 0.296125745 | 0.20630577 | 1.43537305 | 0.15118082 | 0.60905477 |
| AAK1 | 4556.36925 | -0.317257325 | 0.24931516 | -1.2725152 | 0.20319014 | 0.64881821 |
| ANXA4 | 4009.64451 | -0.201583635 | 0.33365383 | -0.60417 | 0.54573062 | 0.84460887 |
| GMCL1 | 678.218999 | -0.347806606 | 0.28045037 | -1.2401717 | 0.2149119 | 0.65925411 |
| MXD1 | 441.441185 | -0.642580059 | 0.36586323 | -1.7563396 | 0.07903045 | 0.51192014 |
| PCBP1 | 624.576842 | -0.40056906 | 0.4261344 | -0.9400064 | 0.34721428 | 0.75209642 |
| C2orf42 | 288.04754 | -0.161551283 | 0.28831875 | -0.5603218 | 0.57525995 | 0.85493746 |
| PCYOX1 | 405.660247 | 0.840536879 | 0.49483922 | 1.69860601 | 0.08939344 | 0.53095823 |
| SNRPG | 270.369279 | -0.404471548 | 0.31228672 | -1.2951929 | 0.19525369 | 0.64558745 |
| FAM136A | 264.473911 | -0.478836945 | 0.30944164 | -1.5474225 | 0.12176141 | 0.5804281 |
| TGFA | 587.870628 | -0.187756543 | 0.28525005 | -0.6582174 | 5.10E-01 | 0.82930674 |
| NAGK | 239.004559 | -0.53804193 | 0.3195758 | -1.6836129 | 0.09225651 | 0.53621484 |
| MCEE | 324.623793 | 0.038633726 | 0.29889758 | 0.12925406 | 0.89715662 | 0.97335173 |
| MPHOSPH10 | 264.198478 | -0.600597886 | 0.36537426 | -1.6437882 | 0.10021997 | 0.54542315 |
| PAIP2B | 566.138781 | -0.435242427 | 0.5759927 | -0.7556388 | 0.4498658 | 0.80321497 |
| ZNF638 | 4405.63614 | 0.044603993 | 0.1895974 | 0.23525635 | 0.81400977 | 0.94881072 |
| DYSF | 392.976347 | -0.79091081 | 0.62042762 | -1.2747834 | 0.20238593 | 0.64881821 |
| EXOC6B | 4271.53182 | -0.391352222 | 0.2940683 | -1.3308208 | 0.18324797 | 0.63276491 |
| SPR | 117.504533 | 0.110426613 | 0.42644145 | 0.25894906 | 0.79567454 | 0.94252227 |

| | | | | | | |
|---|---|---|---|---|---|---|
| EMX1 | 311.395661 | 0.202279234 | 0.31906209 | 0.63398079 | 0.52609336 | 0.83685293 |
| SFXN5 | 2069.9028 | 0.122077654 | 0.49061095 | 0.24882782 | 0.80349397 | 0.94543168 |
| RAB11FIP5 | 380.390415 | -0.177244816 | 0.39474473 | -0.4490112 | 0.65342355 | 0.88977515 |
| SMYD5 | 98.8139585 | -0.39259039 | 0.26982441 | -1.4549847 | 0.1456735 | 0.60203043 |
| PRADC1 | 61.1739277 | 0.366711357 | 0.4502741 | 0.81441806 | 0.41540549 | 0.78632759 |
| CCT7 | 364.921908 | 0.335515326 | 0.2980707 | 1.12562331 | 0.260325 | 0.69280433 |
| ALMS1 | 988.71152 | 0.179011939 | 0.31374673 | 0.57056193 | 0.56829663 | 0.85263558 |
| NAT8 | 1512.20796 | 1.300079209 | 0.97290008 | 1.33629263 | 0.18145364 | 0.6318513 |
| TPRKB | 162.299657 | -0.023800079 | 0.21354769 | -0.1114509 | 0.91125881 | 0.97697493 |
| DUSP11 | 216.961804 | -0.058770752 | 0.25222494 | -0.2330093 | 0.8157542 | 0.94906482 |
| STAMBP | 571.767255 | -0.085578808 | 0.23814187 | -0.3593606 | 0.71932534 | 0.91465123 |
| DGUOK | 445.335251 | -0.50317335 | 0.19910038 | -2.5272345 | 0.01149647 | 0.28627263 |
| DGUOK-AS1 | 52.9114808 | -0.464183189 | 0.37605816 | -1.2343388 | 0.21707668 | 0.66032006 |
| TET3 | 717.574303 | -0.649657519 | 0.2978504 | -2.1811537 | 0.02917205 | 0.39595188 |
| BOLA3 | 95.3247256 | 0.051233479 | 0.45597086 | 0.1123613 | 0.91053694 | 0.97697493 |
| MOB1A | 672.286459 | -0.115694671 | 0.13867199 | -0.8343045 | 0.40410941 | 0.78358009 |
| MTHFD2 | 62.0970416 | -1.473482347 | 0.83992901 | -1.7542939 | 7.94E-02 | 0.51192014 |
| AC006030.1 | 119.340089 | -0.118537589 | 0.54775224 | -0.2164073 | 0.82867027 | 0.95322116 |
| DCTN1 | 130.771555 | -0.325755319 | 0.36710806 | -0.8873554 | 0.37488758 | 0.7697436 |
| WDR54 | 56.0018452 | 0.633393133 | 0.59905377 | 1.05732268 | 0.29036435 | 0.71269438 |
| RTKN | 692.592307 | 0.144498883 | 0.28870214 | 0.50051199 | 0.61671461 | 0.8739075 |
| MOGS | 311.125561 | 0.028786683 | 0.32827781 | 0.08769001 | 0.93012306 | 0.98272895 |
| CCDC142 | 168.020451 | -0.16065399 | 0.34778105 | -0.46194 | 0.64412433 | 0.88546907 |
| TTC31 | 161.399448 | -0.093985974 | 0.36801344 | -0.2553873 | 0.79842394 | 0.94320409 |
| PCGF1 | 44.415134 | -0.900583537 | 0.4162812 | -2.1634019 | 0.03051028 | 0.40445302 |
| AUP1 | 267.595791 | -0.7046248 | 0.37992184 | -1.8546572 | 0.06364519 | 0.48593331 |
| HTRA2 | 84.8379542 | -0.393516736 | 0.37873784 | -1.0390214 | 0.29879476 | 0.71852421 |
| HK2 | 59.6558956 | -1.10821855 | 0.54428151 | -2.0361128 | 4.17E-02 | 4.35E-01 |
| POLE4 | 144.972747 | -0.215135708 | 0.41922656 | -0.5131729 | 0.60783038 | 0.86985416 |
| TACR1 | 72.1881316 | 0.515884047 | 0.64462014 | 0.80029155 | 0.4235419 | 0.78894887 |
| AC007681.1 | 114.240639 | 0.33799528 | 0.44717609 | 0.75584381 | 0.44974285 | 0.8032085 |
| EVA1A | 1962.41683 | 0.156907761 | 0.55830376 | 0.28104371 | 0.77867687 | 0.93646866 |
| AC007099.1 | 319.313515 | 0.515766179 | 0.77316976 | 0.66708012 | 0.50472096 | 0.82715271 |
| MRPL19 | 479.54077 | 0.325597844 | 0.26931195 | 1.20899885 | 0.22666329 | 0.66690202 |
| GCFC2 | 789.839745 | 0.625563279 | 0.2484578 | 2.51778485 | 0.01180954 | 0.29061654 |
| LRRTM4 | 144.885363 | 2.397837243 | 0.77591184 | 3.09034753 | 0.00199922 | 0.14129287 |
| REG1A | 268.01289 | 2.097139843 | 1.49647522 | 1.40138628 | 0.16109859 | 0.61697277 |
| CTNNA2 | 59.7486835 | -0.455144183 | 0.36670231 | -1.2411817 | 0.21453865 | 0.65925411 |
| SUCLG1 | 1517.1642 | 0.148193207 | 0.50308757 | 0.29456742 | 0.76832435 | 0.93198626 |
| DNAH6 | 486.344998 | 1.094803094 | 0.74900077 | 1.46168487 | 0.14382759 | 0.60031813 |
| TRABD2A | 36.2368181 | 0.386104069 | 0.82750062 | 0.46659067 | 0.64079276 | 0.88396933 |
| TMSB10 | 2919.45128 | 0.006858203 | 0.51963535 | 0.01319811 | 9.89E-01 | 0.99725454 |
| KCMF1 | 1591.87373 | -0.644018118 | 0.38121926 | -1.6893641 | 0.09114968 | 0.53537574 |
| TCF7L1 | 87.1767317 | 0.461376552 | 0.58284952 | 0.79158776 | 0.42860108 | 0.79037914 |
| TGOLN2 | 1204.50911 | 0.138500817 | 0.31267123 | 0.4429599 | 0.65779474 | 0.89053559 |
| RETSAT | 425.324341 | -0.097333642 | 0.31860819 | -0.3054964 | 0.75998814 | 0.92952239 |
| ELMOD3 | 512.663474 | 0.641863411 | 0.17687975 | 3.62881239 | 0.00028473 | 0.06116705 |
| CAPG | 127.524974 | 0.206806591 | 0.5071278 | 0.40779975 | 0.6834207 | 0.90047558 |
| SH2D6 | 30.8401744 | 0.311777826 | 0.62586428 | 0.49815564 | 0.61837434 | 0.8747221 |
| MAT2A | 1490.78279 | -0.358954044 | 0.28532331 | -1.2580607 | 0.2083698 | 0.65368257 |
| GGCX | 263.396744 | 0.035570261 | 0.28193619 | 0.12616422 | 0.89960193 | 0.97465308 |
| VAMP8 | 712.407872 | -0.155025752 | 0.29896334 | -0.5185443 | 0.60407853 | 0.86848168 |
| VAMP5 | 154.257434 | -1.097740202 | 0.44394123 | -2.4727152 | 0.0134091 | 0.29875272 |
| RNF181 | 121.701524 | -0.459196789 | 0.45758495 | -1.0035225 | 0.31560883 | 0.72792366 |

| | | | | | | |
|---|---|---|---|---|---|---|
| TMEM150A | 312.264689 | 0.152496219 | 0.32288336 | 0.47229507 | 6.37E-01 | 0.88266168 |
| USP39 | 563.492428 | -0.096278426 | 0.25782447 | -0.3734263 | 0.70883122 | 0.91082291 |
| C2orf68 | 230.253923 | -0.167465077 | 0.23506195 | -0.7124295 | 0.47619883 | 0.81570932 |
| ST3GAL5 | 124.219238 | 0.925955672 | 0.62092621 | 1.49124914 | 0.1358961 | 0.59398208 |
| POLR1A | 460.263772 | -0.537585099 | 0.39642702 | -1.3560758 | 0.17507504 | 0.62699479 |
| PTCD3 | 902.477188 | 0.076501403 | 0.18686469 | 0.40939465 | 0.68225007 | 0.89962501 |
| IMMT | 909.162822 | 0.304553654 | 0.19164683 | 1.58914004 | 0.11202878 | 0.56455091 |
| MRPL35 | 159.489779 | 0.084474451 | 0.27332225 | 0.30906541 | 0.75727177 | 0.92799282 |
| REEP1 | 263.167577 | -0.955955135 | 0.49621192 | -1.9265058 | 0.05404125 | 0.46711025 |
| KDM3A | 1218.29048 | -0.304910892 | 0.37152383 | -0.8207035 | 0.4118152 | 0.78632759 |
| CHMP3 | 187.022382 | 0.093167127 | 0.35856533 | 0.25983306 | 0.79499255 | 0.94252227 |
| RNF103-CHN | 324.852413 | -1.65575262 | 0.83141679 | -1.9914833 | 4.64E-02 | 0.44508419 |
| AC015971.1 | 50.8943891 | 0.239434117 | 0.50797028 | 0.47135458 | 0.63738755 | 0.8827026 |
| RMND5A | 746.193914 | -0.494851428 | 0.25794014 | -1.9184739 | 0.05505095 | 0.46904913 |
| PLGLB1 | 185.804253 | 0.568736542 | 0.41047404 | 1.38556033 | 0.1658812 | 0.62065658 |
| CYTOR | 192.360365 | -0.795698375 | 0.73588556 | -1.0812801 | 2.80E-01 | 7.04E-01 |
| KRCC1 | 798.54192 | 0.494087073 | 0.26044131 | 1.8971148 | 0.05781279 | 0.47684824 |
| FABP1 | 409.813577 | 1.379790982 | 0.97161458 | 1.42010115 | 0.15557824 | 0.61208693 |
| THNSL2 | 415.580607 | 1.128514218 | 0.89439049 | 1.26176902 | 0.20703191 | 0.65251041 |
| EIF2AK3 | 1194.61403 | -0.366121787 | 0.4375897 | -0.8366783 | 4.03E-01 | 0.78260941 |
| RPIA | 211.001472 | -0.763988148 | 0.37667376 | -2.0282489 | 0.04253485 | 0.43836478 |
| AL845331.2 | 287.573761 | 1.470851674 | 0.85511343 | 1.72006616 | 0.08542042 | 0.5226185 |
| ANKRD20A8I | 72.1550427 | -0.218193121 | 0.84214452 | -0.2590923 | 0.79556406 | 0.94252227 |
| MAL | 218.184344 | 0.836872733 | 0.66256597 | 1.26307835 | 0.20656102 | 0.65251041 |
| MRPS5 | 719.027364 | -0.141548015 | 0.25787133 | -0.5489095 | 0.58306758 | 0.85990837 |
| ZNF514 | 75.4664904 | -0.157289755 | 0.36009681 | -0.4367985 | 0.66225747 | 0.89205613 |
| ZNF2 | 88.4783391 | -0.21520991 | 0.40861423 | -0.5266824 | 0.59841418 | 0.86646266 |
| AC092835.1 | 108.435183 | 0.071892408 | 0.36009815 | 0.1996467 | 0.8417569 | 0.95681213 |
| KCNIP3 | 146.218124 | 0.522932 | 0.55544285 | 0.9414686 | 0.34646477 | 0.75166079 |
| FAHD2A | 154.707279 | 0.634919961 | 0.42468237 | 1.49504665 | 0.13490227 | 0.59342278 |
| LINC00342 | 51.5622565 | 0.355930311 | 0.56901891 | 0.62551578 | 0.53163259 | 0.83789294 |
| ANKRD36C | 553.647111 | 0.296134034 | 0.29380811 | 1.00791646 | 0.31349457 | 0.72717675 |
| STARD7 | 567.531962 | -0.15421996 | 0.22997975 | -0.6705806 | 0.50248775 | 0.82638105 |
| TMEM127 | 253.086232 | -0.278352103 | 0.33234668 | -0.8375354 | 0.40229169 | 0.78260179 |
| CIAO1 | 392.103126 | -0.409257469 | 0.29988019 | -1.3647366 | 0.17233585 | 0.62592399 |
| SNRNP200 | 515.729151 | -0.249938511 | 0.32278945 | -0.7743082 | 0.43874856 | 0.79718009 |
| NEURL3 | 91.1921714 | -0.654438409 | 0.65983316 | -0.9918241 | 0.32128336 | 0.73291455 |
| ARID5A | 92.967575 | -1.001504211 | 0.60442805 | -1.6569453 | 9.75E-02 | 5.41E-01 |
| KANSL3 | 724.514139 | -0.291766927 | 0.22285303 | -1.3092347 | 0.19045486 | 0.64281105 |
| LMAN2L | 559.545087 | 0.084022428 | 0.21079528 | 0.3985973 | 6.90E-01 | 0.90417984 |
| CNNM4 | 204.858611 | -0.870863782 | 0.39026787 | -2.2314514 | 0.02565124 | 0.38166806 |
| CNNM3 | 101.063871 | 0.3607434 | 0.29922151 | 1.20560652 | 0.22796925 | 0.66705629 |
| ANKRD23 | 229.744169 | -0.019408732 | 0.383696 | -0.0505836 | 0.95965732 | 0.98912673 |
| SEMA4C | 41.308073 | 0.142999093 | 0.61828922 | 0.23128188 | 0.81709582 | 0.94954998 |
| ANKRD36 | 711.981142 | 0.105958225 | 0.3299061 | 0.32117692 | 0.74807632 | 0.92532868 |
| AC092683.1 | 701.01058 | -0.102221027 | 0.38638143 | -0.2645599 | 0.79134853 | 0.94098694 |
| ANKRD36B | 253.841821 | -0.421453365 | 0.54955162 | -0.766904 | 0.44313857 | 0.7993765 |
| COX5B | 701.038136 | -0.003443361 | 0.49607613 | -0.0069412 | 0.99446177 | 0.998434 |
| ACTR1B | 210.170432 | 0.220022945 | 0.38424375 | 0.57261294 | 0.5669068 | 0.85233132 |
| C2orf92 | 107.396341 | -0.151462378 | 0.3685536 | -0.4109643 | 0.6810987 | 0.89947163 |
| TMEM131 | 1999.3868 | -0.12561374 | 0.14218085 | -0.8834786 | 0.37697772 | 0.77045763 |
| INPP4A | 522.177973 | -0.310497092 | 0.30611734 | -1.0143074 | 0.31043608 | 0.72498238 |
| COA5 | 186.798424 | -0.056302938 | 0.30505888 | -0.1845642 | 0.85357089 | 0.96109041 |
| UNC50 | 241.06705 | 0.209659376 | 0.23613788 | 0.88786848 | 0.3746115 | 0.7697436 |

| | | | | | | |
|---|---|---|---|---|---|---|
| MGAT4A | 1412.71124 | 0.15800713 | 0.46544275 | 0.33947704 | 0.73425039 | 0.91975299 |
| KIAA1211L | 119.266233 | 1.651581094 | 0.83490405 | 1.97816874 | 0.04790967 | 0.45133876 |
| TSGA10 | 1328.85972 | 0.763766802 | 0.28952772 | 2.63797471 | 0.00834028 | 0.25016267 |
| AC079447.1 | 636.046909 | 0.643337689 | 0.23741486 | 2.70976164 | 0.00673316 | 0.23083391 |
| MITD1 | 420.268747 | 0.135281369 | 0.2191121 | 0.61740711 | 0.53696623 | 0.84001325 |
| LYG1 | 338.723116 | 0.742670566 | 0.49771124 | 1.49217158 | 0.13565417 | 0.59381224 |
| TXNDC9 | 350.422177 | -0.130609441 | 0.27883984 | -0.4684031 | 0.63949637 | 0.88321992 |
| EIF5B | 1203.40273 | -0.356832448 | 0.40495171 | -0.8811728 | 0.37822428 | 0.77055888 |
| REV1 | 1274.76664 | -0.123402229 | 0.2617004 | -0.4715401 | 0.6372551 | 0.88267497 |
| AFF3 | 643.195416 | 0.286578532 | 0.65439024 | 0.43793216 | 0.66143546 | 0.89184036 |
| PDCL3 | 49.6958286 | -0.806384517 | 0.40202676 | -2.0057981 | 0.04487779 | 0.44360789 |
| NPAS2 | 3983.87 | -0.154420423 | 0.46814088 | -0.3298589 | 0.7415066 | 0.92235631 |
| RPL31 | 637.632971 | 0.142449343 | 0.34626335 | 0.41139018 | 0.68078645 | 0.89947163 |
| TBC1D8 | 1457.13437 | 0.215662047 | 0.2708555 | 0.79622546 | 4.26E-01 | 0.78894887 |
| CNOT11 | 458.791596 | -0.245472922 | 0.20037149 | -1.225089 | 0.22054167 | 0.66263062 |
| RNF149 | 2372.77318 | -0.28758055 | 0.20651249 | -1.3925577 | 0.16375359 | 0.62065658 |
| MAP4K4 | 2279.03323 | -0.06353431 | 0.54034704 | -0.1175806 | 0.90640001 | 0.97651095 |
| IL1R1 | 1049.70552 | -0.33130364 | 0.41954126 | -0.7896807 | 0.42971426 | 0.79055252 |
| IL1RL2 | 118.909902 | -0.236303565 | 0.59628691 | -0.3962917 | 6.92E-01 | 0.90499641 |
| IL18R1 | 137.808233 | 0.221647839 | 0.58895092 | 0.37634348 | 0.70666156 | 0.91002281 |
| SLC9A2 | 125.259972 | 0.532957673 | 0.69079273 | 0.77151605 | 0.44040111 | 0.79826598 |
| MFSD9 | 164.620499 | -0.196649674 | 0.28435673 | -0.6915598 | 0.48921383 | 0.81970016 |
| TMEM182 | 226.453368 | -0.062530886 | 0.30392863 | -0.205742 | 0.83699244 | 0.95496955 |
| POU3F3 | 654.514452 | -0.203312699 | 0.47445648 | -0.4285171 | 0.66827471 | 0.89532221 |
| AC010884.1 | 63.2719708 | 0.776225158 | 0.76173604 | 1.01902117 | 0.30819291 | 0.72367286 |
| MRPS9 | 484.584958 | 0.201125595 | 0.21000816 | 0.9577037 | 0.33821219 | 0.74507787 |
| TGFBRAP1 | 361.394288 | -0.279754277 | 0.26848676 | -1.0419667 | 0.2974271 | 0.71839551 |
| AC012360.3 | 63.7814507 | -0.95489195 | 0.39356074 | -2.4262886 | 0.01525413 | 0.3105046 |
| C2orf49 | 190.516231 | -0.086180812 | 0.21708107 | -0.3969983 | 0.69136873 | 0.90485927 |
| FHL2 | 307.116464 | 0.08121129 | 0.54167904 | 0.14992511 | 8.81E-01 | 0.96773407 |
| NCK2 | 812.063765 | 0.210856248 | 0.26491792 | 0.79593049 | 0.42607244 | 0.78895108 |
| UXS1 | 552.970765 | -0.694853142 | 0.30162943 | -2.3036649 | 2.12E-02 | 3.57E-01 |
| RGPD3 | 45.9449722 | 0.114668464 | 0.77792901 | 0.14740222 | 0.88281455 | 0.96824408 |
| ST6GAL2 | 57.9363685 | 0.479487234 | 0.66856511 | 0.71718853 | 0.47325777 | 0.81478861 |
| LINC01789 | 284.893126 | 2.383163148 | 0.8908506 | 2.67515469 | 7.47E-03 | 0.23590813 |
| LINC01885 | 89.1647772 | 0.083087327 | 0.70200359 | 0.11835741 | 9.06E-01 | 9.76E-01 |
| LINC01886 | 108.070872 | -0.036003196 | 0.92504559 | -0.0389205 | 9.69E-01 | 9.92E-01 |
| LINC01594 | 203.38469 | 0.637818235 | 0.63356614 | 1.00671137 | 0.3140735 | 0.72719146 |
| SULT1C2 | 1964.94113 | 0.403540469 | 0.44864352 | 0.89946796 | 0.36840345 | 0.76615653 |
| SULT1C4 | 430.374982 | 0.651966302 | 0.64847475 | 1.00538425 | 0.31471186 | 0.72719146 |
| GCC2 | 2405.29494 | -0.134988495 | 0.20908162 | -0.6456258 | 0.5185217 | 0.83325693 |
| GCC2-AS1 | 343.341603 | 0.111269899 | 0.41206441 | 0.27003035 | 0.7871369 | 0.93950968 |
| LIMS1 | 2476.47104 | -0.562704376 | 0.28118357 | -2.0011993 | 0.04537091 | 0.44360789 |
| RANBP2 | 1223.96039 | -0.32947609 | 0.21637748 | -1.5226912 | 0.12783598 | 0.58793678 |
| CCDC138 | 457.416293 | -0.47798797 | 0.30668533 | -1.5585616 | 0.11910018 | 0.57624646 |
| EDAR | 162.775237 | 0.967879651 | 0.60364713 | 1.60338648 | 0.10884935 | 0.55877607 |
| SH3RF3 | 852.178952 | -0.15641914 | 0.7560132 | -0.2069 | 0.83608795 | 0.95468028 |
| 10-Sep | 1612.19244 | -0.464223977 | 0.34734301 | -1.3365001 | 0.18138586 | 0.6318513 |
| SOWAHC | 162.680505 | -0.751344998 | 0.48659353 | -1.5440916 | 0.12256614 | 0.58152096 |
| MALL | 30.8570872 | 0.698558198 | 0.5538667 | 1.26123884 | 0.20722281 | 0.65266614 |
| NPHP1 | 353.116078 | 0.886228097 | 0.56953701 | 1.55605005 | 0.11969619 | 0.57673415 |
| SMIM37 | 294.081053 | -0.458431819 | 0.33517856 | -1.3677242 | 0.17139842 | 0.62520756 |
| LINC01106 | 44.8154295 | -0.5888239 | 0.39981381 | -1.4727453 | 1.41E-01 | 0.59555904 |
| AC112229.3 | 38.1724242 | 0.218895592 | 0.62124414 | 0.35235035 | 0.72457553 | 0.91600702 |

| | | | | | | |
|---|---|---|---|---|---|---|
| MIR4435-2H | 1496.74911 | -0.846707836 | 0.60985355 | -1.388379 | 1.65E-01 | 6.21E-01 |
| BCL2L11 | 385.482673 | -0.77561315 | 0.35420915 | -2.1897039 | 0.02854572 | 0.39280225 |
| AC017002.5 | 101.864592 | -0.627528486 | 0.63179164 | -0.9932523 | 3.21E-01 | 7.32E-01 |
| ANAPC1 | 1056.42954 | -0.321454264 | 0.17948824 | -1.7909489 | 0.0733015 | 0.50459544 |
| MERTK | 414.562622 | 0.360142696 | 0.40074702 | 0.89867842 | 0.36882398 | 0.76615653 |
| TMEM87B | 688.014765 | -0.019555984 | 0.1788825 | -0.1093231 | 9.13E-01 | 9.77E-01 |
| FBLN7 | 58.501123 | 0.372100399 | 0.75535512 | 0.4926165 | 0.62228358 | 0.87714058 |
| ZC3H8 | 556.14464 | -0.124704067 | 0.30765385 | -0.4053389 | 0.68522844 | 0.90112503 |
| ZC3H6 | 599.972016 | 0.983944171 | 0.61171133 | 1.60851061 | 0.10772339 | 0.5565641 |
| RGPD8 | 71.024619 | -0.741053716 | 0.40004136 | -1.8524428 | 0.06396227 | 0.48593331 |
| TTL | 327.552966 | -0.469450159 | 0.61385611 | -0.764756 | 0.44441684 | 0.79994797 |
| POLR1B | 210.058385 | -0.768607599 | 0.44517295 | -1.7265371 | 0.08425084 | 0.52183702 |
| CHCHD5 | 109.535033 | 0.04251212 | 0.39430188 | 0.10781618 | 0.91414151 | 0.9774502 |
| SLC20A1 | 832.146649 | -0.591694109 | 0.61384639 | -0.9639123 | 0.33508988 | 0.74165622 |
| PSD4 | 135.770561 | -0.111332334 | 0.41614267 | -0.2675341 | 0.789058 | 0.94005579 |
| PAX8 | 24108.095 | 0.204900354 | 0.22158084 | 0.92472055 | 3.55E-01 | 0.75757147 |
| CBWD2 | 351.433453 | -0.369154096 | 0.21869744 | -1.6879671 | 0.09141754 | 0.53537574 |
| RABL2A | 261.390755 | 0.028007401 | 0.26295942 | 0.10650845 | 0.91517895 | 0.97782888 |
| AC017074.1 | 175.790873 | 0.665106996 | 0.52589366 | 1.26471766 | 0.20597256 | 0.651931 |
| SLC35F5 | 2785.97505 | -0.064837449 | 0.30787715 | -0.2105952 | 0.83320316 | 0.95355066 |
| ACTR3 | 1696.14575 | -0.056600871 | 0.28969994 | -0.1953776 | 0.84509737 | 0.95760132 |
| LINC01191 | 116.248142 | 0.033929893 | 0.67946377 | 0.04993628 | 9.60E-01 | 9.89E-01 |
| DPP10 | 222.785836 | 1.616357137 | 0.77948373 | 2.07362524 | 0.03811413 | 0.42339465 |
| DDX18 | 460.44092 | -0.58328205 | 0.38367698 | -1.5202425 | 0.12845004 | 0.58793678 |
| AC009404.1 | 88.2854224 | -0.41988262 | 0.50998521 | -0.8233231 | 0.41032426 | 0.78582015 |
| CCDC93 | 1338.32113 | -0.563256845 | 0.25999276 | -2.1664328 | 0.03027813 | 0.40332587 |
| AC009303.2 | 57.5527526 | -0.203543884 | 0.54991502 | -0.370137 | 0.71128043 | 0.91143287 |
| INSIG2 | 765.749623 | -0.543521066 | 0.516003 | -1.0533293 | 2.92E-01 | 0.71399546 |
| C2orf76 | 192.344413 | 0.33146517 | 0.33371598 | 0.99325532 | 0.32058554 | 0.73210121 |
| DBI | 1504.16157 | -0.314256356 | 0.57862467 | -0.5431092 | 0.58705465 | 0.86112855 |
| TMEM37 | 959.396773 | -0.138731992 | 0.62135893 | -0.2232719 | 0.82332389 | 0.95131214 |
| SCTR | 312.903304 | 2.545836445 | 0.96528884 | 2.63738307 | 0.00835484 | 0.25016267 |
| AC013275.1 | 82.7746207 | 0.539945503 | 1.04461832 | 0.51688305 | 0.6052378 | 0.86869053 |
| CFAP221 | 837.800917 | 1.005578054 | 0.56383695 | 1.7834554 | 0.0745122 | 0.5071704 |
| TMEM177 | 87.7516066 | 0.686152563 | 0.45068542 | 1.52246454 | 0.12789272 | 0.58793678 |
| PTPN4 | 2101.86245 | 0.240179629 | 0.2325377 | 1.03286317 | 0.30166794 | 0.71908176 |
| EPB41L5 | 3579.62292 | 0.228235231 | 0.32424428 | 0.70389902 | 0.48149566 | 0.81755903 |
| AC012363.1 | 34.0298321 | 0.627338522 | 0.4946829 | 1.26816293 | 0.20473977 | 0.65068591 |
| RALB | 692.715955 | -0.055119128 | 0.27712528 | -0.1988961 | 0.84234405 | 0.95716002 |
| TFCP2L1 | 1582.54524 | 1.360499714 | 0.54961958 | 2.47534798 | 0.01331064 | 0.29875272 |
| CLASP1 | 3755.24604 | 0.090645447 | 0.27174843 | 0.33356383 | 0.73870872 | 0.9208911 |
| NIFK-AS1 | 229.359849 | 0.414176257 | 0.47803272 | 0.86641822 | 0.38626086 | 0.77325508 |
| NIFK | 130.468077 | -0.566721945 | 0.45550674 | -1.2441571 | 0.21344175 | 0.65818808 |
| TSN | 247.901008 | -0.284166457 | 0.26802195 | -1.0602358 | 0.28903736 | 0.71218341 |
| CNTNAP5 | 81.6720055 | -0.689994491 | 0.77803142 | -0.8868466 | 0.37516152 | 0.7697436 |
| GYPC | 66.0307772 | 0.559475555 | 0.54967215 | 1.01783501 | 3.09E-01 | 0.72418268 |
| BIN1 | 943.688456 | 0.227095288 | 0.41899454 | 0.54200059 | 0.5878181 | 0.8613161 |
| ERCC3 | 393.184117 | -0.614393368 | 0.31806552 | -1.9316566 | 0.0534019 | 0.46711025 |
| MAP3K2 | 2243.09533 | 0.02192505 | 0.10898914 | 0.2011673 | 8.41E-01 | 0.956374 |
| PROC | 278.89292 | 0.270292812 | 0.45705671 | 0.59137697 | 0.55426787 | 0.84727776 |
| IWS1 | 1390.89207 | 0.233200541 | 0.18607468 | 1.25326317 | 0.21010995 | 0.65441616 |
| AC010976.1 | 160.051418 | 0.857009075 | 0.63264136 | 1.3546523 | 0.17552836 | 0.62699479 |
| MYO7B | 1459.52521 | 1.188594936 | 0.61193108 | 1.94236732 | 0.05209265 | 0.46437412 |
| WDR33 | 2102.27387 | -0.302648538 | 0.2916021 | -1.0378819 | 0.29932505 | 0.71852421 |

| | | | | | | |
|---|---|---|---|---|---|---|
| SFT2D3 | 47.2626844 | -0.175706759 | 0.38504689 | -0.4563256 | 0.64815585 | 0.88787189 |
| POLR2D | 173.855158 | -0.733673898 | 0.37205225 | -1.9719647 | 0.04861364 | 0.45329737 |
| AMMECR1L | 247.949589 | -0.206720265 | 0.2984546 | -0.6926355 | 0.4885383 | 0.81933645 |
| SAP130 | 545.989188 | -0.076259732 | 0.2749005 | -0.2774085 | 0.78146646 | 0.93748328 |
| UGGT1 | 662.687382 | -0.051026799 | 0.31228232 | -0.1633996 | 0.87020383 | 0.96564168 |
| HS6ST1 | 266.368723 | -0.223420914 | 0.42317516 | -0.5279632 | 0.59752488 | 0.86614567 |
| SMPD4 | 401.376502 | -0.773478742 | 0.27239786 | -2.8395185 | 0.00451817 | 0.20749797 |
| MZT2B | 521.713181 | -0.192622311 | 0.38230778 | -0.5038409 | 0.61437315 | 0.87225883 |
| CCDC115 | 88.1622448 | 0.328982487 | 0.40477689 | 0.81275017 | 0.41636131 | 0.78632759 |
| IMP4 | 168.966126 | -0.480597053 | 0.46783737 | -1.0272738 | 0.30429158 | 0.72082426 |
| PTPN18 | 412.70712 | -0.040862191 | 0.31207549 | -0.1309369 | 0.89582524 | 0.97309833 |
| FAM168B | 944.115259 | -0.494484204 | 0.28365028 | -1.7432883 | 0.08128327 | 0.51593841 |
| PLEKHB2 | 1092.60059 | -0.404600317 | 0.30178544 | -1.3406887 | 0.18002156 | 0.63124625 |
| MZT2A | 280.243837 | -0.234463261 | 0.21690787 | -1.0809348 | 0.27972613 | 0.70446562 |
| ANKRD30BL | 111.146266 | 0.052973178 | 0.49399413 | 0.10723443 | 0.914603 | 0.9776687 |
| GPR39 | 706.618162 | -0.277587016 | 0.22612617 | -1.2275758 | 0.21960624 | 0.66208719 |
| LYPD1 | 88.8548904 | -0.725390083 | 0.961616 | -0.7543449 | 4.51E-01 | 0.80361623 |
| NCKAP5 | 11487.7455 | 0.900888464 | 0.67272318 | 1.33916668 | 0.18051642 | 0.63166264 |
| AC016909.2 | 57.2690633 | 1.640969268 | 1.06122526 | 1.54629684 | 0.12203289 | 0.58055167 |
| AC010890.1 | 83.3202424 | 1.662109625 | 1.11657203 | 1.48858254 | 0.13659733 | 0.5942049 |
| MGAT5 | 1419.7427 | -0.35197691 | 0.40364978 | -0.8719859 | 0.38321608 | 0.77317162 |
| TMEM163 | 467.420647 | 0.905365524 | 0.57012162 | 1.58802173 | 0.11228143 | 0.56540834 |
| CCNT2-AS1 | 828.002546 | -0.031127451 | 0.51679967 | -0.0602312 | 0.95197152 | 0.98766432 |
| ACMSD | 3562.54782 | 0.519813397 | 0.68290186 | 0.76118316 | 0.44654767 | 0.80205685 |
| CCNT2 | 1207.98011 | -0.024540604 | 0.13502341 | -0.1817507 | 0.85577835 | 0.96211892 |
| MAP3K19 | 48.6856907 | 0.631758951 | 0.5474563 | 1.15398974 | 0.24850438 | 0.6852242 |
| RAB3GAP1 | 2145.39508 | -0.197972675 | 0.16369049 | -1.209433 | 0.22649655 | 0.66690202 |
| ZRANB3 | 826.025297 | -0.179935419 | 0.27039871 | -0.6654448 | 0.50576603 | 0.82738999 |
| R3HDM1 | 1713.61483 | 0.037496572 | 0.22240912 | 0.16859278 | 0.86611696 | 0.96437968 |
| UBXN4 | 1589.91549 | -0.366518113 | 0.28160409 | -1.3015369 | 0.19307474 | 0.64532959 |
| MCM6 | 40.1509879 | 0.596579575 | 0.49795871 | 1.19805029 | 0.23089744 | 0.66878058 |
| DARS | 1636.34661 | -0.149560149 | 0.25746128 | -0.5809035 | 5.61E-01 | 0.84958759 |
| DARS-AS1 | 75.3475654 | -0.854338287 | 0.61448459 | -1.3903331 | 0.16442774 | 0.62065658 |
| THSD7B | 54.6191229 | 0.080201699 | 0.59136544 | 0.13562122 | 0.89212073 | 0.97237481 |
| HNMT | 1018.73453 | 0.736742663 | 0.62718203 | 1.17468714 | 0.2401199 | 0.67862341 |
| AC114763.1 | 370.263024 | 0.260276361 | 0.40175623 | 0.64784648 | 0.51708425 | 0.83276628 |
| SPOPL | 808.018653 | -0.130478101 | 0.21206063 | -0.6152868 | 0.53836536 | 0.84049498 |
| LRP1B | 1061.77258 | -0.525466034 | 0.90834763 | -0.5784856 | 0.56293631 | 0.85057939 |
| KYNU | 1278.31016 | -0.544353282 | 0.40051404 | -1.3591366 | 0.17410331 | 0.62699479 |
| ARHGAP15 | 68.2200971 | 0.333045442 | 0.48029664 | 0.69341614 | 0.48804844 | 0.81921136 |
| GTDC1 | 1558.29032 | 0.05517198 | 0.2164007 | 0.25495287 | 0.7987595 | 0.94327531 |
| ZEB2 | 1913.59314 | 0.072715352 | 0.32412741 | 0.22434188 | 0.82249129 | 0.95114217 |
| TEX41 | 113.883683 | 0.12468743 | 0.7612459 | 0.16379389 | 0.8698934 | 0.96564168 |
| ACVR2A | 790.152837 | 0.124343094 | 0.16193525 | 0.76785685 | 4.43E-01 | 7.99E-01 |
| ORC4 | 1663.85535 | -0.064686631 | 0.30726437 | -0.2105243 | 0.83325845 | 0.95355066 |
| MBD5 | 7014.11487 | 0.50645357 | 0.39793163 | 1.27271504 | 0.20311918 | 0.64881821 |
| EPC2 | 1296.59699 | -0.309068312 | 0.19518104 | -1.5834956 | 0.11330856 | 0.56809124 |
| AC105402.3 | 76.6534889 | -0.637374018 | 1.12922203 | -0.5644364 | 0.57245717 | 0.85408471 |
| MMADHC | 283.480796 | -0.183070217 | 0.29155402 | -0.6279118 | 0.5300617 | 0.83789294 |
| AC007364.1 | 3464.86552 | -0.10139433 | 0.39696796 | -0.2554219 | 0.79839722 | 0.94320409 |
| LINC01931 | 138.880078 | 0.761322395 | 0.78612774 | 0.96844616 | 0.3328216 | 0.73988623 |
| RND3 | 366.001166 | -2.602375352 | 0.88929551 | -2.9263336 | 0.00342983 | 0.1911084 |
| RBM43 | 66.1881635 | 0.057249133 | 0.64036752 | 0.08940043 | 0.92876368 | 0.98272895 |
| NMI | 206.562515 | -0.16778719 | 0.59023631 | -0.2842712 | 0.77620255 | 0.9352344 |

| | | | | | | |
|---|---|---|---|---|---|---|
| RIF1 | 1129.93879 | -0.334820224 | 0.26181692 | -1.2788334 | 0.20095573 | 0.64849129 |
| NEB | 143.288764 | 0.682650348 | 0.48209032 | 1.41602168 | 0.15676916 | 0.61255952 |
| ARL5A | 408.223027 | 0.40463573 | 0.24962598 | 1.62096804 | 0.10502449 | 0.55219437 |
| CACNB4 | 108.353248 | 0.552537846 | 0.32868709 | 1.68104518 | 0.09275414 | 0.53669957 |
| STAM2 | 744.757114 | -0.381771352 | 0.21028891 | -1.8154612 | 0.06945306 | 0.49281529 |
| FMNL2 | 10205.8796 | -0.11338069 | 0.59616468 | -0.1901835 | 0.84916534 | 0.95959925 |
| PRPF40A | 1115.10316 | -0.133293887 | 0.16888381 | -0.7892638 | 0.42995782 | 0.79055252 |
| ARL6IP6 | 357.932473 | 0.037999893 | 0.28497915 | 0.13334271 | 0.89392235 | 0.97272276 |
| GALNT13 | 176.121077 | 1.467822556 | 0.59140503 | 2.48192439 | 0.0130675 | 0.29875272 |
| KCNJ3 | 1916.11924 | -0.107355181 | 0.6938111 | -0.1547326 | 0.87703215 | 0.96643457 |
| LINC01876 | 363.663497 | 1.860393759 | 1.02212236 | 1.82012823 | 0.06873948 | 0.49201674 |
| GPD2 | 572.994493 | -0.050210801 | 0.24486573 | -0.2050544 | 0.83752961 | 0.95496955 |
| ACVR1C | 88.6794044 | 1.113643132 | 0.36908843 | 3.01727996 | 0.00255054 | 0.16568849 |
| AC019186.1 | 120.487357 | 0.674223136 | 0.56237168 | 1.19889241 | 0.23056978 | 0.66865767 |
| ACVR1 | 2530.94788 | -0.334751691 | 0.34347756 | -0.9745955 | 0.32976093 | 0.73849818 |
| UPP2 | 1313.74315 | 2.315133055 | 0.71814917 | 3.22374954 | 0.00126524 | 0.11639359 |
| CCDC148 | 1354.20394 | -0.056498723 | 0.6511749 | -0.0867643 | 0.93085888 | 0.98272895 |
| PKP4 | 7267.5131 | -0.084327941 | 0.28517228 | -0.2957088 | 0.7674525 | 0.93169958 |
| PKP4-AS1 | 171.802574 | 0.006697607 | 0.73994004 | 0.00905155 | 0.992778 | 0.9976934 |
| TANC1 | 1830.55001 | 0.176771768 | 0.20526226 | 0.86119957 | 0.38912814 | 0.7748665 |
| WDSUB1 | 879.913736 | -0.57502547 | 0.31850518 | -1.8053881 | 0.07101397 | 0.49789693 |
| BAZ2B | 4694.38398 | 0.274139422 | 0.32535032 | 0.84259767 | 0.3994535 | 0.7817403 |
| AC009961.1 | 152.143027 | -0.018813222 | 0.33578835 | -0.056027 | 0.95532028 | 0.98840182 |
| 7-Mar | 1958.28628 | -0.311440621 | 0.20352323 | -1.530246 | 0.12595584 | 0.58564339 |
| PLA2R1 | 766.036213 | -0.144752515 | 0.39092823 | -0.370279 | 0.71117462 | 0.91143287 |
| ITGB6 | 1353.26023 | 1.036200481 | 0.72845939 | 1.4224547 | 1.55E-01 | 0.61132626 |
| LINC02478 | 47.9668805 | 0.940913317 | 0.53926027 | 1.74482223 | 0.08101582 | 0.51567042 |
| RBMS1 | 3958.98079 | -0.10071862 | 0.28407122 | -0.3545541 | 0.72292366 | 0.91592159 |
| TANK | 1740.27907 | 0.281183609 | 0.19668531 | 1.42961166 | 1.53E-01 | 6.10E-01 |
| AC009299.3 | 39.4491764 | 0.225572472 | 0.64531373 | 0.34955474 | 0.72667288 | 0.91639348 |
| PSMD14 | 883.176179 | -0.408919018 | 0.3777618 | -1.0824785 | 0.27903998 | 0.70446562 |
| DPP4 | 2268.10711 | -1.119126951 | 0.63453905 | -1.7636849 | 0.07778504 | 0.51185769 |
| IFIH1 | 269.448192 | -0.239166702 | 0.68411434 | -0.3496005 | 0.72663855 | 0.91639348 |
| GCA | 293.465089 | 0.360104598 | 0.44312793 | 0.81264253 | 0.41642304 | 0.78632759 |
| KCNH7 | 59.5780943 | 0.133165157 | 0.65115447 | 0.20450625 | 0.83795791 | 0.95496955 |
| FIGN | 2032.42352 | -0.508894639 | 0.30194822 | -1.6853706 | 0.0919171 | 0.53567268 |
| AC016766.1 | 49.8927568 | -0.054317666 | 0.59283993 | -0.0916228 | 0.92699772 | 0.98184958 |
| GRB14 | 636.914936 | 0.304517964 | 0.47294793 | 0.64387207 | 0.5196584 | 0.8335052 |
| COBLL1 | 3830.00011 | -0.147583266 | 0.38617849 | -0.3821633 | 0.70234021 | 0.90837679 |
| AC019197.1 | 1165.91562 | 1.997581608 | 0.91585792 | 2.18110426 | 0.02917571 | 0.39595188 |
| SLC38A11 | 137.685135 | 0.468337403 | 0.69190036 | 0.67688562 | 0.49847853 | 0.82362007 |
| SCN2A | 153.352766 | 2.734326968 | 0.9627935 | 2.83999317 | 0.00451145 | 0.20749797 |
| CSRNP3 | 763.654199 | 0.701713495 | 0.56498941 | 1.24199406 | 0.21423875 | 0.65906957 |
| GALNT3 | 239.742159 | 0.302897651 | 0.44109866 | 0.68668912 | 0.49227865 | 0.82087587 |
| AC009495.1 | 176.48803 | 0.505787998 | 0.42282276 | 1.19621752 | 2.32E-01 | 6.69E-01 |
| TTC21B | 1407.32421 | 0.214615511 | 0.33399498 | 0.64257107 | 5.21E-01 | 0.833897 |
| AC010127.1 | 266.040719 | -0.272823194 | 0.88210347 | -0.3092871 | 0.75710317 | 0.92799282 |
| SCN9A | 909.133677 | 0.186577026 | 0.78213499 | 0.23854837 | 0.8114558 | 0.94753961 |
| AC073050.1 | 151.679185 | 0.233656697 | 0.86759627 | 0.26931501 | 0.78768728 | 0.93953874 |
| STK39 | 1193.33591 | -0.039179369 | 0.65082713 | -0.0601993 | 9.52E-01 | 9.88E-01 |
| CERS6 | 1434.25322 | 0.13053693 | 0.26981586 | 0.48380007 | 0.62852777 | 0.88009297 |
| NOSTRIN | 1956.92291 | 1.116243447 | 0.76045975 | 1.46785342 | 0.14214404 | 0.59729009 |
| SPC25 | 296.990968 | 1.184300327 | 0.82179693 | 1.44111068 | 0.14955342 | 0.60702655 |
| LRP2 | 28560.0978 | 1.059054015 | 0.87806753 | 1.2061191 | 0.22777157 | 0.66705629 |

| | | | | | | |
|---|---|---|---|---|---|---|
| FASTKD1 | 528.352367 | 0.160653787 | 0.2898492 | 0.55426679 | 0.57939628 | 0.85816191 |
| PPIG | 1274.06825 | -0.146410121 | 0.1633278 | -0.8964189 | 0.37002909 | 0.76669785 |
| CCDC173 | 65.1015603 | 0.246273161 | 0.64059982 | 0.38444151 | 0.70065124 | 0.90766879 |
| KLHL23 | 507.67902 | 0.074545344 | 0.29375092 | 0.25377059 | 0.7996728 | 0.94386593 |
| SSB | 327.563425 | -0.363227855 | 0.24028025 | -1.5116842 | 0.13061423 | 0.58853436 |
| METTL5 | 202.972881 | -0.043212835 | 0.24287758 | -0.1779202 | 0.85878562 | 0.9626966 |
| UBR3 | 2485.86988 | -0.408929213 | 0.18963263 | -2.1564285 | 0.03105022 | 0.40611202 |
| MYO3B | 113.483427 | 0.01818956 | 0.61478629 | 0.0295868 | 0.97639659 | 0.99429832 |
| AC007405.1 | 48.7932414 | 1.061146406 | 0.73507969 | 1.44358008 | 0.14885713 | 0.60622442 |
| GORASP2 | 556.714004 | -0.300277332 | 0.27923101 | -1.0753725 | 0.28220801 | 0.70682473 |
| TLK1 | 1927.3013 | -0.229656766 | 0.16573727 | -1.3856676 | 0.16584841 | 0.62065658 |
| METTL8 | 644.450059 | -0.435403039 | 0.3861162 | -1.1276477 | 0.25946873 | 0.69242517 |
| DCAF17 | 281.068062 | -0.372315628 | 0.27229904 | -1.3673042 | 0.17152997 | 0.62520756 |
| DYNC1I2 | 1155.90779 | -0.09642283 | 0.1602532 | -0.6016905 | 0.54738017 | 0.8448516 |
| SLC25A12 | 1072.188 | 0.67987241 | 0.29678231 | 2.29081175 | 0.0219743 | 0.36066559 |
| HAT1 | 328.192805 | -0.153486405 | 0.21884916 | -0.7013342 | 0.48309445 | 0.8177376 |
| METAP1D | 195.189284 | -0.588474349 | 0.3729369 | -1.5779462 | 0.11457798 | 0.57068327 |
| ITGA6 | 3531.23003 | -0.39875887 | 0.41158776 | -0.9688307 | 0.33262965 | 0.73988623 |
| AC078883.1 | 179.074214 | -0.277915178 | 0.43162445 | -0.6438819 | 5.20E-01 | 8.34E-01 |
| PDK1 | 698.928137 | -0.890713364 | 0.5855565 | -1.5211399 | 1.28E-01 | 5.88E-01 |
| RAPGEF4 | 98.8083293 | -0.016330217 | 0.66016867 | -0.0247364 | 0.9802652 | 0.99494599 |
| MAP3K20 | 410.454082 | -0.7853158 | 0.80490926 | -0.9756576 | 0.32923419 | 0.73829083 |
| SP3 | 1099.04279 | -0.025167378 | 0.19149678 | -0.1314245 | 0.89543947 | 0.97309833 |
| OLA1 | 2114.14478 | -0.060549122 | 0.24589811 | -0.2462366 | 0.80549906 | 0.94585363 |
| CIR1 | 690.00002 | -0.246417865 | 0.30796842 | -0.8001401 | 0.42362966 | 0.78894887 |
| SCRN3 | 232.031387 | 0.145806971 | 0.42826938 | 0.34045622 | 0.73351299 | 0.91962522 |
| GPR155 | 2319.62689 | 0.598927337 | 0.50235622 | 1.19223633 | 0.23316859 | 0.66978724 |
| AC010894.3 | 596.454503 | 0.400667566 | 0.34732243 | 1.15358965 | 0.24866844 | 0.68534254 |
| WIPF1 | 1801.148 | 0.47829459 | 0.2567532 | 1.86285736 | 0.06248234 | 0.4840117 |
| CHN1 | 161.872699 | 0.036998506 | 0.47984703 | 0.07710479 | 0.93854018 | 0.98435387 |
| ATF2 | 1088.3254 | -0.14251969 | 0.19306009 | -0.7382141 | 0.46038434 | 0.80717423 |
| ATP5MC3 | 866.241948 | 0.162085652 | 0.59701321 | 0.27149425 | 0.78601092 | 0.93904951 |
| LNPK | 633.245889 | -0.513520048 | 0.41901776 | -1.2255329 | 0.2203745 | 0.66263062 |
| HOXD-AS2 | 62.9565938 | 0.30655296 | 0.51587424 | 0.59423972 | 0.55235179 | 0.84721285 |
| HOXD8 | 49.2657279 | 0.90777633 | 0.43862711 | 2.06958557 | 0.03849117 | 0.42447711 |
| HOXD3 | 137.073534 | 0.969355891 | 0.48399438 | 2.0028247 | 0.04519611 | 0.44360789 |
| HAGLR | 74.6815064 | 0.743606253 | 0.42051952 | 1.76830379 | 0.07701013 | 0.51013111 |
| MTX2 | 1554.73465 | 0.272683856 | 0.2974817 | 0.91664078 | 0.35933091 | 0.75938155 |
| AC079305.3 | 101.848232 | 0.793170935 | 0.50500388 | 1.57062346 | 0.11627014 | 0.57350419 |
| HNRNPA3 | 1389.24353 | -0.299451031 | 0.25646567 | -1.1676067 | 0.24296546 | 0.68132724 |
| NFE2L2 | 3303.20939 | 0.05350035 | 0.27193093 | 0.19674242 | 0.84402911 | 0.95731603 |
| AGPS | 1619.30718 | -0.242151648 | 0.41763203 | -0.5798206 | 0.56203562 | 0.85033466 |
| TTC30A | 38.426706 | -0.412696235 | 0.50822328 | -0.8120373 | 0.41677026 | 0.78632759 |
| PDE11A | 29.1521355 | 1.939380125 | 0.91228771 | 2.12584266 | 0.03351636 | 0.4131374 |
| RBM45 | 272.644018 | 0.38166372 | 0.33872614 | 1.12676193 | 0.25984315 | 0.69252772 |
| OSBPL6 | 1045.81485 | 0.208440245 | 0.36435865 | 0.57207437 | 0.56727159 | 0.85239988 |
| AC009948.1 | 149.870584 | -0.506010747 | 0.6828705 | -0.7410054 | 0.45869015 | 0.80636987 |
| PRKRA | 581.313063 | -0.050231509 | 0.22302014 | -0.2252331 | 0.82179798 | 0.95105545 |
| PJVK | 147.693823 | 0.470687264 | 0.37619312 | 1.25118519 | 0.21086692 | 0.65540882 |
| PLEKHA3 | 351.933244 | 0.069696528 | 0.26127863 | 0.26675174 | 0.78966032 | 0.94023796 |
| TTN-AS1 | 1655.0378 | 0.61883433 | 0.245436 | 2.52136737 | 0.01168997 | 0.28824618 |
| TTN | 125.520772 | 1.465001062 | 0.71098679 | 2.06051799 | 0.03934905 | 0.42704465 |
| SESTD1 | 3067.31489 | 1.16106662 | 0.38006543 | 3.0549125 | 0.00225126 | 0.15428723 |
| ZNF385B | 1665.66781 | 2.130664498 | 0.62776582 | 3.39404349 | 6.89E-04 | 0.08896424 |

| | | | | | | |
|---|---|---|---|---|---|---|
| CWC22 | 538.867236 | -0.09761179 | 0.20156425 | -0.4842713 | 0.62819331 | 0.88007417 |
| UBE2E3 | 1422.26235 | -0.063072759 | 0.22728829 | -0.2775011 | 0.78139533 | 0.93748328 |
| LINC01934 | 411.394896 | 0.14381976 | 0.84000708 | 0.17121256 | 8.64E-01 | 0.96393376 |
| CERKL | 231.895597 | 0.415102062 | 0.71710453 | 0.57885852 | 0.56268464 | 0.85054239 |
| SSFA2 | 1327.98231 | 0.224643828 | 0.3085589 | 0.72804196 | 0.4665879 | 0.81098649 |
| PDE1A | 622.790192 | 1.383273755 | 0.73506394 | 1.88184141 | 0.05985755 | 0.48004006 |
| DNAJC10 | 586.215373 | -0.113128714 | 0.32146673 | -0.3519142 | 0.72490259 | 0.91600702 |
| NCKAP1 | 2592.5588 | -0.044484597 | 0.22024109 | -0.2019814 | 0.83993129 | 0.95574781 |
| NUP35 | 270.286381 | -0.436884224 | 0.31530449 | -1.3855947 | 0.1658707 | 0.62065658 |
| FSIP2 | 196.818114 | -0.69914704 | 0.49022499 | -1.4261759 | 0.15381759 | 0.6105654 |
| FSIP2-AS1 | 107.737598 | 1.512610375 | 0.6352925 | 2.38096683 | 0.01726727 | 0.32964126 |
| LINC01473 | 283.170384 | 1.440154869 | 0.62388894 | 2.30835134 | 0.0209796 | 0.35422078 |
| AC017071.1 | 46.252871 | 1.199668039 | 0.48274116 | 2.48511654 | 0.0129509 | 0.29875272 |
| ZC3H15 | 617.153388 | -0.297463008 | 0.23732493 | -1.2533998 | 0.21006026 | 0.65441616 |
| ITGAV | 3772.07272 | 0.2534199 | 0.67296483 | 0.37657228 | 0.70649149 | 0.91002281 |
| AC017101.1 | 288.377084 | 0.689869756 | 0.59738073 | 1.15482426 | 0.2481624 | 0.6848323 |
| AC007319.1 | 1617.2341 | 0.456258306 | 0.53700941 | 0.84962814 | 0.39553186 | 0.77931337 |
| CALCRL | 498.366714 | 0.62633897 | 0.52099297 | 1.20220234 | 0.22928514 | 0.66705629 |
| TFPI | 5187.01178 | -0.237182237 | 0.42298477 | -0.5607347 | 5.75E-01 | 0.85474196 |
| LINC01090 | 175.933916 | -0.371282724 | 0.61753138 | -0.601237 | 5.48E-01 | 0.84486687 |
| GULP1 | 4346.16845 | -1.253933671 | 0.5397212 | -2.3232989 | 0.0201631 | 0.34851906 |
| COL5A2 | 351.294997 | 0.543014543 | 0.65652747 | 0.82710102 | 0.40817981 | 0.78481094 |
| WDR75 | 600.617958 | 0.06481123 | 0.26445407 | 0.24507556 | 0.80639792 | 0.94585363 |
| SLC40A1 | 280.493709 | 1.011792398 | 0.36807404 | 2.74888283 | 0.00597988 | 0.22637955 |
| ASNSD1 | 159.271183 | -0.151731095 | 0.28396905 | -0.5343227 | 0.59311832 | 0.86414464 |
| ANKAR | 846.259554 | 0.051887475 | 0.34957847 | 0.14842869 | 0.88200446 | 0.96796225 |
| OSGEPL1 | 93.6513586 | 0.368684997 | 0.47381214 | 0.77812484 | 0.43649542 | 0.79558019 |
| ORMDL1 | 558.941957 | -0.026225074 | 0.31882063 | -0.0822565 | 0.93444273 | 0.98359701 |
| PMS1 | 545.37641 | -0.20924355 | 0.21627604 | -0.9674837 | 0.33330228 | 0.74013767 |
| C2orf88 | 326.51368 | 0.227424084 | 0.38589263 | 0.5893455 | 0.55562953 | 0.84861484 |
| HIBCH | 3882.62367 | -0.139706479 | 0.38614946 | -0.3617938 | 7.18E-01 | 0.91342588 |
| INPP1 | 277.087494 | -0.154422846 | 0.21862049 | -0.7063512 | 4.80E-01 | 8.17E-01 |
| MFSD6 | 276.851971 | -0.161762533 | 0.36503447 | -0.4431432 | 0.65766218 | 0.89053559 |
| NEMP2 | 84.2016427 | 0.251396269 | 0.36142854 | 0.69556286 | 0.48670263 | 0.81885761 |
| AC108047.1 | 68.169644 | 0.208505204 | 0.53131696 | 0.39243092 | 0.69473984 | 0.90589033 |
| NAB1 | 335.851379 | 0.124021547 | 0.41118388 | 0.30162065 | 0.76294127 | 0.93022913 |
| GLS | 9199.61983 | -0.807720391 | 0.33939834 | -2.3798596 | 0.01731924 | 0.33017352 |
| STAT1 | 2187.6981 | 0.327995908 | 1.01306079 | 0.32376725 | 0.74611425 | 0.92443171 |
| STAT4 | 39.0481799 | 0.672784149 | 0.66932223 | 1.00517228 | 0.3148139 | 0.72719146 |
| MYO1B | 582.833499 | -0.260130831 | 0.66593416 | -0.3906255 | 6.96E-01 | 0.90612803 |
| NABP1 | 1664.28652 | -0.972377123 | 0.48685031 | -1.9972815 | 0.04579461 | 0.44430994 |
| AC098617.1 | 119.378889 | 1.009594615 | 0.82882389 | 1.2181051 | 0.22318404 | 0.66408018 |
| TMEFF2 | 36.9993506 | 0.512451771 | 0.56414867 | 0.90836299 | 0.36368647 | 0.76272056 |
| SLC39A10 | 642.10863 | 0.765417213 | 0.24366667 | 3.14124708 | 0.0016823 | 0.13207839 |
| DNAH7 | 1552.73341 | 0.663359653 | 0.4645245 | 1.42804018 | 0.15328029 | 0.61008819 |
| STK17B | 76.6036612 | -0.266223179 | 0.35533862 | -0.7492098 | 0.45373075 | 0.80433814 |
| CCDC150 | 168.068771 | -0.084537074 | 0.56019085 | -0.1509076 | 0.88004858 | 0.96769187 |
| GTF3C3 | 413.072133 | -0.082741676 | 0.1975524 | -0.4188341 | 0.67533741 | 0.89801183 |
| PGAP1 | 225.380338 | 0.356901207 | 0.30829316 | 1.15766825 | 0.24699944 | 0.68482118 |
| ANKRD44 | 654.500371 | 0.903883204 | 0.41030677 | 2.20294488 | 0.02759863 | 0.38942007 |
| SF3B1 | 1348.31616 | -0.084009064 | 0.15601127 | -0.5384807 | 0.5902452 | 0.86307777 |
| COQ10B | 326.30195 | -0.340833673 | 0.38663833 | -0.881531 | 0.37803048 | 0.77055888 |
| HSPD1 | 1898.46251 | -0.986820687 | 0.46039967 | -2.1434001 | 0.03208099 | 0.40988962 |
| MOB4 | 100.131808 | 0.031929592 | 0.33292949 | 0.09590497 | 0.92359605 | 0.98012108 |

| | | | | | | |
|---|---|---|---|---|---|---|
| AC011997.1 | 170.037903 | -0.136820215 | 0.45330337 | -0.3018292 | 0.76278223 | 0.93022076 |
| PLCL1 | 5579.79301 | -0.117614213 | 0.68760932 | -0.171048 | 0.86418601 | 0.96393376 |
| SATB2 | 1519.62954 | 0.440443556 | 0.40564381 | 1.08578891 | 0.27757239 | 0.7036882 |
| SATB2-AS1 | 45.919325 | 0.287540509 | 0.49971966 | 0.57540363 | 0.56501835 | 0.85171209 |
| FTCDNL1 | 559.695126 | 0.543187925 | 0.59810838 | 0.90817642 | 0.36378502 | 0.76272056 |
| C2orf69 | 137.406006 | 0.231394478 | 0.35063329 | 0.65993299 | 0.50929683 | 0.82882082 |
| TYW5 | 408.862401 | 0.088814077 | 0.20391804 | 0.43553811 | 0.66317188 | 0.89267377 |
| MAIP1 | 188.379247 | -0.050841902 | 0.28499836 | -0.1783937 | 0.85841382 | 0.9626966 |
| SPATS2L | 6242.10385 | 0.278465657 | 0.5217358 | 0.53372925 | 0.59352887 | 0.86444505 |
| KCTD18 | 135.585943 | 0.057814569 | 0.32065307 | 0.18030256 | 0.85691505 | 0.96252844 |
| SGO2 | 71.6231384 | -0.72846392 | 0.33523618 | -2.1729872 | 0.02978128 | 0.39981555 |
| AOX1 | 3002.00098 | 1.177318229 | 0.46843939 | 2.51327759 | 0.01196152 | 0.29262893 |
| BZW1 | 248.380624 | -0.824216328 | 0.43502796 | -1.8946284 | 0.05814165 | 0.47775688 |
| CLK1 | 1083.88758 | -0.504646084 | 0.27348071 | -1.8452713 | 0.06499809 | 0.48790819 |
| PPIL3 | 194.878424 | -0.256188618 | 0.32057592 | -0.7991512 | 0.42420277 | 0.78894887 |
| NIF3L1 | 113.87941 | -0.09654581 | 0.30104795 | -0.3206991 | 0.74843843 | 0.92552718 |
| ORC2 | 1282.96186 | -0.217476481 | 0.21564372 | -1.008499 | 0.31321497 | 0.72717675 |
| AC005037.1 | 378.201021 | 0.057644521 | 0.30230774 | 0.19068159 | 0.84877506 | 0.95951833 |
| FAM126B | 1284.88454 | -0.163863334 | 0.30677098 | -0.5341553 | 0.59323411 | 0.86422149 |
| NDUFB3 | 330.040813 | 0.059419025 | 0.47230404 | 0.12580672 | 0.89988492 | 0.97465308 |
| CFLAR | 3459.57526 | 0.116165777 | 0.31147584 | 0.37295277 | 0.7091836 | 0.91086765 |
| CFLAR-AS1 | 294.883534 | 0.533496653 | 0.62618384 | 0.85198087 | 0.39422471 | 0.77823445 |
| CASP10 | 288.414109 | 0.789050196 | 0.4329456 | 1.82251581 | 0.06837676 | 0.49147362 |
| CASP8 | 620.236681 | -0.02086581 | 0.28752548 | -0.0725703 | 0.94214806 | 0.98489601 |
| ALS2CR12 | 84.3215962 | -0.723630391 | 0.75476902 | -0.9587442 | 0.33768765 | 0.74416151 |
| TRAK2 | 943.638099 | -0.264910152 | 0.27725129 | -0.9554876 | 0.3393312 | 0.74644675 |
| STRADB | 680.448949 | 0.323775362 | 0.5000496 | 0.64748649 | 0.51731714 | 0.83284778 |
| C2CD6 | 53.2832245 | 0.859117802 | 0.56617102 | 1.51741749 | 0.12916131 | 0.58793678 |
| TMEM237 | 39.5732906 | 0.195276236 | 0.491718 | 0.39713054 | 0.6912712 | 0.90481802 |
| ALS2 | 1040.86602 | 0.200842955 | 0.2595535 | 0.77380176 | 0.43904803 | 0.7974071 |
| SUMO1 | 493.64422 | -0.279349822 | 0.15073159 | -1.8532931 | 0.06384036 | 0.48593331 |
| NOP58 | 784.976811 | -0.837689962 | 0.29965262 | -2.7955369 | 0.00518136 | 0.21811586 |
| BMPR2 | 2594.60095 | 0.123300991 | 0.19882742 | 0.62014078 | 0.53516511 | 0.83954482 |
| FAM117B | 670.910305 | -0.184776175 | 0.25913358 | -0.7130538 | 4.76E-01 | 0.81570932 |
| ICA1L | 325.615908 | -0.240505449 | 0.42714963 | -0.5630473 | 0.57340267 | 0.8542683 |
| WDR12 | 653.546783 | 0.441983415 | 0.37144141 | 1.18991421 | 0.23408011 | 0.67060722 |
| CARF | 487.753626 | 0.624132568 | 0.46793931 | 1.33378956 | 0.18227284 | 0.63216521 |
| NBEAL1 | 4442.4369 | 0.366084224 | 0.2205766 | 1.65966934 | 0.09698099 | 0.54015377 |
| CYP20A1 | 1004.28155 | -0.481269367 | 0.32541012 | -1.4789625 | 0.13915033 | 0.59474076 |
| ABI2 | 1432.21659 | 0.176898619 | 0.22245219 | 0.79522084 | 0.42648505 | 0.78895108 |
| RAPH1 | 717.718786 | 0.380982504 | 0.22881908 | 1.66499447 | 0.09591386 | 0.53870645 |
| PARD3B | 5738.12534 | 0.22297578 | 0.27657338 | 0.8062084 | 0.4201227 | 0.78798479 |
| NRP2 | 56.6483636 | -1.583236571 | 0.69208087 | -2.2876468 | 0.0221581 | 0.36178312 |
| AC007679.1 | 63.6743324 | 0.240988284 | 0.75422926 | 0.31951596 | 0.74933529 | 0.92565124 |
| INO80D | 2733.78059 | -0.532449606 | 0.64257142 | -0.8286232 | 0.40731764 | 0.78481094 |
| NDUFS1 | 1531.40034 | 0.367986541 | 0.14880768 | 2.47290023 | 0.01340216 | 0.29875272 |
| EEF1B2 | 443.037937 | 0.16771769 | 0.47239899 | 0.35503397 | 0.72256414 | 0.91592159 |
| ZDBF2 | 370.542974 | 0.081390401 | 0.31614129 | 0.25744945 | 0.79683183 | 0.94280281 |
| MDH1B | 67.4451766 | 0.383281917 | 0.49194335 | 0.779118 | 4.36E-01 | 0.79549938 |
| FASTKD2 | 472.634495 | -0.380292973 | 0.24316416 | -1.5639351 | 0.1178328 | 0.57396383 |
| CPO | 89.5052736 | 0.58110185 | 0.39426702 | 1.4738789 | 0.1405142 | 0.59500405 |
| KLF7 | 847.415197 | -1.076218222 | 0.75980038 | -1.4164486 | 0.15664419 | 0.61255952 |
| AC009226.1 | 81.7765897 | -0.278078855 | 0.59211051 | -0.4696401 | 0.63861215 | 0.88284765 |
| CREB1 | 1013.61653 | 0.393596741 | 0.14128928 | 2.78575088 | 0.00534039 | 0.21873395 |

| | | | | | | |
|---|---|---|---|---|---|---|
| METTL21A | 246.821309 | 0.303506856 | 0.30035581 | 1.01049104 | 0.31226009 | 0.7265535 |
| CCNYL1 | 733.580121 | -0.218082723 | 0.26851685 | -0.8121752 | 0.41669112 | 0.78632759 |
| FZD5 | 257.420537 | 0.15058259 | 0.31243896 | 0.48195843 | 0.62983548 | 0.88083664 |
| PLEKHM3 | 985.401506 | 0.005810344 | 0.33553604 | 0.0173166 | 0.98618404 | 0.99609994 |
| IDH1 | 878.378023 | 0.593657444 | 0.56965899 | 1.04212775 | 0.29735245 | 0.71839551 |
| PIKFYVE | 1440.2018 | 0.269740172 | 0.11977174 | 2.25211876 | 0.02431477 | 0.37480999 |
| PTH2R | 26606.0205 | -0.43991363 | 0.48637381 | -0.9044764 | 0.36574285 | 0.76409651 |
| MAP2 | 106.897525 | -0.030661661 | 0.50524218 | -0.0606871 | 0.95160844 | 0.98756033 |
| RPE | 463.170867 | -0.370329789 | 0.2976996 | -1.2439714 | 0.21351009 | 0.65825073 |
| KANSL1L | 6648.08244 | 0.657374444 | 0.4387555 | 1.49827054 | 0.13406298 | 0.59240843 |
| AC007038.2 | 45.4622316 | -0.447048031 | 0.48290069 | -0.9257556 | 0.35457299 | 0.75755608 |
| AC007038.1 | 501.541322 | -0.094298574 | 0.34078883 | -0.2767068 | 0.78200528 | 0.93757221 |
| ACADL | 1023.17891 | 0.473552187 | 0.55936268 | 0.84659239 | 0.39722236 | 0.78031644 |
| LANCL1-AS1 | 101.070216 | 1.201466398 | 0.65985678 | 1.82079875 | 0.06863745 | 0.49154315 |
| LANCL1 | 474.359414 | 0.486161203 | 0.36505463 | 1.33174918 | 0.18294262 | 0.63276491 |
| ERBB4 | 777.222653 | -0.118903646 | 0.54076315 | -0.2198812 | 0.82596368 | 0.95192757 |
| IKZF2 | 280.284842 | 0.593815892 | 0.50181753 | 1.1833303 | 0.23667827 | 0.67375888 |
| SPAG16 | 2987.12365 | 0.658503268 | 0.42244725 | 1.55878224 | 0.11904793 | 0.57619702 |
| AC068051.1 | 140.827787 | 1.303170559 | 0.79981481 | 1.62934037 | 0.10324098 | 0.55022042 |
| BARD1 | 80.7880941 | 1.385112432 | 0.348958 | 3.96928124 | 7.21E-05 | 0.02490203 |
| AC016708.1 | 115.378186 | 0.372832516 | 0.53345442 | 0.69890229 | 0.4846131 | 0.81839266 |
| ATIC | 374.856708 | -0.369548422 | 0.30893038 | -1.1962191 | 0.23161107 | 0.66878058 |
| FN1 | 69.9016414 | -0.174079928 | 0.52197544 | -0.3335021 | 0.73875527 | 0.9208911 |
| MREG | 517.736125 | 0.45054861 | 0.37888952 | 1.18912924 | 0.23438881 | 0.670712 |
| PECR | 950.780885 | 0.804448826 | 0.44055873 | 1.8259741 | 0.06785416 | 0.48963361 |
| TMEM169 | 90.8208189 | 0.91499273 | 0.45958716 | 1.99090141 | 0.04649173 | 0.44508419 |
| XRCC5 | 1922.43652 | -0.137254541 | 0.26018425 | -0.5275282 | 0.59782681 | 0.86636561 |
| SMARCAL1 | 282.447133 | -0.071688915 | 0.39603703 | -0.1810157 | 0.85635526 | 0.96242445 |
| RPL37A | 1095.52163 | -0.177148753 | 0.39475919 | -0.4487514 | 0.65361098 | 0.88985356 |
| IGFBP2 | 65.2053536 | -0.226365421 | 0.60671652 | -0.3730992 | 0.70907465 | 0.91086765 |
| IGFBP5 | 230.520783 | -1.825766618 | 0.51766084 | -3.5269553 | 0.00042037 | 0.07286236 |
| DIRC3 | 1126.12668 | 1.012967961 | 0.74085088 | 1.36730345 | 0.17153021 | 0.62520756 |
| TNS1 | 4332.74175 | -0.005692949 | 0.49011756 | -0.0116155 | 0.9907324 | 0.99737132 |
| ARPC2 | 1028.25992 | -0.649442645 | 0.33949314 | -1.9129772 | 0.05575097 | 0.46997447 |
| AAMP | 241.863761 | -0.175417027 | 0.3495636 | -0.5018172 | 0.61579611 | 0.87323788 |
| PNKD | 725.058665 | 0.025973831 | 0.21861108 | 0.11881296 | 0.90542354 | 0.97613972 |
| TMBIM1 | 680.438922 | -0.217703636 | 0.32657124 | -0.6666344 | 5.05E-01 | 0.82715271 |
| CATIP-AS2 | 101.971192 | 0.39108431 | 0.51963632 | 0.75261157 | 0.45168336 | 0.80384625 |
| CATIP | 85.8031008 | 1.384147943 | 0.72999192 | 1.89611407 | 0.05794496 | 0.47684824 |
| CTDSP1 | 349.356446 | -0.095529434 | 0.30887667 | -0.3092802 | 0.7571084 | 0.92799282 |
| VIL1 | 277.1 | 0.578303526 | 0.70592677 | 0.81921178 | 0.41266559 | 0.78632759 |
| USP37 | 804.802205 | -0.363238719 | 0.19316529 | -1.8804554 | 0.06004604 | 0.48004006 |
| CNOT9 | 419.87617 | -0.079296512 | 0.2001277 | -0.3962296 | 0.69193568 | 0.90499641 |
| PLCD4 | 273.533974 | 1.037282407 | 0.88067567 | 1.17782567 | 0.23886611 | 0.67667639 |
| AC012510.1 | 54.1017779 | -2.234680577 | 0.93951872 | -2.3785376 | 0.01738147 | 0.33064634 |
| ZNF142 | 266.870247 | -0.632499297 | 0.42807704 | -1.4775361 | 0.13953198 | 0.59500405 |
| BCS1L | 100.210354 | -0.19159426 | 0.33503063 | -0.5718709 | 0.56740945 | 0.85251358 |
| RNF25 | 105.30067 | -0.69428067 | 0.36277418 | -1.9138095 | 0.0556445 | 0.46970412 |
| STK36 | 294.869177 | -0.076913371 | 0.29298429 | -0.262517 | 7.93E-01 | 0.94160727 |
| TTLL4 | 237.444187 | 0.006683158 | 0.2992262 | 0.0223348 | 0.98218089 | 0.99522272 |
| CYP27A1 | 687.836756 | 1.04430907 | 0.39015265 | 2.67666792 | 0.00743583 | 0.23590813 |
| IHH | 56.7051815 | 1.107408611 | 0.69196547 | 1.60038132 | 0.10951402 | 0.55969747 |
| CNPPD1 | 361.44639 | -0.176813836 | 0.25559261 | -0.6917799 | 0.48907555 | 0.81970016 |
| RETREG2 | 340.75159 | -0.28624238 | 0.33586108 | -0.8522642 | 0.39406746 | 0.77819948 |

| | | | | | | |
|---|---|---|---|---|---|---|
| ZFAND2B | 290.082477 | -0.120525385 | 0.27157684 | -0.4437985 | 0.6571883 | 0.89053559 |
| ATG9A | 40.1904095 | 0.437814516 | 0.52422912 | 0.83515872 | 0.40362835 | 0.78342308 |
| ANKZF1 | 483.380365 | -0.322683043 | 0.39842671 | -0.8098931 | 0.41800162 | 0.78691775 |
| GLB1L | 210.482321 | -0.4127838 | 0.36275157 | -1.1379242 | 0.25515213 | 0.68976367 |
| STK16 | 104.133913 | -0.289540485 | 0.3333141 | -0.8686716 | 0.38502679 | 0.77317162 |
| TUBA4A | 513.895496 | -0.573411778 | 0.50681274 | -1.1314076 | 0.25788358 | 0.69210282 |
| DNAJB2 | 337.40664 | -1.414358498 | 0.422041 | -3.3512349 | 0.00080452 | 0.09506518 |
| DNPEP | 260.93848 | 0.1475739 | 0.40410677 | 0.36518542 | 0.714973 | 0.91279611 |
| GMPPA | 120.371449 | -0.055070922 | 0.49905341 | -0.1103508 | 0.9121312 | 0.977179 |
| CHPF | 118.34471 | -0.252624121 | 0.67662937 | -0.3733567 | 0.70888297 | 0.91082291 |
| OBSL1 | 210.8021 | 0.498709748 | 0.42284511 | 1.17941471 | 0.23823308 | 0.67642542 |
| STK11IP | 103.955632 | -0.11638801 | 0.32875854 | -0.3540228 | 0.72332181 | 0.91592159 |
| EPHA4 | 275.777787 | -0.452625571 | 0.54384396 | -0.832271 | 0.40525598 | 0.78429302 |
| SGPP2 | 836.58782 | 1.341230573 | 0.67772645 | 1.97901465 | 4.78E-02 | 4.51E-01 |
| FARSB | 488.751882 | -0.364738492 | 0.35191426 | -1.0364414 | 0.29999628 | 0.71879123 |
| MOGAT1 | 1325.71415 | 0.138852977 | 0.63145199 | 0.21989475 | 0.82595313 | 0.95192757 |
| ACSL3 | 894.771095 | -0.38598411 | 0.38334459 | -1.0068855 | 0.3139898 | 0.72719146 |
| AP1S3 | 413.104123 | 0.02596898 | 0.42044695 | 0.06176518 | 0.95074984 | 0.98731502 |
| WDFY1 | 970.769611 | -0.108143432 | 0.29742504 | -0.3635989 | 0.71615751 | 0.91298968 |
| MRPL44 | 164.581323 | 0.002819351 | 0.26660765 | 0.01057491 | 0.9915626 | 0.99748517 |
| SERPINE2 | 2925.31163 | -1.175397318 | 0.6259543 | -1.8777686 | 6.04E-02 | 0.48026727 |
| CUL3 | 2087.38012 | -0.094241581 | 0.19639598 | -0.4798549 | 0.63133054 | 0.88123095 |
| DOCK10 | 56.6798344 | -0.192182134 | 0.45853994 | -0.4191175 | 0.67513023 | 0.89801183 |
| IRS1 | 284.385883 | 0.446225531 | 0.33551577 | 1.32996887 | 0.18352853 | 0.63276491 |
| RHBDD1 | 872.311502 | 0.306928093 | 0.2887208 | 1.06306193 | 0.28775387 | 0.7113446 |
| COL4A4 | 1224.17998 | -0.541760981 | 0.29989089 | -1.806527 | 0.07083606 | 0.49741286 |
| COL4A3 | 1778.57323 | -0.277924389 | 0.37692559 | -0.7373455 | 0.46091226 | 0.80758333 |
| AC097662.1 | 265.933196 | -0.008045112 | 0.4954265 | -0.0162388 | 0.98704391 | 0.9963052 |
| MFF | 807.091761 | 0.06331558 | 0.21806916 | 0.29034633 | 0.7715513 | 0.93316533 |
| AGFG1 | 2302.41428 | 0.014589816 | 0.22586419 | 0.06459553 | 0.94849605 | 0.98687076 |
| SLC19A3 | 332.581738 | 0.629292127 | 0.57314879 | 1.09795596 | 0.27222372 | 0.7005953 |
| DNER | 143.252624 | 0.866109204 | 0.49509136 | 1.74939269 | 0.08022316 | 0.51297858 |
| TRIP12 | 3419.80334 | -0.097951418 | 0.15080719 | -0.6495142 | 0.51600605 | 0.83210529 |
| FBXO36 | 477.457503 | 0.611180598 | 0.26549986 | 2.3019997 | 0.02133519 | 0.35688511 |
| AC009950.1 | 32.091932 | -0.865436862 | 0.9134383 | -0.9474497 | 0.34340967 | 0.75073625 |
| SP110 | 256.577515 | -1.191929931 | 0.61887921 | -1.9259492 | 0.05411072 | 0.46711025 |
| SP140L | 214.040984 | -0.185477957 | 0.47426855 | -0.3910821 | 0.69573653 | 0.90590487 |
| SP100 | 1405.7413 | -0.76498402 | 0.40877931 | -1.8713864 | 0.06129155 | 0.4813972 |
| AC010149.1 | 59.8710491 | -0.70113225 | 0.4169511 | -1.6815695 | 0.09265235 | 0.53669957 |
| CAB39 | 1566.30113 | -0.350745173 | 0.25526152 | -1.3740621 | 0.16942241 | 0.62326131 |
| ITM2C | 217.636416 | 0.762222446 | 0.70984988 | 1.07377978 | 0.2829214 | 0.70756556 |
| C2orf72 | 50.0307977 | 0.697866545 | 0.8873323 | 0.78647712 | 0.43158802 | 0.79135216 |
| PSMD1 | 1424.24863 | -0.40593233 | 0.32086002 | -1.2651384 | 0.20582173 | 0.651931 |
| ARMC9 | 1150.33143 | 0.114796035 | 0.34660147 | 0.3312047 | 0.74048987 | 0.9217959 |
| NCL | 737.677169 | -0.696331695 | 0.43654303 | -1.5951044 | 0.1106889 | 0.56297319 |
| PTMA | 1717.30546 | -0.576721138 | 0.43839621 | -1.3155249 | 0.18833355 | 0.64026838 |
| PDE6D | 351.254776 | -0.22018967 | 0.31896031 | -0.6903356 | 0.48998314 | 0.81973856 |
| COPS7B | 237.891284 | 0.164971938 | 0.21987687 | 0.75029237 | 0.45307864 | 0.80433814 |
| DIS3L2 | 2021.73326 | -0.14732869 | 0.24452722 | -0.6025043 | 0.54683853 | 0.84460887 |
| AC019130.1 | 113.349723 | 0.570111958 | 0.58768939 | 0.97009061 | 0.33200133 | 0.73947712 |
| TIGD1 | 74.0263178 | 0.027253924 | 0.34621459 | 0.07871974 | 0.93725554 | 0.98435387 |
| EIF4E2 | 724.12027 | 0.09193235 | 0.2697325 | 0.34082786 | 0.73323318 | 0.91952673 |
| EFHD1 | 762.736503 | 0.371928902 | 0.34261591 | 1.08555644 | 2.78E-01 | 7.04E-01 |
| GIGYF2 | 3158.06396 | -0.227701001 | 0.14001143 | -1.6263029 | 0.10388523 | 0.55022042 |

| | | | | | | |
|---|---|---|---|---|---|---|
| SNORC | 93.3500574 | 0.823032285 | 0.28073545 | 2.93170061 | 0.00337112 | 0.18932494 |
| NGEF | 729.566642 | 1.009801181 | 0.44064255 | 2.2916561 | 0.0219255 | 0.36066559 |
| INPP5D | 47.4887881 | 0.466052301 | 0.67296531 | 0.6925354 | 0.48860116 | 0.81933645 |
| ATG16L1 | 368.371783 | -0.84455873 | 0.40287906 | -2.0963083 | 0.03605485 | 0.42040764 |
| DGKD | 621.842695 | -0.8609485 | 0.41990117 | -2.0503599 | 0.04032933 | 0.43086056 |
| USP40 | 1806.18559 | 0.133720953 | 0.42073126 | 0.31782985 | 0.75061401 | 0.92590588 |
| AC114812.2 | 1826.7982 | 0.752815003 | 0.96566896 | 0.77957875 | 0.43563887 | 0.79549938 |
| MROH2A | 84.6989784 | 1.521394733 | 0.80885299 | 1.88092862 | 0.05998163 | 0.48004006 |
| ARL4C | 70.3448409 | -0.034265178 | 1.12847674 | -0.0303641 | 0.97577668 | 0.99429832 |
| SH3BP4 | 274.789786 | -1.448499354 | 0.62123324 | -2.3316514 | 0.01971904 | 0.34523522 |
| AGAP1 | 12316.7966 | -0.367744971 | 0.32319266 | -1.1378506 | 0.25518287 | 0.68976367 |
| IQCA1 | 630.282023 | -0.158325223 | 0.41153278 | -0.3847208 | 0.70044428 | 0.90763752 |
| ACKR3 | 76.8233358 | 0.269725691 | 0.66374536 | 0.40636923 | 0.68447133 | 0.9007343 |
| AC105760.2 | 114.582585 | 0.585821502 | 0.4863783 | 1.2044565 | 0.22841319 | 0.66705629 |
| COPS8 | 174.459337 | -0.037099048 | 0.28892036 | -0.1284058 | 0.89782785 | 0.9737247 |
| RAB17 | 1104.44426 | -0.374652563 | 0.41799466 | -0.8963095 | 0.37008751 | 0.76669785 |
| LRRFIP1 | 4287.11084 | -0.610294673 | 0.40966313 | -1.4897476 | 1.36E-01 | 5.94E-01 |
| RBM44 | 41.3444801 | -0.012482053 | 0.73142311 | -0.0170654 | 0.98638441 | 0.99612253 |
| UBE2F-SCLY | 56.2034636 | -0.069677662 | 0.49231435 | -0.1415308 | 0.8874506 | 0.97098615 |
| ILKAP | 672.247718 | -0.222936286 | 0.27141688 | -0.8213796 | 0.41143009 | 0.78632759 |
| PER2 | 262.111792 | -0.411538545 | 0.54161256 | -0.7598394 | 0.44735061 | 0.80223612 |
| TRAF3IP1 | 739.244621 | -0.078831723 | 0.24691445 | -0.3192673 | 0.74952379 | 0.92565124 |
| ASB1 | 147.0338 | -0.376444938 | 0.32518511 | -1.1576328 | 0.24701393 | 0.68482118 |
| HDAC4 | 1378.35299 | -0.257806663 | 0.41119595 | -0.6269679 | 0.53068027 | 0.83789294 |
| AC062017.1 | 38.2815628 | 0.011483275 | 0.83596573 | 0.01373654 | 0.98904017 | 0.99721497 |
| NDUFA10 | 1072.21866 | -0.092575672 | 0.25522566 | -0.3627209 | 0.71681341 | 0.91309429 |
| COPS9 | 263.878109 | -0.570558199 | 0.45114675 | -1.2646843 | 0.20598453 | 0.651931 |
| GPC1 | 62.7225884 | 0.123523819 | 0.47757611 | 0.2586474 | 0.79590731 | 0.94252227 |
| ANKMY1 | 369.522573 | -0.217315529 | 0.33143717 | -0.6556764 | 0.51203236 | 0.83054053 |
| RNPEPL1 | 280.576324 | -0.057150124 | 0.44747256 | -0.1277176 | 0.89837246 | 0.97397701 |
| CAPN10-DT | 45.7777781 | -0.157481479 | 0.48014959 | -0.3279842 | 0.74292361 | 0.92318522 |
| CAPN10 | 137.046484 | -0.031597901 | 0.36712195 | -0.0860692 | 0.93141139 | 0.9827462 |
| GPR35 | 82.2678791 | 0.290564611 | 0.62096092 | 0.46792737 | 0.63983653 | 0.88321992 |
| AGXT | 97.0974199 | -1.551409039 | 0.77552871 | -2.0004534 | 0.04545133 | 0.44360789 |
| C2orf54 | 95.2513219 | -0.432741767 | 1.18338786 | -0.3656804 | 0.71460356 | 0.91252758 |
| SNED1 | 171.337825 | 0.510652734 | 0.46902521 | 1.08875328 | 0.2762627 | 0.70328016 |
| AC005237.1 | 89.6525942 | 1.680896676 | 0.71299362 | 2.35751993 | 0.01839747 | 0.33848871 |
| MTERF4 | 456.152168 | 0.033849814 | 0.29270593 | 0.11564444 | 0.90793434 | 0.97673359 |
| PASK | 82.239881 | 0.456370029 | 0.4062702 | 1.12331652 | 0.26130309 | 0.69358665 |
| PPP1R7 | 302.606951 | 0.276137886 | 0.22649748 | 1.21916541 | 0.22278142 | 0.66398799 |
| ANO7 | 58.6335687 | -0.621807545 | 0.36195831 | -1.7178982 | 8.58E-02 | 5.23E-01 |
| HDLBP | 1703.21243 | -0.664997316 | 0.29224341 | -2.2754912 | 0.02287648 | 0.36631766 |
| 2-Sep | 2047.92483 | -0.031482766 | 0.19340594 | -0.1627808 | 0.87069106 | 0.9657206 |
| FARP2 | 1219.16528 | -0.136856662 | 0.21916897 | -0.6244345 | 0.53234228 | 0.83819165 |
| STK25 | 416.640186 | -0.268698307 | 0.25250359 | -1.0641366 | 0.28726683 | 0.71118494 |
| BOK | 219.971421 | 0.015203101 | 0.48399817 | 0.03141149 | 0.97494138 | 0.99402915 |
| THAP4 | 422.914249 | -0.314701683 | 0.28506475 | -1.1039656 | 0.26960805 | 0.69961625 |
| ATG4B | 1190.14895 | -0.396101069 | 0.2595467 | -1.5261264 | 0.12697839 | 0.58793678 |
| DTYMK | 100.228803 | -0.258866919 | 0.3089547 | -0.8378798 | 0.40209819 | 0.78258224 |
| ING5 | 548.545488 | -0.398705424 | 0.32340714 | -1.2328281 | 0.21763989 | 0.66043612 |
| D2HGDH | 1365.4283 | 1.583842851 | 0.59917967 | 2.64335213 | 8.21E-03 | 0.24877162 |
| CHL1 | 945.679092 | 1.060306641 | 0.84867664 | 1.2493647 | 0.21153171 | 0.65584609 |
| LINC01266 | 522.528736 | 1.835671745 | 0.51804113 | 3.5434865 | 0.00039487 | 0.07121752 |
| CNTN6 | 78.8563161 | 1.636017494 | 0.70783012 | 2.31131377 | 0.02081553 | 0.35313211 |

| | | | | | | |
|---|---|---|---|---|---|---|
| CNTN4 | 7063.50703 | 0.549786898 | 0.48063292 | 1.14388106 | 0.25267297 | 0.68887358 |
| CNTN4-AS2 | 51.1501539 | 1.450010035 | 0.57510399 | 2.52130061 | 0.01169219 | 0.28824618 |
| TRNT1 | 458.762687 | -0.18807138 | 0.32060598 | -0.5866122 | 0.55746419 | 0.84861484 |
| CRBN | 748.48576 | -0.070780637 | 0.39576304 | -0.178846 | 0.85805863 | 0.9626966 |
| AC024060.1 | 46.5195786 | -0.383523428 | 0.38147823 | -1.0053613 | 0.31472293 | 0.72719146 |
| AC034195.1 | 2299.8238 | 0.821128565 | 0.7627644 | 1.07651663 | 0.28169627 | 0.70625976 |
| SUMF1 | 4252.47688 | 0.141626332 | 0.43611368 | 0.32474636 | 0.74537304 | 0.92393094 |
| LRRN1 | 141.655669 | 0.811686977 | 0.41160738 | 1.97199327 | 0.04861037 | 0.45329737 |
| SETMAR | 77.1514541 | 0.287067616 | 0.38613102 | 0.74344614 | 0.45721162 | 0.80552695 |
| ITPR1 | 897.891293 | 0.556046232 | 0.42828616 | 1.29830541 | 0.19418241 | 0.64548758 |
| EGOT | 475.196035 | 0.487486351 | 0.61202084 | 0.7965192 | 0.42573032 | 0.78894887 |
| AC018816.1 | 130.022857 | 0.375960952 | 0.35653248 | 1.05449285 | 0.29165733 | 0.71337138 |
| BHLHE40-AS | 89.9174326 | 0.386252571 | 0.58762994 | 0.65730581 | 0.5109843 | 0.82976684 |
| BHLHE40 | 1074.47463 | -0.609688011 | 0.51145082 | -1.1920755 | 0.23323163 | 0.66978724 |
| ARL8B | 1397.5288 | -0.624805293 | 0.39787091 | -1.5703719 | 0.11632862 | 0.57350419 |
| AC026202.2 | 209.805023 | 0.293325287 | 0.47561024 | 0.61673459 | 0.5374098 | 0.8403387 |
| EDEM1 | 494.03986 | -0.271093359 | 0.43307831 | -0.6259684 | 0.53133564 | 0.83789294 |
| AC069277.1 | 68.9640156 | 1.605171834 | 0.99455842 | 1.6139543 | 0.10653734 | 0.55461729 |
| GRM7-AS3 | 73.4839319 | 0.449335504 | 0.60706005 | 0.74018295 | 0.45918899 | 0.80669065 |
| LMCD1-AS1 | 1220.59016 | -0.617992009 | 0.3777645 | -1.6359187 | 0.1018566 | 0.547724 |
| LMCD1 | 938.982591 | -1.755355875 | 0.88277598 | -1.98845 | 0.04676195 | 0.44635514 |
| RAD18 | 224.842899 | -0.170161551 | 0.3221561 | -0.528196 | 0.59736333 | 0.86614567 |
| SRGAP3 | 860.753894 | 1.246051999 | 0.5143032 | 2.42279649 | 0.01540155 | 0.31183024 |
| THUMPD3-A! | 1008.99406 | -0.143067626 | 0.29334109 | -0.4877177 | 0.62574985 | 0.87854882 |
| THUMPD3 | 213.366718 | -0.270012073 | 0.24312364 | -1.1105957 | 0.26674241 | 0.69812269 |
| SETD5 | 3483.14372 | 0.02264647 | 0.16810782 | 0.13471396 | 0.89283804 | 0.9724379 |
| MTMR14 | 418.665299 | -0.408643183 | 0.27704844 | -1.4749882 | 0.14021571 | 0.59500405 |
| BRPF1 | 82.0211943 | -0.676997164 | 0.46702081 | -1.4496081 | 0.14716782 | 0.60468506 |
| OGG1 | 682.150795 | 0.425797035 | 0.39672911 | 1.07326894 | 0.28315047 | 0.70772128 |
| CAMK1 | 273.518353 | 0.685832107 | 0.42112034 | 1.62858936 | 0.10339998 | 0.55022042 |
| TADA3 | 256.523841 | 0.025524909 | 0.24228847 | 0.10534925 | 0.91609869 | 0.97803292 |
| TTLL3 | 53.8961076 | 0.279618531 | 0.46053284 | 0.60716307 | 0.5437427 | 0.84396451 |
| RPUSD3 | 132.071977 | 0.175988041 | 0.2834482 | 0.62088254 | 0.53467691 | 0.83954482 |
| JAGN1 | 137.985389 | 0.003543543 | 0.38357695 | 0.00923815 | 0.99262912 | 0.9976934 |
| IL17RE | 70.5861419 | 0.611606583 | 0.68954203 | 0.88697507 | 0.37509233 | 0.7697436 |
| IL17RC | 218.202317 | 0.458717915 | 0.43053312 | 1.06546487 | 0.28666562 | 0.71118494 |
| CRELD1 | 215.603156 | 0.599198212 | 0.52117694 | 1.14970208 | 0.2502666 | 0.68617881 |
| PRRT3-AS1 | 53.5517088 | -0.185787026 | 0.39283369 | -0.4729407 | 0.63625551 | 0.88266168 |
| EMC3 | 465.398995 | 0.160679471 | 0.20749919 | 0.77436192 | 0.4387168 | 0.79718009 |
| FANCD2 | 56.8563201 | 0.250388844 | 0.37371949 | 0.6699914 | 0.50286327 | 0.82646845 |
| BRK1 | 405.421457 | -0.109048501 | 0.23889945 | -0.4564619 | 0.64805785 | 0.88787189 |
| VHL | 278.364565 | -0.163857867 | 0.24607832 | -0.6658769 | 0.5054898 | 0.8272145 |
| IRAK2 | 519.113818 | 0.454400321 | 0.4376926 | 1.03817228 | 2.99E-01 | 7.19E-01 |
| GHRL | 44.1451237 | 0.34740928 | 0.55045886 | 0.63112669 | 0.52795769 | 0.83761342 |
| SEC13 | 550.855881 | -0.127065909 | 0.34811017 | -0.3650164 | 0.71509919 | 0.91279611 |
| HRH1 | 598.964793 | 0.676782877 | 0.48853114 | 1.38534234 | 1.66E-01 | 6.21E-01 |
| ATG7 | 1907.99977 | -0.021401497 | 0.25940085 | -0.0825036 | 0.93424628 | 0.98359701 |
| VGLL4 | 1997.24826 | 0.09794901 | 0.20873608 | 0.46924811 | 0.6388923 | 0.88284765 |
| TAMM41 | 636.282317 | 0.051609686 | 0.34327142 | 0.15034659 | 0.88049118 | 0.96773407 |
| SYN2 | 826.322302 | -0.184723371 | 0.42078137 | -0.4390008 | 0.66066093 | 0.89130703 |
| PPARG | 3720.93858 | -0.90731832 | 0.26434604 | -3.4323128 | 0.00059846 | 0.08329748 |
| TSEN2 | 710.71586 | -0.182512805 | 0.42947765 | -0.4249646 | 0.67086247 | 0.89694443 |
| MKRN2OS | 108.338416 | 0.554709912 | 0.48902317 | 1.13432235 | 0.25665936 | 0.69102923 |
| MKRN2 | 187.441871 | -0.232903931 | 0.23276217 | -1.0006091 | 0.31701585 | 0.728983 |

| | | | | | | |
|---|---|---|---|---|---|---|
| RAF1 | 1311.76105 | -0.166981568 | 0.14437528 | -1.1565801 | 0.24744396 | 0.68482118 |
| RPL32 | 1292.43407 | -0.163782954 | 0.52402434 | -0.3125484 | 0.75462381 | 0.92799282 |
| IQSEC1 | 223.480658 | 0.221338276 | 0.34118361 | 0.64873654 | 0.51650868 | 0.83258835 |
| NUP210 | 551.958693 | 0.269156192 | 0.38172506 | 0.70510486 | 4.81E-01 | 0.81715537 |
| HDAC11 | 180.988712 | 0.682712882 | 0.39713769 | 1.71908357 | 0.08559916 | 0.5226185 |
| CHCHD4 | 55.2618548 | 0.281050581 | 0.47811871 | 0.58782594 | 0.55664913 | 0.84861484 |
| TMEM43 | 184.001841 | -0.33004385 | 0.2647515 | -1.2466175 | 0.21253779 | 0.65745584 |
| AC093495.1 | 45.3888765 | 0.26017213 | 0.79246542 | 0.32830723 | 0.74267938 | 0.92309632 |
| XPC | 1144.68364 | 0.010038923 | 0.35843932 | 0.02800732 | 9.78E-01 | 0.99478399 |
| LSM3 | 261.283006 | -0.159709831 | 0.31002702 | -0.5151481 | 0.60644953 | 0.86961018 |
| AC093496.1 | 55.2907784 | -1.739512009 | 0.81736984 | -2.1281823 | 0.03332197 | 0.41254527 |
| SLC6A6 | 2797.41307 | -0.475662995 | 0.31322326 | -1.5186069 | 0.12886148 | 0.58793678 |
| CCDC174 | 367.427933 | -0.161079539 | 0.21823065 | -0.738116 | 0.46044395 | 0.8071755 |
| C3orf20 | 67.2972749 | -2.382100412 | 0.6138679 | -3.8804773 | 0.00010425 | 0.03247677 |
| FGD5 | 979.87623 | 0.063261558 | 0.38234966 | 0.16545472 | 0.86858608 | 0.96536557 |
| FGD5-AS1 | 388.737452 | -0.256503009 | 0.28914078 | -0.8871215 | 0.37501348 | 0.7697436 |
| NR2C2 | 1101.77569 | -0.692324593 | 0.37139325 | -1.8641281 | 0.06230372 | 0.4840117 |
| MRPS25 | 690.556855 | -0.002355991 | 0.17825562 | -0.0132169 | 0.98945473 | 0.99725454 |
| RBSN | 341.358245 | -0.161336452 | 0.19116098 | -0.8439821 | 0.39867939 | 0.78122921 |
| CAPN7 | 1455.11383 | -0.1288805 | 0.16025861 | -0.8042033 | 0.42127959 | 0.78877309 |
| SH3BP5-AS1 | 176.16196 | -0.392390042 | 0.28993738 | -1.3533613 | 0.17594023 | 0.62699479 |
| SH3BP5 | 838.223236 | -0.772848158 | 0.56032722 | -1.37928 | 0.16780843 | 0.62304975 |
| METTL6 | 554.073931 | -0.085480221 | 0.30771763 | -0.2777879 | 0.78117522 | 0.93748328 |
| EAF1 | 220.633405 | -0.050221393 | 0.26704798 | -0.1880613 | 0.85082859 | 0.96006828 |
| COLQ | 252.673884 | 0.173517226 | 0.54190952 | 0.32019593 | 0.74881981 | 0.92554142 |
| HACL1 | 714.606236 | -0.110878238 | 0.27094774 | -0.4092237 | 0.6823755 | 0.89962501 |
| BTD | 361.879433 | 0.145947321 | 0.33930843 | 0.43013172 | 6.67E-01 | 8.94E-01 |
| ANKRD28 | 2039.44722 | -0.013297557 | 0.17456417 | -0.0761758 | 0.93927927 | 0.98435387 |
| DPH3 | 96.6874651 | -0.298679049 | 0.44468188 | -0.671669 | 0.50179442 | 0.82638105 |
| OXNAD1 | 1026.66691 | 0.386913157 | 0.24797042 | 1.56031983 | 0.11868432 | 0.57589001 |
| RFTN1 | 218.936041 | 0.987900024 | 0.34805187 | 2.83837008 | 0.00453446 | 0.20749797 |
| PLCL2 | 2239.83865 | 0.201651311 | 0.28784369 | 0.70055838 | 4.84E-01 | 8.18E-01 |
| TBC1D5 | 5659.0214 | 0.349722572 | 0.29149999 | 1.19973442 | 0.2302425 | 0.66820537 |
| KCNH8 | 88.0402131 | 1.276813738 | 0.75475616 | 1.69169039 | 0.09070502 | 0.5349801 |
| EFHB | 121.175259 | -0.345305672 | 0.57485402 | -0.6006841 | 0.54805041 | 0.84486687 |
| RAB5A | 1106.29181 | -0.246679295 | 0.18798895 | -1.3122011 | 0.1894523 | 0.6416112 |
| KAT2B | 825.993325 | -0.466296256 | 0.35306525 | -1.3207084 | 0.1865986 | 0.6378941 |
| ZNF385D | 603.173393 | -0.480887329 | 0.76355085 | -0.6298039 | 0.52882287 | 0.83765775 |
| UBE2E2 | 10106.5281 | -0.378908002 | 0.3185737 | -1.1893888 | 0.23428669 | 0.670712 |
| UBE2E1 | 1805.82255 | -0.041597907 | 0.26926451 | -0.1544871 | 0.87722564 | 0.96653382 |
| NKIRAS1 | 316.503603 | -0.334399033 | 0.29041414 | -1.1514558 | 0.24954479 | 0.68617881 |
| RPL15 | 1630.30154 | 0.38727902 | 0.45082046 | 0.85905378 | 0.39031085 | 0.77569824 |
| NR1D2 | 546.340971 | -1.004136032 | 0.56068755 | -1.7909012 | 0.07330914 | 0.50459544 |
| THRB | 3858.98972 | 0.201032515 | 0.33874636 | 0.59346029 | 0.55287315 | 0.84721285 |
| RARB | 123.65639 | 0.386858436 | 0.72141393 | 0.5362503 | 0.59178557 | 0.86382106 |
| TOP2B | 1013.4815 | 0.056110656 | 0.18445115 | 0.30420334 | 0.76097298 | 0.92989212 |
| NGLY1 | 1202.07275 | -0.339950396 | 0.28262847 | -1.2028172 | 0.22904705 | 0.66705629 |
| OXSM | 165.421722 | 0.02776675 | 0.38770749 | 0.07161778 | 0.94290609 | 0.98501744 |
| NEK10 | 387.773079 | 0.266138893 | 0.59512405 | 0.44719902 | 0.65473137 | 0.89040608 |
| SLC4A7 | 1206.7573 | 0.018592765 | 0.42875171 | 0.04336488 | 0.96541068 | 0.99074368 |
| LINC01967 | 63.9269808 | 1.479112944 | 0.60312952 | 2.45239688 | 0.0141908 | 0.30468798 |
| CMC1 | 1192.7834 | 0.114412193 | 0.27029613 | 0.42328461 | 0.67208762 | 0.89713835 |
| AZI2 | 508.651488 | -0.184116495 | 0.33891023 | -0.5432604 | 0.58695055 | 0.86112855 |
| RBMS3 | 417.023597 | 1.363229809 | 0.56458482 | 2.41457042 | 0.01575378 | 0.31566878 |

| | | | | | | |
|---|---|---|---|---|---|---|
| RBMS3-AS3 | 167.167612 | 0.426992345 | 0.70736061 | 0.60364167 | 0.5460819 | 0.84460887 |
| TGFBR2 | 810.93191 | -0.38110069 | 0.28903866 | -1.3185111 | 0.1873326 | 0.63858939 |
| STT3B | 1339.79378 | 0.026804076 | 0.23932306 | 0.11199955 | 0.91082376 | 0.97697493 |
| OSBPL10 | 6061.49349 | 0.014239938 | 0.22495733 | 0.06330062 | 0.94952712 | 0.9871942 |
| OSBPL10-AS | 110.961541 | -0.156614208 | 0.47313866 | -0.3310112 | 0.74063601 | 0.92189409 |
| ZNF860 | 113.951807 | 0.020947534 | 0.46109407 | 0.04543006 | 0.96376452 | 0.99012655 |
| GPD1L | 341.031168 | 0.125908074 | 0.26037055 | 0.48357265 | 0.62868919 | 0.88022909 |
| CMTM8 | 1061.60512 | -0.665164084 | 0.33209792 | -2.0029155 | 0.04518636 | 0.44360789 |
| CMTM7 | 722.699104 | -0.206912276 | 0.45915071 | -0.4506413 | 0.65224809 | 0.88935488 |
| CMTM6 | 544.165312 | 0.395702731 | 0.43400917 | 0.91173818 | 0.36190656 | 0.76060307 |
| DYNC1LI1 | 624.645607 | -0.088266784 | 0.19064482 | -0.4629907 | 0.64337102 | 0.8848654 |
| CNOT10 | 789.141512 | -0.221441077 | 0.18642761 | -1.1878127 | 0.23490722 | 0.67119484 |
| TRIM71 | 574.615613 | -0.446899099 | 0.3560438 | -1.2551801 | 0.20941338 | 0.65441616 |
| GLB1 | 828.838603 | -0.024492791 | 0.22373995 | -0.1094699 | 0.91282979 | 0.977179 |
| CRTAP | 452.803368 | 0.261152641 | 0.30928708 | 0.84436971 | 0.39846284 | 0.78122921 |
| FBXL2 | 1336.43392 | -0.047920086 | 0.26924865 | -0.1779771 | 0.85874098 | 0.9626966 |
| UBP1 | 574.570305 | -0.083461293 | 0.2355365 | -0.3543455 | 0.72307999 | 0.91592159 |
| CLASP2 | 2403.67699 | 0.021590773 | 0.20893269 | 0.10333841 | 0.91769439 | 0.97828877 |
| PDCD6IP | 1690.39737 | -0.210807191 | 0.19116581 | -1.1027453 | 0.27013781 | 0.69961625 |
| ARPP21 | 266.418743 | 0.155756493 | 0.59249693 | 0.26288152 | 0.7926419 | 0.94154414 |
| STAC | 133.61047 | -0.634159001 | 0.61905532 | -1.024398 | 0.30564735 | 0.72122 |
| TRANK1 | 159.542517 | -1.018031371 | 0.84675754 | -1.2022702 | 0.22925885 | 0.66705629 |
| EPM2AIP1 | 404.808022 | 0.069618576 | 0.17988908 | 0.38700835 | 0.69875002 | 0.90696517 |
| MLH1 | 242.18711 | -0.103764846 | 0.24636948 | -0.4211757 | 0.67362676 | 0.89799059 |
| LRRFIP2 | 3894.91588 | 0.26082689 | 0.2693172 | 0.9684747 | 0.33280735 | 0.73988623 |
| GOLGA4 | 8425.45236 | -0.739780803 | 0.30519125 | -2.4239909 | 0.01535099 | 0.31126635 |
| ITGA9 | 179.007516 | 0.900020575 | 0.4485775 | 2.00638814 | 0.04481485 | 0.44360789 |
| CTDSPL | 3262.53084 | 0.066902061 | 0.35827657 | 0.18673301 | 0.85186997 | 0.96050023 |
| PLCD1 | 100.647186 | 0.365773251 | 0.33205455 | 1.10154568 | 0.27065923 | 0.69961625 |
| DLEC1 | 40.0150924 | 0.66999872 | 0.52199292 | 1.28353985 | 0.19930301 | 0.64643784 |
| ACAA1 | 1061.26796 | 0.899011646 | 0.47457146 | 1.89436517 | 0.05817656 | 0.47775688 |
| MYD88 | 82.6605236 | -0.461967224 | 0.39816728 | -1.160234 | 0.24595354 | 0.68320879 |
| OXSR1 | 1760.85425 | -0.321265943 | 0.21246077 | -1.5121189 | 0.13050362 | 0.58853436 |
| SLC22A13 | 66.1360814 | 0.834119684 | 0.58003517 | 1.43805019 | 1.50E-01 | 6.08E-01 |
| SLC22A14 | 213.57319 | 1.111151361 | 0.55479079 | 2.00282949 | 4.52E-02 | 4.44E-01 |
| XYLB | 2300.12343 | -0.17639346 | 0.3534776 | -0.499023 | 6.18E-01 | 8.74E-01 |
| ACVR2B | 518.364203 | 0.320007273 | 0.28149177 | 1.13682638 | 0.25561088 | 0.69024 |
| EXOG | 286.314103 | 0.012066054 | 0.2798083 | 0.04312257 | 0.96560382 | 0.99081812 |
| SCN11A | 66.2011917 | 1.389131635 | 0.80404862 | 1.7276712 | 0.08404719 | 0.52183702 |
| WDR48 | 661.808986 | -0.152579066 | 0.45926912 | -0.3322215 | 0.73972203 | 0.92125839 |
| GORASP1 | 149.457965 | 0.029957399 | 0.30316968 | 0.09881397 | 0.92128598 | 0.97939204 |
| TTC21A | 196.008812 | -0.005258927 | 0.43562179 | -0.0120722 | 0.99036799 | 0.99737132 |
| CSRNP1 | 152.153558 | -0.894079173 | 0.44992882 | -1.9871569 | 0.04690501 | 0.44732709 |
| SLC25A38 | 193.915369 | -0.651441364 | 0.24445471 | -2.6648755 | 0.00770168 | 0.24157203 |
| RPSA | 936.581701 | 0.078607401 | 0.41740236 | 0.18832524 | 0.8506217 | 0.95994332 |
| MYRIP | 1151.54884 | -0.931258236 | 0.66428942 | -1.4018863 | 0.1609492 | 0.61675446 |
| EIF1B | 263.461788 | -0.483662737 | 0.42402429 | -1.1406486 | 0.25401617 | 0.68891958 |
| ENTPD3-AS1 | 452.194712 | 0.658165217 | 0.52652505 | 1.25001691 | 0.21129337 | 0.65561118 |
| RPL14 | 1714.62118 | -0.155780386 | 0.57028667 | -0.2731615 | 0.78472904 | 0.93884957 |
| ZNF620 | 53.287897 | -0.478328723 | 0.32849567 | -1.4561188 | 0.14535977 | 0.60165215 |
| ZNF621 | 466.455539 | -0.257615044 | 0.30015043 | -0.8582864 | 0.39073432 | 0.77606489 |
| CTNNB1 | 1901.79326 | -0.127429746 | 0.23705052 | -0.5375637 | 0.59087832 | 0.86333372 |
| ULK4 | 3079.49469 | -0.125049746 | 0.39212251 | -0.3189048 | 0.74979872 | 0.92565124 |
| TRAK1 | 2003.86541 | -0.448445878 | 0.32981014 | -1.3597092 | 0.17392197 | 0.62685996 |

| | | | | | | |
|---|---|---|---|---|---|---|
| SEC22C | 261.118648 | -0.250135685 | 0.33394906 | -0.7490235 | 0.45384306 | 0.80433814 |
| SS18L2 | 162.096542 | 0.073123211 | 0.29792125 | 0.24544476 | 0.80611207 | 0.94585363 |
| NKTR | 2518.17132 | -0.132870275 | 0.19337704 | -0.6871047 | 0.49201675 | 0.82080956 |
| AC006059.1 | 267.223961 | 0.344684344 | 0.53010428 | 0.65021988 | 0.5155502 | 0.83185952 |
| CCDC13 | 76.8365893 | 0.047147881 | 0.58366129 | 0.08077952 | 0.93561729 | 0.98406248 |
| AC099329.3 | 111.662702 | 0.372047313 | 0.44585564 | 0.8344569 | 0.40402356 | 0.78358009 |
| AC092042.3 | 31.4158872 | 0.975652449 | 0.82732344 | 1.17928782 | 0.23828359 | 0.67642542 |
| CYP8B1 | 161.296152 | 2.669128364 | 0.82194407 | 3.24733577 | 0.00116491 | 0.11093704 |
| POMGNT2 | 59.8655442 | -0.425932104 | 0.39960857 | -1.0658733 | 0.28648092 | 0.71118494 |
| SNRK | 1490.6806 | -0.213071878 | 0.31324846 | -0.6802009 | 0.49637727 | 0.82243887 |
| ANO10 | 3240.14664 | -0.7774864 | 0.36830974 | -2.110958 | 0.03477592 | 0.41686761 |
| ABHD5 | 845.765379 | -1.841307124 | 0.59200634 | -3.1102828 | 0.00186908 | 0.13738894 |
| AC006058.1 | 97.1332199 | -0.117048751 | 0.65754386 | -0.178009 | 0.85871588 | 0.9626966 |
| TCAIM | 1384.25488 | 0.681849653 | 0.3933625 | 1.73338754 | 0.08302681 | 0.51935659 |
| ZNF445 | 291.813422 | 0.21306436 | 0.18850187 | 1.13030369 | 0.25834828 | 0.69210282 |
| ZNF852 | 92.6915444 | -0.005739452 | 0.26141729 | -0.0219551 | 0.98248374 | 0.99522272 |
| ZKSCAN7 | 53.2390997 | 0.035373094 | 0.46835621 | 0.07552605 | 0.93979617 | 0.98439448 |
| AC099669.1 | 243.747142 | 0.099897314 | 0.3686677 | 0.27096845 | 0.7864153 | 0.9390702 |
| ZNF197 | 235.882289 | 0.446595675 | 0.3738136 | 1.1947015 | 0.23220367 | 0.66964354 |
| ZNF35 | 71.8968105 | -0.712826038 | 0.46897428 | -1.5199683 | 0.12851894 | 0.58793678 |
| ZNF502 | 58.7159256 | -0.710871599 | 0.30444873 | -2.3349468 | 0.0195462 | 0.34513641 |
| KIAA1143 | 98.8427014 | -0.111541163 | 0.29813754 | -0.3741265 | 0.70831018 | 0.9105991 |
| ZDHHC3 | 365.363296 | 0.186836765 | 0.21538028 | 0.86747387 | 0.38568242 | 0.77317162 |
| EXOSC7 | 191.831027 | -0.786964704 | 0.26032423 | -3.0230176 | 0.00250268 | 0.16335332 |
| CDCP1 | 74.5158699 | 2.562438715 | 0.88526873 | 2.89453205 | 0.00379724 | 0.2001601 |
| LARS2 | 620.136015 | -0.125351543 | 0.15401233 | -0.8139059 | 0.41569887 | 0.78632759 |
| LIMD1 | 442.406897 | -0.720349841 | 0.30749969 | -2.3426035 | 0.01914972 | 0.34084276 |
| SACM1L | 750.371071 | 0.035244891 | 0.15721089 | 0.22418861 | 0.82261055 | 0.95116101 |
| SLC6A20 | 687.59852 | -1.052764306 | 0.64184784 | -1.6402085 | 0.10096181 | 0.54608983 |
| LZTFL1 | 289.101578 | -0.176213498 | 0.23004947 | -0.7659809 | 0.44368767 | 0.7993976 |
| FYCO1 | 357.104451 | 0.027533577 | 0.23121115 | 0.11908412 | 0.90520871 | 0.97598655 |
| LINC02009 | 45.5198346 | 0.322895148 | 0.71838245 | 0.44947527 | 6.53E-01 | 0.88949911 |
| LTF | 3062.65712 | 1.238830544 | 1.49757511 | 0.82722432 | 0.40810993 | 0.78481094 |
| LRRC2 | 68.7867773 | 0.126029321 | 0.58766375 | 0.21445822 | 0.83018975 | 0.95325673 |
| ALS2CL | 201.222131 | -0.450578414 | 0.30036948 | -1.5000805 | 1.34E-01 | 5.92E-01 |
| MYL3 | 80.475868 | 0.61856005 | 0.52908064 | 1.16912245 | 0.2423543 | 0.68080966 |
| PTH1R | 3724.88245 | 1.000178107 | 0.72301931 | 1.38333527 | 1.67E-01 | 0.62216414 |
| CCDC12 | 677.869658 | -0.592165418 | 0.33002164 | -1.794323 | 0.07276163 | 0.50396344 |
| NBEAL2 | 91.488126 | -1.259637222 | 0.48203381 | -2.6131719 | 0.00897062 | 0.2563089 |
| SETD2 | 3831.05771 | -0.357231522 | 0.19613951 | -1.8213134 | 0.06855923 | 0.49154315 |
| KIF9-AS1 | 1305.80586 | 0.420461539 | 0.34482273 | 1.21935563 | 0.22270924 | 0.66398799 |
| KIF9 | 196.97186 | -0.105505002 | 0.27444211 | -0.3844344 | 0.70065647 | 0.90766879 |
| KLHL18 | 395.2063 | -0.551067643 | 0.3827122 | -1.4399009 | 0.14989544 | 0.60758225 |
| AC099778.1 | 46.4971587 | -0.237175274 | 0.51829741 | -0.4576046 | 0.64723654 | 0.88774616 |
| PTPN23 | 305.098067 | -0.614483178 | 0.33628673 | -1.8272597 | 0.06766074 | 0.48963361 |
| SCAP | 732.931918 | -0.20318641 | 0.22688053 | -0.8955657 | 0.37048479 | 0.76698912 |
| ELP6 | 181.910099 | -0.322359719 | 0.37107692 | -0.868714 | 0.38500358 | 0.77317162 |
| SMARCC1 | 3524.72928 | -0.29872284 | 0.21596739 | -1.3831849 | 0.16660817 | 0.62216414 |
| DHX30 | 457.672287 | -0.293802983 | 0.25023833 | -1.1740927 | 0.24035791 | 0.67897455 |
| MAP4 | 4329.67832 | -0.450569448 | 0.33156628 | -1.3589122 | 0.17417442 | 0.62699479 |
| ZNF589 | 144.741079 | -0.188441331 | 0.31374041 | -0.6006282 | 0.54808767 | 0.84486687 |
| NME6 | 123.333518 | -0.477898263 | 0.46766066 | -1.0218911 | 0.30683245 | 0.72190689 |
| PLXNB1 | 803.362527 | -0.027712919 | 0.28336226 | -0.0978003 | 0.92209086 | 0.9796992 |
| CCDC51 | 55.405037 | -0.408042038 | 0.34379084 | -1.1868904 | 0.23527087 | 0.67142574 |

| | | | | | | |
|---|---|---|---|---|---|---|
| TMA7 | 421.76211 | -0.284835916 | 0.40126731 | -0.7098408 | 0.47780285 | 0.81580016 |
| ATRIP | 49.4554297 | 0.237331044 | 0.37724322 | 0.62911943 | 0.52927087 | 0.83782374 |
| SHISA5 | 589.264541 | 0.269950754 | 0.44341643 | 0.60879736 | 0.54265876 | 0.84314483 |
| PFKFB4 | 343.721902 | -0.891912528 | 0.86103898 | -1.0358562 | 3.00E-01 | 7.19E-01 |
| UQCRC1 | 430.859446 | 0.076989678 | 0.34302927 | 0.22444055 | 0.82241452 | 0.95113364 |
| SLC26A6 | 285.883429 | -0.148601317 | 0.68978539 | -0.2154312 | 0.82943112 | 0.95325673 |
| NCKIPSD | 165.865021 | -0.232534869 | 0.36172115 | -0.6428567 | 0.52031708 | 0.833897 |
| IP6K2 | 1126.7147 | -0.59623818 | 0.2539139 | -2.3481904 | 0.01886488 | 0.34084276 |
| PRKAR2A | 1309.6042 | -0.592909121 | 0.28692507 | -2.066425 | 0.03878837 | 0.4256783 |
| SLC25A20 | 387.681763 | 0.157865016 | 0.6136575 | 0.25725265 | 0.79698375 | 0.94280281 |
| ARIH2 | 1140.94957 | -0.325972236 | 0.26474665 | -1.2312611 | 0.21822524 | 0.66064833 |
| P4HTM | 243.404014 | 0.243715253 | 0.32647079 | 0.74651473 | 0.45535654 | 0.80465616 |
| WDR6 | 840.561395 | -0.608919434 | 0.37887061 | -1.6071963 | 0.10801132 | 0.55727735 |
| DALRD3 | 165.591133 | 0.211433433 | 0.42124607 | 0.50192382 | 0.61572111 | 0.87323788 |
| NDUFAF3 | 152.639414 | -0.094405324 | 0.5074331 | -0.1860449 | 0.85240958 | 0.96061646 |
| IMPDH2 | 117.113396 | -0.25623068 | 0.48797313 | -0.5250918 | 0.59951939 | 0.86692818 |
| QRICH1 | 778.819692 | -0.511085274 | 0.24585593 | -2.0787999 | 0.03763575 | 0.42215481 |
| QARS | 175.051867 | 0.102006688 | 0.3510195 | 0.2906012 | 0.77135634 | 0.93316533 |
| USP19 | 126.374835 | 0.109021645 | 0.33349213 | 0.3269092 | 0.74373657 | 0.92350771 |
| LAMB2 | 407.090426 | -0.196002925 | 0.40364377 | -0.4855839 | 0.6272622 | 0.87930896 |
| CCDC71 | 36.9871273 | -0.368787233 | 0.4234797 | -0.8708498 | 0.38383615 | 0.77317162 |
| KLHDC8B | 61.528979 | 0.437645356 | 0.54637991 | 0.80099094 | 0.42313689 | 0.78894887 |
| C3orf62 | 133.820785 | -0.278148953 | 0.44701999 | -0.6222293 | 0.53379107 | 0.83915086 |
| USP4 | 693.561532 | -0.25506576 | 0.31148347 | -0.8188742 | 0.41285822 | 0.78632759 |
| RHOA | 1390.08408 | -0.506668664 | 0.43497383 | -1.1648256 | 0.24408958 | 0.68222743 |
| TCTA | 152.412926 | 0.112831373 | 0.31774332 | 0.35510226 | 0.72251298 | 0.91592159 |
| DAG1 | 552.434131 | -0.420658068 | 0.40175098 | -1.0470617 | 0.29507112 | 0.71646902 |
| BSN | 55.2034659 | 0.24715275 | 0.4294537 | 0.57550499 | 0.56494982 | 0.85171209 |
| APEH | 242.439301 | 0.2679671 | 0.41944849 | 0.6388558 | 0.52291675 | 0.83509494 |
| MST1 | 1002.72867 | 0.187624909 | 0.55415866 | 0.33857616 | 0.73492905 | 0.91997009 |
| RNF123 | 257.804841 | 0.154296804 | 0.31617525 | 0.48801038 | 0.62554249 | 0.87854882 |
| GMPPB | 70.8485772 | 0.2443296 | 0.52657376 | 0.46399881 | 0.6426486 | 0.88441609 |
| IP6K1 | 1063.13619 | -0.636902527 | 0.3945393 | -1.6142943 | 0.10646361 | 0.55444404 |
| UBA7 | 45.4225422 | -0.741030061 | 1.10573544 | -0.6701694 | 0.50274981 | 0.82638105 |
| MON1A | 89.7476356 | -0.410215683 | 0.46808373 | -0.8763724 | 0.3808276 | 0.77210521 |
| RBM6 | 5059.70299 | -0.291264789 | 0.21834677 | -1.3339551 | 0.18221856 | 0.63216521 |
| RBM5 | 1293.41975 | -0.083646693 | 0.13228944 | -0.6323006 | 0.52719047 | 0.83733484 |
| SLC38A3 | 72.5307557 | 2.644856556 | 1.18967078 | 2.22318359 | 0.02620342 | 0.38392873 |
| GNAI2 | 419.611518 | 0.316079563 | 0.29134804 | 1.08488653 | 0.27797191 | 0.7036882 |
| SEMA3B | 757.334586 | 0.81866621 | 0.43833847 | 1.86765768 | 0.0618098 | 0.48267858 |
| IFRD2 | 244.93505 | -0.297676034 | 0.43078609 | -0.6910066 | 0.48956139 | 0.81970016 |
| HYAL1 | 222.029318 | 0.675596726 | 0.4723783 | 1.43020272 | 0.15265885 | 0.61001662 |
| HYAL2 | 97.8707309 | -0.143333065 | 0.42864055 | -0.3343899 | 0.73808537 | 0.92064211 |
| TUSC2 | 134.96183 | -0.199302522 | 0.43620752 | -0.4568984 | 0.64774407 | 0.88774616 |
| RASSF1 | 182.809782 | -0.926993646 | 0.44686511 | -2.0744373 | 0.03803872 | 0.42339465 |
| NPRL2 | 59.8128758 | -0.258507177 | 0.41476666 | -0.6232593 | 0.53311415 | 0.8387736 |
| TMEM115 | 110.184553 | -0.068531264 | 0.47213422 | -0.1451521 | 0.8845908 | 0.9692315 |
| HEMK1 | 203.525732 | 0.111568736 | 0.28150263 | 0.39633284 | 6.92E-01 | 0.90499641 |
| CISH | 283.649189 | -0.16611075 | 0.6040744 | -0.2749839 | 0.78332859 | 0.93839232 |
| MAPKAPK3 | 270.702399 | -0.425796921 | 0.44684851 | -0.9528888 | 0.34064644 | 0.74725037 |
| DOCK3 | 208.776905 | 0.007787258 | 0.47030886 | 0.01655775 | 0.98678943 | 0.9962029 |
| MANF | 259.204188 | 0.838345441 | 0.6833281 | 1.22685638 | 0.21987657 | 0.66238422 |
| RBM15B | 93.676652 | -0.692347879 | 0.48725915 | -1.4209028 | 0.15534503 | 0.6118547 |
| DCAF1 | 643.452098 | -0.361535779 | 0.29831531 | -1.211925 | 0.2255411 | 0.66688879 |

| | | | | | | |
|---|---|---|---|---|---|---|
| RAD54L2 | 661.072681 | -0.194781104 | 0.26222221 | -0.7428093 | 0.45759712 | 0.80561734 |
| TEX264 | 334.679741 | 0.091954055 | 0.36143685 | 0.25441251 | 0.79917689 | 0.94352434 |
| RRP9 | 55.1241532 | -0.701761024 | 0.51872589 | -1.3528552 | 0.17610189 | 0.62699479 |
| PARP3 | 101.154813 | 0.0824658 | 0.59259447 | 0.1391606 | 0.88932324 | 0.97208467 |
| PCBP4 | 68.1642553 | 0.296848072 | 0.49012275 | 0.60566067 | 0.5447401 | 0.84441376 |
| ABHD14B | 65.094088 | 0.296975218 | 0.37216464 | 0.79796732 | 0.42488946 | 0.78894887 |
| ABHD14A | 51.1448819 | 0.792528621 | 0.60202835 | 1.31643073 | 0.18802951 | 0.64002718 |
| ACY1 | 67.9693294 | 0.29489565 | 0.77635775 | 0.37984505 | 7.04E-01 | 9.09E-01 |
| RPL29 | 1849.69239 | 0.337055031 | 0.48569823 | 0.69395977 | 0.48770744 | 0.81893983 |
| POC1A | 59.0202787 | -0.362736736 | 0.49755095 | -0.7290444 | 0.4659745 | 0.81054727 |
| ALAS1 | 134.861656 | 0.018608784 | 0.31174687 | 0.05969197 | 0.95240097 | 0.98777411 |
| WDR82 | 596.107978 | -0.172709929 | 0.23125087 | -0.7468509 | 0.45515354 | 0.80465616 |
| GLYCTK | 210.463758 | 0.246698532 | 0.49098795 | 0.50245334 | 6.15E-01 | 0.87296463 |
| DNAH1 | 292.353543 | 0.438698621 | 0.38803054 | 1.13057756 | 0.25823294 | 0.69210282 |
| BAP1 | 161.492385 | -0.26533583 | 0.40013486 | -0.663116 | 0.50725626 | 0.82832518 |
| PHF7 | 154.049732 | 0.025842803 | 0.34828284 | 0.07420062 | 0.94085075 | 0.98485301 |
| NISCH | 436.287551 | 0.260976782 | 0.25698949 | 1.01551538 | 0.30986022 | 0.72498238 |
| NT5DC2 | 57.0027584 | 0.226770683 | 0.69571956 | 0.32595128 | 0.74446121 | 0.92363594 |
| SMIM4 | 519.448953 | -0.100244304 | 0.36254913 | -0.2764985 | 0.78216518 | 0.93757221 |
| PBRM1 | 1924.21637 | -0.224507923 | 0.24033129 | -0.9341602 | 0.35022127 | 0.75455562 |
| GNL3 | 340.650463 | -0.752221809 | 0.39410536 | -1.908682 | 0.05630312 | 0.47143978 |
| GLT8D1 | 169.666756 | 0.405851181 | 0.27060241 | 1.49980622 | 0.13366461 | 0.59240843 |
| SPCS1 | 417.474787 | 0.211482605 | 0.37792791 | 0.55958451 | 0.57576287 | 0.85537243 |
| NEK4 | 449.993388 | -0.212200352 | 0.22463948 | -0.9446263 | 0.34484971 | 0.75133476 |
| SFMBT1 | 593.344565 | -0.567635003 | 0.26005888 | -2.1827172 | 0.02905664 | 0.39554194 |
| AC096887.1 | 76.7928959 | 0.972437042 | 0.45086019 | 2.15684833 | 0.03101748 | 0.4060713 |
| RFT1 | 80.9341145 | 0.26424128 | 0.44683939 | 0.59135628 | 0.55428173 | 0.84727776 |
| PRKCD | 193.059858 | 0.515792844 | 0.32949938 | 1.56538334 | 0.11749305 | 0.57353179 |
| TKT | 1512.13218 | 0.453038834 | 0.69211199 | 0.65457447 | 0.51274177 | 0.83055441 |
| DCP1A | 1527.98685 | 0.157606567 | 0.1699921 | 0.92714052 | 0.35385358 | 0.75682564 |
| CACNA1D | 3954.09971 | 0.794186594 | 0.61102857 | 1.29975362 | 0.19368543 | 0.64548758 |
| CHDH | 2083.19206 | -0.119381285 | 0.44384967 | -0.2689678 | 0.78795443 | 0.93953874 |
| IL17RB | 3330.53206 | 0.721903526 | 0.70284493 | 1.02711636 | 0.30436567 | 0.72082426 |
| AC012467.2 | 47.7634969 | 0.639087886 | 0.52672817 | 1.21331633 | 0.2250089 | 0.66642114 |
| ACTR8 | 187.951049 | 0.014779559 | 0.25769115 | 0.05735377 | 0.95426339 | 0.98840182 |
| SELENOK | 352.838014 | 0.024658013 | 0.42016789 | 0.05868609 | 0.95320213 | 0.9880091 |
| CACNA2D3 | 1401.27656 | 0.015714196 | 0.41623198 | 0.03775346 | 9.70E-01 | 9.92E-01 |
| WNT5A | 42.9055637 | 2.175395343 | 0.78911037 | 2.75676944 | 0.00583755 | 0.22637955 |
| ERC2 | 747.332276 | 0.788357761 | 0.42092659 | 1.87291035 | 0.06108077 | 0.4813972 |
| CCDC66 | 1063.11847 | 0.150104641 | 0.1887155 | 0.79540177 | 0.42637983 | 0.78895108 |
| FAM208A | 1058.29304 | 0.190500265 | 0.18591805 | 1.02464642 | 0.30553005 | 0.72122 |
| ARHGEF3 | 4952.47195 | 0.427363105 | 0.66229589 | 0.64527519 | 0.51874887 | 0.83329602 |
| SPATA12 | 103.614888 | 0.439090657 | 0.52470258 | 0.83683724 | 0.40268406 | 0.78260941 |
| IL17RD | 429.692432 | 0.240872971 | 0.3876033 | 0.621442 | 0.53430884 | 0.83929051 |
| HESX1 | 42.2329296 | 0.316429772 | 0.43969573 | 0.71965623 | 0.47173668 | 0.81389222 |
| APPL1 | 670.872083 | -0.046454592 | 0.17767668 | -0.2614558 | 0.79374106 | 0.94205634 |
| ASB14 | 126.078071 | 0.386132728 | 0.52245808 | 0.7390693 | 0.45986492 | 0.8069452 |
| DNAH12 | 146.079551 | 0.946177323 | 0.65947664 | 1.43473972 | 0.15136128 | 0.60905477 |
| PDE12 | 184.974179 | -0.235527002 | 0.31982754 | -0.7364188 | 0.46147589 | 0.80795122 |
| ARF4 | 870.582855 | -1.293926162 | 0.52876456 | -2.4470743 | 0.01440212 | 0.30672862 |
| ARF4-AS1 | 36.6221053 | -0.321680838 | 0.48170669 | -0.667794 | 0.5042651 | 0.82715271 |
| DENND6A | 767.706324 | -0.132425609 | 0.22242338 | -0.5953763 | 0.55159197 | 0.84694423 |
| DENND6A-AS | 139.450367 | 0.159637212 | 0.56242344 | 0.28383812 | 0.77653444 | 0.9353214 |
| SLMAP | 3200.40457 | -0.402799923 | 0.28684471 | -1.4042438 | 0.16024625 | 0.61590405 |

| | | | | | | |
|---|---|---|---|---|---|---|
| FLNB | 2126.4039 | -0.425010905 | 0.28912413 | -1.4699946 | 0.14156323 | 0.59611016 |
| ABHD6 | 1312.03066 | 0.208383919 | 0.41764352 | 0.49895164 | 6.18E-01 | 8.74E-01 |
| RPP14 | 125.038086 | 0.301845786 | 0.36888904 | 0.81825632 | 0.41321084 | 0.78632759 |
| PXK | 435.389248 | -0.002546293 | 0.45273714 | -0.0056242 | 0.99551255 | 0.99866448 |
| PDHB | 190.994817 | 0.146613175 | 0.27756253 | 0.52821674 | 0.59734891 | 0.86614567 |
| KCTD6 | 326.399333 | 0.441255925 | 0.45485398 | 0.97010457 | 0.33199437 | 0.73947712 |
| ACOX2 | 390.424533 | 1.142326145 | 0.5885443 | 1.94093485 | 5.23E-02 | 0.46474067 |
| FAM107A | 77.7118154 | 1.89763256 | 0.83318653 | 2.2775603 | 0.02275279 | 0.36561837 |
| FAM3D-AS1 | 83.2038549 | 1.663125747 | 1.01929086 | 1.63164982 | 0.10275328 | 0.55022042 |
| C3orf67 | 151.789729 | 0.557699766 | 0.59596289 | 0.93579613 | 0.34937816 | 0.7542101 |
| C3orf67-AS1 | 42.8486715 | 0.422026604 | 0.60726307 | 0.69496504 | 0.48707721 | 0.81893983 |
| FHIT | 25978.1286 | 0.682593249 | 0.41892322 | 1.62939941 | 0.10322849 | 0.55022042 |
| PTPRG | 15441.0433 | 0.407830145 | 0.21626625 | 1.885778 | 0.05932488 | 0.47855094 |
| C3orf14 | 39.9173877 | -0.052993097 | 0.4604996 | -0.1150774 | 0.90838377 | 0.97679582 |
| THOC7 | 344.006096 | 0.077235187 | 0.16132089 | 0.47876742 | 0.6321041 | 0.88123095 |
| ATXN7 | 322.111928 | -0.081013583 | 0.21924357 | -0.3695141 | 0.7117446 | 0.91159439 |
| PSMD6-AS2 | 147.35834 | 0.248473548 | 0.33290938 | 0.74636993 | 0.45544398 | 0.80465616 |
| PSMD6 | 195.004522 | -0.26495663 | 0.29117936 | -0.909943 | 0.36285256 | 0.76200705 |
| PSMD6-AS1 | 49.3559232 | 0.360983911 | 0.33371293 | 1.08171989 | 0.27937701 | 0.70446562 |
| PRICKLE2 | 2985.29559 | -0.109594105 | 0.34836731 | -0.3145935 | 0.75307028 | 0.92751122 |
| ADAMTS9 | 4568.58979 | 0.466112576 | 0.29372906 | 1.58687936 | 0.11253997 | 0.56608637 |
| ADAMTS9-A! | 10338.4623 | 0.423465893 | 0.82516325 | 0.51319044 | 0.60781811 | 0.86985416 |
| ADAMTS9-A! | 5194.17393 | 0.400581441 | 0.58846305 | 0.68072488 | 0.49604559 | 0.82243887 |
| MAGI1 | 11315.2007 | -0.101799609 | 0.28343041 | -0.3591697 | 0.71946816 | 0.91465123 |
| MAGI1-AS1 | 140.931894 | 0.030733683 | 0.74613692 | 0.04119041 | 0.9671441 | 0.99157802 |
| SLC25A26 | 1214.78975 | -0.884171014 | 0.22863618 | -3.8671527 | 0.00011011 | 0.03354057 |
| LRIG1 | 549.235095 | 0.251372329 | 0.36731481 | 0.68435119 | 0.49375343 | 0.82164359 |
| SUCLG2 | 3487.47785 | -0.024007779 | 0.28108811 | -0.0854102 | 0.93193532 | 0.9829207 |
| SUCLG2-AS1 | 1961.42348 | 0.748164491 | 0.37963705 | 1.97073623 | 0.04875405 | 0.4536808 |
| FAM19A1 | 603.523886 | 0.52981976 | 0.60864046 | 0.87049711 | 0.3840288 | 0.77317162 |
| EOGT | 170.195378 | -0.373106084 | 0.47702734 | -0.7821482 | 0.43412747 | 0.79442004 |
| AC109587.1 | 85.0500474 | 0.50271271 | 0.63849663 | 0.78733808 | 0.43108398 | 0.79133095 |
| TMF1 | 835.666011 | -0.078552011 | 0.14981635 | -0.524322 | 0.60005459 | 0.86733611 |
| UBA3 | 706.105457 | 0.135054193 | 0.1630887 | 0.82810272 | 0.40761233 | 0.78481094 |
| ARL6IP5 | 720.118238 | 0.21876666 | 0.2690028 | 0.81325049 | 0.41607446 | 0.78632759 |
| FRMD4B | 6125.94378 | 1.613711274 | 0.46747222 | 3.45199395 | 0.00055646 | 0.08114255 |
| MITF | 3186.7022 | -0.944257497 | 0.51545727 | -1.8318832 | 0.06696883 | 0.48956893 |
| FOXP1 | 3926.83379 | 0.358238378 | 0.39061634 | 0.91711058 | 0.3590847 | 0.75936492 |
| EIF4E3 | 329.2626 | 0.359807936 | 0.38234695 | 0.94105089 | 0.34667878 | 0.75166079 |
| LINC00870 | 51.3347161 | 0.448087107 | 0.94433768 | 0.47449881 | 0.63514424 | 0.88238823 |
| RYBP | 1067.61371 | -0.478389273 | 0.28512283 | -1.6778357 | 0.09337917 | 0.53688613 |
| SHQ1 | 451.38981 | -0.287248351 | 0.32354538 | -0.8878147 | 0.37464042 | 0.7697436 |
| GXYLT2 | 461.355972 | 0.220065753 | 0.51285659 | 0.42909803 | 6.68E-01 | 0.89501817 |
| PPP4R2 | 1148.86267 | -0.02862378 | 0.28636701 | -0.0999549 | 0.92038015 | 0.97939204 |
| PDZRN3 | 3724.60821 | -0.212100017 | 0.56362294 | -0.3763154 | 0.7066824 | 0.91002281 |
| CNTN3 | 1478.95679 | -0.184169616 | 0.24548119 | -0.7502392 | 4.53E-01 | 8.04E-01 |
| LINC02018 | 99.7767616 | -0.081103164 | 0.65987213 | -0.1229074 | 0.90218043 | 0.9750023 |
| ZNF717 | 494.343503 | 0.44765843 | 0.35842252 | 1.24896849 | 0.2116766 | 0.65584609 |
| ROBO2 | 518.409634 | 0.616578657 | 0.68478231 | 0.90040097 | 0.36790691 | 0.76610076 |
| ROBO1 | 120.043173 | 0.393106094 | 0.52308737 | 0.75151135 | 0.45234498 | 0.80432522 |
| LINC02027 | 4256.85471 | -0.008347088 | 0.73381654 | -0.0113749 | 0.99092434 | 0.99737132 |
| GBE1 | 6323.18678 | -0.850089698 | 0.51786037 | -1.6415423 | 1.01E-01 | 0.54608983 |
| CADM2 | 69.4631503 | 0.248929652 | 0.44919496 | 0.5541684 | 0.5794636 | 0.85816191 |
| CHMP2B | 338.173968 | -0.483991148 | 0.30162803 | -1.604596 | 0.10858273 | 0.55858026 |

| | | | | | | |
|---|---|---|---|---|---|---|
| CGGBP1 | 732.592584 | -0.174330221 | 0.32919693 | -0.5295621 | 0.59641558 | 0.86559503 |
| ZNF654 | 1116.95149 | -0.016296042 | 0.29311904 | -0.0555953 | 0.95566421 | 0.98840182 |
| C3orf38 | 192.429806 | 0.257245433 | 0.21580791 | 1.19201112 | 0.23325688 | 0.66978724 |
| PROS1 | 1494.76901 | 0.540931941 | 0.26304779 | 2.05640177 | 0.03974381 | 0.42869465 |
| ARL13B | 568.696642 | 0.24556591 | 0.30709117 | 0.79965147 | 0.42391276 | 0.78894887 |
| DHFR2 | 61.6350811 | 0.371099476 | 0.42940408 | 0.86421973 | 0.3874672 | 0.77363625 |
| NSUN3 | 441.174161 | -0.114657489 | 0.24256741 | -0.472683 | 0.63643936 | 0.88266168 |
| MTRNR2L12 | 1219.43139 | -1.180429898 | 0.58088827 | -2.0321118 | 0.04214234 | 0.43742111 |
| EPHA6 | 389.571594 | -0.761191453 | 0.9005959 | -0.8452087 | 0.39799435 | 0.78100337 |
| ARL6 | 291.588471 | -0.165659196 | 0.33611913 | -0.4928586 | 0.62211252 | 0.87713418 |
| CRYBG3 | 604.44451 | -0.572106877 | 0.27895227 | -2.0509132 | 0.0402754 | 0.43086056 |
| RIOX2 | 233.844595 | -0.016492826 | 0.29003098 | -0.0568657 | 9.55E-01 | 9.88E-01 |
| CLDND1 | 215.543066 | -0.043179659 | 0.23084721 | -0.1870487 | 0.85162249 | 0.96044014 |
| CPOX | 126.207016 | 0.043741534 | 0.33391634 | 0.13099549 | 0.89577888 | 0.97309833 |
| ST3GAL6-AS1 | 122.527983 | -0.472834201 | 0.5922814 | -0.798327 | 0.42468078 | 0.78894887 |
| ST3GAL6 | 1357.67916 | -0.197356989 | 0.42317287 | -0.4663744 | 0.64094754 | 0.88396933 |
| DCBLD2 | 814.45699 | -0.618041082 | 0.72901613 | -0.8477742 | 0.39656372 | 0.77992032 |
| COL8A1 | 108.865298 | 0.433637468 | 0.55416573 | 0.78250502 | 0.43391783 | 0.79429104 |
| CMSS1 | 1174.2788 | -0.819020355 | 0.40745449 | -2.0100904 | 0.04442162 | 0.44360789 |
| FILIP1L | 262.577827 | -0.132533207 | 0.39191148 | -0.3381713 | 0.73523412 | 0.92009988 |
| TBC1D23 | 904.054535 | -0.390260246 | 0.22185974 | -1.7590404 | 0.07857064 | 0.51192014 |
| NIT2 | 756.691029 | 0.49718541 | 0.41208974 | 1.20649792 | 0.22762556 | 0.66705629 |
| TOMM70 | 800.803938 | 0.397863798 | 0.20828987 | 1.91014476 | 0.05611458 | 0.47126031 |
| LNP1 | 120.038859 | 1.740604269 | 0.52403613 | 3.32153485 | 0.00089524 | 0.10027947 |
| TFG | 1384.0571 | -0.137569989 | 0.21124787 | -0.6512255 | 0.51490096 | 0.83169608 |
| ABI3BP | 767.98569 | 1.436315919 | 0.77328209 | 1.85742816 | 0.06325026 | 0.48572144 |
| SENP7 | 1020.99425 | 0.447832932 | 0.29102058 | 1.53883593 | 0.12384435 | 0.58354574 |
| TRMT10C | 203.666823 | -0.393467042 | 0.31983355 | -1.2302244 | 0.21861307 | 0.66098715 |
| PCNP | 837.008216 | -0.267395337 | 0.1418861 | -1.8845774 | 0.05948691 | 0.47879455 |
| ZBTB11 | 534.802201 | -0.572369759 | 0.37125121 | -1.5417317 | 0.1231388 | 0.5825794 |
| ZBTB11-AS1 | 35.0926165 | -0.365092024 | 0.54634183 | -0.6682483 | 0.50397508 | 0.82715271 |
| RPL24 | 705.874854 | -0.167675236 | 0.42428616 | -0.3951937 | 0.69269991 | 0.90528251 |
| CEP97 | 177.602342 | -0.044799108 | 0.27643004 | -0.1620631 | 0.87125617 | 0.9657206 |
| NFKBIZ | 1492.52806 | 0.592866611 | 0.65332851 | 0.9074556 | 0.36416592 | 0.76273126 |
| ALCAM | 2592.91093 | 0.528005246 | 0.36530017 | 1.44540106 | 0.14834527 | 0.60622442 |
| CBLB | 1894.61294 | -0.335790893 | 0.33184286 | -1.0118973 | 0.31158715 | 0.72600794 |
| LINC00882 | 183.682694 | 1.173916948 | 0.56715951 | 2.06981796 | 0.0384694 | 0.42447711 |
| DUBR | 79.8032053 | 0.36214024 | 0.34501511 | 1.04963589 | 2.94E-01 | 7.16E-01 |
| AC063944.1 | 109.887832 | 0.997871581 | 0.55987615 | 1.78230773 | 0.07469906 | 0.5071704 |
| BBX | 4205.00792 | 0.307474867 | 0.40878478 | 0.75216809 | 0.45194998 | 0.80411194 |
| CD47 | 318.170146 | 0.1746343 | 0.41034338 | 0.42558088 | 0.67041327 | 0.89660989 |
| IFT57 | 541.849862 | 0.431034484 | 0.44245214 | 0.9741946 | 0.32995993 | 0.73849818 |
| HHLA2 | 704.862001 | 0.184894838 | 0.63393143 | 0.29166378 | 0.7705437 | 0.9329619 |
| CIP2A | 86.3230475 | -0.672190249 | 0.60286836 | -1.1149868 | 0.26485609 | 0.69614236 |
| DZIP3 | 787.301517 | 1.097899749 | 0.59818293 | 1.83539131 | 0.06644773 | 0.48798003 |
| C3orf85 | 242.507071 | -0.25005328 | 0.87759252 | -0.284931 | 0.77569704 | 0.9351345 |
| NECTIN3-AS1 | 72.0359152 | -0.127768407 | 0.61356176 | -0.2082405 | 0.83504119 | 0.9543899 |
| NECTIN3 | 1289.59537 | 0.338795406 | 0.12238385 | 2.76830163 | 0.00563493 | 0.22463891 |
| PLCXD2 | 112.364947 | 0.328722135 | 0.47936298 | 0.68574786 | 0.49287212 | 0.82100434 |
| PHLDB2 | 4911.75405 | 0.698932689 | 0.31841178 | 2.19505914 | 0.02815936 | 0.39225639 |
| ABHD10 | 263.768374 | 0.381961493 | 0.44161465 | 0.86492034 | 0.38708251 | 0.77335375 |
| C3orf52 | 356.103511 | -0.035349196 | 0.67066405 | -0.0527078 | 0.95796476 | 0.98876679 |
| AC128688.2 | 58.1006294 | 0.000376434 | 0.67906344 | 0.00055434 | 0.9995577 | 0.99977652 |
| SLC9C1 | 502.774179 | 1.212840829 | 0.54332032 | 2.23227584 | 0.02559674 | 0.38166806 |

| | | | | | | |
|---|---|---|---|---|---|---|
| AC112487.1 | 66.7213287 | 1.975400054 | 0.70205425 | 2.81374275 | 0.00489684 | 0.21582309 |
| CD200 | 166.115176 | 1.380398518 | 0.51518334 | 2.67943159 | 0.00737473 | 0.23590813 |
| AC092894.1 | 443.876643 | 0.179905462 | 0.47531466 | 0.37849761 | 0.70506097 | 0.909554 |
| ATG3 | 302.591643 | -0.21100213 | 0.19724784 | -1.069731 | 0.2847404 | 0.70962486 |
| SLC35A5 | 189.847825 | -0.073392896 | 0.3544819 | -0.2070427 | 0.8359765 | 0.95468028 |
| GTPBP8 | 149.265065 | -0.178790045 | 0.36482988 | -0.4900641 | 0.62408851 | 0.87791268 |
| NEPRO | 345.201409 | 0.026278072 | 0.17595203 | 0.14934793 | 0.8812791 | 0.96774334 |
| AC078785.1 | 87.4438608 | 0.768679626 | 0.52767847 | 1.45671969 | 0.14519377 | 0.60162365 |
| CFAP44 | 639.770851 | 0.320118959 | 0.239736 | 1.33529782 | 0.18177889 | 0.6318513 |
| SPICE1 | 573.357492 | -0.37754734 | 0.17707785 | -2.132098 | 0.03299879 | 0.41231942 |
| USF3 | 619.59658 | 0.136165622 | 0.21205864 | 0.64211307 | 0.52079977 | 0.83404632 |
| NAA50 | 1115.77926 | -0.394378458 | 0.3034093 | -1.2998232 | 0.19366156 | 0.64548758 |
| ATP6V1A | 1643.81248 | -0.044361209 | 0.27507241 | -0.161271 | 0.87187994 | 0.9657206 |
| GRAMD1C | 1142.13624 | 0.63867851 | 0.55895814 | 1.14262315 | 0.25319509 | 0.68887358 |
| ZDHHC23 | 178.759353 | 0.686798571 | 0.48833581 | 1.40640634 | 0.1596035 | 0.61520955 |
| CCDC191 | 564.373575 | 0.392319931 | 0.5680044 | 0.69069875 | 0.48975488 | 0.81970016 |
| QTRT2 | 275.369208 | -0.217383945 | 0.24178184 | -0.8990913 | 0.36860404 | 0.76615653 |
| ZBTB20 | 40075.2076 | 0.196102897 | 0.36061053 | 0.54380802 | 0.58657359 | 0.86102615 |
| ZBTB20-AS1 | 163.446153 | 0.266566095 | 0.81663923 | 0.32641843 | 0.7441078 | 0.92361546 |
| ZBTB20-AS5 | 691.022884 | 0.347136222 | 0.73838601 | 0.47012839 | 0.63826329 | 0.8827026 |
| ZBTB20-AS2 | 199.974058 | -0.205692774 | 0.82719161 | -0.248664 | 0.8036207 | 0.94543168 |
| LSAMP | 737.500445 | 1.142041356 | 0.6481681 | 1.76195241 | 0.07807734 | 0.51192014 |
| IGSF11 | 1148.16777 | 0.679835496 | 0.53222757 | 1.27733987 | 2.01E-01 | 0.6487474 |
| UPK1B | 42.8427545 | 0.71229273 | 0.67036277 | 1.06254817 | 0.2879869 | 0.71170639 |
| B4GALT4 | 643.275353 | 0.621821869 | 0.34596744 | 1.79734218 | 0.07228131 | 0.50215913 |
| B4GALT4-AS | 98.6673133 | 0.46677688 | 0.71296481 | 0.65469835 | 0.51266199 | 0.83055441 |
| ARHGAP31 | 836.980967 | 0.976782594 | 0.37122658 | 2.63123022 | 0.00850764 | 0.25070735 |
| TMEM39A | 363.635562 | -0.067816762 | 0.27550168 | -0.2461573 | 0.80556044 | 0.94585363 |
| POGLUT1 | 198.339962 | -0.079232645 | 0.35733188 | -0.2217341 | 0.82452092 | 0.95180295 |
| TIMMDC1 | 519.881699 | 0.092376337 | 0.16988155 | 0.54376911 | 0.58660037 | 0.86102615 |
| PLA1A | 202.990557 | -0.757283465 | 0.71380532 | -1.0609104 | 0.28873064 | 0.71218341 |
| COX17 | 248.286113 | -0.710048835 | 0.41960835 | -1.6921704 | 0.09061349 | 0.53490056 |
| MAATS1 | 400.391972 | 1.530423535 | 0.49221275 | 3.10927245 | 0.00187549 | 0.13738894 |
| GSK3B | 3599.54437 | 0.288075556 | 0.29310389 | 0.98284452 | 0.32568396 | 0.7372514 |
| AC092910.3 | 140.921485 | 0.631700977 | 0.30270822 | 2.08683125 | 0.03690339 | 0.42215481 |
| GPR156 | 64.839182 | -2.363486179 | 0.82612017 | -2.8609472 | 0.00422377 | 0.20676883 |
| LRRC58 | 582.678711 | -0.693231472 | 0.33143139 | -2.0916289 | 0.03647172 | 0.42151595 |
| FSTL1 | 128.892287 | 0.226801765 | 0.46477922 | 0.48797742 | 0.62556583 | 0.87854882 |
| NDUFB4 | 458.921051 | -0.072616458 | 0.49053864 | -0.1480341 | 0.88231584 | 0.96797688 |
| HGD | 2085.67872 | 0.771158676 | 0.45677377 | 1.68827268 | 0.0913589 | 0.53537574 |
| RABL3 | 335.862036 | -0.172461196 | 0.25886555 | -0.6662192 | 0.50527101 | 0.82715271 |
| GTF2E1 | 152.638532 | 0.2729171 | 0.26276133 | 1.03865016 | 0.29896747 | 0.71852421 |
| STXBP5L | 37.0966587 | -0.160660386 | 0.37996189 | -0.4228329 | 0.67241719 | 0.89727632 |
| GOLGB1 | 3545.04499 | -0.899942716 | 0.35549355 | -2.5315304 | 0.0113566 | 0.28510052 |
| IQCB1 | 1164.2201 | -0.109439492 | 0.19165777 | -0.5710152 | 0.56798936 | 0.85263558 |
| EAF2 | 870.972654 | -0.452836905 | 0.59284598 | -0.7638357 | 4.45E-01 | 8.00E-01 |
| SLC15A2 | 697.664111 | -0.232676494 | 0.69931098 | -0.3327225 | 7.39E-01 | 9.21E-01 |
| ILDR1 | 110.285709 | 1.398616964 | 0.51315465 | 2.72552724 | 0.00641989 | 0.22797255 |
| CASR | 195.122331 | -0.015961778 | 0.64710318 | -0.0246665 | 0.98032097 | 0.99494599 |
| CCDC58 | 388.26603 | -0.54461267 | 0.42350402 | -1.2859681 | 0.19845419 | 0.64582421 |
| FAM162A | 929.990988 | -1.051944631 | 0.52388097 | -2.0079841 | 0.04464499 | 0.44360789 |
| WDR5B | 40.4259677 | -0.305300073 | 0.38414718 | -0.7947477 | 0.4267603 | 0.78913582 |
| AC083798.2 | 260.354454 | 0.21236737 | 0.30795885 | 0.68959658 | 0.49044792 | 0.81989696 |
| KPNA1 | 1398.81799 | -0.317922872 | 0.30294801 | -1.0494305 | 0.29398005 | 0.7156393 |

| | | | | | | |
|---|---|---|---|---|---|---|
| PARP9 | 705.074993 | -0.429229243 | 0.67294064 | -0.6378412 | 0.52357708 | 0.83536399 |
| DTX3L | 285.581061 | -0.338500134 | 0.56198459 | -0.6023299 | 0.54695455 | 0.84460887 |
| PARP14 | 1892.83564 | -1.142892968 | 0.79865629 | -1.4310198 | 0.15242454 | 0.61001662 |
| HSPBAP1 | 527.10744 | -0.145939408 | 0.27081296 | -0.5388937 | 0.58996018 | 0.86292147 |
| DIRC2 | 641.92865 | -0.26782214 | 0.29815617 | -0.8982613 | 0.36904627 | 0.76615653 |
| LINC02035 | 52.9169483 | -0.019436212 | 0.43035769 | -0.0451629 | 0.96397745 | 0.99012655 |
| PDIA5 | 788.865703 | 0.09373814 | 0.36790028 | 0.25479225 | 0.79888356 | 0.94334054 |
| SEC22A | 859.145127 | -0.043463481 | 0.23327136 | -0.1863216 | 0.85219261 | 0.96060889 |
| ADCY5 | 261.01624 | 0.669195454 | 0.76864551 | 0.87061649 | 0.38396359 | 0.77317162 |
| HACD2 | 542.409653 | 0.05964498 | 0.22320047 | 0.26722606 | 0.78929512 | 0.94005579 |
| MYLK-AS1 | 152.607775 | 0.548120391 | 0.23857422 | 2.29748374 | 0.02159119 | 0.35786026 |
| MYLK | 954.221282 | 0.281225942 | 0.44251088 | 0.63552322 | 0.52508723 | 0.835938 |
| CCDC14 | 1103.99626 | 0.092762477 | 0.29809645 | 0.31118276 | 0.75566169 | 0.92799282 |
| KALRN | 784.014221 | 0.433289999 | 0.59793695 | 0.72464162 | 0.46867192 | 0.81217042 |
| UMPS | 251.261582 | -0.143693427 | 0.26268226 | -0.5470237 | 0.58436243 | 0.86025731 |
| ITGB5 | 1068.62578 | -0.334245827 | 0.34407496 | -0.9714332 | 0.33133262 | 0.73916381 |
| ITGB5-AS1 | 49.3245525 | 0.075857454 | 0.81540786 | 0.09303007 | 0.92587967 | 0.98098729 |
| MUC13 | 70.4311024 | 0.756997408 | 0.96409095 | 0.78519294 | 0.43234045 | 0.79246999 |
| HEG1 | 176.877282 | -0.464762163 | 0.72032025 | -0.645216 | 0.51878721 | 0.83329602 |
| SLC12A8 | 411.03809 | 0.107159141 | 0.53634574 | 0.1997949 | 8.42E-01 | 0.95681213 |
| ZNF148 | 3493.90973 | -0.10218783 | 0.16509232 | -0.6189739 | 0.53593357 | 0.83954482 |
| SNX4 | 1582.2001 | -0.082472937 | 0.21417174 | -0.3850785 | 0.70017924 | 0.90737986 |
| OSBPL11 | 490.694987 | 0.310508734 | 0.23244689 | 1.33582655 | 0.18160597 | 0.6318513 |
| AC092902.3 | 65.7683168 | 1.228861723 | 0.53417739 | 2.300475 | 0.02142132 | 0.35743746 |
| AC092903.2 | 53.9398298 | 0.315425602 | 0.53297934 | 0.59181581 | 0.55397394 | 0.84727776 |
| SLC41A3 | 161.402671 | -0.084388182 | 0.29701557 | -0.2841204 | 0.77631811 | 0.9352344 |
| ALDH1L1 | 2370.5691 | 0.709902886 | 0.56498451 | 1.25649974 | 0.20893484 | 0.65401126 |
| ALDH1L1-AS2 | 77.4907328 | 0.806416938 | 0.58602717 | 1.37607433 | 0.16879862 | 0.62304975 |
| AC016924.1 | 67.6337056 | 0.808543128 | 1.00840796 | 0.80180162 | 0.42266772 | 0.78894887 |
| AC063919.1 | 70.4327484 | 1.329116644 | 1.11342401 | 1.19372012 | 0.23258747 | 0.66978724 |
| KLF15 | 306.996006 | 0.52771093 | 0.61815072 | 0.85369298 | 0.39327512 | 0.77753096 |
| ZXDC | 1565.02028 | -0.201510915 | 0.2600249 | -0.7749677 | 0.43835872 | 0.79699999 |
| CHST13 | 55.8357082 | -0.094080675 | 0.4577788 | -0.2055156 | 0.83716933 | 0.95496955 |
| TXNRD3NB | 148.30025 | -0.127201255 | 0.52380678 | -0.24284 | 0.80812932 | 0.94617795 |
| TXNRD3 | 204.203629 | 0.296956776 | 0.33055576 | 0.89835608 | 0.36899574 | 0.76615653 |
| CHCHD6 | 334.548871 | -0.482365929 | 0.35272038 | -1.367559 | 0.17145016 | 0.62520756 |
| PLXNA1 | 131.084622 | -0.714115956 | 0.46946755 | -1.5211189 | 0.12823 | 0.58793678 |
| TPRA1 | 160.668179 | -0.527215866 | 0.40641934 | -1.2972214 | 0.19455502 | 0.64548758 |
| ABTB1 | 109.42903 | -0.33448901 | 0.45047146 | -0.742531 | 0.45776569 | 0.80567467 |
| MGLL | 1960.97662 | -1.324051839 | 0.57899502 | -2.2868104 | 0.02220689 | 0.36178312 |
| SEC61A1 | 813.538222 | -0.631892128 | 0.39395979 | -1.6039508 | 0.10872489 | 0.55858026 |
| RUVBL1 | 594.518195 | 0.243606782 | 0.29467466 | 0.82669742 | 0.40840858 | 0.78481094 |
| EEFSEC | 1294.92874 | 0.267128515 | 0.3272292 | 0.81633459 | 0.41430879 | 0.78632759 |
| RPN1 | 512.372765 | -0.298459942 | 0.36744911 | -0.8122484 | 0.41664913 | 0.78632759 |
| RAB7A | 3089.92216 | -0.906589987 | 0.36217253 | -2.5031992 | 0.01230762 | 0.29629837 |
| ACAD9 | 385.311614 | -0.444985709 | 0.25091606 | -1.7734445 | 0.07615508 | 0.50767193 |
| KIAA1257 | 102.69677 | -0.609628535 | 0.50484757 | -1.2075497 | 0.22722051 | 0.66705629 |
| CNBP | 866.568221 | -0.033291217 | 0.52429801 | -0.0634967 | 0.94937095 | 0.9871942 |
| COPG1 | 376.613633 | -0.482868186 | 0.33651109 | -1.434925 | 0.15130846 | 0.60905477 |
| HMCES | 163.0282 | 0.467825525 | 0.2297876 | 2.03590416 | 0.04175998 | 0.43528825 |
| H1FX | 168.869783 | -0.586072537 | 0.550345 | -1.0649184 | 0.28691284 | 0.71118494 |
| EFCAB12 | 60.0942113 | -0.243235003 | 0.64916351 | -0.3746899 | 0.70789111 | 0.91025029 |
| MBD4 | 722.552929 | 0.025339079 | 0.34647862 | 0.07313317 | 0.94170015 | 0.98489601 |
| IFT122 | 605.754012 | 0.458309348 | 0.33622046 | 1.36312152 | 0.17284422 | 0.62593809 |

| | | | | | | |
|---|---|---|---|---|---|---|
| TMCC1 | 3236.94723 | 0.101878934 | 0.23302849 | 0.4371952 | 0.66196979 | 0.89193781 |
| TMCC1-AS1 | 58.7994658 | 0.131226436 | 0.43382724 | 0.30248547 | 0.76228201 | 0.93022076 |
| ALG1L2 | 44.8234945 | -0.232944794 | 0.63801708 | -0.3651075 | 0.71503119 | 0.91279611 |
| LINC02021 | 64.6909068 | 0.300441894 | 0.70243979 | 0.42771195 | 0.66886085 | 0.89566976 |
| PIK3R4 | 406.626166 | -0.207158474 | 0.17432729 | -1.1883307 | 0.23470315 | 0.67119484 |
| ATP2C1 | 834.274255 | -0.148660253 | 0.25349037 | -0.5864532 | 0.55757097 | 0.84861484 |
| AC055733.2 | 37.3991792 | 0.737863889 | 0.79788201 | 0.9247782 | 0.35508129 | 0.75757147 |
| ASTE1 | 64.1175408 | 0.165400052 | 0.44746803 | 0.36963546 | 0.71165413 | 0.91159439 |
| NEK11 | 1190.01576 | 0.002872826 | 0.35111343 | 0.00818205 | 0.99347174 | 0.9976934 |
| NUDT16 | 213.405388 | 0.374260128 | 0.34891156 | 1.07265041 | 0.283428 | 0.70780772 |
| MRPL3 | 498.769929 | -0.165740818 | 0.31278185 | -0.5298927 | 0.59618634 | 0.86539512 |
| CPNE4 | 500.929668 | 1.717925268 | 0.61837508 | 2.77812823 | 0.0054673 | 0.22038648 |
| DNAJC13 | 1497.66691 | -0.270667246 | 0.23903442 | -1.1323358 | 0.25749327 | 0.691812 |
| NPHP3-ACAD | 83.4506954 | 0.431429259 | 0.56776094 | 0.75987837 | 0.44732729 | 0.80223612 |
| ACKR4 | 148.900244 | 1.230044046 | 1.14834018 | 1.07114954 | 0.2841022 | 0.70897061 |
| UBA5 | 754.957454 | -0.18264651 | 0.62206801 | -0.2936118 | 0.76905456 | 0.93209094 |
| CDV3 | 618.043317 | -0.962564099 | 0.48031113 | -2.0040429 | 0.04506547 | 0.44360789 |
| TOPBP1 | 348.234702 | -0.421381296 | 0.31099964 | -1.3549254 | 0.17544134 | 0.62699479 |
| TF | 61.2530108 | -0.417223408 | 0.5070862 | -0.822786 | 0.41062972 | 0.78592213 |
| SRPRB | 250.898045 | -0.42440788 | 0.46535235 | -0.9120141 | 0.36176132 | 0.76060307 |
| RAB6B | 78.024151 | 0.255037982 | 0.74046476 | 0.3444296 | 7.31E-01 | 0.91890263 |
| SLCO2A1 | 297.576546 | 1.703346847 | 0.77077703 | 2.20990867 | 0.0271115 | 0.38702446 |
| RYK | 1667.27542 | -0.175935216 | 0.23990334 | -0.7333588 | 0.46333965 | 0.8082072 |
| AMOTL2 | 324.970489 | -1.660274988 | 0.55190649 | -3.0082541 | 0.00262753 | 0.16823326 |
| ANAPC13 | 126.900926 | -0.332621387 | 0.32152947 | -1.0344973 | 0.30090372 | 0.71879123 |
| CEP63 | 561.170713 | -0.066581853 | 0.19444842 | -0.342414 | 0.73203938 | 0.91942342 |
| EPHB1 | 168.910916 | 0.096554419 | 0.49362601 | 0.19560237 | 0.8449214 | 0.95760132 |
| PPP2R3A | 11531.2546 | 0.297903829 | 0.22378024 | 1.33123387 | 1.83E-01 | 6.33E-01 |
| MSL2 | 751.935055 | -0.544815252 | 0.41834239 | -1.302319 | 0.19280735 | 0.64518499 |
| PCCB | 359.612774 | 0.247965477 | 0.2222617 | 1.11564644 | 0.2645735 | 0.69567438 |
| STAG1 | 4778.29124 | -0.997640608 | 0.40375156 | -2.4709269 | 0.01347633 | 0.29889988 |
| SLC35G2 | 136.148738 | 0.390564524 | 0.3804298 | 1.02664019 | 3.05E-01 | 7.21E-01 |
| NCK1-DT | 89.7057329 | 0.224483275 | 0.32387484 | 0.69311736 | 0.4882359 | 0.81931938 |
| NCK1 | 838.469512 | -0.042248127 | 0.22313427 | -0.1893395 | 0.84982675 | 0.95959925 |
| IL20RB | 82.7089307 | -0.263026148 | 0.62026583 | -0.4240539 | 0.6715265 | 0.89704376 |
| DZIP1L | 184.945407 | -0.106332433 | 0.27582641 | -0.3855049 | 7.00E-01 | 9.07E-01 |
| DBR1 | 89.6741945 | -0.215453634 | 0.25812513 | -0.8346868 | 0.40389408 | 0.78358009 |
| ARMC8 | 1450.85419 | 0.020804041 | 0.16633748 | 0.12507128 | 0.90046712 | 0.97479173 |
| NME9 | 133.516299 | 0.912581523 | 0.46398767 | 1.96682279 | 0.04920364 | 0.45419182 |
| MRAS | 633.323443 | -0.855383032 | 0.55883876 | -1.5306437 | 1.26E-01 | 0.58564339 |
| ESYT3 | 65.2972535 | -0.180114588 | 0.33276994 | -0.5412586 | 5.88E-01 | 0.86140264 |
| CEP70 | 792.309035 | 0.506574579 | 0.47834034 | 1.05902541 | 0.28958821 | 0.71218341 |
| FAIM | 143.269647 | -0.800211969 | 0.42362731 | -1.8889527 | 0.05889816 | 0.47778423 |
| PIK3CB | 2643.29257 | -0.166920023 | 0.23778304 | -0.7019846 | 4.83E-01 | 8.18E-01 |
| MRPS22 | 141.658677 | 0.019448696 | 0.29305043 | 0.06636638 | 0.94708614 | 0.98646931 |
| COPB2 | 761.336224 | -0.472579179 | 0.20345885 | -2.3227261 | 0.02019387 | 0.34861133 |
| AC097103.2 | 863.55437 | 0.474064918 | 0.44511588 | 1.06503709 | 0.28685915 | 0.71118494 |
| NMNAT3 | 515.220956 | 0.393421391 | 0.42685047 | 0.92168434 | 0.35669325 | 0.75872042 |
| CLSTN2 | 1026.64254 | 1.128082058 | 0.44244802 | 2.54963747 | 0.0107835 | 0.27525031 |
| SLC25A36 | 1782.4663 | -0.520975494 | 0.46458207 | -1.1213853 | 0.2621239 | 0.69415229 |
| PXYLP1 | 326.454739 | 0.410843233 | 0.26841567 | 1.53062311 | 0.12586257 | 0.58564339 |
| AC117383.1 | 373.274337 | 0.053258507 | 0.48428237 | 0.10997408 | 9.12E-01 | 9.77E-01 |
| ZBTB38 | 3649.34611 | -0.373078151 | 0.47897472 | -0.7789099 | 0.43603279 | 0.79549938 |
| RASA2 | 1545.52008 | -0.370812232 | 0.24931382 | -1.4873312 | 1.37E-01 | 0.59474076 |

| | | | | | | |
|---|---|---|---|---|---|---|
| RNF7 | 349.052689 | -0.434371256 | 0.32912337 | -1.3197825 | 1.87E-01 | 0.6378941 |
| ATP1B3 | 799.535823 | -1.353248332 | 0.59949038 | -2.2573312 | 0.02398739 | 0.37387451 |
| TFDP2 | 3089.91708 | 0.157122978 | 0.34274904 | 0.45841989 | 0.6466508 | 0.88751803 |
| GK5 | 473.490841 | 0.614807394 | 0.3044387 | 2.01947843 | 0.04343752 | 0.44147822 |
| XRN1 | 1952.39142 | -0.005683854 | 0.33774965 | -0.0168286 | 0.98657335 | 0.99616655 |
| ATR | 1987.29668 | -0.1897947 | 0.26906909 | -0.7053753 | 0.4805767 | 0.81714875 |
| PLS1 | 1202.52481 | 0.417682434 | 0.39348997 | 1.06148179 | 0.28847101 | 0.71218341 |
| TRPC1 | 556.405975 | 0.636825113 | 0.36211161 | 1.75864318 | 0.07863813 | 0.51192014 |
| PCOLCE2 | 55.3291263 | -0.183417169 | 0.49052324 | -0.3739215 | 0.70846274 | 0.91070982 |
| U2SURP | 1231.31863 | -0.147054625 | 0.23215547 | -0.6334317 | 0.5264518 | 0.83698249 |
| SLC9A9 | 269.264161 | 0.819154396 | 0.49019693 | 1.67107205 | 0.09470745 | 0.53824794 |
| C3orf58 | 155.303001 | -0.705991635 | 0.38162418 | -1.8499657 | 0.0643185 | 0.48593331 |
| AC107021.1 | 265.678053 | -0.704722233 | 0.75178762 | -0.9373954 | 3.49E-01 | 7.54E-01 |
| LNCSRLR | 101.411255 | -1.011470806 | 0.60264388 | -1.6783889 | 9.33E-02 | 5.37E-01 |
| PLOD2 | 2804.68775 | -1.020997695 | 0.62011555 | -1.6464636 | 9.97E-02 | 5.45E-01 |
| PLSCR4 | 1058.76678 | 0.280224536 | 0.52200222 | 0.53682633 | 0.59138758 | 0.86364262 |
| PLSCR2 | 188.078968 | 1.210448073 | 0.60028553 | 2.01645385 | 0.04375254 | 0.44226848 |
| PLSCR1 | 1548.03626 | 0.301567116 | 0.62820095 | 0.48004881 | 0.63119269 | 0.88122598 |
| GYG1 | 148.778583 | -0.382719737 | 0.39950494 | -0.957985 | 0.33807033 | 0.74488507 |
| HLTF | 284.818781 | 0.278368548 | 0.36230786 | 0.76832047 | 0.44229682 | 0.79928982 |
| HPS3 | 670.125315 | 0.367049195 | 0.31480186 | 1.16596896 | 0.24362699 | 0.68164834 |
| CP | 123.467684 | 0.950970573 | 0.79440494 | 1.19708542 | 0.23127326 | 0.66878058 |
| TM4SF18 | 156.277092 | 1.292307037 | 0.75509968 | 1.71143899 | 0.0870001 | 0.52626233 |
| TM4SF1 | 108.187753 | 0.140783412 | 0.75660419 | 0.18607274 | 0.85238772 | 0.96061646 |
| WWTR1 | 4138.79346 | 0.231883487 | 0.28305614 | 0.81921376 | 0.41266447 | 0.78632759 |
| COMMD2 | 249.557271 | -0.087009944 | 0.24773586 | -0.3512206 | 0.72542284 | 0.91618639 |
| ANKUB1 | 690.308311 | 0.742933203 | 0.53716585 | 1.38306111 | 0.16664613 | 0.62216414 |
| RNF13 | 2101.95005 | 0.111181301 | 0.18486498 | 0.60141895 | 0.54756098 | 0.84485526 |
| PFN2 | 38.9131872 | 0.090699391 | 0.38474917 | 0.23573641 | 0.81363721 | 0.94874736 |
| AC117386.2 | 53.1001187 | 1.60658523 | 0.77145015 | 2.08255223 | 0.03729206 | 0.42215481 |
| TSC22D2 | 1598.42591 | -0.277831827 | 0.29594148 | -0.9388066 | 0.34783003 | 0.75307317 |
| SERP1 | 848.054441 | -0.12400984 | 0.2539313 | -0.4883598 | 0.62529502 | 0.87854882 |
| EIF2A | 485.339404 | -0.359922217 | 0.25657268 | -1.4028081 | 0.16067408 | 0.61630095 |
| SELENOT | 453.281012 | -0.181703629 | 0.23944293 | -0.7588599 | 0.44793639 | 0.80223612 |
| ERICH6 | 64.9135769 | -0.419693939 | 0.45204395 | -0.9284361 | 0.35318138 | 0.75673243 |
| ERICH6-AS1 | 129.878717 | 1.289372784 | 0.52109541 | 2.4743507 | 0.01334786 | 0.29875272 |
| SIAH2 | 284.361986 | -0.436212535 | 0.34249938 | -1.2736155 | 0.2027997 | 0.64881821 |
| MED12L | 66.8722899 | 0.124092551 | 0.39794839 | 0.31183077 | 0.75516914 | 0.92799282 |
| SUCNR1 | 283.314973 | 1.25317453 | 0.50878049 | 2.4630947 | 0.01377435 | 0.30354511 |
| MBNL1 | 4963.29055 | -0.06006968 | 0.32413558 | -0.1853227 | 0.85297594 | 0.96073035 |
| RAP2B | 230.991304 | -0.043804866 | 0.4053699 | -0.1080615 | 9.14E-01 | 9.77E-01 |
| ARHGEF26 | 48.0951163 | 0.267961779 | 0.45130354 | 0.59375067 | 0.55267889 | 0.84721285 |
| DHX36 | 1180.14218 | -0.336277564 | 0.18282146 | -1.8393769 | 0.06585977 | 0.48798003 |
| MME | 4119.00671 | 0.139725015 | 0.39832189 | 0.35078417 | 7.26E-01 | 9.16E-01 |
| MME-AS1 | 96.5770793 | 1.136933879 | 0.96542831 | 1.17764713 | 0.23893731 | 0.67667639 |
| DWORF | 98.1209545 | 0.809109384 | 0.99141955 | 0.81611199 | 4.14E-01 | 7.86E-01 |
| PLCH1 | 964.962073 | 1.455748585 | 0.59368394 | 2.4520599 | 0.0142041 | 0.30468798 |
| GMPS | 529.76795 | -0.102212098 | 0.29198118 | -0.350064 | 0.72629067 | 0.91635631 |
| KCNAB1 | 91.3738118 | 0.678018392 | 0.56300176 | 1.20429178 | 0.22847682 | 0.66705629 |
| AC084036.1 | 35.4624085 | -0.57020772 | 0.97185884 | -0.5867187 | 0.55739267 | 0.84861484 |
| SSR3 | 581.637749 | -0.43783379 | 0.39591927 | -1.1058663 | 0.26878438 | 0.69961625 |
| TIPARP | 432.078843 | -0.42812852 | 0.46949016 | -0.9119009 | 0.36182087 | 0.76060307 |
| LINC00886 | 823.835544 | 0.396232747 | 0.55950936 | 0.70817895 | 0.47883414 | 0.81596289 |
| LEKR1 | 623.26714 | 0.179423825 | 0.60026555 | 0.29890741 | 0.76501069 | 0.93083208 |

| | | | | | | |
|---|---|---|---|---|---|---|
| CCNL1 | 2119.62415 | -0.355159784 | 0.3238446 | -1.0966982 | 0.27277335 | 0.70108838 |
| VEPH1 | 332.958338 | 0.894607578 | 0.64508145 | 1.38681337 | 0.16549868 | 0.62065658 |
| PQLC2L | 51.1854174 | 0.425591767 | 0.88747537 | 0.47955333 | 6.32E-01 | 0.88123095 |
| RSRC1 | 1931.84002 | -0.350828886 | 0.21588788 | -1.6250513 | 0.10415163 | 0.55056166 |
| MLF1 | 128.884132 | 0.191657556 | 0.34003797 | 0.56363575 | 0.57300205 | 0.8542679 |
| GFM1 | 451.973902 | -0.40899553 | 0.3324618 | -1.2302031 | 0.21862107 | 0.66098715 |
| LXN | 84.7767084 | -0.002575972 | 0.56714574 | -0.004542 | 0.99637603 | 0.99914596 |
| RARRES1 | 476.176368 | 1.097917448 | 0.69437112 | 1.58116808 | 0.11383961 | 0.56865873 |
| MFSD1 | 321.501142 | -0.391128053 | 0.28460582 | -1.3742799 | 0.16935481 | 0.62326131 |
| IQCJ-SCHIP1 | 2810.08453 | -1.399827259 | 0.50080564 | -2.7951508 | 0.00518755 | 0.21811586 |
| AC079594.2 | 42.9322005 | 0.723010143 | 0.60764563 | 1.18985493 | 0.23410342 | 0.67060722 |
| SMC4 | 253.155635 | -0.280785977 | 0.32385624 | -0.8670081 | 0.38593758 | 0.77317162 |
| KPNA4 | 1313.31142 | -0.478646363 | 0.33495245 | -1.4289979 | 0.15300482 | 0.61001662 |
| PPM1L | 1304.23566 | 0.124240497 | 0.28219111 | 0.44027076 | 0.65974101 | 0.89094706 |
| B3GALNT1 | 244.02512 | 0.304805906 | 0.43645455 | 0.69836803 | 0.48494706 | 0.81839266 |
| NMD3 | 945.057306 | 0.120596056 | 0.24846769 | 0.48535911 | 0.62742163 | 0.87944251 |
| LINC01322 | 816.915967 | 0.194938967 | 0.52896443 | 0.36852944 | 0.7124785 | 0.9117676 |
| PDCD10 | 796.432117 | -0.167745297 | 0.2626117 | -0.6387579 | 0.52298044 | 0.83509936 |
| SERPINI1 | 278.500101 | 0.548554222 | 0.61096718 | 0.89784564 | 0.36926785 | 0.76620563 |
| GOLIM4 | 1016.96883 | -0.274222568 | 0.4980266 | -0.5506183 | 0.58189535 | 0.85938792 |
| LINC02082 | 40.25392 | 0.133849015 | 0.82385116 | 0.16246747 | 0.87093774 | 0.9657206 |
| MECOM | 1447.91401 | -0.419986344 | 0.51109249 | -0.8217424 | 0.41122355 | 0.78632759 |
| MYNN | 213.999063 | -0.078435016 | 0.20717433 | -0.3785943 | 0.70498917 | 0.909554 |
| LRRC34 | 56.0144482 | 0.045013782 | 0.59415572 | 0.07576092 | 0.93960931 | 0.98435387 |
| SEC62 | 1092.19925 | -0.343361881 | 0.18501729 | -1.8558367 | 0.06347684 | 0.48572144 |
| GPR160 | 1300.30354 | 0.106958066 | 0.46959362 | 0.22776729 | 0.81982716 | 0.94990455 |
| PHC3 | 2299.58639 | 0.025999692 | 0.17971215 | 0.14467409 | 0.88496819 | 0.96937383 |
| AC008040.5 | 53.8216538 | 0.765316658 | 0.66549866 | 1.14998978 | 0.25014808 | 0.68617881 |
| PRKCI | 1086.56098 | -0.421777798 | 0.21942387 | -1.9222057 | 0.05457989 | 0.46772575 |
| SKIL | 970.566279 | 0.044307181 | 0.36909786 | 0.12004183 | 0.90445002 | 0.97574711 |
| AC026316.5 | 153.505762 | 0.202240774 | 0.72890345 | 0.27745893 | 7.81E-01 | 9.37E-01 |
| RPL22L1 | 179.301001 | -0.467631403 | 0.53532043 | -0.8735542 | 0.38236108 | 0.77279408 |
| EIF5A2 | 92.3297054 | 0.536231236 | 0.28535413 | 1.87917813 | 0.06022018 | 0.48004006 |
| SLC2A2 | 1073.41613 | 0.989553787 | 0.87927463 | 1.12542061 | 0.26041084 | 0.69280433 |
| TNIK | 5360.82807 | 0.448695311 | 0.42470513 | 1.05648669 | 0.29074592 | 0.71269438 |
| PLD1 | 1375.50301 | 1.292599741 | 0.52563559 | 2.45911763 | 0.0139279 | 0.30363654 |
| FNDC3B | 9269.30461 | 0.000323661 | 0.30381539 | 0.00106532 | 9.99E-01 | 1.00E+00 |
| TNFSF10 | 4238.50631 | 0.861707129 | 0.67442518 | 1.27769122 | 0.20135832 | 0.648653 |
| LINC02068 | 386.2547 | 0.365642952 | 0.63661353 | 0.57435624 | 0.56572676 | 0.85194709 |
| NCEH1 | 2058.72251 | -0.274334749 | 0.43087104 | -0.636698 | 0.52432155 | 0.83589803 |
| ECT2 | 219.220277 | 0.208678413 | 0.39963998 | 0.52216602 | 0.60155475 | 0.86757631 |
| SPATA16 | 51.6449297 | 0.149784522 | 0.70914476 | 0.21121854 | 8.33E-01 | 9.53E-01 |
| NLGN1 | 19470.8264 | 0.708550632 | 0.58067834 | 1.22021192 | 0.22238455 | 0.66373504 |
| NAALADL2 | 4428.11365 | 1.131984532 | 0.54236474 | 2.0871278 | 0.03687658 | 0.42215481 |
| TBL1XR1 | 5836.89595 | -0.941164205 | 0.33537549 | -2.8062999 | 0.0050114 | 0.21686099 |
| LINC00501 | 157.269167 | -1.539920994 | 0.61470502 | -2.5051381 | 0.01224036 | 0.29629837 |
| LINC00578 | 383.836256 | -1.311981976 | 0.87104343 | -1.5062188 | 0.13201102 | 0.59094548 |
| LINC02015 | 305.736803 | -1.188799547 | 0.8155357 | -1.4576916 | 0.14492557 | 0.60109715 |
| KCNMB2-AS1 | 317.723489 | -0.220968841 | 0.83214702 | -0.2655406 | 0.79059302 | 0.94063545 |
| ZMAT3 | 307.871754 | 0.425901652 | 0.36517916 | 1.16628139 | 0.2435007 | 0.68164834 |
| PIK3CA | 1079.15252 | -0.049831627 | 0.23253171 | -0.2143004 | 0.83031285 | 0.95325673 |
| KCNMB3 | 115.283536 | -1.155350608 | 0.43180745 | -2.6756153 | 0.00745922 | 0.23590813 |
| ZNF639 | 201.299318 | -0.234100868 | 0.27014943 | -0.8665607 | 0.38618278 | 0.77321171 |
| MFN1 | 899.924918 | -0.062908348 | 0.28899627 | -0.2176788 | 0.82767942 | 0.95264101 |

| | | | | | | |
|---|---|---|---|---|---|---|
| GNB4 | 39.1292401 | 0.574676959 | 0.4875802 | 1.17863062 | 0.23854529 | 0.67658923 |
| ACTL6A | 159.446187 | -0.289592553 | 0.34703098 | -0.8344862 | 0.40400707 | 0.78358009 |
| MRPL47 | 159.917053 | -0.245105516 | 0.29168525 | -0.8403082 | 0.40073558 | 0.78232083 |
| NDUFB5 | 359.569516 | 0.563235956 | 0.39377611 | 1.43034567 | 0.15261783 | 0.61001662 |
| USP13 | 235.307411 | 1.095644067 | 0.40360724 | 2.7146294 | 0.006635 | 0.22966139 |
| TTC14 | 1070.81556 | -0.241089441 | 0.21683805 | -1.111841 | 0.2662065 | 0.69797523 |
| FXR1 | 1245.02997 | -0.168668294 | 0.24570608 | -0.6864637 | 0.49242077 | 0.82087587 |
| DNAJC19 | 188.604724 | 0.119399986 | 0.41049552 | 0.29086794 | 0.77115232 | 0.93302012 |
| ATP11B | 1496.83329 | -0.070700304 | 0.17438792 | -0.4054197 | 0.68516901 | 0.90112503 |
| DCUN1D1 | 773.343625 | 0.028465721 | 0.18728805 | 0.15198899 | 0.87919562 | 0.9674841 |
| MCCC1 | 1067.4268 | 0.114355833 | 0.29447468 | 0.38833843 | 0.6977656 | 0.90696517 |
| MCF2L2 | 182.593688 | 0.213963732 | 0.45898883 | 0.46616326 | 0.64109864 | 0.88396933 |
| B3GNT5 | 276.712759 | 0.006010999 | 0.35164322 | 0.01709403 | 0.9863616 | 0.99612253 |
| KLHL6 | 37.6113482 | 0.784619296 | 0.68761044 | 1.14108113 | 0.25383616 | 0.68891958 |
| KLHL24 | 1747.94703 | -0.253008627 | 0.36714152 | -0.6891311 | 0.49074079 | 0.81989696 |
| YEATS2 | 874.421091 | -0.553663924 | 0.42454356 | -1.3041392 | 0.19218611 | 0.64497936 |
| ABCC5 | 514.746136 | -0.055199721 | 0.50454965 | -0.1094039 | 0.91288211 | 0.977179 |
| EIF2B5 | 187.398392 | -0.418582855 | 0.27156999 | -1.5413443 | 0.12323301 | 0.5825794 |
| DVL3 | 359.213934 | -0.469856883 | 0.2684162 | -1.7504788 | 0.08003572 | 0.51264427 |
| AP2M1 | 605.280876 | 0.148402151 | 0.37008467 | 0.40099514 | 0.68842371 | 0.90307854 |
| ABCF3 | 128.797011 | -0.59782735 | 0.36028599 | -1.6593133 | 0.09705267 | 0.5401548 |
| ALG3 | 134.095906 | -0.83562064 | 0.48026013 | -1.7399334 | 0.08187071 | 0.51593841 |
| PSMD2 | 364.483751 | -0.570662506 | 0.48063034 | -1.187321 | 0.23510102 | 0.67119484 |
| EIF4G1 | 916.921493 | -0.448424665 | 0.44599403 | -1.0054499 | 0.31468024 | 0.72719146 |
| FAM131A | 101.793445 | 0.258826757 | 0.3934074 | 0.65791024 | 0.51059581 | 0.82943076 |
| CLCN2 | 66.0377383 | 0.514801534 | 0.42336729 | 1.21596907 | 0.2239967 | 0.66543624 |
| POLR2H | 551.449567 | -0.012974023 | 0.28642738 | -0.045296 | 0.96387135 | 0.99012655 |
| THPO | 45.9245799 | 0.283662279 | 0.62017717 | 0.4573891 | 0.6473914 | 0.88774616 |
| CHRD | 158.159993 | 0.402540037 | 0.57552584 | 0.69943 | 0.48428335 | 0.81839266 |
| MAGEF1 | 134.188415 | -0.265635678 | 0.34722565 | -0.7650232 | 0.44425773 | 0.79994797 |
| VPS8 | 1924.8817 | -0.252278975 | 0.19215341 | -1.312904 | 0.18921531 | 0.6413893 |
| EHHADH | 2780.09181 | 0.854700819 | 0.63583776 | 1.34421212 | 0.1788798 | 0.62933919 |
| MAP3K13 | 7193.7714 | 0.85689398 | 0.47104899 | 1.8191186 | 0.06889334 | 0.49260354 |
| TMEM41A | 432.1447 | 0.125474553 | 0.35438555 | 0.35406227 | 0.72329222 | 0.91592159 |
| LIPH | 56.3005232 | -0.102805622 | 0.75585266 | -0.1360128 | 0.89181118 | 0.97234681 |
| SENP2 | 826.350402 | -0.09653026 | 0.19659216 | -0.4910178 | 0.62341383 | 0.8776842 |
| IGF2BP2 | 3416.15559 | -0.317083636 | 0.30042211 | -1.0554604 | 0.29121483 | 0.71317245 |
| IGF2BP2-AS1 | 74.672417 | -0.404805254 | 0.57443648 | -0.7046998 | 0.4809971 | 0.81720798 |
| TRA2B | 851.619929 | -0.109065936 | 0.15343764 | -0.710816 | 0.47719823 | 0.81570932 |
| ETV5 | 524.766264 | 0.679321325 | 0.42117161 | 1.61293239 | 0.1067592 | 0.55488138 |
| DGKG | 502.805227 | -0.081440475 | 0.59331617 | -0.1372632 | 0.89082276 | 0.97230741 |
| CRYGS | 82.3378027 | 1.118502484 | 0.2768059 | 4.04074661 | 5.33E-05 | 0.02213112 |
| TBCCD1 | 142.573955 | 0.397202427 | 0.3581477 | 1.10904644 | 0.26741015 | 0.69874881 |
| DNAJB11 | 148.319728 | -0.548554139 | 0.54147996 | -1.0130645 | 0.31102935 | 0.72558189 |
| AC068631.1 | 45.9260202 | -0.245393226 | 0.51013429 | -0.4810365 | 0.63049054 | 0.8811311 |
| AC068631.2 | 63.9606209 | -0.270249204 | 0.44928326 | -0.6015118 | 0.54749913 | 0.84485526 |
| KNG1 | 63.4956694 | -0.507388906 | 0.68412061 | -0.7416659 | 0.4582898 | 0.80618305 |
| EIF4A2 | 931.816998 | -0.152865694 | 0.27824797 | -0.5493865 | 0.5827402 | 0.85986705 |
| RFC4 | 40.9602639 | -0.350515136 | 0.43947017 | -0.7975857 | 0.42511093 | 0.78894887 |
| ST6GAL1 | 531.879024 | 1.530555331 | 0.87090863 | 1.75742355 | 0.07884564 | 0.51192014 |
| MASP1 | 240.264838 | 0.093249164 | 0.61088208 | 0.15264675 | 0.87867685 | 0.96739146 |
| AC072022.1 | 127.516136 | -0.531692766 | 0.92842579 | -0.572682 | 0.56686001 | 0.85233132 |
| BCL6 | 1965.63258 | -0.469641034 | 0.64692991 | -0.7259535 | 0.46786728 | 0.8118777 |
| LPP | 19857.6756 | 0.265287856 | 0.34098974 | 0.77799367 | 0.43657274 | 0.79558019 |

| | | | | | | |
|---|---|---|---|---|---|---|
| TPRG1 | 444.130846 | -0.110452565 | 0.44562697 | -0.2478588 | 0.80424366 | 0.94551573 |
| P3H2 | 4330.08482 | -0.725907682 | 0.55931823 | -1.2978438 | 0.19434103 | 0.64548758 |
| P3H2-AS1 | 61.3596825 | -0.02404231 | 0.74803661 | -0.0321406 | 0.97435996 | 0.99377398 |
| CLDN1 | 569.469014 | 0.281664955 | 0.50124572 | 0.56192989 | 5.74E-01 | 8.54E-01 |
| CLDN16 | 46.0261815 | 1.045167328 | 0.70039909 | 1.49224542 | 0.13563482 | 0.59381224 |
| IL1RAP | 204.329053 | 0.410264634 | 0.42181907 | 0.97260808 | 0.33074812 | 0.73866225 |
| GMNC | 64.9133262 | 0.904581057 | 0.89794092 | 1.00739485 | 0.31374507 | 0.72717675 |
| CCDC50 | 1710.17455 | -0.10249512 | 0.43932759 | -0.2333 | 0.81552847 | 0.94906482 |
| FGF12 | 80.7340238 | -0.283868171 | 0.38033247 | -0.7463685 | 0.45544485 | 0.80465616 |
| MB21D2 | 932.345784 | -0.185142749 | 0.43887357 | -0.421859 | 0.67312797 | 0.89782405 |
| ATP13A4 | 75.9042682 | 0.188521563 | 0.74669745 | 0.25247383 | 0.80067484 | 0.94407312 |
| OPA1 | 1335.69662 | -0.019578509 | 0.13191144 | -0.1484216 | 0.88201004 | 0.96796225 |
| OPA1-AS1 | 46.5175168 | 0.357734608 | 0.79951478 | 0.44743965 | 0.65455766 | 0.89032312 |
| HES1 | 225.112653 | 0.954661018 | 0.66310368 | 1.43968591 | 0.14995628 | 0.60758225 |
| LINC00887 | 84.885371 | 1.501507629 | 0.93278956 | 1.609696 | 0.10746424 | 0.55606354 |
| CPN2 | 128.897902 | -0.268726948 | 0.6015467 | -0.4467267 | 0.65507243 | 0.89053559 |
| ATP13A3 | 3280.5479 | -0.651816735 | 0.36554526 | -1.7831355 | 0.07456425 | 0.5071704 |
| LINC00884 | 343.2482 | -0.443736802 | 0.42700104 | -1.0391937 | 0.29871464 | 0.71852421 |
| TMEM44-AS | 141.682658 | 0.161561539 | 0.59591317 | 0.27111591 | 0.78630189 | 0.9390702 |
| TMEM44 | 73.4519557 | -0.26986625 | 0.46632797 | -0.5787048 | 5.63E-01 | 8.51E-01 |
| LSG1 | 388.448302 | -0.382465415 | 0.4217507 | -0.9068519 | 0.36448513 | 0.76286421 |
| XXYLT1 | 305.648254 | -0.356744564 | 0.32647407 | -1.0927195 | 0.274517 | 0.7014484 |
| ACAP2 | 1761.1823 | -0.116135883 | 0.19845451 | -0.5852015 | 0.55841222 | 0.84904674 |
| PPP1R2 | 424.756272 | 0.157992835 | 0.20580394 | 0.76768615 | 0.44267367 | 0.79933184 |
| MUC20-OT1 | 5103.92728 | 0.192724699 | 0.49374882 | 0.39032944 | 0.69629296 | 0.90612803 |
| MUC20 | 59.0411979 | 1.086735013 | 0.70170573 | 1.54870477 | 0.12145271 | 0.57985101 |
| MUC4 | 147.763281 | 2.424724024 | 1.04760374 | 2.31454312 | 0.02063795 | 0.35267827 |
| TNK2 | 125.113421 | -0.276273124 | 0.45836325 | -0.6027384 | 0.54668274 | 0.84460887 |
| TFRC | 456.935276 | -0.218800934 | 0.31949583 | -0.6848319 | 0.49345001 | 0.82143785 |
| SLC51A | 182.919057 | 0.719401623 | 0.75904979 | 0.94776605 | 0.34324857 | 0.75073625 |
| PCYT1A | 437.176809 | -0.017038847 | 0.24786508 | -0.0687424 | 0.94519465 | 0.98566355 |
| UBXN7 | 693.407935 | -0.129740093 | 0.21037118 | -0.6167199 | 0.5374195 | 0.8403387 |
| RNF168 | 410.84872 | -0.313163976 | 0.36414354 | -0.8600015 | 0.38978823 | 0.77569824 |
| WDR53 | 109.962703 | -0.314872567 | 0.43942292 | -0.7165593 | 0.47364608 | 0.81479253 |
| FBXO45 | 80.615839 | -0.840805591 | 0.51130293 | -1.6444373 | 0.10008591 | 0.54542315 |
| PIGX | 173.85845 | -0.010935857 | 0.23293449 | -0.0469482 | 0.96255451 | 0.99012655 |
| PAK2 | 1945.66098 | -0.678573333 | 0.33137848 | -2.047729 | 0.04058656 | 0.43086475 |
| SENP5 | 1510.74053 | -0.720966488 | 0.35682763 | -2.0204895 | 0.04333263 | 0.44144764 |
| NCBP2 | 312.164362 | -0.546349229 | 0.30650384 | -1.78252 | 0.07466447 | 0.5071704 |
| NCBP2-AS2 | 129.74506 | -0.272915792 | 0.46746136 | -0.5838253 | 0.55933783 | 0.84936251 |
| PIGZ | 71.7251858 | 0.416842897 | 0.50550437 | 0.8246079 | 0.40959423 | 0.78542936 |
| DLG1 | 4807.264 | 0.058612711 | 0.18742245 | 0.31273046 | 0.75448545 | 0.92799282 |
| BDH1 | 475.438703 | -0.029756515 | 0.44037201 | -0.0675713 | 0.94612689 | 0.98605241 |
| AC024560.1 | 75.5725725 | -2.910098103 | 2.83742501 | -1.0256123 | 0.30507435 | 0.72103276 |
| RUBCN | 381.659936 | -0.146573882 | 0.28585908 | -0.5127487 | 0.60812709 | 0.86985416 |
| FYTTD1 | 423.970127 | -0.128528054 | 0.30665344 | -0.4191313 | 0.67512018 | 0.89801183 |
| LRCH3 | 1699.90817 | -0.454915355 | 0.24692118 | -1.8423505 | 0.0654239 | 0.48790819 |
| IQCG | 295.587536 | -0.050445157 | 0.24638299 | -0.2047429 | 0.83777304 | 0.95496955 |
| RPL35A | 821.949836 | -0.104073992 | 0.48668036 | -0.2138446 | 0.8306682 | 0.95325673 |
| LMLN | 486.772843 | 0.275519407 | 0.30931136 | 0.89075102 | 0.37306276 | 0.76930513 |
| ZNF595 | 323.280907 | 0.029475284 | 0.19901021 | 0.1481094 | 0.88225643 | 0.96797688 |
| ZNF718 | 699.370612 | 0.151709727 | 0.21454272 | 0.70713064 | 0.4794853 | 0.81653684 |
| ZNF141 | 560.932928 | -0.146719959 | 0.17990042 | -0.8155621 | 0.41475066 | 0.78632759 |
| AC092574.2 | 88.9210979 | 0.873961768 | 0.68266912 | 1.28021283 | 0.20047029 | 0.64780303 |

| | | | | | | |
|---|---|---|---|---|---|---|
| ZNF721 | 2268.204 | -0.25855895 | 0.18151919 | -1.4244166 | 0.1543259 | 0.61099212 |
| PIGG | 392.114178 | -0.503603483 | 0.24395259 | -2.0643498 | 0.03898456 | 0.42680622 |
| PDE6B | 36.3726736 | 0.609964754 | 0.67633338 | 0.90186996 | 0.36712595 | 0.76546934 |
| ATP5ME | 327.412474 | -0.496110284 | 0.45093828 | -1.1001734 | 0.2712566 | 0.70002917 |
| MYL5 | 50.0510436 | -0.733905073 | 0.40705465 | -1.8029645 | 0.0713938 | 0.49949133 |
| PCGF3 | 605.83182 | -0.044272286 | 0.25933117 | -0.1707172 | 0.86444615 | 0.96411419 |
| AC139887.2 | 1270.08218 | 0.556589306 | 0.40879816 | 1.36152596 | 0.17334753 | 0.62660197 |
| GAK | 1492.49666 | -0.448685061 | 0.34492545 | -1.3008175 | 0.19332094 | 0.64548758 |
| TMEM175 | 126.816809 | -0.002695677 | 0.25799542 | -0.0104485 | 0.99166342 | 0.99748517 |
| DGKQ | 78.966077 | 0.042013248 | 0.46859913 | 0.08965712 | 0.92855969 | 0.98272895 |
| SLC26A1 | 54.2752312 | -0.316898131 | 0.38869316 | -0.8152912 | 0.41490562 | 0.78632759 |
| IDUA | 96.9238394 | -0.453825321 | 0.42004486 | -1.0804211 | 0.27995471 | 0.70461609 |
| FGFRL1 | 138.637862 | -0.228512825 | 0.36868635 | -0.6198028 | 0.53538764 | 0.83954482 |
| RNF212 | 89.1988757 | -0.896455738 | 0.51146203 | -1.7527318 | 0.07964806 | 0.51231158 |
| AC092535.3 | 235.352202 | -0.58938518 | 0.5407908 | -1.089858 | 0.2757757 | 0.70248235 |
| SPON2 | 612.09032 | -0.16757108 | 0.54330975 | -0.3084264 | 0.75775788 | 0.928294 |
| CTBP1-AS | 163.263319 | -0.374651866 | 0.506205 | -0.7401189 | 0.45922788 | 0.80669065 |
| CTBP1 | 539.829812 | -0.562729814 | 0.28742823 | -1.9578098 | 0.05025233 | 0.45824658 |
| CTBP1-DT | 119.732533 | 0.506992837 | 0.47085596 | 1.0767472 | 0.28159322 | 0.70625976 |
| MAEA | 613.530738 | -0.262951052 | 0.29236379 | -0.8993968 | 0.36844135 | 0.76615653 |
| UVSSA | 765.267735 | -0.144683834 | 0.29955319 | -0.4829988 | 0.62909658 | 0.88057005 |
| FAM53A | 69.1935981 | -0.785212735 | 0.54712493 | -1.4351617 | 0.15124103 | 0.60905477 |
| SLBP | 446.006467 | -0.606363297 | 0.37309065 | -1.6252439 | 0.10411061 | 0.55056166 |
| TMEM129 | 133.871915 | 0.399026109 | 0.39007307 | 1.02295221 | 0.30633044 | 0.72182764 |
| TACC3 | 36.6351891 | -0.153041502 | 0.43175699 | -0.3544621 | 0.72299258 | 0.91592159 |
| FGFR3 | 560.281653 | 1.093358069 | 0.76032559 | 1.43801298 | 0.15043037 | 0.60799621 |
| LETM1 | 430.041472 | -0.381598019 | 0.41132674 | -0.9277248 | 0.35355034 | 0.75673243 |
| NSD2 | 1110.03702 | -0.131415732 | 0.2989158 | -0.4396413 | 0.66019692 | 0.89103182 |
| NELFA | 370.965279 | -0.179181387 | 0.28197253 | -0.6354569 | 0.52513048 | 0.835938 |
| POLN | 236.47619 | -0.366041142 | 0.39770874 | -0.9203749 | 0.35737688 | 0.75903693 |
| MXD4 | 293.518115 | -0.020914638 | 0.38110908 | -0.0548784 | 0.95623538 | 0.9886112 |
| ZFYVE28 | 699.046942 | -0.208933818 | 0.35037038 | -0.5963227 | 0.55095966 | 0.8466552 |
| RNF4 | 794.858693 | -0.418291112 | 0.29951557 | -1.3965588 | 0.16254628 | 0.61912485 |
| FAM193A | 1469.47425 | -0.240135255 | 0.25140914 | -0.9551572 | 0.3394982 | 0.74644675 |
| TNIP2 | 352.964476 | -0.036800575 | 0.41200429 | -0.0893209 | 0.92882692 | 0.98272895 |
| SH3BP2 | 1791.45234 | 0.450351241 | 0.26658463 | 1.68933683 | 0.0911549 | 0.53537574 |
| ADD1 | 2465.77806 | -0.104065657 | 0.21888917 | -0.4754262 | 0.63448319 | 0.88238823 |
| MFSD10 | 311.794672 | -0.257987035 | 0.39108855 | -0.659664 | 0.50946948 | 0.82882082 |
| NOP14-AS1 | 82.3435288 | -0.57011475 | 0.3853806 | -1.4793551 | 0.13904543 | 0.59474076 |
| NOP14 | 256.706066 | -0.8984246 | 0.36899829 | -2.4347663 | 0.01490141 | 0.30894792 |
| GRK4 | 253.221229 | 0.156360892 | 0.5041932 | 0.31012098 | 0.75646896 | 0.92799282 |
| HTT | 1204.83953 | 0.03621521 | 0.15676474 | 0.23101629 | 0.81730215 | 0.94954998 |
| RGS12 | 1268.37252 | -0.204002092 | 0.29070132 | -0.7017584 | 0.48282985 | 0.8177376 |
| LRPAP1 | 579.911543 | -0.095130146 | 0.32904029 | -0.289114 | 0.77249416 | 0.933623 |
| ADRA2C | 62.5359779 | 1.333326502 | 0.84314232 | 1.58137774 | 0.11379169 | 0.56865873 |
| TMEM128 | 135.891897 | 0.205013916 | 0.2742276 | 0.74760496 | 0.45469847 | 0.80465616 |
| LYAR | 53.728218 | -0.690317363 | 0.59527489 | -1.1596615 | 0.24618666 | 0.6834865 |
| ZBTB49 | 113.796658 | -0.242737021 | 0.26347792 | -0.9212803 | 0.3569041 | 0.75893336 |
| STX18 | 1547.11053 | -0.311297501 | 0.20871166 | -1.4915195 | 0.13582517 | 0.59386143 |
| STX18-AS1 | 488.121821 | 0.734346802 | 0.47044402 | 1.56096534 | 0.11853193 | 0.57589001 |
| STK32B | 1597.60838 | -0.084589002 | 0.43080902 | -0.1963492 | 0.84433686 | 0.95742233 |
| EVC2 | 843.002294 | -0.785723953 | 0.17649711 | -4.4517668 | 8.52E-06 | 0.01167379 |
| EVC | 2389.66225 | -1.051648389 | 0.48912366 | -2.1500665 | 0.03154996 | 0.40913458 |
| CRMP1 | 517.139839 | -0.065494512 | 0.64110139 | -0.1021594 | 0.91863018 | 0.97862093 |

| | | | | | | |
|---|---|---|---|---|---|---|
| WFS1 | 56.5003954 | 0.341567512 | 0.42687332 | 0.8001613 | 0.42361735 | 0.78894887 |
| MAN2B2 | 150.350899 | -0.241800315 | 0.39645421 | -0.6099073 | 0.54192322 | 0.84247948 |
| MRFAP1 | 371.985192 | -0.500017383 | 0.38237806 | -1.3076519 | 0.19099142 | 0.6431984 |
| LINC02482 | 64.6039078 | 0.063270158 | 0.47392115 | 0.13350355 | 0.89379515 | 0.97272276 |
| AC093323.1 | 47.2738993 | -0.818685527 | 0.37421051 | -2.1877673 | 0.02868655 | 0.39339119 |
| MRFAP1L1 | 138.466178 | -0.562255262 | 0.25157259 | -2.2349623 | 0.02541982 | 0.38163148 |
| BLOC1S4 | 113.994982 | -0.356516355 | 0.40357214 | -0.8834018 | 0.37701922 | 0.77045763 |
| KIAA0232 | 2067.94949 | -0.135878083 | 0.30194222 | -0.4500135 | 0.65270068 | 0.8894805 |
| TBC1D14 | 2255.61679 | -0.225705017 | 0.3856763 | -0.5852188 | 0.55840061 | 0.84904674 |
| TADA2B | 152.272492 | -0.588422985 | 0.48313259 | -1.2179327 | 0.22324954 | 0.66408018 |
| GRPEL1 | 307.192584 | -0.810381004 | 0.35579308 | -2.277675 | 0.02274595 | 0.36561837 |
| LINC02447 | 42.995618 | 2.248312836 | 0.75137471 | 2.9922658 | 0.00276915 | 0.17020962 |
| SORCS2 | 609.758873 | 1.884868012 | 0.85713647 | 2.19902906 | 0.02787586 | 0.3898922 |
| AFAP1 | 419.084895 | -0.026340101 | 0.37273044 | -0.070668 | 0.94366202 | 0.98522232 |
| ABLIM2 | 553.952309 | 0.46689401 | 0.37948649 | 1.23033106 | 2.19E-01 | 6.61E-01 |
| SH3TC1 | 821.411894 | 0.509310754 | 0.38109876 | 1.3364272 | 0.18140968 | 0.6318513 |
| AC104825.1 | 41.4688355 | 0.470419367 | 0.43439877 | 1.08292057 | 0.27884368 | 0.70446562 |
| ACOX3 | 318.075127 | 0.231497313 | 0.23951513 | 0.96652481 | 0.33378164 | 0.74043452 |
| TRMT44 | 252.295614 | -0.289661406 | 0.29246669 | -0.9904082 | 0.32197465 | 0.73375526 |
| SLC2A9 | 6259.52001 | 0.948338224 | 0.48636763 | 1.9498383 | 0.05119539 | 0.46106129 |
| WDR1 | 569.759496 | 0.06035857 | 0.34782317 | 0.17353234 | 0.86223301 | 0.96368459 |
| ZNF518B | 58.2993923 | -0.330044939 | 0.41317793 | -0.7987961 | 0.42440863 | 0.78894887 |
| CLNK | 39.3952987 | -0.111718043 | 0.4776178 | -0.2339068 | 0.81505735 | 0.94895023 |
| HS3ST1 | 795.49822 | 0.895918229 | 0.87388779 | 1.02520969 | 0.30526426 | 0.7210507 |
| AC005699.1 | 3240.51175 | 1.713959709 | 0.83976753 | 2.04099306 | 0.04125152 | 0.43221049 |
| LINC02360 | 30.7802109 | 1.283433822 | 0.80927242 | 1.58591075 | 0.11275956 | 0.56677494 |
| RAB28 | 1350.73029 | 0.072506113 | 0.2051256 | 0.35347178 | 0.72373478 | 0.91600702 |
| BOD1L1 | 1485.01877 | -1.259596532 | 0.47754162 | -2.6376686 | 0.00834781 | 0.25016267 |
| LINC01182 | 325.23723 | -0.0442765 | 0.53813285 | -0.082278 | 0.93442564 | 0.98359701 |
| LINC00504 | 243.145526 | 0.930427895 | 0.51976335 | 1.79009907 | 0.07343799 | 0.50459544 |
| CPEB2-DT | 372.82197 | 0.534595438 | 0.51537201 | 1.0373001 | 0.29959602 | 0.7186844 |
| CPEB2 | 2770.85158 | -0.404050534 | 0.32536109 | -1.2418527 | 0.21429093 | 0.65906957 |
| AC098829.1 | 2783.03107 | 0.642752014 | 0.61043381 | 1.05294302 | 2.92E-01 | 7.14E-01 |
| C1QTNF7 | 240.532249 | 1.611763821 | 0.84102129 | 1.9164364 | 0.05530957 | 0.46947044 |
| CC2D2A | 799.371917 | 0.499763222 | 0.47163735 | 1.05963452 | 0.2893109 | 0.71218341 |
| FBXL5 | 1949.99556 | 0.302283928 | 0.19995651 | 1.51174834 | 0.1305979 | 0.58853436 |
| FAM200B | 316.016418 | 0.221012055 | 0.43680916 | 0.50596938 | 0.61287815 | 0.87222135 |
| BST1 | 54.6643831 | 0.487532561 | 0.67531815 | 0.7219302 | 0.4703374 | 0.8137284 |
| PROM1 | 704.917036 | 0.68711426 | 0.5567613 | 1.23412718 | 0.21715552 | 0.66032006 |
| TAPT1 | 875.778842 | 0.138909436 | 0.23109404 | 0.60109483 | 0.54777682 | 0.84486687 |
| TAPT1-AS1 | 1951.3141 | 1.31439496 | 0.62563431 | 2.10089974 | 0.03564977 | 0.42040764 |
| AC097515.1 | 371.405269 | 2.096601035 | 0.95252124 | 2.20110687 | 0.02772846 | 0.38942007 |
| LDB2 | 129.602536 | -0.001702715 | 0.49957774 | -0.0034083 | 0.99728057 | 0.99963814 |
| QDPR | 546.132548 | -0.069799268 | 0.35748363 | -0.1952516 | 0.84519595 | 0.95760463 |
| LAP3 | 659.763612 | -0.130299397 | 0.82877366 | -0.1572195 | 0.87507185 | 0.96615698 |
| AC006160.1 | 115.686856 | 0.0249467 | 0.71081687 | 0.03509582 | 0.97200333 | 0.99241376 |
| MED28 | 391.688834 | -0.395316996 | 0.31351321 | -1.2609261 | 0.20733547 | 0.65269814 |
| FAM184B | 84.5379353 | 0.841596805 | 0.52294359 | 1.60934529 | 0.10754086 | 0.55624448 |
| DCAF16 | 278.320739 | -0.284007232 | 0.20487515 | -1.3862454 | 0.16567199 | 0.62065658 |
| LCORL | 1763.5313 | -0.211199897 | 0.20351103 | -1.0377811 | 0.29937198 | 0.71852421 |
| AC024230.1 | 203.702515 | 0.992757483 | 0.62134361 | 1.59775922 | 0.11009657 | 0.56114143 |
| SLIT2 | 5295.40395 | -0.088925561 | 0.58736581 | -0.1513972 | 0.87966237 | 0.96751353 |
| PACRGL | 628.295712 | -0.119155435 | 0.29730832 | -0.4007807 | 0.68858159 | 0.90310859 |
| KCNIP4 | 11765.2597 | 1.170143826 | 0.63570242 | 1.84071003 | 0.06566406 | 0.48790819 |

| | | | | | | |
|---|---|---|---|---|---|---|
| AC096719.1 | 46.904406 | 0.299085523 | 0.67403987 | 0.44372082 | 0.65724444 | 0.89053559 |
| ADGRA3 | 1601.93997 | -0.119717472 | 0.21551523 | -0.5554942 | 0.57855665 | 0.85751254 |
| AC097512.1 | 106.698985 | 0.515703502 | 0.90355944 | 0.57074663 | 0.56817141 | 0.85263558 |
| AC092440.1 | 44.6248822 | 0.918614531 | 0.9329972 | 0.98458445 | 0.32482823 | 0.73628585 |
| AC093607.1 | 421.026726 | 1.379820337 | 0.68287963 | 2.02059087 | 0.04332213 | 0.44144764 |
| PPARGC1A | 2992.55629 | 0.037546441 | 0.4938015 | 0.07603549 | 0.93939086 | 0.98435387 |
| DHX15 | 1173.4304 | -0.47901297 | 0.2326297 | -2.0591221 | 0.03948254 | 0.4275775 |
| SOD3 | 209.243479 | 0.782837919 | 0.72315172 | 1.0825362 | 0.27901434 | 0.70446562 |
| CCDC149 | 293.775052 | -1.180469393 | 0.34147602 | -3.4569613 | 0.0005463 | 0.08106654 |
| SEPSECS | 700.173237 | 0.693596822 | 0.55956008 | 1.2395395 | 0.21514577 | 0.65925411 |
| AC104662.2 | 175.62672 | 1.01210822 | 0.55117212 | 1.8362834 | 0.06631575 | 0.48798003 |
| PI4K2B | 83.3046206 | -0.353151332 | 0.37801508 | -0.9342255 | 0.35018757 | 0.75455562 |
| ZCCHC4 | 266.61575 | -0.06007887 | 0.22186524 | -0.2707899 | 0.78655261 | 0.9391356 |
| ANAPC4 | 544.257767 | -0.476077531 | 0.21403894 | -2.2242566 | 0.02613119 | 0.38392873 |
| SLC34A2 | 1185.42705 | 0.18316937 | 1.00963944 | 0.18142058 | 8.56E-01 | 0.9623313 |
| SEL1L3 | 2347.32757 | -0.2918379 | 0.32999413 | -0.884373 | 3.76E-01 | 0.77035529 |
| AC092436.4 | 35.0876815 | 0.825900668 | 0.54906656 | 1.50419043 | 1.33E-01 | 0.5912647 |
| SMIM20 | 268.556926 | -0.736695391 | 0.41799557 | -1.7624478 | 0.07799366 | 0.51192014 |
| RBPJ | 1801.78128 | -0.349369754 | 0.25976321 | -1.3449547 | 0.17863985 | 0.6288217 |
| TBC1D19 | 2629.63536 | 0.177478941 | 0.39967257 | 0.44406085 | 0.65699859 | 0.89053559 |
| STIM2 | 541.169151 | -0.026933924 | 0.27126139 | -0.0992914 | 0.9209069 | 0.97939204 |
| LINC02506 | 82.7947805 | 0.60930676 | 1.39177963 | 0.43778968 | 0.66153875 | 0.89184036 |
| LINC02484 | 73.5336419 | -0.008916648 | 0.8669579 | -0.010285 | 9.92E-01 | 9.97E-01 |
| ARAP2 | 1161.61486 | 0.096940004 | 0.26185887 | 0.37019943 | 0.7112339 | 0.91143287 |
| LINC02505 | 94.53735 | 4.388144293 | 1.26112081 | 3.47955901 | 0.00050224 | 0.08018476 |
| C4orf19 | 3441.49791 | -0.380882341 | 0.36231459 | -1.0512476 | 0.29314489 | 0.71446249 |
| RELL1 | 272.622364 | 0.383543905 | 0.38531646 | 0.99539975 | 3.20E-01 | 7.31E-01 |
| PGM2 | 136.829752 | -0.248221493 | 0.35630324 | -0.696658 | 0.48601686 | 0.81850757 |
| TBC1D1 | 2319.34538 | 0.344693636 | 0.23465988 | 1.46890739 | 0.14185791 | 0.59663896 |
| KLF3 | 630.98763 | 0.060962817 | 0.31553526 | 0.19320446 | 0.84679885 | 0.95831174 |
| TLR1 | 32.0848293 | 0.391064223 | 0.52640501 | 0.74289609 | 0.45754459 | 0.80561734 |
| FAM114A1 | 332.971638 | -0.562124194 | 0.38610418 | -1.4558873 | 0.14542376 | 0.60166723 |
| KLHL5 | 670.495497 | 0.411227078 | 0.24485363 | 1.67948123 | 0.09305829 | 0.53684765 |
| WDR19 | 1034.53475 | 0.042029912 | 0.17053892 | 0.24645349 | 0.8053312 | 0.94585363 |
| RFC1 | 869.239845 | 0.072977062 | 0.30090122 | 0.2425283 | 0.80837083 | 0.94625542 |
| RPL9 | 151.845317 | -0.334466161 | 0.41846527 | -0.7992686 | 0.42413466 | 0.78894887 |
| LIAS | 293.209145 | 0.714214882 | 0.40492052 | 1.76383968 | 0.07775898 | 0.51185769 |
| UGDH | 1301.8115 | -0.774407277 | 0.62676434 | -1.2355637 | 0.21662078 | 0.66032006 |
| UGDH-AS1 | 521.981037 | 0.253412379 | 0.44208146 | 0.57322553 | 0.566492 | 0.85216263 |
| SMIM14 | 1167.32275 | 0.00807363 | 0.30816622 | 0.02619895 | 0.97909866 | 0.99482978 |
| AC108471.2 | 41.8942781 | 0.89821813 | 0.70772229 | 1.2691675 | 0.20438133 | 0.64998953 |
| UBE2K | 3131.90132 | -0.471346472 | 0.29318819 | -1.6076584 | 0.10791 | 0.55721497 |
| PDS5A | 3218.14831 | -0.386495709 | 0.18672574 | -2.0698577 | 0.03846567 | 0.42447711 |
| N4BP2 | 494.36348 | -0.105811572 | 0.29052149 | -0.3642125 | 0.7156993 | 0.91298152 |
| RBM47 | 12402.0157 | -0.146937702 | 0.23153697 | -0.6346187 | 0.5256771 | 0.83638492 |
| NSUN7 | 723.598343 | 0.617075859 | 0.46875454 | 1.31641574 | 0.18803454 | 0.64002718 |
| APBB2 | 2136.9911 | -0.379719311 | 0.23935913 | -1.5864 | 0.11264861 | 0.56642498 |
| UCHL1 | 142.908161 | -0.758291168 | 0.56266512 | -1.3476776 | 1.78E-01 | 6.28E-01 |
| LIMCH1 | 2837.18871 | 0.276191887 | 0.42230628 | 0.65400847 | 0.51310635 | 0.83073024 |
| TMEM33 | 516.301985 | -0.482512921 | 0.40762202 | -1.1837263 | 0.23652141 | 0.67365231 |
| SLC30A9 | 1971.46987 | -0.101632545 | 0.20158874 | -0.5041579 | 0.61415045 | 0.87225883 |
| ATP8A1 | 346.989391 | 1.021914465 | 0.35114633 | 2.91022393 | 0.0036117 | 0.19706898 |
| KCTD8 | 1233.42158 | 0.016982807 | 0.39202233 | 0.04332102 | 0.96544564 | 0.99074368 |
| GUF1 | 285.441732 | 0.025947711 | 0.26261661 | 0.09880453 | 0.92129347 | 0.97939204 |

| | | | | | | |
|---|---|---|---|---|---|---|
| GNPDA2 | 236.08104 | 0.202180319 | 0.23318614 | 0.86703404 | 0.38592336 | 0.77317162 |
| COMMD8 | 123.164632 | 0.092942914 | 0.42134336 | 0.22058711 | 0.82541394 | 0.95192757 |
| ATP10D | 1006.73935 | -0.276225756 | 0.62488 | -0.4420461 | 0.65845586 | 0.89058983 |
| CORIN | 42.4008055 | -0.016273844 | 0.7052855 | -0.0230741 | 0.98159115 | 0.99516789 |
| AC107068.2 | 102.197452 | 0.572806637 | 0.61960816 | 0.92446594 | 0.35524378 | 0.75757147 |
| NFXL1 | 418.909048 | 0.146122763 | 0.24635595 | 0.59313672 | 0.55308966 | 0.84725078 |
| NIPAL1 | 154.011401 | -0.119291976 | 0.37431547 | -0.3186937 | 0.7499588 | 0.92565124 |
| TXK | 611.627479 | 0.386311021 | 0.40921715 | 0.94402452 | 0.34515712 | 0.75144038 |
| TEC | 908.230776 | -0.122896116 | 0.2395559 | -0.5130164 | 0.60793983 | 0.86985416 |
| SLAIN2 | 896.949156 | -0.275341803 | 0.2191075 | -1.2566516 | 0.20887981 | 0.65401126 |
| FRYL | 3419.06573 | -0.11273745 | 0.21400582 | -0.5267962 | 0.59833513 | 0.86646266 |
| OCIAD1 | 1381.26308 | 0.026896994 | 0.15456491 | 0.17401747 | 0.86185174 | 0.96336945 |
| OCIAD2 | 1125.83086 | -0.704025272 | 0.45544054 | -1.5458116 | 0.12215007 | 0.58055167 |
| CWH43 | 51.1999199 | -2.378821739 | 1.08025933 | -2.202084 | 0.02765938 | 0.38942007 |
| DCUN1D4 | 2245.31548 | -0.026769562 | 0.28295583 | -0.0946069 | 0.9246271 | 0.9804664 |
| LRRC66 | 173.041278 | 0.641134263 | 0.38732952 | 1.65526826 | 0.0978701 | 0.5411478 |
| SGCB | 406.947325 | 0.492416075 | 0.27851084 | 1.76803197 | 0.07705556 | 0.51013111 |
| SPATA18 | 333.908287 | 0.637859546 | 0.55895743 | 1.14115943 | 0.25380358 | 0.68891958 |
| USP46 | 325.037199 | -0.324361328 | 0.32992675 | -0.9831313 | 0.3255428 | 0.73706891 |
| USP46-AS1 | 34.2383204 | 0.220954884 | 0.51020571 | 0.43307019 | 0.66496378 | 0.89315615 |
| SCFD2 | 1900.98593 | -1.046314578 | 0.49315457 | -2.1216767 | 0.03386489 | 0.41378378 |
| AC058822.1 | 575.652969 | 0.195239725 | 0.54488825 | 0.3583115 | 0.72011021 | 0.9148113 |
| LNX1 | 273.569178 | 1.066058874 | 0.54027907 | 1.97316338 | 0.04847695 | 0.45329252 |
| CHIC2 | 1575.82581 | 0.179564584 | 0.24841896 | 0.72282963 | 0.46978456 | 0.81322606 |
| SRD5A3 | 57.910255 | -0.225876285 | 0.52153338 | -0.4331003 | 0.66494187 | 0.89315615 |
| TMEM165 | 1758.89745 | 0.043929244 | 0.20905828 | 0.21012918 | 8.34E-01 | 0.95381091 |
| CLOCK | 1346.42109 | 0.054547132 | 0.16869097 | 0.32335537 | 0.74642613 | 0.92448528 |
| EXOC1 | 1443.58111 | -0.260525047 | 0.21544069 | -1.2092658 | 0.22656076 | 0.66690202 |
| CEP135 | 933.848071 | -0.034301108 | 0.33071609 | -0.1037177 | 9.17E-01 | 9.78E-01 |
| KIAA1211 | 159.51755 | 0.620013264 | 0.75255575 | 0.82387686 | 0.41000953 | 0.7854648 |
| AASDH | 530.893636 | 0.679514429 | 0.36689806 | 1.85205241 | 0.0640183 | 0.48593331 |
| PPAT | 220.291261 | -0.851549045 | 0.45399969 | -1.87566 | 0.06070199 | 0.48026727 |
| PAICS | 258.766874 | -0.201477431 | 0.28740267 | -0.7010284 | 0.4832853 | 0.81782612 |
| SRP72 | 544.496663 | -0.22413521 | 0.27179078 | -0.8246608 | 0.40956417 | 0.78542936 |
| THEGL | 36.8261954 | 0.338225847 | 0.5262564 | 0.64270163 | 5.20E-01 | 0.833897 |
| REST | 665.83206 | -0.180128705 | 0.30414589 | -0.5922444 | 0.55368694 | 0.84727776 |
| NOA1 | 119.639733 | -0.099824965 | 0.37796687 | -0.2641104 | 0.79169488 | 0.94104411 |
| POLR2B | 833.940299 | -0.577055988 | 0.2918024 | -1.9775574 | 0.04797866 | 0.45138969 |
| IGFBP7 | 3061.5653 | 0.299813036 | 0.39024428 | 0.76827016 | 0.4423267 | 0.79928982 |
| AC093725.2 | 131.641391 | 1.081222692 | 0.97378282 | 1.11033248 | 0.26685578 | 0.69812269 |
| ADGRL3 | 77.8469524 | -0.815645718 | 0.54790671 | -1.488658 | 0.13657744 | 0.5942049 |
| AC116049.2 | 420.829237 | 0.373790991 | 0.70381932 | 0.53108942 | 0.59535682 | 0.86510861 |
| AC104806.2 | 217.282778 | 0.784043099 | 0.52295243 | 1.49926276 | 0.13380548 | 0.59240843 |
| CENPC | 630.634926 | -0.322938543 | 0.34056878 | -0.948233 | 0.34301087 | 0.7505827 |
| UBA6 | 1419.17843 | -0.339931871 | 0.22757389 | -1.4937209 | 1.35E-01 | 0.59381224 |
| UBA6-AS1 | 2649.63999 | 0.816213146 | 0.60303449 | 1.3535099 | 0.17589279 | 0.62699479 |
| TMPRSS11D | 38.7569161 | 1.439354365 | 0.98510247 | 1.46112147 | 0.14398211 | 0.60078014 |
| YTHDC1 | 1551.42254 | -0.104936254 | 0.24148741 | -0.4345413 | 0.66389541 | 0.89313476 |
| UGT2A3 | 1314.29841 | 0.223278516 | 0.64495147 | 0.34619429 | 0.72919671 | 0.91799222 |
| UGT2B7 | 9519.49082 | 0.344188427 | 0.56467739 | 0.60953109 | 0.54217247 | 0.84267582 |
| AC111000.4 | 1094.71389 | -1.450213514 | 0.64265257 | -2.2566058 | 0.02403272 | 0.37387451 |
| ENAM | 54.3619782 | 1.057431131 | 0.51696506 | 2.04545958 | 0.04080957 | 0.43095283 |
| UTP3 | 223.618386 | -0.276492175 | 0.33432851 | -0.8270075 | 0.40823283 | 0.78481094 |
| RUFY3 | 8565.02741 | -0.248932852 | 0.29662458 | -0.8392185 | 0.40134668 | 0.7824291 |

| | | | | | | |
|---|---|---|---|---|---|---|
| GRSF1 | 583.243243 | -0.205431649 | 0.20065891 | -1.0237853 | 0.30593668 | 0.72139585 |
| MOB1B | 1472.80915 | -0.253562206 | 0.26628315 | -0.9522277 | 0.34098149 | 0.74757411 |
| SLC4A4 | 11456.5227 | 0.344917089 | 0.60036456 | 0.57451274 | 0.56562088 | 0.85194709 |
| GC | 253.44758 | 1.111486942 | 1.44633369 | 0.76848583 | 0.44219861 | 0.79928982 |
| ADAMTS3 | 206.098305 | 0.642748816 | 0.5823743 | 1.1036696 | 0.26973649 | 0.69961625 |
| COX18 | 116.132127 | 0.529005216 | 0.29168139 | 1.81364061 | 0.06973307 | 0.49371449 |
| ANKRD17 | 6456.05823 | -0.292072673 | 0.27007291 | -1.0814586 | 0.27949317 | 0.70446562 |
| AC053527.1 | 953.345462 | 1.469013588 | 0.60766676 | 2.41746577 | 0.015629 | 0.31503934 |
| ALB | 135.243771 | 2.55190466 | 1.66534497 | 1.53235799 | 0.12543412 | 0.58564339 |
| AFM | 1164.15911 | 0.390436516 | 0.89203265 | 0.43769308 | 6.62E-01 | 8.92E-01 |
| LINC02499 | 995.674344 | 0.108856198 | 1.43143653 | 0.07604682 | 9.39E-01 | 9.84E-01 |
| RASSF6 | 1852.95854 | 0.740723411 | 0.3036419 | 2.43946373 | 1.47E-02 | 3.08E-01 |
| CXCL1 | 93.2469523 | 1.121934815 | 1.10826567 | 1.01233382 | 0.31137847 | 0.72586134 |
| CXCL2 | 194.534403 | 0.642044322 | 0.70179151 | 0.91486476 | 3.60E-01 | 7.60E-01 |
| MTHFD2L | 652.24343 | 0.226547806 | 0.27645174 | 0.81948409 | 0.41251027 | 0.78632759 |
| BTC | 91.2417506 | 0.085329989 | 0.75968642 | 0.11232265 | 0.91056759 | 0.97697493 |
| PARM1 | 1258.65148 | 0.079307766 | 0.47937086 | 0.16544136 | 0.8685966 | 0.96536557 |
| AC110760.1 | 48.2624824 | 0.625005793 | 0.95405937 | 0.65510157 | 0.51240236 | 0.83055441 |
| RCHY1 | 465.099074 | 0.067358874 | 0.21574286 | 0.31221833 | 0.7548746 | 0.92799282 |
| THAP6 | 314.46466 | -0.276273398 | 0.20934907 | -1.3196782 | 0.18694248 | 0.6378941 |
| CDKL2 | 991.664825 | 0.686535276 | 0.55475316 | 1.23755091 | 0.21588263 | 0.65967206 |
| G3BP2 | 837.469225 | -0.129173507 | 0.20688249 | -0.6243811 | 0.53237736 | 0.83819165 |
| USO1 | 1499.98906 | 0.199042586 | 0.24506417 | 0.81220598 | 0.41667345 | 0.78632759 |
| NAAA | 143.874257 | 0.43947704 | 0.28051436 | 1.56668288 | 0.11718883 | 0.57350419 |
| SDAD1 | 567.103231 | -0.086296077 | 0.2892955 | -0.2982973 | 0.76547624 | 0.93091853 |
| NUP54 | 602.085393 | -0.236340471 | 0.16894792 | -1.3988954 | 0.16184435 | 0.61845523 |
| SCARB2 | 1089.40453 | -0.111146994 | 0.2580355 | -0.430743 | 0.66665523 | 0.89420131 |
| FAM47E-STB | 41.8563274 | 1.008469623 | 0.56298641 | 1.79128591 | 0.07324743 | 0.50459544 |
| CCDC158 | 232.76111 | 1.279903627 | 0.72963142 | 1.75417832 | 0.07939996 | 0.51192014 |
| SHROOM3 | 2162.17475 | 0.738645005 | 0.56452871 | 1.30842771 | 0.19072828 | 0.64281105 |
| 11-Sep | 2219.83032 | -0.204321003 | 0.40767727 | -0.5011832 | 0.61624219 | 0.87341864 |
| CCNI | 1418.43146 | 0.225527386 | 0.26295489 | 0.85766568 | 0.3910771 | 0.77606489 |
| CCNG2 | 649.151776 | -0.08376169 | 0.52633425 | -0.1591416 | 0.87355729 | 0.96592775 |
| CXCL13 | 46.8262897 | -0.206225943 | 0.69829881 | -0.2953262 | 0.76774469 | 0.93180413 |
| CNOT6L | 965.004862 | 0.266560219 | 0.25939341 | 1.02762912 | 0.30412432 | 0.72082426 |
| MRPL1 | 593.221703 | 0.002534825 | 0.18690191 | 0.01356233 | 0.98917916 | 0.99721497 |
| FRAS1 | 3560.89671 | 0.494824919 | 0.47055058 | 1.05158709 | 0.29298903 | 0.71433665 |
| ANXA3 | 110.527486 | -0.153765025 | 0.43148036 | -0.3563662 | 0.72156633 | 0.91553361 |
| LINC01094 | 55.480208 | 0.645608057 | 0.92743652 | 0.69612102 | 4.86E-01 | 0.81868653 |
| BMP2K | 2948.10431 | -0.225671366 | 0.32473646 | -0.6949369 | 0.48709482 | 0.81893983 |
| PAQR3 | 183.846864 | -0.30591264 | 0.43941764 | -0.6961774 | 4.86E-01 | 0.81868653 |
| LINC01088 | 108.831537 | -0.200722921 | 0.66588512 | -0.3014378 | 0.7630807 | 0.93023365 |
| ANTXR2 | 302.186618 | 0.941514897 | 0.34540026 | 2.72586618 | 0.0064133 | 0.22797255 |
| FGF5 | 116.17945 | 1.004984243 | 0.78783099 | 1.27563431 | 0.20208482 | 0.64881821 |
| CFAP299 | 539.960972 | 0.970098753 | 0.75887275 | 1.27834179 | 0.20112894 | 0.64849129 |
| PRKG2 | 178.68665 | 1.911705562 | 0.59007823 | 3.23974936 | 1.20E-03 | 1.12E-01 |
| RASGEF1B | 5671.61988 | -0.864142348 | 0.57404309 | -1.5053615 | 0.13223117 | 0.59115873 |
| HNRNPD | 1500.28582 | -0.375734492 | 0.20635493 | -1.8208166 | 0.06863474 | 0.49154315 |
| AC124016.1 | 39.342769 | -0.297559906 | 0.54179446 | -0.5492118 | 0.58286011 | 0.85986705 |
| HNRNPDL | 1406.44702 | -0.247629264 | 0.22054238 | -1.1228194 | 0.2615142 | 0.69374398 |
| ENOPH1 | 275.029267 | -0.562423279 | 0.28105132 | -2.0011409 | 0.04537721 | 0.44360789 |
| TMEM150C | 474.537504 | 0.885598304 | 0.59437858 | 1.48995662 | 0.13623564 | 0.5942049 |
| SCD5 | 575.610474 | 0.878520959 | 0.37295966 | 2.3555388 | 0.01849587 | 0.33938799 |
| SEC31A | 3466.10847 | -0.356003241 | 0.27435891 | -1.2975822 | 0.19443094 | 0.64548758 |

| | | | | | | |
|---|---|---|---|---|---|---|
| THAP9 | 138.829544 | -0.18715212 | 0.72939064 | -0.256587 | 0.79749765 | 0.94308518 |
| LIN54 | 608.072574 | -0.174076083 | 0.42018683 | -0.4142826 | 0.67866717 | 0.8992258 |
| COPS4 | 299.135466 | -0.066191658 | 0.2047408 | -0.3232949 | 0.74647191 | 0.92448528 |
| COQ2 | 39.9293923 | -0.400955884 | 0.41569811 | -0.9645362 | 0.33477716 | 0.74154392 |
| HELQ | 436.946579 | 0.066705455 | 0.29138299 | 0.22892707 | 0.8189256 | 0.94990455 |
| MRPS18C | 181.905096 | -0.130211032 | 0.35076596 | -0.3712191 | 0.71047433 | 0.91123308 |
| ABRAXAS1 | 145.757278 | 0.317981974 | 0.37039241 | 0.85850025 | 0.39061629 | 0.77606489 |
| GPAT3 | 4296.28069 | -1.280666253 | 0.49539374 | -2.5851482 | 0.00973371 | 0.26315585 |
| CDS1 | 526.927483 | -0.196923003 | 0.34877514 | -0.5646131 | 0.57233697 | 0.85408471 |
| WDFY3 | 9226.06813 | -0.0308108 | 0.34836543 | -0.0884439 | 9.30E-01 | 0.98272895 |
| WDFY3-AS1 | 52.4251733 | 0.031933808 | 0.82373321 | 0.03876717 | 0.96907602 | 0.99187457 |
| ARHGAP24 | 4161.25654 | 0.376940738 | 0.49641651 | 0.75932353 | 0.44765904 | 0.80223612 |
| MAPK10 | 5729.03379 | 0.733695608 | 0.62291871 | 1.17783524 | 0.2388623 | 0.67667639 |
| AC104827.1 | 93.534376 | 0.7635695 | 0.72535741 | 1.05268037 | 0.29248751 | 0.71399546 |
| PTPN13 | 2478.83265 | 0.284689938 | 0.38760703 | 0.73448085 | 0.46265573 | 0.8082072 |
| AC093827.5 | 231.705987 | -1.263278957 | 0.57368894 | -2.2020277 | 0.02766335 | 0.38942007 |
| AFF1 | 8850.28798 | -0.900208967 | 0.36562273 | -2.4621253 | 0.01381164 | 0.30363654 |
| KLHL8 | 653.232386 | 0.074365022 | 0.14074866 | 0.52835333 | 5.97E-01 | 8.66E-01 |
| HSD17B11 | 403.375223 | 0.611459677 | 0.36785409 | 1.66223429 | 0.09646581 | 0.53969191 |
| NUDT9 | 631.86611 | -0.31567351 | 0.22817475 | -1.3834726 | 0.16652001 | 0.62216414 |
| SPARCL1 | 114.638545 | 0.405624064 | 0.61583266 | 0.65865955 | 0.51011442 | 0.82923841 |
| SPP1 | 39409.4048 | 0.513716524 | 0.75228379 | 0.68287597 | 0.49468522 | 0.8220963 |
| PKD2 | 2924.60436 | 0.078564988 | 0.2909993 | 0.26998343 | 0.787173 | 0.93950968 |
| ABCG2 | 84.7608591 | 0.195850795 | 0.93028709 | 0.21052726 | 0.83325618 | 0.95355066 |
| PPM1K | 558.748865 | -0.034206692 | 0.48604299 | -0.0703779 | 0.94389287 | 0.98522232 |
| HERC6 | 228.557337 | -2.026518139 | 0.93630088 | -2.1643877 | 0.0304346 | 0.40391981 |
| HERC5 | 280.860956 | -1.145669177 | 0.72966978 | -1.5701201 | 0.11638718 | 0.57350419 |
| PYURF | 135.34643 | -0.316823359 | 0.33801041 | -0.9373183 | 0.34859483 | 0.75377651 |
| AC098582.1 | 47.0440055 | 0.846178373 | 0.90144817 | 0.93868777 | 0.34789107 | 0.75308638 |
| HERC3 | 989.530917 | -0.448712464 | 0.41303454 | -1.08638 | 0.27731091 | 0.7036882 |
| FAM13A-AS1 | 403.027714 | -0.432697973 | 0.25973747 | -1.6659051 | 0.09573232 | 0.538613 |
| FAM13A | 3347.54177 | -0.107424831 | 0.37058506 | -0.289879 | 0.7719088 | 0.93340577 |
| TIGD2 | 72.9569896 | -0.028206869 | 0.33204652 | -0.0849485 | 0.9323023 | 0.98306798 |
| GPRIN3 | 279.203272 | 1.005717671 | 0.70669534 | 1.42312764 | 0.15469916 | 0.61119904 |
| CCSER1 | 6726.41308 | 0.346858476 | 0.37156716 | 0.93350143 | 0.35056113 | 0.75493189 |
| GRID2 | 55.7428825 | -0.252165181 | 0.42214232 | -0.5973464 | 0.55027615 | 0.84620747 |
| SMARCAD1 | 910.446061 | 0.033504453 | 0.20862207 | 0.1605988 | 0.8724094 | 0.96578281 |
| PDLIM5 | 4042.90279 | 0.120177432 | 0.2285336 | 0.52586329 | 0.5989832 | 0.86671329 |
| BMPR1B | 117.420377 | 0.258633431 | 0.63711894 | 0.40594215 | 0.68478511 | 0.90098343 |
| STPG2 | 3280.97694 | 0.805023005 | 0.59033829 | 1.36366388 | 1.73E-01 | 0.62593809 |
| STPG2-AS1 | 58.2804961 | 0.67908466 | 0.97606556 | 0.69573673 | 0.48659372 | 0.81877553 |
| RAP1GDS1 | 1545.8566 | -0.391221495 | 0.28440743 | -1.3755671 | 0.16895571 | 0.62304975 |
| TSPAN5 | 73.414871 | -1.283019447 | 0.38355457 | -3.3450767 | 0.0008226 | 0.09555385 |
| EIF4E | 928.056239 | -0.854405387 | 0.51107329 | -1.6717864 | 0.09456645 | 0.53824794 |
| AC019131.1 | 50.4982658 | 0.913509855 | 0.8881002 | 1.02861125 | 0.30366239 | 0.72082426 |
| METAP1 | 527.126318 | -0.067622354 | 0.21558303 | -0.313672 | 0.75377019 | 0.92783044 |
| ADH5 | 635.305351 | 0.193529965 | 0.26520313 | 0.72974238 | 0.46554767 | 0.81042189 |
| AP002026.1 | 133.560012 | 1.129586877 | 0.61627571 | 1.83292455 | 0.06681379 | 0.48884194 |
| ADH6 | 371.876398 | 2.057190063 | 0.75047316 | 2.74119073 | 0.0061217 | 0.22637955 |
| TRMT10A | 135.530473 | -0.364814334 | 0.37977867 | -0.9605972 | 0.33675473 | 0.74293725 |
| MTTP | 240.232308 | 1.806919011 | 0.76994207 | 2.34682461 | 0.01893416 | 0.34084276 |
| LAMTOR3 | 252.261788 | -0.394787562 | 0.31044977 | -1.2716633 | 0.20349279 | 0.64881965 |
| DNAJB14 | 1198.61093 | 0.122055055 | 0.15664115 | 0.77920171 | 0.4358609 | 0.79549938 |
| H2AFZ | 365.420887 | -0.505686029 | 0.62094855 | -0.8143767 | 0.41542918 | 0.78632759 |

| | | | | | | |
|---|---|---|---|---|---|---|
| DDIT4L | 163.647417 | -2.345865304 | 1.1152079 | -2.1035229 | 0.03542009 | 0.4198859 |
| EMCN | 97.8868851 | -0.34574327 | 0.58002746 | -0.5960809 | 0.55112121 | 0.8466552 |
| PPP3CA | 4044.15392 | -0.087761427 | 0.26786351 | -0.3276349 | 0.74318775 | 0.92333113 |
| BANK1 | 622.005683 | 0.749117994 | 0.80937513 | 0.92555105 | 0.35467934 | 0.75755608 |
| AP002075.1 | 144.091213 | 1.280207273 | 0.65508028 | 1.9542754 | 0.05066866 | 0.45903191 |
| SLC39A8 | 684.521112 | 0.147325412 | 0.52467123 | 0.28079568 | 0.77886712 | 0.93646866 |
| NFKB1 | 1518.6112 | 0.232618504 | 0.30722101 | 0.7571699 | 0.44894808 | 0.80300352 |
| MANBA | 1072.51583 | -0.338622449 | 0.26958291 | -1.2560976 | 0.20908058 | 0.65403148 |
| UBE2D3 | 3126.61103 | -0.280816964 | 0.23204382 | -1.2101893 | 0.22620624 | 0.66688879 |
| AC018797.2 | 92.0627741 | -0.19756528 | 0.38769819 | -0.5095853 | 0.61034206 | 0.87109318 |
| CISD2 | 446.503297 | -0.19427254 | 0.34115423 | -0.5694566 | 0.5690463 | 0.85291608 |
| SLC9B1 | 104.074225 | -0.80097238 | 0.51010372 | -1.5702147 | 0.11636516 | 0.57350419 |
| SLC9B2 | 192.594144 | -0.102179721 | 0.37213635 | -0.274576 | 0.78364198 | 0.93860369 |
| BDH2 | 312.252825 | 0.696514191 | 0.65162687 | 1.06888501 | 0.28512149 | 0.71015169 |
| CXXC4 | 35.7084358 | 1.391399115 | 0.93922382 | 1.48143508 | 0.13849067 | 0.59474076 |
| CXXC4-AS1 | 119.013613 | 1.063755454 | 1.08103506 | 0.98401568 | 0.3251078 | 0.73628585 |
| AC004053.1 | 2800.43719 | 0.699867742 | 0.85374546 | 0.81976159 | 0.41235203 | 0.78632759 |
| AC096577.1 | 6024.61401 | 1.381803997 | 1.04178956 | 1.32637535 | 0.1847154 | 0.63412875 |
| TET2 | 2017.41333 | -0.231762865 | 0.4712204 | -0.4918354 | 0.62283573 | 0.87714058 |
| PPA2 | 1946.16935 | 0.548591525 | 0.24413634 | 2.24707032 | 0.02463553 | 0.37729526 |
| ARHGEF38 | 647.416733 | 1.39074008 | 0.69202094 | 2.0096792 | 0.04446515 | 0.44360789 |
| INTS12 | 375.523389 | -0.013351217 | 0.21385928 | -0.0624299 | 0.95022047 | 0.98724111 |
| GSTCD | 591.678746 | 0.350256513 | 0.23278325 | 1.50464653 | 0.13241498 | 0.5912647 |
| NPNT | 427.985326 | 0.133324514 | 0.84522971 | 0.15773761 | 0.87466356 | 0.96615698 |
| AC109361.1 | 40.1774076 | 0.452282273 | 0.66443829 | 0.68069869 | 0.49606217 | 0.82243887 |
| TBCK | 3072.46916 | 0.78269331 | 0.45281136 | 1.72851958 | 0.08389511 | 0.52128301 |
| AIMP1 | 470.735888 | -0.155911545 | 0.21074396 | -0.739815 | 0.45941225 | 0.80669065 |
| PAPSS1 | 464.439752 | 0.541766232 | 0.28453446 | 1.90404436 | 0.05690441 | 0.4746515 |
| SGMS2 | 1088.88152 | -0.892183011 | 0.4539886 | -1.9652102 | 0.04938992 | 0.45419182 |
| AC096564.1 | 156.650781 | -0.175739106 | 0.63422798 | -0.2770914 | 0.78170994 | 0.93751843 |
| CYP2U1 | 39.8688468 | 0.490566615 | 0.48651085 | 1.00833643 | 0.31329298 | 0.72717675 |
| HADH | 417.852438 | 0.770550011 | 0.43423931 | 1.7744824 | 0.07598339 | 0.50730849 |
| RPL34-AS1 | 318.332864 | 0.394464127 | 0.46604704 | 0.84640411 | 0.39732735 | 0.7803648 |
| RPL34 | 893.500173 | 0.034972401 | 0.47997368 | 0.07286316 | 0.94191501 | 0.98489601 |
| OSTC | 461.791906 | -0.249162903 | 0.37172217 | -0.6702934 | 0.50267079 | 0.82638105 |
| COL25A1 | 59.4136845 | 0.239847515 | 0.40188139 | 0.59681169 | 0.55063311 | 0.84651503 |
| SEC24B-AS1 | 81.0027316 | 1.11181479 | 0.64131191 | 1.73365685 | 0.08297899 | 0.51935659 |
| SEC24B | 1706.9293 | -0.116920691 | 0.17693015 | -0.6608297 | 0.50872156 | 0.82874455 |
| MCUB | 138.041983 | -1.173200232 | 0.66425705 | -1.7661841 | 7.74E-02 | 5.11E-01 |
| CASP6 | 377.809469 | 0.357229229 | 0.4531303 | 0.78835873 | 0.4304869 | 0.79055252 |
| PLA2G12A | 403.656558 | 0.426099634 | 0.4292609 | 0.99263556 | 0.32088759 | 0.73233537 |
| AC126283.2 | 251.032467 | 1.062095429 | 0.73623001 | 1.44261361 | 0.14912935 | 0.60622442 |
| CFI | 35.2176447 | -0.147408073 | 0.77588946 | -0.1899859 | 0.84932016 | 0.95959925 |
| GAR1 | 171.286935 | -1.176320915 | 0.49345426 | -2.38385 | 0.01713259 | 0.32964126 |
| RRH | 34.9301358 | -0.580290934 | 0.79693759 | -0.728151 | 0.46652113 | 0.81098649 |
| EGF | 939.01532 | 0.651355448 | 0.67490212 | 0.96511098 | 0.33448923 | 0.74120095 |
| ELOVL6 | 778.801224 | 1.308972602 | 0.27041734 | 4.84056468 | 1.29E-06 | 0.00591552 |
| ENPEP | 5436.94734 | 0.559494299 | 0.57252446 | 0.97724087 | 0.32844992 | 0.73826734 |
| PANCR | 86.1170591 | 2.268783895 | 1.02336962 | 2.21697405 | 0.02662486 | 0.38563341 |
| PITX2 | 27.961719 | 1.874829081 | 0.91619846 | 2.04631328 | 0.04072556 | 0.43086475 |
| FAM241A | 175.533851 | -0.739837884 | 0.54529999 | -1.3567539 | 0.17485942 | 0.62699479 |
| AP1AR | 922.910444 | -0.227294075 | 0.29220281 | -0.7778641 | 0.43664914 | 0.79558019 |
| TIFA | 66.9766492 | 1.050145421 | 0.60897874 | 1.72443691 | 8.46E-02 | 0.52183702 |
| ALPK1 | 1470.09585 | 0.991988902 | 0.44925361 | 2.20808218 | 0.02723854 | 0.38702446 |

| | | | | | | |
|---|---|---|---|---|---|---|
| ZGRF1 | 74.2542533 | -0.830121375 | 0.42855502 | -1.9370241 | 0.0527424 | 0.46564998 |
| LARP7 | 509.393792 | -0.080507792 | 0.19454975 | -0.413816 | 0.6790089 | 0.89932032 |
| ANK2 | 3462.1345 | -0.70480748 | 0.72739531 | -0.968947 | 0.33257165 | 0.73988623 |
| CAMK2D | 3057.39529 | -0.723777536 | 0.27466635 | -2.6351155 | 0.00841087 | 0.25062554 |
| ARSJ | 855.41168 | 0.020844348 | 0.39251734 | 0.05310427 | 0.95764883 | 0.98861313 |
| UGT8 | 277.98967 | 0.265524191 | 0.45127796 | 0.5883828 | 0.55627538 | 0.84861484 |
| SNHG8 | 268.138212 | -0.050446722 | 0.51226657 | -0.0984775 | 0.92155315 | 0.97939204 |
| PRSS12 | 75.8509212 | -0.022694837 | 0.85602001 | -0.026512 | 0.97884893 | 0.99482978 |
| METTL14 | 377.003499 | -0.092151657 | 0.17731991 | -0.5196916 | 0.60327857 | 0.8678002 |
| SEC24D | 835.082337 | -0.14225916 | 0.39693809 | -0.3583913 | 7.20E-01 | 9.15E-01 |
| USP53 | 1910.52985 | -0.297575622 | 0.21382455 | -1.3916812 | 0.16401894 | 0.62065658 |
| C4orf3 | 633.011489 | -0.270032408 | 0.49900286 | -0.541144 | 0.58840832 | 0.86140264 |
| PDE5A | 462.941452 | 0.946441533 | 0.47454766 | 1.99440774 | 0.04610751 | 0.44508419 |
| AC108866.1 | 318.831347 | 0.738952874 | 0.51527601 | 1.43409137 | 0.15154619 | 0.60934105 |
| PRDM5 | 94.6527862 | 1.37582375 | 0.81860309 | 1.68069699 | 0.09282179 | 0.53669957 |
| QRFPR | 127.58335 | 1.151901273 | 0.82542837 | 1.39551937 | 0.16285928 | 0.6197424 |
| ANXA5 | 1005.24684 | -0.378228202 | 0.55688417 | -0.6791865 | 4.97E-01 | 0.82278993 |
| EXOSC9 | 93.4491621 | -0.340459905 | 0.30114274 | -1.1305599 | 0.25824037 | 0.69210282 |
| BBS7 | 231.896495 | -0.050476284 | 0.30652727 | -0.1646714 | 0.86920261 | 0.96556934 |
| KIAA1109 | 2945.20335 | 0.437120301 | 0.29032679 | 1.50561475 | 0.1321661 | 0.59115873 |
| BBS12 | 117.233344 | 0.599805132 | 0.37702659 | 1.59088286 | 0.11163594 | 0.56443177 |
| FGF2 | 110.16324 | -1.072721716 | 0.86954788 | -1.2336546 | 0.21733165 | 0.66032006 |
| NUDT6 | 104.768271 | 0.405648964 | 0.42480722 | 0.95490129 | 0.33962762 | 0.74644675 |
| SPATA5 | 1509.66249 | -0.749893417 | 0.36640565 | -2.0466208 | 0.04069533 | 0.43086475 |
| SPRY1 | 125.340293 | -0.152015237 | 0.47610558 | -0.3192889 | 0.74950743 | 0.92565124 |
| LINC01091 | 225.006667 | 1.027667506 | 0.7459342 | 1.3776919 | 0.16829843 | 0.62304975 |
| ANKRD50 | 267.183575 | -0.169760978 | 0.44318248 | -0.3830498 | 0.70168281 | 0.90803937 |
| FAT4 | 375.139639 | -0.24102302 | 0.39306545 | -0.613188 | 0.53975203 | 0.84120308 |
| AC093772.1 | 72.9295402 | 3.400971067 | 0.85204924 | 3.9915194 | 6.57E-05 | 0.02490203 |
| INTU | 641.64948 | 0.40339126 | 0.3662929 | 1.10128059 | 0.27077456 | 0.69961625 |
| HSPA4L | 1213.50785 | -0.289188681 | 0.56950458 | -0.5077899 | 0.61160068 | 0.8717981 |
| MFSD8 | 569.988588 | -0.117385369 | 0.17723871 | -0.662301 | 0.50777835 | 0.82868411 |
| ABHD18 | 1507.26799 | -0.141085147 | 0.14452504 | -0.9761987 | 0.32896603 | 0.73829083 |
| LARP1B | 3106.58161 | 0.341705803 | 0.25741204 | 1.32746627 | 0.18435449 | 0.63377801 |
| PGRMC2 | 903.12024 | -0.18026868 | 0.18239058 | -0.9883662 | 0.32297335 | 0.7350762 |
| AC108062.1 | 1049.45611 | 0.308545996 | 0.34637582 | 0.89078388 | 0.37304512 | 0.76930513 |
| AC078850.2 | 279.923835 | 0.842152774 | 0.87632838 | 0.96100137 | 3.37E-01 | 7.43E-01 |
| JADE1 | 912.114095 | -0.350280397 | 0.27189205 | -1.2883069 | 1.98E-01 | 0.64558745 |
| SCLT1 | 796.740237 | 0.045332778 | 0.32757265 | 0.13838999 | 0.8899322 | 0.97213108 |
| C4orf33 | 94.9402994 | 0.181275475 | 0.6838852 | 0.26506711 | 0.79095776 | 0.94078949 |
| SNHG27 | 65.3981764 | 1.907447535 | 0.91691521 | 2.08028781 | 0.03749914 | 0.42215481 |
| AC096711.2 | 665.969718 | 0.110805514 | 0.41216695 | 0.26883649 | 0.78805551 | 0.93953874 |
| SLC7A11 | 188.426371 | -2.333731725 | 1.0024406 | -2.3280499 | 1.99E-02 | 3.47E-01 |
| AC109927.1 | 101.226419 | 0.756394266 | 0.53262924 | 1.42011404 | 0.15557448 | 0.61208693 |
| NOCT | 536.314108 | -0.577239354 | 0.40527985 | -1.4242982 | 0.15436017 | 0.61099212 |
| ELF2 | 2751.77996 | 0.127605848 | 0.17080701 | 0.74707618 | 0.45501757 | 0.80465616 |
| NDUFC1 | 420.797727 | -0.375305514 | 0.36116688 | -1.0391471 | 0.29873633 | 0.71852421 |
| NAA15 | 1181.3968 | -0.208564109 | 0.30626688 | -0.6809881 | 0.495879 | 0.82243887 |
| AC097376.2 | 1692.79454 | -0.125816309 | 0.31461655 | -0.3999037 | 0.68922748 | 0.90367 |
| RAB33B | 120.036232 | 0.337106013 | 0.27173254 | 1.24058023 | 0.21476086 | 0.65925411 |
| SETD7 | 260.377334 | -0.017451682 | 0.54284582 | -0.0321485 | 9.74E-01 | 9.94E-01 |
| MGST2 | 647.818489 | 0.679132753 | 0.42664352 | 1.59180374 | 0.11142881 | 0.56435572 |
| MAML3 | 1628.75552 | 1.326508366 | 0.47043497 | 2.81974866 | 0.00480613 | 0.21388829 |
| SCOC | 587.71849 | 0.222951473 | 0.36077464 | 0.61797989 | 0.5365886 | 0.83980588 |

| | | | | | | |
|---|---|---|---|---|---|---|
| SCOC-AS1 | 117.261285 | 0.41926556 | 0.60634734 | 0.69146104 | 0.48927586 | 0.81970016 |
| ELMOD2 | 370.740686 | 0.22020417 | 0.27606255 | 0.79766041 | 0.42506758 | 0.78894887 |
| TBC1D9 | 257.571517 | -0.202269278 | 0.39464186 | -0.5125388 | 0.60827395 | 0.86985416 |
| RNF150 | 478.087173 | 0.380274813 | 0.92828039 | 0.40965512 | 0.68205896 | 0.89962501 |
| ZNF330 | 330.77925 | 0.057816514 | 0.33122416 | 0.17455403 | 0.86143007 | 0.96302275 |
| AC107220.1 | 154.309589 | -0.630670947 | 1.01447692 | -0.6216711 | 5.34E-01 | 8.39E-01 |
| LINC02432 | 90.3756193 | -0.22819988 | 0.74345845 | -0.3069437 | 0.75888623 | 0.9290866 |
| IL15 | 782.020747 | -0.34044693 | 0.29035895 | -1.1725036 | 0.24099491 | 0.67957469 |
| INPP4B | 602.079774 | 0.700291615 | 0.55728592 | 1.25661099 | 0.20889453 | 0.65401126 |
| AC104596.1 | 137.021531 | 0.811575977 | 0.79397892 | 1.02216313 | 0.3067037 | 0.72190689 |
| USP38 | 396.323872 | -0.249936022 | 0.30597156 | -0.8168603 | 0.41400827 | 0.78632759 |
| GAB1 | 2533.39678 | -0.543451442 | 0.28271399 | -1.9222658 | 0.05457232 | 0.46772575 |
| SMARCA5 | 587.389987 | -0.072109802 | 0.20370253 | -0.3539956 | 0.72334218 | 0.91592159 |
| GYPA | 63.5850798 | 1.0751558 | 0.84294236 | 1.27547962 | 0.20213953 | 0.64881821 |
| ANAPC10 | 1006.05025 | 0.033929436 | 0.1585026 | 0.21406233 | 0.83049845 | 0.95325673 |
| ABCE1 | 367.32604 | -0.356596599 | 0.30031475 | -1.1874095 | 0.23506612 | 0.67119484 |
| OTUD4 | 866.75638 | 0.075366258 | 0.14633293 | 0.51503281 | 0.60653009 | 0.86963473 |
| SMAD1 | 728.503716 | 0.420222662 | 0.30500987 | 1.37773465 | 0.16828523 | 0.62304975 |
| MMAA | 308.585948 | 0.510383291 | 0.35869904 | 1.42287333 | 0.15477288 | 0.61119904 |
| C4orf51 | 59.8874105 | 0.849567519 | 0.81122639 | 1.04726317 | 0.29497822 | 0.71646902 |
| ZNF827 | 2245.43009 | 0.607343944 | 0.41192426 | 1.47440684 | 0.14037208 | 0.59500405 |
| LSM6 | 187.054122 | -0.01160232 | 0.25978082 | -0.044662 | 0.96437676 | 0.99031076 |
| SLC10A7 | 773.210434 | 0.082520405 | 0.2524413 | 0.32688947 | 0.74375148 | 0.92350771 |
| TMEM184C | 364.247443 | -0.186042459 | 0.26842695 | -0.6930841 | 0.48825676 | 0.81931938 |
| PRMT9 | 476.878572 | 0.294132605 | 0.23257705 | 1.26466734 | 0.2059906 | 0.651931 |
| ARHGAP10 | 3430.21794 | -0.410447564 | 0.18586049 | -2.2083637 | 0.02721893 | 0.38702446 |
| NR3C2 | 2020.53878 | 0.205881803 | 0.46108152 | 0.44651932 | 0.65522216 | 0.89053559 |
| AC069272.1 | 46.3066823 | 0.315762132 | 0.52505495 | 0.60138874 | 0.5475811 | 0.84485526 |
| DCLK2 | 257.418404 | -0.407230681 | 0.61063652 | -0.6668954 | 0.50483895 | 0.82715271 |
| LRBA | 9843.65995 | 0.118898554 | 0.20460671 | 0.58110779 | 0.56116781 | 0.84958759 |
| RPS3A | 506.177448 | 0.295652057 | 0.47987684 | 0.61609987 | 0.53782861 | 0.84040975 |
| SH3D19 | 4382.41339 | -1.055987314 | 0.37649219 | -2.8048054 | 0.0050347 | 0.21686099 |
| PRSS48 | 114.480359 | -0.468953067 | 0.60082218 | -0.7805189 | 0.4350855 | 0.79515776 |
| FAM160A1 | 3392.0811 | -0.373409948 | 0.26611181 | -1.403207 | 0.16055513 | 0.61627813 |
| AC092611.2 | 49.5143928 | -0.374846154 | 0.46921564 | -0.7988782 | 0.42436103 | 0.78894887 |
| GATB | 272.00636 | -0.030562374 | 0.29239693 | -0.1045236 | 0.91675386 | 0.97828877 |
| AC079340.1 | 51.3730877 | 0.987884041 | 1.09303137 | 0.9038021 | 0.36610035 | 0.76421682 |
| FBXW7 | 2721.40231 | -0.082990081 | 0.23992894 | -0.3458944 | 0.72942207 | 0.9181916 |
| TIGD4 | 59.2206833 | 0.962192938 | 0.52720758 | 1.82507416 | 0.06798984 | 0.4900219 |
| ARFIP1 | 1291.4898 | 0.146167128 | 0.16251074 | 0.89943057 | 0.36842336 | 0.76615653 |
| FHDC1 | 68.7410482 | 0.059320094 | 0.68758791 | 0.08627274 | 9.31E-01 | 0.98272895 |
| TRIM2 | 926.423775 | 0.348654808 | 0.40570801 | 0.85937375 | 0.39013435 | 0.77569824 |
| TMEM131L | 906.601357 | 0.549134332 | 0.43265594 | 1.26921715 | 0.20436363 | 0.64998953 |
| TLR2 | 162.003238 | 0.599476167 | 0.57224769 | 1.04758164 | 0.2948314 | 0.71646902 |
| PLRG1 | 242.987324 | -0.215620905 | 0.27863193 | -0.7738557 | 0.43901613 | 0.7974071 |
| FGB | 1704.49405 | 0.948669939 | 1.62164873 | 0.58500335 | 0.55854546 | 0.84906096 |
| FGA | 695.921094 | 0.664924813 | 1.43524661 | 0.46328262 | 6.43E-01 | 8.85E-01 |
| LRAT | 441.111268 | -0.276708927 | 0.55008004 | -0.5030339 | 0.61494041 | 0.87260075 |
| MAP9 | 257.536424 | 0.616360064 | 0.32468955 | 1.89830585 | 0.0576558 | 0.47684824 |
| GUCY1A1 | 660.368904 | 0.045554537 | 0.39350663 | 0.11576561 | 0.9078383 | 0.97673359 |
| GUCY1B1 | 333.571572 | 0.477417344 | 0.49380006 | 0.96682318 | 0.33363243 | 0.74022334 |
| CTSO | 181.120491 | -0.113937 | 0.49855897 | -0.2285326 | 0.81923218 | 0.94990455 |
| PDGFC | 4059.38908 | -0.583726167 | 0.46370632 | -1.2588273 | 0.20809272 | 0.65352038 |
| GLRB | 82.3188641 | -0.062328184 | 0.51778546 | -0.1203745 | 0.90418646 | 0.9756941 |

| | | | | | | |
|---|---|---|---|---|---|---|
| TMEM144 | 186.027258 | -0.060281773 | 0.60103167 | -0.1002972 | 0.92010841 | 0.97939204 |
| ETFDH | 887.41252 | -0.444203157 | 0.38537718 | -1.1526452 | 0.24905605 | 0.68557761 |
| PPID | 471.122438 | -0.957492956 | 0.38500484 | -2.4869634 | 0.01288386 | 0.29875272 |
| FNIP2 | 12343.7348 | -0.868162568 | 0.48716761 | -1.7820614 | 0.07473923 | 0.5071704 |
| C4orf45 | 287.238151 | 0.348302168 | 0.58146334 | 0.59900968 | 0.54916643 | 0.84587304 |
| RAPGEF2 | 4994.5022 | -0.165254366 | 0.29832291 | -0.5539446 | 0.57961676 | 0.85816191 |
| FSTL5 | 58.2825082 | -0.396123248 | 0.64679664 | -0.6124386 | 0.54024759 | 0.8414758 |
| NAF1 | 299.803426 | 0.021512353 | 0.1714821 | 0.12544956 | 0.90016766 | 0.97476679 |
| TMA16 | 222.542552 | -0.306571947 | 0.41319105 | -0.7419617 | 0.45811052 | 0.80600032 |
| 1-Mar | 149.184671 | 0.123747278 | 0.59207045 | 0.20900769 | 0.83444223 | 0.95393659 |
| TMEM192 | 1057.65084 | 0.422721686 | 0.40707447 | 1.03843821 | 0.29906609 | 0.71852421 |
| KLHL2 | 953.595589 | 0.25449015 | 0.45145183 | 0.56371496 | 0.57294813 | 0.8542679 |
| GK3P | 91.1643928 | -0.646475846 | 0.53716694 | -1.2034915 | 0.22878618 | 0.66705629 |
| MSMO1 | 615.35334 | 1.338844056 | 0.43902313 | 3.04959799 | 0.00229148 | 0.15549158 |
| CPE | 372.708405 | 0.096950711 | 0.82221616 | 0.1179139 | 0.90613588 | 0.9764469 |
| TLL1 | 720.976282 | 0.202248068 | 0.57874035 | 0.34946253 | 0.7267421 | 0.91639348 |
| DDX60 | 331.162589 | -2.361414449 | 1.13624276 | -2.0782658 | 0.03768489 | 0.42236037 |
| DDX60L | 485.49979 | -1.928431527 | 0.82910262 | -2.3259262 | 0.02002249 | 0.34828456 |
| PALLD | 1030.09399 | -0.235409891 | 0.42189043 | -0.5579882 | 0.57685242 | 0.85619017 |
| CBR4 | 1912.00915 | 0.595897815 | 0.37614031 | 1.58424345 | 0.11313833 | 0.56806517 |
| SH3RF1 | 4044.23446 | -0.570760009 | 0.45301208 | -1.2599223 | 0.2076974 | 0.65309147 |
| AC096741.1 | 73.8101608 | 0.107501275 | 0.62851998 | 0.17103875 | 0.86419329 | 0.96393376 |
| NEK1 | 1496.10619 | -0.113375419 | 0.27391568 | -0.4139063 | 0.67894275 | 0.89932032 |
| CLCN3 | 1039.39366 | 0.276446689 | 0.21085953 | 1.31104672 | 0.18984198 | 0.64203405 |
| HPF1 | 135.309322 | 0.350614558 | 0.3879609 | 0.90373685 | 0.36613495 | 0.76421682 |
| LINC02275 | 948.758611 | 0.58212001 | 0.7675057 | 0.75845692 | 0.44817749 | 0.80229449 |
| AC084866.2 | 54.0554021 | 1.005773963 | 1.12990199 | 0.89014266 | 0.37338929 | 0.76963112 |
| MFAP3L | 823.981353 | -0.0014898 | 0.27970051 | -0.0053264 | 0.99575016 | 0.99866448 |
| AADAT | 48.9474394 | -0.280713896 | 0.65292957 | -0.4299298 | 0.66724669 | 0.8944961 |
| GALNTL6 | 46.3857959 | -0.076347973 | 0.57137412 | -0.1336217 | 0.89370173 | 0.97272276 |
| AC105285.1 | 142.745022 | 0.601468987 | 0.38563863 | 1.55966998 | 0.11883789 | 0.57609025 |
| GALNT7 | 1237.31269 | 0.072111178 | 0.31764891 | 0.22701534 | 8.20E-01 | 0.95036843 |
| HMGB2 | 122.173879 | -0.404737309 | 0.57083719 | -0.7090241 | 0.47830954 | 0.81596289 |
| SAP30 | 83.9991619 | -0.513700053 | 0.49200869 | -1.0440874 | 0.29644497 | 0.71803697 |
| FBXO8 | 213.577237 | -0.136833068 | 0.31509594 | -0.4342584 | 0.66410079 | 0.89313476 |
| CEP44 | 381.834534 | -0.350443712 | 0.3571487 | -0.9812263 | 0.32648114 | 0.73745391 |
| HPGD | 157.940014 | 0.708249441 | 0.63901397 | 1.10834735 | 2.68E-01 | 0.69882424 |
| GPM6A | 1478.25268 | -0.779982228 | 0.58528244 | -1.3326595 | 0.18264356 | 0.63251522 |
| WDR17 | 192.121512 | -0.764169097 | 0.33945834 | -2.2511425 | 0.02437651 | 0.37480999 |
| SPATA4 | 37.883637 | 0.59947973 | 0.74906393 | 0.80030516 | 0.42353401 | 0.78894887 |
| AC019163.1 | 44.1201128 | 0.649649258 | 0.50701933 | 1.28131063 | 0.20008458 | 0.64713529 |
| SPCS3 | 672.546592 | 0.07201622 | 0.40769642 | 0.17664178 | 0.85978978 | 0.96283855 |
| VEGFC | 56.7112258 | 1.193565572 | 0.76477452 | 1.56067643 | 0.11860011 | 0.57589001 |
| AGA | 97.8430616 | 0.368210313 | 0.33938888 | 1.08492157 | 0.27795639 | 0.7036882 |
| AC078881.1 | 485.426502 | 0.635579377 | 0.4309146 | 1.47495439 | 0.14022481 | 0.59500405 |
| AC098864.1 | 118.453574 | 1.531933861 | 0.70004274 | 2.18834332 | 0.0286446 | 0.39339119 |
| TENM3 | 1204.59199 | 1.341792569 | 0.62801209 | 2.13657123 | 0.03263288 | 0.41149854 |
| DCTD | 275.396247 | -0.268136233 | 0.3657546 | -0.7331042 | 0.46349488 | 0.8082072 |
| WWC2 | 3056.06276 | -0.249653409 | 0.16926255 | -1.4749477 | 0.14022661 | 0.59500405 |
| CDKN2AIP | 121.16156 | -0.133635976 | 0.32981624 | -0.4051831 | 0.6853429 | 0.90113631 |
| AC107214.1 | 92.3933068 | 0.555360795 | 0.29765234 | 1.86580359 | 0.06206885 | 0.48347236 |
| ING2 | 97.2604837 | -0.060922326 | 0.32319281 | -0.1885015 | 0.85048355 | 0.95994332 |
| RWDD4 | 328.32006 | 0.121798805 | 0.24065566 | 0.50611236 | 0.61277777 | 0.87222135 |
| TRAPPC11 | 614.698017 | 0.14643178 | 0.27362699 | 0.53515108 | 0.59254539 | 0.86382106 |

| | | | | | | |
|---|---|---|---|---|---|---|
| STOX2 | 7428.82466 | 1.017689912 | 0.48473345 | 2.09948355 | 0.0357743 | 0.42040764 |
| ENPP6 | 926.733165 | 1.911824107 | 0.62928532 | 3.0380879 | 2.38E-03 | 0.15841865 |
| IRF2 | 725.616333 | 0.343035646 | 0.27759372 | 1.23574714 | 0.21655257 | 0.66032006 |
| LINC02427 | 63.1148131 | 1.505048433 | 0.8199511 | 1.83553437 | 0.06642655 | 0.48798003 |
| CASP3 | 239.214105 | 0.155516517 | 0.28084193 | 0.55375107 | 0.57974922 | 0.85816191 |
| PRIMPOL | 192.510022 | 0.612139217 | 0.50397311 | 1.21462675 | 0.22450848 | 0.66551422 |
| ACSL1 | 2805.53983 | -0.205983655 | 0.4418408 | -0.4661943 | 0.64107644 | 0.88396933 |
| SLC25A4 | 204.255903 | 0.486569855 | 0.47996745 | 1.01375594 | 0.31069922 | 0.72537613 |
| CFAP97 | 814.410532 | 0.301313887 | 0.26984226 | 1.11662974 | 0.26415267 | 0.69567438 |
| SNX25 | 1145.90373 | -0.539079627 | 0.30204575 | -1.7847615 | 0.07430001 | 0.5071704 |
| LRP2BP | 133.85291 | 0.363366173 | 0.35888081 | 1.01249819 | 0.31129991 | 0.7258226 |
| AC112722.1 | 182.596099 | 0.046765405 | 0.41895734 | 0.11162331 | 0.91112209 | 0.97697493 |
| ANKRD37 | 294.319284 | -0.675001245 | 0.96267667 | -0.7011713 | 0.48319612 | 0.81777618 |
| UFSP2 | 446.287678 | 0.174325008 | 0.39357093 | 0.44293162 | 0.65781519 | 0.89053559 |
| C4orf47 | 415.240446 | -0.250682454 | 0.67986521 | -0.3687238 | 0.71233364 | 0.91166734 |
| CCDC110 | 43.6854526 | 0.005189766 | 0.61290251 | 0.00846752 | 0.99324397 | 0.9976934 |
| SORBS2 | 5000.55755 | 0.768326661 | 0.50424206 | 1.52372586 | 0.12757721 | 0.58793678 |
| TLR3 | 63.3458988 | -0.048888305 | 0.78264144 | -0.0624658 | 0.95019192 | 0.98724111 |
| FAM149A | 922.698192 | -0.16867096 | 0.35096917 | -0.4805863 | 0.63081058 | 0.88115748 |
| CYP4V2 | 563.804086 | 0.246356955 | 0.42170589 | 0.5841914 | 5.59E-01 | 0.84936251 |
| KLKB1 | 765.968361 | 0.193515705 | 0.46813913 | 0.41337221 | 6.79E-01 | 8.99E-01 |
| F11 | 238.056144 | 0.464473195 | 0.82377819 | 0.56383284 | 5.73E-01 | 0.8542679 |
| F11-AS1 | 313.179133 | 1.133438888 | 0.79226026 | 1.43063958 | 0.15253354 | 0.61001662 |
| AC018709.1 | 284.118789 | 1.021256516 | 0.74706766 | 1.36702011 | 0.171619 | 0.62520756 |
| FAT1 | 3408.68727 | 0.301092051 | 0.2950437 | 1.02049984 | 0.30749147 | 0.72263382 |
| LINC01060 | 3987.43247 | 0.901554758 | 0.72972429 | 1.23547313 | 0.21665447 | 0.66032006 |
| FRG1-DT | 166.722889 | -0.130048208 | 0.56634905 | -0.2296255 | 0.81838276 | 0.94990455 |
| FRG1 | 137.801381 | 0.136084148 | 0.23522993 | 0.57851545 | 0.56291617 | 0.85057939 |
| CCDC127 | 87.5832061 | -0.245924091 | 0.26355819 | -0.9330922 | 0.35077236 | 0.75495648 |
| SDHA | 945.262257 | 0.20392778 | 0.23726056 | 0.85950982 | 0.39005931 | 0.77569824 |
| PDCD6 | 552.27946 | -0.189675 | 0.31694561 | -0.5984465 | 0.54954202 | 0.84612944 |
| AHRR | 30.3315438 | -0.046820805 | 0.98548876 | -0.0475102 | 0.96210657 | 0.99012655 |
| EXOC3-AS1 | 46.5303457 | 0.327602176 | 0.44320042 | 0.73917389 | 0.45980141 | 0.8069452 |
| EXOC3 | 504.551571 | -0.302892531 | 0.22180092 | -1.3656054 | 0.17206285 | 0.62592399 |
| SLC9A3 | 602.062874 | 1.67788047 | 0.59682162 | 2.81136005 | 0.00493325 | 0.21673116 |
| CEP72 | 67.143408 | -0.35936658 | 0.38969492 | -0.9221742 | 0.35643774 | 0.75841231 |
| ZDHHC11B | 110.940428 | 0.784491508 | 0.80232244 | 0.97777585 | 0.3281852 | 0.73826734 |
| ZDHHC11 | 220.747303 | -0.994427311 | 0.49978942 | -1.9896926 | 0.04662481 | 0.44535626 |
| BRD9 | 364.894677 | -0.706694407 | 0.38398936 | -1.840401 | 0.06570938 | 0.48790819 |
| TRIP13 | 42.9073963 | -1.407811679 | 0.67142458 | -2.0967533 | 0.03601542 | 0.42040764 |
| SLC12A7 | 1355.25998 | 0.477786291 | 0.36816354 | 1.29775557 | 0.19437134 | 0.64548758 |
| AC114291.1 | 237.118788 | 2.193802371 | 1.25988575 | 1.74127088 | 8.16E-02 | 5.16E-01 |
| SLC6A19 | 587.331679 | 1.46649117 | 0.74175026 | 1.97706864 | 4.80E-02 | 0.45149028 |
| SLC6A18 | 305.037899 | -0.954408373 | 1.03391426 | -0.923102 | 0.35595403 | 0.75841231 |
| CLPTM1L | 918.24449 | 0.172840382 | 0.46142382 | 0.37458054 | 0.70797245 | 0.91025029 |
| LPCAT1 | 435.92261 | -1.190818794 | 0.62684308 | -1.899708 | 0.05747145 | 0.47684824 |
| MRPL36 | 129.654329 | -0.3594108 | 0.41533966 | -0.8653419 | 0.38685117 | 0.77334938 |
| NDUFS6 | 521.690272 | -0.604534976 | 0.28505285 | -2.1207821 | 0.03394014 | 0.41378378 |
| AC010451.1 | 67.5463287 | 3.435095749 | 1.42181067 | 2.41600082 | 0.01569203 | 0.31566878 |
| LINC01020 | 597.262894 | 4.312404395 | 1.05310251 | 4.09495214 | 4.22E-05 | 0.02067092 |
| ADAMTS16 | 224.164832 | 2.424302857 | 0.91570082 | 2.64748354 | 0.00810933 | 0.24877162 |
| ICE1 | 930.018578 | -0.50918266 | 0.31621532 | -1.6102403 | 0.1073454 | 0.55579103 |
| LINC02145 | 43.4694492 | -0.652529693 | 0.68398207 | -0.9540158 | 0.34007565 | 0.74666297 |
| MED10 | 264.124072 | -0.599764145 | 0.24342171 | -2.4638893 | 0.01374386 | 0.30336076 |

| | | | | | | |
|---|---|---|---|---|---|---|
| NSUN2 | 482.357362 | -0.463544114 | 0.29082653 | -1.5938853 | 0.11096174 | 0.56342465 |
| SRD5A1 | 236.740878 | -0.124428863 | 0.41958384 | -0.296553 | 0.76680777 | 0.93138917 |
| TENT4A | 771.038679 | -0.637653608 | 0.30516744 | -2.0895205 | 0.0366609 | 0.42192353 |
| LINC02196 | 41.6044224 | 0.59726238 | 0.98083203 | 0.60893442 | 0.54256791 | 0.84314228 |
| ADCY2 | 230.891952 | 1.524844983 | 0.69002286 | 2.20984705 | 0.02711578 | 0.38702446 |
| C5orf49 | 44.4670618 | 0.442661405 | 0.51381744 | 0.86151495 | 0.38895449 | 0.7748665 |
| MTRR | 784.61938 | 0.378609559 | 0.27242485 | 1.38977614 | 0.16459687 | 0.62065658 |
| FASTKD3 | 111.488292 | 0.117907361 | 0.33067284 | 0.35656803 | 0.72141522 | 0.91542662 |
| MIR4458HG | 112.791336 | -0.536267197 | 0.45456566 | -1.1797354 | 0.23810548 | 0.67642542 |
| SEMA5A | 3653.6777 | 0.216387309 | 0.60647273 | 0.35679644 | 0.72124421 | 0.91542662 |
| SNHG18 | 189.674623 | 0.535688719 | 0.37870156 | 1.41454058 | 0.15720324 | 0.6128512 |
| LINC02112 | 100.073385 | 0.897554106 | 0.73926362 | 1.21411913 | 0.22470224 | 0.66594455 |
| FAM173B | 234.377001 | 0.401895424 | 0.34576615 | 1.16233306 | 0.2451002 | 0.68281753 |
| CCT5 | 498.197971 | -0.09035264 | 0.3301924 | -0.2736363 | 0.78436411 | 0.93873036 |
| CMBL | 1890.93239 | 0.69892825 | 0.70222808 | 0.99530092 | 0.31958993 | 0.73132207 |
| 6-Mar | 2777.13741 | 0.03500798 | 0.30793475 | 0.11368635 | 0.90948643 | 0.97680747 |
| ANKRD33B | 1829.21823 | -0.300005952 | 0.7279208 | -0.4121409 | 0.68023613 | 0.89941358 |
| DAP | 804.998302 | -0.198356627 | 0.32783198 | -0.6050558 | 0.54514195 | 0.84449215 |
| CTNND2 | 80.4508761 | 0.072763331 | 0.48569342 | 0.14981329 | 8.81E-01 | 9.68E-01 |
| LINC02220 | 484.052546 | 1.312430866 | 0.86555248 | 1.51629266 | 0.12944537 | 0.58817997 |
| DNAH5 | 644.759778 | 1.09649443 | 1.3519667 | 0.81103657 | 0.41734467 | 0.78652162 |
| TRIO | 8390.33215 | -0.485631393 | 0.37441421 | -1.2970432 | 0.19461634 | 0.64548758 |
| OTULINL | 67.686221 | 0.715463128 | 0.55585132 | 1.28714837 | 0.19804258 | 0.64582202 |
| OTULIN | 807.673482 | 0.339777472 | 0.29555753 | 1.14961535 | 0.25030233 | 0.68617881 |
| ANKH | 965.502813 | -0.367662102 | 0.41203924 | -0.8922988 | 0.37223282 | 0.76866079 |
| FBXL7 | 2202.59832 | -0.772900191 | 0.60054095 | -1.2870066 | 0.19809197 | 0.64582202 |
| ZNF622 | 143.670983 | -0.423427213 | 0.34912892 | -1.2128105 | 0.22520228 | 0.66649135 |
| RETREG1 | 5024.21037 | 0.479169405 | 0.67386795 | 0.71107315 | 0.47703891 | 0.81570932 |
| MYO10 | 1403.60054 | 0.193270638 | 0.49756156 | 0.38843563 | 6.98E-01 | 0.90696517 |
| LINC02241 | 300.510186 | 0.603636205 | 0.41267538 | 1.46273858 | 0.14353892 | 0.59966109 |
| AC093274.1 | 34.873755 | 0.636973396 | 0.95034464 | 0.67025516 | 0.50269515 | 0.82638105 |
| CDH12 | 102.546975 | 0.353674204 | 0.53334773 | 0.66312123 | 0.50725292 | 0.82832518 |
| C5orf17 | 351.851642 | -0.942650154 | 0.62753198 | -1.5021548 | 0.13305714 | 0.5919553 |
| CDH9 | 5803.99795 | 1.425126729 | 0.68468208 | 2.081443 | 0.03739338 | 0.42215481 |
| AC113386.1 | 868.261776 | 1.054939252 | 0.74589113 | 1.41433408 | 0.15726383 | 0.6129131 |
| CDH6 | 12711.0776 | 1.097818167 | 0.80455295 | 1.36450705 | 0.17240804 | 0.62593809 |
| DROSHA | 569.651847 | -0.05377456 | 0.2375936 | -0.22633 | 0.82094477 | 0.95072145 |
| C5orf22 | 242.553659 | -0.410309456 | 0.23108777 | -1.7755568 | 0.07580599 | 0.50730849 |
| PDZD2 | 6969.48147 | 1.138131777 | 0.77700193 | 1.46477342 | 0.14298275 | 0.59824924 |
| AC025178.1 | 112.435683 | -0.183118792 | 0.58164377 | -0.3148298 | 0.75289088 | 0.92746251 |
| GOLPH3 | 1691.86518 | -0.303362855 | 0.28175383 | -1.0766947 | 0.28161668 | 0.70625976 |
| MTMR12 | 955.755578 | -0.066547972 | 0.29716232 | -0.2239448 | 0.82280022 | 0.95117842 |
| ZFR | 2494.20158 | -0.06400247 | 0.16712494 | -0.3829618 | 0.70174809 | 0.90803937 |
| SUB1 | 726.700145 | 0.0534896 | 0.23267793 | 0.22988686 | 0.81817968 | 0.94984237 |
| LINC02061 | 411.389506 | 0.597759456 | 1.26613057 | 0.47211518 | 6.37E-01 | 8.83E-01 |
| NPR3 | 2175.47995 | 0.179526059 | 0.44632886 | 0.40222821 | 6.88E-01 | 9.03E-01 |
| AC010343.3 | 51.1977832 | 0.643909402 | 0.48194476 | 1.33606475 | 0.18152811 | 0.6318513 |
| TARS | 391.643407 | 0.008099839 | 0.28413815 | 0.02850669 | 0.97725803 | 0.99473325 |
| ADAMTS12 | 84.5876492 | 0.209209943 | 0.58476886 | 0.35776519 | 0.72051904 | 0.9149807 |
| SLC45A2 | 182.177539 | 1.119355341 | 0.59273464 | 1.88845948 | 0.05896429 | 0.47795598 |
| AMACR | 142.703883 | 1.384965034 | 0.67156693 | 2.06228892 | 0.03918024 | 0.42704465 |
| C1QTNF3-AN | 743.223066 | 0.525492307 | 0.33532919 | 1.56709382 | 0.11709276 | 0.57350419 |
| RAI14 | 2077.28383 | -0.30305398 | 0.34557816 | -0.8769477 | 0.38051503 | 0.7716703 |
| TTC23L | 54.0097281 | 0.543714111 | 0.58473503 | 0.929847 | 0.35245031 | 0.75652927 |

| | | | | | | |
|---|---|---|---|---|---|---|
| RAD1 | 168.192598 | 0.064835757 | 0.24517749 | 0.26444416 | 0.79143769 | 0.94098694 |
| BRIX1 | 163.670673 | -0.445374405 | 0.40593513 | -1.0971566 | 0.27257294 | 0.70083612 |
| DNAJC21 | 435.385018 | -0.016476414 | 0.19507161 | -0.0844634 | 0.93268799 | 0.98310938 |
| AGXT2 | 3551.75536 | 1.62068324 | 0.77319962 | 2.09607351 | 0.03607567 | 0.42040764 |
| PRLR | 2753.13893 | 0.756604674 | 0.54178316 | 1.39650828 | 0.16256149 | 0.61912485 |
| SPEF2 | 1580.68987 | 0.7320499 | 0.58557993 | 1.25012805 | 0.21125278 | 0.65561118 |
| AC137810.1 | 64.5052193 | 0.66505408 | 0.93608693 | 0.71046188 | 0.47741776 | 0.81579656 |
| UGT3A1 | 819.080281 | -0.148007636 | 0.5264795 | -0.2811271 | 0.77861294 | 0.93646866 |
| LMBRD2 | 382.083422 | 0.330491376 | 0.28484227 | 1.16026101 | 0.24594255 | 0.68320879 |
| SKP2 | 94.9604458 | -0.016260513 | 0.32403864 | -0.0501808 | 0.95997832 | 0.98928072 |
| NADK2 | 386.194679 | -0.126366424 | 0.3785193 | -0.3338441 | 0.73849723 | 0.92080006 |
| NIPBL | 4193.759 | -0.307835566 | 0.19448262 | -1.5828435 | 0.11345713 | 0.56809124 |
| CPLANE1 | 1591.08598 | 0.046959295 | 0.15638409 | 0.30028179 | 0.76396222 | 0.93079056 |
| NUP155 | 257.604312 | -0.454473039 | 0.26605254 | -1.7082078 | 0.08759779 | 0.5273546 |
| WDR70 | 2557.02733 | -0.217526182 | 0.18471079 | -1.1776582 | 0.23893289 | 0.67667639 |
| GDNF-AS1 | 211.287357 | 0.305906299 | 0.74556261 | 0.41030263 | 6.82E-01 | 9.00E-01 |
| LINC02107 | 86.9163502 | 1.054372223 | 0.86086494 | 1.2247824 | 2.21E-01 | 6.63E-01 |
| EGFLAM | 39.1264815 | 0.909509279 | 0.82174043 | 1.10680848 | 0.26837674 | 0.69961625 |
| LIFR | 1841.57574 | -0.412285593 | 0.60546995 | -0.6809349 | 0.49591271 | 0.82243887 |
| LIFR-AS1 | 59.0485525 | 1.327531201 | 0.7072531 | 1.87702421 | 0.06051478 | 0.48026727 |
| OSMR-AS1 | 796.752069 | 0.607927291 | 0.4364623 | 1.39285178 | 1.64E-01 | 0.62065658 |
| OSMR | 1668.35639 | 0.191583236 | 0.60835163 | 0.31492188 | 7.53E-01 | 9.27E-01 |
| RICTOR | 2905.32461 | -0.458909541 | 0.15091876 | -3.040772 | 0.00235972 | 0.15777925 |
| FYB1 | 142.716537 | 2.201532719 | 0.73356 | 3.00116245 | 0.00268951 | 0.16909084 |
| C9 | 393.264612 | 0.676191293 | 0.44678254 | 1.5134685 | 0.1301607 | 0.58853436 |
| DAB2 | 1168.75778 | 0.071965347 | 0.36264626 | 0.19844503 | 0.84269689 | 0.95716002 |
| TTC33 | 387.635954 | 0.359524272 | 0.22935516 | 1.56754383 | 0.11698763 | 0.57350419 |
| PRKAA1 | 793.405485 | 0.157676282 | 0.22792737 | 0.691783 | 0.48907361 | 0.81970016 |
| RPL37 | 730.064196 | -0.071255505 | 0.39822348 | -0.1789335 | 0.85798995 | 0.9626966 |
| C6 | 123.212209 | 0.808085985 | 0.96601817 | 0.8365122 | 0.40286681 | 0.78260941 |
| PLCXD3 | 678.3573 | -0.568789551 | 0.86634245 | -0.6565412 | 0.51147594 | 0.83017178 |
| OXCT1 | 291.154137 | -0.684284359 | 0.55825277 | -1.2257608 | 0.2202887 | 0.66263062 |
| C5orf51 | 189.387832 | -0.064185356 | 0.24532188 | -0.2616373 | 0.79360108 | 0.94197176 |
| FBXO4 | 118.576109 | -0.145241671 | 0.46700894 | -0.3110041 | 0.75579754 | 0.92799282 |
| GHR | 2961.64046 | -0.094536313 | 0.47743418 | -0.1980091 | 0.84303794 | 0.95716002 |
| SELENOP | 773.618812 | -0.497168183 | 0.57860033 | -0.8592601 | 0.39019703 | 0.77569824 |
| ZNF131 | 1028.61975 | -0.29412408 | 0.26591595 | -1.1060791 | 0.26869227 | 0.69961625 |
| NIM1K | 57.863161 | -1.162357309 | 0.66192726 | -1.7560197 | 0.07908505 | 0.51192014 |
| HMGCS1 | 1019.85057 | 1.118500626 | 0.34499579 | 3.24207037 | 0.00118665 | 0.11201338 |
| CCL28 | 96.6417246 | 0.082523404 | 0.82719136 | 0.09976338 | 9.21E-01 | 9.79E-01 |
| TMEM267 | 376.682732 | 0.340574005 | 0.29135846 | 1.16891751 | 0.24243687 | 0.68080966 |
| C5orf34 | 40.4968554 | 0.815515679 | 0.60839686 | 1.34043373 | 0.18010438 | 0.63124625 |
| PAIP1 | 430.660779 | -0.19879115 | 0.20075472 | -0.9902191 | 0.32206705 | 0.73375526 |
| NNT-AS1 | 460.975862 | 0.787597795 | 0.29335065 | 2.68483396 | 0.00725658 | 0.23590813 |
| NNT | 1263.24219 | 0.430924398 | 0.27454351 | 1.56960331 | 0.11650743 | 0.57350419 |
| MRPS30-DT | 91.8877268 | 0.411273769 | 0.4622967 | 0.88963164 | 0.37366371 | 0.76973377 |
| MRPS30 | 213.341467 | 0.164185748 | 0.22008177 | 0.74602158 | 0.45565438 | 0.80465616 |
| AC093297.2 | 66.1015724 | 0.127843511 | 0.35583633 | 0.35927616 | 0.71938851 | 0.91465123 |
| PARP8 | 1099.16314 | -0.036188511 | 0.40052786 | -0.090352 | 9.28E-01 | 0.98263409 |
| AC022126.1 | 89.8226931 | 1.060657965 | 0.68614397 | 1.54582421 | 0.12214703 | 0.58055167 |
| ITGA1 | 757.378166 | 0.185998133 | 0.25929914 | 0.71731102 | 4.73E-01 | 0.81478861 |
| ITGA2 | 569.622922 | -0.434231532 | 0.38881183 | -1.1168167 | 0.26407272 | 0.69567438 |
| MOCS2 | 181.013624 | -0.175590486 | 0.45496717 | -0.385941 | 0.69954036 | 0.90721166 |
| NDUFS4 | 1104.39681 | 0.205600417 | 0.2182855 | 0.94188766 | 0.34625016 | 0.75166079 |

| | | | | | | |
|---|---|---|---|---|---|---|
| ARL15 | 4604.99673 | -0.221924068 | 0.38122532 | -0.5821336 | 0.5604767 | 0.84944908 |
| SNX18 | 109.16692 | 0.515266668 | 0.35726042 | 1.44227191 | 0.14922568 | 0.60629349 |
| AC026704.1 | 96.3785619 | 0.897594408 | 0.63319529 | 1.4175633 | 0.1563183 | 0.61255389 |
| DHX29 | 738.04336 | -0.145269661 | 0.22496831 | -0.6457339 | 0.51845172 | 0.83325693 |
| MTREX | 1178.38098 | -0.162866671 | 0.17791275 | -0.91543 | 0.35996592 | 0.75998694 |
| PLPP1 | 2388.0744 | -0.284499517 | 0.41894487 | -0.6790858 | 0.49708349 | 0.82278993 |
| SLC38A9 | 1101.8729 | 0.238428106 | 0.22364819 | 1.06608558 | 0.28638496 | 0.71118494 |
| IL6ST | 1858.63722 | -0.133606993 | 0.30207571 | -0.4422964 | 0.65827475 | 0.89058983 |
| AC008892.1 | 174.846657 | 0.30415424 | 0.48894978 | 0.6220562 | 0.53390492 | 0.83915086 |
| ANKRD55 | 79.4715964 | 0.856331812 | 0.69954005 | 1.22413551 | 0.22090111 | 0.66263062 |
| LINC01948 | 294.680253 | 0.550301972 | 0.84760635 | 0.64924239 | 0.51618171 | 0.83219275 |
| C5orf67 | 1219.51229 | 2.150850517 | 0.9846541 | 2.18437166 | 0.02893495 | 0.39542507 |
| MAP3K1 | 2212.33813 | 0.432397383 | 0.49423407 | 0.8748838 | 0.38163714 | 0.77237191 |
| AC008937.2 | 73.1730621 | 0.647349226 | 0.72928616 | 0.88764775 | 0.37473026 | 0.7697436 |
| SETD9 | 70.7144183 | 0.140554918 | 0.35776187 | 0.39287283 | 0.69441341 | 0.90589033 |
| MIER3 | 263.261527 | 0.003918058 | 0.26374218 | 0.01485564 | 0.98814735 | 0.99671759 |
| GPBP1 | 2446.68368 | -0.33161132 | 0.31483284 | -1.0532933 | 0.29220659 | 0.71399546 |
| PLK2 | 62.5907572 | -0.550699711 | 0.69835081 | -0.7885717 | 0.43036235 | 0.79055252 |
| PDE4D | 32494.8809 | 1.06042374 | 0.55525234 | 1.90980508 | 0.05615832 | 0.47126031 |
| AC092343.1 | 55.1769965 | 1.047431293 | 0.90339729 | 1.15943595 | 0.24627853 | 0.6834865 |
| DEPDC1B | 69.707541 | -1.708871305 | 0.59424985 | -2.8756781 | 0.00403161 | 0.20256898 |
| ELOVL7 | 1354.48072 | -0.337949257 | 0.41887273 | -0.8068065 | 0.41977796 | 0.78798479 |
| ERCC8 | 512.867268 | 0.123501985 | 0.24624276 | 0.50154566 | 0.61598715 | 0.87333999 |
| NDUFAF2 | 1324.94132 | -0.631886689 | 0.34845536 | -1.8133935 | 0.06977115 | 0.49372905 |
| SMIM15 | 154.647914 | -0.551747475 | 0.34414881 | -1.6032236 | 0.10888531 | 0.55877607 |
| SMIM15-AS1 | 77.0394506 | -0.233777683 | 0.41324958 | -0.5657058 | 0.5715938 | 0.85399468 |
| ZSWIM6 | 4551.22929 | -0.462459622 | 0.33669777 | -1.3735156 | 0.16959213 | 0.62326131 |
| KIF2A | 789.894043 | 0.101910709 | 0.34746415 | 0.29329849 | 0.76929401 | 0.93209094 |
| DIMT1 | 156.723846 | -0.335563115 | 0.25633338 | -1.3090886 | 0.19050434 | 0.64281105 |
| IPO11 | 435.389315 | -0.262179255 | 0.25224251 | -1.0393936 | 0.29862171 | 0.71852421 |
| RNF180 | 1847.35045 | 0.712052416 | 0.34222602 | 2.08064955 | 0.03746599 | 0.42215481 |
| SREK1IP1 | 472.188511 | -0.325074489 | 0.20905597 | -1.5549639 | 0.11995467 | 0.57730856 |
| CWC27 | 965.573509 | 0.04094034 | 0.19793886 | 0.20683326 | 0.83614008 | 0.95468028 |
| ADAMTS6 | 64.6437737 | -0.483435368 | 0.69485451 | -0.6957361 | 0.48659411 | 0.81877553 |
| PPWD1 | 425.284279 | 0.224266469 | 0.37332991 | 0.60071926 | 0.54802699 | 0.84486687 |
| TRIM23 | 409.473627 | -0.189832353 | 0.29294292 | -0.6480182 | 0.51697315 | 0.83276628 |
| TRAPPC13 | 463.330107 | 0.361392617 | 0.24202637 | 1.49319518 | 0.1353861 | 0.59381224 |
| SGTB | 118.498192 | -0.178483117 | 0.42492893 | -0.4200305 | 0.67446317 | 0.89801183 |
| NLN | 551.625987 | -0.504087645 | 0.39114388 | -1.2887525 | 1.97E-01 | 6.46E-01 |
| ERBIN | 4604.88631 | -0.132245737 | 0.19091643 | -0.6926892 | 0.48850465 | 0.81933645 |
| SREK1 | 1561.58375 | -0.585719468 | 0.16756419 | -3.495493 | 0.00047319 | 0.07721972 |
| MAST4 | 23826.4955 | -0.669538414 | 0.2553989 | -2.6215399 | 0.00875335 | 0.25397092 |
| AC112206.2 | 67.7920275 | 0.43697221 | 0.53319354 | 0.8195377 | 0.4124797 | 0.78632759 |
| PIK3R1 | 1557.2406 | 1.402122762 | 0.54921111 | 2.55297598 | 0.01068069 | 0.27357367 |
| SLC30A5 | 580.41505 | -0.351605111 | 0.32399002 | -1.0852344 | 0.27781786 | 0.7036882 |
| MRPS36 | 476.809371 | -0.032342103 | 0.36567373 | -0.0884452 | 0.9295228 | 0.98272895 |
| CDK7 | 501.153296 | -0.490509707 | 0.21385184 | -2.2936895 | 0.02180834 | 0.36058739 |
| CCDC125 | 611.769513 | 0.629194977 | 0.42753837 | 1.47166902 | 0.14111028 | 0.59568789 |
| AK6 | 246.112359 | 0.034601713 | 0.26475788 | 0.13069191 | 0.89601903 | 0.97309833 |
| RAD17 | 460.062802 | -0.110296672 | 0.14683772 | -0.7511467 | 0.45256436 | 0.80433814 |
| MARVELD2 | 232.465336 | 0.42797657 | 0.23673124 | 1.80785845 | 0.07062853 | 0.4967563 |
| OCLN | 692.704875 | -0.0358565 | 0.36099287 | -0.0993274 | 0.92087829 | 0.97939204 |
| GTF2H2C | 490.973641 | -0.021256585 | 0.3589032 | -0.0592265 | 0.95277169 | 0.98786081 |
| SMN2 | 35.790054 | -0.959297085 | 1.01201879 | -0.9479044 | 0.34317811 | 0.75073625 |

| | | | | | | |
|---|---|---|---|---|---|---|
| NAIP | 853.709259 | 0.441902856 | 0.31882613 | 1.38603088 | 0.16573747 | 0.62065658 |
| GTF2H2 | 485.39459 | 0.176137969 | 0.53952594 | 0.32646803 | 0.74407028 | 0.92361546 |
| LINC02197 | 232.682604 | 0.661010699 | 0.98906117 | 0.66832135 | 0.50392849 | 0.82715271 |
| BDP1 | 1733.47632 | -0.220894259 | 0.15478671 | -1.4270881 | 0.15355451 | 0.61014911 |
| MCCC2 | 710.466683 | 0.052744162 | 0.21198438 | 0.24881155 | 0.80350655 | 0.94543168 |
| MAP1B | 711.593788 | -2.235895689 | 1.34175578 | -1.6663954 | 9.56E-02 | 0.538613 |
| MRPS27 | 641.408961 | -0.236696535 | 0.26890002 | -0.8802399 | 0.37872935 | 0.77061593 |
| PTCD2 | 151.841292 | 0.090741234 | 0.36589875 | 0.24799547 | 0.80413791 | 0.94551573 |
| TNPO1 | 1743.49942 | -0.366983405 | 0.27164933 | -1.3509454 | 0.17671294 | 0.62709281 |
| FCHO2 | 2051.59055 | 0.190799376 | 0.27080621 | 0.70456055 | 0.48108376 | 0.81720798 |
| TMEM171 | 257.605136 | 0.234422532 | 0.4883126 | 0.48006652 | 0.63118009 | 0.88122598 |
| AC116345.3 | 313.024696 | 0.581640165 | 1.13396938 | 0.51292405 | 0.60800445 | 0.86985416 |
| TMEM174 | 135.528009 | 1.350197774 | 0.78694813 | 1.71573923 | 0.08620978 | 0.52402549 |
| BTF3 | 441.524111 | 0.079893282 | 0.39488424 | 0.20232077 | 0.83966597 | 0.9556011 |
| ANKRA2 | 173.371812 | 0.272992809 | 0.50293232 | 0.54280228 | 0.58726595 | 0.86112855 |
| UTP15 | 64.4488064 | -0.425928339 | 0.39708442 | -1.0726393 | 0.283433 | 0.70780772 |
| ARHGEF28 | 6169.63034 | -0.104542317 | 0.32910191 | -0.3176594 | 0.75074331 | 0.92598205 |
| ENC1 | 40.2407839 | 0.378024374 | 0.69400111 | 0.54470284 | 0.58595791 | 0.8609417 |
| HEXB | 779.144603 | -0.560617912 | 0.44815034 | -1.2509595 | 0.21094926 | 0.6555161 |
| GFM2 | 350.193076 | -0.291989533 | 0.25449657 | -1.1473221 | 0.25124852 | 0.68726073 |
| NSA2 | 258.029638 | -0.075798037 | 0.29969674 | -0.2529158 | 0.80033329 | 0.94403774 |
| FAM169A | 161.667857 | 0.932054965 | 0.84619196 | 1.10146988 | 0.2706922 | 0.69961625 |
| ANKRD31 | 58.9315544 | 0.002729911 | 0.65769175 | 0.00415075 | 0.99668819 | 0.9992561 |
| HMGCR | 373.988299 | 0.740592721 | 0.40951909 | 1.80844492 | 0.07053728 | 0.4967563 |
| COL4A3BP | 2470.43289 | -0.115456254 | 0.22364905 | -0.5162385 | 0.60568783 | 0.86915442 |
| POLK | 1270.51985 | 0.08511467 | 0.21926647 | 0.38817915 | 0.69788346 | 0.90696517 |
| ANKDD1B | 380.725602 | 0.344120689 | 0.60854723 | 0.56547901 | 0.57174799 | 0.85399468 |
| POC5 | 290.740371 | -0.045074256 | 0.18543419 | -0.2430741 | 0.80794797 | 0.94613779 |
| IQGAP2 | 2432.46414 | 0.896510009 | 0.46894015 | 1.91177917 | 0.05590452 | 0.47069 |
| F2R | 34.8656457 | 0.165139223 | 0.5051793 | 0.3268923 | 0.74374935 | 0.92350771 |
| F2RL1 | 229.123784 | 0.197796829 | 0.27958919 | 0.7074552 | 0.47928364 | 0.81649775 |
| AC008581.2 | 95.454631 | 1.038651989 | 0.56670522 | 1.83279059 | 0.06683372 | 0.48884194 |
| ZBED3 | 269.180692 | 0.385589506 | 0.34361979 | 1.12213999 | 0.26180292 | 0.69408034 |
| AC022414.1 | 93.2361393 | 1.277389644 | 0.99472035 | 1.28416961 | 0.19908262 | 0.64606998 |
| WDR41 | 593.039976 | 0.260184963 | 0.26575529 | 0.97903963 | 0.3275604 | 0.73785873 |
| TBCA | 2217.92869 | -0.526145804 | 0.34915302 | -1.5069204 | 0.13183106 | 0.59087581 |
| AP3B1 | 2693.4681 | 0.181347979 | 0.23890227 | 0.75908856 | 0.44779958 | 0.80223612 |
| SCAMP1-AS1 | 58.7666404 | -0.668741122 | 0.54393338 | -1.2294541 | 0.2189016 | 0.66119089 |
| SCAMP1 | 968.021471 | -0.130785846 | 0.20578898 | -0.6355338 | 0.52508035 | 0.835938 |
| LHFPL2 | 1067.40245 | -1.050762099 | 0.60782381 | -1.7287281 | 0.08385777 | 0.52128301 |
| ARSB | 2730.43365 | -0.404965688 | 0.53242807 | -0.7606017 | 0.44689501 | 0.80219878 |
| DMGDH | 2795.03921 | 1.185794336 | 0.55857879 | 2.12287749 | 0.03376412 | 0.41378378 |
| BHMT2 | 4748.02455 | 0.643033089 | 0.47697215 | 1.34815648 | 0.17760806 | 0.6280421 |
| BHMT | 5143.08933 | 1.000675403 | 0.47551769 | 2.10439152 | 3.53E-02 | 4.20E-01 |
| JMY | 886.046233 | -0.370030262 | 0.21797174 | -1.6976066 | 0.08958204 | 0.53155889 |
| HOMER1 | 833.183665 | -1.341040404 | 0.54828446 | -2.4458844 | 0.01444973 | 0.30707362 |
| TENT2 | 3220.50671 | 0.017429345 | 0.20848167 | 0.08360133 | 0.93337341 | 0.98352931 |
| CMYA5 | 951.830446 | -0.229616917 | 0.27856216 | -0.8242933 | 4.10E-01 | 0.78542936 |
| MTX3 | 185.377571 | -0.206540009 | 0.20620828 | -1.0016087 | 0.3165326 | 0.72846078 |
| THBS4 | 178.840006 | 0.177236081 | 0.52541882 | 0.33732344 | 0.73587309 | 0.92019178 |
| SERINC5 | 523.31779 | -0.37678228 | 0.35583758 | -1.0588603 | 0.28966342 | 0.71218341 |
| AC010260.1 | 38.6312796 | 0.48166155 | 0.53770332 | 0.89577566 | 0.37037259 | 0.76698853 |
| ZFYVE16 | 964.341107 | -0.12273934 | 0.14941105 | -0.8214877 | 0.41136854 | 0.78632759 |
| AC008771.1 | 40.0567898 | -0.014459201 | 0.52309858 | -0.0276414 | 0.97794812 | 0.99482978 |

| | | | | | | |
|---|---|---|---|---|---|---|
| FAM151B | 355.37143 | -0.179781031 | 0.32467294 | -0.5537296 | 0.57976389 | 0.85816191 |
| DHFR | 2313.86062 | -1.143912463 | 0.42659345 | -2.681505 | 0.00732918 | 0.23590813 |
| MSH3 | 2216.89296 | 0.53633557 | 0.29525041 | 1.81654472 | 6.93E-02 | 4.93E-01 |
| RASGRF2 | 1194.99172 | -0.090042255 | 0.58173688 | -0.1547818 | 0.87699338 | 0.96643457 |
| CKMT2-AS1 | 760.982895 | -0.476342971 | 0.32219298 | -1.4784399 | 0.13929006 | 0.59478159 |
| CKMT2 | 41.817502 | 0.068826713 | 0.64719401 | 0.10634634 | 0.91530757 | 0.97782888 |
| ZCCHC9 | 290.67651 | -0.096695735 | 0.32577836 | -0.2968145 | 0.76660815 | 0.93138917 |
| ACOT12 | 96.1825126 | 0.200559224 | 0.72153906 | 0.27796031 | 7.81E-01 | 0.93748328 |
| SSBP2 | 1296.10688 | 0.144488026 | 0.41163976 | 0.351006 | 0.72558385 | 0.91630531 |
| ATG10 | 1044.06867 | -0.015592337 | 0.30023971 | -0.051933 | 0.95858211 | 0.98878187 |
| RPS23 | 1539.02861 | -0.110342174 | 0.61077639 | -0.1806589 | 0.85663534 | 0.96245027 |
| ATP6AP1L | 433.549879 | 0.987994051 | 0.52040128 | 1.89852348 | 0.05762716 | 0.47684824 |
| TMEM167A | 372.212993 | -0.291498196 | 0.23983508 | -1.215411 | 0.22420936 | 0.66551422 |
| XRCC4 | 671.08015 | -0.309371272 | 0.28608265 | -1.0814052 | 0.2795169 | 0.70446562 |
| VCAN | 224.757582 | 2.49671345 | 0.86326688 | 2.8921687 | 0.00382592 | 0.2001601 |
| EDIL3 | 397.430999 | 0.247336125 | 0.60036833 | 0.41197397 | 0.6803585 | 0.89941358 |
| AC113383.1 | 107.115483 | 0.591654339 | 0.60027813 | 0.98563367 | 0.32431291 | 0.73614611 |
| COX7C | 882.629765 | -0.001017482 | 0.42778042 | -0.0023785 | 0.99810222 | 0.99977652 |
| RASA1 | 2278.03426 | 0.223416632 | 0.20755606 | 1.07641583 | 0.28174133 | 0.70625976 |
| CCNH | 2067.66611 | 0.69242296 | 0.31719512 | 2.1829559 | 0.02903906 | 0.39554194 |
| TMEM161B | 673.114412 | -0.164936477 | 0.20962543 | -0.7868152 | 0.43139005 | 0.79135216 |
| TMEM161B-/ | 1753.54454 | 0.655702255 | 0.40613028 | 1.6145121 | 0.10641639 | 0.55440878 |
| MEF2C | 64.173642 | -0.552279234 | 0.52313179 | -1.0557172 | 0.29109743 | 0.71302224 |
| MEF2C-AS1 | 63.5052081 | -0.217111954 | 0.72408525 | -0.2998431 | 0.76429685 | 0.93083208 |
| CETN3 | 88.1919286 | 0.430847036 | 0.46200933 | 0.93255051 | 0.3510521 | 0.75515868 |
| MBLAC2 | 78.1630024 | -0.03094911 | 0.34734825 | -0.0891011 | 0.92900156 | 0.98272895 |
| POLR3G | 43.1954978 | -0.554628166 | 0.61042055 | -0.9086001 | 0.36356126 | 0.76267741 |
| LYSMD3 | 280.389451 | -0.267648862 | 0.3168927 | -0.8446041 | 0.39833193 | 0.781221 |
| ADGRV1 | 742.479611 | 0.194724895 | 0.49950173 | 0.38983828 | 6.97E-01 | 9.06E-01 |
| LUCAT1 | 290.165013 | -2.026709978 | 1.02931882 | -1.9689818 | 4.90E-02 | 4.54E-01 |
| ARRDC3 | 1046.14159 | -0.235481448 | 0.41586958 | -0.5662387 | 0.57123153 | 0.85399468 |
| ARRDC3-AS1 | 151.703156 | 0.176206949 | 0.31426377 | 0.56069762 | 0.57500369 | 0.85474196 |
| AC114316.2 | 40.5070308 | 1.166060549 | 0.90805258 | 1.28413329 | 0.19909532 | 0.64606998 |
| AC124854.1 | 526.934173 | 1.466597906 | 1.01207096 | 1.44910581 | 0.14730803 | 0.60507978 |
| NR2F1-AS1 | 247.603187 | 0.383192534 | 0.42558726 | 0.90038535 | 0.36791522 | 0.76610076 |
| NR2F1 | 199.602664 | 0.472859667 | 0.41814334 | 1.13085543 | 2.58E-01 | 0.69210282 |
| FAM172A | 3087.30091 | 0.024915555 | 0.27394588 | 0.09095065 | 0.92753181 | 0.98220631 |
| KIAA0825 | 2158.91171 | 0.6953755 | 0.51890037 | 1.34009444 | 0.18021464 | 0.63124625 |
| SLF1 | 408.72359 | 0.034228354 | 0.35130635 | 0.09743164 | 0.92238362 | 0.97970649 |
| MCTP1 | 126.55212 | 0.357037069 | 0.52919184 | 0.67468362 | 0.49987679 | 0.82482379 |
| TTC37 | 1411.52141 | -0.352621251 | 0.24119156 | -1.4619967 | 0.14374212 | 0.60014416 |
| ARSK | 173.939601 | -0.046380622 | 0.27048327 | -0.1714732 | 0.86385173 | 0.96393376 |
| RHOBTB3 | 436.492371 | 0.609325698 | 0.5613538 | 1.08545751 | 0.27771907 | 0.7036882 |
| GLRX | 1275.49849 | 0.210372519 | 0.62641274 | 0.33583691 | 0.73699385 | 0.92030854 |
| AC008592.4 | 44.0258071 | 1.736865113 | 0.8494261 | 2.04475129 | 0.04087938 | 0.43135775 |
| LINC01554 | 57.2671179 | 2.290970055 | 1.14793591 | 1.99572993 | 0.04596333 | 0.44508419 |
| ELL2 | 2417.18136 | -0.564318864 | 0.33315776 | -1.6938488 | 0.09029402 | 0.5339345 |
| AC104123.1 | 1016.30836 | -0.104099965 | 0.55796119 | -0.1865721 | 0.85199618 | 0.96050023 |
| CAST | 2822.51853 | -0.106133446 | 0.28452679 | -0.3730174 | 7.09E-01 | 9.11E-01 |
| ERAP1 | 470.722787 | 0.404982245 | 0.35885665 | 1.12853489 | 0.25909409 | 0.69241621 |
| AC009126.1 | 72.7600131 | 0.701076678 | 0.6146022 | 1.14069991 | 0.25399482 | 0.68891958 |
| ERAP2 | 158.695212 | 0.339949329 | 0.56891122 | 0.59754372 | 0.55014442 | 0.84620747 |
| LNPEP | 1178.48651 | 0.194101244 | 0.18392026 | 1.0553554 | 0.29126281 | 0.71317245 |
| LIX1-AS1 | 610.278515 | 0.278348797 | 0.43993051 | 0.63271083 | 0.52692249 | 0.83710323 |

| | | | | | | |
|---|---|---|---|---|---|---|
| RIOK2 | 149.907594 | -0.066810676 | 0.34024029 | -0.1963632 | 0.84432589 | 0.95742233 |
| LINC01340 | 57.3676224 | 0.321211137 | 0.85424088 | 0.37601939 | 0.70690248 | 0.91002281 |
| RGMB | 71.0335327 | 0.367935281 | 0.28314588 | 1.29945483 | 0.19378788 | 0.64548758 |
| CHD1 | 2105.61222 | 0.143709036 | 0.21427017 | 0.67069081 | 0.50241752 | 0.82638105 |
| LINC02062 | 337.42887 | 1.048042156 | 0.44645194 | 2.34749156 | 0.0189003 | 0.34084276 |
| AC021086.1 | 58.4770537 | 0.549888545 | 0.69742659 | 0.78845366 | 0.4304314 | 0.79055252 |
| FAM174A | 198.737492 | 0.375881294 | 0.24873356 | 1.51118047 | 0.13074248 | 0.58853436 |
| SLCO4C1 | 1658.34077 | 0.482710162 | 0.471354 | 1.02409264 | 0.30579152 | 0.72122 |
| PAM | 6839.63356 | -0.624812269 | 0.30489832 | -2.049248 | 0.04043787 | 0.43086475 |
| GIN1 | 182.597358 | -0.02434123 | 0.18678286 | -0.1303183 | 0.89631458 | 0.97312927 |
| PPIP5K2 | 1403.43792 | -0.024511066 | 0.2395258 | -0.1023316 | 0.91849345 | 0.97860783 |
| C5orf30 | 68.9302952 | 0.029077438 | 0.3715281 | 0.07826444 | 0.9376177 | 0.98435387 |
| NUDT12 | 230.595177 | 0.365294071 | 0.48436543 | 0.75417041 | 0.4507469 | 0.80364046 |
| AC099520.1 | 86.6484965 | 0.105148687 | 0.58456248 | 0.17987587 | 8.57E-01 | 0.96266803 |
| EFNA5 | 456.462642 | -0.750580995 | 0.70947207 | -1.057943 | 0.29008144 | 0.71260257 |
| FBXL17 | 4967.97571 | 0.222203154 | 0.15679882 | 1.41712263 | 0.15644708 | 0.61255952 |
| FER | 2442.47664 | -0.264565743 | 0.30196665 | -0.8761423 | 0.3809527 | 0.77210521 |
| AC008467.1 | 130.157478 | -0.470010025 | 0.75119357 | -0.6256843 | 0.53152203 | 0.83789294 |
| PJA2 | 1298.3936 | -0.009074907 | 0.16698193 | -0.0543466 | 9.57E-01 | 0.98861313 |
| MAN2A1 | 2318.32861 | 0.422343571 | 0.3023754 | 1.39675243 | 0.16248803 | 0.61912485 |
| TMEM232 | 1446.94143 | 0.839243796 | 0.61301625 | 1.36904005 | 0.17098676 | 0.62504223 |
| SLC25A46 | 701.484349 | -0.100101121 | 0.20553546 | -0.487026 | 0.62623988 | 0.87877458 |
| WDR36 | 397.204162 | -0.373930757 | 0.2194085 | -1.7042674 | 0.08833114 | 0.52894496 |
| STARD4 | 170.676562 | 0.853361331 | 0.50704161 | 1.68302032 | 0.09237116 | 0.53621484 |
| STARD4-AS1 | 136.30134 | 0.511562954 | 0.48948167 | 1.04511156 | 0.29597141 | 0.71727018 |
| NREP | 845.397286 | 0.069433568 | 0.28581591 | 0.24293108 | 8.08E-01 | 0.94617795 |
| EPB41L4A | 4146.77076 | -0.192927069 | 0.39859284 | -0.4840204 | 0.62837139 | 0.88009297 |
| EPB41L4A-A! | 81.3087643 | -0.88363815 | 0.44344521 | -1.9926659 | 0.04629804 | 0.44508419 |
| AC010261.2 | 67.5381633 | 0.140653356 | 0.80280013 | 0.17520345 | 0.86091977 | 0.96302275 |
| AC104126.1 | 120.289923 | 0.082279419 | 0.83001956 | 0.09912949 | 0.92103546 | 0.97939204 |
| APC | 941.762666 | -0.2130249 | 0.19599987 | -1.0868625 | 0.2770976 | 0.7036882 |
| SRP19 | 317.150149 | -0.06361332 | 0.25951539 | -0.2451235 | 0.80636081 | 0.94585363 |
| REEP5 | 983.063558 | -0.399696387 | 0.29153566 | -1.3710034 | 0.17037389 | 0.62424885 |
| DCP2 | 606.059759 | 0.030394551 | 0.16040989 | 0.18948053 | 0.84971621 | 0.95959925 |
| MCC | 500.225931 | 0.312089725 | 0.60025001 | 0.5199329 | 0.60311035 | 0.8678002 |
| YTHDC2 | 1082.08625 | 0.39083842 | 0.22041609 | 1.77318464 | 0.07619812 | 0.50767193 |
| KCNN2 | 58.2759166 | 0.81069675 | 0.94876849 | 0.85447268 | 3.93E-01 | 7.77E-01 |
| PGGT1B | 371.075441 | -0.096391612 | 0.18436655 | -0.5228259 | 0.60109539 | 0.86755264 |
| CCDC112 | 112.667072 | -0.827358706 | 0.42289143 | -1.9564329 | 0.05041418 | 0.45843622 |
| FEM1C | 430.837935 | -0.947337509 | 0.35387955 | -2.6770055 | 7.43E-03 | 0.23590813 |
| TMED7-TICA! | 109.158746 | 0.420467319 | 0.36919136 | 1.13888722 | 0.2547502 | 0.68976367 |
| ATG12 | 453.571411 | -0.117979609 | 0.21339528 | -0.5528689 | 0.5803532 | 0.85831909 |
| AP3S1 | 782.231778 | -0.208592207 | 0.29444274 | -0.7084305 | 0.47867799 | 0.81596289 |
| COMMD10 | 3371.6389 | -0.111707052 | 0.18753456 | -0.5956612 | 0.55140161 | 0.84689779 |
| SEMA6A | 2735.6112 | -0.88311901 | 0.62113925 | -1.421773 | 0.15509216 | 0.611403 |
| SEMA6A-AS1 | 436.468528 | 0.218735072 | 0.39288809 | 0.55673632 | 0.5777076 | 0.85681 |
| LINC00992 | 71.8825571 | 0.975031365 | 0.86460518 | 1.12771863 | 0.25943877 | 0.69242517 |
| LINC02147 | 1969.71062 | 2.133789778 | 0.77217721 | 2.76334209 | 0.00572128 | 0.22501232 |
| AC093206.1 | 101.105281 | -0.011470673 | 0.43258212 | -0.0265168 | 9.79E-01 | 0.99482978 |
| DTWD2 | 932.409848 | -0.071478105 | 0.21216797 | -0.336894 | 7.36E-01 | 9.20E-01 |
| AC008629.1 | 309.65876 | 1.067115436 | 0.53430249 | 1.99721216 | 0.04580214 | 0.44430994 |
| DMXL1 | 4722.88224 | -0.024073359 | 0.34174418 | -0.0704426 | 0.94384136 | 0.98522232 |
| TNFAIP8 | 1110.79186 | 0.226572679 | 0.3356633 | 0.67499986 | 5.00E-01 | 8.25E-01 |
| HSD17B4 | 1019.88981 | 0.29511796 | 0.3322969 | 0.8881153 | 0.37447873 | 0.7697436 |

| | | | | | | |
|---|---|---|---|---|---|---|
| SRFBP1 | 544.074321 | -0.300444158 | 0.33707374 | -0.8913307 | 0.37275181 | 0.769011 |
| AC119150.1 | 67.1693049 | 0.261624656 | 0.54764282 | 0.47772863 | 0.63284336 | 0.88151688 |
| SNX2 | 908.682691 | -0.203085415 | 0.279165 | -0.7274745 | 0.46693534 | 0.81129201 |
| SNX24 | 1542.99929 | -0.312406708 | 0.18719797 | -1.6688573 | 0.09514566 | 0.53824794 |
| PPIC | 203.585073 | -0.000144012 | 0.39839092 | -0.0003615 | 0.99971158 | 0.99985747 |
| PRDM6 | 77.3096404 | 0.568434184 | 0.48180218 | 1.17980826 | 0.23807649 | 0.67642542 |
| CEP120 | 344.522272 | 0.123710391 | 0.28626266 | 0.43215693 | 0.66562736 | 0.89369715 |
| CSNK1G3 | 1072.68845 | 0.299377168 | 0.24006093 | 1.24708826 | 0.21236514 | 0.65708554 |
| ZNF608 | 1898.72711 | 0.047024376 | 0.58401654 | 0.08051891 | 0.93582456 | 0.98406248 |
| AC112196.1 | 62.3235863 | -0.729318721 | 0.92418776 | -0.7891456 | 0.4300269 | 0.79055252 |
| LINC02240 | 77.8540418 | -1.034386255 | 0.52306792 | -1.9775372 | 0.04798094 | 0.45138969 |
| AC010235.1 | 206.452002 | 0.626737217 | 0.80443845 | 0.77909903 | 0.43592138 | 0.79549938 |
| GRAMD2B | 4017.54333 | -0.116341499 | 0.37171644 | -0.3129845 | 0.75429241 | 0.92799282 |
| AC093535.1 | 131.64729 | 0.529500333 | 0.73504032 | 0.7203691 | 0.47129777 | 0.81387913 |
| ALDH7A1 | 969.52808 | 0.435670172 | 0.52551902 | 0.82902837 | 4.07E-01 | 0.78481094 |
| PHAX | 250.624539 | -0.306307908 | 0.21668175 | -1.4136304 | 0.15747045 | 0.61319532 |
| LMNB1 | 57.8986348 | 0.415109866 | 0.57081199 | 0.72722696 | 0.46708694 | 0.81140608 |
| 3-Mar | 141.641 | -0.287149013 | 0.49020719 | -0.5857707 | 0.55802961 | 0.8489192 |
| C5orf63 | 106.345565 | 0.096851929 | 0.38768889 | 0.24981868 | 0.80272757 | 0.94543168 |
| PRRC1 | 436.08106 | -0.454189458 | 0.33227348 | -1.3669146 | 0.17165208 | 0.62520756 |
| CTXN3 | 42.4281391 | 1.0242913 | 0.85058451 | 1.20422049 | 2.29E-01 | 6.67E-01 |
| CCDC192 | 57.4677893 | 1.325159751 | 0.87610722 | 1.51255431 | 0.13039291 | 0.58853436 |
| LINC01184 | 1043.11341 | -0.011783389 | 0.31282157 | -0.0376681 | 0.96995232 | 0.99204339 |
| SLC12A2 | 387.081099 | -0.088206997 | 0.28261853 | -0.3121062 | 0.75495981 | 0.92799282 |
| FBN2 | 51.6765736 | 0.989265407 | 0.69823253 | 1.41681369 | 0.15653741 | 0.61255952 |
| SLC27A6 | 176.946403 | -0.673798826 | 1.60369526 | -0.4201539 | 0.67437303 | 0.89801183 |
| ISOC1 | 188.782285 | 0.149113783 | 0.25310134 | 0.58914656 | 0.55576296 | 0.84861484 |
| CHSY3 | 391.673588 | 1.005498133 | 0.55926679 | 1.79788635 | 0.07219502 | 0.50202435 |
| HINT1 | 1988.33436 | 0.227358384 | 0.41071956 | 0.55356113 | 0.57987923 | 0.85817368 |
| LYRM7 | 225.363128 | 0.537598345 | 0.26626645 | 2.01902398 | 0.04348473 | 0.44147822 |
| CDC42SE2 | 3612.83397 | 0.199986437 | 0.39349769 | 0.50822772 | 0.61129365 | 0.87163238 |
| RAPGEF6 | 36.6916438 | -0.415382758 | 0.3933122 | -1.0561146 | 0.29091586 | 0.71295972 |
| P4HA2-AS1 | 41.018065 | 0.945220447 | 0.78020827 | 1.2114976 | 0.22570475 | 0.66688879 |
| P4HA2 | 1456.5108 | -0.058319015 | 0.48007404 | -0.1214792 | 9.03E-01 | 9.75E-01 |
| SLC22A4 | 502.095141 | 0.187433871 | 0.28675663 | 0.65363396 | 0.51334766 | 0.8308396 |
| SLC22A5 | 2466.74445 | 0.512933712 | 0.53716907 | 0.95488319 | 0.33963677 | 0.74644675 |
| AC116366.3 | 630.123451 | 0.087258338 | 0.56573001 | 0.15424025 | 0.87742031 | 0.96655149 |
| IRF1 | 249.818177 | 0.302963081 | 0.7346732 | 0.41237802 | 0.68006237 | 0.89941358 |
| TH2LCRR | 89.1973561 | 0.124343838 | 0.7082138 | 0.17557387 | 0.86062873 | 0.96302275 |
| AC004039.1 | 42.4343813 | 0.301156311 | 0.76869385 | 0.39177666 | 0.69522324 | 0.90589033 |
| KIF3A | 386.233723 | -0.019098044 | 0.30305643 | -0.0630181 | 0.94975208 | 0.98723887 |
| 8-Sep | 238.81941 | 0.753240295 | 0.44244503 | 1.70244944 | 0.08867115 | 0.52982364 |
| SHROOM1 | 78.290114 | -0.028457152 | 0.50646165 | -0.0561882 | 0.95519191 | 0.98840182 |
| UQCRQ | 436.8733 | -0.234992596 | 0.43284215 | -0.542906 | 0.58719453 | 0.86112855 |
| LEAP2 | 116.287095 | -1.496270651 | 0.94063294 | -1.5907062 | 0.11167571 | 0.56443177 |
| AFF4 | 3429.01346 | -0.569964477 | 0.21564808 | -2.6430306 | 0.00821676 | 0.24877162 |
| ZCCHC10 | 530.232209 | -0.45002578 | 0.19764269 | -2.2769664 | 0.02278823 | 0.36575908 |
| HSPA4 | 932.259084 | -0.686347836 | 0.39124435 | -1.754269 | 0.07938442 | 0.51192014 |
| FSTL4 | 141.551716 | 0.52408402 | 0.67085388 | 0.78121933 | 0.43467351 | 0.79451524 |
| C5orf15 | 856.624406 | 0.398821698 | 0.33093792 | 1.20512542 | 0.22815489 | 0.66705629 |
| VDAC1 | 900.700866 | 0.049643035 | 0.21671053 | 0.22907533 | 0.81881037 | 0.94990455 |
| AC008608.2 | 34.7663062 | -0.249655318 | 0.55899849 | -0.4466118 | 0.65515538 | 0.89053559 |
| TCF7 | 95.7070091 | 0.381093971 | 0.39869603 | 0.95585093 | 3.39E-01 | 0.74644675 |
| SKP1 | 746.493272 | 0.02440262 | 0.27109519 | 0.09001495 | 0.92827533 | 0.98272895 |

| | | | | | | |
|---|---|---|---|---|---|---|
| CDKL3 | 416.692731 | 0.17551393 | 0.31606836 | 0.5553037 | 0.57868695 | 0.8575616 |
| UBE2B | 1266.1839 | -0.424206271 | 0.24027134 | -1.76553 | 0.07747472 | 0.51128842 |
| CDKN2AIPNL | 75.9370128 | -0.007624551 | 0.38129678 | -0.0199964 | 0.98404627 | 0.99523767 |
| LINC01843 | 59.4765065 | 0.697341421 | 0.64467057 | 1.08170197 | 0.27938498 | 0.70446562 |
| JADE2 | 454.948129 | 1.079018544 | 0.43599132 | 2.47486243 | 0.01332875 | 0.29875272 |
| SAR1B | 729.400401 | -0.108432571 | 0.22969041 | -0.4720814 | 0.63686869 | 0.88266168 |
| SEC24A | 729.452095 | -0.088710174 | 0.27676285 | -0.3205277 | 0.7485683 | 0.92552718 |
| CAMLG | 240.255744 | -0.137397099 | 0.30996264 | -0.4432699 | 0.65757055 | 0.89053559 |
| DDX46 | 1186.86383 | -0.560671871 | 0.30527863 | -1.8365906 | 0.06627035 | 0.48798003 |
| C5orf24 | 395.8419 | 0.137132749 | 0.26303554 | 0.52134683 | 0.60212518 | 0.86770668 |
| TXNDC15 | 221.386188 | -0.193632626 | 0.45050454 | -0.4298128 | 0.66733181 | 0.8944961 |
| PCBD2 | 1114.70402 | -0.773939919 | 0.46882499 | -1.6508077 | 0.09877784 | 0.54236074 |
| C5orf66 | 93.5475039 | 0.782204319 | 0.60925709 | 1.28386576 | 0.19918893 | 0.64622075 |
| H2AFY | 697.271319 | -0.432235666 | 0.36833867 | -1.1734735 | 0.24060598 | 0.67925154 |
| AC034206.1 | 51.6064929 | 1.395919099 | 0.76363179 | 1.82800026 | 0.06754951 | 0.48963361 |
| CXCL14 | 2158.56323 | 0.966645379 | 0.42919042 | 2.25225295 | 0.02430629 | 0.37480999 |
| SLC25A48 | 2228.16504 | 0.660706839 | 0.62191806 | 1.06236961 | 0.28806793 | 0.71170639 |
| LECT2 | 62.9645902 | 0.58421669 | 1.07813099 | 0.54187914 | 0.58790177 | 0.8613161 |
| TGFBI | 55.5992961 | -1.246957906 | 0.67560929 | -1.845679 | 0.06493882 | 0.48790819 |
| SMAD5 | 1027.37257 | 0.034195659 | 0.23194262 | 0.14743154 | 0.88279141 | 0.96824408 |
| SMIM32 | 239.260345 | 0.751545222 | 0.49427583 | 1.52049763 | 0.12838595 | 0.58793678 |
| TRPC7 | 168.919155 | 1.517726406 | 0.89940793 | 1.68747278 | 0.09151248 | 0.53537574 |
| TRPC7-AS1 | 589.713399 | 0.832529468 | 0.35552551 | 2.34168703 | 1.92E-02 | 3.41E-01 |
| SPOCK1 | 672.75305 | -1.493959023 | 0.8414422 | -1.7754743 | 0.0758196 | 0.50730849 |
| KLHL3 | 37.9765211 | 0.072955384 | 0.76292288 | 0.09562616 | 0.92381749 | 0.98015759 |
| HNRNPA0 | 287.438316 | -0.478917782 | 0.35832375 | -1.3365505 | 0.18136941 | 0.6318513 |
| AC106791.1 | 76.9170933 | 0.563428808 | 0.82957256 | 0.67917965 | 0.49702404 | 0.82278993 |
| PKD2L2 | 52.0951095 | 0.512039497 | 0.4828712 | 1.06040595 | 0.28895995 | 0.71218341 |
| FAM13B | 1490.68744 | -0.268406984 | 0.18627499 | -1.440918 | 0.14960786 | 0.60706776 |
| NME5 | 457.659686 | 0.804146126 | 0.58860693 | 1.36618529 | 0.17188081 | 0.62575572 |
| BRD8 | 415.184935 | -0.108233058 | 0.32565404 | -0.3323559 | 0.7396205 | 0.92123698 |
| CDC23 | 160.667566 | -0.026237303 | 0.2320843 | -0.1130507 | 0.90999032 | 0.97692341 |
| FAM53C | 214.884505 | -0.267085484 | 0.36097802 | -0.739894 | 0.45936429 | 0.80669065 |
| KDM3B | 1216.90463 | -0.116238644 | 0.18214367 | -0.6381701 | 0.52336295 | 0.83532091 |
| EGR1 | 120.24955 | -0.045186075 | 0.84295516 | -0.0536044 | 0.95725038 | 0.98861313 |
| ETF1 | 645.726846 | -0.718918566 | 0.45231881 | -1.5894068 | 0.11196859 | 0.56455091 |
| HSPA9 | 1562.22632 | -0.858548457 | 0.36866648 | -2.3287945 | 0.01986996 | 0.34651082 |
| CTNNA1 | 4249.51965 | -0.31341261 | 0.1723994 | -1.8179449 | 0.06907256 | 0.49273866 |
| SIL1 | 1732.31219 | 0.126789958 | 0.25868662 | 0.49012956 | 0.62404222 | 0.87791268 |
| AC011405.1 | 53.3319416 | 0.581495132 | 0.66844694 | 0.86991966 | 0.38434431 | 0.77317162 |
| MATR3 | 50.4405332 | -0.604075211 | 0.76416696 | -0.7905016 | 0.42923488 | 0.79037914 |
| PAIP2 | 919.982543 | -0.232194529 | 0.2574408 | -0.9019337 | 0.3670921 | 0.76546934 |
| AC135457.1 | 50.4758213 | 0.145661769 | 0.73579962 | 0.19796391 | 0.8430733 | 0.95716002 |
| SLC23A1 | 224.4675 | -0.024467487 | 0.63283215 | -0.0386635 | 9.69E-01 | 9.92E-01 |
| PROB1 | 40.6903127 | 0.627022798 | 0.56504878 | 1.10967906 | 0.26713735 | 0.69838864 |
| SPATA24 | 66.8297311 | -0.021256225 | 0.45284113 | -0.0469397 | 0.96256129 | 0.99012655 |
| DNAJC18 | 89.0070627 | 0.233147295 | 0.48888211 | 0.4768988 | 0.63343418 | 0.88188803 |
| UBE2D2 | 5802.29542 | -0.352235181 | 0.46344295 | -0.76004 | 0.44723068 | 0.80223612 |
| CXXC5 | 551.919858 | 0.312616342 | 0.37704875 | 0.82911385 | 0.40703999 | 0.78481094 |
| AC008667.1 | 35.1647291 | 0.283811673 | 0.61789037 | 0.45932367 | 0.64600175 | 0.88700494 |
| NRG2 | 316.891582 | 0.964246606 | 0.49264614 | 1.95728033 | 0.05031452 | 0.45824658 |
| PURA | 916.820463 | 0.387996991 | 0.24604782 | 1.57691701 | 0.11481462 | 0.57144663 |
| IGIP | 129.137877 | 0.932316379 | 0.38078226 | 2.44842388 | 0.01434827 | 0.30634236 |
| CYSTM1 | 2498.39207 | -0.177209362 | 0.25813544 | -0.6864976 | 0.49239938 | 0.82087587 |

| | | | | | | |
|---|---|---|---|---|---|---|
| PFDN1 | 630.44304 | -0.195300675 | 0.24030166 | -0.8127313 | 0.41637215 | 0.78632759 |
| AC008438.1 | 102.522648 | -0.499666108 | 0.41039749 | -1.2175175 | 0.22340739 | 0.66426142 |
| ANKHD1-EIF4 | 167.685391 | 0.580807418 | 0.41189651 | 1.41008094 | 0.15851578 | 0.61423843 |
| SRA1 | 237.348694 | 0.057004427 | 0.36498588 | 0.15618255 | 0.87588913 | 0.96617876 |
| APBB3 | 110.841379 | -0.079870051 | 0.36306707 | -0.219987 | 0.82588125 | 0.95192757 |
| SLC35A4 | 171.061645 | -0.205378577 | 0.47554133 | -0.4318838 | 0.6658259 | 0.89378862 |
| NDUFA2 | 290.132733 | -0.219726987 | 0.34975552 | -0.6282302 | 0.52985313 | 0.83789294 |
| TMCO6 | 33.9308017 | 0.365398612 | 0.40327914 | 0.90606872 | 0.36489948 | 0.76338174 |
| IK | 186.721911 | -0.481548783 | 0.35624205 | -1.3517461 | 0.17645656 | 0.62709281 |
| WDR55 | 168.95333 | -0.714102674 | 0.31668886 | -2.2549031 | 0.02413941 | 0.37387451 |
| HARS | 357.906919 | -0.42821998 | 0.2609559 | -1.6409669 | 0.10080429 | 0.54608983 |
| HARS2 | 94.3282716 | 0.121888833 | 0.29570575 | 0.41219635 | 0.68019551 | 0.89941358 |
| ZMAT2 | 148.378268 | -0.336521874 | 0.30284221 | -1.1112119 | 0.26647715 | 0.69812269 |
| TAF7 | 378.335655 | -0.501227614 | 0.20391409 | -2.4580332 | 0.01397003 | 0.30363654 |
| DIAPH1 | 2497.74581 | -0.07486258 | 0.19301102 | -0.3878669 | 0.69811455 | 0.90696517 |
| AC008781.2 | 62.9084365 | 0.033987527 | 0.75132465 | 0.0452368 | 0.96391856 | 0.99012655 |
| HDAC3 | 389.337557 | -0.21044768 | 0.30113626 | -0.6988454 | 0.48464867 | 0.81839266 |
| RELL2 | 75.5160753 | -0.381394369 | 0.63231453 | -0.6031719 | 0.54639432 | 0.84460887 |
| FCHSD1 | 49.0285937 | 0.389502629 | 0.45340697 | 0.85905744 | 0.39030883 | 0.77569824 |
| PCDH1 | 141.460991 | 0.724439013 | 0.4766283 | 1.51992446 | 0.12852996 | 0.58793678 |
| DELE1 | 359.521178 | -0.291872364 | 0.20390321 | -1.4314261 | 0.15230814 | 0.61001662 |
| RNF14 | 578.89764 | 0.044072462 | 0.31181 | 0.14134397 | 0.88759822 | 0.97098615 |
| AC005740.3 | 39.7486025 | -0.266201166 | 0.88599379 | -0.3004549 | 0.7638302 | 0.93079056 |
| GNPDA1 | 580.149024 | -0.612389255 | 0.4752774 | -1.2884881 | 0.19757612 | 0.64558745 |
| NDFIP1 | 1174.98116 | -0.109177376 | 0.19787949 | -0.5517367 | 0.58112877 | 0.85895204 |
| SPRY4 | 185.819646 | 0.92173996 | 0.49965529 | 1.84475172 | 0.06507366 | 0.48790819 |
| SPRY4-AS1 | 287.950183 | 1.370833543 | 0.4927827 | 2.78182157 | 0.00540548 | 0.2192096 |
| FGF1 | 77.1916829 | 0.452496041 | 0.62537651 | 0.72355778 | 0.46933727 | 0.8127866 |
| ARHGAP26 | 6765.74469 | 0.505600681 | 0.47835315 | 1.05696112 | 0.29052933 | 0.71269438 |
| ARHGAP26-A | 60.7653593 | 0.707808803 | 0.65562781 | 1.07958935 | 0.28032508 | 0.70515983 |
| NR3C1 | 2266.51397 | -0.368435688 | 0.25103011 | -1.4676952 | 0.14218703 | 0.59729009 |
| YIPF5 | 222.5753 | -0.654471421 | 0.31106682 | -2.1039577 | 0.03538214 | 0.4198859 |
| KCTD16 | 964.158432 | -0.289452476 | 0.63111749 | -0.4586349 | 0.6464964 | 0.88739497 |
| PRELID2 | 674.603631 | 0.003233956 | 0.60412773 | 0.0053531 | 0.99572886 | 0.99866448 |
| SH3RF2 | 335.545842 | -1.100461136 | 0.78165483 | -1.4078607 | 0.15917231 | 0.61423843 |
| PLAC8L1 | 37.525497 | -0.887953532 | 0.50633619 | -1.7536837 | 0.07948472 | 0.51192014 |
| LARS | 854.55078 | -0.366577306 | 0.21268565 | -1.7235639 | 0.08478661 | 0.52183702 |
| AC091959.3 | 37.2638605 | 0.159628278 | 0.39864955 | 0.40042257 | 0.6888453 | 0.9032816 |
| TCERG1 | 1106.71506 | -0.40528648 | 0.24274143 | -1.6696222 | 0.09499414 | 0.53824794 |
| PPP2R2B | 1717.99296 | 3.374852703 | 0.93809397 | 3.59756359 | 0.00032121 | 0.06474783 |
| STK32A | 294.551024 | 0.878679912 | 0.71154241 | 1.23489465 | 0.21686972 | 0.66032006 |
| AC091948.1 | 297.23549 | 1.433590371 | 0.65437509 | 2.19077772 | 0.02846788 | 0.39280225 |
| FBXO38 | 706.789139 | -0.196883612 | 0.16492115 | -1.1938045 | 0.23255444 | 0.66978724 |
| ABLIM3 | 3092.45053 | -0.537604812 | 0.55256717 | -0.9729221 | 0.33059202 | 0.73849818 |
| AC012613.1 | 79.9515244 | 0.04098661 | 0.81158874 | 0.0505017 | 0.9597226 | 0.98912673 |
| AFAP1L1 | 147.677652 | -0.416577291 | 0.39136028 | -1.0644343 | 2.87E-01 | 7.11E-01 |
| GRPEL2 | 63.4449739 | -0.511395712 | 0.41531109 | -1.2313558 | 0.21818983 | 0.66064833 |
| CSNK1A1 | 3558.00555 | 0.079201667 | 0.19808272 | 0.39984137 | 0.68927335 | 0.90367 |
| ARHGEF37 | 230.33244 | 0.218881487 | 0.45425763 | 0.48184438 | 0.6299165 | 0.88083664 |
| PPARGC1B | 757.086561 | 0.204156382 | 0.51832355 | 0.39387827 | 0.69367091 | 0.90587439 |
| SLC26A2 | 102.38723 | 0.166386649 | 0.56747263 | 0.29320648 | 0.76936433 | 0.93209094 |
| TIGD6 | 41.8060466 | -0.408214296 | 0.3564082 | -1.1453561 | 0.25206169 | 0.68862185 |
| HMGXB3 | 484.27879 | -0.276618142 | 0.31646479 | -0.8740882 | 0.38207022 | 0.77253821 |
| CAMK2A | 142.522827 | 2.309898055 | 1.10571175 | 2.08905988 | 0.03670233 | 0.42204605 |

| | | | | | | |
|---|---|---|---|---|---|---|
| ARSI | 150.109899 | 1.35897488 | 0.81296919 | 1.67161917 | 9.46E-02 | 0.53824794 |
| TCOF1 | 180.029969 | -0.150219556 | 0.4455766 | -0.3371352 | 0.73601499 | 0.92019178 |
| CD74 | 1610.83763 | 0.863037513 | 0.57497898 | 1.50098967 | 0.13335823 | 0.59240843 |
| RPS14 | 1220.36668 | 0.055362263 | 0.46997301 | 0.11779881 | 0.90622707 | 0.97646839 |
| NDST1 | 659.393021 | -0.311346212 | 0.30348198 | -1.0259133 | 0.30493243 | 0.72103276 |
| SYNPO | 116.846132 | 0.138047189 | 0.47032263 | 0.29351594 | 7.69E-01 | 0.93209094 |
| RBM22 | 175.082214 | -0.625632782 | 0.26400121 | -2.3698103 | 0.01779721 | 0.33417314 |
| DCTN4 | 703.513199 | -0.176530773 | 0.19159425 | -0.9213782 | 0.35685299 | 0.75893336 |
| SMIM3 | 54.8396014 | -0.173688334 | 0.47786035 | -0.3634709 | 0.71625314 | 0.91298968 |
| ZNF300 | 101.582983 | -0.925826665 | 0.31305597 | -2.9573838 | 0.00310262 | 0.18216645 |
| GPX3 | 23397.173 | -0.077237592 | 0.61385495 | -0.1258238 | 0.89987136 | 0.97465308 |
| TNIP1 | 1206.7936 | -0.045672708 | 0.28918788 | -0.1579344 | 0.87450851 | 0.96615698 |
| ANXA6 | 363.763153 | 0.262813711 | 0.57866836 | 0.45416983 | 0.64970659 | 0.88850926 |
| GM2A | 54.8091362 | 0.571601713 | 0.44361546 | 1.2885072 | 0.19756946 | 0.64558745 |
| SLC36A2 | 3283.23669 | -0.168514266 | 1.02566486 | -0.1642976 | 0.86949688 | 0.96564168 |
| SLC36A1 | 658.903686 | -0.327817637 | 0.54352163 | -0.6031363 | 0.546418 | 0.84460887 |
| SPARC | 205.88475 | 1.060771048 | 0.62347461 | 1.70138611 | 0.08887051 | 0.53009053 |
| ATOX1 | 525.132951 | -0.110342762 | 0.29164668 | -0.378344 | 0.70517509 | 0.909554 |
| G3BP1 | 1249.21772 | -0.402336713 | 0.22849435 | -1.7608169 | 0.0782694 | 0.51192014 |
| LINC01933 | 70.8646117 | 0.153378384 | 0.35757085 | 0.42894543 | 0.66796294 | 0.89507948 |
| LINC01470 | 56.1799899 | 0.127766018 | 0.39705065 | 0.32178771 | 0.74761353 | 0.92528565 |
| FAM114A2 | 189.612991 | 0.048921393 | 0.24140585 | 0.20265206 | 0.839407 | 0.9556011 |
| MFAP3 | 293.216627 | 0.04126435 | 0.25358404 | 0.16272455 | 0.87073531 | 0.9657206 |
| GALNT10 | 407.839521 | 0.00112923 | 0.40649139 | 0.00277799 | 0.99778349 | 0.99977652 |
| SAP30L-AS1 | 236.853902 | 0.81652059 | 0.30290287 | 2.6956515 | 0.00702511 | 0.23315548 |
| SAP30L | 162.705698 | 0.096972486 | 0.24387284 | 0.39763545 | 0.69089893 | 0.90467631 |
| LARP1 | 2789.09525 | -0.40303873 | 0.27194562 | -1.4820563 | 0.1383253 | 0.59474076 |
| FAXDC2 | 210.788452 | 0.578211224 | 0.56873125 | 1.01666863 | 0.30931109 | 0.72449199 |
| CNOT8 | 290.34271 | -0.247759248 | 0.1784273 | -1.3885725 | 0.16496279 | 0.62065658 |
| GEMIN5 | 203.276069 | -0.456488035 | 0.39754434 | -1.1482695 | 0.25085734 | 0.68648949 |
| MRPL22 | 236.253578 | -0.170248635 | 0.19245174 | -0.8846303 | 0.37635609 | 0.77035529 |
| SGCD | 895.285892 | 2.44070406 | 0.71090697 | 3.43322568 | 0.00059645 | 0.08329748 |
| TIMD4 | 613.869686 | 0.531426369 | 0.445202 | 1.19367472 | 0.23260523 | 0.66978724 |
| HAVCR1 | 1650.90001 | -0.282139768 | 0.4752624 | -0.5936505 | 5.53E-01 | 8.47E-01 |
| HAVCR2 | 1067.86905 | 0.770372834 | 0.38075442 | 2.02328007 | 0.04304428 | 0.44030444 |
| AC011377.1 | 42.6662485 | 1.141793918 | 0.60425617 | 1.88958587 | 5.88E-02 | 0.47778423 |
| CYFIP2 | 1538.46632 | 0.372647364 | 0.56501918 | 0.65953046 | 0.50955519 | 0.82882082 |
| THG1L | 52.6119274 | -0.841679777 | 0.504885 | -1.6670723 | 0.09550004 | 0.53824794 |
| LSM11 | 53.0288344 | -0.652993753 | 0.35937846 | -1.8170086 | 0.06921579 | 0.49273866 |
| CLINT1 | 2016.1904 | -0.07871565 | 0.22394154 | -0.3515009 | 0.72521261 | 0.91600702 |
| EBF1 | 53.015983 | -0.096727327 | 0.62848835 | -0.1539047 | 0.87768486 | 0.96658493 |
| RNF145 | 583.800307 | -0.107617855 | 0.17183054 | -0.6263023 | 5.31E-01 | 8.38E-01 |
| UBLCP1 | 248.535434 | -0.396450562 | 0.30135289 | -1.3155691 | 0.1883187 | 0.64026838 |
| AC008691.1 | 38.8802262 | -1.165287868 | 0.40539673 | -2.8744382 | 0.00404747 | 0.20256898 |
| ADRA1B | 443.746732 | -0.224051042 | 0.64224208 | -0.3488576 | 0.72719621 | 0.91645085 |
| TTC1 | 796.564172 | -0.300138859 | 0.23307282 | -1.2877471 | 0.197834 | 0.64579914 |
| PWWP2A | 555.480477 | 0.081285024 | 0.31408157 | 0.25880227 | 0.7957878 | 0.94252227 |
| ZBED8 | 38.1768035 | 0.764719431 | 0.50484842 | 1.51475056 | 0.12983558 | 0.58853436 |
| SLU7 | 182.359386 | -0.212666002 | 0.28962952 | -0.7342691 | 0.46278475 | 0.8082072 |
| MIR3142HG | 153.892217 | 1.747709686 | 0.79824194 | 2.1894486 | 0.02856425 | 0.39280225 |
| GABRB2 | 217.692089 | 1.097565236 | 1.0686791 | 1.02702975 | 0.30440645 | 0.72082426 |
| AC113414.1 | 2915.11328 | 1.631198714 | 0.76047182 | 2.14498245 | 0.03195425 | 0.40988962 |
| CCNG1 | 465.368417 | 0.752369 | 0.33166462 | 2.26846323 | 0.02330099 | 0.37046453 |
| NUDCD2 | 121.866922 | 0.213317081 | 0.39232265 | 0.54372869 | 0.58662819 | 0.86102615 |

| | | | | | | |
|---|---|---|---|---|---|---|
| MAT2B | 263.137179 | 0.265868687 | 0.30347452 | 0.8760824 | 0.38098524 | 0.77210521 |
| AC109466.1 | 144.173265 | 0.222688211 | 0.95972489 | 0.23203338 | 0.81651209 | 0.94935374 |
| AC008415.1 | 201.740735 | 0.958704387 | 1.1172654 | 0.85808115 | 0.39084765 | 0.77606489 |
| TENM2 | 795.374251 | 1.672699202 | 0.88078868 | 1.89909252 | 0.05755231 | 0.47684824 |
| AC008637.1 | 54.9227007 | 0.391138984 | 0.72673933 | 0.53821084 | 0.5904315 | 0.86315571 |
| WWC1 | 8445.37487 | -0.754986476 | 0.43391138 | -1.7399555 | 0.08186684 | 0.51593841 |
| RARS | 700.988437 | -0.29890087 | 0.29334823 | -1.0189285 | 0.30823692 | 0.72367286 |
| PANK3 | 573.107253 | -0.021264219 | 0.29184036 | -0.0728625 | 0.94191553 | 0.98489601 |
| SLIT3 | 994.448792 | -0.545292745 | 1.00479144 | -0.5426925 | 0.58734156 | 0.86112855 |
| SPDL1 | 56.7013511 | -0.30595631 | 0.41691253 | -0.7338621 | 0.46303278 | 0.8082072 |
| FAM196B | 72.6961378 | 2.378766427 | 0.92506966 | 2.57144572 | 0.01012749 | 0.26582196 |
| KCNIP1 | 85.0011568 | 0.456948817 | 0.53843004 | 0.84866888 | 0.39606557 | 0.7795748 |
| AC008514.1 | 68.9381086 | 0.66369057 | 0.82190894 | 0.80749891 | 0.4193791 | 0.78783502 |
| RANBP17 | 1351.30732 | 0.569589245 | 0.32003144 | 1.77979153 | 0.07511008 | 0.5071704 |
| NPM1 | 632.02161 | -0.504367152 | 0.37556786 | -1.3429455 | 0.17928963 | 0.6302957 |
| FBXW11 | 2233.09362 | -0.209312969 | 0.25563911 | -0.818783 | 0.41291021 | 0.78632759 |
| STK10 | 363.747357 | -1.203168319 | 0.71015203 | -1.6942405 | 0.09021959 | 0.53372463 |
| UBTD2 | 1028.14437 | -0.036652412 | 0.36607221 | -0.1001234 | 0.92024632 | 0.97939204 |
| SH3PXD2B | 265.843321 | 1.108137135 | 0.47786405 | 2.31893807 | 0.02039839 | 0.35081655 |
| DUSP1 | 648.014886 | -0.842792063 | 0.79682876 | -1.0576828 | 0.29020009 | 0.71260706 |
| ERGIC1 | 1508.393 | -0.129213239 | 0.31070822 | -0.4158668 | 0.67750746 | 0.89864474 |
| RPL26L1 | 80.6797901 | -0.221504028 | 0.311382 | -0.7113578 | 0.47686252 | 0.81570932 |
| ATP6V0E1 | 1462.3435 | -0.335767328 | 0.24788281 | -1.3545406 | 0.17556397 | 0.62699479 |
| CREBRF | 1335.51933 | -0.297496543 | 0.3313446 | -0.8978464 | 0.36926747 | 0.76620563 |
| BNIP1 | 97.7461793 | 0.008858458 | 0.2831678 | 0.03128342 | 0.97504351 | 0.99402915 |
| STC2 | 97.2499455 | -1.070216716 | 0.65215787 | -1.6410393 | 0.10078925 | 0.54608983 |
| BOD1 | 185.099066 | -0.287001319 | 0.19104752 | -1.502251 | 0.13303231 | 0.5919553 |
| CPEB4 | 4589.16731 | -0.385118541 | 0.36348937 | -1.0595043 | 2.89E-01 | 0.71218341 |
| SFXN1 | 1405.89144 | -0.127016954 | 0.3579173 | -0.3548779 | 0.72268104 | 0.91592159 |
| THOC3 | 260.955825 | -1.012156382 | 0.52799094 | -1.9169957 | 0.05523847 | 0.46947044 |
| SIMC1 | 356.141732 | 0.021766828 | 0.21019079 | 0.10355748 | 0.91752053 | 0.97828877 |
| KIAA1191 | 792.505465 | 0.081144045 | 0.48733626 | 0.16650525 | 0.86775935 | 0.96490447 |
| AC138956.1 | 66.3990155 | 0.358406097 | 0.8700453 | 0.41193958 | 0.68038371 | 0.89941358 |
| AC138956.2 | 65.4108793 | 0.958694587 | 1.08957147 | 0.87988224 | 0.37892311 | 0.77061593 |
| ARL10 | 206.638752 | -0.092987489 | 0.27100774 | -0.3431175 | 0.73151009 | 0.91917457 |
| NOP16 | 84.9302724 | -1.031209446 | 0.60070579 | -1.7166631 | 0.08604075 | 0.52392738 |
| CLTB | 323.425441 | -0.934536728 | 0.37883824 | -2.466849 | 0.01363078 | 0.30135021 |
| FAF2 | 960.371675 | -0.470592293 | 0.3255944 | -1.4453329 | 0.14836441 | 0.60622442 |
| RNF44 | 201.32679 | 0.004796055 | 0.27606172 | 0.01737313 | 0.98613895 | 0.99609994 |
| CDHR2 | 857.724238 | -0.006586929 | 0.80148768 | -0.0082184 | 0.99344276 | 0.9976934 |
| TSPAN17 | 128.501705 | -0.017801167 | 0.4149676 | -0.0428977 | 0.96578306 | 0.99081812 |
| UIMC1 | 1277.4749 | -0.288930138 | 0.23295088 | -1.240305 | 0.21486262 | 0.65925411 |
| ZNF346 | 510.703381 | 0.0289927 | 0.33390136 | 0.08683013 | 0.93080654 | 0.98272895 |
| FGFR4 | 443.042402 | 0.820847292 | 0.45919081 | 1.78759522 | 0.07384134 | 0.50581869 |
| NSD1 | 2158.89527 | 0.062163361 | 0.14722101 | 0.42224518 | 6.73E-01 | 8.98E-01 |
| RAB24 | 46.3795938 | -0.448403448 | 0.72897186 | -0.6151176 | 0.53847704 | 0.84049498 |
| MXD3 | 50.3419898 | 0.221788414 | 0.46311985 | 0.47890069 | 0.63200928 | 0.88123095 |
| PRELID1 | 554.20951 | -0.410211887 | 0.65934184 | -0.6221536 | 0.53384089 | 0.83915086 |
| LMAN2 | 465.537443 | -0.19417403 | 0.40689729 | -0.4772065 | 0.63321509 | 0.88177189 |
| RGS14 | 406.367267 | 0.210926737 | 0.51554608 | 0.40913265 | 0.68244231 | 0.89962501 |
| SLC34A1 | 1914.91928 | 0.660161627 | 0.86333983 | 0.76466022 | 4.44E-01 | 8.00E-01 |
| F12 | 118.954939 | 0.258587315 | 0.4564183 | 0.56655772 | 5.71E-01 | 0.85399468 |
| GRK6 | 292.711621 | 0.695756581 | 0.3999158 | 1.73975766 | 0.08190158 | 0.51593841 |
| PRR7-AS1 | 42.6053467 | -0.464345508 | 0.56549127 | -0.8211365 | 4.12E-01 | 0.78632759 |

| | | | | | | |
|---|---|---|---|---|---|---|
| PRR7 | 105.670328 | -0.178485207 | 0.47785258 | -0.3735152 | 7.09E-01 | 9.11E-01 |
| DBN1 | 110.958794 | 0.105170376 | 0.36093312 | 0.29138467 | 0.77075714 | 0.93302012 |
| PDLIM7 | 121.00994 | -0.453366876 | 0.58444732 | -0.775719 | 0.43791493 | 0.79682729 |
| DDX41 | 117.215174 | -0.661839679 | 0.40442586 | -1.636492 | 0.10173666 | 0.54729371 |
| FAM193B | 837.621788 | -0.278857238 | 0.25408954 | -1.0974762 | 0.27243326 | 0.70073986 |
| TMED9 | 365.28736 | -0.301942269 | 0.45477005 | -0.6639449 | 0.50672556 | 0.82806179 |
| B4GALT7 | 137.351208 | -0.208971614 | 0.4695853 | -0.4450131 | 0.65631028 | 0.89053559 |
| AC106795.3 | 54.9923837 | -0.691287845 | 0.55348808 | -1.2489661 | 0.21167747 | 0.65584609 |
| RMND5B | 370.668729 | -0.270648202 | 0.32085816 | -0.8435135 | 0.3989413 | 0.78140731 |
| NHP2 | 155.273886 | -0.425573932 | 0.55217926 | -0.770717 | 0.44087471 | 0.79826598 |
| HNRNPAB | 429.156883 | -0.655070011 | 0.41145331 | -1.5920883 | 0.11136486 | 0.56431726 |
| PHYKPL | 2038.70742 | 0.036196463 | 0.30337217 | 0.11931373 | 0.90502681 | 0.97598655 |
| COL23A1 | 680.336603 | -0.312240748 | 0.69466347 | -0.4494849 | 0.65308189 | 0.88949911 |
| CLK4 | 641.312536 | 0.103329806 | 0.38670771 | 0.26720389 | 0.78931218 | 0.94005579 |
| ZNF354A | 330.787875 | 0.18761398 | 0.33186644 | 0.56532978 | 0.57184948 | 0.85403583 |
| ZNF354B | 280.133754 | -0.285795424 | 0.34036969 | -0.8396618 | 0.40109804 | 0.78240897 |
| ZFP2 | 109.559972 | 0.396440672 | 0.39057106 | 1.01502828 | 0.31009235 | 0.72498238 |
| ZNF354C | 89.5549727 | 0.002522969 | 0.44244395 | 0.00570235 | 0.99545021 | 0.99866448 |
| ADAMTS2 | 43.0325295 | -0.52422007 | 0.3916518 | -1.338485 | 0.18073837 | 0.63166264 |
| RUFY1 | 894.518053 | 0.057682661 | 0.2012366 | 0.286641 | 0.77438722 | 0.93453439 |
| AC136604.2 | 86.7538769 | -0.070765744 | 0.32777683 | -0.2158961 | 0.82906873 | 0.95325673 |
| HNRNPH1 | 2973.12477 | -0.295441862 | 0.24302087 | -1.2157057 | 0.22409705 | 0.66544589 |
| CANX | 1867.0221 | 0.018253376 | 0.34430444 | 0.05301522 | 0.95771978 | 0.98861313 |
| MAML1 | 672.995794 | -0.129772642 | 0.34549692 | -0.3756116 | 0.70720568 | 0.91002281 |
| LTC4S | 201.961849 | 0.328801986 | 0.70520648 | 0.46624924 | 0.6410371 | 0.88396933 |
| MGAT4B | 744.666354 | -0.42968463 | 0.47264979 | -0.9090973 | 3.63E-01 | 0.76224345 |
| SQSTM1 | 6947.98662 | -1.207565086 | 0.62687301 | -1.9263313 | 5.41E-02 | 0.46711025 |
| MRNIP | 435.200061 | 0.162553814 | 0.2538639 | 0.64031875 | 0.52196539 | 0.83445062 |
| TBC1D9B | 562.221052 | -0.535372101 | 0.24247784 | -2.2079217 | 0.02724973 | 0.38702446 |
| RNF130 | 1612.72864 | -0.629827139 | 0.37810203 | -1.6657597 | 0.09576128 | 0.538613 |
| MAPK9 | 442.66376 | -0.032393595 | 0.18717073 | -0.1730698 | 0.86259658 | 0.96391903 |
| CNOT6 | 628.298637 | -0.440345512 | 0.26463092 | -1.6639987 | 0.09611268 | 0.53876283 |
| MGAT1 | 396.772337 | -1.02346767 | 0.36550704 | -2.8001312 | 0.00510818 | 0.2174468 |
| LINC00847 | 41.4590742 | -0.458739204 | 0.47134909 | -0.9732473 | 0.33043043 | 0.73849818 |
| ZFP62 | 119.029881 | -0.170286925 | 0.30173155 | -0.5643657 | 0.57250531 | 0.85408471 |
| TRIM41 | 213.869178 | -0.115079407 | 0.3021803 | -0.3808303 | 0.70332919 | 0.90887942 |
| RACK1 | 1122.10965 | -0.063439256 | 0.46580788 | -0.1361919 | 0.89166959 | 0.97234681 |
| TRIM52 | 233.901436 | -0.504029765 | 0.23021843 | -2.1893545 | 0.02857108 | 0.39280225 |
| AC138035.1 | 88.8406736 | -0.059679085 | 0.41833904 | -0.1426572 | 0.8865609 | 0.97033958 |
| DUSP22 | 330.760189 | 0.198545201 | 0.21727462 | 0.9137984 | 3.61E-01 | 0.76009788 |
| EXOC2 | 1219.76014 | -0.287737441 | 0.2676796 | -1.0749323 | 0.28240505 | 0.70682473 |
| AL356130.2 | 59.2070095 | 2.506662956 | 1.13478294 | 2.20893607 | 0.02717909 | 0.38702446 |
| AL033381.1 | 99.8838334 | 1.437494304 | 0.91146602 | 1.5771233 | 0.11476716 | 0.57141787 |
| FOXQ1 | 63.8213261 | 0.520842438 | 0.634235 | 0.82121365 | 0.41152459 | 0.78632759 |
| FOXC1 | 168.565389 | -0.252954139 | 0.60152055 | -0.4205245 | 0.67410232 | 0.89801183 |
| GMDS | 3497.77564 | 0.111742846 | 0.39013511 | 0.28642089 | 0.77455578 | 0.93453439 |
| GMDS-DT | 5648.10637 | 0.399808208 | 0.43503513 | 0.9190251 | 0.35808244 | 0.75936492 |
| MYLK4 | 416.157055 | 0.735532271 | 0.96806016 | 0.75980017 | 0.44737404 | 0.80223612 |
| WRNIP1 | 416.412379 | -0.300549756 | 0.22602185 | -1.3297376 | 0.18360475 | 0.63276491 |
| SERPINB1 | 292.875245 | 0.169800563 | 0.38090987 | 0.44577622 | 0.6557589 | 0.89053559 |
| SERPINB9 | 83.1631166 | -0.262914848 | 0.44714727 | -0.5879827 | 0.55654392 | 0.84861484 |
| SERPINB6 | 800.290051 | -0.050437275 | 0.27161799 | -0.185692 | 0.85268634 | 0.96073035 |
| NQO2 | 2733.8401 | -0.232495052 | 0.49529485 | -0.4694074 | 0.63877848 | 0.88284765 |
| RIPK1 | 835.218371 | -0.193736184 | 0.34439769 | -0.5625362 | 0.57375072 | 0.85447394 |

| | | | | | | |
|---|---|---|---|---|---|---|
| BPHL | 562.655934 | 0.266707454 | 0.46968639 | 0.56784156 | 0.57014255 | 0.85327831 |
| TUBB2A | 121.753546 | 0.155851064 | 0.36893093 | 0.42243968 | 0.67270412 | 0.89757183 |
| PSMG4 | 282.733014 | 0.446879883 | 0.28865277 | 1.54815725 | 0.12158444 | 0.5799266 |
| SLC22A23 | 1500.57976 | 0.322744603 | 0.47103367 | 0.68518372 | 0.49322799 | 0.821168 |
| PXDC1 | 295.864369 | -0.321217009 | 0.35813771 | -0.8969092 | 0.36976737 | 0.76669785 |
| PRPF4B | 1704.44201 | -0.378947 | 0.23346105 | -1.6231701 | 0.10455305 | 0.55077196 |
| FAM217A | 51.0048441 | 0.082240486 | 0.73488196 | 0.11190979 | 0.91089493 | 0.97697493 |
| ECI2 | 684.964079 | -0.226860916 | 0.3843666 | -0.5902202 | 0.55504306 | 0.84801165 |
| CDYL | 1567.34877 | -0.808923843 | 0.42034249 | -1.9244398 | 0.05429947 | 0.46711025 |
| LYRM4 | 990.104008 | -0.536011647 | 0.36651103 | -1.4624707 | 0.14361227 | 0.59978468 |
| FARS2 | 2420.78221 | 0.010678423 | 0.28194853 | 0.03787366 | 0.96978841 | 0.99204339 |
| F13A1 | 165.276595 | 0.507476393 | 0.76979883 | 0.65923248 | 5.10E-01 | 0.82883692 |
| RREB1 | 2680.39036 | -0.177108626 | 0.19089488 | -0.9277809 | 0.35352123 | 0.75673243 |
| SSR1 | 600.167868 | -0.072208195 | 0.39408161 | -0.1832316 | 0.85461632 | 0.96160121 |
| AL139095.4 | 103.790377 | 0.515840482 | 0.54093504 | 0.95360892 | 0.34028164 | 0.74699558 |
| RIOK1 | 171.506967 | -0.478708259 | 0.41912187 | -1.1421696 | 0.25338353 | 0.68891958 |
| DSP | 1254.52525 | -0.751205971 | 0.39223741 | -1.9151818 | 0.05546932 | 0.46947044 |
| SNRNP48 | 248.852175 | -0.017027883 | 0.19229019 | -0.0885531 | 0.92943712 | 0.98272895 |
| BLOC1S5-TXI | 227.944424 | 0.044147304 | 0.58345714 | 0.07566503 | 0.93968559 | 0.98435387 |
| SLC35B3 | 335.26658 | 0.260437427 | 0.28046168 | 0.92860254 | 0.3530951 | 0.75673243 |
| GCNT2 | 2627.01115 | -0.854845769 | 0.40448739 | -2.1134052 | 0.0345661 | 0.41670847 |
| C6orf52 | 35.077885 | 0.32385468 | 0.43335285 | 0.74732329 | 0.45486843 | 0.80465616 |
| PAK1IP1 | 110.773254 | -0.716570399 | 0.48886407 | -1.4657866 | 0.14270644 | 0.59824924 |
| TMEM14C | 252.211372 | 0.183738883 | 0.29201095 | 0.62921916 | 0.52920559 | 0.83781716 |
| AL024498.2 | 42.590427 | 0.864738093 | 0.58523579 | 1.47758921 | 0.13951776 | 0.59500405 |
| MAK | 276.138776 | 0.701042439 | 0.47037986 | 1.49037513 | 0.13612563 | 0.5942049 |
| SMIM13 | 168.482958 | -0.017203039 | 0.24477146 | -0.070282 | 0.94396917 | 0.98522694 |
| AL139807.1 | 378.925439 | -0.652325483 | 0.74002481 | -0.8814914 | 0.37805193 | 0.77055888 |
| NEDD9 | 1339.12343 | -1.113853546 | 0.55847963 | -1.994439 | 0.0461041 | 0.44508419 |
| TMEM170B | 243.770116 | 0.789963125 | 0.24089598 | 3.27927063 | 0.00104076 | 0.1097359 |
| AL157373.2 | 42.1871532 | 0.391725627 | 0.63596672 | 0.61595302 | 0.53792552 | 0.84040975 |
| HIVEP1 | 1296.46675 | 0.827069795 | 0.38493632 | 2.1485886 | 0.03166703 | 0.40930954 |
| EDN1 | 55.01333 | 0.011359806 | 0.63826667 | 0.0177979 | 0.98580008 | 0.99609994 |
| PHACTR1 | 236.04646 | 0.604357504 | 0.53593862 | 1.1276618 | 0.25946278 | 0.69242517 |
| TBC1D7 | 429.84421 | -0.804611976 | 0.66957625 | -1.2016734 | 0.22949008 | 0.66705629 |
| GFOD1 | 118.336247 | -0.589285377 | 0.61868341 | -0.9524829 | 0.34085213 | 0.74741004 |
| SIRT5 | 363.120368 | 0.414233725 | 0.57075271 | 0.72576743 | 0.46798136 | 0.8118777 |
| NOL7 | 298.719231 | -0.541857191 | 0.302263 | -1.7926679 | 0.07302603 | 0.50459544 |
| RANBP9 | 1178.84935 | 0.040623173 | 0.17608224 | 0.23070568 | 0.81754346 | 0.94954998 |
| MCUR1 | 503.47591 | 0.111036482 | 0.29337843 | 0.37847527 | 0.70507756 | 0.909554 |
| CD83 | 143.337044 | 0.518425691 | 0.58023936 | 0.89346867 | 0.37160624 | 0.76820041 |
| JARID2 | 1500.59175 | -0.473366668 | 0.32735163 | -1.4460495 | 0.14816331 | 0.60605027 |
| DTNBP1 | 1970.2853 | -0.011261142 | 0.25349339 | -0.0444238 | 0.96456658 | 0.99031076 |
| MYLIP | 282.663974 | -0.488299705 | 0.3710079 | -1.3161437 | 0.18812581 | 0.64017888 |
| GMPR | 172.105359 | -0.717709363 | 0.48060364 | -1.4933498 | 0.13534564 | 0.59381224 |
| ATXN1 | 5176.14817 | 0.226877845 | 0.2540182 | 0.89315585 | 0.37177372 | 0.76838093 |
| CAP2 | 729.785333 | -0.188515925 | 0.33467114 | -0.5632871 | 0.57323942 | 0.8542683 |
| FAM8A1 | 192.782067 | 0.008173446 | 0.30293073 | 0.02698124 | 0.9784747 | 0.99482978 |
| NUP153 | 1217.02966 | -0.351006277 | 0.22479729 | -1.5614346 | 0.11842124 | 0.57589001 |
| KIF13A | 1656.23698 | -0.38466214 | 0.32131384 | -1.197154 | 0.23124655 | 0.66878058 |
| TPMT | 698.735937 | 0.487016174 | 0.2719901 | 1.7905658 | 0.073363 | 0.50459544 |
| KDM1B | 488.644337 | 0.459958163 | 0.34592176 | 1.32965952 | 0.18363048 | 0.63276491 |
| DEK | 575.658683 | -0.398511107 | 0.31347939 | -1.2712514 | 0.20363924 | 0.64912495 |
| RNF144B | 732.938808 | 1.262164507 | 0.6422244 | 1.96530139 | 0.04937936 | 0.45419182 |

| | | | | | | |
|---|---|---|---|---|---|---|
| MIR548A1HG | 147.381631 | 2.446497938 | 0.74163211 | 3.29880263 | 0.00097098 | 0.10560879 |
| AL022068.1 | 191.495134 | 0.253392392 | 0.64907022 | 0.39039288 | 0.69624605 | 0.90612803 |
| ID4 | 196.729704 | 0.31700636 | 0.43136916 | 0.73488416 | 0.46241005 | 0.8082072 |
| MBOAT1 | 101.161217 | 1.946726688 | 0.75087752 | 2.59260217 | 0.00952529 | 0.26008591 |
| E2F3 | 1731.97213 | -0.011166555 | 0.17054796 | -0.0654746 | 0.94779615 | 0.98659528 |
| CDKAL1 | 5263.6133 | -0.1128507 | 0.29084881 | -0.3880047 | 0.69801257 | 0.90696517 |
| AL513188.1 | 89.1266452 | 0.087034535 | 0.62038969 | 0.14029011 | 0.88843078 | 0.97134248 |
| AL512380.2 | 41.581552 | 0.305014428 | 0.42763952 | 0.71325126 | 0.47569029 | 0.81570932 |
| SOX4 | 725.790961 | -0.278495576 | 0.68613359 | -0.4058912 | 0.68482256 | 0.90098343 |
| CASC15 | 941.791151 | 0.644303743 | 0.68505527 | 0.94051353 | 0.34695422 | 0.75207626 |
| NBAT1 | 38.2815713 | 1.183039366 | 0.63781381 | 1.85483499 | 0.06361979 | 0.48593331 |
| DCDC2 | 3335.83619 | 0.595108055 | 0.40907182 | 1.45477646 | 0.14573116 | 0.60203043 |
| MRS2 | 465.942226 | -0.082636418 | 0.21692053 | -0.3809525 | 0.7032385 | 0.90887942 |
| GPLD1 | 54.4810898 | 0.572952881 | 0.44116391 | 1.29873018 | 0.19403654 | 0.64548758 |
| ALDH5A1 | 187.981733 | 0.666196452 | 0.47014128 | 1.41701331 | 0.15647904 | 0.61255952 |
| KIAA0319 | 158.667402 | -1.308534271 | 0.65911321 | -1.9852952 | 4.71E-02 | 0.44782198 |
| TDP2 | 969.02598 | -0.00911436 | 0.48018997 | -0.0189807 | 0.98485647 | 0.99553301 |
| ACOT13 | 578.45825 | 0.038733356 | 0.55600336 | 0.06966389 | 0.94446118 | 0.9853777 |
| C6orf62 | 1057.89166 | -0.658766875 | 0.34496769 | -1.909648 | 0.05617855 | 0.47126031 |
| GMNN | 101.056861 | -0.349162364 | 0.32955813 | -1.0594864 | 0.28937831 | 0.71218341 |
| RIPOR2 | 878.233231 | 0.341284317 | 0.65904295 | 0.51784837 | 0.60456407 | 0.86865188 |
| CARMIL1 | 2735.0425 | -0.012945 | 0.35689291 | -0.0362714 | 0.97106597 | 0.99220667 |
| SCGN | 52.6664303 | 2.507145479 | 0.95423889 | 2.62737718 | 0.00860459 | 0.25133745 |
| SLC17A4 | 149.546277 | 3.148100994 | 0.86444565 | 3.64175701 | 0.00027078 | 0.05986502 |
| SLC17A1 | 9114.39695 | 0.528181678 | 0.80937104 | 0.65258287 | 0.51402523 | 0.83126535 |
| SLC17A3 | 3636.94126 | 0.800247582 | 0.75453424 | 1.06058485 | 0.28887861 | 0.71218341 |
| TRIM38 | 2785.24234 | 0.955594507 | 0.32608418 | 2.93051479 | 0.00338401 | 0.18932494 |
| U91328.2 | 298.395535 | 1.506356794 | 0.635492 | 2.37037886 | 0.01776987 | 0.33411736 |
| U91328.1 | 43.1424166 | 0.089396764 | 0.48906029 | 0.18279293 | 0.85496049 | 0.96171023 |
| HIST1H1C | 213.262288 | -1.033609079 | 0.43842931 | -2.3575273 | 0.0183971 | 0.33848871 |
| HFE | 90.0852222 | 0.076365086 | 0.703583 | 0.10853742 | 0.9135694 | 0.97731177 |
| HIST1H4C | 215.723248 | 1.157452023 | 0.66447833 | 1.74189581 | 0.08152668 | 0.51593841 |
| HIST1H2BC | 97.6844802 | -2.536999923 | 0.73287736 | -3.4616978 | 0.00053678 | 0.08106654 |
| HIST1H2AC | 812.377896 | -1.600967881 | 0.65701627 | -2.4367249 | 0.01482095 | 0.30873979 |
| HIST1H1E | 59.9639872 | -0.854100534 | 0.74207374 | -1.1509645 | 0.24974684 | 0.68617881 |
| HIST1H2BD | 172.854486 | -1.565625354 | 0.45897111 | -3.4111632 | 0.00064686 | 0.08457928 |
| HIST1H2BE | 55.0507494 | -1.047867872 | 0.37485757 | -2.7953761 | 0.00518394 | 0.21811586 |
| HIST1H4H | 50.0982303 | -1.981780609 | 0.67360805 | -2.9420382 | 0.0032606 | 0.18622085 |
| BTN3A2 | 85.1445983 | 0.42806335 | 0.76959219 | 0.55622101 | 0.57805978 | 0.85714683 |
| BTN2A2 | 76.979068 | 0.881422158 | 0.55689112 | 1.58275492 | 0.11347733 | 0.56809124 |
| BTN3A1 | 76.7045643 | -0.200335255 | 0.94055759 | -0.2129963 | 0.83132988 | 0.95325673 |
| BTN3A3 | 43.7166865 | 0.17936226 | 0.78010129 | 0.22992176 | 0.81815257 | 0.94984237 |
| BTN2A1 | 155.620394 | -0.090087879 | 0.27330078 | -0.3296291 | 0.74168026 | 0.92235631 |
| HCG11 | 62.4172229 | 0.266103679 | 0.54094771 | 0.49192126 | 0.62277501 | 0.87714058 |
| HMGN4 | 169.940829 | -0.324915388 | 0.42616139 | -0.7624233 | 0.44580738 | 0.8012958 |
| ABT1 | 55.4475975 | -0.815753169 | 0.39588432 | -2.0605847 | 0.03934268 | 0.42704465 |
| ZNF322 | 169.373125 | 0.08724399 | 0.23421964 | 0.37248794 | 0.70952959 | 0.91105593 |
| AL513548.1 | 26.7447624 | 1.31013871 | 0.87232518 | 1.50189258 | 0.13312485 | 0.59206437 |
| LINC00240 | 167.651991 | 0.760894957 | 0.55443934 | 1.37236828 | 0.16994882 | 0.6233579 |
| ZNF391 | 66.3760875 | 0.326250203 | 0.45743256 | 0.71322034 | 0.47570942 | 0.81570932 |
| ZNF184 | 109.007871 | -0.245953416 | 0.45902388 | -0.5358183 | 0.5920841 | 0.86382106 |
| HIST1H2BN | 120.961524 | -0.212162741 | 0.34776701 | -0.6100715 | 5.42E-01 | 8.42E-01 |
| AL121944.1 | 135.162619 | -0.954566645 | 0.68247496 | -1.3986838 | 0.16190784 | 0.61845523 |
| ZNF165 | 125.409052 | -2.084058419 | 0.9568769 | -2.1779797 | 0.02940755 | 0.39791639 |

| | | | | | | |
|---|---|---|---|---|---|---|
| ZSCAN16-AS | 182.615972 | 0.085623714 | 0.27490257 | 0.31146931 | 0.75544387 | 0.92799282 |
| ZSCAN16 | 42.4754257 | -0.180270058 | 0.40034594 | -0.4502857 | 0.65250444 | 0.8894167 |
| ZKSCAN8 | 266.227401 | -0.261092922 | 0.18250813 | -1.4305824 | 0.15254993 | 0.61001662 |
| ZSCAN9 | 118.256673 | -0.319770687 | 0.32047607 | -0.9977989 | 0.31837686 | 0.73049743 |
| ZKSCAN4 | 34.4900694 | 0.527575352 | 0.54126114 | 0.974715 | 0.32970165 | 0.73849818 |
| PGBD1 | 124.717527 | 0.558285859 | 0.29715051 | 1.87879825 | 0.06027205 | 0.48004006 |
| ZSCAN31 | 190.292067 | 1.589260749 | 0.8775166 | 1.81108909 | 0.07012706 | 0.49573576 |
| ZKSCAN3 | 78.1372799 | 0.816353632 | 0.42430259 | 1.92398929 | 0.05435593 | 0.46712017 |
| ZSCAN12 | 99.9579943 | -0.126697756 | 0.30961705 | -0.4092079 | 0.68238707 | 0.89962501 |
| TRIM27 | 535.365228 | -0.403058334 | 0.35169254 | -1.1460531 | 0.25177317 | 0.68828377 |
| GABBR1 | 194.63791 | 0.579383009 | 0.59467177 | 0.97429042 | 0.32991236 | 0.73849818 |
| HLA-F | 152.062732 | -0.658857739 | 0.91774191 | -0.7179118 | 0.47281167 | 0.81460134 |
| HLA-A | 2524.81373 | -0.138283989 | 0.51872846 | -0.2665826 | 0.78979054 | 0.9402987 |
| ZNRD1 | 65.9778292 | 0.043613613 | 0.29422935 | 0.14822999 | 0.88216127 | 0.96796225 |
| PPP1R11 | 142.434328 | -0.306014545 | 0.32351328 | -0.9459103 | 0.34419433 | 0.75111515 |
| TRIM10 | 50.6116244 | 0.97203933 | 0.54617165 | 1.77973229 | 0.07511978 | 0.5071704 |
| TRIM15 | 159.62392 | 0.724179904 | 0.46714704 | 1.55021834 | 0.12108912 | 0.57927181 |
| TRIM26 | 641.513886 | 0.133106048 | 0.27640723 | 0.48155776 | 0.63012014 | 0.88094696 |
| HCG17 | 1010.84592 | 0.375175631 | 0.36980796 | 1.01451475 | 0.31033719 | 0.72498238 |
| HLA-E | 1129.69435 | -0.700269701 | 0.46047424 | -1.5207576 | 0.12832067 | 0.58793678 |
| GNL1 | 344.867871 | -0.029731699 | 0.39819458 | -0.0746663 | 0.94048026 | 0.98473477 |
| PRR3 | 147.172264 | 0.401971057 | 0.30065623 | 1.33697898 | 0.1812295 | 0.6318513 |
| ABCF1 | 496.65018 | -0.589302063 | 0.40674376 | -1.4488288 | 0.14738541 | 0.60514005 |
| PPP1R10 | 722.938407 | -0.408645677 | 0.33375558 | -1.224386 | 0.22080663 | 0.66263062 |
| MRPS18B | 299.333793 | 0.261234697 | 0.30796947 | 0.84824867 | 0.3962995 | 0.7795748 |
| ATAT1 | 187.508133 | 0.203960101 | 0.36716334 | 0.55550236 | 0.5785511 | 0.85751254 |
| C6orf136 | 173.999917 | 0.197793868 | 0.26953368 | 0.73383729 | 0.46304791 | 0.8082072 |
| DHX16 | 242.472331 | -0.565186817 | 0.26263814 | -2.1519602 | 0.03140049 | 0.40835531 |
| PPP1R18 | 85.1779819 | -0.451263819 | 0.55668012 | -0.810634 | 0.4175759 | 0.78665652 |
| MDC1 | 202.344703 | 0.390085436 | 0.396451 | 0.98394363 | 0.32514323 | 0.73628585 |
| TUBB | 623.618083 | -0.563582096 | 0.70495675 | -0.7994563 | 0.42402589 | 0.78894887 |
| FLOT1 | 981.449513 | -0.159952072 | 0.40092687 | -0.3989557 | 0.68992582 | 0.90409304 |
| IER3 | 551.258172 | 0.609596723 | 0.71146906 | 0.8568141 | 3.92E-01 | 0.77606943 |
| DDR1 | 266.099061 | 0.650995822 | 0.46098881 | 1.41217273 | 0.15789911 | 0.61369956 |
| VARS2 | 153.250036 | -0.437328562 | 0.45010137 | -0.9716224 | 0.33123844 | 0.73916381 |
| PSORS1C1 | 56.1594481 | 0.106506542 | 1.01958803 | 0.10446037 | 0.91680402 | 0.97828877 |
| CCHCR1 | 159.987941 | 0.379435224 | 0.49255094 | 0.77034717 | 0.44109398 | 0.79837253 |
| POU5F1 | 1303.22896 | 0.368140496 | 0.53988764 | 0.68188354 | 0.4953126 | 0.8222417 |
| HCG27 | 87.580403 | 0.682817347 | 0.49595823 | 1.37676381 | 0.16858528 | 0.62304975 |
| HLA-C | 1039.41639 | -1.211837003 | 0.73174771 | -1.6560858 | 0.09770444 | 0.54093423 |
| HLA-B | 1551.10761 | -0.456923125 | 0.66929652 | -0.6826916 | 0.49480173 | 0.82215397 |
| MICA | 337.376234 | 0.619368677 | 0.40667597 | 1.5230029 | 0.12775798 | 0.58793678 |
| MCCD1 | 37.6520225 | 0.767262111 | 1.14887589 | 0.66783725 | 0.50423749 | 0.82715271 |
| NFKBIL1 | 109.174712 | -0.200977968 | 0.30126017 | -0.6671243 | 0.50469276 | 0.82715271 |
| PRRC2A | 415.556705 | 0.005692697 | 0.26144599 | 0.02177389 | 0.98262832 | 0.99522272 |
| BAG6 | 348.798499 | -0.399390458 | 0.26037731 | -1.5338912 | 0.12505642 | 0.58564339 |
| APOM | 196.410927 | -0.064158128 | 0.78149609 | -0.0820965 | 0.93456994 | 0.98359701 |
| C6orf47 | 88.4247632 | -0.336348576 | 0.40611261 | -0.828215 | 0.40754873 | 0.78481094 |
| GPANK1 | 107.779141 | -0.217484908 | 0.31718942 | -0.6856626 | 0.49292592 | 0.82100434 |
| DDAH2 | 178.171181 | 0.416905802 | 0.48498544 | 0.8596254 | 0.38999557 | 0.77569824 |
| CLIC1 | 1233.35449 | -0.056779449 | 0.47610864 | -0.1192573 | 9.05E-01 | 9.76E-01 |
| VARS | 169.130509 | -0.520263574 | 0.49690853 | -1.0470007 | 0.29509926 | 0.71646902 |
| LSM2 | 136.093334 | 0.17323714 | 0.25043529 | 0.69174411 | 0.48909804 | 0.81970016 |
| HSPA1A | 1223.14096 | -1.887080231 | 0.56394636 | -3.3462052 | 0.00081926 | 0.09555385 |

| | | | | | | |
|---|---|---|---|---|---|---|
| HSPA1B | 521.591473 | -1.776727732 | 0.67484295 | -2.6328018 | 0.00846838 | 0.25070735 |
| C6orf48 | 196.648097 | -0.060146419 | 0.4315154 | -0.1393842 | 0.88914657 | 0.97197002 |
| NEU1 | 384.648544 | -0.44520396 | 0.47145087 | -0.9443274 | 0.34500238 | 0.75134217 |
| SLC44A4 | 991.460705 | 0.63319799 | 0.41956989 | 1.50915974 | 0.13125797 | 0.58930656 |
| EHMT2-AS1 | 80.5361746 | 0.931569977 | 0.57964524 | 1.60713814 | 0.10802407 | 0.55727735 |
| EHMT2 | 182.506755 | -0.265153649 | 0.2747791 | -0.9649702 | 0.33455973 | 0.74120095 |
| C2 | 129.288875 | -0.520046147 | 0.46604484 | -1.1158715 | 0.26447715 | 0.69567438 |
| CFB | 59.7536168 | -0.249577911 | 0.90702103 | -0.2751622 | 0.78319162 | 0.93831025 |
| NELFE | 181.242434 | -0.701818012 | 0.33086149 | -2.1211838 | 0.03390634 | 0.41378378 |
| SKIV2L | 145.157072 | -0.179305517 | 0.29719159 | -0.6033331 | 0.54628713 | 0.84460887 |
| DXO | 54.3070808 | -0.268299172 | 0.47592788 | -0.5637391 | 0.57293168 | 0.8542679 |
| STK19 | 394.753963 | 0.593705265 | 0.41557748 | 1.42862713 | 0.15311143 | 0.61008674 |
| CYP21A2 | 106.844092 | 1.245287118 | 0.77522782 | 1.60634988 | 0.10819706 | 0.55770254 |
| TNXB | 60.7355082 | 0.404597125 | 0.49223881 | 0.82195292 | 0.41110369 | 0.78632759 |
| ATF6B | 356.834034 | -0.185564047 | 0.29069002 | -0.6383571 | 0.52324124 | 0.8352569 |
| AGPAT1 | 146.893952 | 0.428610394 | 0.31983753 | 1.34008787 | 0.18021678 | 0.63124625 |
| RNF5 | 193.459563 | 0.255674136 | 0.48360053 | 0.5286887 | 0.59702142 | 0.86596535 |
| PBX2 | 192.754794 | 0.263357797 | 0.32698077 | 0.80542289 | 0.42057569 | 0.78831722 |
| HLA-DRA | 462.023092 | 0.948554282 | 0.62515458 | 1.51731158 | 0.12918804 | 0.58793678 |
| HLA-DRB5 | 125.695562 | 0.788453025 | 1.00883856 | 0.78154529 | 0.43448185 | 0.79451524 |
| HLA-DRB1 | 443.398801 | 0.958299582 | 0.90050669 | 1.06417819 | 0.28724799 | 0.71118494 |
| HLA-DQB1 | 108.775315 | 1.558346486 | 0.99240694 | 1.57026963 | 0.1163524 | 0.57350419 |
| PSMB8 | 217.895483 | 0.20796228 | 0.72722853 | 0.28596552 | 0.77490453 | 0.93477852 |
| PSMB9 | 136.556215 | -0.134311227 | 0.95064672 | -0.1412841 | 0.88764554 | 0.97098615 |
| TAP1 | 224.222992 | -0.102224302 | 0.76823733 | -0.1330634 | 0.89414321 | 0.97272276 |
| HLA-DMA | 82.2401146 | 0.633041722 | 0.62220974 | 1.01740889 | 0.30895895 | 0.72440992 |
| BRD2 | 835.746331 | -0.745274758 | 0.42081917 | -1.7710095 | 0.07655912 | 0.50867468 |
| HLA-DPA1 | 107.800798 | 0.632830557 | 0.74529716 | 0.84909831 | 0.39582659 | 0.77955046 |
| HLA-DPB1 | 97.2368952 | 0.700436523 | 0.59815461 | 1.17099578 | 0.24160048 | 0.68015474 |
| RXRB | 137.429205 | -0.039686987 | 0.39453517 | -0.1005918 | 0.91987454 | 0.97939204 |
| SLC39A7 | 234.435663 | -0.748751829 | 0.53720536 | -1.3937907 | 0.16338082 | 0.62065658 |
| HSD17B8 | 39.3465979 | 0.194542044 | 0.5532572 | 0.35163039 | 0.72511547 | 0.91600702 |
| RING1 | 237.169069 | -0.249341576 | 0.31482208 | -0.7920079 | 0.42835607 | 0.79037914 |
| HCG25 | 85.2696181 | 0.191288581 | 0.59966893 | 0.31899031 | 0.74973386 | 0.92565124 |
| VPS52 | 261.057165 | -0.22763395 | 0.24630231 | -0.9242055 | 3.55E-01 | 7.58E-01 |
| RPS18 | 784.707384 | 0.093144103 | 0.50984996 | 0.18268924 | 0.85504186 | 0.96171023 |
| WDR46 | 170.845827 | -0.681570008 | 0.48282686 | -1.411624 | 0.15806069 | 0.61369956 |
| PFDN6 | 135.78922 | -0.62298435 | 0.37392621 | -1.6660623 | 0.09570101 | 0.538613 |
| RGL2 | 178.848023 | 0.300275776 | 0.41388355 | 0.72550787 | 0.46814052 | 0.81194509 |
| TAPBP | 946.250225 | 0.263007839 | 0.44205548 | 0.59496568 | 0.55186641 | 0.84701114 |
| ZBTB22 | 43.6124437 | -0.06325458 | 0.41070938 | -0.154013 | 0.87759949 | 0.96658493 |
| DAXX | 96.8646015 | -0.619129237 | 0.436431 | -1.4186188 | 0.15601017 | 0.61255389 |
| PHF1 | 273.477511 | -0.306954389 | 0.27071966 | -1.1338459 | 0.25685917 | 0.69116216 |
| CUTA | 270.759706 | -0.201566043 | 0.40643789 | -0.4959332 | 0.61994155 | 0.87576407 |
| SYNGAP1 | 175.336749 | -0.025955216 | 0.26787939 | -0.0968914 | 0.92281262 | 0.97983687 |
| SYNGAP1-AS | 62.7782541 | 0.725231675 | 0.47988814 | 1.5112515 | 0.13072439 | 0.58853436 |
| BAK1 | 40.5014072 | -0.564152047 | 0.5634376 | -1.001268 | 0.31669725 | 0.7285009 |
| ITPR3 | 227.224101 | 0.44708034 | 0.66375812 | 0.67355912 | 5.01E-01 | 8.26E-01 |
| UQCC2 | 203.898398 | -0.004517818 | 0.21462565 | -0.0210498 | 0.98320596 | 0.99523767 |
| IP6K3 | 100.334775 | 0.04256379 | 0.8348122 | 0.05098607 | 0.95933662 | 0.98912673 |
| LEMD2 | 312.387202 | -0.20673937 | 0.32828836 | -0.6297493 | 0.52885861 | 0.83765775 |
| HMGA1 | 141.992887 | -1.821170465 | 0.92001897 | -1.9794923 | 4.78E-02 | 0.45133876 |
| SMIM29 | 95.9190553 | -0.201019367 | 0.54322338 | -0.3700492 | 0.71134585 | 0.91143287 |
| NUDT3 | 222.162949 | -0.088078778 | 0.57285306 | -0.1537546 | 0.87780325 | 0.96658493 |

| | | | | | | |
|---|---|---|---|---|---|---|
| C6orf106 | 2331.00542 | -0.187730434 | 0.24612163 | -0.7627547 | 0.44560969 | 0.80125568 |
| SNRPC | 201.007683 | -0.566290703 | 0.27059156 | -2.0927878 | 0.0363681 | 0.42151595 |
| UHRF1BP1 | 510.209691 | 0.048174493 | 0.1768477 | 0.27240667 | 0.78530935 | 0.93892226 |
| TAF11 | 300.899202 | -0.178907941 | 0.18879446 | -0.9476334 | 0.34331611 | 0.75073625 |
| ANKS1A | 1951.85951 | -0.00272591 | 0.18135199 | -0.015031 | 0.98800741 | 0.99671759 |
| SCUBE3 | 39.1342441 | 1.326417019 | 0.56349843 | 2.35389655 | 0.01857778 | 0.33968023 |
| ZNF76 | 390.768437 | 0.254710277 | 0.26893885 | 0.94709364 | 0.34359106 | 0.7507483 |
| PPARD | 679.767798 | 0.011379925 | 0.51045838 | 0.02229354 | 0.9822138 | 0.99522272 |
| RPL10A | 828.945309 | 0.150080869 | 0.52331692 | 0.28678772 | 0.77427486 | 0.93453439 |
| TEAD3 | 278.77501 | 0.204693539 | 0.29023549 | 0.7052671 | 0.48064403 | 0.81714875 |
| FKBP5 | 4293.02525 | 0.314677342 | 1.28862144 | 0.24419689 | 0.80707834 | 0.94585363 |
| SRPK1 | 1826.92848 | 0.397553191 | 0.30482561 | 1.30419878 | 0.19216582 | 0.64497936 |
| SLC26A8 | 62.317966 | 1.096107862 | 0.72130226 | 1.51962349 | 0.12860563 | 0.58793678 |
| MAPK14 | 1982.88855 | 0.035040342 | 0.22096966 | 0.15857535 | 0.87400345 | 0.9661168 |
| MAPK13 | 165.198839 | 0.216496788 | 0.41213332 | 0.52530766 | 0.59936934 | 0.86689411 |
| Z84485.1 | 68.4608844 | 0.787607428 | 0.61056645 | 1.28996186 | 0.1970639 | 0.64558745 |
| BRPF3 | 364.154816 | -0.142466848 | 0.2604047 | -0.5470978 | 0.58431152 | 0.86025731 |
| PNPLA1 | 390.983384 | 1.586849668 | 0.66440833 | 2.38836511 | 1.69E-02 | 3.29E-01 |
| ETV7 | 33.8016337 | 0.403617737 | 1.19917924 | 0.33657832 | 0.7364348 | 0.92019178 |
| KCTD20 | 935.562295 | 0.079671348 | 0.15232506 | 0.52303507 | 0.60094985 | 0.86755264 |
| STK38 | 1725.59577 | 0.542523128 | 0.23448766 | 2.31365319 | 0.02068675 | 0.35267827 |
| SRSF3 | 801.724186 | -0.444401431 | 0.23198005 | -1.9156881 | 0.05540481 | 0.46947044 |
| CDKN1A | 339.47933 | -0.879231206 | 0.41221009 | -2.1329687 | 3.29E-02 | 4.12E-01 |
| PPIL1 | 73.6260458 | -0.266177934 | 0.44739886 | -0.5949455 | 0.55187991 | 0.84701114 |
| C6orf89 | 973.408561 | 0.224400313 | 0.25555178 | 0.87810114 | 0.37988883 | 0.77097072 |
| MTCH1 | 706.182692 | -0.145296911 | 0.30186268 | -0.4813345 | 0.6302788 | 0.88101399 |
| PIM1 | 219.824087 | -0.99745708 | 0.5684678 | -1.7546413 | 0.07932068 | 0.51192014 |
| TMEM217 | 233.996769 | -1.205647426 | 0.68483531 | -1.7604925 | 0.07832434 | 0.51192014 |
| TBC1D22B | 833.09497 | -0.017560004 | 0.23905435 | -0.0734561 | 0.94144317 | 0.98489601 |
| RNF8 | 271.400761 | -0.326309248 | 0.24095601 | -1.3542275 | 0.17566381 | 0.62699479 |
| CMTR1 | 228.372818 | -0.335794009 | 0.34121166 | -0.9841223 | 0.32505538 | 0.73628585 |
| CCDC167 | 59.9510918 | -0.061200394 | 0.4724579 | -0.1295362 | 0.89693339 | 0.97324791 |
| ZFAND3 | 10805.3529 | -0.315027454 | 0.23669066 | -1.330967 | 0.18319987 | 0.63276491 |
| BTBD9 | 4727.35465 | -0.721672397 | 0.1674694 | -4.3092791 | 1.64E-05 | 0.01603597 |
| GLO1 | 319.64136 | 0.256286783 | 0.19872038 | 1.28968543 | 0.1971599 | 0.64558745 |
| SAYSD1 | 113.941313 | -0.046222041 | 0.26746267 | -0.1728168 | 0.86279544 | 0.96391903 |
| KCNK5 | 2187.92368 | 1.639045655 | 0.48023511 | 3.4130067 | 0.0006425 | 0.08457928 |
| KIF6 | 160.047805 | 0.303084155 | 0.77380742 | 0.39167905 | 0.69529537 | 0.90589033 |
| AL590999.1 | 98.6663282 | 0.776956612 | 0.85128124 | 0.91269087 | 3.61E-01 | 0.76009788 |
| MOCS1 | 327.006181 | 0.786797763 | 0.37706685 | 2.08662673 | 0.03692189 | 0.42215481 |
| UNC5CL | 1426.75758 | 1.434027334 | 0.67746388 | 2.11675837 | 0.03428036 | 0.41505357 |
| OARD1 | 406.564401 | 0.486877881 | 0.35800013 | 1.35999359 | 0.17383195 | 0.62685996 |
| NFYA | 248.199408 | 0.32136432 | 0.27554732 | 1.16627632 | 0.24350274 | 0.68164834 |
| FOXP4 | 190.806791 | 0.060978237 | 0.43179616 | 0.14121996 | 0.88769618 | 0.97098615 |
| TFEB | 222.080321 | 0.750048783 | 0.42754925 | 1.75429797 | 0.07937946 | 0.51192014 |
| FRS3 | 41.3265687 | 0.303680501 | 0.51577274 | 0.58878742 | 0.55600389 | 0.84861484 |
| USP49 | 386.886697 | 0.309611995 | 0.25762372 | 1.20179927 | 0.22944131 | 0.66705629 |
| MED20 | 87.3965901 | -0.23411961 | 0.31354801 | -0.7466787 | 0.45525755 | 0.80465616 |
| CCND3 | 568.90344 | 0.730484169 | 0.59398401 | 1.22980444 | 0.21877035 | 0.66108579 |
| TAF8 | 340.430019 | -0.049239924 | 0.27351908 | -0.1800237 | 0.85713394 | 0.96266803 |
| C6orf132 | 92.0991055 | 0.283364262 | 0.38805108 | 0.73022412 | 0.4652532 | 0.81042138 |
| MRPS10 | 138.798428 | -0.169190331 | 0.25382326 | -0.6665675 | 0.50504844 | 0.82715271 |
| TRERF1 | 204.904326 | -0.079233925 | 0.71792896 | -0.1103646 | 0.91212025 | 0.977179 |
| UBR2 | 2253.26252 | -0.062656111 | 0.25495421 | -0.2457544 | 0.80587239 | 0.94585363 |

| | | | | | | |
|---|---|---|---|---|---|---|
| TBCC | 58.4577252 | -0.501059673 | 0.56017234 | -0.8944741 | 0.37106825 | 0.76784912 |
| BICRAL | 1112.0258 | 0.301897417 | 0.30067964 | 1.00405007 | 0.31535448 | 0.72782796 |
| RPL7L1 | 498.443184 | -0.430209081 | 0.31697294 | -1.3572423 | 0.17470424 | 0.62699479 |
| C6orf226 | 59.8462437 | -0.938424263 | 0.40729834 | -2.3040218 | 0.02122142 | 0.35680968 |
| CNPY3 | 127.431516 | -0.624253834 | 0.41140779 | -1.5173603 | 0.12917575 | 0.58793678 |
| AL035587.1 | 48.8949776 | -0.975924471 | 0.72562068 | -1.3449513 | 0.17864096 | 0.6288217 |
| PEX6 | 308.158363 | 0.08144032 | 0.30389404 | 0.26798919 | 0.78870764 | 0.93998918 |
| PPP2R5D | 127.758393 | 0.002535007 | 0.2403703 | 0.01054626 | 0.99158546 | 0.99748517 |
| MEA1 | 69.9328841 | -0.20678391 | 0.41757899 | -0.4951971 | 0.62046099 | 0.87604644 |
| KLHDC3 | 224.428486 | -0.303274315 | 0.28389496 | -1.0682624 | 0.28540216 | 0.71029858 |
| RRP36 | 290.796882 | -0.265098349 | 0.31826939 | -0.832937 | 0.40488026 | 0.78422899 |
| CUL7 | 191.660277 | -0.425674709 | 0.33390705 | -1.27483 | 0.20236943 | 0.64881821 |
| KLC4 | 416.875832 | 0.210735524 | 0.3343782 | 0.63023106 | 0.52854342 | 0.83761342 |
| MRPL2 | 155.402227 | -0.005593971 | 0.29781978 | -0.0187831 | 0.98501416 | 0.99561898 |
| PTK7 | 86.6608301 | 0.403865396 | 0.63795624 | 0.63306128 | 0.52669363 | 0.83703067 |
| SRF | 74.5028729 | -0.311477439 | 0.43511193 | -0.7158559 | 0.47408034 | 0.8151865 |
| CUL9 | 282.367195 | 0.162274995 | 0.19991376 | 0.81172499 | 4.17E-01 | 0.78644918 |
| DNPH1 | 418.792747 | 0.151666091 | 0.58743121 | 0.25818528 | 0.79626392 | 0.94260208 |
| SLC22A7 | 2742.33725 | 2.9672066 | 0.87483033 | 3.39175094 | 0.00069448 | 0.08896424 |
| CRIP3 | 92.6730367 | 0.453672882 | 0.90209802 | 0.50290863 | 0.61502851 | 0.87260075 |
| ZNF318 | 217.562469 | 0.234702534 | 0.27648366 | 0.84888392 | 0.39594589 | 0.77955046 |
| ABCC10 | 273.409828 | -0.218547526 | 0.17600878 | -1.2416854 | 0.21435268 | 0.65906957 |
| TJAP1 | 304.83536 | -0.262305509 | 0.2664349 | -0.9845013 | 0.32486909 | 0.73628585 |
| POLR1C | 367.34359 | -0.178165212 | 0.30368151 | -0.5866844 | 0.55741567 | 0.84861484 |
| YIPF3 | 259.670405 | -0.34983419 | 0.37894521 | -0.9231788 | 0.35591402 | 0.75841231 |
| XPO5 | 999.415882 | -0.08017163 | 0.32298795 | -0.2482186 | 0.80396524 | 0.94543168 |
| POLH | 297.177343 | 0.223259109 | 0.31033211 | 0.71941994 | 0.47188221 | 0.81390329 |
| GTPBP2 | 348.332558 | -0.514792425 | 0.36472838 | -1.4114406 | 0.15811475 | 0.61369956 |
| MAD2L1BP | 71.7961573 | -0.252570005 | 0.34366632 | -0.734928 | 0.46238334 | 0.8082072 |
| MRPS18A | 129.488131 | -0.156059111 | 0.34155065 | -0.4569135 | 0.6477332 | 0.88774616 |
| VEGFA | 1409.38033 | -0.66644396 | 0.46836624 | -1.422912 | 0.15476166 | 0.61119904 |
| AL157371.2 | 82.0917685 | -2.267872811 | 1.39323812 | -1.6277711 | 1.04E-01 | 0.55022042 |
| MRPL14 | 333.803719 | -0.353207173 | 0.2608631 | -1.3539944 | 1.76E-01 | 0.62699479 |
| TMEM63B | 441.650687 | 0.326823587 | 0.35292096 | 0.92605321 | 0.35441833 | 0.75755608 |
| SLC29A1 | 44.0518008 | 0.244984744 | 0.40318958 | 0.60761675 | 5.43E-01 | 0.84396451 |
| HSP90AB1 | 1670.27642 | -0.425132599 | 0.45109001 | -0.9424562 | 0.3459591 | 0.75166079 |
| SLC35B2 | 111.188013 | -0.198812086 | 0.48072219 | -0.4135696 | 0.67918934 | 0.89932032 |
| NFKBIE | 67.6998657 | 0.490469326 | 0.4893687 | 1.00224906 | 0.31622332 | 0.72798933 |
| AARS2 | 60.6393498 | -0.173552556 | 0.5592928 | -0.3103072 | 0.75632739 | 0.92799282 |
| CDC5L | 300.292812 | -0.321361608 | 0.24423234 | -1.3158028 | 0.18824022 | 0.64026838 |
| SUPT3H | 2270.60988 | -0.274229252 | 0.38056433 | -0.7205858 | 0.4711644 | 0.81387913 |
| RUNX2 | 193.121022 | 1.012587398 | 0.49576007 | 2.04249485 | 0.04110247 | 0.43143739 |
| CLIC5 | 168.799106 | -0.208010156 | 0.48746157 | -0.4267211 | 0.66958245 | 0.89611079 |
| AL035701.1 | 48.2800549 | 1.135435775 | 0.41639979 | 2.72679235 | 0.00639533 | 0.22797255 |
| ENPP4 | 243.957979 | 0.696114492 | 0.38613501 | 1.80277486 | 0.07142358 | 0.49949133 |
| ENPP5 | 156.160326 | 0.874822893 | 0.46115046 | 1.8970444 | 0.05782208 | 0.47684824 |
| RCAN2 | 184.484894 | 0.211453033 | 0.52732642 | 0.40099078 | 0.68842691 | 0.90307854 |
| CYP39A1 | 112.140877 | 0.883665114 | 0.79698982 | 1.10875332 | 0.26753661 | 0.69874881 |
| SLC25A27 | 688.551583 | 0.050175726 | 0.68357306 | 0.07340214 | 0.94148611 | 0.98489601 |
| TDRD6 | 81.4527938 | 0.786729542 | 0.49679098 | 1.58362284 | 0.11327957 | 0.56809124 |
| ADGRF5 | 61.4456845 | -0.951932149 | 0.51844336 | -1.8361353 | 0.06633765 | 0.48798003 |
| ADGRF1 | 84.0927129 | -1.087274733 | 0.54523286 | -1.9941475 | 0.04613594 | 0.44508419 |
| TNFRSF21 | 1275.54313 | -0.114489747 | 0.52370205 | -0.2186162 | 0.82694904 | 0.95204019 |
| AL355353.1 | 52.3843379 | 0.031034419 | 0.75305465 | 0.04121138 | 0.96712738 | 0.99157802 |

| | | | | | | |
|---|---|---|---|---|---|---|
| CD2AP | 3407.14282 | -0.10207695 | 0.16929089 | -0.6029678 | 0.54653013 | 0.84460887 |
| PTCHD4 | 233.7046 | 2.702865446 | 1.11120209 | 2.43237973 | 0.01499997 | 0.30964551 |
| MUT | 299.841194 | 0.541085242 | 0.4503889 | 1.20137341 | 0.22960639 | 0.66705629 |
| CENPQ | 36.2704655 | 0.002862504 | 0.48280765 | 0.00592887 | 0.99526947 | 0.99866448 |
| AL138826.1 | 87.7295551 | -0.226724845 | 0.80484947 | -0.2816984 | 7.78E-01 | 0.93646866 |
| AL355997.1 | 77.6466858 | 1.514607829 | 0.98346174 | 1.54007805 | 0.12354133 | 0.58332105 |
| PKHD1 | 22581.5413 | 0.769116637 | 0.59120657 | 1.30092708 | 0.19328342 | 0.64548758 |
| MCM3 | 103.440681 | 0.002661635 | 0.41460617 | 0.00641967 | 0.99487788 | 0.99866448 |
| EFHC1 | 439.47466 | 0.39992923 | 0.35478714 | 1.12723712 | 0.25964224 | 0.69242517 |
| TRAM2 | 185.628035 | -0.53274342 | 0.55793053 | -0.9548562 | 0.33965043 | 0.74644675 |
| TRAM2-AS1 | 42.0579371 | 0.256816914 | 0.63019026 | 0.40752282 | 0.68362404 | 0.90058256 |
| TMEM14A | 118.616979 | 0.248372954 | 0.39927195 | 0.62206462 | 0.53389938 | 0.83915086 |
| GSTA2 | 194.758708 | 1.346487304 | 1.24547402 | 1.08110429 | 0.27965073 | 0.70446562 |
| GSTA1 | 320.814215 | 1.721687248 | 0.82368739 | 2.09021927 | 0.03659811 | 0.42192353 |
| GSTA4 | 174.278577 | 1.037869231 | 0.56105007 | 1.84986917 | 0.06433241 | 0.48593331 |
| ICK | 186.739566 | 0.462980516 | 0.37486454 | 1.23506084 | 0.21680786 | 0.66032006 |
| FBXO9 | 876.573609 | 0.185769847 | 0.14909553 | 1.24597862 | 0.21277224 | 0.65745584 |
| GCM1 | 73.7776752 | 1.114301855 | 0.81291726 | 1.3707445 | 0.17045462 | 0.62437773 |
| ELOVL5 | 1398.41483 | -0.431648706 | 0.48976765 | -0.8813336 | 0.37813726 | 0.77055888 |
| GCLC | 2706.65015 | -0.705703479 | 0.43381378 | -1.6267429 | 1.04E-01 | 0.55022042 |
| LINC01564 | 79.424012 | -1.207479361 | 0.75883346 | -1.5912311 | 0.11155759 | 0.56435572 |
| LRRC1 | 839.150832 | 0.682671855 | 0.43701243 | 1.56213373 | 0.11825649 | 0.5756185 |
| MLIP | 127.47927 | 0.901519453 | 0.58748003 | 1.53455335 | 0.12489358 | 0.58564339 |
| TINAG | 8766.74434 | 1.311634316 | 0.49851387 | 2.63108893 | 0.00851118 | 0.25070735 |
| DST | 2722.20741 | 0.010888633 | 0.5049761 | 0.02156267 | 0.98279681 | 0.9952242 |
| KIAA1586 | 135.876202 | 0.114530594 | 0.23282564 | 0.49191573 | 0.62277891 | 0.87714058 |
| ZNF451 | 1382.9969 | -0.23405301 | 0.13071247 | -1.7905943 | 0.07335843 | 0.50459544 |
| ZNF451-AS1 | 136.909529 | 0.383425476 | 0.64831013 | 0.59142293 | 0.55423708 | 0.84727776 |
| BAG2 | 141.756801 | 0.121345893 | 0.34762396 | 0.3490723 | 0.72703503 | 0.91645085 |
| RAB23 | 144.375207 | 0.797098894 | 0.2451986 | 3.25082971 | 0.00115069 | 0.11093704 |
| PRIM2 | 744.090471 | -0.7311707 | 0.32250103 | -2.2671887 | 0.0233787 | 0.37046453 |
| AL021368.2 | 137.754094 | 0.215899712 | 0.41544894 | 0.51967809 | 0.60328796 | 0.8678002 |
| AL445250.1 | 84.0340075 | 0.463319672 | 0.62087414 | 0.74623767 | 0.45552385 | 0.80465616 |
| KHDRBS2 | 2851.61049 | 1.034546967 | 0.59762047 | 1.73111033 | 0.08343209 | 0.51981984 |
| LGSN | 265.661932 | 1.626041517 | 0.9281505 | 1.7519158 | 7.98E-02 | 5.13E-01 |
| PTP4A1 | 836.528888 | -0.206801564 | 0.33712499 | -0.613427 | 0.53959405 | 0.84113285 |
| PHF3 | 2747.62565 | -0.019416541 | 0.16093869 | -0.1206456 | 0.90397177 | 0.9756941 |
| EYS | 288.551353 | 0.52438687 | 0.58676473 | 0.89369187 | 0.37148678 | 0.76820041 |
| ADGRB3 | 107.075307 | 0.503062638 | 0.60433024 | 0.83243003 | 0.40516626 | 0.78429302 |
| LMBRD1 | 1666.87163 | 0.136469792 | 0.34593105 | 0.39449998 | 0.69321194 | 0.90551691 |
| COL19A1 | 1680.66617 | 1.057091093 | 0.42160554 | 2.50729886 | 0.01216578 | 0.29514397 |
| FAM135A | 4147.69625 | 0.756231327 | 0.3399986 | 2.22421897 | 0.02613372 | 0.38392873 |
| SDHAF4 | 129.149964 | 0.497022583 | 0.42520251 | 1.16890792 | 0.24244073 | 0.68080966 |
| SMAP1 | 2085.47546 | -0.386986548 | 0.20407131 | -1.89633 | 0.05791642 | 0.47684824 |
| OGFRL1 | 494.543265 | -0.666276642 | 0.60498909 | -1.1013036 | 2.71E-01 | 7.00E-01 |
| AL136164.2 | 888.947744 | 0.432576191 | 0.47665771 | 0.90751955 | 0.36413212 | 0.76273126 |
| LINC00472 | 355.144301 | 0.290764409 | 0.43367403 | 0.67046765 | 0.50255972 | 0.82638105 |
| KCNQ5 | 123.62221 | 0.36108213 | 0.82867975 | 0.43573181 | 0.66303132 | 0.89263183 |
| AC019205.1 | 60.1081766 | 0.261037023 | 0.53347364 | 0.48931569 | 0.62461821 | 0.87832483 |
| OOEP | 740.725854 | 0.026926952 | 0.45643729 | 0.05899376 | 0.95295708 | 0.98791277 |
| DDX43 | 43.9428921 | 0.605104261 | 0.9233147 | 0.6553608 | 0.51223549 | 0.83055441 |
| MTO1 | 407.517345 | 0.386716211 | 0.1798034 | 2.15077249 | 3.15E-02 | 0.4087978 |
| EEF1A1 | 4224.10332 | 0.279617816 | 0.5036438 | 0.55518963 | 0.57876496 | 0.8575616 |
| AL121972.1 | 113.594256 | 1.001799483 | 0.57130717 | 1.75352163 | 0.07951251 | 0.51192014 |

| | | | | | | |
|---|---|---|---|---|---|---|
| SLC17A5 | 1331.74237 | -0.524884389 | 0.3717954 | -1.411756 | 0.15802182 | 0.61369956 |
| CD109 | 346.770029 | -0.032959122 | 0.30305736 | -0.1087554 | 9.13E-01 | 9.77E-01 |
| AL357507.1 | 67.0424209 | 0.384649333 | 0.42548549 | 0.90402457 | 0.36598237 | 0.76421682 |
| COL12A1 | 80.6976594 | -0.305512827 | 0.61135345 | -0.4997319 | 0.61726386 | 0.87435665 |
| COX7A2 | 475.368824 | -0.223936604 | 0.39170842 | -0.5716921 | 0.56753059 | 0.85260213 |
| TMEM30A | 597.378268 | 0.025884491 | 0.29173466 | 0.08872614 | 0.92929956 | 0.98272895 |
| AL080250.1 | 68.2717089 | 0.624969284 | 0.45008053 | 1.38857215 | 0.16496289 | 0.62065658 |
| FILIP1 | 40.1418142 | 0.699884344 | 0.5781365 | 1.21058668 | 0.22605385 | 0.66688879 |
| SENP6 | 2839.86093 | -0.190483548 | 0.17929813 | -1.0623845 | 0.28806117 | 0.71170639 |
| MYO6 | 7840.35059 | -0.292237561 | 0.21286644 | -1.3728682 | 0.16979332 | 0.62326131 |
| MEI4 | 478.00601 | 0.721747755 | 0.50904369 | 1.41785031 | 0.15623447 | 0.61255389 |
| IRAK1BP1 | 712.946932 | 1.117256422 | 0.52809237 | 2.11564584 | 0.03437494 | 0.41550027 |
| PHIP | 2581.78103 | 0.243386348 | 0.22238084 | 1.09445738 | 0.27375443 | 0.7014484 |
| HMGN3 | 2804.9954 | 0.261894989 | 0.34907214 | 0.75026035 | 0.45309792 | 0.80433814 |
| HMGN3-AS1 | 66.8393876 | 0.057823802 | 0.33789192 | 0.17113106 | 0.86412071 | 0.96393376 |
| LCA5 | 170.281883 | 0.57584314 | 0.38653805 | 1.48974504 | 0.13629129 | 0.5942049 |
| SH3BGRL2 | 353.617408 | 0.416190441 | 0.29720232 | 1.40036067 | 0.16140534 | 0.61746665 |
| BCKDHB | 2283.5848 | 0.606788708 | 0.55889949 | 1.08568484 | 0.27761845 | 0.7036882 |
| AL359715.1 | 109.121139 | 1.432699481 | 0.88087388 | 1.62645245 | 0.10385344 | 0.55022042 |
| TENT5A | 230.346515 | 0.07243956 | 0.5560304 | 0.13027986 | 0.89634502 | 0.97312927 |
| IBTK | 1720.14833 | -0.375603167 | 0.17852191 | -2.1039612 | 0.03538183 | 0.4198859 |
| TPBG | 50.8109287 | 1.110151921 | 0.66560357 | 1.66788757 | 0.09533805 | 0.53824794 |
| UBE3D | 1051.58033 | -0.107717729 | 0.27475754 | -0.3920465 | 0.69502387 | 0.90589033 |
| DOPEY1 | 1638.28821 | 0.117988952 | 0.17334749 | 0.68064989 | 0.49609305 | 0.82243887 |
| PGM3 | 680.15571 | -0.36239484 | 0.34906971 | -1.0381733 | 0.2991894 | 0.71852421 |
| RWDD2A | 97.8185153 | 0.522570536 | 0.37934889 | 1.37754596 | 0.16834352 | 0.62304975 |
| ME1 | 605.391499 | -0.394113594 | 0.55535861 | -0.709656 | 4.78E-01 | 0.81588117 |
| CYB5R4 | 397.722986 | -0.429433372 | 0.30088436 | -1.4272372 | 0.15351152 | 0.61014911 |
| CEP162 | 757.143265 | -0.031774212 | 0.25790965 | -0.123199 | 0.90194952 | 0.97492485 |
| NT5E | 383.097954 | 1.174299335 | 0.64689452 | 1.81528718 | 0.06947979 | 0.49281529 |
| SNX14 | 1524.18526 | 0.087503671 | 0.19368421 | 0.45178526 | 0.65142369 | 0.88935488 |
| AL589666.1 | 89.1581783 | 0.428411553 | 0.39116939 | 1.09520724 | 0.27342585 | 0.70137637 |
| SYNCRIP | 635.295602 | -0.494357982 | 0.36435914 | -1.3567877 | 0.17484869 | 0.62699479 |
| ZNF292 | 5339.19599 | -0.851550979 | 0.28932966 | -2.9431859 | 0.00324853 | 0.18622085 |
| SMIM8 | 427.28418 | 0.334330881 | 0.38370331 | 0.87132655 | 0.38357588 | 0.77317162 |
| SLC35A1 | 66.6350843 | 0.36365306 | 0.44995355 | 0.80820134 | 0.41897469 | 0.78766781 |
| RARS2 | 1004.49446 | -0.237798674 | 0.23913394 | -0.9944162 | 0.32002027 | 0.73157402 |
| ORC3 | 560.461824 | 0.20504166 | 0.36543152 | 0.56109462 | 0.57473304 | 0.85474196 |
| AKIRIN2 | 429.847372 | -0.023708807 | 0.32291133 | -0.073422 | 0.94147028 | 0.98489601 |
| AL139042.1 | 90.7288083 | -0.612906368 | 0.97925961 | -0.6258875 | 0.53138872 | 0.83789294 |
| RNGTT | 1813.03397 | -0.075936811 | 0.24568656 | -0.30908 | 0.75726065 | 0.92799282 |
| PNRC1 | 954.618828 | 0.221789314 | 0.44606256 | 0.49721571 | 0.61903695 | 0.87511751 |
| SRSF12 | 170.069812 | 0.305150087 | 0.38122847 | 0.80043888 | 4.23E-01 | 7.89E-01 |
| PM20D2 | 444.368229 | 0.092677939 | 0.19740282 | 0.4694864 | 6.39E-01 | 8.83E-01 |
| GABRR2 | 44.7261594 | 0.034319585 | 0.62009444 | 0.05534574 | 0.95586302 | 0.98852557 |
| UBE2J1 | 452.695348 | 0.044050214 | 0.21603761 | 0.20390067 | 8.38E-01 | 9.55E-01 |
| RRAGD | 383.414381 | -0.451542692 | 0.33731934 | -1.3386208 | 1.81E-01 | 6.32E-01 |
| ANKRD6 | 1540.64329 | -0.145535644 | 0.3540022 | -0.4111151 | 0.68098815 | 0.89947163 |
| LYRM2 | 332.246023 | -0.067464108 | 0.28439979 | -0.2372157 | 0.81248942 | 0.9478928 |
| MDN1 | 1265.94263 | -0.451623221 | 0.18373652 | -2.4579938 | 0.01397156 | 0.30363654 |
| CASP8AP2 | 425.838839 | -0.130805186 | 0.21399547 | -0.6112521 | 0.54103269 | 0.84192857 |
| BACH2 | 656.539003 | -0.000874549 | 0.59916598 | -0.0014596 | 9.99E-01 | 1.00E+00 |
| MAP3K7 | 1128.13645 | 0.048156115 | 0.2001802 | 0.24056382 | 0.80989319 | 0.94692486 |
| EPHA7 | 2378.40233 | 1.194899438 | 0.58628702 | 2.03807931 | 0.041542 | 0.43367572 |

| | | | | | | |
|---|---|---|---|---|---|---|
| MANEA | 178.02256 | 0.495676881 | 0.29026439 | 1.70767378 | 0.08769689 | 0.5273546 |
| UFL1-AS1 | 344.193203 | 1.099919786 | 0.47443954 | 2.31835608 | 0.02042998 | 0.35091941 |
| UFL1 | 609.768283 | 0.331529402 | 0.38546763 | 0.86007066 | 0.38975009 | 0.77569824 |
| NDUFAF4 | 172.098419 | -0.054416018 | 0.3457384 | -0.1573907 | 0.87493692 | 0.96615698 |
| KLHL32 | 477.820037 | 1.116754468 | 0.64782921 | 1.72384088 | 0.08473657 | 0.52183702 |
| MMS22L | 280.668115 | -0.007768563 | 0.27930949 | -0.0278135 | 0.97781093 | 0.99481657 |
| AL589740.1 | 72.1423928 | -0.180182347 | 0.50040264 | -0.3600747 | 0.71879125 | 0.91438252 |
| FBXL4 | 1167.4085 | -0.145503107 | 0.29373772 | -0.4953504 | 0.62035277 | 0.87598387 |
| COQ3 | 67.5352112 | -0.063775441 | 0.3189322 | -0.1999655 | 8.42E-01 | 9.57E-01 |
| PNISR | 3277.07657 | -0.000586437 | 0.30608881 | -0.0019159 | 0.99847133 | 0.99977652 |
| AL513550.1 | 132.694703 | 2.63E-01 | 0.52680505 | 0.49935476 | 0.61752948 | 0.8744294 |
| USP45 | 880.351599 | 0.419294016 | 0.18441239 | 2.27367598 | 0.02298548 | 0.36677754 |
| TSTD3 | 145.247909 | 0.303177859 | 0.31390643 | 0.96582238 | 0.33413307 | 0.74097427 |
| CCNC | 983.580297 | 0.015136096 | 0.25929927 | 0.05837308 | 0.95345146 | 0.98812635 |
| SIM1 | 346.339575 | 0.299663863 | 0.52372192 | 0.57218125 | 0.56719919 | 0.85239988 |
| ASCC3 | 1785.54906 | -0.179431141 | 0.24392766 | -0.7355916 | 4.62E-01 | 0.8082072 |
| HACE1 | 811.122063 | 0.500884162 | 0.58636145 | 0.85422424 | 0.39298076 | 0.77706105 |
| PREP | 490.173435 | -0.22197403 | 0.27765018 | -0.7994737 | 0.42401576 | 0.78894887 |
| ATG5 | 1106.66402 | -0.222307005 | 0.20160409 | -1.1026909 | 0.2701614 | 0.69961625 |
| CRYBG1 | 701.607665 | 1.283474489 | 0.56726705 | 2.2625578 | 0.02366296 | 0.37281404 |
| RTN4IP1 | 216.835347 | 0.415992459 | 0.40906435 | 1.01693648 | 0.30918365 | 0.72449199 |
| QRSL1 | 459.88964 | 0.035210316 | 0.22252846 | 0.15822837 | 0.87427685 | 0.9661168 |
| LINC02532 | 7470.54892 | 0.13682514 | 0.77661263 | 0.17618197 | 0.86015099 | 0.96283855 |
| CD24 | 2212.26994 | 0.487356489 | 0.4785349 | 1.01843457 | 0.30847148 | 0.72388608 |
| C6orf203 | 306.190547 | 0.226874556 | 0.28359232 | 0.80000247 | 0.42370936 | 0.78894887 |
| BEND3 | 54.0439452 | -0.562085836 | 0.44143927 | -1.2733027 | 0.20291063 | 0.64881821 |
| PDSS2 | 2756.74974 | -0.232148372 | 0.17628167 | -1.3169173 | 0.18786635 | 0.63977245 |
| SOBP | 1504.85028 | 0.383539893 | 0.61797934 | 0.62063546 | 0.5348395 | 0.83954482 |
| SEC63 | 1358.44064 | -0.33986 | 0.2148503 | -1.5818456 | 0.11368483 | 0.56865873 |
| OSTM1 | 611.366297 | -0.105705373 | 0.33655716 | -0.3140785 | 0.75346141 | 0.9277474 |
| SNX3 | 1003.87914 | -0.190827065 | 0.22845852 | -0.835281 | 0.40355949 | 0.78340037 |
| AFG1L | 908.836547 | 0.198994696 | 0.34593972 | 0.57522939 | 0.56513617 | 0.85171209 |
| FOXO3 | 3686.67575 | -0.195618168 | 0.24683389 | -0.7925093 | 0.42806374 | 0.79037914 |
| ARMC2 | 453.835923 | 0.978975886 | 0.37566727 | 2.60596534 | 0.00916157 | 0.25831482 |
| SESN1 | 1017.40001 | 0.310990179 | 0.26767137 | 1.1618358 | 0.24530217 | 0.68281753 |
| CEP57L1 | 473.446604 | 0.187975222 | 0.25865923 | 0.72672922 | 0.46739185 | 0.81166836 |
| CD164 | 1346.79916 | -0.285347326 | 0.26446811 | -1.0789479 | 0.28061093 | 0.70574937 |
| PPIL6 | 224.371637 | 0.947659029 | 0.40334986 | 2.3494716 | 0.01880008 | 0.34084276 |
| SMPD2 | 60.3179534 | -0.004299904 | 0.37984927 | -0.01132 | 0.99096812 | 0.99737132 |
| MICAL1 | 285.639036 | 0.146323922 | 0.25711468 | 0.56909984 | 0.56928838 | 0.85309237 |
| ZBTB24 | 123.165357 | -0.046373123 | 0.31021193 | -0.1494885 | 0.88116817 | 0.96773407 |
| AK9 | 688.409342 | 0.2080975 | 0.28365759 | 0.73362219 | 0.46317903 | 0.8082072 |
| FIG4 | 516.023309 | 0.102115547 | 0.23558339 | 0.43345818 | 0.66468194 | 0.89315615 |
| WASF1 | 864.390852 | -0.205402416 | 0.30214734 | -0.6798088 | 0.49662555 | 0.82252856 |
| CDC40 | 420.604404 | 0.011125668 | 0.21525595 | 0.05168576 | 0.95877908 | 0.98882925 |
| METTL24 | 546.096827 | 0.909873796 | 0.48994595 | 1.85709015 | 0.06329833 | 0.48572144 |
| DDO | 102.251089 | 0.420573645 | 0.86705597 | 0.4850594 | 0.62763421 | 0.87947068 |
| CDK19 | 1596.4868 | 0.143030613 | 0.34079629 | 0.41969533 | 0.67470804 | 0.89801183 |
| AMD1 | 679.713068 | -0.266847065 | 0.27162598 | -0.9824063 | 3.26E-01 | 0.7373175 |
| GTF3C6 | 361.761188 | -0.14992489 | 0.37880107 | -0.3957879 | 0.69226148 | 0.90516342 |
| RPF2 | 359.58498 | -0.510007021 | 0.29601801 | -1.7228919 | 0.08490808 | 0.52183702 |
| SLC16A10 | 2178.2781 | 0.16932983 | 0.5272126 | 0.32117941 | 0.74807443 | 0.92532868 |
| MFSD4B | 863.715776 | -0.138562399 | 0.34520462 | -0.4013921 | 6.88E-01 | 0.90307854 |
| AL080317.1 | 93.9658819 | -0.283447772 | 0.47110171 | -0.60167 | 0.54739382 | 0.8448516 |

| | | | | | | |
|---|---|---|---|---|---|---|
| AL080317.2 | 172.729887 | 0.007838659 | 0.37583602 | 0.02085659 | 0.98336005 | 0.99523767 |
| REV3L | 3027.25763 | -0.055515707 | 0.24051024 | -0.2308247 | 0.81745098 | 0.94954998 |
| AL080317.3 | 326.932338 | 0.093604334 | 0.38915284 | 0.2405336 | 8.10E-01 | 0.94692486 |
| TRAF3IP2-AS | 873.108318 | 0.075987048 | 0.26769933 | 0.28385221 | 0.77652364 | 0.9353214 |
| TRAF3IP2 | 296.042814 | 0.211985725 | 0.23727346 | 0.89342368 | 3.72E-01 | 7.68E-01 |
| FYN | 168.5767 | 0.980083241 | 0.60014474 | 1.63307811 | 0.10245257 | 0.54941994 |
| TUBE1 | 310.568671 | 0.148014896 | 0.40280761 | 0.36745804 | 0.71327738 | 0.91193854 |
| FAM229B | 83.9290342 | 0.791133761 | 0.53551149 | 1.47734226 | 0.13958392 | 0.59500405 |
| LAMA4 | 38.4200739 | 1.897362029 | 0.68164912 | 2.78348783 | 0.00537779 | 0.21873395 |
| MARCKS | 193.373361 | 0.440846548 | 0.65710349 | 0.67089363 | 0.50228829 | 0.82638105 |
| LINC01268 | 124.061201 | 1.353481349 | 0.58908353 | 2.29760515 | 0.02158427 | 0.35786026 |
| HDAC2 | 428.384907 | 0.049988887 | 0.25626046 | 0.19507062 | 0.84533766 | 0.95764192 |
| HDAC2-AS2 | 64.1732247 | 0.152114325 | 0.5778781 | 0.26322909 | 0.79237401 | 0.94141203 |
| FRK | 1786.24063 | 0.569473724 | 0.31113804 | 1.83029283 | 0.06720616 | 0.48963361 |
| NT5DC1 | 3173.49048 | 0.420397497 | 0.34999011 | 1.20116965 | 0.2296854 | 0.66705629 |
| COL10A1 | 122.67719 | 1.344422334 | 0.93720835 | 1.43449675 | 0.15143056 | 0.60905477 |
| TSPYL4 | 88.4164847 | -0.113650876 | 0.36603034 | -0.3104958 | 0.75618397 | 0.92799282 |
| DSE | 889.46262 | 0.511972968 | 0.35352225 | 1.44820577 | 0.14755951 | 0.60556834 |
| TSPYL1 | 279.874764 | 0.234312479 | 0.34079895 | 0.68753873 | 0.4917433 | 0.82065604 |
| RWDD1 | 206.94771 | -0.104453253 | 0.32134273 | -0.3250525 | 0.74514135 | 0.92389439 |
| ZUP1 | 351.843753 | -0.106168134 | 0.23504085 | -0.4517008 | 0.65148456 | 0.88935488 |
| KPNA5 | 640.085072 | 0.054622675 | 0.25434624 | 0.21475715 | 0.82995666 | 0.95325673 |
| DCBLD1 | 311.488563 | -0.332000541 | 0.33336474 | -0.9959078 | 0.31929495 | 0.73132207 |
| GOPC | 419.353343 | 0.23491208 | 0.23596047 | 0.99555694 | 0.31946546 | 0.73132207 |
| NUS1 | 531.102806 | -0.283844683 | 0.31055496 | -0.9139918 | 0.36072116 | 0.76009788 |
| SLC35F1 | 100.873481 | -0.057145464 | 0.37370836 | -0.1529146 | 0.87846562 | 0.96723659 |
| CEP85L | 591.516114 | 0.242495471 | 0.24347558 | 0.99597452 | 0.31926253 | 0.73132207 |
| MCM9 | 1007.7768 | 0.361348866 | 0.318684 | 1.13387829 | 0.2568456 | 0.69116216 |
| ASF1A | 55.907176 | 0.100154056 | 0.43076383 | 0.2325034 | 0.81614705 | 0.94913008 |
| AL365275.1 | 417.779589 | 0.019846508 | 0.63588723 | 0.03121074 | 0.97510148 | 0.99402915 |
| FAM184A | 483.694554 | 0.701323725 | 0.39287301 | 1.78511556 | 0.07424258 | 0.5071704 |
| MAN1A1 | 3052.64868 | 0.261515286 | 0.28075247 | 0.9314799 | 0.35160538 | 0.75548011 |
| TBC1D32 | 619.943065 | -0.400517141 | 0.32674895 | -1.2257641 | 0.22028745 | 0.66263062 |
| HSF2 | 196.32524 | -0.084206483 | 0.26032709 | -0.3234642 | 0.74634374 | 0.92448528 |
| SERINC1 | 888.545552 | -0.10075182 | 0.24601296 | -0.4095387 | 0.6821444 | 0.89962501 |
| PKIB | 274.743246 | -0.454677045 | 0.67796701 | -0.6706477 | 0.50244496 | 0.82638105 |
| SMPDL3A | 246.852329 | -0.109076952 | 0.56354232 | -0.1935559 | 0.84652362 | 0.95807937 |
| NKAIN2 | 136.021195 | -0.018878626 | 0.54611708 | -0.0345688 | 0.97242356 | 0.99248025 |
| RNF217-AS1 | 109.269677 | 0.403010681 | 0.65913597 | 0.61142267 | 0.5409198 | 0.84187438 |
| RNF217 | 1233.7844 | 0.137840108 | 0.25293989 | 0.54495203 | 5.86E-01 | 0.86078213 |
| TPD52L1 | 917.29409 | -0.006913962 | 0.431618 | -0.0160187 | 0.98721947 | 0.9963052 |
| HDDC2 | 231.089132 | 0.290671481 | 0.32839674 | 0.88512293 | 0.37609033 | 0.77010311 |
| AL365259.1 | 270.512482 | 0.837112054 | 0.76514819 | 1.09405219 | 0.27393209 | 0.7014484 |
| HEY2 | 52.6405599 | 1.475570969 | 0.37457395 | 3.93933151 | 8.17E-05 | 0.02604616 |
| NCOA7 | 2973.55668 | 0.646368305 | 0.3072331 | 2.10383679 | 0.03539269 | 0.4198859 |
| NCOA7-AS1 | 48.7249637 | 0.567134792 | 0.71685849 | 0.79113912 | 0.42886281 | 0.79037914 |
| HINT3 | 200.69045 | 0.426674332 | 0.3580923 | 1.19152053 | 0.2334493 | 0.66999363 |
| TRMT11 | 1163.40456 | 0.115498323 | 0.24851647 | 0.46475119 | 0.64210965 | 0.88441609 |
| AC020559.1 | 312.607338 | 0.546688386 | 0.37201213 | 1.46954453 | 0.14168515 | 0.59627829 |
| RNF146 | 578.726538 | 0.67941826 | 0.26942639 | 2.52172129 | 0.01167822 | 0.28824618 |
| ECHDC1 | 1373.47328 | 0.254056972 | 0.28851859 | 0.88055668 | 0.37855781 | 0.77061593 |
| PTPRK | 9428.54767 | 0.075441427 | 0.28621494 | 0.26358312 | 0.79210117 | 0.94132545 |
| AL590006.1 | 55.3213979 | 1.188633971 | 0.78291594 | 1.51821403 | 0.12896045 | 0.58793678 |
| LAMA2 | 82.6733089 | 0.25268091 | 0.61386394 | 0.41162364 | 0.6806153 | 0.89947163 |

| | | | | | | |
|---|---|---|---|---|---|---|
| AL356124.1 | 73.0823236 | 1.031234384 | 0.3549586 | 2.90522441 | 0.0036699 | 0.19726785 |
| ARHGAP18 | 768.400509 | 0.401391473 | 0.26414902 | 1.5195645 | 0.12862047 | 0.58793678 |
| L3MBTL3 | 354.707538 | 0.665814218 | 0.29181592 | 2.28162408 | 0.02251154 | 0.36344609 |
| SAMD3 | 240.619468 | 1.1133504 | 0.59976101 | 1.8563234 | 0.06340748 | 0.48572144 |
| TMEM200A | 641.856261 | 0.487554156 | 0.3706594 | 1.31536974 | 0.18838567 | 0.64026838 |
| SMLR1 | 38.8558743 | -0.921158855 | 0.85569165 | -1.0765079 | 0.28170015 | 0.70625976 |
| EPB41L2 | 1484.70869 | 0.487202345 | 0.38364433 | 1.26993234 | 0.20410873 | 0.64972559 |
| AKAP7 | 409.450232 | 0.484722935 | 0.46576623 | 1.04070005 | 0.29801478 | 0.71852421 |
| MED23 | 730.482381 | -0.086305286 | 0.24549608 | -0.3515546 | 0.72517229 | 0.91600702 |
| ENPP3 | 1289.60249 | 0.171912473 | 0.6622674 | 0.25958166 | 0.79518648 | 0.94252227 |
| ENPP1 | 1321.60336 | -0.725601989 | 0.6714472 | -1.0806538 | 0.27985113 | 0.70448474 |
| CTGF | 125.351976 | -1.18531932 | 0.59273577 | -1.9997432 | 0.045528 | 0.44378987 |
| STX7 | 2126.92058 | 0.00443293 | 0.43211069 | 0.01025878 | 0.99181482 | 0.99749106 |
| VNN1 | 426.972797 | 1.257010304 | 0.76287786 | 1.64772156 | 0.09940982 | 0.54399442 |
| SLC18B1 | 242.203701 | 0.094335564 | 0.4452318 | 0.21187966 | 0.83220092 | 0.95325673 |
| RPS12 | 737.628433 | 0.231048261 | 0.51097066 | 0.4521752 | 0.65114277 | 0.88923125 |
| TBPL1 | 252.787915 | -0.013877623 | 0.27304927 | -0.0508246 | 0.95946528 | 0.98912673 |
| SGK1 | 662.139713 | -1.235653069 | 0.79488273 | -1.5545099 | 0.12006284 | 0.57743907 |
| AL078590.2 | 371.296635 | -2.104472627 | 0.81045353 | -2.5966604 | 0.00941349 | 0.25998727 |
| AL078590.3 | 1295.33061 | -2.077039418 | 0.83416326 | -2.4899675 | 0.01277548 | 0.29875272 |
| ALDH8A1 | 1271.42436 | 0.694611517 | 0.50751757 | 1.36864527 | 0.17111019 | 0.62520756 |
| HBS1L | 608.70207 | 0.014727667 | 0.18019805 | 0.08173045 | 0.93486107 | 0.98373565 |
| AHI1 | 2260.60275 | -0.21435724 | 0.1448434 | -1.4799241 | 0.13889349 | 0.59474076 |
| LINC00271 | 988.006198 | 1.219368235 | 0.69339268 | 1.75855366 | 0.07865335 | 0.51192014 |
| PDE7B | 2053.31445 | -0.575103573 | 0.4023973 | -1.4291934 | 0.15294864 | 0.61001662 |
| AL138828.1 | 4054.989 | -2.005648799 | 0.86658499 | -2.3144283 | 2.06E-02 | 0.35267827 |
| BCLAF1 | 1259.42792 | -0.15057392 | 0.15556881 | -0.9678927 | 0.33309794 | 0.74000014 |
| AL023284.4 | 67.8620887 | -0.375077692 | 0.43263176 | -0.8669675 | 0.3859598 | 0.77317162 |
| MAP7 | 5129.17552 | -0.252576838 | 0.24403485 | -1.0350032 | 0.30066743 | 0.71879123 |
| MAP3K5 | 2247.86527 | 1.221549117 | 0.36792565 | 3.32009769 | 0.00089986 | 0.10027947 |
| PEX7 | 298.426645 | 0.067679471 | 0.29963774 | 0.22587099 | 0.82130176 | 0.95088971 |
| IFNGR1 | 362.125171 | 0.372689484 | 0.24493221 | 1.52160261 | 0.12810868 | 0.58793678 |
| TNFAIP3 | 177.344465 | 0.227436931 | 0.57767905 | 0.39370811 | 6.94E-01 | 0.90587439 |
| PERP | 245.939364 | -0.192223344 | 0.57977321 | -0.3315492 | 7.40E-01 | 0.92173153 |
| ARFGEF3 | 247.0556 | -1.00400785 | 0.40379791 | -2.4864117 | 1.29E-02 | 2.99E-01 |
| HEBP2 | 376.639295 | -0.097696911 | 0.27435875 | -0.3560918 | 0.72177179 | 0.91560945 |
| NHSL1 | 1290.08018 | 0.420291639 | 0.3114848 | 1.34931667 | 0.17723527 | 0.62731853 |
| CCDC28A | 194.84548 | 0.218721203 | 0.42695525 | 0.51228133 | 0.60845413 | 0.86985416 |
| REPS1 | 1587.5206 | 0.077370624 | 0.22231547 | 0.34802177 | 0.72782383 | 0.91665945 |
| ABRACL | 318.066173 | -0.822918606 | 0.46332593 | -1.7761117 | 0.07571451 | 0.50730849 |
| HECA | 164.339529 | -0.091668465 | 0.31712697 | -0.2890592 | 0.77253608 | 0.933623 |
| AL031772.1 | 68.9443498 | 0.079559007 | 0.42651245 | 0.18653384 | 0.85202614 | 0.96050023 |
| CITED2 | 205.461134 | -0.811967828 | 0.49529722 | -1.6393547 | 0.10113941 | 0.54622453 |
| FILNC1 | 68.6723578 | -1.651268326 | 1.10881896 | -1.4892136 | 0.13643112 | 0.5942049 |
| VTA1 | 647.1746 | -0.20628663 | 0.22071275 | -0.9346385 | 0.34997463 | 0.75455562 |
| ADGRG6 | 745.617304 | 0.628294354 | 0.43300976 | 1.45099351 | 0.14678167 | 0.60383734 |
| HIVEP2 | 2616.07984 | -0.383032966 | 0.34899082 | -1.0975445 | 0.27240343 | 0.70073986 |
| LINC01277 | 524.395214 | 0.098823178 | 0.4546935 | 0.21734021 | 0.82794323 | 0.95270465 |
| AL023581.2 | 57.6606394 | -0.408318183 | 0.51543348 | -0.7921841 | 0.42825336 | 0.79037914 |
| AIG1 | 3868.64132 | 0.970370693 | 0.3791924 | 2.55904572 | 0.01049599 | 0.27112183 |
| ADAT2 | 149.947693 | -0.208306995 | 0.38453909 | -0.5417056 | 0.5880213 | 0.86138805 |
| PEX3 | 640.969262 | -0.2758243 | 0.36760537 | -0.7503272 | 0.45305768 | 0.80433814 |
| FUCA2 | 314.236924 | -0.35934069 | 0.27929548 | -1.2865969 | 0.19823483 | 0.64582202 |
| PHACTR2 | 1254.29247 | 0.187670083 | 0.4130075 | 0.45439873 | 0.64954187 | 0.88850926 |

| | | | | | | |
|---|---|---|---|---|---|---|
| LTV1 | 39.5977137 | -0.798073868 | 0.46640012 | -1.7111356 | 0.08705608 | 0.52636862 |
| PLAGL1 | 1376.61616 | -0.344758236 | 0.31898832 | -1.0807864 | 0.27979214 | 0.70446562 |
| SF3B5 | 328.278226 | 0.030511528 | 0.42161703 | 0.07236787 | 0.94230915 | 0.98489601 |
| UTRN | 1120.59387 | 0.772495472 | 0.49899216 | 1.54811145 | 0.12159547 | 0.5799266 |
| EPM2A | 603.947454 | -0.02087518 | 0.33244817 | -0.0627923 | 0.94993191 | 0.98724063 |
| FBXO30 | 239.540168 | -0.436281299 | 0.39948812 | -1.0921008 | 0.27478879 | 0.70166356 |
| AL356599.1 | 530.439537 | 1.140089058 | 0.52894183 | 2.15541481 | 0.03112939 | 0.40675932 |
| SHPRH | 1748.5787 | 0.263887109 | 0.17994547 | 1.46648379 | 0.14251653 | 0.59794127 |
| RAB32 | 211.269003 | 0.399761823 | 0.3100175 | 1.28948149 | 0.19723075 | 0.64558745 |
| STXBP5-AS1 | 91.1132808 | 0.296320715 | 0.25211293 | 1.17534915 | 2.40E-01 | 0.6780147 |
| STXBP5 | 1214.94837 | 0.103627146 | 0.29438309 | 0.3520146 | 0.72482731 | 0.91600702 |
| SAMD5 | 10117.8356 | -0.38852835 | 0.36107035 | -1.0760461 | 0.28190663 | 0.70654493 |
| AL033504.1 | 5656.39176 | -0.170367874 | 0.42754755 | -0.398477 | 0.6902786 | 0.90420955 |
| AL513164.1 | 246.940417 | 0.453946154 | 0.65728719 | 0.69063594 | 0.48979436 | 0.81970016 |
| SASH1 | 6918.8703 | 0.405180689 | 0.61831989 | 0.65529298 | 0.51227914 | 0.83055441 |
| UST | 205.303561 | 0.833926553 | 0.64253294 | 1.29787362 | 0.19433077 | 0.64548758 |
| TAB2 | 1418.67905 | -0.16401418 | 0.25250585 | -0.6495461 | 0.51598549 | 0.83210529 |
| ZC3H12D | 79.357498 | -0.2740565 | 0.61913931 | -0.4426411 | 0.65802536 | 0.89058983 |
| PPIL4 | 561.523748 | -0.061631444 | 0.17516943 | -0.351839 | 0.72495898 | 0.91600702 |
| GINM1 | 843.740034 | 0.011158214 | 0.23364735 | 0.04775665 | 0.96191019 | 0.99000623 |
| AL078581.3 | 56.5513819 | 1.076894745 | 0.79540262 | 1.35389892 | 0.17576863 | 0.62699479 |
| KATNA1 | 362.894656 | -0.039087815 | 0.26524037 | -0.1473675 | 0.88284194 | 0.96824408 |
| LATS1 | 909.320539 | -0.2197853 | 0.24689851 | -0.8901848 | 0.37336667 | 0.76963112 |
| NUP43 | 278.827535 | -0.371941468 | 0.21799583 | -1.7061861 | 0.08797343 | 0.52795613 |
| PCMT1 | 700.314376 | -0.497304154 | 0.32673297 | -1.5220507 | 0.12799637 | 0.58793678 |
| LRP11 | 152.365824 | 0.454504113 | 0.47185806 | 0.9632221 | 0.33543607 | 0.74206299 |
| PPP1R14C | 739.142048 | -0.48151385 | 0.51155001 | -0.941284 | 0.34655933 | 0.75166079 |
| IYD | 495.075132 | 1.840237522 | 0.73610627 | 2.49996176 | 0.01242067 | 0.29816137 |
| PLEKHG1 | 867.233218 | 1.326281925 | 0.50757591 | 2.61297257 | 0.00897585 | 0.2563089 |
| MTHFD1L | 1295.76345 | -0.67363349 | 0.32758753 | -2.0563465 | 3.97E-02 | 4.29E-01 |
| AL138733.1 | 120.800728 | 0.067623513 | 0.69608796 | 0.09714794 | 0.92260891 | 0.97974213 |
| AKAP12 | 3270.96053 | -1.365270752 | 0.48347024 | -2.8238982 | 0.00474434 | 0.21294213 |
| ZBTB2 | 162.363104 | -0.986697345 | 0.31104297 | -3.1722219 | 0.00151277 | 0.13146073 |
| RMND1 | 793.341611 | -0.397413036 | 0.38928857 | -1.02087 | 0.30731602 | 0.72250292 |
| ARMT1 | 259.34226 | 0.071481476 | 0.25727197 | 0.27784401 | 0.7811321 | 0.93748328 |
| CCDC170 | 91.8988638 | 1.517797442 | 0.72992093 | 2.0793998 | 0.03758062 | 0.42215481 |
| ESR1 | 657.368525 | 0.74494201 | 0.61317367 | 1.21489563 | 2.24E-01 | 0.66551422 |
| SYNE1 | 2930.85122 | -0.352685562 | 0.38263443 | -0.9217298 | 0.35666952 | 0.75872042 |
| FBXO5 | 39.8119276 | -0.675574133 | 0.58826274 | -1.1484224 | 0.25079422 | 0.68648949 |
| AL080276.2 | 79.1756664 | 0.486071745 | 0.41839886 | 1.16174252 | 0.24534007 | 0.68281753 |
| MTRF1L | 245.421106 | -0.183761429 | 0.28022447 | -0.6557651 | 0.51197527 | 0.83054053 |
| RGS17 | 91.6340457 | 0.489698814 | 0.98990309 | 0.49469369 | 6.21E-01 | 0.87609696 |
| OPRM1 | 68.3865272 | 0.889189276 | 0.84732963 | 1.04940185 | 0.29399321 | 0.7156393 |
| IPCEF1 | 596.349157 | -0.04881157 | 0.60751042 | -0.0803469 | 0.93596137 | 0.98406248 |
| CNKSR3 | 6002.13599 | -0.031950426 | 0.28527751 | -0.1119977 | 0.91082523 | 0.97697493 |
| SCAF8 | 2067.83288 | -0.296980078 | 0.21499726 | -1.3813203 | 0.1671805 | 0.62242897 |
| TIAM2 | 1018.19409 | 0.273989409 | 0.26362429 | 1.03931777 | 0.29865697 | 0.71852421 |
| TFB1M | 610.721304 | -0.29577778 | 0.29340504 | -1.0080869 | 0.31341274 | 0.72717675 |
| CLDN20 | 40.4249095 | 0.217274642 | 0.61398797 | 0.35387443 | 0.72343299 | 0.91595197 |
| ARID1B | 6215.04062 | 0.002182439 | 0.1897986 | 0.01149871 | 0.99082556 | 0.99737132 |
| TMEM242 | 349.054356 | 0.10696887 | 0.2190763 | 0.48827222 | 0.62535704 | 0.87854882 |
| ZDHHC14 | 2771.92912 | 0.571996105 | 0.50133947 | 1.14093571 | 2.54E-01 | 0.68891958 |
| SNX9 | 2296.90618 | -0.870351458 | 0.31711469 | -2.7445953 | 0.00605856 | 0.22637955 |
| SYNJ2 | 309.723004 | -0.170092901 | 0.61617837 | -0.2760449 | 0.78251357 | 0.93782579 |

| | | | | | | |
|---|---|---|---|---|---|---|
| SERAC1 | 365.545752 | -0.122079168 | 0.31988661 | -0.3816326 | 0.70273387 | 0.90870007 |
| GTF2H5 | 230.594235 | -0.063750172 | 0.33357091 | -0.1911143 | 0.84843604 | 0.95951833 |
| TULP4 | 3001.13881 | -0.41851518 | 0.2991782 | -1.3988826 | 0.16184819 | 0.61845523 |
| TMEM181 | 523.660104 | -0.328468845 | 0.46160956 | -0.7115729 | 0.47672931 | 0.81570932 |
| DYNLT1 | 832.499559 | -0.157748752 | 0.28052323 | -0.5623376 | 5.74E-01 | 8.54E-01 |
| SYTL3 | 649.157873 | -0.643303488 | 0.71450049 | -0.9003542 | 3.68E-01 | 7.66E-01 |
| EZR | 3234.1595 | 0.235694313 | 0.20880658 | 1.12876862 | 0.25899545 | 0.69241621 |
| C6orf99 | 53.2235687 | 0.083492618 | 0.55421304 | 0.15065076 | 0.88025121 | 0.96769187 |
| RSPH3 | 772.38802 | 0.133223004 | 0.20237995 | 0.65828164 | 0.51035717 | 0.82930674 |
| AL035530.2 | 345.785712 | 0.646585223 | 0.487225 | 1.32707727 | 0.18448312 | 0.63377801 |
| SOD2 | 219.017536 | 0.248438609 | 0.5925764 | 0.41925161 | 0.67503226 | 0.89801183 |
| SOD2.1 | 259.978272 | -0.09814583 | 0.38134659 | -0.2573665 | 7.97E-01 | 0.94280281 |
| WTAP | 1952.78105 | -0.161927738 | 0.27149667 | -0.5964262 | 5.51E-01 | 8.47E-01 |
| ACAT2 | 222.195118 | 0.619076221 | 0.46847653 | 1.32146688 | 0.18634574 | 0.63757786 |
| TCP1 | 464.035777 | -0.035771641 | 0.34500678 | -0.1036839 | 0.91742022 | 0.97828877 |
| MRPL18 | 233.853604 | -0.586158163 | 0.28953703 | -2.024467 | 0.04292213 | 0.43971119 |
| IGF2R | 3550.41858 | -0.5292092 | 0.31327209 | -1.6892957 | 0.09116277 | 0.53537574 |
| SLC22A1 | 156.521061 | -0.203707492 | 0.63713661 | -0.3197234 | 0.74917801 | 0.92565124 |
| SLC22A2 | 1943.99109 | 0.712968556 | 0.37109493 | 1.9212565 | 0.05469938 | 0.46772575 |
| SLC22A3 | 773.748278 | -0.693000512 | 0.61019711 | -1.1356994 | 0.25608238 | 0.69083334 |
| LPA | 55.9055522 | 0.515907177 | 1.0131559 | 0.50920809 | 0.61060638 | 0.87128686 |
| PLG | 2192.50727 | 0.639334166 | 0.81088288 | 0.78844206 | 0.43043818 | 0.79055252 |
| MAP3K4 | 866.526816 | -0.151344644 | 0.13839945 | -1.093535 | 0.27415895 | 0.7014484 |
| AGPAT4 | 221.490587 | 0.906978202 | 0.63805423 | 1.4214751 | 0.15517869 | 0.61156822 |
| PRKN | 8046.35494 | 1.091134074 | 0.50293187 | 2.16954649 | 0.03004122 | 0.40173169 |
| PACRG | 628.338809 | 0.768106926 | 0.62805151 | 1.22299989 | 0.22132974 | 0.66263062 |
| QKI | 3127.89285 | -0.014883858 | 0.20069082 | -0.0741631 | 0.94088059 | 0.98485301 |
| PDE10A | 2792.94768 | 0.460493013 | 0.40890794 | 1.12615327 | 0.26010065 | 0.69280433 |
| SFT2D1 | 437.751921 | -0.075818581 | 0.16620032 | -0.4561879 | 0.64825485 | 0.88787189 |
| MPC1 | 1286.21981 | 0.511866873 | 0.30372754 | 1.68528306 | 0.09193397 | 0.53567268 |
| AL022069.1 | 31.8332974 | 0.41205471 | 0.72742557 | 0.56645618 | 0.57108371 | 0.85399468 |
| RPS6KA2 | 682.549049 | 0.207691065 | 0.29546149 | 0.70293785 | 0.48209448 | 0.8177376 |
| AL159163.1 | 40.4810893 | 0.601815165 | 0.52029028 | 1.15669116 | 0.24739856 | 0.68482118 |
| Z94721.1 | 600.862003 | -0.117933605 | 0.42499103 | -0.2774967 | 0.78139874 | 0.93748328 |
| FGFR1OP | 1462.00001 | 0.140146937 | 0.29247125 | 0.47918192 | 0.63180921 | 0.88123095 |
| CCR6 | 38.1541797 | -0.487355755 | 0.64650271 | -0.753834 | 0.45094888 | 0.80384625 |
| LINC01558 | 81.559118 | 0.758243559 | 0.78571097 | 0.96504133 | 0.33452411 | 0.74120095 |
| AFDN | 4289.66591 | -0.076778437 | 0.36296987 | -0.2115284 | 0.83247497 | 0.95334883 |
| FRMD1 | 155.045182 | 0.046425201 | 0.82840123 | 0.05604193 | 0.95530841 | 0.98840182 |
| AL606970.3 | 34.4471572 | 2.628759636 | 0.94898927 | 2.77006254 | 0.00560455 | 0.22462459 |
| DACT2 | 107.621318 | 4.401381933 | 1.11191784 | 3.95836973 | 7.55E-05 | 0.02497299 |
| BX322234.1 | 67.1500179 | 1.568049862 | 0.60729034 | 2.58204316 | 0.00982173 | 0.26385502 |
| THBS2 | 165.647383 | 2.093996139 | 0.93387902 | 2.24225632 | 0.02494481 | 0.37990947 |
| WDR27 | 1104.71604 | -0.079137808 | 0.29354674 | -0.2695918 | 0.78747427 | 0.93953874 |
| C6orf120 | 80.6372505 | -0.093833693 | 0.31407197 | -0.2987649 | 0.7651194 | 0.93083208 |
| PHF10 | 385.149921 | -0.019187423 | 0.2284292 | -0.0839972 | 0.93305862 | 0.9832732 |
| TCTE3 | 59.3305233 | -0.232805852 | 0.43679967 | -0.5329808 | 0.59404685 | 0.86485397 |
| ERMARD | 251.037875 | -0.375491183 | 0.3857546 | -0.9733939 | 0.33035756 | 0.73849818 |
| DLL1 | 45.2308015 | -0.131207311 | 0.54540594 | -0.2405682 | 0.80988982 | 0.94692486 |
| FAM120B | 1363.09916 | -0.309740665 | 0.27630414 | -1.1210135 | 0.26228212 | 0.69443712 |
| PSMB1 | 577.158936 | -0.154053854 | 0.31003914 | -0.4968852 | 0.61927002 | 0.87527555 |
| TBP | 204.82165 | -0.111337823 | 0.23789173 | -0.4680189 | 0.63977108 | 0.88321992 |
| PDCD2 | 227.088584 | -0.16313079 | 0.30968648 | -0.5267611 | 0.5983595 | 0.86646266 |
| AC093627.5 | 51.0831596 | -0.130481449 | 0.43170994 | -0.3022433 | 0.76246658 | 0.93022076 |

| | | | | | | |
|---|---|---|---|---|---|---|
| AC093627.3 | 95.5934845 | 0.726203291 | 0.61146196 | 1.18765081 | 2.35E-01 | 6.71E-01 |
| FAM20C | 2196.1461 | 0.365465162 | 0.4490414 | 0.81387855 | 4.16E-01 | 0.78632759 |
| PDGFA | 168.029586 | 0.430269643 | 0.31622937 | 1.36062519 | 0.17363217 | 0.62663931 |
| PRKAR1B | 808.636302 | -0.713802372 | 0.52698925 | -1.3544913 | 0.17557968 | 0.62699479 |
| DNAAF5 | 559.665381 | -0.868341615 | 0.38009519 | -2.2845372 | 0.02233998 | 0.36273984 |
| SUN1 | 880.553318 | -0.140770705 | 0.22746403 | -0.6188702 | 0.53600189 | 0.83954482 |
| AC073957.3 | 45.7858658 | 1.443557902 | 0.87792709 | 1.64427994 | 0.10011839 | 0.54542315 |
| COX19 | 334.324019 | -0.247309825 | 0.26129775 | -0.9464675 | 0.3439102 | 0.7507483 |
| C7orf50 | 773.910442 | -0.40875138 | 0.29263253 | -1.3968077 | 0.1624714 | 0.61912485 |
| GPR146 | 60.239053 | -0.793774929 | 0.64183441 | -1.2367285 | 0.21618788 | 0.66012192 |
| GPER1 | 93.5023491 | -0.719716639 | 0.59225682 | -1.2152104 | 0.22428586 | 0.66551422 |
| ZFAND2A | 103.983136 | -2.101275097 | 0.61308222 | -3.4273953 | 0.0006094 | 0.08353064 |
| MICALL2 | 142.882183 | -0.439328876 | 0.4902629 | -0.8961088 | 3.70E-01 | 0.76673594 |
| INTS1 | 376.771736 | -0.234336421 | 0.41414228 | -0.5658355 | 0.57150558 | 0.85399468 |
| MAFK | 133.523077 | -0.308889867 | 0.42917975 | -0.7197214 | 0.47169653 | 0.81389222 |
| TMEM184A | 201.925086 | -0.627882518 | 0.35977941 | -1.7451875 | 0.08095224 | 0.51567042 |
| PSMG3 | 119.206598 | -0.324570365 | 0.29675749 | -1.0937226 | 0.27407667 | 0.7014484 |
| PSMG3-AS1 | 96.2795107 | 0.022889968 | 0.43115567 | 0.0530898 | 0.95766036 | 0.98861313 |
| MAD1L1 | 593.702424 | -0.415252712 | 0.38079702 | -1.0904831 | 0.2755004 | 0.70247842 |
| MRM2 | 93.0091271 | -0.709348237 | 0.41014251 | -1.7295165 | 0.0837167 | 0.52069073 |
| SNX8 | 489.387569 | -1.027392909 | 0.42283849 | -2.4297526 | 0.01510913 | 0.31036958 |
| EIF3B | 858.148234 | -0.918508239 | 0.31037007 | -2.9593969 | 0.00308242 | 0.18216645 |
| CHST12 | 303.106317 | -0.708154364 | 0.35114331 | -2.0167104 | 0.04372574 | 0.44226848 |
| BRAT1 | 182.17885 | 0.093262873 | 0.33118385 | 0.28160453 | 0.77824677 | 0.93646866 |
| IQCE | 467.497777 | -0.118618275 | 0.24991104 | -0.474642 | 0.63504216 | 0.88238823 |
| TTYH3 | 1470.83498 | -0.206637161 | 0.39706724 | -0.5204085 | 0.6027789 | 0.8677246 |
| AMZ1 | 236.023389 | 0.971448405 | 0.87826393 | 1.10610077 | 0.26868291 | 0.69961625 |
| GNA12 | 1174.02867 | 0.354973561 | 0.28738631 | 1.23517907 | 0.21676387 | 0.66032006 |
| SDK1 | 6392.38234 | 1.015415478 | 0.45997355 | 2.20755189 | 0.02727553 | 0.38702446 |
| AC017000.1 | 42.9983127 | 1.058657061 | 1.02038394 | 1.03750855 | 0.29949892 | 0.7186844 |
| FOXK1 | 1486.24626 | -0.285871969 | 0.26991693 | -1.0591109 | 0.28954926 | 0.71218341 |
| AP5Z1 | 427.499205 | -1.234481398 | 0.43584292 | -2.8323998 | 0.00462 | 0.20969005 |
| RBAK-RBAKD | 671.196332 | -0.757296638 | 0.30335017 | -2.4964437 | 0.01254456 | 0.29875272 |
| WIPI2 | 879.10402 | -0.322807968 | 0.31545966 | -1.023294 | 0.30616887 | 0.72169506 |
| TNRC18 | 1669.6591 | 0.740125776 | 0.44950317 | 1.64654184 | 0.09965225 | 0.54450139 |
| FBXL18 | 275.042503 | -0.307382588 | 0.48316632 | -0.6361838 | 0.52465663 | 0.83592566 |
| ACTB | 3009.79746 | 0.388715939 | 0.57283602 | 0.67858152 | 0.49740306 | 0.82313869 |
| RNF216 | 1523.19563 | -0.730502337 | 0.27268261 | -2.6789473 | 0.0073854 | 0.23590813 |
| CCZ1 | 223.136392 | -0.192961112 | 0.31845298 | -0.6059328 | 0.54455936 | 0.84441376 |
| PMS2 | 461.445263 | 0.415513409 | 0.35487814 | 1.17086223 | 0.24165416 | 0.68015474 |
| AIMP2 | 252.11327 | -0.209622621 | 0.34111121 | -0.6145287 | 0.53886603 | 0.84070752 |
| EIF2AK1 | 530.66224 | -0.065940161 | 0.18839167 | -0.3500163 | 0.72632645 | 0.91635631 |
| USP42 | 577.22446 | -0.129316316 | 0.24562743 | -0.5264734 | 0.59855931 | 0.86646266 |
| CYTH3 | 371.364885 | -0.208854609 | 0.36080399 | -0.5788589 | 0.56268437 | 0.85054239 |
| FAM220A | 43.5031863 | -0.793580913 | 0.3924526 | -2.0221064 | 0.04316536 | 0.44121371 |
| RAC1 | 1260.13791 | -0.20859815 | 0.2882099 | -0.7237716 | 0.46920595 | 0.81266186 |
| DAGLB | 269.745494 | -1.008603691 | 0.51350006 | -1.9641745 | 0.04950986 | 0.45484699 |
| ZDHHC4 | 125.591427 | -0.187800754 | 0.24113326 | -0.7788256 | 0.43608247 | 0.79549938 |
| C7orf26 | 123.186271 | -0.292683424 | 0.36941143 | -0.7922966 | 0.42818775 | 0.79037914 |
| ZNF316 | 316.86685 | -0.626178144 | 0.46169784 | -1.356251 | 0.17501933 | 0.62699479 |
| AC073343.2 | 165.773038 | 0.406598838 | 0.58288083 | 0.69756769 | 0.48544759 | 0.81839266 |
| ZNF12 | 317.264681 | -0.284428581 | 0.16524849 | -1.7212174 | 0.08521137 | 0.5221244 |
| CCZ1B | 318.125077 | -0.510667103 | 0.30127723 | -1.6950073 | 0.09007403 | 0.53311774 |
| C1GALT1 | 2062.46875 | -0.358913894 | 0.31434989 | -1.1417656 | 0.25355147 | 0.68891958 |

| | | | | | | |
|---|---|---|---|---|---|---|
| COL28A1 | 405.340543 | 0.66617402 | 0.48378493 | 1.3770045 | 0.16851086 | 0.62304975 |
| AC004982.1 | 37.6381091 | -0.252914471 | 0.44401242 | -0.5696113 | 0.56894139 | 0.85291608 |
| AC004982.2 | 109.014732 | -0.073098101 | 0.45114081 | -0.1620295 | 0.87128266 | 0.9657206 |
| MIOS | 456.524021 | -0.033289882 | 0.22313539 | -0.1491914 | 0.8814026 | 0.96774334 |
| RPA3 | 226.922652 | 0.37824321 | 0.57474917 | 0.65810135 | 0.51047301 | 0.82932956 |
| UMAD1 | 2168.01489 | 0.162812051 | 0.22090314 | 0.73702913 | 0.46110463 | 0.80771389 |
| GLCCI1 | 530.167237 | 0.406479045 | 0.24952761 | 1.62899425 | 0.10331424 | 0.55022042 |
| ICA1 | 470.913279 | 1.019731356 | 0.62024548 | 1.64407706 | 0.10016029 | 0.54542315 |
| NDUFA4 | 762.898956 | -0.01838526 | 0.46670645 | -0.0393936 | 0.96857656 | 0.99187457 |
| PHF14 | 1566.46341 | 0.172760255 | 0.37155063 | 0.46497096 | 0.64195225 | 0.88441609 |
| THSD7A | 4343.92825 | -1.096590832 | 0.50665259 | -2.1643841 | 0.03043488 | 0.40391981 |
| TMEM106B | 711.270733 | -0.082988557 | 0.29924017 | -0.2773309 | 0.78152601 | 0.93748328 |
| SCIN | 340.869536 | 0.550888051 | 0.62022701 | 0.8882039 | 0.37443108 | 0.7697436 |
| ARL4A | 49.3797358 | -0.828498778 | 0.60706476 | -1.3647618 | 0.17232794 | 0.62592399 |
| AC011287.1 | 352.205475 | 0.914449611 | 0.48804206 | 1.87371067 | 0.06097031 | 0.48123823 |
| ETV1 | 144.80894 | 1.758914507 | 0.59719999 | 2.94526879 | 0.00322674 | 0.18622085 |
| DGKB | 1371.20676 | 1.182951393 | 0.61464512 | 1.92460878 | 5.43E-02 | 4.67E-01 |
| AGMO | 1864.23358 | 1.568065764 | 0.7418748 | 2.11365282 | 0.03454493 | 0.41670847 |
| ISPD | 1291.10518 | -0.271176332 | 0.36931827 | -0.7342619 | 0.46278914 | 0.8082072 |
| SOSTDC1 | 113.722154 | 0.741349778 | 0.87085558 | 0.85128901 | 0.39460883 | 0.77881976 |
| ANKMY2 | 403.527348 | -0.219065884 | 0.2488055 | -0.8804704 | 0.37860451 | 0.77061593 |
| BZW2 | 277.334863 | -0.262321185 | 0.35966831 | -0.7293419 | 0.46579252 | 0.81043641 |
| AHR | 49.5491255 | 0.325747378 | 0.58359974 | 0.55816916 | 0.57672888 | 0.85615431 |
| AC073332.1 | 221.360286 | 0.069073381 | 0.47098836 | 0.14665624 | 0.88340336 | 0.96846661 |
| AC019117.1 | 179.014605 | 1.718757027 | 0.65181755 | 2.6368683 | 0.00836753 | 0.25016267 |
| SNX13 | 1677.94393 | -0.143464362 | 0.170922 | -0.8393558 | 0.4012697 | 0.78240897 |
| HDAC9 | 300.602421 | 0.296556244 | 0.80983986 | 0.36619122 | 0.71422239 | 0.91221081 |
| TWISTNB | 101.973075 | 0.014730479 | 0.34886456 | 0.04222406 | 0.96632008 | 0.99112162 |
| MACC1 | 3159.23651 | 0.482273706 | 0.51169192 | 0.94250796 | 0.34593264 | 0.75166079 |
| MACC1-AS1 | 61.043518 | 0.700232275 | 0.90843244 | 0.77081381 | 0.4408173 | 0.79826598 |
| ITGB8 | 6966.14898 | 0.657771501 | 0.58364356 | 1.12700892 | 0.25973871 | 0.69242517 |
| SP4 | 157.056795 | -0.344494907 | 0.39403889 | -0.8742663 | 0.38197327 | 0.77252454 |
| DNAH11 | 269.864615 | -2.263459301 | 1.55644702 | -1.4542476 | 0.14587768 | 0.60245416 |
| RAPGEF5 | 1933.81464 | 1.310747089 | 0.92594598 | 1.4155762 | 1.57E-01 | 6.13E-01 |
| STEAP1B | 800.099793 | 0.092595483 | 0.4855059 | 0.19071958 | 0.84874529 | 0.95951833 |
| TOMM7 | 354.952334 | 0.081941199 | 0.40067271 | 0.20450906 | 0.83795571 | 0.95496955 |
| FAM126A | 476.526213 | 0.309370806 | 0.28546283 | 1.08375164 | 0.27847493 | 0.70442645 |
| KLHL7 | 555.227854 | 0.338676764 | 0.20517272 | 1.65069099 | 0.09880169 | 0.54236074 |
| NUPL2 | 559.773218 | 0.258376806 | 0.23433354 | 1.10260273 | 0.27019972 | 0.69961625 |
| MALSU1 | 249.759399 | 0.330388012 | 0.22892056 | 1.4432431 | 0.148952 | 0.60622442 |
| TRA2A | 1636.08216 | -0.110078429 | 0.30464986 | -0.3613277 | 0.71785449 | 0.91369965 |
| CCDC126 | 242.442388 | 0.094061848 | 0.28926139 | 0.32517941 | 0.74504529 | 0.92389439 |
| FAM221A | 157.770107 | -0.201135351 | 0.2843921 | -0.7072466 | 0.47941324 | 0.81653684 |
| AC004485.1 | 2400.09979 | -0.23515939 | 0.98569055 | -0.2385732 | 0.81143652 | 0.94753961 |
| NPY | 117.76815 | -0.711825563 | 1.26574609 | -0.5623763 | 0.57385968 | 0.85447394 |
| MPP6 | 2517.06921 | -0.084362776 | 0.27368715 | -0.3082453 | 0.75789569 | 0.928294 |
| GSDME | 124.524798 | 1.167868972 | 0.41381303 | 2.82221414 | 0.00476933 | 0.21294213 |
| OSBPL3 | 298.710889 | 1.165762659 | 0.64131735 | 1.81776254 | 0.06910043 | 0.49273866 |
| CYCS | 329.741124 | -0.187494645 | 0.42143289 | -0.444898 | 0.65639348 | 0.89053559 |
| NFE2L3 | 201.491485 | 1.400498549 | 0.61843138 | 2.26459813 | 0.02353735 | 0.37138185 |
| HNRNPA2B1 | 4189.23778 | -0.169843477 | 0.18099219 | -0.9384023 | 0.34803773 | 0.75316595 |
| CBX3 | 897.061121 | -0.337638356 | 0.28948461 | -1.166343 | 2.43E-01 | 0.68164834 |
| SNX10 | 3398.73025 | 0.487951781 | 0.42702501 | 1.14267728 | 2.53E-01 | 0.68887358 |
| AC004540.1 | 370.155337 | 1.789621686 | 0.57238207 | 3.12662082 | 0.00176828 | 0.13465439 |

| | | | | | | |
|---|---|---|---|---|---|---|
| SKAP2 | 2464.38998 | 0.815976544 | 0.3379725 | 2.4143282 | 0.01576426 | 0.31566878 |
| HOTAIRM1 | 193.523292 | 0.076170531 | 0.42516417 | 0.17915557 | 0.85781555 | 0.9626966 |
| HOXA3 | 314.919867 | 0.648520569 | 0.43156895 | 1.50270441 | 0.13291529 | 0.59190052 |
| HOXA-AS2 | 172.556343 | 0.59060763 | 0.54703523 | 1.07965191 | 0.28029722 | 0.70515983 |
| HOXA7 | 45.7366127 | 0.18846821 | 0.64216644 | 0.2934881 | 0.76914909 | 0.93209094 |
| HOXA9 | 227.302774 | 0.696421862 | 0.36439423 | 1.91117698 | 0.05598184 | 0.47105163 |
| HOXA10-AS | 59.7044915 | 0.601820406 | 0.39603297 | 1.51962198 | 0.12860601 | 0.58793678 |
| HIBADH | 2727.99384 | 0.362511117 | 0.30259193 | 1.19801976 | 0.23090932 | 0.66878058 |
| TAX1BP1 | 2890.1717 | -0.038906622 | 0.24072672 | -0.1616215 | 0.8716039 | 0.9657206 |
| JAZF1 | 817.541131 | 0.795843977 | 0.4207413 | 1.89152806 | 5.86E-02 | 0.47778423 |
| CREB5 | 3580.16822 | -0.087379255 | 0.63398301 | -0.1378259 | 0.89037804 | 0.97225819 |
| AC005162.3 | 396.292289 | 0.267413425 | 0.52636161 | 0.50804128 | 0.6114244 | 0.87168459 |
| CPVL | 663.331393 | -0.24605974 | 0.43550139 | -0.5650033 | 0.57207149 | 0.85408471 |
| AC007255.1 | 68.4114987 | 0.582305633 | 0.51008827 | 1.14157818 | 0.2536294 | 0.68891958 |
| WIPF3 | 207.277009 | 0.360094534 | 0.45309312 | 0.7947473 | 4.27E-01 | 0.78913582 |
| SCRN1 | 182.997667 | 0.828768757 | 0.5039593 | 1.64451524 | 0.10006982 | 0.54542315 |
| FKBP14 | 142.027812 | 0.073524567 | 0.33475158 | 0.2196392 | 0.82615216 | 0.95192757 |
| PLEKHA8 | 865.40907 | 0.092347939 | 0.28808901 | 0.3205535 | 0.74854878 | 0.92552718 |
| MTURN | 240.183093 | 0.080942249 | 0.36902175 | 0.21934276 | 0.82638305 | 0.95192757 |
| ZNRF2 | 1409.42777 | 0.319895271 | 0.32796845 | 0.97538428 | 0.32936968 | 0.73829083 |
| AC006978.2 | 1349.07091 | 0.307734724 | 0.33292903 | 0.92432529 | 0.35531698 | 0.75757147 |
| NOD1 | 194.391814 | 0.653279881 | 0.5095697 | 1.28202261 | 0.19983471 | 0.64705133 |
| GGCT | 96.7505481 | 0.420725069 | 0.49129283 | 0.85636314 | 0.39179694 | 0.77617585 |
| AC005154.6 | 384.865648 | 0.305003951 | 0.46216896 | 0.65994036 | 0.5092921 | 0.82882082 |
| GARS | 636.612963 | -0.529752426 | 0.42737475 | -1.2395501 | 0.21514183 | 0.65925411 |
| AQP1 | 2348.00421 | 2.011056955 | 0.8239857 | 2.44064546 | 0.01466104 | 0.30832605 |
| PDE1C | 58.553244 | 0.249846411 | 0.64015308 | 0.39029166 | 0.69632089 | 0.90612803 |
| LSM5 | 177.547441 | 0.311507153 | 0.31789073 | 0.97991896 | 0.32712612 | 0.73746094 |
| AVL9 | 2633.55172 | 0.009339666 | 0.17837447 | 0.05235988 | 0.95824194 | 0.98878187 |
| KBTBD2 | 448.087977 | 0.030242778 | 0.20379204 | 0.1484002 | 0.88202694 | 0.96796225 |
| FKBP9 | 263.89254 | 0.147743579 | 0.21711973 | 0.68047054 | 0.49620657 | 0.82243887 |
| NT5C3A | 977.033646 | -0.151039087 | 0.18782139 | -0.8041634 | 0.42130262 | 0.78877309 |
| RP9 | 136.450034 | -0.070417721 | 0.2437387 | -0.2889066 | 0.77265285 | 0.9336818 |
| BBS9 | 1799.50468 | 0.377480926 | 0.30611459 | 1.23313604 | 0.21752501 | 0.66043612 |
| DPY19L1 | 363.535343 | 0.065747105 | 0.41857444 | 0.15707386 | 0.87518664 | 0.96615698 |
| HERPUD2 | 603.068634 | -0.06923005 | 0.19921621 | -0.3475121 | 0.72820661 | 0.91697155 |
| AC018647.2 | 53.4753271 | -0.642718006 | 0.43387631 | -1.4813393 | 0.13851619 | 0.59474076 |
| SEPT7-AS1 | 119.50875 | 0.895329106 | 0.49693822 | 1.80169099 | 0.07159404 | 0.50017302 |
| 7-Sep | 1377.46901 | -0.240890993 | 0.18529763 | -1.300022 | 0.19359344 | 0.64548758 |
| EEPD1 | 285.167624 | -0.837509523 | 0.50701498 | -1.6518438 | 0.09856641 | 0.54215479 |
| KIAA0895 | 539.347483 | 0.189533522 | 0.26883705 | 0.70501265 | 0.48080237 | 0.81715537 |
| AC006960.2 | 86.2520486 | 0.678384526 | 1.03625252 | 0.65465175 | 0.512692 | 0.83055441 |
| AOAH | 596.855643 | 0.793229078 | 1.05899666 | 0.74903832 | 0.45383411 | 0.80433814 |
| AC007349.1 | 45.9073869 | 1.678854981 | 0.78932495 | 2.12695035 | 0.03342421 | 0.41254527 |
| ELMO1 | 133.994402 | -0.53121384 | 0.51436325 | -1.0327601 | 0.30171618 | 0.71908176 |
| EPDR1 | 382.782841 | 0.303549186 | 0.31047881 | 0.97768084 | 0.3282322 | 0.73826734 |
| SFRP4 | 105.708681 | 1.040895338 | 0.80671774 | 1.29028442 | 0.19695192 | 0.64558745 |
| STARD3NL | 373.8148 | 0.291073654 | 0.20471459 | 1.42185107 | 0.15506949 | 0.611403 |
| VPS41 | 1457.47237 | -0.272934908 | 0.21885041 | -1.24713 | 0.21234985 | 0.65708554 |
| YAE1D1 | 176.163222 | 0.348349049 | 0.27865816 | 1.25009456 | 0.21126501 | 0.65561118 |
| RALA | 890.128381 | -0.482165395 | 0.37952529 | -1.2704434 | 0.20392673 | 0.64929703 |
| LINC00265 | 91.9356038 | -0.164783408 | 0.31675021 | -0.5202314 | 0.60290229 | 0.86779185 |
| CDK13 | 2979.14243 | -0.121036584 | 0.19335448 | -0.6259828 | 0.53132621 | 0.83789294 |
| MPLKIP | 310.581487 | 0.157137138 | 0.32167229 | 0.4885007 | 0.62519523 | 0.87854882 |

| | | | | | | |
|---|---|---|---|---|---|---|
| SUGCT | 6832.90618 | 0.865375913 | 0.4995074 | 1.73245864 | 8.32E-02 | 0.51935659 |
| AC004988.1 | 59.9332321 | -1.256029265 | 0.66621624 | -1.8853177 | 0.05938696 | 0.47855094 |
| LINC01449 | 39.6477645 | 2.527081379 | 0.9303735 | 2.71620096 | 0.00660358 | 0.22966139 |
| GLI3 | 327.380453 | 1.617339383 | 0.47312652 | 3.4184078 | 0.00062989 | 0.08457928 |
| MRPL32 | 237.742896 | -0.232840691 | 0.29244889 | -0.7961757 | 0.42592993 | 0.78894887 |
| AC005537.1 | 65.4034577 | -0.08185345 | 0.77566426 | -0.1055269 | 0.91595772 | 0.97801193 |
| HECW1 | 1734.24975 | 2.363081737 | 0.82986158 | 2.84756133 | 0.00440556 | 0.20749797 |
| STK17A | 333.946654 | -0.345666387 | 0.43845247 | -0.7883782 | 0.43047549 | 0.79055252 |
| COA1 | 1680.56026 | -0.212447056 | 0.15724244 | -1.3510796 | 0.17666993 | 0.62709281 |
| BLVRA | 363.301673 | 0.183611085 | 0.26297958 | 0.69819521 | 0.48505512 | 0.81839266 |
| URGCP | 590.098196 | 0.297488544 | 0.36429458 | 0.81661535 | 4.14E-01 | 0.78632759 |
| UBE2D4 | 203.947359 | -0.955686178 | 0.39173463 | -2.4396265 | 0.01470245 | 0.30832605 |
| SPDYE1 | 98.9060398 | 0.292615454 | 0.32229811 | 0.90790311 | 0.36392941 | 0.76272056 |
| AC004951.1 | 117.706694 | -0.224652271 | 0.39549452 | -0.5680288 | 0.57001543 | 0.85327114 |
| DBNL | 336.404916 | -0.30912238 | 0.2044266 | -1.5121436 | 0.13049734 | 0.58853436 |
| POLM | 254.797707 | 0.338486001 | 0.41056573 | 0.82443804 | 0.40969071 | 0.78542936 |
| AEBP1 | 96.4984729 | -0.591161232 | 0.86217302 | -0.6856643 | 0.49292484 | 0.82100434 |
| POLD2 | 178.33652 | 0.278161048 | 0.35166789 | 0.79097652 | 0.42895769 | 0.79037914 |
| YKT6 | 214.31469 | -0.900834902 | 0.51510544 | -1.7488359 | 0.08031938 | 0.51297858 |
| CAMK2B | 42.4166651 | 0.130210417 | 0.4910189 | 0.26518413 | 7.91E-01 | 9.41E-01 |
| NUDCD3 | 1567.5185 | -0.394512698 | 0.25033461 | -1.5759415 | 0.11503929 | 0.57182344 |
| DDX56 | 395.830958 | -0.006915621 | 0.22148761 | -0.0312235 | 0.97509129 | 0.99402915 |
| TMED4 | 564.378514 | 0.197275932 | 0.33729717 | 0.58487277 | 0.55863327 | 0.84910027 |
| OGDH | 2321.34662 | 0.563243635 | 0.29706691 | 1.89601609 | 0.05795792 | 0.47684824 |
| ZMIZ2 | 768.877312 | 0.921574246 | 0.40450997 | 2.27824855 | 0.02271177 | 0.36561837 |
| PPIA | 1494.08596 | -0.221787554 | 0.3733874 | -0.5939878 | 0.55252029 | 0.84721285 |
| H2AFV | 520.744732 | -0.273313802 | 0.24267685 | -1.1262459 | 0.26006146 | 0.69280433 |
| PURB | 354.217259 | -0.129563367 | 0.1693985 | -0.7648436 | 0.44436466 | 0.79994797 |
| SNHG15 | 98.2152866 | -0.357654065 | 0.45898913 | -0.7792212 | 0.43584942 | 0.79549938 |
| CCM2 | 684.772178 | 0.503399514 | 0.2383969 | 2.11160262 | 0.03472055 | 0.41686761 |
| TBRG4 | 248.951293 | -0.158522529 | 0.28959306 | -0.5473975 | 0.58410564 | 0.86025731 |
| IGFBP3 | 94.1439648 | -0.86544425 | 0.69548728 | -1.2443711 | 0.21336303 | 0.65809339 |
| AC004870.4 | 184.563166 | 2.659303829 | 1.19483861 | 2.22565944 | 0.026037 | 0.38392873 |
| AC004870.3 | 118.264342 | 1.499531549 | 1.01282484 | 1.48054381 | 0.13872818 | 0.59474076 |
| TNS3 | 6493.4902 | -0.677834649 | 0.40590792 | -1.6699222 | 0.09493476 | 0.53824794 |
| HUS1 | 283.306457 | -0.446817096 | 0.24465244 | -1.8263341 | 0.06779995 | 0.48963361 |
| UPP1 | 327.091559 | -1.241623576 | 0.66781273 | -1.8592392 | 6.30E-02 | 0.48535606 |
| ABCA13 | 214.088602 | -0.078743163 | 0.89570015 | -0.0879124 | 0.92994629 | 0.98272895 |
| ZPBP | 55.9714485 | -0.267950386 | 0.59257956 | -0.4521762 | 0.65114204 | 0.88923125 |
| FIGNL1 | 45.6978769 | -0.657009002 | 0.5601942 | -1.1728236 | 0.24086653 | 0.67957469 |
| DDC | 195.887495 | -0.60472007 | 0.60740379 | -0.9955817 | 3.19E-01 | 7.31E-01 |
| GRB10 | 2933.1657 | -0.460671154 | 0.32400676 | -1.4217949 | 0.15508581 | 0.611403 |
| COBL | 3664.95625 | -0.470542402 | 0.33414545 | -1.4081963 | 0.15907294 | 0.61423843 |
| SEC61G | 245.270861 | -0.809218531 | 0.44300289 | -1.8266665 | 0.06774993 | 0.48963361 |
| EGFR | 3145.70396 | -0.454030419 | 0.30050489 | -1.5108919 | 0.13081599 | 0.58858527 |
| LANCL2 | 566.918749 | -0.294416478 | 0.38414808 | -0.766414 | 0.44342999 | 0.7993765 |
| VOPP1 | 850.349426 | -0.503278963 | 0.49245654 | -1.0219764 | 3.07E-01 | 0.72190689 |
| NIPSNAP2 | 807.315237 | 0.330311221 | 0.22239064 | 1.48527485 | 0.13747103 | 0.59474076 |
| PSPH | 308.435915 | -0.690621633 | 0.378697 | -1.8236786 | 0.06820067 | 0.49072262 |
| CCT6A | 701.441197 | -0.787592557 | 0.43793725 | -1.7984142 | 0.0721114 | 0.50199643 |
| SUMF2 | 549.507811 | -0.292198375 | 0.21033023 | -1.3892362 | 0.16476094 | 0.62065658 |
| PHKG1 | 127.422214 | 0.118302967 | 0.33766934 | 0.35035152 | 0.7260749 | 0.91635631 |
| CHCHD2 | 720.933697 | -0.305801609 | 0.4519626 | -0.6766082 | 0.49865457 | 0.82369946 |
| ZNF727 | 106.372838 | -0.459631923 | 0.58835127 | -0.7812202 | 0.43467297 | 0.79451524 |

| | | | | | | |
|---|---|---|---|---|---|---|
| ZNF736 | 618.106844 | 0.372264611 | 0.28347404 | 1.31322294 | 0.18910784 | 0.6413893 |
| ZNF680 | 450.76234 | 0.165505481 | 0.40169106 | 0.41202182 | 0.68032343 | 0.89941358 |
| ZNF107 | 146.981446 | 0.015540317 | 0.63546903 | 0.02445488 | 0.98048978 | 0.99500802 |
| ZNF138 | 331.838975 | 0.10128481 | 0.2496214 | 0.40575372 | 0.68492357 | 0.90098343 |
| ZNF273 | 43.817072 | 0.075033872 | 0.53808385 | 0.13944643 | 0.88909739 | 0.97197002 |
| ZNF117 | 174.474805 | 0.789599029 | 0.40438084 | 1.95261235 | 0.05086555 | 0.45990373 |
| ZNF92 | 74.6486455 | -0.148654109 | 0.29928651 | -0.496695 | 0.61940416 | 0.87527555 |
| VKORC1L1 | 904.654813 | -0.892030017 | 0.40805742 | -2.1860404 | 0.02881265 | 0.39414665 |
| GUSB | 233.509394 | -0.225309413 | 0.24366637 | -0.9246636 | 0.35514093 | 0.75757147 |
| ASL | 343.645315 | 0.745459391 | 0.33256113 | 2.24157105 | 0.02498911 | 0.37991912 |
| AC068533.4 | 301.06544 | 0.701720298 | 0.51942039 | 1.35096795 | 0.1767057 | 0.62709281 |
| CRCP | 136.940631 | -0.19654278 | 0.28298286 | -0.6945395 | 0.48734393 | 0.81893983 |
| AC068533.3 | 142.495126 | 0.450047545 | 0.43663915 | 1.03070818 | 0.30267769 | 0.72027831 |
| TPST1 | 1303.19276 | 0.450601688 | 0.3507222 | 1.28478235 | 0.19886835 | 0.64597062 |
| AC008267.5 | 35.7320784 | 0.647740005 | 0.60770991 | 1.0658704 | 0.28648223 | 0.71118494 |
| AC027644.4 | 901.86036 | -0.437375238 | 0.50945666 | -0.8585131 | 0.39060918 | 0.77606489 |
| AC027644.3 | 120.030835 | -0.441942371 | 0.39440312 | -1.1205347 | 0.26248599 | 0.69467275 |
| AC073335.2 | 48.2003172 | 0.484439086 | 0.54704715 | 0.88555271 | 0.3758586 | 0.7697436 |
| TMEM248 | 597.434186 | -0.283465406 | 0.26831487 | -1.0564655 | 0.2907556 | 0.71269438 |
| SBDS | 318.034546 | -0.243010642 | 0.30517746 | -0.7962929 | 0.42586181 | 0.78894887 |
| TYW1 | 1734.65076 | -0.076370715 | 0.2211826 | -0.3452836 | 0.7298812 | 0.91834786 |
| AC006480.2 | 184.43194 | 0.495429983 | 0.38996146 | 1.27045885 | 0.20392123 | 0.64929703 |
| LINC01372 | 121.547574 | 0.455350005 | 0.45508186 | 1.00058923 | 0.31702544 | 0.728983 |
| AUTS2 | 2824.84468 | 0.339787127 | 0.2899614 | 1.17183572 | 0.24126302 | 0.68003129 |
| GALNT17 | 55.3507759 | -0.360179516 | 0.56213256 | -0.6407377 | 0.52169313 | 0.83445062 |
| CALN1 | 113.858787 | 0.181458965 | 0.48747647 | 0.37224148 | 0.70971306 | 0.91112082 |
| TYW1B | 1028.87491 | -0.122184474 | 0.21059081 | -0.5801985 | 5.62E-01 | 8.50E-01 |
| POM121 | 1373.27661 | -0.2316609 | 0.23450957 | -0.9878526 | 0.32322483 | 0.73534319 |
| NSUN5 | 214.976059 | -0.374967431 | 0.31286517 | -1.1984953 | 0.23072426 | 0.66878058 |
| BAZ1B | 931.572787 | -0.229636058 | 0.32940459 | -0.6971246 | 0.48572479 | 0.81839266 |
| BCL7B | 219.274287 | -0.679161664 | 0.32032001 | -2.1202598 | 0.03398414 | 0.41378378 |
| TBL2 | 112.39694 | -0.599773854 | 0.4076478 | -1.471304 | 0.14120891 | 0.59581113 |
| MLXIPL | 2781.17768 | 0.488583901 | 0.66286935 | 0.73707421 | 0.46107722 | 0.80771389 |
| VPS37D | 70.5123591 | 0.273997197 | 0.3941827 | 0.69510204 | 0.48699136 | 0.81893983 |
| DNAJC30 | 75.787939 | 0.180109769 | 0.36861787 | 0.48860834 | 0.625119 | 0.87854882 |
| BUD23 | 332.845449 | -0.458884869 | 0.26730875 | -1.7166848 | 0.08603678 | 0.52392738 |
| STX1A | 48.6871025 | -0.634607365 | 0.46027415 | -1.3787595 | 0.16796891 | 0.62304975 |
| ABHD11 | 307.778878 | -0.367226357 | 0.51753191 | -0.7095724 | 0.47796934 | 0.81588117 |
| CLDN3 | 593.552215 | -0.332624383 | 0.56976991 | -0.5837872 | 5.59E-01 | 0.84936251 |
| CLDN4 | 905.837712 | -0.522525474 | 0.6865165 | -0.7611259 | 0.44658188 | 0.80205685 |
| METTL27 | 82.9617887 | 0.771205288 | 0.65055676 | 1.18545427 | 0.23583789 | 0.67276377 |
| LIMK1 | 101.160302 | 0.199729313 | 0.55341924 | 0.36090056 | 0.71817379 | 0.91393632 |
| EIF4H | 479.21217 | -0.356410277 | 0.17818087 | -2.0002724 | 0.04547086 | 0.44360789 |
| LAT2 | 33.6221214 | 0.019638505 | 0.44354499 | 0.04427624 | 0.96468421 | 0.99033302 |
| RFC2 | 94.341876 | -0.000581859 | 0.42931569 | -0.0013553 | 0.99891861 | 0.99977652 |
| CLIP2 | 217.897517 | -0.100108501 | 0.65024556 | -0.1539549 | 0.87764528 | 0.96658493 |
| GTF2IRD1 | 1777.18485 | -0.170618593 | 0.4217958 | -0.4045052 | 0.68584125 | 0.90141203 |
| GTF2I | 4399.56202 | 0.039574318 | 0.1945836 | 0.20337952 | 0.83883841 | 0.95545195 |
| GTF2IRD2 | 258.054678 | -0.194530976 | 0.51236587 | -0.379672 | 0.70418891 | 0.90922356 |
| CASTOR2 | 510.29023 | -0.398485008 | 0.44009686 | -0.9054484 | 0.36522787 | 0.76362239 |
| RCC1L | 389.777654 | -0.530529583 | 0.30157568 | -1.7591922 | 0.07854487 | 0.51192014 |
| GTF2IRD2B | 469.859541 | -0.278559664 | 0.45024403 | -0.618686 | 0.53612325 | 0.83955689 |
| AC211486.5 | 108.253414 | 0.145999358 | 0.60914976 | 0.23967728 | 0.81058045 | 0.94717173 |
| TRIM73 | 369.041804 | 0.482675844 | 0.47352557 | 1.01932372 | 0.30804931 | 0.7236387 |

| | | | | | | |
|---|---|---|---|---|---|---|
| POM121C | 1094.99161 | -0.375960876 | 0.17168164 | -2.1898724 | 0.02853349 | 0.39280225 |
| SPDYE5 | 270.298688 | 0.690937038 | 0.42546941 | 1.62394057 | 0.10438849 | 0.55066578 |
| HIP1 | 802.47712 | 0.157579791 | 0.37484329 | 0.42038845 | 0.67420171 | 0.89801183 |
| RHBDD2 | 228.706613 | -0.067389044 | 0.31100441 | -0.216682 | 0.82845621 | 0.95305492 |
| POR | 1914.66049 | -0.789648079 | 0.50477832 | -1.5643463 | 0.11773627 | 0.57390152 |
| TMEM120A | 240.271279 | -0.132595671 | 0.27245232 | -0.4866748 | 0.62648882 | 0.87885398 |
| STYXL1 | 551.447719 | 0.314558675 | 0.26495407 | 1.18721965 | 0.23514099 | 0.67119484 |
| MDH2 | 347.317538 | 0.118857724 | 0.29062127 | 0.40897806 | 0.68255576 | 0.89962501 |
| HSPB1 | 1389.31608 | -0.443314059 | 0.37763042 | -1.1739363 | 0.24042054 | 0.67897455 |
| YWHAG | 786.427953 | -0.589019636 | 0.32962252 | -1.786952 | 0.07394524 | 0.50602467 |
| ZP3 | 69.0045774 | -0.090304902 | 0.53424173 | -0.1690338 | 0.86577007 | 0.96437968 |
| DTX2 | 379.404227 | -0.032625353 | 0.34359351 | -0.0949533 | 0.92435188 | 0.9803537 |
| SPDYE16 | 81.7873209 | 0.780254456 | 0.47890943 | 1.62923177 | 0.10326396 | 0.55022042 |
| POMZP3 | 216.946188 | 0.13121552 | 0.37709989 | 0.34795958 | 0.72787054 | 0.91665945 |
| CCDC146 | 910.197888 | 0.803061098 | 0.37438164 | 2.14503333 | 0.03195018 | 0.40988962 |
| FGL2 | 89.0120628 | 1.585319761 | 0.71294104 | 2.22363375 | 0.0261731 | 0.38392873 |
| GSAP | 1144.51681 | 1.007075929 | 0.50752439 | 1.9842907 | 0.04722344 | 0.44857361 |
| PTPN12 | 1971.96673 | 0.095842971 | 0.27320849 | 0.35080525 | 0.72573446 | 0.91634663 |
| APTR | 433.497983 | 0.580468571 | 0.39804639 | 1.45829378 | 0.14475958 | 0.60109715 |
| RSBN1L | 893.367043 | -0.019319713 | 0.17373397 | -0.1112029 | 0.91145549 | 0.97710937 |
| TMEM60 | 102.8382 | -0.011509653 | 0.3313596 | -0.0347346 | 0.97229135 | 0.9924192 |
| PHTF2 | 538.948862 | -0.193906372 | 0.15628486 | -1.240724 | 0.21470773 | 0.65925411 |
| MAGI2 | 8652.56736 | -0.156951174 | 0.30964871 | -0.5068685 | 0.61224709 | 0.87222135 |
| GNAI1 | 1048.40545 | 0.043004259 | 0.51847117 | 0.08294436 | 0.93389578 | 0.98355048 |
| SEMA3C | 305.609328 | -1.019824141 | 0.94169895 | -1.082962 | 2.79E-01 | 0.70446562 |
| CACNA2D1 | 3936.78431 | 0.155595562 | 0.2695059 | 0.57733639 | 0.56371223 | 0.85125081 |
| PCLO | 4551.8173 | 0.593248105 | 0.60212462 | 0.98525802 | 0.32449735 | 0.73614611 |
| SEMA3E | 82.1260536 | -2.206707505 | 0.83144221 | -2.654072 | 0.00795269 | 0.2460665 |
| SEMA3A | 44.719959 | -0.185404828 | 0.48949708 | -0.378766 | 0.70486167 | 0.909554 |
| AC003984.1 | 655.657675 | -1.054700776 | 0.78778186 | -1.3388234 | 1.81E-01 | 0.63166264 |
| SEMA3D | 291.389609 | 1.314774175 | 0.63092142 | 2.08389529 | 0.03716969 | 0.42215481 |
| KIAA1324L | 150.944279 | -0.405573978 | 0.64897088 | -0.6249494 | 0.53200425 | 0.83789294 |
| DMTF1 | 1430.56262 | 0.226718158 | 0.3144986 | 0.72088765 | 0.47097864 | 0.81387913 |
| TMEM243 | 394.612085 | 0.652152709 | 0.3018306 | 2.16065799 | 3.07E-02 | 4.04E-01 |
| TP53TG1 | 51.8418413 | 1.096516684 | 0.47776015 | 2.2951196 | 0.02172627 | 0.35966425 |
| CROT | 229.761578 | 0.675376279 | 0.32138426 | 2.10146037 | 0.03560057 | 0.42040764 |
| ABCB4 | 52.0631391 | 0.327475164 | 0.61850237 | 0.52946468 | 0.59648314 | 0.86559503 |
| ABCB1 | 1626.86938 | -0.862060783 | 0.47129613 | -1.8291276 | 0.06738049 | 0.48963361 |
| RUNDC3B | 793.355153 | -0.183484428 | 0.50591651 | -0.3626773 | 0.71684597 | 0.91309429 |
| SLC25A40 | 286.423563 | 0.405788157 | 0.26839771 | 1.51189128 | 0.13056152 | 0.58853436 |
| DBF4 | 156.398678 | 0.093761561 | 0.36270259 | 0.25850811 | 0.79601479 | 0.94252227 |
| ADAM22 | 510.341428 | 0.108396821 | 0.45598613 | 0.23771956 | 0.81209861 | 0.94784066 |
| SRI | 234.10625 | 0.271174733 | 0.33999725 | 0.79757918 | 0.42511473 | 0.78894887 |
| AC003991.1 | 52.4921855 | 0.772518375 | 0.40225834 | 1.92045336 | 0.05480066 | 0.46800787 |
| ZNF804B | 5293.73859 | 0.392068502 | 0.80804942 | 0.48520362 | 6.28E-01 | 0.87944599 |
| STEAP1 | 98.4174148 | -0.215919059 | 0.50525861 | -0.4273436 | 0.66912905 | 0.89582842 |
| STEAP2 | 380.667775 | 0.86146768 | 0.40794704 | 2.11171449 | 0.03471095 | 0.41686761 |
| CFAP69 | 1333.60947 | 0.514086334 | 0.39520074 | 1.3008233 | 0.19331894 | 0.64548758 |
| AC002064.1 | 85.1063307 | 1.151995603 | 0.47980183 | 2.40098212 | 0.01635114 | 0.32248204 |
| GTPBP10 | 547.590617 | 0.199746826 | 0.25973069 | 0.7690536 | 0.44186149 | 0.79902315 |
| CLDN12 | 843.893863 | 0.323423312 | 0.22529929 | 1.43552742 | 0.15113686 | 0.60905477 |
| CDK14 | 5562.65457 | -0.040074507 | 0.37231372 | -0.1076364 | 0.91428413 | 0.97746607 |
| FZD1 | 179.493097 | 0.371571545 | 0.51641101 | 0.71952677 | 0.47181642 | 0.81389222 |
| AC079760.2 | 207.167759 | 0.498803076 | 0.56792271 | 0.87829394 | 0.37978422 | 0.77095585 |

| | | | | | | |
|---|---|---|---|---|---|---|
| AC079760.1 | 126.382826 | -0.247213935 | 0.62060332 | -0.3983445 | 0.69037624 | 0.90425104 |
| MTERF1 | 745.178871 | -0.103827095 | 0.51017672 | -0.203512 | 0.83873485 | 0.95541749 |
| AKAP9 | 3155.59121 | -0.342386878 | 0.2012024 | -1.7017037 | 0.08881092 | 0.52996572 |
| KRIT1 | 1615.69551 | 0.136163797 | 0.22821535 | 0.59664609 | 0.55074369 | 0.84659008 |
| ANKIB1 | 5158.00926 | 0.010921885 | 0.2251367 | 0.04851223 | 9.61E-01 | 0.98985638 |
| GATAD1 | 404.913799 | 0.676908896 | 0.41564567 | 1.62857198 | 0.10340366 | 0.55022042 |
| AC007566.1 | 364.194149 | 0.825035912 | 0.54277357 | 1.520037 | 0.12850168 | 0.58793678 |
| PEX1 | 454.471466 | 0.375956801 | 0.25395328 | 1.48041718 | 0.13876195 | 0.59474076 |
| RBM48 | 156.428647 | -0.410920925 | 0.34115261 | -1.2045076 | 0.22839343 | 0.66705629 |
| FAM133B | 713.552951 | -0.122069235 | 0.14942001 | -0.8169537 | 0.41395486 | 0.78632759 |
| CDK6 | 1243.38239 | 0.980224233 | 0.46435817 | 2.11092277 | 0.03477895 | 0.41686761 |
| SAMD9 | 67.0945228 | -2.452038465 | 0.78064216 | -3.1410531 | 0.00168342 | 0.13207839 |
| SAMD9L | 202.516088 | -2.289095986 | 1.07070431 | -2.1379348 | 0.03252204 | 0.41149854 |
| VPS50 | 1454.18793 | -0.193564388 | 0.16715373 | -1.1580022 | 0.24686313 | 0.68482118 |
| GNGT1 | 1066.2612 | 0.155560491 | 0.48518538 | 0.32062073 | 0.74849783 | 0.92552718 |
| GNG11 | 67.1420017 | 0.687038627 | 0.48116339 | 1.42786971 | 0.15332936 | 0.61008819 |
| BET1 | 315.249092 | 0.488594969 | 0.32318825 | 1.51179684 | 0.13058555 | 0.58853436 |
| CASD1 | 556.410239 | 0.341231279 | 0.32016285 | 1.06580535 | 0.28651164 | 0.71118494 |
| SGCE | 551.333407 | 0.580537838 | 0.38134598 | 1.52233894 | 0.12792418 | 0.58793678 |
| PEG10 | 39.8505926 | 0.147853952 | 0.47605068 | 0.31058448 | 0.75611653 | 0.92799282 |
| PPP1R9A | 3570.89226 | 0.11392982 | 0.25321053 | 0.44994108 | 0.65275293 | 0.8894805 |
| AC002429.2 | 270.578046 | 0.398217716 | 0.54096119 | 0.73612991 | 0.46165164 | 0.80805249 |
| PON2 | 781.978651 | -0.125323224 | 0.3269155 | -0.3833505 | 0.70145989 | 0.90783786 |
| PDK4 | 2626.9637 | -0.721132115 | 1.4967917 | -0.4817852 | 0.62995853 | 0.88083664 |
| AC002451.1 | 46.2887132 | 0.237048946 | 1.11664965 | 0.21228587 | 0.83188402 | 0.95325673 |
| DYNC1I1 | 415.572275 | 1.038420953 | 0.90866984 | 1.14279236 | 0.25312481 | 0.68887358 |
| SLC25A13 | 2724.94701 | -0.274258317 | 0.25676733 | -1.06812 | 0.28546639 | 0.71029858 |
| SEM1 | 704.891533 | -0.332631546 | 0.20192516 | -1.6473011 | 0.09949617 | 0.54399442 |
| SDHAF3 | 318.207674 | -0.192826156 | 0.2569293 | -0.7505028 | 0.45295194 | 0.80433814 |
| ASNS | 110.574528 | 0.439876736 | 0.52068056 | 0.84481114 | 0.3982163 | 0.78121522 |
| LMTK2 | 2018.42922 | -0.393290264 | 0.18765701 | -2.0957931 | 0.03610055 | 0.42040764 |
| TECPR1 | 232.806675 | -0.179433555 | 0.30755764 | -0.5834144 | 0.55961437 | 0.84941263 |
| BRI3 | 1507.05083 | -0.554702458 | 0.31560165 | -1.7576031 | 0.07881506 | 0.51192014 |
| BAIAP2L1 | 1148.7863 | 0.654010686 | 0.50577575 | 1.2930843 | 0.1959819 | 0.64558745 |
| TRRAP | 1458.31354 | -0.04880051 | 0.17535418 | -0.2782968 | 0.78078452 | 0.93748328 |
| SMURF1 | 2181.80986 | -0.09489901 | 0.32674362 | -0.2904388 | 0.7714806 | 0.93316533 |
| AC004834.1 | 63.7311296 | 0.565571454 | 0.54685723 | 1.03422141 | 0.30103267 | 0.71879123 |
| ARPC1B | 182.311691 | 0.322258333 | 0.40127562 | 0.80308476 | 0.42192575 | 0.78892549 |
| PDAP1 | 242.849615 | -0.587688174 | 0.31309755 | -1.877013 | 0.06051631 | 0.48026727 |
| BUD31 | 158.908791 | -0.794569691 | 0.30564986 | -2.5996075 | 0.00933304 | 0.25998727 |
| PTCD1 | 40.4418984 | -0.366038044 | 0.45183959 | -0.8101062 | 0.41787915 | 0.78679526 |
| CPSF4 | 196.587394 | -0.111106251 | 0.26709761 | -0.4159762 | 0.67742742 | 0.89864474 |
| ZNF789 | 432.534202 | 0.070536112 | 0.28454366 | 0.24789206 | 0.80421793 | 0.94551573 |
| ZNF394 | 224.426615 | 0.105672494 | 0.31381501 | 0.33673498 | 0.73631669 | 0.92019178 |
| ZKSCAN5 | 201.899091 | -0.274718261 | 0.33466769 | -0.8208688 | 0.41172101 | 0.78632759 |
| FAM200A | 146.295161 | 0.256080018 | 0.3130564 | 0.81799963 | 0.4133574 | 0.78632759 |
| ZSCAN25 | 161.684068 | -0.392635168 | 0.29183832 | -1.3453859 | 0.17850064 | 0.6288217 |
| CYP3A5 | 1055.44088 | 1.07098461 | 0.66421265 | 1.61241224 | 0.10687226 | 0.55488138 |
| TRIM4 | 523.73803 | -0.081398456 | 0.33967423 | -0.2396368 | 0.81061181 | 0.94717173 |
| AZGP1 | 725.934277 | -0.602875827 | 0.57866493 | -1.0418392 | 0.29748622 | 0.71839551 |
| ZKSCAN1 | 2562.20043 | 0.754314251 | 0.2921975 | 2.58152194 | 0.00983657 | 0.26385502 |
| ZSCAN21 | 254.057469 | 0.142738225 | 0.26667129 | 0.53525907 | 5.92E-01 | 8.64E-01 |
| ZNF3 | 304.93961 | -0.370537311 | 0.34405149 | -1.0769822 | 0.28148824 | 0.70625976 |
| COPS6 | 273.354249 | -0.03824359 | 0.22942827 | -0.1666908 | 0.86761332 | 0.96487312 |

| | | | | | | |
|---|---|---|---|---|---|---|
| MCM7 | 137.011861 | -0.990585576 | 0.38118734 | -2.5986843 | 0.00935818 | 0.25998727 |
| AP4M1 | 151.886567 | 0.019460016 | 0.27010453 | 0.07204624 | 0.94256511 | 0.98489601 |
| TAF6 | 141.499356 | -0.153406856 | 0.34998677 | -0.4383219 | 0.66115298 | 0.89184036 |
| AC073842.2 | 47.8561499 | -0.231929987 | 0.51460251 | -0.4506973 | 0.65220769 | 0.88935488 |
| CNPY4 | 37.1425663 | -0.057171737 | 0.43900226 | -0.1302311 | 0.89638361 | 0.97312927 |
| LAMTOR4 | 458.27774 | -0.248882841 | 0.29916269 | -0.8319314 | 0.40544766 | 0.78429302 |
| C7orf43 | 77.7779054 | -0.275387683 | 0.46133431 | -0.5969374 | 0.5505492 | 0.84648098 |
| STAG3 | 189.00933 | 0.231891709 | 0.48521847 | 0.47791196 | 0.63271287 | 0.88151688 |
| CASTOR3 | 1069.46664 | -0.200998971 | 0.30849265 | -0.6515519 | 0.51469029 | 0.83169608 |
| SPDYE3 | 399.407634 | 0.931901375 | 0.52094529 | 1.7888661 | 0.07363638 | 0.5056783 |
| PILRB | 1278.04377 | -0.295264253 | 0.44485528 | -0.663731 | 0.50686247 | 0.82806179 |
| PILRA | 242.4405 | -0.456712619 | 0.36945372 | -1.2361836 | 0.21639034 | 0.66032006 |
| ZCWPW1 | 85.4998837 | 0.517765189 | 0.61679583 | 0.8394434 | 0.40122054 | 0.78240897 |
| MEPCE | 109.184042 | -0.952932388 | 0.50070396 | -1.9031853 | 0.05701638 | 0.47509025 |
| PPP1R35 | 230.781464 | -0.274364533 | 0.30482298 | -0.9000782 | 0.36807861 | 0.76615653 |
| TSC22D4 | 181.363558 | 0.379991888 | 0.26782633 | 1.4187996 | 0.15595745 | 0.61255389 |
| AGFG2 | 85.6139831 | 0.070589003 | 0.48705188 | 0.14493118 | 0.88476521 | 0.96934512 |
| LRCH4 | 108.931313 | -0.082154571 | 0.29205579 | -0.2812975 | 0.77848221 | 0.93646866 |
| MOSPD3 | 52.5194856 | 0.460515805 | 0.40586429 | 1.13465466 | 0.25652004 | 0.69092557 |
| GNB2 | 421.808337 | -0.132723501 | 0.42585211 | -0.3116657 | 0.75529459 | 0.92799282 |
| GIGYF1 | 562.533613 | -0.59321296 | 0.4365214 | -1.3589551 | 0.17416083 | 0.62699479 |
| POP7 | 70.2966361 | -0.23814403 | 0.4667406 | -0.5102278 | 0.60989188 | 0.87090196 |
| EPHB4 | 160.17148 | 0.164971415 | 0.37728507 | 0.43725932 | 0.66192329 | 0.89193781 |
| SLC12A9 | 363.283594 | 0.374908073 | 0.34598454 | 1.08359777 | 0.27854318 | 0.70442645 |
| TRIP6 | 326.606838 | -0.733482005 | 0.3780799 | -1.9400185 | 0.05237744 | 0.4649855 |
| SRRT | 294.187725 | -0.919181384 | 0.38949769 | -2.3599148 | 0.01827913 | 0.33797156 |
| TRIM56 | 861.488403 | -0.375553941 | 0.45538744 | -0.824691 | 0.40954702 | 0.78542936 |
| AP1S1 | 264.708958 | 0.797295774 | 0.43484725 | 1.8335077 | 0.0667271 | 0.48884194 |
| MOGAT3 | 99.3705725 | 3.363312484 | 0.87228351 | 3.85575611 | 0.00011537 | 0.03437849 |
| PLOD3 | 408.148577 | -0.056384063 | 0.33244158 | -0.1696059 | 0.86532006 | 0.96428252 |
| ZNHIT1 | 369.479796 | -0.349127871 | 0.29444731 | -1.1857058 | 0.23573852 | 0.67262028 |
| CLDN15 | 92.5625567 | 0.084596561 | 0.46298721 | 0.182719 | 0.85501851 | 0.96171023 |
| FIS1 | 517.402877 | -0.336134163 | 0.32283068 | -1.0412089 | 0.29777863 | 0.71852421 |
| IFT22 | 132.667775 | 0.165642851 | 0.36567702 | 0.45297583 | 0.65056615 | 0.8887773 |
| COL26A1 | 73.5508648 | -0.043232864 | 0.409335 | -0.1056173 | 0.91588599 | 0.97801193 |
| CUX1 | 11541.5162 | -0.136855788 | 0.2397871 | -0.5707387 | 0.56817675 | 0.85263558 |
| SH2B2 | 223.146393 | 0.630702515 | 0.40314298 | 1.56446358 | 0.11770874 | 0.57390152 |
| PRKRIP1 | 2504.98148 | -0.299809125 | 0.22591166 | -1.3271078 | 0.18447302 | 0.63377801 |
| ORAI2 | 344.296317 | -1.238061464 | 0.73777407 | -1.6781038 | 0.09332684 | 0.53688613 |
| ALKBH4 | 108.230409 | 0.104431176 | 0.34932081 | 0.29895493 | 0.76497444 | 0.93083208 |
| LRWD1 | 48.9396034 | -0.636345528 | 0.57794378 | -1.1010509 | 0.2708745 | 0.69961625 |
| POLR2J | 137.478961 | -0.24226962 | 0.32816711 | -0.7382508 | 0.46036208 | 0.80717423 |
| AC093668.2 | 116.847292 | -0.354424843 | 0.56691409 | -0.6251826 | 0.5318512 | 0.83789294 |
| SPDYE2 | 127.126978 | 0.210517477 | 0.68020812 | 0.3094898 | 0.75694897 | 0.92799282 |
| AC105052.1 | 120.920078 | -0.61368118 | 0.50924649 | -1.2050769 | 0.22817361 | 0.66705629 |
| AC105052.3 | 367.461086 | 0.243028246 | 0.48500093 | 0.50108821 | 0.61630904 | 0.87342308 |
| FAM185A | 780.433236 | 0.426160206 | 0.25095703 | 1.69814015 | 0.08948131 | 0.53119115 |
| FBXL13 | 635.96666 | 0.382827217 | 0.31096051 | 1.231112 | 0.21828098 | 0.66064833 |
| LRRC17 | 81.2796492 | 0.925739154 | 0.52064455 | 1.77806366 | 0.0753934 | 0.50728736 |
| ARMC10 | 461.485968 | -0.033888507 | 0.15150835 | -0.2236742 | 0.82301083 | 0.95131214 |
| NAPEPLD | 1492.60047 | 0.357575989 | 0.46687406 | 0.76589388 | 0.44373945 | 0.7993976 |
| PMPCB | 518.239818 | -0.302988624 | 0.26983291 | -1.122875 | 0.26149059 | 0.69374398 |
| DNAJC2 | 482.786548 | -0.743745468 | 0.40123725 | -1.8536301 | 0.06379209 | 0.48593331 |
| PSMC2 | 344.08299 | 0.044958005 | 0.19148182 | 0.23478994 | 0.81437178 | 0.94881072 |

| | | | | | | |
|---|---|---|---|---|---|---|
| SLC26A5 | 69.4285735 | 0.137140179 | 0.54730305 | 0.25057449 | 0.80214311 | 0.94499146 |
| RELN | 95.5504652 | 0.492106788 | 0.59710191 | 0.82415879 | 0.40984934 | 0.78542936 |
| ORC5 | 541.959571 | 0.057474363 | 0.23066377 | 0.24916944 | 0.80322972 | 0.94543168 |
| LHFPL3 | 872.840308 | 0.483536713 | 0.57250262 | 0.84460175 | 0.39833323 | 0.781221 |
| LINC01004 | 1290.22713 | 0.080287854 | 0.24940396 | 0.32191892 | 0.74751412 | 0.92524616 |
| KMT2E-AS1 | 156.147085 | -0.959544448 | 0.43820406 | -2.1897206 | 0.02854451 | 0.39280225 |
| KMT2E | 2565.14907 | -0.031393311 | 0.25461703 | -0.1232962 | 0.90187255 | 0.97492485 |
| SRPK2 | 5705.19266 | -0.310719619 | 0.29410274 | -1.0565002 | 0.29073973 | 0.71269438 |
| PUS7 | 508.329619 | -1.107954599 | 0.38821234 | -2.8539912 | 0.00431737 | 0.20749797 |
| RINT1 | 246.285909 | -0.532778304 | 0.28762272 | -1.8523513 | 0.0639754 | 0.48593331 |
| EFCAB10 | 86.4319213 | -0.51035295 | 0.4521956 | -1.128611 | 0.25906195 | 0.69241621 |
| ATXN7L1 | 2975.71966 | 0.592813942 | 0.35170052 | 1.68556461 | 0.09187969 | 0.53567268 |
| CDHR3 | 3075.6182 | 0.844898329 | 0.50226793 | 1.6821666 | 0.09253654 | 0.53654752 |
| SYPL1 | 696.370337 | 0.133077668 | 0.29685555 | 0.44829098 | 0.65394321 | 0.89009171 |
| NAMPT | 4444.01825 | 1.190435954 | 0.5079514 | 2.34360208 | 1.91E-02 | 3.41E-01 |
| CCDC71L | 85.1162672 | 0.360804551 | 0.41465432 | 0.87013335 | 0.38422753 | 0.77317162 |
| HBP1 | 861.647395 | -0.360731332 | 0.37472147 | -0.9626652 | 0.33571555 | 0.74225095 |
| COG5 | 2892.9834 | -0.524532134 | 0.32672876 | -1.6054055 | 0.1084046 | 0.55819005 |
| DUS4L | 104.405131 | -0.131878174 | 0.28257871 | -0.4666954 | 0.64071785 | 0.88396933 |
| BCAP29 | 427.593321 | 0.061719247 | 0.24128875 | 0.25578999 | 0.798113 | 0.94320409 |
| CBLL1 | 208.179111 | -0.266590348 | 0.30252986 | -0.8812034 | 0.37820773 | 0.77055888 |
| SLC26A3 | 3679.53769 | -0.848253037 | 0.75590615 | -1.1221671 | 0.26179139 | 0.69408034 |
| DLD | 536.675753 | 0.530420214 | 0.28230931 | 1.87886192 | 0.06026335 | 0.48004006 |
| LAMB1 | 3638.93801 | -0.637564138 | 0.59669593 | -1.0684909 | 0.28529915 | 0.71024256 |
| NRCAM | 2680.40787 | -0.294459601 | 0.61326355 | -0.4801518 | 6.31E-01 | 8.81E-01 |
| PNPLA8 | 1003.02233 | -0.303777904 | 0.26841642 | -1.1317411 | 0.25774329 | 0.69204452 |
| DNAJB9 | 292.317594 | 0.208385688 | 0.47239417 | 0.44112672 | 0.65912126 | 0.89058983 |
| IMMP2L | 11940.4183 | 0.507268316 | 0.35172724 | 1.44222074 | 0.14924011 | 0.60629349 |
| LRRN3 | 54.0480564 | 0.607091684 | 0.8379401 | 0.72450488 | 0.46875584 | 0.81217042 |
| DOCK4 | 2884.41745 | -0.788593897 | 0.36525748 | -2.1590082 | 0.03084953 | 0.40503308 |
| ZNF277 | 1521.9484 | -0.041312804 | 0.2437081 | -0.1695176 | 0.86538956 | 0.96428252 |
| AC004112.1 | 66.6775081 | 0.333496786 | 0.41126758 | 0.81089978 | 0.41742323 | 0.78652162 |
| IFRD1 | 1533.17918 | -0.62559902 | 0.5506441 | -1.1361223 | 0.2559054 | 0.69076316 |
| LSMEM1 | 282.931222 | 0.26200402 | 0.60283561 | 0.43461935 | 0.66383875 | 0.89313476 |
| TMEM168 | 418.01898 | 0.125730602 | 0.28770689 | 0.43700936 | 0.66210456 | 0.89193781 |
| BMT2 | 975.189799 | -0.262759297 | 0.36398302 | -0.7218999 | 0.47035603 | 0.8137284 |
| SMIM30 | 120.570167 | -0.276956126 | 0.41879022 | -0.6613242 | 0.5084044 | 0.82874455 |
| FOXP2 | 4464.00798 | 0.145676846 | 0.44617967 | 0.32649817 | 0.74404747 | 0.92361546 |
| MDFIC | 329.290796 | 0.931186384 | 0.53742791 | 1.73267216 | 0.08315395 | 0.51935659 |
| TFEC | 1055.47267 | 0.389977552 | 0.84027975 | 0.46410443 | 0.64257293 | 0.88441609 |
| TES | 513.501957 | -0.184539988 | 0.43301527 | -0.4261743 | 6.70E-01 | 8.96E-01 |
| AC002066.1 | 74.2605016 | 1.104631656 | 0.56101089 | 1.96900217 | 0.04895284 | 0.45368151 |
| CAV2 | 105.567457 | 0.800030847 | 0.31645274 | 2.52812109 | 0.01146748 | 0.28627263 |
| CAV1 | 48.0319972 | 0.694686779 | 0.61206489 | 1.13498877 | 0.25638002 | 0.69092557 |
| AC006159.2 | 360.863614 | 3.756277551 | 0.82134683 | 4.57331475 | 4.80E-06 | 0.00940042 |
| LINC01510 | 603.952263 | 3.420948792 | 0.76271246 | 4.48524049 | 7.28E-06 | 0.01167379 |
| MET | 3692.52168 | -0.461210183 | 0.38668684 | -1.1927227 | 2.33E-01 | 0.66978724 |
| CAPZA2 | 1458.19998 | -0.282881802 | 0.20663464 | -1.368995 | 0.17100083 | 0.62504223 |
| ST7 | 4111.99111 | -0.503983585 | 0.22644136 | -2.2256693 | 0.02603634 | 0.38392873 |
| ST7-AS2 | 193.08772 | -0.251940101 | 0.7162543 | -0.3517467 | 0.72502822 | 0.91600702 |
| CFTR | 52.7107398 | 1.178008344 | 0.68063649 | 1.7307452 | 0.08349722 | 0.51991532 |
| CTTNBP2 | 773.790397 | 0.747708489 | 0.41823307 | 1.78777943 | 0.0738116 | 0.50581869 |
| LSM8 | 382.866258 | -0.070390558 | 0.21706037 | -0.3242902 | 0.74571831 | 0.92410821 |
| ANKRD7 | 112.696076 | 1.360784763 | 0.62619585 | 2.1730977 | 0.02977297 | 0.39981555 |

| | | | | | | |
|---|---|---|---|---|---|---|
| KCND2 | 118.16257 | 0.68788692 | 1.11612382 | 0.61631775 | 0.53768483 | 0.84040975 |
| TSPAN12 | 1042.56115 | 0.674973715 | 0.68981161 | 0.97848993 | 0.32783207 | 0.73799349 |
| ING3 | 234.99424 | 0.107087969 | 0.21643869 | 0.49477277 | 0.62076053 | 0.87609696 |
| FAM3C | 812.111461 | -0.56085758 | 0.39950705 | -1.4038741 | 0.16035636 | 0.61590405 |
| AASS | 1047.54857 | 0.896576335 | 0.66791349 | 1.34235399 | 0.17948124 | 0.63064584 |
| CADPS2 | 2217.45356 | 0.751660433 | 0.42892408 | 1.75243237 | 0.07969949 | 0.51240193 |
| AC004594.1 | 100.951214 | 1.08057141 | 0.72260994 | 1.49537302 | 0.13481712 | 0.59323859 |
| AC073311.1 | 51.1603469 | 3.334765786 | 0.78848616 | 4.22932696 | 2.34E-05 | 0.01647703 |
| SLC13A1 | 5815.62769 | 2.507203425 | 0.82677087 | 3.03252511 | 0.00242517 | 0.15981632 |
| IQUB | 373.12703 | 1.044261086 | 0.49423539 | 2.11288207 | 0.03461086 | 0.41686761 |
| NDUFA5 | 571.712069 | 0.113385024 | 0.36312141 | 0.312251 | 0.75484977 | 0.92799282 |
| ASB15 | 74.0847587 | 1.029830815 | 0.99471612 | 1.03530122 | 3.01E-01 | 0.71879123 |
| WASL | 1501.64197 | 0.124872529 | 0.21677656 | 0.57604259 | 0.56458639 | 0.85163263 |
| AC006333.2 | 37.6355162 | -0.340935246 | 0.48836235 | -0.6981194 | 0.4851025 | 0.81839266 |
| POT1 | 476.571725 | 0.122500631 | 0.32230904 | 0.38007197 | 0.70389199 | 0.90919468 |
| POT1-AS1 | 133.860666 | 0.055276928 | 0.40825363 | 0.1353985 | 0.89229681 | 0.97237481 |
| AC073486.1 | 430.906438 | -0.439985623 | 0.84648019 | -0.5197825 | 0.60321515 | 0.8678002 |
| GRM8 | 188.737874 | -0.108190762 | 0.94827547 | -0.1140921 | 0.90916477 | 0.97680747 |
| ZNF800 | 796.246862 | 0.179937562 | 0.2718536 | 0.66189141 | 0.50804082 | 0.82874455 |
| GCC1 | 50.9148789 | 0.114128682 | 0.40670068 | 0.28062083 | 0.77900124 | 0.93646866 |
| ARF5 | 132.227412 | -0.069405936 | 0.55865363 | -0.1242379 | 0.90112694 | 0.97482505 |
| SND1 | 3562.43786 | -0.480483746 | 0.36916772 | -1.3015324 | 0.19307629 | 0.64532959 |
| LRRC4 | 46.0303825 | 0.08017077 | 0.49339177 | 0.16248907 | 0.87092073 | 0.9657206 |
| RBM28 | 420.707695 | -0.64539072 | 0.23877215 | -2.7029565 | 0.00687257 | 0.23083391 |
| IMPDH1 | 45.9959833 | 0.18282838 | 0.49780584 | 0.36726845 | 0.71341879 | 0.91203426 |
| HILPDA | 844.249631 | -1.443153461 | 0.8851691 | -1.6303704 | 1.03E-01 | 5.50E-01 |
| METTL2B | 379.6742 | -0.14554438 | 0.33514627 | -0.4342712 | 0.6640915 | 0.89313476 |
| AC090114.2 | 57.3439425 | -0.85709266 | 0.52620488 | -1.6288193 | 0.10335128 | 0.55022042 |
| FAM71F2 | 69.674642 | -0.498476119 | 0.57334196 | -0.869422 | 0.38461635 | 0.77317162 |
| CALU | 686.851206 | -0.433465108 | 0.41480816 | -1.0449773 | 0.29603346 | 0.71729374 |
| KCP | 585.905955 | 0.134061447 | 0.48273524 | 0.27771216 | 0.78123332 | 0.93748328 |
| ATP6V1F | 552.356732 | -0.56842064 | 0.41554336 | -1.3678973 | 1.71E-01 | 0.62520756 |
| IRF5 | 146.166998 | 0.726288371 | 0.34046036 | 2.13325383 | 0.03290391 | 0.41180153 |
| TNPO3 | 1676.39366 | -0.188268376 | 0.25760818 | -0.7308323 | 0.4648816 | 0.81002663 |
| TSPAN33 | 488.29114 | 0.869680131 | 0.44984381 | 1.93329355 | 0.05320005 | 0.46624874 |
| SMO | 261.553779 | 0.701972095 | 0.38967662 | 1.80142214 | 7.16E-02 | 0.5001766 |
| AHCYL2 | 3466.4135 | -0.241876221 | 0.28649469 | -0.8442608 | 0.39852371 | 0.78122921 |
| STRIP2 | 424.310617 | 0.289789563 | 0.76846127 | 0.37710367 | 0.70609657 | 0.90988678 |
| AC078846.1 | 130.103419 | -0.427093417 | 0.28960124 | -1.4747638 | 0.14027607 | 0.59500405 |
| NRF1 | 1681.40464 | -0.034463865 | 0.21239354 | -0.1622642 | 0.87109781 | 0.9657206 |
| UBE2H | 5118.15321 | -0.910217017 | 0.37759914 | -2.4105378 | 0.01592902 | 0.31735328 |
| AC073320.1 | 59.9810906 | -0.176532586 | 0.4726635 | -0.3734847 | 0.70878773 | 0.91082291 |
| ZC3HC1 | 680.08998 | -0.422037168 | 0.26158892 | -1.6133602 | 0.10666627 | 0.55486698 |
| KLHDC10 | 2120.20355 | -0.15325353 | 0.19438113 | -0.7884177 | 0.4304524 | 0.79055252 |
| TMEM209 | 198.349966 | -0.402231241 | 0.2714756 | -1.4816479 | 0.13843402 | 0.59474076 |
| AC087071.2 | 41.5121099 | 0.220412517 | 0.584385 | 0.37717005 | 0.70604724 | 0.90988678 |
| CEP41 | 55.409526 | 0.624742883 | 0.49079953 | 1.27290847 | 0.20305052 | 0.64881821 |
| MEST | 92.8852807 | 0.375171821 | 0.52276398 | 0.71766961 | 0.47296103 | 0.81471211 |
| COPG2 | 244.358506 | 0.043051363 | 0.32739804 | 0.13149548 | 0.89538336 | 0.97309833 |
| AC016831.7 | 11482.6546 | -0.634624056 | 0.34365148 | -1.8467083 | 0.06478943 | 0.48782903 |
| LINC00513 | 2101.55629 | -0.753953134 | 0.59251434 | -1.2724639 | 0.20320833 | 0.64881821 |
| AC058791.1 | 205.005773 | -0.379750018 | 0.64596446 | -0.5878807 | 0.55661239 | 0.84861484 |
| MKLN1 | 7210.6206 | -0.26729761 | 0.33977082 | -0.7866997 | 0.43145766 | 0.79135216 |
| PODXL | 115.702366 | -0.256643361 | 0.486084 | -0.5279815 | 0.59751218 | 0.86614567 |

| | | | | | | |
|---|---|---|---|---|---|---|
| PLXNA4 | 77.6855821 | 0.495111472 | 0.5016719 | 0.98692288 | 0.32368045 | 0.73552518 |
| CHCHD3 | 3787.64982 | 0.208968971 | 0.22807145 | 0.91624344 | 0.35953923 | 0.75958757 |
| EXOC4 | 8126.36774 | 0.207581145 | 0.29952788 | 0.69302779 | 0.48829211 | 0.81931938 |
| LRGUK | 436.210545 | 0.336635713 | 0.40491598 | 0.83137177 | 0.40576364 | 0.78429302 |
| SLC35B4 | 645.279796 | 0.529676602 | 0.37409091 | 1.41590342 | 0.15680378 | 0.61255952 |
| AKR1B1 | 709.82031 | -1.191797735 | 0.76538506 | -1.5571218 | 1.19E-01 | 0.57652003 |
| AKR1B10 | 669.531608 | 1.833030905 | 1.78388998 | 1.02754706 | 3.04E-01 | 0.72082426 |
| BPGM | 194.511254 | 0.47843569 | 0.41425363 | 1.15493422 | 0.24811737 | 0.6848323 |
| CALD1 | 2569.07895 | -0.18550595 | 0.27554328 | -0.6732371 | 0.50079648 | 0.82564559 |
| AGBL3 | 918.571848 | 0.423518585 | 0.5315077 | 0.79682493 | 0.42555271 | 0.78894887 |
| CYREN | 428.283131 | -0.301825635 | 0.29521574 | -1.0223901 | 0.30659633 | 0.72190689 |
| TMEM140 | 772.14507 | -0.923861329 | 0.45220469 | -2.0430158 | 0.04105088 | 0.43143739 |
| AC083862.2 | 73.1102495 | -1.396871672 | 0.66210278 | -2.1097505 | 0.03487985 | 0.41718861 |
| WDR91 | 766.242603 | 0.971963031 | 0.54924004 | 1.76965072 | 0.07678534 | 0.50968364 |
| AC009542.1 | 102.187234 | 0.858608056 | 0.76961075 | 1.11563937 | 0.26457653 | 0.69567438 |
| STRA8 | 97.032126 | 1.509686775 | 0.80065536 | 1.88556381 | 0.05935376 | 0.47855094 |
| CNOT4 | 2076.46275 | -0.15099448 | 0.17242374 | -0.8757175 | 0.38118365 | 0.77210521 |
| NUP205 | 400.53695 | -0.220057683 | 0.20280331 | -1.0850793 | 0.27788651 | 0.7036882 |
| STMP1 | 462.83879 | 0.385180836 | 0.24008765 | 1.60433426 | 0.10864039 | 0.55858026 |
| SLC13A4 | 71.8605986 | 1.859923181 | 0.45267043 | 4.10877994 | 3.98E-05 | 0.02019269 |
| MTPN | 549.043425 | -0.127335008 | 0.29447046 | -0.4324203 | 0.66543596 | 0.89369715 |
| AC009264.1 | 39.4169815 | -0.145449749 | 0.59357076 | -0.245042 | 0.80642393 | 0.94585363 |
| PTN | 75.2682888 | 0.876032642 | 0.89941424 | 0.97400353 | 0.33005479 | 0.73849818 |
| DGKI | 201.092884 | -0.725476711 | 0.47316443 | -1.5332444 | 0.12521565 | 0.58564339 |
| CREB3L2 | 939.352728 | -0.092219768 | 0.41583138 | -0.221772 | 0.82449136 | 0.95180295 |
| TRIM24 | 1039.07511 | -0.56086358 | 0.34542185 | -1.6237062 | 0.10443853 | 0.55066578 |
| ATP6V0A4 | 68.5595635 | -0.413042541 | 0.50782014 | -0.8133639 | 0.41600948 | 0.78632759 |
| TMEM213 | 81.9098794 | -0.269898248 | 0.54031707 | -0.4995183 | 0.61741432 | 0.87435665 |
| KIAA1549 | 2257.89747 | -0.014921941 | 0.38277346 | -0.0389837 | 9.69E-01 | 9.92E-01 |
| ZC3HAV1L | 58.5868142 | -0.674619579 | 0.41205496 | -1.6372078 | 1.02E-01 | 0.54713316 |
| ZC3HAV1 | 1237.98201 | -0.024811041 | 0.5039585 | -0.0492323 | 0.96073416 | 0.98961322 |
| TTC26 | 343.637346 | 0.38826555 | 0.36643335 | 1.05958028 | 0.28933559 | 0.71218341 |
| UBN2 | 1393.24408 | 0.138288689 | 0.28701918 | 0.48180992 | 0.62994098 | 0.88083664 |
| LUC7L2 | 259.855349 | -0.243632865 | 0.31112826 | -0.7830625 | 0.43359042 | 0.79391182 |
| HIPK2 | 5610.26912 | -0.266667491 | 0.25137658 | -1.0608287 | 0.28876776 | 0.71218341 |
| TBXAS1 | 212.767548 | 0.007890737 | 0.37913337 | 0.02081256 | 0.98339518 | 0.99523767 |
| PARP12 | 326.02547 | -1.327912101 | 0.65401961 | -2.0303857 | 0.04231734 | 0.43742111 |
| KDM7A | 744.578404 | -0.076604758 | 0.23713186 | -0.3230471 | 0.74665958 | 0.92448528 |
| SLC37A3 | 520.068292 | -0.526586057 | 0.27334212 | -1.9264724 | 0.05404542 | 0.46711025 |
| MKRN1 | 457.843539 | -0.062612827 | 0.22529184 | -0.2779187 | 0.78107473 | 0.93748328 |
| DENND2A | 103.982298 | -0.11313684 | 0.63326624 | -0.178656 | 0.85820779 | 0.9626966 |
| NDUFB2 | 674.722744 | -0.150456263 | 0.36036778 | -0.4175075 | 0.67630722 | 0.89835672 |
| BRAF | 3469.18403 | 0.070647816 | 0.20560328 | 0.3436123 | 0.73113787 | 0.9190831 |
| MRPS33 | 175.729288 | -0.053407042 | 0.25043415 | -0.2132578 | 0.83112587 | 0.95325673 |
| TMEM178B | 2769.63839 | -0.11255232 | 0.49800224 | -0.2260077 | 0.82119546 | 0.95084695 |
| AGK | 427.964041 | 0.264081871 | 0.2190339 | 1.20566663 | 0.22794606 | 0.66705629 |
| AC004918.1 | 249.981127 | 0.909774637 | 0.44184704 | 2.05902621 | 0.03949173 | 0.4275775 |
| KIAA1147 | 642.801135 | 0.901995163 | 0.44198182 | 2.04079699 | 0.04127101 | 0.43221049 |
| WEE2-AS1 | 68.5429388 | -0.101578323 | 0.46026432 | -0.2206956 | 0.82532944 | 0.95192757 |
| SSBP1 | 691.175285 | -0.231170486 | 0.25172729 | -0.918337 | 0.35844246 | 0.75936492 |
| MGAM | 799.094055 | 0.715868705 | 0.73804697 | 0.96995006 | 0.33207139 | 0.739513 |
| MGAM2 | 73.2019611 | 1.271512199 | 0.89165785 | 1.42600909 | 0.15386572 | 0.6105654 |
| GSTK1 | 686.877541 | 0.673627051 | 0.47109973 | 1.42990329 | 0.15274478 | 0.61001662 |
| AC073342.1 | 112.444194 | 0.991325535 | 0.6475485 | 1.53089003 | 0.12579657 | 0.58564339 |

| | | | | | | |
|---|---|---|---|---|---|---|
| TMEM139 | 249.18546 | -0.178893397 | 0.7061383 | -0.2533405 | 0.80000514 | 0.94403774 |
| CASP2 | 131.061915 | -0.404717266 | 0.24767909 | -1.6340389 | 0.10225068 | 0.54898162 |
| ZYX | 172.60435 | -0.369039189 | 0.49860663 | -0.740141 | 0.45921447 | 0.80669065 |
| EPHA1 | 52.952537 | 0.500848843 | 0.54512452 | 0.91877878 | 0.35821129 | 0.75936492 |
| EPHA1-AS1 | 42.3787894 | -0.596918365 | 0.5075328 | -1.1761178 | 2.40E-01 | 6.78E-01 |
| TCAF2 | 362.045351 | -0.418731913 | 0.72904642 | -0.5743556 | 5.66E-01 | 8.52E-01 |
| TCAF1 | 540.068222 | 0.917650547 | 0.3164903 | 2.89945865 | 0.00373808 | 0.1981659 |
| ARHGEF35 | 46.8466974 | -0.07410887 | 0.56756548 | -0.1305733 | 0.8961129 | 0.97309833 |
| AC004889.1 | 356.347111 | 0.268967407 | 0.40593868 | 0.66258138 | 0.5075987 | 0.82858823 |
| AC074386.1 | 740.731709 | -0.151168869 | 0.36262891 | -0.4168693 | 0.67677399 | 0.89843878 |
| ARHGEF5 | 171.900294 | -0.131813218 | 0.39381469 | -0.3347087 | 0.73784481 | 0.92059676 |
| TPK1 | 1124.42962 | -0.277043724 | 0.48600342 | -0.5700448 | 0.56864731 | 0.85269103 |
| CNTNAP2 | 163.177658 | 0.00273755 | 0.33423771 | 0.00819043 | 0.99346506 | 0.9976934 |
| CUL1 | 1320.67294 | -0.220452958 | 0.23607228 | -0.9338367 | 0.35038815 | 0.75467792 |
| EZH2 | 192.572959 | -0.099711532 | 0.42831402 | -0.2328001 | 0.81591666 | 0.94906482 |
| PDIA4 | 390.509488 | -0.416190993 | 0.56755935 | -0.7332995 | 0.46337578 | 0.8082072 |
| ZNF786 | 195.879599 | -0.427443192 | 0.51478874 | -0.8303274 | 0.4063537 | 0.78453757 |
| ZNF425 | 117.584747 | -0.079915146 | 0.40381677 | -0.1978995 | 0.84312368 | 0.95716002 |
| ZNF398 | 488.732392 | -0.220211396 | 0.36993375 | -0.5952725 | 0.55166133 | 0.84695585 |
| ZNF282 | 247.515793 | -0.176926777 | 0.35801365 | -0.49419 | 0.62117198 | 0.87641939 |
| ZNF212 | 201.629677 | -0.884721613 | 0.34768521 | -2.5446052 | 0.01094013 | 0.27821228 |
| ZNF783 | 129.461128 | -0.179759316 | 0.30572491 | -0.5879773 | 0.5565475 | 0.84861484 |
| ZNF777 | 151.267511 | -0.346779375 | 0.40108733 | -0.8645982 | 0.38725938 | 0.77344664 |
| ZNF746 | 386.790912 | -0.502840381 | 0.42124807 | -1.1936918 | 0.23259854 | 0.66978724 |
| KRBA1 | 60.362926 | 0.567135531 | 0.39952489 | 1.4195249 | 0.15574604 | 0.61239559 |
| ZNF467 | 59.967202 | 0.527477157 | 0.73211515 | 0.72048387 | 0.47122713 | 0.81387913 |
| ZNF862 | 377.429933 | 0.61072441 | 0.34881644 | 1.75084758 | 0.07997217 | 0.51264427 |
| ATP6V0E2 | 93.4671305 | 1.320940912 | 0.52664536 | 2.50821715 | 0.01213421 | 0.29489996 |
| AC092666.1 | 114.895414 | 1.082010164 | 0.582672 | 1.85697984 | 0.06331402 | 0.48572144 |
| ACTR3C | 3107.63207 | -0.125238708 | 0.38033102 | -0.3292887 | 0.74193748 | 0.9225925 |
| LRRC61 | 237.68908 | -0.310095388 | 0.61116945 | -0.5073804 | 0.61188795 | 0.87199356 |
| RARRES2 | 412.5641 | -0.479835068 | 0.77665821 | -0.6178201 | 0.53669392 | 0.83987482 |
| AC073111.5 | 40.7907 | -0.706866843 | 0.70084778 | -1.0085883 | 0.31317214 | 0.72717675 |
| TMEM176B | 1775.27924 | 0.457577261 | 0.31963902 | 1.43154383 | 0.15227442 | 0.61001662 |
| TMEM176A | 876.27799 | 0.13888335 | 0.46340233 | 0.29970361 | 0.76440325 | 0.93083208 |
| AOC1 | 306.743057 | -0.020854302 | 0.76304541 | -0.0273304 | 0.97819625 | 0.99482978 |
| ABCB8 | 235.678689 | -0.178086298 | 0.34682517 | -0.5134757 | 0.6076186 | 0.86985416 |
| SLC4A2 | 378.045301 | -0.675706319 | 0.46987961 | -1.4380414 | 0.15042232 | 0.60799621 |
| FASTK | 237.8074 | -0.580119688 | 0.41001416 | -1.4148772 | 0.1571045 | 0.61281486 |
| TMUB1 | 251.657298 | -0.261220031 | 0.42776095 | -0.6106682 | 0.54141923 | 0.84221416 |
| AGAP3 | 450.747671 | -0.084946908 | 0.3704269 | -0.2293217 | 0.81861892 | 0.94990455 |
| CHPF2 | 117.665787 | -0.72382082 | 0.46953818 | -1.541559 | 0.12318079 | 0.5825794 |
| SMARCD3 | 72.7480754 | 0.937503414 | 0.48066278 | 1.95043896 | 0.05112382 | 0.46102252 |
| NUB1 | 438.136162 | -0.15267648 | 0.44989533 | -0.33936 | 0.73433854 | 0.91975299 |
| RHEB | 1137.66957 | -0.555044166 | 0.30008135 | -1.8496456 | 0.06436464 | 0.48593331 |
| PRKAG2 | 2983.58495 | -0.453721036 | 0.56565847 | -0.8021113 | 0.42248859 | 0.78894887 |
| GALNT11 | 2399.87956 | 0.221350882 | 0.29975741 | 0.7384334 | 0.46025112 | 0.80714715 |
| KMT2C | 5430.98428 | -0.310374597 | 0.25371795 | -1.2233056 | 0.22121428 | 0.66263062 |
| ACTR3B | 689.596031 | -0.046674211 | 0.31424563 | -0.1485278 | 0.88192625 | 0.96796225 |
| DPP6 | 3317.8435 | 1.111510176 | 0.46033098 | 2.41458912 | 0.01575297 | 0.31566878 |
| PAXIP1-AS2 | 163.398695 | 0.271325809 | 0.39762735 | 0.68236204 | 0.49501005 | 0.82216554 |
| PAXIP1 | 338.516884 | -0.209600063 | 0.25708251 | -0.8153027 | 0.41489907 | 0.78632759 |
| PAXIP1-AS1 | 122.639779 | 0.201783137 | 0.58403251 | 0.34549984 | 0.72971863 | 0.91834786 |
| INSIG1 | 263.982712 | 0.80073988 | 0.63913733 | 1.2528448 | 0.2102622 | 0.65441616 |

| | | | | | | |
|---|---|---|---|---|---|---|
| AC009403.1 | 517.405239 | 0.9056461 | 0.39538235 | 2.29055775 | 0.02198901 | 0.36066559 |
| RBM33 | 3556.12461 | -0.120836286 | 0.19679267 | -0.6140284 | 0.53919656 | 0.84091105 |
| SHH | 98.2637853 | -0.529962874 | 0.61213062 | -0.8657676 | 0.3866176 | 0.77334938 |
| AC021218.1 | 102.697213 | -0.858806982 | 0.84886794 | -1.0117086 | 0.31167741 | 0.72600794 |
| LINC01006 | 673.676821 | -0.269988142 | 0.32620428 | -0.8276658 | 0.40785976 | 0.78481094 |
| RNF32 | 60.3000545 | 0.427946864 | 0.47570694 | 0.8996019 | 0.36833215 | 0.76615653 |
| LMBR1 | 2263.6565 | 0.106647023 | 0.18148444 | 0.58763729 | 0.55677577 | 0.84861484 |
| NOM1 | 302.193365 | -0.81865536 | 0.46436512 | -1.7629562 | 0.07790788 | 0.51192014 |
| UBE3C | 2483.51712 | -0.438217577 | 0.29005683 | -1.510799 | 0.13083967 | 0.58858527 |
| DNAJB6 | 1092.41521 | -0.672812218 | 0.37744892 | -1.7825252 | 0.07466362 | 0.5071704 |
| PTPRN2 | 51.6639391 | 0.667664659 | 0.58146656 | 1.14824257 | 0.25086844 | 0.68648949 |
| NCAPG2 | 74.1796148 | -0.537124311 | 0.38237995 | -1.4046874 | 0.16011425 | 0.61590405 |
| ESYT2 | 1957.99137 | -0.078917362 | 0.28418886 | -0.2776934 | 0.78124775 | 0.93748328 |
| WDR60 | 2051.7658 | 0.382622738 | 0.20160592 | 1.89787452 | 0.05771261 | 0.47684824 |
| PLCXD1 | 37.9240356 | -0.746265071 | 0.4197528 | -1.777868 | 0.07542554 | 0.50728736 |
| GTPBP6 | 401.672518 | -0.251183536 | 0.37747784 | -0.6654259 | 0.50577812 | 0.82738999 |
| LINC00685 | 212.547079 | 0.237474756 | 0.54060188 | 0.43927845 | 0.66045979 | 0.89124924 |
| PPP2R3B | 124.911573 | -0.416174984 | 0.40511326 | -1.0273053 | 0.30427675 | 0.72082426 |
| IL3RA | 184.063705 | -0.285662839 | 0.60561153 | -0.4716932 | 0.63714579 | 0.88267497 |
| SLC25A6 | 955.609976 | 0.582980842 | 0.43507793 | 1.3399458 | 0.18026297 | 0.63124625 |
| LINC00106 | 290.531182 | -0.33846113 | 0.36307621 | -0.9322041 | 0.35123106 | 0.75518811 |
| ASMTL-AS1 | 191.765069 | -0.246863052 | 0.32773108 | -0.7532488 | 4.51E-01 | 8.04E-01 |
| ASMTL | 241.62204 | -0.609603156 | 0.35034951 | -1.7399858 | 0.08186152 | 0.51593841 |
| AKAP17A | 504.648676 | 0.077804411 | 0.25257581 | 0.3080438 | 0.758049 | 0.92839329 |
| DHRSX | 3130.68762 | -0.491010034 | 0.2537267 | -1.9351926 | 0.05296666 | 0.46599104 |
| CD99 | 520.360671 | 0.00128019 | 0.38946726 | 0.00328703 | 0.99737734 | 0.99963814 |
| GYG2 | 72.415746 | 1.110322509 | 0.7408911 | 1.4986312 | 0.13396933 | 0.59240843 |
| ARSD | 384.235161 | 0.624166337 | 0.5190528 | 1.20251029 | 0.22916588 | 0.66705629 |
| ARSE | 346.201379 | 0.29668089 | 0.30219754 | 0.98174488 | 0.32622555 | 0.7373175 |
| ARSF | 534.965689 | 0.663387202 | 0.6016006 | 1.1027037 | 2.70E-01 | 7.00E-01 |
| PRKX | 676.533919 | 0.476598109 | 0.42529861 | 1.12061996 | 0.26244967 | 0.69467275 |
| BX890604.1 | 172.984345 | -0.480931625 | 0.44003486 | -1.0929398 | 0.27442022 | 0.7014484 |
| AC097347.1 | 41.9339669 | 0.185124973 | 0.67380341 | 0.27474627 | 0.78351118 | 0.93852904 |
| PUDP | 355.611355 | -0.255665528 | 0.39953466 | -0.6399083 | 0.52223224 | 0.83468253 |
| STS | 109.11428 | -1.211746351 | 0.42624487 | -2.8428409 | 0.00447134 | 0.20749797 |
| PNPLA4 | 102.199643 | 0.16192539 | 0.47308692 | 0.34227408 | 0.73214464 | 0.91942342 |
| AC078960.1 | 37.8676199 | -0.269546794 | 0.646444 | -0.4169685 | 0.67670144 | 0.89843878 |
| ANOS1 | 37.3374419 | 0.141186656 | 0.44421036 | 0.31783738 | 0.7506083 | 0.92590588 |
| TBL1X | 2456.49684 | -0.230608337 | 0.27688026 | -0.8328811 | 0.40491178 | 0.78422899 |
| GPR143 | 159.865377 | 0.815911792 | 0.41635288 | 1.95966412 | 0.05003506 | 0.45813666 |
| SHROOM2 | 574.37073 | 0.293323636 | 0.4442145 | 0.66031981 | 0.50904862 | 0.82878364 |
| WWC3 | 559.750899 | 0.126966638 | 0.22453615 | 0.56546191 | 0.57175963 | 0.85399468 |
| CLCN4 | 182.699921 | 0.299359371 | 0.39742599 | 0.75324558 | 0.45130235 | 0.80384625 |
| MID1 | 550.292155 | 0.558836913 | 0.62024787 | 0.90098965 | 0.36759382 | 0.76597879 |
| AC073529.1 | 167.417948 | 0.338137917 | 0.5929202 | 0.57029246 | 0.56847935 | 0.85263558 |
| HCCS | 56.437189 | -0.190276045 | 0.35856792 | -0.5306555 | 0.59565752 | 0.86526372 |
| ARHGAP6 | 7785.66486 | 0.475721637 | 0.51581537 | 0.92227115 | 0.35638716 | 0.75841231 |
| MSL3 | 92.775576 | -0.513579188 | 0.32931699 | -1.5595284 | 0.11887138 | 0.57609025 |
| PRPS2 | 57.5006885 | 0.338139273 | 0.49131159 | 0.68823793 | 0.49130296 | 0.82035445 |
| TMSB4X | 970.757877 | -0.514168873 | 0.54106981 | -0.9502819 | 0.34196901 | 0.74914004 |
| TCEANC | 44.2372036 | 0.518053516 | 0.54214471 | 0.95556316 | 0.33929298 | 0.74644675 |
| RAB9A | 583.47573 | -0.356680118 | 0.34099819 | -1.0459883 | 2.96E-01 | 7.17E-01 |
| TRAPPC2 | 230.913703 | -0.291607793 | 0.34653805 | -0.8414885 | 0.40007431 | 0.7818841 |
| OFD1 | 961.675548 | -0.368783073 | 0.29012278 | -1.2711276 | 0.20368326 | 0.64912495 |

| | | | | | | |
|---|---|---|---|---|---|---|
| GPM6B | 78.5701286 | 0.054245595 | 0.45777134 | 0.11849933 | 0.90567203 | 0.97633014 |
| GEMIN8 | 306.558712 | -0.207095933 | 0.28145942 | -0.7357932 | 0.46185654 | 0.8082072 |
| MOSPD2 | 217.952006 | 0.222427883 | 0.18498221 | 1.20242853 | 0.22919754 | 0.66705629 |
| ASB9 | 115.52637 | 0.365592216 | 0.60189131 | 0.6074057 | 5.44E-01 | 8.44E-01 |
| PIGA | 113.25882 | -0.451386837 | 0.3800982 | -1.1875532 | 0.23500948 | 0.67119484 |
| VEGFD | 110.799048 | 1.488313004 | 0.65167257 | 2.28383559 | 0.0223812 | 0.36273984 |
| PIR | 920.545934 | 0.797507372 | 0.55387344 | 1.43987294 | 0.14990335 | 0.60758225 |
| BMX | 280.954808 | 1.060724497 | 0.73263003 | 1.44783105 | 0.14766431 | 0.60581702 |
| ACE2 | 769.007071 | 0.708805021 | 0.57523288 | 1.23220533 | 0.21787239 | 0.66064833 |
| CLTRN | 2181.73894 | -1.304766835 | 0.57966187 | -2.2509102 | 0.02439122 | 0.37480999 |
| CA5B | 774.799683 | -0.704113215 | 0.29875436 | -2.35683 | 0.01843169 | 0.33866368 |
| ZRSR2 | 351.974027 | -0.543165219 | 0.27929937 | -1.944742 | 0.05180604 | 0.46381799 |
| AP1S2 | 149.786233 | -0.628801443 | 0.29181145 | -2.154821 | 3.12E-02 | 0.40697835 |
| CTPS2 | 372.719321 | 0.131344291 | 0.29092149 | 0.45147676 | 0.65164597 | 0.88935488 |
| SYAP1 | 491.591876 | -0.269973072 | 0.20786798 | -1.2987718 | 0.19402225 | 0.64548758 |
| TXLNG | 415.002966 | -0.277325369 | 0.23776769 | -1.1663711 | 0.24346443 | 0.68164834 |
| RBBP7 | 316.349429 | -0.00037799 | 0.22480558 | -0.0016814 | 0.99865843 | 0.99977652 |
| REPS2 | 2044.43907 | 0.491865423 | 0.77098751 | 0.63796808 | 0.52349446 | 0.83533627 |
| NHS | 15128.0254 | 0.232255966 | 0.45125178 | 0.51469264 | 0.60676782 | 0.86985416 |
| NHS-AS1 | 78.6859977 | 0.509798527 | 0.84031725 | 0.60667388 | 0.54406736 | 0.84408956 |
| SCML1 | 421.525253 | 0.221518238 | 0.30419986 | 0.72819967 | 0.46649137 | 0.81098649 |
| RAI2 | 88.5874367 | 1.219077568 | 0.51940017 | 2.34708736 | 0.01892082 | 0.34084276 |
| SCML2 | 109.654273 | 0.251935245 | 0.33678238 | 0.7480654 | 0.45442071 | 0.80465616 |
| CDKL5 | 1026.26444 | -0.271882273 | 0.32884138 | -0.8267885 | 0.40835695 | 0.78481094 |
| PHKA2 | 769.665397 | 0.05154739 | 0.45058735 | 0.11440044 | 0.90892037 | 0.97680747 |
| ADGRG2 | 76.6227529 | 0.826102798 | 0.67093022 | 1.23127976 | 0.21821825 | 0.66064833 |
| PDHA1 | 553.763266 | 0.462142824 | 0.36395062 | 1.26979541 | 0.20415752 | 0.64972999 |
| MAP3K15 | 140.582151 | -0.497090687 | 0.56356095 | -0.8820531 | 0.37774809 | 0.77055888 |
| SH3KBP1 | 1161.42647 | -1.157857301 | 0.28017629 | -4.1326027 | 3.59E-05 | 0.01966562 |
| BCLAF3 | 628.956787 | -0.251235432 | 0.2867657 | -0.8761 | 0.38097566 | 0.77210521 |
| MAP7D2 | 336.788984 | -0.873772255 | 0.49322836 | -1.7715369 | 0.07647146 | 0.50833864 |
| EIF1AX | 763.764941 | -0.374703452 | 0.33705295 | -1.111705 | 0.26626501 | 0.69797523 |
| RPS6KA3 | 1804.32464 | -0.399262957 | 0.34552937 | -1.1555109 | 0.24788127 | 0.68482118 |
| MBTPS2 | 370.557593 | 0.087367805 | 0.30554634 | 0.28593962 | 0.77492437 | 0.93477852 |
| SMS | 1073.34907 | -0.838485767 | 0.45827264 | -1.8296658 | 0.06729993 | 0.48963361 |
| PHEX | 89.7513925 | -3.077306053 | 0.8884668 | -3.463614 | 0.00053297 | 0.08106654 |
| PTCHD1-AS | 137.751948 | 0.768334529 | 0.61995775 | 1.2393337 | 0.21522194 | 0.65925411 |
| PRDX4 | 140.853646 | -0.43968113 | 0.3790772 | -1.1598723 | 2.46E-01 | 6.83E-01 |
| ACOT9 | 521.039085 | -0.984376898 | 0.46128669 | -2.1339807 | 0.03284436 | 0.41180153 |
| SAT1 | 3363.53962 | -0.826472863 | 0.48162758 | -1.7159999 | 0.08616207 | 0.52402549 |
| APOO | 2080.79485 | 0.157298836 | 0.4318918 | 0.3642089 | 0.71570202 | 0.91298152 |
| KLHL15 | 104.315955 | -0.288630449 | 0.30243986 | -0.95434 | 0.33991157 | 0.74654617 |
| EIF2S3 | 604.818085 | -0.492412192 | 0.34792379 | -1.4152875 | 0.15698421 | 0.61255952 |
| ZFX | 887.41171 | -0.834728663 | 0.26538254 | -3.1453789 | 0.00165872 | 0.13207839 |
| PDK3 | 635.609895 | 0.411344897 | 0.55905622 | 0.73578449 | 0.46186186 | 0.8082072 |
| POLA1 | 554.282486 | -0.397999089 | 0.30955828 | -1.2857 | 0.19854777 | 0.64597062 |
| IL1RAPL1 | 106.408848 | 0.467408099 | 0.62021384 | 0.75362411 | 0.45107496 | 0.80384625 |
| GK | 2571.02636 | -0.530253368 | 0.43649846 | -1.2147886 | 0.22444672 | 0.66551422 |
| GK-AS1 | 146.433668 | 0.043140438 | 0.65559652 | 0.06580334 | 0.9475344 | 0.98646931 |
| TAB3 | 742.738684 | -0.124749551 | 0.2592226 | -0.4812449 | 0.63034247 | 0.88101399 |
| DMD | 257.89896 | -0.012248791 | 0.27058654 | -0.0452676 | 0.96389405 | 0.99012655 |
| TMEM47 | 178.293335 | 0.672992952 | 0.49633141 | 1.35593463 | 0.17511997 | 0.62699479 |
| AF241726.2 | 448.559665 | 0.208899579 | 0.38436887 | 0.54348724 | 0.58679437 | 0.86112855 |
| DYNLT3 | 168.890126 | -0.362492631 | 0.26823868 | -1.3513809 | 0.17657345 | 0.62709281 |

| | | | | | | |
|---|---|---|---|---|---|---|
| RPGR | 293.016993 | -0.167257376 | 0.3738652 | -0.4473735 | 0.65460542 | 0.89032312 |
| MID1IP1 | 141.373065 | -0.447230862 | 0.54095289 | -0.8267464 | 0.40838081 | 0.78481094 |
| BCOR | 790.166143 | -0.140494788 | 0.23619219 | -0.5948325 | 0.55195544 | 0.84702791 |
| ATP6AP2 | 422.631598 | 0.029142616 | 0.34550982 | 0.08434671 | 0.93278078 | 0.98311128 |
| CXorf38 | 369.552794 | 0.173329947 | 0.20614267 | 0.84082517 | 4.00E-01 | 0.78232083 |
| MED14 | 788.480075 | -0.002914895 | 0.2234772 | -0.0130434 | 0.98959319 | 0.99730401 |
| USP9X | 2134.61094 | -0.146628147 | 0.27090236 | -0.5412583 | 0.58832956 | 0.86140264 |
| DDX3X | 1196.85001 | -0.364032428 | 0.32014917 | -1.1370713 | 0.25550849 | 0.6902355 |
| CASK | 1875.5267 | 0.18485014 | 0.26956141 | 0.68574408 | 0.49287451 | 0.82100434 |
| Z93403.1 | 1546.23844 | 0.292025824 | 0.72440699 | 0.40312397 | 0.68685703 | 0.90222801 |
| MAOA | 1666.25696 | 0.31780213 | 0.68563889 | 0.4635124 | 0.64299713 | 0.88473736 |
| MAOB | 1142.17605 | 0.652005676 | 0.51182048 | 1.27389523 | 0.20270055 | 0.64881821 |
| EFHC2 | 61.7632206 | 0.448899208 | 0.94887494 | 0.47308575 | 0.63615201 | 0.88266168 |
| FUNDC1 | 61.0526292 | -0.054758793 | 0.42296297 | -0.1294647 | 0.89698992 | 0.97324791 |
| KDM6A | 2683.41945 | -0.658337021 | 0.38733042 | -1.6996781 | 0.0891915 | 0.53095823 |
| MIR222HG | 176.987196 | -0.93455601 | 0.64178865 | -1.4561741 | 0.1453445 | 0.60165215 |
| KRBOX4 | 367.731725 | 0.008451636 | 0.28062151 | 0.03011756 | 0.97597329 | 0.99429832 |
| ZNF674 | 120.931548 | -0.506573644 | 0.32830299 | -1.5430065 | 0.1228292 | 0.58216759 |
| SLC9A7 | 340.070584 | 0.013285459 | 0.35527857 | 0.03739448 | 0.97017047 | 0.99204339 |
| RP2 | 168.498141 | -0.460534516 | 0.40190603 | -1.1458761 | 0.25184641 | 0.68834672 |
| JADE3 | 509.459493 | -0.141432801 | 0.33661504 | -0.4201619 | 0.67436722 | 0.89801183 |
| RGN | 1256.22768 | 0.568110741 | 0.89120119 | 0.63746632 | 0.52382114 | 0.83556574 |
| NDUFB11 | 307.157166 | 0.079404774 | 0.49874622 | 0.15920877 | 0.87350439 | 0.96592775 |
| RBM10 | 412.534475 | -0.350557628 | 0.23488832 | -1.4924439 | 0.13558282 | 0.59381224 |
| UBA1 | 625.724701 | -0.301481938 | 0.24137654 | -1.249011 | 0.21166107 | 0.65584609 |
| CDK16 | 240.061109 | -0.316527748 | 0.35233128 | -0.8983811 | 0.36898243 | 0.76615653 |
| USP11 | 136.51244 | 0.360638175 | 0.42263127 | 0.85331636 | 0.39348389 | 0.77771935 |
| ZNF41 | 215.98715 | -0.030834214 | 0.25120801 | -0.1227438 | 0.90231002 | 0.9750023 |
| ARAF | 188.72808 | -0.26798667 | 0.34937627 | -0.7670431 | 0.44305587 | 0.7993765 |
| TIMP1 | 291.455508 | 0.183873594 | 0.62698304 | 0.29326725 | 7.69E-01 | 0.93209094 |
| ELK1 | 116.331792 | -0.190524485 | 0.27899307 | -0.6829004 | 0.49466977 | 0.8220963 |
| UXT | 139.707301 | -0.220223707 | 0.23104717 | -0.9531548 | 0.34051166 | 0.74725037 |
| ZNF81 | 657.310432 | 0.314359182 | 0.29553897 | 1.063681 | 0.28747324 | 0.71118494 |
| ZNF182 | 98.7258675 | -0.202868014 | 0.27467971 | -0.7385621 | 0.46017296 | 0.80711334 |
| ZNF630 | 121.192919 | 1.253191088 | 0.68436724 | 1.83116757 | 0.06707554 | 0.48963361 |
| FTSJ1 | 51.784061 | -0.593054215 | 0.43111532 | -1.3756278 | 0.1689369 | 0.62304975 |
| EBP | 232.132702 | 0.618055465 | 0.28317334 | 2.1826047 | 0.02906493 | 0.39554194 |
| TBC1D25 | 162.085929 | -0.31808624 | 0.28717118 | -1.1076538 | 0.26801137 | 0.69933977 |
| RBM3 | 498.471653 | -0.282278703 | 0.37408594 | -0.7545825 | 0.45049953 | 0.80361623 |
| WDR13 | 976.737928 | 0.102341563 | 0.30590086 | 0.33455795 | 7.38E-01 | 0.92059676 |
| WAS | 39.4611475 | 1.415016765 | 0.76801777 | 1.84242712 | 0.0654127 | 0.48790819 |
| GLOD5 | 46.8539201 | 0.48174875 | 0.66596823 | 0.72338098 | 0.46944586 | 0.81287195 |
| HDAC6 | 566.743277 | 0.62890726 | 0.42700616 | 1.47282948 | 0.14079702 | 0.59555904 |
| PCSK1N | 1058.55394 | 0.483534458 | 0.74774043 | 0.64666085 | 0.51785147 | 0.83325693 |
| TIMM17B | 107.334456 | -0.029916299 | 0.42194552 | -0.0709009 | 0.94347666 | 0.98522232 |
| PQBP1 | 130.397865 | -0.670909347 | 0.40666417 | -1.6497872 | 0.09898647 | 0.5425667 |
| SLC35A2 | 81.5082839 | -0.666985485 | 0.44678069 | -1.49287 | 0.13547122 | 0.59381224 |
| OTUD5 | 508.714533 | -0.189003873 | 0.25707996 | -0.7351949 | 0.46222081 | 0.8082072 |
| GRIPAP1 | 437.288764 | 0.178061794 | 0.23502998 | 0.7576131 | 0.44868264 | 0.80267462 |
| TFE3 | 345.726212 | -0.332071219 | 0.35759217 | -0.9286311 | 0.35308029 | 0.75673243 |
| CCDC120 | 243.169408 | 0.705150987 | 0.39108012 | 1.80308574 | 0.07137475 | 0.49949133 |
| PRAF2 | 37.3521858 | -0.223203725 | 0.64081559 | -0.3483119 | 0.72760592 | 0.91657884 |
| WDR45 | 336.660811 | -0.135312853 | 0.25438188 | -0.531928 | 0.59477584 | 0.86499581 |
| GPKOW | 61.0801564 | -0.085958548 | 0.30255329 | -0.2841104 | 0.77632575 | 0.9352344 |

| | | | | | | |
|---|---|---|---|---|---|---|
| MAGIX | 75.6957896 | -0.040848535 | 0.4434995 | -0.092105 | 0.92661459 | 0.9816089 |
| PLP2 | 90.4474604 | -0.011431503 | 0.46427026 | -0.0246225 | 9.80E-01 | 9.95E-01 |
| PRICKLE3 | 37.4133527 | -0.110200187 | 0.41438527 | -0.2659365 | 0.79028808 | 0.94048261 |
| CCDC22 | 57.7280601 | 0.084063984 | 0.42739307 | 0.1966901 | 0.84407006 | 0.95731603 |
| PPP1R3F | 60.6843934 | 0.527005083 | 0.4093046 | 1.28756209 | 0.19789844 | 0.64582202 |
| CLCN5 | 1798.76693 | 0.424845481 | 0.24854713 | 1.70931555 | 8.74E-02 | 0.52706515 |
| SHROOM4 | 2250.24606 | -0.252801368 | 0.36410971 | -0.6943 | 0.48749409 | 0.81893983 |
| GSPT2 | 44.1386183 | 0.520425662 | 0.56547191 | 0.92033867 | 0.35739581 | 0.75903693 |
| MAGED1 | 1387.45531 | -0.042689959 | 0.1964127 | -0.2173483 | 0.82793695 | 0.95270465 |
| FAM156A | 189.485888 | 0.242214839 | 0.27225637 | 0.88965718 | 0.37364999 | 0.76973377 |
| GPR173 | 45.8506908 | 0.62920579 | 0.79056757 | 0.79589122 | 0.42609526 | 0.78895108 |
| TSPYL2 | 95.4234321 | -1.150564999 | 0.46819271 | -2.4574603 | 0.01399233 | 0.30363654 |
| KANTR | 953.019414 | -0.179680909 | 0.27609089 | -0.6508035 | 0.51517335 | 0.83174101 |
| KDM5C | 474.910158 | -0.912157736 | 0.33014577 | -2.7628939 | 0.00572914 | 0.22501232 |
| IQSEC2 | 444.77769 | 0.491799131 | 0.30387542 | 1.61842355 | 0.10557135 | 0.55358321 |
| SMC1A | 393.404421 | -0.429392813 | 0.3149936 | -1.3631795 | 0.17282594 | 0.62593809 |
| HSD17B10 | 183.990856 | 0.36124956 | 0.33358622 | 1.08292711 | 0.27884078 | 0.70446562 |
| HUWE1 | 2007.28954 | -0.252324907 | 0.17499035 | -1.4419361 | 0.1493204 | 0.6064399 |
| PHF8 | 728.43985 | -0.206684532 | 0.31018505 | -0.6663265 | 0.5052024 | 0.82715271 |
| FAM120C | 348.936993 | -0.102901988 | 0.29904883 | -0.3440976 | 0.73077288 | 0.91897609 |
| WNK3 | 116.841135 | 0.115659981 | 0.38280803 | 0.30213573 | 0.7625486 | 0.93022076 |
| TSR2 | 88.212752 | -0.41272002 | 0.32108286 | -1.2854003 | 0.19865242 | 0.64597062 |
| FGD1 | 100.20102 | 0.520180885 | 0.38116037 | 1.36472972 | 0.17233801 | 0.62592399 |
| GNL3L | 140.081529 | -0.867586675 | 0.46989841 | -1.8463282 | 0.06484457 | 0.48782903 |
| MAGED2 | 235.260737 | 0.305721986 | 0.41166869 | 0.74264084 | 0.45769915 | 0.80566101 |
| PFKFB1 | 121.520458 | 1.962409409 | 0.94596083 | 2.07451445 | 0.03803156 | 0.42339465 |
| APEX2 | 78.4591706 | 0.638721493 | 0.31207376 | 2.04670043 | 0.04068751 | 0.43086475 |
| FAM104B | 150.172801 | 0.602132331 | 0.35383414 | 1.70173612 | 0.08880485 | 0.52996572 |
| USP51 | 61.080741 | 1.054211834 | 0.45709374 | 2.30633619 | 0.02109185 | 0.35516714 |
| RRAGB | 850.174851 | 0.183288443 | 0.23940361 | 0.76560433 | 0.44391177 | 0.79956618 |
| AL445472.1 | 920.271365 | 0.645450454 | 0.43036756 | 1.49976559 | 0.13367513 | 0.59240843 |
| KLF8 | 412.805717 | 0.537690532 | 0.62580556 | 0.85919743 | 0.3902316 | 0.77569824 |
| UBQLN2 | 211.263099 | 0.103075292 | 0.30080429 | 0.34266563 | 0.73185003 | 0.91930611 |
| NBDY | 695.628783 | -0.05141263 | 0.24915774 | -0.2063457 | 0.83652088 | 0.95472037 |
| SPIN3 | 261.679282 | 0.280473161 | 0.22781853 | 1.23112532 | 0.218276 | 0.66064833 |
| SPIN2B | 35.3392442 | 0.371823503 | 0.44743128 | 0.83101812 | 0.40596339 | 0.78429302 |
| FAAH2 | 1003.67114 | 0.472057646 | 0.35503476 | 1.32960966 | 0.18364691 | 0.63276491 |
| ZXDB | 34.4424077 | -0.209336943 | 0.46137138 | -0.4537276 | 0.65002488 | 0.88866038 |
| SPIN4 | 39.0212146 | 0.281497834 | 0.43178536 | 0.65193927 | 0.51444035 | 0.83169608 |
| ARHGEF9 | 597.303712 | 0.35510422 | 0.27232347 | 1.3039795 | 0.19224057 | 0.64497936 |
| AMER1 | 50.5469812 | 0.632727238 | 0.45764593 | 1.38256937 | 0.16679695 | 0.6222879 |
| MTMR8 | 300.198721 | 0.036721294 | 0.24897932 | 0.14748732 | 0.88274738 | 0.96824408 |
| ZC4H2 | 536.203723 | -0.190561699 | 0.22792307 | -0.836079 | 0.40311044 | 0.78275037 |
| ZC3H12B | 257.310749 | 0.843552997 | 0.41934046 | 2.0116184 | 0.04426018 | 0.44360789 |
| LAS1L | 146.710962 | -0.405829769 | 0.36855874 | -1.1011264 | 0.27084164 | 0.69961625 |
| MSN | 1989.14284 | -0.234257627 | 0.35908742 | -0.6523693 | 0.51416294 | 0.83138273 |
| AR | 863.365883 | -0.252718486 | 0.39437715 | -0.6408041 | 0.52164999 | 0.83445062 |
| OPHN1 | 1148.98168 | 0.010979904 | 0.26964174 | 0.04072034 | 9.68E-01 | 0.99175746 |
| YIPF6 | 468.566823 | -0.351919173 | 0.24522997 | -1.4350578 | 0.15127064 | 0.60905477 |
| STARD8 | 97.1481158 | -0.026081989 | 0.36207537 | -0.0720347 | 0.9425743 | 0.98489601 |
| EFNB1 | 92.0947165 | 0.51568963 | 0.35001349 | 1.47334215 | 0.1406588 | 0.59524859 |
| EDA | 1983.52573 | 0.019454636 | 0.29811761 | 0.06525826 | 9.48E-01 | 0.98668191 |
| IGBP1 | 429.461697 | -0.036441401 | 0.29244689 | -0.1246086 | 0.90083341 | 0.97479173 |
| PDZD11 | 43.5889115 | 0.175089968 | 0.56380224 | 0.3105521 | 0.75614115 | 0.92799282 |

| | | | | | | |
|---|---|---|---|---|---|---|
| DLG3 | 304.300429 | 0.557345785 | 0.42928261 | 1.29831904 | 0.19417772 | 0.64548758 |
| TEX11 | 46.1741274 | 0.24552194 | 0.43776157 | 0.56085768 | 0.57489456 | 0.85474196 |
| SNX12 | 174.882116 | -0.389252926 | 0.34807092 | -1.118315 | 0.26343248 | 0.69567438 |
| MED12 | 95.4396835 | 0.065217064 | 0.30847931 | 0.21141471 | 0.83256368 | 0.95334883 |
| GJB1 | 182.972153 | 2.075330563 | 0.67474683 | 3.07571741 | 0.00209997 | 0.14685845 |
| ZMYM3 | 235.596071 | 0.717808303 | 0.41966138 | 1.71044642 | 0.08718335 | 0.5268877 |
| NONO | 608.841581 | 0.058486445 | 0.27900589 | 0.20962441 | 0.83396083 | 0.95393659 |
| TAF1 | 1172.62688 | -0.110191445 | 0.20579233 | -0.5354497 | 0.59233891 | 0.86382106 |
| OGT | 1560.93252 | -0.734375555 | 0.34378373 | -2.1361556 | 0.03266673 | 0.41154675 |
| GCNA | 154.861989 | 0.402302088 | 0.40984417 | 0.98159768 | 0.32629809 | 0.7373175 |
| NHSL2 | 149.868538 | -0.072446398 | 0.45942581 | -0.157689 | 0.87470187 | 0.96615698 |
| PIN4 | 432.923165 | -0.008553126 | 0.29109875 | -0.0293822 | 0.97655976 | 0.99433254 |
| RPS4X | 1060.31262 | 0.019225659 | 0.54192371 | 0.03547669 | 0.97169963 | 0.99240251 |
| HDAC8 | 2499.8784 | -0.074261832 | 0.22081131 | -0.3363135 | 0.73663444 | 0.92020873 |
| PHKA1 | 242.712959 | 0.047852828 | 0.65366228 | 0.07320726 | 0.94164119 | 0.98489601 |
| CHIC1 | 722.82629 | -0.091144944 | 0.31280174 | -0.2913825 | 0.77075882 | 0.93302012 |
| TSIX | 76.7787359 | -4.525813505 | 2.79547006 | -1.6189812 | 0.10545131 | 0.55348589 |
| XIST | 2979.19543 | -9.735912861 | 4.18706733 | -2.3252344 | 0.02005943 | 0.34840598 |
| JPX | 2227.692 | -0.342253271 | 0.36867799 | -0.9283257 | 0.35323862 | 0.75673243 |
| FTX | 17051.7946 | 0.290647045 | 0.22954094 | 1.26621005 | 0.20543789 | 0.651931 |
| SLC16A2 | 282.830985 | 1.497929995 | 0.41843637 | 3.57982741 | 0.00034382 | 0.06649962 |
| RLIM | 308.009097 | -0.021383419 | 0.21605043 | -0.0989742 | 0.92115875 | 0.97939204 |
| NEXMIF | 62.1593684 | 0.124168108 | 0.40716151 | 0.30496033 | 0.76039637 | 0.92968986 |
| ABCB7 | 676.724113 | -0.014564421 | 0.24392895 | -0.0597076 | 0.95238849 | 0.98777411 |
| UPRT | 162.510508 | 0.18874413 | 0.22447811 | 0.84081307 | 0.40045266 | 0.78232083 |
| ZDHHC15 | 106.949613 | -0.125170692 | 0.36904556 | -0.3391741 | 0.73447859 | 0.91982622 |
| PBDC1 | 119.713025 | -1.048278634 | 0.51310166 | -2.0430233 | 0.04105013 | 0.43143739 |
| AC233296.1 | 55.4832454 | 0.776329804 | 0.7067588 | 1.0984367 | 0.27201385 | 0.70056282 |
| ATRX | 3361.1121 | -0.114801898 | 0.24135469 | -0.4756564 | 0.6343192 | 0.88238823 |
| MAGT1 | 737.664825 | -0.184223015 | 0.33966899 | -0.5423604 | 0.58757024 | 0.86124068 |
| COX7B | 682.663021 | -0.462934451 | 0.50518362 | -0.9163687 | 0.35947355 | 0.7595659 |
| PGK1 | 2032.16226 | -0.597057267 | 0.48346469 | -1.2349553 | 0.21684715 | 0.66032006 |
| TAF9B | 48.2474143 | 0.089393413 | 0.42115962 | 0.21225542 | 0.83190777 | 0.95325673 |
| BRWD3 | 1035.98598 | 0.051706014 | 0.37210007 | 0.13895728 | 0.88948391 | 0.97208467 |
| HMGN5 | 121.993292 | 0.737402919 | 0.5637133 | 1.30811694 | 0.19083365 | 0.64285004 |
| SH3BGRL | 488.400684 | 0.175500901 | 0.33372812 | 0.52587988 | 0.59897167 | 0.86671329 |
| RPS6KA6 | 409.877545 | 0.336466353 | 0.34486721 | 0.97564032 | 0.32924274 | 0.73829083 |
| HDX | 442.703369 | 0.172367934 | 0.35024436 | 0.49213621 | 0.62262305 | 0.87714058 |
| APOOL | 706.43619 | -0.259086234 | 0.25629821 | -1.0108781 | 0.3120748 | 0.72624945 |
| ZNF711 | 96.8901534 | 0.032607103 | 0.48648318 | 0.06702617 | 0.94656087 | 0.98620476 |
| POF1B | 54.7018536 | 0.036706669 | 0.82532931 | 0.04447518 | 0.96452564 | 0.99031076 |
| CHM | 1029.36053 | -0.231223644 | 0.2382439 | -0.9705333 | 0.33178072 | 0.73939532 |
| DACH2 | 51.7103472 | 0.64152532 | 0.6875553 | 0.93305269 | 0.35079277 | 0.75495648 |
| KLHL4 | 200.60887 | 0.628612058 | 0.47110942 | 1.33432284 | 0.18209808 | 0.63216521 |
| PCDH11X | 100.291267 | -0.081983552 | 0.52577236 | -0.1559297 | 0.8760884 | 0.96617876 |
| DIAPH2 | 2906.02584 | -0.15013052 | 0.25366972 | -0.5918346 | 0.55396135 | 0.84727776 |
| DIAPH2-AS1 | 152.251531 | 0.28951242 | 0.62335754 | 0.46444039 | 0.64233226 | 0.88441609 |
| PCDH19 | 35.24869 | 0.767082992 | 0.51087265 | 1.5015151 | 0.13322239 | 0.59230595 |
| TSPAN6 | 82.7420692 | 0.496811224 | 0.47020592 | 1.05658225 | 0.29070229 | 0.71269438 |
| SYTL4 | 183.620886 | 0.686453288 | 0.46402607 | 1.47934206 | 0.13904892 | 0.59474076 |
| CSTF2 | 43.9537852 | -0.605348608 | 0.39849251 | -1.5190966 | 0.12873819 | 0.58793678 |
| TRMT2B | 322.852893 | 0.171597473 | 0.26350769 | 0.6512048 | 0.51491429 | 0.83169608 |
| TIMM8A | 56.8581548 | -0.393305345 | 0.35241627 | -1.1160249 | 0.26441146 | 0.69567438 |
| GLA | 133.857584 | -1.79110796 | 0.6222856 | -2.8782732 | 0.00399859 | 0.20256898 |

| | | | | | | |
|---|---|---|---|---|---|---|
| HNRNPH2 | 153.692056 | -0.103940587 | 0.33634194 | -0.3090325 | 0.75729682 | 0.92799282 |
| ARMCX4 | 838.950139 | 0.093309491 | 0.51545294 | 0.18102427 | 0.85634853 | 0.96242445 |
| ARMCX1 | 67.6448316 | -1.197781077 | 0.54607189 | -2.1934495 | 0.02827501 | 0.39280225 |
| ARMCX6 | 109.383175 | -0.182129842 | 0.43524561 | -0.418453 | 0.67561592 | 0.89813476 |
| ARMCX3 | 342.370525 | -0.194938752 | 0.27227619 | -0.7159596 | 0.47401629 | 0.8151865 |
| ARMCX2 | 63.7450801 | 0.094967542 | 0.42570546 | 0.22308274 | 0.82347111 | 0.95131214 |
| NXF5 | 42.4942196 | 1.546633327 | 0.86806948 | 1.78169301 | 0.07479931 | 0.5071704 |
| ZMAT1 | 964.490528 | 0.815407048 | 0.67457862 | 1.20876504 | 0.22675313 | 0.66690202 |
| ARMCX5 | 66.3139439 | -0.073259378 | 0.45361621 | -0.1615008 | 0.87169899 | 0.9657206 |
| AL035427.1 | 49.6542309 | 0.671482898 | 0.87408079 | 0.76821605 | 0.44235884 | 0.79928982 |
| GPRASP1 | 74.8315173 | 0.23780159 | 0.34737641 | 0.68456459 | 0.49361872 | 0.82161891 |
| GPRASP2 | 43.404387 | 1.093768068 | 0.52644722 | 2.07764049 | 0.03774248 | 0.42261887 |
| BHLHB9 | 206.599374 | -0.949430156 | 0.48275747 | -1.9666814 | 0.04921995 | 0.45419182 |
| LINC00630 | 548.025358 | -0.254360076 | 0.18575017 | -1.3693666 | 0.17088472 | 0.62504223 |
| Z68871.1 | 80.0777745 | -0.296411651 | 0.28399102 | -1.043736 | 0.29660756 | 0.71805012 |
| BEX4 | 210.814659 | -0.323652867 | 0.33001598 | -0.9807188 | 0.32673144 | 0.73745391 |
| TCEAL8 | 104.682605 | 0.052471031 | 0.29665225 | 0.17687724 | 8.60E-01 | 0.96283855 |
| BEX2 | 207.889391 | -0.758153136 | 0.29269386 | -2.5902598 | 9.59E-03 | 0.26082337 |
| TCEAL9 | 286.367418 | -1.057304336 | 0.29539049 | -3.5793446 | 0.00034446 | 0.06649962 |
| BEX3 | 332.739866 | 0.248528525 | 0.45764275 | 0.54306231 | 0.5870869 | 0.86112855 |
| TCEAL4 | 355.820637 | -0.422045628 | 0.23103173 | -1.8267864 | 0.06773189 | 0.48963361 |
| TCEAL3 | 105.285187 | -0.030606348 | 0.27735258 | -0.1103518 | 9.12E-01 | 0.977179 |
| TCEAL1 | 55.602251 | 0.428604927 | 0.39461117 | 1.08614493 | 0.27741487 | 0.7036882 |
| MORF4L2 | 1309.18122 | -0.457495338 | 0.28310795 | -1.6159749 | 0.10609974 | 0.55440878 |
| SLC25A53 | 447.131781 | -0.350169323 | 0.19440628 | -1.8012243 | 0.07166753 | 0.5001766 |
| FAM199X | 882.564826 | 0.14514467 | 0.31735113 | 0.45736301 | 0.64741015 | 0.88774616 |
| IL1RAPL2 | 671.759116 | 2.091885933 | 0.84072836 | 2.48818289 | 0.01283977 | 0.29875272 |
| NRK | 59.7095574 | 0.04486234 | 0.8136469 | 0.05513736 | 0.95602903 | 0.98852557 |
| MUM1L1 | 125.183887 | -0.104787786 | 0.6273408 | -0.1670349 | 0.86734262 | 0.96486219 |
| CXorf57 | 55.275571 | 0.536858413 | 0.42170384 | 1.27306977 | 0.20299329 | 0.64881821 |
| RNF128 | 541.413281 | 0.231033558 | 0.41813603 | 0.55253205 | 0.58058388 | 0.858475 |
| TBC1D8B | 589.695198 | 0.83503337 | 0.63617608 | 1.31258216 | 0.18932378 | 0.6413893 |
| MORC4 | 438.077898 | 0.356956144 | 0.42508468 | 0.83972949 | 0.40106007 | 0.78240897 |
| CLDN2 | 949.84913 | 1.235781667 | 0.6826136 | 1.81036777 | 0.07023877 | 0.49575843 |
| RBM41 | 646.050649 | 0.128568988 | 0.18971297 | 0.67770267 | 0.49796024 | 0.82326489 |
| NUP62CL | 75.0482203 | 0.291052284 | 0.50627502 | 0.57488968 | 0.56536591 | 0.85194709 |
| PRPS1 | 70.4094595 | -0.10711464 | 0.45935626 | -0.2331842 | 0.81561834 | 0.94906482 |
| TSC22D3 | 328.760041 | -0.033840305 | 0.79838319 | -0.042386 | 0.96619095 | 0.99112162 |
| MID2 | 261.968157 | -0.854401457 | 0.34336432 | -2.4883234 | 0.0128347 | 0.29875272 |
| PSMD10 | 50.1254225 | 0.010229357 | 0.48921155 | 0.02090989 | 0.98331754 | 0.99523767 |
| ATG4A | 132.057214 | -0.089372953 | 0.29299446 | -0.3050329 | 0.76034109 | 0.92968986 |
| COL4A5 | 131.979402 | 0.921366161 | 0.93956401 | 0.9806316 | 0.32677445 | 0.73745391 |
| ACSL4 | 1690.5163 | 0.49265977 | 0.66291972 | 0.74316656 | 0.45738085 | 0.80561734 |
| TMEM164 | 667.654671 | 0.642185543 | 0.30805266 | 2.08466155 | 0.03710003 | 0.42215481 |
| AMMECR1 | 555.339846 | -0.160685942 | 0.21996204 | -0.7305167 | 0.46507442 | 0.81021544 |
| CHRDL1 | 47.771285 | 0.597427953 | 0.67600163 | 0.88376703 | 0.37682199 | 0.77045763 |
| PAK3 | 60.6084751 | 0.293640013 | 0.6515627 | 0.45067039 | 0.65222713 | 0.88935488 |
| ALG13 | 919.653389 | 0.387761677 | 0.36202705 | 1.07108481 | 0.2841313 | 0.70897061 |
| ALG13-AS1 | 65.3326599 | 1.198758923 | 0.7104765 | 1.68726049 | 0.09155327 | 0.53537574 |
| TRPC5 | 123.178879 | 0.860203964 | 0.63385601 | 1.35709679 | 0.17475046 | 0.62699479 |
| AMOT | 358.526754 | 1.109045214 | 0.41188251 | 2.69262516 | 0.00708919 | 0.23330842 |
| PLS3 | 501.712745 | 0.783350693 | 0.32257326 | 2.42844274 | 0.01516382 | 0.3105046 |
| DANT2 | 3962.58052 | 0.373238815 | 0.24664597 | 1.51325732 | 0.13021431 | 0.58853436 |
| KLHL13 | 2395.81182 | 0.995314971 | 0.5864332 | 1.69723504 | 0.08965223 | 0.5317452 |

| | | | | | | |
|---|---|---|---|---|---|---|
| WDR44 | 876.060984 | -0.296559509 | 0.24064665 | -1.2323442 | 0.21782052 | 0.66064833 |
| DOCK11 | 705.152514 | 0.065914413 | 0.40898162 | 0.16116718 | 0.87196173 | 0.9657206 |
| IL13RA1 | 549.737751 | -0.104490947 | 0.44901796 | -0.23271 | 0.81598664 | 0.94906482 |
| LINC01285 | 119.595786 | -0.075955568 | 0.57247986 | -0.1326781 | 0.89444792 | 0.97272276 |
| LONRF3 | 359.68578 | -0.616888391 | 0.90170365 | -0.6841365 | 0.49388897 | 0.82176937 |
| PGRMC1 | 562.236664 | 0.03050131 | 0.37239493 | 0.08190582 | 0.93472161 | 0.98366442 |
| SLC25A43 | 657.296076 | 0.180969516 | 0.26384884 | 0.6858833 | 0.4927867 | 0.82100434 |
| SLC25A5 | 551.605569 | 0.74831329 | 0.53001244 | 1.41187872 | 0.15798567 | 0.61369956 |
| CXorf56 | 91.1455561 | -0.508047962 | 0.29167615 | -1.7418221 | 0.08153958 | 0.51593841 |
| UBE2A | 245.15281 | -0.302583498 | 0.28072125 | -1.0778789 | 0.28108782 | 0.70625976 |
| NKRF | 102.824119 | 0.091401095 | 0.31901616 | 0.2865093 | 0.77448807 | 0.93453439 |
| 6-Sep | 118.475334 | 0.663645018 | 0.63816846 | 1.03992137 | 0.29837643 | 0.71852421 |
| RPL39 | 70.288484 | -0.598242571 | 0.46265779 | -1.2930563 | 0.19599159 | 0.64558745 |
| UPF3B | 565.964386 | -0.506609006 | 0.32334702 | -1.5667656 | 0.11716948 | 0.57350419 |
| NDUFA1 | 441.974275 | -0.274522931 | 0.43517641 | -0.6308314 | 0.52815079 | 0.83761342 |
| NKAP | 191.688107 | -0.38113201 | 0.24084184 | -1.5824991 | 0.11353567 | 0.56817577 |
| RHOXF1-AS1 | 224.472875 | 0.72407033 | 0.56741393 | 1.27608839 | 0.20192427 | 0.64881821 |
| ZBTB33 | 64.9832245 | -0.188330555 | 0.44969 | -0.4188009 | 0.67536168 | 0.89801183 |
| LAMP2 | 1311.55463 | 0.113283789 | 0.3215006 | 0.35235949 | 0.72456868 | 0.91600702 |
| CUL4B | 288.848437 | 0.056160161 | 0.20644554 | 0.27203378 | 0.78559605 | 0.93892226 |
| MCTS1 | 277.178169 | 0.212517385 | 0.20381193 | 1.0427132 | 0.29708114 | 0.71825247 |
| C1GALT1C1 | 88.126458 | -0.241753549 | 0.37526457 | -0.6442216 | 0.51943174 | 0.8335052 |
| GLUD2 | 40.7902476 | -0.060652637 | 0.45378897 | -0.1336582 | 0.89367283 | 0.97272276 |
| GRIA3 | 549.5404 | -0.06847994 | 0.50583878 | -0.135379 | 0.89231224 | 0.97237481 |
| THOC2 | 2360.68313 | -0.224854443 | 0.18576285 | -1.2104381 | 0.22611082 | 0.66688879 |
| XIAP | 1712.61066 | -0.169833522 | 0.21792854 | -0.7793083 | 0.43579812 | 0.79549938 |
| STAG2 | 1861.74637 | 0.074407308 | 0.23566711 | 0.31573056 | 0.75220703 | 0.92686999 |
| TENM1 | 1892.10258 | 1.326975515 | 0.58031145 | 2.28666091 | 0.02221562 | 0.36178312 |
| AL442647.1 | 54.9767202 | 1.856461938 | 0.66693983 | 2.78355237 | 0.00537672 | 0.21873395 |
| SMARCA1 | 790.093066 | 0.32202886 | 0.46396364 | 0.6940821 | 0.48763072 | 0.81893983 |
| OCRL | 281.227258 | 0.001272549 | 0.22676126 | 0.00561185 | 0.99552242 | 0.99866448 |
| XPNPEP2 | 724.97939 | 1.547290246 | 0.86924486 | 1.78003957 | 0.07506948 | 0.5071704 |
| ZDHHC9 | 575.612016 | -0.144304184 | 0.20655685 | -0.6986173 | 0.48479125 | 0.81839266 |
| AL034405.1 | 48.7131984 | 0.033605834 | 0.43154107 | 0.07787401 | 0.93792827 | 0.98435387 |
| UTP14A | 146.322151 | -0.900427997 | 0.43690331 | -2.060932 | 0.03930953 | 0.42704465 |
| BCORL1 | 277.948191 | 0.309651198 | 0.38601561 | 0.80217273 | 0.42245305 | 0.78894887 |
| ELF4 | 62.0762573 | 0.243438725 | 0.52851883 | 0.46060559 | 6.45E-01 | 8.87E-01 |
| AIFM1 | 1037.54302 | 1.015714135 | 0.39431325 | 2.57590668 | 0.00999776 | 0.26511053 |
| ZNF280C | 272.652492 | 0.395892404 | 0.2929306 | 1.35148871 | 0.17653893 | 0.62709281 |
| SLC25A14 | 115.465766 | -0.105624929 | 0.28197389 | -0.3745912 | 0.70796455 | 0.91025029 |
| RBMX2 | 107.18585 | -0.316551077 | 0.32952606 | -0.9606253 | 0.33674059 | 0.74293725 |
| ENOX2 | 1083.75694 | -0.120109748 | 0.3068396 | -0.3914415 | 0.69547093 | 0.90589033 |
| STK26 | 56.3545533 | -0.680693734 | 0.54700615 | -1.2443987 | 0.21335287 | 0.65809339 |
| RAP2C | 298.765036 | -0.262180869 | 0.2623131 | -0.9994959 | 0.31755452 | 0.72939256 |
| RAP2C-AS1 | 1100.44156 | -0.185726537 | 0.36188568 | -0.5132188 | 0.60779827 | 0.86985416 |
| MBNL3 | 242.072807 | 0.568568622 | 0.42092983 | 1.35074444 | 0.17677731 | 0.62709281 |
| HS6ST2 | 57.3281683 | -0.38867157 | 0.5840685 | -0.6654555 | 0.50575922 | 0.82738999 |
| GPC4 | 1337.30381 | -0.923768785 | 0.41315003 | -2.235916 | 0.02535727 | 0.38152813 |
| GPC3 | 455.515605 | -0.911495387 | 0.96655265 | -0.9430375 | 0.34566173 | 0.75166079 |
| PHF6 | 263.71883 | 0.341094356 | 0.22766284 | 1.49824344 | 0.13407001 | 0.59240843 |
| HPRT1 | 209.823638 | 0.162111078 | 0.34190227 | 0.47414449 | 6.35E-01 | 8.82E-01 |
| MIR503HG | 72.5967338 | 0.241117932 | 0.69854236 | 0.34517296 | 0.72996435 | 0.91836818 |
| FAM122B | 152.326769 | -0.187038683 | 0.24753653 | -0.7556003 | 0.44988887 | 0.80321497 |
| FAM122C | 115.547159 | 0.454541617 | 0.61068619 | 0.74431291 | 0.45668719 | 0.80507423 |

| | | | | | | |
|---|---|---|---|---|---|---|
| MOSPD1 | 116.296667 | -0.190630172 | 0.25314339 | -0.7530521 | 0.45141858 | 0.80384625 |
| RTL8C | 294.571634 | -0.14656162 | 0.55606017 | -0.2635715 | 0.79211011 | 0.94132545 |
| RTL8A | 160.508206 | -0.601541641 | 0.4009932 | -1.5001293 | 0.13358092 | 0.59240843 |
| ZNF75D | 1196.76406 | 0.083623672 | 0.269143 | 0.3107035 | 0.75602604 | 0.92799282 |
| ZNF449 | 124.308638 | -0.054464344 | 0.34792671 | -0.1565397 | 0.87560762 | 0.96617876 |
| SMIM10L2A | 111.280798 | 0.194629407 | 0.52417825 | 0.37130386 | 0.71041122 | 0.91123308 |
| INTS6L | 317.074123 | 0.695288685 | 0.35338194 | 1.9675275 | 0.04912243 | 0.45409375 |
| MMGT1 | 98.3513132 | 0.088666162 | 0.32355147 | 0.27404036 | 0.7840536 | 0.93867623 |
| SLC9A6 | 176.828389 | 0.0189276 | 0.21134407 | 0.08955823 | 0.92863828 | 0.98272895 |
| FHL1 | 150.098447 | 0.13834592 | 0.65164242 | 0.21230343 | 0.83187032 | 0.95325673 |
| MAP7D3 | 246.490238 | 0.203423732 | 0.22365394 | 0.90954684 | 0.36306155 | 0.76209565 |
| HTATSF1 | 161.846283 | 0.176776095 | 0.39024159 | 0.45299143 | 0.65055491 | 0.8887773 |
| AL683813.1 | 194.463131 | 0.831888285 | 0.47634583 | 1.74639563 | 0.08074223 | 0.51475987 |
| RBMX | 985.732634 | 0.180754009 | 0.19192645 | 0.94178789 | 0.34630125 | 0.75166079 |
| FGF13 | 2548.99216 | 0.576653982 | 0.69252092 | 0.83268818 | 0.40502061 | 0.78422899 |
| ATP11C | 1145.78598 | 0.215147901 | 0.18317917 | 1.17452162 | 2.40E-01 | 6.79E-01 |
| LDOC1 | 56.8274147 | 0.726266486 | 0.63066863 | 1.15158175 | 0.24949299 | 0.68617881 |
| SLITRK4 | 77.8259073 | 1.01737817 | 0.74821641 | 1.35973785 | 0.1739129 | 0.62685996 |
| AL589669.1 | 4118.48874 | 0.417595612 | 0.60208294 | 0.69358486 | 0.48794259 | 0.81919239 |
| FMR1 | 1072.05908 | 0.236526572 | 0.46150613 | 0.51251014 | 0.60829402 | 0.86985416 |
| AFF2 | 449.231412 | 1.619979162 | 0.76719941 | 2.11154903 | 0.03472515 | 0.41686761 |
| IDS | 360.888099 | 0.095951569 | 0.37292585 | 0.25729396 | 0.79695185 | 0.94280281 |
| LINC00893 | 92.9268531 | 0.323606702 | 0.43034233 | 0.75197507 | 0.45206605 | 0.80415518 |
| TMEM185A | 198.006969 | 0.052474204 | 0.21117022 | 0.24849244 | 0.80375342 | 0.94543168 |
| CXorf40B | 37.7473224 | -0.210785543 | 0.39831817 | -0.5291889 | 0.59667444 | 0.8657369 |
| LINC00894 | 1735.77632 | 0.479659384 | 0.54604098 | 0.87843111 | 0.37970981 | 0.77095585 |
| MAMLD1 | 205.081502 | -0.112155923 | 0.82933578 | -0.1352358 | 0.89242541 | 0.97237481 |
| MTM1 | 930.312247 | 0.225345537 | 0.20024727 | 1.12533635 | 0.26044653 | 0.69280433 |
| MTMR1 | 535.59515 | -0.338124874 | 0.16966425 | -1.9929058 | 0.04627176 | 0.44508419 |
| CD99L2 | 403.555566 | 0.232177332 | 0.19395766 | 1.1970516 | 0.23128644 | 0.66878058 |
| HMGB3 | 70.3588643 | -0.470084583 | 0.5757541 | -0.8164676 | 0.41423273 | 0.78632759 |
| VMA21 | 170.92047 | 0.070285144 | 0.34950514 | 0.20109903 | 0.84062114 | 0.956374 |
| GABRE | 47.3764001 | -0.116592659 | 0.44970878 | -0.2592626 | 0.79543265 | 0.94252227 |
| CETN2 | 89.6595586 | 0.423045132 | 0.49373486 | 0.85682654 | 0.39154075 | 0.77606943 |
| NSDHL | 87.389574 | 0.291198185 | 0.34708946 | 0.83897155 | 0.40148528 | 0.78246874 |
| ZNF185 | 54.036895 | 0.479705986 | 0.78402084 | 0.61185361 | 0.54063461 | 0.8417342 |
| PNMA3 | 59.3771066 | 1.582844451 | 1.1550887 | 1.37032286 | 0.17058614 | 0.62469256 |
| PNMA6A | 56.4873849 | 0.751807857 | 0.44556356 | 1.68731901 | 0.09154203 | 0.53537574 |
| ZNF275 | 60.4017978 | 0.320341662 | 0.56251211 | 0.56948403 | 0.56902771 | 0.85291608 |
| BGN | 50.3731454 | 0.083576749 | 0.61556771 | 0.13577182 | 0.89200167 | 0.97237481 |
| CCNQ | 71.593691 | -0.537412801 | 0.53900146 | -0.9970526 | 0.31873898 | 0.73071671 |
| SLC6A8 | 882.62806 | -0.678114128 | 0.54770544 | -1.2381001 | 0.21567897 | 0.6596384 |
| BCAP31 | 499.465555 | -0.415342642 | 0.40630889 | -1.0222337 | 0.3066703 | 0.72190689 |
| ABCD1 | 67.0563134 | -0.676706306 | 0.36061784 | -1.8765192 | 0.06058402 | 0.48026727 |
| IDH3G | 189.296464 | 0.105723772 | 0.33176401 | 0.31867161 | 0.74997555 | 0.92565124 |
| SSR4 | 231.79031 | -0.25628226 | 0.56754289 | -0.4515646 | 0.6515827 | 0.88935488 |
| RENBP | 391.784042 | -0.790749507 | 0.51479953 | -1.5360339 | 0.12453007 | 0.58516755 |
| HCFC1 | 160.65226 | -0.351841869 | 0.40658797 | -0.8653524 | 0.3868454 | 0.77334938 |
| TMEM187 | 50.4116139 | 0.556665658 | 0.39396421 | 1.41298536 | 0.15766003 | 0.61323667 |
| IRAK1 | 227.215219 | -0.277205217 | 0.60242596 | -0.4601482 | 0.64540986 | 0.88687652 |
| MECP2 | 992.168897 | -0.418023451 | 0.28019672 | -1.4918927 | 0.13572727 | 0.59381224 |
| FLNA | 226.645891 | 1.190459033 | 0.71191414 | 1.67219467 | 0.09448594 | 0.53824794 |
| EMD | 192.233223 | -0.158495592 | 0.33396608 | -0.4745859 | 0.63508216 | 0.88238823 |
| RPL10 | 1827.4246 | 0.456587933 | 0.55707054 | 0.8196232 | 0.41243095 | 0.78632759 |

| | | | | | | |
|---|---|---|---|---|---|---|
| AC245140.2 | 38.9078265 | -0.119496216 | 0.53083226 | -0.2251111 | 0.82189287 | 0.95105545 |
| DNASE1L1 | 154.28049 | -0.221257996 | 0.4464802 | -0.4955606 | 0.62020445 | 0.87586466 |
| TAZ | 281.81284 | -0.018981455 | 0.27430883 | -0.0691974 | 0.9448325 | 0.9853777 |
| AC244090.1 | 48.282648 | -0.201017538 | 0.46719491 | -0.4302648 | 0.66700301 | 0.89449274 |
| ATP6AP1 | 292.459113 | -0.051500235 | 0.40289506 | -0.1278254 | 0.89828712 | 0.97396153 |
| GDI1 | 272.265628 | -0.069403529 | 0.27098784 | -0.2561131 | 0.79786353 | 0.94319236 |
| FAM50A | 349.24435 | -0.906293149 | 0.4686933 | -1.9336593 | 0.05315503 | 0.46624874 |
| PLXNA3 | 170.160722 | -0.298887119 | 0.35992837 | -0.8304072 | 4.06E-01 | 0.78453757 |
| UBL4A | 86.928556 | -0.174422846 | 0.47520488 | -0.3670477 | 0.71358346 | 0.91209742 |
| SLC10A3 | 54.0817536 | -0.596538059 | 0.47782617 | -1.2484416 | 0.21186939 | 0.65614408 |
| FAM3A | 189.739993 | 0.147984522 | 0.22533548 | 0.65672979 | 0.51135467 | 0.83007666 |
| G6PD | 81.794744 | -0.641299933 | 0.6588983 | -0.9732912 | 0.33040859 | 0.73849818 |
| IKBKG | 123.802413 | -0.035797251 | 0.28816419 | -0.1242252 | 0.90113698 | 0.97482505 |
| GAB3 | 45.7342135 | -0.319406203 | 0.59434286 | -0.5374107 | 5.91E-01 | 0.86333372 |
| DKC1 | 185.858518 | -0.737399126 | 0.42970597 | -1.7160551 | 0.08615195 | 0.52402549 |
| MPP1 | 281.201401 | 0.118988251 | 0.28611696 | 0.41587277 | 0.67750311 | 0.89864474 |
| F8 | 164.556097 | 0.257305947 | 0.40868924 | 0.62958826 | 0.52896401 | 0.83772497 |
| F8A1 | 31.1366937 | -0.053202284 | 0.62939428 | -0.0845293 | 0.93263558 | 0.98310938 |
| FUNDC2 | 259.72454 | -0.360317746 | 0.33838493 | -1.0648162 | 0.28695913 | 0.71118494 |
| MTCP1 | 83.8060186 | 0.408686976 | 0.74822649 | 0.54620757 | 0.58492326 | 0.86062622 |
| BRCC3 | 380.951902 | 0.158281749 | 0.16226742 | 0.97543762 | 0.32934323 | 0.73829083 |
| VBP1 | 139.915476 | -0.180560567 | 0.23796868 | -0.7587577 | 0.44799753 | 0.80223612 |
| TMLHE-AS1 | 250.314367 | 0.842113475 | 0.63260915 | 1.33117499 | 0.18313144 | 0.63276491 |
| TMLHE | 657.907807 | 0.4461847 | 0.2754789 | 1.61966922 | 0.10530335 | 0.55323612 |
| VAMP7 | 302.629335 | -0.105036107 | 0.26575426 | -0.3952377 | 0.69266746 | 0.90528251 |
| ZNF596 | 170.057092 | -0.361220225 | 0.2880377 | -1.2540727 | 0.20981557 | 0.65441616 |
| FBXO25 | 503.082835 | -0.348339812 | 0.21565104 | -1.6152939 | 0.10624706 | 0.55440878 |
| TDRP | 702.039351 | -0.741660774 | 0.3085068 | -2.4040338 | 0.01621527 | 0.32071648 |
| ERICH1 | 1613.47765 | -0.795223012 | 0.4711863 | -1.687704 | 0.09146806 | 0.53537574 |
| DLGAP2 | 443.709984 | -1.064540476 | 0.76695839 | -1.3880029 | 0.16513616 | 0.62065658 |
| AC100810.1 | 93.4527392 | 0.378140616 | 0.44260644 | 0.85434956 | 0.39291134 | 0.77703588 |
| AC019257.8 | 36.4324886 | -0.137986025 | 0.61038238 | -0.2260649 | 0.82115095 | 0.95084695 |
| KBTBD11 | 397.905574 | -0.256594631 | 0.37605343 | -0.6823356 | 0.49502679 | 0.82216554 |
| MYOM2 | 191.361397 | 0.282796104 | 0.37172119 | 0.76077478 | 4.47E-01 | 8.02E-01 |
| AC246817.2 | 203.082388 | 1.418572182 | 0.64408832 | 2.20244977 | 0.02763355 | 0.38942007 |
| AC246817.1 | 221.008026 | 0.439477622 | 0.42577459 | 1.03218377 | 0.30198604 | 0.71925676 |
| CSMD1 | 1451.5634 | -0.434909527 | 0.91513547 | -0.4752406 | 0.63461549 | 0.88238823 |
| MCPH1 | 1438.40861 | 0.22199332 | 0.17405841 | 1.27539552 | 0.20216928 | 0.64881821 |
| ANGPT2 | 192.998728 | 0.674037292 | 0.6117849 | 1.10175535 | 0.27056805 | 0.69961625 |
| MCPH1-AS1 | 437.769588 | 1.271062217 | 0.50932216 | 2.49559576 | 0.01257458 | 0.29875272 |
| AGPAT5 | 1073.6451 | -0.175497471 | 0.49032643 | -0.3579197 | 0.72040344 | 0.9149807 |
| AF233439.1 | 127.30488 | -0.511480807 | 0.7291155 | -0.7015086 | 0.48298566 | 0.8177376 |
| DEFB1 | 919.297699 | -0.884727893 | 0.42328235 | -2.0901602 | 3.66E-02 | 0.42192353 |
| FAM85B | 2874.71787 | -0.123600741 | 0.49762328 | -0.2483822 | 0.80383874 | 0.94543168 |
| PRAG1 | 839.803687 | -0.32404986 | 0.52254378 | -0.6201392 | 0.53516617 | 0.83954482 |
| MFHAS1 | 2284.69422 | 0.13751336 | 0.37663853 | 0.36510699 | 0.71503154 | 0.91279611 |
| ERI1 | 620.603967 | 0.035966812 | 0.35519384 | 0.10125967 | 0.91934433 | 0.97913385 |
| PPP1R3B | 295.674969 | -0.05650792 | 0.39543444 | -0.1429009 | 0.88636848 | 0.97024858 |
| AC022784.3 | 86.9878925 | 1.005987717 | 0.78624162 | 1.27948928 | 0.20072481 | 0.64813546 |
| TNKS | 2613.71558 | -0.092400945 | 0.19332477 | -0.4779571 | 0.63268074 | 0.88151688 |
| AC034111.1 | 62.1487844 | 0.457899143 | 0.61031045 | 0.75027249 | 0.4530906 | 0.80433814 |
| MSRA | 8932.64267 | -0.129851427 | 0.27016024 | -0.480646 | 6.31E-01 | 8.81E-01 |
| AC104964.1 | 232.465975 | 0.931640221 | 0.37223889 | 2.50280196 | 0.01232145 | 0.29629837 |
| PRSS51 | 142.783718 | 2.069516661 | 0.86737483 | 2.38595425 | 0.01703487 | 0.32964126 |

| | | | | | | |
|---|---|---|---|---|---|---|
| AC104964.4 | 91.210918 | 0.108762625 | 0.50148441 | 0.21688137 | 0.8283008 | 0.9530361 |
| PRSS55 | 134.044608 | 2.093660501 | 0.74666357 | 2.80402123 | 0.00504696 | 0.21686099 |
| PINX1 | 40.1519397 | -0.72454323 | 0.43741096 | -1.6564359 | 0.09763357 | 0.54093104 |
| XKR6 | 1056.79049 | 0.426928273 | 0.52868369 | 0.80753062 | 0.41936084 | 0.78783502 |
| MTMR9 | 421.44971 | 0.011630129 | 0.25326007 | 0.04592169 | 0.96337267 | 0.99012655 |
| NEIL2 | 85.7085459 | 0.478998118 | 0.43811496 | 1.09331604 | 0.27425506 | 0.7014484 |
| FDFT1 | 884.809589 | 0.930443076 | 0.25645996 | 3.62802467 | 0.0002856 | 0.06116705 |
| CTSB | 3907.7653 | -0.411842446 | 0.57131516 | -0.7208674 | 0.47099113 | 0.81387913 |
| FAM66D | 135.1254 | 0.904177074 | 0.93662011 | 0.96536158 | 0.33436374 | 0.74120095 |
| FAM86B1 | 39.4356014 | -0.209067943 | 0.61735823 | -0.3386493 | 0.73487392 | 0.91997009 |
| AC145124.1 | 310.948325 | 0.203355742 | 0.72395667 | 0.28089491 | 0.77879101 | 0.93646866 |
| LONRF1 | 575.117662 | -0.525767383 | 0.30758934 | -1.709316 | 0.08739243 | 0.52706515 |
| TRMT9B | 1146.43789 | 0.297523313 | 0.64083967 | 0.46427106 | 0.64245355 | 0.88441609 |
| DLC1 | 956.197666 | 0.000789677 | 0.7216012 | 0.00109434 | 0.99912684 | 0.99977652 |
| SGCZ | 42.9068719 | -0.248639721 | 0.46443143 | -0.5353637 | 0.5923984 | 0.86382106 |
| TUSC3 | 1042.72455 | 0.026272692 | 0.46825847 | 0.05610724 | 9.55E-01 | 9.88E-01 |
| MSR1 | 190.92106 | 0.139379417 | 0.76879146 | 0.18129678 | 0.85613463 | 0.9623616 |
| AC068992.1 | 70.9787779 | -0.795876502 | 0.84866975 | -0.9377929 | 0.34835083 | 0.75360555 |
| MICU3 | 337.29903 | 0.798218779 | 0.44562222 | 1.79124547 | 0.07325391 | 0.50459544 |
| ZDHHC2 | 734.167541 | 0.16357181 | 0.25276088 | 0.64714052 | 0.517541 | 0.8330125 |
| CNOT7 | 356.279781 | -0.177354222 | 0.20387557 | -0.8699141 | 0.38434738 | 0.77317162 |
| VPS37A | 726.551604 | -0.408329082 | 0.21771103 | -1.8755553 | 0.06071637 | 0.48026727 |
| SLC7A2 | 251.320089 | -0.217052569 | 0.6500272 | -0.3339131 | 0.73844517 | 0.92080006 |
| MTUS1 | 2096.13885 | 0.197567552 | 0.44532701 | 0.44364601 | 0.65729854 | 0.89053559 |
| AC027117.2 | 470.354601 | 0.367584245 | 0.47521546 | 0.77351071 | 0.43922019 | 0.79750844 |
| PCM1 | 2354.97601 | -0.021972779 | 0.23292827 | -0.0943328 | 0.92484478 | 0.9804664 |
| ASAH1 | 1494.83222 | 0.143994941 | 0.43661179 | 0.32980085 | 0.74155044 | 0.92235631 |
| NAT1 | 337.212316 | 0.719108175 | 0.72313719 | 0.99442843 | 3.20E-01 | 0.73157402 |
| PSD3 | 334.147619 | 1.065235372 | 0.72997128 | 1.45928394 | 0.14448697 | 0.60101297 |
| SH2D4A | 190.758392 | 0.146455125 | 0.34339707 | 0.42648915 | 0.66975145 | 0.89611537 |
| CSGALNACT1 | 80.7544535 | -1.412256845 | 0.50692808 | -2.7859117 | 0.00533774 | 0.21873395 |
| INTS10 | 1137.3823 | 0.335397413 | 0.30949168 | 1.08370415 | 0.278496 | 0.70442645 |
| ATP6V1B2 | 1382.90276 | -0.549788737 | 0.31558334 | -1.7421348 | 0.08148485 | 0.51593841 |
| AC018541.1 | 83.5793408 | 0.744941092 | 0.78078987 | 0.95408653 | 3.40E-01 | 7.47E-01 |
| XPO7 | 1163.15236 | -0.050166602 | 0.20840025 | -0.2407224 | 0.8097703 | 0.94692486 |
| NPM2 | 91.7495523 | -0.015046094 | 0.39008898 | -0.0385709 | 0.96923248 | 0.99187457 |
| DMTN | 488.473126 | 0.355107448 | 0.44210925 | 0.80321198 | 0.42185222 | 0.78892549 |
| FAM160B2 | 208.450786 | 0.128399333 | 0.30858776 | 0.41608692 | 0.6773464 | 0.89864474 |
| NUDT18 | 33.4054822 | -0.181249553 | 0.53176277 | -0.3408466 | 0.73321904 | 0.91952673 |
| BMP1 | 316.086051 | 0.233658312 | 0.46284697 | 0.50482844 | 0.61367934 | 0.87222135 |
| PHYHIP | 59.2656636 | 0.555708191 | 0.6002744 | 0.92575695 | 0.35457231 | 0.75755608 |
| POLR3D | 97.7528277 | -0.429309069 | 0.50606516 | -0.8483277 | 0.39625552 | 0.7795748 |
| PIWIL2 | 66.6479861 | 0.380366517 | 0.57275777 | 0.66409665 | 0.50662846 | 0.82806179 |
| SLC39A14 | 3956.46155 | -0.46275555 | 0.44229051 | -1.0462706 | 0.29543613 | 0.71673327 |
| PPP3CC | 1262.04378 | -0.081215353 | 0.23746318 | -0.3420124 | 0.73234156 | 0.91950222 |
| SORBS3 | 151.075278 | -0.166037268 | 0.29775988 | -0.5576214 | 0.57710297 | 0.85637657 |
| CCAR2 | 190.427958 | -0.286308415 | 0.25209887 | -1.1356989 | 0.2560826 | 0.69083334 |
| AC037459.3 | 36.2904663 | -0.492612611 | 0.60195782 | -0.8183507 | 0.41315696 | 0.78632759 |
| BIN3 | 626.079699 | -0.099054775 | 0.37801492 | -0.2620393 | 0.79329112 | 0.94184854 |
| PEBP4 | 81.8266588 | 0.591915072 | 0.69871021 | 0.84715389 | 0.39690935 | 0.78014596 |
| RHOBTB2 | 178.839852 | 0.614575969 | 0.56657458 | 1.0847221 | 0.27804476 | 0.7036882 |
| TNFRSF10B | 1272.72464 | 0.213025393 | 0.23989794 | 0.8879834 | 3.75E-01 | 7.70E-01 |
| TNFRSF10D | 573.948533 | 0.051239393 | 0.21056986 | 0.24333679 | 8.08E-01 | 0.94613779 |
| TNFRSF10A | 450.573538 | -0.415362119 | 0.341648 | -1.2157604 | 0.22407619 | 0.66544589 |

| | | | | | | |
|---|---|---|---|---|---|---|
| CHMP7 | 175.855264 | -0.488712086 | 0.26026705 | -1.8777332 | 0.06041767 | 0.48026727 |
| R3HCC1 | 79.025108 | -0.18091553 | 0.33413977 | -0.5414367 | 0.58820663 | 0.86140264 |
| LOXL2 | 55.6611553 | 0.178056891 | 0.34804925 | 0.51158534 | 0.60894125 | 0.87017908 |
| ENTPD4 | 394.452961 | -0.119603228 | 0.21234276 | -0.5632555 | 0.57326092 | 0.8542683 |
| SLC25A37 | 1828.48757 | 0.7257926 | 0.41324779 | 1.75631332 | 0.07903492 | 0.51192014 |
| ADAM28 | 293.695928 | 1.695991472 | 0.71895982 | 2.35895168 | 0.01832664 | 0.33809324 |
| AC120193.1 | 101.574124 | 0.569213814 | 0.68008846 | 0.83697026 | 0.40260929 | 0.78260941 |
| DOCK5 | 1868.7757 | 0.0051496 | 0.26054498 | 0.01976473 | 9.84E-01 | 9.95E-01 |
| KCTD9 | 346.319927 | -0.011192117 | 0.35144852 | -0.0318457 | 0.97459512 | 0.99388254 |
| PPP2R2A | 2236.33408 | -0.370814722 | 0.24520055 | -1.5122916 | 0.13045971 | 0.58853436 |
| BNIP3L | 4064.39332 | 0.221817862 | 0.62999205 | 0.35209629 | 0.72476605 | 0.91600702 |
| PNMA2 | 173.28363 | 0.738925567 | 0.59933323 | 1.23291274 | 2.18E-01 | 6.60E-01 |
| DPYSL2 | 1138.06828 | 0.468566909 | 0.35363801 | 1.32499024 | 1.85E-01 | 6.35E-01 |
| ADRA1A | 87.0609826 | 0.103050807 | 0.92122311 | 0.11186303 | 0.91093201 | 0.97697493 |
| AC090150.1 | 251.856291 | -0.265889048 | 0.7414527 | -0.3586055 | 7.20E-01 | 0.9148113 |
| TRIM35 | 273.34435 | -0.062191922 | 0.48496763 | -0.1282393 | 0.89795958 | 0.97374331 |
| PTK2B | 652.178467 | 0.627986943 | 0.38658228 | 1.62445869 | 0.10427795 | 0.5505924 |
| EPHX2 | 1558.336 | 1.567806414 | 0.63138352 | 2.48312851 | 0.01302341 | 0.29875272 |
| CLU | 3427.02292 | 0.266061626 | 0.70596325 | 0.37687744 | 7.06E-01 | 0.90993233 |
| CCDC25 | 213.609322 | 0.089011821 | 0.24509507 | 0.36317263 | 0.71647593 | 0.91298968 |
| NUGGC | 637.314034 | 1.620827986 | 0.70128152 | 2.31123727 | 2.08E-02 | 0.35313211 |
| ELP3 | 854.911235 | -0.02111999 | 0.19768586 | -0.1068361 | 0.91491899 | 0.97782888 |
| ZNF395 | 359.847449 | -0.582967674 | 0.35493723 | -1.6424529 | 0.10049618 | 0.54576115 |
| FBXO16 | 483.598503 | -0.306372125 | 0.45876746 | -0.6678157 | 0.50425123 | 0.82715271 |
| FZD3 | 350.865083 | 0.324119905 | 0.21935029 | 1.47763609 | 0.13950521 | 0.59500405 |
| EXTL3 | 361.50796 | -0.359587566 | 0.47980838 | -0.7494399 | 0.45359208 | 0.80433814 |
| INTS9 | 668.682559 | -0.336115172 | 0.17958577 | -1.8716136 | 0.06126008 | 0.4813972 |
| HMBOX1 | 1792.51013 | 0.03425465 | 0.18090259 | 0.18935411 | 0.84981529 | 0.95959925 |
| KIF13B | 2921.93252 | 0.052454855 | 0.32947198 | 0.15920885 | 0.87350433 | 0.96592775 |
| AC108449.1 | 119.772402 | 0.399727862 | 0.72196803 | 0.55366422 | 0.57980867 | 0.85816191 |
| SARAF | 1053.67885 | -0.059049013 | 0.29975632 | -0.1969901 | 8.44E-01 | 0.95731603 |
| LEPROTL1 | 619.101179 | 0.15704148 | 0.32165146 | 0.48823494 | 0.62538344 | 0.87854882 |
| AC026979.2 | 42.5172392 | -0.955750936 | 0.44285166 | -2.158174 | 0.03091431 | 0.40540296 |
| DCTN6 | 165.686488 | -0.331617001 | 0.28514497 | -1.1629769 | 0.24483889 | 0.68250191 |
| RBPMS-AS1 | 167.16097 | 0.090281196 | 0.48191557 | 0.1873382 | 0.85139547 | 0.96042118 |
| RBPMS | 19595.144 | -0.196548632 | 0.33511351 | -0.5865136 | 0.55753042 | 0.84861484 |
| GTF2E2 | 1132.60457 | -0.376212946 | 0.29687203 | -1.2672563 | 0.20506366 | 0.65140385 |
| SMIM18 | 41.1788982 | 0.26749934 | 0.41712072 | 0.64129957 | 0.52132807 | 0.83445062 |
| GSR | 1123.90827 | -0.536062301 | 0.38219622 | -1.402584 | 0.16074093 | 0.61630095 |
| UBXN8 | 97.1800606 | -0.454668415 | 0.31940372 | -1.4234913 | 0.15459379 | 0.61119904 |
| PPP2CB | 1654.99055 | -0.29023453 | 0.22934041 | -1.2655185 | 0.20568553 | 0.651931 |
| WRN | 781.746119 | -0.29753384 | 0.31769258 | -0.9365464 | 0.34899194 | 0.75415932 |
| NRG1 | 5056.61883 | -0.648149148 | 0.50028001 | -1.2955727 | 0.19512273 | 0.64558745 |
| AC090204.1 | 75.6410271 | 0.194672513 | 0.58757409 | 0.33131569 | 0.74040604 | 0.92177526 |
| FUT10 | 721.916067 | 0.407216641 | 0.25260891 | 1.61204382 | 0.1069524 | 0.55488138 |
| TTI2 | 191.840014 | -0.09927953 | 0.3248253 | -0.3056398 | 0.75987894 | 0.92947176 |
| MAK16 | 83.6839674 | -0.737224204 | 0.46773996 | -1.5761412 | 0.11499327 | 0.57182344 |
| RNF122 | 143.706186 | 0.078097084 | 0.41626366 | 0.18761446 | 0.85117889 | 0.96033492 |
| AF279873.3 | 62.9788336 | 0.480107029 | 0.64758836 | 0.74137687 | 4.58E-01 | 8.06E-01 |
| UNC5D | 2862.89273 | -0.278732908 | 0.6433482 | -0.4332536 | 0.66483056 | 0.89315615 |
| LINC01605 | 140.972973 | 1.016753833 | 0.79538798 | 1.27831179 | 0.20113951 | 0.64849129 |
| ERLIN2 | 274.576889 | -0.017126278 | 0.26024473 | -0.0658084 | 0.9475304 | 0.98646931 |
| PLPBP | 216.951506 | 0.085265658 | 0.19955453 | 0.42728 | 0.6691754 | 0.89582842 |
| RAB11FIP1 | 352.637573 | -0.336433267 | 0.24828561 | -1.3550252 | 0.17540952 | 0.62699479 |

| | | | | | | |
|---|---|---|---|---|---|---|
| EIF4EBP1 | 187.4773 | -0.479137555 | 0.33184148 | -1.4438748 | 0.14877419 | 0.60622442 |
| ASH2L | 473.783196 | -0.210399513 | 0.35832724 | -0.5871714 | 0.55708859 | 0.84861484 |
| LSM1 | 188.106758 | -0.32083944 | 0.26305045 | -1.2196879 | 0.22258322 | 0.66398799 |
| BAG4 | 545.842365 | -0.208268031 | 0.1618875 | -1.2864985 | 1.98E-01 | 0.64582202 |
| DDHD2 | 501.808409 | -0.584226408 | 0.16792437 | -3.4791044 | 0.00050309 | 0.08018476 |
| PLPP5 | 164.037126 | -0.1821427 | 0.5508871 | -0.3306353 | 0.74092001 | 0.92213095 |
| NSD3 | 1550.86606 | -0.061929079 | 0.19025813 | -0.3255003 | 0.74480245 | 0.92380845 |
| FGFR1 | 1429.41258 | -1.32192905 | 0.86333806 | -1.5311836 | 0.12572403 | 0.58564339 |
| TACC1 | 1802.31599 | -0.407676784 | 0.36137059 | -1.1281405 | 0.2592606 | 0.69242517 |
| PLEKHA2 | 1232.31586 | -0.68613112 | 0.44381194 | -1.5459952 | 0.12210573 | 0.58055167 |
| TM2D2 | 141.676276 | -0.145267393 | 0.41123495 | -0.3532467 | 0.7239035 | 0.91600702 |
| ADAM9 | 1878.0024 | -0.423038848 | 0.36959871 | -1.1445896 | 0.25237919 | 0.68874851 |
| ADAM32 | 158.487022 | 0.988823707 | 0.4935602 | 2.00345108 | 0.04512889 | 0.44360789 |
| IDO2 | 225.073026 | 2.216754862 | 1.11364446 | 1.99054091 | 4.65E-02 | 4.45E-01 |
| TCIM | 479.505168 | 0.975250804 | 1.01297601 | 0.96275804 | 0.33566896 | 0.74225095 |
| ZMAT4 | 41.6912347 | -0.011587898 | 0.52105854 | -0.0222391 | 0.98225719 | 0.99522272 |
| SFRP1 | 65.1318685 | -0.249091682 | 0.52332803 | -0.4759762 | 0.63409134 | 0.88238823 |
| GOLGA7 | 310.84301 | -0.24980072 | 0.24901084 | -1.0031721 | 0.31577784 | 0.72798933 |
| AC009630.1 | 69.633646 | -0.022450963 | 0.49037678 | -0.0457831 | 0.96348314 | 0.99012655 |
| GPAT4 | 1072.39784 | -0.548017215 | 0.27778101 | -1.972839 | 0.04851391 | 0.45329252 |
| KAT6A | 2077.44639 | -0.075892444 | 0.20716061 | -0.3663459 | 0.71410697 | 0.91214838 |
| AP3M2 | 276.791753 | 0.053529676 | 0.22098637 | 0.24223066 | 0.80860144 | 0.94630788 |
| PLAT | 49.8795171 | -0.727888455 | 0.61131828 | -1.1906865 | 0.23377666 | 0.67043632 |
| IKBKB | 791.493094 | -0.150961111 | 0.26474895 | -0.5702048 | 0.56853883 | 0.85263558 |
| POLB | 173.996711 | 0.152921463 | 0.2365751 | 0.64639712 | 0.51802221 | 0.83325693 |
| VDAC3 | 564.526422 | 0.068096639 | 0.40986468 | 0.1661442 | 0.86804347 | 0.96498555 |
| SLC20A2 | 1686.68865 | -1.065496081 | 0.33006672 | -3.2281233 | 0.00124605 | 0.11540298 |
| SMIM19 | 277.308565 | -0.384040166 | 0.22899217 | -1.6770886 | 0.09352515 | 0.53688613 |
| THAP1 | 53.8279207 | -0.500132005 | 0.3922236 | -1.2751196 | 0.20226691 | 0.64881821 |
| RNF170 | 594.400461 | 0.325316216 | 0.37339775 | 0.8712324 | 0.38362727 | 0.77317162 |
| HOOK3 | 1186.09759 | -0.009787667 | 0.17510989 | -0.0558944 | 0.95542591 | 0.98840182 |
| FNTA | 293.721439 | -0.254427409 | 0.25909849 | -0.9819718 | 0.32611375 | 0.7373175 |
| POMK | 97.6544345 | 0.002463678 | 0.27856295 | 0.00884424 | 0.99294341 | 0.9976934 |
| HGSNAT | 936.262405 | -0.030828804 | 0.39361874 | -0.0783215 | 0.93757233 | 0.98435387 |
| SPIDR | 3771.06646 | -0.645734919 | 0.27442104 | -2.3530809 | 0.01861858 | 0.33968023 |
| CEBPD | 1871.51568 | -0.26626746 | 0.51429043 | -0.5177375 | 0.60464141 | 0.86865188 |
| PRKDC | 1297.72888 | -0.588286175 | 0.3416312 | -1.7219919 | 0.08507098 | 0.52183702 |
| MCM4 | 55.8661904 | -0.975531839 | 0.50264315 | -1.940804 | 0.05228205 | 0.46474067 |
| UBE2V2 | 578.151447 | -0.358759332 | 0.26318497 | -1.3631452 | 0.17283676 | 0.62593809 |
| SNTG1 | 121.693399 | 0.961331791 | 0.49926179 | 1.92550644 | 0.05416603 | 0.46711025 |
| PXDNL | 218.622639 | 0.093522681 | 0.29275278 | 0.31945959 | 0.74937803 | 0.92565124 |
| PCMTD1 | 2339.84848 | 0.147421793 | 0.25735126 | 0.57284272 | 0.56675119 | 0.85233132 |
| AC064807.1 | 237.227135 | 0.752434219 | 0.37467956 | 2.00820728 | 0.04462127 | 0.44360789 |
| AC064807.2 | 96.5530858 | 0.212031396 | 0.79436528 | 0.26691926 | 0.78953132 | 0.94023509 |
| ST18 | 707.881735 | 0.922945462 | 0.95303879 | 0.96842381 | 0.33283275 | 0.73988623 |
| RB1CC1 | 2336.44427 | -0.541269261 | 0.25194197 | -2.1483886 | 0.03168289 | 0.40930954 |
| ATP6V1H | 1654.55447 | -0.547290111 | 0.28939523 | -1.8911511 | 0.05860418 | 0.47778423 |
| TCEA1 | 608.27576 | -0.490291961 | 0.20584518 | -2.3818481 | 0.017226 | 0.32964126 |
| LYPLA1 | 964.951338 | -0.15029146 | 0.23310046 | -0.6447497 | 0.51908937 | 0.83344946 |
| MRPL15 | 211.881035 | 0.048128477 | 0.30112568 | 0.15982854 | 0.87301614 | 0.96592775 |
| RP1 | 69.8857143 | 0.835166634 | 0.91229472 | 0.91545706 | 0.35995174 | 0.75998694 |
| XKR4 | 1043.57503 | 2.870705778 | 0.62131895 | 4.62034161 | 3.83E-06 | 8.75E-03 |
| TMEM68 | 370.852292 | 0.349053844 | 0.2584595 | 1.35051661 | 0.17685033 | 0.62718951 |
| TGS1 | 403.172445 | -0.275643534 | 0.25320188 | -1.0886315 | 0.27631644 | 0.70328016 |

| | | | | | | |
|---|---|---|---|---|---|---|
| LYN | 368.056662 | -0.164549613 | 0.43502288 | -0.3782551 | 0.70524111 | 0.909554 |
| RPS20 | 711.91207 | 0.019188635 | 0.50209429 | 0.03821719 | 0.96951451 | 0.99187457 |
| PLAG1 | 138.294791 | 0.732759841 | 0.48934124 | 1.49744142 | 0.13427844 | 0.59240843 |
| CHCHD7 | 103.765646 | 0.106091095 | 0.3602136 | 0.29452273 | 0.7683585 | 0.93198626 |
| IMPAD1 | 494.474841 | -0.444412314 | 0.23283699 | -1.9086843 | 0.05630283 | 0.47143978 |
| FAM110B | 735.071524 | 0.473057539 | 0.46633864 | 1.01440778 | 0.31038822 | 0.72498238 |
| AC027698.1 | 45.5302305 | 0.792474609 | 0.56721699 | 1.39712778 | 0.16237515 | 0.61910325 |
| UBXN2B | 593.446209 | 0.041495757 | 0.27401059 | 0.15143852 | 0.87962981 | 0.96751353 |
| SDCBP | 750.106013 | -0.453836634 | 0.46231276 | -0.9816658 | 0.32626451 | 0.7373175 |
| NSMAF | 503.843886 | -0.7540906 | 0.43534631 | -1.7321626 | 0.08324461 | 0.51935659 |
| TOX | 1209.02782 | 2.364799792 | 0.86266221 | 2.7412813 | 0.00612001 | 0.22637955 |
| LINC01301 | 198.520491 | 1.361705557 | 0.65491625 | 2.07920562 | 0.03759845 | 0.42215481 |
| RAB2A | 2114.19103 | -0.311352897 | 0.23788442 | -1.308841 | 0.1905882 | 0.64281105 |
| CHD7 | 1834.73183 | 0.003002279 | 0.20144071 | 0.01490403 | 0.98810874 | 0.99671759 |
| CLVS1 | 74.2762112 | -0.149314307 | 0.37580949 | -0.3973138 | 0.69113605 | 0.90473488 |
| ASPH | 2214.11556 | -0.647702163 | 0.3154618 | -2.0531873 | 0.04005441 | 0.43027104 |
| NKAIN3 | 346.597885 | 1.533074479 | 0.83702738 | 1.83157028 | 0.06701547 | 0.48963361 |
| GGH | 209.529712 | -0.320209203 | 0.46269754 | -0.6920486 | 0.48890679 | 0.81970016 |
| TTPA | 71.1598532 | 1.366765303 | 0.86126368 | 1.58693016 | 0.11252846 | 0.56608637 |
| YTHDF3 | 1024.15182 | -0.194558458 | 0.23739112 | -0.8195692 | 0.41246173 | 0.78632759 |
| AC022639.1 | 40.9278496 | -0.22199249 | 0.47227772 | -0.4700465 | 0.6383218 | 0.8827026 |
| CYP7B1 | 766.119522 | 0.295077644 | 0.43912801 | 0.67196271 | 0.50160744 | 0.82638105 |
| ARMC1 | 404.917007 | -0.433678733 | 0.20733578 | -2.0916734 | 0.03646774 | 0.42151595 |
| MTFR1 | 1659.42535 | -0.070485395 | 0.25457622 | -0.2768734 | 0.78187728 | 0.93757221 |
| PDE7A | 3247.77209 | 0.017860376 | 0.4672019 | 0.03822839 | 0.96950559 | 0.99187457 |
| TRIM55 | 274.064432 | -1.210150075 | 0.42597975 | -2.8408629 | 0.00449916 | 0.20749797 |
| RRS1-AS1 | 57.132738 | 0.695542859 | 0.73807881 | 0.94236936 | 0.34600357 | 0.75166079 |
| ADHFE1 | 638.40067 | 1.131659661 | 0.68163013 | 1.66022542 | 0.09686911 | 0.54003127 |
| VXN | 234.319092 | 0.197378419 | 0.64822107 | 0.30449244 | 7.61E-01 | 0.92989212 |
| VCPIP1 | 396.014605 | -0.108603875 | 0.23761707 | -0.4570542 | 0.6476321 | 0.88774616 |
| SGK3 | 2548.3093 | 0.415156241 | 0.35779194 | 1.16032865 | 0.24591503 | 0.68320879 |
| MCMDC2 | 40.8716912 | 0.860020357 | 0.74779351 | 1.15007731 | 0.25011203 | 0.68617881 |
| PPP1R42 | 95.4701078 | 0.645851941 | 0.56866781 | 1.13572798 | 0.25607043 | 0.69083334 |
| COPS5 | 90.8670755 | 0.200965981 | 0.4532624 | 0.44337668 | 0.6574933 | 0.89053559 |
| CSPP1 | 2417.49731 | -0.185985271 | 0.20828199 | -0.8929494 | 0.37188429 | 0.76845856 |
| ARFGEF1 | 3194.08892 | 0.083456553 | 0.18479574 | 0.45161514 | 0.65154626 | 0.88935488 |
| C8orf34 | 70.4215982 | -0.327743674 | 0.52223022 | -0.6275847 | 0.53027606 | 0.83789294 |
| SULF1 | 44.6205274 | 0.798131347 | 0.64671835 | 1.2341251 | 0.21715629 | 0.66032006 |
| SLCO5A1 | 105.141156 | -1.234755428 | 0.76472073 | -1.6146488 | 0.10638676 | 0.55440878 |
| NCOA2 | 6073.30755 | 0.274176011 | 0.28058826 | 0.97714713 | 0.32849632 | 0.73826734 |
| TRAM1 | 1203.43708 | -0.254393059 | 0.36417182 | -0.6985523 | 0.48483187 | 0.81839266 |
| LACTB2-AS1 | 166.368895 | 1.577766302 | 0.75145301 | 2.09962071 | 0.03576222 | 0.42040764 |
| LACTB2 | 1217.33737 | 0.322824983 | 0.41749366 | 0.77324523 | 0.43937726 | 0.79768796 |
| XKR9 | 452.457916 | -0.333963686 | 0.64499136 | -0.5177801 | 0.60461171 | 0.86865188 |
| MSC-AS1 | 618.810222 | 2.053161354 | 0.62316492 | 3.2947319 | 0.00098516 | 0.10560879 |
| TERF1 | 611.362184 | -0.008103475 | 0.16653903 | -0.0486581 | 0.96119176 | 0.98985638 |
| AC022893.1 | 149.580669 | 0.087813684 | 0.40083895 | 0.21907473 | 0.82659184 | 0.9519561 |
| SBSPON | 104.755469 | 1.723327557 | 0.66572123 | 2.58866247 | 0.00963495 | 0.2615173 |
| RPL7 | 1057.18253 | 0.2365149 | 0.5864616 | 0.40329137 | 0.68673389 | 0.90215271 |
| RDH10 | 663.768722 | -0.864091985 | 0.49897325 | -1.7317401 | 0.08331985 | 0.51935659 |
| RDH10-AS1 | 177.256903 | 0.010190214 | 0.60088125 | 0.01695878 | 0.9864695 | 0.99613507 |
| STAU2 | 2552.02629 | 0.424690193 | 0.22480957 | 1.88911079 | 0.05887699 | 0.47778423 |
| AC027018.1 | 291.198227 | 0.533835503 | 0.59057809 | 0.90392027 | 0.36603768 | 0.76421682 |
| AC022826.2 | 36.266817 | 1.013354965 | 0.51822411 | 1.9554377 | 0.05053143 | 0.45900219 |

| | | | | | | |
|---|---|---|---|---|---|---|
| ELOC | 444.690205 | -0.555160376 | 0.31914577 | -1.7395198 | 0.08194338 | 0.51593841 |
| TMEM70 | 101.177513 | -0.235486536 | 0.40131384 | -0.586789 | 0.55734544 | 0.84861484 |
| JPH1 | 147.128434 | 0.881919338 | 0.51846641 | 1.70101539 | 0.0889401 | 0.53027486 |
| GDAP1 | 743.714542 | 0.239471904 | 0.46123997 | 0.51919157 | 0.60362716 | 0.86810591 |
| MIR2052HG | 172.780328 | 0.546698334 | 0.42545975 | 1.28495901 | 0.19880661 | 0.64597062 |
| CASC9 | 300.726265 | 0.737770706 | 0.82316536 | 0.89626063 | 0.37011358 | 0.76669785 |
| HNF4G | 2477.09919 | 0.248533842 | 0.21087627 | 1.17857664 | 0.2385668 | 0.67658923 |
| PEX2 | 297.43842 | 0.168394878 | 0.21027546 | 0.8008299 | 0.42323013 | 0.78894887 |
| PKIA-AS1 | 49.4925594 | 3.762660031 | 0.95135722 | 3.95504439 | 7.65E-05 | 0.02497299 |
| ZC2HC1A | 330.98787 | 0.139779061 | 0.44790885 | 0.31207032 | 0.75498708 | 0.92799282 |
| IL7 | 742.732253 | 1.579026698 | 0.58286588 | 2.70907384 | 0.00674713 | 0.23083391 |
| AC083837.1 | 81.0791661 | 0.701503238 | 0.84082281 | 0.83430567 | 0.40410876 | 0.78358009 |
| TPD52 | 2472.33103 | -0.663717251 | 0.47731401 | -1.3905254 | 0.16436939 | 0.62065658 |
| AC034114.2 | 345.020676 | -0.010840354 | 0.84275515 | -0.012863 | 9.90E-01 | 9.97E-01 |
| ZBTB10 | 556.176948 | 0.302530937 | 0.42348539 | 0.71438341 | 0.47499013 | 0.81564184 |
| ZNF704 | 1814.60611 | 0.626353155 | 0.28985037 | 2.16095342 | 0.03069894 | 0.40446361 |
| PAG1 | 79.5238606 | -0.052128632 | 0.53016583 | -0.0983251 | 9.22E-01 | 0.97939204 |
| IMPA1 | 412.17854 | -0.267945791 | 0.32644579 | -0.8207972 | 0.41176179 | 0.78632759 |
| ZFAND1 | 429.918869 | 0.145189076 | 0.25162639 | 0.57700257 | 0.56393772 | 0.85149031 |
| CHMP4C | 185.580497 | -0.112004801 | 0.26612826 | -0.4208678 | 0.67385165 | 0.89799059 |
| SNX16 | 283.014324 | -0.528707423 | 0.35625734 | -1.4840604 | 0.13779289 | 0.59474076 |
| RALYL | 4557.65745 | 0.670664866 | 0.73358158 | 0.91423352 | 3.61E-01 | 0.76009788 |
| LRRCC1 | 43.9394372 | 0.199740495 | 0.6130752 | 0.32580097 | 0.74457495 | 0.92369344 |
| E2F5 | 292.437301 | 0.213244082 | 0.29792857 | 0.71575574 | 0.47414218 | 0.8151865 |
| C8orf59 | 262.472797 | -0.278316818 | 0.3731705 | -0.7458168 | 0.4557781 | 0.80465616 |
| CA13 | 649.645079 | 0.130911611 | 0.31343762 | 0.41766401 | 0.67619279 | 0.89835672 |
| CA1 | 204.019271 | 0.552252713 | 0.58746332 | 0.94006331 | 0.34718509 | 0.75209642 |
| CA3 | 930.081806 | 0.792858614 | 0.30418776 | 2.60647766 | 0.00914788 | 0.25831482 |
| CA2 | 486.12606 | 0.054218785 | 0.53444479 | 0.1014488 | 0.9191942 | 0.97905003 |
| AC084128.1 | 141.829805 | 2.271072979 | 1.5397228 | 1.47498821 | 1.40E-01 | 5.95E-01 |
| SLC7A13 | 982.474556 | 3.072779769 | 1.14169404 | 2.69142141 | 7.11E-03 | 2.33E-01 |
| WWP1 | 3092.84256 | 0.358895626 | 0.21907953 | 1.63819791 | 0.10138042 | 0.54666458 |
| RMDN1 | 783.30861 | 0.293982069 | 0.33895742 | 0.86731268 | 0.38577071 | 0.77317162 |
| CPNE3 | 613.23601 | 0.109901143 | 0.19431761 | 0.5655748 | 0.57168286 | 0.85399468 |
| CNGB3 | 77.641552 | 0.514646921 | 0.4453547 | 1.15558884 | 0.24784938 | 0.68482118 |
| CNBD1 | 45.1459322 | -0.506503092 | 0.39240448 | -1.2907679 | 0.19678419 | 0.64558745 |
| AF117829.1 | 1479.26411 | 0.840279341 | 0.4148952 | 2.02528092 | 0.04283853 | 0.43918301 |
| RIPK2 | 287.611193 | -0.006430564 | 0.35658237 | -0.0180339 | 0.98561182 | 0.99607618 |
| OSGIN2 | 413.100355 | -0.506657455 | 0.35557098 | -1.4249123 | 0.15418256 | 0.61099212 |
| NBN | 424.696735 | -0.024180567 | 0.31691995 | -0.0762987 | 0.93918149 | 0.98435387 |
| DECR1 | 1049.99419 | 0.569583118 | 0.34409473 | 1.65530906 | 0.09786182 | 0.5411478 |
| CALB1 | 44.1589696 | 2.088475533 | 0.79855304 | 2.61532476 | 0.00891427 | 0.25615898 |
| LINC00534 | 111.100083 | 0.690965043 | 0.70521833 | 0.97978883 | 0.32719037 | 0.73746094 |
| AB015752.1 | 57.1318015 | -0.485160808 | 0.58579371 | -0.828211 | 0.407551 | 0.78481094 |
| TMEM64 | 482.780566 | 0.00373627 | 0.38408645 | 0.00972768 | 0.99223856 | 0.9976934 |
| NECAB1 | 203.931147 | 0.191240447 | 0.78108693 | 0.24483888 | 0.80658119 | 0.94585363 |
| PIP4P2 | 153.503503 | 0.240433749 | 0.33261803 | 0.72285241 | 0.46977056 | 0.81322606 |
| OTUD6B-AS1 | 207.771376 | 0.576031859 | 0.42267236 | 1.36283303 | 0.17293514 | 0.62610194 |
| OTUD6B | 292.488372 | -0.039810489 | 0.27126383 | -0.1467593 | 0.88332201 | 0.96846661 |
| LRRC69 | 610.182163 | 0.141834881 | 0.22510991 | 0.63006947 | 0.52864913 | 0.83761342 |
| SLC26A7 | 150.073594 | 0.770181019 | 0.41164321 | 1.87099168 | 0.06134623 | 0.4813972 |
| AC117834.1 | 85.8950126 | 0.895425082 | 0.78309156 | 1.14344877 | 0.25285231 | 0.68887358 |
| TRIQK | 510.400311 | 0.336315206 | 0.3010237 | 1.1172383 | 0.26389245 | 0.69567438 |
| LINC00535 | 315.316725 | 0.112917036 | 0.3888303 | 0.29040184 | 0.77150883 | 0.93316533 |

| | | | | | | |
|---|---|---|---|---|---|---|
| FAM92A | 627.101436 | -0.363322195 | 0.31439244 | -1.1556327 | 0.24783142 | 0.68482118 |
| AC120053.1 | 32.809899 | 0.63130995 | 0.51605897 | 1.22332909 | 0.22120543 | 0.66263062 |
| RBM12B | 419.799401 | 0.071067715 | 0.23603128 | 0.30109447 | 0.76334246 | 0.93038725 |
| TMEM67 | 719.646438 | -0.012077875 | 0.29532497 | -0.0408969 | 0.96737809 | 0.99168735 |
| PDP1 | 233.806335 | -0.275654748 | 0.40627926 | -0.6784859 | 0.49746366 | 0.82313869 |
| GEM | 359.199341 | -1.25436254 | 0.73562777 | -1.7051593 | 8.82E-02 | 5.28E-01 |
| RAD54B | 48.134076 | -0.51367986 | 0.62434426 | -0.822751 | 0.41064961 | 0.78592213 |
| VIRMA | 540.875447 | -0.229523855 | 0.17182626 | -1.3357903 | 0.18161783 | 0.6318513 |
| AC023632.2 | 48.2745078 | -0.120094052 | 0.46466881 | -0.2584509 | 0.79605897 | 0.94252227 |
| ESRP1 | 59.9595897 | -0.119227327 | 0.88566162 | -0.1346195 | 0.89291272 | 0.97244197 |
| DPY19L4 | 419.364674 | 0.196166325 | 0.29867156 | 0.65679614 | 0.51131201 | 0.83007666 |
| INTS8 | 568.774087 | 0.274050625 | 0.20555145 | 1.3332459 | 0.18245112 | 0.63216521 |
| NDUFAF6 | 681.9126 | 0.254551709 | 0.28918364 | 0.88024243 | 0.37872799 | 0.77061593 |
| TP53INP1 | 559.383829 | 0.419482731 | 0.44576484 | 0.94104043 | 0.34668414 | 0.75166079 |
| PLEKHF2 | 571.236968 | 0.251445812 | 0.55518224 | 0.4529068 | 0.65061585 | 0.8887773 |
| C8orf37-AS1 | 774.206654 | 1.089061137 | 0.77134176 | 1.4119048 | 0.157978 | 0.61369956 |
| C8orf37 | 58.4875666 | 0.464155188 | 0.5757885 | 0.80612098 | 0.4201731 | 0.78798479 |
| UQCRB | 760.112438 | -0.309410801 | 0.53365287 | -0.5797979 | 0.56205093 | 0.85033466 |
| MTERF3 | 164.593006 | -0.166398696 | 0.26582872 | -0.6259621 | 0.53133982 | 0.83789294 |
| PTDSS1 | 277.860054 | -0.418203754 | 0.37128193 | -1.1263779 | 0.2600056 | 0.69280433 |
| SDC2 | 848.488272 | -0.024820564 | 0.49115804 | -0.0505348 | 0.95969623 | 0.98912673 |
| CPQ | 2310.16834 | 0.367722929 | 0.53226945 | 0.6908586 | 0.4896544 | 0.81970016 |
| MTDH | 2334.73066 | -0.623894489 | 0.37336116 | -1.6710214 | 0.09471745 | 0.53824794 |
| LAPTM4B | 897.716677 | 0.112526469 | 0.40692751 | 0.27652706 | 0.78214327 | 0.93757221 |
| MATN2 | 113.706089 | 0.608820987 | 0.5863853 | 1.03826099 | 0.29914856 | 0.71852421 |
| RPL30 | 858.565288 | -0.036136037 | 0.46152994 | -0.0782962 | 0.93759245 | 0.98435387 |
| ERICH5 | 133.905157 | -0.354943211 | 0.29501582 | -1.2031328 | 0.22892494 | 0.66705629 |
| RIDA | 816.338699 | 0.818100921 | 0.49469836 | 1.65373687 | 0.09818099 | 0.54193402 |
| POP1 | 415.16807 | 0.447034105 | 0.33903586 | 1.31854519 | 0.18732121 | 0.63858939 |
| NIPAL2 | 180.162047 | -0.514685067 | 0.57753543 | -0.8911749 | 0.37283536 | 0.76906762 |
| AP003438.1 | 35.679791 | 0.227168237 | 0.70336808 | 0.32297206 | 0.7467164 | 0.92448528 |
| STK3 | 5068.6543 | -0.477313649 | 0.23775416 | -2.0075933 | 0.04468653 | 0.44360789 |
| AC104986.2 | 289.178671 | 0.236929426 | 0.29300768 | 0.80861166 | 0.41873856 | 0.78765602 |
| VPS13B | 3968.7802 | 0.044083045 | 0.15498185 | 0.28444005 | 0.77607317 | 0.9352344 |
| COX6C | 667.937906 | -0.355283031 | 0.33371395 | -1.0646334 | 0.28704184 | 0.71118494 |
| POLR2K | 157.069364 | -0.260278707 | 0.4228209 | -0.6155767 | 0.53817392 | 0.84049498 |
| SPAG1 | 689.397644 | 0.3774254 | 0.2745447 | 1.37473203 | 0.16921454 | 0.62326131 |
| RNF19A | 4478.21654 | -0.090000779 | 0.39532928 | -0.2276603 | 0.81991034 | 0.94992064 |
| AP000424.1 | 189.245031 | -0.65761942 | 0.89519215 | -0.7346126 | 0.46257547 | 0.8082072 |
| ANKRD46 | 501.601037 | 0.236368173 | 0.39336708 | 0.60088448 | 0.54791693 | 0.84486687 |
| PABPC1 | 1551.88672 | -0.541654613 | 0.46890059 | -1.1551587 | 0.24802543 | 0.68482118 |
| YWHAZ | 1850.93007 | -0.243987758 | 0.26721881 | -0.9130636 | 0.36120912 | 0.76009788 |
| AP003469.4 | 49.6280418 | 0.853681398 | 0.37604503 | 2.27015739 | 0.02319804 | 0.36970978 |
| ZNF706 | 606.669489 | 0.233639934 | 0.23423744 | 0.99744915 | 0.31854654 | 0.73064215 |
| GRHL2 | 93.9949685 | 0.313689421 | 0.91336329 | 0.3434443 | 0.73126423 | 0.9190831 |
| NCALD | 333.269547 | -0.156949651 | 0.51046962 | -0.3074613 | 0.75849228 | 0.9287702 |
| RRM2B | 299.694632 | 0.564820361 | 0.24214711 | 2.33255051 | 0.01967175 | 0.34523522 |
| UBR5-AS1 | 58.3835686 | 0.409945071 | 0.5873126 | 0.6980015 | 0.48517626 | 0.81839266 |
| UBR5 | 2840.04183 | -0.055028025 | 0.21779973 | -0.2526542 | 0.80053542 | 0.94403774 |
| KLF10 | 364.528707 | 0.240582498 | 0.42492446 | 0.5661771 | 0.5712734 | 0.85399468 |
| AZIN1 | 575.643353 | -0.196783569 | 0.25643945 | -0.7673685 | 0.44286243 | 0.7993765 |
| AZIN1-AS1 | 642.081866 | 0.900482394 | 0.56633192 | 1.59002585 | 0.11182898 | 0.56455091 |
| AP003354.2 | 96.6625946 | 0.877000623 | 0.61560627 | 1.42461289 | 0.15426913 | 0.61099212 |
| ATP6V1C1 | 1915.69365 | -0.805000862 | 0.32497934 | -2.4770832 | 0.0132461 | 0.29875272 |

| | | | | | | |
|---|---|---|---|---|---|---|
| BAALC-AS1 | 167.724807 | 1.212051267 | 0.5516188 | 2.19726242 | 0.02800171 | 0.39125325 |
| FZD6 | 538.162146 | -0.095850343 | 0.24757703 | -0.3871536 | 0.69864249 | 0.90696517 |
| SLC25A32 | 373.282488 | 0.432726544 | 0.24975203 | 1.73262472 | 0.08316239 | 0.51935659 |
| DCAF13 | 284.095 | -0.509103024 | 0.39572497 | -1.2865072 | 0.19826611 | 0.64582202 |
| RIMS2 | 1376.92047 | 2.544540892 | 0.72208239 | 3.52389275 | 4.25E-04 | 7.29E-02 |
| DPYS | 6905.69449 | 0.718233051 | 0.72102081 | 0.99613359 | 0.31918524 | 0.73132207 |
| LRP12 | 379.63418 | -0.196084625 | 0.30114578 | -0.6511286 | 0.51496349 | 0.83169608 |
| ZFPM2 | 110.39233 | 0.381464054 | 0.56659329 | 0.67325904 | 0.5007825 | 0.82564559 |
| AC027031.2 | 41.1781871 | -0.812564253 | 0.52554525 | -1.5461357 | 0.12207181 | 0.58055167 |
| OXR1 | 12335.573 | -0.536992376 | 0.36991026 | -1.4516828 | 0.14658981 | 0.60383734 |
| AC090579.1 | 1032.84964 | -0.387037711 | 0.72271776 | -0.5355309 | 0.59228278 | 0.86382106 |
| ANGPT1 | 120.573322 | -0.750096518 | 0.47611261 | -1.5754603 | 1.15E-01 | 5.72E-01 |
| EIF3E | 1512.85871 | 0.300408915 | 0.19290816 | 1.5572639 | 0.11940784 | 0.57652003 |
| EMC2 | 757.64781 | 0.674943573 | 0.42746483 | 1.57894526 | 0.11434861 | 0.56984472 |
| NUDCD1 | 415.915073 | -0.367140245 | 0.32809314 | -1.1190123 | 0.26313491 | 0.69535188 |
| ENY2 | 497.704584 | -0.340073959 | 0.34432811 | -0.9876451 | 0.32332651 | 0.73538915 |
| EBAG9 | 179.595948 | 0.246261336 | 0.33822564 | 0.72809777 | 0.46655374 | 0.81098649 |
| LINC01609 | 23.6686049 | -0.509720457 | 1.0385222 | -0.4908133 | 0.62355854 | 0.87772094 |
| TRPS1 | 8619.06085 | 0.245597855 | 0.31869217 | 0.77064288 | 0.44091864 | 0.79826598 |
| EIF3H | 1061.4195 | -0.201013695 | 0.28725904 | -0.6997645 | 0.48407436 | 0.81839266 |
| UTP23 | 187.349311 | 0.278240036 | 0.34979645 | 0.79543413 | 0.42636102 | 0.78895108 |
| RAD21 | 637.038503 | 0.14885068 | 0.21462778 | 0.69352942 | 0.48797737 | 0.81919239 |
| SLC30A8 | 1350.98031 | 1.46470711 | 0.96630459 | 1.51578201 | 1.30E-01 | 5.88E-01 |
| MED30 | 85.0243183 | 0.022758776 | 0.2681489 | 0.08487365 | 0.93236184 | 0.98306798 |
| EXT1 | 6197.52766 | -0.320922353 | 0.34128132 | -0.9403455 | 0.34704037 | 0.75207626 |
| SAMD12 | 4402.85341 | 0.645957298 | 0.3596387 | 1.79612844 | 0.07247409 | 0.502734 |
| AC023590.1 | 300.905875 | 0.8527332 | 0.54524435 | 1.56394688 | 0.11783004 | 0.57396383 |
| SAMD12-AS1 | 284.499907 | 0.064571102 | 0.51855103 | 0.12452217 | 0.90090184 | 0.97479173 |
| TNFRSF11B | 943.665342 | 1.46286573 | 0.64859526 | 2.25543697 | 0.02410591 | 0.37387451 |
| COLEC10 | 162.164002 | 1.493866975 | 0.62852168 | 2.37679465 | 0.01746381 | 0.33064634 |
| MAL2 | 279.57998 | -0.01349863 | 0.4479621 | -0.0301334 | 0.97596065 | 0.99429832 |
| ENPP2 | 1093.6931 | 0.216633 | 0.47071493 | 0.46022122 | 0.64535744 | 0.88687652 |
| TAF2 | 1219.86325 | -0.362859362 | 0.25636685 | -1.4153911 | 0.15695385 | 0.61255952 |
| DEPTOR | 2810.8666 | 0.369620468 | 0.62741415 | 0.5891172 | 0.55578266 | 0.84861484 |
| AC091563.1 | 149.653451 | 0.644414411 | 0.47829743 | 1.34730895 | 0.17788075 | 0.62808126 |
| COL14A1 | 154.384208 | 0.638882619 | 0.63027562 | 1.01365592 | 0.31074696 | 0.72537613 |
| MRPL13 | 244.930957 | -0.054731998 | 0.32284197 | -0.1695319 | 0.86537832 | 0.96428252 |
| MTBP | 90.2134552 | -0.405205925 | 0.45037566 | -0.8997065 | 0.36827645 | 0.76615653 |
| SNTB1 | 1701.27827 | 0.397252247 | 0.63340604 | 0.62716839 | 0.53054886 | 0.83789294 |
| ZHX2 | 3542.15629 | 0.243522988 | 0.32952564 | 0.73901073 | 0.45990048 | 0.8069452 |
| AC104316.1 | 127.513017 | 0.074490611 | 0.56110319 | 0.13275742 | 0.89438523 | 0.97272276 |
| DERL1 | 379.28463 | -0.103097853 | 0.27801041 | -0.3708417 | 7.11E-01 | 0.91134938 |
| TBC1D31 | 126.69692 | 0.413380762 | 0.25463065 | 1.62345246 | 0.10449272 | 0.55066578 |
| C8orf76 | 86.4987244 | -0.19666493 | 0.32380905 | -0.6073485 | 0.54361968 | 0.84396451 |
| ZHX1-C8orf7( | 42.6677379 | 1.146007629 | 0.46008565 | 2.49085712 | 0.01274354 | 0.29875272 |
| ATAD2 | 88.0906772 | -0.097911751 | 0.32777777 | -0.2987138 | 0.76515841 | 0.93083208 |
| WDYHV1 | 139.99058 | -0.19075055 | 0.46330287 | -0.4117189 | 0.68054547 | 0.89947163 |
| FBXO32 | 375.063927 | -1.353222312 | 0.47050984 | -2.8760765 | 0.00402652 | 0.20256898 |
| FAM91A1 | 980.837734 | 0.126513115 | 0.21431742 | 0.5903072 | 0.55498471 | 0.84801165 |
| FER1L6 | 330.569552 | 2.04226317 | 0.99394043 | 2.05471386 | 0.03990665 | 0.42969399 |
| TMEM65 | 1227.17538 | 0.269648076 | 0.28922568 | 0.93231029 | 0.35117619 | 0.75518811 |
| TRMT12 | 99.5773997 | -0.316196308 | 0.34259349 | -0.922949 | 0.35603379 | 0.75841231 |
| RNF139-AS1 | 47.5350184 | 0.296876109 | 0.425248 | 0.69812465 | 0.48509924 | 0.81839266 |
| RNF139 | 149.25482 | 0.041458896 | 0.23494437 | 0.17646261 | 0.85993052 | 0.96283855 |

| | | | | | | |
|---|---|---|---|---|---|---|
| TATDN1 | 721.669729 | 0.040205268 | 0.18357685 | 0.21901056 | 0.82664182 | 0.9519561 |
| NDUFB9 | 385.167561 | -0.095956767 | 0.32456779 | -0.2956448 | 0.76750139 | 0.93169958 |
| MTSS1 | 1776.21713 | 0.472194845 | 0.43249379 | 1.09179565 | 2.75E-01 | 0.70187532 |
| SQLE | 184.746074 | 1.324668262 | 0.55207502 | 2.39943524 | 0.01642038 | 0.32338248 |
| WASHC5 | 745.694603 | -0.446161137 | 0.28255583 | -1.5790194 | 0.1143316 | 0.56984472 |
| NSMCE2 | 4383.32404 | -0.519537183 | 0.26532383 | -1.9581248 | 0.05021536 | 0.45824658 |
| TRIB1 | 242.343446 | -0.585789645 | 0.41490508 | -1.4118643 | 0.15798994 | 0.61369956 |
| AC091114.1 | 43.372765 | -0.917575352 | 0.64804579 | -1.4159113 | 0.15680148 | 0.61255952 |
| FAM84B | 329.777823 | -0.679606311 | 0.47428025 | -1.4329214 | 0.15188031 | 0.60996876 |
| PCAT1 | 4734.64844 | 0.27015289 | 0.30134623 | 0.8964867 | 0.36999288 | 0.76669785 |
| MYC | 154.629714 | -0.91399603 | 0.58613263 | -1.5593673 | 1.19E-01 | 5.76E-01 |
| PVT1 | 801.237391 | 0.222127357 | 0.54162169 | 0.41011533 | 6.82E-01 | 9.00E-01 |
| CCDC26 | 261.143969 | 1.649696565 | 0.79556586 | 2.07361408 | 0.03811517 | 0.42339465 |
| AC103718.1 | 44.5797015 | 0.709118546 | 0.96239164 | 0.73682949 | 0.46122604 | 0.80772012 |
| FAM49B | 1810.33898 | 0.07857282 | 0.31105412 | 0.25260177 | 0.80057596 | 0.94403774 |
| ASAP1 | 2275.31567 | -0.44712851 | 0.33423204 | -1.3377787 | 0.18096857 | 0.6318513 |
| ADCY8 | 179.412715 | -2.221577199 | 0.7971012 | -2.7870704 | 0.00531869 | 0.21873395 |
| EFR3A | 1880.73741 | 0.102801717 | 0.27508554 | 0.37370818 | 0.70862144 | 0.91082291 |
| KCNQ3 | 1115.1577 | 0.553324472 | 0.45706032 | 1.21061585 | 2.26E-01 | 0.66688879 |
| LRRC6 | 1175.19962 | 0.484395213 | 0.46018506 | 1.0526096 | 0.29251995 | 0.71399546 |
| PHF20L1 | 1078.9634 | 0.005395618 | 0.15324489 | 0.03520912 | 0.97191299 | 0.99240251 |
| TG | 258.019478 | 1.272728198 | 0.47025989 | 2.70643581 | 0.00680097 | 0.23083391 |
| SLA | 61.3481978 | 1.759384385 | 0.8207938 | 2.1435157 | 0.03207171 | 0.40988962 |
| NDRG1 | 6655.33802 | -1.232645109 | 0.66410197 | -1.8561082 | 0.06343814 | 0.48572144 |
| ST3GAL1 | 1576.10521 | -0.344078621 | 0.42058743 | -0.8180906 | 0.41330546 | 0.78632759 |
| AC105180.1 | 71.2126509 | 1.02971461 | 0.69018207 | 1.49194634 | 0.13571321 | 0.59381224 |
| ZFAT | 1537.74844 | 0.467091517 | 0.5290431 | 0.88289879 | 0.37729096 | 0.77048581 |
| AC083843.3 | 249.120531 | -0.153379502 | 0.50784682 | -0.3020192 | 0.76263741 | 0.93022076 |
| KHDRBS3 | 2309.59342 | -0.332144897 | 0.58662751 | -0.5661939 | 0.571262 | 0.85399468 |
| LINC02055 | 106.114394 | 0.267560893 | 1.19850425 | 0.22324568 | 0.8233443 | 0.95131214 |
| AC046195.1 | 483.443161 | -1.362837096 | 1.07144871 | -1.2719574 | 0.20338826 | 0.64881965 |
| FAM135B | 555.767628 | 1.904095453 | 0.81993547 | 2.32225038 | 0.02021946 | 0.34861397 |
| TRAPPC9 | 3583.41582 | 0.178913528 | 0.16380404 | 1.09224121 | 0.27472709 | 0.70166356 |
| AC107375.1 | 68.9768138 | -0.302857169 | 0.39346482 | -0.7697185 | 0.44146686 | 0.79862562 |
| CHRAC1 | 108.069513 | -0.843592215 | 0.34662351 | -2.4337421 | 0.01494364 | 0.30894792 |
| AGO2 | 980.010734 | -0.45636754 | 0.43282969 | -1.0543813 | 0.29170837 | 0.71337138 |
| PTK2 | 6494.74976 | -0.349137899 | 0.21147543 | -1.650962 | 0.09874634 | 0.54236074 |
| DENND3 | 283.834085 | 0.456724719 | 0.4552113 | 1.00332466 | 0.31570424 | 0.72798933 |
| SLC45A4 | 738.968212 | 0.400652015 | 0.49307563 | 0.81255692 | 0.41647214 | 0.78632759 |
| PTP4A3 | 70.0920085 | -0.607092878 | 0.74912515 | -0.8104025 | 0.41770889 | 0.78669082 |
| TSNARE1 | 437.372674 | -0.365074662 | 0.31396761 | -1.1627781 | 0.24491954 | 0.68250191 |
| JRK | 156.525837 | -0.456272876 | 0.40930813 | -1.1147418 | 0.26496108 | 0.69615135 |
| THEM6 | 92.1498667 | -0.092517297 | 0.45747313 | -0.2022355 | 0.83973264 | 0.9556011 |
| LY6E-DT | 252.164804 | 0.990441685 | 0.85121316 | 1.16356482 | 0.24460041 | 0.68250191 |
| LY6E | 812.932338 | -0.37606934 | 0.75776429 | -0.496288 | 0.61969122 | 0.87559092 |
| AC138696.1 | 60.6914751 | -0.036081713 | 0.54662855 | -0.0660077 | 0.94737167 | 0.98646931 |
| GLI4 | 141.193957 | -0.120425672 | 0.36179834 | -0.332853 | 0.73924523 | 0.92108726 |
| ZNF696 | 48.7582261 | -0.295649364 | 0.45925523 | -0.6437583 | 0.51973218 | 0.8335052 |
| TOP1MT | 231.97118 | -0.086580047 | 0.2749993 | -0.3148373 | 0.75288516 | 0.92746251 |
| RHPN1 | 313.091057 | 0.470027481 | 0.53041462 | 0.88615107 | 0.37553613 | 0.7697436 |
| ZC3H3 | 650.020284 | 0.139167597 | 0.463911 | 0.29998771 | 0.76418653 | 0.93083208 |
| AC067930.4 | 59.6868089 | 1.627278591 | 0.75830804 | 2.14593346 | 0.03187829 | 0.40988962 |
| GSDMD | 381.877369 | 0.588488449 | 0.60752475 | 0.96866581 | 0.33271196 | 0.73988623 |
| MROH6 | 57.509643 | 0.425153346 | 0.72193681 | 0.58890659 | 0.55592394 | 0.84861484 |

| | | | | | | |
|---|---|---|---|---|---|---|
| NAPRT | 249.045267 | -0.03722516 | 0.36602718 | -0.1017005 | 0.91899438 | 0.97891327 |
| EEF1D | 1133.12635 | -0.525507768 | 0.46941402 | -1.1194974 | 0.26292801 | 0.69505103 |
| TIGD5 | 44.3407404 | 0.344151682 | 0.34036782 | 1.01111697 | 0.31196045 | 0.72610662 |
| PYCR3 | 41.1475771 | -0.695306811 | 0.67897436 | -1.0240546 | 0.30580949 | 0.72122 |
| TSTA3 | 178.262924 | -0.689039782 | 0.56094562 | -1.228354 | 0.21931412 | 0.6618535 |
| ZNF623 | 250.310317 | -0.569336853 | 0.37860653 | -1.5037692 | 0.1326408 | 0.59142019 |
| ZNF707 | 85.2942149 | -0.347325906 | 0.3879106 | -0.8953762 | 0.37058604 | 0.76708288 |
| FAM83H | 145.464302 | -0.01837456 | 0.46881581 | -0.0391936 | 0.96873607 | 0.99187457 |
| IQANK1 | 117.00664 | 0.172326117 | 0.65545957 | 0.26290884 | 0.79262084 | 0.94154414 |
| SCRIB | 152.157188 | -0.010612605 | 0.23352398 | -0.0454455 | 0.96375225 | 0.99012655 |
| PUF60 | 392.658162 | -0.374213886 | 0.45172716 | -0.8284069 | 0.40744012 | 0.78481094 |
| NRBP2 | 826.060198 | 0.113382609 | 0.6783773 | 0.16713798 | 0.86726149 | 0.96486219 |
| EPPK1 | 149.547099 | 0.398066968 | 0.75599917 | 0.52654419 | 0.59851016 | 0.86646266 |
| PLEC | 561.345786 | -0.055348465 | 0.33015798 | -0.1676424 | 0.86686464 | 0.96486219 |
| PARP10 | 512.965975 | 0.281763419 | 0.5392997 | 0.52246166 | 0.60134893 | 0.86756024 |
| GRINA | 504.037369 | 0.09508954 | 0.39109587 | 0.24313614 | 0.80789994 | 0.94613779 |
| SPATC1 | 113.743325 | 0.731828787 | 0.9672992 | 0.75656921 | 0.449308 | 0.80309271 |
| OPLAH | 117.461087 | 0.579131476 | 0.49770107 | 1.16361308 | 0.24458085 | 0.68250191 |
| EXOSC4 | 66.317074 | -0.62130054 | 0.50782831 | -1.2234461 | 0.22116127 | 0.66263062 |
| GPAA1 | 300.003347 | 0.237604062 | 0.45466123 | 0.52259583 | 0.60125554 | 0.86755264 |
| CYC1 | 347.581551 | 0.129530193 | 0.48702625 | 0.26596142 | 0.79026892 | 0.94048261 |
| SHARPIN | 316.635385 | -0.17324934 | 0.19084028 | -0.9078238 | 0.36397134 | 0.76272056 |
| MAF1 | 249.0201 | -0.482940465 | 0.46531468 | -1.0378793 | 0.29932626 | 0.71852421 |
| HGH1 | 39.42282 | -0.462931989 | 0.57518902 | -0.8048345 | 0.42091517 | 0.78872573 |
| MROH1 | 1188.51041 | -0.975577574 | 0.53060797 | -1.8386033 | 0.06597355 | 0.48798003 |
| BOP1 | 54.9833818 | -0.897862181 | 0.59952925 | -1.497612 | 0.13423409 | 0.59240843 |
| SCX | 85.9243052 | -0.850935117 | 0.69578777 | -1.2229809 | 0.22133693 | 0.66263062 |
| HSF1 | 456.768472 | -0.346157276 | 0.31518459 | -1.0982684 | 0.2720873 | 0.70056282 |
| DGAT1 | 110.688518 | 0.007654709 | 0.39102128 | 0.0195762 | 0.98438145 | 0.99527304 |
| SLC52A2 | 129.185885 | -0.36534102 | 0.54836157 | -0.6662411 | 0.505257 | 0.82715271 |
| FBXL6 | 55.1619011 | -0.40102531 | 0.46216081 | -0.8677181 | 0.38554867 | 0.77317162 |
| ADCK5 | 76.8876255 | -0.212767903 | 0.33350151 | -0.6379818 | 0.52348551 | 0.83533627 |
| CPSF1 | 161.484403 | -0.253589464 | 0.39883997 | -0.6358176 | 0.52489532 | 0.835938 |
| SLC39A4 | 133.800117 | -0.109037221 | 0.47794126 | -0.2281394 | 0.81953789 | 0.94990455 |
| VPS28 | 504.505498 | -0.117039109 | 0.41738987 | -0.2804072 | 0.77916514 | 0.93646866 |
| TONSL | 34.9996981 | 0.117645396 | 0.51405688 | 0.22885677 | 0.81898024 | 0.94990455 |
| CYHR1 | 276.560801 | -0.098524442 | 0.21144204 | -0.4659643 | 6.41E-01 | 0.88403746 |
| KIFC2 | 274.845033 | -0.309183867 | 0.36794274 | -0.8403043 | 0.40073778 | 0.78232083 |
| PPP1R16A | 776.858476 | -0.19888831 | 0.29089313 | -0.6837161 | 0.49415446 | 0.82191181 |
| GPT | 356.53452 | -0.917119547 | 0.64666335 | -1.4182334 | 0.15612264 | 0.61255389 |
| MFSD3 | 105.32283 | -0.244094118 | 0.38642241 | -0.6316769 | 0.52759801 | 0.83749692 |
| LRRC14 | 71.4598788 | -0.01909997 | 0.31827072 | -0.0600117 | 0.95214631 | 0.98766985 |
| ARHGAP39 | 197.524458 | -0.340941044 | 0.44008068 | -0.774724 | 0.4385028 | 0.7970505 |
| ZNF251 | 780.192564 | -0.30449926 | 0.30524432 | -0.9975591 | 0.31849319 | 0.73064202 |
| ZNF34 | 53.5719117 | 0.318355559 | 0.38889202 | 0.818622 | 0.41300211 | 0.78632759 |
| RPL8 | 1528.61949 | -0.023669797 | 0.54507396 | -0.0434249 | 0.96536281 | 0.99074368 |
| ZNF517 | 40.7104769 | -0.021716777 | 0.49853628 | -0.0435611 | 0.96525428 | 0.99074368 |
| ZNF7 | 263.786743 | -0.079331425 | 0.29450746 | -0.2693698 | 0.7876451 | 0.93953874 |
| COMMD5 | 102.317446 | -0.057370751 | 0.36355652 | -0.1578042 | 0.87461108 | 0.96615698 |
| ZNF250 | 316.156609 | 0.946069264 | 0.4997581 | 1.89305438 | 0.05835064 | 0.47778423 |
| ZNF16 | 184.515639 | -0.043752085 | 0.44059415 | -0.0993025 | 0.92089812 | 0.97939204 |
| C8orf33 | 127.919731 | -0.754991741 | 0.45272515 | -1.6676603 | 0.09538319 | 0.53824794 |
| WASHC1 | 1218.62503 | -0.212159388 | 0.15359808 | -1.3812632 | 1.67E-01 | 0.62242897 |
| CBWD1 | 427.84323 | -0.030051755 | 0.36180174 | -0.0830614 | 0.93380273 | 0.98355048 |

| | | | | | | |
|---|---|---|---|---|---|---|
| C9orf66 | 125.333893 | -0.369764901 | 0.85288036 | -0.4335484 | 0.66461642 | 0.89315615 |
| DOCK8 | 3073.93559 | 0.134593541 | 0.47313046 | 0.28447448 | 0.77604679 | 0.9352344 |
| AL161725.1 | 55.4646595 | 0.406848905 | 0.77472124 | 0.52515522 | 0.59947529 | 0.86692818 |
| KANK1 | 6864.97893 | 0.085970663 | 0.18860528 | 0.45582321 | 0.64851711 | 0.88787189 |
| SMARCA2 | 1207.89593 | 0.2173986 | 0.24491561 | 0.88764698 | 0.37473068 | 0.7697436 |
| VLDLR-AS1 | 416.644835 | -0.29357711 | 0.60000879 | -0.489288 | 0.62463781 | 0.87832483 |
| VLDLR | 361.570424 | -0.04507091 | 0.39555693 | -0.1139429 | 0.90928305 | 0.97680747 |
| KCNV2 | 65.0834766 | 0.286779801 | 0.65210867 | 0.43977302 | 0.66010151 | 0.89099078 |
| PUM3 | 3727.95689 | -0.226235139 | 0.42931798 | -0.526964 | 0.59821856 | 0.86646266 |
| RFX3 | 1729.08438 | 0.267986527 | 0.33042779 | 0.81102901 | 0.41734901 | 0.78652162 |
| RFX3-AS1 | 1793.71957 | 0.686888683 | 0.42218074 | 1.62700146 | 0.10373679 | 0.55022042 |
| GLIS3 | 11029.8095 | -0.441499004 | 0.24820413 | -1.7787738 | 0.07527685 | 0.50718602 |
| SLC1A1 | 4135.80457 | 0.350019256 | 0.35233879 | 0.99341674 | 0.32050691 | 0.73210121 |
| SPATA6L | 568.399911 | 0.458265362 | 0.42634108 | 1.07487969 | 0.2824286 | 0.70682473 |
| PLPP6 | 39.1961302 | -0.145281434 | 0.37117189 | -0.3914128 | 0.69549212 | 0.90589033 |
| CDC37L1 | 543.924032 | 0.016366596 | 0.21265572 | 0.07696288 | 0.93865308 | 0.98435387 |
| AK3 | 1540.61908 | 0.046235749 | 0.42869584 | 0.10785211 | 0.91411301 | 0.9774502 |
| RCL1 | 677.608529 | -0.306615247 | 0.29509156 | -1.0390512 | 0.29878092 | 0.71852421 |
| JAK2 | 1375.57499 | 0.256633945 | 0.19007647 | 1.3501616 | 0.17696415 | 0.62726859 |
| PLGRKT | 348.811793 | 0.511374254 | 0.25059526 | 2.0406382 | 0.0412868 | 0.43221049 |
| RIC1 | 1445.9105 | 0.029799441 | 0.21608107 | 0.13790862 | 0.89031264 | 0.97225819 |
| ERMP1 | 384.077203 | 0.145248217 | 0.37592551 | 0.386375 | 0.69921897 | 0.90721166 |
| KIAA2026 | 1969.72703 | -0.142735959 | 0.18496611 | -0.7716871 | 0.44029977 | 0.79826598 |
| RANBP6 | 159.855357 | 0.116239972 | 0.30388871 | 0.38250836 | 0.70208433 | 0.90837679 |
| UHRF2 | 880.17458 | -0.40028543 | 0.26757577 | -1.4959704 | 0.13466137 | 0.59277997 |
| AL133480.1 | 79.2098039 | 0.1832052 | 0.53054378 | 0.34531589 | 0.7298569 | 0.91834786 |
| GLDC | 2406.96976 | 0.174245758 | 0.43269122 | 0.40270232 | 6.87E-01 | 0.90237605 |
| AL162411.1 | 511.810263 | 0.181550609 | 0.28346832 | 0.64046172 | 0.52187247 | 0.83445062 |
| AL354707.1 | 40.2944122 | 0.411707832 | 0.9127573 | 0.45105948 | 0.65194668 | 0.88935488 |
| KDM4C | 4958.76641 | -0.100658662 | 0.17734238 | -0.5675951 | 0.57030996 | 0.85331717 |
| DMAC1 | 151.183704 | 0.341536357 | 0.33034638 | 1.03387347 | 0.30119532 | 0.71879123 |
| PTPRD | 8095.60519 | 0.239985875 | 0.45566624 | 0.52667029 | 0.59842257 | 0.86646266 |
| LURAP1L-AS | 58.6436791 | -0.155923326 | 0.83328357 | -0.1871192 | 0.8515672 | 0.96044014 |
| LURAP1L | 329.704857 | -1.140755135 | 0.76107349 | -1.4988765 | 0.13390568 | 0.59240843 |
| MPDZ | 1311.29474 | 0.108556751 | 0.36348135 | 0.29865838 | 0.76520071 | 0.93083208 |
| NFIB | 9869.48894 | 0.567424675 | 0.37905509 | 1.49694515 | 0.13440753 | 0.59240843 |
| AL136366.1 | 43.0748469 | 0.866214225 | 0.50819885 | 1.70447891 | 0.08829165 | 0.52893954 |
| ZDHHC21 | 1527.3096 | -0.079203428 | 0.2502266 | -0.3165268 | 0.75160268 | 0.92647278 |
| FREM1 | 637.188704 | -0.227576201 | 0.8760252 | -0.2597827 | 7.95E-01 | 0.94252227 |
| TTC39B | 2249.862 | 0.989162848 | 0.54721279 | 1.80763839 | 0.0706628 | 0.4967563 |
| SNAPC3 | 847.364448 | -0.217967665 | 0.26567878 | -0.820418 | 0.41197784 | 0.78632759 |
| PSIP1 | 475.1466 | 0.110217391 | 0.31250519 | 0.35268979 | 0.72432101 | 0.91600702 |
| CCDC171 | 815.261446 | 0.259026154 | 0.568113 | 0.45594126 | 0.64843222 | 0.88787189 |
| BNC2 | 11483.2511 | 0.516751637 | 0.48997353 | 1.05465216 | 0.29158444 | 0.71337138 |
| AL162725.2 | 561.438807 | 3.386616326 | 0.96467783 | 3.51061901 | 0.00044706 | 0.07565327 |
| CNTLN | 980.585464 | 0.263073835 | 0.3519725 | 0.74742723 | 0.45480571 | 0.80465616 |
| SH3GL2 | 1720.73296 | -0.396654633 | 0.51557222 | -0.7693483 | 0.44168655 | 0.79881218 |
| ADAMTSL1 | 126.783115 | 0.10862696 | 0.48891576 | 0.2221793 | 0.82417431 | 0.95180295 |
| SAXO1 | 90.9155684 | 0.284190781 | 0.51307024 | 0.55390229 | 0.57964572 | 0.85816191 |
| RRAGA | 182.730032 | 0.048606963 | 0.36955733 | 0.13152753 | 8.95E-01 | 9.73E-01 |
| HAUS6 | 322.477931 | -0.090411075 | 0.27900214 | -0.3240515 | 0.74589901 | 0.92424857 |
| PLIN2 | 3559.12082 | -1.077953751 | 0.55240292 | -1.9513904 | 0.05101062 | 0.46060777 |
| DENND4C | 2861.43455 | -0.08635159 | 0.22404799 | -0.3854156 | 0.69992952 | 0.90721166 |
| RPS6 | 1710.19055 | 0.11402367 | 0.56227348 | 0.20279041 | 0.83929885 | 0.9556011 |

| | | | | | | |
|---|---|---|---|---|---|---|
| ACER2 | 186.653901 | 0.08120742 | 0.51689256 | 0.15710696 | 0.87516055 | 0.96615698 |
| MLLT3 | 1729.58156 | 0.693277903 | 0.39436586 | 1.75795619 | 0.07875496 | 0.51192014 |
| FOCAD | 1532.85137 | 0.133278206 | 0.3004267 | 0.4436297 | 0.65731033 | 0.89053559 |
| KLHL9 | 210.857112 | 0.01671514 | 0.31546859 | 0.05298512 | 0.95774377 | 0.98861313 |
| DMRTA1 | 133.742343 | 0.973486436 | 0.35542779 | 2.73891479 | 0.00616423 | 0.22637955 |
| LINC01239 | 990.554327 | 2.081835534 | 0.71373857 | 2.91680403 | 0.00353638 | 0.19467126 |
| AL391117.1 | 4151.85723 | 0.935827877 | 0.74652032 | 1.25358661 | 0.2099923 | 0.65441616 |
| TUSC1 | 55.5460305 | -0.420760564 | 0.44853411 | -0.9380793 | 0.34820366 | 0.75340609 |
| CAAP1 | 1175.69741 | 0.698503495 | 0.19691278 | 3.54727349 | 0.00038924 | 0.07113756 |
| PLAA | 402.446922 | -0.197304105 | 0.29852372 | -0.6609328 | 0.50865544 | 0.82874455 |
| IFT74 | 908.471231 | 0.004054152 | 0.20130328 | 0.02013952 | 0.98393207 | 0.99523767 |
| LRRC19 | 690.409294 | 0.434929139 | 0.60037686 | 0.72442688 | 0.4688037 | 0.81217042 |
| MOB3B | 1513.008 | 0.569603998 | 0.44555049 | 1.27842748 | 0.20109874 | 0.64849129 |
| C9orf72 | 335.954291 | 0.027386567 | 0.30667366 | 0.08930199 | 9.29E-01 | 0.98272895 |
| LINGO2 | 955.001384 | 1.01231527 | 0.86539066 | 1.16977836 | 0.24209018 | 0.68080966 |
| ACO1 | 2281.64867 | 0.388460896 | 0.34740539 | 1.11817753 | 0.26349118 | 0.69567438 |
| DDX58 | 1394.29408 | -0.296426939 | 0.54330939 | -0.5455951 | 0.5853443 | 0.86077718 |
| TOPORS | 330.897845 | -0.338222852 | 0.26730618 | -1.2653013 | 0.20576335 | 0.651931 |
| SMIM27 | 54.8454967 | 0.313043359 | 0.46885952 | 0.66766983 | 0.50434437 | 0.82715271 |
| NDUFB6 | 325.730635 | 0.123555383 | 0.16743065 | 0.73794962 | 0.46054507 | 0.80724952 |
| APTX | 991.531704 | 0.389318666 | 0.32404298 | 1.20144144 | 0.22958001 | 0.66705629 |
| DNAJA1 | 583.656227 | -0.543020546 | 0.37754185 | -1.4383056 | 0.15034737 | 0.60799621 |
| SMU1 | 347.297112 | -0.03832597 | 0.17401735 | -0.2202422 | 0.82568251 | 0.95192757 |
| B4GALT1 | 1497.83458 | 0.368438391 | 0.43186769 | 0.85312794 | 0.39358836 | 0.77781367 |
| BAG1 | 293.749706 | -0.405238827 | 0.34403364 | -1.1779047 | 0.23883461 | 0.67667639 |
| CHMP5 | 300.033242 | -0.442692209 | 0.33990306 | -1.3024073 | 0.1927772 | 0.64518499 |
| NFX1 | 1762.80065 | -0.413061851 | 0.29683329 | -1.3915618 | 0.16405515 | 0.62065658 |
| AQP7 | 710.850235 | 0.324785951 | 0.65106779 | 0.49885121 | 0.61788421 | 0.87448001 |
| AQP3 | 484.458713 | -0.341613868 | 0.7010243 | -0.4873067 | 0.62604097 | 0.87876535 |
| NOL6 | 56.4323295 | -0.932712943 | 0.48185184 | -1.9356841 | 0.0529064 | 0.46586548 |
| PRSS3 | 53.831721 | 0.012221857 | 0.70607192 | 0.01730965 | 0.98618959 | 0.99609994 |
| UBE2R2-AS1 | 53.9035795 | -0.706109054 | 0.63945447 | -1.1042366 | 0.2694905 | 0.69961625 |
| UBE2R2 | 2995.49191 | -0.29440294 | 0.3519777 | -0.836425 | 0.40291586 | 0.78260941 |
| UBAP2 | 1651.13917 | -0.699121671 | 0.29420067 | -2.3763429 | 0.01748521 | 0.33064634 |
| DCAF12 | 275.980247 | -0.245482944 | 0.29542823 | -0.8309394 | 0.40600788 | 0.78429302 |
| UBAP1 | 2317.21896 | -0.520746831 | 0.30411934 | -1.7123108 | 0.08683941 | 0.52575431 |
| KIF24 | 98.609248 | 1.007531911 | 0.41895644 | 2.4048608 | 0.01617863 | 0.32046307 |
| NUDT2 | 108.529917 | -0.270328679 | 0.55543802 | -0.4866946 | 0.62647478 | 0.87885398 |
| MYORG | 74.1761612 | 1.17890221 | 0.43168224 | 2.73094907 | 0.00631522 | 0.22719885 |
| C9orf24 | 51.5152258 | -1.218141663 | 0.77791886 | -1.5658981 | 0.11737247 | 0.57350419 |
| FAM219A | 983.152262 | -0.705442854 | 0.43369648 | -1.6265819 | 1.04E-01 | 0.55022042 |
| DNAI1 | 72.6487444 | 0.078888275 | 0.61549952 | 0.12816951 | 0.89801482 | 0.97374331 |
| RPP25L | 63.7205691 | 0.266726138 | 0.51236216 | 0.52058126 | 0.60265851 | 0.8677246 |
| DCTN3 | 103.799369 | 0.376279874 | 0.50407238 | 0.74647985 | 0.4553776 | 0.80465616 |
| SIGMAR1 | 78.0267708 | 0.365519064 | 0.42025541 | 0.86975458 | 0.38443454 | 0.77317162 |
| IL11RA | 78.0543311 | -0.214844439 | 0.68423867 | -0.3139905 | 0.75352826 | 0.9277474 |
| AL162231.2 | 57.1044234 | 0.433607706 | 0.54615737 | 0.79392447 | 0.4272394 | 0.78933193 |
| DNAJB5 | 53.7116247 | -0.811548208 | 0.75486212 | -1.0750946 | 0.28233237 | 0.70682473 |
| VCP | 1010.16157 | -0.525253479 | 0.39298989 | -1.3365572 | 0.18136721 | 0.6318513 |
| PIGO | 72.5071548 | 0.214294402 | 0.42674954 | 0.50215497 | 0.61555852 | 0.87317195 |
| STOML2 | 244.180738 | -0.022296227 | 0.38626362 | -0.0577228 | 0.95396941 | 0.98836423 |
| FAM214B | 150.913565 | -0.304910212 | 0.38635059 | -0.789206 | 0.4299916 | 0.79055252 |
| UNC13B | 1715.95148 | 0.157212454 | 0.20944171 | 0.7506263 | 0.45287759 | 0.80433814 |
| RUSC2 | 1670.92495 | -0.803784672 | 0.43713027 | -1.838776 | 0.06594813 | 0.48798003 |

| | | | | | | |
|---|---|---|---|---|---|---|
| TESK1 | 153.731301 | 0.094207299 | 0.31550309 | 0.2985939 | 0.76524992 | 0.93083208 |
| CD72 | 128.306441 | 0.566532773 | 0.49890355 | 1.13555571 | 0.25614255 | 0.690849 |
| CCDC107 | 215.703379 | -0.13701908 | 0.40814533 | -0.3357115 | 0.73708844 | 0.92030854 |
| TLN1 | 543.804549 | -0.152643349 | 0.27852509 | -0.5480416 | 0.58366331 | 0.86025731 |
| CREB3 | 147.437885 | -0.474727022 | 0.37964659 | -1.2504446 | 0.21113719 | 0.65561118 |
| GBA2 | 452.519635 | 0.558810781 | 0.27947524 | 1.99950013 | 0.04555427 | 0.44378987 |
| RGP1 | 200.792135 | 0.056676949 | 0.23305865 | 0.24318749 | 0.80786016 | 0.94613779 |
| AL133410.1 | 122.46767 | 0.164473079 | 0.4246756 | 0.3872911 | 0.69854071 | 0.90696517 |
| NPR2 | 48.4763173 | -0.518560113 | 0.45633894 | -1.1363486 | 0.2558107 | 0.69064355 |
| HINT2 | 116.832125 | -0.344119346 | 0.39459841 | -0.8720749 | 0.38316754 | 0.77317162 |
| TMEM8B | 107.046848 | 0.994388537 | 0.54182334 | 1.83526337 | 0.06646668 | 0.48798003 |
| HRCT1 | 57.4129935 | 0.615091707 | 0.58206378 | 1.0567428 | 2.91E-01 | 0.71269438 |
| SPAAR | 60.0700801 | 1.707856184 | 0.7479892 | 2.28326316 | 0.02241487 | 0.36273984 |
| RECK | 188.535743 | 0.996466569 | 0.28746752 | 3.46636224 | 0.00052755 | 0.08106654 |
| CLTA | 1139.54039 | -0.440229033 | 0.36933421 | -1.191953 | 0.23327966 | 0.66978724 |
| GNE | 303.843263 | 0.052556304 | 0.31540619 | 0.16663054 | 0.86766076 | 0.96487312 |
| RNF38 | 2486.17713 | 0.057698318 | 0.23507789 | 0.2454434 | 0.80611313 | 0.94585363 |
| MELK | 39.0819021 | -0.07377354 | 0.46147196 | -0.1598657 | 0.87298686 | 0.96592775 |
| EBLN3P | 598.576504 | -0.229781643 | 0.27785585 | -0.8269815 | 0.40824756 | 0.78481094 |
| ZCCHC7 | 4567.14287 | -0.572220067 | 0.24431539 | -2.3421368 | 0.01917369 | 0.34084276 |
| GRHPR | 617.639811 | 0.339889844 | 0.43730206 | 0.77724272 | 0.43701558 | 0.79603623 |
| ZBTB5 | 438.879871 | -0.405655331 | 0.26353945 | -1.5392584 | 0.12374121 | 0.58354574 |
| POLR1E | 48.2255378 | -0.406300221 | 0.59561967 | -0.6821471 | 0.49514595 | 0.82216643 |
| AL138752.2 | 547.275108 | -0.007598741 | 0.2569173 | -0.0295766 | 0.97640473 | 0.99429832 |
| TRMT10B | 413.262697 | 0.3426031 | 0.3783578 | 0.90550029 | 0.36520041 | 0.76362239 |
| DCAF10 | 1343.10193 | -0.027635727 | 0.20322454 | -0.1359862 | 0.89183221 | 0.97234681 |
| AL135785.1 | 135.124861 | 1.260060702 | 1.12840772 | 1.11667147 | 0.26413482 | 0.69567438 |
| ALDH1B1 | 246.72733 | 0.367764354 | 0.56982237 | 0.64540174 | 0.51866687 | 0.83329602 |
| FAM95C | 219.575212 | -0.000600078 | 0.59342152 | -0.0010112 | 0.99919316 | 0.99977652 |
| ANKRD18A | 212.530772 | -0.287181492 | 0.45111777 | -0.6365998 | 0.52438554 | 0.83589803 |
| CNTNAP3 | 648.157417 | 0.684181532 | 0.66474339 | 1.02924158 | 0.30336617 | 0.72079045 |
| GLIDR | 661.269814 | 0.362032035 | 0.66547417 | 0.54402117 | 0.58642691 | 0.86102615 |
| FAM95B1 | 57.8428376 | -0.873623742 | 1.14858855 | -0.7606063 | 0.44689223 | 0.80219878 |
| BMS1P14 | 453.390174 | 0.723512843 | 0.61420545 | 1.17796553 | 0.23881035 | 0.67667639 |
| FP325318.1 | 54.2657075 | 1.681199701 | 0.64438191 | 2.60901134 | 0.00908042 | 0.25769226 |
| CBWD6 | 225.413537 | 0.240293578 | 0.38526977 | 0.62370214 | 0.53282322 | 0.83870036 |
| CNTNAP3B | 8081.29492 | 0.622967389 | 0.65190009 | 0.95561789 | 0.33926532 | 0.74644675 |
| BX088651.4 | 62.1471244 | -0.673029259 | 0.88975239 | -0.7564231 | 0.44939556 | 0.80311147 |
| BX088645.1 | 39.7884771 | 0.666335829 | 0.70651817 | 0.94312625 | 0.34561633 | 0.75166079 |
| FAM27C | 43.2313973 | -0.382606829 | 0.64238192 | -0.5956065 | 0.55143815 | 0.84689779 |
| AL391987.2 | 52.0117276 | 0.621327248 | 0.37263818 | 1.66737408 | 0.09544004 | 0.53824794 |
| AL590399.3 | 37.9673452 | 0.894656186 | 0.81113075 | 1.10297407 | 0.27003843 | 0.69961625 |
| AL590399.1 | 37.9924033 | -1.117914919 | 0.93943114 | -1.1899913 | 0.2340498 | 0.67060722 |
| AL512625.3 | 160.507572 | 0.529194932 | 0.44253248 | 1.19583297 | 0.23176175 | 0.66883909 |
| AL512625.1 | 58.604305 | -0.261542803 | 0.6139094 | -0.4260283 | 0.6700872 | 0.89626124 |
| CBWD5 | 776.13082 | -0.137972568 | 0.31114786 | -0.4434309 | 0.65745411 | 0.89053559 |
| ZNF658 | 168.236889 | 0.46129987 | 0.44938119 | 1.02652244 | 0.30464539 | 0.72083107 |
| CBWD3 | 153.651171 | 0.354805979 | 0.28906013 | 1.22744695 | 0.21965465 | 0.66208719 |
| PGM5 | 234.254676 | 1.132115408 | 0.68229441 | 1.65927699 | 0.09705999 | 0.5401548 |
| AL161457.2 | 107.772508 | 0.828420287 | 0.48749975 | 1.69932455 | 0.08925805 | 0.53095823 |
| TMEM252 | 320.451092 | 1.511916136 | 0.60167872 | 2.51282966 | 0.01197671 | 0.29262893 |
| AL353616.2 | 2663.89834 | 0.87066395 | 0.66740638 | 1.30454844 | 0.19204666 | 0.64497936 |
| PIP5K1B | 265.372465 | 0.466759132 | 0.52840014 | 0.88334408 | 0.37705039 | 0.77045763 |
| APBA1 | 254.14782 | -0.34276752 | 0.39414784 | -0.869642 | 0.38449607 | 0.77317162 |

| | | | | | | |
|---|---|---|---|---|---|---|
| PTAR1 | 1732.25758 | -0.251395027 | 0.29178618 | -0.8615728 | 0.38892266 | 0.7748665 |
| MAMDC2-AS | 379.518364 | 1.143502092 | 0.53657271 | 2.13112236 | 0.03307906 | 0.41254527 |
| MAMDC2 | 773.203153 | 0.104298398 | 0.5315837 | 0.19620315 | 0.84445116 | 0.95742233 |
| SMC5-AS1 | 574.512844 | 0.556615811 | 0.40874826 | 1.36175702 | 0.17327458 | 0.62650347 |
| SMC5 | 1342.05369 | -0.182507953 | 0.20573476 | -0.8871032 | 0.37502337 | 0.7697436 |
| KLF9 | 677.321967 | 0.625560361 | 0.50236264 | 1.24523664 | 0.21304478 | 0.65785194 |
| TRPM3 | 14627.1632 | 0.255492442 | 0.32783349 | 0.77933601 | 0.43578181 | 0.79549938 |
| CEMIP2 | 2281.96181 | 0.042126653 | 0.37165981 | 0.11334735 | 9.10E-01 | 0.97690916 |
| ABHD17B | 701.858028 | -0.039494148 | 0.26848878 | -0.1470979 | 0.88305471 | 0.96839995 |
| C9orf85 | 963.893461 | 0.283275943 | 0.20960241 | 1.35149185 | 0.17653793 | 0.62709281 |
| GDA | 9363.20413 | 1.31408925 | 0.36882347 | 3.56292202 | 0.00036675 | 0.06861791 |
| AL135924.2 | 79.3502583 | 1.269976579 | 0.64548904 | 1.96746422 | 0.04912972 | 0.45409375 |
| LINC01504 | 168.96671 | 0.875390629 | 1.54018129 | 0.56836857 | 0.56978473 | 0.85327114 |
| ZFAND5 | 1794.36692 | -0.084660578 | 0.37053342 | -0.228483 | 0.81927079 | 0.94990455 |
| TMC1 | 293.837462 | 0.774661075 | 0.82442422 | 0.93963891 | 0.34740281 | 0.75238589 |
| ALDH1A1 | 2560.75532 | 0.910542072 | 0.61775866 | 1.47394464 | 0.1404965 | 0.59500405 |
| ANXA1 | 171.361066 | 0.675833624 | 1.07921159 | 0.62622903 | 0.53116473 | 0.83789294 |
| RORB | 147.10068 | 0.294364885 | 0.62403393 | 0.47171295 | 6.37E-01 | 0.88267497 |
| C9orf41-AS1 | 72.4996752 | 0.221974791 | 0.41171405 | 0.53914796 | 0.58978476 | 0.86281961 |
| CARNMT1 | 619.003287 | -0.090727253 | 0.25237588 | -0.3594926 | 0.71922663 | 0.91465123 |
| NMRK1 | 575.479874 | 0.395114071 | 0.46346849 | 0.85251549 | 0.39392804 | 0.77814839 |
| OSTF1 | 323.302808 | -0.104675752 | 0.21453386 | -0.4879218 | 0.62560521 | 0.87854882 |
| PCSK5 | 4967.71538 | -1.047560592 | 0.46388745 | -2.2582215 | 0.02393186 | 0.37387451 |
| RFK | 227.596171 | 0.042447938 | 0.30581489 | 0.13880272 | 0.88960605 | 0.97208467 |
| GCNT1 | 219.553952 | -0.510500815 | 0.4908685 | -1.0399951 | 0.2983422 | 0.71852421 |
| PRUNE2 | 10183.1905 | -0.554105899 | 0.3217515 | -1.7221548 | 0.08504147 | 0.52183702 |
| PCA3 | 979.265284 | -0.172407932 | 0.75485685 | -0.2283982 | 0.8193367 | 0.94990455 |
| VPS13A | 5069.42945 | -0.248132601 | 0.25484509 | -0.9736605 | 0.33022513 | 0.73849818 |
| GNA14 | 224.992341 | 0.131335826 | 0.43488614 | 0.30200049 | 7.63E-01 | 9.30E-01 |
| GNAQ | 5408.60988 | -0.109710665 | 0.20596161 | -0.5326753 | 0.59425835 | 0.86488065 |
| CEP78 | 270.912946 | -0.059703717 | 0.32498733 | -0.1837109 | 0.85424025 | 0.96133589 |
| PSAT1 | 924.64581 | 0.565854857 | 0.62003587 | 0.91261632 | 0.36144437 | 0.76009788 |
| TLE4 | 853.705509 | 0.237562951 | 0.34085266 | 0.6969667 | 0.48582363 | 0.81839266 |
| AL161782.1 | 32.5147311 | 0.250948583 | 0.49109695 | 0.51099602 | 6.09E-01 | 0.87048326 |
| LINC01507 | 46.2374288 | 0.80965562 | 0.56434 | 1.43469473 | 0.15137411 | 0.60905477 |
| TLE1 | 572.631202 | 0.046975381 | 0.41622759 | 0.11285984 | 9.10E-01 | 9.77E-01 |
| AL591368.1 | 91.8034953 | -0.025295595 | 1.04080536 | -0.0243039 | 0.98061023 | 0.99505659 |
| RASEF | 847.368247 | -0.156337836 | 0.50548384 | -0.3092835 | 0.75710585 | 0.92799282 |
| AL137847.1 | 57.5226449 | 0.055865868 | 0.50938585 | 0.10967299 | 0.91266872 | 0.977179 |
| FRMD3 | 2630.98286 | -0.958867433 | 0.39140842 | -2.4497875 | 0.01429406 | 0.30566089 |
| IDNK | 340.909585 | 0.555370929 | 0.54753229 | 1.01431631 | 0.31043184 | 0.72498238 |
| UBQLN1 | 1051.40976 | -0.083779345 | 0.18615592 | -0.4500493 | 0.65267488 | 0.8894805 |
| GKAP1 | 555.576094 | 0.36442886 | 0.3054276 | 1.19317595 | 0.23280047 | 0.66978724 |
| KIF27 | 695.975792 | 0.250987313 | 0.23533817 | 1.06649639 | 0.28619931 | 0.71118494 |
| C9orf64 | 112.314062 | 0.106307303 | 0.25793772 | 0.4121433 | 6.80E-01 | 8.99E-01 |
| HNRNPK | 1316.81221 | -0.133472114 | 0.20375366 | -0.6550661 | 0.51242519 | 0.83055441 |
| AL354733.3 | 91.1750105 | 0.870721431 | 0.75729049 | 1.14978524 | 0.25023233 | 0.68617881 |
| RMI1 | 156.515275 | 0.059521892 | 0.36436238 | 0.16335905 | 0.87023574 | 0.96564168 |
| NTRK2 | 84.2315193 | 0.589717353 | 0.55671236 | 1.05928554 | 0.28946976 | 0.71218341 |
| AGTPBP1 | 1150.22366 | -0.160363151 | 0.25653276 | -0.6251176 | 0.53189385 | 0.83789294 |
| NAA35 | 973.414039 | -0.219729022 | 0.1763189 | -1.2462023 | 0.21269014 | 0.65745584 |
| GOLM1 | 422.780098 | -1.095946855 | 0.45776966 | -2.3941011 | 0.01666116 | 0.32578383 |
| C9orf153 | 95.8174612 | 1.415308399 | 0.75199128 | 1.88208085 | 0.05982504 | 0.48004006 |
| ISCA1 | 320.966111 | -0.352729397 | 0.34023802 | -1.0367136 | 0.29986933 | 0.71879123 |

| | | | | | | |
|---|---|---|---|---|---|---|
| TUT7 | 1091.65113 | -0.397432308 | 0.31889153 | -1.2462931 | 0.21265681 | 0.65745584 |
| AL513318.2 | 87.151293 | 0.970001573 | 0.48379451 | 2.00498671 | 0.04496447 | 0.44360789 |
| DAPK1 | 2891.79545 | -0.852626127 | 0.34384771 | -2.479662 | 0.0131507 | 0.29875272 |
| CTSL | 1597.32539 | -1.23269869 | 0.70476475 | -1.7490924 | 0.08027504 | 0.51297858 |
| AL772337.2 | 37.9465276 | 0.728656717 | 0.91273349 | 0.79832363 | 0.42468271 | 0.78894887 |
| CDK20 | 51.4103736 | -0.126748313 | 0.47135173 | -0.2689039 | 0.78800365 | 0.93953874 |
| SPIN1 | 1859.17965 | 0.169562476 | 0.23619385 | 0.7178954 | 0.47282179 | 0.81460134 |
| NXNL2 | 476.862058 | 1.257697595 | 0.65345147 | 1.92469931 | 0.05426699 | 0.46711025 |
| SHC3 | 234.745745 | -0.379472717 | 0.53676621 | -0.7069609 | 0.47959079 | 0.81661502 |
| CKS2 | 139.176948 | -0.499117304 | 0.62804565 | -0.794715 | 0.4267793 | 0.78913582 |
| SECISBP2 | 1132.26519 | 0.100666407 | 0.3073569 | 0.32752284 | 0.74327247 | 0.92333113 |
| SEMA4D | 2594.14943 | -0.326133049 | 0.37548276 | -0.86857 | 0.38508239 | 0.77317162 |
| GADD45G | 135.03212 | -0.492876485 | 0.61830862 | -0.7971367 | 0.42537165 | 0.78894887 |
| LINC01508 | 302.637061 | 0.804436362 | 0.78345219 | 1.02678424 | 0.30452207 | 0.72082426 |
| SYK | 61.058452 | 1.87132197 | 0.98622562 | 1.89745828 | 0.05776748 | 0.47684824 |
| AL158071.1 | 73.7343811 | -0.292375657 | 0.74925016 | -0.3902243 | 0.69637066 | 0.90612803 |
| AL158071.4 | 280.975003 | -0.177176389 | 0.58493279 | -0.3029004 | 0.76196575 | 0.93016063 |
| AUH | 3040.14549 | 0.042038136 | 0.38604962 | 0.10889309 | 0.91328729 | 0.97720308 |
| NFIL3 | 838.552579 | -0.479432019 | 0.48966714 | -0.9790978 | 3.28E-01 | 0.73785873 |
| SPTLC1 | 857.132631 | -0.323654919 | 0.24785282 | -1.3058351 | 0.19160863 | 0.64451029 |
| IARS | 1023.61092 | -1.146377661 | 0.50748292 | -2.2589483 | 0.0238866 | 0.37387451 |
| NOL8 | 405.369013 | -0.379257982 | 0.31575084 | -1.2011305 | 0.22970057 | 0.66705629 |
| CENPP | 672.291464 | -0.553737051 | 0.35326729 | -1.5674733 | 0.1170041 | 0.57350419 |
| ECM2 | 126.644516 | 0.560931101 | 0.75370563 | 0.74423101 | 0.45673673 | 0.80507423 |
| IPPK | 281.196873 | -0.359426005 | 0.27902618 | -1.2881444 | 0.1976957 | 0.64558745 |
| BICD2 | 255.33142 | -0.113322862 | 0.31221367 | -0.3629657 | 0.71663049 | 0.91298968 |
| ZNF484 | 159.20643 | -0.107895476 | 0.29036661 | -0.3715836 | 0.71020289 | 0.91123308 |
| SUSD3 | 191.918418 | 1.06726489 | 0.62484357 | 1.70805132 | 0.08762682 | 0.5273546 |
| CARD19 | 249.87853 | 0.481655064 | 0.29816087 | 1.61542011 | 0.10621975 | 0.55440878 |
| NINJ1 | 282.888494 | 0.785479743 | 0.18612955 | 4.22007003 | 2.44E-05 | 0.01647703 |
| FAM120AOS | 148.125083 | -0.263090565 | 0.31046119 | -0.8474185 | 0.39676188 | 0.78014596 |
| FAM120A | 1517.43398 | 0.125403185 | 0.1576828 | 0.79528764 | 0.4264462 | 0.78895108 |
| PHF2 | 302.22456 | -0.064584605 | 0.35081971 | -0.1840963 | 0.85393792 | 0.96119142 |
| PTPDC1 | 670.209189 | 0.165321399 | 0.28527649 | 0.57951287 | 0.56224316 | 0.85043776 |
| AL158152.2 | 660.91662 | 0.13084202 | 0.22501713 | 0.58147582 | 0.56091981 | 0.84953736 |
| ZNF169 | 214.849765 | -0.16045863 | 0.27111241 | -0.5918528 | 0.55394919 | 0.84727776 |
| MFSD14B | 637.25874 | -0.112614336 | 0.30705506 | -0.3667562 | 0.71380091 | 0.91209742 |
| PCAT7 | 100.664646 | 0.557075545 | 0.41072832 | 1.35631149 | 0.17500008 | 0.62699479 |
| FBP1 | 1424.40738 | 0.62073857 | 0.57061315 | 1.08784484 | 2.77E-01 | 0.70345459 |
| C9orf3 | 7292.8079 | -0.133275652 | 0.25502628 | -0.5225957 | 6.01E-01 | 0.86755264 |
| AL353768.1 | 68.9826019 | 0.373064974 | 0.6941956 | 0.53740613 | 0.59098711 | 0.86333372 |
| FANCC | 3097.7116 | 0.458063947 | 0.32755244 | 1.39844462 | 0.16197959 | 0.61845523 |
| PTCH1 | 189.539807 | 0.499760469 | 0.58394924 | 0.85582861 | 0.39209259 | 0.77642489 |
| LINC00476 | 626.198258 | 0.07587368 | 0.48723934 | 0.15572158 | 0.87625249 | 0.96617876 |
| ERCC6L2 | 968.596755 | -0.066823997 | 0.23729825 | -0.2816034 | 0.77824763 | 0.93646866 |
| AL160269.1 | 149.004058 | 0.937739659 | 0.29300648 | 3.20040586 | 0.00137234 | 0.12294569 |
| SLC35D2 | 786.40334 | 0.244282239 | 0.26230502 | 0.93129073 | 0.35170319 | 0.75548011 |
| ZNF367 | 84.9020535 | -0.760376694 | 0.4410996 | -1.7238209 | 0.08474019 | 0.52183702 |
| HABP4 | 882.720988 | -0.503805365 | 0.42811339 | -1.1768036 | 0.23927392 | 0.67720991 |
| CDC14B | 2363.18463 | 0.196289788 | 0.34489313 | 0.5691322 | 0.56926642 | 0.85309237 |
| AAED1 | 125.187114 | -0.783190806 | 0.36020856 | -2.1742704 | 0.02968483 | 0.39981555 |
| ZNF510 | 127.177164 | 0.127495996 | 0.26615135 | 0.47903569 | 0.63191324 | 0.88123095 |
| ZNF782 | 759.191192 | 0.193557041 | 0.23715061 | 0.81617771 | 0.41439851 | 0.78632759 |
| MFSD14C | 1408.12227 | -0.078504551 | 0.2115725 | -0.3710527 | 0.71059826 | 0.91123308 |

| | | | | | | |
|---|---|---|---|---|---|---|
| AL590705.1 | 245.414908 | 1.252946334 | 0.59106389 | 2.11981538 | 0.03402162 | 0.41378378 |
| CCDC180 | 197.873021 | 1.301347568 | 0.59513937 | 2.18662658 | 0.0287698 | 0.39395363 |
| TDRD7 | 416.295602 | 0.026202338 | 0.35415465 | 0.07398558 | 0.94102186 | 0.98489601 |
| TSTD2 | 353.469361 | -0.089134713 | 0.17632312 | -0.5055191 | 0.61319426 | 0.87222135 |
| NCBP1 | 318.47941 | -0.073400099 | 0.25477442 | -0.2880984 | 0.77327142 | 0.93400456 |
| XPA | 342.619917 | 0.180545876 | 0.21928544 | 0.82333729 | 0.41031622 | 0.78582015 |
| TRMO | 112.542443 | -0.760667508 | 0.39582345 | -1.9217343 | 0.05463919 | 0.46772575 |
| ANP32B | 436.828488 | -0.5183668 | 0.38053224 | -1.3622152 | 0.17312998 | 0.6264249 |
| NANS | 219.334025 | -0.247345636 | 0.35199693 | -0.7026926 | 0.48224735 | 0.8177376 |
| TRIM14 | 883.937193 | 0.110442314 | 0.52964506 | 0.20852137 | 0.8348219 | 0.95429103 |
| CORO2A | 151.490693 | 0.975291061 | 0.51676788 | 1.88729042 | 0.05912127 | 0.47855094 |
| TBC1D2 | 128.33554 | 0.375677943 | 0.42936897 | 0.87495364 | 0.38159913 | 0.77237191 |
| ANKS6 | 390.565223 | 0.010587378 | 0.27135798 | 0.03901628 | 0.96887741 | 0.99187457 |
| GALNT12 | 71.6527156 | -0.483798168 | 0.67118785 | -0.7208089 | 0.47102711 | 0.81387913 |
| TGFBR1 | 615.151283 | -0.177520448 | 0.26639765 | -0.6663739 | 0.50517212 | 0.82715271 |
| ALG2 | 237.059851 | -0.423282162 | 0.38610246 | -1.0962949 | 0.27294974 | 0.70118792 |
| SEC61B | 616.740545 | -0.475114231 | 0.3384665 | -1.403726 | 0.16040045 | 0.61590405 |
| AL359710.1 | 344.18154 | 1.034713392 | 0.56898863 | 1.81851332 | 0.06898571 | 0.49273866 |
| STX17-AS1 | 32.861897 | 0.321440193 | 0.67884838 | 0.47350808 | 0.63585074 | 0.88257807 |
| STX17 | 516.421552 | 0.324966207 | 0.36077434 | 0.90074646 | 0.36772314 | 0.76604024 |
| ERP44 | 1326.11882 | -0.297569515 | 0.27225033 | -1.0929997 | 0.27439395 | 0.7014484 |
| INVS | 3458.40222 | -0.092334147 | 0.27441291 | -0.3364789 | 0.73650978 | 0.92019178 |
| TEX10 | 356.133602 | -0.304496836 | 0.17916751 | -1.6995092 | 0.08922329 | 0.53095823 |
| MSANTD3 | 83.1669289 | -1.272713643 | 0.67515815 | -1.88506 | 0.05942173 | 0.47855094 |
| PLPPR1 | 1650.89083 | 0.107380258 | 0.89141711 | 0.12046017 | 9.04E-01 | 9.76E-01 |
| MRPL50 | 145.61362 | 0.164243007 | 0.40930328 | 0.4012746 | 0.68821797 | 0.90307854 |
| ZNF189 | 656.579189 | 0.572831926 | 0.41602466 | 1.37691819 | 0.16853754 | 0.62304975 |
| ALDOB | 6114.26512 | 0.258564589 | 0.63086503 | 0.40985722 | 0.68191069 | 0.89962501 |
| TMEM246-A! | 1473.96328 | 0.741983378 | 0.84875301 | 0.87420412 | 3.82E-01 | 7.73E-01 |
| TMEM246 | 217.582888 | 0.929621124 | 0.59200781 | 1.57028523 | 0.11634877 | 0.57350419 |
| RNF20 | 300.226651 | 0.456376237 | 0.34801057 | 1.31138615 | 0.18972733 | 0.64196311 |
| LINC01492 | 36.6442489 | -0.000436065 | 0.52520193 | -0.0008303 | 0.99933753 | 0.99977652 |
| SMC2 | 94.3842154 | -0.142657182 | 0.32673205 | -0.4366183 | 0.66238821 | 0.89214456 |
| NIPSNAP3A | 125.336292 | -0.144315639 | 0.47594126 | -0.3032215 | 0.76172104 | 0.93014791 |
| ABCA1 | 1095.66449 | -2.099245504 | 0.50088133 | -4.1911035 | 2.78E-05 | 0.01647703 |
| SLC44A1 | 1301.81773 | -0.119148317 | 0.18978867 | -0.6277947 | 0.53013846 | 0.83789294 |
| FSD1L | 983.163117 | -0.230214143 | 0.23088699 | -0.9970858 | 3.19E-01 | 7.31E-01 |
| FKTN | 435.919263 | -0.063105968 | 0.29301068 | -0.2153709 | 0.82947817 | 0.95325673 |
| TMEM38B | 762.650273 | -0.746185235 | 0.44610102 | -1.6726822 | 0.09438988 | 0.5381872 |
| LINC01505 | 726.376567 | 0.045017677 | 0.43455635 | 0.10359457 | 0.9174911 | 0.97828877 |
| ZNF462 | 2507.94681 | -0.205441332 | 0.32371364 | -0.6346391 | 0.52566382 | 0.83638492 |
| AL807761.4 | 65.1151987 | 0.53370569 | 0.52016617 | 1.02602923 | 0.30487781 | 0.72103276 |
| RAD23B | 1646.59732 | -0.239017408 | 0.19569912 | -1.2213515 | 0.22195297 | 0.66334898 |
| ELP1 | 369.456994 | 0.01758052 | 0.19830966 | 0.08865186 | 0.92935859 | 0.98272895 |
| FAM206A | 140.208003 | 0.078113601 | 0.21787716 | 0.35852129 | 0.71995324 | 0.9148113 |
| CTNNAL1 | 489.670781 | -0.804130986 | 0.41544571 | -1.9355862 | 5.29E-02 | 0.46586548 |
| TMEM245 | 2367.12639 | -0.168678988 | 0.37986693 | -0.4440476 | 0.65700819 | 0.89053559 |
| EPB41L4B | 61.0500979 | 0.119061958 | 0.45885348 | 0.25947707 | 0.79526717 | 0.94252227 |
| PTPN3 | 3049.54471 | -0.228205708 | 0.25823056 | -0.8837285 | 0.37684278 | 0.77045763 |
| PALM2 | 721.186178 | 0.253278986 | 0.56578841 | 0.44765672 | 0.65440096 | 0.89032312 |
| PALM2-AKAF | 218.164578 | 0.443617539 | 0.48277956 | 0.91888218 | 0.3581572 | 0.75936492 |
| TXN | 1368.54493 | -0.678198537 | 0.46885838 | -1.4464891 | 0.14804006 | 0.60605027 |
| LPAR1 | 197.552313 | -0.198567147 | 0.82590479 | -0.2404238 | 8.10E-01 | 9.47E-01 |
| ECPAS | 2268.38768 | -0.503510993 | 0.23497907 | -2.1427908 | 0.0321299 | 0.40988962 |

| | | | | | | |
|---|---|---|---|---|---|---|
| ZNF483 | 362.069208 | 0.032025862 | 0.46240355 | 0.06925955 | 0.94478302 | 0.9853777 |
| PTGR1 | 6060.75995 | -1.451682627 | 0.55805416 | -2.6013293 | 0.00928633 | 0.25998727 |
| C9orf84 | 218.034511 | 1.409759427 | 0.67783425 | 2.07979962 | 0.03754392 | 0.42215481 |
| UGCG | 5572.54848 | -0.213537286 | 0.48095963 | -0.4439817 | 0.6570558 | 0.89053559 |
| SUSD1 | 572.814068 | -0.419756276 | 0.29804506 | -1.4083651 | 0.15902298 | 0.61423843 |
| PTBP3 | 2290.35144 | -0.514899607 | 0.30596579 | -1.6828666 | 0.09240092 | 0.53621484 |
| HSDL2 | 2608.13237 | 0.0851456 | 0.34708719 | 0.24531473 | 0.80621275 | 0.94585363 |
| C9orf147 | 288.2993 | 0.288169153 | 0.58787042 | 0.49019162 | 0.62399831 | 0.87791268 |
| KIAA1958 | 3563.07465 | 0.220627156 | 0.29376938 | 0.75102163 | 0.45263964 | 0.80433814 |
| INIP | 353.003251 | 0.133821191 | 0.31940318 | 0.41897263 | 0.67523614 | 0.89801183 |
| SNX30 | 1643.22668 | -0.456407267 | 0.27420177 | -1.6644942 | 9.60E-02 | 0.53870645 |
| SLC31A2 | 106.672363 | -0.281593642 | 0.454854 | -0.6190858 | 0.53585986 | 0.83954482 |
| FKBP15 | 400.178016 | -0.197370445 | 0.18323292 | -1.077156 | 0.28141057 | 0.70625976 |
| SLC31A1 | 563.231652 | -0.179596529 | 0.22493504 | -0.7984373 | 0.42461675 | 0.78894887 |
| CDC26 | 188.278751 | 0.141592997 | 0.30266325 | 0.46782355 | 0.63991078 | 0.88321992 |
| PRPF4 | 340.583343 | -0.620068071 | 0.48742741 | -1.2721239 | 0.20332909 | 0.64881965 |
| RNF183 | 341.626351 | -0.750570264 | 0.50655876 | -1.4817042 | 0.13841901 | 0.59474076 |
| WDR31 | 223.502079 | 0.491943669 | 0.46901846 | 1.04887912 | 0.29423376 | 0.71584347 |
| BSPRY | 227.603343 | -0.310112856 | 0.36981065 | -0.838572 | 0.40170953 | 0.78246874 |
| HDHD3 | 1337.66745 | -0.198362748 | 0.32588761 | -0.6086845 | 0.54273356 | 0.84316546 |
| ALAD | 349.089993 | 0.464331001 | 0.51330291 | 0.90459452 | 3.66E-01 | 7.64E-01 |
| POLE3 | 75.1756511 | -1.343747269 | 0.64028778 | -2.0986614 | 0.03584676 | 0.42040764 |
| RGS3 | 1032.05286 | 1.725324459 | 0.49781811 | 3.46577278 | 0.00052871 | 0.08106654 |
| ZNF618 | 1073.81875 | 0.586345061 | 0.48770883 | 1.20224409 | 0.22926897 | 0.66705629 |
| AMBP | 42.3177493 | 0.394168902 | 0.66132299 | 0.59603085 | 0.55115462 | 0.8466552 |
| KIF12 | 1563.95002 | 0.290723674 | 0.46401186 | 0.62654363 | 0.53095843 | 0.83789294 |
| COL27A1 | 2992.14505 | -0.016592038 | 0.45346952 | -0.0365891 | 0.97081264 | 0.99220667 |
| AKNA | 43.4260787 | -0.424010236 | 0.55893104 | -0.7586092 | 0.44808636 | 0.80223612 |
| WHRN | 229.834471 | -0.209841504 | 0.4148232 | -0.5058577 | 0.61295656 | 0.87222135 |
| ATP6V1G1 | 566.375149 | -0.342369188 | 0.246379 | -1.3896038 | 1.65E-01 | 0.62065658 |
| TMEM268 | 136.159738 | 0.100910884 | 0.61602839 | 0.16380882 | 0.86988165 | 0.96564168 |
| TNFSF15 | 558.492263 | 1.492088602 | 0.89427308 | 1.66849326 | 0.09521785 | 0.53824794 |
| TNC | 346.184214 | 0.966273714 | 0.94052685 | 1.02737493 | 0.30424395 | 0.72082426 |
| PAPPA | 3002.48646 | 0.645188935 | 0.65181479 | 0.98983476 | 0.32225489 | 0.73387566 |
| AL137024.1 | 138.338358 | 0.820539722 | 0.59441601 | 1.38041323 | 0.16745945 | 0.62296534 |
| ASTN2 | 5979.49587 | -0.240320085 | 0.43142464 | -0.5570384 | 0.5775012 | 0.85659658 |
| ASTN2-AS1 | 121.141261 | 0.179919049 | 0.84939867 | 0.21181932 | 0.83224799 | 0.95325673 |
| AL133284.1 | 43.8216948 | 0.945612908 | 0.78177053 | 1.2095786 | 0.22644063 | 0.66690202 |
| TRIM32 | 77.9558134 | 0.13552997 | 0.31685394 | 0.42773643 | 0.66884303 | 0.89566976 |
| BRINP1 | 245.505101 | 0.979730668 | 0.64907375 | 1.50942889 | 1.31E-01 | 0.58930656 |
| CDK5RAP2 | 630.994831 | -0.465773049 | 0.28828128 | -1.6156895 | 0.10616145 | 0.55440878 |
| MEGF9 | 800.516014 | 0.5216717 | 0.27356654 | 1.90692799 | 0.05652991 | 0.47276115 |
| FBXW2 | 524.549375 | -0.313254035 | 0.27929652 | -1.1215823 | 0.26204007 | 0.69408034 |
| PSMD5 | 375.433476 | 0.277452874 | 0.24017765 | 1.15519857 | 0.24800912 | 0.68482118 |
| TRAF1 | 77.6329834 | 0.281423588 | 0.48183565 | 0.58406551 | 0.55917624 | 0.84936251 |
| C5 | 111.097307 | 0.161756371 | 0.52793806 | 0.3063927 | 0.75930566 | 0.92930182 |
| CNTRL | 155.983221 | -1.058984874 | 0.48513806 | -2.1828526 | 0.02904667 | 0.39554194 |
| RAB14 | 631.799675 | -0.058177813 | 0.19022193 | -0.3058418 | 0.75972512 | 0.92936655 |
| GSN | 404.245867 | 0.060752245 | 0.40243173 | 0.15096286 | 0.88000501 | 0.96769187 |
| STOM | 1237.85764 | -0.127423703 | 0.34478683 | -0.3695724 | 0.7117011 | 0.91159439 |
| AL359644.1 | 272.612736 | 0.999897623 | 0.54469485 | 1.83570237 | 0.06640169 | 0.48798003 |
| DAB2IP | 1256.41124 | -0.308525878 | 0.4322045 | -0.7138424 | 0.47532467 | 0.81564184 |
| TTLL11 | 321.523383 | 0.161367274 | 0.29320823 | 0.55035042 | 0.58207904 | 0.85938792 |
| NDUFA8 | 204.870147 | 0.103121079 | 0.24666183 | 0.41806663 | 0.6758984 | 0.89833381 |

| | | | | | | |
|---|---|---|---|---|---|---|
| RBM18 | 122.220794 | 0.118021821 | 0.25694695 | 0.45932369 | 0.64600174 | 0.88700494 |
| MRRF | 870.24926 | 0.073068573 | 0.21241153 | 0.34399532 | 0.73084981 | 0.91897609 |
| PDCL | 127.210496 | -0.012025442 | 0.24019003 | -0.0500664 | 0.9600695 | 0.9893003 |
| RC3H2 | 869.085064 | -0.23782806 | 0.18290836 | -1.300258 | 0.19351256 | 0.64548758 |
| ZBTB6 | 68.6014278 | -0.06021037 | 0.34838743 | -0.1728259 | 0.86278828 | 0.96391903 |
| ZBTB26 | 143.883162 | -0.06289923 | 0.22801319 | -0.2758579 | 0.78265724 | 0.93791597 |
| RABGAP1 | 2262.95279 | -0.213273772 | 0.1719014 | -1.240675 | 0.21472582 | 0.65925411 |
| STRBP | 1690.83799 | -0.349700113 | 0.20300681 | -1.7226028 | 0.08496037 | 0.52183702 |
| DENND1A | 4421.68923 | 0.237464603 | 0.32920705 | 0.72132297 | 0.47071083 | 0.81387913 |
| AL390774.2 | 53.3502068 | 0.842468728 | 0.96821759 | 0.87012335 | 0.384233 | 0.77317162 |
| NEK6 | 1332.76371 | -0.460262546 | 0.35826265 | -1.284707 | 1.99E-01 | 0.64597062 |
| PSMB7 | 1030.3634 | -0.655366871 | 0.30747081 | -2.1314767 | 0.03304989 | 0.41254527 |
| NR6A1 | 4889.17955 | -0.107335448 | 0.33088319 | -0.3243908 | 0.74564222 | 0.92410821 |
| MIR181A2HG | 321.859854 | 0.023875183 | 0.52106171 | 0.04582026 | 0.96345351 | 0.99012655 |
| RPL35 | 822.107387 | -0.378649672 | 0.50977442 | -0.7427789 | 0.45761557 | 0.80561734 |
| ARPC5L | 261.890196 | -0.872684001 | 0.43070314 | -2.0261845 | 0.04274589 | 0.43882523 |
| GOLGA1 | 382.469152 | 0.120234874 | 0.46319569 | 0.25957684 | 0.7951902 | 0.94252227 |
| SCAI | 406.935326 | -0.091578861 | 0.1944287 | -0.4710151 | 0.63762993 | 0.8827026 |
| PPP6C | 576.043908 | -0.187455979 | 0.23506013 | -0.797481 | 0.42517175 | 0.78894887 |
| RABEPK | 246.145542 | 0.075576031 | 0.45205577 | 0.16718298 | 0.86722608 | 0.96486219 |
| HSPA5 | 998.299777 | -0.21775491 | 0.63003392 | -0.3456241 | 0.72962523 | 0.91834786 |
| GAPVD1 | 2002.85319 | -0.244041167 | 0.20261472 | -1.2044592 | 0.22841213 | 0.66705629 |
| MAPKAP1 | 2361.80634 | -0.499429724 | 0.30563357 | -1.6340801 | 0.10224204 | 0.54898162 |
| PBX3 | 411.069606 | -0.53263439 | 0.79948085 | -0.6662253 | 0.50526709 | 0.82715271 |
| MVB12B | 117.761948 | 0.314150343 | 0.50711018 | 0.61949129 | 0.53559276 | 0.83954482 |
| ZBTB43 | 792.06997 | -0.707443487 | 0.48670193 | -1.4535457 | 0.14607231 | 0.60307624 |
| ZBTB34 | 205.267267 | -0.319133969 | 0.44591501 | -0.7156834 | 0.47418687 | 0.8151865 |
| RALGPS1 | 876.49964 | 0.024886728 | 0.28847429 | 0.08627018 | 0.93125165 | 0.98272895 |
| GARNL3 | 627.714788 | 0.36515577 | 0.29941863 | 1.21954928 | 0.22263578 | 0.66398799 |
| SLC2A8 | 108.616473 | 0.022008059 | 0.33762402 | 0.06518511 | 0.94802662 | 0.98668191 |
| ZNF79 | 85.7962132 | -0.265650082 | 0.41992696 | -0.6326102 | 0.52698822 | 0.83711061 |
| RPL12 | 375.344095 | -0.314424642 | 0.53276219 | -0.5901782 | 0.55507117 | 0.84801165 |
| LRSAM1 | 646.682826 | -0.194752805 | 0.28252369 | -0.6893327 | 0.49061395 | 0.81989696 |
| FAM129B | 415.306657 | -0.551569242 | 0.57389492 | -0.961098 | 3.37E-01 | 7.43E-01 |
| STXBP1 | 339.89063 | 0.167088762 | 0.42661874 | 0.39165828 | 0.69531072 | 0.90589033 |
| PTRH1 | 138.392791 | 0.049633494 | 0.28075052 | 0.17678861 | 0.85967444 | 0.96283855 |
| TOR2A | 32.8877757 | -0.021020914 | 0.47350709 | -0.0443941 | 0.96459028 | 0.99031076 |
| CDK9 | 249.481898 | -0.357386754 | 0.24017504 | -1.4880262 | 0.13674399 | 0.59465413 |
| FPGS | 304.108479 | -0.476510616 | 0.47236297 | -1.0087806 | 0.31307985 | 0.72717675 |
| ENG | 65.0073926 | -0.060182418 | 0.52579073 | -0.1144608 | 0.90887253 | 0.97680747 |
| ST6GALNAC4 | 113.07688 | -0.270671582 | 0.75805221 | -0.3570619 | 0.72104545 | 0.91529635 |
| DPM2 | 201.260165 | -1.557283738 | 0.56961075 | -2.7339438 | 0.00625807 | 0.22644975 |
| FAM102A | 912.396003 | -1.150658999 | 0.83821741 | -1.3727453 | 0.16983155 | 0.62326131 |
| NAIF1 | 33.0564484 | -0.0293574 | 0.51221359 | -0.0573148 | 0.95429446 | 0.98840182 |
| SLC25A25 | 253.861213 | -0.030374848 | 0.62253273 | -0.0487924 | 0.96108476 | 0.98982559 |
| SLC25A25-AS | 146.585613 | -0.904301559 | 0.36510911 | -2.4767981 | 0.01325669 | 0.29875272 |
| PTGES2 | 281.744948 | 0.092286466 | 0.31100916 | 0.29673231 | 0.76667089 | 0.93138917 |
| LCN2 | 318.599256 | -0.686421792 | 0.84440056 | -0.8129102 | 4.16E-01 | 7.86E-01 |
| C9orf16 | 204.58211 | 0.507762781 | 0.36799057 | 1.37982551 | 0.16764038 | 0.62304975 |
| CIZ1 | 295.283216 | -0.353196261 | 0.29548789 | -1.1952986 | 0.23197039 | 0.66925238 |
| DNM1 | 1065.5596 | 0.002291085 | 0.33947482 | 0.00674891 | 0.99461519 | 0.99847594 |
| GOLGA2 | 649.917673 | -0.482322833 | 0.37468229 | -1.2872848 | 0.19799502 | 0.64582202 |
| SWI5 | 113.217261 | 0.030717636 | 0.25264237 | 0.12158545 | 0.90322734 | 0.97536781 |
| TRUB2 | 179.898638 | -0.57510927 | 0.29885852 | -1.9243529 | 0.05431036 | 0.46711025 |

| | | | | | | |
|---|---|---|---|---|---|---|
| COQ4 | 626.065274 | 0.012055471 | 0.2514877 | 0.04793662 | 9.62E-01 | 0.99000623 |
| SLC27A4 | 374.149352 | 0.128025009 | 0.37978433 | 0.33709924 | 0.73604209 | 0.92019178 |
| URM1 | 239.810153 | -0.564914999 | 0.23452254 | -2.4087877 | 0.01600561 | 0.31841631 |
| ODF2 | 223.940045 | -0.653071537 | 0.43723449 | -1.4936414 | 0.13526937 | 0.59381224 |
| GLE1 | 357.024905 | -0.124585438 | 0.32110124 | -0.3879943 | 0.69802029 | 0.90696517 |
| SPTAN1 | 1060.114 | -0.376794708 | 0.1762639 | -2.1376737 | 0.03254323 | 0.41149854 |
| WDR34 | 221.29865 | 0.453116458 | 0.46829898 | 0.96757942 | 0.33325446 | 0.74013767 |
| SET | 707.821414 | -0.442792645 | 0.38632086 | -1.1461785 | 0.25172132 | 0.6882793 |
| PKN3 | 53.808514 | -0.218866377 | 0.49628977 | -0.4410052 | 0.65920922 | 0.89058983 |
| ZDHHC12 | 101.388109 | -0.01678721 | 0.6476628 | -0.0259197 | 0.97932141 | 0.99482978 |
| ZER1 | 1100.42735 | 0.44563117 | 0.3622625 | 1.23013331 | 0.21864719 | 0.66098715 |
| TBC1D13 | 450.503288 | -0.212807099 | 0.26808748 | -0.7937972 | 4.27E-01 | 7.89E-01 |
| ENDOG | 155.44274 | -0.114221647 | 0.41284532 | -0.2766694 | 0.78203401 | 0.93757221 |
| SPOUT1 | 117.310602 | -0.457818878 | 0.33616866 | -1.3618726 | 0.1732381 | 0.62650347 |
| KYAT1 | 447.855139 | -0.421168649 | 0.67159902 | -0.6271133 | 0.530585 | 0.83789294 |
| LRRC8A | 274.063395 | -0.727342579 | 0.49031267 | -1.483426 | 0.13796125 | 0.59474076 |
| PHYHD1 | 233.882007 | 0.837803488 | 0.66238855 | 1.26482182 | 0.20593521 | 0.651931 |
| DOLK | 34.4537229 | -0.324111479 | 0.64833974 | -0.4999099 | 0.61713849 | 0.87432737 |
| NUP188 | 268.72839 | -0.343711338 | 0.33635222 | -1.0218792 | 0.30683808 | 0.72190689 |
| SH3GLB2 | 203.305703 | -0.025545437 | 0.44676574 | -0.0571786 | 0.95440293 | 0.98840182 |
| MIGA2 | 235.766679 | -0.092997638 | 0.22862998 | -0.4067605 | 0.68418392 | 0.9007343 |
| DOLPP1 | 35.06599 | -0.320016703 | 0.48220634 | -0.663651 | 0.50691373 | 0.82806179 |
| CRAT | 199.482487 | 0.60886688 | 0.30137017 | 2.02032897 | 0.04334928 | 0.44144764 |
| PTPA | 379.92238 | 0.609810739 | 0.30423331 | 2.00441808 | 0.04502529 | 0.44360789 |
| IER5L | 434.052395 | -0.254407872 | 0.49568165 | -0.5132485 | 6.08E-01 | 8.70E-01 |
| LINC01503 | 74.8215729 | 0.192712681 | 0.45947405 | 0.41942016 | 6.75E-01 | 0.89801183 |
| NTMT1 | 202.520696 | -0.005466002 | 0.2484135 | -0.0220036 | 0.98244505 | 0.99522272 |
| ASB6 | 86.4254791 | -0.170548355 | 0.40130954 | -0.4249796 | 0.67085157 | 0.89694443 |
| TOR1B | 121.700628 | 0.026033203 | 0.33504389 | 0.07770087 | 0.938066 | 0.98435387 |
| TOR1A | 97.8772405 | -0.217511384 | 0.35918955 | -0.6055616 | 0.54480593 | 0.84441376 |
| C9orf78 | 321.889721 | -0.463836259 | 0.34919237 | -1.3283116 | 0.18407521 | 0.63359771 |
| USP20 | 300.661143 | 0.185940059 | 0.29692304 | 0.62622308 | 0.53116863 | 0.83789294 |
| FNBP1 | 2031.53147 | -0.124303665 | 0.27472585 | -0.4524644 | 0.65093447 | 0.88912394 |
| GPR107 | 754.643761 | -0.378489563 | 0.229023 | -1.6526269 | 0.09840683 | 0.54193402 |
| NCS1 | 607.774851 | -0.808217479 | 0.47744418 | -1.6928 | 0.09049355 | 0.53465307 |
| HMCN2 | 51.3367104 | -0.715005743 | 0.66659965 | -1.0726164 | 2.83E-01 | 7.08E-01 |
| ASS1 | 1965.72479 | -0.244165651 | 0.43622282 | -0.5597269 | 5.76E-01 | 8.55E-01 |
| FUBP3 | 873.277365 | -0.559686453 | 0.26277019 | -2.1299465 | 0.03317603 | 0.41254527 |
| EXOSC2 | 72.9627868 | -0.567206956 | 0.43415734 | -1.3064548 | 0.19139793 | 0.6439596 |
| ABL1 | 3286.11753 | -0.34763277 | 0.24563296 | -1.4152529 | 0.15699435 | 0.61255952 |
| AIF1L | 718.140957 | 0.209293346 | 0.38286685 | 0.54664787 | 0.58462068 | 0.86045266 |
| NUP214 | 1291.51337 | -0.29533874 | 0.22152176 | -1.3332268 | 0.18245739 | 0.63216521 |
| AL157938.2 | 285.835846 | 0.493779963 | 0.76832746 | 0.64266864 | 0.52043913 | 0.833897 |
| AL157938.3 | 74.435211 | -0.310963745 | 0.53215935 | -0.5843433 | 0.55898937 | 0.84936251 |
| PLPP7 | 52.3764447 | 0.222299078 | 0.42018488 | 0.52905064 | 0.59677032 | 0.86578438 |
| PRRC2B | 2646.90604 | -0.22591673 | 0.31605966 | -0.7147914 | 0.47473795 | 0.81544274 |
| POMT1 | 243.433026 | 0.121211591 | 0.2271326 | 0.53366003 | 0.59357677 | 0.86444505 |
| UCK1 | 134.051277 | -0.148951752 | 0.28418207 | -0.524142 | 0.60017979 | 0.86740066 |
| RAPGEF1 | 2518.14643 | -0.799025619 | 0.36011367 | -2.218815 | 0.02649931 | 0.38507144 |
| MED27 | 1008.48255 | -0.095285632 | 0.23938285 | -0.398047 | 0.69059552 | 0.90445182 |
| SETX | 2427.19771 | -0.367786078 | 0.21819199 | -1.6856076 | 0.0918714 | 0.53567268 |
| TTF1 | 305.585634 | -0.400892524 | 0.25866657 | -1.5498428 | 0.12117925 | 0.57935261 |
| DDX31 | 602.286427 | -0.370536981 | 0.35139378 | -1.0544779 | 0.29166417 | 0.71337138 |
| GTF3C4 | 196.55389 | -0.138006999 | 0.35783761 | -0.3856694 | 0.69974156 | 0.90721166 |

| | | | | | | |
|---|---|---|---|---|---|---|
| AK8 | 194.769988 | 0.475424341 | 0.61555302 | 0.7723532 | 0.43990527 | 0.79826598 |
| SPACA9 | 37.8337737 | 0.256015362 | 0.47147784 | 0.54300614 | 0.58712557 | 0.86112855 |
| TSC1 | 751.897972 | -0.176930003 | 0.21747046 | -0.8135818 | 0.41588458 | 0.78632759 |
| GTF3C5 | 223.788938 | -0.84092982 | 0.41949888 | -2.0046056 | 0.04500522 | 0.44360789 |
| RALGDS | 118.9911 | -0.965169779 | 0.39851702 | -2.4219035 | 0.01543945 | 0.31213646 |
| ABO | 854.833952 | 0.895956754 | 0.49693818 | 1.80295416 | 0.07139541 | 0.49949133 |
| SURF6 | 208.157566 | -0.622777182 | 0.46160103 | -1.3491677 | 0.17728312 | 0.62731853 |
| MED22 | 96.8679203 | -0.349673727 | 0.403834 | -0.8658848 | 0.38655334 | 0.77334938 |
| RPL7A | 877.688548 | 0.076609053 | 0.52374971 | 0.14627035 | 0.88370797 | 0.9685739 |
| SURF1 | 141.914992 | 0.000458303 | 0.27795442 | 0.00164884 | 0.99868441 | 0.99977652 |
| SURF2 | 70.5026497 | -0.626529114 | 0.3855032 | -1.6252242 | 0.1041148 | 0.55056166 |
| SURF4 | 675.599759 | -0.697644219 | 0.37327188 | -1.8689975 | 0.06162316 | 0.48266777 |
| REXO4 | 178.629605 | -0.263074041 | 0.41464753 | -0.6344522 | 0.52578574 | 0.83646068 |
| ADAMTS13 | 69.9052991 | 0.181014555 | 0.61535158 | 0.29416444 | 0.76863225 | 0.93198626 |
| CACFD1 | 145.143575 | 0.195982685 | 0.4019823 | 0.48754058 | 0.62587529 | 0.87862276 |
| ADAMTSL2 | 53.0202591 | 0.134043893 | 0.84313813 | 0.15898213 | 8.74E-01 | 9.66E-01 |
| DBH-AS1 | 84.5854709 | 1.345071799 | 0.95995988 | 1.40117502 | 0.16116174 | 0.61697277 |
| SARDH | 245.547205 | 1.085877699 | 0.75690788 | 1.43462332 | 1.51E-01 | 0.60905477 |
| VAV2 | 1372.03088 | -0.184171285 | 0.27072072 | -0.6802999 | 0.49631458 | 0.82243887 |
| BRD3OS | 80.5555597 | 0.351369712 | 0.36582205 | 0.96049352 | 0.33680689 | 0.74293725 |
| BRD3 | 222.844874 | 0.133344032 | 0.44604615 | 0.29894672 | 0.7649807 | 0.93083208 |
| WDR5 | 290.771158 | -0.203311812 | 0.33224757 | -0.6119287 | 0.54058496 | 0.8417342 |
| RXRA | 476.294929 | 0.32728069 | 0.3662308 | 0.89364601 | 0.37151132 | 0.76820041 |
| PPP1R26-AS1 | 396.742705 | 0.268719877 | 0.52341585 | 0.51339652 | 0.60767398 | 0.86985416 |
| PPP1R26 | 192.811409 | 0.279207125 | 0.28056627 | 0.99515569 | 0.31966055 | 0.73136156 |
| C9orf116 | 53.9633374 | 0.904746126 | 0.6134806 | 1.47477545 | 0.14027292 | 0.59500405 |
| MRPS2 | 153.276702 | -0.139961181 | 0.46228023 | -0.3027626 | 0.76207076 | 0.93016063 |
| CAMSAP1 | 594.508221 | -0.246550721 | 0.37334982 | -0.6603746 | 0.5090135 | 0.82878364 |
| AL355574.1 | 53.6272095 | -0.52142028 | 0.37071001 | -1.4065449 | 0.15956238 | 0.61520955 |
| UBAC1 | 277.918338 | 0.544189667 | 0.41079619 | 1.32471936 | 0.18526425 | 0.63533076 |
| NACC2 | 603.060557 | -0.257896568 | 0.40356152 | -0.6390514 | 0.52278947 | 0.83504325 |
| TMEM250 | 113.443822 | -0.854763968 | 0.50863032 | -1.6805211 | 0.09285598 | 0.53669957 |
| QSOX2 | 149.46271 | -0.450854883 | 0.2436869 | -1.85014 | 0.06429337 | 0.48593331 |
| SNAPC4 | 289.971784 | -0.857694792 | 0.40525421 | -2.1164365 | 0.0343077 | 0.41505357 |
| ENTR1 | 271.322169 | -0.556425861 | 0.3275349 | -1.6988292 | 0.08935137 | 0.53095823 |
| PMPCA | 268.852743 | -0.547996684 | 0.30454393 | -1.7994011 | 0.07195525 | 0.50167376 |
| INPP5E | 111.643879 | -0.034910109 | 0.32806338 | -0.1064127 | 0.91525492 | 0.97782888 |
| SEC16A | 556.755427 | -0.47474513 | 0.39274255 | -1.2087947 | 0.22674171 | 0.66690202 |
| NOTCH1 | 343.828278 | -0.056186944 | 0.38813271 | -0.1447622 | 8.85E-01 | 0.96937383 |
| EGFL7 | 95.4269555 | -0.471496702 | 0.56221415 | -0.8386425 | 0.40166993 | 0.78246874 |
| AGPAT2 | 216.240498 | -0.531226542 | 0.43427064 | -1.2232615 | 0.22123096 | 0.66263062 |
| SNHG7 | 256.789616 | -0.310944244 | 0.31596657 | -0.9841049 | 0.32506395 | 0.73628585 |
| RABL6 | 696.013545 | 0.00209682 | 0.38116862 | 0.00550103 | 0.99561084 | 0.99866448 |
| AJM1 | 84.8238129 | -0.081266029 | 0.774724 | -0.1048967 | 0.91645774 | 0.97822899 |
| PHPT1 | 532.453514 | 0.39628989 | 0.42824484 | 0.92538158 | 0.35476746 | 0.75755608 |
| MAMDC4 | 33.3479925 | -0.04575981 | 0.70490009 | -0.0649167 | 0.9482403 | 0.98675446 |
| EDF1 | 794.165944 | -0.460293235 | 0.41248138 | -1.1159128 | 0.26445948 | 0.69567438 |
| TRAF2 | 161.23424 | 0.143326825 | 0.37740464 | 0.37976965 | 0.70411642 | 0.90921561 |
| FBXW5 | 541.474984 | -0.145120119 | 0.46145151 | -0.3144862 | 0.75315182 | 0.92751122 |
| LCN12 | 61.0319491 | 0.880881818 | 0.60646238 | 1.4524921 | 0.14636482 | 0.60383734 |
| PAXX | 133.241615 | 0.023594108 | 0.42765324 | 0.05517112 | 0.95600214 | 0.98852557 |
| ABCA2 | 50.3908223 | 1.811704881 | 0.62286583 | 2.90865994 | 0.00362981 | 0.19706898 |
| ENTPD2 | 94.3757322 | 1.841483003 | 0.67766871 | 2.71737941 | 0.00658011 | 0.22966139 |
| UAP1L1 | 156.110969 | -0.758415623 | 0.43983719 | -1.7243099 | 0.08465191 | 0.52183702 |

| | | | | | | |
|---|---|---|---|---|---|---|
| MAN1B1 | 283.379606 | -0.327441582 | 0.33654709 | -0.9729443 | 0.33058098 | 0.73849818 |
| DPP7 | 278.335216 | -0.282577013 | 0.38454562 | -0.7348335 | 0.46244089 | 0.8082072 |
| ANAPC2 | 52.0340624 | -0.173433905 | 0.38098566 | -0.4552242 | 0.64894793 | 0.88800332 |
| SSNA1 | 323.843061 | -0.247009459 | 0.38223461 | -0.6462247 | 0.51813382 | 0.83325693 |
| TPRN | 296.684444 | -0.011996928 | 0.3196485 | -0.0375316 | 0.97006112 | 0.99204339 |
| TMEM203 | 90.2575751 | 0.015675432 | 0.38746196 | 0.0404567 | 0.96772903 | 0.99177684 |
| NDOR1 | 98.9803478 | -0.335248206 | 0.32771943 | -1.0229732 | 0.3063205 | 0.72182764 |
| BX255925.3 | 85.2179289 | -0.620740356 | 0.49929862 | -1.2432247 | 0.21378506 | 0.65849767 |
| RNF208 | 57.4729212 | -0.543470428 | 0.60337065 | -0.900724 | 0.36773509 | 0.76604024 |
| SLC34A3 | 109.13737 | -0.233439643 | 0.63628239 | -0.3668806 | 7.14E-01 | 9.12E-01 |
| TUBB4B | 862.993708 | 0.05136151 | 0.35512527 | 0.14462927 | 0.88500358 | 0.96937383 |
| STPG3-AS1 | 92.5324375 | 0.54357006 | 1.11778915 | 0.48629033 | 0.62676133 | 0.87890915 |
| NELFB | 379.960979 | -0.086323677 | 0.25143753 | -0.3433206 | 0.7313573 | 0.91910833 |
| EXD3 | 753.449139 | 0.0243515 | 0.22712211 | 0.10721766 | 0.91461631 | 0.9776687 |
| NOXA1 | 214.935153 | 0.317701158 | 0.35504497 | 0.89481948 | 0.3708836 | 0.76758289 |
| ENTPD8 | 37.8549162 | 0.634852806 | 0.65244201 | 0.97304097 | 3.31E-01 | 0.73849818 |
| NSMF | 408.302539 | 0.518458723 | 0.40697653 | 1.27392781 | 0.202689 | 0.64881821 |
| PNPLA7 | 579.016955 | 0.605296266 | 0.63997004 | 0.94581969 | 0.34424056 | 0.75111515 |
| MRPL41 | 614.701517 | -0.464711316 | 0.42149494 | -1.1025312 | 0.27023081 | 0.69961625 |
| DPH7 | 217.644614 | -0.421912075 | 0.27019274 | -1.5615226 | 0.1184005 | 0.57589001 |
| ZMYND19 | 77.1616822 | -0.730766536 | 0.56367914 | -1.2964229 | 0.19482983 | 0.64558745 |
| ARRDC1 | 290.156275 | -0.408257884 | 0.28539359 | -1.4305082 | 0.15257121 | 0.61001662 |
| ARRDC1-AS1 | 73.195377 | -0.138148519 | 0.29394832 | -0.4699755 | 0.6383725 | 0.8827026 |
| EHMT1 | 2426.6436 | -0.219704879 | 0.28093092 | -0.7820601 | 0.43417923 | 0.79442004 |
| LINC01001 | 68.113813 | -1.013031663 | 0.47863296 | -2.1165105 | 0.03430142 | 0.41505357 |
| AC069287.3 | 60.873162 | -0.007401114 | 0.70438613 | -0.0105072 | 0.99161663 | 0.99748517 |
| BET1L | 166.848717 | -0.324670093 | 0.34965033 | -0.9285565 | 0.35311896 | 0.75673243 |
| RIC8A | 154.228621 | -0.649045974 | 0.43114339 | -1.5054063 | 0.13221965 | 0.59115873 |
| SIRT3 | 217.703441 | 0.332694733 | 0.41531642 | 0.80106328 | 0.42309502 | 0.78894887 |
| PSMD13 | 374.085498 | -0.298370315 | 0.29764452 | -1.0024384 | 0.31613188 | 0.72798933 |
| PGGHG | 322.558106 | 0.624098721 | 0.67265111 | 0.92781936 | 0.35350128 | 0.75673243 |
| IFITM2 | 489.615024 | 0.096175138 | 0.5320345 | 0.18076861 | 8.57E-01 | 0.96245027 |
| IFITM3 | 1734.79042 | -0.632616801 | 0.72256387 | -0.8755168 | 3.81E-01 | 0.77210521 |
| SIGIRR | 664.161007 | -0.021784165 | 0.46806985 | -0.0465404 | 0.96287952 | 0.99012655 |
| ANO9 | 329.469034 | 0.405082202 | 0.59349542 | 0.68253635 | 0.49489986 | 0.82215397 |
| PTDSS2 | 379.336846 | -0.092573225 | 0.25792282 | -0.3589183 | 0.7196562 | 0.9147188 |
| RNH1 | 446.526366 | 0.217735968 | 0.31836007 | 0.6839299 | 0.49401943 | 0.82179535 |
| HRAS | 192.665938 | -0.480569452 | 0.46557847 | -1.0321986 | 0.30197909 | 0.71925676 |
| LMNTD2 | 67.1580016 | -0.151859169 | 0.50928666 | -0.2981801 | 7.66E-01 | 0.9309447 |
| AP006284.1 | 64.0650848 | -0.65278202 | 0.77807895 | -0.8389663 | 0.40148822 | 0.78246874 |
| RASSF7 | 427.699685 | -0.010422809 | 0.44353792 | -0.0234993 | 0.98125204 | 0.9951185 |
| MIR210HG | 200.662461 | -0.292368312 | 0.76642764 | -0.3814689 | 7.03E-01 | 9.09E-01 |
| PHRF1 | 445.563481 | -0.33152507 | 0.29558782 | -1.1215789 | 0.26204151 | 0.69408034 |
| IRF7 | 142.248192 | -1.880677493 | 0.74515655 | -2.523869 | 0.01160712 | 0.28822241 |
| CDHR5 | 833.012885 | 0.326529794 | 0.52686053 | 0.61976515 | 0.53541242 | 0.83954482 |
| DEAF1 | 971.376484 | -0.416530123 | 0.31124756 | -1.3382599 | 0.18081173 | 0.63175794 |
| AC131934.1 | 134.387562 | -0.654181509 | 0.59967188 | -1.0908991 | 0.27531728 | 0.70235883 |
| EPS8L2 | 1671.61648 | -0.025374686 | 0.379725 | -0.0668238 | 0.94672194 | 0.9862976 |
| AP006621.4 | 247.017322 | 0.306237613 | 0.70032916 | 0.43727669 | 0.6619107 | 0.89193781 |
| TALDO1 | 869.185418 | -0.177426788 | 0.45428399 | -0.3905636 | 6.96E-01 | 0.90612803 |
| GATD1 | 282.986363 | -0.74092404 | 0.39162463 | -1.891924 | 0.05850111 | 0.47778423 |
| AP006621.3 | 569.537002 | -0.03142658 | 0.39088744 | -0.080398 | 0.93592069 | 0.98406248 |
| SLC25A22 | 85.2560503 | -0.66340607 | 0.40920033 | -1.6212256 | 0.10496926 | 0.55219437 |
| PIDD1 | 134.108908 | -0.198990286 | 0.2792634 | -0.7125541 | 0.47612171 | 0.81570932 |

| | | | | | | |
|---|---|---|---|---|---|---|
| RPLP2 | 976.433692 | -0.479239224 | 0.51368387 | -0.9329458 | 0.35084794 | 0.75495648 |
| PNPLA2 | 398.803033 | -0.224249822 | 0.21186391 | -1.0584616 | 0.28984503 | 0.71239004 |
| CD151 | 795.95258 | -0.090500862 | 0.42529449 | -0.2127958 | 0.83148628 | 0.95325673 |
| POLR2L | 577.753112 | -0.14258856 | 0.46747739 | -0.305017 | 0.76035319 | 0.92968986 |
| TSPAN4 | 265.876786 | -0.001683621 | 0.31319762 | -0.0053756 | 0.99571092 | 0.99866448 |
| AP006623.1 | 35.1885661 | -0.058064487 | 0.41769205 | -0.1390127 | 0.88944014 | 0.97208467 |
| CHID1 | 173.690265 | -0.191482799 | 0.35956223 | -0.5325443 | 0.5943491 | 0.86488065 |
| AP2A2 | 1703.48734 | -0.11005462 | 0.29812441 | -0.3691567 | 0.71201094 | 0.91165921 |
| TOLLIP | 507.373729 | -0.381289712 | 0.31929811 | -1.1941496 | 0.23241945 | 0.66978724 |
| MOB2 | 249.326619 | -0.505190505 | 0.28612319 | -1.7656399 | 0.07745629 | 0.51128842 |
| DUSP8 | 92.6214755 | -0.273751602 | 0.7439776 | -0.3679568 | 0.71290546 | 0.91188831 |
| KRTAP5-AS1 | 94.200644 | -0.198394436 | 1.0175591 | -0.1949709 | 0.84541571 | 0.95764192 |
| MRPL23 | 140.445043 | 1.230829318 | 0.44180448 | 2.78591406 | 0.0053377 | 0.21873395 |
| CD81-AS1 | 81.1517891 | 0.812193954 | 0.4562053 | 1.78032557 | 0.07502269 | 0.5071704 |
| CD81 | 705.622721 | 0.202229591 | 0.41293844 | 0.48973303 | 0.62432283 | 0.87815218 |
| TSSC4 | 129.829872 | -0.837916508 | 0.52013284 | -1.6109664 | 0.10718705 | 0.55579103 |
| KCNQ1 | 753.526609 | 1.061580326 | 0.38085469 | 2.78736315 | 0.00531389 | 0.21873395 |
| KCNQ1OT1 | 4552.53356 | 0.987222163 | 0.39773983 | 2.48208021 | 0.01306179 | 0.29875272 |
| CDKN1C | 416.76303 | 1.169281613 | 0.62163502 | 1.88097769 | 0.05997495 | 0.48004006 |
| SLC22A18AS | 94.1964345 | 0.752583485 | 0.62525282 | 1.20364669 | 0.22872616 | 0.66705629 |
| SLC22A18 | 742.324124 | 0.74896791 | 0.70310457 | 1.06522976 | 0.28677197 | 0.71118494 |
| NAP1L4 | 2006.04817 | 0.064880508 | 0.18644328 | 0.3479906 | 0.72784724 | 0.91665945 |
| CARS | 637.771879 | -0.286345975 | 0.35471858 | -0.8072483 | 0.41952344 | 0.78783502 |
| OSBPL5 | 133.355226 | 0.857677063 | 0.45165741 | 1.89895493 | 0.0575704 | 0.47684824 |
| ZNF195 | 536.688847 | 0.083758828 | 0.24892595 | 0.3364809 | 0.73650825 | 0.92019178 |
| AC127526.1 | 548.15805 | 0.198444153 | 0.41480766 | 0.47840041 | 0.63236524 | 0.88150416 |
| NUP98 | 1179.65408 | -0.209183684 | 0.32535724 | -0.6429354 | 0.52026602 | 0.833897 |
| PGAP2 | 174.527877 | 0.394587981 | 0.37988818 | 1.03869507 | 0.29894658 | 0.71852421 |
| RHOG | 101.105287 | -0.595186447 | 0.38495508 | -1.5461192 | 0.12207577 | 0.58055167 |
| STIM1 | 1482.21815 | 0.376423257 | 0.33674194 | 1.11783897 | 0.26363578 | 0.69567438 |
| RRM1 | 99.991728 | -0.057680608 | 0.43426828 | -0.1328225 | 0.89433374 | 0.97272276 |
| TRIM21 | 51.0661938 | -0.4430882 | 0.65984891 | -0.6714995 | 0.50190239 | 0.82638105 |
| AC104389.4 | 496.135716 | 0.252071512 | 0.70887575 | 0.35559336 | 0.72214511 | 0.9157845 |
| TRIM5 | 110.787842 | -0.92207141 | 0.70815707 | -1.3020719 | 0.19289181 | 0.64518499 |
| TRIM22 | 264.219808 | -2.132060777 | 1.18726677 | -1.7957723 | 0.07253074 | 0.50287245 |
| FAM160A2 | 437.015251 | 0.209838763 | 0.2487595 | 0.8435407 | 0.39892611 | 0.78140731 |
| CAVIN3 | 63.1079376 | 0.254670485 | 0.62531732 | 0.407266 | 6.84E-01 | 0.90059706 |
| SMPD1 | 355.719147 | -0.16337475 | 0.48467184 | -0.3370832 | 0.73605416 | 0.92019178 |
| APBB1 | 222.896721 | 0.037723204 | 0.36568083 | 0.10315882 | 0.91783692 | 0.97832208 |
| TRIM3 | 77.4502978 | 0.072318542 | 0.44091734 | 0.16401837 | 0.86971668 | 0.96564168 |
| ARFIP2 | 176.482236 | -0.429619352 | 0.31620725 | -1.3586638 | 0.17425314 | 0.62699479 |
| AC084337.2 | 314.810186 | 0.147932242 | 0.57535997 | 0.2571125 | 0.79709193 | 0.94280402 |
| DNHD1 | 482.904277 | 0.195534509 | 0.46542085 | 0.42012409 | 0.67439481 | 0.89801183 |
| RRP8 | 107.504143 | 0.043447442 | 0.31342829 | 0.13862004 | 0.88975041 | 0.97212576 |
| ILK | 376.004627 | 0.075277475 | 0.30232693 | 0.24899362 | 0.80336572 | 0.94543168 |
| TAF10 | 162.523437 | -0.09490208 | 0.39469238 | -0.2404457 | 0.80998476 | 0.94692486 |
| TPP1 | 486.137403 | -0.540023562 | 0.35229782 | -1.5328609 | 0.12531012 | 0.58564339 |
| MRPL17 | 163.583347 | -0.597446776 | 0.41511107 | -1.4392456 | 0.15008095 | 0.6079077 |
| AC087280.2 | 325.439699 | -0.3342314 | 0.53430416 | -0.6255452 | 0.5316133 | 0.83789294 |
| ZNF215 | 168.960842 | 0.026572001 | 0.49723575 | 0.05343944 | 0.95738178 | 0.98861313 |
| ZNF214 | 119.029312 | 0.510041652 | 0.56005951 | 0.91069189 | 0.36245774 | 0.76129455 |
| NLRP14 | 42.0271318 | 0.259401059 | 0.51005707 | 0.50857261 | 0.61105183 | 0.87163238 |
| SYT9 | 532.999418 | 0.594159935 | 0.45570994 | 1.30381166 | 0.1922978 | 0.64497936 |
| AC107884.1 | 266.734271 | 0.310587812 | 0.40475717 | 0.76734356 | 0.44287727 | 0.7993765 |

| | | | | | | |
|---|---|---|---|---|---|---|
| PPFIBP2 | 1043.36807 | -0.040856146 | 0.42582457 | -0.095946 | 0.92356349 | 0.98012108 |
| EIF3F | 295.471601 | 0.023347002 | 0.29552619 | 0.07900146 | 0.93703146 | 0.98435387 |
| TUB | 685.030698 | 1.432699716 | 0.77802719 | 1.84145198 | 0.06555535 | 0.48790819 |
| RIC3 | 639.450082 | -0.40903977 | 0.41185979 | -0.993153 | 0.32063542 | 0.73210121 |
| STK33 | 2731.86517 | -0.035362246 | 0.33499648 | -0.1055601 | 0.91593142 | 0.97801193 |
| TRIM66 | 936.10225 | -0.599920528 | 0.58538387 | -1.0248327 | 0.30544214 | 0.72122 |
| RPL27A | 915.591799 | -0.173751581 | 0.4737505 | -0.3667576 | 0.71379986 | 0.91209742 |
| ST5 | 268.972535 | 1.042739663 | 0.55997559 | 1.86211627 | 0.0625867 | 0.48421747 |
| AKIP1 | 71.8652989 | 0.274325874 | 0.35113625 | 0.78125193 | 0.43465434 | 0.79451524 |
| TMEM9B | 180.728357 | 0.094161302 | 0.21843028 | 0.43108172 | 0.66640895 | 0.89420131 |
| TMEM9B-AS | 160.190394 | 0.690684421 | 0.6706341 | 1.02989756 | 0.3030581 | 0.72043311 |
| DENND5A | 1928.92905 | -0.267866259 | 0.33458039 | -0.8006036 | 4.23E-01 | 0.78894887 |
| TMEM41B | 1082.50286 | -0.675711884 | 0.32759567 | -2.0626398 | 0.03914686 | 0.42704465 |
| IPO7 | 1031.73324 | -0.279566335 | 0.38590435 | -0.7244446 | 0.46879281 | 0.81217042 |
| AC132192.2 | 37.1900977 | -0.089984636 | 0.68256893 | -0.1318323 | 0.89511693 | 0.97309833 |
| ZNF143 | 406.140996 | -0.220694088 | 0.51218011 | -0.4308916 | 0.66654722 | 0.89420131 |
| WEE1 | 129.743034 | -0.22115385 | 0.32962238 | -0.6709309 | 0.50226454 | 0.82638105 |
| SWAP70 | 654.349462 | -0.27334184 | 0.28010984 | -0.975838 | 0.32914473 | 0.73829083 |
| SBF2-AS1 | 272.680466 | -0.173596791 | 0.31440672 | -0.5521408 | 5.81E-01 | 8.59E-01 |
| SBF2 | 5483.62978 | 0.274428484 | 0.29461842 | 0.93147088 | 0.35161004 | 0.75548011 |
| AC011092.2 | 172.786124 | 0.643343039 | 0.58286391 | 1.10376201 | 0.26969639 | 0.69961625 |
| AC100763.1 | 51.296928 | 0.482279731 | 0.75898948 | 0.63542347 | 0.52515226 | 0.835938 |
| ADM | 572.32879 | -1.276362984 | 1.00813199 | -1.2660673 | 0.20548898 | 0.651931 |
| AMPD3 | 151.1313 | -0.30206912 | 0.59805912 | -0.5050824 | 6.14E-01 | 8.72E-01 |
| MTRNR2L8 | 644.370252 | -0.698009473 | 0.80836375 | -0.8634844 | 0.3878712 | 0.77373447 |
| RNF141 | 132.0346 | 0.071602378 | 0.21136819 | 0.33875665 | 0.73479306 | 0.9199679 |
| CTR9 | 374.80347 | -0.262399283 | 0.27040387 | -0.9703977 | 0.33184831 | 0.73939532 |
| EIF4G2 | 1103.1344 | -0.381745684 | 0.27014533 | -1.4131123 | 0.15762271 | 0.61323667 |
| ZBED5 | 452.093679 | 0.131428634 | 0.23300559 | 0.56405786 | 0.57271475 | 0.8542679 |
| ZBED5-AS1 | 177.226645 | 0.667033133 | 0.64964263 | 1.02676934 | 0.30452909 | 0.72082426 |
| GALNT18 | 2922.28258 | 0.896206369 | 0.50168863 | 1.78637967 | 0.0740378 | 0.50640528 |
| USP47 | 2640.37226 | 0.041337813 | 0.20439306 | 0.20224666 | 0.8397239 | 0.9556011 |
| MICAL2 | 177.394658 | -0.880179934 | 0.60389792 | -1.4574979 | 0.144979 | 0.60109715 |
| PARVA | 883.501516 | -0.103414663 | 0.35854773 | -0.2884265 | 0.77302029 | 0.93396114 |
| TEAD1 | 5628.78968 | -0.181154336 | 0.34692597 | -0.52217 | 0.60155197 | 0.86757631 |
| LINC00958 | 622.320019 | -0.995739946 | 0.51467971 | -1.9346788 | 5.30E-02 | 0.46624655 |
| ARNTL | 485.855332 | 0.01238527 | 0.50041362 | 0.02475007 | 0.98025432 | 0.99494599 |
| BTBD10 | 603.273366 | -0.328237547 | 0.34452167 | -0.9527341 | 0.3407248 | 0.74725037 |
| FAR1 | 960.913081 | -0.021966915 | 0.25999777 | -0.0844889 | 0.93266776 | 0.98310938 |
| SPON1 | 1043.40986 | 0.42824491 | 0.5751565 | 0.7445711 | 0.45653104 | 0.80507423 |
| SPON1-AS1 | 364.752407 | -2.246295421 | 0.7889134 | -2.8473283 | 0.00440879 | 0.20749797 |
| RRAS2 | 4706.78437 | -1.962611423 | 0.52957046 | -3.706044 | 0.00021052 | 0.05350954 |
| COPB1 | 881.782474 | -0.337801149 | 0.2267791 | -1.4895603 | 0.13633988 | 0.5942049 |
| AC018523.2 | 40.7372568 | -1.131974749 | 0.67310048 | -1.6817322 | 0.09262078 | 0.53669957 |
| PSMA1 | 278.561494 | 0.153617506 | 0.43348052 | 0.35438157 | 0.72305294 | 0.91592159 |
| PDE3B | 833.375621 | -0.156928931 | 0.33424458 | -0.4695033 | 0.63870993 | 0.88284765 |
| CYP2R1 | 155.532038 | 0.098492161 | 0.31031948 | 0.31738955 | 0.75094804 | 0.9260679 |
| CALCB | 300.11736 | 0.475144723 | 0.62133116 | 0.76472058 | 0.44443795 | 0.79994797 |
| INSC | 75.6103776 | 0.26943198 | 0.94258149 | 0.28584476 | 0.77499702 | 0.9347839 |
| SOX6 | 6953.51457 | 0.361520396 | 0.50893182 | 0.71035134 | 0.47748629 | 0.81579656 |
| C11orf58 | 927.6463 | -0.213218058 | 0.17634876 | -1.2090703 | 0.22663583 | 0.66690202 |
| PLEKHA7 | 9116.40571 | 0.053310899 | 0.24025397 | 0.22189393 | 8.24E-01 | 9.52E-01 |
| RPS13 | 737.307011 | -0.030374099 | 0.54148688 | -0.0560939 | 0.95526702 | 0.98840182 |
| PIK3C2A | 2204.93729 | -0.025261752 | 0.14886672 | -0.1696938 | 0.86525099 | 0.96428252 |

| | | | | | | |
|---|---|---|---|---|---|---|
| NUCB2 | 325.64351 | 0.285686451 | 0.41112996 | 0.69488114 | 0.4871298 | 0.81893983 |
| NCR3LG1 | 194.338897 | 0.065076351 | 0.56338265 | 0.11551004 | 0.90804086 | 0.97673359 |
| AC124798.1 | 88.3220098 | 0.06114103 | 0.52555314 | 0.11633653 | 0.90738583 | 0.97673359 |
| USH1C | 1467.71416 | 0.5854628 | 0.38181869 | 1.5333529 | 0.12518893 | 0.58564339 |
| KCNC1 | 76.3582729 | 0.778615227 | 0.68461264 | 1.13730771 | 0.25540967 | 0.69010455 |
| SERGEF | 1031.46285 | 0.371320884 | 0.20065189 | 1.8505726 | 0.06423106 | 0.48593331 |
| SAAL1 | 91.2154687 | -0.238082122 | 0.2899375 | -0.8211498 | 0.41156094 | 0.78632759 |
| SAA1 | 2344.04605 | 1.904748904 | 1.74265082 | 1.09301811 | 2.74E-01 | 0.7014484 |
| HPS5 | 260.285311 | -0.036134138 | 0.21742929 | -0.166188 | 0.86800901 | 0.96498555 |
| GTF2H1 | 693.563744 | -0.561645307 | 0.26245327 | -2.1399821 | 0.03235621 | 0.41065429 |
| LDHA | 1844.11059 | -0.236641927 | 0.54142659 | -0.4370711 | 6.62E-01 | 8.92E-01 |
| LDHC | 45.0331712 | 2.047935731 | 0.91331959 | 2.24229915 | 0.02494204 | 0.37990947 |
| TSG101 | 907.123546 | -0.423156488 | 0.30363734 | -1.3936247 | 0.16343098 | 0.62065658 |
| UEVLD | 675.885441 | 0.027721295 | 0.185028 | 0.14982216 | 0.88090493 | 0.96773407 |
| SPTY2D1OS | 159.52607 | 0.553063367 | 0.27394985 | 2.01884897 | 0.04350292 | 0.44147822 |
| SPTY2D1 | 224.768042 | -0.882369014 | 0.43467231 | -2.0299637 | 0.04236023 | 0.43742111 |
| ZDHHC13 | 372.529973 | -0.130749566 | 0.27420086 | -0.4768386 | 0.63347702 | 0.88188803 |
| NAV2 | 1059.81906 | -2.271744183 | 1.08399433 | -2.0957159 | 0.0361074 | 0.42040764 |
| HTATIP2 | 273.923979 | 0.237071335 | 0.45354548 | 0.52270686 | 6.01E-01 | 8.68E-01 |
| PRMT3 | 757.276872 | -0.14086319 | 0.30205283 | -0.4663528 | 0.64096298 | 0.88396933 |
| NELL1 | 233.099172 | -0.307327075 | 0.57174876 | -0.5375212 | 5.91E-01 | 8.63E-01 |
| ANO5 | 463.590819 | 0.668950815 | 0.38026422 | 1.75917371 | 0.07854801 | 0.51192014 |
| FANCF | 50.3177534 | 0.8588296 | 0.48230494 | 1.78067761 | 0.07496513 | 0.5071704 |
| GAS2 | 406.480114 | 0.537314652 | 0.47511198 | 1.13092213 | 0.25808787 | 0.69210282 |
| SVIP | 253.541949 | 0.436110736 | 0.29214527 | 1.49278726 | 0.13549288 | 0.59381224 |
| AC006299.1 | 144.513109 | 0.725322987 | 0.63358483 | 1.14479223 | 0.25229524 | 0.68874851 |
| AC090092.1 | 31.0066912 | 0.536861874 | 0.69650021 | 0.77079929 | 0.44082591 | 0.79826598 |
| LUZP2 | 191.363913 | -1.915522522 | 0.61872845 | -3.0959018 | 0.00196215 | 0.14007934 |
| ANO3 | 32.0692392 | 0.671035076 | 0.51241456 | 1.30955504 | 0.19034641 | 0.64278842 |
| SLC5A12 | 7706.45061 | -0.231133132 | 1.07711657 | -0.2145851 | 0.83009084 | 0.95325673 |
| BBOX1 | 3351.26472 | 0.849227102 | 0.80300426 | 1.05756239 | 0.290255 | 0.71261423 |
| BBOX1-AS1 | 784.774643 | 0.137235321 | 0.54057189 | 0.25387062 | 0.79959551 | 0.94385599 |
| CCDC34 | 53.7897761 | 0.785241775 | 0.40636136 | 1.93237315 | 0.05331347 | 0.46664603 |
| LGR4 | 4604.60104 | -0.159603705 | 0.30981982 | -0.5151501 | 0.60644815 | 0.86961018 |
| LIN7C | 292.666742 | -0.281527097 | 0.3010997 | -0.9349963 | 0.34979022 | 0.75455562 |
| BDNF-AS | 914.68378 | 0.472475094 | 0.26549312 | 1.7796133 | 0.07513927 | 0.5071704 |
| KIF18A | 36.7613607 | 1.150494308 | 0.51845473 | 2.21908344 | 0.02648105 | 0.38507144 |
| METTL15 | 2380.68796 | 0.376280453 | 0.32417522 | 1.1607317 | 0.24575103 | 0.68320879 |
| AC090833.1 | 61.0521265 | 1.564403944 | 0.71394647 | 2.19120622 | 0.02843687 | 0.39280225 |
| ARL14EP | 249.040456 | 0.176660761 | 0.21998729 | 0.80304985 | 0.42194592 | 0.78892549 |
| MPPED2 | 370.871954 | 1.606302168 | 0.77691695 | 2.06753395 | 0.03868387 | 0.42523931 |
| DCDC1 | 1246.79967 | 0.313418955 | 0.44443319 | 0.70521051 | 0.48067924 | 0.81714875 |
| DNAJC24 | 513.259742 | -0.745056057 | 0.35730112 | -2.0852329 | 0.03704816 | 0.42215481 |
| IMMP1L | 1220.74383 | 0.215556652 | 0.33353639 | 0.64627626 | 0.51810046 | 0.83325693 |
| ELP4 | 2177.72399 | 0.215794307 | 0.28159594 | 0.76632605 | 0.44348232 | 0.7993765 |
| AC131571.1 | 199.437594 | 0.831326088 | 0.78203032 | 1.06303562 | 0.2877658 | 0.7113446 |
| RCN1 | 204.885815 | -0.716974943 | 0.49572146 | -1.4463262 | 0.14808572 | 0.60605027 |
| AL078612.1 | 41.2349013 | -0.025412264 | 0.52414782 | -0.048483 | 0.9613313 | 0.98985638 |
| WT1 | 46.4926616 | -0.049861812 | 0.57117226 | -0.0872973 | 0.93043518 | 0.98272895 |
| EIF3M | 467.807329 | -0.133286798 | 0.29321328 | -0.4545729 | 0.64941657 | 0.88850926 |
| CCDC73 | 471.299512 | 0.156006513 | 0.5046042 | 0.3091661 | 0.75719518 | 0.92799282 |
| PRRG4 | 554.821264 | -0.282659718 | 0.53147744 | -0.5318377 | 0.59483844 | 0.86499581 |
| QSER1 | 891.395333 | 0.106561613 | 0.28083222 | 0.37944939 | 0.70435418 | 0.9093513 |
| DEPDC7 | 377.908293 | 0.108005364 | 0.67095307 | 0.16097305 | 0.87211463 | 0.9657206 |

| | | | | | | |
|---|---|---|---|---|---|---|
| TCP11L1 | 994.606396 | 0.207518441 | 0.35682107 | 0.58157564 | 0.56085256 | 0.84953736 |
| CSTF3 | 908.939604 | 0.216736327 | 0.1916301 | 1.13101399 | 0.2580492 | 0.69210282 |
| HIPK3 | 2446.02387 | -0.369219732 | 0.25211149 | -1.4645097 | 0.14305473 | 0.59836777 |
| KIAA1549L | 158.846387 | 2.330029462 | 0.660815 | 3.52599357 | 0.0004219 | 0.07286236 |
| AL049629.1 | 453.238614 | 1.326259406 | 0.54640526 | 2.42724494 | 0.01521398 | 0.3105046 |
| CD59 | 167.663631 | 0.32991817 | 0.30469262 | 1.08279014 | 0.27890158 | 0.70446562 |
| FBXO3 | 632.501766 | 0.033194251 | 0.28359506 | 0.11704806 | 0.90682197 | 0.97664815 |
| LMO2 | 71.245885 | -0.092350499 | 0.41775842 | -0.221062 | 0.82504418 | 0.95192757 |
| CAPRIN1 | 1051.44648 | -0.04832799 | 0.26165777 | -0.1846992 | 0.85346495 | 0.96109041 |
| NAT10 | 361.045648 | -0.421536938 | 0.28347783 | -1.4870191 | 0.13700977 | 0.59474076 |
| ABTB2 | 3080.32094 | -0.916159201 | 0.48305873 | -1.8965793 | 5.79E-02 | 4.77E-01 |
| CAT | 1411.31003 | 0.013861992 | 0.5414996 | 0.02559927 | 0.97957697 | 0.99484461 |
| APIP | 251.632757 | -0.661339427 | 0.34251482 | -1.9308345 | 0.05350353 | 0.46711025 |
| PDHX | 726.204589 | 0.199836958 | 0.22754816 | 0.8782183 | 0.37982526 | 0.77095585 |
| CD44 | 171.540738 | 2.057591827 | 0.9140755 | 2.25100861 | 2.44E-02 | 0.37480999 |
| FJX1 | 55.5648003 | -0.540040409 | 0.84429195 | -0.639637 | 0.52240859 | 0.83477439 |
| TRIM44 | 3133.33101 | 0.176074807 | 0.23645806 | 0.74463441 | 0.45649276 | 0.80507423 |
| LDLRAD3 | 619.505072 | -0.107456498 | 0.41480375 | -0.2590538 | 7.96E-01 | 0.94252227 |
| COMMD9 | 177.78023 | -0.344244072 | 0.2304586 | -1.493735 | 0.1352449 | 0.59381224 |
| PRR5L | 64.9582041 | 0.479592002 | 0.4892764 | 0.98020669 | 0.3269841 | 0.73745391 |
| TRAF6 | 385.478725 | 0.444093765 | 0.22480988 | 1.97541921 | 0.04822059 | 0.45192142 |
| RAG1 | 86.0403491 | 1.26945813 | 0.48908221 | 2.59559255 | 0.0094428 | 0.25998727 |
| C11orf74 | 144.71231 | 0.007869224 | 0.3265406 | 0.02409876 | 0.98077383 | 0.99510207 |
| LRRC4C | 308.152108 | -0.546326433 | 0.78436622 | -0.6965196 | 0.4861035 | 0.8185529 |
| API5 | 735.327663 | -0.476240748 | 0.23396653 | -2.035508 | 0.04179978 | 0.43537207 |
| TTC17 | 3795.67336 | 0.04657423 | 0.16473264 | 0.28272617 | 0.77738675 | 0.93618346 |
| AC068205.1 | 94.7649511 | -0.033889621 | 0.33180799 | -0.1021362 | 0.91864854 | 0.97862093 |
| HSD17B12 | 2125.35988 | 0.259199126 | 0.19496076 | 1.32949385 | 0.18368509 | 0.63276491 |
| AC087521.2 | 43.9652132 | 1.181943108 | 0.60846118 | 1.94251195 | 0.05207516 | 0.46437412 |
| ALKBH3 | 221.275249 | -0.190748231 | 0.20576882 | -0.9270026 | 0.35392517 | 0.75682564 |
| C11orf96 | 122.424404 | -1.426101002 | 1.051465 | -1.3562991 | 0.17500403 | 0.62699479 |
| ACCS | 213.028644 | 1.087986796 | 0.69217674 | 1.5718338 | 0.1159891 | 0.57350419 |
| EXT2 | 691.098409 | 0.034756066 | 0.22504653 | 0.15443947 | 0.87726324 | 0.96653382 |
| CD82 | 53.6673483 | 0.292733943 | 0.58730036 | 0.49843992 | 0.61817401 | 0.87470952 |
| TSPAN18 | 970.576345 | 0.14803044 | 0.51173993 | 0.28926889 | 0.77237562 | 0.933623 |
| TP53I11 | 67.0888288 | 0.023686761 | 0.52800134 | 0.04486118 | 0.96421796 | 0.99029939 |
| PRDM11 | 146.194862 | -0.016896311 | 0.40249056 | -0.0419794 | 0.96651512 | 0.99122723 |
| SLC35C1 | 78.2569735 | 0.234920775 | 0.49960104 | 0.47021675 | 0.63820017 | 0.8827026 |
| CRY2 | 532.230679 | 0.58334559 | 0.39250292 | 1.48621974 | 0.13722101 | 0.59474076 |
| MAPK8IP1 | 65.4201086 | 1.115002057 | 0.405595 | 2.7490528 | 0.00597678 | 0.22637955 |
| PEX16 | 113.436882 | -0.254012627 | 0.31510822 | -0.8061123 | 0.42017807 | 0.78798479 |
| LARGE2 | 149.612591 | 0.651900386 | 0.61231173 | 1.06465441 | 0.28703235 | 0.71118494 |
| PHF21A | 2173.03425 | 0.369679413 | 0.39946505 | 0.9254362 | 0.35473906 | 0.75755608 |
| DGKZ | 530.21695 | -0.156964692 | 0.20933255 | -0.7498342 | 0.4533546 | 0.80433814 |
| MDK | 98.1823859 | -1.008942957 | 0.91650833 | -1.1008552 | 0.27095968 | 0.69961625 |
| AMBRA1 | 2964.27607 | 0.068283256 | 0.29109827 | 0.23457115 | 0.8145416 | 0.94881072 |
| HARBI1 | 55.4745417 | -0.417866617 | 0.53717591 | -0.7778953 | 0.43663074 | 0.79558019 |
| ATG13 | 568.45308 | -0.23374706 | 0.26696817 | -0.8755615 | 0.38126845 | 0.77210521 |
| ARHGAP1 | 336.386447 | 0.067682319 | 0.35623106 | 0.18999556 | 0.84931261 | 0.95959925 |
| ZNF408 | 38.1355022 | -0.518216933 | 0.39160774 | -1.3233062 | 0.18573358 | 0.63661432 |
| CKAP5 | 636.714433 | -0.307289058 | 0.19694304 | -1.5602941 | 0.1186904 | 0.57589001 |
| LRP4 | 473.72074 | 0.36397908 | 0.52823064 | 0.68905333 | 0.49078971 | 0.81989696 |
| C11orf49 | 2557.57796 | -0.271676747 | 0.20423952 | -1.330187 | 0.18345668 | 0.63276491 |
| ARFGAP2 | 297.419392 | -0.168386377 | 0.26473392 | -0.6360589 | 0.524738 | 0.83593508 |

| | | | | | | |
|---|---|---|---|---|---|---|
| PACSIN3 | 85.0241715 | -0.225848868 | 0.33586974 | -0.6724299 | 0.50131003 | 0.8260948 |
| DDB2 | 236.866625 | 0.342847961 | 0.61026591 | 0.56180094 | 0.57425165 | 0.85448213 |
| ACP2 | 145.883369 | -0.085603877 | 0.32127073 | -0.266454 | 0.78988957 | 0.94031124 |
| NR1H3 | 234.473278 | 0.591649461 | 0.29470138 | 2.00762368 | 0.0446833 | 0.44360789 |
| MADD | 366.445601 | -0.137481437 | 0.17637087 | -0.7795019 | 4.36E-01 | 0.79549938 |
| SLC39A13 | 207.359797 | -0.512415055 | 0.40716198 | -1.2585042 | 0.20820947 | 0.65352123 |
| PSMC3 | 146.015427 | -0.453645212 | 0.42399452 | -1.0699318 | 0.28465002 | 0.70952861 |
| CELF1 | 3394.59489 | -0.568121271 | 0.33262004 | -1.7080188 | 0.08763286 | 0.5273546 |
| NDUFS3 | 215.973305 | -0.062107876 | 0.30032744 | -0.2068005 | 0.83616564 | 0.95468028 |
| KBTBD4 | 43.4008723 | -0.232528221 | 0.38871209 | -0.5982017 | 0.54970538 | 0.84612944 |
| MTCH2 | 1243.30986 | 0.029476957 | 0.19863549 | 0.14839723 | 0.88202929 | 0.96796225 |
| AGBL2 | 49.6411955 | 0.019613523 | 0.61158307 | 0.03207009 | 0.97441616 | 0.99377398 |
| FNBP4 | 2088.48482 | -0.434493893 | 0.26587909 | -1.6341785 | 0.10222137 | 0.54898162 |
| NUP160 | 959.89341 | -0.008645536 | 0.22454727 | -0.0385021 | 0.96928738 | 0.99187457 |
| PTPRJ | 6656.38778 | 0.247220882 | 0.34803954 | 0.71032413 | 0.47750316 | 0.81579656 |
| FOLH1 | 288.546767 | -0.021533515 | 0.71311463 | -0.0301964 | 9.76E-01 | 9.94E-01 |
| TNKS1BP1 | 349.925117 | 0.193864264 | 0.38910084 | 0.49823656 | 6.18E-01 | 8.75E-01 |
| SSRP1 | 174.305686 | -0.175933038 | 0.37796827 | -0.4654704 | 0.64159464 | 0.8841196 |
| TIMM10 | 104.066228 | -0.227981267 | 0.55483746 | -0.4108974 | 0.68114777 | 0.89947163 |
| UBE2L6 | 85.7045469 | -0.680481616 | 1.00533081 | -0.6768733 | 0.49848633 | 0.82362007 |
| SERPING1 | 467.022232 | 0.274339855 | 0.58693659 | 0.4674097 | 0.64020679 | 0.88353951 |
| CLP1 | 36.0730843 | -0.029113606 | 0.4623019 | -0.0629753 | 0.94978616 | 0.98723887 |
| ZDHHC5 | 671.031559 | -0.122673959 | 0.27724149 | -0.4424805 | 0.65814152 | 0.89058983 |
| MED19 | 119.404853 | -0.267200238 | 0.30048955 | -0.8892164 | 0.37388679 | 0.7697436 |
| CTNND1 | 1724.37053 | 0.508567202 | 0.29366046 | 1.73182052 | 0.08330552 | 0.51935659 |
| OR9Q1 | 418.287191 | 2.206436642 | 0.95437929 | 2.3119075 | 0.02078278 | 0.35313211 |
| LPXN | 403.470237 | 0.295072412 | 0.39513331 | 0.74676673 | 0.45520438 | 0.80465616 |
| GLYAT | 3866.30172 | 0.199465395 | 0.81423348 | 0.24497322 | 0.80647717 | 0.94585363 |
| GLYATL2 | 89.7497538 | 1.070204841 | 0.49999592 | 2.14042717 | 3.23E-02 | 0.41057814 |
| GLYATL1 | 2231.83138 | 1.57014418 | 0.76631148 | 2.04896341 | 0.04046569 | 0.43086475 |
| AP001636.3 | 169.466653 | 1.456605897 | 0.84387877 | 1.7260843 | 0.08433225 | 0.52183702 |
| FAM111A-D1 | 53.3736272 | 0.303541922 | 0.48559256 | 0.62509591 | 0.53190812 | 0.83789294 |
| FAM111A | 275.735097 | 0.583764588 | 0.45134463 | 1.29338991 | 0.19587624 | 0.64558745 |
| DTX4 | 296.527857 | 0.281833739 | 0.64164034 | 0.43923943 | 0.66048806 | 0.89124924 |
| OSBP | 898.414382 | -0.447636497 | 0.20704723 | -2.1620018 | 0.03061803 | 0.40446361 |
| AP000442.2 | 47.9520771 | 0.329137785 | 0.51186769 | 0.6430134 | 0.52021541 | 0.833897 |
| PATL1 | 616.704826 | -0.336871663 | 0.30327967 | -1.1107624 | 0.26667062 | 0.69812269 |
| STX3 | 2570.90303 | -0.663936069 | 0.3944611 | -1.6831471 | 0.09234661 | 0.53621484 |
| MRPL16 | 164.160858 | -0.020110665 | 0.48905582 | -0.0411214 | 0.96719911 | 0.99157802 |
| GIF | 36.2889918 | 1.838923867 | 0.71511128 | 2.57152128 | 0.01012528 | 0.26582196 |
| AP002347.1 | 83.2960226 | 2.108217891 | 0.6246115 | 3.3752467 | 0.0007375 | 0.09083761 |
| AP000790.3 | 60.8806435 | 2.721279287 | 0.84618555 | 3.21593684 | 0.00130019 | 0.11802493 |
| PRPF19 | 252.142559 | -0.829306184 | 0.51505769 | -1.6101229 | 0.10737103 | 0.55579103 |
| TMEM109 | 89.2633983 | 0.335958158 | 0.38891017 | 0.86384513 | 0.38767298 | 0.77373447 |
| TMEM132A | 46.6933929 | 0.229189438 | 0.75361583 | 0.30411972 | 7.61E-01 | 0.92989212 |
| VPS37C | 285.049107 | -0.330050648 | 0.32077892 | -1.0289038 | 0.30352489 | 0.72082426 |
| VWCE | 40.5722667 | 1.308818747 | 0.71308994 | 1.83541888 | 0.06644365 | 0.48798003 |
| DDB1 | 596.123731 | -0.268891129 | 0.31357451 | -0.8575032 | 0.39116687 | 0.77606489 |
| TKFC | 258.015563 | 0.607800674 | 0.54446073 | 1.1163352 | 2.64E-01 | 6.96E-01 |
| CYB561A3 | 93.7856822 | -0.152725072 | 0.27616238 | -0.5530263 | 0.58024537 | 0.85831909 |
| TMEM138 | 160.582539 | -0.258607739 | 0.27678652 | -0.934322 | 0.35013781 | 0.75455562 |
| TMEM216 | 48.1310628 | -0.362370865 | 0.46254035 | -0.7834362 | 0.43337099 | 0.79380258 |
| CPSF7 | 709.197627 | -0.424985521 | 0.23739393 | -1.7902122 | 0.0734198 | 0.50459544 |
| AP003108.2 | 448.347571 | 0.490314298 | 0.38804242 | 1.26355848 | 0.20638854 | 0.65243721 |

| | | | | | | |
|---|---|---|---|---|---|---|
| DAGLA | 61.6146559 | 0.178327699 | 0.57622649 | 0.30947501 | 0.75696022 | 0.92799282 |
| MYRF | 176.452704 | 1.501005453 | 0.52515342 | 2.85822273 | 0.00426021 | 0.20749797 |
| TMEM258 | 193.462794 | 0.072650235 | 0.31286173 | 0.23221196 | 0.81637339 | 0.94927299 |
| FADS2 | 139.735045 | 0.921628159 | 0.84545737 | 1.09009419 | 0.27567166 | 0.70247842 |
| FADS1 | 178.184143 | -0.468226884 | 0.77632391 | -0.6031334 | 5.46E-01 | 8.45E-01 |
| FADS3 | 463.483424 | -0.308170806 | 0.48899609 | -0.6302112 | 5.29E-01 | 0.83761342 |
| RAB3IL1 | 105.068989 | -1.320953091 | 0.49828455 | -2.6510015 | 0.00802535 | 0.24775553 |
| FTH1 | 15342.8017 | 0.04818225 | 0.70162491 | 0.06867238 | 0.94525041 | 0.98566355 |
| AP003733.3 | 32.8081771 | 0.228654116 | 1.06610062 | 0.21447705 | 0.83017506 | 0.95325673 |
| INCENP | 69.3802384 | 0.055642093 | 0.43193599 | 0.12882023 | 0.8974999 | 0.97357005 |
| ASRGL1 | 1140.91285 | 0.145614996 | 0.4795159 | 0.30367084 | 0.76137868 | 0.93000994 |
| AHNAK | 1378.90304 | -1.21736403 | 0.35692238 | -3.4107249 | 0.0006479 | 0.08457928 |
| MTA2 | 66.4566751 | -0.687292173 | 0.46692857 | -1.4719429 | 0.14103631 | 0.59555904 |
| EML3 | 127.225628 | -0.301374785 | 0.36541091 | -0.8247558 | 0.40951022 | 0.78542936 |
| B3GAT3 | 206.113577 | -0.257571007 | 0.33883642 | -0.7601633 | 0.44715698 | 0.80223612 |
| GANAB | 697.341463 | -0.117823567 | 0.22747294 | -0.5179674 | 0.60448102 | 0.86865188 |
| INTS5 | 52.2594372 | 0.052835601 | 0.41971431 | 0.12588468 | 0.89982321 | 0.97465308 |
| CSKMT | 66.9075398 | -0.12282232 | 0.41834415 | -0.2935916 | 0.76907001 | 0.93209094 |
| UQCC3 | 181.006836 | -0.636972158 | 0.35403855 | -1.7991605 | 0.0719933 | 0.50168387 |
| UBXN1 | 223.805852 | -0.299751147 | 0.36773399 | -0.8151304 | 0.41499768 | 0.78632759 |
| TTC9C | 268.186155 | 0.102263161 | 0.28080781 | 0.36417491 | 0.7157274 | 0.91298152 |
| POLR2G | 79.203531 | -0.32909134 | 0.38790427 | -0.8483829 | 0.39622478 | 0.7795748 |
| TAF6L | 170.433231 | -0.374044579 | 0.39501515 | -0.946912 | 0.34368363 | 0.7507483 |
| TMEM223 | 165.512155 | -0.114833524 | 0.27851989 | -0.4122992 | 0.68012015 | 0.89941358 |
| TMEM179B | 239.525132 | 0.050187545 | 0.36146743 | 0.13884389 | 0.88957351 | 0.97208467 |
| NXF1 | 311.046073 | -0.576447474 | 0.40928232 | -1.4084348 | 0.15900235 | 0.61423843 |
| STX5 | 526.220319 | -0.164386099 | 0.30159918 | -0.5450482 | 0.58572035 | 0.86077718 |
| AP001160.1 | 42.8246393 | 0.147782765 | 0.43010302 | 0.34359853 | 0.73114823 | 0.9190831 |
| WDR74 | 108.922235 | -0.856207383 | 0.41216402 | -2.0773462 | 0.03776962 | 0.42261887 |
| SLC3A2 | 1276.16021 | -1.317763968 | 0.44258315 | -2.9774382 | 0.00290668 | 0.17474518 |
| SLC22A6 | 4506.83278 | 1.624160767 | 0.97561813 | 1.6647505 | 0.09596254 | 0.53870645 |
| SLC22A8 | 1422.81261 | 2.538880909 | 0.9042011 | 2.80787196 | 0.00498701 | 0.21686099 |
| SLC22A24 | 789.514516 | 2.342124327 | 0.56000093 | 4.1823579 | 2.89E-05 | 0.01647703 |
| SLC22A10 | 134.471515 | 0.608987978 | 0.76385518 | 0.79725581 | 0.42530248 | 0.78894887 |
| HRASLS5 | 21.8123328 | 1.29884275 | 1.46666126 | 0.88557786 | 0.37584504 | 0.7697436 |
| RARRES3 | 830.49524 | 0.049636295 | 0.85941264 | 0.05775607 | 0.95394293 | 0.98836423 |
| HRASLS2 | 35.0841468 | -0.96331014 | 0.85605192 | -1.1252941 | 0.26046445 | 0.69280433 |
| PLA2G16 | 1079.66941 | 0.023611997 | 0.26747186 | 0.08827843 | 0.92965538 | 0.98272895 |
| ATL3 | 475.06032 | -0.016122547 | 0.26758712 | -0.0602516 | 0.95195526 | 0.98766432 |
| RTN3 | 1077.99863 | -0.486994695 | 0.36697811 | -1.3270402 | 0.1844954 | 0.63377801 |
| C11orf95 | 88.3345298 | 0.531689664 | 0.29885923 | 1.77906386 | 0.07522929 | 0.50718602 |
| SPINDOC | 82.0252383 | 0.338199807 | 0.39428612 | 0.85775226 | 0.39102928 | 0.77606489 |
| MARK2 | 842.255005 | -0.226493718 | 0.31982056 | -0.70819 | 0.47882729 | 0.81596289 |
| NAA40 | 204.998015 | -0.06257231 | 0.32208723 | -0.1942713 | 0.84596344 | 0.95784082 |
| OTUB1 | 261.414162 | -0.249582346 | 0.22181605 | -1.1251771 | 0.26051399 | 0.69280433 |
| MACROD1 | 880.880747 | 0.101973665 | 0.40693592 | 0.250589 | 0.80213189 | 0.94499146 |
| AP000721.2 | 137.022909 | 0.539105545 | 0.6937296 | 0.77711193 | 0.43709274 | 0.79607097 |
| FLRT1 | 43.9393594 | 0.827635326 | 0.83839964 | 0.98716088 | 0.32356378 | 0.7354443 |
| STIP1 | 377.056297 | -1.017699687 | 0.41004105 | -2.4819458 | 0.01306671 | 0.29875272 |
| TRPT1 | 107.934522 | -0.007592844 | 0.34938516 | -0.021732 | 0.98266172 | 0.99522272 |
| NUDT22 | 209.334141 | 0.130343872 | 0.24194799 | 0.53872683 | 0.59007536 | 0.86292147 |
| AP001453.1 | 78.7566357 | 0.600887657 | 0.48931298 | 1.22802313 | 0.21943829 | 0.66186058 |
| DNAJC4 | 109.374295 | 0.030127441 | 0.41949391 | 0.07181854 | 0.94274632 | 0.98492559 |
| VEGFB | 424.732892 | 0.143518762 | 0.3738667 | 0.38387683 | 0.70106974 | 0.90783786 |

| | | | | | | |
|---|---|---|---|---|---|---|
| FKBP2 | 448.597859 | 0.055137199 | 0.40418873 | 0.13641449 | 0.89149362 | 0.97234681 |
| PPP1R14B | 244.068023 | -0.37661809 | 0.43029516 | -0.8752552 | 3.81E-01 | 0.7722792 |
| PLCB3 | 79.148777 | 0.000761534 | 0.37749783 | 0.00201732 | 0.99839041 | 0.99977652 |
| BAD | 217.996965 | 0.170761324 | 0.33839017 | 0.50462849 | 0.61381979 | 0.87225883 |
| GPR137 | 203.04154 | -0.155217745 | 0.29462577 | -0.5268302 | 0.59831152 | 0.86646266 |
| ESRRA | 773.659343 | 0.385296763 | 0.15558834 | 2.47638586 | 0.01327201 | 0.29875272 |
| TRMT112 | 276.466265 | -0.704467783 | 0.50748942 | -1.3881428 | 0.16509356 | 0.62065658 |
| PRDX5 | 406.399114 | -0.445566169 | 0.41348657 | -1.0775832 | 0.28121982 | 0.70625976 |
| CCDC88B | 39.7289538 | 0.24679973 | 0.57277699 | 0.43088276 | 0.66655362 | 0.89420131 |
| RPS6KA4 | 107.059608 | -0.409505936 | 0.63414417 | -0.6457616 | 0.51843378 | 0.83325693 |
| SLC22A11 | 1191.50263 | 0.378966193 | 0.32626727 | 1.16152071 | 0.24543021 | 0.68292973 |
| SLC22A12 | 1310.83071 | -0.149858768 | 0.41664273 | -0.3596817 | 0.71908518 | 0.91465123 |
| NRXN2 | 110.326715 | 1.291299004 | 0.4796596 | 2.69211542 | 0.00710004 | 0.23330842 |
| SF1 | 1463.51524 | -0.377681125 | 0.22785479 | -1.6575519 | 0.09740795 | 0.54093104 |
| MAP4K2 | 57.6380239 | 0.815294419 | 0.5300631 | 1.53810824 | 0.12402215 | 0.58398201 |
| MEN1 | 132.617335 | 0.048427426 | 0.29004716 | 0.16696398 | 0.86739839 | 0.96486219 |
| CDC42BPG | 33.2959389 | 0.768702877 | 0.45165054 | 1.70198598 | 0.088758 | 0.52996572 |
| EHD1 | 359.240084 | -0.950252623 | 0.32153654 | -2.9553488 | 0.00312316 | 0.18216645 |
| MIR194-2HG | 142.738751 | 1.428896127 | 0.68886432 | 2.07427805 | 0.0380535 | 0.42339465 |
| ATG2A | 296.24587 | -0.39397647 | 0.40891447 | -0.9634691 | 0.33531214 | 0.74190856 |
| PPP2R5B | 176.764398 | 0.202797204 | 0.38261373 | 0.53003118 | 0.59609031 | 0.86539512 |
| MAJIN | 155.434491 | -0.357766134 | 0.73836747 | -0.4845367 | 0.62800503 | 0.87990033 |
| BATF2 | 60.1527129 | -0.221437562 | 0.69819181 | -0.3171586 | 0.75112323 | 0.92620063 |
| SAC3D1 | 41.6624745 | -0.224088184 | 0.5374623 | -0.4169375 | 0.67672413 | 0.89843878 |
| ZFPL1 | 103.32521 | -0.314634364 | 0.34097815 | -0.9227406 | 0.35614243 | 0.75841231 |
| VPS51 | 501.845766 | 0.191002846 | 0.33793692 | 0.56520265 | 0.57193593 | 0.85407188 |
| TM7SF2 | 116.871407 | 0.453401139 | 0.41221884 | 1.09990398 | 0.27137396 | 0.70002917 |
| ZNHIT2 | 49.5223137 | -0.17002136 | 0.5263716 | -0.3230063 | 0.74669045 | 0.92448528 |
| FAU | 821.847892 | -0.243181291 | 0.46994608 | -0.5174664 | 0.60483064 | 0.86865188 |
| SYVN1 | 174.883224 | -0.16691781 | 0.41488193 | -0.4023261 | 0.68744408 | 0.90265313 |
| MRPL49 | 92.9740576 | -0.031188684 | 0.43004099 | -0.0725249 | 0.94218418 | 0.98489601 |
| AP003068.2 | 33.6194177 | 0.607089101 | 0.49462364 | 1.22737582 | 0.21968137 | 0.66208719 |
| CAPN1 | 463.535461 | 0.126786836 | 0.29171037 | 0.43463259 | 6.64E-01 | 0.89313476 |
| POLA2 | 98.040375 | -0.462023076 | 0.29965669 | -1.5418413 | 0.12311215 | 0.5825794 |
| DPF2 | 271.072442 | -0.132302338 | 0.1720854 | -0.7688179 | 0.44200142 | 0.79917075 |
| SLC25A45 | 222.720926 | 0.48379307 | 0.39650222 | 1.22015224 | 0.22240717 | 0.66373504 |
| FRMD8 | 230.858078 | -0.072743809 | 0.35048648 | -0.207551 | 0.83557959 | 0.95467946 |
| NEAT1 | 226385.829 | 0.660034337 | 0.35424238 | 1.86322806 | 6.24E-02 | 0.4840117 |
| MALAT1 | 756554.886 | 0.335080065 | 0.43315319 | 0.77358327 | 0.43917727 | 0.79750844 |
| SCYL1 | 292.917751 | -0.175056233 | 0.37419165 | -0.4678251 | 0.6399097 | 0.88321992 |
| LTBP3 | 357.152883 | 0.41740276 | 0.39871147 | 1.04687924 | 0.29515528 | 0.71646902 |
| SSSCA1 | 106.744814 | -0.787696781 | 0.58364774 | -1.3496099 | 0.17714114 | 0.62731853 |
| FAM89B | 83.6286612 | 0.152397233 | 0.46528658 | 0.32753413 | 0.74326394 | 0.92333113 |
| EHBP1L1 | 70.0865995 | 0.48897674 | 0.54596645 | 0.89561682 | 0.37045745 | 0.76698912 |
| MAP3K11 | 269.636968 | -0.20874628 | 0.3395821 | -0.6147152 | 0.53874281 | 0.84070752 |
| PCNX3 | 181.530035 | -0.343669596 | 0.3960709 | -0.8676972 | 0.38556014 | 0.77317162 |
| SIPA1 | 60.8630061 | -0.17295615 | 0.49411351 | -0.3500332 | 0.72631375 | 0.91635631 |
| RELA | 197.458911 | -0.349775157 | 0.27363893 | -1.2782361 | 2.01E-01 | 0.64849129 |
| KAT5 | 204.446713 | -0.384769446 | 0.31177283 | -1.234134 | 0.21715299 | 0.66032006 |
| RNASEH2C | 180.865099 | -0.295082355 | 0.45792617 | -0.6443885 | 0.51932354 | 0.8335052 |
| AP001266.2 | 58.3632364 | 0.603977941 | 0.90971321 | 0.66392126 | 0.50674071 | 0.82806179 |
| AP5B1 | 103.957413 | -0.040899551 | 0.29514243 | -0.1385756 | 0.88978549 | 0.97212576 |
| OVOL1 | 40.3401826 | 0.585510705 | 0.65619989 | 0.89227492 | 0.37224559 | 0.76866079 |
| CFL1 | 807.49068 | -0.389984877 | 0.44274895 | -0.8808262 | 0.3784119 | 0.77061593 |

| | | | | | | |
|---|---|---|---|---|---|---|
| SNX32 | 37.6454327 | 0.312432547 | 0.50387529 | 0.62005927 | 0.53521877 | 0.83954482 |
| MUS81 | 145.492451 | 0.003127802 | 0.28434261 | 0.01100012 | 0.99122335 | 0.99748517 |
| EFEMP2 | 72.9605886 | 0.740737512 | 0.61148455 | 1.21137568 | 0.22575145 | 0.66688879 |
| FIBP | 71.4912453 | 0.153520944 | 0.44940633 | 0.34160833 | 0.73264567 | 0.91952673 |
| CCDC85B | 66.941997 | 0.19746704 | 0.5243594 | 0.37658721 | 0.7064804 | 0.91002281 |
| FOSL1 | 153.59293 | -2.685685403 | 1.52456492 | -1.7616078 | 7.81E-02 | 5.12E-01 |
| C11orf68 | 66.5531512 | -0.612363448 | 0.54234405 | -1.1291051 | 0.25885349 | 0.69241621 |
| DRAP1 | 167.623653 | -0.8345118 | 0.31866434 | -2.61878 | 0.00882448 | 0.25464675 |
| SART1 | 161.369445 | -0.838371728 | 0.46305849 | -1.8105093 | 0.07021684 | 0.49575843 |
| EIF1AD | 75.3996286 | -0.434471073 | 0.32163916 | -1.3508028 | 0.17675861 | 0.62709281 |
| BANF1 | 177.205605 | -0.460421164 | 0.33429847 | -1.3772757 | 0.16842702 | 0.62304975 |
| SF3B2 | 528.752129 | -0.429277748 | 0.1688784 | -2.541934 | 0.0110241 | 0.27879579 |
| PACS1 | 3359.83771 | 0.040096449 | 0.29207285 | 0.13728236 | 0.89080761 | 0.97230741 |
| KLC2 | 143.437994 | -0.235212789 | 0.40073448 | -0.5869542 | 0.55723446 | 0.84861484 |
| RAB1B | 249.342353 | -0.173725281 | 0.39085503 | -0.444475 | 0.65669922 | 0.89053559 |
| YIF1A | 174.800729 | -0.926453483 | 0.54249801 | -1.7077546 | 0.08768188 | 0.5273546 |
| BRMS1 | 132.882328 | -0.812821523 | 0.407942 | -1.9924928 | 0.04631701 | 0.44508419 |
| B4GAT1 | 70.9908958 | 0.729805464 | 0.58146125 | 1.25512313 | 0.20943406 | 0.65441616 |
| AP001107.9 | 51.1130175 | 0.684246936 | 0.64846795 | 1.05517464 | 2.91E-01 | 7.13E-01 |
| SLC29A2 | 51.1470086 | 0.594292334 | 0.51717698 | 1.14910825 | 0.25051135 | 0.68627228 |
| MRPL11 | 88.6662507 | -0.074282217 | 0.32943664 | -0.2254826 | 0.82160389 | 0.95105545 |
| AP002748.3 | 43.0253172 | 0.517397697 | 0.51600447 | 1.00270002 | 0.31600562 | 0.72798933 |
| DPP3 | 92.2502674 | -0.568709094 | 0.50157615 | -1.133844 | 0.25686 | 0.69116216 |
| ZDHHC24 | 158.334592 | 0.205403599 | 0.30698229 | 0.66910569 | 0.50342806 | 0.82660378 |
| CTSF | 117.815165 | 0.537134576 | 0.44846988 | 1.19770491 | 0.23103192 | 0.66878058 |
| CCS | 449.239595 | 0.504581318 | 0.34925839 | 1.44472209 | 0.14853596 | 0.60622442 |
| RBM14-RBM | 390.338951 | -0.171808638 | 0.36340474 | -0.4727749 | 0.63637379 | 0.88266168 |
| RBM4 | 222.936664 | -0.232901914 | 0.44521392 | -0.5231236 | 0.60088824 | 0.86755264 |
| RBM4B | 185.367604 | 0.13053674 | 0.4375739 | 0.2983193 | 0.76545947 | 0.93091853 |
| AP001157.1 | 46.8358757 | 0.035200135 | 0.57568045 | 0.06114527 | 0.95124352 | 0.9874059 |
| SPTBN2 | 69.0804161 | 0.285385062 | 0.5022541 | 0.56820853 | 0.56989338 | 0.85327114 |
| C11orf80 | 755.125567 | 0.213235379 | 0.25470163 | 0.83719676 | 4.02E-01 | 7.83E-01 |
| RCE1 | 53.2760292 | -0.248211635 | 0.37646839 | -0.659316 | 0.50969287 | 0.82883692 |
| PC | 1311.606 | -0.546077223 | 0.52520597 | -1.0397392 | 0.2984611 | 0.71852421 |
| RHOD | 107.433983 | -0.617128564 | 0.39772638 | -1.551641 | 0.12074815 | 0.5789069 |
| KDM2A | 3559.59798 | -0.458507209 | 0.19867649 | -2.3078081 | 0.02100981 | 0.35422078 |
| GRK2 | 190.265775 | -0.305119151 | 0.32204166 | -0.9474524 | 0.34340829 | 0.75073625 |
| ANKRD13D | 282.94549 | -0.13572586 | 0.40168842 | -0.3378884 | 0.73544728 | 0.92019178 |
| SSH3 | 165.07122 | 0.494804994 | 0.24909296 | 1.98642707 | 0.04698592 | 0.44732709 |
| RAD9A | 1417.28198 | 0.114067692 | 0.24019677 | 0.4748927 | 0.63486345 | 0.88238823 |
| POLD4 | 62.1713141 | -0.447972068 | 0.48864626 | -0.9167615 | 0.35926764 | 0.75936492 |
| CLCF1 | 124.523116 | -0.493398776 | 0.35215273 | -1.4010932 | 0.16118622 | 0.61697277 |
| PPP1CA | 376.623979 | -0.014646464 | 0.37723353 | -0.038826 | 0.96902913 | 0.99187457 |
| RPS6KB2 | 184.398023 | -0.244346956 | 0.35641242 | -0.6855736 | 0.49298202 | 0.82100434 |
| CORO1B | 187.023772 | 0.082921912 | 0.41521018 | 0.19971069 | 0.84170685 | 0.95681213 |
| TMEM134 | 220.55617 | 0.455216416 | 0.38824881 | 1.17248631 | 0.24100187 | 0.67957469 |
| AIP | 107.949705 | -0.394390343 | 0.40594549 | -0.9715352 | 0.33128182 | 0.73916381 |
| PITPNM1 | 180.011074 | 0.122614577 | 0.57393797 | 0.21363733 | 0.83082988 | 0.95325673 |
| CDK2AP2 | 351.608854 | -0.304170229 | 0.57132693 | -0.5323926 | 0.59445411 | 0.86488065 |
| GSTP1 | 1547.46912 | 0.376628401 | 0.69229782 | 0.54402655 | 0.5864232 | 0.86102615 |
| NDUFV1 | 539.989518 | 0.417200584 | 0.42553568 | 0.9804127 | 0.32688245 | 0.73745391 |
| NUDT8 | 162.957238 | 0.21931584 | 0.45456816 | 0.48247075 | 0.62947157 | 0.88083664 |
| ACY3 | 257.948993 | 1.016206056 | 0.72525378 | 1.40117306 | 0.16116233 | 0.61697277 |
| AP003716.1 | 534.020739 | 0.918931241 | 0.55974221 | 1.64170438 | 0.10065128 | 0.54608983 |

| | | | | | | |
|---|---|---|---|---|---|---|
| UNC93B1 | 169.332106 | -0.847667882 | 0.43844633 | -1.9333447 | 0.05319375 | 0.46624874 |
| ALDH3B1 | 140.127762 | 0.160835456 | 0.61914176 | 0.25977162 | 0.79503995 | 0.94252227 |
| NDUFS8 | 514.045407 | -0.000647933 | 0.31946894 | -0.0020282 | 0.99838177 | 0.99977652 |
| TCIRG1 | 39.2250951 | 0.24387003 | 0.73381169 | 0.33233326 | 0.73963763 | 0.92123698 |
| CHKA | 1751.67571 | -0.912921677 | 0.49931932 | -1.8283324 | 0.06749968 | 0.48963361 |
| KMT5B | 1257.7086 | -0.19405603 | 0.22836725 | -0.8497542 | 0.39546176 | 0.77931337 |
| C11orf24 | 116.150024 | -0.172103276 | 0.45760484 | -0.3760958 | 0.70684565 | 0.91002281 |
| LRP5 | 999.390773 | 0.182356797 | 0.37001389 | 0.4928377 | 0.62212727 | 0.87713418 |
| PPP6R3 | 3976.76055 | -0.209462489 | 0.20639564 | -1.0148591 | 0.31017301 | 0.72498238 |
| TESMIN | 99.1621902 | 0.508136207 | 0.47640558 | 1.06660423 | 0.28615059 | 0.71118494 |
| CPT1A | 1403.77867 | -0.320968424 | 0.48450082 | -0.6624724 | 0.50766851 | 0.82860351 |
| MRPL21 | 128.410091 | -0.357476543 | 0.26266935 | -1.3609374 | 0.17353348 | 0.62663931 |
| IGHMBP2 | 302.106281 | -0.593526516 | 0.30688666 | -1.9340252 | 0.05311004 | 0.46624874 |
| TPCN2 | 770.226752 | 0.363838313 | 0.49632196 | 0.73306914 | 0.46351626 | 0.8082072 |
| AP003071.2 | 47.5936677 | 2.296002706 | 1.1603641 | 1.9786916 | 0.04785074 | 0.45133876 |
| CCND1 | 289.553952 | -1.152005841 | 0.63093729 | -1.8258643 | 0.06787071 | 0.48963361 |
| ORAOV1 | 339.151602 | -0.463254263 | 0.3331557 | -1.3905038 | 0.16437595 | 0.62065658 |
| ANO1 | 85.1356531 | 0.224036501 | 0.98035243 | 0.22852649 | 0.81923696 | 0.94990455 |
| FADD | 86.6106494 | -0.234243628 | 0.29109281 | -0.8047043 | 0.42099036 | 0.78872573 |
| PPFIA1 | 2139.92749 | -0.664564052 | 0.35675515 | -1.8628016 | 0.06249019 | 0.4840117 |
| AP002336.2 | 718.415504 | -0.184443035 | 0.49997299 | -0.368906 | 0.71219779 | 0.91165921 |
| AP000487.1 | 208.209048 | -0.622981701 | 0.58031761 | -1.0735185 | 0.28303854 | 0.70757053 |
| CTTN | 2818.56939 | -0.632914272 | 0.28434985 | -2.2258294 | 2.60E-02 | 0.38392873 |
| SHANK2 | 5720.10986 | 0.573125011 | 0.37363913 | 1.53389985 | 0.12505429 | 0.58564339 |
| SHANK2-AS1 | 41.6732525 | 1.409879701 | 0.7486622 | 1.88319872 | 0.05967344 | 0.47945128 |
| DHCR7 | 61.7309105 | 0.70854874 | 0.48357631 | 1.46522632 | 0.14285918 | 0.59824924 |
| NADSYN1 | 588.740484 | 0.060978007 | 0.29823785 | 0.204461 | 0.83799327 | 0.95496955 |
| AP003498.1 | 83.7902762 | 0.179263417 | 0.89266317 | 0.20081865 | 0.84084038 | 0.95654403 |
| FAM86C1 | 52.8641534 | -0.418115877 | 0.54516914 | -0.7669471 | 0.44311299 | 0.7993765 |
| AP002495.2 | 411.990886 | 0.402489891 | 0.3210804 | 1.25354861 | 0.21000612 | 0.65441616 |
| RNF121 | 933.885194 | 0.143171859 | 0.20963054 | 0.68297234 | 0.49462432 | 0.8220963 |
| IL18BP | 88.4689386 | -0.318652146 | 0.36546766 | -0.8719024 | 0.38326161 | 0.77317162 |
| NUMA1 | 807.473208 | -0.069631786 | 0.35023248 | -0.1988159 | 0.84240676 | 0.95716002 |
| AP002490.1 | 37.3273115 | -0.266192523 | 0.55244471 | -0.4818446 | 0.62991631 | 0.88083664 |
| LAMTOR1 | 454.409841 | -0.299943334 | 0.32806515 | -0.9142798 | 0.36056985 | 0.76009788 |
| ANAPC15 | 69.5744675 | -0.122214755 | 0.42276873 | -0.2890818 | 0.77251877 | 0.933623 |
| FOLR1 | 170.588787 | -0.58770862 | 0.60311736 | -0.9744515 | 0.3298324 | 0.73849818 |
| INPPL1 | 478.370288 | 0.11948566 | 0.26596699 | 0.44924996 | 0.65325135 | 0.88962904 |
| CLPB | 246.971682 | -0.822237748 | 0.39553112 | -2.0788193 | 0.03763396 | 0.42215481 |
| ARAP1 | 428.832187 | -0.43210369 | 0.29925253 | -1.4439433 | 0.14875492 | 0.60622442 |
| ATG16L2 | 152.409741 | 0.827525948 | 0.49362166 | 1.67643768 | 0.09365249 | 0.53688613 |
| FCHSD2 | 1503.20632 | 0.619086991 | 0.35546826 | 1.74160977 | 0.08157675 | 0.51593841 |
| P2RY6 | 209.07746 | 1.506318541 | 1.14963154 | 1.31026202 | 0.19010721 | 0.64261395 |
| ARHGEF17 | 401.723369 | 0.315237265 | 0.27578554 | 1.1430522 | 0.25301692 | 0.68887358 |
| FAM168A | 1285.33989 | -0.067559924 | 0.29022251 | -0.2327866 | 0.81592709 | 0.94906482 |
| RAB6A | 2053.45735 | -0.480600671 | 0.21627565 | -2.2221673 | 0.026272 | 0.38417182 |
| MRPL48 | 315.132615 | -0.305767855 | 0.19808965 | -1.5435832 | 0.12268933 | 0.58190405 |
| COA4 | 324.315488 | -0.159514016 | 0.2495131 | -0.6393012 | 0.52262704 | 0.8349241 |
| PAAF1 | 650.781866 | 0.640760938 | 0.44549815 | 1.43830213 | 0.15034835 | 0.60799621 |
| C2CD3 | 491.054604 | -0.092395786 | 0.19905154 | -0.4641802 | 0.64251864 | 0.88441609 |
| PPME1 | 866.597452 | -0.249759651 | 0.30052392 | -0.8310808 | 0.405928 | 0.78429302 |
| P4HA3 | 123.406747 | -0.073943305 | 0.80763885 | -0.0915549 | 0.92705168 | 0.98184958 |
| PGM2L1 | 223.50364 | 0.420539091 | 0.46734827 | 0.8998409 | 3.68E-01 | 0.76615653 |
| KCNE3 | 134.095227 | 1.557691319 | 0.66359144 | 2.34736501 | 0.01890672 | 0.34084276 |

| | | | | | | |
|---|---|---|---|---|---|---|
| POLD3 | 570.514328 | 0.404623897 | 0.44757353 | 0.90403894 | 0.36597475 | 0.76421682 |
| RNF169 | 1079.66314 | -0.777263495 | 0.45975653 | -1.690598 | 0.0909136 | 0.53537574 |
| XRRA1 | 947.060849 | -0.9977851 | 0.46177959 | -2.1607388 | 0.03071552 | 0.40446361 |
| SPCS2 | 694.591222 | 0.013886048 | 0.31734648 | 0.04375674 | 0.96509831 | 0.99068393 |
| NEU3 | 320.587461 | 0.236419833 | 0.22885759 | 1.03304346 | 0.30158356 | 0.71908176 |
| SLCO2B1 | 301.511487 | 1.450282109 | 0.78720812 | 1.84231092 | 0.06542968 | 0.48790819 |
| ARRB1 | 80.0994734 | -0.487728794 | 0.53637924 | -0.9092984 | 0.36319262 | 0.76213737 |
| RPS3 | 1304.36916 | 0.074046701 | 0.4371931 | 0.16936841 | 0.86550687 | 0.96428252 |
| SERPINH1 | 259.624296 | -0.676180302 | 0.52277656 | -1.2934404 | 0.1958588 | 0.64558745 |
| UVRAG | 1930.4968 | -0.26666969 | 0.22622018 | -1.1788059 | 0.23847546 | 0.67658923 |
| THAP12 | 255.115948 | -0.116582542 | 0.24566795 | -0.4745533 | 0.63510538 | 0.88238823 |
| GVQW3 | 142.792371 | 0.51875901 | 0.62894199 | 0.82481217 | 0.40947823 | 0.78542936 |
| AP002360.1 | 53.7974363 | 0.39445849 | 0.62309197 | 0.63306624 | 0.52669039 | 0.83703067 |
| EMSY | 1059.29629 | -0.099543307 | 0.17869654 | -0.5570522 | 0.57749174 | 0.85659658 |
| TSKU | 418.269099 | -0.098306594 | 0.75919006 | -0.1294888 | 0.8969709 | 0.97324791 |
| AP003119.2 | 61.9769746 | 0.190753138 | 0.79423291 | 0.24017279 | 0.8101963 | 0.94707153 |
| ACER3 | 1532.29738 | 0.296899111 | 0.25661072 | 1.15700198 | 0.24727155 | 0.68482118 |
| CAPN5 | 198.743903 | 1.093495879 | 0.6347641 | 1.72268073 | 0.08494628 | 0.52183702 |
| MYO7A | 287.301264 | -0.238673279 | 0.74916728 | -0.3185848 | 0.75004141 | 0.92565124 |
| PAK1 | 1079.30684 | 0.301608411 | 0.22798355 | 1.32293934 | 0.18585555 | 0.63661432 |
| CLNS1A | 766.562078 | 0.160881731 | 0.28054156 | 0.57346845 | 0.56632756 | 0.85215881 |
| AQP11 | 153.184074 | 1.181235018 | 0.54678783 | 2.16031697 | 0.03074814 | 0.40446361 |
| RSF1 | 1982.31791 | -0.009204333 | 0.23385398 | -0.0393593 | 0.96860391 | 0.99187457 |
| AAMDC | 495.802999 | 0.326140315 | 0.40205856 | 0.81117615 | 0.41726452 | 0.78652162 |
| AP002812.2 | 212.177962 | -0.058580803 | 0.34236818 | -0.1711047 | 0.86414145 | 0.96393376 |
| INTS4 | 528.379753 | -0.312116058 | 0.23113413 | -1.3503677 | 0.17689808 | 0.62719659 |
| ALG8 | 318.854035 | -0.577949405 | 0.34158643 | -1.6919566 | 0.09065426 | 0.53491084 |
| KCTD21-AS1 | 68.9190834 | -0.140106292 | 0.82726417 | -0.169361 | 0.86551269 | 0.96428252 |
| KCTD21 | 126.118156 | -0.219958483 | 0.37741327 | -0.5828054 | 0.56002433 | 0.84941263 |
| USP35 | 111.070105 | 0.252100268 | 0.58523507 | 0.43076753 | 0.66663741 | 0.89420131 |
| GAB2 | 1956.71644 | -0.550718024 | 0.45673964 | -1.2057592 | 0.22791035 | 0.66705629 |
| AP003086.2 | 484.040876 | -0.013036809 | 0.60500866 | -0.0215481 | 0.98280841 | 0.9952242 |
| NARS2 | 796.980275 | -0.324558581 | 0.2118882 | -1.5317445 | 0.1255855 | 0.58564339 |
| AP003110.1 | 46.3356346 | 0.118261089 | 0.40657752 | 0.29086972 | 0.77115096 | 0.93302012 |
| TENM4 | 1512.77708 | 1.078065452 | 0.38428319 | 2.80539318 | 0.00502552 | 0.21686099 |
| AP001978.1 | 130.150257 | 1.010141516 | 0.59988275 | 1.68389826 | 9.22E-02 | 5.36E-01 |
| MIR4300HG | 112.524716 | -0.363331105 | 0.68553258 | -0.5299983 | 0.59611311 | 0.86539512 |
| PRCP | 517.072089 | 0.834251436 | 0.39309106 | 2.12228543 | 0.03381378 | 0.41378378 |
| RAB30 | 385.601689 | -0.149545455 | 0.36699457 | -0.4074868 | 0.68365046 | 0.90058256 |
| RAB30-AS1 | 1272.08486 | 0.150877788 | 0.24851827 | 0.60710944 | 0.54377829 | 0.84396451 |
| AP001767.3 | 98.9859352 | 0.6971797 | 0.71774498 | 0.97134737 | 0.33137533 | 0.73916381 |
| PCF11 | 761.61436 | 0.256393134 | 0.26644451 | 0.96227592 | 0.33591102 | 0.74239478 |
| PCF11-AS1 | 65.239637 | -0.508474833 | 0.35168673 | -1.4458175 | 0.1482284 | 0.60613563 |
| ANKRD42 | 550.513109 | 0.13696987 | 0.3376583 | 0.40564639 | 0.68500244 | 0.90100071 |
| CCDC90B | 280.533555 | -0.052785086 | 0.31734237 | -0.1663348 | 0.86789347 | 0.96497532 |
| AP000446.1 | 289.527793 | -0.053857115 | 0.27757395 | -0.194028 | 0.84615397 | 0.95789828 |
| DLG2 | 9777.77217 | 1.347138497 | 0.54835402 | 2.45669486 | 0.01402217 | 0.30363654 |
| AP001825.1 | 75.1681594 | 1.043534461 | 0.78018294 | 1.33755098 | 0.18104285 | 0.6318513 |
| TMEM126B | 231.060594 | -0.100405489 | 0.23740752 | -0.4229246 | 0.67235025 | 0.89727632 |
| TMEM126A | 87.8514386 | 0.008794232 | 0.31308478 | 0.02808898 | 0.97759118 | 0.99478399 |
| CREBZF | 906.42982 | -0.080622617 | 0.25472256 | -0.3165115 | 0.75161431 | 0.92647278 |
| SYTL2 | 897.254522 | 1.135295719 | 0.31087723 | 3.65191013 | 0.0002603 | 0.05940045 |
| PICALM | 7195.32535 | 0.028400673 | 0.24458216 | 0.11611915 | 0.90755811 | 0.97673359 |
| EED | 324.994621 | 0.034219314 | 0.20072996 | 0.17047437 | 0.86463709 | 0.96417024 |

| | | | | | | |
|---|---|---|---|---|---|---|
| HIKESHI | 389.951816 | 0.055275387 | 0.26027789 | 0.21237066 | 0.83181787 | 0.95325673 |
| AP001831.1 | 101.701509 | 0.646440068 | 0.41276003 | 1.56614018 | 0.1173158 | 0.57350419 |
| ME3 | 311.211074 | 0.031554991 | 0.36890041 | 0.08553797 | 0.93183371 | 0.98290134 |
| PRSS23 | 373.459088 | 0.766761977 | 0.7990189 | 0.95962934 | 0.3372418 | 0.74353761 |
| AP001528.2 | 523.429148 | 2.263040999 | 0.86125524 | 2.62760783 | 0.00859876 | 0.25133745 |
| FZD4 | 444.783333 | 0.41414037 | 0.50505196 | 0.81999557 | 0.41221863 | 0.78632759 |
| AP001528.1 | 231.400012 | 0.817824758 | 0.59599577 | 1.37219893 | 0.17000152 | 0.62338438 |
| TMEM135 | 2299.34766 | 0.010265366 | 0.39655962 | 0.02588606 | 0.97934822 | 0.99482978 |
| RAB38 | 178.442259 | -0.737804036 | 0.28534949 | -2.5856154 | 0.00972053 | 0.26315585 |
| CTSC | 823.11204 | 0.339341728 | 0.4118229 | 0.82399917 | 0.40994003 | 0.78544143 |
| GRM5-AS1 | 49.5426551 | 1.272693915 | 0.72857003 | 1.74683813 | 0.08066542 | 0.51474903 |
| GRM5 | 342.592239 | 1.139620656 | 0.58829067 | 1.93717274 | 0.05272423 | 0.46564998 |
| NOX4 | 8286.65092 | 0.906199398 | 0.44621693 | 2.03084945 | 4.23E-02 | 4.37E-01 |
| AP003400.1 | 1551.23636 | 0.688251134 | 0.85043989 | 0.80928839 | 0.41834928 | 0.78724788 |
| CHORDC1 | 509.675855 | -1.002462798 | 0.54201527 | -1.8495102 | 0.06438417 | 0.48593331 |
| SLC36A4 | 669.48356 | -0.617482181 | 0.26483089 | -2.3316094 | 1.97E-02 | 0.34523522 |
| SMCO4 | 161.860443 | -0.375807243 | 0.42698567 | -0.8801402 | 0.37878338 | 0.77061593 |
| CEP295 | 591.578127 | -0.239106591 | 0.17259925 | -1.3853281 | 0.16595217 | 0.62065658 |
| TAF1D | 2277.11566 | -0.132466033 | 0.34098401 | -0.3884817 | 0.69765962 | 0.90696517 |
| AP001273.2 | 190.19133 | 0.291365111 | 0.40793178 | 0.7142496 | 0.47507285 | 0.81564184 |
| VSTM5 | 98.4890874 | 0.163991335 | 0.58278863 | 0.28139076 | 0.77841071 | 0.93646866 |
| PANX1 | 155.100813 | -0.290592436 | 0.48637605 | -0.5974645 | 0.55019728 | 0.84620747 |
| MRE11 | 417.516896 | -0.304615036 | 0.18960639 | -1.6065652 | 0.10814977 | 0.55770254 |
| ANKRD49 | 125.722162 | -0.440586075 | 0.30595468 | -1.4400371 | 0.14985691 | 0.60758225 |
| PIWIL4 | 342.165829 | 2.025167994 | 0.64363145 | 3.14647146 | 0.00165253 | 0.13207839 |
| AP000943.3 | 85.0067923 | 2.053582695 | 0.74664018 | 2.75043155 | 0.00595168 | 0.22637955 |
| AMOTL1 | 351.517799 | 1.32061201 | 0.4777674 | 2.76413168 | 0.00570745 | 0.22501232 |
| CWC15 | 274.986986 | -0.336375212 | 0.19548209 | -1.7207469 | 0.08529676 | 0.52241404 |
| SRSF8 | 268.594402 | -0.526004379 | 0.39808101 | -1.3213501 | 0.18638466 | 0.63757786 |
| ENDOD1 | 766.144151 | -0.984309062 | 0.46880053 | -2.099633 | 0.03576114 | 0.42040764 |
| AP000787.1 | 281.790253 | -1.134035297 | 0.40179981 | -2.8223889 | 0.00476673 | 0.21294213 |
| SESN3 | 559.776369 | 0.498863066 | 0.48145536 | 1.03615642 | 0.30012917 | 0.71879123 |
| FAM76B | 245.190265 | 0.188145287 | 0.19221342 | 0.97883531 | 0.32766136 | 0.73795117 |
| CEP57 | 626.399455 | 0.724343168 | 0.32539085 | 2.22607111 | 0.02600942 | 0.38392873 |
| MTMR2 | 1200.17575 | 0.201762108 | 0.24016577 | 0.8400952 | 0.40085501 | 0.78236076 |
| MAML2 | 17426.5191 | 0.543136303 | 0.48493982 | 1.12000764 | 0.26271051 | 0.69467275 |
| CCDC82 | 619.785101 | -0.248824084 | 0.30571944 | -0.8138968 | 0.41570405 | 0.78632759 |
| JRKL | 164.444952 | 0.055609151 | 0.29025069 | 0.19159008 | 0.84806332 | 0.95926753 |
| CNTN5 | 289.103382 | -0.136167335 | 0.43952889 | -0.3098029 | 0.75671083 | 0.92799282 |
| ARHGAP42 | 3399.94643 | 0.146303024 | 0.34544512 | 0.4235203 | 0.67191569 | 0.89704376 |
| TRPC6 | 156.313474 | 0.056832511 | 0.72338827 | 0.07856433 | 0.93737916 | 0.98435387 |
| CEP126 | 510.925945 | 0.857869395 | 0.45059601 | 1.90385484 | 0.05692909 | 0.4746515 |
| YAP1 | 3197.20352 | -0.024078122 | 0.27170997 | -0.088617 | 0.9293863 | 0.98272895 |
| BIRC3 | 1035.12984 | 1.617570763 | 0.53947653 | 2.99840802 | 2.71E-03 | 1.69E-01 |
| BIRC2 | 831.276639 | 0.119579731 | 0.21425641 | 0.55811508 | 0.5767658 | 0.85615431 |
| TMEM123 | 783.55562 | -0.10062645 | 0.27794119 | -0.3620422 | 7.17E-01 | 0.91327434 |
| MMP7 | 607.410288 | 0.300547046 | 0.87333196 | 0.34413838 | 0.73074222 | 0.91897609 |
| DCUN1D5 | 335.701802 | -0.064345628 | 0.23216506 | -0.2771547 | 0.78166135 | 0.93751843 |
| DYNC2H1 | 1390.0051 | 0.470585902 | 0.4857469 | 0.96878828 | 0.33265083 | 0.73988623 |
| PDGFD | 91.388428 | 0.683072801 | 0.59971004 | 1.1390051 | 0.25470103 | 0.68976367 |
| CASP4 | 582.310632 | -0.080833008 | 0.53673897 | -0.1506002 | 0.88029109 | 0.96769187 |
| GRIA4 | 565.009121 | 0.983646197 | 1.10606641 | 0.88931929 | 0.37383151 | 0.7697436 |
| MSANTD4 | 223.584262 | 0.271417196 | 0.2701836 | 1.00456575 | 0.31510599 | 0.72774353 |
| KBTBD3 | 175.803326 | 0.290648232 | 0.41032346 | 0.7083393 | 0.47873458 | 0.81596289 |

| | | | | | | |
|---|---|---|---|---|---|---|
| AASDHPPT | 452.520325 | 0.092633258 | 0.54947682 | 0.16858447 | 0.8661235 | 0.96437968 |
| GUCY1A2 | 106.856904 | 0.742703107 | 0.42165684 | 1.76139228 | 0.07817203 | 0.51192014 |
| AP000766.1 | 117.650105 | 0.050937787 | 0.29601924 | 0.17207593 | 8.63E-01 | 0.96393376 |
| CWF19L2 | 1211.45256 | -0.394519275 | 0.14711098 | -2.68178 | 7.32E-03 | 0.23590813 |
| ALKBH8 | 180.375663 | -0.397181488 | 0.2606805 | -1.5236333 | 0.12760035 | 0.58793678 |
| ELMOD1 | 288.003764 | -0.877164877 | 0.56772122 | -1.5450627 | 0.1223311 | 0.58080789 |
| SLC35F2 | 464.403295 | 0.32352354 | 0.37821548 | 0.85539475 | 0.39233265 | 0.77678804 |
| RAB39A | 53.9681902 | 0.519188422 | 0.51612435 | 1.00593669 | 0.31444603 | 0.72719146 |
| CUL5 | 2277.01431 | -0.130035451 | 0.18718323 | -0.694696 | 0.48724581 | 0.81893983 |
| AP002433.1 | 51.2688414 | 0.698634998 | 0.65953964 | 1.05927673 | 0.28947377 | 0.71218341 |
| ACAT1 | 1814.68721 | 0.041321191 | 0.34578405 | 0.11949999 | 0.90487925 | 0.97593672 |
| NPAT | 488.912309 | 0.149850041 | 0.22353919 | 0.67035244 | 0.50263314 | 0.82638105 |
| ATM | 1607.09665 | -0.144271122 | 0.2045206 | -0.7054112 | 0.48055438 | 0.81714875 |
| C11orf65 | 348.30893 | 1.320578312 | 0.55051628 | 2.39879974 | 0.01644891 | 0.32347946 |
| KDELC2 | 173.123434 | 0.143486877 | 0.36671654 | 0.39127463 | 0.69559425 | 0.90589033 |
| EXPH5 | 434.032938 | 0.357487795 | 0.38932212 | 0.91823139 | 0.35849774 | 0.75936492 |
| DDX10 | 1603.05831 | -0.344341971 | 0.32988208 | -1.0438335 | 0.29656242 | 0.71805012 |
| AP003049.2 | 395.020867 | 0.636641352 | 1.06067655 | 0.60022196 | 5.48E-01 | 8.45E-01 |
| AP002444.1 | 142.301069 | 1.284144494 | 1.83688976 | 0.69908631 | 4.84E-01 | 8.18E-01 |
| RDX | 3917.22058 | 0.536493409 | 0.70230118 | 0.76390788 | 0.44492214 | 0.80041177 |
| ZC3H12C | 2249.67372 | 0.910131896 | 0.22823322 | 3.98772752 | 6.67E-05 | 0.02490203 |
| AP000924.1 | 120.811311 | 1.556159423 | 0.69713154 | 2.23223214 | 2.56E-02 | 3.82E-01 |
| FDX1 | 516.12995 | 0.174691548 | 0.26530265 | 0.65846138 | 0.51024171 | 0.82930674 |
| ARHGAP20 | 43.7119292 | -0.151601951 | 0.63207359 | -0.2398486 | 0.81044765 | 0.94717173 |
| COLCA1 | 42.1301405 | 0.607843133 | 0.86402173 | 0.70350445 | 0.48174143 | 0.8177376 |
| COLCA2 | 70.9482368 | 0.313773337 | 0.60036374 | 0.52263872 | 0.60122569 | 0.86755264 |
| SIK2 | 3997.15952 | 0.51112804 | 0.34629923 | 1.47597219 | 0.13995136 | 0.59500405 |
| PPP2R1B | 927.440645 | 0.460601445 | 0.50032964 | 0.92059597 | 0.35726141 | 0.75903693 |
| C11orf1 | 83.1327147 | 0.618701182 | 0.53839241 | 1.14916401 | 0.25048836 | 0.68627228 |
| CRYAB | 10227.3117 | -0.291265832 | 0.69684277 | -0.4179793 | 0.67596228 | 0.89833381 |
| DIXDC1 | 961.976479 | 0.365198174 | 0.64994955 | 0.56188695 | 0.57419304 | 0.85448213 |
| DLAT | 276.004444 | 0.350133148 | 0.21227104 | 1.64946264 | 0.09905289 | 0.54265306 |
| PIH1D2 | 62.5536854 | -0.419689807 | 0.49183229 | -0.8533189 | 0.39348246 | 0.77771935 |
| NKAPD1 | 182.555534 | -0.265015146 | 0.22836705 | -1.1604789 | 0.24585386 | 0.68320879 |
| TIMM8B | 206.444212 | 0.00648672 | 0.30954719 | 0.02095551 | 0.98328115 | 0.99523767 |
| AP002884.3 | 176.737439 | -0.225559889 | 0.41459838 | -0.5440443 | 0.58641099 | 0.86102615 |
| IL18 | 294.408778 | -0.055160244 | 0.40916588 | -0.1348114 | 8.93E-01 | 0.9724379 |
| BCO2 | 93.7647563 | -0.115994564 | 0.49368641 | -0.234956 | 0.81424292 | 0.94881072 |
| PTS | 463.96721 | 0.092342359 | 0.30159657 | 0.30617841 | 0.75946881 | 0.92930182 |
| AP002884.1 | 293.535827 | 0.779323568 | 0.61093034 | 1.27563409 | 0.2020849 | 0.64881821 |
| AP003063.1 | 72.1216356 | 2.049007738 | 0.92119006 | 2.22430509 | 0.02612792 | 0.38392873 |
| TTC12 | 466.198167 | 0.197010585 | 0.2883452 | 0.68324558 | 0.49445168 | 0.8220963 |
| ZW10 | 210.987983 | -0.152902655 | 0.2702591 | -0.5657632 | 0.57155476 | 0.85399468 |
| USP28 | 299.643676 | -0.241758629 | 0.35166129 | -0.6874758 | 0.49178294 | 0.82065604 |
| ZBTB16 | 3224.86093 | -0.035772301 | 0.7855702 | -0.0455367 | 0.9636795 | 0.99012655 |
| AP002518.2 | 302.015636 | 0.986203431 | 0.87508242 | 1.12698348 | 0.25974946 | 0.69242517 |
| NNMT | 2023.53946 | 0.777590025 | 0.91951003 | 0.84565693 | 3.98E-01 | 7.81E-01 |
| C11orf71 | 92.0603899 | 0.206454139 | 0.56988832 | 0.36227122 | 0.71714936 | 0.91317971 |
| AP002373.1 | 63.4158494 | -0.64015681 | 0.71765265 | -0.8920148 | 0.37238498 | 0.76879661 |
| REXO2 | 354.492256 | -0.02949161 | 0.3943397 | -0.0747873 | 0.94038393 | 0.98470913 |
| CADM1 | 1487.98821 | -0.524417144 | 0.44109771 | -1.1888911 | 0.23448252 | 0.670712 |
| AP000462.1 | 50.5270586 | -0.296419908 | 0.62321403 | -0.475631 | 0.63433729 | 0.88238823 |
| BUD13 | 117.965299 | -0.13730095 | 0.2632828 | -0.5214961 | 0.60202123 | 0.86770668 |
| ZPR1 | 100.64847 | -0.768386565 | 0.43337376 | -1.7730344 | 0.07622301 | 0.50767193 |

| | | | | | | |
|---|---|---|---|---|---|---|
| SIK3 | 6860.79689 | -0.616351933 | 0.3144515 | -1.9600859 | 0.04998576 | 0.45799114 |
| PAFAH1B2 | 708.623621 | -0.380175447 | 0.33684208 | -1.1286459 | 0.25904722 | 0.69241621 |
| SIDT2 | 154.828107 | -0.22592069 | 0.36148473 | -0.6249799 | 0.53198425 | 0.83789294 |
| TAGLN | 89.6369479 | -0.378789373 | 0.60524196 | -0.6258478 | 0.53141475 | 0.83789294 |
| PCSK7 | 522.953461 | -0.066410029 | 0.27936347 | -0.2377191 | 0.81209898 | 0.94784066 |
| RNF214 | 473.620669 | -0.202764067 | 0.17249904 | -1.1754504 | 0.23981456 | 0.6780147 |
| BACE1 | 223.555478 | 0.5203143 | 0.26523819 | 1.96168695 | 0.04979894 | 0.45679453 |
| CEP164 | 577.692077 | 0.280057728 | 0.35950469 | 0.77900993 | 4.36E-01 | 0.79549938 |
| AP000892.1 | 92.6758229 | 1.10943689 | 0.83667677 | 1.32600418 | 0.18483831 | 0.63418742 |
| DSCAML1 | 4320.77526 | 0.937294028 | 0.76643895 | 1.22292066 | 0.22135967 | 0.66263062 |
| FXYD2 | 840.930332 | -0.214543294 | 0.69069395 | -0.3106199 | 0.75608959 | 0.92799282 |
| AP000757.2 | 61.0197046 | -0.020125302 | 0.75137588 | -0.0267846 | 0.97863154 | 0.99482978 |
| AP000757.1 | 36.1679612 | 0.007809747 | 0.89050087 | 0.00877006 | 0.99300259 | 0.9976934 |
| SMIM35 | 397.5428 | 0.060214815 | 0.79526464 | 0.0757167 | 9.40E-01 | 9.84E-01 |
| TMPRSS4 | 80.8186312 | -0.771514348 | 0.6539434 | -1.1797876 | 2.38E-01 | 6.76E-01 |
| MPZL3 | 241.631659 | -0.611108269 | 0.3543349 | -1.7246629 | 0.08458823 | 0.52183702 |
| MPZL2 | 127.823473 | 0.170588317 | 0.5083911 | 0.33554544 | 0.73721367 | 0.92030854 |
| UBE4A | 748.481706 | -0.253743939 | 0.1781056 | -1.4246825 | 0.15424899 | 0.61099212 |
| ATP5MG | 830.063414 | -0.165517754 | 0.37528309 | -0.4410477 | 0.65917845 | 0.89058983 |
| KMT2A | 1934.71385 | -0.07996813 | 0.21004844 | -0.3807128 | 0.70341637 | 0.90887942 |
| AP001267.3 | 96.8071038 | 0.696400481 | 0.36318184 | 1.91749807 | 0.05517469 | 0.46944719 |
| TMEM25 | 151.081348 | 0.152237028 | 0.4307753 | 0.35340241 | 0.72378678 | 0.91600702 |
| IFT46 | 162.742896 | 0.095082933 | 0.45342833 | 0.20969782 | 0.83390353 | 0.95393659 |
| ARCN1 | 412.675831 | -0.305115675 | 0.26373881 | -1.1568858 | 0.24731903 | 0.68482118 |
| PHLDB1 | 552.045764 | -0.208258099 | 0.50482101 | -0.4125385 | 0.67994477 | 0.89941358 |
| TREH | 156.387842 | 0.494223253 | 0.69883157 | 0.70721368 | 4.79E-01 | 0.81653684 |
| AP002954.1 | 104.017551 | -0.006342853 | 0.41107403 | -0.01543 | 0.98768917 | 0.99648575 |
| DDX6 | 1041.17448 | 0.037517766 | 0.17750673 | 0.21135968 | 0.83260662 | 0.95334883 |
| BCL9L | 338.415964 | -0.632028232 | 0.56246163 | -1.1236824 | 0.26114779 | 0.69330869 |
| UPK2 | 37.6064777 | 0.104751674 | 0.58499175 | 0.17906522 | 0.85788649 | 0.9626966 |
| CCDC84 | 715.301259 | -0.675155847 | 0.21398032 | -3.155224 | 0.00160375 | 0.13207839 |
| RPS25 | 977.670335 | -0.13515467 | 0.46968617 | -0.2877553 | 0.77353408 | 0.93415955 |
| TRAPPC4 | 256.199293 | 0.3896519 | 0.36418373 | 1.06993221 | 0.28464983 | 0.70952861 |
| SLC37A4 | 288.494132 | 0.632707516 | 0.62942367 | 1.00521722 | 0.31479226 | 0.72719146 |
| HYOU1 | 193.120082 | -0.456894968 | 0.61295496 | -0.7453973 | 0.45603158 | 0.80499998 |
| VPS11 | 175.004575 | -0.129898625 | 0.206062 | -0.6303861 | 0.52844198 | 0.83761342 |
| HMBS | 65.6242798 | 0.136823802 | 0.36789461 | 0.37191032 | 0.70995962 | 0.91118132 |
| H2AFX | 63.4863326 | -0.272420268 | 0.43737181 | -0.6228574 | 0.53337824 | 0.8389965 |
| DPAGT1 | 88.1864055 | -0.423050328 | 0.36157976 | -1.1700056 | 0.24199873 | 0.68080966 |
| C2CD2L | 124.709387 | 0.237111816 | 0.41742964 | 0.56802822 | 0.57001581 | 0.85327114 |
| HINFP | 223.204071 | -0.337276838 | 0.28604408 | -1.1791079 | 0.2383552 | 0.67642542 |
| NLRX1 | 138.861156 | 0.353644685 | 0.41999916 | 0.84201285 | 0.39978076 | 0.78186732 |
| PDZD3 | 552.304947 | 1.429338562 | 0.93945616 | 1.52145319 | 0.12814615 | 0.58793678 |
| CBL | 813.799347 | -0.429770704 | 0.35621978 | -1.2064762 | 0.22763392 | 0.66705629 |
| MCAM | 85.7923617 | -0.597311432 | 0.84119105 | -0.7100782 | 0.47765564 | 0.81580016 |
| RNF26 | 52.6422728 | 0.158007231 | 0.49328359 | 0.32031723 | 0.74872787 | 0.92554142 |
| USP2 | 344.860118 | 0.138619023 | 0.64295375 | 0.21559719 | 8.29E-01 | 0.95325673 |
| USP2-AS1 | 270.267405 | 1.425238903 | 0.87090479 | 1.63650369 | 0.10173421 | 0.54729371 |
| THY1 | 209.408101 | 0.935964867 | 0.67724208 | 1.3820241 | 1.67E-01 | 6.22E-01 |
| NECTIN1 | 76.1783123 | -0.054468685 | 0.43936427 | -0.1239716 | 0.90133777 | 0.97487863 |
| TRIM29 | 80.2288485 | 0.243889291 | 0.71296118 | 0.34207934 | 0.73229118 | 0.91950222 |
| OAF | 1329.23977 | 0.164675446 | 0.29965126 | 0.54955699 | 0.58262327 | 0.85982677 |
| TMEM136 | 39.944672 | 1.212889829 | 0.5313935 | 2.28247022 | 0.02246159 | 0.3630673 |
| ARHGEF12 | 6459.53032 | 0.258563598 | 0.25721167 | 1.00525608 | 0.31477356 | 0.72719146 |

| | | | | | | |
|---|---|---|---|---|---|---|
| TECTA | 41.2060756 | -0.087024473 | 0.47664775 | -0.1825761 | 0.85513066 | 0.96171023 |
| SC5D | 191.9122 | 0.573579504 | 0.26683985 | 2.14952719 | 0.03159263 | 0.40930077 |
| SORL1 | 150.010084 | 0.764867693 | 0.5938097 | 1.2880687 | 0.19772205 | 0.64558745 |
| AP000755.1 | 43.1650239 | 1.689414113 | 0.75673617 | 2.23250082 | 0.02558188 | 0.38166806 |
| UBASH3B | 611.697915 | -0.407085927 | 0.80674788 | -0.5046012 | 6.14E-01 | 8.72E-01 |
| JHY | 744.98955 | 0.388896235 | 0.40750446 | 0.95433613 | 0.33991352 | 0.74654617 |
| HSPA8 | 879.261903 | 0.240595285 | 0.49697401 | 0.48412046 | 0.62830038 | 0.88009297 |
| GRAMD1B | 3144.07685 | 1.779348869 | 0.83169595 | 2.1394223 | 0.03240148 | 0.4107005 |
| ZNF202 | 120.598081 | -0.48644887 | 0.32341625 | -1.5040953 | 0.13255683 | 0.5912647 |
| VWA5A | 130.167289 | 0.526827672 | 0.5033315 | 1.04668131 | 0.29524659 | 0.71646902 |
| TBRG1 | 680.50025 | 0.050430446 | 0.21154578 | 0.23839023 | 0.81157845 | 0.94755586 |
| SIAE | 308.738815 | 0.660034096 | 0.36362164 | 1.81516726 | 0.06949821 | 0.49281529 |
| SPA17 | 77.2062059 | -0.198097652 | 0.3035626 | -0.6525759 | 0.5140297 | 0.83126535 |
| MSANTD2 | 555.023579 | 0.071719407 | 0.2429556 | 0.29519553 | 0.76784452 | 0.9318145 |
| AP000866.1 | 55.0801733 | 0.075721045 | 0.54952353 | 0.137794 | 0.89040323 | 0.97225819 |
| TMEM218 | 64.1702394 | 0.347187403 | 0.49760883 | 0.6977115 | 0.48535763 | 0.81839266 |
| PKNOX2 | 109.195088 | 1.902267159 | 0.97649289 | 1.94806043 | 0.05140773 | 0.46176002 |
| EI24 | 269.586059 | -0.127921182 | 0.41046722 | -0.3116477 | 0.75530825 | 0.92799282 |
| STT3A-AS1 | 69.3462323 | 0.317099101 | 0.52590633 | 0.60295737 | 0.54653705 | 0.84460887 |
| STT3A | 329.181968 | -0.34092899 | 0.32949645 | -1.034697 | 0.30081043 | 0.71879123 |
| CHEK1 | 90.8274495 | 0.040371028 | 0.34427235 | 0.1172648 | 0.90665022 | 0.97661505 |
| HYLS1 | 91.5233449 | 0.114502602 | 0.3273083 | 0.34983103 | 0.72646551 | 0.91635631 |
| PUS3 | 79.8239845 | 0.161614339 | 0.41073582 | 0.39347515 | 0.69396858 | 0.90589033 |
| CDON | 277.102957 | -0.799615835 | 0.69180324 | -1.1558429 | 0.24774544 | 0.68482118 |
| RPUSD4 | 208.385784 | -0.535711473 | 0.31054022 | -1.7250953 | 0.0845103 | 0.52183702 |
| FAM118B | 415.545318 | -0.214595888 | 0.24249442 | -0.8849519 | 0.37618259 | 0.77017697 |
| SRPRA | 255.932341 | -0.855540015 | 0.44165206 | -1.9371358 | 0.05272875 | 0.46564998 |
| FOXRED1 | 250.616876 | 0.081844834 | 0.46718919 | 0.17518563 | 0.86093377 | 0.96302275 |
| TIRAP | 118.820577 | -0.302715383 | 0.35886168 | -0.8435434 | 0.39892463 | 0.78140731 |
| DCPS | 108.87046 | -1.298190258 | 0.35253769 | -3.6824155 | 0.00023103 | 0.05581517 |
| ST3GAL4 | 85.4036532 | -1.076022078 | 0.55513572 | -1.9383045 | 0.05258608 | 0.4656314 |
| ETS1 | 1072.87922 | -0.95434591 | 0.57673606 | -1.654736 | 9.80E-02 | 5.41E-01 |
| KCNJ1 | 53.2559838 | -0.331491805 | 0.57486889 | -0.576639 | 0.56418336 | 0.85163263 |
| ARHGAP32 | 3515.32206 | 0.175200491 | 0.29084847 | 0.60237722 | 0.54692307 | 0.84460887 |
| BARX2 | 181.486965 | -0.020871901 | 0.64727792 | -0.0322457 | 0.97427615 | 0.99377398 |
| TMEM45B | 105.594113 | 0.237218923 | 0.62044854 | 0.38233456 | 0.70221322 | 0.90837679 |
| NFRKB | 406.016889 | -0.073824622 | 0.1920415 | -0.3844202 | 0.70066707 | 0.90766879 |
| PRDM10 | 514.700663 | -0.362018046 | 0.32939016 | -1.0990554 | 0.27174388 | 0.70054417 |
| APLP2 | 1678.91669 | 0.115431092 | 0.23213688 | 0.49725444 | 0.61900964 | 0.87511751 |
| ST14 | 672.608691 | -0.320070214 | 0.48350195 | -0.6619833 | 0.50798193 | 0.82874455 |
| ZBTB44 | 1658.9041 | -0.738104596 | 0.29829046 | -2.4744492 | 0.01334418 | 0.29875272 |
| AP002986.1 | 297.730475 | 0.207545768 | 0.75659081 | 0.27431706 | 0.78384097 | 0.93864304 |
| ADAMTS15 | 142.379024 | -1.299374828 | 0.76290549 | -1.7031924 | 0.08853207 | 0.5293868 |
| AP003486.1 | 40.4444916 | -0.009579144 | 0.4614111 | -0.0207605 | 0.98343668 | 0.99523767 |
| SNX19 | 674.590358 | -0.97240977 | 0.49189624 | -1.9768595 | 0.04805751 | 0.45149028 |
| NTM | 75.1206538 | 0.290557985 | 0.49372025 | 0.58850733 | 0.55619182 | 0.84861484 |
| OPCML | 248.925576 | 0.096264215 | 0.7174313 | 0.134179 | 8.93E-01 | 9.73E-01 |
| NCAPD3 | 160.311364 | 0.056133166 | 0.33887648 | 0.16564492 | 0.8684364 | 0.96534407 |
| VPS26B | 173.081565 | -0.290178586 | 0.27859991 | -1.0415602 | 0.29761562 | 0.71839551 |
| THYN1 | 55.9532095 | 0.129949312 | 0.42139767 | 0.30837691 | 0.75779555 | 0.928294 |
| ACAD8 | 192.691728 | 0.416019513 | 0.28105004 | 1.48023291 | 0.1388111 | 0.59474076 |
| GLB1L3 | 37.0852148 | -0.878161593 | 0.93802366 | -0.9361828 | 0.34917909 | 0.7542101 |
| GLB1L2 | 136.561018 | 0.231356272 | 0.36810607 | 0.62850436 | 0.52967359 | 0.83789294 |
| B3GAT1 | 99.7170133 | 0.642412375 | 0.89605586 | 0.7169334 | 0.47341519 | 0.81478861 |

| | | | | | | |
|---|---|---|---|---|---|---|
| AP004550.1 | 67.3735689 | 1.050921701 | 0.92116287 | 1.14086415 | 0.25392646 | 0.68891958 |
| ZMYND11 | 1476.07007 | -0.152522624 | 0.21216906 | -0.7188731 | 0.47221914 | 0.81407467 |
| DIP2C | 7505.82089 | -0.039157682 | 0.20891899 | -0.18743 | 0.85132352 | 0.96041905 |
| PRR26 | 135.959165 | -0.39901325 | 0.55477453 | -0.719235 | 0.47199615 | 0.81395965 |
| LARP4B | 2845.11362 | -0.396122923 | 0.18822823 | -2.104482 | 0.03533642 | 0.4198859 |
| GTPBP4 | 395.447342 | -0.830272159 | 0.48066097 | -1.7273551 | 0.08410391 | 0.52183702 |
| IDI1 | 246.139601 | 0.924421976 | 0.37290956 | 2.47894414 | 0.01317719 | 0.29875272 |
| WDR37 | 1004.91156 | -0.199914967 | 0.22652477 | -0.8825303 | 0.37749012 | 0.77055888 |
| ADARB2 | 47.7394343 | 0.74714286 | 0.65908646 | 1.13360372 | 0.25696081 | 0.69129771 |
| PFKP | 931.212185 | -0.314633319 | 0.66466308 | -0.4733726 | 6.36E-01 | 8.83E-01 |
| PITRM1 | 508.366551 | -0.671490383 | 0.40579903 | -1.6547363 | 0.097978 | 0.54130807 |
| PITRM1-AS1 | 54.4470854 | 0.275385339 | 0.60713831 | 0.45357925 | 0.65013169 | 0.88866038 |
| KLF6 | 4050.81408 | -0.55991945 | 0.44299614 | -1.2639375 | 2.06E-01 | 6.52E-01 |
| AKR1E2 | 55.7836412 | 0.089602448 | 0.54819467 | 0.16345005 | 0.8701641 | 0.96564168 |
| AKR1C1 | 2299.19227 | -1.326541948 | 0.8195843 | -1.6185546 | 1.06E-01 | 5.54E-01 |
| AKR1C2 | 443.447589 | -2.487226499 | 0.90431761 | -2.7503904 | 5.95E-03 | 0.22637955 |
| AKR1C3 | 1038.21619 | -0.076564179 | 0.44647457 | -0.1714861 | 0.86384157 | 0.96393376 |
| AL355303.1 | 57.550117 | -1.411469982 | 0.86733888 | -1.627357 | 0.1036613 | 0.55022042 |
| TUBAL3 | 86.7666229 | 0.377643729 | 0.62447042 | 0.60474238 | 0.54535019 | 0.84449215 |
| NET1 | 745.825146 | 0.20286209 | 0.4560976 | 0.44477781 | 0.65648034 | 0.89053559 |
| ASB13 | 671.830989 | 0.66143022 | 0.43299509 | 1.52756977 | 0.12661939 | 0.58713532 |
| FAM208B | 2360.2601 | -0.431997936 | 0.2336015 | -1.8492943 | 0.06441532 | 0.48593331 |
| AL365356.1 | 119.884739 | 0.064384162 | 0.6403434 | 0.1005463 | 0.91991062 | 0.97939204 |
| GDI2 | 1457.04506 | -0.218026983 | 0.30474429 | -0.7154424 | 0.47433573 | 0.81526268 |
| ANKRD16 | 194.599332 | 0.038780206 | 0.30684928 | 0.12638193 | 0.8994296 | 0.97458352 |
| FBH1 | 861.33551 | 0.016487025 | 0.28488069 | 0.05787344 | 0.95384944 | 0.98836423 |
| IL15RA | 64.8626814 | 1.205209221 | 0.57211678 | 2.10657906 | 3.52E-02 | 4.20E-01 |
| RBM17 | 608.564855 | -0.298539549 | 0.19597027 | -1.523392 | 0.12766065 | 0.58793678 |
| PFKFB3 | 635.263989 | 0.314922347 | 0.69073321 | 0.45592472 | 0.64844411 | 0.88787189 |
| AL137145.2 | 71.2012334 | 0.761066932 | 0.6980775 | 1.09023271 | 0.27561065 | 0.70247842 |
| PRKCQ | 153.454352 | 0.935822647 | 0.69001232 | 1.35624048 | 0.17502267 | 0.62699479 |
| SFMBT2 | 248.6789 | 1.512511309 | 0.64521005 | 2.34421536 | 0.01906716 | 0.34084276 |
| ITIH5 | 252.336442 | 2.16805748 | 0.74604597 | 2.90606418 | 0.00366006 | 0.19726785 |
| KIN | 272.534956 | -0.209179383 | 0.21633481 | -0.9669243 | 0.33358188 | 0.74022334 |
| ATP5F1C | 1042.25949 | 0.354076647 | 0.30270878 | 1.16969399 | 0.24212414 | 0.68080966 |
| TAF3 | 1162.0481 | -0.225901678 | 0.23202862 | -0.973594 | 0.33025815 | 0.73849818 |
| CELF2 | 197.704329 | 0.4311938 | 0.8190413 | 0.5264616 | 0.59856752 | 0.86646266 |
| USP6NL | 1081.30639 | 0.155425765 | 0.23511004 | 0.66107668 | 0.50856314 | 0.82874455 |
| ECHDC3 | 293.986579 | 0.721277603 | 0.37174068 | 1.94027084 | 0.05234678 | 0.4649855 |
| PROSER2 | 369.82713 | -0.884960951 | 0.3750494 | -2.359585 | 1.83E-02 | 3.38E-01 |
| UPF2 | 1238.52677 | -0.491784271 | 0.22025115 | -2.2328341 | 0.02555989 | 0.38166806 |
| DHTKD1 | 1510.64944 | 0.287639222 | 0.33724565 | 0.85290714 | 3.94E-01 | 0.77794348 |
| SEC61A2 | 225.24127 | 0.067730805 | 0.24748131 | 0.27368048 | 0.78433018 | 0.93873036 |
| NUDT5 | 628.211439 | -0.40629016 | 0.25462512 | -1.5956405 | 0.11056908 | 0.56264122 |
| CDC123 | 514.73745 | -0.469236467 | 0.29905659 | -1.5690558 | 0.11663494 | 0.57350419 |
| CAMK1D | 179.023339 | 0.200596272 | 0.55930981 | 0.35864966 | 0.71985719 | 0.9148113 |
| CCDC3 | 73.5992395 | -0.61200747 | 0.52162709 | -1.1732663 | 0.24068905 | 0.67925154 |
| OPTN | 1311.87806 | -0.156060582 | 0.33547659 | -0.4651907 | 0.64179491 | 0.88430668 |
| PHYH | 356.918532 | 0.346214598 | 0.46043305 | 0.75193255 | 0.45209162 | 0.80415518 |
| SEPHS1 | 369.843007 | -0.127656976 | 0.31219101 | -0.4089066 | 0.68260818 | 0.89962501 |
| BEND7 | 2355.82619 | -0.248308161 | 0.20347884 | -1.2203144 | 0.2223457 | 0.66373504 |
| PRPF18 | 207.810322 | -0.330187252 | 0.26509152 | -1.2455595 | 0.21292617 | 0.65763385 |
| AL157392.5 | 430.439229 | -0.050858464 | 0.2304554 | -0.2206868 | 0.82533631 | 0.95192757 |
| FRMD4A | 1904.64843 | 0.417684038 | 0.59924766 | 0.69701405 | 0.485794 | 0.81839266 |

| | | | | | | |
|---|---|---|---|---|---|---|
| AL157392.2 | 61.5205518 | 1.791282878 | 1.09433242 | 1.63687272 | 0.10165707 | 0.54729371 |
| FAM107B | 2485.24277 | -0.30474138 | 0.4231261 | -0.7202141 | 0.47139319 | 0.81387913 |
| CDNF | 91.6993162 | 0.524917368 | 0.67586044 | 0.77666533 | 0.43735625 | 0.79633928 |
| HSPA14.1 | 136.796765 | 0.324826989 | 0.28457108 | 1.1414617 | 0.25367784 | 0.68891958 |
| SUV39H2 | 77.5088925 | 0.037382441 | 0.46725318 | 0.08000468 | 0.93623353 | 0.98415216 |
| DCLRE1C | 259.059421 | 0.119040254 | 0.22994024 | 0.51770083 | 0.60466702 | 0.86865188 |
| RPP38 | 455.692779 | -0.128859167 | 0.19059992 | -0.6760715 | 0.49899527 | 0.82396436 |
| NMT2 | 477.856541 | -0.326716875 | 0.23662907 | -1.3807132 | 0.16736716 | 0.62288941 |
| FAM171A1 | 979.669369 | 0.769379115 | 0.63053608 | 1.2201984 | 0.22238967 | 0.66373504 |
| MINDY3 | 710.730078 | -0.316642837 | 0.18972878 | -1.6689236 | 0.09513253 | 0.53824794 |
| PTER | 2642.87787 | -0.192841734 | 0.3217648 | -0.5993251 | 5.49E-01 | 8.46E-01 |
| C1QL3 | 164.504682 | 0.86186484 | 0.80330369 | 1.07290039 | 0.28331581 | 0.70780772 |
| RSU1 | 1807.32281 | -0.457287542 | 0.33825324 | -1.3519088 | 0.17640449 | 0.62709281 |
| CUBN | 26731.6181 | 2.139706759 | 0.7850268 | 2.72564804 | 0.00641754 | 0.22797255 |
| TRDMT1 | 137.887535 | 0.16747803 | 0.33298853 | 0.50295435 | 0.61499637 | 0.87260075 |
| VIM-AS1 | 46.7155367 | -1.082050469 | 0.47010202 | -2.3017354 | 0.0213501 | 0.35688511 |
| VIM | 1726.14357 | -0.576787425 | 0.67673624 | -0.8523076 | 3.94E-01 | 0.77819948 |
| HACD1 | 82.3734991 | 0.309997782 | 0.48410098 | 0.64035768 | 5.22E-01 | 0.83445062 |
| STAM | 658.341439 | -0.68279177 | 0.42963187 | -1.5892484 | 0.11200432 | 0.56455091 |
| CACNB2 | 920.45281 | 1.330481587 | 0.7117871 | 1.86921284 | 6.16E-02 | 0.48266777 |
| AL450384.2 | 271.386496 | 0.320642283 | 0.31268975 | 1.02543267 | 0.30515908 | 0.72103276 |
| NSUN6 | 801.580495 | 0.214172151 | 0.23062841 | 0.928646 | 0.35307258 | 0.75673243 |
| ARL5B | 357.919785 | -0.227280831 | 0.4045747 | -0.5617772 | 0.57426785 | 0.85448213 |
| MALRD1 | 61.6194593 | -0.445069574 | 0.51418926 | -0.8655754 | 0.38672305 | 0.77334938 |
| PLXDC2 | 6766.20629 | 0.440493656 | 0.39926618 | 1.10325811 | 0.26991509 | 0.69961625 |
| NEBL | 2009.13016 | 0.190783118 | 0.46001039 | 0.41473654 | 0.67833478 | 0.89913305 |
| CASC10 | 43.611589 | -0.54331917 | 0.66304373 | -0.8194319 | 0.41254005 | 0.78632759 |
| MLLT10 | 4126.159 | -0.239569032 | 0.20923228 | -1.1449908 | 0.25221298 | 0.68874851 |
| DNAJC1 | 3336.00644 | 0.089251016 | 0.30342215 | 0.29414799 | 0.76864482 | 0.93198626 |
| BMI1 | 88.6135568 | -0.388740021 | 0.32336206 | -1.2021819 | 0.22929306 | 0.66705629 |
| PIP4K2A | 1991.56579 | 0.219953021 | 0.39878856 | 0.55155298 | 0.58125466 | 0.85895204 |
| ARMC3 | 108.48932 | 1.094952596 | 0.90265782 | 1.21303176 | 0.22511768 | 0.6664553 |
| MSRB2 | 350.554805 | 0.456587037 | 0.44095015 | 1.03546181 | 0.30045329 | 0.71879123 |
| C10orf67 | 425.464449 | 1.638872467 | 0.74055664 | 2.21302786 | 0.02689573 | 0.38702446 |
| OTUD1 | 117.273035 | -0.357622014 | 0.34026036 | -1.0510246 | 0.29324728 | 0.71458498 |
| KIAA1217 | 4487.88353 | -0.065716281 | 0.33757065 | -0.1946742 | 0.84564805 | 0.95764192 |
| ARHGAP21 | 1574.87367 | -0.400237841 | 0.19833661 | -2.0179726 | 0.04359412 | 0.44197082 |
| THNSL1 | 50.63888 | 1.068696456 | 0.66362994 | 1.61038012 | 0.1073149 | 0.55579103 |
| MYO3A | 232.822145 | 2.16972978 | 0.7340107 | 2.95599202 | 0.00311665 | 0.18216645 |
| APBB1IP | 738.301586 | 0.495459857 | 0.47275491 | 1.04802689 | 0.29462622 | 0.71628974 |
| PDSS1 | 572.14369 | 0.189610076 | 0.31156355 | 0.60857592 | 0.54280557 | 0.84318177 |
| ABI1 | 1881.78045 | -0.10107222 | 0.24311224 | -0.415743 | 0.67759805 | 0.89867794 |
| ANKRD26 | 1482.63396 | 0.174889227 | 0.32485104 | 0.53836746 | 0.59032339 | 0.86310002 |
| YME1L1 | 937.050312 | -0.234719243 | 0.16491345 | -1.4232874 | 0.15465285 | 0.61119904 |
| MASTL | 203.728832 | -0.064937927 | 0.300736 | -0.21593 | 0.8290423 | 0.95325673 |
| ACBD5 | 550.79748 | 0.044911254 | 0.21781985 | 0.20618531 | 0.83664616 | 0.95478386 |
| RAB18 | 1202.43908 | -0.616804411 | 0.30689633 | -2.0098136 | 0.04445092 | 0.44360789 |
| MPP7 | 856.423235 | 0.446985037 | 0.37458155 | 1.19329166 | 0.23275517 | 0.66978724 |
| AL358234.1 | 55.7170268 | 1.601221944 | 0.92879401 | 1.72397961 | 0.08471152 | 0.52183702 |
| WAC-AS1 | 339.294271 | -0.133534453 | 0.45379669 | -0.2942605 | 0.76855883 | 0.93198626 |
| WAC | 3551.90003 | -0.271124109 | 0.22338345 | -1.2137162 | 0.22485612 | 0.66625656 |
| BAMBI | 104.081913 | 1.120985427 | 0.49028573 | 2.28639212 | 0.02223133 | 0.36178312 |
| SVIL | 4400.04594 | -0.26130533 | 0.53547215 | -0.4879905 | 0.62555656 | 0.87854882 |
| MTPAP | 268.430815 | -0.220860918 | 0.21090564 | -1.0472025 | 0.29500618 | 0.71646902 |

| | | | | | | |
|---|---|---|---|---|---|---|
| MAP3K8 | 289.597266 | 0.318677133 | 0.41757308 | 0.76316493 | 4.45E-01 | 0.80102593 |
| ZNF438 | 985.388772 | 0.135484902 | 0.24653733 | 0.54955126 | 0.5826272 | 0.85982677 |
| ZEB1-AS1 | 52.6201164 | 0.091772583 | 0.4664496 | 0.19674705 | 0.84402549 | 0.95731603 |
| ZEB1 | 1854.04769 | -0.802977415 | 0.62310832 | -1.2886643 | 0.19751484 | 0.64558745 |
| ARHGAP12 | 1769.81763 | -0.213262842 | 0.23344199 | -0.9135582 | 0.36094904 | 0.76009788 |
| KIF5B | 1145.19494 | -0.0169484 | 0.22297321 | -0.0760109 | 0.93941041 | 0.98435387 |
| EPC1 | 1716.40635 | -0.16482259 | 0.19927261 | -0.8271211 | 0.40816841 | 0.78481094 |
| AL391839.1 | 50.4397818 | 0.801542146 | 0.64740397 | 1.23808655 | 0.21568397 | 0.6596384 |
| CCDC7 | 1121.14694 | 0.816039394 | 0.56538308 | 1.4433389 | 0.14892503 | 0.60622442 |
| ITGB1 | 1495.75692 | -0.486997582 | 0.37742967 | -1.2903002 | 0.19694645 | 0.64558745 |
| ITGB1-DT | 63.040933 | -1.771769441 | 0.74309563 | -2.3843088 | 0.01711124 | 0.32964126 |
| NRP1 | 9428.85451 | 0.18032256 | 0.54687847 | 0.32973059 | 0.74160354 | 0.92235631 |
| PARD3 | 20127.0892 | -0.345293659 | 0.29330351 | -1.1772572 | 0.23909288 | 0.6769771 |
| CUL2 | 633.727916 | -0.100100382 | 0.19967957 | -0.5013051 | 0.61615643 | 0.8733874 |
| CREM | 981.189372 | -0.384169815 | 0.39641183 | -0.9691179 | 0.33248635 | 0.73988623 |
| CCNY | 3884.38921 | -0.582457667 | 0.2121492 | -2.7455096 | 0.0060417 | 0.22637955 |
| AL117336.3 | 388.2425 | -0.162813257 | 0.33513227 | -0.4858179 | 0.62709628 | 0.87916628 |
| FZD8 | 58.41376 | 1.384664101 | 0.55791099 | 2.4818728 | 0.01306939 | 0.29875272 |
| ZNF248 | 956.746614 | 0.745985345 | 0.37851884 | 1.97080111 | 0.04874663 | 0.4536808 |
| ZNF25 | 95.5766726 | 0.664075684 | 0.45264364 | 1.46710487 | 0.14234753 | 0.59759804 |
| ZNF33A | 1935.07993 | -0.123497394 | 0.22961902 | -0.5378361 | 0.59069021 | 0.86326802 |
| AL117339.5 | 95.6847358 | 1.695810606 | 0.53550439 | 3.16675388 | 0.00154151 | 0.13146073 |
| ZNF33B | 1242.25257 | 0.031182016 | 0.35343557 | 0.08822546 | 0.92969748 | 0.98272895 |
| BMS1 | 617.210273 | -0.287137844 | 0.22774003 | -1.2608141 | 0.20737582 | 0.65269814 |
| CSGALNACT2 | 417.847889 | -0.412848826 | 0.41695003 | -0.9901638 | 0.32209407 | 0.73375526 |
| RASGEF1A | 42.3623777 | 1.092566298 | 0.61498043 | 1.77658709 | 0.0756362 | 0.50730849 |
| FXYD4 | 64.1962239 | -0.109001497 | 0.83423947 | -0.1306597 | 0.8960445 | 0.97309833 |
| HNRNPF | 953.333231 | -0.512469946 | 0.38385174 | -1.3350726 | 0.18185258 | 0.6318513 |
| ZNF487 | 276.083247 | 0.186098239 | 0.23293945 | 0.7989125 | 0.42434115 | 0.78894887 |
| ZNF239 | 43.5506223 | -0.973149152 | 0.36541143 | -2.6631601 | 0.00774106 | 0.24225268 |
| ZNF32 | 168.729518 | -0.082163817 | 0.24443261 | -0.336141 | 0.73676456 | 0.92020873 |
| CXCL12 | 95.4034369 | 1.140940847 | 0.98108759 | 1.16293475 | 0.24485598 | 0.68250191 |
| TMEM72-AS1 | 746.12226 | 1.453333214 | 0.53526427 | 2.7151695 | 0.00662419 | 0.22966139 |
| RASSF4 | 3296.46383 | 1.620082928 | 0.51319894 | 3.15683218 | 0.00159493 | 0.13207839 |
| DEPP1 | 224.022211 | -0.576980263 | 0.43470858 | -1.3272806 | 1.84E-01 | 6.34E-01 |
| ZNF22 | 184.022503 | -0.371203317 | 0.29631222 | -1.2527439 | 2.10E-01 | 0.65441616 |
| ALOX5 | 111.893103 | 1.770435091 | 0.7390291 | 2.39562297 | 1.66E-02 | 0.32489795 |
| 8-Mar | 908.203261 | -0.418404856 | 0.21732426 | -1.9252561 | 0.05419733 | 0.46711025 |
| ZFAND4 | 356.782423 | -0.242175116 | 0.2469915 | -0.9804998 | 0.32683948 | 0.73745391 |
| WASHC2C | 568.08278 | -0.631624765 | 0.27851267 | -2.2678493 | 0.02333839 | 0.37046453 |
| AGAP4 | 625.75773 | 0.314912645 | 0.30671113 | 1.0267402 | 0.30454281 | 0.72082426 |
| TIMM23 | 421.100843 | -0.455654781 | 0.27521569 | -1.6556279 | 0.09779719 | 0.5411478 |
| NCOA4 | 503.495383 | 0.036039979 | 0.19450537 | 0.1852904 | 0.85300126 | 0.96073035 |
| AL356056.2 | 61.4957509 | 0.472898172 | 0.48451468 | 0.97602445 | 0.32905235 | 0.73829083 |
| PTPN20 | 887.919417 | 0.916023623 | 0.62461397 | 1.46654361 | 0.14250025 | 0.59794127 |
| AGAP9 | 513.454925 | 0.928429751 | 0.44308004 | 2.09539963 | 0.03613548 | 0.42040764 |
| BX547991.1 | 155.377641 | 0.2547311 | 0.82192703 | 0.30991936 | 0.75662228 | 0.92799282 |
| MAPK8 | 3383.54934 | -0.228307208 | 0.17859323 | -1.2783643 | 0.201121 | 0.64849129 |
| ARHGAP22 | 334.899951 | -0.15294481 | 0.53405891 | -0.2863819 | 7.75E-01 | 9.35E-01 |
| WDFY4 | 213.437229 | 0.84961675 | 1.1697182 | 0.72634311 | 0.46762846 | 0.81184083 |
| VSTM4 | 352.232874 | 1.024934794 | 0.57558256 | 1.7806912 | 0.07496291 | 0.5071704 |
| AL138760.1 | 47.9490812 | 0.832021259 | 0.71275405 | 1.16733292 | 0.24307595 | 0.68149766 |
| ERCC6 | 535.341865 | -0.315689863 | 0.1570656 | -2.0099237 | 0.04443927 | 0.44360789 |
| OGDHL | 559.766315 | 1.379894375 | 0.50856695 | 2.71329933 | 0.00666169 | 0.2300045 |

| | | | | | | |
|---|---|---|---|---|---|---|
| PARG | 1721.51029 | 0.003226761 | 0.23718641 | 0.01360432 | 0.98914565 | 0.99721497 |
| TIMM23B | 1573.40839 | -0.806861279 | 0.55157327 | -1.4628361 | 0.14351224 | 0.59966109 |
| LINC00843 | 359.112009 | 0.011800734 | 0.25848878 | 0.04565279 | 0.96358699 | 0.99012655 |
| AGAP6 | 522.993442 | -0.10501693 | 0.26140818 | -0.4017354 | 0.68787874 | 0.90296436 |
| WASHC2A | 538.292201 | -0.981531603 | 0.3101135 | -3.1650722 | 0.00155044 | 0.13146073 |
| SGMS1 | 2567.51818 | 0.029811912 | 0.18954411 | 0.15728219 | 0.87502246 | 0.96615698 |
| SGMS1-AS1 | 67.9325654 | -0.130880935 | 0.49499969 | -0.2644061 | 0.79146702 | 0.94098694 |
| ASAH2B | 140.98003 | -0.077979544 | 0.46521355 | -0.167621 | 0.86688148 | 0.96486219 |
| A1CF | 1435.92261 | 1.333462018 | 0.66600878 | 2.00216883 | 0.04526658 | 0.44360789 |
| PRKG1 | 684.732596 | 0.071008051 | 0.65057369 | 0.10914682 | 0.91308604 | 0.97720308 |
| CSTF2T | 45.0824175 | -0.176890237 | 0.42268494 | -0.4184919 | 0.67558749 | 0.89813476 |
| PCDH15 | 2408.0398 | 2.326555286 | 0.90053947 | 2.58351286 | 9.78E-03 | 0.26385502 |
| IPMK | 639.566419 | -0.513360527 | 0.24217106 | -2.1198261 | 3.40E-02 | 0.41378378 |
| CISD1 | 844.066221 | 0.150865221 | 0.28341267 | 0.53231643 | 0.59450685 | 0.86488065 |
| UBE2D1 | 289.881874 | -0.409486496 | 0.36557578 | -1.1201139 | 0.26266525 | 0.69467275 |
| TFAM | 207.263846 | -0.224527782 | 0.25800826 | -0.8702348 | 0.38417208 | 0.77317162 |
| BICC1 | 16000.8726 | -0.827112228 | 0.32405065 | -2.5524165 | 0.01069785 | 0.27357367 |
| PHYHIPL | 2013.91517 | 0.034988262 | 0.49591447 | 0.07055302 | 0.9437535 | 0.98522232 |
| FAM13C | 42.0283611 | 1.192431633 | 0.78872513 | 1.51184689 | 0.13057282 | 0.58853436 |
| SLC16A9 | 5878.08188 | 1.001354198 | 0.92190302 | 1.0861817 | 0.27739861 | 0.7036882 |
| CCDC6 | 1324.24352 | -0.179831013 | 0.25046918 | -0.7179766 | 0.4727717 | 0.81460134 |
| ANK3 | 14429.0274 | 1.092665435 | 0.53836733 | 2.0295909 | 0.04239814 | 0.43742111 |
| AL592430.1 | 66.5418822 | 1.536433045 | 0.96679719 | 1.58919892 | 0.11201549 | 0.56455091 |
| RHOBTB1 | 5413.04013 | 0.170761839 | 0.68718574 | 0.24849444 | 0.80375187 | 0.94543168 |
| TMEM26 | 148.302034 | 1.449266335 | 0.57271314 | 2.53052748 | 1.14E-02 | 0.28539418 |
| AC022398.1 | 43.3710859 | 0.068759376 | 0.71144889 | 0.09664697 | 0.92300676 | 0.97983687 |
| ARID5B | 3401.98538 | -0.665056116 | 0.42841591 | -1.5523609 | 0.12057589 | 0.57856254 |
| RTKN2 | 69.9086858 | -0.40105308 | 0.60746751 | -0.660205 | 0.50912229 | 0.82880513 |
| AC024597.1 | 130.519978 | -2.847461246 | 1.74845795 | -1.6285558 | 0.1034071 | 0.55022042 |
| ZNF365 | 333.362726 | -0.157827226 | 1.53101674 | -0.1030865 | 0.91789428 | 0.97832208 |
| ADO | 57.3439239 | 0.310297347 | 0.43189804 | 0.71845047 | 0.47247958 | 0.81442122 |
| NRBF2 | 183.530332 | 0.178524535 | 0.34099733 | 0.52353646 | 0.60060099 | 0.86755264 |
| JMJD1C | 7635.53377 | -0.495587267 | 0.20849066 | -2.3770238 | 0.01745296 | 0.33064634 |
| REEP3 | 892.863523 | -0.424615547 | 0.34896244 | -1.2167944 | 0.22368245 | 0.66479083 |
| LINC01515 | 1817.53663 | 1.205954236 | 0.67626818 | 1.78324852 | 0.07454586 | 0.5071704 |
| CTNNA3 | 199.267311 | -0.178961159 | 0.50913698 | -0.351499 | 7.25E-01 | 0.91600702 |
| DNAJC12 | 200.459196 | 0.313826932 | 0.45533612 | 0.68922038 | 0.49068459 | 0.81989696 |
| SIRT1 | 531.436341 | -0.396637135 | 0.24772062 | -1.601147 | 0.10934436 | 0.55948242 |
| HERC4 | 4870.22281 | -0.55164065 | 0.25923021 | -2.1279952 | 0.03333748 | 0.41254527 |
| AC016395.1 | 253.827818 | 1.103394238 | 0.73918145 | 1.49272446 | 1.36E-01 | 0.59381224 |
| PBLD | 1794.24304 | 0.606428189 | 0.47311934 | 1.28176579 | 2.00E-01 | 6.47E-01 |
| HNRNPH3 | 1222.60256 | -0.325938159 | 0.25339782 | -1.2862706 | 0.19834865 | 0.64582202 |
| RUFY2 | 870.256382 | -0.099096409 | 0.22992497 | -0.4309946 | 0.66647233 | 0.89420131 |
| DNA2 | 363.221151 | -0.167987772 | 0.47596828 | -0.352939 | 0.72413417 | 0.91600702 |
| SLC25A16 | 1988.21532 | -0.223452463 | 0.33098334 | -0.6751169 | 0.49960147 | 0.82476476 |
| TET1 | 161.13643 | 0.350147906 | 0.53365701 | 0.65612912 | 0.51174105 | 0.83042711 |
| CCAR1 | 1638.68205 | -0.263081103 | 0.28854324 | -0.9117562 | 0.36189705 | 0.76060307 |
| STOX1 | 85.5159164 | 0.208757423 | 0.65455624 | 0.3189297 | 0.74977982 | 0.92565124 |
| DDX50 | 587.789725 | -0.515684659 | 0.24340064 | -2.118666 | 0.03411871 | 0.41441585 |
| DDX21 | 846.503937 | -0.703673185 | 0.54391488 | -1.2937193 | 0.19576239 | 0.64558745 |
| KIF1BP | 248.427405 | -0.109399231 | 0.29378501 | -0.3723785 | 0.70961104 | 0.91107517 |
| VPS26A | 657.111187 | -0.324595953 | 0.21390375 | -1.517486 | 0.12914403 | 0.58793678 |
| SUPV3L1 | 376.215107 | -0.604642981 | 0.24741885 | -2.4438033 | 0.01453335 | 0.30802371 |
| AL596223.1 | 94.8192311 | -0.808572477 | 0.86311439 | -0.936808 | 3.49E-01 | 0.75410619 |

| | | | | | | |
|---|---|---|---|---|---|---|
| HKDC1 | 3185.55428 | -1.350663155 | 0.73417104 | -1.8397118 | 6.58E-02 | 0.48798003 |
| HK1 | 421.323911 | -0.64849733 | 0.4265483 | -1.5203374 | 1.28E-01 | 5.88E-01 |
| TSPAN15 | 132.288862 | 1.517166512 | 0.68568158 | 2.21264005 | 0.02692247 | 0.38702446 |
| FAM241B | 57.043768 | 0.195770866 | 0.5175536 | 0.37826201 | 0.70523596 | 0.909554 |
| H2AFY2 | 225.812074 | 1.433904032 | 0.53505131 | 2.67993744 | 0.00736359 | 0.23590813 |
| AIFM2 | 283.97517 | -0.196578521 | 0.29793753 | -0.6597978 | 0.50938361 | 0.82882082 |
| TYSND1 | 113.541358 | 0.188093621 | 0.34257945 | 0.5490511 | 0.58297038 | 0.85987034 |
| SAR1A | 453.772711 | -0.363044701 | 0.20336306 | -1.7852047 | 0.07422811 | 0.5071704 |
| PPA1 | 1027.07085 | -0.067008538 | 0.3535896 | -0.1895094 | 8.50E-01 | 0.95959925 |
| LRRC20 | 53.4011224 | 0.234031416 | 0.36539809 | 0.64048342 | 0.52185837 | 0.83445062 |
| EIF4EBP2 | 861.939303 | -0.171519876 | 0.25463153 | -0.6736003 | 0.50056546 | 0.82560579 |
| SGPL1 | 580.194268 | -0.271643163 | 0.2028019 | -1.3394508 | 0.18042398 | 0.63152999 |
| PCBD1 | 473.419693 | -0.843667159 | 0.5186812 | -1.6265621 | 0.10383014 | 0.55022042 |
| SLC29A3 | 61.8764404 | -0.746873838 | 0.59216612 | -1.2612573 | 0.20721616 | 0.65266614 |
| CDH23 | 38.1920675 | -0.338526462 | 0.72694314 | -0.4656849 | 0.64144105 | 0.88403746 |
| PSAP | 1534.03572 | -0.924735825 | 0.39941442 | -2.315229 | 0.02060041 | 0.35267827 |
| CHST3 | 163.295242 | -0.014186987 | 0.28060711 | -0.0505582 | 0.95967758 | 0.98912673 |
| ASCC1 | 564.21686 | -0.433491427 | 0.15947493 | -2.7182418 | 0.00656299 | 0.22966139 |
| ANAPC16 | 1186.80408 | 0.213287217 | 0.27969097 | 0.76258172 | 0.44571289 | 0.8012958 |
| DDIT4 | 456.295453 | -0.774774792 | 0.7522186 | -1.0299862 | 3.03E-01 | 0.72043311 |
| DNAJB12 | 290.194304 | -0.397514115 | 0.39595515 | -1.0039372 | 0.31540888 | 0.72782988 |
| MICU1 | 3007.92158 | -0.412866366 | 0.23926765 | -1.725542 | 0.08442985 | 0.52183702 |
| MCU | 2395.66614 | -0.850541708 | 0.48401446 | -1.7572651 | 0.07887264 | 0.51192014 |
| PLA2G12B | 173.581829 | 0.730507081 | 0.77910109 | 0.93762811 | 0.34843556 | 0.75366991 |
| P4HA1 | 3017.72125 | -0.957600532 | 0.61580457 | -1.5550397 | 0.11993662 | 0.57730856 |
| NUDT13 | 147.625298 | -0.068518176 | 0.57016668 | -0.1201722 | 0.90434675 | 0.97574629 |
| ECD | 382.727527 | -0.126421377 | 0.16919054 | -0.747213 | 0.45493497 | 0.80465616 |
| FAM149B1 | 618.441025 | 0.494644838 | 0.35893216 | 1.37810119 | 0.16817205 | 0.62304975 |
| DNAJC9 | 458.08701 | 0.356452058 | 0.25555337 | 1.39482436 | 0.16306881 | 0.62002336 |
| DNAJC9-AS1 | 34.0742663 | 0.029857239 | 0.58780557 | 0.05079441 | 0.95948934 | 0.98912673 |
| MRPS16 | 296.876451 | -0.302306828 | 0.25960218 | -1.1645003 | 0.24422131 | 0.68222743 |
| CFAP70 | 616.724061 | 0.725025749 | 0.54915273 | 1.3202625 | 0.18674739 | 0.6378941 |
| ANXA7 | 1016.74823 | 0.048207578 | 0.17760296 | 0.27143455 | 0.78605684 | 0.93904951 |
| MSS51 | 49.716906 | 0.472648667 | 0.35810776 | 1.31985041 | 0.18688497 | 0.6378941 |
| PPP3CB | 1034.25425 | -0.023869279 | 0.22887259 | -0.1042907 | 0.91693867 | 0.97828877 |
| USP54 | 885.374514 | -0.241932706 | 0.29618555 | -0.8168282 | 0.41402662 | 0.78632759 |
| AGAP5 | 198.225807 | 0.404739073 | 0.22135502 | 1.82846128 | 0.06748035 | 0.48963361 |
| AC022400.6 | 90.1059484 | 0.518669511 | 0.30043237 | 1.72641019 | 0.08427364 | 0.52183702 |
| SEC24C | 403.94134 | -0.368555727 | 0.23516054 | -1.5672516 | 0.1170559 | 0.57350419 |
| FUT11 | 291.844575 | -0.502125217 | 0.41664462 | -1.2051643 | 2.28E-01 | 6.67E-01 |
| CHCHD1 | 197.990611 | -0.366857912 | 0.4276992 | -0.8577475 | 0.39103192 | 0.77606489 |
| ZSWIM8 | 937.486525 | -0.106541885 | 0.23018217 | -0.462859 | 0.64346545 | 0.8848654 |
| CAMK2G | 1064.7538 | 0.43528348 | 0.38974096 | 1.11685331 | 2.64E-01 | 0.69567438 |
| VCL | 1065.28031 | -0.209426078 | 0.34185719 | -0.6126128 | 0.54013242 | 0.8414758 |
| AP3M1 | 298.432099 | -0.467721776 | 0.30059061 | -1.5560093 | 0.11970589 | 0.57673415 |
| ADK | 4146.04288 | -0.380600089 | 0.3911492 | -0.9730305 | 0.33053816 | 0.73849818 |
| KAT6B | 1940.22971 | -0.643659227 | 0.26516372 | -2.4274031 | 0.01520735 | 0.3105046 |
| SAMD8 | 636.990433 | -0.222845058 | 0.28717296 | -0.775996 | 0.43775137 | 0.7967412 |
| VDAC2 | 409.628375 | 0.019243012 | 0.27581058 | 0.06976894 | 0.94437757 | 0.9853777 |
| COMTD1 | 230.878589 | -0.403054017 | 0.41406272 | -0.973413 | 0.3303481 | 0.73849818 |
| AC010997.3 | 161.027709 | 0.116639142 | 0.60211297 | 0.19371637 | 0.84639797 | 0.95801627 |
| ZNF503-AS2 | 33.6829649 | -0.239450923 | 0.40653932 | -0.5889982 | 0.55586248 | 0.84861484 |
| AC010997.5 | 61.2938661 | -0.107251678 | 0.49739993 | -0.2156246 | 0.82928035 | 0.95325673 |
| LRMDA | 20315.2866 | 0.165738601 | 0.32626086 | 0.50799413 | 0.61145746 | 0.87168459 |

| | | | | | | |
|---|---|---|---|---|---|---|
| KCNMA1 | 882.072567 | -0.059056835 | 1.05364161 | -0.0560502 | 0.95530181 | 0.98840182 |
| DLG5 | 615.204602 | -0.178939015 | 0.35189929 | -0.508495 | 0.61110627 | 0.87163238 |
| POLR3A | 510.683618 | -0.286869679 | 0.30770959 | -0.9322741 | 0.35119489 | 0.75518811 |
| RPS24 | 1717.31698 | -0.02647126 | 0.48348686 | -0.0547507 | 0.95633705 | 0.9886112 |
| ZMIZ1-AS1 | 82.6613536 | 0.702152565 | 0.4407204 | 1.5931928 | 0.11111695 | 0.56347761 |
| ZMIZ1 | 1163.74034 | 0.118185352 | 0.43188537 | 0.27364982 | 0.78435375 | 0.93873036 |
| PPIF | 174.0791 | -0.772198245 | 0.49259961 | -1.5675982 | 0.11697494 | 0.57350419 |
| ZCCHC24 | 98.1593075 | 0.466295887 | 0.5912571 | 0.78865165 | 0.43031563 | 0.79055252 |
| EIF5AL1 | 56.6035352 | -0.884596827 | 0.53036998 | -1.6678863 | 0.0953383 | 0.53824794 |
| NUTM2B-AS | 2238.99674 | 0.170947594 | 0.3360602 | 0.50868146 | 0.61097552 | 0.87163212 |
| AL135925.1 | 42.2350032 | 0.036934933 | 0.58225169 | 0.06343465 | 0.94942039 | 0.9871942 |
| TMEM254-A! | 112.427057 | 0.117776151 | 0.50665262 | 0.23245938 | 8.16E-01 | 0.94913008 |
| TMEM254 | 214.535956 | 0.414663629 | 0.33825556 | 1.22588858 | 0.22024061 | 0.66263062 |
| ANXA11 | 1084.72493 | 0.046852127 | 0.28356742 | 0.16522394 | 8.69E-01 | 9.65E-01 |
| FAM213A | 540.833609 | 0.505039787 | 0.30285051 | 1.66762073 | 0.09539104 | 0.53824794 |
| TSPAN14 | 598.439581 | 0.204844355 | 0.35785854 | 0.57241712 | 0.56703942 | 0.85233132 |
| NRG3 | 5693.89764 | 0.090151901 | 0.73132246 | 0.12327244 | 0.90189137 | 0.97492485 |
| GHITM | 606.156683 | 0.116587612 | 0.34056475 | 0.34233611 | 0.73209797 | 0.91942342 |
| CCSER2 | 2122.48923 | -0.011774041 | 0.22555975 | -0.0521992 | 0.95836996 | 0.98878187 |
| GRID1 | 144.190491 | 1.645509248 | 0.6393778 | 2.57361025 | 0.01006436 | 0.26580376 |
| WAPL | 1970.73428 | -0.342681474 | 0.2045221 | -1.675523 | 0.09383166 | 0.53723919 |
| BMPR1A | 2328.77947 | -0.082414113 | 0.20860164 | -0.3950789 | 0.69278464 | 0.90528251 |
| MMRN2 | 66.1801421 | -0.043602268 | 0.61303208 | -0.0711256 | 0.9432978 | 0.98520138 |
| ADIRF | 643.204507 | 0.831531864 | 0.8254304 | 1.00739186 | 0.3137465 | 0.72717675 |
| GLUD1 | 1726.99113 | 0.503107116 | 0.34784162 | 1.44636836 | 0.14807391 | 0.60605027 |
| SHLD2 | 2754.67434 | -0.443127841 | 0.21832221 | -2.0296965 | 0.04238739 | 0.43742111 |
| NUTM2A-AS | 2335.58074 | 0.00709544 | 0.2701472 | 0.02626509 | 0.9790459 | 0.99482978 |
| LINC00864 | 178.24176 | 0.948597037 | 0.89184375 | 1.06363591 | 0.28749367 | 0.71118494 |
| MINPP1 | 400.432593 | 0.051323926 | 0.45396984 | 0.11305581 | 0.90998631 | 0.97692341 |
| PAPSS2 | 594.147371 | 0.437930286 | 0.39699875 | 1.10310243 | 0.26998269 | 0.69961625 |
| ATAD1 | 942.722509 | 0.026634752 | 0.22772753 | 0.11695886 | 0.90689266 | 0.97664815 |
| PTEN | 2093.99254 | 0.151022868 | 0.18555148 | 0.81391356 | 0.41569447 | 0.78632759 |
| RNLS | 938.867064 | 0.030835533 | 0.40624187 | 0.07590437 | 0.93949518 | 0.98435387 |
| STAMBPL1 | 55.6081507 | 0.361938749 | 0.59095402 | 0.61246516 | 0.54023004 | 0.8414758 |
| ACTA2 | 85.3843037 | 1.097894472 | 0.48422996 | 2.26729976 | 0.02337192 | 0.37046453 |
| FAS | 231.760319 | 1.507614963 | 0.48015882 | 3.13982561 | 1.69E-03 | 0.13207839 |
| LIPA | 1088.07015 | 0.293759762 | 0.53322222 | 0.55091433 | 0.5816924 | 0.85918726 |
| IFIT2 | 297.018533 | -2.077399263 | 0.87148571 | -2.3837445 | 0.0171375 | 0.32964126 |
| IFIT3 | 485.130392 | -3.12587979 | 0.90393488 | -3.4580807 | 0.00054404 | 0.08106654 |
| IFIT1 | 466.807986 | -3.78037456 | 1.09413337 | -3.4551314 | 0.00055002 | 0.08106654 |
| IFIT5 | 115.19387 | -0.481045374 | 0.48786413 | -0.9860233 | 0.3241217 | 0.73603979 |
| SLC16A12 | 3582.5593 | 0.083285862 | 0.6408047 | 0.12997074 | 8.97E-01 | 0.97319871 |
| SLC16A12-AS | 129.442211 | 0.298773035 | 1.01394772 | 0.29466316 | 0.76825121 | 0.93198626 |
| PANK1 | 1183.24663 | 0.307970376 | 0.80905079 | 0.38065642 | 0.70345821 | 0.90887942 |
| AL157400.3 | 49.7493206 | 1.280277843 | 1.02990602 | 1.24310162 | 0.21383039 | 0.65849767 |
| KIF20B | 148.776885 | -0.393675537 | 0.48827612 | -0.806256 | 0.42009527 | 0.78798479 |
| RPP30 | 186.822272 | -0.234745631 | 0.27844952 | -0.8430456 | 0.39920297 | 0.78161291 |
| PCGF5 | 2239.96079 | -0.514705794 | 0.24879764 | -2.0687728 | 0.03856741 | 0.42479978 |
| HECTD2 | 361.068854 | 0.52630715 | 0.3031236 | 1.736279 | 0.08251451 | 0.5183475 |
| PPP1R3C | 130.171306 | -0.756267127 | 0.97333592 | -0.7769847 | 4.37E-01 | 0.79610188 |
| TNKS2 | 1579.50695 | -0.460341659 | 0.15028697 | -3.0630842 | 0.00219068 | 0.15242488 |
| BTAF1 | 1283.68796 | -0.315723682 | 0.22198718 | -1.4222609 | 0.15495053 | 0.61137216 |
| CPEB3 | 1251.9135 | -0.110962758 | 0.34163254 | -0.3248015 | 0.74533132 | 0.92393094 |
| 5-Mar | 636.868811 | -0.280569201 | 0.34459707 | -0.814195 | 0.41553326 | 0.78632759 |

| | | | | | | |
|---|---|---|---|---|---|---|
| IDE | 778.695826 | -0.254298329 | 0.25048951 | -1.0152055 | 0.31000789 | 0.72498238 |
| EXOC6 | 554.933025 | -0.421603252 | 0.28316803 | -1.4888801 | 0.13651893 | 0.5942049 |
| MYOF | 423.027001 | -0.94159104 | 0.56993819 | -1.6520933 | 9.85E-02 | 0.54209253 |
| RBP4 | 705.526669 | -1.123910882 | 0.89925267 | -1.2498277 | 0.2113625 | 0.65561118 |
| FRA10AC1 | 532.874077 | 0.174062769 | 0.19368492 | 0.89869037 | 0.36881761 | 0.76615653 |
| PLCE1 | 159.599218 | 1.555569876 | 0.50856636 | 3.05873529 | 0.00222273 | 0.15310061 |
| NOC3L | 261.749884 | -0.514205926 | 0.38202086 | -1.3460153 | 0.17829758 | 0.6285261 |
| TBC1D12 | 787.678664 | -0.053756621 | 0.20153345 | -0.266738 | 0.78967093 | 0.94023796 |
| HELLS | 103.137702 | 0.204049126 | 0.38267324 | 0.53322027 | 0.59388111 | 0.86472188 |
| CYP2C9 | 89.7447396 | -0.348060824 | 0.98437822 | -0.3535844 | 0.72365034 | 0.91600702 |
| AL157834.1 | 48.256949 | 0.241342278 | 0.58412109 | 0.41317166 | 0.67948085 | 0.89941358 |
| ACSM6 | 41.7008595 | 1.027339222 | 0.80441404 | 1.27712741 | 0.20155727 | 0.64879726 |
| PDLIM1 | 227.708927 | 0.329638329 | 0.47737572 | 0.69052177 | 0.48986613 | 0.81970016 |
| SORBS1 | 1877.47851 | -0.117740634 | 0.56208752 | -0.2094703 | 0.83408113 | 0.95393659 |
| ALDH18A1 | 506.544117 | -0.26095196 | 0.29457622 | -0.8858555 | 3.76E-01 | 0.7697436 |
| TCTN3 | 419.440934 | 0.111043081 | 0.21134952 | 0.52540021 | 0.59930501 | 0.86689411 |
| ENTPD1 | 55.0540709 | 1.28054525 | 0.69946428 | 1.83075146 | 0.06713765 | 0.48963361 |
| ENTPD1-AS1 | 2213.4146 | 0.514191175 | 0.44429185 | 1.15732749 | 0.24713858 | 0.68482118 |
| AL365273.2 | 260.837882 | 0.428736098 | 0.67412652 | 0.63598758 | 0.52478452 | 0.83593508 |
| CC2D2B | 134.241425 | 0.521422668 | 0.73901099 | 0.70556822 | 0.48045669 | 0.81714875 |
| CCNJ | 67.0102415 | 6.73E-05 | 0.32105468 | 0.00020976 | 0.99983264 | 0.99985901 |
| ZNF518A | 1651.21112 | 0.341914655 | 0.42142888 | 0.81132231 | 0.4171806 | 0.78652162 |
| BLNK | 449.107813 | 0.942182451 | 0.76526353 | 1.23118693 | 0.21825296 | 0.66064833 |
| TLL2 | 49.1324531 | -0.114253915 | 0.51147563 | -0.223381 | 8.23E-01 | 9.51E-01 |
| TM9SF3 | 1732.01565 | -0.150797515 | 0.19594817 | -0.7695786 | 0.4415499 | 0.79867043 |
| PIK3AP1 | 863.59313 | 0.343649184 | 0.38429352 | 0.89423622 | 0.37119552 | 0.76799652 |
| LCOR | 3181.49037 | -0.119364893 | 0.2352278 | -0.5074438 | 0.61184344 | 0.87199356 |
| RRP12 | 382.149339 | -0.763918444 | 0.45557076 | -1.6768382 | 0.09357413 | 0.53688613 |
| PGAM1 | 646.236954 | -0.058204571 | 0.44857695 | -0.1297538 | 8.97E-01 | 0.97324791 |
| EXOSC1 | 256.808776 | -0.113887946 | 0.30162268 | -0.3775842 | 0.70573954 | 0.90980406 |
| ZDHHC16 | 234.919029 | -0.242837011 | 0.24478223 | -0.9920533 | 0.32117155 | 0.73286139 |
| MMS19 | 606.532467 | -0.351824535 | 0.16881349 | -2.0841021 | 0.03715088 | 0.42215481 |
| UBTD1 | 371.991852 | -0.549755692 | 0.37370927 | -1.4710786 | 0.14126987 | 0.59581113 |
| AL355315.1 | 198.237205 | 1.070837687 | 0.53645816 | 1.99612525 | 0.04592029 | 0.44508419 |
| MORN4 | 185.544877 | 0.807641167 | 0.35812618 | 2.25518605 | 0.02412165 | 0.37387451 |
| PI4K2A | 70.1487133 | -0.298240298 | 0.55511354 | -0.53726 | 0.59108804 | 0.86336496 |
| ZFYVE27 | 477.564788 | 0.23995135 | 0.32527137 | 0.73769588 | 0.46069928 | 0.80741657 |
| R3HCC1L | 1330.52959 | -0.212478814 | 0.19387051 | -1.0959832 | 0.27308614 | 0.70123486 |
| LOXL4 | 39.1751974 | 1.253189284 | 0.47399134 | 2.64390754 | 0.0081955 | 0.24877162 |
| PYROXD2 | 125.972191 | 0.531641961 | 0.32089419 | 1.65675158 | 0.09756971 | 0.54093104 |
| HPS1 | 394.690258 | -0.350609206 | 0.25065702 | -1.3987608 | 0.16188473 | 0.61845523 |
| HPSE2 | 47.1954354 | -0.335239661 | 0.39461935 | -0.8495267 | 0.3955883 | 0.77931337 |
| CNNM1 | 166.058301 | -1.639808445 | 0.74803094 | -2.1921666 | 0.02836748 | 0.39280225 |
| GOT1 | 1177.80788 | 0.186624056 | 0.2297177 | 0.81240608 | 0.41655866 | 0.78632759 |
| AL391684.1 | 58.5619341 | 0.369095187 | 0.61314077 | 0.60197463 | 0.54719103 | 0.84482399 |
| SLC25A28 | 281.198103 | 0.146457201 | 0.41978556 | 0.34888575 | 0.72717508 | 0.91645085 |
| ENTPD7 | 219.707652 | -0.262470817 | 0.51196049 | -0.5126779 | 0.60817667 | 0.86985416 |
| CUTC | 512.719859 | -0.142645931 | 0.24242524 | -0.588412 | 5.56E-01 | 0.84861484 |
| COX15 | 324.975057 | 0.058379066 | 0.21429915 | 0.27241856 | 0.7853002 | 0.93892226 |
| ABCC2 | 5082.84063 | -1.072977801 | 0.69893032 | -1.5351714 | 0.12474174 | 0.58564339 |
| DNMBP | 887.843697 | -0.450972179 | 0.32980559 | -1.3673879 | 0.17150374 | 0.62520756 |
| DNMBP-AS1 | 59.9607224 | 0.600989934 | 0.63634438 | 0.94444134 | 0.34494417 | 0.75133476 |
| ERLIN1 | 259.900979 | 0.189901207 | 0.31107138 | 0.6104747 | 0.5415474 | 0.84221416 |
| CHUK | 305.866202 | -0.224764215 | 0.32048001 | -0.7013362 | 0.48309326 | 0.8177376 |

| | | | | | | |
|---|---|---|---|---|---|---|
| CWF19L1 | 321.373899 | -0.36548801 | 0.20091397 | -1.8191269 | 0.06889207 | 0.49260354 |
| BLOC1S2 | 168.658716 | -0.575824216 | 0.3450194 | -1.6689618 | 9.51E-02 | 0.53824794 |
| SCD | 188.400655 | -0.249023792 | 0.69814962 | -0.3566912 | 7.21E-01 | 0.91542662 |
| OLMALINC | 104.25867 | 0.190141412 | 0.51435962 | 0.36966629 | 0.71163115 | 0.91159439 |
| SEC31B | 158.709246 | 0.928796509 | 0.51069996 | 1.81867355 | 0.06896125 | 0.49273866 |
| HIF1AN | 316.318543 | 0.237267987 | 0.22737783 | 1.04349656 | 0.29671837 | 0.71819154 |
| PAX2 | 2919.99677 | 0.150076634 | 0.62741861 | 0.23919697 | 0.81095285 | 0.94730959 |
| SLF2 | 810.973541 | 0.312402251 | 0.22033953 | 1.41782208 | 0.15624272 | 0.61255389 |
| MRPL43 | 187.166812 | 0.214083086 | 0.3612956 | 0.59254274 | 0.55348721 | 0.84727776 |
| SEMA4G | 221.203055 | 0.984470142 | 0.57001067 | 1.72710827 | 0.08414822 | 0.52183702 |
| TWNK | 34.0183497 | -1.129293986 | 0.64319171 | -1.7557658 | 0.07912841 | 0.51192014 |
| LZTS2 | 405.396249 | -0.405461205 | 0.27022624 | -1.5004509 | 0.13349765 | 0.59240843 |
| PDZD7 | 50.4001689 | 0.551411377 | 0.81450669 | 0.67698815 | 0.49841347 | 0.82362007 |
| SFXN3 | 196.821619 | -0.192486285 | 0.42824958 | -0.4494722 | 0.65309105 | 0.88949911 |
| BTRC | 1948.99953 | -0.100292664 | 0.2284414 | -0.4390302 | 0.66063968 | 0.89130703 |
| DPCD | 195.243567 | 0.154271614 | 0.37943196 | 0.40658571 | 0.6843123 | 0.9007343 |
| POLL | 249.213165 | 0.128915748 | 0.28979567 | 0.44485049 | 0.6564278 | 0.89053559 |
| FBXW4 | 1033.50213 | 0.274450345 | 0.31237263 | 0.87859919 | 0.37961863 | 0.77095585 |
| AC010789.1 | 47.6506735 | -0.488068268 | 0.73992615 | -0.6596175 | 0.50949929 | 0.82882082 |
| NPM3 | 44.3459471 | -0.295719638 | 0.56479486 | -0.5235877 | 0.60056535 | 0.86755264 |
| OGA | 3510.89944 | -0.079297391 | 0.31049697 | -0.2553886 | 0.79842296 | 0.94320409 |
| KCNIP2 | 34.3955008 | 0.380765159 | 0.3668381 | 1.03796512 | 0.29928629 | 0.71852421 |
| ARMH3 | 1646.63362 | -0.108033942 | 0.17282338 | -0.6251118 | 0.53189765 | 0.83789294 |
| HPS6 | 35.4582653 | 0.413197471 | 0.588555 | 0.70205413 | 0.48264541 | 0.8177376 |
| LDB1 | 116.951558 | -0.021595192 | 0.39473944 | -0.0547075 | 0.95637153 | 0.9886112 |
| PPRC1 | 224.419532 | -1.240858575 | 0.7125582 | -1.7414136 | 0.0816111 | 0.51593841 |
| NOLC1 | 166.857749 | -0.873841401 | 0.60843249 | -1.4362175 | 0.15094046 | 0.60905477 |
| GBF1 | 2456.3777 | -0.216004678 | 0.27458745 | -0.7866517 | 0.4314858 | 0.79135216 |
| NFKB2 | 221.063953 | 0.361958086 | 0.5896809 | 0.61382026 | 0.53933411 | 0.84102987 |
| FBXL15 | 221.748344 | 0.121881214 | 0.41170164 | 0.29604258 | 0.76719756 | 0.93168929 |
| CUEDC2 | 185.731453 | 0.1946101 | 0.26762522 | 0.72717398 | 0.46711938 | 0.81140608 |
| MFSD13A | 35.4015718 | 0.807426696 | 0.47804587 | 1.68901512 | 0.09121653 | 0.53537574 |
| ACTR1A | 288.495622 | -0.278243899 | 0.35893462 | -0.7751938 | 0.43822514 | 0.79696434 |
| SUFU | 1050.53329 | 0.681231453 | 0.30828375 | 2.20975469 | 0.02712219 | 0.38702446 |
| TRIM8 | 467.866323 | 0.202079275 | 0.33370175 | 0.60556851 | 0.54480132 | 0.84441376 |
| ARL3 | 431.091761 | 0.663525608 | 0.35739413 | 1.85656552 | 0.06337299 | 0.48572144 |
| SFXN2 | 962.666715 | 0.827567358 | 0.46897215 | 1.76464075 | 7.76E-02 | 5.11E-01 |
| WBP1L | 1282.54772 | 0.065600087 | 0.22359158 | 0.29339248 | 7.69E-01 | 9.32E-01 |
| CYP17A1 | 118.211218 | -1.334832435 | 0.74159257 | -1.7999539 | 0.07186792 | 0.50131988 |
| BORCS7-ASN | 104.914477 | 0.290126798 | 0.53227746 | 0.54506685 | 0.58570754 | 0.86077718 |
| CNNM2 | 1749.6912 | -0.310327735 | 0.31794784 | -0.9760335 | 3.29E-01 | 0.73829083 |
| NT5C2 | 3996.71176 | -0.173500706 | 0.2121351 | -0.8178783 | 0.41342666 | 0.78632759 |
| PCGF6 | 307.993413 | -0.180507531 | 0.30906976 | -0.5840349 | 0.55919683 | 0.84936251 |
| TAF5 | 53.2626372 | -0.497726102 | 0.50249994 | -0.9904998 | 0.32192988 | 0.73375526 |
| ATP5MD | 422.560021 | -0.452353596 | 0.33338272 | -1.3568598 | 0.17482577 | 0.62699479 |
| PDCD11 | 365.477042 | -0.736636193 | 0.42660112 | -1.7267563 | 0.08421143 | 0.52183702 |
| SH3PXD2A | 234.627681 | 0.495513176 | 0.4765125 | 1.03987445 | 0.29839824 | 0.71852421 |
| STN1 | 131.217451 | 0.922256885 | 0.39074037 | 2.36028049 | 0.01826112 | 0.33797156 |
| SLK | 862.15732 | 0.223586811 | 0.14870306 | 1.50357913 | 0.13268977 | 0.59142019 |
| CFAP43 | 599.82446 | 0.988013355 | 0.55850778 | 1.76902345 | 0.07688996 | 0.50988423 |
| GSTO1 | 576.175314 | -0.48537562 | 0.42097694 | -1.1529744 | 0.2489209 | 0.68557761 |
| GSTO2 | 296.594121 | 0.816810095 | 0.47518877 | 1.71891709 | 0.08562947 | 0.5226185 |
| ITPRIP | 39.3909375 | -0.103592943 | 0.63937818 | -0.1620214 | 0.87128901 | 0.9657206 |
| SORCS3 | 96.8073819 | 0.455693882 | 0.72668937 | 0.62708208 | 0.53060543 | 0.83789294 |

| | | | | | | |
|---|---|---|---|---|---|---|
| SORCS1 | 2607.66016 | -0.157971708 | 0.69189118 | -0.2283187 | 0.81939847 | 0.94990455 |
| LINC01435 | 160.181692 | -0.322613987 | 1.01250022 | -0.318631 | 0.75000632 | 0.92565124 |
| XPNPEP1 | 650.460335 | -0.026773561 | 0.27929949 | -0.0958597 | 9.24E-01 | 0.98012108 |
| ADD3 | 2987.39952 | 0.318962718 | 0.38290365 | 0.83301038 | 0.40483887 | 0.78422899 |
| MXI1 | 1124.72012 | -0.100259778 | 0.45903658 | -0.2184135 | 0.82710696 | 0.95214203 |
| SMNDC1 | 311.505429 | -0.317974561 | 0.24591082 | -1.2930483 | 0.19599436 | 0.64558745 |
| DUSP5 | 117.82813 | -0.523165605 | 0.92911877 | -0.5630772 | 5.73E-01 | 0.8542683 |
| SMC3 | 427.712538 | -0.355826738 | 0.18532186 | -1.9200473 | 0.05485192 | 0.46815399 |
| RBM20 | 706.475062 | -0.489311601 | 0.38442002 | -1.2728567 | 0.2030689 | 0.64881821 |
| PDCD4 | 482.428848 | -0.085767402 | 0.44807419 | -0.1914134 | 0.84820172 | 0.95934492 |
| BBIP1 | 268.51342 | 0.369028037 | 0.21870193 | 1.68735607 | 0.09153491 | 0.53537574 |
| SHOC2 | 1423.68328 | -0.173333743 | 0.20074198 | -0.8634653 | 0.38788168 | 0.77373447 |
| GPAM | 82.6800416 | 0.175981776 | 0.3069088 | 0.57340088 | 0.56637329 | 0.85215881 |
| ZDHHC6 | 220.791317 | -0.149247876 | 0.19973279 | -0.7472377 | 0.45492008 | 0.80465616 |
| VTI1A | 3762.09216 | -0.13672924 | 0.20587774 | -0.6641283 | 0.50660818 | 0.82806179 |
| TCF7L2 | 2949.72115 | 0.226683884 | 0.3249749 | 0.69754274 | 0.4854632 | 0.81839266 |
| HABP2 | 138.614284 | 0.882491135 | 0.86875712 | 1.01580881 | 0.30972044 | 0.72498238 |
| CASP7 | 282.973018 | -0.523199217 | 0.36879984 | -1.4186536 | 0.15600004 | 0.61255389 |
| DCLRE1A | 48.1580852 | 0.051772495 | 0.41505793 | 0.12473559 | 0.90073288 | 0.97479173 |
| NHLRC2 | 697.587423 | 0.297210035 | 0.19855602 | 1.49685736 | 0.13443038 | 0.59240843 |
| CCDC186 | 1020.10736 | 0.207052919 | 0.2347536 | 0.88200106 | 0.37777624 | 0.77055888 |
| ABLIM1 | 6473.53285 | -0.232196803 | 0.25569316 | -0.9081072 | 0.36382158 | 0.76272056 |
| FAM160B1 | 574.877657 | -0.090871428 | 0.22906126 | -0.3967123 | 0.6915796 | 0.90491747 |
| TRUB1 | 293.844247 | 0.067970848 | 0.26283502 | 0.25860651 | 0.79593886 | 0.94252227 |
| ATRNL1 | 767.054254 | -0.361128101 | 0.76384455 | -0.472777 | 6.36E-01 | 8.83E-01 |
| GFRA1 | 102.425644 | 1.218655925 | 0.66246338 | 1.83958233 | 0.06582958 | 0.48798003 |
| HSPA12A | 1371.9963 | 0.536536703 | 0.38680347 | 1.38710418 | 0.16541 | 0.62065658 |
| ENO4 | 175.518694 | 0.363079497 | 0.41286712 | 0.87941006 | 0.37917898 | 0.77076106 |
| SHTN1 | 2431.88507 | -0.488198111 | 0.26599944 | -1.8353351 | 0.06645605 | 0.48798003 |
| PDZD8 | 2429.44456 | -0.242379184 | 0.27637424 | -0.8769963 | 0.38048864 | 0.7716703 |
| AC005871.2 | 105.860187 | 0.984858003 | 0.83171121 | 1.18413458 | 0.23635979 | 0.6735965 |
| EMX2OS | 2048.02756 | 0.031999424 | 0.42223126 | 0.07578649 | 0.93958897 | 0.98435387 |
| EMX2 | 718.253721 | -0.166707487 | 0.33427061 | -0.4987201 | 0.61797655 | 0.87452039 |
| RAB11FIP2 | 824.053683 | 0.116926923 | 0.26472159 | 0.44169772 | 0.65870796 | 0.89058983 |
| AC022395.1 | 58.909423 | 0.610813091 | 0.7690066 | 0.79428849 | 0.4270275 | 0.78933193 |
| CASC2 | 737.670961 | 0.7110143 | 0.64989882 | 1.09403846 | 0.27393811 | 0.7014484 |
| FAM204A | 654.105516 | -0.259906354 | 0.19817669 | -1.311488 | 0.18969294 | 0.64196311 |
| CACUL1 | 1162.07397 | -0.191950219 | 0.1755664 | -1.0933198 | 0.27425343 | 0.7014484 |
| EIF3A | 1735.29531 | -0.557711081 | 0.44618682 | -1.2499497 | 0.21131792 | 0.65561118 |
| FAM45A | 301.609216 | -0.10098428 | 0.21134516 | -0.4778168 | 0.63278057 | 0.88151688 |
| SFXN4 | 471.710965 | 0.165667085 | 0.35250603 | 0.4699695 | 0.63837681 | 0.8827026 |
| PRDX3 | 388.808992 | 0.520707173 | 0.40297691 | 1.29215139 | 0.19630472 | 0.64558745 |
| GRK5 | 182.745154 | -0.595997488 | 0.4094601 | -1.4555691 | 0.14551177 | 0.60184967 |
| TIAL1 | 895.205558 | -0.200178654 | 0.2529214 | -0.7914659 | 0.42867218 | 0.79037914 |
| BAG3 | 638.39538 | -0.976809302 | 0.41500771 | -2.3537136 | 0.01858692 | 0.33968023 |
| INPP5F | 1122.4165 | 0.25956421 | 0.38931147 | 0.66672634 | 0.50494695 | 0.82715271 |
| MCMBP | 403.024766 | 0.049229643 | 0.2175171 | 0.22632539 | 0.82094835 | 0.95072145 |
| SEC23IP | 751.616048 | -0.56871024 | 0.34421669 | -1.6521867 | 0.09849649 | 0.54209253 |
| PLPP4 | 367.488959 | 0.469831429 | 0.50834007 | 0.92424629 | 0.3553581 | 0.75757147 |
| WDR11-AS1 | 162.27554 | 0.265728237 | 0.30337052 | 0.87591978 | 0.38107365 | 0.77210521 |
| WDR11 | 1057.55122 | 0.171935716 | 0.28773136 | 0.59755641 | 0.55013596 | 0.84620747 |
| FGFR2 | 1376.9248 | -0.000878694 | 0.75868886 | -0.0011582 | 0.99907591 | 0.99977652 |
| ATE1 | 2624.83931 | 0.113548167 | 0.18815991 | 0.60346632 | 0.54619851 | 0.84460887 |
| NSMCE4A | 179.865246 | 0.22457033 | 0.32190615 | 0.69762672 | 0.48541067 | 0.81839266 |

| | | | | | | |
|---|---|---|---|---|---|---|
| TACC2 | 1267.78517 | -0.306739826 | 0.38319719 | -0.8004751 | 0.42343556 | 0.78894887 |
| PLEKHA1 | 9048.84854 | 0.091786615 | 0.40181125 | 0.22843217 | 8.19E-01 | 9.50E-01 |
| HTRA1 | 157.448501 | 0.834854855 | 0.32481269 | 2.57026548 | 0.01016206 | 0.26582321 |
| C10orf88 | 222.154352 | -0.214396404 | 0.23635596 | -0.9070912 | 0.36435858 | 0.76283229 |
| PSTK | 114.122809 | -0.169860574 | 0.4114406 | -0.4128435 | 0.67972129 | 0.89941358 |
| IKZF5 | 307.284647 | -0.398155668 | 0.36255188 | -1.0982033 | 0.27211573 | 0.70056282 |
| ACADSB | 1354.62475 | 0.273720004 | 0.84855511 | 0.32257186 | 0.74701951 | 0.92471745 |
| BUB3 | 364.296509 | -0.278688147 | 0.2838948 | -0.9816599 | 0.32626742 | 0.7373175 |
| CHST15 | 167.490914 | -0.193781434 | 0.5287405 | -0.3664963 | 0.71399478 | 0.91214838 |
| OAT | 314.395866 | 0.024058637 | 0.37019445 | 0.06498919 | 0.94818261 | 0.98675446 |
| LHPP | 545.0663 | 0.167054532 | 0.38570382 | 0.4331161 | 0.66493043 | 0.89315615 |
| ABRAXAS2 | 312.492809 | -0.279690284 | 0.18686375 | -1.4967605 | 0.13445558 | 0.59240843 |
| ZRANB1 | 745.234476 | -0.415139851 | 0.34268967 | -1.2114163 | 0.2257359 | 0.66688879 |
| CTBP2 | 4166.77319 | -0.444067427 | 0.29151429 | -1.5233127 | 1.28E-01 | 5.88E-01 |
| EDRF1 | 421.835066 | -0.412266835 | 0.19531596 | -2.1107688 | 0.03479219 | 0.41686761 |
| UROS | 283.838947 | -0.139952152 | 0.3371093 | -0.4151536 | 0.67802944 | 0.89898912 |
| BCCIP | 428.827644 | -0.63747489 | 0.23573266 | -2.7042281 | 0.00684633 | 0.23083391 |
| DHX32 | 1281.51456 | -0.591755491 | 0.23017188 | -2.5709287 | 0.01014262 | 0.26582196 |
| FANK1 | 751.145664 | 0.143822157 | 0.2755547 | 0.52193686 | 0.60171429 | 0.86763074 |
| ADAM12 | 72.5632285 | 0.232916997 | 0.28052648 | 0.83028525 | 4.06E-01 | 7.85E-01 |
| DOCK1 | 3821.49734 | 0.05433867 | 0.24846048 | 0.21870146 | 0.82688261 | 0.95204019 |
| AL359094.1 | 48.7003324 | 0.242832427 | 0.64190079 | 0.37830211 | 0.70520618 | 0.909554 |
| FAM196A | 139.690709 | 0.321694901 | 0.57139965 | 0.56299457 | 0.57343858 | 0.8542683 |
| CLRN3 | 350.784755 | 1.421325731 | 0.63838028 | 2.22645618 | 0.02598364 | 0.38392873 |
| PTPRE | 326.762651 | 0.744288459 | 0.37849249 | 1.96645504 | 4.92E-02 | 0.45419182 |
| MGMT | 1748.83434 | 0.596510426 | 0.35391728 | 1.68545155 | 0.09190149 | 0.53567268 |
| AL157832.3 | 187.825834 | 0.437405455 | 0.47672928 | 0.91751329 | 0.35887373 | 0.75936492 |
| C10orf143 | 145.024947 | 0.597784724 | 0.40026316 | 1.49347925 | 0.13531178 | 0.59381224 |
| GLRX3 | 820.018175 | -0.540441231 | 0.39623386 | -1.3639451 | 0.17258485 | 0.62593809 |
| PPP2R2D | 1872.47685 | -0.632141502 | 0.3277267 | -1.9288679 | 0.05374727 | 0.46711025 |
| BNIP3 | 1385.91487 | -0.815963697 | 0.59456483 | -1.3723713 | 0.16994789 | 0.6233579 |
| AL162274.2 | 34.8581091 | 0.153036296 | 0.61620928 | 0.24835117 | 0.80386271 | 0.94543168 |
| STK32C | 160.121569 | -0.046900326 | 0.35928952 | -0.1305363 | 0.89614213 | 0.97309833 |
| LRRC27 | 219.110452 | -0.382988776 | 0.52178257 | -0.7340007 | 0.4629483 | 0.8082072 |
| PWWP2B | 48.0977224 | -0.498330395 | 0.56224182 | -0.8863275 | 0.37544107 | 0.7697436 |
| INPP5A | 636.705614 | -0.063092486 | 0.2568668 | -0.2456234 | 0.80597381 | 0.94585363 |
| CFAP46 | 253.947774 | 1.025139567 | 0.51891273 | 1.97555292 | 0.04820544 | 0.45192142 |
| ADAM8 | 48.9622562 | 0.169389522 | 0.41576068 | 0.40742074 | 6.84E-01 | 9.01E-01 |
| TUBGCP2 | 1018.46707 | 0.062142746 | 0.23046461 | 0.26964117 | 0.78743633 | 0.93953874 |
| AL360181.1 | 58.7850469 | 1.212549201 | 0.72611746 | 1.66990779 | 0.09493761 | 0.53824794 |
| PRAP1 | 41.0233969 | -0.071163321 | 0.56585773 | -0.1257619 | 0.89992043 | 0.97465308 |
| FUOM | 148.924106 | 0.292169815 | 0.52213826 | 0.55956408 | 0.57577681 | 0.85537243 |
| ECHS1 | 1065.73071 | 0.241215917 | 0.43925663 | 0.54914576 | 0.58290542 | 0.85986705 |
| PAOX | 110.099784 | 0.590620209 | 0.35131976 | 1.68114712 | 0.09273434 | 0.53669957 |
| SCART1 | 97.3878269 | 0.096321249 | 0.56283177 | 0.17113684 | 0.86411617 | 0.96393376 |
| IQSEC3 | 370.071878 | 1.155880549 | 0.99114366 | 1.16620889 | 0.24353 | 0.68164834 |
| AC026369.1 | 76.0857582 | -0.144533895 | 1.28613606 | -0.1123784 | 0.91052339 | 0.97697493 |
| SLC6A12 | 1441.67524 | -0.217507082 | 0.64465516 | -0.3374007 | 0.73581488 | 0.92019178 |
| SLC6A13 | 3642.26087 | 2.035903285 | 0.5717274 | 3.56096852 | 0.00036949 | 0.06861791 |
| KDM5A | 1429.25259 | -0.143173057 | 0.13478823 | -1.0622074 | 0.28814153 | 0.71175994 |
| CCDC77 | 97.9690508 | -0.550075359 | 0.36737022 | -1.4973325 | 0.13430677 | 0.59240843 |
| B4GALNT3 | 69.1871558 | 0.58647451 | 0.43883388 | 1.33643855 | 0.18140597 | 0.6318513 |
| WNK1 | 2131.97635 | 0.123771126 | 0.16298683 | 0.75939341 | 0.44761726 | 0.80223612 |
| RAD52 | 374.599134 | 0.351280456 | 0.29034381 | 1.20987754 | 0.22632589 | 0.66690202 |

| | | | | | | |
|---|---|---|---|---|---|---|
| ERC1 | 5172.51626 | -0.349672623 | 0.22114779 | -1.5811717 | 0.11383879 | 0.56865873 |
| WNT5B | 105.534464 | -0.107940566 | 0.61223768 | -0.176305 | 0.86005433 | 0.96283855 |
| FBXL14 | 112.365517 | -0.206854408 | 0.35957843 | -0.5752692 | 0.56510922 | 0.85171209 |
| ADIPOR2 | 1023.42203 | -0.157236885 | 0.35640598 | -0.4411735 | 0.65908738 | 0.89058983 |
| DCP1B | 456.813666 | 0.813506086 | 0.35154227 | 2.31410602 | 0.02066191 | 0.35267827 |
| CACNA1C | 221.656505 | 0.119692725 | 0.74281677 | 0.16113358 | 0.87198819 | 0.9657206 |
| FKBP4 | 366.626071 | -1.170840996 | 0.49167332 | -2.3813393 | 0.01724982 | 0.32964126 |
| ITFG2-AS1 | 65.4482417 | 0.303740506 | 0.42595445 | 0.71308213 | 0.47579494 | 0.81570932 |
| ITFG2 | 852.805403 | -0.514825586 | 0.3960613 | -1.2998634 | 0.19364779 | 0.64548758 |
| TULP3 | 1887.44115 | 0.135857485 | 0.29273543 | 0.46409649 | 0.64257862 | 0.88441609 |
| TEAD4 | 593.105195 | -0.687526673 | 0.54327426 | -1.2655241 | 0.20568352 | 0.651931 |
| TSPAN9 | 3798.83614 | 0.071956071 | 0.35105093 | 0.20497331 | 0.83759298 | 0.95496955 |
| CRACR2A | 41.8058594 | -1.161310936 | 0.5154367 | -2.2530622 | 0.02425523 | 0.37480999 |
| PARP11 | 291.805923 | -0.047656897 | 0.25716918 | -0.1853134 | 0.85298322 | 0.96073035 |
| CCND2 | 70.7434565 | 2.305824293 | 0.8125166 | 2.83787962 | 0.00454143 | 0.20749797 |
| TIGAR | 188.418636 | -0.320325076 | 0.55471439 | -0.5774595 | 0.56362911 | 0.85121907 |
| C12orf4 | 276.400705 | -0.158555775 | 0.15855328 | -1.0000158 | 0.31730288 | 0.72925395 |
| AKAP3 | 36.8931918 | -0.001944986 | 0.58921847 | -0.003301 | 0.99736622 | 0.99963814 |
| AC005833.2 | 46.2553362 | 0.398787239 | 0.43243884 | 0.92218184 | 0.35643374 | 0.75841231 |
| CD9 | 344.775203 | -0.720275894 | 0.54896605 | -1.3120591 | 0.1895002 | 0.6416112 |
| PLEKHG6 | 52.8247093 | 0.200779854 | 0.52529091 | 0.38222602 | 0.70229372 | 0.90837679 |
| TNFRSF1A | 1190.68667 | 0.319903111 | 0.43835414 | 0.72978235 | 0.46552324 | 0.81042189 |
| SCNN1A | 807.388314 | 0.121934873 | 0.50685049 | 0.24057365 | 0.80988557 | 0.94692486 |
| LTBR | 927.293301 | -0.518959299 | 0.38928527 | -1.333108 | 0.18249637 | 0.63216521 |
| AC005840.2 | 51.6780929 | -0.706621397 | 0.71780533 | -0.9844193 | 0.32490941 | 0.73628585 |
| TAPBPL | 122.984678 | 0.667741496 | 0.54333223 | 1.22897457 | 0.21908134 | 0.661588 |
| MRPL51 | 259.749104 | -0.407225816 | 0.36834608 | -1.1055522 | 0.26892039 | 0.69961625 |
| NCAPD2 | 207.910329 | -0.057261756 | 0.31938693 | -0.1792865 | 0.85771277 | 0.9626966 |
| GAPDH | 3368.42093 | 0.226778777 | 0.50997814 | 0.44468333 | 6.57E-01 | 0.89053559 |
| IFFO1 | 73.1987519 | -0.10100031 | 0.32374383 | -0.311976 | 0.75505875 | 0.92799282 |
| CHD4 | 323.00774 | 0.293176212 | 0.41224136 | 0.71117612 | 0.4769751 | 0.81570932 |
| AC006064.2 | 93.2440131 | 0.570723788 | 0.4749028 | 1.20176968 | 0.22945278 | 0.66705629 |
| ING4 | 256.299258 | -0.348383506 | 0.33237332 | -1.0481693 | 0.29456062 | 0.71625731 |
| ZNF384 | 287.035206 | -0.113636778 | 0.19365238 | -0.5868081 | 0.55733262 | 0.84861484 |
| COPS7A | 201.858192 | -0.234608934 | 0.26611427 | -0.8816097 | 0.37798789 | 0.77055888 |
| MLF2 | 428.669167 | -0.178908094 | 0.30900645 | -0.5789785 | 0.56260367 | 0.85054239 |
| PTMS | 619.180195 | 0.191538537 | 0.35036722 | 0.54667939 | 0.58459901 | 0.86045266 |
| USP5 | 187.053156 | 0.047739555 | 0.43273977 | 0.11031932 | 0.91215614 | 0.977179 |
| TPI1 | 994.437151 | -0.156599649 | 0.48103035 | -0.3255505 | 0.7447645 | 0.92380845 |
| SPSB2 | 117.577413 | -0.414622427 | 0.37158833 | -1.1158112 | 0.26450295 | 0.69567438 |
| LRRC23 | 485.233823 | -0.952635401 | 0.40997789 | -2.3236263 | 0.02014553 | 0.34851906 |
| ENO2 | 81.9527808 | -1.216683204 | 0.54897087 | -2.2162983 | 2.67E-02 | 3.86E-01 |
| ATN1 | 387.147028 | -0.382479435 | 0.29190408 | -1.3102915 | 0.19009725 | 0.64261395 |
| C12orf57 | 143.992137 | 0.125247456 | 0.60922256 | 0.20558572 | 0.83711454 | 0.95496955 |
| PTPN6 | 153.727556 | -0.156882031 | 0.40877569 | -0.3837851 | 0.70113771 | 0.90783786 |
| PHB2 | 374.082293 | -0.036404355 | 0.36389615 | -0.1000405 | 0.92031217 | 0.97939204 |
| EMG1.1 | 121.318742 | -0.136238894 | 0.47426309 | -0.2872644 | 0.77390989 | 0.93429478 |
| LPCAT3 | 1843.66104 | -0.046354454 | 0.29524454 | -0.1570036 | 0.87524202 | 0.96615698 |
| C1S | 581.383155 | 0.168526271 | 0.44857555 | 0.37569206 | 0.70714584 | 0.91002281 |
| C1R | 503.624502 | 0.881516339 | 0.60752066 | 1.45100637 | 1.47E-01 | 6.04E-01 |
| C1RL | 849.207085 | 0.79436602 | 0.46116584 | 1.72251705 | 0.0849759 | 0.52183702 |
| C1RL-AS1 | 170.135443 | 0.496305074 | 0.41803178 | 1.18724244 | 0.235132 | 0.67119484 |
| AC233309.1 | 50.1441686 | 1.900105228 | 0.666737 | 2.8498572 | 0.00437389 | 0.20749797 |
| RBP5 | 929.284099 | 1.607846483 | 0.39435449 | 4.07716029 | 4.56E-05 | 0.02082964 |

| | | | | | | |
|---|---|---|---|---|---|---|
| AC018653.3 | 45.5156532 | 1.212206878 | 0.81931328 | 1.47954013 | 0.13899601 | 0.59474076 |
| CLSTN3 | 768.008686 | 1.139125752 | 0.40127848 | 2.83874116 | 0.00452919 | 0.20749797 |
| PEX5 | 322.046635 | 0.495555588 | 0.27861509 | 1.77863872 | 0.07529901 | 0.50718602 |
| CD163L1 | 78.2380641 | 0.307016434 | 0.32876796 | 0.93383929 | 0.3503868 | 0.75467792 |
| SLC2A14 | 57.5880618 | 0.706458722 | 0.84382866 | 0.83720636 | 0.40247658 | 0.78260941 |
| FOXJ2 | 97.3222862 | 0.289801893 | 0.45213992 | 0.64095622 | 0.52155113 | 0.83445062 |
| NECAP1 | 152.827972 | -0.354644978 | 0.23871545 | -1.485639 | 0.13737464 | 0.59474076 |
| LINC00937 | 678.4464 | 0.532761459 | 0.34656043 | 1.53728303 | 0.12422402 | 0.58412988 |
| RIMKLB | 755.947366 | -0.069478911 | 0.38769064 | -0.1792122 | 0.85777105 | 0.9626966 |
| A2ML1-AS1 | 95.5676735 | -0.139454573 | 0.69624628 | -0.2002949 | 0.84124996 | 0.95677175 |
| AC006581.2 | 61.0815664 | 0.237506306 | 0.48172998 | 0.49302787 | 0.6219929 | 0.87713418 |
| PHC1 | 183.709087 | 0.529480452 | 0.41944051 | 1.26234935 | 0.2068231 | 0.65251041 |
| M6PR | 459.488046 | -0.341435244 | 0.33707753 | -1.0129279 | 0.31109458 | 0.72558189 |
| KLRG1 | 150.810869 | -0.352089864 | 0.35362298 | -0.9956646 | 0.31941316 | 0.73132207 |
| A2M | 74.4381194 | 0.144344676 | 0.47543127 | 0.30360787 | 0.76142665 | 0.93000994 |
| CLEC2D | 158.012069 | 0.390011344 | 0.54386797 | 0.71710666 | 0.47330829 | 0.81478861 |
| CLEC7A | 90.6712727 | -2.791561884 | 1.21450452 | -2.2985191 | 0.02153226 | 0.35786026 |
| GABARAPL1 | 1302.68198 | -0.708914346 | 0.58627155 | -1.2091911 | 0.22658943 | 0.66690202 |
| KLRD1 | 704.112933 | 0.549973016 | 0.79205266 | 0.6943642 | 0.48745384 | 0.81893983 |
| AC022075.1 | 66.3167776 | 1.230791381 | 0.49347444 | 2.49413402 | 0.01262649 | 0.29875272 |
| EIF2S3B | 78.055464 | -0.390546094 | 0.38082277 | -1.0255324 | 0.30511204 | 0.72103276 |
| MAGOHB | 95.5735019 | -0.270185182 | 0.41896877 | -0.6448814 | 0.51900402 | 0.83341004 |
| YBX3 | 1990.75702 | -0.331591816 | 0.38179862 | -0.8684992 | 0.38512109 | 0.77317162 |
| PRH1 | 122.225166 | 1.182876604 | 0.32132585 | 3.68123701 | 0.00023211 | 0.05581517 |
| SMIM10L1 | 152.351518 | -0.141957769 | 0.32190646 | -0.4409907 | 0.6592197 | 0.89058983 |
| AC244131.2 | 362.481226 | 0.606328836 | 0.6832518 | 0.88741638 | 0.37485477 | 0.7697436 |
| PRB2 | 53.9589146 | 1.613272701 | 1.03846246 | 1.55352048 | 0.12029884 | 0.57816838 |
| LINC01252 | 158.845772 | 1.327089736 | 0.85706102 | 1.5484192 | 0.1215214 | 0.5799266 |
| ETV6 | 4139.98575 | 0.523597501 | 0.45511664 | 1.15046882 | 0.24995083 | 0.68617881 |
| BCL2L14 | 295.006475 | 0.198631847 | 0.47198549 | 0.42084312 | 0.67386964 | 0.89799059 |
| LRP6 | 1371.38888 | 0.193380969 | 0.32061021 | 0.60316534 | 0.5463987 | 0.84460887 |
| MANSC1 | 71.5048423 | 0.140481715 | 0.35679992 | 0.39372686 | 0.69378271 | 0.90587439 |
| BORCS5 | 454.933934 | 0.095958017 | 0.22500102 | 0.42647814 | 0.66975947 | 0.89611537 |
| DUSP16 | 1161.27986 | -0.10680494 | 0.35735112 | -0.2988795 | 0.76503195 | 0.93083208 |
| CREBL2 | 314.158534 | -0.050363579 | 0.245122 | -0.2054633 | 0.83721015 | 0.95496955 |
| CDKN1B | 141.609108 | 0.577507482 | 0.30051405 | 1.92173203 | 0.05463948 | 0.46772575 |
| APOLD1 | 68.4920544 | -0.466433346 | 0.59266739 | -0.7870069 | 0.4312778 | 0.79135216 |
| GPRC5D-AS1 | 234.570911 | -0.492063779 | 0.38244821 | -1.2866154 | 0.19822836 | 0.64582202 |
| HEBP1 | 187.528843 | 0.164789435 | 0.21456114 | 0.76803018 | 0.44246925 | 0.79933184 |
| AC023790.2 | 34.967279 | 0.622434119 | 0.57530896 | 1.08191278 | 0.27929129 | 0.70446562 |
| FAM234B | 41.4739093 | 0.066345899 | 0.39194632 | 0.16927292 | 0.86558197 | 0.96428252 |
| EMP1 | 110.428743 | -0.139406078 | 0.79540271 | -0.1752648 | 0.86087159 | 0.96302275 |
| ATF7IP | 2111.87401 | -0.034347236 | 0.34063793 | -0.1008321 | 0.91968375 | 0.97934316 |
| PLBD1 | 123.417169 | 0.534741844 | 0.35193915 | 1.51941562 | 0.12865791 | 0.58793678 |
| PLBD1-AS1 | 140.028626 | 0.443222605 | 0.4855171 | 0.91288773 | 0.36130159 | 0.76009788 |
| AC010168.1 | 320.345953 | 0.440741017 | 0.59571083 | 0.73985731 | 0.45938658 | 0.80669065 |
| HIST4H4 | 34.3125455 | -0.061722605 | 0.51571272 | -0.1196841 | 0.90473341 | 0.97585622 |
| H2AFJ | 455.438564 | -0.118878421 | 0.46123915 | -0.2577371 | 0.79660984 | 0.94276732 |
| WBP11 | 270.711651 | 0.17336661 | 0.23622289 | 0.73391113 | 0.46300291 | 0.8082072 |
| C12orf60 | 129.298085 | 0.7280502 | 0.52438424 | 1.38839071 | 0.1650181 | 0.62065658 |
| MGP | 144.196706 | -0.853829487 | 0.42611248 | -2.0037655 | 0.04509518 | 0.44360789 |
| RERG | 1628.80644 | -0.116754502 | 0.54939085 | -0.2125163 | 0.83170428 | 0.95325673 |
| RERG-AS1 | 49.7441161 | 1.503612739 | 1.00060035 | 1.50271059 | 0.13291369 | 0.59190052 |
| PTPRO | 156.842781 | 0.220469628 | 0.75458388 | 0.29217379 | 0.77015375 | 0.93266075 |

| | | | | | | |
|---|---|---|---|---|---|---|
| EPS8 | 6430.75123 | -0.047812426 | 0.29020189 | -0.1647557 | 0.86913624 | 0.96556934 |
| STRAP | 543.86833 | -0.148777825 | 0.23943806 | -0.6213625 | 0.53436115 | 0.83929051 |
| DERA | 2278.12059 | -0.249710562 | 0.39577382 | -0.6309426 | 0.52807806 | 0.83761342 |
| MGST1 | 5284.81452 | -0.484182694 | 0.40210749 | -1.2041126 | 0.22854606 | 0.66705629 |
| AC007529.2 | 392.361283 | 0.375240952 | 0.61814248 | 0.60704605 | 0.54382035 | 0.84396451 |
| RERGL | 216.705229 | 1.326813543 | 0.64585164 | 2.05436273 | 0.0399406 | 0.42972196 |
| PIK3C2G | 111.32543 | 0.973661125 | 0.56639259 | 1.71905696 | 0.085604 | 0.5226185 |
| PLCZ1 | 145.280318 | 1.562743914 | 0.80049962 | 1.95221069 | 5.09E-02 | 0.4600311 |
| PLEKHA5 | 17480.2 | -0.032702332 | 0.36962708 | -0.0884739 | 9.30E-01 | 0.98272895 |
| AEBP2 | 4350.6856 | 0.224280063 | 0.24751559 | 0.90612502 | 0.36486969 | 0.76338174 |
| AC024901.1 | 377.945317 | 0.695236888 | 0.51625206 | 1.34670046 | 0.17807672 | 0.62849772 |
| LINC02398 | 106.795003 | 0.891876007 | 0.6108034 | 1.46016871 | 0.14424371 | 0.60086567 |
| PDE3A | 1183.55412 | -0.08642147 | 0.74652006 | -0.1157658 | 0.90783818 | 0.97673359 |
| PYROXD1 | 329.732174 | -0.347057357 | 0.3466266 | -1.0012427 | 0.31670948 | 0.7285009 |
| RECQL | 327.271352 | -0.072154741 | 0.29617894 | -0.2436187 | 0.80752611 | 0.94604106 |
| GOLT1B | 302.016864 | -0.193308728 | 0.22933698 | -0.8429026 | 3.99E-01 | 0.78161291 |
| SPX | 227.911676 | 0.882794713 | 0.65096647 | 1.35612932 | 0.17505803 | 0.62699479 |
| LDHB | 1907.31666 | 0.713726218 | 0.38582747 | 1.84985846 | 0.06433395 | 0.48593331 |
| CMAS | 327.459126 | 0.300632432 | 0.27290496 | 1.10160122 | 0.27063508 | 0.69961625 |
| C2CD5 | 1177.80542 | -0.25069053 | 0.24198992 | -1.0359544 | 0.30022341 | 0.71879123 |
| ETNK1 | 2171.62185 | -0.015351747 | 0.36630387 | -0.0419099 | 0.96657055 | 0.99122723 |
| AC084816.1 | 203.667869 | -0.60166799 | 0.75631979 | -0.7955206 | 0.42631072 | 0.78895108 |
| SOX5 | 1991.62143 | -0.353002037 | 0.81673483 | -0.4322113 | 0.66558784 | 0.89369715 |
| BCAT1 | 104.884182 | -3.120670315 | 1.19688398 | -2.607329 | 0.00912516 | 0.25831482 |
| LRMP | 447.530802 | 0.51464854 | 0.84952108 | 0.60581021 | 0.54464079 | 0.84441376 |
| CASC1 | 356.276509 | 1.460146448 | 0.7075799 | 2.06357818 | 0.03905773 | 0.42692527 |
| ETFRF1 | 316.50195 | 0.641968239 | 0.41897468 | 1.53223636 | 0.12546412 | 0.58564339 |
| KRAS | 640.692603 | -0.117290177 | 0.312404 | -0.3754439 | 0.70733036 | 0.91002281 |
| RASSF8-AS1 | 1193.41525 | -0.472990244 | 0.39728377 | -1.1905602 | 0.23382629 | 0.67043632 |
| RASSF8 | 2516.25921 | -0.679043521 | 0.40638553 | -1.6709343 | 0.09473465 | 0.53824794 |
| BHLHE41 | 150.320913 | -1.389656947 | 0.54220444 | -2.562976 | 1.04E-02 | 2.69E-01 |
| SSPN | 63.4201692 | 0.954440285 | 0.57237137 | 1.66751927 | 0.09541119 | 0.53824794 |
| AC022509.2 | 42.4001945 | 0.848134849 | 0.55358116 | 1.53208762 | 0.12550081 | 0.58564339 |
| AC024145.1 | 123.629244 | 0.601897907 | 0.72856353 | 0.82614334 | 0.40872279 | 0.78511023 |
| ITPR2 | 4762.59677 | 0.014177413 | 0.25339587 | 0.05594966 | 0.95538191 | 0.98840182 |
| AC023051.1 | 93.2081429 | 0.777770615 | 0.7436785 | 1.04584254 | 0.29563374 | 0.71683206 |
| INTS13 | 247.940882 | -0.411035606 | 0.30656838 | -1.3407632 | 0.17999734 | 0.63124625 |
| FGFR1OP2 | 798.62459 | -0.107418979 | 0.35688862 | -0.3009874 | 0.7634241 | 0.93040403 |
| TM7SF3 | 2030.19 | 0.476247355 | 0.50742115 | 0.93856425 | 0.34795451 | 0.75310477 |
| MED21 | 255.073465 | -0.061138526 | 0.21872806 | -0.2795184 | 0.77984699 | 0.93684161 |
| STK38L | 726.267534 | 0.179342495 | 0.22675258 | 0.79091711 | 4.29E-01 | 7.90E-01 |
| ARNTL2 | 174.725942 | -0.46450729 | 0.62275721 | -0.7458883 | 4.56E-01 | 8.05E-01 |
| PPFIBP1 | 4515.53879 | -0.36957533 | 0.36704813 | -1.0068852 | 0.31398994 | 0.72719146 |
| REP15 | 39.7877537 | -0.207402966 | 0.61640721 | -0.3364707 | 0.73651594 | 0.92019178 |
| MRPS35 | 443.631333 | -0.124890507 | 0.21419714 | -0.5830634 | 0.55985064 | 0.84941263 |
| KLHL42 | 642.787289 | -0.276702819 | 0.28957352 | -0.9555529 | 0.33929817 | 0.74644675 |
| CCDC91 | 3470.55253 | -0.19479465 | 0.23931601 | -0.8139641 | 0.41566551 | 0.78632759 |
| AC022364.1 | 59.3408895 | 0.029676547 | 0.42819592 | 0.069306 | 0.94474605 | 0.9853777 |
| FAR2 | 149.461783 | 1.296592156 | 0.38897349 | 3.33336898 | 0.00085801 | 0.09882986 |
| AC009318.1 | 117.737078 | -0.077522816 | 0.59195227 | -0.1309613 | 0.89580595 | 0.97309833 |
| ERGIC2 | 500.836722 | -0.111006782 | 0.18423867 | -0.6025162 | 0.54683058 | 0.84460887 |
| TMTC1 | 256.252082 | -0.298790624 | 0.60048732 | -0.4975802 | 0.61877994 | 0.87502492 |
| IPO8 | 656.256346 | 0.281951176 | 0.21352955 | 1.32043165 | 0.18669094 | 0.6378941 |
| CAPRIN2 | 378.82749 | 0.124313746 | 0.26215244 | 0.47420404 | 0.63535442 | 0.88246953 |

| | | | | | | |
|---|---|---|---|---|---|---|
| AC010198.1 | 61.9828068 | 0.482641676 | 0.54575273 | 0.88435962 | 0.37650211 | 0.77035529 |
| DDX11-AS1 | 38.1426633 | 0.750987563 | 0.41770257 | 1.79790029 | 0.07219281 | 0.50202435 |
| DDX11 | 157.530948 | -0.542923875 | 0.55260744 | -0.9824766 | 0.32586511 | 0.7373175 |
| SINHCAF | 777.802126 | -0.331953121 | 0.26418929 | -1.2564973 | 0.20893574 | 0.65401126 |
| DENND5B | 4426.48122 | 0.034058827 | 0.24923437 | 0.13665381 | 8.91E-01 | 9.72E-01 |
| DENND5B-AS | 110.842567 | 0.468532426 | 0.80383241 | 0.58287327 | 0.55997862 | 0.84941263 |
| ETFBKMT | 115.756752 | 0.848188882 | 0.50002049 | 1.69630824 | 0.08982752 | 0.53232417 |
| AMN1 | 309.593117 | -0.09629011 | 0.39511961 | -0.2436986 | 0.80746423 | 0.94604106 |
| KIAA1551 | 548.284793 | -0.001272754 | 0.28740455 | -0.0044284 | 0.99646663 | 0.99916372 |
| BICD1 | 1217.94686 | 0.615908952 | 0.4475478 | 1.37618585 | 1.69E-01 | 0.62304975 |
| FGD4 | 7429.49725 | -0.019254337 | 0.32689314 | -0.058901 | 0.95303095 | 0.98791464 |
| DNM1L | 924.063469 | 0.206491135 | 0.14792571 | 1.39591111 | 0.16274126 | 0.61960665 |
| YARS2 | 384.072965 | -0.117246644 | 0.19937702 | -0.588065 | 0.55648868 | 0.84861484 |
| PKP2 | 379.004877 | 0.378617305 | 0.35428278 | 1.06868674 | 0.28521085 | 0.71015169 |
| ALG10B | 87.66147 | -0.087571157 | 0.41143606 | -0.2128427 | 0.83144967 | 0.95325673 |
| CPNE8 | 2539.24077 | -0.226967908 | 0.21586676 | -1.0514259 | 0.29306301 | 0.71438996 |
| KIF21A | 3516.54525 | -0.03413205 | 0.28366994 | -0.1203231 | 0.9042272 | 0.9756941 |
| C12orf40 | 148.825461 | 0.996484973 | 0.53346963 | 1.86793197 | 0.06177155 | 0.48267858 |
| SLC2A13 | 2391.10411 | 0.383457006 | 0.3462816 | 1.10735599 | 0.26814005 | 0.69954237 |
| LINC02471 | 1784.18744 | 2.010887752 | 1.35484142 | 1.48422371 | 0.13774958 | 0.59474076 |
| AC079630.1 | 344.8417 | 1.502729085 | 0.84776452 | 1.77257842 | 0.07629859 | 0.50768193 |
| LRRK2 | 4365.74714 | 0.663237834 | 0.6451075 | 1.02810436 | 0.30390074 | 0.72082426 |
| AC090630.1 | 67.4808164 | -0.174645045 | 0.92595551 | -0.1886106 | 0.85039801 | 0.95992798 |
| GXYLT1 | 635.283413 | -0.349009392 | 0.24182665 | -1.4432214 | 0.14895812 | 0.60622442 |
| YAF2 | 1559.73061 | -0.343055151 | 0.2705545 | -1.2679706 | 0.20480846 | 0.65074398 |
| PPHLN1 | 1811.67835 | -0.150760174 | 0.15095425 | -0.9987143 | 0.31793311 | 0.72996802 |
| ZCRB1 | 191.91396 | -0.033107147 | 0.27286019 | -0.1213337 | 0.9034267 | 0.97536781 |
| PRICKLE1 | 675.11175 | 0.947109174 | 0.56652234 | 1.67179492 | 0.09456477 | 0.53824794 |
| PUS7L | 309.064963 | -0.008704117 | 0.33336209 | -0.0261101 | 0.97916952 | 0.99482978 |
| IRAK4 | 293.395186 | 0.329246405 | 0.4035227 | 0.81593032 | 0.41453999 | 0.78632759 |
| TWF1 | 202.919906 | -0.158235322 | 0.31332739 | -0.5050159 | 0.61354765 | 0.87222135 |
| TMEM117 | 1621.03981 | 0.360542083 | 0.28555722 | 1.26259139 | 0.20673606 | 0.65251041 |
| ANO6 | 2493.23917 | -0.674378959 | 0.48691625 | -1.3849999 | 0.1660525 | 0.62079891 |
| AC008124.1 | 122.007816 | 0.429814568 | 0.27473147 | 1.56448976 | 0.1177026 | 0.57390152 |
| ARID2 | 2261.33253 | 0.030748457 | 0.18687864 | 0.16453704 | 0.86930839 | 0.9656086 |
| SCAF11 | 2833.33202 | -0.040241352 | 0.20593889 | -0.1954043 | 0.84507642 | 0.95760132 |
| SLC38A1 | 609.786643 | 0.091651986 | 0.61310171 | 0.14948904 | 8.81E-01 | 9.68E-01 |
| SLC38A2 | 1958.09088 | -0.606951424 | 0.45392591 | -1.3371156 | 0.1811849 | 0.6318513 |
| AC008014.1 | 5458.9177 | 0.905787289 | 0.55274455 | 1.63870867 | 0.10127395 | 0.54652047 |
| SLC38A4 | 364.714721 | -0.119644882 | 0.99713803 | -0.1199883 | 0.90449243 | 0.97574711 |
| PCED1B | 109.82826 | 1.223578933 | 0.81644555 | 1.49866568 | 0.13396038 | 0.59240843 |
| RPAP3 | 236.598361 | -0.411740784 | 0.29138787 | -1.4130334 | 0.15764591 | 0.61323667 |
| AC004241.1 | 41.0619646 | 0.750878214 | 0.54062818 | 1.3888995 | 0.16486331 | 0.62065658 |
| RAPGEF3 | 830.649028 | -0.249330343 | 0.39345554 | -0.6336938 | 0.52628066 | 0.83695661 |
| SLC48A1 | 317.782196 | 0.138611562 | 0.44367719 | 0.31241534 | 0.75472489 | 0.92799282 |
| HDAC7 | 332.282569 | -0.548974537 | 0.58324912 | -0.9412351 | 0.3465844 | 0.75166079 |
| VDR | 798.050877 | 0.350733495 | 0.58576986 | 0.59875647 | 0.54933529 | 0.84594302 |
| TMEM106C | 195.829879 | 0.613769312 | 0.36816359 | 1.66711031 | 0.09549247 | 0.53824794 |
| SENP1 | 285.715079 | -0.241163811 | 0.27406729 | -0.8799438 | 0.37888977 | 0.77061593 |
| PFKM | 140.916455 | 0.490366333 | 0.41239503 | 1.18906946 | 0.23441234 | 0.670712 |
| ASB8 | 210.640339 | 0.04606856 | 0.29092217 | 0.15835355 | 0.87417821 | 0.9661168 |
| ZNF641 | 134.601571 | 0.240274108 | 0.3153319 | 0.76197209 | 0.44607666 | 0.80146452 |
| AC090115.1 | 68.4517581 | 0.974044431 | 0.57839037 | 1.68406061 | 0.09216996 | 0.53619162 |
| KANSL2 | 378.325492 | -0.331991958 | 0.31831588 | -1.0429639 | 0.29696503 | 0.71825247 |

| | | | | | | |
|---|---|---|---|---|---|---|
| CCNT1 | 484.346574 | -0.296889249 | 0.21362112 | -1.3897935 | 0.16459158 | 0.62065658 |
| ADCY6 | 336.864419 | 0.22755269 | 0.25072529 | 0.90757772 | 0.36410137 | 0.76273126 |
| DDX23 | 224.494515 | -0.350478381 | 0.26789085 | -1.308288 | 0.19077565 | 0.64281265 |
| AC073610.3 | 112.022465 | 0.204504563 | 0.49209553 | 0.41557899 | 0.6777181 | 0.8987502 |
| CCDC65 | 237.245569 | 0.447689252 | 0.38154324 | 1.1733644 | 0.24064971 | 0.67925154 |
| PRKAG1 | 324.779821 | -0.571180642 | 0.41188024 | -1.3867639 | 0.16551377 | 0.62065658 |
| KMT2D | 409.061959 | -0.312642424 | 0.40800485 | -0.7662713 | 0.44351487 | 0.7993765 |
| LMBR1L | 352.614049 | 0.707520209 | 0.30934695 | 2.28714134 | 0.02218757 | 0.36178312 |
| AC011603.2 | 57.7109298 | 0.009857551 | 0.37872626 | 0.02602817 | 0.97923487 | 0.99482978 |
| TUBA1B | 688.983854 | 0.083032619 | 0.77638493 | 0.10694775 | 0.91483043 | 0.97782133 |
| TUBA1A | 262.40946 | 1.456809481 | 0.72587018 | 2.00698352 | 0.04475142 | 0.44360789 |
| TUBA1C | 751.692944 | -0.410786864 | 0.59072589 | -0.6953934 | 0.48680881 | 0.81889885 |
| AC010173.1 | 59.4532781 | 1.355054256 | 0.49439468 | 2.74083504 | 0.00612833 | 0.22637955 |
| DNAJC22 | 402.146401 | 1.29547569 | 0.35767148 | 3.62197094 | 0.00029237 | 0.06165345 |
| SPATS2 | 1230.7759 | 0.371801567 | 0.29569064 | 1.25740052 | 0.20860864 | 0.65401126 |
| MCRS1 | 158.967704 | -0.493917753 | 0.38296312 | -1.2897267 | 0.19714557 | 0.64558745 |
| PRPF40B | 649.518932 | -0.052779648 | 0.30647258 | -0.1722165 | 0.86326729 | 0.96393376 |
| FMNL3 | 53.5645486 | 1.483415053 | 0.87299881 | 1.69921772 | 0.08927817 | 0.53095823 |
| TMBIM6 | 2903.49682 | -0.151076936 | 0.39171025 | -0.3856854 | 0.69972965 | 0.90721166 |
| NCKAP5L | 224.562254 | 0.17623374 | 0.28461521 | 0.6192 | 0.53578461 | 0.83954482 |
| BCDIN3D | 119.893768 | -0.289044485 | 0.36497363 | -0.7919599 | 0.42838405 | 0.79037914 |
| AQP2 | 277.821911 | -1.767164373 | 0.72058395 | -2.452406 | 0.01419045 | 0.30468798 |
| RACGAP1 | 91.3346142 | 0.05505319 | 0.31521122 | 0.17465492 | 0.86135079 | 0.96302275 |
| SMARCD1 | 266.311634 | -0.178486287 | 0.31654546 | -0.5638567 | 0.57285164 | 0.8542679 |
| GPD1 | 497.449079 | 1.754726344 | 0.5450149 | 3.21959336 | 0.00128373 | 0.11730684 |
| AC074032.1 | 829.886197 | 0.316919345 | 0.20167637 | 1.5714253 | 0.11608389 | 0.57350419 |
| CERS5 | 462.741618 | -0.048412007 | 0.225341 | -0.2148389 | 0.82989294 | 0.95325673 |
| LIMA1 | 1372.02901 | 0.101650757 | 0.30103941 | 0.33766595 | 0.73561494 | 0.92019178 |
| FAM186A | 35.8915797 | 0.301707984 | 0.83931816 | 0.35946796 | 0.71924504 | 0.91465123 |
| LARP4 | 3024.01407 | -0.27624738 | 0.19399599 | -1.423985 | 0.15445081 | 0.6111597 |
| DIP2B | 5117.9522 | -0.649479147 | 0.37027651 | -1.7540382 | 0.07942396 | 0.51192014 |
| ATF1 | 651.416652 | -0.24025516 | 0.26026547 | -0.9231158 | 0.35594688 | 0.75841231 |
| METTL7A | 712.538419 | 0.759556715 | 0.5170246 | 1.46909202 | 0.14180783 | 0.59661141 |
| HIGD1C | 34.7541172 | 1.789834619 | 0.66804253 | 2.67922255 | 0.00737933 | 0.23590813 |
| SLC11A2 | 1320.69954 | -0.375963219 | 0.25658213 | -1.4652744 | 0.14284608 | 0.59824924 |
| LETMD1 | 637.038243 | 0.118221016 | 0.24648289 | 0.47963174 | 0.63148927 | 0.88123095 |
| CSRNP2 | 234.24471 | -0.033787075 | 0.29735515 | -0.1136253 | 0.90953481 | 0.97680747 |
| TFCP2 | 1323.56074 | -0.027784001 | 0.2240448 | -0.1240109 | 0.90130663 | 0.97487863 |
| POU6F1 | 157.591335 | -0.153279334 | 0.31260559 | -0.4903282 | 0.62390168 | 0.87791268 |
| DAZAP2 | 762.852949 | -0.317953045 | 0.31727032 | -1.0021519 | 0.31627025 | 0.72798933 |
| SMAGP | 122.105299 | -0.011032458 | 0.43792158 | -0.0251928 | 0.9799012 | 0.99487538 |
| GALNT6 | 46.1607296 | 0.963474422 | 0.75763684 | 1.27168371 | 0.20348552 | 0.64881965 |
| SLC4A8 | 48.4559435 | 0.121085135 | 0.53920258 | 0.22456335 | 0.82231898 | 0.95112619 |
| SCN8A | 478.668492 | 0.203537663 | 0.43222079 | 0.47091132 | 0.63770407 | 0.8827026 |
| AC068987.2 | 59.8517098 | 0.611385015 | 0.75796647 | 0.80661222 | 0.41988993 | 0.78798479 |
| AC068987.3 | 78.6220313 | 0.763340851 | 0.48567908 | 1.57169803 | 0.1160206 | 0.57350419 |
| FIGNL2 | 613.10501 | -0.010112148 | 0.49812046 | -0.0203006 | 0.98380357 | 0.99523767 |
| AC068987.1 | 30.2674146 | 0.678793007 | 0.55376856 | 1.22577022 | 0.22028515 | 0.66263062 |
| ACVR1B | 1487.12342 | 0.233564428 | 0.27174354 | 0.85950315 | 0.39006298 | 0.77569824 |
| NR4A1 | 80.2688947 | -0.863982056 | 0.80161816 | -1.0777975 | 0.28112413 | 0.70625976 |
| ATG101 | 143.634723 | -0.792435309 | 0.42513862 | -1.8639457 | 0.06232933 | 0.4840117 |
| AC078864.2 | 190.985247 | 0.172529228 | 0.90702234 | 0.19021497 | 0.84914068 | 0.95959925 |
| AC078864.1 | 497.895365 | -0.043427583 | 0.56308723 | -0.0771241 | 0.93852484 | 0.98435387 |
| KRT7 | 196.889243 | 1.104951517 | 0.63395947 | 1.74293717 | 0.08134459 | 0.51593841 |

| | | | | | | |
|---|---|---|---|---|---|---|
| KRT8 | 1239.42274 | -0.034877114 | 0.51155722 | -0.0681783 | 0.94564368 | 0.98584713 |
| KRT18 | 1203.79182 | -0.09327918 | 0.67988497 | -0.1371985 | 0.89087392 | 0.97230741 |
| EIF4B | 1137.37053 | -0.451685857 | 0.3676257 | -1.2286569 | 0.21920047 | 0.66165621 |
| AC068888.1 | 370.329545 | 0.292092019 | 0.45083512 | 0.64789101 | 0.51705544 | 0.83276628 |
| TNS2 | 190.303694 | 0.856849592 | 0.56519147 | 1.51603419 | 0.12951071 | 0.58820519 |
| SPRYD3 | 82.5278084 | -0.233520394 | 0.49617615 | -0.4706401 | 0.63789776 | 0.8827026 |
| IGFBP6 | 48.1440295 | 1.362652679 | 0.72693212 | 1.87452534 | 0.06085805 | 0.48107343 |
| CSAD | 730.183053 | 0.281621415 | 0.30838237 | 0.91322151 | 0.36112605 | 0.76009788 |
| ZNF740 | 142.149393 | -0.013202152 | 0.27306417 | -0.0483482 | 0.96143877 | 0.98989268 |
| MFSD5 | 85.8672199 | -0.777114755 | 0.40016562 | -1.9419828 | 0.05213918 | 0.46437412 |
| ESPL1 | 76.0185205 | -0.112749851 | 0.97602453 | -0.1155195 | 0.90803338 | 0.97673359 |
| PFDN5 | 653.672054 | -0.072616765 | 0.41340146 | -0.1756568 | 0.8605636 | 0.96302275 |
| C12orf10 | 88.8412999 | -0.12413834 | 0.45638009 | -0.2720065 | 0.78561704 | 0.93892226 |
| AAAS | 119.167986 | -0.240969533 | 0.29452393 | -0.8181662 | 0.41326227 | 0.78632759 |
| SP1 | 982.30113 | 0.394981283 | 0.33333146 | 1.18495049 | 0.23603702 | 0.67305169 |
| PRR13 | 740.247508 | 0.295737722 | 0.34488811 | 0.85748889 | 0.39117475 | 0.77606489 |
| PCBP2 | 2080.64419 | -0.365592707 | 0.2001043 | -1.8270107 | 0.06769815 | 0.48963361 |
| MAP3K12 | 163.401789 | 0.887674165 | 0.30517611 | 2.90872761 | 0.00362903 | 0.19706898 |
| TARBP2 | 78.4756223 | -0.60637615 | 0.36691746 | -1.6526228 | 0.09840766 | 0.54193402 |
| ATP5MC2 | 771.343663 | 0.005415272 | 0.26329693 | 0.02056717 | 0.98359093 | 0.99523767 |
| CALCOCO1 | 406.641111 | -0.002590273 | 0.37580706 | -0.0068926 | 0.99450058 | 0.998434 |
| HOXC4 | 151.045437 | 0.651461687 | 0.52917354 | 1.2310927 | 0.21828819 | 0.66064833 |
| AC023794.1 | 89.7769466 | 0.517651373 | 0.52273317 | 0.99027841 | 0.32203806 | 0.73375526 |
| SMUG1 | 209.268084 | 0.525595935 | 0.31727803 | 1.65657842 | 0.09760474 | 0.54093104 |
| LINC02381 | 70.1095863 | 0.396915915 | 0.50874631 | 0.78018436 | 0.43528237 | 0.79530997 |
| CBX5 | 432.921968 | 0.71066678 | 0.299626 | 2.37184615 | 0.01769946 | 0.33325066 |
| AC078778.1 | 36.5546143 | 1.447627206 | 0.63139687 | 2.29273738 | 0.02186313 | 0.36066559 |
| HNRNPA1 | 475.196765 | -0.232796392 | 0.35061062 | -0.6639742 | 0.50670684 | 0.82806179 |
| COPZ1 | 424.52494 | -0.397467132 | 0.25519754 | -1.5574881 | 0.11935464 | 0.57652003 |
| PPP1R1A | 332.851509 | -0.726201998 | 0.57615897 | -1.2604195 | 0.20751807 | 0.65289883 |
| METTL7B | 115.17685 | 1.902892032 | 0.68928539 | 2.76067369 | 0.00576823 | 0.22590027 |
| ITGA7 | 166.873419 | 1.460801776 | 0.62706647 | 2.32958044 | 0.01982834 | 0.3462255 |
| CD63 | 1072.65783 | -0.483363577 | 0.50505518 | -0.957051 | 0.3385415 | 0.74568349 |
| AC009779.2 | 30.2441295 | 0.470948109 | 0.57520192 | 0.81875266 | 0.41292754 | 0.78632759 |
| ORMDL2 | 131.388866 | -0.20023658 | 0.37745022 | -0.530498 | 5.96E-01 | 8.65E-01 |
| PYM1 | 161.947228 | -0.286880162 | 0.34659107 | -0.8277194 | 0.40782941 | 0.78481094 |
| CDK2 | 39.3734871 | 0.289889054 | 0.51228839 | 0.56587082 | 0.5714816 | 0.85399468 |
| RAB5B | 432.982612 | -0.016138173 | 0.22222914 | -0.0726195 | 0.94210889 | 0.98489601 |
| SUOX | 227.616491 | 1.17040135 | 0.68090697 | 1.71888584 | 0.08563516 | 0.5226185 |
| IKZF4 | 171.758384 | 0.931104596 | 0.34003723 | 2.73824307 | 0.00617684 | 0.22637955 |
| RPS26 | 207.635004 | -0.514230429 | 0.36301118 | -1.4165691 | 0.15660894 | 0.61255952 |
| ERBB3 | 819.760806 | 0.672351861 | 0.50003035 | 1.3446221 | 0.1787473 | 0.62903448 |
| PA2G4 | 227.545562 | -0.534256728 | 0.49089447 | -1.0883332 | 0.27644805 | 0.70328016 |
| RPL41 | 2298.80001 | 0.319967989 | 0.42272648 | 0.75691494 | 0.44910082 | 0.80300352 |
| ESYT1 | 275.864522 | -0.228750033 | 0.29185044 | -0.783792 | 0.43316218 | 0.79355172 |
| AC034102.5 | 46.1663937 | 0.462907162 | 0.62129315 | 0.74507045 | 0.45622913 | 0.80507423 |
| MYL6B | 105.748893 | -0.065562255 | 0.39302087 | -0.1668162 | 0.86751466 | 0.9648672 |
| MYL6 | 1208.48617 | -0.456116967 | 0.44677813 | -1.0209026 | 0.30730057 | 0.72250292 |
| SMARCC2 | 656.657976 | 0.380543998 | 0.27427736 | 1.38744223 | 0.16530696 | 0.62065658 |
| RNF41 | 224.204001 | 0.172496611 | 0.27666723 | 0.62348046 | 0.53296884 | 0.83872659 |
| NABP2 | 77.7063333 | -0.154660542 | 0.35614043 | -0.4342684 | 0.66409353 | 0.89313476 |
| SLC39A5 | 678.283301 | 0.715869149 | 0.50744939 | 1.41072028 | 1.58E-01 | 6.14E-01 |
| ANKRD52 | 257.929981 | -0.131230794 | 0.3321579 | -0.3950856 | 0.69277973 | 0.90528251 |
| COQ10A | 76.4455431 | -0.111676193 | 0.34848614 | -0.3204609 | 0.74861895 | 0.92552718 |

| | | | | | | |
|---|---|---|---|---|---|---|
| CS | 406.42627 | -0.267366147 | 0.30470273 | -0.8774655 | 0.38023383 | 0.77144243 |
| AC073896.1 | 108.248221 | -0.575240882 | 0.52139874 | -1.1032648 | 0.26991218 | 0.69961625 |
| AC073896.2 | 106.730564 | 0.09928212 | 0.33079925 | 0.30012801 | 0.76407952 | 0.93079056 |
| PAN2 | 311.453133 | 0.65146149 | 0.39376434 | 1.6544451 | 0.09803711 | 0.54141605 |
| STAT2 | 644.911832 | -0.443743394 | 0.55684518 | -0.7968883 | 0.42551592 | 0.78894887 |
| TIMELESS | 66.9955594 | -0.53046678 | 0.43025467 | -1.2329135 | 0.21760805 | 0.66043612 |
| SPRYD4 | 92.6342985 | -0.098026567 | 0.40939473 | -0.2394427 | 0.81076235 | 0.94717173 |
| GLS2 | 50.5768607 | 0.490080934 | 0.42979662 | 1.14026242 | 0.25417699 | 0.68908306 |
| RBMS2 | 525.094451 | 0.040519241 | 0.35572385 | 0.11390645 | 0.90931195 | 0.97680747 |
| BAZ2A | 830.208875 | 0.05906765 | 0.20709189 | 0.28522435 | 0.77547227 | 0.9351209 |
| ATP5F1B | 1058.32436 | 0.457944065 | 0.40403226 | 1.13343441 | 0.25703187 | 0.69135319 |
| PTGES3 | 1625.40488 | -0.448423657 | 0.25072309 | -1.7885216 | 0.0736919 | 0.50573515 |
| NACA | 720.918928 | -0.039888387 | 0.41747717 | -0.0955463 | 0.92388094 | 0.98015759 |
| NEMP1 | 148.860579 | 0.11333469 | 0.30989245 | 0.36572266 | 0.71457204 | 0.91252758 |
| NAB2 | 116.319779 | -0.216377534 | 0.33706977 | -0.6419369 | 0.52091413 | 0.83413201 |
| STAT6 | 958.015858 | -0.014788005 | 0.22826478 | -0.0647844 | 0.94834563 | 0.98678916 |
| LRP1 | 85.799188 | 0.819054346 | 0.50729817 | 1.61454228 | 0.10640985 | 0.55440878 |
| SHMT2 | 144.865647 | 0.250671845 | 0.28813679 | 0.86997514 | 0.38431399 | 0.77317162 |
| R3HDM2 | 2953.60811 | 0.287458257 | 0.15638753 | 1.83811494 | 6.60E-02 | 4.88E-01 |
| MARS | 289.244283 | -0.632233297 | 0.38375607 | -1.6474874 | 0.0994579 | 0.54399442 |
| DDIT3 | 424.540287 | -1.131870356 | 0.45006568 | -2.5149004 | 0.0119066 | 0.29195667 |
| MBD6 | 209.825315 | -0.732970149 | 0.39729363 | -1.8449079 | 0.06505094 | 0.48790819 |
| DCTN2 | 631.189554 | -0.427092477 | 0.25995531 | -1.6429458 | 0.10039415 | 0.54542315 |
| PIP4K2C | 173.979052 | -0.141366441 | 0.3243692 | -0.4358196 | 0.66296765 | 0.89263183 |
| DTX3 | 58.1957488 | 0.822932614 | 0.73088791 | 1.12593546 | 0.26019284 | 0.69280433 |
| OS9 | 529.313266 | -0.065706541 | 0.25408377 | -0.2586019 | 0.79594243 | 0.94252227 |
| AC025165.2 | 47.6791186 | 0.727753446 | 0.55589407 | 1.3091585 | 0.19048067 | 0.64281105 |
| TSPAN31 | 130.293569 | 0.383167071 | 0.30007749 | 1.27689373 | 0.20163977 | 0.64879726 |
| CDK4 | 225.322415 | -0.867644947 | 0.44520753 | -1.9488551 | 0.05131274 | 0.4615116 |
| 9-Mar | 268.242968 | 0.292978196 | 0.31597306 | 0.92722523 | 0.35380961 | 0.75682564 |
| CYP27B1 | 123.159647 | 0.262910309 | 0.74365376 | 0.3535386 | 0.7236847 | 0.91600702 |
| TSFM | 250.316518 | 0.417507497 | 0.33536207 | 1.24494548 | 0.21315179 | 0.65801421 |
| AVIL | 227.563325 | 0.155637163 | 0.498314 | 0.31232749 | 7.55E-01 | 0.92799282 |
| CTDSP2 | 403.12391 | -0.055823301 | 0.22078958 | -0.2528349 | 0.80039583 | 0.94403774 |
| AC084033.3 | 43.2077898 | 0.001970052 | 0.35068864 | 0.00561767 | 0.99551777 | 0.99866448 |
| ATP23 | 52.476887 | 0.644684282 | 0.47196166 | 1.3659675 | 0.17194917 | 0.62583835 |
| LRIG3 | 543.410772 | 0.206281181 | 0.49496498 | 0.41675914 | 0.67685459 | 0.89843878 |
| AC068305.2 | 105.467712 | 0.200603845 | 0.82138019 | 0.24422776 | 0.80705443 | 0.94585363 |
| SLC16A7 | 430.098348 | 0.399157449 | 0.57086149 | 0.69921943 | 0.48441491 | 0.81839266 |
| FAM19A2 | 485.889549 | 0.508065008 | 0.57334692 | 0.8861389 | 0.37554268 | 0.7697436 |
| USP15 | 2160.25055 | -0.256665192 | 0.22410911 | -1.1452689 | 0.25209779 | 0.68862185 |
| MON2 | 2570.90296 | -0.246579892 | 0.13943882 | -1.7683733 | 0.07699852 | 0.51013111 |
| AC048341.1 | 192.901192 | -0.029924673 | 0.38851879 | -0.0770225 | 0.93860568 | 0.98435387 |
| PPM1H | 1322.45854 | -0.4926966 | 0.3933025 | -1.2527167 | 0.21030883 | 0.65441616 |
| DPY19L2 | 908.249718 | 0.037304248 | 0.43126948 | 0.0864987 | 0.93106999 | 0.98272895 |
| RXYLT1 | 205.308149 | -0.323953991 | 0.46998785 | -0.6892816 | 0.49064606 | 0.81989696 |
| SRGAP1 | 3069.4751 | -0.683334261 | 0.4004975 | -1.7062136 | 0.08796832 | 0.52795613 |
| AC020611.2 | 98.332744 | 0.460350225 | 0.56375056 | 0.81658496 | 0.41416565 | 0.78632759 |
| C12orf66 | 120.578946 | 0.203799661 | 0.37753108 | 0.53982221 | 0.58931965 | 0.86246043 |
| C12orf56 | 50.9643902 | -0.227597385 | 0.44388875 | -0.5127352 | 0.60813657 | 0.86985416 |
| XPOT | 357.860345 | -0.834259058 | 0.42531894 | -1.9614905 | 0.04982183 | 0.45679453 |
| TBK1 | 902.888732 | -0.363906167 | 0.20665238 | -1.760958 | 0.07824551 | 0.51192014 |
| RASSF3 | 225.739022 | -0.263470195 | 0.52805726 | -0.4989425 | 0.61781991 | 0.8744793 |
| GNS | 1128.37559 | -0.308733597 | 0.21787207 | -1.4170407 | 0.15647103 | 0.61255952 |

| | | | | | | |
|---|---|---|---|---|---|---|
| TBC1D30 | 266.801034 | -0.292763211 | 0.45905339 | -0.6377542 | 0.52363372 | 0.83536399 |
| LINC02389 | 41.6267889 | 0.225067691 | 0.58179467 | 0.38685072 | 0.69886672 | 0.90696517 |
| LEMD3 | 835.457966 | 0.2106922 | 0.20744678 | 1.0156446 | 0.30979866 | 0.72498238 |
| MSRB3 | 57.6930441 | 0.634390927 | 0.67881105 | 0.93456188 | 0.35001413 | 0.75455562 |
| HMGA2 | 217.24325 | -2.222303297 | 1.05112064 | -2.1142229 | 3.45E-02 | 0.41659905 |
| LLPH | 100.159308 | -0.506952734 | 0.34896734 | -1.4527226 | 0.1463008 | 0.60383734 |
| HELB | 412.780064 | 0.767598556 | 0.48045248 | 1.59765761 | 0.1101192 | 0.56114143 |
| GRIP1 | 6605.47529 | 0.00608574 | 0.36369402 | 0.01673313 | 9.87E-01 | 9.96E-01 |
| CAND1 | 857.417465 | 0.103105564 | 0.15901709 | 0.64839299 | 0.5167308 | 0.83258835 |
| DYRK2 | 205.513676 | 0.355011679 | 0.2822108 | 1.2579663 | 0.20840394 | 0.65368257 |
| MDM1 | 176.13802 | 0.2627735 | 0.26855198 | 0.97848282 | 3.28E-01 | 7.38E-01 |
| LINC02384 | 196.163089 | 3.300267865 | 0.73857174 | 4.46844593 | 7.88E-06 | 0.01167379 |
| RAP1B | 2759.86176 | -0.067201846 | 0.24783295 | -0.2711578 | 0.78626965 | 0.9390702 |
| NUP107 | 610.808546 | -0.320676838 | 0.3013913 | -1.0639884 | 0.28733397 | 0.71118494 |
| SLC35E3 | 613.217698 | -0.312781812 | 0.44720322 | -0.6994176 | 0.48429108 | 0.81839266 |
| MDM2 | 1149.40765 | 0.162793382 | 0.35634926 | 0.45683659 | 0.64778851 | 0.88774616 |
| CPM | 2990.45755 | -0.683931934 | 0.52990529 | -1.2906682 | 0.19681874 | 0.64558745 |
| CPSF6 | 1199.05989 | -0.230133315 | 0.19821951 | -1.1610023 | 0.24564095 | 0.68310013 |
| YEATS4 | 92.9477885 | 0.302977393 | 0.26764325 | 1.13201957 | 0.25762621 | 0.69186568 |
| FRS2 | 1418.85277 | -0.281197195 | 0.19015234 | -1.4787996 | 0.13919389 | 0.59474148 |
| CCT2 | 495.910376 | -0.3623773 | 0.30515503 | -1.1875187 | 0.23502309 | 0.67119484 |
| BEST3 | 43.0229789 | 0.292757857 | 0.63764909 | 0.45912064 | 0.64614754 | 0.88700494 |
| RAB3IP | 1701.37259 | 0.328425248 | 0.25208506 | 1.30283506 | 0.19263108 | 0.64507333 |
| AC025263.2 | 40.3728931 | 1.628223686 | 0.67336139 | 2.41805324 | 0.01560379 | 0.3149944 |
| MYRFL | 382.393386 | 1.087634546 | 0.73178007 | 1.48628611 | 0.13720346 | 0.59474076 |
| AC025159.1 | 543.240224 | 1.681005398 | 0.71544146 | 2.34960581 | 0.0187933 | 0.34084276 |
| CNOT2 | 2083.92975 | -0.114350446 | 0.24855552 | -0.46006 | 0.64547317 | 0.88687652 |
| KCNMB4 | 318.781638 | 1.11809106 | 0.53378521 | 2.094646 | 0.03620247 | 0.42040764 |
| TSPAN8 | 54.8992965 | -0.122554332 | 0.67410471 | -0.1818031 | 0.85573724 | 0.96211892 |
| ZFC3H1 | 2120.03745 | -0.367821728 | 0.24820788 | -1.48191 | 0.13836425 | 0.59474076 |
| THAP2 | 87.9342543 | 0.60310554 | 0.47367998 | 1.27323418 | 0.20293496 | 0.64881821 |
| TMEM19 | 170.693196 | 0.773859051 | 0.37554892 | 2.06060786 | 0.03934047 | 0.42704465 |
| RAB21 | 1436.36243 | -0.103812738 | 0.22663074 | -0.45807 | 0.64690216 | 0.88768525 |
| TBC1D15 | 1404.08737 | -0.619824196 | 0.16888429 | -3.6701117 | 0.00024244 | 0.05632519 |
| TRHDE | 2719.95966 | 0.256586451 | 0.53018994 | 0.48395194 | 0.62841998 | 0.88009297 |
| AC090502.1 | 56.1677339 | -1.255150368 | 1.03967825 | -1.2072488 | 0.22733633 | 0.66705629 |
| ATXN7L3B | 372.926413 | -0.126369231 | 0.31553114 | -0.4004969 | 0.6887906 | 0.9032816 |
| CAPS2 | 949.743202 | 0.871556973 | 0.48214097 | 1.8076808 | 0.07065619 | 0.4967563 |
| GLIPR1L1 | 69.7246559 | 0.995463055 | 0.90082514 | 1.10505691 | 0.26913493 | 0.69961625 |
| KRR1 | 344.037042 | -0.186268892 | 0.26364198 | -0.7065221 | 0.47986349 | 0.81677498 |
| AC078923.1 | 436.164697 | -0.719715601 | 0.53680195 | -1.340747 | 0.18000261 | 0.63124625 |
| PHLDA1 | 118.706242 | -0.763912501 | 0.62310785 | -1.2259716 | 0.22020938 | 0.66263062 |
| NAP1L1 | 873.330847 | -0.074497943 | 0.32249153 | -0.2310074 | 0.81730902 | 0.94954998 |
| BBS10 | 56.8577019 | 0.008903368 | 0.37289569 | 0.0238763 | 0.98095128 | 0.9951185 |
| OSBPL8 | 3814.29571 | -0.323997884 | 0.2810782 | -1.1526966 | 0.24903494 | 0.68557761 |
| AC107032.2 | 89.0687009 | -0.322661452 | 0.47232232 | -0.6831383 | 0.49451947 | 0.8220963 |
| ZDHHC17 | 993.101378 | -0.223757071 | 0.18271187 | -1.2246444 | 0.22070923 | 0.66263062 |
| CSRP2 | 232.998303 | -0.48023899 | 0.64635254 | -0.7429985 | 0.45748257 | 0.80561734 |
| NAV3 | 545.525932 | -1.595253052 | 0.87607126 | -1.820917 | 0.06861947 | 0.49154315 |
| AC128707.1 | 1017.11755 | 3.652902255 | 0.88766681 | 4.11517275 | 3.87E-05 | 0.02019269 |
| AC079362.1 | 58.7596323 | 0.907096701 | 1.22477003 | 0.74062614 | 0.45892015 | 0.80656732 |
| SYT1 | 116.939904 | 0.565271843 | 0.87287785 | 0.64759559 | 0.51724655 | 0.83283196 |
| AC027288.3 | 2966.45303 | -0.021979049 | 0.60348026 | -0.0364205 | 9.71E-01 | 9.92E-01 |
| PAWR | 2797.61163 | -0.517832512 | 0.3141417 | -1.6484042 | 0.09926975 | 0.54362383 |

| | | | | | | |
|---|---|---|---|---|---|---|
| AC073569.2 | 288.431854 | 0.12395964 | 0.46690904 | 0.26548991 | 0.79063208 | 0.94063545 |
| PPP1R12A | 2266.66069 | 0.043846694 | 0.21220411 | 0.20662509 | 0.83630267 | 0.95470981 |
| OTOGL | 646.602331 | 1.122202899 | 0.82555001 | 1.35933969 | 1.74E-01 | 6.27E-01 |
| PTPRQ | 51.2250562 | -0.37601144 | 0.44654621 | -0.8420437 | 0.39976347 | 0.78186732 |
| LIN7A | 883.866463 | 1.178028991 | 0.41451065 | 2.84197519 | 0.0044835 | 0.20749797 |
| ACSS3 | 1836.10726 | 1.150545359 | 0.49009507 | 2.34759625 | 0.01889499 | 0.34084276 |
| PPFIA2 | 57.2861665 | 0.700504959 | 0.64748453 | 1.08188679 | 0.27930283 | 0.70446562 |
| CCDC59 | 190.220203 | -0.021090931 | 0.24735485 | -0.0852659 | 0.93205001 | 0.98296603 |
| METTL25 | 764.580343 | -0.528356862 | 0.23892119 | -2.2114274 | 0.02700625 | 0.38702446 |
| TMTC2 | 3637.06556 | 1.245782611 | 0.50208388 | 2.4812241 | 0.0130932 | 0.29875272 |
| LRRIQ1 | 371.48722 | 0.996711329 | 0.61520512 | 1.62012848 | 0.10520468 | 0.55292966 |
| MGAT4C | 40.1770723 | -0.102527307 | 0.48070293 | -0.2132862 | 0.83110373 | 0.95325673 |
| C12orf29 | 153.669326 | -0.20246169 | 0.32394702 | -0.624984 | 0.53198159 | 0.83789294 |
| CEP290 | 1047.18769 | -0.046196057 | 0.21204133 | -0.2178635 | 0.8275355 | 0.95255535 |
| TMTC3 | 599.501658 | 0.294638659 | 0.23353614 | 1.26164055 | 0.20707816 | 0.65251041 |
| KITLG | 208.372478 | 0.760718348 | 0.57504121 | 1.32289361 | 0.18587076 | 0.63661432 |
| DUSP6 | 331.623674 | 0.107982318 | 0.45936158 | 0.23507041 | 0.81415408 | 0.94881072 |
| POC1B | 777.325046 | 0.388825434 | 0.27570587 | 1.41029071 | 0.15845386 | 0.61423843 |
| POC1B-AS1 | 298.097044 | 0.067704787 | 0.30338829 | 0.22316216 | 0.8234093 | 0.95131214 |
| AC025034.1 | 141.482397 | 0.161567346 | 0.41059355 | 0.39349704 | 0.69395241 | 0.90589033 |
| ATP2B1 | 1190.95032 | -0.251155705 | 0.13564975 | -1.8515014 | 0.06409746 | 0.48593331 |
| DCN | 41.8644208 | -0.56175019 | 0.51294075 | -1.0951561 | 0.27344826 | 0.70137637 |
| BTG1 | 816.812368 | -0.047079171 | 0.4167196 | -0.1129757 | 9.10E-01 | 9.77E-01 |
| AC025164.1 | 48.4566849 | 0.497708463 | 0.65533984 | 0.75946621 | 0.44757372 | 0.80223612 |
| EEA1 | 1692.36076 | -0.458594936 | 0.34207414 | -1.3406302 | 0.18004053 | 0.63124625 |
| AC138123.1 | 2495.26586 | 0.817387578 | 0.61386907 | 1.33153406 | 0.18301335 | 0.63276491 |
| AC138123.2 | 96.6749927 | 1.20252924 | 0.83146513 | 1.44627742 | 0.14809941 | 0.60605027 |
| AC124947.1 | 208.185891 | 0.006460812 | 0.59942723 | 0.01077831 | 0.99140032 | 0.99748517 |
| NUDT4 | 826.831634 | 0.366690669 | 0.40424738 | 0.90709474 | 0.3643567 | 0.76283229 |
| UBE2N | 512.405437 | -0.209464596 | 0.26151407 | -0.8009687 | 0.42314974 | 0.78894887 |
| MRPL42 | 424.413972 | -0.168156103 | 0.22937539 | -0.7331044 | 0.46349476 | 0.8082072 |
| SOCS2 | 64.1150814 | 0.586395472 | 0.58616908 | 1.00038623 | 0.31712363 | 0.72908648 |
| CRADD | 3693.48942 | 0.335685185 | 0.37838397 | 0.88715488 | 0.37499552 | 0.7697436 |
| AC012085.2 | 224.044342 | 0.898602386 | 0.7556827 | 1.18912659 | 0.23438986 | 0.670712 |
| PLXNC1 | 154.368229 | 0.761274139 | 0.44687324 | 1.70355722 | 0.08846384 | 0.52927711 |
| CEP83 | 662.544291 | 0.016627125 | 0.28660167 | 0.05801475 | 0.95373688 | 0.98834742 |
| TMCC3 | 716.286376 | -0.062124336 | 0.54510247 | -0.1139682 | 0.90926302 | 0.97680747 |
| NDUFA12 | 280.848738 | -0.0849002 | 0.2860924 | -0.296758 | 0.76665129 | 0.93138917 |
| NR2C1 | 793.518073 | 0.10335627 | 0.19522515 | 0.52942089 | 0.59651351 | 0.86559503 |
| FGD6 | 1218.2341 | 0.093891186 | 0.22660219 | 0.41434368 | 0.67862243 | 0.8992258 |
| VEZT | 1740.26347 | 0.024222574 | 0.17666829 | 0.13710765 | 0.8909457 | 0.97230834 |
| METAP2 | 557.740002 | -0.112706773 | 0.26596156 | -0.4237709 | 0.67173289 | 0.89704376 |
| NTN4 | 2478.40974 | -0.345601956 | 0.41278055 | -0.8372535 | 4.02E-01 | 7.83E-01 |
| SNRPF | 149.921373 | -0.215711513 | 0.2207808 | -0.9770393 | 0.3285497 | 0.73826734 |
| CCDC38 | 45.4264117 | 0.155800537 | 0.52159673 | 0.29869922 | 0.76516955 | 0.93083208 |
| AMDHD1 | 185.439761 | 0.32190128 | 0.46192949 | 0.69686238 | 0.48588892 | 0.81839266 |
| LTA4H | 305.502347 | 0.242546886 | 0.2591833 | 0.93581217 | 0.3493699 | 0.7542101 |
| ELK3 | 131.40186 | -0.374371932 | 0.68918909 | -0.5432064 | 0.58698769 | 0.86112855 |
| CDK17 | 1120.93955 | 0.003070402 | 0.19694864 | 0.01558986 | 0.98756159 | 0.99648575 |
| CFAP54 | 136.322767 | 1.197084456 | 0.8387581 | 1.42721061 | 0.1535192 | 0.61014911 |
| NEDD1 | 355.129646 | -0.046438161 | 0.41351528 | -0.112301 | 0.91058479 | 0.97697493 |
| TMPO | 523.570684 | 0.688694575 | 0.40672194 | 1.69328111 | 0.09040197 | 0.53434231 |
| SLC25A3 | 1111.53806 | 0.356948422 | 0.4045628 | 0.88230659 | 0.37761103 | 0.77055888 |
| IKBIP | 255.28213 | -0.640004278 | 0.34489216 | -1.8556649 | 6.35E-02 | 4.86E-01 |

| | | | | | | |
|---|---|---|---|---|---|---|
| APAF1 | 231.686196 | 0.049896611 | 0.27198756 | 0.18345182 | 0.85444352 | 0.9614857 |
| ANKS1B | 6012.56994 | 1.598644617 | 0.67716787 | 2.36078038 | 0.01823653 | 0.33797156 |
| AC069437.1 | 91.3353371 | 1.262768053 | 0.85529592 | 1.4764107 | 0.13983367 | 0.59500405 |
| UHRF1BP1L | 1146.83208 | -0.159845914 | 0.26596368 | -0.6010066 | 0.54783562 | 0.84486687 |
| ACTR6 | 366.035373 | 0.350975326 | 0.30544912 | 1.14904677 | 0.2505367 | 0.68627228 |
| DEPDC4 | 227.471717 | 0.72955773 | 0.59775507 | 1.22049609 | 0.22227687 | 0.66373504 |
| SCYL2 | 1274.52246 | -0.133267448 | 0.23600522 | -0.5646801 | 0.57229138 | 0.85408471 |
| NR1H4 | 4936.31053 | -0.179561502 | 0.37875446 | -0.4740842 | 0.63543987 | 0.88246953 |
| GAS2L3 | 981.976321 | -0.011007148 | 0.48921407 | -0.0224997 | 0.98204939 | 0.99522272 |
| ANO4 | 77.8166655 | 0.16796891 | 0.57717242 | 0.29102034 | 0.77103577 | 0.93302012 |
| SLC5A8 | 2279.37358 | -0.904366961 | 1.04230192 | -0.8676631 | 0.38557877 | 0.77317162 |
| UTP20 | 415.325833 | -0.523887291 | 0.31489683 | -1.6636791 | 0.09617657 | 0.53876283 |
| ARL1 | 336.6473 | -0.102842149 | 0.32484041 | -0.3165928 | 0.75155257 | 0.92647278 |
| CHPT1 | 2337.59045 | 0.564833189 | 0.4487899 | 1.2585693 | 0.20818594 | 0.65352123 |
| SYCP3 | 86.0117386 | 1.415544664 | 0.73768449 | 1.91890256 | 0.05499667 | 0.46904913 |
| GNPTAB | 550.054765 | 0.025633479 | 0.29972458 | 0.08552345 | 0.93184526 | 0.98290134 |
| DRAM1 | 1318.27101 | -0.822634076 | 0.31988544 | -2.5716522 | 0.01012145 | 0.26582196 |
| WASHC3 | 553.075458 | 0.159887941 | 0.39175618 | 0.40813126 | 0.68317731 | 0.90024144 |
| NUP37 | 153.681027 | -0.052721565 | 0.37444762 | -0.1407982 | 0.88802934 | 0.97120196 |
| PARPBP | 40.4153578 | -0.871350927 | 0.58905128 | -1.4792446 | 0.13907495 | 0.59474076 |
| PAH | 2578.46979 | 0.953910762 | 0.52745307 | 1.80852254 | 7.05E-02 | 4.97E-01 |
| NT5DC3 | 268.154365 | 0.347246623 | 0.30482285 | 1.13917517 | 2.55E-01 | 6.90E-01 |
| HSP90B1 | 2005.52606 | -0.098420267 | 0.57346387 | -0.1716242 | 0.863733 | 0.96393376 |
| C12orf73 | 166.424414 | 0.479647756 | 0.31616593 | 1.51707602 | 0.12924749 | 0.58798386 |
| TDG | 547.627166 | -0.298750311 | 0.3257264 | -0.9171817 | 0.35904741 | 0.75936492 |
| HCFC2 | 340.336258 | 0.084841621 | 0.26375607 | 0.32166698 | 0.74770499 | 0.92531168 |
| NFYB | 205.853244 | 0.220963652 | 0.24074763 | 0.91782276 | 3.59E-01 | 0.75936492 |
| TXNRD1 | 6274.4615 | -1.04141102 | 0.53162665 | -1.9589143 | 5.01E-02 | 4.58E-01 |
| CHST11 | 1959.11936 | -0.699605123 | 0.6766232 | -1.0339656 | 0.30115224 | 0.71879123 |
| SLC41A2 | 1690.61964 | -0.653633058 | 0.35399799 | -1.8464316 | 6.48E-02 | 0.48782903 |
| C12orf45 | 191.352337 | -0.43886156 | 0.39714492 | -1.1050414 | 0.26914167 | 0.69961625 |
| WASHC4 | 2286.16589 | -0.088261127 | 0.23383031 | -0.377458 | 0.70583326 | 0.90980406 |
| APPL2 | 984.712138 | 0.419028835 | 0.18148585 | 2.30887881 | 0.02095031 | 0.35422078 |
| C12orf75 | 1497.28305 | 0.851034588 | 0.3581392 | 2.37626765 | 0.01748877 | 0.33064634 |
| AC011313.1 | 205.154394 | 0.080989661 | 0.57045005 | 0.14197503 | 0.88709973 | 0.97073894 |
| NUAK1 | 138.733318 | 0.527469526 | 0.59542437 | 0.88587158 | 0.37568673 | 0.7697436 |
| CKAP4 | 130.348997 | 0.631934 | 0.46015009 | 1.37332149 | 0.16965243 | 0.62326131 |
| TCP11L2 | 289.867472 | 0.364599812 | 0.41294512 | 0.88292559 | 0.37727648 | 0.77048581 |
| POLR3B | 313.790998 | -0.094290433 | 0.28249534 | -0.3337769 | 7.39E-01 | 0.92080006 |
| RIC8B | 946.843673 | 0.220882788 | 0.19230392 | 1.14861303 | 0.25071558 | 0.68648949 |
| TMEM263 | 560.83721 | -0.161520997 | 0.34354652 | -0.4701576 | 0.63824245 | 0.8827026 |
| MTERF2 | 206.808296 | 0.383929615 | 0.31422356 | 1.22183588 | 0.2217697 | 0.66313203 |
| CRY1 | 865.153966 | -0.800456776 | 0.67700944 | -1.1823421 | 0.23706998 | 0.67445376 |
| BTBD11 | 1293.09112 | -1.361167765 | 0.81046665 | -1.6794865 | 9.31E-02 | 0.53684765 |
| PWP1 | 356.530104 | -0.397257062 | 0.29125836 | -1.3639336 | 0.17258846 | 0.62593809 |
| PRDM4 | 586.377888 | -0.435561206 | 0.35761654 | -1.217956 | 0.2232407 | 0.66408018 |
| AC007622.2 | 47.1601164 | 0.736302803 | 0.68029783 | 1.0823242 | 0.2791085 | 0.70446562 |
| FICD | 59.526588 | -0.145699835 | 0.57532667 | -0.2532471 | 0.80007724 | 0.94403774 |
| SART3 | 391.781921 | 0.089990177 | 0.19182124 | 0.46913563 | 0.63897269 | 0.88284765 |
| ISCU | 661.814874 | 0.30613806 | 0.40297501 | 0.75969491 | 0.44743697 | 0.80223612 |
| AC007569.1 | 31.6714438 | 1.835137582 | 0.54210749 | 3.38519134 | 0.00071129 | 0.08944591 |
| CORO1C | 1293.06085 | -0.714534312 | 0.35302421 | -2.0240377 | 0.04296627 | 0.4398347 |
| SSH1 | 944.527398 | -0.076413177 | 0.31521403 | -0.2424168 | 0.80845722 | 0.94625542 |
| DAO | 266.731937 | 1.607447965 | 0.56668392 | 2.83658651 | 4.56E-03 | 2.08E-01 |

| | | | | | | |
|---|---|---|---|---|---|---|
| USP30 | 318.847686 | 0.353951795 | 0.42903601 | 0.82499321 | 0.40937544 | 0.78542936 |
| ALKBH2 | 54.3047881 | -0.454605208 | 0.46054395 | -0.9871049 | 0.3235912 | 0.7354443 |
| UNG | 122.003433 | 0.496147425 | 0.38085484 | 1.30272055 | 0.19267018 | 0.64507333 |
| ACACB | 1672.97488 | 0.710304174 | 0.57586911 | 1.23344725 | 0.21740895 | 0.66032006 |
| KCTD10 | 203.68687 | 0.008194195 | 0.32131869 | 0.02550177 | 0.97965474 | 0.99484461 |
| UBE3B | 393.046115 | 0.392774385 | 0.31336686 | 1.25340116 | 0.21005975 | 0.65441616 |
| MMAB | 340.39645 | 0.579374171 | 0.28881292 | 2.0060535 | 0.04485054 | 0.44360789 |
| MVK | 440.632619 | 0.319577914 | 0.34829822 | 0.91754106 | 0.35885919 | 0.75936492 |
| FAM222A | 52.5504444 | -0.585585851 | 0.74461205 | -0.7864308 | 0.43161514 | 0.79135216 |
| TRPV4 | 1305.86276 | 1.600664314 | 0.60934918 | 2.62684247 | 8.62E-03 | 0.25133745 |
| GLTP | 199.60502 | -0.230482275 | 0.38843682 | -0.5933585 | 0.55294128 | 0.84721285 |
| TCHP | 417.511616 | 0.151428814 | 0.31765242 | 0.4767123 | 0.63356701 | 0.88192373 |
| GIT2 | 1049.21302 | -0.057854945 | 0.23954714 | -0.241518 | 0.80915367 | 0.94673968 |
| ANKRD13A | 656.872787 | -0.583520437 | 0.3647525 | -1.5997709 | 0.10964942 | 0.56010179 |
| C12orf76 | 196.255888 | 0.05848587 | 0.34842166 | 0.16785946 | 0.86669385 | 0.96481544 |
| IFT81 | 599.963881 | 0.104283725 | 0.31173476 | 0.3345271 | 0.73798185 | 0.92059676 |
| ATP2A2 | 1798.08908 | -0.064622516 | 0.36450221 | -0.1772898 | 0.85928078 | 0.96283855 |
| ANAPC7 | 520.920714 | -0.688949759 | 0.41507618 | -1.6598152 | 0.09695163 | 0.54015377 |
| ARPC3 | 999.585527 | -0.214453394 | 0.24185569 | -0.8866998 | 3.75E-01 | 0.7697436 |
| GPN3 | 99.8167349 | 0.054816646 | 0.26513828 | 0.20674738 | 0.83620715 | 0.95468028 |
| FAM216A | 53.1524951 | -0.027105951 | 0.3983201 | -0.0680507 | 0.9457453 | 0.98584713 |
| VPS29 | 550.60217 | 0.032733288 | 0.19611175 | 0.16691141 | 0.86743976 | 0.96486219 |
| PPTC7 | 676.985907 | -0.24935804 | 0.26874916 | -0.9278468 | 0.35348705 | 0.75673243 |
| TCTN1 | 416.560641 | 0.612161556 | 0.44007192 | 1.3910489 | 0.1642106 | 0.62065658 |
| HVCN1 | 84.7946719 | 0.921900291 | 0.69509556 | 1.32629288 | 0.1847427 | 0.63412875 |
| PPP1CC | 295.124382 | -0.022259425 | 0.24673841 | -0.0902147 | 0.92811662 | 0.98267377 |
| PHETA1 | 87.7297446 | 1.051669453 | 0.4930138 | 2.13314403 | 0.03291291 | 0.41180153 |
| SH2B3 | 136.408811 | 0.158843808 | 0.46345896 | 0.34273544 | 0.7317975 | 0.91930611 |
| ATXN2 | 4502.42405 | -0.342589424 | 0.21831422 | -1.5692493 | 0.11658987 | 0.57350419 |
| BRAP | 420.10534 | -0.340712561 | 0.20145546 | -1.6912551 | 0.0907881 | 0.53523977 |
| ACAD10 | 547.434517 | -0.034396419 | 0.20110334 | -0.1710385 | 8.64E-01 | 0.96393376 |
| ALDH2 | 1352.03006 | 0.21636603 | 0.47722912 | 0.45337977 | 0.6502753 | 0.8887244 |
| MAPKAPK5-A | 107.767181 | -0.122308277 | 0.36035641 | -0.3394092 | 0.73430149 | 0.91975299 |
| MAPKAPK5 | 442.982252 | -0.209522518 | 0.26511922 | -0.7902955 | 0.42935524 | 0.79037914 |
| TMEM116 | 399.582346 | 0.113567727 | 0.33391656 | 0.3401081 | 0.73377512 | 0.91972212 |
| ERP29 | 551.540718 | -0.078400947 | 0.35675431 | -0.2197617 | 0.82605672 | 0.95192757 |
| NAA25 | 1032.56125 | -0.684679625 | 0.33497047 | -2.044 | 0.04095355 | 0.43143739 |
| TRAFD1 | 251.982901 | 0.384212709 | 0.37050728 | 1.03699097 | 0.29974006 | 0.71879123 |
| HECTD4 | 2357.54081 | -0.196958335 | 0.1637999 | -1.2024325 | 0.22919599 | 0.66705629 |
| RPL6 | 1222.83798 | -0.02322798 | 0.47021876 | -0.0493982 | 0.96060193 | 0.98955138 |
| PTPN11 | 1016.15253 | -0.407524432 | 0.3930428 | -1.0368449 | 0.29980814 | 0.71879123 |
| OAS1 | 239.191691 | -2.44555597 | 1.08329504 | -2.2575161 | 0.02397585 | 0.37387451 |
| AC004551.1 | 39.9019536 | -1.86178413 | 1.11054276 | -1.6764633 | 0.09364749 | 0.53688613 |
| OAS3 | 67.6494297 | -3.19192522 | 1.1052817 | -2.8878839 | 0.00387843 | 0.20213552 |
| RASAL1 | 361.314895 | -0.042766284 | 0.4104626 | -0.1041905 | 9.17E-01 | 9.78E-01 |
| DDX54 | 252.664617 | -0.409107849 | 0.41745794 | -0.9799978 | 0.32708722 | 0.73746094 |
| RITA1 | 84.4659537 | -0.295494904 | 0.29909977 | -0.9879476 | 0.32317831 | 0.73534319 |
| IQCD | 95.5043286 | 0.020017173 | 0.55964033 | 0.03576792 | 0.97146741 | 0.99240251 |
| TPCN1 | 3367.17947 | 1.656735235 | 0.60118623 | 2.75577711 | 0.00585529 | 0.22637955 |
| SLC8B1 | 219.630259 | 0.515225385 | 0.2432047 | 2.11848451 | 0.03413406 | 0.41441585 |
| PLBD2 | 228.872224 | -0.550903873 | 0.41086692 | -1.3408329 | 0.17997472 | 0.63124625 |
| SDSL | 173.801347 | 0.530804034 | 0.44554814 | 1.19135058 | 0.23351599 | 0.67004472 |
| RBM19 | 459.934931 | -0.694020854 | 0.4223579 | -1.6432056 | 0.10034041 | 0.54542315 |
| MED13L | 6728.81563 | 0.096307184 | 0.21653028 | 0.44477468 | 0.65648259 | 0.89053559 |

| | | | | | | |
|---|---|---|---|---|---|---|
| MAP1LC3B2 | 55.9978245 | -0.224153291 | 0.36009232 | -0.6224884 | 5.34E-01 | 8.39E-01 |
| C12orf49 | 441.479169 | -1.193238437 | 0.48495265 | -2.4605256 | 0.01387337 | 0.30363654 |
| RNFT2 | 68.5029697 | -0.688632407 | 0.69912041 | -0.9849983 | 0.32462491 | 0.73614611 |
| FBXW8 | 698.330805 | -0.416577967 | 0.35996028 | -1.1572887 | 0.24715443 | 0.68482118 |
| TESC | 267.211498 | 1.839133748 | 0.88735092 | 2.07261153 | 3.82E-02 | 0.42406735 |
| FBXO21 | 1197.23945 | 0.09261383 | 0.38618822 | 0.23981527 | 0.81047347 | 0.94717173 |
| KSR2 | 73.2770398 | 0.150712867 | 0.53807235 | 0.28009777 | 0.7794025 | 0.93655387 |
| RFC5 | 72.0518066 | -0.777534371 | 0.46796516 | -1.6615219 | 0.09660867 | 0.53969191 |
| WSB2 | 364.308377 | -0.263677698 | 0.31323706 | -0.8417832 | 0.39990931 | 0.78186732 |
| VSIG10 | 471.316284 | 0.141966438 | 0.3972126 | 0.35740669 | 0.72078737 | 0.91513824 |
| PEBP1 | 2717.36478 | 0.26040156 | 0.44807052 | 0.581162 | 0.56113128 | 0.84958759 |
| TAOK3 | 2328.07099 | 0.148090676 | 0.21638025 | 0.68440015 | 0.49372252 | 0.82164359 |
| SUDS3 | 477.357207 | -0.095603956 | 0.17518464 | -0.5457325 | 0.58524981 | 0.86077718 |
| HSPB8 | 537.88419 | 0.489258847 | 0.34855322 | 1.40368476 | 1.60E-01 | 6.16E-01 |
| CCDC60 | 56.9469285 | 1.067415629 | 0.61678962 | 1.7305992 | 0.08352328 | 0.51991532 |
| AC002070.1 | 411.974944 | 1.050987941 | 0.49241465 | 2.13435553 | 0.03281369 | 0.41180153 |
| PRKAB1 | 725.019919 | 0.582488226 | 0.45186261 | 1.28908259 | 0.19736938 | 0.64558745 |
| CIT | 755.557272 | 0.635957209 | 0.78023301 | 0.81508627 | 0.41502293 | 0.78632759 |
| BICDL1 | 6504.15053 | -0.17426269 | 0.44016125 | -0.3959065 | 0.69217402 | 0.90516342 |
| RAB35 | 258.212499 | -0.865280181 | 0.44964065 | -1.9243816 | 0.05430677 | 0.46711025 |
| GCN1 | 512.742919 | -0.58096476 | 0.4122926 | -1.4091079 | 0.15880327 | 0.61423843 |
| RPLP0 | 1810.59811 | 0.195630868 | 0.57565591 | 0.33983994 | 0.73397707 | 0.91975299 |
| PXN-AS1 | 80.0401621 | -0.208393571 | 0.27783183 | -0.7500709 | 0.453212 | 0.80433814 |
| PXN | 536.496848 | -0.520644082 | 0.34433803 | -1.5120145 | 0.13053018 | 0.58853436 |
| SIRT4 | 85.285249 | -0.055042417 | 0.43423189 | -0.1267581 | 0.89913185 | 0.97458352 |
| TRIAP1 | 41.6615779 | -0.596439594 | 0.44050184 | -1.3540002 | 0.17573631 | 0.62699479 |
| GATC | 148.540897 | -0.12115854 | 0.31427173 | -0.3855216 | 0.69985101 | 0.90721166 |
| SRSF9 | 393.384445 | -0.139589113 | 0.32562607 | -0.4286792 | 0.66815673 | 0.89525164 |
| DYNLL1 | 622.804024 | 0.633289671 | 0.42274931 | 1.49802651 | 0.13412636 | 0.59240843 |
| NRAV | 44.1256867 | -0.045732218 | 0.3592955 | -0.127283 | 0.8987164 | 0.97427283 |
| COQ5 | 215.588097 | 0.228090385 | 0.27390326 | 0.83274067 | 4.05E-01 | 7.84E-01 |
| RNF10 | 1060.8794 | -0.349815599 | 0.23607421 | -1.4818035 | 0.13839258 | 0.59474076 |
| POP5 | 167.463404 | 0.14092304 | 0.33938112 | 0.41523535 | 0.67796962 | 0.89898912 |
| CABP1 | 287.638514 | -0.659303749 | 0.92039848 | -0.7163242 | 0.47379116 | 0.81493982 |
| MLEC | 1147.54199 | 0.139832406 | 0.45689668 | 0.3060482 | 0.75956795 | 0.92934017 |
| UNC119B | 202.247799 | 0.273333901 | 0.25048857 | 1.0912031 | 0.27518352 | 0.702279 |
| ACADS | 281.539938 | 0.069664572 | 0.32693075 | 0.21308663 | 0.83125939 | 0.95325673 |
| SPPL3 | 2897.93625 | -0.272795824 | 0.17838377 | -1.5292637 | 0.1261991 | 0.5860467 |
| HNF1A | 431.524156 | 1.002235664 | 0.53329872 | 1.87931385 | 0.06020165 | 0.48004006 |
| C12orf43 | 89.5784865 | -0.941540686 | 0.30200425 | -3.1176405 | 0.00182305 | 0.13687118 |
| OASL | 43.7835562 | -3.365312869 | 1.4478393 | -2.3243691 | 0.02010572 | 0.34840598 |
| P2RX4 | 632.149953 | -0.616435966 | 0.36196079 | -1.7030462 | 0.08855942 | 0.5293868 |
| CAMKK2 | 528.872469 | 0.046429164 | 0.30628637 | 0.15158743 | 0.87951235 | 0.96751353 |
| ANAPC5 | 934.393502 | -0.245263176 | 0.25073003 | -0.9781963 | 0.32797726 | 0.73806934 |
| RNF34 | 422.792203 | -0.608009275 | 0.45803814 | -1.3274206 | 0.18436958 | 0.63377801 |
| KDM2B | 311.902506 | 0.041873225 | 0.34492224 | 0.12139903 | 0.90337499 | 0.97536781 |
| ORAI1 | 126.086108 | -0.347813391 | 0.43184848 | -0.8054061 | 0.42058538 | 0.78831722 |
| TMEM120B | 846.854864 | -0.432533102 | 0.4963218 | -0.8714771 | 0.38349369 | 0.77317162 |
| RHOF | 215.108819 | -1.425750816 | 0.68870286 | -2.0701973 | 0.03843387 | 0.42447711 |
| SETD1B | 235.311801 | -0.421993755 | 0.49249328 | -0.8568518 | 0.39152679 | 0.77606943 |
| HPD | 675.862575 | 0.25748604 | 0.8889555 | 0.28965009 | 0.77208394 | 0.93340577 |
| WDR66 | 335.197075 | -0.754442215 | 0.44732311 | -1.6865711 | 0.09168586 | 0.53567268 |
| BCL7A | 676.13996 | -0.396848348 | 0.37215466 | -1.0663533 | 0.28626397 | 0.71118494 |
| MLXIP | 1177.89442 | -0.167808532 | 0.43434711 | -0.3863466 | 0.69923999 | 0.90721166 |

| | | | | | | |
|---|---|---|---|---|---|---|
| LRRC43 | 77.1928619 | -0.493672199 | 0.38568748 | -1.2799798 | 0.20055222 | 0.64788341 |
| B3GNT4 | 38.0710333 | 0.288039397 | 0.8262541 | 0.34860874 | 0.72738307 | 0.91646656 |
| CLIP1 | 2550.72778 | -0.667878474 | 0.42507338 | -1.5712075 | 0.11613447 | 0.57350419 |
| ZCCHC8 | 478.802321 | -0.298365695 | 0.25861586 | -1.1537023 | 0.24862226 | 0.68534254 |
| RSRC2 | 793.755456 | -0.313619873 | 0.17077662 | -1.8364333 | 0.06629359 | 0.48798003 |
| KNTC1 | 126.59704 | -0.417279981 | 0.33981513 | -1.2279618 | 0.21946132 | 0.66186058 |
| DENR | 801.592464 | -0.200549884 | 0.28697143 | -0.6988496 | 0.484646 | 0.81839266 |
| CCDC62 | 47.0561893 | 0.200388168 | 0.40289087 | 0.4973758 | 0.61892407 | 0.87511751 |
| HIP1R | 977.246652 | 0.40624168 | 0.39920178 | 1.01763495 | 0.30885147 | 0.7242818 |
| VPS37B | 686.160467 | -0.255402491 | 0.33418005 | -0.7642661 | 0.44470867 | 0.80026543 |
| ABCB9 | 100.454438 | 0.702522453 | 0.43467612 | 1.61619748 | 0.10605162 | 0.55440878 |
| OGFOD2 | 76.6085758 | -0.207964254 | 0.31931138 | -0.6512898 | 0.51485942 | 0.83169608 |
| ARL6IP4 | 686.575537 | -0.104462938 | 0.26292803 | -0.3973062 | 0.69114167 | 0.90473488 |
| PITPNM2 | 1066.92782 | -0.518314401 | 0.40130703 | -1.2915657 | 0.19650758 | 0.64558745 |
| MPHOSPH9 | 744.466554 | -0.006046944 | 0.18489001 | -0.0327056 | 9.74E-01 | 0.9937007 |
| C12orf65 | 355.042899 | -0.478833905 | 0.28836608 | -1.660507 | 0.0968125 | 0.54003127 |
| AC068768.1 | 43.8765831 | -0.064785007 | 0.33024746 | -0.1961711 | 0.84447625 | 0.95742233 |
| SBNO1 | 1129.05262 | -0.345717414 | 0.24212405 | -1.4278524 | 0.15333434 | 0.61008819 |
| AC137767.1 | 79.3915626 | 0.262550366 | 0.61508262 | 0.42685382 | 0.6694858 | 0.89611079 |
| KMT5A | 192.292302 | -0.443129952 | 0.34730854 | -1.2758971 | 0.20199191 | 0.64881821 |
| RILPL2 | 294.000049 | -0.239271058 | 0.38336033 | -0.6241414 | 0.53253471 | 0.83834309 |
| SNRNP35 | 279.946325 | 0.059951109 | 0.20258913 | 0.29592461 | 7.67E-01 | 0.93169958 |
| RILPL1 | 805.759025 | -0.25058655 | 0.20021199 | -1.2516061 | 0.21071343 | 0.65524759 |
| AC055713.1 | 64.5259572 | -0.269763188 | 0.3802486 | -0.709439 | 0.47805209 | 0.81592081 |
| TMED2 | 386.523861 | -0.480064986 | 0.35660785 | -1.3461986 | 0.17823848 | 0.6285261 |
| DDX55 | 239.442957 | -0.414110323 | 0.34367371 | -1.204952 | 0.22822184 | 0.66705629 |
| EIF2B1 | 241.017377 | -0.385122428 | 0.20744923 | -1.856466 | 0.06338717 | 0.48572144 |
| GTF2H3 | 256.594479 | -0.281013847 | 0.20463171 | -1.3732664 | 0.16966955 | 0.62326131 |
| TCTN2 | 162.030758 | -0.228779717 | 0.39714083 | -0.576067 | 0.56456991 | 0.85163263 |
| ATP6V0A2 | 263.548476 | -0.445622874 | 0.3990375 | -1.1167444 | 0.26410364 | 0.69567438 |
| DNAH10 | 103.838649 | 0.731059018 | 0.81497371 | 0.89703387 | 0.36970084 | 0.76669785 |
| CCDC92 | 981.645019 | -0.000957117 | 0.23173462 | -0.0041302 | 0.99670456 | 0.9992561 |
| RFLNA | 79.9062137 | 0.925857525 | 0.70534994 | 1.31262155 | 0.18931051 | 0.6413893 |
| NCOR2 | 1026.67123 | -0.379462193 | 0.36612119 | -1.0364388 | 0.29999749 | 0.71879123 |
| SCARB1 | 674.166304 | 0.607728862 | 0.70440301 | 0.86275734 | 0.3882709 | 0.77411334 |
| UBC | 3030.32256 | -0.842924348 | 0.27048814 | -3.116308 | 1.83E-03 | 1.37E-01 |
| DHX37 | 149.98151 | -0.906233386 | 0.47013281 | -1.9276115 | 0.05390347 | 0.46711025 |
| BRI3BP | 176.44957 | 0.598851305 | 0.44955361 | 1.33210209 | 0.18282664 | 0.63276491 |
| AACS | 288.396594 | 0.351328256 | 0.23809071 | 1.47560676 | 1.40E-01 | 0.59500405 |
| TMEM132B | 49.2427822 | 0.622959112 | 0.48674729 | 1.27984096 | 0.20060108 | 0.64788853 |
| LINC00939 | 175.860534 | 1.328896652 | 0.72090925 | 1.8433619 | 0.06527619 | 0.48790819 |
| LINC02405 | 37.3938267 | 0.414892439 | 0.78899164 | 0.5258515 | 0.59899139 | 0.86671329 |
| SLC15A4 | 668.893947 | -0.269460824 | 0.37177943 | -0.7247868 | 4.69E-01 | 8.12E-01 |
| GLT1D1 | 95.6678832 | 0.603454761 | 0.83729416 | 0.72072014 | 0.47108172 | 0.81387913 |
| TMEM132D | 411.686644 | -1.555944992 | 0.78646461 | -1.9784043 | 4.79E-02 | 0.45133876 |
| STX2 | 451.537396 | -0.382088682 | 0.31530187 | -1.2118186 | 0.22558182 | 0.66688879 |
| RAN | 416.20518 | -0.210554931 | 0.47854728 | -0.4399877 | 0.659946 | 0.89094706 |
| SFSWAP | 979.734171 | -0.367714807 | 0.28810369 | -1.276328 | 0.20183959 | 0.64881821 |
| ULK1 | 353.966798 | -0.21707854 | 0.32035806 | -0.6776122 | 0.49801758 | 0.82326489 |
| PUS1 | 244.356584 | -0.881558868 | 0.467251 | -1.8866923 | 0.05920172 | 0.47855094 |
| EP400 | 904.707824 | -0.311681212 | 0.30866443 | -1.0097736 | 0.31260375 | 0.72674009 |
| DDX51 | 112.496199 | -0.677925354 | 0.45676664 | -1.4841832 | 0.13776034 | 0.59474076 |
| NOC4L | 43.4644136 | -0.469742838 | 0.55965827 | -0.8393387 | 0.40127928 | 0.78240897 |
| GALNT9 | 76.2200355 | 0.701011186 | 0.82755361 | 0.84708855 | 0.39694577 | 0.78014596 |

| | | | | | | |
|---|---|---|---|---|---|---|
| FBRSL1 | 527.712655 | -0.304494583 | 0.32249795 | -0.9441753 | 0.3450801 | 0.75139204 |
| POLE | 160.964982 | -0.549867052 | 0.31982753 | -1.7192612 | 0.08556683 | 0.5226185 |
| AC135586.2 | 52.8367942 | 0.509422438 | 0.54874359 | 0.9283433 | 0.35322952 | 0.75673243 |
| PGAM5 | 88.5402967 | -0.989796424 | 0.48740083 | -2.0307647 | 0.04227887 | 0.43742111 |
| ANKLE2 | 956.793546 | -1.103964725 | 0.43143658 | -2.5588112 | 0.01050308 | 0.27112183 |
| GOLGA3 | 805.777672 | -1.129867546 | 0.48171447 | -2.345513 | 0.01900092 | 0.34084276 |
| CHFR | 878.498049 | -0.44051019 | 0.32312418 | -1.3632845 | 0.17279286 | 0.62593809 |
| ZNF26 | 545.723437 | -0.461069418 | 0.21980279 | -2.0976504 | 0.03593604 | 0.42040764 |
| ZNF84 | 582.789969 | -0.552238292 | 0.24561964 | -2.2483475 | 0.02455404 | 0.37682963 |
| ZNF140 | 311.386561 | -0.535178337 | 0.1996809 | -2.6801678 | 0.00735853 | 0.23590813 |
| ZNF891 | 224.578616 | -0.210255753 | 0.20934223 | -1.0043638 | 0.31520329 | 0.72782155 |
| ZNF268 | 95.9186592 | -0.416530597 | 0.28289561 | -1.4723827 | 0.14091756 | 0.59555904 |
| MPHOSPH8 | 1349.54378 | 0.346837652 | 0.27978676 | 1.23964997 | 0.21510489 | 0.65925411 |
| PSPC1 | 2006.51128 | -0.066447686 | 0.24746423 | -0.2685143 | 0.78830347 | 0.93975262 |
| ZMYM5 | 763.396733 | 0.034419444 | 0.27846312 | 0.12360504 | 0.901628 | 0.97491785 |
| AL355001.1 | 48.0548159 | 0.31364114 | 0.41803415 | 0.75027635 | 0.45308828 | 0.80433814 |
| ZMYM2 | 3679.81397 | -0.06018443 | 0.25820685 | -0.2330861 | 0.81569454 | 0.94906482 |
| GJB2 | 67.1011074 | 1.025501966 | 0.50993942 | 2.01102704 | 0.0443226 | 0.44360789 |
| CRYL1 | 4113.6519 | 0.913941514 | 0.4714284 | 1.93866453 | 0.0525422 | 0.46554359 |
| IFT88 | 3105.47865 | 0.009474196 | 0.31915332 | 0.02968541 | 0.97631795 | 0.99429832 |
| EEF1AKMT1 | 199.695776 | 0.370447201 | 0.43248113 | 0.8565627 | 0.39168661 | 0.77606943 |
| XPO4 | 1080.1728 | -0.254478027 | 0.21946231 | -1.1595523 | 0.24623113 | 0.6834865 |
| LATS2 | 910.961367 | -0.987597459 | 0.39083497 | -2.5268913 | 0.01150771 | 0.28627263 |
| SAP18 | 457.602542 | -0.272474439 | 0.32863895 | -0.8290996 | 0.40704803 | 0.78481094 |
| MRPL57 | 331.03475 | -0.140194623 | 0.34094872 | -0.4111898 | 0.6809334 | 0.89947163 |
| ZDHHC20 | 954.582591 | -0.205484784 | 0.22873718 | -0.8983445 | 0.36900193 | 0.76615653 |
| MICU2 | 1814.22081 | -0.113770333 | 0.19218998 | -0.5919681 | 0.55387196 | 0.84727776 |
| FGF9 | 59.7750294 | -0.205212638 | 0.79198279 | -0.2591125 | 0.79554844 | 0.94252227 |
| AL136962.1 | 954.741018 | 0.228478679 | 0.77435901 | 0.29505523 | 0.76795169 | 0.93186206 |
| LINC00621 | 2060.13853 | 0.643197591 | 0.84517116 | 0.76102643 | 0.44664128 | 0.80205844 |
| AL157931.1 | 840.405101 | 1.37077594 | 0.75576892 | 1.81375007 | 0.06971621 | 0.49371449 |
| SACS | 204.766738 | -0.844635558 | 0.6550758 | -1.2893707 | 0.19726923 | 0.64558745 |
| TNFRSF19 | 682.62117 | 1.186219503 | 0.67664922 | 1.75307895 | 0.07958846 | 0.51216853 |
| MIPEP | 1120.86726 | -0.220191263 | 0.25259808 | -0.871706 | 0.38336878 | 0.77317162 |
| PARP4 | 2448.42612 | -0.057985079 | 0.25940664 | -0.2235297 | 8.23E-01 | 0.95131214 |
| CENPJ | 109.459288 | -0.580511325 | 0.38402832 | -1.5116368 | 0.13062629 | 0.58853436 |
| PABPC3 | 47.7856562 | -0.85372473 | 0.55356615 | -1.5422271 | 0.12301841 | 0.58245711 |
| MTMR6 | 480.757034 | 0.063991658 | 0.26423861 | 0.24217376 | 0.80864553 | 0.94630788 |
| NUP58 | 1385.20395 | -0.31098753 | 0.29095378 | -1.0688554 | 0.28513482 | 0.71015169 |
| ATP8A2 | 36.9668263 | 0.109346567 | 0.53973883 | 0.20259163 | 0.83945424 | 0.9556011 |
| RNF6 | 201.033077 | -0.226130734 | 0.43040864 | -0.5253861 | 0.59931478 | 0.86689411 |
| CDK8 | 1917.66729 | -0.084943622 | 0.18823155 | -0.451272 | 0.65179355 | 0.88935488 |
| WASF3 | 250.125966 | -0.314093019 | 0.46122566 | -0.6809964 | 0.49587376 | 0.82243887 |
| USP12 | 1234.42373 | -0.105054701 | 0.33841066 | -0.3104356 | 0.75622976 | 0.92799282 |
| LINC02340 | 393.212865 | 0.359944656 | 0.50434012 | 0.71369427 | 0.47541625 | 0.81564184 |
| RPL21 | 557.386136 | 0.206895767 | 0.46362992 | 0.44625198 | 0.65541524 | 0.89053559 |
| GTF3A | 270.366845 | -0.120862619 | 0.27426289 | -0.4406816 | 0.6594435 | 0.89071659 |
| MTIF3 | 499.425335 | 0.244627343 | 0.1952648 | 1.25279799 | 0.21027923 | 0.65441616 |
| LNX2 | 964.337602 | -0.255813232 | 0.27994893 | -0.9137854 | 0.36082964 | 0.76009788 |
| POLR1D | 585.751145 | -0.187094794 | 0.21642171 | -0.8644918 | 0.38731779 | 0.7734506 |
| FLT3 | 102.268401 | 0.305582905 | 0.79177013 | 0.38594902 | 0.69953443 | 0.90721166 |
| PAN3 | 4542.40439 | -0.062624133 | 0.20073441 | -0.3119751 | 0.75505946 | 0.92799282 |
| FLT1 | 79.759731 | -0.036652921 | 0.49259775 | -0.0744074 | 0.94068622 | 0.98484365 |
| POMP | 375.450062 | -0.413667638 | 0.26584298 | -1.55606 | 0.11969382 | 0.57673415 |

| | | | | | | |
|---|---|---|---|---|---|---|
| SLC46A3 | 93.8143852 | 1.571510431 | 0.39459986 | 3.9825418 | 6.82E-05 | 0.02490203 |
| MTUS2 | 39.3471829 | -0.016691825 | 0.45776806 | -0.0364635 | 0.97091278 | 0.99220667 |
| SLC7A1 | 94.8725478 | -0.871863738 | 0.68294484 | -1.2766239 | 0.20173504 | 0.64881721 |
| UBL3 | 751.006576 | 0.011609833 | 0.18535954 | 0.06263413 | 0.95005785 | 0.98724111 |
| KATNAL1 | 724.995202 | 0.186809489 | 0.28007926 | 0.66698793 | 0.50477984 | 0.82715271 |
| HMGB1 | 1888.37567 | 0.354227733 | 0.30362896 | 1.16664672 | 0.24335307 | 0.68164834 |
| USPL1 | 592.726875 | -0.389765453 | 0.34425885 | -1.1321872 | 0.25755575 | 0.691812 |
| HSPH1 | 1238.83278 | -2.305641491 | 0.7430673 | -3.1028703 | 0.00191654 | 0.13753904 |
| B3GLCT | 318.78934 | -0.353178902 | 0.21304476 | -1.6577685 | 0.0973642 | 0.54093104 |
| FRY | 446.582587 | 0.524763383 | 0.66715969 | 0.78656339 | 0.4315375 | 0.79135216 |
| BRCA2 | 55.3573018 | -1.542478879 | 0.63231243 | -2.4394252 | 0.01471065 | 0.30832605 |
| N4BP2L1 | 718.540426 | 1.147766686 | 0.42500454 | 2.70059865 | 0.00692148 | 0.23083391 |
| N4BP2L2 | 8281.9215 | -0.374134782 | 0.18118889 | -2.0648881 | 0.03893359 | 0.42680351 |
| PDS5B | 1508.42119 | -0.055640775 | 0.22100505 | -0.2517625 | 0.80122468 | 0.9444777 |
| KL | 1350.79293 | 0.361215304 | 0.42114431 | 0.85769959 | 3.91E-01 | 7.76E-01 |
| STARD13 | 2528.86761 | -0.390073328 | 0.30258903 | -1.2891192 | 0.19735665 | 0.64558745 |
| AL139383.1 | 398.246531 | 1.702529059 | 0.75475385 | 2.2557408 | 0.02408687 | 0.37387451 |
| RFC3 | 182.10248 | -0.080542619 | 0.53428761 | -0.1507477 | 0.88017476 | 0.96769187 |
| LINC02343 | 68.740201 | 1.012670039 | 0.73480066 | 1.37815613 | 0.16815509 | 0.62304975 |
| NBEA | 3564.18584 | -0.008018336 | 0.2677446 | -0.0299477 | 0.97610876 | 0.99429832 |
| DCLK1 | 206.559073 | -1.414700355 | 0.9232334 | -1.5323323 | 0.12544046 | 0.58564339 |
| SPART | 1680.64722 | 0.073689465 | 0.17495867 | 0.42118214 | 0.67362209 | 0.89799059 |
| SPART-AS1 | 155.371739 | 0.646632255 | 0.46341698 | 1.39535727 | 0.16290813 | 0.61975625 |
| RFXAP | 134.972097 | -0.110561178 | 0.21223481 | -0.520938 | 0.60240996 | 0.8677246 |
| SMAD9 | 674.086434 | -0.278735046 | 0.75719798 | -0.3681138 | 0.71278836 | 0.91188831 |
| ALG5 | 346.387722 | -0.031447911 | 0.19482076 | -0.1614197 | 0.87176285 | 0.9657206 |
| EXOSC8 | 147.346843 | -0.122090833 | 0.33208572 | -0.3676486 | 0.71313531 | 0.91191793 |
| SUPT20H | 1197.19613 | 0.32679825 | 0.264104 | 1.23738469 | 0.2159443 | 0.65967206 |
| LINC02334 | 381.665294 | 0.835264751 | 0.51396571 | 1.6251371 | 1.04E-01 | 5.51E-01 |
| LINC00571 | 494.752968 | -4.80E-05 | 0.27186215 | -0.0001767 | 0.99985901 | 0.99985901 |
| UFM1 | 661.069226 | -0.492779321 | 0.21540203 | -2.287719 | 2.22E-02 | 0.36178312 |
| FREM2 | 2824.7952 | 0.379384017 | 0.65105109 | 0.58272541 | 0.56007816 | 0.84941263 |
| FREM2-AS1 | 50.7512988 | 0.302788745 | 1.00258978 | 0.30200661 | 0.76264702 | 0.93022076 |
| PROSER1 | 328.235022 | 0.013595225 | 0.17404167 | 0.07811477 | 0.93773676 | 0.98435387 |
| NHLRC3 | 628.477237 | 0.782473525 | 0.50452213 | 1.55092013 | 0.12092083 | 0.5792686 |
| AL354809.1 | 98.9486012 | 0.820473697 | 0.29817298 | 2.75167019 | 0.00592922 | 0.22637955 |
| LHFPL6 | 458.067885 | -0.24494242 | 0.78177313 | -0.3133165 | 0.75404022 | 0.92796097 |
| COG6 | 955.226929 | -0.224929301 | 0.2267921 | -0.9917863 | 0.32130178 | 0.73291455 |
| LINC00598 | 312.810698 | 1.036591431 | 0.62416182 | 1.66077353 | 0.09675894 | 0.54001417 |
| FOXO1 | 3415.3339 | 0.332308909 | 0.28832453 | 1.15255164 | 0.24909446 | 0.68557761 |
| MRPS31 | 744.015208 | 0.140456399 | 0.35597511 | 0.39456803 | 0.69316171 | 0.90551691 |
| SLC25A15 | 82.584135 | -0.753107269 | 0.44338154 | -1.6985535 | 0.08940334 | 0.53095823 |
| ELF1 | 2473.69742 | -0.393139184 | 0.26196966 | -1.500705 | 0.13343187 | 0.59240843 |
| WBP4 | 488.300266 | -0.245237598 | 0.18666614 | -1.3137765 | 0.18892142 | 0.64129418 |
| KBTBD6 | 64.1961932 | 0.011803743 | 0.40546299 | 0.02911176 | 0.97677545 | 0.99446862 |
| AL354696.2 | 232.871417 | 0.270343314 | 0.45547325 | 0.59354378 | 0.55281729 | 0.84721285 |
| KBTBD7 | 74.3867473 | 0.064051002 | 0.5133099 | 0.12478038 | 0.90069742 | 0.97479173 |
| MTRF1 | 268.376829 | -0.627630527 | 0.34575834 | -1.8152289 | 0.06948874 | 0.49281529 |
| NAA16 | 740.820233 | 0.12236767 | 0.2488594 | 0.49171408 | 0.62292148 | 0.87717123 |
| VWA8 | 5250.3988 | 0.104991248 | 0.32371231 | 0.32433505 | 0.74568439 | 0.92410821 |
| DGKH | 4382.04871 | -0.573781574 | 0.36306561 | -1.5803798 | 0.11401991 | 0.56918057 |
| AKAP11 | 853.471416 | 0.235078239 | 0.17674797 | 1.33001948 | 0.18351185 | 0.63276491 |
| LINC02341 | 68.8966434 | 0.111987488 | 0.33645181 | 0.33284852 | 7.39E-01 | 9.21E-01 |
| EPSTI1 | 1749.62796 | -1.174747776 | 0.93056499 | -1.2624027 | 0.20680391 | 0.65251041 |

| | | | | | | |
|---|---|---|---|---|---|---|
| DNAJC15 | 1059.5324 | 0.155676075 | 0.34752728 | 0.44795354 | 0.65418673 | 0.89019532 |
| ENOX1 | 466.657923 | 0.417962766 | 0.49177562 | 0.84990543 | 0.39537767 | 0.77931337 |
| CCDC122 | 673.643966 | 1.433665596 | 0.44020274 | 3.25683025 | 0.00112664 | 0.11093704 |
| LINC00390 | 338.755161 | 0.51573263 | 0.8406849 | 0.61346722 | 0.53956745 | 0.84113285 |
| SMIM2-AS1 | 8782.19225 | 0.219895176 | 0.34053931 | 0.64572626 | 0.51845665 | 0.83325693 |
| SMIM2 | 81.3316526 | 1.047372748 | 0.76504437 | 1.3690353 | 0.17098824 | 0.62504223 |
| SERP2 | 85.615118 | -0.343568457 | 0.42757958 | -0.8035193 | 0.42167463 | 0.78892549 |
| TSC22D1 | 3757.53919 | 0.152067021 | 0.14845173 | 1.02435334 | 0.30566841 | 0.72122 |
| TSC22D1-AS1 | 33.1158803 | -0.262214094 | 0.57110956 | -0.459131 | 0.64614011 | 0.88700494 |
| NUFIP1 | 198.097739 | -0.154496808 | 0.31363357 | -0.4926029 | 0.62229319 | 0.87714058 |
| AL359706.1 | 113.155824 | 0.472339913 | 0.65786499 | 0.71798914 | 0.47276399 | 0.81460134 |
| GPALPP1 | 468.235081 | 0.040446002 | 0.16937015 | 0.23880241 | 0.8112588 | 0.94753961 |
| GTF2F2 | 1040.99605 | -0.591669947 | 0.4233917 | -1.3974529 | 0.16227743 | 0.61907506 |
| TPT1 | 3986.19992 | 0.1440004 | 0.53146444 | 0.27095021 | 0.78642933 | 0.9390702 |
| AL138963.1 | 89.567377 | -0.328189406 | 0.40924523 | -0.8019383 | 0.42258867 | 0.78894887 |
| SLC25A30 | 649.454086 | -0.34469823 | 0.5212113 | -0.6613407 | 0.50839386 | 0.82874455 |
| SLC25A30-AS | 40.0147491 | -0.468296411 | 0.77521662 | -0.6040846 | 0.54578741 | 0.84460887 |
| COG3 | 768.979026 | -0.092868065 | 0.18440343 | -0.5036135 | 0.61453298 | 0.87225883 |
| ZC3H13 | 914.758103 | -0.037159834 | 0.23105059 | -0.1608299 | 0.87222741 | 0.9657206 |
| AL138686.1 | 206.988561 | -1.260540429 | 0.71191127 | -1.7706426 | 0.07662015 | 0.50883353 |
| LRCH1 | 3919.68243 | 0.392934115 | 0.34375552 | 1.14306272 | 0.25301255 | 0.68887358 |
| ESD | 435.626001 | 0.141891441 | 0.28062297 | 0.50563018 | 0.61311629 | 0.87222135 |
| SUCLA2 | 1168.27247 | -0.520582431 | 0.36480219 | -1.4270266 | 0.15357222 | 0.61014911 |
| NUDT15 | 54.5311047 | -0.629538038 | 0.38317102 | -1.6429688 | 0.10038939 | 0.54542315 |
| MED4 | 616.484561 | -0.132495885 | 0.26069425 | -0.5082425 | 0.61128332 | 0.87163238 |
| ITM2B | 4987.08996 | 0.31682199 | 0.38350613 | 0.82611975 | 0.40873616 | 0.78511023 |
| RB1 | 2189.61817 | 0.293068053 | 0.18343859 | 1.59763578 | 0.11012406 | 0.56114143 |
| LPAR6 | 339.460564 | 0.218172362 | 0.72693507 | 0.30012634 | 0.76408079 | 0.93079056 |
| RCBTB2 | 398.749753 | 0.180021898 | 0.33762353 | 0.53320305 | 0.59389303 | 0.86472188 |
| FNDC3A | 3941.24123 | -0.169094506 | 0.22491941 | -0.7518004 | 0.45217109 | 0.80419218 |
| CDADC1 | 406.243944 | 0.026700049 | 0.34960732 | 0.07637154 | 9.39E-01 | 9.84E-01 |
| CAB39L | 289.818907 | -0.751344 | 0.39574795 | -1.8985417 | 0.05762475 | 0.47684824 |
| SETDB2 | 614.966146 | -0.079107136 | 0.29956157 | -0.2640764 | 0.79172106 | 0.94104411 |
| PHF11 | 357.585271 | -0.682350271 | 0.37350078 | -1.8269046 | 0.06771412 | 0.48963361 |
| RCBTB1 | 341.333991 | -0.612382041 | 0.27884614 | -2.1961288 | 0.02808273 | 0.39158689 |
| EBPL | 188.928314 | -0.032359449 | 0.40862365 | -0.0791913 | 0.93688044 | 0.98435387 |
| KPNA3 | 1048.84084 | -0.195322497 | 0.31036077 | -0.6293402 | 0.52912639 | 0.83778854 |
| SPRYD7 | 256.844403 | -0.034929282 | 0.27195558 | -0.1284375 | 0.89780279 | 0.9737247 |
| DLEU2 | 3191.18444 | 0.188007378 | 0.38001334 | 0.49473889 | 0.62078444 | 0.87609696 |
| TRIM13 | 806.209138 | 0.121212079 | 0.46179346 | 0.26248115 | 0.79295051 | 0.94160727 |
| RNASEH2B | 247.768875 | 0.099033672 | 0.37036765 | 0.26739288 | 0.78916668 | 0.94005579 |
| FAM124A | 469.83333 | 0.260744615 | 0.41430209 | 0.62935868 | 0.52911427 | 0.83778854 |
| INTS6 | 1496.56913 | 0.328374163 | 0.25667239 | 1.27935131 | 0.20077337 | 0.64813957 |
| INTS6-AS1 | 176.879751 | 0.207156224 | 0.49130803 | 0.42164225 | 0.67328616 | 0.89782405 |
| WDFY2 | 1050.94593 | 0.143995223 | 0.17726622 | 0.81231054 | 0.41661347 | 0.78632759 |
| ATP7B | 358.081313 | 0.159642067 | 0.26690985 | 0.59811231 | 0.54976499 | 0.84612944 |
| ALG11 | 134.162073 | -0.133535193 | 0.3144312 | -0.4246881 | 0.67106405 | 0.89694443 |
| UTP14C | 159.491243 | -0.110807245 | 0.38252453 | -0.2896736 | 0.77206598 | 0.93340577 |
| NEK5 | 135.006171 | 0.541303275 | 0.67361755 | 0.80357656 | 0.42164157 | 0.78892549 |
| NEK3 | 291.606729 | 0.126919188 | 0.53429578 | 0.23754481 | 0.81223416 | 0.94787133 |
| VPS36 | 765.32849 | -0.035923236 | 0.15580347 | -0.2305676 | 0.81765072 | 0.94954998 |
| CKAP2 | 104.190553 | 0.503634375 | 0.32442424 | 1.5523944 | 0.12056788 | 0.57856254 |
| SUGT1 | 560.196761 | -0.054973247 | 0.16762927 | -0.3279454 | 0.74295295 | 0.92318522 |
| DIAPH3 | 297.026774 | 0.050092768 | 0.50035425 | 0.10011461 | 0.92025334 | 0.97939204 |

| | | | | | | |
|---|---|---|---|---|---|---|
| TDRD3 | 1503.3097 | -0.068616153 | 0.25459314 | -0.269513 | 0.78753496 | 0.93953874 |
| PCDH9 | 200.230754 | 0.60730176 | 0.53517309 | 1.13477633 | 0.25646905 | 0.69092557 |
| KLHL1 | 55.9092068 | 0.581858307 | 0.58592154 | 0.99306523 | 3.21E-01 | 0.73210121 |
| DACH1 | 1659.55399 | 0.744231799 | 0.59600458 | 1.24870147 | 2.12E-01 | 6.56E-01 |
| MZT1 | 92.4399324 | -0.153534809 | 0.37488394 | -0.4095529 | 0.68213395 | 0.89962501 |
| BORA | 69.5413704 | -0.043291479 | 0.32915323 | -0.1315238 | 0.89536097 | 0.97309833 |
| DIS3 | 415.489116 | -0.062708127 | 0.32184618 | -0.1948388 | 0.84551914 | 0.95764192 |
| PIBF1 | 2369.57916 | -0.120465303 | 0.21698759 | -0.5551714 | 0.57877744 | 0.8575616 |
| KLF5 | 228.968202 | 0.097433041 | 0.52803077 | 0.18452152 | 8.54E-01 | 9.61E-01 |
| KLF12 | 2137.02961 | 0.270504704 | 0.28693552 | 0.94273692 | 3.46E-01 | 0.75166079 |
| TBC1D4 | 2324.54956 | 0.615283787 | 0.45746056 | 1.34499855 | 0.1786257 | 0.6288217 |
| COMMD6 | 328.4879 | -0.134055168 | 0.44267216 | -0.3028317 | 0.76201812 | 0.93016063 |
| UCHL3 | 503.641154 | -0.54408307 | 0.416218 | -1.307207 | 0.19114243 | 0.64341583 |
| LMO7-AS1 | 63.3118801 | 0.828869947 | 0.52705872 | 1.57263301 | 0.11580382 | 0.57350419 |
| LMO7 | 1546.59699 | 0.243737994 | 0.45189276 | 0.53937132 | 0.58963067 | 0.86281961 |
| KCTD12 | 79.6657499 | 1.262425662 | 0.68260322 | 1.84942822 | 0.064396 | 0.48593331 |
| AC001226.2 | 177.6733 | 0.820758664 | 0.59472464 | 1.380065 | 0.16756663 | 0.62296534 |
| FBXL3 | 504.406605 | -0.149469105 | 0.2340455 | -0.6386327 | 0.52306191 | 0.83513217 |
| MYCBP2 | 2573.22122 | -0.079635897 | 0.17488337 | -0.4553658 | 0.64884613 | 0.88800332 |
| SLAIN1 | 271.022462 | 0.733906728 | 0.33961463 | 2.16099857 | 0.03069545 | 0.40446361 |
| RNF219 | 277.686507 | -0.127440958 | 0.38052252 | -0.3349104 | 0.73769266 | 0.92057113 |
| RBM26 | 2364.37057 | 0.145812702 | 0.18793625 | 0.77586256 | 0.43783014 | 0.79677878 |
| RBM26-AS1 | 69.1580614 | 0.587954231 | 0.53239871 | 1.10434946 | 0.26944158 | 0.69961625 |
| NDFIP2 | 1673.85357 | -0.288858555 | 0.27591475 | -1.0469123 | 0.29514002 | 0.71646902 |
| SPRY2 | 102.435824 | -0.349199363 | 0.42000669 | -0.8314138 | 0.4057399 | 0.78429302 |
| MIR4500HG | 103.548971 | 2.852003381 | 0.89898405 | 3.17247382 | 0.00151146 | 0.13146073 |
| LINC01040 | 90.0927886 | 1.345335694 | 0.99797197 | 1.34806962 | 0.17763599 | 0.6280421 |
| LINC00410 | 28.3215836 | 0.51307457 | 1.15531757 | 0.4440983 | 0.65697151 | 0.89053559 |
| GPC5 | 242.2481 | 0.498188152 | 0.76189112 | 0.65388366 | 0.51318676 | 0.83073024 |
| GPC6 | 6341.02923 | -0.192088688 | 0.51051003 | -0.3762682 | 0.70671752 | 0.91002281 |
| TGDS | 256.842877 | -0.436174603 | 0.20924444 | -2.0845218 | 0.03711273 | 0.42215481 |
| GPR180 | 306.115857 | -0.723890416 | 0.30748526 | -2.354228 | 0.01856122 | 0.33968023 |
| ABCC4 | 5392.9986 | -0.548139171 | 0.28167562 | -1.9459944 | 0.05165541 | 0.4629611 |
| CLDN10 | 11288.8556 | 2.506821996 | 0.65844248 | 3.80719968 | 0.00014055 | 0.0393165 |
| CLDN10-AS1 | 1632.29892 | 1.833420582 | 0.63684747 | 2.87890061 | 0.00399064 | 0.20256898 |
| DZIP1 | 1159.56658 | 1.19759168 | 0.43419156 | 2.75821042 | 0.00581188 | 0.22631649 |
| DNAJC3-DT | 188.877654 | 0.451896793 | 0.5336088 | 0.84686909 | 0.39706809 | 0.78019099 |
| DNAJC3 | 1595.22686 | -0.527006322 | 0.43874702 | -1.2011622 | 0.2296883 | 0.66705629 |
| UGGT2 | 2304.26964 | -0.039776153 | 0.31901667 | -0.1246836 | 0.90077403 | 0.97479173 |
| MBNL2 | 3395.24846 | -0.515688421 | 0.54572486 | -0.9449605 | 0.34467906 | 0.75124763 |
| RAP2A | 276.741055 | 0.266167976 | 0.20626983 | 1.2903873 | 0.19691622 | 0.64558745 |
| IPO5 | 514.085484 | -0.198484184 | 0.31342365 | -0.6332776 | 0.52655237 | 0.83700027 |
| FARP1 | 12883.1384 | 0.098537209 | 0.38085229 | 0.25872815 | 0.795845 | 0.94252227 |
| STK24 | 1925.94805 | 0.113933621 | 0.26021723 | 0.43784042 | 0.66150197 | 0.89184036 |
| SLC15A1 | 1428.16852 | 1.791179952 | 0.61365776 | 2.91885814 | 0.00351316 | 0.19417295 |
| DOCK9 | 1080.10237 | -0.327367315 | 0.33336064 | -0.9820215 | 0.32608925 | 0.7373175 |
| UBAC2 | 2543.5212 | -0.137560468 | 0.21621675 | -0.6362156 | 0.52463592 | 0.83592566 |
| TM9SF2 | 1205.48428 | -0.161314827 | 0.19435654 | -0.8299943 | 0.40654199 | 0.78474455 |
| CLYBL | 2490.95906 | 0.669174137 | 0.42000878 | 1.59323844 | 0.11110671 | 0.56347761 |
| AL355338.1 | 137.340094 | 0.08304911 | 0.49259709 | 0.1685944 | 0.86611569 | 0.96437968 |
| PCCA | 9011.80891 | 0.162940895 | 0.24667618 | 0.66054572 | 0.50890369 | 0.82874455 |
| PCCA-AS1 | 70.1688338 | 0.541038562 | 0.71928139 | 0.75219319 | 0.45193489 | 0.80411194 |
| GGACT | 407.688848 | -0.470919791 | 0.43099184 | -1.092642 | 0.27455101 | 0.7014484 |
| TMTC4 | 451.327764 | 0.227483062 | 0.37769844 | 0.60228754 | 0.54698275 | 0.84460887 |

| | | | | | | |
|---|---|---|---|---|---|---|
| FGF14 | 616.664624 | 1.451880588 | 0.66950038 | 2.16860307 | 0.03011283 | 0.40229687 |
| TPP2 | 909.304686 | 0.114386888 | 0.16299968 | 0.7017614 | 0.48282797 | 0.8177376 |
| TEX30 | 180.662249 | -0.28874641 | 0.33365622 | -0.865401 | 0.38681875 | 0.77334938 |
| KDELC1 | 55.139045 | 0.379512493 | 0.41213752 | 0.92083947 | 0.35713424 | 0.75903693 |
| SLC10A2 | 75.2829662 | -0.420899762 | 0.80545705 | -0.5225602 | 0.60128037 | 0.86755264 |
| ARGLU1 | 1322.71018 | -0.161171814 | 0.22641594 | -0.7118395 | 0.47656416 | 0.81570932 |
| FAM155A | 97.6784045 | 0.018729138 | 0.72430773 | 0.02585798 | 0.97937061 | 0.99482978 |
| LIG4 | 188.080242 | -0.007899204 | 0.2720796 | -0.0290327 | 0.97683852 | 0.99446862 |
| ABHD13 | 254.466256 | 0.336958206 | 0.24609794 | 1.36920366 | 0.17093562 | 0.62504223 |
| TNFSF13B | 300.640868 | 0.974842618 | 0.34795808 | 2.80160938 | 0.00508484 | 0.21719901 |
| MYO16 | 58.5289713 | 1.001880626 | 0.75032165 | 1.33526818 | 0.18178859 | 0.6318513 |
| IRS2 | 368.960032 | -1.054147417 | 0.51667536 | -2.040251 | 0.04132533 | 0.43221049 |
| COL4A1 | 3123.45513 | -0.780598858 | 0.31501608 | -2.4779651 | 0.01321341 | 0.29875272 |
| COL4A2 | 4051.24932 | -0.454470784 | 0.27659256 | -1.6431056 | 0.10036109 | 0.54542315 |
| RAB20 | 938.035455 | 0.227119547 | 0.4497181 | 0.50502648 | 0.61354024 | 0.87222135 |
| NAXD | 325.81016 | 0.073257428 | 0.17123642 | 0.42781452 | 0.66878616 | 0.89566976 |
| CARS2 | 720.438474 | -0.352205672 | 0.34503318 | -1.0207878 | 0.30735497 | 0.72250292 |
| ING1 | 79.3995103 | 0.111664259 | 0.25630421 | 0.43567079 | 0.6630756 | 0.89263183 |
| ANKRD10 | 2551.5775 | 0.145951291 | 0.24663276 | 0.59177576 | 0.55400076 | 0.84727776 |
| ARHGEF7 | 2433.00383 | -0.229941391 | 0.27612886 | -0.8327322 | 0.40499579 | 0.78422899 |
| TUBGCP3 | 1114.18752 | 0.349683257 | 0.44758603 | 0.78126491 | 0.4346467 | 0.79451524 |
| ATP11AUN | 64.0039897 | 0.783784079 | 0.65021194 | 1.20542861 | 2.28E-01 | 6.67E-01 |
| ATP11A | 3647.7655 | -0.138127274 | 0.26541877 | -0.5204126 | 0.60277602 | 0.8677246 |
| MCF2L | 1945.21726 | 0.582428454 | 0.4884186 | 1.19247804 | 0.23307386 | 0.66978724 |
| F10 | 34.3590305 | -0.635257834 | 0.60226545 | -1.0547805 | 0.29152574 | 0.71337138 |
| PROZ | 161.487937 | 0.790453894 | 0.70935808 | 1.11432282 | 2.65E-01 | 6.96E-01 |
| PCID2 | 484.748029 | -0.11390531 | 0.22467153 | -0.506986 | 0.61216467 | 0.87222135 |
| CUL4A | 1483.20443 | -0.608464628 | 0.3145521 | -1.9343843 | 0.0530659 | 0.46624874 |
| LAMP1 | 1088.72926 | -0.564299042 | 0.21116078 | -2.6723667 | 0.00753183 | 0.23733051 |
| GRTP1 | 1084.93558 | 0.227130013 | 0.4360184 | 0.52091842 | 0.60242361 | 0.8677246 |
| GRTP1-AS1 | 73.3305293 | 0.148225096 | 0.69636656 | 0.21285499 | 0.83144007 | 0.95325673 |
| ADPRHL1 | 46.4843981 | -1.581594609 | 0.66176169 | -2.3899761 | 0.01684947 | 0.32852872 |
| DCUN1D2 | 406.724011 | -0.482089107 | 0.27090418 | -1.7795558 | 0.07514869 | 0.5071704 |
| TMCO3 | 662.881647 | -0.435410156 | 0.33710278 | -1.2916244 | 1.96E-01 | 6.46E-01 |
| TFDP1 | 334.872054 | -0.849161574 | 0.42094779 | -2.017261 | 0.04366828 | 0.44226848 |
| GAS6 | 67.8948824 | 0.856033637 | 0.70476297 | 1.21464049 | 0.22450324 | 0.66551422 |
| RASA3 | 149.941221 | -0.328800974 | 0.7760459 | -0.4236875 | 6.72E-01 | 8.97E-01 |
| CDC16 | 690.379508 | -0.361583237 | 0.18163339 | -1.9907311 | 0.04651046 | 0.44508419 |
| UPF3A | 764.177274 | -0.63711738 | 0.29338953 | -2.171575 | 0.02988773 | 0.4000695 |
| CHAMP1 | 170.036302 | 0.034746793 | 0.33854078 | 0.10263695 | 0.91825112 | 0.97857783 |
| TTC5 | 207.278474 | -0.013342112 | 0.31901556 | -0.0418228 | 0.96663999 | 0.99122723 |
| CCNB1IP1 | 333.776216 | -0.179671868 | 0.28724759 | -0.6254948 | 0.53164637 | 0.83789294 |
| PARP2 | 73.9775757 | -0.461918408 | 0.34175546 | -1.3516051 | 0.17650169 | 0.62709281 |
| TEP1 | 377.336284 | -0.383422846 | 0.35699964 | -1.0740147 | 0.28281611 | 0.70756556 |
| OSGEP | 210.009451 | -0.855648392 | 0.41713225 | -2.0512641 | 0.04024124 | 0.43086056 |
| APEX1 | 237.344068 | -0.454144816 | 0.40282732 | -1.1273933 | 0.25957624 | 0.69242517 |
| PIP4P1 | 128.066036 | -0.530205019 | 0.34026169 | -1.5582272 | 0.1191794 | 0.57642628 |
| PNP | 329.021051 | 0.011623048 | 0.4461623 | 0.02605117 | 0.97921653 | 0.99482978 |
| METTL17 | 310.733048 | -0.44063703 | 0.24016553 | -1.8347222 | 0.06654686 | 0.48830715 |
| NDRG2 | 121.616686 | 1.064775333 | 0.4379824 | 2.43109159 | 0.01505341 | 0.30991786 |
| ARHGEF40 | 208.00028 | 0.105929442 | 0.42239297 | 0.2507841 | 0.80198103 | 0.94496295 |
| ZNF219 | 99.8353616 | 0.775158679 | 0.42883507 | 1.80759164 | 0.07067008 | 0.4967563 |
| HNRNPC | 5430.35001 | -0.308747122 | 0.26233328 | -1.176927 | 0.23922464 | 0.67720991 |
| SUPT16H | 548.368894 | -0.1000587 | 0.2996491 | -0.3339196 | 0.73844025 | 0.92080006 |

| | | | | | | |
|---|---|---|---|---|---|---|
| CHD8 | 992.599314 | -0.279331698 | 0.27014819 | -1.0339943 | 0.30113881 | 0.71879123 |
| RAB2B | 332.61511 | -0.316968264 | 0.30556311 | -1.037325 | 0.29958441 | 0.7186844 |
| TOX4 | 463.549305 | -0.179516221 | 0.22705168 | -0.7906403 | 0.4291539 | 0.79037914 |
| METTL3 | 210.634484 | -0.397907303 | 0.27578485 | -1.4428179 | 0.14907179 | 0.60622442 |
| DAD1 | 687.245464 | -0.641333285 | 0.36933747 | -1.7364425 | 0.08248562 | 0.5183475 |
| ABHD4 | 156.83824 | 0.022541507 | 0.39637729 | 0.05686882 | 0.9546497 | 0.98840182 |
| OXA1L | 352.898687 | -0.166095763 | 0.25530166 | -0.6505863 | 0.51531358 | 0.83181198 |
| SLC7A7 | 1355.93171 | 0.104446888 | 0.47632465 | 0.21927668 | 8.26E-01 | 9.52E-01 |
| MRPL52 | 244.681154 | -0.148416987 | 0.32654238 | -0.4545106 | 0.64946134 | 0.88850926 |
| MMP14 | 233.07767 | 0.207177079 | 0.5366899 | 0.38602754 | 0.69947628 | 0.90721166 |
| LRP10 | 583.938383 | 0.224305211 | 0.41232693 | 0.54399845 | 0.58644254 | 0.86102615 |
| RBM23 | 569.664691 | -0.028911507 | 0.26364417 | -0.1096611 | 0.91267816 | 0.977179 |
| PRMT5 | 118.219893 | 0.434902737 | 0.34855255 | 1.24773938 | 0.21212652 | 0.65673943 |
| AL132780.1 | 96.1052386 | 0.75875579 | 0.69650665 | 1.08937337 | 0.27598927 | 0.70289575 |
| AL132780.3 | 129.772408 | -1.021957798 | 0.52385266 | -1.9508497 | 0.05107493 | 0.46088481 |
| AL132780.2 | 99.9583118 | -0.526754028 | 0.39153452 | -1.3453578 | 1.79E-01 | 0.6288217 |
| C14orf93 | 89.1810981 | 0.346904047 | 0.32774825 | 1.05844668 | 0.28985185 | 0.71239004 |
| PSMB5 | 417.072361 | -0.162093273 | 0.33235899 | -0.4877054 | 0.62575853 | 0.87854882 |
| ACIN1 | 1351.08674 | -0.330747107 | 0.18413838 | -1.7961878 | 0.07246466 | 0.502734 |
| C14orf119 | 200.247095 | -0.170222606 | 0.32338733 | -0.5263738 | 0.59862854 | 0.86646266 |
| SLC7A8 | 2595.46987 | 0.13321932 | 0.57195247 | 0.23292026 | 8.16E-01 | 9.49E-01 |
| RNF212B | 2093.50705 | 0.577581953 | 0.86634992 | 0.66668437 | 5.05E-01 | 8.27E-01 |
| HOMEZ | 165.67121 | 0.200546715 | 0.29444041 | 0.68111138 | 0.49580102 | 0.82243887 |
| PPP1R3E | 185.518086 | 0.211707657 | 0.51608979 | 0.41021477 | 0.6816484 | 0.89962501 |
| BCL2L2 | 50.0676902 | 0.366304887 | 0.68954441 | 0.5312274 | 0.5952612 | 0.86510861 |
| BCL2L2-PABI | 360.787456 | 0.002814977 | 0.28943215 | 0.00972586 | 0.99224001 | 0.9976934 |
| PABPN1 | 831.579347 | -0.298437635 | 0.36260128 | -0.8230463 | 0.41048167 | 0.78582015 |
| SLC22A17 | 137.120599 | 0.35233061 | 0.60499584 | 0.58236865 | 0.5603184 | 0.84944908 |
| NGDN | 232.294179 | -0.547570248 | 0.27486115 | -1.9921704 | 0.04635236 | 0.44508419 |
| ZFHX2 | 181.390348 | 0.999391016 | 0.52165013 | 1.91582627 | 0.05538721 | 0.46947044 |
| THTPA | 42.7871746 | 0.144349277 | 0.48834871 | 0.29558648 | 0.7675459 | 0.93169958 |
| AP1G2 | 265.849926 | -0.587504646 | 0.35585532 | -1.6509649 | 0.09874574 | 0.54236074 |
| AL135999.1 | 102.113093 | -0.359655915 | 0.41184598 | -0.8732777 | 0.38251172 | 0.77279408 |
| LINC00596 | 81.5831007 | 0.128608603 | 0.92905938 | 0.13842883 | 0.88990151 | 0.97213108 |
| DHRS4-AS1 | 377.962957 | 0.341614652 | 0.52270744 | 0.65354848 | 0.51340275 | 0.8308396 |
| DHRS4 | 198.697137 | 0.376927146 | 0.57334312 | 0.65741985 | 5.11E-01 | 0.82974608 |
| DHRS4L2 | 294.400022 | 0.935386871 | 0.55731298 | 1.67838702 | 0.09327157 | 0.53688613 |
| NRL | 144.156346 | 0.696868307 | 0.54077213 | 1.28865425 | 0.19751831 | 0.64558745 |
| PCK2 | 358.174736 | 0.5581109 | 0.36178622 | 1.54265383 | 1.23E-01 | 5.82E-01 |
| DCAF11 | 65.4323766 | 0.886563675 | 0.46157033 | 1.92075534 | 0.05476256 | 0.46797407 |
| PSME1 | 316.382608 | 0.060519439 | 0.51932301 | 0.11653525 | 0.90722835 | 0.97673359 |
| EMC9 | 76.5836556 | 0.244420752 | 0.52596154 | 0.46471222 | 0.64213756 | 0.88441609 |
| PSME2 | 87.9528354 | 0.129539663 | 0.55225704 | 0.23456408 | 0.8145471 | 0.94881072 |
| REC8 | 127.874352 | 0.527214169 | 0.73819494 | 0.71419369 | 0.47510742 | 0.81564184 |
| IPO4 | 106.494386 | -0.094026273 | 0.58953373 | -0.1594926 | 0.87328078 | 0.96592775 |
| AL136295.1 | 240.41058 | 0.183613306 | 0.23198643 | 0.79148296 | 0.42866222 | 0.79037914 |
| GMPR2 | 119.262835 | 0.488064883 | 0.35198814 | 1.38659467 | 0.1655654 | 0.62065658 |
| TINF2 | 81.4910318 | -0.35330711 | 0.37493612 | -0.9423128 | 0.34603251 | 0.75166079 |
| RABGGTA | 51.1447354 | -0.557282475 | 0.50272097 | -1.1085324 | 0.26763196 | 0.69874881 |
| DHRS1 | 238.128261 | 0.026411583 | 0.25484653 | 0.10363721 | 0.91745726 | 0.97828877 |
| CIDEB | 124.709464 | -0.285371899 | 0.57515854 | -0.4961621 | 0.61978001 | 0.87562612 |
| LTB4R | 176.314995 | -0.641923196 | 0.56229825 | -1.1416062 | 0.25361773 | 0.68891958 |
| KHNYN | 189.33842 | -0.259803477 | 0.2231762 | -1.1641182 | 0.2443761 | 0.68249048 |
| SDR39U1 | 201.30915 | -0.482713321 | 0.32106883 | -1.5034574 | 0.13272113 | 0.59142019 |

| | | | | | | |
|---|---|---|---|---|---|---|
| STXBP6 | 209.585735 | 0.741905423 | 0.45551865 | 1.62870483 | 0.10337552 | 0.55022042 |
| NOVA1 | 1252.92669 | 1.049135284 | 0.55277889 | 1.89792936 | 0.05770539 | 0.47684824 |
| LINC02588 | 61.4472629 | 0.825764056 | 0.62906607 | 1.31268256 | 0.18928994 | 0.6413893 |
| LINC02294 | 471.842844 | 0.718668325 | 0.8973278 | 0.80089832 | 0.42319051 | 0.78894887 |
| AL110292.1 | 3251.57548 | 1.616433008 | 0.86177802 | 1.87569533 | 0.06069713 | 0.48026727 |
| PRKD1 | 9092.21027 | 0.127798215 | 0.54488634 | 0.23454105 | 0.81456497 | 0.94881072 |
| G2E3-AS1 | 43.8712023 | 0.117948938 | 0.52407179 | 0.22506256 | 0.82193061 | 0.95105545 |
| G2E3 | 344.332287 | -0.242007991 | 0.18409484 | -1.3145832 | 0.18865001 | 0.640849 |
| SCFD1 | 1557.11344 | -0.251420551 | 0.17266794 | -1.4560928 | 0.14536697 | 0.60165215 |
| COCH | 181.370823 | -0.00889181 | 0.3533444 | -0.0251647 | 0.97992358 | 0.99487538 |
| AL049830.3 | 41.1087088 | -0.300457219 | 0.64013548 | -0.469365 | 0.63880873 | 0.88284765 |
| STRN3 | 5256.86035 | 0.040811944 | 0.24398192 | 0.16727446 | 0.8671541 | 0.96486219 |
| AP4S1 | 192.407526 | -0.040467045 | 0.48644052 | -0.0831901 | 0.93370037 | 0.98355048 |
| HECTD1 | 2515.23968 | -0.583780046 | 0.28797939 | -2.0271591 | 0.04264614 | 0.43852266 |
| HEATR5A | 348.439144 | 0.094966834 | 0.27713814 | 0.34266966 | 0.73184699 | 0.91930611 |
| AL139353.1 | 112.703193 | 0.269415018 | 0.40338989 | 0.66787746 | 0.50421182 | 0.82715271 |
| NUBPL | 1420.06411 | 0.343502174 | 0.32387506 | 1.06060089 | 0.28887132 | 0.71218341 |
| ARHGAP5 | 3191.76356 | -0.416262343 | 0.32278234 | -1.2896069 | 0.19718717 | 0.64558745 |
| AKAP6 | 1018.89582 | 0.096982945 | 0.46986543 | 0.20640579 | 0.83647395 | 0.95472037 |
| NPAS3 | 907.835409 | 1.163646829 | 0.56359638 | 2.06468117 | 0.03895318 | 0.42680351 |
| EGLN3 | 405.434803 | 0.691301892 | 0.47654165 | 1.45066416 | 1.47E-01 | 0.60383734 |
| EAPP | 430.776569 | -0.089124334 | 0.33047184 | -0.2696881 | 0.78740019 | 0.93953874 |
| SNX6 | 1162.14785 | -0.347131183 | 0.28929728 | -1.1999117 | 0.23017364 | 0.66814699 |
| CFL2 | 147.526505 | -0.377933595 | 0.34633056 | -1.0912511 | 0.27516242 | 0.702279 |
| BAZ1A | 1187.10306 | -0.318341057 | 0.32984243 | -0.9651307 | 0.33447935 | 0.74120095 |
| SRP54 | 1298.84919 | -0.57428633 | 0.21349305 | -2.6899533 | 0.0071462 | 0.23377804 |
| FAM177A1 | 1336.65595 | 0.043264194 | 0.29801238 | 0.14517583 | 0.88457205 | 0.9692315 |
| PPP2R3C | 489.876772 | -0.196106358 | 0.24811497 | -0.790385 | 0.42930294 | 0.79037914 |
| NFKBIA | 697.123642 | -0.093618056 | 0.43567443 | -0.2148808 | 0.82986028 | 0.95325673 |
| RALGAPA1 | 2857.33217 | -0.126941536 | 0.20202267 | -0.6283529 | 0.52977277 | 0.83789294 |
| BRMS1L | 208.736276 | 0.297334487 | 0.22056548 | 1.3480554 | 0.17764056 | 0.6280421 |
| PTCSC3 | 57.9881131 | 0.224051219 | 0.72527608 | 0.30891853 | 7.57E-01 | 9.28E-01 |
| MBIP | 365.726636 | 0.260778412 | 0.25690206 | 1.01508882 | 0.31006349 | 0.72498238 |
| SLC25A21 | 737.073284 | -0.380645571 | 0.52027814 | -0.7316194 | 0.46440092 | 0.80945363 |
| MIPOL1 | 1058.92817 | 0.109857365 | 0.35640278 | 0.30823936 | 0.75790021 | 0.928294 |
| SEC23A | 1607.47663 | 0.22849661 | 0.23346028 | 0.97873869 | 0.32770911 | 0.73795117 |
| GEMIN2 | 111.032089 | 0.349762351 | 0.38697518 | 0.90383664 | 0.36608203 | 0.76421682 |
| TRAPPC6B | 524.447026 | -0.122017806 | 0.19694896 | -0.6195402 | 0.53556053 | 0.83954482 |
| PNN | 1028.41671 | -0.493273956 | 0.3208329 | -1.5374793 | 0.12417597 | 0.58410435 |
| MIA2 | 5706.79762 | 0.293041878 | 0.25937608 | 1.12979529 | 0.25856249 | 0.6923454 |
| FBXO33 | 534.721739 | -0.050561166 | 0.18690918 | -0.270512 | 0.78676642 | 0.93930907 |
| AL049875.1 | 212.321472 | 0.161578145 | 0.31190899 | 0.51802979 | 0.60443749 | 0.86865188 |
| LINC02307 | 134.211855 | -0.486323659 | 0.7775526 | -0.6254544 | 5.32E-01 | 0.83789294 |
| C14orf28 | 78.6317875 | -0.404158461 | 0.34608037 | -1.1678168 | 0.24288069 | 0.68132724 |
| KLHL28 | 636.08581 | -0.05456187 | 0.24916666 | -0.2189774 | 0.82666764 | 0.9519561 |
| TOGARAM1 | 1362.53499 | -0.241345871 | 0.20245257 | -1.1921107 | 0.23321786 | 0.66978724 |
| PRPF39 | 672.307812 | -0.296664525 | 0.27099579 | -1.09472 | 0.27363931 | 0.70140695 |
| FKBP3 | 309.553176 | -0.176727002 | 0.32061502 | -0.5512125 | 0.58148802 | 0.85915432 |
| FANCM | 289.594772 | -0.536288825 | 0.38938334 | -1.3772773 | 0.16842654 | 0.62304975 |
| MIS18BP1 | 263.331711 | -0.023405535 | 0.28516694 | -0.0820766 | 0.9345858 | 0.98359701 |
| LINC02303 | 168.821145 | -0.464814321 | 0.9054962 | -0.5133255 | 0.60772363 | 0.86985416 |
| LINC00871 | 5447.27901 | 0.892066505 | 0.80737175 | 1.1049018 | 0.26920214 | 0.69961625 |
| MDGA2 | 47.3599462 | 0.248820498 | 0.47365773 | 0.52531709 | 0.59936278 | 0.86689411 |
| RPS29 | 691.094886 | -0.06307279 | 0.29506331 | -0.2137602 | 0.83073407 | 0.95325673 |

| | | | | | | |
|---|---|---|---|---|---|---|
| LRR1 | 86.6829218 | -0.51758779 | 0.4966678 | -1.0421207 | 0.29735573 | 0.71839551 |
| RPL36AL | 777.510484 | -0.503737317 | 0.46015666 | -1.0947083 | 0.27364445 | 0.70140695 |
| MGAT2 | 68.0875399 | -0.284140354 | 0.54291331 | -0.5233623 | 0.60072216 | 0.86755264 |
| DNAAF2 | 199.13324 | 0.033638465 | 0.2095983 | 0.16049016 | 0.87249497 | 0.96578281 |
| POLE2 | 140.421332 | 0.445746601 | 0.47460014 | 0.93920453 | 0.34762575 | 0.75274978 |
| KLHDC1 | 395.849715 | 1.047446429 | 0.49073234 | 2.1344557 | 0.0328055 | 0.41180153 |
| KLHDC2 | 873.211632 | 0.073496069 | 0.4285546 | 0.17149756 | 0.86383255 | 0.96393376 |
| NEMF | 1792.31973 | -0.104493263 | 0.14577381 | -0.7168178 | 0.47348653 | 0.81478861 |
| AL627171.2 | 51.0429817 | -1.358983622 | 0.65294089 | -2.0813272 | 0.03740397 | 0.42215481 |
| ARF6 | 254.25841 | -0.216459416 | 0.37743358 | -0.5735033 | 0.56630394 | 0.85215881 |
| LINC01588 | 174.135565 | 0.049150473 | 0.46399262 | 0.10592943 | 0.91563834 | 0.9779223 |
| VCPKMT | 244.947376 | 0.076186102 | 0.32397444 | 0.23516084 | 0.8140839 | 0.94881072 |
| SOS2 | 1714.69744 | 0.096292871 | 0.20045488 | 0.48037179 | 0.63096305 | 0.88122598 |
| L2HGDH | 1371.03635 | 0.685903104 | 0.3233823 | 2.12102856 | 3.39E-02 | 0.41378378 |
| ATP5S | 316.451502 | 0.104991651 | 0.24285126 | 0.43232903 | 0.66550229 | 0.89369715 |
| AL359397.2 | 53.998829 | -0.128945085 | 0.42569992 | -0.3029014 | 7.62E-01 | 9.30E-01 |
| CDKL1 | 1199.48676 | -0.356526742 | 0.32679813 | -1.0909693 | 2.75E-01 | 7.02E-01 |
| MAP4K5 | 1356.7127 | -0.330125394 | 0.23081485 | -1.4302606 | 0.15264223 | 0.61001662 |
| ATL1 | 207.159376 | 0.292454891 | 0.50751923 | 0.57624396 | 0.5644503 | 0.85163263 |
| SAV1 | 439.185512 | -0.26269111 | 0.22688089 | -1.1578371 | 2.47E-01 | 0.68482118 |
| NIN | 506.129993 | 0.026442636 | 0.31985048 | 0.08267187 | 0.93411246 | 0.98359701 |
| PYGL | 194.461986 | 0.549135252 | 0.39862891 | 1.37756001 | 0.16833917 | 0.62304975 |
| TRIM9 | 148.617864 | -0.327402054 | 0.77771905 | -0.4209773 | 6.74E-01 | 0.89799059 |
| TMX1 | 261.051887 | -0.29275104 | 0.2506361 | -1.1680322 | 0.24279376 | 0.68126388 |
| FRMD6 | 78.0065243 | -0.824550216 | 0.54734238 | -1.5064615 | 0.13194874 | 0.59087581 |
| RTRAF | 284.832162 | -0.303181905 | 0.29929297 | -1.0129937 | 0.31106315 | 0.72558189 |
| NID2 | 173.7876 | -0.086463589 | 0.87233801 | -0.0991171 | 0.92104532 | 0.97939204 |
| TXNDC16 | 184.873279 | 0.145145291 | 0.34776584 | 0.41736501 | 0.67641145 | 0.89840811 |
| GPR137C | 53.9082437 | 0.346918135 | 0.63919808 | 0.54273964 | 0.58730908 | 0.86112855 |
| ERO1A | 1243.77386 | -0.770930929 | 0.48214195 | -1.5989709 | 1.10E-01 | 5.60E-01 |
| AL133453.1 | 103.248443 | 0.147945417 | 0.73146271 | 0.20225969 | 8.40E-01 | 9.56E-01 |
| PSMC6 | 741.936001 | -0.315352816 | 0.19637607 | -1.6058617 | 0.1083043 | 0.55788313 |
| STYX | 205.390457 | 0.208657515 | 0.24408304 | 0.85486281 | 0.3926271 | 0.77703588 |
| GNPNAT1 | 142.936804 | -0.536439938 | 0.39626083 | -1.3537547 | 0.17581467 | 0.62699479 |
| FERMT2 | 1834.69219 | -0.45115785 | 0.32432211 | -1.3910795 | 0.16420131 | 0.62065658 |
| DDHD1 | 517.745438 | 0.088677954 | 0.28203547 | 0.31442128 | 0.75320109 | 0.92751122 |
| CNIH1 | 244.908678 | 0.329014887 | 0.2865223 | 1.14830463 | 0.25084283 | 0.68648949 |
| GMFB | 365.407623 | -0.009915022 | 0.33223673 | -0.0298432 | 0.97619207 | 0.99429832 |
| CGRRF1 | 242.763736 | -0.177500686 | 0.33106716 | -0.5361471 | 0.59185691 | 0.86382106 |
| SAMD4A | 6067.86374 | -0.421823686 | 0.50538073 | -0.8346652 | 0.40390627 | 0.78358009 |
| GCH1 | 710.230193 | -0.208194406 | 0.28095529 | -0.7410233 | 4.59E-01 | 0.80636987 |
| WDHD1 | 120.343568 | -0.869100921 | 0.51122952 | -1.700021 | 0.08912698 | 0.53095823 |
| SOCS4 | 285.959626 | 0.027692346 | 0.24814846 | 0.11159588 | 0.91114384 | 0.97697493 |
| MAPK1IP1L | 1199.19279 | 0.044935324 | 0.30297046 | 0.14831586 | 0.8820935 | 0.96796225 |
| LGALS3 | 210.435932 | 0.096282882 | 0.45431659 | 0.21192905 | 0.83216239 | 0.95325673 |
| FBXO34 | 2926.34247 | -0.313179013 | 0.28861981 | -1.0850919 | 0.27788096 | 0.7036882 |
| AL158801.3 | 156.258995 | -0.157849861 | 0.60939437 | -0.2590274 | 0.79561407 | 0.94252227 |
| ATG14 | 532.692257 | -0.315555135 | 0.30289469 | -1.0417982 | 0.29750526 | 0.71839551 |
| KTN1-AS1 | 123.23685 | -0.303301352 | 0.32571195 | -0.931195 | 0.35175268 | 0.75548011 |
| KTN1 | 2754.06686 | -0.143328588 | 0.19650377 | -0.7293936 | 0.46576095 | 0.81043641 |
| AL355773.1 | 44.0193767 | -0.239974804 | 0.70398993 | -0.3408782 | 0.73319529 | 0.91952673 |
| PELI2 | 2468.00292 | 0.34300777 | 0.26103482 | 1.31403071 | 0.18883587 | 0.64116256 |
| LINC02284 | 45.3038622 | 0.294596413 | 0.56734713 | 0.5192525 | 0.60358468 | 0.86810591 |
| TMEM260 | 768.73864 | 0.346808746 | 0.24364829 | 1.42339906 | 0.15462051 | 0.61119904 |

| | | | | | | |
|---|---|---|---|---|---|---|
| EXOC5 | 1119.23124 | 0.196203053 | 0.16494696 | 1.1894918 | 0.2342462 | 0.670712 |
| AP5M1 | 728.200452 | 0.548467424 | 0.18919678 | 2.89892582 | 0.00374443 | 0.1981659 |
| NAA30 | 318.306392 | 0.098896196 | 0.16991144 | 0.58204556 | 0.560536 | 0.84944908 |
| CCDC198 | 3324.70208 | 0.644509977 | 0.50171398 | 1.28461635 | 0.19892638 | 0.64597062 |
| SLC35F4 | 68.5777868 | 1.131254123 | 0.61097359 | 1.85155977 | 0.06408906 | 0.48593331 |
| ARMH4 | 846.631333 | 0.296591924 | 0.27768278 | 1.0680962 | 0.28547712 | 0.71029858 |
| ACTR10 | 834.733768 | -0.202519702 | 0.19349881 | -1.0466199 | 0.29527493 | 0.71646902 |
| PSMA3 | 724.38466 | -0.187392728 | 0.3279283 | -0.5714442 | 0.56769857 | 0.85263558 |
| ARID4A | 1239.27479 | 0.19270397 | 0.31648268 | 0.60889263 | 0.54259561 | 0.84314228 |
| AL139021.2 | 114.800934 | 1.006009268 | 0.59887968 | 1.67981866 | 0.0929926 | 0.53684765 |
| AL139021.1 | 184.387044 | -0.106419521 | 0.48111886 | -0.2211917 | 0.82494314 | 0.95192757 |
| TIMM9 | 301.68597 | -0.013462762 | 0.19967715 | -0.0674226 | 0.94624524 | 0.98610078 |
| KIAA0586 | 684.08039 | 0.365608337 | 0.29640924 | 1.23345796 | 0.21740495 | 0.66032006 |
| DAAM1 | 907.144722 | -0.483782539 | 0.34233411 | -1.4131882 | 0.15760039 | 0.61323667 |
| GPR135 | 873.367736 | 0.600341554 | 0.39809262 | 1.50804491 | 0.13154303 | 0.59000666 |
| L3HYPDH | 700.835878 | 0.003688684 | 0.38876448 | 0.00948822 | 0.99242961 | 0.9976934 |
| JKAMP | 473.64966 | 0.34067226 | 0.29629894 | 1.14975864 | 0.2502433 | 0.68617881 |
| RTN1 | 113.775595 | 2.17201513 | 0.5861278 | 3.70570227 | 0.00021081 | 0.05350954 |
| LRRC9 | 48.8740599 | 0.566623754 | 0.68674143 | 0.82509039 | 0.40932028 | 0.78542936 |
| AL157911.1 | 299.930943 | 0.313317256 | 0.42108115 | 0.74407809 | 0.45682923 | 0.80507423 |
| PCNX4 | 2118.05137 | -0.449542856 | 0.36214413 | -1.2413369 | 0.21448133 | 0.65925411 |
| DHRS7 | 206.012855 | 0.307731943 | 0.26880286 | 1.14482393 | 0.2522821 | 0.68874851 |
| AL157756.1 | 45.4740165 | -0.434068536 | 0.5396964 | -0.8042828 | 4.21E-01 | 0.78877309 |
| PPM1A | 1242.814 | -0.278395178 | 0.32675215 | -0.8520072 | 0.39421011 | 0.77823445 |
| MNAT1 | 2444.19632 | 0.123943204 | 0.2607596 | 0.47531598 | 0.63456177 | 0.88238823 |
| TRMT5 | 88.796547 | -0.0734703 | 0.29293633 | -0.2508064 | 0.80196381 | 0.94496295 |
| SLC38A6 | 758.131492 | -0.030175008 | 0.30654339 | -0.0984363 | 0.92158582 | 0.97939204 |
| PRKCH | 153.367006 | 0.218903255 | 0.5213827 | 0.4198514 | 0.67459402 | 0.89801183 |
| LINC01303 | 53.2036591 | -0.19749741 | 0.83951875 | -0.2352507 | 0.81401412 | 0.94881072 |
| HIF1A | 10042.6488 | 0.51947602 | 0.59385192 | 0.87475683 | 0.38170623 | 0.77237191 |
| HIF1A-AS2 | 1280.62121 | 0.376488448 | 0.54714787 | 0.68809269 | 4.91E-01 | 8.20E-01 |
| SNAPC1 | 555.118516 | -0.094401809 | 0.42718558 | -0.2209855 | 0.82510374 | 0.95192757 |
| PPP2R5E | 2826.39044 | -0.131818523 | 0.36184783 | -0.3642927 | 0.71563945 | 0.91298152 |
| WDR89 | 340.047003 | -0.149682741 | 0.22813723 | -0.6561083 | 5.12E-01 | 0.83042711 |
| SGPP1 | 1357.52949 | -0.649208753 | 0.23173588 | -2.801503 | 0.00508652 | 0.21719901 |
| SYNE2 | 29226.1449 | 0.224451666 | 0.40920774 | 0.54850298 | 0.58334658 | 0.85996253 |
| ESR2 | 1316.73007 | 0.368136288 | 0.65649298 | 0.56076196 | 0.57495982 | 0.85474196 |
| MTHFD1 | 572.96885 | -0.06558772 | 0.29354035 | -0.2234368 | 0.82319556 | 0.95131214 |
| AL122035.1 | 141.502424 | 0.059210546 | 0.39264728 | 0.15079831 | 0.88013482 | 0.96769187 |
| ZBTB25 | 383.395526 | -0.019016247 | 0.31947407 | -0.0595236 | 0.95253507 | 0.98777411 |
| ZBTB1 | 776.03098 | -0.337075424 | 0.23767044 | -1.4182471 | 0.15611862 | 0.61255389 |
| AL049869.3 | 72.0055336 | 0.381448058 | 0.48727172 | 0.78282412 | 4.34E-01 | 0.79406205 |
| PLEKHG3 | 652.186392 | -0.163717282 | 0.42301655 | -0.3870234 | 0.69873892 | 0.90696517 |
| CHURC1-FNT | 230.23488 | 0.65399192 | 0.40066483 | 1.63226685 | 0.10262328 | 0.54996682 |
| GPX2 | 186.495828 | 4.466460952 | 1.59246906 | 2.80473955 | 0.00503573 | 0.21686099 |
| RAB15 | 188.667959 | 0.743908675 | 0.60023427 | 1.23936389 | 0.21521077 | 0.65925411 |
| FNTB | 38.9044827 | 0.493129986 | 0.51603155 | 0.95561984 | 0.33926433 | 0.74644675 |
| MAX | 986.414366 | -0.035942072 | 0.28745009 | -0.1250376 | 0.90049378 | 0.97479173 |
| AL139022.1 | 56.2321659 | 0.633176796 | 0.77788647 | 0.81397071 | 0.41566173 | 0.78632759 |
| FUT8 | 2343.45 | 0.282265757 | 0.3197398 | 0.88279833 | 3.77E-01 | 0.77048581 |
| AL391261.2 | 45.9157524 | 1.827151859 | 0.8161202 | 2.23882689 | 0.02516718 | 0.37991912 |
| AL359232.1 | 1226.71971 | 0.71183299 | 0.56930702 | 1.25034992 | 0.21117175 | 0.65561118 |
| CCDC196 | 54.1377637 | 0.760876444 | 0.8710503 | 0.87351608 | 0.38238185 | 0.77279408 |
| GPHN | 11741.4727 | 0.320286859 | 0.30792372 | 1.04014999 | 2.98E-01 | 7.19E-01 |

| | | | | | | |
|---|---|---|---|---|---|---|
| FAM71D | 35.0617373 | 0.269766933 | 0.42181474 | 0.63953889 | 5.22E-01 | 0.83477439 |
| MPP5 | 1644.48889 | -0.189197255 | 0.2430949 | -0.7782856 | 0.43640067 | 0.79555046 |
| ATP6V1D | 624.665944 | -0.679825695 | 0.36521468 | -1.8614413 | 0.06268188 | 0.48421747 |
| EIF2S1 | 400.157701 | -0.356313373 | 0.31864805 | -1.1182035 | 0.26348008 | 0.69567438 |
| PLEKHH1 | 498.609017 | -0.221025796 | 0.26428512 | -0.8363157 | 0.40297733 | 0.78260941 |
| PIGH | 191.984035 | -0.471313184 | 0.30405551 | -1.5500893 | 0.12112009 | 0.57927181 |
| ARG2 | 655.798049 | -0.092100461 | 0.82572316 | -0.1115392 | 0.91118882 | 0.97697493 |
| VTI1B | 238.859508 | 0.052715674 | 0.26614632 | 0.19807027 | 0.84299008 | 0.95716002 |
| RDH11 | 125.821189 | 0.218579927 | 0.34940449 | 0.62557847 | 0.53159146 | 0.83789294 |
| RDH12 | 73.0442937 | -0.330691929 | 0.64580378 | -0.5120625 | 6.09E-01 | 8.70E-01 |
| ZFYVE26 | 473.52775 | -0.551623635 | 0.23581797 | -2.3391925 | 0.01932547 | 0.34268338 |
| RAD51B | 2609.23428 | 0.134872716 | 0.58228265 | 0.23162757 | 0.81682729 | 0.94946888 |
| ZFP36L1 | 2148.17295 | -0.482154725 | 0.49962868 | -0.9650261 | 0.33453173 | 0.74120095 |
| ACTN1 | 525.63292 | 0.230963893 | 0.41562953 | 0.55569654 | 0.57841832 | 0.85751254 |
| DCAF5 | 1972.33075 | 0.287099913 | 0.22003899 | 1.30476836 | 0.19197174 | 0.64497936 |
| AL359317.2 | 120.517711 | 0.041558518 | 0.46456954 | 0.08945597 | 0.92871955 | 0.98272895 |
| EXD2 | 371.844786 | -0.280185191 | 0.17074627 | -1.6409447 | 0.10080889 | 0.54608983 |
| ERH | 290.521875 | -0.338960183 | 0.24539259 | -1.3812975 | 0.1671875 | 0.62242897 |
| SLC39A9 | 602.20981 | -0.157793241 | 0.22966955 | -0.6870447 | 0.49205458 | 0.82080956 |
| SUSD6 | 1269.48808 | 0.262342352 | 0.21115332 | 1.24242588 | 0.21407947 | 0.65896863 |
| SYNJ2BP-CO | 31.2404581 | 0.599743303 | 0.46374853 | 1.2932511 | 0.19592423 | 0.64558745 |
| AL357153.2 | 409.928119 | 0.78589301 | 0.43345119 | 1.81310614 | 0.06981545 | 0.49378762 |
| MED6 | 319.532058 | -0.176456386 | 0.27214413 | -0.6483931 | 0.51673071 | 0.83258835 |
| TTC9 | 60.5532418 | -0.362021701 | 0.72880478 | -0.4967334 | 6.19E-01 | 8.75E-01 |
| MAP3K9 | 513.342952 | -0.314768483 | 0.38571106 | -0.8160733 | 0.41445823 | 0.78632759 |
| PCNX1 | 3131.11011 | -0.164547593 | 0.2869773 | -0.5733819 | 0.56638613 | 0.85215881 |
| AC005476.2 | 97.2388025 | 1.055511339 | 0.52227243 | 2.02099762 | 0.04328001 | 0.44144764 |
| SIPA1L1 | 7608.08673 | 0.077521025 | 0.32664003 | 0.23732861 | 0.81240187 | 0.94787133 |
| RGS6 | 153.140353 | -0.516293425 | 0.63984843 | -0.8068996 | 0.41972434 | 0.78798479 |
| DPF3 | 1563.16378 | 0.794599648 | 0.7434551 | 1.06879305 | 0.28516293 | 0.71015169 |
| DCAF4 | 201.739594 | -0.631009566 | 0.28426162 | -2.2198197 | 0.02643101 | 0.38507144 |
| ZFYVE1 | 796.716635 | -0.297921619 | 0.30195689 | -0.9866363 | 0.32382098 | 0.73572256 |
| RBM25 | 1393.54539 | -0.378402298 | 0.26157388 | -1.4466364 | 0.14799879 | 0.60605027 |
| PSEN1 | 1528.20456 | -0.111819959 | 0.18474136 | -0.6052784 | 0.54499402 | 0.84447581 |
| PAPLN | 1236.06082 | 0.478020891 | 0.48519682 | 0.98521028 | 0.32452079 | 0.73614611 |
| AC004846.2 | 41.2512103 | 0.643965157 | 0.45751765 | 1.4075198 | 0.15927331 | 0.61445519 |
| NUMB | 6051.6674 | 0.033375991 | 0.24412925 | 0.13671443 | 0.89125652 | 0.97234284 |
| HEATR4 | 218.101562 | 0.374156202 | 0.61945705 | 0.60400669 | 0.54583919 | 0.84460887 |
| AC005225.3 | 54.6114977 | 0.812822356 | 0.42154176 | 1.92821314 | 0.05382862 | 0.46711025 |
| ACOT2 | 73.4521155 | 0.289003571 | 0.6325014 | 0.45692163 | 0.64772738 | 0.88774616 |
| ACOT4 | 63.0432451 | 0.382861896 | 0.66218639 | 0.57817844 | 0.56314365 | 0.85057962 |
| ACOT6 | 94.5742963 | 0.566954438 | 0.79685364 | 0.71149131 | 0.47677983 | 0.81570932 |
| AC005225.2 | 131.067068 | -0.11770483 | 0.54303541 | -0.2167535 | 0.82840045 | 0.95305492 |
| DNAL1 | 445.468957 | -0.00620501 | 0.31236657 | -0.0198645 | 0.98415145 | 0.99523767 |
| PNMA1 | 104.445256 | 0.082538027 | 0.36850474 | 0.22398091 | 0.82277216 | 0.95117842 |
| ELMSAN1 | 636.90841 | -0.301339806 | 0.45334848 | -0.664698 | 0.5062437 | 0.82785522 |
| AC005520.2 | 48.0215558 | 0.726270983 | 0.86146183 | 0.8430681 | 0.39919036 | 0.78161291 |
| LINC02274 | 357.865495 | 0.047457967 | 0.65645871 | 0.07229391 | 0.94236801 | 0.98489601 |
| AC005480.1 | 254.008457 | 0.4742037 | 0.44283514 | 1.07083575 | 0.28424329 | 0.70898277 |
| FAM161B | 48.6670087 | 0.349136327 | 0.39317855 | 0.88798418 | 0.37454926 | 0.7697436 |
| COQ6 | 161.626406 | -0.004471938 | 0.32788002 | -0.0136389 | 0.98911803 | 0.99721497 |
| ENTPD5 | 1709.58905 | 0.358308085 | 0.49747161 | 0.72025836 | 0.47136594 | 0.81387913 |
| BBOF1 | 659.742644 | 0.51657472 | 0.40205639 | 1.28483151 | 0.19885117 | 0.64597062 |

| | | | | | | |
|---|---|---|---|---|---|---|
| ALDH6A1 | 1571.13859 | 0.278669635 | 0.47867829 | 0.58216477 | 5.60E-01 | 0.84944908 |
| LIN52 | 2570.91294 | 0.094576101 | 0.34708802 | 0.27248449 | 7.85E-01 | 9.39E-01 |
| ABCD4 | 998.862784 | 0.38787653 | 0.17957674 | 2.15994859 | 0.03077665 | 0.40446361 |
| NPC2 | 545.776669 | 0.297988782 | 0.32864447 | 0.90672083 | 0.36455445 | 0.76289279 |
| ISCA2 | 90.7743966 | 0.037750781 | 0.388705 | 0.09711936 | 0.92263161 | 0.97974213 |
| AREL1 | 490.681631 | -0.237776641 | 0.25153191 | -0.945314 | 0.34449858 | 0.75124763 |
| FCF1 | 227.726628 | -0.358544489 | 0.24082184 | -1.4888371 | 0.13653028 | 0.5942049 |
| YLPM1 | 1031.71594 | 0.149674453 | 0.16303987 | 0.91802362 | 0.3586065 | 0.75936492 |
| DLST | 671.020328 | 0.029788427 | 0.3192205 | 0.09331615 | 0.9256524 | 0.98097398 |
| RPS6KL1 | 41.8961641 | -0.934998649 | 0.94674553 | -0.9875924 | 0.32335233 | 0.73538915 |
| EIF2B2 | 162.685684 | 0.078876103 | 0.28879878 | 0.27311785 | 0.78476262 | 0.93884957 |
| AL049780.1 | 45.9408692 | 0.361993775 | 0.41335897 | 0.87573708 | 0.38117298 | 0.77210521 |
| MLH3 | 1021.38473 | 0.353438061 | 0.27196979 | 1.29954898 | 0.1937556 | 0.64548758 |
| ACYP1 | 299.560271 | 0.445922112 | 0.36799165 | 1.21177238 | 0.22559953 | 0.66688879 |
| ZC2HC1C | 125.001353 | 0.483240666 | 0.64186166 | 0.75287355 | 0.4515259 | 0.80384625 |
| NEK9 | 688.187619 | 0.141805923 | 0.20267189 | 0.69968223 | 0.48412577 | 0.81839266 |
| AL049780.2 | 91.1903065 | 0.793709671 | 0.43682642 | 1.81699101 | 0.06921849 | 0.49273866 |
| TMED10 | 751.165061 | 0.393747003 | 0.27264215 | 1.44418975 | 0.14868561 | 0.60622442 |
| FOS | 100.334284 | -1.132196605 | 0.83363047 | -1.3581517 | 0.17441556 | 0.62699479 |
| JDP2 | 233.377594 | 0.845523727 | 0.56185523 | 1.50487826 | 1.32E-01 | 5.91E-01 |
| FLVCR2 | 648.084344 | -0.177914938 | 0.3801313 | -0.4680355 | 0.63975921 | 0.88321992 |
| TTLL5 | 1341.62018 | 0.101993987 | 0.1733631 | 0.58832583 | 0.55631361 | 0.84861484 |
| ERG28 | 108.999483 | 0.771504616 | 0.32893415 | 2.34546827 | 0.0190032 | 0.34084276 |
| AF107885.2 | 60.8896849 | 0.472697874 | 0.63514195 | 0.74423973 | 0.45673146 | 0.80507423 |
| IFT43 | 1605.23844 | 0.650468232 | 0.25224409 | 2.5787254 | 0.00991656 | 0.26496347 |
| GPATCH2L | 2429.43883 | -0.151696713 | 0.34019848 | -0.4459065 | 0.65566479 | 0.89053559 |
| ESRRB | 192.412666 | -0.700999353 | 0.76730608 | -0.913585 | 0.36093493 | 0.76009788 |
| VASH1-AS1 | 43.8082463 | -0.130860027 | 0.45424922 | -0.2880798 | 0.77328565 | 0.93400456 |
| ANGEL1 | 190.824078 | -0.05841502 | 0.33187457 | -0.1760154 | 0.86028188 | 0.96283855 |
| IRF2BPL | 166.42571 | 0.744401892 | 0.45358122 | 1.6411656 | 0.10076304 | 0.54608983 |
| POMT2 | 334.857313 | -0.434075871 | 0.40807898 | -1.0637055 | 0.28746212 | 0.71118494 |
| GSTZ1 | 99.0953161 | 0.157216335 | 0.29998778 | 0.5240758 | 0.60022582 | 0.86740066 |
| TMED8 | 621.388375 | 0.243024393 | 0.18353266 | 1.32414796 | 0.18545391 | 0.6358221 |
| SAMD15 | 52.8022632 | -0.308605034 | 0.46569954 | -0.6626698 | 0.50754204 | 0.82858823 |
| NOXRED1 | 146.730004 | -0.302726522 | 0.42589441 | -0.7108018 | 0.47720706 | 0.81570932 |
| VIPAS39 | 249.887032 | 0.276553397 | 0.33869438 | 0.81652785 | 0.4141983 | 0.78632759 |
| AHSA1 | 241.15774 | -0.614462688 | 0.33707132 | -1.8229456 | 0.06831163 | 0.4912631 |
| SPTLC2 | 1267.58198 | -0.367844464 | 0.24260532 | -1.5162259 | 0.12946224 | 0.58817997 |
| ALKBH1 | 372.168373 | -0.431499337 | 0.21464659 | -2.0102781 | 0.04440177 | 0.44360789 |
| SLIRP | 380.405137 | 0.11184526 | 0.33562537 | 0.33324436 | 0.73894984 | 0.92099078 |
| SNW1 | 441.279256 | -0.368068977 | 0.22859229 | -1.6101548 | 0.10736406 | 0.55579103 |
| ADCK1 | 294.32934 | -0.775818545 | 0.33556426 | -2.3119821 | 0.02077867 | 0.35313211 |
| NRXN3 | 198.672606 | 0.135042983 | 0.40131242 | 0.33650338 | 0.73649131 | 0.92019178 |
| CEP128 | 393.367664 | -0.833933508 | 0.32718391 | -2.5488219 | 0.01080875 | 0.27538203 |
| TSHR | 68.1548141 | 1.003808442 | 0.65860321 | 1.52414752 | 0.12747186 | 0.58793678 |
| AC007262.2 | 339.485757 | 0.10965691 | 0.46404317 | 0.23630756 | 0.81319402 | 0.94839194 |
| GTF2A1 | 373.153183 | 0.245232899 | 0.3459783 | 0.70881006 | 0.47844235 | 0.81596289 |
| STON2 | 3030.62556 | -0.862955729 | 0.65369025 | -1.3201294 | 0.18679181 | 0.6378941 |
| SEL1L | 755.914576 | -0.180418923 | 0.31780329 | -0.5677063 | 0.57023443 | 0.85329731 |
| GALC | 174.765502 | -0.092375789 | 0.41936643 | -0.2202746 | 0.82565727 | 0.95192757 |
| KCNK10 | 361.614326 | -0.365417521 | 0.50436021 | -0.7245169 | 4.69E-01 | 8.12E-01 |
| SPATA7 | 826.735608 | 0.421111399 | 0.27502372 | 1.53118211 | 0.12572439 | 0.58564339 |
| PTPN21 | 543.228546 | -0.234774457 | 0.31188852 | -0.7527512 | 0.45159944 | 0.80384625 |
| ZC3H14 | 1279.83759 | -0.022897557 | 0.16668088 | -0.1373736 | 0.89073548 | 0.97230741 |

| | | | | | | |
|---|---|---|---|---|---|---|
| EML5 | 526.244337 | -2.157447801 | 0.97247449 | -2.2185135 | 0.02651984 | 0.38507144 |
| TTC8 | 528.811151 | 0.167315973 | 0.33856677 | 0.494189 | 0.62117271 | 0.87641939 |
| FOXN3 | 1601.44661 | -0.210778369 | 0.23138953 | -0.9109244 | 0.36233521 | 0.76115383 |
| EFCAB11 | 601.605352 | -0.041943418 | 0.53086081 | -0.0790102 | 0.93702451 | 0.98435387 |
| TDP1 | 315.947836 | -0.321247234 | 0.22448997 | -1.4310093 | 0.15242756 | 0.61001662 |
| KCNK13 | 73.7388753 | -0.196931136 | 0.60354011 | -0.3262934 | 0.74420241 | 0.92362993 |
| PSMC1 | 370.610555 | -0.433215346 | 0.34012596 | -1.2736909 | 0.20277297 | 0.64881821 |
| NRDE2 | 469.145312 | -0.028253346 | 0.24831758 | -0.1137791 | 0.90941292 | 0.97680747 |
| CALM1 | 1419.9246 | -0.231940017 | 0.29352773 | -0.7901809 | 0.42942211 | 0.79037914 |
| TTC7B | 2136.67736 | 0.452477575 | 0.34576693 | 1.30862016 | 0.19066305 | 0.64281105 |
| RPS6KA5 | 778.619632 | -0.004044911 | 0.26077975 | -0.0155108 | 9.88E-01 | 0.99648575 |
| DGLUCY | 1811.32719 | 0.356996933 | 0.41066596 | 0.86931221 | 0.38467639 | 0.77317162 |
| GPR68 | 31.6114291 | 0.540012074 | 0.81259511 | 0.66455245 | 0.50633679 | 0.82790867 |
| CCDC88C | 688.95688 | -0.141838815 | 0.26876474 | -0.5277434 | 0.59767746 | 0.86627523 |
| PPP4R3A | 1520.47188 | -0.246597257 | 0.16601799 | -1.4853647 | 0.13744725 | 0.59474076 |
| CATSPERB | 488.832303 | 0.043980522 | 0.46528268 | 0.0945243 | 0.92469268 | 0.9804664 |
| TC2N | 444.092142 | -0.201236427 | 0.3983874 | -0.5051275 | 0.61346929 | 0.87222135 |
| FBLN5 | 314.083398 | 0.021537867 | 0.28600831 | 0.07530504 | 0.93997201 | 0.98439739 |
| TRIP11 | 1289.77301 | 0.168452656 | 0.13656617 | 1.23348743 | 0.21739397 | 0.66032006 |
| ATXN3 | 701.142179 | -0.448344411 | 0.23858905 | -1.8791492 | 0.06022413 | 0.48004006 |
| NDUFB1 | 394.393558 | -0.121058486 | 0.26885396 | -0.450276 | 0.65251144 | 0.8894167 |
| CPSF2 | 460.257435 | 0.014511475 | 0.23641972 | 0.06138014 | 0.95105647 | 0.98731502 |
| RIN3 | 454.048846 | -0.041708285 | 0.48450091 | -0.086085 | 0.93139881 | 0.9827462 |
| LGMN | 1462.15578 | -0.147868073 | 0.63428439 | -0.2331258 | 0.8156637 | 0.94906482 |
| GOLGA5 | 450.372568 | -0.375289496 | 0.27876944 | -1.3462361 | 0.17822638 | 0.6285261 |
| ITPK1 | 729.315547 | -0.178007622 | 0.37096485 | -0.4798504 | 0.63133379 | 0.88123095 |
| MOAP1 | 155.37637 | -0.39380227 | 0.43374451 | -0.907913 | 0.3639242 | 0.76272056 |
| AL110118.2 | 70.1538273 | -0.434155844 | 0.45965721 | -0.9445209 | 0.34490352 | 0.75133476 |
| GON7 | 78.9924916 | -0.260748835 | 0.27702614 | -0.9412427 | 0.34658049 | 0.75166079 |
| UBR7 | 85.9155215 | 0.125754101 | 0.38907142 | 0.32321598 | 0.74653168 | 0.92448528 |
| BTBD7 | 3609.81349 | 0.041587594 | 0.3051719 | 0.13627596 | 0.89160313 | 0.97234681 |
| UNC79 | 1127.53282 | 2.80463245 | 0.66790471 | 4.19915058 | 2.68E-05 | 0.01647703 |
| DDX24 | 885.381101 | -0.049056177 | 0.24563613 | -0.1997108 | 0.8417068 | 0.95681213 |
| IFI27L1 | 108.786193 | -0.127806855 | 0.37468396 | -0.3411058 | 0.73302397 | 0.91952673 |
| IFI27 | 631.52743 | -3.844112067 | 1.1408773 | -3.3694352 | 0.00075322 | 0.09084673 |
| IFI27L2 | 60.1510806 | 0.152777439 | 0.75240937 | 0.20305095 | 0.83909521 | 0.95558973 |
| PPP4R4 | 76.9131948 | 1.618854668 | 0.66407476 | 2.43775966 | 0.0147786 | 0.30832605 |
| SERPINA6 | 51.1464132 | -0.139728938 | 0.84704011 | -0.1649614 | 0.86897435 | 0.96555054 |
| SERPINA1 | 3810.63588 | -0.549276817 | 0.92765424 | -0.5921137 | 0.55377443 | 0.84727776 |
| DICER1 | 1629.23956 | 0.022861304 | 0.18710511 | 0.12218428 | 0.90275308 | 0.97517822 |
| DICER1-AS1 | 191.548718 | 0.629317318 | 0.50061853 | 1.25707956 | 0.20872483 | 0.65401126 |
| CLMN | 2994.16927 | -0.077017074 | 0.47261403 | -0.1629598 | 0.87055011 | 0.9657206 |
| SYNE3 | 50.0728879 | 0.118204606 | 1.01622067 | 0.11631785 | 0.90740064 | 0.97673359 |
| SNHG10 | 48.6657044 | -0.580801307 | 0.34892342 | -1.6645524 | 0.09600209 | 0.53870645 |
| GLRX5 | 245.172336 | -0.240439364 | 0.36390392 | -0.6607221 | 0.50879054 | 0.82874455 |
| ATG2B | 1066.65349 | 0.135804358 | 0.20770228 | 0.65384145 | 0.51321396 | 0.83073024 |
| GSKIP | 198.254012 | -0.21559243 | 0.2757291 | -0.7818994 | 0.43427368 | 0.79442004 |
| AK7 | 135.460372 | 1.104230321 | 0.53209172 | 2.07526312 | 0.03796216 | 0.42339465 |
| PAPOLA | 1580.41634 | -0.049700973 | 0.23715087 | -0.2095753 | 0.83399913 | 0.95393659 |
| VRK1 | 181.810955 | -0.031442735 | 0.37157254 | -0.0846207 | 0.93256293 | 0.98310938 |
| LINC01550 | 311.502059 | 1.19458605 | 0.6496892 | 1.83870386 | 0.06595875 | 0.48798003 |
| SETD3 | 3115.6951 | 0.143215132 | 0.33154681 | 0.43196052 | 0.66577011 | 0.89378862 |
| CCNK | 451.505785 | -0.306949651 | 0.20688932 | -1.4836418 | 0.13790397 | 0.59474076 |
| CCDC85C | 417.091827 | 0.396198319 | 0.3347725 | 1.18348527 | 0.23661688 | 0.67372405 |

| | | | | | | |
|---|---|---|---|---|---|---|
| AL160313.1 | 47.2067684 | -0.145107102 | 0.6675562 | -0.2173706 | 0.82791953 | 0.95270465 |
| CYP46A1 | 111.024878 | 1.30959307 | 0.65663607 | 1.9943971 | 0.04610867 | 0.44508419 |
| EML1 | 207.295553 | 0.223560848 | 0.44233173 | 0.50541445 | 0.61326777 | 0.87222135 |
| EVL | 1490.97705 | 0.265684647 | 0.39580581 | 0.67124999 | 0.50206129 | 0.82638105 |
| AL133523.1 | 78.0989979 | -0.098151438 | 0.28768634 | -0.3411752 | 0.73297171 | 0.91952673 |
| YY1 | 1225.30406 | -0.199220186 | 0.25834912 | -0.7711278 | 0.44063118 | 0.79826598 |
| SLC25A29 | 1062.51233 | 0.549488353 | 0.41958886 | 1.30958755 | 0.1903354 | 0.64278842 |
| WARS | 662.192578 | 0.239534952 | 0.62655014 | 0.38230771 | 0.70223313 | 0.90837679 |
| AL157871.4 | 129.028215 | 0.832704299 | 0.74023272 | 1.12492231 | 0.26062196 | 0.69280433 |
| WDR25 | 520.015738 | -0.404700552 | 0.41970316 | -0.9642542 | 0.33491848 | 0.74156916 |
| MEG3 | 162.926399 | 1.380868257 | 1.08666992 | 1.27073385 | 0.20382335 | 0.64926949 |
| MEG8 | 117.522372 | 0.951786449 | 1.13081116 | 0.8416847 | 0.39996447 | 0.78186732 |
| PPP2R5C | 1444.9259 | 0.424260077 | 0.20690153 | 2.05054101 | 0.04031167 | 0.43086056 |
| AL118558.3 | 33.3537952 | -0.077647989 | 0.67639531 | -0.1147968 | 0.90860622 | 0.97680747 |
| DYNC1H1 | 1517.48975 | -0.391335779 | 0.2803691 | -1.3957878 | 0.1627784 | 0.61960665 |
| HSP90AA1 | 3222.33418 | -0.618010716 | 0.41599117 | -1.4856342 | 0.1373759 | 0.59474076 |
| WDR20 | 1536.21266 | -0.236183095 | 0.25468385 | -0.9273579 | 0.35374072 | 0.75682564 |
| MOK | 1051.75746 | -0.121240358 | 0.35108735 | -0.3453282 | 0.72984766 | 0.91834786 |
| ZNF839 | 315.208437 | 0.141966595 | 0.30629727 | 0.46349286 | 0.64301113 | 0.88473736 |
| CINP | 129.410262 | -0.361560016 | 0.27908759 | -1.2955073 | 0.19514528 | 0.64558745 |
| TECPR2 | 633.569586 | -0.140005025 | 0.32454163 | -0.4313931 | 0.66618256 | 0.89417982 |
| ANKRD9 | 131.949911 | 0.021511576 | 0.40101022 | 0.05364346 | 0.95721923 | 0.98861313 |
| RCOR1 | 1128.80239 | -0.394874719 | 0.223385 | -1.7676868 | 0.07711327 | 0.51013111 |
| TRAF3 | 1244.90304 | -0.161590178 | 0.36344982 | -0.4446011 | 0.65660808 | 0.89053559 |
| AMN | 1254.78135 | -0.399312138 | 0.48982988 | -0.8152058 | 0.41495454 | 0.78632759 |
| CDC42BPB | 1575.95514 | -0.612928817 | 0.3133182 | -1.9562503 | 0.05043568 | 0.45843622 |
| AL117209.1 | 50.7284778 | 0.105980611 | 0.49999152 | 0.21196482 | 0.83213448 | 0.95325673 |
| EXOC3L4 | 348.646426 | 0.922782291 | 0.68776507 | 1.34171149 | 0.17968956 | 0.63089262 |
| TNFAIP2 | 649.691943 | 0.467572411 | 0.6524503 | 0.71664065 | 0.47359586 | 0.81479253 |
| AL161669.3 | 108.232603 | 0.457843491 | 0.60351762 | 0.7586249 | 0.44807697 | 0.80223612 |
| EIF5 | 1029.84488 | -0.543239379 | 0.40189857 | -1.3516828 | 0.17647681 | 0.62709281 |
| MARK3 | 3223.09474 | 0.065250556 | 0.20498303 | 0.31832174 | 0.7502409 | 0.92569556 |
| CKB | 562.834787 | 0.664582045 | 0.55958782 | 1.18762779 | 0.23498009 | 0.67119484 |
| TRMT61A | 62.5676423 | -0.765316234 | 0.65379187 | -1.1705808 | 0.2417673 | 0.68033349 |
| BAG5 | 320.017664 | -0.230090062 | 0.27079608 | -0.8496802 | 3.96E-01 | 7.79E-01 |
| KLC1 | 723.938025 | -0.191820882 | 0.30904332 | -0.6206925 | 0.53480195 | 0.83954482 |
| AL049840.1 | 81.1619088 | -0.146535792 | 0.74101402 | -0.1977504 | 0.84324038 | 0.95716002 |
| ZFYVE21 | 238.88069 | 0.358300495 | 0.24673936 | 1.45214161 | 0.14646223 | 0.60383734 |
| PPP1R13B | 2468.46581 | -0.331591689 | 0.25108147 | -1.3206538 | 0.18661684 | 0.6378941 |
| AL132712.1 | 92.597408 | -0.473121208 | 0.6453079 | -0.7331713 | 0.46345398 | 0.8082072 |
| ATP5MPL | 473.851834 | -0.179684011 | 0.3194818 | -0.5624233 | 0.57382765 | 0.85447394 |
| TDRD9 | 262.949531 | -0.596026584 | 0.59154623 | -1.007574 | 0.31365903 | 0.72717675 |
| ASPG | 211.897432 | 0.042448156 | 0.59544682 | 0.0712879 | 9.43E-01 | 9.85E-01 |
| INF2 | 325.837435 | -0.336504088 | 0.28804751 | -1.1682243 | 2.43E-01 | 0.68118603 |
| ADSSL1 | 60.9173009 | 0.113606576 | 0.76002348 | 0.14947772 | 0.88117669 | 0.96773407 |
| SIVA1 | 221.300672 | -0.258118786 | 0.31320872 | -0.8241111 | 0.40987643 | 0.78542936 |
| AKT1 | 269.391649 | 0.052200343 | 0.2929878 | 0.17816558 | 0.85859294 | 0.9626966 |
| AL583810.1 | 60.075253 | -0.11853446 | 0.36913034 | -0.3211182 | 0.74812084 | 0.92532868 |
| CEP170B | 243.162514 | -0.539364313 | 0.35057596 | -1.5385091 | 0.12392418 | 0.58372123 |
| AHNAK2 | 80.0945731 | -0.628695631 | 0.94343877 | -0.6663873 | 5.05E-01 | 0.82715271 |
| CLBA1 | 133.469766 | -0.466295535 | 0.23190587 | -2.0107104 | 0.04435606 | 0.44360789 |
| NUDT14 | 116.236141 | 0.387617516 | 0.53415301 | 0.72566756 | 0.46804259 | 0.81187798 |
| BRF1 | 407.65407 | -0.146878231 | 0.29402848 | -0.4995374 | 0.61740083 | 0.87435665 |
| BTBD6 | 71.0629083 | -0.318056653 | 0.47516798 | -0.6693562 | 0.50326826 | 0.82652042 |

| | | | | | | |
|---|---|---|---|---|---|---|
| PACS2 | 728.440781 | -0.230599971 | 0.30426906 | -0.7578818 | 0.44852177 | 0.80259633 |
| TEX22 | 95.8267085 | 0.257084661 | 0.45728978 | 0.56219202 | 0.5739852 | 0.85448213 |
| MTA1 | 282.868248 | -0.145044826 | 0.38163711 | -0.3800595 | 0.70390122 | 0.90919468 |
| CRIP2 | 108.300308 | 0.959686551 | 0.72864805 | 1.31707832 | 0.18781237 | 0.63974756 |
| AC087386.1 | 152.480964 | -0.660229942 | 0.42796706 | -1.5427121 | 0.12290065 | 0.58216759 |
| AC026495.1 | 89.3075214 | 0.869411846 | 0.9689471 | 0.89727483 | 3.70E-01 | 7.67E-01 |
| NIPA1 | 466.592591 | -0.108621335 | 0.2464181 | -0.440801 | 0.6593571 | 0.89068767 |
| NIPA2 | 442.852919 | -0.344922336 | 0.405497 | -0.8506162 | 0.39498256 | 0.77922078 |
| CYFIP1 | 1067.37423 | -0.295490779 | 0.32220944 | -0.9170767 | 0.35910243 | 0.75936492 |
| TUBGCP5 | 451.492329 | -0.632229298 | 0.29348834 | -2.1541888 | 0.03122537 | 0.407237 |
| PWRN1 | 46.813334 | 0.324281446 | 0.77924837 | 0.41614645 | 0.67730284 | 0.89864474 |
| AC124312.1 | 2058.61192 | 0.533737771 | 0.23347396 | 2.28606978 | 0.02225018 | 0.36178312 |
| PWAR6 | 429.578663 | 0.801570886 | 0.36820955 | 2.17694217 | 0.02948488 | 0.39817664 |
| UBE3A | 2266.11686 | -0.189010106 | 0.15081662 | -1.2532446 | 0.21011672 | 0.65441616 |
| ATP10A | 121.830238 | -0.426686601 | 0.89935326 | -0.4744372 | 0.6351882 | 0.88238823 |
| GABRB3 | 873.276014 | 0.927533051 | 0.48629534 | 1.90734512 | 0.05647591 | 0.47259787 |
| GABRG3 | 75.3058465 | -0.313678575 | 0.63026246 | -0.4976952 | 0.61869892 | 0.87502492 |
| HERC2 | 2330.28149 | -0.071909218 | 0.19164475 | -0.3752214 | 0.70749579 | 0.91012042 |
| GOLGA8M | 219.655755 | -0.205285893 | 1.16466138 | -0.1762623 | 0.86008788 | 0.96283855 |
| APBA2 | 73.7602826 | 0.928460723 | 0.84508964 | 1.09865354 | 0.27191921 | 0.70056282 |
| FAM189A1 | 1356.45728 | 1.185501097 | 0.35068119 | 3.38056651 | 0.00072337 | 0.09013794 |
| AC022613.1 | 43.956903 | 0.253598275 | 0.54433436 | 0.46588695 | 0.64129642 | 0.88403746 |
| TJP1 | 2264.05989 | -0.232393959 | 0.31728141 | -0.7324538 | 0.46389166 | 0.80866883 |
| AC091057.2 | 43.7943325 | 0.463430407 | 0.45064506 | 1.02837121 | 0.30377525 | 0.72082426 |
| AC091057.6 | 65.933933 | -0.036061561 | 0.3486336 | -0.1034368 | 0.91761627 | 0.97828877 |
| FAN1 | 437.035413 | 0.173180799 | 0.33171188 | 0.52208198 | 0.60161325 | 0.86757631 |
| MTMR10 | 1836.67248 | -0.043725041 | 0.26681519 | -0.1638776 | 0.86982748 | 0.96564168 |
| LINC02352 | 38.0599499 | -0.633031106 | 0.57237133 | -1.1059798 | 0.26873528 | 0.69961625 |
| KLF13 | 815.506514 | 0.102383529 | 0.39957116 | 0.25623353 | 0.79777052 | 0.94316375 |
| OTUD7A | 1536.67821 | -0.203139348 | 0.42342257 | -0.4797556 | 0.63140119 | 0.88123095 |
| CHRNA7 | 82.1994472 | -0.402176535 | 0.39327575 | -1.0226324 | 0.30648168 | 0.72190689 |
| LINC02256 | 1369.08323 | 0.796266009 | 0.65200345 | 1.22126042 | 0.22198744 | 0.66334898 |
| AC123768.3 | 207.070494 | 0.088131628 | 0.65342286 | 0.13487687 | 0.89270923 | 0.9724379 |
| GOLGA8N | 147.347542 | 0.261294708 | 0.71934985 | 0.36323731 | 0.71642762 | 0.91298968 |
| FMN1 | 484.255051 | 0.171988116 | 0.56512132 | 0.3043384 | 0.76087009 | 0.92989212 |
| RYR3 | 75.0409893 | -0.067431246 | 0.43411834 | -0.1553292 | 0.87656181 | 0.966227 |
| AC010809.2 | 58.2341333 | -0.513448201 | 0.52486931 | -0.9782401 | 0.3279556 | 0.73806934 |
| AC010809.1 | 107.770887 | -0.835128024 | 0.41451928 | -2.0146904 | 0.0439371 | 0.44347996 |
| AVEN | 1272.40349 | -0.782098993 | 0.296645 | -2.6364813 | 0.00837708 | 0.25016267 |
| CHRM5 | 142.762211 | -0.099149346 | 0.54307107 | -0.1825716 | 0.85513418 | 0.96171023 |
| EMC7 | 236.430892 | 0.106073695 | 0.31999203 | 0.33148856 | 0.74027548 | 0.92173153 |
| PGBD4 | 48.2655318 | -0.745390581 | 0.34044414 | -2.1894652 | 0.02856305 | 0.39280225 |
| KATNBL1 | 1198.49257 | -0.196590085 | 0.24328338 | -0.8080703 | 0.41905009 | 0.78770153 |
| EMC4 | 111.513078 | -0.604759587 | 0.34727621 | -1.7414369 | 0.08160703 | 0.51593841 |
| SLC12A6 | 2415.43344 | 0.229318632 | 0.35198377 | 0.65150343 | 5.15E-01 | 8.32E-01 |
| NOP10 | 383.459122 | -0.582873893 | 0.46108993 | -1.2641219 | 0.20618627 | 0.65224906 |
| LPCAT4 | 110.160245 | -0.151940117 | 0.41139014 | -0.3693334 | 0.71187924 | 0.91161746 |
| GOLGA8A | 464.598913 | -1.420245405 | 0.67089554 | -2.1169397 | 0.03426496 | 0.41505357 |
| GOLGA8B | 896.011056 | 0.405288255 | 0.35952822 | 1.12727801 | 0.25962496 | 0.69242517 |
| AQR | 786.177521 | -0.087120587 | 0.21003545 | -0.4147899 | 0.6782957 | 0.89913305 |
| ZNF770 | 466.73721 | 0.145656837 | 0.29957339 | 0.4862142 | 0.62681531 | 0.87890915 |
| AC021231.1 | 633.407329 | 1.408206996 | 0.63631921 | 2.2130512 | 0.02689412 | 0.38702446 |
| DPH6 | 2465.19745 | 0.331329411 | 0.55589591 | 0.59602779 | 0.55115666 | 0.8466552 |
| DPH6-DT | 1699.3874 | 0.805814345 | 0.67532995 | 1.19321577 | 0.23278488 | 0.66978724 |

| | | | | | | |
|---|---|---|---|---|---|---|
| C15orf41 | 408.422395 | -0.381782317 | 0.26289284 | -1.4522355 | 0.14643613 | 0.60383734 |
| MEIS2 | 143.233682 | 0.422886094 | 0.60080595 | 0.70386469 | 0.48151704 | 0.81755903 |
| SPRED1 | 1304.98641 | 0.737756594 | 0.32418358 | 2.2757371 | 0.02286175 | 0.36631766 |
| FAM98B | 266.363145 | 0.006806283 | 0.22644599 | 0.03005698 | 0.97602161 | 0.99429832 |
| RASGRP1 | 471.908559 | 0.192444842 | 0.27029471 | 0.71198152 | 0.47647622 | 0.81570932 |
| C15orf53 | 232.0553 | 0.174818416 | 0.81319457 | 0.21497735 | 0.82978497 | 0.95325673 |
| AC013652.1 | 731.194052 | 0.861511879 | 0.65592176 | 1.31343695 | 0.18903575 | 0.6413893 |
| THBS1 | 484.958462 | -0.745274678 | 0.55058379 | -1.3536081 | 1.76E-01 | 0.62699479 |
| FSIP1 | 205.27267 | 0.286567234 | 0.44572025 | 0.6429307 | 0.52026907 | 0.833897 |
| GPR176 | 226.110332 | -0.321521735 | 0.53016529 | -0.6064557 | 0.54421222 | 0.84416152 |
| EIF2AK4 | 707.299822 | -0.319071134 | 0.28252033 | -1.1293741 | 0.25874007 | 0.69241621 |
| SRP14 | 981.745714 | -0.285598889 | 0.39378966 | -0.7252575 | 4.68E-01 | 0.8121087 |
| SRP14-AS1 | 168.297708 | 0.392343907 | 0.42723044 | 0.91834259 | 0.35843954 | 0.75936492 |
| BMF | 59.9378141 | 2.322340904 | 0.95531379 | 2.43097182 | 0.01505839 | 0.30991786 |
| CCDC9B | 58.9436498 | 0.354182811 | 0.56181733 | 0.63042343 | 0.52841758 | 0.83761342 |
| KNSTRN | 69.9983654 | 0.38055919 | 0.4429906 | 0.85906831 | 0.39030283 | 0.77569824 |
| IVD | 474.57345 | 0.671206237 | 0.42796679 | 1.56836056 | 0.11679701 | 0.57350419 |
| BAHD1 | 211.218688 | 0.048180811 | 0.30856853 | 0.15614299 | 0.87592032 | 0.96617876 |
| AC091045.1 | 63.7070784 | 0.609863936 | 0.61437541 | 0.99265681 | 0.32087723 | 0.73233537 |
| CCDC32 | 413.871515 | -0.211775572 | 0.40635547 | -0.5211584 | 0.60225642 | 0.8677246 |
| RPUSD2 | 42.9910622 | -0.370736896 | 0.365507 | -1.0143086 | 0.31043552 | 0.72498238 |
| KNL1 | 164.531749 | 0.381325624 | 0.64467755 | 0.59149823 | 0.55418664 | 0.84727776 |
| RMDN3 | 319.45111 | -0.279213265 | 0.29603455 | -0.943178 | 0.34558988 | 0.75166079 |
| GCHFR | 289.152563 | 0.463612028 | 0.62715786 | 0.73922701 | 0.45976916 | 0.8069452 |
| DNAJC17 | 498.188196 | 0.124959308 | 0.22551064 | 0.55411713 | 0.57949869 | 0.85816191 |
| ZFYVE19 | 263.975243 | -0.136699068 | 0.34397198 | -0.3974134 | 0.69106265 | 0.90473488 |
| SPINT1-AS1 | 112.23622 | 0.841078474 | 0.4050075 | 2.07669849 | 0.0378294 | 0.42294252 |
| SPINT1 | 510.763813 | 0.237886601 | 0.33421014 | 0.71178751 | 4.77E-01 | 0.81570932 |
| VPS18 | 155.261635 | -0.488234458 | 0.438512 | -1.113389 | 0.26554138 | 0.69700797 |
| INO80 | 1196.36707 | -0.637116184 | 0.29612757 | -2.1514923 | 0.03143736 | 0.40844733 |
| AC020661.3 | 38.8015373 | -0.521574924 | 0.56083987 | -0.929989 | 0.35237676 | 0.75652927 |
| INO80-AS1 | 38.8990621 | 0.225847098 | 0.56351579 | 0.4007822 | 0.68858048 | 0.90310859 |
| CHP1 | 763.055623 | 0.033518357 | 0.40229873 | 0.08331709 | 0.93359942 | 0.98355048 |
| OIP5 | 44.0402506 | 0.155337696 | 0.48105785 | 0.32290855 | 0.7467645 | 0.92448528 |
| NUSAP1 | 131.225655 | -0.635508116 | 0.61378419 | -1.0353934 | 0.30048521 | 0.71879123 |
| NDUFAF1 | 148.576076 | -0.100641017 | 0.26206658 | -0.3840284 | 0.70095737 | 0.90782041 |
| RTF1 | 692.05586 | -0.389908529 | 0.32581604 | -1.1967137 | 0.23141816 | 0.66878058 |
| RPAP1 | 144.021283 | -0.036388765 | 0.29448736 | -0.1235665 | 0.90165854 | 0.97491785 |
| MGA | 1500.49729 | -0.57108817 | 0.20563358 | -2.7772126 | 0.00548273 | 0.22038648 |
| MAPKBP1 | 474.046807 | -0.790408602 | 0.48332228 | -1.6353655 | 0.10197244 | 0.54813186 |
| AC020659.1 | 70.8355079 | 0.625864196 | 0.92740062 | 0.6748585 | 0.49976567 | 0.82476476 |
| EHD4 | 202.745127 | 0.071356731 | 0.29155238 | 0.24474755 | 8.07E-01 | 9.46E-01 |
| VPS39 | 686.120829 | -0.153059986 | 0.11588791 | -1.3207589 | 0.18658176 | 0.6378941 |
| TMEM87A | 1199.01451 | -0.206388157 | 0.2675135 | -0.7715056 | 0.44040732 | 0.79826598 |
| GANC | 929.6239 | 0.575518889 | 0.45868067 | 1.2547267 | 0.20957799 | 0.65441616 |
| ZNF106 | 798.937189 | 0.007145912 | 0.2222804 | 0.03214819 | 0.97435388 | 0.99377398 |
| SNAP23 | 843.033459 | -0.288126706 | 0.4304189 | -0.66941 | 0.50323399 | 0.82652042 |
| LRRC57 | 99.0846381 | -0.120091844 | 0.31129055 | -0.385787 | 0.69965444 | 0.90721166 |
| HAUS2 | 168.449388 | -0.338322742 | 0.26873014 | -1.2589683 | 0.20804177 | 0.65352038 |
| STARD9 | 167.55029 | -0.21249604 | 0.30822266 | -0.6894238 | 0.49055661 | 0.81989696 |
| CDAN1 | 95.4160469 | -0.112790692 | 0.33599532 | -0.3356913 | 0.7371037 | 0.92030854 |
| TTBK2 | 1276.7141 | 0.131322387 | 0.25451803 | 0.51596497 | 0.60587887 | 0.86915559 |
| UBR1 | 1252.84708 | 0.001958075 | 0.16555073 | 0.01182765 | 0.99056312 | 0.99737132 |
| AC068724.3 | 202.539658 | 0.204043075 | 0.3679736 | 0.55450465 | 0.57923352 | 0.85805186 |

| | | | | | | |
|---|---|---|---|---|---|---|
| TMEM62 | 330.122659 | -0.210293471 | 0.27119854 | -0.7754226 | 0.43809 | 0.79696434 |
| CCNDBP1 | 143.296982 | 0.273447748 | 0.27434959 | 0.9967128 | 0.31890393 | 0.73097261 |
| ADAL | 265.045628 | 0.078161957 | 0.41347805 | 0.18903532 | 0.85006513 | 0.9597103 |
| ZSCAN29 | 115.862965 | -0.335059363 | 0.33137137 | -1.0111295 | 0.31195446 | 0.72610662 |
| TUBGCP4 | 271.110495 | -0.194753688 | 0.2680204 | -0.7266375 | 0.46744802 | 0.81166836 |
| TP53BP1 | 953.998385 | -0.039694485 | 0.27730953 | -0.1431414 | 0.88617848 | 0.97019556 |
| PPIP5K1 | 240.58254 | 0.275846617 | 0.2851303 | 0.96744057 | 0.33332384 | 0.74013767 |
| AC011330.3 | 43.4649305 | 0.323262687 | 0.65591134 | 0.4928451 | 0.62212204 | 0.87713418 |
| CATSPER2 | 297.524691 | 0.523318602 | 0.55924892 | 0.93575255 | 0.34940061 | 0.7542101 |
| PDIA3 | 899.074482 | 0.041921978 | 0.3800075 | 0.11031882 | 0.91215654 | 0.977179 |
| AC018512.1 | 48.0709718 | -0.284246298 | 0.53421663 | -0.5320806 | 0.59467018 | 0.86499581 |
| SERF2 | 2639.52162 | -0.531562709 | 0.38589717 | -1.3774724 | 0.16836624 | 0.62304975 |
| MFAP1 | 184.577796 | -0.205981331 | 0.23609072 | -0.8724669 | 0.38295373 | 0.77317162 |
| FRMD5 | 1223.89978 | 1.117238648 | 0.53868387 | 2.0740154 | 0.03807788 | 0.42339465 |
| CASC4 | 2429.14538 | 0.033129987 | 0.24478775 | 0.13534168 | 0.89234173 | 0.97237481 |
| AC025043.1 | 83.4352055 | 0.37899661 | 0.44543235 | 0.85085112 | 0.39485206 | 0.77907546 |
| CTDSPL2 | 1416.47519 | -0.25910305 | 0.21564945 | -1.2015011 | 0.22955689 | 0.66705629 |
| EIF3J-DT | 300.301526 | 0.301565028 | 0.33234226 | 0.90739296 | 0.36419903 | 0.76273126 |
| EIF3J | 672.520903 | -0.595447882 | 0.39291581 | -1.5154592 | 0.12965615 | 0.58838111 |
| SPG11 | 2096.16799 | -0.05466772 | 0.17902637 | -0.3053613 | 0.76009102 | 0.92956527 |
| PATL2 | 72.6886919 | -0.492062132 | 0.68651314 | -0.7167556 | 0.47352492 | 0.81478861 |
| B2M | 4508.29682 | -0.300827049 | 0.53289415 | -0.5645156 | 0.5724033 | 0.85408471 |
| TRIM69 | 125.595411 | 0.257317912 | 0.47176441 | 0.54543732 | 0.58545278 | 0.86077718 |
| SORD | 1217.28468 | 1.048835287 | 0.66986982 | 1.56573001 | 0.11741184 | 0.57350419 |
| DUOX1 | 138.718288 | 0.069728819 | 0.52236548 | 0.13348665 | 0.89380852 | 0.97272276 |
| SHF | 33.8251199 | 0.473452045 | 0.51439928 | 0.92039795 | 0.35736484 | 0.75903693 |
| AC051619.5 | 62.6413644 | 1.885912056 | 0.88898174 | 2.12142947 | 0.03388568 | 0.41378378 |
| SLC28A2 | 44.6669606 | 2.156559028 | 0.92405738 | 2.33379341 | 0.01960654 | 0.34523522 |
| GATM | 2023.82093 | 1.217641259 | 0.46987032 | 2.59144109 | 9.56E-03 | 0.26044636 |
| AC025580.3 | 34.7404994 | 0.810521694 | 0.90434985 | 0.89624794 | 0.37012035 | 0.76669785 |
| SPATA5L1 | 206.964216 | -0.112495632 | 0.33537874 | -0.3354286 | 0.73730177 | 0.92033471 |
| AC025580.2 | 98.2420421 | 0.442831487 | 0.50360191 | 0.87932845 | 0.37922321 | 0.77076106 |
| SLC30A4 | 418.537842 | 0.32359295 | 0.34616531 | 0.93479311 | 0.34989493 | 0.75455562 |
| SQOR | 356.584496 | 0.134461127 | 0.46761539 | 0.28754641 | 7.74E-01 | 0.93419867 |
| AC068722.1 | 49.8349928 | 1.012900331 | 0.59063271 | 1.71494114 | 0.08635602 | 0.52421699 |
| SLC24A5 | 47.3229876 | 1.066322243 | 0.74654946 | 1.42833437 | 0.15319564 | 0.61008819 |
| MYEF2 | 1283.24713 | 0.367068373 | 0.30375598 | 1.20843175 | 0.22688123 | 0.66692281 |
| SLC12A1 | 429.875388 | 0.415174536 | 0.80248623 | 0.51736033 | 6.05E-01 | 0.86866716 |
| DUT | 285.498565 | 0.231765541 | 0.33776534 | 0.68617325 | 0.49260387 | 0.82100434 |
| FBN1 | 151.167249 | 0.585917486 | 0.24748635 | 2.36747398 | 0.01790998 | 0.33537171 |
| CEP152 | 533.308957 | 0.145471026 | 0.28137433 | 0.51700177 | 0.60515492 | 0.86869053 |
| EID1 | 368.235057 | 0.05198202 | 0.39499272 | 0.13160248 | 0.89529873 | 0.97309833 |
| SECISBP2L | 1748.62143 | 0.18224721 | 0.26956792 | 0.67607159 | 0.4989952 | 0.82396436 |
| COPS2 | 489.344714 | -0.058875326 | 0.20972659 | -0.2807242 | 0.77892196 | 0.93646866 |
| GALK2 | 2117.95956 | 0.394040473 | 0.26836774 | 1.46828556 | 0.14202667 | 0.5969824 |
| FAM227B | 679.171536 | 1.322182987 | 0.63420785 | 2.08477866 | 0.0370894 | 0.42215481 |
| FGF7 | 47.4084701 | 1.006506625 | 0.54272517 | 1.854542 | 0.06366166 | 0.48593331 |
| DTWD1 | 433.614059 | -0.060354196 | 0.31428605 | -0.1920359 | 0.8477141 | 0.95908381 |
| ATP8B4 | 1548.48789 | 0.746224637 | 0.64029894 | 1.16543163 | 0.24384432 | 0.68197798 |
| SLC27A2 | 2491.30948 | 0.736075796 | 0.54689341 | 1.34592187 | 0.17832772 | 0.6285261 |
| GABPB1 | 635.528709 | -0.490062463 | 0.26912194 | -1.8209681 | 0.06861171 | 0.49154315 |
| GABPB1-AS1 | 900.439037 | -0.005279635 | 0.40784348 | -0.0129452 | 0.98967147 | 0.99730401 |
| USP8 | 1596.96471 | -0.017929532 | 0.13966992 | -0.1283707 | 0.89785558 | 0.9737247 |
| USP50 | 188.93192 | -0.781383935 | 0.51392601 | -1.5204211 | 0.12840518 | 0.58793678 |

| | | | | | | |
|---|---|---|---|---|---|---|
| TRPM7 | 3245.59846 | -0.452350934 | 0.32193017 | -1.4051213 | 0.15998522 | 0.61564218 |
| SPPL2A | 1288.4891 | 0.037637968 | 0.14283758 | 0.26350186 | 0.79216379 | 0.94132545 |
| AC021752.1 | 83.538112 | 0.145973465 | 0.49630533 | 0.29412029 | 0.76866599 | 0.93198626 |
| AP4E1 | 645.878169 | -0.053092466 | 0.17303433 | -0.306832 | 0.75897127 | 0.92910772 |
| AC066613.2 | 48.8355417 | 0.533811528 | 0.79235872 | 0.67369931 | 0.5005025 | 0.82560579 |
| DMXL2 | 1753.55207 | 0.301332062 | 0.26481723 | 1.13788693 | 2.55E-01 | 6.90E-01 |
| LYSMD2 | 42.6470652 | 0.20146002 | 0.50913326 | 0.39569212 | 0.69233217 | 0.9051695 |
| TMOD2 | 53.6845798 | 1.242330618 | 0.55478125 | 2.23931614 | 0.02513535 | 0.37991912 |
| TMOD3 | 1644.0365 | -0.39505516 | 0.27560975 | -1.433386 | 0.15174755 | 0.60961422 |
| AC090971.2 | 74.3082787 | 0.442207015 | 0.57298805 | 0.77175609 | 0.4402589 | 0.79826598 |
| LEO1 | 210.652769 | -0.281011082 | 0.25279567 | -1.1116135 | 0.26630436 | 0.69797523 |
| MAPK6 | 1437.62832 | -0.554781678 | 0.38924575 | -1.4252736 | 0.15407814 | 0.61099212 |
| AC023906.5 | 40.0612263 | -0.395777124 | 0.60178983 | -0.6576667 | 0.51075233 | 0.82958671 |
| GNB5 | 679.011936 | -0.030281445 | 0.32907033 | -0.0920212 | 0.9266812 | 0.9816089 |
| MYO5C | 256.465105 | 1.007708242 | 0.53271724 | 1.89163811 | 0.05853921 | 0.47778423 |
| MYO5A | 751.650478 | -0.189515109 | 0.23461768 | -0.8077614 | 0.41922795 | 0.78783502 |
| ARPP19 | 779.832859 | -0.500390708 | 0.26474146 | -1.8901109 | 0.05874313 | 0.47778423 |
| FAM214A | 1412.28524 | 0.200100148 | 0.27685622 | 0.72275837 | 0.46982835 | 0.81322606 |
| WDR72 | 13887.7322 | 0.462150444 | 0.75197016 | 0.61458615 | 0.53882806 | 0.84070752 |
| UNC13C | 49.0455802 | 0.099984117 | 0.79956594 | 0.12504799 | 0.90048556 | 0.97479173 |
| RSL24D1 | 205.026096 | -0.072515901 | 0.29630625 | -0.2447329 | 0.80666322 | 0.94585363 |
| RAB27A | 429.682424 | 0.215995632 | 0.35255006 | 0.61266657 | 0.54009683 | 0.8414758 |
| PIGB | 403.762703 | -0.002209239 | 0.25037013 | -0.0088239 | 0.99295965 | 0.9976934 |
| CCPG1 | 1004.22987 | 0.22418803 | 0.15781101 | 1.42061082 | 0.15542993 | 0.6118547 |
| C15orf65 | 48.2891457 | 0.853755975 | 0.47030498 | 1.81532412 | 0.06947411 | 0.49281529 |
| DNAAF4 | 501.165165 | 0.075849626 | 0.31851637 | 0.23813415 | 0.81177705 | 0.94770701 |
| PRTG | 186.544191 | 0.354404588 | 0.56090304 | 0.63184644 | 0.52748723 | 0.83749692 |
| NEDD4 | 2323.75974 | 0.391367453 | 0.26804357 | 1.46008897 | 0.14426562 | 0.60086567 |
| RFX7 | 2013.04152 | 0.5684422 | 0.20835655 | 2.72821856 | 0.00636774 | 0.22797255 |
| TEX9 | 794.805702 | 0.6487651 | 0.37288386 | 1.73985835 | 0.08188389 | 0.51593841 |
| MNS1 | 76.0030944 | 1.473072599 | 0.52912154 | 2.78399665 | 0.00536936 | 0.21873395 |
| AC090518.1 | 109.455961 | 1.455072771 | 0.52649727 | 2.76368533 | 0.00571526 | 0.22501232 |
| ZNF280D | 602.10318 | 0.983084697 | 0.37198167 | 2.64283103 | 0.00822161 | 0.24877162 |
| AC090517.4 | 450.647448 | 0.840275917 | 0.43143261 | 1.94764119 | 0.05145791 | 0.46190805 |
| TCF12 | 6778.63492 | -0.100034354 | 0.20182589 | -0.4956468 | 0.62014364 | 0.87586466 |
| CGNL1 | 10185.7216 | 0.290139365 | 0.58829868 | 0.49318378 | 0.62188274 | 0.87713418 |
| AC025271.4 | 32.3558874 | 0.090504503 | 0.76001728 | 0.11908216 | 0.90521026 | 0.97598655 |
| POLR2M | 105.889501 | -0.446452328 | 0.3389483 | -1.3171694 | 0.18778185 | 0.63974756 |
| ALDH1A2 | 615.233575 | -0.695338469 | 0.54605953 | -1.2733748 | 2.03E-01 | 6.49E-01 |
| LIPC | 413.791314 | -0.599913532 | 0.26790718 | -2.2392589 | 0.02513907 | 0.37991912 |
| ADAM10 | 2290.26013 | -0.196350605 | 0.2042669 | -0.9612454 | 0.33642881 | 0.7429163 |
| MINDY2 | 1035.25042 | -0.430637015 | 0.21256228 | -2.0259334 | 4.28E-02 | 0.43882523 |
| AC090515.4 | 48.497314 | -0.347708505 | 0.35473613 | -0.9801891 | 0.32699276 | 0.73745391 |
| RNF111 | 1872.94745 | -0.228978051 | 0.17372087 | -1.3180803 | 0.18747677 | 0.63892195 |
| SLTM | 2146.09083 | -0.249338505 | 0.19984573 | -1.2476549 | 0.21215745 | 0.65673943 |
| MYO1E | 2201.54965 | -0.502104123 | 0.34483769 | -1.4560593 | 1.45E-01 | 6.02E-01 |
| FAM81A | 94.5752218 | 0.887541036 | 0.46345494 | 1.91505357 | 0.05548567 | 0.46947044 |
| GCNT3 | 282.8605 | 0.394911659 | 0.48669183 | 0.81142035 | 4.17E-01 | 7.87E-01 |
| GTF2A2 | 334.730227 | -0.122974916 | 0.21538945 | -0.5709421 | 0.56803886 | 0.85263558 |
| BNIP2 | 629.621772 | -0.076202494 | 0.18579543 | -0.4101419 | 0.68170184 | 0.89962501 |
| ANXA2 | 1361.03122 | -0.381493383 | 0.59570464 | -0.6404069 | 5.22E-01 | 8.34E-01 |
| ICE2 | 677.37293 | 0.556932339 | 0.31746268 | 1.75432384 | 0.07937503 | 0.51192014 |
| RORA-AS1 | 537.391926 | 0.123461599 | 0.82799502 | 0.14910911 | 0.88146754 | 0.96774334 |
| RORA | 12493.627 | 0.18445858 | 0.52018237 | 0.35460367 | 0.72288652 | 0.91592159 |

| | | | | | | |
|---|---|---|---|---|---|---|
| AC012404.1 | 173.740435 | 1.081980344 | 1.03974781 | 1.04061806 | 0.29805285 | 0.71852421 |
| AC009554.1 | 266.607806 | 0.319310953 | 0.55639304 | 0.57389458 | 0.56603914 | 0.85213602 |
| VPS13C | 2698.89666 | 0.137138582 | 0.18795872 | 0.72962076 | 4.66E-01 | 0.81043641 |
| TLN2 | 3015.23894 | 0.166222383 | 0.44264386 | 0.37552172 | 0.7072725 | 0.91002281 |
| AC103740.1 | 57.4192613 | 0.526323362 | 0.74623692 | 0.7053033 | 0.48062151 | 0.81714875 |
| TPM1 | 2398.30365 | 0.889543456 | 0.3933172 | 2.2616439 | 0.02371941 | 0.37327439 |
| LACTB | 438.940432 | -0.212493626 | 0.34180564 | -0.6216797 | 0.5341525 | 0.83916403 |
| RPS27L | 800.136667 | -0.016960378 | 0.44181915 | -0.0383876 | 0.96937865 | 0.99187457 |
| RAB8B | 869.267828 | 0.013399376 | 0.24050837 | 0.05571272 | 0.95557066 | 0.98840182 |
| APH1B | 188.524623 | 0.419397676 | 0.50953087 | 0.82310553 | 0.41044798 | 0.78582015 |
| CA12 | 3669.42564 | 0.604813113 | 0.39711063 | 1.52303431 | 0.12775012 | 0.58793678 |
| LINC02568 | 44.252757 | 0.499994256 | 0.7468114 | 0.66950539 | 0.50317315 | 0.82652042 |
| USP3 | 1492.27682 | -0.125167443 | 0.14612256 | -0.8565922 | 0.3916703 | 0.77606943 |
| USP3-AS1 | 318.410331 | 0.177303976 | 0.50638047 | 0.35013984 | 0.72623375 | 0.91635631 |
| HERC1 | 1878.75584 | -0.171921823 | 0.13139016 | -1.3084832 | 0.19070947 | 0.64281105 |
| AC073167.1 | 60.3627699 | 0.815472912 | 0.73052213 | 1.11628776 | 0.26429898 | 0.69567438 |
| DAPK2 | 60.1021984 | -0.456570605 | 0.69045842 | -0.6612572 | 0.50844738 | 0.82874455 |
| FAM96A | 423.040036 | 0.058433231 | 0.32587293 | 0.17931293 | 0.85769199 | 0.9626966 |
| SNX1 | 1749.84533 | 0.073845537 | 0.21188928 | 0.34851002 | 0.72745719 | 0.91647571 |
| PPIB | 838.526446 | 0.075581261 | 0.47529387 | 0.15902006 | 0.87365307 | 0.96592775 |
| CSNK1G1 | 299.039175 | 0.099681541 | 0.26206831 | 0.38036472 | 0.7036747 | 0.90907343 |
| TRIP4 | 1415.52554 | -0.081492043 | 0.21527857 | -0.3785423 | 7.05E-01 | 9.10E-01 |
| ZNF609 | 3492.78454 | -0.318911696 | 0.27836127 | -1.1456755 | 0.25192943 | 0.68843633 |
| OAZ2 | 255.515129 | 0.286414548 | 0.26277782 | 1.08994947 | 0.2757354 | 0.70248235 |
| AC100830.1 | 74.8994807 | 0.737550005 | 0.51774613 | 1.42453987 | 0.15429025 | 0.61099212 |
| RBPMS2 | 227.48378 | 0.458272265 | 0.41117633 | 1.11453951 | 0.2650478 | 0.69622296 |
| AC069368.1 | 51.5290421 | 1.191815446 | 0.6386461 | 1.86615943 | 0.06201906 | 0.48347236 |
| ANKDD1A | 56.5548603 | 0.748362841 | 0.5397466 | 1.38650773 | 0.16559192 | 0.62065658 |
| SPG21 | 594.44776 | -0.64836677 | 0.26386387 | -2.4572018 | 0.0140024 | 0.30363654 |
| MTFMT | 477.34833 | 0.094057439 | 0.26252067 | 0.35828584 | 0.72012942 | 0.9148113 |
| SLC51B | 104.285758 | -0.274480124 | 0.76961072 | -0.356648 | 0.72135534 | 0.91542662 |
| UBAP1L | 228.182782 | -0.136443948 | 0.49484173 | -0.2757325 | 0.78275353 | 0.93794935 |
| PDCD7 | 331.883986 | -0.289325242 | 0.28132022 | -1.0284552 | 0.30373575 | 0.72082426 |
| CLPX | 703.690447 | -0.427507373 | 0.30756931 | -1.3899546 | 0.16454267 | 0.62065658 |
| PARP16 | 199.057003 | -0.070026659 | 0.36968165 | -0.1894242 | 0.84976034 | 0.95959925 |
| DPP8 | 970.741883 | -0.213216371 | 0.18459044 | -1.1550781 | 0.24805845 | 0.68482118 |
| HACD3 | 538.358448 | 0.063000981 | 0.23536562 | 0.26767282 | 0.78895117 | 0.94005579 |
| INTS14 | 230.394808 | -0.350294322 | 0.23411678 | -1.4962376 | 0.13459176 | 0.59277997 |
| SLC24A1 | 158.664376 | 0.308187918 | 0.3824472 | 0.80583128 | 0.42034014 | 0.78818088 |
| DENND4A | 3788.50774 | 0.116361995 | 0.35280881 | 0.32981601 | 0.74153899 | 0.92235631 |
| RAB11A | 1517.98387 | 0.10225108 | 0.46061488 | 0.22198823 | 0.82432305 | 0.95180295 |
| MEGF11 | 1749.02386 | 1.236601327 | 0.93113296 | 1.32806096 | 1.84E-01 | 6.34E-01 |
| DIS3L | 287.699208 | 0.277972808 | 0.35618155 | 0.78042451 | 0.43514105 | 0.79515776 |
| TIPIN | 180.03008 | 0.780916478 | 0.61103989 | 1.27801228 | 0.2012451 | 0.6485595 |
| MAP2K1 | 1100.19398 | -0.89313979 | 0.32368721 | -2.7592681 | 0.0057931 | 0.22622796 |
| SNAPC5 | 120.117447 | 0.047260654 | 0.33062639 | 0.14294277 | 0.88633538 | 0.97024858 |
| RPL4 | 977.127899 | -0.02783334 | 0.52163182 | -0.0533582 | 0.9574465 | 0.98861313 |
| ZWILCH | 60.8730861 | -0.075681811 | 0.49793222 | -0.1519922 | 0.87919309 | 0.9674841 |
| SMAD6 | 121.961828 | 0.235654679 | 0.63664387 | 0.3701515 | 0.71126962 | 0.91143287 |
| AC093334.1 | 37.7477643 | -0.476790018 | 0.70365469 | -0.6775909 | 0.49803112 | 0.82326489 |
| AC087482.1 | 3312.63144 | -0.161404169 | 0.72985696 | -0.2211449 | 0.82497958 | 0.95192757 |
| SMAD3 | 4877.93916 | -1.010216412 | 0.33513766 | -3.0143327 | 0.00257545 | 0.16651755 |
| AAGAB | 809.378593 | -0.158828866 | 0.23693226 | -0.6703556 | 0.50263113 | 0.82638105 |
| IQCH | 721.530377 | 0.553262779 | 0.45256713 | 1.22249881 | 0.22151905 | 0.66267168 |

| | | | | | | |
|---|---|---|---|---|---|---|
| IQCH-AS1 | 502.407748 | 0.106219634 | 0.38390893 | 0.27667925 | 0.78202641 | 0.93757221 |
| C15orf61 | 82.3559656 | -0.337953448 | 0.4401841 | -0.7677548 | 0.44263288 | 0.79933184 |
| MAP2K5 | 1643.26217 | 0.214302535 | 0.27074126 | 0.79153999 | 0.42862895 | 0.79037914 |
| PIAS1 | 2559.18937 | -0.068289271 | 0.20014356 | -0.3412014 | 0.73295195 | 0.91952673 |
| FEM1B | 347.614879 | 0.068261963 | 0.21238278 | 0.32141006 | 0.74789966 | 0.92531168 |
| CORO2B | 128.923287 | 0.413907082 | 0.76754586 | 0.5392604 | 5.90E-01 | 0.86281961 |
| ANP32A | 757.047334 | 0.300111907 | 0.31632839 | 0.94873529 | 0.34275526 | 0.7505827 |
| SPESP1 | 185.344798 | 0.893676779 | 0.45847435 | 1.94924051 | 0.05126671 | 0.46140039 |
| NOX5 | 45.565607 | 0.963631672 | 0.93953832 | 1.02564382 | 0.30505951 | 0.72103276 |
| GLCE | 1228.53238 | 0.184244579 | 0.38381357 | 0.48003665 | 0.63120133 | 0.88122598 |
| AC026992.2 | 280.786052 | 0.255597384 | 0.62397671 | 0.40962648 | 0.68207997 | 0.89962501 |
| AC026992.1 | 181.676585 | 0.969646597 | 0.79256344 | 1.2234309 | 0.22116699 | 0.66263062 |
| PAQR5 | 3756.18689 | -0.692568776 | 0.52124696 | -1.3286769 | 0.1839546 | 0.63338605 |
| AC027237.3 | 80.6855331 | 0.558732506 | 0.70919307 | 0.78784259 | 0.43078879 | 0.79100093 |
| KIF23 | 371.110235 | 0.112311099 | 0.70414326 | 0.15950035 | 0.87327468 | 0.96592775 |
| RPLP1 | 1998.7212 | 0.054072898 | 0.51025196 | 0.10597294 | 0.91560383 | 0.9779223 |
| DRAIC | 1541.84973 | 0.772829093 | 0.73669495 | 1.04904899 | 0.29415557 | 0.71584347 |
| TLE3 | 403.73753 | 0.11885817 | 0.36562507 | 0.32508212 | 0.74511892 | 0.92389439 |
| UACA | 1837.69547 | 0.820718012 | 0.20294721 | 4.04399752 | 5.25E-05 | 0.02213112 |
| AC009269.2 | 42.2372021 | 1.056181924 | 0.50234989 | 2.10248266 | 0.03551102 | 0.41997369 |
| LARP6 | 119.062143 | -0.156723438 | 0.29733521 | -0.5270934 | 0.5981287 | 0.86646266 |
| LRRC49 | 158.289186 | 0.705693513 | 0.58405107 | 1.20827365 | 0.22694202 | 0.66695847 |
| THSD4 | 309.777775 | 0.505654488 | 0.53509014 | 0.94498936 | 0.34466431 | 0.75124763 |
| THSD4-AS1 | 349.474203 | -0.407246036 | 0.28225269 | -1.442842 | 0.14906499 | 0.60622442 |
| MYO9A | 6505.51831 | -0.028309227 | 0.31528969 | -0.089788 | 0.92845569 | 0.98272895 |
| AC022872.1 | 142.593861 | 0.042244845 | 0.72083276 | 0.05860561 | 0.95326624 | 0.9880091 |
| SENP8 | 171.81723 | 0.523520808 | 0.41805911 | 1.25226503 | 0.21047331 | 0.6547793 |
| GRAMD2A | 146.062068 | 1.728276545 | 0.53020558 | 3.25963475 | 0.00111556 | 0.11093704 |
| PKM | 2594.28565 | -0.090526837 | 0.34267768 | -0.2641749 | 7.92E-01 | 9.41E-01 |
| PARP6 | 529.92872 | -0.384669143 | 0.35177791 | -1.0935 | 0.27417433 | 0.7014484 |
| AC009690.3 | 38.7011845 | -0.56436331 | 0.79303471 | -0.7116502 | 0.47668141 | 0.81570932 |
| AC009690.2 | 294.298071 | 0.1611435 | 0.61536614 | 0.26186605 | 0.79342472 | 0.94189286 |
| AC009690.1 | 1392.74083 | -0.563784893 | 0.58606469 | -0.9619841 | 0.33605761 | 0.74259901 |
| TMEM202-A: | 167.788767 | 0.207259275 | 0.59442532 | 0.34867168 | 0.72733581 | 0.91646656 |
| ARIH1 | 3451.80089 | -0.383873761 | 0.23153066 | -1.6579824 | 0.09732102 | 0.54093104 |
| BBS4 | 503.333109 | 0.267307155 | 0.32166701 | 0.83100582 | 0.40597034 | 0.78429302 |
| ADPGK | 535.202247 | -0.488006317 | 0.27878508 | -1.750475 | 0.08003639 | 0.51264427 |
| NEO1 | 2538.99678 | 0.583273361 | 0.36803412 | 1.58483503 | 0.11300382 | 0.56779449 |
| NPTN | 912.953867 | -0.152614091 | 0.20446017 | -0.7464246 | 0.45541099 | 0.80465616 |
| CD276 | 171.188979 | -0.626006811 | 0.5198213 | -1.2042731 | 0.22848404 | 0.66705629 |
| LOXL1-AS1 | 44.4870786 | 1.308321267 | 0.58627098 | 2.23159821 | 0.02564153 | 0.38166806 |
| LOXL1 | 68.7340846 | 0.31723884 | 0.55612447 | 0.57044574 | 0.56837541 | 0.85263558 |
| STOML1 | 50.1925085 | 0.165190667 | 0.44042218 | 0.37507345 | 0.70760584 | 0.91012042 |
| PML | 355.026663 | 0.313938725 | 0.30373041 | 1.03360978 | 3.01E-01 | 7.19E-01 |
| STRA6 | 97.2275096 | 2.311842392 | 1.04701039 | 2.2080415 | 2.72E-02 | 3.87E-01 |
| UBL7 | 50.5113851 | -0.176826539 | 0.34203099 | -0.5169898 | 0.60516327 | 0.86869053 |
| UBL7-AS1 | 42.0739266 | -0.000340837 | 0.50962839 | -0.0006688 | 0.99946638 | 0.99977652 |
| AC012435.2 | 65.90043 | 0.193501697 | 0.39800153 | 0.48618329 | 0.62683721 | 0.87890915 |
| ARID3B | 295.877193 | -0.981182797 | 0.31487126 | -3.1161396 | 0.00183235 | 0.13687118 |
| CLK3 | 245.308213 | -0.256738474 | 0.20805394 | -1.2339996 | 0.21720306 | 0.66032006 |
| EDC3 | 350.613666 | -0.431685412 | 0.23196008 | -1.8610332 | 0.06273949 | 0.48421747 |
| CSK | 109.185157 | -0.603016685 | 0.48011364 | -1.2559874 | 0.20912053 | 0.65403148 |
| ULK3 | 288.487153 | -0.397829887 | 0.33245189 | -1.196654 | 2.31E-01 | 0.66878058 |
| SCAMP2 | 595.179244 | -0.337051251 | 0.25011331 | -1.3475942 | 0.17778892 | 0.62808062 |

| | | | | | | |
|---|---|---|---|---|---|---|
| MPI | 115.970469 | 0.169152726 | 0.36135199 | 0.46811069 | 0.63970544 | 0.88321992 |
| FAM219B | 199.967655 | 0.010385222 | 0.26264259 | 0.03954127 | 0.96845885 | 0.99187457 |
| COX5A | 454.1419 | -0.263184314 | 0.28321434 | -0.9292761 | 0.35274603 | 0.75673243 |
| SCAMP5 | 344.719867 | 0.759808932 | 0.34179727 | 2.22298126 | 0.02621706 | 0.38392873 |
| PPCDC | 778.449906 | -0.454621795 | 0.45359209 | -1.0022701 | 0.31621315 | 0.72798933 |
| C15orf39 | 39.5070272 | -0.599849749 | 0.47568007 | -1.2610361 | 0.20729584 | 0.65269814 |
| COMMD4 | 213.849866 | -0.377257344 | 0.31929863 | -1.1815188 | 0.23739667 | 0.67510294 |
| NEIL1 | 310.83396 | 0.195800161 | 0.42194472 | 0.46404222 | 0.6426175 | 0.88441609 |
| MAN2C1 | 520.547061 | -0.158151393 | 0.25588652 | -0.6180528 | 0.53654051 | 0.83980588 |
| SIN3A | 655.621097 | 0.231550418 | 0.24509906 | 0.94472178 | 0.34480094 | 0.75133476 |
| PTPN9 | 1118.89013 | 0.107269549 | 0.27684718 | 0.38746845 | 0.69840944 | 0.90696517 |
| SNUPN | 188.830579 | -0.070670488 | 0.38828037 | -0.1820089 | 0.85557573 | 0.96204894 |
| IMP3 | 154.343168 | -0.524307019 | 0.38487868 | -1.3622657 | 0.17311406 | 0.6264249 |
| SNX33 | 105.660998 | 0.039622496 | 0.32267573 | 0.12279354 | 0.90227059 | 0.9750023 |
| UBE2Q2 | 493.149424 | -0.143344468 | 0.24360127 | -0.5884389 | 5.56E-01 | 8.49E-01 |
| FBXO22 | 636.40129 | 0.36029211 | 0.24347979 | 1.47976187 | 0.13893681 | 0.59474076 |
| NRG4 | 45.6241412 | 0.646967246 | 0.54648787 | 1.18386388 | 0.23646695 | 0.67365231 |
| TMEM266 | 90.2404502 | 1.024570434 | 0.57148776 | 1.79281255 | 0.0730029 | 0.50459544 |
| ETFA | 1681.41722 | 0.058492564 | 0.26353417 | 0.22195438 | 0.8243494 | 0.95180295 |
| SCAPER | 2911.62174 | 0.612448548 | 0.28757705 | 2.12968504 | 0.03319762 | 0.41254527 |
| RCN2 | 370.092602 | 0.434941233 | 0.2331522 | 1.86548202 | 0.06211387 | 0.48347236 |
| TSPAN3 | 1015.88084 | -0.129537711 | 0.27092682 | -0.478128 | 0.63255908 | 0.88151688 |
| PEAK1 | 1612.32006 | -0.420768035 | 0.28106692 | -1.4970386 | 0.13438321 | 0.59240843 |
| HMG20A | 764.71727 | 0.221201067 | 0.17989539 | 1.22960944 | 0.21884339 | 0.66116077 |
| LINGO1 | 6132.91213 | -0.958893764 | 0.67071332 | -1.4296626 | 0.15281387 | 0.61001662 |
| TBC1D2B | 531.237077 | -0.083984456 | 0.33468185 | -0.2509382 | 0.8018619 | 0.94496295 |
| CIB2 | 60.259546 | 0.333127742 | 0.6221363 | 0.53545782 | 0.59233332 | 0.86382106 |
| IDH3A | 608.007147 | 0.478102045 | 0.23085442 | 2.071011 | 0.03835777 | 0.42447711 |
| DNAJA4 | 298.445295 | -1.80601785 | 0.81696971 | -2.2106301 | 0.02706146 | 0.38702446 |
| WDR61 | 343.145017 | 0.200572606 | 0.19236808 | 1.04265017 | 0.29711035 | 0.71825247 |
| IREB2 | 1004.53783 | -0.164000845 | 0.1434625 | -1.1431618 | 0.25297143 | 0.68887358 |
| HYKK | 288.471516 | 1.213264515 | 0.49805493 | 2.43600542 | 0.01485046 | 0.3088851 |
| PSMA4 | 405.927937 | -0.188244651 | 0.32504538 | -0.5791334 | 0.56249914 | 0.85054239 |
| MORF4L1 | 1100.08931 | -0.059776451 | 0.14705076 | -0.4065021 | 0.68437368 | 0.9007343 |
| CTSH | 1456.12903 | 0.807383577 | 0.3299687 | 2.44684899 | 0.01441112 | 0.30672862 |
| RASGRF1 | 142.468315 | 1.19846156 | 0.62485814 | 1.91797383 | 0.05511433 | 0.46922491 |
| TMED3 | 269.789428 | -0.901167002 | 0.60200457 | -1.4969438 | 0.13440788 | 0.59240843 |
| MTHFS | 143.060841 | -0.559679353 | 0.46300675 | -1.2087931 | 0.22674235 | 0.66690202 |
| ZFAND6 | 2820.61675 | -0.114383089 | 0.21135567 | -0.5411877 | 0.58837821 | 0.86140264 |
| FAH | 451.145904 | 0.456844278 | 0.33305121 | 1.37169379 | 0.17015878 | 0.62379417 |
| CTXND1 | 223.787483 | 1.492838694 | 0.78614606 | 1.898933 | 0.05757329 | 0.47684824 |
| AC016705.2 | 139.783571 | 1.642333099 | 0.66931784 | 2.4537417 | 0.01413785 | 0.30468798 |
| ARNT2 | 409.918537 | 1.044121551 | 0.77998462 | 1.33864377 | 0.18068667 | 0.63166264 |
| ABHD17C | 159.258878 | -0.041331402 | 0.5071799 | -0.0814926 | 0.93505022 | 0.98379126 |
| CEMIP | 147.904094 | 1.587617074 | 0.85694398 | 1.85264979 | 0.06393257 | 0.48593331 |
| AC027808.1 | 265.070838 | 1.085076645 | 0.32769789 | 3.31121031 | 0.00092893 | 0.10268462 |
| MESD | 589.736857 | 0.184744692 | 0.25660848 | 0.71994773 | 0.47155718 | 0.81389222 |
| TLNRD1 | 140.876346 | -0.420247711 | 0.61971038 | -0.6781357 | 0.49768568 | 0.82326489 |
| CFAP161 | 119.148113 | -0.996994782 | 1.0922816 | -0.9127635 | 0.36136694 | 0.76009788 |
| AC104041.1 | 89.386059 | 0.795912066 | 0.70881211 | 1.12288159 | 0.26148779 | 0.69374398 |
| EFL1 | 1008.40806 | -0.538754964 | 0.42867364 | -1.2567952 | 0.20882781 | 0.65401126 |
| SAXO2 | 77.3644717 | 1.361762476 | 0.69170115 | 1.96871507 | 0.04898582 | 0.45368151 |
| GOLGA6L10 | 45.2639688 | 0.061373439 | 0.86552138 | 0.07090921 | 0.94347001 | 0.98522232 |
| GOLGA6L9 | 55.4733619 | 0.363179083 | 0.36013356 | 1.00845663 | 0.3132353 | 0.72717675 |

| | | | | | | |
|---|---|---|---|---|---|---|
| RPS17 | 99.0336222 | -0.414120764 | 0.65504934 | -0.6321978 | 0.52725762 | 0.83734448 |
| CPEB1 | 106.896302 | -0.898952673 | 0.58895503 | -1.526352 | 0.12692223 | 0.58793678 |
| AP3B2 | 63.5023768 | -0.543173349 | 0.62276178 | -0.8722008 | 0.38309882 | 0.77317162 |
| AC105339.3 | 42.7979802 | 0.543802859 | 0.73606744 | 0.73879489 | 0.46003156 | 0.80696859 |
| SNHG21 | 67.4688629 | -0.004590219 | 0.37667949 | -0.012186 | 0.99027721 | 0.99737132 |
| WHAMM | 885.615431 | -0.224872484 | 0.27793739 | -0.809076 | 0.41847141 | 0.78736961 |
| HOMER2 | 510.97858 | -1.141334394 | 0.6574237 | -1.7360713 | 0.08255123 | 0.5183475 |
| AC022558.2 | 125.481424 | 0.032257792 | 0.61758644 | 0.05223203 | 0.95834381 | 0.98878187 |
| C15orf40 | 347.757791 | 0.069077441 | 0.2325846 | 0.29699921 | 0.76646712 | 0.93138917 |
| BTBD1 | 658.37475 | -0.162909219 | 0.25745958 | -0.6327565 | 0.52689267 | 0.83710323 |
| HDGFL3 | 46.5605333 | 0.788135248 | 0.78478014 | 1.00427522 | 0.31524597 | 0.72782155 |
| ADAMTSL3 | 384.316682 | -1.14934848 | 0.77631581 | -1.4805166 | 0.13873542 | 0.59474076 |
| GOLGA6L4 | 64.2060317 | -0.899379632 | 0.36941056 | -2.4346343 | 0.01490685 | 0.30894792 |
| AC048382.5 | 197.050609 | 0.26394854 | 0.36155699 | 0.73003301 | 0.46537001 | 0.81042138 |
| ZSCAN2 | 187.976822 | 0.409939742 | 0.29426239 | 1.39310951 | 0.16358668 | 0.62065658 |
| WDR73 | 701.281179 | -0.039506303 | 0.16202598 | -0.243827 | 0.80736484 | 0.94604106 |
| NMB | 68.6611267 | -0.343831788 | 0.56946967 | -0.6037754 | 0.54599297 | 0.84460887 |
| SEC11A | 1137.04867 | -0.105046267 | 0.25547318 | -0.4111832 | 0.68093824 | 0.89947163 |
| ZNF592 | 936.870428 | 0.072072469 | 0.22110724 | 0.3259616 | 0.74445341 | 0.92363594 |
| ALPK3 | 605.278203 | -0.874162446 | 0.77760483 | -1.1241731 | 0.26093959 | 0.69289016 |
| SLC28A1 | 5646.85107 | 1.344336287 | 0.61202128 | 2.19655154 | 0.02805249 | 0.39156363 |
| PDE8A | 4644.04416 | 0.34327046 | 0.30986035 | 1.10782311 | 0.26793822 | 0.69928201 |
| AKAP13 | 5753.3566 | -0.307494572 | 0.25390708 | -1.2110516 | 0.22587563 | 0.66688879 |
| AC087286.2 | 180.202557 | 0.155776231 | 0.5705813 | 0.27301321 | 0.78484306 | 0.93884957 |
| AC021739.2 | 39.1279572 | -0.577780472 | 0.55816261 | -1.0351472 | 0.30060016 | 0.71879123 |
| KLHL25 | 170.182862 | -0.66363241 | 0.49285159 | -1.3465157 | 0.17813626 | 0.62849772 |
| AGBL1 | 58.3879388 | 0.411868211 | 0.57595626 | 0.71510327 | 0.47454523 | 0.81531007 |
| MRPL46 | 144.855909 | 0.021898365 | 0.38619845 | 0.05670236 | 0.95478229 | 0.98840182 |
| MRPS11 | 249.572408 | -0.219457812 | 0.34740967 | -0.6316975 | 0.52758457 | 0.83749692 |
| DET1 | 186.446052 | 0.23275099 | 0.44533318 | 0.52264462 | 0.60122158 | 0.86755264 |
| AEN | 103.843599 | -1.116692224 | 0.72986955 | -1.5299888 | 0.12601951 | 0.58574071 |
| ISG20 | 204.05735 | -2.324195055 | 0.73701482 | -3.1535255 | 1.61E-03 | 1.32E-01 |
| ABHD2 | 584.2243 | 0.877208016 | 0.438394 | 2.00095809 | 0.04539691 | 0.44360789 |
| FANCI | 72.1151986 | -0.433014027 | 0.35783067 | -1.2101087 | 0.22623717 | 0.66688879 |
| POLG | 314.445399 | -0.201271572 | 0.26938233 | -0.7471595 | 0.45496727 | 0.80465616 |
| PEX11A | 165.345149 | 0.591814861 | 0.43850344 | 1.3496242 | 0.17713656 | 0.62731853 |
| WDR93 | 100.944776 | 1.030858465 | 0.54551354 | 1.88970281 | 0.05879772 | 0.47778423 |
| MESP1 | 41.6817923 | -0.407262092 | 0.41313508 | -0.9857843 | 0.32423895 | 0.73614611 |
| ANPEP | 6432.01137 | -0.009543272 | 0.493312 | -0.0193453 | 0.98456564 | 0.99538584 |
| AP3S2 | 73.9280379 | -0.224069923 | 0.38748283 | -0.5782706 | 0.56308145 | 0.85057939 |
| ZNF710 | 198.95113 | 0.186251491 | 0.27690907 | 0.67260885 | 5.01E-01 | 0.8260948 |
| ZNF710-AS1 | 69.655697 | 1.558050368 | 0.49756718 | 3.13133666 | 0.00174013 | 0.13403387 |
| IDH2 | 558.738898 | 0.560068794 | 0.41354309 | 1.35431787 | 0.17563499 | 0.62699479 |
| SEMA4B | 1317.48402 | 0.398028681 | 0.43594775 | 0.91301924 | 0.36123242 | 0.76009788 |
| CIB1 | 777.809627 | 0.081984318 | 0.41612778 | 0.19701717 | 0.8438141 | 0.95731603 |
| AC091167.6 | 72.749895 | -0.015446764 | 0.38577329 | -0.040041 | 0.96806041 | 0.99187457 |
| ZNF774 | 36.3532025 | -0.162499942 | 0.36350473 | -0.4470367 | 0.65484858 | 0.89047714 |
| IQGAP1 | 1599.18959 | -0.337107939 | 0.28882804 | -1.1671579 | 2.43E-01 | 6.82E-01 |
| CRTC3 | 1463.25149 | 0.297129121 | 0.16666796 | 1.78276091 | 0.07462523 | 0.5071704 |
| CRTC3-AS1 | 177.191737 | 0.224173565 | 0.41730563 | 0.53719277 | 0.59113447 | 0.86336496 |
| FURIN | 151.274426 | -0.246951424 | 0.40405382 | -0.6111845 | 0.54107746 | 0.84192857 |
| MAN2A2 | 212.65093 | 0.22003888 | 0.29576396 | 0.74396785 | 0.45689593 | 0.80507423 |
| HDDC3 | 84.94468 | -0.048496735 | 0.37417457 | -0.1296099 | 0.89687506 | 0.97324791 |
| UNC45A | 313.785439 | 0.146218101 | 0.28858644 | 0.50667003 | 0.61238636 | 0.87222135 |

| | | | | | | |
|---|---|---|---|---|---|---|
| RCCD1 | 102.573511 | 0.312661538 | 0.47746834 | 0.65483198 | 0.51257594 | 0.83055441 |
| AC104035.1 | 301.389551 | 1.194342429 | 0.91718802 | 1.3021784 | 0.19285541 | 0.64518499 |
| SLCO3A1 | 1271.04124 | 1.012963822 | 0.45136446 | 2.24422593 | 2.48E-02 | 3.79E-01 |
| FAM174B | 303.599476 | -0.66527254 | 0.3448552 | -1.9291359 | 0.05371399 | 0.46711025 |
| AC013394.1 | 1238.7126 | 0.0649847 | 0.26600126 | 0.24430222 | 0.80699676 | 0.94585363 |
| CHD2 | 2445.56792 | 0.016768446 | 0.31610319 | 0.05304738 | 0.95769416 | 0.98861313 |
| MCTP2 | 1711.31473 | 0.221042477 | 0.25003839 | 0.88403415 | 0.37667778 | 0.77038531 |
| AC012409.2 | 270.526039 | 0.941770764 | 0.77976151 | 1.20776769 | 0.22713663 | 0.66705629 |
| NR2F2-AS1 | 2158.95166 | 1.083181269 | 0.67598572 | 1.602373 | 0.10907315 | 0.55932125 |
| NR2F2 | 640.862494 | 0.295423493 | 0.26194066 | 1.12782601 | 0.2593934 | 0.69242517 |
| AC087477.2 | 155.969654 | 0.985967362 | 0.59746625 | 1.65024779 | 0.09889227 | 0.5425667 |
| LINC00923 | 155.003451 | 0.363420303 | 1.03219949 | 0.3520834 | 0.72477572 | 0.91600702 |
| ARRDC4 | 107.275344 | 1.102596278 | 0.45835222 | 2.40556548 | 0.01614746 | 0.32030855 |
| FAM169B | 182.789231 | -0.501117421 | 0.72201986 | -0.6940494 | 0.48765125 | 0.81893983 |
| IGF1R | 11548.4031 | -0.643161732 | 0.29400569 | -2.1875826 | 0.02870002 | 0.39339119 |
| AC036108.1 | 90.3334844 | 0.248338709 | 0.57777125 | 0.42982185 | 0.66732524 | 0.8944961 |
| SYNM | 43.9640904 | -1.035324286 | 0.59113798 | -1.7514089 | 0.07987551 | 0.51257191 |
| TTC23 | 724.097548 | 0.242178628 | 0.21057723 | 1.15007033 | 0.2501149 | 0.68617881 |
| LRRC28 | 1702.36179 | 0.644397664 | 0.38239327 | 1.68517001 | 0.09195578 | 0.53567268 |
| MEF2A | 3590.56718 | -0.11773263 | 0.2929844 | -0.4018392 | 0.68780234 | 0.90295054 |
| LYSMD4 | 139.609612 | 0.984754974 | 0.31119202 | 3.16446091 | 0.00155371 | 0.13146073 |
| AC090825.1 | 186.298401 | 0.600149092 | 0.62992157 | 0.95273622 | 0.34072374 | 0.74725037 |
| ADAMTS17 | 212.662509 | -1.718863758 | 0.66290292 | -2.5929344 | 0.00951609 | 0.26008591 |
| AC027020.2 | 44.6556192 | -0.523957515 | 0.51156029 | -1.0242341 | 0.3057247 | 0.72122 |
| LINS1 | 261.162641 | 0.011760893 | 0.21863191 | 0.05379312 | 0.95709999 | 0.98861313 |
| ASB7 | 481.626869 | 0.21782057 | 0.17380226 | 1.25326663 | 0.21010869 | 0.65441616 |
| AC087762.1 | 6166.09991 | 0.146491603 | 0.73512349 | 0.19927482 | 0.84204777 | 0.95704385 |
| LRRK1 | 85.0855802 | 0.600039461 | 0.71543507 | 0.83870568 | 0.40163449 | 0.78246874 |
| CHSY1 | 557.573936 | -0.454755266 | 0.25384105 | -1.7914961 | 0.07321371 | 0.50459544 |
| SELENOS | 526.327075 | -0.396691658 | 0.40115594 | -0.9888714 | 3.23E-01 | 0.7346962 |
| SNRPA1 | 319.794306 | -0.128870538 | 0.20833154 | -0.6185839 | 0.53619051 | 0.83956629 |
| PCSK6 | 44.0522895 | -0.215027993 | 0.49613967 | -0.4334021 | 0.66472265 | 0.89315615 |
| TM2D3 | 357.175039 | -0.329708573 | 0.27651982 | -1.1923506 | 0.23312381 | 0.66978724 |
| TARSL2 | 732.287058 | -0.136105849 | 0.34060903 | -0.3995955 | 0.68945444 | 0.90373453 |
| OR4F15 | 27.992529 | 1.246565578 | 0.92889384 | 1.34198928 | 0.17959947 | 0.63073788 |
| POLR3K | 38.3364201 | -0.497366601 | 0.36171188 | -1.3750353 | 0.1691205 | 0.62326131 |
| SNRNP25 | 65.7208523 | -0.076622825 | 0.57659861 | -0.1328876 | 0.89428225 | 0.97272276 |
| RHBDF1 | 63.1494442 | -0.422334923 | 0.36373837 | -1.1610953 | 2.46E-01 | 6.83E-01 |
| MPG | 231.05559 | 0.280499305 | 0.21697406 | 1.29277805 | 1.96E-01 | 6.46E-01 |
| NPRL3 | 619.888068 | 0.189442796 | 0.23479025 | 0.80685973 | 0.4197473 | 0.78798479 |
| LUC7L | 1234.29109 | 0.002291231 | 0.2611654 | 0.0087731 | 0.99300017 | 0.9976934 |
| FAM234A | 198.43821 | 0.47290818 | 0.42049365 | 1.12465 | 0.26073738 | 0.69280433 |
| AXIN1 | 401.091122 | -0.726957172 | 0.49658038 | -1.4639265 | 0.14321404 | 0.59866875 |
| MRPL28 | 151.486121 | -0.025459276 | 0.47882038 | -0.0531708 | 0.9575958 | 0.98861313 |
| TMEM8A | 284.1446 | -0.075233407 | 0.39047956 | -0.1926693 | 0.847218 | 0.9586278 |
| NME4 | 99.8248027 | 0.038241656 | 0.58115149 | 0.06580325 | 0.94753447 | 0.98646931 |
| DECR2 | 278.194102 | 0.236346953 | 0.41135669 | 0.57455479 | 0.56559244 | 0.85194709 |
| RAB11FIP3 | 9281.46405 | -0.225036732 | 0.54295643 | -0.4144655 | 0.6785332 | 0.8992258 |
| CAPN15 | 505.161212 | -0.689298648 | 0.35281124 | -1.9537321 | 0.05073291 | 0.45923264 |
| PIGQ | 240.808586 | 0.093087004 | 0.36660644 | 0.25391535 | 0.79956095 | 0.94385599 |
| RAB40C | 583.231598 | -0.302253196 | 0.33898598 | -0.8916392 | 0.37258634 | 0.76891189 |
| METTL26 | 154.006893 | -0.483647215 | 0.43122409 | -1.1215682 | 0.26204609 | 0.69408034 |
| MCRIP2 | 571.135892 | -0.126250386 | 0.42405466 | -0.297722 | 0.76591538 | 0.93112214 |
| WDR90 | 168.586769 | -0.345263091 | 0.34133807 | -1.0114989 | 0.31177769 | 0.72604649 |

| | | | | | | |
|---|---|---|---|---|---|---|
| RHOT2 | 316.482298 | 0.074602409 | 0.27306427 | 0.27320458 | 0.78469596 | 0.93884957 |
| RHBDL1 | 36.0577917 | 0.817152168 | 0.58029661 | 1.40816293 | 0.15908283 | 0.61423843 |
| STUB1 | 403.729067 | -0.218658383 | 0.33295196 | -0.6567265 | 0.51135677 | 0.83007666 |
| JMJD8 | 174.998119 | 0.000594818 | 0.45432038 | 0.00130925 | 0.99895537 | 0.99977652 |
| WDR24 | 44.3883982 | -0.562060713 | 0.37952077 | -1.4809748 | 0.13861328 | 0.59474076 |
| METRN | 207.253407 | 1.095097639 | 0.34213802 | 3.20074815 | 0.00137071 | 0.12294569 |
| FAM173A | 250.36366 | 0.257256322 | 0.35351966 | 0.72770019 | 0.46679713 | 0.8111547 |
| HAGHL | 67.1191758 | 0.389763968 | 0.70443099 | 0.55330326 | 0.58005577 | 0.85824961 |
| NARFL | 178.041335 | -0.179608836 | 0.25200658 | -0.7127149 | 0.47602222 | 0.81570932 |
| RPUSD1 | 44.8438218 | -0.581754872 | 0.59033315 | -0.9854688 | 0.32439387 | 0.73614611 |
| LMF1 | 522.031504 | 0.125186393 | 0.39926104 | 0.31354523 | 0.75386646 | 0.92783044 |
| CACNA1H | 96.9651743 | 0.768662597 | 1.06627321 | 0.72088709 | 0.47097898 | 0.81387913 |
| AC120498.4 | 88.36923 | -0.233655059 | 1.0897463 | -0.2144123 | 0.83022552 | 0.95325673 |
| UBE2I | 797.498483 | -0.417951028 | 0.3444577 | -1.2133595 | 0.2249924 | 0.66642114 |
| TSR3 | 181.575764 | -0.097075662 | 0.39409504 | -0.2463255 | 0.80543027 | 0.94585363 |
| GNPTG | 411.883149 | -0.550319505 | 0.38498675 | -1.4294505 | 0.15287479 | 0.61001662 |
| UNKL | 264.213354 | -0.213825409 | 0.3185878 | -0.6711664 | 0.50211456 | 0.82638105 |
| C16orf91 | 47.0437143 | -0.055096621 | 0.54496191 | -0.1011018 | 0.91946966 | 0.97919126 |
| CCDC154 | 26.5404311 | 0.710893546 | 0.73095267 | 0.97255755 | 0.33077325 | 0.73866225 |
| CLCN7 | 689.022698 | -0.707626473 | 0.38796532 | -1.8239426 | 0.06816075 | 0.49069295 |
| TELO2 | 113.575035 | -0.129894193 | 0.57569964 | -0.2256284 | 0.82149044 | 0.95102783 |
| IFT140 | 435.108409 | 0.285585981 | 0.41802217 | 0.68318382 | 0.4944907 | 0.8220963 |
| AL133297.2 | 35.0286639 | 0.115454977 | 0.66141657 | 0.17455713 | 0.86142764 | 0.96302275 |
| CRAMP1 | 74.3540012 | -0.583251003 | 0.3403287 | -1.7137873 | 0.0865678 | 0.52503751 |
| JPT2 | 379.268678 | -0.706645939 | 0.22584554 | -3.1288903 | 1.75E-03 | 0.1343652 |
| MAPK8IP3 | 824.146906 | -0.366226886 | 0.24820376 | -1.475509 | 0.14007574 | 0.59500405 |
| NME3 | 488.464965 | 0.745589479 | 0.4687849 | 1.59047246 | 0.11172835 | 0.56445264 |
| MRPS34 | 280.961536 | -0.464980671 | 0.33144403 | -1.4028935 | 0.1606486 | 0.61630095 |
| EME2 | 143.077852 | -0.245785224 | 0.32913202 | -0.7467679 | 0.45520369 | 0.80465616 |
| SPSB3 | 355.251939 | -0.322379904 | 0.23830001 | -1.3528321 | 0.17610928 | 0.62699479 |
| NUBP2 | 115.973359 | -0.325846968 | 0.32350018 | -1.0072544 | 0.31381256 | 0.72719146 |
| HAGH | 1150.68682 | 0.417807057 | 0.40981304 | 1.01950649 | 0.30796257 | 0.72355897 |
| FAHD1 | 441.769746 | -0.091969417 | 0.47235289 | -0.1947049 | 0.84562398 | 0.95764192 |
| MEIOB | 50.5951512 | 0.515877982 | 0.66950953 | 0.7705312 | 0.44098486 | 0.79828043 |
| MSRB1 | 341.410516 | -0.217128387 | 0.36656783 | -0.592328 | 0.55363098 | 0.84727776 |
| NDUFB10 | 358.43501 | -0.226230813 | 0.44812675 | -0.5048367 | 0.61367357 | 0.87222135 |
| RPS2 | 909.993092 | 0.018594818 | 0.50041301 | 0.03715894 | 0.97035828 | 0.99205112 |
| SNHG9 | 120.770084 | -0.76357359 | 0.40888106 | -1.8674712 | 0.06183582 | 0.48267858 |
| TBL3 | 55.139896 | -0.447327276 | 0.56467272 | -0.7921886 | 0.42825073 | 0.79037914 |
| GFER | 72.0868253 | -0.067508834 | 0.31346408 | -0.2153639 | 0.82948365 | 0.95325673 |
| ZNF598 | 131.534851 | -0.382424287 | 0.53674318 | -0.7124903 | 0.47616125 | 0.81570932 |
| SLC9A3R2 | 58.7439037 | -0.236800327 | 0.34775781 | -0.6809346 | 0.49591287 | 0.82243887 |
| TSC2 | 427.937255 | -0.202784803 | 0.24496596 | -0.8278081 | 0.40777919 | 0.78481094 |
| PKD1 | 338.78679 | -0.479347544 | 0.36251168 | -1.3222954 | 0.18606979 | 0.6371368 |
| SNHG19 | 155.74785 | -0.121551867 | 0.45791092 | -0.2654487 | 0.79066381 | 0.94063545 |
| TRAF7 | 182.664714 | 0.006992949 | 0.3547728 | 0.01971106 | 0.98427387 | 0.99523767 |
| MLST8 | 81.1580834 | -0.199029167 | 0.53547527 | -0.3716869 | 0.71012594 | 0.91123308 |
| BRICD5 | 36.2864212 | 0.096039089 | 0.66664962 | 0.14406232 | 0.88545126 | 0.96957206 |
| PGP | 299.013826 | 0.522855496 | 0.40790342 | 1.28181199 | 0.19990861 | 0.64705133 |
| E4F1 | 169.318787 | -0.227361766 | 0.34683473 | -0.6555334 | 0.51212436 | 0.83054053 |
| ECI1 | 738.489369 | 0.192532135 | 0.15550893 | 1.23807764 | 0.21568727 | 0.6596384 |
| RNPS1 | 473.295851 | -0.49316618 | 0.27531506 | -1.7912793 | 0.07324848 | 0.50459544 |
| ABCA3 | 255.572198 | -0.103625188 | 0.47129012 | -0.2198756 | 0.82596806 | 0.95192757 |
| TBC1D24 | 41.9332126 | -0.130424568 | 0.52882335 | -0.2466316 | 0.80519332 | 0.94585363 |

| | | | | | | |
|---|---|---|---|---|---|---|
| AMDHD2 | 370.048971 | -1.198121983 | 0.38263239 | -3.1312612 | 0.00174057 | 0.13403387 |
| PDPK1 | 1895.60559 | 0.220981401 | 0.22435949 | 0.9849434 | 0.32465187 | 0.73614611 |
| AC093525.6 | 200.702606 | 0.444902692 | 0.42964359 | 1.03551571 | 0.30042813 | 0.71879123 |
| AC093525.7 | 157.05409 | 0.281900438 | 0.42270141 | 0.66690205 | 0.5048347 | 0.82715271 |
| AC093525.4 | 556.182563 | 0.058512344 | 0.44062317 | 0.13279452 | 0.89435589 | 0.97272276 |
| KCTD5 | 144.232319 | -0.196640808 | 0.43378665 | -0.4533123 | 0.65032391 | 0.8887244 |
| SRRM2-AS1 | 45.4000426 | -0.710139633 | 0.40692389 | -1.7451412 | 0.0809603 | 0.51567042 |
| SRRM2 | 4093.09226 | -0.385079651 | 0.20642008 | -1.8655145 | 0.06210932 | 0.48347236 |
| ELOB | 446.714361 | -0.270887747 | 0.43453426 | -0.6233979 | 0.53302308 | 0.83872659 |
| FLYWCH2 | 98.3269008 | 0.601844635 | 0.31952046 | 1.88358718 | 0.05962084 | 0.47942559 |
| FLYWCH1 | 318.854934 | 0.490253484 | 0.35518261 | 1.38028574 | 0.16749869 | 0.62296534 |
| TNFRSF12A | 557.422622 | -0.752816157 | 0.66345093 | -1.1346976 | 2.57E-01 | 6.91E-01 |
| HCFC1R1 | 236.717957 | -0.591845308 | 0.38045308 | -1.5556328 | 0.11979543 | 0.57696272 |
| THOC6 | 66.9426693 | -0.944768649 | 0.55724558 | -1.695426 | 0.08999464 | 0.53308405 |
| MMP25-AS1 | 130.057394 | 1.235463822 | 0.47307476 | 2.61156148 | 0.00901298 | 0.2563089 |
| IL32 | 1760.60923 | 0.096458323 | 0.53397357 | 0.18064251 | 0.85664819 | 0.96245027 |
| ZNF213-AS1 | 195.729856 | 0.030757452 | 0.19927849 | 0.15434406 | 0.87733846 | 0.966539 |
| ZNF205 | 127.392775 | -0.068611663 | 0.42215383 | -0.1625276 | 0.87089037 | 0.9657206 |
| ZNF213 | 268.88516 | 0.325000289 | 0.40945367 | 0.79374131 | 0.42734604 | 0.78933193 |
| ZNF200 | 58.6102941 | 0.360567489 | 0.30950321 | 1.16498788 | 0.2440239 | 0.68220184 |
| ZNF263 | 136.626004 | -0.52215371 | 0.26966003 | -1.9363407 | 0.05282598 | 0.46564998 |
| TIGD7 | 111.96233 | -0.146920026 | 0.33241358 | -0.4419796 | 0.65850396 | 0.89058983 |
| ZNF75A | 444.374607 | -1.058712346 | 0.48609143 | -2.1780107 | 0.02940524 | 0.39791639 |
| AC025283.3 | 121.911521 | -0.012462128 | 0.46832839 | -0.0266098 | 0.97877095 | 0.99482978 |
| ZSCAN32 | 125.638869 | -0.261558051 | 0.30086948 | -0.8693406 | 0.38466087 | 0.77317162 |
| NAA60 | 204.003426 | -0.48953005 | 0.28197731 | -1.7360619 | 0.0825529 | 0.5183475 |
| CLUAP1 | 485.410034 | 0.423616506 | 0.34223242 | 1.23780356 | 0.21578891 | 0.6596384 |
| SLX4 | 60.7809813 | -0.343774348 | 0.43831915 | -0.7843015 | 0.43286324 | 0.79332216 |
| DNASE1 | 580.781143 | 0.341738366 | 0.29328428 | 1.16521201 | 0.24393318 | 0.68208734 |
| TRAP1 | 870.610522 | -0.222689556 | 0.20931466 | -1.0638985 | 0.28737467 | 0.71118494 |
| CREBBP | 1836.2939 | -0.175068417 | 0.18866595 | -0.927928 | 0.35344492 | 0.75673243 |
| ADCY9 | 3132.56358 | -0.06718895 | 0.29336854 | -0.2290258 | 0.8188489 | 0.94990455 |
| LINC01569 | 70.9054426 | -0.296513849 | 0.71905671 | -0.412365 | 0.68007188 | 0.89941358 |
| TFAP4 | 53.3359876 | -0.758675561 | 0.50885422 | -1.4909487 | 0.13597495 | 0.59413728 |
| GLIS2 | 377.627014 | 0.411634513 | 0.46702198 | 0.88140287 | 0.37809981 | 0.77055888 |
| GLIS2-AS1 | 40.9743237 | 0.45899199 | 0.62949692 | 0.72914097 | 0.46591544 | 0.81054727 |
| PAM16 | 53.882526 | 0.312718741 | 0.51803113 | 0.60366785 | 0.54606449 | 0.84460887 |
| VASN | 176.105157 | 1.17303794 | 0.61607971 | 1.904036 | 0.0569055 | 0.4746515 |
| DNAJA3 | 412.70461 | -0.031473037 | 0.20084863 | -0.1567003 | 0.87548106 | 0.96617876 |
| NMRAL1 | 369.003966 | 0.360283322 | 0.21984416 | 1.63881233 | 0.10125235 | 0.54652047 |
| HMOX2 | 742.124151 | 0.064807351 | 0.36261826 | 0.17872059 | 0.8581571 | 0.9626966 |
| CDIP1 | 396.28936 | -0.197115163 | 0.20233 | -0.9742261 | 0.32994431 | 0.73849818 |
| UBALD1 | 188.020339 | -0.590975987 | 0.26557199 | -2.2252949 | 0.02606145 | 0.38392873 |
| MGRN1 | 813.224283 | -0.176187885 | 0.24959482 | -0.7058956 | 0.48025306 | 0.81714875 |
| NUDT16L1 | 145.623312 | 0.342820726 | 0.38613802 | 0.88781914 | 0.37463805 | 0.7697436 |
| ANKS3 | 571.646425 | -0.544959457 | 0.16746352 | -3.254198 | 0.00113713 | 0.11093704 |
| ZNF500 | 180.497523 | -0.305258627 | 0.35911803 | -0.8500231 | 0.39531223 | 0.77931337 |
| ROGDI | 136.885494 | 0.251945145 | 0.5317868 | 0.47377097 | 0.63566324 | 0.88257807 |
| GLYR1 | 1085.77142 | -0.365427651 | 0.29838553 | -1.2246829 | 0.22069472 | 0.66263062 |
| UBN1 | 652.349425 | -0.462328475 | 0.39668172 | -1.1654897 | 0.24382082 | 0.68197798 |
| PPL | 280.221646 | 0.184695995 | 0.31742757 | 0.5818524 | 0.56066611 | 0.84945842 |
| NAGPA | 136.00168 | 0.240118164 | 0.3375033 | 0.71145427 | 0.47680278 | 0.81570932 |
| ALG1 | 179.637724 | 0.033021973 | 0.40976842 | 0.08058692 | 0.93577047 | 0.98406248 |
| EEF2KMT | 47.0351242 | -0.333745296 | 0.57297226 | -0.5824807 | 0.56024292 | 0.84944908 |

| | | | | | | |
|---|---|---|---|---|---|---|
| AC026458.2 | 80.8532995 | -0.279031039 | 0.45228184 | -0.6169406 | 5.37E-01 | 0.84030274 |
| RBFOX1 | 430.059598 | 1.054242922 | 0.47050763 | 2.24065001 | 0.02504875 | 0.37991912 |
| METTL22 | 208.987082 | -0.380490625 | 0.33516307 | -1.1352403 | 0.25627465 | 0.69092557 |
| ABAT | 1284.56387 | 0.888171939 | 0.46375697 | 1.91516675 | 0.05547124 | 0.46947044 |
| TMEM186 | 71.9173161 | -0.172662285 | 0.31227587 | -0.5529159 | 0.58032102 | 0.85831909 |
| PMM2 | 1209.73718 | -0.391635043 | 0.38918873 | -1.0062857 | 0.31427817 | 0.72719146 |
| AC022167.2 | 76.0236817 | -0.566876126 | 0.43746819 | -1.2958111 | 0.19504058 | 0.64558745 |
| CARHSP1 | 1515.33702 | -0.300483755 | 0.44753728 | -0.6714162 | 0.50195546 | 0.82638105 |
| USP7 | 576.017471 | -0.217501787 | 0.20433347 | -1.0644452 | 0.28712705 | 0.71118494 |
| C16orf72 | 642.573392 | -0.363685684 | 0.2814036 | -1.2923988 | 0.19621906 | 0.64558745 |
| AC087190.1 | 69.3713337 | -0.093955821 | 0.66514205 | -0.1412568 | 0.8876671 | 0.97098615 |
| ATF7IP2 | 432.091478 | 0.908085975 | 0.48750404 | 1.86272502 | 0.06250096 | 0.4840117 |
| NUBP1 | 160.226139 | -0.436498472 | 0.31929779 | -1.3670576 | 0.17160725 | 0.62520756 |
| AC133065.1 | 93.0232129 | 1.613730477 | 0.83221304 | 1.93908339 | 0.05249119 | 0.46539242 |
| CIITA | 266.322237 | 2.387894732 | 0.66653838 | 3.58253147 | 0.00034028 | 0.06649962 |
| DEXI | 206.291289 | 0.146865208 | 0.30405408 | 0.48302332 | 0.62907918 | 0.88057005 |
| CLEC16A | 1375.08288 | 0.164939072 | 0.23427995 | 0.70402555 | 0.48141686 | 0.81755903 |
| AC007014.1 | 58.1946974 | 0.515780413 | 0.53781086 | 0.95903681 | 0.3375402 | 0.74404633 |
| AC007220.1 | 86.8319622 | -0.121388095 | 0.43023095 | -0.2821464 | 0.77783129 | 0.93646866 |
| SOCS1 | 74.9113704 | 0.250438962 | 0.65643377 | 0.38151444 | 0.70282156 | 0.90870007 |
| AC099489.1 | 59.6239287 | -0.071847311 | 0.52505711 | -0.1368371 | 0.89115953 | 0.97234284 |
| LITAF | 814.132879 | -0.39441656 | 0.36237323 | -1.0884263 | 2.76E-01 | 0.70328016 |
| TXNDC11 | 1275.9373 | -0.119108475 | 0.18019737 | -0.660989 | 0.50861938 | 0.82874455 |
| ZC3H7A | 1169.53526 | 0.137104463 | 0.19637231 | 0.69818635 | 0.48506066 | 0.81839266 |
| RSL1D1 | 250.533328 | -0.75243736 | 0.40290662 | -1.8675229 | 0.0618286 | 0.48267858 |
| AC007216.4 | 387.644291 | -0.064778191 | 0.3092458 | -0.2094715 | 0.83408016 | 0.95393659 |
| GSPT1 | 1077.64886 | -0.438970869 | 0.26380838 | -1.6639762 | 0.09611719 | 0.53876283 |
| NPIPB2 | 233.572485 | -0.315874977 | 0.81509857 | -0.3875298 | 0.69836403 | 0.90696517 |
| SNX29 | 17676.6552 | 0.082928863 | 0.3404829 | 0.24356249 | 0.80756968 | 0.94604106 |
| AC010333.2 | 130.153576 | 0.920331618 | 0.75401133 | 1.22058062 | 0.22224485 | 0.66373504 |
| CPPED1 | 1443.51487 | -0.705069433 | 0.42738395 | -1.6497331 | 0.09899754 | 0.5425667 |
| AC109597.1 | 77.3291659 | 0.099510764 | 0.73861276 | 0.13472657 | 0.89282807 | 0.9724379 |
| SHISA9 | 934.432354 | 1.450211374 | 0.67692096 | 2.14236441 | 0.03216417 | 0.40988962 |
| U91319.1 | 427.827615 | 1.86904266 | 0.75046627 | 2.49050855 | 0.01275604 | 0.29875272 |
| ERCC4 | 225.454511 | 0.470011055 | 0.2117636 | 2.2195082 | 0.02645217 | 0.38507144 |
| MKL2 | 2510.25732 | 0.133752802 | 0.21372441 | 0.62581904 | 0.53143364 | 0.83789294 |
| AC130650.1 | 48.3642537 | 0.083289089 | 0.75099586 | 0.11090486 | 0.91169179 | 0.977179 |
| MIR193BHG | 359.169983 | -0.94039056 | 0.59786309 | -1.5729196 | 0.11573744 | 0.57350419 |
| PARN | 1035.86892 | -0.164473914 | 0.2585238 | -0.6362041 | 0.52464338 | 0.83592566 |
| BFAR | 495.728009 | -0.398976066 | 0.2867363 | -1.391439 | 0.16409235 | 0.62065658 |
| NOMO1 | 200.273771 | -0.256518293 | 0.3800309 | -0.6749933 | 0.49968005 | 0.82476476 |
| NPIPA1 | 99.7876864 | -0.533229812 | 0.40602805 | -1.3132832 | 0.18908754 | 0.6413893 |
| PDXDC1 | 2709.04468 | -0.160741494 | 0.21550157 | -0.7458948 | 0.45573098 | 0.80465616 |
| NTAN1 | 154.842331 | -0.570823546 | 0.3262594 | -1.7496003 | 0.08018731 | 0.51297858 |
| RRN3 | 484.589525 | -0.323944897 | 0.31307378 | -1.0347238 | 0.3007979 | 0.71879123 |
| NPIPA5 | 32.7511581 | -0.521643027 | 0.68651835 | -0.7598384 | 0.44735116 | 0.80223612 |
| C16orf45 | 434.728704 | 0.891867173 | 0.52057424 | 1.71323724 | 0.0866689 | 0.52518597 |
| MARF1 | 669.405741 | -0.368577404 | 0.1859214 | -1.9824367 | 0.04743039 | 0.44991578 |
| NDE1 | 170.170857 | 0.055262381 | 0.47022063 | 0.11752437 | 0.90644454 | 0.97651095 |
| MYH11 | 39.5205356 | 0.71890801 | 0.39277787 | 1.83031701 | 0.06720255 | 0.48963361 |
| FOPNL | 292.979782 | -0.32483108 | 0.23765622 | -1.3668108 | 0.17168463 | 0.62520756 |
| ABCC1 | 545.727068 | -0.577106993 | 0.41545427 | -1.3890987 | 0.16480274 | 0.62065658 |
| ABCC6 | 1830.93793 | 0.260619252 | 0.41289381 | 0.63120164 | 5.28E-01 | 0.83761342 |
| NOMO3 | 72.0583577 | 0.07084462 | 0.28703254 | 0.24681738 | 0.80504956 | 0.94585363 |

| | | | | | | |
|---|---|---|---|---|---|---|
| XYLT1 | 94.6168603 | 0.463252022 | 0.90849202 | 0.50991314 | 0.61011231 | 0.87103525 |
| NOMO2 | 118.831259 | -0.33863127 | 0.46998673 | -0.7205124 | 0.47120956 | 0.81387913 |
| RPS15A | 235.324775 | 0.027955489 | 0.5836809 | 0.04789516 | 0.9617998 | 0.99000623 |
| SMG1 | 3422.625 | -0.413123618 | 0.18723737 | -2.2064164 | 0.02735486 | 0.38774871 |
| AC092287.1 | 132.853159 | -0.792802749 | 0.55600504 | -1.4258913 | 0.15389972 | 0.6105654 |
| TMC7 | 63.0146854 | 0.018943603 | 0.40921123 | 0.04629297 | 0.96307674 | 0.99012655 |
| COQ7 | 131.517486 | -0.086969642 | 0.31881842 | -0.2727874 | 0.78501666 | 0.93892226 |
| SYT17 | 52.5157775 | 1.704664165 | 0.99144691 | 1.7193701 | 0.085547 | 0.5226185 |
| GDE1 | 178.14038 | 0.003835716 | 0.36227784 | 0.01058777 | 0.99155234 | 0.99748517 |
| CCP110 | 227.873432 | 0.067791292 | 0.26050888 | 0.26022641 | 0.79468913 | 0.94252227 |
| VPS35L | 1264.85931 | -0.096194972 | 0.27010327 | -0.3561415 | 0.72173463 | 0.91560945 |
| KNOP1 | 191.258875 | -1.224124659 | 0.50859449 | -2.4068776 | 0.01608956 | 0.31962267 |
| IQCK | 773.421272 | 0.146278943 | 0.42100537 | 0.34745149 | 0.72825216 | 0.91697155 |
| UMOD | 733.450335 | -0.525022066 | 0.63453093 | -0.8274176 | 0.4080004 | 0.78481094 |
| ACSM5 | 461.794675 | -0.044914898 | 0.86983864 | -0.0516359 | 0.95881882 | 0.98882925 |
| AC137056.1 | 64.443814 | -0.981029573 | 0.98191582 | -0.9990974 | 0.31774749 | 0.72966408 |
| ACSM2A | 10082.0474 | 0.499615198 | 0.85133569 | 0.58686039 | 0.55729747 | 0.84861484 |
| ACSM2B | 10403.044 | 0.678394243 | 0.85698982 | 0.79160129 | 0.4285932 | 0.79037914 |
| ACSM3 | 1142.26674 | 0.984246656 | 0.62171166 | 1.58312401 | 0.1133932 | 0.56809124 |
| ACSM1 | 84.8119458 | 0.568676983 | 0.6789877 | 0.83753651 | 0.40229107 | 0.78260179 |
| THUMPD1 | 511.44976 | -0.026098313 | 0.20779287 | -0.1255977 | 0.90005036 | 0.97471678 |
| ERI2 | 213.09975 | 0.374464437 | 0.42264816 | 0.88599567 | 0.37561986 | 0.7697436 |
| REXO5 | 37.1370034 | 0.57602838 | 0.65026669 | 0.88583405 | 0.37570696 | 0.7697436 |
| LYRM1 | 336.558723 | -0.198022362 | 0.30722291 | -0.644556 | 0.51921498 | 0.8335052 |
| DNAH3 | 53.0676061 | 0.140223422 | 0.55833071 | 0.25114761 | 0.80169999 | 0.94487548 |
| TMEM159 | 266.352599 | -0.476914113 | 0.37641643 | -1.2669854 | 0.20516052 | 0.65140961 |
| ANKS4B | 294.760999 | 0.691123471 | 0.50927553 | 1.35707182 | 0.17475839 | 0.62699479 |
| CRYM | 704.938055 | 0.257806517 | 0.74347077 | 0.3467608 | 0.72877103 | 0.9175406 |
| NPIPB3 | 118.858191 | -0.760682942 | 0.61738998 | -1.2320947 | 0.2179137 | 0.66064833 |
| METTL9 | 1503.12135 | 0.087409331 | 0.33753476 | 0.25896394 | 0.79566306 | 0.94252227 |
| IGSF6 | 66.0790891 | 0.0599764 | 0.68389559 | 0.08769818 | 0.93011656 | 0.98272895 |
| NPIPB4 | 421.821356 | -0.895985833 | 0.42930087 | -2.0870813 | 0.03688079 | 0.42215481 |
| UQCRC2 | 997.490888 | 0.117279256 | 0.18596313 | 0.63065865 | 0.52826374 | 0.83761342 |
| MOSMO | 969.404217 | -0.352558197 | 0.27353958 | -1.2888745 | 0.19744171 | 0.64558745 |
| EEF2K | 537.378642 | -1.169818472 | 0.53142953 | -2.2012674 | 0.0277171 | 0.38942007 |
| POLR3E | 329.50378 | -0.163966829 | 0.27369035 | -0.5990961 | 0.54910879 | 0.84587304 |
| CDR2 | 2882.12719 | -0.304043363 | 0.29034674 | -1.0471733 | 0.29501964 | 0.71646902 |
| AC092338.2 | 80.5253148 | 0.422237855 | 0.5668415 | 0.7448958 | 0.45633472 | 0.80507423 |
| NPIPB5 | 1100.89857 | -0.394459308 | 0.33281118 | -1.1852345 | 0.23592473 | 0.67287145 |
| USP31 | 1263.96331 | 0.026076292 | 0.34031422 | 0.07662416 | 0.93892254 | 0.98435387 |
| COG7 | 672.688343 | -0.174636796 | 0.1985887 | -0.8793894 | 0.37919018 | 0.77076106 |
| AC008915.2 | 46.7516877 | -0.473859291 | 0.35678118 | -1.328151 | 0.18412822 | 0.63359771 |
| GGA2 | 1186.56931 | -1.075773973 | 0.31355614 | -3.4308815 | 0.00060162 | 0.08329748 |
| EARS2 | 199.276975 | -0.752696855 | 0.36654716 | -2.0534789 | 0.04002615 | 0.43027104 |
| UBFD1 | 222.352688 | -0.678664203 | 0.30144183 | -2.2513936 | 0.02436062 | 0.37480999 |
| NDUFAB1 | 290.488348 | -0.234316059 | 0.22339289 | -1.0488967 | 0.29422568 | 0.71584347 |
| PALB2 | 239.451218 | -0.206537021 | 0.39857795 | -0.5181848 | 0.60432937 | 0.86865188 |
| DCTN5 | 412.039961 | -0.089651824 | 0.2800837 | -0.3200894 | 0.74890056 | 0.92554142 |
| PRKCB | 58.5445895 | 0.90011056 | 0.87413255 | 1.02971862 | 0.30314211 | 0.72044214 |
| RBBP6 | 835.674807 | -0.074195002 | 0.27851942 | -0.2663908 | 0.78993827 | 0.94031124 |
| TNRC6A | 6488.77254 | -0.463928939 | 0.26130139 | -1.7754553 | 0.07582273 | 0.50730849 |
| AC008731.1 | 150.618948 | -0.03671834 | 0.66660006 | -0.055083 | 0.95607233 | 0.98852557 |
| SLC5A11 | 1499.93759 | 0.79472193 | 0.89231282 | 0.89063153 | 0.37312688 | 0.76932162 |
| ARHGAP17 | 932.213894 | -0.238693473 | 0.1790075 | -1.3334273 | 0.18239163 | 0.63216521 |

| | | | | | | |
|---|---|---|---|---|---|---|
| LCMT1 | 559.022329 | 0.096641367 | 0.16370345 | 0.59034411 | 0.55495997 | 0.84801165 |
| ZKSCAN2 | 47.0984465 | -0.025321189 | 0.4113081 | -0.0615626 | 0.95091118 | 0.98731502 |
| AC008741.2 | 75.5288884 | -0.261161837 | 0.44099713 | -0.5922076 | 0.55371161 | 0.84727776 |
| HS3ST4 | 39.855669 | 0.03228399 | 0.64911942 | 0.04973506 | 0.96033352 | 0.98936382 |
| KDM8 | 51.2675958 | -0.22443308 | 0.41726247 | -0.5378703 | 0.59066661 | 0.86326802 |
| NSMCE1 | 263.722409 | 0.566515226 | 0.28524614 | 1.98605745 | 0.04702695 | 0.44732709 |
| AC106739.1 | 135.211098 | 1.71181018 | 0.77219583 | 2.21680838 | 0.02663619 | 0.38563341 |
| IL4R | 787.656176 | 0.321737007 | 0.48160348 | 0.66805374 | 5.04E-01 | 0.82715271 |
| GTF3C1 | 746.285267 | -0.353575956 | 0.25157403 | -1.4054549 | 0.15988605 | 0.61543334 |
| KIAA0556 | 1010.44706 | -0.266809332 | 0.26203164 | -1.0182332 | 0.30856712 | 0.72398657 |
| XPO6 | 925.467045 | -0.904978582 | 0.39791617 | -2.2742945 | 0.02294828 | 0.36677754 |
| NPIPB6 | 25.7786369 | -1.092738655 | 1.14277883 | -0.9562118 | 0.33896522 | 0.7463769 |
| NUPR1 | 594.979643 | -1.222045952 | 0.75117552 | -1.6268448 | 0.10377007 | 0.55022042 |
| SGF29 | 201.50777 | 0.039165517 | 0.42019756 | 0.09320739 | 0.9257388 | 0.98098729 |
| SULT1A2 | 49.3758978 | 1.407178344 | 0.90400142 | 1.55661077 | 0.11956293 | 0.57673415 |
| SULT1A1 | 195.101185 | 0.424380255 | 0.67299381 | 0.63058568 | 0.52831147 | 0.83761342 |
| ATXN2L | 549.676011 | -0.272737167 | 0.22767042 | -1.1979473 | 0.23093753 | 0.66878058 |
| TUFM | 580.522005 | 0.167907228 | 0.4635365 | 0.36223087 | 0.71717951 | 0.91317971 |
| SH2B1 | 463.261123 | 0.483581965 | 0.27395387 | 1.76519485 | 0.07753102 | 0.51128995 |
| AC133550.2 | 84.4169845 | -0.1647357 | 0.3215168 | -0.5123704 | 0.60839178 | 0.86985416 |
| ATP2A1 | 38.837713 | -0.489963274 | 0.34983596 | -1.4005515 | 0.16134824 | 0.61742053 |
| RABEP2 | 251.742575 | -0.122018673 | 0.40432563 | -0.3017832 | 0.76281736 | 0.93022076 |
| NFATC2IP | 887.024997 | -0.221884291 | 0.35233906 | -0.6297465 | 0.52886045 | 0.83765775 |
| AC109460.2 | 70.5482352 | -0.262821931 | 0.52413105 | -0.5014432 | 0.61605927 | 0.87333999 |
| SPNS1 | 305.375802 | -1.001562979 | 0.42930167 | -2.3330051 | 0.01964788 | 0.34523522 |
| LAT | 87.4366932 | -0.669329226 | 0.50065335 | -1.3369115 | 0.18125152 | 0.6318513 |
| AC009093.2 | 101.987982 | -2.169831461 | 0.90777439 | -2.3902761 | 0.01683571 | 0.32852872 |
| NPIPB11 | 91.7916112 | -0.201363404 | 0.38817771 | -0.5187403 | 6.04E-01 | 0.86837631 |
| NPIPB12 | 29.9437403 | -1.100494448 | 0.61752094 | -1.7821168 | 0.07473018 | 0.5071704 |
| QPRT | 698.524524 | 0.454695412 | 0.34861316 | 1.30429792 | 0.19213203 | 0.64497936 |
| KIF22 | 161.265777 | 0.122713685 | 0.29826663 | 0.41142278 | 0.68076255 | 0.89947163 |
| MAZ | 206.476603 | -0.359077983 | 0.43417769 | -0.82703 | 0.40822005 | 0.78481094 |
| AC009133.1 | 846.755505 | -0.748408128 | 0.46742675 | -1.6011239 | 0.10934949 | 0.55948242 |
| MVP | 1657.26701 | 0.245321277 | 0.32817589 | 0.74752986 | 0.45474379 | 0.80465616 |
| CDIPT | 232.266814 | -0.031491356 | 0.28735818 | -0.1095892 | 0.91273517 | 0.977179 |
| SEZ6L2 | 86.8830396 | 1.491542651 | 0.6661356 | 2.23909765 | 0.02514956 | 0.37991912 |
| ASPHD1 | 173.131179 | 1.094522165 | 0.46132639 | 2.37255483 | 0.01766554 | 0.33306956 |
| KCTD13 | 424.672713 | -0.203940613 | 0.35439903 | -0.5754548 | 0.56498377 | 0.85171209 |
| TMEM219 | 347.266933 | 0.37922319 | 0.37429135 | 1.01317646 | 0.31097588 | 0.72558189 |
| TAOK2 | 171.51953 | -0.074339035 | 0.40096277 | -0.1854013 | 0.85291426 | 0.96073035 |
| HIRIP3 | 39.3174444 | 0.439543645 | 0.40094881 | 1.09625877 | 0.27296555 | 0.70118792 |
| INO80E | 271.71542 | -0.03840057 | 0.33229231 | -0.1155626 | 0.90799919 | 0.97673359 |
| PPP4C | 370.326311 | -0.751010304 | 0.32431769 | -2.3156625 | 0.0205767 | 0.35267827 |
| YPEL3 | 326.061955 | -0.271295226 | 0.39885882 | -0.6801786 | 0.49639139 | 0.82243887 |
| MAPK3 | 103.94163 | 0.637597509 | 0.35083993 | 1.81734591 | 0.06916416 | 0.49273866 |
| NPIPB13 | 282.250511 | -0.550096222 | 0.36184672 | -1.5202465 | 0.12844902 | 0.58793678 |
| CD2BP2 | 245.950803 | -0.401813792 | 0.41514461 | -0.9678887 | 0.33309994 | 0.74000014 |
| TBC1D10B | 176.885244 | -0.221482747 | 0.41332331 | -0.5358583 | 0.59205645 | 0.86382106 |
| ZNF48 | 63.9088051 | -0.317106823 | 0.35944321 | -0.8822167 | 0.37765961 | 0.77055888 |
| 1-Sep | 58.7569937 | 0.095321807 | 0.37825259 | 0.2520057 | 0.80103666 | 0.9444185 |
| ZNF771 | 166.385295 | -0.422532692 | 0.51061413 | -0.827499 | 0.40795428 | 0.78481094 |
| DCTPP1 | 80.151496 | -0.031774564 | 0.41793064 | -0.0760283 | 0.93939657 | 0.98435387 |
| SEPHS2 | 851.8433 | -0.124161922 | 0.44128996 | -0.2813613 | 0.77843329 | 0.93646866 |
| ITGAL | 116.225426 | 0.700230456 | 0.80550515 | 0.869306 | 0.38467978 | 0.77317162 |

| | | | | | | |
|---|---|---|---|---|---|---|
| ZNF768 | 153.101136 | -0.319951173 | 0.23952588 | -1.3357687 | 1.82E-01 | 0.6318513 |
| AC002310.4 | 83.2792021 | -0.201832112 | 0.38158414 | -0.5289321 | 0.59685253 | 0.86581201 |
| AC002310.1 | 38.169492 | -0.177140477 | 0.45653255 | -0.3880128 | 0.69800655 | 0.90696517 |
| ZNF688 | 84.3736365 | -0.121226992 | 0.35597951 | -0.3405449 | 0.73344625 | 0.91962522 |
| ZNF785 | 209.617089 | 0.050277718 | 0.38364353 | 0.13105321 | 0.89573321 | 0.97309833 |
| AC093249.2 | 98.6755367 | 0.906467721 | 0.67540895 | 1.34210203 | 0.17956291 | 0.63073788 |
| ZNF689 | 61.6305445 | 0.788674388 | 0.67884785 | 1.16178374 | 0.24532332 | 0.68281753 |
| PRR14 | 159.85929 | -0.432756587 | 0.39355581 | -1.0996067 | 0.27150354 | 0.70005626 |
| FBRS | 238.425974 | 0.040748596 | 0.23328929 | 0.17466981 | 0.86133909 | 0.96302275 |
| SRCAP | 59.7372801 | -0.283210276 | 0.35308874 | -0.8020938 | 0.42249873 | 0.78894887 |
| PHKG2 | 213.041829 | -0.158106146 | 0.35337188 | -0.4474214 | 0.65457081 | 0.89032312 |
| CCDC189 | 55.7863735 | 0.057789475 | 0.41999824 | 0.13759457 | 0.89056085 | 0.97230741 |
| RNF40 | 283.861843 | -0.231833537 | 0.25389271 | -0.9131162 | 0.36118145 | 0.76009788 |
| ZNF629 | 72.9857082 | -0.186603121 | 0.49478581 | -0.3771392 | 0.70607017 | 0.90988678 |
| BCL7C | 361.155785 | 0.27499216 | 0.37506487 | 0.73318559 | 0.46344524 | 0.8082072 |
| MIR762HG | 111.879837 | 0.257083923 | 0.36634169 | 0.70175995 | 0.48282888 | 0.8177376 |
| FBXL19-AS1 | 76.0219262 | -1.485635546 | 0.67150015 | -2.2124128 | 0.02693815 | 0.38702446 |
| FBXL19 | 170.519151 | 0.134748278 | 0.46298478 | 0.29104257 | 0.77101876 | 0.93302012 |
| ORAI3 | 130.758391 | 0.353559014 | 0.35927988 | 0.98407687 | 0.32507772 | 0.73628585 |
| SETD1A | 231.938844 | -0.95627878 | 0.4172193 | -2.2920291 | 0.02190396 | 0.36066559 |
| HSD3B7 | 237.143251 | -0.699207067 | 0.64991272 | -1.0758476 | 0.2819954 | 0.70663819 |
| STX1B | 39.354035 | 0.228106703 | 0.56796172 | 0.40162338 | 0.68796122 | 0.90298616 |
| STX4 | 290.274288 | -0.595690356 | 0.29821532 | -1.9975176 | 0.04576899 | 0.44430994 |
| AC135050.3 | 96.1772196 | -0.689047787 | 0.59051362 | -1.1668618 | 0.24326618 | 0.68164834 |
| ZNF668 | 77.4328767 | -0.406086195 | 0.62422229 | -0.6505474 | 0.51533868 | 0.83181198 |
| ZNF646 | 56.3955608 | -0.710071602 | 0.40006881 | -1.7748737 | 0.07591875 | 0.50730849 |
| AC135050.2 | 33.3955655 | -0.345651989 | 0.65859188 | -0.5248349 | 0.59969798 | 0.86700351 |
| BCKDK | 203.131739 | -0.042498036 | 0.392218 | -0.1083531 | 0.9137156 | 0.9773919 |
| KAT8 | 328.237087 | -0.383083759 | 0.39042471 | -0.9811975 | 0.32649534 | 0.73745391 |
| PRSS8 | 390.412158 | -0.513922655 | 0.47053792 | -1.0922024 | 0.27474414 | 0.70166356 |
| FUS | 1433.86313 | -0.182789671 | 0.24360657 | -0.7503479 | 0.45304523 | 0.80433814 |
| AC026471.4 | 147.174186 | 0.1542714 | 0.41970747 | 0.36756887 | 0.71319473 | 0.91191793 |
| AC026471.1 | 47.3684935 | 0.059415324 | 0.47833114 | 0.12421379 | 0.901146 | 0.97482505 |
| ARMC5 | 79.0742914 | -0.633637523 | 0.49407211 | -1.2824799 | 0.19967437 | 0.64687699 |
| SLC5A2 | 1105.74342 | 0.739496437 | 0.7121846 | 1.03834938 | 2.99E-01 | 7.19E-01 |
| AC026471.3 | 30.3930699 | 1.119934401 | 1.2257793 | 0.91365093 | 0.36090029 | 0.76009788 |
| C16orf58 | 295.054824 | 0.004925081 | 0.30545268 | 0.01612387 | 0.98713557 | 0.9963052 |
| ZNF720 | 546.198928 | 0.659857289 | 0.3211193 | 2.05486651 | 0.0398919 | 0.42969399 |
| ZNF267 | 399.570643 | 0.449467573 | 0.24468739 | 1.83690537 | 6.62E-02 | 0.48798003 |
| AC136944.1 | 34.8499526 | -0.101978323 | 0.4444626 | -0.2294419 | 0.81852551 | 0.94990455 |
| VPS35 | 621.482546 | -0.067681821 | 0.22668778 | -0.2985685 | 0.76526933 | 0.93083208 |
| MYLK3 | 36.5204607 | 0.838857058 | 0.44410348 | 1.88887748 | 0.05890825 | 0.47778423 |
| C16orf87 | 645.263142 | 0.544195278 | 0.34545716 | 1.57529019 | 0.11518948 | 0.57182344 |
| GPT2 | 1047.04882 | 0.14007548 | 0.60178822 | 0.23276541 | 0.81594358 | 0.94906482 |
| DNAJA2 | 263.828141 | 0.126181728 | 0.25256726 | 0.49959654 | 0.6173592 | 0.87435665 |
| ITFG1 | 3174.62958 | 0.003274958 | 0.26099967 | 0.01254775 | 0.98998861 | 0.99737132 |
| PHKB | 2264.28474 | 0.127700037 | 0.21690793 | 0.58872922 | 0.55604294 | 0.84861484 |
| LINC02133 | 48.1994826 | 0.842993527 | 0.59403412 | 1.4190995 | 0.15587001 | 0.6125316 |
| LONP2 | 1682.45567 | 0.013545682 | 0.15453453 | 0.08765473 | 0.9301511 | 0.98272895 |
| SIAH1 | 467.194981 | -0.263100184 | 0.20609315 | -1.2766081 | 0.20174064 | 0.64881721 |
| N4BP1 | 1124.45211 | -0.241989098 | 0.24639055 | -0.9821363 | 0.32603272 | 0.7373175 |
| AC026470.2 | 43.2192247 | 0.33186325 | 0.81530788 | 0.4070404 | 0.68397831 | 0.90069082 |
| AC007611.1 | 75.6210256 | -0.822093795 | 0.51559916 | -1.5944436 | 0.11083671 | 0.56317482 |
| CNEP1R1 | 114.245658 | 0.136668613 | 0.23135618 | 0.59072818 | 0.55470256 | 0.8478265 |

| | | | | | | |
|---|---|---|---|---|---|---|
| HEATR3 | 225.177625 | -0.314969218 | 0.35060293 | -0.8983645 | 0.36899127 | 0.76615653 |
| TENT4B | 1295.45634 | 0.081057766 | 0.33166545 | 0.24439617 | 0.80692401 | 0.94585363 |
| BRD7 | 356.245676 | -0.086677099 | 0.22404002 | -0.3868822 | 0.69884341 | 0.90696517 |
| CYLD | 498.700214 | -0.879138297 | 0.40959436 | -2.1463633 | 0.03184401 | 0.40988962 |
| AC025810.1 | 499.927208 | 2.001772282 | 0.76034339 | 2.63272134 | 0.00847038 | 0.25070735 |
| SALL1 | 776.564543 | 0.145627799 | 0.57475629 | 0.25337313 | 0.79997989 | 0.94403774 |
| AC087564.1 | 1741.5529 | 1.510440696 | 0.46639043 | 3.23857566 | 0.00120128 | 0.11201338 |
| AC007344.1 | 235.265994 | 0.433287327 | 0.67630408 | 0.64066939 | 0.5217375 | 0.83445062 |
| HNRNPA1P4 | 61.187868 | -0.385989281 | 0.44502694 | -0.8673391 | 0.38575622 | 0.77317162 |
| TOX3 | 1477.59608 | 0.795366488 | 0.53848623 | 1.47704147 | 0.13966452 | 0.59500405 |
| AC007906.2 | 87.4263636 | 0.274243089 | 0.74752757 | 0.36686686 | 0.71371833 | 0.91209742 |
| CHD9 | 6370.43603 | 0.147387712 | 0.22931098 | 0.64274164 | 0.52039176 | 0.833897 |
| AC007342.4 | 45.955225 | 0.9723619 | 0.79755667 | 1.21917594 | 0.22277742 | 0.66398799 |
| RBL2 | 1132.66811 | 0.257378602 | 0.37549702 | 0.68543447 | 0.4930698 | 0.82100434 |
| AKTIP | 165.006729 | 0.013115494 | 0.2886159 | 0.04544273 | 0.96375442 | 0.99012655 |
| RPGRIP1L | 288.010223 | -0.038866913 | 0.34157276 | -0.1137881 | 0.90940578 | 0.97680747 |
| FTO | 4762.58537 | -0.04658589 | 0.2606705 | -0.1787156 | 0.858161 | 0.9626966 |
| IRX3 | 131.851079 | 0.16522762 | 0.65395954 | 0.25265725 | 0.80053309 | 0.94403774 |
| CRNDE | 265.744413 | 0.412405821 | 0.47627216 | 0.86590369 | 0.38654298 | 0.77334938 |
| IRX5 | 68.5523579 | 0.538444409 | 0.62255938 | 0.86488843 | 0.38710003 | 0.77335375 |
| LPCAT2 | 115.084431 | 0.515022412 | 0.39782257 | 1.2946033 | 0.19545712 | 0.64558745 |
| AMFR | 791.64393 | -0.044588789 | 0.28212011 | -0.158049 | 0.87441822 | 0.96615698 |
| NUDT21 | 311.414489 | -0.13248144 | 0.18848686 | -0.7028683 | 0.48213782 | 0.8177376 |
| OGFOD1 | 270.992707 | -0.228087301 | 0.29912678 | -0.7625105 | 0.4457554 | 0.8012958 |
| BBS2 | 567.684838 | -0.023274919 | 0.3089215 | -0.0753425 | 0.93994221 | 0.98439739 |
| MT2A | 7396.0749 | -1.691667557 | 0.75676649 | -2.2353891 | 0.02539181 | 0.38162895 |
| MT1E | 2891.13849 | -2.917036324 | 0.6760812 | -4.3146242 | 1.60E-05 | 0.01603597 |
| MT1M | 449.639 | -3.680020263 | 1.07209472 | -3.4325514 | 0.00059793 | 0.08329748 |
| MT1F | 782.116318 | -2.04008852 | 0.5664905 | -3.6012758 | 0.00031666 | 0.06474783 |
| MT1G | 12204.5379 | -2.520821524 | 0.75678031 | -3.3309819 | 0.0008654 | 0.09885057 |
| MT1H | 4341.43822 | -3.00714563 | 0.84463538 | -3.5602885 | 0.00037045 | 0.06861791 |
| MT1X | 2884.98461 | -1.908369882 | 0.69877982 | -2.7310031 | 0.00631419 | 0.22719885 |
| NUP93 | 376.541378 | -0.679154765 | 0.4121906 | -1.6476717 | 0.09942007 | 0.54399442 |
| SLC12A3 | 87.2118433 | 0.143452968 | 0.56159847 | 0.25543689 | 7.98E-01 | 0.94320409 |
| HERPUD1 | 1031.91132 | -0.502593127 | 0.5222005 | -0.9624524 | 0.3358224 | 0.74231859 |
| NLRC5 | 243.312826 | -1.489748594 | 0.61982031 | -2.4035169 | 1.62E-02 | 0.32071648 |
| CPNE2 | 50.1796028 | -0.170614264 | 0.37153141 | -0.4592189 | 6.46E-01 | 8.87E-01 |
| FAM192A | 578.50957 | -0.52578842 | 0.26530355 | -1.9818371 | 0.04749747 | 0.44992941 |
| RSPRY1 | 1012.93903 | -0.178851713 | 0.19541318 | -0.915249 | 0.36006093 | 0.75998694 |
| ARL2BP | 372.170795 | 0.005388845 | 0.26178 | 0.0205854 | 0.98357639 | 0.99523767 |
| PLLP | 334.979927 | 0.700711157 | 0.41772157 | 1.67745985 | 0.09345259 | 0.53688613 |
| CX3CL1 | 258.739156 | 1.053941133 | 0.57266506 | 1.8404146 | 0.06570739 | 0.48790819 |
| CIAPIN1 | 377.937335 | -0.056780941 | 0.17958839 | -0.3161727 | 0.75187145 | 0.9267064 |
| COQ9 | 593.907778 | 0.338757437 | 0.27804691 | 1.21834634 | 0.22309239 | 0.66408018 |
| POLR2C | 194.415592 | -0.112005097 | 0.3195757 | -0.3504806 | 0.72597802 | 0.91635631 |
| DOK4 | 396.178115 | 0.227838692 | 0.25345436 | 0.89893379 | 0.36868793 | 0.76615653 |
| CCDC102A | 36.274841 | 0.688134487 | 0.46430326 | 1.48207982 | 0.13831906 | 0.59474076 |
| ADGRG5 | 301.586087 | 1.800153172 | 0.62219392 | 2.8932349 | 3.81E-03 | 0.2001601 |
| ADGRG1 | 2009.33578 | -0.269877451 | 0.68852781 | -0.391963 | 0.69508553 | 0.90589033 |
| KATNB1 | 44.6144747 | 0.040623633 | 0.43792672 | 0.09276354 | 0.92609142 | 0.9811358 |
| KIFC3 | 3316.69993 | -0.485094134 | 0.32748102 | -1.4812893 | 1.39E-01 | 5.95E-01 |
| AC092118.1 | 60.2694614 | 0.194986243 | 0.41860101 | 0.46580452 | 0.64135543 | 0.88403746 |
| USB1 | 145.553438 | -0.192283472 | 0.33022732 | -0.5822761 | 0.56038074 | 0.84944908 |
| MMP15 | 89.0083901 | 0.209637419 | 0.47216272 | 0.44399401 | 0.65704691 | 0.89053559 |

| | | | | | | |
|---|---|---|---|---|---|---|
| CFAP20 | 102.471965 | 0.044520553 | 0.21638968 | 0.2057425 | 0.83699206 | 0.95496955 |
| CSNK2A2 | 520.555367 | -0.128077612 | 0.26945137 | -0.4753274 | 0.63455365 | 0.88238823 |
| CCDC113 | 43.4666789 | 0.400567335 | 0.55153127 | 0.72628218 | 0.4676658 | 0.81184083 |
| SETD6 | 136.061 | -0.709264825 | 0.30581962 | -2.319226 | 0.02038278 | 0.35081655 |
| CNOT1 | 1601.85166 | -0.21979736 | 0.21103821 | -1.0415051 | 0.29764121 | 0.71839551 |
| AC009118.3 | 37.4920418 | -0.860523426 | 0.51724808 | -1.6636571 | 0.09618098 | 0.53876283 |
| SLC38A7 | 320.046299 | -0.967998562 | 0.44858695 | -2.1578839 | 0.03093686 | 0.40540296 |
| GOT2 | 430.147948 | -0.045755454 | 0.20865505 | -0.2192875 | 0.82642606 | 0.95192757 |
| AC092378.1 | 102.211192 | 0.011274752 | 0.41315502 | 0.0272894 | 0.97822891 | 0.99482978 |
| AC138305.1 | 307.65244 | 1.178955801 | 0.48074935 | 2.45232947 | 0.01419346 | 0.30468798 |
| CDH11 | 85.5915398 | 0.734642111 | 0.6784101 | 1.08288794 | 0.27885817 | 0.70446562 |
| BEAN1 | 103.744505 | 1.991831536 | 0.63250207 | 3.14913048 | 0.00163757 | 0.13207839 |
| AC010542.3 | 233.231101 | 1.612647283 | 0.51169374 | 3.15158692 | 0.00162386 | 0.13207839 |
| TK2 | 207.711772 | 0.744704633 | 0.41491757 | 1.79482551 | 0.07268151 | 0.50366299 |
| CMTM3 | 59.3381514 | 0.412150436 | 0.49190955 | 0.83785817 | 4.02E-01 | 7.83E-01 |
| CMTM4 | 1377.0835 | -0.024623532 | 0.28984358 | -0.0849546 | 0.93229752 | 0.98306798 |
| DYNC1LI2 | 1163.40671 | -0.250700745 | 0.24820583 | -1.0100518 | 0.31247048 | 0.7265535 |
| AC018557.1 | 51.0809818 | 0.826586219 | 0.55873506 | 1.47938848 | 0.13903652 | 0.59474076 |
| NAE1 | 285.475713 | -0.038995017 | 0.2043572 | -0.1908179 | 0.84866824 | 0.95951833 |
| PDP2 | 222.420271 | 0.156854187 | 0.39706106 | 0.39503795 | 0.69281488 | 0.90528251 |
| CDH16 | 932.508471 | 0.184198246 | 0.46981006 | 0.39206961 | 0.69500678 | 0.90589033 |
| RRAD | 383.438895 | 0.910080982 | 0.67753677 | 1.34322005 | 0.17920073 | 0.63014479 |
| FAM96B | 487.403835 | -0.296711879 | 0.38670743 | -0.7672774 | 0.4429166 | 0.7993765 |
| CES2 | 731.259165 | 0.370551932 | 0.3959578 | 0.93583693 | 0.34935715 | 0.7542101 |
| CES3 | 215.284936 | 1.572260151 | 0.8500082 | 1.84969999 | 0.0643568 | 0.48593331 |
| CES4A | 135.928402 | 1.795873496 | 0.76992544 | 2.33252909 | 0.01967287 | 0.34523522 |
| CBFB | 1086.26554 | -0.135895578 | 0.18990162 | -0.7156104 | 0.47423194 | 0.8151865 |
| C16orf70 | 719.825952 | -0.33382676 | 0.33400998 | -0.9994515 | 0.31757604 | 0.72939256 |
| TRADD | 190.986772 | 0.593696782 | 0.39627634 | 1.49818882 | 0.1340842 | 0.59240843 |
| FBXL8 | 33.8396923 | 0.497523123 | 0.62643611 | 0.79421207 | 0.42707198 | 0.78933193 |
| HSF4 | 132.322004 | 0.561809369 | 0.73958844 | 0.75962433 | 0.44747917 | 0.80223612 |
| NOL3 | 291.622407 | 0.775950787 | 0.50814371 | 1.52703019 | 0.1267535 | 0.58755843 |
| KIAA0895L | 144.141417 | 1.080851123 | 0.44560335 | 2.42559022 | 0.01528352 | 0.3105046 |
| E2F4 | 216.916548 | -0.413233405 | 0.36487267 | -1.1325414 | 0.25740688 | 0.691812 |
| ELMO3 | 146.803095 | 0.234414451 | 0.35568841 | 0.6590444 | 0.50986726 | 0.82893495 |
| LRRC29 | 135.526646 | 0.008644809 | 0.50040363 | 0.01727567 | 0.98621669 | 0.99609994 |
| TMEM208 | 228.150166 | -0.292637237 | 0.42554043 | -0.6876838 | 0.49165195 | 0.82063728 |
| FHOD1 | 84.0229745 | 0.198465161 | 0.34664888 | 0.57252504 | 0.56696633 | 0.85233132 |
| LRRC36 | 163.790507 | 0.072415828 | 0.75168799 | 0.09633761 | 0.92325244 | 0.98002177 |
| ZDHHC1 | 204.996986 | 0.348919567 | 0.35340047 | 0.98732062 | 0.32348549 | 0.7354443 |
| HSD11B2 | 198.945831 | 0.793828217 | 0.78580857 | 1.0102056 | 3.12E-01 | 0.7265535 |
| ATP6V0D1 | 600.168152 | -0.563911064 | 0.41158469 | -1.3700973 | 0.17065654 | 0.62478344 |
| AC009061.2 | 133.870426 | 0.821870631 | 0.50105317 | 1.64028627 | 0.10094566 | 0.54608983 |
| AC027682.6 | 70.970767 | -0.920321082 | 0.64263826 | -1.4320982 | 0.15211573 | 0.61001662 |
| RIPOR1 | 738.958131 | -0.917820798 | 0.35128036 | -2.6127872 | 0.00898072 | 0.2563089 |
| AC027682.4 | 75.7828226 | -0.074903308 | 0.39767052 | -0.1883552 | 0.85059822 | 0.95994332 |
| CTCF | 713.102848 | -0.622031539 | 0.39703677 | -1.566685 | 0.11718834 | 0.57350419 |
| ENKD1 | 146.902932 | 0.170646159 | 0.44284217 | 0.38534307 | 0.69998326 | 0.90721166 |
| C16orf86 | 35.8692429 | 1.012297075 | 0.60595588 | 1.67057885 | 0.09480489 | 0.53824794 |
| GFOD2 | 417.080133 | -0.193623473 | 0.37084562 | -0.5221134 | 0.60159137 | 0.86757631 |
| RANBP10 | 775.501658 | 0.135187582 | 0.27548287 | 0.49072955 | 0.62361775 | 0.87772094 |
| TSNAXIP1 | 58.4262854 | 0.413932127 | 0.51931942 | 0.79706653 | 0.42541239 | 0.78894887 |
| CENPT | 150.036118 | -1.251921566 | 0.3855386 | -3.2472016 | 0.00116546 | 0.11093704 |
| THAP11 | 37.3041232 | -0.256309088 | 0.41576346 | -0.6164781 | 0.53757899 | 0.84039635 |

| | | | | | | |
|---|---|---|---|---|---|---|
| NUTF2 | 169.200616 | -0.67233199 | 0.41341265 | -1.6262976 | 0.10388637 | 0.55022042 |
| PSKH1 | 238.183688 | -0.010999524 | 0.29357821 | -0.0374671 | 0.97011257 | 0.99204339 |
| LCAT | 71.8535142 | -0.52459621 | 0.57572835 | -0.911187 | 0.36219683 | 0.76104518 |
| SLC12A4 | 96.0329895 | 0.02287955 | 0.37078315 | 0.06170601 | 0.95079695 | 0.98731502 |
| DUS2 | 191.347879 | -0.328668838 | 0.23287314 | -1.4113643 | 0.15813722 | 0.61369956 |
| DDX28 | 34.1595623 | -0.361106851 | 0.51932936 | -0.695333 | 0.48684663 | 0.81889885 |
| NFATC3 | 2909.72291 | 0.051786572 | 0.34109585 | 0.15182411 | 0.87932567 | 0.9674841 |
| AC020978.5 | 57.7357561 | -0.311395007 | 0.48327424 | -0.6443443 | 0.51935218 | 0.8335052 |
| ESRP2 | 117.189354 | -0.335396664 | 0.4117906 | -0.8144835 | 0.415368 | 0.78632759 |
| PLA2G15 | 410.305694 | 0.453680101 | 0.45619793 | 0.99448085 | 0.31998883 | 0.73157402 |
| SLC7A6 | 200.338629 | -0.47625265 | 0.42453491 | -1.1218221 | 0.26193808 | 0.69408034 |
| SLC7A6OS | 245.882736 | -0.580360768 | 0.36703491 | -1.5812141 | 0.1138291 | 0.56865873 |
| PRMT7 | 330.748804 | -0.10936134 | 0.39340906 | -0.2779838 | 0.78102481 | 0.93748328 |
| ZFP90 | 294.567848 | 0.561160252 | 0.43632252 | 1.28611342 | 0.19840348 | 0.64582202 |
| CDH1 | 390.322478 | 0.519393174 | 0.61986037 | 0.83791964 | 0.40207584 | 0.78258224 |
| TANGO6 | 1328.00103 | 0.033346979 | 0.32049864 | 0.10404718 | 9.17E-01 | 0.97828877 |
| AC009137.1 | 38.0760497 | 0.291574099 | 0.65905731 | 0.44241084 | 0.65819194 | 0.89058983 |
| CHTF8 | 478.396065 | -0.254598313 | 0.38504523 | -0.6612166 | 0.50847339 | 0.82874455 |
| UTP4 | 894.27061 | -0.128132864 | 0.23350408 | -0.5487393 | 0.58318438 | 0.85990837 |
| SNTB2 | 1371.96653 | -0.179491279 | 0.4583964 | -0.3915635 | 0.69538079 | 0.90589033 |
| VPS4A | 351.411425 | 0.294845459 | 0.30367933 | 0.97091051 | 0.33159284 | 0.73939532 |
| COG8 | 228.403491 | 0.436503835 | 0.37525211 | 1.1632282 | 0.24473693 | 0.68250191 |
| AC026464.3 | 37.2808148 | 0.006558096 | 0.44379442 | 0.01477733 | 0.98820983 | 0.99671759 |
| NIP7 | 92.9599854 | -0.532679771 | 0.37305478 | -1.4278862 | 0.15332462 | 0.61008819 |
| TERF2 | 310.367721 | -0.121049404 | 0.22115991 | -0.5473388 | 0.58414598 | 0.86025731 |
| CYB5B | 711.848109 | 0.134970974 | 0.23577657 | 0.57245288 | 0.5670152 | 0.85233132 |
| NFAT5 | 5166.67675 | -0.094930009 | 0.15839617 | -0.5993201 | 0.54895944 | 0.84583938 |
| NQO1 | 539.690905 | 0.668262872 | 1.01158026 | 0.6606128 | 0.50886066 | 0.82874455 |
| AC092115.3 | 74.7283936 | 0.10708866 | 0.67587461 | 0.15844457 | 0.8741065 | 0.9661168 |
| NOB1 | 123.356126 | -0.488081781 | 0.34428887 | -1.4176519 | 0.15629241 | 0.61255389 |
| WWP2 | 1467.59861 | -0.501889043 | 0.31220468 | -1.6075641 | 0.10793067 | 0.55721497 |
| CLEC18A | 87.7911623 | 1.404269325 | 0.96755523 | 1.45135832 | 0.14668011 | 0.60383734 |
| PDPR | 970.729933 | -0.657255131 | 0.37586309 | -1.7486557 | 0.08035055 | 0.51297858 |
| AC009060.1 | 839.411974 | -0.018112183 | 0.58780733 | -0.0308131 | 0.97541857 | 0.99420467 |
| CLEC18C | 87.7321142 | 2.038853224 | 0.95787685 | 2.1285129 | 0.03329458 | 0.41254527 |
| EXOSC6 | 238.053059 | -0.751128967 | 0.37591966 | -1.9981104 | 0.0457047 | 0.44430994 |
| AARS | 475.144358 | -0.487592839 | 0.37585479 | -1.2972905 | 0.19453126 | 0.64548758 |
| AC012184.2 | 62.9382561 | 0.010292802 | 0.43710766 | 0.02354752 | 0.98121353 | 0.9951185 |
| DDX19A | 85.9665841 | -0.251509351 | 0.35372611 | -0.7110285 | 0.47706657 | 0.81570932 |
| ST3GAL2 | 413.876609 | 0.182216389 | 0.40866229 | 0.44588501 | 0.65568031 | 0.89053559 |
| FUK | 209.271992 | 0.080628101 | 0.24392181 | 0.33054896 | 0.7409852 | 0.92213095 |
| COG4 | 420.099941 | 0.005505782 | 0.23182892 | 0.02374933 | 0.98105256 | 0.9951185 |
| SF3B3 | 589.818806 | -0.165594689 | 0.22682419 | -0.7300574 | 0.46535507 | 0.81042138 |
| IL34 | 52.3076088 | 0.747231617 | 0.65981552 | 1.13248567 | 0.25743031 | 0.691812 |
| MTSS1L | 315.738838 | -0.54925196 | 0.49835425 | -1.1021316 | 0.27040447 | 0.69961625 |
| VAC14 | 672.488996 | -0.63916023 | 0.47697333 | -1.3400335 | 0.18023446 | 0.63124625 |
| HYDIN | 1309.59804 | 0.669785696 | 0.53166085 | 1.25979878 | 0.20774196 | 0.65309147 |
| CMTR2 | 369.076832 | -0.403111049 | 0.35842138 | -1.1246847 | 0.26072266 | 0.69280433 |
| ZNF19 | 337.607248 | 1.00088369 | 0.468414 | 2.13675017 | 0.03261832 | 0.41149854 |
| MARVELD3 | 552.166495 | 0.323945683 | 0.42331307 | 0.76526265 | 0.44411516 | 0.79982742 |
| PHLPP2 | 525.411811 | 0.218446983 | 0.2130489 | 1.0253373 | 3.05E-01 | 0.72103276 |
| AP1G1 | 1881.34454 | 0.073456406 | 0.22470757 | 0.32689777 | 0.7437452 | 0.92350771 |
| IST1 | 1119.29626 | -0.241353076 | 0.14080582 | -1.7140845 | 0.08651321 | 0.52493871 |
| ZNF821 | 318.096906 | 0.473017042 | 0.40615249 | 1.16462918 | 0.24416913 | 0.68222743 |

| | | | | | | |
|---|---|---|---|---|---|---|
| PKD1L3 | 56.8203612 | 1.938026787 | 0.66836362 | 2.89965928 | 0.00373568 | 0.1981659 |
| DHODH | 123.651455 | 0.163894425 | 0.36870376 | 0.44451519 | 0.65667015 | 0.89053559 |
| TXNL4B | 311.677998 | 0.503552828 | 0.33209907 | 1.51627293 | 0.12945035 | 0.58817997 |
| HP | 467.364406 | -2.637808856 | 1.73850362 | -1.517287 | 1.29E-01 | 0.58793678 |
| DHX38 | 241.909062 | -0.393171858 | 0.32665638 | -1.2036252 | 0.22873446 | 0.66705629 |
| PMFBP1 | 52.1715499 | 1.274065474 | 0.70926103 | 1.79632804 | 0.07244236 | 0.502734 |
| LINC01572 | 757.356245 | 0.684211357 | 0.57804087 | 1.18367298 | 0.23654254 | 0.67365231 |
| AC004943.2 | 304.348742 | 0.350866246 | 0.38030549 | 0.92259054 | 0.35622063 | 0.75841231 |
| ZFHX3 | 1329.46149 | -0.328034943 | 0.26507859 | -1.2375007 | 0.21590126 | 0.65967206 |
| AC138627.1 | 405.678729 | 1.300280523 | 0.65212972 | 1.99389857 | 0.04616314 | 0.44508419 |
| PSMD7 | 463.524434 | -0.455441087 | 0.34947469 | -1.3032162 | 0.19250095 | 0.64497936 |
| NPIPB15 | 231.682418 | 0.079481537 | 0.99390677 | 0.07996881 | 0.93626207 | 0.98415216 |
| CLEC18B | 105.631307 | 1.971043976 | 1.07307587 | 1.83681697 | 0.06623692 | 0.48798003 |
| AC009053.3 | 639.08693 | -0.935612295 | 0.61115635 | -1.5308886 | 0.12579693 | 0.58564339 |
| GLG1 | 3922.60348 | -0.32331222 | 0.17092871 | -1.8915033 | 0.05855719 | 0.47778423 |
| RFWD3 | 344.948852 | -0.258414858 | 0.32732278 | -0.7894802 | 0.42983139 | 0.79055252 |
| MLKL | 65.9087022 | -0.403410066 | 0.52639322 | -0.7663664 | 4.43E-01 | 7.99E-01 |
| FA2H | 63.617937 | 0.513869825 | 0.54526091 | 0.94242925 | 0.34597292 | 0.75166079 |
| WDR59 | 1809.11435 | -0.171346251 | 0.24357706 | -0.7034581 | 0.48177032 | 0.8177376 |
| ZNRF1 | 762.81559 | -0.079347926 | 0.35336686 | -0.2245483 | 0.82233069 | 0.95112619 |
| LDHD | 115.52811 | 0.582469393 | 0.6530344 | 0.8919429 | 0.37242354 | 0.76879661 |
| ZFP1 | 102.274924 | -0.184809404 | 0.27307933 | -0.6767609 | 0.49855768 | 0.82363868 |
| BCAR1 | 1173.36621 | -0.731796176 | 0.43502965 | -1.6821754 | 0.09253483 | 0.53654752 |
| CFDP1 | 2906.98239 | -0.427369354 | 0.22128369 | -1.9313188 | 0.05344364 | 0.46711025 |
| AC009163.4 | 42.5332214 | -0.36792526 | 0.4617625 | -0.7967846 | 0.42557613 | 0.78894887 |
| TMEM231 | 37.7950929 | 0.421316198 | 0.55855945 | 0.7542907 | 0.45067469 | 0.80361623 |
| GABARAPL2 | 282.119335 | 0.141894164 | 0.38421278 | 0.36931141 | 0.71189562 | 0.91161746 |
| ADAT1 | 148.643309 | -0.115858407 | 0.3412284 | -0.3395333 | 0.73420801 | 0.91975299 |
| KARS | 279.824225 | -0.551155911 | 0.42457224 | -1.298144 | 0.19423785 | 0.64548758 |
| TERF2IP | 760.558972 | 0.564985046 | 0.45061085 | 1.25382032 | 0.20990732 | 0.65441616 |
| AC010528.1 | 44.1561718 | 0.440545896 | 0.71513216 | 0.61603424 | 0.53787192 | 0.84040975 |
| MON1B | 94.657578 | -0.302970508 | 0.49256926 | -0.615082 | 0.53850054 | 0.84049498 |
| SYCE1L | 51.7760142 | 0.29785734 | 0.61152711 | 0.48707136 | 0.62620776 | 0.87877458 |
| NUDT7 | 102.016083 | 0.408621432 | 0.58002942 | 0.70448398 | 0.48113142 | 0.81720798 |
| VAT1L | 49.1073938 | -0.151075 | 0.50057417 | -0.3018034 | 0.76280192 | 0.93022076 |
| WWOX | 14136.0206 | 0.336462793 | 0.29305385 | 1.14812615 | 0.25091649 | 0.68648949 |
| AC046158.1 | 120.866108 | 0.243360634 | 0.77601183 | 0.31360428 | 0.75382161 | 0.92783044 |
| MAF | 1670.7581 | 0.03370183 | 0.45719755 | 0.07371393 | 0.94123801 | 0.98489601 |
| AC009159.3 | 27.7693967 | -0.812152355 | 0.74192086 | -1.0946617 | 0.27366487 | 0.70140695 |
| LINC01229 | 59.7168462 | 1.08054134 | 1.00887564 | 1.07103522 | 0.28415359 | 0.70897061 |
| MAFTRR | 136.286709 | 0.426090139 | 0.80410613 | 0.52989291 | 0.59618618 | 0.86539512 |
| AC105411.1 | 94.3954321 | 1.056126086 | 0.67650663 | 1.5611467 | 0.11848914 | 0.57589001 |
| DYNLRB2 | 67.9039978 | 0.453237246 | 0.58868652 | 0.76991273 | 0.44135166 | 0.79862562 |
| CDYL2 | 1139.00694 | 0.465005984 | 0.33382129 | 1.39297881 | 0.1636262 | 0.62065658 |
| CMC2 | 734.477846 | 0.381977909 | 0.19541016 | 1.95474944 | 0.05061265 | 0.45903191 |
| CENPN | 58.8051883 | -0.443561744 | 0.46158626 | -0.9609509 | 0.33657684 | 0.7429163 |
| ATMIN | 189.272133 | -0.425838854 | 0.37680069 | -1.1301435 | 0.25841575 | 0.69210282 |
| BCO1 | 181.552438 | 0.515829057 | 0.36577095 | 1.41025158 | 0.15846541 | 0.61423843 |
| GAN | 525.823645 | -0.107773555 | 0.29725236 | -0.3625658 | 0.71692922 | 0.91311549 |
| CMIP | 5426.20022 | -0.709605065 | 0.34734874 | -2.0429182 | 0.04106053 | 0.43143739 |
| PLCG2 | 1411.13828 | -0.184019987 | 0.36533832 | -0.5036975 | 0.61447399 | 0.87225883 |
| SDR42E1 | 38.9819074 | -0.099525856 | 0.40866803 | -0.2435372 | 0.80758929 | 0.94604106 |
| AC092142.1 | 50.2048757 | -0.496951607 | 0.42739059 | -1.1627575 | 0.24492791 | 0.68250191 |
| MPHOSPH6 | 210.667015 | -0.395067448 | 0.26219557 | -1.5067663 | 0.13187056 | 0.59087581 |

| | | | | | | |
|---|---|---|---|---|---|---|
| CDH13 | 58.010117 | -0.3682443 | 0.46190771 | -0.7972248 | 0.42532046 | 0.78894887 |
| HSBP1 | 385.651874 | -0.330176476 | 0.32356035 | -1.0204479 | 0.30751609 | 0.72263382 |
| MLYCD | 122.250028 | 0.006283472 | 0.49256477 | 0.01275664 | 0.98982195 | 0.99731619 |
| AC009119.2 | 165.390799 | 0.240130138 | 0.59409016 | 0.40419814 | 0.68606702 | 0.90153587 |
| OSGIN1 | 381.056983 | -1.551501542 | 0.69334459 | -2.2377063 | 0.02524022 | 0.38060247 |
| MBTPS1 | 680.579017 | -0.096649873 | 0.2208249 | -0.4376765 | 0.6616208 | 0.89184036 |
| HSDL1 | 177.646066 | -0.653413833 | 0.46207341 | -1.414091 | 0.1573352 | 0.61301692 |
| DNAAF1 | 135.460618 | 0.937865901 | 0.77592731 | 1.2087033 | 0.22677685 | 0.66690202 |
| TAF1C | 220.39908 | -0.349741203 | 0.23525613 | -1.4866401 | 0.13710988 | 0.59474076 |
| ATP2C2 | 248.035105 | 0.516738313 | 0.52983648 | 0.97527886 | 0.32942195 | 0.73829083 |
| ATP2C2-AS1 | 38.7206601 | 0.498938687 | 0.49984297 | 0.99819087 | 0.31818681 | 0.73030586 |
| TLDC1 | 365.224204 | 0.242326125 | 0.33389088 | 0.72576442 | 0.4679832 | 0.8118777 |
| COTL1 | 792.545586 | -0.69260372 | 0.56522655 | -1.225356 | 0.22044113 | 0.66263062 |
| KLHL36 | 813.364598 | 0.217954944 | 0.20835933 | 1.04605319 | 0.29553647 | 0.71679565 |
| USP10 | 1219.55546 | -0.335440594 | 0.29354077 | -1.1427394 | 0.25314681 | 0.68887358 |
| ZDHHC7 | 584.169739 | -0.353811739 | 0.23767777 | -1.4886194 | 0.13658762 | 0.5942049 |
| KIAA0513 | 149.819391 | -0.491567345 | 0.5426931 | -0.9057925 | 0.36504569 | 0.76357107 |
| GSE1 | 1906.21595 | -0.250876817 | 0.30417231 | -0.8247852 | 0.40949355 | 0.78542936 |
| EMC8 | 197.145293 | -0.34352754 | 0.36762435 | -0.9344526 | 0.35007046 | 0.75455562 |
| COX4I1 | 1344.77056 | 0.059635681 | 0.45831834 | 0.13011847 | 0.8964727 | 0.97314891 |
| IRF8 | 108.452649 | 0.387200337 | 0.91173901 | 0.4246833 | 0.67106756 | 0.89694443 |
| MTHFSD | 132.506536 | -0.139486155 | 0.37923652 | -0.3678078 | 0.71301654 | 0.91191793 |
| C16orf95 | 46.4535466 | 0.115326306 | 0.44861286 | 0.25707312 | 0.79712233 | 0.94280402 |
| AC010531.1 | 46.1928761 | -0.239925958 | 0.42681932 | -0.5621253 | 0.57403062 | 0.85448213 |
| FBXO31 | 461.927355 | -0.897804865 | 0.4050654 | -2.2164442 | 0.02666109 | 0.38563341 |
| MAP1LC3B | 604.993381 | -0.837452361 | 0.42507313 | -1.9701371 | 0.04882266 | 0.45368151 |
| ZCCHC14 | 1877.57243 | -0.965760457 | 0.25363327 | -3.8077042 | 0.00014026 | 0.0393165 |
| KLHDC4 | 624.666359 | -0.595360204 | 0.51485043 | -1.1563751 | 0.24752777 | 0.68482118 |
| SLC7A5 | 108.919021 | -0.524114919 | 0.65659349 | -0.7982335 | 0.424735 | 0.78894887 |
| BANP | 893.222299 | -0.738005207 | 0.48502016 | -1.521597 | 0.1281101 | 0.58793678 |
| ZFPM1 | 74.9534006 | -0.123441781 | 0.42773874 | -0.2885915 | 0.77289397 | 0.93389084 |
| ZC3H18 | 424.538658 | -0.601999613 | 0.3976921 | -1.5137329 | 0.1300936 | 0.58853436 |
| CYBA | 1165.97861 | 0.135154978 | 0.5457838 | 0.24763465 | 0.8044171 | 0.94555743 |
| MVD | 241.360381 | 0.842829605 | 0.35382942 | 2.38202238 | 0.01721785 | 0.32964126 |
| SNAI3-AS1 | 83.9403999 | 0.675325034 | 0.3064926 | 2.20339749 | 0.02756674 | 0.38942007 |
| SNAI3 | 35.5512131 | 1.532954563 | 0.7760657 | 1.97528967 | 0.04823528 | 0.45192142 |
| RNF166 | 73.6917267 | 0.00320564 | 0.38694278 | 0.00828453 | 0.99338997 | 0.9976934 |
| PIEZO1 | 433.846736 | -1.070131948 | 0.53293416 | -2.0080003 | 0.04464326 | 0.44360789 |
| APRT | 370.416583 | -0.010471034 | 0.45274339 | -0.023128 | 0.9815482 | 0.99516789 |
| GALNS | 350.436303 | 0.140810727 | 0.40365104 | 0.34884272 | 0.72720739 | 0.91645085 |
| TRAPPC2L | 123.947315 | -0.593614043 | 0.3803873 | -1.5605517 | 0.11862956 | 0.57589001 |
| ACSF3 | 318.611464 | -0.093262546 | 0.37861424 | -0.246326 | 0.80542985 | 0.94585363 |
| AC009113.1 | 35.4103522 | -0.067687507 | 0.58647873 | -0.1154134 | 0.90811745 | 0.97673932 |
| ZNF778 | 143.177889 | -0.577268176 | 0.36174753 | -1.5957764 | 0.11053872 | 0.56264122 |
| ANKRD11 | 3535.33183 | -0.795243568 | 0.43579856 | -1.8247962 | 0.06803179 | 0.4900219 |
| AC137932.3 | 36.9609665 | -0.742007912 | 0.55730079 | -1.3314317 | 0.18304702 | 0.63276491 |
| SPG7 | 1838.94939 | -0.957677577 | 0.39723415 | -2.4108642 | 0.01591477 | 0.31735328 |
| RPL13 | 2942.10796 | -0.061784905 | 0.47146376 | -0.1310491 | 0.89573646 | 0.97309833 |
| CPNE7 | 129.726875 | 1.03868245 | 0.55661024 | 1.86608578 | 0.06202937 | 0.48347236 |
| DPEP1 | 1122.38846 | 0.87755841 | 0.44099985 | 1.98992906 | 0.04659875 | 0.44535626 |
| CHMP1A | 357.777697 | -0.302440523 | 0.3739397 | -0.8087949 | 0.41863313 | 0.78756578 |
| SPATA33 | 175.09842 | 0.521945179 | 0.25854883 | 2.01874895 | 0.04351332 | 0.44147822 |
| CDK10 | 851.401238 | 0.240126167 | 0.2099961 | 1.14347917 | 0.2528397 | 0.68887358 |
| LINC02166 | 96.7370275 | 0.39915857 | 0.51436277 | 0.77602539 | 0.437734 | 0.7967412 |

| | | | | | | |
|---|---|---|---|---|---|---|
| SPATA2L | 130.904737 | 0.2430392 | 0.29041878 | 0.83685772 | 0.40267255 | 0.78260941 |
| VPS9D1 | 281.089249 | 0.003753067 | 0.32967983 | 0.01138397 | 0.9909171 | 0.99737132 |
| ZNF276 | 235.225164 | -0.042649857 | 0.26381064 | -0.1616684 | 0.87156695 | 0.9657206 |
| FANCA | 55.0170461 | -0.57411201 | 0.50203656 | -1.1435661 | 0.25280361 | 0.68887358 |
| SPIRE2 | 61.4232178 | 0.326453093 | 0.50148604 | 0.65097145 | 0.51506492 | 0.8317258 |
| TCF25 | 1856.1636 | -0.222764229 | 0.36203706 | -0.6153078 | 0.53835146 | 0.84049498 |
| MC1R | 53.416976 | -0.709274983 | 0.73091499 | -0.9703933 | 0.3318505 | 0.73939532 |
| DEF8 | 220.77869 | -0.509473853 | 0.28988315 | -1.7575145 | 0.07883015 | 0.51192014 |
| DBNDD1 | 94.5226909 | 0.134173172 | 0.34196906 | 0.39235471 | 0.69479614 | 0.90589033 |
| GAS8 | 668.970271 | 0.100035272 | 0.27607107 | 0.36235333 | 0.71708801 | 0.91317971 |
| FAM157C | 357.472215 | -0.629096918 | 0.35135692 | -1.7904782 | 0.07337707 | 0.50459544 |
| AC133919.2 | 41.9301775 | 0.424430968 | 0.93493647 | 0.45396771 | 0.64985207 | 0.88857633 |
| LINC02193 | 135.756843 | -0.099755524 | 0.42130049 | -0.23678 | 0.81282748 | 0.94804512 |
| AC240565.1 | 31.9920801 | -0.097570396 | 0.54860905 | -0.1778505 | 0.85884038 | 0.9626966 |
| RPH3AL | 605.098156 | -0.035485923 | 0.37224617 | -0.0953292 | 0.92405336 | 0.98018878 |
| AC129507.2 | 84.2079489 | 0.515665226 | 0.49929727 | 1.03278199 | 0.30170593 | 0.71908176 |
| C17orf97 | 69.505705 | 0.510770894 | 0.56059185 | 0.91112794 | 0.36222797 | 0.76104518 |
| VPS53 | 1962.35743 | -0.500032609 | 0.29434612 | -1.6987912 | 0.08935853 | 0.53095823 |
| AC015853.3 | 82.7829325 | 0.258981003 | 0.53213304 | 0.48668469 | 0.6264818 | 0.87885398 |
| FAM57A | 80.5508147 | -0.788932687 | 0.39988861 | -1.9728811 | 4.85E-02 | 4.53E-01 |
| GEMIN4 | 35.9118815 | -0.785257813 | 0.53420209 | -1.4699639 | 0.14157152 | 0.59611016 |
| GLOD4 | 276.005018 | 0.104650275 | 0.18015258 | 0.580898 | 0.5613092 | 0.84958759 |
| MRM3 | 67.073451 | -0.136511541 | 0.30910194 | -0.4416392 | 0.6587503 | 0.89058983 |
| NXN | 183.360582 | 0.769440884 | 0.37539324 | 2.04969298 | 0.0403944 | 0.43086475 |
| TIMM22 | 105.296866 | -0.513447543 | 0.29090261 | -1.7650152 | 0.0775612 | 0.51128995 |
| ABR | 354.122786 | 0.579863197 | 0.64214682 | 0.90300719 | 0.36652208 | 0.76479192 |
| YWHAE | 1955.9532 | -0.395008714 | 0.20984654 | -1.8823694 | 0.05978588 | 0.48004006 |
| CRK | 1059.30029 | -0.640144135 | 0.50584196 | -1.2655022 | 0.20569135 | 0.651931 |
| MYO1C | 473.781727 | -0.106253468 | 0.35830569 | -0.2965442 | 7.67E-01 | 0.93138917 |
| INPP5K | 135.905841 | -0.679305011 | 0.40459665 | -1.6789684 | 0.09315819 | 0.53688613 |
| PITPNA | 1012.02 | -0.002937423 | 0.35447835 | -0.0082866 | 0.99338832 | 0.9976934 |
| SLC43A2 | 994.641325 | 1.200863855 | 0.5733631 | 2.09442125 | 0.03622247 | 0.42040764 |
| SCARF1 | 62.394289 | 0.115224333 | 0.41551075 | 0.27730771 | 7.82E-01 | 0.93748328 |
| RILP | 65.8659526 | -0.205900118 | 0.48558337 | -0.4240263 | 0.67154664 | 0.89704376 |
| PRPF8 | 724.898068 | -0.118008513 | 0.26037183 | -0.4532307 | 0.65038261 | 0.8887244 |
| TLCD2 | 51.3554156 | -0.828538082 | 0.53881736 | -1.5376974 | 0.12412261 | 0.58405375 |
| MIR22HG | 444.069659 | -1.541761805 | 0.75134685 | -2.0519974 | 0.04016991 | 0.4304996 |
| WDR81 | 393.153679 | -0.855086118 | 0.58657217 | -1.457768 | 0.14490451 | 0.60109715 |
| SERPINF2 | 1557.41118 | 1.618365557 | 0.37046623 | 4.36845634 | 1.25E-05 | 0.01429273 |
| SMYD4 | 230.250252 | 0.104453794 | 0.35176855 | 0.29693898 | 0.76651309 | 0.93138917 |
| RPA1 | 371.631725 | -0.053239747 | 0.23978266 | -0.2220333 | 0.82428792 | 0.95180295 |
| RTN4RL1 | 76.5821061 | 1.33419701 | 0.48658601 | 2.74195516 | 0.00610747 | 0.22637955 |
| DPH1 | 103.808 | -0.390294439 | 0.40544965 | -0.9626212 | 0.33573765 | 0.74225095 |
| SMG6 | 1994.03857 | -0.707861118 | 0.34173293 | -2.0713869 | 0.03832265 | 0.42447711 |
| AC090617.4 | 43.9354273 | -0.205891011 | 0.33933141 | -0.6067549 | 0.54401356 | 0.84408956 |
| SRR | 90.0838374 | -0.17884285 | 0.4541619 | -0.3937866 | 0.69373863 | 0.90587439 |
| TSR1 | 242.198254 | -0.96407131 | 0.41472177 | -2.3246219 | 0.02009219 | 0.34840598 |
| SGSM2 | 290.722336 | 0.1888063 | 0.2893104 | 0.65260806 | 0.51400899 | 0.83126535 |
| MNT | 179.442644 | -0.745219337 | 0.5050072 | -1.4756608 | 0.14003497 | 0.59500405 |
| METTL16 | 753.596311 | -0.453817501 | 0.22790594 | -1.9912491 | 0.04645351 | 0.44508419 |
| PAFAH1B1 | 3168.66362 | -0.366499854 | 0.1720604 | -2.1300652 | 0.03316623 | 0.41254527 |
| CLUH | 1069.90109 | -0.482901632 | 0.3269215 | -1.477118 | 0.13964401 | 0.59500405 |
| RAP1GAP2 | 308.613957 | 0.996222821 | 0.56878704 | 1.75148649 | 0.07986215 | 0.51257191 |
| SPATA22 | 411.399499 | -0.209726861 | 1.00205622 | -0.2092965 | 0.83421678 | 0.95393659 |

| | | | | | | |
|---|---|---|---|---|---|---|
| ASPA | 1465.54452 | -0.289587849 | 1.01710418 | -0.284718 | 0.77586021 | 0.9352344 |
| TRPV1 | 313.786754 | 0.370819836 | 0.4305259 | 0.86131829 | 0.38906276 | 0.7748665 |
| CTNS | 319.683675 | -0.41410734 | 0.24635312 | -1.6809503 | 0.09277258 | 0.53669957 |
| TAX1BP3 | 160.9543 | -0.988147727 | 0.61930313 | -1.5955801 | 0.11058258 | 0.56264122 |
| EMC6 | 174.921709 | -0.301764345 | 0.48575739 | -0.6212244 | 0.53445198 | 0.839337 |
| ITGAE | 165.309229 | -0.111003115 | 0.30194592 | -0.3676258 | 0.71315227 | 0.91191793 |
| NCBP3 | 529.900308 | -0.096403028 | 0.20119825 | -0.4791445 | 0.63183586 | 0.88123095 |
| CAMKK1 | 52.0406686 | -0.729029792 | 0.60911417 | -1.1968689 | 0.23135767 | 0.66878058 |
| ZZEF1 | 1187.13058 | -0.254988789 | 0.25184591 | -1.0124794 | 0.3113089 | 0.7258226 |
| CYB5D2 | 300.527689 | 0.134790725 | 0.35970487 | 0.37472589 | 0.70786434 | 0.91025029 |
| ANKFY1 | 852.346338 | -0.274112791 | 0.22395811 | -1.2239467 | 0.22097233 | 0.66263062 |
| UBE2G1 | 1336.18965 | -0.360346021 | 0.31736125 | -1.1354443 | 0.25618921 | 0.690849 |
| SPNS3 | 136.92427 | 0.012635286 | 0.52503422 | 0.02406564 | 0.98080025 | 0.99510207 |
| AC127521.1 | 156.92058 | -0.370431489 | 0.46068561 | -0.8040874 | 4.21E-01 | 7.89E-01 |
| SPNS2 | 967.704801 | -0.54600072 | 0.43320568 | -1.260373 | 0.20753485 | 0.65289883 |
| MYBBP1A | 132.929825 | -1.011043463 | 0.49711044 | -2.0338407 | 0.04196765 | 0.43678864 |
| GGT6 | 111.191335 | 0.199804366 | 0.50893634 | 0.39259206 | 6.95E-01 | 9.06E-01 |
| SMTNL2 | 719.336903 | -0.450228305 | 0.38399192 | -1.1724942 | 0.24099869 | 0.67957469 |
| PELP1 | 231.713469 | -0.535403945 | 0.38596697 | -1.3871756 | 0.16538823 | 0.62065658 |
| AC091153.2 | 36.9794287 | -0.544997814 | 0.4987268 | -1.0927783 | 0.27449116 | 0.7014484 |
| ARRB2 | 82.6492219 | 0.717822443 | 0.42894196 | 1.67347221 | 0.09423437 | 0.53783536 |
| MED11 | 120.163041 | -0.119567701 | 0.38854951 | -0.3077283 | 0.75828905 | 0.9286043 |
| CXCL16 | 123.590638 | -0.19534149 | 0.49910428 | -0.3913841 | 0.69551332 | 0.90589033 |
| TM4SF5 | 68.1376804 | -0.788772923 | 0.70464024 | -1.1193981 | 0.26297035 | 0.69505103 |
| GLTPD2 | 75.8621615 | -0.522594059 | 0.84325989 | -0.6197307 | 0.53543509 | 0.83954482 |
| PSMB6 | 248.1047 | -0.070480701 | 0.46994683 | -0.1499759 | 0.88078363 | 0.96773407 |
| PLD2 | 90.574717 | 0.483880924 | 0.47335299 | 1.02224119 | 0.30666676 | 0.72190689 |
| MINK1 | 761.889196 | 0.100686841 | 0.29600758 | 0.34014953 | 0.73374392 | 0.91972212 |
| SLC25A11 | 200.844929 | 0.48112394 | 0.31705515 | 1.51747716 | 0.12914626 | 0.58793678 |
| RNF167 | 184.797577 | -0.306068855 | 0.26507588 | -1.1546462 | 0.24823535 | 0.6848328 |
| PFN1 | 998.702688 | -0.710450121 | 0.45627036 | -1.5570815 | 0.11945115 | 0.57652003 |
| SPAG7 | 337.547362 | -1.013305376 | 0.472863 | -2.1429153 | 0.0321199 | 0.40988962 |
| CAMTA2 | 240.135134 | 0.080487827 | 0.22103934 | 0.3641335 | 0.71575833 | 0.91298152 |
| KIF1C | 518.003412 | 0.194895941 | 0.39059795 | 0.49896816 | 0.61780181 | 0.8744793 |
| ZFP3 | 81.2623118 | -0.259141621 | 0.36300591 | -0.7138771 | 0.47530316 | 0.81564184 |
| ZNF232 | 85.6374153 | -0.646815224 | 0.77788709 | -0.8315027 | 0.4056897 | 0.78429302 |
| USP6 | 37.6393599 | -0.140280284 | 0.609648 | -0.2301005 | 0.81801371 | 0.94981058 |
| ZNF594 | 234.600604 | -0.132036165 | 0.23935761 | -0.5516272 | 0.5812038 | 0.85895204 |
| AC087500.1 | 358.218178 | 0.33256563 | 0.37483914 | 0.88722225 | 0.37495925 | 0.7697436 |
| SCIMP | 62.7010303 | 0.832648056 | 0.56487709 | 1.47403404 | 0.14047243 | 0.59500405 |
| RABEP1 | 1582.26792 | -0.414615156 | 0.27491363 | -1.5081651 | 0.13151227 | 0.59000666 |
| NUP88 | 726.419732 | 0.058130895 | 0.21590747 | 0.26923985 | 0.78774512 | 0.93953874 |
| RPAIN | 388.400343 | -0.217403475 | 0.26620524 | -0.8166762 | 0.41411352 | 0.78632759 |
| C1QBP | 245.070551 | -0.102337756 | 0.36327096 | -0.2817119 | 0.77816442 | 0.93646866 |
| DHX33 | 94.4813358 | -0.534799745 | 0.42352775 | -1.2627266 | 0.20668746 | 0.65251041 |
| DERL2 | 244.398287 | -0.062220298 | 0.32673809 | -0.1904287 | 0.84897324 | 0.95959925 |
| MIS12 | 149.52225 | -0.425169526 | 0.31732821 | -1.3398416 | 0.18029686 | 0.63124625 |
| NLRP1 | 42.7006553 | 0.301308599 | 0.55074816 | 0.54708962 | 0.58431717 | 0.86025731 |
| WSCD1 | 87.7493372 | -0.462418898 | 0.75294781 | -0.6141447 | 0.53911972 | 0.84091105 |
| PITPNM3 | 60.3766305 | 0.398464122 | 0.72819967 | 0.54719075 | 0.58424769 | 0.86025731 |
| KIAA0753 | 603.74059 | 0.04046854 | 0.26625308 | 0.15199276 | 0.87919265 | 0.9674841 |
| TXNDC17 | 165.227451 | -0.472807174 | 0.47766632 | -0.9898273 | 0.32225852 | 0.73387566 |
| MED31 | 129.545978 | 0.15397083 | 0.30405119 | 0.50639773 | 0.61257747 | 0.87222135 |
| XAF1 | 246.955528 | -5.467560388 | 1.65963645 | -3.2944326 | 0.00098621 | 0.10560879 |

| | | | | | | |
|---|---|---|---|---|---|---|
| ALOX12-AS1 | 459.494849 | 1.670102887 | 0.61091546 | 2.73377089 | 0.00626136 | 0.22644975 |
| SLC16A13 | 272.614924 | 0.503441586 | 0.57944584 | 0.86883287 | 0.38493855 | 0.77317162 |
| ACADVL | 2185.32489 | -0.364889217 | 0.49013366 | -0.7444688 | 0.45659291 | 0.80507423 |
| DVL2 | 201.722434 | -0.267598599 | 0.24361859 | -1.0984326 | 0.27201563 | 0.70056282 |
| PHF23 | 132.892469 | -0.417876095 | 0.34813561 | -1.2003256 | 0.23001291 | 0.66782184 |
| ELP5 | 159.403567 | 0.150354211 | 0.23618883 | 0.63658475 | 0.52439536 | 0.83589803 |
| EIF5A | 336.2276 | -0.948158244 | 0.58363849 | -1.6245643 | 0.10425543 | 0.5505924 |
| GPS2 | 264.028743 | -0.693438225 | 0.41740745 | -1.6612981 | 0.09665359 | 0.53969191 |
| NEURL4 | 38.0787395 | 0.325103261 | 0.67519795 | 0.48149326 | 0.63016597 | 0.88094696 |
| KCTD11 | 38.5714271 | -0.662884942 | 0.37759083 | -1.7555642 | 0.07916285 | 0.51192014 |
| TNK1 | 42.3619368 | 0.111363442 | 0.36040369 | 0.3089964 | 0.75732427 | 0.92799282 |
| NLGN2 | 85.8859944 | -0.177042829 | 0.69433915 | -0.2549803 | 0.79873829 | 0.94327531 |
| TMEM102 | 33.5958287 | 0.319225755 | 0.67840335 | 0.47055451 | 0.6379589 | 0.8827026 |
| CHRNB1 | 45.1420096 | 0.05758717 | 0.55262179 | 0.1042072 | 0.91700492 | 0.97828877 |
| ZBTB4 | 433.942949 | 0.231022559 | 0.34509801 | 0.66944042 | 0.50321457 | 0.82652042 |
| POLR2A | 759.861613 | 0.041105778 | 0.38150984 | 0.107745 | 0.91419797 | 0.9774502 |
| EIF4A1 | 382.037947 | -0.373713465 | 0.51973522 | -0.7190459 | 0.47211266 | 0.81399348 |
| CD68 | 116.805311 | -1.176863582 | 0.67773421 | -1.7364677 | 0.08248117 | 0.5183475 |
| MPDU1 | 246.688595 | -0.618399024 | 0.29108482 | -2.1244633 | 0.03363142 | 0.41378378 |
| FXR2 | 260.971573 | -0.352015438 | 0.2960818 | -1.1889128 | 0.23447399 | 0.670712 |
| SHBG | 69.8165196 | -0.156573868 | 0.59479271 | -0.2632411 | 0.79236478 | 0.94141203 |
| SAT2 | 1469.77389 | 0.12061165 | 0.59539487 | 0.20257422 | 0.83946784 | 0.9556011 |
| TP53 | 249.97279 | 0.288147577 | 0.42340546 | 0.68054762 | 0.49615778 | 0.82243887 |
| WRAP53 | 59.6873924 | -0.962869734 | 0.60506156 | -1.5913583 | 0.11152897 | 0.56435572 |
| DNAH2 | 37.4615833 | 0.439287428 | 0.55243523 | 0.79518359 | 0.42650671 | 0.78895108 |
| KDM6B | 540.65256 | -1.441441506 | 0.60992146 | -2.3633232 | 0.01811187 | 0.33730895 |
| TMEM88 | 43.8088685 | -1.514944579 | 0.7921373 | -1.9124773 | 0.055815 | 0.47022512 |
| NAA38 | 391.070131 | 0.262230113 | 0.32436197 | 0.80844902 | 0.41883214 | 0.78766781 |
| CYB5D1 | 124.209985 | -0.479704236 | 0.30382939 | -1.5788605 | 0.11436805 | 0.56984472 |
| CHD3 | 147.788889 | 1.134370179 | 0.48374087 | 2.34499553 | 0.01902731 | 0.34084276 |
| TRAPPC1 | 99.3803651 | -0.320687889 | 0.53980653 | -0.5940793 | 0.55245907 | 0.84721285 |
| CNTROB | 169.146205 | -0.023275778 | 0.3649087 | -0.0637852 | 0.94914125 | 0.9871942 |
| PER1 | 499.200244 | -0.361406349 | 0.62644355 | -0.5769177 | 0.56399508 | 0.85149031 |
| TMEM107 | 79.7598712 | 0.314742675 | 0.36130412 | 0.8711295 | 0.38368345 | 0.77317162 |
| BORCS6 | 37.4969205 | -0.394479196 | 0.51970102 | -0.7590503 | 0.44782249 | 0.80223612 |
| CTC1 | 222.876714 | -0.018210024 | 0.21606396 | -0.0842807 | 0.93283325 | 0.98311128 |
| PFAS | 69.5561962 | -0.143186884 | 0.426029 | -0.3360966 | 0.73679805 | 0.92020873 |
| SLC25A35 | 27.4755031 | 0.56954776 | 0.60272859 | 0.94494896 | 0.34468493 | 0.75124763 |
| RANGRF | 70.0602031 | 0.068818022 | 0.36039103 | 0.19095376 | 0.84856182 | 0.95951833 |
| AC135178.3 | 36.6936129 | 0.001002417 | 0.48471051 | 0.00206807 | 0.99834992 | 0.99977652 |
| NDEL1 | 725.633914 | -0.779941296 | 0.3442121 | -2.2658742 | 0.02345909 | 0.37088088 |
| MYH10 | 1080.35311 | 0.409615953 | 0.27787644 | 1.47409386 | 0.14045632 | 0.59500405 |
| STX8 | 1709.084 | -0.578660894 | 0.21334864 | -2.7122784 | 0.00668225 | 0.23013451 |
| AC005695.2 | 69.5509031 | -0.28878884 | 0.60653567 | -0.4761284 | 0.63398293 | 0.88238823 |
| AC118755.2 | 45.6017984 | 0.025868643 | 0.4255611 | 0.06078714 | 0.95152873 | 0.98756033 |
| USP43 | 316.857399 | 0.124728516 | 0.34176638 | 0.36495256 | 0.71514682 | 0.91279611 |
| GAS7 | 42.177951 | -0.583115809 | 0.36055964 | -1.617252 | 0.10582389 | 0.55405959 |
| MYHAS | 182.262464 | 2.156474623 | 0.86704823 | 2.48714494 | 0.01287729 | 0.29875272 |
| MYH3 | 70.9675537 | -0.915511359 | 0.4102012 | -2.2318593 | 0.02562427 | 0.38166806 |
| SCO1 | 271.217508 | -0.209747362 | 0.32053787 | -0.6543606 | 0.51287949 | 0.83067933 |
| ADPRM | 65.0520122 | -0.254659754 | 0.31876771 | -0.7988882 | 0.42435526 | 0.78894887 |
| TMEM220 | 347.130746 | -0.359553925 | 0.37416762 | -0.9609435 | 0.3365806 | 0.7429163 |
| TMEM220-A! | 390.569263 | 0.247124058 | 0.46525386 | 0.53115961 | 0.59530818 | 0.86510861 |
| AC015908.3 | 39.90272 | 1.378846402 | 0.74192449 | 1.8584727 | 0.06310192 | 0.48572144 |

| | | | | | | |
|---|---|---|---|---|---|---|
| AC015908.2 | 66.7613348 | 1.113234443 | 0.81404648 | 1.36753179 | 0.17145868 | 0.62520756 |
| SHISA6 | 123.701132 | 0.774546851 | 0.6607198 | 1.17227735 | 0.24108572 | 0.67967133 |
| DNAH9 | 90.0047916 | 0.894963798 | 0.64622649 | 1.38490733 | 0.1660808 | 0.62079891 |
| AC005410.2 | 33.6888166 | 0.575194785 | 0.62512863 | 0.92012229 | 0.35750886 | 0.7591594 |
| ZNF18 | 87.5069547 | -0.114913854 | 0.43391589 | -0.2648298 | 0.79114059 | 0.9409253 |
| MAP2K4 | 2245.18482 | -0.189336131 | 0.17786765 | -1.0644776 | 0.2871124 | 0.71118494 |
| ARHGAP44 | 142.768493 | 0.518264695 | 0.40627319 | 1.27565568 | 0.20207726 | 0.64881821 |
| ELAC2 | 253.422879 | -0.291800224 | 0.28365422 | -1.0287181 | 0.30361217 | 0.72082426 |
| COX10 | 525.229612 | 0.108319742 | 0.20990303 | 0.51604659 | 0.60582187 | 0.86915559 |
| PMP22 | 139.047898 | -3.388485619 | 1.06139801 | -3.1924741 | 0.0014106 | 0.1247422 |
| TRIM16 | 239.191479 | -0.137572219 | 0.37032572 | -0.3714898 | 0.71027277 | 0.91123308 |
| ZSWIM7 | 548.293989 | 0.092345682 | 0.21761432 | 0.4243548 | 0.67130708 | 0.89704376 |
| TTC19 | 1220.06028 | 0.119682932 | 0.22371174 | 0.53498726 | 0.59265867 | 0.86382106 |
| NCOR1 | 3808.12477 | -0.08012031 | 0.14325901 | -0.5592689 | 0.57597822 | 0.85554112 |
| PIGL | 1242.77782 | -0.037359533 | 0.42993453 | -0.0868959 | 0.93075429 | 0.98272895 |
| CENPV | 156.542543 | 0.473453898 | 0.43046542 | 1.09986511 | 0.2713909 | 0.70002917 |
| UBB | 1107.16256 | -0.310705405 | 0.46415437 | -0.669401 | 0.5032397 | 0.82652042 |
| LRRC75A | 98.9446096 | 0.266576489 | 0.38189191 | 0.69804173 | 0.4851511 | 0.81839266 |
| ZNF287 | 30.2566753 | 0.076196823 | 0.47277218 | 0.16117028 | 0.87195929 | 0.9657206 |
| ZNF624 | 74.0368049 | 0.684331069 | 0.3189231 | 2.14575576 | 0.03189247 | 0.40988962 |
| AC098850.3 | 485.093754 | -0.007496676 | 0.76181733 | -0.0098405 | 0.99214853 | 0.9976934 |
| MPRIP | 1114.12621 | -0.416326109 | 0.29430842 | -1.4145912 | 0.15718838 | 0.6128512 |
| COPS3 | 179.430841 | -0.5080041 | 0.27775099 | -1.8289911 | 0.06740094 | 0.48963361 |
| NT5M | 41.0172838 | -0.127719733 | 0.56791193 | -0.2248936 | 0.82206208 | 0.95112619 |
| MED9 | 211.535506 | 0.227792097 | 0.32577519 | 0.69923095 | 0.48440771 | 0.81839266 |
| RASD1 | 425.678684 | 0.028327526 | 0.79268883 | 0.035736 | 0.97149287 | 0.99240251 |
| PEMT | 147.338414 | -0.0838644 | 0.52590547 | -0.1594667 | 0.87330121 | 0.96592775 |
| RAI1 | 563.346999 | 0.078807837 | 0.38396052 | 0.20524984 | 0.83737693 | 0.95496955 |
| SMCR5 | 51.1760443 | 0.506211098 | 0.5023066 | 1.00777314 | 0.31356338 | 0.72717675 |
| SREBF1 | 513.703231 | -0.824751984 | 0.409675 | -2.013186 | 0.04409506 | 0.44360789 |
| TOM1L2 | 1335.88712 | -0.43877943 | 0.25628884 | -1.7120505 | 0.08688737 | 0.52581243 |
| DRC3 | 197.863679 | 0.581074769 | 0.45694058 | 1.27166375 | 0.20349261 | 0.64881965 |
| ATPAF2 | 204.913022 | -0.171391904 | 0.20620669 | -0.8311656 | 0.40588009 | 0.78429302 |
| GID4 | 156.429797 | -0.358166461 | 0.2508902 | -1.4275825 | 0.15341206 | 0.61014911 |
| DRG2 | 177.918697 | -0.363877061 | 0.32914458 | -1.1055235 | 0.26893283 | 0.69961625 |
| MYO15A | 38.569526 | 0.18792678 | 0.75154602 | 0.25005359 | 0.80254591 | 0.94538473 |
| ALKBH5 | 727.451457 | -0.608548212 | 0.33131663 | -1.8367572 | 0.06624574 | 0.48798003 |
| FLII | 457.981829 | -0.544274578 | 0.35889461 | -1.5165304 | 0.1293853 | 0.58817997 |
| MIEF2 | 99.5522208 | -0.665256865 | 0.3528216 | -1.8855333 | 0.05935788 | 0.47855094 |
| TOP3A | 291.254543 | -0.306688344 | 0.36417675 | -0.8421415 | 0.39970878 | 0.78186732 |
| SMCR8 | 161.874954 | -0.882701456 | 0.37521224 | -2.3525391 | 0.01864573 | 0.33968023 |
| SHMT1 | 2441.45866 | 0.765969408 | 0.29516918 | 2.59501828 | 0.00945859 | 0.25998727 |
| TRIM16L | 226.250551 | -1.078434293 | 0.49773989 | -2.1666624 | 0.03026061 | 0.40332587 |
| TVP23B | 305.34552 | -0.072844504 | 0.25535842 | -0.2852638 | 0.77544207 | 0.9351209 |
| PRPSAP2 | 1222.38988 | -0.220214809 | 0.26194582 | -0.8406884 | 4.01E-01 | 0.78232083 |
| SLC5A10 | 2005.74551 | 0.00729138 | 0.33305015 | 0.02189274 | 9.83E-01 | 9.95E-01 |
| FAM83G | 195.131891 | -0.872081628 | 0.4413937 | -1.9757455 | 0.0481836 | 0.45192142 |
| AC007952.6 | 48.7525403 | 0.114940225 | 0.70254538 | 0.16360541 | 8.70E-01 | 0.96564168 |
| EPN2 | 1074.42654 | 0.001993679 | 0.2358248 | 0.00845407 | 0.99325471 | 0.9976934 |
| EPN2-AS1 | 37.3821698 | 0.150600385 | 0.66075148 | 0.22792289 | 0.81970619 | 0.94990455 |
| B9D1 | 247.987519 | 0.169624368 | 0.33034394 | 0.51347807 | 0.60761695 | 0.86985416 |
| MAPK7 | 63.295559 | -0.731380352 | 0.40586788 | -1.8020159 | 0.07154291 | 0.50007072 |
| AC004448.2 | 94.3148579 | -0.245053195 | 0.34514541 | -0.7099999 | 0.47770422 | 0.81580016 |
| SLC47A1 | 3042.66162 | 1.260315223 | 0.44786526 | 2.81404997 | 4.89E-03 | 0.21582309 |

| | | | | | | |
|---|---|---|---|---|---|---|
| AC025627.1 | 314.118745 | 1.449570989 | 0.79072272 | 1.83322289 | 0.06676943 | 0.48884194 |
| ALDH3A2 | 881.980409 | -0.030740772 | 0.42453598 | -0.0724103 | 0.9422754 | 0.98489601 |
| SLC47A2 | 4577.18637 | 1.467652094 | 0.55791471 | 2.63060295 | 0.00852335 | 0.25070735 |
| AC005722.2 | 44.0567854 | 0.29595845 | 0.57298437 | 0.51652098 | 0.60549059 | 0.86896236 |
| ULK2 | 409.114818 | 0.381407637 | 0.38059211 | 1.00214278 | 0.31627464 | 0.72798933 |
| AKAP10 | 1322.29373 | -0.282594513 | 0.20965616 | -1.3478951 | 0.17769212 | 0.62806237 |
| SPECC1 | 100.52196 | -0.457031268 | 0.46678345 | -0.9791077 | 3.28E-01 | 7.38E-01 |
| CCDC144NL-/ | 205.489435 | 0.727081135 | 0.41705619 | 1.74336493 | 0.08126989 | 0.51593841 |
| USP22 | 1197.82926 | 0.005906075 | 0.26195092 | 0.0225465 | 0.98201202 | 0.99522272 |
| DHRS7B | 919.82682 | 0.18793862 | 0.35428129 | 0.53047854 | 0.59578019 | 0.86526372 |
| TMEM11 | 125.251814 | -0.388150448 | 0.35004132 | -1.1088704 | 0.26748611 | 0.69874881 |
| AC087294.1 | 130.005811 | -0.2519626 | 0.37646824 | -0.6692798 | 0.50331699 | 0.82652042 |
| NATD1 | 61.4338038 | -0.478148666 | 0.48475935 | -0.986363 | 0.32395503 | 0.73590515 |
| MAP2K3 | 708.44348 | -0.989121446 | 0.62145685 | -1.5916173 | 0.11147073 | 0.56435572 |
| C17orf51 | 48.1936817 | -0.165248329 | 0.60883305 | -0.2714181 | 0.78606946 | 0.93904951 |
| WSB1 | 3599.39403 | -0.307397171 | 0.51886489 | -0.5924416 | 0.55355489 | 0.84727776 |
| KSR1 | 896.325369 | 0.46216962 | 0.3166365 | 1.45962205 | 0.14439397 | 0.60101297 |
| LGALS9 | 36.4532077 | -1.131037818 | 0.85978305 | -1.3154921 | 0.18834457 | 0.64026838 |
| AC005697.1 | 63.2093425 | 1.167520842 | 0.80798915 | 1.44497094 | 0.14846605 | 0.60622442 |
| NLK | 2164.74802 | 0.418221211 | 0.13990038 | 2.98942162 | 0.00279506 | 0.1710353 |
| IFT20 | 143.069876 | -0.737250912 | 0.26652164 | -2.7661953 | 0.00567146 | 0.22501232 |
| TNFAIP1 | 374.272213 | -0.16735355 | 0.30245892 | -0.55331 | 0.58005114 | 0.85824961 |
| POLDIP2 | 306.055429 | 0.002931548 | 0.21681635 | 0.01352088 | 0.98921223 | 0.99721497 |
| TMEM199 | 101.314543 | -0.308116348 | 0.24979465 | -1.2334786 | 0.21739727 | 0.66032006 |
| SARM1 | 202.876553 | -0.21354441 | 0.33593546 | -0.6356709 | 0.52499093 | 0.835938 |
| SLC46A1 | 46.8208141 | -0.079804883 | 0.43359656 | -0.1840533 | 0.85397163 | 0.96119142 |
| AC005726.1 | 1779.56735 | 0.922109938 | 0.60499005 | 1.52417374 | 0.12746532 | 0.58793678 |
| SLC13A2 | 258.039873 | 2.816189574 | 0.88146642 | 3.19489151 | 0.00139883 | 0.12450532 |
| AC005726.5 | 123.364914 | 1.142648387 | 0.73867754 | 1.54688389 | 0.12189125 | 0.5805293 |
| KIAA0100 | 1227.90486 | 0.219602022 | 0.22500053 | 0.97600667 | 0.32906116 | 0.73829083 |
| SDF2 | 292.296534 | -0.350286988 | 0.26237108 | -1.3350823 | 0.18184941 | 0.6318513 |
| SUPT6H | 991.17857 | -0.609360499 | 0.30746044 | -1.9819152 | 0.04748874 | 0.44992941 |
| PROCA1 | 34.2550549 | 0.966113944 | 0.80775629 | 1.19604633 | 0.23167848 | 0.66883255 |
| RAB34 | 125.022494 | 0.282718614 | 0.56737327 | 0.49829386 | 0.61827694 | 0.8747221 |
| RPL23A | 983.632006 | -0.195354701 | 0.47654065 | -0.4099434 | 0.68184746 | 0.89962501 |
| TLCD1 | 49.3804728 | -0.865879648 | 0.51527633 | -1.6804181 | 0.092876 | 0.53669957 |
| NEK8 | 107.769832 | -0.277381661 | 0.45093723 | -0.6151226 | 0.53847379 | 0.84049498 |
| TRAF4 | 564.235753 | -0.193315716 | 0.32048574 | -0.603196 | 0.54637831 | 0.84460887 |
| FAM222B | 2199.66418 | -0.208740716 | 0.29899229 | -0.6981475 | 0.48508497 | 0.81839266 |
| ERAL1 | 117.712296 | -0.166790633 | 0.35152016 | -0.4744838 | 0.63515492 | 0.88238823 |
| FLOT2 | 919.411799 | 0.481765254 | 0.43578159 | 1.10551998 | 0.26893434 | 0.69961625 |
| DHRS13 | 88.5961178 | 0.50100875 | 0.59063361 | 0.84825641 | 0.39629519 | 0.7795748 |
| PHF12 | 486.376239 | 0.162659314 | 0.25762594 | 0.63137785 | 0.5277935 | 0.83761342 |
| PIPOX | 389.656961 | 0.031536002 | 0.64344503 | 0.04901118 | 0.96091038 | 0.98972037 |
| MYO18A | 560.284142 | -0.669831628 | 0.49620109 | -1.3499197 | 0.17704174 | 0.62731853 |
| AC024619.3 | 59.4643843 | 0.55959366 | 0.7710394 | 0.72576532 | 0.46798265 | 0.8118777 |
| NUFIP2 | 1722.94858 | -0.287027973 | 0.35871691 | -0.8001518 | 0.42362287 | 0.78894887 |
| TAOK1 | 2544.40286 | -0.13492044 | 0.22554995 | -0.5981843 | 0.54971695 | 0.84612944 |
| ABHD15-AS1 | 344.12343 | 0.467476035 | 0.57117297 | 0.81844915 | 0.41310077 | 0.78632759 |
| ABHD15 | 79.6249711 | 0.519403439 | 0.383858 | 1.35311349 | 0.17601938 | 0.62699479 |
| GIT1 | 204.453408 | -0.190554967 | 0.44212242 | -0.4310005 | 0.66646803 | 0.89420131 |
| SSH2 | 5660.60514 | -0.014182219 | 0.14549105 | -0.0974783 | 0.92234657 | 0.97970649 |
| AC023389.1 | 158.049329 | 0.137524295 | 0.35311273 | 0.38946286 | 0.69693378 | 0.90649868 |
| EFCAB5 | 177.465772 | 0.469946976 | 0.42052649 | 1.1175205 | 2.64E-01 | 6.96E-01 |

| | | | | | | |
|---|---|---|---|---|---|---|
| NSRP1 | 1677.5216 | 0.025670562 | 0.19270638 | 0.13321075 | 0.89402671 | 0.97272276 |
| AC104984.3 | 64.6733736 | 0.237093479 | 0.72100596 | 0.32883706 | 0.74227885 | 0.9229205 |
| BLMH | 602.082126 | 0.117651156 | 0.15588671 | 0.75472217 | 0.4504157 | 0.80361623 |
| CPD | 2246.83329 | -0.044441795 | 0.41741443 | -0.1064692 | 9.15E-01 | 9.78E-01 |
| GOSR1 | 954.149971 | -0.412399987 | 0.25472298 | -1.6190137 | 0.10544432 | 0.55348589 |
| CRLF3 | 612.315036 | -0.055707292 | 0.27863159 | -0.1999317 | 0.84153399 | 0.95681213 |
| ATAD5 | 165.899608 | 0.107802126 | 0.45969688 | 0.23450698 | 0.81459142 | 0.94881072 |
| AC130324.2 | 97.1243906 | 0.599837354 | 0.61156968 | 0.98081604 | 0.32668347 | 0.73745391 |
| TEFM | 320.872322 | 0.205829416 | 0.2163419 | 0.95140799 | 0.34139731 | 0.74809832 |
| ADAP2 | 331.415911 | 0.074295291 | 0.31151417 | 0.23849731 | 0.8114954 | 0.94753961 |
| RNF135 | 341.071634 | 0.332873079 | 0.27611486 | 1.20556019 | 0.22798712 | 0.66705629 |
| NF1 | 5096.45813 | -0.160688104 | 0.17043404 | -0.942817 | 0.34577453 | 0.75166079 |
| OMG | 91.6002333 | 0.075471905 | 0.59701529 | 0.12641536 | 0.89940314 | 0.97458352 |
| RAB11FIP4 | 208.719854 | 0.389735446 | 0.37740313 | 1.03267678 | 0.30175518 | 0.71908176 |
| COPRS | 151.331486 | -0.387014876 | 0.40128237 | -0.9644453 | 0.33482274 | 0.74154392 |
| UTP6 | 392.227821 | -0.513053213 | 0.28242759 | -1.8165832 | 0.06928096 | 0.49273866 |
| AC004253.1 | 94.7596212 | 0.396352839 | 0.52499641 | 0.75496295 | 0.45027121 | 0.80360602 |
| SUZ12 | 1072.26809 | 0.073476224 | 0.18782907 | 0.39118665 | 0.69565927 | 0.90589033 |
| LRRC37B | 424.127199 | -0.213652216 | 0.19754774 | -1.0815219 | 0.27946501 | 0.70446562 |
| RHOT1 | 917.749767 | -0.029535415 | 0.16743177 | -0.1764027 | 0.85997759 | 0.96283855 |
| C17orf75 | 153.075591 | 0.575246258 | 0.44467043 | 1.29364629 | 0.19578763 | 0.64558745 |
| ZNF207 | 2606.11137 | -0.092526506 | 0.17897878 | -0.5169691 | 0.60517771 | 0.86869053 |
| PSMD11 | 937.469281 | -0.42068938 | 0.30349719 | -1.3861393 | 0.16570438 | 0.62065658 |
| MYO1D | 1275.61259 | 0.329187489 | 0.3811656 | 0.86363379 | 3.88E-01 | 0.77373447 |
| TMEM98 | 134.619041 | 1.123105991 | 0.44753447 | 2.50954075 | 0.01208883 | 0.29431891 |
| ASIC2 | 79.4232455 | 0.110008574 | 0.55475656 | 0.19830063 | 0.84280986 | 0.95716002 |
| CCL2 | 197.922087 | -0.127511333 | 0.8370633 | -0.1523318 | 0.87892527 | 0.9674841 |
| TMEM132E | 86.2756693 | 0.04983052 | 0.44472275 | 0.11204851 | 9.11E-01 | 9.77E-01 |
| CCT6B | 116.523054 | -0.276658159 | 0.54028542 | -0.5120593 | 0.60860953 | 0.86985416 |
| ZNF830 | 124.757143 | -0.687676657 | 0.38760077 | -1.774188 | 0.07603205 | 0.50738624 |
| LIG3 | 128.532473 | -0.24801881 | 0.20345231 | -1.2190513 | 0.22282472 | 0.66398799 |
| RFFL | 407.670696 | -0.214752999 | 0.50689062 | -0.4236673 | 0.67180844 | 0.89704376 |
| SLFN5 | 184.275912 | 0.608880264 | 0.61306953 | 0.99316673 | 3.21E-01 | 0.73210121 |
| SLFN12 | 43.7443062 | 0.25206173 | 0.49351604 | 0.51074679 | 0.60952837 | 0.87056428 |
| SLFN13 | 74.1719567 | 1.092797733 | 0.77745091 | 1.40561638 | 0.15983807 | 0.61542146 |
| LINC02001 | 172.278893 | 0.118131364 | 0.24168749 | 0.48877732 | 0.62499936 | 0.87854882 |
| AP2B1 | 1192.42579 | 0.107758302 | 0.17770033 | 0.60640464 | 0.54424609 | 0.84416152 |
| TAF15 | 2139.74296 | -0.169816941 | 0.16714803 | -1.0159673 | 0.30964495 | 0.72498238 |
| AC015849.3 | 118.844041 | -0.202092855 | 0.54874426 | -0.3682824 | 0.71266267 | 0.91183303 |
| CCL15-CCL14 | 49.6843934 | 1.429320304 | 0.8243048 | 1.73397063 | 0.0829233 | 0.51935659 |
| ZNHIT3 | 122.923428 | -0.287637559 | 0.29871508 | -0.9629161 | 0.33558963 | 0.74225095 |
| MYO19 | 554.649487 | -0.464078157 | 0.4125528 | -1.124894 | 0.26063397 | 0.69280433 |
| GGNBP2 | 1279.29032 | -0.197995362 | 0.23623162 | -0.8381408 | 0.40195163 | 0.78258224 |
| DHRS11 | 58.5480998 | 0.223042805 | 0.4459808 | 0.5001175 | 6.17E-01 | 0.87421068 |
| MRM1 | 71.4008072 | 0.188694837 | 0.35544543 | 0.53086865 | 0.5955098 | 0.86519223 |
| LHX1-DT | 212.789974 | 1.602322155 | 0.55875613 | 2.86765922 | 0.00413521 | 0.20365996 |
| LHX1 | 141.856464 | 1.726544186 | 0.52785404 | 3.27087423 | 0.00107216 | 0.11082848 |
| AATF | 1169.23903 | -0.410824094 | 0.28199946 | -1.4568258 | 0.14516446 | 0.60162365 |
| ACACA | 1648 | -0.166633125 | 0.29064537 | -0.5733211 | 0.5664273 | 0.85215881 |
| TADA2A | 389.61154 | 0.003418507 | 0.27879923 | 0.01226154 | 0.99021695 | 0.99737132 |
| DUSP14 | 91.8808954 | -0.582210365 | 0.48799314 | -1.1930708 | 0.23284165 | 0.66978724 |
| SYNRG | 1511.79847 | 0.121166433 | 0.25883863 | 0.46811573 | 0.63970183 | 0.88321992 |
| DDX52 | 839.907648 | -0.253354771 | 0.32912317 | -0.7697871 | 0.44142619 | 0.79862562 |
| AC243585.2 | 40.8153612 | 0.084510435 | 0.73146556 | 0.11553577 | 0.90802047 | 0.97673359 |

| | | | | | | |
|---|---|---|---|---|---|---|
| HNF1B | 2180.0556 | 0.494864222 | 0.4548248 | 1.08803262 | 0.27658071 | 0.70345459 |
| MRPL45 | 335.689885 | -0.273980642 | 0.18446781 | -1.485249 | 0.13747786 | 0.59474076 |
| SOCS7 | 426.755143 | -0.072757691 | 0.29311913 | -0.2482188 | 8.04E-01 | 0.94543168 |
| MLLT6 | 816.023301 | 0.240959099 | 0.30486457 | 0.79038079 | 0.42930541 | 0.79037914 |
| CISD3 | 217.929964 | 0.048357662 | 0.29608595 | 0.16332306 | 0.87026408 | 0.96564168 |
| PCGF2 | 247.636062 | 0.287106356 | 0.36506413 | 0.78645457 | 0.43160123 | 0.79135216 |
| PSMB3 | 351.240724 | -0.451655529 | 0.32436751 | -1.3924192 | 0.16379551 | 0.62065658 |
| PIP4K2B | 357.168788 | 0.074092015 | 0.33231457 | 0.22295747 | 0.82356861 | 0.95134459 |
| CWC25 | 472.757411 | -0.681183269 | 0.39147071 | -1.7400619 | 0.08184815 | 0.51593841 |
| RPL23 | 776.234169 | 0.176979837 | 0.52103371 | 0.33967061 | 0.7341046 | 0.91975299 |
| LASP1 | 835.813423 | -0.255797452 | 0.39183999 | -0.652811 | 0.51387814 | 0.83126535 |
| RPL19 | 1335.69659 | -0.112920016 | 0.55544923 | -0.2032949 | 0.83890451 | 0.95545195 |
| FBXL20 | 2347.70973 | -0.421720216 | 0.26706543 | -1.5790895 | 0.11431553 | 0.56984472 |
| MED1 | 387.88355 | -0.309157807 | 0.27400173 | -1.128306 | 0.2591907 | 0.69242517 |
| CDK12 | 1217.17447 | -0.051545064 | 0.21757511 | -0.236907 | 0.81272895 | 0.94801888 |
| STARD3 | 278.629722 | -0.2172113 | 0.33874048 | -0.6412322 | 0.52137184 | 0.83445062 |
| PGAP3 | 90.7340036 | 0.476102979 | 0.42124418 | 1.13023041 | 0.25837915 | 0.69210282 |
| ERBB2 | 671.636682 | 0.663440294 | 0.45431413 | 1.46031182 | 0.1442044 | 0.60086567 |
| MIEN1 | 165.338242 | -0.193258756 | 0.39293357 | -0.4918357 | 0.6228355 | 0.87714058 |
| GRB7 | 585.010206 | 0.340451487 | 0.43740312 | 0.77834719 | 0.43636437 | 0.79555046 |
| GSDMB | 112.741677 | 0.576434255 | 0.50482335 | 1.14185339 | 0.25351497 | 0.68891958 |
| ORMDL3 | 99.6458026 | -0.189992855 | 0.34449428 | -0.5515124 | 0.58128249 | 0.85895204 |
| AC090844.2 | 80.071232 | 0.859637636 | 0.66208906 | 1.2983716 | 0.19415967 | 0.64548758 |
| PSMD3 | 285.683489 | -0.618762159 | 0.33304033 | -1.8579196 | 0.06318043 | 0.48572144 |
| MED24 | 343.762204 | 0.07430465 | 0.16772771 | 0.44300759 | 0.65776024 | 0.89053559 |
| THRA | 218.187931 | 0.11758526 | 0.38307185 | 0.30695353 | 0.75887874 | 0.9290866 |
| NR1D1 | 129.52235 | -1.400543715 | 0.67751468 | -2.0671784 | 0.03871735 | 0.42523931 |
| MSL1 | 459.240078 | 0.197290851 | 0.25465693 | 0.77473191 | 0.43849809 | 0.7970505 |
| CASC3 | 513.612928 | -0.387734918 | 0.20407102 | -1.8999999 | 0.05743314 | 0.47684824 |
| RAPGEFL1 | 54.7406602 | -0.879106546 | 0.40435817 | -2.1740789 | 0.02969921 | 0.39981555 |
| WIPF2 | 1385.92837 | -0.458271259 | 0.30634362 | -1.4959386 | 0.13466964 | 0.59277997 |
| RARA | 214.479573 | -1.026843307 | 0.43422354 | -2.3647804 | 0.01804076 | 0.33690022 |
| IGFBP4 | 1553.46925 | 0.064083567 | 0.22950869 | 0.27922066 | 0.78007549 | 0.93703398 |
| KRT10 | 309.391261 | -0.632679605 | 0.45960505 | -1.3765723 | 0.16864451 | 0.62304975 |
| TMEM99 | 98.524381 | 0.003634096 | 0.43440047 | 0.00836577 | 0.99332516 | 0.9976934 |
| KRT19 | 317.138667 | 0.415746883 | 0.66996309 | 0.62055192 | 0.53489448 | 0.83954482 |
| EIF1 | 1443.74457 | -0.370520611 | 0.3961011 | -0.9354193 | 0.34957226 | 0.75434303 |
| JUP | 828.619347 | 0.215944759 | 0.30317238 | 0.71228374 | 0.47628909 | 0.81570932 |
| P3H4 | 105.871255 | -0.010810704 | 0.36627455 | -0.0295153 | 0.97645362 | 0.99429832 |
| NT5C3B | 204.120586 | -0.061206492 | 0.30805384 | -0.1986876 | 0.84250709 | 0.95716002 |
| KLHL11 | 43.4691393 | -0.398008427 | 0.52515527 | -0.7578871 | 0.44851854 | 0.80259633 |
| ACLY | 1539.83557 | -0.218892633 | 0.37626992 | -0.5817436 | 0.56073937 | 0.84947553 |
| TTC25 | 178.422005 | 0.248353308 | 0.48748452 | 0.50945886 | 0.61043063 | 0.87112677 |
| CNP | 269.182118 | -0.327770763 | 0.54796476 | -0.5981603 | 0.54973298 | 0.84612944 |
| DNAJC7 | 971.904882 | -0.436966543 | 0.18701125 | -2.336579 | 0.01946109 | 0.34419759 |
| NKIRAS2 | 123.383149 | -0.29858855 | 0.26244666 | -1.1377114 | 0.25524102 | 0.68978482 |
| DHX58 | 72.0395767 | -1.162632794 | 1.01144208 | -1.1494803 | 0.25035797 | 0.68619409 |
| KAT2A | 172.597713 | -0.280945212 | 0.44141475 | -0.6364654 | 0.52447313 | 0.8359248 |
| GHDC | 134.113418 | 0.245114459 | 0.43039067 | 0.5695162 | 0.56900588 | 0.85291608 |
| STAT5B | 1557.0962 | 0.356257894 | 0.21235966 | 1.67761571 | 0.09342214 | 0.53688613 |
| STAT5A | 163.198856 | 0.730595751 | 0.3409221 | 2.14299911 | 0.03211317 | 0.40988962 |
| STAT3 | 3180.76963 | 0.123950539 | 0.34515328 | 0.35911737 | 0.71950728 | 0.91465123 |
| CAVIN1 | 81.9989455 | 0.006997957 | 0.59898962 | 0.01168294 | 0.99067858 | 0.99737132 |
| ATP6V0A1 | 2618.41244 | 0.172647746 | 0.22026144 | 0.783831 | 0.43313928 | 0.79355172 |

| | | | | | | |
|---|---|---|---|---|---|---|
| NAGLU | 490.86124 | -0.440866141 | 0.4726931 | -0.9326689 | 0.35099096 | 0.75514568 |
| AC067852.2 | 34.3895387 | 0.227777691 | 0.48073927 | 0.47380713 | 0.63563745 | 0.88257807 |
| COASY | 276.980953 | -0.064607457 | 0.33008119 | -0.195732 | 0.84481992 | 0.95760132 |
| MLX | 260.889958 | 0.301294794 | 0.41276903 | 0.72993557 | 0.46542957 | 0.81042189 |
| RETREG3 | 552.232598 | -0.382785681 | 0.28044449 | -1.3649249 | 0.17227665 | 0.62592399 |
| TUBG1 | 105.076916 | 0.424860348 | 0.29282534 | 1.45090024 | 0.14680764 | 0.60383734 |
| TUBG2 | 352.373669 | 0.050937249 | 0.27210564 | 0.1871966 | 0.85150649 | 0.96044014 |
| PLEKHH3 | 180.905171 | 0.838836171 | 0.50494061 | 1.66125708 | 0.09666183 | 0.53969191 |
| EZH1 | 771.667368 | -0.233682726 | 0.25093059 | -0.9312644 | 0.35171681 | 0.75548011 |
| VPS25 | 173.581674 | 0.056323461 | 0.35582607 | 0.1582893 | 0.87422884 | 0.9661168 |
| WNK4 | 54.1793573 | 0.785916862 | 0.67624329 | 1.16218064 | 0.2451621 | 0.68281753 |
| COA3 | 342.789303 | 0.067190629 | 0.40202008 | 0.16713252 | 0.86726578 | 0.96486219 |
| BECN1 | 278.483471 | -0.064334685 | 0.2453514 | -0.2622145 | 0.7931561 | 0.94176981 |
| PSME3 | 216.173749 | -0.483552718 | 0.33507565 | -1.4431151 | 0.14898806 | 0.60622442 |
| LINC00671 | 3173.97867 | 0.830875576 | 1.00641256 | 0.82558148 | 0.40904154 | 0.78542936 |
| G6PC | 991.956576 | -0.623271759 | 1.61632287 | -0.3856109 | 0.69978484 | 0.90721166 |
| RUNDC1 | 282.119372 | -0.59191316 | 0.33726418 | -1.7550431 | 0.07925194 | 0.51192014 |
| RPL27 | 560.81908 | -0.190949349 | 0.45356545 | -0.4209962 | 0.67375783 | 0.89799059 |
| IFI35 | 217.598673 | -1.365101101 | 0.85619762 | -1.5943762 | 0.1108518 | 0.56317482 |
| VAT1 | 312.450271 | -0.395848723 | 0.47484318 | -0.833641 | 0.40448334 | 0.78408331 |
| BRCA1 | 244.594576 | 0.244077112 | 0.47592288 | 0.51285013 | 0.60805616 | 0.86985416 |
| NBR1 | 1109.70281 | 0.58431211 | 0.44059274 | 1.32619551 | 0.18477495 | 0.63412875 |
| TMEM106A | 919.302544 | 0.519863808 | 0.58003452 | 0.89626356 | 0.37011202 | 0.76669785 |
| LINC00854 | 1059.72943 | 0.344705668 | 0.60707215 | 0.56781664 | 0.57015948 | 0.85327831 |
| LINC00910 | 37.185328 | 0.364575505 | 0.6495242 | 0.56129626 | 0.57459559 | 0.8547017 |
| ARL4D | 74.3214885 | 1.017765535 | 0.55890452 | 1.82100072 | 0.06860675 | 0.49154315 |
| DHX8 | 339.498283 | 0.023097624 | 0.28346385 | 0.08148349 | 0.93505745 | 0.98379126 |
| DUSP3 | 502.571308 | -0.574265679 | 0.33305768 | -1.7242229 | 0.08466762 | 0.52183702 |
| MPP3 | 109.655506 | -0.742661833 | 0.42522592 | -1.7465112 | 0.08072216 | 0.51475987 |
| PYY | 72.61378 | 1.260986252 | 0.70935754 | 1.77764551 | 0.07546209 | 0.50728736 |
| NAGS | 93.9956079 | 0.958893392 | 0.72080044 | 1.33031743 | 0.18341371 | 0.63276491 |
| TMEM101 | 139.30211 | -0.013786256 | 0.27028653 | -0.0510061 | 0.95932068 | 0.98912673 |
| LSM12 | 377.736492 | -0.832920539 | 0.32335766 | -2.5758491 | 0.00999943 | 0.26511053 |
| G6PC3 | 117.002422 | 0.006914399 | 0.47511434 | 0.01455313 | 0.98838869 | 0.99682465 |
| HDAC5 | 227.550555 | -0.262506756 | 0.3694274 | -0.7105774 | 0.47734616 | 0.81579656 |
| ASB16-AS1 | 85.0577132 | -0.214769624 | 0.30254198 | -0.7098837 | 0.47777624 | 0.81580016 |
| TMUB2 | 256.443956 | 0.064524757 | 0.29839233 | 0.21624134 | 0.82879963 | 0.95325673 |
| ATXN7L3 | 124.921234 | -0.455493495 | 0.39426418 | -1.1553002 | 0.24796751 | 0.68482118 |
| AC004596.1 | 35.8003358 | 0.830539762 | 0.4139802 | 2.00623062 | 0.04483165 | 0.44360789 |
| UBTF | 358.244475 | -0.333289025 | 0.33660304 | -0.9901545 | 0.32209859 | 0.73375526 |
| AC003102.1 | 48.0335691 | -0.095207033 | 0.63620461 | -0.1496484 | 0.88104198 | 0.96773407 |
| SLC25A39 | 411.195023 | -0.338957513 | 0.45777937 | -0.7404386 | 0.45903391 | 0.80666382 |
| GRN | 485.653376 | -0.551946353 | 0.49658678 | -1.1114802 | 0.26636173 | 0.69797523 |
| GPATCH8 | 2806.10891 | -0.058674342 | 0.21813251 | -0.2689849 | 0.78794134 | 0.93953874 |
| CCDC43 | 107.586065 | -0.541459969 | 0.36534386 | -1.4820558 | 0.13832546 | 0.59474076 |
| DBF4B | 102.671908 | -0.015875504 | 0.56092616 | -0.0283023 | 0.97742104 | 0.99473325 |
| EFTUD2 | 449.459014 | -0.315844509 | 0.36650025 | -0.8617853 | 0.3888057 | 0.7748665 |
| DCAKD | 467.573307 | 0.399601446 | 0.3356572 | 1.19050463 | 0.23384811 | 0.67043632 |
| NMT1 | 880.750296 | -0.198708269 | 0.23864546 | -0.8326505 | 0.40504185 | 0.78422899 |
| PLCD3 | 59.9673717 | 0.768256521 | 0.50590705 | 1.51857249 | 0.12887014 | 0.58793678 |
| ACBD4 | 360.591724 | 0.393812206 | 0.57086911 | 0.68984676 | 4.90E-01 | 0.81989696 |
| HEXIM1 | 270.441174 | -0.298439238 | 0.35357776 | -0.8440555 | 0.39863842 | 0.78122921 |
| AC138150.1 | 399.684345 | 0.647449689 | 0.44760904 | 1.44646249 | 0.14804752 | 0.60605027 |
| HEXIM2 | 78.4709662 | 0.028971677 | 0.4372448 | 0.06625963 | 9.47E-01 | 9.86E-01 |

| | | | | | | |
|---|---|---|---|---|---|---|
| MAP3K14 | 1468.64204 | 0.345555692 | 0.33509098 | 1.03122948 | 0.30243322 | 0.71994653 |
| ARHGAP27 | 453.584658 | 0.359706179 | 0.23792779 | 1.51182922 | 0.13057731 | 0.58853436 |
| PLEKHM1 | 700.51749 | -1.385425561 | 0.61363617 | -2.2577312 | 0.02396242 | 0.37387451 |
| AC091132.2 | 41.1174889 | -0.457574722 | 0.6980237 | -0.6555289 | 0.51212727 | 0.83054053 |
| LINC02210-C | 181.940124 | 0.226459146 | 0.48536722 | 0.46657281 | 0.64080555 | 0.88396933 |
| MAPT-AS1 | 87.7115908 | 0.992462065 | 1.04654865 | 0.94831909 | 3.43E-01 | 0.7505827 |
| MAPT | 1567.26226 | -0.038092101 | 0.49883364 | -0.0763623 | 0.93913084 | 0.98435387 |
| KANSL1 | 6397.28642 | -0.433702278 | 0.31555684 | -1.374403 | 0.16931659 | 0.62326131 |
| KANSL1-AS1 | 45.5420892 | 0.357253988 | 0.61215416 | 0.58360134 | 0.55948856 | 0.84941263 |
| ARL17B | 1210.32615 | 0.760035059 | 1.07166286 | 0.70921097 | 0.47819356 | 0.81596289 |
| LRRC37A | 37.2518276 | 0.40409416 | 0.56130447 | 0.71991973 | 0.47157442 | 0.81389222 |
| LRRC37A2 | 237.746049 | -0.088233305 | 0.3096298 | -0.2849639 | 0.77567183 | 0.9351345 |
| ARL17A | 318.82773 | -0.890074691 | 0.58864082 | -1.5120846 | 0.13051235 | 0.58853436 |
| NSF | 1207.39875 | -0.14566327 | 0.41274425 | -0.3529141 | 0.72415283 | 0.91600702 |
| WNT3 | 42.0012397 | 0.433574961 | 0.49711681 | 0.87217923 | 0.38311061 | 0.77317162 |
| AC005670.2 | 143.22945 | 0.872888178 | 0.52029437 | 1.67768137 | 0.09340932 | 0.53688613 |
| CDC27 | 1277.74646 | -0.076037821 | 0.20482616 | -0.371231 | 0.71046549 | 0.91123308 |
| AC068234.1 | 1586.88973 | 1.012632944 | 0.70423611 | 1.43791681 | 1.50E-01 | 0.60799621 |
| AC068234.2 | 244.905486 | 1.326826407 | 0.37491808 | 3.53897684 | 4.02E-04 | 7.15E-02 |
| EFCAB13 | 933.920796 | 1.334209139 | 0.438527 | 3.04247891 | 2.35E-03 | 0.15766817 |
| AC040934.1 | 122.298265 | 0.053199735 | 0.46183595 | 0.11519185 | 0.90829306 | 0.97677491 |
| NPEPPS | 3114.86947 | -0.372865537 | 0.17163962 | -2.1723746 | 0.02982742 | 0.39981555 |
| AC025682.1 | 228.337485 | -0.025556916 | 0.41460467 | -0.0616416 | 0.95084821 | 0.98731502 |
| KPNB1 | 1340.23378 | -0.40305768 | 0.35913284 | -1.1223081 | 0.26173148 | 0.69408034 |
| TBKBP1 | 105.562709 | -1.13084835 | 0.55266235 | -2.0461831 | 0.04073836 | 0.43086475 |
| OSBPL7 | 44.3709585 | 0.017410439 | 0.49326294 | 0.03529647 | 0.97184334 | 0.99240251 |
| MRPL10 | 97.807902 | -0.425952932 | 0.31271426 | -1.3621155 | 0.17316145 | 0.6264249 |
| SCRN2 | 584.017347 | 0.569788641 | 0.6266009 | 0.90933262 | 0.36317457 | 0.76213737 |
| SP2 | 151.686925 | -0.379588016 | 0.25336667 | -1.4981766 | 0.13408737 | 0.59240843 |
| SP2-AS1 | 103.47997 | 0.411914361 | 0.36130747 | 1.14006598 | 0.25425881 | 0.68916859 |
| PNPO | 264.748264 | 0.926665506 | 0.37825862 | 2.44981991 | 0.01429277 | 0.30566089 |
| AC018521.2 | 68.8477499 | 1.372739104 | 0.79716923 | 1.72201718 | 0.08506641 | 0.52183702 |
| PRR15L | 82.7023326 | 1.647654655 | 1.05100549 | 1.56769366 | 0.11695264 | 0.57350419 |
| CDK5RAP3 | 460.341265 | 0.283740417 | 0.36773912 | 0.77158072 | 0.4403628 | 0.79826598 |
| COPZ2 | 51.617347 | 0.347550082 | 0.59121945 | 0.58785292 | 0.55663102 | 0.84861484 |
| NFE2L1 | 938.459176 | -0.028041563 | 0.24018158 | -0.1167515 | 0.90705697 | 0.97668111 |
| CBX1 | 287.643499 | -0.502367487 | 0.30669615 | -1.6379974 | 0.10142224 | 0.54667503 |
| SNX11 | 104.643308 | -0.585528407 | 0.29796727 | -1.9650762 | 0.04940541 | 0.45419182 |
| SKAP1 | 247.893027 | 1.091741239 | 0.7921455 | 1.37820796 | 0.16813909 | 0.62304975 |
| HOXB2 | 101.538076 | 0.435153204 | 0.55011199 | 0.79102658 | 0.42892848 | 0.79037914 |
| HOXB-AS1 | 83.7493785 | 0.497548777 | 0.55332697 | 0.89919488 | 0.36854886 | 0.76615653 |
| HOXB3 | 209.096283 | 1.288205331 | 0.5766318 | 2.23401717 | 0.02548194 | 0.38166806 |
| HOXB-AS3 | 46.7507971 | 0.461223686 | 0.61583919 | 0.74893526 | 0.45389622 | 0.80433814 |
| HOXB7 | 92.1553394 | 0.304871851 | 0.49706098 | 0.613349 | 0.5396456 | 0.84113285 |
| TTLL6 | 113.728093 | -0.501190091 | 0.54649857 | -0.9170931 | 3.59E-01 | 0.75936492 |
| CALCOCO2 | 705.673858 | -0.057754489 | 0.24677132 | -0.2340405 | 0.81495352 | 0.94895023 |
| ATP5MC1 | 390.205316 | -0.049098601 | 0.42057932 | -0.1167404 | 0.90706577 | 0.97668111 |
| UBE2Z | 804.452183 | -0.343315154 | 0.33027991 | -1.0394673 | 0.29858747 | 0.71852421 |
| SNF8 | 323.439427 | -0.501708292 | 0.22531641 | -2.2266833 | 0.02596845 | 0.38392873 |
| AC091133.1 | 55.1122591 | 0.108643404 | 0.39425067 | 0.27556936 | 0.78287885 | 0.93801751 |
| IGF2BP1 | 118.447918 | 0.563737933 | 0.73509575 | 0.76689048 | 0.44314664 | 0.7993765 |
| ZNF652 | 1015.64585 | -0.266807881 | 0.21724405 | -1.2281482 | 0.21939135 | 0.66186058 |
| AC091180.5 | 64.3274086 | -0.120840074 | 0.54569845 | -0.2214411 | 0.82474898 | 0.95192757 |
| PHB | 494.807117 | 0.473647937 | 0.40448213 | 1.17099843 | 0.24159941 | 0.68015474 |

| | | | | | | |
|---|---|---|---|---|---|---|
| SPOP | 923.811296 | -0.261646099 | 0.14152539 | -1.8487573 | 0.06449287 | 0.48598338 |
| SLC35B1 | 146.770044 | -0.218117212 | 0.35920321 | -0.6072251 | 0.54370152 | 0.84396451 |
| FAM117A | 225.109953 | -0.103691589 | 0.39834655 | -0.260305 | 0.79462853 | 0.94252227 |
| KAT7 | 319.684809 | -0.048405156 | 0.18037427 | -0.2683595 | 0.78842258 | 0.93976423 |
| TAC4 | 55.948862 | 0.118525457 | 0.44795527 | 0.26459217 | 0.79132365 | 0.94098694 |
| ITGA3 | 2130.94635 | -0.250339976 | 0.68688969 | -0.3644544 | 7.16E-01 | 9.13E-01 |
| PDK2 | 248.28961 | 0.876490734 | 0.4550543 | 1.92612342 | 0.05408897 | 0.46711025 |
| PPP1R9B | 130.174363 | -0.186419006 | 0.5132652 | -0.3632021 | 0.71645391 | 0.91298968 |
| AC002401.1 | 122.798684 | 0.109257514 | 0.79863963 | 0.13680452 | 8.91E-01 | 9.72E-01 |
| XYLT2 | 133.200002 | 0.226198159 | 0.30053051 | 0.75266287 | 0.45165253 | 0.80384625 |
| MRPL27 | 285.190211 | -0.315232847 | 0.32485479 | -0.9703808 | 0.33185672 | 0.73939532 |
| LRRC59 | 233.191241 | -0.679262873 | 0.5226148 | -1.2997391 | 0.19369042 | 0.64548758 |
| ACSF2 | 3895.23294 | 1.124821072 | 0.55038036 | 2.04371585 | 4.10E-02 | 4.31E-01 |
| CHAD | 286.56268 | 1.229476717 | 0.95320195 | 1.28983866 | 0.19710668 | 0.64558745 |
| AC021491.4 | 90.5416383 | 1.43253218 | 0.81174139 | 1.76476424 | 7.76E-02 | 5.11E-01 |
| RSAD1 | 521.692179 | -0.607382846 | 0.29451385 | -2.0623236 | 3.92E-02 | 4.27E-01 |
| SPATA20 | 234.668936 | 0.429711888 | 0.49199283 | 0.87341087 | 0.38243917 | 0.77279408 |
| ABCC3 | 3965.2186 | 0.742917948 | 0.3882517 | 1.91349567 | 5.57E-02 | 4.70E-01 |
| ANKRD40 | 223.357757 | -0.568211305 | 0.26401738 | -2.1521738 | 0.03138367 | 0.40835531 |
| AC005921.2 | 49.1678699 | -0.014527596 | 0.41127477 | -0.0353233 | 0.97182192 | 0.99240251 |
| LUC7L3 | 2322.90379 | -0.474495073 | 0.19120638 | -2.481586 | 0.01307991 | 0.29875272 |
| TOB1 | 540.719099 | 0.624881286 | 0.5159068 | 1.21122901 | 0.22580765 | 0.66688879 |
| SPAG9 | 6858.42601 | -0.974954634 | 0.3599473 | -2.7086038 | 0.0067567 | 0.23083391 |
| AC005920.2 | 450.405158 | -0.354468112 | 0.69134229 | -0.5127245 | 0.60814407 | 0.86985416 |
| MBTD1 | 1266.61172 | -0.454368982 | 0.18340998 | -2.4773406 | 0.01323655 | 0.29875272 |
| UTP18 | 316.883949 | -0.46645118 | 0.28650289 | -1.6280854 | 0.10350677 | 0.55022042 |
| TOM1L1 | 2012.8205 | 0.314002502 | 0.30113704 | 1.04272294 | 0.29707663 | 0.71825247 |
| COX11 | 339.900584 | 0.241279598 | 0.22761216 | 1.06004706 | 0.28912319 | 0.71218341 |
| STXBP4 | 1104.49622 | 0.256730142 | 0.30697979 | 0.8363096 | 0.40298075 | 0.78260941 |
| HLF | 586.844604 | 0.157250595 | 0.4099075 | 0.38362458 | 0.70125672 | 0.90783786 |
| MMD | 133.588887 | 0.702186786 | 0.52459582 | 1.33852912 | 0.18072401 | 0.63166264 |
| PCTP | 472.537959 | 0.577470992 | 0.53139189 | 1.08671398 | 0.27716323 | 0.7036882 |
| ANKFN1 | 129.225156 | -0.695008598 | 1.00632361 | -0.6906413 | 0.48979102 | 0.81970016 |
| C17orf67 | 44.7138222 | 0.324922124 | 0.74790696 | 0.43444191 | 0.66396757 | 0.89313476 |
| DGKE | 72.1316176 | -0.070611659 | 0.58453882 | -0.1207989 | 9.04E-01 | 0.97567145 |
| TRIM25 | 341.786858 | -1.238099246 | 0.42204656 | -2.9335608 | 0.00335098 | 0.18902009 |
| COIL | 184.132154 | -0.326625935 | 0.18475393 | -1.7678971 | 0.07707811 | 0.51013111 |
| SCPEP1 | 199.449309 | -0.171657333 | 0.27456929 | -0.6251877 | 0.5318479 | 0.83789294 |
| AC007114.1 | 48.857182 | 0.875502586 | 0.58200286 | 1.50429258 | 0.13250605 | 0.5912647 |
| AKAP1 | 522.735604 | 0.231995169 | 0.28877311 | 0.80338218 | 0.42175387 | 0.78892549 |
| AC007114.2 | 70.7318635 | 0.566233611 | 0.64420901 | 0.87895948 | 0.37942325 | 0.77095585 |
| MSI2 | 4894.64816 | 0.473406049 | 0.41586624 | 1.13836133 | 0.25496963 | 0.68976367 |
| AC015845.2 | 535.60524 | -1.766262664 | 0.99383806 | -1.7772137 | 0.07553308 | 0.50730849 |
| MRPS23 | 298.863677 | -0.156689859 | 0.25995894 | -0.6027485 | 0.54667601 | 0.84460887 |
| CUEDC1 | 998.619773 | -0.471504257 | 0.49882962 | -0.9452211 | 0.34454603 | 0.75124763 |
| VEZF1 | 435.241098 | 0.268547325 | 0.25554805 | 1.05086821 | 0.29331912 | 0.71463299 |
| AC015813.2 | 324.883012 | 0.121268693 | 0.27794982 | 0.43629708 | 0.6626212 | 0.89232896 |
| SRSF1 | 145.90362 | -0.385130052 | 0.31129725 | -1.2371778 | 0.21602108 | 0.65975957 |
| DYNLL2 | 365.799459 | -0.254548758 | 0.31160557 | -0.8168941 | 0.41398894 | 0.78632759 |
| MKS1 | 149.440318 | 0.000461068 | 0.41632517 | 0.00110747 | 0.99911637 | 0.99977652 |
| TSPOAP1 | 187.667586 | 1.534838225 | 0.76731202 | 2.00027914 | 0.04547013 | 0.44360789 |
| TSPOAP1-AS | 80.9973108 | 0.352765438 | 0.576655 | 0.61174435 | 0.54070691 | 0.8417342 |
| SUPT4H1 | 122.709179 | -0.458573235 | 0.32081865 | -1.4293846 | 0.15289373 | 0.61001662 |
| RNF43 | 274.440949 | 0.784979711 | 0.49800062 | 1.57626253 | 0.11496531 | 0.57182344 |

| | | | | | | |
|---|---|---|---|---|---|---|
| MTMR4 | 446.193767 | 0.683654152 | 0.44746956 | 1.52782269 | 0.12655657 | 0.58713532 |
| SEPT4-AS1 | 56.9530506 | -0.196814205 | 0.62298824 | -0.3159196 | 0.75206352 | 0.92679087 |
| TEX14 | 42.9631465 | -0.112453544 | 0.45594764 | -0.246637 | 0.80518919 | 0.94585363 |
| RAD51C | 88.5813811 | 0.110519564 | 0.24964829 | 0.44270107 | 0.65798197 | 0.89058983 |
| PPM1E | 187.942279 | -1.059846065 | 0.62085844 | -1.7070656 | 0.08780986 | 0.52761721 |
| TRIM37 | 573.643328 | -0.534479662 | 0.28310184 | -1.8879413 | 0.05903383 | 0.47823683 |
| SKA2 | 139.397532 | -0.263400161 | 0.34169877 | -0.7708549 | 0.44079296 | 0.79826598 |
| GDPD1 | 84.5775001 | 0.737386098 | 0.49792953 | 1.48090454 | 0.13863201 | 0.59474076 |
| YPEL2 | 2050.46983 | -0.041339304 | 0.5493467 | -0.0752518 | 0.94001441 | 0.98439739 |
| LINC01476 | 37.925048 | 0.067974107 | 0.70258834 | 0.09674813 | 0.92292642 | 0.97983687 |
| DHX40 | 433.061954 | 0.148810303 | 0.38438443 | 0.38713925 | 0.69865312 | 0.90696517 |
| CLTC | 2676.70866 | 0.06632984 | 0.21833252 | 0.30380193 | 0.7612788 | 0.93000994 |
| PTRH2 | 389.897034 | -0.312287558 | 0.37940797 | -0.8230917 | 0.41045584 | 0.78582015 |
| VMP1 | 7690.24768 | 0.202507936 | 0.3800372 | 0.53286345 | 0.59412811 | 0.86488041 |
| TUBD1 | 306.127736 | 0.257879077 | 0.29699363 | 0.86829834 | 0.38523103 | 0.77317162 |
| RPS6KB1 | 561.827965 | -0.180870837 | 0.23484727 | -0.7701637 | 0.44120281 | 0.79846407 |
| AC005702.1 | 42.4115984 | 0.642410991 | 0.65368774 | 0.98274902 | 0.32573098 | 0.7372514 |
| RNFT1 | 186.848013 | 0.096799712 | 0.2683272 | 0.36075251 | 0.71828447 | 0.91399231 |
| RNFT1-DT | 40.8279811 | 0.596287471 | 0.63254662 | 0.9426775 | 0.34584588 | 0.75166079 |
| HEATR6 | 132.192036 | -0.103774075 | 0.27173072 | -0.3819004 | 0.70253523 | 0.9085433 |
| AC025048.4 | 38.0972876 | 0.982773638 | 0.50948375 | 1.92895975 | 0.05373586 | 0.46711025 |
| CA4 | 77.4248484 | 0.122416607 | 0.78007146 | 0.15692999 | 0.87530002 | 0.96615698 |
| USP32 | 2714.20228 | -0.269903277 | 0.24494871 | -1.1018767 | 0.27051528 | 0.69961625 |
| APPBP2 | 2131.17902 | -0.075424597 | 0.1401606 | -0.5381298 | 0.59048742 | 0.86315571 |
| AC011921.1 | 252.792706 | 0.72061499 | 0.44926986 | 1.60396915 | 0.10872085 | 0.55858026 |
| PPM1D | 514.016398 | -0.212784529 | 0.24595819 | -0.8651248 | 0.38697031 | 0.77335375 |
| BCAS3 | 5789.56387 | -0.322783569 | 0.34263693 | -0.9420572 | 0.34616338 | 0.75166079 |
| AC005746.2 | 75.0453985 | 0.666111257 | 0.60225439 | 1.10602972 | 0.26871366 | 0.69961625 |
| TBX2 | 365.049269 | 0.700651505 | 0.4581802 | 1.52920511 | 0.1262136 | 0.5860467 |
| NACA2 | 85.903464 | 0.437214297 | 0.47624896 | 0.91803728 | 0.35859935 | 0.75936492 |
| BRIP1 | 78.2091194 | -0.137913675 | 0.75389575 | -0.1829347 | 0.85484927 | 0.96171023 |
| INTS2 | 298.751666 | 0.034959881 | 0.29157132 | 0.11990164 | 0.90456106 | 0.97574711 |
| MED13 | 3962.01797 | -0.473719874 | 0.29538672 | -1.6037277 | 0.10877408 | 0.55862357 |
| EFCAB3 | 85.3916023 | -0.107554649 | 0.41080769 | -0.2618127 | 7.93E-01 | 0.94189286 |
| METTL2A | 156.709115 | -0.373677274 | 0.29127466 | -1.2829035 | 0.1995259 | 0.64671356 |
| TLK2 | 1601.11606 | -0.376137054 | 0.20257691 | -1.8567617 | 0.06334506 | 0.48572144 |
| TANC2 | 1710.37038 | -0.53432665 | 0.2636464 | -2.0266791 | 4.27E-02 | 0.43869842 |
| AC015923.1 | 94.0198176 | -0.266027257 | 0.73861946 | -0.3601682 | 0.71872134 | 0.91437845 |
| CYB561 | 202.711828 | 0.641895553 | 0.35508377 | 1.80772992 | 0.07064855 | 0.4967563 |
| ACE | 64.1160079 | 2.638764997 | 0.99010586 | 2.6651342 | 0.00769576 | 0.24157203 |
| KCNH6 | 296.23321 | 2.561788145 | 0.81592224 | 3.13974544 | 0.00169095 | 0.13207839 |
| DCAF7 | 1178.77438 | 0.047943291 | 0.23213855 | 0.20652878 | 0.83637789 | 0.95471617 |
| TACO1 | 122.135701 | 0.307994638 | 0.37665108 | 0.81771871 | 0.41351783 | 0.78632759 |
| MAP3K3 | 466.889649 | -0.266329694 | 0.30409799 | -0.8758022 | 0.38113756 | 0.77210521 |
| CCDC47 | 295.410818 | -0.550521941 | 0.23620731 | -2.3306728 | 0.01977062 | 0.34565805 |
| DDX42 | 825.747769 | -0.364654554 | 0.28077553 | -1.2987405 | 0.19403301 | 0.64548758 |
| FTSJ3 | 237.906758 | -0.083798702 | 0.38186267 | -0.2194472 | 0.82630169 | 0.95192757 |
| PSMC5 | 207.880165 | -0.36893304 | 0.38893246 | -0.9485787 | 0.34283493 | 0.7505827 |
| SMARCD2 | 259.663847 | 0.395488494 | 0.2362466 | 1.67404943 | 0.09412088 | 0.53783536 |
| ERN1 | 617.878729 | 0.880100454 | 0.2928259 | 3.00554172 | 0.00265108 | 0.16823326 |
| SNHG25 | 441.421678 | -0.375329008 | 0.49742713 | -0.7545407 | 0.45052462 | 0.80361623 |
| TEX2 | 1016.78585 | 0.376823202 | 0.23287697 | 1.61812134 | 0.10563645 | 0.55370151 |
| PECAM1 | 69.0272302 | -0.776364252 | 0.43591531 | -1.7809979 | 0.07491279 | 0.5071704 |
| POLG2 | 316.282535 | -0.467151406 | 0.19191325 | -2.4341801 | 0.01492557 | 0.30894792 |

| | | | | | | |
|---|---|---|---|---|---|---|
| DDX5 | 3231.58966 | 0.804784454 | 0.27043715 | 2.97586498 | 0.00292163 | 0.17487706 |
| CEP95 | 1107.63126 | -0.132728549 | 0.2194847 | -0.604728 | 0.54535973 | 0.84449215 |
| SMURF2 | 1326.47464 | 0.131503746 | 0.19779857 | 0.66483669 | 0.50615495 | 0.82782978 |
| LRRC37A3 | 1501.54651 | -0.497779062 | 0.25163172 | -1.9782047 | 0.04790562 | 0.45133876 |
| AC103810.2 | 46.4887954 | -1.084288211 | 0.71632421 | -1.5136836 | 0.1301061 | 0.58853436 |
| GNA13 | 1353.56934 | -1.057673068 | 0.37376045 | -2.8298154 | 0.00465749 | 0.21069366 |
| RGS9 | 213.271428 | -0.286404789 | 0.40636172 | -0.7048026 | 4.81E-01 | 8.17E-01 |
| AC004805.1 | 52.380106 | 1.302121977 | 0.40028327 | 3.25300126 | 0.00114193 | 0.11093704 |
| CEP112 | 1998.6103 | 0.816226092 | 0.44701233 | 1.82595877 | 0.06785647 | 0.48963361 |
| PRKCA | 5732.19558 | -0.164870397 | 0.32295582 | -0.5105045 | 0.60969807 | 0.87071593 |
| PRKCA-AS1 | 200.217428 | 0.026786438 | 0.46826585 | 0.05720348 | 0.9543831 | 0.98840182 |
| HELZ | 3400.16321 | -0.273482067 | 0.25268883 | -1.0822879 | 0.27912461 | 0.70446562 |
| AC007448.3 | 76.0169523 | -0.524673937 | 0.57104894 | -0.9187898 | 3.58E-01 | 7.59E-01 |
| PSMD12 | 615.965534 | -0.542628363 | 0.34829239 | -1.5579679 | 0.11924085 | 0.57652003 |
| PITPNC1 | 5310.04325 | -0.460331337 | 0.64380867 | -0.7150126 | 0.47460123 | 0.81531007 |
| NOL11 | 371.545693 | -0.289987543 | 0.23299475 | -1.2446098 | 0.21327523 | 0.65809339 |
| BPTF | 2694.06895 | -0.21365126 | 0.24086955 | -0.8869999 | 0.37507897 | 0.7697436 |
| KPNA2 | 111.702418 | -0.387355615 | 0.53979444 | -0.7175984 | 0.47300496 | 0.81471211 |
| AC005332.7 | 114.145526 | -0.223915078 | 0.23446798 | -0.9549921 | 0.33958167 | 0.74644675 |
| AC005332.6 | 41.8657177 | -0.061569968 | 0.39415456 | -0.1562077 | 0.87586933 | 0.96617876 |
| AMZ2 | 92.1685882 | 0.264338698 | 0.36679843 | 0.72066475 | 0.4711158 | 0.81387913 |
| ARSG | 272.797853 | -0.357407187 | 0.45919039 | -0.7783421 | 0.43636739 | 0.79555046 |
| WIPI1 | 791.482982 | 0.863081389 | 0.37798151 | 2.28339577 | 0.02240707 | 0.36273984 |
| PRKAR1A | 1199.56579 | 0.528253018 | 0.2823459 | 1.87094279 | 0.06135301 | 0.4813972 |
| FAM20A | 1917.99379 | 0.814948863 | 0.86591907 | 0.94113745 | 0.34663443 | 0.75166079 |
| AC079210.1 | 93.9799914 | 0.945931358 | 0.73122172 | 1.29363137 | 0.19579278 | 0.64558745 |
| LINC01482 | 281.881191 | 1.18536417 | 0.75532755 | 1.56933792 | 0.11656922 | 0.57350419 |
| ABCA9 | 43.0819671 | 0.75691508 | 0.55022756 | 1.37564008 | 0.16893309 | 0.62304975 |
| ABCA6 | 63.7633807 | 0.601167022 | 0.63497963 | 0.9467501 | 0.34376614 | 0.7507483 |
| ABCA10 | 248.303654 | 0.764309126 | 0.38334264 | 1.99380148 | 0.04617375 | 0.44508419 |
| ABCA5 | 959.104818 | 0.440790584 | 0.33819933 | 1.30334552 | 0.19245683 | 0.64497936 |
| MAP2K6 | 109.12878 | 0.513663239 | 0.76577946 | 0.67077176 | 0.50236594 | 0.82638105 |
| AC005208.1 | 1714.27071 | 0.957785614 | 0.80061521 | 1.19631204 | 0.23157481 | 0.66878058 |
| KCNJ16 | 3604.70965 | 0.069477466 | 0.46299642 | 0.15006048 | 0.8807169 | 0.96773407 |
| SOX9 | 685.364073 | 2.003231158 | 0.80549208 | 2.48696568 | 0.01288378 | 0.29875272 |
| LINC00511 | 515.590489 | 2.839997743 | 0.70114416 | 4.050519 | 5.11E-05 | 0.02213112 |
| SLC39A11 | 2915.34344 | 0.190533232 | 0.22933343 | 0.83081314 | 0.4060792 | 0.78429302 |
| SSTR2 | 327.373667 | -0.305895059 | 0.89946316 | -0.3400863 | 0.73379157 | 0.91972212 |
| COG1 | 217.976789 | -0.154570179 | 0.19755551 | -0.7824139 | 0.43397137 | 0.79429104 |
| FAM104A | 264.165119 | -0.049258917 | 0.22853741 | -0.2155398 | 0.82934645 | 0.95325673 |
| C17orf80 | 148.001903 | 0.155226416 | 0.31540113 | 0.49215555 | 0.62260939 | 0.87714058 |
| CDC42EP4 | 222.479658 | 0.17617315 | 0.35600435 | 0.49486235 | 0.62069729 | 0.87609696 |
| SDK2 | 64.0041298 | -0.113016263 | 0.81926477 | -0.1379484 | 0.8902812 | 0.97225819 |
| RPL38 | 458.929932 | -0.263210356 | 0.46432362 | -0.5668683 | 0.57080364 | 0.85386942 |
| GPRC5C | 860.950006 | 0.472917957 | 0.19594373 | 2.41353967 | 0.01579841 | 0.31566878 |
| SLC9A3R1 | 1026.9807 | 0.020782382 | 0.22736404 | 0.09140576 | 0.92717019 | 0.98189923 |
| NAT9 | 223.025014 | -0.630502359 | 0.41996539 | -1.5013198 | 0.13327286 | 0.59233824 |
| TMEM104 | 145.34013 | -0.84630068 | 0.55885797 | -1.5143395 | 0.12993977 | 0.58853436 |
| FDXR | 53.3899821 | 0.955803441 | 0.47398803 | 2.01651388 | 0.04374627 | 0.44226848 |
| CDR2L | 70.7926832 | 0.323104129 | 0.56787463 | 0.56897088 | 0.5693759 | 0.85313024 |
| MRPL58 | 95.1098999 | -0.082744312 | 0.3645367 | -0.2269849 | 0.82043551 | 0.95036843 |
| KCTD2 | 196.016044 | -0.271816404 | 0.28753664 | -0.9453279 | 0.3444915 | 0.75124763 |
| ATP5PD | 290.683921 | -0.324222269 | 0.43576053 | -0.7440377 | 0.45685366 | 0.80507423 |
| SLC16A5 | 107.99816 | 0.183984946 | 0.61299975 | 0.3001387 | 0.76407136 | 0.93079056 |

| | | | | | | |
|---|---|---|---|---|---|---|
| ARMC7 | 136.515707 | 0.465459624 | 0.53685075 | 0.86701867 | 0.38593178 | 0.77317162 |
| NT5C | 306.425493 | -0.389895386 | 0.29496048 | -1.3218564 | 0.18621598 | 0.63745505 |
| JPT1 | 279.319088 | 0.180914061 | 0.23616884 | 0.76603695 | 4.44E-01 | 7.99E-01 |
| SUMO2 | 225.465337 | -0.38595305 | 0.3732373 | -1.0340688 | 0.301104 | 0.71879123 |
| NUP85 | 263.747282 | -0.205089357 | 0.25056929 | -0.8184936 | 0.4130754 | 0.78632759 |
| GGA3 | 371.437115 | -0.346343801 | 0.30327384 | -1.1420167 | 0.25344707 | 0.68891958 |
| MRPS7 | 197.84874 | 0.075124855 | 0.30332308 | 0.24767273 | 0.80438763 | 0.94555743 |
| MIF4GD | 40.2059575 | 0.220244372 | 0.41491131 | 0.53082277 | 0.5955416 | 0.86519223 |
| SLC25A19 | 50.8512385 | -0.074356133 | 0.3509396 | -0.2118773 | 0.83220277 | 0.95325673 |
| GRB2 | 1256.89343 | -0.265545051 | 0.26331987 | -1.0084505 | 0.31323824 | 0.72717675 |
| AC011933.4 | 150.092786 | 0.060176408 | 0.52622176 | 0.11435561 | 0.9089559 | 0.97680747 |
| TMEM94 | 288.175948 | -0.164756764 | 0.20135612 | -0.8182357 | 0.41322261 | 0.78632759 |
| CASKIN2 | 70.0285071 | 0.273893551 | 0.30876648 | 0.88705728 | 0.37504806 | 0.7697436 |
| TSEN54 | 177.184217 | 0.133093399 | 0.28029769 | 0.47482874 | 0.63490904 | 0.88238823 |
| LLGL2 | 1853.38113 | 0.670667601 | 0.35317034 | 1.89899187 | 0.05756555 | 0.47684824 |
| MYO15B | 548.437545 | 0.40449296 | 0.26418399 | 1.53110322 | 0.12574388 | 0.58564339 |
| RECQL5 | 232.039696 | -0.02926335 | 0.26866294 | -0.1089222 | 9.13E-01 | 0.97720308 |
| SMIM5 | 74.4564775 | -1.16901711 | 0.72211256 | -1.6188849 | 0.10547203 | 0.55348589 |
| SAP30BP | 470.271528 | -0.681004171 | 0.39257672 | -1.7347034 | 0.08279335 | 0.51935659 |
| ITGB4 | 134.995906 | 1.191908729 | 0.41772705 | 2.85331951 | 0.00432651 | 0.20749797 |
| GALK1 | 104.616391 | -0.110821553 | 0.48467692 | -0.2286504 | 0.81914068 | 0.94990455 |
| H3F3B | 1270.93773 | -0.309442972 | 0.36592256 | -0.8456515 | 0.39774717 | 0.78063007 |
| UNK | 611.47868 | -0.127512347 | 0.28549904 | -0.4466297 | 0.65514245 | 0.89053559 |
| UNC13D | 35.3626163 | -0.487487413 | 0.57179457 | -0.8525569 | 0.39390506 | 0.77814839 |
| WBP2 | 525.379995 | -0.306397333 | 0.25258592 | -1.213042 | 0.22511375 | 0.6664553 |
| TRIM47 | 288.656625 | 0.058102 | 0.38674989 | 0.15023146 | 0.88058201 | 0.96773407 |
| TRIM65 | 157.028428 | 0.307982903 | 0.35053847 | 0.87859945 | 0.3796185 | 0.77095585 |
| FBF1 | 48.5657611 | 0.156065397 | 0.47792586 | 0.32654729 | 0.74401031 | 0.92361546 |
| ACOX1 | 1466.42198 | 0.324294006 | 0.39424386 | 0.82257212 | 0.41075136 | 0.7860071 |
| TEN1 | 166.829014 | -0.379477507 | 0.3606838 | -1.0521058 | 0.29275102 | 0.71426455 |
| CDK3 | 109.29965 | 0.058730288 | 0.46463542 | 0.1264008 | 0.89941467 | 0.97458352 |
| EVPL | 296.521245 | 0.412300398 | 0.44474622 | 0.92704643 | 0.35390243 | 0.75682564 |
| SRP68 | 294.085516 | -0.158100318 | 0.19985855 | -0.7910611 | 0.42890836 | 0.79037914 |
| EXOC7 | 566.450839 | -0.168658918 | 0.32352542 | -0.5213158 | 0.60214678 | 0.86770668 |
| RNF157-AS1 | 46.5118646 | 0.428506634 | 0.67485882 | 0.63495745 | 0.52545617 | 0.83628497 |
| RNF157 | 795.411189 | 0.433456176 | 0.39379998 | 1.10070137 | 0.27102665 | 0.69961625 |
| UBALD2 | 280.32663 | -0.082350903 | 0.37657042 | -0.2186866 | 0.82689419 | 0.95204019 |
| QRICH2 | 46.5891955 | -0.804434817 | 0.58992134 | -1.3636306 | 0.17268385 | 0.62593809 |
| PRPSAP1 | 335.123076 | 0.0039101 | 0.25031836 | 0.01562051 | 0.98753714 | 0.99648575 |
| SPHK1 | 84.7283367 | -2.124704916 | 1.0208566 | -2.0812962 | 3.74E-02 | 4.22E-01 |
| UBE2O | 817.897013 | -0.57372303 | 0.32568276 | -1.7616009 | 0.07813676 | 0.51192014 |
| RHBDF2 | 172.720796 | 0.406073281 | 0.41980328 | 0.96729421 | 0.33339698 | 0.74018017 |
| AC015802.4 | 54.6357089 | 0.485155806 | 0.48153231 | 1.00752494 | 0.31368258 | 0.72717675 |
| ST6GALNAC2 | 208.878268 | 1.665489762 | 0.58573641 | 2.84341172 | 0.00446334 | 0.20749797 |
| MXRA7 | 183.283602 | 0.288423153 | 0.52461582 | 0.54977975 | 0.58247045 | 0.85981507 |
| JMJD6 | 241.433523 | -0.736114017 | 0.34332366 | -2.1440818 | 0.03202633 | 0.40988962 |
| METTL23 | 122.871321 | 0.203426957 | 0.37754932 | 0.53880896 | 0.59001868 | 0.86292147 |
| AC005837.2 | 81.2711817 | 0.342052398 | 0.86572575 | 0.3951048 | 0.69276555 | 0.90528251 |
| SRSF2 | 869.935478 | -0.340906077 | 0.2861981 | -1.1911542 | 0.23359304 | 0.67012553 |
| MFSD11 | 507.303166 | -0.797436553 | 0.34797282 | -2.2916633 | 0.02192508 | 0.36066559 |
| SEC14L1 | 684.421787 | -0.404246283 | 0.28331438 | -1.426847 | 0.15362399 | 0.61017793 |
| 9-Sep | 1605.1949 | 0.052511414 | 0.49394886 | 0.10630941 | 0.91533686 | 0.97782888 |
| TNRC6C | 824.891309 | -0.621063486 | 0.40383784 | -1.5379031 | 0.1240723 | 0.58401752 |
| TMC6 | 80.2708213 | 0.436375579 | 0.52402732 | 0.83273441 | 0.40499453 | 0.78422899 |

| | | | | | | |
|---|---|---|---|---|---|---|
| SYNGR2 | 269.155488 | -0.024639926 | 0.45327905 | -0.0543593 | 0.95664892 | 0.98861313 |
| TK1 | 45.2712253 | -0.218321396 | 0.44350262 | -0.4922663 | 0.6225311 | 0.87714058 |
| AFMID | 1595.91408 | 0.42327722 | 0.39517587 | 1.071111 | 2.84E-01 | 7.09E-01 |
| SOCS3 | 243.924121 | -0.182972486 | 0.75842907 | -0.2412519 | 8.09E-01 | 9.47E-01 |
| PGS1 | 516.19049 | -0.446823401 | 0.3586127 | -1.2459776 | 0.21277261 | 0.65745584 |
| DNAH17 | 45.5573944 | -1.14446229 | 0.49021853 | -2.3345961 | 0.01956453 | 0.34513641 |
| CYTH1 | 1052.98572 | -0.278523639 | 0.30840717 | -0.9031037 | 0.36647089 | 0.76479192 |
| USP36 | 752.527074 | -1.069450181 | 0.53399554 | -2.0027324 | 0.04520602 | 0.44360789 |
| TIMP2 | 81.1933288 | 0.541563059 | 0.64903201 | 0.83441656 | 0.40404629 | 0.78358009 |
| LGALS3BP | 268.459852 | -0.430534158 | 0.79076793 | -0.5444507 | 0.58613136 | 0.86101196 |
| CANT1 | 108.956225 | -0.27020166 | 0.43196212 | -0.6255217 | 0.53162874 | 0.83789294 |
| ENGASE | 151.134397 | -0.601085708 | 0.56538835 | -1.0631378 | 0.28771948 | 0.7113446 |
| CBX4 | 105.423875 | -0.009215809 | 0.34717055 | -0.0265455 | 0.97882226 | 0.99482978 |
| TBC1D16 | 1048.73056 | 0.137428121 | 0.2281811 | 0.60227654 | 0.54699007 | 0.84460887 |
| CCDC40 | 406.160261 | 0.685886762 | 0.42466565 | 1.61512184 | 0.10628431 | 0.55440878 |
| GAA | 245.592093 | 0.287689942 | 0.33257954 | 0.86502598 | 0.38702453 | 0.77335375 |
| EIF4A3 | 199.221005 | -0.44438778 | 0.30799802 | -1.4428268 | 1.49E-01 | 0.60622442 |
| CARD14 | 95.6318462 | 0.4853334 | 0.50567541 | 0.9597726 | 0.33716967 | 0.74349819 |
| AC087741.1 | 271.549514 | -0.057161254 | 0.3331555 | -0.1715753 | 0.86377143 | 0.96393376 |
| SGSH | 162.022345 | -0.183775758 | 0.25221741 | -0.7286402 | 0.46622176 | 0.81077158 |
| SLC26A11 | 561.136177 | -0.175135256 | 0.33699779 | -0.5196926 | 0.60327785 | 0.8678002 |
| RNF213 | 9020.87145 | 0.501224063 | 0.86462846 | 0.57969878 | 0.56211777 | 0.85034193 |
| AC124319.2 | 1764.03623 | 0.256137086 | 0.45366673 | 0.56459305 | 0.57235059 | 0.85408471 |
| ENDOV | 416.298288 | 0.123451991 | 0.19175246 | 0.64380916 | 0.51969919 | 0.8335052 |
| RPTOR | 1463.23897 | -0.235915094 | 0.24417168 | -0.9661853 | 0.33395145 | 0.74069134 |
| CHMP6 | 103.240731 | -0.243525807 | 0.42864752 | -0.568126 | 0.56994941 | 0.85327114 |
| BAIAP2-DT | 58.3444658 | 1.225299631 | 0.55965845 | 2.18937037 | 0.02856993 | 0.39280225 |
| BAIAP2 | 566.060986 | 0.295088853 | 0.66791529 | 0.44180581 | 0.65862974 | 0.89058983 |
| CEP131 | 32.8200076 | 0.513135344 | 0.48513572 | 1.05771503 | 0.29018538 | 0.71260706 |
| TEPSIN | 130.938776 | -0.340119526 | 0.31585796 | -1.0768116 | 0.28156442 | 0.70625976 |
| NDUFAF8 | 223.962196 | 0.065917059 | 0.31687718 | 0.20802085 | 0.83521269 | 0.95449894 |
| SLC38A10 | 598.083661 | 0.297991591 | 0.33856949 | 0.88014898 | 0.37877861 | 0.77061593 |
| LINC00482 | 64.4492809 | -0.347917998 | 1.03796258 | -0.3351932 | 0.73747935 | 0.92047254 |
| AC110285.6 | 68.5813398 | -0.295619162 | 0.72731107 | -0.4064549 | 0.68440837 | 0.9007343 |
| AC110285.2 | 33.2796141 | 0.03717968 | 0.81297342 | 0.04573296 | 0.96352309 | 0.99012655 |
| BAHCC1 | 79.7153417 | 0.18624215 | 0.5322534 | 0.34991256 | 0.72640432 | 0.91635631 |
| ACTG1 | 1638.41629 | 0.39194551 | 0.50164883 | 0.78131451 | 0.43461754 | 0.79451524 |
| FAAP100 | 121.129008 | -0.393946848 | 0.49145937 | -0.8015858 | 0.4227926 | 0.78894887 |
| NPLOC4 | 2564.33663 | -0.425828382 | 0.21312207 | -1.9980492 | 0.04571133 | 0.44430994 |
| TSPAN10 | 38.9207957 | -0.757062265 | 0.48234541 | -1.5695438 | 0.11652128 | 0.57350419 |
| OXLD1 | 134.739903 | -0.091209916 | 0.26903792 | -0.3390225 | 0.73459276 | 0.91988516 |
| CCDC137 | 239.384297 | -0.464784337 | 0.38749183 | -1.1994688 | 0.23034573 | 0.66836344 |
| ARL16 | 368.260026 | -0.402125363 | 0.27807818 | -1.4460875 | 0.14815267 | 0.60605027 |
| HGS | 561.475739 | -1.053554579 | 0.40085722 | -2.6282539 | 0.00858244 | 0.25133745 |
| AC139530.1 | 42.7904923 | -0.199167878 | 0.48346019 | -0.4119633 | 0.68036629 | 0.89941358 |
| SLC25A10 | 198.837392 | 0.335380125 | 0.38695337 | 0.86671973 | 3.86E-01 | 0.77317162 |
| MCRIP1 | 274.496544 | -0.486982246 | 0.52025733 | -0.9360411 | 0.34925202 | 0.7542101 |
| P4HB | 1341.47088 | -0.827322411 | 0.50987249 | -1.6226065 | 0.10467355 | 0.55098325 |
| ARHGDIA | 652.234078 | -0.820446973 | 0.65122124 | -1.259859 | 0.20772022 | 0.65309147 |
| ALYREF | 38.0520034 | -0.550818423 | 0.48404702 | -1.137944 | 0.25514385 | 0.68976367 |
| ANAPC11 | 304.272331 | -0.194397679 | 0.31884141 | -0.6097002 | 0.5420604 | 0.84259718 |
| PCYT2 | 179.586402 | 0.393877874 | 0.43033996 | 0.91527144 | 0.36004915 | 0.75998694 |
| SIRT7 | 324.828022 | -0.58475226 | 0.45758569 | -1.2779077 | 0.20128198 | 0.6485595 |
| MAFG | 382.066197 | -0.570068465 | 0.38404057 | -1.4843965 | 1.38E-01 | 5.95E-01 |

| | | | | | | |
|---|---|---|---|---|---|---|
| ASPSCR1 | 440.594152 | -0.666501279 | 0.4662675 | -1.4294397 | 0.15287789 | 0.61001662 |
| CENPX | 126.979318 | -0.221617867 | 0.42586938 | -0.5203893 | 0.60279227 | 0.8677246 |
| LRRC45 | 57.6744694 | 0.236487426 | 0.45738245 | 0.51704526 | 0.60512456 | 0.86869053 |
| RAC3 | 63.917514 | 0.017497838 | 0.43432806 | 0.04028715 | 0.9678642 | 0.99181479 |
| DCXR | 457.055268 | -0.560789355 | 0.48778308 | -1.1496696 | 0.25028 | 0.68617881 |
| RFNG | 319.357227 | -0.023091346 | 0.22555368 | -0.1023763 | 0.91845801 | 0.97860783 |
| GPS1 | 288.490208 | -0.07225501 | 0.36988481 | -0.1953446 | 0.84512317 | 0.95760132 |
| DUS1L | 293.036684 | -0.28936716 | 0.34810739 | -0.8312583 | 0.40582772 | 0.78429302 |
| FASN | 122.672342 | -0.101404616 | 0.48043027 | -0.2110704 | 0.83283232 | 0.95344798 |
| CCDC57 | 1177.10338 | 0.105128642 | 0.31029446 | 0.33880284 | 0.73475826 | 0.9199679 |
| SLC16A3 | 329.743793 | -0.179292781 | 0.53181044 | -0.3371366 | 0.7360139 | 0.92019178 |
| CSNK1D | 1182.65667 | -0.445184145 | 0.34959891 | -1.273414 | 0.20287118 | 0.64881821 |
| AC132872.1 | 58.5296445 | -0.741116534 | 0.44332525 | -1.6717219 | 0.09457917 | 0.53824794 |
| SECTM1 | 116.463089 | -1.205241187 | 0.55768854 | -2.1611367 | 0.03068478 | 0.40446361 |
| OGFOD3 | 192.370126 | -0.157800267 | 0.35253771 | -0.4476124 | 0.65443292 | 0.89032312 |
| HEXDC | 501.097549 | -0.099045569 | 0.29025745 | -0.3412335 | 0.73292779 | 0.91952673 |
| CYBC1 | 244.278421 | 0.396320126 | 0.45801279 | 0.86530363 | 0.38687216 | 0.77334938 |
| NARF | 406.930847 | -0.269559171 | 0.39115201 | -0.6891417 | 0.49073409 | 0.81989696 |
| FOXK2 | 1258.84846 | -0.469097921 | 0.25472794 | -1.8415645 | 0.06553888 | 0.48790819 |
| WDR45B | 805.223551 | -0.967011849 | 0.22783377 | -4.2443745 | 2.19E-05 | 0.01647703 |
| RAB40B | 718.367635 | -0.320696182 | 0.43604582 | -0.7354644 | 0.4620567 | 0.8082072 |
| FN3KRP | 191.867318 | -0.434019465 | 0.23561465 | -1.8420733 | 0.06546443 | 0.48790819 |
| FN3K | 968.102125 | 0.333961227 | 0.64214142 | 0.52007427 | 0.60301181 | 0.8678002 |
| TBCD | 900.987665 | -0.454161475 | 0.20705456 | -2.1934387 | 0.02827579 | 0.39280225 |
| B3GNTL1 | 60.8716401 | -0.49564566 | 0.38694913 | -1.2809065 | 0.2002265 | 0.64744153 |
| METRNL | 98.6448979 | -0.708320211 | 0.3216968 | -2.2018255 | 0.02767764 | 0.38942007 |
| USP14 | 975.380692 | -0.348293848 | 0.27766907 | -1.2543487 | 0.20971528 | 0.65441616 |
| THOC1 | 638.757199 | -0.382212417 | 0.24503371 | -1.5598361 | 0.11879862 | 0.57609025 |
| TYMS | 154.453104 | 0.730091674 | 0.49487424 | 1.4753075 | 0.14012989 | 0.59500405 |
| ENOSF1 | 5237.46818 | -0.133313941 | 0.31057301 | -0.4292515 | 0.66774018 | 0.89495597 |
| YES1 | 1624.21453 | -0.133113007 | 0.22734519 | -0.5855106 | 0.55820447 | 0.8489192 |
| LINC00470 | 108.772706 | 1.698529406 | 1.21758642 | 1.394997 | 0.16301674 | 0.61999736 |
| AP005230.1 | 91.5109251 | -1.43246132 | 1.10206231 | -1.2998007 | 0.1936693 | 0.64548758 |
| METTL4 | 210.729732 | 0.027934908 | 0.23645157 | 0.11814219 | 0.90595499 | 0.97637317 |
| SMCHD1 | 1782.59172 | 0.152959204 | 0.18947639 | 0.80727318 | 0.41950911 | 0.78783502 |
| AP001011.1 | 237.173732 | 0.656932044 | 0.5770407 | 1.1384501 | 0.25493258 | 0.68976367 |
| LPIN2 | 2355.17714 | -0.318741313 | 0.25719234 | -1.239311 | 0.21523033 | 0.65925411 |
| AP000919.2 | 161.729233 | 0.193409173 | 0.58829619 | 0.32876156 | 0.74233592 | 0.9229205 |
| AP000919.3 | 55.054458 | -0.211893435 | 0.72516712 | -0.2921995 | 0.77013413 | 0.93266075 |
| MYOM1 | 108.699894 | 1.358227762 | 0.71873674 | 1.88974306 | 0.05879233 | 0.47778423 |
| MYL12A | 675.085121 | -0.148505276 | 0.40914133 | -0.3629682 | 7.17E-01 | 0.91298968 |
| MYL12B | 1405.99908 | -0.205115987 | 0.33205491 | -0.6177171 | 5.37E-01 | 0.83988525 |
| TGIF1 | 475.038595 | -0.576144169 | 0.41785914 | -1.3788 | 0.16795644 | 0.62304975 |
| DLGAP1 | 4245.74962 | 1.056957918 | 0.7507293 | 1.40790817 | 0.15915826 | 0.61423843 |
| DLGAP1-AS1 | 91.3280992 | -0.503630652 | 0.47892017 | -1.0515963 | 0.29298483 | 0.71433665 |
| DLGAP1-AS2 | 43.2849116 | -0.926480118 | 0.63562216 | -1.4575957 | 1.45E-01 | 0.60109715 |
| DLGAP1-AS3 | 59.2652172 | 0.389864628 | 1.04629055 | 0.37261603 | 0.70943424 | 0.91101885 |
| LINC00667 | 216.984693 | 0.152583802 | 0.29297564 | 0.52080713 | 0.60250114 | 0.8677246 |
| ZBTB14 | 36.4617365 | 0.310334857 | 0.37016252 | 0.83837461 | 0.40182034 | 0.78257338 |
| EPB41L3 | 7223.81996 | -0.020519049 | 0.34422321 | -0.0596097 | 0.95246648 | 0.98777411 |
| AP005059.1 | 40.9723501 | -0.012110565 | 1.03714996 | -0.0116768 | 0.99068349 | 0.99737132 |
| L3MBTL4 | 9525.91991 | 0.121415435 | 0.32333765 | 0.37550664 | 0.70728371 | 0.91002281 |
| AP001021.2 | 419.824934 | 0.453838908 | 0.72219736 | 0.62841397 | 0.52973279 | 0.83789294 |
| ARHGAP28 | 518.376266 | 0.076289757 | 0.43849594 | 0.17398053 | 8.62E-01 | 9.63E-01 |

| | | | | | | |
|---|---|---|---|---|---|---|
| LAMA1 | 1302.68669 | -0.428067052 | 0.41880782 | -1.0221085 | 3.07E-01 | 7.22E-01 |
| PTPRM | 13720.0895 | -0.501000714 | 0.31256704 | -1.6028584 | 0.10896592 | 0.55898048 |
| RAB12 | 620.5318 | 0.017527612 | 0.20243605 | 0.08658345 | 0.93100262 | 0.98272895 |
| MTCL1 | 549.587611 | -0.811905725 | 0.5366331 | -1.5129624 | 0.1302892 | 0.58853436 |
| NDUFV2 | 878.650602 | -0.525815171 | 0.21939885 | -2.3966177 | 0.01654718 | 0.32448091 |
| AP005899.1 | 57.9392318 | -0.228933685 | 0.61840182 | -0.3702022 | 0.71123187 | 0.91143287 |
| NDUFV2-AS1 | 91.7777637 | -0.283129008 | 0.44667033 | -0.6338657 | 0.52616847 | 0.83687528 |
| ANKRD12 | 4614.26242 | -0.753394559 | 0.27216615 | -2.7681421 | 0.00563769 | 0.22463891 |
| TWSG1 | 406.030051 | -0.160845708 | 0.31448736 | -0.5114536 | 0.60903344 | 0.87022009 |
| RALBP1 | 549.476917 | -0.138987617 | 0.19626624 | -0.7081586 | 0.4788468 | 0.81596289 |
| PPP4R1 | 884.54323 | -0.240515048 | 0.23623778 | -1.0181058 | 0.3086277 | 0.72400477 |
| RAB31 | 246.521818 | -1.437922488 | 0.76996618 | -1.8675138 | 0.06182987 | 0.48267858 |
| VAPA | 1530.45205 | -0.164785455 | 0.26571315 | -0.620163 | 0.5351505 | 0.83954482 |
| NAPG | 656.042356 | -0.113734174 | 0.32347668 | -0.3515993 | 0.72513879 | 0.91600702 |
| PIEZO2 | 35.8788056 | 0.035977239 | 0.51841672 | 0.0693983 | 0.94467258 | 0.9853777 |
| LINC01928 | 76.0935205 | 3.315829635 | 2.07188188 | 1.60039511 | 0.10951096 | 0.55969747 |
| LINC01255 | 58.9608923 | 6.325795029 | 2.78700935 | 2.26974302 | 0.02322318 | 0.36970978 |
| GNAL | 233.319848 | 0.370785238 | 0.28958697 | 1.28039337 | 0.20040682 | 0.64780303 |
| MPPE1 | 479.162678 | -0.181255244 | 0.28048277 | -0.6462259 | 0.51813304 | 0.83325693 |
| IMPA2 | 1777.89783 | 0.934350317 | 0.43631259 | 2.14146997 | 0.03223616 | 0.40988962 |
| TUBB6 | 158.946389 | -0.910193373 | 0.56585168 | -1.608537 | 1.08E-01 | 0.5565641 |
| AFG3L2 | 1200.55504 | -0.357852504 | 0.27915487 | -1.2819139 | 0.19987285 | 0.64705133 |
| SPIRE1 | 882.249782 | -0.239013087 | 0.52696435 | -0.4535659 | 6.50E-01 | 0.88866038 |
| PSMG2 | 315.967594 | -0.115007648 | 0.20703105 | -0.5555092 | 0.57854643 | 0.85751254 |
| CEP76 | 108.807058 | 0.013558353 | 0.37563184 | 0.03609479 | 0.97120678 | 0.99227275 |
| PTPN2 | 1216.60982 | -0.01308993 | 0.25233789 | -0.0518746 | 0.95862861 | 0.98878187 |
| SEH1L | 384.0194 | -0.584926494 | 0.29147238 | -2.006799 | 0.04477107 | 0.44360789 |
| CEP192 | 1053.96696 | -0.330590328 | 0.24028378 | -1.3758329 | 0.16887338 | 0.62304975 |
| LDLRAD4 | 805.321433 | 0.786629534 | 0.65652286 | 1.1981754 | 0.23084874 | 0.66878058 |
| FAM210A | 658.734216 | -0.858924376 | 0.36180113 | -2.3740235 | 0.01759543 | 0.33220465 |
| RNMT | 1029.78348 | -0.825485916 | 0.24841124 | -3.3230619 | 0.00089035 | 0.10027947 |
| ZNF519 | 69.9148315 | -0.035015432 | 0.52946215 | -0.066134 | 0.94727117 | 0.98646931 |
| ANKRD30B | 61.6228078 | -0.264754469 | 0.67864231 | -0.3901237 | 0.69644506 | 0.90613881 |
| ROCK1 | 2022.59855 | -0.018846239 | 0.13351829 | -0.141151 | 0.88775067 | 0.97098615 |
| GREB1L | 286.215179 | 0.251799841 | 0.57507853 | 0.43785297 | 0.66149287 | 0.89184036 |
| ESCO1 | 937.660094 | -0.429819529 | 0.2054689 | -2.0918958 | 0.03644784 | 0.42151595 |
| SNRPD1 | 206.693516 | -0.810339308 | 0.42163844 | -1.9218819 | 0.05462061 | 0.46772575 |
| ABHD3 | 654.974628 | -0.672967065 | 0.43918948 | -1.5322932 | 0.12545009 | 0.58564339 |
| MIB1 | 1620.53102 | -0.197865622 | 0.2872918 | -0.688727 | 0.49099507 | 0.82014007 |
| CTAGE1 | 81.9061769 | -0.334145328 | 0.35472862 | -0.9419745 | 3.46E-01 | 0.75166079 |
| AC090912.1 | 123.694784 | 0.182346003 | 0.49002718 | 0.37211406 | 0.70980793 | 0.91115727 |
| RBBP8 | 1249.00235 | -0.269124706 | 0.33678063 | -0.7991098 | 4.24E-01 | 0.78894887 |
| CABLES1 | 1437.43438 | -0.460462267 | 0.37130298 | -1.2401254 | 0.21492901 | 0.65925411 |
| TMEM241 | 911.646209 | 0.085456475 | 0.28809183 | 0.29662929 | 0.76674954 | 0.93138917 |
| RIOK3 | 2084.79345 | -0.268782261 | 0.3015335 | -0.8913844 | 0.37272297 | 0.769011 |
| RMC1 | 687.887056 | -0.207642606 | 0.24254239 | -0.8561085 | 0.39193776 | 0.77634262 |
| NPC1 | 3231.08884 | -1.385927058 | 0.69764454 | -1.9865805 | 0.0469689 | 0.44732709 |
| ANKRD29 | 94.6748567 | 0.41892346 | 0.67663121 | 0.61913115 | 0.53582997 | 0.83954482 |
| LAMA3 | 126.930105 | -0.044686935 | 0.46209225 | -0.0967057 | 9.23E-01 | 9.80E-01 |
| TTC39C | 232.543491 | 0.585304543 | 0.52335975 | 1.11835987 | 2.63E-01 | 6.96E-01 |
| AC090772.3 | 34.0109733 | -0.189659202 | 0.47916705 | -0.3958102 | 6.92E-01 | 0.90516342 |
| OSBPL1A | 1802.73292 | -0.250293815 | 0.28456533 | -0.8795654 | 0.37909479 | 0.77076106 |
| IMPACT | 162.420538 | 0.021007431 | 0.44526675 | 0.04717943 | 0.96237022 | 0.99012655 |
| ZNF521 | 886.403657 | 0.42646159 | 0.29942145 | 1.42428537 | 1.54E-01 | 0.61099212 |

| | | | | | | |
|---|---|---|---|---|---|---|
| SS18 | 1331.21046 | -0.017574921 | 0.18015583 | -0.097554 | 0.92228646 | 0.97970649 |
| TAF4B | 480.322769 | -0.396654777 | 0.35827595 | -1.1071208 | 0.26824168 | 0.69961625 |
| KCTD1 | 692.622305 | 0.859796023 | 0.44595101 | 1.92800555 | 0.05385444 | 0.46711025 |
| AQP4-AS1 | 570.815423 | 1.434946191 | 1.05442984 | 1.36087404 | 0.1735535 | 0.62663931 |
| CHST9 | 172.550206 | 0.819319398 | 0.69179918 | 1.18433126 | 0.23628196 | 0.6735965 |
| CDH2 | 5097.60835 | 0.047658395 | 0.27174725 | 0.17537765 | 0.8607829 | 0.96302275 |
| DSC2 | 311.967248 | 0.083690896 | 0.19773688 | 0.42324374 | 6.72E-01 | 8.97E-01 |
| DSG2 | 448.298612 | -0.056662719 | 0.45974006 | -0.1232495 | 0.90190955 | 0.97492485 |
| B4GALT6 | 242.074318 | 0.007817736 | 0.39302539 | 0.01989117 | 0.98413019 | 0.99523767 |
| TRAPPC8 | 1545.00994 | -0.040195098 | 0.18608628 | -0.2160025 | 0.82898582 | 0.95325673 |
| RNF125 | 51.3118855 | 0.527328679 | 0.76816938 | 0.68647448 | 0.49241395 | 0.82087587 |
| RNF138 | 509.478512 | 0.114903112 | 0.20920883 | 0.54922688 | 0.58284976 | 0.85986705 |
| GAREM1 | 4579.99477 | -0.274713467 | 0.50344276 | -0.5456697 | 0.58529299 | 0.86077718 |
| AC025887.2 | 79.1296741 | 0.028660248 | 0.47217492 | 0.06069837 | 0.95159943 | 0.98756033 |
| KLHL14 | 105.935999 | 0.83427001 | 0.61173652 | 1.36377343 | 0.17263888 | 0.62593809 |
| CCDC178 | 350.193217 | 1.311784476 | 0.71227747 | 1.84167622 | 0.06552252 | 0.48790819 |
| ASXL3 | 407.139968 | 0.597779353 | 0.54348648 | 1.09989737 | 0.27137684 | 0.70002917 |
| NOL4 | 510.597162 | 2.099638386 | 0.60900647 | 3.44764545 | 5.65E-04 | 0.08159212 |
| DTNA | 2252.14023 | 0.624580353 | 0.76524759 | 0.81618075 | 0.41439676 | 0.78632759 |
| MAPRE2 | 561.051285 | 0.105696468 | 0.36875042 | 0.28663416 | 7.74E-01 | 0.93453439 |
| ZNF397 | 390.414708 | 0.100862377 | 0.34748257 | 0.29026601 | 0.77161274 | 0.93316533 |
| ZSCAN30 | 429.200523 | 0.303479694 | 0.35159971 | 0.86313976 | 0.38806063 | 0.77389948 |
| ZNF24 | 555.913699 | -0.212474611 | 0.20204779 | -1.0516057 | 0.29298049 | 0.71433665 |
| ZNF396 | 59.6531042 | 0.249907317 | 0.39150461 | 0.63832536 | 0.52326191 | 0.8352569 |
| GALNT1 | 697.610586 | -0.102563852 | 0.27637619 | -0.3711023 | 0.71056131 | 0.91123308 |
| C18orf21 | 61.993047 | -0.430090548 | 0.34911287 | -1.2319527 | 0.21796674 | 0.66064833 |
| RPRD1A | 1235.91035 | -0.189902606 | 0.25101517 | -0.7565384 | 0.44932647 | 0.80309271 |
| SLC39A6 | 194.010478 | -0.296095491 | 0.34828241 | -0.8501592 | 0.39523659 | 0.77931337 |
| ELP2 | 750.280304 | -0.263627604 | 0.13520046 | -1.9499016 | 0.05118785 | 0.46106129 |
| MOCOS | 82.2038769 | 0.423769666 | 0.49267672 | 0.86013739 | 3.90E-01 | 7.76E-01 |
| FHOD3 | 4200.41623 | -0.122214079 | 0.29235988 | -0.4180262 | 0.67592799 | 0.89833381 |
| TPGS2 | 401.028609 | 0.059440312 | 0.26092133 | 0.22780933 | 8.20E-01 | 9.50E-01 |
| KIAA1328 | 1386.67343 | -0.118551417 | 0.23264391 | -0.5095831 | 0.61034354 | 0.87109318 |
| AC016493.1 | 35.7491453 | 1.178651072 | 0.75856152 | 1.55379759 | 0.1202327 | 0.5780532 |
| CELF4 | 130.859119 | 0.972269209 | 0.79809691 | 1.21823452 | 0.22313487 | 0.66408018 |
| AC016205.1 | 281.79787 | 0.232301677 | 0.42611524 | 0.54516162 | 0.58564236 | 0.86077718 |
| PIK3C3 | 1701.20436 | -0.196381493 | 0.27659636 | -0.709993 | 0.47770845 | 0.81580016 |
| LINC00907 | 1444.12225 | 0.808811234 | 0.60807558 | 1.33011629 | 0.18347996 | 0.63276491 |
| SETBP1 | 6095.69566 | -0.436159044 | 0.29446592 | -1.4811869 | 0.13855679 | 0.59474076 |
| SLC14A2 | 158.170258 | 0.394690201 | 0.85942221 | 0.45925064 | 0.64605419 | 0.88700494 |
| EPG5 | 1482.5993 | -0.178345618 | 0.26826285 | -0.6648167 | 0.50616775 | 0.82782978 |
| PSTPIP2 | 264.95877 | 0.368063401 | 0.82755446 | 0.44476034 | 0.65649296 | 0.89053559 |
| ATP5F1A | 880.19133 | 0.272907374 | 0.34674839 | 0.78704727 | 0.4312542 | 0.79135216 |
| HAUS1 | 105.94389 | -0.454088125 | 0.40885106 | -1.1106444 | 0.26672146 | 0.69812269 |
| C18orf25 | 675.932844 | -0.604665228 | 0.27801637 | -2.1749267 | 0.02963561 | 0.39981555 |
| PIAS2 | 1936.18558 | -0.622781549 | 0.36511761 | -1.7057012 | 0.08806373 | 0.52826674 |
| KATNAL2 | 993.479196 | 0.384347219 | 0.59047487 | 0.65091206 | 0.51510325 | 0.8317258 |
| AC012254.1 | 106.499115 | -1.320916928 | 0.67568809 | -1.9549211 | 0.05059238 | 0.45903191 |
| HDHD2 | 70.053146 | 0.009627174 | 0.47834255 | 0.02012611 | 0.98394277 | 0.99523767 |
| SMAD2 | 1785.28872 | -0.054494201 | 0.16113204 | -0.3381959 | 0.73521554 | 0.92009988 |
| ZBTB7C | 59.9041089 | 1.060736943 | 0.77229079 | 1.37349422 | 0.16959876 | 0.62326131 |
| CTIF | 806.260477 | -0.317502084 | 0.28065975 | -1.1312705 | 0.25794127 | 0.69210282 |
| SMAD7 | 184.651088 | -0.845363762 | 0.57423445 | -1.4721579 | 0.14097825 | 0.59555904 |
| DYM | 3386.78219 | -0.229325679 | 0.20832769 | -1.1007931 | 0.27098673 | 0.69961625 |

| | | | | | | |
|---|---|---|---|---|---|---|
| C18orf32 | 54.5526191 | 0.298045599 | 0.30826622 | 0.96684481 | 0.33362162 | 0.74022334 |
| LIPG | 241.580643 | 0.658963557 | 0.5265013 | 1.25158961 | 0.21071944 | 0.65524759 |
| ACAA2 | 748.138182 | 0.330585755 | 0.53292696 | 0.62032095 | 0.53504651 | 0.83954482 |
| SNHG22 | 464.288945 | 0.837148557 | 0.45440026 | 1.84231531 | 0.06542904 | 0.48790819 |
| MYO5B | 3825.07289 | -0.037799701 | 0.36688321 | -0.1030292 | 0.91793976 | 0.97832208 |
| CFAP53 | 139.430997 | -0.110739793 | 0.29926419 | -0.3700402 | 0.71135251 | 0.91143287 |
| MBD1 | 281.809574 | -0.077826782 | 0.28694287 | -0.2712274 | 0.78621611 | 0.9390702 |
| CXXC1 | 202.129565 | -0.707179032 | 0.32906425 | -2.1490607 | 0.03162959 | 0.40930954 |
| MRO | 171.291053 | 0.080050293 | 0.90628565 | 0.08832788 | 9.30E-01 | 9.83E-01 |
| ME2 | 358.998095 | -0.529622377 | 0.46347466 | -1.1427213 | 0.25315431 | 0.68887358 |
| AC091551.1 | 121.585718 | 0.288793052 | 0.51009006 | 0.56616091 | 0.5712844 | 0.85399468 |
| SMAD4 | 539.827774 | -0.058513526 | 0.18929692 | -0.3091098 | 0.75723803 | 0.92799282 |
| MEX3C | 195.308244 | -0.027102828 | 0.21739271 | -0.1246722 | 0.90078307 | 0.97479173 |
| DCC | 1129.26994 | 1.854465803 | 0.84154856 | 2.20363494 | 0.02755003 | 0.38942007 |
| MBD2 | 1567.63555 | -0.160651323 | 0.24353376 | -0.6596676 | 0.50946719 | 0.82882082 |
| POLI | 559.746293 | 0.508722571 | 0.33769686 | 1.50644744 | 0.13195235 | 0.59087581 |
| CCDC68 | 171.449209 | 0.977850233 | 0.43651045 | 2.24015311 | 0.02508098 | 0.37991912 |
| TCF4 | 284.40221 | 0.162282948 | 0.38062649 | 0.42635747 | 0.66984739 | 0.89611537 |
| AC009271.1 | 92.0246618 | 2.39466788 | 0.73716063 | 3.24850213 | 0.00116014 | 0.11093704 |
| AC006305.1 | 694.429228 | 1.348170293 | 1.10241996 | 1.22291898 | 0.2213603 | 0.66263062 |
| LINC01905 | 39.8874085 | 1.933113022 | 1.10832537 | 1.74417465 | 0.08112864 | 0.51593841 |
| TXNL1 | 562.169755 | -0.159294015 | 0.19455385 | -0.8187657 | 0.41292012 | 0.78632759 |
| WDR7 | 1556.8137 | 0.063231595 | 0.26881266 | 0.23522551 | 0.81403371 | 0.94881072 |
| FECH | 159.92479 | -0.142285428 | 0.31873144 | -0.4464117 | 0.65529992 | 0.89053559 |
| NARS | 803.457581 | 0.087327548 | 0.29214276 | 0.29892081 | 0.76500047 | 0.93083208 |
| AC027097.2 | 524.037773 | -0.538123598 | 0.28073131 | -1.9168635 | 0.05525528 | 0.46947044 |
| ATP8B1 | 416.715045 | -0.072200168 | 0.30774583 | -0.2346097 | 0.81451165 | 0.94881072 |
| NEDD4L | 12782.6818 | -0.607690303 | 0.39561243 | -1.5360749 | 0.12452002 | 0.58516755 |
| ALPK2 | 1696.19264 | 0.198734732 | 0.47255525 | 0.42055343 | 0.67408121 | 0.89801183 |
| MALT1 | 2745.80443 | 0.118689098 | 0.42189643 | 0.28132283 | 0.7784628 | 0.93646866 |
| AC104365.1 | 303.523023 | 0.369740384 | 0.73301538 | 0.50441013 | 0.6139732 | 0.87225883 |
| ZNF532 | 311.856002 | -0.341732509 | 0.39395824 | -0.8674333 | 0.38570463 | 0.77317162 |
| SEC11C | 194.413799 | -0.214137799 | 0.48297689 | -0.4433707 | 0.65749763 | 0.89053559 |
| LMAN1 | 693.223894 | -0.22068934 | 0.27935346 | -0.7900004 | 0.42952755 | 0.79037914 |
| AC113137.1 | 55.2061104 | 1.542909349 | 0.65834203 | 2.34362883 | 0.01909717 | 0.34084276 |
| CDH20 | 70.9920184 | 2.743572038 | 0.93402381 | 2.93736843 | 0.00331011 | 0.188264 |
| AC105094.2 | 308.224798 | 2.217665259 | 0.83413022 | 2.65865592 | 0.0078453 | 0.24384485 |
| RNF152 | 7315.30171 | 0.793200033 | 0.82714988 | 0.95895563 | 0.3375811 | 0.74404633 |
| PIGN | 1074.89791 | 0.157787108 | 0.15846667 | 0.99571167 | 0.31939026 | 0.73132207 |
| KIAA1468 | 1222.63693 | 0.015674 | 0.15952471 | 0.09825437 | 0.92173031 | 0.97939204 |
| ZCCHC2 | 928.791682 | 0.11352843 | 0.20599471 | 0.55112304 | 0.58154934 | 0.85915432 |
| PHLPP1 | 1413.80574 | -0.296263594 | 0.19513175 | -1.5182747 | 0.12894517 | 0.58793678 |
| BCL2 | 3109.04751 | -0.089874965 | 0.50538419 | -0.1778349 | 0.85885261 | 0.9626966 |
| KDSR | 777.662803 | -0.421641325 | 0.34181528 | -1.2335356 | 0.217376 | 0.66032006 |
| AC036176.1 | 44.2841912 | 0.792039742 | 0.5036849 | 1.57249053 | 0.11583683 | 0.57350419 |
| VPS4B | 741.281388 | -0.105311927 | 0.21461162 | -0.4907094 | 0.62363203 | 0.87772094 |
| SERPINB8 | 70.2395014 | -1.161102221 | 0.59411712 | -1.9543322 | 5.07E-02 | 0.45903191 |
| DSEL | 263.986301 | 0.616910467 | 0.44458836 | 1.38759924 | 0.16525911 | 0.62065658 |
| AC110597.3 | 36.8322949 | 0.904883722 | 0.89446463 | 1.01164842 | 0.31170619 | 0.72600794 |
| AC114689.3 | 337.243556 | 0.70760316 | 0.63714032 | 1.11059235 | 0.26674386 | 0.69812269 |
| TMX3 | 750.259968 | -0.252068966 | 0.28364144 | -0.8886888 | 0.37417036 | 0.7697436 |
| CCDC102B | 243.25968 | 0.092720336 | 0.59511425 | 0.15580258 | 0.87618864 | 0.96617876 |
| DOK6 | 3855.53158 | 0.38218261 | 0.42007451 | 0.90979719 | 0.36292947 | 0.76205182 |
| AC119868.2 | 136.949978 | 0.836115949 | 0.56130289 | 1.4895985 | 0.13632984 | 0.5942049 |

| | | | | | | |
|---|---|---|---|---|---|---|
| CD226 | 241.901091 | 0.192700774 | 0.52059564 | 0.37015441 | 7.11E-01 | 9.11E-01 |
| RTTN | 1450.97238 | 0.280296803 | 0.39371309 | 0.71193164 | 4.77E-01 | 8.16E-01 |
| SOCS6 | 641.009996 | 0.14575861 | 0.27216434 | 0.53555366 | 0.59226706 | 0.86382106 |
| LINC01909 | 44.1160292 | -0.03914014 | 0.46885695 | -0.0834799 | 0.93346994 | 0.98355048 |
| AC027458.1 | 111.956777 | 2.540415142 | 1.12507922 | 2.25798779 | 0.02394642 | 0.37387451 |
| FBXO15 | 102.872196 | 0.284459325 | 0.70349108 | 0.40435385 | 0.68595253 | 0.90147185 |
| TIMM21 | 132.654614 | -0.111059989 | 0.23511247 | -0.4723696 | 0.63666298 | 0.88266168 |
| CYB5A | 3552.30109 | 0.467982595 | 0.56322994 | 0.83089083 | 0.4060353 | 0.78429302 |
| CNDP2 | 2577.38162 | 0.45404216 | 0.4037769 | 1.12448771 | 0.26080618 | 0.69280433 |
| CNDP1 | 54.9575382 | 0.391796574 | 0.57834761 | 0.67744133 | 0.49812599 | 0.82332243 |
| LINC00909 | 159.769084 | 0.239239626 | 0.39179885 | 0.6106185 | 0.54145217 | 0.84221416 |
| ZNF407 | 3462.10394 | -0.390546036 | 0.27844546 | -1.4025944 | 0.16073783 | 0.61630095 |
| ZADH2 | 227.023066 | 0.455144599 | 0.3525025 | 1.29118121 | 0.19664085 | 0.64558745 |
| TSHZ1 | 1046.82966 | 0.679111833 | 0.47521642 | 1.429058 | 0.15298756 | 0.61001662 |
| ZNF516 | 1827.02938 | -0.230264492 | 0.37720867 | -0.6104433 | 0.54156822 | 0.84221416 |
| AC009716.1 | 51.7306381 | 0.42041522 | 0.72309958 | 0.58140709 | 0.56096612 | 0.84953736 |
| ZNF236-DT | 95.3624036 | 0.147870271 | 0.28942048 | 0.51091848 | 0.60940814 | 0.87048326 |
| ZNF236 | 666.194918 | -0.441486371 | 0.25668566 | -1.7199495 | 0.08544162 | 0.5226185 |
| MBP | 2866.38508 | -0.627829773 | 0.3846496 | -1.6322122 | 1.03E-01 | 0.54996682 |
| AC124254.1 | 84.5438773 | -0.640484486 | 1.36859683 | -0.4679862 | 0.63979443 | 0.88321992 |
| ATP9B | 2055.88913 | 0.049780403 | 0.23647553 | 0.21050975 | 0.83326985 | 0.95355066 |
| NFATC1 | 43.4831121 | 0.775122597 | 0.76324681 | 1.01555957 | 0.30983917 | 0.72498238 |
| CTDP1 | 435.302025 | -0.893413825 | 0.28194263 | -3.1687788 | 0.00153081 | 0.13146073 |
| AC021594.2 | 121.499921 | 2.307189016 | 1.21474055 | 1.89932658 | 0.05752155 | 0.47684824 |
| KCNG2 | 69.8851334 | 1.096036371 | 1.07879528 | 1.0159818 | 0.30963805 | 0.72498238 |
| PQLC1 | 1255.3447 | 0.282159181 | 0.40458885 | 0.69739732 | 0.48555418 | 0.81839266 |
| HSBP1L1 | 527.581191 | -0.607210127 | 0.40709998 | -1.4915504 | 0.13581706 | 0.59386143 |
| TXNL4A | 498.614111 | -0.385838963 | 0.26885407 | -1.4351241 | 0.15125173 | 0.60905477 |
| ADNP2 | 882.363937 | -0.837354272 | 0.40883516 | -2.0481464 | 0.04054565 | 0.43086475 |
| PARD6G-AS1 | 232.352603 | -0.558936015 | 0.64704463 | -0.8638291 | 0.38768175 | 0.77373447 |
| PARD6G | 146.264214 | 0.379634105 | 0.37004515 | 1.02591294 | 3.05E-01 | 0.72103276 |
| C20orf96 | 157.583498 | 0.042795288 | 0.4566541 | 0.09371489 | 0.92533564 | 0.9808658 |
| ZCCHC3 | 133.089465 | -0.304823192 | 0.30124518 | -1.0118774 | 0.31159668 | 0.72600794 |
| NRSN2-AS1 | 465.456568 | 0.10237669 | 0.28917789 | 0.35402669 | 0.72331888 | 0.91592159 |
| SOX12 | 97.4316856 | 0.043464904 | 0.44554105 | 0.09755533 | 0.9222854 | 0.97970649 |
| NRSN2 | 133.477216 | -0.530843608 | 0.42306708 | -1.2547505 | 0.20956936 | 0.65441616 |
| RBCK1 | 525.301616 | -0.398421862 | 0.30879341 | -1.2902538 | 0.19696256 | 0.64558745 |
| TBC1D20 | 390.114637 | -0.506776913 | 0.32847413 | -1.5428214 | 0.12287412 | 0.58216759 |
| CSNK2A1 | 3102.11228 | -0.524050829 | 0.2698346 | -1.9421187 | 0.05212273 | 0.46437412 |
| RSPO4 | 45.625492 | 0.426887765 | 0.72965159 | 0.585057 | 0.55850939 | 0.84906096 |
| PSMF1 | 705.721217 | -0.378159758 | 0.33626685 | -1.1245823 | 0.26076607 | 0.69280433 |
| AL136531.2 | 304.135525 | -0.029809406 | 0.47663319 | -0.0625416 | 0.95013153 | 0.98724111 |
| SDCBP2-AS1 | 646.613104 | 0.310685937 | 0.39454972 | 0.78744432 | 0.43102181 | 0.79132279 |
| NSFL1C | 345.842424 | -0.586357462 | 0.26993552 | -2.1722131 | 0.02983959 | 0.39981555 |
| SIRPA | 307.848883 | -0.213063866 | 0.41291276 | -0.5160021 | 0.60585292 | 0.86915559 |
| STK35 | 313.204626 | -0.459820112 | 0.34250917 | -1.3425045 | 0.17943246 | 0.63063608 |
| SNRPB | 276.169729 | -0.652320631 | 0.40461785 | -1.6121895 | 0.10692072 | 0.55488138 |
| AL049650.1 | 69.9597352 | 0.487695605 | 0.43788358 | 1.11375633 | 0.26538374 | 0.69672762 |
| NOP56 | 418.868369 | -1.365453827 | 0.65080026 | -2.098115 | 0.03589498 | 0.42040764 |
| IDH3B | 176.567789 | 0.103562955 | 0.36783098 | 0.28155039 | 0.77828829 | 0.93646866 |
| EBF4 | 68.0378594 | 1.212225752 | 0.73747887 | 1.64374302 | 0.1002293 | 0.54542315 |
| PCED1A | 201.805617 | 0.177101647 | 0.3232382 | 0.54789825 | 5.84E-01 | 0.86025731 |
| VPS16 | 95.2390367 | -0.435215394 | 0.48743534 | -0.8928679 | 0.37192789 | 0.76845856 |
| PTPRA | 1490.79724 | -0.052829268 | 0.18083164 | -0.2921462 | 0.77017488 | 0.93266075 |

| | | | | | | |
|---|---|---|---|---|---|---|
| MRPS26 | 252.623582 | 0.202850837 | 0.38643549 | 0.52492807 | 0.59963319 | 0.86700128 |
| UBOX5-AS1 | 161.281958 | 1.022649405 | 0.7257513 | 1.4090907 | 0.15880835 | 0.61423843 |
| UBOX5 | 237.158039 | 0.641440174 | 0.54930934 | 1.16772122 | 0.24291923 | 0.68132724 |
| LZTS3 | 285.698292 | -0.36822149 | 0.46248124 | -0.7961869 | 0.42592343 | 0.78894887 |
| DDRGK1 | 308.733496 | -0.203828725 | 0.24696171 | -0.8253455 | 4.09E-01 | 0.78542936 |
| ITPA | 480.36332 | -0.356178785 | 0.32094165 | -1.109793 | 0.26708826 | 0.69838864 |
| C20orf194 | 4081.55112 | 0.226195538 | 0.31757668 | 0.71225487 | 0.47630696 | 0.81570932 |
| ATRN | 1476.18898 | -0.337094486 | 0.13815644 | -2.4399477 | 0.01468939 | 0.30832605 |
| ADAM33 | 122.862914 | -1.744563577 | 1.67775935 | -1.0398175 | 0.29842469 | 0.71852421 |
| C20orf27 | 59.0274025 | 0.06410055 | 0.47108133 | 0.13607109 | 0.89176508 | 0.97234681 |
| CENPB | 197.447156 | -0.294093579 | 0.55330341 | -0.5315232 | 0.59505629 | 0.86510861 |
| CDC25B | 256.173397 | 0.581753735 | 0.64568099 | 0.90099251 | 0.3675923 | 0.76597879 |
| AP5S1 | 52.8969483 | -0.390386177 | 0.4790033 | -0.8149968 | 0.41507412 | 0.78632759 |
| MAVS | 818.166866 | 0.196453109 | 0.28294046 | 0.69432668 | 0.48747736 | 0.81893983 |
| PANK2 | 521.341611 | -0.286298673 | 0.25820683 | -1.1087959 | 0.26751824 | 0.69874881 |
| RNF24 | 686.993827 | -0.889228409 | 0.39106351 | -2.2738721 | 2.30E-02 | 3.67E-01 |
| SMOX | 158.030337 | 0.214645085 | 0.49534981 | 0.43332021 | 6.65E-01 | 8.93E-01 |
| PRNP | 575.512977 | -0.398442276 | 0.40231725 | -0.9903684 | 0.3219941 | 0.73375526 |
| SLC23A2 | 243.463681 | 0.197207578 | 0.33247352 | 0.59315274 | 0.55307894 | 0.84725078 |
| TMEM230 | 369.936578 | -0.164454606 | 0.27296996 | -0.6024641 | 0.54686524 | 0.84460887 |
| PCNA | 84.9865657 | 0.256585056 | 0.29106578 | 0.8815363 | 0.37802761 | 0.77055888 |
| CDS2 | 835.593617 | -0.133623779 | 0.24425154 | -0.5470745 | 0.58432758 | 0.86025731 |
| GPCPD1 | 833.369374 | 0.410334661 | 0.57023812 | 0.71958476 | 0.4717807 | 0.81389222 |
| SHLD1 | 325.516051 | -0.67728823 | 0.34916715 | -1.939725 | 0.05241313 | 0.4650011 |
| TRMT6 | 95.941301 | -0.955030833 | 0.43008771 | -2.220549 | 0.02638152 | 0.38507144 |
| MCM8 | 228.919376 | -0.680411909 | 0.33154334 | -2.0522563 | 0.04014476 | 0.4304996 |
| MCM8-AS1 | 36.4532344 | 0.123647187 | 0.68711331 | 0.17995167 | 0.85719051 | 0.96266803 |
| CRLS1 | 1680.45432 | -0.031412732 | 0.36298869 | -0.0865391 | 0.93103784 | 0.98272895 |
| LRRN4 | 58.3785385 | 0.041150537 | 0.57754419 | 0.07125089 | 0.94319808 | 0.9851723 |
| FERMT1 | 209.909987 | -0.111800141 | 0.34784973 | -0.3214036 | 0.74790458 | 0.92531168 |
| BMP2 | 47.4787857 | -0.519355355 | 0.62679646 | -0.8285869 | 0.40733819 | 0.78481094 |
| AL096799.1 | 79.9910292 | 1.02400416 | 0.83754229 | 1.2226298 | 0.22146955 | 0.66267168 |
| LINC01428 | 393.913298 | 2.315241392 | 0.75635883 | 3.06103571 | 2.21E-03 | 0.15269652 |
| TMX4 | 294.242244 | 0.53689505 | 0.33082505 | 1.6228972 | 0.10461138 | 0.55086754 |
| PLCB1 | 1335.08 | 0.033927454 | 0.53237175 | 0.06372888 | 0.94918611 | 0.9871942 |
| PLCB4 | 1544.91229 | -0.505529706 | 0.60030349 | -0.8421235 | 0.3997188 | 0.78186732 |
| PAK5 | 34.533857 | 0.442043822 | 0.45371659 | 0.97427299 | 0.32992101 | 0.73849818 |
| ANKEF1 | 67.9924634 | 0.286943967 | 0.30947017 | 0.92721041 | 0.3538173 | 0.75682564 |
| SNAP25-AS1 | 63.7472993 | 1.805523275 | 0.65929195 | 2.73857927 | 0.00617053 | 0.22637955 |
| SNAP25 | 121.915326 | 1.671604035 | 0.72982577 | 2.2904152 | 0.02199726 | 0.36066559 |
| MKKS | 158.949582 | 0.43375404 | 0.18872991 | 2.29827926 | 0.0215459 | 0.35786026 |
| SLX4IP | 1134.97114 | 0.197827886 | 0.31605694 | 0.62592483 | 0.53136425 | 0.83789294 |
| JAG1 | 268.139003 | 0.078993748 | 0.50192212 | 0.15738248 | 0.87494342 | 0.96615698 |
| AL050403.2 | 247.889048 | -0.286169914 | 0.62439597 | -0.4583148 | 0.64672629 | 0.88753277 |
| BTBD3 | 399.846604 | 0.256225031 | 0.30442589 | 0.84166637 | 0.39997473 | 0.78186732 |
| SPTLC3 | 2805.41195 | 1.521032338 | 0.49814021 | 3.05342215 | 0.00226247 | 0.15428723 |
| ISM1 | 127.824332 | 1.398620426 | 0.33338616 | 4.19519639 | 2.73E-05 | 0.01647703 |
| TASP1 | 2313.29228 | -0.079349882 | 0.21802178 | -0.3639539 | 0.71589241 | 0.91298263 |
| ESF1 | 886.62099 | -0.453519276 | 0.3203254 | -1.415808 | 0.15683172 | 0.61255952 |
| NDUFAF5 | 547.234465 | 0.397512326 | 0.30648551 | 1.29700202 | 0.19463049 | 0.64548758 |
| MACROD2 | 4520.42128 | 0.012543818 | 0.49484699 | 0.02534888 | 0.97977668 | 0.99487362 |
| FLRT3 | 772.933407 | 0.659250468 | 0.39310777 | 1.67702221 | 9.35E-02 | 0.53688613 |
| KIF16B | 2184.90545 | 0.044257356 | 0.30188343 | 0.14660412 | 0.8834445 | 0.96846661 |
| SNRPB2 | 312.270365 | -0.202548553 | 0.3469002 | -0.5838813 | 0.55930015 | 0.84936251 |

| | | | | | | |
|---|---|---|---|---|---|---|
| DSTN | 1283.1903 | 0.244380943 | 0.21362147 | 1.14399052 | 0.25262757 | 0.68887358 |
| RRBP1 | 1615.72566 | -0.103286476 | 0.39299701 | -0.2628175 | 0.79269128 | 0.94154414 |
| SNX5 | 745.600684 | 0.142848463 | 0.2670422 | 0.53492843 | 0.59269935 | 0.86382106 |
| OVOL2 | 243.764275 | 0.458330864 | 0.46977573 | 0.9756376 | 0.32924408 | 0.73829083 |
| MGME1 | 353.576219 | 0.341240767 | 0.36705763 | 0.92966538 | 0.35254437 | 0.7565877 |
| KAT14 | 168.03013 | 0.629110634 | 0.378587 | 1.66173333 | 0.09656626 | 0.53969191 |
| ZNF133 | 428.036751 | 0.10872823 | 0.26097915 | 0.41661654 | 0.67695891 | 0.89843878 |
| DZANK1 | 323.088403 | 0.280629698 | 0.30610142 | 0.91678665 | 0.35925445 | 0.75936492 |
| POLR3F | 138.149087 | -0.21417938 | 0.30084368 | -0.7119291 | 0.47650867 | 0.81570932 |
| RBBP9 | 96.9337161 | 0.690304551 | 0.37286663 | 1.85134442 | 0.06412002 | 0.48593331 |
| SEC23B | 697.101961 | 0.18713285 | 0.18765595 | 0.99721245 | 0.3186614 | 0.73071671 |
| DTD1 | 137.23883 | -0.024570256 | 0.43998912 | -0.0558429 | 0.95546698 | 0.98840182 |
| SLC24A3 | 64.670033 | 0.897253285 | 0.70922679 | 1.26511477 | 0.20583019 | 0.651931 |
| AL121761.1 | 402.970068 | 0.569172083 | 0.35744329 | 1.59234236 | 0.1113078 | 0.5642367 |
| RIN2 | 1630.82205 | 0.779842389 | 0.3573497 | 2.18229476 | 0.02908779 | 0.39554194 |
| NAA20 | 543.401623 | -0.397479988 | 0.28840088 | -1.3782205 | 0.16813523 | 0.62304975 |
| CRNKL1 | 163.756155 | -0.270327345 | 0.25922085 | -1.0428457 | 0.29701978 | 0.71825247 |
| CFAP61 | 128.431971 | 0.16616438 | 0.45149698 | 0.36802988 | 0.71285095 | 0.91188831 |
| RALGAPA2 | 5153.34247 | 0.450900864 | 0.26925996 | 1.67459305 | 0.0940141 | 0.53761006 |
| KIZ | 545.061023 | 0.316989853 | 0.38072533 | 0.83259459 | 0.40507341 | 0.78422899 |
| KIZ-AS1 | 126.690648 | 0.189915486 | 0.48411368 | 0.39229523 | 0.69484009 | 0.90589033 |
| XRN2 | 1105.11982 | -0.204234373 | 0.26462102 | -0.7717995 | 0.44023319 | 0.79826598 |
| NXT1 | 58.0673988 | -0.769262552 | 0.5029145 | -1.529609 | 0.12611354 | 0.58597906 |
| GZF1 | 86.4413156 | -0.584262055 | 0.42703866 | -1.3681713 | 0.17125846 | 0.62520756 |
| NAPB | 518.73847 | 0.249276116 | 0.25650489 | 0.97181819 | 0.33114101 | 0.73912226 |
| CST3 | 1442.66319 | -0.259070669 | 0.44432492 | -0.5830658 | 0.559849 | 0.84941263 |
| SYNDIG1 | 630.283846 | 1.433921208 | 0.93140954 | 1.53951742 | 0.12367803 | 0.58354574 |
| APMAP | 178.735975 | -0.159016252 | 0.32996572 | -0.4819175 | 0.62986456 | 0.88083664 |
| ACSS1 | 627.229074 | -0.001820486 | 0.36036788 | -0.0050517 | 0.99596931 | 0.99881119 |
| ENTPD6 | 324.65689 | 0.294628343 | 0.31946994 | 0.9222412 | 0.35640278 | 0.75841231 |
| PYGB | 354.270525 | -0.142904121 | 0.50819706 | -0.2811982 | 0.77855835 | 0.93646866 |
| ABHD12 | 618.528763 | -0.444354895 | 0.34310421 | -1.2951018 | 0.1952851 | 0.64558745 |
| GINS1 | 57.0806204 | -0.72120064 | 0.3767667 | -1.9141836 | 0.0555967 | 0.46970412 |
| NINL | 3240.50522 | -0.055491531 | 0.38524616 | -0.1440417 | 0.8854675 | 0.96957206 |
| NANP | 94.3677004 | -0.075154179 | 0.36792877 | -0.2042629 | 0.8381481 | 0.95506659 |
| ZNF337-AS1 | 386.555083 | -0.585402215 | 0.48139328 | -1.2160581 | 0.22396278 | 0.66543624 |
| ZNF337 | 290.74405 | 0.002826577 | 0.26482926 | 0.01067321 | 0.99148418 | 0.99748517 |
| FAM182B | 327.368066 | 0.59572901 | 0.39863561 | 1.49441995 | 0.13506589 | 0.59381224 |
| HM13 | 1331.34095 | -0.256699975 | 0.35452436 | -0.7240687 | 0.46902359 | 0.81244866 |
| ID1 | 137.675872 | 0.299531091 | 0.6956653 | 0.43056782 | 0.66678264 | 0.89428471 |
| BCL2L1 | 2808.19157 | -0.895342541 | 0.50674465 | -1.7668515 | 0.0772531 | 0.51080959 |
| AL117381.1 | 479.274666 | -0.031180046 | 0.65268669 | -0.0477718 | 0.96189808 | 0.99000623 |
| DUSP15 | 142.147666 | 0.521790238 | 0.76439589 | 0.6826178 | 0.49484839 | 0.82215397 |
| TTLL9 | 78.7294827 | 0.800125109 | 0.52489632 | 1.52434888 | 0.12742158 | 0.58793678 |
| PDRG1 | 75.2327493 | -0.58298747 | 0.33633285 | -1.7333646 | 0.08303088 | 0.51935659 |
| TM9SF4 | 868.310373 | -0.117378668 | 0.19389184 | -0.6053822 | 0.54492508 | 0.84446444 |
| PLAGL2 | 153.54973 | 0.235604831 | 0.32223758 | 0.73115255 | 0.46468598 | 0.80984752 |
| POFUT1 | 257.487808 | -0.082243707 | 0.30176498 | -0.2725422 | 0.78520511 | 0.93892226 |
| KIF3B | 907.177467 | -0.041907688 | 0.2486345 | -0.1685514 | 0.86614953 | 0.96437968 |
| ASXL1 | 2274.0427 | -0.4678594 | 0.1918512 | -2.4386577 | 0.01474193 | 0.30832605 |
| NOL4L | 237.86694 | 0.908697344 | 0.34133236 | 2.66220684 | 0.00776302 | 0.24233539 |
| FO393400.1 | 88.6734401 | -0.427826717 | 0.56588722 | -0.7560282 | 0.44963226 | 0.80311564 |
| DNMT3B | 201.965706 | -0.180006754 | 0.41408202 | -0.4347128 | 0.6637709 | 0.89313476 |
| MAPRE1 | 247.418987 | -0.563770534 | 0.3790911 | -1.4871637 | 0.13697158 | 0.59474076 |

| | | | | | | |
|---|---|---|---|---|---|---|
| AL035071.2 | 66.3659967 | 0.36096876 | 0.41907598 | 0.86134442 | 3.89E-01 | 0.7748665 |
| AL035071.1 | 75.345413 | -0.501192549 | 0.54831749 | -0.9140554 | 0.36068775 | 0.76009788 |
| CDK5RAP1 | 414.263043 | -0.160675811 | 0.26300745 | -0.6109173 | 0.54125431 | 0.84210815 |
| SNTA1 | 166.07968 | -0.187145447 | 0.38939399 | -0.4806069 | 0.63079589 | 0.88115748 |
| CBFA2T2 | 2842.99281 | -0.095194855 | 0.2193865 | -0.4339139 | 0.66435096 | 0.89315615 |
| NECAB3 | 75.4871479 | 0.257217105 | 0.37134632 | 0.69266098 | 0.48852233 | 0.81933645 |
| PXMP4 | 58.3511172 | 0.519174483 | 0.49070238 | 1.05802316 | 0.29004488 | 0.71260257 |
| ZNF341 | 92.8178937 | -0.398258354 | 0.290894 | -1.3690841 | 0.17097298 | 0.62504223 |
| CHMP4B | 590.312526 | -0.525452801 | 0.40120466 | -1.3096877 | 0.19030152 | 0.64278842 |
| RALY | 944.017308 | -0.101407169 | 0.2872374 | -0.3530431 | 0.72405616 | 0.91600702 |
| EIF2S2 | 671.1355 | -0.486309589 | 0.34624485 | -1.4045251 | 0.16016253 | 0.61590405 |
| ASIP | 200.972337 | -0.284402119 | 0.75343374 | -0.3774746 | 0.70582092 | 0.90980406 |
| AHCY | 518.293084 | 0.016120292 | 0.34118585 | 0.04724783 | 0.9623157 | 0.99012655 |
| ITCH | 3153.91694 | -0.443811428 | 0.18062671 | -2.4570643 | 0.01400776 | 0.30363654 |
| DYNLRB1 | 803.380773 | -0.3041509 | 0.19432785 | -1.5651432 | 0.11754935 | 0.57360231 |
| MAP1LC3A | 193.213755 | -0.33341566 | 0.59221321 | -0.5629994 | 0.57343532 | 0.8542683 |
| PIGU | 429.785451 | -0.085552805 | 0.27903211 | -0.3066056 | 0.75914359 | 0.92923569 |
| NCOA6 | 1739.51105 | -0.191381007 | 0.18787429 | -1.0186652 | 0.30836193 | 0.72375291 |
| TP53INP2 | 126.494003 | 0.440842471 | 0.35501651 | 1.24175202 | 0.21432807 | 0.65906957 |
| GGT7 | 503.930949 | 0.590279675 | 0.1977053 | 2.98565427 | 0.00282972 | 0.17238676 |
| ACSS2 | 2471.99725 | 1.065037383 | 0.49638575 | 2.14558412 | 0.03190617 | 0.40988962 |
| GSS | 427.964742 | 0.717266599 | 0.38486074 | 1.86370427 | 0.06236324 | 0.4840117 |
| TRPC4AP | 1479.41487 | -0.356173325 | 0.1654837 | -2.1523167 | 0.03137242 | 0.40835531 |
| EDEM2 | 316.774009 | 0.17979007 | 0.26522489 | 0.67787783 | 0.49784916 | 0.82326489 |
| PROCR | 94.5605256 | 0.349532959 | 0.43523603 | 0.80308829 | 0.4219237 | 0.78892549 |
| MMP24OS | 250.552734 | 0.320641434 | 0.2966123 | 1.08101191 | 0.27969181 | 0.70446562 |
| MMP24 | 419.502469 | -0.677432388 | 0.48785384 | -1.388597 | 0.16495533 | 0.62065658 |
| EIF6 | 66.7216324 | -1.068912493 | 0.68561653 | -1.559053 | 0.11898384 | 0.57609025 |
| UQCC1 | 975.761044 | -0.257767072 | 0.1539226 | -1.6746538 | 0.09400217 | 0.53761006 |
| CEP250 | 387.691086 | -2.212801793 | 0.67352479 | -3.2854051 | 0.00101836 | 0.1082065 |
| ERGIC3 | 446.561947 | -0.362574423 | 0.36921953 | -0.9820023 | 0.32609872 | 0.7373175 |
| CPNE1 | 389.132739 | 0.078081463 | 0.26414069 | 0.29560559 | 0.7675313 | 0.93169958 |
| NFS1 | 101.341526 | 0.237031348 | 0.33178353 | 0.71441565 | 4.75E-01 | 0.81564184 |
| ROMO1 | 273.652725 | -0.665796435 | 0.51643031 | -1.289228 | 0.19731882 | 0.64558745 |
| RBM39 | 3312.74819 | -0.232913843 | 0.21453384 | -1.0856741 | 0.27762319 | 0.7036882 |
| PHF20 | 1424.1101 | -0.447927962 | 0.30718607 | -1.458165 | 0.14479508 | 0.60109715 |
| SCAND1 | 474.925647 | -0.707628313 | 0.44415604 | -1.5931975 | 0.11111588 | 0.56347761 |
| CNBD2 | 82.8928913 | -0.088850756 | 0.58418387 | -0.1520938 | 0.87911294 | 0.9674841 |
| NORAD | 995.524921 | -0.085090935 | 0.24661294 | -0.3450384 | 0.7300655 | 0.91841114 |
| EPB41L1 | 1820.2493 | -0.152247389 | 0.40277287 | -0.3779981 | 0.70543199 | 0.90971458 |
| AAR2 | 137.854985 | -0.284885741 | 0.33008987 | -0.8630551 | 0.38810717 | 0.77389948 |
| DLGAP4 | 1209.10262 | 0.14834249 | 0.35013442 | 0.42367297 | 0.67180433 | 0.89704376 |
| DLGAP4-AS1 | 415.338053 | 0.039586685 | 0.32566931 | 0.12155485 | 0.90325157 | 0.97536781 |
| MYL9 | 113.91541 | 0.077129007 | 0.45760429 | 0.16854957 | 0.86615095 | 0.96437968 |
| RAB5IF | 353.244081 | -0.655957215 | 0.68843066 | -0.9528297 | 0.34067634 | 0.74725037 |
| NDRG3 | 409.593656 | 0.022720502 | 0.3265425 | 0.069579 | 0.94452875 | 0.9853777 |
| DSN1 | 77.8089737 | 0.015547938 | 0.36220298 | 0.04292604 | 0.96576049 | 0.99081812 |
| SOGA1 | 273.651875 | 0.334173416 | 0.47745472 | 0.69990599 | 0.48398602 | 0.81839266 |
| SAMHD1 | 143.734717 | -0.699895102 | 0.67622641 | -1.0350011 | 0.30066838 | 0.71879123 |
| RBL1 | 141.6811 | -0.482887884 | 0.39640991 | -1.2181529 | 0.22316588 | 0.66408018 |
| MROH8 | 216.555936 | 0.824245726 | 0.50268268 | 1.6396939 | 0.10106883 | 0.54608983 |
| RPN2 | 680.534372 | -0.283878228 | 0.25883622 | -1.0967485 | 0.27275135 | 0.70108838 |
| MANBAL | 212.029949 | -0.269634584 | 0.2700565 | -0.9984377 | 0.31806717 | 0.73015352 |
| SRC | 362.326829 | -0.156931091 | 0.34899994 | -0.4496594 | 0.65295607 | 0.88949911 |

| | | | | | | |
|---|---|---|---|---|---|---|
| BLCAP | 875.617818 | -0.226750543 | 0.35826381 | -0.632915 | 0.52678915 | 0.83708543 |
| CTNNBL1 | 660.274201 | -0.327086247 | 0.20556737 | -1.5911389 | 0.11157832 | 0.56435572 |
| TTI1 | 341.775182 | -0.146429172 | 0.24675125 | -0.5934283 | 0.55289456 | 0.84721285 |
| RPRD1B | 657.4295 | -0.408245134 | 0.23003103 | -1.7747394 | 0.07594092 | 0.50730849 |
| TGM2 | 1211.49377 | 1.332203648 | 0.74502988 | 1.78812111 | 7.38E-02 | 5.06E-01 |
| KIAA1755 | 350.866117 | 4.83354024 | 0.99492996 | 4.85817136 | 1.18E-06 | 0.00591552 |
| LBP | 562.037071 | -0.075867654 | 0.56647736 | -0.1339288 | 0.89345885 | 0.97272276 |
| AL391095.2 | 58.4383292 | -0.719201948 | 0.70736116 | -1.0167394 | 0.30927743 | 0.72449199 |
| RALGAPB | 1159.08823 | -0.279310337 | 0.1855354 | -1.5054288 | 0.13221386 | 0.59115873 |
| ARHGAP40 | 99.5322075 | -0.721796401 | 0.6230937 | -1.1584075 | 0.24669779 | 0.68451145 |
| ACTR5 | 128.450995 | -0.528804809 | 0.34905657 | -1.5149545 | 0.12978394 | 0.58853436 |
| PPP1R16B | 1129.91334 | 1.001831989 | 0.9023321 | 1.1102697 | 0.26688283 | 0.69812269 |
| FAM83D | 55.4417425 | -1.114639306 | 0.67435069 | -1.6529075 | 0.09834969 | 0.54193402 |
| DHX35 | 378.436265 | -0.146252691 | 0.24485786 | -0.5972963 | 0.55030958 | 0.84620747 |
| TOP1 | 1682.40505 | 0.203359201 | 0.19999828 | 1.01680475 | 0.30924632 | 0.72449199 |
| PLCG1-AS1 | 50.8615413 | 0.930713002 | 0.53723414 | 1.73241595 | 0.08319953 | 0.51935659 |
| PLCG1 | 371.30291 | 0.250094988 | 0.27303514 | 0.91598097 | 0.35967688 | 0.75976128 |
| ZHX3 | 3251.526 | 0.38675293 | 0.28781182 | 1.34377012 | 0.17902273 | 0.62968042 |
| LPIN3 | 1089.60284 | 0.495445646 | 0.57377798 | 0.86347971 | 0.38787377 | 0.77373447 |
| CHD6 | 1610.12631 | 0.275367724 | 0.31622144 | 0.87080662 | 0.38385975 | 0.77317162 |
| PTPRT | 135.206247 | 1.103881205 | 0.61080332 | 1.8072613 | 0.07072155 | 0.49686328 |
| SRSF6 | 669.564135 | -0.125485663 | 0.28415613 | -0.4416081 | 6.59E-01 | 0.89058983 |
| L3MBTL1 | 301.435552 | 0.160242901 | 0.34040355 | 0.47074392 | 0.63782362 | 0.8827026 |
| Z98752.3 | 110.033223 | 0.751031185 | 0.50649254 | 1.482808 | 0.13812543 | 0.59474076 |
| SGK2 | 719.552145 | -0.02594415 | 0.38700401 | -0.0670385 | 0.94655109 | 0.98620476 |
| IFT52 | 349.856407 | 0.137580667 | 0.29065766 | 0.47334264 | 0.63596875 | 0.88257807 |
| OSER1 | 336.948164 | -0.603525289 | 0.38746836 | -1.5576118 | 0.11932531 | 0.57652003 |
| OSER1-DT | 53.1619505 | 0.443554161 | 0.61380683 | 0.72262826 | 0.4699083 | 0.81326175 |
| GDAP1L1 | 43.0254449 | 2.172682511 | 0.80413 | 2.70190454 | 0.00689436 | 0.23083391 |
| FITM2 | 150.416471 | 0.166551199 | 0.28900453 | 0.5762927 | 0.56441736 | 0.85163263 |
| R3HDML | 101.502894 | 0.949399737 | 1.25767247 | 0.75488631 | 0.4503172 | 0.80360602 |
| AL117382.1 | 437.764364 | 3.252307153 | 1.72464478 | 1.88578378 | 0.0593241 | 0.47855094 |
| HNF4A | 1942.5735 | 1.263448476 | 0.85916616 | 1.47055196 | 0.14141232 | 0.59604512 |
| HNF4A-AS1 | 1103.69397 | 3.549491124 | 1.34501537 | 2.63899671 | 0.00831518 | 0.25016267 |
| TTPAL | 129.834887 | -0.418394252 | 0.28681651 | -1.4587523 | 0.14463329 | 0.60109715 |
| SERINC3 | 487.665666 | -0.180476227 | 0.17027739 | -1.0598954 | 0.28919218 | 0.71218341 |
| PKIG | 934.32234 | 0.116513212 | 0.27323641 | 0.42641906 | 0.66980252 | 0.89611537 |
| YWHAB | 811.388638 | -0.000212106 | 0.36226683 | -0.0005855 | 0.99953284 | 0.99977652 |
| PABPC1L | 216.061524 | -0.080819862 | 0.29500337 | -0.2739625 | 0.78411344 | 0.93867623 |
| TOMM34 | 138.300884 | -0.189706313 | 0.3720256 | -0.5099281 | 0.61010182 | 0.87103525 |
| STK4 | 633.091901 | -0.329592504 | 0.32780797 | -1.0054438 | 0.31468318 | 0.72719146 |
| SLPI | 910.173191 | 0.190437217 | 0.96841751 | 0.19664785 | 8.44E-01 | 9.57E-01 |
| SDC4 | 1832.85554 | 0.028857397 | 0.37343352 | 0.07727586 | 0.93840409 | 0.98435387 |
| SYS1-DBNDD | 133.666464 | -0.361744722 | 0.49826719 | -0.7260055 | 0.46783539 | 0.8118777 |
| TP53TG5 | 42.1435191 | 0.413553349 | 0.51533194 | 0.80249896 | 0.42226439 | 0.78894887 |
| PIGT | 169.984469 | 0.001491228 | 0.37688757 | 0.00395669 | 0.99684303 | 0.99932183 |
| WFDC2 | 2362.90836 | 0.774966495 | 0.7029023 | 1.10252377 | 2.70E-01 | 7.00E-01 |
| DNTTIP1 | 180.85602 | 0.085147771 | 0.27803956 | 0.30624336 | 0.75941936 | 0.92930182 |
| SNX21 | 90.4193469 | 0.797569379 | 0.42852331 | 1.86120419 | 0.06271535 | 0.48421747 |
| ACOT8 | 172.744698 | 0.106317355 | 0.2311841 | 0.45988178 | 0.64560108 | 0.88687652 |
| ZSWIM3 | 87.4511977 | 0.769288028 | 0.37342782 | 2.06007154 | 0.0393917 | 0.42716935 |
| NEURL2 | 95.7491594 | 0.445665126 | 0.79128009 | 0.56322045 | 0.57328478 | 0.8542683 |
| CTSA | 1000.17366 | -0.226405539 | 0.35916063 | -0.630374 | 0.52844989 | 0.83761342 |
| PLTP | 304.479478 | -0.745888051 | 1.09188895 | -0.6831171 | 0.49453283 | 0.8220963 |

| | | | | | | |
|---|---|---|---|---|---|---|
| PCIF1 | 64.8084312 | -0.006325675 | 0.35717739 | -0.0177102 | 0.98587006 | 0.99609994 |
| ZNF335 | 224.442453 | -0.40615943 | 0.29937717 | -1.3566814 | 0.17488247 | 0.62699479 |
| NCOA5 | 512.572118 | -0.344792406 | 0.35347068 | -0.9754484 | 0.32933789 | 0.73829083 |
| CD40 | 62.6693535 | 0.877840006 | 0.61706502 | 1.42260537 | 0.15485059 | 0.61132626 |
| SLC35C2 | 291.284766 | -0.044456609 | 0.30533874 | -0.1455977 | 0.88423901 | 0.96900096 |
| AL133227.1 | 41.9208672 | 1.256123456 | 0.62782995 | 2.00073836 | 0.04542059 | 0.44360789 |
| ELMO2 | 251.437043 | 0.10897978 | 0.25838182 | 0.42177806 | 0.67318702 | 0.89782405 |
| ZNF334 | 239.303933 | 0.308703598 | 0.58936161 | 0.52379319 | 0.60042239 | 0.86755264 |
| SLC13A3 | 13309.4771 | 1.157932251 | 0.374703 | 3.09026679 | 0.00199977 | 0.14129287 |
| TP53RK | 82.4199782 | -0.251289706 | 0.35451404 | -0.7088286 | 0.47843082 | 0.81596289 |
| EYA2 | 1274.56323 | 0.324847624 | 0.40830501 | 0.79560039 | 4.26E-01 | 7.89E-01 |
| ZMYND8 | 1692.60367 | -0.013261791 | 0.26282382 | -0.0504589 | 0.95975673 | 0.98912673 |
| NCOA3 | 1355.38903 | -0.200206439 | 0.18246096 | -1.0972563 | 0.27252936 | 0.70083612 |
| SULF2 | 1048.17174 | 0.946029251 | 0.39651784 | 2.38584284 | 0.01704003 | 0.32964126 |
| LINC00494 | 54.4922338 | 0.394352399 | 0.77573525 | 0.50835952 | 0.61120124 | 0.87163238 |
| ARFGEF2 | 1821.62751 | -0.344862694 | 0.21389628 | -1.6122893 | 0.10689899 | 0.55488138 |
| CSE1L | 447.911167 | -0.046418593 | 0.2597712 | -0.1786903 | 0.85818089 | 0.9626966 |
| STAU1 | 1515.60221 | -0.210216541 | 0.26377771 | -0.7969458 | 0.42548251 | 0.78894887 |
| DDX27 | 264.525146 | -0.352615273 | 0.31015712 | -1.1368924 | 0.25558327 | 0.69024 |
| ZNFX1 | 463.278604 | -0.746062395 | 0.42492599 | -1.7557467 | 0.07913167 | 0.51192014 |
| ZFAS1 | 440.810464 | -0.493724816 | 0.3563888 | -1.3853545 | 1.66E-01 | 0.62065658 |
| B4GALT5 | 4980.15657 | -0.102620816 | 0.36101779 | -0.2842542 | 0.7762156 | 0.9352344 |
| SLC9A8 | 637.244776 | -0.116890529 | 0.43910851 | -0.2661996 | 0.79008546 | 0.94040477 |
| SPATA2 | 73.0819913 | -0.096286976 | 0.38084868 | -0.2528221 | 0.80040567 | 0.94403774 |
| RNF114 | 860.492954 | 0.383732351 | 0.27837458 | 1.37847481 | 0.16805674 | 0.62304975 |
| CEBPB | 552.597635 | -0.189138491 | 0.38403222 | -0.4925068 | 6.22E-01 | 8.77E-01 |
| PTPN1 | 5140.61042 | -0.712342705 | 0.54255918 | -1.3129309 | 0.18920623 | 0.6413893 |
| RIPOR3 | 656.650552 | 0.101827588 | 0.43685663 | 0.23309155 | 0.81569032 | 0.94906482 |
| PARD6B | 2030.78741 | -0.581474896 | 0.31079553 | -1.8709243 | 0.06135558 | 0.4813972 |
| BCAS4 | 87.4872846 | -0.840315531 | 0.43902317 | -1.9140574 | 0.05561282 | 0.46970412 |
| ADNP | 1626.47001 | 0.02086516 | 0.28922974 | 0.07214044 | 0.94249014 | 0.98489601 |
| DPM1 | 480.304712 | -0.371250352 | 0.22867023 | -1.6235185 | 0.1044786 | 0.55066578 |
| NFATC2 | 95.7444451 | 1.139697767 | 0.70575868 | 1.61485476 | 0.10634215 | 0.55440878 |
| ATP9A | 364.003411 | 1.080320051 | 0.54884588 | 1.96834867 | 0.04902793 | 0.45376495 |
| ZFP64 | 552.931543 | 0.144363927 | 0.2585348 | 0.55839262 | 0.57657631 | 0.85615117 |
| TSHZ2 | 149.839175 | 1.131981027 | 0.66187867 | 1.71025458 | 0.08721881 | 0.5268877 |
| AL157838.1 | 42.3628495 | 1.01775233 | 0.56778184 | 1.79250594 | 0.07305196 | 0.50459544 |
| ZNF217 | 696.769593 | 0.204455565 | 0.30825529 | 0.66326701 | 0.50715956 | 0.82832518 |
| CYP24A1 | 4399.57887 | 1.643418528 | 0.96957657 | 1.69498581 | 0.09007811 | 0.53311774 |
| PFDN4 | 140.722392 | -0.189181249 | 0.31576685 | -0.5991169 | 0.54909495 | 0.84587304 |
| DOK5 | 48.1547825 | 1.471430453 | 0.63898289 | 2.30276974 | 0.0212918 | 0.35688511 |
| FAM210B | 111.476158 | 0.412484344 | 0.35801209 | 1.15215199 | 0.24925862 | 0.68578642 |
| AURKA | 37.8631416 | 0.408501639 | 0.52546784 | 0.7774056 | 0.43691951 | 0.795967 |
| CSTF1 | 106.085842 | -0.164443845 | 0.23308066 | -0.7055233 | 0.48048462 | 0.81714875 |
| RTF2 | 331.031708 | -0.510231841 | 0.25489661 | -2.0017208 | 0.04531477 | 0.44360789 |
| BMP7 | 74.3065397 | -1.920917966 | 0.61197419 | -3.1388872 | 0.00169591 | 0.13207839 |
| RAE1 | 210.558296 | -0.283688711 | 0.28208705 | -1.0056779 | 0.31457055 | 0.72719146 |
| RBM38 | 98.5521947 | 0.809647873 | 0.38053889 | 2.12763503 | 0.03336736 | 0.41254527 |
| PCK1 | 6748.272 | 0.265167333 | 1.02877639 | 0.25775021 | 0.79659969 | 0.94276732 |
| PMEPA1 | 137.087676 | -0.147738065 | 0.90462835 | -0.1633135 | 0.87027157 | 0.96564168 |
| RAB22A | 811.449333 | -0.245255908 | 0.2635006 | -0.9307603 | 0.35197755 | 0.75584463 |
| VAPB | 963.652915 | -0.240113981 | 0.15650753 | -1.5342008 | 0.12498026 | 0.58564339 |
| STX16-NPEPI | 65.5033411 | -0.148801328 | 0.60737714 | -0.24499 | 0.80646417 | 0.94585363 |
| NPEPL1 | 182.116987 | 0.339128202 | 0.47528812 | 0.71352131 | 0.47552323 | 0.81566724 |

| | | | | | | |
|---|---|---|---|---|---|---|
| GNAS | 1255.15488 | 0.24770286 | 0.26913087 | 0.92038072 | 3.57E-01 | 7.59E-01 |
| NELFCD | 375.786105 | -0.188037617 | 0.231107 | -0.8136388 | 0.41585192 | 0.78632759 |
| CTSZ | 547.103908 | -0.878799454 | 0.56711364 | -1.5496003 | 0.12123749 | 0.57942895 |
| ATP5F1E | 1135.76264 | -0.488649485 | 0.46121706 | -1.0594783 | 0.28938199 | 0.71218341 |
| PRELID3B | 129.53436 | -0.156206959 | 0.34634729 | -0.4510125 | 0.65198054 | 0.88935488 |
| SYCP2 | 124.617371 | 0.604942441 | 0.64566734 | 0.93692587 | 0.34879669 | 0.75409404 |
| FAM217B | 154.207046 | -0.035388352 | 0.37510538 | -0.0943424 | 0.92483715 | 0.9804664 |
| PPP1R3D | 38.3334864 | 0.584814571 | 0.62038461 | 0.94266453 | 0.34585252 | 0.75166079 |
| CDH26 | 93.2751976 | 0.560356902 | 0.57838628 | 0.96882813 | 0.33263095 | 0.73988623 |
| MIR646HG | 928.438677 | 0.048951327 | 0.91697697 | 0.05338338 | 0.95742645 | 0.98861313 |
| CDH4 | 85.9038376 | 0.902432078 | 0.77496999 | 1.16447358 | 0.24423214 | 0.68222743 |
| TAF4 | 738.198901 | 0.046790003 | 0.32541125 | 0.1437873 | 0.88566843 | 0.96971461 |
| LSM14B | 204.799952 | 0.167735866 | 0.35516712 | 0.47227307 | 0.63673189 | 0.88266168 |
| PSMA7 | 567.411327 | -0.257088016 | 0.33209002 | -0.7741516 | 0.43884115 | 0.79724264 |
| SS18L1 | 705.485758 | -0.365747555 | 0.39738967 | -0.9203751 | 0.35737678 | 0.75903693 |
| MTG2 | 441.205287 | -0.144834276 | 0.25016607 | -0.5789525 | 0.56262122 | 0.85054239 |
| OSBPL2 | 454.765915 | -0.289502212 | 0.2228361 | -1.299171 | 0.19388524 | 0.64548758 |
| ADRM1 | 136.720635 | -0.909346851 | 0.51296802 | -1.7727165 | 0.0762757 | 0.50768193 |
| AL354836.1 | 116.896656 | 0.668249436 | 0.62454006 | 1.06998651 | 0.28462538 | 0.70952861 |
| LAMA5 | 926.622804 | 0.256475363 | 0.32495946 | 0.78925342 | 0.42996391 | 0.79055252 |
| RPS21 | 134.394704 | -0.538654687 | 0.50011014 | -1.0770721 | 0.28144805 | 0.70625976 |
| CABLES2 | 42.2053352 | -0.739009367 | 0.46171389 | -1.6005786 | 0.10947029 | 0.55969747 |
| SLCO4A1 | 1365.05602 | -0.403709139 | 0.43504179 | -0.9279778 | 3.53E-01 | 7.57E-01 |
| MRGBP | 132.077581 | -0.85389295 | 0.3478498 | -2.4547749 | 0.01409728 | 0.30468798 |
| OGFR | 482.474384 | 0.222551758 | 0.34220921 | 0.65033831 | 0.51547372 | 0.83185952 |
| TCFL5 | 99.7174367 | 0.174661908 | 0.3712315 | 0.47049323 | 0.63800267 | 0.8827026 |
| DIDO1 | 1062.51624 | -0.339439295 | 0.17213223 | -1.9719682 | 0.04861323 | 0.45329737 |
| GID8 | 277.399444 | -0.496058529 | 0.20550124 | -2.4138956 | 0.01578298 | 0.31566878 |
| YTHDF1 | 424.891173 | -0.603272761 | 0.32257112 | -1.8702008 | 0.06145594 | 0.48190882 |
| NKAIN4 | 734.8433 | 0.326729844 | 0.91330132 | 0.35774594 | 0.72053345 | 0.9149807 |
| AL121827.2 | 60.8398933 | 0.475856141 | 0.89009004 | 0.53461573 | 0.5929156 | 0.86394112 |
| ARFGAP1 | 589.374212 | -0.495928276 | 0.53232292 | -0.9316305 | 0.35152751 | 0.75548011 |
| CHRNA4 | 105.261686 | 1.992209617 | 1.07267043 | 1.85724298 | 6.33E-02 | 0.48572144 |
| PPDPF | 664.549437 | 0.13430414 | 0.45832832 | 0.29303042 | 0.7694989 | 0.93217158 |
| HELZ2 | 117.243131 | -1.433538542 | 0.98787442 | -1.4511344 | 0.14674244 | 0.60383734 |
| GMEB2 | 228.973617 | -0.410036484 | 0.19822005 | -2.0685924 | 0.03858435 | 0.42479978 |
| MHENCR | 121.129763 | -0.110252236 | 0.51960608 | -0.2121843 | 0.83196328 | 0.95325673 |
| STMN3 | 80.7632985 | 0.928400536 | 0.3087523 | 3.00694286 | 0.00263889 | 0.16823326 |
| ARFRP1 | 195.556655 | 0.051768313 | 0.27729369 | 0.18669128 | 0.85190269 | 0.96050023 |
| SLC2A4RG | 497.643001 | 0.989970957 | 0.53218074 | 1.86021569 | 0.06285502 | 0.48483608 |
| ZBTB46 | 493.282293 | 0.561021317 | 0.39814004 | 1.40910551 | 0.15880397 | 0.61423843 |
| TPD52L2 | 461.820112 | -0.77464398 | 0.3630218 | -2.1338773 | 0.03285283 | 0.41180153 |
| DNAJC5 | 1096.09446 | -0.664728099 | 0.40744486 | -1.6314553 | 0.10279428 | 0.55022042 |
| UCKL1 | 228.215998 | -0.364604942 | 0.31188529 | -1.1690354 | 0.24238938 | 0.68080966 |
| ZNF512B | 290.919517 | 0.356380962 | 0.3918087 | 0.90957899 | 0.36304458 | 0.76209565 |
| SAMD10 | 37.6032041 | 0.222369159 | 0.57255386 | 0.3883812 | 0.69773395 | 0.90696517 |
| PRPF6 | 459.263101 | -0.289902197 | 0.23792283 | -1.2184715 | 0.22304484 | 0.66408018 |
| TCEA2 | 112.70485 | -0.134190778 | 0.37884141 | -0.3542136 | 0.72317881 | 0.91592159 |
| PCMTD2 | 499.541425 | 0.033385631 | 0.2672542 | 0.12492088 | 0.90058619 | 0.97479173 |
| LINC00266-1 | 48.2294079 | -0.093881108 | 0.32190434 | -0.2916429 | 0.77055969 | 0.9329619 |
| PLPP2 | 54.4766496 | 0.720491755 | 0.8881813 | 0.81119897 | 4.17E-01 | 0.78652162 |
| MIER2 | 97.3514159 | 0.101677066 | 0.58099475 | 0.17500514 | 0.8610756 | 0.96302275 |
| SHC2 | 130.56683 | -0.633582827 | 0.74505602 | -0.8503828 | 0.39511229 | 0.77931337 |
| MADCAM1 | 52.6763685 | 0.17702935 | 0.45736587 | 0.38706288 | 0.69870966 | 0.90696517 |

| | | | | | | |
|---|---|---|---|---|---|---|
| TPGS1 | 219.053179 | -0.594937245 | 0.55175556 | -1.0782623 | 0.2809167 | 0.70625976 |
| CDC34 | 178.700312 | -0.494152646 | 0.31004888 | -1.5937895 | 0.11098319 | 0.56342465 |
| BSG | 906.825527 | -0.182378425 | 0.38091558 | -0.4787896 | 0.6320883 | 0.88123095 |
| POLRMT | 86.440799 | -0.470393075 | 0.28941306 | -1.6253346 | 0.10409127 | 0.55056166 |
| RNF126 | 315.007063 | -0.629580544 | 0.22953085 | -2.7429016 | 0.00608989 | 0.22637955 |
| FSTL3 | 230.404726 | -1.274322131 | 0.79925255 | -1.5943923 | 0.11084819 | 0.56317482 |
| PALM | 343.846587 | 1.43403725 | 0.57589853 | 2.49008668 | 1.28E-02 | 2.99E-01 |
| MISP | 162.458621 | 0.4334394 | 0.39532933 | 1.09640081 | 0.27290341 | 0.70118792 |
| PTBP1 | 790.006207 | -0.61796789 | 0.27977841 | -2.2087762 | 0.02719021 | 0.38702446 |
| MED16 | 155.339251 | -0.072752588 | 0.42803229 | -0.1699699 | 0.86503384 | 0.96428252 |
| R3HDM4 | 126.784639 | -0.133359427 | 0.48036962 | -0.2776184 | 0.78130533 | 0.93748328 |
| ARID3A | 387.897694 | -0.33713428 | 0.36321784 | -0.9281876 | 0.35331028 | 0.75673243 |
| WDR18 | 171.31115 | -0.462393875 | 0.47275285 | -0.978088 | 0.32803082 | 0.73806934 |
| TMEM259 | 649.780518 | -0.081898806 | 0.27144096 | -0.3017187 | 0.76286653 | 0.93022076 |
| CNN2 | 55.5865101 | 0.225438379 | 0.6149392 | 0.36660271 | 0.71391538 | 0.91214838 |
| ABCA7 | 131.35682 | -0.178710395 | 0.60006107 | -0.2978203 | 0.76584028 | 0.93111343 |
| POLR2E | 347.406671 | 0.061707957 | 0.29514667 | 0.20907557 | 0.83438925 | 0.95393659 |
| GPX4 | 1318.17806 | -0.127408592 | 0.44671409 | -0.2852128 | 0.77548109 | 0.9351209 |
| SBNO2 | 557.199258 | -0.089414998 | 0.38845548 | -0.2301808 | 0.81795127 | 0.94981058 |
| STK11 | 340.041948 | -0.290323686 | 0.3681271 | -0.7886507 | 0.4303162 | 0.79055252 |
| ATP5F1D | 756.665044 | -0.264852633 | 0.41063937 | -0.6449762 | 0.51894259 | 0.83340902 |
| MIDN | 998.480509 | -0.416146004 | 0.50158354 | -0.8296644 | 0.40672855 | 0.78481094 |
| CIRBP | 909.88555 | 0.194820243 | 0.42795755 | 0.45523263 | 0.64894189 | 0.88800332 |
| C19orf24 | 228.590511 | -0.622787753 | 0.50416131 | -1.2352946 | 0.21672088 | 0.66032006 |
| MUM1 | 655.135547 | 0.48563895 | 0.30149256 | 1.61078256 | 0.10722712 | 0.55579103 |
| NDUFS7 | 264.595968 | 0.044733989 | 0.33585762 | 0.13319331 | 0.8940405 | 0.97272276 |
| GAMT | 280.383079 | 0.70306375 | 0.44896586 | 1.5659626 | 0.11735737 | 0.57350419 |
| DAZAP1 | 851.614191 | -0.253445791 | 0.2376342 | -1.0665375 | 0.28618073 | 0.71118494 |
| RPS15 | 1827.271 | -0.214819303 | 0.4785473 | -0.4488988 | 0.65350467 | 0.88979721 |
| C19orf25 | 536.460085 | -0.447734287 | 0.39214336 | -1.1417617 | 0.25355307 | 0.68891958 |
| PCSK4 | 56.6560043 | 0.542318677 | 0.60321683 | 0.89904434 | 0.36862904 | 0.76615653 |
| REEP6 | 56.398414 | 1.247063041 | 0.5235205 | 2.38207107 | 0.01721557 | 0.32964126 |
| MBD3 | 222.946799 | -0.527290494 | 0.49042771 | -1.0751646 | 0.28230106 | 0.70682473 |
| UQCR11 | 26.8183948 | 0.018786069 | 0.86330714 | 0.02176059 | 0.98263893 | 0.99522272 |
| TCF3 | 238.302501 | -0.390178067 | 0.49051707 | -0.7954424 | 0.42635622 | 0.78895108 |
| REXO1 | 448.06127 | -0.439565694 | 0.3549397 | -1.2384236 | 0.21555904 | 0.6596384 |
| KLF16 | 79.6213133 | -0.470639363 | 0.62394807 | -0.7542925 | 0.45067359 | 0.80361623 |
| AC012615.1 | 32.3522274 | -0.332868623 | 0.48224364 | -0.6902499 | 0.49003705 | 0.81973856 |
| ABHD17A | 234.517949 | -0.270039953 | 0.41656169 | -0.6482592 | 0.51681732 | 0.83262987 |
| SCAMP4 | 282.813891 | -0.42435781 | 0.37716485 | -1.1251256 | 0.26053583 | 0.69280433 |
| CSNK1G2 | 505.514202 | -0.223414575 | 0.41198752 | -0.5422848 | 0.58762231 | 0.86124068 |
| BTBD2 | 462.917857 | -0.31192235 | 0.37058807 | -0.8416956 | 0.39995836 | 0.78186732 |
| MKNK2 | 260.594381 | -0.267408572 | 0.47039707 | -0.5684741 | 0.56971306 | 0.85327114 |
| MOB3A | 300.252245 | -0.218398184 | 0.43548681 | -0.5015036 | 0.61601678 | 0.87333999 |
| AP3D1 | 1150.07553 | -0.598344492 | 0.38717278 | -1.5454198 | 0.12224475 | 0.58059901 |
| DOT1L | 692.173783 | -0.702821387 | 0.48512692 | -1.4487371 | 0.147411 | 0.60514005 |
| PLEKHJ1 | 425.217278 | -0.165172438 | 0.30882621 | -0.5348394 | 0.59276088 | 0.86382106 |
| SF3A2 | 148.548419 | -0.609108566 | 0.41710403 | -1.4603277 | 0.14420003 | 0.60086567 |
| AC005258.1 | 44.5633708 | -0.906244357 | 1.02497146 | -0.8841655 | 0.3766069 | 0.77035529 |
| LSM7 | 347.776426 | -0.665522571 | 0.40047498 | -1.6618331 | 0.09654625 | 0.53969191 |
| SPPL2B | 513.811443 | 0.280877782 | 0.2605221 | 1.07813418 | 0.28097388 | 0.70625976 |
| TMPRSS9 | 43.5528512 | 0.095060634 | 0.56012013 | 0.16971473 | 0.86523449 | 0.96428252 |
| TIMM13 | 285.591028 | -0.296074408 | 0.441757 | -0.6702201 | 0.50271751 | 0.82638105 |
| LMNB2 | 84.0739723 | -0.337602211 | 0.49137174 | -0.6870607 | 0.49204448 | 0.82080956 |

| | | | | | | |
|---|---|---|---|---|---|---|
| GADD45B | 681.925709 | -0.725501307 | 0.68137539 | -1.0647601 | 0.28698452 | 0.71118494 |
| GNG7 | 575.555035 | -0.98157771 | 0.66997101 | -1.4651047 | 0.14289234 | 0.59824924 |
| SGTA | 273.21329 | -0.431746488 | 0.42549016 | -1.0147038 | 0.31024703 | 0.72498238 |
| THOP1 | 135.960312 | -0.047303658 | 0.32807996 | -0.1441833 | 0.88535571 | 0.96957206 |
| ZNF554 | 82.7455472 | 0.338715205 | 0.30840095 | 1.09829496 | 0.27207571 | 0.70056282 |
| ZNF555 | 110.450288 | 0.402298617 | 0.34675796 | 1.16017125 | 0.24597909 | 0.68320879 |
| ZNF57 | 78.6780684 | -0.239173769 | 0.45850355 | -0.52164 | 0.60192104 | 0.86765503 |
| AC119403.1 | 33.2697599 | 0.564262144 | 0.51746252 | 1.09044062 | 0.2755191 | 0.70247842 |
| ZNF77 | 49.5450449 | 0.032194346 | 0.43298182 | 0.07435496 | 0.94072795 | 0.98484365 |
| TLE2 | 50.3129959 | 0.471196939 | 0.73977341 | 0.63694766 | 0.52415894 | 0.83589803 |
| AES | 657.466931 | 0.262758594 | 0.42960388 | 0.61162993 | 0.54078263 | 0.84175647 |
| GNA11 | 363.346782 | -0.254037684 | 0.3772936 | -0.6733156 | 0.5007465 | 0.82564559 |
| NCLN | 227.138985 | -0.610479711 | 0.3856912 | -1.5828199 | 0.11346251 | 0.56809124 |
| NFIC | 1157.34925 | -0.055533662 | 0.33913315 | -0.1637518 | 0.86992654 | 0.96564168 |
| SMIM24 | 506.245038 | 0.818168812 | 0.63509408 | 1.28826396 | 0.19765409 | 0.64558745 |
| DOHH | 97.0963454 | -0.574765581 | 0.46354824 | -1.2399261 | 0.21500271 | 0.65925411 |
| FZR1 | 175.691685 | -0.056755367 | 0.33693411 | -0.1684465 | 0.86623204 | 0.96437968 |
| MFSD12 | 838.149726 | -0.309509833 | 0.35119312 | -0.8813095 | 0.37815033 | 0.77055888 |
| C19orf71 | 56.2155709 | 0.276782887 | 0.48075409 | 0.57572654 | 0.56480003 | 0.85167372 |
| AC005786.3 | 131.332255 | 0.207489433 | 0.42009802 | 0.4939072 | 0.62137172 | 0.87660994 |
| HMG20B | 229.668692 | 0.086502288 | 0.27678522 | 0.31252496 | 0.7546416 | 0.92799282 |
| CACTIN | 70.4457001 | -0.681751906 | 0.40833928 | -1.6695722 | 0.09500404 | 0.53824794 |
| PIP5K1C | 235.180082 | -0.24456565 | 0.38611081 | -0.6334079 | 0.52646729 | 0.83698249 |
| TJP3 | 191.489904 | 0.583236201 | 0.57976638 | 1.00598486 | 0.31442285 | 0.72719146 |
| APBA3 | 91.8131764 | -0.316542241 | 0.30742237 | -1.0296656 | 0.30316701 | 0.72044214 |
| MRPL54 | 190.571129 | -0.581922955 | 0.51870228 | -1.1218824 | 0.26191244 | 0.69408034 |
| DAPK3 | 217.015862 | -0.515200808 | 0.32855954 | -1.5680592 | 0.11686731 | 0.57350419 |
| EEF2 | 1809.87834 | 0.279518727 | 0.48928988 | 0.57127429 | 0.56781373 | 0.85263558 |
| PIAS4 | 201.240504 | -0.355039742 | 0.39721893 | -0.8938138 | 0.37142155 | 0.76820041 |
| ZBTB7A | 569.397117 | -0.226646122 | 0.49322932 | -0.4595147 | 0.6458646 | 0.88700494 |
| MAP2K2 | 444.846089 | 0.055102988 | 0.31290556 | 0.17610102 | 0.86021458 | 0.96283855 |
| SIRT6 | 132.946898 | -0.678827095 | 0.32736726 | -2.0735949 | 0.03811695 | 0.42339465 |
| ANKRD24 | 115.011505 | 0.283365631 | 0.57788695 | 0.49034787 | 6.24E-01 | 0.87791268 |
| YJU2 | 101.464661 | -0.669643298 | 0.36983464 | -1.8106559 | 0.07019413 | 0.49575843 |
| STAP2 | 245.511027 | 0.817831606 | 0.42367111 | 1.93034546 | 0.05356405 | 0.46711025 |
| MPND | 83.4263031 | 0.676680993 | 0.36690719 | 1.84428382 | 0.06514179 | 0.48790819 |
| AC007292.3 | 46.1714289 | -0.207974327 | 0.55085038 | -0.3775514 | 0.70576388 | 0.90980406 |
| SH3GL1 | 890.219282 | -0.750577086 | 0.25672416 | -2.9236714 | 0.0034593 | 0.19196998 |
| CHAF1A | 52.1121778 | -0.433610314 | 0.50611808 | -0.8567374 | 0.39159 | 0.77606943 |
| UBXN6 | 348.312735 | 0.014200332 | 0.40556485 | 0.03501372 | 0.9720688 | 0.99241376 |
| HDGFL2 | 191.490066 | 0.269244061 | 0.29098607 | 0.92528162 | 0.35481944 | 0.75755608 |
| PLIN4 | 99.9296965 | 1.74452709 | 0.51788205 | 3.36857993 | 0.00075556 | 0.09084673 |
| PLIN5 | 1091.30103 | 1.009246682 | 0.61958658 | 1.6289034 | 0.10333347 | 0.55022042 |
| TNFAIP8L1 | 204.665398 | 0.211079957 | 0.32904176 | 0.64149899 | 5.21E-01 | 0.83445062 |
| MYDGF | 462.409396 | -0.177190696 | 0.35094316 | -0.5048986 | 0.61363009 | 0.87222135 |
| DPP9 | 417.956353 | -0.211418297 | 0.4187348 | -0.5048978 | 0.61363059 | 0.87222135 |
| FEM1A | 52.352068 | -0.796802296 | 0.51373811 | -1.5509893 | 0.12090426 | 0.5792686 |
| TICAM1 | 408.42464 | 0.250973971 | 0.38443365 | 0.6528408 | 0.51385892 | 0.83126535 |
| PLIN3 | 267.425918 | 0.256819901 | 0.42965275 | 0.59773829 | 5.50E-01 | 0.84620747 |
| KDM4B | 1271.12916 | -0.257830914 | 0.2374747 | -1.0857195 | 0.27760311 | 0.7036882 |
| PTPRS | 781.687232 | 0.405320057 | 0.40298277 | 1.00579996 | 0.3145118 | 0.72719146 |
| TINCR | 94.6112564 | -0.757862754 | 0.33967744 | -2.2311248 | 0.02567286 | 0.38166806 |
| SAFB2 | 1499.79201 | 0.045633006 | 0.24151544 | 0.18894446 | 8.50E-01 | 0.95971165 |
| SAFB | 1038.60592 | -0.300112334 | 0.19875707 | -1.5099454 | 0.13105735 | 0.58917777 |

| | | | | | | |
|---|---|---|---|---|---|---|
| RPL36 | 1067.29409 | -0.225005115 | 0.49882482 | -0.4510704 | 0.6519388 | 0.88935488 |
| C19orf70 | 346.409523 | -0.187181226 | 0.29851684 | -0.6270374 | 0.53063471 | 0.83789294 |
| LONP1 | 605.14703 | -1.114413246 | 0.38871996 | -2.8668794 | 0.00414541 | 0.20365996 |
| PRR22 | 56.0783635 | -0.280995287 | 0.63837909 | -0.4401699 | 0.65981403 | 0.89094706 |
| DUS3L | 76.1933359 | 0.140709836 | 0.46710093 | 0.30124075 | 0.76323092 | 0.93033403 |
| NRTN | 90.3417286 | 1.103876987 | 0.62207405 | 1.77451058 | 0.07597873 | 0.50730849 |
| FUT6 | 1516.29576 | 1.543665015 | 0.57144243 | 2.70134826 | 6.91E-03 | 0.23083391 |
| FUT3 | 175.314833 | 0.565317614 | 0.6151155 | 0.91904303 | 3.58E-01 | 7.59E-01 |
| AC024592.2 | 37.7405207 | 1.178540725 | 0.69920141 | 1.68555255 | 9.19E-02 | 5.36E-01 |
| AC104532.2 | 62.4937239 | 0.424720679 | 0.46442501 | 0.91450863 | 0.36044964 | 0.76009788 |
| CAPS | 109.659717 | 0.495445166 | 0.56013788 | 0.88450574 | 0.37642327 | 0.77035529 |
| RANBP3 | 254.011503 | -0.333861906 | 0.23684965 | -1.4095942 | 0.15865953 | 0.61423843 |
| RFX2 | 375.948722 | 0.026668811 | 0.49221275 | 0.05418147 | 0.95679058 | 0.98861313 |
| ACSBG2 | 42.6247744 | 1.050706555 | 0.73088641 | 1.43757845 | 0.1505537 | 0.60820499 |
| MLLT1 | 259.296218 | -0.324063201 | 0.39322134 | -0.8241242 | 0.40986902 | 0.78542936 |
| CLPP | 157.025417 | 0.030301883 | 0.26602708 | 0.11390526 | 0.9093129 | 0.97680747 |
| ALKBH7 | 163.873551 | 0.426016974 | 0.55170166 | 0.77218723 | 0.44000355 | 0.79826598 |
| GTF2F1 | 225.252465 | -0.495823623 | 0.30441449 | -1.628778 | 0.10336003 | 0.55022042 |
| KHSRP | 350.845212 | -0.542504089 | 0.38646969 | -1.4037429 | 0.16039542 | 0.61590405 |
| SLC25A23 | 246.04701 | 0.432041412 | 0.36940064 | 1.16957408 | 0.24217242 | 0.68080966 |
| CRB3 | 76.1012053 | -0.617002615 | 0.47701402 | -1.2934685 | 0.19584907 | 0.64558745 |
| DENND1C | 46.0629385 | -0.013647373 | 0.57757268 | -0.0236288 | 0.98114867 | 0.9951185 |
| AC010503.4 | 113.475575 | 0.075525357 | 0.48557407 | 0.15553828 | 0.87639698 | 0.96617876 |
| C3 | 1550.47717 | 1.960656416 | 0.82948506 | 2.36370312 | 1.81E-02 | 0.33730895 |
| GPR108 | 86.6457693 | 0.404880118 | 0.3939386 | 1.02777467 | 0.30405584 | 0.72082426 |
| TRIP10 | 504.301216 | 0.239144216 | 0.36474249 | 0.65565219 | 0.51204793 | 0.83054053 |
| SH2D3A | 252.715616 | -0.098414657 | 0.29416449 | -0.3345565 | 0.73795963 | 0.92059676 |
| ZNF557 | 129.428795 | -0.540464161 | 0.32698905 | -1.652851 | 0.0983612 | 0.54193402 |
| INSR | 9557.54471 | -0.243623166 | 0.36798847 | -0.6620402 | 5.08E-01 | 0.82874455 |
| PEX11G | 53.8256537 | 0.598894974 | 0.57930518 | 1.03381601 | 0.30122219 | 0.71879123 |
| ZNF358 | 183.346582 | 0.267958917 | 0.48830252 | 0.54875596 | 0.58317294 | 0.85990837 |
| MCOLN1 | 176.81789 | -0.912200687 | 0.48199257 | -1.8925617 | 0.05841618 | 0.47778423 |
| PNPLA6 | 407.025161 | -0.305385845 | 0.37868616 | -0.8064352 | 0.41999196 | 0.78798479 |
| CAMSAP3 | 572.847086 | 0.298626583 | 0.30991994 | 0.9635604 | 0.33526637 | 0.74190856 |
| XAB2 | 129.168211 | -0.12898284 | 0.27002798 | -0.4776647 | 0.63288886 | 0.88151688 |
| STXBP2 | 241.971107 | -0.504513565 | 0.41805401 | -1.2068143 | 0.22750366 | 0.66705629 |
| EVI5L | 181.802525 | 0.235170181 | 0.2838652 | 0.82845725 | 0.40741159 | 0.78481094 |
| MAP2K7 | 458.251007 | -0.742528623 | 0.48820512 | -1.5209358 | 0.12827596 | 0.58793678 |
| SNAPC2 | 43.0516504 | -0.020446051 | 0.39131817 | -0.0522492 | 0.95833015 | 0.98878187 |
| TIMM44 | 248.668973 | -0.3732631 | 0.34239809 | -1.0901436 | 0.27564987 | 0.70247842 |
| ELAVL1 | 724.086162 | -0.199254584 | 0.29370671 | -0.6784134 | 0.49750958 | 0.82313869 |
| AC008946.1 | 38.3116771 | -0.02864477 | 0.45149984 | -0.0634436 | 0.94941328 | 0.9871942 |
| CERS4 | 1342.17369 | -0.150159868 | 0.32936134 | -0.4559122 | 6.48E-01 | 0.88787189 |
| AC022146.2 | 280.324851 | 0.316923348 | 0.52409475 | 0.6047062 | 0.54537423 | 0.84449215 |
| CD320 | 125.678876 | -0.278217483 | 0.41395079 | -0.6721028 | 0.50151826 | 0.8263386 |
| RPS28 | 492.755486 | -0.143523411 | 0.39529217 | -0.3630818 | 0.71654375 | 0.91298968 |
| KANK3 | 41.1367956 | -0.004167798 | 0.70766839 | -0.0058895 | 0.9953009 | 0.99866448 |
| ANGPTL4 | 1470.25749 | -1.017939876 | 1.01420535 | -1.0036822 | 0.31553181 | 0.72786848 |
| RAB11B-AS1 | 138.945188 | -0.552454028 | 0.35991518 | -1.5349562 | 0.12479459 | 0.58564339 |
| RAB11B | 316.278294 | -0.035664053 | 0.33957111 | -0.1050268 | 0.91635457 | 0.97822899 |
| 2-Mar | 194.356352 | 0.14660724 | 0.31676767 | 0.46282261 | 0.64349152 | 0.8848654 |
| HNRNPM | 1396.97666 | -0.361539828 | 0.14627279 | -2.4716821 | 0.0134479 | 0.29875272 |
| ZNF414 | 58.7549707 | -1.238030047 | 0.43072483 | -2.8742946 | 0.00404931 | 0.20256898 |
| ADAMTS10 | 42.8750264 | 1.775535575 | 0.70929121 | 2.50325331 | 0.01230574 | 0.29629837 |

| | | | | | | |
|---|---|---|---|---|---|---|
| ZNF558 | 98.1793019 | 0.226914179 | 0.35404979 | 0.64091036 | 0.52158092 | 0.83445062 |
| ZNF317 | 179.213891 | -0.37182365 | 0.214992 | -1.7294767 | 0.08372382 | 0.52069073 |
| ZNF699 | 57.0972389 | 0.266718235 | 0.41053064 | 0.64969142 | 0.51589157 | 0.83210529 |
| ZNF559-ZNF | 71.0505524 | 0.393582364 | 0.64272364 | 0.6123664 | 0.54029536 | 0.8414758 |
| ZNF266 | 190.485972 | 0.092508326 | 0.38753161 | 0.23871169 | 0.81132915 | 0.94753961 |
| ZNF426 | 100.523931 | 0.001085212 | 0.39528897 | 0.00274536 | 0.99780952 | 0.99977652 |
| ZNF121 | 282.742008 | -0.454420306 | 0.3250903 | -1.397828 | 0.16216474 | 0.61881739 |
| ZNF561 | 192.446724 | 0.043903307 | 0.28353412 | 0.15484311 | 0.876945 | 0.96643457 |
| ZNF562 | 420.744435 | -0.231276112 | 0.31527046 | -0.73358 | 0.46320474 | 0.8082072 |
| ZNF846 | 368.318775 | 0.459658889 | 0.38098148 | 1.20651242 | 0.22761997 | 0.66705629 |
| FBXL12 | 245.573778 | 0.032371921 | 0.23004191 | 0.14072184 | 0.88808969 | 0.97120196 |
| UBL5 | 218.516874 | -0.507898595 | 0.46352482 | -1.0957312 | 0.27319644 | 0.70126562 |
| PIN1 | 180.934413 | 0.257335939 | 0.31877408 | 0.80726744 | 0.41951242 | 0.78783502 |
| C19orf66 | 254.966408 | -0.399550955 | 0.75672224 | -0.5280021 | 0.59749786 | 0.86614567 |
| EIF3G | 200.584825 | -0.597197541 | 0.49004304 | -1.2186635 | 0.22297195 | 0.66408018 |
| DNMT1 | 166.485091 | -0.559009051 | 0.3706519 | -1.508178 | 0.13150896 | 0.59000666 |
| MRPL4 | 221.968342 | 0.107346672 | 0.29534132 | 0.36346649 | 0.71625644 | 0.91298968 |
| AC011511.2 | 141.143144 | 1.291754866 | 0.45385917 | 2.84615794 | 4.43E-03 | 2.07E-01 |
| ICAM1 | 531.216988 | 0.939022474 | 0.86859392 | 1.08108341 | 2.80E-01 | 0.70446562 |
| RAVER1 | 72.3008413 | -0.580536328 | 0.46841211 | -1.2393709 | 0.21520818 | 0.65925411 |
| TYK2 | 295.747646 | 0.067619491 | 0.29320647 | 0.23062073 | 0.81760946 | 0.94954998 |
| CDC37 | 530.216575 | -0.606881655 | 0.35890681 | -1.6909171 | 0.09085264 | 0.53537574 |
| KEAP1 | 257.037412 | -0.626689758 | 0.46279465 | -1.3541422 | 0.17569102 | 0.62699479 |
| ATG4D | 214.125836 | 0.440256465 | 0.2786139 | 1.58016689 | 0.11406865 | 0.56918057 |
| KRI1 | 200.592252 | -0.73598038 | 0.35197201 | -2.0910196 | 0.03652631 | 0.42179115 |
| CDKN2D | 62.0307836 | -0.132967312 | 0.52140453 | -0.2550176 | 0.79870954 | 0.94327531 |
| AP1M2 | 286.348547 | -0.019199317 | 0.53904864 | -0.035617 | 0.97158772 | 0.99240251 |
| SLC44A2 | 128.233855 | 0.137110761 | 0.74575243 | 0.1838556 | 0.85412674 | 0.96128707 |
| ILF3-DT | 224.885435 | -0.200826829 | 0.40429933 | -0.4967281 | 0.61938082 | 0.87527555 |
| ILF3 | 712.139096 | -0.548376104 | 0.39379036 | -1.3925585 | 0.16375335 | 0.62065658 |
| QTRT1 | 354.460212 | -0.370346509 | 0.3550564 | -1.0430639 | 2.97E-01 | 0.71825247 |
| DNM2 | 2414.89571 | 0.063825949 | 0.30840727 | 0.20695345 | 0.83604621 | 0.95468028 |
| TMED1 | 110.384131 | -0.310374006 | 0.3589529 | -0.864665 | 0.38722269 | 0.77344664 |
| CARM1 | 317.2495 | -0.267676818 | 0.32175064 | -0.8319387 | 0.40544353 | 0.78429302 |
| YIPF2 | 192.467405 | -0.779397589 | 0.36011371 | -2.1643097 | 0.03044059 | 0.40391981 |
| TIMM29 | 132.322247 | -0.249079387 | 0.26398372 | -0.9435407 | 0.34540439 | 0.75166079 |
| SMARCA4 | 1175.13514 | -0.291797953 | 0.39619382 | -0.736503 | 0.46142462 | 0.80795122 |
| LDLR | 293.602538 | 0.311856596 | 0.69193947 | 0.45069925 | 0.65220633 | 0.88935488 |
| KANK2 | 196.06177 | 0.302959191 | 0.52216568 | 0.58019744 | 0.56178148 | 0.85011467 |
| DOCK6 | 341.063438 | 0.440731838 | 0.47600474 | 0.925898 | 0.35449899 | 0.75755608 |
| RAB3D | 100.181955 | -0.038973159 | 0.53801562 | -0.0724387 | 0.94225278 | 0.98489601 |
| CCDC159 | 97.2162908 | 0.928569822 | 0.55389456 | 1.67643788 | 0.09365245 | 0.53688613 |
| PLPPR2 | 47.6904029 | 0.026406218 | 0.34822682 | 0.07583051 | 0.93955394 | 0.98435387 |
| SWSAP1 | 67.0739944 | 0.374043202 | 0.52576069 | 0.71143243 | 0.47681631 | 0.81570932 |
| EPOR | 116.605335 | -0.138085598 | 0.5348179 | -0.2581918 | 0.79625888 | 0.94260208 |
| RGL3 | 869.072711 | 0.67920314 | 0.38536394 | 1.76249789 | 0.07798521 | 0.51192014 |
| PRKCSH | 355.608403 | -0.216883785 | 0.33556104 | -0.6463318 | 0.51806447 | 0.83325693 |
| ECSIT | 308.252885 | -0.326162906 | 0.28209308 | -1.1562244 | 0.24758937 | 0.68482118 |
| ELOF1 | 105.94057 | -0.618871681 | 0.42432325 | -1.4584911 | 0.14470523 | 0.60109715 |
| ZNF627 | 426.980516 | 0.125261959 | 0.3297747 | 0.37984102 | 0.70406343 | 0.90921561 |
| ACP5 | 206.163379 | -0.096679438 | 0.4662731 | -0.2073451 | 0.83574036 | 0.95468028 |
| AC008543.1 | 68.7066326 | -0.020869002 | 0.52278861 | -0.0399186 | 0.968158 | 0.99187457 |
| ZNF823 | 385.101515 | -0.344054022 | 0.27326237 | -1.2590611 | 0.20800827 | 0.65352038 |
| ZNF441 | 214.853089 | 0.188250546 | 0.42462097 | 0.44333785 | 0.65752139 | 0.89053559 |

| | | | | | | |
|---|---|---|---|---|---|---|
| ZNF491 | 42.2838713 | 0.543083954 | 0.3798875 | 1.42959155 | 0.15283428 | 0.61001662 |
| ZNF440 | 438.31968 | -0.084110768 | 0.48016241 | -0.1751715 | 0.86094488 | 0.96302275 |
| ZNF439 | 176.003678 | 0.208832151 | 0.34133282 | 0.61181387 | 0.54066091 | 0.8417342 |
| ZNF69 | 933.332617 | -0.250324381 | 0.27656604 | -0.9051161 | 3.65E-01 | 0.76379289 |
| AC008770.2 | 88.3504922 | 0.492437801 | 0.51720599 | 0.95211156 | 0.34104041 | 0.7475837 |
| ZNF763 | 31.8410849 | 0.007130343 | 0.7900053 | 0.00902569 | 0.99279864 | 0.9976934 |
| ZNF433 | 67.407261 | 0.319927076 | 0.55680129 | 0.57458034 | 0.56557515 | 0.85194709 |
| ZNF844 | 271.711591 | -0.023446105 | 0.38077412 | -0.0615748 | 0.95090141 | 0.98731502 |
| ZNF788P | 169.035491 | 0.35547882 | 0.47735725 | 0.74468089 | 0.45646466 | 0.80507423 |
| ZNF20 | 70.9137482 | 0.836584051 | 0.56261762 | 1.48694961 | 0.13702812 | 0.59474076 |
| ZNF136 | 153.091231 | -0.412910028 | 0.34482466 | -1.1974493 | 0.2311315 | 0.66878058 |
| ZNF44 | 1243.81245 | -0.002600609 | 0.34296258 | -0.0075828 | 0.99394988 | 0.99802732 |
| ZNF563 | 157.440408 | -0.032811478 | 0.49627791 | -0.0661151 | 0.94728617 | 0.98646931 |
| ZNF442 | 86.4223638 | 0.423900706 | 0.461136 | 0.91925313 | 0.35796319 | 0.75936492 |
| AC008758.4 | 54.9560245 | 0.702052894 | 0.45955576 | 1.52767729 | 0.12659268 | 0.58713532 |
| AC008758.5 | 182.97356 | 0.079339039 | 0.57523363 | 0.1379249 | 0.89029977 | 0.97225819 |
| AC008758.6 | 73.0693923 | 0.497889363 | 0.4284797 | 1.16199054 | 0.24523931 | 0.68281753 |
| AC008758.1 | 128.854015 | -1.008577159 | 0.54487457 | -1.8510263 | 0.06416577 | 0.48593331 |
| AC010422.6 | 41.3435568 | 0.195541052 | 0.38111054 | 0.51308224 | 0.60789379 | 0.86985416 |
| ZNF490 | 339.436867 | 0.199344123 | 0.31538908 | 0.63205778 | 0.52734912 | 0.83739277 |
| ZNF791 | 565.093499 | -0.354586576 | 0.26316125 | -1.3474118 | 0.17784765 | 0.62808126 |
| MAN2B1 | 136.954401 | -0.02631828 | 0.33685595 | -0.0781292 | 0.9377253 | 0.98435387 |
| WDR83 | 102.509434 | -0.037888049 | 0.38506092 | -0.0983949 | 0.92161869 | 0.97939204 |
| DHPS | 134.982073 | 0.090634108 | 0.26962316 | 0.33615104 | 0.73675697 | 0.92020873 |
| FBXW9 | 74.9560127 | -0.494454291 | 0.48814885 | -1.012917 | 0.31109978 | 0.72558189 |
| TNPO2 | 486.862933 | -0.245845095 | 0.3342341 | -0.7355476 | 0.46200606 | 0.8082072 |
| TRIR | 536.132846 | -0.324354291 | 0.25092631 | -1.2926277 | 0.19613987 | 0.64558745 |
| ASNA1 | 179.687375 | -0.026788578 | 0.33887199 | -0.0790522 | 0.9369911 | 0.98435387 |
| HOOK2 | 1262.67202 | 0.379600991 | 0.34062804 | 1.11441499 | 0.2651012 | 0.69622296 |
| JUNB | 417.584586 | 0.146243104 | 0.58675435 | 0.24924077 | 0.80317455 | 0.94543168 |
| DNASE2 | 35.0268574 | 0.569648008 | 0.54734631 | 1.04074512 | 0.29799386 | 0.71852421 |
| GCDH | 335.249636 | 0.388853288 | 0.54959856 | 0.70752239 | 0.4792419 | 0.81649775 |
| SYCE2 | 47.4940887 | 0.58542065 | 0.84099274 | 0.69610666 | 0.48636204 | 0.81868653 |
| FARSA | 230.068465 | -0.289246682 | 0.4156509 | -0.6958885 | 0.48649866 | 0.81877553 |
| CALR | 801.885965 | 0.24765686 | 0.55276362 | 0.44803393 | 0.65412871 | 0.89019532 |
| AC092069.1 | 147.56887 | -0.061678946 | 0.70896302 | -0.0869988 | 0.93067245 | 0.98272895 |
| RAD23A | 360.277688 | -0.669390488 | 0.3534308 | -1.8939789 | 0.05822781 | 0.47775688 |
| GADD45GIP1 | 406.770771 | -0.481611798 | 0.39266643 | -1.2265164 | 0.22000443 | 0.66262374 |
| NFIX | 306.01783 | 0.863468734 | 0.42324452 | 2.04011791 | 0.04133858 | 0.43221049 |
| TRMT1 | 177.489953 | -0.854095556 | 0.41058717 | -2.0801808 | 0.03750895 | 0.42215481 |
| NACC1 | 255.112427 | -0.544561804 | 0.52549036 | -1.0362927 | 0.30006563 | 0.71879123 |
| STX10 | 153.346706 | -0.571259887 | 0.36148742 | -1.5803036 | 0.11403735 | 0.56918057 |
| IER2 | 391.631649 | -0.195817515 | 0.27865897 | -0.7027138 | 0.4822341 | 0.8177376 |
| CACNA1A | 64.4097507 | -0.088490422 | 0.46679718 | -0.1895693 | 0.84964663 | 0.95959925 |
| CCDC130 | 578.497268 | -0.146921695 | 0.19519799 | -0.7526804 | 0.451642 | 0.80384625 |
| MRI1 | 385.411533 | 0.280408112 | 0.24867517 | 1.12760799 | 0.25948551 | 0.69242517 |
| AC008686.1 | 40.859331 | 0.237359986 | 0.74651458 | 0.3179576 | 0.7505171 | 0.92590588 |
| C19orf53 | 371.207972 | -0.433004376 | 0.48011698 | -0.9018727 | 0.36712452 | 0.76546934 |
| ZSWIM4 | 257.856142 | -0.327038206 | 0.52851189 | -0.6187906 | 0.5360543 | 0.83954482 |
| CC2D1A | 345.388284 | 0.010288269 | 0.31965598 | 0.03218544 | 0.97432417 | 0.99377398 |
| DCAF15 | 95.0340657 | 0.03567188 | 0.44381795 | 0.08037503 | 0.93593899 | 0.98406248 |
| RFX1 | 358.377348 | -0.106979422 | 0.32419491 | -0.3299849 | 0.74141138 | 0.92235631 |
| IL27RA | 42.3914109 | 1.093700625 | 0.62198032 | 1.75841678 | 0.07867662 | 0.51192014 |
| PALM3 | 319.405899 | 0.666743143 | 0.51140113 | 1.30375767 | 0.19231622 | 0.64497936 |

| | | | | | | |
|---|---|---|---|---|---|---|
| MISP3 | 120.165706 | 0.404370491 | 0.49465474 | 0.81748028 | 0.41365402 | 0.78632759 |
| SAMD1 | 96.1755198 | -0.33561036 | 0.52093498 | -0.6442462 | 0.51941582 | 0.8335052 |
| PRKACA | 212.453934 | 0.317567617 | 0.27292821 | 1.16355734 | 0.24460345 | 0.68250191 |
| AC022098.1 | 68.2950161 | -0.281934746 | 0.54483645 | -0.5174668 | 0.60483037 | 0.86865188 |
| ADGRL1 | 65.1739279 | 0.855325118 | 0.52159521 | 1.63982549 | 0.10104146 | 0.54608983 |
| ADGRE5 | 60.5417175 | -1.078139893 | 0.8968668 | -1.2021182 | 0.22931774 | 0.66705629 |
| DDX39A | 209.761061 | -1.321579735 | 0.51666098 | -2.5579244 | 0.0105299 | 0.27130321 |
| PKN1 | 283.298337 | -0.080553513 | 0.3014512 | -0.2672191 | 0.78930049 | 0.94005579 |
| GIPC1 | 447.813731 | 0.129388795 | 0.34706035 | 0.37281353 | 0.70928723 | 0.9109154 |
| DNAJB1 | 500.286857 | -1.893874413 | 0.46388903 | -4.0826023 | 4.45E-05 | 0.02082964 |
| TECR | 375.195984 | -0.212256968 | 0.23856841 | -0.8897111 | 3.74E-01 | 0.76973377 |
| NDUFB7 | 466.143665 | -0.442687928 | 0.34414494 | -1.2863415 | 0.1983239 | 0.64582202 |
| ADGRE3 | 43.1488712 | -0.129704406 | 0.51970772 | -0.2495718 | 0.80291849 | 0.94543168 |
| ZNF333 | 132.651737 | -1.252319483 | 0.4628708 | -2.7055487 | 0.00681917 | 0.23083391 |
| SYDE1 | 43.9436573 | 0.294633314 | 0.54338685 | 0.54221649 | 0.58766937 | 0.86124068 |
| ILVBL | 276.819361 | 0.396256178 | 0.35601865 | 1.11302086 | 0.26569947 | 0.69728943 |
| BRD4 | 1966.51148 | -0.389467685 | 0.2305365 | -1.689397 | 0.09114337 | 0.53537574 |
| AKAP8 | 527.293274 | -0.354751761 | 0.25836723 | -1.3730524 | 0.16973604 | 0.62326131 |
| AKAP8L | 1235.48024 | -0.114674355 | 0.25704235 | -0.4461302 | 0.65550321 | 0.89053559 |
| WIZ | 117.686866 | -0.283792904 | 0.32911198 | -0.8622989 | 0.38852305 | 0.7745034 |
| CYP4F3 | 2104.8933 | 0.201312376 | 0.71802827 | 0.28036832 | 0.77919494 | 0.93646866 |
| CYP4F12 | 171.1087 | 0.734650768 | 0.96391042 | 0.76215668 | 0.4459665 | 0.80137163 |
| CYP4F2 | 159.415247 | 2.43673781 | 0.98378869 | 2.47689147 | 0.01325322 | 0.29875272 |
| CYP4F11 | 116.801915 | 0.240899577 | 0.98438735 | 0.24472031 | 0.806673 | 0.94585363 |
| TPM4 | 217.835797 | 1.125562868 | 0.57774339 | 1.94820552 | 0.05139038 | 0.46176002 |
| RAB8A | 171.095393 | -0.320218271 | 0.26166035 | -1.2237937 | 0.22103008 | 0.66263062 |
| FAM32A | 148.902044 | -0.539046086 | 0.30724851 | -1.7544303 | 0.0793568 | 0.51192014 |
| AP1M1 | 490.295203 | -0.121066574 | 0.27437602 | -0.4412433 | 0.65903689 | 0.89058983 |
| KLF2 | 56.2298162 | 0.873866236 | 0.90258942 | 0.96817692 | 0.33295602 | 0.74000014 |
| EPS15L1 | 1280.46483 | -0.195921678 | 0.36890625 | -0.531088 | 0.59535781 | 0.86510861 |
| AC008764.1 | 51.6894133 | -0.336929652 | 0.46682389 | -0.721749 | 0.47044884 | 0.81378625 |
| C19orf44 | 193.812692 | -0.209944302 | 0.31493218 | -0.6666334 | 0.50500634 | 0.82715271 |
| AC008764.8 | 58.4705459 | -0.203438321 | 0.3827163 | -0.5315643 | 0.5950278 | 0.86510861 |
| TMEM38A | 147.639312 | 0.080594667 | 0.51823447 | 0.15551777 | 0.87641315 | 0.96617876 |
| SIN3B | 580.030787 | -0.449067916 | 0.32403587 | -1.3858587 | 0.16579007 | 0.62065658 |
| HAUS8 | 58.456667 | -0.456120842 | 0.47455466 | -0.9611555 | 0.33647397 | 0.7429163 |
| MYO9B | 709.986192 | -0.769671154 | 0.42065414 | -1.8297006 | 0.06729471 | 0.48963361 |
| USE1 | 130.194005 | -0.506350915 | 0.33262201 | -1.5223013 | 0.12793361 | 0.58793678 |
| OCEL1 | 153.193094 | 0.082292294 | 0.41517866 | 0.19820935 | 0.84288127 | 0.95716002 |
| NR2F6 | 321.525387 | 0.603286618 | 0.35650392 | 1.69222998 | 0.09060213 | 0.53490056 |
| USHBP1 | 96.60135 | 0.455160052 | 0.51389126 | 0.88571278 | 0.37577232 | 0.7697436 |
| ABHD8 | 126.802367 | 0.43788622 | 0.23810651 | 1.83903506 | 0.06591003 | 0.48798003 |
| MRPL34 | 259.553216 | -0.132697182 | 0.4286472 | -0.309572 | 0.75688643 | 0.92799282 |
| DDA1 | 158.012153 | -0.24617999 | 0.28193503 | -0.8731799 | 0.38256506 | 0.77279408 |
| ANO8 | 61.6826191 | 0.427605121 | 0.410448 | 1.04180097 | 0.29750396 | 0.71839551 |
| GTPBP3 | 85.067291 | 0.046984802 | 0.3598206 | 0.13057841 | 0.89610882 | 0.97309833 |
| PLVAP | 46.9487688 | -1.322673437 | 0.58838381 | -2.2479773 | 0.02457764 | 0.37682963 |
| BST2 | 67.8271724 | -2.201931583 | 0.91835444 | -2.3976925 | 0.01649871 | 0.32399397 |
| MVB12A | 145.025578 | 0.383634019 | 0.37977179 | 1.01016988 | 0.31241391 | 0.7265535 |
| SLC27A1 | 217.224318 | 0.468503335 | 0.30998226 | 1.51138757 | 0.13068974 | 0.58853436 |
| AC010618.3 | 243.367302 | -0.26225942 | 0.37379751 | -0.7016082 | 0.48292351 | 0.8177376 |
| PGLS | 354.763706 | 0.002060387 | 0.3465697 | 0.00594509 | 0.99525653 | 0.99866448 |
| COLGALT1 | 207.53554 | -0.58233831 | 0.45161738 | -1.2894506 | 0.19724147 | 0.64558745 |
| MAP1S | 81.9373988 | -0.346734545 | 0.47631319 | -0.7279549 | 0.46664116 | 0.81098649 |

| | | | | | | |
|---|---|---|---|---|---|---|
| B3GNT3 | 143.724427 | -1.020744629 | 0.57022456 | -1.7900748 | 0.07344188 | 0.50459544 |
| RPL18A | 100.383429 | -0.178873435 | 0.583918 | -0.3063331 | 0.75935101 | 0.92930182 |
| CCDC124 | 143.588151 | -0.68172571 | 0.38325334 | -1.7787861 | 0.07527484 | 0.50718602 |
| ARRDC2 | 56.6716993 | -0.240688837 | 0.55454785 | -0.4340272 | 0.66426871 | 0.89315615 |
| MAST3 | 483.014724 | 0.84518528 | 0.40635147 | 2.07993656 | 0.03753135 | 0.42215481 |
| AC007192.2 | 76.1283369 | 1.068760813 | 0.80127767 | 1.33382079 | 0.1822626 | 0.63216521 |
| JUND | 1120.27966 | -0.948928978 | 0.5038167 | -1.8834806 | 0.05963527 | 0.47942559 |
| LSM4 | 365.404012 | 0.019637126 | 0.27163971 | 0.07229107 | 0.94237027 | 0.98489601 |
| PGPEP1 | 1887.81322 | 0.371475435 | 0.38047872 | 0.97633696 | 0.32889751 | 0.73829083 |
| GDF15 | 826.769991 | -0.463371217 | 0.35546248 | -1.3035728 | 1.92E-01 | 6.45E-01 |
| SSBP4 | 228.190822 | 0.336023469 | 0.32617682 | 1.03018808 | 0.30292173 | 0.72043311 |
| ISYNA1 | 68.2749617 | -0.154000783 | 0.52892265 | -0.2911594 | 0.77092944 | 0.93302012 |
| ELL | 802.977986 | -0.1494612 | 0.18243649 | -0.8192506 | 0.41264347 | 0.78632759 |
| FKBP8 | 328.237884 | 0.051637666 | 0.33007488 | 0.15644228 | 0.87568442 | 0.96617876 |
| KXD1 | 205.519461 | -0.337894336 | 0.24602142 | -1.3734346 | 0.16961727 | 0.62326131 |
| AC005253.1 | 40.1172907 | -0.050426526 | 0.57873369 | -0.0871325 | 0.93056617 | 0.98272895 |
| UBA52 | 638.042456 | -0.325754303 | 0.59047009 | -0.5516864 | 0.58116324 | 0.85895204 |
| CRLF1 | 57.867095 | 0.151437824 | 0.57057337 | 0.26541341 | 0.79069101 | 0.94063545 |
| REX1BD | 189.518963 | -0.165551314 | 0.49805637 | -0.3323947 | 0.73959122 | 0.92123698 |
| KLHL26 | 43.2533063 | 0.166294617 | 0.4954976 | 0.33561135 | 0.73716397 | 0.92030854 |
| CRTC1 | 536.712624 | -0.44151505 | 0.26738548 | -1.6512305 | 0.09869152 | 0.54236074 |
| UPF1 | 432.06391 | -0.549412889 | 0.30247202 | -1.816409 | 0.06930766 | 0.49273866 |
| COPE | 305.507921 | -0.504156735 | 0.39950821 | -1.2619434 | 0.20696917 | 0.65251041 |
| DDX49 | 172.487855 | -0.600363005 | 0.40983819 | -1.4648781 | 0.14295417 | 0.59824924 |
| HOMER3 | 33.1342887 | -0.293640958 | 0.75649336 | -0.3881607 | 0.69789715 | 0.90696517 |
| SUGP2 | 1518.8241 | 0.196624704 | 0.23172723 | 0.84851792 | 0.3961496 | 0.7795748 |
| ARMC6 | 123.083867 | -0.517159681 | 0.40091966 | -1.2899334 | 0.19707377 | 0.64558745 |
| SLC25A42 | 1507.84027 | 0.225623261 | 0.49530626 | 0.45552273 | 0.64873322 | 0.88797545 |
| TMEM161A | 224.83497 | 0.175471937 | 0.32125821 | 0.54620219 | 0.58492696 | 0.86062622 |
| BORCS8-MEI | 71.3641858 | -0.094191063 | 0.57294918 | -0.1643969 | 0.86941871 | 0.96564168 |
| RFXANK | 89.5558467 | -0.312259772 | 0.3303087 | -0.9453574 | 0.34447645 | 0.75124763 |
| NR2C2AP | 38.0706972 | -0.782127526 | 0.41791773 | -1.8714868 | 0.06127764 | 0.4813972 |
| TM6SF2 | 119.298208 | 1.430296905 | 0.97569581 | 1.46592503 | 0.14266871 | 0.59824924 |
| SUGP1 | 289.463858 | -0.087026072 | 0.22699672 | -0.3833803 | 0.7014378 | 0.90783786 |
| MAU2 | 637.936614 | -0.161781472 | 0.23155144 | -0.6986848 | 0.48474903 | 0.81839266 |
| GATAD2A | 1925.89872 | -0.29985341 | 0.40171525 | -0.7464327 | 0.45540606 | 0.80465616 |
| YJEFN3 | 37.2948185 | -0.904751497 | 0.62959883 | -1.4370286 | 0.15070988 | 0.60865655 |
| ATP13A1 | 198.582159 | -0.532565629 | 0.40521704 | -1.3142725 | 0.18875451 | 0.6410451 |
| ZNF101 | 72.6909658 | -0.091960678 | 0.45181633 | -0.2035355 | 0.83871647 | 0.95541749 |
| ZNF14 | 181.114971 | 0.071392091 | 0.36057733 | 0.19799384 | 0.84304988 | 0.95716002 |
| AC011477.4 | 96.8261666 | 0.612219965 | 0.3788554 | 1.61597266 | 0.10610022 | 0.55440878 |
| ZNF253 | 317.775768 | 0.913852469 | 0.34733534 | 2.63103798 | 0.00851245 | 0.25070735 |
| ZNF93 | 81.1585906 | 0.117841035 | 0.41337703 | 0.28506914 | 0.77559117 | 0.9351345 |
| ZNF682 | 231.633067 | 0.459890996 | 0.36944007 | 1.24483247 | 0.21319334 | 0.65801421 |
| ZNF90 | 649.306032 | 0.232579312 | 0.47583347 | 0.488783 | 0.62499534 | 0.87854882 |
| AC011447.3 | 189.343959 | 0.830384615 | 0.43119978 | 1.9257538 | 0.05413512 | 0.46711025 |
| ZNF486 | 83.8114657 | 1.741925025 | 0.58092269 | 2.99854877 | 0.00271269 | 0.16909084 |
| AC008554.1 | 1095.77721 | 0.689823379 | 0.80587363 | 0.85599448 | 0.39200083 | 0.77635535 |
| ZNF737 | 282.39331 | 0.353041426 | 0.36623104 | 0.96398553 | 0.33505318 | 0.74165622 |
| AC010636.2 | 77.5892433 | 1.697635499 | 0.68069951 | 2.49395729 | 0.01263277 | 0.29875272 |
| ZNF66 | 53.8312145 | 0.150074308 | 0.49341385 | 0.30415504 | 0.76100978 | 0.92989212 |
| ZNF85 | 58.8302482 | 0.987948512 | 0.60249622 | 1.63975886 | 0.10105532 | 0.54608983 |
| ZNF430 | 220.885289 | -0.156517637 | 0.32781832 | -0.4774524 | 0.63304002 | 0.88163783 |
| ZNF714 | 135.0337 | 0.212829524 | 0.4685313 | 0.45424825 | 0.64965016 | 0.88850926 |

| | | | | | | |
|---|---|---|---|---|---|---|
| ZNF431 | 457.035904 | 0.04033424 | 0.32883127 | 0.12265938 | 0.90237683 | 0.9750023 |
| ZNF708 | 453.2648 | 0.099455558 | 0.28909964 | 0.34401827 | 0.73083254 | 0.91897609 |
| ZNF738 | 40.3427596 | 0.227355912 | 0.49825112 | 0.45630788 | 0.6481686 | 0.88787189 |
| ZNF493 | 203.848418 | 0.252808902 | 0.2721197 | 0.92903566 | 0.35287061 | 0.75673243 |
| AC010615.4 | 311.624599 | 1.971354719 | 0.69392522 | 2.84087489 | 0.004499 | 0.20749797 |
| ZNF429 | 461.054945 | 0.830959983 | 0.29393953 | 2.82697596 | 0.00469899 | 0.21187169 |
| AC123912.1 | 175.790083 | 0.768663784 | 0.82524754 | 0.93143421 | 0.351629 | 0.75548011 |
| AC123912.4 | 38.735166 | 1.625532599 | 0.61970204 | 2.62308738 | 0.00871369 | 0.25358509 |
| ZNF100 | 244.922355 | 0.645716878 | 0.53325444 | 1.21089826 | 0.2259344 | 0.66688879 |
| ZNF43 | 400.564742 | 0.522945703 | 0.43704795 | 1.19654081 | 0.23148558 | 0.66878058 |
| ZNF208 | 62.2297718 | 1.124037245 | 0.81684377 | 1.37607372 | 0.16879881 | 0.62304975 |
| ZNF98 | 75.2479362 | 1.905066268 | 0.70731138 | 2.69339124 | 0.00707292 | 0.23330842 |
| AC092329.3 | 141.768579 | -0.199814005 | 0.43854799 | -0.4556263 | 6.49E-01 | 0.88796216 |
| ZNF91 | 2173.76844 | -0.011845023 | 0.30896164 | -0.0383382 | 0.96941806 | 0.99187457 |
| ZNF675 | 182.105748 | -0.223406569 | 0.30928931 | -0.7223223 | 0.47009633 | 0.81348446 |
| ZNF254 | 1254.57265 | 0.067023874 | 0.23976743 | 0.27953703 | 0.77983273 | 0.93684161 |
| LINC00662 | 480.529115 | -0.122710243 | 0.2875241 | -0.4267825 | 0.66953779 | 0.89611079 |
| AC006504.5 | 457.968448 | 1.557363178 | 0.50183112 | 3.10336112 | 0.00191336 | 0.13753904 |
| LINC01532 | 30.3555585 | 1.671472528 | 0.96491233 | 1.73225326 | 0.08322848 | 0.51935659 |
| UQCRFS1 | 442.67091 | 0.373471911 | 0.38097657 | 0.98030152 | 0.32693731 | 0.73745391 |
| POP4 | 73.5281882 | -0.563226291 | 0.38449965 | -1.4648291 | 0.14296754 | 0.59824924 |
| PLEKHF1 | 44.5875666 | 0.494356054 | 0.52988082 | 0.93295707 | 0.35084213 | 0.75495648 |
| C19orf12 | 741.455374 | 0.210350309 | 0.23724953 | 0.88662056 | 0.37528322 | 0.7697436 |
| URI1 | 1726.53844 | -0.076665261 | 0.17384616 | -0.4409948 | 0.65921673 | 0.89058983 |
| ZNF507 | 538.107022 | -0.112531989 | 0.1587626 | -0.7088067 | 0.47844446 | 0.81596289 |
| DPY19L3 | 448.766004 | -0.495830215 | 0.39627857 | -1.2512163 | 0.21085556 | 0.65540882 |
| PDCD5 | 225.267219 | -0.281080199 | 0.23139757 | -1.2147068 | 0.22447794 | 0.66551422 |
| ANKRD27 | 551.751756 | -0.23702283 | 0.26136026 | -0.9068817 | 0.36446937 | 0.76286421 |
| NUDT19 | 206.062175 | -0.219201412 | 0.26953459 | -0.8132589 | 0.41606962 | 0.78632759 |
| SLC7A9 | 789.549036 | 0.884306009 | 0.49816463 | 1.77512805 | 7.59E-02 | 0.50730849 |
| CEP89 | 518.43599 | -0.188408265 | 0.25569255 | -0.7368547 | 0.4612107 | 0.80772012 |
| RHPN2 | 2426.22247 | 0.160933304 | 0.44995963 | 0.35766165 | 0.72059653 | 0.9149807 |
| GPATCH1 | 191.645634 | -0.124885598 | 0.29481552 | -0.4236059 | 0.67185324 | 0.89704376 |
| WDR88 | 43.8228171 | -0.049879911 | 0.65271106 | -0.0764196 | 0.93908529 | 0.98435387 |
| CEBPG | 176.51804 | -0.774282469 | 0.42031761 | -1.8421366 | 0.06545517 | 0.48790819 |
| PEPD | 2273.33189 | 0.124385399 | 0.31627024 | 0.39328835 | 0.69410652 | 0.90589033 |
| LSM14A | 2439.1976 | -0.227265683 | 0.14513286 | -1.5659147 | 0.11736858 | 0.57350419 |
| KIAA0355 | 1241.02166 | -0.183418338 | 0.2623222 | -0.6992101 | 0.48442073 | 0.81839266 |
| GPI | 1568.34601 | -0.380694879 | 0.30091218 | -1.2651362 | 2.06E-01 | 0.651931 |
| UBA2 | 830.715623 | -0.279790005 | 0.16658049 | -1.6796085 | 0.09303352 | 0.53684765 |
| WTIP | 42.7951632 | 0.338671369 | 0.4980744 | 0.67996141 | 0.4965289 | 0.82252856 |
| SCGB2B2 | 529.137954 | -0.104350377 | 0.50989261 | -0.2046517 | 0.83784428 | 0.95496955 |
| ZNF302 | 499.665432 | -0.26236425 | 0.41778034 | -0.6279957 | 0.53000677 | 0.83789294 |
| ZNF181 | 84.7522795 | 0.209720407 | 0.46550751 | 0.45051992 | 0.65233559 | 0.88935488 |
| ZNF30 | 76.528064 | 0.314145525 | 0.40879429 | 0.76846847 | 0.44220892 | 0.79928982 |
| GRAMD1A | 424.650494 | 0.544823129 | 0.42293371 | 1.28819982 | 0.19767641 | 0.64558745 |
| HPN | 3906.60888 | 1.257899081 | 0.4948236 | 2.54211618 | 0.01101835 | 0.27879579 |
| HPN-AS1 | 51.3397567 | 1.163845729 | 0.84255553 | 1.38132822 | 0.16717807 | 0.62242897 |
| LSR | 317.720487 | -0.369813543 | 0.46569718 | -0.7941073 | 0.42713295 | 0.78933193 |
| USF2 | 273.244623 | -0.55519137 | 0.35317976 | -1.5719796 | 0.11595528 | 0.57350419 |
| DMKN | 121.961589 | 0.835191562 | 0.61376217 | 1.36077393 | 0.17358514 | 0.62663931 |
| TMEM147-A! | 38.0087246 | 1.070564313 | 0.55804867 | 1.9184067 | 0.05505946 | 0.46904913 |
| TMEM147 | 175.825058 | 0.033816385 | 0.46346002 | 0.07296505 | 0.94183393 | 0.98489601 |
| HAUS5 | 41.5512123 | 0.367934892 | 0.71952511 | 0.51135796 | 0.60910043 | 0.87022509 |

| | | | | | | |
|---|---|---|---|---|---|---|
| RBM42 | 111.528129 | -0.508331172 | 0.36561486 | -1.390346 | 0.16442383 | 0.62065658 |
| COX6B1 | 593.543616 | -0.199171913 | 0.36229827 | -0.5497457 | 0.58249382 | 0.85981507 |
| KMT2B | 360.086759 | -0.018671586 | 0.31072432 | -0.0600905 | 0.95208354 | 0.98766985 |
| PROSER3 | 162.186777 | -0.192035763 | 0.37852809 | -0.5073224 | 0.61192866 | 0.87199356 |
| PRODH2 | 1362.27847 | 0.727719357 | 0.62922495 | 1.1565329 | 2.47E-01 | 6.85E-01 |
| NPHS1 | 43.0711954 | 0.482496305 | 0.64363595 | 0.74964164 | 0.45347057 | 0.80433814 |
| NFKBID | 154.553426 | 0.333100669 | 0.50403158 | 0.66087262 | 0.508694 | 0.82874455 |
| LRFN3 | 34.370543 | 0.053991346 | 0.49243335 | 0.10964193 | 0.91269335 | 0.977179 |
| SDHAF1 | 69.2506895 | -0.165552092 | 0.43490391 | -0.3806636 | 0.70345287 | 0.90887942 |
| SYNE4 | 82.5147915 | 0.471184654 | 0.41316867 | 1.1404172 | 0.25411253 | 0.6890446 |
| ALKBH6 | 50.8281481 | -0.143232387 | 0.34760943 | -0.4120498 | 0.68030294 | 0.89941358 |
| THAP8 | 65.028511 | -0.418362409 | 0.3819148 | -1.0954339 | 0.2733266 | 0.70137637 |
| POLR2I | 51.8425951 | -0.774330725 | 0.5383922 | -1.438228 | 0.15036938 | 0.60799621 |
| TBCB | 167.09708 | -0.100265419 | 0.37250444 | -0.2691657 | 0.78780217 | 0.93953874 |
| CAPNS1 | 563.53894 | -0.220709356 | 0.46464217 | -0.4750093 | 0.63478034 | 0.88238823 |
| COX7A1 | 36.1378617 | -1.956903843 | 0.92118958 | -2.1243226 | 0.03364318 | 0.41378378 |
| ZNF565 | 550.256518 | -0.20400802 | 0.36124629 | -0.5647339 | 0.57225479 | 0.85408471 |
| ZNF146 | 690.285155 | -0.529381079 | 0.24950426 | -2.1217316 | 0.03386028 | 0.41378378 |
| AC012617.1 | 33.898361 | 0.462768073 | 0.53738608 | 0.86114636 | 0.38915744 | 0.7748665 |
| LINC00665 | 337.095715 | -0.108108529 | 0.25779963 | -0.419351 | 0.67495964 | 0.89801183 |
| ZFP14 | 658.434771 | 0.667904535 | 0.35826272 | 1.86428701 | 0.06228141 | 0.4840117 |
| ZFP82 | 200.602849 | -0.515339653 | 0.25785164 | -1.9985898 | 0.04565275 | 0.44430994 |
| ZNF566 | 351.774512 | -0.535253703 | 0.18618277 | -2.8748832 | 0.00404177 | 0.20256898 |
| ZNF260 | 230.108771 | 0.004423259 | 0.2137045 | 0.02069801 | 0.98348655 | 0.99523767 |
| ZNF529 | 389.954492 | -0.284159888 | 0.22403303 | -1.2683839 | 0.2046609 | 0.65068591 |
| ZNF529-AS1 | 310.205219 | 0.152045044 | 0.28623246 | 0.53119428 | 0.59528415 | 0.86510861 |
| ZNF382 | 109.918599 | 0.168059645 | 0.3279056 | 0.51252447 | 0.608284 | 0.86985416 |
| ZNF461 | 193.851414 | -0.798266245 | 0.27450665 | -2.9080033 | 0.00363744 | 0.19706898 |
| ZNF567 | 202.595768 | -0.310955148 | 0.26902533 | -1.1558583 | 0.24773914 | 0.68482118 |
| ZNF850 | 49.4835029 | -0.230869113 | 0.42327093 | -0.5454405 | 0.58545059 | 0.86077718 |
| AC020928.1 | 317.301609 | -0.184869127 | 0.31519631 | -0.5865206 | 0.55752573 | 0.84861484 |
| ZNF790 | 351.541582 | 0.369191169 | 0.3440604 | 1.07304172 | 0.2832524 | 0.70780772 |
| ZNF345 | 364.350066 | 0.32910891 | 0.44384018 | 0.74150319 | 0.4583884 | 0.80625302 |
| ZNF829 | 38.2768033 | -0.43380604 | 0.43124974 | -1.0059277 | 0.31445037 | 0.72719146 |
| ZNF568 | 186.380819 | -0.426629435 | 0.28619856 | -1.4906764 | 0.13604647 | 0.5942049 |
| ZNF420 | 256.184646 | -0.034277252 | 0.22648756 | -0.1513428 | 0.87970534 | 0.96751353 |
| AC012309.1 | 39.6964153 | 0.437007284 | 0.70690535 | 0.61819773 | 0.536445 | 0.83977292 |
| ZNF383 | 176.10116 | 0.063905198 | 0.24678971 | 0.25894596 | 0.79567693 | 0.94252227 |
| LINC01535 | 29.5295757 | 1.368422807 | 0.70419207 | 1.94325222 | 0.05198569 | 0.46437412 |
| HKR1 | 592.178735 | 0.233712535 | 0.22155763 | 1.05486115 | 0.29148883 | 0.71337138 |
| ZNF527 | 59.8185275 | -0.300689568 | 0.37582716 | -0.8000741 | 0.42366788 | 0.78894887 |
| ZNF569 | 198.396888 | 0.171641245 | 0.30791839 | 0.55742447 | 0.57723745 | 0.85648341 |
| ZNF570 | 64.1026462 | -0.245035119 | 0.27280547 | -0.8982046 | 0.3690765 | 0.76615653 |
| ZNF793 | 176.92681 | 0.723140106 | 0.37017257 | 1.95352159 | 0.05075782 | 0.45923264 |
| ZNF540 | 178.265151 | 0.865606956 | 0.53502882 | 1.61786978 | 0.10569066 | 0.55370151 |
| ZNF571 | 194.63655 | 0.371232643 | 0.31913637 | 1.16324142 | 0.24473157 | 0.68250191 |
| ZFP30 | 331.390759 | 0.114629701 | 0.34771356 | 0.32966704 | 0.74165156 | 0.92235631 |
| AC093227.1 | 38.2893618 | -0.348602885 | 0.36251421 | -0.9616254 | 0.33623779 | 0.74287741 |
| ZNF573 | 297.913713 | 0.134495534 | 0.25644732 | 0.52445676 | 0.59996089 | 0.86729212 |
| SIPA1L3 | 4542.18502 | -1.151114535 | 0.45742855 | -2.5164904 | 0.01185301 | 0.29116356 |
| SPINT2 | 298.280165 | -0.024592143 | 0.30896673 | -0.0795948 | 0.93655953 | 0.98435387 |
| AC011479.1 | 57.4940387 | 0.836980528 | 0.51385608 | 1.62882285 | 0.10335053 | 0.55022042 |
| YIF1B | 63.1397456 | -0.217337294 | 0.36991693 | -0.58753 | 0.55684781 | 0.84861484 |
| CATSPERG | 233.012996 | -0.085261195 | 0.55035463 | -0.1549205 | 0.87688402 | 0.96643457 |

| | | | | | | |
|---|---|---|---|---|---|---|
| PSMD8 | 394.310955 | -0.1960626 | 0.3744584 | -0.5235898 | 0.60056388 | 0.86755264 |
| FAM98C | 58.5347363 | 0.357327196 | 0.30398691 | 1.17546902 | 0.23980713 | 0.6780147 |
| EIF3K | 464.579404 | -0.017618351 | 0.25453672 | -0.0692173 | 0.94481663 | 0.9853777 |
| ACTN4 | 3743.49653 | -0.533781566 | 0.36682283 | -1.4551482 | 0.14562825 | 0.60203043 |
| CAPN12 | 1568.5647 | 1.383814235 | 0.48644547 | 2.84474689 | 0.00444467 | 0.20749797 |
| AC022144.1 | 100.582908 | 0.800610125 | 0.44984866 | 1.77973214 | 0.07511981 | 0.5071704 |
| ECH1 | 497.892779 | 0.529742661 | 0.5324877 | 0.99484488 | 0.31981171 | 0.73157402 |
| HNRNPL | 705.850086 | -0.438682835 | 0.26008474 | -1.6866919 | 0.09166261 | 0.53567268 |
| RINL | 34.3069256 | 0.186443503 | 0.47000497 | 0.39668411 | 0.69160042 | 0.90491747 |
| SIRT2 | 350.629612 | 0.283572716 | 0.39726653 | 0.71380974 | 0.47534484 | 0.81564184 |
| NFKBIB | 142.420126 | -0.347146104 | 0.32569162 | -1.0658736 | 0.28648078 | 0.71118494 |
| SARS2 | 33.6524536 | -0.443164884 | 0.56379316 | -0.7860416 | 0.43184311 | 0.79166423 |
| AC011455.2 | 67.8535436 | 0.373775787 | 0.68890299 | 0.54256665 | 0.58742821 | 0.86116347 |
| MRPS12 | 174.763399 | -0.319321159 | 0.39380624 | -0.8108585 | 0.41744691 | 0.78652162 |
| FBXO17 | 1704.60772 | -0.035538766 | 0.37184786 | -0.0955734 | 0.92385939 | 0.98015759 |
| FBXO27 | 115.555858 | 0.000265962 | 0.42235336 | 0.00062971 | 0.99949756 | 0.99977652 |
| PAK4 | 439.649384 | -0.070788906 | 0.318879 | -0.221993 | 0.82431933 | 0.95180295 |
| AC011443.1 | 54.11079 | 0.481804757 | 0.58894978 | 0.81807443 | 0.41331469 | 0.78632759 |
| SAMD4B | 626.563938 | -0.364968712 | 0.32334465 | -1.1287297 | 0.25901187 | 0.69241621 |
| PAF1 | 138.18139 | -0.007716456 | 0.35108278 | -0.021979 | 0.98246469 | 0.99522272 |
| MED29 | 155.663315 | -0.088753686 | 0.44460269 | -0.1996247 | 0.84177409 | 0.95681213 |
| ZFP36 | 733.202269 | -0.000788224 | 0.51720916 | -0.001524 | 9.99E-01 | 1.00E+00 |
| RPS16 | 806.380792 | -0.092345713 | 0.512007 | -0.1803603 | 0.85686975 | 0.96252844 |
| SUPT5H | 721.974096 | -0.670432614 | 0.25836715 | -2.5948834 | 0.0094623 | 0.25998727 |
| TIMM50 | 145.584659 | 0.102853559 | 0.30128809 | 0.34137944 | 0.73281795 | 0.91952673 |
| EID2B | 34.1682986 | -0.072513316 | 0.42498575 | -0.1706253 | 0.86451841 | 0.96411634 |
| EID2 | 69.4565131 | -0.085737664 | 0.34963998 | -0.245217 | 0.80628842 | 0.94585363 |
| DYRK1B | 127.440552 | -0.13923101 | 0.39425229 | -0.3531521 | 0.72397445 | 0.91600702 |
| FBL | 114.767037 | -0.373052477 | 0.33726177 | -1.1061214 | 0.26867396 | 0.69961625 |
| FCGBP | 66.3440554 | -0.274917969 | 0.58060382 | -0.4735036 | 6.36E-01 | 0.88257807 |
| PSMC4 | 276.075436 | -0.426126532 | 0.40964502 | -1.0402336 | 0.29823137 | 0.71852421 |
| ZNF546 | 213.451257 | 0.663126373 | 0.42775559 | 1.55024596 | 0.12108249 | 0.57927181 |
| ZNF780B | 775.127568 | 0.041809555 | 0.33016778 | 0.12663124 | 0.89923227 | 0.97458352 |
| ZNF780A | 164.829598 | -0.23606984 | 0.29603038 | -0.7974514 | 0.42518892 | 0.78894887 |
| MAP3K10 | 224.209088 | -0.094284612 | 0.39724762 | -0.2373447 | 0.8123894 | 0.94787133 |
| AKT2 | 934.466225 | 0.45627509 | 0.34565928 | 1.32001402 | 0.18683034 | 0.6378941 |
| C19orf47 | 61.1764276 | -0.978987388 | 0.49996865 | -1.9580975 | 0.05021857 | 0.45824658 |
| PLD3 | 929.332736 | -0.208667061 | 0.32215134 | -0.6477299 | 0.51715965 | 0.83278987 |
| SERTAD1 | 62.5071187 | -0.582418692 | 0.41840026 | -1.3920132 | 0.1639184 | 0.62065658 |
| SERTAD3 | 68.7187022 | -0.378856675 | 0.35995755 | -1.0525038 | 0.29256848 | 0.71399546 |
| BLVRB | 688.965819 | 0.052378077 | 0.39900851 | 0.13127058 | 0.89556127 | 0.97309833 |
| SPTBN4 | 87.1870254 | -0.342760599 | 0.57040808 | -0.6009042 | 0.54790381 | 0.84486687 |
| SHKBP1 | 124.956202 | -0.153434818 | 0.39177316 | -0.391642 | 0.69532276 | 0.90589033 |
| NUMBL | 77.0747475 | 0.319198264 | 0.36243111 | 0.8807143 | 0.37847247 | 0.77061593 |
| COQ8B | 226.067342 | 0.111391941 | 0.35756212 | 0.31153171 | 0.75539644 | 0.92799282 |
| ITPKC | 450.456172 | -0.044706836 | 0.25295385 | -0.1767391 | 8.60E-01 | 9.63E-01 |
| C19orf54 | 65.3750472 | -0.209796588 | 0.31318295 | -0.6698851 | 0.50293104 | 0.82648074 |
| SNRPA | 331.604321 | -0.381002862 | 0.34279334 | -1.1114652 | 0.26636816 | 0.69797523 |
| AC008537.4 | 36.3109061 | 1.200344843 | 0.40663785 | 2.95187679 | 0.00315849 | 0.18344667 |
| AC008537.1 | 361.438762 | -0.011357136 | 0.36068684 | -0.0314875 | 0.97488074 | 0.99402915 |
| CYP2B6 | 180.248469 | -0.254957274 | 0.69127553 | -0.3688215 | 0.71226078 | 0.91165921 |
| HNRNPUL1 | 457.147979 | -0.237399698 | 0.31752455 | -0.7476578 | 0.45466661 | 0.80465616 |
| TGFB1 | 227.237083 | 0.512636215 | 0.36334875 | 1.41086548 | 0.15828428 | 0.61408877 |
| CCDC97 | 123.996514 | -0.447084274 | 0.35492037 | -1.2596749 | 0.20778667 | 0.65309147 |

| | | | | | | |
|---|---|---|---|---|---|---|
| B9D2 | 39.9851405 | -0.288060726 | 0.46469032 | -0.6198983 | 0.53532476 | 0.83954482 |
| EXOSC5 | 45.7097836 | -0.755430835 | 0.55893111 | -1.3515634 | 0.17651502 | 0.62709281 |
| DMAC2 | 154.781272 | -0.145431772 | 0.23371423 | -0.6222632 | 0.5337688 | 0.83915086 |
| RPS19 | 2284.89428 | 0.048177925 | 0.41115349 | 0.11717747 | 9.07E-01 | 0.97661505 |
| ARHGEF1 | 256.815373 | -0.227333867 | 0.39851987 | -0.5704455 | 0.56837558 | 0.85263558 |
| RABAC1 | 448.509881 | -0.570139692 | 0.49588144 | -1.14975 | 0.25024685 | 0.68617881 |
| ZNF574 | 37.1209027 | -0.361649808 | 0.48011932 | -0.7532499 | 0.45129978 | 0.80384625 |
| POU2F2 | 58.0751823 | -0.447769979 | 0.63848252 | -0.7013034 | 0.48311367 | 0.8177376 |
| DEDD2 | 132.34133 | -1.224118259 | 0.42525345 | -2.8785616 | 0.00399493 | 0.20256898 |
| ZNF526 | 81.0132302 | -0.665994156 | 0.48325589 | -1.3781398 | 0.16816014 | 0.62304975 |
| GSK3A | 133.244478 | -0.306218237 | 0.44216079 | -0.6925495 | 0.48859231 | 0.81933645 |
| CIC | 189.057619 | 0.017078175 | 0.27569022 | 0.06194697 | 0.95060506 | 0.98731502 |
| PAFAH1B3 | 80.8690408 | 0.55286946 | 0.52493533 | 1.05321442 | 0.29224273 | 0.71399546 |
| MEGF8 | 251.752451 | 0.655355702 | 0.36888059 | 1.77660662 | 0.07563299 | 0.50730849 |
| LIPE-AS1 | 217.594326 | 0.886796786 | 0.47331217 | 1.87359811 | 0.06098584 | 0.48123823 |
| CEACAM1 | 389.010492 | 1.248393905 | 0.60574506 | 2.06092297 | 0.03931039 | 0.42704465 |
| PHLDB3 | 231.216779 | 0.910235765 | 0.30570963 | 2.9774521 | 0.00290655 | 0.17474518 |
| ETHE1 | 294.147525 | 1.122051403 | 0.35549358 | 3.15631972 | 0.00159774 | 0.13207839 |
| XRCC1 | 80.6699048 | 0.355972095 | 0.32339374 | 1.10073897 | 0.27101028 | 0.69961625 |
| PINLYP | 38.4173149 | -0.466254607 | 0.58324242 | -0.7994182 | 0.42404796 | 0.78894887 |
| ZNF428 | 241.24426 | -0.078371027 | 0.34392357 | -0.2278734 | 0.81974468 | 0.94990455 |
| CADM4 | 124.363521 | 0.788976842 | 0.47371522 | 1.66550875 | 0.0958113 | 0.53867329 |
| SMG9 | 166.928834 | -0.004527219 | 0.26139024 | -0.0173198 | 0.98618151 | 0.99609994 |
| ZNF283 | 139.548395 | -0.244769356 | 0.40422722 | -0.6055242 | 0.54483076 | 0.84441376 |
| ZNF404 | 33.1919759 | -0.730496655 | 0.79608831 | -0.9176076 | 0.35882435 | 0.75936492 |
| ZNF45 | 98.6208782 | 0.355309296 | 0.38424699 | 0.92468985 | 0.35512726 | 0.75757147 |
| ZNF155 | 152.507843 | 0.025467863 | 0.37185959 | 0.06848785 | 0.94539729 | 0.98574172 |
| ZNF230 | 44.0497803 | -0.004628298 | 0.40558444 | -0.0114114 | 0.99089519 | 0.99737132 |
| ZNF224 | 316.407043 | -0.28497886 | 0.32276331 | -0.8829345 | 0.37727166 | 0.77048581 |
| ZNF225 | 53.597265 | -0.169879961 | 0.33094545 | -0.5133171 | 0.60772952 | 0.86985416 |
| ZNF234 | 132.523594 | -0.053852771 | 0.31785445 | -0.1694259 | 0.86546167 | 0.96428252 |
| ZNF226 | 162.695489 | -0.076042461 | 0.34561748 | -0.2200191 | 0.82585625 | 0.95192757 |
| ZNF227 | 122.399174 | -0.628972817 | 0.31362772 | -2.0054759 | 0.0449122 | 0.44360789 |
| ZNF235 | 646.097204 | -0.008346031 | 0.4795512 | -0.0174038 | 0.98611445 | 0.99609994 |
| ZNF229 | 87.5884078 | -0.70073478 | 0.51225055 | -1.3679532 | 0.17132672 | 0.62520756 |
| ZNF180 | 87.4467272 | -0.182150214 | 0.35785441 | -0.5090065 | 0.61074768 | 0.87139778 |
| AC243964.2 | 349.684396 | -0.287974778 | 0.36997584 | -0.778361 | 0.4363562 | 0.79555046 |
| PVR | 181.96772 | -1.611094512 | 0.57834167 | -2.785714 | 0.005341 | 0.21873395 |
| CEACAM19 | 408.542701 | -1.830811753 | 0.63827148 | -2.8683904 | 0.00412566 | 0.20365996 |
| BCL3 | 218.523632 | 0.292141567 | 0.74631559 | 0.39144508 | 0.69546828 | 0.90589033 |
| CBLC | 272.274368 | 0.181666697 | 0.53158726 | 0.34174389 | 0.73254364 | 0.91952673 |
| BCAM | 598.600398 | 0.131055913 | 0.51139291 | 0.25627245 | 0.79774047 | 0.94316375 |
| NECTIN2 | 551.977437 | 0.294201146 | 0.33544588 | 0.87704503 | 0.38046218 | 0.7716703 |
| TOMM40 | 231.171997 | -0.746734706 | 0.45048046 | -1.6576406 | 0.09739003 | 0.54093104 |
| APOE | 5094.07775 | -1.103027007 | 0.63297153 | -1.7426171 | 0.08140052 | 0.51593841 |
| CLPTM1 | 564.6994 | -0.194017875 | 0.22241498 | -0.8723238 | 0.38303176 | 0.77317162 |
| RELB | 395.647081 | 0.472319113 | 0.65453177 | 0.72161373 | 0.470532 | 0.81382739 |
| CLASRP | 646.864046 | -0.35920719 | 0.30039309 | -1.1957905 | 0.23177834 | 0.66883909 |
| GEMIN7-AS1 | 253.810109 | -0.15953206 | 0.42451943 | -0.3757945 | 0.70706969 | 0.91002281 |
| MARK4 | 443.202181 | 0.426474064 | 0.34760088 | 1.22690734 | 0.21985742 | 0.66238422 |
| TRAPPC6A | 202.23237 | 0.374423772 | 0.34097361 | 1.09810191 | 0.27215999 | 0.70056282 |
| ERCC2 | 136.43624 | 0.520007863 | 0.35605846 | 1.46045643 | 0.14416467 | 0.60086567 |
| PPP1R13L | 398.146579 | -0.389918675 | 0.37034507 | -1.0528523 | 0.29240867 | 0.71399546 |
| CD3EAP | 49.8837354 | -1.451732164 | 0.7930027 | -1.8306775 | 0.0671487 | 0.48963361 |

| | | | | | | |
|---|---|---|---|---|---|---|
| ERCC1 | 191.375541 | -1.716027809 | 0.61148114 | -2.8063463 | 0.00501068 | 0.21686099 |
| VASP | 315.673902 | -0.269369996 | 0.37664931 | -0.7151745 | 0.47450121 | 0.81531007 |
| OPA3 | 671.236374 | -0.4798232 | 0.38992383 | -1.2305562 | 0.21848888 | 0.66098715 |
| EML2 | 167.204667 | 0.659163239 | 0.34534383 | 1.90871586 | 0.05629876 | 0.47143978 |
| SNRPD2 | 327.017045 | -0.449159241 | 0.35787806 | -1.2550622 | 0.20945617 | 0.65441616 |
| QPCTL | 102.68968 | -0.257887152 | 0.47713407 | -0.540492 | 0.58885777 | 0.86196855 |
| FBXO46 | 211.275788 | -0.216635116 | 0.30835198 | -0.7025579 | 0.48233132 | 0.8177376 |
| SIX5 | 34.9754703 | 0.431795466 | 0.53100907 | 0.81316024 | 0.41612619 | 0.78632759 |
| DM1-AS | 81.1589985 | 0.500513092 | 0.50643263 | 0.98831129 | 0.32300022 | 0.7350762 |
| DMPK | 237.967738 | -0.228378894 | 0.45305993 | -0.504081 | 0.61420447 | 0.87225883 |
| SYMPK | 797.906256 | -0.253061164 | 0.37703624 | -0.6711852 | 0.50210252 | 0.82638105 |
| IRF2BP1 | 32.0508619 | -0.051274347 | 0.58708266 | -0.0873375 | 0.93040323 | 0.98272895 |
| MYPOP | 229.238588 | 0.104027218 | 0.3050096 | 0.34106211 | 7.33E-01 | 0.91952673 |
| CCDC61 | 64.3257098 | 0.189682003 | 0.33596288 | 0.56459215 | 0.57235121 | 0.85408471 |
| PPP5C | 383.298152 | -0.384751529 | 0.22115938 | -1.7397025 | 0.08191127 | 0.51593841 |
| PPP5D1 | 145.402594 | -0.129227319 | 0.39623775 | -0.3261358 | 0.7443216 | 0.92362993 |
| CALM3 | 321.737679 | 0.124500717 | 0.39204436 | 0.31756793 | 0.7508127 | 0.92598432 |
| PRKD2 | 405.814635 | 0.130133504 | 0.38314674 | 0.33964403 | 0.73412462 | 0.91975299 |
| STRN4 | 193.692338 | -0.631187165 | 0.27445184 | -2.2998103 | 0.02145897 | 0.35743746 |
| FKRP | 135.467436 | 0.027611486 | 0.3355955 | 0.08227609 | 0.93442717 | 0.98359701 |
| SLC1A5 | 228.500685 | -0.08452523 | 0.55721215 | -0.1516931 | 0.87942901 | 0.96751353 |
| AP2S1 | 335.436963 | -0.185685485 | 0.31082718 | -0.5973914 | 0.5502461 | 0.84620747 |
| ARHGAP35 | 1247.54361 | -0.190198653 | 0.19725923 | -0.9642066 | 0.33494234 | 0.74156916 |
| TMEM160 | 218.136305 | 0.384270317 | 0.4905969 | 0.78327098 | 0.433468 | 0.79380258 |
| ZC3H4 | 254.708067 | -0.381911094 | 0.34617263 | -1.1032388 | 0.26992346 | 0.69961625 |
| SAE1 | 342.260356 | -1.067596971 | 0.48535146 | -2.1996369 | 0.02783267 | 0.38968579 |
| BBC3 | 52.481485 | 0.341611488 | 0.58589697 | 0.58305727 | 5.60E-01 | 8.49E-01 |
| CCDC9 | 102.731353 | -0.619409192 | 0.42661112 | -1.4519293 | 0.14652126 | 0.60383734 |
| INAFM1 | 65.494904 | -0.487865236 | 0.49796194 | -0.979724 | 0.3272224 | 0.73746094 |
| C5AR1 | 513.581448 | 1.052286392 | 0.75760139 | 1.38897105 | 0.16484155 | 0.62065658 |
| DHX34 | 375.616042 | -0.764382753 | 0.39553912 | -1.9325086 | 0.05329676 | 0.46664603 |
| KPTN | 40.1524581 | -0.331405493 | 0.44930469 | -0.7375963 | 0.46075978 | 0.80741938 |
| NAPA-AS1 | 82.4191723 | 0.285409394 | 0.49763693 | 0.57352937 | 0.56628632 | 0.85215881 |
| NAPA | 534.096789 | -0.779646431 | 0.28655763 | -2.7207317 | 0.00651376 | 0.22952213 |
| BICRA-AS1 | 63.4892318 | 0.288264113 | 0.49194505 | 0.58596811 | 0.55789694 | 0.84889425 |
| NOP53 | 152.488231 | -0.221124463 | 0.4581777 | -0.4826173 | 0.62936753 | 0.88081894 |
| SELENOW | 458.06504 | 0.086699896 | 0.31874722 | 0.27200205 | 0.78562045 | 0.93892226 |
| PLA2G4C | 650.022794 | -0.432853012 | 0.53456305 | -0.8097324 | 4.18E-01 | 0.78697737 |
| LIG1 | 112.520805 | -0.621316683 | 0.46566114 | -1.3342678 | 0.18211609 | 0.63216521 |
| ZSWIM9 | 161.819553 | -0.221049972 | 0.27379545 | -0.8073544 | 0.41946231 | 0.78783502 |
| CARD8 | 389.745471 | 0.187485017 | 0.31447043 | 0.59619284 | 0.55104641 | 0.8466552 |
| EMP3 | 68.0451368 | 0.438163881 | 0.63385802 | 0.69126503 | 0.489399 | 0.81970016 |
| TMEM143 | 210.281832 | 0.883949835 | 0.45936882 | 1.92427044 | 0.05432069 | 0.46711025 |
| KDELR1 | 386.062147 | 0.204760693 | 0.29830604 | 0.68641149 | 0.49245366 | 0.82087587 |
| GRWD1 | 67.0565816 | -0.38416298 | 0.60067818 | -0.6395487 | 0.52246601 | 0.83477439 |
| KCNJ14 | 59.4857455 | -0.551045771 | 0.64911215 | -0.8489223 | 0.39592453 | 0.77955046 |
| CYTH2 | 236.240085 | 0.149966673 | 0.38513033 | 0.38939201 | 0.69698618 | 0.90649868 |
| LMTK3 | 57.5323728 | 0.137181281 | 0.60197076 | 0.22788695 | 0.81973413 | 0.94990455 |
| SULT2B1 | 51.8887854 | 0.165868004 | 0.80930416 | 0.20495138 | 0.83761011 | 0.95496955 |
| RPL18 | 858.839597 | 0.055311664 | 0.52785965 | 0.10478479 | 0.91654658 | 0.97822899 |
| SPHK2 | 263.638743 | 0.273123774 | 0.38550601 | 0.70848123 | 0.47864646 | 0.81596289 |
| DBP | 50.993057 | 0.265253762 | 0.56438326 | 0.46998871 | 0.63836308 | 0.8827026 |
| CA11 | 38.6521656 | 0.288330726 | 0.44382361 | 0.64965162 | 0.51591728 | 0.83210529 |
| BCAT2 | 395.280779 | -0.049225593 | 0.36741809 | -0.1339771 | 0.89342072 | 0.97272276 |

| | | | | | | |
|---|---|---|---|---|---|---|
| HSD17B14 | 830.379287 | -0.252496529 | 0.45165659 | -0.5590454 | 0.57613075 | 0.85567496 |
| PLEKHA4 | 139.812238 | 0.265437464 | 0.75417589 | 0.35195697 | 0.72487053 | 0.91600702 |
| PPP1R15A | 258.365976 | -1.499994093 | 0.47338813 | -3.1686348 | 1.53E-03 | 0.13146073 |
| NUCB1 | 1052.57461 | -0.626318363 | 0.45508562 | -1.3762649 | 0.16873964 | 0.62304975 |
| NUCB1-AS1 | 150.20099 | -0.930162162 | 0.81523312 | -1.1409769 | 0.25387954 | 0.68891958 |
| DHDH | 239.062126 | 0.890220599 | 0.61006693 | 1.45921792 | 0.14450514 | 0.60101297 |
| BAX | 399.826461 | -0.229218167 | 0.2401181 | -0.9546059 | 0.33977701 | 0.74648557 |
| FTL | 24154.4632 | -0.171246488 | 0.70917846 | -0.2414716 | 0.80918959 | 0.94673968 |
| GYS1 | 654.594477 | -0.209663654 | 0.30340586 | -0.6910336 | 0.4895444 | 0.81970016 |
| RUVBL2 | 155.59422 | -0.668030698 | 0.29581913 | -2.2582404 | 0.02393068 | 0.37387451 |
| SNRNP70 | 2322.74154 | -0.27966778 | 0.4114592 | -0.6796975 | 0.49669603 | 0.82254591 |
| LIN7B | 65.3989829 | 0.908289299 | 0.40262781 | 2.25590301 | 0.02407671 | 0.37387451 |
| PPFIA3 | 38.5477815 | 1.432801484 | 0.55187809 | 2.59622825 | 0.00942534 | 0.25998727 |
| TRPM4 | 249.924836 | 0.820692693 | 0.70986553 | 1.15612417 | 0.24763037 | 0.68482118 |
| SLC6A16 | 64.4516717 | 0.501613963 | 0.85667463 | 0.58553615 | 0.55818727 | 0.8489192 |
| PIH1D1 | 232.066682 | -0.298553866 | 0.27642841 | -1.0800405 | 0.28012416 | 0.70491314 |
| RPL13A | 1593.75521 | 0.092042129 | 0.51876246 | 0.17742635 | 0.85917351 | 0.96283855 |
| RPS11 | 1051.51665 | -0.146928603 | 0.51001312 | -0.2880879 | 0.77327946 | 0.93400456 |
| FCGRT | 481.732074 | 0.695679679 | 0.40113275 | 1.73428791 | 0.08286701 | 0.51935659 |
| RCN3 | 142.86877 | -0.008969023 | 0.42641277 | -0.0210337 | 9.83E-01 | 9.95E-01 |
| NOSIP | 294.232925 | -0.17101625 | 0.19359673 | -0.8833633 | 0.37704001 | 0.77045763 |
| PRRG2 | 86.6949083 | 0.026594117 | 0.33713611 | 0.07888243 | 0.93712614 | 0.98435387 |
| PRR12 | 228.048789 | 0.278708025 | 0.38848551 | 0.71742193 | 0.47311379 | 0.81478861 |
| RRAS | 93.4145592 | 0.813542046 | 0.53584798 | 1.51823291 | 0.12895569 | 0.58793678 |
| SCAF1 | 189.861275 | -0.360867981 | 0.29963685 | -1.2043511 | 0.22845389 | 0.66705629 |
| IRF3 | 402.409 | 0.300717501 | 0.34385941 | 0.87453619 | 0.38182632 | 0.77250086 |
| PRMT1 | 294.788074 | -0.016122501 | 0.33717043 | -0.0478171 | 0.96186203 | 0.99000623 |
| AP2A1 | 547.061352 | -0.083793395 | 0.3104182 | -0.2699371 | 0.78720863 | 0.93950968 |
| FUZ | 58.5847785 | 0.47240111 | 0.55515889 | 0.85092957 | 0.39480848 | 0.77907546 |
| MED25 | 339.684911 | 0.09573436 | 0.34019657 | 0.28140895 | 0.77839675 | 0.93646866 |
| PTOV1-AS1 | 59.8555353 | 0.352303563 | 0.4123297 | 0.854422 | 0.39287121 | 0.77703588 |
| PTOV1 | 350.393033 | 0.016353042 | 0.25915233 | 0.06310204 | 0.94968525 | 0.98723887 |
| PNKP | 415.620496 | -0.693484577 | 0.43810618 | -1.5829144 | 0.11344098 | 0.56809124 |
| TBC1D17 | 267.190345 | -0.067910243 | 0.26242528 | -0.2587794 | 0.79580549 | 0.94252227 |
| IL4I1 | 55.084936 | 0.963628498 | 0.70976635 | 1.35767003 | 1.75E-01 | 0.62699479 |
| ATF5 | 149.893472 | -0.124041977 | 0.39744992 | -0.3120946 | 0.75496863 | 0.92799282 |
| VRK3 | 881.565074 | -0.048209583 | 0.284049 | -0.1697228 | 0.86522816 | 0.96428252 |
| ZNF473 | 105.393578 | -0.456820797 | 0.42421269 | -1.0768674 | 0.28153952 | 0.70625976 |
| MYH14 | 484.235899 | 0.85239516 | 0.26176901 | 3.25628756 | 0.00112879 | 0.11093704 |
| KCNC3 | 540.088443 | 0.055647179 | 0.30500582 | 0.18244629 | 0.8552325 | 0.96174189 |
| NR1H2 | 490.051257 | -0.170500967 | 0.42660865 | -0.399666 | 0.68940254 | 0.90373453 |
| NAPSA | 1075.41331 | -0.62686103 | 0.57624958 | -1.087829 | 0.27667059 | 0.70345459 |
| POLD1 | 38.8320044 | -0.446120343 | 0.41073203 | -1.0861591 | 0.2774086 | 0.7036882 |
| EMC10 | 421.502885 | 0.044554944 | 0.43088829 | 0.10340254 | 0.9176435 | 0.97828877 |
| JOSD2 | 100.310152 | -0.059101891 | 0.48323167 | -0.1223055 | 0.90265708 | 0.97515137 |
| ASPDH | 95.99244 | -0.109692818 | 0.62727552 | -0.1748718 | 8.61E-01 | 9.63E-01 |
| C19orf48 | 99.7791625 | -1.424768626 | 0.60305774 | -2.3625741 | 0.01814851 | 0.33753276 |
| CTU1 | 91.1468686 | -0.645043775 | 0.40323898 | -1.5996563 | 0.10967485 | 0.56010179 |
| ETFB | 671.404143 | 0.535278202 | 0.38806468 | 1.37935307 | 0.16778592 | 0.62304975 |
| ZNF175 | 168.177282 | -0.358037947 | 0.22730544 | -1.5751403 | 0.11522407 | 0.57182344 |
| SPACA6 | 277.51297 | -0.38272711 | 0.48441798 | -0.7900762 | 0.42948328 | 0.79037914 |
| ZNF577 | 257.596961 | 0.743953993 | 0.51974303 | 1.43138811 | 0.15231901 | 0.61001662 |
| ZNF649 | 50.4718831 | 0.457776752 | 0.53282072 | 0.85915718 | 0.3902538 | 0.77569824 |
| ZNF613 | 55.940235 | -0.357611936 | 0.34495272 | -1.0366984 | 0.29987644 | 0.71879123 |

| | | | | | | |
|---|---|---|---|---|---|---|
| ZNF350-AS1 | 44.5322718 | 0.207189846 | 0.46920565 | 0.44157576 | 0.65879622 | 0.89058983 |
| ZNF350 | 89.9509051 | -0.530465463 | 0.36089708 | -1.4698525 | 0.14160171 | 0.59611016 |
| ZNF615 | 106.617678 | 0.231023154 | 0.31230553 | 0.73973444 | 0.45946115 | 0.80669065 |
| ZNF614 | 49.7225903 | -0.250328048 | 0.36697192 | -0.682145 | 0.49514729 | 0.82216643 |
| ZNF432 | 272.380273 | -0.745985385 | 0.43438627 | -1.7173319 | 0.08591855 | 0.52364856 |
| ZNF616 | 97.0966095 | -0.011317943 | 0.29239681 | -0.0387075 | 0.96912361 | 0.99187457 |
| AC011468.3 | 95.5669164 | 0.100897172 | 0.64421413 | 0.15662055 | 0.8755439 | 0.96617876 |
| ZNF836 | 73.0103235 | -0.168262096 | 0.31211584 | -0.5391014 | 0.58981687 | 0.86281961 |
| PPP2R1A | 840.111207 | -0.067351455 | 0.30048489 | -0.2241426 | 0.82264637 | 0.95116101 |
| ZNF766 | 256.422412 | 0.153097981 | 0.33111264 | 0.46237431 | 0.64381292 | 0.88521854 |
| ZNF480 | 260.971243 | -0.635749422 | 0.36837282 | -1.7258315 | 0.08437773 | 0.52183702 |
| AC010320.4 | 52.453364 | 0.019688015 | 0.66156699 | 0.02975967 | 0.97625872 | 0.99429832 |
| ZNF610 | 42.6345239 | 0.724776474 | 0.6174185 | 1.17388201 | 0.24044229 | 0.67897455 |
| ZNF880 | 45.9908169 | -0.120703394 | 0.48563573 | -0.2485472 | 0.80371106 | 0.94543168 |
| ZNF528 | 98.6812099 | -0.764644707 | 0.45813251 | -1.669047 | 0.09510806 | 0.53824794 |
| ZNF808 | 241.130586 | 0.008611666 | 0.32274371 | 0.02668268 | 0.97871283 | 0.99482978 |
| ZNF701 | 313.750283 | -0.154501403 | 0.26447643 | -0.5841783 | 0.55910033 | 0.84936251 |
| ZNF83 | 2616.51754 | -0.761240262 | 0.46278479 | -1.644912 | 0.09998796 | 0.54542315 |
| ZNF611 | 1365.25525 | -0.932573349 | 0.40550759 | -2.2997679 | 0.02146137 | 0.35743746 |
| ZNF600 | 144.727279 | -0.505172055 | 0.35018292 | -1.4425948 | 0.14913466 | 0.60622442 |
| ZNF28 | 89.542149 | -0.115817182 | 0.30726383 | -0.3769307 | 0.70622508 | 0.90993233 |
| AC010487.3 | 62.0204976 | 0.757328254 | 0.68581689 | 1.10427181 | 0.26947525 | 0.69961625 |
| ZNF160 | 675.709866 | -0.07056422 | 0.321219 | -0.2196764 | 0.82612322 | 0.95192757 |
| ZNF347 | 116.174693 | 0.06854927 | 0.32919626 | 0.20823222 | 0.83504765 | 0.9543899 |
| ZNF677 | 44.376866 | 0.066526872 | 0.61021637 | 0.10902178 | 0.91318522 | 0.97720308 |
| ZNF845 | 62.9267075 | -0.545731749 | 0.38823658 | -1.405668 | 0.15982273 | 0.61542146 |
| ZNF525 | 138.799106 | 0.046204394 | 0.34600287 | 0.1335376 | 0.89376823 | 0.97272276 |
| ZNF765 | 272.590364 | -0.06509152 | 0.31336379 | -0.2077187 | 0.83544861 | 0.95460938 |
| ZNF761 | 338.079638 | -0.764722159 | 0.3136348 | -2.4382567 | 0.01475829 | 0.30832605 |
| ZNF813 | 77.0367431 | -0.449203416 | 0.3398584 | -1.321737 | 0.18625574 | 0.63745505 |
| ZNF331 | 242.87408 | -0.334645692 | 0.29991792 | -1.1157909 | 0.26451164 | 0.69567438 |
| NDUFA3 | 165.848844 | -0.26597408 | 0.47622505 | -0.558505 | 0.57649958 | 0.85612999 |
| TFPT | 68.3889903 | -0.25883964 | 0.34768323 | -0.7444697 | 0.45659235 | 0.80507423 |
| PRPF31 | 332.137489 | -0.426232812 | 0.23399116 | -1.8215765 | 0.06851927 | 0.49154315 |
| CNOT3 | 186.525839 | -0.459417044 | 0.29829053 | -1.5401664 | 0.1235198 | 0.58332105 |
| LENG1 | 52.407293 | -0.330019156 | 0.38868833 | -0.8490586 | 0.39584871 | 0.77955046 |
| TMC4 | 401.303563 | 0.741372854 | 0.54320482 | 1.36481273 | 0.17231192 | 0.62592399 |
| MBOAT7 | 308.409227 | -0.182434209 | 0.38737235 | -0.4709531 | 0.63767423 | 0.8827026 |
| TSEN34 | 170.759829 | 0.358520786 | 0.34320388 | 1.04462919 | 0.29619438 | 0.71755681 |
| RPS9 | 1946.93861 | 0.067832076 | 0.42772576 | 0.15858777 | 0.87399366 | 0.9661168 |
| TTYH1 | 35.7884133 | 0.666916972 | 0.63367453 | 1.05245981 | 0.29258864 | 0.71399546 |
| LENG8-AS1 | 238.583117 | 0.246651255 | 0.3553689 | 0.69407103 | 0.48763767 | 0.81893983 |
| LENG8 | 1217.04883 | -0.514093497 | 0.33566175 | -1.531582 | 0.12562562 | 0.58564339 |
| LENG9 | 47.0044025 | 0.140571534 | 0.45959669 | 0.30585845 | 0.75971242 | 0.92936655 |
| AC011476.3 | 82.5335058 | 0.253927085 | 0.60386745 | 0.42050136 | 0.67411923 | 0.89801183 |
| RDH13 | 182.245874 | -0.080659735 | 0.35808056 | -0.2252558 | 0.82178026 | 0.95105545 |
| PPP1R12C | 383.698676 | -0.453133942 | 0.35731756 | -1.2681547 | 0.20474271 | 0.65068591 |
| TMEM86B | 43.5102615 | -0.841096093 | 0.62326999 | -1.3494891 | 0.1771799 | 0.62731853 |
| PPP6R1 | 385.646175 | -0.616741177 | 0.47993715 | -1.2850457 | 0.19877631 | 0.64597062 |
| HSPBP1 | 125.702446 | -0.534526583 | 0.45318668 | -1.1794843 | 0.23820537 | 0.67642542 |
| TMEM150B | 213.271751 | -0.024505717 | 0.70332546 | -0.0348426 | 0.97220522 | 0.9924192 |
| KMT5C | 106.30931 | 1.071487706 | 0.40911922 | 2.61901095 | 0.00881851 | 0.25464675 |
| TMEM238 | 112.783731 | 0.220870101 | 0.46833911 | 0.47160294 | 0.63721023 | 0.88267497 |
| RPL28 | 1391.35314 | -0.19562408 | 0.47588606 | -0.4110734 | 0.68101875 | 0.89947163 |

| | | | | | | |
|---|---|---|---|---|---|---|
| UBE2S | 75.2042447 | -0.672621168 | 0.5193606 | -1.2950947 | 0.19528756 | 0.64558745 |
| ISOC2 | 381.995332 | 0.467898113 | 0.48992439 | 0.95504148 | 0.33955672 | 0.74644675 |
| ZNF628 | 35.9807244 | -1.1214951 | 0.4695931 | -2.3882274 | 0.01692986 | 0.32896109 |
| NAT14 | 57.1943942 | 0.504956493 | 0.56291473 | 0.89703905 | 0.36969808 | 0.76669785 |
| ZNF579 | 94.549107 | 0.119518142 | 0.41495977 | 0.28802345 | 0.77332879 | 0.93400456 |
| FIZ1 | 52.7516878 | -0.264628128 | 0.40795466 | -0.6486704 | 0.51655141 | 0.83258835 |
| ZNF524 | 74.5005703 | -0.148485741 | 0.43874956 | -0.3384294 | 0.73503964 | 0.9200245 |
| ZNF865 | 95.5069501 | -0.564678221 | 0.40527404 | -1.3933244 | 0.16352171 | 0.62065658 |
| ZNF784 | 33.9463657 | 0.087170008 | 0.48920707 | 0.17818632 | 0.85857665 | 0.9626966 |
| CCDC106 | 125.870477 | -0.139363753 | 0.39745712 | -0.3506385 | 0.7258596 | 0.91635631 |
| U2AF2 | 294.252451 | -0.655378955 | 0.23485933 | -2.790517 | 0.00526239 | 0.21873395 |
| AC008735.2 | 120.919086 | 0.183112199 | 0.58194499 | 0.31465551 | 0.75302322 | 0.92751122 |
| EPN1 | 502.044422 | -0.351779272 | 0.36614739 | -0.9607586 | 0.33667354 | 0.74293725 |
| ZNF787 | 308.60365 | -0.985830255 | 0.46469647 | -2.1214499 | 0.03388396 | 0.41378378 |
| ZNF444 | 361.225598 | 0.026009032 | 0.2990751 | 0.08696489 | 0.93069942 | 0.98272895 |
| ZSCAN5A | 748.99957 | -0.264631403 | 0.28768913 | -0.9198519 | 0.35765015 | 0.75934179 |
| EDDM13 | 72.8014849 | 0.6094729 | 0.58386385 | 1.04386133 | 0.29654955 | 0.71805012 |
| ZNF582 | 56.3795075 | -0.043463954 | 0.39141714 | -0.1110425 | 0.91158261 | 0.97716922 |
| ZNF583 | 86.102965 | -0.793009271 | 0.35416759 | -2.2390791 | 0.02515077 | 0.37991912 |
| ZNF667 | 132.593039 | -1.038781339 | 0.45539533 | -2.281054 | 0.02254525 | 0.36356209 |
| ZNF667-AS1 | 128.799203 | -0.387293806 | 0.30894601 | -1.253597 | 0.20998851 | 0.65441616 |
| ZNF471 | 166.499818 | -0.39578069 | 0.35877229 | -1.1031529 | 0.26996077 | 0.69961625 |
| ZFP28 | 58.2121559 | -0.226291752 | 0.39835683 | -0.5680629 | 0.56999223 | 0.85327114 |
| AC005498.3 | 89.102812 | -0.309877968 | 0.60080092 | -0.5157748 | 0.60601171 | 0.86925518 |
| ZNF470 | 122.764895 | -0.985140751 | 0.58321206 | -1.6891639 | 0.09118803 | 0.53537574 |
| ZNF71 | 74.1972284 | -0.488035958 | 0.27060953 | -1.8034692 | 0.07131456 | 0.49949133 |
| ZNF264 | 357.144429 | -0.161986158 | 0.18894226 | -0.8573315 | 0.39126169 | 0.77606943 |
| ZNF805 | 75.5858382 | -0.332324142 | 0.35115885 | -0.9463641 | 0.34396289 | 0.7507483 |
| AC005261.3 | 38.8375579 | -0.224145095 | 0.45927199 | -0.4880443 | 0.62551843 | 0.87854882 |
| ZNF460-AS1 | 100.126956 | 0.190986661 | 0.32178269 | 0.59352683 | 0.55282863 | 0.84721285 |
| ZNF460 | 161.350126 | -0.339445293 | 0.371868 | -0.9128113 | 0.36134182 | 0.76009788 |
| AC005261.1 | 494.972997 | -0.085649764 | 0.31469987 | -0.2721633 | 0.78549644 | 0.93892226 |
| ZNF304 | 37.0207349 | -0.194476094 | 0.38484361 | -0.505338 | 0.61332147 | 0.87222135 |
| AC003002.2 | 49.6088051 | 0.128928398 | 0.39522942 | 0.32621154 | 0.74426431 | 0.92362993 |
| AC004076.1 | 158.289281 | 0.041133511 | 0.44000518 | 0.09348415 | 0.92551894 | 0.98090838 |
| ZNF773 | 34.6911094 | -0.414900599 | 0.61693547 | -0.6725186 | 0.50125358 | 0.8260948 |
| ZNF549 | 103.09249 | -0.277176679 | 0.3668568 | -0.7555446 | 0.44992227 | 0.80321497 |
| ZNF550 | 61.6112054 | 0.203185921 | 0.32677405 | 0.62179331 | 0.53407778 | 0.83916403 |
| ZNF134 | 60.2324257 | -0.024959567 | 0.35886892 | -0.0695507 | 0.94455131 | 0.9853777 |
| ZNF211 | 45.0597555 | 0.423986775 | 0.34250164 | 1.23791168 | 0.21574881 | 0.6596384 |
| AC003006.1 | 64.9456179 | 0.174194211 | 0.49474472 | 0.35208908 | 0.72477146 | 0.91600702 |
| ZNF671 | 39.3571821 | 0.1589651 | 0.40603815 | 0.39150286 | 0.69542557 | 0.90589033 |
| ZNF776 | 80.3526196 | -0.478534172 | 0.27996498 | -1.7092644 | 0.08740199 | 0.52706515 |
| ZNF586 | 1030.97447 | -0.336857867 | 0.27356744 | -1.2313522 | 0.21819116 | 0.66064833 |
| ZNF552 | 266.409119 | 0.185769362 | 0.30009751 | 0.61903 | 0.5358966 | 0.83954482 |
| AC010522.1 | 63.8208738 | -0.384691407 | 0.31964872 | -1.2034818 | 0.22878995 | 0.66705629 |
| ZNF814 | 654.689836 | -0.270167926 | 0.28462465 | -0.9492078 | 0.34251495 | 0.75021612 |
| ZNF587 | 343.976309 | -0.572916033 | 0.28366749 | -2.0196746 | 0.04341715 | 0.44147822 |
| AC010326.2 | 37.7040099 | -0.527821867 | 0.49892663 | -1.0579148 | 0.29009428 | 0.71260257 |
| ZNF418 | 78.2041355 | -0.922983542 | 0.4412558 | -2.0917199 | 0.03646358 | 0.42151595 |
| C19orf18 | 74.6825956 | -0.120590815 | 0.39641951 | -0.3042 | 0.76097552 | 0.92989212 |
| ZNF606 | 128.750262 | 0.346036924 | 0.41813826 | 0.8275658 | 0.40791644 | 0.78481094 |
| ZNF135 | 108.661758 | 1.620309437 | 0.582936 | 2.77956658 | 0.00544315 | 0.22008628 |
| ZSCAN18 | 262.420193 | 0.746610571 | 0.45891461 | 1.6269052 | 0.10375723 | 0.55022042 |

| | | | | | | |
|---|---|---|---|---|---|---|
| ZNF329 | 282.087422 | -0.252675881 | 0.27561161 | -0.9167824 | 0.35925666 | 0.75936492 |
| ZNF274 | 472.829865 | -0.3328859 | 0.27543857 | -1.2085668 | 0.22682933 | 0.66691328 |
| AC020915.3 | 138.839059 | -0.369179403 | 0.36769059 | -1.0040491 | 0.31535495 | 0.72782796 |
| AC010642.2 | 70.2672987 | -0.076878931 | 0.41181126 | -0.1866849 | 0.85190772 | 0.96050023 |
| ZNF497 | 53.5786315 | -0.011016821 | 0.46254713 | -0.0238177 | 0.980998 | 0.9951185 |
| ZNF837 | 45.2375402 | -0.304700606 | 0.36031223 | -0.8456571 | 0.39774405 | 0.78063007 |
| RPS5 | 695.704198 | -0.082711591 | 0.5124443 | -0.161406 | 0.87177363 | 0.9657206 |
| ZNF584 | 140.587601 | 1.25686445 | 0.2861694 | 4.39202954 | 1.12E-05 | 0.01399328 |
| ZNF132 | 47.4152445 | 0.053271029 | 0.40263831 | 0.13230492 | 0.89474312 | 0.97296659 |
| ZNF324B | 35.9943394 | 0.116746173 | 0.45657878 | 0.25569776 | 0.79818422 | 0.94320409 |
| ZNF324 | 93.9243966 | -0.12610353 | 0.44846994 | -0.2811861 | 0.77856764 | 0.93646866 |
| ZNF446 | 81.0226757 | -0.031471444 | 0.37802 | -0.0832534 | 0.93365007 | 0.98355048 |
| SLC27A5 | 105.109513 | 0.255916562 | 0.32095692 | 0.79735488 | 0.42524496 | 0.78894887 |
| ZBTB45 | 46.1888536 | -0.048918682 | 0.51887001 | -0.0942793 | 0.92488732 | 0.9804664 |
| TRIM28 | 331.690391 | -0.567742595 | 0.45863016 | -1.2379094 | 0.21574965 | 0.6596384 |
| CHMP2A | 217.789136 | -0.202897109 | 0.40272452 | -0.5038112 | 0.61439408 | 0.87225883 |
| UBE2M | 218.759988 | -0.397593506 | 0.35589964 | -1.1171506 | 0.26392992 | 0.69567438 |
| MZF1-AS1 | 133.211259 | 0.084516366 | 0.3825067 | 0.22095395 | 0.82512829 | 0.95192757 |
| MZF1 | 549.839047 | -0.195758687 | 0.3459543 | -0.5658513 | 0.57149487 | 0.85399468 |
| RPS4Y1 | 832.391021 | 0.351139958 | 2.00789154 | 0.17487994 | 0.86117397 | 0.96302275 |
| ZFY | 361.696548 | 0.116927342 | 2.18761783 | 0.05344962 | 0.95737367 | 0.98861313 |
| ZFY-AS1 | 151.856156 | 2.001321519 | 1.82904894 | 1.09418697 | 0.27387298 | 0.7014484 |
| LINC00278 | 6644.04286 | 2.050633937 | 2.44055244 | 0.84023351 | 0.40077747 | 0.78232083 |
| PCDH11Y | 50.7952807 | 0.305222005 | 0.37448519 | 0.81504427 | 0.41504697 | 0.78632759 |
| TBL1Y | 327.762854 | -0.988095816 | 1.96887674 | -0.5018576 | 0.61576767 | 0.87323788 |
| TTTY15 | 437.403451 | 1.277240611 | 1.97966972 | 0.64517864 | 0.51881143 | 0.83329602 |
| USP9Y | 2036.07806 | 1.553956385 | 2.19907682 | 0.70664034 | 0.47979002 | 0.81675133 |
| DDX3Y | 354.795098 | 1.178901341 | 2.44455467 | 0.48225607 | 0.62962405 | 0.88083664 |
| UTY | 2635.23335 | 1.548194717 | 2.27723477 | 0.67985731 | 0.49659481 | 0.82252856 |
| NLGN4Y | 997.43144 | 1.990404256 | 1.85400671 | 1.07356907 | 0.28301587 | 0.70757053 |
| TTTY14 | 4874.60387 | 1.732099277 | 1.58155418 | 1.09518807 | 0.27343425 | 0.70137637 |
| KDM5D | 382.445143 | 1.132319058 | 2.30366616 | 0.49152914 | 0.62305224 | 0.87726524 |
| TTTY10 | 171.750467 | 1.583702271 | 1.53552005 | 1.03137844 | 0.30236339 | 0.71990532 |
| EIF1AY | 128.246016 | 1.292365143 | 2.51378352 | 0.51411155 | 0.607174 | 0.86985416 |
| IL17RA | 249.427543 | -0.03402985 | 0.32080192 | -0.1060775 | 0.9155209 | 0.9779223 |
| HDHD5 | 304.416604 | -0.639316109 | 0.32586994 | -1.961875 | 0.04977703 | 0.45679453 |
| HDHD5-AS1 | 92.1020516 | 0.601118671 | 0.68504442 | 0.8774886 | 0.38022131 | 0.77144243 |
| ADA2 | 48.0215196 | -0.760642676 | 1.01409566 | -0.7500699 | 0.45321258 | 0.80433814 |
| CECR3 | 46.266017 | -0.890558797 | 1.23754489 | -0.7196174 | 0.47176062 | 0.81389222 |
| CECR2 | 540.287876 | 0.133543593 | 0.97748959 | 0.13661894 | 0.891332 | 0.97234284 |
| SLC25A18 | 57.9678327 | -0.168183081 | 0.98345576 | -0.1710124 | 0.86421405 | 0.96393376 |
| ATP6V1E1 | 777.445605 | -0.361504893 | 0.27736238 | -1.3033667 | 0.19244959 | 0.64497936 |
| BCL2L13 | 1844.06508 | 0.00956645 | 0.23900605 | 0.04002598 | 0.96807242 | 0.99187457 |
| BID | 138.171882 | -0.90419975 | 0.4081833 | -2.2151807 | 0.02674767 | 0.38633327 |
| MICAL3 | 3893.81882 | -0.390600035 | 0.32369682 | -1.2066848 | 0.22755356 | 0.66705629 |
| PEX26 | 487.852527 | 0.197821816 | 0.17427575 | 1.13510811 | 0.25633003 | 0.69092557 |
| USP18 | 114.659371 | -1.595771692 | 0.99914989 | -1.5971294 | 0.11023686 | 0.56150749 |
| AC023490.3 | 38.9644206 | 1.203378088 | 0.80395368 | 1.49682516 | 0.13443876 | 0.59240843 |
| AC007326.4 | 63.5599207 | -0.192675755 | 0.87768921 | -0.2195262 | 0.82624019 | 0.95192757 |
| DGCR2 | 845.481855 | -0.141086952 | 0.31452128 | -0.4485768 | 0.65373697 | 0.8899367 |
| ESS2 | 47.6476893 | -0.460799601 | 0.37206041 | -1.2385075 | 0.21552795 | 0.6596384 |
| SLC25A1 | 385.875965 | 0.374882026 | 0.28619606 | 1.30987836 | 0.19023699 | 0.64278842 |
| CLTCL1 | 283.712503 | -0.054702425 | 0.29651445 | -0.1844849 | 0.85363311 | 0.96109041 |
| HIRA | 407.679416 | -0.270614592 | 0.22134186 | -1.2226092 | 0.22147734 | 0.66267168 |

| | | | | | | |
|---|---|---|---|---|---|---|
| MRPL40 | 145.348497 | -0.080471081 | 0.40430911 | -0.1990336 | 0.84223649 | 0.95716002 |
| UFD1 | 335.750558 | -0.756982218 | 0.35341037 | -2.1419355 | 0.03219867 | 0.40988962 |
| GNB1L | 70.9003609 | -0.09081792 | 0.54997474 | -0.1651311 | 0.86884082 | 0.96548043 |
| TXNRD2 | 278.09072 | -0.577569552 | 0.34671782 | -1.6658202 | 0.09574923 | 0.538613 |
| COMT | 570.084445 | 0.289493122 | 0.1949521 | 1.48494491 | 0.13755842 | 0.59474076 |
| ARVCF | 213.49031 | -0.34284665 | 0.23311196 | -1.4707382 | 0.14136194 | 0.59601605 |
| TANGO2 | 357.913365 | -0.497402734 | 0.40025085 | -1.2427275 | 0.21396827 | 0.65877428 |
| AC006547.1 | 58.1664503 | 0.319192581 | 0.66418739 | 0.48057609 | 0.63081781 | 0.88115748 |
| DGCR8 | 286.037099 | -0.225445399 | 0.39127456 | -0.5761821 | 0.5644921 | 0.85163263 |
| TRMT2A | 108.913416 | -0.798637852 | 0.37288643 | -2.1417724 | 0.0322118 | 0.40988962 |
| RANBP1 | 116.672372 | 0.123696324 | 0.28524392 | 0.43365105 | 0.66454186 | 0.89315615 |
| ZDHHC8 | 212.607821 | -0.29853329 | 0.45283194 | -0.6592585 | 0.50972981 | 0.82883692 |
| DGCR6L | 95.7952855 | 0.23630757 | 0.46733137 | 0.50565313 | 0.61310017 | 0.87222135 |
| ZNF74 | 39.085169 | 0.382370484 | 0.47310056 | 0.80822243 | 0.41896255 | 0.78766781 |
| KLHL22 | 634.630726 | 0.34277962 | 0.31171888 | 1.09964344 | 0.27148751 | 0.70005626 |
| MED15 | 1225.27895 | -0.41894128 | 0.25550403 | -1.639666 | 0.10107463 | 0.54608983 |
| PI4KA | 1509.85107 | -0.34208973 | 0.24739843 | -1.3827482 | 0.16674208 | 0.62225259 |
| SNAP29 | 237.714032 | -0.424926912 | 0.3364235 | -1.2630714 | 0.20656351 | 0.65251041 |
| CRKL | 375.364704 | -0.479841035 | 0.24968235 | -1.921806 | 0.05463017 | 0.46772575 |
| LZTR1 | 170.06783 | -0.20195685 | 0.24082021 | -0.8386208 | 0.40168211 | 0.78246874 |
| THAP7 | 88.4225567 | -0.567573057 | 0.40061415 | -1.4167574 | 0.15655387 | 0.61255952 |
| THAP7-AS1 | 43.5923761 | -0.666610653 | 0.58508804 | -1.1393339 | 0.25456391 | 0.68976367 |
| GGT2 | 3162.00503 | 1.458455497 | 0.71230374 | 2.04751908 | 0.04060715 | 0.43086475 |
| HIC2 | 291.779291 | -0.400124937 | 0.42562409 | -0.94009 | 0.34717141 | 0.75209642 |
| UBE2L3 | 850.98737 | -0.557806148 | 0.29865982 | -1.8676973 | 0.06180427 | 0.48267858 |
| YDJC | 85.3559426 | -0.21656824 | 0.53661789 | -0.40358 | 0.68652161 | 0.90196028 |
| SDF2L1 | 188.019609 | -0.285533742 | 0.6768324 | -0.4218677 | 0.67312157 | 0.89782405 |
| PPIL2 | 603.422606 | -0.455644298 | 0.49856352 | -0.9139142 | 0.36076192 | 0.76009788 |
| YPEL1 | 143.555601 | -0.355746125 | 0.62322978 | -0.5708105 | 0.56812808 | 0.85263558 |
| MAPK1 | 1702.2882 | -0.44929573 | 0.21549478 | -2.0849495 | 0.03707388 | 0.42215481 |
| PPM1F | 168.140125 | -0.149407017 | 0.46481086 | -0.3214362 | 0.74787988 | 0.92531168 |
| AC245452.1 | 54.1713707 | 0.152051918 | 0.52913835 | 0.28735759 | 0.77383853 | 0.93429092 |
| TOP3B | 228.936664 | -0.02723781 | 0.33046782 | -0.082422 | 0.93431116 | 0.98359701 |
| AC245060.6 | 53.8125536 | -0.427724783 | 0.62719731 | -0.6819621 | 0.49526293 | 0.8222417 |
| AC245060.5 | 420.10951 | -0.649928381 | 0.50715007 | -1.2815307 | 0.20000733 | 0.64705133 |
| ZNF280B | 51.9580755 | -0.248218314 | 0.42244708 | -0.5875726 | 0.55681923 | 0.84861484 |
| RSPH14 | 104.497397 | 1.237795593 | 0.64136646 | 1.92993503 | 0.05361489 | 0.46711025 |
| RAB36 | 118.892686 | 0.741710866 | 0.5014764 | 1.47905438 | 0.13912578 | 0.59474076 |
| BCR | 1689.29201 | 0.185826076 | 0.39319257 | 0.4726083 | 0.63649266 | 0.88266168 |
| PCAT14 | 38.0521857 | 0.777398827 | 0.6756982 | 1.1505119 | 0.24993309 | 0.68617881 |
| AP000345.2 | 166.509474 | -1.778223989 | 0.65671704 | -2.7077476 | 0.00677415 | 0.23083391 |
| DRICH1 | 419.445406 | -0.80957814 | 0.52529397 | -1.5411906 | 1.23E-01 | 0.5825794 |
| RGL4 | 62.0995547 | 0.278495807 | 0.77961585 | 0.35722184 | 0.72092574 | 0.91522915 |
| ZNF70 | 57.0296595 | -0.780802232 | 0.53563685 | -1.4577082 | 0.14492099 | 0.60109715 |
| CHCHD10 | 927.100501 | -0.098668524 | 0.50037798 | -0.197188 | 0.84368044 | 0.95731603 |
| SMARCB1 | 483.308783 | 0.152745856 | 0.24331131 | 0.6277795 | 0.5301484 | 0.83789294 |
| SLC2A11 | 66.5893955 | 0.982503646 | 0.68458499 | 1.43518141 | 0.15123541 | 0.60905477 |
| AP000350.4 | 71.8120376 | 0.461880735 | 0.35999748 | 1.28301102 | 1.99E-01 | 0.64671356 |
| AP000350.6 | 33.3236387 | 0.134627768 | 1.18796811 | 0.11332608 | 0.90977204 | 0.97690916 |
| GSTT2B | 45.9128301 | 0.561187711 | 0.64308575 | 0.87264834 | 0.38285478 | 0.77317162 |
| DDT | 663.601967 | -0.059512874 | 0.48572967 | -0.1225226 | 0.90248513 | 0.97504246 |
| CABIN1 | 1215.55154 | -0.244113124 | 0.3012053 | -0.8104543 | 0.41767913 | 0.78669082 |
| SUSD2 | 106.830368 | -1.630918633 | 1.0086632 | -1.616911 | 0.10589749 | 0.55423326 |
| ADORA2A-A! | 1030.90655 | -0.417031319 | 0.49761475 | -0.8380606 | 0.40199667 | 0.78258224 |

| | | | | | | |
|---|---|---|---|---|---|---|
| UPB1 | 1414.04757 | 0.782336793 | 0.69095735 | 1.13225048 | 0.25752914 | 0.691812 |
| GUCD1 | 576.054655 | -0.217439754 | 0.29296612 | -0.742201 | 0.45796555 | 0.80592294 |
| SNRPD3 | 409.048048 | -0.424751931 | 0.214624 | -1.9790514 | 0.04781022 | 0.45133876 |
| GGT1 | 1716.43023 | 0.179643444 | 0.32801451 | 0.54766919 | 0.58391907 | 0.86025731 |
| LRRC75B | 420.421052 | 1.551257371 | 0.60988216 | 2.54353625 | 0.01097367 | 0.27854824 |
| SGSM1 | 88.4621999 | 0.465226268 | 0.5709619 | 0.8148114 | 0.41518027 | 0.78632759 |
| KIAA1671 | 3899.49104 | -0.740388157 | 0.4779909 | -1.5489587 | 0.12139165 | 0.57985101 |
| AL022323.4 | 268.957048 | -0.391995046 | 0.66268709 | -0.5915236 | 0.55416965 | 0.84727776 |
| AL022323.1 | 60.9134659 | -0.364836859 | 0.709155 | -0.514467 | 0.60692551 | 0.86985416 |
| LRP5L | 337.59475 | -0.090697718 | 0.43626719 | -0.2078949 | 0.83531104 | 0.95453176 |
| GRK3 | 397.532887 | 0.889805897 | 0.46132839 | 1.92879067 | 0.05375685 | 0.46711025 |
| HPS4 | 451.058002 | 0.197191083 | 0.41474268 | 0.47545405 | 0.63446338 | 0.88238823 |
| SRRD | 76.7442757 | -0.48094893 | 0.26515209 | -1.8138606 | 0.06969919 | 0.49371449 |
| TFIP11 | 167.417392 | -0.226557887 | 0.36134344 | -0.6269877 | 0.53066731 | 0.83789294 |
| TPST2 | 197.675854 | -0.497787291 | 0.36471092 | -1.3648818 | 0.17229021 | 0.62592399 |
| MIATNB | 264.898689 | -0.416146806 | 0.29435518 | -1.4137574 | 0.15743315 | 0.61319532 |
| PITPNB | 1226.73382 | -0.271627965 | 0.25861799 | -1.0503057 | 0.29357756 | 0.71502415 |
| TTC28 | 7452.62446 | -0.196496417 | 0.34082293 | -0.5765352 | 0.56425348 | 0.85163263 |
| CHEK2 | 162.020196 | 0.479441661 | 0.28466082 | 1.68425588 | 0.09213223 | 0.53619162 |
| HSCB | 356.24449 | -0.008389876 | 0.29511794 | -0.0284289 | 0.97732008 | 0.99473325 |
| CCDC117 | 184.619433 | 0.210910756 | 0.22241643 | 0.94826967 | 0.34299219 | 0.7505827 |
| XBP1 | 532.982903 | 0.311798016 | 0.41076265 | 0.759071 | 0.44781008 | 0.80223612 |
| Z93930.2 | 386.362514 | 0.91140285 | 0.48594639 | 1.87552138 | 0.06072103 | 0.48026727 |
| ZNRF3 | 4279.03872 | -0.115135437 | 0.26342284 | -0.4370746 | 0.66205724 | 0.89193781 |
| ZNRF3-AS1 | 64.4741279 | 0.273441915 | 0.5269683 | 0.51889633 | 0.60383304 | 0.86831089 |
| KREMEN1 | 462.663588 | 0.869553584 | 0.44167034 | 1.96878419 | 0.04897788 | 0.45368151 |
| EMID1 | 37.2806517 | 0.517374718 | 0.75962684 | 0.68109062 | 0.49581415 | 0.82243887 |
| RHBDD3 | 215.760487 | -1.00911178 | 0.4660817 | -2.1650963 | 0.03038031 | 0.40391981 |
| EWSR1 | 1356.80949 | -0.662028424 | 0.20245358 | -3.2700258 | 0.00107538 | 0.11082848 |
| GAS2L1 | 72.7731232 | 0.223449628 | 0.50378369 | 0.4435428 | 0.65737317 | 0.89053559 |
| AP1B1 | 559.692554 | 0.342551784 | 0.19508777 | 1.75588546 | 0.07910797 | 0.51192014 |
| THOC5 | 272.815157 | -0.259873322 | 0.2569461 | -1.0113924 | 0.31182868 | 0.72604649 |
| NIPSNAP1 | 513.884972 | 1.127771489 | 0.43286568 | 2.60536129 | 0.00917774 | 0.25831482 |
| NF2 | 612.896707 | -0.582873132 | 0.39422417 | -1.4785322 | 0.13926538 | 0.59478159 |
| ZMAT5 | 106.137984 | -0.257168903 | 0.29227884 | -0.8798752 | 0.37892692 | 0.77061593 |
| UQCR10 | 713.835883 | -0.5080159 | 0.53626572 | -0.9473212 | 0.34347512 | 0.7507468 |
| ASCC2 | 425.82413 | -0.582037346 | 0.27783855 | -2.0948761 | 0.03618201 | 0.42040764 |
| MTMR3 | 1914.50804 | -0.54122208 | 0.19911037 | -2.7182013 | 0.00656379 | 0.22966139 |
| LIF | 107.388359 | -1.605759617 | 0.78502565 | -2.0454868 | 0.04080689 | 0.43095283 |
| TBC1D10A | 332.44505 | -0.043132768 | 0.37025635 | -0.1164943 | 0.90726077 | 0.97673359 |
| SF3A1 | 164.283433 | -0.146309122 | 0.23517388 | -0.6221317 | 0.53385528 | 0.83915086 |
| CCDC157 | 71.8727556 | -0.100630303 | 0.28265447 | -0.3560188 | 0.72182649 | 0.91560945 |
| RNF215 | 497.076784 | -0.777177371 | 0.40800393 | -1.9048281 | 0.05680241 | 0.4746515 |
| SEC14L2 | 119.672649 | 0.292342204 | 0.48126345 | 0.60744735 | 0.54355408 | 0.84396451 |
| SEC14L6 | 234.474434 | 0.157720846 | 0.78731533 | 0.20032742 | 0.84122452 | 0.95677175 |
| GAL3ST1 | 278.486274 | -0.05212684 | 0.74464217 | -0.0700025 | 0.94419164 | 0.9853777 |
| PES1 | 386.965779 | -0.443442012 | 0.36898993 | -1.2017727 | 0.22945162 | 0.66705629 |
| TCN2 | 506.963744 | 0.095767137 | 0.54258546 | 0.17650148 | 0.85989999 | 0.96283855 |
| SLC35E4 | 79.1374761 | 0.584346243 | 0.42547926 | 1.37338361 | 0.16963312 | 0.62326131 |
| DUSP18 | 176.491721 | 0.103620423 | 0.32540638 | 0.3184339 | 0.75015582 | 0.92567392 |
| OSBP2 | 470.223778 | 0.025084709 | 0.29202705 | 0.08589858 | 0.93154704 | 0.98281368 |
| MORC2-AS1 | 39.9987104 | -0.093055186 | 0.51937878 | -0.1791663 | 0.85780711 | 0.9626966 |
| MORC2 | 346.356958 | -0.430251784 | 0.30038814 | -1.4323195 | 0.15205241 | 0.61001662 |
| SMTN | 395.027507 | -0.127898844 | 0.71649478 | -0.1785063 | 0.85832536 | 0.9626966 |

| | | | | | | |
|---|---|---|---|---|---|---|
| SELENOM | 256.386398 | -0.455779732 | 0.49865112 | -0.9140253 | 0.36070356 | 0.76009788 |
| RNF185 | 701.14698 | -0.77049168 | 0.30157296 | -2.5549097 | 0.01062154 | 0.2726393 |
| LIMK2 | 760.124514 | 0.390892666 | 0.33169582 | 1.17846728 | 0.23861037 | 0.67658923 |
| PIK3IP1 | 103.170559 | 1.592855511 | 0.61829661 | 2.57619967 | 0.00998929 | 0.26511053 |
| PIK3IP1-AS1 | 77.6999463 | 0.72284284 | 0.40209416 | 1.79769542 | 0.07222529 | 0.50202435 |
| PATZ1 | 122.03404 | 0.763223446 | 0.38342419 | 1.99054589 | 0.04653083 | 0.44508419 |
| DRG1 | 531.88991 | -0.197014924 | 0.24817709 | -0.7938481 | 0.42728384 | 0.78933193 |
| EIF4ENIF1 | 639.257502 | -0.807643983 | 0.31272883 | -2.5825696 | 0.00980676 | 0.26385502 |
| SFI1 | 563.249512 | -1.072207604 | 0.27025639 | -3.967372 | 7.27E-05 | 0.02490203 |
| PISD | 1272.86237 | -0.938815954 | 0.30207644 | -3.1078755 | 0.00188437 | 0.13738894 |
| PRR14L | 1147.12721 | -0.349878353 | 0.3367539 | -1.0389734 | 0.29881709 | 0.71852421 |
| YWHAH | 150.601193 | -0.156958322 | 0.53323561 | -0.2943508 | 0.76848987 | 0.93198626 |
| SLC5A1 | 375.978286 | 1.516387424 | 0.95547138 | 1.58705688 | 0.11249976 | 0.56608637 |
| SLC5A4-AS1 | 63.1582758 | -1.067022072 | 0.81818654 | -1.3041306 | 0.19218907 | 0.64497936 |
| RTCB | 933.973924 | 0.379803087 | 0.23190277 | 1.63776862 | 0.10146997 | 0.54671733 |
| FBXO7 | 715.902732 | 0.081515385 | 0.2292547 | 0.35556691 | 0.72216493 | 0.9157845 |
| SYN3 | 66.7096063 | -0.017440619 | 0.69745354 | -0.0250061 | 0.98005007 | 0.99493011 |
| TIMP3 | 295.2162 | -0.556511227 | 0.52919851 | -1.0516115 | 0.29297784 | 0.71433665 |
| LARGE1 | 2557.92117 | -0.083271672 | 0.29635869 | -0.2809827 | 0.77872365 | 0.93646866 |
| HMGXB4 | 347.871691 | -0.222489182 | 0.29427297 | -0.7560639 | 0.44961087 | 0.80311564 |
| TOM1 | 765.7662 | -0.323852548 | 0.29314377 | -1.1047567 | 0.26926502 | 0.69961625 |
| HMOX1 | 2980.40684 | -0.709659725 | 0.66276059 | -1.0707633 | 0.28427587 | 0.70898277 |
| MCM5 | 35.1435331 | -0.019992014 | 0.53490615 | -0.0373748 | 0.97018616 | 0.99204339 |
| APOL6 | 664.652675 | 0.338989676 | 0.59680542 | 0.56800703 | 0.57003019 | 0.85327114 |
| RBFOX2 | 2118.06215 | -0.171824363 | 0.21710635 | -0.7914295 | 0.42869342 | 0.79037914 |
| APOL2 | 397.585998 | -0.777006183 | 0.59682816 | -1.3018926 | 0.19295309 | 0.64523251 |
| APOL1 | 88.5776332 | -1.207344337 | 0.59112654 | -2.0424465 | 0.04110726 | 0.43143739 |
| MYH9 | 1984.43665 | -1.028820566 | 0.54757168 | -1.8788783 | 0.06026111 | 0.48004006 |
| TXN2 | 216.600673 | 0.196513348 | 0.32410526 | 0.6063257 | 0.5442985 | 0.84416152 |
| FOXRED2 | 60.8858774 | -0.276621102 | 0.40056776 | -0.6905726 | 0.4898342 | 0.81970016 |
| EIF3D | 415.590316 | -0.439634052 | 0.39247495 | -1.1201582 | 0.26264634 | 0.69467275 |
| IFT27 | 150.145083 | 0.127061872 | 0.38048703 | 0.33394534 | 0.73842081 | 0.92080006 |
| TST | 483.50905 | 0.445569645 | 0.30917175 | 1.44117193 | 0.14953612 | 0.60702655 |
| MPST | 770.850801 | 0.063541316 | 0.39879938 | 0.15933153 | 0.87340768 | 0.96592775 |
| C1QTNF6 | 85.0157579 | -0.094125132 | 0.41054318 | -0.2292697 | 0.81865926 | 0.94990455 |
| ELFN2 | 149.256546 | 1.82109561 | 0.87582859 | 2.07928314 | 0.03759133 | 0.42215481 |
| CARD10 | 291.892423 | 0.673561088 | 0.246312 | 2.73458492 | 0.0062459 | 0.22644975 |
| CDC42EP1 | 130.602851 | -0.446892086 | 0.52158484 | -0.8567965 | 3.92E-01 | 7.76E-01 |
| LGALS2 | 476.710677 | 0.592928841 | 0.49453099 | 1.19897207 | 0.23053881 | 0.66865767 |
| GGA1 | 542.874547 | -0.151680584 | 0.25550724 | -0.593645 | 0.55274961 | 0.84721285 |
| LGALS1 | 84.0734237 | 0.102836069 | 0.46457964 | 0.22135294 | 0.82481763 | 0.95192757 |
| TRIOBP | 444.412934 | 0.449288161 | 0.34324 | 1.30896213 | 0.19054718 | 0.64281105 |
| H1F0 | 465.339598 | -1.841819121 | 0.47919299 | -3.8435853 | 0.00012125 | 0.03536109 |
| GCAT | 256.120938 | 0.180349107 | 0.49546127 | 0.36400243 | 0.71585619 | 0.91298263 |
| ANKRD54 | 176.892506 | -0.230451542 | 0.25161979 | -0.9158721 | 0.359734 | 0.75976485 |
| EIF3L | 596.155743 | -0.4107837 | 0.40692277 | -1.0094881 | 0.3127406 | 0.72693495 |
| MICALL1 | 155.46177 | -0.851734382 | 0.58001713 | -1.4684642 | 0.14197818 | 0.59696161 |
| POLR2F | 165.771358 | -0.42003568 | 0.28242308 | -1.4872569 | 0.13694697 | 0.59474076 |
| PICK1 | 233.616003 | -0.400168189 | 0.21090489 | -1.897387 | 0.05777688 | 0.47684824 |
| AL031587.2 | 84.4366961 | 0.222998856 | 0.58148591 | 0.3834983 | 0.70135033 | 0.90783786 |
| BAIAP2L2 | 869.119962 | 0.548441115 | 0.35636857 | 1.53897163 | 0.12381122 | 0.58354574 |
| PLA2G6 | 240.287815 | 0.57838441 | 0.35142363 | 1.64583243 | 9.98E-02 | 0.54499395 |
| MAFF | 189.39852 | -1.099641576 | 0.32238378 | -3.4109705 | 6.47E-04 | 8.46E-02 |
| TMEM184B | 382.642739 | -0.327110957 | 0.4583258 | -0.7137084 | 0.47540753 | 0.81564184 |

| | | | | | | |
|---|---|---|---|---|---|---|
| CSNK1E | 104.383445 | -0.160181169 | 0.5103141 | -0.3138874 | 0.75360656 | 0.92776047 |
| TPTEP2-CSNI | 940.629743 | 0.41721405 | 0.30019095 | 1.38982889 | 0.16458084 | 0.62065658 |
| DDX17 | 5475.03378 | 0.119881343 | 0.27885512 | 0.42990548 | 0.6672644 | 0.8944961 |
| FAM227A | 270.542965 | 0.319788746 | 0.37037835 | 0.86341102 | 0.38791152 | 0.77373447 |
| CBY1 | 291.464942 | 0.212093949 | 0.23531505 | 0.9013191 | 0.36741868 | 0.76584669 |
| AL021707.2 | 44.9736056 | 0.100645896 | 0.44127687 | 0.22807879 | 0.81958499 | 0.94990455 |
| TOMM22 | 87.6571588 | -0.52049594 | 0.50353729 | -1.033679 | 0.30128623 | 0.71879123 |
| JOSD1 | 332.963721 | -0.29728673 | 0.35154017 | -0.8456693 | 0.39773726 | 0.78063007 |
| GTPBP1 | 254.668757 | 0.091469647 | 0.25200642 | 0.36296555 | 0.71663062 | 0.91298968 |
| SUN2 | 442.686829 | 0.120069731 | 0.37124059 | 0.32342834 | 0.74637087 | 0.92448528 |
| AL021707.1 | 154.827671 | 0.020509979 | 0.33414904 | 0.06137973 | 0.95105679 | 0.98731502 |
| NPTXR | 47.7301727 | -0.17622275 | 0.78190784 | -0.2253753 | 0.82168729 | 0.95105545 |
| CBX6 | 665.962875 | 0.54322825 | 0.62126491 | 0.87439069 | 0.38190553 | 0.77252454 |
| CBX7 | 254.176675 | 0.890170776 | 0.44379714 | 2.00580559 | 0.04487699 | 0.44360789 |
| AL031846.2 | 83.0693682 | 1.085670667 | 0.55896059 | 1.94230272 | 0.05210046 | 0.46437412 |
| PDGFB | 88.230935 | -0.974168257 | 0.71704584 | -1.3585858 | 0.17427789 | 0.62699479 |
| AL031590.1 | 62.129355 | -0.066419755 | 0.46572334 | -0.1426163 | 0.8865932 | 0.97033958 |
| RPL3 | 1021.99215 | 0.137011482 | 0.57222041 | 0.2394383 | 0.81076573 | 0.94717173 |
| SYNGR1 | 99.2675698 | 0.927369464 | 0.59711187 | 1.55309167 | 0.12040124 | 0.57845767 |
| TAB1 | 259.405199 | -0.020427854 | 0.26720565 | -0.0764499 | 0.93906115 | 0.98435387 |
| MGAT3 | 48.7962277 | 1.251853139 | 0.78756686 | 1.58951983 | 0.11194308 | 0.56455091 |
| MIEF1 | 223.832204 | 0.044196952 | 0.2842721 | 0.15547411 | 0.87644756 | 0.96617876 |
| ATF4 | 722.98424 | -1.083138592 | 0.43120232 | -2.5119035 | 0.01200819 | 0.29287599 |
| RPS19BP1 | 391.812862 | -0.346772501 | 0.40073085 | -0.8653502 | 0.38684663 | 0.77334938 |
| FAM83F | 191.611625 | 0.883672707 | 0.46986089 | 1.88071134 | 0.0600112 | 0.48004006 |
| TNRC6B | 5715.59578 | 0.103317529 | 0.23401399 | 0.44150151 | 0.65884997 | 0.89058983 |
| SGSM3 | 36.3697078 | -0.448013416 | 0.47297455 | -0.9472252 | 0.34352403 | 0.7507468 |
| AL022238.2 | 53.7638551 | 0.444182246 | 0.57468338 | 0.77291646 | 0.43957182 | 0.7979355 |
| MKL1 | 2057.32805 | -0.857081105 | 0.42671266 | -2.0085673 | 0.04458304 | 0.44360789 |
| SLC25A17 | 577.500408 | -0.290475911 | 0.27482126 | -1.056963 | 0.29052847 | 0.71269438 |
| ST13 | 963.299865 | -0.119038609 | 0.18892354 | -0.6300888 | 0.52863647 | 0.83761342 |
| XPNPEP3 | 784.442027 | 0.286191322 | 0.28775555 | 0.99456404 | 0.31994834 | 0.73157402 |
| RBX1 | 465.94914 | -0.472493282 | 0.22713147 | -2.0802634 | 0.03750138 | 0.42215481 |
| EP300 | 1603.49049 | -0.189469388 | 0.17593709 | -1.0769156 | 0.281518 | 0.70625976 |
| L3MBTL2 | 157.647752 | -0.565500155 | 0.3379957 | -1.6730987 | 0.09430787 | 0.53794341 |
| AL035681.1 | 48.6086064 | 0.000616624 | 0.47554405 | 0.00129667 | 0.99896541 | 0.99977652 |
| RANGAP1 | 433.966855 | -0.852422223 | 0.4659428 | -1.8294568 | 0.0673312 | 0.48963361 |
| ZC3H7B | 566.775597 | 0.091500971 | 0.39646466 | 0.23079225 | 0.8174762 | 0.94954998 |
| TEF | 165.870957 | -0.336184269 | 0.74552407 | -0.4509368 | 0.65203507 | 0.88935488 |
| TOB2 | 577.203203 | 0.323116786 | 0.32791866 | 0.9853565 | 3.24E-01 | 0.73614611 |
| PHF5A | 97.3540586 | -0.77189863 | 0.39854907 | -1.9367719 | 0.05277323 | 0.46564998 |
| ACO2 | 1710.59311 | 0.180471922 | 0.20669501 | 0.87313147 | 3.83E-01 | 0.77279408 |
| POLR3H | 227.288548 | -0.71648048 | 0.42815998 | -1.6733943 | 0.09424969 | 0.53783536 |
| CSDC2 | 543.578376 | 0.001264217 | 0.48423632 | 0.00261074 | 0.99791693 | 0.99977652 |
| PMM1 | 292.671413 | 0.367683694 | 0.41425701 | 0.88757385 | 0.37477002 | 0.7697436 |
| DESI1 | 390.553136 | 0.136865129 | 0.19933344 | 0.68661399 | 4.92E-01 | 8.21E-01 |
| XRCC6 | 841.240947 | -0.267581216 | 0.23257973 | -1.1504924 | 0.24994111 | 0.68617881 |
| SNU13 | 533.36762 | -0.544075809 | 0.28965534 | -1.8783559 | 0.06033249 | 0.48024245 |
| C22orf46 | 61.7315912 | -0.019875704 | 0.58163024 | -0.0341724 | 0.97273967 | 0.9926729 |
| CCDC134 | 101.259853 | -0.103982474 | 0.42383439 | -0.2453375 | 0.80619511 | 0.94585363 |
| SREBF2 | 1348.16239 | 0.253503674 | 0.24531429 | 1.03338324 | 0.30142459 | 0.71891888 |
| WBP2NL | 238.282201 | 0.391529919 | 0.52952923 | 0.73939246 | 0.45966872 | 0.8069452 |
| NAGA | 69.6413912 | 0.414043042 | 0.42034394 | 0.98501014 | 0.32461909 | 0.73614611 |
| SMDT1 | 236.570459 | -0.092319413 | 0.47615851 | -0.1938838 | 0.84626688 | 0.957947 |

| | | | | | | |
|---|---|---|---|---|---|---|
| NDUFA6 | 253.003743 | -0.748929962 | 0.45207905 | -1.656635 | 0.0975933 | 0.54093104 |
| TCF20 | 2241.93291 | -0.623223241 | 0.39228872 | -1.5886851 | 0.11213151 | 0.56486093 |
| RRP7A | 211.540828 | -0.610737596 | 0.50688267 | -1.2048895 | 0.22824597 | 0.66705629 |
| SERHL2 | 58.7609658 | 0.459289801 | 0.38525926 | 1.19215772 | 0.23319941 | 0.66978724 |
| POLDIP3 | 308.72446 | -0.336058337 | 0.18835157 | -1.7842078 | 0.07438991 | 0.5071704 |
| CYB5R3 | 887.421732 | 0.473255034 | 0.29872807 | 1.58423356 | 0.11314058 | 0.56806517 |
| A4GALT | 694.512708 | 0.788216254 | 0.44947762 | 1.75362738 | 0.07949438 | 0.51192014 |
| ARFGAP3 | 1139.6147 | -0.309241031 | 0.31103458 | -0.9942336 | 3.20E-01 | 7.32E-01 |
| PACSIN2 | 2694.22762 | -0.712238105 | 0.28568423 | -2.4930956 | 0.01266347 | 0.29875272 |
| TTLL1 | 93.0457127 | 0.306366537 | 0.52193256 | 0.58698491 | 0.55721384 | 0.84861484 |
| BIK | 57.2110335 | -0.289668517 | 0.50994066 | -0.5680436 | 0.57000538 | 0.85327114 |
| MCAT | 153.263269 | -0.066758514 | 0.34973324 | -0.1908841 | 0.8486164 | 0.95951833 |
| TSPO | 421.979382 | -0.021114149 | 0.37186385 | -0.0567792 | 9.55E-01 | 9.88E-01 |
| TTLL12 | 114.618692 | -0.035184374 | 0.30775189 | -0.1143271 | 0.90897852 | 0.97680747 |
| SCUBE1 | 56.8724412 | 2.087188187 | 1.10951388 | 1.88117356 | 0.05994831 | 0.48004006 |
| EFCAB6 | 1076.12896 | 0.589990174 | 0.40298329 | 1.46405618 | 0.1431786 | 0.59866875 |
| PNPLA3 | 963.453498 | 1.813449115 | 0.67594688 | 2.68282785 | 7.30E-03 | 0.23590813 |
| SAMM50 | 265.560932 | 0.114199143 | 0.24645942 | 0.46335881 | 6.43E-01 | 0.88476704 |
| PARVB | 257.217351 | 0.546813973 | 0.51785403 | 1.05592298 | 0.29100341 | 0.71302224 |
| RTL6 | 54.4790124 | -0.113135199 | 0.47643632 | -0.2374613 | 0.81229892 | 0.94787133 |
| PRR5 | 443.361484 | -0.346808231 | 0.59603693 | -0.581857 | 0.56066304 | 0.84945842 |
| PRR5-ARHG/ | 140.379631 | -0.060897708 | 0.69907656 | -0.0871116 | 0.93058277 | 0.98272895 |
| NUP50-DT | 850.342518 | 0.091102805 | 0.40529759 | 0.22478003 | 0.8221504 | 0.95112619 |
| NUP50 | 307.705603 | -0.247206588 | 0.32278318 | -0.7658596 | 0.44375986 | 0.7993976 |
| KIAA0930 | 410.451853 | -0.954760761 | 0.60723847 | -1.5722995 | 0.1158811 | 0.57350419 |
| FAM118A | 239.276315 | -0.65605959 | 0.5933667 | -1.1056562 | 0.26887534 | 0.69961625 |
| ATXN10 | 1951.66924 | 0.067244623 | 0.17169528 | 0.39165098 | 0.69531611 | 0.90589033 |
| MIRLET7BHG | 676.020628 | 0.189601622 | 0.31409654 | 0.60364123 | 0.5460822 | 0.84460887 |
| PPARA | 4877.02453 | 0.414262174 | 0.44890641 | 0.92282526 | 0.35609827 | 0.75841231 |
| TTC38 | 1970.1304 | 1.001675288 | 0.33387382 | 3.00016122 | 2.70E-03 | 0.16909084 |
| TRMU | 450.939688 | -0.105807346 | 0.2534194 | -0.4175187 | 0.67629903 | 0.89835672 |
| CELSR1 | 1905.75459 | -0.116114966 | 0.41449582 | -0.2801354 | 0.7793736 | 0.93655387 |
| AL021392.1 | 54.807228 | 0.295184189 | 0.69061055 | 0.42742496 | 0.66906983 | 0.89582842 |
| GRAMD4 | 489.259422 | 0.149780687 | 0.27213194 | 0.55039731 | 0.5820469 | 0.85938792 |
| CERK | 362.658626 | -0.187815287 | 0.29581771 | -0.6349021 | 5.25E-01 | 0.83628497 |
| AL118516.1 | 71.4082401 | -0.999381215 | 0.67315298 | -1.4846272 | 0.13764261 | 0.59474076 |
| TBC1D22A | 3038.08852 | -0.327244339 | 0.25317616 | -1.2925559 | 0.1961647 | 0.64558745 |
| FAM19A5 | 119.873798 | -1.289876723 | 0.85855048 | -1.5023889 | 0.1329967 | 0.5919553 |
| BRD1 | 668.419984 | -0.677683315 | 0.32309917 | -2.0974468 | 0.03595404 | 0.42040764 |
| ZBED4 | 476.046751 | -0.854116184 | 0.43894957 | -1.9458185 | 0.05167655 | 0.4629611 |
| ALG12 | 121.056783 | -0.013773301 | 0.30477287 | -0.045192 | 0.96395426 | 0.99012655 |
| CRELD2 | 104.841279 | 0.419690043 | 0.54450186 | 0.77077798 | 0.44083854 | 0.79826598 |
| PIM3 | 1001.66833 | -0.538361814 | 0.4214604 | -1.2773722 | 0.20147086 | 0.6487474 |
| PANX2 | 87.8636644 | 0.439910491 | 0.59103023 | 0.74431132 | 0.45668815 | 0.80507423 |
| TRABD | 322.309708 | -0.03483062 | 0.31751065 | -0.1096991 | 0.91264805 | 0.977179 |
| SELENOO | 923.162665 | 0.001336732 | 0.22461811 | 0.00595113 | 0.99525171 | 0.99866448 |
| TUBGCP6 | 188.92692 | -0.277314687 | 0.33344627 | -0.8316623 | 0.40559959 | 0.78429302 |
| HDAC10 | 82.7317102 | -0.603971778 | 0.37712716 | -1.601507 | 0.10926468 | 0.55946616 |
| MAPK12 | 172.018601 | 0.037231788 | 0.65272449 | 0.05704059 | 0.95451286 | 0.98840182 |
| MAPK11 | 56.2894494 | 1.219687733 | 0.48831035 | 2.4977716 | 0.01249767 | 0.29875272 |
| PLXNB2 | 1032.17021 | 0.19326324 | 0.36022134 | 0.53651247 | 0.59160442 | 0.86382106 |
| DENND6B | 199.928748 | 0.689510017 | 0.49007017 | 1.40696182 | 0.15943871 | 0.6149202 |
| PPP6R2 | 2461.68764 | -0.168688228 | 0.21070476 | -0.8005905 | 0.42336876 | 0.78894887 |
| SBF1 | 303.269947 | -0.421595423 | 0.37818444 | -1.1147879 | 0.26494134 | 0.69615135 |

| | | | | | | |
|---|---|---|---|---|---|---|
| ADM2 | 122.528938 | -0.042439418 | 0.44982685 | -0.0943461 | 0.92483421 | 0.9804664 |
| MIOX | 3391.55949 | -1.226539826 | 0.83322726 | -1.4720352 | 0.14101138 | 0.59555904 |
| LMF2 | 230.095518 | -0.1728225 | 0.37818521 | -0.4569785 | 0.64768652 | 0.88774616 |
| NCAPH2 | 67.0771217 | -0.590623842 | 0.51315752 | -1.1509601 | 0.24974864 | 0.68617881 |
| TYMP | 166.734419 | 0.591024354 | 0.40205177 | 1.47002052 | 1.42E-01 | 5.96E-01 |
| ODF3B | 77.7350953 | -0.213105712 | 0.78262275 | -0.2722969 | 0.78539378 | 0.93892226 |
| ARSA | 185.900272 | 0.094877036 | 0.31316476 | 0.30296204 | 0.76191879 | 0.93016063 |
| SHANK3 | 439.064918 | 0.175094682 | 0.53483039 | 0.32738357 | 0.74337779 | 0.92337829 |
| RABL2B | 288.895481 | -0.391205398 | 0.25022401 | -1.5634207 | 0.11795368 | 0.57434852 |
| CU633967.1 | 286.274655 | 1.817068795 | 0.45792036 | 3.96808909 | 7.25E-05 | 0.02490203 |
| U2AF1L5 | 50.0191038 | -0.001627947 | 1.12477539 | -0.0014474 | 0.99884518 | 0.99977652 |
| FP671120.1 | 5835.36822 | -1.574421465 | 0.63097648 | -2.4952142 | 0.01258811 | 0.29875272 |
| FP236383.1 | 3223.40581 | -1.6016452 | 0.62408712 | -2.5663808 | 0.01027659 | 0.26767103 |
| CR381653.1 | 421.004248 | -0.423625037 | 0.46330497 | -0.9143546 | 0.36053055 | 0.76009788 |
| CR382287.2 | 34.9386066 | -0.901103165 | 0.50522345 | -1.7835735 | 0.07449299 | 0.5071704 |
| HSPA13 | 125.88605 | -0.26182823 | 0.39317634 | -0.6659308 | 0.50545534 | 0.8272145 |
| AF127577.4 | 281.374369 | 0.299714901 | 0.36915656 | 0.81189104 | 0.41685416 | 0.78637764 |
| NRIP1 | 2484.47346 | -0.525895833 | 0.29946055 | -1.7561439 | 0.07906384 | 0.51192014 |
| AF127577.2 | 71.9490381 | 0.09682162 | 0.6376161 | 0.1518494 | 0.87930572 | 0.9674841 |
| AJ009632.2 | 259.363949 | 0.413066107 | 0.77552826 | 0.53262547 | 5.94E-01 | 8.65E-01 |
| USP25 | 2619.04195 | -0.555066584 | 0.2259117 | -2.4570068 | 0.01401 | 0.30363654 |
| MIR99AHG | 4422.21542 | 0.686391889 | 0.60259964 | 1.13905127 | 2.55E-01 | 0.68976367 |
| CXADR | 386.173947 | 0.479590515 | 0.61548837 | 0.77920321 | 0.43586002 | 0.79549938 |
| BTG3 | 94.0432781 | 1.027307762 | 0.61183347 | 1.67906434 | 0.09313951 | 0.53688613 |
| C21orf91 | 139.146167 | 0.333209711 | 0.29816528 | 1.11753357 | 0.26376625 | 0.69567438 |
| CHODL | 153.111083 | 0.538124178 | 0.71841419 | 0.74904447 | 0.4538304 | 0.80433814 |
| NCAM2 | 152.147074 | 1.401377407 | 0.66527018 | 2.10647862 | 0.0351628 | 0.41984014 |
| LINC01687 | 153.561404 | -1.760519445 | 1.006089 | -1.7498645 | 0.08014169 | 0.51297858 |
| MRPL39 | 148.174476 | -0.090696762 | 0.3281937 | -0.2763513 | 0.78227825 | 0.93762574 |
| ATP5PF | 609.577852 | -0.099419395 | 0.41780704 | -0.2379553 | 0.81191577 | 0.94778824 |
| GABPA | 278.186585 | -0.297299302 | 0.20506306 | -1.4497945 | 0.14711583 | 0.60465267 |
| APP | 7190.8744 | -0.09483748 | 0.47966515 | -0.197716 | 0.84326726 | 0.95716002 |
| AP001439.1 | 135.845406 | -0.233414221 | 0.60230266 | -0.3875364 | 6.98E-01 | 0.90696517 |
| CYYR1 | 48.3995072 | 0.794215249 | 0.75991635 | 1.04513509 | 0.29596054 | 0.71727018 |
| ADAMTS1 | 1217.40443 | 0.701845701 | 0.51599647 | 1.36017539 | 1.74E-01 | 6.27E-01 |
| ADAMTS5 | 85.0637962 | 0.775867526 | 0.60605891 | 1.280185 | 0.20048008 | 0.64780303 |
| LINC00113 | 95.3586505 | 0.067862373 | 1.1200891 | 0.06058659 | 0.95168846 | 0.9875686 |
| LINC01695 | 46.7058845 | 0.867027546 | 0.67599612 | 1.28259249 | 0.19963488 | 0.64687699 |
| AF165147.1 | 222.249968 | 0.426113009 | 0.45670846 | 0.9330088 | 0.35081542 | 0.75495648 |
| N6AMT1 | 93.7018329 | 0.614032333 | 0.38793391 | 1.58282719 | 0.11346086 | 0.56809124 |
| LTN1 | 785.888734 | -0.336885443 | 0.21723376 | -1.5507969 | 0.12095037 | 0.5792686 |
| RWDD2B | 103.555716 | 0.608028745 | 0.38731806 | 1.56984353 | 0.11645152 | 0.57350419 |
| USP16 | 437.392807 | -0.5493976 | 0.273455 | -2.009097 | 0.04452685 | 0.44360789 |
| CCT8 | 363.051323 | -0.068784238 | 0.22792903 | -0.3017792 | 0.7628204 | 0.93022076 |
| MAP3K7CL | 177.119037 | 0.538571307 | 0.53652583 | 1.00381244 | 0.31546903 | 0.72784614 |
| BACH1 | 4197.68086 | -0.118288216 | 0.3976954 | -0.2974342 | 0.76613503 | 0.93130657 |
| GRIK1 | 157.235037 | 0.685395631 | 0.66527561 | 1.03024314 | 0.30289588 | 0.72043311 |
| TIAM1 | 535.574668 | 0.713861279 | 0.64220645 | 1.111576 | 0.2663205 | 0.69797523 |
| AP000253.1 | 42.7286025 | 1.511745459 | 0.76348095 | 1.98006965 | 0.0476957 | 0.45133876 |
| SOD1 | 983.627116 | 0.033138699 | 0.33561471 | 0.09874031 | 0.92134446 | 0.97939204 |
| SCAF4 | 972.048378 | 0.013014134 | 0.20428768 | 0.06370494 | 0.94920517 | 0.9871942 |
| HUNK | 698.103707 | -0.18045695 | 0.45230145 | -0.398975 | 0.68991166 | 0.90409304 |
| MIS18A | 48.2155347 | -0.871527852 | 0.50738677 | -1.7176795 | 0.08585509 | 0.52349455 |
| URB1 | 376.462584 | -0.613612301 | 0.42655724 | -1.4385228 | 0.15028578 | 0.60799621 |

| | | | | | | |
|---|---|---|---|---|---|---|
| URB1-AS1 | 37.5918699 | -0.181131667 | 0.66099822 | -0.2740275 | 0.78406352 | 0.93867623 |
| EVA1C | 719.571838 | 1.347712148 | 0.45010804 | 2.99419702 | 0.00275168 | 0.16989777 |
| SYNJ1 | 670.378772 | 0.368224527 | 0.16409397 | 2.24398578 | 0.02483332 | 0.37905375 |
| PAXBP1-AS1 | 56.8895757 | 0.834943354 | 0.67950283 | 1.22875625 | 0.21916321 | 0.66165621 |
| PAXBP1 | 1121.89013 | -0.289061062 | 0.15129696 | -1.9105543 | 0.05606188 | 0.47114668 |
| C21orf62-AS1 | 269.317691 | 0.381234099 | 0.65416293 | 0.58278157 | 0.56004035 | 0.84941263 |
| C21orf62 | 74.2356868 | 1.470051971 | 0.77959936 | 1.88565056 | 0.05934206 | 0.47855094 |
| IFNAR1 | 547.815065 | -0.100618831 | 0.27279778 | -0.3688404 | 0.71224672 | 0.91165921 |
| IFNGR2 | 989.108894 | 0.302151436 | 0.26735987 | 1.1301301 | 0.25842141 | 0.69210282 |
| TMEM50B | 611.577802 | -0.034354729 | 0.27800416 | -0.1235763 | 0.90165075 | 0.97491785 |
| DNAJC28 | 45.5343837 | -0.205290858 | 0.35647122 | -0.5758974 | 5.65E-01 | 0.85167372 |
| GART | 445.087511 | -0.118497463 | 0.29135409 | -0.4067129 | 0.68421887 | 0.9007343 |
| SON | 2053.96059 | -0.518334866 | 0.3747974 | -1.3829735 | 0.16667299 | 0.62216414 |
| DONSON | 89.3766054 | 0.127632083 | 0.36487789 | 0.34979397 | 0.72649333 | 0.91635631 |
| AP000311.1 | 619.021023 | 0.221060247 | 0.61235239 | 0.36100169 | 0.71809818 | 0.91392496 |
| ITSN1 | 2662.42287 | -0.160180076 | 0.31157981 | -0.51409 | 0.60718904 | 0.86985416 |
| LINC00649 | 82.8937839 | 0.464774108 | 0.65617633 | 0.70830673 | 0.4787548 | 0.81596289 |
| MRPS6 | 1155.87629 | -0.322005932 | 0.42535527 | -0.7570282 | 0.44903297 | 0.80300352 |
| LINC00310 | 103.89608 | 0.223069206 | 0.57252873 | 0.38962098 | 0.69681683 | 0.90645045 |
| AP000317.1 | 93.3412074 | 0.081687332 | 0.54741062 | 0.14922497 | 0.88137612 | 0.96774334 |
| KCNE1 | 76.6141215 | -0.292943537 | 0.64016431 | -0.4576068 | 0.64723496 | 0.88774616 |
| RCAN1 | 784.864956 | 0.28705736 | 0.32194693 | 0.89162945 | 0.37259158 | 0.76891189 |
| CLIC6 | 61.9578032 | 1.180110482 | 0.76417787 | 1.54428769 | 0.12251865 | 0.58149695 |
| RUNX1 | 1196.8145 | -0.721537837 | 0.58016341 | -1.2436803 | 0.21361723 | 0.65834762 |
| SETD4 | 423.947465 | 0.017627148 | 0.21722096 | 0.08114847 | 0.93532388 | 0.98399604 |
| CBR1 | 499.375599 | 0.405033163 | 0.50370039 | 0.80411524 | 0.42133042 | 0.78877309 |
| AP000688.2 | 47.2493475 | 0.768078351 | 0.61662242 | 1.24562185 | 0.21290326 | 0.65763385 |
| CBR3 | 110.948567 | -0.543914277 | 0.62491857 | -0.8703762 | 0.38409483 | 0.77317162 |
| DOPEY2 | 349.648287 | -0.322972203 | 0.54016709 | -0.5979117 | 0.54989888 | 0.84620747 |
| MORC3 | 912.933992 | -0.396802919 | 0.21560524 | -1.8404141 | 0.06570747 | 0.48790819 |
| CLDN14 | 51.4112077 | -0.2645782 | 0.64700774 | -0.4089259 | 0.68259407 | 0.89962501 |
| SIM2 | 64.9388201 | -0.68071479 | 0.98788818 | -0.6890606 | 0.49078515 | 0.81989696 |
| HLCS | 1338.01962 | 0.241256924 | 0.31255361 | 0.77188974 | 0.44017973 | 0.79826598 |
| PIGP | 119.402099 | 0.602142315 | 0.33606584 | 1.79173911 | 0.07317477 | 0.50459544 |
| TTC3 | 2490.98394 | -0.478439439 | 0.22320739 | -2.1434749 | 0.03207498 | 0.40988962 |
| VPS26C | 536.869987 | -0.084599041 | 0.28484105 | -0.2970044 | 0.76646316 | 0.93138917 |
| DYRK1A | 3128.00094 | -0.446207787 | 0.23454395 | -1.9024485 | 0.05711254 | 0.47560246 |
| KCNJ6 | 69.1299991 | 0.89363633 | 0.74542281 | 1.19883148 | 0.23059348 | 0.66865767 |
| KCNJ15 | 9530.09046 | 0.904839038 | 0.5460004 | 1.65721314 | 0.09747639 | 0.54093104 |
| ETS2 | 912.887345 | -0.407283552 | 0.42812196 | -0.951326 | 3.41E-01 | 0.74809832 |
| PSMG1 | 121.705652 | -0.643301056 | 0.24987595 | -2.5744817 | 0.01003904 | 0.26564693 |
| BRWD1 | 2756.66415 | 0.112910479 | 0.16697566 | 0.6762092 | 0.49890783 | 0.82396436 |
| HMGN1 | 638.472341 | -0.051709144 | 0.24182744 | -0.2138266 | 0.83068227 | 0.95325673 |
| WRB | 338.010826 | 0.24150461 | 0.24795792 | 0.97397415 | 0.33006937 | 0.73849818 |
| LCA5L | 146.510805 | 0.19665882 | 0.33704406 | 0.58348105 | 0.55956951 | 0.84941263 |
| SH3BGR | 93.1413608 | -0.687821326 | 0.54757688 | -1.2561183 | 0.20907309 | 0.65403148 |
| B3GALT5 | 57.0192002 | 0.023028375 | 0.98873153 | 0.02329083 | 0.98141829 | 0.99513985 |
| PCP4 | 174.562254 | 1.121138737 | 0.80203785 | 1.39786263 | 0.16215432 | 0.61881739 |
| DSCAM | 50.1330329 | -0.64733409 | 0.34978902 | -1.8506415 | 0.06422114 | 0.48593331 |
| BACE2 | 527.860152 | 0.097337592 | 0.49754446 | 0.19563597 | 8.45E-01 | 9.58E-01 |
| MX1 | 744.244405 | -4.126775609 | 1.36230469 | -3.0292604 | 0.00245153 | 0.16078068 |
| AP001610.1 | 46.6989646 | -3.826938704 | 1.47366395 | -2.596887 | 0.00940729 | 0.25998727 |
| TMPRSS2 | 345.787199 | -0.385255658 | 0.41433157 | -0.9298245 | 0.35246195 | 0.75652927 |
| RIPK4 | 405.17425 | -0.522445328 | 0.60131357 | -0.8688401 | 0.38493461 | 0.77317162 |

| | | | | | | |
|---|---|---|---|---|---|---|
| PRDM15 | 176.480465 | -0.650027932 | 0.50574737 | -1.2852819 | 0.1986938 | 0.64597062 |
| C2CD2 | 320.44051 | 0.01601332 | 0.28204721 | 0.05677532 | 0.95472418 | 0.98840182 |
| ZBTB21 | 291.049581 | -0.289392184 | 0.319747 | -0.9050661 | 3.65E-01 | 0.76379289 |
| ABCG1 | 119.422661 | -1.895769632 | 0.61564656 | -3.0793149 | 0.00207477 | 0.14584054 |
| RSPH1 | 90.760539 | 2.314426411 | 0.62721244 | 3.69001996 | 0.00022424 | 0.05581517 |
| SLC37A1 | 309.513335 | 1.335345428 | 0.48574579 | 2.74906228 | 0.0059766 | 0.22637955 |
| LINC01671 | 134.289129 | 0.077844162 | 0.67140874 | 0.11594154 | 0.90769887 | 0.97673359 |
| PDE9A | 703.77746 | 0.7647697 | 0.52848482 | 1.4470987 | 0.14786928 | 0.60605027 |
| AP001628.1 | 135.305581 | -1.573665338 | 0.91158574 | -1.7262944 | 0.08429447 | 0.52183702 |
| WDR4 | 264.545861 | -1.082154666 | 0.65452426 | -1.6533454 | 0.09826059 | 0.54193402 |
| NDUFV3 | 671.691738 | -0.218940106 | 0.23127333 | -0.9466725 | 0.34380568 | 0.7507483 |
| PKNOX1 | 617.698988 | -0.508608963 | 0.30954513 | -1.643085 | 0.10036534 | 0.54542315 |
| U2AF1 | 63.0963047 | -0.28826588 | 1.3626704 | -0.2115448 | 0.83246215 | 0.95334883 |
| RRP1B | 249.063706 | -0.814565198 | 0.44054409 | -1.8489981 | 0.06445808 | 0.48598338 |
| PDXK | 1243.64761 | -0.483471072 | 0.33726092 | -1.4335224 | 0.15170862 | 0.60961422 |
| CSTB | 748.171555 | -0.434215313 | 0.38421192 | -1.1301453 | 2.58E-01 | 6.92E-01 |
| RRP1 | 148.831489 | -0.84374943 | 0.42731084 | -1.9745566 | 0.04831849 | 0.45239176 |
| AGPAT3 | 2016.44788 | -0.0818558 | 0.35295629 | -0.2319148 | 0.81660416 | 0.94938026 |
| TRAPPC10 | 1071.22207 | -0.178109438 | 0.22530834 | -0.7905142 | 0.42922754 | 0.79037914 |
| GATD3A | 68.8780229 | 0.321092531 | 1.31493322 | 0.24418923 | 0.80708427 | 0.94585363 |
| PFKL | 1044.1163 | 0.076119475 | 0.19846366 | 0.38354365 | 0.70131671 | 0.90783786 |
| C21orf2 | 132.383891 | 0.427082837 | 0.55402392 | 0.77087437 | 0.4407814 | 0.79826598 |
| UBE2G2 | 1246.37839 | -0.214275536 | 0.20989702 | -1.0208603 | 0.30732063 | 0.72250292 |
| SUMO3 | 231.268425 | -0.787834783 | 0.43806684 | -1.7984351 | 0.07210809 | 0.50199643 |
| PTTG1IP | 1268.24859 | -0.441220185 | 0.28300182 | -1.5590719 | 0.11897936 | 0.57609025 |
| FAM207A | 136.137253 | -0.572410747 | 0.32007636 | -1.7883569 | 0.07371845 | 0.50573515 |
| ADARB1 | 448.736406 | -0.308756327 | 0.36134161 | -0.8544721 | 0.39284346 | 0.77703588 |
| POFUT2 | 289.296814 | -0.715646631 | 0.27918764 | -2.5633177 | 0.01036771 | 0.26890396 |
| COL18A1 | 1896.89333 | -0.398617378 | 0.41522114 | -0.9600122 | 0.33704906 | 0.7433518 |
| COL18A1-AS: | 40.4593242 | -0.470654699 | 0.81383783 | -0.5783151 | 0.56305141 | 0.85057939 |
| SLC19A1 | 112.115212 | 0.776325522 | 0.47526284 | 1.63346565 | 0.1023711 | 0.54919791 |
| COL6A1 | 219.427525 | 0.363385174 | 0.33386609 | 1.08841593 | 0.27641153 | 0.70328016 |
| COL6A2 | 69.1028083 | 0.560819252 | 0.70933695 | 0.79062461 | 0.42916308 | 0.79037914 |
| FTCD | 3225.73346 | -0.034693669 | 0.47740949 | -0.0726707 | 0.94206819 | 0.98489601 |
| SPATC1L | 37.4312076 | 0.033154496 | 0.48993743 | 0.06767088 | 0.94604763 | 0.98604478 |
| LSS | 344.240007 | 0.171879765 | 0.33566562 | 0.51205651 | 0.60861146 | 0.86985416 |
| MCM3AP-AS | 103.692519 | 0.263775087 | 0.34883111 | 0.75616847 | 0.4495482 | 0.80311564 |
| MCM3AP | 665.949363 | -0.163978198 | 0.138484 | -1.1840949 | 0.23637551 | 0.6735965 |
| YBEY | 140.369874 | -0.494146597 | 0.38838197 | -1.2723212 | 0.20325903 | 0.64881965 |
| C21orf58 | 44.3043918 | -0.065066143 | 0.50488829 | -0.1288724 | 0.89745865 | 0.97357005 |
| PCNT | 415.882541 | -0.689327 | 0.3959237 | -1.7410602 | 0.08167303 | 0.51593841 |
| DIP2A | 938.532113 | -0.074499958 | 0.22926836 | -0.3249465 | 0.74522154 | 0.92391025 |
| PRMT2 | 923.947498 | 0.102394976 | 0.33644725 | 0.30434184 | 0.76086747 | 0.92989212 |
| AC011043.1 | 200.662082 | 1.3867377 | 0.61729694 | 2.24646781 | 0.02467406 | 0.37746354 |
| AL592183.1 | 1308.53242 | 0.487408056 | 0.57020061 | 0.85480101 | 0.39266132 | 0.77703588 |
| AL354822.1 | 98.6324877 | -0.696449208 | 0.5330236 | -1.3066011 | 0.19134823 | 0.64395047 |
| AC240274.1 | 63.6942317 | 0.77172229 | 0.4977004 | 1.55057598 | 0.12100333 | 0.57927181 |

**Thin limbs**

*This means that positive fold changes imply higher mean value in non-COVID AKI than in COVID AKI and vice versa.

| | baseMean | log2 fold change | lfcSE | stat | pvalue | padj |
|---|---|---|---|---|---|---|
| AL669831.5 | 52.2154787 | 0.228420232 | 0.28613791 | 0.79828719 | 0.42470385 | 0.90167091 |
| NOC2L | 55.91855 | -0.454304608 | 0.31783929 | -1.4293532 | 0.15290275 | 0.72832193 |
| HES4 | 57.0154683 | -1.663534516 | 0.61463645 | -2.7065341 | 0.00679896 | 0.25593763 |
| ISG15 | 305.258589 | -5.927513644 | 1.57972888 | -3.7522348 | 0.00017527 | 0.03426256 |
| AGRN | 248.033876 | 0.073973927 | 0.54177142 | 0.13654084 | 0.89139374 | 0.987861 |
| C1orf159 | 58.7516737 | -0.16413038 | 0.367619 | -0.4464687 | 0.65525871 | 0.953185 |
| AL390719.2 | 26.314542 | -0.385268058 | 0.65341831 | -0.5896193 | 0.5554459 | 0.93523904 |
| SDF4 | 94.2080825 | 0.123256655 | 0.31443142 | 0.39199853 | 0.6950593 | 0.96220115 |
| UBE2J2 | 90.0143783 | 0.057708094 | 0.29199844 | 0.19763152 | 0.84333337 | 0.98517901 |
| INTS11 | 71.0480294 | -0.008943887 | 0.27335684 | -0.0327187 | 0.97389889 | 0.99681784 |
| DVL1 | 29.0869628 | -0.108752238 | 0.33857814 | -0.3212028 | 0.74805673 | 0.97028355 |
| AURKAIP1 | 103.585098 | -0.258161991 | 0.48498135 | -0.5323132 | 0.59450907 | 0.94660943 |
| CCNL2 | 182.325605 | 0.171500682 | 0.26468039 | 0.64795388 | 0.51701478 | 0.93230176 |
| MRPL20 | 113.341296 | -0.52646933 | 0.36761073 | -1.4321381 | 0.15210431 | 0.72632461 |
| VWA1 | 41.9851934 | 0.686800068 | 0.69906518 | 0.98245498 | 0.32587575 | 0.85509839 |
| ATAD3B | 41.2815756 | -0.34897322 | 0.30505447 | -1.1439702 | 0.25263599 | 0.80798674 |
| ATAD3A | 25.5240541 | -0.150128649 | 0.44628859 | -0.3363937 | 0.73657403 | 0.9698508 |
| SSU72 | 109.015086 | -0.093215322 | 0.31482542 | -0.2960858 | 0.76716459 | 0.97221495 |
| MIB2 | 30.654244 | 0.466418318 | 0.37524167 | 1.24298113 | 0.21387479 | 0.78135263 |
| CDK11B | 39.7338448 | -0.267813127 | 0.31603877 | -0.8474059 | 0.3967689 | 0.88863925 |
| SLC35E2B | 50.1812219 | -0.084452894 | 0.354493 | -0.2382357 | 0.81169827 | 0.97930627 |
| SLC35E2A | 50.0978225 | 0.840623039 | 0.33928488 | 2.47763188 | 0.01322575 | 0.31992293 |
| NADK | 47.6562208 | -0.368166179 | 0.29407292 | -1.2519554 | 0.21058612 | 0.77730294 |
| GNB1 | 513.707706 | -0.307975245 | 0.25722625 | -1.1972932 | 0.23119229 | 0.79432962 |
| PRKCZ | 99.1976964 | -0.171793055 | 0.28199313 | -0.6092101 | 0.5423852 | 0.93523904 |
| FAAP20 | 80.0415192 | -0.202372538 | 0.32754626 | -0.6178441 | 0.53667808 | 0.93469807 |
| SKI | 180.491474 | 0.532125136 | 0.27771787 | 1.91606369 | 0.05535699 | 0.53915335 |
| MORN1 | 60.8365544 | 0.120187885 | 0.33810734 | 0.35547257 | 0.72223559 | 0.96779196 |
| RER1 | 93.0631492 | -0.188561857 | 0.2600879 | -0.7249928 | 0.46845645 | 0.91462428 |
| PEX10 | 27.2832986 | -0.569751302 | 0.3040862 | -1.8736507 | 0.06097859 | 0.56071154 |
| TNFRSF14 | 82.7536551 | -0.077296216 | 0.42181809 | -0.1832454 | 0.85460549 | 0.9864199 |
| ARHGEF16 | 39.2844351 | -0.147027103 | 0.44729672 | -0.3287015 | 0.74238132 | 0.9698508 |
| TPRG1L | 30.4018568 | 0.065285945 | 0.33718776 | 0.19361896 | 0.84647425 | 0.98525378 |
| LRRC47 | 30.8531533 | -0.182489359 | 0.3072892 | -0.5938685 | 0.5526001 | 0.93523904 |
| CEP104 | 78.7240175 | -0.247800884 | 0.24027676 | -1.0313144 | 0.3023934 | 0.84375949 |
| AJAP1 | 49.3217402 | -0.269671875 | 0.80085903 | -0.3367283 | 0.73632175 | 0.9698508 |
| NPHP4 | 52.7220754 | 0.345714391 | 0.29786283 | 1.16064967 | 0.2457844 | 0.80488052 |
| RPL22 | 97.4727796 | 0.103384987 | 0.50650952 | 0.20411262 | 0.83826549 | 0.98462033 |
| RNF207 | 35.4724407 | 0.007651251 | 0.34396469 | 0.02224429 | 0.98225309 | 0.99681784 |
| ICMT | 42.4148785 | -0.12399533 | 0.36579364 | -0.3389762 | 0.73462768 | 0.9698508 |
| NOL9 | 66.3724511 | -0.416659706 | 0.2903264 | -1.4351423 | 0.15124655 | 0.72403332 |
| ZBTB48 | 27.2207303 | 0.513648791 | 0.30690955 | 1.67361617 | 0.09420606 | 0.6377384 |
| KLHL21 | 35.1219449 | 0.275799287 | 0.48022292 | 0.57431513 | 0.56575458 | 0.93809297 |
| DNAJC11 | 139.659154 | -0.039908503 | 0.31046885 | -0.1285427 | 0.89771951 | 0.98876109 |
| CAMTA1 | 528.805391 | 0.092505632 | 0.19054305 | 0.48548414 | 0.62733296 | 0.94857496 |
| VAMP3 | 48.0905403 | -0.056554298 | 0.27838974 | -0.2031479 | 0.83901941 | 0.98462033 |
| PER3 | 136.569854 | -0.271947252 | 0.61464287 | -0.4424476 | 0.65816536 | 0.95399905 |
| PARK7 | 186.269627 | -0.006766862 | 0.43745195 | -0.0154688 | 0.98765817 | 0.99804429 |
| ERRFI1 | 688.8972 | 0.447232889 | 0.79922913 | 0.55958032 | 0.57576573 | 0.94034284 |
| RERE | 1733.27225 | -0.075573792 | 0.17876953 | -0.4227443 | 6.72E-01 | 0.95887225 |
| ENO1 | 810.640303 | -0.292284772 | 0.41021517 | -0.7125158 | 4.76E-01 | 0.91809716 |
| MIR34AHG | 65.1425322 | 1.146287484 | 0.57559615 | 1.99147873 | 0.04642828 | 0.50827029 |

| | | | | | | |
|---|---|---|---|---|---|---|
| H6PD | 122.004374 | 0.48543898 | 0.38643272 | 1.25620569 | 0.2090414 | 0.77583638 |
| SPSB1 | 147.701827 | 0.291764787 | 0.85395643 | 0.34166238 | 0.73260499 | 0.96980299 |
| SLC25A33 | 61.5537764 | -0.879629968 | 0.57490276 | -1.53005 | 0.12600435 | 0.69492095 |
| PIK3CD | 33.2398565 | 0.291249391 | 0.32601496 | 0.89336203 | 0.37166333 | 0.87605028 |
| CLSTN1 | 236.701946 | -0.07455662 | 0.34522681 | -0.2159642 | 0.82901568 | 0.98142144 |
| CTNNBIP1 | 45.0622262 | -0.613284106 | 0.31771358 | -1.930305 | 0.05356906 | 0.53396253 |
| LZIC | 52.2020534 | -0.296044766 | 0.28614845 | -1.0345846 | 0.30086297 | 0.8426686 |
| NMNAT1 | 57.5549054 | 0.325152986 | 0.2544058 | 1.27808793 | 0.20121842 | 0.77110248 |
| UBE4B | 268.72592 | 0.156522274 | 0.22137762 | 0.70703746 | 0.4795432 | 0.91971919 |
| KIF1B | 408.840987 | -0.001131847 | 0.20195988 | -0.0056043 | 0.99552843 | 0.99866813 |
| PGD | 36.5927084 | -0.42913144 | 0.33444295 | -1.283123 | 0.19944899 | 0.77110248 |
| DFFA | 68.1828516 | -0.026597432 | 0.24716914 | -0.1076082 | 0.91430647 | 0.99159829 |
| PEX14 | 169.071488 | -0.0731834 | 0.25515332 | -0.2868213 | 0.77424917 | 0.97411309 |
| CASZ1 | 135.081393 | 0.202545945 | 0.59451372 | 0.34069179 | 0.73333562 | 0.9698508 |
| SRM | 39.0533713 | -1.050407113 | 0.53570513 | -1.9607935 | 0.04990312 | 0.52415717 |
| EXOSC10 | 51.7419569 | -0.726254185 | 0.26774294 | -2.7125055 | 0.00667767 | 0.25396993 |
| MTOR | 106.717495 | -0.088737251 | 0.26090064 | -0.340119 | 0.73376695 | 0.9698508 |
| AGTRAP | 62.4399309 | 0.289318131 | 0.53118404 | 0.54466646 | 0.58598293 | 0.9423304 |
| MTHFR | 51.100282 | 0.707523721 | 0.50661368 | 1.39657444 | 0.16254158 | 0.74071637 |
| CLCN6 | 47.7383561 | -0.101208361 | 0.28257065 | -0.3581701 | 0.72021602 | 0.96779196 |
| KIAA2013 | 42.0539237 | 0.285575673 | 0.3870649 | 0.7377979 | 0.46063727 | 0.91368087 |
| MFN2 | 53.8389851 | 0.158952148 | 0.27540592 | 0.57715588 | 0.56383416 | 0.93809297 |
| VPS13D | 357.56489 | 0.074916608 | 0.15866147 | 0.47217895 | 0.63679907 | 0.94944486 |
| DHRS3 | 114.57007 | -0.776472802 | 0.69091138 | -1.1238385 | 0.26108155 | 0.81281978 |
| PRDM2 | 211.762686 | -0.425009687 | 0.22312039 | -1.9048447 | 0.05680026 | 0.54566463 |
| KAZN | 4685.04528 | 0.339321812 | 0.39918554 | 0.85003533 | 0.39530545 | 0.88760715 |
| TMEM51-AS | 32.2407885 | -0.848420928 | 0.78465631 | -1.0812644 | 0.27957952 | 0.82428201 |
| TMEM51 | 263.448769 | -0.305198459 | 0.43754003 | -0.6975327 | 0.48546951 | 0.92034617 |
| FHAD1 | 61.7483917 | 1.242530608 | 0.47021227 | 2.64248867 | 0.00822992 | 0.27246484 |
| CASP9 | 42.1720736 | -0.139560511 | 0.52435867 | -0.2661547 | 0.79012009 | 0.97423208 |
| DNAJC16 | 123.54841 | -0.124524932 | 0.17156741 | -0.7258076 | 0.46795673 | 0.91462428 |
| DDI2 | 145.821708 | -0.058470201 | 0.17417977 | -0.3356888 | 0.73710553 | 0.9698508 |
| PLEKHM2 | 92.8596555 | -0.243227359 | 0.23299649 | -1.04391 | 0.29652705 | 0.83973489 |
| FBLIM1 | 65.6164787 | -1.149658453 | 0.34287353 | -3.3530103 | 0.00079938 | 0.09115758 |
| SPEN | 320.282587 | -0.044495834 | 0.140595 | -0.3164823 | 0.75163642 | 0.97058015 |
| EPHA2 | 66.5083576 | -0.334389928 | 0.36239861 | -0.922713 | 0.35615677 | 0.87316144 |
| FBXO42 | 258.89644 | 0.018429592 | 0.27306366 | 0.06749193 | 0.94619009 | 0.9952642 |
| SZRD1 | 83.6736509 | -0.194059342 | 0.30243557 | -0.6416552 | 0.52109711 | 0.93272217 |
| NECAP2 | 44.8584866 | 0.094744449 | 0.33879994 | 0.27964718 | 0.77974821 | 0.97423208 |
| NBPF1 | 63.0307114 | 0.203099466 | 0.2502832 | 0.81147863 | 0.41709086 | 0.89862171 |
| CROCC | 59.9099106 | 0.181849004 | 0.33836505 | 0.53743436 | 0.59096762 | 0.94527451 |
| ATP13A2 | 27.7784801 | -0.321861239 | 0.50719392 | -0.6345921 | 0.5256945 | 0.93272384 |
| SDHB | 80.9296177 | 0.148679266 | 0.27867041 | 0.53353086 | 0.59366615 | 0.94649617 |
| RCC2 | 51.3879274 | -0.33694345 | 0.27360482 | -1.2314968 | 0.21813713 | 0.78523615 |
| ARHGEF10L | 558.977623 | 0.315543957 | 0.50231859 | 0.62817496 | 0.52988933 | 0.93366189 |
| UBR4 | 261.720216 | -0.01602248 | 0.18054556 | -0.0887448 | 0.92928473 | 0.99350462 |
| AL035413.1 | 50.0126856 | 0.240568799 | 0.30843597 | 0.77996349 | 0.43541237 | 0.90514499 |
| EMC1 | 39.0455323 | -0.294186022 | 0.31740088 | -0.9268595 | 0.35399949 | 0.87241657 |
| MRTO4 | 38.9048284 | -0.715885854 | 0.45127718 | -1.5863551 | 0.11265878 | 0.66409749 |
| AKR7A2 | 37.1624927 | 0.510910403 | 0.46095553 | 1.10837243 | 0.26770101 | 0.81510106 |
| CAPZB | 283.114069 | -0.067191271 | 0.18134949 | -0.3705071 | 0.7110047 | 0.96587538 |
| NBL1 | 42.0508772 | 0.493043589 | 0.49105754 | 1.00404444 | 0.31535719 | 0.8490543 |
| TMCO4 | 160.359504 | -0.381033833 | 0.44536036 | -0.855563 | 0.39223956 | 0.88760715 |
| OTUD3 | 41.083136 | -0.089301948 | 0.31994411 | -0.2791173 | 0.78015478 | 0.97423208 |

| | | | | | | |
|---|---|---|---|---|---|---|
| CAMK2N1 | 107.855002 | -0.136471825 | 0.42295894 | -0.3226597 | 0.74695294 | 0.97028355 |
| PINK1 | 48.0149176 | 0.298603248 | 0.30291976 | 0.98575031 | 0.32425565 | 0.85448278 |
| DDOST | 51.2360372 | -0.168813291 | 0.36017004 | -0.4687044 | 0.63928093 | 0.94944486 |
| HP1BP3 | 329.046103 | -0.021992145 | 0.2202815 | -0.0998365 | 0.92047409 | 0.99214128 |
| EIF4G3 | 1004.29916 | -0.020665505 | 0.20852413 | -0.0991037 | 0.92105597 | 0.99214128 |
| ECE1 | 117.171501 | -0.044310359 | 0.28235959 | -0.1569288 | 0.87530094 | 0.98766275 |
| NBPF3 | 27.6841536 | 0.290266277 | 0.35180742 | 0.82507149 | 0.409331 | 0.8959746 |
| RAP1GAP | 125.978253 | 1.447128907 | 0.66718656 | 2.16900189 | 0.03008254 | 0.43492739 |
| USP48 | 207.50747 | 0.115656866 | 0.17945482 | 0.64449017 | 0.51925763 | 0.93259336 |
| HSPG2 | 241.633486 | -0.5495608 | 0.44028681 | -1.2481882 | 0.21196214 | 0.77732978 |
| CDC42 | 233.877957 | -0.345698092 | 0.22818009 | -1.515023 | 0.12976658 | 0.69904834 |
| ZBTB40 | 60.0207708 | 0.308521614 | 0.3584838 | 0.86062918 | 0.38944231 | 0.88586416 |
| EPHB2 | 175.101967 | -1.139595878 | 0.6592163 | -1.7287131 | 0.08386045 | 0.61656297 |
| KDM1A | 147.974466 | 0.045908787 | 0.21442637 | 0.21410047 | 0.83046871 | 0.9823695 |
| AL031428.1 | 38.9406991 | 0.327908221 | 0.38064112 | 0.86146295 | 0.38898312 | 0.88586416 |
| LUZP1 | 138.914955 | 0.575551286 | 0.33791907 | 1.70322227 | 0.08852648 | 0.62958606 |
| LINC01355 | 41.8559851 | 0.057077957 | 0.36993855 | 0.15429037 | 0.87738079 | 0.98766275 |
| HNRNPR | 297.485801 | -0.345518971 | 0.14650405 | -2.3584261 | 0.01835262 | 0.36034684 |
| TCEA3 | 83.5415436 | 0.651841383 | 0.59509978 | 1.09534804 | 0.27336419 | 0.82047446 |
| ASAP3 | 40.2871574 | 1.127729408 | 0.56262277 | 2.0044148 | 0.04502564 | 0.50340725 |
| ID3 | 103.071869 | -1.2779925 | 0.91863763 | -1.3911824 | 0.16417013 | 0.74306594 |
| RPL11 | 632.823467 | 0.018829466 | 0.63759322 | 0.0295321 | 0.97644022 | 0.99681784 |
| ELOA | 50.7478309 | 0.180957449 | 0.31398603 | 0.57632325 | 0.56439671 | 0.93809297 |
| ELOA-AS1 | 48.3682283 | 0.046677873 | 0.32702704 | 0.14273398 | 0.88650028 | 0.98766275 |
| PITHD1 | 44.1391474 | -0.31602524 | 0.26293083 | -1.201933 | 0.22938948 | 0.79370986 |
| LYPLA2 | 48.9175373 | -0.103306634 | 0.46008179 | -0.2245397 | 0.82233737 | 0.98094663 |
| GALE | 35.5043585 | -0.339149053 | 0.42124424 | -0.8051126 | 4.21E-01 | 9.00E-01 |
| HMGCL | 68.098303 | -0.295194076 | 0.2310221 | -1.2777742 | 0.20132905 | 0.77110248 |
| FUCA1 | 47.979151 | 0.393149668 | 0.45216613 | 0.86948057 | 0.38458433 | 0.88328503 |
| PNRC2 | 85.6690606 | -0.261193398 | 0.30135746 | -0.8667229 | 0.38609387 | 0.88353166 |
| SRSF10 | 210.722978 | -0.095904841 | 0.23360275 | -0.4105467 | 0.68140495 | 0.96060087 |
| IFNLR1 | 30.3786586 | 0.087154064 | 0.35824753 | 0.2432789 | 0.80778935 | 0.97930627 |
| STPG1 | 38.0320007 | 0.383446985 | 0.30624428 | 1.25209518 | 0.21053518 | 0.77730294 |
| NIPAL3 | 156.683614 | -0.051412997 | 0.51251943 | -0.1003142 | 0.92009485 | 0.99214128 |
| RCAN3 | 40.1987952 | -0.118943222 | 0.42078305 | -0.2826711 | 0.77742894 | 0.97423208 |
| SRRM1 | 192.969848 | -0.175069845 | 0.18970467 | -0.9228547 | 0.35608294 | 0.87316144 |
| CLIC4 | 879.929095 | 0.41802411 | 0.23417986 | 1.78505575 | 0.07425227 | 0.59321312 |
| SYF2 | 69.5679224 | 0.344697803 | 0.40830152 | 0.84422366 | 0.39854443 | 0.8890678 |
| RSRP1 | 383.247482 | 0.389945058 | 0.28916039 | 1.34854242 | 0.17748398 | 0.75126783 |
| TMEM50A | 128.227296 | 0.02117328 | 0.26061701 | 0.08124289 | 0.93524879 | 0.99464285 |
| MACO1 | 115.811048 | -0.103194778 | 0.20445171 | -0.5047391 | 0.61374207 | 0.94791646 |
| LDLRAP1 | 28.3535324 | 0.601288722 | 0.44709084 | 1.34489161 | 0.17866024 | 0.7513166 |
| MAN1C1 | 56.0910358 | 1.43799103 | 0.97714328 | 1.47162762 | 0.14112146 | 0.7148274 |
| MTFR1L | 36.9994463 | 0.216255741 | 0.38060854 | 0.56818415 | 0.56990993 | 0.93998433 |
| PAQR7 | 40.2037752 | -0.295691164 | 0.47845274 | -0.6180154 | 0.53656518 | 0.93469807 |
| STMN1 | 58.9064339 | -0.718621452 | 0.72753669 | -0.987746 | 0.32327706 | 0.85354768 |
| PAFAH2 | 37.9725317 | 0.277625952 | 0.25272207 | 1.09854257 | 0.27196764 | 0.81898083 |
| ZNF593 | 37.4417451 | -0.142724613 | 0.53662179 | -0.2659687 | 0.79026329 | 0.97423208 |
| SH3BGRL3 | 98.2284848 | -0.96040616 | 0.64818052 | -1.4816955 | 0.13842134 | 0.71091391 |
| UBXN11 | 41.580528 | 0.122736148 | 0.32564886 | 0.37689722 | 0.70624999 | 0.96587538 |
| DHDDS | 53.3789636 | -0.295431642 | 0.36190936 | -0.8163139 | 0.41432063 | 0.89750731 |
| HMGN2 | 138.643183 | -0.40801006 | 0.33599576 | -1.214331 | 0.22462137 | 0.78901652 |
| RPS6KA1 | 82.6313641 | -0.140988543 | 0.22564538 | -0.6248235 | 0.53208688 | 0.93366189 |
| ARID1A | 238.832712 | -0.019666699 | 0.24757959 | -0.0794359 | 0.93668594 | 0.99526299 |

| | | | | | | |
|---|---|---|---|---|---|---|
| GPN2 | 31.7608977 | -0.171330843 | 0.34985341 | -0.4897218 | 0.62433077 | 0.94808496 |
| NUDC | 100.625358 | -0.180431632 | 0.27213064 | -0.6630331 | 0.50730934 | 0.92666294 |
| TRNP1 | 38.8680188 | -0.683502601 | 0.68811651 | -0.9932949 | 0.32056628 | 0.85084632 |
| SLC9A1 | 34.94285 | -0.065553357 | 0.31182784 | -0.2102229 | 0.8334937 | 0.98351801 |
| WDTC1 | 96.7161072 | 0.304745454 | 0.22767988 | 1.33848217 | 0.18073931 | 0.75448394 |
| TMEM222 | 37.3974699 | -0.373275389 | 0.28292576 | -1.3193404 | 0.18705533 | 0.76207437 |
| WASF2 | 265.004584 | 0.139661134 | 0.24531312 | 0.56931783 | 0.56914047 | 0.93998433 |
| AHDC1 | 103.912831 | 0.438866354 | 0.43801906 | 1.00193439 | 0.31637528 | 0.8490543 |
| IFI6 | 182.616365 | -5.090919453 | 1.52818688 | -3.3313461 | 0.00086427 | 0.09407783 |
| FAM76A | 49.5930421 | 0.063171957 | 0.33233001 | 0.19008803 | 0.84924015 | 0.98572641 |
| STX12 | 145.801098 | -0.076604975 | 0.25325887 | -0.302477 | 0.76228849 | 0.9714717 |
| PPP1R8 | 37.8854955 | -0.422621546 | 0.26570828 | -1.5905472 | 0.11171152 | 0.66300165 |
| RPA2 | 47.7312989 | -0.002556546 | 0.3770183 | -0.006781 | 0.99458962 | 0.99866813 |
| EYA3 | 210.115327 | 0.000180095 | 0.24608963 | 0.00073183 | 0.99941609 | 0.99971834 |
| DNAJC8 | 77.4147257 | -0.038507525 | 0.27992012 | -0.1375661 | 0.89058333 | 0.98766275 |
| ATP5IF1 | 219.74406 | 0.258653516 | 0.39878436 | 0.64860496 | 0.51659375 | 0.9320507 |
| SESN2 | 38.6041552 | 0.727669928 | 0.24807834 | 2.93322639 | 0.00335459 | 0.18902139 |
| PHACTR4 | 365.776054 | 0.103808163 | 0.20931015 | 0.49595379 | 0.61992701 | 0.9480368 |
| RCC1 | 84.8765137 | -0.487937565 | 0.42561174 | -1.1464382 | 0.25161388 | 0.80744031 |
| TRNAU1AP | 80.7563474 | -0.015495482 | 0.26349268 | -0.058808 | 0.95310502 | 0.99531843 |
| TAF12 | 47.2918339 | 0.046041642 | 0.36382895 | 0.12654749 | 0.89929856 | 0.98956835 |
| GMEB1 | 47.2787702 | -0.612331152 | 0.3270527 | -1.8722706 | 0.06116918 | 0.56071154 |
| YTHDF2 | 122.121792 | 0.014880679 | 0.20033998 | 0.07427714 | 0.94078987 | 0.99526299 |
| EPB41 | 265.217138 | 0.482431023 | 0.24968089 | 1.93219038 | 0.05333601 | 0.53386172 |
| SRSF4 | 334.309014 | 0.141644121 | 0.27133413 | 0.52202839 | 0.60165056 | 0.94683415 |
| MECR | 47.27038 | -0.399716995 | 0.26167365 | -1.5275401 | 0.12662677 | 0.69494309 |
| PTPRU | 34.5712101 | -0.51850409 | 0.45827794 | -1.1314184 | 0.25787903 | 0.81223967 |
| PUM1 | 653.674723 | -0.033326606 | 0.13805646 | -0.2413984 | 0.80924636 | 0.97930627 |
| SNRNP40 | 62.9104188 | -0.34307633 | 0.36336037 | -0.9441765 | 0.34507945 | 0.86990996 |
| ZCCHC17 | 71.9321851 | -0.353610127 | 0.29540929 | -1.1970176 | 0.23129969 | 0.79441368 |
| SERINC2 | 148.252558 | -0.454743397 | 0.4962812 | -0.9163019 | 0.35950859 | 0.87316144 |
| TINAGL1 | 120.684174 | -0.343108108 | 0.46386871 | -0.7396664 | 0.45950243 | 0.91368087 |
| PEF1 | 42.5798954 | -0.51786716 | 0.30481671 | -1.6989461 | 0.08932934 | 0.63290368 |
| PTP4A2 | 52.9752532 | 0.246586471 | 0.43433979 | 0.5677271 | 0.57022029 | 0.93998433 |
| KHDRBS1 | 288.166095 | -0.389618772 | 0.22313623 | -1.7461027 | 0.08079311 | 0.6074703 |
| TMEM39B | 26.9837697 | -0.859939022 | 0.33057295 | -2.6013593 | 0.00928551 | 0.28622448 |
| KPNA6 | 214.812582 | -0.097013115 | 0.22676683 | -0.42781 | 0.66878947 | 0.95764829 |
| TXLNA | 63.4858878 | -0.249170055 | 0.2394553 | -1.0405702 | 0.29807506 | 0.83973489 |
| EIF3I | 65.5454085 | -0.156058766 | 0.49096452 | -0.3178616 | 0.75058992 | 0.97058015 |
| HDAC1 | 89.8106362 | 0.391800401 | 0.28391772 | 1.37997867 | 0.16759321 | 0.74445571 |
| MARCKSL1 | 38.4905563 | -0.123875623 | 0.56400057 | -0.2196374 | 0.82615356 | 0.98099974 |
| BSDC1 | 97.1925494 | 0.284612755 | 0.24459839 | 1.16359208 | 0.24458936 | 0.80476923 |
| ZBTB8OS | 132.198746 | 0.04657723 | 0.16306084 | 0.28564326 | 0.77515137 | 0.97423208 |
| RBBP4 | 164.497548 | -0.001502494 | 0.15415111 | -0.0097469 | 0.99222323 | 0.99866813 |
| SYNC | 38.4745747 | 0.634373149 | 0.44135626 | 1.43732672 | 0.15062518 | 0.72322737 |
| KIAA1522 | 151.739252 | -0.107191557 | 0.21650422 | -0.4951015 | 0.6205285 | 0.9480368 |
| YARS | 80.4625303 | -0.457320049 | 0.33753631 | -1.3548766 | 0.17545687 | 0.74934286 |
| S100PBP | 136.253703 | 0.28178115 | 0.19754422 | 1.42642061 | 0.15374697 | 0.7288378 |
| TMEM54 | 48.052993 | 1.102354997 | 0.33603146 | 3.28051124 | 0.00103619 | 0.10514713 |
| RNF19B | 77.7652461 | -0.810543836 | 0.56785835 | -1.4273698 | 0.15347334 | 0.7288378 |
| AK2 | 119.519118 | 0.087935203 | 0.23083389 | 0.3809458 | 0.70324346 | 0.9645815 |
| ZNF362 | 51.0457704 | 0.139945518 | 0.43923174 | 0.31861431 | 0.750019 | 0.97054289 |
| AL513327.1 | 44.9041214 | 0.034036959 | 0.50762567 | 0.0670513 | 0.94654087 | 0.9952642 |
| PHC2 | 246.58113 | -0.965165499 | 0.31257902 | -3.0877488 | 0.00201679 | 0.14702642 |

| | | | | | | |
|---|---|---|---|---|---|---|
| SMIM12 | 43.9424992 | -0.583138813 | 0.28121709 | -2.0736251 | 0.03811414 | 0.47153643 |
| ZMYM1 | 79.295495 | 0.373779879 | 0.23745747 | 1.57409191 | 0.1154662 | 0.67260842 |
| SFPQ | 361.384965 | -0.367313034 | 0.21973355 | -1.6716293 | 0.09459744 | 0.63858274 |
| ZMYM4 | 620.965211 | 0.154596915 | 0.13764873 | 1.12312637 | 0.26138383 | 0.81281978 |
| KIAA0319L | 305.428847 | -0.231798688 | 0.19200493 | -1.2072538 | 0.2273344 | 0.79244405 |
| PSMB2 | 86.254854 | -0.315412566 | 0.35581414 | -0.8864532 | 0.37537339 | 0.87742842 |
| AGO4 | 118.83606 | 0.343844808 | 0.25750583 | 1.33528939 | 0.18178165 | 0.75448394 |
| AGO1 | 52.6151596 | -0.288921874 | 0.28430913 | -1.0162244 | 0.30952254 | 0.84554063 |
| AGO3 | 350.542744 | -0.192149797 | 0.13676255 | -1.4049884 | 0.16002472 | 0.73816762 |
| ADPRHL2 | 43.5739511 | 0.214992304 | 0.45844853 | 0.46895625 | 0.63910091 | 0.94944486 |
| TRAPPC3 | 40.7714197 | 0.348736943 | 0.33628476 | 1.0370287 | 0.29972248 | 0.84120763 |
| MAP7D1 | 36.6983021 | -0.079136253 | 0.43487017 | -0.1819767 | 0.85560099 | 0.9864199 |
| THRAP3 | 280.776608 | -0.22726125 | 0.17101072 | -1.3289298 | 0.18387112 | 0.75689793 |
| EVA1B | 56.2370791 | 0.887287603 | 0.69762975 | 1.27186032 | 0.20342275 | 0.77155952 |
| STK40 | 71.5197207 | 0.031148727 | 0.42869971 | 0.07265861 | 0.94207778 | 0.99526299 |
| OSCP1 | 32.3555162 | 0.1432226 | 0.38273654 | 0.37420676 | 0.70825049 | 0.96587538 |
| MRPS15 | 32.4483304 | -0.226558617 | 0.40244996 | -0.5629485 | 0.57346992 | 0.93998433 |
| MEAF6 | 70.0796879 | -0.43872051 | 0.28306246 | -1.549907 | 0.12116383 | 0.69072302 |
| SNIP1 | 34.3853412 | -0.328192713 | 0.31510158 | -1.0415458 | 0.29762233 | 0.83973489 |
| DNALI1 | 43.5151118 | 0.566511226 | 0.34100473 | 1.6613002 | 0.09665317 | 0.64295596 |
| GNL2 | 27.7594564 | -0.672859616 | 0.4540955 | -1.4817579 | 0.13840471 | 0.71091391 |
| C1orf122 | 49.6950006 | -0.005350405 | 0.50271687 | -0.010643 | 0.99150829 | 0.99866813 |
| MTF1 | 54.5472862 | -0.09264237 | 0.26778594 | -0.3459568 | 0.72937518 | 0.96862399 |
| INPP5B | 80.9674806 | 0.484637281 | 0.23805576 | 2.03581413 | 0.04176902 | 0.48852927 |
| SF3A3 | 53.1899258 | 0.199278914 | 0.35344508 | 0.56381861 | 0.57287758 | 0.93998433 |
| RRAGC | 44.648645 | -0.188427111 | 0.31659617 | -0.5951655 | 0.55173287 | 0.93523904 |
| AL139260.1 | 76.4795603 | -0.453117451 | 0.46788017 | -0.9684476 | 0.33282085 | 0.86067542 |
| AKIRIN1 | 123.512197 | 0.039423879 | 0.38539797 | 0.10229395 | 0.91852336 | 0.99214128 |
| NDUFS5 | 201.33101 | -0.284926751 | 0.50736774 | -0.5615784 | 0.57440331 | 0.94034284 |
| MACF1 | 1560.1094 | 0.001949068 | 0.31985066 | 0.00609368 | 0.99513798 | 0.99866813 |
| PABPC4 | 68.5163453 | -0.619240665 | 0.26137284 | -2.3691852 | 0.01782732 | 0.35777027 |
| PPIE | 42.315566 | 0.047902533 | 0.3253077 | 0.14725299 | 0.88293233 | 0.98766275 |
| TRIT1 | 104.526439 | -0.356483461 | 0.21736126 | -1.6400506 | 0.10099465 | 0.64808588 |
| CAP1 | 155.76781 | -0.626603244 | 0.40508681 | -1.5468369 | 0.12190257 | 0.69196376 |
| PPT1 | 50.7614585 | -0.606298747 | 0.50100872 | -1.2101561 | 0.226219 | 0.79085834 |
| RLF | 247.609756 | -0.031647327 | 0.30473556 | -0.1038518 | 0.91728698 | 0.99214128 |
| ZMPSTE24 | 69.5766846 | -0.443081526 | 0.24178013 | -1.8325804 | 0.06686499 | 0.57853902 |
| SMAP2 | 46.3374308 | -0.209887032 | 0.29885986 | -0.7022925 | 0.48249678 | 0.91971919 |
| AL603839.3 | 42.0843172 | 0.373464996 | 0.33856753 | 1.10307386 | 0.26999509 | 0.81705157 |
| ZNF684 | 26.6050638 | 0.654047047 | 0.46004644 | 1.42169788 | 0.15511398 | 0.72978231 |
| NFYC | 140.190681 | 0.207964372 | 0.26353356 | 0.78913811 | 0.43003129 | 0.90364119 |
| CITED4 | 84.8032316 | -0.974692092 | 0.46885089 | -2.0788957 | 3.76E-02 | 0.46930788 |
| CTPS1 | 50.0940164 | -1.332672109 | 0.46103556 | -2.8906059 | 0.003845 | 0.19908785 |
| SLFNL1-AS1 | 36.0845288 | -0.077371064 | 0.41645422 | -0.1857853 | 0.85261315 | 0.9864199 |
| SCMH1 | 840.897399 | -0.1588881 | 0.22259132 | -0.7138109 | 0.47534414 | 0.91809716 |
| AC093151.3 | 35.0011844 | 0.816844605 | 0.81840082 | 0.99809846 | 0.31823162 | 0.84919767 |
| HIVEP3 | 159.566924 | -0.573540565 | 0.31894052 | -1.7982681 | 0.07213453 | 0.59090333 |
| FOXJ3 | 340.60575 | -0.050111352 | 0.18805011 | -0.2664787 | 0.78987055 | 0.97423208 |
| PPCS | 40.5723322 | 0.158277753 | 0.36570955 | 0.43279633 | 0.66516274 | 0.95511825 |
| CCDC30 | 332.827055 | 0.846901206 | 0.32919891 | 2.57261241 | 0.01009342 | 0.29271368 |
| PPIH | 35.37524 | -0.577931146 | 0.30790251 | -1.8769939 | 0.06051892 | 0.56071154 |
| YBX1 | 229.702122 | -0.682604809 | 0.56209134 | -1.2144019 | 0.22459428 | 0.78901652 |
| ERMAP | 36.6433142 | -0.238991709 | 0.36693951 | -0.6513109 | 0.5148458 | 0.93113905 |
| SLC2A1 | 52.1225598 | -0.964424941 | 0.80336345 | -1.200484 | 0.22995144 | 0.79370986 |

| | | | | | | |
|---|---|---|---|---|---|---|
| EBNA1BP2 | 59.7831549 | 0.075508289 | 0.30759493 | 0.24547963 | 0.80608508 | 0.97930627 |
| ELOVL1 | 34.8369559 | -0.102368909 | 0.39800388 | -0.2572058 | 0.7970199 | 0.97534702 |
| SZT2 | 68.0968502 | -0.281770383 | 0.26277384 | -1.0722924 | 0.28358873 | 0.82918183 |
| PTPRF | 272.249249 | 0.218526525 | 0.28712051 | 0.76109689 | 0.4465992 | 0.90827467 |
| ATP6V0B | 157.652154 | 0.192938414 | 0.32713461 | 0.58978295 | 0.55533618 | 0.93523904 |
| B4GALT2 | 34.2916159 | -0.582168015 | 0.39062095 | -1.4903656 | 0.13612814 | 0.70752658 |
| CCDC24 | 45.828969 | 0.012657365 | 0.43670429 | 0.02898383 | 0.97687748 | 0.99681784 |
| DMAP1 | 27.2130501 | -0.650870625 | 0.45021176 | -1.4456989 | 0.14826169 | 0.72246729 |
| ERI3 | 221.534343 | 0.17110864 | 0.23386685 | 0.73164984 | 0.46438232 | 0.91462428 |
| RNF220 | 50.8315638 | 0.101833283 | 0.30910699 | 0.32944348 | 0.74182051 | 0.9698508 |
| ARMH1 | 38.6277048 | 1.49547714 | 0.45626702 | 3.27763582 | 0.0010468 | 0.10514713 |
| RPS8 | 537.876307 | 0.264928678 | 0.50159417 | 0.52817336 | 0.59737902 | 0.94683415 |
| EIF2B3 | 92.2996744 | -0.267982752 | 0.31431734 | -0.8525866 | 0.3938886 | 0.88760715 |
| UROD | 24.8422876 | -0.246212919 | 0.35094944 | -0.7015624 | 0.48295213 | 0.91971919 |
| ZSWIM5 | 239.017759 | 0.196451673 | 0.34150512 | 0.5752525 | 0.56512054 | 0.93809297 |
| TESK2 | 68.2700951 | 0.638061622 | 0.30873441 | 2.06670069 | 0.03876237 | 0.47547104 |
| PRDX1 | 481.051176 | -0.338626144 | 0.55888004 | -0.6059013 | 0.54458029 | 0.93523904 |
| AKR1A1 | 70.486806 | -0.049634422 | 0.33610797 | -0.147674 | 0.88260002 | 0.98766275 |
| NASP | 114.171192 | -0.252044585 | 0.28535953 | -0.8832527 | 0.37709974 | 0.87842274 |
| GPBP1L1 | 334.882771 | 0.041956054 | 0.18204376 | 0.23047235 | 0.81772475 | 0.98010849 |
| IPP | 70.2513553 | 0.020515486 | 0.3274692 | 0.0626486 | 0.95004633 | 0.99531843 |
| MAST2 | 104.634049 | 0.155042237 | 0.26848465 | 0.57747152 | 0.56362097 | 0.93809297 |
| TSPAN1 | 92.2676015 | 1.482980934 | 0.42753844 | 3.46864933 | 0.00052308 | 0.07057183 |
| POMGNT1 | 50.8320882 | 0.203594801 | 0.25515349 | 0.79793068 | 0.42491072 | 0.90167091 |
| LURAP1 | 28.8574237 | 0.870510595 | 0.52925851 | 1.64477391 | 0.10001644 | 0.64666342 |
| LRRC41 | 94.3747275 | -0.153107972 | 0.25759038 | -0.5943854 | 0.55225435 | 0.93523904 |
| UQCRH | 152.366579 | 0.374454295 | 0.42330935 | 0.88458783 | 0.37637898 | 0.87842274 |
| NSUN4 | 61.9846814 | -0.330965938 | 0.21474483 | -1.5412056 | 0.12326676 | 0.69314363 |
| FAAH | 28.4650849 | 0.664687793 | 0.52641385 | 1.26267156 | 0.20670724 | 0.77401635 |
| MKNK1 | 57.1740307 | 0.029009447 | 0.26695756 | 0.10866689 | 0.91346671 | 0.99140013 |
| ATPAF1 | 67.7311263 | 0.401054935 | 0.24242518 | 1.65434524 | 0.09805739 | 0.64312419 |
| EFCAB14 | 126.309826 | -0.266557753 | 0.22551418 | -1.182 | 0.23720572 | 0.80063199 |
| PDZK1IP1 | 642.444455 | 1.439708453 | 0.63599907 | 2.26369583 | 0.02359283 | 0.40141334 |
| CMPK1 | 153.986984 | 0.3059189 | 0.26021948 | 1.17561874 | 0.23974726 | 0.80193227 |
| TRABD2B | 1274.79463 | -0.698872079 | 0.42304861 | -1.65199 | 0.09853659 | 0.64312419 |
| SPATA6 | 286.633888 | 0.440067346 | 0.24413206 | 1.80257909 | 0.07145434 | 0.59090333 |
| AGBL4 | 678.062899 | 1.064894246 | 0.52326178 | 2.03510803 | 0.04184 | 0.48876268 |
| FAF1 | 786.789592 | -0.208532917 | 0.15982475 | -1.3047598 | 0.19197464 | 0.7646258 |
| RNF11 | 151.126515 | 0.09637645 | 0.19496286 | 0.49433236 | 0.62107147 | 0.9480368 |
| TTC39A | 95.0368541 | -0.127174848 | 0.41634798 | -0.3054533 | 7.60E-01 | 9.71E-01 |
| EPS15 | 417.600548 | -0.182189808 | 0.18153391 | -1.0036131 | 0.31556514 | 0.8490543 |
| OSBPL9 | 370.267157 | 0.356966296 | 0.17703082 | 2.01640765 | 0.04375737 | 0.49836909 |
| NRDC | 314.08265 | -0.023314854 | 0.17514156 | -0.1331201 | 0.89409844 | 0.98842187 |
| RAB3B | 35.8329903 | -0.878807058 | 0.54620583 | -1.6089302 | 0.1076316 | 0.65699665 |
| TXNDC12 | 70.5235281 | -0.124192797 | 0.27165639 | -0.4571687 | 0.6475498 | 0.95027667 |
| BTF3L4 | 129.185802 | -0.353255769 | 0.18423901 | -1.9173777 | 0.05518997 | 0.53849427 |
| ZFYVE9 | 224.835795 | 0.012202291 | 0.24240659 | 0.05033812 | 0.95985295 | 0.99572129 |
| CC2D1B | 47.4666625 | -0.288572805 | 0.41857904 | -0.6894105 | 0.49056495 | 0.92175788 |
| PRPF38A | 47.061888 | -0.100946755 | 0.33834826 | -0.2983516 | 0.7654348 | 0.97177982 |
| TUT4 | 576.873357 | 0.099384913 | 0.16913397 | 0.5876106 | 0.55679369 | 0.93533595 |
| ZYG11B | 172.507192 | 0.137923441 | 0.26412671 | 0.52218665 | 0.60154038 | 0.94683415 |
| ECHDC2 | 308.216912 | 0.628127643 | 0.30588292 | 2.05349038 | 0.04002504 | 0.482264 |
| SCP2 | 201.67368 | 0.228825671 | 0.17216453 | 1.3291104 | 0.18381155 | 0.75689793 |
| CPT2 | 25.7152376 | 0.635607798 | 0.32477835 | 1.95705101 | 0.05034147 | 0.52415717 |

| | | | | | | |
|---|---|---|---|---|---|---|
| C1orf123 | 26.2131279 | -0.036709487 | 0.41688932 | -0.0880557 | 0.92983239 | 0.99362798 |
| MAGOH | 44.387942 | -0.358991439 | 0.29619567 | -1.2120077 | 0.22550942 | 0.78977225 |
| GLIS1 | 87.147892 | 0.336096165 | 0.44826638 | 0.74976884 | 0.45339394 | 0.91077355 |
| NDC1 | 59.1370707 | -0.723609429 | 0.32560634 | -2.2223444 | 0.02626004 | 0.42051885 |
| YIPF1 | 39.234008 | 0.204461687 | 0.26168193 | 0.78133667 | 0.43460451 | 0.90514499 |
| HSPB11 | 82.6362223 | 0.002127106 | 0.36848527 | 0.00577257 | 0.99539418 | 0.99866813 |
| LRRC42 | 47.6603854 | -0.245090337 | 0.26400322 | -0.9283612 | 0.35322026 | 0.8720098 |
| TMEM59 | 287.559064 | 0.498981957 | 0.35686993 | 1.39821797 | 0.16204762 | 0.74071637 |
| TCEANC2 | 30.6066357 | 0.191458311 | 0.39778641 | 0.48130933 | 0.63029667 | 0.94916079 |
| MRPL37 | 25.0506842 | -0.146610172 | 0.3599157 | -0.4073459 | 0.68375398 | 0.96161098 |
| SSBP3 | 270.840316 | 0.073820651 | 0.50875554 | 0.14510044 | 0.88463157 | 0.98766275 |
| ACOT11 | 151.233569 | 0.347547167 | 0.4945603 | 0.70273972 | 0.48221797 | 0.91971919 |
| DHCR24 | 108.105885 | -0.533088414 | 0.33031233 | -1.613892 | 0.10655086 | 0.65699665 |
| USP24 | 381.561784 | -0.222563026 | 0.13661356 | -1.629143 | 0.10328275 | 0.65288882 |
| PLPP3 | 161.856305 | 0.334911485 | 0.61876488 | 0.54125807 | 0.58832971 | 0.94444601 |
| PRKAA2 | 383.951175 | -0.05890501 | 0.16493761 | -0.3571351 | 7.21E-01 | 9.68E-01 |
| FYB2 | 73.3070328 | 1.364702616 | 0.84529689 | 1.61446543 | 0.10642651 | 0.65699665 |
| DAB1 | 296.628484 | -0.590100246 | 1.01188805 | -0.5831675 | 0.55978053 | 0.93744835 |
| TACSTD2 | 340.331377 | -1.566904427 | 0.79736473 | -1.9651038 | 0.04940223 | 0.52308378 |
| MYSM1 | 170.414229 | -0.147090361 | 0.24254769 | -0.6064389 | 0.54422332 | 0.93523904 |
| AL136985.3 | 48.5729415 | -0.452079748 | 0.81937131 | -0.5517398 | 0.58112665 | 0.94046497 |
| JUN | 287.830035 | 0.526080174 | 0.33023121 | 1.5930662 | 0.11114534 | 0.66251476 |
| LINC01135 | 61.7535965 | 0.644892479 | 0.50774081 | 1.27012142 | 0.20404138 | 0.77155952 |
| FGGY | 355.068937 | -0.125561148 | 0.38060241 | -0.3299011 | 0.7414747 | 0.9698508 |
| HOOK1 | 231.908085 | 0.424436806 | 0.25667199 | 1.65361561 | 0.09820564 | 0.64312419 |
| NFIA | 1121.83967 | 0.477685114 | 0.30176408 | 1.58297539 | 0.11342707 | 0.66657543 |
| TM2D1 | 151.589241 | -0.159798228 | 0.16989836 | -0.9405519 | 0.34693453 | 0.87015462 |
| PATJ | 1727.61455 | -0.118127305 | 0.24684693 | -0.4785448 | 0.63226252 | 0.94932781 |
| USP1 | 43.5607379 | -0.281551087 | 0.28057812 | -1.0034677 | 0.31563525 | 0.8490543 |
| DOCK7 | 238.719559 | 0.095156983 | 0.24725673 | 0.38485093 | 0.70034786 | 0.96360703 |
| ATG4C | 70.312279 | 0.101204371 | 0.33385903 | 0.30313504 | 0.76178695 | 0.9714717 |
| ALG6 | 37.3408644 | -0.366343782 | 0.24951089 | -1.4682477 | 0.14203696 | 0.7148274 |
| ITGB3BP | 119.88349 | 0.628413236 | 0.45028747 | 1.39558232 | 0.16284031 | 0.74167981 |
| EFCAB7 | 70.6677 | 0.539675804 | 0.38756839 | 1.392466 | 0.16378133 | 0.74283209 |
| PGM1 | 79.5944965 | 0.58167145 | 0.37534465 | 1.5496996 | 0.12121363 | 0.69072302 |
| ROR1 | 161.366784 | -0.117995408 | 0.64819267 | -0.1820376 | 0.85555325 | 0.9864199 |
| CACHD1 | 84.5697007 | 0.467423415 | 0.68567234 | 0.68170084 | 0.49542813 | 0.92310953 |
| RAVER2 | 139.383949 | -0.4052838 | 0.2784332 | -1.4555872 | 0.14550678 | 0.71957119 |
| JAK1 | 561.086845 | -0.041663946 | 0.29903366 | -0.1393286 | 0.88919048 | 0.98766275 |
| AK4 | 187.46412 | 0.758671593 | 0.42827351 | 1.77146513 | 0.07648339 | 0.59726395 |
| DNAJC6 | 188.125981 | 0.469332444 | 0.36880121 | 1.27258921 | 0.20316385 | 0.77155952 |
| LEPR | 310.318668 | 0.425923312 | 0.35148031 | 1.21179849 | 0.22558952 | 0.78977225 |
| PDE4B | 43.288202 | -1.029835741 | 0.63426796 | -1.6236604 | 0.10444832 | 0.65551333 |
| SGIP1 | 591.888444 | -1.153470106 | 0.72305436 | -1.5952744 | 0.11065089 | 0.66038931 |
| WDR78 | 239.977455 | 0.939577772 | 0.31854171 | 2.94962238 | 0.00318163 | 0.18249574 |
| MIER1 | 224.040024 | -0.186648422 | 0.22643699 | -0.8242841 | 0.40977813 | 0.8959746 |
| SLC35D1 | 62.9448272 | 0.087875069 | 0.24651916 | 0.35646345 | 0.72149352 | 0.96779196 |
| SERBP1 | 243.377981 | -0.268329988 | 0.22335477 | -1.2013622 | 0.22961071 | 0.79370986 |
| GADD45A | 90.2397422 | 0.461677476 | 0.49660519 | 0.92966704 | 0.3525435 | 0.8720098 |
| GNG12 | 375.540838 | 0.343529435 | 0.3711265 | 0.92563974 | 0.35463323 | 0.87258503 |
| GNG12-AS1 | 127.451841 | 0.456645599 | 0.39537818 | 1.15495904 | 0.2481072 | 0.80672756 |
| WLS | 174.081886 | 0.311009484 | 0.46732618 | 0.66550837 | 0.50572539 | 0.9257297 |
| LRRC40 | 69.6071607 | -0.032990219 | 0.23104543 | -0.1427867 | 0.88645863 | 0.98766275 |
| SRSF11 | 423.851525 | 0.199471732 | 0.19148896 | 1.04168788 | 0.29755641 | 0.83973489 |

| | | | | | | |
|---|---|---|---|---|---|---|
| ANKRD13C | 201.920311 | -0.100250956 | 0.17436041 | -0.574964 | 0.56531566 | 0.93809297 |
| ZRANB2 | 356.111548 | -0.027731869 | 0.21387769 | -0.1296623 | 0.89683363 | 0.98842187 |
| NEGR1 | 62.1489493 | -0.098261161 | 0.49229047 | -0.1996 | 0.84179345 | 0.9851606 |
| CRYZ | 176.792058 | -0.047095171 | 0.29598604 | -0.1591128 | 0.87357999 | 0.98766275 |
| TYW3 | 56.9921571 | -0.632941254 | 0.30723712 | -2.0601067 | 0.03938834 | 0.479886 |
| SLC44A5 | 831.684975 | -0.53466488 | 0.69165563 | -0.7730218 | 0.43950948 | 0.90561284 |
| ACADM | 121.431057 | 0.180086363 | 0.23869209 | 0.75447144 | 0.45056619 | 0.91054293 |
| RABGGTB | 57.2005005 | -0.222777733 | 0.46853735 | -0.4754749 | 0.63444854 | 0.94932781 |
| ST6GALNAC3 | 49.7162088 | 0.432932953 | 0.7109849 | 0.60892004 | 0.54257744 | 0.93523904 |
| PIGK | 93.1472474 | 0.004086613 | 0.23273667 | 0.01755896 | 0.9859907 | 0.99764442 |
| AC118549.1 | 507.637521 | -0.205062548 | 0.19383154 | -1.0579421 | 0.29008183 | 0.83506029 |
| USP33 | 312.641294 | 0.123083691 | 0.17071764 | 0.72097815 | 0.47092296 | 0.91611922 |
| MIGA1 | 205.64314 | -0.084124531 | 0.17014217 | -0.4944367 | 0.6209978 | 0.9480368 |
| FUBP1 | 261.878485 | -0.282555838 | 0.20772541 | -1.3602373 | 0.17375486 | 0.74910274 |
| GIPC2 | 102.617484 | 1.486146398 | 0.35170705 | 4.22552347 | 2.38E-05 | 0.00945023 |
| PTGFR | 202.646101 | 0.934986475 | 1.13885201 | 0.82099032 | 0.41165178 | 0.89695293 |
| IFI44L | 244.089937 | -4.027810066 | 1.42733584 | -2.8219078 | 0.00477389 | 0.21962686 |
| IFI44 | 91.3519066 | -3.092277428 | 1.1689356 | -2.6453788 | 0.00815995 | 0.27246484 |
| AL136234.1 | 60.8622399 | 1.901542843 | 0.35454519 | 5.36333001 | 8.17E-08 | 0.00019565 |
| LINC01781 | 460.558568 | 2.264079555 | 0.48376155 | 4.68015609 | 2.87E-06 | 0.00200368 |
| ADGRL2 | 336.999929 | 0.729839111 | 0.37800095 | 1.93078646 | 0.05350946 | 0.53392413 |
| LINC01725 | 207.645142 | -0.17628578 | 0.43775906 | -0.4027005 | 0.68716859 | 0.96161098 |
| TTLL7 | 44.9072568 | 1.968218682 | 0.64268526 | 3.06249232 | 0.00219502 | 0.15271309 |
| PRKACB | 190.381204 | 0.235401574 | 0.23040353 | 1.02169257 | 0.30692643 | 0.84488735 |
| RPF1 | 27.969632 | 0.105208029 | 0.31513789 | 0.3338476 | 0.73849456 | 0.9698508 |
| GNG5 | 114.886757 | -0.159774692 | 0.34334143 | -0.4653522 | 0.64167926 | 0.94944486 |
| SPATA1 | 42.4936269 | 0.65580782 | 0.64848984 | 1.01128465 | 0.31188021 | 0.84811918 |
| CTBS | 65.4001211 | -0.084385892 | 0.28922996 | -0.2917605 | 0.77046971 | 0.97313769 |
| SSX2IP | 58.6363305 | -0.28636603 | 0.27553558 | -1.0393069 | 0.29866202 | 0.83973489 |
| SYDE2 | 61.0320524 | 0.299205016 | 0.38983717 | 0.76751279 | 4.43E-01 | 0.90615787 |
| C1orf52 | 55.9035652 | 0.080195533 | 0.24983396 | 0.32099532 | 0.74821394 | 0.97028355 |
| BCL10 | 30.8220022 | -0.084932736 | 0.308523 | -0.2752882 | 0.78309484 | 0.97423208 |
| AL078459.1 | 95.1907948 | 0.558930923 | 0.40606302 | 1.37646348 | 0.16867819 | 0.74445571 |
| DDAH1 | 487.053884 | 0.296319075 | 0.30178017 | 0.98190373 | 0.32614728 | 0.85523262 |
| CYR61 | 143.10354 | -0.882038966 | 0.58969417 | -1.4957566 | 1.35E-01 | 7.06E-01 |
| ZNHIT6 | 138.01247 | -0.298790975 | 0.22732222 | -1.3143941 | 0.18871363 | 0.76207437 |
| ODF2L | 269.49649 | -0.006637338 | 0.2902862 | -0.0228648 | 0.98175811 | 0.99681784 |
| SH3GLB1 | 183.187612 | -0.25886805 | 0.2131382 | -1.2145549 | 0.22453589 | 0.78901652 |
| SELENOF | 196.006657 | -0.120262671 | 0.18543109 | -0.6485572 | 0.51662461 | 0.9320507 |
| HS2ST1 | 156.177815 | 0.216025634 | 0.25389299 | 0.85085307 | 0.39485098 | 0.88760715 |
| LMO4 | 67.0630918 | -0.217384303 | 0.25852148 | -0.8408752 | 0.40041786 | 0.8907577 |
| PKN2-AS1 | 41.965207 | 0.407451445 | 0.49529748 | 0.82263985 | 0.41071283 | 0.89649693 |
| PKN2 | 476.938574 | 0.034781441 | 0.15536313 | 0.22387191 | 0.82285697 | 0.98094663 |
| GTF2B | 70.9863221 | 0.017029904 | 0.25197334 | 0.06758614 | 0.94611509 | 0.9952642 |
| KYAT3 | 97.9257214 | -0.121405445 | 0.25094283 | -0.4837972 | 0.62852979 | 0.94858781 |
| GBP3 | 47.2994934 | 0.219915026 | 0.59829226 | 0.36757123 | 0.71319297 | 0.96587538 |
| GBP2 | 90.3349679 | 0.629062669 | 0.66503062 | 0.94591535 | 0.34419176 | 0.86969477 |
| LRRC8B | 76.2639539 | -0.549696703 | 0.37310652 | -1.473297 | 0.14067098 | 0.7148274 |
| ZNF326 | 142.146336 | 0.202155943 | 0.16137603 | 1.25270121 | 0.21031447 | 0.77730294 |
| ZNF644 | 422.96022 | -0.052665109 | 0.18567957 | -0.2836344 | 0.77669058 | 0.97423208 |
| HFM1 | 57.7653698 | -0.048691527 | 0.40236806 | -0.1210124 | 0.90368121 | 0.99050161 |
| TGFBR3 | 162.543066 | 0.157246614 | 0.55068457 | 0.28554752 | 0.7752247 | 0.97423208 |
| BTBD8 | 73.3946296 | 0.542217813 | 0.38651324 | 1.40284409 | 0.16066334 | 0.73816762 |
| GLMN | 80.1591984 | -0.10928522 | 0.20808455 | -0.5251962 | 0.59944679 | 0.94683415 |

| | | | | | | |
|---|---|---|---|---|---|---|
| RPAP2 | 245.701606 | -0.156847728 | 0.17088082 | -0.9178779 | 0.35868278 | 0.87316144 |
| EVI5 | 409.801073 | 0.168638969 | 0.17527485 | 0.96214016 | 0.3359792 | 0.86391445 |
| RPL5 | 315.503196 | 0.095928173 | 0.44433499 | 0.21589156 | 0.82907228 | 0.98142144 |
| FAM69A | 79.9071348 | 0.603973284 | 0.30510305 | 1.97957143 | 0.04775171 | 0.51394786 |
| MTF2 | 268.497943 | -0.104138395 | 0.23887488 | -0.4359537 | 0.6628703 | 0.95477098 |
| TMED5 | 143.771117 | 0.421708241 | 0.27697389 | 1.52255595 | 0.12786984 | 0.69635098 |
| CCDC18 | 68.7966222 | 0.276111149 | 0.342634 | 0.80584865 | 0.42033012 | 0.90012434 |
| DR1 | 53.7817126 | -0.067298999 | 0.32230388 | -0.208806 | 0.83459965 | 0.98367279 |
| FNBP1L | 539.2885 | 0.323431924 | 0.28429902 | 1.13764701 | 0.25526791 | 0.80897837 |
| BCAR3 | 284.931633 | -1.320137597 | 0.34530981 | -3.8230527 | 0.00013181 | 0.02918429 |
| DNTTIP2 | 27.6564744 | -0.718327652 | 0.46469739 | -1.5457966 | 0.1221537 | 0.69242609 |
| GCLM | 89.3269095 | -0.239465192 | 0.34435116 | -0.6954099 | 4.87E-01 | 9.20E-01 |
| ABCA4 | 83.418315 | 0.85199233 | 1.36427154 | 0.62450349 | 0.53229698 | 0.93368848 |
| ARHGAP29 | 2190.23433 | 0.140040713 | 0.33280352 | 0.42079096 | 0.67390774 | 0.95910677 |
| ABCD3 | 258.953052 | 0.411056189 | 0.22572152 | 1.82107663 | 0.06859521 | 0.58250536 |
| SLC44A3 | 187.869211 | 0.042093575 | 0.27421768 | 0.15350424 | 0.87800065 | 0.98766275 |
| CNN3 | 385.333202 | -0.04696592 | 0.35803334 | -0.1311775 | 0.89563489 | 0.98842187 |
| AC105942.1 | 35.4375047 | -0.387364934 | 0.28254029 | -1.3710078 | 0.17037254 | 0.74445571 |
| ALG14 | 68.1697793 | -0.202212209 | 0.40422519 | -0.5002464 | 0.61690157 | 0.94791646 |
| PTBP2 | 487.122763 | 0.084249868 | 0.1190107 | 0.7079184 | 0.47899593 | 0.91971919 |
| DPYD | 432.949683 | 0.086244923 | 0.48407141 | 0.1781657 | 0.85859284 | 0.9864199 |
| SNX7 | 84.4753449 | 0.27057014 | 0.29700221 | 0.9110038 | 0.36229337 | 0.8752623 |
| AGL | 101.673619 | -0.220944507 | 0.21520472 | -1.0266713 | 0.30457528 | 0.84488735 |
| TRMT13 | 80.0986668 | -0.130678404 | 0.18938341 | -0.6900203 | 0.4901814 | 0.92157694 |
| DBT | 76.9701072 | -0.108057277 | 0.24957919 | -0.4329579 | 0.66504537 | 0.9550929 |
| RTCA | 52.9546304 | -0.15363691 | 0.24670168 | -0.6227639 | 0.53343966 | 0.93436907 |
| CDC14A | 150.168213 | 0.412257149 | 0.3001131 | 1.37367264 | 0.16954333 | 0.74445571 |
| VCAM1 | 1287.50893 | 1.248918766 | 0.85985382 | 1.45247802 | 0.14636874 | 0.71974648 |
| EXTL2 | 38.1504041 | 0.391901393 | 0.38077348 | 1.02922449 | 0.3033742 | 0.84470484 |
| SLC30A7 | 166.623425 | -0.195901558 | 0.21324935 | -0.9186502 | 0.35827856 | 0.87316144 |
| DPH5 | 44.6765899 | 0.0250001 | 0.2841343 | 0.08798691 | 0.92988708 | 0.99362798 |
| AC093157.1 | 37.5161444 | 0.202813125 | 0.39751167 | 0.51020671 | 0.60990665 | 0.94791646 |
| RNPC3 | 142.309998 | 0.018997114 | 0.21297899 | 0.08919713 | 0.92892525 | 0.99345382 |
| AMY2B | 49.5460648 | 0.233176457 | 0.35641716 | 0.65422343 | 0.51296787 | 0.92957231 |
| VAV3 | 479.119314 | 0.08971796 | 0.74891955 | 0.11979653 | 0.90464433 | 0.99061193 |
| SLC25A24 | 100.453515 | -0.086010311 | 0.27417063 | -0.3137109 | 0.75374063 | 0.9710175 |
| PRPF38B | 110.559686 | -0.224112884 | 0.25174912 | -0.8902231 | 0.37334609 | 0.87666478 |
| STXBP3 | 154.093918 | 0.004701394 | 0.20446229 | 0.02299394 | 0.98165511 | 0.99681784 |
| GPSM2 | 33.3237679 | 0.223391204 | 0.32377078 | 0.68996715 | 0.49021484 | 0.92157694 |
| WDR47 | 33.0758161 | 0.525568659 | 0.28955711 | 1.81507773 | 0.06951197 | 0.58408344 |
| TAF13 | 29.1951112 | -0.096247484 | 0.3730393 | -0.258009 | 0.79639999 | 0.97524554 |
| TMEM167B | 41.2280698 | 0.043081364 | 0.25462186 | 0.16919743 | 0.86564135 | 0.98670067 |
| SARS | 104.408831 | 0.086120751 | 0.23785693 | 0.36206955 | 0.71730005 | 0.96742001 |
| CELSR2 | 62.2270261 | 0.093967158 | 0.40899395 | 0.22975195 | 8.18E-01 | 0.98012053 |
| SORT1 | 226.88837 | -0.048824941 | 0.34625649 | -0.141008 | 0.88786361 | 0.98766275 |
| PSMA5 | 108.84471 | -0.408774046 | 0.33427824 | -1.2228557 | 0.22138421 | 0.78687916 |
| SYPL2 | 44.3268864 | 0.285787566 | 0.48931223 | 0.58405973 | 0.55918013 | 0.93732665 |
| GNAI3 | 129.697301 | -0.125416075 | 0.19401283 | -0.6464319 | 0.5179997 | 0.93257822 |
| AMPD2 | 31.5996147 | -0.23786173 | 0.39638009 | -0.600085 | 0.54844961 | 0.93523904 |
| GSTM3 | 69.6715534 | -0.625655378 | 0.81677457 | -0.7660074 | 0.44367189 | 0.90667728 |
| CSF1 | 87.3995719 | 0.961792328 | 1.42574769 | 0.67458803 | 5.00E-01 | 0.92398825 |
| AHCYL1 | 304.954046 | 0.343773976 | 0.37314625 | 0.92128482 | 0.35690174 | 0.87316144 |
| STRIP1 | 48.3546841 | 0.165424649 | 0.25377015 | 0.65186803 | 0.51448631 | 0.93091506 |
| RBM15 | 49.9214581 | 0.300004739 | 0.28351179 | 1.05817377 | 0.28997623 | 0.83506029 |

| | | | | | | |
|---|---|---|---|---|---|---|
| SLC16A4 | 87.0378387 | 1.279677723 | 0.26208335 | 4.88271273 | 1.05E-06 | 0.00091119 |
| LAMTOR5 | 158.831776 | 0.371487258 | 0.38959928 | 0.95351114 | 0.34033115 | 0.86777921 |
| LRIF1 | 33.8088149 | 0.170045238 | 0.47813566 | 0.35564224 | 0.72210851 | 0.96779196 |
| DRAM2 | 89.1269842 | 0.886632249 | 0.30113397 | 2.94431167 | 0.00323674 | 0.18455191 |
| CEPT1 | 123.512763 | 0.668668463 | 0.22394115 | 2.98591155 | 0.00282734 | 0.17277569 |
| DENND2D | 43.5714029 | 0.127780769 | 0.32132577 | 0.39766735 | 0.69087541 | 0.96220115 |
| WDR77 | 28.3812129 | -0.282946168 | 0.35253003 | -0.8026158 | 0.42219684 | 0.90066187 |
| ATP5PB | 110.010513 | 0.084728083 | 0.28157113 | 0.30091182 | 0.76348173 | 0.97149197 |
| RAP1A | 311.046283 | -0.183277619 | 0.24133342 | -0.7594374 | 0.44759097 | 0.90951928 |
| KCND3 | 66.5548702 | 0.77988948 | 0.48396397 | 1.61146186 | 0.10707909 | 0.65699665 |
| CTTNBP2NL | 158.576409 | -0.061740586 | 0.24721506 | -0.2497444 | 0.80278499 | 0.97835591 |
| WNT2B | 36.8037243 | 1.20402058 | 0.46451328 | 2.59200464 | 0.00954185 | 0.28916146 |
| ST7L | 141.202704 | -0.003734509 | 0.2569839 | -0.0145321 | 0.98840549 | 0.99820747 |
| CAPZA1 | 195.295114 | -0.222577956 | 0.14689712 | -1.5151962 | 0.12972273 | 0.69904834 |
| MOV10 | 60.6543683 | -0.03196681 | 0.48986424 | -0.0652565 | 0.9479698 | 0.9952642 |
| RHOC | 115.827277 | -0.164087527 | 0.54758716 | -0.2996555 | 0.76443991 | 0.97177982 |
| SLC16A1-AS1 | 120.224329 | 0.546295312 | 0.88386042 | 0.61807872 | 0.53652345 | 0.93469807 |
| LRIG2 | 117.250042 | -0.080035243 | 0.23462482 | -0.3411201 | 0.73301317 | 0.9698508 |
| MAGI3 | 623.970229 | 0.143199256 | 0.41537129 | 0.34475001 | 0.73028232 | 0.96888841 |
| PHTF1 | 73.8402686 | 0.251165132 | 0.33869675 | 0.74156347 | 0.45835187 | 0.91336645 |
| RSBN1 | 98.04728 | 0.136236481 | 0.2285692 | 0.59604041 | 0.55114823 | 0.93523904 |
| HIPK1 | 118.653949 | 0.355840209 | 0.24812717 | 1.43410415 | 0.15154254 | 0.72436429 |
| TRIM33 | 292.423206 | -0.164650534 | 0.18822407 | -0.8747581 | 0.38170553 | 0.88128509 |
| BCAS2 | 37.5732052 | 0.171432027 | 0.41026372 | 0.41785812 | 0.67605086 | 0.9591071 |
| NRAS | 40.4916067 | -0.501167085 | 0.40758533 | -1.2296004 | 0.21884679 | 0.78523615 |
| CSDE1 | 313.898177 | -0.094085128 | 0.19191362 | -0.4902473 | 0.62395893 | 0.94808496 |
| SIKE1 | 39.4917813 | -0.361643881 | 0.39343627 | -0.919193 | 0.35799461 | 0.87316144 |
| VANGL1 | 25.4532647 | -0.215512049 | 0.41483779 | -0.5195092 | 0.6034057 | 0.94683415 |
| SLC22A15 | 48.6409141 | 0.480541925 | 0.43807005 | 1.09695224 | 0.27266227 | 0.82004141 |
| ATP1A1 | 705.836242 | -0.065285103 | 0.31236882 | -0.2090001 | 0.83444819 | 0.98367279 |
| LINC01762 | 50.1118131 | 1.97972966 | 0.56834448 | 3.48332698 | 0.00049522 | 0.06874989 |
| CD58 | 91.6575921 | -0.000979122 | 0.27866452 | -0.0035136 | 0.99719654 | 0.99917842 |
| IGSF3 | 96.7555786 | 0.029603284 | 0.42596158 | 0.06949755 | 0.94459359 | 0.9952642 |
| PTGFRN | 54.9170069 | 0.260610443 | 0.33378554 | 0.78077213 | 0.43493653 | 0.90514499 |
| TTF2 | 41.9045358 | -0.526401003 | 0.28289028 | -1.8607956 | 0.06277304 | 0.56713487 |
| VTCN1 | 35.1198223 | -0.849492227 | 0.97668901 | -0.8697674 | 0.38442755 | 0.88328503 |
| MAN1A2 | 388.595544 | -0.04447422 | 0.18812485 | -0.236408 | 0.81311609 | 0.97930627 |
| GDAP2 | 98.5217183 | -0.083561462 | 0.23892681 | -0.3497366 | 0.72653635 | 0.96794043 |
| WDR3 | 34.5615816 | -0.753522081 | 0.30122961 | -2.5014874 | 0.01236728 | 0.30967852 |
| SPAG17 | 184.918157 | 0.490354843 | 0.56137176 | 0.87349396 | 0.3823939 | 0.88177874 |
| WARS2 | 77.3368962 | -0.413042567 | 0.22341591 | -1.8487608 | 0.06449237 | 0.57467198 |
| AL359915.2 | 232.788662 | -0.476633982 | 0.38684702 | -1.2320994 | 0.21791195 | 0.7847288 |
| ZNF697 | 47.1703428 | -0.115857772 | 0.38865741 | -0.2980974 | 0.76562881 | 0.97177982 |
| PHGDH | 56.7944243 | -0.417065485 | 0.43155266 | -0.9664301 | 0.33382901 | 0.86191987 |
| NOTCH2 | 241.995894 | 0.052793512 | 0.35045859 | 0.15064123 | 0.88025874 | 0.98766275 |
| SEC22B | 129.735938 | 0.089294825 | 0.24705962 | 0.36143026 | 0.71777783 | 0.96743968 |
| NBPF26 | 45.467905 | -0.461377381 | 0.5460971 | -0.8448633 | 0.39818718 | 0.8887831 |
| LINC00623 | 254.287762 | -0.254659175 | 0.37146911 | -0.685546 | 0.49299942 | 0.92289011 |
| SRGAP2C | 59.0813473 | -0.47170025 | 0.39382503 | -1.1977407 | 0.231018 | 0.79430058 |
| AC239800.3 | 35.2789346 | 0.149045559 | 0.36910672 | 0.40380072 | 0.68635927 | 0.96161098 |
| NBPF15 | 86.4104161 | 0.020896798 | 0.35185478 | 0.0593904 | 0.95264116 | 0.99531843 |
| FP700111.1 | 58.6484726 | -0.248910841 | 0.31453142 | -0.7913704 | 0.4287279 | 0.90278843 |
| SRGAP2B | 71.3925343 | 0.130042615 | 0.35643677 | 0.36484063 | 0.71523037 | 0.9667635 |
| GPR89A | 39.0284331 | -0.422313942 | 0.30734834 | -1.3740564 | 0.16942418 | 0.74445571 |

| | | | | | | |
|---|---|---|---|---|---|---|
| PDZK1 | 342.843915 | 1.540272425 | 0.34657834 | 4.44422583 | 8.82E-06 | 0.00421757 |
| RNF115 | 298.147076 | -0.259842863 | 0.17056705 | -1.523406 | 0.12765717 | 0.69635098 |
| POLR3C | 43.4258767 | -0.022331747 | 0.25043437 | -0.0891721 | 0.92894517 | 0.99345382 |
| PIAS3 | 24.8941426 | -0.889002974 | 0.40763159 | -2.1808981 | 0.02919095 | 0.43284852 |
| PEX11B | 31.4026471 | -0.600050539 | 0.33517705 | -1.7902495 | 0.07341381 | 0.59320069 |
| RBM8A | 79.085764 | -0.464414896 | 0.43347043 | -1.0713877 | 0.28399513 | 0.82918183 |
| LIX1L | 30.2903987 | 0.488606451 | 0.42425803 | 1.15167284 | 0.24945554 | 0.80672756 |
| TXNIP | 1439.85098 | 0.699442373 | 0.51677177 | 1.35348409 | 0.17590103 | 0.74950983 |
| NBPF12 | 69.2394617 | -0.060000168 | 0.2777671 | -0.2160089 | 0.8289808 | 0.98142144 |
| PRKAB2 | 30.4477462 | 0.32477294 | 0.33208976 | 0.97796733 | 0.32809048 | 0.8573817 |
| AC242426.2 | 92.6884196 | -0.441478025 | 0.28961199 | -1.5243776 | 0.12741442 | 0.69623657 |
| FMO5 | 37.8659737 | 1.229866557 | 0.44927381 | 2.73745435 | 0.00619167 | 0.24799848 |
| CHD1L | 81.1018486 | -0.262449734 | 0.28076205 | -0.9347764 | 0.34990356 | 0.8720098 |
| BCL9 | 88.9036319 | 0.082356833 | 0.43453404 | 0.18952907 | 0.84967817 | 0.9858744 |
| ACP6 | 105.613158 | -0.508372742 | 0.3095256 | -1.6424255 | 0.10050185 | 0.64698063 |
| GPR89B | 42.63211 | -0.186818625 | 0.33157732 | -0.563424 | 0.57314619 | 0.93998433 |
| NBPF11 | 67.4940936 | -0.017533013 | 0.2650383 | -0.0661528 | 0.94725621 | 0.9952642 |
| AC239809.3 | 66.853158 | -0.356206319 | 0.29690578 | -1.1997285 | 0.23024481 | 0.79391882 |
| NBPF14 | 52.1217715 | -0.117021199 | 0.25518142 | -0.4585804 | 0.64653551 | 0.95008515 |
| PDE4DIP | 315.546667 | 0.20995891 | 0.51305322 | 0.40923417 | 0.68236782 | 0.96101988 |
| NBPF9 | 47.5422155 | -0.177092343 | 0.37754334 | -0.4690649 | 0.63902322 | 0.94944486 |
| AC245297.3 | 465.154514 | -0.205144459 | 0.35537816 | -0.5772568 | 0.56376599 | 0.93809297 |
| NBPF19 | 75.9093062 | 0.027159908 | 0.71312296 | 0.03808587 | 0.96961922 | 0.99681784 |
| SF3B4 | 27.8420085 | -0.394560671 | 0.33801428 | -1.1672899 | 0.2430933 | 0.80476923 |
| OTUD7B | 198.096754 | 0.339501506 | 0.42820177 | 0.79285404 | 0.42786287 | 0.90272191 |
| VPS45 | 138.186118 | 0.122854638 | 0.15720447 | 0.78149582 | 0.43451094 | 0.90514499 |
| ANP32E | 105.613158 | -0.27425142 | 0.46811974 | -0.5858574 | 0.55797134 | 0.93620477 |
| APH1A | 73.9238959 | -0.35611582 | 0.32435826 | -1.0979089 | 0.27224427 | 0.81929873 |
| MRPS21 | 128.218654 | -0.045562196 | 0.27260712 | -0.167135 | 0.86726381 | 0.98670067 |
| PRPF3 | 120.267896 | -0.371609251 | 0.24284806 | -1.5302129 | 0.12596403 | 0.69492095 |
| RPRD2 | 382.899337 | -0.144661879 | 0.24664189 | -0.586526 | 0.55752209 | 0.93594535 |
| TARS2 | 29.4771183 | -0.182202469 | 0.32649666 | -0.5580531 | 0.57680813 | 0.94034284 |
| MCL1 | 193.74444 | -0.339459045 | 0.37772761 | -0.8986874 | 0.36881919 | 0.87605028 |
| ENSA | 334.336638 | -0.056785866 | 0.19883964 | -0.2855862 | 0.77519504 | 0.97423208 |
| GOLPH3L | 95.1943952 | 0.136369436 | 0.27730277 | 0.49177091 | 0.6228813 | 0.94808496 |
| ARNT | 185.548999 | 0.02612041 | 0.19955387 | 0.13089403 | 8.96E-01 | 0.98842187 |
| SETDB1 | 102.059659 | -0.283290854 | 0.23772224 | -1.1916885 | 0.23338343 | 0.79622042 |
| CERS2 | 117.108983 | 0.22424657 | 0.22112147 | 1.01413296 | 0.31051931 | 0.84670211 |
| MINDY1 | 29.8953959 | 0.835178079 | 0.37013031 | 2.25644334 | 0.02404288 | 0.40690238 |
| PRUNE1 | 38.0010817 | -0.008312654 | 0.36879205 | -0.0225402 | 0.98201703 | 0.99681784 |
| C1orf56 | 29.1799089 | 0.049242023 | 0.54998544 | 0.08953332 | 0.92865807 | 0.99336859 |
| CDC42SE1 | 77.5719253 | -0.04402573 | 0.22514814 | -0.1955412 | 0.84496932 | 0.98520136 |
| GABPB2 | 112.102221 | 0.857319586 | 0.4282342 | 2.00198765 | 0.04528606 | 0.50441298 |
| VPS72 | 26.3199992 | 0.127039302 | 0.33271637 | 0.38182462 | 0.70259145 | 0.9645815 |
| PIP5K1A | 328.953739 | 0.319379003 | 0.19356576 | 1.64997674 | 0.09894769 | 0.6434623 |
| PSMD4 | 81.4466445 | -0.030074521 | 0.2668318 | -0.1127097 | 0.91026075 | 0.99100813 |
| PI4KB | 77.5739772 | 0.130081151 | 0.2001207 | 0.65001346 | 0.51568353 | 0.93140514 |
| PSMB4 | 58.5246777 | -0.394885832 | 0.39113055 | -1.0096011 | 0.31268644 | 0.84826686 |
| POGZ | 506.261782 | 0.14481031 | 0.22818663 | 0.63461348 | 0.52568054 | 0.93272384 |
| CGN | 68.1511562 | 0.464777533 | 0.40648806 | 1.14339776 | 0.25287348 | 0.80798674 |
| TUFT1 | 86.5624345 | 0.348692921 | 0.40861465 | 0.85335394 | 0.39346306 | 0.88760715 |
| SNX27 | 138.748435 | 0.398874982 | 0.17759671 | 2.24595928 | 0.02470662 | 0.41230428 |
| MRPL9 | 28.5448447 | -0.031335001 | 0.43297091 | -0.0723721 | 0.94230581 | 0.99526299 |
| RORC | 36.6396473 | 0.062132695 | 0.36044617 | 0.17237718 | 0.86314101 | 0.98670067 |

| | | | | | | |
|---|---|---|---|---|---|---|
| THEM4 | 58.5782506 | 0.023564607 | 0.28212617 | 0.08352507 | 0.93343405 | 0.99425828 |
| S100A10 | 526.650077 | -0.833343049 | 0.67750178 | -1.2300234 | 0.21868834 | 0.78523615 |
| S100A11 | 867.88199 | -1.020094976 | 0.61976331 | -1.6459428 | 0.09977553 | 0.64621354 |
| FLG-AS1 | 153.021938 | 0.101763042 | 0.36344409 | 0.27999641 | 7.79E-01 | 0.97423208 |
| S100A9 | 88.2615934 | -1.113752907 | 1.13904999 | -0.9777911 | 0.32817767 | 0.8573817 |
| S100A6 | 1265.35628 | -1.952726864 | 0.7000067 | -2.7895831 | 0.0052776 | 0.22669993 |
| S100A4 | 31.0132761 | -1.529525173 | 1.27178463 | -1.2026605 | 0.22910771 | 0.7931636 |
| S100A2 | 64.2981637 | -3.15321595 | 1.1794194 | -2.6735324 | 0.0075057 | 0.26951269 |
| S100A16 | 162.898165 | -0.856412595 | 0.5825583 | -1.4700891 | 0.14153763 | 0.7148274 |
| S100A14 | 116.08328 | -1.204977316 | 0.82763385 | -1.4559304 | 0.14541185 | 0.71947319 |
| S100A13 | 278.682658 | -0.730129967 | 0.56450358 | -1.2934018 | 0.19587212 | 0.77017438 |
| S100A1 | 116.868491 | -0.322922218 | 0.80077181 | -0.4032637 | 0.68675423 | 0.96161098 |
| CHTOP | 90.1086678 | -0.308775876 | 0.2965298 | -1.041298 | 0.29773728 | 0.83973489 |
| ILF2 | 87.5570504 | -0.025561643 | 0.37008153 | -0.0690703 | 0.94493366 | 0.9952642 |
| NPR1 | 200.382882 | 0.600405225 | 0.59193729 | 1.01430547 | 0.31043702 | 0.84670211 |
| INTS3 | 195.592959 | -0.267802709 | 0.27694052 | -0.9670044 | 0.33354183 | 0.8614174 |
| GATAD2B | 400.878199 | 0.083961893 | 0.20250966 | 0.41460685 | 0.67842973 | 0.95924874 |
| DENND4B | 52.5368406 | 0.110920121 | 0.26132104 | 0.42445921 | 0.67123095 | 0.95800185 |
| CRTC2 | 30.5591919 | -0.088136565 | 0.36671596 | -0.2403401 | 0.81006658 | 0.97930627 |
| SLC39A1 | 72.9419345 | -0.242644838 | 0.45597888 | -0.5321405 | 0.59462868 | 0.94664253 |
| JTB | 67.4858168 | -0.025502946 | 0.43385651 | -0.058782 | 0.95312576 | 0.99531843 |
| RAB13 | 128.720475 | -0.258235352 | 0.45444071 | -0.5682487 | 0.5698661 | 0.93998433 |
| RPS27 | 140.616161 | 0.274767269 | 0.45377572 | 0.60551338 | 0.54483794 | 0.93523904 |
| TPM3 | 139.103246 | -0.504652182 | 0.32552419 | -1.5502755 | 0.12107541 | 0.69072302 |
| C1orf43 | 109.827998 | -0.061191888 | 0.20112413 | -0.3042494 | 0.76093792 | 0.9714717 |
| UBAP2L | 126.971414 | -0.1394898 | 0.18913049 | -0.737532 | 0.46079886 | 0.91368087 |
| HAX1 | 30.0915316 | 0.031363574 | 0.42649625 | 0.07353775 | 0.94137821 | 0.99526299 |
| IL6R | 88.8353038 | 0.363627994 | 0.7075825 | 0.5139019 | 0.60732058 | 0.94708529 |
| UBE2Q1 | 40.8782523 | -0.31079995 | 0.37180126 | -0.8359303 | 0.4031941 | 0.89319988 |
| ADAR | 199.757472 | -0.54853843 | 0.285022 | -1.9245477 | 0.05428597 | 0.53752502 |
| PMVK | 45.3074058 | 0.594755611 | 0.38902454 | 1.52883831 | 0.12630453 | 0.69492095 |
| PBXIP1 | 42.4586482 | 1.036328099 | 0.50294106 | 2.06053588 | 0.03934734 | 0.479886 |
| ZBTB7B | 23.7026663 | 0.076151987 | 0.33010059 | 0.23069328 | 0.8175531 | 0.98010849 |
| EFNA1 | 180.265876 | 0.24312499 | 0.46005125 | 0.52847371 | 0.59717059 | 0.94683415 |
| DPM3 | 50.5074823 | -0.370251053 | 0.36949937 | -1.0020343 | 0.31632701 | 0.8490543 |
| THBS3 | 35.8557143 | 0.636281535 | 0.31671855 | 2.00898097 | 0.04453915 | 0.50134023 |
| GBA | 29.750251 | -0.302687176 | 0.35565192 | -0.851077 | 0.39472657 | 0.88760715 |
| SCAMP3 | 48.73878 | -0.239580906 | 0.2504828 | -0.9564765 | 0.33883156 | 0.86680923 |
| CLK2 | 40.5872221 | 0.095018663 | 0.28658414 | 0.3315559 | 0.74022462 | 0.9698508 |
| FDPS | 116.905848 | 0.220874304 | 0.31337464 | 0.70482508 | 0.4809191 | 0.91971919 |
| ASH1L | 805.409959 | 0.142808479 | 0.14430436 | 0.98963383 | 0.32235313 | 0.85246644 |
| YY1AP1 | 159.851588 | 0.262601904 | 0.19357579 | 1.35658444 | 0.17491329 | 0.74910274 |
| DAP3 | 135.164326 | -0.404440885 | 0.26739811 | -1.5125047 | 0.13040552 | 0.69957738 |
| GON4L | 185.627031 | -0.016593152 | 0.21781007 | -0.0761818 | 0.9392745 | 0.99526299 |
| RIT1 | 55.3145895 | 0.282975692 | 0.32040222 | 0.88318893 | 0.3771342 | 0.87842274 |
| KHDC4 | 152.451969 | -0.447776793 | 0.3027857 | -1.4788571 | 0.1391785 | 0.71184729 |
| ARHGEF2 | 67.5164803 | 0.05064608 | 0.26348279 | 0.19221779 | 0.84757161 | 0.98545133 |
| SSR2 | 103.794301 | 0.293314246 | 0.43629149 | 0.67228963 | 0.50139933 | 0.92484445 |
| UBQLN4 | 38.869744 | 0.244717448 | 0.35587719 | 0.68764578 | 0.49167587 | 0.92275924 |
| LAMTOR2 | 29.0283209 | -0.039552215 | 0.43471742 | -0.0909837 | 0.92750552 | 0.99336859 |
| RAB25 | 42.0336426 | -0.103868482 | 0.66230633 | -0.1568285 | 0.87538004 | 0.98766275 |
| LMNA | 218.010214 | -0.300934573 | 0.34142092 | -0.8814181 | 0.37809158 | 0.87964763 |
| SEMA4A | 94.605269 | -0.397019857 | 0.57914582 | -0.6855266 | 0.49301169 | 0.92289011 |
| SLC25A44 | 26.9465275 | -0.087177451 | 0.32953193 | -0.2645493 | 0.79135666 | 0.97423208 |

| | | | | | | |
|---|---|---|---|---|---|---|
| SMG5 | 102.651489 | -0.14118622 | 0.26797794 | -0.5268576 | 0.59829246 | 0.94683415 |
| CCT3 | 125.00387 | -0.164962575 | 0.39387728 | -0.4188172 | 0.67534974 | 0.95910677 |
| MEF2D | 49.7966076 | -0.037562015 | 0.24345314 | -0.1542885 | 0.87738228 | 0.98766275 |
| NAXE | 32.9501651 | -0.374095309 | 0.60166638 | -0.6217654 | 0.53409617 | 0.93446646 |
| GPATCH4 | 27.2280269 | -0.854735571 | 0.56546153 | -1.5115716 | 0.13064289 | 0.69990393 |
| ISG20L2 | 29.3271994 | -0.269352222 | 0.30593003 | -0.8804373 | 0.37862243 | 0.87970706 |
| RRNAD1 | 31.2764864 | 0.168758913 | 0.28882493 | 0.58429482 | 0.55902198 | 0.93731341 |
| MRPL24 | 28.6082303 | -0.425401343 | 0.45192586 | -0.9413078 | 0.34654714 | 0.86990996 |
| HDGF | 170.690606 | -0.150934802 | 0.40473392 | -0.3729235 | 0.70920536 | 0.96587538 |
| PRCC | 74.3626406 | -0.331941885 | 0.25385675 | -1.3075952 | 0.19101063 | 0.76328615 |
| ARHGEF11 | 174.84091 | 0.045670734 | 0.25677373 | 0.17786373 | 0.85882999 | 0.9864199 |
| KIRREL1 | 381.60514 | -0.343234794 | 0.46025061 | -0.7457563 | 0.45581464 | 0.91077355 |
| IFI16 | 137.527298 | -0.222563906 | 1.48943245 | -0.1494287 | 0.88121539 | 0.98766275 |
| DUSP23 | 197.137491 | 0.510898545 | 0.52160602 | 0.9794721 | 3.27E-01 | 8.57E-01 |
| TAGLN2 | 150.999853 | -0.11260897 | 0.47714044 | -0.236008 | 0.81342645 | 0.97930627 |
| IGSF8 | 31.0914199 | 0.129693052 | 0.32065749 | 0.40445976 | 0.68587466 | 0.96161098 |
| PEA15 | 50.2105617 | 0.207617736 | 0.52189892 | 0.39781216 | 0.69076865 | 0.96220115 |
| DCAF8 | 53.62542 | 0.134543488 | 0.38316039 | 0.35114143 | 0.72548225 | 0.96781837 |
| COPA | 214.726639 | 0.247456019 | 0.18653163 | 1.32661696 | 0.18463542 | 0.75873991 |
| NCSTN | 57.0059161 | 0.159507505 | 0.42758779 | 0.37304036 | 0.7091184 | 0.96587538 |
| F11R | 79.8713519 | -0.546619437 | 0.40371866 | -1.3539613 | 0.17574873 | 0.74934286 |
| AL591806.3 | 45.3766009 | 0.238154048 | 0.46406247 | 0.51319394 | 0.60781567 | 0.94708529 |
| PFDN2 | 83.8718303 | -0.389147498 | 0.43302392 | -0.8986744 | 0.36882613 | 0.87605028 |
| NIT1 | 60.8379428 | 0.548229506 | 0.36256212 | 1.51209812 | 0.1305089 | 0.69957738 |
| DEDD | 30.1159262 | 0.580433271 | 0.35072154 | 1.65496898 | 0.09793079 | 0.64312419 |
| UFC1 | 61.7138025 | 0.426435667 | 0.44874317 | 0.95028893 | 0.34196546 | 0.86846099 |
| B4GALT3 | 26.1364413 | 0.219637909 | 0.31837601 | 0.68986953 | 0.49027624 | 0.92157694 |
| NDUFS2 | 64.8243761 | -0.09058742 | 0.28643703 | -0.316256 | 0.75180823 | 0.97058015 |
| SDHC | 152.159669 | 0.286989854 | 0.19440235 | 1.47626742 | 0.13987212 | 0.71251643 |
| AL592295.4 | 28.8212992 | -0.195733139 | 0.46517641 | -0.4207718 | 0.67392169 | 0.95910677 |
| ATF6 | 270.53157 | -0.0200756 | 0.16647856 | -0.1205897 | 0.90401603 | 0.99057075 |
| NOS1AP | 115.255128 | -0.380599489 | 0.38361908 | -0.9921287 | 0.32113477 | 0.85131972 |
| UHMK1 | 118.885388 | -0.182025114 | 0.2356452 | -0.7724541 | 0.4398455 | 0.90561284 |
| UAP1 | 29.9848858 | 0.040375214 | 0.40021012 | 0.10088504 | 0.91964172 | 0.99214128 |
| HSD17B7 | 31.5845433 | 0.425528407 | 0.34629718 | 1.22879546 | 2.19E-01 | 0.78523615 |
| PBX1 | 873.091748 | 0.352605608 | 0.27927421 | 1.26257847 | 0.20674071 | 0.77401635 |
| MGST3 | 326.187885 | -0.028248511 | 0.26668826 | -0.1059233 | 0.91564318 | 0.99159829 |
| ALDH9A1 | 67.9799759 | 0.446199319 | 0.39638037 | 1.12568469 | 0.260299 | 0.81263292 |
| TMCO1 | 112.139125 | 0.022476258 | 0.2359116 | 0.09527407 | 0.92409714 | 0.99258328 |
| UCK2 | 41.3210431 | -0.085801547 | 0.52285369 | -0.1641024 | 0.86965053 | 0.98748014 |
| POGK | 48.4005897 | -0.15325786 | 0.32322111 | -0.474158 | 0.63538727 | 0.94932781 |
| ILDR2 | 56.2001577 | 1.710688096 | 0.89661062 | 1.90794985 | 0.0563977 | 0.54349452 |
| POU2F1 | 407.433838 | -0.085258733 | 0.24578055 | -0.3468897 | 0.72867421 | 0.96836435 |
| CREG1 | 66.9781469 | -0.078848658 | 0.47194193 | -0.1670728 | 0.86731277 | 0.98670067 |
| MPZL1 | 205.346422 | -0.294456396 | 0.20651101 | -1.4258629 | 1.54E-01 | 0.72912162 |
| MPC2 | 127.342005 | 0.34786674 | 0.42264774 | 0.82306542 | 0.41047079 | 0.8964268 |
| DCAF6 | 540.303733 | 0.06148289 | 0.17800191 | 0.34540578 | 0.72978933 | 0.96877106 |
| TIPRL | 53.0032259 | 0.161447269 | 0.24969534 | 0.64657702 | 0.51790574 | 0.93257822 |
| SFT2D2 | 125.32898 | 0.410088899 | 0.31314288 | 1.30959037 | 0.19033445 | 0.76219391 |
| LINC00970 | 50.5881898 | 0.423643508 | 0.91653364 | 0.46222363 | 6.44E-01 | 9.49E-01 |
| ATP1B1 | 725.401123 | 0.291082904 | 0.45350969 | 0.64184495 | 0.52097386 | 0.93272217 |
| NME7 | 310.257499 | 0.095763493 | 0.20700184 | 0.46262145 | 0.64363573 | 0.94944486 |
| BLZF1 | 88.4685614 | -0.028735897 | 0.25768718 | -0.1115147 | 0.91120824 | 0.99100813 |
| SLC19A2 | 84.6679862 | -0.324293432 | 0.56942296 | -0.5695124 | 0.56900846 | 0.93998433 |

| | | | | | | |
|---|---|---|---|---|---|---|
| F5 | 51.2525461 | 2.578893925 | 0.8092096 | 3.18692948 | 0.00143792 | 0.12082298 |
| SCYL3 | 76.7161929 | 0.193569801 | 0.21337939 | 0.90716258 | 0.36432083 | 0.87605028 |
| KIFAP3 | 161.097478 | -0.116726613 | 0.26591877 | -0.4389559 | 0.66069349 | 0.95407427 |
| FMO4 | 39.7955336 | 0.865217218 | 0.52984336 | 1.63296794 | 0.10247574 | 0.65288882 |
| PRRC2C | 495.086348 | -0.156495947 | 0.17320101 | -0.903551 | 0.36623354 | 0.87605028 |
| VAMP4 | 46.5508853 | 0.070424883 | 0.34976896 | 0.20134687 | 0.84042736 | 0.98464453 |
| DNM3 | 648.695723 | -0.50117143 | 0.67804788 | -0.7391387 | 0.45982278 | 0.91368087 |
| PIGC | 66.6966546 | -0.017877682 | 0.34895854 | -0.0512315 | 0.95914102 | 0.99572129 |
| SUCO | 209.70529 | 0.106701064 | 0.18239708 | 0.58499326 | 0.55855225 | 0.93668977 |
| AL645568.1 | 101.29298 | 0.198350666 | 0.31999606 | 0.61985345 | 5.35E-01 | 0.93460153 |
| PRDX6 | 212.285887 | 0.380488166 | 0.3446713 | 1.10391602 | 0.26962957 | 0.8167745 |
| KLHL20 | 104.923998 | 0.05487553 | 0.19494105 | 0.28149808 | 0.7783284 | 0.97423208 |
| ZBTB37 | 79.1519686 | -0.103658522 | 0.33893135 | -0.3058393 | 0.75972701 | 0.9714717 |
| RC3H1 | 283.164095 | 0.027371755 | 0.19386521 | 0.14118962 | 0.88772015 | 0.98766275 |
| RABGAP1L | 909.599015 | -0.091262556 | 0.21799996 | -0.4186357 | 0.67548242 | 0.9591071 |
| CACYBP | 30.860084 | -0.184442087 | 0.31291684 | -0.5894284 | 0.5555739 | 0.93523904 |
| MRPS14 | 33.7260721 | 0.168881383 | 0.37158764 | 0.45448601 | 0.64947906 | 0.95027667 |
| KIAA0040 | 46.7837586 | -0.450961448 | 0.24591437 | -1.833815 | 0.06668146 | 0.5780981 |
| COP1 | 454.742845 | -0.120839512 | 0.1643797 | -0.7351243 | 0.46226379 | 0.91462428 |
| PAPPA2 | 115.223073 | 2.779917501 | 1.56017494 | 1.78179858 | 0.07478208 | 0.59444897 |
| SEC16B | 43.2244842 | 1.417330942 | 0.49967403 | 2.83651112 | 0.00456094 | 0.21628334 |
| RASAL2 | 723.535382 | 0.000508347 | 0.37422945 | 0.00135838 | 0.99891617 | 0.99971834 |
| RALGPS2 | 505.73022 | -0.16111532 | 0.36457879 | -0.4419218 | 0.65854578 | 0.95402584 |
| ANGPTL1 | 35.5239783 | -0.203360593 | 0.58781629 | -0.3459594 | 0.7293732 | 0.96862399 |
| FAM20B | 72.2769511 | -0.520992485 | 0.2740323 | -1.9012083 | 0.05727473 | 0.54616991 |
| ABL2 | 280.646559 | 0.120340695 | 0.20946059 | 0.57452666 | 0.56561146 | 0.93809297 |
| SOAT1 | 119.711596 | 0.471648916 | 0.18857907 | 2.50106719 | 0.01238197 | 0.30967852 |
| TOR1AIP2 | 190.946393 | 0.191143031 | 0.30571503 | 0.6252327 | 0.53181834 | 0.93366189 |
| TOR1AIP1 | 77.7788779 | 0.051732532 | 0.27787983 | 0.18616871 | 0.85231246 | 0.9864199 |
| CEP350 | 335.558695 | 0.073818351 | 0.16283031 | 0.45334528 | 0.65030013 | 0.95061176 |
| QSOX1 | 42.4450956 | 0.084990951 | 0.44240043 | 0.19211318 | 0.84765355 | 0.98545133 |
| ACBD6 | 305.566784 | -0.194883207 | 0.19107736 | -1.0199178 | 0.30776744 | 0.84488735 |
| XPR1 | 417.446079 | 0.123337288 | 0.25185776 | 0.48971009 | 0.62433906 | 0.94808496 |
| STX6 | 67.4631146 | 0.24508045 | 0.29560533 | 0.82907994 | 0.40705917 | 0.89585737 |
| MR1 | 29.8063435 | 0.293026461 | 0.42111408 | 0.69583629 | 0.48653136 | 0.92034617 |
| GLUL | 132.639092 | -0.135280916 | 0.48672417 | -0.2779416 | 0.78105715 | 0.97423208 |
| DHX9 | 276.370192 | -0.278777466 | 0.25609127 | -1.0885864 | 0.27633632 | 0.82231303 |
| LAMC1 | 448.751599 | -0.184251112 | 0.27412104 | -0.6721524 | 0.50148669 | 0.92484445 |
| LAMC2 | 89.6910734 | -1.292809336 | 0.53619618 | -2.4110752 | 0.01590557 | 0.34086104 |
| SMG7 | 252.859384 | -0.027953705 | 0.15154609 | -0.1844568 | 0.85365512 | 0.9864199 |
| ARPC5 | 104.633593 | -0.298977006 | 0.37518441 | -0.7968802 | 0.42552064 | 0.90167091 |
| RGL1 | 139.590053 | 1.710174436 | 0.50509541 | 3.38584436 | 0.0007096 | 0.08496527 |
| TSEN15 | 73.0845144 | -0.218018226 | 0.24869721 | -0.8766412 | 0.38068155 | 0.8807629 |
| C1orf21 | 46.0656453 | 0.700789222 | 0.80437676 | 0.87122012 | 0.38363398 | 0.88328503 |
| EDEM3 | 137.747901 | 0.14657333 | 0.24936395 | 0.58778877 | 0.55667408 | 0.93533595 |
| FAM129A | 107.715297 | 0.771437141 | 0.83915162 | 0.91930603 | 0.35793553 | 0.87316144 |
| RNF2 | 61.2747543 | -0.481085871 | 0.22848855 | -2.1055141 | 0.03524658 | 0.4562527 |
| TRMT1L | 68.207907 | 0.338228404 | 0.40541357 | 0.83427993 | 0.40412326 | 0.89360496 |
| SWT1 | 82.6662692 | 0.118891621 | 0.23851642 | 0.49846304 | 0.61815771 | 0.9480368 |
| IVNS1ABP | 412.987478 | -0.815736558 | 0.74640743 | -1.0928838 | 0.27444484 | 0.82102033 |
| HMCN1 | 44.1291415 | -0.058318316 | 1.2569537 | -0.0463966 | 0.96299419 | 0.99572129 |
| TPR | 217.693156 | -0.091445795 | 0.15257271 | -0.5993588 | 0.54893364 | 0.93523904 |
| ODR4 | 118.10008 | -0.116937754 | 0.20063344 | -0.5828428 | 0.55999914 | 0.93744835 |
| PLA2G4A | 40.2640091 | -1.10184111 | 0.99374338 | -1.1087783 | 0.26752583 | 0.81510106 |

| | | | | | | |
|---|---|---|---|---|---|---|
| ERVMER61-1 | 54.1164184 | 0.105190605 | 0.34486732 | 0.30501761 | 0.76035274 | 0.9714717 |
| AL136454.1 | 150.000554 | -0.518457983 | 0.53988902 | -0.9603047 | 0.33690186 | 0.86431445 |
| UCHL5 | 76.839715 | 0.058325454 | 0.26874126 | 0.217032 | 0.82818341 | 0.98142144 |
| TROVE2 | 158.689329 | 0.082482415 | 0.19851606 | 0.41549492 | 0.67777964 | 0.95924874 |
| CDC73 | 277.942786 | -0.156865475 | 0.19228636 | -0.815791 | 0.4146197 | 0.89757799 |
| KCNT2 | 859.244089 | 0.356024425 | 1.19053641 | 0.29904539 | 0.76490541 | 0.97177982 |
| CFH | 816.300827 | 0.659456924 | 1.58481968 | 0.41610849 | 0.67733062 | 0.95924874 |
| ZBTB41 | 49.890668 | 0.01803894 | 0.26469473 | 0.06814998 | 0.94566625 | 0.9952642 |
| CRB1 | 39.8803441 | 0.374815586 | 0.68134495 | 0.55011134 | 0.58224301 | 0.94091128 |
| DENND1B | 331.136321 | -0.061544366 | 0.20047128 | -0.3069984 | 7.59E-01 | 0.9714717 |
| NEK7 | 452.123484 | 0.425520384 | 0.24219366 | 1.75694271 | 0.07892758 | 0.60435431 |
| MIR181A1HG | 73.5065807 | -0.193694462 | 0.52702279 | -0.3675258 | 0.71322686 | 0.96587538 |
| ZNF281 | 31.3078202 | 0.333148039 | 0.35287388 | 0.94409947 | 0.34511882 | 0.86990996 |
| DDX59 | 31.1381874 | -0.454362367 | 0.32252433 | -1.4087693 | 0.1589034 | 0.73686885 |
| CAMSAP2 | 303.010767 | -0.202233033 | 0.25269405 | -0.8003079 | 0.42353246 | 0.90167091 |
| INAVA | 28.370069 | -0.180345895 | 0.41075649 | -0.4390579 | 0.66061956 | 0.95407427 |
| TMEM9 | 60.2517639 | 0.570442046 | 0.36077396 | 1.58116193 | 0.11384102 | 0.66818818 |
| TNNT2 | 62.9413437 | 1.400699299 | 1.34348854 | 1.04258374 | 0.29714113 | 0.83973489 |
| LAD1 | 70.0152306 | 0.085770951 | 0.58306998 | 0.14710233 | 0.88305125 | 0.98766275 |
| CSRP1 | 50.1019497 | -0.21619518 | 0.43233118 | -0.5000684 | 0.61702689 | 0.94791646 |
| IPO9 | 151.174973 | -0.378173206 | 0.19897155 | -1.9006396 | 0.05734924 | 0.54616991 |
| TIMM17A | 45.8583223 | 0.132477176 | 0.32963037 | 0.40189615 | 0.68776045 | 0.96161098 |
| RNPEP | 82.350276 | 0.009220809 | 0.2768769 | 0.03330292 | 0.97343303 | 0.99681784 |
| ELF3 | 352.295957 | 0.580451554 | 0.38568195 | 1.50500058 | 0.13232393 | 0.70223319 |
| ARL8A | 57.6882564 | -0.148393605 | 0.25105881 | -0.5910711 | 0.55447279 | 0.93523904 |
| PPP1R12B | 319.575717 | 0.210661979 | 0.21545888 | 0.97773633 | 0.32820475 | 0.8573817 |
| KDM5B | 314.545716 | 0.301621244 | 0.1656639 | 1.8206818 | 0.06865524 | 0.58250536 |
| KLHL12 | 38.3610763 | -0.363427272 | 0.36574967 | -0.9936503 | 0.32039314 | 0.85075884 |
| ADIPOR1 | 34.4660705 | -0.192552919 | 0.3213239 | -0.5992487 | 0.54900708 | 0.93523904 |
| CYB5R1 | 31.3325024 | -0.200155201 | 0.38019407 | -0.5264553 | 0.59857187 | 0.94683415 |
| BTG2 | 60.538137 | 0.610945944 | 0.79178677 | 0.77160413 | 0.44034892 | 0.90561284 |
| ATP2B4 | 183.461905 | 0.132504929 | 0.20884349 | 0.63447 | 0.52577414 | 0.93272384 |
| SNRPE | 39.0221695 | -0.233681266 | 0.38408506 | -0.6084102 | 0.54291546 | 0.93523904 |
| SOX13 | 72.1889329 | 0.619449122 | 0.4045045 | 1.53137756 | 0.12567611 | 0.69492095 |
| PLEKHA6 | 261.11317 | 0.219843848 | 0.36764993 | 0.59797059 | 0.54985956 | 0.93523904 |
| PPP1R15B | 52.7921602 | 0.105926621 | 0.32890657 | 0.32205687 | 0.74740961 | 0.97028355 |
| PIK3C2B | 66.1093343 | 0.387531643 | 0.49325286 | 0.78566527 | 0.43206361 | 0.90503769 |
| MDM4 | 282.310289 | -0.066627196 | 0.22502268 | -0.296091 | 0.76716056 | 0.97221495 |
| LRRN2 | 122.570222 | -1.734202017 | 0.75848124 | -2.2864139 | 0.02223006 | 0.39000316 |
| RBBP5 | 54.578242 | -0.183868507 | 0.30695532 | -0.5990074 | 0.54916795 | 0.93523904 |
| DSTYK | 98.5422858 | 0.007001407 | 0.22655533 | 0.03090374 | 0.97534631 | 0.99681784 |
| NUAK2 | 190.203881 | -0.767425696 | 0.4707543 | -1.6302043 | 1.03E-01 | 0.65288882 |
| CDK18 | 49.8622142 | 0.565728484 | 0.39728907 | 1.42397193 | 0.1544546 | 0.72978231 |
| ELK4 | 145.965091 | -0.029882349 | 0.3195411 | -0.0935164 | 0.92549328 | 0.99316012 |
| NUCKS1 | 489.037952 | -0.090505282 | 0.23068781 | -0.392328 | 0.6948159 | 0.96220115 |
| RAB29 | 55.737291 | 0.340420647 | 0.34086451 | 0.99869782 | 0.3179411 | 0.8490543 |
| SLC41A1 | 34.4550382 | 0.119254189 | 0.31394305 | 0.37985931 | 0.70404985 | 0.96461079 |
| RHEX | 2007.64222 | 0.278758233 | 0.65576564 | 0.4250882 | 0.67077238 | 0.95771779 |
| SRGAP2 | 141.21943 | -0.107899726 | 0.34577001 | -0.3120563 | 0.7549977 | 0.97125609 |
| EIF2D | 41.1041673 | 0.31611308 | 0.26795685 | 1.17971634 | 0.23811305 | 0.80145582 |
| MAPKAPK2 | 138.504244 | -0.195704682 | 0.30459035 | -0.6425177 | 0.52053712 | 0.93272217 |
| PIGR | 412.0148 | 2.562090802 | 0.67089969 | 3.81888802 | 1.34E-04 | 0.02918429 |
| PFKFB2 | 28.7660671 | 0.042442737 | 0.35956789 | 0.11803817 | 0.90603741 | 0.99100813 |
| CD55 | 60.2552532 | -0.934242763 | 0.55231937 | -1.6914901 | 0.09074323 | 0.6330877 |

| | | | | | | |
|---|---|---|---|---|---|---|
| CD46 | 331.606316 | -0.030153079 | 0.26106814 | -0.1154989 | 0.9080497 | 0.99100813 |
| MIR29B2CHC | 182.650371 | 0.206675221 | 0.48121884 | 0.42948282 | 0.6675719 | 0.95728611 |
| PLXNA2 | 80.5905119 | -0.386681454 | 0.32072977 | -1.2056301 | 0.22796016 | 0.79284833 |
| IRF6 | 24.4836099 | 1.006847911 | 0.52289103 | 1.9255406 | 0.05416176 | 0.53752502 |
| UTP25 | 28.3427564 | -0.50115975 | 0.34807419 | -1.4398073 | 0.14992193 | 0.72246729 |
| HHAT | 90.6194864 | 0.507535943 | 0.51602466 | 0.98354978 | 0.32533693 | 0.855037 |
| RCOR3 | 351.833024 | 0.126442274 | 0.22978267 | 0.55026898 | 0.5821349 | 0.94091128 |
| TRAF5 | 75.7761402 | -0.048309703 | 0.36078713 | -0.1339009 | 0.89348098 | 0.98842187 |
| LINC00467 | 61.8884535 | -0.164093573 | 0.34537553 | -0.4751164 | 0.63470402 | 0.94932781 |
| SLC30A1 | 62.8816595 | 0.227449352 | 0.42876703 | 0.53047305 | 0.59578399 | 0.94683415 |
| LPGAT1 | 426.187417 | 0.322000795 | 0.44224581 | 0.72810367 | 0.46655013 | 0.91462428 |
| INTS7 | 46.8314707 | -0.425825564 | 0.5400398 | -0.7885077 | 0.43039977 | 0.90364119 |
| PPP2R5A | 341.085641 | 0.225356143 | 0.18313635 | 1.23053748 | 0.2184959 | 0.78523615 |
| AL360091.3 | 50.1818695 | 0.589219483 | 0.49637043 | 1.18705597 | 0.23520554 | 0.79894819 |
| NENF | 71.6490721 | 0.313878877 | 0.33554297 | 0.9354357 | 0.34956381 | 0.8720098 |
| ATF3 | 67.2471391 | 0.417832876 | 0.72545794 | 0.57595741 | 0.56464397 | 0.93809297 |
| NSL1 | 102.944165 | 0.015152303 | 0.25007026 | 0.06059218 | 0.951684 | 0.99531843 |
| TATDN3 | 44.6461044 | -0.266959858 | 0.41267141 | -0.6469066 | 0.5176924 | 0.93257822 |
| FLVCR1 | 69.8357443 | 0.152068703 | 0.28228058 | 0.53871472 | 0.59008371 | 0.94516215 |
| ANGEL2 | 76.1470946 | 0.179319162 | 0.23924765 | 0.74951274 | 0.45354823 | 0.91077355 |
| RPS6KC1 | 257.8316 | -0.031404419 | 0.32687069 | -0.096076 | 0.92346024 | 0.99214128 |
| SMYD2 | 100.013174 | -0.372470437 | 0.29458714 | -1.2643812 | 0.20609324 | 0.7735572 |
| PTPN14 | 970.357347 | -0.490075787 | 0.28957406 | -1.6924023 | 0.0905693 | 0.6330877 |
| KCTD3 | 203.737928 | 0.107313868 | 0.25105874 | 0.42744526 | 0.66905505 | 0.95764829 |
| ESRRG | 1095.46176 | 0.180174918 | 0.38827875 | 0.46403497 | 0.64262269 | 0.94944486 |
| GPATCH2 | 310.463182 | -0.043115593 | 0.28000705 | -0.1539804 | 0.87762521 | 0.98766275 |
| SPATA17 | 182.560915 | 0.741953816 | 0.34624339 | 2.14286779 | 0.03212372 | 0.44049392 |
| RRP15 | 53.900945 | -0.502751715 | 0.31235167 | -1.6095695 | 0.10749188 | 0.65699665 |
| TGFB2 | 80.6394056 | 0.100049688 | 0.51091993 | 0.19582264 | 0.84474899 | 0.98520136 |
| LYPLAL1 | 330.282977 | 0.296968987 | 0.31606063 | 0.93959501 | 0.34742534 | 0.87017396 |
| EPRS | 119.445595 | -0.381017827 | 0.19696005 | -1.9344929 | 0.05305255 | 0.53381345 |
| BPNT1 | 38.0230799 | 0.483511341 | 0.29180667 | 1.65695779 | 0.09752801 | 0.64312419 |
| IARS2 | 102.993898 | -0.274608776 | 0.24656963 | -1.113717 | 0.26540063 | 0.81378765 |
| RAB3GAP2 | 294.87147 | -0.275733443 | 0.12141806 | -2.2709427 | 0.02315045 | 0.39839181 |
| MARK1 | 146.350552 | -0.823264707 | 0.32538283 | -2.5301418 | 0.01140164 | 0.30834879 |
| C1orf115 | 33.5828224 | 0.403091431 | 0.36156759 | 1.11484391 | 0.26491732 | 0.81378765 |
| 2-Mar | 104.868834 | 0.884383326 | 0.19162027 | 4.61529118 | 3.93E-06 | 0.00221187 |
| AL513314.2 | 30.678143 | 0.040935696 | 0.5153932 | 0.07942615 | 0.93669367 | 0.99526299 |
| MIA3 | 104.435553 | 0.023279098 | 0.18446641 | 0.12619695 | 0.89957602 | 0.98956835 |
| AIDA | 51.0463669 | 0.035424546 | 0.25858621 | 0.13699317 | 0.89103619 | 0.98766275 |
| BROX | 69.5621269 | -0.045992412 | 0.21741581 | -0.2115412 | 0.83246495 | 0.98306468 |
| FAM177B | 64.0064154 | 0.627993457 | 0.62404223 | 1.00633166 | 0.31425606 | 0.8490543 |
| AC093152.1 | 229.496968 | -0.14874955 | 0.47626022 | -0.3123283 | 0.75479103 | 0.97125609 |
| DISP1 | 29.3439517 | -0.377327998 | 0.56244414 | -0.6708719 | 0.50230211 | 0.92484445 |
| SUSD4 | 43.3312845 | -1.052004495 | 0.91538056 | -1.1492537 | 0.25045138 | 0.80680142 |
| CAPN2 | 268.993043 | -0.462868537 | 0.3296397 | -1.404165 | 0.16026972 | 0.73816762 |
| TP53BP2 | 81.1166779 | -0.044072895 | 0.28628402 | -0.1539481 | 0.87765062 | 0.98766275 |
| FBXO28 | 145.719141 | 0.017585105 | 0.1790963 | 0.09818799 | 0.92178302 | 0.99214128 |
| DEGS1 | 73.5034181 | 0.602580611 | 0.28159407 | 2.139891 | 0.03236358 | 0.44049392 |
| NVL | 127.569997 | -0.347584102 | 0.27824726 | -1.2491915 | 0.21159505 | 0.77730294 |
| CNIH4 | 71.9027123 | -0.093928386 | 0.31437868 | -0.2987747 | 0.76511198 | 0.97177982 |
| WDR26 | 269.123116 | -0.009811826 | 0.19775608 | -0.0496158 | 0.96042855 | 0.99572129 |
| CNIH3 | 142.722516 | 0.219412109 | 0.4872672 | 0.45029116 | 0.65250052 | 0.95176155 |
| DNAH14 | 687.733335 | -0.2798531 | 0.2093373 | -1.3368525 | 0.18127078 | 0.75448394 |

| | | | | | | |
|---|---|---|---|---|---|---|
| LBR | 267.058823 | -0.282748294 | 0.33697297 | -0.839083 | 0.40142273 | 0.89216435 |
| ENAH | 179.168179 | -0.184672165 | 0.28989821 | -0.6370242 | 0.5241091 | 0.93272384 |
| SRP9 | 132.426749 | 0.230977557 | 0.29498898 | 0.78300401 | 0.43362476 | 0.90514499 |
| EPHX1 | 43.6231748 | 0.117877592 | 0.54136325 | 0.21774214 | 0.82763003 | 0.98142144 |
| AL591895.1 | 41.2007585 | 0.606540858 | 0.43800693 | 1.38477459 | 0.1661214 | 0.74445571 |
| TMEM63A | 99.8661323 | 0.027323904 | 0.33982546 | 0.0804057 | 0.93591459 | 0.99501952 |
| H3F3A | 33.1537512 | 0.008638313 | 0.35780202 | 0.02414272 | 0.98073877 | 0.99681784 |
| ACBD3 | 169.943243 | 0.415216363 | 0.26496772 | 1.56704511 | 0.11710415 | 0.6781987 |
| PARP1 | 62.9948239 | 0.012200946 | 0.35604603 | 0.03426789 | 0.97266353 | 0.99681784 |
| ITPKB | 61.0207009 | -0.228606969 | 0.35690984 | -0.6405174 | 0.52183626 | 0.93272217 |
| COQ8A | 46.2243626 | 0.244615158 | 0.33662876 | 0.7266615 | 0.46743334 | 0.91462428 |
| CDC42BPA | 789.017001 | 0.060964414 | 0.14284239 | 0.42679497 | 0.66952867 | 0.95764829 |
| ZNF678 | 72.6396772 | 0.105644209 | 0.21646802 | 0.4880361 | 0.62552427 | 0.94808496 |
| SNAP47 | 37.989115 | 0.109505435 | 0.33554986 | 0.32634624 | 0.74416241 | 0.97012548 |
| ARF1 | 188.064808 | -0.382202124 | 0.27701032 | -1.3797397 | 0.16766682 | 0.74445571 |
| MRPL55 | 30.5043363 | -0.277974554 | 0.40924423 | -0.6792388 | 0.49698657 | 0.92398825 |
| GUK1 | 116.227063 | -0.316252021 | 0.38862743 | -0.8137666 | 0.41577868 | 0.89825994 |
| OBSCN | 164.522637 | 0.190746997 | 0.30988257 | 0.61554607 | 0.53819415 | 0.93516669 |
| RNF187 | 56.5791492 | -0.544769731 | 0.36761675 | -1.4818958 | 0.13836801 | 0.71091391 |
| RHOU | 70.3970895 | -0.587133229 | 0.51637829 | -1.1370215 | 0.25552929 | 0.80897837 |
| RAB4A | 61.9590299 | 0.394752938 | 0.20727852 | 1.90445658 | 0.05685075 | 0.54566463 |
| NUP133 | 99.5209109 | -0.088910301 | 0.18727944 | -0.4747467 | 0.63496751 | 0.94932781 |
| ABCB10 | 45.4589583 | -0.063196044 | 0.27473394 | -0.2300263 | 0.8180713 | 0.98010849 |
| GALNT2 | 272.018365 | -0.016487392 | 0.37204719 | -0.0443153 | 0.96465306 | 0.99586809 |
| COG2 | 49.989004 | -0.063515471 | 0.23667247 | -0.2683686 | 0.78841557 | 0.97423208 |
| C1orf198 | 38.2336626 | -0.330921993 | 0.46338509 | -0.7141404 | 0.47514039 | 0.91809716 |
| TTC13 | 68.4056525 | -0.578640347 | 0.25242391 | -2.2923357 | 0.02188628 | 0.38823827 |
| ARV1 | 28.6225796 | -0.349350345 | 0.31897722 | -1.0952204 | 0.2734201 | 0.82047446 |
| C1orf131 | 26.5740598 | -0.372900218 | 0.32707309 | -1.1401128 | 0.25423931 | 0.80897837 |
| GNPAT | 87.3020448 | -0.1402371 | 0.21727712 | -0.6454297 | 0.51864878 | 0.93257822 |
| EGLN1 | 251.340318 | -0.265619177 | 0.3527851 | -0.7529206 | 0.45149761 | 0.91077355 |
| DISC1 | 93.41229 | 0.13102895 | 0.53429545 | 0.24523688 | 0.80627302 | 0.97930627 |
| SIPA1L2 | 262.284594 | 0.820338118 | 1.04625391 | 0.78407173 | 0.43299802 | 0.90514499 |
| NTPCR | 53.3223807 | -0.036513149 | 0.25452967 | -0.1434534 | 0.88593211 | 0.98766275 |
| PCNX2 | 396.929242 | -0.259895151 | 0.26670223 | -0.9744769 | 0.32981983 | 0.85851742 |
| MAP3K21 | 299.277345 | 0.165535461 | 0.33312032 | 0.49692393 | 0.61924269 | 0.9480368 |
| KCNK1 | 52.2086409 | 1.302710129 | 0.51440569 | 2.53245668 | 0.01132664 | 0.30823253 |
| SLC35F3 | 208.325451 | -0.92232493 | 0.72487376 | -1.2723939 | 0.20323321 | 0.77155952 |
| TARBP1 | 82.5483381 | -0.206989185 | 0.32508285 | -0.6367275 | 0.52430237 | 0.93272384 |
| IRF2BP2 | 132.504952 | 0.471676773 | 0.24576525 | 1.91921669 | 0.05495692 | 0.53804407 |
| TOMM20 | 160.150278 | 0.241264726 | 0.4490856 | 0.5372355 | 0.59110495 | 0.94527451 |
| ARID4B | 629.38089 | 0.165564871 | 0.17681606 | 0.93636783 | 0.34908383 | 0.87178925 |
| B3GALNT2 | 87.3438073 | 0.099426414 | 0.26226958 | 0.37910006 | 0.70461356 | 0.96461079 |
| LYST | 155.455732 | 0.122088805 | 0.21024423 | 0.5806999 | 0.56144273 | 0.93747805 |
| GPR137B | 78.2247005 | 0.468278834 | 0.27763953 | 1.68664322 | 0.09167198 | 0.63464873 |
| ERO1B | 79.3047899 | 0.580305791 | 0.46435387 | 1.24970594 | 0.21140699 | 0.77730294 |
| LGALS8 | 104.417512 | -0.274217523 | 0.23999184 | -1.1426118 | 0.25319978 | 0.80798674 |
| HEATR1 | 85.6574263 | -0.41999511 | 0.25508762 | -1.6464739 | 0.09966622 | 0.64594233 |
| MTR | 421.501502 | 0.218142743 | 0.338085 | 0.64523047 | 0.51877784 | 0.93257822 |
| LINC01139 | 42.1176774 | -0.019297602 | 0.70954188 | -0.0271973 | 0.97830239 | 0.99681784 |
| FMN2 | 74.9264397 | 2.333877538 | 1.26154023 | 1.85002228 | 0.06431034 | 0.57358354 |
| FH | 28.9019231 | 0.064719882 | 0.33837659 | 0.19126584 | 0.84831732 | 0.98545133 |
| OPN3 | 33.8516802 | 0.163305535 | 0.46554247 | 0.35078547 | 0.7257493 | 0.96781837 |
| CEP170 | 200.700074 | -0.359270565 | 0.26460664 | -1.3577534 | 0.17454194 | 0.74910274 |

| | | | | | | |
|---|---|---|---|---|---|---|
| SDCCAG8 | 446.21734 | -0.281189467 | 0.13978525 | -2.0115819 | 0.04426403 | 0.49952159 |
| AKT3 | 787.212304 | 0.283033084 | 0.37262014 | 0.75957538 | 0.44750844 | 0.90951928 |
| ZBTB18 | 45.2703705 | 0.240850058 | 0.34706029 | 0.69397182 | 0.48769988 | 0.92034617 |
| AL645465.1 | 33.7130334 | -0.871364292 | 0.38983538 | -2.2352109 | 0.02540351 | 0.41555526 |
| ADSS | 156.913786 | -0.660594212 | 0.41177077 | -1.6042766 | 0.10865309 | 0.65764016 |
| CATSPERE | 111.687736 | -0.223228785 | 0.38292038 | -0.582964 | 0.55991755 | 0.93744835 |
| DESI2 | 104.68154 | 0.186705754 | 0.1959786 | 0.9526844 | 0.34075 | 0.86809688 |
| COX20 | 113.740735 | 0.081959323 | 0.27108718 | 0.30233566 | 0.7623962 | 0.9714717 |
| HNRNPU | 579.774603 | -0.173511756 | 0.18474217 | -0.9392104 | 0.34762275 | 0.87032889 |
| EFCAB2 | 235.773719 | 0.066731323 | 0.40727208 | 0.16384949 | 0.86984963 | 0.98758914 |
| KIF26B | 428.294976 | 0.112123616 | 0.65453871 | 0.17130173 | 0.86398652 | 0.98670067 |
| SMYD3 | 1025.82503 | -0.259228922 | 0.17314182 | -1.4972057 | 0.13433973 | 0.70619627 |
| CNST | 114.725669 | -0.048353867 | 0.19800824 | -0.2442013 | 0.80707493 | 0.97930627 |
| SCCPDH | 270.339851 | -0.542568397 | 0.57250385 | -0.9477113 | 0.34327642 | 0.86909179 |
| AHCTF1 | 198.955778 | -0.017464002 | 0.19126066 | -0.09131 | 0.92724631 | 0.99336859 |
| ZNF124 | 126.24333 | -0.091029727 | 0.3942059 | -0.2309192 | 0.81737755 | 0.98010849 |
| AL390728.6 | 39.5968142 | -0.708279835 | 0.41828428 | -1.6932978 | 0.0903988 | 0.6330877 |
| ZNF496 | 75.5703652 | -0.204584702 | 0.36218492 | -0.5648626 | 0.57216725 | 0.93998433 |
| ZNF672 | 25.8131499 | -0.234749263 | 0.30294193 | -0.7748986 | 0.43839961 | 0.90561284 |
| ZNF692 | 31.1454975 | 0.062847832 | 0.37212816 | 0.1688876 | 0.86588505 | 0.98670067 |
| FAM110C | 98.512504 | -0.996777267 | 0.42147606 | -2.3649677 | 0.01803164 | 0.35858401 |
| SH3YL1 | 190.780108 | -0.062654321 | 0.22928222 | -0.2732629 | 0.78465114 | 0.97423208 |
| ACP1 | 75.6339456 | -0.324936758 | 0.31626395 | -1.0274227 | 0.30422147 | 0.84488735 |
| SNTG2 | 63.6722942 | 0.85777008 | 0.67419055 | 1.2722962 | 0.2032679 | 0.77155952 |
| PXDN | 108.279446 | 0.618590009 | 0.49385813 | 1.25256622 | 0.21036361 | 0.77730294 |
| EIPR1 | 101.724014 | -0.395797414 | 0.39598815 | -0.9995183 | 0.31754366 | 0.8490543 |
| TRAPPC12 | 179.678173 | 0.066289988 | 0.32955708 | 0.20114873 | 0.84058228 | 0.9847056 |
| ADI1 | 58.9435441 | 0.347820371 | 0.31948606 | 1.08868717 | 0.27629186 | 0.82231303 |
| RPS7 | 386.070281 | -0.060758581 | 0.61095523 | -0.0994485 | 0.92078218 | 0.99214128 |
| RNF144A | 63.2565754 | -0.440187173 | 0.45555981 | -0.9662555 | 0.33391634 | 0.86191987 |
| ID2 | 139.516879 | 0.484319357 | 0.43905728 | 1.10308923 | 0.26998842 | 0.81705157 |
| KIDINS220 | 260.875594 | -0.120742442 | 0.1783259 | -0.6770887 | 0.4983497 | 0.92398825 |
| MBOAT2 | 151.898368 | -0.241281367 | 0.57606461 | -0.4188443 | 0.67532994 | 0.95910677 |
| ASAP2 | 370.032646 | -0.579484858 | 0.50287311 | -1.1523481 | 0.24917807 | 0.80672756 |
| ITGB1BP1 | 68.9810522 | -0.05469474 | 0.31082537 | -0.1759661 | 0.86032054 | 0.9864199 |
| CPSF3 | 37.1968102 | -0.493667069 | 0.2873642 | -1.7179143 | 0.08581225 | 0.62178179 |
| IAH1 | 65.1269796 | 0.056094906 | 0.26487685 | 0.21177731 | 0.83228077 | 0.98306468 |
| ADAM17 | 171.01924 | 0.02941124 | 0.17610252 | 0.16701203 | 0.86736058 | 0.98670067 |
| YWHAQ | 204.153561 | 0.041041769 | 0.17441022 | 0.23531745 | 0.81396235 | 0.97937868 |
| TAF1B | 108.20542 | -0.226655075 | 0.33414191 | -0.6783198 | 0.49756895 | 0.92398825 |
| GRHL1 | 34.8664857 | 0.315267639 | 0.42704627 | 0.73825171 | 0.4603615 | 0.91368087 |
| KLF11 | 33.1602752 | 0.241917783 | 0.43609724 | 0.55473358 | 0.57907691 | 0.94034284 |
| CYS1 | 172.159239 | 0.307559186 | 0.72317064 | 0.42529269 | 0.67062332 | 0.95771779 |
| HPCAL1 | 196.389853 | 0.191669629 | 0.31505734 | 0.60836426 | 0.54294591 | 0.93523904 |
| ODC1 | 233.109862 | -0.519863995 | 0.63036602 | -0.8247018 | 0.40954091 | 0.8959746 |
| NOL10 | 181.120181 | -0.370593528 | 0.24476535 | -1.5140768 | 0.13000637 | 0.69904834 |
| PDIA6 | 184.011442 | 0.339135067 | 0.35499511 | 0.9553232 | 0.33941428 | 0.8672311 |
| PQLC3 | 27.9598015 | 0.876880102 | 0.44002489 | 1.9927966 | 0.04628372 | 0.50827029 |
| ROCK2 | 549.289465 | -0.180349629 | 0.21205106 | -0.850501 | 3.95E-01 | 0.88760715 |
| E2F6 | 34.2441228 | -0.248295883 | 0.28962881 | -0.85729 | 0.39128463 | 0.88670817 |
| LPIN1 | 249.086544 | -0.939225285 | 0.56957173 | -1.6490026 | 0.0991471 | 0.64388477 |
| MIR3681HG | 148.745744 | -1.337574347 | 1.26448792 | -1.0577992 | 0.29014699 | 0.83506029 |
| TRIB2 | 52.2104841 | -1.43339292 | 0.48807408 | -2.9368347 | 0.00331581 | 0.18794154 |
| FAM84A | 24.7002863 | 0.089266403 | 0.6160588 | 0.14489916 | 0.88479049 | 0.98766275 |

| | | | | | | |
|---|---|---|---|---|---|---|
| NBAS | 385.52088 | -0.189392183 | 0.16081606 | -1.1776944 | 0.23891844 | 0.80174646 |
| DDX1 | 119.921732 | -0.294253854 | 0.16634355 | -1.7689526 | 0.07690178 | 0.59726395 |
| SMC6 | 94.4386682 | -0.082411866 | 0.30469742 | -0.2704712 | 0.7867978 | 0.97423208 |
| KCNS3 | 45.1252808 | 0.372229612 | 0.63945998 | 0.58209994 | 0.56049937 | 0.93747805 |
| RDH14 | 32.5682178 | 0.192011492 | 0.40861158 | 0.46991202 | 0.63841787 | 0.94944486 |
| LINC01376 | 186.995295 | 0.384361588 | 0.33028056 | 1.16374268 | 0.24452831 | 0.80476923 |
| TTC32 | 35.2573869 | -0.339493396 | 0.3463263 | -0.9802703 | 0.32695269 | 0.85641576 |
| WDR35 | 171.399867 | 0.139516527 | 0.17024596 | 0.81949977 | 0.41250133 | 0.89706788 |
| LAPTM4A | 466.847748 | 0.340088143 | 0.35909409 | 0.94707253 | 0.34360182 | 0.86909179 |
| SDC1 | 33.5011573 | -0.970332971 | 0.54233036 | -1.7891917 | 0.07358395 | 0.59320069 |
| PUM2 | 576.15312 | -0.175498482 | 0.11085594 | -1.583122 | 0.11339366 | 0.66657543 |
| RHOB | 138.448793 | 0.072554921 | 0.58431182 | 0.12417158 | 0.90117943 | 0.9900784 |
| HS1BP3 | 30.6959136 | 0.11974905 | 0.41019177 | 0.29193431 | 0.77033685 | 0.97313769 |
| GDF7 | 49.5445456 | 0.417943713 | 0.44987892 | 0.92901376 | 0.35288195 | 0.8720098 |
| LDAH | 183.461687 | -0.217152414 | 0.25469511 | -0.8525975 | 0.39388255 | 0.88760715 |
| KLHL29 | 354.489846 | -0.849363529 | 0.26183581 | -3.2438784 | 0.00117914 | 0.10860566 |
| ATAD2B | 290.752801 | -0.113711546 | 0.22720568 | -0.5004784 | 0.61673823 | 0.94791646 |
| UBXN2A | 118.941182 | 0.118859842 | 0.3283705 | 0.3619687 | 0.71737542 | 0.96742001 |
| SF3B6 | 81.3196679 | -0.077548345 | 0.54075932 | -0.1434064 | 0.88596923 | 0.98766275 |
| FAM228B | 55.718928 | 0.740131875 | 0.48817483 | 1.51612052 | 0.12948888 | 0.69904834 |
| TP53I3 | 28.5725664 | -0.369415092 | 0.38394231 | -0.962163 | 0.33596772 | 0.86391445 |
| ITSN2 | 374.260344 | 0.34352248 | 0.17189792 | 1.99840972 | 0.04567226 | 0.50577407 |
| NCOA1 | 641.014633 | 0.360387488 | 0.17151125 | 2.10124687 | 0.0356193 | 0.45736901 |
| DNAJC27 | 44.2116593 | 0.161713625 | 0.33719523 | 0.47958456 | 0.63152283 | 0.94932781 |
| DNAJC27-AS | 39.6369206 | -0.026351675 | 0.72110245 | -0.0365436 | 0.97084892 | 0.99681784 |
| DNMT3A | 140.838093 | 0.457461191 | 0.32696389 | 1.39911839 | 0.16177748 | 0.74071637 |
| DTNB | 284.153819 | -0.105439767 | 0.34856406 | -0.3024975 | 0.76227282 | 0.9714717 |
| ASXL2 | 194.059017 | -0.292272545 | 0.15823554 | -1.8470727 | 0.0647366 | 0.57495764 |
| RAB10 | 438.140814 | -0.117259268 | 0.19912702 | -0.5888667 | 0.5559507 | 0.93523904 |
| HADHA | 147.01358 | 0.266054232 | 0.18378776 | 1.44761673 | 0.14772427 | 0.72246583 |
| HADHB | 203.986987 | 0.188912995 | 0.24476254 | 0.7718215 | 0.44022015 | 0.90561284 |
| SELENOI | 109.566528 | -0.54951162 | 0.28378207 | -1.936386 | 0.05282045 | 0.5325969 |
| MAPRE3 | 30.5657623 | 0.298428522 | 0.29644894 | 1.00667763 | 0.31408971 | 0.8490543 |
| TMEM214 | 32.248413 | 0.249564967 | 0.39982149 | 0.62419097 | 0.53250217 | 0.93387739 |
| AGBL5 | 48.5888651 | 0.424781475 | 0.33095085 | 1.28351831 | 0.19931055 | 0.77110248 |
| OST4 | 90.9423291 | 0.381486365 | 0.43873597 | 0.8695124 | 0.38456693 | 0.88328503 |
| ATRAID | 73.3847418 | -0.313632669 | 0.49950478 | -0.6278872 | 0.53007782 | 0.93366189 |
| CAD | 30.4069318 | -0.337265801 | 0.40418343 | -0.8344375 | 0.40403451 | 0.89360496 |
| MPV17 | 47.2975133 | -0.275064914 | 0.26432685 | -1.0406242 | 0.29804999 | 0.83973489 |
| GTF3C2 | 95.2137176 | -0.299183437 | 0.20790234 | -1.4390576 | 0.15013421 | 0.72246729 |
| SNX17 | 41.1268101 | 0.113165536 | 0.45916362 | 0.24646015 | 0.80532605 | 0.97930627 |
| PPM1G | 162.583681 | 0.177066646 | 0.37862356 | 0.46765881 | 0.6400286 | 0.94944486 |
| NRBP1 | 68.1822168 | -0.576979541 | 0.27706735 | -2.0824523 | 0.03730117 | 0.46890807 |
| IFT172 | 71.3787261 | 0.929527955 | 0.33557622 | 2.76994585 | 0.00560656 | 0.23658701 |
| ZNF512 | 101.295632 | -0.034410388 | 0.21100087 | -0.1630817 | 0.87045408 | 0.98766275 |
| SUPT7L | 34.4922785 | -0.006264395 | 0.29452797 | -0.0212693 | 0.98303086 | 0.99721376 |
| SLC4A1AP | 43.6501037 | -0.226258606 | 0.27576284 | -0.8204826 | 0.41194106 | 0.89701828 |
| MRPL33 | 218.188128 | -0.245949771 | 0.31815918 | -0.77304 | 0.43949871 | 0.90561284 |
| RBKS | 158.147927 | 0.077575111 | 0.43916434 | 0.17664256 | 0.85978917 | 0.9864199 |
| BABAM2 | 836.2662 | -0.07787432 | 0.23811049 | -0.3270512 | 0.74362916 | 0.97012548 |
| FOSL2 | 353.547812 | -0.14256216 | 0.52648633 | -0.2707804 | 0.78655996 | 0.97423208 |
| PPP1CB | 179.006189 | 0.096505763 | 0.2881573 | 0.33490653 | 0.73769559 | 0.9698508 |
| TRMT61B | 61.3884235 | -0.154868521 | 0.27449929 | -0.5641855 | 0.57262789 | 0.93998433 |
| WDR43 | 156.055387 | -0.763093314 | 0.2706703 | -2.8192724 | 0.00481326 | 0.21962686 |

| | | | | | | |
|---|---|---|---|---|---|---|
| CLIP4 | 199.587917 | 0.646939541 | 0.53173284 | 1.21666275 | 0.22373256 | 0.78791697 |
| YPEL5 | 87.6478339 | 0.274337777 | 0.28792422 | 0.95281244 | 0.34068511 | 0.86809688 |
| LBH | 70.5624264 | 0.154568007 | 0.69006206 | 0.22399146 | 0.82276395 | 0.98094663 |
| LCLAT1 | 223.07687 | 0.124206691 | 0.19871767 | 0.62504099 | 0.53194416 | 0.93366189 |
| GALNT14 | 1314.28776 | 0.084949356 | 0.26973732 | 0.31493364 | 0.75281204 | 0.97096495 |
| DPY30 | 70.313831 | -0.105324254 | 0.30632246 | -0.3438346 | 0.73097069 | 0.96926471 |
| SPAST | 91.2371521 | 0.209448934 | 0.21024456 | 0.99621573 | 0.31914534 | 0.84966459 |
| SLC30A6 | 100.66942 | -0.250969936 | 0.18809105 | -1.3343003 | 0.18210548 | 0.75456023 |
| YIPF4 | 175.355871 | 0.192297618 | 0.14939022 | 1.28721692 | 0.19801869 | 0.77110248 |
| BIRC6 | 883.209934 | -0.076035538 | 0.12775589 | -0.5951627 | 0.55173474 | 0.93523904 |
| TTC27 | 137.674997 | -0.477669147 | 0.25569241 | -1.8681397 | 0.0617426 | 0.56326893 |
| LINC00486 | 165.167448 | -0.262606628 | 0.21222448 | -1.2374002 | 0.21593853 | 0.78293142 |
| LTBP1 | 124.317699 | -1.886198915 | 1.37458841 | -1.3721918 | 0.17000375 | 0.74445571 |
| RASGRP3 | 79.2244789 | -0.975177353 | 0.65574334 | -1.4871327 | 0.13697977 | 0.71041106 |
| FAM98A | 64.3734776 | 0.232475466 | 0.34911238 | 0.66590439 | 0.50547222 | 0.9257297 |
| LINC01317 | 35.6431598 | 0.491719143 | 0.67413683 | 0.72940555 | 0.46575363 | 0.91462428 |
| AC068599.1 | 64.5653653 | 0.795492659 | 0.55236308 | 1.44016262 | 0.1498214 | 0.72246729 |
| LINC01320 | 2598.7516 | -0.017994158 | 0.44599967 | -0.0403457 | 0.96781754 | 0.9965306 |
| AC073218.2 | 30.4778409 | 0.257926155 | 0.83771363 | 0.30789299 | 0.75816376 | 0.9714717 |
| AC012593.1 | 1743.07318 | 1.314841677 | 1.04247798 | 1.26126566 | 0.20721315 | 0.77401635 |
| CRIM1 | 1211.93078 | 0.132446372 | 0.39772203 | 0.33301241 | 0.73912492 | 0.9698508 |
| FEZ2 | 225.686278 | -0.091363902 | 0.3425026 | -0.2667539 | 0.78965865 | 0.97423208 |
| STRN | 370.09179 | -0.353545679 | 0.32760449 | -1.0791845 | 0.2805055 | 0.825559 |
| HEATR5B | 136.217821 | -0.06006172 | 0.31350467 | -0.1915816 | 0.84806997 | 0.98545133 |
| EIF2AK2 | 202.799693 | -1.433083111 | 0.5851724 | -2.448993 | 0.01432562 | 0.32762834 |
| CEBPZOS | 99.6680945 | 0.148500112 | 0.34138359 | 0.43499488 | 0.66356615 | 0.95482952 |
| CEBPZ | 105.21008 | -0.139832676 | 0.32830029 | -0.4259292 | 0.67015945 | 0.95764829 |
| NDUFAF7 | 62.9881528 | 0.36841899 | 0.22912285 | 1.60795396 | 0.10784526 | 0.65699665 |
| PRKD3 | 182.501206 | 0.118268831 | 0.2323866 | 0.50893136 | 0.61080033 | 0.94791646 |
| QPCT | 28.0214944 | 0.421009508 | 0.40443326 | 1.04098637 | 0.29788188 | 0.83973489 |
| CDC42EP3 | 49.2338348 | -0.68393177 | 0.40667999 | -1.6817443 | 0.09261842 | 0.63577454 |
| RMDN2 | 108.404317 | 0.471278831 | 0.34768668 | 1.35546991 | 0.17526789 | 0.74934286 |
| CYP1B1 | 217.879905 | 0.801182372 | 0.64609454 | 1.24003891 | 0.214961 | 0.78293142 |
| CYP1B1-AS1 | 149.502611 | 1.008597376 | 0.50336986 | 2.00369046 | 0.04510323 | 0.50354759 |
| ATL2 | 202.662106 | 0.345531146 | 0.2143278 | 1.61216208 | 0.10692667 | 0.65699665 |
| HNRNPLL | 134.118085 | 0.171160914 | 0.2737277 | 0.62529629 | 5.32E-01 | 0.93366189 |
| GALM | 95.3734757 | 0.607707001 | 0.27570129 | 2.20422253 | 0.0275087 | 0.42500938 |
| SRSF7 | 136.278417 | -0.100302075 | 0.25759172 | -0.3893839 | 0.69699216 | 0.96220115 |
| DHX57 | 117.615032 | 0.141582901 | 0.21204091 | 0.66771502 | 0.50431552 | 0.92552859 |
| MORN2 | 39.0606734 | 1.365921372 | 0.38762068 | 3.52386097 | 0.00042531 | 0.06267722 |
| SOS1 | 519.939012 | 0.117949352 | 0.19353514 | 0.60944671 | 0.54222839 | 0.93523904 |
| MAP4K3 | 534.897289 | 0.284537027 | 0.22149489 | 1.28462117 | 0.1989247 | 0.77110248 |
| AC007388.1 | 238.050178 | 0.662569701 | 0.28396503 | 2.33327925 | 0.01963349 | 0.37455228 |
| TMEM178A | 40.9580677 | 1.425325011 | 0.79178604 | 1.8001391 | 7.18E-02 | 0.59090333 |
| THUMPD2 | 103.641972 | -0.264102296 | 0.16416909 | -1.6087212 | 0.10767732 | 0.65699665 |
| SLC8A1 | 82.1260764 | -0.926839507 | 1.25344435 | -0.7394341 | 0.45964344 | 0.91368087 |
| EML4 | 486.771635 | 0.417145143 | 0.30480617 | 1.36855874 | 0.17113725 | 0.74445571 |
| COX7A2L | 121.006277 | 0.113459109 | 0.28904014 | 0.39253755 | 0.69466107 | 0.96220115 |
| MTA3 | 264.073046 | 0.275571482 | 0.21691929 | 1.27038719 | 0.20394675 | 0.77155952 |
| ZFP36L2 | 296.309125 | 0.010941116 | 0.36610034 | 0.02988557 | 0.97615832 | 0.99681784 |
| THADA | 447.229573 | -0.230814997 | 0.1830362 | -1.2610347 | 0.20729635 | 0.77401635 |
| PLEKHH2 | 573.456228 | -0.034218731 | 0.30575062 | -0.1119171 | 9.11E-01 | 0.99100813 |
| DYNC2LI1 | 358.824121 | -0.024518232 | 0.37873842 | -0.0647366 | 9.48E-01 | 0.9952642 |
| LRPPRC | 260.36409 | -0.365042658 | 0.20130957 | -1.8133398 | 0.06977943 | 0.58513328 |

| | | | | | | |
|---|---|---|---|---|---|---|
| PPM1B | 251.478262 | 0.435059989 | 0.31579332 | 1.37767317 | 0.16830422 | 0.74445571 |
| SLC3A1 | 274.830358 | 2.086291776 | 0.50873755 | 4.10091955 | 4.12E-05 | 0.01313957 |
| PREPL | 287.064235 | 0.362933745 | 0.26051107 | 1.39316056 | 0.16357124 | 0.74227469 |
| CAMKMT | 944.507601 | 0.017719203 | 0.24322851 | 0.07285003 | 0.94192546 | 0.99526299 |
| SRBD1 | 153.212727 | -0.273247084 | 0.22226035 | -1.2294009 | 0.21892152 | 0.78523615 |
| PRKCE | 583.991281 | -1.29553283 | 0.39430739 | -3.2855911 | 0.00101769 | 0.1048217 |
| EPAS1 | 69.7614512 | 1.187332079 | 0.50557318 | 2.34848709 | 0.01884985 | 0.36488708 |
| RHOQ | 138.080549 | 0.067054062 | 0.30936794 | 0.21674535 | 0.82840681 | 0.98142144 |
| PIGF | 67.4063397 | -0.23905334 | 0.32649034 | -0.7321912 | 0.4640519 | 0.91462428 |
| CRIPT | 29.1829976 | 0.97806622 | 0.48702776 | 2.00823503 | 0.04461832 | 0.50164191 |
| SOCS5 | 113.022721 | -0.054969687 | 0.26832214 | -0.2048645 | 0.83767798 | 0.98462033 |
| MCFD2 | 86.0059036 | -0.171063857 | 0.26028638 | -0.657214 | 0.51104332 | 0.92819651 |
| TTC7A | 98.8341514 | 0.036718713 | 0.29908016 | 0.12277215 | 0.90228753 | 0.9900784 |
| AC106869.1 | 93.9145663 | 0.212655254 | 0.5493359 | 0.38711334 | 0.6986723 | 0.96256506 |
| EPCAM | 309.164634 | -0.377936614 | 0.60813954 | -0.6214636 | 0.53429461 | 0.93457315 |
| MSH2 | 114.196505 | 0.266638931 | 0.21800792 | 1.22306994 | 0.22130328 | 0.78687916 |
| MSH6 | 50.5756722 | 0.154326974 | 0.2554046 | 0.60424508 | 0.54568071 | 0.93523904 |
| FBXO11 | 462.300534 | -0.003349384 | 0.16190614 | -0.0206872 | 0.98349518 | 0.99731258 |
| FOXN2 | 61.7134745 | -0.297133981 | 0.34741834 | -0.8552628 | 0.39240568 | 0.88760715 |
| PPP1R21 | 289.43187 | 0.39079391 | 0.26737226 | 1.46160978 | 0.14384817 | 0.71631428 |
| GPR75-ASB3 | 80.9392287 | 0.112272088 | 0.28227578 | 0.397739 | 0.69082259 | 0.96220115 |
| ERLEC1 | 60.1400994 | -0.001251679 | 0.24083526 | -0.0051972 | 0.99585322 | 0.99866813 |
| PSME4 | 264.206368 | -0.063175376 | 0.20846167 | -0.3030551 | 0.76184786 | 0.9714717 |
| ACYP2 | 407.483791 | 0.042656839 | 0.27401065 | 0.15567584 | 0.87628854 | 0.98766275 |
| SPTBN1 | 647.020299 | 0.035997875 | 0.23373171 | 0.15401365 | 0.87759897 | 0.98766275 |
| EML6 | 182.753954 | 0.007951676 | 0.76351 | 0.01041463 | 0.99169048 | 0.99866813 |
| RTN4 | 812.590984 | -0.332612627 | 0.35508307 | -0.9367178 | 0.34890374 | 0.87178925 |
| CLHC1 | 93.5173658 | 0.70985595 | 0.28640711 | 2.47848581 | 0.01319414 | 0.31992293 |
| RPS27A | 691.819877 | -0.076816553 | 0.56033907 | -0.1370894 | 0.89096012 | 0.98766275 |
| MTIF2 | 42.0571225 | -0.059852512 | 0.3225094 | -0.1855838 | 0.85277118 | 0.9864199 |
| AC012358.3 | 48.0162611 | -0.262931032 | 0.41217577 | -0.63791 | 0.52353229 | 0.93272384 |
| CCDC88A | 112.420028 | 0.380481847 | 0.18045097 | 2.10850536 | 0.0349873 | 0.45597142 |
| CFAP36 | 124.51991 | 0.276912666 | 0.17782014 | 1.55726268 | 0.11940813 | 0.68603665 |
| PPP4R3B | 258.749524 | -0.026912509 | 0.19734294 | -0.1363743 | 0.89152537 | 0.987861 |
| PNPT1 | 109.191329 | -1.6502453 | 0.44497884 | -3.7085928 | 0.00020841 | 0.03842424 |
| EFEMP1 | 86.7913569 | -0.258117232 | 0.63033357 | -0.4094931 | 0.68217785 | 0.9609694 |
| CCDC85A | 172.479608 | 0.822855724 | 1.02724358 | 0.80103272 | 0.4231127 | 0.90126675 |
| VRK2 | 170.180724 | -0.313136826 | 0.24250343 | -1.2912676 | 0.1966109 | 0.77059568 |
| FANCL | 119.871471 | -0.30852665 | 0.32011771 | -0.9637913 | 0.33515059 | 0.863477 |
| BCL11A | 33.3597731 | 0.737249942 | 0.56760296 | 1.29888318 | 0.19398402 | 0.7687432 |
| PAPOLG | 65.5894205 | 0.180936806 | 0.2892592 | 0.62551789 | 0.53163121 | 0.93366189 |
| REL | 82.6198061 | -0.041193533 | 0.36192823 | -0.1138169 | 0.90938298 | 0.99100813 |
| PUS10 | 42.8651243 | 1.031370537 | 0.36787068 | 2.80362259 | 0.0050532 | 0.22203954 |
| PEX13 | 63.9578224 | 0.096828484 | 0.26264691 | 0.36866409 | 0.71237812 | 0.96587538 |
| KIAA1841 | 46.7702728 | 0.005788439 | 0.32795089 | 0.01765032 | 0.98591781 | 0.99764442 |
| C2orf74 | 32.7764258 | -0.71872632 | 0.60721702 | -1.18364 | 0.23655562 | 0.79958336 |
| USP34 | 961.066236 | -0.022062785 | 0.14325202 | -0.1540138 | 0.87759887 | 0.98766275 |
| XPO1 | 471.820652 | -0.353319225 | 0.36153524 | -0.9772746 | 0.3284332 | 0.8574012 |
| FAM161A | 29.941112 | 0.764851491 | 0.4270683 | 1.79093482 | 0.07330375 | 0.59320069 |
| CCT4 | 109.313087 | 0.075607055 | 0.30435541 | 0.24841699 | 0.80381179 | 0.97880236 |
| COMMD1 | 373.076423 | -0.083776498 | 0.14143435 | -0.5923349 | 0.55362637 | 0.93523904 |
| B3GNT2 | 52.4239157 | 0.728107824 | 0.27497719 | 2.64788447 | 0.00809972 | 0.27246484 |
| EHBP1 | 679.127713 | 0.134265793 | 0.25195023 | 0.53290601 | 0.59409864 | 0.94649617 |
| WDPCP | 473.544213 | -0.011518129 | 0.20412961 | -0.0564256 | 0.95500279 | 0.99531843 |

| | | | | | | |
|---|---|---|---|---|---|---|
| MDH1 | 110.954693 | 0.623910055 | 0.33141983 | 1.88253689 | 0.05976316 | 0.55922073 |
| UGP2 | 164.475364 | 0.084837508 | 0.31350922 | 0.27060611 | 0.786694 | 0.97423208 |
| VPS54 | 342.025157 | 0.102735952 | 0.19181741 | 0.53559244 | 0.59224026 | 0.94649617 |
| PELI1 | 188.749923 | -0.049847657 | 0.28796896 | -0.1731008 | 0.8625722 | 0.98669442 |
| AC008074.1 | 47.4838651 | 0.141810653 | 0.50964006 | 0.27825649 | 0.78081548 | 0.97423208 |
| AFTPH | 221.121539 | 0.039297693 | 0.21118835 | 0.18607888 | 0.8523829 | 0.9864199 |
| SERTAD2 | 87.2893335 | -0.036686901 | 0.30633953 | -0.119759 | 0.9046741 | 0.99061193 |
| CEP68 | 45.9468201 | 0.397343992 | 0.27785698 | 1.43003065 | 0.15270822 | 0.72775725 |
| RAB1A | 394.031905 | 0.236973296 | 0.22112019 | 1.07169454 | 0.28385725 | 0.82918183 |
| ACTR2 | 263.656083 | -0.183094461 | 0.23257334 | -0.7872547 | 0.43113276 | 0.90447235 |
| SPRED2 | 253.832918 | -0.100795323 | 0.16007012 | -0.6296948 | 0.5288943 | 0.93366189 |
| MEIS1 | 88.4027131 | 1.485571784 | 0.81375842 | 1.82556856 | 0.06791528 | 0.5809835 |
| ETAA1 | 43.2075689 | -0.005247051 | 0.2547845 | -0.0205941 | 0.98356947 | 0.99731258 |
| C1D | 72.5343545 | 0.200627148 | 0.29429731 | 0.68171588 | 0.49541862 | 0.92310953 |
| PNO1 | 26.1167936 | -0.697067907 | 0.43318419 | -1.6091721 | 0.10757872 | 0.65699665 |
| APLF | 103.678919 | 0.564073071 | 0.35733486 | 1.57855594 | 0.11443795 | 0.66978434 |
| ANTXR1 | 73.6189424 | 1.422042097 | 0.60298909 | 2.35832145 | 0.01835779 | 0.36034684 |
| GFPT1 | 198.360346 | 0.437323027 | 0.2432963 | 1.79749151 | 0.07225762 | 0.59090333 |
| NFU1 | 59.9212035 | 0.402490809 | 0.25279264 | 1.59217769 | 0.11134478 | 0.66300165 |
| AAK1 | 469.595723 | 0.150659216 | 0.242517 | 0.62123156 | 0.53444727 | 0.93457315 |
| ANXA4 | 453.504106 | 0.236750197 | 0.33231769 | 0.71242129 | 0.47620394 | 0.91809716 |
| GMCL1 | 98.4288339 | -0.021286181 | 0.24867993 | -0.0855967 | 0.93178702 | 0.99411667 |
| MXD1 | 40.0797048 | 0.00435128 | 0.33202067 | 0.01310545 | 0.98954366 | 0.99839055 |
| PCBP1 | 151.427474 | -0.218122918 | 0.42001176 | -0.5193257 | 0.6035336 | 0.94683415 |
| C2orf42 | 30.7173335 | 0.229310667 | 0.36652821 | 0.62562897 | 0.53155833 | 0.93366189 |
| PCYOX1 | 96.5077885 | 0.907655197 | 0.49198914 | 1.84486836 | 0.06505669 | 0.57495764 |
| SNRPG | 53.9850066 | -0.284796299 | 0.39327338 | -0.7241688 | 0.46896213 | 0.91462428 |
| FAM136A | 43.2537797 | -0.207390474 | 0.33526278 | -0.6185908 | 0.53618595 | 0.93469807 |
| TGFA | 113.562748 | -0.338351235 | 0.52922845 | -0.6393293 | 0.52260875 | 0.93272217 |
| NAGK | 41.5606604 | -0.085425999 | 0.34962007 | -0.2443395 | 0.80696788 | 0.97930627 |
| MCEE | 58.7887365 | 0.466401529 | 0.33698237 | 1.38405321 | 0.16634216 | 0.74445571 |
| MPHOSPH10 | 47.3217818 | -0.692797958 | 0.2579863 | -2.685406 | 0.00724418 | 0.26418897 |
| PAIP2B | 33.3018784 | 0.597260718 | 0.52380731 | 1.14022982 | 0.25419057 | 0.80897837 |
| ZNF638 | 751.189825 | 0.231057502 | 0.16993928 | 1.35964743 | 0.17394152 | 0.74910274 |
| EXOC6B | 746.10235 | -0.204632747 | 0.18793166 | -1.0888678 | 0.27621217 | 0.82231303 |
| SFXN5 | 65.7421842 | -0.198585345 | 0.26709712 | -0.7434949 | 0.45718211 | 0.91188421 |
| RAB11FIP5 | 35.5342209 | -0.171910886 | 0.36416275 | -0.4720716 | 0.6368757 | 0.94944486 |
| CCT7 | 99.8667908 | -0.139551065 | 0.39925496 | -0.3495287 | 0.72669243 | 0.96801374 |
| ALMS1 | 196.280563 | -0.191072419 | 0.20488883 | -0.9325663 | 0.35104394 | 0.8720098 |
| DUSP11 | 33.6507938 | 0.347466355 | 0.29352499 | 1.18377094 | 0.23650375 | 0.79958336 |
| STAMBP | 76.2504373 | -0.322139751 | 0.22335321 | -1.4422884 | 0.14922103 | 0.72246729 |
| DGUOK | 78.4870276 | -0.73894094 | 0.34043329 | -2.1705895 | 0.02996221 | 0.43492739 |
| TET3 | 116.966773 | -0.800497341 | 0.25781607 | -3.1049164 | 0.00190333 | 0.14133317 |
| MOB1A | 145.01518 | -0.05341632 | 0.22600511 | -0.23635 | 0.81316105 | 0.97930627 |
| AC006030.1 | 34.4870226 | -0.426538883 | 0.36246144 | -1.1767842 | 0.23928167 | 0.80174646 |
| RTKN | 58.7700268 | -0.012541798 | 0.34089397 | -0.0367909 | 0.97065173 | 0.99681784 |
| MOGS | 51.0976372 | 0.193353628 | 0.26666894 | 0.72506991 | 0.46840914 | 0.91462428 |
| AUP1 | 39.3014051 | -0.569431715 | 0.42135227 | -1.3514386 | 0.17655498 | 0.75067397 |
| HK2 | 55.9892884 | -0.931066861 | 0.91050196 | -1.0225863 | 0.30650347 | 0.84488735 |
| POLE4 | 34.6279447 | -0.456983645 | 0.43648863 | -1.0469543 | 0.29512066 | 0.83911606 |
| EVA1A | 89.4516773 | 0.216076122 | 0.50670287 | 0.42643556 | 0.66979049 | 0.95764829 |
| MRPL19 | 119.832542 | -0.130259671 | 0.28908348 | -0.4505954 | 0.65228122 | 0.95159942 |
| GCFC2 | 156.869597 | 0.017368372 | 0.35255355 | 0.04926449 | 0.96070852 | 0.99572129 |
| SUCLG1 | 106.905364 | 0.70884945 | 0.2194568 | 3.23001819 | 0.00123782 | 0.11242415 |

| | | | | | | |
|---|---|---|---|---|---|---|
| DNAH6 | 165.883892 | 0.460742 | 0.55442807 | 0.83102214 | 0.40596112 | 0.89569633 |
| TMSB10 | 920.484419 | -0.238297428 | 0.57759658 | -0.4125672 | 0.67992371 | 0.95948574 |
| KCMF1 | 238.746427 | -0.167824462 | 0.21201262 | -0.7915777 | 0.42860696 | 0.90278843 |
| TCF7L1 | 285.138275 | -0.409015712 | 0.60583434 | -0.675128 | 0.49959446 | 0.92398825 |
| TGOLN2 | 228.861424 | 0.060971685 | 0.30063227 | 0.20281151 | 0.83928236 | 0.98462777 |
| RETSAT | 30.1729431 | 0.64757998 | 0.29845872 | 2.16974723 | 0.030026 | 0.43492739 |
| ELMOD3 | 78.4718189 | 0.173118911 | 0.22492643 | 0.76966904 | 0.44149624 | 0.90561284 |
| CAPG | 49.861974 | -0.231270118 | 0.33273238 | -0.6950635 | 0.48701554 | 0.92034617 |
| MAT2A | 149.621877 | -0.302791206 | 0.36794178 | -0.8229324 | 0.41054644 | 0.8964268 |
| GGCX | 50.8248757 | 0.283730311 | 0.24051654 | 1.17967067 | 0.23813122 | 0.80145582 |
| VAMP8 | 182.820924 | -0.147444658 | 0.43854999 | -0.3362095 | 0.73671291 | 0.9698508 |
| VAMP5 | 39.8986487 | -0.088531637 | 0.48268356 | -0.1834155 | 0.85447203 | 0.9864199 |
| USP39 | 100.701666 | 0.00489105 | 0.29372537 | 0.01665178 | 0.98671442 | 0.99764442 |
| C2orf68 | 39.8351073 | 0.037951996 | 0.40317642 | 0.09413248 | 0.92500392 | 0.99311954 |
| ATOH8 | 79.7170297 | -0.99183667 | 1.20725996 | -0.8215601 | 0.41132728 | 0.89649693 |
| ST3GAL5 | 49.4949824 | -0.100642921 | 0.55003839 | -0.1829744 | 0.85481814 | 0.9864199 |
| POLR1A | 64.3175103 | -0.635742694 | 0.25791392 | -2.4649414 | 0.01370357 | 0.32173155 |
| PTCD3 | 128.428 | -0.182974289 | 0.17328574 | -1.0559108 | 0.29100897 | 0.83575635 |
| IMMT | 150.26583 | -0.229265979 | 0.19863998 | -1.1541784 | 0.24842703 | 0.80672756 |
| MRPL35 | 26.5881989 | -0.407824373 | 0.38083781 | -1.070861 | 0.28423191 | 0.82918183 |
| KDM3A | 141.471721 | -0.027247971 | 0.47968154 | -0.0568043 | 0.95470109 | 0.99531843 |
| CHMP3 | 54.7534787 | -0.56620163 | 0.46774474 | -1.2104928 | 0.22608987 | 0.79069544 |
| RMND5A | 103.210292 | 0.167783854 | 0.26411708 | 0.63526316 | 0.52525679 | 0.93272384 |
| PLGLB1 | 30.4019689 | 0.707628942 | 0.50867523 | 1.39112128 | 0.16418865 | 0.74306594 |
| CYTOR | 64.8679309 | -0.766406338 | 0.45157767 | -1.697175 | 0.08966358 | 0.6330877 |
| KRCC1 | 122.958859 | -0.044226686 | 0.22078306 | -0.2003174 | 0.84123236 | 0.98515707 |
| THNSL2 | 40.9561663 | -0.631650148 | 0.95209667 | -0.6634307 | 0.50705476 | 0.92666294 |
| EIF2AK3 | 156.577531 | -0.095133648 | 0.68127029 | -0.1396416 | 0.8889432 | 0.98766275 |
| RPIA | 44.989206 | -0.327570311 | 0.32721396 | -1.001089 | 0.31678376 | 0.8490543 |
| MAL | 274.04545 | -0.352811213 | 0.81391216 | -0.4334758 | 0.66466915 | 0.95505623 |
| MRPS5 | 121.355873 | 0.118394654 | 0.20681811 | 0.57245787 | 0.56701182 | 0.9385359 |
| ANKRD36C | 129.672709 | -0.102115398 | 0.3214275 | -0.3176934 | 0.75071751 | 0.97058015 |
| STARD7 | 92.6835283 | 0.013233828 | 0.1982004 | 0.06676994 | 0.94676486 | 0.9952642 |
| TMEM127 | 49.5754917 | -0.038557112 | 0.24259797 | -0.1589342 | 0.87372072 | 0.98766275 |
| CIAO1 | 55.4718052 | -0.534377173 | 0.25465536 | -2.0984328 | 0.03586693 | 0.45984105 |
| SNRNP200 | 131.424616 | -0.396947087 | 0.26891456 | -1.4761086 | 0.13991475 | 0.71251643 |
| NEURL3 | 30.9345517 | -1.686257043 | 0.70026005 | -2.408044 | 0.01603825 | 0.3414008 |
| KANSL3 | 147.450098 | -0.428903087 | 0.19651405 | -2.1825569 | 0.02906846 | 0.43178298 |
| FER1L5 | 24.3700369 | -0.434076656 | 0.71680857 | -0.6055685 | 0.54480136 | 0.93523904 |
| LMAN2L | 64.9597822 | 0.221001516 | 0.25360018 | 0.87145645 | 0.38350498 | 0.88328503 |
| CNNM4 | 55.7453706 | -0.794326252 | 0.52856418 | -1.5028 | 0.13289062 | 0.70368119 |
| ANKRD23 | 43.9444281 | 0.274877031 | 0.3055161 | 0.89971375 | 0.36827261 | 0.87605028 |
| ANKRD36 | 198.995664 | -0.139514836 | 0.29238063 | -0.4771685 | 0.63324211 | 0.94932781 |
| AC092683.1 | 110.537101 | 0.606775646 | 0.41867578 | 1.44927333 | 0.14726127 | 0.72167017 |
| ANKRD36B | 59.7381516 | 0.073406128 | 0.44412835 | 0.16528134 | 0.86872255 | 0.98676155 |
| COX5B | 134.988118 | 0.161386137 | 0.46439725 | 0.34751742 | 0.72820263 | 0.96827499 |
| ACTR1B | 48.1470323 | -0.301251431 | 0.28152968 | -1.0700521 | 0.28459584 | 0.82918183 |
| TMEM131 | 355.24955 | -0.316755622 | 0.27274144 | -1.1613769 | 0.24548864 | 0.80476923 |
| INPP4A | 106.309214 | 0.068219046 | 0.22054659 | 0.30931808 | 0.75707958 | 0.9714717 |
| COA5 | 44.8912203 | 0.243967298 | 0.33368039 | 0.73114065 | 0.46469325 | 0.91462428 |
| UNC50 | 52.2025399 | -0.17220677 | 0.38010511 | -0.4530504 | 0.65051245 | 0.95075661 |
| MGAT4A | 412.715302 | 0.369362645 | 0.27016727 | 1.36716281 | 0.17157428 | 0.74445571 |
| TSGA10 | 200.878451 | 0.606214881 | 0.32576384 | 1.86090291 | 0.06275789 | 0.56713487 |
| AC079447.1 | 112.227539 | 0.02530085 | 0.31781236 | 0.0796094 | 0.93654792 | 0.99526299 |

| | | | | | | |
|---|---|---|---|---|---|---|
| MITD1 | 55.2515974 | 0.533052801 | 0.32684837 | 1.63088711 | 0.10291414 | 0.65288882 |
| LYG1 | 44.144881 | 1.222451422 | 0.54161754 | 2.2570381 | 0.02400569 | 0.40690238 |
| TXNDC9 | 54.7177852 | -0.092162588 | 0.38810739 | -0.2374667 | 0.81229474 | 0.97930627 |
| EIF5B | 248.88661 | -0.368645953 | 0.23025763 | -1.6010151 | 0.10937357 | 0.65764016 |
| REV1 | 245.070107 | 0.061091699 | 0.17812512 | 0.34297072 | 7.32E-01 | 0.96972363 |
| AFF3 | 155.298921 | 0.686928823 | 0.60992237 | 1.12625615 | 0.26005711 | 0.81263292 |
| LONRF2 | 53.8172204 | -0.240971825 | 0.73336449 | -0.328584 | 0.74247016 | 0.9698508 |
| NPAS2 | 362.093257 | 0.092864476 | 0.55415087 | 0.16757977 | 0.86691389 | 0.98670067 |
| RPL31 | 180.799584 | 0.462634661 | 0.3705975 | 1.24834804 | 0.21190363 | 0.77732978 |
| TBC1D8 | 241.088977 | -0.005284333 | 0.36005431 | -0.0146765 | 0.98829028 | 0.99820747 |
| CNOT11 | 71.5790909 | -0.403528894 | 0.19499521 | -2.0694297 | 0.03850578 | 0.4740962 |
| RNF149 | 430.952372 | -0.426786084 | 0.20947408 | -2.0374171 | 0.04160826 | 0.48843812 |
| MAP4K4 | 442.664902 | -0.211457691 | 0.46227623 | -0.4574271 | 0.64736407 | 0.95027667 |
| IL1R1 | 491.290156 | -0.256457035 | 0.533964 | -0.480289 | 0.63102191 | 0.94932781 |
| IL18R1 | 31.8033217 | -0.272523937 | 0.66031045 | -0.4127209 | 0.67981109 | 0.95948574 |
| TMEM182 | 42.2801658 | 0.432176495 | 0.29851612 | 1.44774925 | 0.14768719 | 0.72246583 |
| POU3F3 | 267.482428 | -0.797509118 | 0.71408724 | -1.1168231 | 0.26406997 | 0.81378765 |
| MRPS9 | 100.493594 | -0.154402722 | 0.3472717 | -0.4446165 | 0.65659694 | 0.95399905 |
| TGFBRAP1 | 74.4688975 | -0.164481704 | 0.24183801 | -0.6801317 | 0.49642106 | 0.92353293 |
| C2orf49 | 32.4964288 | 0.548765395 | 0.30079915 | 1.82435817 | 0.06809794 | 0.58138162 |
| FHL2 | 170.923971 | -1.443565249 | 0.55731447 | -2.5902167 | 0.00959155 | 0.28916146 |
| NCK2 | 183.575499 | 0.053458986 | 0.27449931 | 0.19475089 | 0.84558797 | 0.98520136 |
| UXS1 | 176.784956 | -0.741566481 | 0.17241947 | -4.3009441 | 1.70E-05 | 0.00708313 |
| SULT1C2 | 148.556787 | 0.392788526 | 0.78070136 | 0.50312263 | 0.61487805 | 0.94791646 |
| SULT1C4 | 70.5224103 | 0.522496468 | 0.52261608 | 0.99977114 | 0.31742128 | 0.8490543 |
| GCC2 | 383.170024 | 0.561902551 | 0.18664238 | 3.01058395 | 0.00260746 | 0.16762984 |
| GCC2-AS1 | 49.0033654 | 0.435212696 | 0.27933562 | 1.55802791 | 0.11922663 | 0.68597919 |
| LIMS1 | 371.863243 | 0.141510464 | 0.29392125 | 0.48145707 | 0.63019168 | 0.94916079 |
| RANBP2 | 219.991938 | 0.038139427 | 0.24155364 | 0.15789217 | 0.87454177 | 0.98766275 |
| CCDC138 | 61.1965461 | -0.178794535 | 0.35087796 | -0.5095633 | 0.61035743 | 0.94791646 |
| EDAR | 31.6470573 | -0.301590692 | 0.43369049 | -0.6954054 | 0.4868013 | 0.92034617 |
| SH3RF3 | 109.422684 | 0.694515261 | 1.13046314 | 0.61436347 | 0.53897517 | 0.93523904 |
| 10-Sep | 269.389954 | -0.390224209 | 0.13070954 | -2.9854303 | 0.0028318 | 0.17277569 |
| SOWAHC | 31.1342421 | -0.526297979 | 0.50007583 | -1.0524363 | 0.2925994 | 0.83697123 |
| NPHP1 | 103.202409 | 0.536908478 | 0.29858604 | 1.79817005 | 0.07215006 | 0.59090333 |
| SMIM37 | 52.4408464 | -0.337113885 | 0.45198224 | -0.7458565 | 0.45575411 | 0.91077355 |
| MIR4435-2H | 482.439914 | -0.866132442 | 0.26320919 | -3.2906619 | 1.00E-03 | 0.10425141 |
| BCL2L11 | 48.1097588 | 0.863664507 | 0.32421376 | 2.66387371 | 0.00772466 | 0.27068528 |
| ANAPC1 | 170.699407 | -0.410401716 | 0.21404916 | -1.9173246 | 0.05519671 | 0.53849427 |
| MERTK | 50.4970547 | 0.525506824 | 0.4077454 | 1.28881115 | 0.19746375 | 0.77110248 |
| TMEM87B | 140.152089 | -0.25105172 | 0.21013506 | -1.194716 | 0.23219801 | 0.79493378 |
| ZC3H8 | 62.2624168 | -0.251519549 | 0.28646978 | -0.8779968 | 0.37994546 | 0.88059823 |
| ZC3H6 | 165.498141 | -0.327536336 | 0.39505598 | -0.8290884 | 0.40705438 | 0.89585737 |
| TTL | 39.5060815 | -0.520973099 | 0.39246865 | -1.327426 | 0.1843678 | 0.7579653 |
| POLR1B | 36.9211327 | -0.430469421 | 0.31193576 | -1.3799938 | 0.16758854 | 0.74445571 |
| CHCHD5 | 29.5811209 | 0.272278245 | 0.41173464 | 0.66129545 | 0.50842286 | 0.92666294 |
| SLC20A1 | 75.9327416 | -0.631951226 | 0.48911368 | -1.2920334 | 0.19634557 | 0.77050152 |
| PSD4 | 47.095609 | -0.873294358 | 0.39809126 | -2.1937039 | 0.0282567 | 0.4250936 |
| PAX8 | 4716.51084 | -0.34274398 | 0.27680675 | -1.2382067 | 0.21563942 | 0.78293142 |
| CBWD2 | 73.6818128 | -0.349181451 | 0.32666077 | -1.0689421 | 0.28509576 | 0.82925512 |
| RABL2A | 59.1812133 | 0.344727572 | 0.31786659 | 1.08450394 | 0.27814142 | 0.8229426 |
| SLC35F5 | 851.077743 | -0.061809579 | 0.21099688 | -0.2929407 | 0.76956746 | 0.97261718 |
| ACTR3 | 404.846129 | -0.135841793 | 0.22520913 | -0.6031807 | 0.54638852 | 0.93523904 |
| DDX18 | 98.5697671 | -0.260442452 | 0.30910049 | -0.8425818 | 0.39946236 | 0.88987208 |

| | | | | | | |
|---|---|---|---|---|---|---|
| CCDC93 | 158.593796 | -0.299895261 | 0.17330818 | -1.7304161 | 0.08355597 | 0.61656297 |
| INSIG2 | 101.10922 | 0.410138477 | 0.44074591 | 0.93055539 | 0.3520836 | 0.8720098 |
| C2orf76 | 30.9483216 | -0.463490173 | 0.37782476 | -1.2267332 | 0.21992289 | 0.78584902 |
| DBI | 164.278071 | 0.010009017 | 0.41959399 | 0.02385405 | 0.98096902 | 0.99681784 |
| TMEM37 | 71.6460036 | 0.54373033 | 0.60248083 | 0.90248569 | 0.36679892 | 0.87605028 |
| SCTR | 153.729568 | -0.162355877 | 1.43512022 | -0.1131305 | 0.90992709 | 0.99100813 |
| CFAP221 | 428.053139 | 0.0606377 | 0.60485541 | 0.10025156 | 0.92014461 | 0.99214128 |
| PTPN4 | 222.187171 | 0.148373972 | 0.19753979 | 0.75110931 | 0.45258688 | 0.91077355 |
| EPB41L5 | 598.785803 | 0.61244265 | 0.54746924 | 1.11867957 | 0.26327686 | 0.81378765 |
| RALB | 65.1030358 | -0.124474246 | 0.28127069 | -0.4425425 | 0.65809665 | 0.95399905 |
| TFCP2L1 | 187.629924 | 0.404556513 | 0.66003431 | 0.61293255 | 0.53992094 | 0.93523904 |
| CLASP1 | 669.410923 | 0.020880963 | 0.24811529 | 0.08415831 | 0.93293057 | 0.99425828 |
| NIFK-AS1 | 70.7317685 | 0.250477723 | 0.39079741 | 0.64094008 | 0.52156162 | 0.93272217 |
| NIFK | 33.9097206 | -0.367234114 | 0.4056504 | -0.9052971 | 0.36530803 | 0.87605028 |
| TSN | 52.5235649 | -0.270059284 | 0.28461825 | -0.9488474 | 0.34269824 | 0.86846099 |
| BIN1 | 132.785218 | 0.372509712 | 0.42653795 | 0.87333311 | 0.38248154 | 0.88177874 |
| ERCC3 | 61.8744904 | -0.278695818 | 0.27884601 | -0.9994614 | 0.31757124 | 0.8490543 |
| MAP3K2 | 404.425099 | 0.120505242 | 0.1839728 | 0.65501662 | 0.51245705 | 0.92919856 |
| IWS1 | 186.055822 | 0.190365367 | 0.26287624 | 0.72416346 | 0.46896539 | 0.91462428 |
| MYO7B | 39.0180153 | 1.47389432 | 0.69732814 | 2.11363093 | 0.0345468 | 0.45223457 |
| LIMS2 | 41.0279917 | 1.204779376 | 0.67770627 | 1.77773091 | 0.07544806 | 0.59465671 |
| WDR33 | 303.045762 | 0.068556479 | 0.19501193 | 0.35155018 | 0.72517563 | 0.96781837 |
| POLR2D | 31.2870185 | -0.279962272 | 0.30158372 | -0.928307 | 0.35324835 | 0.8720098 |
| AMMECR1L | 34.9533571 | -0.03948016 | 0.28354764 | -0.1392364 | 0.88926333 | 0.98766275 |
| SAP130 | 87.4403929 | -0.045808932 | 0.20489393 | -0.2235739 | 0.82308889 | 0.98094663 |
| UGGT1 | 99.300612 | -0.316661181 | 0.26275935 | -1.2051376 | 0.22815018 | 0.79284833 |
| HS6ST1 | 235.858754 | -0.376733772 | 0.47491448 | -0.7932666 | 0.42762254 | 0.90272191 |
| SMPD4 | 82.4214127 | -0.572660257 | 0.26751213 | -2.140689 | 0.03229913 | 0.44049392 |
| MZT2B | 154.223906 | 0.140352486 | 0.50954743 | 0.27544538 | 0.78297408 | 0.97423208 |
| CCDC115 | 29.37784 | 0.031541563 | 0.37134031 | 0.08493978 | 0.93230927 | 0.99416721 |
| IMP4 | 34.5436943 | -0.260039902 | 0.43163568 | -0.6024523 | 0.54687312 | 0.93523904 |
| PTPN18 | 55.418473 | -0.184909606 | 0.32289285 | -0.5726655 | 0.56687119 | 0.9385359 |
| FAM168B | 165.477271 | -0.149424104 | 0.22015787 | -0.6787134 | 0.49731944 | 0.92398825 |
| PLEKHB2 | 144.731053 | -0.428443617 | 0.20487465 | -2.0912476 | 0.03650588 | 0.46377956 |
| MZT2A | 78.997288 | -0.167051994 | 0.3007893 | -0.5553788 | 0.5786356 | 0.94034284 |
| GPR39 | 99.6113783 | -0.682747626 | 0.48882387 | -1.396715 | 0.1624993 | 0.74071637 |
| LYPD1 | 37.1707268 | 0.405100452 | 0.67250341 | 0.6023768 | 0.54692335 | 0.93523904 |
| NCKAP5 | 177.375182 | 1.099230876 | 0.71946811 | 1.52783822 | 0.12655271 | 0.69494309 |
| MGAT5 | 239.553054 | 0.281056215 | 0.35353169 | 0.79499582 | 0.42661594 | 0.90250752 |
| TMEM163 | 476.688517 | 0.503012628 | 0.80310769 | 0.62633273 | 0.53109672 | 0.93366189 |
| CCNT2-AS1 | 77.3279193 | 0.072550967 | 0.37604535 | 0.19293143 | 0.84701267 | 0.98533374 |
| ACMSD | 108.852033 | 1.717977786 | 0.46456556 | 3.69803084 | 0.00021728 | 0.03854279 |
| CCNT2 | 200.419916 | -0.143117551 | 0.21629384 | -0.6616811 | 0.50817561 | 0.92666294 |
| RAB3GAP1 | 348.608602 | -0.087860215 | 0.14278963 | -0.6153123 | 0.5383485 | 0.93516669 |
| ZRANB3 | 164.727721 | -0.528714853 | 0.24912122 | -2.1223196 | 0.03381091 | 0.44937373 |
| R3HDM1 | 303.389929 | 0.052881005 | 0.15512588 | 0.34089093 | 0.73318569 | 0.9698508 |
| UBXN4 | 278.750163 | -0.156858589 | 0.18386295 | -0.8531278 | 0.39358846 | 0.88760715 |
| DARS | 262.498909 | -0.224368222 | 0.23450026 | -0.956793 | 0.33867172 | 0.86680923 |
| HNMT | 191.760111 | 0.453417614 | 0.52634705 | 0.86144231 | 0.38899448 | 0.88586416 |
| AC114763.1 | 111.789434 | -0.704622838 | 0.42965645 | -1.639968 | 0.10101182 | 0.64808588 |
| SPOPL | 188.532508 | 0.062448206 | 0.19604147 | 0.31854589 | 0.75007089 | 0.97054289 |
| LRP1B | 46.1447912 | 0.023502229 | 0.4011645 | 0.05858502 | 0.95328264 | 0.99531843 |
| GTDC1 | 293.824956 | 0.294099595 | 0.37790336 | 0.77824023 | 0.43642741 | 0.90525098 |
| ZEB2 | 145.155549 | 0.590482902 | 0.86381986 | 0.68357181 | 0.49424559 | 0.92289011 |

| | | | | | | |
|---|---|---|---|---|---|---|
| TEX41 | 128.137206 | 0.135510885 | 0.51371523 | 0.263786 | 0.79194482 | 0.97423208 |
| ACVR2A | 133.796104 | 0.112464371 | 0.27667858 | 0.40648023 | 0.68438978 | 0.96161098 |
| ORC4 | 263.355607 | 0.196476849 | 0.24428268 | 0.80430118 | 0.42122306 | 0.90012434 |
| MBD5 | 1309.31795 | 0.147009505 | 0.25642594 | 0.57330201 | 0.56644022 | 0.93845576 |
| EPC2 | 272.653311 | -0.060215774 | 0.20433451 | -0.2946921 | 0.76822907 | 0.97239879 |
| MMADHC | 58.9222056 | 0.560397494 | 0.40515246 | 1.38317681 | 0.16661066 | 0.74445571 |
| AC007364.1 | 70.921598 | 1.126843413 | 0.5052331 | 2.23034359 | 0.02572464 | 0.4183639 |
| RND3 | 100.284534 | -0.387467649 | 0.63370455 | -0.6114326 | 0.5409132 | 0.93523904 |
| NMI | 48.5445062 | -0.710130967 | 0.59363511 | -1.1962415 | 0.23160232 | 0.79459836 |
| RIF1 | 265.43002 | -0.094781883 | 0.28667275 | -0.3306275 | 0.7409259 | 0.9698508 |
| ARL5A | 70.011349 | 0.481546071 | 0.3514635 | 1.37011688 | 0.17065042 | 0.74445571 |
| STAM2 | 128.213925 | -0.16914296 | 0.18406997 | -0.9189058 | 0.35814485 | 0.87316144 |
| FMNL2 | 967.269152 | -0.381890203 | 0.51262496 | -0.74497 | 0.45628987 | 0.91134292 |
| PRPF40A | 189.131577 | -0.455984905 | 0.16729033 | -2.7257099 | 0.00641634 | 0.25167097 |
| ARL6IP6 | 91.8332489 | -0.02139891 | 0.27105948 | -0.0789454 | 0.93707602 | 0.99526299 |
| KCNJ3 | 425.09304 | -0.856724434 | 1.0393063 | -0.8243233 | 0.40975586 | 0.8959746 |
| LINC01876 | 157.831886 | 0.389673758 | 0.75669473 | 0.51496825 | 0.60657521 | 0.94708529 |
| GPD2 | 201.197376 | -0.213503598 | 0.23818657 | -0.8963713 | 0.37005451 | 0.87605028 |
| ACVR1 | 253.744354 | -0.221034802 | 0.3943261 | -0.5605381 | 0.57511247 | 0.94034284 |
| UPP2 | 150.335023 | 1.127769337 | 0.56057326 | 2.01181437 | 0.04423951 | 0.49952159 |
| CCDC148 | 424.910679 | 0.183615206 | 0.59248667 | 0.30990605 | 0.7566324 | 0.9714717 |
| PKP4 | 1520.22208 | -0.183746363 | 0.35594901 | -0.5162154 | 0.60570397 | 0.94708529 |
| PKP4-AS1 | 50.1972217 | 0.555809868 | 0.58059986 | 0.9573028 | 0.33841444 | 0.86672435 |
| TANC1 | 479.683274 | 0.096252615 | 0.26974559 | 0.3568274 | 0.72122103 | 0.96779196 |
| WDSUB1 | 154.489944 | -0.32096135 | 0.31871465 | -1.0070492 | 0.31391111 | 0.8490543 |
| BAZ2B | 862.154154 | 0.081491527 | 0.20471693 | 0.39806931 | 0.69057909 | 0.96220115 |
| 7-Mar | 325.524664 | 0.049732461 | 0.1698706 | 0.29276673 | 0.76970046 | 0.97261718 |
| PLA2R1 | 171.261625 | 0.759434572 | 1.02897221 | 0.73805158 | 0.4604831 | 0.91368087 |
| ITGB6 | 1080.9893 | -1.694810939 | 0.69400893 | -2.4420593 | 0.01460375 | 0.32915131 |
| LINC02478 | 39.3373202 | -0.660017223 | 0.68064344 | -0.969696 | 0.33219804 | 0.85980141 |
| RBMS1 | 1047.39058 | -0.401013717 | 0.30504904 | -1.3145877 | 0.18864851 | 0.76207437 |
| TANK | 406.836965 | 0.145323315 | 0.24511736 | 0.5928724 | 0.55326655 | 0.93523904 |
| PSMD14 | 125.98916 | -0.603452634 | 0.25633721 | -2.3541359 | 0.01856582 | 0.36368509 |
| DPP4 | 118.784297 | -0.093296544 | 0.55902388 | -0.1668919 | 0.86745512 | 0.98670067 |
| IFIH1 | 65.7592 | -1.921594511 | 0.87486844 | -2.1964382 | 0.02806059 | 0.4250936 |
| GCA | 84.0954407 | -0.340384065 | 0.34988384 | -0.9728488 | 0.33062847 | 0.85923151 |
| FIGN | 423.289115 | 0.111652751 | 0.28074693 | 0.39769892 | 6.91E-01 | 0.96220115 |
| GRB14 | 232.762354 | -0.2455763 | 0.30075895 | -0.816522 | 0.41420164 | 0.89745251 |
| COBLL1 | 1803.09891 | -0.574508318 | 0.60750015 | -0.9456925 | 0.34430546 | 0.86975264 |
| AC019197.1 | 811.380298 | 1.054701748 | 0.59715789 | 1.76620248 | 0.07736188 | 0.5981029 |
| SLC38A11 | 248.233916 | -2.053227494 | 1.30275009 | -1.5760717 | 0.11500929 | 0.67134308 |
| SCN3A | 36.9726197 | -0.198022853 | 0.32648996 | -0.6065205 | 0.54416917 | 0.93523904 |
| SCN2A | 111.572759 | 1.402419298 | 0.68182661 | 2.05685621 | 0.03970006 | 0.48094465 |
| CSRNP3 | 132.543297 | 0.297912089 | 0.2202526 | 1.35259282 | 1.76E-01 | 7.50E-01 |
| GALNT3 | 58.6140532 | -0.479768215 | 0.46207625 | -1.038288 | 0.29913601 | 0.84005388 |
| TTC21B | 152.538341 | -0.013335978 | 0.22361532 | -0.059638 | 9.52E-01 | 0.99531843 |
| SCN9A | 60.0265784 | 0.770513127 | 0.94289894 | 0.81717467 | 0.41382862 | 0.89732895 |
| STK39 | 334.267253 | -1.140398141 | 0.30134951 | -3.784304 | 0.00015414 | 0.03134677 |
| CERS6 | 195.450269 | -0.101102391 | 0.27573321 | -0.3666674 | 0.71386709 | 0.96611089 |
| NOSTRIN | 190.888495 | -0.393381554 | 0.63056095 | -0.6238597 | 0.53271974 | 0.93391698 |
| SPC25 | 37.6285144 | -0.670207251 | 0.56633312 | -1.1834152 | 0.23664462 | 0.79958336 |
| LRP2 | 773.256741 | 1.197134052 | 0.71192026 | 1.68155637 | 0.0926549 | 0.63577454 |
| FASTKD1 | 69.6749352 | -0.129795856 | 0.33155157 | -0.3914801 | 0.69544236 | 0.96220115 |
| PPIG | 223.694271 | -0.11081181 | 0.16928921 | -0.654571 | 0.51274399 | 0.92934242 |

| | | | | | | |
|---|---|---|---|---|---|---|
| CCDC173 | 28.8885213 | 0.849770715 | 0.41927806 | 2.02674741 | 0.04268825 | 0.49385357 |
| KLHL23 | 88.0622627 | 0.00374188 | 0.24244038 | 0.01543423 | 0.98768576 | 0.99804429 |
| SSB | 100.501501 | -0.057525665 | 0.29995955 | -0.1917781 | 0.84791605 | 0.98545133 |
| METTL5 | 38.6361408 | -0.174018287 | 0.27471196 | -0.6334573 | 0.52643508 | 0.93331883 |
| UBR3 | 442.827299 | 0.032524657 | 0.17990062 | 0.18079236 | 0.85653056 | 0.9864199 |
| MYO3B | 162.819358 | -0.810332817 | 0.82719611 | -0.9796139 | 0.32727674 | 0.85650224 |
| GORASP2 | 93.0810338 | -0.124958452 | 0.27180289 | -0.4597392 | 0.6457034 | 0.94981463 |
| TLK1 | 359.364042 | -0.005619371 | 0.20866218 | -0.0269305 | 0.97851519 | 0.99681784 |
| METTL8 | 68.5304389 | -0.928413484 | 0.31372918 | -2.9592832 | 0.00308356 | 0.18079751 |
| DCAF17 | 58.7584717 | -0.072554557 | 0.24784354 | -0.2927434 | 0.76971831 | 0.97261718 |
| CYBRD1 | 48.9172324 | 1.429765135 | 0.86546388 | 1.65202173 | 0.09853012 | 0.64312419 |
| DYNC1I2 | 192.946468 | -0.021388819 | 0.19219969 | -0.1112844 | 0.91139086 | 0.99100813 |
| SLC25A12 | 170.577576 | 0.088764844 | 0.22334947 | 0.39742582 | 0.69105348 | 0.96220115 |
| HAT1 | 81.528038 | -0.310309541 | 0.24827339 | -1.2498703 | 0.21134692 | 0.77730294 |
| METAP1D | 43.163764 | -0.418156291 | 0.2633659 | -1.587739 | 0.11234538 | 0.66401239 |
| ITGA6 | 77.4194909 | 0.281950181 | 0.51640329 | 0.54598835 | 0.58507394 | 0.94229635 |
| PDK1 | 128.425521 | -0.375051374 | 0.66118315 | -0.5672428 | 0.57054924 | 0.93998433 |
| RAPGEF4 | 33.8096974 | 0.123245707 | 0.47046446 | 0.26196603 | 0.79334763 | 0.9745418 |
| MAP3K20 | 194.58474 | -0.509387008 | 0.42811049 | -1.1898494 | 0.23410559 | 0.79634141 |
| SP3 | 207.6331 | -0.233154838 | 0.18138499 | -1.2854142 | 0.19864759 | 0.77110248 |
| OLA1 | 323.939579 | -0.250312519 | 0.25078325 | -0.9981229 | 0.31821974 | 0.84919767 |
| CIR1 | 118.70269 | -0.054860013 | 0.30692052 | -0.1787434 | 0.8581392 | 0.9864199 |
| SCRN3 | 45.4067679 | 0.163936336 | 0.37681594 | 0.4350568 | 0.6635212 | 0.95482952 |
| GPR155 | 130.165063 | -0.169718315 | 0.40808719 | -0.4158874 | 0.67749241 | 0.95924874 |
| AC010894.3 | 227.214373 | 0.133356107 | 0.41703978 | 0.31976832 | 0.74914397 | 0.97028355 |
| WIPF1 | 172.020709 | 0.339231106 | 0.29540422 | 1.14836243 | 0.25081898 | 0.80680142 |
| CHN1 | 109.996945 | 0.342303089 | 1.07025566 | 0.31983301 | 0.74909492 | 0.97028355 |
| ATF2 | 217.52338 | 0.216776705 | 0.19695339 | 1.10064977 | 0.27104912 | 0.81775732 |
| ATP5MC3 | 161.24873 | 0.392093288 | 0.56481249 | 0.69420081 | 0.48755628 | 0.92034617 |
| LNPK | 80.9596711 | -0.071945368 | 0.25606204 | -0.2809685 | 0.77873456 | 0.97423208 |
| HOXD10 | 60.3712017 | 0.375233347 | 0.67073369 | 0.55943716 | 0.57586341 | 0.94034284 |
| HOXD8 | 28.3028196 | -1.040670047 | 0.69021869 | -1.5077396 | 0.13162119 | 0.700349 |
| HOXD3 | 61.5782337 | -0.35441037 | 0.70267547 | -0.5043728 | 0.61399945 | 0.94791646 |
| MTX2 | 225.555732 | -0.008924719 | 0.18519571 | -0.0481907 | 0.96156423 | 0.99572129 |
| HNRNPA3 | 367.907524 | -0.550772746 | 0.2406931 | -2.2882781 | 0.02212133 | 0.39000316 |
| NFE2L2 | 600.298887 | -0.084864014 | 0.21053539 | -0.4030867 | 0.68688445 | 0.96161098 |
| AGPS | 183.844468 | 0.659383991 | 0.33789907 | 1.95142295 | 0.05100675 | 0.52593509 |
| RBM45 | 38.9842601 | -0.116859469 | 0.32192434 | -0.3630029 | 0.71660272 | 0.96742001 |
| AC009948.1 | 41.7963626 | -0.834809229 | 0.65949757 | -1.2658261 | 0.20557535 | 0.77352973 |
| PRKRA | 110.674219 | -0.202558672 | 0.2887243 | -0.7015643 | 0.4829509 | 0.91971919 |
| PJVK | 27.445105 | 0.380386789 | 0.40041308 | 0.94998593 | 0.3421194 | 0.86846099 |
| PLEKHA3 | 65.8481598 | -0.113418908 | 0.27947235 | -0.4058323 | 0.68486582 | 0.96161098 |
| TTN-AS1 | 267.822384 | 0.337237955 | 0.23126634 | 1.45822327 | 0.14477901 | 0.71738996 |
| SESTD1 | 279.441786 | 0.485719101 | 0.37491993 | 1.29552756 | 0.1951383 | 0.76968045 |
| ZNF385B | 259.681939 | 1.615079459 | 0.49088014 | 3.29017069 | 0.00100127 | 0.10425141 |
| CWC22 | 87.634271 | 0.004348397 | 0.317107 | 0.01371271 | 0.98905918 | 0.99833487 |
| UBE2E3 | 333.621347 | -0.165531778 | 0.22097594 | -0.7490941 | 0.45380049 | 0.91077355 |
| SSFA2 | 200.855394 | 0.000573257 | 0.28937437 | 0.00198102 | 0.99841937 | 0.99971834 |
| PDE1A | 1432.23757 | 1.024516898 | 1.41308605 | 0.72502088 | 0.46843922 | 0.91462428 |
| DNAJC10 | 135.756178 | -0.154634354 | 0.23511562 | -0.6576949 | 0.51073418 | 0.92819651 |
| NCKAP1 | 523.910478 | -0.113806294 | 0.22600367 | -0.5035595 | 0.61457097 | 0.94791646 |
| NUP35 | 62.2711915 | -0.62264453 | 0.27725989 | -2.2457072 | 0.02472277 | 0.41230428 |
| FSIP2 | 42.1287981 | -0.359290446 | 0.57355691 | -0.6264251 | 0.53103615 | 0.93366189 |
| FSIP2-AS1 | 35.8830447 | -0.238353446 | 0.48784985 | -0.4885795 | 0.62513941 | 0.94808496 |

| | | | | | | |
|---|---|---|---|---|---|---|
| LINC01473 | 78.4936881 | -0.069867016 | 0.43186794 | -0.1617787 | 0.87148016 | 0.98766275 |
| ZC3H15 | 125.805932 | -0.054887659 | 0.25096299 | -0.2187082 | 0.82687737 | 0.98142144 |
| ITGAV | 574.245069 | 0.656521155 | 0.32483364 | 2.02109967 | 0.04326945 | 0.49484046 |
| AC017101.1 | 60.2284561 | 1.632408577 | 0.67201884 | 2.42911133 | 0.01513588 | 0.33639589 |
| AC007319.1 | 381.216631 | 1.140765433 | 0.65992666 | 1.72862455 | 0.08387631 | 0.61656297 |
| TFPI | 878.194402 | 0.597485601 | 0.46030882 | 1.29801034 | 1.94E-01 | 0.7687432 |
| LINC01090 | 31.3793956 | 0.54155584 | 0.58737339 | 0.92199588 | 0.35653073 | 0.87316144 |
| GULP1 | 723.277189 | -0.654816536 | 0.55901347 | -1.1713788 | 0.24144655 | 0.803412 |
| WDR75 | 67.1635942 | -0.370695412 | 0.24144344 | -1.5353302 | 0.12470274 | 0.69439514 |
| SLC40A1 | 85.276334 | 1.280587653 | 0.40034788 | 3.19868724 | 0.00138055 | 0.11702898 |
| ASNSD1 | 31.5658295 | 0.11508213 | 0.33845852 | 0.34001841 | 0.73384266 | 0.9698508 |
| ANKAR | 161.930971 | -0.236910335 | 0.33376696 | -0.7098076 | 0.47782344 | 0.91890599 |
| ORMDL1 | 71.4261124 | 0.198924648 | 0.33701439 | 0.59025566 | 0.55501926 | 0.93523904 |
| PMS1 | 102.697556 | -0.542867262 | 0.19573847 | -2.7734316 | 0.00554685 | 0.23510302 |
| C2orf88 | 43.0117827 | 0.633957702 | 0.35665975 | 1.77748599 | 0.07548831 | 0.59465671 |
| HIBCH | 282.009988 | 0.290432697 | 0.26940858 | 1.07803805 | 0.28101678 | 0.8264844 |
| INPP1 | 33.0874481 | 0.056004638 | 0.2685017 | 0.20858206 | 0.83477451 | 0.98367279 |
| MFSD6 | 77.3506531 | 0.209735835 | 0.2388998 | 0.87792386 | 0.37998504 | 0.88059823 |
| NAB1 | 108.788285 | 0.364583161 | 0.25591933 | 1.4246019 | 0.15427231 | 0.72976516 |
| GLS | 1323.9712 | -0.287608507 | 0.32634844 | -0.8812927 | 0.37815941 | 0.87964763 |
| STAT1 | 567.634367 | -0.712020477 | 0.79694564 | -0.8934367 | 0.37162336 | 0.87605028 |
| MYO1B | 147.462668 | -0.271567309 | 0.63638569 | -0.4267338 | 0.6695732 | 0.95764829 |
| NABP1 | 87.1063691 | -0.018439878 | 0.7356695 | -0.0250654 | 0.98000277 | 0.99681784 |
| AC098617.1 | 136.018712 | 0.377421055 | 0.64258157 | 0.58735119 | 0.55696787 | 0.93533595 |
| SLC39A10 | 122.68021 | 0.571669525 | 0.36983124 | 1.54575779 | 0.12216307 | 0.69242609 |
| DNAH7 | 414.555287 | 0.665432792 | 0.37776155 | 1.76151541 | 0.07815121 | 0.60226099 |
| STK17B | 30.1543289 | -0.239784375 | 0.35188677 | -0.6814248 | 0.49560273 | 0.92325527 |
| GTF3C3 | 58.88712 | 0.234806702 | 0.25871077 | 0.90760313 | 0.36408794 | 0.87605028 |
| PGAP1 | 78.1799927 | -0.331604659 | 0.37999087 | -0.8726648 | 0.38284581 | 0.88240616 |
| ANKRD44 | 314.576153 | -0.434454966 | 0.33408773 | -1.3004218 | 0.19345645 | 0.76797319 |
| SF3B1 | 247.6228 | 0.133324466 | 0.22890323 | 0.58244904 | 0.56026426 | 0.93744835 |
| COQ10B | 45.2255839 | -0.213594129 | 0.3060599 | -0.6978834 | 0.4852501 | 0.92034617 |
| HSPD1 | 207.24381 | 0.052724354 | 0.35305808 | 0.14933621 | 0.88128835 | 0.98766275 |
| MOB4 | 32.2733546 | -0.126365065 | 0.47825951 | -0.2642186 | 0.79161147 | 0.97423208 |
| PLCL1 | 443.218092 | 2.480614475 | 0.70916911 | 3.49791672 | 0.00046891 | 0.06755525 |
| SATB2 | 513.670005 | -0.364489953 | 0.65610351 | -0.5555373 | 0.57852722 | 0.94034284 |
| FTCDNL1 | 36.0903049 | 0.229870968 | 0.37556542 | 0.61206638 | 0.54049384 | 0.93523904 |
| C2orf69 | 28.4822794 | 0.031721321 | 0.43404034 | 0.07308381 | 0.94173943 | 0.99526299 |
| TYW5 | 59.1467985 | 0.062480426 | 0.21260676 | 0.29387789 | 0.76885121 | 0.972511 |
| MAIP1 | 29.1001384 | -0.076704346 | 0.30370316 | -0.2525635 | 0.80060551 | 0.97723068 |
| SPATS2L | 577.020476 | 0.116671916 | 0.59486527 | 0.19613167 | 0.84450711 | 0.98520136 |
| AOX1 | 47.691673 | 2.04400803 | 0.71564812 | 2.85616349 | 0.00428794 | 0.20962265 |
| BZW1 | 48.7515292 | -0.795768811 | 0.38823429 | -2.0497128 | 0.04039246 | 0.48486138 |
| CLK1 | 163.947804 | 0.038752955 | 0.22814206 | 0.16986327 | 0.86511767 | 0.98670067 |
| PPIL3 | 37.7118149 | -0.635317526 | 0.37657453 | -1.6870964 | 0.09158482 | 0.63464873 |
| ORC2 | 174.497015 | -0.404249302 | 0.16031371 | -2.5216141 | 0.01168178 | 0.30872898 |
| AC005037.1 | 49.1284114 | 0.007802865 | 0.34499457 | 0.02261736 | 0.9819555 | 0.99681784 |
| FAM126B | 153.642363 | -0.084281577 | 0.2266125 | -0.3719194 | 0.70995289 | 0.96587538 |
| NDUFB3 | 83.3879725 | 0.321599395 | 0.53208331 | 0.60441549 | 0.54556744 | 0.93523904 |
| CFLAR | 537.543121 | -0.399819126 | 0.40890383 | -0.9777828 | 0.32818177 | 0.8573817 |
| CFLAR-AS1 | 46.0053086 | 0.260014969 | 0.65116604 | 0.39930671 | 0.68966722 | 0.96220115 |
| CASP10 | 42.5734582 | -0.270412573 | 0.45576538 | -0.5933153 | 0.55297016 | 0.93523904 |
| CASP8 | 79.2149175 | 0.019719872 | 0.37764658 | 0.0522178 | 0.95835515 | 0.99572129 |
| TRAK2 | 118.443004 | 0.182790691 | 0.27144971 | 0.67338695 | 0.50070115 | 0.92414927 |

| | | | | | | |
|---|---|---|---|---|---|---|
| STRADB | 69.7703762 | 0.827028341 | 0.40583505 | 2.03784358 | 0.04156558 | 0.48843812 |
| ALS2 | 118.679867 | -0.042850745 | 0.25180605 | -0.1701736 | 0.8648736 | 0.98670067 |
| SUMO1 | 115.464753 | -0.111713864 | 0.27180842 | -0.4110022 | 0.68107091 | 0.9603972 |
| NOP58 | 143.779291 | -0.530230215 | 0.24363351 | -2.1763435 | 0.02952958 | 0.43492739 |
| BMPR2 | 596.818789 | 0.172270903 | 0.20672758 | 0.83332327 | 0.40466244 | 0.89373953 |
| FAM117B | 165.757212 | 0.023100396 | 0.31737445 | 0.07278593 | 0.94197646 | 0.99526299 |
| ICA1L | 100.831093 | -0.277712506 | 0.38149831 | -0.7279521 | 0.46664289 | 0.91462428 |
| WDR12 | 77.1169009 | -0.001394447 | 0.33603162 | -0.0041498 | 0.99668899 | 0.99898334 |
| CARF | 116.210753 | 0.169811964 | 0.37499533 | 0.45283754 | 0.65066573 | 0.95083555 |
| NBEAL1 | 783.135243 | 0.196068887 | 0.21290704 | 0.92091314 | 0.35709578 | 0.87316144 |
| CYP20A1 | 166.641667 | 0.028960522 | 0.2172599 | 0.13329898 | 0.89395694 | 0.98842187 |
| ABI2 | 353.103353 | 0.331324953 | 0.194106 | 1.70692794 | 0.08783545 | 0.62705598 |
| RAPH1 | 186.487286 | -0.088115105 | 0.28157579 | -0.3129357 | 0.75432955 | 0.9712249 |
| PARD3B | 1832.9897 | 0.098563096 | 0.26192592 | 0.37630142 | 0.70669282 | 0.96587538 |
| NRP2 | 62.8433998 | 1.062078546 | 1.1904036 | 0.89220038 | 0.37228554 | 0.87605028 |
| INO80D | 418.694806 | -0.146129063 | 0.58353219 | -0.2504216 | 0.80226133 | 0.97835591 |
| NDUFS1 | 201.089102 | 0.246953825 | 0.16018146 | 1.54171292 | 0.12314337 | 0.69314363 |
| EEF1B2 | 149.572702 | -0.285931025 | 0.6521077 | -0.4384721 | 0.6610441 | 0.95407427 |
| ZDBF2 | 69.6057099 | 0.400514136 | 0.38091621 | 1.05144945 | 0.29305222 | 0.83733658 |
| FASTKD2 | 68.6852952 | -0.347898217 | 0.23244359 | -1.4966996 | 0.13447145 | 0.70619627 |
| KLF7 | 219.697656 | -0.147073442 | 0.49675246 | -0.2960699 | 0.76717671 | 0.97221495 |
| CREB1 | 212.958498 | 0.129372641 | 0.16378406 | 0.78989762 | 0.42958756 | 0.90333313 |
| METTL21A | 68.1911154 | 0.04266923 | 0.22578998 | 0.18897752 | 0.85011043 | 0.9858744 |
| CCNYL1 | 106.364918 | -0.386667425 | 0.30585829 | -1.2642045 | 0.20615664 | 0.7735572 |
| FZD5 | 34.9821996 | -0.240614817 | 0.53778982 | -0.4474142 | 6.55E-01 | 0.95280477 |
| PLEKHM3 | 125.698666 | 0.044236809 | 0.33754168 | 0.13105584 | 0.89573113 | 0.98842187 |
| IDH1 | 31.8211876 | 0.229093503 | 0.39814285 | 0.57540529 | 0.56501723 | 0.93809297 |
| PIKFYVE | 170.139051 | -0.008581677 | 0.16626051 | -0.0516158 | 0.95883479 | 0.99572129 |
| PTH2R | 1061.59442 | 1.630366802 | 0.64547367 | 2.52584555 | 0.01154202 | 0.30872898 |
| MAP2 | 179.287673 | 0.153178602 | 0.78503406 | 0.19512351 | 0.84529626 | 0.98520136 |
| RPE | 73.4197142 | 0.007961716 | 0.21685561 | 0.03671436 | 0.97071276 | 0.99681784 |
| KANSL1L | 765.212911 | 0.683346033 | 0.22724018 | 3.00715324 | 0.00263707 | 0.16840315 |
| AC007038.1 | 95.7918355 | 0.022460903 | 0.264717 | 0.08484873 | 0.93238165 | 0.99416721 |
| ACADL | 176.199146 | -0.165753042 | 0.54909877 | -0.3018638 | 0.76275589 | 0.9714717 |
| LANCL1 | 99.7899471 | 0.178749309 | 0.20706496 | 0.86325231 | 0.38799876 | 0.8853359 |
| ERBB4 | 957.567994 | 0.990984639 | 0.64815709 | 1.52892663 | 0.12628263 | 0.69492095 |
| IKZF2 | 133.487215 | 0.38640043 | 0.22793282 | 1.69523822 | 0.09003024 | 0.6330877 |
| SPAG16 | 495.071642 | 0.678511888 | 0.30009053 | 2.26102401 | 0.02375777 | 0.40350298 |
| AC068051.1 | 84.3444851 | 2.144802036 | 0.71657781 | 2.99311811 | 0.00276143 | 0.17081506 |
| BARD1 | 44.9333145 | 0.129576812 | 0.27035345 | 0.4792867 | 0.63173468 | 0.94932781 |
| AC016708.1 | 57.9907542 | 0.340012698 | 0.48851136 | 0.696018 | 0.48641756 | 0.92034617 |
| ATIC | 60.7938186 | -0.43422873 | 0.3115887 | -1.3935959 | 0.16343967 | 0.74227469 |
| MREG | 99.7412997 | -0.406799199 | 0.35001383 | -1.1622375 | 0.245139 | 0.80476923 |
| PECR | 124.218723 | -0.024697411 | 0.36731314 | -0.067238 | 0.94639221 | 0.9952642 |
| TMEM169 | 32.2420466 | -0.35376488 | 0.46275232 | -0.76448 | 0.44458126 | 0.90694419 |
| XRCC5 | 485.167744 | -0.436799856 | 0.21449274 | -2.0364319 | 0.04170701 | 0.4884665 |
| SMARCAL1 | 52.8264384 | 0.046757998 | 0.25992234 | 0.17989218 | 0.85723721 | 0.9864199 |
| RPL37A | 354.547205 | 0.033695164 | 0.37978893 | 0.08872077 | 0.92930383 | 0.99350462 |
| IGFBP2 | 238.902756 | -0.540075106 | 0.62951347 | -0.8579246 | 0.39093409 | 0.88670817 |
| IGFBP5 | 440.633421 | -2.335337199 | 1.13516083 | -2.0572743 | 0.03965985 | 0.48094465 |
| AC007563.2 | 43.5126005 | -0.367534301 | 1.19819264 | -0.3067406 | 0.75904084 | 0.9714717 |
| DIRC3 | 138.814422 | 1.198350494 | 0.52551643 | 2.28032926 | 0.02258817 | 0.39240055 |
| TNS1 | 675.717288 | 0.761981159 | 0.49107327 | 1.55166491 | 0.12074242 | 0.68926798 |
| ARPC2 | 187.598438 | 0.008537855 | 0.28855436 | 0.02958838 | 0.97639534 | 0.99681784 |

| | | | | | | |
|---|---|---|---|---|---|---|
| AAMP | 54.781928 | 0.193256344 | 0.2989245 | 0.64650553 | 0.51795202 | 0.93257822 |
| PNKD | 151.379068 | 0.260969786 | 0.33922485 | 0.76931211 | 0.44170805 | 0.90561284 |
| TMBIM1 | 93.5034816 | -0.11544072 | 0.21806551 | -0.5293855 | 0.59653806 | 0.94683415 |
| CTDSP1 | 91.0215465 | -0.402140763 | 0.26216425 | -1.5339268 | 0.12504767 | 0.69439514 |
| USP37 | 157.630866 | -0.181642508 | 0.28761956 | -0.6315374 | 0.5276892 | 0.93366189 |
| CNOT9 | 71.6736913 | -0.429509603 | 0.37135108 | -1.1566133 | 0.24743038 | 0.80672756 |
| ZNF142 | 28.3830383 | -0.393991836 | 0.30678743 | -1.2842503 | 0.1990544 | 0.77110248 |
| STK36 | 35.7807511 | 0.355877787 | 0.28915939 | 1.23073224 | 0.21842302 | 0.78523615 |
| TTLL4 | 28.436393 | 0.190274473 | 0.44696464 | 0.42570364 | 0.67032381 | 0.95764829 |
| CYP27A1 | 57.0703058 | 0.789021275 | 0.40213994 | 1.96205648 | 0.04975591 | 0.52415717 |
| CNPPD1 | 36.6894678 | -0.021975638 | 0.29322251 | -0.0749453 | 0.94025826 | 0.99526299 |
| RETREG2 | 59.5175791 | 0.197329502 | 0.40582994 | 0.48623692 | 0.6267992 | 0.94836669 |
| ZFAND2B | 34.3377835 | 0.162564028 | 0.36472576 | 0.44571578 | 0.65580256 | 0.95368646 |
| ANKZF1 | 57.1077984 | 0.344956908 | 0.29118807 | 1.18465331 | 0.23615455 | 0.79958336 |
| TUBA4A | 63.4982485 | -0.378875945 | 0.64965437 | -0.5831962 | 0.55976124 | 0.93744835 |
| DNAJB2 | 47.8960818 | -0.269766122 | 0.38645862 | -0.6980466 | 0.48514806 | 0.92034617 |
| DNPEP | 39.7044681 | -0.315921418 | 0.39548013 | -0.7988301 | 0.42438895 | 0.90167091 |
| OBSL1 | 62.3801799 | 0.448181499 | 0.27986868 | 1.60139926 | 0.10928852 | 0.65764016 |
| EPHA4 | 154.099568 | -0.057407484 | 0.77647543 | -0.0739334 | 0.94106336 | 0.99526299 |
| SGPP2 | 360.692512 | 0.624870822 | 0.65848189 | 0.94895673 | 0.34264262 | 0.86846099 |
| FARSB | 90.1522677 | -1.042317733 | 0.28471351 | -3.6609352 | 0.0002513 | 0.042231 |
| ACSL3 | 152.181743 | -0.131512169 | 0.27015222 | -0.4868077 | 0.62639464 | 0.94831213 |
| AP1S3 | 231.840451 | -1.458155756 | 0.49019979 | -2.9746152 | 0.00293356 | 0.17565323 |
| WDFY1 | 135.676061 | -0.188851679 | 0.31538408 | -0.598799 | 0.54930691 | 0.93523904 |
| SERPINE2 | 99.1940178 | -0.819829965 | 0.8517047 | -0.9625754 | 3.36E-01 | 0.86391445 |
| CUL3 | 405.35996 | -0.018807381 | 0.14113904 | -0.1332543 | 0.89399229 | 0.98842187 |
| IRS1 | 97.9688471 | 0.099223003 | 0.31785521 | 0.31216415 | 0.75491577 | 0.97125609 |
| RHBDD1 | 187.937781 | -0.575343165 | 0.25655007 | -2.2426155 | 0.02492162 | 0.41230428 |
| COL4A4 | 411.895403 | -0.734022682 | 0.48601489 | -1.5102885 | 0.13096983 | 0.700349 |
| COL4A3 | 596.418071 | -0.488309094 | 0.56316685 | -0.8670771 | 0.38589976 | 0.88353166 |
| AC097662.1 | 92.6308874 | 0.301111963 | 0.71222592 | 0.42277591 | 0.67245876 | 0.95887225 |
| MFF | 173.207112 | 0.240576014 | 0.25491495 | 0.94375013 | 0.34529735 | 0.86990996 |
| AGFG1 | 315.921293 | 0.00172921 | 0.1973664 | 0.00876142 | 0.99300949 | 0.99866813 |
| TRIP12 | 477.106266 | -0.193878157 | 0.22126734 | -0.8762168 | 0.38091219 | 0.8807629 |
| FBXO36 | 63.2416366 | -0.26577585 | 0.30223882 | -0.8793571 | 0.37920768 | 0.88016244 |
| SP110 | 79.853943 | -1.828491955 | 0.71205264 | -2.5679168 | 0.01023117 | 0.29271368 |
| SP140L | 62.8629344 | -0.503806708 | 0.4094903 | -1.2303264 | 0.21857492 | 0.78523615 |
| SP100 | 292.507286 | -0.744396611 | 0.64085916 | -1.1615604 | 0.24541408 | 0.80476923 |
| CAB39 | 293.145593 | -0.104398561 | 0.23745384 | -0.4396583 | 0.66018458 | 0.95407427 |
| ITM2C | 98.7752984 | 1.137670916 | 0.57123462 | 1.99160007 | 0.04641496 | 0.50827029 |
| PSMD1 | 210.378767 | -0.543296223 | 0.24387823 | -2.2277356 | 0.02589815 | 0.4190513 |
| ARMC9 | 325.671452 | 0.102958808 | 0.22242948 | 0.46288293 | 0.64344828 | 0.94944486 |
| NCL | 214.443017 | -0.688023748 | 0.4295583 | -1.6017005 | 0.10922186 | 0.65764016 |
| PTMA | 936.012656 | -0.647764477 | 0.44290981 | -1.4625201 | 0.14359875 | 0.71631428 |
| PDE6D | 58.8716657 | 0.172658475 | 0.28737243 | 0.60081782 | 0.54796134 | 0.93523904 |
| COPS7B | 38.2473958 | 0.196730809 | 0.26914631 | 0.73094373 | 0.46481352 | 0.91462428 |
| DIS3L2 | 338.778456 | -0.212573638 | 0.22992302 | -0.9245427 | 0.35520386 | 0.87261977 |
| EIF4E2 | 78.25627 | 0.545614125 | 0.26927648 | 2.02622274 | 0.04274196 | 0.4938785 |
| EFHD1 | 199.479546 | 0.242367276 | 0.40944602 | 0.59193951 | 0.55389109 | 0.93523904 |
| GIGYF2 | 475.494299 | -0.134827026 | 0.1768967 | -0.7621794 | 0.44595292 | 0.90734558 |
| ATG16L1 | 41.2392028 | -0.857630806 | 0.27565872 | -3.1112051 | 0.00186325 | 0.14133317 |
| DGKD | 99.0053357 | -0.378060357 | 0.34866803 | -1.0842989 | 0.2782323 | 0.8229426 |
| USP40 | 263.127424 | -0.061410324 | 0.3094788 | -0.1984314 | 0.84270752 | 0.98517901 |
| ARL4C | 66.3228927 | -1.575057571 | 0.70162805 | -2.2448612 | 0.02477704 | 0.41230428 |

| | | | | | | |
|---|---|---|---|---|---|---|
| SH3BP4 | 172.062534 | -0.027066198 | 0.61070989 | -0.0443192 | 0.96464994 | 0.99586809 |
| AGAP1 | 1367.25758 | -0.40718865 | 0.3029707 | -1.3439869 | 0.17895263 | 0.75150688 |
| IQCA1 | 112.73161 | 0.347290068 | 0.32084503 | 1.08242307 | 0.27906458 | 0.82352422 |
| COPS8 | 40.7063924 | -0.294540219 | 0.25848612 | -1.1394818 | 0.25450227 | 0.80897837 |
| MLPH | 33.5759705 | -1.100327178 | 0.75059768 | -1.4659347 | 0.14266608 | 0.71517697 |
| RAB17 | 107.275882 | 0.538384841 | 0.39653179 | 1.35773439 | 0.17454798 | 0.74910274 |
| LRRFIP1 | 779.772457 | -0.520884413 | 0.2398768 | -2.1714664 | 0.02989594 | 0.43492739 |
| ILKAP | 127.828274 | -0.277304377 | 0.20436798 | -1.3568876 | 0.17481692 | 0.74910274 |
| PER2 | 72.9480868 | -0.295367226 | 0.41481719 | -0.7120419 | 0.47643882 | 0.91819563 |
| TRAF3IP1 | 132.867135 | 0.355812288 | 0.22468021 | 1.58363878 | 0.11327594 | 0.66657543 |
| HDAC4 | 252.038761 | -0.343507996 | 0.26881954 | -1.2778386 | 0.20130633 | 0.77110248 |
| NDUFA10 | 164.739652 | -0.217501202 | 0.17686711 | -1.2297437 | 0.21879311 | 0.78523615 |
| COPS9 | 50.617597 | -0.202456525 | 0.45071423 | -0.4491904 | 0.65329428 | 0.95220723 |
| AC124861.1 | 39.7032354 | -0.244938145 | 0.85696364 | -0.2858209 | 0.77501528 | 0.97423208 |
| GPC1 | 57.9798585 | -1.742197414 | 0.31089347 | -5.6038405 | 2.10E-08 | 0.00014889 |
| ANKMY1 | 72.9745527 | -0.12998516 | 0.33374437 | -0.3894752 | 0.69692464 | 0.96220115 |
| RNPEPL1 | 50.7835354 | -0.3912355 | 0.34334515 | -1.1394817 | 0.25450232 | 0.80897837 |
| SNED1 | 180.159496 | -0.084692124 | 0.54284054 | -0.1560166 | 0.87601995 | 0.98766275 |
| AC005237.1 | 42.8874461 | 0.425567305 | 0.67617515 | 0.62937437 | 0.529104 | 0.93366189 |
| MTERF4 | 100.659088 | 0.31117859 | 0.41101326 | 0.7571011 | 0.4489893 | 0.90994378 |
| PPP1R7 | 61.4772631 | 0.029496836 | 0.23549032 | 0.1252571 | 0.90032001 | 0.98983972 |
| HDLBP | 274.685035 | -0.170542524 | 0.17939004 | -0.95068 | 0.34176684 | 0.86846099 |
| 2-Sep | 439.262676 | -0.052990618 | 0.23267548 | -0.2277447 | 0.81984468 | 0.98080333 |
| FARP2 | 158.13893 | 0.171960896 | 0.17185359 | 1.0006244 | 0.31700843 | 0.8490543 |
| STK25 | 74.3822565 | -0.046341105 | 0.27500202 | -0.1685119 | 0.86618061 | 0.98670067 |
| BOK | 39.4465596 | 0.807984476 | 0.57740615 | 1.39933471 | 0.16171263 | 0.74071637 |
| THAP4 | 74.9713901 | 0.133715268 | 0.24062988 | 0.55568854 | 0.5784238 | 0.94034284 |
| ATG4B | 116.067928 | -0.213809456 | 0.22214387 | -0.9624819 | 0.3358076 | 0.86391445 |
| DTYMK | 25.0467037 | -0.213036086 | 0.37725819 | -0.5646957 | 0.57228075 | 0.93998433 |
| ING5 | 86.571866 | -0.406525214 | 0.22124459 | -1.837447 | 0.06614393 | 0.57792071 |
| D2HGDH | 129.51922 | 0.447080895 | 0.3238988 | 1.38031042 | 0.16749109 | 0.74445571 |
| CHL1 | 344.733613 | -0.490071952 | 0.73220022 | -0.6693141 | 0.50329512 | 0.92517058 |
| LINC01266 | 184.03426 | 0.315036871 | 0.56383369 | 0.55874077 | 0.57633865 | 0.94034284 |
| CNTN4 | 1181.87309 | 0.016460396 | 0.39355215 | 0.0418252 | 0.96663805 | 0.99601626 |
| TRNT1 | 80.1747575 | -0.387102641 | 0.25123799 | -1.5407807 | 0.12337017 | 0.69314363 |
| CRBN | 112.1734 | 0.130696465 | 0.37467774 | 0.34882367 | 0.72722169 | 0.9681562 |
| AC034195.1 | 142.382727 | 0.234728551 | 0.46491174 | 0.50488841 | 0.61363721 | 0.94791646 |
| SUMF1 | 673.429141 | -0.399945124 | 0.33006911 | -1.2117012 | 0.22562679 | 0.78977225 |
| ITPR1 | 270.545493 | 0.343179947 | 0.32444413 | 1.05774744 | 0.2901706 | 0.83506029 |
| EGOT | 87.1189714 | 1.035284651 | 0.86956584 | 1.1905765 | 0.23381988 | 0.79633808 |
| AC018816.1 | 42.2146899 | -0.183934808 | 0.40115222 | -0.4585162 | 0.6465816 | 0.95008515 |
| BHLHE40-AS | 34.7441043 | 0.058954 | 0.65607722 | 0.08985832 | 0.9283998 | 0.99336859 |
| BHLHE40 | 297.644836 | -0.943583322 | 0.76397988 | -1.2350892 | 0.2167973 | 0.78319543 |
| ARL8B | 205.807164 | -0.203289668 | 0.26202318 | -0.7758461 | 0.43783985 | 0.90561284 |
| AC026202.2 | 36.9548872 | 0.162259475 | 0.47786001 | 0.33955441 | 0.73419212 | 0.9698508 |
| EDEM1 | 52.3568076 | -0.014383977 | 0.30960871 | -0.0464586 | 0.96294476 | 0.99572129 |
| LMCD1-AS1 | 286.351435 | 0.453758015 | 0.29220437 | 1.55287895 | 0.12045206 | 0.68884194 |
| LMCD1 | 39.5110287 | -0.34508302 | 0.58976395 | -0.5851206 | 0.55846665 | 0.93668977 |
| RAD18 | 51.6664083 | -0.456479519 | 0.39018971 | -1.1698912 | 0.24204474 | 0.80393432 |
| SRGAP3 | 300.130371 | -0.736904209 | 0.62476471 | -1.1794908 | 0.23820281 | 0.80145582 |
| THUMPD3-A! | 165.349496 | -0.114182522 | 0.20860452 | -0.5473636 | 0.58412895 | 0.94184538 |
| THUMPD3 | 41.4316241 | -0.172453091 | 0.33659535 | -0.5123454 | 0.60840927 | 0.94727544 |
| SETD5 | 607.435416 | 0.08826323 | 0.17858071 | 0.49424841 | 0.62113075 | 0.9480368 |
| MTMR14 | 64.4498949 | -0.305806722 | 0.24419863 | -1.2522868 | 0.21046537 | 0.77730294 |

| | | | | | | |
|---|---|---|---|---|---|---|
| OGG1 | 43.9854212 | 0.421681562 | 0.36084453 | 1.16859626 | 0.24256634 | 0.80476923 |
| TADA3 | 42.3128216 | 0.153598507 | 0.25536993 | 0.60147452 | 0.54752398 | 0.93523904 |
| RPUSD3 | 27.6573309 | -0.199376688 | 0.37388648 | -0.5332546 | 0.59385735 | 0.94649617 |
| JAGN1 | 33.5651677 | 0.101680466 | 0.45879402 | 0.22162553 | 0.82460541 | 0.98094663 |
| IL17RC | 30.7700746 | 0.214328693 | 0.32381484 | 0.66188656 | 0.50804392 | 0.92666294 |
| CRELD1 | 48.687805 | 0.829157233 | 0.36968939 | 2.24284836 | 0.02490659 | 0.41230428 |
| EMC3 | 83.3179655 | 0.021734385 | 0.26846522 | 0.08095791 | 0.93547542 | 0.99466301 |
| BRK1 | 88.3432066 | -0.012236105 | 0.45983973 | -0.0266095 | 0.9787712 | 0.99681784 |
| VHL | 59.7091907 | 0.510251481 | 0.24142553 | 2.1134943 | 0.03455848 | 0.45223457 |
| IRAK2 | 88.8645055 | 0.168678708 | 0.57008203 | 0.29588498 | 0.76731792 | 0.97221728 |
| SEC13 | 73.1763894 | -0.116711226 | 0.2872488 | -0.4063071 | 0.68451698 | 0.96161098 |
| HRH1 | 481.020614 | -0.468430187 | 0.68652607 | -0.6823196 | 0.49503689 | 0.92307934 |
| ATG7 | 451.94581 | -0.274643451 | 0.26691168 | -1.0289675 | 0.30349493 | 0.84470484 |
| VGLL4 | 438.727207 | 0.290999196 | 0.29922865 | 0.97249779 | 0.33080297 | 0.85923151 |
| TAMM41 | 114.176662 | -0.440077436 | 0.22257704 | -1.9771915 | 0.04801998 | 0.51547342 |
| PPARG | 280.55073 | 0.560384146 | 0.38847283 | 1.44253113 | 0.1491526 | 0.72246729 |
| TSEN2 | 156.478398 | -0.399718149 | 0.22486844 | -1.7775645 | 0.07547541 | 0.59465671 |
| MKRN2OS | 26.8000085 | -0.378037846 | 0.47958862 | -0.7882544 | 0.43054791 | 0.90364119 |
| MKRN2 | 32.9892453 | -0.164355471 | 0.28193359 | -0.5829581 | 0.55992151 | 0.93744835 |
| RAF1 | 264.973274 | -0.137364639 | 0.11501766 | -1.1942917 | 0.23236388 | 0.79498513 |
| RPL32 | 543.983019 | -0.084624419 | 0.58891208 | -0.1436962 | 0.88574038 | 0.98766275 |
| IQSEC1 | 47.7621439 | 0.422831872 | 0.3624476 | 1.16660139 | 0.24337138 | 0.80476923 |
| NUP210 | 55.4678014 | -0.507096024 | 0.41357854 | -1.2261178 | 0.22015434 | 0.78584902 |
| HDAC11 | 36.0236832 | -0.127820273 | 0.3538245 | -0.3612533 | 0.71791009 | 0.9674421 |
| TMEM43 | 26.7363642 | -0.147379518 | 0.42338576 | -0.3480975 | 0.72776697 | 0.96823331 |
| XPC | 176.300724 | 0.548502152 | 0.28530684 | 1.92249916 | 0.05454298 | 0.53752502 |
| LSM3 | 73.6622234 | -0.051848603 | 0.52339658 | -0.0990618 | 0.92108921 | 0.99214128 |
| SLC6A6 | 523.191462 | -0.541944371 | 0.41104363 | -1.3184595 | 0.18734988 | 0.76207437 |
| CCDC174 | 51.3431524 | 0.183857348 | 0.27724056 | 0.663169 | 0.50722232 | 0.92666294 |
| FGD5-AS1 | 104.069424 | -0.288141019 | 0.33496132 | -0.8602218 | 0.3896668 | 0.88586416 |
| NR2C2 | 190.2272 | 0.038998054 | 0.19708484 | 0.19787445 | 0.8431433 | 0.98517901 |
| MRPS25 | 109.576274 | -0.043904535 | 0.20281374 | -0.2164771 | 0.82861585 | 0.98142144 |
| RBSN | 62.5698402 | 0.114035312 | 0.24591434 | 0.46371966 | 0.64284861 | 0.94944486 |
| CAPN7 | 254.26231 | 0.010013238 | 0.19426652 | 0.05154382 | 0.95889219 | 0.99572129 |
| SH3BP5 | 43.4407262 | 0.404784859 | 0.76201505 | 0.53120323 | 0.59527795 | 0.94664456 |
| METTL6 | 54.9333839 | -0.116756712 | 0.32597221 | -0.35818 | 0.72020865 | 0.96779196 |
| EAF1 | 28.3459213 | 0.028216516 | 0.30732263 | 0.09181399 | 0.92684583 | 0.99336859 |
| COLQ | 33.8797358 | 0.159432025 | 0.40544667 | 0.39322564 | 0.69415283 | 0.96220115 |
| HACL1 | 84.7302608 | 0.058313938 | 0.22199059 | 0.26268653 | 0.7927922 | 0.97443119 |
| BTD | 27.201576 | 0.693269442 | 0.32361399 | 2.14227275 | 0.03217154 | 0.44049392 |
| ANKRD28 | 434.445064 | -0.034431325 | 0.20129801 | -0.1710465 | 0.86418718 | 0.98670067 |
| OXNAD1 | 106.366955 | -0.013040425 | 0.2807154 | -0.0464543 | 0.96294819 | 0.99572129 |
| RFTN1 | 205.471064 | 0.719496587 | 1.07376509 | 0.6700689 | 0.50281387 | 0.92492012 |
| PLCL2 | 89.9795023 | 1.590412336 | 0.41814581 | 3.80348748 | 0.00014267 | 0.03037037 |
| TBC1D5 | 1189.92114 | -0.153664388 | 0.2156372 | -0.7126061 | 0.47608953 | 0.91809716 |
| KCNH8 | 36.4513574 | -0.393616828 | 0.46538672 | -0.8457844 | 0.39767303 | 0.88863925 |
| RAB5A | 160.635979 | -0.150182028 | 0.19238932 | -0.7806152 | 0.43502885 | 0.90514499 |
| KAT2B | 113.521767 | 1.14806455 | 0.39829841 | 2.88242314 | 0.00394629 | 0.2001215 |
| ZNF385D | 50.0684605 | 1.165969038 | 0.9561116 | 1.21949053 | 0.22265807 | 0.78710314 |
| UBE2E2 | 869.22202 | 0.353236186 | 0.31356041 | 1.1265331 | 0.25993994 | 0.81263292 |
| UBE2E1 | 423.612153 | -0.236484017 | 0.31218131 | -0.7575214 | 0.44873755 | 0.90994378 |
| NKIRAS1 | 36.4615458 | 0.406687943 | 0.39095383 | 1.04024546 | 0.29822588 | 0.83973489 |
| RPL15 | 757.142489 | 0.400233282 | 0.56142406 | 0.71288944 | 0.47591417 | 0.91809716 |
| NR1D2 | 105.693667 | -0.383106855 | 0.41621614 | -0.9204517 | 0.35733677 | 0.87316144 |

| | | | | | | |
|---|---|---|---|---|---|---|
| THRB | 858.084591 | 0.729350471 | 0.57527527 | 1.26782865 | 0.20485915 | 0.77352973 |
| TOP2B | 246.068902 | -0.080579207 | 0.18683186 | -0.4312926 | 0.6662556 | 0.95613858 |
| NGLY1 | 124.698717 | -0.05829407 | 0.28878729 | -0.2018582 | 0.84002761 | 0.98464453 |
| NEK10 | 91.5496527 | 1.238471016 | 0.37047893 | 3.34289187 | 0.0008291 | 0.09295873 |
| SLC4A7 | 255.622215 | 0.274890241 | 0.56576644 | 0.4858723 | 0.62705771 | 0.94856269 |
| CMC1 | 156.746486 | 0.018592584 | 0.19227126 | 0.09669975 | 0.92296484 | 0.99214128 |
| AZI2 | 96.6434488 | -0.036244813 | 0.33935162 | -0.1068061 | 0.91494283 | 0.99159829 |
| RBMS3 | 226.202057 | 0.680697059 | 0.78260124 | 0.86978786 | 0.38441635 | 0.88328503 |
| RBMS3-AS3 | 85.6088847 | 0.343423872 | 0.88909236 | 0.38626344 | 0.69930157 | 0.96280283 |
| TGFBR2 | 668.217071 | -0.665478352 | 0.8068333 | -0.8248028 | 0.40948357 | 0.8959746 |
| STT3B | 244.790508 | 0.257928734 | 0.25008456 | 1.0313661 | 0.30236917 | 0.84375949 |
| OSBPL10 | 1165.91587 | -0.055665847 | 0.37325287 | -0.1491371 | 0.88144546 | 0.98766275 |
| CMTM8 | 67.5900083 | 0.181801818 | 0.26906334 | 0.67568408 | 0.49924125 | 0.92398825 |
| CMTM7 | 77.024192 | 0.408676117 | 0.54563125 | 0.7489969 | 0.45385907 | 0.91077355 |
| CMTM6 | 107.039894 | 0.341727947 | 0.31552065 | 1.08306048 | 0.27878158 | 0.8229426 |
| DYNC1LI1 | 74.9775466 | 0.319607667 | 0.28601049 | 1.11746834 | 0.26379413 | 0.81378765 |
| CNOT10 | 148.277656 | -0.004553285 | 0.20541751 | -0.022166 | 0.98231554 | 0.99681784 |
| GLB1 | 125.811524 | 0.04288648 | 0.21115352 | 0.20310568 | 0.83905243 | 0.98462033 |
| CRTAP | 87.7462306 | -0.186937796 | 0.31538014 | -0.592738 | 0.5533565 | 0.93523904 |
| FBXL2 | 311.093585 | 0.219493727 | 0.24342144 | 0.90170252 | 0.36721491 | 0.87605028 |
| UBP1 | 107.456787 | -0.313797643 | 0.20490753 | -1.531411 | 0.12566786 | 0.69492095 |
| CLASP2 | 342.513579 | 0.026921785 | 0.18722168 | 0.1437963 | 0.88566133 | 0.98766275 |
| PDCD6IP | 255.834391 | -0.132489391 | 0.16723443 | -0.7922375 | 0.4282222 | 0.90272191 |
| LINC01811 | 104.391366 | -0.068911319 | 1.50147541 | -0.0458957 | 0.96339335 | 0.99572129 |
| TRANK1 | 72.0512221 | -0.723745458 | 0.8174391 | -0.8853815 | 0.3759509 | 0.877999 |
| EPM2AIP1 | 97.9449414 | 0.193101721 | 0.24080093 | 0.80191435 | 0.42260251 | 0.90066187 |
| MLH1 | 50.5481193 | -0.646445355 | 0.355986 | -1.8159292 | 0.06938124 | 0.58408344 |
| LRRFIP2 | 501.039477 | 0.074103029 | 0.27914621 | 0.26546314 | 0.7906527 | 0.97423208 |
| GOLGA4 | 728.87975 | -0.132462537 | 0.21962472 | -0.6031313 | 0.54642137 | 0.93523904 |
| ITGA9 | 65.9105195 | 0.027386134 | 0.56307835 | 0.04863645 | 0.96120902 | 0.99572129 |
| CTDSPL | 588.947294 | -0.238293699 | 0.18198665 | -1.3094021 | 0.19039817 | 0.76219391 |
| ACAA1 | 76.2025078 | 0.857517354 | 0.21918392 | 3.91231875 | 9.14E-05 | 0.02304358 |
| OXSR1 | 222.902749 | 0.181082216 | 0.19845397 | 0.91246455 | 0.36152422 | 0.8752623 |
| XYLB | 41.0616854 | 0.636896657 | 0.54123162 | 1.17675434 | 0.23929357 | 0.80174646 |
| ACVR2B | 57.3857772 | 0.216424315 | 0.3647878 | 0.59328825 | 0.55298826 | 0.93523904 |
| EXOG | 60.0794854 | -0.087795996 | 0.26861021 | -0.3268528 | 0.74377923 | 0.97012548 |
| WDR48 | 109.812142 | 0.310350167 | 0.28447043 | 1.09097514 | 0.27528382 | 0.82164863 |
| GORASP1 | 29.7424167 | -0.009503709 | 0.30447217 | -0.0312137 | 0.9750991 | 0.99681784 |
| TTC21A | 39.1642408 | 0.657680411 | 0.50232293 | 1.30927809 | 0.19044017 | 0.76219391 |
| CSRNP1 | 26.2519436 | -0.638760115 | 0.52540925 | -1.2157382 | 0.22408466 | 0.78828752 |
| RPSA | 383.788112 | -0.136885329 | 0.42798921 | -0.3198336 | 0.74909448 | 0.97028355 |
| MYRIP | 180.263639 | 0.589685642 | 0.66848678 | 0.88212012 | 0.37771185 | 0.87924711 |
| EIF1B | 73.3484011 | -0.125520408 | 0.43861332 | -0.2861755 | 0.77474367 | 0.97423208 |
| ENTPD3-AS1 | 49.7714801 | 0.389493058 | 0.65994734 | 0.59018808 | 0.55506456 | 0.93523904 |
| RPL14 | 641.397048 | 0.094342319 | 0.63958363 | 0.14750584 | 0.88273277 | 0.98766275 |
| ZNF621 | 88.7548266 | -0.52247434 | 0.24193933 | -2.1595263 | 0.03080936 | 0.43851837 |
| CTNNB1 | 364.649572 | 0.233475232 | 0.27311706 | 0.85485408 | 0.39263193 | 0.88760715 |
| ULK4 | 383.534292 | 0.20909528 | 0.29150755 | 0.71728942 | 0.47319553 | 0.91793034 |
| TRAK1 | 241.126959 | 0.691552408 | 0.47090589 | 1.46855756 | 0.14195283 | 0.7148274 |
| SEC22C | 50.5127664 | -0.403623249 | 0.27894163 | -1.446981 | 0.14790223 | 0.72246583 |
| SS18L2 | 30.4594593 | -0.457096571 | 0.39403401 | -1.1600434 | 0.24603112 | 0.80516983 |
| NKTR | 491.068697 | 0.064169008 | 0.13191864 | 0.48642865 | 0.62666328 | 0.94831213 |
| AC006059.1 | 70.2728613 | 0.326251915 | 0.40928338 | 0.79712964 | 0.42537574 | 0.90167091 |
| SNRK | 535.736738 | -1.109094482 | 0.67002223 | -1.65531 | 0.09786163 | 0.64312419 |

| | | | | | | |
|---|---|---|---|---|---|---|
| ANO10 | 355.311121 | 0.236123781 | 0.2714959 | 0.86971397 | 0.38445673 | 0.88328503 |
| ABHD5 | 60.2376345 | -0.300265709 | 0.51127212 | -0.5872914 | 0.55700803 | 0.93533595 |
| TCAIM | 220.019326 | -0.19792988 | 0.28790751 | -0.6874773 | 0.49178199 | 0.9227776 |
| ZNF445 | 52.6067168 | -0.116775311 | 0.31690643 | -0.3684851 | 0.71251152 | 0.96587538 |
| AC099669.1 | 100.134645 | 0.013986469 | 0.39035588 | 0.03583004 | 0.97141788 | 0.99681784 |
| ZNF197 | 78.0538681 | 0.081325915 | 0.2570585 | 0.31637124 | 0.75172075 | 0.97058015 |
| KIAA1143 | 34.2238957 | -0.179215596 | 0.29643915 | -0.6045612 | 0.54547062 | 0.93523904 |
| ZDHHC3 | 85.9374325 | 0.192256516 | 0.24216106 | 0.79392002 | 0.42724199 | 0.90272191 |
| EXOSC7 | 44.2018493 | -0.211003868 | 0.31842343 | -0.6626518 | 0.50755357 | 0.92666294 |
| LARS2 | 101.654011 | -0.240909275 | 0.26967507 | -0.8933317 | 0.37167959 | 0.87605028 |
| LIMD1 | 77.1473104 | 0.126832516 | 0.32989716 | 0.38446076 | 0.70063697 | 0.96386637 |
| SACM1L | 129.474191 | -0.153746917 | 0.17397424 | -0.8837338 | 0.37683993 | 0.87842274 |
| LZTFL1 | 48.232001 | 0.196640137 | 0.25532561 | 0.77015437 | 0.44120833 | 0.90561284 |
| FYCO1 | 66.4749032 | -0.104967054 | 0.20808188 | -0.5044507 | 0.61394469 | 0.94791646 |
| LTF | 212.78516 | 2.251767789 | 1.40996635 | 1.59703654 | 0.11025756 | 0.65927415 |
| ALS2CL | 35.0264903 | -0.757834526 | 0.42100467 | -1.8000621 | 0.07185084 | 0.59090333 |
| PTH1R | 57.16233 | 1.597974745 | 0.65941431 | 2.42332432 | 0.01537919 | 0.33889992 |
| CCDC12 | 123.346837 | -0.364442444 | 0.23107441 | -1.5771649 | 0.1147576 | 0.67080211 |
| NBEAL2 | 29.0829853 | -1.243483928 | 0.54779799 | -2.269968 | 0.02320952 | 0.39839181 |
| SETD2 | 573.369897 | 0.144526208 | 0.15934824 | 0.90698342 | 0.36441557 | 0.87605028 |
| KIF9-AS1 | 144.368305 | 0.592661151 | 0.34445835 | 1.72055967 | 0.08533076 | 0.61974555 |
| KIF9 | 47.5649578 | 0.326649007 | 0.3154809 | 1.03540025 | 0.30048203 | 0.84221847 |
| KLHL18 | 64.8927652 | 0.087501386 | 0.37602025 | 0.23270392 | 0.81599133 | 0.98007572 |
| PTPN23 | 46.8320953 | -0.203781752 | 0.33976175 | -0.5997784 | 0.54865394 | 0.93523904 |
| SCAP | 98.1937212 | 0.003585842 | 0.25872065 | 0.0138599 | 0.98894175 | 0.99832154 |
| ELP6 | 32.8926201 | 0.086104398 | 0.3263058 | 0.2638764 | 0.79187516 | 0.97423208 |
| SMARCC1 | 432.885873 | -0.199156106 | 0.22296478 | -0.8932178 | 0.37174056 | 0.87605028 |
| DHX30 | 85.3003799 | -0.129992156 | 0.20066118 | -0.6478192 | 0.51710192 | 0.93230176 |
| MAP4 | 491.43321 | -0.08607712 | 0.23783182 | -0.3619243 | 0.71740858 | 0.96742001 |
| ZNF589 | 27.9057716 | 0.186727423 | 0.49822022 | 0.37478893 | 0.70781745 | 0.96587538 |
| PLXNB1 | 185.482861 | 0.050403671 | 0.29843587 | 0.16889281 | 0.86588096 | 0.98670067 |
| TMA7 | 118.234014 | -0.325823631 | 0.5832226 | -0.5586608 | 0.57639321 | 0.94034284 |
| SHISA5 | 120.50293 | -0.435864945 | 0.33970245 | -1.2830786 | 1.99E-01 | 0.77110248 |
| PFKFB4 | 60.736574 | -0.505163667 | 0.89612638 | -0.5637192 | 0.57294523 | 0.93998433 |
| UQCRC1 | 56.4606016 | 0.311055258 | 0.27475804 | 1.13210611 | 0.25758983 | 0.8120529 |
| NCKIPSD | 27.7855574 | 0.481769642 | 0.3803629 | 1.26660524 | 0.20529648 | 0.77352973 |
| IP6K2 | 140.732262 | -0.331486879 | 0.27502484 | -1.205298 | 0.22808829 | 0.79284833 |
| PRKAR2A | 197.567589 | -0.112196447 | 0.2178115 | -0.515108 | 0.60647756 | 0.94708529 |
| ARIH2 | 180.029381 | -0.08153295 | 0.16275332 | -0.5009603 | 0.61639906 | 0.94791646 |
| P4HTM | 64.8318419 | 0.337393392 | 0.24464238 | 1.37912898 | 0.16785499 | 0.74445571 |
| WDR6 | 118.652715 | -0.239144285 | 0.26893431 | -0.8892294 | 0.37387982 | 0.87666478 |
| DALRD3 | 26.1686095 | 0.641821615 | 0.32802888 | 1.95660094 | 0.0503944 | 0.52415717 |
| NDUFAF3 | 57.0276096 | 0.134731255 | 0.57168608 | 0.23567349 | 0.81368604 | 0.97930627 |
| IMPDH2 | 45.7118595 | -0.221691827 | 0.45018011 | -0.4924514 | 0.62240027 | 0.94808496 |
| QRICH1 | 116.898155 | 0.024756619 | 0.23441777 | 0.10560897 | 0.91589261 | 0.99159829 |
| QARS | 36.1610234 | 0.471615825 | 0.30553957 | 1.54355073 | 0.12269721 | 0.69314363 |
| USP19 | 24.475377 | -0.142246002 | 0.35147244 | -0.4047145 | 0.68568737 | 0.96161098 |
| LAMB2 | 76.0941487 | 0.39375912 | 0.29083949 | 1.35387087 | 0.17577758 | 0.74934286 |
| USP4 | 106.182853 | -0.306798071 | 0.23302321 | -1.3165988 | 0.18797315 | 0.76207437 |
| RHOA | 287.031752 | 0.157026704 | 0.23862079 | 0.65805962 | 0.51049982 | 0.92819651 |
| TCTA | 30.4359581 | 0.077432637 | 0.44381168 | 0.17447183 | 0.86149467 | 0.98669442 |
| DAG1 | 94.3046957 | -0.306704365 | 0.42293864 | -0.7251746 | 0.46834491 | 0.91462428 |
| APEH | 40.6968501 | 0.668374782 | 0.45124759 | 1.48117087 | 0.13856105 | 0.71091391 |
| MST1 | 54.9262888 | 1.219679538 | 0.48707288 | 2.50410069 | 0.01227631 | 0.30967852 |

| | | | | | | |
|---|---|---|---|---|---|---|
| RNF123 | 27.1624725 | 0.165546102 | 0.35932348 | 0.46071607 | 0.64500233 | 0.94954229 |
| IP6K1 | 167.613673 | -0.181485497 | 0.24162162 | -0.7511145 | 0.45258377 | 0.91077355 |
| RBM6 | 622.319889 | -0.075040398 | 0.19606743 | -0.3827275 | 0.70192181 | 0.9645815 |
| RBM5 | 240.794284 | -0.012949394 | 0.17013393 | -0.0761129 | 9.39E-01 | 0.99526299 |
| SEMA3F | 53.0853857 | -1.292324998 | 0.52641377 | -2.4549605 | 0.01409001 | 0.32375679 |
| GNAI2 | 122.025325 | -0.047084506 | 0.40601477 | -0.1159675 | 0.90767832 | 0.99100813 |
| SEMA3B | 93.3395475 | -0.17841031 | 0.29892769 | -0.5968343 | 0.55061799 | 0.93523904 |
| IFRD2 | 31.3927004 | -0.242598003 | 0.38546421 | -0.6293658 | 0.52910958 | 0.93366189 |
| HYAL2 | 30.5998851 | -0.276415286 | 0.37538384 | -0.7363537 | 0.46151546 | 0.91462428 |
| TUSC2 | 26.7381918 | -0.322076355 | 0.3468279 | -0.9286345 | 0.35307853 | 0.8720098 |
| HEMK1 | 26.5647757 | 0.372414945 | 0.36941685 | 1.00811574 | 0.3133989 | 0.8490543 |
| MAPKAPK3 | 42.5162587 | -0.729610344 | 0.39074134 | -1.8672464 | 0.06186719 | 0.56386853 |
| DOCK3 | 107.464515 | 0.397532364 | 0.6920563 | 0.574422 | 0.56568227 | 0.93809297 |
| MANF | 48.445097 | 0.425619066 | 0.69072611 | 0.61619079 | 0.53776861 | 0.93516669 |
| RBM15B | 24.7377599 | -0.518636154 | 0.54965106 | -0.9435735 | 0.34538766 | 0.86990996 |
| DCAF1 | 99.5119368 | -0.320543479 | 0.23567995 | -1.3600795 | 0.17380475 | 0.74910274 |
| RAD54L2 | 110.543894 | -0.411593261 | 0.3018035 | -1.3637789 | 0.17263715 | 0.74625056 |
| TEX264 | 59.3391599 | 0.265857944 | 0.21786142 | 1.22030759 | 0.22234829 | 0.78710314 |
| RPL29 | 710.017589 | 0.29650778 | 0.59545032 | 0.49795553 | 0.61851539 | 0.9480368 |
| WDR82 | 122.047924 | 0.172412743 | 0.16437038 | 1.04892829 | 0.29421113 | 0.8387644 |
| DNAH1 | 27.3528463 | 0.119586557 | 0.46742543 | 0.25584093 | 0.79807367 | 0.97584218 |
| BAP1 | 30.6142642 | 0.099480668 | 0.3110429 | 0.31982941 | 0.74909765 | 0.97028355 |
| NISCH | 110.20664 | 0.287446008 | 0.30503586 | 0.94233513 | 0.34602109 | 0.86990996 |
| SMIM4 | 60.5440625 | -0.395570562 | 0.28148951 | -1.4052764 | 0.1599391 | 0.73816762 |
| PBRM1 | 285.663331 | -0.095055423 | 0.2019294 | -0.4707359 | 0.63782932 | 0.94944486 |
| GNL3 | 71.1900246 | -0.402514722 | 0.45905605 | -0.8768313 | 0.38057826 | 0.8807629 |
| GLT8D1 | 39.8000771 | 0.21909883 | 0.28397791 | 0.77153478 | 0.44039002 | 0.90561284 |
| SPCS1 | 145.015253 | 0.046635704 | 0.45622424 | 0.10222101 | 0.91858125 | 0.99214128 |
| NEK4 | 87.3405305 | 0.34855572 | 0.21685551 | 1.60731779 | 0.10798467 | 0.65707482 |
| SFMBT1 | 71.3901376 | -0.334286092 | 0.33400048 | -1.0008551 | 0.31689685 | 0.8490543 |
| TKT | 168.595065 | 0.023546036 | 0.51814424 | 0.04544301 | 9.64E-01 | 0.99581912 |
| DCP1A | 173.440256 | -0.279988791 | 0.20166373 | -1.3883944 | 0.16501698 | 0.74445571 |
| CACNA1D | 138.166265 | 1.24500884 | 0.53183535 | 2.34096667 | 0.01923388 | 0.36922119 |
| CHDH | 78.5050502 | 0.557050638 | 0.29876823 | 1.8644909 | 0.0622528 | 0.56576808 |
| SELENOK | 72.531365 | 0.042136129 | 0.5068007 | 0.08314142 | 0.93373909 | 0.9943329 |
| CACNA2D3 | 794.792034 | -1.120077048 | 0.69780691 | -1.6051389 | 0.10846323 | 0.65764016 |
| ERC2 | 160.74306 | -0.204219177 | 0.27815241 | -0.7341989 | 0.46282755 | 0.91462428 |
| CCDC66 | 190.039086 | 0.237562612 | 0.33876712 | 0.7012564 | 0.48314302 | 0.91971919 |
| FAM208A | 225.801247 | 0.091753863 | 0.18545124 | 0.49476004 | 0.62076951 | 0.9480368 |
| ARHGEF3 | 318.038593 | 0.461590652 | 0.429752 | 1.0740861 | 2.83E-01 | 8.29E-01 |
| IL17RD | 46.1343795 | 1.541794097 | 0.75356363 | 2.04600387 | 0.04075599 | 0.48617886 |
| APPL1 | 125.698546 | -0.134506385 | 0.20449928 | -0.6577352 | 0.51070827 | 0.92819651 |
| PDE12 | 29.4001968 | -0.451102218 | 0.33450166 | -1.3485799 | 0.17747193 | 0.75126783 |
| ARF4 | 126.595045 | 0.26160594 | 0.36310028 | 0.72047849 | 0.47123044 | 0.91634658 |
| DENND6A | 168.960342 | -0.258035762 | 0.20542638 | -1.2560984 | 0.20908028 | 0.77583638 |
| DENND6A-AS | 37.5635505 | -0.359675619 | 0.34105739 | -1.0545897 | 0.291613 | 0.83658609 |
| SLMAP | 462.088093 | 0.017076216 | 0.17728427 | 0.0963211 | 0.92326556 | 0.99214128 |
| FLNB | 452.218134 | 0.117731689 | 0.23816754 | 0.49432298 | 0.6210781 | 0.9480368 |
| ABHD6 | 36.0315764 | 1.206307794 | 0.39377834 | 3.06341834 | 0.00218824 | 0.15271309 |
| PXK | 141.312967 | 0.215028843 | 0.37153539 | 0.57875736 | 0.5627529 | 0.93805441 |
| PDHB | 42.812537 | 0.503861264 | 0.37168251 | 1.35562276 | 0.17521923 | 0.74934286 |
| C3orf67 | 42.0426066 | 0.301486533 | 0.56107309 | 0.53733915 | 0.59103337 | 0.94527451 |
| FHIT | 2074.26584 | 0.365415703 | 0.29614645 | 1.23390202 | 0.21723942 | 0.78353439 |
| PTPRG | 2456.90838 | -0.008942332 | 0.38660672 | -0.0231303 | 0.98154633 | 0.99681784 |

| | | | | | | |
|---|---|---|---|---|---|---|
| THOC7 | 69.1197649 | -0.072907541 | 0.25909259 | -0.2813957 | 0.77840691 | 0.97423208 |
| ATXN7 | 45.9934821 | 0.343083418 | 0.37023115 | 0.92667356 | 0.35409605 | 0.87241657 |
| PSMD6 | 30.8053261 | -0.309978588 | 0.4229414 | -0.7329114 | 0.46361245 | 0.91462428 |
| PRICKLE2 | 299.111637 | 0.693778644 | 0.67437213 | 1.02877716 | 0.3035844 | 0.84470484 |
| ADAMTS9 | 1010.0218 | -1.131459277 | 0.44561749 | -2.5390818 | 0.01111438 | 0.30331814 |
| ADAMTS9-AS | 661.159552 | 1.214660513 | 0.53543206 | 2.2685614 | 0.02329501 | 0.39839181 |
| ADAMTS9-AS | 2256.61488 | 1.14459757 | 0.51554792 | 2.2201575 | 0.02640808 | 0.42051885 |
| MAGI1 | 2380.80088 | 0.120830876 | 0.37779219 | 0.31983424 | 0.74909399 | 0.97028355 |
| MAGI1-AS1 | 36.0099085 | 0.225368059 | 0.54732417 | 0.41176339 | 0.68051285 | 0.95989244 |
| SLC25A26 | 166.833826 | -0.544458311 | 0.25887459 | -2.103174 | 0.03545056 | 0.45686942 |
| LRIG1 | 98.4028634 | -0.187570482 | 0.38623738 | -0.4856352 | 0.62722581 | 0.94856269 |
| SUCLG2 | 373.057908 | -0.175027551 | 0.1951987 | -0.8966635 | 0.36989849 | 0.87605028 |
| SUCLG2-AS1 | 196.094979 | 0.448486006 | 0.42606562 | 1.05262192 | 0.29251431 | 0.83697123 |
| EOGT | 34.2899908 | -0.350773728 | 0.33538948 | -1.0458698 | 0.29562114 | 0.83911606 |
| TMF1 | 177.172762 | 0.226779156 | 0.15843079 | 1.43140839 | 0.15231321 | 0.72695974 |
| UBA3 | 112.816395 | -0.031177821 | 0.1860362 | -0.1675901 | 0.86690578 | 0.98670067 |
| ARL6IP5 | 116.320887 | 0.119839981 | 0.27153601 | 0.44134103 | 0.65896613 | 0.95402584 |
| FRMD4B | 503.52141 | 0.675815106 | 0.27831812 | 2.42821092 | 0.01517352 | 0.33645164 |
| MITF | 483.201337 | 0.181728138 | 0.25293741 | 0.71847078 | 0.47246706 | 0.91737234 |
| FOXP1 | 1139.91691 | 0.329095031 | 0.2252733 | 1.46087012 | 0.14405109 | 0.71634214 |
| EIF4E3 | 50.1779096 | -0.062798395 | 0.41275044 | -0.1521462 | 0.87907166 | 0.98766275 |
| RYBP | 216.501413 | 0.080858336 | 0.34534984 | 0.23413457 | 0.81488051 | 0.97975906 |
| SHQ1 | 78.1925775 | -0.196338546 | 0.3308399 | -0.5934549 | 0.55287679 | 0.93523904 |
| GXYLT2 | 67.9145183 | 1.2614007 | 0.58458281 | 2.15777933 | 0.03094499 | 0.43905874 |
| PPP4R2 | 214.946006 | 0.119377906 | 0.20558542 | 0.58067301 | 0.56146085 | 0.93747805 |
| PDZRN3 | 343.166037 | 0.579510157 | 0.75977716 | 0.76273701 | 0.44562026 | 0.90724685 |
| CNTN3 | 150.591029 | 0.052712569 | 0.74495602 | 0.0707593 | 0.94358932 | 0.9952642 |
| ZNF717 | 98.6843673 | -0.333121213 | 0.32973562 | -1.0102676 | 0.3123671 | 0.84811918 |
| ROBO2 | 172.91309 | 2.17113997 | 0.86337489 | 2.5147129 | 0.01191294 | 0.30967852 |
| ROBO1 | 241.988129 | 0.941160838 | 1.08666046 | 0.86610387 | 0.3864332 | 0.88365806 |
| LINC02027 | 87.6977283 | 1.757173587 | 0.73665455 | 2.38534273 | 0.01706322 | 0.35074796 |
| GBE1 | 614.854457 | -0.242046977 | 0.42468845 | -0.5699401 | 0.56871833 | 0.93991595 |
| CADM2 | 199.677188 | 1.175389829 | 1.30309496 | 0.9019986 | 0.36705761 | 0.87605028 |
| CHMP2B | 66.2214552 | -0.275519485 | 0.24850248 | -1.1087192 | 0.26755132 | 0.81510106 |
| CGGBP1 | 176.904925 | 0.055274281 | 0.29084954 | 0.19004425 | 0.84927446 | 0.98572641 |
| ZNF654 | 197.85669 | 0.046134199 | 0.27814989 | 0.16586093 | 0.8682664 | 0.98671532 |
| C3orf38 | 51.2880349 | 0.006033448 | 0.26251697 | 0.02298308 | 0.98166377 | 0.99681784 |
| PROS1 | 167.783748 | 0.699193136 | 0.51895852 | 1.34730063 | 0.17788343 | 0.7513166 |
| ARL13B | 151.373181 | 0.296334378 | 0.29998187 | 0.98784096 | 0.32323055 | 0.85354768 |
| NSUN3 | 72.95416 | 0.137196556 | 0.25944639 | 0.52880502 | 0.59694071 | 0.94683415 |
| MTRNR2L12 | 264.611502 | -0.330152218 | 0.60350579 | -0.5470573 | 0.5843394 | 0.94184538 |
| EPHA6 | 64.0401383 | -0.352480847 | 0.79534187 | -0.4431816 | 0.65763442 | 0.95399905 |
| ARL6 | 56.6355771 | 0.674429235 | 0.28747004 | 2.34608527 | 0.01897177 | 0.36565503 |
| CRYBG3 | 110.290284 | -0.396286845 | 0.30987979 | -1.2788406 | 0.2009532 | 0.77110248 |
| RIOX2 | 48.957168 | -0.054357807 | 0.3800269 | -0.1430367 | 0.88626117 | 0.98766275 |
| CLDND1 | 53.0451924 | -0.165434682 | 0.38763163 | -0.4267832 | 0.66953722 | 0.95764829 |
| ST3GAL6 | 217.446789 | -0.574101262 | 0.54927013 | -1.0452075 | 0.29592707 | 0.83955237 |
| DCBLD2 | 305.255187 | -0.542494684 | 0.50049037 | -1.0839263 | 0.27839747 | 0.8229426 |
| COL8A1 | 190.293826 | 0.112501765 | 0.62193168 | 0.18089087 | 0.85645323 | 0.9864199 |
| CMSS1 | 338.083443 | -0.544628502 | 0.22394442 | -2.4319807 | 0.01501651 | 0.3345189 |
| FILIP1L | 111.209445 | 0.500103684 | 0.36657783 | 1.36424965 | 0.17248901 | 0.74625056 |
| TBC1D23 | 115.851827 | -0.024212673 | 0.21349102 | -0.1134131 | 0.90970307 | 0.99100813 |
| NIT2 | 52.4477127 | 0.334751321 | 0.24434419 | 1.36999912 | 0.17068718 | 0.74445571 |
| TOMM70 | 107.171702 | 0.230140305 | 0.18737626 | 1.22822552 | 0.21936232 | 0.7854948 |

| | | | | | | |
|---|---|---|---|---|---|---|
| TFG | 215.82046 | 0.213137435 | 0.21575173 | 0.98788287 | 0.32321002 | 0.85354768 |
| ABI3BP | 234.644055 | 0.621783002 | 0.54209463 | 1.14700085 | 0.25138127 | 0.80744031 |
| SENP7 | 225.364133 | 0.006628543 | 0.27945499 | 0.02371954 | 0.98107632 | 0.99681784 |
| TRMT10C | 35.512994 | -0.160823556 | 0.45618244 | -0.3525422 | 0.72443168 | 0.96781837 |
| PCNP | 177.956154 | -0.220751697 | 0.1865178 | -1.1835422 | 0.23659432 | 0.79958336 |
| ZBTB11 | 60.4290818 | -0.278840613 | 0.28102527 | -0.9922261 | 0.32108723 | 0.85131972 |
| RPL24 | 270.848271 | -0.259670743 | 0.54767844 | -0.4741299 | 0.63540725 | 0.94932781 |
| CEP97 | 40.1128968 | 0.348007315 | 0.30600085 | 1.13727566 | 0.25542307 | 0.80897837 |
| NFKBIZ | 556.5116 | -0.548066821 | 0.49110883 | -1.1159784 | 0.2644314 | 0.81378765 |
| ALCAM | 288.588264 | 0.374255264 | 0.26159731 | 1.4306541 | 0.15252938 | 0.72726674 |
| CBLB | 475.218245 | 0.41435115 | 0.53299455 | 0.77740222 | 0.4369215 | 0.90561284 |
| LINC00882 | 96.6035987 | -0.144232942 | 0.52309361 | -0.2757307 | 0.78275495 | 0.97423208 |
| BBX | 720.467791 | 0.334231694 | 0.33424396 | 0.99996329 | 0.31732827 | 0.8490543 |
| CD47 | 131.099839 | 0.111939566 | 0.50838086 | 0.2201884 | 0.82572443 | 0.98094663 |
| IFT57 | 107.353943 | 0.564367261 | 0.20764047 | 2.71800228 | 0.00656774 | 0.2536789 |
| DZIP3 | 168.664625 | 0.350747853 | 0.53279205 | 0.65832036 | 0.5103323 | 0.92819651 |
| NECTIN3 | 130.042182 | 0.067041806 | 0.36436811 | 0.18399472 | 0.8540176 | 0.9864199 |
| PHLDB2 | 739.123605 | -0.268393294 | 0.15934014 | -1.6844048 | 0.09210347 | 0.63464873 |
| ABHD10 | 37.8126873 | 0.811924672 | 0.33792186 | 2.40269947 | 0.01627456 | 0.34282836 |
| SLC9C1 | 80.0549978 | 0.775567055 | 0.37078199 | 2.09170636 | 0.03646479 | 0.46377956 |
| CD200 | 178.031258 | -0.032755434 | 0.39795771 | -0.0823088 | 0.93440113 | 0.99442609 |
| AC092894.1 | 220.98456 | -0.229374472 | 0.32241521 | -0.7114257 | 4.77E-01 | 0.91845229 |
| ATG3 | 48.1446958 | -0.239792131 | 0.27583525 | -0.869331 | 0.38466611 | 0.88328503 |
| SLC35A5 | 37.5244908 | -0.15470527 | 0.28400367 | -0.5447298 | 0.58593935 | 0.9423304 |
| NEPRO | 61.3359428 | -0.279009652 | 0.2119195 | -1.3165832 | 0.18797838 | 0.76207437 |
| BOC | 128.788691 | -0.389815175 | 0.61456071 | -0.6342989 | 0.52588578 | 0.93272384 |
| CFAP44 | 153.296898 | 1.001999624 | 0.32209926 | 3.11084109 | 0.00186555 | 0.14133317 |
| SPICE1 | 95.6752215 | 0.259145759 | 0.1919804 | 1.34985529 | 0.17706241 | 0.75126783 |
| USF3 | 93.5135415 | -0.065168978 | 0.20714041 | -0.3146126 | 0.75305583 | 0.97096495 |
| NAA50 | 194.56855 | -0.241637735 | 0.17426519 | -1.3866093 | 0.16556093 | 0.74445571 |
| ATP6V1A | 245.522254 | 0.140925927 | 0.30941529 | 0.45545883 | 0.64877918 | 0.95027667 |
| GRAMD1C | 125.439248 | 0.53774159 | 0.26339443 | 2.04158299 | 0.04119291 | 0.48752866 |
| ZDHHC23 | 26.8041744 | 0.31843553 | 0.32793313 | 0.97103801 | 0.33152935 | 0.85923151 |
| CCDC191 | 138.896001 | 0.492455841 | 0.43378182 | 1.1352616 | 0.25626573 | 0.80988075 |
| QTRT2 | 58.4643311 | -0.337041352 | 0.32378113 | -1.0409543 | 0.29789677 | 0.83973489 |
| ZBTB20 | 6580.32414 | 0.290813371 | 0.20642137 | 1.40883367 | 0.15888436 | 0.73686885 |
| ZBTB20-AS1 | 30.386651 | 0.568720726 | 0.48693071 | 1.16797055 | 0.24281865 | 0.80476923 |
| ZBTB20-AS5 | 144.568866 | 0.961274501 | 0.54296782 | 1.77040786 | 0.07665922 | 0.59726395 |
| ZBTB20-AS2 | 35.5786702 | 0.754381084 | 0.73124129 | 1.03164454 | 0.30223867 | 0.84375949 |
| LSAMP | 122.161109 | -0.527424978 | 1.05084366 | -0.5019062 | 0.61573348 | 0.94791646 |
| IGSF11 | 44.6107406 | 0.766176573 | 0.34510848 | 2.22010359 | 0.02641174 | 0.42051885 |
| B4GALT4 | 53.2383902 | 0.754887259 | 0.3635347 | 2.07652052 | 0.03784584 | 0.47081202 |
| ARHGAP31 | 136.48547 | -0.26458617 | 0.27241891 | -0.9712475 | 0.33142507 | 0.85923151 |
| TMEM39A | 52.8219225 | 0.295201783 | 0.30840646 | 0.95718417 | 0.3384743 | 0.86672435 |
| POGLUT1 | 42.7864708 | -0.146126381 | 0.36242469 | -0.403191 | 0.68680771 | 0.96161098 |
| TIMMDC1 | 80.7836883 | 0.27983539 | 0.20075231 | 1.3939336 | 0.16333766 | 0.74222552 |
| COX17 | 49.2010946 | 0.248832339 | 0.39786702 | 0.62541584 | 0.53169817 | 0.93366189 |
| MAATS1 | 128.097734 | 0.77061635 | 0.36987275 | 2.08346344 | 0.037209 | 0.46890807 |
| GSK3B | 554.98759 | -0.226702968 | 0.23407338 | -0.9685124 | 0.33278854 | 0.86067542 |
| LRRC58 | 84.582589 | -0.198572381 | 0.30620927 | -0.6484858 | 0.51667076 | 0.9320507 |
| FSTL1 | 163.66664 | 0.379336623 | 0.40790085 | 0.92997263 | 0.35238526 | 0.8720098 |
| NDUFB4 | 83.4566378 | -0.286397341 | 0.39655943 | -0.7222053 | 0.47016824 | 0.91502267 |
| HGD | 86.502557 | 0.870173011 | 0.48457669 | 1.79573848 | 0.07253612 | 0.59133913 |
| RABL3 | 51.8021048 | -0.216766595 | 0.34853746 | -0.621932 | 0.53398659 | 0.93446646 |

| | | | | | | |
|---|---|---|---|---|---|---|
| GOLGB1 | 385.660673 | 0.129218675 | 0.14095838 | 0.9167151 | 0.35929195 | 0.87316144 |
| IQCB1 | 161.464242 | -0.120593602 | 0.17345433 | -0.695247 | 0.48690052 | 0.92034617 |
| SLC15A2 | 70.3988893 | 1.100899115 | 0.73825909 | 1.49120969 | 0.13590645 | 0.70722407 |
| ILDR1 | 26.8284555 | -0.462390116 | 0.57647065 | -0.8021052 | 0.42249209 | 0.90066187 |
| CASR | 60.049231 | 0.529478438 | 1.47729861 | 0.35840989 | 0.72003659 | 0.96779196 |
| CCDC58 | 28.8273112 | -0.244755202 | 0.32420166 | -0.7549474 | 0.45028055 | 0.91037995 |
| FAM162A | 132.076535 | -0.266661859 | 0.53236773 | -0.5008979 | 0.616443 | 0.94791646 |
| AC083798.2 | 77.9630474 | 0.545931662 | 0.53674414 | 1.01711713 | 0.30909771 | 0.84554063 |
| KPNA1 | 193.527281 | 0.06332275 | 0.27490351 | 0.23034537 | 0.81782341 | 0.98010849 |
| PARP9 | 142.804301 | -1.364330303 | 0.75081988 | -1.8171206 | 0.06919864 | 0.58408344 |
| DTX3L | 65.8228727 | -0.822946512 | 0.6165363 | -1.33479 | 0.18194507 | 0.75448394 |
| PARP14 | 465.438033 | -1.949880683 | 0.79026147 | -2.4673868 | 0.01361033 | 0.32032755 |
| HSPBAP1 | 66.3066073 | -0.050927502 | 0.34995886 | -0.1455243 | 0.88429697 | 0.98766275 |
| DIRC2 | 104.546183 | 0.151380484 | 0.32557346 | 0.46496567 | 0.64195604 | 0.94944486 |
| SEMA5B | 63.6772922 | 1.366195098 | 0.7598941 | 1.79787565 | 0.07219672 | 0.59090333 |
| PDIA5 | 59.1973622 | 0.184278461 | 0.34791826 | 0.52966022 | 0.59634753 | 0.94683415 |
| SEC22A | 169.328657 | -0.008329557 | 0.17430873 | -0.0477862 | 0.96188661 | 0.99572129 |
| ADCY5 | 182.653543 | 0.160060879 | 0.49222337 | 0.32517935 | 0.74504533 | 0.97028355 |
| HACD2 | 88.9700315 | -0.591441058 | 0.33772238 | -1.751264 | 0.07990044 | 0.60643125 |
| MYLK-AS1 | 30.2709301 | -0.022189537 | 0.50108704 | -0.0442828 | 0.96467898 | 0.99586809 |
| MYLK | 220.205929 | 0.859843725 | 1.01197771 | 0.84966666 | 0.39551044 | 0.88760715 |
| CCDC14 | 189.299585 | 0.218727501 | 0.16678754 | 1.31141394 | 0.18971795 | 0.76207437 |
| KALRN | 190.082762 | -0.041450096 | 0.44701232 | -0.092727 | 0.92612047 | 0.99316012 |
| UMPS | 35.087436 | 0.029534298 | 0.37715058 | 0.07830904 | 0.93758223 | 0.99526299 |
| ITGB5 | 111.47045 | 0.602439531 | 0.42883502 | 1.40482821 | 0.16007237 | 0.73816762 |
| HEG1 | 114.430013 | -0.88654088 | 0.50266168 | -1.763693 | 0.07778368 | 0.59991135 |
| SLC12A8 | 168.472366 | -0.486261093 | 0.36379407 | -1.3366383 | 0.18134073 | 0.75448394 |
| ZNF148 | 827.954937 | -0.052271624 | 0.26510577 | -0.1971727 | 0.84369238 | 0.98517901 |
| SNX4 | 205.056996 | 0.070753453 | 0.19748335 | 0.35827553 | 0.72013713 | 0.96779196 |
| OSBPL11 | 75.2930119 | -0.045399101 | 0.30373324 | -0.1494703 | 0.88118254 | 0.98766275 |
| SLC41A3 | 38.2679155 | -0.068555824 | 0.26486964 | -0.2588285 | 0.79576753 | 0.97524554 |
| ALDH1L1 | 89.5448062 | 0.995343023 | 0.43578129 | 2.2840426 | 0.02236903 | 0.39172379 |
| ZXDC | 192.812784 | -0.179230529 | 0.17983485 | -0.9966396 | 0.31893949 | 0.84935263 |
| TXNRD3NB | 26.6477394 | -0.793751831 | 0.43278087 | -1.8340733 | 0.06664311 | 0.5780981 |
| TXNRD3 | 30.2403193 | -0.182492442 | 0.32895438 | -0.5547652 | 0.57905527 | 0.94034284 |
| CHCHD6 | 71.1590328 | -0.662617886 | 0.22605594 | -2.931212 | 0.00337642 | 0.18913892 |
| PLXNA1 | 30.2040939 | -0.336083426 | 0.39614213 | -0.848391 | 0.39622024 | 0.88822693 |
| TPRA1 | 35.2322071 | 0.030817128 | 0.35637381 | 0.08647417 | 0.93108949 | 0.99408228 |
| MGLL | 225.809391 | -0.458686671 | 0.40276963 | -1.1388313 | 0.25477352 | 0.80897837 |
| SEC61A1 | 116.855031 | 0.171922822 | 0.39222456 | 0.43832753 | 0.66114887 | 0.95407427 |
| RUVBL1 | 75.5523778 | 0.084541399 | 0.21742137 | 0.38883665 | 0.69739698 | 0.96220115 |
| EEFSEC | 116.062854 | 0.056472509 | 0.18047555 | 0.31290948 | 0.75434944 | 0.9712249 |
| RPN1 | 92.0869849 | -0.163293182 | 0.35072347 | -0.4655895 | 0.64150933 | 0.94944486 |
| RAB7A | 362.856751 | -0.330274214 | 0.20537847 | -1.6081248 | 0.10780784 | 0.65699665 |
| ACAD9 | 67.7493438 | -0.225169735 | 0.24647497 | -0.9135603 | 0.36094795 | 0.87465733 |
| CNBP | 200.808014 | 0.043677782 | 0.39931692 | 0.10938124 | 0.91290011 | 0.99113939 |
| COPG1 | 49.6160305 | -0.219556782 | 0.29414789 | -0.7464163 | 0.45541598 | 0.91077355 |
| HMCES | 44.8511116 | -0.424321591 | 0.34106268 | -1.2441161 | 0.21345682 | 0.78012319 |
| H1FX | 127.474942 | 0.012034501 | 0.7184684 | 0.01675022 | 0.98663589 | 0.99764442 |
| MBD4 | 85.9325735 | -0.050445605 | 0.22504216 | -0.2241607 | 0.82263227 | 0.98094663 |
| IFT122 | 202.309266 | 0.269957615 | 0.29987153 | 0.90024424 | 0.36799029 | 0.87605028 |
| TMCC1 | 581.543959 | 0.094890584 | 0.2292652 | 0.41389005 | 0.67895463 | 0.95924874 |
| PIK3R4 | 73.0474851 | -0.374486801 | 0.22883477 | -1.6364943 | 0.10173618 | 0.65098923 |
| ATP2C1 | 227.748685 | 0.02898216 | 0.25184354 | 0.11508002 | 0.90838169 | 0.99100813 |

| | | | | | | |
|---|---|---|---|---|---|---|
| NEK11 | 341.608341 | 0.39321713 | 0.32941209 | 1.1936937 | 0.23259781 | 0.79519257 |
| NUDT16 | 32.8126062 | 0.301497121 | 0.3224385 | 0.9350531 | 0.34976094 | 0.8720098 |
| MRPL3 | 94.9748718 | -0.249703053 | 0.23906501 | -1.0444985 | 0.29625479 | 0.83973489 |
| DNAJC13 | 185.823335 | -0.019438126 | 0.29604752 | -0.0656588 | 0.94764947 | 0.9952642 |
| UBA5 | 97.5507724 | 0.242698083 | 0.2990367 | 0.81159965 | 0.41702139 | 0.89862171 |
| CDV3 | 125.002328 | -0.172665903 | 0.37582326 | -0.4594338 | 0.64592269 | 0.94991115 |
| TOPBP1 | 56.7586271 | -0.504400099 | 0.26353293 | -1.9139927 | 0.05562109 | 0.53981202 |
| SRPRB | 33.9252488 | -0.046069459 | 0.33275198 | -0.1384498 | 0.88988491 | 0.98766275 |
| RAB6B | 32.5877505 | 1.223374046 | 0.45115821 | 2.71162978 | 0.00669533 | 0.25396993 |
| SLCO2A1 | 160.043572 | -0.58853438 | 0.80463752 | -0.731428 | 0.46451779 | 0.91462428 |
| RYK | 302.233353 | 0.033457075 | 0.15113244 | 0.22137586 | 0.82479978 | 0.98094663 |
| AMOTL2 | 110.13685 | 0.08551935 | 0.59342117 | 0.1441124 | 0.88541171 | 0.98766275 |
| ANAPC13 | 41.9894263 | -0.010706816 | 0.35051102 | -0.0305463 | 0.97563136 | 0.99681784 |
| CEP63 | 131.882571 | -0.072226936 | 0.19967635 | -0.36172 | 0.71756125 | 0.96742001 |
| EPHB1 | 36.8102861 | -1.151954311 | 0.52036137 | -2.2137583 | 0.02684541 | 0.4234302 |
| PPP2R3A | 1084.4156 | -0.068926002 | 0.28250888 | -0.2439782 | 0.80724772 | 0.97930627 |
| MSL2 | 102.535096 | 0.184279137 | 0.21553671 | 0.85497796 | 0.39256335 | 0.88760715 |
| PCCB | 53.8429381 | -0.283468078 | 0.36750462 | -0.771332 | 0.44051017 | 0.90561284 |
| STAG1 | 1011.51809 | -0.193572252 | 0.14720061 | -1.3150235 | 0.18850202 | 0.76207437 |
| NCK1 | 108.519111 | 0.237729963 | 0.37442797 | 0.63491507 | 0.52548381 | 0.93272384 |
| DZIP1L | 82.5371342 | 0.14144964 | 0.32707966 | 0.43246236 | 6.65E-01 | 0.95532349 |
| ARMC8 | 234.348674 | -0.089997762 | 0.20655784 | -0.4357025 | 0.66305261 | 0.95477098 |
| NME9 | 41.9350557 | 0.453819497 | 0.36054157 | 1.25871616 | 0.20813287 | 0.77455509 |
| MRAS | 58.9801686 | 0.688534817 | 0.50435164 | 1.36518801 | 0.17219396 | 0.74601809 |
| CEP70 | 258.690553 | -0.205853691 | 0.57596632 | -0.3574058 | 0.72078805 | 0.96779196 |
| FAIM | 38.9193357 | -0.239269178 | 0.33589065 | -0.7123425 | 0.47625272 | 0.91809716 |
| PIK3CB | 540.265061 | -0.034636936 | 0.21952946 | -0.1577781 | 8.75E-01 | 0.98766275 |
| COPB2 | 113.75368 | -0.024480864 | 0.24010683 | -0.1019582 | 0.91878984 | 0.99214128 |
| AC097103.2 | 226.29732 | -0.341334955 | 0.32753456 | -1.0421342 | 0.29734948 | 0.83973489 |
| NMNAT3 | 170.152836 | -0.592557382 | 0.54315335 | -1.0909578 | 2.75E-01 | 0.82164863 |
| CLSTN2 | 274.348188 | 0.822619394 | 0.56008542 | 1.46873916 | 0.14190355 | 0.7148274 |
| SLC25A36 | 291.544508 | -0.11126351 | 0.33678174 | -0.3303728 | 0.74111832 | 0.9698508 |
| PXYLP1 | 29.3578677 | 0.388154913 | 0.36977733 | 1.04969905 | 0.2938565 | 0.83825235 |
| ZBTB38 | 229.116648 | 0.197621589 | 0.30366545 | 0.6507872 | 0.51518387 | 0.93140514 |
| RASA2 | 371.911168 | -0.529871994 | 0.17487957 | -3.0299251 | 0.00244614 | 0.16049049 |
| RNF7 | 77.993049 | -0.009468214 | 0.34960622 | -0.0270825 | 0.97839392 | 0.99681784 |
| ATP1B3 | 167.344327 | -1.007773721 | 0.70639071 | -1.426652 | 0.15368023 | 0.7288378 |
| TFDP2 | 375.111145 | 0.289092821 | 0.23371575 | 1.23694198 | 0.21610862 | 0.78293142 |
| GK5 | 211.184296 | -0.02250696 | 0.30063005 | -0.074866 | 0.94032135 | 0.99526299 |
| XRN1 | 354.297764 | -0.402958667 | 0.37765511 | -1.0670018 | 0.28597104 | 0.83003998 |
| ATR | 280.682185 | -0.40227785 | 0.20226716 | -1.9888441 | 0.04671841 | 0.5102801 |
| PLS1 | 120.770876 | 0.452562917 | 0.4552394 | 0.99412071 | 0.3201641 | 0.85075884 |
| TRPC1 | 194.363291 | 0.164139944 | 0.29985218 | 0.54740287 | 0.58410198 | 0.94184538 |
| U2SURP | 247.366859 | -0.058745303 | 0.21823603 | -0.2691824 | 0.78778931 | 0.97423208 |
| SLC9A9 | 125.559989 | -0.392788324 | 0.62056901 | -0.6329487 | 0.52676717 | 0.93356202 |
| C3orf58 | 62.9294291 | -0.274144181 | 0.43051888 | -0.6367762 | 0.52427062 | 0.93272384 |
| AC107021.1 | 86.6984296 | 0.046389157 | 0.81203317 | 0.05712717 | 0.95444389 | 0.99531843 |
| LNCSRLR | 30.72241 | -0.750084458 | 0.52858318 | -1.4190472 | 0.15588527 | 0.73197303 |
| PLOD2 | 723.799613 | -0.103357741 | 0.50357715 | -0.2052471 | 0.83737909 | 0.98462033 |
| PLSCR4 | 118.359282 | 0.703781663 | 0.38919173 | 1.80831607 | 0.07055732 | 0.58797037 |
| PLSCR2 | 50.4730357 | 1.030643807 | 0.46917858 | 2.19669835 | 0.028042 | 0.4250936 |
| PLSCR1 | 400.535591 | -0.351057169 | 0.5851563 | -0.5999374 | 0.54854793 | 0.93523904 |
| GYG1 | 34.2077575 | -0.712368734 | 0.27019027 | -2.6365447 | 0.00837552 | 0.272888 |
| HLTF | 86.111638 | 0.131606496 | 0.27942036 | 0.47099823 | 0.637642 | 0.94944486 |

| | | | | | | |
|---|---|---|---|---|---|---|
| HPS3 | 80.253156 | 0.188352753 | 0.25212415 | 0.74706352 | 0.45502522 | 0.91077355 |
| CP | 82.6972644 | 0.727385189 | 1.29802406 | 0.56037882 | 5.75E-01 | 0.94034284 |
| TM4SF1 | 78.2672612 | -0.707391096 | 0.76454039 | -0.9252501 | 0.35483581 | 0.87258503 |
| WWTR1 | 427.100971 | 0.207711462 | 0.27143728 | 0.76522821 | 0.44413566 | 0.90667728 |
| COMMD2 | 64.7666248 | -0.051621018 | 0.35612122 | -0.1449535 | 0.88474758 | 0.98766275 |
| ANKUB1 | 123.812579 | -0.343902367 | 0.49465064 | -0.6952429 | 0.48690307 | 0.92034617 |
| RNF13 | 276.974538 | -0.117889498 | 0.17282962 | -0.682114 | 0.49516688 | 0.92307934 |
| TSC22D2 | 291.81042 | -0.138217602 | 0.20514721 | -0.6737484 | 0.5004713 | 0.92405832 |
| SERP1 | 201.132439 | 0.08585546 | 0.38393873 | 0.2236176 | 0.82305486 | 0.98094663 |
| EIF2A | 83.2278201 | 0.271162835 | 0.27535736 | 0.98476699 | 0.32473854 | 0.855037 |
| SELENOT | 99.9786947 | 0.032807106 | 0.27735085 | 0.11828738 | 0.90583996 | 0.99100813 |
| SIAH2 | 28.0096961 | -0.256481772 | 0.3380983 | -0.7586012 | 0.44809116 | 0.90976372 |
| MBNL1 | 701.694437 | -0.125518699 | 0.1682107 | -0.7461992 | 0.45554707 | 0.91077355 |
| DHX36 | 233.426921 | -0.187437628 | 0.16284229 | -1.1510378 | 0.24971669 | 0.80672756 |
| PLCH1 | 203.249582 | 0.349263376 | 0.444486 | 0.78576913 | 0.43200276 | 0.90503769 |
| GMPS | 100.089683 | -0.039963452 | 0.21650772 | -0.1845821 | 0.85355682 | 0.9864199 |
| SSR3 | 80.3496373 | 0.384844351 | 0.23997152 | 1.60370847 | 0.10877833 | 0.65764016 |
| TIPARP | 40.3319055 | 0.258928614 | 0.64800844 | 0.39957599 | 0.68946884 | 0.96220115 |
| LINC00886 | 89.9557175 | 0.49505949 | 0.44901647 | 1.10254194 | 0.27022614 | 0.8173338 |
| LEKR1 | 125.211172 | 0.493565374 | 0.52025861 | 0.94869236 | 0.3427771 | 0.86846099 |
| CCNL1 | 225.078344 | 0.275778228 | 0.35314596 | 0.78091853 | 0.43485041 | 0.90514499 |
| VEPH1 | 154.535939 | -0.480059014 | 0.54637057 | -0.8786326 | 0.3796005 | 0.88042813 |
| RSRC1 | 311.906285 | -0.274919122 | 0.20902444 | -1.3152487 | 0.18842633 | 0.76207437 |
| MLF1 | 86.9656595 | 0.686904102 | 0.28621777 | 2.39993521 | 0.01639797 | 0.34446534 |
| GFM1 | 74.2427621 | -0.402262515 | 0.32432176 | -1.2403192 | 0.21485734 | 0.7828522 |
| RARRES1 | 77.6706735 | 0.893245934 | 1.16690003 | 0.76548626 | 4.44E-01 | 0.90667728 |
| MFSD1 | 39.5712974 | 0.149205922 | 0.33263178 | 0.44856183 | 0.65374777 | 0.95243345 |
| IQCJ-SCHIP1 | 258.417294 | 0.298039705 | 0.21133045 | 1.41030175 | 1.58E-01 | 7.36E-01 |
| SMC4 | 37.6803235 | -0.574371188 | 0.31097727 | -1.8469877 | 0.06474893 | 0.57495764 |
| KPNA4 | 219.61236 | -0.011363373 | 0.24004465 | -0.0473386 | 0.96224338 | 0.99572129 |
| PPM1L | 211.254231 | 0.399096888 | 0.51243542 | 0.77882378 | 0.43608353 | 0.90514499 |
| NMD3 | 170.739055 | 0.015806342 | 0.36216438 | 0.04364411 | 0.96518809 | 0.99601598 |
| LINC01322 | 43.8577679 | 0.402189605 | 0.60576085 | 0.66394123 | 0.50672792 | 0.92666294 |
| PDCD10 | 111.744188 | -0.396726241 | 0.24188596 | -1.6401376 | 0.10097657 | 0.64808588 |
| GOLIM4 | 89.2762332 | 0.047418632 | 0.23929387 | 0.19816066 | 0.84291936 | 0.98517901 |
| MECOM | 1607.41137 | -1.083790789 | 0.81873138 | -1.323744 | 0.18558806 | 0.76046631 |
| MYNN | 33.8020553 | 0.059893879 | 0.36089959 | 0.16595718 | 0.86819065 | 0.98671532 |
| SEC62 | 269.239377 | 0.187176176 | 0.18831007 | 0.9939786 | 0.32023329 | 0.85075884 |
| GPR160 | 107.274547 | -0.524474372 | 0.55286187 | -0.9486535 | 0.34279685 | 0.86846099 |
| PHC3 | 460.819949 | 0.085870973 | 0.21883942 | 0.39239262 | 0.69476813 | 0.96220115 |
| PRKCI | 451.41916 | -0.240861173 | 0.27093634 | -0.8889954 | 3.74E-01 | 0.87666478 |
| SKIL | 190.852724 | 0.038539398 | 0.29217253 | 0.1319063 | 0.89505841 | 0.98842187 |
| RPL22L1 | 43.033452 | -0.166507316 | 0.54842135 | -0.303612 | 0.76142349 | 0.9714717 |
| TNIK | 588.531093 | 0.501891162 | 0.39658192 | 1.26554221 | 0.20567703 | 0.77352973 |
| PLD1 | 81.1656203 | 1.15095097 | 0.45767326 | 2.51478744 | 0.01191042 | 0.30967852 |
| FNDC3B | 1777.08811 | 0.186350702 | 0.29654142 | 0.62841374 | 0.52973294 | 0.93366189 |
| TNFSF10 | 1141.95745 | 0.340949853 | 0.61906079 | 0.55075343 | 0.58180272 | 0.94091128 |
| NCEH1 | 439.597139 | -0.152105859 | 0.48781562 | -0.3118101 | 0.75518482 | 0.97125609 |
| ECT2 | 43.2256494 | -0.138519888 | 0.50151679 | -0.2762019 | 0.782393 | 0.97423208 |
| NLGN1 | 1876.85215 | 0.051242979 | 0.21392386 | 0.2395384 | 0.81068812 | 0.97930627 |
| NAALADL2 | 955.619163 | 0.19315052 | 0.2776509 | 0.69565963 | 0.48664201 | 0.92034617 |
| TBL1XR1 | 1099.34721 | 0.041688462 | 0.36978329 | 0.11273755 | 0.91023863 | 0.99100813 |
| KCNMB2-AS: | 281.197082 | 1.615605772 | 0.73446492 | 2.19970449 | 0.02782787 | 0.4250936 |
| ZMAT3 | 89.8296608 | 0.673281438 | 0.2701852 | 2.49192566 | 0.01270526 | 0.31439454 |

| | | | | | | |
|---|---|---|---|---|---|---|
| PIK3CA | 211.821033 | 0.013934955 | 0.20972245 | 0.06644475 | 0.94702374 | 0.9952642 |
| KCNMB3 | 28.0397408 | -0.362860324 | 0.4267291 | -0.8503295 | 0.39514194 | 0.88760715 |
| ZNF639 | 38.0864227 | -0.00117795 | 0.33389022 | -0.003528 | 0.9971851 | 0.99917842 |
| MFN1 | 155.877263 | -0.118733888 | 0.20525131 | -0.5784805 | 0.56293974 | 0.93805441 |
| ACTL6A | 48.1646873 | -0.669726245 | 0.25589243 | -2.6172179 | 0.00886497 | 0.28118401 |
| MRPL47 | 30.4602492 | -0.51385295 | 0.39560651 | -1.2988991 | 0.19397855 | 0.7687432 |
| NDUFB5 | 62.7630008 | 0.889473163 | 0.39848719 | 2.23212483 | 0.02560671 | 0.41715427 |
| USP13 | 38.2258264 | 0.027984984 | 0.4114654 | 0.06801297 | 0.94577531 | 0.9952642 |
| TTC14 | 222.790886 | -0.014454281 | 0.22630179 | -0.0638717 | 0.94907238 | 0.9952642 |
| FXR1 | 228.005689 | 0.147678829 | 0.1661083 | 0.88905149 | 0.37397541 | 0.87666478 |
| DNAJC19 | 37.2633076 | 0.184885069 | 0.35618606 | 0.5190688 | 0.60371277 | 0.94683415 |
| ATP11B | 335.738152 | -0.073473305 | 0.23633651 | -0.3108843 | 0.7558886 | 0.9714717 |
| DCUN1D1 | 161.643741 | 0.278872836 | 0.24321993 | 1.14658709 | 0.25155231 | 0.80744031 |
| MCCC1 | 198.537464 | -0.114722776 | 0.27274776 | -0.4206186 | 0.67403363 | 0.95910677 |
| B3GNT5 | 33.7654362 | 0.685201617 | 0.34897437 | 1.96347261 | 0.04959128 | 0.523743 |
| KLHL24 | 250.510829 | 0.51071209 | 0.30451525 | 1.67713138 | 0.09351679 | 0.63721577 |
| YEATS2 | 169.355757 | -0.432767802 | 0.33423034 | -1.294819 | 0.19538267 | 0.76968045 |
| ABCC5 | 134.04237 | -0.626003202 | 0.39919219 | -1.568175 | 0.1168403 | 0.67748987 |
| EIF2B5 | 39.9444615 | -0.432078604 | 0.30123034 | -1.4343795 | 0.15146401 | 0.72436429 |
| DVL3 | 68.8051315 | -0.424054475 | 0.27892227 | -1.5203321 | 0.12842754 | 0.69700134 |
| AP2M1 | 139.023046 | -0.209243717 | 0.42765545 | -0.4892811 | 0.62464272 | 0.94808496 |
| PSMD2 | 69.3752686 | -0.599094711 | 0.4716954 | -1.2700881 | 0.20405325 | 0.77155952 |
| EIF4G1 | 138.143207 | -0.259843405 | 0.27575984 | -0.9422815 | 0.34604851 | 0.86990996 |
| POLR2H | 74.1384482 | -0.172857222 | 0.31474851 | -0.5491915 | 0.58287401 | 0.94121612 |
| MAGEF1 | 37.5219406 | -0.13767804 | 0.37033755 | -0.3717637 | 0.71006883 | 0.96587538 |
| VPS8 | 227.48757 | -0.130930307 | 0.19129708 | -0.6844344 | 0.49370089 | 0.92289011 |
| EHHADH | 100.448472 | 0.162440306 | 0.34335216 | 0.47310116 | 0.63614101 | 0.94932781 |
| MAP3K13 | 329.774859 | 0.438982172 | 0.18463433 | 2.37757617 | 0.01742684 | 0.35441982 |
| TMEM41A | 46.1673426 | -0.261142093 | 0.40454724 | -0.6455169 | 0.51859225 | 0.93257822 |
| LIPH | 93.0901695 | -2.061071897 | 0.81548264 | -2.5274258 | 0.01149021 | 0.30872898 |
| SENP2 | 115.448553 | 0.375660973 | 0.24145819 | 1.55580134 | 0.11975534 | 0.68649695 |
| IGF2BP2 | 467.046501 | 0.17202687 | 0.223132 | 0.77096457 | 0.44072793 | 0.90561284 |
| TRA2B | 197.038954 | 0.019887395 | 0.20202239 | 0.09844154 | 0.92158169 | 0.99214128 |
| ETV5 | 31.4958433 | 0.566760797 | 0.38634713 | 1.46697296 | 0.14238341 | 0.7148274 |
| EIF4A2 | 235.84026 | 0.172028223 | 0.33810074 | 0.50880759 | 0.6108871 | 0.94791646 |
| ST6GAL1 | 224.699282 | 1.164177653 | 1.09694273 | 1.06129301 | 0.28855676 | 0.83356008 |
| AC072022.1 | 27.9809195 | 0.905351133 | 0.62161538 | 1.45644905 | 0.14526853 | 0.71913551 |
| BCL6 | 367.318436 | 0.341743856 | 0.39384178 | 0.86771865 | 0.38554837 | 0.88353166 |
| LPP | 2354.68376 | 0.303994353 | 0.36799989 | 0.82607186 | 0.40876333 | 0.8959746 |
| TPRG1 | 89.0065664 | 0.953307965 | 0.90674752 | 1.05134885 | 0.2930984 | 0.83733658 |
| P3H2 | 1154.85594 | -0.324797231 | 0.52353933 | -0.6203874 | 0.53500273 | 0.93460153 |
| P3H2-AS1 | 25.4507221 | -0.022977896 | 0.72769638 | -0.0315762 | 0.97481001 | 0.99681784 |
| CLDN1 | 572.144665 | 0.50360148 | 0.74718207 | 0.67400102 | 0.50031066 | 0.92398825 |
| CLDN16 | 47.3053926 | 0.595983511 | 0.46672551 | 1.27694651 | 0.20162113 | 0.77110248 |
| IL1RAP | 68.8409517 | 0.629439407 | 0.91441803 | 0.68834973 | 0.49123257 | 0.92246947 |
| CCDC50 | 258.98517 | 0.439136482 | 0.22073499 | 1.98942847 | 0.04665393 | 0.51015752 |
| MB21D2 | 73.0337301 | 0.58858495 | 0.50672975 | 1.1615362 | 0.24542391 | 0.80476923 |
| ATP13A4 | 48.645452 | 0.599874236 | 0.79678647 | 0.75286699 | 0.45152984 | 0.91077355 |
| OPA1 | 205.14901 | -0.276018666 | 0.13616401 | -2.0271045 | 0.04265173 | 0.49385357 |
| HES1 | 188.903297 | -0.358848108 | 0.67360391 | -0.5327287 | 0.59422143 | 0.94649617 |
| ATP13A3 | 637.265001 | 0.053839137 | 0.24075287 | 0.22362823 | 0.8230466 | 0.98094663 |
| LINC00884 | 33.7855516 | -0.009016583 | 0.54986666 | -0.0163978 | 0.98691706 | 0.99764442 |
| LSG1 | 57.7378172 | -0.511843009 | 0.2991399 | -1.7110489 | 0.08707208 | 0.62570402 |
| XXYLT1 | 51.9971003 | -0.322503269 | 0.33146504 | -0.9729631 | 0.33057162 | 0.85923151 |

| | | | | | | |
|---|---|---|---|---|---|---|
| ACAP2 | 341.050925 | -0.051992733 | 0.12911406 | -0.4026884 | 0.68717747 | 0.96161098 |
| PPP1R2 | 85.0999051 | 0.272396561 | 0.25775677 | 1.05679693 | 0.29060427 | 0.83531608 |
| MUC20-OT1 | 514.401366 | 0.335254125 | 0.44523787 | 0.75297756 | 0.4514634 | 0.91077355 |
| TFRC | 79.6782754 | -0.117544659 | 0.46128602 | -0.2548195 | 0.79886254 | 0.9761581 |
| PCYT1A | 53.2752373 | -0.279254759 | 0.24018807 | -1.1626504 | 0.24497137 | 0.80476923 |
| UBXN7 | 143.358729 | 0.245131035 | 0.1976326 | 1.24033706 | 0.21485075 | 0.7828522 |
| RNF168 | 100.89399 | -0.233302176 | 0.24181426 | -0.9647991 | 0.33464545 | 0.86285304 |
| PIGX | 40.6428514 | -0.268949204 | 0.26616813 | -1.0104486 | 0.31228043 | 0.84811918 |
| PAK2 | 237.851389 | -0.01928782 | 0.16722687 | -0.1153392 | 0.90817624 | 0.99100813 |
| SENP5 | 241.941708 | -0.391761402 | 0.15351627 | -2.551921 | 0.01071308 | 0.29948783 |
| NCBP2 | 49.256563 | -0.109303078 | 0.31071068 | -0.3517841 | 0.72500018 | 0.96781837 |
| NCBP2-AS2 | 29.4177533 | -0.332476163 | 0.47844488 | -0.6949101 | 0.48711166 | 0.92034617 |
| DLG1 | 731.537584 | -0.023584295 | 0.20950588 | -0.112571 | 0.91037065 | 0.99100813 |
| BDH1 | 46.3553397 | -0.580206579 | 0.4307362 | -1.3470114 | 0.17797655 | 0.7513166 |
| RUBCN | 51.3905814 | -0.237970584 | 0.3505102 | -0.6789263 | 0.49718457 | 0.92398825 |
| FYTTD1 | 90.1317312 | -0.221251146 | 0.3131871 | -0.7064504 | 0.47990809 | 0.91971919 |
| LRCH3 | 325.911748 | 0.021210524 | 0.20748169 | 0.10222841 | 0.91857538 | 0.99214128 |
| IQCG | 67.0955493 | 0.193880229 | 0.37750809 | 0.51357901 | 0.60754636 | 0.94708529 |
| RPL35A | 312.647342 | 0.318898344 | 0.50158988 | 0.63577508 | 0.52492303 | 0.93272384 |
| LMLN | 81.1439164 | 0.655921953 | 0.28464644 | 2.3043392 | 0.02120361 | 0.38394968 |
| ZNF595 | 50.6118619 | -0.36186551 | 0.2577282 | -1.4040587 | 0.16030138 | 0.73816762 |
| ZNF718 | 151.93688 | -0.270499134 | 0.18482173 | -1.4635678 | 0.14331207 | 0.71631428 |
| ZNF141 | 122.54826 | -0.011019843 | 0.28913497 | -0.0381131 | 0.96959747 | 0.99681784 |
| ZNF721 | 339.025931 | 0.033035334 | 0.20737052 | 0.15930584 | 0.87342792 | 0.98766275 |
| PIGG | 83.7526544 | -0.173603544 | 0.25696628 | -0.6755888 | 0.49930175 | 0.92398825 |
| ATP5ME | 49.0714578 | -0.029321455 | 0.41278478 | -0.0710333 | 0.94337127 | 0.9952642 |
| PCGF3 | 113.45616 | -0.353924582 | 0.2770703 | -1.2773819 | 0.20146745 | 0.77110248 |
| AC139887.2 | 190.256853 | 0.54707085 | 0.29841742 | 1.83324034 | 0.06676684 | 0.57826359 |
| GAK | 124.755737 | -0.414153749 | 0.2181867 | -1.8981622 | 0.05767472 | 0.54753827 |
| FGFRL1 | 27.9487812 | -0.372165039 | 0.41057781 | -0.9064422 | 0.36470187 | 0.87605028 |
| RNF212 | 32.7834835 | 0.056575113 | 0.49489852 | 0.11431659 | 0.90898683 | 0.99100813 |
| AC092535.3 | 82.2680709 | -0.326095599 | 0.39474387 | -0.8260941 | 0.40875069 | 0.8959746 |
| SPON2 | 94.4134312 | 1.627582821 | 0.70721795 | 2.30138788 | 0.02136972 | 0.38550002 |
| CTBP1-AS | 33.3951181 | 0.433542367 | 0.54845761 | 0.79047561 | 0.42925006 | 0.90333313 |
| CTBP1 | 91.3262469 | -0.059748949 | 0.26585752 | -0.2247405 | 0.82218117 | 0.98094663 |
| CTBP1-DT | 27.1463707 | -0.111825994 | 0.31666858 | -0.3531326 | 0.72398905 | 0.96779196 |
| MAEA | 86.2305464 | 0.004917654 | 0.1805752 | 0.02723328 | 0.97827367 | 0.99681784 |
| UVSSA | 106.919485 | -0.603644094 | 0.26544935 | -2.2740462 | 0.02296321 | 0.39776593 |
| SLBP | 91.6137515 | -0.396680605 | 0.28804634 | -1.3771416 | 0.16846846 | 0.74445571 |
| TMEM129 | 30.067174 | 0.202406436 | 0.38500534 | 0.52572371 | 0.59908018 | 0.94683415 |
| FGFR3 | 96.1430002 | 0.666722252 | 0.72654141 | 0.91766587 | 0.35879382 | 0.87316144 |
| LETM1 | 53.7755453 | -0.512416803 | 0.32376883 | -1.5826626 | 0.11349839 | 0.6665856 |
| NSD2 | 225.615643 | 0.027647947 | 0.29472888 | 0.09380807 | 0.92526162 | 0.99316012 |
| NELFA | 88.4794121 | -0.087289304 | 0.23482863 | -0.3717149 | 0.71010512 | 0.96587538 |
| MXD4 | 80.9679825 | 0.260112051 | 0.45926143 | 0.56637033 | 0.57114206 | 0.93998433 |
| ZFYVE28 | 73.6041536 | 0.769913797 | 0.50657212 | 1.51985032 | 0.1285486 | 0.69707674 |
| RNF4 | 99.4876736 | 0.018942402 | 0.21137807 | 0.08961385 | 0.92859408 | 0.99336859 |
| FAM193A | 211.995289 | 0.053577323 | 0.27156171 | 0.19729336 | 0.84359797 | 0.98517901 |
| TNIP2 | 35.2975983 | -0.256025585 | 0.3506211 | -0.7302059 | 0.46526435 | 0.91462428 |
| SH3BP2 | 92.9612107 | -0.148483581 | 0.32701151 | -0.4540622 | 6.50E-01 | 0.95031069 |
| ADD1 | 313.979473 | -0.012416267 | 0.19363415 | -0.0641223 | 0.94887284 | 0.9952642 |
| MFSD10 | 71.9759079 | 0.154914152 | 0.39974546 | 0.38753198 | 0.69836241 | 0.96256506 |
| NOP14 | 32.8570332 | -0.464791493 | 0.38886105 | -1.1952637 | 0.23198403 | 0.79471003 |
| GRK4 | 32.0157713 | 0.075471487 | 0.39708212 | 0.19006519 | 0.84925805 | 0.98572641 |

| | | | | | | |
|---|---|---|---|---|---|---|
| HTT | 205.782522 | -0.216354103 | 0.19742325 | -1.0958897 | 0.27312706 | 0.82047446 |
| RGS12 | 242.611294 | -0.358258437 | 0.27023599 | -1.3257244 | 0.18493101 | 0.75962872 |
| LRPAP1 | 121.34909 | 0.458523913 | 0.33046959 | 1.38749199 | 0.16529179 | 0.74445571 |
| TMEM128 | 34.9245843 | -0.019724026 | 0.32205164 | -0.0612449 | 0.95116416 | 0.99531843 |
| STX18 | 216.181759 | 0.042594988 | 0.26307391 | 0.16191262 | 0.87137466 | 0.98766275 |
| STX18-AS1 | 114.189294 | -0.371524312 | 0.34761038 | -1.0687952 | 0.28516195 | 0.82925512 |
| STK32B | 399.915799 | 0.01525431 | 0.60297738 | 0.02529831 | 0.97981702 | 0.99681784 |
| EVC2 | 185.984272 | 0.067078214 | 0.41614449 | 0.16118972 | 0.87194398 | 0.98766275 |
| EVC | 716.71641 | 0.08941305 | 0.70814354 | 0.12626402 | 0.89952294 | 0.98956835 |
| CRMP1 | 184.578314 | 0.449577983 | 0.72730705 | 0.61814056 | 0.53648269 | 0.93469807 |
| MRFAP1 | 134.858949 | -0.260721917 | 0.51415173 | -0.5070914 | 0.61209069 | 0.94791646 |
| MRFAP1L1 | 34.4155624 | -0.269923307 | 0.32882832 | -0.8208639 | 0.41172377 | 0.89695293 |
| KIAA0232 | 291.254657 | 0.072754855 | 0.24515229 | 0.29677412 | 0.76663897 | 0.97221495 |
| TBC1D14 | 135.242863 | 0.163573585 | 0.2929099 | 0.55844334 | 0.57654168 | 0.94034284 |
| TADA2B | 28.4786053 | -0.330134363 | 0.30555399 | -1.0804453 | 0.27994394 | 0.82464035 |
| GRPEL1 | 37.1346271 | -0.045525717 | 0.53713 | -0.0847574 | 0.9324543 | 0.99416721 |
| AFAP1 | 170.753926 | 0.313066806 | 0.33268897 | 0.9410195 | 0.34669487 | 0.87005243 |
| ABLIM2 | 156.111096 | 1.068319658 | 1.42925167 | 0.74746784 | 0.45478121 | 0.91077355 |
| SH3TC1 | 74.6288951 | 0.513277243 | 0.45746393 | 1.12200593 | 0.26185992 | 0.81308142 |
| ACOX3 | 60.1326092 | 0.14352574 | 0.25215266 | 0.56920178 | 0.56921921 | 0.93998433 |
| TRMT44 | 60.2738738 | 0.201475689 | 0.30108752 | 0.66915988 | 0.5033935 | 0.92517389 |
| SLC2A9 | 192.637248 | 0.972907528 | 0.48655795 | 1.99957175 | 0.04554653 | 0.50496549 |
| WDR1 | 117.852953 | -0.080951487 | 0.29020382 | -0.278947 | 0.7802855 | 0.97423208 |
| HS3ST1 | 95.3170362 | -0.736646228 | 0.73327459 | -1.0045981 | 3.15E-01 | 8.49E-01 |
| AC005699.1 | 134.366915 | 0.199029168 | 0.57589549 | 0.34559945 | 7.30E-01 | 0.96871206 |
| RAB28 | 240.065969 | -0.139309208 | 0.17848665 | -0.7805021 | 0.43509538 | 0.90514499 |
| BOD1L1 | 198.495189 | -0.055257072 | 0.1775106 | -0.3112889 | 0.75558103 | 0.9714717 |
| CPEB2 | 258.217852 | -0.457001497 | 0.35460094 | -1.2887769 | 0.19747566 | 0.77110248 |
| AC098829.1 | 149.751478 | -0.916533235 | 0.41537951 | -2.206496 | 0.02734929 | 0.42391395 |
| CC2D2A | 147.144854 | 0.214213095 | 0.22749814 | 0.94160374 | 0.34639556 | 0.86990996 |
| FBXL5 | 288.413056 | 0.017873026 | 0.18963721 | 0.09424852 | 0.92491174 | 0.99311954 |
| FAM200B | 66.7446468 | 0.576213613 | 0.34033279 | 1.69308874 | 0.09043858 | 0.6330877 |
| PROM1 | 876.375033 | -0.431976976 | 0.31245606 | -1.3825207 | 0.16681187 | 0.74445571 |
| TAPT1 | 161.761047 | 0.13417947 | 0.16095811 | 0.83362975 | 0.40448966 | 0.89373953 |
| TAPT1-AS1 | 270.373055 | 1.005617319 | 0.33593136 | 2.99352027 | 0.00275779 | 0.17081506 |
| LDB2 | 91.0252846 | 0.502012768 | 0.90955696 | 0.55193109 | 0.58099556 | 0.94041171 |
| QDPR | 34.0457892 | -0.019681811 | 0.35549322 | -0.0553648 | 0.95584784 | 0.99572129 |
| LAP3 | 74.202511 | -1.16667072 | 0.82592766 | -1.412558 | 0.15778573 | 0.73580079 |
| MED28 | 79.9241098 | -0.149475856 | 0.25701853 | -0.5815762 | 0.56085218 | 0.93747805 |
| DCAF16 | 82.3745115 | -0.346762109 | 0.29967319 | -1.1571343 | 0.24721751 | 0.80672756 |
| LCORL | 225.652704 | 0.031708113 | 0.16740859 | 0.18940553 | 0.84977499 | 0.9858744 |
| AC024230.1 | 79.6712079 | 0.511244503 | 0.60529655 | 0.84461823 | 0.39832403 | 0.8887831 |
| SLIT2 | 79.103815 | 0.560818662 | 0.89716423 | 0.62510145 | 0.53190447 | 0.93366189 |
| PACRGL | 138.116872 | -0.305922705 | 0.18374944 | -1.6648906 | 0.09593459 | 0.64267209 |
| KCNIP4 | 2356.95919 | 0.686045548 | 0.80731387 | 0.84978789 | 0.39544302 | 0.88760715 |
| ADGRA3 | 312.8613 | -0.674690296 | 0.33400287 | -2.0200135 | 0.04338199 | 0.49484046 |
| PPARGC1A | 389.986458 | 0.268895143 | 0.48606331 | 0.55321013 | 0.58011953 | 0.94034284 |
| DHX15 | 279.509172 | -0.309789544 | 0.21090951 | -1.4688268 | 0.14187977 | 0.7148274 |
| SOD3 | 185.267771 | 0.778871332 | 1.29635957 | 0.60081427 | 0.5479637 | 0.93523904 |
| CCDC149 | 81.3138599 | -0.688918789 | 0.40902109 | -1.6843112 | 0.09212155 | 0.63464873 |
| LGI2 | 54.0853175 | -0.389333246 | 1.38382217 | -0.2813463 | 0.7784448 | 0.97423208 |
| SEPSECS | 74.7117675 | -0.109313493 | 0.3751717 | -0.2913692 | 0.77076894 | 0.97313769 |
| ZCCHC4 | 46.5849439 | -0.552940152 | 0.32073372 | -1.7239851 | 0.08471053 | 0.6175359 |
| ANAPC4 | 102.991085 | -0.104223632 | 0.18461418 | -0.5645484 | 0.572381 | 0.93998433 |

| | | | | | | |
|---|---|---|---|---|---|---|
| SLC34A2 | 364.690055 | 0.650943007 | 0.69793916 | 0.9326644 | 3.51E-01 | 8.72E-01 |
| SEL1L3 | 429.277956 | -0.03146977 | 0.47493039 | -0.0662619 | 0.94716935 | 0.9952642 |
| SMIM20 | 49.1297718 | -0.313660213 | 0.41972266 | -0.7473035 | 0.45488038 | 0.91077355 |
| RBPJ | 295.78378 | -0.037158355 | 0.25435581 | -0.1460881 | 0.88385185 | 0.98766275 |
| TBC1D19 | 224.720851 | 0.462266765 | 0.27550115 | 1.67791231 | 0.09336422 | 0.63721577 |
| STIM2 | 110.788159 | -0.050671095 | 0.19875344 | -0.2549445 | 0.79876597 | 0.9761581 |
| ARAP2 | 331.801663 | 0.003365434 | 0.32240417 | 0.01043856 | 0.99167139 | 0.99866813 |
| C4orf19 | 272.552453 | -0.113689498 | 0.47334205 | -0.2401847 | 0.81018711 | 0.97930627 |
| RELL1 | 82.6559509 | 0.020346861 | 0.26809646 | 0.07589381 | 0.93950358 | 0.99526299 |
| PGM2 | 24.647964 | -0.132717677 | 0.39812734 | -0.3333548 | 0.73886645 | 0.9698508 |
| TBC1D1 | 461.599981 | -0.339558025 | 0.2712555 | -1.2518014 | 0.21064223 | 0.77730294 |
| KLF3 | 169.592661 | -0.643519209 | 0.4259429 | -1.510811 | 0.13083661 | 0.700349 |
| FAM114A1 | 78.3756878 | 0.056036518 | 0.36509855 | 0.15348327 | 0.87801718 | 0.98766275 |
| KLHL5 | 101.268378 | 0.217847115 | 0.18418716 | 1.18274866 | 0.23690877 | 0.79991158 |
| WDR19 | 182.078293 | 0.137798668 | 0.21349562 | 0.64544026 | 0.51864192 | 0.93257822 |
| RFC1 | 140.165052 | 0.081027489 | 0.20155698 | 0.40200785 | 0.68767825 | 0.96161098 |
| RPL9 | 35.1400925 | -0.247363295 | 0.38496756 | -0.6425562 | 0.52051211 | 0.93272217 |
| LIAS | 37.9762962 | 0.456596614 | 0.35799683 | 1.27542083 | 0.20216033 | 0.77110248 |
| UGDH | 67.3399333 | -0.27114873 | 0.56633993 | -0.4787738 | 0.63209955 | 0.94932781 |
| UGDH-AS1 | 84.8065799 | 0.637991389 | 0.25081771 | 2.54364567 | 0.01097023 | 0.30196505 |
| SMIM14 | 220.460712 | 0.116111413 | 0.32465547 | 0.35764502 | 0.72060898 | 0.96779196 |
| UBE2K | 493.921494 | -0.157834273 | 0.17853772 | -0.8840388 | 0.37667526 | 0.87842274 |
| PDS5A | 455.880096 | 0.095445527 | 0.15910936 | 0.59987373 | 0.54859039 | 0.93523904 |
| N4BP2 | 112.052817 | 0.090185134 | 0.24064287 | 0.37476754 | 0.70783336 | 0.96587538 |
| RBM47 | 1235.38272 | -0.192283976 | 0.30428312 | -0.6319246 | 0.52743618 | 0.93366189 |
| NSUN7 | 132.229378 | 0.253075146 | 0.29784463 | 0.84968847 | 0.39549831 | 0.88760715 |
| APBB2 | 502.619364 | -0.191961054 | 0.38005676 | -0.5050852 | 0.61349898 | 0.94791646 |
| UCHL1 | 51.4987254 | -1.720444604 | 0.63825249 | -2.6955549 | 0.00702715 | 0.25989602 |
| LIMCH1 | 330.302062 | -0.054315989 | 0.52064772 | -0.1043239 | 0.91691233 | 0.99207822 |
| TMEM33 | 80.2571753 | -0.256210926 | 0.30592718 | -0.8374899 | 0.40231725 | 0.89239743 |
| SLC30A9 | 255.957724 | 0.065653303 | 0.13621851 | 0.48197051 | 0.6298269 | 0.94916079 |
| ATP8A1 | 124.76629 | -0.76677913 | 0.4201023 | -1.82522 | 0.06796783 | 0.5809835 |
| KCTD8 | 78.3763568 | 1.149020778 | 0.49368924 | 2.32741709 | 0.01994308 | 0.3747883 |
| GUF1 | 45.8276951 | -0.631798268 | 0.27398322 | -2.3059744 | 0.02111207 | 0.38374283 |
| GNPDA2 | 37.3614313 | -0.468211492 | 0.2819492 | -1.6606236 | 0.09678907 | 0.64295596 |
| GABRA2 | 112.559101 | -1.632281083 | 0.77138638 | -2.1160357 | 0.03434178 | 0.45186795 |
| COMMD8 | 29.5590484 | 0.110695438 | 0.45982785 | 0.24073235 | 0.80976257 | 0.97930627 |
| ATP10D | 190.710589 | 0.294837259 | 0.29788388 | 0.98977245 | 0.32228536 | 0.85246644 |
| AC107068.2 | 26.7003075 | 0.024080652 | 0.54209433 | 0.04442152 | 0.96456841 | 0.99586809 |
| NFXL1 | 91.7037798 | -0.294639458 | 0.2112232 | -1.3949199 | 0.16303998 | 0.74167981 |
| NIPAL1 | 32.2828701 | -0.823865613 | 0.46482033 | -1.7724389 | 0.07632174 | 0.59726395 |
| TXK | 43.0247864 | 0.530873851 | 0.61948856 | 0.85695505 | 0.39146973 | 0.88691782 |
| TEC | 84.886554 | 0.196688206 | 0.34440605 | 0.57109393 | 0.56793597 | 0.93910057 |
| SLAIN2 | 189.124018 | 0.080489878 | 0.15987604 | 0.5034518 | 0.61464666 | 0.94791646 |
| FRYL | 476.510286 | -0.021219253 | 0.14778337 | -0.1435835 | 0.88582938 | 0.98766275 |
| OCIAD1 | 256.825179 | 0.021997653 | 0.14515043 | 0.15155072 | 0.8795413 | 0.98766275 |
| OCIAD2 | 100.480323 | 0.005841936 | 0.48353614 | 0.01208169 | 0.99036044 | 0.99859607 |
| DCUN1D4 | 246.047118 | 0.183038836 | 0.26703729 | 0.68544297 | 0.49306444 | 0.92289011 |
| SGCB | 46.5690718 | 0.30854138 | 0.35806176 | 0.86169877 | 0.3888533 | 0.88586416 |
| USP46 | 67.3634546 | -0.564913183 | 0.26528974 | -2.1294196 | 0.03321956 | 0.44692436 |
| SCFD2 | 261.987255 | -1.003794565 | 0.38731799 | -2.5916549 | 0.00955155 | 0.28916146 |
| AC058822.1 | 140.554524 | 0.369689892 | 0.50211508 | 0.73626527 | 0.46156928 | 0.91462428 |
| LNX1 | 101.977545 | 0.000764233 | 0.76451135 | 0.00099964 | 0.99920241 | 0.99971834 |
| CHIC2 | 218.922305 | 0.42293748 | 0.2772232 | 1.52562079 | 0.12710433 | 0.69570738 |

| | | | | | | |
|---|---|---|---|---|---|---|
| TMEM165 | 330.00612 | 0.081491552 | 0.36569054 | 0.22284293 | 0.82365775 | 0.98094663 |
| CLOCK | 233.353396 | 0.031976416 | 0.21299936 | 0.15012447 | 0.88066642 | 0.98766275 |
| EXOC1 | 178.348563 | -0.213138124 | 0.24590725 | -0.8667419 | 0.38608342 | 0.88353166 |
| CEP135 | 87.0484831 | -0.004639118 | 0.24932532 | -0.0186067 | 0.98515487 | 0.99752755 |
| AASDH | 83.6012433 | 0.381657224 | 0.20835254 | 1.8317858 | 0.06698334 | 0.57856935 |
| PPAT | 39.1948866 | -0.750857824 | 0.31982918 | -2.347684 | 0.01889054 | 0.36488708 |
| PAICS | 65.7999163 | -0.853192074 | 0.28214712 | -3.0239262 | 0.00249517 | 0.16259361 |
| SRP72 | 99.7603133 | -0.158470742 | 0.28661765 | -0.5528994 | 0.58033227 | 0.94034284 |
| REST | 135.820437 | 0.058342143 | 0.23610066 | 0.24710708 | 0.80482536 | 0.97930627 |
| NOA1 | 26.1448285 | 0.205147581 | 0.44510173 | 0.46090043 | 0.64487004 | 0.94954229 |
| POLR2B | 131.204947 | -0.185929264 | 0.19179427 | -0.9694203 | 0.33233552 | 0.8599249 |
| IGFBP7 | 2505.91301 | 0.287827338 | 0.55594677 | 0.51772464 | 0.60465041 | 0.94683415 |
| ADGRL3 | 508.464794 | -1.514698625 | 1.13110294 | -1.3391342 | 0.18052699 | 0.75447995 |
| AC116049.2 | 46.0028193 | 0.144705894 | 0.62118478 | 0.23295145 | 0.81579911 | 0.98007572 |
| CENPC | 123.897355 | -0.63374459 | 0.34086492 | -1.859225 | 0.06299525 | 0.56713487 |
| UBA6 | 221.242625 | -0.518553853 | 0.3245306 | -1.5978581 | 0.11007455 | 0.65905549 |
| UBA6-AS1 | 337.735902 | -0.684935356 | 0.38805204 | -1.7650606 | 0.07755359 | 0.59861872 |
| YTHDC1 | 397.313851 | 0.117540362 | 0.24345674 | 0.48279774 | 0.62923936 | 0.94891118 |
| UGT2A3 | 170.172477 | 0.795591335 | 0.63599011 | 1.25094923 | 0.210953 | 0.77730294 |
| UGT2B7 | 1651.78485 | 0.118092532 | 0.60806243 | 0.1942112 | 0.84601052 | 0.98520136 |
| AC111000.4 | 124.60864 | 1.120819695 | 0.85891009 | 1.30493251 | 0.19191583 | 0.7646258 |
| UTP3 | 35.1516751 | 0.248458213 | 0.29968023 | 0.82907774 | 0.40706042 | 0.89585737 |
| RUFY3 | 472.749217 | -0.25048426 | 0.27807224 | -0.9007884 | 0.36770081 | 0.87605028 |
| GRSF1 | 88.8241464 | -0.13488692 | 0.17452541 | -0.7728784 | 0.43959435 | 0.90561284 |
| MOB1B | 296.415721 | 0.104582603 | 0.28979288 | 0.36088742 | 0.71818361 | 0.9674421 |
| SLC4A4 | 134.143867 | -0.145470242 | 0.51086702 | -0.2847517 | 0.7758344 | 0.97423208 |
| ADAMTS3 | 276.31026 | -0.847509678 | 0.51097478 | -1.6586135 | 0.0971937 | 0.64312419 |
| ANKRD17 | 869.957419 | 0.006042796 | 0.15108463 | 0.0399961 | 0.96809624 | 0.99671043 |
| AC053527.1 | 152.946165 | 0.410146235 | 0.43291216 | 0.94741213 | 0.34342881 | 0.86909179 |
| RASSF6 | 139.332709 | 0.093601294 | 0.48825016 | 0.19170766 | 0.84797121 | 0.98545133 |
| CXCL1 | 188.886331 | 2.150363247 | 1.87920953 | 1.14429137 | 0.25250282 | 0.80798674 |
| MTHFD2L | 123.870668 | 0.096047633 | 0.29940722 | 0.32079264 | 0.74836754 | 0.97028355 |
| BTC | 31.8836942 | -1.190452283 | 0.65227805 | -1.8250687 | 0.06799066 | 0.5809835 |
| RCHY1 | 106.601788 | 0.176152864 | 0.23960371 | 0.73518422 | 0.46222731 | 0.91462428 |
| THAP6 | 62.715721 | 0.059932551 | 0.22053239 | 0.27176304 | 0.78580423 | 0.97423208 |
| CDKL2 | 205.853941 | -0.359941168 | 0.47750503 | -0.7537956 | 4.51E-01 | 0.91077355 |
| G3BP2 | 156.689018 | -0.078994726 | 0.20831136 | -0.3792147 | 0.70452846 | 0.96461079 |
| USO1 | 273.436957 | 0.118330301 | 0.23914609 | 0.4948034 | 0.6207389 | 0.9480368 |
| NAAA | 43.0104246 | 0.729769133 | 0.47340918 | 1.54151877 | 0.12319058 | 0.69314363 |
| SDAD1 | 95.216319 | -0.521687563 | 0.21801441 | -2.392904 | 0.01671561 | 0.34884284 |
| NUP54 | 117.703655 | -0.229801678 | 0.24844748 | -0.9249507 | 0.35499153 | 0.87258503 |
| SCARB2 | 302.294104 | -0.133388707 | 0.21825593 | -0.6111573 | 0.54109545 | 0.93523904 |
| SHROOM3 | 534.901335 | -0.128486185 | 0.40317254 | -0.3186878 | 0.74996324 | 0.97054289 |
| 11-Sep | 412.104219 | -0.056107126 | 0.36277108 | -0.1546626 | 0.8770873 | 0.98766275 |
| CCNI | 430.03852 | 0.469089467 | 0.30407029 | 1.54270074 | 0.1229034 | 0.69314363 |
| CCNG2 | 107.561971 | 0.200222066 | 0.38870601 | 0.51509897 | 0.60648386 | 0.94708529 |
| CNOT6L | 202.625905 | 0.080058989 | 0.21040089 | 0.3805069 | 0.70356918 | 0.9645815 |
| MRPL1 | 108.610027 | -0.452723126 | 0.18380115 | -2.4631137 | 0.01377362 | 0.32179883 |
| FRAS1 | 729.600155 | -0.525483485 | 0.44608729 | -1.1779835 | 2.39E-01 | 0.80174646 |
| ANXA3 | 134.630715 | -1.119303022 | 0.47190902 | -2.3718619 | 0.0176987 | 0.35767064 |
| BMP2K | 183.31146 | -0.069030865 | 0.26843887 | -0.2571567 | 0.79705778 | 0.97534702 |
| PAQR3 | 30.5759109 | -1.00061468 | 0.445952 | -2.2437721 | 0.02484707 | 0.41230428 |
| CFAP299 | 101.381858 | 0.439589197 | 0.5843201 | 0.75230887 | 0.45186533 | 0.91077355 |
| RASGEF1B | 869.978313 | -0.410086359 | 0.54681365 | -0.7499563 | 0.45328101 | 0.91077355 |

| | | | | | | |
|---|---|---|---|---|---|---|
| HNRNPD | 305.411995 | -0.279712211 | 0.12553733 | -2.2281198 | 0.02587253 | 0.4190513 |
| HNRNPDL | 252.392302 | -0.474666024 | 0.20396776 | -2.3271621 | 0.01995664 | 0.3747883 |
| ENOPH1 | 51.9824731 | -0.644888002 | 0.25735574 | -2.5058232 | 0.01221667 | 0.30967852 |
| TMEM150C | 69.8424134 | 0.374768169 | 0.37520766 | 0.99882868 | 0.31787769 | 0.8490543 |
| SCD5 | 108.535505 | 0.36764811 | 0.66531841 | 0.55258971 | 0.58054439 | 0.94041171 |
| SEC31A | 410.546322 | 0.12902777 | 0.15560457 | 0.82920296 | 0.40698957 | 0.89585737 |
| LIN54 | 69.5564061 | 0.030131055 | 0.23258089 | 0.12955086 | 0.89692178 | 0.98842187 |
| COPS4 | 60.7191314 | 0.010978912 | 0.25559108 | 0.04295499 | 0.96573741 | 0.99601598 |
| HELQ | 65.0651739 | 0.044992376 | 0.25492679 | 0.17649136 | 0.85990794 | 0.9864199 |
| MRPS18C | 38.2789383 | 0.360352254 | 0.43147339 | 0.83516681 | 0.4036238 | 0.89344441 |
| ABRAXAS1 | 44.3717933 | -0.0613999 | 0.28295526 | -0.2169951 | 0.82821218 | 0.98142144 |
| GPAT3 | 223.610697 | -0.252370228 | 0.68911325 | -0.3662246 | 0.71419748 | 0.96628499 |
| CDS1 | 218.366018 | -0.018302085 | 0.28446681 | -0.0643382 | 0.94870093 | 0.9952642 |
| WDFY3 | 1347.84129 | 0.012389866 | 0.22879794 | 0.054152 | 9.57E-01 | 9.96E-01 |
| ARHGAP24 | 745.378794 | 0.225435602 | 0.36484164 | 0.61789988 | 0.53664134 | 0.93469807 |
| MAPK10 | 1436.04317 | 0.438414656 | 0.31974801 | 1.37112551 | 0.17033583 | 0.74445571 |
| PTPN13 | 967.515051 | -0.064536463 | 0.31921389 | -0.2021731 | 0.8397814 | 0.98464453 |
| AC093827.5 | 37.0842064 | -0.299638394 | 0.47686635 | -0.6283488 | 0.52977547 | 0.93366189 |
| AFF1 | 1545.39591 | -0.152138236 | 0.15577746 | -0.9766383 | 0.32874825 | 0.8574012 |
| KLHL8 | 100.162577 | 0.16023184 | 0.30735795 | 0.52131997 | 0.60214389 | 0.94683415 |
| NUDT9 | 69.28072 | 0.051313924 | 0.26855292 | 0.19107565 | 0.84846633 | 0.98548777 |
| SPP1 | 9555.38623 | 0.171180766 | 0.57339546 | 0.29853875 | 7.65E-01 | 9.72E-01 |
| PKD2 | 521.94612 | 0.059167724 | 0.21416748 | 0.27626848 | 0.78234187 | 0.97423208 |
| PPM1K | 88.201403 | -1.083532385 | 0.61383582 | -1.7651827 | 0.07753306 | 0.59861872 |
| HERC6 | 56.1630034 | -2.891390011 | 0.95008756 | -3.0432879 | 0.00234008 | 0.15891203 |
| HERC5 | 55.5827789 | -1.925284754 | 0.95046849 | -2.0256166 | 0.04280409 | 0.49400044 |
| PYURF | 35.4426583 | -0.216278739 | 0.45532723 | -0.4749963 | 0.63478963 | 0.94932781 |
| HERC3 | 108.304197 | -0.198680549 | 0.38791313 | -0.5121779 | 0.60852648 | 0.94727544 |
| FAM13A-AS1 | 40.7765044 | -0.924648542 | 0.39029854 | -2.3690802 | 0.01783239 | 0.35777027 |
| FAM13A | 359.460531 | 0.364561964 | 0.27608879 | 1.32045191 | 0.18668418 | 0.76192918 |
| CCSER1 | 1568.45607 | -0.478635482 | 0.61385607 | -0.7797194 | 0.43555606 | 0.90514499 |
| SMARCAD1 | 172.840642 | -0.041371829 | 0.21015481 | -0.1968636 | 0.84393429 | 0.98517901 |
| PDLIM5 | 478.426067 | 0.403823055 | 0.44111011 | 0.91546996 | 0.35994497 | 0.87367102 |
| BMPR1B | 280.846058 | -1.240380565 | 0.97507052 | -1.2720932 | 0.20334001 | 0.77155952 |
| UNC5C | 192.346286 | 1.045581867 | 1.47863764 | 0.70712515 | 0.47948871 | 0.91971919 |
| STPG2 | 289.551505 | -0.490147719 | 0.45762409 | -1.0710706 | 0.28413768 | 0.82918183 |
| RAP1GDS1 | 212.967536 | -0.162321793 | 0.22000137 | -0.7378217 | 0.46062277 | 0.91368087 |
| TSPAN5 | 437.954323 | 0.454299889 | 1.34071232 | 0.33884964 | 0.734723 | 0.9698508 |
| EIF4E | 135.583829 | -0.109745262 | 0.23907306 | -0.4590449 | 0.64620194 | 0.94996446 |
| METAP1 | 69.730342 | -0.533886347 | 0.20248537 | -2.6366663 | 0.00837251 | 0.272888 |
| ADH5 | 96.0551814 | 0.333877256 | 0.29670249 | 1.12529305 | 0.26046487 | 0.81263292 |
| AP002026.1 | 48.1730907 | 0.036575513 | 0.66708877 | 0.05482855 | 0.95627505 | 0.99572129 |
| LAMTOR3 | 46.349778 | -0.382152491 | 0.29251193 | -1.306451 | 0.19139924 | 0.76328615 |
| DNAJB14 | 224.40297 | -0.084415887 | 0.17748285 | -0.4756284 | 0.63433913 | 0.94932781 |
| H2AFZ | 87.2059115 | -0.247496397 | 0.72917319 | -0.3394206 | 0.73429291 | 0.9698508 |
| DDIT4L | 83.3890976 | -0.73622631 | 0.58106605 | -1.2670269 | 2.05E-01 | 0.77352973 |
| PPP3CA | 459.449725 | 0.450626185 | 0.31890758 | 1.41303064 | 0.15764671 | 0.73580079 |
| BANK1 | 75.2346521 | -0.019844254 | 0.48815559 | -0.0406515 | 0.96757373 | 0.99648221 |
| SLC39A8 | 132.466285 | 1.624921128 | 0.33102361 | 4.90877712 | 9.16E-07 | 0.00091119 |
| NFKB1 | 278.047989 | 0.069031111 | 0.33898463 | 0.20364083 | 0.83863418 | 0.98462033 |
| MANBA | 112.45627 | 0.268413964 | 0.35884103 | 0.74800244 | 0.45445869 | 0.91077355 |
| UBE2D3 | 538.805465 | 0.099240165 | 0.11571293 | 0.85764113 | 0.39109066 | 0.88670817 |
| CISD2 | 58.2811725 | 0.271097482 | 0.27303951 | 0.99288739 | 0.32076484 | 0.8511375 |
| SLC9B1 | 31.8972835 | 0.298440439 | 0.76833809 | 0.38842333 | 0.69770278 | 0.96231749 |

| | | | | | | |
|---|---|---|---|---|---|---|
| SLC9B2 | 28.0166265 | 0.261226737 | 0.43369888 | 0.60232283 | 0.54695926 | 0.93523904 |
| TET2 | 284.886154 | 0.116597235 | 0.27490953 | 0.42412948 | 0.67147139 | 0.95800185 |
| PPA2 | 328.402718 | 0.21744067 | 0.17810707 | 1.22084246 | 0.22214567 | 0.78710314 |
| ARHGEF38 | 211.052167 | -0.23148336 | 0.59772655 | -0.387273 | 0.6985541 | 0.96256506 |
| INTS12 | 99.7923471 | -0.860697745 | 0.31314375 | -2.7485708 | 0.00598557 | 0.24502474 |
| GSTCD | 125.565647 | -0.577461752 | 0.28084167 | -2.0561826 | 0.03976492 | 0.48094465 |
| NPNT | 318.370214 | -0.405014791 | 0.51658365 | -0.7840256 | 0.43302511 | 0.90514499 |
| TBCK | 559.78509 | 0.373318584 | 0.32333725 | 1.15457959 | 0.24826263 | 0.80672756 |
| AIMP1 | 88.1915664 | -0.151991465 | 0.26213415 | -0.5798232 | 0.56203384 | 0.93762898 |
| PAPSS1 | 112.848404 | 0.360829282 | 0.35341292 | 1.02098497 | 0.30726156 | 0.84488735 |
| SGMS2 | 129.186992 | -0.444901534 | 0.46243619 | -0.962082 | 0.33600841 | 0.86391445 |
| HADH | 44.6307463 | -0.060875293 | 0.37603232 | -0.1618885 | 0.87139369 | 0.98766275 |
| RPL34-AS1 | 29.8602032 | 0.659869578 | 0.67082295 | 0.98367173 | 0.32527694 | 0.855037 |
| RPL34 | 304.495871 | 0.538526037 | 0.48180319 | 1.11773032 | 0.26368219 | 0.81378765 |
| OSTC | 77.2039 | 0.160046107 | 0.40944398 | 0.39088646 | 0.69588117 | 0.96220115 |
| SEC24B | 317.455404 | -0.112022843 | 0.18859503 | -0.5939862 | 0.55252135 | 0.93523904 |
| MCUB | 47.2650502 | -0.922920427 | 0.63428856 | -1.4550482 | 0.14565592 | 0.71974648 |
| CASP6 | 43.5052769 | 0.653851585 | 0.28808198 | 2.26967194 | 0.02322749 | 0.39839181 |
| PLA2G12A | 53.8744059 | 0.518333567 | 0.27372594 | 1.89362238 | 0.05827515 | 0.55214408 |
| GAR1 | 30.5961497 | -0.701547456 | 0.31599087 | -2.2201511 | 0.02640852 | 0.42051885 |
| EGF | 49.7878213 | 1.747760803 | 0.57238423 | 3.05347478 | 0.00226208 | 0.15588805 |
| ELOVL6 | 119.806823 | -0.165069236 | 0.36060762 | -0.457753 | 0.64712988 | 0.95027667 |
| ENPEP | 139.073162 | 0.845817369 | 0.6421192 | 1.31722797 | 0.18776222 | 0.76207437 |
| FAM241A | 33.2555489 | -0.844423167 | 0.47776698 | -1.7674373 | 0.07715503 | 0.59795146 |
| AP1AR | 78.6826212 | -0.464346733 | 0.30704448 | -1.5123109 | 0.13045478 | 0.69957738 |
| ALPK1 | 298.075054 | 0.094695229 | 0.33932094 | 0.27907276 | 7.80E-01 | 0.97423208 |
| LARP7 | 104.481515 | -0.338010325 | 0.22543962 | -1.4993386 | 0.13378581 | 0.70530229 |
| ANK2 | 273.932193 | -1.415636507 | 0.42892723 | -3.3004119 | 0.00096543 | 0.10275397 |
| CAMK2D | 554.128702 | -0.075050313 | 0.16135856 | -0.4651152 | 0.64184899 | 0.94944486 |
| ARSJ | 149.906618 | 0.091234645 | 0.30140907 | 0.30269376 | 0.76212325 | 0.9714717 |
| UGT8 | 160.652958 | -0.467101556 | 0.68688644 | -0.6800273 | 0.49648715 | 0.92353293 |
| SNHG8 | 67.7261259 | -0.288664768 | 0.4870055 | -0.5927341 | 0.55335911 | 0.93523904 |
| METTL14 | 63.641508 | -0.181320502 | 0.25425864 | -0.7131341 | 0.47576279 | 0.91809716 |
| SEC24D | 132.970733 | 0.467620567 | 0.37821554 | 1.23638646 | 0.21631495 | 0.78293142 |
| USP53 | 345.258519 | -0.16351815 | 0.22567208 | -0.724583 | 0.46870791 | 0.91462428 |
| C4orf3 | 141.701679 | 0.275679203 | 0.4341574 | 0.63497525 | 0.52544456 | 0.93272384 |
| PDE5A | 150.219709 | 0.315807919 | 0.34156596 | 0.92458838 | 0.35518006 | 0.87261977 |
| AC108866.1 | 77.5837153 | -0.866666846 | 0.30494634 | -2.8420306 | 0.00448272 | 0.2151382 |
| PRDM5 | 101.23197 | -0.795782141 | 0.32386364 | -2.4571518 | 0.01400435 | 0.32375679 |
| ANXA5 | 249.25635 | 0.0535522 | 0.42767694 | 0.12521648 | 0.90035217 | 0.98983972 |
| BBS7 | 48.9012672 | 0.210824277 | 0.26422878 | 0.79788537 | 0.42493702 | 0.90167091 |
| KIAA1109 | 467.959661 | 0.220100834 | 0.16125472 | 1.36492646 | 0.17227616 | 0.74603679 |
| FGF2 | 90.5204744 | 0.16606137 | 0.45089225 | 0.36829502 | 0.71265327 | 0.96587538 |
| NUDT6 | 28.9949298 | 0.276281583 | 0.4117287 | 0.67102824 | 0.50220254 | 0.92484445 |
| SPATA5 | 307.514139 | -0.273301945 | 0.18288549 | -1.4943884 | 0.13507414 | 0.70620568 |
| SPRY1 | 37.9909002 | 0.030438696 | 0.62195858 | 0.04894007 | 0.96096706 | 0.99572129 |
| LINC01091 | 51.3050408 | 1.692666 | 0.36050688 | 4.69523907 | 2.66E-06 | 0.00200368 |
| ANKRD50 | 53.2429289 | 0.057119003 | 0.50220497 | 0.11373643 | 0.90944673 | 0.99100813 |
| FAT4 | 160.289753 | 0.219395796 | 0.42941478 | 0.51091813 | 0.60940839 | 0.94791646 |
| INTU | 198.238426 | 0.090984518 | 0.30634361 | 0.29700152 | 0.76646535 | 0.97221495 |
| HSPA4L | 73.8064795 | 1.022825455 | 0.61270335 | 1.66936489 | 0.09504509 | 0.63980105 |
| MFSD8 | 91.1788774 | -0.105281757 | 0.23218461 | -0.4534399 | 0.65023204 | 0.95061176 |
| ABHD18 | 232.249871 | -0.278948629 | 0.15426337 | -1.8082622 | 0.0705657 | 0.58797037 |
| LARP1B | 325.185501 | 0.459375644 | 0.20956841 | 2.19200809 | 0.02837892 | 0.42608417 |

| | | | | | | |
|---|---|---|---|---|---|---|
| PGRMC2 | 126.463807 | 0.282481833 | 0.22618933 | 1.24887338 | 0.21171139 | 0.77730294 |
| AC108062.1 | 183.636253 | 0.321826856 | 0.44396056 | 0.72489966 | 0.46851359 | 0.91462428 |
| JADE1 | 184.997474 | 0.04577542 | 0.35742957 | 0.12806836 | 0.89809487 | 0.98894709 |
| SCLT1 | 171.933014 | -0.588280937 | 0.31484327 | -1.8684882 | 0.06169405 | 0.56326893 |
| AC096711.2 | 92.1894046 | -0.134683943 | 0.38030261 | -0.3541494 | 0.72322692 | 0.96779196 |
| AC079380.1 | 45.0458611 | 0.665401039 | 0.44787429 | 1.48568705 | 0.13736192 | 0.71046965 |
| ELF2 | 505.011728 | -0.194412704 | 0.27150659 | -0.7160515 | 0.47395954 | 0.91809716 |
| NDUFC1 | 93.0862294 | -0.121092391 | 0.33759178 | -0.3586947 | 0.71982348 | 0.96779196 |
| NAA15 | 237.17581 | -0.347303806 | 0.19422895 | -1.7881156 | 0.07375737 | 0.59320069 |
| AC097376.2 | 253.231755 | -0.071586258 | 0.26515333 | -0.2699806 | 0.78717517 | 0.97423208 |
| SETD7 | 56.5169262 | 0.234448395 | 0.37202995 | 0.63018688 | 0.52857232 | 0.93366189 |
| MGST2 | 69.8173355 | 0.342601405 | 0.23743943 | 1.44290023 | 0.14904858 | 0.72246729 |
| MAML3 | 1101.30619 | 0.011092042 | 0.53296628 | 0.0208119 | 0.9833957 | 0.99731258 |
| SCOC | 148.191662 | -0.249467826 | 0.3360159 | -0.7424286 | 0.45782768 | 0.91270163 |
| ELMOD2 | 63.3511488 | 0.038806232 | 0.21427622 | 0.18110377 | 0.85628613 | 0.9864199 |
| TBC1D9 | 87.7970972 | 0.156866658 | 0.30298435 | 0.51773849 | 0.60464074 | 0.94683415 |
| RNF150 | 156.610374 | 1.003645435 | 0.76794731 | 1.30691965 | 0.19124 | 0.76328615 |
| ZNF330 | 49.1097837 | -0.046865477 | 0.29237335 | -0.1602933 | 0.87265007 | 0.98766275 |
| IL15 | 161.200101 | -0.082080461 | 0.5971215 | -0.1374602 | 0.89066702 | 0.98766275 |
| INPP4B | 70.6289475 | 0.50646276 | 0.52327654 | 0.96786827 | 0.33311017 | 0.86067542 |
| AC104596.1 | 74.5889124 | -0.086199682 | 0.53866262 | -0.1600254 | 0.87286109 | 0.98766275 |
| USP38 | 50.9069962 | -0.071272284 | 0.26502169 | -0.26893 | 0.78798354 | 0.97423208 |
| GAB1 | 534.860997 | -0.632996005 | 0.3542587 | -1.7868185 | 0.07396683 | 0.59320069 |
| SMARCA5 | 148.737305 | 0.20233324 | 0.23618188 | 0.85668402 | 0.39161954 | 0.88704743 |
| ANAPC10 | 148.590769 | -0.197797046 | 0.23168641 | -0.8537274 | 0.39325603 | 0.88760715 |
| ABCE1 | 83.1358549 | -0.39502502 | 0.26921967 | -1.4672963 | 0.14229547 | 0.7148274 |
| OTUD4 | 155.582568 | 0.030918553 | 0.18254511 | 0.16937486 | 0.8655018 | 0.98670067 |
| SMAD1 | 102.9221 | -0.277176956 | 0.29071178 | -0.9534425 | 0.34036594 | 0.86777921 |
| MMAA | 43.0477557 | 0.05864611 | 0.34670646 | 0.16915205 | 0.86567704 | 0.98670067 |
| ZNF827 | 276.132089 | 0.81106879 | 0.42154807 | 1.92402444 | 0.05435152 | 0.53752502 |
| LSM6 | 42.8330365 | -0.63362261 | 0.46291111 | -1.3687782 | 0.17106863 | 0.74445571 |
| SLC10A7 | 125.040379 | 0.020003367 | 0.24434658 | 0.08186473 | 0.93475428 | 0.99444816 |
| TMEM184C | 76.8198966 | 0.155095373 | 0.42558963 | 0.3644247 | 0.71554089 | 0.96687349 |
| PRMT9 | 81.8823928 | 0.020319216 | 0.38708691 | 0.05249265 | 0.95813616 | 0.99572129 |
| ARHGAP10 | 588.935067 | -0.026548179 | 0.37542452 | -0.0707151 | 0.94362451 | 0.9952642 |
| NR3C2 | 630.355017 | 0.374432318 | 0.48009084 | 0.77991974 | 0.43543812 | 0.90514499 |
| DCLK2 | 85.3069582 | 0.666839108 | 0.83138058 | 0.80208646 | 0.42250294 | 0.90066187 |
| LRBA | 1505.10126 | -0.159825982 | 0.22292869 | -0.7169377 | 0.47341254 | 0.91804805 |
| RPS3A | 217.249287 | 0.199450799 | 0.55929229 | 0.35661282 | 0.72138168 | 0.96779196 |
| SH3D19 | 384.032461 | -0.260661851 | 0.33010426 | -0.7896349 | 0.429741 | 0.90333313 |
| FAM160A1 | 461.242597 | -0.258276157 | 0.31779299 | -0.8127182 | 0.41637962 | 0.89862171 |
| GATB | 32.4808224 | -0.448432991 | 0.37239745 | -1.2041785 | 0.22852061 | 0.79311555 |
| FBXW7 | 500.433518 | 0.045368146 | 0.29075994 | 0.156033 | 0.87600701 | 0.98766275 |
| ARFIP1 | 234.221428 | 0.044648168 | 0.17062974 | 0.26166697 | 0.79357821 | 0.97464093 |
| TRIM2 | 305.911938 | 0.029843768 | 0.43978702 | 0.06785959 | 0.9458974 | 0.9952642 |
| TMEM131L | 92.6045489 | 0.013009316 | 0.54697001 | 0.02378433 | 0.98102464 | 0.99681784 |
| PLRG1 | 49.3501374 | 0.010736661 | 0.29904805 | 0.0359028 | 0.97135987 | 0.99681784 |
| FGB | 300.253085 | 3.41973087 | 1.37996979 | 2.47812009 | 1.32E-02 | 0.31992293 |
| FGA | 115.508155 | 3.453693912 | 1.41298575 | 2.44425247 | 1.45E-02 | 0.32792859 |
| LRAT | 47.9547319 | 0.474134372 | 0.69457203 | 0.68262809 | 0.49484188 | 0.92307934 |
| MAP9 | 75.365823 | 0.650524647 | 0.29459873 | 2.20817196 | 0.02723229 | 0.42391395 |
| GUCY1A1 | 222.528125 | 0.276541899 | 0.35737093 | 0.77382314 | 0.43903538 | 0.90561284 |
| GUCY1B1 | 125.653117 | 0.478726465 | 0.83001846 | 0.57676605 | 0.56409751 | 0.93809297 |
| TDO2 | 27.6831835 | -1.420460254 | 1.31332883 | -1.0815724 | 0.27944256 | 0.82413186 |

| | | | | | | |
|---|---|---|---|---|---|---|
| CTSO | 82.0394837 | -1.14051591 | 0.31397882 | -3.6324612 | 0.00028073 | 0.04481867 |
| PDGFC | 385.075405 | 0.213843954 | 0.41859731 | 0.51085841 | 0.60945021 | 0.94791646 |
| TMEM144 | 31.0754398 | -0.847038969 | 0.38810821 | -2.1824815 | 0.02907402 | 0.43178298 |
| ETFDH | 103.496533 | 0.524822743 | 0.3057732 | 1.71637916 | 0.08609267 | 0.62276717 |
| PPID | 45.72793 | -0.10761942 | 0.32040986 | -0.3358805 | 0.73696099 | 0.9698508 |
| FNIP2 | 705.557616 | -0.52689061 | 0.43578378 | -1.2090643 | 0.22663814 | 0.79116864 |
| C4orf45 | 28.1799007 | 0.137348096 | 0.50588638 | 0.27149989 | 0.78600659 | 0.97423208 |
| RAPGEF2 | 648.397847 | 0.351718282 | 0.22854054 | 1.53897547 | 0.12381029 | 0.69436693 |
| NAF1 | 81.0689076 | -0.139294725 | 0.2327555 | -0.5984594 | 0.54953342 | 0.93523904 |
| TMA16 | 48.9865309 | -0.480292614 | 0.3038282 | -1.5808033 | 0.11392302 | 0.66826004 |
| 1-Mar | 832.584885 | -1.343423292 | 1.21148896 | -1.1089026 | 0.26747219 | 0.81510106 |
| TMEM192 | 101.928594 | -0.119947768 | 0.27888699 | -0.4300945 | 0.66712688 | 0.95691417 |
| KLHL2 | 255.210697 | -0.738641155 | 0.43870729 | -1.6836765 | 0.0922442 | 0.63464873 |
| MSMO1 | 120.298841 | 0.405022384 | 0.5794098 | 0.69902577 | 0.48453592 | 0.92034617 |
| CPE | 602.384037 | -0.222786417 | 1.08788009 | -0.2047895 | 0.83773659 | 0.98462033 |
| DDX60 | 130.934906 | -2.386723083 | 0.87224668 | -2.7362937 | 0.00621355 | 0.24799848 |
| DDX60L | 131.826071 | -2.589684932 | 0.88431348 | -2.9284694 | 0.00340635 | 0.18951432 |
| PALLD | 290.864698 | 0.760815941 | 0.76177375 | 0.99874267 | 0.31791937 | 0.8490543 |
| CBR4 | 294.55259 | -0.192391499 | 0.23761137 | -0.8096898 | 0.41811848 | 0.89862171 |
| SH3RF1 | 433.457396 | -0.448016911 | 0.24452205 | -1.8322148 | 0.06691943 | 0.57853902 |
| NEK1 | 240.87207 | -0.022374792 | 0.22838822 | -0.0979682 | 0.92195751 | 0.99214128 |
| CLCN3 | 233.662857 | 0.087887827 | 0.19628824 | 0.44774881 | 0.65433449 | 0.95280477 |
| HPF1 | 36.2813267 | 0.094288107 | 0.32651404 | 0.28877198 | 0.77275587 | 0.97346509 |
| MFAP3L | 103.949945 | 0.224851679 | 0.60238891 | 0.37326663 | 0.70895001 | 0.96587538 |
| GALNT7 | 158.971736 | 0.336969424 | 0.22786396 | 1.47881843 | 0.13918885 | 0.71184729 |
| HMGB2 | 56.3851959 | -0.560204503 | 0.59497992 | -0.941552 | 0.34642208 | 0.86990996 |
| FBXO8 | 40.709199 | 0.267086908 | 0.31489012 | 0.84819083 | 0.39633171 | 0.88826893 |
| CEP44 | 71.4966467 | 0.02924568 | 0.25194343 | 0.11608034 | 0.90758887 | 0.99100813 |
| GPM6A | 134.239703 | -0.391491216 | 0.56748081 | -0.6898757 | 0.49027236 | 0.92157694 |
| WDR17 | 40.3874941 | 0.271129362 | 0.40621991 | 0.66744478 | 0.50448807 | 0.92552859 |
| SPCS3 | 93.4812448 | -0.000418472 | 0.32692543 | -0.00128 | 0.99897869 | 0.99971834 |
| AC078881.1 | 75.0374536 | -0.261186558 | 0.46042338 | -0.5672747 | 0.57052753 | 0.93998433 |
| TENM3 | 677.061415 | 1.027875642 | 0.72832487 | 1.41128731 | 0.15815991 | 0.73580079 |
| DCTD | 71.6903022 | -0.365711302 | 0.20356225 | -1.7965576 | 0.07240589 | 0.59090333 |
| WWC2 | 731.896987 | -0.237172059 | 0.17643765 | -1.3442259 | 1.79E-01 | 7.52E-01 |
| CDKN2AIP | 28.9642482 | 0.756152861 | 0.74481689 | 1.01521981 | 0.31000106 | 0.84601144 |
| RWDD4 | 40.3505307 | -0.46330271 | 0.30194638 | -1.5343874 | 0.12493439 | 0.69439514 |
| TRAPPC11 | 120.997168 | -0.259573993 | 0.25954446 | -1.0001138 | 0.31725545 | 0.8490543 |
| STOX2 | 1617.24577 | 0.502712395 | 0.26278612 | 1.91300968 | 0.05574681 | 0.54048454 |
| IRF2 | 135.498159 | 0.07111129 | 0.30284291 | 0.23481246 | 0.81435429 | 0.97937868 |
| CASP3 | 38.0376419 | 0.175495679 | 0.29706399 | 0.59076726 | 0.55467638 | 0.93523904 |
| PRIMPOL | 59.037448 | 0.799594597 | 0.3956196 | 2.02111979 | 0.04326737 | 0.49484046 |
| ACSL1 | 164.081043 | 0.617675114 | 0.34736856 | 1.77815493 | 0.07537841 | 0.59465671 |
| SLC25A4 | 49.0217984 | 0.694854255 | 0.37348116 | 1.86048009 | 0.06281763 | 0.56713487 |
| CFAP97 | 118.222619 | 0.016753096 | 0.29489803 | 0.05680979 | 0.95469671 | 0.99531843 |
| SNX25 | 267.742936 | -0.161196043 | 0.27521041 | -0.5857193 | 0.55806418 | 0.93620477 |
| LRP2BP | 37.2539985 | 0.979613576 | 0.52657186 | 1.86036067 | 0.06283452 | 0.56713487 |
| AC112722.1 | 27.8678586 | 0.036945618 | 0.42069972 | 0.08781945 | 0.93002018 | 0.99362798 |
| ANKRD37 | 52.5435162 | 0.683111574 | 1.24305644 | 0.54954188 | 0.58263363 | 0.94115473 |
| UFSP2 | 61.9478696 | 0.363632546 | 0.35463993 | 1.02535702 | 0.30519476 | 0.84488735 |
| C4orf47 | 111.306592 | 0.780273232 | 0.71305649 | 1.09426566 | 0.27383848 | 0.82047446 |
| SORBS2 | 289.175438 | 1.11775451 | 0.47298491 | 2.36319273 | 0.01811824 | 0.35858401 |
| FAM149A | 153.25561 | 0.174967542 | 0.41246914 | 0.42419547 | 0.67142326 | 0.95800185 |
| CYP4V2 | 56.5316584 | -0.120826892 | 0.36487833 | -0.331143 | 0.7405365 | 0.9698508 |

| | | | | | | |
|---|---|---|---|---|---|---|
| KLKB1 | 40.387073 | 0.444233817 | 0.31129105 | 1.42706905 | 0.15355999 | 0.7288378 |
| FAT1 | 662.979162 | 0.212634753 | 0.27563217 | 0.77144389 | 4.40E-01 | 0.90561284 |
| FRG1-DT | 67.1605817 | -1.023948232 | 0.51690292 | -1.9809295 | 0.04759918 | 0.51346008 |
| FRG1 | 30.111144 | -0.138679748 | 0.39765745 | -0.3487417 | 0.72728321 | 0.9681562 |
| SDHA | 124.670909 | 0.460101327 | 0.20847235 | 2.20701365 | 0.02731311 | 0.42391395 |
| PDCD6 | 79.8999512 | -0.335353356 | 0.30926825 | -1.0843446 | 0.27821203 | 0.8229426 |
| EXOC3 | 97.8168964 | -0.419031449 | 0.27411346 | -1.5286788 | 0.12634408 | 0.69492095 |
| SLC9A3 | 120.000418 | 0.207679734 | 0.84859832 | 0.24473267 | 0.80666343 | 0.97930627 |
| BRD9 | 45.8373034 | -0.561840991 | 0.25692042 | -2.1868289 | 0.02875502 | 0.43038175 |
| SLC12A7 | 91.3546394 | 0.425866498 | 0.40976835 | 1.03928597 | 0.29867176 | 0.83973489 |
| CLPTM1L | 73.1541737 | 0.349435603 | 0.33285221 | 1.04982209 | 0.29379992 | 0.83825235 |
| LPCAT1 | 59.3962726 | -0.779703451 | 0.44848323 | -1.7385342 | 0.08211672 | 0.61261376 |
| MRPL36 | 30.3265665 | -0.21877706 | 0.41643922 | -0.5253517 | 0.59933871 | 0.94683415 |
| NDUFS6 | 88.4947104 | -0.292579163 | 0.47344977 | -0.617973 | 0.53659316 | 0.93469807 |
| IRX2 | 25.6007887 | -2.18304384 | 0.98632381 | -2.2133135 | 0.02687604 | 0.4234302 |
| IRX1 | 32.8712976 | -1.907813721 | 1.25361191 | -1.5218535 | 0.12804578 | 0.69635098 |
| ADAMTS16 | 124.029491 | 2.379361978 | 0.90071304 | 2.64164264 | 0.00825051 | 0.27246484 |
| ICE1 | 163.131451 | -0.396898563 | 0.17101431 | -2.32085 | 0.02029494 | 0.37800094 |
| MED10 | 51.4687296 | -0.383091635 | 0.26828866 | -1.4279084 | 0.15331822 | 0.7288378 |
| NSUN2 | 78.9760723 | -0.277984976 | 0.32079834 | -0.8665412 | 0.38619345 | 0.88353166 |
| TENT4A | 122.289926 | -0.522089762 | 0.23801253 | -2.193539 | 0.02826857 | 0.4250936 |
| MTRR | 84.5185251 | 0.219236015 | 0.29721765 | 0.73762785 | 0.46074063 | 0.91368087 |
| SEMA5A | 240.057621 | 1.339324557 | 0.69855985 | 1.9172653 | 0.05520424 | 0.53849427 |
| SNHG18 | 30.0542198 | 1.302052945 | 0.48373409 | 2.69167085 | 0.00710951 | 0.26193066 |
| FAM173B | 46.018448 | 0.141380659 | 0.35673769 | 0.39631545 | 0.69187233 | 0.96220115 |
| CCT5 | 101.656861 | -0.29077325 | 0.42202012 | -0.6890033 | 0.4908212 | 0.92187769 |
| CMBL | 53.6344569 | 1.43072521 | 0.66394443 | 2.15488698 | 0.03117068 | 0.43905874 |
| 6-Mar | 399.443292 | 0.031064487 | 0.18797894 | 0.16525514 | 0.86874316 | 0.98676155 |
| ANKRD33B | 231.35714 | -0.185318089 | 0.4672773 | -0.3965912 | 0.69166891 | 0.96220115 |
| DAP | 186.914768 | 0.159202068 | 0.30289795 | 0.52559638 | 0.59916867 | 0.94683415 |
| DNAH5 | 220.328425 | 0.750240333 | 0.54824204 | 1.3684473 | 0.17117211 | 0.74445571 |
| TRIO | 1444.86124 | -0.207347612 | 0.19283251 | -1.0752731 | 0.28225248 | 0.82808469 |
| OTULIN | 65.4539475 | 0.405166616 | 0.23156454 | 1.749692 | 0.08017147 | 0.60643125 |
| ANKH | 117.26069 | -0.213713455 | 0.58169371 | -0.3673986 | 0.71332172 | 0.96587538 |
| LINC02149 | 37.4346317 | 0.52442433 | 1.38311358 | 0.37916216 | 0.70456745 | 0.96461079 |
| FBXL7 | 600.947148 | 0.523874143 | 0.61877876 | 0.84662593 | 0.39720366 | 0.88863925 |
| RETREG1 | 525.485538 | 0.631949318 | 0.46317488 | 1.36438599 | 0.17244611 | 0.74625056 |
| MYO10 | 524.656668 | -0.153484798 | 0.37561514 | -0.4086225 | 0.68281671 | 0.96101988 |
| LINC02241 | 81.278346 | 0.385626441 | 0.31159703 | 1.23758059 | 0.21587161 | 0.78293142 |
| C5orf17 | 66.6477688 | -1.317898569 | 0.54426567 | -2.4214251 | 0.01545979 | 0.33965433 |
| CDH9 | 89.5984685 | 1.709897363 | 1.24565313 | 1.37269142 | 0.1698483 | 0.74445571 |
| AC113386.1 | 76.0586411 | 1.012988474 | 0.71098013 | 1.42477747 | 0.15422154 | 0.72976516 |
| CDH6 | 934.624212 | 0.700596898 | 0.8078472 | 0.86723938 | 0.38581087 | 0.88353166 |
| DROSHA | 120.378836 | -0.333257222 | 0.20403982 | -1.633295 | 0.10240696 | 0.65288882 |
| C5orf22 | 32.1718647 | -0.238968608 | 0.36765435 | -0.6499817 | 0.51570403 | 0.93140514 |
| PDZD2 | 313.463331 | -0.134851327 | 0.4939418 | -0.2730106 | 0.7848451 | 0.97423208 |
| GOLPH3 | 313.877376 | -0.046865032 | 0.23801729 | -0.1968976 | 0.84390767 | 0.98517901 |
| MTMR12 | 165.027209 | 0.531171844 | 0.25574032 | 2.07699688 | 0.03780185 | 0.47081202 |
| ZFR | 430.204743 | -0.031129042 | 0.17382977 | -0.1790777 | 0.85787666 | 0.9864199 |
| SUB1 | 204.44477 | -0.19147217 | 0.37741042 | -0.5073314 | 0.61192228 | 0.94791646 |
| NPR3 | 240.477611 | 1.182421662 | 0.56776161 | 2.08260233 | 0.03728749 | 0.46890807 |
| TARS | 66.4490665 | -0.142359229 | 0.30798639 | -0.4622257 | 0.64391946 | 0.94944486 |
| C1QTNF3-AN | 209.474609 | -0.227966086 | 0.2098541 | -1.0863075 | 0.27734297 | 0.8229426 |
| RAI14 | 468.495205 | -0.314213366 | 0.27474676 | -1.1436472 | 0.25276997 | 0.80798674 |

| | | | | | | |
|---|---|---|---|---|---|---|
| TTC23L | 31.0681668 | 0.672292968 | 0.64197031 | 1.04723374 | 0.29499179 | 0.83911606 |
| RAD1 | 28.7172351 | -0.831955442 | 0.35049265 | -2.3736744 | 0.01761207 | 0.35667241 |
| BRIX1 | 42.5204 | -0.309259349 | 0.30013014 | -1.0304175 | 0.30281406 | 0.84398487 |
| DNAJC21 | 85.0659299 | -0.175194132 | 0.22613789 | -0.7747226 | 0.4385036 | 0.90561284 |
| PRLR | 63.1732376 | 1.379057227 | 0.4400908 | 3.13357433 | 0.00172691 | 0.13559088 |
| SPEF2 | 156.465932 | 0.570391067 | 0.47047784 | 1.21236542 | 0.22537253 | 0.78977225 |
| LMBRD2 | 60.5983942 | 0.074417945 | 0.2208224 | 0.3370036 | 0.73611418 | 0.9698508 |
| NADK2 | 94.6147548 | -0.395243919 | 0.44324112 | -0.8917131 | 0.37254672 | 0.87605028 |
| RANBP3L | 112.530602 | -0.436186994 | 1.59365867 | -0.2737016 | 7.84E-01 | 0.97423208 |
| NIPBL | 861.78271 | -0.070418758 | 0.14462766 | -0.4868969 | 0.62633141 | 0.94831213 |
| CPLANE1 | 369.858887 | -0.374255618 | 0.26937497 | -1.3893482 | 0.16472691 | 0.74430144 |
| NUP155 | 65.4500709 | -0.632924533 | 0.27609009 | -2.2924566 | 0.02187931 | 0.38823827 |
| WDR70 | 419.082559 | -0.078937445 | 0.15084848 | -0.5232896 | 0.60077272 | 0.94683415 |
| LIFR | 514.686228 | -0.20680484 | 0.39721709 | -0.5206343 | 0.60262155 | 0.94683415 |
| LIFR-AS1 | 40.6295976 | -0.729508514 | 0.73405348 | -0.9938084 | 0.32031616 | 0.85075884 |
| OSMR-AS1 | 257.45247 | 0.19221848 | 0.54361711 | 0.35359166 | 0.72364492 | 0.96779196 |
| OSMR | 305.44699 | -0.137688445 | 0.48817705 | -0.2820461 | 0.77790814 | 0.97423208 |
| RICTOR | 443.279435 | -0.037352894 | 0.25908674 | -0.1441714 | 0.88536514 | 0.98766275 |
| DAB2 | 46.0076101 | 0.23066965 | 0.53140271 | 0.43407692 | 6.64E-01 | 9.55E-01 |
| TTC33 | 84.7776716 | 0.363288923 | 0.24245107 | 1.49840096 | 0.13402911 | 0.70542022 |
| PRKAA1 | 136.017106 | 0.110709576 | 0.26403618 | 0.419297 | 0.67499909 | 0.95910677 |
| RPL37 | 241.578301 | 0.105453508 | 0.31333292 | 0.3365542 | 0.73645299 | 0.9698508 |
| OXCT1 | 98.6550788 | -0.504380082 | 0.22960672 | -2.1967131 | 0.02804094 | 0.4250936 |
| C5orf51 | 29.4462532 | 0.152910203 | 0.32197521 | 0.47491297 | 0.634849 | 0.94932781 |
| GHR | 91.4387523 | 1.623467959 | 0.91570878 | 1.7729086 | 0.07624385 | 0.59726395 |
| SELENOP | 96.660551 | 0.831338684 | 0.36322013 | 2.28880125 | 0.0220909 | 0.39000316 |
| ZNF131 | 230.563578 | -0.062942335 | 0.17372229 | -0.3623158 | 0.71711604 | 0.96742001 |
| HMGCS1 | 255.781694 | 1.370662539 | 0.63899076 | 2.14504281 | 0.03194942 | 0.44049392 |
| CCL28 | 101.829091 | 0.379906504 | 0.44713194 | 0.84965189 | 0.39551865 | 0.88760715 |
| TMEM267 | 154.078147 | 0.289624733 | 0.22715925 | 1.27498542 | 0.2023144 | 0.77110248 |
| PAIP1 | 93.3662284 | 0.047672845 | 0.27438073 | 0.17374706 | 0.86206425 | 0.98669442 |
| NNT-AS1 | 82.9414164 | 0.14201947 | 0.25227733 | 0.56294978 | 0.57346908 | 0.93998433 |
| NNT | 165.802708 | 0.187089145 | 0.26576933 | 0.7039531 | 0.48146198 | 0.91971919 |
| MRPS30-DT | 38.5939905 | 0.598161032 | 0.60784993 | 0.98406037 | 0.32508583 | 0.855037 |
| MRPS30 | 37.9485015 | 0.307536519 | 0.3503394 | 0.87782453 | 0.38003895 | 0.88059823 |
| PARP8 | 299.219714 | -0.042546219 | 0.40402807 | -0.1053051 | 0.91613371 | 0.99159829 |
| ITGA1 | 374.487528 | 0.54594303 | 0.4861934 | 1.12289273 | 0.26148306 | 0.81281978 |
| ITGA2 | 453.566004 | -2.747560439 | 0.52454633 | -5.2379748 | 1.62E-07 | 0.0002976 |
| MOCS2 | 33.1431582 | 0.202755525 | 0.43726333 | 0.46369204 | 0.6428684 | 0.94944486 |
| NDUFS4 | 185.415302 | 0.050314637 | 0.18408262 | 0.27332639 | 0.78460233 | 0.97423208 |
| ARL15 | 1057.37465 | 0.265685369 | 0.21514503 | 1.23491288 | 2.17E-01 | 7.83E-01 |
| SNX18 | 34.780515 | 0.236354606 | 0.34988356 | 0.67552361 | 0.49934315 | 0.92398825 |
| AC026704.1 | 29.2629505 | 0.540257068 | 0.66648576 | 0.81060557 | 0.41759222 | 0.89862171 |
| DHX29 | 118.552193 | -0.013994998 | 0.18252647 | -0.0766738 | 0.93888305 | 0.99526299 |
| MTREX | 241.533275 | 0.054190295 | 0.16928489 | 0.32011302 | 0.74888266 | 0.97028355 |
| PLPP1 | 211.679224 | 0.443870862 | 0.211229 | 2.10137273 | 0.03560826 | 0.45736901 |
| SLC38A9 | 149.750548 | 0.11231147 | 0.26268156 | 0.42755749 | 0.66897332 | 0.95764829 |
| IL6ST | 478.596591 | 0.041218106 | 0.32285479 | 0.12766763 | 0.89841201 | 0.9891826 |
| LINC01948 | 86.6360893 | 0.230529645 | 0.55086418 | 0.41848727 | 0.67559089 | 0.9591071 |
| MAP3K1 | 868.945136 | -0.059393146 | 0.33476313 | -0.1774184 | 0.85917974 | 0.9864199 |
| AC008937.2 | 38.3564326 | -0.240880791 | 0.58195974 | -0.4139132 | 0.67893771 | 0.95924874 |
| MIER3 | 46.8981829 | 0.204858342 | 0.24922723 | 0.82197414 | 0.41109161 | 0.89649693 |
| GPBP1 | 454.00761 | -0.030149058 | 0.22347112 | -0.1349125 | 0.89268102 | 0.9882865 |
| PDE4D | 3248.84549 | 0.551521802 | 0.3073585 | 1.79439256 | 0.07275053 | 0.59192805 |

| | | | | | | |
|---|---|---|---|---|---|---|
| ELOVL7 | 208.03613 | -0.581476227 | 0.39256851 | -1.4812096 | 0.13855074 | 0.71091391 |
| ERCC8 | 92.5228749 | -0.337926477 | 0.19558733 | -1.7277524 | 0.08403263 | 0.61656297 |
| NDUFAF2 | 206.115784 | -0.324446557 | 0.35393437 | -0.9166857 | 0.35930738 | 0.87316144 |
| SMIM15 | 37.4709793 | -0.259668417 | 0.4180136 | -0.6211961 | 0.5344706 | 0.93457315 |
| ZSWIM6 | 775.10996 | -0.014067291 | 0.2684802 | -0.052396 | 0.95821315 | 0.99572129 |
| KIF2A | 177.527741 | 0.063082536 | 0.23037033 | 0.27383099 | 0.78421451 | 0.97423208 |
| DIMT1 | 31.703595 | -0.514504763 | 0.38227217 | -1.3459122 | 0.17833085 | 0.7513166 |
| IPO11 | 90.0226474 | -0.216764513 | 0.20138504 | -1.0763685 | 0.28176249 | 0.82749734 |
| RNF180 | 378.123359 | -0.034463381 | 0.24027638 | -0.1434322 | 0.88594882 | 0.98766275 |
| SREK1IP1 | 86.7548882 | -0.200032403 | 0.19591423 | -1.0210203 | 0.30724482 | 0.84488735 |
| CWC27 | 206.980464 | -0.216658455 | 0.21712103 | -0.9978695 | 0.31834264 | 0.84919767 |
| PPWD1 | 85.2506264 | 0.207637237 | 0.29798524 | 0.69680377 | 0.48592561 | 0.92034617 |
| TRIM23 | 68.9759095 | 0.644794066 | 0.37382504 | 1.7248552 | 0.08455357 | 0.61686112 |
| TRAPPC13 | 74.4973211 | -0.256537713 | 0.24191615 | -1.0604406 | 0.28894419 | 0.83399936 |
| NLN | 74.3606903 | -0.872185865 | 0.42794005 | -2.0381029 | 0.04153964 | 0.48843812 |
| ERBIN | 730.754019 | 0.054502247 | 0.17450797 | 0.31231953 | 0.7547977 | 0.97125609 |
| SREK1 | 239.445662 | 0.053134724 | 0.13615382 | 0.3902551 | 6.96E-01 | 0.96220115 |
| MAST4 | 429.558482 | 0.416240037 | 0.28164414 | 1.47789347 | 0.13943629 | 0.71191199 |
| PIK3R1 | 374.069863 | -0.344907019 | 0.67193914 | -0.513301 | 0.6077408 | 0.94708529 |
| SLC30A5 | 97.2404227 | -0.296668063 | 0.29052601 | -1.0211411 | 0.30718757 | 0.84488735 |
| MRPS36 | 63.1571905 | 0.110056309 | 0.37516151 | 0.29335714 | 0.76924918 | 0.97261718 |
| CDK7 | 65.1109256 | -0.030407166 | 0.23274291 | -0.130647 | 0.89605455 | 0.98842187 |
| CCDC125 | 94.2516392 | -0.178018236 | 0.41595458 | -0.4279752 | 0.66866919 | 0.95764829 |
| AK6 | 45.8670158 | -0.127779567 | 0.39243229 | -0.3256092 | 0.74472004 | 0.97028355 |
| RAD17 | 83.1437107 | -0.098208328 | 0.22201602 | -0.4423479 | 0.65823745 | 0.95399905 |
| MARVELD2 | 66.3766354 | 0.12590012 | 0.4786744 | 0.26301828 | 0.79253649 | 0.97429505 |
| OCLN | 204.868549 | 0.3577113 | 0.32682323 | 1.09451001 | 0.27373136 | 0.82047446 |
| GTF2H2C | 82.0957676 | 0.071019072 | 0.23932803 | 0.29674364 | 0.76666224 | 0.97221495 |
| NAIP | 171.220488 | 0.174439512 | 0.34378686 | 0.50740599 | 0.61186998 | 0.94791646 |
| GTF2H2 | 70.2535039 | 0.076478491 | 0.55404427 | 0.13803679 | 0.89021134 | 0.98766275 |
| BDP1 | 317.106352 | -0.378854205 | 0.197064 | -1.9224932 | 0.05454372 | 0.53752502 |
| MCCC2 | 133.740742 | -0.014295743 | 0.37246706 | -0.0383812 | 0.96938373 | 0.99681784 |
| MRPS27 | 132.013999 | -0.323048279 | 0.18877712 | -1.7112682 | 0.08703162 | 0.62570402 |
| TNPO1 | 297.69961 | -0.180386669 | 0.17516112 | -1.0298328 | 0.3030885 | 0.84446908 |
| FCHO2 | 377.99333 | 0.070293319 | 0.18753938 | 0.37481898 | 0.7077951 | 0.96587538 |
| TMEM171 | 34.8442855 | 0.388895174 | 0.35149749 | 1.10639532 | 0.26855545 | 0.8158873 |
| BTF3 | 173.872514 | 0.078910901 | 0.51027299 | 0.15464448 | 0.8771016 | 0.98766275 |
| ANKRA2 | 44.0050559 | 0.509660378 | 0.3476546 | 1.46599634 | 0.14264928 | 0.71517697 |
| ARHGEF28 | 868.746507 | 0.170869409 | 0.33242901 | 0.51400271 | 0.6072501 | 0.94708529 |
| HEXB | 71.3754302 | -0.405586242 | 0.31801948 | -1.2753503 | 0.20218527 | 0.77110248 |
| GFM2 | 59.7717179 | -0.03186835 | 0.2659152 | -0.119844 | 0.9046067 | 0.99061193 |
| NSA2 | 72.976378 | -0.023352732 | 0.32936916 | -0.0709014 | 0.94347624 | 0.9952642 |
| FAM169A | 63.8846368 | -0.710210105 | 0.54356525 | -1.3065775 | 0.19135625 | 0.76328615 |
| HMGCR | 82.2317085 | 0.106777221 | 0.41131855 | 0.25959739 | 0.79517435 | 0.97524554 |
| COL4A3BP | 286.243168 | 0.118544177 | 0.25754073 | 0.46029293 | 0.64530597 | 0.94954229 |
| POLK | 220.43502 | 0.100275274 | 0.29458754 | 0.34039211 | 0.73356126 | 0.9698508 |
| ANKDD1B | 50.8422743 | -0.422504781 | 0.44781143 | -0.9434882 | 0.34543127 | 0.86990996 |
| POC5 | 44.8110133 | -0.173459531 | 0.35410947 | -0.4898472 | 0.62424203 | 0.94808496 |
| IQGAP2 | 68.1697766 | 1.615855769 | 0.73135796 | 2.20939111 | 0.02714745 | 0.42391395 |
| F2RL1 | 42.525241 | 0.248776899 | 0.68822705 | 0.36147504 | 0.71774436 | 0.96743968 |
| ZBED3 | 44.1212749 | 0.119803471 | 0.25456615 | 0.47061822 | 0.63791339 | 0.94944486 |
| WDR41 | 137.046039 | 0.056715051 | 0.21475014 | 0.26409786 | 0.79170451 | 0.97423208 |
| TBCA | 338.269192 | -0.347842275 | 0.21336279 | -1.6302856 | 0.10304115 | 0.65288882 |
| AP3B1 | 446.968779 | 0.01286185 | 0.25752563 | 0.04994396 | 0.96016705 | 0.99572129 |

| | | | | | | |
|---|---|---|---|---|---|---|
| SCAMP1 | 166.572638 | 3.37E-01 | 0.22237654 | 1.51643663 | 0.12940898 | 0.69904834 |
| LHFPL2 | 83.2563207 | -0.711109017 | 0.3282601 | -2.1662975 | 0.03028846 | 0.43694756 |
| ARSB | 148.808208 | -0.476749824 | 0.47810534 | -0.9971648 | 0.31868451 | 0.84935263 |
| DMGDH | 148.569777 | 0.754123633 | 0.26997236 | 2.79333647 | 0.00521674 | 0.22611381 |
| BHMT2 | 205.261606 | 1.154875566 | 0.36493088 | 3.16464192 | 0.00155274 | 0.12712554 |
| BHMT | 76.4332253 | 1.786592707 | 0.67821965 | 2.63423909 | 0.00843261 | 0.2738168 |
| JMY | 156.39633 | 0.12957822 | 0.28144623 | 0.46040134 | 0.64522818 | 0.94954229 |
| HOMER1 | 494.248102 | -0.731249906 | 0.63069057 | -1.1594432 | 2.46E-01 | 0.80556213 |
| TENT2 | 478.074418 | -0.080475901 | 0.18627248 | -0.4320332 | 0.66571726 | 0.95562801 |
| CMYA5 | 40.8267027 | 1.428585605 | 0.53327633 | 2.67888431 | 0.00738679 | 0.26701159 |
| MTX3 | 39.7143566 | -0.294543139 | 0.3069896 | -0.9594564 | 0.33732887 | 0.86467039 |
| THBS4 | 36.9100983 | -0.291309013 | 0.5041466 | -0.577826 | 0.5633816 | 0.93809297 |
| SERINC5 | 265.685949 | -0.236006993 | 0.3093372 | -0.7629441 | 0.44549673 | 0.90718817 |
| ZFYVE16 | 212.070875 | -0.104223772 | 0.21646055 | -0.4814908 | 0.63016769 | 0.94916079 |
| FAM151B | 50.6209919 | 0.501865291 | 0.33586008 | 1.49426895 | 0.13510534 | 0.70620568 |
| DHFR | 382.35303 | -0.321209865 | 0.38713444 | -0.8297114 | 0.40670197 | 0.89585737 |
| MSH3 | 260.377228 | -0.12042285 | 0.22077158 | -0.5454636 | 0.58543473 | 0.9423304 |
| RASGRF2 | 89.7059179 | -0.926627591 | 0.75954969 | -1.2199697 | 0.22247637 | 0.78710314 |
| CKMT2-AS1 | 118.823258 | -0.143590504 | 0.24142537 | -0.5947615 | 0.55200294 | 0.93523904 |
| ZCCHC9 | 32.3326176 | 0.145505423 | 0.28555762 | 0.50954837 | 0.6103679 | 0.94791646 |
| SSBP2 | 556.277144 | 0.526763897 | 0.34380132 | 1.53217533 | 0.12547917 | 0.69492095 |
| ATG10 | 258.525668 | -0.240531405 | 0.29855358 | -0.8056557 | 0.42044138 | 0.90012434 |
| RPS23 | 602.0299 | 0.097010736 | 0.59092181 | 0.16416848 | 0.86959851 | 0.98748014 |
| ATP6AP1L | 103.625963 | 0.226615672 | 0.48429019 | 0.46793364 | 0.63983205 | 0.94944486 |
| TMEM167A | 66.1493917 | -0.208275208 | 0.23404182 | -0.8899059 | 0.37351639 | 0.87666478 |
| XRCC4 | 113.097258 | -0.188878637 | 0.31180132 | -0.605766 | 0.54467015 | 0.93523904 |
| VCAN | 177.060576 | 1.686060415 | 1.09160024 | 1.5445768 | 0.12244866 | 0.69314363 |
| EDIL3 | 709.504502 | -0.089381448 | 0.26593927 | -0.3360972 | 0.73679756 | 0.9698508 |
| AC113383.1 | 179.502094 | -0.047003073 | 0.33648791 | -0.1396873 | 0.8889071 | 0.98766275 |
| COX7C | 209.729567 | 0.327510457 | 0.40345001 | 0.81177457 | 0.416921 | 0.89862171 |
| RASA1 | 405.849713 | -0.019911436 | 0.15374059 | -0.1295132 | 0.89695158 | 0.98842187 |
| CCNH | 513.138938 | 0.128421225 | 0.30504677 | 0.42098864 | 0.67376338 | 0.95910677 |
| TMEM161B | 100.368251 | -0.316686207 | 0.26289915 | -1.204592 | 0.22836085 | 0.79284833 |
| TMEM161B-/ | 289.78077 | 0.292264241 | 0.26789264 | 1.09097524 | 0.27528378 | 0.82164863 |
| MEF2C | 50.1909273 | 0.307996395 | 0.92013107 | 0.334731 | 0.73782801 | 0.9698508 |
| CETN3 | 35.3862203 | 0.288246642 | 0.36236204 | 0.79546588 | 0.42634255 | 0.90237497 |
| LYSMD3 | 38.0902571 | 0.475440344 | 0.30534817 | 1.55704338 | 0.11946019 | 0.68603665 |
| ADGRV1 | 176.447252 | -0.725276855 | 0.7118472 | -1.0188659 | 0.30826663 | 0.84488735 |
| ARRDC3 | 327.87926 | -0.6049579 | 0.52593243 | -1.1502578 | 0.25003769 | 0.80672756 |
| ARRDC3-AS1 | 31.3058505 | -0.208510573 | 0.40869019 | -0.5101923 | 0.60991677 | 0.94791646 |
| AC124854.1 | 194.308132 | -0.091970187 | 1.69583424 | -0.054233 | 9.57E-01 | 0.99572129 |
| NR2F1-AS1 | 49.3105778 | 0.788812279 | 0.95502402 | 0.82596067 | 0.4088264 | 0.8959746 |
| FAM172A | 628.492184 | -0.108056786 | 0.15355798 | -0.7036872 | 0.4816276 | 0.91971919 |
| KIAA0825 | 359.790835 | 0.340069082 | 0.27997451 | 1.21464301 | 0.22450228 | 0.78901652 |
| SLF1 | 46.6901684 | -0.482630896 | 0.36013716 | -1.3401308 | 0.18020283 | 0.75378294 |
| MCTP1 | 154.962822 | -0.802804697 | 0.57807125 | -1.3887643 | 0.16490444 | 0.74445571 |
| TTC37 | 247.051451 | -0.310136345 | 0.19191499 | -1.616009 | 0.10609237 | 0.65699665 |
| ARSK | 35.1060184 | 0.204120672 | 0.35647379 | 0.5726106 | 0.56690838 | 0.9385359 |
| RHOBTB3 | 361.323125 | -0.220259181 | 0.47063296 | -0.4680063 | 0.6397801 | 0.94944486 |
| GLRX | 111.668584 | 0.013864407 | 0.55562485 | 0.02495282 | 0.9800926 | 0.99681784 |
| ELL2 | 212.027257 | -0.462639687 | 0.53297384 | -0.8680345 | 0.38537544 | 0.88353166 |
| AC104123.1 | 101.477166 | -0.416081029 | 0.28441831 | -1.4629193 | 0.14348947 | 0.71631428 |
| CAST | 463.935551 | -0.243693583 | 0.1842036 | -1.3229578 | 0.18584942 | 0.76046631 |
| ERAP1 | 107.516953 | -0.441333568 | 0.25547223 | -1.7275207 | 0.08407419 | 0.61656297 |

| | | | | | | |
|---|---|---|---|---|---|---|
| AC009126.1 | 58.2728492 | 0.109930847 | 0.55965813 | 0.196425 | 0.84427753 | 0.98517901 |
| ERAP2 | 58.6982468 | 0.084376693 | 0.47688986 | 0.1769312 | 0.85956243 | 0.9864199 |
| LNPEP | 363.989003 | -0.025271039 | 0.19975831 | -0.1265081 | 0.89932976 | 0.98956835 |
| LIX1-AS1 | 78.8412346 | 0.405651891 | 0.31368817 | 1.29316923 | 0.19595253 | 0.77017438 |
| LIX1 | 40.9182845 | 0.219697733 | 1.26845245 | 0.17320139 | 0.86249313 | 0.98669442 |
| CHD1 | 319.605418 | 0.124095242 | 0.15603522 | 0.79530275 | 0.42643742 | 0.90237497 |
| LINC02062 | 97.4856384 | 0.283703587 | 0.50316763 | 0.56383514 | 0.57286634 | 0.93998433 |
| FAM174A | 36.3260483 | 0.395648581 | 0.32317065 | 1.2242714 | 0.22084986 | 0.78614672 |
| SLCO4C1 | 165.676042 | -0.009218149 | 0.47623859 | -0.0193562 | 0.98455699 | 0.99752755 |
| PAM | 720.668549 | 0.227425633 | 0.31091382 | 0.73147483 | 0.46448917 | 0.91462428 |
| GIN1 | 31.9984679 | 0.029298307 | 0.42813692 | 0.0684321 | 0.94544167 | 0.9952642 |
| PPIP5K2 | 200.073892 | -0.189732623 | 0.18554024 | -1.0225956 | 0.30649911 | 0.84488735 |
| NUDT12 | 29.9766002 | -0.160143264 | 0.33778983 | -0.4740914 | 0.63543471 | 0.94932781 |
| EFNA5 | 995.48974 | -0.905911069 | 0.89746159 | -1.0094149 | 0.31277571 | 0.84826686 |
| FBXL17 | 708.464607 | -0.032819279 | 0.1550117 | -0.2117213 | 0.83232447 | 0.98306468 |
| FER | 786.727351 | -0.045415508 | 0.18224599 | -0.2491989 | 8.03E-01 | 0.97874556 |
| PJA2 | 257.877957 | 0.286498724 | 0.1900459 | 1.50752386 | 0.13167643 | 0.700349 |
| MAN2A1 | 298.857381 | 0.120073401 | 0.41249445 | 0.29109095 | 0.77098176 | 0.97327811 |
| TMEM232 | 308.496278 | 0.80657343 | 0.44630201 | 1.80723682 | 0.07072536 | 0.58815298 |
| SLC25A46 | 103.720427 | -0.030150684 | 0.25849836 | -0.1166378 | 0.90714707 | 0.99100813 |
| WDR36 | 75.669084 | -0.572609217 | 0.32007768 | -1.7889695 | 0.07361972 | 0.59320069 |
| STARD4 | 46.1385867 | -0.48066329 | 0.63117977 | -0.7615315 | 0.44633966 | 0.9079396 |
| NREP | 108.493285 | -0.818551275 | 0.58107424 | -1.4086862 | 0.15892797 | 0.73686885 |
| EPB41L4A | 440.221859 | -0.088329719 | 0.22377392 | -0.3947275 | 0.69304401 | 0.96220115 |
| EPB41L4A-AS | 27.828658 | -0.806973789 | 0.58453223 | -1.3805463 | 0.16741851 | 0.74445571 |
| APC | 128.614502 | -0.303676922 | 0.20512599 | -1.480441 | 0.13875559 | 0.7113761 |
| SRP19 | 62.8904529 | 0.137852281 | 0.29059317 | 0.47438239 | 0.63522725 | 0.94932781 |
| REEP5 | 276.652106 | -0.029843412 | 0.24459802 | -0.12201 | 0.90289107 | 0.99034419 |
| DCP2 | 117.417401 | 0.218449354 | 0.21360423 | 1.0226827 | 0.30645789 | 0.84488735 |
| MCC | 70.3339852 | 1.574727255 | 0.8379695 | 1.87921786 | 0.06021475 | 0.56071154 |
| YTHDC2 | 182.810686 | 0.158965519 | 0.21247569 | 0.74815863 | 0.45436448 | 0.91077355 |
| PGGT1B | 70.1373486 | 0.130092919 | 0.22647345 | 0.574429 | 0.56567753 | 0.93809297 |
| FEM1C | 50.3641573 | -0.233403232 | 0.36636819 | -0.6370729 | 0.52407739 | 0.93272384 |
| TMED7-TICAI | 28.4395444 | 0.170641666 | 0.33967894 | 0.50236164 | 0.61541316 | 0.94791646 |
| ATG12 | 64.7790137 | -0.216875507 | 0.31290222 | -0.6931095 | 0.48824082 | 0.92081962 |
| AP3S1 | 101.845549 | -0.169916747 | 0.34874544 | -0.4872229 | 0.62610041 | 0.94831213 |
| COMMD10 | 480.92559 | -0.600528597 | 0.22040563 | -2.7246518 | 0.00643693 | 0.25167097 |
| SEMA6A | 184.978893 | 0.783861561 | 0.48677924 | 1.61030194 | 1.07E-01 | 0.65699665 |
| SEMA6A-AS1 | 78.4903191 | -0.57282831 | 0.32205672 | -1.7786566 | 0.07529608 | 0.59465671 |
| DTWD2 | 111.175387 | 0.040526517 | 0.22690423 | 0.17860627 | 0.85824687 | 0.9864199 |
| AC008629.1 | 60.1075133 | 0.357143749 | 0.59695536 | 0.59827547 | 0.54965614 | 0.93523904 |
| DMXL1 | 404.593726 | 0.207074839 | 0.17176933 | 1.20554025 | 0.22799481 | 0.79284833 |
| TNFAIP8 | 380.8644 | -0.273899874 | 0.35352839 | -0.7747606 | 0.43848113 | 0.90561284 |
| HSD17B4 | 141.785285 | 0.703547314 | 0.17280832 | 4.07125824 | 4.68E-05 | 0.01399728 |
| SRFBP1 | 94.9675923 | -0.526085706 | 0.25645355 | -2.0513879 | 0.04022919 | 0.4841148 |
| SNX2 | 192.223231 | 0.174807431 | 0.30948103 | 0.56484053 | 0.57218224 | 0.93998433 |
| SNX24 | 226.355322 | -0.198272233 | 0.17907806 | -1.1071833 | 0.26821468 | 0.81584693 |
| PPIC | 46.0536206 | -0.018562921 | 0.40221122 | -0.0461522 | 0.96318896 | 0.99572129 |
| CEP120 | 68.2657963 | -0.499413697 | 0.3463053 | -1.4421197 | 0.14926861 | 0.72246729 |
| CSNK1G3 | 173.726367 | 0.104293583 | 0.18837378 | 0.55365233 | 0.5798168 | 0.94034284 |
| ZNF608 | 329.495154 | 0.560414686 | 0.65848855 | 0.85106216 | 0.39473483 | 0.88760715 |
| GRAMD2B | 485.74216 | -0.450749242 | 0.30172874 | -1.493889 | 0.13520464 | 0.70620568 |
| ALDH7A1 | 93.1244351 | 0.048558196 | 0.28577251 | 0.16991906 | 0.86507379 | 0.98670067 |
| PHAX | 43.5892902 | 0.162298988 | 0.26750413 | 0.60671581 | 0.54403953 | 0.93523904 |

| | | | | | | |
|---|---|---|---|---|---|---|
| C5orf63 | 32.139416 | -0.138579512 | 0.33908557 | -0.408686 | 0.68277014 | 0.96101988 |
| PRRC1 | 76.1781234 | -0.231864664 | 0.34034552 | -0.6812626 | 0.49570536 | 0.92326689 |
| LINC01184 | 186.369039 | -0.437183937 | 0.25608833 | -1.7071607 | 0.08779218 | 0.62705598 |
| SLC12A2 | 64.1303693 | -0.387895004 | 0.35792175 | -1.0837425 | 0.278479 | 0.8229426 |
| FBN2 | 94.6640791 | -0.302521208 | 0.46454227 | -0.6512243 | 5.15E-01 | 0.93113905 |
| SLC27A6 | 59.8588172 | -0.545657987 | 1.14278776 | -0.4774797 | 0.63302055 | 0.94932781 |
| HINT1 | 536.086282 | 0.386509527 | 0.52588902 | 0.73496406 | 0.46236138 | 0.91462428 |
| LYRM7 | 48.7888708 | 0.098882038 | 0.2975372 | 0.33233504 | 0.73963628 | 0.9698508 |
| CDC42SE2 | 850.329627 | -0.116316349 | 0.43800742 | -0.2655579 | 0.79057968 | 0.97423208 |
| P4HA2 | 33.9475311 | 0.771203593 | 0.37970683 | 2.03105011 | 0.04224991 | 0.49175201 |
| SLC22A4 | 41.8732414 | -0.299906947 | 0.38024046 | -0.7887297 | 0.43027 | 0.90364119 |
| SLC22A5 | 101.646185 | 0.154041901 | 0.22142742 | 0.69567673 | 0.4866313 | 0.92034617 |
| AC116366.3 | 182.101624 | -0.904601934 | 0.61812069 | -1.4634714 | 0.14333844 | 0.71631428 |
| IRF1 | 75.75095 | 0.042052786 | 0.68414277 | 0.06146785 | 0.95098662 | 0.99531843 |
| KIF3A | 108.386047 | 0.551664171 | 0.20922469 | 2.63670682 | 0.00837151 | 0.272888 |
| 8-Sep | 107.561893 | -0.114901562 | 0.46276051 | -0.248296 | 0.80390543 | 0.97880236 |
| UQCRQ | 75.5821885 | 0.418096409 | 0.42161557 | 0.99165315 | 0.32136675 | 0.85131972 |
| AFF4 | 487.523361 | -0.028751266 | 0.1810175 | -0.1588314 | 0.87380169 | 0.98766275 |
| ZCCHC10 | 68.2009293 | -0.106610772 | 0.26147757 | -0.4077244 | 0.68347606 | 0.96144376 |
| HSPA4 | 135.85107 | -0.152617825 | 0.23465901 | -0.6503813 | 0.51544596 | 0.93140514 |
| C5orf15 | 98.0489657 | 0.502085451 | 0.35793482 | 1.40272871 | 0.16069775 | 0.73816762 |
| VDAC1 | 135.930694 | 0.025221638 | 0.36619201 | 0.06887545 | 0.94508876 | 0.9952642 |
| SKP1 | 269.677213 | 0.135243289 | 0.47642835 | 0.2838691 | 0.7765107 | 0.97423208 |
| CDKL3 | 92.3939757 | -0.217830773 | 0.32183371 | -0.6768426 | 0.49850581 | 0.92398825 |
| UBE2B | 200.674224 | -0.250294023 | 0.22345596 | -1.1201045 | 0.26266923 | 0.81377303 |
| JADE2 | 90.2041167 | 0.117828394 | 0.28750582 | 0.40982959 | 0.68193096 | 0.96095398 |
| SAR1B | 90.9002907 | 0.325306879 | 0.26553687 | 1.22509119 | 0.22054087 | 0.78592297 |
| SEC24A | 100.069245 | 0.091040057 | 0.28788501 | 0.31623758 | 0.75182219 | 0.97058015 |
| CAMLG | 50.5058357 | 0.703063385 | 0.32883199 | 2.13806262 | 0.03251166 | 0.44049392 |
| DDX46 | 190.941249 | -0.26005647 | 0.23158783 | -1.1229281 | 0.26146805 | 0.81281978 |
| C5orf24 | 72.0270578 | -0.049748288 | 0.35778276 | -0.1390461 | 0.88941374 | 0.98766275 |
| TXNDC15 | 51.1725918 | -0.048265955 | 0.26241579 | -0.1839293 | 0.85406891 | 0.9864199 |
| PCBD2 | 140.192858 | -0.614838297 | 0.32836349 | -1.872432 | 0.06114687 | 0.56071154 |
| H2AFY | 177.079391 | -0.62715721 | 0.29939837 | -2.0947249 | 0.03619546 | 0.46105889 |
| CXCL14 | 174.036325 | 0.928667798 | 0.42968975 | 2.1612519 | 0.03067589 | 0.43815485 |
| SLC25A48 | 62.1872102 | 0.718428994 | 0.61820054 | 1.16212935 | 0.24518293 | 0.80476923 |
| SMAD5 | 188.470334 | 0.012515561 | 0.16825217 | 0.07438573 | 0.94070346 | 0.99526299 |
| SPOCK1 | 169.635362 | 0.254673849 | 0.64276909 | 0.39621359 | 0.69194747 | 0.96220115 |
| HNRNPA0 | 87.1809461 | -0.583993348 | 0.40054462 | -1.4579982 | 0.14484103 | 0.71738996 |
| FAM13B | 334.5004 | 0.055725443 | 0.250927 | 0.22207831 | 0.82425292 | 0.98094663 |
| NME5 | 84.5390582 | 0.704383217 | 0.39327473 | 1.79107164 | 0.0732818 | 0.59320069 |
| BRD8 | 72.9282436 | -0.030936122 | 0.20851073 | -0.1483671 | 0.8820531 | 0.98766275 |
| CDC23 | 37.8399091 | 0.395883478 | 0.26514885 | 1.49306124 | 0.13542116 | 0.7063854 |
| FAM53C | 33.3240479 | 0.261894091 | 0.3814246 | 0.68662087 | 0.49232167 | 0.92289011 |
| KDM3B | 202.65299 | 0.041688461 | 0.18914212 | 0.22040813 | 0.82555331 | 0.98094663 |
| EGR1 | 252.357162 | 0.218125809 | 0.79767237 | 0.27345288 | 0.78450511 | 0.97423208 |
| ETF1 | 119.307538 | -0.599977289 | 0.29636972 | -2.0244217 | 0.04292678 | 0.49422552 |
| HSPA9 | 207.080053 | -0.308102903 | 0.2811839 | -1.0957345 | 0.27319498 | 0.82047446 |
| CTNNA1 | 728.858003 | -0.214187042 | 0.18423118 | -1.1625993 | 0.24499211 | 0.80476923 |
| SIL1 | 302.113131 | 0.084098288 | 0.24568592 | 0.3423 | 0.73212514 | 0.96980299 |
| PAIP2 | 149.474953 | 0.441612121 | 0.22262334 | 1.983674 | 0.04729219 | 0.51284792 |
| UBE2D2 | 440.442426 | 0.07173169 | 0.29710869 | 0.24143249 | 0.80921994 | 0.97930627 |
| CXXC5 | 214.860714 | -0.202561031 | 0.45328013 | -0.4468782 | 0.65496297 | 0.95294428 |
| PURA | 195.377642 | 0.307738284 | 0.27011909 | 1.13926893 | 0.254591 | 0.80897837 |

| | | | | | | |
|---|---|---|---|---|---|---|
| IGIP | 32.148248 | 0.2684093 | 0.42042215 | 0.63842807 | 0.52319506 | 0.93272384 |
| CYSTM1 | 524.352624 | 0.029159974 | 0.48769659 | 0.05979122 | 0.95232192 | 0.99531843 |
| PFDN1 | 97.5723224 | -0.417389062 | 0.25221505 | -1.6548936 | 0.09794609 | 0.64312419 |
| SRA1 | 40.0985148 | 0.159305277 | 0.37790944 | 0.42154352 | 0.67335824 | 0.95910677 |
| SLC35A4 | 41.9368217 | -0.525811745 | 0.36179356 | -1.4533474 | 0.14612731 | 0.71974648 |
| NDUFA2 | 71.6378362 | 0.29798969 | 0.44384584 | 0.67138107 | 0.5019778 | 0.92484445 |
| IK | 45.547807 | 0.136937282 | 0.36862114 | 0.37148515 | 0.71027621 | 0.96587538 |
| HARS | 59.8738836 | -0.333644557 | 0.28465134 | -1.1721166 | 0.24115025 | 0.80319133 |
| ZMAT2 | 36.7256328 | 0.03452281 | 0.39508538 | 0.08738063 | 0.93036896 | 0.99375606 |
| TAF7 | 91.7216862 | -0.232027015 | 0.32211573 | -0.7203219 | 0.47132682 | 0.91634658 |
| DIAPH1 | 445.248476 | 0.089155059 | 0.37644185 | 0.2368362 | 0.81278387 | 0.97930627 |
| HDAC3 | 61.0738057 | -0.037215681 | 0.33879578 | -0.1098469 | 0.91253076 | 0.99113939 |
| DELE1 | 57.302293 | -0.177316894 | 0.32005974 | -0.5540119 | 0.57957073 | 0.94034284 |
| RNF14 | 57.0290812 | 0.083663888 | 0.33148631 | 0.25239018 | 0.80073949 | 0.97723068 |
| GNPDA1 | 43.783963 | -0.501347187 | 0.5078569 | -0.987182 | 0.32355343 | 0.85404196 |
| NDFIP1 | 284.486906 | 0.075324439 | 0.18870075 | 0.39917404 | 0.68976497 | 0.96220115 |
| SPRY4 | 36.9461429 | -0.730507759 | 0.43645905 | -1.6737143 | 0.09418676 | 0.6377384 |
| SPRY4-AS1 | 160.535549 | -0.499278362 | 0.68121792 | -0.7329202 | 0.46360711 | 0.91462428 |
| FGF1 | 30.378492 | 0.688736081 | 0.37353307 | 1.84384232 | 0.06520612 | 0.57495764 |
| ARHGAP26 | 1250.67734 | 0.157940942 | 0.46802083 | 0.33746562 | 0.73576592 | 0.9698508 |
| NR3C1 | 372.108384 | 0.312894011 | 0.5425632 | 0.57669597 | 0.56414486 | 0.93809297 |
| YIPF5 | 39.4514477 | 0.079332072 | 0.27072243 | 0.29303842 | 0.76949279 | 0.97261718 |
| KCTD16 | 448.53238 | 0.279731909 | 0.55863743 | 0.50073964 | 0.61655437 | 0.94791646 |
| PRELID2 | 240.665679 | -0.43407102 | 0.42342532 | -1.0251419 | 0.30529626 | 0.84488735 |
| SH3RF2 | 46.2392157 | -0.229742147 | 0.83938009 | -0.2737045 | 0.78431168 | 0.97423208 |
| LARS | 181.377089 | -0.109500533 | 0.21382244 | -0.5121096 | 0.60857428 | 0.94727544 |
| TCERG1 | 208.262278 | -0.523450961 | 0.19080164 | -2.7434301 | 0.0060801 | 0.24646394 |
| PPP2R2B | 184.697144 | 0.092976857 | 0.78125803 | 0.11900915 | 0.9052681 | 0.99092254 |
| DPYSL3 | 84.6528832 | 0.830371773 | 1.38593771 | 0.59914076 | 0.54907902 | 0.93523904 |
| AC091948.1 | 56.459383 | 0.2111296 | 0.78429706 | 0.26919596 | 0.78777889 | 0.97423208 |
| FBXO38 | 92.8976076 | -0.009189327 | 0.19414212 | -0.047333 | 0.96224784 | 0.99572129 |
| CSNK1A1 | 669.000982 | -0.142772871 | 0.14478201 | -0.986123 | 0.32407274 | 0.85444355 |
| ARHGEF37 | 48.3657727 | -0.392966685 | 0.57025259 | -0.6891099 | 0.49075413 | 0.92187769 |
| SLC26A2 | 45.9440241 | -1.097386232 | 0.3794222 | -2.8922562 | 0.00382486 | 0.19908785 |
| HMGXB3 | 78.6618025 | -0.627368933 | 0.20926096 | -2.9980219 | 0.00271738 | 0.17081506 |
| PDGFRB | 62.6104188 | 0.1099566 | 1.69365015 | 0.06492285 | 0.94823542 | 0.9952642 |
| TCOF1 | 34.4458409 | -1.057416713 | 0.39004515 | -2.711011 | 0.00670784 | 0.25396993 |
| CD74 | 223.14648 | 0.551182224 | 0.6239059 | 0.88343807 | 0.37699963 | 0.87842274 |
| RPS14 | 485.29131 | -0.006588693 | 0.50724041 | -0.0129893 | 0.98963634 | 0.99839055 |
| NDST1 | 99.0794664 | -0.072072836 | 0.31947183 | -0.2256 | 0.82151256 | 0.98094663 |
| SYNPO | 37.6440914 | 0.239976608 | 0.69548025 | 0.34505165 | 0.73005554 | 0.96885592 |
| RBM22 | 33.2643796 | -0.259133508 | 0.28132246 | -0.9211263 | 0.35698451 | 0.87316144 |
| DCTN4 | 154.405611 | -0.290078574 | 0.15595798 | -1.8599791 | 0.06288848 | 0.56713487 |
| SMIM3 | 34.1612641 | -0.962522101 | 0.66365053 | -1.4503448 | 0.14696239 | 0.72098466 |
| GPX3 | 1413.61785 | 0.769814966 | 0.84457392 | 0.91148323 | 0.36204082 | 0.8752623 |
| TNIP1 | 199.117646 | 0.377236697 | 0.48771626 | 0.77347575 | 0.43924088 | 0.90561284 |
| SLC36A1 | 28.7287273 | 0.012162424 | 0.45587886 | 0.02667907 | 0.97871571 | 0.99681784 |
| SPARC | 134.709833 | 1.408947969 | 1.28198363 | 1.09903741 | 0.27175175 | 0.81858805 |
| ATOX1 | 87.5958171 | -0.147784463 | 0.29061262 | -0.5085273 | 0.61108358 | 0.94791646 |
| G3BP1 | 228.451389 | -0.264559254 | 0.18855659 | -1.4030762 | 0.16059413 | 0.73816762 |
| FAM114A2 | 36.0423072 | -0.21714354 | 0.25802107 | -0.8415729 | 0.40002707 | 0.89050878 |
| MFAP3 | 50.2001196 | 0.19431803 | 0.3533667 | 0.54990477 | 0.58238469 | 0.94091128 |
| GALNT10 | 198.633443 | -0.235266157 | 0.41961589 | -0.5606703 | 0.57502233 | 0.94034284 |
| SAP30L-AS1 | 44.1847036 | 0.126468438 | 0.35145433 | 0.35984317 | 0.71896442 | 0.96779196 |

| | | | | | | |
|---|---|---|---|---|---|---|
| SAP30L | 25.2855158 | 0.077669167 | 0.42297957 | 0.18362392 | 0.8543085 | 0.9864199 |
| LARP1 | 320.616811 | -0.348003949 | 0.26411484 | -1.3176236 | 0.18762968 | 0.76207437 |
| FAXDC2 | 45.2977725 | 1.40554942 | 0.59957089 | 2.34425894 | 0.01906493 | 0.36671278 |
| CNOT8 | 56.9808129 | 0.433361756 | 0.22327731 | 1.94091263 | 0.05226888 | 0.53151121 |
| GEMIN5 | 47.2611763 | -0.291661008 | 0.2478291 | -1.1768635 | 0.23925001 | 0.80174646 |
| MRPL22 | 47.2260203 | -0.313361826 | 0.2682033 | -1.1683742 | 0.24265584 | 0.80476923 |
| SGCD | 100.074248 | 1.870252412 | 1.16072827 | 1.61127497 | 0.1071198 | 0.65699665 |
| TIMD4 | 47.0943776 | 0.333777629 | 0.70707825 | 0.47205189 | 0.63688975 | 0.94944486 |
| HAVCR1 | 157.421324 | 0.473810533 | 1.06649257 | 0.44426989 | 0.65684747 | 0.95399905 |
| CYFIP2 | 199.311642 | -0.508776325 | 0.62168869 | -0.818378 | 0.4131414 | 0.89724628 |
| CLINT1 | 279.101766 | 0.276625406 | 0.29106845 | 0.95037922 | 0.3419196 | 0.86846099 |
| EBF1 | 231.1484 | -0.408233276 | 1.52925696 | -0.2669488 | 0.7895086 | 0.97423208 |
| RNF145 | 135.27202 | 0.22479041 | 0.25098312 | 0.89563954 | 0.37044531 | 0.87605028 |
| UBLCP1 | 45.04094 | -0.555507008 | 0.29522077 | -1.8816664 | 0.05988132 | 0.55922073 |
| TTC1 | 112.151508 | -0.276438632 | 0.23715534 | -1.1656437 | 0.24375853 | 0.80476923 |
| PWWP2A | 155.298041 | -0.068180873 | 0.25576098 | -0.2665804 | 0.78979222 | 0.97423208 |
| CCNJL | 69.0769829 | -1.035601811 | 0.77535018 | -1.3356569 | 0.18166144 | 0.75448394 |
| SLU7 | 35.0483533 | -0.080780247 | 0.30482781 | -0.2650029 | 0.79100725 | 0.97423208 |
| AC113414.1 | 92.724953 | 0.51928214 | 0.39883545 | 1.30199596 | 0.19291777 | 0.76647004 |
| CCNG1 | 78.1497937 | 0.884454504 | 0.2989939 | 2.95810215 | 0.0030954 | 0.18079751 |
| NUDCD2 | 31.697554 | -0.220443943 | 0.35253119 | -0.6253176 | 0.53176265 | 0.93366189 |
| MAT2B | 66.802831 | 0.040416408 | 0.34239775 | 0.11803935 | 0.90603648 | 0.99100813 |
| TENM2 | 56.1934909 | 1.269621216 | 0.69944914 | 1.81517304 | 0.06949732 | 0.58408344 |
| WWC1 | 1722.61598 | -0.548617929 | 0.42015885 | -1.3057393 | 0.19164122 | 0.76393312 |
| RARS | 113.493659 | -0.270593264 | 0.19853091 | -1.362978 | 0.17288944 | 0.74700403 |
| PANK3 | 106.457504 | -0.143321049 | 0.20570731 | -0.6967232 | 0.48597607 | 0.92034617 |
| SLIT3 | 857.557353 | 0.872127485 | 0.82090767 | 1.06239413 | 0.2880568 | 0.83300685 |
| RANBP17 | 156.929628 | 0.196476902 | 0.33726407 | 0.58256102 | 0.56018885 | 0.93744835 |
| NPM1 | 177.084114 | -0.282497647 | 0.35337795 | -0.7994207 | 0.42404651 | 0.90167091 |
| FBXW11 | 383.962615 | -0.147619761 | 0.18619832 | -0.7928093 | 0.42788893 | 0.90272191 |
| UBTD2 | 113.170248 | 0.273772822 | 0.26479992 | 1.03388558 | 0.30118966 | 0.84277781 |
| SH3PXD2B | 44.3802729 | 0.038756909 | 0.53399254 | 0.07257949 | 0.94214074 | 0.99526299 |
| DUSP1 | 320.396169 | -0.035617359 | 0.71559464 | -0.0497731 | 0.96030321 | 0.99572129 |
| ERGIC1 | 237.531013 | -0.077127312 | 0.33489954 | -0.2302998 | 0.81785878 | 0.98010849 |
| ATP6V0E1 | 272.873726 | -0.016212509 | 0.22609393 | -0.071707 | 0.94283511 | 0.9952642 |
| CREBRF | 231.513615 | 0.086894509 | 0.27593028 | 0.31491473 | 0.7528264 | 0.97096495 |
| BOD1 | 32.7765009 | -0.278090666 | 0.33122684 | -0.8395777 | 0.40114521 | 0.89196147 |
| CPEB4 | 698.54262 | 0.490270253 | 0.35598767 | 1.37721134 | 0.16844692 | 0.74445571 |
| SFXN1 | 129.424687 | 0.346061487 | 0.21754888 | 1.59072978 | 0.1116704 | 0.66300165 |
| THOC3 | 47.8553312 | -0.267280899 | 0.29959409 | -0.8921434 | 0.37231606 | 0.87605028 |
| SIMC1 | 133.389265 | 0.333288089 | 0.35838992 | 0.92995945 | 0.35239208 | 0.8720098 |
| KIAA1191 | 74.3914594 | 0.111515764 | 0.27737359 | 0.40204175 | 0.6876533 | 0.96161098 |
| ARL10 | 43.1037572 | 0.462105434 | 0.37974848 | 1.21687237 | 0.22365278 | 0.78791697 |
| CLTB | 62.7581338 | -0.937206761 | 0.36730606 | -2.551569 | 0.01072391 | 0.29948783 |
| FAF2 | 147.893012 | -0.164985607 | 0.18503726 | -0.8916345 | 0.37258888 | 0.87605028 |
| RNF44 | 38.1440775 | 0.234922265 | 0.26022535 | 0.90276473 | 0.36665078 | 0.87605028 |
| TSPAN17 | 30.0292612 | -0.137652408 | 0.38735729 | -0.3553629 | 0.72231774 | 0.96779196 |
| UIMC1 | 210.326552 | -0.12870538 | 0.21417103 | -0.6009467 | 0.54787547 | 0.93523904 |
| ZNF346 | 94.7244087 | 0.381278778 | 0.26113483 | 1.46008394 | 0.144267 | 0.71634214 |
| NSD1 | 457.691403 | -0.082262942 | 0.20504019 | -0.401204 | 0.68826995 | 0.96161098 |
| PRELID1 | 64.2223505 | -0.245928754 | 0.5063345 | -0.4857041 | 0.62717696 | 0.94856269 |
| LMAN2 | 87.9210171 | -0.146753778 | 0.35319028 | -0.4155091 | 0.67776925 | 0.95924874 |
| DBN1 | 35.5006766 | 0.120375666 | 0.52329251 | 0.23003514 | 0.81806446 | 0.98010849 |
| PDLIM7 | 31.8342777 | -0.255934274 | 0.67212885 | -0.3807816 | 0.70336534 | 0.9645815 |

| | | | | | | |
|---|---|---|---|---|---|---|
| FAM193B | 116.477793 | 0.086087928 | 0.31810707 | 0.27062563 | 0.78667898 | 0.97423208 |
| TMED9 | 108.012711 | 0.141585711 | 0.43238369 | 0.32745386 | 0.74332463 | 0.97012548 |
| B4GALT7 | 27.5646957 | -0.553530908 | 0.36253377 | -1.5268396 | 0.1268009 | 0.69494309 |
| RMND5B | 64.1147652 | -0.427697451 | 0.25169478 | -1.6992702 | 0.08926828 | 0.63290368 |
| NHP2 | 62.3793727 | -0.573281026 | 0.52270902 | -1.0967498 | 0.27275077 | 0.82005009 |
| HNRNPAB | 82.966649 | -0.482056375 | 0.40917256 | -1.1781249 | 2.39E-01 | 0.80174646 |
| PHYKPL | 222.621338 | 0.482327395 | 0.27484695 | 1.7548945 | 0.07927735 | 0.60556378 |
| COL23A1 | 64.9388414 | 0.803265155 | 0.83535927 | 0.96158046 | 0.33626039 | 0.86401242 |
| CLK4 | 96.3770236 | 0.216464555 | 0.39770524 | 0.54428388 | 0.58624614 | 0.9423304 |
| ZNF354A | 48.7085627 | 0.192877547 | 0.29169193 | 0.66123718 | 0.50846022 | 0.92666294 |
| ZNF354B | 36.2962683 | -0.265035147 | 0.32481767 | -0.8159505 | 0.41452843 | 0.89757799 |
| RUFY1 | 173.159282 | 0.012920166 | 0.23523316 | 0.05492493 | 0.95619827 | 0.99572129 |
| HNRNPH1 | 418.942379 | 0.152465286 | 0.22066899 | 0.69092301 | 4.90E-01 | 0.92157694 |
| CANX | 375.290057 | -0.020813223 | 0.31565772 | -0.065936 | 0.94742874 | 0.9952642 |
| MAML1 | 63.9239325 | -0.385782637 | 0.25491997 | -1.513348 | 0.13019128 | 0.69904834 |
| MGAT4B | 90.3961563 | -0.559257408 | 0.45615763 | -1.2260179 | 0.22019195 | 0.78584902 |
| SQSTM1 | 446.278027 | -0.528525782 | 0.29004941 | -1.8221922 | 0.06842582 | 0.58250536 |
| MRNIP | 75.5876983 | 0.293547923 | 0.28864566 | 1.01698366 | 0.3091612 | 0.84554063 |
| TBC1D9B | 95.3896825 | -0.265408053 | 0.27729024 | -0.9571489 | 0.3384921 | 0.86672435 |
| RNF130 | 227.397083 | -0.098281578 | 0.29261444 | -0.335874 | 0.7369659 | 0.9698508 |
| MAPK9 | 77.6573489 | -0.263109062 | 0.23806495 | -1.1051986 | 0.26907353 | 0.81655182 |
| CNOT6 | 128.360529 | -0.022808735 | 0.17506692 | -0.1302858 | 0.89634031 | 0.98842187 |
| MGAT1 | 56.4912539 | -0.478869781 | 0.26074809 | -1.8365227 | 0.06628039 | 0.57792071 |
| ZFP62 | 25.889056 | 0.019962278 | 0.3118241 | 0.06401775 | 0.94895609 | 0.9952642 |
| TRIM41 | 29.4276641 | 0.190228756 | 0.31286555 | 0.60802079 | 0.54317369 | 0.93523904 |
| RACK1 | 354.095043 | -0.007473482 | 0.46942392 | -0.0159205 | 0.98729778 | 0.99792397 |
| TRIM52 | 43.5725602 | -0.203408089 | 0.30814841 | -0.6600978 | 0.50919107 | 0.92764192 |
| DUSP22 | 58.4916954 | -0.100011544 | 0.30636424 | -0.3264465 | 0.74408653 | 0.97012548 |
| EXOC2 | 169.853893 | -0.414950825 | 0.23360362 | -1.7763031 | 0.07568298 | 0.59499139 |
| FOXC1 | 264.368437 | -0.204270352 | 0.59503683 | -0.3432903 | 0.7313801 | 0.96953917 |
| GMDS | 1049.15277 | -0.418152876 | 0.22797808 | -1.8341802 | 0.06662725 | 0.5780981 |
| GMDS-DT | 891.589564 | 0.00562616 | 0.24746459 | 0.02273521 | 0.98186149 | 0.99681784 |
| WRNIP1 | 70.3830498 | 0.240634076 | 0.22913215 | 1.05019778 | 0.29362719 | 0.83825235 |
| SERPINB1 | 58.9400818 | -0.237686517 | 0.26738456 | -0.8889313 | 0.37403998 | 0.87666478 |
| SERPINB6 | 113.60774 | -0.081243445 | 0.19613784 | -0.4142161 | 0.67871587 | 0.95924874 |
| NQO2 | 55.2387446 | 0.31424931 | 0.34177995 | 0.91944923 | 0.35786065 | 0.87316144 |
| RIPK1 | 103.399221 | -0.376474669 | 0.28306309 | -1.3300027 | 0.18351739 | 0.75689793 |
| BPHL | 35.630105 | 0.343629935 | 0.30228991 | 1.13675623 | 0.2556402 | 0.80897837 |
| PSMG4 | 62.2497001 | -0.150841253 | 0.29101762 | -0.5183234 | 0.60423263 | 0.94683415 |
| SLC22A23 | 145.005572 | -0.377377259 | 0.60866146 | -0.6200117 | 0.53525005 | 0.93460153 |
| PXDC1 | 55.1231318 | -0.170961639 | 0.51736116 | -0.3304493 | 0.74106049 | 0.9698508 |
| PRPF4B | 226.393325 | -0.1924616 | 0.21056589 | -0.9140208 | 0.36070592 | 0.87429201 |
| ECI2 | 57.4644828 | 0.592181397 | 0.30544423 | 1.9387546 | 0.05253123 | 0.53192032 |
| CDYL | 255.058918 | -0.165930878 | 0.22001889 | -0.7541665 | 0.45074924 | 0.91072073 |
| LYRM4 | 148.233616 | -0.272602137 | 0.32240251 | -0.8455335 | 0.39781301 | 0.88867791 |
| FARS2 | 465.513945 | -0.058593449 | 0.20652367 | -0.283713 | 0.77663034 | 0.97423208 |
| RREB1 | 380.576666 | 0.008084451 | 0.32909905 | 0.0245654 | 0.98040162 | 0.99681784 |
| SSR1 | 141.413951 | 0.043329825 | 0.31936641 | 0.13567434 | 0.89207874 | 0.98811543 |
| AL139095.4 | 26.4750593 | 0.778626511 | 0.58585229 | 1.3290492 | 0.18383174 | 0.75689793 |
| RIOK1 | 34.0827164 | -0.314596895 | 0.3952959 | -0.7958517 | 0.42611826 | 0.90237497 |
| DSP | 185.730655 | -0.418914937 | 0.20571596 | -2.0363755 | 0.04171266 | 0.4884665 |
| SNRNP48 | 43.1321624 | -0.362147899 | 0.26487076 | -1.3672627 | 0.17154299 | 0.74445571 |
| BMP6 | 149.321697 | -1.036688871 | 0.41974197 | -2.4698242 | 0.01351795 | 0.32032755 |
| BLOC1S5-TXI | 54.2586145 | 0.291033385 | 0.48347599 | 0.60196037 | 0.54720052 | 0.93523904 |

| | | | | | | |
|---|---|---|---|---|---|---|
| SLC35B3 | 81.4774682 | 0.229992569 | 0.38379836 | 0.59925366 | 0.54900375 | 0.93523904 |
| GCNT2 | 187.99 | -0.77464692 | 0.57181605 | -1.3547135 | 0.17550886 | 0.74934286 |
| TMEM14C | 71.3927019 | -0.037920507 | 0.41973614 | -0.0903437 | 0.92801411 | 0.99336859 |
| MAK | 107.117136 | 0.425214191 | 0.33564147 | 1.26687026 | 0.20520169 | 0.77352973 |
| AL139807.1 | 178.98264 | 0.313292762 | 1.08671806 | 0.28829259 | 0.77312278 | 0.9735432 |
| NEDD9 | 1070.5792 | -1.166531205 | 0.61967601 | -1.8824857 | 0.0597701 | 0.55922073 |
| TMEM170B | 37.2822644 | 0.513770146 | 0.28208095 | 1.82135711 | 0.06855259 | 0.58250536 |
| HIVEP1 | 274.018059 | 0.103643879 | 0.23892324 | 0.43379572 | 6.64E-01 | 0.95505623 |
| PHACTR1 | 313.067713 | 0.425648416 | 0.5074593 | 0.83878336 | 0.40159089 | 0.89233104 |
| TBC1D7 | 25.5689398 | -0.665417087 | 0.46398551 | -1.4341333 | 0.15153422 | 0.72436429 |
| GFOD1 | 42.3779095 | 0.009139332 | 0.72168494 | 0.01266388 | 0.98989596 | 0.99839055 |
| SIRT5 | 67.8817371 | 0.295010573 | 0.27120482 | 1.08777778 | 0.27669322 | 0.82286383 |
| NOL7 | 84.5816162 | -0.262860458 | 0.42398405 | -0.6199772 | 0.53527277 | 0.93460153 |
| RANBP9 | 222.231941 | 0.118510255 | 0.16989971 | 0.69753065 | 0.48547076 | 0.92034617 |
| MCUR1 | 75.93539 | -0.003401309 | 0.43687891 | -0.0077855 | 0.99378815 | 0.99866813 |
| JARID2 | 210.536191 | 0.088551929 | 0.3010542 | 0.29413949 | 7.69E-01 | 0.972511 |
| DTNBP1 | 201.631084 | 0.14911509 | 0.28682263 | 0.51988606 | 0.60314299 | 0.94683415 |
| MYLIP | 64.903854 | 0.29799305 | 0.3623883 | 0.82230318 | 0.41090437 | 0.89649693 |
| GMPR | 81.8220746 | -1.186159426 | 0.75521469 | -1.5706255 | 0.11626967 | 0.67559092 |
| ATXN1 | 699.160936 | 0.644225356 | 0.34810657 | 1.85065557 | 0.06421912 | 0.57330377 |
| CAP2 | 118.238392 | 0.038174154 | 0.27431335 | 0.13916258 | 8.89E-01 | 0.98766275 |
| FAM8A1 | 36.2785351 | 0.573507775 | 0.31157636 | 1.84066523 | 0.06567063 | 0.57606134 |
| NUP153 | 220.608165 | -0.28998504 | 0.19133096 | -1.51562 | 0.12961546 | 0.69904834 |
| KIF13A | 357.013716 | -0.003534682 | 0.3351808 | -0.0105456 | 0.99158599 | 0.99866813 |
| TPMT | 55.608966 | 0.540948944 | 0.33943961 | 1.59365295 | 0.11101379 | 0.66214263 |
| KDM1B | 63.3982596 | 0.136523344 | 0.38050796 | 0.35879234 | 0.71975044 | 0.96779196 |
| DEK | 185.321407 | -0.457713308 | 0.19813807 | -2.3100725 | 0.02088414 | 0.38104604 |
| RNF144B | 943.326276 | -0.096630665 | 0.39606617 | -0.2439761 | 0.80724936 | 0.97930627 |
| MIR548A1HG | 97.7096682 | 1.977038942 | 0.74479291 | 2.65448141 | 0.00794304 | 0.27173712 |
| AL022068.1 | 297.47134 | -0.62996378 | 0.46778346 | -1.3466996 | 0.17807701 | 0.7513166 |
| ID4 | 260.795666 | -1.67277877 | 0.65253585 | -2.5635048 | 0.01036213 | 0.29453656 |
| MBOAT1 | 50.6968459 | 0.240994173 | 0.72067082 | 0.33440257 | 0.7380758 | 0.9698508 |
| E2F3 | 116.848873 | -0.341923941 | 0.37382366 | -0.9146664 | 0.36036679 | 0.87413358 |
| CDKAL1 | 1006.14234 | -0.357436429 | 0.23198701 | -1.5407605 | 0.12337508 | 0.69314363 |
| SOX4 | 240.658763 | -0.430469412 | 0.53901614 | -0.7986206 | 0.42451042 | 0.90167091 |
| CASC15 | 1218.88608 | 0.035444894 | 0.36372009 | 0.09745102 | 0.92236823 | 0.99214128 |
| NBAT1 | 47.9261224 | -0.219868608 | 0.40749705 | -0.5395588 | 0.58950135 | 0.94491859 |
| DCDC2 | 1349.33793 | 0.172194303 | 0.43416864 | 0.39660696 | 0.69165732 | 0.96220115 |
| MRS2 | 61.9109058 | -0.175618585 | 0.27397321 | -0.6410064 | 0.52151852 | 0.93272217 |
| ALDH5A1 | 33.2451395 | 0.679711717 | 0.40142277 | 1.6932565 | 0.09040665 | 0.6330877 |
| TDP2 | 63.9520187 | 0.176903227 | 0.25282556 | 0.69970467 | 0.48411176 | 0.92034617 |
| ACOT13 | 44.9344507 | 0.186294263 | 0.35445052 | 0.52558608 | 0.59917583 | 0.94683415 |
| C6orf62 | 160.60613 | -0.369906633 | 0.31602251 | -1.1705072 | 0.2417969 | 0.803412 |
| RIPOR2 | 455.64496 | -0.742519922 | 0.54045056 | -1.3738906 | 0.16947566 | 0.74445571 |
| CARMIL1 | 530.646966 | -0.071411257 | 0.23463146 | -0.304355 | 0.76085745 | 0.9714717 |
| SLC17A1 | 105.152276 | 2.077174679 | 1.0631241 | 1.95384027 | 0.05072011 | 0.52415717 |
| TRIM38 | 208.490777 | 0.054988538 | 0.49854595 | 0.11029783 | 0.91217318 | 0.9911192 |
| HIST1H1C | 82.9933834 | -0.353526605 | 0.69295579 | -0.5101719 | 0.60993101 | 0.94791646 |
| HIST1H4C | 85.1114101 | 0.418530464 | 0.61274382 | 0.68304314 | 0.49457958 | 0.92289011 |
| HIST1H2AC | 327.175047 | 0.53776012 | 0.78189047 | 0.68776912 | 0.49159818 | 0.92275924 |
| HIST1H1E | 42.4347813 | 0.054497205 | 0.88589574 | 0.0615165 | 0.95094787 | 0.99531843 |
| HIST1H2BD | 63.2544113 | 1.290335708 | 0.78410623 | 1.64561339 | 0.09984338 | 0.64621604 |
| HIST1H2BG | 38.3174082 | 1.171592961 | 1.25716542 | 0.93193222 | 0.35137155 | 0.8720098 |
| BTN2A1 | 33.8059406 | 0.204416693 | 0.33497964 | 0.61023616 | 0.54170538 | 0.93523904 |

| | | | | | | |
|---|---|---|---|---|---|---|
| HCG11 | 35.7087438 | -0.444267803 | 0.58186355 | -0.7635257 | 0.44514991 | 0.90703449 |
| HMGN4 | 29.6442253 | 0.135117296 | 0.40406524 | 0.33439475 | 0.7380817 | 0.9698508 |
| LINC00240 | 102.169262 | -0.106981859 | 0.61137967 | -0.1749843 | 0.86109195 | 0.98658849 |
| ZSCAN16-AS | 34.8340681 | 0.32704315 | 0.39214615 | 0.83398282 | 0.40429066 | 0.89373953 |
| ZKSCAN8 | 42.8268446 | 0.361112242 | 0.28409942 | 1.271077 | 0.20370126 | 0.77155952 |
| TRIM27 | 112.35491 | -0.254606547 | 0.25706325 | -0.9904432 | 0.32195753 | 0.85217773 |
| GABBR1 | 89.5631506 | 1.037738417 | 0.30385408 | 3.41525249 | 0.00063723 | 0.08031602 |
| HLA-F | 42.3820037 | -0.988214062 | 0.83764358 | -1.1797548 | 0.23809774 | 0.80145582 |
| HLA-A | 837.706712 | -0.312813756 | 0.43283621 | -0.722707 | 0.46985992 | 0.91502267 |
| PPP1R11 | 33.8750122 | 0.391525145 | 0.35373881 | 1.10681986 | 0.26837182 | 0.81584693 |
| TRIM26 | 97.8977934 | -0.414311071 | 0.31573684 | -1.3122038 | 0.18945139 | 0.76207437 |
| HCG17 | 191.724276 | -0.173630185 | 0.24175027 | -0.7182213 | 0.47262087 | 0.91737234 |
| HLA-E | 230.045158 | -0.805506844 | 0.46582862 | -1.7291914 | 8.38E-02 | 0.61656297 |
| GNL1 | 78.3199892 | -0.033019177 | 0.25631396 | -0.1288232 | 0.89749757 | 0.98863032 |
| PRR3 | 29.932418 | 0.173008211 | 0.36577806 | 0.47298684 | 0.63622256 | 0.94932781 |
| ABCF1 | 73.358229 | -0.314662972 | 0.23873028 | -1.318069 | 0.18748056 | 0.76207437 |
| PPP1R10 | 88.4502232 | -0.329953181 | 0.28672713 | -1.1507567 | 0.24983233 | 0.80672756 |
| MRPS18B | 62.4784084 | 0.105084784 | 0.26583926 | 0.39529445 | 6.93E-01 | 0.96220115 |
| ATAT1 | 30.8382511 | -0.119590012 | 0.28177384 | -0.4244184 | 0.67126068 | 0.95800185 |
| DHX16 | 31.4142214 | 0.034642681 | 0.31621244 | 0.10955509 | 0.91276223 | 0.99113939 |
| MDC1 | 48.452967 | -0.05031858 | 0.28124868 | -0.1789113 | 0.85800731 | 0.9864199 |
| TUBB | 220.76236 | -0.874062646 | 0.69407887 | -1.2593131 | 0.20791725 | 0.77435432 |
| FLOT1 | 192.426928 | 0.28858318 | 0.27858279 | 1.03589737 | 0.30025003 | 0.84194819 |
| IER3 | 277.218578 | 0.605714469 | 1.11303557 | 0.54420046 | 0.58630353 | 0.9423304 |
| DDR1 | 135.568181 | 0.299540235 | 0.36142184 | 0.82878289 | 0.40722727 | 0.89585737 |
| CCHCR1 | 41.9631413 | 0.741379698 | 0.33530183 | 2.21108159 | 0.02703019 | 0.42391395 |
| POU5F1 | 165.825053 | -0.354693518 | 0.39403593 | -0.9001553 | 0.36803763 | 0.87605028 |
| HLA-C | 336.1179 | -1.187142773 | 0.61839085 | -1.9197289 | 0.05489215 | 0.53804407 |
| HLA-B | 546.436371 | -0.784022526 | 0.55533466 | -1.4118019 | 0.1580083 | 0.73580079 |
| MICA | 49.2245869 | 0.09690047 | 0.37728616 | 0.25683548 | 0.79730578 | 0.97552588 |
| NFKBIL1 | 26.3936454 | 0.297173754 | 0.36660259 | 0.81061553 | 0.4175865 | 0.89862171 |
| PRRC2A | 108.916487 | -0.140971905 | 0.29078718 | -0.4847941 | 0.62782242 | 0.94858781 |
| BAG6 | 54.887353 | -0.43006495 | 0.32666347 | -1.3165382 | 0.18799345 | 0.76207437 |
| DDAH2 | 72.5166033 | 0.207828559 | 0.46434734 | 0.44757133 | 0.6544626 | 0.95280477 |
| CLIC1 | 307.539366 | 0.020543486 | 0.44748223 | 0.04590906 | 9.63E-01 | 0.99572129 |
| VARS | 34.7044262 | -0.970579107 | 0.46670644 | -2.0796351 | 0.03755901 | 0.46930788 |
| LSM2 | 47.0015543 | -0.362657471 | 0.36966096 | -0.9810543 | 0.32656597 | 0.85588725 |
| HSPA1A | 229.553822 | 1.653591371 | 0.74179856 | 2.22916498 | 0.02580293 | 0.41892588 |
| HSPA1B | 114.53796 | 1.490949174 | 0.73634053 | 2.02480933 | 0.04288695 | 0.49422552 |
| C6orf48 | 70.471127 | 0.467995492 | 0.50759173 | 0.92199195 | 0.35653277 | 0.87316144 |
| NEU1 | 58.477918 | 0.851623293 | 0.46338805 | 1.83781885 | 0.0660891 | 0.57792071 |
| SLC44A4 | 117.689255 | 1.310219359 | 0.35335532 | 3.70793732 | 0.00020895 | 0.03842424 |
| EHMT2 | 40.2160536 | -0.220443743 | 0.41917598 | -0.5258978 | 0.59895918 | 0.94683415 |
| NELFE | 28.3834457 | -0.478873814 | 0.46357752 | -1.0329962 | 0.30160568 | 0.84328103 |
| STK19 | 43.1656853 | 0.934855347 | 0.36653303 | 2.55053508 | 0.01075577 | 0.29950442 |
| ATF6B | 66.3590222 | 0.01689669 | 0.26539604 | 0.06366594 | 0.94923622 | 0.9952642 |
| AGPAT1 | 32.7243008 | 0.298512956 | 0.39499439 | 0.75573973 | 0.44980526 | 0.91034958 |
| RNF5 | 45.8472678 | 0.082716428 | 0.59877411 | 0.13814296 | 0.89012743 | 0.98766275 |
| PBX2 | 52.1282099 | -0.28208008 | 0.23169791 | -1.2174477 | 0.22343393 | 0.78791697 |
| AL662796.1 | 65.7684007 | -0.560847831 | 0.67783538 | -0.8274101 | 0.40800467 | 0.8959746 |
| HLA-DRA | 47.7163122 | 0.690844306 | 0.84965519 | 0.81308785 | 0.41616769 | 0.89846075 |
| HLA-DRB1 | 43.4488905 | -1.387060513 | 1.32406857 | -1.0475745 | 0.29483467 | 0.83911606 |
| PSMB8 | 69.2958444 | -0.223250743 | 0.44879819 | -0.4974413 | 0.61887791 | 0.9480368 |
| PSMB9 | 42.7819892 | -1.773637267 | 0.84284965 | -2.1043341 | 0.03534931 | 0.45634914 |

| | | | | | | |
|---|---|---|---|---|---|---|
| TAP1 | 57.2970886 | -1.559729449 | 0.77970104 | -2.0004199 | 0.04545495 | 0.50469451 |
| BRD2 | 143.768844 | -0.325406956 | 0.27114984 | -1.2001001 | 0.23010046 | 0.79370986 |
| RXRB | 26.4157007 | 0.012525403 | 0.41551 | 0.03014465 | 0.97595169 | 0.99681784 |
| SLC39A7 | 45.0911371 | 0.170663677 | 0.61032236 | 0.27962875 | 0.77976235 | 0.97423208 |
| RING1 | 53.6651206 | 0.137931594 | 0.31560629 | 0.4370369 | 0.66208459 | 0.9545911 |
| VPS52 | 44.9227431 | 0.026271388 | 0.25580871 | 0.10269935 | 0.91820159 | 0.99214128 |
| RPS18 | 332.947849 | 0.003238545 | 0.56510508 | 0.00573087 | 0.99542745 | 0.99866813 |
| PFDN6 | 39.2343338 | -0.566338014 | 0.3598274 | -1.5739158 | 0.11550693 | 0.67260842 |
| RGL2 | 65.0897614 | 0.401994217 | 0.29800778 | 1.34893868 | 0.17735666 | 0.75126783 |
| TAPBP | 223.70391 | -0.141368539 | 0.38281638 | -0.3692855 | 0.71191493 | 0.96587538 |
| PHF1 | 50.2813847 | 0.099394381 | 0.32919157 | 0.30193477 | 0.76270179 | 0.9714717 |
| CUTA | 75.571318 | 0.012463002 | 0.48763209 | 0.02555821 | 0.97960972 | 0.99681784 |
| SYNGAP1 | 27.0604509 | 0.612900948 | 0.30882585 | 1.98461674 | 0.04718713 | 0.51284792 |
| ITPR3 | 149.363511 | -0.606376486 | 0.3609342 | -1.6800195 | 0.09295353 | 0.63691121 |
| UQCC2 | 40.3762109 | -0.355430686 | 0.39600215 | -0.8975474 | 0.36942691 | 0.87605028 |
| LEMD2 | 55.1278239 | -0.087130051 | 0.24862786 | -0.3504436 | 0.72600578 | 0.96781837 |
| HMGA1 | 45.8331926 | -1.747626527 | 0.70641972 | -2.4739209 | 0.01336393 | 0.32022149 |
| NUDT3 | 90.7802989 | -0.404437101 | 0.45841569 | -0.8822497 | 0.37764179 | 0.87924711 |
| C6orf106 | 213.987178 | 0.280082172 | 0.16696842 | 1.677456 | 0.09345335 | 0.63721577 |
| SNRPC | 45.9219877 | -0.450630286 | 0.47827997 | -0.9421893 | 0.34609571 | 0.86990996 |
| UHRF1BP1 | 98.5837325 | -0.068071097 | 0.2280999 | -0.2984267 | 0.76537752 | 0.97177982 |
| TAF11 | 55.8687578 | 0.049237161 | 0.3257721 | 0.1511399 | 0.87986536 | 0.98766275 |
| ANKS1A | 298.369924 | 0.140328298 | 0.19225254 | 0.7299165 | 0.46544123 | 0.91462428 |
| ZNF76 | 61.2281155 | -0.012378222 | 0.25916966 | -0.0477611 | 0.96190666 | 0.99572129 |
| PPARD | 102.505588 | 0.752494384 | 0.337011 | 2.23284812 | 0.02555896 | 0.41708568 |
| RPL10A | 366.048712 | 0.302192374 | 0.55785359 | 0.54170552 | 0.58802138 | 0.94428446 |
| TEAD3 | 68.4372596 | -0.123372323 | 0.32375705 | -0.3810645 | 0.70315537 | 0.9645815 |
| FKBP5 | 463.951222 | -0.337374003 | 1.23217673 | -0.2738033 | 0.78423582 | 0.97423208 |
| SRPK1 | 155.098875 | -0.234573518 | 0.36600684 | -0.6408993 | 0.52158812 | 0.93272217 |
| MAPK14 | 248.842741 | 0.245734763 | 0.32453175 | 0.7571979 | 0.44893131 | 0.90994378 |
| MAPK13 | 29.949429 | -0.648066033 | 0.57290597 | -1.1311909 | 0.25797474 | 0.81223967 |
| BRPF3 | 35.1082082 | 0.283490918 | 0.37933768 | 0.74733129 | 0.45486361 | 0.91077355 |
| KCTD20 | 140.101772 | 0.172073657 | 0.15936302 | 1.07975904 | 0.28024949 | 0.82524128 |
| STK38 | 283.612152 | 0.010153795 | 0.29705949 | 0.03418102 | 0.9727328 | 0.99681784 |
| SRSF3 | 187.723201 | -0.487024132 | 0.30163671 | -1.614605 | 0.10639627 | 0.65699665 |
| CDKN1A | 69.2135544 | 0.541853007 | 0.69249758 | 0.78246195 | 0.43394314 | 0.90514499 |
| C6orf89 | 124.052819 | 0.230849206 | 0.23719614 | 0.97324183 | 0.33043312 | 0.85923151 |
| MTCH1 | 132.331639 | -0.256842164 | 0.27630781 | -0.9295509 | 0.35260368 | 0.8720098 |
| PIM1 | 35.4549085 | 0.142106252 | 0.52865996 | 0.26880464 | 0.78808002 | 0.97423208 |
| TBC1D22B | 98.0457901 | 0.166082354 | 0.21990728 | 0.75523809 | 0.45010614 | 0.91037995 |
| RNF8 | 63.4141018 | -0.404318405 | 0.37244319 | -1.0855841 | 0.27766303 | 0.8229426 |
| CMTR1 | 51.6817406 | -0.619945353 | 0.41654777 | -1.4882935 | 0.1366735 | 0.70997583 |
| ZFAND3 | 1561.23984 | -0.122317552 | 0.21902109 | -0.5584738 | 0.57652086 | 0.94034284 |
| BTBD9 | 628.304327 | -0.100906933 | 0.27733248 | -0.3638482 | 0.71597135 | 0.96709741 |
| GLO1 | 93.478953 | 0.014787888 | 0.25220334 | 0.05863479 | 0.953243 | 0.99531843 |
| KCNK5 | 75.3266352 | 0.126284315 | 0.22118013 | 0.57095686 | 0.56802889 | 0.93910057 |
| KIF6 | 64.2328012 | 0.359591832 | 0.59322574 | 0.60616357 | 0.54440614 | 0.93523904 |
| UNC5CL | 123.009644 | 0.824495366 | 0.60166676 | 1.37035219 | 0.17057699 | 0.74445571 |
| OARD1 | 64.5615538 | 0.107998663 | 0.33260919 | 0.32470138 | 0.74540709 | 0.97028355 |
| NFYA | 55.0510178 | 0.030916325 | 0.27724085 | 0.11151432 | 0.91120851 | 0.99100813 |
| FOXP4 | 38.045192 | -0.047147506 | 0.70748176 | -0.0666413 | 0.94686726 | 0.9952642 |
| TFEB | 39.5473831 | 0.665555591 | 0.49112761 | 1.35515815 | 0.17536717 | 0.74934286 |
| USP49 | 66.083892 | 0.53918223 | 0.30349275 | 1.77659014 | 0.0756357 | 0.59499139 |
| CCND3 | 375.870364 | -0.323114134 | 0.62366627 | -0.5180882 | 0.60439674 | 0.94683415 |

| | | | | | | |
|---|---|---|---|---|---|---|
| TAF8 | 52.1624227 | 0.310997402 | 0.25626636 | 1.2135709 | 0.22491162 | 0.78917125 |
| C6orf132 | 58.6310719 | -0.475948274 | 0.57289291 | -0.8307805 | 0.40609762 | 0.89569633 |
| MRPS10 | 27.3303282 | -0.086388099 | 0.31238442 | -0.2765442 | 0.78213011 | 0.97423208 |
| TRERF1 | 33.051554 | 0.190186693 | 0.69788901 | 0.27251711 | 0.78522444 | 0.97423208 |
| UBR2 | 392.785497 | 0.000861628 | 0.16545402 | 0.00520766 | 0.99584491 | 0.99866813 |
| BICRAL | 254.404003 | -0.136133237 | 0.30172169 | -0.4511881 | 0.65185398 | 0.95159942 |
| RPL7L1 | 124.100183 | -0.316563685 | 0.25852419 | -1.2245032 | 0.22076247 | 0.78612778 |
| CNPY3 | 32.6119318 | -0.633257929 | 0.35368755 | -1.7904445 | 0.07338248 | 0.59320069 |
| PEX6 | 52.7314216 | 1.017090767 | 0.32239176 | 3.15482866 | 0.00160592 | 0.13036563 |
| PPP2R5D | 27.3581765 | 0.332630718 | 0.30254453 | 1.09944384 | 0.27157452 | 0.81831151 |
| MEA1 | 22.760679 | -0.471265149 | 0.53559774 | -0.8798864 | 0.37892087 | 0.87973265 |
| KLHDC3 | 38.7194796 | -0.107163859 | 0.39152798 | -0.2737068 | 0.78430998 | 0.97423208 |
| RRP36 | 52.3535995 | -0.440514108 | 0.24738596 | -1.7806755 | 0.07496548 | 0.59444897 |
| CUL7 | 32.0258591 | -0.423810281 | 0.32053116 | -1.3222124 | 0.18609742 | 0.76083106 |
| KLC4 | 42.670076 | 0.213614575 | 0.32046009 | 0.66658713 | 0.50503589 | 0.9257297 |
| MRPL2 | 32.1228741 | 0.205490194 | 0.41257786 | 0.49806403 | 0.61843891 | 0.9480368 |
| PTK7 | 89.3760935 | 0.023747812 | 0.36788094 | 0.06455298 | 0.94852992 | 0.9952642 |
| CUL9 | 41.3009962 | -0.050519359 | 0.29179055 | -0.1731357 | 0.86254477 | 0.98669442 |
| DNPH1 | 74.4455185 | 0.098057862 | 0.49166009 | 0.19944239 | 0.84191671 | 0.98517901 |
| ZNF318 | 37.2457171 | 0.293964504 | 0.34324828 | 0.85641946 | 0.39176581 | 0.88709734 |
| ABCC10 | 35.2841992 | -0.348662858 | 0.31909516 | -1.0926611 | 0.27454264 | 0.82105649 |
| TJAP1 | 40.3963099 | -0.118949805 | 0.25093277 | -0.4740306 | 0.6354781 | 0.94932781 |
| POLR1C | 84.7577267 | -0.039675967 | 0.25175306 | -0.1575987 | 0.87477299 | 0.98766275 |
| YIPF3 | 70.2358126 | 0.342582182 | 0.39143545 | 0.87519457 | 0.38146805 | 0.88128509 |
| XPO5 | 261.370461 | -0.163245959 | 0.34171117 | -0.4777308 | 0.6328418 | 0.94932781 |
| POLH | 45.3589713 | 0.994017427 | 0.32229939 | 3.08414307 | 0.00204139 | 0.14702642 |
| GTPBP2 | 50.0363656 | -0.108266751 | 0.27777488 | -0.3897644 | 0.6967108 | 0.96220115 |
| MRPS18A | 31.3839565 | -0.031347941 | 0.30466522 | -0.1028931 | 0.91804784 | 0.99214128 |
| VEGFA | 138.767129 | 0.152431518 | 0.50858689 | 0.29971578 | 0.76439396 | 0.97177982 |
| MRPL14 | 71.0518576 | -0.060624678 | 0.48751249 | -0.1243551 | 0.9010341 | 0.9900784 |
| TMEM63B | 46.1045821 | -0.111539123 | 0.26933062 | -0.4141346 | 0.67877555 | 0.95924874 |
| SLC29A1 | 43.1393262 | -0.324356485 | 0.41251054 | -0.7862987 | 0.43169254 | 0.90498 |
| HSP90AB1 | 488.56661 | -0.393009283 | 0.41109413 | -0.956008 | 0.33906819 | 0.86680923 |
| SLC35B2 | 31.2413388 | -0.378786363 | 0.4934407 | -0.7676431 | 0.44269923 | 0.90615787 |
| CDC5L | 73.6833114 | 0.007859684 | 0.23119606 | 0.03399575 | 0.97288054 | 0.99681784 |
| SUPT3H | 598.412796 | -0.062487042 | 0.29883239 | -0.209104 | 0.83436707 | 0.98367279 |
| RUNX2 | 62.5515913 | 1.424053864 | 0.81695599 | 1.74312188 | 0.08131233 | 0.60850843 |
| CLIC5 | 150.264216 | 0.89790026 | 0.84207244 | 1.06629811 | 0.2862889 | 0.83026382 |
| ENPP4 | 45.3018685 | 0.030598077 | 0.36587827 | 0.08362912 | 0.93335131 | 0.99425828 |
| ENPP5 | 46.3209081 | 0.45808154 | 0.34145355 | 1.34156327 | 0.17973764 | 0.75265164 |
| RCAN2 | 143.656609 | 0.379042538 | 0.47824141 | 0.79257574 | 0.42802505 | 0.90272191 |
| CYP39A1 | 33.9590705 | 0.996279786 | 0.58112808 | 1.71438933 | 0.08645725 | 0.62381659 |
| SLC25A27 | 136.480833 | 0.645670078 | 0.57395736 | 1.12494433 | 0.26061263 | 0.81263292 |
| ADGRF1 | 162.535832 | -1.085064039 | 0.58527398 | -1.853942 | 0.06374745 | 0.57068863 |
| TNFRSF21 | 44.4597439 | 0.919076064 | 0.51973925 | 1.7683407 | 0.07700396 | 0.59726395 |
| CD2AP | 554.027686 | -0.02465317 | 0.21274461 | -0.1158815 | 0.90774644 | 0.99100813 |
| PTCHD4 | 124.237102 | 1.344410885 | 0.61904416 | 2.1717528 | 0.02987431 | 0.43492739 |
| MUT | 54.0029092 | 0.092109471 | 0.35937592 | 0.25630396 | 0.79771614 | 0.9756541 |
| PKHD1 | 4334.9091 | 0.001732193 | 0.6569834 | 0.00263659 | 0.99789631 | 0.99956591 |
| MCM3 | 47.9548138 | -1.283024531 | 0.27523269 | -4.6615994 | 3.14E-06 | 0.00200368 |
| EFHC1 | 146.738725 | 0.582116088 | 0.24165762 | 2.40884637 | 0.01600303 | 0.3414008 |
| TRAM2 | 83.2538741 | -0.286712083 | 0.38730229 | -0.7402798 | 0.45913021 | 0.91362347 |
| TMEM14A | 24.5474834 | 0.525113535 | 0.50803191 | 1.03362313 | 0.30131238 | 0.84277781 |
| GSTA4 | 47.4111513 | 0.137231889 | 0.45976516 | 0.29848257 | 0.76533487 | 0.97177982 |

| | | | | | | |
|---|---|---|---|---|---|---|
| ICK | 44.3823073 | 0.056450845 | 0.34664344 | 0.16284989 | 0.87063663 | 0.98766275 |
| FBXO9 | 130.552415 | 0.212472254 | 0.24287856 | 0.8748086 | 0.38167806 | 0.88128509 |
| ELOVL5 | 130.835268 | 0.231609483 | 0.37768955 | 0.61322714 | 0.53972616 | 0.93523904 |
| GCLC | 94.5839546 | 0.052046035 | 0.37337119 | 0.13939489 | 8.89E-01 | 9.88E-01 |
| LRRC1 | 169.08026 | 0.323236065 | 0.3097516 | 1.04353316 | 0.29670143 | 0.83973489 |
| TINAG | 113.289505 | 1.422257317 | 0.53278249 | 2.66948962 | 0.00759666 | 0.26951269 |
| FAM83B | 68.1733454 | -0.973998304 | 0.53357992 | -1.8254028 | 0.06794026 | 0.5809835 |
| DST | 607.720553 | 0.286213903 | 0.59123738 | 0.48409304 | 0.62831984 | 0.94858781 |
| ZNF451 | 190.565985 | 0.259827848 | 0.1776974 | 1.46219272 | 0.14368841 | 0.71631428 |
| PRIM2 | 179.727391 | -0.460738905 | 0.28644674 | -1.6084627 | 0.10773387 | 0.65699665 |
| AL021368.2 | 32.2310398 | 0.155496982 | 0.32398278 | 0.47995447 | 0.63125977 | 0.94932781 |
| PTP4A1 | 204.058403 | -0.276003594 | 0.65059865 | -0.4242302 | 0.67139794 | 0.95800185 |
| PHF3 | 363.322882 | 0.305702379 | 0.19123612 | 1.59855982 | 0.10991844 | 0.65905549 |
| LMBRD1 | 283.596832 | 0.094131221 | 0.20116064 | 0.46794054 | 0.63982711 | 0.94944486 |
| COL19A1 | 109.483733 | 0.538279029 | 0.30691982 | 1.75380992 | 0.07946308 | 0.60556378 |
| FAM135A | 408.477036 | -0.227086485 | 0.19865801 | -1.1431026 | 2.53E-01 | 0.80798674 |
| SMAP1 | 228.915155 | -0.262696421 | 0.20068389 | -1.309006 | 0.19053231 | 0.76219391 |
| OGFRL1 | 175.5462 | -0.694289689 | 0.45474023 | -1.5267831 | 0.12681496 | 0.69494309 |
| AL136164.2 | 438.015055 | -0.978751351 | 0.55686305 | -1.7576159 | 0.07881288 | 0.60395889 |
| LINC00472 | 109.189869 | -0.641953848 | 0.55318575 | -1.1604671 | 0.24585869 | 0.80488052 |
| AC019205.1 | 45.0836364 | 0.455554154 | 0.40048603 | 1.13750323 | 0.25532797 | 0.80897837 |
| OOEP | 316.562548 | -0.034067428 | 0.49709933 | -0.0685324 | 0.9453618 | 0.9952642 |
| MTO1 | 52.9290345 | 0.014110511 | 0.22889833 | 0.06164532 | 0.95084529 | 0.99531843 |
| EEF1A1 | 1677.0619 | 0.638446855 | 0.55433665 | 1.15173126 | 0.24943152 | 0.80672756 |
| SLC17A5 | 75.9056175 | 0.002299588 | 0.27961402 | 0.00822415 | 0.99343815 | 0.99866813 |
| COL12A1 | 71.901333 | 0.424485241 | 0.48053498 | 0.88335971 | 3.77E-01 | 8.78E-01 |
| COX7A2 | 98.1141921 | -0.001527373 | 0.48042264 | -0.0031792 | 0.99746335 | 0.99934122 |
| TMEM30A | 127.879651 | 0.100303414 | 0.35047914 | 0.2861894 | 0.77473306 | 0.97423208 |
| SENP6 | 673.830103 | -0.192586753 | 0.16008587 | -1.2030216 | 0.22896797 | 0.7931636 |
| MYO6 | 611.073676 | 0.191163403 | 0.33096593 | 0.5775924 | 0.56353934 | 0.93809297 |
| IRAK1BP1 | 145.906289 | 0.408331143 | 0.30942731 | 1.31963509 | 0.18695688 | 0.76207437 |
| PHIP | 537.109448 | -0.140154894 | 0.16896278 | -0.8295016 | 0.40682063 | 0.89585737 |
| HMGN3 | 230.336187 | 0.166082309 | 0.239305 | 0.6940194 | 0.48767004 | 0.92034617 |
| LCA5 | 57.5400827 | 0.656708999 | 0.37741033 | 1.74003982 | 0.08185203 | 0.61159169 |
| SH3BGRL2 | 84.3063908 | -0.04610051 | 0.42213143 | -0.1092089 | 0.9130368 | 0.99113939 |
| BCKDHB | 265.455969 | -0.410949265 | 0.29936139 | -1.372753 | 0.16982913 | 0.74445571 |
| TENT5A | 58.0138775 | -1.572089301 | 0.59146929 | -2.657939 | 0.00786201 | 0.27165021 |
| IBTK | 239.984436 | -0.653649236 | 0.22058465 | -2.963258 | 0.00304401 | 0.17999135 |
| TPBG | 34.8750525 | 0.003652568 | 0.38672136 | 0.00944496 | 0.99246412 | 0.99866813 |
| UBE3D | 186.321895 | -0.277673958 | 0.31773373 | -0.8739203 | 0.38216165 | 0.88146556 |
| DOPEY1 | 237.925493 | 0.007223657 | 0.1980391 | 0.03647591 | 0.97090289 | 0.99681784 |
| PGM3 | 58.3531188 | 0.836368913 | 0.37369262 | 2.23811994 | 0.02521323 | 0.4135769 |
| ME1 | 54.1242176 | -0.068868045 | 0.28736758 | -0.2396514 | 0.81060051 | 0.97930627 |
| CYB5R4 | 65.7371814 | -0.362406028 | 0.26921703 | -1.3461482 | 0.17825472 | 0.7513166 |
| CEP162 | 136.634913 | 0.204132014 | 0.26982685 | 0.75652965 | 0.4493317 | 0.90994378 |
| SNX14 | 204.238371 | 0.123332958 | 0.16342658 | 0.75466889 | 0.45044768 | 0.91049553 |
| SYNCRIP | 100.512728 | -0.583737099 | 0.28021369 | -2.0831855 | 0.03723432 | 0.46890807 |
| ZNF292 | 770.015828 | -0.332133009 | 0.33170712 | -1.0012839 | 0.31668956 | 0.8490543 |
| SMIM8 | 101.597433 | -0.134728489 | 0.36015478 | -0.374085 | 0.7083411 | 0.96587538 |
| RARS2 | 159.277214 | -0.01661537 | 0.20000107 | -0.0830764 | 0.93379079 | 0.9943329 |
| ORC3 | 134.449318 | -0.404627144 | 0.23917999 | -1.6917266 | 0.09069812 | 0.6330877 |
| AKIRIN2 | 46.6806732 | 0.195224847 | 0.31793415 | 0.61404177 | 0.53918773 | 0.93523904 |
| RNGTT | 281.776359 | -0.39834915 | 0.14836238 | -2.684974 | 0.00725354 | 0.26418897 |
| PNRC1 | 146.67081 | 0.507380106 | 0.46144892 | 1.09953689 | 0.27153395 | 0.81831151 |

| | | | | | | |
|---|---|---|---|---|---|---|
| SRSF12 | 33.4033227 | 0.436648956 | 0.35913978 | 1.21581898 | 0.22405389 | 0.78828752 |
| PM20D2 | 76.9199013 | 0.062862795 | 0.23707085 | 0.26516459 | 0.79088267 | 0.97423208 |
| UBE2J1 | 75.8368834 | -0.044656098 | 0.25090356 | -0.1779811 | 0.8587378 | 0.9864199 |
| RRAGD | 47.3549583 | 0.221452489 | 0.37151192 | 0.59608448 | 5.51E-01 | 0.93523904 |
| ANKRD6 | 250.880142 | 0.491238782 | 0.41902063 | 1.17234988 | 0.24105661 | 0.80319133 |
| LYRM2 | 74.6886564 | 0.398974164 | 0.29108663 | 1.37063723 | 0.17048807 | 0.74445571 |
| MDN1 | 202.686918 | -0.444775663 | 0.24136447 | -1.8427553 | 0.06536474 | 0.57495764 |
| CASP8AP2 | 83.648464 | -0.375351428 | 0.23144465 | -1.6217762 | 0.10485127 | 0.6568805 |
| BACH2 | 208.160736 | 0.584187178 | 0.55952347 | 1.04407984 | 0.29644845 | 0.83973489 |
| MAP3K7 | 187.497496 | -0.001824522 | 0.24001278 | -0.0076018 | 0.99393472 | 0.99866813 |
| EPHA7 | 576.041632 | 0.245168251 | 0.36021806 | 0.68061065 | 0.49611788 | 0.92334178 |
| MANEA | 41.3618681 | 0.135108473 | 0.33646006 | 0.40155873 | 0.68800881 | 0.96161098 |
| FUT9 | 96.0936242 | -0.763679867 | 1.14478613 | -0.6670939 | 0.50471214 | 0.92564381 |
| UFL1-AS1 | 66.9634034 | 0.109942579 | 0.4781359 | 0.22994002 | 0.81813838 | 0.98010849 |
| UFL1 | 120.107773 | -0.26553578 | 0.22096615 | -1.2017034 | 0.22947846 | 0.79370986 |
| NDUFAF4 | 28.9045543 | 0.234420499 | 0.46930861 | 0.49950181 | 0.61742591 | 0.94791646 |
| KLHL32 | 69.0169404 | 0.468788191 | 0.54600402 | 0.85858011 | 0.39057222 | 0.88670817 |
| MMS22L | 72.0418978 | -0.74885514 | 0.30690936 | -2.4399879 | 0.01468775 | 0.33026756 |
| FBXL4 | 227.955397 | -0.124223126 | 0.2919253 | -0.4255305 | 0.67044997 | 0.9576857 |
| PNISR | 537.611139 | 0.248216264 | 0.1936532 | 1.28175661 | 2.00E-01 | 0.77110248 |
| AL513550.1 | 28.577104 | -0.129443574 | 0.51186969 | -0.2528838 | 0.80035798 | 0.97723068 |
| USP45 | 108.085863 | 0.152252455 | 0.29468318 | 0.51666491 | 0.6053901 | 0.94699994 |
| CCNC | 178.325293 | -0.079869116 | 0.26148471 | -0.3054447 | 0.76002749 | 0.9714717 |
| SIM1 | 236.827746 | -0.392472726 | 0.79882754 | -0.491311 | 0.62320653 | 0.94808496 |
| ASCC3 | 416.790711 | 0.071423748 | 0.20600213 | 0.34671364 | 0.72880646 | 0.96840575 |
| HACE1 | 108.240022 | 0.176624985 | 0.2879536 | 0.61338002 | 0.53962509 | 0.93523904 |
| PREP | 69.679379 | -0.347651805 | 0.22136522 | -1.5704897 | 0.11630122 | 0.67559092 |
| ATG5 | 186.857765 | -0.064481433 | 0.20047823 | -0.3216381 | 0.7477269 | 0.97028355 |
| CRYBG1 | 93.7189409 | 0.096907661 | 0.36453808 | 0.26583686 | 0.79036485 | 0.97423208 |
| RTN4IP1 | 43.5771321 | -0.407268363 | 0.522699 | -0.7791642 | 0.43588298 | 0.90514499 |
| QRSL1 | 64.9436631 | -0.474535429 | 0.3288711 | -1.4429223 | 0.14904237 | 0.72246729 |
| LINC02532 | 942.84189 | 0.630974895 | 0.49189329 | 1.28274751 | 0.19958055 | 0.77110248 |
| CD24 | 1140.62738 | 0.311252437 | 0.34046843 | 0.91418884 | 0.36061763 | 0.87429201 |
| C6orf203 | 32.7790636 | 0.113890554 | 0.30629745 | 0.37182991 | 0.71001949 | 0.96587538 |
| PDSS2 | 337.606018 | -0.03713242 | 0.1842233 | -0.201562 | 0.84025914 | 0.98464453 |
| SOBP | 91.2658972 | 0.546594299 | 0.91660769 | 0.59632306 | 0.55095943 | 0.93523904 |
| SEC63 | 250.276663 | 0.144714642 | 0.23221164 | 0.6232015 | 0.53315212 | 0.93433299 |
| OSTM1 | 66.6885651 | -0.165201628 | 0.26193659 | -0.6306932 | 0.52824113 | 0.93366189 |
| SNX3 | 151.2504 | 0.369951428 | 0.24357321 | 1.51885104 | 0.12879999 | 0.69783662 |
| AFG1L | 111.721082 | -0.320617961 | 0.31372847 | -1.02196 | 0.30679982 | 0.84488735 |
| FOXO3 | 615.586533 | 0.036134966 | 0.36614095 | 0.09869141 | 0.92138329 | 0.99214128 |
| ARMC2 | 130.676607 | 0.825244208 | 0.29043779 | 2.84138025 | 0.00449187 | 0.2151382 |
| SESN1 | 158.870611 | 0.23539045 | 0.2688952 | 0.87539847 | 0.38135713 | 0.88128509 |
| CEP57L1 | 55.3978701 | -0.044067337 | 0.35811155 | -0.1230548 | 0.90206372 | 0.9900784 |
| CD164 | 250.392316 | -0.077022579 | 0.21672472 | -0.3553936 | 0.72229474 | 0.96779196 |
| PPIL6 | 103.23514 | 0.647690542 | 0.42372828 | 1.52855159 | 0.12637564 | 0.69492095 |
| AK9 | 97.0745165 | 0.658981465 | 0.42337455 | 1.5564976 | 0.11958981 | 0.68618625 |
| FIG4 | 98.3849926 | -0.053175551 | 0.23781456 | -0.2236009 | 0.82306786 | 0.98094663 |
| WASF1 | 80.7522342 | 0.740883423 | 0.23843619 | 3.10726076 | 0.0018883 | 0.14133317 |
| CDC40 | 66.3151652 | 0.51357315 | 0.28219983 | 1.81989178 | 0.06877549 | 0.5830092 |
| CDK19 | 238.443323 | 0.702605158 | 0.28544031 | 2.46147838 | 0.01383657 | 0.32248306 |
| AMD1 | 119.751539 | -0.374414438 | 0.33714054 | -1.1105589 | 0.26675825 | 0.81462808 |
| GTF3C6 | 80.744456 | -0.235961434 | 0.51868467 | -0.4549227 | 0.64916485 | 0.95027667 |
| RPF2 | 54.9917987 | -0.690270569 | 0.25424803 | -2.7149495 | 0.00662859 | 0.25396993 |

| | | | | | | |
|---|---|---|---|---|---|---|
| SLC16A10 | 140.395372 | 1.143138804 | 0.51437525 | 2.22238298 | 0.02625743 | 0.42051885 |
| MFSD4B | 46.4537259 | 0.319673846 | 0.30203213 | 1.05841008 | 0.28986853 | 0.83506029 |
| REV3L | 430.333091 | 0.338905866 | 0.24533079 | 1.38142412 | 0.16714859 | 0.74445571 |
| TRAF3IP2-AS | 228.670439 | 0.267139542 | 0.3156792 | 0.84623739 | 0.39742033 | 0.88863925 |
| TRAF3IP2 | 49.0768319 | 0.743197642 | 0.38488685 | 1.93095097 | 0.05348912 | 0.53392413 |
| FYN | 160.514734 | -0.042328388 | 0.44843114 | -0.0943922 | 0.92479763 | 0.99311954 |
| TUBE1 | 59.9764848 | -0.246056883 | 0.39895408 | -0.6167549 | 0.53739641 | 0.93504069 |
| FAM229B | 32.3180556 | 0.574544658 | 0.30207929 | 1.90196637 | 0.05717555 | 0.54616991 |
| MARCKS | 133.632527 | 0.046443787 | 0.30460412 | 0.15247261 | 0.87881419 | 0.98766275 |
| HDAC2 | 99.4865115 | -0.13295484 | 0.23587239 | -0.5636728 | 0.57297686 | 0.93998433 |
| FRK | 140.473035 | 0.745733488 | 0.3586807 | 2.07910124 | 0.03760805 | 0.46930788 |
| NT5DC1 | 227.576566 | 0.062013777 | 0.26305467 | 0.23574482 | 0.81363069 | 0.97930627 |
| TSPYL4 | 32.2822868 | 0.246629376 | 0.37294873 | 0.66129566 | 0.50842273 | 0.92666294 |
| DSE | 191.784007 | 0.283844845 | 0.38591925 | 0.7355032 | 0.46203309 | 0.91462428 |
| TSPYL1 | 88.5692668 | 0.092358747 | 0.33093868 | 0.27908115 | 0.78018255 | 0.97423208 |
| RWDD1 | 57.2751144 | -0.29381378 | 0.4163069 | -0.7057625 | 0.48033587 | 0.91971919 |
| ZUP1 | 38.8389854 | -0.596874771 | 0.35340709 | -1.6889157 | 0.09123559 | 0.63421316 |
| KPNA5 | 141.370304 | 0.490705357 | 0.26810636 | 1.83026374 | 0.06721051 | 0.57884645 |
| DCBLD1 | 87.4535931 | -0.405812654 | 0.39371432 | -1.0307287 | 0.30266805 | 0.84398487 |
| GOPC | 91.7950659 | -0.018831765 | 0.24943222 | -0.0754985 | 0.93981807 | 0.99526299 |
| NUS1 | 94.9020848 | -0.147134441 | 0.27796164 | -0.5293336 | 0.59657404 | 0.94683415 |
| SLC35F1 | 31.4922715 | 0.10068377 | 0.2992991 | 0.3363985 | 0.73657038 | 0.9698508 |
| CEP85L | 129.332603 | 0.263056366 | 0.35206995 | 0.74717074 | 0.45496051 | 0.91077355 |
| MCM9 | 147.752201 | -0.243291392 | 0.21852765 | -1.1133209 | 0.26557065 | 0.81387543 |
| AL365275.1 | 60.9213885 | -0.010968808 | 0.6527662 | -0.0168036 | 0.98659332 | 0.99764442 |
| FAM184A | 51.7787545 | 1.452798613 | 0.3461511 | 4.19700702 | 2.70E-05 | 0.00996458 |
| MAN1A1 | 239.167496 | 0.940904513 | 0.2817168 | 3.33989493 | 0.0008381 | 0.09295873 |
| TBC1D32 | 195.414352 | -0.260564192 | 0.26791891 | -0.9725487 | 0.33077765 | 0.85923151 |
| GJA1 | 35.7872941 | -1.537315655 | 0.88577861 | -1.7355529 | 0.08264292 | 0.6141478 |
| HSF2 | 41.4584393 | -0.05506041 | 0.28748299 | -0.1915258 | 0.84811367 | 0.98545133 |
| SERINC1 | 184.849679 | 0.166644293 | 0.26501451 | 0.62881195 | 0.52947217 | 0.93366189 |
| RNF217 | 247.195943 | 0.080015586 | 0.23055887 | 0.34705056 | 0.72855333 | 0.96833805 |
| TPD52L1 | 106.502868 | 0.213634237 | 0.61283967 | 0.34859727 | 0.72739168 | 0.9681562 |
| HDDC2 | 60.1577427 | -0.083541158 | 0.45318093 | -0.1843439 | 0.85374365 | 0.9864199 |
| NCOA7 | 345.108467 | 0.543631564 | 0.33954801 | 1.60104475 | 0.10936701 | 0.65764016 |
| HINT3 | 48.3806751 | 0.242196892 | 0.35779651 | 0.67691239 | 0.49846154 | 0.92398825 |
| TRMT11 | 117.351646 | -0.181952785 | 0.24360392 | -0.7469206 | 0.45511149 | 0.91077355 |
| RNF146 | 67.5559878 | 0.188271843 | 0.31827179 | 0.59154425 | 0.55415582 | 0.93523904 |
| ECHDC1 | 132.677352 | 0.109947467 | 0.18427249 | 0.59665697 | 0.55073643 | 0.93523904 |
| PTPRK | 961.184556 | -0.100362061 | 0.29616582 | -0.3388712 | 0.73470678 | 0.9698508 |
| ARHGAP18 | 134.88607 | -0.196376533 | 0.33527704 | -0.5857142 | 0.55806757 | 0.93620477 |
| L3MBTL3 | 63.8896696 | 0.058073872 | 0.32293605 | 0.17983088 | 0.85728534 | 0.9864199 |
| EPB41L2 | 242.196689 | 0.329772074 | 0.44487546 | 0.7412683 | 0.45853078 | 0.91350475 |
| AKAP7 | 89.1537197 | -0.757190249 | 0.53629256 | -1.4118977 | 0.15798007 | 0.73580079 |
| MED23 | 107.251367 | -0.075820508 | 0.20112088 | -0.3769897 | 0.70618123 | 0.96587538 |
| ENPP1 | 84.7876132 | 0.399997938 | 0.61557864 | 0.64979178 | 5.16E-01 | 0.93140514 |
| CTGF | 555.907138 | 0.889248976 | 1.18346757 | 0.75139277 | 0.45241631 | 0.91077355 |
| STX7 | 254.088147 | 0.13603609 | 0.29241366 | 0.46521797 | 0.64177537 | 0.94944486 |
| SLC18B1 | 47.6685928 | 0.316898261 | 0.39291117 | 0.80653921 | 0.41993201 | 0.90012434 |
| RPS12 | 301.674313 | 0.400535272 | 0.51876447 | 0.77209465 | 0.44005837 | 0.90561284 |
| TBPL1 | 44.674094 | 0.275315583 | 0.26469293 | 1.04013197 | 0.29827859 | 0.83973489 |
| SGK1 | 71.0748797 | -1.283976185 | 0.75060836 | -1.7105807 | 0.08715854 | 0.62585583 |
| ALDH8A1 | 39.1811764 | 0.841500166 | 0.42405177 | 1.9844279 | 0.04720816 | 0.51284792 |
| HBS1L | 116.390705 | 0.054740585 | 0.17867369 | 0.30637182 | 0.75932156 | 0.9714717 |

| | | | | | | |
|---|---|---|---|---|---|---|
| AHI1 | 488.65373 | 0.02562786 | 0.23274828 | 0.11010977 | 0.91232232 | 0.99113939 |
| LINC00271 | 285.454416 | 0.323027679 | 0.41143706 | 0.78512052 | 0.43238291 | 0.90514499 |
| PDE7B | 900.119723 | 0.253576304 | 0.33942058 | 0.74708582 | 0.45501176 | 0.91077355 |
| AL138828.1 | 149.895482 | -0.281110631 | 0.56905082 | -0.4939992 | 0.62130677 | 0.9480368 |
| BCLAF1 | 281.337207 | 0.145117647 | 0.13337684 | 1.08802733 | 0.27658304 | 0.8227916 |
| MAP7 | 577.490255 | -0.281695761 | 0.39571927 | -0.7118576 | 0.47655297 | 0.91819563 |
| MAP3K5 | 957.214952 | 0.470295894 | 0.58135135 | 0.80897016 | 0.41853231 | 0.89910765 |
| PEX7 | 66.8228888 | -0.175936107 | 0.24476438 | -0.7187978 | 0.47226549 | 0.91737234 |
| IFNGR1 | 115.43112 | 0.235636216 | 0.52670056 | 0.44738174 | 0.65459945 | 0.95280477 |
| TNFAIP3 | 61.8894273 | 0.590228832 | 1.33305066 | 0.44276549 | 0.65793536 | 0.95399905 |
| PERP | 221.013518 | -0.200035233 | 0.53769864 | -0.3720211 | 0.70987715 | 0.96587538 |
| ARFGEF3 | 92.4251092 | -0.332980574 | 0.54727176 | -0.6084373 | 0.54289746 | 0.93523904 |
| HEBP2 | 84.7232636 | -0.339831659 | 0.29552177 | -1.1499378 | 0.25016949 | 0.80672756 |
| NHSL1 | 87.1535639 | -0.655852112 | 0.37084434 | -1.7685375 | 0.07697109 | 0.59726395 |
| CCDC28A | 41.9153394 | 0.438743703 | 0.45020987 | 0.9745315 | 0.32979271 | 0.85851742 |
| REPS1 | 223.998465 | -0.140520944 | 0.17868449 | -0.7864194 | 0.43162184 | 0.90498 |
| HECA | 33.1500215 | 0.073637746 | 0.39039273 | 0.18862479 | 0.85038691 | 0.9858744 |
| CITED2 | 132.609203 | -0.28096397 | 0.5006172 | -0.5612352 | 0.57463724 | 0.94034284 |
| VTA1 | 98.677858 | -0.328666061 | 0.21769994 | -1.5097205 | 0.13111476 | 0.700349 |
| ADGRG6 | 241.011038 | -0.044990728 | 0.5946553 | -0.0756585 | 0.93969079 | 0.99526299 |
| HIVEP2 | 549.662655 | -0.453813311 | 0.35735023 | -1.2699399 | 0.20410605 | 0.77155952 |
| AIG1 | 344.697143 | -0.160359769 | 0.22827452 | -0.7024865 | 0.48237581 | 0.91971919 |
| ADAT2 | 34.6713642 | -0.539149359 | 0.43251534 | -1.2465439 | 0.21256478 | 0.77822114 |
| PEX3 | 128.95946 | -0.048980273 | 0.30912822 | -0.1584465 | 0.874105 | 0.98766275 |
| FUCA2 | 44.1307774 | -0.53067284 | 0.24948504 | -2.1270728 | 0.03341403 | 0.44828157 |
| PHACTR2 | 244.737127 | -0.105951752 | 0.28302114 | -0.3743598 | 0.70813661 | 0.96587538 |
| PLAGL1 | 251.506336 | 0.023607592 | 0.30716668 | 0.07685597 | 0.93873813 | 0.99526299 |
| SF3B5 | 94.1840948 | -0.004969013 | 0.43977901 | -0.0112989 | 0.99098499 | 0.99866813 |
| UTRN | 1735.79905 | -0.163478788 | 0.26578937 | -0.615069 | 5.39E-01 | 0.93516669 |
| EPM2A | 65.8712959 | -0.922875909 | 0.36648491 | -2.5181826 | 0.01179622 | 0.30957794 |
| AL356599.1 | 92.0381086 | 0.69662588 | 0.33609584 | 2.07270009 | 0.0382002 | 0.47154601 |
| SHPRH | 336.853583 | 0.215523684 | 0.18734863 | 1.15038837 | 0.24998395 | 0.80672756 |
| STXBP5 | 297.116697 | -0.0174192 | 0.21948689 | -0.0793633 | 0.93674367 | 0.99526299 |
| SAMD5 | 268.920458 | -1.2409861 | 0.57197054 | -2.1696679 | 0.03003202 | 0.43492739 |
| AL033504.1 | 167.607857 | -1.428438382 | 0.66877382 | -2.1359066 | 0.03268703 | 0.44162068 |
| SASH1 | 1546.81349 | -0.338751945 | 0.21959448 | -1.5426251 | 0.12292177 | 0.69314363 |
| UST | 44.8741388 | 0.994133823 | 0.78887947 | 1.26018468 | 0.20760275 | 0.77435432 |
| TAB2 | 249.080533 | 0.309481992 | 0.2298177 | 1.34664124 | 0.17809581 | 0.7513166 |
| PPIL4 | 103.447419 | 0.140019308 | 0.28681357 | 0.48818926 | 0.62541579 | 0.94808496 |
| GINM1 | 125.71796 | 0.205536362 | 0.17671273 | 1.16311011 | 0.24478483 | 0.80476923 |
| KATNA1 | 51.7250901 | -0.081474029 | 0.30841576 | -0.2641695 | 0.79164933 | 0.97423208 |
| LATS1 | 168.961491 | 0.030563201 | 0.17368321 | 0.17597096 | 0.86031676 | 0.9864199 |
| NUP43 | 52.6859958 | -0.712667476 | 0.27758561 | -2.5673791 | 0.01024705 | 0.29271368 |
| PCMT1 | 116.42825 | -0.045601939 | 0.22772778 | -0.2002476 | 0.84128695 | 0.98515707 |
| PPP1R14C | 190.301258 | -0.731892242 | 0.42027359 | -1.7414662 | 0.0816019 | 0.61019873 |
| PLEKHG1 | 66.2314998 | 0.722299627 | 0.46352371 | 1.55827977 | 0.11916694 | 0.68597919 |
| MTHFD1L | 280.823859 | -0.574296148 | 0.39898478 | -1.4393936 | 1.50E-01 | 0.72246729 |
| AKAP12 | 470.936924 | -0.27852138 | 0.56262729 | -0.4950371 | 0.62057394 | 0.9480368 |
| RMND1 | 70.1472569 | -0.477004355 | 0.19966247 | -2.3890537 | 0.01689183 | 0.35074796 |
| ARMT1 | 38.7987749 | -0.024824672 | 0.29316223 | -0.084679 | 0.93251662 | 0.99416721 |
| SYNE1 | 2403.17371 | 0.664404477 | 0.54727733 | 1.21401791 | 0.22474089 | 0.78914698 |
| MTRF1L | 36.2738538 | -0.401051727 | 0.33416757 | -1.2001516 | 0.23008048 | 0.79370986 |
| IPCEF1 | 183.31303 | -0.099040668 | 0.68456332 | -0.1446771 | 0.88496578 | 0.98766275 |
| CNKSR3 | 694.251655 | 0.2461001 | 0.33794442 | 0.72822656 | 0.46647491 | 0.91462428 |

| | | | | | | |
|---|---|---|---|---|---|---|
| SCAF8 | 436.085041 | -0.103489285 | 0.17555299 | -0.5895046 | 0.55552285 | 0.93523904 |
| TIAM2 | 240.58573 | 0.056225325 | 0.18964485 | 0.29647694 | 0.76686588 | 0.97221495 |
| TFB1M | 93.9412535 | -0.577409203 | 0.23400982 | -2.4674572 | 0.01360765 | 0.32032755 |
| ARID1B | 1314.02023 | 0.011161178 | 0.21761408 | 0.05128886 | 0.95909535 | 0.99572129 |
| TMEM242 | 53.4140151 | -0.18262875 | 0.27341382 | -0.6679573 | 0.50416087 | 0.92552859 |
| ZDHHC14 | 1212.6484 | -0.335195157 | 0.3484445 | -0.9619757 | 0.33606179 | 0.86391445 |
| SNX9 | 306.78488 | -0.526908859 | 0.22084912 | -2.3858318 | 0.01704054 | 0.35074796 |
| SYNJ2 | 133.046009 | 0.014067421 | 0.56926464 | 0.02471157 | 0.98028503 | 0.99681784 |
| SERAC1 | 54.4609757 | 0.043993187 | 0.3560118 | 0.12357227 | 0.90165395 | 0.9900784 |
| GTF2H5 | 73.243301 | -0.24769355 | 0.3067897 | -0.8073724 | 0.41945194 | 0.90007396 |
| TULP4 | 377.998214 | -0.35637464 | 0.21313553 | -1.6720565 | 0.09451318 | 0.63858274 |
| TMEM181 | 83.6520723 | -0.141867882 | 0.29591902 | -0.4794145 | 0.63164375 | 0.94932781 |
| DYNLT1 | 171.672914 | -0.014401526 | 0.43194599 | -0.033341 | 0.97340263 | 0.99681784 |
| SYTL3 | 56.4315943 | -1.117158696 | 0.54662341 | -2.0437447 | 0.04097877 | 0.48641347 |
| EZR | 484.44969 | 0.139966606 | 0.30288116 | 0.46211724 | 0.64399725 | 0.94944486 |
| RSPH3 | 115.693062 | 0.715829627 | 0.29994854 | 2.38650813 | 0.01700923 | 0.35074796 |
| AL035530.2 | 59.6866534 | 0.655105027 | 0.40568541 | 1.6148104 | 0.10635176 | 0.65699665 |
| SOD2.1 | 46.0356925 | -0.136055622 | 0.42033993 | -0.32368 | 0.74618032 | 0.97028355 |
| WTAP | 281.528388 | 0.161997453 | 0.36076646 | 0.4490369 | 0.65340503 | 0.95222377 |
| TCP1 | 77.905622 | 0.13105949 | 0.37368451 | 0.3507223 | 0.72579669 | 0.96781837 |
| MRPL18 | 42.9886206 | -0.244342952 | 0.35228855 | -0.6935876 | 0.4879409 | 0.92043834 |
| IGF2R | 421.577459 | 0.064866259 | 0.20139464 | 0.32208532 | 0.74738806 | 0.97028355 |
| SLC22A2 | 153.785782 | 1.149017769 | 0.36884875 | 3.11514616 | 0.00183854 | 0.14133317 |
| SLC22A3 | 65.9643206 | 0.520200171 | 0.63479352 | 0.81947933 | 0.41251299 | 0.89706788 |
| MAP3K4 | 179.706374 | -0.234899613 | 0.21411553 | -1.0970695 | 0.27261102 | 0.82004141 |
| AGPAT4 | 118.607798 | -0.150940611 | 0.42289402 | -0.356923 | 0.72114943 | 0.96779196 |
| PRKN | 882.071589 | -0.040996702 | 0.18298568 | -0.2240432 | 0.82272367 | 0.98094663 |
| PACRG | 302.092816 | 0.325564998 | 0.45805098 | 0.71076149 | 0.47723205 | 0.91864961 |
| QKI | 777.24251 | 0.244340465 | 0.19091113 | 1.27986497 | 0.20059263 | 0.77110248 |
| PDE10A | 202.968399 | -0.611444796 | 0.44350301 | -1.3786711 | 0.16799617 | 0.74445571 |
| SFT2D1 | 95.1463699 | -0.397840037 | 0.21751002 | -1.8290654 | 0.06738982 | 0.57946773 |
| MPC1 | 168.357713 | 0.487772174 | 0.35666121 | 1.36760645 | 0.1714353 | 0.74445571 |
| RPS6KA2 | 285.528695 | -0.523186303 | 0.3143164 | -1.6645212 | 0.09600833 | 0.64267209 |
| Z94721.1 | 51.3646676 | -0.484250251 | 0.28332479 | -1.7091701 | 0.08741945 | 0.62705598 |
| FGFR1OP | 184.073451 | -0.050001826 | 0.225725 | -0.2215166 | 0.82469024 | 0.98094663 |
| AFDN | 786.241355 | 0.251029533 | 0.33550152 | 0.74822174 | 0.45432642 | 0.91077355 |
| SMOC2 | 111.519673 | -1.436151735 | 1.59413047 | -0.9008997 | 0.36764163 | 0.87605028 |
| WDR27 | 223.392916 | 0.16488391 | 0.32164084 | 0.51263362 | 0.60820763 | 0.94716646 |
| PHF10 | 83.1674268 | 0.02014638 | 0.24437731 | 0.08243965 | 0.93429711 | 0.99442609 |
| ERMARD | 50.8137086 | -0.548078366 | 0.42730037 | -1.2826536 | 0.19961345 | 0.77110248 |
| DLL1 | 35.3828503 | -0.277754206 | 0.78312603 | -0.3546737 | 0.72283406 | 0.96779196 |
| FAM120B | 282.626493 | -0.186227967 | 0.31974289 | -0.5824304 | 0.56027683 | 0.93744835 |
| PSMB1 | 153.185681 | -0.407245895 | 0.33227623 | -1.2256245 | 0.22034001 | 0.78584902 |
| TBP | 40.0767992 | -0.44652785 | 0.28948022 | -1.5425159 | 0.12294827 | 0.69314363 |
| PDCD2 | 48.2856241 | -0.243172082 | 0.3445624 | -0.7057418 | 0.48034873 | 0.91971919 |
| FAM20C | 106.559945 | 1.160809498 | 0.4631386 | 2.50639765 | 0.01219684 | 0.30967852 |
| PDGFA | 70.3388044 | -0.672237652 | 0.56062271 | -1.199091 | 0.23049256 | 0.79391882 |
| PRKAR1B | 71.6017001 | -0.274053065 | 0.34981962 | -0.7834125 | 0.43338491 | 0.90514499 |
| DNAAF5 | 73.8838614 | -0.370755942 | 0.30359661 | -1.2212124 | 0.22200562 | 0.78710314 |
| SUN1 | 178.786161 | -0.089085613 | 0.19102282 | -0.4663611 | 0.64095705 | 0.94944486 |
| COX19 | 51.6688804 | 0.019370848 | 0.29242955 | 0.06624108 | 0.94718589 | 0.9952642 |
| C7orf50 | 140.039651 | -0.353774259 | 0.22075508 | -1.6025645 | 0.10903084 | 0.65764016 |
| MICALL2 | 60.7619736 | 0.186606514 | 0.28625739 | 0.65188366 | 0.51447623 | 0.93091506 |
| INTS1 | 74.4423354 | -0.40859762 | 0.34811273 | -1.1737509 | 0.24049483 | 0.8030067 |

| | | | | | | |
|---|---|---|---|---|---|---|
| MAFK | 53.0484508 | -0.750748967 | 0.42158867 | -1.7807617 | 0.07495139 | 0.59444897 |
| PSMG3 | 30.0208465 | -0.472778685 | 0.40619057 | -1.1639332 | 0.24445109 | 0.80476923 |
| MAD1L1 | 160.558129 | -0.550453548 | 0.29974885 | -1.8363825 | 0.0663011 | 0.57792071 |
| SNX8 | 78.1555922 | -0.058663634 | 0.38994248 | -0.1504418 | 0.88041609 | 0.98766275 |
| EIF3B | 140.947279 | -0.592757198 | 0.31894906 | -1.8584698 | 0.06310233 | 0.56733665 |
| CHST12 | 36.7334243 | 0.415834611 | 0.37138475 | 1.11968681 | 0.26284725 | 0.81377303 |
| BRAT1 | 35.0927308 | -0.054776513 | 0.39782296 | -0.1376907 | 0.89048489 | 0.98766275 |
| IQCE | 78.5323967 | 0.316780218 | 0.28769508 | 1.10109709 | 0.2708544 | 0.81775732 |
| TTYH3 | 80.0832387 | 0.495267658 | 0.46228301 | 1.07135162 | 0.28401136 | 0.82918183 |
| GNA12 | 133.800478 | -0.133311533 | 0.23260531 | -0.5731233 | 0.56656119 | 0.93845576 |
| SDK1 | 708.717108 | 0.928508427 | 0.61164743 | 1.51804516 | 0.12900301 | 0.69814682 |
| FOXK1 | 212.081556 | -0.122989994 | 0.29538259 | -0.4163752 | 0.67713545 | 0.95924874 |
| AP5Z1 | 27.9981193 | -0.141145777 | 0.38382617 | -0.3677336 | 0.71307189 | 0.96587538 |
| RBAK-RBAKD | 110.635988 | -0.168090785 | 0.2546082 | -0.6601939 | 0.5091294 | 0.92764192 |
| WIPI2 | 92.573928 | -0.149893719 | 0.24082732 | -0.6224116 | 0.53367125 | 0.93446646 |
| TNRC18 | 378.646193 | -0.000389415 | 0.34849459 | -0.0011174 | 0.99910843 | 0.99971834 |
| FBXL18 | 33.1110312 | -0.172471785 | 0.34436315 | -0.5008427 | 0.6164818 | 0.94791646 |
| ACTB | 1167.03282 | -0.197525714 | 0.62639464 | -0.3153375 | 0.75250542 | 0.97096495 |
| RNF216 | 250.924454 | -0.317398609 | 0.27342439 | -1.1608277 | 0.24571196 | 0.80488052 |
| CCZ1 | 37.691414 | 0.084854738 | 0.4046967 | 0.20967489 | 0.83392142 | 0.98351801 |
| PMS2 | 57.5110547 | -0.364148973 | 0.35734805 | -1.0190316 | 0.30818794 | 0.84488735 |
| AIMP2 | 37.0955762 | -0.790102136 | 0.35151266 | -2.2477203 | 0.02459403 | 0.41230428 |
| EIF2AK1 | 109.625206 | 0.163087262 | 0.18944984 | 0.86084665 | 0.38932251 | 0.88586416 |
| USP42 | 96.7728463 | 0.043940729 | 0.24888075 | 0.17655335 | 0.85985924 | 0.9864199 |
| CYTH3 | 65.4809736 | 0.524760155 | 0.4917171 | 1.06719932 | 0.28588184 | 0.83003998 |
| RAC1 | 335.799347 | -0.25827169 | 0.21222891 | -1.2169486 | 0.22362375 | 0.78791697 |
| DAGLB | 32.4226986 | 0.121741326 | 0.29781854 | 0.40877686 | 0.68270342 | 0.96101988 |
| ZDHHC4 | 25.360159 | 0.321771867 | 0.33711753 | 0.95447978 | 0.33984084 | 0.86742596 |
| ZNF316 | 40.1665498 | -0.014500074 | 0.34163574 | -0.0424431 | 0.96614548 | 0.99601626 |
| AC073343.2 | 35.2625289 | 0.404473801 | 0.37518208 | 1.07807336 | 0.28100102 | 0.8264844 |
| ZNF12 | 56.0985308 | 0.270329566 | 0.26897252 | 1.00504529 | 0.31487504 | 0.8490543 |
| CCZ1B | 42.3253862 | -0.199220041 | 0.29900438 | -0.666278 | 0.50523342 | 0.9257297 |
| C1GALT1 | 202.733844 | -0.177941222 | 0.37368113 | -0.4761847 | 0.63394282 | 0.94932781 |
| COL28A1 | 96.8347538 | -1.09607677 | 0.66674182 | -1.6439298 | 0.1001907 | 0.64671608 |
| MIOS | 108.957114 | -0.851121628 | 0.32164164 | -2.6461799 | 0.00814065 | 0.27246484 |
| RPA3 | 56.4125578 | 0.32307398 | 0.51691487 | 0.62500423 | 0.53196828 | 0.93366189 |
| UMAD1 | 462.493786 | 0.01615253 | 0.29018665 | 0.05566255 | 0.95561063 | 0.99567464 |
| GLCCI1 | 289.04901 | -0.219347999 | 0.49446501 | -0.4436067 | 0.65732695 | 0.95399905 |
| ICA1 | 153.375858 | -0.471444645 | 0.47263309 | -0.9974855 | 0.31852892 | 0.84920593 |
| NDUFA4 | 235.158689 | 0.149699762 | 0.55049156 | 0.27193834 | 0.78566943 | 0.97423208 |
| PHF14 | 323.065721 | -0.189623811 | 0.19118944 | -0.9918111 | 0.32128968 | 0.85131972 |
| THSD7A | 1635.0481 | -1.669535641 | 0.57911678 | -2.8828998 | 0.00394033 | 0.2001215 |
| TMEM106B | 168.25159 | -0.705471548 | 0.30031892 | -2.3490746 | 0.01882014 | 0.36488708 |
| SCIN | 579.227992 | -0.644685161 | 0.53944234 | -1.1950956 | 0.23204969 | 0.79471003 |
| ETV1 | 41.2408239 | 0.150061398 | 0.60317494 | 0.24878586 | 0.80352643 | 0.97880236 |
| ISPD | 128.95551 | -0.382314729 | 0.32635605 | -1.1714651 | 0.24141187 | 0.803412 |
| ANKMY2 | 71.4706394 | 0.150375632 | 0.31053375 | 0.48424892 | 0.62820922 | 0.94858781 |
| BZW2 | 83.2905141 | -0.976363411 | 0.45758262 | -2.1337424 | 0.03286388 | 0.44338462 |
| AC073332.1 | 53.1694773 | -0.181778508 | 0.44360025 | -0.40978 | 0.68196734 | 0.96095398 |
| AC019117.1 | 120.408672 | 1.883663797 | 1.3401768 | 1.4055338 | 0.1598626 | 0.73816762 |
| SNX13 | 290.364016 | 0.117478652 | 0.12285905 | 0.95620678 | 0.33896778 | 0.86680923 |
| HDAC9 | 139.647102 | -0.315119727 | 1.33465431 | -0.2361059 | 0.81335051 | 0.97930627 |
| TWISTNB | 30.0363341 | -0.175161708 | 0.47638686 | -0.3676879 | 0.71310593 | 0.96587538 |
| MACC1 | 719.733561 | -0.253021096 | 0.61914352 | -0.4086631 | 0.68278693 | 0.96101988 |

| | | | | | | |
|---|---|---|---|---|---|---|
| ITGB8 | 1593.44624 | 0.793425077 | 0.36724647 | 2.16047027 | 0.03073628 | 0.43815485 |
| SP4 | 72.4416813 | 0.248118171 | 0.23846637 | 1.04047445 | 2.98E-01 | 0.83973489 |
| DNAH11 | 175.464294 | 0.021151925 | 1.05359517 | 0.02007595 | 9.84E-01 | 0.99752755 |
| CDCA7L | 30.5019508 | -0.810482586 | 0.35532796 | -2.2809423 | 0.02255186 | 0.39240055 |
| RAPGEF5 | 128.138358 | 0.875089275 | 0.56686014 | 1.54374812 | 0.12264936 | 0.69314363 |
| STEAP1B | 302.254316 | 0.607906951 | 0.45170945 | 1.34579196 | 0.17836962 | 0.7513166 |
| TOMM7 | 126.105534 | 0.548654082 | 0.46293456 | 1.18516554 | 0.235952 | 0.79952815 |
| FAM126A | 43.6859925 | 0.092728995 | 0.38727893 | 0.23943723 | 0.81076656 | 0.97930627 |
| KLHL7 | 93.1601122 | 0.017507991 | 0.28960112 | 0.06045554 | 0.95179283 | 0.99531843 |
| NUPL2 | 74.369495 | -0.34538876 | 0.23003957 | -1.5014319 | 0.13324388 | 0.70399508 |
| AC005082.1 | 28.7351062 | -1.409933542 | 0.93576958 | -1.5067102 | 0.13188496 | 0.7008294 |
| GPNMB | 62.413217 | -1.859576604 | 0.95861764 | -1.9398523 | 0.05239765 | 0.53169179 |
| MALSU1 | 63.2835478 | 0.10331905 | 0.27685135 | 0.37319322 | 0.70900464 | 0.96587538 |
| TRA2A | 409.733535 | 0.216569004 | 0.25349087 | 0.85434636 | 0.39291311 | 0.88760715 |
| CCDC126 | 38.4057441 | 0.210662409 | 0.2882806 | 0.73075473 | 0.46492898 | 0.91462428 |
| FAM221A | 29.4773046 | -0.501910742 | 0.39390306 | -1.2741986 | 0.20259303 | 0.77110248 |
| MPP6 | 343.715462 | -0.302472944 | 0.32593014 | -0.92803 | 0.35339201 | 0.8720098 |
| OSBPL3 | 234.591139 | -1.024229106 | 0.71228485 | -1.4379487 | 0.1504486 | 0.72274179 |
| CYCS | 76.7859632 | -0.018322835 | 0.54629515 | -0.0335402 | 0.97324383 | 0.99681784 |
| NFE2L3 | 40.6648035 | -0.022273664 | 0.28328086 | -0.0786275 | 0.93732892 | 0.99526299 |
| HNRNPA2B1 | 765.741423 | -0.264244555 | 0.21352773 | -1.2375187 | 0.21589458 | 0.78293142 |
| CBX3 | 144.228204 | -0.136394326 | 0.20701232 | -0.6588706 | 5.10E-01 | 0.92819651 |
| SNX10 | 259.13621 | 0.049060728 | 0.41828838 | 0.11728924 | 9.07E-01 | 9.91E-01 |
| AC004540.1 | 152.262196 | 0.304542919 | 0.5297298 | 0.57490237 | 0.56535733 | 0.93809297 |
| SKAP2 | 669.227858 | 0.128353326 | 0.29301462 | 0.43804411 | 0.66135431 | 0.95422698 |
| HOTAIRM1 | 89.5030723 | -0.204790172 | 0.3660982 | -0.5593859 | 0.57589838 | 0.94034284 |
| HOXA3 | 143.728015 | -0.189495155 | 0.39590975 | -0.4786322 | 0.63220031 | 0.94932781 |
| HOXA-AS2 | 66.1704833 | -0.367508302 | 0.52181862 | -0.7042836 | 0.48125617 | 0.91971919 |
| HOXA7 | 35.55364 | -0.470379068 | 0.56167458 | -0.8374584 | 0.40233498 | 0.89239743 |
| HOXA9 | 77.3921155 | -0.06831225 | 0.29653984 | -0.2303645 | 0.81780855 | 0.98010849 |
| HOXA10-AS | 28.2245106 | -0.438745229 | 0.57046166 | -0.7691056 | 0.44183065 | 0.90561284 |
| HIBADH | 343.53762 | 0.421988773 | 0.20747454 | 2.03393037 | 0.04195862 | 0.4889557 |
| TAX1BP1 | 434.411779 | 0.344962921 | 0.19935785 | 1.73037037 | 0.08356412 | 0.61656297 |
| JAZF1 | 363.869958 | 0.528548436 | 0.33547138 | 1.57553958 | 0.11513195 | 0.67164978 |
| CREB5 | 224.207354 | 0.732240802 | 0.57357706 | 1.2766215 | 0.20173591 | 0.77110248 |
| AC005162.3 | 121.538966 | -0.144253579 | 0.59142549 | -0.2439083 | 0.80730185 | 0.97930627 |
| CPVL | 406.819571 | -0.273100011 | 0.5399136 | -0.5058217 | 0.61298183 | 0.94791646 |
| WIPF3 | 54.7725272 | 0.19405432 | 0.5334135 | 0.36379716 | 0.71600948 | 0.96709741 |
| SCRN1 | 110.42467 | 0.140630085 | 0.28470041 | 0.49395814 | 0.62133574 | 0.9480368 |
| FKBP14 | 25.2823721 | -0.250484249 | 0.39805995 | -0.6292626 | 0.52917714 | 0.93366189 |
| PLEKHA8 | 80.9538553 | -0.267064212 | 0.29888084 | -0.8935474 | 0.37156408 | 0.87605028 |
| MTURN | 110.017218 | -0.356364935 | 0.70693426 | -0.5040991 | 0.61419173 | 0.94791646 |
| ZNRF2 | 110.797982 | 0.173352635 | 0.19514616 | 0.88832205 | 0.37436754 | 0.87678297 |
| AC006978.2 | 89.7474268 | 0.309411667 | 0.2802491 | 1.10405946 | 0.26956734 | 0.8167745 |
| AC005154.6 | 69.7167025 | -0.011631992 | 0.32574236 | -0.0357092 | 0.97151425 | 0.99681784 |
| GARS | 91.0222506 | -0.000821903 | 0.36469483 | -0.0022537 | 0.99820183 | 0.99971834 |
| AQP1 | 603.516193 | 1.226146074 | 0.66104442 | 1.85486186 | 0.06361596 | 0.57043925 |
| PDE1C | 57.808629 | 0.225571392 | 0.83089655 | 0.27147952 | 0.78602225 | 0.97423208 |
| LSM5 | 32.3510718 | -0.139601976 | 0.45141254 | -0.3092559 | 0.7571269 | 0.9714717 |
| AVL9 | 319.901889 | -0.141617152 | 0.1853874 | -0.7638985 | 0.44492774 | 0.90699357 |
| KBTBD2 | 78.0877032 | 0.125328361 | 0.26020131 | 0.48165922 | 0.63004805 | 0.94916079 |
| FKBP9 | 56.6057257 | 0.129141353 | 0.33408203 | 0.38655582 | 0.69908507 | 0.96280283 |
| NT5C3A | 122.357275 | -0.699684787 | 0.39126908 | -1.7882445 | 0.07373657 | 0.59320069 |
| RP9 | 26.8687564 | 0.180795122 | 0.34996733 | 0.51660572 | 0.60543143 | 0.94699994 |

| | | | | | | |
|---|---|---|---|---|---|---|
| BBS9 | 345.134458 | 0.500984047 | 0.26268658 | 1.90715508 | 0.05650051 | 0.54393806 |
| DPY19L1 | 89.7576906 | -0.347199488 | 0.32417653 | -1.0710198 | 0.28416051 | 0.82918183 |
| HERPUD2 | 107.650158 | 0.050910206 | 0.18235121 | 0.27918765 | 0.78010082 | 0.97423208 |
| 7-Sep | 301.691888 | -0.00685928 | 0.14563106 | -0.0471004 | 0.96243322 | 0.99572129 |
| KIAA0895 | 67.2166688 | 0.665354007 | 0.23931425 | 2.7802523 | 0.00543167 | 0.23227655 |
| ELMO1 | 119.98452 | 0.827053528 | 1.34984462 | 0.61270276 | 0.5400729 | 0.93523904 |
| STARD3NL | 76.785246 | 0.084193974 | 0.24179516 | 0.34820371 | 0.7276872 | 0.96823331 |
| VPS41 | 457.82527 | 0.010127962 | 0.28694354 | 0.03529601 | 0.97184371 | 0.99681784 |
| POU6F2 | 96.2550242 | 2.836752034 | 1.1532671 | 2.45975284 | 0.01390327 | 0.32325109 |
| YAE1D1 | 37.0205056 | 0.212711264 | 0.27384972 | 0.7767445 | 0.43730953 | 0.90561284 |
| RALA | 150.51041 | -0.038464349 | 0.22687927 | -0.1695366 | 0.86537456 | 0.98670067 |
| CDK13 | 545.001777 | 0.085688931 | 0.17260584 | 0.49644282 | 0.61958202 | 0.9480368 |
| MPLKIP | 62.5046444 | 0.075207745 | 0.26577485 | 0.28297541 | 0.77719569 | 0.97423208 |
| SUGCT | 398.823666 | 0.530954042 | 0.41654788 | 1.27465309 | 0.20243206 | 0.77110248 |
| GLI3 | 152.322248 | -0.403885848 | 0.48012504 | -0.8412097 | 0.40023046 | 0.89075456 |
| MRPL32 | 43.943662 | 0.124895038 | 0.40884802 | 0.30548036 | 0.76000033 | 0.9714717 |
| STK17A | 56.2922959 | 0.412286956 | 0.31658862 | 1.30227978 | 0.19282076 | 0.76640252 |
| COA1 | 357.077488 | -0.077762993 | 0.19841123 | -0.3919284 | 0.69511113 | 0.96220115 |
| BLVRA | 61.3072695 | 0.36730095 | 0.29938638 | 1.22684591 | 0.21988051 | 0.78584902 |
| URGCP | 68.0088529 | 0.083324906 | 0.35037299 | 0.23781772 | 0.81202247 | 0.97930627 |
| UBE2D4 | 27.2463232 | -0.509479309 | 0.33481399 | -1.5216787 | 0.12808962 | 0.69635098 |
| DBNL | 58.0935219 | -0.197770682 | 0.30721206 | -0.6437595 | 0.51973141 | 0.93272217 |
| POLM | 60.5871212 | -0.054303251 | 0.40264998 | -0.1348647 | 0.89271888 | 0.9882865 |
| AEBP1 | 184.934326 | 0.160838542 | 0.58705194 | 0.27397668 | 0.78410254 | 0.97423208 |
| POLD2 | 50.1855142 | -0.184433952 | 0.41757076 | -0.4416831 | 0.65871853 | 0.95402584 |
| YKT6 | 27.7723089 | -0.502288335 | 0.36970796 | -1.3586084 | 0.17427072 | 0.74910274 |
| NUDCD3 | 195.489964 | -0.137154822 | 0.18910208 | -0.7252952 | 0.46827097 | 0.91462428 |
| DDX56 | 40.6775481 | -0.465689089 | 0.30999979 | -1.5022239 | 0.13303929 | 0.70369044 |
| TMED4 | 89.9911332 | -0.156799206 | 0.30062601 | -0.5215756 | 0.60196582 | 0.94683415 |
| OGDH | 226.371222 | 0.048495537 | 0.17712503 | 0.27379268 | 0.78424395 | 0.97423208 |
| ZMIZ2 | 93.675081 | 0.121065068 | 0.35249098 | 0.34345579 | 0.7312556 | 0.96950828 |
| PPIA | 402.782456 | -0.110753819 | 0.50029758 | -0.2213759 | 0.82479977 | 0.98094663 |
| H2AFV | 128.105704 | -0.044535385 | 0.33451125 | -0.1331357 | 0.89408607 | 0.98842187 |
| PURB | 112.922012 | 0.081188487 | 0.22437629 | 0.36184077 | 0.71747103 | 0.96742001 |
| CCM2 | 119.988758 | -0.099810943 | 0.28946703 | -0.3448094 | 0.73023769 | 0.96888841 |
| TBRG4 | 29.9403075 | -0.002916492 | 0.33300372 | -0.0087581 | 0.99301211 | 0.99866813 |
| IGFBP3 | 287.08391 | -2.832490291 | 1.10026395 | -2.5743734 | 0.01004218 | 0.29238317 |
| AC004870.4 | 261.264807 | 2.841930673 | 1.70145948 | 1.67028995 | 0.09486201 | 0.63901772 |
| TNS3 | 1159.50983 | -0.285194835 | 0.23664906 | -1.2051383 | 0.22814993 | 0.79284833 |
| HUS1 | 46.5804498 | -0.323864239 | 0.30966079 | -1.0458678 | 0.29562209 | 0.83911606 |
| UPP1 | 56.5944107 | -1.210459523 | 0.49955667 | -2.4230675 | 1.54E-02 | 0.33889992 |
| ABCA13 | 387.105436 | -1.060117162 | 0.70672799 | -1.5000356 | 0.13360517 | 0.70512614 |
| GRB10 | 176.810755 | -0.113702795 | 0.4894006 | -0.2323307 | 0.81628115 | 0.98008989 |
| COBL | 359.043024 | -0.113850241 | 0.32233559 | -0.3532041 | 0.72393546 | 0.96779196 |
| SEC61G | 61.9743678 | -0.132077858 | 0.30291763 | -0.4360191 | 0.66282291 | 0.95477098 |
| EGFR | 804.903479 | -0.382676407 | 0.28365996 | -1.3490674 | 0.17731532 | 0.75126783 |
| LANCL2 | 64.7054326 | -0.214417091 | 0.27272726 | -0.786196 | 0.43175264 | 0.90498 |
| VOPP1 | 175.225071 | 0.146339052 | 0.26831913 | 0.54539179 | 0.58548409 | 0.9423304 |
| NIPSNAP2 | 207.21246 | 0.01997844 | 0.26639331 | 0.07499603 | 0.94021787 | 0.99526299 |
| PSPH | 39.0038141 | 0.203868429 | 0.49018395 | 0.41590188 | 0.6774818 | 0.95924874 |
| CCT6A | 91.7477657 | -0.174024116 | 0.43364335 | -0.401307 | 0.6881941 | 0.96161098 |
| SUMF2 | 101.892875 | 0.172099628 | 0.20179199 | 0.85285661 | 0.39373883 | 0.88760715 |
| CHCHD2 | 213.120575 | 0.028608557 | 0.60564402 | 0.04723659 | 0.96232467 | 0.99572129 |
| ZNF727 | 36.0346519 | -0.266630891 | 0.5254083 | -0.5074737 | 0.61182249 | 0.94791646 |

| | | | | | | |
|---|---|---|---|---|---|---|
| ZNF736 | 119.039879 | 0.286931721 | 0.30389465 | 0.94418153 | 0.34507689 | 0.86990996 |
| ZNF680 | 77.6006463 | 0.01229672 | 0.41129898 | 0.02989728 | 0.97614898 | 0.99681784 |
| ZNF107 | 40.9890282 | -0.956794029 | 0.43066734 | -2.2216545 | 0.02630666 | 0.42051885 |
| ZNF138 | 68.9734401 | 0.015532178 | 0.38093998 | 0.04077329 | 0.96747663 | 0.99648221 |
| ZNF117 | 62.580707 | 0.365787242 | 0.24165477 | 1.5136769 | 0.13010781 | 0.69904834 |
| VKORC1L1 | 179.494317 | -0.287915204 | 0.23610452 | -1.2194396 | 0.22267738 | 0.78710314 |
| GUSB | 38.6876255 | 0.468019301 | 0.2753277 | 1.69986277 | 0.08915674 | 0.63290368 |
| AC068533.4 | 32.9662129 | 0.327086081 | 0.48760295 | 0.67080415 | 0.50234531 | 0.92484445 |
| CRCP | 31.3174483 | -0.250459838 | 0.36191236 | -0.6920455 | 0.48890873 | 0.92121587 |
| AC068533.3 | 28.9475934 | 0.489355359 | 0.51487498 | 0.95043531 | 0.34189111 | 0.86846099 |
| TPST1 | 720.818799 | -0.273757902 | 0.37807014 | -0.7240929 | 0.46900871 | 0.91462428 |
| AC027644.4 | 124.102699 | -0.065263162 | 0.27260191 | -0.2394083 | 0.81078899 | 0.97930627 |
| AC027644.3 | 28.4036838 | 0.417601704 | 0.4478678 | 0.93242181 | 0.35111858 | 0.8720098 |
| TMEM248 | 119.879742 | -0.156678436 | 0.2361096 | -0.6635835 | 0.50695692 | 0.92666294 |
| SBDS | 96.207053 | -0.15190737 | 0.39737618 | -0.382276 | 0.70225667 | 0.9645815 |
| TYW1 | 235.92846 | -0.202137572 | 0.20303401 | -0.9955848 | 0.31945193 | 0.85024454 |
| AUTS2 | 1139.21983 | -0.08980817 | 0.6339296 | -0.141669 | 0.88734147 | 0.98766275 |
| TYW1B | 166.212507 | 0.136484185 | 0.21255194 | 0.64212156 | 0.52079426 | 0.93272217 |
| POM121 | 175.516795 | -0.180337961 | 0.23565812 | -0.7652525 | 0.44412122 | 0.90667728 |
| NSUN5 | 38.4752908 | -0.499006004 | 0.32653894 | -1.5281669 | 0.1264711 | 0.69494309 |
| BAZ1B | 242.327104 | 0.150248861 | 0.18798086 | 0.79927745 | 4.24E-01 | 0.90167091 |
| BCL7B | 38.0563363 | -0.212118417 | 0.34277958 | -0.6188187 | 0.53603579 | 0.93469807 |
| MLXIPL | 81.3206564 | 1.602396151 | 0.55046933 | 2.91096357 | 0.00360316 | 0.19174818 |
| BUD23 | 69.9711029 | -0.694211595 | 0.32231726 | -2.1538146 | 0.03125472 | 0.43905874 |
| ABHD11 | 41.7648649 | -0.015033795 | 0.24411449 | -0.061585 | 9.51E-01 | 9.95E-01 |
| CLDN3 | 158.391659 | 0.066697153 | 0.45461885 | 0.14671005 | 0.88336089 | 0.98766275 |
| CLDN4 | 207.126239 | -0.27741707 | 0.52171911 | -0.5317365 | 0.59490854 | 0.94664456 |
| LIMK1 | 32.7724614 | -0.099996446 | 0.48999367 | -0.204077 | 0.83829332 | 0.98462033 |
| EIF4H | 120.594432 | -0.274984278 | 0.30801192 | -0.8927715 | 0.37197953 | 0.87605028 |
| CLIP2 | 144.960463 | -0.492475238 | 0.38889803 | -1.2663351 | 0.20539312 | 0.77352973 |
| GTF2IRD1 | 495.403035 | -0.003896872 | 0.39130412 | -0.0099587 | 0.99205425 | 0.99866813 |
| GTF2I | 844.069408 | 0.271144273 | 0.21881954 | 1.23912278 | 0.21530003 | 0.78293142 |
| GTF2IRD2 | 65.5043342 | -0.773617817 | 0.40397531 | -1.9150126 | 0.05549089 | 0.53915335 |
| CASTOR2 | 86.8350136 | 0.810403682 | 0.62199903 | 1.30290183 | 0.19260828 | 0.76619381 |
| RCC1L | 76.3492375 | -0.05881907 | 0.41261627 | -0.1425515 | 0.8866444 | 0.98766275 |
| GTF2IRD2B | 102.771227 | -0.376111175 | 0.25953378 | -1.44918 | 0.14728731 | 0.72167017 |
| TRIM73 | 50.6730221 | 0.341902648 | 0.46859579 | 0.72963234 | 0.46561495 | 0.91462428 |
| POM121C | 151.476886 | -0.155141775 | 0.1993865 | -0.7780957 | 0.43651261 | 0.90525098 |
| SPDYE5 | 41.0553916 | 0.87933532 | 0.45782681 | 1.92067239 | 0.05477302 | 0.53804407 |
| HIP1 | 294.675572 | 0.84748634 | 0.36565225 | 2.31773862 | 0.02046353 | 0.3784173 |
| RHBDD2 | 44.933952 | -0.111313004 | 0.32245968 | -0.3451998 | 0.7299442 | 0.96884238 |
| POR | 152.142074 | 0.444992777 | 0.22830123 | 1.94914755 | 0.05127781 | 0.52609271 |
| TMEM120A | 26.2953323 | 0.681809666 | 0.32291558 | 2.11141767 | 0.03473643 | 0.45394303 |
| STYXL1 | 81.7444526 | 0.408164048 | 0.3341813 | 1.22138507 | 0.22194026 | 0.78710314 |
| MDH2 | 75.3437815 | -0.010343102 | 0.44105277 | -0.0234509 | 0.98129057 | 0.99681784 |
| HSPB1 | 233.880798 | 0.597025025 | 0.41578647 | 1.43589333 | 0.1510327 | 0.72403332 |
| YWHAG | 119.119178 | -0.294615251 | 0.30320605 | -0.9716668 | 0.33121634 | 0.85923151 |
| DTX2 | 61.6103463 | 0.374449946 | 0.27278203 | 1.3727075 | 0.1698433 | 0.74445571 |
| POMZP3 | 40.6921904 | 0.394263432 | 0.4365293 | 0.90317747 | 0.36643171 | 0.87605028 |
| CCDC146 | 205.374305 | 0.664565432 | 0.42272986 | 1.57208064 | 0.11593185 | 0.67467265 |
| GSAP | 137.247058 | 0.002166648 | 0.37834746 | 0.00572661 | 0.99543085 | 0.99866813 |
| PTPN12 | 417.691745 | -0.110913011 | 0.21626817 | -0.5128494 | 0.60805664 | 0.94708529 |
| APTR | 62.0617641 | 0.623834577 | 0.36739923 | 1.69797466 | 0.08951255 | 0.6330877 |
| RSBN1L | 213.231642 | 0.140477782 | 0.19291689 | 0.72817771 | 0.46650481 | 0.91462428 |

| | | | | | | |
|---|---|---|---|---|---|---|
| PHTF2 | 148.055979 | 0.081018597 | 0.25830459 | 0.31365527 | 0.75378287 | 0.9710175 |
| MAGI2 | 2154.22563 | 0.254517237 | 0.45554271 | 0.55871213 | 0.5763582 | 0.94034284 |
| GNAI1 | 440.459812 | -0.073465182 | 0.3432811 | -0.2140088 | 0.83054018 | 0.9823695 |
| SEMA3C | 82.3365732 | -1.056325235 | 0.53279024 | -1.9826287 | 0.04740892 | 0.51284792 |
| CACNA2D1 | 248.174669 | 0.001763849 | 0.29898845 | 0.00589939 | 0.995293 | 0.99866813 |
| PCLO | 424.111807 | -0.771136954 | 0.52733128 | -1.4623387 | 0.14364841 | 0.71631428 |
| AC003984.1 | 323.201754 | 1.847288994 | 0.56708926 | 3.25749245 | 0.00112401 | 0.10732865 |
| KIAA1324L | 43.9339827 | 0.012087885 | 0.65337334 | 0.01850073 | 0.98523939 | 0.99752755 |
| DMTF1 | 283.153305 | 0.19606287 | 0.2981661 | 0.65756257 | 0.51081925 | 0.92819651 |
| TMEM243 | 92.1973579 | 0.005028237 | 0.27886379 | 0.01803116 | 0.985614 | 0.99764442 |
| CROT | 56.6083727 | -0.348765951 | 0.33139641 | -1.0524132 | 0.29261002 | 0.83697123 |
| ABCB1 | 228.128977 | -0.079540184 | 0.32673389 | -0.2434403 | 0.80766436 | 0.97930627 |
| RUNDC3B | 31.1322505 | 0.461722072 | 0.31970373 | 1.4442186 | 0.1486775 | 0.72246729 |
| SLC25A40 | 54.450097 | 0.332697771 | 0.24301486 | 1.36904291 | 0.17098586 | 0.74445571 |
| ADAM22 | 40.4080925 | 0.523425752 | 0.69335027 | 0.75492254 | 0.45029546 | 0.91037995 |
| SRI | 61.9922321 | 0.16137578 | 0.43922663 | 0.36740892 | 0.71331402 | 0.96587538 |
| STEAP2 | 45.9682839 | 0.046759903 | 0.63775846 | 0.07331914 | 0.94155216 | 0.99526299 |
| CFAP69 | 153.49507 | 0.688464859 | 0.3711858 | 1.85477154 | 0.06362886 | 0.57043925 |
| GTPBP10 | 70.1867166 | -0.170990429 | 0.27955899 | -0.6116435 | 0.54077368 | 0.93523904 |
| CLDN12 | 71.1910018 | 0.515128721 | 0.23843617 | 2.16044704 | 0.03073808 | 0.43815485 |
| CDK14 | 378.781984 | 0.471714483 | 0.51533065 | 0.91536276 | 0.36000122 | 0.87367102 |
| FZD1 | 33.224457 | 1.022209649 | 0.41318133 | 2.47399769 | 0.01336106 | 0.32022149 |
| AC079760.1 | 35.9862943 | -1.388018762 | 0.36723913 | -3.7796048 | 0.00015708 | 0.03134677 |
| MTERF1 | 166.855912 | -0.642799463 | 0.30207511 | -2.1279458 | 0.03334158 | 0.44793682 |
| AKAP9 | 608.547471 | 0.028660922 | 0.15910601 | 0.18013726 | 0.85704481 | 0.9864199 |
| KRIT1 | 193.555524 | 0.271061304 | 0.19762347 | 1.37160479 | 0.1701865 | 0.74445571 |
| ANKIB1 | 485.116994 | 0.031323417 | 0.20727431 | 0.15112059 | 0.87988059 | 0.98766275 |
| GATAD1 | 91.0069791 | 0.026490918 | 0.27464294 | 0.09645585 | 0.92315854 | 0.99214128 |
| AC007566.1 | 74.7594863 | -0.043129424 | 0.41775806 | -0.1032402 | 0.91777234 | 0.99214128 |
| PEX1 | 80.3146312 | -0.073227587 | 0.22818077 | -0.3209192 | 0.74827163 | 0.97028355 |
| FAM133B | 103.92547 | -0.145258689 | 0.23447912 | -0.6194952 | 0.53559019 | 0.93469807 |
| CDK6 | 264.278207 | 0.267532634 | 0.49790271 | 0.53731909 | 0.59104722 | 0.94527451 |
| SAMD9L | 79.881083 | -3.053044337 | 1.28905808 | -2.3684304 | 0.01786374 | 0.35777027 |
| VPS50 | 159.743907 | 0.251716092 | 0.19177552 | 1.31255594 | 0.18933262 | 0.76207437 |
| CALCR | 54.2501271 | -0.256281666 | 1.38033749 | -0.1856659 | 0.85270674 | 0.9864199 |
| GNGT1 | 218.438466 | 0.357111893 | 0.47761156 | 0.74770362 | 0.45463895 | 0.91077355 |
| TFPI2 | 72.0789125 | -2.257805557 | 0.78042492 | -2.8930465 | 0.00381525 | 0.19908785 |
| GNG11 | 51.638276 | -1.24706543 | 0.52769947 | -2.3632114 | 0.01811733 | 0.35858401 |
| BET1 | 70.0938168 | -0.146728917 | 0.32079767 | -0.4573877 | 0.64739238 | 0.95027667 |
| CASD1 | 138.919216 | -0.236907975 | 0.19124094 | -1.2387932 | 0.2154221 | 0.78293142 |
| SGCE | 141.863212 | 0.73666989 | 0.37458939 | 1.9666064 | 0.0492286 | 0.52279468 |
| PPP1R9A | 653.583036 | 0.164877275 | 0.20458252 | 0.80592063 | 0.42028862 | 0.90012434 |
| AC002429.2 | 49.8096052 | 0.419428571 | 0.36543956 | 1.14773717 | 0.25107708 | 0.80724549 |
| PON2 | 287.610021 | -0.013762094 | 0.41543088 | -0.0331273 | 0.97357309 | 0.99681784 |
| PDK4 | 140.584531 | 0.06605231 | 1.16991669 | 0.05645899 | 0.95497617 | 0.99531843 |
| SLC25A13 | 296.581643 | -0.085086723 | 0.28780205 | -0.2956432 | 0.76750257 | 0.97221728 |
| SEM1 | 153.434725 | -0.267780249 | 0.23812967 | -1.1245144 | 0.26079485 | 0.81267205 |
| SDHAF3 | 39.6518982 | -0.15133742 | 0.28280097 | -0.5351376 | 0.59255474 | 0.94649617 |
| ASNS | 28.5365651 | -0.230802795 | 0.44042191 | -0.5240493 | 0.60024424 | 0.94683415 |
| LMTK2 | 230.913057 | 0.381517159 | 0.21303832 | 1.7908382 | 0.07331926 | 0.59320069 |
| TECPR1 | 29.9881662 | -0.64261395 | 0.40587373 | -1.5832854 | 0.11335643 | 0.66657543 |
| BRI3 | 248.224683 | -0.305286242 | 0.3609487 | -0.8457885 | 0.39767076 | 0.88863925 |
| BAIAP2L1 | 575.097361 | 0.276418096 | 0.55699209 | 0.49626934 | 0.6197044 | 0.9480368 |
| TMEM130 | 60.0386352 | 0.455132785 | 1.4012939 | 0.32479467 | 0.74533648 | 0.97028355 |

| | | | | | | |
|---|---|---|---|---|---|---|
| TRRAP | 269.295245 | 0.161620092 | 0.21076328 | 0.76683229 | 0.44318124 | 0.90651998 |
| SMURF1 | 351.973598 | 0.076585504 | 0.28916862 | 0.26484721 | 0.79112716 | 0.97423208 |
| ARPC1B | 43.8906882 | -0.211381246 | 0.5463603 | -0.3868898 | 0.69883777 | 0.96263546 |
| PDAP1 | 60.5582349 | -0.441590294 | 0.33028795 | -1.3369858 | 0.18122728 | 0.75448394 |
| BUD31 | 46.4744435 | -0.401688535 | 0.26712092 | -1.5037704 | 0.1326405 | 0.7031341 |
| CPSF4 | 29.1776897 | 0.027654151 | 0.38864785 | 0.07115478 | 0.94327457 | 0.9952642 |
| ZNF789 | 58.8122041 | -0.023741257 | 0.22249769 | -0.1067034 | 0.9150243 | 0.99159829 |
| ZNF394 | 41.8538264 | 0.045688717 | 0.29211161 | 0.15640842 | 0.8757111 | 0.98766275 |
| CYP3A5 | 158.20404 | 2.394251761 | 1.14876023 | 2.08420495 | 0.03714153 | 0.46890807 |
| TRIM4 | 77.6166157 | -0.011940529 | 0.25916558 | -0.046073 | 0.96325209 | 0.99572129 |
| ZKSCAN1 | 226.721916 | 0.210673368 | 0.17616544 | 1.19588362 | 0.23174198 | 0.79471003 |
| ZNF3 | 43.9914929 | -0.726371321 | 0.32434126 | -2.2395279 | 0.02512159 | 0.4135769 |
| COPS6 | 61.0912338 | 0.127848454 | 0.30786528 | 0.41527403 | 0.67794131 | 0.95924874 |
| MCM7 | 32.7025336 | -0.746877428 | 0.36356137 | -2.0543366 | 0.03994312 | 0.48194503 |
| AP4M1 | 34.9477246 | -0.211519648 | 0.32136664 | -0.658188 | 0.51041737 | 0.92819651 |
| TAF6 | 32.1022244 | -0.219687647 | 0.30214931 | -0.7270831 | 0.46717507 | 0.91462428 |
| LAMTOR4 | 125.871243 | -0.094247455 | 0.40462891 | -0.2329232 | 0.81582105 | 0.98007572 |
| STAG3 | 32.3746219 | 0.755266217 | 0.5818894 | 1.29795493 | 0.19430282 | 0.7687432 |
| CASTOR3 | 200.737832 | 0.332742983 | 0.26340301 | 1.2632467 | 0.20650053 | 0.77401635 |
| SPDYE3 | 110.334086 | 0.392767248 | 0.45788494 | 0.8577859 | 0.3910107 | 0.88670817 |
| PILRB | 159.968567 | -0.380456119 | 0.30053607 | -1.265925 | 0.20553993 | 0.77352973 |
| PILRA | 36.1381303 | -0.180479508 | 0.30451092 | -0.5926865 | 0.55339099 | 0.93523904 |
| MEPCE | 25.4892665 | -0.370464366 | 0.44462814 | -0.8332005 | 0.40473169 | 0.89373953 |
| PPP1R35 | 40.5671669 | -0.442799997 | 0.36818062 | -1.2026706 | 0.22910382 | 0.7931636 |
| TSC22D4 | 65.3341916 | -0.409448481 | 0.38164783 | -1.0728437 | 0.28334124 | 0.82918183 |
| AGFG2 | 42.580805 | -0.774601003 | 0.60228988 | -1.2860933 | 0.19841049 | 0.77110248 |
| LRCH4 | 26.8468324 | -0.645954114 | 0.37195075 | -1.7366657 | 0.08244619 | 0.6141478 |
| GNB2 | 135.336194 | -0.027460282 | 0.41324262 | -0.0664507 | 0.94701897 | 0.9952642 |
| GIGYF1 | 85.709363 | 0.030469451 | 0.24558384 | 0.12406944 | 0.90126029 | 0.9900784 |
| EPHB4 | 36.8903439 | 0.051709727 | 0.4368831 | 0.11836056 | 0.90578198 | 0.99100813 |
| SLC12A9 | 74.6823994 | -0.172207727 | 0.4621734 | -0.3726042 | 0.70944308 | 0.96587538 |
| TRIP6 | 67.5828564 | -0.040259449 | 0.4144124 | -0.0971483 | 0.92260865 | 0.99214128 |
| SRRT | 42.9881558 | -0.421045204 | 0.2598626 | -1.6202609 | 0.10517625 | 0.65699665 |
| TRIM56 | 198.7164 | -0.327082402 | 0.34780888 | -0.9404084 | 0.3470081 | 0.87015462 |
| AP1S1 | 46.6897909 | 0.182135513 | 0.43378037 | 0.41987956 | 0.67457344 | 0.95910677 |
| PLOD3 | 46.3369454 | -0.181650416 | 0.35046571 | -0.5183115 | 0.60424093 | 0.94683415 |
| ZNHIT1 | 79.797594 | 0.092726168 | 0.40573994 | 0.22853596 | 0.8192296 | 0.98079697 |
| FIS1 | 143.555043 | -0.069654592 | 0.40755195 | -0.1709097 | 0.86429475 | 0.98670067 |
| IFT22 | 45.1603884 | 0.806320665 | 0.37461179 | 2.15241667 | 0.03136455 | 0.43905874 |
| COL26A1 | 229.118917 | -0.649423694 | 0.62833127 | -1.033569 | 3.01E-01 | 0.84277781 |
| CUX1 | 1047.37164 | 0.477307299 | 0.26646311 | 1.79126975 | 0.07325002 | 0.59320069 |
| PRKRIP1 | 337.640232 | -0.191595602 | 0.27478932 | -0.6972454 | 0.4856492 | 0.92034617 |
| ORAI2 | 122.617425 | -0.583076606 | 0.4414923 | -1.3206948 | 0.18660313 | 0.76192302 |
| POLR2J | 34.624869 | -0.216656176 | 0.43676939 | -0.4960425 | 0.61986443 | 0.9480368 |
| SPDYE2 | 27.4867813 | 0.704114713 | 0.39382645 | 1.7878807 | 0.07379526 | 0.59320069 |
| AC105052.3 | 72.4463726 | 0.484943568 | 0.48728445 | 0.99519606 | 0.31964092 | 0.85051121 |
| FAM185A | 124.461458 | 0.151203403 | 0.31771012 | 0.47591623 | 0.63413406 | 0.94932781 |
| FBXL13 | 158.15764 | 0.363013441 | 0.32159894 | 1.12877687 | 0.25899197 | 0.81263292 |
| ARMC10 | 93.7138655 | -0.113788463 | 0.1959947 | -0.5805691 | 0.56153092 | 0.93747805 |
| NAPEPLD | 226.990561 | -0.051961331 | 0.38924797 | -0.1334916 | 8.94E-01 | 0.98842187 |
| PMPCB | 74.8521871 | 0.105824298 | 0.24004477 | 0.44085233 | 0.65931991 | 0.95407427 |
| DNAJC2 | 70.6577316 | -0.119249988 | 0.31193746 | -0.3822881 | 0.70224766 | 0.9645815 |
| PSMC2 | 61.3936167 | -0.291318661 | 0.2358654 | -1.2351055 | 0.21679124 | 0.78319543 |
| ORC5 | 96.839984 | -0.355785741 | 0.22203864 | -1.6023595 | 0.10907613 | 0.65764016 |

| | | | | | | |
|---|---|---|---|---|---|---|
| LHFPL3 | 247.689311 | 0.4179713 | 0.58442526 | 0.7151835 | 0.47449567 | 0.91809716 |
| LINC01004 | 205.497771 | -0.001948441 | 0.24825029 | -0.0078487 | 0.99373771 | 0.99866813 |
| KMT2E | 617.70012 | 0.123085078 | 0.24247041 | 0.50762926 | 0.61171336 | 0.94791646 |
| SRPK2 | 688.639292 | 0.048316211 | 0.23178311 | 0.20845441 | 0.83487418 | 0.98367279 |
| PUS7 | 80.3684132 | -0.727549719 | 0.48210606 | -1.5091072 | 0.13127139 | 0.700349 |
| RINT1 | 41.5371477 | -0.08124195 | 0.3142338 | -0.2585398 | 0.79599032 | 0.97524554 |
| ATXN7L1 | 242.018567 | -0.043736035 | 0.2488747 | -0.1757352 | 0.86050201 | 0.98644672 |
| CDHR3 | 103.618216 | 1.568371172 | 0.57979811 | 2.70502983 | 0.00682983 | 0.25593763 |
| SYPL1 | 170.505573 | -0.135033338 | 0.2411301 | -0.560002 | 0.57547808 | 0.94034284 |
| NAMPT | 1505.86016 | 0.493644768 | 0.78223879 | 0.63106659 | 5.28E-01 | 9.34E-01 |
| PRKAR2B | 72.6163886 | -1.24302732 | 0.71133087 | -1.7474671 | 0.08055634 | 0.60722226 |
| HBP1 | 143.106991 | 0.217275068 | 0.24276765 | 0.89499185 | 0.37079144 | 0.87605028 |
| COG5 | 388.050372 | -0.108849605 | 0.15473796 | -0.7034448 | 0.48177862 | 0.91971919 |
| BCAP29 | 74.1572525 | 0.201855519 | 0.35959837 | 0.56133603 | 0.57456849 | 0.94034284 |
| CBLL1 | 29.7625273 | -0.079059061 | 0.27503303 | -0.287453 | 0.77376551 | 0.97396843 |
| SLC26A3 | 621.333007 | -0.552845878 | 0.62158821 | -0.8894086 | 0.37378354 | 0.87666478 |
| DLD | 102.465339 | 0.302193104 | 0.25386979 | 1.19034685 | 0.2339101 | 0.79633808 |
| LAMB1 | 870.691895 | -1.076200398 | 0.46908665 | -2.2942465 | 0.02177634 | 0.38823827 |
| NRCAM | 107.844023 | 0.404240357 | 0.80855272 | 0.49995547 | 0.61710643 | 0.94791646 |
| PNPLA8 | 130.657693 | -0.067783728 | 0.2759987 | -0.2455944 | 0.80599625 | 0.97930627 |
| DNAJB9 | 36.7707028 | 0.801925808 | 0.54370254 | 1.47493482 | 0.14023007 | 0.7133658 |
| IMMP2L | 3479.72174 | -0.110557839 | 0.37450323 | -0.295212 | 0.76783195 | 0.97236012 |
| DOCK4 | 809.379342 | -1.035046398 | 0.50604487 | -2.0453648 | 0.0408189 | 0.48618451 |
| ZNF277 | 197.633884 | 0.025811175 | 0.24482236 | 0.10542818 | 0.91603606 | 0.99159829 |
| IFRD1 | 131.177979 | 0.176272401 | 0.43526646 | 0.40497585 | 0.68549527 | 0.96161098 |
| TMEM168 | 138.812422 | -0.287075718 | 0.15363493 | -1.8685576 | 0.06168439 | 0.56326893 |
| BMT2 | 163.276597 | 0.101956349 | 0.24415724 | 0.41758479 | 0.67625072 | 0.9591378 |
| SMIM30 | 29.1526231 | -0.081423008 | 0.48265572 | -0.1686979 | 0.86603427 | 0.98670067 |
| FOXP2 | 1918.99365 | -0.176596434 | 0.64931859 | -0.2719719 | 0.7856436 | 0.97423208 |
| MDFIC | 187.029561 | 0.090836145 | 0.3273869 | 0.27745809 | 0.78142838 | 0.97423208 |
| TES | 110.859991 | -0.378827673 | 0.43018878 | -0.880608 | 0.37853004 | 0.87970706 |
| CAV2 | 64.2538148 | 0.294290319 | 0.48461841 | 0.60726195 | 0.54367709 | 0.93523904 |
| MET | 606.48818 | -0.841096608 | 0.33605743 | -2.5028359 | 0.01232026 | 0.30967852 |
| CAPZA2 | 271.776591 | -0.200053922 | 0.23943414 | -0.835528 | 0.40342051 | 0.89337058 |
| ST7 | 293.705378 | -0.213623743 | 0.27081083 | -0.7888301 | 0.4302113 | 0.90364119 |
| CFTR | 66.2059966 | 2.273841003 | 0.74733523 | 3.04259843 | 0.00234545 | 0.15891203 |
| CTTNBP2 | 696.028709 | 0.390779718 | 0.60488732 | 0.64603722 | 0.51825525 | 0.93257822 |
| LSM8 | 74.4172651 | -0.14680487 | 0.24719563 | -0.5938813 | 0.55259149 | 0.93523904 |
| TSPAN12 | 344.569044 | 0.996158384 | 0.54879042 | 1.81518909 | 0.06949486 | 0.58408344 |
| ING3 | 46.0556506 | -0.127215891 | 0.25752188 | -0.4940003 | 0.62130596 | 0.9480368 |
| FAM3C | 206.250211 | -0.275206747 | 0.33339656 | -0.8254637 | 0.4091084 | 0.8959746 |
| AASS | 99.9008356 | -0.23324901 | 0.37398842 | -0.6236798 | 0.53283792 | 0.93395323 |
| CADPS2 | 429.679765 | 0.119892021 | 0.36483748 | 0.32861761 | 0.74244474 | 0.9698508 |
| AC004594.1 | 27.97976 | -0.033144144 | 0.66343206 | -0.0499586 | 0.96015537 | 0.99572129 |
| SLC13A1 | 105.476656 | 2.035088374 | 1.24086729 | 1.64005321 | 0.1009941 | 0.64808588 |
| IQUB | 90.7058103 | 0.747743575 | 0.41923792 | 1.78357812 | 0.07449224 | 0.59413921 |
| NDUFA5 | 128.252552 | 0.206870994 | 0.42834672 | 0.48295221 | 0.62912967 | 0.94891118 |
| WASL | 237.879814 | 0.148462968 | 0.19704014 | 0.75346561 | 0.45117017 | 0.91077355 |
| POT1 | 99.6528281 | -0.187751451 | 0.23348408 | -0.8041296 | 0.42132216 | 0.90012434 |
| POT1-AS1 | 39.3706775 | 0.558501107 | 0.4498356 | 1.24156716 | 0.2143963 | 0.78237483 |
| GRM8 | 105.204234 | -0.878113386 | 1.28356388 | -0.6841213 | 0.49389857 | 0.92289011 |
| ZNF800 | 136.915263 | 0.280132301 | 0.25863192 | 1.08313118 | 0.2787502 | 0.8229426 |
| ARF5 | 40.3418045 | -0.213963168 | 0.46927943 | -0.4559398 | 0.64843327 | 0.95027667 |
| SND1 | 472.556434 | -0.007376944 | 0.32856036 | -0.0224523 | 0.98208714 | 0.99681784 |

| | | | | | | |
|---|---|---|---|---|---|---|
| RBM28 | 64.0983524 | -0.263966102 | 0.29753502 | -0.8871766 | 0.37498384 | 0.8769473 |
| HILPDA | 122.137738 | 0.762257018 | 1.45957046 | 0.5222475 | 0.60149802 | 0.94683415 |
| METTL2B | 70.1539735 | -0.102519799 | 0.2751209 | -0.3726354 | 0.70941979 | 0.96587538 |
| CALU | 115.730743 | 0.127756849 | 0.31674366 | 0.40334461 | 0.68669472 | 0.96161098 |
| KCP | 111.235613 | -0.088198985 | 0.53027116 | -0.1663281 | 0.86789877 | 0.98670067 |
| ATP6V1F | 120.165757 | -0.142484277 | 0.53949584 | -0.2641064 | 0.79169797 | 0.97423208 |
| TNPO3 | 239.000452 | -0.091570151 | 0.18395113 | -0.4977961 | 0.61862777 | 0.9480368 |
| TSPAN33 | 48.9828755 | 0.280955068 | 0.70742892 | 0.39714953 | 0.69125719 | 0.96220115 |
| SMO | 53.4659845 | 1.274354519 | 0.48743094 | 2.61443091 | 0.00893762 | 0.28162334 |
| AHCYL2 | 327.236852 | 0.115174681 | 0.2385382 | 0.48283537 | 0.62921264 | 0.94891118 |
| NRF1 | 271.463827 | 0.119748522 | 0.1850189 | 0.64722319 | 0.51748751 | 0.93247044 |
| UBE2H | 587.485691 | -0.088489132 | 0.19675087 | -0.4497522 | 0.65288915 | 0.95176155 |
| ZC3HC1 | 66.7354413 | 0.239025569 | 0.27787887 | 0.860179 | 0.38969038 | 0.88586416 |
| KLHDC10 | 269.921577 | -0.036344365 | 0.20109971 | -0.1807281 | 0.85658102 | 0.9864199 |
| TMEM209 | 47.622061 | -0.407882868 | 0.37588698 | -1.085121 | 0.27786806 | 0.8229426 |
| COPG2 | 88.7411084 | -0.763138061 | 0.27014417 | -2.8249289 | 0.00472911 | 0.21962686 |
| AC016831.7 | 1933.30542 | -0.202768145 | 0.33570595 | -0.6040052 | 0.54584017 | 0.93523904 |
| LINC00513 | 223.474592 | -0.091848439 | 0.55044915 | -0.1668609 | 0.8674795 | 0.98670067 |
| AC058791.1 | 41.0161858 | 0.202350816 | 0.51892418 | 0.38994293 | 0.69657875 | 0.96220115 |
| MKLN1 | 1461.81955 | -0.376736636 | 0.2739953 | -1.3749748 | 0.16913926 | 0.74445571 |
| PODXL | 33.2525347 | 0.663308126 | 0.6198031 | 1.07019169 | 0.28453303 | 0.82918183 |
| PLXNA4 | 46.7855692 | -1.682062918 | 1.39104492 | -1.2092082 | 0.22658287 | 0.79116864 |
| CHCHD3 | 698.744169 | 0.233823093 | 0.27383841 | 0.85387252 | 0.39317562 | 0.88760715 |
| EXOC4 | 1455.14643 | -0.016311517 | 0.17142473 | -0.0951527 | 0.92419358 | 0.99258328 |
| LRGUK | 76.3758678 | 0.351295812 | 0.41858213 | 0.83925181 | 0.40132802 | 0.89216085 |
| SLC35B4 | 42.9444618 | -0.304030951 | 0.291874 | -1.0416514 | 0.29757335 | 0.83973489 |
| AKR1B1 | 84.6309634 | -0.023168152 | 0.72776438 | -0.0318347 | 0.97460388 | 0.99681784 |
| BPGM | 27.3815781 | 1.09076883 | 0.58586984 | 1.86179379 | 0.06263216 | 0.56713487 |
| CALD1 | 357.174774 | -0.073952104 | 0.31917667 | -0.2316965 | 0.81677378 | 0.98010849 |
| AGBL3 | 132.1447 | 0.872103688 | 0.37148431 | 2.34761915 | 0.01889383 | 0.36488708 |
| CYREN | 79.5604702 | -0.556055443 | 0.30753296 | -1.8081165 | 0.07058836 | 0.58797037 |
| TMEM140 | 65.2699947 | -0.3837569 | 0.86462381 | -0.4438426 | 0.65715637 | 0.95399905 |
| WDR91 | 126.487998 | 0.133901591 | 0.26964271 | 0.49658896 | 0.61947894 | 0.9480368 |
| CNOT4 | 371.740493 | 0.11010612 | 0.172175 | 0.6395012 | 0.52249693 | 0.93272217 |
| NUP205 | 92.2692163 | -0.712821467 | 0.29111064 | -2.4486273 | 0.01434018 | 0.32762834 |
| STMP1 | 148.702222 | 0.136810107 | 0.3753944 | 0.36444366 | 0.71552674 | 0.96687349 |
| MTPN | 122.28227 | -0.07644272 | 0.27590119 | -0.2770656 | 0.78172976 | 0.97423208 |
| CREB3L2 | 159.80987 | 0.45680713 | 0.39587017 | 1.15393168 | 0.24852818 | 0.80672756 |
| TRIM24 | 177.920731 | -0.230507107 | 0.32164816 | -0.7166436 | 0.47359401 | 0.91804805 |
| ATP6V0A4 | 27.5373332 | -1.102985611 | 0.67866049 | -1.6252392 | 0.10411161 | 0.65481622 |
| KIAA1549 | 87.0881278 | 1.165174939 | 0.34185789 | 3.4083605 | 0.00065354 | 0.08130268 |
| ZC3HAV1 | 156.255918 | -0.797494355 | 0.4831759 | -1.6505259 | 0.09883541 | 0.6434623 |
| TTC26 | 70.854342 | 0.575381009 | 0.26517463 | 2.16981919 | 0.03002055 | 0.43492739 |
| UBN2 | 196.961874 | 0.301288287 | 0.22986029 | 1.31074527 | 0.18994384 | 0.76207437 |
| LUC7L2 | 65.645701 | -0.199483822 | 0.24753565 | -0.8058792 | 0.42031252 | 0.90012434 |
| KLRG2 | 206.728452 | 0.529121836 | 0.41116992 | 1.28686901 | 1.98E-01 | 0.77110248 |
| HIPK2 | 1690.14003 | -0.634424602 | 0.22539729 | -2.8146949 | 0.00488236 | 0.21962686 |
| TBXAS1 | 123.370705 | -0.9521439 | 0.36176683 | -2.631927 | 0.00849021 | 0.27475583 |
| PARP12 | 94.8538876 | -1.645783314 | 0.72541632 | -2.2687432 | 0.02328395 | 0.39839181 |
| KDM7A | 178.317934 | 0.223175264 | 0.33548521 | 0.6652313 | 0.50590256 | 0.92587708 |
| SLC37A3 | 120.570384 | 0.08308845 | 0.2803081 | 0.2964183 | 0.76691065 | 0.97221495 |
| MKRN1 | 93.9396989 | 0.241679687 | 0.21380054 | 1.13039792 | 0.25830859 | 0.8123237 |
| DENND2A | 121.005742 | 0.242191408 | 0.55660715 | 0.4351209 | 0.66347467 | 0.95482952 |
| NDUFB2 | 128.368862 | 0.134745826 | 0.30824707 | 0.43713578 | 0.66201288 | 0.9545911 |

| | | | | | | |
|---|---|---|---|---|---|---|
| BRAF | 763.310506 | 0.02507552 | 0.16225 | 0.15454866 | 8.77E-01 | 9.88E-01 |
| MRPS33 | 45.1815394 | 0.100085491 | 0.34898326 | 0.28679167 | 0.77427184 | 0.97411309 |
| TMEM178B | 451.55319 | -0.023149235 | 0.80364996 | -0.0288051 | 0.97702002 | 0.99681784 |
| AGK | 94.9121882 | -0.306157345 | 0.20803229 | -1.4716819 | 0.14110681 | 0.7148274 |
| AC004918.1 | 27.3927387 | 0.208956036 | 0.4148334 | 0.50371074 | 0.61446466 | 0.94791646 |
| KIAA1147 | 98.2109546 | 0.341870998 | 0.29626954 | 1.15391881 | 0.24853346 | 0.80672756 |
| SSBP1 | 122.65443 | -0.323610408 | 0.1990703 | -1.6256087 | 0.10403293 | 0.65475128 |
| GSTK1 | 94.3265911 | 0.459787971 | 0.29922318 | 1.53660547 | 0.12438995 | 0.69439514 |
| TMEM139 | 30.85729 | -0.254536416 | 0.53579021 | -0.4750673 | 0.634739 | 0.94932781 |
| CASP2 | 46.5968628 | -0.178276847 | 0.31663647 | -0.5630332 | 0.57341227 | 0.93998433 |
| ZYX | 56.3816702 | -0.174978479 | 0.37868396 | -0.46207 | 0.64403116 | 0.94944486 |
| EPHA1 | 23.9639061 | -0.117064661 | 0.51370447 | -0.2278833 | 0.81973698 | 0.98079697 |
| TCAF2 | 70.6029267 | -0.753862532 | 0.71985593 | -1.0472408 | 0.29498851 | 0.83911606 |
| TCAF1 | 133.935748 | -0.129607768 | 0.22062156 | -0.5874665 | 0.55689047 | 0.93533595 |
| AC004889.1 | 189.346663 | 0.06833148 | 0.52470193 | 0.13022914 | 8.96E-01 | 0.98842187 |
| AC074386.1 | 281.28236 | -0.493880397 | 0.40511966 | -1.2190976 | 0.22280716 | 0.78710314 |
| ARHGEF5 | 65.5917981 | -0.553561376 | 0.5293132 | -1.0458106 | 0.29564847 | 0.83911606 |
| TPK1 | 352.710833 | 0.129804443 | 0.31106207 | 0.41729434 | 0.67646313 | 0.9591378 |
| CNTNAP2 | 32.5524272 | 0.347799958 | 0.47843353 | 0.72695565 | 0.46725312 | 0.91462428 |
| CUL1 | 212.56677 | -0.003673559 | 0.1539806 | -0.0238573 | 0.98096645 | 0.99681784 |
| EZH2 | 32.7489693 | -0.181841987 | 0.3752563 | -0.4845808 | 0.62797376 | 0.94858781 |
| PDIA4 | 71.0897118 | -0.114212376 | 0.60887875 | -0.1875782 | 0.85120732 | 0.98617742 |
| ZNF398 | 65.7590719 | -0.102389793 | 0.27513655 | -0.3721417 | 0.70978732 | 0.96587538 |
| ZNF282 | 42.391429 | -0.009982333 | 0.32967645 | -0.0302792 | 0.9758444 | 0.99681784 |
| ZNF783 | 34.5190485 | -0.52498493 | 0.28040059 | -1.8722676 | 0.0611696 | 0.56071154 |
| ZNF746 | 27.3295798 | 0.070899874 | 0.38844137 | 0.18252401 | 0.85517152 | 0.9864199 |
| ZNF862 | 78.3142777 | 0.452163042 | 0.25686783 | 1.76029452 | 0.07835788 | 0.60239981 |
| ACTR3C | 274.757454 | -0.63770434 | 0.42950219 | -1.4847523 | 0.13760946 | 0.71091391 |
| LRRC61 | 39.4260204 | 0.358295398 | 0.29622869 | 1.20952294 | 0.226462 | 0.79113039 |
| RARRES2 | 77.4767382 | 0.441988475 | 0.53020507 | 0.83361798 | 0.40449629 | 0.89373953 |
| TMEM176B | 168.685332 | 1.69641603 | 0.50823576 | 3.33785256 | 0.00084429 | 0.09295873 |
| TMEM176A | 95.0253895 | 1.108059607 | 0.49373586 | 2.24423563 | 0.02481724 | 0.41230428 |
| ABCB8 | 26.0699857 | 0.352494192 | 0.4599328 | 0.76640368 | 0.44343615 | 0.90658382 |
| SLC4A2 | 48.9754702 | -0.029904639 | 0.36693053 | -0.0814995 | 0.93504476 | 0.99464285 |
| FASTK | 34.5544285 | 0.114281312 | 0.40448508 | 0.28253529 | 0.77753309 | 0.97423208 |
| TMUB1 | 41.432882 | -0.031938169 | 0.47949233 | -0.0666083 | 0.94689354 | 0.9952642 |
| AGAP3 | 56.2705624 | 0.864179787 | 0.35782975 | 2.41505851 | 0.01573268 | 0.34086104 |
| SMARCD3 | 78.0830315 | 2.629729953 | 1.22336933 | 2.14957976 | 0.03158847 | 0.43905874 |
| NUB1 | 104.368754 | -0.370151651 | 0.48864757 | -0.7575023 | 0.448749 | 0.90994378 |
| RHEB | 214.971145 | -0.003913211 | 0.23582586 | -0.0165936 | 0.98676079 | 0.99764442 |
| PRKAG2 | 178.991988 | -0.126247796 | 0.47376022 | -0.2664804 | 0.78986928 | 0.97423208 |
| GALNT11 | 268.572816 | 0.392141691 | 0.21965299 | 1.78527819 | 0.0742162 | 0.59321312 |
| KMT2C | 1057.5349 | 0.001994992 | 0.16001146 | 0.01246781 | 0.99005239 | 0.99839055 |
| ACTR3B | 100.246199 | -0.332246593 | 0.33082966 | -1.004283 | 0.31524224 | 0.8490543 |
| DPP6 | 453.197273 | 0.647127261 | 0.74953876 | 0.86336731 | 0.38793555 | 0.8853359 |
| PAXIP1-AS2 | 40.2707294 | 0.875258297 | 0.57380892 | 1.52534802 | 0.12717232 | 0.69570738 |
| PAXIP1 | 71.779307 | 0.01410641 | 0.24377368 | 0.05786683 | 0.9538547 | 0.99531843 |
| PAXIP1-AS1 | 31.0165627 | -0.490867773 | 0.36817719 | -1.3332379 | 0.18245376 | 0.75495661 |
| INSIG1 | 84.4855418 | 0.329090497 | 0.75891182 | 0.4336347 | 0.66455373 | 0.95505623 |
| AC009403.1 | 62.1363392 | 0.216237906 | 0.23688467 | 0.91284045 | 3.61E-01 | 8.75E-01 |
| RBM33 | 427.474119 | -0.003020886 | 0.1995384 | -0.0151394 | 0.98792099 | 0.99804429 |
| LINC01006 | 96.0231596 | -0.77257425 | 0.35729266 | -2.1623009 | 0.03059499 | 0.43815485 |
| LMBR1 | 320.760729 | 0.019106892 | 0.21268369 | 0.08983713 | 0.92841664 | 0.99336859 |
| NOM1 | 35.3842825 | -0.19212599 | 0.38395376 | -0.5003884 | 0.61680165 | 0.94791646 |

| | | | | | | |
|---|---|---|---|---|---|---|
| UBE3C | 357.561755 | -0.145314591 | 0.25209061 | -0.5764379 | 0.5643192 | 0.93809297 |
| DNAJB6 | 186.166299 | 0.227741135 | 0.23345293 | 0.9755334 | 0.32929574 | 0.85808594 |
| ESYT2 | 418.897283 | -0.314200332 | 0.41734728 | -0.752851 | 0.45153944 | 0.91077355 |
| WDR60 | 292.427656 | 0.174105447 | 0.19978977 | 0.87144323 | 0.38351219 | 0.88328503 |
| GTPBP6 | 76.4413195 | -0.377844552 | 0.25889216 | -1.459467 | 0.14443663 | 0.71634214 |
| IL3RA | 27.0448306 | 1.008884338 | 0.44497731 | 2.26727142 | 0.02337365 | 0.39839181 |
| SLC25A6 | 291.336396 | 0.502380223 | 0.42735112 | 1.17556783 | 0.23976762 | 0.80193227 |
| LINC00106 | 37.0770464 | -0.203783264 | 0.41346889 | -0.4928624 | 0.62210982 | 0.94808496 |
| ASMTL | 52.2770796 | -0.047423245 | 0.25971418 | -0.1825978 | 0.85511359 | 0.9864199 |
| AKAP17A | 50.8943265 | 0.05400611 | 0.3151931 | 0.17134293 | 0.86395412 | 0.98670067 |
| DHRSX | 371.815962 | -0.154657877 | 0.27487286 | -0.5626524 | 0.57367159 | 0.94015401 |
| CD99 | 50.4163315 | 0.855453958 | 0.69037641 | 1.23911238 | 0.21530388 | 0.78293142 |
| ARSD | 127.100657 | -0.151368273 | 0.47814376 | -0.3165748 | 0.75156624 | 0.97058015 |
| ARSE | 37.9216559 | 2.476554181 | 0.50495345 | 4.90451977 | 9.37E-07 | 0.00091119 |
| PRKX | 241.543178 | -0.372264102 | 0.39884078 | -0.9333652 | 0.35063145 | 0.8720098 |
| BX890604.1 | 37.3609918 | -0.958123051 | 0.40076436 | -2.3907392 | 0.01681449 | 0.35014356 |
| PUDP | 75.7294099 | -0.261558828 | 0.37813624 | -0.6917053 | 0.48912244 | 0.92121587 |
| TBL1X | 918.277645 | 0.100736924 | 0.32608679 | 0.30892672 | 0.75737727 | 0.9714717 |
| GPR143 | 32.1379629 | 0.683607971 | 0.44175623 | 1.54747782 | 0.12174807 | 0.69196376 |
| SHROOM2 | 116.614125 | 0.477063293 | 0.52642631 | 0.90622995 | 0.36481415 | 0.87605028 |
| WWC3 | 83.4001313 | 0.232732813 | 0.27607497 | 0.84300585 | 0.39922517 | 0.88955057 |
| MID1 | 399.063682 | 0.373163737 | 0.50939985 | 0.73255565 | 0.46382949 | 0.91462428 |
| AC073529.1 | 43.6742888 | -0.70636457 | 0.32006661 | -2.2069299 | 0.02731896 | 0.42391395 |
| ARHGAP6 | 898.865734 | 0.230393054 | 0.29086507 | 0.79209597 | 0.42830472 | 0.90272191 |
| TMSB4X | 372.152513 | -0.534833001 | 0.5718269 | -0.9353058 | 0.34963074 | 0.8720098 |
| RAB9A | 76.2007075 | 0.229969103 | 0.32838466 | 0.70030404 | 0.48373745 | 0.92004859 |
| TRAPPC2 | 32.3208843 | -0.354028872 | 0.2756419 | -1.2843798 | 0.19900911 | 0.77110248 |
| OFD1 | 158.707951 | -0.067436772 | 0.29197063 | -0.2309711 | 0.81733727 | 0.98010849 |
| GEMIN8 | 51.6322323 | 0.375242241 | 0.25709122 | 1.4595685 | 0.1444087 | 0.71634214 |
| MOSPD2 | 42.661709 | -0.032247486 | 0.25405116 | -0.126933 | 0.89899339 | 0.98948153 |
| PIR | 60.1118 | 0.031314009 | 0.34337118 | 0.0911958 | 0.92733701 | 0.99336859 |
| CLTRN | 151.713947 | 0.975234505 | 0.51094433 | 1.90869032 | 0.05630205 | 0.5431192 |
| CA5B | 118.536443 | -0.840102894 | 0.34942234 | -2.4042621 | 0.01620515 | 0.34266913 |
| ZRSR2 | 48.4454398 | -0.077518641 | 0.35179608 | -0.2203511 | 0.82559775 | 0.98094663 |
| AP1S2 | 34.633903 | -0.307437514 | 0.34692839 | -0.88617 | 0.37552595 | 0.87757089 |
| CTPS2 | 108.376859 | 0.087956626 | 0.20539422 | 0.42823321 | 0.66848134 | 0.95764829 |
| SYAP1 | 99.5791177 | 0.042901375 | 0.30923515 | 0.13873382 | 0.88966049 | 0.98766275 |
| TXLNG | 73.3733492 | -0.10927765 | 0.33007737 | -0.3310668 | 0.74059405 | 0.9698508 |
| RBBP7 | 88.0835452 | 0.052556607 | 0.2889022 | 0.18191833 | 0.85564682 | 0.9864199 |
| REPS2 | 223.561161 | 0.082295549 | 0.67054922 | 0.12272857 | 0.90232204 | 0.9900784 |
| NHS | 1064.87641 | -0.05528031 | 0.3876262 | -0.1426124 | 0.88659629 | 0.98766275 |
| SCML1 | 72.6898169 | -0.157535202 | 0.46091122 | -0.3417908 | 0.73250836 | 0.96980299 |
| RAI2 | 67.0615739 | 0.273906477 | 0.35880363 | 0.76338826 | 0.44523188 | 0.90703449 |
| CDKL5 | 160.239936 | 0.148535121 | 0.23316572 | 0.63703671 | 0.52410093 | 0.93272384 |
| PHKA2 | 81.2214146 | -0.05232419 | 0.27437448 | -0.1907036 | 0.84875785 | 0.98548777 |
| PDHA1 | 95.2593038 | 0.667393894 | 0.35285961 | 1.89138646 | 0.05857277 | 0.55386832 |
| MAP3K15 | 66.3021573 | 0.083569585 | 0.60536769 | 0.13804765 | 0.89020276 | 0.98766275 |
| SH3KBP1 | 95.0599434 | 0.711897941 | 0.8715848 | 0.81678564 | 0.41405094 | 0.89732895 |
| BCLAF3 | 91.1066147 | -0.081880961 | 0.20307272 | -0.40321 | 0.68679371 | 0.96161098 |
| EIF1AX | 168.899894 | -0.54398681 | 0.2376501 | -2.2890242 | 0.02207795 | 0.39000316 |
| RPS6KA3 | 260.765176 | -0.089958068 | 0.20613123 | -0.4364116 | 0.66253809 | 0.95477098 |
| MBTPS2 | 66.133517 | -0.229778193 | 0.21164287 | -1.0856883 | 0.2776169 | 0.8229426 |
| SMS | 172.439828 | -0.446324758 | 0.24232837 | -1.841818 | 0.06550178 | 0.57563442 |
| PRDX4 | 26.4726697 | -0.240553155 | 0.42041237 | -0.5721838 | 0.56719745 | 0.9385359 |

| | | | | | | |
|---|---|---|---|---|---|---|
| ACOT9 | 86.0121148 | -0.356961419 | 0.35038732 | -1.0187624 | 0.30831579 | 0.84488735 |
| SAT1 | 1016.81145 | 0.535387602 | 0.70750968 | 0.75672124 | 0.44921689 | 0.90994378 |
| APOO | 818.561987 | -0.057693915 | 0.57052928 | -0.1011235 | 0.91945243 | 0.99214128 |
| EIF2S3 | 113.424498 | -0.063826127 | 0.24175159 | -0.2640153 | 0.7917681 | 0.97423208 |
| ZFX | 157.446114 | -0.1054965 | 0.36309963 | -0.2905442 | 0.77139992 | 0.97329019 |
| PDK3 | 126.006179 | 0.148321253 | 0.51283379 | 0.28921896 | 0.77241383 | 0.97329019 |
| POLA1 | 140.89082 | -0.533952811 | 0.25037205 | -2.1326374 | 0.03295448 | 0.44398169 |
| GK | 52.1689706 | 0.647437309 | 0.29356156 | 2.20545669 | 0.02742206 | 0.42435533 |
| TAB3 | 109.069138 | 0.083855384 | 0.19941427 | 0.42050844 | 0.67411407 | 0.95910677 |
| DMD | 93.2404116 | 1.090251116 | 0.51848082 | 2.10278002 | 0.035485 | 0.45686942 |
| TMEM47 | 39.749199 | 0.861568099 | 0.42748471 | 2.01543608 | 0.04385898 | 0.49836909 |
| AF241726.2 | 105.449531 | -0.049724662 | 0.41118395 | -0.1209305 | 0.90374612 | 0.99050161 |
| DYNLT3 | 46.8659737 | -0.543118135 | 0.38866915 | -1.3973791 | 0.16229961 | 0.74071637 |
| RPGR | 104.094336 | 0.291928887 | 0.36010328 | 0.81068099 | 0.41754889 | 0.89862171 |
| MID1IP1 | 37.3973659 | 0.294920999 | 0.40830517 | 0.72230533 | 0.47010678 | 0.91502267 |
| BCOR | 172.257037 | -0.184905612 | 0.3182478 | -0.5810114 | 0.56123275 | 0.93747805 |
| ATP6AP2 | 147.587548 | 0.271675559 | 0.35514403 | 0.7649729 | 0.44428768 | 0.90667728 |
| CXorf38 | 67.7697684 | -0.201164283 | 0.26970074 | -0.7458796 | 0.45574015 | 0.91077355 |
| MED14 | 163.228246 | -0.216179238 | 0.24304884 | -0.8894477 | 0.37376251 | 0.87666478 |
| USP9X | 453.896549 | 0.355172568 | 0.23962015 | 1.48223163 | 0.13827867 | 0.71091391 |
| DDX3X | 200.952937 | -0.204415175 | 0.29110402 | -0.7022066 | 0.48255031 | 0.91971919 |
| CASK | 394.681432 | -0.363753438 | 0.31585204 | -1.1516577 | 0.24946176 | 0.80672756 |
| Z93403.1 | 117.004032 | 0.884512176 | 0.67910545 | 1.30246662 | 0.19275692 | 0.76640252 |
| MAOA | 311.620084 | 0.011430552 | 0.33884949 | 0.03373342 | 0.97308973 | 0.99681784 |
| MAOB | 126.106973 | 0.625246239 | 0.69803855 | 0.89571878 | 0.37040298 | 0.87605028 |
| EFHC2 | 71.121945 | 1.18215444 | 0.57698303 | 2.04885478 | 0.04047632 | 0.48525988 |
| KDM6A | 467.821367 | 0.031446271 | 0.37598996 | 0.08363593 | 0.9333459 | 0.99425828 |
| MIR222HG | 56.0477308 | -0.916558032 | 0.55285653 | -1.6578587 | 0.09734598 | 0.64312419 |
| KRBOX4 | 56.7993001 | 0.219694393 | 0.24092335 | 0.91188502 | 0.36182925 | 0.8752623 |
| SLC9A7 | 128.703396 | -0.727348827 | 0.40820933 | -1.7818035 | 0.07478127 | 0.59444897 |
| RP2 | 30.5560927 | 0.070149071 | 0.35773933 | 0.1960899 | 0.8445398 | 0.98520136 |
| JADE3 | 95.6012618 | 0.012863736 | 0.28709101 | 0.04480717 | 0.96426101 | 0.99586809 |
| RGN | 37.1134619 | 1.652765749 | 0.77278367 | 2.13871722 | 0.03245858 | 0.44049392 |
| NDUFB11 | 84.9042278 | 0.138235179 | 0.50889725 | 0.27163672 | 0.78590136 | 0.97423208 |
| RBM10 | 67.0461062 | -0.015828374 | 0.22223902 | -0.0712223 | 0.94322083 | 0.9952642 |
| UBA1 | 127.509335 | 0.023359051 | 0.24337449 | 0.09597987 | 0.92353656 | 0.99214128 |
| CDK16 | 39.6334009 | 0.099648765 | 0.35289879 | 0.28237208 | 0.77765822 | 0.97423208 |
| USP11 | 60.3256647 | 0.259253155 | 0.35217087 | 0.7361573 | 0.46163498 | 0.91462428 |
| ZNF41 | 33.1944868 | -0.070599097 | 0.32014958 | -0.2205191 | 0.8254669 | 0.98094663 |
| ARAF | 40.1307736 | -0.06710633 | 0.35762585 | -0.187644 | 0.85115577 | 0.98617742 |
| TIMP1 | 212.681379 | -0.233283899 | 0.76564494 | -0.3046894 | 0.76060272 | 0.9714717 |
| UXT | 26.0211179 | -0.164608691 | 0.36117806 | -0.455755 | 0.6485662 | 0.95027667 |
| ZNF81 | 85.2656757 | 0.097874771 | 0.33541214 | 0.2918045 | 0.77043611 | 0.97313769 |
| ZNF630 | 31.5856766 | 0.260554028 | 0.52672883 | 0.49466445 | 0.620837 | 0.9480368 |
| EBP | 33.6812303 | 0.117672931 | 0.36804315 | 0.31972591 | 0.74917612 | 0.97028355 |
| RBM3 | 102.539807 | -0.259938615 | 0.46709532 | -0.5565001 | 0.577869 | 0.94034284 |
| WDR13 | 98.575198 | 0.435464553 | 0.26171457 | 1.66389115 | 0.09613418 | 0.64295596 |
| HDAC6 | 54.1124691 | 0.267176382 | 0.27656907 | 0.96603854 | 0.3340249 | 0.8619678 |
| PCSK1N | 68.0831236 | 0.404738475 | 0.64469027 | 0.62780298 | 0.53013301 | 0.93366189 |
| TIMM17B | 32.063132 | -0.272394997 | 0.51088188 | -0.5331859 | 0.59390493 | 0.94649617 |
| OTUD5 | 86.2224414 | 0.057494684 | 0.21768349 | 0.26412055 | 0.79168703 | 0.97423208 |
| GRIPAP1 | 75.7498616 | 0.052028998 | 0.18768025 | 0.27722149 | 0.78161004 | 0.97423208 |
| TFE3 | 47.8427324 | -0.412590572 | 0.42945497 | -0.9607307 | 0.33668759 | 0.86431445 |
| CCDC120 | 28.1486282 | -0.126713588 | 0.31554811 | -0.4015666 | 0.688003 | 0.96161098 |

| | | | | | | |
|---|---|---|---|---|---|---|
| WDR45 | 46.2320373 | -0.090923069 | 0.30314274 | -0.2999348 | 0.76422686 | 0.97177982 |
| PLP2 | 43.8763135 | -0.794285122 | 0.44476683 | -1.7858461 | 0.07412418 | 0.59321312 |
| CLCN5 | 168.873189 | 0.420156632 | 0.4446015 | 0.94501847 | 0.34464945 | 0.86990996 |
| SHROOM4 | 191.039271 | -0.33060853 | 0.29438805 | -1.1230365 | 0.26142198 | 0.81281978 |
| MAGED1 | 226.21774 | 0.127894588 | 0.2053495 | 0.6228142 | 0.53340662 | 0.93436907 |
| FAM156A | 30.776596 | -0.611597491 | 0.4657049 | -1.3132726 | 0.1890911 | 0.76207437 |
| GPR173 | 49.3784949 | 0.561585624 | 0.48606238 | 1.15537767 | 0.2479358 | 0.80672756 |
| KANTR | 165.047146 | -0.173041974 | 0.30426996 | -0.568712 | 0.56955161 | 0.93998433 |
| KDM5C | 75.8990926 | -0.515619303 | 0.36736154 | -1.4035745 | 0.16044558 | 0.73816762 |
| IQSEC2 | 40.8093317 | 0.573404109 | 0.35782555 | 1.60246831 | 0.10905209 | 0.65764016 |
| SMC1A | 83.4223954 | -0.426586133 | 0.30903155 | -1.3803967 | 0.16746453 | 0.74445571 |
| HSD17B10 | 48.6662423 | -0.293511484 | 0.52633534 | -0.5576511 | 0.57708265 | 0.94034284 |
| HUWE1 | 361.769606 | 0.081096673 | 0.15485886 | 0.52368121 | 0.60050029 | 0.94683415 |
| PHF8 | 95.5770156 | 0.101451655 | 0.23864002 | 0.42512424 | 0.67074611 | 0.95771779 |
| FAM120C | 62.9910729 | -0.194304522 | 0.28768751 | -0.6754013 | 0.49942083 | 0.92398825 |
| WNK3 | 31.1475533 | 0.300646687 | 0.37862746 | 0.79404353 | 0.42717009 | 0.90272191 |
| TSR2 | 31.8745523 | -0.455913792 | 0.38250567 | -1.1919138 | 0.23329504 | 0.79622042 |
| GNL3L | 26.3734163 | -0.588392374 | 0.31751995 | -1.8530879 | 0.06386977 | 0.57099806 |
| MAGED2 | 71.1252515 | -0.029842814 | 0.40179556 | -0.0742736 | 0.94079266 | 0.99526299 |
| FAM104B | 30.3903275 | -0.085752891 | 0.37595059 | -0.2280962 | 0.81957148 | 0.98079697 |
| RRAGB | 117.205346 | 0.240828872 | 0.20942001 | 1.14998023 | 0.25015201 | 0.80672756 |
| AL445472.1 | 182.774354 | 0.134475916 | 0.20107739 | 0.66877692 | 0.50363779 | 0.92537933 |
| KLF8 | 47.5211161 | 0.74065333 | 0.39014655 | 1.89839772 | 0.05764371 | 0.54753827 |
| UBQLN2 | 42.0308695 | -0.011469239 | 0.36207163 | -0.0316767 | 0.97472987 | 0.99681784 |
| NBDY | 151.845845 | 0.046374537 | 0.19754176 | 0.23475815 | 0.81439645 | 0.97937868 |
| SPIN3 | 40.6543693 | -0.04195073 | 0.29110777 | -0.1441072 | 0.88541581 | 0.98766275 |
| FAAH2 | 220.275654 | 0.221883032 | 0.31683129 | 0.70031919 | 0.48372799 | 0.92004859 |
| ARHGEF9 | 72.0314253 | 0.114594527 | 0.29561699 | 0.38764527 | 0.69827856 | 0.96256506 |
| MTMR8 | 28.8744566 | -0.219819592 | 0.36239139 | -0.6065806 | 0.54412927 | 0.93523904 |
| ZC4H2 | 87.9048593 | 0.427920211 | 0.23197817 | 1.84465725 | 0.06508741 | 0.57495764 |
| ZC3H12B | 57.7839213 | 0.667480662 | 0.26124318 | 2.55501656 | 0.01061827 | 0.29899856 |
| LAS1L | 35.3682807 | -0.445242173 | 0.30443899 | -1.4625005 | 0.14360411 | 0.71631428 |
| MSN | 386.059447 | -0.002610911 | 0.26078284 | -0.0100118 | 0.99201186 | 0.99866813 |
| AR | 48.5668177 | 1.471507608 | 0.56005502 | 2.62743403 | 0.00860315 | 0.27620989 |
| OPHN1 | 399.849823 | -0.345263971 | 0.45989086 | -0.750752 | 4.53E-01 | 0.91077355 |
| YIPF6 | 90.1839903 | -0.057210699 | 0.20633487 | -0.2772711 | 0.78157194 | 0.97423208 |
| EDA | 174.065726 | 0.096036093 | 0.39856759 | 0.24095309 | 0.80959147 | 0.97930627 |
| IGBP1 | 69.707423 | 0.225464736 | 0.27920323 | 0.80752912 | 0.41936171 | 0.90007396 |
| DLG3 | 69.0686819 | 0.151529437 | 0.32460187 | 0.46681628 | 0.64063133 | 0.94944486 |
| SNX12 | 33.4959777 | 0.093918016 | 0.30828389 | 0.30464782 | 0.76063439 | 0.9714717 |
| ZMYM3 | 40.5351062 | -0.063213426 | 0.36390128 | -0.1737104 | 0.86209309 | 0.98669442 |
| NONO | 117.716538 | -0.083245688 | 0.2576866 | -0.3230501 | 0.74665728 | 0.97028355 |
| TAF1 | 191.916739 | -0.248126334 | 0.21159804 | -1.1726306 | 0.24094397 | 0.80319133 |
| OGT | 256.629538 | -0.057077812 | 0.23247608 | -0.2455212 | 0.80605288 | 0.97930627 |
| NHSL2 | 123.021992 | 0.454545651 | 0.74643363 | 0.60895655 | 0.54255324 | 0.93523904 |
| PIN4 | 80.6125152 | -0.471281613 | 0.24741454 | -1.9048258 | 0.05680272 | 0.54566463 |
| RPS4X | 494.564723 | 0.655021916 | 0.51568466 | 1.27019857 | 0.20401391 | 0.77155952 |
| HDAC8 | 394.219204 | 0.030887609 | 0.21256033 | 0.1453122 | 0.88446439 | 0.98766275 |
| CHIC1 | 137.156153 | 0.162271499 | 0.1789381 | 0.90685827 | 0.36448175 | 0.87605028 |
| XIST | 687.484043 | -0.607044913 | 4.13841247 | -0.1466855 | 0.8833803 | 0.98766275 |
| JPX | 519.19882 | -0.032195256 | 0.44979267 | -0.071578 | 0.94293776 | 0.9952642 |
| FTX | 2687.96429 | 0.239873325 | 0.26929295 | 0.89075235 | 0.37306204 | 0.87651737 |
| SLC16A2 | 34.8649663 | 1.117190551 | 0.50585817 | 2.20850551 | 0.02720905 | 0.42391395 |
| RLIM | 71.7815338 | 0.031696021 | 0.2806442 | 0.11294023 | 0.91007794 | 0.99100813 |

| | | | | | | |
|---|---|---|---|---|---|---|
| NEXMIF | 86.4814283 | 1.167231647 | 0.77303838 | 1.50992715 | 0.13106201 | 0.700349 |
| ABCB7 | 129.187801 | -0.350603743 | 0.1800912 | -1.9468121 | 0.05155726 | 0.52722328 |
| ZDHHC15 | 31.6600035 | -0.265007237 | 0.30567279 | -0.8669638 | 0.38596186 | 0.88353166 |
| PBDC1 | 25.1056724 | -0.220435353 | 0.43799321 | -0.5032849 | 0.61476401 | 0.94791646 |
| ATRX | 679.355955 | -0.116745681 | 0.16098418 | -0.7251997 | 0.46832951 | 0.91462428 |
| MAGT1 | 112.814117 | -0.064449076 | 0.2692963 | -0.239324 | 0.81085434 | 0.97930627 |
| COX7B | 132.458559 | 0.007334426 | 0.42741882 | 0.01715981 | 0.98630912 | 0.99764442 |
| PGK1 | 285.328958 | 0.23020731 | 0.41702186 | 0.55202696 | 0.58092989 | 0.94041171 |
| BRWD3 | 189.55983 | -0.040603114 | 0.28322769 | -0.1433586 | 0.88600701 | 0.98766275 |
| HMGN5 | 35.991606 | 0.214290156 | 0.45783982 | 0.46804613 | 0.6397516 | 0.94944486 |
| SH3BGRL | 134.361688 | -0.108696483 | 0.27022481 | -0.4022446 | 0.68750399 | 0.96161098 |
| RPS6KA6 | 159.281364 | 0.02627907 | 0.44669325 | 0.05883024 | 0.95308732 | 0.99531843 |
| HDX | 71.7539031 | -0.26497981 | 0.47133542 | -0.5621895 | 0.57398693 | 0.94034284 |
| APOOL | 83.3072256 | 0.026858126 | 0.22002886 | 0.12206638 | 0.90284645 | 0.99034419 |
| ZNF711 | 30.2878328 | 0.295622624 | 0.58180596 | 0.50811206 | 0.61137476 | 0.94791646 |
| POF1B | 47.0226492 | -0.632118785 | 0.84638829 | -0.7468425 | 0.45515861 | 0.91077355 |
| CHM | 181.545748 | 0.132244957 | 0.23792153 | 0.55583433 | 0.57832411 | 0.94034284 |
| DIAPH2 | 416.713609 | 0.231617195 | 0.36110383 | 0.64141439 | 0.52125348 | 0.93272217 |
| TSPAN6 | 28.3977985 | 0.604466488 | 0.52997432 | 1.14055808 | 0.25405387 | 0.80897837 |
| SYTL4 | 45.9191547 | 0.891773794 | 0.44751383 | 1.99272903 | 0.04629113 | 0.50827029 |
| TRMT2B | 57.7581948 | 0.353580981 | 0.36126845 | 0.97872091 | 0.3277179 | 0.85723914 |
| HNRNPH2 | 50.722249 | -0.171224002 | 0.44746218 | -0.3826558 | 0.70197498 | 0.9645815 |
| ARMCX4 | 239.608213 | -0.537969421 | 0.39004272 | -1.3792577 | 0.16781532 | 0.74445571 |
| ARMCX3 | 74.1588955 | 0.314164678 | 0.30820725 | 1.0193293 | 0.30804666 | 0.84488735 |
| ARMCX2 | 31.5273515 | 0.063236146 | 0.39500434 | 0.16008975 | 0.87281038 | 0.98766275 |
| ZMAT1 | 156.974349 | 0.348359454 | 0.47655377 | 0.73099716 | 0.46478089 | 0.91462428 |
| GPRASP1 | 28.5402552 | 0.89279524 | 0.60324106 | 1.47999747 | 0.13887392 | 0.7113761 |
| BHLHB9 | 33.1752964 | 0.412800056 | 0.30876103 | 1.33695648 | 0.18123684 | 0.75448394 |
| LINC00630 | 114.305723 | 0.057479756 | 0.28134927 | 0.20430036 | 0.83811879 | 0.98462033 |
| BEX4 | 90.7552127 | -0.299992103 | 0.40388527 | -0.7427657 | 0.45762357 | 0.91248464 |
| TCEAL8 | 42.5570641 | -0.015226476 | 0.38299762 | -0.0397561 | 0.96828761 | 0.99680032 |
| BEX2 | 69.7161073 | -0.108693706 | 0.34646689 | -0.3137203 | 0.75373346 | 0.9710175 |
| TCEAL9 | 117.780106 | 0.02362382 | 0.40090424 | 0.05892634 | 0.95301078 | 0.99531843 |
| BEX3 | 219.464519 | 0.251214512 | 0.44550701 | 0.56388453 | 0.57283272 | 0.93998433 |
| TCEAL4 | 114.249036 | -0.194611545 | 0.22456582 | -0.8666125 | 0.38615437 | 0.88353166 |
| TCEAL3 | 43.6986134 | 0.01889236 | 0.39340232 | 0.048023 | 0.96169791 | 0.99572129 |
| MORF4L2 | 173.758454 | 0.034738014 | 0.30865995 | 0.11254461 | 0.9103916 | 0.99100813 |
| SLC25A53 | 76.510685 | 0.231253876 | 0.25016691 | 0.92439835 | 0.35527895 | 0.87261977 |
| FAM199X | 127.983888 | 0.114813337 | 0.25263056 | 0.45447129 | 0.64948965 | 0.95027667 |
| NRK | 165.647513 | 0.406266845 | 0.4817663 | 0.84328614 | 0.39906843 | 0.88955057 |
| RNF128 | 89.9769426 | 0.248757151 | 0.23742861 | 1.04771345 | 0.29477065 | 0.83911606 |
| TBC1D8B | 60.4235354 | 0.792375403 | 0.30178233 | 2.62565212 | 0.00864832 | 0.27620989 |
| MORC4 | 70.6682077 | 0.699668703 | 0.20030809 | 3.49296283 | 0.00047769 | 0.06755525 |
| CLDN2 | 160.458545 | 1.846480035 | 0.83153795 | 2.22056015 | 0.02638077 | 0.42051885 |
| RBM41 | 106.929986 | 0.47353907 | 0.21890715 | 2.16319599 | 0.03052611 | 0.43815485 |
| TSC22D3 | 74.1079452 | 0.720420886 | 0.62541867 | 1.15190179 | 0.24936143 | 0.80672756 |
| MID2 | 42.4579078 | -0.069238367 | 0.32537348 | -0.2127966 | 0.83148562 | 0.98282427 |
| COL4A6 | 76.3625524 | 0.640191139 | 0.50258256 | 1.27380292 | 0.20273327 | 0.77110248 |
| COL4A5 | 491.368562 | 0.293034358 | 0.31061261 | 0.9434078 | 3.45E-01 | 0.86990996 |
| ACSL4 | 590.372785 | 1.111724503 | 0.58906702 | 1.88726318 | 0.05912494 | 0.55686386 |
| TMEM164 | 102.084204 | -0.135673133 | 0.32509813 | -0.4173298 | 0.6764372 | 0.9591378 |
| AMMECR1 | 154.78625 | -0.484085454 | 0.2913724 | -1.6613978 | 0.09663359 | 0.64295596 |
| PAK3 | 54.8707177 | 1.125122801 | 0.91778747 | 1.22590778 | 0.22023338 | 0.78584902 |
| ALG13 | 197.36174 | 0.092484504 | 0.30007488 | 0.30820475 | 0.75792654 | 0.9714717 |

| | | | | | | |
|---|---|---|---|---|---|---|
| AMOT | 73.0011326 | 0.383743761 | 0.2752627 | 1.39410013 | 0.16328737 | 0.74222552 |
| PLS3 | 74.8013555 | 0.740560304 | 0.37847608 | 1.95668987 | 0.05038394 | 0.52415717 |
| DANT2 | 627.84256 | -0.180775888 | 0.26925553 | -0.6713915 | 0.50197113 | 0.92484445 |
| KLHL13 | 400.520151 | -0.267154131 | 0.61639849 | -0.4334114 | 0.66471592 | 0.95505623 |
| WDR44 | 147.532362 | -0.074803669 | 0.20095145 | -0.3722475 | 0.7097086 | 0.96587538 |
| DOCK11 | 556.160133 | -0.261578611 | 0.38328078 | -0.6824725 | 4.95E-01 | 9.23E-01 |
| IL13RA1 | 114.068515 | -0.03013073 | 0.4515777 | -0.0667232 | 0.94680203 | 0.9952642 |
| LONRF3 | 50.0967134 | -0.632208046 | 0.72092194 | -0.8769438 | 0.38051715 | 0.8807629 |
| PGRMC1 | 142.550546 | 0.249291102 | 0.45763737 | 0.54473502 | 0.58593577 | 0.9423304 |
| SLC25A43 | 77.1023927 | 0.478603632 | 0.40885258 | 1.17060197 | 0.2417588 | 0.803412 |
| SLC25A5 | 100.302435 | 0.337662391 | 0.56776342 | 0.59472375 | 0.55202814 | 0.93523904 |
| UBE2A | 51.0480656 | -0.302157295 | 0.33585911 | -0.8996549 | 0.36830394 | 0.87605028 |
| 6-Sep | 64.3737339 | -0.611736097 | 0.6054826 | -1.0103281 | 0.31233811 | 0.84811918 |
| UPF3B | 88.6110832 | -0.556640538 | 0.24103911 | -2.309337 | 0.02092489 | 0.38106367 |
| NDUFA1 | 100.685918 | 0.128855131 | 0.33964827 | 0.37937814 | 0.70440708 | 0.96461079 |
| NKAP | 34.7241638 | -0.309995832 | 0.29432607 | -1.0532394 | 0.29223126 | 0.83697123 |
| RHOXF1-AS1 | 59.7918076 | 0.295454071 | 0.38546033 | 0.76649671 | 0.44338081 | 0.90658382 |
| LAMP2 | 167.215391 | 0.239608295 | 0.26117716 | 0.91741673 | 0.35892431 | 0.87316144 |
| CUL4B | 50.6398529 | -0.382032665 | 0.23849722 | -1.6018328 | 0.10919259 | 0.65764016 |
| MCTS1 | 64.4621287 | 0.060280418 | 0.2530057 | 0.23825716 | 0.81168165 | 0.97930627 |
| THOC2 | 332.012555 | -0.197088558 | 0.15511996 | -1.2705558 | 0.20388673 | 0.77155952 |
| XIAP | 240.129102 | -0.02374006 | 0.17964109 | -0.1321527 | 0.89486349 | 0.98842187 |
| STAG2 | 374.989708 | -0.06177135 | 0.20434088 | -0.3022956 | 0.76242673 | 0.9714717 |
| TENM1 | 119.100035 | -0.330817533 | 0.7027312 | -0.4707597 | 0.63781234 | 0.94944486 |
| SMARCA1 | 166.444432 | 0.51814289 | 0.23364532 | 2.21764724 | 0.02657889 | 0.42222094 |
| OCRL | 49.8499563 | -0.088185045 | 0.23643868 | -0.3729722 | 0.70916917 | 0.96587538 |
| ZDHHC9 | 37.7359674 | -0.012233493 | 0.37243154 | -0.0328476 | 0.9737961 | 0.99681784 |
| UTP14A | 29.567941 | -0.882916725 | 0.34260165 | -2.5770942 | 0.00996348 | 0.29238317 |
| BCORL1 | 50.9043365 | 0.707964616 | 0.39817819 | 1.77800954 | 0.07540229 | 0.59465671 |
| AIFM1 | 64.5067705 | 0.922017693 | 0.41174145 | 2.23931228 | 0.0251356 | 0.4135769 |
| ZNF280C | 38.5622244 | -0.393844274 | 0.28785736 | -1.3681925 | 0.17125184 | 0.74445571 |
| ENOX2 | 194.75746 | -0.418320986 | 0.27990133 | -1.4945302 | 0.1350371 | 0.70620568 |
| RAP2C | 35.1362643 | 0.096400304 | 0.36065277 | 0.26729395 | 0.78924284 | 0.97423208 |
| RAP2C-AS1 | 156.81731 | -0.116140801 | 0.36691442 | -0.3165338 | 0.75159741 | 0.97058015 |
| MBNL3 | 77.724248 | -0.266650641 | 0.28099165 | -0.9489629 | 0.34263949 | 0.86846099 |
| GPC4 | 666.9858 | -0.326286305 | 0.59073621 | -0.5523384 | 0.58071651 | 0.94041171 |
| GPC3 | 110.790497 | 0.526706851 | 0.53490574 | 0.98467226 | 0.32478508 | 0.855037 |
| PHF6 | 98.6958255 | -0.357578034 | 0.31837569 | -1.1231324 | 0.26138129 | 0.81281978 |
| HPRT1 | 92.9161469 | -0.213395399 | 0.30738488 | -0.6942287 | 0.4875388 | 0.92034617 |
| FAM122B | 39.8237118 | -0.356885726 | 0.3999023 | -0.8924323 | 0.37216127 | 0.87605028 |
| RTL8C | 43.8763831 | -0.111218376 | 0.39583595 | -0.2809709 | 0.77873274 | 0.97423208 |
| RTL8A | 36.0900628 | -0.531033589 | 0.43077294 | -1.2327459 | 0.21767057 | 0.78415436 |
| ZNF75D | 88.5921961 | -0.513371389 | 0.26639462 | -1.9271087 | 0.05396609 | 0.53680286 |
| INTS6L | 63.7906462 | -0.10511959 | 0.32232313 | -0.3261311 | 0.74432518 | 0.97018518 |
| MMGT1 | 25.8531123 | -0.065664139 | 0.47081159 | -0.1394701 | 0.88907868 | 0.98766275 |
| SLC9A6 | 38.118671 | -0.415637376 | 0.33287624 | -1.2486244 | 0.2118025 | 0.77732978 |
| FHL1 | 125.958213 | 0.590928959 | 0.61284919 | 0.96423226 | 0.33492949 | 0.86313953 |
| MAP7D3 | 45.5082961 | 0.05997842 | 0.35248474 | 0.17015891 | 0.86488516 | 0.98670067 |
| HTATSF1 | 66.3260655 | -0.43742542 | 0.43241457 | -1.0115881 | 0.31173506 | 0.84811918 |
| AL683813.1 | 69.1043731 | 0.356935343 | 0.36774485 | 0.97060596 | 0.33174454 | 0.85955665 |
| RBMX | 254.371384 | 0.087146121 | 0.13231149 | 0.65864365 | 0.51012463 | 0.92819651 |
| FGF13 | 264.042966 | 0.124212949 | 0.65917492 | 0.18843701 | 0.85053409 | 0.9858744 |
| ATP11C | 132.210521 | 0.128506667 | 0.2721163 | 0.47224907 | 0.63674902 | 0.94944486 |
| SLITRK4 | 35.8163728 | 1.81876236 | 0.4986429 | 3.6474246 | 0.00026488 | 0.04347958 |

| | | | | | | |
|---|---|---|---|---|---|---|
| AL589669.1 | 104.751407 | 0.718337424 | 0.61261945 | 1.17256713 | 0.24096944 | 0.80319133 |
| FMR1 | 134.812041 | 0.177444246 | 0.33287917 | 0.53305903 | 0.59399272 | 0.94649617 |
| IDS | 75.9894676 | 0.079838352 | 0.3612359 | 0.22101444 | 0.82508119 | 0.98094663 |
| TMEM185A | 37.7949532 | 0.314264101 | 0.29713083 | 1.0576624 | 0.29020938 | 0.83506029 |
| LINC00894 | 299.386092 | 0.511112561 | 0.42467418 | 1.20354046 | 0.22876724 | 0.7931636 |
| MAMLD1 | 82.3807005 | -0.794840096 | 0.40660423 | -1.954825 | 0.05060373 | 0.52415717 |
| MTM1 | 121.842925 | 0.759919101 | 0.18491444 | 4.10957149 | 3.96E-05 | 0.01313957 |
| MTMR1 | 115.145775 | -0.303648571 | 0.26090069 | -1.1638473 | 0.2444859 | 0.80476923 |
| CD99L2 | 65.5731982 | -0.010538762 | 0.22083059 | -0.0477233 | 0.96193677 | 0.99572129 |
| VMA21 | 39.5464784 | -0.206338627 | 0.38131042 | -0.5411303 | 0.58841777 | 0.94444601 |
| CETN2 | 55.7937705 | 0.498424692 | 0.476351 | 1.04633912 | 0.29540449 | 0.83911606 |
| ZNF185 | 39.5236313 | -0.149787978 | 0.53556025 | -0.2796846 | 0.77971946 | 0.97423208 |
| BGN | 190.195305 | 1.491159178 | 1.1388365 | 1.30937073 | 0.1904088 | 0.76219391 |
| SLC6A8 | 67.7668932 | 0.277903479 | 0.38388941 | 0.72391546 | 4.69E-01 | 0.91465049 |
| BCAP31 | 133.979122 | -0.265654747 | 0.44517052 | -0.5967483 | 0.55067545 | 0.93523904 |
| IDH3G | 30.8444558 | -0.085280244 | 0.44719456 | -0.1907005 | 0.84876022 | 0.98548777 |
| SSR4 | 77.4605711 | -0.225210037 | 0.54225757 | -0.4153193 | 0.67790817 | 0.95924874 |
| HCFC1 | 36.5133027 | -0.320918121 | 0.31477338 | -1.0195212 | 0.30795561 | 0.84488735 |
| IRAK1 | 43.8928073 | -0.759760033 | 0.68740373 | -1.1052603 | 0.26904683 | 0.81655182 |
| MECP2 | 200.603779 | -0.100115204 | 0.21928359 | -0.4565558 | 0.64799033 | 0.95027667 |
| FLNA | 312.009083 | -0.206383349 | 0.60434304 | -0.3415003 | 0.73272695 | 0.96980299 |
| EMD | 63.4269939 | -0.268662063 | 0.42014592 | -0.6394494 | 0.52253061 | 0.93272217 |
| RPL10 | 740.442882 | 0.33016308 | 0.6214383 | 0.5312886 | 0.59521881 | 0.94664456 |
| TAZ | 52.1661185 | -0.003351943 | 0.26230108 | -0.012779 | 0.98980412 | 0.99839055 |
| ATP6AP1 | 69.2663238 | -0.039124613 | 0.45032467 | -0.0868809 | 0.93076618 | 0.99399379 |
| GDI1 | 58.9684836 | 0.468106795 | 0.34134063 | 1.37137732 | 0.17025736 | 0.74445571 |
| FAM50A | 47.0311849 | -0.546757665 | 0.36423981 | -1.5010925 | 0.13333163 | 0.70407038 |
| PLXNA3 | 41.5078859 | -0.201206904 | 0.43083883 | -0.467012 | 0.64049129 | 0.94944486 |
| FAM3A | 32.3655968 | -0.090941387 | 0.32850632 | -0.276833 | 0.78190835 | 0.97423208 |
| DKC1 | 48.0921016 | -0.489311855 | 0.37670489 | -1.2989262 | 0.19396925 | 0.7687432 |
| MPP1 | 28.0570957 | 1.11266957 | 0.3462188 | 3.21377571 | 0.00131002 | 0.11421765 |
| FUNDC2 | 55.8332595 | -0.12489706 | 0.48458803 | -0.2577386 | 0.79660862 | 0.97524554 |
| BRCC3 | 66.117992 | -0.029786023 | 0.22094108 | -0.1348143 | 0.89275868 | 0.9882865 |
| VBP1 | 26.2674868 | -0.401139553 | 0.42666652 | -0.9401712 | 0.34712978 | 0.87017396 |
| TMLHE-AS1 | 48.5673474 | 0.387409198 | 0.43653033 | 0.88747372 | 0.37482391 | 0.8769473 |
| TMLHE | 120.736553 | 0.283999454 | 0.30556321 | 0.92942947 | 0.35266656 | 0.8720098 |
| VAMP7 | 49.7777623 | 0.23546177 | 0.25321765 | 0.92987899 | 0.35243374 | 0.8720098 |
| ZNF596 | 27.3645402 | -0.272599063 | 0.32994625 | -0.8261923 | 0.40869499 | 0.8959746 |
| FBXO25 | 89.6917116 | -0.063198825 | 0.30466474 | -0.2074373 | 0.83566837 | 0.98423382 |
| TDRP | 166.80616 | -0.30726468 | 0.37736816 | -0.8142305 | 0.41551289 | 0.89807127 |
| ERICH1 | 182.671418 | -0.247508592 | 0.36848277 | -0.6716965 | 0.50177692 | 0.92484445 |
| KBTBD11 | 75.0543077 | -0.738208813 | 0.52100381 | -1.4168972 | 0.156513 | 0.7335972 |
| AC246817.1 | 64.4856791 | -0.700513732 | 0.54614446 | -1.2826528 | 0.19961374 | 0.77110248 |
| CSMD1 | 123.677407 | 0.437590467 | 1.04637217 | 0.41819773 | 0.67580255 | 0.9591071 |
| MCPH1 | 284.496135 | -0.024681066 | 0.23224797 | -0.1062703 | 0.91536788 | 0.99159829 |
| ANGPT2 | 67.4378048 | -0.06956778 | 0.37412039 | -0.1859503 | 0.85248377 | 0.9864199 |
| MCPH1-AS1 | 69.111081 | 0.129500105 | 0.29868358 | 0.43356955 | 0.66460105 | 0.95505623 |
| AGPAT5 | 157.180435 | -0.078294037 | 0.36691277 | -0.213386 | 0.83102592 | 0.98264379 |
| AF233439.1 | 63.4183811 | -0.194280254 | 0.54742507 | -0.3548984 | 0.72266574 | 0.96779196 |
| DEFB1 | 520.590174 | -0.384734755 | 0.4663722 | -0.8249522 | 4.09E-01 | 0.8959746 |
| FAM85B | 256.412059 | 0.788121942 | 0.52749883 | 1.49407336 | 0.13515645 | 0.70620568 |
| PRAG1 | 113.387155 | -0.894954327 | 0.41353477 | -2.1641574 | 0.03045227 | 0.43815485 |
| MFHAS1 | 553.088905 | -0.091241415 | 0.3193905 | -0.2856735 | 0.77512817 | 0.97423208 |
| ERI1 | 71.2120876 | 0.096969383 | 0.36679428 | 0.26436995 | 0.79149487 | 0.97423208 |

| | | | | | | |
|---|---|---|---|---|---|---|
| PPP1R3B | 37.8790211 | 0.567772769 | 0.56899293 | 0.99785558 | 0.31834939 | 0.84919767 |
| TNKS | 551.015378 | -0.145097528 | 0.18320729 | -0.7919855 | 0.42836911 | 0.90272191 |
| MSRA | 194.831417 | 0.132351688 | 0.40306582 | 0.32836247 | 0.74263762 | 0.9698508 |
| XKR6 | 260.062226 | 0.744169886 | 0.58015256 | 1.28271413 | 0.19959225 | 0.77110248 |
| MTMR9 | 75.180683 | 0.02484845 | 0.2480336 | 0.10018179 | 0.9202 | 0.99214128 |
| FDFT1 | 153.108816 | 0.81273208 | 0.46591914 | 1.74436292 | 0.08109583 | 0.60850843 |
| CTSB | 248.228602 | -0.13982692 | 0.45317886 | -0.3085469 | 0.75766624 | 0.9714717 |
| FAM66D | 82.2693954 | -0.326035806 | 0.78005614 | -0.4179645 | 0.67597304 | 0.9591071 |
| AC145124.1 | 32.3093671 | 1.265958807 | 0.71949852 | 1.75950161 | 0.07849235 | 0.6024665 |
| LONRF1 | 123.928226 | -0.450255452 | 0.38286581 | -1.1760137 | 0.23958939 | 0.80193227 |
| TRMT9B | 73.2155191 | 0.562094852 | 0.36523927 | 1.53897704 | 0.1238099 | 0.69436693 |
| DLC1 | 173.750457 | -0.35957319 | 0.69094117 | -0.5204107 | 0.60277734 | 0.94683415 |
| TUSC3 | 295.860768 | 0.015943992 | 0.31356848 | 0.05084692 | 0.9594475 | 0.99572129 |
| MSR1 | 82.7121759 | -1.657882606 | 0.88485728 | -1.8736158 | 0.06098339 | 0.56071154 |
| MICU3 | 123.500918 | 0.375723077 | 0.24288905 | 1.54689178 | 0.12188934 | 0.69196376 |
| ZDHHC2 | 137.854941 | 0.246847195 | 0.34047988 | 0.7249979 | 0.46845332 | 0.91462428 |
| CNOT7 | 66.3352771 | -0.048491221 | 0.22596564 | -0.2145956 | 0.83008266 | 0.98213461 |
| VPS37A | 95.9145144 | -0.365328469 | 0.23013339 | -1.587464 | 0.11240759 | 0.66401239 |
| SLC7A2 | 54.2505437 | 0.163751852 | 0.67937669 | 0.24103248 | 0.80952994 | 0.97930627 |
| MTUS1 | 152.570283 | 0.586258117 | 0.4124937 | 1.42125351 | 0.15524307 | 0.73003113 |
| PCM1 | 529.302898 | 0.144719906 | 0.18826957 | 0.76868451 | 0.44208062 | 0.90562239 |
| ASAH1 | 246.313005 | -0.059728134 | 0.32641277 | -0.1829835 | 0.854811 | 0.9864199 |
| PSD3 | 198.21076 | -0.537109969 | 0.61961006 | -0.8668516 | 0.38602333 | 0.88353166 |
| SH2D4A | 76.7013419 | -0.013020794 | 0.43208727 | -0.0301346 | 0.97595967 | 0.99681784 |
| CSGALNACT1 | 139.532127 | -2.268703278 | 0.87295081 | -2.5988902 | 0.00935257 | 0.28622448 |
| INTS10 | 215.622756 | -0.147151748 | 0.26509477 | -0.5550911 | 0.57883234 | 0.94034284 |
| ATP6V1B2 | 98.1697627 | -0.24054215 | 0.21283758 | -1.1301677 | 0.25840557 | 0.81236199 |
| XPO7 | 156.915981 | 0.011774078 | 0.1672232 | 0.07040935 | 0.94386785 | 0.9952642 |
| DMTN | 64.9643025 | -0.705399182 | 0.35365697 | -1.9945858 | 0.04608807 | 0.50827029 |
| FAM160B2 | 48.0105881 | 0.356299851 | 0.26016239 | 1.36952866 | 0.17083408 | 0.74445571 |
| BMP1 | 91.8514801 | 0.502311821 | 0.78864874 | 0.63692719 | 0.52417227 | 0.93272384 |
| SLC39A14 | 339.562719 | 1.100026253 | 1.05941006 | 1.0383385 | 0.29911249 | 0.84005388 |
| PPP3CC | 263.093281 | -0.512810863 | 0.25083024 | -2.0444539 | 0.04090873 | 0.48618451 |
| SORBS3 | 34.5909374 | -0.009606802 | 0.34241178 | -0.0280563 | 0.97761726 | 0.99681784 |
| CCAR2 | 41.2668277 | -0.464383347 | 0.2861715 | -1.6227449 | 0.10464395 | 0.65601071 |
| BIN3 | 73.7500473 | 0.021962952 | 0.28871733 | 0.07607078 | 0.93936279 | 0.99526299 |
| TNFRSF10B | 159.819674 | 0.120898238 | 0.29302274 | 0.41258996 | 0.67990706 | 0.95948574 |
| TNFRSF10D | 65.4909925 | -0.540164699 | 0.51920076 | -1.0403773 | 0.29816463 | 0.83973489 |
| TNFRSF10A | 36.2714488 | 0.018890137 | 0.39615041 | 0.04768426 | 9.62E-01 | 9.96E-01 |
| CHMP7 | 26.4014016 | 0.000170261 | 0.37050898 | 0.00045953 | 0.99963335 | 0.99971834 |
| LOXL2 | 33.4719461 | 0.665914938 | 0.74755506 | 0.89079049 | 0.37304158 | 0.87651737 |
| ENTPD4 | 66.386261 | -0.254760981 | 0.27592922 | -0.9232838 | 0.35585933 | 0.87316144 |
| SLC25A37 | 561.123493 | 0.479248701 | 0.55027733 | 0.87092212 | 3.84E-01 | 0.88328503 |
| DOCK5 | 346.883222 | 0.36261676 | 0.33995511 | 1.06666073 | 2.86E-01 | 0.83003998 |
| KCTD9 | 89.1205949 | -0.105298698 | 0.27596658 | -0.3815632 | 0.70278536 | 0.9645815 |
| PPP2R2A | 298.057738 | -0.028838991 | 0.19665616 | -0.1466468 | 0.88341084 | 0.98766275 |
| BNIP3L | 418.819503 | 0.954142421 | 0.52528791 | 1.81641801 | 0.06930627 | 0.58408344 |
| DPYSL2 | 148.368598 | 0.472452698 | 0.37316079 | 1.26608345 | 0.20548321 | 0.77352973 |
| PTK2B | 125.307016 | 0.0095516 | 0.39926948 | 0.02392269 | 0.98091428 | 0.99681784 |
| EPHX2 | 63.1207312 | 0.968842998 | 0.49998398 | 1.9377481 | 0.05265396 | 0.53195745 |
| CLU | 1218.31833 | 0.551792077 | 0.59754081 | 0.92343831 | 0.35577883 | 0.87316144 |
| SCARA3 | 93.3567015 | -0.516110046 | 0.62598065 | -0.8244824 | 0.4096655 | 0.8959746 |
| CCDC25 | 55.3524324 | -0.333091329 | 0.32825651 | -1.0147288 | 0.31023513 | 0.84640909 |
| ELP3 | 148.016924 | -0.586055766 | 0.22135131 | -2.6476273 | 0.00810588 | 0.27246484 |

| | | | | | | |
|---|---|---|---|---|---|---|
| ZNF395 | 57.0420603 | -0.24452446 | 0.4212586 | -0.5804616 | 0.56160335 | 0.93747805 |
| FBXO16 | 66.2764671 | 0.217596613 | 0.4330202 | 0.50250916 | 0.61530941 | 0.94791646 |
| FZD3 | 52.0660833 | 0.107200674 | 0.23166463 | 0.4627408 | 0.64355017 | 0.94944486 |
| EXTL3 | 60.0122651 | 0.343903722 | 0.31912099 | 1.07765938 | 0.28118579 | 0.82672766 |
| INTS9 | 78.2381756 | -0.498983342 | 0.26451337 | -1.8864201 | 0.05923836 | 0.55686386 |
| HMBOX1 | 339.279182 | -0.055970423 | 0.16158844 | -0.3463764 | 0.72905986 | 0.9686081 |
| KIF13B | 401.942266 | -0.492865237 | 0.36285486 | -1.3582986 | 0.17436897 | 0.74910274 |
| SARAF | 184.465217 | 0.545995337 | 0.33374881 | 1.63594691 | 0.1018507 | 0.65103232 |
| LEPROTL1 | 68.5002299 | 0.695085331 | 0.29988853 | 2.31781233 | 0.02045952 | 0.3784173 |
| DCTN6 | 38.2995168 | -0.278197382 | 0.33622536 | -0.8274134 | 0.40800276 | 0.8959746 |
| RBPMS-AS1 | 27.5601426 | -0.122522163 | 0.55366692 | -0.2212922 | 0.82486493 | 0.98094663 |
| RBPMS | 3840.12025 | -0.255959873 | 0.28547658 | -0.8966055 | 0.36992946 | 0.87605028 |
| GTF2E2 | 128.071791 | -0.000592384 | 0.19887405 | -0.0029787 | 0.99762335 | 0.999397 |
| GSR | 84.8517133 | -0.212832742 | 0.23760614 | -0.8957375 | 0.37039296 | 0.87605028 |
| PPP2CB | 279.048356 | -0.370896777 | 0.17871774 | -2.0753215 | 0.03795674 | 0.47106462 |
| PURG | 52.4005924 | 0.931720079 | 0.60170632 | 1.54846317 | 0.12151082 | 0.69161235 |
| WRN | 316.654069 | -0.321379386 | 0.19200457 | -1.6738111 | 0.09416772 | 0.6377384 |
| NRG1 | 435.732555 | 1.021812908 | 0.55347997 | 1.84616061 | 0.06486889 | 0.57495764 |
| FUT10 | 76.3849524 | 0.278373007 | 0.35624147 | 0.78141663 | 0.43455749 | 0.90514499 |
| TTI2 | 29.1566955 | 0.280439294 | 0.48021071 | 0.58399217 | 0.55922558 | 0.93732665 |
| UNC5D | 835.441428 | -0.862572375 | 0.66712416 | -1.2929713 | 0.196021 | 0.77017438 |
| ERLIN2 | 68.1104908 | -0.237346747 | 0.32846508 | -0.7225935 | 0.46992963 | 0.91502267 |
| PLPBP | 45.6211381 | -0.19768014 | 0.36348343 | -0.5438491 | 0.58654531 | 0.94254614 |
| RAB11FIP1 | 135.02171 | -0.365886423 | 0.39796485 | -0.9193938 | 0.35788962 | 0.87316144 |
| EIF4EBP1 | 45.2946905 | -0.53438221 | 0.40244286 | -1.3278462 | 1.84E-01 | 0.75788529 |
| ASH2L | 66.5499771 | -0.247559077 | 0.22850813 | -1.083371 | 0.2786438 | 0.8229426 |
| LSM1 | 41.9718297 | -0.2158369 | 0.43885481 | -0.4918185 | 0.62284768 | 0.94808496 |
| BAG4 | 68.8483186 | 0.390124854 | 0.26226216 | 1.48753769 | 0.13687286 | 0.71041106 |
| DDHD2 | 106.669717 | -0.347338061 | 0.20532137 | -1.69168 | 0.090707 | 0.6330877 |
| PLPP5 | 43.3255076 | -0.037033894 | 0.42368284 | -0.0874095 | 0.93034604 | 0.99375606 |
| NSD3 | 400.921053 | 0.178380817 | 0.19283135 | 0.92506131 | 0.35493402 | 0.87258503 |
| FGFR1 | 263.660429 | -0.18985303 | 0.28648443 | -0.6626993 | 0.50752316 | 0.92666294 |
| AC069120.1 | 38.7894484 | -0.067974042 | 0.67491232 | -0.1007154 | 0.91977641 | 0.99214128 |
| TACC1 | 270.806189 | 0.502523962 | 0.36295819 | 1.38452299 | 0.16619837 | 0.74445571 |
| PLEKHA2 | 264.910333 | -0.317017182 | 0.33400043 | -0.949152 | 0.34254331 | 0.86846099 |
| ADAM9 | 192.621302 | -0.016395131 | 0.25620729 | -0.0639917 | 0.94897686 | 0.9952642 |
| TCIM | 284.610522 | -0.404644401 | 1.15581854 | -0.3500934 | 0.72626863 | 0.96781837 |
| GOLGA7 | 53.5596178 | -0.089766871 | 0.27015172 | -0.3322832 | 0.73967544 | 0.9698508 |
| GPAT4 | 75.738166 | -0.211325529 | 0.33317922 | -0.6342698 | 0.52590473 | 0.93272384 |
| KAT6A | 390.339106 | 0.119811707 | 0.15954194 | 0.75097312 | 0.45266883 | 0.91077355 |
| AP3M2 | 40.5754072 | 0.035212155 | 0.29935404 | 0.11762712 | 0.90636311 | 0.99100813 |
| IKBKB | 126.461538 | -0.10607072 | 0.27283117 | -0.3887779 | 0.69744047 | 0.96220115 |
| POLB | 38.4441713 | 0.075991723 | 0.31366787 | 0.24226812 | 0.80857242 | 0.97930627 |
| VDAC3 | 89.5460477 | -0.145251462 | 0.24682233 | -0.5884859 | 0.55620619 | 0.93523904 |
| SLC20A2 | 415.466351 | -0.386329369 | 0.24200121 | -1.5963944 | 0.11040077 | 0.65930736 |
| SMIM19 | 69.5073476 | -0.284532487 | 0.27730748 | -1.0260541 | 0.30486607 | 0.84488735 |
| RNF170 | 108.313963 | -0.013393773 | 0.28495907 | -0.0470024 | 0.96251128 | 0.99572129 |
| HOOK3 | 206.664623 | 0.153382566 | 0.13931742 | 1.10095756 | 0.27091513 | 0.81775732 |
| FNTA | 64.8288303 | -0.246577821 | 0.272839 | -0.9037484 | 0.36612881 | 0.87605028 |
| HGSNAT | 192.277168 | -0.106367374 | 0.18511937 | -0.574588 | 0.56556996 | 0.93809297 |
| SPIDR | 1076.78786 | -0.059866001 | 0.3413368 | -0.1753869 | 0.86077564 | 0.98652427 |
| CEBPD | 696.197597 | -0.256923201 | 0.55317724 | -0.4644501 | 0.64232533 | 0.94944486 |
| PRKDC | 283.685019 | -0.6890549 | 0.22178401 | -3.1068737 | 0.00189077 | 0.14133317 |
| UBE2V2 | 104.21223 | -0.133736776 | 0.24847031 | -0.5382405 | 0.59041104 | 0.94516215 |

| | | | | | | |
|---|---|---|---|---|---|---|
| PCMTD1 | 356.09301 | 0.321831854 | 0.22271476 | 1.44504054 | 0.1484465 | 0.72246729 |
| AC064807.1 | 41.9637494 | -0.363567992 | 0.36198896 | -1.0043621 | 0.31520412 | 0.8490543 |
| RB1CC1 | 283.470939 | 0.245971957 | 0.14449587 | 1.70227676 | 0.0887035 | 0.63033443 |
| ATP6V1H | 195.116035 | -0.110755111 | 0.27624926 | -0.4009245 | 0.68847567 | 0.96163728 |
| TCEA1 | 131.292449 | -0.511524164 | 0.16970324 | -3.0142275 | 0.00257635 | 0.16674875 |
| LYPLA1 | 135.851218 | -0.65592081 | 0.25546449 | -2.5675616 | 0.01024166 | 0.29271368 |
| MRPL15 | 35.3356303 | -0.562020056 | 0.41246437 | -1.3625906 | 0.17301159 | 0.74719017 |
| RP1 | 121.126313 | -0.302053249 | 1.17740305 | -0.2565419 | 0.79753241 | 0.97555967 |
| XKR4 | 562.48634 | -0.183573967 | 0.48223726 | -0.3806715 | 7.03E-01 | 0.9645815 |
| TMEM68 | 99.0535063 | 0.012424656 | 0.29676211 | 0.04186739 | 0.96660441 | 0.99601626 |
| TGS1 | 85.8096306 | -0.730731885 | 0.35738455 | -2.0446656 | 0.04088784 | 0.48618451 |
| LYN | 79.7305663 | -0.354547321 | 0.41664684 | -0.8509541 | 0.39479487 | 0.88760715 |
| RPS20 | 246.208998 | 0.251066179 | 0.43606862 | 0.57574924 | 0.56478468 | 0.93809297 |
| PLAG1 | 82.2095539 | -0.341793152 | 0.52991572 | -0.6449953 | 0.51893023 | 0.93257822 |
| CHCHD7 | 32.2858552 | -0.337891244 | 0.39120541 | -0.8637182 | 0.38774269 | 0.8853359 |
| IMPAD1 | 95.3471624 | -0.056498312 | 0.36061672 | -0.1566714 | 0.87550386 | 0.98766275 |
| LINC01606 | 70.4220615 | -0.975547454 | 1.32171337 | -0.738093 | 0.46045791 | 0.91368087 |
| FAM110B | 370.039087 | -0.222976191 | 0.33011683 | -0.6754463 | 0.49939225 | 0.92398825 |
| UBXN2B | 106.023719 | 0.009576146 | 0.26386638 | 0.03629165 | 0.97104981 | 0.99681784 |
| SDCBP | 129.428691 | -0.243860759 | 0.31991942 | -0.7622568 | 0.44590673 | 0.90734558 |
| NSMAF | 116.488257 | -0.580500751 | 0.21211441 | -2.7367342 | 0.00620524 | 0.24799848 |
| TOX | 67.5546905 | 0.347251044 | 0.75074351 | 0.46254285 | 0.64369208 | 0.94944486 |
| LINC01301 | 40.9107389 | 1.286227982 | 0.59090871 | 2.1766949 | 0.02950334 | 0.43492739 |
| RAB2A | 372.886916 | -0.013337882 | 0.20024902 | -0.0666065 | 0.94689499 | 0.9952642 |
| CHD7 | 341.871672 | 0.057257245 | 0.26031013 | 0.2199578 | 0.82590402 | 0.98094663 |
| ASPH | 336.338137 | -0.541025119 | 0.18188332 | -2.9745724 | 0.00293397 | 0.17565323 |
| YTHDF3 | 149.816492 | 0.031978482 | 0.21014682 | 0.1521721 | 0.8790512 | 0.98766275 |
| CYP7B1 | 141.347487 | 0.775185076 | 0.45075389 | 1.71975239 | 0.08547746 | 0.62029438 |
| ARMC1 | 64.85024 | -0.376418266 | 0.21962802 | -1.71389 | 0.08654893 | 0.62381659 |
| MTFR1 | 109.411382 | 0.3207181 | 0.2886256 | 1.11119078 | 0.26648624 | 0.81444418 |
| PDE7A | 190.036936 | 1.07358446 | 0.3110488 | 3.4514985 | 0.00055748 | 0.07416845 |
| TRIM55 | 38.9118767 | 0.528116768 | 0.75983576 | 0.69504069 | 0.4870298 | 0.92034617 |
| ADHFE1 | 71.2895204 | 0.152014632 | 0.60142366 | 0.25275798 | 0.80045524 | 0.97723068 |
| VCPIP1 | 58.6601462 | 0.009443027 | 0.2594678 | 0.03639383 | 0.97096833 | 0.99681784 |
| SGK3 | 125.021123 | 0.320741008 | 0.26333375 | 1.21800191 | 0.22322325 | 0.78766444 |
| PPP1R42 | 38.3936521 | 0.931139798 | 0.57090161 | 1.63099873 | 0.10289059 | 0.65288882 |
| CSPP1 | 321.534065 | 0.30996744 | 0.22574884 | 1.37306327 | 0.16973268 | 0.74445571 |
| ARFGEF1 | 559.854992 | 0.144081686 | 0.14554811 | 0.9899248 | 0.32221088 | 0.85246644 |
| PREX2 | 49.6651463 | -2.645717089 | 1.29680595 | -2.0401796 | 0.04133244 | 0.48819163 |
| C8orf34 | 267.044341 | -1.137113093 | 0.6385436 | -1.7807916 | 0.0749465 | 0.59444897 |
| SULF1 | 59.021345 | 0.306488989 | 0.66598742 | 0.46020237 | 0.64537097 | 0.94954229 |
| NCOA2 | 867.71206 | 0.180126892 | 0.24642992 | 0.7309457 | 0.46481232 | 0.91462428 |
| TRAM1 | 224.465724 | 0.368075038 | 0.31748776 | 1.15933617 | 0.24631918 | 0.80556213 |
| LACTB2 | 66.0523336 | 0.571366823 | 0.36528472 | 1.56416844 | 0.11777802 | 0.68127756 |
| EYA1 | 114.988554 | 0.278637407 | 0.87179977 | 0.3196117 | 0.7492627 | 0.97028355 |
| MSC-AS1 | 304.411333 | 1.671241122 | 1.2385502 | 1.34935275 | 0.17722369 | 0.75126783 |
| TERF1 | 97.8608481 | 0.452754528 | 0.18593922 | 2.43495981 | 0.01489344 | 0.3338856 |
| RPL7 | 452.629382 | 0.312255444 | 0.65248054 | 0.47856668 | 0.63224693 | 0.94932781 |
| RDH10 | 118.89145 | -0.872557116 | 0.75675986 | -1.1530172 | 0.24890334 | 0.80672756 |
| RDH10-AS1 | 45.9354408 | -0.230744226 | 0.86423829 | -0.2669914 | 0.78947575 | 0.97423208 |
| STAU2 | 329.65487 | 0.327857821 | 0.17151597 | 1.91152943 | 0.05593658 | 0.54122877 |
| AC027018.1 | 49.9187944 | 1.164949787 | 0.57266018 | 2.03427761 | 0.04192361 | 0.4889557 |
| ELOC | 74.9699554 | -0.106255268 | 0.26710973 | -0.3977963 | 0.69078033 | 0.96220115 |
| GDAP1 | 70.9765667 | 0.04403566 | 0.38050812 | 0.11572857 | 0.90786766 | 0.99100813 |

| | | | | | | |
|---|---|---|---|---|---|---|
| HNF4G | 64.1573412 | 0.501031504 | 0.50899202 | 0.98436023 | 0.32493842 | 0.855037 |
| LINC01111 | 44.6709183 | -0.191795095 | 0.84473537 | -0.2270475 | 8.20E-01 | 0.98094663 |
| ZFHX4 | 42.8106171 | -0.186657459 | 0.80310535 | -0.2324196 | 0.81621209 | 0.98008989 |
| PEX2 | 69.0846521 | -0.006936253 | 0.24841139 | -0.0279224 | 0.97772401 | 0.99681784 |
| ZC2HC1A | 96.8371308 | -0.1146111 | 0.43834123 | -0.2614655 | 0.79373357 | 0.97464093 |
| IL7 | 59.9555385 | 0.472573606 | 0.4490635 | 1.05235364 | 0.29263733 | 0.83697123 |
| TPD52 | 184.403382 | -0.040591823 | 0.37166192 | -0.1092171 | 0.91303033 | 0.99113939 |
| ZBTB10 | 101.00751 | 0.566370765 | 0.2865131 | 1.9767709 | 0.04806753 | 0.51547342 |
| ZNF704 | 203.388434 | 0.463884945 | 0.57254502 | 0.81021567 | 0.41781623 | 0.89862171 |
| PAG1 | 248.123548 | -0.198872838 | 1.04737439 | -0.1898775 | 0.84940512 | 0.98575862 |
| IMPA1 | 53.6192759 | -0.333202235 | 0.40304816 | -0.8267058 | 0.40840386 | 0.8959746 |
| ZFAND1 | 88.4745076 | 0.032476315 | 0.2976741 | 0.10910024 | 0.91312299 | 0.99113939 |
| CHMP4C | 32.7871041 | -0.086867564 | 0.43830176 | -0.1981912 | 0.84289545 | 0.98517901 |
| SNX16 | 49.4325059 | -0.549422624 | 0.31106743 | -1.7662493 | 0.07735403 | 0.5981029 |
| E2F5 | 49.7297282 | 0.10888869 | 0.31011526 | 0.35112328 | 0.72549586 | 0.96781837 |
| C8orf59 | 53.597773 | -0.600522578 | 0.27626813 | -2.1736948 | 0.02972807 | 0.43492739 |
| CA13 | 66.8626845 | -0.083067236 | 0.28117543 | -0.2954285 | 0.76766656 | 0.97229645 |
| CA3 | 88.2012034 | 0.316151998 | 0.41431088 | 0.76307916 | 0.44541618 | 0.90718817 |
| CA2 | 61.5877995 | 0.114925174 | 0.49923226 | 0.23020382 | 0.81793339 | 0.98010849 |
| WWP1 | 394.886433 | 0.253797218 | 0.18488903 | 1.37270022 | 0.16984556 | 0.74445571 |
| RMDN1 | 115.892765 | -0.305611933 | 0.17911201 | -1.7062615 | 0.0879594 | 0.62737387 |
| CPNE3 | 102.287159 | -0.273292205 | 0.28459932 | -0.9602701 | 0.33691932 | 0.86431445 |
| AF117829.1 | 309.633747 | 0.56421135 | 0.46093829 | 1.22404964 | 0.2209335 | 0.78615231 |
| RIPK2 | 41.0023928 | 0.086931352 | 0.53173815 | 0.16348526 | 0.87013638 | 0.98766275 |
| OSGIN2 | 54.7788373 | -0.001828768 | 0.32852753 | -0.0055666 | 0.99555855 | 0.99866813 |
| NBN | 105.814874 | -0.582228951 | 0.34560262 | -1.6846775 | 0.09205081 | 0.63464873 |
| DECR1 | 143.558057 | 0.331417421 | 0.26267028 | 1.26172407 | 0.20704809 | 0.77401635 |
| TMEM64 | 130.130696 | 0.478273493 | 0.43630644 | 1.0961871 | 0.2729969 | 0.82047446 |
| NECAB1 | 106.362332 | 0.514116769 | 0.66093052 | 0.7778681 | 0.43664677 | 0.90533321 |
| PIP4P2 | 37.181111 | -0.10445608 | 0.4548218 | -0.2296637 | 0.81835307 | 0.98012053 |
| OTUD6B-AS1 | 41.9088475 | 0.063555912 | 0.38215775 | 0.16630806 | 0.86791453 | 0.98670067 |
| OTUD6B | 45.0712635 | 0.141038107 | 0.33665772 | 0.4189362 | 0.67526276 | 0.95910677 |
| LRRC69 | 65.0643237 | 0.293578286 | 0.28874159 | 1.01675094 | 0.30927193 | 0.84554063 |
| TRIQK | 108.114807 | 0.096628814 | 0.20629093 | 0.46841037 | 0.63949115 | 0.94944486 |
| FAM92A | 41.5033989 | 0.718514114 | 0.33839249 | 2.12331578 | 0.0337274 | 0.44933903 |
| RBM12B | 76.313161 | 0.212324286 | 0.20583575 | 1.03152289 | 0.30229568 | 0.84375949 |
| TMEM67 | 140.343357 | 0.38892515 | 0.27430017 | 1.41788156 | 0.15622535 | 0.73307914 |
| VIRMA | 112.029486 | -0.088818989 | 0.19627972 | -0.4525123 | 0.65089996 | 0.95103275 |
| ESRP1 | 97.747885 | -0.211595519 | 0.71703741 | -0.2950969 | 0.76791985 | 0.97236012 |
| DPY19L4 | 102.299596 | -0.062793048 | 0.22581031 | -0.2780787 | 0.78095191 | 0.97423208 |
| INTS8 | 101.916183 | -0.076533503 | 0.25323039 | -0.3022287 | 0.7624777 | 0.9714717 |
| NDUFAF6 | 120.200863 | -0.178026077 | 0.22847294 | -0.7791998 | 0.43586201 | 0.90514499 |
| TP53INP1 | 42.0979278 | 0.984284174 | 0.42228231 | 2.33086766 | 0.01976034 | 0.3747883 |
| PLEKHF2 | 52.4900938 | 0.378022886 | 0.521523 | 0.72484414 | 0.46854766 | 0.91462428 |
| C8orf37-AS1 | 91.2777194 | 0.945116132 | 0.52843446 | 1.78852101 | 0.07369199 | 0.59320069 |
| UQCRB | 188.287202 | 0.491201481 | 0.45928359 | 1.06949494 | 0.28484671 | 0.82918183 |
| PTDSS1 | 34.9923581 | -0.624878714 | 0.40018449 | -1.5614766 | 0.11841134 | 0.68370239 |
| SDC2 | 446.772653 | -0.163884657 | 0.35026322 | -0.46789 | 0.63986325 | 0.94944486 |
| CPQ | 345.555681 | 0.069067761 | 0.33235621 | 0.20781246 | 0.83537541 | 0.9841423 |
| MTDH | 405.418211 | -0.048217355 | 0.14912506 | -0.323335 | 0.74644153 | 0.97028355 |
| LAPTM4B | 171.980328 | -0.593931219 | 0.39237087 | -1.5136985 | 0.13010231 | 0.69904834 |
| RPL30 | 324.276424 | -0.018099817 | 0.52226018 | -0.0346567 | 0.97235348 | 0.99681784 |
| RIDA | 58.4880907 | 0.913793442 | 0.35850858 | 2.54887467 | 0.01080711 | 0.29965266 |
| NIPAL2 | 44.3561022 | -0.254867582 | 0.36271617 | -0.702664 | 0.48226519 | 0.91971919 |

| | | | | | | |
|---|---|---|---|---|---|---|
| STK3 | 795.33065 | -0.300154949 | 0.17362704 | -1.7287339 | 0.08385673 | 0.61656297 |
| AC104986.2 | 61.6021269 | 0.397952194 | 0.37127507 | 1.07185273 | 0.28378619 | 0.82918183 |
| VPS13B | 799.086454 | -0.102050534 | 0.16162719 | -0.6313946 | 0.52778254 | 0.93366189 |
| COX6C | 130.376338 | -0.354857781 | 0.30360772 | -1.1688036 | 0.24248279 | 0.80476923 |
| POLR2K | 53.9094612 | -0.263014221 | 0.54923806 | -0.4788711 | 0.63203035 | 0.94932781 |
| SPAG1 | 141.934481 | 0.462138237 | 0.40976761 | 1.12780568 | 0.259402 | 0.81263292 |
| RNF19A | 318.67474 | 0.228945679 | 0.31402206 | 0.72907515 | 0.4659557 | 0.91462428 |
| ANKRD46 | 62.9734836 | -0.125972855 | 0.24832816 | -0.5072838 | 0.6119557 | 0.94791646 |
| PABPC1 | 329.06014 | -0.123581989 | 0.3932691 | -0.3142428 | 0.75333663 | 0.97096495 |
| YWHAZ | 419.888328 | -0.246820665 | 0.26065002 | -0.9469428 | 0.34366792 | 0.86909179 |
| ZNF706 | 141.651979 | 0.182999059 | 0.28597448 | 0.63991395 | 0.52222855 | 0.93272217 |
| AP001207.3 | 29.7372225 | -0.799345441 | 1.4128061 | -0.5657857 | 0.57153949 | 0.93998433 |
| GRHL2 | 373.387277 | -0.584228814 | 0.82555036 | -0.707684 | 0.47914149 | 0.91971919 |
| NCALD | 56.7074294 | 0.368440239 | 0.66642926 | 0.55285723 | 0.58036117 | 0.94034284 |
| RRM2B | 96.2714809 | 0.352135538 | 0.46331751 | 0.76003072 | 0.44723622 | 0.90918416 |
| UBR5 | 557.562105 | 0.020233856 | 0.17248004 | 0.11731129 | 0.90661338 | 0.99100813 |
| KLF10 | 122.752416 | 0.346961895 | 0.53340003 | 0.6504722 | 0.51538725 | 0.93140514 |
| AZIN1 | 118.663941 | 0.185352479 | 0.30057593 | 0.61665776 | 0.53746049 | 0.93504069 |
| AZIN1-AS1 | 255.823303 | -0.268652458 | 0.33992726 | -0.7903234 | 0.42933895 | 0.90333313 |
| AP003354.2 | 70.448432 | 0.042323491 | 0.39961913 | 0.10590957 | 0.9156541 | 0.99159829 |
| ATP6V1C1 | 176.675253 | 0.169240955 | 0.31675102 | 0.53430279 | 0.59313207 | 0.94649617 |
| BAALC-AS1 | 72.9394886 | -0.124232187 | 0.60416209 | -0.2056272 | 0.83708209 | 0.98462033 |
| FZD6 | 174.395157 | 0.168118125 | 0.2064054 | 0.8145045 | 0.415356 | 0.89807127 |
| SLC25A32 | 53.8133284 | -0.074141028 | 0.26838068 | -0.2762532 | 0.78235358 | 0.97423208 |
| DCAF13 | 38.2747228 | -0.56287512 | 0.42201808 | -1.3337702 | 0.18227919 | 0.75456023 |
| RIMS2 | 135.194385 | 1.754522804 | 1.08387139 | 1.61875552 | 0.10549988 | 0.65699665 |
| DPYS | 43.736251 | 1.897621937 | 0.80454672 | 2.35862242 | 0.01834291 | 0.36034684 |
| LRP12 | 51.8656477 | 0.126797074 | 0.3175153 | 0.39934162 | 0.68964151 | 0.96220115 |
| ZFPM2 | 685.641815 | 0.363079746 | 1.19846497 | 0.30295399 | 0.76192492 | 0.9714717 |
| OXR1 | 2630.16534 | -0.535296549 | 0.23913979 | -2.2384252 | 0.02519334 | 0.4135769 |
| AC090579.1 | 268.981111 | 0.179937965 | 0.5796383 | 0.31043146 | 0.75623287 | 0.9714717 |
| EIF3E | 237.522677 | 0.181865721 | 0.15202704 | 1.19627219 | 0.23159036 | 0.79459836 |
| EMC2 | 135.662303 | 0.063996228 | 0.33233395 | 0.19256603 | 0.84729885 | 0.98534366 |
| NUDCD1 | 88.7962607 | -0.589414998 | 0.43114883 | -1.3670801 | 0.17160019 | 0.74445571 |
| ENY2 | 89.6785177 | -0.069768316 | 0.30982426 | -0.2251867 | 0.821834 | 0.98094663 |
| EBAG9 | 51.2961766 | -0.30002082 | 0.29250668 | -1.0256888 | 0.3050383 | 0.84488735 |
| TRPS1 | 629.099024 | 0.52965481 | 0.21017086 | 2.52011539 | 0.01173164 | 0.30872898 |
| EIF3H | 253.019962 | -0.033588373 | 0.22429885 | -0.1497483 | 0.8809632 | 0.98766275 |
| UTP23 | 43.9369197 | -0.108375108 | 0.27729247 | -0.3908332 | 0.69592052 | 0.96220115 |
| RAD21 | 133.418989 | 0.085377472 | 0.21966396 | 0.38867309 | 0.69751799 | 0.96220115 |
| MED30 | 26.0315107 | -0.185544321 | 0.39684429 | -0.4675494 | 0.64010684 | 0.94944486 |
| EXT1 | 1191.32832 | 0.001054769 | 0.53004647 | 0.00198996 | 0.99841225 | 0.99971834 |
| SAMD12 | 883.199804 | 0.044323696 | 0.22161143 | 0.20000637 | 0.8414756 | 0.98515707 |
| AC023590.1 | 85.2803121 | 0.579618131 | 0.5385599 | 1.07623708 | 0.28182124 | 0.82749734 |
| SAMD12-AS1 | 58.9113429 | -0.437630878 | 0.33787065 | -1.2952616 | 0.19523 | 0.76968045 |
| TNFRSF11B | 531.993313 | 0.701533917 | 0.39896688 | 1.75837631 | 0.0786835 | 0.60345017 |
| COLEC10 | 116.926336 | 1.04500486 | 0.53169457 | 1.96542323 | 0.04936527 | 0.52308378 |
| MAL2 | 198.480905 | -0.921370947 | 0.47033651 | -1.9589612 | 0.05011734 | 0.52415717 |
| TAF2 | 159.952518 | -0.220173859 | 0.22519442 | -0.9777056 | 0.32821993 | 0.8573817 |
| DEPTOR | 917.474469 | -0.993898458 | 0.62626516 | -1.587025 | 0.11250699 | 0.66401239 |
| AC091563.1 | 69.0337969 | -0.247645651 | 0.62006305 | -0.3993879 | 0.68960744 | 0.96220115 |
| COL14A1 | 83.7658522 | -0.243702321 | 0.60578185 | -0.4022939 | 0.68746777 | 0.96161098 |
| MRPL13 | 48.3986323 | 0.120462686 | 0.28195116 | 0.42724664 | 0.66919969 | 0.95764829 |
| MTBP | 30.7695402 | -0.540175959 | 0.42692145 | -1.2652818 | 0.20577031 | 0.7735572 |

| | | | | | | |
|---|---|---|---|---|---|---|
| SNTB1 | 663.164981 | -0.31576968 | 0.20743748 | -1.5222403 | 0.12794889 | 0.69635098 |
| ZHX2 | 640.931537 | 0.092076821 | 0.28964682 | 0.31789343 | 0.75056578 | 0.97058015 |
| AC104316.1 | 38.4429514 | -0.25054536 | 0.54265678 | -0.4617013 | 0.64429551 | 0.94944486 |
| DERL1 | 126.118686 | -0.410509678 | 0.28545838 | -1.4380719 | 0.15041365 | 0.72274179 |
| WDYHV1 | 28.5063331 | 0.133367776 | 0.47067254 | 0.28335576 | 0.77690414 | 0.97423208 |
| FAM91A1 | 119.527291 | 0.137625808 | 0.2753157 | 0.49988362 | 0.61715703 | 0.94791646 |
| TMEM65 | 184.095744 | -0.080372598 | 0.22484301 | -0.357461 | 0.72074675 | 0.96779196 |
| RNF139 | 26.6592776 | -0.156880608 | 0.29747458 | -0.5273748 | 0.59793331 | 0.94683415 |
| TATDN1 | 113.149149 | -0.056040435 | 0.23073669 | -0.2428761 | 0.80810135 | 0.97930627 |
| NDUFB9 | 84.4089389 | -0.075669692 | 0.31100634 | -0.2433059 | 0.8077684 | 0.97930627 |
| MTSS1 | 546.617677 | -0.089067243 | 0.53134371 | -0.1676264 | 0.86687719 | 0.98670067 |
| SQLE | 88.4450851 | 0.660870968 | 0.66446815 | 0.99458638 | 0.31993748 | 0.85075884 |
| WASHC5 | 69.7374468 | -0.273262892 | 0.21010441 | -1.3006052 | 0.19339362 | 0.76797319 |
| NSMCE2 | 573.513466 | -0.346542955 | 0.21610892 | -1.6035569 | 0.10881175 | 0.65764016 |
| TRIB1 | 69.4288593 | 0.221806883 | 0.3382478 | 0.65575263 | 0.5119833 | 0.9287766 |
| FAM84B | 121.867898 | -1.311784161 | 0.63051902 | -2.0804831 | 0.03748125 | 0.46930788 |
| PCAT1 | 996.87955 | -0.5790072 | 0.57721066 | -1.0031124 | 0.31580661 | 0.8490543 |
| PVT1 | 240.330931 | 0.122143262 | 0.7506672 | 0.16271293 | 0.87074446 | 0.98766275 |
| FAM49B | 377.420681 | -0.334604282 | 0.25326773 | -1.3211485 | 0.18645185 | 0.76162997 |
| ASAP1 | 400.965952 | -0.018252136 | 0.49639388 | -0.0367695 | 0.97066882 | 0.99681784 |
| EFR3A | 236.851045 | -0.245217765 | 0.20765404 | -1.1808957 | 0.23764415 | 0.80098286 |
| KCNQ3 | 142.50388 | 0.697178411 | 1.0604123 | 0.65745975 | 0.51088534 | 0.92819651 |
| LRRC6 | 172.095743 | 0.56806033 | 0.30954854 | 1.83512523 | 0.06648714 | 0.5780981 |
| PHF20L1 | 135.914461 | -0.014942601 | 0.2319383 | -0.0644249 | 0.94863191 | 0.9952642 |
| NDRG1 | 383.112095 | 0.149996917 | 0.81634153 | 0.18374285 | 0.85421519 | 0.9864199 |
| ST3GAL1 | 81.7239547 | 0.652702925 | 0.82796958 | 0.78831752 | 0.430511 | 0.90364119 |
| ZFAT | 62.7414973 | 0.003802329 | 0.30481048 | 0.0124744 | 0.99004713 | 0.99839055 |
| AC083843.3 | 41.534817 | -0.122619931 | 0.43196135 | -0.2838678 | 0.77651167 | 0.97423208 |
| KHDRBS3 | 99.929385 | 0.341498394 | 0.41449132 | 0.82389757 | 0.40999776 | 0.89625024 |
| TRAPPC9 | 549.78407 | -0.207697518 | 0.256482 | -0.8097938 | 0.41805872 | 0.89862171 |
| CHRAC1 | 28.6229144 | -0.680275755 | 0.32908149 | -2.0671955 | 0.03871574 | 0.47547104 |
| AGO2 | 218.951636 | -0.177649709 | 0.18662026 | -0.9519315 | 0.34113171 | 0.86837647 |
| PTK2 | 1370.16147 | -0.217773196 | 0.20913565 | -1.0413011 | 0.29773581 | 0.83973489 |
| DENND3 | 112.275813 | -0.288663626 | 0.35560143 | -0.8117617 | 0.41692837 | 0.89862171 |
| SLC45A4 | 225.796214 | -0.360428968 | 0.62660251 | -0.5752115 | 0.56514827 | 0.93809297 |
| TSNARE1 | 59.9792485 | -0.093215829 | 0.30808868 | -0.3025617 | 0.76222392 | 0.9714717 |
| JRK | 31.6439404 | 0.549175786 | 0.42869043 | 1.28105447 | 0.20017453 | 0.77110248 |
| LY6E | 239.229268 | -0.542695113 | 1.26869441 | -0.4277587 | 0.66882679 | 0.95764829 |
| RHPN1 | 42.7664723 | 0.030217301 | 0.38478973 | 0.07852938 | 0.93740696 | 0.99526299 |
| ZC3H3 | 52.4653937 | -0.142624588 | 0.34048744 | -0.4188836 | 0.67530124 | 0.95910677 |
| GSDMD | 67.1948772 | 0.214438046 | 0.55566664 | 0.38591132 | 0.69956235 | 0.96280283 |
| EEF1D | 314.778861 | 0.030582132 | 0.45852159 | 0.06669726 | 0.94682272 | 0.9952642 |
| TSTA3 | 37.0728257 | 0.282809536 | 0.53260056 | 0.53099744 | 0.59542055 | 0.94664456 |
| ZNF623 | 36.5954667 | -0.34572192 | 0.39261195 | -0.880569 | 0.37855113 | 0.87970706 |
| MAPK15 | 31.2562807 | 0.606646948 | 0.603955 | 1.0044572 | 0.31515829 | 0.8490543 |
| FAM83H | 41.3981341 | -0.585311482 | 0.35428283 | -1.6521023 | 0.09851369 | 0.64312419 |
| IQANK1 | 31.1796069 | -0.236877041 | 0.53929379 | -0.4392356 | 0.66049082 | 0.95407427 |
| SCRIB | 27.6122167 | 0.035312986 | 0.39433176 | 0.08955146 | 0.92864366 | 0.99336859 |
| PUF60 | 68.557405 | -0.23000242 | 0.39035006 | -0.5892209 | 0.55571308 | 0.93523904 |
| NRBP2 | 79.8308008 | 0.504028098 | 0.65045583 | 0.77488444 | 0.43840795 | 0.90561284 |
| PLEC | 92.3476685 | -0.179649496 | 0.35219889 | -0.5100797 | 0.60999564 | 0.94791646 |
| PARP10 | 49.9989235 | -1.067809359 | 0.74129855 | -1.4404579 | 0.14973789 | 0.72246729 |
| GRINA | 83.3788567 | -0.149231217 | 0.47086993 | -0.3169266 | 0.75129928 | 0.97058015 |
| GPAA1 | 57.1698179 | 0.245257156 | 0.47135376 | 0.52032502 | 0.60283706 | 0.94683415 |

| | | | | | | |
|---|---|---|---|---|---|---|
| CYC1 | 54.145749 | 0.058958632 | 0.46808406 | 0.12595736 | 0.89976568 | 0.98956835 |
| SHARPIN | 40.7307194 | 0.233058094 | 0.24711375 | 0.94312071 | 0.34561916 | 0.86990996 |
| MAF1 | 50.9438764 | 0.116832576 | 0.39193899 | 0.29808868 | 0.76563548 | 0.97177982 |
| MROH1 | 152.776327 | -0.193380089 | 0.39968468 | -0.4838316 | 0.62850537 | 0.94858781 |
| HSF1 | 89.9968959 | -0.18144615 | 0.2565099 | -0.7073651 | 0.47933962 | 0.91971919 |
| CPSF1 | 25.1049429 | 0.401766021 | 0.55227176 | 0.72747884 | 0.46693267 | 0.91462428 |
| VPS28 | 120.313006 | 0.02611494 | 0.43063872 | 0.06064234 | 0.95164405 | 0.99531843 |
| CYHR1 | 62.3309039 | 0.092717836 | 0.26359472 | 0.35174391 | 0.72503033 | 0.96781837 |
| PPP1R16A | 79.1276542 | 0.273893239 | 0.38018243 | 0.72042583 | 0.47126285 | 0.91634658 |
| ARHGAP39 | 39.4786574 | 0.395930655 | 0.32859584 | 1.20491682 | 0.22823542 | 0.79284833 |
| ZNF251 | 115.660308 | 0.1051105 | 0.24629875 | 0.42676018 | 0.66955401 | 0.95764829 |
| RPL8 | 650.67521 | -0.131046856 | 0.57717036 | -0.2270506 | 0.82038442 | 0.98094663 |
| ZNF7 | 38.5869833 | -0.678674532 | 0.27436285 | -2.4736386 | 0.01337449 | 0.32022149 |
| ZNF250 | 36.8386592 | -0.083709085 | 0.32293062 | -0.2592169 | 0.79546787 | 0.97524554 |
| C8orf33 | 29.5196648 | -0.576587713 | 0.37907844 | -1.5210248 | 0.12825361 | 0.69645198 |
| WASHC1 | 154.500073 | 0.037430896 | 0.28888215 | 0.12957151 | 0.89690545 | 0.98842187 |
| CBWD1 | 49.8739167 | -0.000538978 | 0.37588552 | -0.0014339 | 0.99885592 | 0.99971834 |
| DOCK8 | 163.169727 | 0.025089246 | 0.5265899 | 0.04764475 | 0.96199936 | 0.99572129 |
| KANK1 | 1174.41694 | 0.013193238 | 0.16111321 | 0.081888 | 9.35E-01 | 9.94E-01 |
| SMARCA2 | 235.184231 | -0.002998495 | 0.36137003 | -0.0082976 | 0.99337957 | 0.99866813 |
| VLDLR-AS1 | 41.9718508 | -0.480654017 | 0.41731504 | -1.1517774 | 0.24941257 | 0.80672756 |
| VLDLR | 38.7678349 | 0.07087832 | 0.42406958 | 0.16713842 | 0.86726114 | 0.98670067 |
| PUM3 | 352.39612 | -0.709017448 | 0.31265086 | -2.2677611 | 0.02334377 | 0.39839181 |
| RFX3 | 439.272609 | 0.528834857 | 0.27684632 | 1.91021086 | 0.05610607 | 0.54232092 |
| RFX3-AS1 | 389.11428 | -0.018526554 | 0.38323031 | -0.0483431 | 0.96144278 | 0.99572129 |
| GLIS3 | 2072.41547 | 0.365243934 | 0.33364189 | 1.09471845 | 0.27364 | 0.82047446 |
| SLC1A1 | 248.431034 | -0.296345482 | 0.36009613 | -0.8229621 | 0.41052952 | 0.8964268 |
| SPATA6L | 280.320884 | 0.46959753 | 0.41847009 | 1.12217704 | 0.26178717 | 0.81308142 |
| CDC37L1 | 80.7821686 | 0.081873124 | 0.26636934 | 0.30736692 | 0.7585641 | 0.9714717 |
| AK3 | 287.497277 | 0.576561845 | 0.21826403 | 2.64157979 | 0.00825204 | 0.27246484 |
| RCL1 | 113.36457 | -0.767398924 | 0.31033378 | -2.4728179 | 0.01340524 | 0.32022149 |
| JAK2 | 130.150801 | -0.067908302 | 0.28744739 | -0.236246 | 0.81324177 | 0.97930627 |
| PLGRKT | 73.3171745 | -0.05026052 | 0.29478264 | -0.1705003 | 0.86461672 | 0.98670067 |
| RIC1 | 360.635918 | -0.310499696 | 0.2539776 | -1.2225476 | 0.22150062 | 0.78700092 |
| ERMP1 | 531.704329 | -1.463786918 | 0.68117294 | -2.1489211 | 0.03164065 | 0.43905874 |
| KIAA2026 | 344.108507 | -0.102935525 | 0.18856615 | -0.5458855 | 0.58514465 | 0.94229635 |
| RANBP6 | 29.1643685 | -0.287845706 | 0.38104095 | -0.7554194 | 0.4499974 | 0.91035378 |
| UHRF2 | 170.770984 | -0.294393055 | 0.20432995 | -1.4407729 | 0.14964886 | 0.72246729 |
| GLDC | 92.8396632 | -0.91087858 | 0.66626773 | -1.367136 | 0.17158269 | 0.74445571 |
| AL162411.1 | 49.6252367 | 0.410109158 | 0.3494726 | 1.17350877 | 0.24059184 | 0.8030067 |
| KDM4C | 559.204465 | 0.113547888 | 0.21144652 | 0.53700523 | 0.591264 | 0.94537103 |
| DMAC1 | 47.273841 | -0.020066782 | 0.40997636 | -0.0489462 | 0.96096218 | 0.99572129 |
| PTPRD | 470.591558 | 1.057559767 | 0.38209443 | 2.76779685 | 0.00564366 | 0.2367511 |
| MPDZ | 162.104007 | 0.828306101 | 0.30178139 | 2.74472226 | 0.00605621 | 0.24646394 |
| NFIB | 745.772945 | 0.416536511 | 0.3463506 | 1.20264411 | 0.22911407 | 0.7931636 |
| ZDHHC21 | 277.433938 | -0.06382331 | 0.22200775 | -0.2874824 | 7.74E-01 | 0.97396843 |
| TTC39B | 242.213369 | 0.429806698 | 0.53281961 | 0.80666456 | 0.41985977 | 0.90012434 |
| SNAPC3 | 111.422014 | -0.080641975 | 0.21266323 | -0.3792004 | 0.70453908 | 0.96461079 |
| PSIP1 | 103.829921 | 0.203773494 | 0.28034338 | 0.72687108 | 0.46730494 | 0.91462428 |
| CCDC171 | 125.112704 | 0.054437436 | 0.39223968 | 0.13878615 | 0.88961914 | 0.98766275 |
| BNC2 | 833.809455 | 0.482080268 | 0.5540985 | 0.8700263 | 0.38428603 | 0.88328503 |
| CNTLN | 169.220179 | 0.511621013 | 0.26594223 | 1.92380506 | 0.05437902 | 0.53752502 |
| RRAGA | 43.3831898 | 0.07172065 | 0.48008268 | 0.14939229 | 0.8812441 | 0.98766275 |
| HAUS6 | 42.4393176 | -0.062395154 | 0.26796075 | -0.2328518 | 0.81587645 | 0.98007572 |

| | | | | | | |
|---|---|---|---|---|---|---|
| PLIN2 | 128.81303 | 0.22837722 | 0.65104857 | 0.35078369 | 7.26E-01 | 0.96781837 |
| DENND4C | 435.473548 | 0.008525968 | 0.21033539 | 0.04053511 | 0.96766652 | 0.99648221 |
| RPS6 | 705.935248 | 0.145635989 | 0.64610437 | 0.22540629 | 0.82166322 | 0.98094663 |
| MLLT3 | 742.492071 | 0.143170172 | 0.31975776 | 0.44774574 | 0.65433671 | 0.95280477 |
| FOCAD | 471.533514 | -0.085381494 | 0.23988506 | -0.3559267 | 0.72189547 | 0.96779196 |
| KLHL9 | 45.5030629 | -0.194801767 | 0.38438447 | -0.5067889 | 0.61230298 | 0.94791646 |
| LINC01239 | 43.8439463 | 0.937867587 | 0.38977592 | 2.40617117 | 0.01612071 | 0.34229417 |
| CAAP1 | 183.383755 | -0.103769889 | 0.29859216 | -0.3475305 | 0.7281928 | 0.96827499 |
| PLAA | 70.7819882 | -0.439516562 | 0.21999339 | -1.9978625 | 0.04573156 | 0.50584602 |
| IFT74 | 227.931964 | 0.196078259 | 0.26748745 | 0.73303723 | 0.46353572 | 0.91462428 |
| LRRC19 | 71.2749357 | 0.664498745 | 0.53818598 | 1.23470095 | 0.21694182 | 0.78319543 |
| MOB3B | 425.937894 | 0.326625758 | 0.6524633 | 0.50060403 | 0.61664983 | 0.94791646 |
| C9orf72 | 75.4119332 | -0.153412482 | 0.42864341 | -0.3579023 | 0.7204164 | 0.96779196 |
| ACO1 | 114.82045 | 0.794786795 | 0.36232237 | 2.19359018 | 0.02826488 | 0.4250936 |
| DDX58 | 167.857414 | -2.050755463 | 0.86063502 | -2.3828399 | 0.01717966 | 0.35165689 |
| TOPORS | 44.926508 | 0.162180327 | 0.26417695 | 0.61390795 | 0.53927616 | 0.93523904 |
| NDUFB6 | 65.0613391 | -0.197586758 | 0.3412346 | -0.5790349 | 0.5625656 | 0.93800102 |
| APTX | 121.272088 | -0.021594856 | 0.37417358 | -0.0577135 | 0.95397686 | 0.99531843 |
| DNAJA1 | 112.198793 | 0.05727038 | 0.33717619 | 0.16985298 | 0.86512576 | 0.98670067 |
| SMU1 | 76.2401567 | 0.105619408 | 0.2857579 | 0.36961151 | 0.71167197 | 0.96587538 |
| B4GALT1 | 314.967426 | 0.24836317 | 0.41520687 | 0.5981673 | 0.54972831 | 0.93523904 |
| BAG1 | 84.9520215 | -0.48079461 | 0.52337582 | -0.9186412 | 0.35828326 | 0.87316144 |
| CHMP5 | 78.7464674 | -0.218430973 | 0.42479052 | -0.5142087 | 0.6071061 | 0.94708529 |
| NFX1 | 221.303068 | -0.55102853 | 0.2344644 | -2.3501587 | 0.01876541 | 0.36488708 |
| AQP3 | 73.1309448 | 1.881641033 | 0.75103874 | 2.50538479 | 0.01223182 | 0.30967852 |
| UBE2R2 | 359.887834 | 0.032206194 | 0.28101777 | 0.11460554 | 0.90875779 | 0.99100813 |
| UBAP2 | 421.467932 | -0.235283831 | 0.19586903 | -1.2012304 | 0.22966186 | 0.79370986 |
| DCAF12 | 55.6651988 | -0.167379657 | 0.2812425 | -0.5951435 | 0.55174753 | 0.93523904 |
| UBAP1 | 327.473571 | 0.056713494 | 0.23866497 | 0.23762806 | 0.81216958 | 0.97930627 |
| FAM219A | 72.6177134 | 0.115812735 | 0.37849581 | 0.30598155 | 0.75961869 | 0.9714717 |
| DNAI1 | 44.3721474 | 1.569236592 | 0.52633233 | 2.9814558 | 0.00286881 | 0.17392642 |
| DCTN3 | 39.2779223 | 0.639706584 | 0.50225153 | 1.27367772 | 0.20277765 | 0.77110248 |
| VCP | 147.588216 | -0.20108449 | 0.32329637 | -0.6219819 | 0.53395377 | 0.93446646 |
| STOML2 | 51.6991892 | 0.031823878 | 0.50664172 | 0.06281338 | 0.94991511 | 0.99531843 |
| UNC13B | 185.082611 | 0.268235233 | 0.26247336 | 1.02195224 | 0.30680351 | 0.84488735 |
| RUSC2 | 205.138505 | 0.057579601 | 0.46615392 | 0.12352058 | 0.90169488 | 0.9900784 |
| CCDC107 | 40.0895728 | 0.188312796 | 0.37706752 | 0.49941399 | 0.61748776 | 0.94791646 |
| TLN1 | 115.997896 | 0.21593227 | 0.34476753 | 0.62631267 | 0.53110988 | 0.93366189 |
| CREB3 | 28.3502623 | 0.492579578 | 0.51162352 | 0.96277744 | 0.33565923 | 0.86391445 |
| GBA2 | 59.8041127 | -0.148524557 | 0.23031498 | -0.6448758 | 0.51900767 | 0.93257822 |
| RGP1 | 42.6257553 | -0.068609002 | 0.33418412 | -0.205303 | 0.8373354 | 0.98462033 |
| AL133410.1 | 54.0185229 | -0.093071234 | 0.40611953 | -0.229172 | 0.81873522 | 0.98033308 |
| TMEM8B | 43.662208 | 0.135755596 | 0.44577351 | 0.3045394 | 0.76071698 | 0.9714717 |
| RECK | 98.0070522 | 0.647095613 | 0.39470723 | 1.63943186 | 0.10112335 | 0.64836718 |
| CLTA | 274.581487 | -0.442179357 | 0.2760902 | -1.6015757 | 0.10924947 | 0.65764016 |
| GNE | 176.099227 | 0.060716041 | 0.34259325 | 0.17722486 | 0.85933177 | 0.9864199 |
| RNF38 | 334.915461 | -0.11856712 | 0.17181402 | -0.6900899 | 0.49013764 | 0.92157694 |
| EBLN3P | 93.8700726 | 0.012166084 | 0.24126646 | 0.05042592 | 0.95978298 | 0.99572129 |
| ZCCHC7 | 766.539709 | -0.084211267 | 0.21448513 | -0.3926205 | 0.69459976 | 0.96220115 |
| GRHPR | 62.9433155 | 0.589700121 | 0.31450695 | 1.87499871 | 0.0607929 | 0.56071154 |
| ZBTB5 | 53.1442412 | 0.07161942 | 0.22343145 | 0.32054315 | 0.74855663 | 0.97028355 |
| AL138752.2 | 93.3369309 | 0.075187456 | 0.35552564 | 0.21148252 | 0.83251077 | 0.98306468 |
| TRMT10B | 76.4345226 | 0.314615494 | 0.28865149 | 1.0899493 | 0.27573548 | 0.82164863 |
| DCAF10 | 246.609871 | 0.096311129 | 0.19928137 | 0.48329218 | 0.62888829 | 0.94891118 |

| | | | | | | |
|---|---|---|---|---|---|---|
| FAM95C | 38.7488943 | -0.573329605 | 0.36770889 | -1.5591943 | 0.1189504 | 0.68597919 |
| ANKRD18A | 37.0357987 | -0.719825052 | 0.33863445 | -2.1256699 | 0.03353076 | 0.44921837 |
| GLIDR | 50.5473114 | -0.265566751 | 0.39714967 | -0.6686818 | 0.50369849 | 0.92537933 |
| CNTNAP3B | 191.353751 | 1.184062125 | 0.84439818 | 1.40225566 | 0.16083891 | 0.73822519 |
| CBWD5 | 88.553469 | -0.867558252 | 0.34358236 | -2.5250373 | 0.0115686 | 0.30872898 |
| PGM5 | 95.0443582 | 0.371353308 | 0.7962671 | 0.46636776 | 0.64095228 | 0.94944486 |
| AL353616.2 | 108.636035 | 1.131769868 | 0.69981996 | 1.61723005 | 0.10582863 | 0.65699665 |
| PIP5K1B | 82.2970502 | 0.258302895 | 0.47977158 | 0.53838724 | 0.59030974 | 0.94516215 |
| APBA1 | 145.763808 | 0.255521433 | 0.76094183 | 0.33579628 | 0.7370245 | 0.9698508 |
| PTAR1 | 259.080853 | 0.154927515 | 0.15310962 | 1.01187313 | 0.31159872 | 0.84811918 |
| MAMDC2-AS | 105.023595 | -0.290851971 | 0.43697873 | -0.6655976 | 0.50566837 | 0.9257297 |
| MAMDC2 | 393.366916 | 0.198574405 | 0.48925487 | 0.40587108 | 0.68483733 | 0.96161098 |
| SMC5-AS1 | 99.0921915 | -0.353882937 | 0.45690625 | -0.7745198 | 0.43862347 | 0.90561284 |
| SMC5 | 310.318584 | -0.138617919 | 0.12062638 | -1.1491509 | 0.25049375 | 0.80680142 |
| KLF9 | 120.260726 | 0.278467675 | 0.50230432 | 0.55438041 | 0.57931853 | 0.94034284 |
| TRPM3 | 307.670026 | 0.881370344 | 0.56631307 | 1.55633057 | 0.11962951 | 0.68618625 |
| CEMIP2 | 134.263707 | 0.173187764 | 0.42943514 | 0.40329202 | 0.68673341 | 0.96161098 |
| ABHD17B | 52.7436881 | 0.261624178 | 0.30766297 | 0.85035966 | 0.39512516 | 0.88760715 |
| C9orf85 | 87.3762688 | 0.049330305 | 0.24437299 | 0.2018648 | 0.84002242 | 0.98464453 |
| ZFAND5 | 295.351669 | 0.05701024 | 0.44043051 | 0.12944208 | 0.89700786 | 0.98842187 |
| ALDH1A1 | 213.499664 | 1.141378381 | 0.44285038 | 2.5773454 | 0.00995624 | 0.29238317 |
| ANXA1 | 100.424245 | 1.368713249 | 1.13331713 | 1.20770543 | 0.22716059 | 0.79241488 |
| CARNMT1 | 115.45141 | -0.060167264 | 0.23838205 | -0.2523985 | 0.80073308 | 0.97723068 |
| NMRK1 | 77.9537111 | 0.393714608 | 0.33881817 | 1.1620233 | 0.245226 | 0.80476923 |
| OSTF1 | 42.9064332 | -0.323195781 | 0.29685257 | -1.0887417 | 0.27626779 | 0.82231303 |
| PCSK5 | 66.5077676 | -0.060700803 | 0.38745035 | -0.1566673 | 0.87550706 | 0.98766275 |
| RFK | 46.3807941 | -0.262875329 | 0.38273933 | -0.686826 | 0.49219238 | 0.92289011 |
| PRUNE2 | 905.32671 | -0.077258939 | 0.4287601 | -0.1801915 | 0.85700221 | 0.9864199 |
| PCA3 | 109.163107 | 0.363543207 | 0.79651701 | 0.45641613 | 0.64809077 | 0.95027667 |
| VPS13A | 444.965456 | 0.062476292 | 0.18723436 | 0.33367963 | 0.73862133 | 0.9698508 |
| GNAQ | 1141.91224 | -0.189407636 | 0.13904779 | -1.3621765 | 0.17314219 | 0.74719017 |
| CEP78 | 41.9506573 | -0.472154866 | 0.37216155 | -1.2686825 | 0.20455431 | 0.77294901 |
| TLE4 | 229.73143 | 0.90674501 | 0.53098638 | 1.70766152 | 0.08769917 | 0.62705598 |
| TLE1 | 158.992845 | -0.003492169 | 0.31685955 | -0.0110212 | 0.99120654 | 0.99866813 |
| RASEF | 83.6283378 | -0.639624067 | 0.57368203 | -1.1149453 | 0.26487387 | 0.81378765 |
| FRMD3 | 481.391915 | -0.694223678 | 0.40919213 | -1.6965714 | 0.08977772 | 0.6330877 |
| IDNK | 26.9104361 | -0.339634739 | 0.4477083 | -0.7586072 | 0.44808756 | 0.90976372 |
| UBQLN1 | 169.421121 | -0.127166352 | 0.17170233 | -0.740621 | 0.45892327 | 0.91362347 |
| GKAP1 | 105.852707 | 0.346699314 | 0.30663372 | 1.13066272 | 0.25819708 | 0.81223967 |
| KIF27 | 193.172118 | 0.356006943 | 0.3025172 | 1.17681553 | 0.23926914 | 0.80174646 |
| HNRNPK | 377.50979 | -0.271435308 | 0.31044674 | -0.8743378 | 0.38193434 | 0.88128509 |
| RMI1 | 39.2978067 | 0.075539243 | 0.32627388 | 0.23152096 | 0.8169101 | 0.98010849 |
| NTRK2 | 173.213172 | -1.449709458 | 1.12513759 | -1.288473 | 0.19758135 | 0.77110248 |
| AGTPBP1 | 170.125498 | -0.186679436 | 0.21910972 | -0.8519907 | 0.39421926 | 0.88760715 |
| NAA35 | 176.004963 | -0.2986996 | 0.15833067 | -1.8865556 | 0.05922013 | 0.55686386 |
| GOLM1 | 84.3549805 | -0.600021496 | 0.4061329 | -1.4774018 | 0.13956796 | 0.71191199 |
| ISCA1 | 41.7645825 | -0.223632278 | 0.25121967 | -0.8901862 | 0.37336592 | 0.87666478 |
| TUT7 | 131.030598 | 0.093710423 | 0.20823895 | 0.45001392 | 0.65270041 | 0.95176155 |
| DAPK1 | 1218.33958 | -0.194696555 | 0.28345134 | -0.6868782 | 4.92E-01 | 0.92289011 |
| CTSL | 173.568797 | -1.614788262 | 0.63055404 | -2.5609039 | 0.01044002 | 0.29587276 |
| SPIN1 | 399.208252 | 0.120343407 | 0.27375617 | 0.4396007 | 0.66022633 | 0.95407427 |
| SECISBP2 | 264.806924 | -0.285623778 | 0.16081553 | -1.7760957 | 0.07571714 | 0.59499139 |
| SEMA4D | 673.708241 | -0.430676057 | 0.38526378 | -1.1178732 | 0.26362115 | 0.81378765 |
| LINC01508 | 71.3880151 | 0.396352784 | 1.13996318 | 0.34768911 | 0.72807368 | 0.96827499 |

| | | | | | | |
|---|---|---|---|---|---|---|
| SYK | 155.455868 | -0.118933511 | 0.33049515 | -0.3598646 | 0.71894838 | 0.96779196 |
| AL158071.4 | 27.9605041 | 0.047433976 | 0.41082367 | 0.11546067 | 0.90807999 | 0.99100813 |
| AUH | 390.03071 | 0.322557366 | 0.21383894 | 1.50841269 | 0.13144894 | 0.700349 |
| NFIL3 | 49.9990568 | 0.469743832 | 0.73488929 | 0.63920353 | 0.52269053 | 0.93272217 |
| ROR2 | 42.9420087 | 0.563738701 | 0.95095614 | 0.59281252 | 0.55330663 | 0.93523904 |
| SPTLC1 | 148.948608 | -0.237127272 | 0.1783004 | -1.3299312 | 0.18354093 | 0.75689793 |
| IARS | 154.125651 | -0.571741625 | 0.27128684 | -2.107517 | 0.03507279 | 0.45597142 |
| NOL8 | 52.3400888 | -0.38958998 | 0.24651328 | -1.5804016 | 0.11401493 | 0.66838982 |
| CENPP | 91.3691997 | -1.280038989 | 0.45681458 | -2.8020975 | 0.00507715 | 0.22207326 |
| IPPK | 35.8867721 | -0.048484688 | 0.27515701 | -0.1762073 | 0.86013105 | 0.9864199 |
| BICD2 | 45.9581116 | -0.042166694 | 0.37380091 | -0.1128052 | 0.91018498 | 0.99100813 |
| NINJ1 | 42.352511 | 0.941699 | 0.37925661 | 2.48301273 | 0.01302764 | 0.31992293 |
| FAM120AOS | 37.9050649 | -0.544630769 | 0.39153468 | -1.3910154 | 0.16422075 | 0.74306594 |
| FAM120A | 295.013253 | -0.200371587 | 0.2237985 | -0.8953214 | 0.37061529 | 0.87605028 |
| PHF2 | 80.7179099 | -0.00202032 | 0.27634586 | -0.0073108 | 0.99416685 | 0.99866813 |
| PTPDC1 | 77.1439423 | 0.321216687 | 0.31507359 | 1.01949735 | 0.30796691 | 0.84488735 |
| AL158152.2 | 118.122328 | 0.610069252 | 0.31697595 | 1.92465468 | 0.05427257 | 0.53752502 |
| ZNF169 | 37.6823294 | -0.124155002 | 0.27226401 | -0.4560096 | 0.64838307 | 0.95027667 |
| MFSD14B | 89.0860698 | -0.31915344 | 0.18192538 | -1.7543096 | 0.07937747 | 0.60556378 |
| FBP1 | 32.6447594 | 1.539039183 | 0.31457335 | 4.89246519 | 9.96E-07 | 0.00091119 |
| C9orf3 | 1089.4998 | 0.163983355 | 0.32808515 | 0.4998195 | 0.61720218 | 0.94791646 |
| FANCC | 561.19125 | -0.153324524 | 0.25869315 | -0.5926888 | 0.55338947 | 0.93523904 |
| PTCH1 | 87.2063357 | 0.886501054 | 0.5055808 | 1.753431 | 0.07952805 | 0.60556378 |
| LINC00476 | 156.0344 | 0.003772306 | 0.43409145 | 0.00869012 | 0.99306638 | 0.99866813 |
| ERCC6L2 | 215.828724 | -0.019757327 | 0.20595219 | -0.0959316 | 0.92357488 | 0.99214128 |
| SLC35D2 | 41.3124707 | 0.350888178 | 0.31790452 | 1.10375334 | 0.26970015 | 0.8167745 |
| HABP4 | 130.700099 | -0.072791477 | 0.3575339 | -0.2035932 | 0.83867139 | 0.98462033 |
| CDC14B | 182.798009 | 0.920618512 | 0.28172928 | 3.2677417 | 0.00108409 | 0.10596452 |
| ZNF510 | 25.8331071 | -0.483268541 | 0.34955002 | -1.3825447 | 0.16680451 | 0.74445571 |
| ZNF782 | 84.7532495 | 0.135695028 | 0.26707196 | 0.50808414 | 0.61139434 | 0.94791646 |
| MFSD14C | 243.835809 | -0.193735164 | 0.21582487 | -0.8976499 | 0.36937224 | 0.87605028 |
| TDRD7 | 61.1949766 | -0.364363822 | 0.59574094 | -0.6116145 | 0.54079281 | 0.93523904 |
| TSTD2 | 60.9056365 | -0.227476628 | 0.2901514 | -0.7839929 | 0.43304429 | 0.90514499 |
| NCBP1 | 56.0974892 | -0.082278701 | 0.27193397 | -0.3025687 | 0.76221859 | 0.9714717 |
| XPA | 58.7281807 | 0.092706246 | 0.26821898 | 0.34563641 | 0.72961598 | 0.96871206 |
| ANP32B | 119.864088 | -0.251600294 | 0.24603488 | -1.0226204 | 0.30648734 | 0.84488735 |
| TRIM14 | 51.9770599 | -1.354613338 | 0.92678678 | -1.4616235 | 0.14384441 | 0.71631428 |
| ANKS6 | 99.1718588 | -0.333098035 | 0.24375932 | -1.3665038 | 0.1717809 | 0.74490231 |
| TGFBR1 | 74.2585431 | 0.018230033 | 0.37169723 | 0.04904538 | 0.96088313 | 0.99572129 |
| ALG2 | 36.6157875 | 0.192933255 | 0.40259421 | 0.47922511 | 0.63177849 | 0.94932781 |
| SEC61B | 127.49887 | -0.098276446 | 0.39501307 | -0.2487929 | 0.80352098 | 0.97880236 |
| AL359710.1 | 75.8497302 | 0.222263107 | 0.55519811 | 0.40033117 | 0.68891261 | 0.96190309 |
| STX17 | 123.166948 | -0.0792125 | 0.32674491 | -0.2424292 | 0.80844763 | 0.97930627 |
| ERP44 | 153.077515 | 0.023243731 | 0.21731824 | 0.10695711 | 0.914823 | 0.99159829 |
| INVS | 341.446701 | 0.195887409 | 0.19917261 | 0.98350576 | 0.32535858 | 0.855037 |
| TEX10 | 62.0608718 | -0.088394173 | 0.23585512 | -0.3747817 | 0.70782286 | 0.96587538 |
| MRPL50 | 26.5858484 | 0.172592224 | 0.44586575 | 0.3870946 | 0.69868617 | 0.96256506 |
| ZNF189 | 36.9436861 | 0.487493596 | 0.34868576 | 1.39808865 | 0.16208645 | 0.74071637 |
| ALDOB | 96.9613354 | 0.363792585 | 1.1771521 | 0.30904467 | 0.75728755 | 0.9714717 |
| TMEM246 | 30.8142932 | 0.456511611 | 0.32467129 | 1.40607325 | 0.15970237 | 0.73816762 |
| RNF20 | 40.1489145 | 0.03688003 | 0.30956009 | 0.1191369 | 0.9051669 | 0.99092254 |
| NIPSNAP3A | 30.730939 | -0.013079844 | 0.35344717 | -0.0370065 | 0.97047982 | 0.99681784 |
| ABCA1 | 48.4316753 | 0.35892033 | 0.56429637 | 0.63604934 | 0.52474427 | 0.93272384 |
| SLC44A1 | 167.488895 | -0.163137712 | 0.23881239 | -0.6831208 | 0.49453051 | 0.92289011 |

| | | | | | | |
|---|---|---|---|---|---|---|
| FSD1L | 114.918444 | 0.235618535 | 0.26296614 | 0.89600334 | 0.37025098 | 0.87605028 |
| FKTN | 112.448742 | -0.180791824 | 0.20680052 | -0.8742329 | 0.38199141 | 0.88128509 |
| TMEM38B | 165.687065 | -0.03902837 | 0.43402625 | -0.0899217 | 0.92834945 | 0.99336859 |
| LINC01505 | 115.906131 | -0.372118405 | 0.43397962 | -0.8574559 | 0.39119295 | 0.88670817 |
| ZNF462 | 401.723998 | -0.085212308 | 0.24012252 | -0.3548701 | 0.7226869 | 0.96779196 |
| RAD23B | 287.189586 | 0.037276743 | 0.21418491 | 0.17404001 | 0.86183402 | 0.98669442 |
| ELP1 | 73.5100388 | 0.121875241 | 0.21070843 | 0.57840705 | 0.56298933 | 0.93805441 |
| FAM206A | 31.315471 | 0.323582037 | 0.28434421 | 1.13799411 | 0.25512294 | 0.80897837 |
| CTNNAL1 | 131.70741 | -0.007947997 | 0.55660464 | -0.0142794 | 0.98860705 | 0.99820747 |
| TMEM245 | 388.004448 | 0.179011416 | 0.22496791 | 0.79571977 | 0.42619493 | 0.90237497 |
| EPB41L4B | 44.9499466 | -0.388535091 | 0.92284684 | -0.421018 | 0.67374197 | 0.95910677 |
| PTPN3 | 365.310831 | -0.477364914 | 0.29261269 | -1.6313883 | 0.10280841 | 0.65288882 |
| PALM2 | 49.8022538 | -0.155927233 | 0.6510001 | -0.2395195 | 0.81070276 | 0.97930627 |
| TXN | 131.616718 | -0.40559595 | 0.52565121 | -0.7716066 | 0.44034745 | 0.90561284 |
| LPAR1 | 74.4865214 | -0.105821601 | 0.49832942 | -0.2123527 | 0.83183188 | 0.98290906 |
| ECPAS | 307.014861 | -0.382421189 | 0.23153503 | -1.6516775 | 0.09860032 | 0.64312419 |
| ZNF483 | 72.1874984 | 0.625764907 | 0.50001892 | 1.25148245 | 0.21075851 | 0.77730294 |
| PTGR1 | 241.265663 | 0.19326281 | 0.28392085 | 0.68069256 | 0.49606604 | 0.92334178 |
| C9orf84 | 76.7034376 | 1.06676432 | 0.61735452 | 1.72796065 | 0.08399528 | 0.61656297 |
| UGCG | 865.765113 | 0.54117113 | 0.48891623 | 1.10687904 | 0.26834623 | 0.81584693 |
| SUSD1 | 294.245121 | -1.079335453 | 0.3212075 | -3.3602436 | 0.00077874 | 0.08987383 |
| PTBP3 | 922.159092 | -1.045506319 | 0.30823797 | -3.3918803 | 0.00069415 | 0.08416759 |
| HSDL2 | 168.813283 | 0.284448382 | 0.21490288 | 1.32361366 | 0.18563139 | 0.76046631 |
| C9orf147 | 29.6922595 | -0.050247346 | 0.54109241 | -0.0928628 | 0.92601257 | 0.99316012 |
| KIAA1958 | 331.170863 | -0.053722669 | 0.22156225 | -0.2424721 | 0.80841436 | 0.97930627 |
| INIP | 59.7598222 | -0.048108059 | 0.22607634 | -0.2127956 | 0.83148637 | 0.98282427 |
| SNX30 | 152.013115 | 0.268661748 | 0.35819652 | 0.75004008 | 0.45323057 | 0.91077355 |
| FKBP15 | 59.9024964 | -0.286553415 | 0.23381008 | -1.2255819 | 0.22035604 | 0.78584902 |
| SLC31A1 | 42.9586377 | 0.33796348 | 0.38061483 | 0.8879409 | 0.37457254 | 0.87678297 |
| CDC26 | 33.4924868 | -0.01074992 | 0.46499979 | -0.0231181 | 0.98155605 | 0.99681784 |
| PRPF4 | 35.9425518 | -0.992594276 | 0.35255741 | -2.8154117 | 0.00487148 | 0.21962686 |
| WDR31 | 39.8183318 | 0.816141075 | 0.41414462 | 1.97066685 | 0.04876199 | 0.52130708 |
| BSPRY | 34.9086934 | -0.161538336 | 0.48443291 | -0.3334586 | 0.73878811 | 0.9698508 |
| HDHD3 | 62.1787576 | 0.217053254 | 0.25916196 | 0.83751974 | 0.40230049 | 0.89239743 |
| RGS3 | 92.4670353 | -0.362515928 | 0.34868198 | -1.039675 | 2.98E-01 | 0.83973489 |
| ZNF618 | 297.563159 | -0.006141266 | 0.37414344 | -0.0164142 | 0.98690395 | 0.99764442 |
| KIF12 | 235.162958 | 0.175618291 | 0.42491115 | 0.41330592 | 0.67938249 | 0.95936177 |
| COL27A1 | 334.521566 | 0.028474337 | 0.35507638 | 0.08019215 | 0.93608443 | 0.99508965 |
| WHRN | 53.1177175 | -0.251494307 | 0.3362391 | -0.7479627 | 0.45448265 | 0.91077355 |
| ATP6V1G1 | 166.928571 | -0.117942338 | 0.45334677 | -0.2601592 | 0.79474096 | 0.97524554 |
| TNFSF15 | 42.9585067 | -1.118123541 | 0.85569924 | -1.3066782 | 0.19132203 | 0.76328615 |
| TNC | 474.168942 | -0.662092459 | 0.52644936 | -1.2576565 | 0.20851601 | 0.77477691 |
| PAPPA | 177.822022 | -0.284954089 | 0.46772766 | -0.6092308 | 0.54237148 | 0.93523904 |
| ASTN2 | 802.998073 | -0.941116217 | 0.53090699 | -1.7726574 | 0.07628549 | 0.59726395 |
| CDK5RAP2 | 123.515049 | 0.159269649 | 0.25379373 | 0.6275555 | 0.53029517 | 0.93366189 |
| MEGF9 | 261.550466 | -0.041887928 | 0.37754365 | -0.1109486 | 0.91165713 | 0.99100813 |
| FBXW2 | 83.8458082 | -0.447703675 | 0.23158812 | -1.9331893 | 0.05321288 | 0.53386172 |
| PSMD5 | 85.6242126 | 0.190480997 | 0.31302551 | 0.6085159 | 0.54284536 | 0.93523904 |
| TRAF1 | 68.3997857 | -1.4240441 | 1.1347543 | -1.2549361 | 0.20950197 | 0.77686608 |
| CNTRL | 71.8127412 | -0.405391995 | 0.35617345 | -1.138187 | 0.2550424 | 0.80897837 |
| RAB14 | 152.296886 | 0.104227626 | 0.21327087 | 0.48871008 | 0.62504696 | 0.94808496 |
| GSN | 111.759355 | -0.401617298 | 0.29284909 | -1.3714139 | 0.17024598 | 0.74445571 |
| STOM | 157.272075 | 0.408636682 | 0.29332304 | 1.39312847 | 1.64E-01 | 0.74227469 |
| DAB2IP | 180.589659 | 0.556040336 | 0.54690136 | 1.01671047 | 0.30929118 | 0.84554063 |

| | | | | | | |
|---|---|---|---|---|---|---|
| TTLL11 | 48.9940827 | 0.289791351 | 0.35356049 | 0.81963726 | 0.41242292 | 0.89706788 |
| NDUFA8 | 35.5947141 | 0.02095276 | 0.41146282 | 0.05092261 | 0.95938719 | 0.99572129 |
| RBM18 | 28.0662202 | -0.174653998 | 0.33930381 | -0.5147422 | 0.60673317 | 0.94708529 |
| MRRF | 112.16129 | -0.118060563 | 0.230041 | -0.5132153 | 0.60780072 | 0.94708529 |
| PDCL | 31.2348085 | -0.21945293 | 0.31700208 | -0.692276 | 0.48876402 | 0.92121587 |
| RC3H2 | 153.419221 | 0.102606807 | 0.23252169 | 0.44127843 | 0.65901144 | 0.95402584 |
| RABGAP1 | 406.570148 | -0.083711687 | 0.20219889 | -0.4140067 | 0.67886923 | 0.95924874 |
| STRBP | 256.330076 | 0.406386399 | 0.2444574 | 1.66240167 | 0.09643226 | 0.64295596 |
| CRB2 | 66.2989502 | 1.135601583 | 1.43064057 | 0.79377141 | 0.42732852 | 0.90272191 |
| DENND1A | 439.454873 | 0.308963629 | 0.27013097 | 1.14375494 | 0.25272528 | 0.80798674 |
| NEK6 | 122.392417 | 0.992407735 | 0.8069807 | 1.22977877 | 0.21877996 | 0.78523615 |
| PSMB7 | 136.052974 | -0.225965242 | 0.29769324 | -0.759054 | 0.44782026 | 0.90976372 |
| NR6A1 | 325.692155 | 0.284925025 | 0.29330516 | 0.97142863 | 0.33133488 | 0.85923151 |
| MIR181A2HG | 91.3824731 | 0.037435744 | 0.34767502 | 0.10767453 | 0.91425387 | 0.99159829 |
| RPL35 | 270.506138 | -0.285466067 | 0.47429439 | -0.6018753 | 0.54725717 | 0.93523904 |
| ARPC5L | 44.7812563 | -0.519346579 | 0.42035897 | -1.2354835 | 0.2166506 | 0.78319543 |
| GOLGA1 | 59.1150416 | -0.483090213 | 0.3108296 | -1.5541963 | 0.1201376 | 0.68786496 |
| SCAI | 127.901735 | -0.049061706 | 0.22516086 | -0.2178962 | 0.82750995 | 0.98142144 |
| PPP6C | 117.22908 | -0.080272843 | 0.21828897 | -0.3677366 | 0.71306966 | 0.96587538 |
| RABEPK | 33.2792893 | -1.044517893 | 0.41929897 | -2.4911053 | 0.01273464 | 0.31439454 |
| HSPA5 | 225.228651 | 0.612199062 | 0.65607945 | 0.93311727 | 0.35075943 | 0.8720098 |
| GAPVD1 | 200.727251 | 0.248893243 | 0.17992788 | 1.38329451 | 0.16657458 | 0.74445571 |
| MAPKAP1 | 302.147316 | -0.426891997 | 0.18432431 | -2.3159831 | 0.02055919 | 0.37872401 |
| PBX3 | 299.999942 | -0.813466179 | 0.36205532 | -2.2468008 | 0.02465276 | 0.41230428 |
| MVB12B | 58.9745681 | -0.244767282 | 0.35395962 | -0.6915119 | 0.48924392 | 0.92126351 |
| ZBTB43 | 68.4663097 | -0.291164372 | 0.26113576 | -1.1149923 | 0.26485371 | 0.81378765 |
| RALGPS1 | 190.459238 | 0.071176992 | 0.51147805 | 0.13915943 | 0.88932417 | 0.98766275 |
| GARNL3 | 77.442377 | 0.192001232 | 0.30506981 | 0.62936818 | 0.52910804 | 0.93366189 |
| RPL12 | 117.420855 | 0.094095437 | 0.46149344 | 0.20389334 | 0.83843685 | 0.98462033 |
| LRSAM1 | 105.618712 | -0.015258766 | 0.35544118 | -0.0429291 | 0.96575806 | 0.99601598 |
| FAM129B | 235.012415 | -0.810950819 | 0.31490912 | -2.57519 | 0.0100185 | 0.29238317 |
| STXBP1 | 81.4789038 | 0.708718379 | 0.50417835 | 1.40568982 | 0.15981625 | 0.73816762 |
| CDK9 | 43.277212 | 0.120406407 | 0.41462722 | 0.29039677 | 0.77151271 | 0.97329019 |
| FPGS | 54.6557361 | -0.681918572 | 0.31591995 | -2.158517 | 0.03088766 | 0.43898056 |
| DPM2 | 28.2564008 | -1.147562784 | 0.36430524 | -3.1500035 | 0.00163269 | 0.13097113 |
| FAM102A | 35.3084414 | -0.565787244 | 0.41885167 | -1.3508057 | 0.17675767 | 0.75098991 |
| SLC25A25 | 30.1002429 | 0.310221336 | 0.44423029 | 0.69833449 | 0.48496803 | 0.92034617 |
| PTGES2 | 53.388387 | -0.228157849 | 0.46842669 | -0.4870727 | 0.62620681 | 0.94831213 |
| LCN2 | 461.424183 | 0.805154506 | 1.26080849 | 0.63860175 | 5.23E-01 | 9.33E-01 |
| C9orf16 | 94.1288751 | -0.538947531 | 0.5809272 | -0.9277368 | 0.35354412 | 0.87216048 |
| CIZ1 | 59.002358 | -0.480201287 | 0.24644284 | -1.9485301 | 0.05135157 | 0.52609271 |
| DNM1 | 256.021035 | 0.460890909 | 0.4890587 | 0.94240406 | 0.34598581 | 0.86990996 |
| GOLGA2 | 109.816969 | 0.254911221 | 0.2742528 | 0.92947537 | 0.35264278 | 0.8720098 |
| SWI5 | 31.456303 | -0.11398465 | 0.38447217 | -0.2964705 | 0.76687081 | 0.97221495 |
| TRUB2 | 31.4906289 | -0.066229264 | 0.3245775 | -0.2040476 | 0.83831629 | 0.98462033 |
| COQ4 | 112.604903 | 0.226499721 | 0.19827269 | 1.14236471 | 0.25330245 | 0.80798674 |
| SLC27A4 | 38.1496448 | -0.568594546 | 0.35222324 | -1.6143016 | 0.10646202 | 0.65699665 |
| URM1 | 38.2647877 | -0.811304979 | 0.2873925 | -2.8229859 | 0.00475787 | 0.21962686 |
| CERCAM | 51.3825965 | -0.441765626 | 0.7523105 | -0.5872118 | 0.55706145 | 0.93533595 |
| ODF2 | 50.9076602 | 0.049862238 | 0.27672614 | 0.18018622 | 0.85700638 | 0.9864199 |
| GLE1 | 56.0480643 | -0.393734049 | 0.23049229 | -1.7082309 | 0.0875935 | 0.62705598 |
| SPTAN1 | 242.562287 | -0.196835231 | 0.3217779 | -0.6117115 | 0.54072868 | 0.93523904 |
| WDR34 | 59.5596336 | 0.502266045 | 0.43517346 | 1.15417435 | 0.2484287 | 0.80672756 |
| SET | 232.091254 | -0.447602828 | 0.40505935 | -1.1050302 | 0.26914649 | 0.81655182 |

| | | | | | | |
|---|---|---|---|---|---|---|
| ZDHHC12 | 29.6991877 | -0.347049599 | 0.57087952 | -0.6079209 | 0.54323993 | 0.93523904 |
| ZER1 | 150.809426 | 0.605738539 | 0.26700667 | 2.26862698 | 0.02329102 | 0.39839181 |
| TBC1D13 | 37.7405809 | -0.605441071 | 0.33630473 | -1.8002752 | 0.0718172 | 0.59090333 |
| KYAT1 | 46.2484975 | 0.999536801 | 0.42963636 | 2.32647162 | 0.0199934 | 0.3747883 |
| LRRC8A | 31.4193806 | 0.226174674 | 0.37802952 | 0.59829898 | 0.54964046 | 0.93523904 |
| NUP188 | 38.2909164 | -0.714026478 | 0.25905076 | -2.7563188 | 0.0058456 | 0.24135775 |
| SH3GLB2 | 57.0486122 | 0.058132775 | 0.37251929 | 0.15605306 | 0.8759912 | 0.98766275 |
| MIGA2 | 29.0592309 | -0.038829265 | 0.34986352 | -0.110984 | 0.91162901 | 0.99100813 |
| PTPA | 66.6640955 | 0.058241361 | 0.30145079 | 0.19320354 | 0.84679956 | 0.98533374 |
| IER5L | 97.6758146 | -0.321546587 | 0.40942132 | -0.7853684 | 4.32E-01 | 9.05E-01 |
| LINC01503 | 47.2893635 | -0.648448695 | 0.44235699 | -1.4658945 | 0.14267702 | 0.71517697 |
| NTMT1 | 30.9128514 | -0.067833198 | 0.33410114 | -0.2030319 | 0.83911008 | 0.98462033 |
| C9orf78 | 52.7387451 | -0.028750737 | 0.42087041 | -0.0683126 | 0.94553682 | 0.9952642 |
| USP20 | 47.1146741 | 0.275147869 | 0.3577441 | 0.76911924 | 0.44182253 | 0.90561284 |
| FNBP1 | 481.80163 | 0.442733201 | 0.28422436 | 1.55768914 | 0.11930696 | 0.68597919 |
| GPR107 | 168.678735 | 0.069778892 | 0.19666683 | 0.35480764 | 0.72273371 | 0.96779196 |
| NCS1 | 74.407543 | -0.76534364 | 0.38428673 | -1.9915953 | 0.04641548 | 0.50827029 |
| ASS1 | 80.1605723 | 1.338637302 | 0.75766219 | 1.76679966 | 0.07726179 | 0.5981029 |
| FUBP3 | 144.109571 | -0.320783067 | 0.20193015 | -1.5885843 | 0.11215428 | 0.66401239 |
| ABL1 | 353.513532 | -0.054652371 | 0.2484092 | -0.2200094 | 0.8258638 | 0.98094663 |
| AIF1L | 545.412759 | -0.17104109 | 0.56701573 | -0.3016514 | 0.76291782 | 0.97149197 |
| NUP214 | 352.35615 | -0.208371092 | 0.33223895 | -0.6271724 | 5.31E-01 | 0.93366189 |
| AL157938.2 | 89.6401825 | 0.228906939 | 0.67780795 | 0.33771651 | 0.73557683 | 0.9698508 |
| AL157938.3 | 27.4367276 | -0.719180308 | 0.54707883 | -1.3145826 | 0.18865023 | 0.76207437 |
| PRRC2B | 420.643102 | -0.191292187 | 0.18454209 | -1.0365776 | 0.29993278 | 0.84130486 |
| POMT1 | 36.1269616 | 0.410822384 | 0.2683084 | 1.53115736 | 0.12573051 | 0.69492095 |
| RAPGEF1 | 233.062547 | -0.099575147 | 0.21849571 | -0.4557304 | 0.64858383 | 0.95027667 |
| MED27 | 190.847773 | -0.124966982 | 0.24176052 | -0.516904 | 0.60522317 | 0.94698346 |
| SETX | 351.692226 | -0.111603662 | 0.2533807 | -0.4404584 | 0.65960513 | 0.95407427 |
| TTF1 | 66.9664693 | -0.313029407 | 0.2561362 | -1.2221209 | 0.22166191 | 0.78710314 |
| DDX31 | 169.769812 | 0.011807242 | 0.26767438 | 0.04411047 | 0.96481635 | 0.99590256 |
| GTF3C4 | 50.6449645 | 0.310291485 | 0.36524047 | 0.84955396 | 0.39557312 | 0.88760715 |
| AK8 | 46.5567767 | 0.730823458 | 0.45575099 | 1.60355868 | 0.10881137 | 0.65764016 |
| TSC1 | 113.959758 | -0.083431439 | 0.21897445 | -0.3810099 | 0.70319588 | 0.9645815 |
| GTF3C5 | 33.419854 | -0.40904982 | 0.33644573 | -1.2157973 | 0.22406214 | 0.78828752 |
| ABO | 178.911358 | 0.137951136 | 0.43055675 | 0.32040175 | 0.7486638 | 0.97028355 |
| SURF6 | 33.6980257 | -0.541336356 | 0.29370247 | -1.8431454 | 0.06530778 | 0.57495764 |
| RPL7A | 362.809561 | -0.172669172 | 0.53355262 | -0.3236216 | 0.7462245 | 0.97028355 |
| SURF1 | 28.8068535 | 0.075481482 | 0.33094931 | 0.22807566 | 0.81958742 | 0.98079697 |
| SURF4 | 103.300084 | -0.085624379 | 0.44840005 | -0.1909553 | 0.84856059 | 0.98548777 |
| REXO4 | 28.0826785 | -0.07651702 | 0.38903273 | -0.1966853 | 0.84407381 | 0.98517901 |
| VAV2 | 234.485092 | -0.348211489 | 0.37605738 | -0.9259531 | 0.35447037 | 0.87258503 |
| BRD3OS | 25.0063528 | 0.596773938 | 0.44265045 | 1.34818328 | 0.17759944 | 0.7513166 |
| BRD3 | 68.3635376 | -0.008377535 | 0.2929367 | -0.0285984 | 0.97718485 | 0.99681784 |
| WDR5 | 42.1836854 | -0.440032513 | 0.3503514 | -1.2559748 | 0.20912512 | 0.77583638 |
| RXRA | 52.2102543 | 0.054726536 | 0.54840458 | 0.09979227 | 0.92050925 | 0.99214128 |
| COL5A1 | 94.1771733 | 1.465565773 | 1.1773949 | 1.24475295 | 0.21322257 | 0.77986218 |
| PPP1R26-AS1 | 32.7339217 | 0.312738535 | 0.44103199 | 0.70910625 | 0.47825855 | 0.9193736 |
| PPP1R26 | 28.500396 | 0.249169442 | 0.33750507 | 0.73826874 | 0.46035116 | 0.91368087 |
| MRPS2 | 29.7686523 | 0.221866569 | 0.36135858 | 0.61397896 | 0.53922923 | 0.93523904 |
| CAMSAP1 | 89.9476918 | -0.309952751 | 0.29278626 | -1.0586315 | 0.28976765 | 0.83506029 |
| UBAC1 | 44.0673667 | 0.173271501 | 0.24570216 | 0.7052095 | 0.48067987 | 0.91971919 |
| NACC2 | 71.0391778 | 0.339313335 | 0.39065032 | 0.86858583 | 0.38507371 | 0.88353166 |
| QSOX2 | 26.1348403 | -0.198068947 | 0.32476678 | -0.6098806 | 0.54194093 | 0.93523904 |

| | | | | | | |
|---|---|---|---|---|---|---|
| SNAPC4 | 37.3605696 | -0.245344409 | 0.26421898 | -0.9285647 | 0.35311475 | 0.8720098 |
| ENTR1 | 38.0308307 | -0.820543744 | 0.40871038 | -2.0076411 | 0.04468145 | 0.50176269 |
| PMPCA | 36.0162353 | -0.191332909 | 0.28670825 | -0.6673436 | 0.5045527 | 0.92552859 |
| SEC16A | 81.6635972 | -0.309932954 | 0.23648905 | -1.3105594 | 0.19000666 | 0.76207437 |
| NOTCH1 | 67.6961122 | -0.365410122 | 0.40560207 | -0.9009079 | 0.36763727 | 0.87605028 |
| AGPAT2 | 47.4768156 | 0.221864148 | 0.40520499 | 0.54753558 | 0.58401084 | 0.94184538 |
| SNHG7 | 67.9492888 | 0.113801237 | 0.52612812 | 0.21629947 | 0.82875432 | 0.98142144 |
| RABL6 | 121.187566 | 0.017975451 | 0.28777695 | 0.06246314 | 0.95019402 | 0.99531843 |
| PHPT1 | 123.503182 | 0.557648052 | 0.45744874 | 1.21903942 | 0.22282923 | 0.78710314 |
| EDF1 | 174.417008 | -0.297745327 | 0.4491211 | -0.6629511 | 0.50736187 | 0.92666294 |
| TRAF2 | 32.6152045 | 0.074461696 | 0.43421011 | 0.17148771 | 0.8638403 | 0.98670067 |
| FBXW5 | 67.9038146 | 0.045843598 | 0.46560785 | 0.09845968 | 0.92156729 | 0.99214128 |
| PAXX | 37.9098078 | 0.515859613 | 0.54054654 | 0.9543297 | 0.33991678 | 0.86742596 |
| MAN1B1 | 56.9354011 | 0.056738323 | 0.22642684 | 0.25058126 | 0.80213787 | 0.97835591 |
| DPP7 | 49.3162723 | 0.058229926 | 0.3629084 | 0.16045351 | 0.87252384 | 0.98766275 |
| SSNA1 | 53.5956423 | -0.091482517 | 0.43411124 | -0.2107352 | 0.83309391 | 0.98338959 |
| TPRN | 42.5064294 | -0.174294591 | 0.31566996 | -0.5521418 | 0.58085118 | 0.94041171 |
| TMEM203 | 28.2986399 | 0.193302486 | 0.41242783 | 0.46869408 | 0.63928832 | 0.94944486 |
| TUBB4B | 193.828728 | -0.442703016 | 0.5744338 | -0.7706772 | 0.4408983 | 0.90561284 |
| NELFB | 74.6869901 | -0.155313205 | 0.3027946 | -0.5129325 | 0.60799851 | 0.94708529 |
| EXD3 | 102.840975 | 0.282377372 | 0.31805835 | 0.88781625 | 0.3746396 | 0.87678297 |
| NOXA1 | 37.1649993 | -0.047419148 | 0.42865732 | -0.1106225 | 0.9119157 | 0.99106427 |
| NSMF | 69.981553 | 0.671372767 | 0.60224546 | 1.11478261 | 0.26494359 | 0.81378765 |
| PNPLA7 | 69.7947076 | 1.310377952 | 0.73515058 | 1.78246197 | 0.07467393 | 0.59444897 |
| MRPL41 | 100.378889 | 0.053346322 | 0.35100208 | 0.15198292 | 0.87920041 | 0.98766275 |
| DPH7 | 42.9455158 | -0.362819751 | 0.28342342 | -1.2801333 | 0.20049827 | 0.77110248 |
| ARRDC1 | 54.8176845 | -0.381626246 | 0.26252592 | -1.4536707 | 0.14603761 | 0.71974648 |
| EHMT1 | 400.523319 | -0.3329083 | 0.19983929 | -1.6658801 | 0.0957373 | 0.64264648 |
| BET1L | 31.8601279 | -0.300884363 | 0.35437149 | -0.8490648 | 0.39584523 | 0.88785003 |
| RIC8A | 37.3477943 | -0.263089416 | 0.31606057 | -0.8324019 | 0.40518214 | 0.89450098 |
| SIRT3 | 37.9197546 | -0.022662834 | 0.39919919 | -0.0567707 | 0.95472782 | 0.99531843 |
| PSMD13 | 79.4582963 | -0.433301169 | 0.30020562 | -1.443348 | 0.14892248 | 0.72246729 |
| AC136475.3 | 27.8259868 | -1.639223425 | 1.45261885 | -1.1284608 | 0.25912536 | 0.81263292 |
| PGGHG | 170.293851 | -1.40902327 | 0.78934124 | -1.7850623 | 0.07425121 | 0.59321312 |
| IFITM2 | 462.453098 | -0.116503205 | 0.70536134 | -0.1651681 | 0.86881166 | 0.98676155 |
| IFITM3 | 851.231518 | -0.177835899 | 0.73080735 | -0.2433417 | 0.80774071 | 0.97930627 |
| SIGIRR | 46.326669 | 0.497205105 | 0.28350271 | 1.75379316 | 0.07946596 | 0.60556378 |
| PTDSS2 | 48.7640499 | -0.036408839 | 0.25864681 | -0.1407666 | 0.88805431 | 0.98766275 |
| RNH1 | 97.8913923 | -0.103772701 | 0.28318028 | -0.3664545 | 0.71402592 | 0.96618933 |
| HRAS | 41.7479284 | -1.037848406 | 0.4927681 | -2.1061599 | 0.03519047 | 0.45614273 |
| RASSF7 | 128.059866 | 0.039669264 | 0.32852927 | 0.12074804 | 0.90389061 | 0.99054664 |
| MIR210HG | 29.9300123 | -0.17219838 | 0.72371235 | -0.2379376 | 0.81192949 | 0.97930627 |
| PHRF1 | 62.9654871 | -0.366333381 | 0.24319834 | -1.5063153 | 0.13198624 | 0.7008294 |
| DEAF1 | 144.259592 | 0.03619749 | 0.19606691 | 0.18461805 | 0.85352863 | 0.9864199 |
| EPS8L2 | 287.59054 | 0.295855957 | 0.36336088 | 0.81422072 | 0.41551852 | 0.89807127 |
| AP006621.4 | 52.9473095 | 1.117416703 | 0.71723033 | 1.55796076 | 0.11924255 | 0.68597919 |
| TALDO1 | 151.304538 | -0.407060247 | 0.38949466 | -1.0450984 | 0.29597749 | 0.83955237 |
| GATD1 | 35.9184018 | -0.090173208 | 0.33204224 | -0.2715715 | 0.78595152 | 0.97423208 |
| AP006621.3 | 45.3449666 | 0.426774288 | 0.45221957 | 0.94373246 | 0.34530638 | 0.86990996 |
| RPLP2 | 302.735262 | -0.260131407 | 0.49713863 | -0.5232573 | 0.60079523 | 0.94683415 |
| PNPLA2 | 82.2668459 | 0.008345898 | 0.27035841 | 0.03086975 | 0.97537341 | 0.99681784 |
| CD151 | 231.732849 | 0.594294312 | 0.53020144 | 1.120884 | 0.26233724 | 0.81377303 |
| POLR2L | 154.960036 | -0.249443204 | 0.52166712 | -0.4781655 | 0.63253244 | 0.94932781 |
| TSPAN4 | 89.8433285 | 0.05177894 | 0.36772388 | 0.14080929 | 0.8880206 | 0.98766275 |

| | | | | | | |
|---|---|---|---|---|---|---|
| CHID1 | 45.5507092 | -0.103983251 | 0.28044746 | -0.3707762 | 0.71080421 | 0.96587538 |
| AP2A2 | 220.337483 | 0.395648534 | 0.28027127 | 1.41166284 | 0.15804926 | 0.73580079 |
| TOLLIP | 68.0126306 | -0.158319973 | 0.30160624 | -0.5249227 | 0.59963691 | 0.94683415 |
| MOB2 | 40.7640861 | -0.049797916 | 0.25619751 | -0.1943731 | 0.84588372 | 0.98520136 |
| CD81-AS1 | 34.5980759 | 0.251250174 | 0.34016799 | 0.73860616 | 0.46014618 | 0.91368087 |
| CD81 | 434.097178 | -0.021781428 | 0.37659691 | -0.0578375 | 0.95387806 | 0.99531843 |
| TSSC4 | 30.3279933 | -0.376288288 | 0.36900743 | -1.0197309 | 0.30785609 | 0.84488735 |
| KCNQ1 | 139.253024 | 1.499512722 | 0.32279157 | 4.64545199 | 3.39E-06 | 0.00203154 |
| KCNQ1OT1 | 541.096251 | 1.742397449 | 0.4178771 | 4.16964091 | 3.05E-05 | 0.01082356 |
| CDKN1C | 118.903819 | 2.915018456 | 0.53352416 | 5.46370466 | 4.66E-08 | 0.00014889 |
| SLC22A18 | 35.4290946 | 0.243185363 | 0.54885846 | 0.44307482 | 0.65771162 | 0.95399905 |
| NAP1L4 | 257.743757 | -0.023265287 | 0.23882591 | -0.0974153 | 0.92239663 | 0.99214128 |
| CARS | 68.7936919 | 0.03305218 | 0.30508253 | 0.10833849 | 0.91372719 | 0.99154413 |
| OSBPL5 | 47.7589304 | 0.146477936 | 0.29446426 | 0.49743875 | 0.61887968 | 0.9480368 |
| ZNF195 | 66.2744073 | -0.408996397 | 0.36700035 | -1.1144305 | 0.26509453 | 0.81378765 |
| AC127526.1 | 46.6906146 | 0.6481288 | 0.4620949 | 1.40258809 | 0.1607397 | 0.73816762 |
| NUP98 | 214.924435 | -0.330079618 | 0.21471273 | -1.5373081 | 0.12421787 | 0.69439514 |
| PGAP2 | 43.9283331 | -0.33152668 | 0.35648467 | -0.9299886 | 0.35237698 | 0.8720098 |
| STIM1 | 222.331896 | -0.294862451 | 0.36376079 | -0.8105944 | 0.41759865 | 0.89862171 |
| RRM1 | 36.996205 | -0.620869021 | 0.36254431 | -1.7125328 | 0.08679853 | 0.62478747 |
| AC104389.4 | 128.639466 | -0.817073224 | 0.65859569 | -1.2406295 | 0.21474266 | 0.7828522 |
| TRIM5 | 39.1357347 | -0.300951815 | 0.81716557 | -0.3682874 | 0.71265892 | 0.96587538 |
| TRIM22 | 133.682246 | -1.311404574 | 0.83081864 | -1.5784487 | 0.11446257 | 0.66978434 |
| FAM160A2 | 52.3147834 | -0.141533904 | 0.26221096 | -0.5397711 | 0.58935489 | 0.94491859 |
| SMPD1 | 43.705857 | 0.529028286 | 0.31831869 | 1.66194539 | 0.09652372 | 0.64295596 |
| APBB1 | 45.8067358 | 0.270038209 | 0.30577932 | 0.88311468 | 0.37717431 | 0.87842274 |
| ARFIP2 | 28.2962663 | -0.220814321 | 0.36737781 | -0.6010551 | 0.54780326 | 0.93523904 |
| AC084337.2 | 45.8858953 | 0.093559405 | 0.35981066 | 0.26002399 | 0.79484527 | 0.97524554 |
| DNHD1 | 65.830875 | -0.103118641 | 0.38526761 | -0.2676546 | 0.78896521 | 0.97423208 |
| ILK | 93.5835784 | 0.305690304 | 0.33545571 | 0.91126875 | 0.36215379 | 0.8752623 |
| TAF10 | 39.2283781 | -0.550746593 | 0.40101627 | -1.3733772 | 0.16963513 | 0.74445571 |
| TPP1 | 46.0214573 | 0.204624554 | 0.31911057 | 0.64123402 | 0.52137065 | 0.93272217 |
| MRPL17 | 46.7418417 | -0.234302663 | 0.45553065 | -0.514351 | 0.60700658 | 0.94708529 |
| ZNF214 | 26.9460888 | 0.99683805 | 0.37126156 | 2.68500202 | 0.00725294 | 0.26418897 |
| AC107884.1 | 30.178796 | 0.579625462 | 0.44359744 | 1.30664743 | 0.19133248 | 0.76328615 |
| PPFIBP2 | 85.6855236 | 0.984389164 | 0.43799637 | 2.24748246 | 0.02460921 | 0.41230428 |
| EIF3F | 92.6001294 | 0.109136036 | 0.32738484 | 0.33335703 | 0.7388648 | 0.9698508 |
| TUB | 124.333821 | 0.589390086 | 0.8415014 | 0.70040298 | 0.48367568 | 0.92004859 |
| RIC3 | 101.295087 | -0.37492112 | 0.54794557 | -0.6842306 | 0.49382957 | 0.92289011 |
| STK33 | 483.165211 | 0.657483699 | 0.29931022 | 2.19666303 | 0.02804452 | 0.4250936 |
| TRIM66 | 103.891269 | -0.526738396 | 0.40242843 | -1.3088996 | 0.19056837 | 0.76219391 |
| RPL27A | 337.887991 | -0.130401071 | 0.48883907 | -0.2667566 | 0.78965653 | 0.97423208 |
| ST5 | 204.096187 | 0.042200931 | 0.4600614 | 0.09172891 | 0.92691343 | 0.99336859 |
| TMEM9B | 51.2011957 | -0.538016475 | 0.31872865 | -1.6880079 | 0.09140972 | 0.63464873 |
| TMEM9B-AS | 48.1210063 | 0.522450969 | 0.50848204 | 1.02747183 | 0.30419834 | 0.84488735 |
| DENND5A | 250.356907 | -0.018959988 | 0.42389309 | -0.0447282 | 0.96432393 | 0.99586809 |
| TMEM41B | 148.688266 | 0.049476915 | 0.24762559 | 0.19980534 | 0.84163282 | 0.98515707 |
| IPO7 | 237.124921 | -0.285204632 | 0.32543811 | -0.8763713 | 0.38082819 | 0.8807629 |
| ZNF143 | 63.5214915 | -0.05552839 | 0.30394597 | -0.1826916 | 0.85503997 | 0.9864199 |
| WEE1 | 66.7987832 | -0.435597087 | 0.58351225 | -0.7465089 | 0.45536006 | 0.91077355 |
| SWAP70 | 204.769399 | -0.399539355 | 0.27543544 | -1.4505735 | 0.14689866 | 0.72098466 |
| SBF2-AS1 | 55.1706049 | -0.103028982 | 0.36482934 | -0.2824032 | 0.77763434 | 0.97423208 |
| SBF2 | 1120.02939 | 0.125293843 | 0.23739749 | 0.52778084 | 0.59765146 | 0.94683415 |
| AC011092.2 | 45.358629 | 0.310761622 | 0.36133463 | 0.8600383 | 0.38976793 | 0.88586416 |

| | | | | | | |
|---|---|---|---|---|---|---|
| ADM | 96.9284066 | -1.023861163 | 1.22695401 | -0.834474 | 0.40401396 | 0.89360496 |
| MTRNR2L8 | 101.336151 | 0.342954586 | 0.6766484 | 0.50684312 | 0.6122649 | 0.94791646 |
| RNF141 | 42.3633644 | -0.402915819 | 0.35609196 | -1.1314937 | 0.25784735 | 0.81223967 |
| CTR9 | 70.3654303 | -0.163294317 | 0.27695202 | -0.5896123 | 0.5554506 | 0.93523904 |
| EIF4G2 | 249.364989 | -0.284700304 | 0.27814773 | -1.0235579 | 0.30604414 | 0.84488735 |
| ZBED5 | 98.5179163 | 0.130698922 | 0.25224955 | 0.51813342 | 0.60436519 | 0.94683415 |
| ZBED5-AS1 | 36.1971689 | 0.05669776 | 0.60901155 | 0.093098 | 0.9258257 | 0.99316012 |
| GALNT18 | 631.548133 | 0.07518108 | 0.44579775 | 0.16864392 | 0.86607673 | 0.98670067 |
| USP47 | 405.813243 | 0.08531613 | 0.13775576 | 0.61932892 | 0.53569969 | 0.93469807 |
| MICAL2 | 77.9753753 | 0.175165225 | 0.80768581 | 0.21687298 | 0.82830734 | 0.98142144 |
| PARVA | 216.159371 | 0.160027047 | 0.34014504 | 0.47046709 | 0.63802134 | 0.94944486 |
| TEAD1 | 924.106068 | -0.569352707 | 0.19527609 | -2.9156295 | 0.00354972 | 0.18995949 |
| LINC00958 | 109.047746 | -0.570677193 | 0.39975151 | -1.4275798 | 0.15341284 | 0.7288378 |
| ARNTL | 47.326169 | 0.851003725 | 0.41378749 | 2.05662024 | 0.03972277 | 0.48094465 |
| BTBD10 | 108.514186 | -0.329322921 | 0.27180563 | -1.2116119 | 0.22566099 | 0.78977225 |
| FAR1 | 218.75342 | 0.087374192 | 0.18138292 | 0.48171123 | 0.63001109 | 0.94916079 |
| SPON1 | 210.049 | 1.0908844 | 0.67689445 | 1.61160194 | 0.10704859 | 0.65699665 |
| RRAS2 | 290.742646 | -0.190179377 | 0.31072828 | -0.612044 | 0.54050866 | 0.93523904 |
| COPB1 | 141.739256 | 0.129546528 | 0.23679867 | 0.54707456 | 0.58432751 | 0.94184538 |
| SOX6 | 1311.12093 | 0.41673966 | 0.26424723 | 1.57708243 | 0.11477656 | 0.67080211 |
| C11orf58 | 224.02132 | 0.021441445 | 0.21250727 | 0.10089746 | 0.91963186 | 0.99214128 |
| PLEKHA7 | 1401.39949 | 0.23541307 | 0.23968208 | 0.98218885 | 0.32600682 | 0.85509839 |
| RPS13 | 297.624053 | 0.042569431 | 0.6227958 | 0.06835215 | 0.94550531 | 0.9952642 |
| PIK3C2A | 399.931725 | 0.035161113 | 0.17268042 | 0.20361957 | 0.8386508 | 0.98462033 |
| NUCB2 | 89.0754023 | 0.448417841 | 0.35842298 | 1.25108563 | 0.21090324 | 0.77730294 |
| USH1C | 80.2659914 | 0.997253976 | 0.61783861 | 1.61410109 | 0.1065055 | 0.65699665 |
| SERGEF | 155.833035 | -0.279735823 | 0.16691563 | -1.6759115 | 0.09375552 | 0.63753181 |
| SAA1 | 895.760918 | 2.405432961 | 1.66193716 | 1.44736698 | 1.48E-01 | 0.72246583 |
| HPS5 | 57.7112933 | -0.302555047 | 0.39581198 | -0.7643908 | 0.44463438 | 0.90694419 |
| GTF2H1 | 96.6418253 | -0.27134472 | 0.20821792 | -1.3031766 | 0.19251449 | 0.76613887 |
| LDHA | 396.871454 | -0.274594648 | 0.54240867 | -0.5062505 | 0.61268083 | 0.94791646 |
| TSG101 | 136.870716 | 0.08880968 | 0.21444727 | 0.41413295 | 0.67877674 | 0.95924874 |
| UEVLD | 93.6834096 | 0.065295187 | 0.19544462 | 0.33408536 | 0.73831515 | 0.9698508 |
| SPTY2D1OS | 31.513 | 0.065593101 | 0.29548132 | 0.2219873 | 0.82432377 | 0.98094663 |
| SPTY2D1 | 35.7796453 | -0.383215536 | 0.38516862 | -0.9949293 | 0.31977067 | 0.85062016 |
| ZDHHC13 | 85.8162679 | -0.312994152 | 0.21055387 | -1.4865276 | 0.13713961 | 0.71044919 |
| NAV2 | 435.845245 | -1.173204178 | 0.5807237 | -2.020245 | 0.04335798 | 0.49484046 |
| HTATIP2 | 32.1930538 | 0.457394903 | 0.4307332 | 1.06189843 | 0.2882818 | 0.8330772 |
| PRMT3 | 130.536296 | -0.747914628 | 0.27386961 | -2.730915 | 0.00631588 | 0.25103639 |
| ANO5 | 51.0059033 | 0.422419438 | 0.41944027 | 1.00710272 | 0.31388542 | 0.8490543 |
| GAS2 | 48.9598726 | -0.200785087 | 0.52971589 | -0.379043 | 0.70465595 | 0.96461079 |
| SVIP | 80.1328293 | 0.022250187 | 0.45644411 | 0.04874679 | 0.96112108 | 0.99572129 |
| MUC15 | 24.1398711 | -0.547057213 | 1.65979889 | -0.3295925 | 0.74170791 | 0.9698508 |
| BBOX1 | 51.2630531 | 1.466819238 | 0.74666984 | 1.9644817 | 0.04947426 | 0.52308378 |
| LGR4 | 979.899968 | -0.2034736 | 0.3805877 | -0.53463 | 0.59290574 | 0.94649617 |
| LIN7C | 52.4581659 | 0.246121552 | 0.38852415 | 0.63347814 | 0.52642145 | 0.93331883 |
| BDNF-AS | 248.558653 | -0.008913886 | 0.28352005 | -0.0314401 | 0.9749186 | 0.99681784 |
| METTL15 | 471.310027 | -0.079840124 | 0.29109533 | -0.2742748 | 0.78387342 | 0.97423208 |
| ARL14EP | 44.2710436 | 0.13934299 | 0.24607154 | 0.56627024 | 0.57121009 | 0.93998433 |
| MPPED2 | 148.322313 | 0.966539358 | 0.49556713 | 1.9503702 | 0.05113201 | 0.52609271 |
| DCDC1 | 296.992016 | 0.79124172 | 0.35745869 | 2.2135193 | 0.02686186 | 0.4234302 |
| DNAJC24 | 67.0179656 | -0.610818545 | 0.25264733 | -2.4176727 | 0.01562012 | 0.34086104 |
| IMMP1L | 171.270805 | -0.004105085 | 0.26921663 | -0.0152483 | 0.98783412 | 0.99804429 |
| ELP4 | 266.710558 | 0.320899233 | 0.20939706 | 1.53249159 | 0.12540117 | 0.69492095 |

| | | | | | | |
|---|---|---|---|---|---|---|
| AC131571.1 | 35.333898 | 0.645960999 | 0.65897757 | 0.98024732 | 0.32696405 | 0.85641576 |
| RCN1 | 88.884528 | -0.264226517 | 0.47519439 | -0.5560388 | 0.57818433 | 0.94034284 |
| WT1 | 148.85842 | 1.011544036 | 1.45718608 | 0.6941763 | 0.48757165 | 0.92034617 |
| EIF3M | 121.922905 | -0.062483747 | 0.49621867 | -0.1259198 | 0.89979542 | 0.98956835 |
| PRRG4 | 86.0096102 | 0.102519964 | 0.33835811 | 0.30299249 | 0.76189559 | 0.9714717 |
| QSER1 | 181.579173 | -0.084133672 | 0.16853283 | -0.4992124 | 0.61762978 | 0.94796918 |
| TCP11L1 | 65.6921678 | 0.37974781 | 0.2957978 | 1.28380878 | 0.19920887 | 0.77110248 |
| CSTF3 | 153.122769 | -0.144217882 | 0.23874197 | -0.6040743 | 0.54579427 | 0.93523904 |
| HIPK3 | 483.231274 | 0.052202559 | 0.31898832 | 0.16365038 | 0.87000638 | 0.98765005 |
| FBXO3 | 95.4481537 | 0.360685348 | 0.22072893 | 1.63406469 | 0.10224527 | 0.65288882 |
| CAPRIN1 | 208.676509 | -0.106530846 | 0.15903239 | -0.6698689 | 0.50294139 | 0.92492012 |
| NAT10 | 69.2940466 | -0.748110792 | 0.2663131 | -2.80914 | 0.0049674 | 0.22029054 |
| ABTB2 | 171.147585 | -0.048407346 | 0.32589132 | -0.1485383 | 0.88191795 | 0.98766275 |
| CAT | 218.601842 | 0.283913606 | 0.62324977 | 0.45553744 | 0.64872264 | 0.95027667 |
| ELF5 | 35.9166064 | 1.821790552 | 2.60766411 | 0.6986293 | 0.48478373 | 0.92034617 |
| EHF | 201.475349 | -0.983171 | 0.66375699 | -1.4812213 | 0.13854761 | 0.71091391 |
| APIP | 74.1328131 | -0.600939393 | 0.27386391 | -2.1942993 | 0.0282139 | 0.4250936 |
| PDHX | 127.171523 | -0.155124532 | 0.25714197 | -0.6032641 | 0.54633298 | 0.93523904 |
| TRIM44 | 411.894163 | -0.006384951 | 0.1926293 | -0.0331463 | 0.97355791 | 0.99681784 |
| LDLRAD3 | 97.6207847 | 0.565570297 | 0.50252592 | 1.12545498 | 0.26039628 | 0.81263292 |
| TRAF6 | 40.9486051 | 0.129457029 | 0.33077233 | 0.39137805 | 0.69551782 | 0.96220115 |
| C11orf74 | 42.1912808 | 0.484553076 | 0.38776789 | 1.24959566 | 0.21144729 | 0.77730294 |
| LRRC4C | 472.753719 | -0.540638349 | 0.53478879 | -1.0109381 | 0.31204607 | 0.84811918 |
| API5 | 111.904239 | -0.295499453 | 0.20969654 | -1.4091766 | 0.15878297 | 0.73686885 |
| TTC17 | 635.291191 | -0.184498077 | 0.17004117 | -1.08502 | 0.27791279 | 0.8229426 |
| HSD17B12 | 318.975511 | -0.112698671 | 0.24036643 | -0.4688619 | 0.63916832 | 0.94944486 |
| ALKBH3 | 40.4251541 | 0.051631287 | 0.32882118 | 0.15701935 | 0.8752296 | 0.98766275 |
| EXT2 | 124.964985 | -0.123116965 | 0.2484305 | -0.4955791 | 0.62019139 | 0.9480368 |
| CD82 | 39.056488 | -0.327765093 | 0.62934984 | -0.5207995 | 0.60250644 | 0.94683415 |
| TSPAN18 | 42.8632949 | 1.214700974 | 0.68986507 | 1.76078052 | 0.07827556 | 0.60239981 |
| TP53I11 | 29.0976642 | -0.025666085 | 0.45178687 | -0.0568102 | 0.95469642 | 0.99531843 |
| PRDM11 | 50.4343442 | -0.034229053 | 0.30576781 | -0.1119446 | 0.91086734 | 0.99100813 |
| CRY2 | 79.7117837 | 0.057694517 | 0.34015205 | 0.1696139 | 0.86531379 | 0.98670067 |
| LARGE2 | 26.5816281 | 0.345941426 | 0.51663391 | 0.6696065 | 0.50310867 | 0.92500536 |
| PHF21A | 585.095435 | 0.120732881 | 0.25184134 | 0.47940056 | 0.63165369 | 0.94932781 |
| DGKZ | 27.2219767 | -0.626611148 | 0.36591537 | -1.7124483 | 0.08681409 | 0.62478747 |
| MDK | 50.7728294 | -0.643535025 | 1.17545951 | -0.5474753 | 0.58405225 | 0.94184538 |
| AMBRA1 | 748.804787 | 0.217870643 | 0.30541325 | 0.71336342 | 0.4756209 | 0.91809716 |
| ATG13 | 78.785728 | 0.30093831 | 0.24894622 | 1.20884868 | 0.22672099 | 0.79116952 |
| ARHGAP1 | 59.6988117 | -0.641726811 | 0.27986813 | -2.2929614 | 0.02185023 | 0.38823827 |
| CKAP5 | 109.877538 | -0.188650938 | 0.19732166 | -0.9560579 | 0.33904296 | 0.86680923 |
| LRP4 | 114.812643 | -0.05979238 | 0.4707565 | -0.1270134 | 0.8989298 | 0.98948153 |
| C11orf49 | 437.003894 | 0.198649596 | 0.18877163 | 1.05232758 | 0.29264928 | 0.83697123 |
| ARFGAP2 | 60.3949904 | -0.26065355 | 0.365074 | -0.7139746 | 0.4752429 | 0.91809716 |
| PACSIN3 | 28.9670597 | -0.176541176 | 0.42405332 | -0.4163183 | 0.67717707 | 0.95924874 |
| DDB2 | 67.9422753 | 0.679941884 | 0.52570627 | 1.29338743 | 0.1958771 | 0.77017438 |
| MADD | 46.159749 | -0.439983284 | 0.31034671 | -1.4177153 | 0.15627389 | 0.73307914 |
| SLC39A13 | 40.944151 | -0.305122607 | 0.33538832 | -0.9097592 | 0.36294952 | 0.87551319 |
| PSMC3 | 28.234122 | -0.750130718 | 0.43446028 | -1.7265806 | 0.084243 | 0.61656297 |
| CELF1 | 498.596017 | -0.040074381 | 0.20245881 | -0.1979384 | 0.84309322 | 0.98517901 |
| NDUFS3 | 47.2654677 | 0.152303536 | 0.31459685 | 0.4841229 | 0.62829865 | 0.94858781 |
| MTCH2 | 106.68361 | -0.041327771 | 0.28857061 | -0.1432155 | 0.88612002 | 0.98766275 |
| FNBP4 | 305.225742 | -0.072799303 | 0.17253937 | -0.4219286 | 0.6730771 | 0.95910677 |
| NUP160 | 200.730343 | -0.121758545 | 0.1759731 | -0.6919157 | 0.4889903 | 0.92121587 |

| | | | | | | |
|---|---|---|---|---|---|---|
| PTPRJ | 887.443975 | -0.555383547 | 0.1926364 | -2.8830665 | 0.00393824 | 0.2001215 |
| TNKS1BP1 | 83.4716332 | -0.307604778 | 0.39532754 | -0.7781011 | 0.43650944 | 0.90525098 |
| SSRP1 | 56.8348224 | -0.49636681 | 0.3291049 | -1.5082328 | 0.13149495 | 0.700349 |
| TIMM10 | 33.7840962 | -0.138656299 | 0.46620385 | -0.2974156 | 0.76614923 | 0.97221495 |
| UBE2L6 | 50.9606477 | -0.838773268 | 0.652273 | -1.2859236 | 0.19846972 | 0.77110248 |
| SERPING1 | 353.778172 | -0.177319415 | 0.32033671 | -0.5535407 | 5.80E-01 | 9.40E-01 |
| ZDHHC5 | 109.531633 | -0.468225183 | 0.23386802 | -2.0020915 | 0.04527489 | 0.50441298 |
| CTNND1 | 276.071478 | 0.102927298 | 0.1991179 | 0.51691635 | 0.60521455 | 0.94698346 |
| LPXN | 63.3368036 | -0.219481124 | 0.43358093 | -0.5062057 | 0.61271228 | 0.94791646 |
| GLYAT | 97.632588 | 1.815721803 | 0.72001621 | 2.52177906 | 0.0116763 | 0.30872898 |
| FAM111A | 73.4141055 | 0.389369264 | 0.37538025 | 1.03726624 | 0.2996118 | 0.84114343 |
| OSBP | 140.501355 | -0.1965868 | 0.20131996 | -0.9764894 | 0.32882201 | 0.8574012 |
| PATL1 | 98.0677376 | -0.418828073 | 0.35890807 | -1.1669508 | 0.24323022 | 0.80476923 |
| STX3 | 120.093276 | 0.278694788 | 0.34660382 | 0.80407304 | 0.42135479 | 0.90012434 |
| MRPL16 | 33.133513 | 0.262485649 | 0.3726576 | 0.70436146 | 0.4812077 | 0.91971919 |
| PRPF19 | 47.3263462 | -0.560314839 | 0.38146158 | -1.4688631 | 0.14186993 | 0.7148274 |
| TMEM109 | 40.4251756 | 0.081913632 | 0.48145532 | 0.17013756 | 0.86490196 | 0.98670067 |
| VPS37C | 47.8812256 | -0.291196614 | 0.3209224 | -0.9073739 | 0.36420911 | 0.87605028 |
| DDB1 | 129.510291 | -0.03327349 | 0.25776697 | -0.1290836 | 0.89729149 | 0.98851699 |
| CPSF7 | 101.621336 | 0.013518799 | 0.18472804 | 0.07318217 | 0.94166115 | 0.99526299 |
| AP003108.2 | 72.5022202 | -0.023511932 | 0.36101222 | -0.0651278 | 0.94807225 | 0.9952642 |
| MYRF | 68.9700627 | 1.323258654 | 0.62479603 | 2.11790502 | 0.03418311 | 0.45111962 |
| TMEM258 | 73.2689839 | 0.189218659 | 0.35835916 | 0.52801402 | 0.59748961 | 0.94683415 |
| FADS1 | 50.8136849 | -0.031554646 | 0.56747754 | -0.0556051 | 0.9556564 | 0.99567464 |
| FADS3 | 48.3156714 | 0.233787923 | 0.49669084 | 0.47069103 | 0.63786139 | 0.94944486 |
| FTH1 | 2132.14107 | 0.893460875 | 0.57192673 | 1.56219465 | 0.11824214 | 0.68313719 |
| ASRGL1 | 126.744554 | 0.401074073 | 0.41075773 | 0.97642489 | 0.32885395 | 0.8574012 |
| AHNAK | 626.93241 | -0.179252848 | 0.34170284 | -0.5245869 | 0.59987037 | 0.94683415 |
| EML3 | 25.9832086 | -0.306889354 | 0.39242141 | -0.7820403 | 0.4341909 | 0.90514499 |
| B3GAT3 | 44.338629 | -0.958347629 | 0.35976935 | -2.6637834 | 0.00772673 | 0.27068528 |
| GANAB | 149.740191 | 0.445674005 | 0.25788643 | 1.72817935 | 0.08395607 | 0.61656297 |
| UQCC3 | 40.8980113 | -0.620554734 | 0.4871382 | -1.2738782 | 0.20270659 | 0.77110248 |
| UBXN1 | 66.2141545 | 0.107968009 | 0.46416011 | 0.23260941 | 0.81606472 | 0.98007572 |
| TTC9C | 52.6714161 | 0.319319182 | 0.24282373 | 1.31502459 | 0.18850164 | 0.76207437 |
| TMEM223 | 34.2759247 | 0.08996005 | 0.33379084 | 0.26951024 | 0.78753706 | 0.97423208 |
| TMEM179B | 58.1842153 | 0.233786284 | 0.42213487 | 0.55381893 | 0.57970277 | 0.94034284 |
| NXF1 | 33.1641569 | -0.128048061 | 0.29042052 | -0.4409057 | 0.65928127 | 0.95407427 |
| STX5 | 96.667005 | 0.329132273 | 0.24875462 | 1.32312024 | 0.1857954 | 0.76046631 |
| SLC3A2 | 108.489212 | -0.708903565 | 0.24820927 | -2.856072 | 0.00428918 | 0.20962265 |
| RARRES3 | 128.394518 | 0.152861425 | 1.08561064 | 0.14080686 | 0.88802252 | 0.98766275 |
| PLA2G16 | 201.692336 | 0.837479756 | 0.38971722 | 2.14894215 | 0.03163899 | 0.43905874 |
| ATL3 | 113.866146 | 0.117894445 | 0.25542462 | 0.46156257 | 0.64439504 | 0.94944486 |
| RTN3 | 236.519593 | -0.08415897 | 0.17864153 | -0.4711053 | 0.63756556 | 0.94944486 |
| MARK2 | 172.05171 | -0.066990343 | 0.18652514 | -0.3591491 | 0.71948353 | 0.96779196 |
| NAA40 | 52.4715733 | 0.034111925 | 0.30735561 | 0.1109852 | 0.91162808 | 0.99100813 |
| OTUB1 | 53.7376721 | -0.046212138 | 0.2900791 | -0.1593087 | 0.87342563 | 0.98766275 |
| MACROD1 | 88.6869356 | -0.214296169 | 0.44249655 | -0.4842889 | 0.62818085 | 0.94858781 |
| STIP1 | 62.2094585 | -0.201732907 | 0.28284988 | -0.7132155 | 0.47571245 | 0.91809716 |
| VEGFB | 67.5602236 | 0.070791685 | 0.2647645 | 0.26737605 | 0.78917964 | 0.97423208 |
| FKBP2 | 83.4965445 | 0.299288502 | 0.43199478 | 0.69280584 | 0.48843141 | 0.92081962 |
| PPP1R14B | 55.0099 | -0.796472679 | 0.60282485 | -1.321234 | 0.18642336 | 0.76162997 |
| BAD | 46.1665416 | 0.326929613 | 0.31417124 | 1.04060961 | 0.29805677 | 0.83973489 |
| GPR137 | 29.7594339 | -0.003835549 | 0.32583381 | -0.0117715 | 0.99060793 | 0.99866813 |
| ESRRA | 79.1804956 | 0.291681765 | 0.44548602 | 0.65474954 | 0.51262902 | 0.92930988 |

| | | | | | | |
|---|---|---|---|---|---|---|
| TRMT112 | 94.743414 | -0.390136364 | 0.54810169 | -0.7117956 | 0.47659136 | 0.91819563 |
| PRDX5 | 95.2308755 | 0.189514525 | 0.53148134 | 0.35657795 | 0.72140779 | 0.96779196 |
| SF1 | 269.553348 | -0.161634608 | 0.20250573 | -0.798173 | 0.4247701 | 0.90167091 |
| EHD1 | 53.1102879 | -0.00920908 | 0.55127719 | -0.016705 | 0.98667197 | 0.99764442 |
| ATG2A | 31.0784801 | -0.36716019 | 0.27314994 | -1.3441709 | 0.17889314 | 0.75150688 |
| VPS51 | 93.8580634 | 0.234573928 | 0.24795104 | 0.94604937 | 0.34412341 | 0.86969477 |
| TM7SF2 | 23.4863004 | -0.013289182 | 0.50208406 | -0.026468 | 0.97888402 | 0.99681784 |
| FAU | 320.487861 | -0.358276799 | 0.52296502 | -0.6850875 | 0.49328871 | 0.92289011 |
| SYVN1 | 31.8631501 | 0.142613265 | 0.36797391 | 0.38756352 | 0.69833907 | 0.96256506 |
| MRPL49 | 29.312858 | 0.612647228 | 0.35109244 | 1.74497412 | 0.08098937 | 0.6084684 |
| CAPN1 | 116.626291 | -0.67662756 | 0.23188014 | -2.9180056 | 0.00352278 | 0.18995949 |
| DPF2 | 60.5853829 | 0.003531409 | 0.26119523 | 0.01352019 | 0.98921278 | 0.9983847 |
| FRMD8 | 38.3388325 | -0.288199155 | 0.28159903 | -1.023438 | 0.3061008 | 0.84488735 |
| NEAT1 | 7503.38749 | 0.316512549 | 0.37829245 | 0.83668745 | 0.40276827 | 0.89266944 |
| MALAT1 | 87344.7714 | -0.238536206 | 0.39190732 | -0.6086546 | 0.54275339 | 0.93523904 |
| SCYL1 | 56.4183066 | 0.027680714 | 0.24806068 | 0.11158848 | 0.91114971 | 0.99100813 |
| LTBP3 | 147.007006 | -0.315373794 | 0.35369878 | -0.8916451 | 0.37258317 | 0.87605028 |
| SSSCA1 | 31.5996363 | -0.306221826 | 0.7376334 | -0.4151409 | 0.67803873 | 0.95924874 |
| EHBP1L1 | 43.9726737 | 0.500581099 | 0.57983226 | 0.86332054 | 0.38796125 | 0.8853359 |
| MAP3K11 | 43.3239151 | 0.100355391 | 0.35356979 | 0.28383475 | 0.77653702 | 0.97423208 |
| PCNX3 | 30.7125437 | -0.159256081 | 0.40891262 | -0.3894624 | 0.69693414 | 0.96220115 |
| SIPA1 | 28.8102373 | -0.382505756 | 0.49399994 | -0.7743032 | 0.43875149 | 0.90561284 |
| RELA | 46.7259393 | -0.076405017 | 0.32104915 | -0.2379854 | 0.8118924 | 0.97930627 |
| KAT5 | 45.0906449 | -0.287604043 | 0.28642193 | -1.0041272 | 0.31531731 | 0.8490543 |
| RNASEH2C | 39.9157116 | 0.234775818 | 0.48056637 | 0.48853984 | 0.62516751 | 0.94808496 |
| CFL1 | 228.339753 | -0.375960892 | 0.55416427 | -0.6784286 | 0.49749997 | 0.92398825 |
| MUS81 | 30.4961348 | -0.110155916 | 0.40878051 | -0.2694745 | 0.78756458 | 0.97423208 |
| EFEMP2 | 51.6093308 | 0.20113202 | 0.39122426 | 0.51410928 | 0.60717559 | 0.94708529 |
| DRAP1 | 47.6202462 | -0.547751123 | 0.57526382 | -0.9521738 | 0.34100886 | 0.8682945 |
| SART1 | 31.6644619 | -0.611746332 | 0.34049662 | -1.7966297 | 0.07239443 | 0.59090333 |
| BANF1 | 72.3228159 | -0.857271025 | 0.45422624 | -1.8873217 | 0.05911707 | 0.55686386 |
| SF3B2 | 107.343213 | -0.340331487 | 0.19476576 | -1.7473887 | 0.08056994 | 0.60722226 |
| PACS1 | 493.493063 | 0.032271339 | 0.27751568 | 0.11628655 | 0.90742545 | 0.99100813 |
| KLC2 | 26.7448867 | -0.160523725 | 0.42118304 | -0.3811258 | 0.7031099 | 0.9645815 |
| RAB1B | 56.9919613 | -0.014751394 | 0.33964859 | -0.0434313 | 0.96535769 | 0.99601598 |
| YIF1A | 25.1973449 | 0.022149668 | 0.35997479 | 0.06153116 | 0.9509362 | 0.99531843 |
| BRMS1 | 27.5196378 | -0.276442426 | 0.38493953 | -0.7181451 | 0.47266784 | 0.91737234 |
| SLC29A2 | 24.2287447 | -0.188740713 | 0.56291808 | -0.3352898 | 0.73740646 | 0.9698508 |
| MRPL11 | 26.9171497 | -0.028653332 | 0.46119472 | -0.0621285 | 0.95046051 | 0.99531843 |
| ZDHHC24 | 30.152718 | -0.279052054 | 0.38481982 | -0.7251499 | 0.46836007 | 0.91462428 |
| CCS | 71.5002844 | 0.458081429 | 0.3180202 | 1.44041616 | 0.14974969 | 0.72246729 |
| RBM14-RBM | 55.1340688 | -0.021750288 | 0.2673136 | -0.0813662 | 0.93515074 | 0.99464285 |
| RBM4 | 48.5169167 | 0.0835568 | 0.33051525 | 0.2528077 | 0.80041681 | 0.97723068 |
| RBM4B | 47.0073585 | 0.273225102 | 0.39952619 | 0.68387282 | 0.49405548 | 0.92289011 |
| C11orf80 | 195.331113 | 0.231932677 | 0.3068657 | 0.75581166 | 0.44976212 | 0.91034958 |
| PC | 38.636502 | 0.205224314 | 0.40471121 | 0.50708829 | 0.61209287 | 0.94791646 |
| RHOD | 37.5096497 | -0.528312136 | 0.45434016 | -1.1628119 | 0.24490583 | 0.80476923 |
| KDM2A | 592.468362 | -0.096147645 | 0.20325522 | -0.473039 | 0.63618535 | 0.94932781 |
| GRK2 | 57.8111179 | -0.646954243 | 0.35176334 | -1.8391747 | 0.06588949 | 0.57692455 |
| ANKRD13D | 63.5380483 | -0.290397703 | 0.24927759 | -1.1649571 | 0.24403635 | 0.80476923 |
| SSH3 | 38.7213038 | -0.178842891 | 0.28176366 | -0.6347266 | 0.52560673 | 0.93272384 |
| RAD9A | 259.348641 | 0.542408923 | 0.2500678 | 2.16904745 | 0.03007908 | 0.43492739 |
| CLCF1 | 23.6813881 | -1.787715046 | 0.48690752 | -3.67157 | 0.00024107 | 0.04172264 |
| PPP1CA | 82.9454779 | -0.070706602 | 0.47604707 | -0.1485286 | 0.88192562 | 0.98766275 |

| | | | | | | |
|---|---|---|---|---|---|---|
| RPS6KB2 | 29.7515626 | -0.205140867 | 0.40813394 | -0.5026312 | 0.61522356 | 0.94791646 |
| CORO1B | 41.2206647 | -0.333552702 | 0.3326287 | -1.0027779 | 0.31596804 | 0.8490543 |
| TMEM134 | 55.9437865 | -0.446577836 | 0.37946893 | -1.1768495 | 0.23925557 | 0.80174646 |
| AIP | 29.557982 | -0.048822612 | 0.49683136 | -0.098268 | 0.92171951 | 0.99214128 |
| PITPNM1 | 40.1640904 | -0.483729152 | 0.39664735 | -1.2195447 | 0.22263754 | 0.78710314 |
| CDK2AP2 | 62.7468431 | 0.051772715 | 0.56086601 | 0.09230853 | 0.92645291 | 0.99322803 |
| GSTP1 | 806.958886 | -0.607892947 | 0.65055782 | -0.934418 | 0.35008831 | 0.8720098 |
| NDUFV1 | 95.9928445 | -0.276422504 | 0.26505021 | -1.0429062 | 0.29699174 | 0.83973489 |
| UNC93B1 | 44.5174732 | -0.262357969 | 0.76207144 | -0.3442695 | 0.7306436 | 0.96896512 |
| NDUFS8 | 86.4673923 | 0.209217456 | 0.32921606 | 0.63550198 | 0.52510108 | 0.93272384 |
| CHKA | 158.040277 | -0.140681739 | 0.40655471 | -0.346034 | 0.72931719 | 0.96862399 |
| KMT5B | 245.434349 | 0.283527562 | 0.22113098 | 1.28217026 | 0.19978292 | 0.77110248 |
| LRP5 | 116.61524 | -0.268615118 | 0.34796833 | -0.7719528 | 0.4401424 | 0.90561284 |
| PPP6R3 | 689.451698 | 0.072221556 | 0.14015435 | 0.51530016 | 6.06E-01 | 0.94708529 |
| CPT1A | 121.690138 | 0.917889446 | 0.46841711 | 1.95955576 | 0.05004774 | 0.52415717 |
| MRPL21 | 27.7995829 | 0.130103704 | 0.32932341 | 0.39506364 | 0.69279593 | 0.96220115 |
| IGHMBP2 | 31.5949488 | -0.295078532 | 0.29866295 | -0.9879984 | 0.32315341 | 0.85354768 |
| TPCN2 | 42.1825338 | 0.755645654 | 0.36318931 | 2.08058342 | 0.03747205 | 0.46930788 |
| CCND1 | 168.09621 | -2.416177648 | 0.61185936 | -3.9489102 | 7.85E-05 | 0.02080127 |
| ORAOV1 | 43.0569469 | -0.848585561 | 0.31425015 | -2.7003505 | 0.00692665 | 0.25717187 |
| PPFIA1 | 280.678483 | 0.190122547 | 0.25636045 | 0.74162198 | 0.4583164 | 0.91336645 |
| AP002336.2 | 103.638812 | 0.759368316 | 0.48453137 | 1.56722218 | 0.11706277 | 0.6781987 |
| CTTN | 301.950275 | -0.298024803 | 0.2678552 | -1.112634 | 0.26586565 | 0.81387543 |
| SHANK2 | 1025.76937 | -0.093173358 | 0.3391178 | -0.2747522 | 0.78350663 | 0.97423208 |
| NADSYN1 | 48.3579685 | 0.165599603 | 0.30367695 | 0.54531503 | 0.58553687 | 0.9423304 |
| AP002495.2 | 121.223957 | 0.450462502 | 0.29937952 | 1.50465369 | 0.13241313 | 0.7023175 |
| RNF121 | 170.740324 | -0.030600539 | 0.20804461 | -0.1470864 | 0.8830638 | 0.98766275 |
| NUMA1 | 175.496983 | -0.356907518 | 0.29620653 | -1.2049279 | 0.22823114 | 0.79284833 |
| LAMTOR1 | 95.6911073 | -0.169145282 | 0.37312074 | -0.4533259 | 0.65031411 | 0.95061176 |
| INPPL1 | 61.2826873 | 0.497124413 | 0.29847477 | 1.66554922 | 0.09580323 | 0.64264648 |
| CLPB | 45.3112122 | -0.117226206 | 0.31402985 | -0.3732964 | 0.70892787 | 0.96587538 |
| ARAP1 | 52.9924725 | 0.355990882 | 0.32943029 | 1.08062584 | 0.27986358 | 0.82464035 |
| ATG16L2 | 30.724751 | 0.453579087 | 0.41053626 | 1.10484538 | 0.26922659 | 0.81655182 |
| FCHSD2 | 531.865935 | 0.224641421 | 0.43784427 | 0.51306237 | 0.60790769 | 0.94708529 |
| P2RY6 | 33.8886624 | -0.764680685 | 0.49603795 | -1.541577 | 0.12317642 | 0.69314363 |
| ARHGEF17 | 90.9155246 | 0.005016843 | 0.38908941 | 0.0128938 | 0.98971252 | 0.99839055 |
| FAM168A | 213.07255 | 0.252021293 | 0.19092407 | 1.32000794 | 0.18683237 | 0.76207437 |
| RAB6A | 285.089336 | -0.137247645 | 0.16856025 | -0.8142349 | 0.41551038 | 0.89807127 |
| MRPL48 | 44.904387 | -0.147099862 | 0.26403642 | -0.5571196 | 0.57744573 | 0.94034284 |
| COA4 | 61.8612446 | -0.07994807 | 0.44749225 | -0.178658 | 0.85820625 | 0.9864199 |
| PAAF1 | 81.4243568 | 0.325806903 | 0.27082549 | 1.20301416 | 0.22897084 | 0.7931636 |
| C2CD3 | 108.946766 | -0.181345967 | 0.22022101 | -0.8234726 | 0.4102393 | 0.8964268 |
| PPME1 | 155.887059 | -0.06227133 | 0.31581597 | -0.197176 | 0.84368981 | 0.98517901 |
| P4HA3 | 29.25487 | -0.192999673 | 0.48660983 | -0.396621 | 0.69164697 | 0.96220115 |
| PGM2L1 | 73.8756257 | 0.00280784 | 0.52152773 | 0.00538387 | 0.99570431 | 0.99866813 |
| POLD3 | 80.4133577 | -0.009722456 | 0.22942564 | -0.0423774 | 0.96619787 | 0.99601626 |
| RNF169 | 186.91404 | -0.366227684 | 0.22457221 | -1.6307792 | 0.10293692 | 0.65288882 |
| XRRA1 | 167.923139 | -0.43801697 | 0.56474764 | -0.7755977 | 0.43798656 | 0.90561284 |
| SPCS2 | 156.435264 | 0.040137803 | 0.32699183 | 0.12274864 | 0.90230615 | 0.9900784 |
| NEU3 | 32.2031921 | -0.350243654 | 0.30857569 | -1.1350332 | 0.25636141 | 0.80988075 |
| ARRB1 | 37.6796726 | -0.924159121 | 0.53221716 | -1.7364324 | 0.08248741 | 0.6141478 |
| RPS3 | 587.081194 | 0.203498952 | 0.51929817 | 0.39187304 | 0.69515203 | 0.96220115 |
| SERPINH1 | 47.5334622 | 0.16116836 | 0.28142769 | 0.57268125 | 0.56686053 | 0.9385359 |
| UVRAG | 369.915932 | -0.010562361 | 0.22177244 | -0.047627 | 0.96201349 | 0.99572129 |

| | | | | | | |
|---|---|---|---|---|---|---|
| THAP12 | 38.2492816 | -0.239000022 | 0.31993017 | -0.7470381 | 0.45504057 | 0.91077355 |
| EMSY | 183.026028 | -0.038547758 | 0.16425626 | -0.2346806 | 0.81445665 | 0.97937868 |
| ACER3 | 175.097119 | 0.308811736 | 0.27271593 | 1.13235678 | 0.25748447 | 0.8120529 |
| CAPN5 | 25.3048296 | 0.217020201 | 0.57625938 | 0.37660159 | 0.70646971 | 0.96587538 |
| PAK1 | 130.66532 | 0.737176495 | 0.29867311 | 2.46817161 | 0.01358052 | 0.32032755 |
| CLNS1A | 131.076381 | -0.040466388 | 0.18334021 | -0.2207175 | 0.82531243 | 0.98094663 |
| RSF1 | 409.862626 | 0.203305395 | 0.16258385 | 1.25046486 | 0.21112978 | 0.77730294 |
| AAMDC | 110.339196 | 0.214975212 | 0.32928715 | 0.6528503 | 0.51385279 | 0.93064774 |
| AP002812.2 | 46.1282914 | 0.013090373 | 0.38353726 | 0.03413064 | 0.97277297 | 0.99681784 |
| INTS4 | 102.375046 | -0.227544505 | 0.20406002 | -1.1150862 | 0.2648135 | 0.81378765 |
| ALG8 | 59.0521939 | -0.45614657 | 0.33976986 | -1.3425163 | 0.17942865 | 0.75251624 |
| GAB2 | 170.404382 | 0.131145505 | 0.58985943 | 0.22233349 | 0.82405428 | 0.98094663 |
| AP003086.2 | 50.8778039 | 0.651783098 | 0.55953524 | 1.16486514 | 0.24407359 | 0.80476923 |
| NARS2 | 154.873762 | -0.174677573 | 0.24011067 | -0.7274878 | 0.46692721 | 0.91462428 |
| TENM4 | 170.029059 | 0.472999574 | 0.76199413 | 0.62073913 | 0.53477128 | 0.93460153 |
| PRCP | 62.417987 | 0.181331616 | 0.34381391 | 0.5274121 | 0.59790744 | 0.94683415 |
| RAB30 | 33.4784702 | 0.537074828 | 0.41869525 | 1.28273446 | 0.19958512 | 0.77110248 |
| RAB30-AS1 | 174.215233 | 0.27607737 | 0.34408855 | 0.80234396 | 0.42235402 | 0.90066187 |
| PCF11 | 133.139673 | 0.056890937 | 0.22733152 | 0.25025538 | 0.80238986 | 0.97835591 |
| ANKRD42 | 73.2128242 | 0.631427645 | 0.29335215 | 2.15245615 | 0.03136145 | 0.43905874 |
| CCDC90B | 49.4970297 | -0.133808929 | 0.33965878 | -0.393951 | 0.69361719 | 0.96220115 |
| DLG2 | 1041.87906 | 0.426209058 | 0.42525171 | 1.00225124 | 0.31622226 | 0.8490543 |
| TMEM126B | 42.6526752 | 0.008405585 | 0.31244029 | 0.02690301 | 0.97853709 | 0.99681784 |
| CREBZF | 132.407923 | 0.497531914 | 0.24960074 | 1.99331109 | 0.04622739 | 0.50827029 |
| SYTL2 | 410.024361 | -0.379379423 | 0.30151451 | -1.258246 | 0.20830281 | 0.77462844 |
| PICALM | 1218.99384 | -0.012196105 | 0.19350023 | -0.0630289 | 9.50E-01 | 9.95E-01 |
| EED | 53.6586639 | 0.028672771 | 0.33356509 | 0.08595855 | 0.93149937 | 0.99411667 |
| HIKESHI | 69.9557584 | 0.028646729 | 0.23526343 | 0.12176448 | 0.90308555 | 0.99034419 |
| AP001831.1 | 46.8965246 | 0.589182369 | 0.5227081 | 1.12717283 | 0.25966942 | 0.81263292 |
| ME3 | 112.552185 | -0.548593911 | 0.34278762 | -1.6003901 | 0.10951208 | 0.65769039 |
| PRSS23 | 137.011968 | 0.369248957 | 0.54421559 | 0.67849757 | 0.49745626 | 0.92398825 |
| AP001528.2 | 92.6218006 | 0.448044637 | 0.68761618 | 0.65159118 | 0.51466494 | 0.93106241 |
| FZD4 | 49.5467986 | 0.377482918 | 0.46876539 | 0.80527045 | 0.42066363 | 0.90012434 |
| TMEM135 | 260.837389 | 0.588265604 | 0.20129395 | 2.92242068 | 0.00347322 | 0.18995949 |
| CTSC | 310.019092 | 1.265236546 | 0.67259885 | 1.88111614 | 0.05995612 | 0.55922073 |
| GRM5-AS1 | 61.0231764 | 1.737544981 | 0.46548843 | 3.73273504 | 0.00018941 | 0.03628751 |
| GRM5 | 178.771485 | 1.783101746 | 0.55228696 | 3.22857842 | 0.00124407 | 0.11242415 |
| NOX4 | 156.234695 | 0.801355422 | 0.73289228 | 1.09341501 | 0.27421163 | 0.82083537 |
| CHORDC1 | 60.0584233 | 0.3353781 | 0.28805283 | 1.16429373 | 0.244305 | 0.80476923 |
| FAT3 | 129.380625 | -2.782133637 | 1.08010871 | -2.5757904 | 0.01000113 | 0.29238317 |
| SLC36A4 | 152.257878 | -0.21568746 | 0.39595584 | -0.544726 | 0.58594195 | 0.9423304 |
| SMCO4 | 44.448281 | 0.158229395 | 0.30577764 | 0.51746555 | 0.60483121 | 0.94683415 |
| CEP295 | 81.2999831 | -0.348414803 | 0.26335971 | -1.3229617 | 0.18584812 | 0.76046631 |
| TAF1D | 296.24599 | 0.041569591 | 0.29287928 | 0.14193421 | 0.88713197 | 0.98766275 |
| MRE11 | 89.7582785 | -0.074845162 | 0.26810918 | -0.2791593 | 0.78012261 | 0.97423208 |
| PIWIL4 | 49.5727429 | 1.095607915 | 0.60889924 | 1.79932546 | 0.07196721 | 0.59090333 |
| AMOTL1 | 98.6044683 | 0.458867743 | 0.50078402 | 0.91629869 | 0.35951025 | 0.87316144 |
| CWC15 | 47.333942 | -0.337008944 | 0.30283643 | -1.1128415 | 0.26577651 | 0.81387543 |
| SRSF8 | 47.5811657 | -0.409173776 | 0.3802353 | -1.0761068 | 0.28187951 | 0.82749734 |
| ENDOD1 | 98.5676437 | -0.239934843 | 0.54251619 | -0.442263 | 0.65829889 | 0.95399905 |
| AP000787.1 | 64.6129446 | -0.429363679 | 0.65783039 | -0.6526966 | 0.51395187 | 0.93065123 |
| SESN3 | 302.003073 | 0.122548695 | 0.55959621 | 0.21899486 | 0.82665405 | 0.98135074 |
| FAM76B | 50.237196 | 0.032993253 | 0.24941295 | 0.13228364 | 0.89475995 | 0.98842187 |
| CEP57 | 126.38082 | 0.638601237 | 0.19809332 | 3.22373936 | 0.00126529 | 0.11327257 |

| | | | | | | |
|---|---|---|---|---|---|---|
| MTMR2 | 167.640912 | 0.004976706 | 0.25065705 | 0.01985464 | 0.98415933 | 0.99752755 |
| MAML2 | 3501.08544 | -0.056563734 | 0.25633843 | -0.2206604 | 0.82535689 | 0.98094663 |
| CCDC82 | 85.6000524 | 0.07693078 | 0.24736023 | 0.31100706 | 0.75579525 | 0.9714717 |
| CNTN5 | 187.663716 | -0.420814764 | 0.68459689 | -0.6146899 | 0.53875956 | 0.93523904 |
| ARHGAP42 | 198.673993 | 1.153923027 | 0.49344671 | 2.33849574 | 0.01936155 | 0.37074405 |
| CEP126 | 119.217622 | 0.345444135 | 0.42673837 | 0.80949865 | 0.41822838 | 0.89865627 |
| YAP1 | 677.989505 | -0.175900069 | 0.22712861 | -0.7744514 | 0.4386639 | 0.90561284 |
| BIRC3 | 263.985746 | 0.160884261 | 0.99694102 | 0.16137791 | 8.72E-01 | 0.98766275 |
| BIRC2 | 117.221271 | 0.065988379 | 0.31977719 | 0.20635737 | 0.83651177 | 0.98462033 |
| TMEM123 | 291.681209 | 0.104284113 | 0.23633185 | 0.44126135 | 0.6590238 | 0.95402584 |
| MMP7 | 463.100793 | 0.886676409 | 0.72737328 | 1.21901152 | 2.23E-01 | 0.78710314 |
| DCUN1D5 | 55.0223877 | -0.310896662 | 0.25805651 | -1.204762 | 0.22829521 | 0.79284833 |
| DYNC2H1 | 376.101439 | 0.286582568 | 0.22454282 | 1.27629361 | 0.20185175 | 0.77110248 |
| PDGFD | 150.184866 | 0.078011895 | 0.55168021 | 0.14140782 | 0.88754778 | 0.98766275 |
| CASP4 | 87.9317188 | 0.525881783 | 0.42328852 | 1.24237196 | 0.21409936 | 0.78187485 |
| MSANTD4 | 32.3603273 | -0.598894496 | 0.29076734 | -2.0597034 | 0.0394269 | 0.479886 |
| KBTBD3 | 29.4736512 | 0.381429392 | 0.32919086 | 1.15868766 | 0.24658352 | 0.80571615 |
| AASDHPPT | 70.2181918 | 0.209736822 | 0.28917726 | 0.72528808 | 0.46827532 | 0.91462428 |
| CWF19L2 | 156.921981 | -0.403807839 | 0.24274759 | -1.6634886 | 0.09621468 | 0.64295596 |
| ALKBH8 | 28.6992363 | -0.758960362 | 0.28742528 | -2.6405484 | 0.0082772 | 0.27246484 |
| SLC35F2 | 58.6298118 | -0.584035994 | 0.3625739 | -1.6108054 | 0.10722214 | 0.65699665 |
| CUL5 | 344.441464 | -0.123464159 | 0.19194406 | -0.6432299 | 0.52007494 | 0.93272217 |
| ACAT1 | 126.095616 | 0.632628039 | 0.20647211 | 3.06398789 | 0.00218408 | 0.15271309 |
| NPAT | 91.7727083 | 0.366398729 | 0.26652469 | 1.37472717 | 0.16921604 | 0.74445571 |
| ATM | 394.568052 | 0.299403098 | 0.34197087 | 0.87552222 | 0.38128983 | 0.88128509 |
| C11orf65 | 87.3375569 | 0.574122805 | 0.53451102 | 1.07410845 | 0.28277408 | 0.82913646 |
| EXPH5 | 126.887156 | -0.749773591 | 0.50932722 | -1.4720862 | 0.1409976 | 0.7148274 |
| DDX10 | 199.246088 | -0.29996669 | 0.3019156 | -0.9935449 | 0.3204445 | 0.85075884 |
| RDX | 286.76039 | 0.330160273 | 0.19099539 | 1.72862956 | 0.08387541 | 0.61656297 |
| ZC3H12C | 137.533654 | 0.738122877 | 0.31290367 | 2.35894608 | 0.01832692 | 0.36034684 |
| FDX1 | 70.4345309 | 0.181071799 | 0.30454913 | 0.59455694 | 0.55213967 | 0.93523904 |
| SIK2 | 598.711595 | 0.212029397 | 0.40482566 | 0.52375483 | 0.60044908 | 0.94683415 |
| PPP2R1B | 115.955181 | 0.168581943 | 0.27811994 | 0.60614835 | 0.54441625 | 0.93523904 |
| C11orf1 | 26.1361615 | 0.071290575 | 0.62650476 | 0.11379095 | 0.90940351 | 0.99100813 |
| CRYAB | 1938.03333 | -0.355468463 | 0.67866636 | -0.523775 | 0.60043505 | 0.94683415 |
| DIXDC1 | 206.180594 | -0.192944295 | 0.45015023 | -0.428622 | 0.66819832 | 0.95764829 |
| DLAT | 51.8062583 | -0.281911307 | 0.29061566 | -0.9700486 | 0.33202228 | 0.85980141 |
| NKAPD1 | 32.1896983 | 0.259620062 | 0.31383029 | 0.8272626 | 0.40808824 | 0.8959746 |
| TIMM8B | 47.7875354 | -0.365532879 | 0.50131481 | -0.7291484 | 0.46591091 | 0.91462428 |
| IL18 | 55.8175925 | 1.203697241 | 0.962356 | 1.25078167 | 0.21101415 | 0.77730294 |
| PTS | 106.444938 | -0.522720797 | 0.31596261 | -1.6543755 | 0.09805124 | 0.64312419 |
| AP002884.1 | 69.2317632 | 0.199112007 | 0.54208714 | 0.36730628 | 0.71339057 | 0.96587538 |
| TTC12 | 94.2642635 | 0.306466622 | 0.25990931 | 1.17912905 | 0.23834679 | 0.80165869 |
| ZW10 | 32.169248 | 0.134791661 | 0.30301316 | 0.44483764 | 0.65643709 | 0.95399905 |
| USP28 | 57.5629151 | 0.258729633 | 0.31489184 | 0.82164603 | 0.41127838 | 0.89649693 |
| ZBTB16 | 773.332373 | -0.649114074 | 0.61388476 | -1.0573875 | 0.29033478 | 0.83517022 |
| AP002518.2 | 74.2614838 | 1.514639555 | 0.7892589 | 1.91906553 | 0.05497604 | 0.53804407 |
| NNMT | 410.68323 | 1.827778566 | 1.25182865 | 1.46008686 | 1.44E-01 | 0.71634214 |
| REXO2 | 67.7536067 | 0.288364186 | 0.34060963 | 0.84661195 | 0.39721145 | 0.88863925 |
| CADM1 | 425.854004 | -0.623402601 | 0.42871263 | -1.454127 | 0.14591111 | 0.71974648 |
| SIK3 | 1188.11642 | -0.089973006 | 0.25101415 | -0.358438 | 0.72001557 | 0.96779196 |
| PAFAH1B2 | 144.681969 | 0.021764124 | 0.18144143 | 0.11995124 | 0.90452177 | 0.99061193 |
| SIDT2 | 28.7646326 | 0.044781262 | 0.41044372 | 0.10910451 | 0.9131196 | 0.99113939 |
| TAGLN | 55.4445323 | 0.766495115 | 0.84045989 | 0.91199488 | 0.36177141 | 0.8752623 |

| | | | | | | |
|---|---|---|---|---|---|---|
| PCSK7 | 91.0382481 | -0.447238022 | 0.20147071 | -2.2198663 | 0.02642785 | 0.42051885 |
| RNF214 | 75.6571683 | -0.205559665 | 0.28079719 | -0.7320574 | 0.46413352 | 0.91462428 |
| BACE1 | 58.9460453 | 0.600819991 | 0.35049254 | 1.71421622 | 0.08648902 | 0.62381659 |
| CEP164 | 95.9972821 | -0.173943147 | 0.37579362 | -0.4628688 | 0.6434584 | 0.94944486 |
| DSCAML1 | 1160.06851 | 0.659323906 | 0.6762886 | 0.97491501 | 0.32960242 | 0.85841805 |
| FXYD2 | 237.937563 | -0.013319037 | 0.69243701 | -0.019235 | 0.98465362 | 0.99752755 |
| MPZL3 | 41.6500488 | -0.513983054 | 0.52718295 | -0.9749614 | 0.32957938 | 0.85841805 |
| UBE4A | 103.116086 | -0.224011313 | 0.22010465 | -1.0177491 | 0.30879719 | 0.84554063 |
| ATP5MG | 180.221616 | 0.089661553 | 0.4861821 | 0.18441969 | 0.85368422 | 0.9864199 |
| KMT2A | 468.382629 | -0.094741298 | 0.15534625 | -0.6098718 | 0.54194673 | 0.93523904 |
| IFT46 | 34.8842453 | 0.165009872 | 0.29115949 | 0.56673363 | 0.57089517 | 0.93998433 |
| ARCN1 | 77.7708456 | -0.186781899 | 0.28019951 | -0.6666032 | 0.5050256 | 0.9257297 |
| PHLDB1 | 252.096258 | 0.122783125 | 0.32173066 | 0.38163327 | 0.7027334 | 0.9645815 |
| AP002954.1 | 37.4180241 | 0.470362906 | 0.53352636 | 0.88161136 | 0.37798701 | 0.87964763 |
| DDX6 | 231.563523 | 0.156317736 | 0.21113341 | 0.74037423 | 0.45907295 | 0.91362347 |
| BCL9L | 61.2450985 | 0.357095884 | 0.41843742 | 0.85340331 | 0.39343568 | 0.88760715 |
| CCDC84 | 136.607294 | -0.182384178 | 0.28513215 | -0.6396479 | 0.52240155 | 0.93272217 |
| RPS25 | 387.81766 | 0.003385528 | 0.48831074 | 0.00693314 | 0.9944682 | 0.99866813 |
| TRAPPC4 | 69.0858832 | 0.345423131 | 0.38234733 | 0.90342761 | 0.36629899 | 0.87605028 |
| SLC37A4 | 40.0974971 | 0.336842209 | 0.525934 | 0.6404648 | 0.52187047 | 0.93272217 |
| HYOU1 | 41.0394194 | -0.288533841 | 0.56869901 | -0.5073577 | 0.61190384 | 0.94791646 |
| VPS11 | 26.2684732 | 0.157829137 | 0.38948412 | 0.40522612 | 0.68531131 | 0.96161098 |
| HINFP | 25.8163327 | 0.045319167 | 0.29116799 | 0.15564612 | 0.87631197 | 0.98766275 |
| NLRX1 | 29.076563 | -0.315705091 | 0.40730511 | -0.7751071 | 0.43827637 | 0.90561284 |
| CBL | 121.738004 | -0.01653042 | 0.27156715 | -0.0608705 | 0.95146237 | 0.99531843 |
| THY1 | 90.3690377 | 2.161568412 | 1.30953849 | 1.65063374 | 0.09881339 | 0.6434623 |
| OAF | 49.4202175 | 0.455526894 | 0.43514686 | 1.04683485 | 0.29517576 | 0.83911606 |
| ARHGEF12 | 807.911766 | 0.285905522 | 0.16442444 | 1.73882623 | 0.08206533 | 0.61261376 |
| SC5D | 37.5912454 | 0.70175104 | 0.43334292 | 1.61938966 | 0.10536345 | 0.65699665 |
| SORL1 | 166.603809 | -0.122121649 | 0.37170989 | -0.3285402 | 0.74250324 | 0.9698508 |
| UBASH3B | 268.793615 | -0.840857001 | 0.49651202 | -1.693528 | 0.09035501 | 0.6330877 |
| JHY | 137.086526 | 0.744300962 | 0.36922585 | 2.01584194 | 0.04381651 | 0.49836909 |
| HSPA8 | 195.011797 | 0.216031036 | 0.53115942 | 0.406716 | 0.68421658 | 0.96161098 |
| GRAMD1B | 38.4073202 | -0.974156564 | 0.59302605 | -1.6426877 | 0.10044757 | 0.64698063 |
| VWA5A | 36.6780984 | -0.013388538 | 0.3213648 | -0.0416615 | 0.96676855 | 0.99601626 |
| TBRG1 | 92.8995721 | 0.077978308 | 0.24968535 | 0.31230631 | 0.75480775 | 0.97125609 |
| SIAE | 72.8148037 | 0.491124457 | 0.46232646 | 1.06228931 | 0.28810436 | 0.83300685 |
| ESAM | 43.6769322 | -0.219445204 | 0.52239006 | -0.4200792 | 0.67442759 | 0.95910677 |
| MSANTD2 | 139.580951 | -0.094788684 | 0.2423711 | -0.3910891 | 0.69573141 | 0.96220115 |
| PKNOX2 | 73.4937608 | 1.385889564 | 0.79197576 | 1.74991412 | 0.08013313 | 0.60643125 |
| EI24 | 76.7876991 | -0.542065801 | 0.42441892 | -1.2771952 | 0.20153335 | 0.77110248 |
| STT3A | 59.2962429 | 0.084831973 | 0.27553324 | 0.30788289 | 0.75817144 | 0.9714717 |
| CDON | 54.1591582 | 0.712797659 | 1.13626141 | 0.62731837 | 0.53045056 | 0.93366189 |
| RPUSD4 | 27.4793696 | -0.339009019 | 0.35629712 | -0.9514784 | 0.34136157 | 0.86846099 |
| FAM118B | 41.554651 | 0.077781956 | 0.24076818 | 0.32305745 | 0.74665173 | 0.97028355 |
| SRPRA | 56.2059427 | -0.514164725 | 0.41412858 | -1.2415582 | 0.21439962 | 0.78237483 |
| FOXRED1 | 38.5999916 | -0.300509797 | 0.27469214 | -1.0939876 | 0.27396042 | 0.82047446 |
| DCPS | 35.470608 | -0.616487865 | 0.55170573 | -1.1174215 | 0.26381413 | 0.81378765 |
| ST3GAL4 | 71.8209724 | -0.41347141 | 0.63796142 | -0.6481135 | 5.17E-01 | 0.93230176 |
| KIRREL3 | 136.511242 | 0.68471308 | 1.14465631 | 0.59818224 | 0.54971834 | 0.93523904 |
| ETS1 | 186.202812 | -0.100028182 | 0.45985866 | -0.2175194 | 0.82780359 | 0.98142144 |
| ARHGAP32 | 296.555982 | 0.551307845 | 0.27400518 | 2.01203439 | 0.04421631 | 0.49952159 |
| TMEM45B | 30.71147 | -0.074211937 | 0.53858969 | -0.1377894 | 0.89040688 | 0.98766275 |
| NFRKB | 57.1262912 | 0.289234282 | 0.2568177 | 1.12622407 | 0.26007069 | 0.81263292 |

| | | | | | | |
|---|---|---|---|---|---|---|
| PRDM10 | 67.4705386 | -0.331972115 | 0.31409934 | -1.0569017 | 0.29055646 | 0.83531608 |
| APLP2 | 463.041453 | 0.270906655 | 0.29748528 | 0.91065567 | 0.36247683 | 0.8752623 |
| ST14 | 152.692951 | -0.553429517 | 0.48445194 | -1.1423827 | 0.25329498 | 0.80798674 |
| ZBTB44 | 338.88871 | 0.133580802 | 0.24826758 | 0.53805173 | 5.91E-01 | 0.94516215 |
| AP002986.1 | 41.6314385 | 0.431713221 | 0.68944774 | 0.62617251 | 0.53120179 | 0.93366189 |
| ADAMTS15 | 108.908597 | -0.912253631 | 0.73160367 | -1.2469232 | 0.21242564 | 0.77822114 |
| SNX19 | 94.3321814 | -0.1597967 | 0.36546842 | -0.437238 | 0.66193872 | 0.9545911 |
| NTM | 274.675481 | -1.52950419 | 0.60470304 | -2.5293476 | 0.01142748 | 0.30834879 |
| NCAPD3 | 34.5546547 | -0.403768901 | 0.34734407 | -1.1624465 | 0.24505415 | 0.80476923 |
| VPS26B | 37.5257414 | -0.008044793 | 0.33007291 | -0.0243728 | 0.98055526 | 0.99681784 |
| ZMYND11 | 316.837651 | -0.045182629 | 0.16709275 | -0.2704045 | 0.78684909 | 0.97423208 |
| DIP2C | 689.577121 | -0.296828686 | 0.24930119 | -1.1906429 | 0.23379382 | 0.79633808 |
| LARP4B | 472.454118 | -0.306812734 | 0.18301071 | -1.6764742 | 0.09364534 | 0.63753181 |
| GTPBP4 | 57.12288 | -0.818364011 | 0.31347376 | -2.6106301 | 0.00903756 | 0.28291106 |
| IDI1 | 68.7536094 | 0.395337706 | 0.47266287 | 0.83640525 | 0.40292696 | 0.89281456 |
| WDR37 | 131.739367 | -0.037966106 | 0.17495805 | -0.2170012 | 0.82820742 | 0.98142144 |
| PFKP | 184.562092 | -0.079769164 | 0.57018443 | -0.1399006 | 0.8887385 | 0.98766275 |
| PITRM1 | 73.8378297 | -0.356327897 | 0.28989007 | -1.2291829 | 0.21900326 | 0.78523615 |
| KLF6 | 416.257916 | -0.744719891 | 0.52662588 | -1.4141346 | 0.15732238 | 0.73580079 |
| AKR1C1 | 143.979428 | -0.707329596 | 1.03928928 | -0.6805897 | 0.49613113 | 0.92334178 |
| AKR1C2 | 82.9001125 | -0.914403873 | 0.91037857 | -1.0044216 | 0.31517546 | 0.8490543 |
| AKR1C3 | 60.795935 | 0.425954144 | 0.37775735 | 1.12758664 | 0.25949453 | 0.81263292 |
| NET1 | 56.1999578 | 0.35496183 | 0.32650583 | 1.08715311 | 0.27696915 | 0.8229426 |
| ASB13 | 43.7092654 | 0.002032436 | 0.39028543 | 0.00520756 | 0.99584499 | 0.99866813 |
| FAM208B | 314.639327 | -0.13377816 | 0.22329205 | -0.5991174 | 0.54909457 | 0.93523904 |
| GDI2 | 196.184258 | -0.203644278 | 0.22209699 | -0.916916 | 0.35918668 | 0.87316144 |
| ANKRD16 | 31.5401074 | 0.045844868 | 0.4744272 | 0.09663204 | 0.92301861 | 0.99214128 |
| FBH1 | 104.35391 | 0.177824187 | 0.22380117 | 0.79456326 | 0.42686759 | 0.90264121 |
| RBM17 | 124.209329 | -0.293245804 | 0.22650423 | -1.2946593 | 0.19543779 | 0.76968045 |
| PFKFB3 | 854.716961 | -1.697485612 | 0.66826127 | -2.5401526 | 0.01108041 | 0.30325501 |
| AL137145.2 | 146.176806 | -1.171792159 | 0.87061784 | -1.3459317 | 0.17832454 | 0.7513166 |
| PRKCQ | 235.944133 | -0.500134353 | 0.63015407 | -0.79367 | 0.42738759 | 0.90272191 |
| SFMBT2 | 234.152081 | -0.215839009 | 0.73250617 | -0.2946583 | 0.76825494 | 0.97239879 |
| ITIH5 | 176.308123 | 2.340301008 | 0.52779724 | 4.434091 | 9.25E-06 | 0.00421757 |
| KIN | 44.5712166 | -0.079252202 | 0.2596801 | -0.3051917 | 0.76022018 | 0.9714717 |
| ATP5F1C | 146.69505 | 0.234039573 | 0.17566525 | 1.33230429 | 0.18276022 | 0.75589815 |
| TAF3 | 269.783319 | 0.236765187 | 0.23626982 | 1.00209663 | 0.31629692 | 0.8490543 |
| CELF2 | 97.7268357 | -0.650317626 | 0.38824058 | -1.6750378 | 0.09392683 | 0.63765064 |
| USP6NL | 174.156207 | 0.054759702 | 0.19246775 | 0.28451365 | 0.77601678 | 0.97423208 |
| ECHDC3 | 40.2762718 | 1.137304851 | 0.35392277 | 3.21342662 | 0.00131161 | 0.11421765 |
| PROSER2 | 251.844254 | -0.583572554 | 0.5467374 | -1.0673727 | 0.28580359 | 0.83003998 |
| UPF2 | 197.28519 | -0.080652304 | 0.16652079 | -0.4843378 | 0.62814619 | 0.94858781 |
| DHTKD1 | 91.1301664 | 0.223624609 | 0.21269947 | 1.05136422 | 0.29309134 | 0.83733658 |
| SEC61A2 | 35.5832371 | -0.297729286 | 0.31789501 | -0.9365648 | 0.34898245 | 0.87178925 |
| NUDT5 | 73.4732959 | 0.188171796 | 0.24038219 | 0.78280257 | 0.43374306 | 0.90514499 |
| CDC123 | 73.5617096 | -0.258204609 | 0.22513572 | -1.1468842 | 0.25142947 | 0.80744031 |
| CAMK1D | 299.351939 | -0.316154056 | 0.56036517 | -0.5641929 | 0.57262285 | 0.93998433 |
| OPTN | 130.247975 | 0.091860397 | 0.58546812 | 0.15690077 | 0.87532306 | 0.98766275 |
| PHYH | 31.6607092 | -0.359235856 | 0.45596371 | -0.7878606 | 0.43077823 | 0.90392654 |
| SEPHS1 | 53.4897872 | -0.055267934 | 0.39110795 | -0.1413112 | 0.8876241 | 0.98766275 |
| BEND7 | 231.350697 | -0.076287692 | 0.18092569 | -0.4216521 | 0.67327899 | 0.95910677 |
| PRPF18 | 28.6773642 | 0.756626109 | 0.37295009 | 2.02875968 | 0.04248277 | 0.49326362 |
| AL157392.5 | 53.9503891 | 0.428575401 | 0.32523208 | 1.31775253 | 0.1875865 | 0.76207437 |
| FRMD4A | 1275.45524 | 0.291053861 | 0.67845162 | 0.42899722 | 0.66792525 | 0.95764829 |

| | | | | | | |
|---|---|---|---|---|---|---|
| FAM107B | 262.029545 | -0.355662408 | 0.23148366 | -1.5364471 | 0.12442876 | 0.69439514 |
| DCLRE1C | 52.2862601 | -0.193395082 | 0.25196574 | -0.7675452 | 0.44275746 | 0.90615787 |
| RPP38 | 86.8348102 | 7.45E-01 | 0.32344838 | 2.30257605 | 0.02130271 | 0.38501631 |
| NMT2 | 83.6835973 | 0.361176672 | 0.39682273 | 0.91017132 | 0.36273216 | 0.87551319 |
| FAM171A1 | 461.294579 | -0.282266032 | 0.53464186 | -0.5279535 | 0.59753162 | 0.94683415 |
| MINDY3 | 122.528701 | -0.132793103 | 0.18257631 | -0.7273293 | 0.46702423 | 0.91462428 |
| PTER | 89.6508567 | 0.180455719 | 0.24084939 | 0.74924715 | 0.45370826 | 0.91077355 |
| RSU1 | 162.126019 | -0.057401119 | 0.27346142 | -0.2099057 | 0.83374125 | 0.98351801 |
| CUBN | 167.605507 | 1.887254983 | 0.41564564 | 4.5405384 | 5.61E-06 | 0.00298603 |
| TRDMT1 | 44.5916114 | 0.477513243 | 0.32459541 | 1.47110287 | 0.1412633 | 0.7148274 |
| VIM | 320.872052 | 1.24395544 | 0.77853366 | 1.59781844 | 0.11008339 | 0.65905549 |
| HACD1 | 46.9706689 | 0.627469519 | 0.50667036 | 1.23841765 | 0.21556124 | 0.78293142 |
| STAM | 103.958882 | 0.055449909 | 0.32156951 | 0.17243522 | 0.86309539 | 0.98670067 |
| CACNB2 | 155.926122 | 0.718209511 | 0.47156668 | 1.52302853 | 0.12775157 | 0.69635098 |
| AL450384.2 | 55.2804199 | -0.464809137 | 0.36979277 | -1.2569449 | 0.20877358 | 0.77543317 |
| NSUN6 | 132.42512 | -0.36385739 | 0.25505757 | -1.4265696 | 0.15370398 | 0.7288378 |
| ARL5B | 73.3193812 | 0.274058199 | 0.40304419 | 0.67997059 | 0.49652308 | 0.92353293 |
| MALRD1 | 48.7472465 | -0.267971106 | 0.78351375 | -0.342012 | 0.73234186 | 0.96980299 |
| PLXDC2 | 776.252797 | 1.037225872 | 0.48494418 | 2.1388562 | 0.03244732 | 0.44049392 |
| NEBL | 1374.09315 | -0.309362435 | 0.519678 | -0.5952964 | 0.55164537 | 0.93523904 |
| MLLT10 | 623.297738 | -0.007797443 | 0.17180396 | -0.0453857 | 0.96379988 | 0.99581912 |
| DNAJC1 | 457.911756 | 0.112709442 | 0.23593366 | 0.47771666 | 0.63285188 | 0.94932781 |
| BMI1 | 29.2737256 | -0.543616091 | 0.31342424 | -1.7344418 | 0.08283973 | 0.61513318 |
| PIP4K2A | 128.857019 | 0.533694026 | 0.50857921 | 1.0493823 | 0.2940022 | 0.8384183 |
| ARMC3 | 70.6885959 | 1.292497318 | 0.74818643 | 1.72750702 | 0.08407665 | 0.61656297 |
| MSRB2 | 69.2199292 | 0.727081439 | 0.33010776 | 2.20255787 | 0.02762593 | 0.4250936 |
| OTUD1 | 33.3442974 | -0.436623666 | 0.52261349 | -0.8354619 | 0.40345768 | 0.89337058 |
| KIAA1217 | 857.060615 | 0.330953484 | 0.36536952 | 0.90580485 | 0.36503915 | 0.87605028 |
| ARHGAP21 | 274.454729 | -0.096034477 | 0.22834011 | -0.4205765 | 0.67406439 | 0.95910677 |
| MYO3A | 71.19025 | 0.377952273 | 0.60131288 | 0.62854511 | 5.30E-01 | 0.93366189 |
| APBB1IP | 75.2399062 | 1.235563072 | 0.81181155 | 1.52198262 | 0.12801344 | 0.69635098 |
| ABI1 | 237.135064 | -0.095409359 | 0.19104735 | -0.4994016 | 0.61749647 | 0.94791646 |
| ANKRD26 | 210.237643 | 0.279083874 | 0.26977828 | 1.03449348 | 0.30090552 | 0.8426686 |
| YME1L1 | 195.174079 | -0.163208195 | 0.15993386 | -1.0204731 | 0.30750415 | 0.84488735 |
| MASTL | 28.7055266 | -0.274597518 | 0.39035701 | -0.7034522 | 0.48177395 | 0.91971919 |
| ACBD5 | 95.2491099 | 0.093216567 | 0.27953366 | 0.3334717 | 0.73877824 | 0.9698508 |
| RAB18 | 177.807383 | 0.080873605 | 0.14861358 | 0.54418716 | 0.58631268 | 0.9423304 |
| MPP7 | 163.023507 | 0.05640087 | 0.51107146 | 0.11035809 | 0.91212539 | 0.9911192 |
| AL358234.1 | 102.692442 | -0.158696082 | 1.39970879 | -0.1133779 | 0.90973094 | 0.99100813 |
| WAC-AS1 | 108.941398 | -0.461700226 | 0.54317881 | -0.8499968 | 0.39532689 | 0.88760715 |
| WAC | 745.340711 | -0.029507249 | 0.16277494 | -0.1812764 | 0.85615066 | 0.9864199 |
| BAMBI | 28.0073323 | 1.042160321 | 0.58793832 | 1.77256744 | 0.07630041 | 0.59726395 |
| SVIL | 631.53911 | -0.537034558 | 0.33080113 | -1.6234363 | 0.10449617 | 0.65551333 |
| MTPAP | 45.0171181 | -0.343403354 | 0.24737701 | -1.3881781 | 0.16508281 | 0.74445571 |
| MAP3K8 | 66.4250261 | 0.713686417 | 0.57672344 | 1.23748468 | 0.2159072 | 0.78293142 |
| ZNF438 | 159.940965 | -0.210386938 | 0.23122605 | -0.9098756 | 0.36288812 | 0.87551319 |
| ZEB1 | 364.521802 | -0.793981195 | 0.63338963 | -1.2535431 | 0.21000813 | 0.77730294 |
| ARHGAP12 | 250.194391 | 0.0645925 | 0.20780821 | 0.31082748 | 0.75593178 | 0.9714717 |
| KIF5B | 223.747489 | -0.069225005 | 0.18917938 | -0.3659226 | 0.71442286 | 0.96631694 |
| EPC1 | 363.821961 | -0.168857807 | 0.15738695 | -1.0728831 | 0.28332356 | 0.82918183 |
| CCDC7 | 197.377445 | -0.209943752 | 0.38174867 | -0.5499528 | 0.58235174 | 0.94091128 |
| ITGB1 | 403.090507 | -0.696845883 | 0.40604182 | -1.7161924 | 0.08612684 | 0.62276717 |
| NRP1 | 1665.43176 | 0.275921771 | 0.56450049 | 0.48878925 | 0.62499091 | 0.94808496 |
| PARD3 | 2684.96717 | -0.168914211 | 0.30452144 | -0.5546874 | 0.57910849 | 0.94034284 |

| | | | | | | |
|---|---|---|---|---|---|---|
| CUL2 | 106.613885 | -0.246739859 | 0.24133751 | -1.0223851 | 0.30659869 | 0.84488735 |
| CREM | 183.256487 | -0.597617171 | 0.3932474 | -1.5196977 | 0.12858697 | 0.69707674 |
| CCNY | 654.137722 | -0.229295533 | 0.17987118 | -1.2747764 | 0.20238839 | 0.77110248 |
| AL117336.3 | 74.6883195 | 0.357848059 | 0.42548784 | 0.84103006 | 0.4003311 | 0.8907577 |
| ZNF248 | 148.491081 | 0.155931018 | 0.28907431 | 0.53941499 | 0.58960054 | 0.94491945 |
| ZNF25 | 31.8862512 | 0.093535795 | 0.48205464 | 0.19403567 | 0.84614795 | 0.98520136 |
| ZNF33A | 308.241339 | 0.068956815 | 0.180223 | 0.3826194 | 0.70200199 | 0.9645815 |
| ZNF33B | 249.817277 | 0.020403355 | 0.20833731 | 0.09793424 | 0.92198451 | 0.99214128 |
| BMS1 | 96.4009575 | -0.553285397 | 0.22627817 | -2.4451558 | 0.01447896 | 0.32788161 |
| CSGALNACT2 | 64.186637 | 0.142442801 | 0.29038038 | 0.49053865 | 0.62375279 | 0.94808496 |
| HNRNPF | 203.946425 | -0.330648576 | 0.28553658 | -1.1579903 | 0.24686801 | 0.80598115 |
| ZNF487 | 50.278364 | 0.309361192 | 0.29311346 | 1.05543155 | 0.291228 | 0.83594927 |
| ZNF32 | 41.6038562 | -0.195467522 | 0.31666469 | -0.6172697 | 0.53705685 | 0.93489692 |
| TMEM72-AS1 | 34.1083708 | 1.021402698 | 0.47365746 | 2.15641636 | 0.03105117 | 0.43905874 |
| RASSF4 | 102.311512 | 0.777087768 | 0.58671891 | 1.32446348 | 0.18534916 | 0.76046631 |
| ZNF22 | 47.6031433 | -0.241341843 | 0.36480768 | -0.6615591 | 0.50825382 | 0.92666294 |
| 8-Mar | 107.757916 | 0.000183299 | 0.30661334 | 0.00059782 | 0.99952301 | 0.99971834 |
| ZFAND4 | 57.9781159 | 0.591736427 | 0.21576776 | 2.74246915 | 0.00609792 | 0.24646394 |
| WASHC2C | 57.0332765 | -0.058208096 | 0.25010773 | -0.2327321 | 0.81596945 | 0.98007572 |
| AGAP4 | 64.0507918 | 0.5719621 | 0.35097034 | 1.62965938 | 0.10317351 | 0.65288882 |
| TIMM23 | 65.0189332 | -0.162781839 | 0.27988774 | -0.5815969 | 0.56083824 | 0.93747805 |
| NCOA4 | 84.6659716 | 0.253408056 | 0.21920822 | 1.15601529 | 0.2476749 | 0.80672756 |
| AL356056.2 | 49.6948841 | -0.128481665 | 0.69716635 | -0.1842913 | 0.85378497 | 0.9864199 |
| PTPN20 | 38.9480995 | 0.901026063 | 0.48017398 | 1.87645751 | 0.06059248 | 0.56071154 |
| AGAP9 | 52.1795731 | 0.715823501 | 0.2899289 | 2.46896223 | 0.01355055 | 0.32032755 |
| MAPK8 | 511.877765 | -0.372851483 | 0.16096196 | -2.316395 | 0.02053671 | 0.37872401 |
| ARHGAP22 | 60.3007092 | -1.892769005 | 0.62418304 | -3.0323942 | 0.00242622 | 0.16028122 |
| ERCC6 | 76.8555443 | 0.008809165 | 0.20948407 | 0.04205172 | 0.96645747 | 0.99601626 |
| PARG | 202.858575 | -0.22124484 | 0.17068188 | -1.296241 | 0.19489248 | 0.76968045 |
| TIMM23B | 155.75631 | -0.39794662 | 0.3693864 | -1.077318 | 0.28133822 | 0.82692199 |
| LINC00843 | 44.709963 | -0.178582631 | 0.29574824 | -0.6038333 | 0.54595449 | 0.93523904 |
| AGAP6 | 61.4504308 | -0.057315381 | 0.28814617 | -0.1989108 | 0.84233253 | 0.98517901 |
| WASHC2A | 64.0512059 | -0.253107775 | 0.42892537 | -0.5900975 | 0.55512531 | 0.93523904 |
| SGMS1 | 705.69971 | -0.108717226 | 0.1895118 | -0.5736699 | 0.56619117 | 0.93832962 |
| PRKG1 | 551.349902 | -0.406787452 | 0.55107911 | -0.7381652 | 0.46041404 | 0.91368087 |
| IPMK | 64.8823865 | -0.179042543 | 0.36704296 | -0.4877972 | 0.62569347 | 0.94819138 |
| CISD1 | 119.206907 | 0.045321531 | 0.28996563 | 0.15629967 | 0.87579682 | 0.98766275 |
| UBE2D1 | 50.5071196 | -0.224112123 | 0.34619939 | -0.6473499 | 0.51740554 | 0.93247044 |
| TFAM | 40.7385081 | -0.459690686 | 0.27751158 | -1.6564739 | 0.09762589 | 0.64312419 |
| BICC1 | 5875.93154 | -0.257207464 | 0.30947469 | -0.8311099 | 0.40591157 | 0.89569633 |
| PHYHIPL | 530.593774 | 0.354024577 | 0.60963879 | 0.58071202 | 0.56143456 | 0.93747805 |
| SLC16A9 | 67.5406947 | 0.54661483 | 0.90720028 | 0.60252939 | 0.54682181 | 0.93523904 |
| CCDC6 | 168.818899 | 0.054648836 | 0.22521975 | 0.24264673 | 0.80827908 | 0.97930627 |
| ANK3 | 1271.41261 | 0.066895994 | 0.3150878 | 0.21230906 | 0.83186593 | 0.98290906 |
| RHOBTB1 | 43.8644482 | 0.761689158 | 0.43684756 | 1.74360399 | 0.08122817 | 0.60850843 |
| CABCOCO1 | 62.9785238 | 0.35146766 | 0.56178509 | 0.62562654 | 0.53155993 | 0.93366189 |
| ARID5B | 564.403237 | 0.393536254 | 0.24093237 | 1.6333889 | 0.10238723 | 0.65288882 |
| JMJD1C | 1180.51466 | 0.040382891 | 0.18023664 | 0.22405483 | 0.82271464 | 0.98094663 |
| REEP3 | 169.495566 | -0.325983845 | 0.19011191 | -1.7146945 | 0.08640125 | 0.62381659 |
| LINC01515 | 142.690958 | 0.202850948 | 0.3096982 | 0.65499556 | 0.51247061 | 0.92919856 |
| SIRT1 | 78.986396 | 0.206880816 | 0.29008817 | 0.71316529 | 0.47574348 | 0.91809716 |
| HERC4 | 602.874033 | 0.109456261 | 0.22309981 | 0.49061566 | 0.62369831 | 0.94808496 |
| PBLD | 51.1112585 | 0.790497806 | 0.40237477 | 1.96458093 | 0.04946276 | 0.52308378 |
| HNRNPH3 | 186.444815 | -0.049169522 | 0.24339077 | -0.2020188 | 0.83990199 | 0.98464453 |

| | | | | | | |
|---|---|---|---|---|---|---|
| RUFY2 | 163.987185 | 0.315543995 | 0.16154094 | 1.95333758 | 0.05077961 | 0.52415717 |
| SLC25A16 | 161.84571 | -0.234247556 | 0.27131875 | -0.8633666 | 0.38793592 | 0.8853359 |
| TET1 | 34.2075211 | 1.088344017 | 0.45689799 | 2.38202844 | 0.01721757 | 0.35165689 |
| CCAR1 | 257.819445 | -0.247531509 | 0.22117806 | -1.1191504 | 0.26307599 | 0.81378765 |
| DDX50 | 104.258404 | -0.200470189 | 0.22407754 | -0.8946465 | 0.37097607 | 0.87605028 |
| DDX21 | 143.678799 | -0.483571989 | 0.47561247 | -1.0167353 | 0.30927937 | 0.84554063 |
| KIF1BP | 49.4850472 | 0.42027066 | 0.25821913 | 1.62757366 | 0.10361532 | 0.65384132 |
| VPS26A | 123.124308 | -0.482348827 | 0.20787874 | -2.3203374 | 0.02032263 | 0.37800094 |
| SUPV3L1 | 51.2504632 | 0.016416231 | 0.22272758 | 0.07370543 | 0.94124478 | 0.99526299 |
| HKDC1 | 156.222259 | 0.252411603 | 0.40167555 | 0.62839673 | 0.52974408 | 0.93366189 |
| HK1 | 215.915155 | -0.793061561 | 0.36913814 | -2.148414 | 0.03168088 | 0.43905874 |
| TSPAN15 | 38.6659366 | 0.925415286 | 0.53733959 | 1.72221682 | 0.08503025 | 0.61939525 |
| H2AFY2 | 77.6580385 | -0.146536405 | 0.30125028 | -0.4864274 | 0.62666414 | 0.94831213 |
| AIFM2 | 35.4931573 | -0.197478259 | 0.29709217 | -0.6647037 | 0.50624003 | 0.92631772 |
| SAR1A | 82.4156178 | 0.048339006 | 0.23679397 | 0.20413952 | 0.83824448 | 0.98462033 |
| PPA1 | 148.862103 | 0.17418031 | 0.52371733 | 0.33258458 | 0.73944788 | 0.9698508 |
| EIF4EBP2 | 77.5832919 | 0.056646934 | 0.2546125 | 0.22248293 | 0.82393796 | 0.98094663 |
| SGPL1 | 139.797031 | 0.199658674 | 0.20802918 | 0.95976283 | 0.33717459 | 0.86450666 |
| PCBD1 | 79.7809741 | 0.213983299 | 0.48400591 | 0.44210886 | 0.65841044 | 0.95400296 |
| PSAP | 365.233044 | -0.045228114 | 0.32211103 | -0.1404116 | 0.88833482 | 0.98766275 |
| CHST3 | 51.5718688 | -0.008903288 | 0.47355443 | -0.018801 | 9.85E-01 | 0.99752755 |
| SPOCK2 | 173.161872 | -0.64579414 | 0.71477666 | -0.9034908 | 0.36626546 | 0.87605028 |
| ASCC1 | 136.995175 | -0.614016407 | 0.22234085 | -2.7615996 | 0.0057519 | 0.23851696 |
| ANAPC16 | 338.033416 | -0.065919585 | 0.2533491 | -0.2601927 | 0.79471514 | 0.97524554 |
| DDIT4 | 138.470481 | 0.532145343 | 1.27610643 | 0.41700702 | 0.67667327 | 0.9591378 |
| DNAJB12 | 42.264508 | -0.288196299 | 0.31965676 | -0.9015805 | 0.36727975 | 0.87605028 |
| MICU1 | 328.729588 | -0.201118339 | 0.21297464 | -0.94433 | 0.34500104 | 0.86990996 |
| MCU | 369.073219 | -0.488133646 | 0.27728087 | -1.7604303 | 0.07833487 | 0.60239981 |
| P4HA1 | 251.820523 | -0.139631898 | 0.58617534 | -0.2382084 | 0.81171945 | 0.97930627 |
| ECD | 65.4304876 | -0.112619662 | 0.29795521 | -0.3779751 | 0.70544906 | 0.96535665 |
| FAM149B1 | 103.012688 | 0.560320741 | 0.25710006 | 2.17938785 | 0.02930287 | 0.43383643 |
| DNAJC9 | 93.7763433 | 0.198241705 | 0.30758038 | 0.64452 | 0.51923829 | 0.93259336 |
| MRPS16 | 59.5187538 | -0.214214402 | 0.29228058 | -0.7329067 | 0.46361532 | 0.91462428 |
| CFAP70 | 146.318521 | 0.65418821 | 0.65059325 | 1.00552566 | 0.3146438 | 0.8490543 |
| ANXA7 | 210.655343 | -0.219839058 | 0.21754219 | -1.0105583 | 0.3122279 | 0.84811918 |
| PPP3CB | 227.491121 | -0.026823702 | 0.18347689 | -0.1461966 | 0.88376617 | 0.98766275 |
| USP54 | 264.639399 | 0.070520029 | 0.23287004 | 0.30282997 | 0.76201944 | 0.9714717 |
| AGAP5 | 25.7659617 | 0.52941045 | 0.33131116 | 1.59792518 | 0.11005963 | 0.65905549 |
| SEC24C | 58.5647686 | 0.052462937 | 0.23475874 | 0.22347597 | 0.82316508 | 0.98094663 |
| FUT11 | 60.5111226 | -0.112436643 | 0.4342052 | -0.2589482 | 0.79567523 | 0.97524554 |
| CHCHD1 | 45.1354947 | -0.104232559 | 0.45946663 | -0.2268556 | 0.82053606 | 0.98094663 |
| ZSWIM8 | 87.9157475 | 0.262610751 | 0.23914536 | 1.0981219 | 0.27215127 | 0.81927623 |
| CAMK2G | 76.8535259 | -0.242496596 | 0.40631718 | -0.596816 | 0.55063023 | 0.93523904 |
| VCL | 436.006162 | -0.518688656 | 0.4758391 | -1.0900505 | 0.27569089 | 0.82164863 |
| AP3M1 | 43.5590954 | -0.348172209 | 0.27283455 | -1.2761295 | 0.20190976 | 0.77110248 |
| ADK | 739.102633 | -0.72478133 | 0.311189 | -2.3290712 | 0.0198553 | 0.3747883 |
| KAT6B | 263.207579 | 0.037344963 | 0.32971636 | 0.11326391 | 0.90982133 | 0.99100813 |
| SAMD8 | 106.280839 | 0.228227135 | 0.17630554 | 1.29449779 | 0.19549353 | 0.76968045 |
| VDAC2 | 79.7346301 | 0.339621314 | 0.37538291 | 0.90473301 | 0.36560685 | 0.87605028 |
| COMTD1 | 29.2987575 | 0.111658585 | 0.43984703 | 0.25385777 | 0.79960544 | 0.97659321 |
| LRMDA | 1561.24377 | 0.841525813 | 0.82345614 | 1.0219437 | 0.30680755 | 0.84488735 |
| KCNMA1 | 77.5592302 | -2.108376656 | 0.80154352 | -2.6303957 | 0.00852855 | 0.27506737 |
| DLG5 | 93.7782114 | -0.109696725 | 0.2664249 | -0.411736 | 0.68053293 | 0.95989244 |
| POLR3A | 55.5469885 | -0.206298239 | 0.22647415 | -0.910913 | 0.36234123 | 0.8752623 |

| | | | | | | |
|---|---|---|---|---|---|---|
| RPS24 | 716.842668 | 0.084119478 | 0.52798584 | 0.15932147 | 0.87341561 | 0.98766275 |
| ZMIZ1 | 373.290851 | 0.202679115 | 0.38795406 | 0.52243071 | 0.60137048 | 0.94683415 |
| NUTM2B-AS | 317.32996 | 0.169295665 | 0.25091639 | 0.67470946 | 0.49986037 | 0.92398825 |
| TMEM254 | 33.2717632 | 0.493423837 | 0.29475317 | 1.67402387 | 0.09412591 | 0.6377384 |
| ANXA11 | 237.188808 | -0.208105754 | 0.19540629 | -1.0649901 | 0.28688043 | 0.83122432 |
| FAM213A | 81.6440011 | 0.604698122 | 0.28056055 | 2.15532125 | 0.0311367 | 0.43905874 |
| TSPAN14 | 160.815281 | 0.269550636 | 0.2638377 | 1.02165323 | 3.07E-01 | 8.45E-01 |
| NRG3 | 2081.61391 | 0.760707007 | 1.32704948 | 0.57323183 | 0.56648773 | 0.93845576 |
| GHITM | 94.697044 | 0.037197665 | 0.28480364 | 0.13060811 | 0.89608533 | 0.98842187 |
| CCSER2 | 258.478308 | -0.021346672 | 0.20481485 | -0.1042242 | 0.9169914 | 0.99207822 |
| WAPL | 248.738502 | -0.047311086 | 0.17390041 | -0.2720585 | 0.78557703 | 0.97423208 |
| BMPR1A | 516.216948 | -0.006056023 | 0.19649737 | -0.0308199 | 0.97541319 | 0.99681784 |
| ADIRF | 181.271855 | -0.195082283 | 1.09445066 | -0.1782468 | 0.85852919 | 0.9864199 |
| GLUD1 | 179.014789 | 0.476951877 | 0.28420463 | 1.67819885 | 0.09330829 | 0.63721577 |
| SHLD2 | 292.540104 | -0.039427926 | 0.32043515 | -0.1230449 | 0.90207151 | 0.9900784 |
| NUTM2A-AS | 348.068364 | -0.482004063 | 0.23143664 | -2.082661 | 0.03728214 | 0.46890807 |
| MINPP1 | 38.3405756 | 0.860925236 | 0.34260817 | 2.51285668 | 0.0119758 | 0.30967852 |
| PAPSS2 | 96.519561 | 0.508736336 | 0.43003331 | 1.18301611 | 0.23680276 | 0.79983556 |
| ATAD1 | 148.944357 | -0.021371426 | 0.20044797 | -0.1066183 | 0.91509179 | 0.99159829 |
| PTEN | 431.994995 | 0.202632458 | 0.21389774 | 0.94733332 | 0.34346896 | 0.86909179 |
| RNLS | 180.28053 | -0.332792626 | 0.48494033 | -0.6862548 | 0.49255245 | 0.92289011 |
| FAS | 42.5507382 | 0.187662795 | 0.36680534 | 0.51161413 | 0.6089211 | 0.94766128 |
| LIPA | 117.809052 | 0.474307067 | 0.41672098 | 1.13818859 | 0.25504174 | 0.80897837 |
| IFIT3 | 129.871263 | -3.104781527 | 1.20446174 | -2.5777336 | 0.00994506 | 0.29238317 |
| IFIT5 | 30.0296944 | -0.444039254 | 0.49122911 | -0.9039351 | 0.3660298 | 0.87605028 |
| SLC16A12 | 251.845601 | 1.540365321 | 0.72026894 | 2.13859745 | 0.03246828 | 0.44049392 |
| PANK1 | 33.469479 | 0.14428564 | 0.47678166 | 0.30262414 | 0.76217631 | 0.9714717 |
| RPP30 | 30.9530457 | -0.071639375 | 0.28634568 | -0.2501849 | 0.80244433 | 0.97835591 |
| PCGF5 | 313.675049 | -0.223634224 | 0.2041764 | -1.0952991 | 0.27338562 | 0.82047446 |
| HECTD2 | 146.578912 | 0.134142199 | 0.27934236 | 0.48020715 | 0.63108011 | 0.94932781 |
| PPP1R3C | 61.4616954 | 1.311920694 | 0.89735084 | 1.46199304 | 0.14374312 | 0.71631428 |
| TNKS2 | 252.309834 | -0.041245926 | 0.20721827 | -0.1990458 | 0.84222692 | 0.98517901 |
| BTAF1 | 233.191215 | -0.096394197 | 0.27247319 | -0.353775 | 0.72350752 | 0.96779196 |
| CPEB3 | 239.512528 | -0.039227048 | 0.40618595 | -0.0965741 | 0.92306461 | 0.99214128 |
| 5-Mar | 115.155294 | -0.021119737 | 0.18629184 | -0.1133691 | 0.90973794 | 0.99100813 |
| IDE | 120.551206 | 0.046952206 | 0.17783474 | 0.26402156 | 0.79176331 | 0.97423208 |
| EXOC6 | 175.208245 | -0.346409506 | 0.25116471 | -1.3792125 | 0.16782924 | 0.74445571 |
| MYOF | 366.632296 | -0.148030905 | 0.35337123 | -0.4189105 | 0.67528157 | 0.95910677 |
| RBP4 | 127.603233 | 1.012080101 | 0.6595451 | 1.53451236 | 0.12490366 | 0.69439514 |
| FRA10AC1 | 104.506432 | -0.005416066 | 0.24890189 | -0.0217598 | 0.98263953 | 0.99700286 |
| PLCE1 | 140.710854 | 1.554612994 | 1.18603811 | 1.31076142 | 0.18993838 | 0.76207437 |
| NOC3L | 51.401684 | -0.359344041 | 0.3515073 | -1.0222947 | 0.30664145 | 0.84488735 |
| TBC1D12 | 103.720584 | 0.291307833 | 0.24602066 | 1.18407873 | 0.2363819 | 0.79958336 |
| HELLS | 48.8940027 | -0.56268099 | 0.41209296 | -1.3654225 | 0.1721203 | 0.74601809 |
| PDLIM1 | 48.1092814 | 1.223856404 | 0.81237659 | 1.50651363 | 0.13193537 | 0.7008294 |
| SORBS1 | 151.118788 | 0.131248041 | 0.32151835 | 0.40821322 | 0.68311714 | 0.96116027 |
| ALDH18A1 | 69.888052 | -0.071612533 | 0.31569248 | -0.2268427 | 0.82054606 | 0.98094663 |
| TCTN3 | 75.8336795 | -0.223685023 | 0.25031456 | -0.8936157 | 0.37152754 | 0.87605028 |
| ENTPD1-AS1 | 339.803101 | 0.338888096 | 0.36907021 | 0.9182212 | 0.35850307 | 0.87316144 |
| AL365273.2 | 27.4886157 | 0.385732039 | 0.47729852 | 0.80815679 | 0.41900033 | 0.8998918 |
| ZNF518A | 291.37577 | 0.081110809 | 0.3304571 | 0.24545035 | 0.80610775 | 0.97930627 |
| BLNK | 98.3990016 | 0.255660178 | 0.57693969 | 0.44313155 | 0.65767058 | 0.95399905 |
| TLL2 | 38.7333596 | -1.668915995 | 0.74211884 | -2.2488527 | 0.02452187 | 0.41230428 |
| TM9SF3 | 319.526792 | -0.00443961 | 0.15009328 | -0.029579 | 0.97640281 | 0.99681784 |

| | | | | | | |
|---|---|---|---|---|---|---|
| PIK3AP1 | 217.039976 | 0.45783227 | 0.62705747 | 0.73012808 | 0.4653119 | 0.91462428 |
| LCOR | 603.448219 | 0.040916174 | 0.20688602 | 0.19777158 | 0.84322379 | 0.98517901 |
| RRP12 | 59.9063504 | -0.192009976 | 0.35913147 | -0.5346509 | 0.59289126 | 0.94649617 |
| PGAM1 | 127.372286 | -0.130018199 | 0.38514553 | -0.337582 | 0.73567821 | 0.9698508 |
| EXOSC1 | 47.2692079 | -0.028146828 | 0.25116494 | -0.1120651 | 0.91077178 | 0.99100813 |
| ZDHHC16 | 28.4472324 | -0.220614684 | 0.32580693 | -0.6771332 | 0.49832146 | 0.92398825 |
| MMS19 | 122.20761 | -0.369320879 | 0.17087435 | -2.1613594 | 0.03066759 | 0.43815485 |
| UBTD1 | 115.791141 | -0.583236371 | 0.32976763 | -1.7686283 | 0.07695592 | 0.59726395 |
| MORN4 | 42.9269155 | 0.278288173 | 0.34299591 | 0.81134545 | 0.41716732 | 0.89862171 |
| ZFYVE27 | 54.216189 | 0.124297645 | 0.29197444 | 0.42571414 | 0.67031616 | 0.95764829 |
| CRTAC1 | 124.583194 | 0.432819662 | 1.78205444 | 0.24287679 | 0.80810085 | 0.97930627 |
| R3HCC1L | 146.061181 | 0.085450251 | 0.23068426 | 0.37042081 | 0.71106897 | 0.96587538 |
| LOXL4 | 48.0798353 | 0.419803611 | 0.52395948 | 0.80121388 | 0.42300783 | 0.90124378 |
| HPS1 | 44.8047886 | 0.301687327 | 0.29827784 | 1.01143058 | 0.31181039 | 0.84811918 |
| CNNM1 | 35.8428781 | -0.989246983 | 0.50253481 | -1.9685144 | 0.04900889 | 0.52219814 |
| GOT1 | 132.900733 | 0.399131918 | 0.34267123 | 1.16476634 | 0.24411359 | 0.80476923 |
| SLC25A28 | 53.5052934 | -0.733224398 | 0.56587734 | -1.2957303 | 0.19506843 | 0.76968045 |
| CUTC | 78.4844831 | -0.163536995 | 0.28864788 | -0.5665623 | 0.57101162 | 0.93998433 |
| COX15 | 53.6350415 | 0.089023519 | 0.30103704 | 0.29572281 | 0.76744178 | 0.97221728 |
| DNMBP | 140.04997 | -1.006091222 | 0.41972242 | -2.3970395 | 0.01652814 | 0.34643995 |
| ERLIN1 | 44.1138272 | -0.271713205 | 0.32538318 | -0.8350561 | 0.40368612 | 0.89344441 |
| CHUK | 49.6143597 | -0.25970705 | 0.42590333 | -0.6097793 | 0.54200799 | 0.93523904 |
| CWF19L1 | 62.0240898 | -0.090259054 | 0.28940402 | -0.3118791 | 0.75513244 | 0.97125609 |
| BLOC1S2 | 30.5125412 | -0.05232022 | 0.26813462 | -0.1951267 | 0.84529377 | 0.98520136 |
| HIF1AN | 53.3677171 | 0.302734154 | 0.26354461 | 1.14870175 | 0.25067898 | 0.80680142 |
| PAX2 | 707.891234 | 0.117679921 | 0.36766253 | 0.32007592 | 0.74891078 | 0.97028355 |
| SLF2 | 126.418553 | 0.091801372 | 0.33971315 | 0.27023203 | 0.78698176 | 0.97423208 |
| MRPL43 | 42.1525574 | 0.276598941 | 0.35973682 | 0.76889251 | 0.44195713 | 0.90561284 |
| LZTS2 | 64.0021674 | -0.448632758 | 0.33556239 | -1.3369578 | 0.18123641 | 0.75448394 |
| BTRC | 286.963952 | 0.162259564 | 0.23722289 | 0.68399622 | 0.49397755 | 0.92289011 |
| DPCD | 42.0227835 | 0.686276489 | 0.36615034 | 1.87430247 | 0.06088875 | 0.56071154 |
| POLL | 40.7859889 | 0.591301067 | 0.27019415 | 2.18843028 | 0.02863827 | 0.42930522 |
| FBXW4 | 250.926507 | 0.060467484 | 0.21931145 | 0.27571513 | 0.78276688 | 0.97423208 |
| OGA | 438.497607 | 0.049288097 | 0.29025312 | 0.16981074 | 0.86515898 | 0.98670067 |
| ARMH3 | 261.508672 | 0.119453509 | 0.16393208 | 0.72867681 | 0.46619938 | 0.91462428 |
| LDB1 | 32.2562241 | 0.729359429 | 0.42277579 | 1.72516837 | 0.08449713 | 0.61686112 |
| PPRC1 | 39.4109805 | -1.13357409 | 0.44116573 | -2.5694971 | 0.01018462 | 0.29271368 |
| NOLC1 | 45.9800846 | -0.726833111 | 0.41060939 | -1.7701327 | 0.07670504 | 0.59726395 |
| GBF1 | 356.509787 | -0.037105871 | 0.15639654 | -0.2372551 | 0.81245892 | 0.97930627 |
| NFKB2 | 42.2157818 | 0.172380072 | 0.65521545 | 0.26308915 | 0.79248187 | 0.97429505 |
| FBXL15 | 30.447218 | -0.017884821 | 0.505984 | -0.0353466 | 0.97180335 | 0.99681784 |
| CUEDC2 | 45.745118 | 0.129248299 | 0.28734974 | 0.44979438 | 0.65285871 | 0.95176155 |
| ACTR1A | 49.2975527 | -0.023925479 | 0.30571797 | -0.07826 | 0.93762126 | 0.99526299 |
| SUFU | 150.791846 | 0.883808512 | 0.31569375 | 2.79957559 | 0.00511698 | 0.22279811 |
| TRIM8 | 95.1469455 | 0.278866967 | 0.34113392 | 0.81747065 | 0.41365952 | 0.89732895 |
| ARL3 | 127.982513 | 0.536717752 | 0.19239976 | 2.78959683 | 0.00527737 | 0.22669993 |
| SFXN2 | 38.957545 | -0.027870884 | 0.48121222 | -0.0579181 | 0.95381389 | 0.99531843 |
| WBP1L | 162.491494 | 0.247393328 | 0.35138735 | 0.70404734 | 0.48140329 | 0.91971919 |
| CNNM2 | 187.268626 | 0.564379469 | 0.41053193 | 1.3747517 | 0.16920844 | 0.74445571 |
| NT5C2 | 443.029618 | 0.021031357 | 0.22893789 | 0.0918649 | 0.92680538 | 0.99336859 |
| PCGF6 | 51.0270131 | -0.010215681 | 0.2753196 | -0.0371048 | 0.97040144 | 0.99681784 |
| ATP5MD | 56.7300152 | 0.058689076 | 0.31710361 | 0.18507855 | 0.85316743 | 0.9864199 |
| PDCD11 | 48.8430023 | -0.484156551 | 0.31660987 | -1.5291897 | 0.12621742 | 0.69492095 |
| SH3PXD2A | 61.6005599 | 0.335804478 | 0.26596098 | 1.26260805 | 0.20673007 | 0.77401635 |

| | | | | | | |
|---|---|---|---|---|---|---|
| STN1 | 35.9595075 | -0.393681911 | 0.38579703 | -1.0204379 | 0.30752083 | 0.84488735 |
| SLK | 148.336283 | -0.314135846 | 0.17106472 | -1.8363567 | 0.06630492 | 0.57792071 |
| CFAP43 | 173.604131 | 1.038858224 | 0.58372841 | 1.77969447 | 0.07512598 | 0.59465671 |
| GSTO1 | 92.8842925 | -0.641812824 | 0.38480648 | -1.6678847 | 0.09533863 | 0.64087627 |
| GSTO2 | 110.780456 | -0.203345716 | 0.35107182 | -0.579214 | 0.56244478 | 0.93796284 |
| SORCS1 | 66.0723946 | 0.862440873 | 0.58965659 | 1.4626155 | 0.14357262 | 0.71631428 |
| LINC01435 | 1600.89114 | 0.941892568 | 0.59211279 | 1.59073167 | 0.11166998 | 0.66300165 |
| AL353740.1 | 56.8686587 | 2.124074537 | 0.81495761 | 2.60636198 | 0.00915097 | 0.2846011 |
| XPNPEP1 | 83.9659681 | -0.152082227 | 0.31137813 | -0.4884165 | 0.62525483 | 0.94808496 |
| ADD3 | 309.484424 | 0.654465997 | 0.28057098 | 2.33262182 | 0.019668 | 0.37455228 |
| MXI1 | 170.486619 | 0.225027717 | 0.38723472 | 0.58111452 | 0.56116328 | 0.93747805 |
| SMNDC1 | 50.0770297 | 0.11797882 | 0.3017956 | 0.39092293 | 0.69585421 | 0.96220115 |
| SMC3 | 99.9863314 | -0.198114429 | 0.27412031 | -0.722728 | 0.469847 | 0.91502267 |
| PDCD4 | 135.744053 | 0.248072934 | 0.33006815 | 0.75158096 | 0.4523031 | 0.91077355 |
| BBIP1 | 63.9070504 | -0.01216147 | 0.25827462 | -0.0470874 | 0.9624436 | 0.99572129 |
| SHOC2 | 195.343354 | 0.112770028 | 0.2109922 | 0.53447486 | 0.59301304 | 0.94649617 |
| ZDHHC6 | 52.3092631 | -0.210174791 | 0.27049741 | -0.7769937 | 0.43716247 | 0.90561284 |
| VTI1A | 561.193798 | -0.239173542 | 0.16466487 | -1.4524867 | 0.14636632 | 0.71974648 |
| TCF7L2 | 537.799535 | 0.178427336 | 0.32658106 | 0.54634931 | 0.58482585 | 0.94229635 |
| CASP7 | 52.1262294 | -0.158621602 | 0.4487169 | -0.3535004 | 0.72371333 | 0.96779196 |
| NHLRC2 | 151.157836 | 0.11616283 | 0.15234186 | 0.76251417 | 0.44575319 | 0.90732465 |
| CCDC186 | 150.717347 | -0.019762426 | 0.23797159 | -0.0830453 | 0.93381551 | 0.9943329 |
| AFAP1L2 | 41.2927173 | 0.446988545 | 0.56151494 | 0.79604034 | 0.42600859 | 0.90237497 |
| ABLIM1 | 602.112696 | 0.613690941 | 0.28225919 | 2.17421066 | 0.02968932 | 0.43492739 |
| FAM160B1 | 90.6529565 | 0.289381155 | 0.34090201 | 0.84886901 | 0.39595418 | 0.88785003 |
| TRUB1 | 49.3728211 | -0.278063021 | 0.2486563 | -1.1182625 | 0.26345488 | 0.81378765 |
| ATRNL1 | 242.246522 | 0.938465704 | 0.33056866 | 2.83894334 | 0.00452632 | 0.21570952 |
| HSPA12A | 291.312744 | -0.203744966 | 0.48132231 | -0.4233026 | 0.67207453 | 0.95871957 |
| ENO4 | 37.7061202 | 0.654525417 | 0.4504989 | 1.45289016 | 0.14625425 | 0.71974648 |
| SHTN1 | 299.309736 | 0.133987205 | 0.20343748 | 0.65861612 | 0.51014231 | 0.92819651 |
| PDZD8 | 253.970279 | -0.247094126 | 0.1712102 | -1.4432208 | 0.14895827 | 0.72246729 |
| EMX2OS | 270.814199 | 0.438619191 | 0.4722892 | 0.92870892 | 0.35303995 | 0.8720098 |
| EMX2 | 108.353707 | 0.020596723 | 0.49050664 | 0.04199071 | 0.9665061 | 0.99601626 |
| RAB11FIP2 | 246.22113 | 0.007882403 | 0.33213264 | 0.0237327 | 0.98106583 | 0.99681784 |
| CASC2 | 172.498563 | 0.204949842 | 0.47318292 | 0.43313026 | 0.66492014 | 0.95505623 |
| FAM204A | 117.543626 | -0.176068054 | 0.21283664 | -0.827245 | 0.4080982 | 0.8959746 |
| CACUL1 | 202.368936 | -0.314361684 | 0.25614456 | -1.2272823 | 0.21971652 | 0.78584902 |
| EIF3A | 271.591706 | -0.249867042 | 0.31941859 | -0.7822558 | 0.43406426 | 0.90514499 |
| FAM45A | 48.0457183 | -0.045704615 | 0.26128518 | -0.1749223 | 0.86114066 | 0.98658849 |
| SFXN4 | 52.4150021 | -0.463999853 | 0.28290196 | -1.6401436 | 0.10097531 | 0.64808588 |
| PRDX3 | 104.496236 | 0.08367301 | 0.48148141 | 0.17378243 | 0.86203645 | 0.98669442 |
| TIAL1 | 145.392828 | -0.153050263 | 0.18469455 | -0.8286669 | 0.40729292 | 0.89585737 |
| BAG3 | 43.3029658 | 0.641203484 | 0.24778415 | 2.58775019 | 0.0096605 | 0.28918107 |
| INPP5F | 111.312066 | 0.311992166 | 0.20494764 | 1.52230185 | 0.12793347 | 0.69635098 |
| MCMBP | 84.7587822 | -0.300093134 | 0.21909277 | -1.369708 | 0.17077806 | 0.74445571 |
| SEC23IP | 104.09529 | 0.073242842 | 0.23904616 | 0.30639623 | 0.75930298 | 0.9714717 |
| PLPP4 | 43.4291332 | 0.123117675 | 0.46413217 | 0.26526426 | 0.7908059 | 0.97423208 |
| WDR11-AS1 | 31.0981361 | -0.496143241 | 0.55617204 | -0.8920679 | 0.37235652 | 0.87605028 |
| WDR11 | 166.385933 | 0.104325368 | 0.16566178 | 0.62974917 | 0.52885871 | 0.93366189 |
| FGFR2 | 364.636836 | -0.23407452 | 0.39512658 | -0.5924039 | 0.55358017 | 0.93523904 |
| ATE1 | 337.598766 | -0.018929301 | 0.13092158 | -0.144585 | 0.88503852 | 0.98766275 |
| NSMCE4A | 31.6366387 | -0.918644892 | 0.35532506 | -2.5853648 | 0.0097276 | 0.29028254 |
| TACC2 | 201.181483 | -0.166408666 | 0.426385 | -0.390278 | 0.69633102 | 0.96220115 |
| PLEKHA1 | 706.238768 | -0.257602783 | 0.33616334 | -0.7663024 | 0.44349637 | 0.90658382 |

| | | | | | | |
|---|---|---|---|---|---|---|
| HTRA1 | 30.4718597 | 0.387355253 | 0.54169574 | 0.71507901 | 0.47456023 | 0.91809716 |
| C10orf88 | 29.8686123 | -0.129835013 | 0.31662815 | -0.4100552 | 0.68176547 | 0.96095224 |
| IKZF5 | 37.517905 | 0.290737129 | 0.31170157 | 0.93274194 | 0.35095322 | 0.8720098 |
| ACADSB | 62.8091545 | 0.209488053 | 0.41742144 | 0.50186223 | 0.61576444 | 0.94791646 |
| BUB3 | 57.4861562 | 0.173763293 | 0.33178677 | 0.52371978 | 0.60047345 | 0.94683415 |
| CHST15 | 82.1368964 | -0.192823695 | 0.60718948 | -0.3175676 | 0.75081297 | 0.97058015 |
| OAT | 28.4994086 | 0.679205284 | 0.42010364 | 1.61675649 | 0.10593086 | 0.65699665 |
| LHPP | 33.5894876 | 0.791636398 | 0.42080809 | 1.88122904 | 0.05994077 | 0.55922073 |
| ABRAXAS2 | 44.1025053 | -0.191314554 | 0.3575786 | -0.535028 | 0.59263051 | 0.94649617 |
| ZRANB1 | 116.76113 | 0.078045747 | 0.1801472 | 0.43323318 | 0.66484537 | 0.95505623 |
| CTBP2 | 641.007007 | -0.156425666 | 0.20705076 | -0.7554943 | 0.44995247 | 0.91035378 |
| EDRF1 | 70.1836972 | -0.112471569 | 0.25654683 | -0.4384056 | 0.66109229 | 0.95407427 |
| UROS | 41.8818224 | -0.410718221 | 0.30391541 | -1.3514228 | 0.17656002 | 0.75067397 |
| BCCIP | 77.9029819 | -0.412826463 | 0.25236784 | -1.6358125 | 0.10187884 | 0.65103232 |
| DHX32 | 187.521829 | -0.245323074 | 0.20487896 | -1.1974049 | 0.23114876 | 0.79432962 |
| FANK1 | 155.908132 | 0.528047644 | 0.27232626 | 1.93902577 | 0.0524982 | 0.53192032 |
| DOCK1 | 979.79362 | -0.456658371 | 0.21669175 | -2.1074101 | 0.03508205 | 0.45597142 |
| FAM196A | 46.4927463 | 0.043555263 | 0.56005483 | 0.07776964 | 0.9380113 | 0.99526299 |
| PTPRE | 52.4981862 | 0.977286758 | 0.69056115 | 1.4152067 | 0.1570079 | 0.73472335 |
| MGMT | 280.640193 | 0.141426184 | 0.26351627 | 0.53668862 | 0.59148272 | 0.94556291 |
| GLRX3 | 107.211963 | -0.479532084 | 0.21657071 | -2.2142056 | 0.02681464 | 0.4234302 |
| PPP2R2D | 195.884886 | 0.044084968 | 0.22410055 | 0.19671959 | 0.84404698 | 0.98517901 |
| BNIP3 | 176.736499 | 0.445382369 | 0.59869134 | 0.74392653 | 0.45692092 | 0.91187931 |
| STK32C | 37.3244621 | -0.694494619 | 0.29329818 | -2.3678791 | 0.01789038 | 0.35777027 |
| LRRC27 | 38.7308887 | 0.023430632 | 0.44061672 | 0.0531769 | 0.95759096 | 0.99572129 |
| PWWP2B | 22.9469306 | -0.890544635 | 0.72676708 | -1.2253508 | 0.22044307 | 0.78586683 |
| INPP5A | 112.116523 | 0.082485448 | 0.33593334 | 0.24554112 | 0.80603748 | 0.97930627 |
| CFAP46 | 42.4425987 | 1.332476985 | 0.43916804 | 3.03409371 | 0.00241259 | 0.16028122 |
| TUBGCP2 | 120.619144 | -0.013258866 | 0.21690435 | -0.0611277 | 0.9512575 | 0.99531843 |
| ECHS1 | 107.025514 | 0.69733716 | 0.47523987 | 1.46733725 | 0.14228433 | 0.7148274 |
| SLC6A12 | 155.441485 | -1.059835606 | 0.67131574 | -1.5787439 | 0.11439482 | 0.66978434 |
| SLC6A13 | 82.189672 | 1.999828462 | 0.52290544 | 3.82445526 | 0.00013106 | 0.02918429 |
| KDM5A | 257.362027 | 0.023992771 | 0.19817971 | 0.12106573 | 0.90363898 | 0.99050161 |
| B4GALNT3 | 50.9680636 | -0.536501283 | 0.56995324 | -0.9413075 | 0.34654728 | 0.86990996 |
| WNK1 | 389.441583 | 0.507019705 | 0.31187261 | 1.62572694 | 0.10400776 | 0.65475128 |
| RAD52 | 67.1557487 | 0.027920976 | 0.32868588 | 0.0849473 | 0.93230329 | 0.99416721 |
| ERC1 | 796.822905 | 0.015025011 | 0.20437774 | 0.07351589 | 0.9413956 | 0.99526299 |
| ADIPOR2 | 139.762209 | -0.109330504 | 0.25192949 | -0.4339726 | 0.66430832 | 0.95505623 |
| DCP1B | 77.2129795 | -0.107695105 | 0.32849201 | -0.327847 | 0.74302739 | 0.9699454 |
| FKBP4 | 70.0667558 | 0.241712178 | 0.43061838 | 0.56131412 | 0.57458341 | 0.94034284 |
| ITFG2 | 104.465005 | -0.071615871 | 0.24759821 | -0.2892423 | 0.77239598 | 0.97329019 |
| TULP3 | 170.476172 | 0.712183947 | 0.17414671 | 4.08956297 | 4.32E-05 | 0.01335457 |
| TEAD4 | 100.598885 | -0.115266083 | 0.41986936 | -0.2745284 | 0.78367855 | 0.97423208 |
| TSPAN9 | 314.693198 | 0.32075962 | 0.4043634 | 0.79324593 | 0.42763455 | 0.90272191 |
| PARP11 | 72.7750472 | 0.657481246 | 0.39911107 | 1.64736409 | 0.09948323 | 0.64519288 |
| CCND2 | 50.8070067 | 1.419584938 | 1.23451685 | 1.14991135 | 0.25018039 | 0.80672756 |
| TIGAR | 28.7141066 | -0.332865688 | 0.44936501 | -0.7407468 | 0.45884698 | 0.91362347 |
| C12orf4 | 54.1214041 | 0.248006629 | 0.23521595 | 1.05437846 | 0.29170968 | 0.83661288 |
| CD9 | 329.49816 | -0.261939422 | 0.46970965 | -0.5576624 | 0.57707492 | 0.94034284 |
| TNFRSF1A | 188.038678 | 0.153984711 | 0.45102418 | 0.34141121 | 7.33E-01 | 0.96980299 |
| SCNN1A | 238.612153 | 0.020483612 | 0.57415596 | 0.03567604 | 0.97154067 | 0.99681784 |
| LTBR | 105.945097 | -0.119953906 | 0.28600231 | -0.4194159 | 6.75E-01 | 0.95910677 |
| MRPL51 | 80.1834659 | -0.361799886 | 0.61289517 | -0.5903128 | 0.55498093 | 0.93523904 |
| NCAPD2 | 40.23528 | 0.036329068 | 0.41151124 | 0.08828208 | 0.92965248 | 0.99362798 |

| | | | | | | |
|---|---|---|---|---|---|---|
| GAPDH | 713.283486 | -0.394273355 | 0.63834277 | -0.6176515 | 0.5368051 | 0.93474933 |
| CHD4 | 64.0413169 | 0.281708535 | 0.2503268 | 1.12536306 | 0.26043522 | 0.81263292 |
| AC006064.2 | 27.1784431 | 0.166389253 | 0.35574391 | 0.467722 | 0.63998341 | 0.94944486 |
| ING4 | 41.7192193 | 0.250161733 | 0.27259757 | 0.91769612 | 0.35877798 | 0.87316144 |
| ZNF384 | 42.6519743 | -0.254356959 | 0.25268904 | -1.0066007 | 0.31412671 | 0.8490543 |
| COPS7A | 39.5526683 | -0.247826042 | 0.3567899 | -0.6945994 | 0.4873064 | 0.92034617 |
| MLF2 | 90.5539715 | -0.269368212 | 0.41845256 | -0.6437246 | 0.51975403 | 0.93272217 |
| PTMS | 153.281827 | -0.325727746 | 0.51824354 | -0.6285225 | 0.52966169 | 0.93366189 |
| USP5 | 34.6767308 | 0.021613061 | 0.37029459 | 0.05836721 | 0.95345613 | 0.99531843 |
| TPI1 | 204.359451 | 0.030488165 | 0.55843924 | 0.05459531 | 0.95646087 | 0.99572129 |
| LRRC23 | 65.5107027 | -0.071227428 | 0.32871564 | -0.216684 | 0.8284546 | 0.98142144 |
| ENO2 | 27.4324187 | -1.197794968 | 0.58770859 | -2.0380763 | 0.0415423 | 0.48843812 |
| ATN1 | 119.92736 | -0.18991469 | 0.19787675 | -0.9597625 | 0.33717474 | 0.86450666 |
| C12orf57 | 51.3346141 | 0.1028642 | 0.51972181 | 0.19792165 | 0.84310636 | 0.98517901 |
| PTPN6 | 32.2510706 | -0.427774612 | 0.32034544 | -1.3353542 | 0.18176046 | 0.75448394 |
| PHB2 | 87.5149061 | 0.074534485 | 0.33873962 | 0.22003474 | 0.8258441 | 0.98094663 |
| LPCAT3 | 204.732791 | 0.288725503 | 0.25158969 | 1.14760466 | 0.25113181 | 0.80724549 |
| C1S | 155.089437 | 0.363433337 | 0.73832481 | 0.49224045 | 0.62254937 | 0.94808496 |
| C1R | 171.407099 | 1.603681265 | 1.4338119 | 1.11847395 | 0.26336462 | 0.81378765 |
| C1RL | 230.323346 | 0.567586347 | 0.58647352 | 0.96779535 | 0.33314659 | 0.86067542 |
| C1RL-AS1 | 30.0904598 | -0.247663903 | 0.58646682 | -0.4222982 | 0.67280735 | 0.95910677 |
| RBP5 | 41.9169676 | 2.031736699 | 0.43494069 | 4.67129598 | 2.99E-06 | 0.00200368 |
| CLSTN3 | 54.6875292 | 0.901757212 | 0.33772116 | 2.67012353 | 0.00758233 | 0.26951269 |
| PEX5 | 37.8917479 | -0.147636226 | 0.281729 | -0.5240363 | 0.60025328 | 0.94683415 |
| NECAP1 | 32.4606869 | -0.154173036 | 0.55363562 | -0.2784738 | 0.78064864 | 0.97423208 |
| LINC00937 | 111.889375 | 0.410196728 | 0.37003188 | 1.10854428 | 0.26762682 | 0.81510106 |
| RIMKLB | 178.073122 | -0.184654093 | 0.38390247 | -0.4809922 | 0.63052204 | 0.94932781 |
| PHC1 | 77.0905514 | 0.004670943 | 0.25741583 | 0.01814552 | 0.98552277 | 0.99764442 |
| M6PR | 106.18117 | -0.191001019 | 0.21707911 | -0.8798683 | 0.37893067 | 0.87973265 |
| CLEC2D | 90.883923 | -0.056227645 | 0.27121419 | -0.2073182 | 0.83576134 | 0.98423382 |
| GABARAPL1 | 233.927079 | -0.046924014 | 0.532389 | -0.0881386 | 0.92976653 | 0.99362798 |
| KLRD1 | 152.768064 | 1.32010616 | 0.63628549 | 2.0747073 | 0.03801367 | 0.47106462 |
| MAGOHB | 25.3692222 | -0.398496003 | 0.33598082 | -1.1860677 | 0.23559556 | 0.79942255 |
| YBX3 | 365.183611 | -0.292715337 | 0.35488511 | -0.8248172 | 0.40947539 | 0.8959746 |
| SMIM10L1 | 28.5410411 | -0.516433865 | 0.34580755 | -1.4934141 | 0.13532883 | 0.7063854 |
| ETV6 | 665.820205 | 0.113274373 | 0.44332919 | 0.2555085 | 0.79833038 | 0.97603149 |
| BCL2L14 | 76.8932398 | -0.188275526 | 0.28667415 | -0.656758 | 0.51133657 | 0.92819651 |
| LRP6 | 264.659173 | 0.153973633 | 0.2212358 | 0.69597071 | 0.48644717 | 0.92034617 |
| BORCS5 | 65.4583942 | -0.007616636 | 0.2602661 | -0.0292648 | 0.9766534 | 0.99681784 |
| DUSP16 | 190.277775 | 0.873439489 | 0.33738492 | 2.58885157 | 0.00962966 | 0.28916146 |
| CREBL2 | 67.7273909 | 0.472605704 | 0.22652372 | 2.08634094 | 0.03694775 | 0.46815146 |
| CDKN1B | 44.7511133 | 0.49782725 | 0.44128192 | 1.12813878 | 0.25926131 | 0.81263292 |
| GPRC5D-AS1 | 49.0657125 | -0.514078294 | 0.45866905 | -1.1208044 | 0.26237113 | 0.81377303 |
| HEBP1 | 74.8701292 | -0.443453379 | 0.36541275 | -1.2135684 | 0.22491257 | 0.78917125 |
| EMP1 | 152.814462 | -1.830769111 | 0.85514716 | -2.1408819 | 0.03228356 | 0.44049392 |
| ATF7IP | 466.770415 | 0.210002592 | 0.26889001 | 0.78099813 | 0.4348036 | 0.90514499 |
| H2AFJ | 120.008732 | 0.154928847 | 0.29212256 | 0.53035563 | 0.59586538 | 0.94683415 |
| WBP11 | 74.3718028 | -0.062698306 | 0.27866449 | -0.2249957 | 0.82198263 | 0.98094663 |
| MGP | 100.868483 | 0.904453326 | 0.82987187 | 1.08987105 | 0.27576995 | 0.82164863 |
| ERP27 | 75.8033854 | 2.669107688 | 1.1144862 | 2.39492216 | 0.01662389 | 0.3476862 |
| RERG | 536.670512 | 0.613924417 | 1.03667995 | 0.59220246 | 0.55371502 | 0.93523904 |
| PTPRO | 95.597913 | 0.847287192 | 0.73737824 | 1.1490537 | 0.25053384 | 0.80680142 |
| EPS8 | 423.541465 | 0.624331281 | 0.59093621 | 1.05651215 | 0.29073429 | 0.83531608 |
| STRAP | 96.5214871 | -0.143569226 | 0.30229828 | -0.4749257 | 0.63483992 | 0.94932781 |

| | | | | | | |
|---|---|---|---|---|---|---|
| DERA | 171.949277 | -0.111887053 | 0.16100513 | -0.6949285 | 0.48710012 | 0.92034617 |
| MGST1 | 201.365069 | 0.93159454 | 0.64597033 | 1.44216306 | 0.14925638 | 0.72246729 |
| PLEKHA5 | 794.751077 | 0.777733714 | 0.39307679 | 1.97857958 | 0.04786336 | 0.51457141 |
| AEBP2 | 350.367931 | 0.439048783 | 0.24033593 | 1.82681296 | 0.0677279 | 0.58081067 |
| PDE3A | 161.064282 | 0.416940458 | 1.43903022 | 0.28973711 | 0.77201736 | 0.97329019 |
| SLCO1A2 | 126.702262 | -1.385177869 | 0.9404428 | -1.4728997 | 1.41E-01 | 7.15E-01 |
| PYROXD1 | 55.8891318 | -0.250710253 | 0.39931735 | -0.6278471 | 0.53010408 | 0.93366189 |
| RECQL | 50.408912 | -0.13074935 | 0.29402552 | -0.4446871 | 0.65654591 | 0.95399905 |
| GOLT1B | 44.8026344 | -0.192831337 | 0.37240532 | -0.5177996 | 0.60459808 | 0.94683415 |
| SPX | 120.214195 | 0.888507281 | 0.61603533 | 1.44229923 | 0.14921798 | 0.72246729 |
| LDHB | 401.700592 | 0.330548781 | 0.37627868 | 0.87846799 | 0.3796898 | 0.88042813 |
| CMAS | 51.8973128 | 0.093900702 | 0.33431081 | 0.28087845 | 0.77880363 | 0.97423208 |
| C2CD5 | 184.226307 | -0.24299032 | 0.24733909 | -0.9824178 | 0.32589407 | 0.85509839 |
| ETNK1 | 334.121577 | 0.010419915 | 0.25608876 | 0.04068869 | 0.96754408 | 0.99648221 |
| AC084816.1 | 43.6858421 | 1.1473498 | 0.74748463 | 1.53494769 | 0.12479668 | 0.69439514 |
| CASC1 | 125.118339 | 1.238457245 | 0.50427365 | 2.45592298 | 0.01405233 | 0.32375679 |
| ETFRF1 | 56.429705 | 0.266732014 | 0.36764181 | 0.72552143 | 0.4681322 | 0.91462428 |
| KRAS | 105.838979 | -0.015091123 | 0.26234418 | -0.0575241 | 0.95412768 | 0.99531843 |
| RASSF8-AS1 | 81.5647847 | -0.296606666 | 0.32298874 | -0.9183189 | 0.35845193 | 0.87316144 |
| RASSF8 | 284.308458 | -0.041010814 | 0.31540625 | -0.1300254 | 0.89654636 | 0.98842187 |
| BHLHE41 | 66.1183219 | 0.408379405 | 0.70439592 | 0.57975833 | 0.5620776 | 0.93762898 |
| SSPN | 49.9504873 | 0.749778126 | 0.7313877 | 1.02514457 | 0.30529498 | 0.84488735 |
| ITPR2 | 305.418616 | -0.109714351 | 0.24709048 | -0.444025 | 0.6570245 | 0.95399905 |
| INTS13 | 32.8475685 | -0.11092366 | 0.31668909 | -0.3502604 | 0.72614326 | 0.96781837 |
| FGFR1OP2 | 110.031021 | 0.30655041 | 0.2982051 | 1.02798514 | 0.30395682 | 0.84488735 |
| TM7SF3 | 129.558497 | 0.231167742 | 0.267989 | 0.86260161 | 0.38835655 | 0.88586416 |
| MED21 | 52.1499237 | 0.15666166 | 0.36114273 | 0.4337943 | 0.66443782 | 0.95505623 |
| STK38L | 77.6397768 | 0.32802674 | 0.23693929 | 1.3844337 | 0.1662257 | 0.74445571 |
| ARNTL2 | 51.2690867 | -0.913423035 | 0.53078766 | -1.7208822 | 0.0852722 | 0.61974555 |
| PPFIBP1 | 1003.73677 | -0.484538432 | 0.21806565 | -2.2219842 | 0.02628438 | 0.42051885 |
| MRPS35 | 80.4779302 | 0.148635853 | 0.26274983 | 0.56569342 | 0.57160221 | 0.93998433 |
| KLHL42 | 70.2001491 | 0.476087697 | 0.31822203 | 1.49608654 | 0.1346311 | 0.70620568 |
| CCDC91 | 500.073779 | -0.017262274 | 0.16695639 | -0.1033939 | 0.91765035 | 0.99214128 |
| FAR2 | 39.0675686 | -0.726425851 | 0.76535689 | -0.9491335 | 0.34255273 | 0.86846099 |
| AC009318.1 | 28.3690184 | -1.195396028 | 0.61176723 | -1.9540047 | 0.05070066 | 0.52415717 |
| ERGIC2 | 96.3208398 | -0.101916186 | 0.19760784 | -0.5157497 | 0.60602924 | 0.94708529 |
| IPO8 | 118.664452 | 0.062195809 | 0.28785319 | 0.21606781 | 0.82893489 | 0.98142144 |
| CAPRIN2 | 62.775318 | -0.102133653 | 0.30878585 | -0.3307588 | 0.74082665 | 0.9698508 |
| SINHCAF | 197.285726 | -0.237531816 | 0.21952811 | -1.0820109 | 0.27924768 | 0.82381074 |
| DENND5B | 231.512268 | 0.696728323 | 0.27769244 | 2.50899275 | 0.0121076 | 0.30967852 |
| AMN1 | 47.5202194 | 0.379799512 | 0.35823641 | 1.06019237 | 0.28905709 | 0.83399936 |
| KIAA1551 | 95.6849932 | 0.22924392 | 0.29728437 | 0.77112671 | 0.44063183 | 0.90561284 |
| BICD1 | 764.8229 | 0.299033733 | 0.53289793 | 0.56114636 | 0.57469777 | 0.94034284 |
| FGD4 | 895.369845 | 0.231650568 | 0.38453959 | 0.60241019 | 0.54690113 | 0.93523904 |
| DNM1L | 197.023065 | -0.390912976 | 0.18645073 | -2.096602 | 0.03602882 | 0.46016006 |
| YARS2 | 73.3088888 | 0.000912559 | 0.24731188 | 0.00368991 | 0.99705588 | 0.99917842 |
| PKP2 | 52.3249586 | 0.159243501 | 0.36755168 | 0.43325472 | 0.66482973 | 0.95505623 |
| SYT10 | 38.870982 | -0.864056211 | 1.67432659 | -0.5160619 | 0.60581116 | 0.94708529 |
| ALG10B | 30.1108721 | -0.476009735 | 0.39994403 | -1.1901909 | 0.23397137 | 0.79633808 |
| CPNE8 | 384.105734 | -0.132286989 | 0.18315808 | -0.7222558 | 0.47013722 | 0.91502267 |
| KIF21A | 268.09286 | 0.396200645 | 0.31985071 | 1.23870492 | 0.2154548 | 0.78293142 |
| C12orf40 | 54.7115452 | 0.209935298 | 0.5794342 | 0.36231085 | 0.71711975 | 0.96742001 |
| SLC2A13 | 741.153977 | -0.053007896 | 0.54947412 | -0.0964702 | 0.92314712 | 0.99214128 |
| LINC02471 | 350.987275 | 1.453075824 | 1.09278432 | 1.32970047 | 0.18361698 | 0.75689793 |

| | | | | | | |
|---|---|---|---|---|---|---|
| AC079630.1 | 141.531303 | 0.475877711 | 0.76975526 | 0.6182195 | 0.53643066 | 0.93469807 |
| LRRK2 | 1039.70673 | 0.248417933 | 0.62858353 | 0.39520274 | 0.69269327 | 0.96220115 |
| GXYLT1 | 103.956349 | 0.133543155 | 0.25039198 | 0.5333364 | 0.59380073 | 0.94649617 |
| YAF2 | 199.899202 | -0.087359751 | 0.19654517 | -0.4444767 | 0.65669796 | 0.95399905 |
| PPHLN1 | 274.550404 | -0.009943102 | 0.15274065 | -0.0650979 | 0.94809602 | 0.9952642 |
| ZCRB1 | 53.1321661 | -0.514582328 | 0.34250633 | -1.5024024 | 0.13299321 | 0.70369044 |
| PRICKLE1 | 180.240265 | 0.453649305 | 0.71323066 | 0.63604852 | 0.5247448 | 0.93272384 |
| PUS7L | 61.2055765 | -0.487312927 | 0.29357197 | -1.6599437 | 0.09692579 | 0.64297238 |
| IRAK4 | 53.2092642 | -0.057728914 | 0.23240993 | -0.2483926 | 0.80383064 | 0.97880236 |
| TWF1 | 33.4162809 | -0.207432097 | 0.39220019 | -0.5288934 | 0.5968794 | 0.94683415 |
| TMEM117 | 148.297576 | 0.59833854 | 0.41377745 | 1.44603952 | 0.14816612 | 0.72246729 |
| ANO6 | 283.636858 | -0.331511767 | 0.22243627 | -1.4903674 | 0.13612765 | 0.70752658 |
| AC008124.1 | 35.5282374 | 0.216952937 | 0.46772697 | 0.46384526 | 0.64275862 | 0.94944486 |
| ARID2 | 484.852572 | -0.055993845 | 0.15236931 | -0.3674877 | 0.71325527 | 0.96587538 |
| SCAF11 | 550.627366 | -0.078544177 | 0.13337341 | -0.5889043 | 0.55592545 | 0.93523904 |
| SLC38A1 | 364.561533 | -0.823407799 | 0.42496555 | -1.9375872 | 0.05267361 | 0.53195745 |
| SLC38A2 | 307.524378 | -0.287269197 | 0.43576025 | -0.6592368 | 0.50974371 | 0.92819651 |
| AC008014.1 | 651.871142 | 0.10144373 | 0.38466269 | 0.26372126 | 0.79199471 | 0.97423208 |
| PCED1B | 199.370842 | -0.161714639 | 0.48470214 | -0.3336371 | 0.73865338 | 0.9698508 |
| RPAP3 | 46.144975 | -0.361725038 | 0.37029885 | -0.9768462 | 0.32864529 | 0.8574012 |
| RAPGEF3 | 299.141346 | 0.039829078 | 0.70515406 | 0.0564828 | 0.9549572 | 0.99531843 |
| SLC48A1 | 152.071113 | -0.1262609 | 0.70866604 | -0.178167 | 0.85859183 | 0.9864199 |
| HDAC7 | 182.892461 | -0.439343204 | 0.26519692 | -1.6566678 | 0.09758666 | 0.64312419 |
| VDR | 26.7860154 | -0.052884544 | 0.58715854 | -0.0900686 | 0.92823271 | 0.99336859 |
| TMEM106C | 42.6186123 | 0.013329884 | 0.42793659 | 0.0311492 | 0.97515055 | 0.99681784 |
| SENP1 | 63.9145426 | -0.303507031 | 0.24813776 | -1.2231393 | 0.2212771 | 0.78687916 |
| PFKM | 26.864556 | 0.324160589 | 0.30616248 | 1.05878615 | 0.28969718 | 0.83506029 |
| ASB8 | 33.8771442 | -0.111494558 | 0.26558874 | -0.4198015 | 0.67463045 | 0.95910677 |
| ZNF641 | 28.2261923 | -0.052435519 | 0.36985766 | -0.1417722 | 0.88725997 | 0.98766275 |
| KANSL2 | 39.7420357 | -0.339068697 | 0.30203782 | -1.1226035 | 0.26160595 | 0.81281978 |
| CCNT1 | 76.9340052 | -0.014217236 | 0.201993 | -0.0703848 | 0.94388739 | 0.9952642 |
| ADCY6 | 75.0491338 | 0.176748162 | 0.25132687 | 0.7032601 | 0.48189367 | 0.91971919 |
| DDX23 | 47.8371912 | -0.131336254 | 0.24183919 | -0.5430727 | 0.58707977 | 0.94308857 |
| CCDC65 | 55.0662616 | 0.136374005 | 0.34729394 | 0.39267603 | 0.69455877 | 0.96220115 |
| PRKAG1 | 61.7622572 | -0.448146237 | 0.23402318 | -1.9149652 | 0.05549694 | 0.53915335 |
| KMT2D | 57.2443977 | -0.383075424 | 0.34102235 | -1.1233147 | 0.26130386 | 0.81281978 |
| LMBR1L | 48.8466021 | 0.333298282 | 0.26972745 | 1.23568545 | 0.21657551 | 0.78319543 |
| TUBA1B | 222.379247 | -1.117004571 | 0.61411359 | -1.8188892 | 0.06892834 | 0.58378828 |
| TUBA1A | 46.1501814 | 0.598570073 | 0.77795843 | 0.76941139 | 0.44164913 | 0.90561284 |
| TUBA1C | 116.178321 | -0.656095461 | 0.61879223 | -1.0602839 | 0.28901545 | 0.83399936 |
| DNAJC22 | 49.1906134 | 1.363735505 | 0.35663305 | 3.82391793 | 0.00013135 | 0.02918429 |
| SPATS2 | 242.099863 | 0.086879817 | 0.28656988 | 0.30317149 | 0.76175918 | 0.9714717 |
| MCRS1 | 28.723857 | -0.386444789 | 0.30336036 | -1.2738803 | 0.20270584 | 0.77110248 |
| PRPF40B | 141.148557 | 0.161504386 | 0.27156318 | 0.59472123 | 0.55202983 | 0.93523904 |
| FMNL3 | 40.2411768 | 1.035125468 | 0.48932269 | 2.11542503 | 0.03439374 | 0.45193089 |
| TMBIM6 | 446.446697 | -0.168172643 | 0.37490965 | -0.4485685 | 0.65374299 | 0.95243345 |
| NCKAP5L | 46.6501566 | 0.104613804 | 0.30067676 | 0.3479278 | 0.72789441 | 0.96826837 |
| AQP2 | 45.4510232 | -0.18643896 | 0.65351321 | -0.2852872 | 0.77542411 | 0.97423208 |
| SMARCD1 | 49.8328962 | 0.122726304 | 0.40740251 | 0.30124091 | 0.76323079 | 0.97149197 |
| AC074032.1 | 142.72007 | 0.115240581 | 0.30083478 | 0.38306934 | 0.70166835 | 0.9645815 |
| CERS5 | 112.427763 | 0.293334083 | 0.21985842 | 1.33419534 | 0.18213985 | 0.75456023 |
| LIMA1 | 219.149655 | 0.056910799 | 0.20047789 | 0.28387569 | 0.77650564 | 0.97423208 |
| LARP4 | 222.355009 | 0.12125646 | 0.23025321 | 0.52662224 | 0.59845595 | 0.94683415 |
| DIP2B | 376.093059 | 0.020285417 | 0.31577598 | 0.0642399 | 0.94877921 | 0.9952642 |

| | | | | | | |
|---|---|---|---|---|---|---|
| ATF1 | 115.040285 | -0.183116269 | 0.2251821 | -0.8131919 | 0.41610802 | 0.89846075 |
| METTL7A | 111.99948 | 1.34296211 | 0.5252232 | 2.55693601 | 0.01055986 | 0.29838625 |
| SLC11A2 | 203.851349 | 0.269102452 | 0.17689077 | 1.52129169 | 0.12818665 | 0.69645198 |
| LETMD1 | 102.397516 | -0.127868591 | 0.19054665 | -0.6710619 | 0.50218112 | 0.92484445 |
| CSRNP2 | 34.8504201 | 0.153511955 | 0.30052713 | 0.51080898 | 0.60948482 | 0.94791646 |
| TFCP2 | 251.331455 | 0.327461198 | 0.23193388 | 1.41187309 | 0.15798733 | 0.73580079 |
| POU6F1 | 40.1550596 | -0.432390581 | 0.43259485 | -0.9995278 | 0.31753907 | 0.8490543 |
| DAZAP2 | 192.795131 | -0.139304197 | 0.24675623 | -0.5645418 | 0.57238549 | 0.93998433 |
| SMAGP | 25.7896132 | -0.328506319 | 0.54715697 | -0.6003877 | 0.54824788 | 0.93523904 |
| FIGNL2 | 67.6419236 | -0.30003522 | 0.53033861 | -0.5657427 | 0.57156868 | 0.93998433 |
| ACVR1B | 227.537797 | -0.459483209 | 0.32899521 | -1.3966258 | 0.16252612 | 0.74071637 |
| NR4A1 | 149.260451 | -0.262356685 | 1.22627294 | -0.2139464 | 8.31E-01 | 9.82E-01 |
| ATG101 | 27.7025529 | 0.376492753 | 0.52540973 | 0.71656982 | 0.47363957 | 0.91804805 |
| KRT7 | 251.526259 | -0.987817982 | 0.62200644 | -1.5881154 | 0.11226025 | 0.66401239 |
| KRT8 | 402.665819 | -0.656184003 | 0.52088042 | -1.2597594 | 2.08E-01 | 0.77435432 |
| KRT18 | 475.597721 | -1.081121434 | 0.59479532 | -1.8176361 | 6.91E-02 | 0.58408344 |
| EIF4B | 235.903745 | -0.058714079 | 0.29153098 | -0.2013991 | 0.84038651 | 0.98464453 |
| AC068888.1 | 84.8142592 | 0.007047001 | 0.21666985 | 0.03252414 | 0.97405406 | 0.99681784 |
| TNS2 | 87.4911216 | 0.881742391 | 0.65097868 | 1.35448735 | 0.17558095 | 0.74934286 |
| SPRYD3 | 25.538408 | -0.255369122 | 0.35155924 | -0.72639 | 0.46759971 | 0.91462428 |
| CSAD | 97.3899423 | 0.070596407 | 0.29548768 | 0.23891489 | 0.81117158 | 0.97930627 |
| ZNF740 | 25.7796037 | -0.11657472 | 0.29414606 | -0.3963158 | 0.6918721 | 0.96220115 |
| RARG | 40.32707 | -0.346173414 | 0.49317333 | -0.7019305 | 0.48272249 | 0.91971919 |
| PFDN5 | 196.618801 | 0.229799674 | 0.46593719 | 0.49319882 | 0.62187211 | 0.94808496 |
| SP1 | 130.749266 | 0.234799398 | 0.29720576 | 0.79002305 | 0.42951431 | 0.90333313 |
| PRR13 | 107.181001 | 0.069052774 | 0.394199 | 0.17517237 | 0.86094419 | 0.98658849 |
| PCBP2 | 317.351281 | 0.018796959 | 0.14800013 | 0.12700637 | 0.89893536 | 0.98948153 |
| MAP3K12 | 36.0661839 | 0.94034519 | 0.31835437 | 2.95376876 | 0.00313919 | 0.18154635 |
| ATP5MC2 | 237.961883 | -0.153677038 | 0.41634439 | -0.3691104 | 0.71204545 | 0.96587538 |
| CALCOCO1 | 74.507462 | 0.855875627 | 0.40659942 | 2.10496029 | 0.03529476 | 0.45625985 |
| HOXC4 | 51.2939408 | -0.237387354 | 0.38533114 | -0.6160607 | 0.53785449 | 0.93516669 |
| AC023794.1 | 29.4299471 | -0.515113603 | 0.55763655 | -0.9237443 | 0.35561944 | 0.87316144 |
| SMUG1 | 30.3704091 | 0.273048448 | 0.35252543 | 0.77454966 | 0.43860581 | 0.90561284 |
| LINC02381 | 63.1653517 | -0.46368579 | 0.58552895 | -0.7919093 | 0.42841359 | 0.90272191 |
| CBX5 | 215.197676 | -0.116220682 | 0.26572882 | -0.4373657 | 0.66184612 | 0.9545911 |
| HNRNPA1 | 179.395139 | -0.398084329 | 0.37833569 | -1.0521987 | 0.29270838 | 0.83697123 |
| COPZ1 | 101.952069 | -0.237967234 | 0.21412656 | -1.1113392 | 0.26642239 | 0.81444418 |
| PPP1R1A | 45.388496 | 0.71547693 | 0.53421324 | 1.33930961 | 0.1804699 | 0.75447995 |
| CD63 | 291.351817 | -0.343125093 | 0.54019905 | -0.6351827 | 0.52530927 | 0.93272384 |
| ORMDL2 | 32.2998208 | -0.209958129 | 0.41781903 | -0.5025097 | 0.61530901 | 0.94791646 |
| RAB5B | 85.1054406 | 0.073951301 | 0.23947552 | 0.30880526 | 0.75746966 | 0.9714717 |
| SUOX | 25.0896927 | 0.098557109 | 0.43687161 | 0.22559742 | 0.82151455 | 0.98094663 |
| RPS26 | 52.6461512 | -1.043430558 | 0.32803113 | -3.1808888 | 0.00146824 | 0.12148388 |
| ERBB3 | 45.0680194 | 0.863955623 | 0.4310514 | 2.0042984 | 0.04503811 | 0.50340725 |
| PA2G4 | 51.2015622 | -0.295192665 | 0.58467474 | -0.5048836 | 0.61364062 | 0.94791646 |
| RPL41 | 993.180066 | 0.159004839 | 0.47954841 | 0.33157203 | 0.74021244 | 0.9698508 |
| ESYT1 | 65.4053221 | -0.282271331 | 0.29347142 | -0.9618358 | 0.33613208 | 0.86391445 |
| MYL6B | 24.9231645 | 0.003447472 | 0.38772267 | 0.00889159 | 0.99290563 | 0.99866813 |
| MYL6 | 422.860956 | -0.131498099 | 0.49320636 | -0.2666188 | 0.78976266 | 0.97423208 |
| SMARCC2 | 151.590489 | -0.155023475 | 0.18082586 | -0.8573081 | 0.39127464 | 0.88670817 |
| RNF41 | 48.3705449 | -0.141610191 | 0.35924342 | -0.3941901 | 0.69344071 | 0.96220115 |
| ANKRD52 | 41.5291779 | 0.094962094 | 0.2620085 | 0.36243898 | 0.71702401 | 0.96742001 |
| CS | 58.3829778 | -0.079955568 | 0.34758346 | -0.2300327 | 0.81806635 | 0.98010849 |
| PAN2 | 49.0170401 | 0.033083688 | 0.28385789 | 0.11655018 | 0.90721652 | 0.99100813 |

| | | | | | | |
|---|---|---|---|---|---|---|
| STAT2 | 125.310221 | -0.473994336 | 0.57340454 | -0.8266316 | 0.40844588 | 0.8959746 |
| RBMS2 | 224.278652 | -0.375396745 | 0.27895513 | -1.3457245 | 1.78E-01 | 0.7513166 |
| BAZ2A | 152.953763 | -0.132194898 | 0.22044401 | -0.5996756 | 0.54872245 | 0.93523904 |
| ATP5F1B | 198.736726 | 0.098813179 | 0.34159175 | 0.28927273 | 0.77237268 | 0.97329019 |
| PTGES3 | 311.603619 | 0.051194275 | 0.15661891 | 0.3268716 | 0.743765 | 0.97012548 |
| NACA | 243.590474 | 0.100556598 | 0.55273641 | 0.18192505 | 0.85564155 | 0.9864199 |
| NEMP1 | 36.2666652 | -0.136712335 | 0.34324629 | -0.3982922 | 0.69041478 | 0.96220115 |
| NAB2 | 26.8407627 | -0.579011546 | 0.43954059 | -1.3173108 | 0.18773448 | 0.76207437 |
| STAT6 | 150.846927 | -0.141381928 | 0.23147605 | -0.6107843 | 5.41E-01 | 0.93523904 |
| LRP1 | 40.4850213 | 1.274317789 | 0.44667772 | 2.8528797 | 0.0043325 | 0.21066515 |
| R3HDM2 | 384.783013 | 0.101546624 | 0.24202629 | 0.41956857 | 0.67480065 | 0.95910677 |
| MARS | 44.556304 | -0.468958822 | 0.37531253 | -1.2495155 | 0.21147658 | 0.77730294 |
| DDIT3 | 47.7304201 | -0.566321538 | 0.66904619 | -0.846461 | 0.3972956 | 0.88863925 |
| MBD6 | 43.7855977 | -0.304830614 | 0.27479516 | -1.1093012 | 0.26730025 | 0.81510106 |
| DCTN2 | 77.828706 | 0.138364633 | 0.26271129 | 0.52667944 | 0.59841622 | 0.94683415 |
| PIP4K2C | 33.2709066 | -0.139621768 | 0.35399535 | -0.394417 | 0.69327323 | 0.96220115 |
| DTX3 | 35.6388438 | 0.279949598 | 0.3589971 | 0.77981019 | 0.43550261 | 0.90514499 |
| OS9 | 108.456318 | 0.286807239 | 0.23248596 | 1.23365402 | 0.21733185 | 0.78353439 |
| TSPAN31 | 36.7542708 | -0.027863422 | 0.26823614 | -0.1038765 | 0.91726738 | 0.99214128 |
| CDK4 | 38.5436722 | -0.640916016 | 0.39670779 | -1.6155872 | 0.1061836 | 0.65699665 |
| 9-Mar | 35.1427481 | -0.225657908 | 0.33453598 | -0.67454 | 0.49996805 | 0.92398825 |
| TSFM | 36.8454419 | 0.421498526 | 0.29626188 | 1.42272277 | 0.15481654 | 0.72978231 |
| CTDSP2 | 112.740492 | 0.08147974 | 0.26309173 | 0.30970088 | 0.75678843 | 0.9714717 |
| LRIG3 | 116.528097 | 0.190698844 | 0.45254033 | 0.42139635 | 0.67346568 | 0.95910677 |
| SLC16A7 | 250.954134 | -0.515824811 | 0.3998604 | -1.2900122 | 0.19704641 | 0.77110248 |
| FAM19A2 | 126.673846 | -0.018921649 | 0.49129449 | -0.0385139 | 0.96927798 | 0.99681784 |
| USP15 | 347.042997 | -0.310208656 | 0.22238535 | -1.394915 | 0.16304148 | 0.74167981 |
| MON2 | 497.627468 | 0.002044542 | 0.14399514 | 0.01419868 | 0.98867147 | 0.99820747 |
| AC048341.1 | 97.3976565 | -1.478897049 | 0.33148218 | -4.4614677 | 8.14E-06 | 0.00410386 |
| PPM1H | 542.609431 | -0.928424339 | 0.48381677 | -1.9189586 | 0.05498958 | 0.53804407 |
| DPY19L2 | 184.264938 | 0.300968562 | 0.42388562 | 0.71002305 | 0.47768984 | 0.91883353 |
| RXYLT1 | 40.4405551 | -0.387999729 | 0.3022839 | -1.2835607 | 0.19929572 | 0.77110248 |
| SRGAP1 | 604.611057 | 0.469095769 | 0.58501806 | 0.80184836 | 0.42264068 | 0.90066187 |
| XPOT | 62.3122936 | -0.106970876 | 0.34730185 | -0.3080055 | 0.75807816 | 0.9714717 |
| TBK1 | 141.998013 | -0.024700183 | 0.2429791 | -0.1016556 | 0.91903006 | 0.99214128 |
| RASSF3 | 51.6186397 | -0.38942425 | 0.50742733 | -0.7674483 | 0.44281501 | 0.90615787 |
| GNS | 159.861314 | 0.139636417 | 0.27680724 | 0.50445363 | 0.61394264 | 0.94791646 |
| LEMD3 | 176.095626 | 0.020336613 | 0.17078009 | 0.1190807 | 0.90521142 | 0.99092254 |
| MSRB3 | 118.804932 | 0.618741182 | 0.75027497 | 0.82468589 | 0.40954994 | 0.8959746 |
| HMGA2 | 82.4490501 | -1.140265827 | 0.45618018 | -2.4995953 | 0.01243353 | 0.31015819 |
| AC090673.1 | 166.318643 | -0.227309866 | 0.57815359 | -0.3931652 | 0.69419748 | 0.96220115 |
| HELB | 57.3612579 | 0.153842174 | 0.43885819 | 0.35055099 | 0.72592523 | 0.96781837 |
| GRIP1 | 1109.13472 | 0.412575655 | 0.28084592 | 1.46904626 | 0.14182024 | 0.7148274 |
| CAND1 | 199.683837 | -0.072490805 | 0.22919051 | -0.3162906 | 0.75178195 | 0.97058015 |
| DYRK2 | 43.6895522 | 0.081183639 | 0.29969353 | 0.27088887 | 0.78647651 | 0.97423208 |
| MDM1 | 31.6865984 | 0.20990858 | 0.31060034 | 0.67581568 | 4.99E-01 | 0.92398825 |
| RAP1B | 428.404736 | 0.207561325 | 0.31492068 | 0.6590908 | 0.50983746 | 0.92819651 |
| NUP107 | 102.850919 | -0.1038117 | 0.2988674 | -0.3473504 | 0.72832812 | 0.96829915 |
| SLC35E3 | 64.5401746 | -0.207611363 | 0.36108883 | -0.5749592 | 0.56531886 | 0.93809297 |
| MDM2 | 192.090753 | 0.198771865 | 0.35999986 | 0.55214429 | 0.5808495 | 0.94041171 |
| CPM | 342.85712 | -0.514078675 | 0.73265137 | -0.7016689 | 0.48288568 | 0.91971919 |
| CPSF6 | 309.058034 | -0.300762921 | 0.27083369 | -1.1105078 | 0.26678028 | 0.81462808 |
| FRS2 | 208.226144 | 0.092103754 | 0.14268845 | 0.6454885 | 0.51861067 | 0.93257822 |
| CCT2 | 89.5571373 | -0.084824388 | 0.3120684 | -0.2718135 | 0.78576546 | 0.97423208 |

| | | | | | | |
|---|---|---|---|---|---|---|
| RAB3IP | 193.75061 | -0.035203763 | 0.34274436 | -0.1027114 | 0.918192 | 0.99214128 |
| MYRFL | 52.100092 | 0.884600953 | 1.16889588 | 0.75678336 | 0.44917966 | 0.90994378 |
| AC025159.1 | 165.491681 | -0.084042997 | 0.52158612 | -0.1611297 | 0.87199128 | 0.98766275 |
| CNOT2 | 326.931609 | 0.171853953 | 0.17337793 | 0.99121006 | 0.32158302 | 0.85142171 |
| ZFC3H1 | 232.230045 | 0.029694696 | 0.16675212 | 0.17807687 | 0.85866261 | 0.9864199 |
| RAB21 | 232.408571 | -0.142729928 | 0.20315497 | -0.7025668 | 0.48232578 | 0.91971919 |
| TBC1D15 | 190.561114 | -0.032794204 | 0.21270759 | -0.1541751 | 0.87747171 | 0.98766275 |
| TRHDE | 598.267819 | -0.245842357 | 0.41775108 | -0.5884901 | 0.5562034 | 0.93523904 |
| ATXN7L3B | 89.6664228 | -0.177868642 | 0.29294015 | -0.6071842 | 0.54372865 | 0.93523904 |
| CAPS2 | 158.445195 | 0.683519225 | 0.40589929 | 1.68396259 | 0.0921889 | 0.63464873 |
| KRR1 | 67.6914353 | -0.516559863 | 0.24726759 | -2.0890723 | 0.03670122 | 0.46564369 |
| PHLDA1 | 49.0493832 | 0.114375436 | 0.80326825 | 0.1423876 | 0.88677386 | 0.98766275 |
| NAP1L1 | 255.483127 | -0.013131262 | 0.19729234 | -0.0665574 | 0.94693407 | 0.9952642 |
| OSBPL8 | 284.337709 | -0.089693868 | 0.18748669 | -0.4784013 | 0.63236464 | 0.94932781 |
| ZDHHC17 | 189.632951 | 0.067216272 | 0.17395507 | 0.38640019 | 0.69920031 | 0.96280283 |
| NAV3 | 110.206614 | -0.283814308 | 1.09190175 | -0.2599266 | 0.79492039 | 0.97524554 |
| AC027288.3 | 152.32949 | 1.277862198 | 1.11740905 | 1.14359392 | 0.25279208 | 0.80798674 |
| PAWR | 951.408916 | 0.541750104 | 0.62495342 | 0.86686476 | 0.38601611 | 0.88353166 |
| AC073569.2 | 87.6260003 | 2.035037332 | 0.84401544 | 2.41113758 | 0.01590285 | 0.34086104 |
| PPP1R12A | 410.025472 | 0.191943626 | 0.24962533 | 0.76892688 | 0.44193672 | 0.90561284 |
| LIN7A | 66.5621369 | 2.242224074 | 0.76825507 | 2.91859327 | 0.00351615 | 0.18995949 |
| ACSS3 | 175.84158 | 1.191550915 | 0.62317264 | 1.91207192 | 0.05586697 | 0.54110182 |
| PPFIA2 | 70.1772916 | 3.047201613 | 1.48609242 | 2.05047921 | 0.04031769 | 0.48457109 |
| CCDC59 | 47.0702474 | -0.433788412 | 0.32993318 | -1.3147765 | 0.18858501 | 0.76207437 |
| METTL25 | 119.856188 | 0.119988322 | 0.19864917 | 0.60402126 | 5.46E-01 | 0.93523904 |
| TMTC2 | 519.927009 | -0.345778349 | 0.36724703 | -0.9415416 | 0.34642739 | 0.86990996 |
| LRRIQ1 | 218.95416 | 0.465183104 | 0.32189886 | 1.44512194 | 0.14842364 | 0.72246729 |
| MGAT4C | 176.45495 | 2.578769413 | 0.8479217 | 3.04128249 | 0.00235573 | 0.15891203 |
| CEP290 | 208.727797 | -0.249711307 | 0.20955381 | -1.1916333 | 0.23340505 | 0.79622042 |
| TMTC3 | 99.9006695 | -0.232919927 | 0.32085915 | -0.7259258 | 0.46788428 | 0.91462428 |
| KITLG | 193.91501 | -1.028777781 | 0.62273424 | -1.6520334 | 0.09852774 | 0.64312419 |
| DUSP6 | 100.697008 | -0.354317311 | 0.58443168 | -0.6062596 | 0.54434239 | 0.93523904 |
| POC1B | 109.601094 | 0.248252226 | 0.27790255 | 0.89330676 | 0.37169292 | 0.87605028 |
| POC1B-AS1 | 58.840572 | 0.332837001 | 0.27484295 | 1.21100796 | 0.22589235 | 0.79029323 |
| AC025034.1 | 31.8015094 | 0.100538867 | 0.42341592 | 0.23744706 | 0.81230998 | 0.97930627 |
| ATP2B1 | 269.879019 | 0.31446749 | 0.27950351 | 1.12509316 | 0.26054956 | 0.81263292 |
| BTG1 | 235.242324 | 0.571834868 | 0.50648959 | 1.12901603 | 0.25889107 | 0.81263292 |
| EEA1 | 209.499406 | 0.035813339 | 0.21537544 | 0.1662833 | 0.86793401 | 0.98670067 |
| AC138123.1 | 169.105065 | 1.178648305 | 0.5094428 | 2.31360284 | 0.02068952 | 0.3802773 |
| NUDT4 | 108.722947 | -0.075374803 | 0.38727895 | -0.1946266 | 0.84568524 | 0.98520136 |
| UBE2N | 100.480877 | -0.282428316 | 0.26515429 | -1.0651471 | 0.28680938 | 0.83122432 |
| MRPL42 | 94.6473388 | -0.359453531 | 0.25319264 | -1.419684 | 0.15569971 | 0.7314603 |
| CRADD | 392.055056 | 0.100722347 | 0.20007062 | 0.50343398 | 0.61465919 | 0.94791646 |
| AC012085.2 | 34.0869885 | 1.064822908 | 0.62823115 | 1.69495402 | 0.09008414 | 0.6330877 |
| CEP83 | 146.99096 | -0.241501666 | 0.19326817 | -1.2495677 | 0.21145751 | 0.77730294 |
| TMCC3 | 55.672659 | -0.316356284 | 0.44318777 | -0.71382 | 4.75E-01 | 0.91809716 |
| NDUFA12 | 68.3751382 | -0.169826232 | 0.35368334 | -0.4801646 | 0.63111034 | 0.94932781 |
| NR2C1 | 117.884144 | 0.208198142 | 0.1819162 | 1.14447277 | 0.25242763 | 0.80798674 |
| FGD6 | 115.3003 | -0.345806957 | 0.44578879 | -0.7757193 | 0.43791476 | 0.90561284 |
| VEZT | 343.292583 | -0.154877748 | 0.13285652 | -1.1657519 | 0.24371476 | 0.80476923 |
| METAP2 | 142.86778 | -0.15515382 | 0.17708184 | -0.8761702 | 0.38093754 | 0.8807629 |
| NTN4 | 913.045143 | 0.249162989 | 0.38807877 | 0.64204231 | 5.21E-01 | 9.33E-01 |
| SNRPF | 43.4460038 | -0.766411771 | 0.32993668 | -2.3229056 | 0.02018422 | 0.37762629 |
| LTA4H | 66.5645618 | 0.458855666 | 0.22811683 | 2.01149414 | 0.04427329 | 0.49952159 |

| | | | | | | |
|---|---|---|---|---|---|---|
| ELK3 | 53.9889723 | -1.014594222 | 0.65371569 | -1.552042 | 0.12065217 | 0.68916345 |
| CDK17 | 199.549593 | -0.094612827 | 0.25673571 | -0.3685223 | 0.71248384 | 0.96587538 |
| CFAP54 | 215.817235 | 0.896190742 | 0.69550129 | 1.28855368 | 0.19755329 | 0.77110248 |
| NEDD1 | 62.763361 | -0.065529413 | 0.29809885 | -0.2198244 | 0.82600788 | 0.98094837 |
| TMPO | 151.214438 | 0.259455904 | 0.3423378 | 0.7578944 | 0.4485142 | 0.90994378 |
| SLC25A3 | 199.526842 | 0.229970518 | 0.48124503 | 0.47786575 | 0.63274576 | 0.94932781 |
| IKBIP | 44.7671715 | 0.132821316 | 0.5529674 | 0.24019737 | 0.81017725 | 0.97930627 |
| APAF1 | 75.3944685 | 0.186066792 | 0.25487245 | 0.73003886 | 0.46536643 | 0.91462428 |
| ANKS1B | 113.751905 | 0.843417782 | 0.7521024 | 1.12141349 | 0.2621119 | 0.81360008 |
| UHRF1BP1L | 118.2296 | 0.154461138 | 0.20414697 | 0.75661734 | 0.44927915 | 0.90994378 |
| ACTR6 | 47.8283865 | 0.288417962 | 0.27508317 | 1.04847551 | 0.29441958 | 0.83885935 |
| SCYL2 | 153.49749 | 0.205183572 | 0.17916104 | 1.14524657 | 0.25210703 | 0.80798674 |
| NR1H4 | 377.091697 | 0.363256154 | 0.35698836 | 1.01755741 | 0.30888834 | 0.84554063 |
| SLC5A8 | 56.9172342 | 1.442208412 | 0.62423169 | 2.31037358 | 0.02086748 | 0.38104604 |
| UTP20 | 85.3090073 | -0.707757725 | 0.32578691 | -2.172456 | 0.02982128 | 0.43492739 |
| ARL1 | 87.070883 | -0.007151108 | 0.24572856 | -0.0291017 | 0.97678351 | 0.99681784 |
| CHPT1 | 133.285942 | 0.978664235 | 0.3187265 | 3.07054552 | 0.00213668 | 0.15271309 |
| GNPTAB | 99.3823889 | -0.165716858 | 0.21182528 | -0.782328 | 0.43402181 | 0.90514499 |
| DRAM1 | 175.202205 | -0.224557574 | 0.23156362 | -0.9697446 | 0.3321738 | 0.85980141 |
| WASHC3 | 93.1607399 | 0.128189168 | 0.31687444 | 0.40454246 | 0.68581386 | 0.96161098 |
| NUP37 | 29.932365 | -0.403146037 | 0.29770309 | -1.3541883 | 0.17567632 | 0.74934286 |
| HSP90B1 | 338.406943 | 0.019567744 | 0.52904297 | 0.03698706 | 0.97049532 | 0.99681784 |
| C12orf73 | 27.1372244 | 0.161387766 | 0.36471808 | 0.44250004 | 0.6581274 | 0.95399905 |
| TDG | 70.5377227 | -0.215940385 | 0.23303098 | -0.9266596 | 0.35410331 | 0.87241657 |
| HCFC2 | 54.8833757 | -0.043241117 | 0.27989649 | -0.1544897 | 0.87722366 | 0.98766275 |
| NFYB | 40.9031417 | 0.137210871 | 0.2889633 | 0.4748384 | 0.63490216 | 0.94932781 |
| TXNRD1 | 865.203359 | -0.112045722 | 0.35641342 | -0.3143701 | 0.75323997 | 0.97096495 |
| CHST11 | 79.969549 | 0.523756828 | 0.63406657 | 0.82602814 | 4.09E-01 | 8.96E-01 |
| SLC41A2 | 142.226975 | 0.512914405 | 0.30673028 | 1.67220011 | 0.09448487 | 0.63858274 |
| C12orf45 | 36.7004242 | -0.902027912 | 0.37813174 | -2.3854859 | 0.01705658 | 0.35074796 |
| WASHC4 | 312.686106 | 0.062547283 | 0.26595296 | 0.23518175 | 0.81406767 | 0.97937868 |
| APPL2 | 153.214451 | 0.327022182 | 0.23140378 | 1.41321018 | 0.15759393 | 0.73580079 |
| C12orf75 | 133.783921 | 0.832287595 | 0.28893039 | 2.88058165 | 0.00396942 | 0.2001215 |
| NUAK1 | 68.7008784 | -0.091842839 | 0.41540188 | -0.2210939 | 0.82501929 | 0.98094663 |
| CKAP4 | 48.2263594 | 0.424439008 | 0.5449608 | 0.7788432 | 0.43607209 | 0.90514499 |
| TCP11L2 | 65.0152926 | 0.680896336 | 0.52098148 | 1.30694921 | 0.19122996 | 0.76328615 |
| POLR3B | 64.9643174 | -0.535903664 | 0.3299663 | -1.6241163 | 0.10435098 | 0.65551333 |
| RIC8B | 262.047265 | 1.032928029 | 0.66707245 | 1.54844954 | 0.1215141 | 0.69161235 |
| TMEM263 | 115.422202 | 0.418909596 | 0.31014229 | 1.35070131 | 0.17679113 | 0.75098991 |
| MTERF2 | 42.7873082 | 0.816063306 | 0.43889914 | 1.85934131 | 0.06297878 | 0.56713487 |
| CRY1 | 103.032137 | -0.602385836 | 0.4078151 | -1.4771053 | 0.13964742 | 0.71191199 |
| BTBD11 | 55.3564404 | -3.558370307 | 0.81858604 | -4.3469716 | 1.38E-05 | 0.00600995 |
| PWP1 | 56.3178121 | -0.250021033 | 0.33447132 | -0.7475111 | 0.45475509 | 0.91077355 |
| PRDM4 | 62.3222782 | -0.376335747 | 0.27743085 | -1.3565029 | 0.17493922 | 0.74910274 |
| SART3 | 65.2284721 | -0.077667676 | 0.20782661 | -0.3737138 | 0.70861724 | 0.96587538 |
| ISCU | 131.017114 | 0.592946689 | 0.51088076 | 1.16063618 | 0.24578989 | 0.80488052 |
| CORO1C | 97.9476703 | 0.442175167 | 0.50911918 | 0.86851013 | 0.38511513 | 0.88353166 |
| SSH1 | 104.87423 | 0.517823233 | 0.39482432 | 1.31152821 | 0.18967937 | 0.76207437 |
| USP30 | 35.2434862 | 0.823629019 | 0.28775295 | 2.86227831 | 0.00420607 | 0.20768032 |
| ACACB | 142.270252 | 0.677293515 | 0.4932377 | 1.37315845 | 0.16970309 | 0.74445571 |
| KCTD10 | 39.887408 | -0.356140703 | 0.30013375 | -1.1866066 | 0.23538281 | 0.79926692 |
| UBE3B | 74.26012 | 0.061914427 | 0.22797963 | 0.27157877 | 0.78594593 | 0.97423208 |
| MMAB | 74.4975748 | -0.272917608 | 0.27517023 | -0.9918137 | 0.32128841 | 0.85131972 |
| MVK | 45.6576195 | -0.428390794 | 0.40265248 | -1.0639219 | 0.28736408 | 0.83136832 |

| | | | | | | |
|---|---|---|---|---|---|---|
| TRPV4 | 33.2816978 | 2.000601172 | 0.4743174 | 4.21785318 | 2.47E-05 | 0.00945023 |
| GLTP | 75.3843104 | -0.158905016 | 0.43323387 | -0.3667881 | 0.71377711 | 0.96611089 |
| TCHP | 57.8319082 | 0.091287194 | 0.30537219 | 0.29893748 | 0.76498775 | 0.97177982 |
| GIT2 | 116.701133 | -0.027726757 | 0.24765922 | -0.1119553 | 0.91085887 | 0.99100813 |
| ANKRD13A | 140.476065 | 0.098152113 | 0.3609456 | 0.27193049 | 0.78567547 | 0.97423208 |
| C12orf76 | 35.3606873 | -0.008816132 | 0.29048066 | -0.0303502 | 0.9757878 | 0.99681784 |
| IFT81 | 127.691508 | 0.274077352 | 0.25594052 | 1.07086346 | 0.28423083 | 0.82918183 |
| ATP2A2 | 280.042756 | -0.182346847 | 0.26162373 | -0.6969813 | 0.48581451 | 0.92034617 |
| ANAPC7 | 89.2979577 | -0.181236237 | 0.3902977 | -0.4643538 | 0.64239425 | 0.94944486 |
| ARPC3 | 221.845684 | -0.04935542 | 0.37681721 | -0.1309797 | 0.89579133 | 0.98842187 |
| GPN3 | 27.1799425 | -0.135853921 | 0.51535443 | -0.2636126 | 0.79207845 | 0.97423208 |
| VPS29 | 104.635421 | 0.025768305 | 0.20960474 | 0.1229376 | 0.90215651 | 0.9900784 |
| PPTC7 | 74.710656 | 0.017471677 | 0.30609925 | 0.05707847 | 0.95448269 | 0.99531843 |
| TCTN1 | 78.4210194 | 0.646045827 | 0.23750958 | 2.72008324 | 0.00652655 | 0.25310853 |
| PPP1CC | 64.684023 | -0.058943061 | 0.31977969 | -0.184324 | 0.85375931 | 0.9864199 |
| SH2B3 | 48.232105 | -0.020011624 | 0.66053172 | -0.0302962 | 0.9758308 | 0.99681784 |
| ATXN2 | 506.599524 | 0.043575707 | 0.2172539 | 0.20057503 | 0.84103089 | 0.98511065 |
| BRAP | 63.9814577 | -0.080220984 | 0.2367772 | -0.3388037 | 0.73475764 | 0.9698508 |
| ACAD10 | 63.4361314 | 0.13231965 | 0.3477511 | 0.38050102 | 0.70357354 | 0.9645815 |
| ALDH2 | 88.7639344 | 0.940640509 | 0.34553456 | 2.72227622 | 0.00648339 | 0.25245699 |
| MAPKAPK5 | 85.8545884 | -0.272175582 | 0.24540323 | -1.1090953 | 0.26738905 | 0.81510106 |
| TMEM116 | 87.7573506 | -0.56404006 | 0.56577376 | -0.9969357 | 0.31879572 | 0.84935263 |
| ERP29 | 123.585589 | 0.698782624 | 0.34934137 | 2.00028593 | 0.0454694 | 0.50469451 |
| NAA25 | 162.490332 | -0.479984736 | 0.25830188 | -1.8582317 | 0.06313612 | 0.56733665 |
| TRAFD1 | 60.5150998 | 0.333496494 | 0.28295407 | 1.17862414 | 0.23854787 | 0.80174646 |
| HECTD4 | 406.613585 | -0.109402685 | 0.1686905 | -0.6485409 | 0.51663519 | 0.9320507 |
| RPL6 | 431.790188 | 0.122400151 | 0.6349064 | 0.19278456 | 0.84712769 | 0.98533374 |
| PTPN11 | 159.730582 | -0.412916748 | 0.27913744 | -1.4792596 | 0.13907094 | 0.71184729 |
| RASAL1 | 116.430673 | -0.463723076 | 0.39860955 | -1.1633517 | 0.24468686 | 0.80476923 |
| DDX54 | 50.0595277 | -0.695758407 | 0.43704478 | -1.5919614 | 0.11139339 | 0.66300165 |
| TPCN1 | 268.845716 | 0.617648246 | 0.28132032 | 2.19553371 | 0.02812534 | 0.4250936 |
| SLC8B1 | 39.6948663 | 0.071777826 | 0.29014728 | 0.24738411 | 0.80461097 | 0.97930627 |
| PLBD2 | 34.211816 | 0.029959437 | 0.26501394 | 0.11304853 | 0.90999208 | 0.99100813 |
| SDSL | 39.5865182 | -0.431770442 | 0.52761805 | -0.818339 | 0.41316363 | 0.89724628 |
| RBM19 | 69.9091212 | -0.771072829 | 0.25985924 | -2.9672711 | 0.00300456 | 0.17876193 |
| MED13L | 1964.73439 | 0.149544445 | 0.20520987 | 0.72873904 | 0.46616131 | 0.91462428 |
| C12orf49 | 80.8153722 | -0.090752898 | 0.31494714 | -0.2881577 | 0.773226 | 0.9735452 |
| FBXW8 | 133.053164 | -0.11674797 | 0.23149007 | -0.5043325 | 0.61402773 | 0.94791646 |
| TESC | 45.6739715 | 0.145334618 | 0.55370064 | 0.26247869 | 0.79295241 | 0.97443119 |
| FBXO21 | 150.310207 | 0.142499795 | 0.33141287 | 0.42997665 | 0.66721263 | 0.95691417 |
| KSR2 | 34.1618508 | -0.192240216 | 0.85675929 | -0.2243807 | 0.82246112 | 0.98094663 |
| WSB2 | 63.5856128 | -0.848256237 | 0.21452771 | -3.9540637 | 7.68E-05 | 0.02080127 |
| VSIG10 | 83.4735904 | -0.064004279 | 0.57615767 | -0.1110881 | 0.91154646 | 0.99100813 |
| PEBP1 | 663.156954 | 0.447193216 | 0.62100258 | 0.72011491 | 0.47145425 | 0.91640833 |
| TAOK3 | 355.592256 | -0.528464449 | 0.34399585 | -1.5362524 | 0.12447649 | 0.69439514 |
| SUDS3 | 88.1577173 | 0.143054123 | 0.22891217 | 0.62493019 | 0.53201688 | 0.93366189 |
| HSPB8 | 94.5451885 | 0.643589082 | 0.57472508 | 1.11982077 | 0.26279015 | 0.81377303 |
| AC002070.1 | 40.8859165 | 0.864492781 | 0.4783637 | 1.80718726 | 0.07073309 | 0.58815298 |
| PRKAB1 | 99.4883023 | 0.105627302 | 0.44630145 | 0.23667255 | 0.81291083 | 0.97930627 |
| CIT | 124.046256 | 0.64538302 | 0.58999164 | 1.09388503 | 0.27400541 | 0.82047446 |
| BICDL1 | 365.62744 | -0.491567186 | 0.4193478 | -1.1722183 | 0.24110941 | 0.80319133 |
| RAB35 | 29.9191389 | -0.234712988 | 0.38400519 | -0.6112235 | 0.54105164 | 0.93523904 |
| GCN1 | 83.1447447 | -0.356730052 | 0.24441593 | -1.4595205 | 0.14442191 | 0.71634214 |
| RPLP0 | 611.77128 | 0.155525905 | 0.57701315 | 0.26953615 | 0.78751713 | 0.97423208 |

| | | | | | | |
|---|---|---|---|---|---|---|
| PXN | 97.7060667 | -0.785912342 | 0.29685863 | -2.6474296 | 0.00811062 | 0.27246484 |
| GATC | 33.5028757 | -0.066915934 | 0.39273948 | -0.1703825 | 0.86470933 | 0.98670067 |
| SRSF9 | 121.757811 | -0.377258516 | 0.4474335 | -0.8431611 | 0.39913836 | 0.88955057 |
| DYNLL1 | 210.868856 | 0.744239584 | 0.46788528 | 1.59064542 | 0.1116894 | 0.66300165 |
| COQ5 | 36.0320529 | 0.45769823 | 0.32431066 | 1.41129566 | 0.15815745 | 0.73580079 |
| RNF10 | 140.216441 | 0.101089489 | 0.15768886 | 0.6410693 | 0.52147766 | 0.93272217 |
| POP5 | 34.9729782 | -0.215042655 | 0.37610823 | -0.5717574 | 0.56748636 | 0.93885178 |
| MLEC | 176.442022 | 0.190568267 | 0.26986529 | 0.70616071 | 4.80E-01 | 0.91971919 |
| UNC119B | 44.367204 | 0.405277941 | 0.30319968 | 1.33667007 | 0.18133036 | 0.75448394 |
| SPPL3 | 477.532329 | -0.083525799 | 0.18099811 | -0.4614733 | 0.64445905 | 0.94944486 |
| P2RX4 | 35.0192943 | 0.147601804 | 0.54640913 | 0.27013056 | 0.78705981 | 0.97423208 |
| CAMKK2 | 52.1672505 | -0.071378242 | 0.36980658 | -0.1930151 | 0.84694717 | 0.98533374 |
| ANAPC5 | 149.985564 | -0.302489355 | 0.20348064 | -1.4865756 | 0.13712693 | 0.71044919 |
| RNF34 | 57.0688518 | -0.468132471 | 0.26431707 | -1.7711019 | 0.07654375 | 0.59726395 |
| KDM2B | 69.1377399 | 0.218876951 | 0.45473183 | 0.48133193 | 0.63028061 | 0.94916079 |
| TMEM120B | 93.0315264 | -0.213569994 | 0.28181082 | -0.7578488 | 0.4485415 | 0.90994378 |
| RHOF | 25.3415703 | -0.113761483 | 0.58484416 | -0.1945159 | 0.84577195 | 0.98520136 |
| SETD1B | 58.3125183 | -0.116201736 | 0.2551711 | -0.4553875 | 0.64883047 | 0.95027667 |
| WDR66 | 143.042956 | 0.552695992 | 0.39406343 | 1.40255589 | 0.16074931 | 0.73816762 |
| BCL7A | 135.764561 | -0.022935636 | 0.36236107 | -0.063295 | 0.94953161 | 0.99531843 |
| MLXIP | 279.162104 | 0.166693567 | 0.32084037 | 0.51955297 | 0.60337519 | 0.94683415 |
| CLIP1 | 270.383143 | -0.315947013 | 0.29533783 | -1.0697817 | 0.28471758 | 0.82918183 |
| ZCCHC8 | 92.2927474 | -0.355966264 | 0.19669239 | -1.8097613 | 0.07033282 | 0.58797037 |
| RSRC2 | 152.153931 | -0.173540621 | 0.21620413 | -0.8026702 | 0.42216536 | 0.90066187 |
| DENR | 149.244275 | -0.216707296 | 0.23985001 | -0.9035117 | 0.36625437 | 0.87605028 |
| HIP1R | 162.026287 | 0.244061923 | 0.380844 | 0.64084487 | 0.52162348 | 0.93272217 |
| VPS37B | 154.231142 | -0.345578738 | 0.4316728 | -0.8005571 | 0.42338808 | 0.90165283 |
| ARL6IP4 | 132.001978 | -0.261978081 | 0.40824042 | -0.641725 | 0.52105176 | 0.93272217 |
| PITPNM2 | 154.555758 | -0.423461892 | 0.30641825 | -1.3819735 | 0.16697984 | 0.74445571 |
| MPHOSPH9 | 111.295361 | 0.232357846 | 0.29522615 | 0.78705035 | 0.43125239 | 0.90452521 |
| C12orf65 | 56.4438122 | 0.341505578 | 0.27281991 | 1.25176193 | 0.21065663 | 0.77730294 |
| SBNO1 | 168.916894 | -0.012094845 | 0.18729601 | -0.0645761 | 0.94851152 | 0.9952642 |
| KMT5A | 42.4789528 | -0.428080583 | 0.24509376 | -1.7465993 | 0.08070687 | 0.60729857 |
| RILPL2 | 61.1578042 | 0.019374887 | 0.25232904 | 0.07678421 | 0.93879521 | 0.99526299 |
| SNRNP35 | 62.6242016 | -0.058894979 | 0.26285599 | -0.224058 | 0.8227122 | 0.98094663 |
| RILPL1 | 163.037636 | -0.223335083 | 0.23418755 | -0.9536591 | 0.34025624 | 0.86777921 |
| TMED2 | 82.8989028 | 0.001422549 | 0.30647097 | 0.00464171 | 0.99629647 | 0.99890348 |
| DDX55 | 41.1795798 | -0.441192803 | 0.33035285 | -1.3355199 | 0.18170625 | 0.75448394 |
| EIF2B1 | 43.018495 | -0.214467427 | 0.27391619 | -0.7829673 | 0.43364629 | 0.90514499 |
| GTF2H3 | 46.2983982 | -0.443237483 | 0.26316084 | -1.6842836 | 0.09212688 | 0.63464873 |
| TCTN2 | 44.7520723 | 0.323141771 | 0.35015351 | 0.92285743 | 0.35608151 | 0.87316144 |
| ATP6V0A2 | 64.9087031 | -0.570315461 | 0.44257318 | -1.2886354 | 0.19752488 | 0.77110248 |
| CCDC92 | 151.65486 | 0.196423517 | 0.16289427 | 1.20583443 | 0.22788134 | 0.79284833 |
| NCOR2 | 289.870051 | -0.150038462 | 0.35078139 | -0.4277264 | 0.66885033 | 0.95764829 |
| SCARB1 | 42.8176702 | -0.15352603 | 0.33134894 | -0.4633364 | 0.64312324 | 0.94944486 |
| UBC | 633.946794 | -0.346220768 | 0.31679153 | -1.0928978 | 0.27443867 | 0.82102033 |
| DHX37 | 27.5089705 | -0.764966047 | 0.37095393 | -2.0621592 | 0.03919259 | 0.47885941 |
| BRI3BP | 26.6727951 | -0.412474098 | 0.39705913 | -1.0388228 | 0.29888714 | 0.83989343 |
| AACS | 48.924009 | 0.459107825 | 0.27798671 | 1.65154595 | 0.09862715 | 0.64312419 |
| SLC15A4 | 95.5315774 | -0.271797425 | 0.34000718 | -0.7993873 | 0.42406587 | 0.90167091 |
| TMEM132D | 235.357034 | -1.041477433 | 0.53898606 | -1.9322901 | 0.05332371 | 0.53386172 |
| RIMBP2 | 38.4102147 | -0.970849976 | 0.7333131 | -1.3239229 | 0.18552866 | 0.76046631 |
| STX2 | 129.755134 | 0.028641903 | 0.28812901 | 0.09940652 | 9.21E-01 | 0.99214128 |
| RAN | 147.550471 | -0.135835078 | 0.52377841 | -0.2593369 | 0.7953753 | 0.97524554 |

| | | | | | | |
|---|---|---|---|---|---|---|
| SFSWAP | 164.903169 | -0.347901391 | 0.1575975 | -2.2075311 | 0.02727697 | 0.42391395 |
| ULK1 | 68.6449813 | 0.308693222 | 0.26217164 | 1.17744703 | 0.23901713 | 0.80174646 |
| PUS1 | 31.3014486 | -0.978485915 | 0.42865425 | -2.2826927 | 0.02244848 | 0.39239777 |
| EP400 | 166.036038 | -0.137754007 | 0.20146182 | -0.6837723 | 0.49411898 | 0.92289011 |
| DDX51 | 25.7734675 | -0.540907613 | 0.30155387 | -1.7937346 | 0.07285553 | 0.59192805 |
| FBRSL1 | 75.6015767 | -0.186399777 | 0.26470343 | -0.7041835 | 0.48131853 | 0.91971919 |
| ANKLE2 | 161.702147 | -0.471641917 | 0.34330237 | -1.3738382 | 0.16949191 | 0.74445571 |
| GOLGA3 | 103.504954 | -0.283872549 | 0.30945696 | -0.9173248 | 0.35897245 | 0.87316144 |
| CHFR | 105.488131 | -0.139880753 | 0.23555481 | -0.5938353 | 0.55262231 | 0.93523904 |
| ZNF26 | 68.8765959 | -0.169407724 | 0.23855531 | -0.7101402 | 0.47761718 | 0.91883353 |
| ZNF84 | 104.208635 | 0.048109196 | 0.31418558 | 0.1531235 | 0.87830088 | 0.98766275 |
| ZNF140 | 47.7976323 | 0.093038957 | 0.32124813 | 0.28961712 | 0.77210917 | 0.97329019 |
| ZNF891 | 54.7695043 | 0.14436859 | 0.255726 | 0.56454404 | 5.72E-01 | 0.93998433 |
| MPHOSPH8 | 211.282511 | 0.283101158 | 0.23628184 | 1.19815028 | 0.23085852 | 0.79403724 |
| PSPC1 | 341.784095 | -0.035827324 | 0.23163002 | -0.1546748 | 0.87707771 | 0.98766275 |
| ZMYM5 | 104.955619 | 0.319568002 | 0.25084154 | 1.27398358 | 0.20266923 | 0.77110248 |
| ZMYM2 | 641.95888 | 0.07420278 | 0.20311891 | 0.36531695 | 0.71487483 | 0.96671397 |
| CRYL1 | 171.182496 | 0.493271125 | 0.25654932 | 1.92271461 | 0.0545159 | 0.53752502 |
| IFT88 | 410.099862 | 0.160359801 | 0.25492281 | 0.62905238 | 0.52931476 | 0.93366189 |
| EEF1AKMT1 | 39.6869878 | -0.298133108 | 0.2648681 | -1.1255908 | 0.26033875 | 0.81263292 |
| XPO4 | 196.536671 | -0.230418913 | 0.20915803 | -1.1016498 | 0.27061393 | 0.81775732 |
| LATS2 | 196.831584 | -0.097516945 | 0.31858082 | -0.306098 | 0.75953005 | 0.9714717 |
| SAP18 | 137.208559 | -0.302037836 | 0.4924238 | -0.6133697 | 0.53963192 | 0.93523904 |
| MRPL57 | 82.3709979 | -0.063609021 | 0.4806581 | -0.1323374 | 0.89471746 | 0.98842187 |
| ZDHHC20 | 177.86709 | -0.069381898 | 0.247064 | -0.2808256 | 0.77884416 | 0.97423208 |
| MICU2 | 290.365458 | -0.128612144 | 0.15861116 | -0.8108644 | 0.41744353 | 0.89862171 |
| LINC00621 | 122.975155 | 0.886561079 | 0.65513088 | 1.35325796 | 0.17597323 | 0.74950983 |
| AL157931.1 | 50.7495763 | 0.605341749 | 0.65898518 | 0.91859691 | 0.35830645 | 0.87316144 |
| SACS | 66.2849515 | -0.38435015 | 0.49797914 | -0.7718198 | 0.44022118 | 0.90561284 |
| TNFRSF19 | 114.093034 | 0.776941096 | 0.33231938 | 2.33793497 | 0.01939062 | 0.37074405 |
| MIPEP | 192.61824 | -0.128887879 | 0.2835732 | -0.4545136 | 0.6494592 | 0.95027667 |
| PARP4 | 451.803825 | -0.398877856 | 0.40455687 | -0.9859624 | 0.32415157 | 0.85444355 |
| MTMR6 | 69.3684194 | -0.089657101 | 0.29397475 | -0.3049823 | 0.76037962 | 0.9714717 |
| NUP58 | 232.166622 | 0.050657576 | 0.25782419 | 0.19648108 | 0.84423364 | 0.98517901 |
| RNF6 | 30.2392179 | 0.020156387 | 0.38335513 | 0.05257889 | 0.95806744 | 0.99572129 |
| CDK8 | 247.654419 | 0.180177249 | 0.23576746 | 0.76421594 | 0.44473857 | 0.90694419 |
| WASF3 | 97.0213189 | -0.158277302 | 0.29406478 | -0.5382396 | 0.59041166 | 0.94516215 |
| USP12 | 153.705089 | 0.189212054 | 0.21756506 | 0.86968032 | 0.38447513 | 0.88328503 |
| LINC02340 | 55.3501906 | 0.375695058 | 0.7055326 | 0.53249851 | 0.59438077 | 0.9465625 |
| RPL21 | 204.114735 | 0.153884249 | 0.53078186 | 0.28991995 | 0.77187748 | 0.97329019 |
| GTF3A | 50.7060052 | -0.588575627 | 0.39252051 | -1.4994774 | 0.13374983 | 0.70530229 |
| MTIF3 | 90.072374 | 0.121669396 | 0.30128336 | 0.40383709 | 0.68633252 | 0.96161098 |
| LNX2 | 119.650449 | -0.034947433 | 0.41790589 | -0.0836251 | 0.93335449 | 0.99425828 |
| POLR1D | 93.6835032 | -0.196275257 | 0.18317258 | -1.0715318 | 0.28393036 | 0.82918183 |
| PAN3 | 718.962733 | 0.087374597 | 0.15181667 | 0.57552705 | 0.56493491 | 0.93809297 |
| POMP | 82.3404897 | -0.657977744 | 0.3956996 | -1.6628214 | 0.0963482 | 0.64295596 |
| SLC7A1 | 91.6299112 | -0.746851228 | 0.45854789 | -1.6287311 | 0.10336995 | 0.65288882 |
| UBL3 | 203.106863 | 0.218320885 | 0.30575707 | 0.7140338 | 0.47520628 | 0.91809716 |
| KATNAL1 | 134.427276 | 0.157428066 | 0.27115778 | 0.58057736 | 0.56152533 | 0.93747805 |
| HMGB1 | 496.728511 | -0.549050434 | 0.20563177 | -2.6700662 | 0.00758363 | 0.26951269 |
| USPL1 | 80.4390806 | -0.155250538 | 0.27901474 | -0.5564241 | 0.57792094 | 0.94034284 |
| HSPH1 | 92.3701532 | 0.425380108 | 0.4094967 | 1.03878764 | 0.29890351 | 0.83989343 |
| B3GLCT | 59.8467964 | 0.324102269 | 0.3608602 | 0.89813804 | 0.36911196 | 0.87605028 |
| FRY | 758.479517 | 0.508241168 | 0.5001247 | 1.01622888 | 0.30952041 | 0.84554063 |

| | | | | | | |
|---|---|---|---|---|---|---|
| N4BP2L1 | 135.568283 | 0.577408904 | 0.27829839 | 2.0747835 | 0.0380066 | 0.47106462 |
| N4BP2L2 | 1027.4287 | 0.267753096 | 0.19580434 | 1.36745228 | 0.17148359 | 0.74445571 |
| PDS5B | 340.519798 | 0.143211305 | 0.20859001 | 0.68656839 | 0.49235476 | 0.92289011 |
| KL | 48.4290116 | 0.177965052 | 0.51071998 | 0.34845916 | 0.72749538 | 0.9681562 |
| STARD13 | 271.778744 | -0.247890465 | 0.21536396 | -1.1510304 | 0.24971974 | 0.80672756 |
| AL139383.1 | 48.1288019 | 1.280040228 | 0.59779076 | 2.14128475 | 0.03225108 | 0.44049392 |
| RFC3 | 57.8849797 | -0.330701895 | 0.36823183 | -0.8980807 | 0.36914254 | 0.87605028 |
| NBEA | 426.604284 | 0.191947323 | 0.11058805 | 1.73569672 | 0.08261747 | 0.6141478 |
| DCLK1 | 35.5824304 | -2.712435432 | 0.73937232 | -3.668565 | 0.00024392 | 0.04172264 |
| SPART | 180.492813 | 0.384822792 | 0.18870176 | 2.03931744 | 0.04141836 | 0.48843812 |
| RFXAP | 29.8324441 | -0.4404635 | 0.32994712 | -1.3349518 | 0.18189213 | 0.75448394 |
| SMAD9 | 329.710045 | -0.900904111 | 0.70701026 | -1.2742448 | 0.20257669 | 0.77110248 |
| ALG5 | 69.5984517 | -0.259985241 | 0.21698881 | -1.1981504 | 0.23085846 | 0.79403724 |
| EXOSC8 | 43.7081742 | -0.503910505 | 0.35432436 | -1.4221729 | 0.15497606 | 0.72978231 |
| SUPT20H | 185.245933 | 0.046718042 | 0.25758525 | 0.18136924 | 0.85607776 | 0.9864199 |
| LINC00571 | 53.5261347 | -0.449268137 | 0.45614572 | -0.9849224 | 0.32466219 | 0.855037 |
| UFM1 | 117.900898 | -0.040707518 | 0.25605518 | -0.1589795 | 0.87368505 | 0.98766275 |
| FREM2 | 293.171516 | 0.176106747 | 0.63479591 | 0.27742262 | 0.78145561 | 0.97423208 |
| PROSER1 | 50.7136679 | -0.52281966 | 0.28369873 | -1.8428692 | 0.06534811 | 0.57495764 |
| NHLRC3 | 93.2481396 | 0.216444204 | 0.36352533 | 0.59540337 | 0.55157388 | 0.93523904 |
| COG6 | 128.491589 | 0.227263314 | 0.21576575 | 1.05328724 | 0.29220937 | 0.83697123 |
| FOXO1 | 409.689086 | 0.401549836 | 0.20778139 | 1.9325592 | 0.05329052 | 0.53386172 |
| MRPS31 | 101.693874 | 0.099915216 | 0.21549132 | 0.46366238 | 0.64288966 | 0.94944486 |
| ELF1 | 464.446936 | -0.193124782 | 0.1964244 | -0.9832016 | 0.32550822 | 0.85509839 |
| WBP4 | 82.875808 | -0.034910152 | 0.23796905 | -0.1467004 | 0.88336851 | 0.98766275 |
| AL354696.2 | 46.3242346 | -0.029468558 | 0.54940774 | -0.053637 | 0.95722441 | 0.99572129 |
| MTRF1 | 31.3277353 | -0.23141191 | 0.34167281 | -0.677291 | 0.49822135 | 0.92398825 |
| NAA16 | 180.666739 | -0.02805166 | 0.19449592 | -0.1442275 | 0.88532082 | 0.98766275 |
| VWA8 | 440.298668 | 0.14122662 | 0.22898299 | 0.61675593 | 0.53739572 | 0.93504069 |
| DGKH | 653.364194 | -0.199047418 | 0.31717901 | -0.6275555 | 0.5302952 | 0.93366189 |
| AKAP11 | 154.040575 | -0.102712491 | 0.2326944 | -0.4414051 | 0.65891976 | 0.95402584 |
| EPSTI1 | 206.911315 | -1.861671477 | 1.21284202 | -1.5349662 | 0.12479214 | 0.69439514 |
| DNAJC15 | 135.0462 | 0.071571423 | 0.29819277 | 0.2400173 | 0.81031685 | 0.97930627 |
| ENOX1 | 210.63142 | 0.244033357 | 1.13078543 | 0.21580872 | 0.82913685 | 0.98142144 |
| CCDC122 | 135.081277 | 0.355322784 | 0.31054666 | 1.14418485 | 0.25254698 | 0.80798674 |
| SMIM2-AS1 | 193.963722 | 0.857683716 | 0.77456577 | 1.10730909 | 0.26816032 | 0.81584693 |
| SERP2 | 33.0055836 | -0.395778262 | 0.30834512 | -1.2835561 | 0.19929732 | 0.77110248 |
| TSC22D1 | 923.200358 | 0.134747372 | 0.36405488 | 0.37012929 | 0.71128616 | 0.96587538 |
| NUFIP1 | 31.4753226 | -0.686343673 | 0.31001028 | -2.2139384 | 0.02683302 | 0.4234302 |
| AL359706.1 | 28.8161236 | 0.157084187 | 0.64580738 | 0.2432369 | 0.80782188 | 0.97930627 |
| GPALPP1 | 89.9639633 | -0.250754112 | 0.25528771 | -0.9822412 | 0.32598103 | 0.85509839 |
| GTF2F2 | 161.768515 | -0.232745278 | 0.24724005 | -0.9413737 | 0.34651339 | 0.86990996 |
| TPT1 | 1033.84969 | 0.674086295 | 0.57104526 | 1.18044285 | 0.23782413 | 0.80130754 |
| SLC25A30 | 66.5867028 | 0.279108606 | 0.34470427 | 0.80970452 | 0.41811002 | 0.89862171 |
| COG3 | 123.357507 | 0.033094074 | 0.22299916 | 0.14840448 | 0.88202357 | 0.98766275 |
| ZC3H13 | 193.751833 | -0.375738241 | 0.20347209 | -1.8466328 | 0.06480038 | 0.57495764 |
| LRRC63 | 29.5483969 | -0.65215461 | 1.30536822 | -0.4995944 | 0.61736073 | 0.94791646 |
| AL138686.1 | 40.7401581 | -0.235428076 | 0.62321304 | -0.377765 | 0.70560518 | 0.96543236 |
| LRCH1 | 455.854095 | 0.289574715 | 0.29363199 | 0.98618246 | 0.32404358 | 0.85444355 |
| ESD | 96.7181953 | -0.116262294 | 0.24497256 | -0.4745931 | 0.63507701 | 0.94932781 |
| SUCLA2 | 144.737968 | -0.512060531 | 0.21992483 | -2.3283435 | 0.01989387 | 0.3747883 |
| MED4 | 119.706354 | -0.112276848 | 0.21064965 | -0.5330028 | 0.59403166 | 0.94649617 |
| ITM2B | 1494.62826 | 0.679393529 | 0.38405697 | 1.76899154 | 0.07689529 | 0.59726395 |
| RB1 | 442.660317 | 0.378238885 | 0.20626241 | 1.83377515 | 0.06668738 | 0.5780981 |

| | | | | | | |
|---|---|---|---|---|---|---|
| LPAR6 | 70.7489214 | 0.367080837 | 0.54466283 | 0.67395977 | 0.50033689 | 0.92398825 |
| RCBTB2 | 98.4882848 | 0.003133872 | 0.27661014 | 0.01132956 | 0.99096051 | 0.99866813 |
| FNDC3A | 625.133242 | -0.027288391 | 0.23995198 | -0.1137244 | 0.90945628 | 0.99100813 |
| CDADC1 | 43.6383378 | -0.296646425 | 0.36610075 | -0.8102863 | 0.41777564 | 0.89862171 |
| CAB39L | 77.6949564 | 0.19079302 | 0.5416911 | 0.35221738 | 0.72467524 | 0.96781837 |
| SETDB2 | 87.8818628 | 0.036453539 | 0.36123476 | 0.10091371 | 0.91961896 | 0.99214128 |
| PHF11 | 81.6190579 | -1.010438428 | 0.37406074 | -2.7012683 | 0.00690756 | 0.25717187 |
| RCBTB1 | 37.6791241 | -0.17646091 | 0.391253 | -0.4510148 | 0.65197885 | 0.95159942 |
| EBPL | 50.9123859 | -0.574460081 | 0.26744648 | -2.147944 | 0.0317182 | 0.43905874 |
| KPNA3 | 151.485724 | -0.133134422 | 0.21991281 | -0.6053964 | 0.54491564 | 0.93523904 |
| SPRYD7 | 33.7089738 | -0.119418225 | 0.28449804 | -0.4197506 | 0.67466765 | 0.95910677 |
| DLEU2 | 284.954018 | 0.504142953 | 0.41991209 | 1.20059165 | 0.22990964 | 0.79370986 |
| TRIM13 | 73.5701544 | 0.105637495 | 0.27615081 | 0.38253552 | 0.70206419 | 0.9645815 |
| RNASEH2B | 80.3007597 | -0.596692959 | 0.30099641 | -1.9823923 | 0.04743535 | 0.51284792 |
| INTS6 | 268.36052 | -0.018708535 | 0.35084443 | -0.0533243 | 0.95747352 | 0.99572129 |
| INTS6-AS1 | 37.5227197 | 0.92770136 | 0.33529014 | 2.7668614 | 0.00565988 | 0.2367511 |
| WDFY2 | 155.470406 | -0.310088726 | 0.27797284 | -1.115536 | 0.26462078 | 0.81378765 |
| ATP7B | 37.2161377 | 0.168148494 | 0.36627901 | 0.45907216 | 0.64618235 | 0.94996446 |
| UTP14C | 42.4480086 | -0.494134807 | 0.38569233 | -1.2811632 | 0.20013636 | 0.77110248 |
| NEK5 | 46.3015601 | -0.183694666 | 0.66498174 | -0.2762402 | 0.78236361 | 0.97423208 |
| NEK3 | 49.4919433 | -0.428683843 | 0.42732069 | -1.00319 | 0.3157692 | 0.8490543 |
| VPS36 | 131.894497 | -0.427563238 | 0.19886322 | -2.1500368 | 0.0315523 | 0.43905874 |
| SUGT1 | 118.641446 | 0.060972589 | 0.21633439 | 0.28184418 | 0.77806299 | 0.97423208 |
| DIAPH3 | 151.788777 | -0.057198607 | 0.40233344 | -0.1421672 | 8.87E-01 | 0.98766275 |
| TDRD3 | 299.890756 | 0.008882714 | 0.20585928 | 0.04314945 | 0.9655824 | 0.99601598 |
| PCDH9 | 268.55162 | 0.598375071 | 1.25622086 | 0.47632951 | 0.63383964 | 0.94932781 |
| DIS3 | 71.3745676 | -0.294549179 | 0.2172943 | -1.3555311 | 0.17524841 | 0.74934286 |
| PIBF1 | 396.432291 | -0.074469459 | 0.17049585 | -0.4367816 | 0.66226971 | 0.95468496 |
| KLF5 | 59.2682829 | -0.687658449 | 0.56407808 | -1.2190838 | 0.22281239 | 0.78710314 |
| KLF12 | 332.949758 | 0.117984912 | 0.31529416 | 0.37420583 | 0.70825119 | 0.96587538 |
| TBC1D4 | 1080.04018 | -0.257114576 | 0.52393085 | -0.4907414 | 0.62360934 | 0.94808496 |
| COMMD6 | 92.2527746 | 0.42991667 | 0.41667586 | 1.03177724 | 0.30217648 | 0.84375949 |
| UCHL3 | 84.4858531 | -0.564019938 | 0.37741991 | -1.4944096 | 0.13506859 | 0.70620568 |
| LMO7-AS1 | 45.6246776 | 0.299940848 | 0.5793445 | 0.51772451 | 0.60465049 | 0.94683415 |
| LMO7 | 595.398912 | -0.260413624 | 0.35280768 | -0.7381178 | 4.60E-01 | 0.91368087 |
| AC001226.2 | 57.4129377 | 0.376519892 | 0.63532422 | 0.59264212 | 0.55342069 | 0.93523904 |
| FBXL3 | 112.918222 | 0.12027707 | 0.32340917 | 0.37190371 | 0.70996454 | 0.96587538 |
| MYCBP2 | 455.747453 | -0.38366236 | 0.21474504 | -1.7865948 | 0.07400301 | 0.59320069 |
| SLAIN1 | 53.4923111 | 0.386705545 | 0.36499134 | 1.05949237 | 0.2893756 | 0.83466693 |
| RNF219 | 43.8008941 | -0.393911865 | 0.33785471 | -1.1659209 | 0.24364643 | 0.80476923 |
| RBM26 | 407.358589 | -0.168106611 | 0.16867314 | -0.9966413 | 0.31893866 | 0.84935263 |
| NDFIP2 | 146.999186 | -0.200214514 | 0.37764099 | -0.5301716 | 0.59599299 | 0.94683415 |
| SPRY2 | 34.5323718 | -0.572084243 | 0.51063968 | -1.1203286 | 0.26257375 | 0.81377303 |
| GPC5 | 77.9486433 | 1.133616443 | 0.68552996 | 1.65363515 | 0.09820167 | 0.64312419 |
| GPC6 | 343.08103 | -1.740048971 | 0.64852479 | -2.6830878 | 0.00729458 | 0.26467731 |
| TGDS | 39.0191755 | -0.067710655 | 0.30706289 | -0.2205107 | 0.82547343 | 0.98094663 |
| GPR180 | 61.2538911 | -0.667777017 | 0.22893766 | -2.9168509 | 0.00353585 | 0.18995949 |
| ABCC4 | 832.051291 | -0.481187545 | 0.51621225 | -0.9321506 | 0.35125872 | 0.8720098 |
| CLDN10 | 164.288719 | 1.401225357 | 0.4012767 | 3.49191804 | 0.00047957 | 0.06755525 |
| DZIP1 | 88.1796119 | 1.583832169 | 0.87341543 | 1.81337781 | 0.06977357 | 0.58513328 |
| DNAJC3 | 196.991257 | 0.622611096 | 0.32707537 | 1.90357073 | 0.05696611 | 0.54616991 |
| UGGT2 | 469.748623 | 0.0077462 | 0.17379088 | 0.04457196 | 0.96444849 | 0.99586809 |
| MBNL2 | 501.628488 | 0.131976308 | 0.30685651 | 0.43009127 | 0.66712925 | 0.95691417 |
| RAP2A | 32.8861155 | 0.84307642 | 0.33881055 | 2.48834168 | 0.01283404 | 0.31603403 |

| | | | | | | |
|---|---|---|---|---|---|---|
| IPO5 | 113.009425 | -0.010277686 | 0.2815665 | -0.0365018 | 0.97088224 | 0.99681784 |
| FARP1 | 1941.45582 | 0.172678579 | 0.25611852 | 0.67421356 | 0.50017555 | 0.92398825 |
| STK24 | 314.548517 | 0.067696821 | 0.15414159 | 0.43918597 | 0.66052679 | 0.95407427 |
| SLC15A1 | 39.6125191 | 2.191673861 | 0.53403714 | 4.10397277 | 4.06E-05 | 0.01313957 |
| DOCK9 | 342.810127 | 0.036118853 | 0.29637131 | 0.12187027 | 0.90300176 | 0.99034419 |
| UBAC2 | 314.467293 | -0.226134225 | 0.1590422 | -1.4218504 | 0.15506968 | 0.72978231 |
| TM9SF2 | 219.308982 | 0.18613848 | 0.13713605 | 1.35732711 | 0.1746773 | 0.74910274 |
| CLYBL | 224.462927 | 0.077290354 | 0.32758969 | 0.23593646 | 0.81348197 | 0.97930627 |
| PCCA | 822.357618 | 0.11305851 | 0.25954646 | 0.43560028 | 0.66312677 | 0.95477098 |
| GGACT | 66.2750524 | -0.67434788 | 0.34119889 | -1.976407 | 0.0481087 | 0.51547342 |
| TMTC4 | 49.9028736 | -0.084734743 | 0.34912526 | -0.2427058 | 0.80823328 | 0.97930627 |
| FGF14 | 315.757767 | -0.48629734 | 0.76025097 | -0.6396537 | 0.52239777 | 0.93272217 |
| TPP2 | 178.432219 | 0.012844305 | 0.165923 | 0.07741124 | 0.9382964 | 0.99526299 |
| EFNB2 | 64.26215 | 0.32992134 | 0.59353295 | 0.55586019 | 0.57830643 | 0.94034284 |
| ARGLU1 | 346.970839 | 0.138109393 | 0.17313881 | 0.79768014 | 0.42505613 | 0.90167091 |
| FAM155A | 465.790912 | -0.45322875 | 1.24417026 | -0.3642819 | 0.71564749 | 0.96688114 |
| ABHD13 | 39.633289 | 0.102780957 | 0.25292203 | 0.40637407 | 0.68446777 | 0.96161098 |
| TNFSF13B | 40.03946 | 0.477264452 | 0.42319407 | 1.12776734 | 0.25941819 | 0.81263292 |
| IRS2 | 64.4360576 | 0.249535176 | 0.55210741 | 0.45196853 | 0.65129165 | 0.95145992 |
| COL4A1 | 769.346766 | -0.123084828 | 0.24708984 | -0.498138 | 6.18E-01 | 9.48E-01 |
| COL4A2 | 908.878777 | -0.209379304 | 0.23459849 | -0.8925006 | 0.37212464 | 0.87605028 |
| RAB20 | 66.5011791 | 0.825933563 | 0.46025553 | 1.7945109 | 0.07273166 | 0.59192805 |
| NAXD | 35.1320623 | 0.131505707 | 0.33526725 | 0.39224144 | 0.69487983 | 0.96220115 |
| CARS2 | 123.568313 | -0.102526534 | 0.21482411 | -0.477258 | 0.63317839 | 0.94932781 |
| ANKRD10 | 419.984268 | 0.094763331 | 0.15664158 | 0.6049692 | 0.54519946 | 0.93523904 |
| ARHGEF7 | 254.730865 | 0.047143726 | 0.26625734 | 0.17706076 | 0.85946066 | 0.9864199 |
| TUBGCP3 | 129.865286 | -0.36926558 | 0.30804313 | -1.1987463 | 0.23062659 | 0.79403724 |
| ATP11A | 426.473835 | -0.139807069 | 0.32055141 | -0.4361455 | 0.66273113 | 0.95477098 |
| MCF2L | 216.325035 | -0.127624732 | 0.32185083 | -0.3965338 | 0.69171128 | 0.96220115 |
| PCID2 | 86.5315724 | -0.020611092 | 0.19180678 | -0.1074576 | 0.91442598 | 0.99159829 |
| CUL4A | 198.888067 | -0.178458716 | 0.23976559 | -0.744305 | 0.456692 | 0.91187931 |
| LAMP1 | 189.118467 | 0.156025542 | 0.31933786 | 0.4885908 | 0.62513143 | 0.94808496 |
| GRTP1 | 44.7416194 | 1.06322213 | 0.54407006 | 1.95420076 | 0.05067748 | 0.52415717 |
| DCUN1D2 | 73.2403409 | 0.096417486 | 0.24703681 | 0.39029603 | 0.69631766 | 0.96220115 |
| TMCO3 | 148.237734 | 0.182068531 | 0.33108616 | 0.54991284 | 0.58237916 | 0.94091128 |
| TFDP1 | 62.9505828 | -0.468064763 | 0.32614345 | -1.43515 | 0.15124437 | 0.72403332 |
| GAS6 | 46.5898397 | 2.084735031 | 0.52866949 | 3.94336172 | 8.03E-05 | 0.02080127 |
| RASA3 | 69.378382 | -0.778330278 | 0.37154798 | -2.0948311 | 0.03618601 | 0.46105889 |
| CDC16 | 142.50856 | -0.208960887 | 0.22964882 | -0.9099149 | 0.36286738 | 0.87551319 |
| UPF3A | 134.704293 | -0.007977003 | 0.21712471 | -0.0367393 | 0.9706929 | 0.99681784 |
| CHAMP1 | 26.5059016 | 0.258594274 | 0.36271149 | 0.71294758 | 0.4758782 | 0.91809716 |
| TTC5 | 45.2531001 | -0.194321572 | 0.32049293 | -0.6063209 | 0.54430166 | 0.93523904 |
| CCNB1IP1 | 58.8416517 | -0.178205944 | 0.3341656 | -0.5332863 | 0.59383539 | 0.94649617 |
| TEP1 | 37.5486809 | -0.272264025 | 0.32231723 | -0.8447083 | 0.39827375 | 0.8887831 |
| OSGEP | 34.521082 | -0.170791995 | 0.30976139 | -0.5513663 | 0.58138259 | 0.94072024 |
| APEX1 | 59.9426051 | -0.390267284 | 0.53161452 | -0.734117 | 0.4628774 | 0.91462428 |
| PNP | 30.3043505 | 0.019970771 | 0.4765193 | 0.04190968 | 9.67E-01 | 0.99601626 |
| METTL17 | 33.1268065 | -0.208060449 | 0.29179234 | -0.7130429 | 0.47581922 | 0.91809716 |
| NDRG2 | 33.1487991 | 1.38311355 | 0.48075573 | 2.87695695 | 0.0040153 | 0.20137484 |
| ARHGEF40 | 39.9589021 | 1.035562684 | 0.61497728 | 1.68390398 | 0.09220023 | 0.63464873 |
| ZNF219 | 30.8593906 | -0.136411619 | 0.32627996 | -0.4180815 | 0.67588752 | 0.9591071 |
| HNRNPC | 706.525176 | 0.044510481 | 0.182514 | 0.24387434 | 0.80732814 | 0.97930627 |
| SUPT16H | 126.82203 | -0.294708884 | 0.21702616 | -1.3579417 | 0.17448217 | 0.74910274 |
| CHD8 | 156.762842 | -0.257662376 | 0.18824537 | -1.3687581 | 0.17107491 | 0.74445571 |

| | | | | | | |
|---|---|---|---|---|---|---|
| RAB2B | 61.833398 | -0.301993835 | 0.23836829 | -1.2669212 | 0.20518347 | 0.77352973 |
| TOX4 | 86.0475751 | 0.30208167 | 0.30052986 | 1.00516359 | 0.31481808 | 0.8490543 |
| METTL3 | 38.9973746 | -0.538851872 | 0.34337879 | -1.5692637 | 0.11658651 | 0.67683766 |
| DAD1 | 173.978689 | -0.142158442 | 0.44438969 | -0.3198959 | 0.74904724 | 0.97028355 |
| ABHD4 | 26.4931318 | 0.133614096 | 0.38099751 | 0.35069546 | 0.72581683 | 0.96781837 |
| OXA1L | 41.1559089 | 0.205649815 | 0.31362786 | 0.65571285 | 0.5120089 | 0.9287766 |
| MRPL52 | 44.5554117 | -0.1501665 | 0.47065327 | -0.3190597 | 0.74968123 | 0.97046004 |
| MMP14 | 45.2052731 | 0.112862825 | 0.28086361 | 0.40184211 | 0.68780023 | 0.96161098 |
| LRP10 | 151.301385 | -0.070746448 | 0.22270626 | -0.3176671 | 7.51E-01 | 0.97058015 |
| RBM23 | 99.3268851 | 0.103528573 | 0.20892888 | 0.49552066 | 0.62023265 | 0.9480368 |
| PRMT5 | 29.7808694 | 0.005718135 | 0.29592058 | 0.01932321 | 0.98458327 | 0.99752755 |
| PSMB5 | 90.6014039 | 0.050712968 | 0.38883277 | 0.1304236 | 0.8962313 | 0.98842187 |
| ACIN1 | 191.863608 | -0.208136323 | 0.18540352 | -1.1226126 | 0.26160208 | 0.81281978 |
| C14orf119 | 28.2270247 | -0.44701828 | 0.4590003 | -0.9738954 | 0.33010848 | 0.85880205 |
| BCL2L2-PABI | 46.683226 | -0.072532569 | 0.29808839 | -0.2433257 | 0.80775309 | 0.97930627 |
| PABPN1 | 117.791547 | -0.37587213 | 0.31603098 | -1.1893522 | 0.23430112 | 0.79672361 |
| SLC22A17 | 39.1580051 | 0.62593189 | 0.55062702 | 1.13676203 | 0.25563778 | 0.80897837 |
| NGDN | 41.8465249 | -0.126122121 | 0.25806618 | -0.4887201 | 0.62503989 | 0.94808496 |
| ZFHX2 | 45.6518969 | 1.084230796 | 0.51270815 | 2.11471342 | 0.03445438 | 0.45210754 |
| AP1G2 | 34.1471257 | -0.195610277 | 0.38517319 | -0.5078502 | 0.61155841 | 0.94791646 |
| DHRS4-AS1 | 33.5216611 | -0.013054212 | 0.37989186 | -0.034363 | 0.97258771 | 0.99681784 |
| DHRS4L2 | 38.4351734 | 0.224180691 | 0.48276373 | 0.46436937 | 0.64238313 | 0.94944486 |
| PSME1 | 118.054197 | 0.023404428 | 0.49420696 | 0.04735754 | 0.96222827 | 0.99572129 |
| AL136295.1 | 35.3975802 | -0.440825783 | 0.29034077 | -1.5183048 | 0.12893758 | 0.69814682 |
| GMPR2 | 28.4709012 | 0.374211645 | 0.36684216 | 1.02008899 | 0.30768626 | 0.84488735 |
| KHNYN | 37.930923 | 0.094473467 | 0.31446261 | 0.3004283 | 0.76385048 | 0.97174831 |
| SDR39U1 | 34.8627163 | -0.363581746 | 0.33031422 | -1.1007148 | 0.2710208 | 0.81775732 |
| STXBP6 | 81.6414651 | -0.177348328 | 0.40582321 | -0.4370088 | 6.62E-01 | 9.55E-01 |
| NOVA1 | 101.923122 | 1.15274022 | 0.59306213 | 1.94370903 | 0.05193055 | 0.52863201 |
| PRKD1 | 1858.73668 | -0.413656467 | 0.26042806 | -1.5883713 | 0.1122024 | 0.66401239 |
| G2E3 | 82.3888815 | 0.068451768 | 0.20283294 | 0.33747856 | 0.73575617 | 0.9698508 |
| SCFD1 | 289.772221 | -0.26016088 | 0.17304093 | -1.5034644 | 0.13271933 | 0.70316286 |
| COCH | 22.6702848 | 0.297711994 | 0.44127869 | 0.67465754 | 0.49989337 | 0.92398825 |
| STRN3 | 748.043357 | 0.314753353 | 0.27244539 | 1.15528971 | 0.24797181 | 0.80672756 |
| AP4S1 | 68.5457819 | -0.036181672 | 0.37389135 | -0.0967706 | 0.92290861 | 0.99214128 |
| HECTD1 | 374.786454 | -0.01037132 | 0.1682229 | -0.0616522 | 0.95083977 | 0.99531843 |
| HEATR5A | 56.1198077 | 0.177454332 | 0.35184796 | 0.50434947 | 0.61401582 | 0.94791646 |
| AL139353.1 | 32.1514834 | 0.007553485 | 0.42591926 | 0.01773455 | 0.98585062 | 0.99764442 |
| NUBPL | 306.430463 | -0.105783026 | 0.23655209 | -0.447187 | 0.65474003 | 0.95286459 |
| ARHGAP5 | 489.609555 | -0.022263945 | 0.16705606 | -0.1332723 | 0.89397803 | 0.98842187 |
| AKAP6 | 117.976044 | 0.222586283 | 0.44817806 | 0.49664698 | 0.61943802 | 0.9480368 |
| NPAS3 | 581.704818 | 0.859038028 | 0.66999159 | 1.28216241 | 0.19978568 | 0.77110248 |
| EGLN3 | 168.341954 | -0.243693276 | 0.77602783 | -0.3140265 | 0.75350094 | 0.9710175 |
| EAPP | 89.494837 | 0.244394253 | 0.32733656 | 0.7466146 | 0.45529623 | 0.91077355 |
| SNX6 | 230.523557 | 0.038551206 | 0.23165904 | 0.16641356 | 0.8678315 | 0.98670067 |
| CFL2 | 35.3891579 | 0.522706764 | 0.42917556 | 1.21793228 | 0.22324971 | 0.78766444 |
| BAZ1A | 195.180968 | -0.197497482 | 0.34782489 | -0.5678072 | 0.57016589 | 0.93998433 |
| SRP54 | 189.271773 | 0.034880824 | 0.19677843 | 0.17725939 | 0.85930465 | 0.9864199 |
| FAM177A1 | 115.809161 | 0.041819023 | 0.27997117 | 0.14936903 | 0.88126245 | 0.98766275 |
| PPP2R3C | 61.5420017 | -0.017794846 | 0.31237334 | -0.0569666 | 0.9545718 | 0.99531843 |
| NFKBIA | 218.714206 | 0.435852044 | 0.72543859 | 0.60081178 | 0.54796536 | 0.93523904 |
| RALGAPA1 | 540.097074 | -0.071566323 | 0.188185 | -0.3802977 | 0.70372444 | 0.96461079 |
| BRMS1L | 36.7615258 | 0.219157791 | 0.3495669 | 0.62694092 | 0.53069796 | 0.93366189 |
| MBIP | 69.0624399 | 0.195992211 | 0.29243927 | 0.670198 | 0.50273158 | 0.92492012 |

| | | | | | | |
|---|---|---|---|---|---|---|
| SLC25A21 | 79.1722987 | -0.179393711 | 0.32553153 | -0.5510794 | 0.58157927 | 0.94087956 |
| MIPOL1 | 309.522929 | -0.359799843 | 0.17158632 | -2.0969028 | 0.03600218 | 0.46016006 |
| SEC23A | 192.659517 | 0.296795786 | 0.19867623 | 1.49386663 | 0.13521048 | 0.70620568 |
| TRAPPC6B | 82.2560034 | -0.030119475 | 0.21989205 | -0.1369739 | 0.89105141 | 0.98766275 |
| PNN | 202.147423 | -0.233640659 | 0.19009613 | -1.2290659 | 0.21904711 | 0.78523615 |
| MIA2 | 460.546914 | 0.479967565 | 0.20863457 | 2.30051786 | 0.0214189 | 0.38566096 |
| FBXO33 | 90.5326755 | 0.024758187 | 0.25158642 | 0.09840828 | 0.9216081 | 0.99214128 |
| AL049875.1 | 35.9511593 | -0.575700963 | 0.37050844 | -1.5538134 | 0.12022894 | 0.68797673 |
| LRFN5 | 133.943373 | 1.326657401 | 1.41690723 | 0.93630505 | 0.34911615 | 0.87178925 |
| KLHL28 | 103.154958 | 0.293072813 | 0.29902878 | 0.98008229 | 0.3270455 | 0.85641576 |
| TOGARAM1 | 214.341143 | 0.260059471 | 0.19002787 | 1.36853331 | 0.1711452 | 0.74445571 |
| PRPF39 | 78.4839998 | 0.062375378 | 0.21069609 | 0.29604431 | 0.76719624 | 0.97221495 |
| FKBP3 | 60.4188665 | 0.214016395 | 0.33667144 | 0.63568326 | 0.52498289 | 0.93272384 |
| FANCM | 45.7901561 | -0.787031156 | 0.29566956 | -2.6618606 | 0.007771 | 0.27068528 |
| MIS18BP1 | 71.9887452 | -0.381480363 | 0.29650142 | -1.2866055 | 0.19823182 | 0.77110248 |
| MDGA2 | 124.631879 | 1.186135427 | 1.65095881 | 0.71845247 | 0.47247835 | 0.91737234 |
| RPS29 | 191.628892 | 0.035199775 | 0.26990754 | 0.1304142 | 0.89623873 | 0.98842187 |
| RPL36AL | 233.370061 | -0.291714477 | 0.55417513 | -0.526394 | 0.59861447 | 0.94683415 |
| DNAAF2 | 27.6780522 | 0.280260625 | 0.33110852 | 0.84643134 | 0.39731216 | 0.88863925 |
| POLE2 | 35.4260931 | -0.182303143 | 0.44519385 | -0.4094916 | 0.68217893 | 0.9609694 |
| KLHDC1 | 101.64441 | 0.556099522 | 0.38698575 | 1.43700259 | 0.15071726 | 0.72330692 |
| KLHDC2 | 119.296461 | 0.387785842 | 0.28827573 | 1.34519075 | 0.17856364 | 0.7513166 |
| NEMF | 316.536462 | -0.048600604 | 0.16095086 | -0.3019593 | 0.76268311 | 0.9714717 |
| ARF6 | 78.9722342 | -0.260070878 | 0.41949408 | -0.6199632 | 0.53528203 | 0.93460153 |
| LINC01588 | 29.0463922 | 0.156084272 | 0.53110633 | 0.29388516 | 0.76884566 | 0.972511 |
| VCPKMT | 39.2620787 | -0.09576067 | 0.25984426 | -0.368531 | 0.71247732 | 0.96587538 |
| SOS2 | 328.756273 | 0.254773975 | 0.2380262 | 1.07036107 | 0.28445682 | 0.82918183 |
| L2HGDH | 121.051972 | 0.120043482 | 0.3429879 | 0.34999334 | 0.72634369 | 0.96781837 |
| ATP5S | 68.5774545 | -0.177888329 | 0.22511347 | -0.7902163 | 0.42940147 | 0.90333313 |
| CDKL1 | 306.857742 | -0.459741111 | 0.45376431 | -1.0131716 | 0.3109782 | 0.8474709 |
| MAP4K5 | 341.291111 | -0.052962523 | 0.23107728 | -0.2291983 | 0.81871478 | 0.98033308 |
| SAV1 | 86.1144914 | 0.378980086 | 0.23542704 | 1.609756 | 0.10745113 | 0.65699665 |
| NIN | 101.783392 | 0.507145779 | 0.29684406 | 1.70845856 | 0.08755129 | 0.62705598 |
| TMX1 | 54.8325705 | -0.436417376 | 0.32167081 | -1.3567205 | 0.17487004 | 0.74910274 |
| RTRAF | 84.0015567 | 0.108327393 | 0.35639521 | 0.303953 | 0.7611637 | 0.9714717 |
| NID2 | 257.605551 | 1.176135713 | 0.5537396 | 2.12398699 | 0.03367123 | 0.44933903 |
| PTGER2 | 34.6710297 | -1.165288167 | 0.44107379 | -2.6419347 | 0.00824339 | 0.27246484 |
| TXNDC16 | 58.0097356 | -0.058297524 | 0.3360698 | -0.1734685 | 0.86228319 | 0.98669442 |
| ERO1A | 154.421159 | -0.323584203 | 0.56333206 | -0.5744111 | 0.56568963 | 0.93809297 |
| PSMC6 | 119.686859 | -0.269029091 | 0.20000117 | -1.3451376 | 0.17858081 | 0.7513166 |
| STYX | 37.006966 | 0.217362782 | 0.26486548 | 0.82065351 | 0.41184366 | 0.8970101 |
| FERMT2 | 235.415392 | 0.027148629 | 0.28097928 | 0.09662146 | 0.92302701 | 0.99214128 |
| DDHD1 | 85.0275475 | 0.323499744 | 0.21837098 | 1.48142276 | 0.13849395 | 0.71091391 |
| CNIH1 | 47.158157 | 0.160911219 | 0.30609599 | 0.52568876 | 0.59910448 | 0.94683415 |
| GMFB | 56.5221569 | -0.226185559 | 0.262836 | -0.8605577 | 0.38948167 | 0.88586416 |
| CGRRF1 | 30.1215725 | 0.137320626 | 0.35302111 | 0.38898701 | 0.69728576 | 0.96220115 |
| SAMD4A | 645.139683 | 0.182181727 | 0.32255633 | 0.56480593 | 0.57220577 | 0.93998433 |
| GCH1 | 36.7024123 | -0.220228989 | 0.68720142 | -0.3204723 | 0.74861036 | 0.97028355 |
| SOCS4 | 45.2911011 | 0.198858486 | 0.29687875 | 0.66983065 | 0.50296575 | 0.92492012 |
| MAPK1IP1L | 191.695078 | -0.140089986 | 0.23996628 | -0.5837903 | 0.55936141 | 0.9373903 |
| LGALS3 | 123.709287 | -0.943500315 | 0.61001094 | -1.5466941 | 0.12193703 | 0.69196376 |
| FBXO34 | 290.551901 | 0.074909663 | 0.20016299 | 0.37424332 | 0.7082233 | 0.96587538 |
| ATG14 | 65.7124552 | -0.108343525 | 0.35747505 | -0.30308 | 0.76182892 | 0.9714717 |
| KTN1-AS1 | 36.5011719 | -0.570779067 | 0.49064217 | -1.1633306 | 0.24469538 | 0.80476923 |

| | | | | | | |
|---|---|---|---|---|---|---|
| KTN1 | 615.935865 | -0.078631555 | 0.18853536 | -0.4170653 | 0.67663066 | 0.9591378 |
| PELI2 | 417.193593 | 0.355797019 | 0.36639343 | 0.97107913 | 0.33150887 | 0.85923151 |
| TMEM260 | 139.911799 | -0.086124675 | 0.24210211 | -0.355737 | 0.72203755 | 0.96779196 |
| EXOC5 | 157.312089 | 0.131637883 | 0.19723211 | 0.66742624 | 0.50449991 | 0.92552859 |
| AP5M1 | 78.1969859 | 0.115061367 | 0.23825876 | 0.48292608 | 0.62914822 | 0.94891118 |
| NAA30 | 58.9341762 | -0.006900438 | 0.29140737 | -0.0236797 | 0.9811081 | 0.99681784 |
| CCDC198 | 809.415615 | 0.160533239 | 0.40957464 | 0.39195112 | 6.95E-01 | 0.96220115 |
| SLC35F4 | 94.8117993 | 0.253722147 | 0.90699507 | 0.27973928 | 0.77967754 | 0.97423208 |
| ARMH4 | 232.125184 | -0.182521055 | 0.3594665 | -0.5077554 | 0.61162489 | 0.94791646 |
| ACTR10 | 145.388833 | -0.073736211 | 0.17297229 | -0.4262892 | 0.66989716 | 0.95764829 |
| PSMA3 | 114.959953 | -0.112939933 | 0.26249267 | -0.4302594 | 0.66700698 | 0.95691417 |
| ARID4A | 217.791137 | 0.602743376 | 0.25246266 | 2.38745557 | 0.01696545 | 0.35074796 |
| TIMM9 | 51.4778196 | -0.08713581 | 0.26054972 | -0.3344306 | 0.73805462 | 0.9698508 |
| KIAA0586 | 110.22448 | 0.035936889 | 0.22480993 | 0.15985455 | 0.87299565 | 0.98766275 |
| DAAM1 | 171.951559 | 0.175972725 | 0.37585328 | 0.46819526 | 0.63964496 | 0.94944486 |
| GPR135 | 59.6744096 | 0.258710764 | 0.54642444 | 0.47346119 | 0.63588418 | 0.94932781 |
| L3HYPDH | 54.6178492 | -0.381531472 | 0.58391123 | -0.6534066 | 0.51349416 | 0.93035003 |
| JKAMP | 67.404181 | 0.276312778 | 0.24302873 | 1.13695521 | 0.25555701 | 0.80897837 |
| PCNX4 | 314.319097 | -0.533746615 | 0.28992148 | -1.841004 | 0.06562097 | 0.57606134 |
| DHRS7 | 76.5062885 | -0.696128939 | 0.44661985 | -1.558661 | 0.11907664 | 0.68597919 |
| PPM1A | 136.693681 | 0.098468512 | 0.23404571 | 0.42072342 | 0.67395706 | 0.95910677 |
| MNAT1 | 397.712252 | 0.180377636 | 0.23384165 | 0.7713666 | 0.44048967 | 0.90561284 |
| SLC38A6 | 87.9241852 | 0.083121682 | 0.31584352 | 0.26317362 | 0.79241677 | 0.97429505 |
| PRKCH | 94.9770566 | -0.533742428 | 0.66969604 | -0.7969921 | 0.42545564 | 0.90167091 |
| TMEM30B | 33.8754586 | -0.323898025 | 0.69686941 | -0.4647901 | 0.64208176 | 0.94944486 |
| HIF1A | 2456.24318 | -0.216558834 | 0.4896749 | -0.4422502 | 0.65830815 | 0.95399905 |
| HIF1A-AS2 | 403.222929 | 0.201424702 | 0.613258 | 0.32845018 | 0.7425713 | 0.9698508 |
| SNAPC1 | 113.694296 | 0.302173694 | 0.24281678 | 1.24445146 | 0.21333345 | 0.77996991 |
| PPP2R5E | 409.934003 | 0.012675618 | 0.20757481 | 0.0610653 | 0.9513072 | 0.99531843 |
| WDR89 | 52.9479826 | -0.135731544 | 0.25246207 | -0.5376314 | 0.59083152 | 0.94527451 |
| SGPP1 | 141.522052 | -0.003269782 | 0.29913298 | -0.0109309 | 0.9912786 | 0.99866813 |
| SYNE2 | 3411.4046 | -0.040850699 | 0.23766308 | -0.1718849 | 0.863528 | 0.98670067 |
| ESR2 | 268.90352 | -0.181140078 | 0.67012765 | -0.2703068 | 0.78692421 | 0.97423208 |
| MTHFD1 | 73.2783775 | -0.812175528 | 0.2827078 | -2.8728444 | 0.00406794 | 0.20295225 |
| AL122035.1 | 62.5271523 | -0.187423068 | 0.52975976 | -0.3537888 | 0.72349717 | 0.96779196 |
| ZBTB25 | 96.8205989 | 0.145501566 | 0.38874773 | 0.37428274 | 0.70819397 | 0.96587538 |
| AKAP5 | 30.1255385 | -1.015186234 | 0.68582996 | -1.4802302 | 1.39E-01 | 0.7113761 |
| ZBTB1 | 192.843611 | 0.261121138 | 0.24930962 | 1.0473769 | 0.29492578 | 0.83911606 |
| HSPA2 | 30.1424228 | -0.399477819 | 0.81437063 | -0.4905356 | 0.62375492 | 0.94808496 |
| PLEKHG3 | 210.856391 | -0.372587768 | 0.39822621 | -0.9356184 | 0.3494697 | 0.8720098 |
| CHURC1-FNT | 41.2009999 | -0.025134578 | 0.26173266 | -0.0960315 | 9.23E-01 | 0.99214128 |
| MAX | 160.580106 | -0.277270234 | 0.40925102 | -0.6775065 | 0.49808463 | 0.92398825 |
| FUT8 | 403.497839 | -0.005629941 | 0.21143571 | -0.0266272 | 0.97875708 | 0.99681784 |
| AL359232.1 | 133.249542 | -0.139980676 | 0.58668608 | -0.2385955 | 0.81141923 | 0.97930627 |
| GPHN | 1459.0511 | 0.074784446 | 0.37837502 | 0.19764636 | 0.84332176 | 0.98517901 |
| MPP5 | 204.829033 | 0.034328107 | 0.1456582 | 0.23567576 | 0.81368428 | 0.97930627 |
| ATP6V1D | 118.305039 | -0.465922978 | 0.25823919 | -1.8042303 | 0.07119521 | 0.59054593 |
| EIF2S1 | 73.9340328 | -0.552360414 | 0.23187639 | -2.382133 | 0.01721268 | 0.35165689 |
| PLEKHH1 | 134.799313 | -0.662366284 | 0.40938298 | -1.6179625 | 0.10567069 | 0.65699665 |
| PIGH | 51.3392038 | 0.176385392 | 0.47859714 | 0.36854669 | 0.71246564 | 0.96587538 |
| ARG2 | 240.766972 | -1.190038548 | 1.21939325 | -0.9759268 | 0.32910074 | 0.85781116 |
| VTI1B | 57.5445658 | -0.055675147 | 0.41280702 | -0.1348697 | 0.89271492 | 0.9882865 |
| ZFYVE26 | 55.6677103 | -0.401906413 | 0.2487292 | -1.6158393 | 0.10612906 | 0.65699665 |
| RAD51B | 765.736197 | -0.214503729 | 0.48921506 | -0.4384651 | 0.66104917 | 0.95407427 |

| | | | | | | |
|---|---|---|---|---|---|---|
| ZFP36L1 | 816.174686 | -0.425909051 | 0.26842026 | -1.5867247 | 0.11257502 | 0.66401239 |
| ACTN1 | 215.759104 | -9.38E-05 | 0.25733661 | -0.0003645 | 0.99970918 | 0.99971834 |
| DCAF5 | 307.494483 | 0.282143196 | 0.20048207 | 1.40732385 | 0.15933138 | 0.7380248 |
| EXD2 | 61.5179458 | -0.037168096 | 0.24431115 | -0.1521343 | 0.87908104 | 0.98766275 |
| ERH | 111.256112 | -0.572865136 | 0.34411407 | -1.6647536 | 0.09596192 | 0.64267209 |
| SLC39A9 | 100.129734 | 0.049713039 | 0.2008012 | 0.24757342 | 0.80446448 | 0.97928139 |
| SUSD6 | 322.234524 | 0.276997884 | 0.50002877 | 0.55396389 | 0.57960356 | 0.94034284 |
| SRSF5 | 341.804547 | -0.013818807 | 0.31831215 | -0.0434128 | 0.96537251 | 0.99601598 |
| AL357153.2 | 63.5176894 | 0.887404054 | 0.36453358 | 2.43435472 | 0.01491837 | 0.3338856 |
| MED6 | 64.0890025 | -0.027064657 | 0.25789979 | -0.1049425 | 0.91642141 | 0.99179761 |
| MAP3K9 | 84.4602411 | -0.122378674 | 0.30407513 | -0.402462 | 0.68734407 | 0.96161098 |
| PCNX1 | 402.481044 | 0.418839758 | 0.18264817 | 2.29315059 | 0.02183934 | 0.38823827 |
| SIPA1L1 | 765.796608 | 0.257973762 | 0.29933934 | 0.86181043 | 0.38879185 | 0.88586416 |
| RGS6 | 235.315961 | -3.824330672 | 1.11305996 | -3.4358712 | 0.00059065 | 0.07556838 |
| ZFYVE1 | 60.2966519 | 0.69709353 | 0.31146641 | 2.23810179 | 0.02521442 | 0.4135769 |
| RBM25 | 282.915485 | -0.441397341 | 0.14310211 | -3.0844921 | 0.002039 | 0.14702642 |
| PSEN1 | 208.135732 | 0.179729248 | 0.17919809 | 1.00296407 | 0.3158782 | 0.8490543 |
| PAPLN | 184.736834 | 1.578401613 | 0.61796915 | 2.55417542 | 0.01064396 | 0.29899856 |
| NUMB | 706.721991 | 0.377749005 | 0.17555317 | 2.15176407 | 0.03141594 | 0.43905874 |
| DNAL1 | 95.130306 | 0.552309946 | 0.20421608 | 2.70453696 | 0.00683997 | 0.25593763 |
| PNMA1 | 36.2340242 | 0.371197245 | 0.43071276 | 0.86182086 | 0.3887861 | 0.88586416 |
| ELMSAN1 | 115.296009 | -0.459547667 | 0.26295534 | -1.7476263 | 0.08052875 | 0.60722226 |
| LINC02274 | 42.0696873 | 0.134210085 | 0.69167756 | 0.19403562 | 0.84614799 | 0.98520136 |
| AC005480.1 | 53.2576838 | 0.145299857 | 0.43070868 | 0.33735066 | 0.73585258 | 0.9698508 |
| ENTPD5 | 63.4110246 | -0.216530403 | 0.37205422 | -0.5819862 | 0.56057601 | 0.93747805 |
| BBOF1 | 89.464479 | 0.866716107 | 0.32669515 | 2.65298129 | 0.00797843 | 0.27197642 |
| ALDH6A1 | 59.0944279 | 1.175191501 | 0.46795064 | 2.51135782 | 0.01202677 | 0.30967852 |
| LIN52 | 108.439557 | -0.304007589 | 0.28171543 | -1.07913 | 0.28052976 | 0.825559 |
| ABCD4 | 117.511428 | 0.247164971 | 0.1886022 | 1.31050948 | 0.19002354 | 0.76207437 |
| NPC2 | 122.506776 | 0.489662243 | 0.46001756 | 1.06444251 | 0.28712828 | 0.83129986 |
| ISCA2 | 31.6343387 | 0.079276455 | 0.55033485 | 0.14405131 | 0.88545995 | 0.98766275 |
| AREL1 | 82.301483 | -0.114433675 | 0.21844472 | -0.5238564 | 0.60037839 | 0.94683415 |
| FCF1 | 46.4407943 | -0.083499637 | 0.2848681 | -0.2931168 | 0.76943285 | 0.97261718 |
| YLPM1 | 222.140899 | 0.186296775 | 0.16941049 | 1.09967674 | 0.27147299 | 0.81831151 |
| DLST | 80.201512 | 0.091214319 | 0.27022528 | 0.33754917 | 0.73570295 | 0.9698508 |
| MLH3 | 148.67549 | 0.060154711 | 0.23603568 | 0.25485432 | 0.79883562 | 0.9761581 |
| ACYP1 | 66.2389705 | 0.298322587 | 0.36843973 | 0.80969169 | 0.4181174 | 0.89862171 |
| NEK9 | 121.652651 | -0.064745974 | 0.16951863 | -0.3819402 | 0.70250574 | 0.9645815 |
| TMED10 | 194.097132 | 0.237042542 | 0.2174326 | 1.09018858 | 0.27563008 | 0.82164863 |
| FOS | 116.055313 | 0.733043991 | 0.8652261 | 0.84722825 | 0.39686791 | 0.88863925 |
| JDP2 | 30.1472175 | -0.438863645 | 0.59781007 | -0.7341189 | 0.4628763 | 0.91462428 |
| FLVCR2 | 110.928271 | 0.645399482 | 0.57385624 | 1.12467102 | 0.26072847 | 0.81267205 |
| TTLL5 | 299.750187 | -0.024027747 | 0.19484593 | -0.1233166 | 0.90185636 | 0.9900784 |
| ERG28 | 34.2455183 | 0.050801579 | 0.38955185 | 0.13041031 | 0.89624181 | 0.98842187 |
| IFT43 | 213.889909 | 0.640453513 | 0.27090506 | 2.3641253 | 0.0180727 | 0.35858401 |
| GPATCH2L | 217.185722 | -0.281270098 | 0.2734191 | -1.0287141 | 0.30361403 | 0.84470484 |
| ANGEL1 | 26.9007202 | -0.299634284 | 0.31861525 | -0.9404267 | 0.34699874 | 0.87015462 |
| IRF2BPL | 87.4816284 | -0.062728415 | 0.47356154 | -0.132461 | 0.8946197 | 0.98842187 |
| POMT2 | 45.9164329 | -0.262352606 | 0.29412523 | -0.8919759 | 0.37240587 | 0.87605028 |
| TMED8 | 115.399229 | 0.043497462 | 0.19443687 | 0.22370995 | 0.822983 | 0.98094663 |
| VIPAS39 | 32.505102 | -0.16652041 | 0.27429136 | -0.6070932 | 0.54378909 | 0.93523904 |
| AHSA1 | 49.2866278 | -0.054065254 | 0.34145277 | -0.1583389 | 0.87418976 | 0.98766275 |
| SPTLC2 | 345.183543 | -0.322663311 | 0.30908252 | -1.043939 | 0.2965136 | 0.83973489 |
| ALKBH1 | 51.9341687 | -0.174751804 | 0.2440308 | -0.7161055 | 0.47392619 | 0.91809716 |

| | | | | | | |
|---|---|---|---|---|---|---|
| SLIRP | 81.7472792 | -0.271438067 | 0.24680265 | -1.0998183 | 0.27141131 | 0.81831151 |
| SNW1 | 76.1295587 | -0.275417413 | 0.23599712 | -1.1670372 | 0.24319536 | 0.80476923 |
| NRXN3 | 164.08022 | 0.584097292 | 1.05152133 | 0.55547831 | 0.57856754 | 0.94034284 |
| CEP128 | 103.521477 | -0.77129812 | 0.28883642 | -2.6703631 | 0.00757693 | 0.26951269 |
| AC007262.2 | 53.6998079 | 1.215138721 | 0.60927711 | 1.99439418 | 0.04610899 | 0.50827029 |
| GTF2A1 | 84.4593108 | 0.246026341 | 0.2173353 | 1.13201278 | 0.25762906 | 0.8120529 |
| STON2 | 239.445626 | 0.598011928 | 0.60842285 | 0.98288868 | 0.32566223 | 0.85509839 |
| SEL1L | 168.719234 | 0.103285439 | 0.33122896 | 0.31182491 | 0.75517359 | 0.97125609 |
| GALC | 51.9074528 | -0.251478974 | 0.28318534 | -0.8880367 | 0.374521 | 0.87678297 |
| SPATA7 | 122.981227 | -0.018684578 | 0.32652785 | -0.057222 | 0.95436835 | 0.99531843 |
| PTPN21 | 134.945426 | -0.056595738 | 0.26984182 | -0.2097367 | 0.83387317 | 0.98351801 |
| ZC3H14 | 198.134611 | 0.154131362 | 0.1993503 | 0.77316845 | 0.4394227 | 0.90561284 |
| EML5 | 101.243057 | -1.517276831 | 0.38028528 | -3.9898384 | 6.61E-05 | 0.01895844 |
| TTC8 | 110.429414 | 0.170432499 | 0.26543142 | 0.64209617 | 0.52081074 | 0.93272217 |
| FOXN3 | 190.74953 | 0.603874534 | 0.29255281 | 2.06415566 | 0.03900296 | 0.47776132 |
| EFCAB11 | 80.5450353 | 0.320633564 | 0.40165698 | 0.79827708 | 0.42470971 | 0.90167091 |
| TDP1 | 50.9603014 | -0.103211016 | 0.25875083 | -0.3988819 | 0.68998024 | 0.96220115 |
| PSMC1 | 68.6434967 | -0.650441536 | 0.23067918 | -2.8196802 | 0.00480715 | 0.21962686 |
| NRDE2 | 68.0434047 | 0.185363053 | 0.39990954 | 0.46351246 | 0.64299709 | 0.94944486 |
| CALM1 | 270.313253 | -0.198591835 | 0.3725193 | -0.5331048 | 0.59396102 | 0.94649617 |
| TTC7B | 240.437565 | -0.632961597 | 0.37926512 | -1.6689159 | 0.09513404 | 0.63995013 |
| RPS6KA5 | 104.779172 | 0.664051567 | 0.30619344 | 2.16873217 | 0.03010302 | 0.43492739 |
| DGLUCY | 171.997001 | -0.297715828 | 0.32930524 | -0.9040726 | 0.36595693 | 0.87605028 |
| PPP4R3A | 229.07628 | -0.061481824 | 0.1493629 | -0.4116271 | 0.68061274 | 0.95989244 |
| CATSPERB | 83.9875903 | 0.352423829 | 0.41650785 | 0.8461397 | 0.39747482 | 0.88863925 |
| TC2N | 94.2735665 | 0.384675415 | 0.32715026 | 1.17583711 | 0.23965997 | 0.80193227 |
| TRIP11 | 246.738823 | 0.314518773 | 0.17504412 | 1.79679718 | 0.07236782 | 0.59090333 |
| ATXN3 | 155.511016 | 0.323333341 | 0.27760018 | 1.16474471 | 0.24412235 | 0.80476923 |
| NDUFB1 | 65.6197362 | -0.218424663 | 0.3511072 | -0.6221025 | 0.53387447 | 0.93446646 |
| CPSF2 | 88.0753598 | -0.036558185 | 0.25120885 | -0.1455291 | 0.88429318 | 0.98766275 |
| LGMN | 75.9603084 | -0.211154214 | 0.45965885 | -0.4593716 | 0.64596735 | 0.94991115 |
| GOLGA5 | 59.5484905 | -0.133984979 | 0.20609632 | -0.6501085 | 0.51562212 | 0.93140514 |
| ITPK1 | 98.7926733 | 0.06147655 | 0.26047215 | 0.23601967 | 0.81341741 | 0.97930627 |
| MOAP1 | 36.7578512 | -0.110500935 | 0.43382239 | -0.2547147 | 0.79894347 | 0.9761581 |
| BTBD7 | 529.022631 | -0.207759114 | 0.24643166 | -0.8430699 | 0.39918934 | 0.88955057 |
| UNC79 | 51.9649716 | 1.68710612 | 0.58912225 | 2.86376234 | 0.00418642 | 0.20768032 |
| DDX24 | 192.627944 | 0.073642304 | 0.22892784 | 0.32168349 | 0.74769249 | 0.97028355 |
| IFI27L2 | 33.9434593 | -0.030451405 | 0.46615214 | -0.065325 | 0.94791521 | 0.9952642 |
| SERPINA1 | 1320.89685 | -0.110530512 | 0.77190527 | -0.1431918 | 8.86E-01 | 0.98766275 |
| DICER1 | 368.621186 | 0.066894999 | 0.2071429 | 0.3229413 | 0.74673969 | 0.97028355 |
| DICER1-AS1 | 44.5962248 | -0.079025719 | 0.55086254 | -0.1434581 | 0.88592836 | 0.98766275 |
| CLMN | 506.192476 | 0.44632784 | 0.32839985 | 1.35909879 | 0.17411529 | 0.74910274 |
| GLRX5 | 55.7004637 | -0.038813112 | 0.5180922 | -0.0749155 | 0.94028198 | 0.99526299 |
| ATG2B | 195.248402 | -0.129923932 | 0.2606687 | -0.4984255 | 0.61818416 | 0.9480368 |
| GSKIP | 35.4127229 | -0.099565825 | 0.32750821 | -0.3040102 | 0.76112015 | 0.9714717 |
| AK7 | 57.4585792 | 0.490489858 | 0.61848515 | 0.79305034 | 0.4277485 | 0.90272191 |
| PAPOLA | 322.328047 | 0.112381593 | 0.14567843 | 0.77143605 | 0.44044851 | 0.90561284 |
| VRK1 | 43.3656498 | -0.272002205 | 0.32478317 | -0.8374886 | 0.40231798 | 0.89239743 |
| LINC01550 | 41.414702 | 0.845088765 | 0.52440887 | 1.61150737 | 0.10706918 | 0.65699665 |
| SETD3 | 256.292668 | 0.142668886 | 0.19036833 | 0.74943605 | 0.45359443 | 0.91077355 |
| CCNK | 62.5124944 | 0.190668803 | 0.2372971 | 0.80350245 | 0.42168438 | 0.90042681 |
| CCDC85C | 38.6168146 | 0.09214391 | 0.40673219 | 0.22654688 | 0.8207761 | 0.98094663 |
| EML1 | 46.1762308 | 0.83005498 | 0.51657294 | 1.60684951 | 0.10808738 | 0.65707482 |
| EVL | 182.757152 | -0.054547103 | 0.44263882 | -0.1232316 | 0.90192368 | 0.9900784 |

| | | | | | | |
|---|---|---|---|---|---|---|
| YY1 | 224.056423 | -0.217795087 | 0.13746631 | -1.5843525 | 0.11311353 | 0.66636809 |
| SLC25A29 | 141.290042 | 0.252187032 | 0.28257206 | 0.89246979 | 0.37214118 | 0.87605028 |
| WARS | 85.1339942 | -0.840000023 | 0.48181026 | -1.7434249 | 0.08125941 | 0.60850843 |
| WDR25 | 66.9618882 | -0.137175579 | 0.26083824 | -0.5259029 | 0.5989557 | 0.94683415 |
| PPP2R5C | 239.559695 | -0.072420068 | 0.22252208 | -0.3254512 | 0.74483964 | 0.97028355 |
| DYNC1H1 | 251.708862 | -0.217251554 | 0.19297399 | -1.1258074 | 0.26024704 | 0.81263292 |
| HSP90AA1 | 858.72466 | 0.213302137 | 0.39596415 | 0.53869053 | 0.59010041 | 0.94516215 |
| WDR20 | 251.392712 | -0.141279777 | 0.20840914 | -0.6778962 | 0.49783748 | 0.92398825 |
| MOK | 201.512812 | 0.514124184 | 0.33458622 | 1.53659703 | 0.12439202 | 0.69439514 |
| ZNF839 | 41.9131094 | -0.314410468 | 0.3654916 | -0.8602399 | 0.3896568 | 0.88586416 |
| CINP | 24.2035772 | -0.033317114 | 0.36649679 | -0.090907 | 0.92756651 | 0.99336859 |
| TECPR2 | 92.6223558 | -0.211640603 | 0.24287828 | -0.8713855 | 0.38354372 | 0.88328503 |
| RCOR1 | 172.099259 | -0.030762267 | 0.20221957 | -0.1521231 | 0.87908985 | 0.98766275 |
| TRAF3 | 207.132425 | 0.086106413 | 0.3548301 | 0.24266941 | 0.8082615 | 0.97930627 |
| AMN | 45.0849068 | 1.024429571 | 0.63100133 | 1.62349828 | 0.10448293 | 0.65551333 |
| CDC42BPB | 290.180483 | -0.432707945 | 0.19670281 | -2.1998056 | 0.02782069 | 0.4250936 |
| TNFAIP2 | 231.998417 | 0.80155938 | 0.79398994 | 1.00953342 | 0.31271888 | 0.84826686 |
| EIF5 | 138.960747 | -0.100904044 | 0.34769834 | -0.2902057 | 0.77165887 | 0.97329019 |
| MARK3 | 559.277429 | -0.182486947 | 0.21677909 | -0.8418107 | 0.39989395 | 0.89041937 |
| CKB | 107.527072 | 0.926254721 | 0.37837202 | 2.44800005 | 0.01436516 | 0.32762834 |
| BAG5 | 68.6224736 | 0.094380666 | 0.19932268 | 0.4735069 | 6.36E-01 | 0.94932781 |
| KLC1 | 138.938709 | -0.02761481 | 0.24221264 | -0.1140106 | 0.90922938 | 0.99100813 |
| ZFYVE21 | 52.8781963 | 0.088345963 | 0.32856285 | 0.26888604 | 0.78801738 | 0.97423208 |
| PPP1R13B | 458.781531 | -0.050781133 | 0.28377139 | -0.1789509 | 0.85797629 | 0.9864199 |
| ATP5MPL | 86.2900221 | 0.181604706 | 0.3401503 | 0.53389547 | 0.59341386 | 0.94649617 |
| INF2 | 60.4503341 | -0.086629664 | 0.73684927 | -0.1175677 | 0.90641021 | 0.99100813 |
| SIVA1 | 47.6048408 | -0.242552773 | 0.40778036 | -0.5948123 | 0.55196895 | 0.93523904 |
| AKT1 | 61.8471611 | -0.42424818 | 0.29533849 | -1.4364812 | 0.15086547 | 0.72365564 |
| CEP170B | 39.2392969 | -0.436737665 | 0.39471165 | -1.1064727 | 0.26852198 | 0.8158873 |
| CLBA1 | 29.4661834 | 0.036673294 | 0.35755209 | 0.10256769 | 0.91830608 | 0.99214128 |
| BRF1 | 60.9869595 | -0.211572719 | 0.29038364 | -0.7285972 | 0.46624806 | 0.91462428 |
| PACS2 | 116.263319 | -0.30939322 | 0.30805538 | -1.0043428 | 0.31521339 | 0.8490543 |
| TEX22 | 27.9088739 | -0.623194033 | 0.41322035 | -1.5081397 | 0.13151877 | 0.700349 |
| MTA1 | 76.7214636 | -0.598488032 | 0.3151304 | -1.8991758 | 0.05754136 | 0.54735721 |
| CRIP2 | 73.5435509 | 0.295417914 | 0.56919958 | 0.51900585 | 0.60375666 | 0.94683415 |
| NIPA1 | 91.0745987 | -0.351620862 | 0.34016895 | -1.0336654 | 0.30129263 | 0.84277781 |
| NIPA2 | 75.6952502 | -0.195779137 | 0.37385735 | -0.5236734 | 0.60050574 | 0.94683415 |
| CYFIP1 | 169.302077 | -0.358042194 | 0.25345586 | -1.4126412 | 0.15776124 | 0.73580079 |
| TUBGCP5 | 53.6100836 | -0.672481009 | 0.34658879 | -1.9402849 | 0.05234507 | 0.53169179 |
| AC124312.1 | 431.470317 | 0.382778633 | 0.35047382 | 1.09217468 | 2.75E-01 | 0.82143911 |
| PWAR6 | 98.0241924 | 0.747023694 | 0.37898778 | 1.97110232 | 0.04871217 | 0.52130708 |
| UBE3A | 341.569539 | -0.029099229 | 0.12933531 | -0.2249906 | 0.82198659 | 0.98094663 |
| ATP10A | 125.846765 | -1.810821856 | 0.59650417 | -3.0357237 | 0.00239959 | 0.16028122 |
| OCA2 | 63.0091073 | 0.752276321 | 0.81152885 | 0.92698654 | 0.35393352 | 0.87241657 |
| HERC2 | 352.77442 | -0.163916142 | 0.21030467 | -0.7794223 | 0.43573102 | 0.90514499 |
| GOLGA8M | 33.9800262 | -0.657986113 | 0.81849963 | -0.803893 | 0.42145878 | 0.90014574 |
| FAM189A1 | 367.153361 | 1.258911271 | 1.1089511 | 1.13522704 | 0.2562802 | 0.80988075 |
| TJP1 | 390.450988 | 0.038139274 | 0.27002587 | 0.14124304 | 0.88767795 | 0.98766275 |
| FAN1 | 84.1675898 | -0.19853286 | 0.2859858 | -0.6942053 | 0.48755346 | 0.92034617 |
| MTMR10 | 414.345422 | -0.24696567 | 0.41816432 | -0.5905948 | 0.55479195 | 0.93523904 |
| KLF13 | 117.101502 | 0.03293447 | 0.38265307 | 0.08606875 | 0.93141177 | 0.99411667 |
| OTUD7A | 101.452218 | 0.255459965 | 0.23840694 | 1.07152906 | 0.28393161 | 0.82918183 |
| LINC02256 | 219.591747 | 0.190411426 | 0.2640742 | 0.72105274 | 0.47087706 | 0.91611922 |
| AC123768.3 | 50.2024893 | 0.056398923 | 0.52581685 | 0.10725963 | 0.91458301 | 0.99159829 |

| | | | | | | |
|---|---|---|---|---|---|---|
| FMN1 | 76.3460566 | -0.285353822 | 0.5433876 | -0.5251386 | 0.59948682 | 0.94683415 |
| RYR3 | 61.9846513 | 0.331180409 | 0.62700767 | 0.52819196 | 0.59736611 | 0.94683415 |
| AVEN | 134.020446 | -0.213488727 | 0.48573828 | -0.4395139 | 0.66028921 | 0.95407427 |
| EMC7 | 57.5325632 | 0.064264557 | 0.3653554 | 0.17589601 | 0.86037564 | 0.9864199 |
| KATNBL1 | 215.570143 | -0.53055203 | 0.19954684 | -2.6587844 | 0.00784231 | 0.27165021 |
| EMC4 | 32.4870891 | -0.370634794 | 0.36641555 | -1.0115149 | 0.31177005 | 0.84811918 |
| SLC12A6 | 126.928964 | 0.289970053 | 0.36369172 | 0.79729626 | 0.42527899 | 0.90167091 |
| NOP10 | 88.4946841 | -0.128696222 | 0.58965775 | -0.2182558 | 0.82722981 | 0.98142144 |
| GOLGA8A | 134.955904 | -0.748214294 | 0.41492355 | -1.8032582 | 0.07134767 | 0.59090333 |
| GOLGA8B | 141.641984 | 0.714628512 | 0.47589906 | 1.50163885 | 0.1331904 | 0.70399508 |
| AQR | 157.352448 | -0.167005918 | 0.18863907 | -0.8853199 | 0.37598412 | 0.877999 |
| ZNF770 | 94.1204655 | 0.109166706 | 0.23851177 | 0.45769946 | 0.64716838 | 0.95027667 |
| AC021231.1 | 197.160396 | 0.25409878 | 0.19661227 | 1.29238514 | 0.1962238 | 0.77033926 |
| DPH6 | 536.195963 | -0.443166997 | 0.15770328 | -2.8101318 | 0.00495212 | 0.22029054 |
| DPH6-DT | 228.266349 | 1.222574091 | 0.62095806 | 1.96885133 | 0.04897017 | 0.52219814 |
| C15orf41 | 93.8305767 | -0.547609232 | 0.26816235 | -2.042081 | 0.0411435 | 0.48752866 |
| MEIS2 | 275.673635 | 1.068872311 | 0.98688024 | 1.08308209 | 0.27877199 | 0.8229426 |
| SPRED1 | 272.321907 | 0.014492864 | 0.22873243 | 0.06336165 | 0.94947853 | 0.99531843 |
| FAM98B | 57.1055493 | -0.083194471 | 0.27221176 | -0.3056241 | 0.75989088 | 0.9714717 |
| RASGRP1 | 117.807663 | 0.556232449 | 0.46949227 | 1.18475316 | 0.23611506 | 0.79958336 |
| C15orf53 | 77.2545459 | 1.412159671 | 0.75517558 | 1.86997529 | 0.06148725 | 0.56254667 |
| AC013652.1 | 235.404931 | 0.089314929 | 0.43339621 | 0.20608147 | 0.83672727 | 0.98462033 |
| THBS1 | 295.32727 | -0.092983513 | 0.67257862 | -0.1382493 | 0.89004341 | 0.98766275 |
| FSIP1 | 66.5466005 | 0.584880753 | 0.29926614 | 1.95438332 | 5.07E-02 | 0.52415717 |
| GPR176 | 101.681646 | 0.735251317 | 0.90532026 | 0.812145 | 0.41670844 | 0.89862171 |
| EIF2AK4 | 141.120253 | -0.398749921 | 0.30320385 | -1.3151216 | 0.18846905 | 0.76207437 |
| SRP14 | 374.096081 | -0.000314081 | 0.52854308 | -0.0005942 | 0.99952587 | 0.99971834 |
| SRP14-AS1 | 26.4341632 | 0.056693637 | 0.57781505 | 0.09811727 | 0.92183917 | 0.99214128 |
| IVD | 82.9595335 | 0.167657516 | 0.33887465 | 0.49474788 | 0.6207781 | 0.9480368 |
| BAHD1 | 29.192383 | 0.462773398 | 0.45261418 | 1.02244565 | 0.30657002 | 0.84488735 |
| CCDC32 | 86.4882544 | -0.648038767 | 0.37017978 | -1.7506055 | 0.08001389 | 0.60643125 |
| RMDN3 | 47.3446076 | -0.493250508 | 0.32152424 | -1.5341005 | 0.12500492 | 0.69439514 |
| GCHFR | 48.2276844 | 0.449913781 | 0.4741443 | 0.94889631 | 0.34267335 | 0.86846099 |
| DNAJC17 | 69.5193533 | -0.234286625 | 0.23428961 | -0.9999873 | 0.31731667 | 0.8490543 |
| ZFYVE19 | 29.0047018 | 0.277664044 | 0.37330718 | 0.74379507 | 0.45700046 | 0.91187931 |
| SPINT1-AS1 | 35.4227262 | 0.012720934 | 0.52525327 | 0.02421867 | 0.98067819 | 0.99681784 |
| SPINT1 | 97.4887068 | 0.347576226 | 0.31842187 | 1.09155891 | 0.27502702 | 0.82164863 |
| INO80 | 229.920127 | -0.421356319 | 0.32911425 | -1.2802737 | 0.20044889 | 0.77110248 |
| CHP1 | 112.471332 | 0.239724465 | 0.65037887 | 0.36859203 | 0.71243184 | 0.96587538 |
| RTF1 | 140.938235 | -0.232402669 | 0.1755535 | -1.3238282 | 0.18556012 | 0.76046631 |
| RPAP1 | 28.0432824 | -0.300418463 | 0.32979826 | -0.9109158 | 0.36233972 | 0.8752623 |
| MGA | 195.118263 | 0.164155154 | 0.24807619 | 0.66171266 | 0.50815539 | 0.92666294 |
| MAPKBP1 | 75.1943142 | -0.04174633 | 0.30373365 | -0.1374439 | 0.89067995 | 0.98766275 |
| EHD4 | 65.6170042 | -0.405067235 | 0.39370593 | -1.0288573 | 0.30354672 | 0.84470484 |
| VPS39 | 90.145339 | 0.059982394 | 0.18039392 | 0.33250784 | 0.73950582 | 0.9698508 |
| TMEM87A | 199.768273 | -0.312707428 | 0.19221586 | -1.6268555 | 0.10376778 | 0.65437236 |
| GANC | 77.7103259 | -0.358895031 | 0.52044611 | -0.6895911 | 0.49045134 | 0.9217252 |
| ZNF106 | 125.783605 | 0.093325586 | 0.19612668 | 0.47584341 | 0.63418594 | 0.94932781 |
| SNAP23 | 101.414405 | 0.2047271 | 0.24452414 | 0.83724699 | 0.40245375 | 0.89239743 |
| HAUS2 | 31.3931415 | -0.293629853 | 0.32927883 | -0.8917362 | 0.37253434 | 0.87605028 |
| TTBK2 | 221.95941 | 0.390936469 | 0.19868721 | 1.96759753 | 0.04911436 | 0.52272749 |
| UBR1 | 323.658375 | 0.075017164 | 0.1905333 | 0.39372205 | 0.69378626 | 0.96220115 |
| AC068724.3 | 72.0388281 | 0.478895972 | 0.41347642 | 1.15821835 | 0.24677494 | 0.80595199 |
| TMEM62 | 77.7787173 | -0.427346697 | 0.37589606 | -1.1368746 | 0.25559069 | 0.80897837 |

| | | | | | | |
|---|---|---|---|---|---|---|
| CCNDBP1 | 39.2686272 | 0.360774792 | 0.4404903 | 0.81903005 | 0.41276927 | 0.89719011 |
| ADAL | 31.115425 | 0.057518237 | 0.42356431 | 0.13579576 | 0.89198274 | 0.98811543 |
| TUBGCP4 | 47.1254595 | -0.431376815 | 0.30629523 | -1.4083693 | 0.15902173 | 0.73694687 |
| TP53BP1 | 213.592341 | 0.337026391 | 0.19833726 | 1.69925909 | 0.08927038 | 0.63290368 |
| PPIP5K1 | 49.7003736 | 0.139198046 | 0.3732453 | 0.37293985 | 0.70919321 | 0.96587538 |
| CATSPER2 | 66.8267752 | 0.261002693 | 0.52729214 | 0.49498688 | 0.62060938 | 0.9480368 |
| PDIA3 | 210.164014 | -0.067966295 | 0.38975897 | -0.1743803 | 0.86156658 | 0.98669442 |
| SERF2 | 546.145951 | 0.030154776 | 0.48203237 | 0.06255758 | 0.95011881 | 0.99531843 |
| MFAP1 | 41.6835602 | -0.272022289 | 0.33262375 | -0.8178078 | 0.41346696 | 0.89732895 |
| FRMD5 | 44.620092 | 0.594187402 | 0.53295852 | 1.11488488 | 0.26489975 | 0.81378765 |
| CASC4 | 474.02782 | 0.131993352 | 0.16936633 | 0.77933641 | 0.43578157 | 0.90514499 |
| CTDSPL2 | 275.912939 | -0.162878893 | 0.15892639 | -1.02487 | 0.30542452 | 0.84488735 |
| EIF3J-DT | 72.9515082 | 0.259670577 | 0.40700769 | 0.63799919 | 0.5234742 | 0.93272384 |
| EIF3J | 144.813174 | 0.125212531 | 0.33811326 | 0.37032719 | 0.71113872 | 0.96587538 |
| SPG11 | 299.032543 | -0.050484454 | 0.16768974 | -0.3010587 | 0.76336973 | 0.97149197 |
| B2M | 1361.1058 | -0.337415738 | 0.5480112 | -0.6157096 | 0.53808622 | 0.93516669 |
| DUOX1 | 32.9295886 | 0.331907244 | 0.45950764 | 0.7223106 | 0.47010354 | 0.91502267 |
| GATM | 93.8082823 | 1.51937841 | 0.4228723 | 3.59299583 | 0.0003269 | 0.04892741 |
| SPATA5L1 | 35.3834122 | 0.10950976 | 0.33491004 | 0.32698261 | 0.74368104 | 0.97012548 |
| SQOR | 60.196184 | -0.024825703 | 0.59407067 | -0.0417891 | 0.96666679 | 0.99601626 |
| SEMA6D | 65.3104028 | 0.418185721 | 0.80522546 | 0.51933991 | 0.60352373 | 0.94683415 |
| MYEF2 | 312.137214 | 0.214982463 | 0.30352441 | 0.70828722 | 0.47876691 | 0.91971919 |
| SLC12A1 | 305.873592 | 0.307058084 | 1.41277904 | 0.21734332 | 0.8279408 | 0.98142144 |
| DUT | 81.1579537 | 0.353808621 | 0.29657731 | 1.19297267 | 0.23288008 | 0.79556286 |
| CEP152 | 36.6412754 | -0.388762611 | 0.36821801 | -1.0557947 | 0.29106203 | 0.83575635 |
| EID1 | 172.625389 | 0.102619196 | 0.41769916 | 0.24567729 | 0.80593205 | 0.97930627 |
| SECISBP2L | 264.140776 | 0.067837208 | 0.24272229 | 0.27948488 | 0.77987274 | 0.97423208 |
| COPS2 | 85.1392079 | 0.176280087 | 0.26800218 | 0.65775617 | 0.51069483 | 0.92819651 |
| GALK2 | 202.331797 | 0.48836898 | 0.23012764 | 2.12216566 | 0.03382383 | 0.44937373 |
| FAM227B | 114.327705 | 0.510528883 | 0.52533961 | 0.97180734 | 0.33114641 | 0.85923151 |
| DTWD1 | 117.394182 | -0.176973735 | 0.40339425 | -0.4387116 | 0.66087052 | 0.95407427 |
| ATP8B4 | 41.6969961 | 1.261590539 | 0.62682165 | 2.01267862 | 0.04414845 | 0.49952159 |
| SLC27A2 | 106.489169 | 0.704111028 | 0.47515512 | 1.48185508 | 0.13837886 | 0.71091391 |
| GABPB1 | 109.889085 | -0.560587283 | 0.18730983 | -2.9928343 | 0.002764 | 0.17081506 |
| GABPB1-AS1 | 290.668266 | 0.280394038 | 0.4004429 | 0.70020979 | 0.4837963 | 0.92004859 |
| USP8 | 195.837526 | 0.188472812 | 0.14551929 | 1.29517411 | 0.19526017 | 0.76968045 |
| TRPM7 | 381.606113 | -0.170832254 | 0.20278882 | -0.8424145 | 0.39955595 | 0.88987363 |
| SPPL2A | 213.370731 | -0.123927488 | 0.21872096 | -0.5666009 | 0.57098538 | 0.93998433 |
| AC021752.1 | 40.8436485 | -0.479665479 | 0.72996956 | -0.6571034 | 0.51111444 | 0.92819651 |
| AP4E1 | 98.1644869 | -0.165529878 | 0.26382789 | -0.6274162 | 0.53038648 | 0.93366189 |
| DMXL2 | 111.698815 | 0.386165788 | 0.24079716 | 1.60369744 | 0.10878076 | 0.65764016 |
| TMOD3 | 165.775252 | -0.34898804 | 0.23887924 | -1.4609392 | 0.14403213 | 0.71634214 |
| MAPK6 | 124.578122 | -0.061370598 | 0.24724867 | -0.2482141 | 0.80396878 | 0.97880236 |
| GNB5 | 84.2427324 | 0.740842447 | 0.26311827 | 2.81562527 | 0.00486824 | 0.21962686 |
| MYO5C | 163.574743 | -0.398536135 | 0.27100005 | -1.4706128 | 0.14139587 | 0.7148274 |
| MYO5A | 91.0738644 | 0.294957643 | 0.45827841 | 0.64362108 | 0.51982118 | 0.93272217 |
| ARPP19 | 141.502148 | -0.470639837 | 0.25217626 | -1.866313 | 0.06199759 | 0.56445068 |
| FAM214A | 205.125578 | 0.571631701 | 0.23367477 | 2.44627052 | 0.01443427 | 0.32788161 |
| WDR72 | 224.322414 | 1.109993839 | 0.71408367 | 1.55443107 | 0.12008163 | 0.68786496 |
| UNC13C | 174.580129 | -1.160775013 | 1.36157469 | -0.8525239 | 0.39392336 | 0.88760715 |
| RSL24D1 | 67.7032204 | 0.387340415 | 0.45157003 | 0.85776378 | 0.39102292 | 0.88670817 |
| RAB27A | 129.564815 | -0.320609614 | 0.1833964 | -1.7481783 | 0.08043316 | 0.60722226 |
| PIGB | 60.6859758 | 0.311258843 | 0.31152365 | 0.99914996 | 0.31772205 | 0.8490543 |
| CCPG1 | 220.959534 | 0.148613144 | 0.28344893 | 0.52430308 | 0.60006776 | 0.94683415 |

| | | | | | | |
|---|---|---|---|---|---|---|
| DNAAF4 | 125.812841 | 0.378906957 | 0.2552505 | 1.48445136 | 0.13768922 | 0.71091391 |
| PYGO1 | 38.2824949 | -0.294051393 | 0.56478009 | -0.5206476 | 0.60261229 | 0.94683415 |
| PRTG | 100.095088 | -0.311013311 | 0.68361923 | -0.4549511 | 0.64914443 | 0.95027667 |
| NEDD4 | 146.17993 | 0.246396478 | 0.52016209 | 0.47369172 | 0.63571976 | 0.94932781 |
| RFX7 | 282.250065 | 0.243703677 | 0.21068408 | 1.15672565 | 0.24738447 | 0.80672756 |
| TEX9 | 246.14483 | 0.381146898 | 0.25717155 | 1.48207254 | 0.13832099 | 0.71091391 |
| MNS1 | 64.483411 | 0.709602717 | 0.47500559 | 1.49388287 | 0.13520624 | 0.70620568 |
| ZNF280D | 145.206975 | 0.334888536 | 0.26082255 | 1.2839708 | 0.19915217 | 0.77110248 |
| AC090517.4 | 104.445569 | 0.172421605 | 0.20980586 | 0.821815 | 0.41118219 | 0.89649693 |
| TCF12 | 1206.06129 | -0.101221311 | 0.14060058 | -0.719921 | 0.47157364 | 0.91645443 |
| CGNL1 | 303.431615 | 0.653062166 | 0.54619526 | 1.19565696 | 0.23183045 | 0.79471003 |
| POLR2M | 24.4486571 | -0.138304206 | 0.33429215 | -0.4137226 | 0.6790773 | 0.95928056 |
| ALDH1A2 | 2956.88114 | 0.177411138 | 0.53577573 | 0.33112948 | 0.74054669 | 0.9698508 |
| AQP9 | 50.7099821 | 1.338275145 | 0.77782972 | 1.72052457 | 0.08533713 | 0.61974555 |
| LIPC | 271.715259 | -0.083483999 | 0.28810959 | -0.2897647 | 0.77199623 | 0.97329019 |
| ADAM10 | 1349.05624 | -0.313262051 | 0.2036909 | -1.5379286 | 0.12406608 | 0.69439514 |
| MINDY2 | 528.166628 | -0.48332116 | 0.18539261 | -2.6070142 | 0.00913356 | 0.2846011 |
| RNF111 | 332.752996 | -0.070406358 | 0.21388622 | -0.3291767 | 0.74202214 | 0.9698508 |
| SLTM | 448.754415 | -0.165603043 | 0.15910563 | -1.0408371 | 0.29795116 | 0.83973489 |
| MYO1E | 485.740248 | -0.546957413 | 0.2877374 | -1.9008909 | 0.0573163 | 0.54616991 |
| FAM81A | 41.8775428 | 0.725452905 | 0.52714729 | 1.37618636 | 0.16876395 | 0.74445571 |
| GCNT3 | 119.954531 | -0.670482357 | 0.7551268 | -0.887907 | 0.37459079 | 0.87678297 |
| GTF2A2 | 86.7643676 | -0.14144903 | 0.48029485 | -0.2945046 | 0.76837237 | 0.97241894 |
| BNIP2 | 111.797018 | -0.076542236 | 0.15952649 | -0.4798089 | 0.63136326 | 0.94932781 |
| ANXA2 | 420.799381 | -0.53429352 | 0.39829628 | -1.3414474 | 0.17977523 | 0.75265164 |
| ICE2 | 163.302401 | 0.02288516 | 0.26716092 | 0.08566058 | 0.93173624 | 0.99411667 |
| RORA-AS1 | 88.3558698 | 0.111370706 | 0.62159057 | 0.17917052 | 0.85780381 | 0.9864199 |
| RORA | 2093.7286 | 0.356932814 | 0.46660729 | 0.76495336 | 0.44429932 | 0.90667728 |
| AC012404.1 | 58.0487324 | 0.307512153 | 0.57149432 | 0.53808435 | 0.59051882 | 0.94516215 |
| AC009554.1 | 109.562724 | 0.6898021 | 0.60719834 | 1.13604082 | 0.25593948 | 0.80939065 |
| VPS13C | 780.288724 | -0.111651852 | 0.19459557 | -0.5737636 | 0.56612779 | 0.93832962 |
| TLN2 | 372.11478 | 0.388827539 | 0.26371822 | 1.4744053 | 0.1403725 | 0.71370921 |
| TPM1 | 1269.28513 | 0.285355918 | 0.27658316 | 1.03171832 | 0.30220409 | 0.84375949 |
| LACTB | 54.2128869 | 0.17372767 | 0.26093931 | 0.66577808 | 5.06E-01 | 0.9257297 |
| RPS27L | 117.730426 | 0.430320889 | 0.31316429 | 1.37410588 | 0.16940881 | 0.74445571 |
| RAB8B | 105.022146 | 0.186099247 | 0.36634906 | 0.50798341 | 0.61146498 | 0.94791646 |
| APH1B | 69.9195475 | 0.503603172 | 0.38802921 | 1.29784861 | 0.19433936 | 0.7687432 |
| CA12 | 141.365154 | 0.44359703 | 0.54480802 | 0.81422633 | 0.41551531 | 0.89807127 |
| USP3 | 308.567573 | -0.083394958 | 0.21253618 | -0.3923801 | 0.69477741 | 0.96220115 |
| USP3-AS1 | 62.9034252 | 0.000114811 | 0.32523481 | 0.00035301 | 0.99971834 | 0.99971834 |
| HERC1 | 312.624662 | 0.016380081 | 0.16257888 | 0.10075159 | 0.91974765 | 0.99214128 |
| FAM96A | 52.1346936 | 0.226826225 | 0.25079475 | 0.90442971 | 0.36576759 | 0.87605028 |
| SNX1 | 165.781414 | 0.2590506 | 0.17831222 | 1.45279222 | 0.14628145 | 0.71974648 |
| PPIB | 247.148993 | 0.077714643 | 0.53103866 | 0.1463446 | 0.88364936 | 0.98766275 |
| CSNK1G1 | 49.2649973 | 0.294603116 | 0.34460194 | 0.85490846 | 0.39260183 | 0.88760715 |
| TRIP4 | 105.572997 | 0.152175654 | 0.27525514 | 0.55285309 | 0.58036401 | 0.94034284 |
| ZNF609 | 575.475746 | -0.055881007 | 0.17371532 | -0.3216815 | 0.74769399 | 0.97028355 |
| OAZ2 | 57.0270977 | 0.296576469 | 0.26599646 | 1.11496398 | 0.26486586 | 0.81378765 |
| SPG21 | 89.3718925 | -0.020435716 | 0.1847126 | -0.1106352 | 0.91190563 | 0.99106427 |
| MTFMT | 49.2811962 | -0.347332779 | 0.24274098 | -1.4308782 | 0.15246512 | 0.72726674 |
| UBAP1L | 32.4036556 | 0.164659408 | 0.33773027 | 0.48754709 | 0.62587068 | 0.94830991 |
| PDCD7 | 53.3721181 | -0.10997358 | 0.2368761 | -0.4642663 | 0.64245699 | 0.94944486 |
| CLPX | 103.358914 | 0.01201117 | 0.1639751 | 0.07324997 | 0.94160721 | 0.99526299 |
| PARP16 | 37.4396111 | -0.00389178 | 0.37481228 | -0.0103833 | 0.99171549 | 0.99866813 |

| | | | | | | |
|---|---|---|---|---|---|---|
| DPP8 | 164.913905 | -0.300319936 | 0.17866117 | -1.6809469 | 0.09277324 | 0.63613087 |
| HACD3 | 117.201426 | -0.04796976 | 0.26005104 | -0.1844629 | 0.85365036 | 0.9864199 |
| INTS14 | 45.8063765 | -0.442767817 | 0.29198418 | -1.5164103 | 0.12941564 | 0.69904834 |
| SLC24A1 | 37.2075456 | -0.449803806 | 0.37953518 | -1.1851439 | 0.23596055 | 0.79952815 |
| DENND4A | 693.937903 | 0.429557434 | 0.30753875 | 1.39675872 | 0.16248614 | 0.74071637 |
| RAB11A | 397.371821 | 0.209314505 | 0.42803573 | 0.48901176 | 0.62483337 | 0.94808496 |
| MEGF11 | 80.2079697 | 0.387611416 | 1.13230569 | 0.34232047 | 7.32E-01 | 9.70E-01 |
| DIS3L | 52.6132974 | -0.189606266 | 0.31353811 | -0.6047312 | 0.54535764 | 0.93523904 |
| TIPIN | 47.5768266 | -0.051286388 | 0.55293644 | -0.0927528 | 0.92609997 | 0.99316012 |
| MAP2K1 | 154.144155 | -0.260887715 | 0.25663195 | -1.0165832 | 0.30935177 | 0.84554063 |
| SNAPC5 | 27.5373578 | 0.044086344 | 0.38690385 | 0.11394651 | 0.90928019 | 0.99100813 |
| RPL4 | 375.319146 | -0.018573496 | 0.58960285 | -0.0315017 | 0.97486943 | 0.99681784 |
| SMAD6 | 142.617512 | -0.683015596 | 0.31897824 | -2.1412608 | 0.03225302 | 0.44049392 |
| SMAD3 | 797.252281 | -0.965473912 | 0.4364501 | -2.212106 | 0.02695934 | 0.42391395 |
| AAGAB | 148.334323 | -0.155231378 | 0.22491405 | -0.6901809 | 0.49008043 | 0.92157694 |
| IQCH | 205.189752 | 0.037922316 | 0.30410391 | 0.12470183 | 0.9007596 | 0.9900784 |
| IQCH-AS1 | 68.35252 | -0.351980033 | 0.39979683 | -0.8803973 | 0.37864414 | 0.87970706 |
| MAP2K5 | 287.276297 | -0.176221442 | 0.1588852 | -1.1091118 | 0.26738197 | 0.81510106 |
| PIAS1 | 588.849901 | -0.084326731 | 0.30863676 | -0.2732232 | 0.78468163 | 0.97423208 |
| FEM1B | 64.8566494 | 0.062685177 | 0.25896647 | 0.24205905 | 0.80873441 | 0.97930627 |
| CORO2B | 60.2832146 | -0.248190901 | 1.23020509 | -0.2017476 | 0.84011406 | 0.98464453 |
| ANP32A | 128.925681 | -0.284223827 | 0.20341228 | -1.3972796 | 0.16232952 | 0.74071637 |
| SPESP1 | 33.0228927 | 0.739025175 | 0.49541315 | 1.49173509 | 0.13576861 | 0.70719276 |
| GLCE | 93.0980673 | 0.389774444 | 0.38868622 | 1.00279975 | 0.31595749 | 0.8490543 |
| PAQR5 | 302.08847 | -0.890092537 | 0.60994084 | -1.4593096 | 0.14447991 | 0.71634214 |
| RPLP1 | 676.060041 | -0.008866842 | 0.51619691 | -0.0171772 | 0.98629521 | 0.99764442 |
| TLE3 | 72.985505 | -0.36046573 | 0.42676407 | -0.8446487 | 0.39830699 | 0.8887831 |
| UACA | 394.782442 | 0.151523501 | 0.31913189 | 0.47479899 | 0.63493025 | 0.94932781 |
| LARP6 | 29.4003803 | 0.20742872 | 0.51716707 | 0.40108648 | 0.68835646 | 0.96161098 |
| LRRC49 | 89.745946 | 0.40823778 | 0.52544511 | 0.77693706 | 0.43719591 | 0.90561284 |
| THSD4 | 274.346066 | -0.454368388 | 0.51953012 | -0.8745756 | 0.38180485 | 0.88128509 |
| THSD4-AS1 | 140.256307 | -0.96652183 | 0.43869851 | -2.2031573 | 0.02758366 | 0.4250936 |
| MYO9A | 2646.83663 | 0.025155754 | 0.34339859 | 0.07325526 | 0.94160299 | 0.99526299 |
| AC022872.1 | 77.3423331 | 0.19783996 | 0.58956083 | 0.33557175 | 0.73719383 | 0.9698508 |
| SENP8 | 35.1515309 | -0.02531037 | 0.43537176 | -0.0581351 | 0.95364104 | 0.99531843 |
| GRAMD2A | 26.4661265 | -0.358928605 | 0.69564871 | -0.5159624 | 0.60588064 | 0.94708529 |
| PKM | 722.666506 | -0.475446403 | 0.32172913 | -1.4777847 | 1.39E-01 | 0.71191199 |
| PARP6 | 92.9245718 | -0.128925927 | 0.19208154 | -0.6712041 | 0.50209049 | 0.92484445 |
| AC009690.2 | 33.7213165 | 1.062963372 | 0.65783052 | 1.61586205 | 0.10612414 | 0.65699665 |
| AC009690.1 | 124.833855 | 0.621943612 | 0.49448902 | 1.25775008 | 0.20848215 | 0.77477691 |
| ARIH1 | 616.230293 | -0.044278683 | 0.21573315 | -0.2052475 | 0.83737879 | 0.98462033 |
| BBS4 | 135.225807 | 0.078800743 | 0.23891651 | 0.32982545 | 0.74153186 | 0.9698508 |
| ADPGK | 71.1938013 | -0.248777035 | 0.27610383 | -0.9010271 | 0.36757391 | 0.87605028 |
| NEO1 | 290.995028 | 0.450385237 | 0.20152075 | 2.23493227 | 0.0254218 | 0.41555526 |
| NPTN | 271.769441 | -0.130769077 | 0.23047576 | -0.5673876 | 0.5704509 | 0.93998433 |
| CD276 | 31.2152816 | 0.217847735 | 0.52394989 | 0.41577971 | 0.67757121 | 0.95924874 |
| LOXL1 | 50.2166103 | 0.266638785 | 0.68150929 | 0.39124747 | 0.69561432 | 0.96220115 |
| PML | 91.9038958 | -0.199956922 | 0.66626631 | -0.3001156 | 0.76408897 | 0.97174831 |
| ARID3B | 29.0656895 | 0.153972304 | 0.39022035 | 0.39457784 | 0.69315447 | 0.96220115 |
| CLK3 | 43.8915333 | -0.097322708 | 0.30701909 | -0.3169924 | 0.75124939 | 0.97058015 |
| EDC3 | 51.8623832 | 0.135916952 | 0.24284803 | 0.55967903 | 0.57569839 | 0.94034284 |
| SCAMP2 | 69.0865247 | -0.444083295 | 0.30937586 | -1.4354168 | 0.15116837 | 0.72403332 |
| FAM219B | 35.000326 | 0.364580383 | 0.29708377 | 1.22719722 | 0.21974847 | 0.78584902 |
| COX5A | 67.0789468 | -0.245199401 | 0.41136468 | -0.5960633 | 0.55113292 | 0.93523904 |

| | | | | | | |
|---|---|---|---|---|---|---|
| PPCDC | 94.8876613 | -0.445564029 | 0.30479412 | -1.4618525 | 0.14378165 | 0.71631428 |
| COMMD4 | 48.171226 | -0.265682955 | 0.3875015 | -0.6856308 | 0.49294597 | 0.92289011 |
| NEIL1 | 25.6693082 | -0.122574228 | 0.39116363 | -0.3133579 | 0.75400874 | 0.97117786 |
| MAN2C1 | 80.5532865 | -0.014521592 | 0.2914915 | -0.0498182 | 0.96026724 | 0.99572129 |
| SIN3A | 159.450257 | -0.206748904 | 0.17437677 | -1.1856448 | 0.23576262 | 0.79952815 |
| PTPN9 | 242.110607 | -0.102183768 | 0.291825 | -0.3501543 | 0.72622293 | 0.96781837 |
| SNUPN | 26.377725 | -0.700053753 | 0.4587019 | -1.5261627 | 0.12696934 | 0.69539125 |
| IMP3 | 32.4523867 | -0.203671485 | 0.39355793 | -0.5175134 | 0.60479785 | 0.94683415 |
| UBE2Q2 | 96.1259581 | 0.453386441 | 0.31513257 | 1.43871654 | 0.15023085 | 0.72246729 |
| FBXO22 | 109.20093 | 0.209661937 | 0.23021633 | 0.91071707 | 0.36244447 | 0.8752623 |
| ETFA | 190.817891 | 0.155937354 | 0.18628044 | 0.83711072 | 0.40253034 | 0.89239743 |
| SCAPER | 605.894338 | 0.385907873 | 0.1907609 | 2.02299256 | 0.04307391 | 0.49473025 |
| RCN2 | 97.3627728 | -0.196991941 | 0.29692187 | -0.6634471 | 0.50704428 | 0.92666294 |
| TSPAN3 | 164.555603 | 0.124499894 | 0.25261947 | 0.49283571 | 0.62212867 | 0.94808496 |
| PEAK1 | 264.826293 | 0.571646673 | 0.23498388 | 2.43270593 | 0.01498647 | 0.3345189 |
| HMG20A | 121.358352 | -0.098629474 | 0.15900726 | -0.6202829 | 0.53507158 | 0.93460153 |
| LINGO1 | 1017.04684 | -0.533257725 | 0.59046236 | -0.9031189 | 0.36646279 | 0.87605028 |
| TBC1D2B | 78.0775497 | 0.049650178 | 0.43817055 | 0.11331245 | 0.90978285 | 0.99100813 |
| IDH3A | 79.1796724 | -0.284495668 | 0.32678799 | -0.8705818 | 0.38398256 | 0.88328503 |
| WDR61 | 70.0244159 | 0.055166103 | 0.21433248 | 0.25738565 | 0.79688108 | 0.97534702 |
| IREB2 | 179.872549 | -0.241483804 | 0.1422735 | -1.6973211 | 0.08963598 | 0.6330877 |
| PSMA4 | 83.8856379 | -0.075074375 | 0.3662612 | -0.204975 | 0.83759169 | 0.98462033 |
| MORF4L1 | 297.081093 | 0.127934668 | 0.20793686 | 0.6152573 | 0.53838481 | 0.93516669 |
| CTSH | 111.548228 | -0.278006591 | 0.30861981 | -0.900806 | 0.36769146 | 0.87605028 |
| TMED3 | 51.373438 | -1.074413433 | 0.29670945 | -3.6210961 | 0.00029336 | 0.04532374 |
| ZFAND6 | 376.792943 | 0.105050159 | 0.16752043 | 0.62708865 | 0.53060112 | 0.93366189 |
| AC016705.2 | 112.036255 | 1.047248154 | 0.61469062 | 1.70369959 | 0.08843722 | 0.62958606 |
| ARNT2 | 363.648977 | 0.768588531 | 0.57774709 | 1.33032004 | 0.18341285 | 0.75689793 |
| ABHD17C | 40.2836134 | 0.000244428 | 0.50087166 | 0.000488 | 0.99961063 | 0.99971834 |
| AC027808.1 | 27.0189933 | 0.731830064 | 0.37888682 | 1.93152686 | 0.05341793 | 0.53392413 |
| MESD | 72.9950808 | 0.114724138 | 0.21258397 | 0.53966504 | 0.58942805 | 0.94491859 |
| AC104041.1 | 42.6992074 | 0.517202132 | 0.88044716 | 0.5874312 | 0.55691414 | 0.93533595 |
| EFL1 | 84.9459431 | -0.41149523 | 0.32634811 | -1.2609089 | 0.20734168 | 0.77401635 |
| RPS17 | 32.8642658 | -0.205741758 | 0.51872216 | -0.3966319 | 0.69163892 | 0.96220115 |
| WHAMM | 137.527453 | -0.125519008 | 0.3273458 | -0.3834447 | 0.70139007 | 0.9645815 |
| HOMER2 | 75.884685 | -0.427835049 | 0.30227821 | -1.4153685 | 0.15696049 | 0.73472335 |
| C15orf40 | 52.5698019 | 0.469710277 | 0.26034808 | 1.8041626 | 0.07120582 | 0.59054593 |
| BTBD1 | 112.041933 | -0.22662166 | 0.2525018 | -0.8975051 | 0.36944944 | 0.87605028 |
| HDGFL3 | 116.749626 | -0.084040342 | 0.25301745 | -0.3321524 | 0.73977422 | 0.9698508 |
| WDR73 | 110.558773 | 0.08634818 | 0.28762795 | 0.30020789 | 0.76401859 | 0.97174831 |
| SEC11A | 213.657017 | 0.227418135 | 0.20655891 | 1.10098439 | 0.27090345 | 0.81775732 |
| ZNF592 | 125.717746 | 0.116975928 | 0.21255173 | 0.55034098 | 0.58208552 | 0.94091128 |
| ALPK3 | 116.279936 | -1.130316788 | 0.47523062 | -2.3784595 | 0.01738515 | 0.3543241 |
| SLC28A1 | 75.7237237 | 0.92392004 | 0.73965619 | 1.24912095 | 0.21162084 | 0.77730294 |
| PDE8A | 326.348304 | 0.211678057 | 0.33573167 | 0.63049773 | 0.52836899 | 0.93366189 |
| AKAP13 | 823.777805 | -0.188594605 | 0.32103646 | -0.5874554 | 0.55689788 | 0.93533595 |
| MRPS11 | 41.519487 | -0.034377022 | 0.40664354 | -0.0845385 | 0.93262832 | 0.9941739 |
| DET1 | 49.3198534 | 0.033307563 | 0.47145828 | 0.07064795 | 0.94367794 | 0.9952642 |
| ABHD2 | 98.8019685 | -0.196504591 | 0.29515443 | -0.6657687 | 0.50555893 | 0.9257297 |
| POLG | 39.3942544 | -0.076070915 | 0.31746188 | -0.2396222 | 0.81062315 | 0.97930627 |
| ANPEP | 52.596426 | 0.786151444 | 0.79913671 | 0.98375088 | 0.32523802 | 0.855037 |
| ZNF710 | 63.776976 | 0.047547484 | 0.36146643 | 0.13154053 | 0.89534773 | 0.98842187 |
| IDH2 | 72.0878443 | 0.524589566 | 0.2282642 | 2.29816835 | 0.02155221 | 0.38588526 |
| SEMA4B | 364.33215 | -0.81420566 | 0.40290943 | -2.0208156 | 4.33E-02 | 0.49484046 |

| | | | | | | |
|---|---|---|---|---|---|---|
| CIB1 | 192.531523 | -0.233023875 | 0.48155593 | -0.4838978 | 6.28E-01 | 0.94858781 |
| IQGAP1 | 397.038726 | -0.042671263 | 0.21363905 | -0.1997353 | 0.84168759 | 0.98515707 |
| CRTC3 | 221.228452 | 0.173590673 | 0.21842883 | 0.79472417 | 0.42677397 | 0.90264121 |
| FURIN | 50.7207518 | -0.554692117 | 0.43408233 | -1.27785 | 0.20130231 | 0.77110248 |
| MAN2A2 | 31.430883 | 0.054322174 | 0.30385159 | 0.17877864 | 0.85811152 | 0.9864199 |
| UNC45A | 46.2613172 | 0.059193086 | 0.25386387 | 0.23316861 | 0.81563048 | 0.98007572 |
| AC104035.1 | 78.1620098 | -0.089068537 | 1.09958866 | -0.0810017 | 0.93544061 | 0.99466301 |
| SLCO3A1 | 834.042026 | 0.080889028 | 0.24650538 | 0.32814305 | 0.7428035 | 0.96991749 |
| FAM174B | 63.9777181 | -0.294588414 | 0.39494349 | -0.7459002 | 0.45572773 | 0.91077355 |
| AC013394.1 | 257.530125 | -0.002061067 | 0.25910648 | -0.0079545 | 0.99365328 | 0.99866813 |
| CHD2 | 330.219782 | -0.274607224 | 0.25898753 | -1.0603106 | 0.28900332 | 0.83399936 |
| MCTP2 | 454.419106 | 0.232125382 | 0.29789204 | 0.77922652 | 0.43584629 | 0.90514499 |
| LINC01197 | 102.146659 | 1.409468661 | 1.41302429 | 0.99748367 | 0.31852979 | 0.84920593 |
| AC012409.2 | 133.004611 | 1.264835119 | 0.54867552 | 2.30525162 | 0.02115249 | 0.38374935 |
| NR2F2-AS1 | 939.210903 | 0.791078553 | 0.41623422 | 1.90056107 | 0.05735953 | 0.54616991 |
| NR2F2 | 347.303621 | 0.358051454 | 0.51547918 | 0.69459925 | 0.48730648 | 0.92034617 |
| AC087477.2 | 81.376689 | 0.311458287 | 0.73108203 | 0.42602372 | 0.67009056 | 0.95764829 |
| ARRDC4 | 77.5800416 | 0.947675212 | 0.46291813 | 2.04717671 | 0.04064074 | 0.48617886 |
| IGF1R | 1242.05343 | 0.173441475 | 0.2845392 | 0.60955212 | 0.54215854 | 0.93523904 |
| TTC23 | 103.143019 | 0.103692322 | 0.24518415 | 0.42291608 | 0.67235649 | 0.95887225 |
| LRRC28 | 138.848346 | -0.100415139 | 0.22878527 | -0.4389056 | 0.66072993 | 0.95407427 |
| MEF2A | 716.873666 | 0.167234192 | 0.23843223 | 0.70139088 | 0.48305911 | 0.91971919 |
| AC090825.1 | 102.279424 | 1.290214225 | 0.5355188 | 2.40927907 | 0.01598407 | 0.3414008 |
| LINS1 | 36.2211996 | -0.324609445 | 0.31381267 | -1.0344052 | 0.30094679 | 0.8426686 |
| ASB7 | 57.3721774 | 0.255176864 | 0.23857139 | 1.06960378 | 0.28479769 | 0.82918183 |
| ALDH1A3 | 80.8198215 | -1.714736089 | 1.36283326 | -1.2582141 | 0.20831432 | 0.77462844 |
| LRRK1 | 71.364261 | -1.160768336 | 0.72514593 | -1.6007376 | 0.10943506 | 0.65764016 |
| CHSY1 | 77.9930518 | -0.144175054 | 0.45824952 | -0.3146213 | 0.75304922 | 0.97096495 |
| SELENOS | 85.7833054 | 0.601335394 | 0.3435977 | 1.75011475 | 0.08009852 | 0.60643125 |
| SNRPA1 | 63.983921 | -0.219512143 | 0.23503534 | -0.9339538 | 0.35032774 | 0.8720098 |
| TM2D3 | 67.130042 | 0.145631937 | 0.36384607 | 0.400257 | 0.68896723 | 0.96190309 |
| TARSL2 | 122.735822 | -0.006660316 | 0.35971143 | -0.0185157 | 0.98522744 | 0.99752755 |
| RHBDF1 | 27.7074983 | -0.384832099 | 0.41601175 | -0.925051 | 0.35493937 | 0.87258503 |
| MPG | 55.502013 | -0.059759024 | 0.27696612 | -0.2157629 | 8.29E-01 | 0.98142144 |
| NPRL3 | 103.368112 | -0.236741198 | 0.26777886 | -0.8840922 | 0.37664645 | 0.87842274 |
| LUC7L | 188.036749 | -0.043722606 | 0.1926368 | -0.2269691 | 0.82044775 | 0.98094663 |
| AXIN1 | 53.3432326 | -0.496878793 | 0.29874729 | -1.6632077 | 0.09627086 | 0.64295596 |
| MRPL28 | 38.4167609 | 0.259419406 | 0.53094733 | 0.48859725 | 0.62512686 | 0.94808496 |
| TMEM8A | 48.6236849 | -0.140971873 | 0.39619737 | -0.3558122 | 0.72198118 | 0.96779196 |
| NME4 | 28.787467 | 0.438496143 | 0.34141623 | 1.28434475 | 0.19902135 | 0.77110248 |
| RAB11FIP3 | 257.625891 | 0.299818708 | 0.43864648 | 0.68350875 | 0.49428542 | 0.92289011 |
| CAPN15 | 46.6653024 | -1.048077113 | 0.41162085 | -2.5462197 | 0.01088966 | 0.30061121 |
| PIGQ | 39.211038 | -0.095016252 | 0.31632597 | -0.3003745 | 0.76389152 | 0.97174831 |
| RAB40C | 84.1063269 | 0.036216333 | 0.39914072 | 0.09073575 | 0.92770256 | 0.99336859 |
| METTL26 | 31.6573654 | 0.503024099 | 0.52816919 | 0.95239197 | 0.34089823 | 0.86824359 |
| MCRIP2 | 43.5636942 | 0.205066355 | 0.36837364 | 0.55668031 | 0.57774587 | 0.94034284 |
| WDR90 | 30.5211952 | 0.127761389 | 0.3182442 | 0.40145708 | 0.68808363 | 0.96161098 |
| RHOT2 | 53.1372892 | -0.092534492 | 0.30487463 | -0.3035165 | 0.76149625 | 0.9714717 |
| STUB1 | 80.4345581 | -0.104310633 | 0.35478389 | -0.2940118 | 0.76874892 | 0.972511 |
| JMJD8 | 46.1722233 | -0.267843575 | 0.37916906 | -0.7063962 | 0.4799418 | 0.91971919 |
| METRN | 33.4799938 | 0.864011085 | 0.45762421 | 1.88803622 | 0.05902109 | 0.55686386 |
| FAM173A | 48.1624981 | 0.370878237 | 0.52002381 | 0.71319473 | 0.47572527 | 0.91809716 |
| LMF1 | 82.2266601 | 0.214005818 | 0.34190181 | 0.62592772 | 0.53136235 | 0.93366189 |
| CACNA1H | 34.4223868 | 0.945069307 | 0.65689933 | 1.4386821 | 0.15024061 | 0.72246729 |

| | | | | | | |
|---|---|---|---|---|---|---|
| UBE2I | 136.533948 | -0.31028381 | 0.2454446 | -1.2641704 | 0.20616886 | 0.7735572 |
| TSR3 | 40.2671198 | 0.070886117 | 0.4358706 | 0.16263111 | 0.87080889 | 0.98766275 |
| GNPTG | 52.2705632 | 0.310090137 | 0.30529855 | 1.01569477 | 0.30977476 | 0.84587583 |
| UNKL | 45.0798789 | 0.210000971 | 0.30016855 | 0.69961017 | 0.48417079 | 0.92034617 |
| CLCN7 | 74.9729863 | -0.629421876 | 0.22435959 | -2.8054155 | 0.00502518 | 0.22182562 |
| TELO2 | 26.992513 | -0.095935295 | 0.4068357 | -0.2358084 | 0.81358131 | 0.97930627 |
| IFT140 | 84.9743238 | 0.602413757 | 0.26339697 | 2.28709447 | 0.02219031 | 0.39000316 |
| JPT2 | 117.498677 | -0.437160315 | 0.20395733 | -2.1433911 | 0.03208171 | 0.44049392 |
| MAPK8IP3 | 139.386285 | 0.044474599 | 0.24011067 | 0.18522542 | 0.85305223 | 0.9864199 |
| NME3 | 107.310633 | 0.493768015 | 0.37524687 | 1.31584844 | 0.18822492 | 0.76207437 |
| MRPS34 | 55.7013565 | -0.208525573 | 0.43456771 | -0.479846 | 0.63133691 | 0.94932781 |
| SPSB3 | 74.1648525 | 0.152147575 | 0.36598827 | 0.41571708 | 0.67761704 | 0.95924874 |
| NUBP2 | 27.921106 | -0.086197755 | 0.36730876 | -0.2346738 | 0.8144619 | 0.97937868 |
| HAGH | 98.3600265 | 0.391496927 | 0.19562855 | 2.00122593 | 0.04536805 | 0.50469451 |
| FAHD1 | 35.4921047 | 0.450712743 | 0.32865245 | 1.37139629 | 1.70E-01 | 0.74445571 |
| MSRB1 | 32.6484984 | 0.183484078 | 0.468934 | 0.39127911 | 0.69559094 | 0.96220115 |
| NDUFB10 | 76.6088886 | -0.02436196 | 0.56253077 | -0.0433078 | 0.96545619 | 0.99601598 |
| RPS2 | 401.24367 | -0.277076598 | 0.51220383 | -0.5409499 | 0.58854213 | 0.94448736 |
| TSC2 | 86.697245 | -0.196028597 | 0.34441523 | -0.5691636 | 0.56924515 | 0.93998433 |
| PKD1 | 67.007749 | 0.041510717 | 0.3052798 | 0.13597597 | 0.89184028 | 0.98807981 |
| SNHG19 | 34.7121081 | -0.487259062 | 0.32986086 | -1.4771654 | 0.1396313 | 0.71191199 |
| TRAF7 | 32.6221912 | 0.016143242 | 0.44635834 | 0.03616655 | 0.97114956 | 0.99681784 |
| PGP | 46.6191614 | -0.183326254 | 0.33091635 | -0.5539958 | 0.57958175 | 0.94034284 |
| ECI1 | 96.0358969 | 0.232061535 | 0.21783802 | 1.06529399 | 0.28674291 | 0.83122432 |
| RNPS1 | 99.5049289 | -0.410740124 | 0.21394186 | -1.919868 | 0.05487457 | 0.53804407 |
| ABCA3 | 43.6038911 | 0.319588971 | 0.47987752 | 0.66598029 | 0.5054237 | 0.9257297 |
| PDPK1 | 184.718408 | 0.343619571 | 0.23823568 | 1.44235143 | 0.14920326 | 0.72246729 |
| AC093525.4 | 27.3482434 | -0.01190093 | 0.35486664 | -0.0335363 | 0.97324688 | 0.99681784 |
| SRRM2 | 493.79883 | -0.19441691 | 0.15374657 | -1.2645284 | 0.20604042 | 0.7735572 |
| ELOB | 127.121044 | 0.180380262 | 0.46726755 | 0.38603208 | 0.69947291 | 0.96280283 |
| PRSS22 | 31.1477304 | -1.787327132 | 0.81776726 | -2.1856184 | 0.02884354 | 0.43103315 |
| FLYWCH1 | 51.9130514 | 0.061037358 | 0.39145608 | 0.1559239 | 0.87609301 | 0.98766275 |
| TNFRSF12A | 143.361889 | -1.757705054 | 0.64427853 | -2.7281757 | 0.00636857 | 0.25167097 |
| HCFC1R1 | 100.897048 | -1.105278492 | 0.6131724 | -1.8025575 | 0.07145774 | 0.59090333 |
| IL32 | 116.690408 | 0.999679021 | 0.77707523 | 1.28646363 | 0.1982813 | 0.77110248 |
| ZNF213-AS1 | 37.3458161 | -0.558264353 | 0.40153985 | -1.3903087 | 0.16443515 | 0.74368476 |
| ZNF213 | 64.8089451 | 0.124186262 | 0.42839491 | 0.28988735 | 0.77190242 | 0.97329019 |
| ZNF263 | 33.331403 | 0.004043894 | 0.26748276 | 0.01511834 | 0.98793777 | 0.99804429 |
| ZNF75A | 73.7767844 | -0.521694618 | 0.26520536 | -1.9671345 | 0.04916771 | 0.52272749 |
| AC025283.3 | 35.8201024 | 0.221232225 | 0.56002712 | 0.39503842 | 0.69281454 | 0.96220115 |
| NAA60 | 30.4677805 | 0.088946202 | 0.34519729 | 0.25766773 | 0.79666335 | 0.97524554 |
| CLUAP1 | 99.7946199 | 0.420130088 | 0.22284484 | 1.88530322 | 0.05938891 | 0.55718549 |
| DNASE1 | 115.133392 | 0.353788808 | 0.29067323 | 1.21713586 | 0.22355253 | 0.78791697 |
| TRAP1 | 101.211194 | -0.037766971 | 0.26093978 | -0.1447344 | 0.88492055 | 0.98766275 |
| CREBBP | 336.29755 | 0.100140134 | 0.16785306 | 0.59659405 | 0.55077844 | 0.93523904 |
| ADCY9 | 288.964479 | 0.303920367 | 0.29991879 | 1.0133422 | 0.31089674 | 0.8474709 |
| GLIS2 | 56.6542695 | 0.170413572 | 0.28631841 | 0.59518901 | 0.55171713 | 0.93523904 |
| VASN | 72.1819104 | 1.229714448 | 0.5797334 | 2.12117232 | 0.0339073 | 0.44985881 |
| DNAJA3 | 65.2651992 | 0.037892069 | 0.20411343 | 0.18564221 | 0.85272535 | 0.9864199 |
| NMRAL1 | 46.898905 | -0.301336759 | 0.26894827 | -1.1204264 | 0.26253208 | 0.81377303 |
| HMOX2 | 47.1394233 | -0.070828977 | 0.30447171 | -0.2326291 | 0.81604943 | 0.98007572 |
| CDIP1 | 39.4341401 | 0.389438014 | 0.42098656 | 0.92506044 | 0.35493447 | 0.87258503 |
| MGRN1 | 115.833529 | -0.019400897 | 0.22374726 | -0.086709 | 0.93090284 | 0.99399379 |
| NUDT16L1 | 32.3115308 | 0.690150044 | 0.46511888 | 1.48381429 | 0.13785819 | 0.71091391 |

| | | | | | | |
|---|---|---|---|---|---|---|
| ANKS3 | 83.3182016 | -0.124464828 | 0.18780723 | -0.6627265 | 0.50750573 | 0.92666294 |
| ZNF500 | 29.2974003 | -0.195274004 | 0.33491979 | -0.5830471 | 0.55986161 | 0.93744835 |
| GLYR1 | 201.341392 | -0.121805307 | 0.25754826 | -0.4729417 | 0.63625478 | 0.94932781 |
| UBN1 | 102.999803 | -0.178840115 | 0.26266689 | -0.6808628 | 0.49595831 | 0.92334178 |
| PPL | 124.407307 | -0.678215651 | 0.51176655 | -1.3252442 | 0.18509017 | 0.75995661 |
| RBFOX1 | 2377.46128 | 1.205635425 | 1.50125072 | 0.80308732 | 0.42192426 | 0.90066187 |
| METTL22 | 27.4886687 | -0.221130098 | 0.30488398 | -0.7252926 | 0.46827254 | 0.91462428 |
| ABAT | 55.8806288 | 1.570174117 | 0.28718075 | 5.46754657 | 4.56E-08 | 0.00014889 |
| PMM2 | 134.569245 | -0.205010744 | 0.28186697 | -0.7273316 | 0.46702287 | 0.91462428 |
| CARHSP1 | 305.860006 | 0.826458197 | 0.53531477 | 1.54387333 | 0.12261902 | 0.69314363 |
| USP7 | 121.345507 | -0.277222133 | 0.26301038 | -1.054035 | 0.29186691 | 0.83681327 |
| C16orf72 | 118.193937 | -0.267408868 | 0.18957286 | -1.4105862 | 0.15836666 | 0.73640497 |
| GRIN2A | 48.5799389 | -0.969036872 | 1.20105378 | -0.8068222 | 0.41976892 | 0.90012434 |
| ATF7IP2 | 162.393305 | 0.879047996 | 0.34136139 | 2.57512429 | 0.01002041 | 0.29238317 |
| EMP2 | 45.0829487 | 0.499967651 | 0.47605239 | 1.05023661 | 0.29360934 | 0.83825235 |
| NUBP1 | 29.0584567 | -0.20522521 | 0.27397783 | -0.7490577 | 0.45382241 | 0.91077355 |
| DEXI | 33.5833807 | 0.129496527 | 0.33255777 | 0.38939558 | 0.69698354 | 0.96220115 |
| CLEC16A | 373.570644 | 0.016416195 | 0.31183895 | 0.05264318 | 0.95801621 | 0.99572129 |
| LITAF | 214.551428 | 0.212252121 | 0.23825396 | 0.89086503 | 0.37300158 | 0.87651737 |
| TXNDC11 | 160.900743 | 0.323887808 | 0.16646562 | 1.94567385 | 0.05169393 | 0.52734415 |
| ZC3H7A | 184.101126 | 0.309921827 | 0.20188527 | 1.53513836 | 0.12474984 | 0.69439514 |
| RSL1D1 | 70.2435891 | -0.480618809 | 0.35859926 | -1.3402671 | 0.18015851 | 0.75378294 |
| AC007216.4 | 101.513779 | 0.063246102 | 0.36321884 | 0.17412671 | 0.86176588 | 0.98669442 |
| GSPT1 | 250.963353 | -0.382612735 | 0.26484455 | -1.4446691 | 0.14855086 | 0.72246729 |
| SNX29 | 1299.27815 | -0.309045348 | 0.40195709 | -0.7688516 | 0.44198142 | 0.90561284 |
| CPPED1 | 98.7117059 | -0.167252482 | 0.48168456 | -0.3472241 | 0.72842297 | 0.96829915 |
| SHISA9 | 306.519416 | -0.991194752 | 0.60785958 | -1.6306311 | 0.10296818 | 0.65288882 |
| U91319.1 | 158.047474 | -0.860003089 | 0.64563603 | -1.3320246 | 0.1828521 | 0.75595176 |
| ERCC4 | 39.8141779 | 0.042234003 | 0.34318858 | 0.12306354 | 0.90205678 | 0.9900784 |
| MKL2 | 443.938511 | 0.284280362 | 0.19364862 | 1.46802162 | 0.14209835 | 0.7148274 |
| PARN | 182.458233 | -0.337004985 | 0.23641315 | -1.4254917 | 0.15401511 | 0.72926879 |
| BFAR | 59.2589828 | -0.251658429 | 0.24694938 | -1.0190689 | 0.30817027 | 0.84488735 |
| PDXDC1 | 352.67248 | -0.06318095 | 0.17919345 | -0.3525852 | 0.72439947 | 0.96781837 |
| RRN3 | 62.580876 | 0.209627237 | 0.2757618 | 0.76017504 | 0.44714996 | 0.90918416 |
| C16orf45 | 68.2689204 | 0.394496017 | 0.70863186 | 0.55670093 | 0.57773178 | 0.94034284 |
| MARF1 | 103.488301 | -0.17864799 | 0.21342023 | -0.8370715 | 0.40255238 | 0.89239743 |
| NDE1 | 43.609775 | -0.093339577 | 0.29226037 | -0.3193713 | 0.74944496 | 0.97038839 |
| FOPNL | 44.105947 | -0.165294701 | 0.24523456 | -0.6740269 | 0.50029418 | 0.92398825 |
| ABCC1 | 160.890858 | -0.098030874 | 0.26292265 | -0.3728506 | 0.70925962 | 0.96587538 |
| XYLT1 | 102.853628 | 0.581440488 | 0.74125368 | 0.78440148 | 0.43280457 | 0.90514499 |
| RPS15A | 90.5316597 | 0.544216036 | 0.55369212 | 0.98288564 | 0.32566373 | 0.85509839 |
| SMG1 | 453.152985 | -0.176514702 | 0.21880719 | -0.8067134 | 0.4198316 | 0.90012434 |
| COQ7 | 29.8689831 | 0.183601219 | 0.34793058 | 0.52769498 | 0.59771106 | 0.94683415 |
| GDE1 | 30.7707415 | -0.35090088 | 0.36699152 | -0.9561553 | 0.33899379 | 0.86680923 |
| CCP110 | 61.1713302 | 0.654481347 | 0.26660658 | 2.45485822 | 0.01409402 | 0.32375679 |
| VPS35L | 187.192738 | -0.086165791 | 0.24312015 | -0.3544165 | 0.72302677 | 0.96779196 |
| KNOP1 | 44.3862094 | -0.470191197 | 0.34658531 | -1.3566392 | 0.1748959 | 0.74910274 |
| IQCK | 249.300775 | 0.262043524 | 0.30145431 | 0.86926447 | 0.38470249 | 0.88328503 |
| GPRC5B | 78.145986 | -0.078277795 | 0.48729323 | -0.160638 | 0.87237855 | 0.98766275 |
| UMOD | 148.251134 | 0.051038288 | 0.72726226 | 0.07017866 | 0.94405146 | 0.9952642 |
| ACSM2A | 162.05198 | 2.41336853 | 0.78122578 | 3.08920748 | 0.00200691 | 0.14702642 |
| ACSM2B | 102.696092 | 2.171216595 | 0.82696726 | 2.62551701 | 0.00865175 | 0.27620989 |
| ACSM3 | 238.830624 | 3.420678059 | 0.65625751 | 5.21240218 | 1.86E-07 | 0.0002976 |
| THUMPD1 | 89.7873454 | 0.551117132 | 0.23314393 | 2.36384942 | 0.01808616 | 0.35858401 |

| | | | | | | |
|---|---|---|---|---|---|---|
| ERI2 | 39.576634 | 0.542069466 | 0.54130398 | 1.00141416 | 0.31662662 | 0.8490543 |
| LYRM1 | 47.34765 | 0.017295255 | 0.23934489 | 0.07226081 | 0.94239436 | 0.99526299 |
| TMEM159 | 41.5128613 | 0.164681059 | 0.35756692 | 0.46056011 | 0.64511424 | 0.94954229 |
| CRYM | 32.0384323 | 0.166424456 | 0.51313499 | 0.3243288 | 0.74568912 | 0.97028355 |
| METTL9 | 194.9761 | 0.030870722 | 0.1650629 | 0.18702399 | 0.85164182 | 0.98636998 |
| NPIPB4 | 52.6260604 | -0.626498683 | 0.31948613 | -1.9609574 | 0.049884 | 0.52415717 |
| UQCRC2 | 135.594694 | 0.271825251 | 0.2372304 | 1.14582807 | 0.2518663 | 0.80770916 |
| MOSMO | 144.708367 | -0.161002239 | 0.15804968 | -1.0186812 | 0.30835433 | 0.84488735 |
| EEF2K | 45.6043499 | -0.320302686 | 0.36233979 | -0.8839843 | 0.37670468 | 0.87842274 |
| POLR3E | 57.6879786 | -0.7724554 | 0.24097091 | -3.2055961 | 0.00134783 | 0.11631411 |
| CDR2 | 677.892375 | -0.090879726 | 0.37846506 | -0.2401271 | 0.81023171 | 0.97930627 |
| NPIPB5 | 154.949002 | 0.132192564 | 0.29303477 | 0.45111563 | 0.65190621 | 0.95159942 |
| USP31 | 150.226629 | -0.059883298 | 0.40618963 | -0.1474269 | 0.88279504 | 0.98766275 |
| COG7 | 124.87962 | 0.200393764 | 0.21566178 | 0.92920387 | 0.35278344 | 0.8720098 |
| GGA2 | 126.256028 | -0.070409416 | 0.3179483 | -0.2214493 | 0.82474264 | 0.98094663 |
| EARS2 | 25.5257457 | -0.464566155 | 0.37261842 | -1.2467611 | 0.21248511 | 0.77822114 |
| UBFD1 | 48.1002536 | -0.264066051 | 0.29037335 | -0.9094018 | 0.36313807 | 0.87553474 |
| NDUFAB1 | 55.2173338 | -0.011116018 | 0.3597039 | -0.0309032 | 0.9753467 | 0.99681784 |
| PALB2 | 32.9565367 | -0.296947967 | 0.32828757 | -0.9045361 | 0.36571119 | 0.87605028 |
| DCTN5 | 76.3952854 | -0.272079827 | 0.22439587 | -1.2124992 | 0.22532133 | 0.78977225 |
| RBBP6 | 158.129719 | -0.0421105 | 0.22416855 | -0.187852 | 0.85099271 | 0.98617742 |
| TNRC6A | 704.195928 | 0.087453575 | 0.29334806 | 0.29812222 | 0.76560988 | 0.97177982 |
| ARHGAP17 | 132.848469 | 0.061803366 | 0.23831678 | 0.25933284 | 0.79537845 | 0.97524554 |
| LCMT1 | 124.011079 | -0.171012744 | 0.21889205 | -0.7812652 | 0.43464653 | 0.90514499 |
| NSMCE1 | 64.7994779 | 0.0336068 | 0.2674935 | 0.12563595 | 0.9000201 | 0.98970182 |
| IL4R | 213.731827 | -0.097557344 | 0.49564264 | -0.19683 | 0.84396057 | 0.98517901 |
| GTF3C1 | 138.507765 | 0.034125749 | 0.30280882 | 0.11269734 | 0.91027051 | 0.99100813 |
| KIAA0556 | 210.707698 | 0.019331923 | 0.27672195 | 0.06986046 | 0.94430472 | 0.9952642 |
| XPO6 | 152.74066 | -0.674708453 | 0.27334368 | -2.4683521 | 0.01357367 | 0.32032755 |
| NUPR1 | 574.584036 | -0.567425039 | 0.69467901 | -0.8168162 | 0.4140335 | 0.89732895 |
| SGF29 | 36.6211605 | -0.399113036 | 0.28818189 | -1.3849345 | 0.1660725 | 0.74445571 |
| SULT1A1 | 28.1481961 | 0.079800777 | 0.46492889 | 0.17164082 | 0.86371991 | 0.98670067 |
| ATXN2L | 85.0126305 | -0.404230959 | 0.2439417 | -1.6570802 | 0.09750326 | 0.64312419 |
| TUFM | 113.10652 | 0.473625915 | 0.43031036 | 1.10066119 | 0.27104414 | 0.81775732 |
| SH2B1 | 71.3594231 | -0.023660328 | 0.27666496 | -0.0855198 | 0.93184817 | 0.99411667 |
| RABEP2 | 38.2922613 | -0.406537566 | 0.27355808 | -1.4861106 | 0.13724986 | 0.71044919 |
| NFATC2IP | 147.714331 | 0.060928362 | 0.21447848 | 0.28407681 | 0.77635152 | 0.97423208 |
| SPNS1 | 36.5412015 | -0.134816882 | 0.29687453 | -0.4541207 | 0.64974192 | 0.95031069 |
| MAZ | 57.2227221 | -0.193899705 | 0.38549558 | -0.5029881 | 0.61497261 | 0.94791646 |
| AC009133.1 | 34.7064268 | 0.219542643 | 0.47434315 | 0.46283507 | 0.64348259 | 0.94944486 |
| MVP | 168.048043 | 0.255588876 | 0.50553633 | 0.50557964 | 0.61315178 | 0.94791646 |
| CDIPT | 54.3637157 | 0.327394749 | 0.30986955 | 1.05655669 | 0.29071396 | 0.83531608 |
| KCTD13 | 47.4283477 | -0.264831174 | 0.35367384 | -0.7488006 | 0.45397741 | 0.91077355 |
| TMEM219 | 85.5287542 | -0.26289573 | 0.38476227 | -0.683268 | 0.49443754 | 0.92289011 |
| TAOK2 | 29.8143937 | -0.172011828 | 0.42699596 | -0.4028418 | 0.6870646 | 0.96161098 |
| INO80E | 42.2509507 | -0.399054969 | 0.34001183 | -1.1736502 | 0.24053515 | 0.8030067 |
| PPP4C | 64.4767314 | -0.159242525 | 0.39062068 | -0.4076654 | 0.68351937 | 0.96144376 |
| YPEL3 | 75.2054954 | 0.103292645 | 0.37560385 | 0.27500422 | 0.78331299 | 0.97423208 |
| MAPK3 | 33.0752777 | 0.480271651 | 0.40843799 | 1.1758741 | 0.23964519 | 0.80193227 |
| NPIPB13 | 38.5815193 | -0.624984029 | 0.30617625 | -2.0412558 | 0.04122541 | 0.48752866 |
| CD2BP2 | 41.658138 | -0.013270225 | 0.39534069 | -0.0335666 | 0.97322279 | 0.99681784 |
| TBC1D10B | 25.9655428 | -0.16274679 | 0.32854377 | -0.495358 | 0.62034743 | 0.9480368 |
| DCTPP1 | 26.8097972 | -0.292405595 | 0.51597108 | -0.5667093 | 0.57091172 | 0.93998433 |
| SEPHS2 | 90.5882804 | 0.467426847 | 0.2980613 | 1.56822389 | 0.11682889 | 0.67748987 |

| | | | | | | |
|---|---|---|---|---|---|---|
| ZNF768 | 32.5796986 | -0.338928639 | 0.40985196 | -0.8269538 | 0.40826325 | 0.8959746 |
| ZNF785 | 40.1182418 | -0.184513072 | 0.25790244 | -0.7154375 | 0.47433875 | 0.91809716 |
| PRR14 | 24.5209527 | -0.432654533 | 0.42943477 | -1.0074977 | 0.31369567 | 0.8490543 |
| FBRS | 45.7011496 | -0.456200511 | 0.26071644 | -1.7497957 | 0.08015357 | 0.60643125 |
| PHKG2 | 33.3547611 | -0.124124854 | 0.26230986 | -0.4731993 | 0.63607097 | 0.94932781 |
| RNF40 | 51.2842845 | 0.175269424 | 0.34425143 | 0.50913202 | 0.61065969 | 0.94791646 |
| BCL7C | 148.972145 | -0.266675738 | 0.32243003 | -0.827081 | 0.40819117 | 0.8959746 |
| FBXL19 | 41.8325656 | 0.046591308 | 0.3147511 | 0.14802588 | 0.88232235 | 0.98766275 |
| SETD1A | 34.0807627 | -0.362043419 | 0.39977442 | -0.9056193 | 0.36513741 | 0.87605028 |
| STX4 | 33.6408944 | -0.064219275 | 0.41669231 | -0.1541168 | 0.87751766 | 0.98766275 |
| BCKDK | 32.3725054 | 0.026875735 | 0.2882606 | 0.09323416 | 0.92571754 | 0.99316012 |
| KAT8 | 63.5101417 | -0.412001979 | 0.32040624 | -1.2858738 | 0.19848712 | 0.77110248 |
| PRSS8 | 58.5607369 | -0.520399008 | 0.72906193 | -0.7137926 | 0.47535544 | 0.91809716 |
| FUS | 317.486808 | -0.305772036 | 0.15207712 | -2.010638 | 0.04436371 | 0.49995287 |
| AC026471.4 | 38.1254736 | -0.339411403 | 0.38177685 | -0.8890309 | 0.37398649 | 0.87666478 |
| C16orf58 | 46.5745243 | -0.083615945 | 0.26917972 | -0.3106324 | 0.75608009 | 0.9714717 |
| ZNF720 | 85.0031066 | 0.653644249 | 0.19001989 | 3.43987283 | 0.00058199 | 0.07556838 |
| ZNF267 | 36.8605536 | 0.402107656 | 0.49425927 | 0.81355613 | 0.41589928 | 0.89828618 |
| VPS35 | 95.1138536 | 0.095210757 | 0.22825992 | 0.41711553 | 0.6765939 | 0.9591378 |
| C16orf87 | 37.0316686 | 0.435676125 | 0.34128517 | 1.27657504 | 0.20175232 | 0.77110248 |
| DNAJA2 | 55.8909821 | 0.308831456 | 0.34557495 | 0.89367432 | 0.37149617 | 0.87605028 |
| ITFG1 | 552.62958 | -0.318020174 | 0.20200155 | -1.5743452 | 0.11540766 | 0.67260842 |
| PHKB | 340.088363 | 0.112289069 | 0.16580603 | 0.67723153 | 0.49825907 | 0.92398825 |
| LONP2 | 272.366544 | -0.058673554 | 0.19377569 | -0.3027911 | 0.76204907 | 0.9714717 |
| SIAH1 | 72.3038906 | -0.127250936 | 0.25117239 | -0.5066279 | 0.61241594 | 0.94791646 |
| N4BP1 | 140.548629 | -0.468083349 | 0.37737938 | -1.2403522 | 0.21484515 | 0.7828522 |
| ZNF423 | 67.9036689 | 2.142983197 | 1.15217718 | 1.85994241 | 0.06289367 | 0.56713487 |
| HEATR3 | 26.6573377 | -0.525901559 | 0.43958041 | -1.1963717 | 0.23155154 | 0.79459836 |
| TENT4B | 129.714108 | 0.11000454 | 0.24443899 | 0.45002862 | 0.6526898 | 0.95176155 |
| BRD7 | 106.629774 | -0.104606498 | 0.23585958 | -0.4435118 | 0.65739561 | 0.95399905 |
| CYLD | 94.8924099 | 0.153294769 | 0.31214397 | 0.49110276 | 0.62335377 | 0.94808496 |
| SALL1 | 110.378757 | -0.138265534 | 0.46175916 | -0.2994321 | 0.76461034 | 0.97177982 |
| AC087564.1 | 319.311874 | 0.343753375 | 0.31670739 | 1.08539739 | 0.27774569 | 0.8229426 |
| HNRNPA1P4 | 28.8082032 | -0.382919552 | 0.62798683 | -0.6097573 | 0.54202259 | 0.93523904 |
| TOX3 | 198.371992 | 0.027813024 | 0.59752087 | 0.04654737 | 0.96287398 | 0.99572129 |
| AC007906.2 | 69.8292408 | 0.748109221 | 0.54498982 | 1.3727031 | 0.16984466 | 0.74445571 |
| CHD9 | 826.569645 | 0.114733559 | 0.14248233 | 0.80524761 | 0.42067681 | 0.90012434 |
| RBL2 | 174.427867 | 0.114508598 | 0.28356168 | 0.40382254 | 0.68634322 | 0.96161098 |
| AKTIP | 34.4958489 | -0.153874206 | 0.47572244 | -0.3234537 | 0.74635163 | 0.97028355 |
| RPGRIP1L | 76.2853164 | 0.367417536 | 0.32986808 | 1.11383172 | 0.26535138 | 0.81378765 |
| FTO | 984.326511 | 0.18336644 | 0.1977696 | 0.92717203 | 0.35383722 | 0.87241657 |
| IRX3 | 251.58434 | -0.695567174 | 0.64379885 | -1.0804107 | 0.27995933 | 0.82464035 |
| CRNDE | 218.377842 | 0.092727602 | 0.62070107 | 0.14939172 | 0.88124454 | 0.98766275 |
| IRX5 | 78.5979448 | -0.312752174 | 0.73409986 | -0.4260349 | 0.6700824 | 0.95764829 |
| LPCAT2 | 155.518382 | 0.174486866 | 0.45986436 | 0.37943115 | 0.70436772 | 0.96461079 |
| AMFR | 121.741878 | 0.337954611 | 0.27723917 | 1.21900022 | 0.22284411 | 0.78710314 |
| NUDT21 | 96.7710737 | -0.135280167 | 0.24786062 | -0.5457913 | 0.58520941 | 0.94229635 |
| OGFOD1 | 36.6433878 | -0.333676409 | 0.26171726 | -1.2749499 | 0.20232696 | 0.77110248 |
| BBS2 | 81.7695431 | -0.496730843 | 0.19825196 | -2.5055532 | 0.012226 | 0.30967852 |
| MT2A | 925.364806 | -1.35516521 | 0.74168873 | -1.8271347 | 6.77E-02 | 0.58081067 |
| MT1E | 361.454742 | -1.856726925 | 1.00624215 | -1.8452089 | 0.06500716 | 0.57495764 |
| MT1F | 68.4223982 | -0.999717986 | 0.90496037 | -1.1047091 | 0.26928565 | 0.81655182 |
| MT1G | 338.025193 | -1.899555635 | 1.09883557 | -1.7286987 | 0.08386303 | 0.61656297 |
| MT1H | 86.2521483 | -3.007018401 | 1.12732532 | -2.6673919 | 0.00764425 | 0.2702002 |

| | | | | | | |
|---|---|---|---|---|---|---|
| MT1X | 268.774187 | -1.274511482 | 0.75188841 | -1.6950806 | 0.09006012 | 0.6330877 |
| NUP93 | 76.2683626 | -0.72942726 | 0.33228526 | -2.1951839 | 0.02815041 | 0.4250936 |
| SLC12A3 | 37.3914233 | 0.242883201 | 0.66496196 | 0.36525879 | 7.15E-01 | 9.67E-01 |
| HERPUD1 | 127.75763 | 0.754257319 | 0.46313031 | 1.62860713 | 0.10339621 | 0.65288882 |
| NLRC5 | 38.4638119 | -1.477487537 | 0.66349847 | -2.2268138 | 0.02595972 | 0.41933927 |
| FAM192A | 99.6007675 | -0.238828408 | 0.18962319 | -1.2594895 | 0.20785359 | 0.77435432 |
| RSPRY1 | 141.248687 | 0.152883862 | 0.21499607 | 0.71110071 | 0.47702183 | 0.91864961 |
| ARL2BP | 57.7513029 | 0.083524371 | 0.2895514 | 0.28846129 | 0.77299365 | 0.97350952 |
| PLLP | 46.2996922 | 0.259663964 | 0.69560322 | 0.37329322 | 0.70893022 | 0.96587538 |
| CX3CL1 | 60.6482893 | 0.662452818 | 0.39726867 | 1.66751841 | 0.09541137 | 0.64091548 |
| CIAPIN1 | 42.0160878 | -0.253283011 | 0.29873979 | -0.8478382 | 0.39652808 | 0.88850116 |
| COQ9 | 73.269715 | 0.451307159 | 0.20790159 | 2.17077302 | 0.02994833 | 0.43492739 |
| POLR2C | 26.369122 | 0.146570845 | 0.40821181 | 0.35905586 | 0.7195533 | 0.96779196 |
| DOK4 | 28.5800102 | 1.01799602 | 0.42870325 | 2.37459365 | 0.01756828 | 0.35653929 |
| ADGRG1 | 306.195152 | -0.065293576 | 0.63171422 | -0.1033594 | 0.91767777 | 0.99214128 |
| KIFC3 | 424.794021 | 0.016809179 | 0.29427857 | 0.05711996 | 0.95444964 | 0.99531843 |
| AC092118.1 | 31.4265778 | -0.011902785 | 0.53812719 | -0.0221189 | 0.9823531 | 0.99681784 |
| CSNK2A2 | 86.2957655 | -0.61748194 | 0.23809 | -2.5934812 | 0.00950097 | 0.28916146 |
| CNOT1 | 234.06669 | -0.026102699 | 0.11562934 | -0.2257446 | 0.82140006 | 0.98094663 |
| GOT2 | 61.3198997 | -0.073588782 | 0.40451411 | -0.181919 | 0.85564633 | 0.9864199 |
| AC138305.1 | 44.3058095 | 0.399406622 | 0.35880491 | 1.11315819 | 0.26564049 | 0.81387543 |
| TK2 | 28.1301634 | -0.671733501 | 0.46169946 | -1.4549151 | 0.14569277 | 0.71974648 |
| CMTM4 | 185.305542 | 0.286062795 | 0.27281691 | 1.04855228 | 0.29438423 | 0.83885935 |
| DYNC1LI2 | 166.261817 | 0.081109858 | 0.21335262 | 0.38016809 | 0.70382065 | 0.96461079 |
| NAE1 | 67.5060994 | 0.024644854 | 0.21349277 | 0.11543648 | 0.90809916 | 0.99100813 |
| CDH16 | 167.959509 | 0.033420464 | 0.46112976 | 0.07247518 | 0.94222375 | 0.99526299 |
| RRAD | 117.619693 | -0.732135455 | 0.50094518 | -1.4615081 | 1.44E-01 | 0.71631428 |
| FAM96B | 94.9331639 | -0.360738277 | 0.47210995 | -0.764098 | 0.44480883 | 0.90694419 |
| CES2 | 65.9075163 | -0.324919179 | 0.37059186 | -0.8767575 | 0.38061838 | 0.8807629 |
| CBFB | 253.941874 | 0.148169404 | 0.21138847 | 0.70093417 | 0.4833441 | 0.91991915 |
| C16orf70 | 43.1633874 | 0.153275624 | 0.32175773 | 0.47636967 | 0.63381104 | 0.94932781 |
| NOL3 | 36.5058872 | 0.031070103 | 0.45418878 | 0.0684079 | 0.94546093 | 0.9952642 |
| E2F4 | 25.6431055 | -0.662652204 | 0.37130775 | -1.7846441 | 0.07431906 | 0.59325188 |
| TMEM208 | 51.6567952 | 0.142587132 | 0.44416109 | 0.32102572 | 0.7481909 | 0.97028355 |
| ZDHHC1 | 40.9877132 | 0.021353832 | 0.33494684 | 0.0637529 | 0.94916698 | 0.9952642 |
| ATP6V0D1 | 72.0168136 | 0.183737461 | 0.25982748 | 0.70715175 | 0.47947218 | 0.91971919 |
| AC009061.2 | 34.5885071 | -0.009422487 | 0.4157526 | -0.0226637 | 0.98191854 | 0.99681784 |
| RIPOR1 | 75.9229965 | 0.038161863 | 0.41221589 | 0.09257737 | 0.92623932 | 0.99316012 |
| CTCF | 128.835457 | -0.531350477 | 0.19128092 | -2.777854 | 0.00547192 | 0.23295784 |
| ENKD1 | 35.1530681 | 0.33575311 | 0.37864942 | 0.88671234 | 0.37523379 | 0.87731621 |
| GFOD2 | 45.9718108 | 0.151660746 | 0.32415998 | 0.46785771 | 0.63988635 | 0.94944486 |
| RANBP10 | 114.888235 | 0.004091154 | 0.18269705 | 0.0223931 | 0.98213438 | 0.99681784 |
| NUTF2 | 33.4788571 | -0.352190245 | 0.37721752 | -0.9336529 | 0.35048295 | 0.8720098 |
| PSKH1 | 35.5693629 | 0.159418999 | 0.27466155 | 0.58041978 | 0.56163157 | 0.93747805 |
| SLC12A4 | 35.2285478 | 0.270620105 | 0.3386833 | 0.79903587 | 0.42426961 | 0.90167091 |
| DUS2 | 29.7415546 | -0.262484012 | 0.41336231 | -0.6349974 | 0.52543008 | 0.93272384 |
| NFATC3 | 304.093583 | 0.195712572 | 0.25874522 | 0.75639107 | 0.44941476 | 0.90994378 |
| ESRP2 | 24.354992 | -0.43909932 | 0.48414934 | -0.9069502 | 0.36443316 | 0.87605028 |
| PLA2G15 | 39.6200913 | 0.360583574 | 0.46004521 | 0.78380031 | 0.43315729 | 0.90514499 |
| SLC7A6 | 94.9050259 | 0.210817394 | 0.36324617 | 0.58037058 | 0.56166474 | 0.93747805 |
| SLC7A6OS | 55.3180862 | -0.418746741 | 0.38648285 | -1.0834808 | 0.27859508 | 0.8229426 |
| PRMT7 | 54.2340714 | 0.097285089 | 0.36901267 | 0.26363618 | 0.79206028 | 0.97423208 |
| ZFP90 | 71.0827254 | 0.464959306 | 0.25583519 | 1.81741735 | 0.06915323 | 0.58408344 |
| CDH1 | 273.300221 | 0.205319622 | 0.36346245 | 0.56489913 | 0.57214237 | 0.93998433 |

| | | | | | | |
|---|---|---|---|---|---|---|
| TANGO6 | 134.697364 | -0.311230113 | 0.29493053 | -1.0552659 | 0.29130375 | 0.83594927 |
| CHTF8 | 59.9054757 | 0.328851135 | 0.3283382 | 1.00156221 | 0.31655508 | 0.8490543 |
| UTP4 | 48.7227995 | -0.113438743 | 0.2975551 | -0.3812361 | 0.70302807 | 0.9645815 |
| SNTB2 | 92.1842687 | -0.119982135 | 0.36806975 | -0.3259766 | 0.74444204 | 0.97020548 |
| VPS4A | 60.2735037 | -0.216886121 | 0.2746301 | -0.7897391 | 0.42968017 | 0.90333313 |
| COG8 | 33.1460476 | 0.126752949 | 0.36797534 | 0.34446045 | 0.73050004 | 0.96891435 |
| TERF2 | 69.7711505 | -0.287953479 | 0.29047148 | -0.9913313 | 0.32152381 | 0.85142171 |
| CYB5B | 217.834031 | -0.537892546 | 0.14907021 | -3.6083169 | 0.00030819 | 0.04685954 |
| NFAT5 | 2561.65778 | -0.83263088 | 0.43176845 | -1.9284199 | 0.05380291 | 0.53573608 |
| NQO1 | 37.4232424 | 0.070587435 | 0.66796282 | 0.10567569 | 0.91583967 | 0.99159829 |
| NOB1 | 31.8886656 | -0.472957335 | 0.3264868 | -1.4486262 | 0.147442 | 0.72205873 |
| WWP2 | 235.4973 | -0.252611724 | 0.29250241 | -0.8636227 | 0.38779518 | 0.8853359 |
| PDPR | 103.527379 | 0.057374879 | 0.26350239 | 0.2177395 | 0.82763209 | 0.98142144 |
| AC009060.1 | 37.9030983 | 0.710833408 | 0.42184298 | 1.68506633 | 0.09197577 | 0.63464873 |
| EXOSC6 | 54.0350696 | -0.603519846 | 0.32260235 | -1.8707857 | 0.0613748 | 0.56205465 |
| AARS | 66.9874083 | -0.358695003 | 0.26333982 | -1.3620994 | 0.17316653 | 0.74719017 |
| ST3GAL2 | 74.7327947 | 1.275764072 | 0.55200673 | 2.31113861 | 0.0208252 | 0.38104604 |
| FUK | 32.3573066 | 0.461937831 | 0.39105441 | 1.18126231 | 0.23749853 | 0.80077383 |
| COG4 | 64.5853518 | 0.063949336 | 0.25169375 | 0.25407597 | 0.79943686 | 0.97651182 |
| SF3B3 | 144.59319 | -0.274357557 | 0.20196355 | -1.3584508 | 0.17432067 | 0.74910274 |
| MTSS1L | 34.3957508 | 0.491872979 | 0.71707995 | 0.68593883 | 0.49275169 | 0.92289011 |
| VAC14 | 98.0863359 | -0.087053165 | 0.43104154 | -0.20196 | 0.83994797 | 0.98464453 |
| HYDIN | 517.456324 | 0.35603589 | 0.4406334 | 0.80800932 | 0.41908522 | 0.8998918 |
| CMTR2 | 47.2687215 | -0.292551542 | 0.35452358 | -0.8251963 | 0.40926016 | 0.8959746 |
| ZNF19 | 70.2173133 | 0.29755575 | 0.35896398 | 0.82892926 | 0.40714444 | 0.89585737 |
| MARVELD3 | 70.5711883 | -0.316821843 | 0.57747755 | -0.5486306 | 0.58325899 | 0.94152981 |
| PHLPP2 | 87.9160544 | -0.180664572 | 0.28311458 | -0.6381323 | 0.52338753 | 0.93272384 |
| AP1G1 | 254.759063 | -0.027149647 | 0.19641481 | -0.1382261 | 0.89006175 | 0.98766275 |
| IST1 | 178.342664 | 0.04932244 | 0.15531258 | 0.31756886 | 0.750812 | 0.97058015 |
| ZNF821 | 57.6970022 | 0.472098943 | 0.39313309 | 1.2008629 | 0.22980439 | 0.79370986 |
| TXNL4B | 34.8246121 | 0.459417074 | 0.30345486 | 1.51395525 | 0.13003719 | 0.69904834 |
| DHX38 | 38.8063342 | -0.682608252 | 0.28306154 | -2.4115189 | 0.01588623 | 0.34086104 |
| LINC01572 | 119.698612 | 0.380623864 | 0.38466601 | 0.98949179 | 0.32242258 | 0.85246644 |
| AC004943.2 | 88.5982462 | -0.009034254 | 0.33241741 | -0.0271774 | 0.97831821 | 0.99681784 |
| ZFHX3 | 475.143548 | -0.908946756 | 0.39156525 | -2.3213162 | 0.02026978 | 0.37800094 |
| AC138627.1 | 45.17839 | 0.262145318 | 0.79091447 | 0.33144585 | 0.74030774 | 0.9698508 |
| PSMD7 | 73.6105602 | -0.394392259 | 0.39028488 | -1.0105241 | 0.31224428 | 0.84811918 |
| GLG1 | 509.652468 | 0.144601964 | 0.21411395 | 0.67535051 | 0.4994531 | 0.92398825 |
| RFWD3 | 56.2034783 | -0.579928669 | 0.23008101 | -2.5205412 | 0.01171745 | 0.30872898 |
| WDR59 | 220.001833 | 0.2301214 | 0.16761673 | 1.37290232 | 0.16978271 | 0.74445571 |
| ZNRF1 | 102.064698 | 0.006108712 | 0.29031751 | 0.02104149 | 0.98321256 | 0.99726738 |
| BCAR1 | 148.420994 | -0.528708427 | 0.47567385 | -1.1114936 | 0.26635595 | 0.81444418 |
| CFDP1 | 461.421591 | -0.376253593 | 0.20164227 | -1.865946 | 0.06204892 | 0.56445068 |
| GABARAPL2 | 108.108307 | 0.297234207 | 0.53283214 | 0.55783836 | 0.57695476 | 0.94034284 |
| ADAT1 | 37.4159305 | -0.617066842 | 0.33985053 | -1.8157007 | 0.0694163 | 0.58408344 |
| KARS | 59.6078904 | -0.328066058 | 0.32116756 | -1.0214795 | 0.30702734 | 0.84488735 |
| TERF2IP | 262.375524 | 0.388677839 | 0.50114954 | 0.77557257 | 0.4380014 | 0.90561284 |
| ADAMTS18 | 44.2526467 | -1.847914013 | 0.87020937 | -2.1235281 | 0.03370963 | 0.44933903 |
| VAT1L | 63.6497237 | 1.118026486 | 1.34188071 | 0.83317875 | 0.40474393 | 0.89373953 |
| WWOX | 2008.1057 | 0.457615935 | 0.42842972 | 1.0681237 | 0.28546472 | 0.82963183 |
| MAF | 81.4041261 | 1.451074949 | 0.8969144 | 1.61785221 | 0.10569445 | 0.65699665 |
| CDYL2 | 37.6295259 | 1.517623706 | 0.55167842 | 2.7509209 | 0.0059428 | 0.24431791 |
| CMC2 | 84.8055757 | 0.133699076 | 0.30380668 | 0.44007945 | 0.65987956 | 0.95407427 |
| ATMIN | 44.238345 | -0.203524487 | 0.28455421 | -0.7152398 | 0.47446089 | 0.91809716 |

| | | | | | | |
|---|---|---|---|---|---|---|
| GAN | 95.8290697 | -0.100248234 | 0.27883788 | -0.3595216 | 0.71920494 | 0.96779196 |
| CMIP | 424.997343 | -0.478073891 | 0.34957801 | -1.3675743 | 0.17144538 | 0.74445571 |
| PLCG2 | 152.033413 | 0.293367806 | 0.5491692 | 0.53420295 | 0.59320113 | 0.94649617 |
| MPHOSPH6 | 36.329865 | -0.388258493 | 0.33159372 | -1.1708862 | 0.24164454 | 0.803412 |
| HSBP1 | 117.623885 | 0.019731147 | 0.38072346 | 0.0518254 | 0.95866781 | 0.99572129 |
| MBTPS1 | 151.949802 | 0.058153178 | 0.20098714 | 0.2893378 | 0.77232289 | 0.97329019 |
| HSDL1 | 28.4201877 | -0.092145165 | 0.47667163 | -0.1933095 | 0.84671657 | 0.98533374 |
| TAF1C | 30.5781536 | -0.138038513 | 0.30347675 | -0.454857 | 0.64921215 | 0.95027667 |
| TLDC1 | 79.8937639 | 0.021001312 | 0.43572576 | 0.04819846 | 0.96155807 | 0.99572129 |
| COTL1 | 69.871267 | -0.562280891 | 0.40722615 | -1.3807583 | 0.16735328 | 0.74445571 |
| KLHL36 | 108.265491 | -0.158109626 | 0.34230121 | -0.461902 | 0.64415158 | 0.94944486 |
| USP10 | 186.952922 | -0.436706989 | 0.24695741 | -1.7683494 | 0.07700251 | 0.59726395 |
| ZDHHC7 | 101.240605 | -0.416150348 | 0.2097639 | -1.9838988 | 0.04726712 | 0.51284792 |
| GSE1 | 223.407778 | -0.056847208 | 0.38682989 | -0.1469566 | 0.88316627 | 0.98766275 |
| EMC8 | 31.4205335 | -0.603665187 | 0.29975872 | -2.0138369 | 0.04402666 | 0.49952159 |
| COX4I1 | 294.460331 | 0.235080618 | 0.47138336 | 0.49870368 | 0.61798815 | 0.9480368 |
| MTHFSD | 25.3327529 | 0.018072237 | 0.35911301 | 0.05032465 | 0.95986368 | 0.99572129 |
| FBXO31 | 37.7719867 | -0.086224268 | 0.32134663 | -0.2683217 | 0.78845172 | 0.97423208 |
| MAP1LC3B | 94.4877442 | -0.047648104 | 0.34943834 | -0.1363563 | 0.89153965 | 0.987861 |
| ZCCHC14 | 237.297279 | -0.146536699 | 0.41709602 | -0.3513261 | 0.72534375 | 0.96781837 |
| KLHDC4 | 87.7359814 | 0.157816227 | 0.44194802 | 0.35709228 | 0.72102272 | 0.96779196 |
| BANP | 103.831474 | -0.262271849 | 0.31564148 | -0.8309169 | 0.40602056 | 0.89569633 |
| ZFPM1 | 42.3865663 | -0.579302761 | 0.25193827 | -2.2993837 | 0.02148316 | 0.38588526 |
| ZC3H18 | 69.416633 | -0.288083184 | 0.2598784 | -1.1085307 | 0.2676327 | 0.81510106 |
| CYBA | 351.964771 | 0.738084703 | 0.54724401 | 1.34873052 | 0.17742354 | 0.75126783 |
| MVD | 40.9370097 | 0.465302276 | 0.46508102 | 1.00047574 | 0.31708033 | 0.8490543 |
| PIEZO1 | 72.785298 | -0.052654338 | 0.51994095 | -0.1012698 | 0.91933626 | 0.99214128 |
| APRT | 100.0527 | -0.109968866 | 0.5411927 | -0.2031972 | 0.83898086 | 0.98462033 |
| GALNS | 45.776609 | 0.566531269 | 0.35008797 | 1.61825405 | 0.10560786 | 0.65699665 |
| TRAPPC2L | 26.6486565 | -0.001871725 | 0.38612665 | -0.0048474 | 0.99613232 | 0.99884345 |
| ACSF3 | 51.9983541 | -0.063308372 | 0.30655882 | -0.206513 | 0.83639024 | 0.98462033 |
| ANKRD11 | 582.89754 | -0.279629095 | 0.21284805 | -1.3137498 | 0.18893041 | 0.76207437 |
| SPG7 | 210.2454 | -0.344950741 | 0.30923038 | -1.1155137 | 0.26463034 | 0.81378765 |
| RPL13 | 1093.947 | 0.192302727 | 0.48030309 | 0.40037787 | 0.68887822 | 0.96190309 |
| CHMP1A | 59.8171596 | -0.127058513 | 0.370859 | -0.342606 | 0.7318949 | 0.96980299 |
| CDK10 | 89.7437237 | 0.337329088 | 0.29036689 | 1.161734 | 0.24534353 | 0.80476923 |
| VPS9D1 | 39.0904873 | 0.169002469 | 0.37486141 | 0.45083987 | 0.65210496 | 0.95159942 |
| ZNF276 | 39.9170708 | -0.21057344 | 0.31152962 | -0.6759339 | 0.4990826 | 0.92398825 |
| TCF25 | 305.955192 | -0.149062889 | 0.2179144 | -0.6840433 | 0.49394782 | 0.92289011 |
| DEF8 | 37.9544676 | -0.612094254 | 0.30893606 | -1.9812975 | 0.04755792 | 0.51346008 |
| GAS8 | 84.0192572 | -0.070509676 | 0.31585692 | -0.223233 | 0.8233542 | 0.98094663 |
| FAM157C | 82.625773 | -0.083189807 | 0.56795945 | -0.1464714 | 0.88354928 | 0.98766275 |
| RPH3AL | 169.422965 | -0.418877375 | 0.51475517 | -0.813741 | 0.41579336 | 0.89825994 |
| VPS53 | 281.392687 | -0.234343683 | 0.23004586 | -1.0186825 | 0.30835373 | 0.84488735 |
| GLOD4 | 50.5777896 | -0.214858726 | 0.29523677 | -0.7277506 | 0.46676629 | 0.91462428 |
| NXN | 167.321337 | 0.821240688 | 0.37321951 | 2.20042274 | 0.02777692 | 0.4250936 |
| ABR | 141.944314 | 0.175386183 | 0.51715275 | 0.33913806 | 0.73450573 | 0.9698508 |
| YWHAE | 447.517139 | -0.08947243 | 0.17100129 | -0.5232266 | 0.60081654 | 0.94683415 |
| CRK | 143.278694 | 0.003229462 | 0.29385769 | 0.01098988 | 0.99123152 | 0.99866813 |
| MYO1C | 168.12425 | -0.055229096 | 0.31473663 | -0.1754772 | 0.8607047 | 0.98652427 |
| INPP5K | 24.2803222 | -0.275812664 | 0.3421702 | -0.8060686 | 0.42020327 | 0.90012434 |
| PITPNA | 173.807787 | 0.034735749 | 0.17050624 | 0.20372128 | 0.83857131 | 0.98462033 |
| SLC43A2 | 75.2412636 | 0.34029599 | 0.65727382 | 0.51773854 | 0.6046407 | 0.94683415 |
| PRPF8 | 157.830863 | 0.069492254 | 0.18741963 | 0.37078428 | 0.71079821 | 0.96587538 |

| | | | | | | |
|---|---|---|---|---|---|---|
| WDR81 | 31.6624593 | -0.192660339 | 0.38274418 | -0.5033658 | 0.61470709 | 0.94791646 |
| SERPINF2 | 86.2266641 | 2.036337186 | 0.52786483 | 3.8576868 | 0.00011447 | 0.02741155 |
| SMYD4 | 43.4397802 | -0.097420699 | 0.33921116 | -0.2871978 | 0.77396087 | 0.97408634 |
| RPA1 | 95.910607 | 0.034858834 | 0.20732666 | 0.16813483 | 0.86647721 | 0.98670067 |
| SMG6 | 337.224797 | -0.040427947 | 0.23998231 | -0.1684622 | 0.86621968 | 0.98670067 |
| TSR1 | 38.3405251 | -0.667831295 | 0.35496558 | -1.8813973 | 0.05991789 | 0.55922073 |
| SGSM2 | 55.9335662 | 0.432767512 | 0.25707499 | 1.68342905 | 0.09229206 | 0.63464873 |
| MNT | 29.5468957 | 0.019441784 | 0.41354775 | 0.04701219 | 0.96250351 | 0.99572129 |
| METTL16 | 147.35756 | -0.392692517 | 0.2274364 | -1.7266037 | 0.08423887 | 0.61656297 |
| PAFAH1B1 | 498.925984 | -0.008151321 | 0.13082363 | -0.0623077 | 0.95031779 | 0.99531843 |
| CLUH | 65.4066117 | 0.149289095 | 0.42120651 | 0.35443207 | 0.7230151 | 0.96779196 |
| RAP1GAP2 | 155.334554 | -0.593542086 | 0.41700857 | -1.4233331 | 0.15463963 | 0.72978231 |
| CTNS | 35.6790046 | -0.387150644 | 0.41187047 | -0.9399816 | 0.34722702 | 0.87017396 |
| TAX1BP3 | 36.1184679 | -0.049982422 | 0.5188741 | -0.0963286 | 0.92325959 | 0.99214128 |
| EMC6 | 38.7593049 | -0.498255724 | 0.5465277 | -0.9116752 | 0.36193975 | 0.8752623 |
| ITGAE | 28.2356987 | 0.014257301 | 0.38807637 | 0.03673839 | 0.9706936 | 0.99681784 |
| NCBP3 | 105.5958 | -0.109135099 | 0.19642867 | -0.5555966 | 0.57848665 | 0.94034284 |
| ZZEF1 | 150.093058 | -0.109044537 | 0.22414662 | -0.4864875 | 0.62662155 | 0.94831213 |
| CYB5D2 | 52.2780616 | 0.041141713 | 0.38543811 | 0.10674013 | 0.91499515 | 0.99159829 |
| ANKFY1 | 145.486795 | -0.321779689 | 0.35045812 | -0.9181687 | 0.35853053 | 0.87316144 |
| UBE2G1 | 183.246909 | 0.067773469 | 0.2380335 | 0.2847224 | 0.77585683 | 0.97423208 |
| SPNS2 | 187.656246 | -0.125262153 | 0.41620662 | -0.3009615 | 0.76344388 | 0.97149197 |
| MYBBP1A | 27.4237937 | -0.763990688 | 0.31095994 | -2.4568782 | 0.01401502 | 0.32375679 |
| SMTNL2 | 54.906916 | 0.552328251 | 0.43879042 | 1.25875184 | 0.20811998 | 0.77455509 |
| PELP1 | 46.6786236 | -0.425494472 | 0.31958594 | -1.331393 | 0.18305974 | 0.75648372 |
| CXCL16 | 41.2319766 | -0.757294057 | 0.50592508 | -1.4968502 | 0.13443224 | 0.70619627 |
| PSMB6 | 66.7078029 | -0.131740817 | 0.55410508 | -0.2377542 | 0.81207173 | 0.97930627 |
| MINK1 | 120.37132 | -0.40518284 | 0.28076138 | -1.4431573 | 0.14897617 | 0.72246729 |
| SLC25A11 | 47.506221 | 0.404332139 | 0.40226 | 1.00515124 | 0.31482403 | 0.8490543 |
| RNF167 | 43.2269305 | -0.205740325 | 0.41008608 | -0.5017003 | 0.61587832 | 0.94791646 |
| PFN1 | 295.123712 | -0.737540697 | 0.58557877 | -1.2595072 | 0.20784718 | 0.77435432 |
| SPAG7 | 69.9928668 | -0.378097696 | 0.39621172 | -0.954282 | 0.33994094 | 0.86742596 |
| CAMTA2 | 41.980305 | -0.243797405 | 0.28416077 | -0.8579559 | 0.39091682 | 0.88670817 |
| KIF1C | 96.7307343 | -0.102241785 | 0.32428721 | -0.3152816 | 0.75254787 | 0.97096495 |
| ZNF594 | 38.2999573 | 0.227960275 | 0.31777493 | 0.71736394 | 0.47314956 | 0.91793034 |
| AC087500.1 | 78.4674822 | 0.638866915 | 0.3070378 | 2.08074356 | 0.03745738 | 0.46930788 |
| RABEP1 | 231.669964 | -0.181032034 | 0.22328607 | -0.8107628 | 0.41750192 | 0.89862171 |
| NUP88 | 146.020742 | 0.05338453 | 0.29416111 | 0.18148059 | 0.85599037 | 0.9864199 |
| RPAIN | 74.6264329 | -0.369383477 | 0.22469231 | -1.6439524 | 0.10018603 | 0.64671608 |
| C1QBP | 58.1453211 | 0.12643688 | 0.55443909 | 0.22804467 | 0.81961151 | 0.98079697 |
| DERL2 | 53.3721222 | -0.139960177 | 0.4092201 | -0.3420169 | 0.7323382 | 0.96980299 |
| KIAA0753 | 115.082334 | -0.011618884 | 0.24973721 | -0.0465244 | 0.96289226 | 0.99572129 |
| TXNDC17 | 29.5107742 | -0.416129145 | 0.50851128 | -0.8183282 | 0.41316978 | 0.89724628 |
| MED31 | 30.0418716 | 0.241103976 | 0.40318899 | 0.59799245 | 0.54984497 | 0.93523904 |
| XAF1 | 126.393915 | -3.768879094 | 1.86219766 | -2.0238878 | 0.0429817 | 0.49426374 |
| ALOX12-AS1 | 41.2419901 | 1.450096182 | 0.57517575 | 2.52113581 | 0.01169767 | 0.30872898 |
| ACADVL | 187.147527 | 0.362097524 | 0.29657116 | 1.22094652 | 0.22210627 | 0.78710314 |
| DVL2 | 29.7920653 | 0.440342595 | 0.36497516 | 1.20650018 | 0.22762469 | 0.79284833 |
| PHF23 | 27.7386438 | 0.163215982 | 0.31789781 | 0.5134228 | 0.6076556 | 0.94708529 |
| ELP5 | 31.4626201 | 0.064027052 | 0.3053375 | 0.20969272 | 0.83390751 | 0.98351801 |
| EIF5A | 80.3482079 | -1.078106296 | 0.55762336 | -1.9333951 | 0.05318754 | 0.53386172 |
| GPS2 | 56.8273663 | -0.453698 | 0.36400183 | -1.2464168 | 0.21261141 | 0.77822114 |
| ZBTB4 | 162.713267 | 0.633186748 | 0.35010316 | 1.80857192 | 0.07051753 | 0.58797037 |
| POLR2A | 134.057082 | -0.157956618 | 0.35233336 | -0.4483158 | 0.65392529 | 0.9525472 |

| | | | | | | |
|---|---|---|---|---|---|---|
| EIF4A1 | 96.2680522 | -0.481166832 | 0.50422121 | -0.9542772 | 0.33994332 | 0.86742596 |
| MPDU1 | 32.7382987 | -0.300121399 | 0.5769461 | -0.5201897 | 0.60293138 | 0.94683415 |
| FXR2 | 42.1579485 | -0.478783831 | 0.24452131 | -1.9580454 | 0.05022468 | 0.52415717 |
| SAT2 | 88.9732438 | 0.164619396 | 0.31522159 | 0.52223388 | 0.6015075 | 0.94683415 |
| TP53 | 51.053446 | 0.161274424 | 0.36662029 | 0.43989497 | 0.66001318 | 0.95407427 |
| KDM6B | 59.0394651 | -0.101644475 | 0.3848013 | -0.264148 | 0.79166592 | 0.97423208 |
| NAA38 | 142.146422 | 0.347094192 | 0.28057708 | 1.23707253 | 0.21606016 | 0.78293142 |
| CHD3 | 112.914994 | 0.273531936 | 0.33390276 | 0.81919639 | 0.41267437 | 0.89718743 |
| TRAPPC1 | 28.6469039 | -0.254750014 | 0.59572804 | -0.427628 | 0.66892195 | 0.95764829 |
| PER1 | 86.9514665 | -0.343174341 | 0.63557426 | -0.5399437 | 0.58923583 | 0.94491859 |
| CTC1 | 36.8108858 | 0.151283708 | 0.3670351 | 0.41217777 | 0.68020913 | 0.95974713 |
| NDEL1 | 106.735875 | -0.442587939 | 0.30441188 | -1.4539115 | 0.14597084 | 0.71974648 |
| MYH10 | 193.541068 | 0.269042015 | 0.34682827 | 0.77572113 | 0.43791366 | 0.90561284 |
| STX8 | 302.160603 | -0.317640111 | 0.22979626 | -1.3822684 | 0.1668893 | 0.74445571 |
| USP43 | 78.1046298 | -0.390829956 | 0.34722081 | -1.1255948 | 0.26033705 | 0.81263292 |
| SCO1 | 51.6860003 | -0.190840046 | 0.29846207 | -0.6394114 | 0.52255534 | 0.93272217 |
| TMEM220 | 61.487317 | -0.009874345 | 0.34070656 | -0.028982 | 0.97687898 | 0.99681784 |
| TMEM220-A! | 38.1028138 | -0.250524267 | 0.53622752 | -0.4671977 | 0.64035843 | 0.94944486 |
| SHISA6 | 37.4186628 | 1.399930509 | 0.60379122 | 2.31856717 | 0.02041852 | 0.3784173 |
| DNAH9 | 46.0308862 | 1.460926143 | 0.44236892 | 3.30250633 | 0.00095825 | 0.10275397 |
| MAP2K4 | 372.332117 | -0.010938047 | 0.14871669 | -0.0735496 | 0.94136881 | 0.99526299 |
| ARHGAP44 | 59.3864216 | 0.211924191 | 0.52431807 | 0.40419013 | 0.68607291 | 0.96161098 |
| ELAC2 | 36.0047292 | -0.394317397 | 0.34653507 | -1.137886 | 0.2551681 | 0.80897837 |
| COX10 | 74.2163223 | -0.309280513 | 0.32195919 | -0.9606202 | 0.33674314 | 0.86431445 |
| TRIM16 | 37.6082142 | -0.716533796 | 0.31567529 | -2.2698444 | 0.02321703 | 0.39839181 |
| ZSWIM7 | 101.271017 | 0.043178237 | 0.25277167 | 0.17081913 | 0.86436599 | 0.98670067 |
| TTC19 | 228.233405 | -0.083819826 | 0.2380247 | -0.3521476 | 0.72472758 | 0.96781837 |
| NCOR1 | 570.642079 | -0.146345575 | 0.15227382 | -0.9610685 | 0.33651773 | 0.86431445 |
| PIGL | 169.920414 | -0.758313616 | 0.19579804 | -3.8729377 | 0.00010753 | 0.02641135 |
| UBB | 320.962126 | 0.512753685 | 0.43541317 | 1.17762559 | 0.2389459 | 0.80174646 |
| LRRC75A | 30.6047172 | 0.527497888 | 0.46585051 | 1.13233296 | 0.25749448 | 0.8120529 |
| AC098850.3 | 78.8936311 | 0.721848199 | 0.61833165 | 1.16741266 | 0.24304376 | 0.80476923 |
| MPRIP | 320.432496 | -0.564419528 | 0.3984597 | -1.4165034 | 0.15662817 | 0.73366319 |
| COPS3 | 38.5962864 | -0.142603693 | 0.30901995 | -0.4614708 | 0.64446085 | 0.94944486 |
| MED9 | 37.5595167 | -0.195927163 | 0.38683266 | -0.5064907 | 0.61251218 | 0.94791646 |
| RASD1 | 260.443632 | 0.136176504 | 1.15625961 | 0.11777329 | 0.90624729 | 0.99100813 |
| PEMT | 47.5479799 | 0.003658118 | 0.54668787 | 0.00669142 | 0.99466106 | 0.99866813 |
| RAI1 | 224.335418 | -0.452329983 | 0.40447749 | -1.1183069 | 0.26343593 | 0.81378765 |
| SREBF1 | 42.3730374 | -0.540823838 | 0.42721374 | -1.2659327 | 0.20553718 | 0.77352973 |
| TOM1L2 | 218.813332 | 0.204357898 | 0.21029552 | 0.97176533 | 0.33116731 | 0.85923151 |
| DRC3 | 46.3648647 | 1.219412638 | 0.33458031 | 3.64460375 | 0.0002678 | 0.04347958 |
| ATPAF2 | 34.4823165 | -0.182817723 | 0.32001745 | -0.5712742 | 0.56781376 | 0.93906906 |
| DRG2 | 27.0879427 | -0.288231075 | 0.39341231 | -0.7326438 | 0.46377573 | 0.91462428 |
| ALKBH5 | 94.7482548 | -0.034903902 | 0.26801847 | -0.1302295 | 0.89638488 | 0.98842187 |
| FLII | 72.9299483 | -0.480417677 | 0.25928323 | -1.8528683 | 0.06390123 | 0.57099806 |
| TOP3A | 50.3420164 | -0.158541755 | 0.27802651 | -0.5702397 | 0.56851513 | 0.93974227 |
| SHMT1 | 203.413068 | 0.856597762 | 0.17049111 | 5.02429575 | 5.05E-07 | 0.00069144 |
| TVP23B | 46.4591767 | 0.131538256 | 0.3971409 | 0.33121307 | 0.74048355 | 0.9698508 |
| PRPSAP2 | 140.730034 | -0.342447694 | 0.28890391 | -1.1853342 | 0.23588532 | 0.79952815 |
| SLC5A10 | 43.2686669 | 0.102163297 | 0.29120158 | 0.35083359 | 0.72571319 | 0.96781837 |
| EPN2 | 310.597027 | -0.129190023 | 0.1818051 | -0.7105963 | 0.47733446 | 0.91864961 |
| B9D1 | 63.8133008 | 0.119613016 | 0.25614611 | 0.46697182 | 0.64052004 | 0.94944486 |
| AC004448.2 | 38.4885301 | -1.048054445 | 0.49666809 | -2.1101707 | 0.03484366 | 0.45472394 |
| SLC47A1 | 73.1840731 | 0.888954365 | 0.27775449 | 3.20050407 | 0.00137187 | 0.11702898 |

| | | | | | | |
|---|---|---|---|---|---|---|
| ALDH3A2 | 101.255005 | 0.133348998 | 0.33644842 | 0.39634306 | 0.69185196 | 0.96220115 |
| ULK2 | 79.2445676 | 0.175103703 | 0.28326357 | 0.61816527 | 0.5364664 | 0.93469807 |
| AKAP10 | 230.43548 | 0.062523458 | 0.19796021 | 0.31583851 | 0.75212509 | 0.97084035 |
| CCDC144NL-/ | 43.2150969 | -0.108357036 | 0.47964034 | -0.2259131 | 0.82126901 | 0.98094663 |
| USP22 | 198.916585 | -0.011741977 | 0.19957668 | -0.0588344 | 0.953084 | 0.99531843 |
| DHRS7B | 161.377561 | -0.106218723 | 0.27168091 | -0.3909687 | 0.6958204 | 0.96220115 |
| TMEM11 | 25.9348079 | -0.272942401 | 0.37687744 | -0.7242206 | 0.46893033 | 0.91462428 |
| AC087294.1 | 27.585456 | 0.203980517 | 0.35440745 | 0.5755537 | 0.56491689 | 0.93809297 |
| MAP2K3 | 54.7664262 | -0.224947678 | 0.40440718 | -0.5562406 | 0.57804642 | 0.94034284 |
| WSB1 | 774.71289 | -0.241462812 | 0.49554189 | -0.4872702 | 0.62606684 | 0.94831213 |
| KSR1 | 135.314634 | 0.049498658 | 0.34743113 | 0.14247042 | 0.88670844 | 0.98766275 |
| NLK | 505.705681 | 0.047623344 | 0.18205753 | 0.26158404 | 7.94E-01 | 0.97464093 |
| IFT20 | 28.5431549 | 0.430016603 | 0.3062609 | 1.40408585 | 0.16029329 | 0.73816762 |
| TNFAIP1 | 55.0501276 | 0.100662094 | 0.38233647 | 0.26328143 | 0.79233367 | 0.97429505 |
| POLDIP2 | 47.8641633 | -0.092700306 | 0.26180962 | -0.3540753 | 0.72328249 | 0.96779196 |
| SARM1 | 35.7432533 | 0.135165216 | 0.58472333 | 0.23116098 | 0.81718975 | 0.98010849 |
| KIAA0100 | 88.8059621 | 0.061657176 | 0.28421014 | 0.21694221 | 0.82825338 | 0.98142144 |
| SDF2 | 47.6808537 | 0.219696733 | 0.36783059 | 0.59727695 | 0.55032249 | 0.93523904 |
| SUPT6H | 105.503585 | -0.239783463 | 0.20485724 | -1.1704906 | 0.24180361 | 0.803412 |
| RAB34 | 65.5568626 | -0.35556105 | 0.42410146 | -0.8383868 | 0.40181353 | 0.89239743 |
| RPL23A | 337.182259 | -0.081107138 | 0.55913803 | -0.1450575 | 0.88466551 | 0.98766275 |
| NEK8 | 39.1292822 | -0.176922298 | 0.63463384 | -0.2787785 | 0.78041478 | 0.97423208 |
| TRAF4 | 81.7168056 | -0.27651608 | 0.3523606 | -0.7847531 | 0.43259833 | 0.90514499 |
| FAM222B | 309.496577 | 0.044929994 | 0.26565818 | 0.16912709 | 0.86569668 | 0.98670067 |
| FLOT2 | 157.763344 | 0.26181139 | 0.28523774 | 0.91787079 | 0.35868652 | 0.87316144 |
| PHF12 | 92.7302677 | 0.035855062 | 0.27343188 | 0.13112978 | 0.89567264 | 0.98842187 |
| MYO18A | 57.8803669 | 1.013826767 | 0.50002705 | 2.02754383 | 0.04260683 | 0.49385357 |
| NUFIP2 | 244.856331 | -0.041941662 | 0.21072978 | -0.1990305 | 0.84223886 | 0.98517901 |
| TAOK1 | 424.092813 | 0.12240121 | 0.23337511 | 0.5244827 | 0.59994285 | 0.94683415 |
| ABHD15-AS1 | 58.9286018 | 0.647515339 | 0.39028904 | 1.65906615 | 0.09710246 | 0.64312419 |
| GIT1 | 37.5775932 | 0.247548081 | 0.43961054 | 0.56310771 | 0.57336155 | 0.93998433 |
| SSH2 | 796.420656 | 0.029751197 | 0.22674515 | 0.13120985 | 0.8956093 | 0.98842187 |
| AC023389.1 | 30.5286526 | 0.354604554 | 0.32740158 | 1.08308747 | 0.2787696 | 0.8229426 |
| NSRP1 | 296.287813 | -0.220735821 | 0.17304888 | -1.2755692 | 0.20210784 | 0.77110248 |
| BLMH | 66.5863947 | 0.208088543 | 0.3090267 | 0.67336752 | 0.50071351 | 0.92414927 |
| CPD | 400.255937 | 0.558273862 | 0.42526685 | 1.3127613 | 0.18926339 | 0.76207437 |
| GOSR1 | 150.94124 | -0.19042966 | 0.21703853 | -0.8774003 | 0.38026927 | 0.8807629 |
| CRLF3 | 136.846029 | 0.033611191 | 0.26179029 | 0.12838975 | 0.89784054 | 0.9887807 |
| ATAD5 | 31.7365426 | -0.488757151 | 0.4241982 | -1.1521905 | 0.24924279 | 0.80672756 |
| TEFM | 55.8365319 | -0.371732039 | 0.32200213 | -1.1544397 | 0.24831995 | 0.80672756 |
| RNF135 | 39.5320076 | -0.453225245 | 0.35498303 | -1.2767519 | 0.20168986 | 0.77110248 |
| NF1 | 928.175751 | -0.055191199 | 0.1825463 | -0.3023408 | 0.76239225 | 0.9714717 |
| RAB11FIP4 | 34.015361 | -0.462756895 | 0.38974594 | -1.1873296 | 0.23509763 | 0.79886493 |
| COPRS | 37.0317925 | -0.054011267 | 0.49109197 | -0.109982 | 0.91242367 | 0.99113939 |
| UTP6 | 58.4870557 | -0.280329148 | 0.28694224 | -0.9769532 | 0.32859231 | 0.8574012 |
| SUZ12 | 243.530037 | -0.182751783 | 0.21202624 | -0.86193 | 0.38872602 | 0.88586416 |
| LRRC37B | 63.5290217 | -0.053327669 | 0.27864154 | -0.1913845 | 0.84822437 | 0.98545133 |
| RHOT1 | 143.711981 | 0.02502183 | 0.20388855 | 0.12272308 | 0.90232639 | 0.9900784 |
| C17orf75 | 25.5223272 | -0.275871121 | 0.33817461 | -0.8157653 | 0.41463437 | 0.89757799 |
| ZNF207 | 366.078824 | 0.015652577 | 0.20653805 | 0.07578544 | 0.9395898 | 0.99526299 |
| PSMD11 | 91.1965618 | -0.432560003 | 0.29395948 | -1.4714954 | 0.1411572 | 0.7148274 |
| MYO1D | 657.444687 | -0.23735912 | 0.2700071 | -0.8790847 | 0.37935534 | 0.88018641 |
| TMEM98 | 51.6614149 | 1.294114584 | 0.52187532 | 2.47973899 | 0.01314786 | 0.31992293 |
| CCL2 | 148.473391 | 0.251258258 | 1.13209811 | 0.22194036 | 0.82436031 | 0.98094663 |

| | | | | | | |
|---|---|---|---|---|---|---|
| LIG3 | 32.0232695 | -0.544212942 | 0.27145877 | -2.004772 | 0.04498742 | 0.50340725 |
| RFFL | 34.330975 | 0.37339281 | 0.4132136 | 0.90363146 | 0.36619085 | 0.87605028 |
| SLFN5 | 100.245711 | -0.141194044 | 0.55372291 | -0.2549904 | 7.99E-01 | 0.9761581 |
| SLFN13 | 28.5160906 | 0.282548158 | 0.62549414 | 0.45171991 | 0.65147077 | 0.95157647 |
| LINC02001 | 33.9410204 | -0.032731513 | 0.35190446 | -0.0930125 | 0.92589363 | 0.99316012 |
| AP2B1 | 197.895317 | 0.033993238 | 0.21493041 | 0.15815927 | 0.87433129 | 0.98766275 |
| TAF15 | 407.13604 | -0.2477992 | 0.21937506 | -1.1295687 | 0.258658 | 0.81263292 |
| ZNHIT3 | 30.4191624 | 0.264696581 | 0.48489179 | 0.54588795 | 0.58514297 | 0.94229635 |
| MYO19 | 82.6039197 | -0.386249737 | 0.29353723 | -1.3158458 | 0.18822579 | 0.76207437 |
| GGNBP2 | 177.634385 | 0.124075608 | 0.18114071 | 0.68496811 | 0.49336404 | 0.92289011 |
| AATF | 142.656338 | -0.254012986 | 0.19284223 | -1.3172062 | 0.18776951 | 0.76207437 |
| ACACA | 299.608239 | -0.451459791 | 0.20676046 | -2.1834919 | 0.0289996 | 0.43178298 |
| TADA2A | 88.8643572 | -0.74089833 | 0.25605773 | -2.8934816 | 0.00380997 | 0.19908785 |
| SYNRG | 205.203652 | 0.177477206 | 0.19120933 | 0.92818279 | 0.35331276 | 0.8720098 |
| DDX52 | 142.688351 | -0.387832886 | 0.2588194 | -1.4984691 | 0.1340114 | 0.70542022 |
| HNF1B | 564.046406 | -0.923233795 | 0.33857997 | -2.7267821 | 0.00639553 | 0.25167097 |
| MRPL45 | 46.9795135 | -0.324120399 | 0.24385458 | -1.3291544 | 0.18379703 | 0.75689793 |
| SOCS7 | 74.7283926 | 0.309075572 | 0.45055332 | 0.68599111 | 0.49271871 | 0.92289011 |
| ARHGAP23 | 60.5872328 | 0.222821735 | 0.7200899 | 0.309436 | 0.75698989 | 0.9714717 |
| MLLT6 | 235.488515 | 0.084979394 | 0.24023254 | 0.35373806 | 0.72353519 | 0.96779196 |
| CISD3 | 37.7643532 | -0.062287344 | 0.43368548 | -0.1436233 | 0.88579793 | 0.98766275 |
| PCGF2 | 50.8265995 | -0.103932876 | 0.30350283 | -0.3424445 | 0.7320164 | 0.96980299 |
| PSMB3 | 78.127645 | -0.672516252 | 0.38573691 | -1.7434584 | 0.08125358 | 0.60850843 |
| PIP4K2B | 72.3881329 | 0.111149794 | 0.35134053 | 0.31635916 | 0.75172991 | 0.97058015 |
| CWC25 | 44.0016385 | -0.0880543 | 0.33383998 | -0.263762 | 0.79196332 | 0.97423208 |
| RPL23 | 262.622266 | 0.178088611 | 0.48350784 | 0.36832621 | 0.71263001 | 0.96587538 |
| LASP1 | 129.433165 | -0.02555255 | 0.34614343 | -0.0738207 | 0.94115306 | 0.99526299 |
| RPL19 | 536.536723 | -0.044869522 | 0.60562371 | -0.0740881 | 0.94094027 | 0.99526299 |
| FBXL20 | 326.498425 | 0.56555358 | 0.26294263 | 2.15086301 | 0.03148701 | 0.43905874 |
| MED1 | 71.4036477 | -0.225495896 | 0.21121809 | -1.0675975 | 0.28570213 | 0.83003998 |
| CDK12 | 198.314644 | 0.064481591 | 0.19385096 | 0.33263488 | 0.73940991 | 0.9698508 |
| STARD3 | 41.8013379 | -0.222059632 | 0.23745318 | -0.9351723 | 0.34969953 | 0.8720098 |
| PGAP3 | 26.2765554 | 0.599939044 | 0.51580753 | 1.16310642 | 0.24478633 | 0.80476923 |
| ERBB2 | 171.824765 | 0.152718043 | 0.46363196 | 0.32939498 | 0.74185716 | 0.9698508 |
| MIEN1 | 36.0355071 | -0.424650546 | 0.44446761 | -0.9554139 | 0.33936841 | 0.8672311 |
| GRB7 | 111.126989 | -0.060054638 | 0.31782207 | -0.1889568 | 0.85012669 | 0.9858744 |
| PSMD3 | 59.5427339 | -0.517346291 | 0.30757951 | -1.681992 | 0.09257038 | 0.63577454 |
| MED24 | 57.2763947 | -0.308744428 | 0.27172845 | -1.1362241 | 0.25586277 | 0.80939065 |
| THRA | 68.1425643 | 0.624182093 | 0.31839916 | 1.96037606 | 0.04995185 | 0.52415717 |
| MSL1 | 105.394775 | -0.328609724 | 0.27219416 | -1.2072622 | 0.22733118 | 0.79244405 |
| CASC3 | 104.093974 | 0.104592802 | 0.23407445 | 0.44683562 | 0.65499375 | 0.95294428 |
| WIPF2 | 197.739479 | -0.110745819 | 0.18650146 | -0.5938067 | 0.55264141 | 0.93523904 |
| RARA | 39.4698322 | 0.041341725 | 0.48822905 | 0.0846769 | 0.93251826 | 0.99416721 |
| IGFBP4 | 411.533252 | 0.399544517 | 0.72854358 | 0.54841539 | 0.58340671 | 0.94160958 |
| KRT10 | 89.9937708 | -0.461577395 | 0.49214855 | -0.9378823 | 0.34830492 | 0.87112607 |
| KRT19 | 395.367646 | -0.090075436 | 0.46525219 | -0.1936056 | 0.8464847 | 0.98525378 |
| EIF1 | 408.405884 | 0.138503422 | 0.43093942 | 0.32139882 | 0.74790818 | 0.97028355 |
| JUP | 291.361116 | 0.091298645 | 0.27509935 | 0.33187517 | 0.73998352 | 0.9698508 |
| P3H4 | 38.5805985 | 0.205920658 | 0.45246431 | 0.45510918 | 0.6490307 | 0.95027667 |
| FKBP10 | 34.1411769 | 0.029095152 | 0.92166107 | 0.03156817 | 0.97481643 | 0.99681784 |
| NT5C3B | 44.1883447 | -0.109658155 | 0.2599804 | -0.4217939 | 0.67317543 | 0.95910677 |
| ACLY | 161.852556 | 0.587039901 | 0.55154399 | 1.06435734 | 0.28716685 | 0.83129986 |
| TTC25 | 36.4223819 | 0.915417225 | 0.42785765 | 2.13953691 | 0.03239221 | 0.44049392 |
| CNP | 90.1502264 | 0.008172961 | 0.44040606 | 0.01855779 | 0.98519388 | 0.99752755 |

| | | | | | | |
|---|---|---|---|---|---|---|
| DNAJC7 | 169.936269 | -0.030794789 | 0.16045109 | -0.1919263 | 0.84779991 | 0.98545133 |
| NKIRAS2 | 26.4102507 | -0.642256208 | 0.29844982 | -2.1519739 | 0.03139941 | 0.43905874 |
| KAT2A | 35.2142665 | -0.277710414 | 0.29559135 | -0.9395079 | 0.34747003 | 0.87017396 |
| GHDC | 23.4080309 | 0.322405053 | 0.39316466 | 0.8200255 | 0.41220157 | 0.89706788 |
| STAT5B | 175.54028 | 0.350546468 | 0.1836425 | 1.90885262 | 0.05628111 | 0.5431192 |
| STAT5A | 27.149819 | 0.686531764 | 0.47437181 | 1.44724403 | 0.14782859 | 0.72246583 |
| STAT3 | 677.881146 | 0.20170067 | 0.30706963 | 0.65685646 | 0.51127322 | 0.92819651 |
| CAVIN1 | 109.949717 | -0.454899871 | 0.60894446 | -0.7470302 | 0.45504536 | 0.91077355 |
| ATP6V0A1 | 192.126096 | 0.213117759 | 0.24997993 | 0.85253947 | 0.39391474 | 0.88760715 |
| NAGLU | 38.3608299 | -0.060737723 | 0.60282463 | -0.1007552 | 0.91974478 | 0.99214128 |
| COASY | 36.2660697 | -0.172798111 | 0.29862645 | -0.578643 | 0.56283007 | 0.93805441 |
| MLX | 31.5762312 | -0.269472365 | 0.31831603 | -0.8465561 | 0.39724262 | 0.88863925 |
| RETREG3 | 102.732017 | -0.0442282 | 0.18743481 | -0.2359658 | 0.81345923 | 0.97930627 |
| TUBG2 | 31.4215345 | -0.21785219 | 0.27720435 | -0.7858902 | 0.4319318 | 0.90503769 |
| EZH1 | 100.24283 | -0.19488962 | 0.21318665 | -0.9141737 | 0.36062561 | 0.87429201 |
| VPS25 | 49.4968314 | -0.371149068 | 0.34256388 | -1.0834448 | 0.27861102 | 0.8229426 |
| COA3 | 64.8488371 | 0.160348133 | 0.48687077 | 0.32934434 | 0.74189543 | 0.9698508 |
| BECN1 | 42.4393183 | -0.02362445 | 0.28548744 | -0.0827513 | 0.93404931 | 0.99442609 |
| PSME3 | 33.9402064 | -0.181703738 | 0.31339247 | -0.5797961 | 0.56205211 | 0.93762898 |
| RUNDC1 | 39.7078739 | -0.116956767 | 0.34602429 | -0.3380016 | 0.73536198 | 0.9698508 |
| RPL27 | 172.65527 | -0.313906371 | 0.5085753 | -0.6172269 | 0.53708505 | 0.93489692 |
| IFI35 | 42.7752329 | -1.583606668 | 0.80421246 | -1.9691397 | 0.04893705 | 0.52219814 |
| VAT1 | 47.6632111 | -0.101487513 | 0.32411641 | -0.3131206 | 0.75418907 | 0.9712249 |
| BRCA1 | 53.9883819 | 0.09605941 | 0.37444826 | 0.25653587 | 0.79753709 | 0.97555967 |
| NBR1 | 143.739697 | 0.660292014 | 0.23900228 | 2.76270172 | 0.00573251 | 0.23851696 |
| LINC00854 | 219.45542 | -0.861816323 | 0.57164625 | -1.5076043 | 0.13165584 | 0.700349 |
| DHX8 | 53.7657979 | 0.120320512 | 0.23224008 | 0.51808676 | 0.60439774 | 0.94683415 |
| DUSP3 | 80.9811245 | -0.200164158 | 0.41517236 | -0.482123 | 0.62971855 | 0.94916079 |
| MPP3 | 48.7124489 | -0.349846254 | 0.51294755 | -0.6820312 | 0.4952192 | 0.92307934 |
| TMEM101 | 31.2650221 | 0.483124909 | 0.28915312 | 1.67082723 | 0.09475581 | 0.63901772 |
| LSM12 | 71.5844347 | -0.402862783 | 0.24899369 | -1.6179638 | 0.10567039 | 0.65699665 |
| G6PC3 | 35.1119569 | -0.361643809 | 0.454746 | -0.7952655 | 0.42645907 | 0.90237497 |
| HDAC5 | 45.4409491 | 0.744661555 | 0.31991965 | 2.3276518 | 0.0199306 | 0.3747883 |
| TMUB2 | 37.2572072 | 0.035328677 | 0.29780784 | 0.1186291 | 0.90556921 | 0.99100813 |
| ATXN7L3 | 32.3621948 | -0.921101209 | 0.38173707 | -2.4129205 | 0.01582527 | 0.34086104 |
| UBTF | 71.1568148 | -0.43738665 | 0.39051733 | -1.1200185 | 0.26270587 | 0.81377303 |
| SLC25A39 | 51.3962315 | -0.865473257 | 0.35136905 | -2.4631459 | 0.01377238 | 0.32179883 |
| GRN | 159.961802 | 0.011444881 | 0.3573429 | 0.03202773 | 0.97444994 | 0.99681784 |
| GPATCH8 | 576.379471 | 0.051417093 | 0.20504123 | 0.25076466 | 0.80199607 | 0.97835591 |
| EFTUD2 | 77.4025589 | -0.705319106 | 0.21669423 | -3.2549048 | 0.0011343 | 0.10732865 |
| DCAKD | 96.4054781 | 0.06503172 | 0.41232939 | 0.15771789 | 0.8746791 | 0.98766275 |
| NMT1 | 123.952629 | -0.219451754 | 0.20131492 | -1.0900918 | 0.27567269 | 0.82164863 |
| PLCD3 | 57.5594297 | 0.110574139 | 0.42364106 | 0.26100902 | 0.79408555 | 0.97494815 |
| ACBD4 | 37.2806575 | 0.349955024 | 0.38183395 | 0.91651103 | 0.35939893 | 0.87316144 |
| HEXIM1 | 51.1374738 | -0.10169651 | 0.35101497 | -0.2897213 | 0.77202946 | 0.97329019 |
| AC138150.1 | 91.2493331 | 0.114548613 | 0.44457296 | 0.25765987 | 0.79666942 | 0.97524554 |
| MAP3K14 | 148.302658 | 0.015569992 | 0.3500587 | 0.04447823 | 0.9645232 | 0.99586809 |
| ARHGAP27 | 35.9165435 | -0.268764962 | 0.30977285 | -0.8676195 | 0.38560268 | 0.88353166 |
| PLEKHM1 | 59.7770632 | -0.453987932 | 0.2736027 | -1.6592962 | 0.09705611 | 0.64312419 |
| LINC02210-C | 52.8196683 | 0.570591651 | 0.40182741 | 1.41999187 | 0.15561005 | 0.73139777 |
| KANSL1 | 1040.52773 | 0.408072657 | 0.330929 | 1.23311243 | 0.21753382 | 0.78395654 |
| ARL17B | 244.428857 | 2.322835467 | 1.06624389 | 2.17852171 | 0.02936722 | 0.43411818 |
| LRRC37A2 | 61.9658395 | 0.277496817 | 0.38048691 | 0.72932027 | 0.46580578 | 0.91462428 |
| ARL17A | 69.70518 | -0.282184811 | 0.4176548 | -0.6756412 | 0.49926845 | 0.92398825 |

| | | | | | | |
|---|---|---|---|---|---|---|
| NSF | 216.213903 | -0.184261718 | 0.42245506 | -0.4361688 | 0.66271426 | 0.95477098 |
| AC005670.2 | 32.0073723 | 0.411363315 | 0.50454782 | 0.81531085 | 0.4148944 | 0.89793797 |
| CDC27 | 270.526361 | -0.197138537 | 0.23513809 | -0.8383947 | 0.40180905 | 0.89239743 |
| AC068234.1 | 763.697634 | -0.123090717 | 0.57282316 | -0.2148843 | 0.82985751 | 0.98198951 |
| AC068234.2 | 142.225806 | 0.613954944 | 0.56566018 | 1.08537769 | 0.27775441 | 0.8229426 |
| EFCAB13 | 372.494762 | 0.502408317 | 0.37574884 | 1.33708549 | 0.18119473 | 0.75448394 |
| NPEPPS | 525.344669 | -0.119594113 | 0.21628656 | -0.5529429 | 0.58030253 | 0.94034284 |
| AC025682.1 | 63.0305784 | 0.313909488 | 0.33266563 | 0.94361863 | 0.34536457 | 0.86990996 |
| KPNB1 | 230.977182 | -0.499893252 | 0.19136992 | -2.612183 | 0.00899661 | 0.2825525 |
| SCRN2 | 36.7856587 | 0.052635069 | 0.29631513 | 0.17763206 | 0.85901194 | 0.9864199 |
| SP2 | 24.9138658 | -0.609296111 | 0.33295745 | -1.8299519 | 0.06725714 | 0.57884645 |
| SP2-AS1 | 29.7138379 | -0.183887858 | 0.31248926 | -0.5884614 | 0.55622265 | 0.93523904 |
| PNPO | 29.2570393 | 0.110817853 | 0.37859878 | 0.29270526 | 0.76974745 | 0.97261718 |
| CDK5RAP3 | 58.8195666 | 0.381780002 | 0.27435026 | 1.39157877 | 0.16404999 | 0.74306594 |
| NFE2L1 | 188.544554 | 0.177843573 | 0.21768471 | 0.8169778 | 0.41394112 | 0.89732895 |
| CBX1 | 81.8749149 | 0.100011153 | 0.30443617 | 0.32851272 | 0.74252403 | 0.9698508 |
| SKAP1 | 482.639995 | -1.092771574 | 0.55803643 | -1.9582441 | 0.05020137 | 0.52415717 |
| HOXB2 | 46.3902699 | -0.641004554 | 0.71506756 | -0.8964252 | 0.37002573 | 0.87605028 |
| HOXB3 | 224.441653 | -0.650334506 | 0.63733259 | -1.0204005 | 0.30753855 | 0.84488735 |
| HOXB-AS3 | 53.8575987 | 0.283054275 | 0.78377501 | 0.36114225 | 0.71799311 | 0.9674421 |
| HOXB7 | 62.0945986 | -0.598583714 | 0.55972349 | -1.0694275 | 0.28487707 | 0.82918183 |
| AC103702.2 | 33.8326204 | -1.314810151 | 0.50567538 | -2.6001071 | 0.00931947 | 0.28622448 |
| LINC02086 | 66.4364513 | -1.12593536 | 0.5548017 | -2.0294375 | 0.04241375 | 0.49305988 |
| CALCOCO2 | 103.14548 | 0.193781557 | 0.30613903 | 0.63298547 | 0.52674313 | 0.93356202 |
| ATP5MC1 | 85.1026838 | -0.383457754 | 0.48893505 | -0.7842713 | 0.43288091 | 0.90514499 |
| UBE2Z | 149.170433 | -0.035908418 | 0.33987084 | -0.1056531 | 0.91585757 | 0.99159829 |
| SNF8 | 61.9481769 | -0.05369832 | 0.25132579 | -0.2136602 | 0.83081205 | 0.98251217 |
| ZNF652 | 221.420987 | -0.301054566 | 0.21741613 | -1.3846928 | 0.16614641 | 0.74445571 |
| PHB | 108.335898 | -0.202864266 | 0.46586664 | -0.4354557 | 0.66323171 | 0.95477856 |
| SPOP | 176.013014 | 0.042738644 | 0.28179023 | 0.1516683 | 0.87944856 | 0.98766275 |
| SLC35B1 | 25.1305365 | -0.173223489 | 0.5243611 | -0.3303515 | 0.74113437 | 0.9698508 |
| FAM117A | 27.4231097 | -0.255318031 | 0.42811157 | -0.596382 | 0.55092006 | 0.93523904 |
| KAT7 | 81.7082595 | -0.153864485 | 0.26058506 | -0.5904578 | 0.55488376 | 0.93523904 |
| ITGA3 | 644.09584 | -1.158602841 | 0.65848254 | -1.7595043 | 0.0784919 | 0.6024665 |
| PDK2 | 28.3917456 | 1.409996339 | 0.54239826 | 2.5995591 | 0.00933436 | 0.28622448 |
| PPP1R9B | 24.4323981 | -0.77669295 | 0.41184613 | -1.8858814 | 0.05931094 | 0.5569995 |
| XYLT2 | 38.2564237 | 0.190056731 | 0.30094875 | 0.63152525 | 0.52769715 | 0.93366189 |
| MRPL27 | 53.5206613 | -0.224181594 | 0.39932268 | -0.5614046 | 0.57452174 | 0.94034284 |
| LRRC59 | 35.1459577 | -0.400829418 | 0.44716195 | -0.8963853 | 0.37004699 | 0.87605028 |
| ACSF2 | 68.3923555 | 1.264967559 | 0.43374 | 2.91641898 | 0.00354075 | 0.18995949 |
| RSAD1 | 37.402849 | -0.338789208 | 0.27403068 | -1.2363186 | 0.21634018 | 0.78293142 |
| SPATA20 | 35.3452307 | -0.194181969 | 0.38122983 | -0.5093567 | 0.61050222 | 0.94791646 |
| ABCC3 | 556.672426 | 0.288134588 | 0.67294259 | 0.42817113 | 0.66852654 | 0.95764829 |
| ANKRD40 | 41.8144778 | -0.489668867 | 0.27898364 | -1.7551885 | 0.07922708 | 0.60556378 |
| LUC7L3 | 449.72405 | -0.021745132 | 0.11523216 | -0.1887071 | 8.50E-01 | 9.86E-01 |
| TOB1 | 86.6132586 | 0.348473868 | 0.48826156 | 0.71370326 | 0.47541069 | 0.91809716 |
| SPAG9 | 766.9623 | -0.007270073 | 0.29136258 | -0.024952 | 0.98009327 | 0.99681784 |
| AC005920.2 | 75.0542363 | -0.027760263 | 0.66692282 | -0.0416244 | 0.96679812 | 0.99601626 |
| MBTD1 | 230.203689 | 0.030201836 | 0.22046057 | 0.13699427 | 0.89103532 | 0.98766275 |
| UTP18 | 55.4286665 | -0.356348903 | 0.257778 | -1.3823868 | 0.16685297 | 0.74445571 |
| TOM1L1 | 318.80774 | 0.063577096 | 0.2527161 | 0.25157517 | 0.80136945 | 0.97775035 |
| COX11 | 56.1017272 | -0.079581509 | 0.25311095 | -0.3144135 | 0.75320698 | 0.97096495 |
| STXBP4 | 210.233607 | 0.492935089 | 0.25292281 | 1.94895467 | 0.05130084 | 0.52609271 |
| HLF | 81.3111192 | 0.261111116 | 0.6507741 | 0.40123157 | 0.68824964 | 0.96161098 |

| | | | | | | |
|---|---|---|---|---|---|---|
| DGKE | 29.8978572 | 0.170766581 | 0.28277355 | 0.60389871 | 0.54591098 | 0.93523904 |
| TRIM25 | 48.5053363 | -0.852100406 | 0.38694389 | -2.2021291 | 0.02765619 | 0.4250936 |
| COIL | 39.9021147 | -0.009348866 | 0.28260647 | -0.0330809 | 0.9736101 | 0.99681784 |
| SCPEP1 | 26.3828633 | 0.515232217 | 0.48417054 | 1.06415441 | 0.28725876 | 0.83131469 |
| AKAP1 | 48.2712621 | 0.093957542 | 0.28352084 | 0.33139554 | 0.74034574 | 0.9698508 |
| MSI2 | 1252.50528 | 0.12950357 | 0.23319981 | 0.55533309 | 0.57866685 | 0.94034284 |
| MRPS23 | 43.5275374 | -0.165467439 | 0.33847202 | -0.4888659 | 0.62493662 | 0.94808496 |
| CUEDC1 | 152.621336 | -0.317817352 | 0.27431116 | -1.1586016 | 0.24661862 | 0.80571615 |
| VEZF1 | 108.599101 | 0.140420025 | 0.21221151 | 0.66169846 | 0.50816449 | 0.92666294 |
| AC015813.2 | 51.010101 | -0.239863598 | 0.26894137 | -0.8918806 | 0.37245691 | 0.87605028 |
| SRSF1 | 25.9925256 | -0.069151198 | 0.39017209 | -0.1772326 | 0.85932572 | 0.9864199 |
| DYNLL2 | 76.6125475 | -0.330386219 | 0.23881091 | -1.3834637 | 0.16652274 | 0.74445571 |
| MKS1 | 27.319259 | 0.225029639 | 0.32039227 | 0.70235663 | 0.48245678 | 0.91971919 |
| TSPOAP1 | 63.5038919 | 0.860399991 | 0.39974204 | 2.15238807 | 0.0313668 | 0.43905874 |
| SUPT4H1 | 36.0514191 | -0.169840883 | 0.36518012 | -0.465088 | 0.64186845 | 0.94944486 |
| RNF43 | 89.4351484 | -0.312772369 | 0.55620711 | -0.5623308 | 0.57389068 | 0.94034284 |
| MTMR4 | 37.5018791 | 0.199150781 | 0.32111458 | 0.62018604 | 0.53513531 | 0.93460153 |
| RAD51C | 31.2925016 | 0.110221916 | 0.29753738 | 0.37044729 | 0.71104924 | 0.96587538 |
| PPM1E | 59.1071323 | 1.009250525 | 0.52170003 | 1.93454182 | 0.05304655 | 0.53381345 |
| TRIM37 | 108.302126 | -0.496485637 | 0.19881606 | -2.497211 | 0.01251745 | 0.31109683 |
| SKA2 | 65.9935514 | -0.448810071 | 0.31193835 | -1.4387781 | 0.1502134 | 0.72246729 |
| YPEL2 | 369.023233 | 0.624755561 | 0.33284098 | 1.87703916 | 0.06051273 | 0.56071154 |
| DHX40 | 90.1850714 | 0.519990753 | 0.25254457 | 2.05900591 | 0.03949367 | 0.48008869 |
| CLTC | 409.189573 | -0.023061306 | 0.24926085 | -0.0925188 | 0.92628588 | 0.99316012 |
| PTRH2 | 68.4917332 | -0.114521105 | 0.3215395 | -0.356165 | 0.72171703 | 0.96779196 |
| VMP1 | 1072.30922 | -0.467560377 | 0.45586727 | -1.0256502 | 0.30505648 | 0.84488735 |
| TUBD1 | 61.6281312 | 0.498298456 | 0.39517374 | 1.26096045 | 0.2073231 | 0.77401635 |
| RPS6KB1 | 97.229731 | -0.06915577 | 0.25272031 | -0.2736455 | 0.78435709 | 0.97423208 |
| HEATR6 | 30.4538672 | -0.140440785 | 0.29933375 | -0.4691779 | 0.63894247 | 0.94944486 |
| USP32 | 380.293262 | -0.023940361 | 0.25805957 | -0.0927707 | 0.92608575 | 0.99316012 |
| APPBP2 | 259.074481 | 0.302598796 | 0.17431934 | 1.73588771 | 0.08258369 | 0.6141478 |
| PPM1D | 89.3081402 | 0.463494436 | 0.26142649 | 1.77294362 | 0.07623805 | 0.59726395 |
| BCAS3 | 820.004882 | 0.066430151 | 0.23029217 | 0.28846031 | 0.77299441 | 0.97350952 |
| NACA2 | 36.7670312 | 0.46958912 | 0.7355836 | 0.63838988 | 0.52321992 | 0.93272384 |
| INTS2 | 53.3889638 | -0.353568227 | 0.2740106 | -1.2903451 | 0.19693087 | 0.77110248 |
| MED13 | 503.701585 | -0.011047433 | 0.15311489 | -0.0721513 | 0.94248153 | 0.99526299 |
| METTL2A | 26.2304809 | -0.57373579 | 0.36658685 | -1.5650746 | 0.11756541 | 0.68045863 |
| TLK2 | 301.485344 | -0.037397847 | 0.21675841 | -0.1725324 | 0.863019 | 0.98670067 |
| MRC2 | 161.449728 | -0.797123842 | 0.66746644 | -1.1942531 | 0.23237899 | 0.79498513 |
| TANC2 | 475.35954 | -0.276455694 | 0.42570078 | -0.6494132 | 0.51607136 | 0.93167123 |
| AC015923.1 | 34.6722384 | 0.065601956 | 0.60608636 | 0.10823863 | 0.9138064 | 0.99154413 |
| CYB561 | 44.7112665 | 0.033070868 | 0.36869734 | 0.08969652 | 0.92852838 | 0.99336859 |
| DCAF7 | 207.514494 | 0.035767092 | 0.22262353 | 0.16066177 | 0.8723598 | 0.98766275 |
| MAP3K3 | 110.767343 | -0.095944163 | 0.28723817 | -0.334023 | 0.73836218 | 0.9698508 |
| CCDC47 | 52.9602497 | -0.259349773 | 0.37156335 | -0.6979961 | 0.48517962 | 0.92034617 |
| DDX42 | 146.890788 | -0.316904658 | 0.19173759 | -1.652804 | 0.09837075 | 0.64312419 |
| FTSJ3 | 29.5482679 | 0.258255911 | 0.38139208 | 0.6771402 | 0.49831701 | 0.92398825 |
| PSMC5 | 39.2953998 | -0.39039464 | 0.44165659 | -0.8839326 | 0.37673262 | 0.87842274 |
| SMARCD2 | 44.4734996 | 0.259751559 | 0.24145001 | 1.07579852 | 0.28201738 | 0.82764842 |
| ERN1 | 131.105323 | 0.117431655 | 0.65406724 | 0.17954065 | 0.85751321 | 0.9864199 |
| SNHG25 | 41.0655164 | -0.143191271 | 0.55957807 | -0.2558915 | 0.79803462 | 0.97584218 |
| TEX2 | 71.8403135 | 0.120125146 | 0.29260865 | 0.41053177 | 0.6814159 | 0.96060087 |
| POLG2 | 50.2606606 | -0.542808429 | 0.31449631 | -1.7259612 | 0.08435439 | 0.61680068 |
| DDX5 | 468.619596 | -0.066964795 | 0.23784681 | -0.2815459 | 0.77829172 | 0.97423208 |

| | | | | | | |
|---|---|---|---|---|---|---|
| CEP95 | 199.337693 | -0.24488499 | 0.30710624 | -0.797395 | 0.42522165 | 0.90167091 |
| SMURF2 | 295.958996 | -0.014092111 | 0.18267052 | -0.077145 | 0.93850822 | 0.99526299 |
| LRRC37A3 | 227.648496 | 0.412726928 | 0.31209556 | 1.32243766 | 0.18602245 | 0.76083106 |
| GNA13 | 143.532967 | -0.075572467 | 0.21559512 | -0.3505296 | 0.72594129 | 0.96781837 |
| RGS9 | 45.2912862 | 0.313878234 | 0.63954268 | 0.49078544 | 0.62357821 | 0.94808496 |
| CEP112 | 519.933436 | 0.778594052 | 0.30982587 | 2.51300529 | 0.01197075 | 0.30967852 |
| PRKCA | 517.313623 | 0.842739235 | 0.49655905 | 1.69715813 | 0.08966677 | 0.6330877 |
| HELZ | 461.448555 | -0.072214196 | 0.20407305 | -0.3538644 | 0.72344048 | 0.96779196 |
| PSMD12 | 92.958473 | -0.470990387 | 0.19579974 | -2.4054699 | 0.01615168 | 0.34229417 |
| PITPNC1 | 838.565622 | -0.613912442 | 0.79736489 | -0.7699266 | 0.44134343 | 0.90561284 |
| NOL11 | 54.7559575 | -0.42885811 | 0.32300239 | -1.3277243 | 0.18426922 | 0.75788529 |
| BPTF | 581.118462 | -0.134504814 | 0.21383064 | -0.629025 | 0.52933271 | 0.93366189 |
| ARSG | 62.4429704 | -0.502101457 | 0.45987832 | -1.0918137 | 0.27491498 | 0.82164863 |
| WIPI1 | 78.5018295 | 0.446465397 | 0.3967169 | 1.1254005 | 0.26041936 | 0.81263292 |
| PRKAR1A | 227.491236 | -0.03731292 | 0.2527903 | -0.1476042 | 0.88265511 | 0.98766275 |
| ABCA10 | 34.6426772 | 0.681261974 | 0.40271764 | 1.69166161 | 0.09071051 | 0.6330877 |
| ABCA5 | 141.642661 | 0.450488065 | 0.28155046 | 1.60002606 | 0.1095928 | 0.65776282 |
| AC005208.1 | 188.660775 | 0.614593111 | 0.48741852 | 1.26091457 | 0.20733963 | 0.77401635 |
| KCNJ16 | 684.186324 | -0.037122164 | 0.33339859 | -0.1113447 | 0.91134301 | 0.99100813 |
| SOX9 | 88.0347336 | 1.158156078 | 0.68324841 | 1.69507321 | 0.09006153 | 0.6330877 |
| LINC00511 | 285.116372 | -1.122328591 | 1.00290011 | -1.1190831 | 0.26310468 | 0.81378765 |
| SLC39A11 | 198.086638 | -0.16451758 | 0.25195293 | -0.6529695 | 0.51377594 | 0.93064774 |
| COG1 | 28.0092673 | 0.030701944 | 0.30156823 | 0.10180762 | 0.91890938 | 0.99214128 |
| FAM104A | 45.7625916 | -0.141565688 | 0.25310795 | -0.5593095 | 0.57595049 | 0.94034284 |
| C17orf80 | 28.345405 | -0.395419085 | 0.28166129 | -1.4038816 | 0.16035412 | 0.73816762 |
| CDC42EP4 | 55.0784192 | -0.404178334 | 0.42926335 | -0.9415626 | 0.34641661 | 0.86990996 |
| RPL38 | 159.788308 | 0.134201928 | 0.40422917 | 0.33199467 | 0.73989329 | 0.9698508 |
| GPRC5C | 201.52955 | 0.167844383 | 0.25731584 | 0.65228936 | 5.14E-01 | 0.93091506 |
| SLC9A3R1 | 49.6896734 | 1.174640139 | 0.65094431 | 1.8045171 | 0.07115028 | 0.59054593 |
| NAT9 | 29.7470669 | -0.358732326 | 0.28696765 | -1.2500793 | 0.21127058 | 0.77730294 |
| KCTD2 | 39.5196942 | 0.001857143 | 0.29850307 | 0.00622152 | 0.99503598 | 0.99866813 |
| ATP5PD | 63.7035007 | 0.049226376 | 0.42050412 | 0.11706515 | 0.90680843 | 0.99100813 |
| SLC16A5 | 204.05057 | -1.210763527 | 0.84090136 | -1.4398401 | 0.14991263 | 0.72246729 |
| ARMC7 | 72.1991007 | -0.370752705 | 0.62639354 | -0.5918846 | 0.55392785 | 0.93523904 |
| NT5C | 77.2656217 | -0.417425832 | 0.3678204 | -1.1348632 | 0.25643265 | 0.80988075 |
| JPT1 | 80.9005806 | -0.814396377 | 0.63760811 | -1.2772679 | 0.20150768 | 0.77110248 |
| SUMO2 | 61.4389579 | -0.14725277 | 0.29873643 | -0.4929187 | 0.62207004 | 0.94808496 |
| NUP85 | 50.1657891 | -0.191938746 | 0.31408454 | -0.6111054 | 0.54112983 | 0.93523904 |
| GGA3 | 59.0123666 | -0.131205923 | 0.22867012 | -0.5737782 | 0.56611792 | 0.93832962 |
| MRPS7 | 45.5777793 | -0.1267158 | 0.3488995 | -0.3631871 | 0.71646512 | 0.96742001 |
| GRB2 | 180.316865 | -0.134908775 | 0.17098574 | -0.789006 | 0.43010851 | 0.90364119 |
| AC011933.4 | 30.5003729 | 0.992268456 | 0.51217588 | 1.93735883 | 0.0527015 | 0.53195745 |
| TMEM94 | 52.1880228 | -0.105738966 | 0.30304069 | -0.3489266 | 0.72714439 | 0.9681562 |
| CASKIN2 | 26.9821906 | -0.218435813 | 0.3793672 | -0.5757899 | 0.56475718 | 0.93809297 |
| TSEN54 | 36.6680952 | -0.120076036 | 0.25889498 | -0.4638021 | 0.64278954 | 0.94944486 |
| LLGL2 | 301.90857 | 0.087832889 | 0.28565357 | 0.30748045 | 0.7584777 | 0.9714717 |
| MYO15B | 92.8725754 | 0.085129225 | 0.31691908 | 0.26861502 | 0.78822595 | 0.97423208 |
| RECQL5 | 48.4678619 | -0.682184149 | 0.36382993 | -1.8750083 | 0.06079157 | 0.56071154 |
| SAP30BP | 71.7855944 | -0.640099084 | 0.27027655 | -2.3683116 | 0.01786948 | 0.35777027 |
| H3F3B | 714.631385 | -0.13780773 | 0.48291426 | -0.2853669 | 0.77536309 | 0.97423208 |
| UNK | 156.813636 | -0.127988902 | 0.30979754 | -0.4131372 | 0.67950607 | 0.95936177 |
| WBP2 | 100.138986 | -0.355906348 | 0.21721674 | -1.6384849 | 1.01E-01 | 0.64876329 |
| TRIM47 | 35.3671113 | -0.052849427 | 0.50003694 | -0.105691 | 9.16E-01 | 9.92E-01 |
| TRIM65 | 24.6239062 | -0.577026871 | 0.33308828 | -1.7323542 | 0.08321052 | 0.61656297 |

| | | | | | | |
|---|---|---|---|---|---|---|
| ACOX1 | 91.4417945 | 0.284862397 | 0.20274684 | 1.40501524 | 0.16001675 | 0.73816762 |
| EVPL | 44.2042056 | -0.491351311 | 0.44900789 | -1.0943044 | 0.2738215 | 0.82047446 |
| SRP68 | 57.8485214 | -0.36480395 | 0.22580065 | -1.6156018 | 0.10618043 | 0.65699665 |
| EXOC7 | 115.480047 | -0.195732337 | 0.26799823 | -0.7303494 | 0.46517665 | 0.91462428 |
| FOXJ1 | 30.4065855 | 0.493339218 | 0.53823602 | 0.91658529 | 0.35936 | 0.87316144 |
| RNF157 | 88.8415781 | 0.888044667 | 0.47408595 | 1.87317228 | 0.0610446 | 0.56071154 |
| UBALD2 | 66.6836426 | -0.008918488 | 0.28710032 | -0.031064 | 0.97521849 | 0.99681784 |
| PRPSAP1 | 76.8358837 | 0.005335267 | 0.21484589 | 0.02483299 | 0.98018817 | 0.99681784 |
| UBE2O | 85.6580324 | -0.412469472 | 0.31967892 | -1.2902617 | 0.19695981 | 0.77110248 |
| RHBDF2 | 46.3359074 | 0.003094192 | 0.44210538 | 0.00699877 | 0.99441584 | 0.99866813 |
| MXRA7 | 97.8212426 | -0.009919536 | 0.52107057 | -0.0190368 | 0.98481172 | 0.99752755 |
| JMJD6 | 28.730883 | -0.220586331 | 0.42462685 | -0.5194828 | 0.60342413 | 0.94683415 |
| SRSF2 | 146.174583 | -0.167035924 | 0.32123394 | -0.5199822 | 0.60307599 | 0.94683415 |
| MFSD11 | 79.7204168 | 0.000156775 | 0.27996851 | 0.00055997 | 0.99955321 | 0.99971834 |
| SEC14L1 | 163.281327 | -0.084502309 | 0.21890842 | -0.3860167 | 0.69948429 | 0.96280283 |
| 9-Sep | 333.938762 | 0.002446057 | 0.43251061 | 0.00565548 | 0.9954876 | 0.99866813 |
| TNRC6C | 146.229815 | -0.174292242 | 0.24315451 | -0.7167963 | 0.47349981 | 0.91804805 |
| TMC6 | 35.4856543 | -0.262390686 | 0.53478404 | -0.4906479 | 0.62367547 | 0.94808496 |
| SYNGR2 | 83.2502866 | -0.640208758 | 0.45538887 | -1.4058507 | 0.15976846 | 0.73816762 |
| AFMID | 88.8720804 | -0.268991187 | 0.45388113 | -0.5926468 | 0.55341757 | 0.93523904 |
| SOCS3 | 86.9753637 | -0.157537263 | 0.5920208 | -0.2661009 | 7.90E-01 | 0.97423208 |
| PGS1 | 76.9549756 | -0.561109816 | 0.29713438 | -1.8884042 | 0.05897171 | 0.55686386 |
| DNAH17 | 28.009301 | -1.145969807 | 0.92630748 | -1.2371376 | 0.21603601 | 0.78293142 |
| CYTH1 | 168.295018 | -0.441900673 | 0.34590802 | -1.2775092 | 0.20142254 | 0.77110248 |
| USP36 | 109.991605 | -0.630139079 | 0.32831339 | -1.9193219 | 0.05494361 | 0.53804407 |
| TIMP2 | 82.9785676 | 0.326369537 | 0.3712968 | 0.87899906 | 0.37940178 | 0.88018641 |
| LGALS3BP | 113.140053 | -1.036795755 | 0.75000614 | -1.382383 | 0.16685413 | 0.74445571 |
| CANT1 | 28.7683613 | -0.168655915 | 0.31628402 | -0.533242 | 0.59386609 | 0.94649617 |
| TBC1D16 | 400.292769 | -0.141427289 | 0.28162668 | -0.50218 | 0.6155409 | 0.94791646 |
| CCDC40 | 146.966224 | 0.17591972 | 0.35753507 | 0.49203487 | 0.62269469 | 0.94808496 |
| GAA | 72.7418598 | 0.202498361 | 0.41723314 | 0.48533624 | 0.62743785 | 0.94858384 |
| EIF4A3 | 53.9382364 | -0.137080449 | 0.44617513 | -0.3072346 | 0.75866479 | 0.9714717 |
| AC087741.1 | 29.7135532 | 0.728296326 | 0.44970932 | 1.6194824 | 0.10534351 | 0.65699665 |
| SLC26A11 | 65.7281142 | 0.445262311 | 0.33898313 | 1.3135235 | 0.18900661 | 0.76207437 |
| RNF213 | 680.016433 | -1.061623785 | 0.55839791 | -1.9011959 | 0.05727636 | 0.54616991 |
| AC124319.2 | 222.356535 | 0.579025053 | 0.37164638 | 1.55799998 | 0.11923325 | 0.68597919 |
| ENDOV | 37.5175066 | -0.216261151 | 0.31665555 | -0.6829539 | 0.49463596 | 0.92289011 |
| RPTOR | 217.132269 | -0.320356117 | 0.25357289 | -1.263369 | 0.20645661 | 0.77401635 |
| BAIAP2-DT | 24.0988791 | 0.434498231 | 0.4608549 | 0.94280919 | 0.34577851 | 0.86990996 |
| BAIAP2 | 82.512507 | 0.024328153 | 0.37783461 | 0.06438836 | 0.94866099 | 0.9952642 |
| NDUFAF8 | 47.2526934 | -0.19694559 | 0.31195813 | -0.6313206 | 0.52783093 | 0.93366189 |
| SLC38A10 | 111.472368 | 0.016375174 | 0.32233138 | 0.0508023 | 0.95948306 | 0.99572129 |
| BAHCC1 | 39.468038 | 0.233496455 | 0.43270851 | 0.53961604 | 0.58946185 | 0.94491859 |
| ACTG1 | 1033.51381 | -0.312246661 | 0.632916 | -0.4933461 | 0.62176804 | 0.94808496 |
| FAAP100 | 25.3568304 | -0.4707246 | 0.40938524 | -1.1498329 | 0.25021271 | 0.80672756 |
| NPLOC4 | 190.127706 | -0.001885597 | 0.24736099 | -0.0076229 | 0.9939179 | 0.99866813 |
| OXLD1 | 26.8094393 | -0.377939727 | 0.4159096 | -0.9087064 | 0.36350511 | 0.87605028 |
| CCDC137 | 33.0405103 | -0.45112707 | 0.49983909 | -0.9025446 | 0.36676764 | 0.87605028 |
| ARL16 | 39.7890821 | 0.179640032 | 0.35810653 | 0.50163853 | 0.61592181 | 0.94791646 |
| HGS | 67.2480718 | -0.431973912 | 0.30380218 | -1.4218921 | 0.15505758 | 0.72978231 |
| MCRIP1 | 62.9109721 | -0.174082319 | 0.53018187 | -0.3283445 | 0.74265117 | 0.9698508 |
| P4HB | 219.647268 | 0.084907725 | 0.39234119 | 0.21641298 | 0.82866585 | 0.98142144 |
| ARHGDIA | 136.078418 | -0.647006256 | 0.48079154 | -1.3457106 | 0.17839588 | 0.7513166 |
| ANAPC11 | 76.9467799 | -0.088481882 | 0.47213725 | -0.1874071 | 0.85134144 | 0.98621353 |

| | | | | | | |
|---|---|---|---|---|---|---|
| SIRT7 | 26.7224166 | -0.125865995 | 0.31371228 | -0.4012147 | 0.68826202 | 0.96161098 |
| ASPSCR1 | 40.8619819 | -0.298102305 | 0.45866627 | -0.6499329 | 0.51573556 | 0.93140514 |
| CENPX | 29.7406135 | -0.547532153 | 0.56451519 | -0.9699157 | 0.33208851 | 0.85980141 |
| DCXR | 42.102376 | -0.80278723 | 0.31855073 | -2.5201236 | 0.01173136 | 0.30872898 |
| RFNG | 38.3645994 | 0.167740935 | 0.29496712 | 0.56867673 | 0.56957554 | 0.93998433 |
| GPS1 | 38.5515308 | -0.464940705 | 0.49640459 | -0.9366165 | 0.34895588 | 0.87178925 |
| DUS1L | 47.1775919 | -0.644987012 | 0.39089956 | -1.650007 | 0.0989415 | 0.6434623 |
| CCDC57 | 219.503093 | -0.019941728 | 0.26733655 | -0.0745941 | 0.94053767 | 0.99526299 |
| SLC16A3 | 44.8818936 | -0.090551852 | 0.46776502 | -0.1935841 | 0.84650158 | 0.98525378 |
| CSNK1D | 148.288902 | -0.135909736 | 0.17540786 | -0.7748212 | 0.4384453 | 0.90561284 |
| OGFOD3 | 43.9296679 | -0.47559462 | 0.28123562 | -1.6910896 | 0.0908197 | 0.63316002 |
| HEXDC | 77.946831 | -0.429528092 | 0.28880595 | -1.487255 | 0.13694748 | 0.71041106 |
| CYBC1 | 40.8734242 | 0.382946764 | 0.41835384 | 0.91536572 | 0.35999967 | 0.87367102 |
| NARF | 58.9585476 | 0.164925825 | 0.24681866 | 0.66820646 | 0.50400181 | 0.92552859 |
| FOXK2 | 202.351018 | -0.290255314 | 0.21724778 | -1.3360565 | 0.1815308 | 0.75448394 |
| WDR45B | 147.967181 | -0.31180309 | 0.34931411 | -0.8926152 | 0.37206327 | 0.87605028 |
| RAB40B | 34.9842926 | 0.445053583 | 0.30119142 | 1.47764361 | 0.13950319 | 0.71191199 |
| FN3KRP | 27.3785321 | -0.144098023 | 0.44694266 | -0.3224083 | 0.7471434 | 0.97028355 |
| FN3K | 27.714537 | 0.26242404 | 0.73523993 | 0.356923 | 0.72114945 | 0.96779196 |
| TBCD | 104.245573 | -0.26592002 | 0.3078931 | -0.8636765 | 0.38776565 | 0.8853359 |
| METRNL | 46.3366763 | 0.443047794 | 0.39780559 | 1.11372942 | 0.26539528 | 0.81378765 |
| USP14 | 161.444546 | -0.227996771 | 0.2341072 | -0.973899 | 0.33010668 | 0.85880205 |
| THOC1 | 145.277694 | -0.579464527 | 0.23209378 | -2.4966827 | 0.01253611 | 0.31109683 |
| ENOSF1 | 466.776518 | -0.214388633 | 0.36065472 | -0.5944429 | 0.55221593 | 0.93523904 |
| YES1 | 194.151076 | -0.018129328 | 0.22074222 | -0.082129 | 0.93454417 | 0.99444552 |
| METTL4 | 44.3576211 | 0.160695521 | 0.29833944 | 0.53863318 | 0.59013999 | 0.94516215 |
| SMCHD1 | 393.602423 | -0.213893802 | 0.25893204 | -0.8260615 | 0.40876918 | 0.8959746 |
| AP001011.1 | 62.4270193 | 0.74378887 | 0.48369776 | 1.5377141 | 0.12411854 | 0.69439514 |
| LPIN2 | 248.510238 | -0.284608742 | 0.46844239 | -0.607564 | 0.54347667 | 0.93523904 |
| MYOM1 | 41.2750585 | -0.375979619 | 0.56654685 | -0.6636338 | 0.50692474 | 0.92666294 |
| MYL12A | 232.319371 | -0.081178317 | 0.45575845 | -0.178117 | 0.85863111 | 0.9864199 |
| MYL12B | 447.151909 | -0.259570589 | 0.497071 | -0.5222002 | 0.60153093 | 0.94683415 |
| TGIF1 | 108.143994 | -0.817569028 | 0.45509938 | -1.7964626 | 0.07242097 | 0.59090333 |
| DLGAP1 | 254.072589 | 0.796756614 | 0.79673414 | 1.00002821 | 0.31729686 | 0.8490543 |
| LINC00667 | 47.4364959 | -0.249250498 | 0.29107536 | -0.8563092 | 0.39182679 | 0.88709734 |
| EPB41L3 | 127.037464 | 0.446616844 | 0.3912607 | 1.14148148 | 0.25366961 | 0.80861938 |
| L3MBTL4 | 676.278974 | 0.327320654 | 0.28629436 | 1.1433011 | 0.2529136 | 0.80798674 |
| AP001021.2 | 40.8930378 | 0.847426647 | 0.36870172 | 2.29840711 | 0.02153863 | 0.38588526 |
| ARHGAP28 | 62.0517482 | 0.631543375 | 0.63653789 | 0.99215362 | 0.32112259 | 0.85131972 |
| LAMA1 | 216.983432 | -1.371807966 | 0.67032212 | -2.0464907 | 0.04070812 | 0.48617886 |
| PTPRM | 1577.22921 | 0.139003074 | 0.25313234 | 0.54913203 | 0.58291485 | 0.94121612 |
| RAB12 | 116.115174 | -0.315359786 | 0.20837728 | -1.5134077 | 0.13017613 | 0.69904834 |
| MTCL1 | 87.760986 | -0.631705202 | 0.37462178 | -1.6862479 | 0.09174805 | 0.63464873 |
| NDUFV2 | 128.152713 | -0.304605907 | 0.26902842 | -1.1322443 | 0.25753172 | 0.8120529 |
| ANKRD12 | 686.408598 | 0.04215899 | 0.21084849 | 0.19994921 | 0.8415203 | 0.98515707 |
| TWSG1 | 86.728143 | -0.115667447 | 0.25669064 | -0.4506103 | 0.65227044 | 0.95159942 |
| RALBP1 | 126.584224 | 0.032467907 | 0.24764936 | 0.13110435 | 0.89569276 | 0.98842187 |
| PPP4R1 | 177.899746 | -0.291616746 | 0.28698541 | -1.0161379 | 0.30956373 | 0.84554063 |
| RAB31 | 53.5221218 | -0.29312986 | 0.44649881 | -0.6565076 | 0.51149759 | 0.92831288 |
| VAPA | 315.937619 | -0.104816139 | 0.2292336 | -0.457246 | 0.64749426 | 0.95027667 |
| NAPG | 92.056208 | -0.001079198 | 0.25909181 | -0.0041653 | 0.99667657 | 0.99898334 |
| MPPE1 | 91.2146895 | -0.03772454 | 0.31059586 | -0.1214586 | 0.9033278 | 0.99038308 |
| IMPA2 | 183.685679 | 1.075467717 | 0.3712699 | 2.89672744 | 0.00377077 | 0.19908785 |
| TUBB6 | 90.4367814 | -1.121653597 | 0.36633731 | -3.0618055 | 0.00220006 | 0.15271309 |

| | | | | | | |
|---|---|---|---|---|---|---|
| AFG3L2 | 130.25102 | 0.02223956 | 0.22567638 | 0.09854625 | 0.92149855 | 0.99214128 |
| SPIRE1 | 168.50083 | 0.095682478 | 0.29321178 | 0.3263255 | 0.7441781 | 0.97012548 |
| PSMG2 | 80.9836733 | -0.021464502 | 0.20260051 | -0.105945 | 0.91562603 | 0.99159829 |
| PTPN2 | 236.126923 | -0.303173748 | 0.31945298 | -0.9490403 | 0.34260012 | 0.86846099 |
| SEH1L | 76.8950967 | -0.318180523 | 0.29780901 | -1.0684046 | 0.28533803 | 0.82951531 |
| CEP192 | 176.985298 | -0.279260818 | 0.26506145 | -1.0535701 | 0.29207979 | 0.83697123 |
| LDLRAD4 | 272.576156 | 0.785987587 | 0.76593792 | 1.02617662 | 0.30480834 | 0.84488735 |
| FAM210A | 70.494888 | -0.035579289 | 0.40526568 | -0.0877925 | 0.93004159 | 0.99362798 |
| RNMT | 162.293912 | -0.17163332 | 0.34038466 | -0.5042334 | 0.61409737 | 0.94791646 |
| ZNF519 | 29.3518058 | -0.595414479 | 0.41539364 | -1.4333741 | 0.15175096 | 0.72499872 |
| ROCK1 | 398.965874 | -0.026639724 | 0.16381198 | -0.1626238 | 0.87081466 | 0.98766275 |
| GREB1L | 156.47491 | 0.268051964 | 0.43547789 | 0.61553519 | 0.53820134 | 0.93516669 |
| ESCO1 | 131.251262 | -0.141055103 | 0.29870471 | -0.4722226 | 0.63676793 | 0.94944486 |
| SNRPD1 | 56.7062376 | -0.718287023 | 0.36645162 | -1.9601142 | 0.04998244 | 0.52415717 |
| ABHD3 | 70.4640186 | -0.537851752 | 0.36069609 | -1.4911494 | 0.13592228 | 0.70722407 |
| MIB1 | 311.577226 | -0.205744221 | 0.23166261 | -0.8881201 | 0.37447615 | 0.87678297 |
| AC090912.1 | 40.2926541 | 0.307358471 | 0.60203245 | 0.51053473 | 0.60967689 | 0.94791646 |
| RBBP8 | 336.364297 | -0.22410852 | 0.37292456 | -0.6009487 | 0.54787417 | 0.93523904 |
| CABLES1 | 235.917006 | -0.572493442 | 0.35577242 | -1.6091563 | 0.10758216 | 0.65699665 |
| TMEM241 | 155.649355 | 0.014133396 | 0.28002593 | 0.05047174 | 0.95974647 | 0.99572129 |
| RIOK3 | 216.838974 | 0.15030931 | 0.31486719 | 0.47737368 | 0.63309605 | 0.94932781 |
| RMC1 | 61.0745136 | -0.190239899 | 0.25972527 | -0.7324659 | 0.46388426 | 0.91462428 |
| NPC1 | 195.869928 | -0.935024544 | 0.38780607 | -2.4110622 | 0.01590614 | 0.34086104 |
| LAMA3 | 76.1889793 | -1.104963218 | 0.7759391 | -1.4240334 | 0.15443679 | 0.72978231 |
| TTC39C | 150.016444 | 0.251574401 | 0.41604537 | 0.6046802 | 0.5453915 | 0.93523904 |
| OSBPL1A | 118.307489 | 0.382187378 | 0.39580194 | 0.9656026 | 0.33424308 | 0.86229853 |
| IMPACT | 34.1907299 | -0.22637899 | 0.33588276 | -0.6739822 | 0.50032266 | 0.92398825 |
| SS18 | 227.582358 | -0.018117634 | 0.21007036 | -0.0862456 | 0.93127122 | 0.99411667 |
| TAF4B | 47.5603787 | -0.310994864 | 0.2595113 | -1.1983866 | 0.23076655 | 0.79403724 |
| KCTD1 | 281.504568 | 0.299940463 | 0.36577565 | 0.82001211 | 0.4122092 | 0.89706788 |
| AQP4-AS1 | 83.0248976 | 1.433882509 | 0.45086173 | 3.18031539 | 0.00147115 | 0.12148388 |
| CHST9 | 246.497491 | 0.420961065 | 0.45996452 | 0.91520334 | 0.36008489 | 0.87367102 |
| CDH2 | 751.402593 | 0.608781642 | 0.24326971 | 2.50249664 | 0.01233208 | 0.30967852 |
| DSC2 | 106.50066 | -0.009845534 | 0.25762852 | -0.038216 | 0.96951546 | 0.99681784 |
| DSG2 | 145.827282 | 0.417853402 | 0.33402814 | 1.25095271 | 0.21095173 | 0.77730294 |
| TRAPPC8 | 229.484429 | 0.090334656 | 0.20542892 | 0.43973681 | 0.66012774 | 0.95407427 |
| RNF138 | 102.631126 | -0.362274947 | 0.23550923 | -1.5382622 | 0.12398451 | 0.69439514 |
| GAREM1 | 597.857949 | 0.411930892 | 0.29966767 | 1.37462573 | 0.16924751 | 0.74445571 |
| KLHL14 | 92.5104156 | -0.038821951 | 0.59100206 | -0.0656884 | 0.94762595 | 0.9952642 |
| CCDC178 | 104.016658 | 0.796454643 | 0.64971511 | 1.22585212 | 0.22025433 | 0.78584902 |
| ASXL3 | 127.334008 | 0.702407988 | 0.390139 | 1.80040442 | 0.0717968 | 0.59090333 |
| NOL4 | 81.2829238 | 0.649271206 | 0.65996442 | 0.98379729 | 0.32521519 | 0.855037 |
| DTNA | 201.819597 | 0.727507678 | 1.04085435 | 0.69895243 | 0.48458176 | 0.92034617 |
| MAPRE2 | 66.3630861 | 0.866528519 | 0.35842824 | 2.4175788 | 0.01562415 | 0.34086104 |
| ZNF397 | 81.8965342 | 0.162403586 | 0.20815313 | 0.78021206 | 0.43526606 | 0.90514499 |
| ZSCAN30 | 71.4042109 | 0.244343416 | 0.27018162 | 0.90436728 | 0.36580068 | 0.87605028 |
| ZNF24 | 109.94428 | 0.052754799 | 0.20254183 | 0.26046372 | 0.7945061 | 0.97524554 |
| GALNT1 | 126.323216 | 0.285646121 | 0.41177037 | 0.69370246 | 0.48786882 | 0.92043834 |
| RPRD1A | 194.639046 | -0.155442591 | 0.18730372 | -0.8298959 | 0.40659764 | 0.89585737 |
| SLC39A6 | 64.4772324 | 0.280675468 | 0.34983365 | 0.80231123 | 0.42237295 | 0.90066187 |
| ELP2 | 156.71758 | -0.260855518 | 0.21052108 | -1.2390946 | 0.21531048 | 0.78293142 |
| FHOD3 | 724.034918 | 0.545427722 | 0.3705855 | 1.47179995 | 0.14107491 | 0.7148274 |
| TPGS2 | 93.7650852 | 0.068583469 | 0.26602755 | 0.25780589 | 0.79655672 | 0.97524554 |
| KIAA1328 | 443.12258 | -0.479438717 | 0.26644867 | -1.7993661 | 0.07196079 | 0.59090333 |

| | | | | | | |
|---|---|---|---|---|---|---|
| AC016205.1 | 91.9679919 | -0.046424934 | 0.3821588 | -0.1214807 | 0.90331027 | 0.99038308 |
| PIK3C3 | 264.905064 | -0.078384913 | 0.20982202 | -0.3735781 | 0.70871822 | 0.96587538 |
| LINC00907 | 46.069562 | 0.222426369 | 0.47817887 | 0.46515307 | 0.64182184 | 0.94944486 |
| SETBP1 | 1163.81303 | -0.040036587 | 0.28751849 | -0.1392487 | 0.8892536 | 0.98766275 |
| SLC14A2 | 138.442323 | -0.045697425 | 1.24934978 | -0.036577 | 9.71E-01 | 0.99681784 |
| EPG5 | 197.172354 | 0.117482865 | 0.19926287 | 0.58958735 | 0.55546733 | 0.93523904 |
| PSTPIP2 | 32.6230177 | 0.204205687 | 0.43757637 | 0.4666744 | 0.64073285 | 0.94944486 |
| ATP5F1A | 163.721482 | 0.250548132 | 0.2807548 | 0.89240908 | 0.37217371 | 0.87605028 |
| C18orf25 | 110.505593 | 0.102700521 | 0.27275903 | 0.37652473 | 0.70652684 | 0.96587538 |
| PIAS2 | 224.542895 | 0.168927925 | 0.22025981 | 0.76694847 | 0.44311216 | 0.90651998 |
| KATNAL2 | 66.2664722 | 0.159779582 | 0.54827551 | 0.29142206 | 0.77072854 | 0.97313769 |
| SMAD2 | 445.343808 | -0.041668156 | 0.26699394 | -0.156064 | 0.87598254 | 0.98766275 |
| ZBTB7C | 195.569855 | 0.653725201 | 1.23434354 | 0.52961366 | 0.59637982 | 0.94683415 |
| CTIF | 208.773518 | 0.81571448 | 0.62914995 | 1.29653429 | 0.19479148 | 0.76968045 |
| SMAD7 | 95.1575392 | -1.084367814 | 0.6652822 | -1.6299366 | 0.10311489 | 0.65288882 |
| DYM | 503.672675 | -0.00783909 | 0.21060658 | -0.0372215 | 0.97030841 | 0.99681784 |
| LIPG | 134.722136 | -1.652524692 | 0.7224972 | -2.2872403 | 0.0221818 | 0.39000316 |
| ACAA2 | 116.392838 | 0.702503694 | 0.38848664 | 1.8083085 | 0.0705585 | 0.58797037 |
| SNHG22 | 97.2804144 | 0.315423027 | 0.40995858 | 0.76940219 | 0.44165459 | 0.90561284 |
| MYO5B | 336.093405 | 0.074733644 | 0.35507756 | 0.21047132 | 0.83329983 | 0.98351147 |
| MBD1 | 45.1794734 | 0.148780781 | 0.27541913 | 0.54019771 | 0.58906069 | 0.94491859 |
| CXXC1 | 34.6580471 | 0.197576581 | 0.31009133 | 0.63715609 | 0.52402317 | 0.93272384 |
| ME2 | 73.5589411 | -0.43095037 | 0.37464369 | -1.1502939 | 0.25002283 | 0.80672756 |
| AC091551.1 | 28.8049115 | -0.514272399 | 0.55353976 | -0.9290614 | 0.35285729 | 0.8720098 |
| SMAD4 | 128.819355 | 0.208644168 | 0.22416616 | 0.93075677 | 0.35197939 | 0.8720098 |
| MEX3C | 31.3504417 | 0.060471501 | 0.28995301 | 0.20855621 | 0.8347947 | 0.98367279 |
| MBD2 | 313.566809 | -0.378231273 | 0.23679963 | -1.5972629 | 0.11020711 | 0.65927415 |
| POLI | 110.40109 | 0.253014974 | 0.24710555 | 1.02391456 | 0.30587563 | 0.84488735 |
| TCF4 | 83.6911944 | -0.113809225 | 0.6177388 | -0.1842352 | 0.85382895 | 0.9864199 |
| TXNL1 | 105.412251 | -0.012596173 | 0.23740369 | -0.053058 | 0.95768567 | 0.99572129 |
| WDR7 | 239.250309 | -0.288787201 | 0.16160969 | -1.7869424 | 0.0739468 | 0.59320069 |
| FECH | 29.8283958 | 0.136642018 | 0.32928937 | 0.4149603 | 0.67817097 | 0.95924874 |
| NARS | 112.406645 | -0.24607117 | 0.2757729 | -0.8922964 | 0.37223408 | 0.87605028 |
| AC027097.2 | 87.711266 | -0.396086713 | 0.41044637 | -0.9650145 | 0.33453753 | 0.86282581 |
| ATP8B1 | 181.373241 | -0.16778705 | 0.27697978 | -0.6057736 | 0.54466507 | 0.93523904 |
| NEDD4L | 723.278787 | 0.012031512 | 0.24254422 | 0.04960544 | 9.60E-01 | 9.96E-01 |
| ALPK2 | 206.065775 | 0.742739101 | 0.459273 | 1.61720611 | 0.1058338 | 0.65699665 |
| MALT1 | 305.549245 | -0.166107822 | 0.33534424 | -0.4953352 | 0.6203635 | 0.9480368 |
| AC104365.1 | 47.8051541 | 0.695432372 | 0.53587478 | 1.29775164 | 0.19437269 | 0.7687432 |
| ZNF532 | 221.594755 | -0.631475596 | 0.45442133 | -1.3896258 | 0.16464255 | 0.74427133 |
| SEC11C | 38.6370631 | -0.714320538 | 0.48205435 | -1.4818257 | 0.13838668 | 0.71091391 |
| LMAN1 | 187.456023 | -0.120379126 | 0.29681123 | -0.4055747 | 0.68505512 | 0.96161098 |
| RNF152 | 54.2560012 | 0.668584 | 0.5569233 | 1.20049565 | 0.2299469 | 0.79370986 |
| PIGN | 188.205269 | -0.197306967 | 0.16797002 | -1.1746559 | 0.24013242 | 0.80287207 |
| KIAA1468 | 213.147228 | -0.162650642 | 0.15013674 | -1.0833501 | 0.27865308 | 0.8229426 |
| ZCCHC2 | 125.624224 | -0.177605572 | 0.2992598 | -0.5934829 | 0.55285802 | 0.93523904 |
| PHLPP1 | 251.682294 | 0.250867235 | 0.19714218 | 1.27251933 | 0.20318866 | 0.77155952 |
| BCL2 | 1170.98871 | -0.386667946 | 0.24906126 | -1.5525014 | 1.21E-01 | 6.89E-01 |
| KDSR | 139.984287 | -0.248234613 | 0.28865305 | -0.8599757 | 3.90E-01 | 0.88586416 |
| VPS4B | 124.609756 | 0.188458606 | 0.22937689 | 0.82161112 | 0.41129826 | 0.89649693 |
| TMX3 | 112.244672 | -0.583616816 | 0.27557494 | -2.1178153 | 0.03419071 | 0.45111962 |
| CCDC102B | 51.432135 | 1.1937815 | 0.6502859 | 1.83577947 | 0.06639028 | 0.5780981 |
| DOK6 | 50.6661879 | 1.7589621 | 0.61059298 | 2.88074404 | 0.00396738 | 0.2001215 |
| RTTN | 138.648383 | -0.028405394 | 0.3397434 | -0.0836084 | 0.93336781 | 0.99425828 |

| | | | | | | |
|---|---|---|---|---|---|---|
| SOCS6 | 54.746111 | 0.265654506 | 0.29705308 | 0.89429978 | 0.37116152 | 0.87605028 |
| CYB5A | 310.885098 | 0.950335136 | 0.33764369 | 2.81460948 | 0.00488365 | 0.21962686 |
| CNDP2 | 406.025748 | 0.164874892 | 0.29978345 | 0.54997997 | 0.58233311 | 0.94091128 |
| LINC00909 | 31.2414985 | -0.099420833 | 0.35559721 | -0.2795883 | 0.77979335 | 0.97423208 |
| ZNF407 | 444.286356 | -0.151359546 | 0.16683127 | -0.9072612 | 0.36426866 | 0.87605028 |
| ZADH2 | 46.729468 | -0.120296588 | 0.32174469 | -0.3738883 | 0.70848739 | 0.96587538 |
| TSHZ1 | 89.4661683 | -0.247017392 | 0.27515935 | -0.8977249 | 0.36933225 | 0.87605028 |
| ZNF516 | 115.688687 | 0.859620976 | 0.5972067 | 1.43940276 | 0.15003644 | 0.72246729 |
| ZNF236 | 116.713084 | -0.210042102 | 0.29078299 | -0.7223328 | 0.47008987 | 0.91502267 |
| MBP | 227.637335 | -0.219390405 | 0.24296961 | -0.9029541 | 0.36655025 | 0.87605028 |
| ATP9B | 358.758855 | 0.048683097 | 0.24144752 | 0.20163014 | 0.84020588 | 0.98464453 |
| CTDP1 | 38.2495069 | -0.197775652 | 0.31759716 | -0.6227249 | 0.53346533 | 0.93436907 |
| PQLC1 | 70.0616319 | 0.531558075 | 0.27619311 | 1.92458847 | 0.05428086 | 0.53752502 |
| HSBP1L1 | 69.4502888 | -0.041423536 | 0.278203 | -0.1488968 | 0.88163507 | 0.98766275 |
| TXNL4A | 91.4318311 | -0.268493172 | 0.33366574 | -0.8046771 | 0.42100605 | 0.90012434 |
| ADNP2 | 80.3920809 | -0.01546103 | 0.26817916 | -0.0576519 | 0.95402593 | 0.99531843 |
| NRSN2-AS1 | 66.9121985 | 0.059453198 | 0.39604642 | 0.15011674 | 0.88067251 | 0.98766275 |
| NRSN2 | 32.6557467 | -0.327923351 | 0.48946566 | -0.6699619 | 0.50288207 | 0.92492012 |
| RBCK1 | 99.8668413 | -0.301900431 | 0.28172914 | -1.0715982 | 0.28390055 | 0.82918183 |
| TBC1D20 | 54.4451787 | 0.127804334 | 0.228643 | 0.55896894 | 0.57618292 | 0.94034284 |
| CSNK2A1 | 298.449085 | 0.109519179 | 0.20153708 | 0.5434195 | 0.586841 | 0.94286313 |
| PSMF1 | 151.312002 | -0.520398651 | 0.25231389 | -2.062505 | 0.03915969 | 0.47885941 |
| AL136531.2 | 54.3800806 | -0.520830602 | 0.40633042 | -1.2817908 | 0.19991603 | 0.77110248 |
| SDCBP2-AS1 | 86.4291283 | -0.171028843 | 0.31149567 | -0.5490569 | 0.58296641 | 0.94121612 |
| NSFL1C | 60.1428252 | -0.497482665 | 0.30895137 | -1.6102297 | 0.10734772 | 0.65699665 |
| SIRPA | 108.370579 | -0.266997351 | 0.25866114 | -1.0322283 | 0.30196518 | 0.84375949 |
| STK35 | 47.6839654 | -0.30332201 | 0.28168394 | -1.0768169 | 0.28156209 | 0.82732616 |
| SNRPB | 99.7905607 | -0.812983111 | 0.48020789 | -1.6929816 | 0.09045898 | 0.6330877 |
| NOP56 | 54.436641 | -1.223795766 | 0.4595669 | -2.6629328 | 0.00774629 | 0.27068528 |
| IDH3B | 30.603539 | -0.200468653 | 0.36669089 | -0.5466966 | 0.58458721 | 0.94208628 |
| EBF4 | 67.3796763 | 0.612537177 | 0.31396982 | 1.95094283 | 0.05106385 | 0.52595762 |
| PCED1A | 41.2008494 | 0.310230344 | 0.30173978 | 1.02813869 | 0.30388459 | 0.84488735 |
| PTPRA | 337.696875 | -0.092717542 | 0.18053184 | -0.51358 | 0.60754566 | 0.94708529 |
| MRPS26 | 58.9597342 | 0.003196871 | 0.53929964 | 0.00592782 | 0.99527031 | 0.99866813 |
| UBOX5 | 30.3891488 | -0.101925387 | 0.36123099 | -0.2821612 | 0.77781987 | 0.97423208 |
| LZTS3 | 44.9692954 | 0.859161108 | 0.38246803 | 2.24636056 | 0.02468092 | 0.41230428 |
| DDRGK1 | 61.8351358 | -0.335641358 | 0.35288606 | -0.9511324 | 0.34153718 | 0.86846099 |
| ITPA | 95.0362058 | -0.380496462 | 0.31450829 | -1.2098138 | 0.22635036 | 0.79102885 |
| SLC4A11 | 209.503988 | 1.456388907 | 0.51636365 | 2.82047141 | 4.80E-03 | 0.21962686 |
| C20orf194 | 586.771086 | 0.410448655 | 0.37144628 | 1.10500139 | 0.26915899 | 0.81655182 |
| ATRN | 219.311541 | 0.032903922 | 0.22355686 | 0.14718368 | 0.88298704 | 0.98766275 |
| CENPB | 52.3794765 | -0.573439773 | 0.37553144 | -1.5270087 | 0.12675885 | 0.69494309 |
| CDC25B | 40.4216365 | -0.158402879 | 0.40071173 | -0.3953038 | 0.69261868 | 0.96220115 |
| MAVS | 130.173568 | -0.033400174 | 0.2019573 | -0.1653824 | 0.86864304 | 0.98676155 |
| PANK2 | 143.27432 | -0.369801713 | 0.31124203 | -1.1881484 | 0.23477495 | 0.79805155 |
| RNF24 | 324.446028 | -0.569004788 | 0.62754425 | -0.9067166 | 0.36455668 | 0.87605028 |
| SMOX | 31.5874637 | -1.682192339 | 0.72257821 | -2.3280419 | 1.99E-02 | 0.3747883 |
| PRNP | 98.0907898 | 0.078630853 | 0.3757133 | 0.20928419 | 0.83422639 | 0.98367279 |
| SLC23A2 | 97.7000867 | -0.112393492 | 0.25238858 | -0.4453192 | 0.65608906 | 0.95395828 |
| TMEM230 | 99.3532102 | -0.066504389 | 0.24106482 | -0.2758776 | 0.78264206 | 0.97423208 |
| PCNA | 30.9414441 | -0.393066545 | 0.42051914 | -0.9347174 | 0.34993397 | 0.8720098 |
| CDS2 | 132.932507 | 0.04169643 | 0.1939542 | 0.21498081 | 0.82978228 | 0.98198951 |
| GPCPD1 | 321.538907 | -0.185193889 | 0.46016568 | -0.4024505 | 0.68735254 | 0.96161098 |
| SHLD1 | 91.6255197 | -0.578107301 | 0.30868511 | -1.8728059 | 0.06109519 | 0.56071154 |

| | | | | | | |
|---|---|---|---|---|---|---|
| MCM8 | 41.1655 | -0.933270528 | 0.42257398 | -2.2085376 | 0.02720682 | 0.42391395 |
| CRLS1 | 221.466763 | -0.225752299 | 0.3203316 | -0.7047456 | 0.48096855 | 0.91971919 |
| TMX4 | 174.134899 | 0.009425092 | 0.29057994 | 0.03243545 | 0.97412479 | 0.99681784 |
| PLCB1 | 1161.85936 | -1.558627574 | 0.92926227 | -1.6772741 | 0.09348889 | 0.63721577 |
| PLCB4 | 436.394658 | 0.52293086 | 0.80927446 | 0.64617245 | 0.51816768 | 0.93257822 |
| SNAP25 | 44.6095889 | 1.703371537 | 0.70500216 | 2.41612243 | 0.01568679 | 0.34086104 |
| SLX4IP | 177.155205 | 0.082611739 | 0.2696301 | 0.30638915 | 0.75930837 | 0.9714717 |
| JAG1 | 365.469412 | -1.194798507 | 0.50848671 | -2.3497143 | 0.01878783 | 0.36488708 |
| AL050403.2 | 61.3461463 | -1.009991469 | 0.39633292 | -2.5483411 | 0.01082366 | 0.29965266 |
| BTBD3 | 117.680417 | -0.379362789 | 0.28810855 | -1.3167356 | 0.18792726 | 0.76207437 |
| SPTLC3 | 252.012317 | 1.081338755 | 0.34343833 | 3.1485675 | 0.00164073 | 0.13097113 |
| ISM1 | 102.109971 | 1.247259289 | 1.38026557 | 0.90363718 | 0.36618782 | 0.87605028 |
| TASP1 | 362.796639 | -0.098263199 | 0.17441138 | -0.563399 | 0.57316321 | 0.93998433 |
| ESF1 | 121.55405 | -0.394753802 | 0.26458352 | -1.4919818 | 0.13570391 | 0.70719276 |
| NDUFAF5 | 52.4753931 | -0.044787469 | 0.27441242 | -0.1632123 | 0.87035132 | 0.98766275 |
| MACROD2 | 808.342173 | 0.781139698 | 0.30877014 | 2.52984213 | 0.01141139 | 0.30834879 |
| FLRT3 | 161.9254 | 0.199873285 | 0.50966134 | 0.39216882 | 0.69493348 | 0.96220115 |
| KIF16B | 224.255713 | 0.118100956 | 0.2574234 | 0.45878096 | 0.64639147 | 0.95008515 |
| SNRPB2 | 53.1825665 | -0.095475532 | 0.2925064 | -0.3264049 | 0.74411801 | 0.97012548 |
| DSTN | 571.532639 | 0.202335165 | 0.38570529 | 0.52458488 | 0.5998718 | 0.94683415 |
| RRBP1 | 197.743433 | 0.291108876 | 0.44399477 | 0.65565835 | 5.12E-01 | 0.9287766 |
| SNX5 | 123.193648 | -0.094255786 | 0.24233903 | -0.3889418 | 0.69731918 | 0.96220115 |
| OVOL2 | 54.9964541 | -0.146896591 | 0.45475652 | -0.3230225 | 0.7466782 | 0.97028355 |
| MGME1 | 62.151044 | -0.131359555 | 0.35989872 | -0.3649903 | 0.71511862 | 0.9667635 |
| KAT14 | 25.3995293 | -0.02837636 | 0.37682015 | -0.0753048 | 0.93997221 | 0.99526299 |
| ZNF133 | 80.854393 | 0.213696818 | 0.26135195 | 0.81765917 | 0.41355183 | 0.89732895 |
| DZANK1 | 71.0971211 | -0.228879015 | 0.2947115 | -0.7766206 | 0.43738265 | 0.90561284 |
| POLR3F | 27.1443679 | -0.369243306 | 0.38356579 | -0.9626596 | 0.33571836 | 0.86391445 |
| SEC23B | 89.7237459 | 0.156347812 | 0.31566252 | 0.49530052 | 0.620388 | 0.9480368 |
| DTD1 | 35.1326381 | -0.546060081 | 0.37022697 | -1.4749333 | 0.14023048 | 0.7133658 |
| SLC24A3 | 70.8517576 | 0.520635787 | 1.14288371 | 0.45554573 | 0.64871668 | 0.95027667 |
| AL121761.1 | 96.3864553 | 0.336634744 | 0.36096351 | 0.93260048 | 0.35102629 | 0.8720098 |
| RIN2 | 239.52236 | 0.468581079 | 0.31035438 | 1.50982589 | 0.13108786 | 0.700349 |
| NAA20 | 97.5994832 | -0.04080586 | 0.27090506 | -0.1506279 | 0.88026925 | 0.98766275 |
| CRNKL1 | 30.5359988 | -0.12399615 | 0.38227692 | -0.3243621 | 0.7456639 | 0.97028355 |
| CFAP61 | 44.8133548 | 0.351453127 | 0.66137115 | 0.53140075 | 0.5951411 | 0.94664456 |
| RALGAPA2 | 532.520258 | 0.080526531 | 0.23800913 | 0.3383338 | 0.73511166 | 0.9698508 |
| KIZ | 131.619812 | 0.606773327 | 0.31129499 | 1.94919078 | 0.05127265 | 0.52609271 |
| KIZ-AS1 | 44.903632 | 0.579288589 | 0.50893999 | 1.13822572 | 0.25502624 | 0.80897837 |
| XRN2 | 215.22336 | -0.161475723 | 0.21329052 | -0.7570694 | 0.44900829 | 0.90994378 |
| NAPB | 86.6340681 | 0.272065517 | 0.29603783 | 0.91902281 | 0.35808364 | 0.87316144 |
| CST3 | 311.876216 | 0.399088757 | 0.53061058 | 0.75213117 | 0.45197218 | 0.91077355 |
| APMAP | 55.7203593 | 0.154416118 | 0.39097237 | 0.39495404 | 0.6928768 | 0.96220115 |
| ACSS1 | 119.19664 | 0.932221297 | 0.60539809 | 1.53984844 | 0.1235973 | 0.69398509 |
| ENTPD6 | 63.0059569 | -0.19370182 | 0.26792754 | -0.7229635 | 0.46970234 | 0.91502267 |
| PYGB | 98.380217 | -0.206410896 | 0.32437845 | -0.6363274 | 0.52456305 | 0.93272384 |
| ABHD12 | 166.980349 | -0.317596024 | 0.49476704 | -0.6419102 | 0.52093147 | 0.93272217 |
| NINL | 246.447015 | 0.015503211 | 0.30517328 | 0.05080134 | 0.95948382 | 0.99572129 |
| ZNF337-AS1 | 63.2259858 | -0.302275535 | 0.33580069 | -0.9001635 | 0.36803324 | 0.87605028 |
| ZNF337 | 66.5320326 | 0.503234841 | 0.27626821 | 1.82154451 | 0.06852413 | 0.58250536 |
| FAM182B | 66.612878 | 0.95272304 | 0.37118743 | 2.5666899 | 0.01026744 | 0.29271368 |
| HM13 | 225.751409 | 0.178350582 | 0.32243985 | 0.55312823 | 0.58017561 | 0.94034284 |
| ID1 | 373.339458 | -1.846307184 | 0.76170187 | -2.4239236 | 0.01535383 | 0.33889992 |
| BCL2L1 | 291.420736 | -1.090396789 | 0.34762509 | -3.1367034 | 0.00170859 | 0.13526092 |

| | | | | | | |
|---|---|---|---|---|---|---|
| AL117381.1 | 51.5595966 | -0.431497743 | 0.50583786 | -0.8530357 | 0.39363952 | 0.88760715 |
| TM9SF4 | 120.127512 | -0.230032877 | 0.19619522 | -1.1724693 | 0.24100867 | 0.80319133 |
| POFUT1 | 45.6403375 | -0.27235485 | 0.24087075 | -1.1307095 | 0.25817738 | 0.81223967 |
| KIF3B | 131.060051 | 0.31045723 | 0.20716006 | 1.49863461 | 0.13396845 | 0.70542022 |
| ASXL1 | 409.815863 | 0.023925508 | 0.20965524 | 0.11411834 | 0.90914398 | 0.99100813 |
| NOL4L | 55.727939 | 0.46993129 | 0.45744683 | 1.0272916 | 0.30428318 | 0.84488735 |
| MAPRE1 | 53.9943138 | -0.194561902 | 0.43135907 | -0.451044 | 0.65195787 | 0.95159942 |
| CDK5RAP1 | 58.0947668 | -0.039028167 | 0.23100149 | -0.168952 | 0.86583438 | 0.98670067 |
| SNTA1 | 33.1144297 | 0.691373769 | 0.4302897 | 1.60676346 | 0.10810627 | 0.65707482 |
| CBFA2T2 | 531.724834 | 0.155801378 | 0.2310294 | 0.67437901 | 0.50007038 | 0.92398825 |
| CHMP4B | 107.318533 | -0.215583645 | 0.29421457 | -0.7327429 | 0.46371523 | 0.91462428 |
| RALY | 150.941682 | -0.155758126 | 0.18655776 | -0.8349056 | 0.40377084 | 0.89344441 |
| EIF2S2 | 122.782629 | -0.140901335 | 0.3619643 | -0.3892686 | 0.69707746 | 0.96220115 |
| AHCY | 51.797557 | 0.102931091 | 0.28893904 | 0.35623808 | 7.22E-01 | 0.96779196 |
| ITCH | 415.53547 | 0.14253916 | 0.15776626 | 0.90348318 | 0.36626951 | 0.87605028 |
| DYNLRB1 | 209.121744 | -0.092481403 | 0.38845642 | -0.2380741 | 0.81182364 | 0.97930627 |
| MAP1LC3A | 36.0912788 | 0.871628974 | 0.40651573 | 2.14414573 | 0.03202121 | 0.44049392 |
| PIGU | 95.6359753 | -0.39809222 | 0.19852759 | -2.0052237 | 0.04493914 | 0.50340725 |
| NCOA6 | 250.025992 | -0.036001709 | 0.20360346 | -0.1768227 | 0.85964768 | 0.9864199 |
| GGT7 | 87.7765593 | 0.378250203 | 0.31887573 | 1.18619942 | 0.23554356 | 0.79942255 |
| ACSS2 | 156.210193 | 0.469370596 | 0.33207018 | 1.41346807 | 0.15751814 | 0.73580079 |
| GSS | 56.2497256 | -0.108832527 | 0.27561501 | -0.3948715 | 0.69293769 | 0.96220115 |
| TRPC4AP | 242.243788 | -0.178335094 | 0.19655731 | -0.9072931 | 0.36425182 | 0.87605028 |
| EDEM2 | 42.456767 | -0.062465362 | 0.27088649 | -0.2305961 | 0.81762862 | 0.98010849 |
| PROCR | 34.9154484 | 0.810166691 | 0.46949417 | 1.72561608 | 0.0844165 | 0.61680068 |
| MMP24OS | 89.4354705 | 0.15511885 | 0.29824542 | 0.52010471 | 0.60299059 | 0.94683415 |
| MMP24 | 59.669785 | 0.246013199 | 0.2737894 | 0.89854902 | 0.36889293 | 0.87605028 |
| UQCC1 | 142.314862 | -0.338659642 | 0.17676749 | -1.915848 | 0.05538444 | 0.53915335 |
| CEP250 | 40.6243037 | -1.107825253 | 0.33837732 | -3.2739347 | 0.00106061 | 0.10514713 |
| ERGIC3 | 74.2428433 | 0.11230052 | 0.24312527 | 0.46190395 | 0.6441502 | 0.94944486 |
| CPNE1 | 75.7766151 | -0.409970822 | 0.31456139 | -1.3033094 | 0.19246914 | 0.76613887 |
| ROMO1 | 63.1721979 | -0.721974938 | 0.40750544 | -1.771694 | 0.07644537 | 0.59726395 |
| RBM39 | 565.79247 | -0.004001769 | 0.2270864 | -0.0176222 | 0.98594022 | 0.99764442 |
| PHF20 | 248.21377 | -0.154345497 | 0.14439367 | -1.0689215 | 0.28510504 | 0.82925512 |
| SCAND1 | 103.573926 | -0.232542827 | 0.41299767 | -0.5630609 | 0.57339343 | 0.93998433 |
| NORAD | 281.941338 | -0.031381412 | 0.21104391 | -0.1486961 | 0.88179341 | 0.98766275 |
| EPB41L1 | 466.197444 | 0.262554341 | 0.44787535 | 0.5862219 | 0.5577264 | 0.93612427 |
| AAR2 | 32.9427893 | -0.193303778 | 0.27484134 | -0.7033286 | 0.48185098 | 0.91971919 |
| DLGAP4 | 191.480595 | -0.163280535 | 0.24430278 | -0.6683531 | 0.50390819 | 0.92552859 |
| DLGAP4-AS1 | 83.7375995 | -0.368268183 | 0.49014972 | -0.7513381 | 0.45244918 | 0.91077355 |
| MYL9 | 123.937338 | 0.770569186 | 0.59960014 | 1.28513843 | 0.19874391 | 0.77110248 |
| RAB5IF | 33.9633089 | -0.178828854 | 0.54900691 | -0.3257315 | 0.7446275 | 0.97028355 |
| NDRG3 | 54.1751036 | -0.002867478 | 0.27310603 | -0.0104995 | 0.99162276 | 0.99866813 |
| SOGA1 | 227.015498 | -0.903043516 | 0.64091503 | -1.408991 | 0.15883783 | 0.73686885 |
| SAMHD1 | 49.4058048 | -2.1878878 | 0.86089926 | -2.5413982 | 0.01104101 | 0.30304253 |
| RBL1 | 41.3970165 | -0.973821553 | 0.37105968 | -2.6244338 | 0.00867932 | 0.27620989 |
| RPN2 | 125.833288 | 0.013064804 | 0.23724425 | 0.055069 | 0.95608349 | 0.99572129 |
| MANBAL | 39.644787 | -0.357577681 | 0.28252636 | -1.2656436 | 0.20564071 | 0.77352973 |
| SRC | 54.8877642 | -0.667351529 | 0.43778586 | -1.5243789 | 0.12741408 | 0.69623657 |
| BLCAP | 96.1420536 | -0.207444863 | 0.19447495 | -1.066692 | 0.28611096 | 0.83003998 |
| CTNNBL1 | 82.5968771 | -0.029207342 | 0.20598353 | -0.1417945 | 0.88724229 | 0.98766275 |
| TTI1 | 55.931885 | -0.355846214 | 0.25014245 | -1.4225743 | 0.15485961 | 0.72978231 |
| RPRD1B | 100.106303 | -0.395164299 | 0.2388295 | -1.6545874 | 0.09800821 | 0.64312419 |
| TGM2 | 231.370803 | 0.211376885 | 0.57078844 | 0.3703244 | 7.11E-01 | 9.66E-01 |

| | | | | | | |
|---|---|---|---|---|---|---|
| RALGAPB | 180.101279 | -0.034262026 | 0.17538282 | -0.1953557 | 0.84511453 | 0.98520136 |
| DHX35 | 46.661922 | -0.247071702 | 0.26341894 | -0.9379421 | 0.34827419 | 0.87112607 |
| TOP1 | 299.074951 | 0.070748475 | 0.25188395 | 0.28087727 | 0.77880454 | 0.97423208 |
| PLCG1 | 63.7950676 | -0.374902362 | 0.27158982 | -1.3803992 | 0.16746378 | 0.74445571 |
| ZHX3 | 408.245799 | -0.226618973 | 0.31235327 | -0.7255214 | 0.46813224 | 0.91462428 |
| LPIN3 | 63.1127126 | 0.054827474 | 0.30707681 | 0.17854645 | 0.85829385 | 0.9864199 |
| CHD6 | 384.313625 | -0.023129011 | 0.16248451 | -0.1423459 | 0.88680676 | 0.98766275 |
| SRSF6 | 101.978364 | -0.079774644 | 0.39307317 | -0.2029511 | 0.83917323 | 0.98462033 |
| L3MBTL1 | 52.502769 | 0.51398687 | 0.25958559 | 1.98002848 | 0.04770033 | 0.51394786 |
| SGK2 | 26.8215104 | 0.354496579 | 0.41349986 | 0.85730762 | 0.3912749 | 0.88670817 |
| IFT52 | 80.6974039 | 0.434951547 | 0.22264576 | 1.95355867 | 0.05075343 | 0.52415717 |
| OSER1 | 41.1834105 | -0.345019503 | 0.37285128 | -0.9253542 | 0.35478169 | 0.87258503 |
| SERINC3 | 94.9566124 | -0.105207055 | 0.29060252 | -0.3620308 | 0.71732903 | 0.96742001 |
| PKIG | 136.700768 | 0.539401216 | 0.21546951 | 2.50337604 | 0.01230148 | 0.30967852 |
| YWHAB | 265.524535 | -0.465315 | 0.41766174 | -1.1140954 | 0.26523828 | 0.81378765 |
| PABPC1L | 40.5348733 | 0.117865476 | 0.36025754 | 0.32717004 | 0.74353928 | 0.97012548 |
| STK4 | 107.208118 | -0.401674922 | 0.20243864 | -1.9841811 | 0.04723565 | 0.51284792 |
| SLPI | 1566.61228 | -0.17529608 | 0.67950545 | -0.257976 | 0.79642545 | 0.97524554 |
| SDC4 | 413.046007 | 0.321255547 | 0.45539885 | 0.70543777 | 0.48053785 | 0.91971919 |
| PIGT | 43.5396866 | 0.01561557 | 0.3114297 | 0.05014156 | 0.96000958 | 0.99572129 |
| WFDC2 | 1191.58307 | 1.351589392 | 0.5167854 | 2.61537845 | 8.91E-03 | 2.82E-01 |
| DNTTIP1 | 34.337925 | -0.206680136 | 0.30503552 | -0.6775609 | 0.49805016 | 0.92398825 |
| CTSA | 74.0208003 | 0.733348301 | 0.44217649 | 1.65849681 | 0.09721723 | 0.64312419 |
| ZNF335 | 37.2557435 | -0.078370227 | 0.32599969 | -0.2403997 | 0.81002041 | 0.97930627 |
| NCOA5 | 74.8823364 | -0.328260617 | 0.2145518 | -1.529983 | 0.12602093 | 0.69492095 |
| CD40 | 51.6650644 | 0.154007431 | 0.61935972 | 0.24865587 | 0.80362699 | 0.97880236 |
| SLC35C2 | 54.0628903 | 0.141816013 | 0.29608746 | 0.47896663 | 0.63196237 | 0.94932781 |
| ELMO2 | 41.2911805 | 0.331851847 | 0.28898212 | 1.14834734 | 0.2508252 | 0.80680142 |
| ZNF334 | 42.7325033 | 0.261705416 | 0.6446034 | 0.40599447 | 0.68474666 | 0.96161098 |
| SLC13A3 | 61.8917997 | -0.494484189 | 0.58838294 | -0.8404122 | 0.40067732 | 0.89112794 |
| EYA2 | 139.980353 | 2.100318464 | 1.170714 | 1.79404915 | 0.07280532 | 0.59192805 |
| ZMYND8 | 380.54406 | -0.621490285 | 0.37176365 | -1.6717349 | 0.0945766 | 0.63858274 |
| NCOA3 | 269.361728 | -0.14120511 | 0.13497512 | -1.0461566 | 0.29548874 | 0.83911606 |
| SULF2 | 94.4434476 | 1.181950393 | 0.77582224 | 1.52348093 | 0.12763843 | 0.69635098 |
| ARFGEF2 | 239.314946 | 0.002615265 | 0.18505791 | 0.01413215 | 0.98872455 | 0.99820747 |
| CSE1L | 91.849808 | -0.519557375 | 0.24764771 | -2.0979696 | 0.03590783 | 0.45984105 |
| STAU1 | 229.751037 | 0.031005141 | 0.1976733 | 0.15685043 | 0.87536273 | 0.98766275 |
| DDX27 | 43.1468913 | -0.212938695 | 0.26148317 | -0.8143495 | 0.41544474 | 0.89807127 |
| ZNFX1 | 100.169449 | -0.83223149 | 0.50603168 | -1.6446233 | 0.10004752 | 0.64666342 |
| ZFAS1 | 106.002195 | -0.272188249 | 0.34155628 | -0.796906 | 0.42550563 | 0.90167091 |
| B4GALT5 | 326.343503 | 0.187543477 | 0.49199025 | 0.38119348 | 0.70305968 | 0.9645815 |
| SLC9A8 | 95.3979682 | -0.066744396 | 0.27563788 | -0.2421452 | 0.80866764 | 0.97930627 |
| RNF114 | 157.402135 | -0.365767511 | 0.4469308 | -0.8183985 | 0.41312966 | 0.89724628 |
| CEBPB | 123.464313 | 1.468701033 | 0.7084095 | 2.07323734 | 3.82E-02 | 0.47153643 |
| PTPN1 | 389.314981 | -0.52750918 | 0.479086 | -1.1010741 | 0.27086441 | 0.81775732 |
| RIPOR3 | 155.702926 | -0.207364315 | 0.46476632 | -0.446169 | 0.65547519 | 0.95335512 |
| PARD6B | 408.780644 | 0.15338307 | 0.36987333 | 0.41469081 | 0.67836826 | 0.95924874 |
| BCAS4 | 48.0260572 | -0.467189014 | 0.57585864 | -0.8112911 | 0.41719849 | 0.89862171 |
| ADNP | 385.686722 | -0.183794615 | 0.19254819 | -0.9545382 | 0.33981126 | 0.86742596 |
| DPM1 | 78.7784998 | 0.024379222 | 0.27160461 | 0.08975997 | 0.92847796 | 0.99336859 |
| ATP9A | 162.146245 | 0.117561777 | 0.26986687 | 0.43562879 | 0.66310608 | 0.95477098 |
| ZFP64 | 67.1334364 | -0.370826383 | 0.25240925 | -1.4691473 | 0.14179283 | 0.7148274 |
| TSHZ2 | 969.116068 | 0.101140823 | 0.69311333 | 0.14592249 | 0.88398258 | 0.98766275 |
| AL109930.1 | 83.5043489 | 0.57358594 | 0.64744459 | 0.88592283 | 0.37565911 | 0.87766796 |

| | | | | | | |
|---|---|---|---|---|---|---|
| ZNF217 | 118.873962 | 0.110906997 | 0.22563963 | 0.49152268 | 0.62305681 | 0.94808496 |
| BCAS1 | 98.7063222 | -1.911955094 | 0.90001717 | -2.124354 | 0.03364055 | 0.44933903 |
| CYP24A1 | 469.357325 | -0.088921317 | 0.82649023 | -0.1075891 | 9.14E-01 | 9.92E-01 |
| PFDN4 | 31.3037746 | 0.147046251 | 0.40544462 | 0.36267901 | 0.71684468 | 0.96742001 |
| DOK5 | 46.2527885 | 0.129305653 | 0.57938036 | 0.22317921 | 0.82339603 | 0.98094663 |
| RTF2 | 51.4175368 | -0.33706264 | 0.30534614 | -1.1038707 | 0.26964925 | 0.8167745 |
| BMP7 | 119.090124 | 1.162884476 | 0.83197584 | 1.39773827 | 0.16219168 | 0.74071637 |
| RAE1 | 32.9199773 | -0.105412107 | 0.27814679 | -0.3789801 | 0.70470261 | 0.96461079 |
| PCK1 | 120.614003 | 0.070958604 | 1.20883053 | 0.05870021 | 0.95319089 | 0.99531843 |
| PMEPA1 | 66.5522084 | 0.438596198 | 1.12368897 | 0.39031815 | 0.69630131 | 0.96220115 |
| RAB22A | 120.834388 | -0.281226038 | 0.20935579 | -1.3432924 | 0.17917731 | 0.7521207 |
| VAPB | 143.628892 | -0.039213602 | 0.19021744 | -0.2061515 | 0.83667261 | 0.98462033 |
| NPEPL1 | 29.0956025 | -0.250916934 | 0.38188463 | -0.6570491 | 0.51114937 | 0.92819651 |
| GNAS | 533.454197 | -0.065699503 | 0.27856735 | -0.2358478 | 0.81355075 | 0.97930627 |
| NELFCD | 76.429898 | -0.271277926 | 0.2134644 | -1.2708345 | 0.20378754 | 0.77155952 |
| CTSZ | 121.817008 | 0.400477352 | 0.36958149 | 1.08359688 | 0.27854358 | 0.8229426 |
| ATP5F1E | 250.739543 | -0.181383995 | 0.39950184 | -0.4540254 | 0.64981052 | 0.95031069 |
| PRELID3B | 25.5496482 | -0.290341049 | 0.37310588 | -0.7781733 | 0.43646683 | 0.90525098 |
| PHACTR3 | 73.7477335 | 1.727807378 | 1.28475351 | 1.34485516 | 0.17867201 | 0.7513166 |
| MIR646HG | 67.4736836 | 0.412497597 | 0.86726983 | 0.47562775 | 0.6343396 | 0.94932781 |
| TAF4 | 81.9401993 | -0.173536521 | 0.31654161 | -0.5482266 | 0.58353635 | 0.94166015 |
| LSM14B | 29.4964387 | -0.154833004 | 0.30300916 | -0.5109846 | 0.60936187 | 0.94791646 |
| PSMA7 | 140.503769 | -0.267972686 | 0.42581554 | -0.6293164 | 0.52914197 | 0.93366189 |
| SS18L1 | 59.258377 | -0.300813228 | 0.24479439 | -1.2288404 | 0.21913166 | 0.78523615 |
| MTG2 | 57.0170056 | 0.147304067 | 0.24609447 | 0.59856715 | 0.54946157 | 0.93523904 |
| OSBPL2 | 87.8895799 | -0.297922487 | 0.26774858 | -1.1126949 | 0.26583947 | 0.81387543 |
| ADRM1 | 26.3524358 | -0.423163431 | 0.61664036 | -0.6862402 | 0.49256163 | 0.92289011 |
| LAMA5 | 180.846829 | 0.266526776 | 0.34046709 | 0.78282683 | 0.43372881 | 0.90514499 |
| RPS21 | 35.7065362 | -0.531477412 | 0.42143202 | -1.2611225 | 0.20726471 | 0.77401635 |
| SLCO4A1 | 91.7866442 | -1.288552469 | 0.737602 | -1.7469482 | 0.08064633 | 0.60729857 |
| OGFR | 75.2974047 | -0.52181094 | 0.39093166 | -1.3347881 | 0.1819457 | 0.75448394 |
| TCFL5 | 28.9176424 | 0.668086631 | 0.36506673 | 1.83003975 | 0.06724399 | 0.57884645 |
| DIDO1 | 180.263495 | 0.201114615 | 0.19513855 | 1.03062472 | 0.30271684 | 0.84398487 |
| GID8 | 63.1396149 | 0.302921097 | 0.32714443 | 0.92595524 | 0.35446924 | 0.87258503 |
| YTHDF1 | 70.0287931 | 0.010397498 | 0.21156794 | 0.04914496 | 0.96080377 | 0.99572129 |
| ARFGAP1 | 53.6608118 | -0.221760968 | 0.44562909 | -0.4976358 | 0.6187408 | 0.9480368 |
| PPDPF | 279.788548 | -0.136391574 | 0.4806395 | -0.283771 | 0.77658584 | 0.97423208 |
| GMEB2 | 48.8650113 | 0.006727306 | 0.23740765 | 0.02833652 | 0.97739375 | 0.99681784 |
| STMN3 | 29.8853152 | -0.863885465 | 0.53700666 | -1.6087053 | 1.08E-01 | 0.65699665 |
| ARFRP1 | 43.9865508 | -0.400743032 | 0.33609713 | -1.1923429 | 0.23312682 | 0.79612186 |
| SLC2A4RG | 69.6824352 | -0.075783775 | 0.30338124 | -0.2497972 | 0.80274421 | 0.97835591 |
| ZBTB46 | 122.76631 | 1.261035146 | 0.48692376 | 2.5898 | 0.00960317 | 0.28916146 |
| TPD52L2 | 77.9173221 | -0.43541468 | 0.24067915 | -1.8091085 | 0.07043416 | 0.58797037 |
| DNAJC5 | 147.049901 | 0.145771737 | 0.29789178 | 0.48934461 | 0.62459774 | 0.94808496 |
| UCKL1 | 38.3032898 | -0.056130339 | 0.35438879 | -0.1583863 | 0.87415239 | 0.98766275 |
| ZNF512B | 29.583315 | -0.092869133 | 0.50357949 | -0.184418 | 0.85368554 | 0.9864199 |
| PRPF6 | 101.466556 | -0.023266799 | 0.2259752 | -0.1029617 | 0.91799334 | 0.99214128 |
| PCMTD2 | 140.330435 | 0.327292698 | 0.28357619 | 1.15416144 | 0.24843399 | 0.80672756 |
| MIER2 | 30.5004418 | 0.127838635 | 0.35702035 | 0.358071 | 0.72029018 | 0.96779196 |
| SHC2 | 34.8242719 | 1.031992911 | 0.36714721 | 2.81084234 | 0.0049412 | 0.22029054 |
| TPGS1 | 34.624582 | -0.220637937 | 0.4644402 | -0.4750621 | 0.63474271 | 0.94932781 |
| CDC34 | 34.6196108 | -0.060458756 | 0.35931211 | -0.1682625 | 0.86637677 | 0.98670067 |
| BSG | 163.656148 | 0.126728532 | 0.43232765 | 0.29313076 | 0.76942221 | 0.97261718 |
| RNF126 | 42.2116622 | -0.498826998 | 0.34298476 | -1.4543707 | 0.14584355 | 0.71974648 |

| | | | | | | |
|---|---|---|---|---|---|---|
| FSTL3 | 70.0641391 | -0.258905984 | 0.82339086 | -0.3144387 | 0.75318784 | 0.97096495 |
| PTBP1 | 146.956465 | -0.471820694 | 0.30501594 | -1.5468722 | 0.12189405 | 0.69196376 |
| MED16 | 42.2066701 | 0.040842929 | 0.39788116 | 0.10265108 | 0.9182399 | 0.99214128 |
| R3HDM4 | 36.1466726 | 0.114375927 | 0.43565365 | 0.26253866 | 0.79290618 | 0.97443119 |
| ARID3A | 94.2119808 | 0.034165223 | 0.40845183 | 0.08364566 | 0.93333816 | 0.99425828 |
| WDR18 | 28.279958 | -0.571094655 | 0.49975587 | -1.1427473 | 0.25314354 | 0.80798674 |
| TMEM259 | 106.701453 | -0.524949015 | 0.31149844 | -1.685238 | 0.09194266 | 0.63464873 |
| CNN2 | 39.2611381 | -0.255504791 | 0.50752126 | -0.5034366 | 0.61465732 | 0.94791646 |
| POLR2E | 61.2008831 | 0.060606417 | 0.26042953 | 0.23271714 | 0.81598106 | 0.98007572 |
| GPX4 | 195.323036 | 0.118105733 | 0.46423789 | 0.25440778 | 0.79918054 | 0.97632322 |
| SBNO2 | 119.099675 | -0.236090018 | 0.38379101 | -0.6151525 | 5.38E-01 | 0.93516669 |
| STK11 | 66.0751024 | -0.020953285 | 0.28969763 | -0.0723281 | 0.94234079 | 0.99526299 |
| ATP5F1D | 170.019519 | -0.019746657 | 0.41124967 | -0.0480162 | 0.96170331 | 0.99572129 |
| MIDN | 212.581873 | -0.519880044 | 0.37128743 | -1.4002091 | 0.16145071 | 0.74032378 |
| CIRBP | 280.520523 | -0.097799128 | 0.20773957 | -0.4707776 | 0.63779958 | 0.94944486 |
| C19orf24 | 39.8294557 | -0.688710741 | 0.40349739 | -1.706853 | 0.08784937 | 0.62705598 |
| MUM1 | 111.982362 | 0.275439169 | 0.37034387 | 0.74373897 | 0.45703441 | 0.91187931 |
| NDUFS7 | 51.0926177 | 0.039041318 | 0.4313312 | 0.09051355 | 0.92787913 | 0.99336859 |
| DAZAP1 | 119.66712 | -0.354468771 | 0.24411909 | -1.4520322 | 0.14649266 | 0.71998622 |
| RPS15 | 691.097192 | -0.194127273 | 0.49196015 | -0.3945996 | 0.69313842 | 0.96220115 |
| C19orf25 | 67.0035884 | -0.243254678 | 0.21155468 | -1.149843 | 0.25020852 | 0.80672756 |
| MBD3 | 42.5723676 | -0.18687261 | 0.48803376 | -0.3829092 | 0.7017871 | 0.9645815 |
| TCF3 | 51.8252077 | 0.125625132 | 0.4172221 | 0.30109894 | 0.76333905 | 0.97149197 |
| REXO1 | 40.0540637 | -0.194632421 | 0.36618252 | -0.5315175 | 0.59506021 | 0.94664456 |
| ABHD17A | 47.0167759 | -0.039320697 | 0.31946524 | -0.1230829 | 0.90204148 | 0.9900784 |
| SCAMP4 | 65.6562145 | -0.091712944 | 0.27440601 | -0.3342235 | 0.73821089 | 0.9698508 |
| CSNK1G2 | 91.4089957 | -0.238823862 | 0.25181299 | -0.9484175 | 0.34291693 | 0.8685355 |
| BTBD2 | 106.252703 | -0.384967142 | 0.3941615 | -0.9766736 | 0.32873075 | 0.8574012 |
| MKNK2 | 54.3487432 | -0.517203428 | 0.28003974 | -1.846893 | 0.06476266 | 0.57495764 |
| MOB3A | 58.7593048 | 0.259708648 | 0.36131224 | 0.71879283 | 0.47226858 | 0.91737234 |
| AP3D1 | 123.798426 | -0.25304528 | 0.34142971 | -0.7411343 | 0.458612 | 0.91350475 |
| DOT1L | 75.7005122 | -0.569306502 | 0.36111618 | -1.5765189 | 0.11490626 | 0.67115064 |
| PLEKHJ1 | 53.6132083 | 0.077768612 | 0.37494307 | 0.20741445 | 0.83568619 | 0.98423382 |
| LSM7 | 95.2561804 | -0.5872104 | 0.43758734 | -1.3419273 | 0.17961956 | 0.75265164 |
| SPPL2B | 76.217872 | 0.157615463 | 0.31309381 | 0.5034129 | 0.614674 | 0.94791646 |
| TIMM13 | 65.2513383 | 0.143337868 | 0.44574342 | 0.32157035 | 0.74777821 | 0.97028355 |
| GADD45B | 113.082402 | 0.300013157 | 0.74200607 | 0.40432709 | 0.6859722 | 0.96161098 |
| GNG7 | 81.1156123 | 0.614867362 | 0.37805463 | 1.62639817 | 0.10386498 | 0.65455437 |
| SGTA | 44.5609075 | -0.397964697 | 0.32259233 | -1.2336459 | 0.21733489 | 0.78353439 |
| THOP1 | 28.0192173 | -0.517380567 | 0.29033195 | -1.7820311 | 0.07474415 | 0.59444897 |
| TLE2 | 73.7988477 | 0.638524943 | 0.46427591 | 1.37531353 | 0.16903426 | 0.74445571 |
| AES | 233.91401 | -0.183152376 | 0.31076488 | -0.58936 | 0.55561983 | 0.93523904 |
| GNA11 | 85.2001332 | -0.037485227 | 0.26056271 | -0.1438626 | 0.88560898 | 0.98766275 |
| NCLN | 45.9970891 | -0.908226145 | 0.31883108 | -2.8486123 | 0.00439104 | 0.21243296 |
| NFIC | 274.934238 | 0.208885005 | 0.29657451 | 0.70432555 | 0.48123006 | 0.91971919 |
| SMIM24 | 39.9028162 | 1.815520111 | 0.55814617 | 3.25276818 | 0.00114287 | 0.10732865 |
| FZR1 | 36.5936447 | 0.160995658 | 0.33381193 | 0.4822945 | 0.62959676 | 0.94916079 |
| MFSD12 | 66.0706967 | -0.370315083 | 0.33244484 | -1.1139144 | 0.26531593 | 0.81378765 |
| HMG20B | 38.8555455 | 0.459033867 | 0.3327115 | 1.37967536 | 0.16768662 | 0.74445571 |
| PIP5K1C | 40.0255647 | 0.220241514 | 0.35778566 | 0.61556832 | 0.53817947 | 0.93516669 |
| MRPL54 | 45.5262561 | -0.450479307 | 0.50305612 | -0.8954852 | 0.37052778 | 0.87605028 |
| DAPK3 | 38.1794062 | -0.110226089 | 0.32974916 | -0.3342725 | 0.73817391 | 0.9698508 |
| EEF2 | 452.263387 | 0.549989139 | 0.40076202 | 1.37235843 | 0.16995188 | 0.74445571 |
| PIAS4 | 32.8363829 | 0.148813301 | 0.40289537 | 0.36935967 | 0.71185965 | 0.96587538 |

| | | | | | | |
|---|---|---|---|---|---|---|
| ZBTB7A | 101.971971 | -0.192777618 | 0.25003108 | -0.7710146 | 0.44069826 | 0.90561284 |
| MAP2K2 | 99.5462486 | 0.147472755 | 0.25441432 | 0.57965588 | 0.5621467 | 0.93762898 |
| STAP2 | 36.519904 | 0.091163578 | 0.33235194 | 0.27429832 | 0.78385538 | 0.97423208 |
| SH3GL1 | 96.5088997 | -0.342243144 | 0.35206095 | -0.9721133 | 0.33099417 | 0.85923151 |
| UBXN6 | 70.4487969 | 0.173351878 | 0.43890896 | 0.3949609 | 0.69287175 | 0.96220115 |
| HDGFL2 | 41.6249551 | 0.222046975 | 0.34457289 | 0.6444122 | 0.51930817 | 0.93259336 |
| PLIN5 | 49.5086408 | 1.643477789 | 0.76044409 | 2.16120791 | 0.03067928 | 0.43815485 |
| MYDGF | 83.703023 | 0.355293453 | 0.36216196 | 0.98103471 | 0.32657562 | 0.85588725 |
| DPP9 | 56.2672932 | -0.345290583 | 0.29227044 | -1.1814078 | 0.23744076 | 0.80077383 |
| PLIN3 | 80.0511136 | -0.024496224 | 0.40088989 | -0.0611046 | 0.95127589 | 0.99531843 |
| KDM4B | 195.584187 | 0.008912466 | 0.33003273 | 0.02700479 | 0.97845591 | 0.99681784 |
| PTPRS | 165.827403 | 0.430488605 | 0.23350175 | 1.84362044 | 0.06523848 | 0.57495764 |
| TINCR | 30.2995745 | -0.423374524 | 0.49724963 | -0.8514325 | 0.39452912 | 0.88760715 |
| SAFB2 | 131.07381 | -0.044904758 | 0.21776832 | -0.2062043 | 0.83663135 | 0.98462033 |
| SAFB | 158.235913 | -0.115540683 | 0.17191958 | -0.6720624 | 0.50154399 | 0.92484445 |
| RPL36 | 343.613203 | -0.313170729 | 0.45757988 | -0.6844067 | 0.4937184 | 0.92289011 |
| C19orf70 | 73.3715623 | -0.084010566 | 0.40793387 | -0.2059416 | 0.83683651 | 0.98462033 |
| LONP1 | 77.4563221 | -0.219141843 | 0.31450833 | -0.696776 | 0.48594301 | 0.92034617 |
| RANBP3 | 56.7060503 | -0.016170646 | 0.24551341 | -0.0658646 | 0.94748561 | 0.9952642 |
| RFX2 | 158.233045 | 0.552751371 | 0.59446 | 0.92983778 | 0.35245508 | 0.8720098 |
| MLLT1 | 74.117279 | 0.035060231 | 0.27792624 | 0.12614941 | 0.89961366 | 0.98956835 |
| CLPP | 38.1872996 | -0.071428059 | 0.42549568 | -0.1678702 | 0.86668537 | 0.98670067 |
| ALKBH7 | 50.3650701 | 0.409081751 | 0.53578586 | 0.76351726 | 0.44515497 | 0.90703449 |
| GTF2F1 | 53.5021388 | 0.057381321 | 0.29547357 | 0.1942012 | 0.84601835 | 0.98520136 |
| KHSRP | 71.6158734 | -0.209100893 | 0.40083379 | -0.5216648 | 0.60190372 | 0.94683415 |
| SLC25A23 | 47.9315671 | 0.576921356 | 0.34148124 | 1.6894672 | 0.09112993 | 0.63421316 |
| C3 | 587.657405 | 1.641953103 | 1.20380507 | 1.36396925 | 1.73E-01 | 0.74625056 |
| TRIP10 | 94.460534 | -0.365148692 | 0.34887137 | -1.0466571 | 0.29525776 | 0.83911606 |
| SH2D3A | 58.3124736 | -1.282182116 | 0.6264362 | -2.046788 | 0.0406789 | 0.48617886 |
| INSR | 380.501648 | 0.674707652 | 0.31851781 | 2.11827294 | 0.03415196 | 0.45111962 |
| ZNF358 | 41.261253 | 0.365935187 | 0.44801123 | 0.81679914 | 0.41404322 | 0.89732895 |
| PNPLA6 | 54.3446612 | 0.202889931 | 0.31684219 | 0.6403501 | 0.52194502 | 0.93272217 |
| CAMSAP3 | 29.1928162 | 0.91732527 | 0.33773289 | 2.71612656 | 0.00660507 | 0.25396993 |
| STXBP2 | 27.6593941 | 0.608809529 | 0.47353079 | 1.28568097 | 0.19855442 | 0.77110248 |
| EVI5L | 26.3236341 | -0.161306111 | 0.3450973 | -0.4674221 | 0.6401979 | 0.94944486 |
| MAP2K7 | 70.3091812 | -0.026752349 | 0.32512817 | -0.0822825 | 0.93442209 | 0.99442609 |
| TIMM44 | 35.3933919 | -0.402802924 | 0.26287056 | -1.5323242 | 0.12544246 | 0.69492095 |
| ELAVL1 | 112.484908 | -0.182192607 | 0.21736573 | -0.8381846 | 0.40192704 | 0.89239743 |
| CERS4 | 110.17901 | 1.029552939 | 0.39548718 | 2.60325237 | 0.0092344 | 0.28622448 |
| AC022146.2 | 33.6593585 | 0.88595681 | 0.3623136 | 2.44527614 | 0.01447413 | 0.32788161 |
| CD320 | 35.7780386 | -0.327503237 | 0.46484575 | -0.7045418 | 0.48109546 | 0.91971919 |
| RPS28 | 127.29697 | 0.248970018 | 0.43978076 | 0.56612303 | 0.57131015 | 0.93998433 |
| ANGPTL4 | 80.2905984 | 0.578129769 | 1.11495686 | 0.5185221 | 0.60409405 | 0.94683415 |
| RAB11B | 72.09022 | 0.372535208 | 0.40082147 | 0.92942927 | 0.35266666 | 0.8720098 |
| 2-Mar | 35.5086435 | 0.228954435 | 0.40448952 | 0.56603305 | 0.57137132 | 0.93998433 |
| HNRNPM | 294.550583 | -0.305423839 | 0.18534933 | -1.6478281 | 0.09938794 | 0.64501158 |
| ZNF558 | 28.1878212 | -0.22466086 | 0.32036486 | -0.7012656 | 0.48313727 | 0.91971919 |
| ZNF317 | 29.6484399 | -0.130241722 | 0.28577694 | -0.4557461 | 0.64857258 | 0.95027667 |
| ZNF559-ZNF | 28.6519132 | 0.828430316 | 0.53615435 | 1.54513401 | 0.12231385 | 0.69287072 |
| ZNF266 | 49.0470644 | -0.171297676 | 0.24800562 | -0.6907008 | 0.48975359 | 0.92157694 |
| ZNF426 | 29.2842186 | 0.00125167 | 0.27882118 | 0.00448915 | 0.99641819 | 0.99892097 |
| ZNF121 | 68.0350434 | -0.783910251 | 0.24081527 | -3.2552349 | 0.00113299 | 0.10732865 |
| ZNF561 | 35.9044737 | -0.685934864 | 0.3209102 | -2.1374667 | 0.03256005 | 0.44052646 |
| ZNF562 | 87.2447095 | -0.251955833 | 0.22012299 | -1.1446139 | 0.25236913 | 0.80798674 |

| | | | | | | |
|---|---|---|---|---|---|---|
| ZNF846 | 155.16746 | 0.627465435 | 0.50015598 | 1.2545395 | 0.20964598 | 0.77686608 |
| FBXL12 | 36.1839566 | 0.234878794 | 0.32947651 | 0.71288481 | 0.47591704 | 0.91809716 |
| UBL5 | 68.0706362 | -0.118567265 | 0.49721525 | -0.2384626 | 0.81152228 | 0.97930627 |
| PIN1 | 54.7876105 | -0.345311796 | 0.32432111 | -1.0647219 | 0.28700179 | 0.83129986 |
| C19orf66 | 68.9566836 | -0.931122711 | 0.69807169 | -1.3338497 | 0.18225312 | 0.75456023 |
| EIF3G | 53.7412474 | -0.069535425 | 0.52024612 | -0.1336587 | 0.89367246 | 0.98842187 |
| DNMT1 | 42.0490253 | -1.115765378 | 0.33115429 | -3.3693219 | 0.00075353 | 0.08802561 |
| MRPL4 | 39.6440166 | -0.410821596 | 0.4782531 | -0.8590046 | 0.390338 | 0.88670817 |
| ICAM1 | 86.9602783 | 1.052528586 | 1.2227836 | 0.86076439 | 0.38936782 | 0.88586416 |
| TYK2 | 42.1858737 | -0.158009387 | 0.28452849 | -0.5553377 | 0.57866372 | 0.94034284 |
| CDC37 | 119.014644 | -0.884443204 | 0.26232126 | -3.3716032 | 0.00074732 | 0.08802561 |
| KEAP1 | 36.5818454 | -0.375353846 | 0.4086804 | -0.9184533 | 0.35838162 | 0.87316144 |
| KRI1 | 36.8078045 | -0.093587863 | 0.35050588 | -0.267008 | 0.78946302 | 0.97423208 |
| AP1M2 | 48.4352019 | -0.140848338 | 0.37732795 | -0.3732783 | 0.70894132 | 0.96587538 |
| SLC44A2 | 74.7276043 | -0.673242306 | 0.52130242 | -1.2914621 | 0.1965435 | 0.77059568 |
| ILF3-DT | 60.8113632 | -0.642368864 | 0.56721954 | -1.1324872 | 0.25742966 | 0.8120529 |
| ILF3 | 192.554328 | -0.281458453 | 0.25189905 | -1.1173462 | 0.26384632 | 0.81378765 |
| QTRT1 | 49.0643347 | -0.436821235 | 0.29475026 | -1.4820046 | 0.13833908 | 0.71091391 |
| DNM2 | 385.685479 | -0.247204016 | 0.26356141 | -0.9379371 | 0.34827675 | 0.87112607 |
| TMED1 | 25.4890952 | 0.362667915 | 0.39508586 | 0.91794709 | 0.35864657 | 0.87316144 |
| CARM1 | 51.1389416 | 0.040749165 | 0.26694902 | 0.15264774 | 0.87867607 | 0.98766275 |
| YIPF2 | 34.8856497 | 0.100399614 | 0.30550978 | 0.32862979 | 0.74243553 | 0.9698508 |
| SMARCA4 | 247.267523 | -0.270541386 | 0.32016482 | -0.8450066 | 0.39810715 | 0.8887831 |
| LDLR | 86.4044509 | 0.110416431 | 0.82054018 | 0.13456554 | 0.89295539 | 0.9882865 |
| KANK2 | 88.7644056 | 0.280914933 | 0.46122387 | 0.60906417 | 0.54248191 | 0.93523904 |
| DOCK6 | 102.187967 | -0.209780679 | 0.38639136 | -0.5429228 | 0.58718296 | 0.94309617 |
| RAB3D | 51.8516624 | -0.09801497 | 0.37891007 | -0.2586761 | 0.79588518 | 0.97524554 |
| EPOR | 23.3000782 | -0.064371878 | 0.64102935 | -0.1004195 | 0.92001125 | 0.99214128 |
| RGL3 | 218.996538 | 0.087795332 | 0.59169923 | 0.14837831 | 0.88204422 | 0.98766275 |
| PRKCSH | 77.3992392 | 0.109990623 | 0.43645848 | 0.25200707 | 0.8010356 | 0.97746751 |
| ECSIT | 44.6260076 | -0.273261349 | 0.32377846 | -0.8439763 | 0.39868265 | 0.88916906 |
| ELOF1 | 26.3053059 | -0.614391044 | 0.40489606 | -1.5174044 | 0.12916462 | 0.69862672 |
| ZNF627 | 55.4771053 | 0.010831439 | 0.58027688 | 0.01866598 | 0.98510756 | 0.99752755 |
| ZNF823 | 59.1060773 | 0.038438331 | 0.31930491 | 0.12038127 | 0.90418113 | 0.99061193 |
| ZNF441 | 44.7680803 | -0.120806173 | 0.40094851 | -0.301301 | 0.76318501 | 0.97149197 |
| ZNF440 | 49.905938 | 0.309813073 | 0.26399645 | 1.17355016 | 0.24057525 | 0.8030067 |
| ZNF439 | 68.1872099 | 0.191501844 | 0.34514806 | 0.55483969 | 0.57900432 | 0.94034284 |
| ZNF69 | 151.682438 | -0.61789404 | 0.29335318 | -2.1063145 | 0.03517705 | 0.45614273 |
| ZNF844 | 40.3590646 | 0.368061988 | 0.49192686 | 0.7482047 | 0.4543367 | 0.91077355 |
| ZNF136 | 33.6821905 | 0.087082999 | 0.3094597 | 0.28140336 | 0.77840104 | 0.97423208 |
| ZNF44 | 233.072873 | -0.034908395 | 0.2667268 | -0.130877 | 0.89587263 | 0.98842187 |
| AC008758.5 | 28.037833 | -0.516440495 | 0.44299439 | -1.1657947 | 0.24369747 | 0.80476923 |
| ZNF490 | 40.2955259 | 0.279373521 | 0.32326979 | 0.86421167 | 0.38747162 | 0.8853359 |
| ZNF791 | 83.2314973 | -0.04762231 | 0.2186813 | -0.2177704 | 0.82760802 | 0.98142144 |
| DHPS | 31.3210222 | -0.144457413 | 0.35491842 | -0.4070158 | 0.68399637 | 0.96161098 |
| TNPO2 | 81.6995621 | 0.029033352 | 0.29250872 | 0.09925636 | 0.92093472 | 0.99214128 |
| TRIR | 132.360163 | -0.204798345 | 0.41952516 | -0.488167 | 0.62543155 | 0.94808496 |
| ASNA1 | 38.3520444 | -0.062273037 | 0.45208386 | -0.1377467 | 0.89044065 | 0.98766275 |
| HOOK2 | 156.676146 | -0.102628272 | 0.37729782 | -0.2720087 | 0.78561536 | 0.97423208 |
| JUNB | 226.726232 | 0.084498945 | 0.7028027 | 0.12023139 | 0.90429985 | 0.99061193 |
| GCDH | 36.9840164 | -0.216693448 | 0.28575348 | -0.758323 | 0.44825762 | 0.90990882 |
| FARSA | 36.3299371 | -0.315985872 | 0.33369955 | -0.9469173 | 3.44E-01 | 0.86909179 |
| CALR | 220.960231 | 0.137225176 | 0.54908664 | 0.24991534 | 0.80265282 | 0.97835591 |
| AC092069.1 | 41.0322156 | 0.241694293 | 0.66031057 | 0.36603123 | 0.71434177 | 0.96631694 |

| | | | | | | |
|---|---|---|---|---|---|---|
| RAD23A | 66.3116723 | -0.87867395 | 0.25324525 | -3.4696562 | 0.00052112 | 0.07057183 |
| GADD45GIP1 | 110.888931 | -0.189771752 | 0.57168318 | -0.3319527 | 0.73992501 | 0.9698508 |
| NFIX | 51.8373846 | 0.265812118 | 0.8266327 | 0.32156013 | 0.74778595 | 0.97028355 |
| NACC1 | 25.414581 | -0.678347654 | 0.54067288 | -1.254636 | 0.20961093 | 0.77686608 |
| IER2 | 182.825816 | -0.214189903 | 0.67137622 | -0.3190311 | 0.74970292 | 0.97046004 |
| CCDC130 | 87.5585528 | 0.146312717 | 0.40107202 | 0.3648041 | 0.71525764 | 0.9667635 |
| MRI1 | 48.6321714 | 1.077030757 | 0.42894124 | 2.5109051 | 0.01204221 | 0.30967852 |
| C19orf53 | 109.98787 | -0.211951391 | 0.45288674 | -0.4680009 | 0.63978396 | 0.94944486 |
| ZSWIM4 | 46.757662 | -0.100102677 | 0.29061181 | -0.344455 | 0.73050417 | 0.96891435 |
| AC020916.1 | 28.7192425 | -0.935741935 | 0.79703338 | -1.1740311 | 0.24038259 | 0.8030067 |
| CC2D1A | 56.2238792 | -0.061567456 | 0.28974003 | -0.2124921 | 0.83172317 | 0.98290906 |
| RFX1 | 46.6845642 | 0.127644443 | 0.34405843 | 0.37099641 | 0.7106402 | 0.96587538 |
| PALM3 | 56.0438879 | -0.321115718 | 0.65653869 | -0.489104 | 0.62476805 | 0.94808496 |
| PRKACA | 38.7484235 | 0.077300983 | 0.32103145 | 0.24078944 | 0.80971831 | 0.97930627 |
| ADGRL1 | 46.8887895 | 0.052215501 | 0.32676093 | 0.15979726 | 0.87304079 | 0.98766275 |
| PKN1 | 69.369772 | -0.106277711 | 0.35506119 | -0.2993222 | 0.76469418 | 0.97177982 |
| GIPC1 | 126.452737 | -0.533725627 | 0.28056319 | -1.9023366 | 0.05712716 | 0.54616991 |
| DNAJB1 | 50.7508709 | 0.62566811 | 0.43962129 | 1.42319794 | 0.15467878 | 0.72978231 |
| TECR | 56.5116286 | 0.079615681 | 0.37773019 | 0.21077394 | 0.83306367 | 0.98338959 |
| NDUFB7 | 93.1324745 | -0.120977897 | 0.43066711 | -0.2809081 | 0.77878086 | 0.97423208 |
| ZNF333 | 30.0083516 | -0.656719437 | 0.36749826 | -1.7870001 | 0.07393747 | 0.59320069 |
| ILVBL | 40.6398018 | 0.546721476 | 0.31229291 | 1.75066886 | 0.08000297 | 0.60643125 |
| NOTCH3 | 25.7469311 | 0.327191135 | 0.7016594 | 0.46631049 | 0.64099327 | 0.94944486 |
| BRD4 | 297.590944 | -0.007043584 | 0.26381859 | -0.0266986 | 0.97870014 | 0.99681784 |
| AKAP8 | 74.861413 | 0.167941644 | 0.23634486 | 0.7105788 | 0.47734529 | 0.91864961 |
| AKAP8L | 184.032172 | -0.076418796 | 0.1817932 | -0.4203611 | 0.67422166 | 0.95910677 |
| TPM4 | 116.189382 | 0.868808285 | 0.5152404 | 1.68621926 | 0.09175358 | 0.63464873 |
| RAB8A | 35.1530441 | 0.145124209 | 0.37628861 | 0.3856726 | 0.69973915 | 0.96290782 |
| FAM32A | 32.7550208 | 0.139915421 | 0.401339 | 0.34862154 | 0.72737345 | 0.9681562 |
| AP1M1 | 39.1566397 | 0.163276221 | 0.35655321 | 0.45792947 | 0.64700312 | 0.95027667 |
| EPS15L1 | 109.568355 | -0.007921735 | 0.27244244 | -0.0290767 | 0.9768034 | 0.99681784 |
| C19orf44 | 31.8726412 | 0.609287956 | 0.3016422 | 2.01990287 | 0.04339346 | 0.49484046 |
| SIN3B | 114.015708 | 0.271447333 | 0.31799484 | 0.85362182 | 0.39331456 | 0.88760715 |
| CPAMD8 | 98.7630268 | 1.149458864 | 0.80588331 | 1.42633413 | 0.15377192 | 0.7288378 |
| MYO9B | 107.334204 | -0.28760322 | 0.33095951 | -0.8689982 | 0.38484811 | 0.88340763 |
| NR2F6 | 57.7106926 | -0.066604651 | 0.31915023 | -0.2086937 | 0.83468734 | 0.98367279 |
| ABHD8 | 28.6702712 | 0.325914137 | 0.40887319 | 0.79710323 | 0.42539108 | 0.90167091 |
| MRPL34 | 44.4153096 | 0.532967081 | 0.31813279 | 1.67529754 | 0.09387587 | 0.63765064 |
| DDA1 | 27.1797557 | -0.174294865 | 0.32719057 | -0.5327014 | 0.59424032 | 0.94649617 |
| BST2 | 84.1757023 | -2.358614543 | 1.37074404 | -1.720682 | 0.08530855 | 0.61974555 |
| MVB12A | 41.9124398 | 0.35237387 | 0.28694301 | 1.22802737 | 0.21943669 | 0.7854948 |
| SLC27A1 | 49.682598 | 0.902317728 | 0.45974193 | 1.96266139 | 0.04968553 | 0.52415717 |
| AC010618.3 | 36.2804645 | 0.486277202 | 0.37649959 | 1.2915743 | 0.19650461 | 0.77059568 |
| PGLS | 73.0125459 | 0.077244473 | 0.34894707 | 0.22136444 | 0.82480868 | 0.98094663 |
| COLGALT1 | 34.8248358 | -0.170626308 | 0.28980351 | -0.5887655 | 0.55601859 | 0.93523904 |
| RPL18A | 49.2182671 | -0.355254204 | 0.52988209 | -0.6704401 | 0.50257727 | 0.92492012 |
| CCDC124 | 33.502765 | -0.397808481 | 0.56356209 | -0.7058823 | 0.48026136 | 0.91971919 |
| JUND | 214.177727 | 0.024735919 | 0.35751574 | 0.06918833 | 0.94483971 | 0.9952642 |
| LSM4 | 76.1999375 | -0.168036058 | 0.3896314 | -0.4312693 | 0.66627258 | 0.95613858 |
| PGPEP1 | 79.9262459 | 0.183638568 | 0.39537966 | 0.46446134 | 0.64231725 | 0.94944486 |
| GDF15 | 270.178388 | -0.234504469 | 1.0690023 | -0.2193676 | 0.82636371 | 0.98112766 |
| SSBP4 | 89.4109763 | -0.067339989 | 0.21861614 | -0.3080284 | 0.75806069 | 0.9714717 |
| ISYNA1 | 32.7170383 | -0.175874224 | 0.60493531 | -0.2907323 | 0.77125608 | 0.97329019 |
| ELL | 108.701792 | 0.02347406 | 0.36357349 | 0.06456483 | 0.94852049 | 0.9952642 |

| | | | | | | |
|---|---|---|---|---|---|---|
| FKBP8 | 86.0213531 | 0.071340311 | 0.29151319 | 0.24472413 | 0.80667005 | 0.97930627 |
| KXD1 | 34.7957773 | -0.340490454 | 0.27864332 | -1.2219581 | 0.22172348 | 0.78710314 |
| UBA52 | 201.015507 | 0.064808486 | 0.55444615 | 0.11688869 | 0.90694826 | 0.99100813 |
| REX1BD | 33.9049225 | -0.334376915 | 0.39265875 | -0.8515713 | 0.39445208 | 0.88760715 |
| CRTC1 | 108.625189 | 0.061098134 | 0.35420264 | 0.17249486 | 0.86304851 | 0.98670067 |
| UPF1 | 74.047087 | -0.136960515 | 0.24633917 | -0.5559835 | 0.57822213 | 0.94034284 |
| COPE | 50.7720708 | -0.158210018 | 0.47004919 | -0.3365818 | 0.73643215 | 0.9698508 |
| DDX49 | 26.2462158 | -0.329639147 | 0.32842475 | -1.0036976 | 0.31552437 | 0.8490543 |
| SUGP2 | 161.418259 | 0.096086757 | 0.20540817 | 0.46778449 | 0.63993872 | 0.94944486 |
| SLC25A42 | 52.245012 | 0.381304081 | 0.30257294 | 1.26020548 | 0.20759525 | 0.77435432 |
| TMEM161A | 29.3033594 | 0.225958369 | 0.32660025 | 0.69184995 | 0.48903156 | 0.92121587 |
| SUGP1 | 46.0970949 | -0.186807008 | 0.26837131 | -0.6960767 | 0.48638081 | 0.92034617 |
| MAU2 | 120.301825 | -0.297311287 | 0.22044083 | -1.3487124 | 0.17742935 | 0.75126783 |
| GATAD2A | 246.62095 | -0.167691706 | 0.2297257 | -0.729965 | 0.46541161 | 0.91462428 |
| PBX4 | 39.7445325 | 1.509509415 | 0.439398 | 3.43540346 | 0.00059167 | 0.07556838 |
| ATP13A1 | 36.1651273 | -0.303866148 | 0.41066583 | -0.7399353 | 0.45933925 | 0.91362347 |
| ZNF14 | 34.1846888 | 0.127740175 | 0.38363201 | 0.3329758 | 0.73915255 | 0.9698508 |
| ZNF253 | 63.831228 | 0.421578524 | 0.33788442 | 1.24770038 | 0.2121408 | 0.77768723 |
| ZNF682 | 45.3038396 | -0.024796637 | 0.48417741 | -0.0512139 | 0.95915504 | 0.99572129 |
| ZNF90 | 302.210876 | 0.171835113 | 0.36690041 | 0.46834266 | 0.63953957 | 0.94944486 |
| AC011447.3 | 121.871965 | -0.312720084 | 0.48811186 | -0.640673 | 5.22E-01 | 0.93272217 |
| ZNF486 | 95.0314146 | -0.326067873 | 0.57530021 | -0.5667786 | 0.57086458 | 0.93998433 |
| AC008554.1 | 149.924566 | 1.284874373 | 0.83782922 | 1.53357551 | 0.12513412 | 0.69447259 |
| ZNF737 | 50.2974185 | 0.222128044 | 0.34615875 | 0.64169414 | 0.52107179 | 0.93272217 |
| ZNF430 | 58.1457527 | -0.161816728 | 0.21748804 | -0.7440259 | 0.45686083 | 0.91187931 |
| ZNF714 | 33.8962896 | -1.243008527 | 0.37979588 | -3.2728331 | 0.00106475 | 0.10514713 |
| ZNF431 | 133.281224 | -0.392932595 | 0.27602854 | -1.4235216 | 0.154585 | 0.72978231 |
| ZNF708 | 97.3836458 | 0.176145035 | 0.22212563 | 0.79299736 | 0.42777936 | 0.90272191 |
| ZNF493 | 36.8092671 | 0.457994378 | 0.31220004 | 1.46699015 | 0.14237874 | 0.7148274 |
| AC010615.4 | 55.5664675 | 0.914211554 | 0.65684326 | 1.39182604 | 0.16397508 | 0.74306594 |
| ZNF429 | 104.292331 | 0.269099177 | 0.23846426 | 1.12846755 | 0.25912251 | 0.81263292 |
| AC123912.1 | 35.8333503 | 0.987708507 | 0.65460514 | 1.50886153 | 0.13133417 | 0.700349 |
| ZNF100 | 55.7700739 | 0.008628621 | 0.43343249 | 0.01990765 | 0.98411705 | 0.99752755 |
| ZNF43 | 110.034825 | 0.079320759 | 0.25611547 | 0.30970702 | 0.75678376 | 0.9714717 |
| ZNF208 | 90.0920552 | 0.331719236 | 0.29746217 | 1.11516444 | 0.26477996 | 0.81378765 |
| ZNF98 | 49.7988732 | 0.36216695 | 0.56135061 | 0.64517067 | 0.51881659 | 0.93257822 |
| ZNF91 | 315.757879 | 0.077398025 | 0.21478538 | 0.36035053 | 0.71858502 | 0.96779196 |
| ZNF675 | 33.1600071 | -0.220507132 | 0.29293129 | -0.7527606 | 0.45159379 | 0.91077355 |
| ZNF254 | 256.890707 | -0.250420365 | 0.25873897 | -0.9678494 | 0.33311957 | 0.86067542 |
| LINC00662 | 90.4694364 | -0.299985348 | 0.30847491 | -0.9724789 | 0.33081234 | 0.85923151 |
| AC006504.5 | 99.1138113 | 0.422539208 | 0.37833678 | 1.11683356 | 0.2640655 | 0.81378765 |
| UQCRFS1 | 73.4383395 | 0.566092963 | 0.46228985 | 1.2245412 | 0.22074814 | 0.78612778 |
| C19orf12 | 53.1036358 | 0.370049331 | 0.45971797 | 0.8049486 | 0.42084934 | 0.90012434 |
| URI1 | 326.116859 | -0.044213751 | 0.16277432 | -0.2716261 | 0.78590954 | 0.97423208 |
| ZNF536 | 152.704982 | 0.72388386 | 1.58687137 | 0.45617047 | 0.6482674 | 0.95027667 |
| ZNF507 | 101.950489 | -0.147899795 | 0.27382098 | -0.5401332 | 0.58910519 | 0.94491859 |
| DPY19L3 | 156.387093 | -0.558446134 | 0.19792492 | -2.821505 | 0.00477989 | 0.21962686 |
| PDCD5 | 40.4734529 | -0.73954079 | 0.35718446 | -2.0704731 | 0.03840807 | 0.47350176 |
| ANKRD27 | 81.7219009 | -0.134746089 | 0.25841896 | -0.5214249 | 0.60207078 | 0.94683415 |
| CEP89 | 107.701848 | 0.368788301 | 0.21792193 | 1.69229551 | 0.09058964 | 0.6330877 |
| RHPN2 | 262.879263 | -0.439338606 | 0.33432661 | -1.3141 | 0.18881255 | 0.76207437 |
| GPATCH1 | 27.6310098 | -0.330031041 | 0.35233641 | -0.936693 | 0.34891651 | 0.87178925 |
| PEPD | 83.4855962 | 0.461304394 | 0.34096758 | 1.35292746 | 0.17607881 | 0.74962617 |
| LSM14A | 525.827576 | -0.014114247 | 0.15515963 | -0.090966 | 9.28E-01 | 0.99336859 |

| | | | | | | |
|---|---|---|---|---|---|---|
| KIAA0355 | 191.885735 | 0.084850192 | 0.23906044 | 0.35493197 | 0.72264056 | 0.96779196 |
| GPI | 164.163118 | -0.569318801 | 0.4780376 | -1.1909498 | 0.23367328 | 0.79633808 |
| UBA2 | 177.975276 | -0.156075724 | 0.24315713 | -0.6418719 | 0.52095638 | 0.93272217 |
| WTIP | 40.4189364 | 0.164774916 | 0.57468594 | 0.28672167 | 0.77432544 | 0.97411309 |
| SCGB2B2 | 94.3629242 | -0.033683879 | 0.45900618 | -0.0733844 | 0.94150026 | 0.99526299 |
| ZNF302 | 111.295858 | -0.326277863 | 0.32134933 | -1.015337 | 0.30994522 | 0.84601144 |
| GRAMD1A | 94.7064262 | 0.554958147 | 0.5397509 | 1.02817457 | 0.30386772 | 0.84488735 |
| HPN | 256.625642 | 0.539627368 | 0.444765 | 1.2132865 | 0.2250203 | 0.78926016 |
| LSR | 71.7292565 | -0.254812832 | 0.28256108 | -0.9017973 | 0.36716453 | 0.87605028 |
| USF2 | 60.6383614 | -0.593286539 | 0.30326721 | -1.9563161 | 0.05042793 | 0.52415717 |
| DMKN | 161.93928 | -0.222985167 | 0.35152086 | -0.6343441 | 0.52585631 | 0.93272384 |
| TMEM147 | 45.9860459 | -0.224908288 | 0.58087466 | -0.387189 | 0.69861628 | 0.96256506 |
| RBM42 | 31.1547784 | -0.553795203 | 0.44857823 | -1.2345566 | 0.21699558 | 0.78319543 |
| COX6B1 | 146.661986 | -0.130018912 | 0.4247318 | -0.30612 | 0.75951326 | 0.9714717 |
| KMT2B | 52.5099986 | -0.232145894 | 0.24352124 | -0.9532881 | 0.34044413 | 0.86777921 |
| PROSER3 | 31.2650099 | -0.402516782 | 0.34879179 | -1.1540317 | 0.24848717 | 0.80672756 |
| NPHS1 | 53.6256209 | 1.345112263 | 1.40526917 | 0.9571919 | 0.33847041 | 0.86672435 |
| NFKBID | 34.015101 | 0.120140448 | 0.37304549 | 0.32205308 | 0.74741248 | 0.97028355 |
| TBCB | 41.0104556 | -0.192224171 | 0.41289424 | -0.465553 | 0.64153546 | 0.94944486 |
| CAPNS1 | 179.958988 | -0.550879776 | 0.41270706 | -1.3347961 | 0.18194309 | 0.75448394 |
| ZNF565 | 104.503544 | -0.2173619 | 0.32554532 | -0.6676855 | 0.50433435 | 0.92552859 |
| ZNF146 | 131.674947 | -0.307313228 | 0.20097415 | -1.5291182 | 0.12623514 | 0.69492095 |
| LINC00665 | 52.9193766 | -0.216242057 | 0.38665611 | -0.559262 | 0.57598294 | 0.94034284 |
| ZFP14 | 72.1949104 | -0.028613928 | 0.29037608 | -0.0985409 | 0.92150278 | 0.99214128 |
| ZFP82 | 31.9997621 | 0.169401446 | 0.49594182 | 0.34157524 | 0.73267057 | 0.96980299 |
| ZNF566 | 55.3097134 | -0.314841487 | 0.2146144 | -1.4670101 | 0.14237332 | 0.7148274 |
| ZNF260 | 42.1680519 | 0.233780235 | 0.30556289 | 0.76508059 | 0.44422356 | 0.90667728 |
| ZNF529 | 70.3779759 | 0.036796832 | 0.28453403 | 0.12932313 | 0.89710197 | 0.98842187 |
| ZNF529-AS1 | 62.6594578 | 0.431927178 | 0.32194599 | 1.34161379 | 0.17972125 | 0.75265164 |
| ZNF461 | 29.7710799 | -0.254143834 | 0.3159671 | -0.8043364 | 0.42120273 | 0.90012434 |
| ZNF567 | 33.8302401 | 0.301372217 | 0.36286508 | 0.83053518 | 0.40623626 | 0.89579585 |
| AC020928.1 | 72.9151488 | 0.411638863 | 0.28039934 | 1.46804505 | 0.14209199 | 0.7148274 |
| ZNF790 | 46.2579087 | 0.460523023 | 0.30103294 | 1.52980939 | 0.12606392 | 0.69492095 |
| ZNF345 | 68.2979729 | 0.290737536 | 0.33725576 | 0.86206839 | 0.38864988 | 0.88586416 |
| ZNF568 | 47.3855093 | 0.244472396 | 0.34804811 | 0.70240978 | 0.48242365 | 0.91971919 |
| ZNF420 | 71.3413014 | 0.711612199 | 0.42838332 | 1.66115756 | 0.09668181 | 0.64295596 |
| ZNF383 | 26.8185482 | -0.221014802 | 0.41456052 | -0.5331304 | 0.59394334 | 0.94649617 |
| HKR1 | 107.650562 | 0.140943118 | 0.22693815 | 0.62106402 | 0.5345575 | 0.93457315 |
| ZNF569 | 27.5269209 | -0.190660308 | 0.31552594 | -0.6042619 | 0.54566952 | 0.93523904 |
| ZNF793 | 51.1963131 | 0.188216803 | 0.30165066 | 0.62395622 | 0.53265634 | 0.93391698 |
| ZNF540 | 35.8090599 | 1.233325409 | 0.75267396 | 1.63859184 | 0.10129829 | 0.64876329 |
| ZNF571 | 29.0371194 | 0.564092523 | 0.32243831 | 1.74945875 | 0.08021175 | 0.60643125 |
| ZFP30 | 49.8688158 | 0.628076205 | 0.34133234 | 1.84007236 | 0.06575762 | 0.57629661 |
| ZNF573 | 59.2491161 | 0.017920695 | 0.38559258 | 0.04647572 | 0.96293108 | 0.99572129 |
| SIPA1L3 | 618.316315 | -0.638848392 | 0.30180249 | -2.1167764 | 0.03427883 | 0.45166013 |
| SPINT2 | 41.5844904 | -0.020639681 | 0.41695588 | -0.0495009 | 0.96052014 | 0.99572129 |
| CATSPERG | 34.0776371 | 0.059694772 | 0.57138932 | 0.10447303 | 0.91679397 | 0.99207822 |
| PSMD8 | 89.2637418 | 0.039324606 | 0.42144841 | 0.09330823 | 0.92565869 | 0.99316012 |
| EIF3K | 88.2036474 | 0.122500048 | 0.33698531 | 0.36351747 | 0.71621837 | 0.96724316 |
| ACTN4 | 640.384863 | -0.115862326 | 0.27723612 | -0.4179193 | 0.67600612 | 0.9591071 |
| CAPN12 | 41.8733995 | 1.505451823 | 0.4827298 | 3.11862211 | 0.00181699 | 0.14133317 |
| ECH1 | 75.2289404 | 0.965402391 | 0.44188425 | 2.18474042 | 0.02890788 | 0.43132185 |
| HNRNPL | 92.8912905 | -0.262864451 | 0.22772058 | -1.1543289 | 0.24836534 | 0.80672756 |
| SIRT2 | 40.642831 | 0.303048824 | 0.34796131 | 0.87092679 | 0.38379413 | 0.88328503 |

| | | | | | | |
|---|---|---|---|---|---|---|
| MRPS12 | 36.6031482 | -0.157329596 | 0.4851414 | -0.3242964 | 0.74571367 | 0.97028355 |
| FBXO17 | 208.932127 | 0.512041576 | 0.54916546 | 0.93239945 | 0.35113013 | 0.8720098 |
| PAK4 | 73.1160339 | -0.009977243 | 0.36349415 | -0.0274482 | 0.97810229 | 0.99681784 |
| SAMD4B | 162.168398 | -0.120277374 | 0.24100344 | -0.4990691 | 0.61773069 | 0.94797217 |
| PAF1 | 27.4273568 | 0.260316727 | 0.37567577 | 0.69292924 | 0.48835396 | 0.92081962 |
| MED29 | 39.1928211 | 0.245831078 | 0.39354333 | 0.62466077 | 0.53219373 | 0.93367834 |
| ZFP36 | 126.379086 | 0.236909228 | 0.67990645 | 0.34844386 | 0.72750687 | 0.9681562 |
| RPS16 | 306.056714 | 0.03973184 | 0.51906351 | 0.07654524 | 0.93898533 | 0.99526299 |
| SUPT5H | 108.108996 | -0.251311914 | 0.24163764 | -1.0400363 | 0.29832304 | 0.83973489 |
| TIMM50 | 25.7228318 | -0.393287856 | 0.37037897 | -1.0618525 | 0.28830263 | 0.8330772 |
| FBL | 42.7571479 | -0.551382317 | 0.3404335 | -1.6196477 | 0.10530799 | 0.65699665 |
| FCGBP | 56.293843 | -1.261319005 | 1.16298967 | -1.0845488 | 0.27812156 | 0.8229426 |
| PSMC4 | 36.3128024 | -0.075044022 | 0.4776208 | -0.1571205 | 0.87514988 | 0.98766275 |
| ZNF546 | 29.1266078 | 0.228571232 | 0.55275001 | 0.41351647 | 0.67922826 | 0.95935233 |
| ZNF780B | 135.346251 | 0.525420032 | 0.30341001 | 1.73171621 | 0.0833241 | 0.61656297 |
| ZNF780A | 36.4432572 | 0.295282393 | 0.38559923 | 0.76577537 | 0.44380997 | 0.90667728 |
| MAP3K10 | 28.8852358 | -0.13782676 | 0.29931304 | -0.460477 | 0.6451739 | 0.94954229 |
| CNTD2 | 70.0124561 | -3.149966109 | 1.95922596 | -1.6077605 | 0.10788764 | 0.65699665 |
| AKT2 | 269.956343 | -0.730886919 | 0.440233 | -1.6602275 | 0.0968687 | 0.64297238 |
| PLD3 | 168.688194 | -0.12594473 | 0.28438892 | -0.4428609 | 0.65786635 | 0.95399905 |
| BLVRB | 98.1720755 | -0.093132998 | 0.35546706 | -0.2620018 | 0.79332008 | 0.9745418 |
| COQ8B | 37.3123035 | -0.079489859 | 0.28601329 | -0.2779237 | 0.78107096 | 0.97423208 |
| ITPKC | 53.7441251 | -0.026230641 | 0.30541106 | -0.0858863 | 0.93155677 | 0.99411667 |
| SNRPA | 49.9474747 | -0.348650185 | 0.2798954 | -1.2456446 | 0.21289492 | 0.7789612 |
| AC008537.1 | 56.0228697 | 0.493674429 | 0.3587089 | 1.37625364 | 0.16874312 | 0.74445571 |
| AXL | 67.3722477 | -0.478735552 | 0.53247293 | -0.8990796 | 0.36861026 | 0.87605028 |
| HNRNPUL1 | 102.084882 | -0.514244324 | 0.26422579 | -1.9462306 | 0.05162704 | 0.52722328 |
| TGFB1 | 69.168069 | 0.124852486 | 0.34024669 | 0.36694695 | 0.71365859 | 0.9661017 |
| DMAC2 | 26.7951963 | 0.191355607 | 0.40860162 | 0.46831827 | 0.639557 | 0.94944486 |
| RPS19 | 858.50481 | -0.217616786 | 0.39760285 | -0.547322 | 0.58415754 | 0.94184538 |
| ARHGEF1 | 46.1252776 | -0.331794181 | 0.33213453 | -0.9989753 | 0.31780668 | 0.8490543 |
| RABAC1 | 106.077756 | 0.062008897 | 0.5810351 | 0.10672143 | 0.91500998 | 0.99159829 |
| GRIK5 | 64.5122398 | -0.065044329 | 0.3351469 | -0.1940771 | 0.84611553 | 0.98520136 |
| GSK3A | 23.7247958 | -0.258258996 | 0.46160088 | -0.5594855 | 0.57583043 | 0.94034284 |
| CIC | 40.4163902 | 0.29497869 | 0.35863684 | 0.82249968 | 0.41079257 | 0.89649693 |
| MEGF8 | 49.9439662 | 0.123913184 | 0.28894141 | 0.42885228 | 0.66803073 | 0.95764829 |
| LIPE-AS1 | 50.2429758 | 0.775927345 | 0.49396947 | 1.57080022 | 0.11622906 | 0.67559092 |
| PHLDB3 | 39.1284809 | 0.799851322 | 0.31858211 | 2.51065985 | 0.01205058 | 0.30967852 |
| ETHE1 | 39.7324109 | -0.487660994 | 0.40977732 | -1.1900634 | 0.23402147 | 0.79633808 |
| ZNF428 | 60.5859204 | -0.104896073 | 0.38860364 | -0.2699308 | 0.78721353 | 0.97423208 |
| SMG9 | 27.1208294 | -0.169639242 | 0.34631332 | -0.4898433 | 0.62424482 | 0.94808496 |
| ZNF283 | 27.973834 | 0.1814391 | 0.4136827 | 0.43859485 | 0.66095513 | 0.95407427 |
| ZNF155 | 40.1129462 | 1.087487194 | 0.37153881 | 2.92698143 | 0.00342269 | 0.18951432 |
| ZNF224 | 49.7995596 | 0.135969568 | 0.33293331 | 0.40839881 | 0.68298091 | 0.96110976 |
| ZNF234 | 27.8612021 | -0.414743197 | 0.3540924 | -1.1712852 | 0.24148415 | 0.803412 |
| ZNF226 | 36.8587623 | 0.190462919 | 0.36322672 | 0.52436373 | 0.60002558 | 0.94683415 |
| ZNF235 | 99.5306237 | 0.715298615 | 0.38791771 | 1.84394421 | 0.06519127 | 0.57495764 |
| AC243964.2 | 78.2327411 | 0.09669901 | 0.51529802 | 0.18765648 | 0.85114595 | 0.98617742 |
| PVR | 37.2007349 | -0.87558697 | 0.36233515 | -2.4165113 | 0.01567004 | 0.34086104 |
| CEACAM19 | 86.0443373 | -0.667243766 | 0.36622504 | -1.8219501 | 0.06846255 | 0.58250536 |
| BCL3 | 42.7154904 | 0.022507146 | 0.51474032 | 0.04372525 | 9.65E-01 | 9.96E-01 |
| CBLC | 45.42175 | 0.035345845 | 0.58546336 | 0.06037243 | 0.95185902 | 0.99531843 |
| BCAM | 570.914137 | 0.634560432 | 0.62760413 | 1.0110839 | 0.31197627 | 0.84811918 |
| NECTIN2 | 210.954386 | -0.028823554 | 0.20879324 | -0.1380483 | 0.89020223 | 0.98766275 |

| | | | | | | |
|---|---|---|---|---|---|---|
| TOMM40 | 37.1766011 | -0.769115053 | 0.31068299 | -2.4755621 | 0.01330266 | 0.32022149 |
| APOE | 139.579945 | -0.654570326 | 0.77397007 | -0.8457308 | 0.39770294 | 0.88863925 |
| CLPTM1 | 83.8837969 | -0.071032793 | 0.34737318 | -0.2044855 | 0.83797409 | 0.98462033 |
| RELB | 80.5802843 | 0.511585488 | 0.60042277 | 0.85204212 | 0.39419071 | 0.88760715 |
| CLASRP | 90.8464316 | -0.113387905 | 0.28666979 | -0.3955349 | 0.69244817 | 0.96220115 |
| GEMIN7-AS1 | 29.8333355 | 0.094718633 | 0.36689287 | 0.25816427 | 0.79628013 | 0.97524554 |
| MARK4 | 100.705727 | 0.175256293 | 0.21425015 | 0.81799847 | 0.41335806 | 0.89732895 |
| TRAPPC6A | 49.990674 | 0.68975812 | 0.43464198 | 1.58695696 | 0.11252239 | 0.66401239 |
| ERCC2 | 27.4290523 | 0.04897349 | 0.36385085 | 0.13459771 | 0.89292995 | 0.9882865 |
| PPP1R13L | 95.8921375 | -0.820472533 | 0.21660295 | -3.7879102 | 0.00015192 | 0.03134677 |
| ERCC1 | 43.7298124 | -1.027284962 | 0.30284167 | -3.3921519 | 0.00069346 | 0.08416759 |
| VASP | 54.3087159 | -0.238136669 | 0.32032575 | -0.7434203 | 0.45722726 | 0.91188421 |
| OPA3 | 72.3851631 | -0.199142206 | 0.38780218 | -0.5135149 | 0.60759118 | 0.94708529 |
| EML2 | 55.437417 | 0.613167397 | 0.30782905 | 1.99190881 | 0.04638106 | 0.50827029 |
| SNRPD2 | 108.021031 | -0.30609825 | 0.45609071 | -0.6711346 | 0.5021348 | 0.92484445 |
| FBXO46 | 36.9514278 | -0.069955842 | 0.30166645 | -0.231898 | 0.81661725 | 0.98010849 |
| DMPK | 33.801594 | 0.680432366 | 0.38276944 | 1.77765593 | 0.07546038 | 0.59465671 |
| SYMPK | 154.817618 | -0.289431 | 0.17256962 | -1.6771839 | 0.09350652 | 0.63721577 |
| HIF3A | 55.8731603 | 0.065395154 | 0.86052336 | 0.07599463 | 0.93942337 | 0.99526299 |
| PPP5C | 87.1308727 | -0.111996339 | 0.1886189 | -0.5937705 | 0.55266562 | 0.93523904 |
| CCDC8 | 23.1744251 | -0.48631473 | 0.62267022 | -0.7810149 | 0.43479372 | 0.90514499 |
| PPP5D1 | 41.533813 | -0.159208282 | 0.52476196 | -0.3033914 | 0.76159158 | 0.9714717 |
| CALM3 | 86.1225353 | 0.360538405 | 0.40040891 | 0.90042552 | 0.36789384 | 0.87605028 |
| PRKD2 | 94.98949 | -0.222986964 | 0.33721879 | -0.6612531 | 0.50845002 | 0.92666294 |
| STRN4 | 34.3021913 | -0.345474893 | 0.32909186 | -1.0497826 | 0.2938181 | 0.83825235 |
| FKRP | 25.8116224 | 0.434737481 | 0.29956374 | 1.45123534 | 0.14671434 | 0.72070597 |
| AP2S1 | 62.972955 | 0.078333934 | 0.45122375 | 0.1736033 | 0.86217724 | 0.98669442 |
| ARHGAP35 | 172.593143 | 0.001709178 | 0.27670295 | 0.00617694 | 0.99507154 | 0.99866813 |
| TMEM160 | 64.1391956 | -0.237932885 | 0.61113785 | -0.3893277 | 0.69703376 | 0.96220115 |
| ZC3H4 | 51.0184087 | -0.562025387 | 0.35345869 | -1.5900737 | 0.11181818 | 0.66322377 |
| SAE1 | 46.3670907 | -0.514295739 | 0.36802998 | -1.3974289 | 0.16228463 | 0.74071637 |
| C5AR1 | 35.2519481 | 1.918711198 | 0.74412267 | 2.57848776 | 0.00992338 | 0.29238317 |
| DHX34 | 50.9967965 | 0.047011624 | 0.33918366 | 0.13860226 | 0.88976445 | 0.98766275 |
| NAPA | 100.472853 | -0.366117647 | 0.31828208 | -1.150293 | 0.25002322 | 0.80672756 |
| NOP53 | 48.9535903 | 0.273130146 | 0.55459977 | 0.49248154 | 0.62237897 | 0.94808496 |
| SELENOW | 79.5765679 | 0.07265917 | 0.43632152 | 0.16652667 | 0.86774249 | 0.98670067 |
| PLA2G4C | 37.1914985 | 0.417780093 | 0.785553 | 0.53182929 | 0.59484423 | 0.94664456 |
| LIG1 | 27.9411203 | -0.260750956 | 0.39132742 | -0.6663243 | 0.50520386 | 0.9257297 |
| ZSWIM9 | 31.2637715 | -0.322184193 | 0.28966009 | -1.1122837 | 0.26601619 | 0.81387543 |
| CARD8 | 59.3468041 | -0.179122641 | 0.33729709 | -0.531053 | 0.59538204 | 0.94664456 |
| EMP3 | 37.8505077 | -0.503828626 | 0.90181194 | -0.5586848 | 0.57637685 | 0.94034284 |
| KDELR1 | 101.57788 | 0.12973596 | 0.39107812 | 0.33173924 | 0.74008617 | 0.9698508 |
| CYTH2 | 47.8910649 | -0.16321949 | 0.26251626 | -0.62175 | 0.53410626 | 0.93446646 |
| LMTK3 | 35.6391936 | -1.042700037 | 0.55088291 | -1.8927798 | 0.05838716 | 0.55265871 |
| SULT2B1 | 45.968213 | -1.453509432 | 1.21782181 | -1.1935321 | 0.23266104 | 0.79519257 |
| RPL18 | 363.497116 | -0.119814436 | 0.49011597 | -0.2444614 | 0.80687349 | 0.97930627 |
| BCAT2 | 45.9520889 | 0.370115487 | 0.33823752 | 1.09424727 | 0.27384654 | 0.82047446 |
| PLEKHA4 | 73.1167419 | -0.522458326 | 0.42522721 | -1.2286569 | 0.21920049 | 0.78523615 |
| PPP1R15A | 37.2681588 | -0.121408884 | 0.47044032 | -0.258075 | 0.79634903 | 0.97524554 |
| NUCB1 | 112.188311 | 0.401812895 | 0.3297749 | 1.21844597 | 0.22305455 | 0.78755603 |
| BAX | 69.3602041 | 0.467761734 | 0.33015056 | 1.41681338 | 0.1565375 | 0.7335972 |
| FTL | 1610.01798 | 0.100840085 | 0.6083066 | 0.16577181 | 0.86833653 | 0.98671532 |
| GYS1 | 64.33397 | -0.085096506 | 0.40549177 | -0.20986 | 0.83377694 | 0.98351801 |
| RUVBL2 | 25.9138885 | -0.644361605 | 0.35250664 | -1.8279417 | 0.0675583 | 0.58039545 |

| | | | | | | |
|---|---|---|---|---|---|---|
| SNRNP70 | 322.42021 | -0.223167373 | 0.22118037 | -1.0089836 | 0.31298249 | 0.84858739 |
| TRPM4 | 33.3126449 | -0.081683878 | 0.5958453 | -0.1370891 | 0.89096039 | 0.98766275 |
| SLC6A16 | 59.4713694 | 0.378379163 | 0.53785992 | 0.70349016 | 0.48175034 | 0.91971919 |
| TEAD2 | 25.6657362 | -0.82819508 | 0.33395631 | -2.4799504 | 0.01314007 | 0.31992293 |
| PIH1D1 | 35.1745485 | 0.475980316 | 0.36281665 | 1.31190317 | 0.18955281 | 0.76207437 |
| RPL13A | 522.336399 | 0.269917608 | 0.44283056 | 0.60952796 | 0.54217455 | 0.93523904 |
| RPS11 | 395.422695 | 0.160430248 | 0.48401702 | 0.3314558 | 0.74030022 | 0.9698508 |
| FCGRT | 96.3611738 | 0.742625186 | 0.39935116 | 1.85957937 | 0.06294506 | 0.56713487 |
| NOSIP | 42.3799113 | -0.202285871 | 0.27527657 | -0.7348459 | 0.46243332 | 0.91462428 |
| PRR12 | 49.6263597 | -0.256802297 | 0.39087763 | -0.656989 | 0.511188 | 0.92819651 |
| RRAS | 29.7470891 | 0.828930079 | 0.53059495 | 1.56226529 | 0.11822551 | 0.68313719 |
| SCAF1 | 38.5235577 | -0.198904202 | 0.30075343 | -0.6613531 | 0.50838591 | 0.92666294 |
| IRF3 | 86.5810091 | -0.004552711 | 0.24375372 | -0.0186775 | 0.98509838 | 0.99752755 |
| PRMT1 | 92.8253261 | -0.325844194 | 0.39261598 | -0.829931 | 0.40657777 | 0.89585737 |
| AP2A1 | 63.1557684 | -0.166487843 | 0.26931115 | -0.6181988 | 0.53644427 | 0.93469807 |
| MED25 | 73.1922357 | 0.173555728 | 0.35497068 | 0.48892975 | 0.62489143 | 0.94808496 |
| PTOV1 | 61.5028898 | 0.31953353 | 0.31590441 | 1.01148803 | 0.31178291 | 0.84811918 |
| PNKP | 52.2742455 | -0.59417934 | 0.31676042 | -1.8758004 | 0.06068269 | 0.56071154 |
| TBC1D17 | 46.2321991 | 0.060522844 | 0.28401013 | 0.21310101 | 0.83124818 | 0.98278528 |
| VRK3 | 52.6246574 | 0.450040906 | 0.32129731 | 1.40069928 | 0.16130402 | 0.74000536 |
| MYH14 | 85.7087894 | -0.595077855 | 0.58414535 | -1.0187154 | 0.30833811 | 0.84488735 |
| KCNC3 | 118.581632 | 0.191738 | 0.38355725 | 0.49989409 | 0.61714966 | 0.94791646 |
| NR1H2 | 109.053747 | -0.10755047 | 0.40450406 | -0.2658823 | 0.79032986 | 0.97423208 |
| NAPSA | 190.227143 | 0.722381712 | 0.36433048 | 1.98276496 | 0.04739369 | 0.51284792 |
| EMC10 | 86.6499394 | -0.194102667 | 0.36923058 | -0.525695 | 0.59910014 | 0.94683415 |
| ETFB | 91.8931637 | 0.878196292 | 0.39968259 | 2.1972343 | 0.02800372 | 0.4250936 |
| ZNF175 | 27.3404686 | 0.347324848 | 0.33413306 | 1.03948065 | 0.29858125 | 0.83973489 |
| SPACA6 | 45.4428721 | -0.698164323 | 0.38507182 | -1.8130756 | 0.06982016 | 0.58513328 |
| ZNF577 | 52.62772 | 0.217330349 | 0.36455692 | 0.59614929 | 0.5510755 | 0.93523904 |
| ZNF350-AS1 | 47.9495891 | 0.438196601 | 0.61487557 | 0.71265899 | 0.4760568 | 0.91809716 |
| ZNF432 | 60.9357461 | 0.414783811 | 0.30887618 | 1.34288052 | 0.17931066 | 0.75235079 |
| AC011468.3 | 35.5736693 | 0.121953785 | 0.60032865 | 0.20314504 | 0.83902167 | 0.98462033 |
| PPP2R1A | 211.149038 | 0.063967099 | 0.33940364 | 0.1884691 | 0.85050893 | 0.9858744 |
| ZNF766 | 53.1539435 | 0.025474818 | 0.24164047 | 0.10542447 | 0.916039 | 0.99159829 |
| ZNF480 | 34.8148652 | -0.353325235 | 0.34421704 | -1.0264606 | 0.30467452 | 0.84488735 |
| ZNF528 | 32.4966046 | -0.05810611 | 0.4309374 | -0.1348365 | 0.89274111 | 0.9882865 |
| ZNF701 | 39.8515372 | 0.210636894 | 0.30746302 | 0.68508042 | 0.49329317 | 0.92289011 |
| ZNF83 | 407.787045 | -0.019929079 | 0.30548953 | -0.0652365 | 0.94798567 | 0.9952642 |
| ZNF611 | 196.138116 | -0.028532586 | 0.36835059 | -0.0774604 | 0.93825729 | 0.99526299 |
| ZNF28 | 27.1049922 | -0.555213033 | 0.32858819 | -1.6896926 | 0.09108678 | 0.63421316 |
| AC010487.3 | 57.4573498 | -0.928462009 | 0.56197561 | -1.6521393 | 0.09850615 | 0.64312419 |
| ZNF160 | 127.660014 | -0.099601191 | 0.24110837 | -0.4130972 | 0.6795354 | 0.95936177 |
| ZNF347 | 46.8025726 | -0.316981399 | 0.31564196 | -1.0042435 | 0.31526124 | 0.8490543 |
| ZNF677 | 31.6233174 | -0.363200744 | 0.41544985 | -0.8742349 | 0.38199036 | 0.88128509 |
| ZNF765 | 40.5723972 | 0.254854328 | 0.30813071 | 0.8270981 | 0.40818146 | 0.8959746 |
| ZNF761 | 48.5448966 | -0.481216323 | 0.30299984 | -1.5881735 | 0.11224711 | 0.66401239 |
| ZNF331 | 96.8393708 | 0.320132204 | 0.80387827 | 0.39823468 | 0.69045721 | 0.96220115 |
| MYADM | 40.1960597 | -0.050325402 | 0.5801161 | -0.0867506 | 0.93086978 | 0.99399379 |
| NDUFA3 | 32.1621094 | 0.17893587 | 0.53559535 | 0.33408779 | 0.73831331 | 0.9698508 |
| PRPF31 | 60.6197175 | 0.189556103 | 0.35008936 | 0.54145062 | 0.58819702 | 0.94440819 |
| CNOT3 | 39.4167926 | -0.338290234 | 0.36275101 | -0.9325687 | 0.3510427 | 0.8720098 |
| TMC4 | 81.7849799 | 0.169384429 | 0.41734285 | 0.40586398 | 0.68484255 | 0.96161098 |
| MBOAT7 | 62.5522699 | 0.067376439 | 0.21420842 | 0.31453684 | 0.75311334 | 0.97096495 |
| TSEN34 | 40.8293543 | 0.369824767 | 0.49195548 | 0.75174438 | 0.4522048 | 0.91077355 |

| | | | | | | |
|---|---|---|---|---|---|---|
| RPS9 | 754.745107 | 0.141565053 | 0.53981329 | 0.26224818 | 0.79313011 | 0.97452454 |
| LENG8-AS1 | 24.3715673 | 0.305222794 | 0.35875064 | 0.85079373 | 0.39488394 | 0.88760715 |
| LENG8 | 182.084608 | -0.470033991 | 0.36363113 | -1.2926121 | 0.19614526 | 0.77033926 |
| RDH13 | 55.9084961 | -0.454675665 | 0.26494668 | -1.7161025 | 0.08614328 | 0.62276717 |
| EPS8L1 | 59.7991052 | 0.26631555 | 0.38525327 | 0.69127396 | 0.48939339 | 0.92136386 |
| PPP1R12C | 60.6242168 | -0.265553528 | 0.33327095 | -0.7968097 | 0.42556156 | 0.90167091 |
| PPP6R1 | 52.0434945 | -0.411495911 | 0.30757208 | -1.3378845 | 0.18093409 | 0.75448394 |
| HSPBP1 | 27.4291273 | -0.084962441 | 0.32849256 | -0.2586434 | 0.79591038 | 0.97524554 |
| TMEM238 | 32.7209259 | -0.52318971 | 0.64990737 | -0.805022 | 0.42080701 | 0.90012434 |
| RPL28 | 513.680055 | -0.154755741 | 0.4969787 | -0.3113931 | 0.7555018 | 0.9714717 |
| ISOC2 | 42.1823007 | 0.401072765 | 0.56802005 | 0.70608909 | 0.48013273 | 0.91971919 |
| NAT14 | 38.1639562 | 0.846890676 | 0.52759469 | 1.60519182 | 0.1084516 | 0.65764016 |
| ZNF579 | 26.3287856 | -0.189242939 | 0.39021826 | -0.4849669 | 0.62769983 | 0.94858781 |
| U2AF2 | 48.5016545 | -0.002268163 | 0.30231741 | -0.0075026 | 0.99401386 | 0.99866813 |
| EPN1 | 86.5067531 | -0.252378377 | 0.30655462 | -0.8232738 | 0.41035232 | 0.8964268 |
| ZNF787 | 38.045045 | -0.424962154 | 0.38529485 | -1.1029531 | 0.27004753 | 0.81705157 |
| ZNF444 | 42.9442874 | 0.187877013 | 0.33769907 | 0.55634448 | 0.57797539 | 0.94034284 |
| ZSCAN5A | 107.065104 | 0.040842188 | 0.33395304 | 0.12229919 | 0.90266207 | 0.99033329 |
| ZNF667 | 38.4527255 | 0.079412455 | 0.4401302 | 0.18042946 | 0.85681543 | 0.9864199 |
| ZNF667-AS1 | 45.6255004 | 0.319671887 | 0.4451843 | 0.7180664 | 0.47271635 | 0.91737234 |
| ZNF471 | 45.7896522 | 0.433619925 | 0.45580568 | 0.95132628 | 0.34143877 | 0.86846099 |
| ZNF264 | 66.3436747 | 0.161628811 | 0.21302176 | 0.75874318 | 0.4480062 | 0.90976372 |
| AC005261.1 | 92.6405069 | 0.092102736 | 0.30998604 | 0.29711898 | 0.76637567 | 0.97221495 |
| AC004076.1 | 52.0177235 | 0.119892159 | 0.32087847 | 0.37363728 | 0.7086742 | 0.96587538 |
| ZNF549 | 32.2778095 | 0.497807855 | 0.30532734 | 1.6304071 | 0.10301549 | 0.65288882 |
| ZNF586 | 167.47482 | 0.269267617 | 0.1924494 | 1.39916056 | 0.16176484 | 0.74071637 |
| ZNF552 | 39.4163893 | 0.334905225 | 0.34718379 | 0.96463383 | 0.33472825 | 0.86285304 |
| ZNF814 | 119.167248 | 0.200601096 | 0.29161506 | 0.68789689 | 0.49151771 | 0.92275924 |
| ZNF587 | 62.2384828 | 0.257754664 | 0.28288815 | 0.91115397 | 0.36221425 | 0.8752623 |
| ZNF606 | 34.1000564 | -0.284341932 | 0.38424957 | -0.7399929 | 0.45930433 | 0.91362347 |
| ZNF135 | 46.8293282 | 0.706498092 | 0.37616384 | 1.87816588 | 0.06035848 | 0.56071154 |
| ZSCAN18 | 104.410939 | 0.407009892 | 0.38961542 | 1.04464524 | 0.29618696 | 0.83973489 |
| ZNF329 | 38.6665732 | 0.466079686 | 0.35924462 | 1.29738809 | 0.19449769 | 0.76892009 |
| ZNF274 | 95.8101426 | 0.094725455 | 0.42936182 | 0.22061918 | 0.82538896 | 0.98094663 |
| RPS5 | 255.425514 | -0.015736534 | 0.58507768 | -0.0268965 | 0.9785423 | 0.99681784 |
| TRIM28 | 86.1324718 | -0.351773363 | 0.44072039 | -0.7981781 | 0.42476714 | 0.90167091 |
| CHMP2A | 70.6259986 | -0.242816059 | 0.42479882 | -0.5716025 | 0.56759131 | 0.93886327 |
| UBE2M | 49.4309132 | -0.055154925 | 0.37820851 | -0.1458321 | 0.88405397 | 0.98766275 |
| MZF1-AS1 | 25.918355 | 0.405167281 | 0.50158128 | 0.80777991 | 0.41921729 | 0.89997365 |
| MZF1 | 91.2315333 | -0.019083706 | 0.26243026 | -0.0727192 | 0.94202961 | 0.99526299 |
| RPS4Y1 | 135.85729 | 0.088803433 | 2.80706087 | 0.03163573 | 0.97476255 | 0.99681784 |
| ZFY | 48.0595532 | 0.498881898 | 2.1646377 | 0.23046901 | 0.81772735 | 0.98010849 |
| LINC00278 | 405.930981 | 1.208650165 | 2.99452445 | 0.40362007 | 0.68649213 | 0.96161098 |
| TBL1Y | 68.5713483 | -0.159542871 | 1.79600131 | -0.0888323 | 0.92921521 | 0.99350462 |
| TTTY15 | 73.6199851 | 0.403071017 | 2.23675304 | 0.18020363 | 0.85699271 | 0.9864199 |
| USP9Y | 313.905035 | 0.363737188 | 2.60796504 | 0.13947165 | 0.88907746 | 0.98766275 |
| DDX3Y | 78.2322202 | 0.485530497 | 2.6230411 | 0.18510213 | 0.85314893 | 0.9864199 |
| UTY | 499.010239 | 0.422677514 | 2.26064379 | 0.18697219 | 0.85168244 | 0.98636998 |
| NLGN4Y | 216.6342 | 0.68974182 | 2.24300013 | 0.3075086 | 0.75845628 | 0.9714717 |
| TTTY14 | 563.072967 | 0.206013215 | 1.24296517 | 0.16574335 | 0.86835893 | 0.98671532 |
| KDM5D | 54.4140824 | 0.425330104 | 2.50783622 | 0.16960043 | 0.86532438 | 0.98670067 |
| TTTY10 | 34.9879718 | -0.55661676 | 1.71082301 | -0.3253503 | 0.74491597 | 0.97028355 |
| EIF1AY | 27.7465065 | 0.48868326 | 2.31515716 | 0.21107995 | 0.83282488 | 0.98331438 |
| IL17RA | 47.2390268 | 0.285649355 | 0.29516575 | 0.96775913 | 0.33316468 | 0.86067542 |

| | | | | | | |
|---|---|---|---|---|---|---|
| HDHD5 | 40.1109061 | -0.210350035 | 0.40545551 | -0.5187993 | 0.60390071 | 0.94683415 |
| CECR2 | 54.5606413 | 1.959850386 | 0.83402701 | 2.34986441 | 0.01878025 | 0.36488708 |
| ATP6V1E1 | 95.9050023 | -0.012651409 | 0.2773043 | -0.0456228 | 0.96361087 | 0.99581912 |
| BCL2L13 | 181.274907 | -0.160264451 | 0.14964182 | -1.070987 | 0.28417525 | 0.82918183 |
| MICAL3 | 582.951424 | -0.007980437 | 0.31877919 | -0.0250344 | 0.98002755 | 0.99681784 |
| PEX26 | 72.1611883 | -0.108124767 | 0.26459506 | -0.4086424 | 0.6828021 | 0.96101988 |
| DGCR2 | 140.758403 | -0.339653081 | 0.23784915 | -1.4280189 | 0.15328642 | 0.7288378 |
| SLC25A1 | 55.0099914 | 0.621770449 | 0.38945769 | 1.59650321 | 0.11037649 | 0.65930736 |
| CLTCL1 | 43.1229191 | 0.355438741 | 0.27116193 | 1.31079882 | 0.18992574 | 0.76207437 |
| HIRA | 56.1018156 | -0.231728035 | 0.36924515 | -0.6275723 | 0.53028414 | 0.93366189 |
| MRPL40 | 47.8751764 | -0.291151469 | 0.6554752 | -0.4441838 | 0.6569097 | 0.95399905 |
| UFD1 | 34.9176825 | -0.448849317 | 0.40354512 | -1.1122655 | 0.26602401 | 0.81387543 |
| TXNRD2 | 46.3393739 | -0.222923734 | 0.28616166 | -0.7790133 | 0.43597189 | 0.90514499 |
| COMT | 162.025012 | 0.119605063 | 0.26973615 | 0.44341503 | 0.65746557 | 0.95399905 |
| TANGO2 | 32.1944919 | -0.286891749 | 0.31540708 | -0.909592 | 0.36303773 | 0.87551319 |
| DGCR8 | 53.8483746 | -0.302948272 | 0.3063946 | -0.988752 | 0.32278449 | 0.85318781 |
| RANBP1 | 42.3794302 | -1.006301202 | 0.44659785 | -2.2532603 | 0.02424274 | 0.40956125 |
| ZDHHC8 | 39.4762509 | -0.481104994 | 0.48786191 | -0.9861499 | 0.32405954 | 0.85444355 |
| KLHL22 | 103.835516 | 0.63581479 | 0.3724978 | 1.70689544 | 0.08784149 | 0.62705598 |
| MED15 | 199.935305 | -0.386080359 | 0.29462826 | -1.3103983 | 0.19006114 | 0.76207437 |
| PI4KA | 211.631165 | 0.208506207 | 0.23120907 | 0.90180807 | 0.36715883 | 0.87605028 |
| SNAP29 | 47.4489604 | -0.607842503 | 0.2516945 | -2.4150011 | 0.01573516 | 0.34086104 |
| CRKL | 51.0239207 | 0.010783153 | 0.29852264 | 0.03612172 | 0.9711853 | 0.99681784 |
| LZTR1 | 27.3339994 | -0.453356259 | 0.34522374 | -1.3132245 | 0.18910733 | 0.76207437 |
| GGT2 | 228.733539 | 0.255623492 | 1.05091015 | 0.2432401 | 0.8078194 | 0.97930627 |
| HIC2 | 45.546181 | -0.237419124 | 0.26280122 | -0.9034171 | 0.36630455 | 0.87605028 |
| UBE2L3 | 145.539342 | -0.113401198 | 0.23760923 | -0.4772592 | 0.63317754 | 0.94932781 |
| SDF2L1 | 48.2724845 | -0.014241632 | 0.67056504 | -0.0212383 | 0.9830556 | 0.99721376 |
| PPIL2 | 84.166291 | -0.392933191 | 0.24412046 | -1.6095873 | 0.10748799 | 0.65699665 |
| YPEL1 | 36.4372976 | 1.135300068 | 0.4274841 | 2.65577148 | 0.00791272 | 0.27167011 |
| MAPK1 | 308.699625 | -0.240425013 | 0.23193467 | -1.0366066 | 0.29991924 | 0.84130486 |
| PPM1F | 40.9193844 | 0.149393558 | 0.52828584 | 0.28278925 | 0.77733839 | 0.97423208 |
| TOP3B | 31.0515231 | -0.165740505 | 0.32672672 | -0.5072756 | 0.61196143 | 0.94791646 |
| AC245060.5 | 55.3440221 | 0.132052807 | 0.4665792 | 0.28302335 | 0.77715894 | 0.97423208 |
| RAB36 | 40.6911004 | 0.319378133 | 0.33572267 | 0.95131536 | 0.34144431 | 0.86846099 |
| BCR | 197.219269 | -0.493549663 | 0.3860914 | -1.2783234 | 0.20113543 | 0.77110248 |
| PCAT14 | 31.3603221 | 0.296823931 | 1.23208915 | 0.24091108 | 0.80962403 | 0.97930627 |
| CHCHD10 | 160.545899 | -0.074110738 | 0.37377569 | -0.198276 | 0.84282915 | 0.98517901 |
| SMARCB1 | 78.0671912 | -0.181508581 | 0.22931769 | -0.7915158 | 0.42864304 | 0.90278843 |
| DDT | 107.505 | 0.460170449 | 0.36400592 | 1.26418398 | 0.206164 | 0.7735572 |
| CABIN1 | 188.549017 | 0.021745865 | 0.2862442 | 0.07596963 | 0.93944326 | 0.99526299 |
| GUCD1 | 61.3944693 | 0.100728436 | 0.22858613 | 0.44065857 | 6.59E-01 | 9.54E-01 |
| SNRPD3 | 123.551658 | -0.331711467 | 0.36686903 | -0.9041686 | 0.36590599 | 0.87605028 |
| GGT1 | 92.3374134 | 0.29515479 | 0.42345109 | 0.69702216 | 0.48578893 | 0.92034617 |
| LRRC75B | 34.0176025 | 0.432557696 | 0.41780336 | 1.03531407 | 0.30052226 | 0.84221847 |
| SGSM1 | 70.7538688 | 0.686068153 | 0.66583297 | 1.03039078 | 0.3028266 | 0.84398487 |
| KIAA1671 | 267.498324 | -0.343593615 | 0.22955372 | -1.4967896 | 0.13444802 | 0.70619627 |
| LRP5L | 48.9852121 | -0.028881481 | 0.5154896 | -0.0560273 | 0.95532007 | 0.99554079 |
| GRK3 | 81.5236591 | 0.293977224 | 0.40247111 | 0.73043061 | 0.46512701 | 0.91462428 |
| HPS4 | 83.99794 | -0.660308364 | 0.30732681 | -2.1485544 | 0.03166974 | 0.43905874 |
| TPST2 | 64.9740825 | -0.367621359 | 0.56981655 | -0.6451574 | 0.5188252 | 0.93257822 |
| MIATNB | 67.9992533 | -0.477152001 | 0.36330422 | -1.3133676 | 0.1890591 | 0.76207437 |
| PITPNB | 222.628421 | -0.086086445 | 0.31097629 | -0.2768264 | 0.78191342 | 0.97423208 |
| TTC28 | 845.643982 | 0.531099705 | 0.45819576 | 1.15911091 | 0.24641098 | 0.8055873 |

| | | | | | | |
|---|---|---|---|---|---|---|
| HSCB | 62.2669901 | -0.047912472 | 0.24629392 | -0.1945337 | 0.845758 | 0.98520136 |
| CCDC117 | 33.865159 | 0.246748493 | 0.3011122 | 0.81945699 | 0.41252573 | 0.89706788 |
| XBP1 | 108.442901 | 0.21003573 | 0.46181227 | 0.4548076 | 0.64924766 | 0.95027667 |
| Z93930.2 | 102.355879 | 0.205681425 | 0.55895252 | 0.36797656 | 0.71289071 | 0.96587538 |
| ZNRF3 | 691.754073 | -0.72655532 | 0.43747628 | -1.6607879 | 0.09675605 | 0.64295596 |
| KREMEN1 | 70.2268774 | 0.544003441 | 0.37741412 | 1.44139663 | 0.14947266 | 0.72246729 |
| EMID1 | 40.1947671 | 1.67637664 | 1.00361396 | 1.6703401 | 0.09485209 | 0.63901772 |
| RHBDD3 | 30.3958554 | -0.154733329 | 0.41020105 | -0.3772134 | 0.70601503 | 0.96585518 |
| EWSR1 | 211.838454 | -0.221828392 | 0.17088864 | -1.2980874 | 0.1942573 | 0.7687432 |
| AP1B1 | 72.1002986 | -0.31902881 | 0.3394511 | -0.9398373 | 0.34730102 | 0.87017396 |
| THOC5 | 25.970047 | 0.217173708 | 0.37796142 | 0.57459227 | 0.56556708 | 0.93809297 |
| NIPSNAP1 | 41.6863912 | 0.230699806 | 0.31171104 | 0.74010791 | 0.45923452 | 0.91362347 |
| NF2 | 91.5654314 | -0.20228835 | 0.23352105 | -0.8662532 | 0.38635134 | 0.88365806 |
| UQCR10 | 150.049619 | 0.004744435 | 0.49483111 | 0.00958799 | 0.99235001 | 0.99866813 |
| ASCC2 | 57.0745707 | -0.376065839 | 0.23631372 | -1.5913839 | 0.11152322 | 0.66300165 |
| MTMR3 | 336.427403 | 0.207302252 | 0.17548351 | 1.18132039 | 0.23747547 | 0.80077383 |
| SF3A1 | 43.5308624 | 0.302466421 | 0.28224841 | 1.07163198 | 0.28388536 | 0.82918183 |
| RNF215 | 39.497141 | -0.117213402 | 0.4089237 | -0.2866388 | 0.77438889 | 0.97411309 |
| PES1 | 46.7634277 | -0.740791479 | 0.29882059 | -2.479051 | 0.01317325 | 0.31992293 |
| TCN2 | 36.2621062 | 0.019094655 | 0.56113037 | 0.03402891 | 0.9728541 | 0.99681784 |
| DUSP18 | 40.2946306 | 0.560867673 | 0.34139594 | 1.64286566 | 0.10041073 | 0.64698063 |
| OSBP2 | 68.7914706 | 0.48637882 | 0.34039714 | 1.42885696 | 0.15304535 | 0.72863887 |
| MORC2 | 54.2476844 | -0.305851579 | 0.29889163 | -1.0232859 | 0.30617271 | 0.84488735 |
| SELENOM | 94.5468011 | 0.595944411 | 0.47255558 | 1.26110967 | 0.20726934 | 0.77401635 |
| RNF185 | 81.2284392 | 0.082575279 | 0.19366635 | 0.42637908 | 0.66983164 | 0.95764829 |
| LIMK2 | 201.708025 | 0.454411017 | 0.41017652 | 1.10784257 | 0.26792981 | 0.8155385 |
| PATZ1 | 41.3944397 | -0.278335862 | 0.42993651 | -0.6473883 | 0.51738067 | 0.93247044 |
| DRG1 | 98.0013177 | -0.374400775 | 0.21640592 | -1.7300857 | 0.08361497 | 0.61656297 |
| EIF4ENIF1 | 105.414032 | -0.289497209 | 0.25600181 | -1.1308405 | 0.25812223 | 0.81223967 |
| SFI1 | 104.00298 | -0.364236658 | 0.31898812 | -1.1418502 | 0.25351628 | 0.80839961 |
| PISD | 212.293918 | -0.19822766 | 0.21534153 | -0.9205269 | 0.3572975 | 0.87316144 |
| PRR14L | 165.431762 | -0.219707028 | 0.19161405 | -1.1466123 | 0.25154188 | 0.80744031 |
| YWHAH | 36.9121284 | -0.243022285 | 0.38239867 | -0.6355207 | 0.52508885 | 0.93272384 |
| RTCB | 90.6015319 | 0.193182779 | 0.21024399 | 0.91885044 | 0.35817381 | 0.87316144 |
| FBXO7 | 116.378388 | 0.198334683 | 0.2510266 | 0.7900943 | 0.4294727 | 0.90333313 |
| TIMP3 | 156.239627 | 0.204023913 | 0.53409129 | 0.38200195 | 0.70245992 | 0.9645815 |
| LARGE1 | 704.235471 | -0.469343862 | 0.27788036 | -1.6890142 | 0.09121671 | 0.63421316 |
| HMGXB4 | 74.2148177 | -0.327803677 | 0.28207553 | -1.1621131 | 0.24518952 | 0.80476923 |
| TOM1 | 67.7143381 | 0.219066604 | 0.25807166 | 0.84885959 | 0.39595942 | 0.88785003 |
| HMOX1 | 80.0687256 | 1.068193278 | 0.62646446 | 1.70511394 | 8.82E-02 | 6.28E-01 |
| APOL6 | 119.09762 | -0.521329894 | 0.81637216 | -0.6385934 | 0.52308746 | 0.93272384 |
| RBFOX2 | 519.309546 | -0.12221603 | 0.21356837 | -0.5722572 | 0.56714776 | 0.9385359 |
| APOL2 | 55.8992278 | -0.52435065 | 0.68072987 | -0.7702771 | 0.44113552 | 0.90561284 |
| APOL1 | 39.4548966 | -0.987638427 | 0.64086937 | -1.5410916 | 0.12329448 | 0.69314363 |
| MYH9 | 325.441279 | -0.199208801 | 0.3182009 | -0.6260473 | 0.53128395 | 0.93366189 |
| TXN2 | 46.833598 | 0.366219233 | 0.39225232 | 0.93363179 | 0.35049386 | 0.8720098 |
| EIF3D | 97.2178435 | 0.062999874 | 0.3269184 | 0.19270826 | 0.84718746 | 0.98533374 |
| IFT27 | 34.7285326 | -0.142147916 | 0.31529202 | -0.4508453 | 0.65210108 | 0.95159942 |
| TST | 49.5162933 | 0.641446611 | 0.39947668 | 1.60571728 | 0.10833604 | 0.65764016 |
| MPST | 75.3940191 | 0.439485044 | 0.33961049 | 1.29408559 | 0.19563586 | 0.76992435 |
| CDC42EP1 | 36.8413635 | 0.580050681 | 0.8174534 | 0.70958256 | 0.47796304 | 0.91898995 |
| GGA1 | 84.440166 | 0.017554504 | 0.21240327 | 0.08264705 | 0.93413219 | 0.99442609 |
| TRIOBP | 142.318347 | -0.066288024 | 0.19899013 | -0.3331222 | 0.73904206 | 0.9698508 |
| H1F0 | 172.82134 | -0.211884003 | 0.72670277 | -0.291569 | 0.77061618 | 0.97313769 |

| | | | | | | |
|---|---|---|---|---|---|---|
| ANKRD54 | 25.6242212 | 0.002316207 | 0.38787657 | 0.00597151 | 0.99523546 | 0.99866813 |
| EIF3L | 127.740897 | 0.148994399 | 0.28605442 | 0.52086034 | 0.60246407 | 0.94683415 |
| POLR2F | 48.3810137 | -0.42162014 | 0.30549058 | -1.3801412 | 0.16754317 | 0.74445571 |
| PICK1 | 39.5635552 | -0.488889918 | 0.3225405 | -1.5157474 | 0.12958324 | 0.69904834 |
| BAIAP2L2 | 37.8628973 | 0.326358669 | 0.35041317 | 0.93135388 | 0.35167054 | 0.8720098 |
| PLA2G6 | 28.1974084 | 0.771721376 | 0.4605076 | 1.67580594 | 0.09377621 | 0.63753181 |
| MAFF | 30.5000321 | 0.086449754 | 0.49030785 | 0.1763173 | 0.86004467 | 0.9864199 |
| TMEM184B | 78.5256527 | -0.063043368 | 0.252168 | -0.2500054 | 0.80258315 | 0.97835591 |
| CSNK1E | 36.9292904 | -0.704789109 | 0.34102835 | -2.0666584 | 0.03876635 | 0.47547104 |
| TPTEP2-CSNI | 261.473894 | 0.342808848 | 0.28586956 | 1.19917926 | 0.23045825 | 0.79391882 |
| DDX17 | 1246.4215 | 0.184671616 | 0.28325451 | 0.65196355 | 0.51442469 | 0.93091506 |
| FAM227A | 68.5641081 | 1.074742625 | 0.29019214 | 3.7035552 | 0.0002126 | 0.03842424 |
| CBY1 | 50.2921029 | 0.297719409 | 0.23834837 | 1.24909353 | 0.21163087 | 0.77730294 |
| TOMM22 | 24.855222 | -0.199670655 | 0.51470593 | -0.3879315 | 0.69806671 | 0.96256506 |
| JOSD1 | 64.8092312 | -0.24854488 | 0.31166542 | -0.7974734 | 0.42517615 | 0.90167091 |
| GTPBP1 | 51.2195383 | -0.583585659 | 0.47204542 | -1.2362913 | 0.2163503 | 0.78293142 |
| SUN2 | 55.8173104 | 0.135905947 | 0.29536085 | 0.46013527 | 0.64541913 | 0.94954229 |
| AL021707.1 | 29.7198025 | -0.026031828 | 0.34470876 | -0.0755183 | 0.93980232 | 0.99526299 |
| CBX6 | 116.950831 | 0.131929533 | 0.49928087 | 0.26423911 | 0.79159568 | 0.97423208 |
| CBX7 | 68.2005979 | 0.864500189 | 0.3737579 | 2.31299508 | 0.02072291 | 0.3802773 |
| PDGFB | 29.8579024 | -1.216166488 | 0.66456694 | -1.8300135 | 0.06724792 | 0.57884645 |
| RPL3 | 446.582417 | 0.430618693 | 0.50161038 | 0.85847246 | 0.39063163 | 0.88670817 |
| SYNGR1 | 119.674019 | 0.131379186 | 0.36691019 | 0.35806905 | 0.72029164 | 0.96779196 |
| TAB1 | 52.5696768 | -0.296501042 | 0.31607566 | -0.9380698 | 0.34820853 | 0.87112607 |
| MIEF1 | 30.9219831 | -0.551789434 | 0.3066583 | -1.7993625 | 0.07196136 | 0.59090333 |
| ATF4 | 103.943897 | -0.471992789 | 0.32546264 | -1.4502211 | 0.14699687 | 0.72098466 |
| RPS19BP1 | 86.0455295 | -0.162694055 | 0.49089001 | -0.3314267 | 0.7403222 | 0.9698508 |
| TNRC6B | 1059.58348 | 0.205865439 | 0.19797161 | 1.03987357 | 0.29839864 | 0.83973489 |
| MKL1 | 349.118097 | -0.049953276 | 0.18341154 | -0.2723562 | 0.78534813 | 0.97423208 |
| SLC25A17 | 79.8427659 | -0.145217277 | 0.20439129 | -0.7104866 | 0.47740242 | 0.91864961 |
| ST13 | 189.241825 | 0.349338011 | 0.2933781 | 1.19074331 | 0.23375437 | 0.79633808 |
| XPNPEP3 | 137.870761 | -0.095747798 | 0.22071675 | -0.4338039 | 0.66443081 | 0.95505623 |
| RBX1 | 107.646163 | -0.420227439 | 0.1842987 | -2.2801433 | 0.02259919 | 0.39240055 |
| EP300 | 290.063105 | 0.036241536 | 0.15438281 | 0.23475111 | 0.81440191 | 0.97937868 |
| L3MBTL2 | 29.9487192 | -0.285208672 | 0.39256137 | -0.7265327 | 0.46751227 | 0.91462428 |
| RANGAP1 | 56.8714841 | -0.183115514 | 0.463633 | -0.3949579 | 0.69287396 | 0.96220115 |
| ZC3H7B | 138.804801 | -0.11776493 | 0.3493196 | -0.3371266 | 0.73602146 | 0.9698508 |
| TEF | 35.7355117 | -0.246108256 | 0.6496473 | -0.3788336 | 0.7048114 | 0.96462186 |
| TOB2 | 77.8619743 | 0.045121765 | 0.22356533 | 0.2018281 | 0.84005111 | 0.98464453 |
| ACO2 | 124.826299 | -0.03991721 | 0.21111397 | -0.189079 | 0.85003093 | 0.9858744 |
| POLR3H | 30.1408612 | -0.195900895 | 0.33056971 | -0.592616 | 0.55343818 | 0.93523904 |
| DESI1 | 63.6013597 | -0.596173725 | 0.29992099 | -1.9877692 | 0.04683722 | 0.51099511 |
| XRCC6 | 197.951527 | -0.309828325 | 0.30997408 | -0.9995298 | 0.31753812 | 0.8490543 |
| SNU13 | 161.260502 | -0.398641103 | 0.3504631 | -1.1374696 | 0.25534204 | 0.80897837 |
| SREBF2 | 353.207461 | -0.069692099 | 0.39499245 | -0.1764391 | 0.85994901 | 0.9864199 |
| WBP2NL | 63.0594311 | 0.326048504 | 0.55201644 | 0.59064998 | 0.55475497 | 0.93523904 |
| SMDT1 | 81.0410894 | 0.1850477 | 0.52403956 | 0.35311781 | 0.72400012 | 0.96779196 |
| NDUFA6 | 53.4898522 | -0.31540175 | 0.35823774 | -0.8804258 | 0.37862868 | 0.87970706 |
| TCF20 | 254.257632 | -0.002027089 | 0.26905883 | -0.007534 | 0.9939888 | 0.99866813 |
| RRP7A | 48.5108602 | -0.379553302 | 0.41332796 | -0.9182861 | 0.35846912 | 0.87316144 |
| POLDIP3 | 78.1464148 | -0.171859759 | 0.25555479 | -0.6724967 | 0.50126752 | 0.92484445 |
| CYB5R3 | 167.628213 | 0.777570998 | 0.4146155 | 1.87540263 | 0.06073735 | 0.56071154 |
| A4GALT | 156.84557 | -0.062297771 | 0.39732534 | -0.1567928 | 8.75E-01 | 0.98766275 |
| ARFGAP3 | 168.487411 | -0.129945417 | 0.28503669 | -0.4558901 | 0.64846898 | 0.95027667 |

| | | | | | | |
|---|---|---|---|---|---|---|
| PACSIN2 | 335.543766 | -0.326903707 | 0.21259211 | -1.5377039 | 0.12412104 | 0.69439514 |
| TSPO | 123.008546 | 0.298514789 | 0.54822005 | 0.54451637 | 0.58608619 | 0.9423304 |
| EFCAB6 | 220.954703 | 0.614580766 | 0.3509657 | 1.75111345 | 0.07992637 | 0.60643125 |
| NUP50-DT | 98.4727694 | 0.514083894 | 0.35696224 | 1.44016323 | 0.14982122 | 0.72246729 |
| NUP50 | 61.6607904 | -0.287873311 | 0.23914157 | -1.2037778 | 0.22867547 | 0.7931636 |
| KIAA0930 | 57.5589372 | -0.273027233 | 0.47265506 | -0.5776458 | 0.56350325 | 0.93809297 |
| FAM118A | 40.3132548 | -0.486204432 | 0.48927442 | -0.9937254 | 0.32035656 | 0.85075884 |
| ATXN10 | 564.366464 | -0.303239299 | 0.17563689 | -1.7265126 | 0.08425524 | 0.61656297 |
| WNT7B | 37.7054013 | -1.688722514 | 1.24291342 | -1.3586807 | 0.17424778 | 0.74910274 |
| MIRLET7BHG | 182.462787 | -0.235844879 | 0.29311391 | -0.8046185 | 0.42103985 | 0.90012434 |
| PPARA | 160.014128 | 0.518935903 | 0.17299511 | 2.99971431 | 0.00270233 | 0.17081506 |
| TTC38 | 72.6579129 | 1.239968611 | 0.31110716 | 3.98566404 | 6.73E-05 | 0.01895844 |
| TRMU | 53.3729494 | -0.280514582 | 0.2346884 | -1.195264 | 0.23198392 | 0.79471003 |
| CELSR1 | 225.575223 | -0.854070524 | 0.37460565 | -2.2799189 | 0.0226125 | 0.39240055 |
| GRAMD4 | 61.5276136 | 0.353374949 | 0.29609245 | 1.19346153 | 0.23268867 | 0.79519257 |
| CERK | 46.1677347 | 0.54224129 | 0.29238377 | 1.85455329 | 0.06366004 | 0.57043925 |
| TBC1D22A | 410.588544 | -0.069956399 | 0.28419442 | -0.2461568 | 0.80556083 | 0.97930627 |
| BRD1 | 99.5757917 | -0.290222047 | 0.31508381 | -0.9210948 | 0.35700095 | 0.87316144 |
| ZBED4 | 66.7248669 | -0.562507083 | 0.44256384 | -1.2710191 | 0.20372186 | 0.77155952 |
| PIM3 | 127.623482 | -0.39802264 | 0.4338069 | -0.9175111 | 0.35887489 | 0.87316144 |
| TRABD | 41.5718669 | 0.013197476 | 0.34989548 | 0.03771834 | 0.96991225 | 0.99681784 |
| SELENOO | 80.405314 | -0.287102971 | 0.25209891 | -1.1388505 | 0.25476551 | 0.80897837 |
| PLXNB2 | 336.627382 | 0.330304454 | 0.32708244 | 1.00985076 | 0.3125668 | 0.84826686 |
| DENND6B | 56.6449244 | 0.771090047 | 0.44873257 | 1.71837327 | 0.08572855 | 0.62164558 |
| PPP6R2 | 357.181458 | 0.179856917 | 0.20434315 | 0.88017103 | 0.37876667 | 0.87973265 |
| SBF1 | 27.1895233 | 0.340431713 | 0.33334573 | 1.02125716 | 0.30713262 | 0.84488735 |
| LMF2 | 40.2077186 | 0.353767312 | 0.41578779 | 0.85083623 | 0.39486034 | 0.88760715 |
| RABL2B | 61.1644212 | 0.457525864 | 0.23526503 | 1.94472531 | 0.05180805 | 0.52794606 |
| CU633967.1 | 48.5892436 | 0.789584371 | 0.44982623 | 1.75530977 | 0.07920634 | 0.60556378 |
| FP671120.1 | 853.865247 | -0.894029565 | 0.3477288 | -2.5710541 | 0.01013895 | 0.29271368 |
| FP236383.1 | 489.157491 | -0.903799665 | 0.47644228 | -1.8969762 | 0.05783108 | 0.5484791 |
| CR381653.1 | 65.4974238 | -0.595416772 | 0.54957526 | -1.0834126 | 0.27862531 | 0.8229426 |
| HSPA13 | 30.1745696 | 0.335550845 | 0.38868127 | 0.86330593 | 0.38796928 | 0.8853359 |
| NRIP1 | 286.735004 | -0.489593398 | 0.30053248 | -1.6290865 | 0.10329471 | 0.65288882 |
| USP25 | 579.548012 | -0.534049721 | 0.3153329 | -1.6936061 | 0.09034016 | 0.6330877 |
| MIR99AHG | 723.254106 | 0.266954092 | 0.48069349 | 0.555352 | 0.57865392 | 0.94034284 |
| CXADR | 158.544171 | 0.167249743 | 0.29470028 | 0.56752488 | 0.57035763 | 0.93998433 |
| BTG3 | 54.929253 | 0.371583945 | 0.32345939 | 1.14878082 | 0.25064637 | 0.80680142 |
| C21orf91 | 30.6947864 | 0.047771074 | 0.35969661 | 0.13280936 | 0.89434415 | 0.98842187 |
| CHODL | 68.2196684 | 1.578930327 | 0.5942515 | 2.6570069 | 0.00788378 | 0.27165021 |
| MRPL39 | 34.3927823 | 0.225759089 | 0.33259268 | 0.67878551 | 0.49727378 | 0.92398825 |
| ATP5PF | 114.557259 | 0.004280632 | 0.29584343 | 0.01446925 | 0.98845561 | 0.99820747 |
| GABPA | 56.3098633 | 0.056808627 | 0.23043973 | 0.24652271 | 0.80527763 | 0.97930627 |
| APP | 1451.78866 | 0.045774818 | 0.26405231 | 0.17335511 | 0.8623723 | 0.98669442 |
| ADAMTS1 | 654.52106 | 0.076190006 | 0.65566832 | 0.11620205 | 0.90749241 | 0.99100813 |
| LINC00113 | 40.5666297 | -0.495255245 | 1.25687313 | -0.3940376 | 0.6935533 | 0.96220115 |
| AF165147.1 | 59.4094846 | 0.229602281 | 0.49813396 | 0.46092477 | 0.64485258 | 0.94954229 |
| LTN1 | 121.830825 | 0.065647439 | 0.20701599 | 0.31711289 | 0.75115794 | 0.97058015 |
| USP16 | 87.9770826 | -0.162113148 | 0.22969619 | -0.705772 | 0.48032991 | 0.91971919 |
| CCT8 | 72.5359763 | 0.054180485 | 0.28661708 | 0.18903439 | 0.85006586 | 0.9858744 |
| MAP3K7CL | 36.1540032 | 0.386915864 | 0.52438069 | 0.73785299 | 0.46060379 | 0.91368087 |
| BACH1 | 478.600617 | -0.52281504 | 0.44174982 | -1.1835093 | 0.23660735 | 0.79958336 |
| CLDN8 | 21.5952631 | 0.035157521 | 0.89445869 | 0.03930592 | 0.96864649 | 0.99681784 |
| SOD1 | 166.542402 | 0.340455913 | 0.48025025 | 0.70891355 | 0.47837812 | 0.91941895 |

| | | | | | | |
|---|---|---|---|---|---|---|
| SCAF4 | 133.072303 | -0.070039175 | 0.23746021 | -0.2949512 | 0.76803116 | 0.97237252 |
| HUNK | 245.972464 | -0.332299374 | 0.63793476 | -0.5208987 | 0.60243736 | 0.94683415 |
| URB1 | 66.2586081 | -0.3966026 | 0.4128964 | -0.9605378 | 0.33678462 | 0.86431445 |
| EVA1C | 390.64979 | 0.023353804 | 0.69688569 | 0.03351167 | 0.97326656 | 0.99681784 |
| SYNJ1 | 88.5457211 | 0.478831646 | 0.23308936 | 2.05428354 | 0.03994826 | 0.48194503 |
| PAXBP1 | 217.429122 | 0.193374477 | 0.17183051 | 1.12537919 | 0.26042839 | 0.81263292 |
| C21orf62-AS | 77.9188946 | 0.585793492 | 0.59355649 | 0.98692121 | 0.32368126 | 0.85414403 |
| IFNAR1 | 133.293613 | 0.303626435 | 0.26493719 | 1.14603178 | 0.251782 | 0.80770916 |
| IFNGR2 | 203.135903 | 0.463777961 | 0.25448449 | 1.82242131 | 0.06839108 | 0.58250536 |
| TMEM50B | 119.739953 | 0.004001952 | 0.3927155 | 0.01019046 | 0.99186933 | 0.99866813 |
| GART | 86.9764986 | -0.35481661 | 0.27697298 | -1.2810513 | 0.20017563 | 0.77110248 |
| SON | 416.316948 | 0.064544982 | 0.1740839 | 0.37076939 | 0.7108093 | 0.96587538 |
| AP000311.1 | 77.4877662 | 0.856183993 | 0.57617373 | 1.48598234 | 0.13728379 | 0.71044919 |
| ITSN1 | 262.207727 | 0.295926774 | 0.30211864 | 0.97950518 | 0.32733043 | 0.85650224 |
| LINC00649 | 30.8924642 | 0.185375674 | 0.77985503 | 0.2377053 | 0.81210967 | 0.97930627 |
| MRPS6 | 485.459122 | -1.125394627 | 0.75383653 | -1.4928895 | 0.13546612 | 0.7063854 |
| LINC00310 | 39.8777684 | -1.400197507 | 0.66106584 | -2.1180908 | 3.42E-02 | 0.45111962 |
| AP000317.1 | 56.1244973 | -1.460557017 | 0.45425244 | -3.2152981 | 0.00130309 | 0.11421765 |
| RCAN1 | 71.5153194 | 0.122013297 | 0.53519714 | 0.22797823 | 0.81966316 | 0.98079697 |
| CLIC6 | 51.8148634 | 0.885790677 | 0.76966151 | 1.15088342 | 0.2497802 | 0.80672756 |
| RUNX1 | 244.724896 | 0.221611349 | 0.43005829 | 0.51530538 | 6.06E-01 | 0.94708529 |
| SETD4 | 50.6794634 | -0.149285795 | 0.2833235 | -0.5269093 | 0.59825656 | 0.94683415 |
| CBR1 | 63.6389958 | -0.282060661 | 0.28184679 | -1.0007588 | 3.17E-01 | 0.8490543 |
| DOPEY2 | 63.8331068 | -0.642381947 | 0.38942478 | -1.6495662 | 0.0990317 | 0.64357166 |
| MORC3 | 181.091068 | 0.032306141 | 0.20716148 | 0.15594666 | 0.87607507 | 0.98766275 |
| CLDN14 | 111.275014 | -0.812415269 | 0.75952895 | -1.0696304 | 0.2847857 | 0.82918183 |
| SIM2 | 361.675629 | -2.235069069 | 1.33270841 | -1.6770878 | 0.09352531 | 0.63721577 |
| HLCS | 314.802015 | -0.26153457 | 0.34044974 | -0.7682032 | 0.44236649 | 0.90601423 |
| PIGP | 25.1995296 | 0.0882644 | 0.45312975 | 0.19478836 | 0.84555864 | 0.98520136 |
| TTC3 | 478.391297 | 0.121195859 | 0.23509646 | 0.51551546 | 0.60619287 | 0.94708529 |
| VPS26C | 81.034811 | -0.189290075 | 0.21576262 | -0.8773071 | 0.38031987 | 0.8807629 |
| DYRK1A | 658.521565 | -0.034075626 | 0.1606646 | -0.2120917 | 0.8320355 | 0.98298817 |
| KCNJ15 | 587.902353 | -0.497045364 | 0.54556703 | -0.911062 | 0.36226273 | 0.8752623 |
| ETS2 | 222.59247 | -1.024072942 | 0.42625632 | -2.4024815 | 0.01628426 | 0.34282836 |
| BRWD1 | 528.297452 | 0.188661991 | 0.11713329 | 1.61066078 | 0.10725368 | 0.65699665 |
| HMGN1 | 219.935592 | -0.322925212 | 0.24290613 | -1.3294239 | 0.18370817 | 0.75689793 |
| WRB | 56.378624 | 0.082994075 | 0.22992889 | 0.3609554 | 0.71813279 | 0.9674421 |
| PCP4 | 66.1668509 | 3.769571853 | 1.2764847 | 2.95308815 | 0.00314612 | 0.18154635 |
| BACE2 | 281.768737 | -0.625572423 | 0.38075621 | -1.6429737 | 0.10038838 | 0.64698063 |
| MX1 | 191.062352 | -4.93900888 | 1.3613209 | -3.6281004 | 0.00028551 | 0.0448351 |
| TMPRSS2 | 96.4455575 | -0.643062894 | 0.69114404 | -0.9304325 | 0.35214718 | 0.8720098 |
| RIPK4 | 53.1179557 | -0.00533886 | 0.41811756 | -0.0127688 | 0.98981225 | 0.99839055 |
| PRDM15 | 24.5398459 | -0.854720787 | 0.43913425 | -1.9463769 | 0.05160948 | 0.52722328 |
| C2CD2 | 68.3941435 | -0.215167858 | 0.27419919 | -0.7847137 | 0.43262145 | 0.90514499 |
| ZBTB21 | 46.2045176 | 0.417940433 | 0.37616958 | 1.11104262 | 0.26655001 | 0.81444418 |
| ABCG1 | 260.280515 | 1.397296986 | 1.08924617 | 1.28281102 | 0.19955829 | 0.77110248 |
| RSPH1 | 44.9243794 | 1.100396093 | 0.33864901 | 3.2493705 | 0.00115661 | 0.10756448 |
| SLC37A1 | 93.9884509 | 0.145021795 | 0.28033615 | 0.51731394 | 0.60493703 | 0.94684506 |
| LINC01671 | 27.7010716 | -0.154334983 | 0.62441694 | -0.2471666 | 0.80477933 | 0.97930627 |
| PDE9A | 330.715702 | 0.147255232 | 0.41319448 | 0.35638238 | 7.22E-01 | 0.96779196 |
| WDR4 | 30.0580344 | -0.975581565 | 0.54251951 | -1.7982424 | 0.07213861 | 0.59090333 |
| NDUFV3 | 102.952715 | 0.111200743 | 0.31577977 | 0.3521465 | 0.72472839 | 0.96781837 |
| PKNOX1 | 109.567509 | 0.146577384 | 0.35375804 | 0.41434361 | 0.67862248 | 0.95924874 |
| RRP1B | 64.3520387 | -0.238561705 | 0.30110964 | -0.7922752 | 0.42820021 | 0.90272191 |

| | | | | | | |
|---|---|---|---|---|---|---|
| PDXK | 555.886474 | -0.55208411 | 0.42601095 | -1.2959388 | 1.95E-01 | 0.76968045 |
| CSTB | 233.780968 | -0.586010158 | 0.51906675 | -1.1289688 | 0.258911 | 0.81263292 |
| AGPAT3 | 177.218864 | -0.288453501 | 0.1719992 | -1.677063 | 0.09353016 | 0.63721577 |
| TRAPPC10 | 129.222104 | -0.251719453 | 0.19390783 | -1.2981397 | 0.19423934 | 0.7687432 |
| PFKL | 135.681635 | 0.38118262 | 0.27327963 | 1.39484459 | 0.16306271 | 0.74167981 |
| C21orf2 | 35.6212601 | 0.099134381 | 0.448745 | 0.22091473 | 0.82515883 | 0.98094663 |
| UBE2G2 | 159.157536 | 0.191433681 | 0.27094933 | 0.70652945 | 0.47985894 | 0.91971919 |
| SUMO3 | 70.1511012 | -0.310014691 | 0.41300819 | -0.750626 | 0.45287776 | 0.91077355 |
| PTTG1IP | 291.702898 | 0.319459726 | 0.41483624 | 0.77008636 | 0.44124867 | 0.90561284 |
| FAM207A | 31.3184829 | -0.443547595 | 0.40692735 | -1.0899921 | 0.27571663 | 0.82164863 |
| ADARB1 | 145.16179 | 0.550337124 | 0.75933192 | 0.7247649 | 0.46859627 | 0.91462428 |
| POFUT2 | 42.293724 | -0.019145628 | 0.2564784 | -0.0746481 | 0.94049469 | 0.99526299 |
| COL18A1 | 397.043151 | -0.093656532 | 0.28553368 | -0.3280052 | 0.74290773 | 0.96992138 |
| COL6A1 | 53.5490166 | 0.761600899 | 0.63509414 | 1.19919372 | 0.23045263 | 0.79391882 |
| COL6A2 | 60.9355472 | 1.060097622 | 1.42187297 | 0.74556423 | 0.4559307 | 0.91081547 |
| LSS | 52.8123196 | 0.824121982 | 0.48386958 | 1.70319031 | 0.08853246 | 0.62958606 |
| MCM3AP-AS | 38.2194706 | 0.295111047 | 0.33264114 | 0.88717543 | 0.37498446 | 0.8769473 |
| MCM3AP | 119.251163 | -0.029052791 | 0.22284975 | -0.1303694 | 0.89627416 | 0.98842187 |
| PCNT | 59.3229666 | 0.075657023 | 0.29248834 | 0.2586668 | 0.79589234 | 0.97524554 |
| DIP2A | 152.328819 | -0.17457106 | 0.18448991 | -0.9462364 | 0.34402804 | 0.86969477 |
| PRMT2 | 198.263055 | 0.158122383 | 0.2990674 | 0.52871822 | 0.59700094 | 0.94683415 |
| AC011043.1 | 55.9380946 | 0.648723939 | 0.3457704 | 1.87616966 | 6.06E-02 | 5.61E-01 |
| AL592183.1 | 146.293065 | 0.145279448 | 0.50255975 | 0.28907896 | 0.77252096 | 0.97329715 |

**Thick ascending limbs**

| | baseMean | log2 fold cha | lfcSE | stat | pvalue | padj |
|---|---|---|---|---|---|---|
| AL627309.1 | 46.4832515 | -1.0333059 | 0.5163351 | -2.0012313 | 0.04536747 | 0.3886245 |
| AL669831.5 | 439.99619 | -0.2103515 | 0.22730744 | -0.9254054 | 0.35475506 | 0.77252365 |
| SAMD11 | 43.3508878 | -0.3726436 | 0.92167179 | -0.4043127 | 0.68598277 | 0.92053037 |
| NOC2L | 246.032503 | 0.03413814 | 0.37083843 | 0.09205664 | 0.92665304 | 0.98243518 |
| KLHL17 | 42.5399791 | -0.2117019 | 0.43586293 | -0.4857074 | 0.62717462 | 0.89919293 |
| HES4 | 107.829312 | -0.0507462 | 0.5474385 | -0.0926976 | 0.92614378 | 0.98234748 |
| ISG15 | 1233.31999 | 0.3110828 | 1.35933064 | 0.22884999 | 0.81898551 | 0.96006637 |
| AGRN | 623.565741 | 0.59380893 | 0.48782393 | 1.21726078 | 0.22350501 | 0.67624946 |
| C1orf159 | 358.43042 | -0.1980774 | 0.28980082 | -0.683495 | 0.49429409 | 0.84882719 |
| LINC01342 | 92.9066276 | 0.26918023 | 0.56101221 | 0.47981171 | 0.63136128 | 0.90011746 |
| AL390719.2 | 226.481438 | -0.2422605 | 0.46561791 | -0.5202989 | 0.60285527 | 0.88971776 |
| TTLL10 | 46.6818208 | -1.4160014 | 0.34006047 | -4.1639695 | 3.13E-05 | 0.00860682 |
| SDF4 | 373.668797 | 0.17492772 | 0.23420352 | 0.74690474 | 0.45512106 | 0.83121393 |
| UBE2J2 | 376.948358 | -0.2526698 | 0.18752254 | -1.3474101 | 0.1778482 | 0.63179521 |
| ACAP3 | 170.899697 | -0.2200588 | 0.29669599 | -0.741698 | 0.45827031 | 0.8331421 |
| INTS11 | 425.128513 | -0.0816001 | 0.22765292 | -0.358441 | 0.7200133 | 0.93001079 |
| CPTP | 67.4225987 | 0.30610106 | 0.40783036 | 0.75055975 | 0.45291765 | 0.82976273 |
| DVL1 | 146.978285 | 0.2054307 | 0.29780513 | 0.68981586 | 0.49031 | 0.84614483 |
| AURKAIP1 | 378.462361 | 0.47577015 | 0.46591822 | 1.02114519 | 0.30718566 | 0.74341546 |
| CCNL2 | 1074.37096 | -0.2558999 | 0.22655567 | -1.1295234 | 0.25867709 | 0.7090404 |
| MRPL20 | 464.634064 | 0.25733656 | 0.24474083 | 1.0514656 | 0.2930448 | 0.73527014 |
| VWA1 | 143.614962 | 1.65063423 | 0.64445351 | 2.56129294 | 0.01042834 | 0.20713973 |
| ATAD3C | 93.6429029 | 0.91133815 | 0.4433237 | 2.05569464 | 0.03981196 | 0.37279391 |
| ATAD3B | 281.766483 | -0.2239106 | 0.25661121 | -0.8725673 | 0.38289896 | 0.79039105 |
| ATAD3A | 140.836864 | 0.01645051 | 0.30453128 | 0.05401911 | 0.95691994 | 0.99065996 |
| SSU72 | 472.675214 | 0.01071198 | 0.21079645 | 0.05081668 | 0.9594716 | 0.99121386 |
| MIB2 | 194.819587 | -0.0735994 | 0.3047452 | -0.2415113 | 0.80915887 | 0.95617995 |
| CDK11B | 263.712491 | -0.357918 | 0.28294335 | -1.2649812 | 0.20587805 | 0.65813563 |
| FO704657.1 | 41.2857048 | -0.0246161 | 0.61138173 | -0.0402631 | 0.96788341 | 0.99248831 |
| SLC35E2B | 407.982983 | -0.7645788 | 0.28676545 | -2.6662164 | 0.00767103 | 0.17544898 |
| CDK11A | 155.127438 | -0.4010666 | 0.37278322 | -1.0758708 | 0.28198504 | 0.72604837 |
| SLC35E2A | 423.599377 | -0.1622986 | 0.29730824 | -0.5458933 | 0.5851393 | 0.88487193 |
| NADK | 324.730353 | 0.00934591 | 0.19652832 | 0.04755501 | 0.96207089 | 0.99223192 |
| GNB1 | 1981.08381 | -0.1973965 | 0.17796755 | -1.1091713 | 2.67E-01 | 0.71749473 |
| PRKCZ | 664.806574 | -0.4655067 | 0.31006278 | -1.5013305 | 0.13327011 | 0.57682553 |
| FAAP20 | 265.69668 | 0.2417197 | 0.3135839 | 0.77082943 | 0.44080804 | 0.82265742 |
| SKI | 709.808908 | 0.23757617 | 0.31192227 | 0.76165182 | 0.44626784 | 0.82571115 |
| MORN1 | 337.484844 | -0.1187118 | 0.26698694 | -0.4446351 | 0.65658347 | 0.90974386 |
| RER1 | 536.393505 | 0.44901392 | 0.28010539 | 1.60301776 | 0.10893073 | 0.53262539 |
| PEX10 | 137.27454 | -0.044546 | 0.31904973 | -0.1396208 | 0.88895964 | 0.97729111 |
| PANK4 | 84.762803 | -0.1511641 | 0.28467062 | -0.5310139 | 0.59540912 | 0.88859628 |
| TNFRSF14 | 152.281098 | 0.63600235 | 0.55325976 | 1.1495547 | 0.25032733 | 0.70099977 |
| FAM213B | 66.6313488 | 0.61890016 | 0.45413387 | 1.36281436 | 0.17294103 | 0.6286116 |
| LINC00982 | 1140.99666 | -0.6008462 | 0.81317394 | -0.7388902 | 0.4599737 | 0.83401408 |
| PRDM16 | 3088.24604 | -0.542251 | 0.39678823 | -1.3666006 | 0.17175053 | 0.6286116 |
| ARHGEF16 | 265.815045 | -0.153095 | 0.29472376 | -0.5194525 | 0.60344524 | 0.89001883 |
| TPRG1L | 104.351169 | 0.41866961 | 0.34193698 | 1.22440576 | 0.2207992 | 0.67356503 |
| WRAP73 | 125.658482 | -0.5282683 | 0.24250694 | -2.1783638 | 0.02937896 | 0.33012415 |
| LRRC47 | 161.011643 | 0.06547291 | 0.27267443 | 0.24011387 | 0.81024198 | 0.95679908 |
| CEP104 | 549.366667 | -0.4428215 | 0.23709675 | -1.8676825 | 0.06180634 | 0.43264436 |
| DFFB | 105.67844 | 0.00304801 | 0.29205521 | 0.01043642 | 0.99167309 | 0.99772135 |
| C1orf174 | 123.204754 | -0.427526 | 0.37169831 | -1.1501962 | 0.25006307 | 0.70069645 |

*This means that positive fold changes imply higher mean value in non-COVID AKI than in COVID AKI and vice versa.

| | | | | | | |
|---|---|---|---|---|---|---|
| AJAP1 | 407.943299 | 0.59877068 | 0.64776953 | 0.92435758 | 0.35530017 | 0.77296991 |
| NPHP4 | 362.142763 | 0.25189972 | 0.2470308 | 1.01970977 | 0.30786613 | 0.74370953 |
| RPL22 | 281.28664 | 0.04960319 | 0.24561744 | 0.20195306 | 0.83995342 | 0.96617873 |
| RNF207 | 271.340846 | -0.0868511 | 0.22646579 | -0.3835063 | 0.7013444 | 0.92446444 |
| ICMT | 162.41816 | 0.46254045 | 0.33857648 | 1.36613285 | 0.17189727 | 0.6286116 |
| ACOT7 | 60.7604388 | -0.1675228 | 0.49130742 | -0.3409735 | 0.73312355 | 0.93552616 |
| ESPN | 44.3894082 | -0.6694543 | 0.78606452 | -0.8516532 | 0.39440663 | 0.79876591 |
| PLEKHG5 | 65.4619324 | 0.20473623 | 0.52034805 | 0.39346017 | 0.69397964 | 0.92229578 |
| NOL9 | 406.977548 | -0.3597768 | 0.25590054 | -1.4059244 | 0.15974658 | 0.60996285 |
| ZBTB48 | 194.593165 | 0.00906134 | 0.26162194 | 0.03463523 | 0.97237061 | 0.99327086 |
| KLHL21 | 376.215506 | -0.1009309 | 0.34521125 | -0.2923743 | 0.77000049 | 0.94387751 |
| PHF13 | 103.887466 | 0.41411816 | 0.36995294 | 1.11938064 | 0.26297779 | 0.71328 |
| THAP3 | 106.412643 | 0.1364064 | 0.22303173 | 0.61160089 | 0.54080185 | 0.86582694 |
| DNAJC11 | 1691.04368 | -0.087184 | 0.3219704 | -0.2707826 | 7.87E-01 | 9.50E-01 |
| CAMTA1 | 2858.2314 | -0.5145727 | 0.19120122 | -2.6912627 | 0.00711821 | 0.17279872 |
| VAMP3 | 183.397645 | 0.28477156 | 0.23133688 | 1.2309821 | 0.21832956 | 0.67177152 |
| PER3 | 855.527492 | -0.8794902 | 0.84391312 | -1.0421572 | 0.29733879 | 0.73673761 |
| PARK7 | 637.475919 | 0.52480048 | 0.30237276 | 1.73560769 | 0.08263323 | 0.48594425 |
| ERRFI1 | 2866.89484 | 0.20761142 | 0.76488107 | 0.27142967 | 0.78606058 | 0.94978829 |
| AL034417.2 | 60.4835317 | -0.4109702 | 0.87663605 | -0.4688037 | 0.63920997 | 0.90330608 |
| RERE | 12891.9394 | -0.2086228 | 0.23030068 | -0.9058713 | 0.36500398 | 0.78095617 |
| AL096855.1 | 46.7644027 | -0.0212695 | 0.46400122 | -0.0458394 | 0.96343826 | 0.99224805 |
| ENO1 | 3209.537 | 0.2612206 | 0.40533219 | 0.64446053 | 0.51927685 | 0.85924115 |
| GPR157 | 213.193704 | 0.21372791 | 0.33612195 | 0.63586419 | 0.52486495 | 0.86059348 |
| MIR34AHG | 229.700908 | 1.26534073 | 0.54399081 | 2.32603327 | 0.02001678 | 0.2836927 |
| H6PD | 403.914126 | 0.19243792 | 0.21432449 | 0.89788114 | 0.36924892 | 0.78095617 |
| SPSB1 | 361.481046 | -0.57352 | 0.58175531 | -0.9858441 | 3.24E-01 | 7.54E-01 |
| SLC25A33 | 682.156208 | 0.00917913 | 0.40325668 | 0.02276251 | 0.98183971 | 0.99518016 |
| TMEM201 | 67.6395888 | 0.54924678 | 0.38179365 | 1.43859589 | 0.15026505 | 0.59663305 |
| PIK3CD | 202.007305 | -0.7290275 | 0.32337145 | -2.2544583 | 0.02416735 | 0.30500338 |
| CLSTN1 | 1079.04305 | 0.03510777 | 0.2453841 | 0.14307272 | 0.88623275 | 0.97690465 |
| CTNNBIP1 | 350.832358 | -0.2282259 | 0.31755033 | -0.7187077 | 0.47232101 | 0.83921324 |
| LZIC | 220.844897 | -0.1635984 | 0.21237424 | -0.7703309 | 0.44110364 | 0.82285053 |
| NMNAT1 | 586.655703 | 0.18900698 | 0.23949646 | 0.78918488 | 0.43000396 | 0.81675008 |
| UBE4B | 1710.44353 | -0.2634182 | 0.20421515 | -1.2899053 | 0.19708353 | 0.64964978 |
| KIF1B | 2279.97157 | -0.08105 | 0.20803382 | -0.3896003 | 0.69683216 | 0.92322348 |
| PGD | 235.509084 | -0.3639288 | 0.30869584 | -1.1789237 | 0.23842855 | 0.68982886 |
| DFFA | 339.495305 | 0.41690374 | 0.25115632 | 1.65993729 | 0.09692707 | 0.51327103 |
| PEX14 | 1127.06303 | 0.00844422 | 0.13234626 | 0.06380402 | 0.94912628 | 0.98919519 |
| CASZ1 | 1649.75102 | 0.93205696 | 0.5164558 | 1.80471776 | 0.07111886 | 0.46413489 |
| AL139423.1 | 96.2556696 | 0.53750971 | 0.55729116 | 0.96450428 | 0.33479317 | 0.76242642 |
| SRM | 122.461013 | -0.3806126 | 0.40903141 | -0.9305217 | 0.35210106 | 0.77151834 |
| EXOSC10 | 311.850959 | -0.4887641 | 0.23806471 | -2.0530725 | 0.04006555 | 0.37342745 |
| AL109811.1 | 54.2917914 | -0.5682732 | 0.27090604 | -2.0976762 | 0.03593376 | 0.35661333 |
| MTOR | 1082.43368 | -0.5330036 | 0.24194695 | -2.2029774 | 0.02759634 | 0.32099424 |
| UBIAD1 | 106.05204 | -0.5519715 | 0.26925754 | -2.0499762 | 0.04036676 | 0.37344844 |
| FBXO2 | 196.848861 | -0.3618557 | 0.49324956 | -0.7336159 | 0.46318288 | 0.83531269 |
| FBXO44 | 208.428961 | 0.0372851 | 0.30251551 | 0.12325021 | 0.90190897 | 0.97941893 |
| MAD2L2 | 81.7613741 | 1.09241383 | 0.39701121 | 2.75159444 | 0.00593059 | 0.15829975 |
| AGTRAP | 249.053822 | 0.12525588 | 0.37057099 | 0.33800779 | 7.35E-01 | 9.36E-01 |
| MTHFR | 251.91192 | 0.24306993 | 0.3350372 | 0.72550132 | 0.46814453 | 0.83825637 |
| CLCN6 | 346.559879 | -0.3518535 | 0.22674145 | -1.5517827 | 0.12071422 | 0.55468726 |
| KIAA2013 | 156.736317 | 0.50783999 | 0.24654163 | 2.0598549 | 0.03941241 | 0.37234059 |
| PLOD1 | 112.833193 | 0.26728109 | 0.31008134 | 0.8619709 | 0.38870353 | 0.79634939 |

| | | | | | | |
|---|---|---|---|---|---|---|
| MFN2 | 387.731519 | 0.07650092 | 0.2589636 | 0.29541188 | 0.76767926 | 0.94316227 |
| MIIP | 53.2930201 | 0.30072086 | 0.36260568 | 0.829333 | 0.406916 | 0.80535713 |
| VPS13D | 3632.92704 | -0.1117799 | 0.13722432 | -0.8145778 | 0.41531403 | 0.80966301 |
| DHRS3 | 622.590568 | -0.7826709 | 0.69393097 | -1.1278801 | 0.25937056 | 0.70921184 |
| PRDM2 | 1436.14374 | -0.1166779 | 0.28804712 | -0.4050653 | 0.68542955 | 0.92053037 |
| KAZN | 29880.1677 | -0.7097185 | 0.24313742 | -2.9190016 | 0.00351154 | 0.11452837 |
| KAZN-AS1 | 223.742557 | -0.9947034 | 0.80415546 | -1.2369541 | 0.21610412 | 0.66944638 |
| TMEM51-AS | 277.313584 | -0.2903541 | 0.62587045 | -0.4639205 | 0.64270473 | 0.90477318 |
| TMEM51 | 1622.63107 | -0.2987565 | 0.32113176 | -0.9303238 | 0.35220347 | 0.77151834 |
| C1orf195 | 78.1801608 | 0.07674551 | 0.41178602 | 0.18637231 | 0.85215281 | 0.97076831 |
| FHAD1 | 126.001906 | 0.77238244 | 0.32320534 | 2.3897577 | 0.01685949 | 0.25961947 |
| EFHD2 | 79.6567939 | -0.0652533 | 0.37433001 | -0.1743204 | 0.8616137 | 0.97249042 |
| CASP9 | 200.064495 | -0.1789803 | 0.22802475 | -0.784916 | 0.43250279 | 0.8177683 |
| DNAJC16 | 651.515765 | -0.4083011 | 0.18056823 | -2.2612013 | 0.02374679 | 0.30189738 |
| DDI2 | 648.021648 | -0.3591082 | 0.19030225 | -1.8870412 | 0.05915479 | 0.42419956 |
| PLEKHM2 | 479.247076 | -0.2145341 | 0.17321577 | -1.2385365 | 0.21551718 | 0.66885336 |
| FBLIM1 | 56.7718002 | 0.73691797 | 0.32949346 | 2.23651778 | 2.53E-02 | 3.11E-01 |
| UQCRHL | 83.667853 | 0.86148451 | 0.39045888 | 2.20633863 | 0.0273603 | 0.31979365 |
| AL450998.2 | 146.419548 | 0.05368626 | 0.39144013 | 0.13715064 | 0.89091173 | 0.97729111 |
| SPEN | 1497.30304 | -0.0261373 | 0.20176109 | -0.129546 | 0.89692565 | 0.97823597 |
| ZBTB17 | 174.040975 | -0.4856076 | 0.25286328 | -1.9204356 | 0.0548029 | 0.41684503 |
| HSPB7 | 132.849329 | -0.5630501 | 0.81601927 | -0.6899961 | 0.49019666 | 0.84614483 |
| CLCNKA | 1964.20145 | -0.0643322 | 1.09529173 | -0.0587352 | 0.95316299 | 0.98949465 |
| CLCNKB | 1268.41096 | 2.20033948 | 0.82150782 | 2.67841576 | 0.00739713 | 0.17324245 |
| FAM131C | 119.100522 | -0.2522326 | 0.75982144 | -0.331963 | 0.73991718 | 0.93552616 |
| EPHA2 | 147.419924 | 0.01376674 | 0.44022861 | 0.03127179 | 0.97505279 | 0.99408793 |
| FBXO42 | 1360.37562 | -0.3052007 | 0.26719223 | -1.1422515 | 0.25334951 | 0.70408029 |
| SZRD1 | 397.336621 | -0.053459 | 0.19604649 | -0.2726854 | 0.78509507 | 0.94978829 |
| SPATA21 | 49.1036907 | -0.0498361 | 0.43068582 | -0.1157134 | 0.90787972 | 0.98047754 |
| NECAP2 | 184.535241 | 0.05329559 | 0.22714706 | 0.23463031 | 0.81449568 | 0.95845643 |
| NBPF1 | 273.557513 | -0.6224548 | 0.27911412 | -2.2301086 | 0.02574023 | 0.31211455 |
| CROCC | 477.125842 | -0.6207341 | 0.36049625 | -1.7218878 | 0.08508984 | 0.49231347 |
| ATP13A2 | 160.601784 | -0.1337691 | 0.30217688 | -0.4426849 | 0.65799365 | 0.91037018 |
| SDHB | 729.477143 | 0.08319147 | 0.19094322 | 0.43568697 | 0.66306386 | 0.91138507 |
| PADI2 | 283.855099 | 0.72995296 | 0.6954609 | 1.04959598 | 0.29390391 | 0.73527014 |
| RCC2 | 272.544087 | -0.4062502 | 0.30682732 | -1.3240354 | 0.18549128 | 0.64030106 |
| ARHGEF10L | 2082.06486 | -0.0401039 | 0.42335729 | -0.0947283 | 0.92453061 | 0.98215577 |
| KLHDC7A | 81.0290699 | 1.12911391 | 0.89604081 | 1.26011439 | 0.2076281 | 0.6607067 |
| UBR4 | 1227.24946 | -0.2136633 | 0.18500408 | -1.1549114 | 0.24812669 | 0.69838949 |
| AL035413.1 | 292.85581 | 0.13245098 | 0.27519977 | 0.4812903 | 0.63031019 | 0.90005881 |
| EMC1 | 164.414003 | -0.1918585 | 0.23713143 | -0.8090807 | 0.41846876 | 0.81120543 |
| MRTO4 | 123.322192 | -0.3507572 | 0.34302679 | -1.0225358 | 0.30652737 | 0.74295884 |
| AKR7A2 | 163.357576 | 0.66764235 | 0.34609653 | 1.92906397 | 0.05372292 | 0.41326474 |
| PQLC2 | 63.0404902 | -0.2527984 | 0.34924779 | -0.7238369 | 0.46916588 | 0.83870782 |
| CAPZB | 1441.34879 | -0.2062445 | 0.24788201 | -0.8320271 | 0.40539365 | 0.80485885 |
| TMCO4 | 1131.63758 | 0.01827524 | 0.39896055 | 0.04580713 | 0.96346398 | 0.99224805 |
| OTUD3 | 248.217377 | -0.0639848 | 0.25734511 | -0.2486343 | 0.80364369 | 0.95450237 |
| UBXN10 | 50.7198729 | 0.60836704 | 0.43663509 | 1.39330771 | 0.16352676 | 0.61416689 |
| CAMK2N1 | 610.404493 | 0.31861652 | 0.47682267 | 0.66820758 | 0.5040011 | 0.85525037 |
| MUL1 | 110.272123 | 0.18345074 | 0.26344034 | 0.69636542 | 0.48620001 | 0.8447301 |
| CDA | 58.5483082 | 1.35582985 | 0.68045381 | 1.99253766 | 0.0463121 | 0.3904347 |
| PINK1 | 294.621842 | 0.14264639 | 0.27511751 | 0.51849258 | 0.60411464 | 0.89001883 |
| PINK1-AS | 44.1159384 | -0.4424524 | 0.38303254 | -1.1551301 | 0.24803715 | 0.69838949 |
| DDOST | 207.270636 | 0.53215638 | 0.27026931 | 1.96898565 | 0.04895474 | 0.39768952 |

| | | | | | | |
|---|---|---|---|---|---|---|
| HP1BP3 | 2076.22625 | 0.09563122 | 0.16379049 | 0.58386307 | 0.55931244 | 0.87295536 |
| EIF4G3 | 7477.46763 | -0.3166498 | 0.40291334 | -0.7859005 | 0.43192577 | 0.81761357 |
| ECE1 | 849.25907 | -0.2149252 | 0.36299201 | -0.5920936 | 0.55378791 | 0.87121136 |
| NBPF3 | 158.56914 | -0.3177303 | 0.31278882 | -1.0157983 | 0.30972544 | 0.74450725 |
| RAP1GAP | 1436.02404 | -0.0438339 | 0.64029052 | -0.0684594 | 0.94541994 | 0.98860449 |
| USP48 | 1317.10882 | -0.1104362 | 0.16014752 | -0.6895904 | 0.49045179 | 0.84614483 |
| HSPG2 | 539.985898 | 0.77367449 | 0.48929516 | 1.58120202 | 0.11383185 | 0.54008799 |
| CDC42 | 1028.48967 | 0.02989599 | 0.21523178 | 0.13890137 | 0.88952809 | 0.97729111 |
| ZBTB40 | 398.858329 | 0.02478322 | 0.32345222 | 0.07662096 | 0.93892508 | 0.98715467 |
| EPHB2 | 584.425577 | -1.3933817 | 0.36607867 | -3.8062357 | 0.0001411 | 0.01998715 |
| KDM1A | 833.547622 | 0.01379581 | 0.13111545 | 0.1052188 | 0.91620219 | 0.98195017 |
| AL031428.1 | 222.771759 | 0.15231933 | 0.46358634 | 0.32856733 | 0.74248274 | 0.93598004 |
| LUZP1 | 772.944826 | 0.16758044 | 0.17625644 | 0.95077623 | 0.34171798 | 0.76545664 |
| LINC01355 | 207.502083 | 0.44987327 | 0.22333773 | 2.01431824 | 0.04397613 | 0.38494416 |
| AL109936.6 | 66.2258145 | 0.36073381 | 0.45745605 | 0.78856496 | 0.43036632 | 0.81700326 |
| HNRNPR | 1189.34655 | 0.13472418 | 0.1237447 | 1.08872681 | 0.27627437 | 0.72241028 |
| ZNF436 | 51.4359527 | 0.27199153 | 0.53971352 | 0.50395537 | 0.61429274 | 0.89488165 |
| ZNF436-AS1 | 39.3549025 | 0.09082406 | 0.4770888 | 0.1903714 | 0.84901811 | 0.97017477 |
| TCEA3 | 561.557468 | 0.07226722 | 0.52550573 | 0.13751937 | 0.89062028 | 0.97729111 |
| ASAP3 | 535.433436 | 0.29849632 | 0.37355694 | 0.79906511 | 0.42425266 | 0.81452811 |
| ID3 | 49.7183759 | -0.4451064 | 0.6572387 | -0.677237 | 0.49825559 | 0.85092921 |
| RPL11 | 1333.02905 | 0.44989202 | 0.42640193 | 1.05508909 | 0.29138458 | 0.73349819 |
| ELOA | 209.376038 | -0.0718863 | 0.21879657 | -0.328553 | 0.74249357 | 0.93598004 |
| ELOA-AS1 | 286.884546 | -0.2530629 | 0.3647299 | -0.6938364 | 0.4877848 | 0.8451268 |
| PITHD1 | 196.922161 | 0.12183161 | 0.19863201 | 0.61335333 | 0.53964274 | 0.86566791 |
| LYPLA2 | 197.715037 | 0.3363515 | 0.33058868 | 1.017432 | 0.30894797 | 0.74399658 |
| GALE | 263.890284 | -0.5821234 | 0.2835297 | -2.0531303 | 4.01E-02 | 0.37342745 |
| HMGCL | 344.129233 | -0.2112195 | 0.32628938 | -0.6473379 | 0.51741325 | 0.85893097 |
| FUCA1 | 338.172373 | 0.25081433 | 0.35241277 | 0.71170615 | 0.47664676 | 0.84150767 |
| PNRC2 | 360.550642 | 0.16054235 | 0.28912915 | 0.55526173 | 0.57871565 | 0.8821758 |
| SRSF10 | 1431.48887 | -0.0658972 | 0.25369302 | -0.2597516 | 0.79505539 | 0.95272759 |
| IFNLR1 | 317.818634 | -0.019561 | 0.27828158 | -0.0702923 | 0.94396103 | 0.98793925 |
| STPG1 | 184.276652 | -0.0828549 | 0.25077497 | -0.3303954 | 0.74110119 | 0.93598004 |
| NIPAL3 | 633.34168 | -0.115772 | 0.30109942 | -0.3844976 | 0.70060971 | 0.92377257 |
| RCAN3 | 173.78702 | 0.05940124 | 0.40617432 | 0.14624569 | 0.88372744 | 0.97664137 |
| NCMAP | 102.325343 | 0.7840471 | 0.57727929 | 1.35817637 | 1.74E-01 | 6.29E-01 |
| SRRM1 | 946.855721 | -0.1938523 | 0.17818948 | -1.0879 | 0.27663927 | 0.72258811 |
| CLIC4 | 1062.13505 | -0.0058248 | 0.56253824 | -0.0103545 | 0.99173846 | 0.99772135 |
| SYF2 | 272.86708 | 0.27421956 | 0.19909442 | 1.37733427 | 0.16840892 | 0.62253512 |
| RSRP1 | 3062.31768 | 0.31613355 | 0.20063311 | 1.57567985 | 0.1150996 | 0.54371891 |
| RHD | 99.72733 | 0.11754807 | 1.02186069 | 0.11503336 | 0.90841868 | 0.98047754 |
| TMEM50A | 401.776813 | 0.2126599 | 0.25279326 | 0.84124039 | 0.40021328 | 0.80238662 |
| MACO1 | 786.026737 | -0.4907361 | 0.16061431 | -3.0553699 | 0.00224783 | 0.09083568 |
| LDLRAP1 | 156.428727 | 0.28377773 | 0.2772587 | 1.02351244 | 0.30606562 | 0.74290739 |
| MAN1C1 | 490.534412 | 1.01858028 | 0.68723005 | 1.48215329 | 0.13829951 | 0.58206378 |
| AL031280.1 | 43.5316474 | 0.91622254 | 0.62991609 | 1.45451522 | 0.14580352 | 0.59218144 |
| SELENON | 62.0630075 | 0.07206572 | 0.39631139 | 0.18184114 | 0.85570739 | 0.97118784 |
| AL020996.1 | 50.5492431 | 0.59078134 | 0.45217846 | 1.30652252 | 0.19137493 | 0.64571429 |
| MTFR1L | 308.88551 | 0.29772984 | 0.4811364 | 0.61880549 | 0.5360445 | 0.8642718 |
| AL033528.2 | 55.1405465 | 0.4331056 | 0.46265647 | 0.93612784 | 0.34920737 | 0.76982416 |
| PAQR7 | 157.900574 | 0.18591959 | 0.51111367 | 0.3637539 | 0.71604179 | 0.92890921 |
| STMN1 | 178.29508 | -0.2346714 | 0.43671362 | -0.5373577 | 0.59102059 | 0.8867913 |
| PAFAH2 | 220.648269 | 0.00549137 | 0.2086465 | 0.02631903 | 0.97900288 | 0.99455895 |
| PDIK1L | 99.8563535 | -0.0879797 | 0.2604404 | -0.3378114 | 0.73550532 | 0.93552616 |

| | | | | | | |
|---|---|---|---|---|---|---|
| ZNF593 | 115.218981 | 0.51748615 | 0.48350243 | 1.07028654 | 0.28449035 | 0.72738535 |
| CNKSR1 | 164.307658 | -0.4725465 | 0.28718788 | -1.6454263 | 0.09988193 | 0.51665771 |
| CEP85 | 165.721319 | 0.01064125 | 0.32388744 | 0.03285477 | 0.9737904 | 0.99345929 |
| SH3BGRL3 | 316.58541 | -0.0206214 | 0.47731818 | -0.0432027 | 9.66E-01 | 9.92E-01 |
| UBXN11 | 128.261238 | -0.0212135 | 0.29576019 | -0.0717254 | 0.94282041 | 0.98772472 |
| DHDDS | 389.333766 | -0.2365564 | 0.29396001 | -0.804723 | 0.42097956 | 0.81260593 |
| HMGN2 | 415.81012 | 0.24230081 | 0.25941131 | 0.93404104 | 0.35028272 | 0.77044053 |
| RPS6KA1 | 247.200303 | 0.21184581 | 0.25564576 | 0.82866937 | 0.40729152 | 0.80561751 |
| ARID1A | 1742.00268 | -0.1618805 | 0.26438071 | -0.6123009 | 0.5403387 | 0.86569031 |
| PIGV | 56.0597952 | 0.34488847 | 0.28743849 | 1.19986875 | 0.23019032 | 0.68251041 |
| ZDHHC18 | 129.293933 | -0.0794262 | 0.29086194 | -0.2730719 | 0.78479794 | 0.94978829 |
| SFN | 101.007161 | 0.1630064 | 0.88033474 | 0.18516412 | 8.53E-01 | 9.71E-01 |
| GPN2 | 203.222827 | -0.0967349 | 0.23002285 | -0.4205448 | 0.6740875 | 0.91636338 |
| GPATCH3 | 69.8203798 | -0.0360162 | 0.24254128 | -0.148495 | 0.8819521 | 0.97664137 |
| NUDC | 482.633392 | 0.01654814 | 0.1213498 | 0.13636722 | 0.89153098 | 0.97736183 |
| KDF1 | 86.9573896 | -0.0762818 | 0.23531513 | -0.3241688 | 0.74581021 | 0.93675395 |
| TRNP1 | 282.03366 | 0.89973967 | 0.56790293 | 1.58431949 | 0.11312103 | 0.53923339 |
| TENT5B | 43.7563377 | 0.6365914 | 0.86131176 | 0.73909522 | 0.45984918 | 0.83390278 |
| SLC9A1 | 281.201021 | -0.1773017 | 0.23026936 | -0.7699754 | 0.44131449 | 0.82285053 |
| WDTC1 | 638.068313 | -0.0545052 | 0.146123 | -0.3730093 | 0.7091415 | 0.92577218 |
| TMEM222 | 187.718233 | 0.08959774 | 0.22187526 | 0.40382035 | 0.68634483 | 0.92067928 |
| MAP3K6 | 76.2425555 | 0.29088296 | 0.76969136 | 0.37792157 | 0.70548886 | 0.92556842 |
| WASF2 | 1591.36499 | 0.03492369 | 0.17744722 | 0.1968117 | 0.8439749 | 0.96831902 |
| FO393419.2 | 41.8053953 | -0.0307559 | 0.59601765 | -0.0516023 | 0.95884556 | 0.99103216 |
| AHDC1 | 795.412983 | 0.16200625 | 0.28593125 | 0.56659161 | 0.57099167 | 0.87863805 |
| IFI6 | 971.449025 | 1.0379377 | 1.18268651 | 0.87761016 | 0.38015531 | 0.7880311 |
| FAM76A | 231.782771 | -0.1964209 | 0.24248714 | -0.8100261 | 0.41792515 | 0.81062751 |
| STX12 | 855.287486 | -0.1099383 | 0.24234854 | -0.4536372 | 0.65008998 | 0.90825486 |
| PPP1R8 | 212.382771 | -0.0862909 | 0.23391988 | -0.3688908 | 0.7122091 | 0.92748721 |
| RPA2 | 230.326667 | -0.0782532 | 0.19780946 | -0.3955987 | 0.69240106 | 0.92224682 |
| SMPDL3B | 47.2117088 | -0.0296033 | 0.50251805 | -0.0589099 | 9.53E-01 | 9.89E-01 |
| EYA3 | 1309.78674 | 0.05246526 | 0.14965872 | 0.350566 | 0.72591397 | 0.93301523 |
| PTAFR | 89.3919035 | 1.18128918 | 0.55320346 | 2.13536113 | 0.03273152 | 0.34532802 |
| DNAJC8 | 329.236997 | 0.03415766 | 0.16972617 | 0.20125159 | 0.84050186 | 0.96617873 |
| ATP5IF1 | 668.055922 | 0.56434551 | 0.36232695 | 1.55755874 | 0.11933789 | 0.55193772 |
| AL353622.1 | 73.559887 | -0.1424318 | 0.56438802 | -0.2523651 | 0.80075889 | 0.95370871 |
| SESN2 | 176.421255 | 0.52094777 | 0.31343029 | 1.66208496 | 0.09649574 | 0.51229242 |
| PHACTR4 | 2796.83078 | -0.0411411 | 0.21311354 | -0.1930477 | 0.84692163 | 0.96891207 |
| RCC1 | 329.430386 | 0.16644157 | 0.42311309 | 0.39337373 | 0.69404347 | 0.92229578 |
| TRNAU1AP | 493.228782 | -0.1313482 | 0.22283049 | -0.5894534 | 0.55555715 | 0.87184917 |
| SNHG12 | 114.995773 | -0.1461814 | 0.36801019 | -0.3972211 | 0.69120439 | 0.92206635 |
| TAF12 | 300.941201 | -0.2555352 | 0.22938715 | -1.1139908 | 0.26528314 | 0.71512755 |
| GMEB1 | 293.216948 | -0.5430055 | 0.27954034 | -1.9424942 | 0.0520773 | 0.40866415 |
| YTHDF2 | 729.443104 | 0.10993843 | 0.16720962 | 0.65748863 | 0.51086678 | 0.85762435 |
| EPB41 | 1990.75635 | 0.37428006 | 0.14807201 | 2.52768953 | 0.01148158 | 0.21884594 |
| SRSF4 | 2660.87287 | -0.1693703 | 0.15938504 | -1.0626489 | 0.2879412 | 0.7307234 |
| MECR | 301.944065 | -0.1286731 | 0.30768237 | -0.4182011 | 0.67580012 | 0.91649321 |
| PTPRU | 102.575507 | 0.28412942 | 0.50703335 | 0.5603762 | 5.75E-01 | 8.81E-01 |
| PUM1 | 3372.90429 | -0.1627445 | 0.1658722 | -0.9811437 | 0.32652187 | 0.75554697 |
| SNRNP40 | 326.95538 | -0.3393051 | 0.17621162 | -1.9255548 | 0.05415998 | 0.41520559 |
| ZCCHC17 | 665.058167 | -0.2269315 | 0.2016368 | -1.125447 | 0.26039967 | 0.71022491 |
| AL451070.1 | 99.4419507 | -0.0318251 | 0.41487925 | -0.0767093 | 0.93885478 | 0.98715467 |
| FABP3 | 110.878464 | 0.47781298 | 0.52154683 | 0.91614588 | 0.35959039 | 0.77668667 |
| SERINC2 | 909.479388 | -0.1788726 | 0.2943865 | -0.6076114 | 0.54344525 | 0.86622038 |

| | | | | | | |
|---|---|---|---|---|---|---|
| TINAGL1 | 249.176954 | 0.56123191 | 0.43553206 | 1.28861218 | 0.19753295 | 0.64964978 |
| PEF1 | 191.828678 | 0.13533929 | 0.28671545 | 0.47203349 | 0.63690289 | 0.90255504 |
| PTP4A2 | 305.064941 | -0.0777741 | 0.28182784 | -0.2759632 | 0.78257631 | 0.94919728 |
| KHDRBS1 | 1215.58138 | -0.4228105 | 0.21381939 | -1.9774187 | 0.04799432 | 0.39511605 |
| AL445248.1 | 55.8436976 | -0.1458738 | 0.4302669 | -0.339031 | 0.73458639 | 0.93552616 |
| TMEM39B | 132.423516 | -0.2926653 | 0.29407118 | -0.9952193 | 0.31962963 | 0.75094569 |
| KPNA6 | 1155.21906 | -0.0338651 | 0.17345285 | -0.1952409 | 0.84520434 | 0.96859165 |
| TXLNA | 325.234804 | 0.33778507 | 0.27150112 | 1.24413879 | 0.21344849 | 0.66763937 |
| DCDC2B | 55.0849034 | -0.2360117 | 0.40551335 | -0.5820072 | 0.56056183 | 0.873671 |
| TMEM234 | 118.533077 | -0.1533524 | 0.21843591 | -0.7020476 | 0.4826495 | 0.84407748 |
| EIF3I | 224.96332 | 0.53741241 | 0.34949549 | 1.53768054 | 0.12412675 | 0.56105491 |
| HDAC1 | 441.833448 | 0.23069875 | 0.20315709 | 1.13556831 | 0.25613728 | 0.70709713 |
| TSSK3 | 89.7270849 | 0.38931177 | 0.48145836 | 0.80860943 | 0.41873984 | 0.81125469 |
| FAM229A | 47.3276105 | 0.10336173 | 0.40210254 | 0.25705316 | 0.79713774 | 0.95281098 |
| BSDC1 | 545.767542 | -0.0676327 | 0.20149003 | -0.3356628 | 0.73712519 | 0.93552616 |
| ZBTB8A | 57.7399839 | -0.356454 | 0.41405525 | -0.8608851 | 0.38930132 | 0.79663535 |
| ZBTB8OS | 894.303585 | -0.061486 | 0.19085682 | -0.3221576 | 0.74733327 | 0.93737776 |
| RBBP4 | 862.39732 | 0.03989002 | 0.14507521 | 0.274961 | 0.7833462 | 0.94954758 |
| SYNC | 189.737444 | 0.16352175 | 0.41771968 | 0.39146287 | 0.69545513 | 0.92229578 |
| KIAA1522 | 890.479649 | 0.06179133 | 0.30282608 | 0.20404889 | 0.83831529 | 0.96566728 |
| YARS | 408.860007 | -0.550997 | 0.27254364 | -2.0216834 | 0.04320907 | 0.38262319 |
| S100PBP | 953.954052 | -0.2158289 | 0.12333124 | -1.7499937 | 0.08011939 | 0.48302012 |
| TMEM54 | 92.2089728 | 0.42834702 | 0.49819167 | 0.85980364 | 3.90E-01 | 0.79698174 |
| RNF19B | 398.968543 | -0.2890765 | 0.38759161 | -0.7458274 | 0.45577166 | 0.83130183 |
| AL020995.1 | 379.47875 | -0.2704963 | 0.71377426 | -0.3789662 | 0.70471298 | 0.92528913 |
| AK2 | 1058.26458 | 0.07765357 | 0.34621083 | 0.22429562 | 0.82252729 | 0.9608882 |
| AZIN2 | 169.497989 | -0.3433559 | 0.3867593 | -0.8877768 | 0.37466084 | 0.78426576 |
| TRIM62 | 141.906156 | -0.5137787 | 0.3487958 | -1.4730072 | 0.14074909 | 0.58629918 |
| ZNF362 | 278.570401 | -0.125953 | 0.30150977 | -0.417741 | 0.67613651 | 0.91650575 |
| AL513327.1 | 617.718928 | 0.00063663 | 0.61377508 | 0.00103724 | 0.9991724 | 0.99985365 |
| PHC2 | 2352.10526 | -0.348886 | 0.27516936 | -1.2678955 | 0.20483528 | 0.65806513 |
| AL513327.2 | 54.3917637 | 0.10842892 | 0.47744697 | 0.2271015 | 0.82034482 | 0.96020688 |
| ZSCAN20 | 60.8630913 | -0.3305322 | 0.26685984 | -1.2385986 | 0.21549418 | 0.66885336 |
| CSMD2 | 122.554511 | -2.664952 | 1.02304962 | -2.6049098 | 0.00918985 | 0.19453312 |
| SMIM12 | 286.029186 | -0.0280578 | 0.17862244 | -0.1570788 | 0.87518271 | 0.97604597 |
| AC114490.3 | 41.7399458 | -0.2216762 | 0.36340278 | -0.6100015 | 0.54186082 | 0.86582694 |
| ZMYM1 | 475.320861 | 0.13502162 | 0.14294964 | 0.94453977 | 0.34489389 | 0.76724655 |
| SFPQ | 2055.68217 | 0.03364169 | 0.22179787 | 0.15167725 | 0.8794415 | 0.9765717 |
| ZMYM4 | 3384.28615 | -0.0683906 | 0.14368414 | -0.4759785 | 0.63408968 | 0.90136076 |
| ZMYM4-AS1 | 80.0287945 | 0.0782442 | 0.67997062 | 0.11506998 | 0.90838966 | 0.98047754 |
| KIAA0319L | 1845.83219 | -0.2208564 | 0.13460583 | -1.6407639 | 0.10084644 | 0.51819907 |
| NCDN | 49.3037794 | 0.4224185 | 0.42385331 | 0.99661484 | 0.3189515 | 0.75094569 |
| PSMB2 | 334.55972 | 0.28622982 | 0.18579959 | 1.54052985 | 0.12343125 | 0.56085429 |
| AGO4 | 896.739708 | -0.1735181 | 0.24228027 | -0.7161874 | 0.47387566 | 0.84075535 |
| AGO1 | 280.213784 | -0.23994 | 0.26739303 | -0.8973308 | 0.3695424 | 0.78095617 |
| AGO3 | 2467.31387 | -0.1730024 | 0.17035098 | -1.0155646 | 0.30983677 | 0.74450725 |
| AL138787.2 | 141.289392 | -0.0959511 | 0.467771 | -0.2051242 | 0.83747512 | 0.96527001 |
| ADPRHL2 | 171.689987 | 0.18242751 | 0.41551525 | 0.43903926 | 0.66063309 | 0.91099119 |
| TRAPPC3 | 222.795914 | 0.09541736 | 0.19606452 | 0.48666305 | 0.62649713 | 0.89873262 |
| MAP7D1 | 164.830276 | 0.14408261 | 0.40093504 | 0.35936647 | 0.71932095 | 0.93001079 |
| THRAP3 | 1563.92863 | -0.0994588 | 0.16020294 | -0.6208299 | 0.53471154 | 0.86419602 |
| SH3D21 | 157.963162 | 1.24590578 | 0.48021426 | 2.59447892 | 0.00947344 | 0.19737334 |
| EVA1B | 170.333146 | 1.10353399 | 0.51720242 | 2.13365973 | 0.03287064 | 0.34596681 |
| AL591845.1 | 123.715231 | 1.49942198 | 0.46988014 | 3.1910733 | 0.00141745 | 0.06883507 |

| | | | | | | |
|---|---|---|---|---|---|---|
| STK40 | 427.789151 | 0.07213689 | 0.26659956 | 0.27058142 | 0.78671299 | 0.94978829 |
| LSM10 | 77.8773664 | 0.88558475 | 0.36323725 | 2.43803395 | 0.01476739 | 0.24869534 |
| OSCP1 | 127.395989 | 0.27250164 | 0.27064568 | 1.0068575 | 0.31400326 | 0.74777437 |
| MRPS15 | 168.762305 | 0.12243094 | 0.31699673 | 0.38622146 | 0.69933266 | 0.92367966 |
| LINC01137 | 146.63904 | 0.30346174 | 0.34606495 | 0.87689246 | 0.38054505 | 0.78824206 |
| ZC3H12A | 98.137825 | 0.83486362 | 0.41953128 | 1.98999136 | 4.66E-02 | 0.39097111 |
| MEAF6 | 338.287258 | -0.0322011 | 0.21875588 | -0.1472013 | 0.88297314 | 0.97664137 |
| SNIP1 | 160.226736 | -0.0285479 | 0.25709918 | -0.1110383 | 0.91158595 | 0.98125148 |
| DNALI1 | 116.942243 | 0.59318164 | 0.31149064 | 1.90433216 | 0.05686694 | 0.41872623 |
| GNL2 | 139.178436 | -0.3300712 | 0.21682159 | -1.5223171 | 0.12792963 | 0.56762585 |
| C1orf109 | 97.6596137 | -0.3619633 | 0.33748994 | -1.0725158 | 0.28348841 | 0.72668316 |
| YRDC | 50.9347582 | 0.08124287 | 0.35588467 | 0.22828426 | 0.81942526 | 0.96006637 |
| C1orf122 | 290.085457 | 0.59677738 | 0.48650446 | 1.22666373 | 0.219949 | 0.67299198 |
| MTF1 | 252.29115 | 0.21794965 | 0.19038625 | 1.14477624 | 0.25230186 | 0.70278705 |
| AL929472.2 | 106.118002 | 0.76747375 | 0.36568067 | 2.09875393 | 0.0358386 | 0.35661333 |
| INPP5B | 815.047291 | 0.35968156 | 0.30380347 | 1.18392839 | 0.23644141 | 0.68867797 |
| SF3A3 | 305.693652 | -0.0360073 | 0.23935248 | -0.1504364 | 0.88042031 | 0.97664137 |
| UTP11 | 80.7704052 | 0.02143222 | 0.31440701 | 0.06816713 | 0.9456526 | 0.98861299 |
| RRAGC | 181.882174 | -0.2196391 | 0.27741159 | -0.7917445 | 0.42850967 | 0.81511869 |
| AL139260.1 | 453.509526 | -0.5263325 | 0.3834504 | -1.372622 | 0.16986989 | 0.62571428 |
| RHBDL2 | 93.6604907 | -0.1977787 | 0.49756858 | -0.3974904 | 0.69100589 | 0.92206635 |
| AKIRIN1 | 1018.93917 | 0.18307712 | 0.29916034 | 0.61196987 | 0.54055769 | 0.86579684 |
| NDUFS5 | 635.398288 | 0.60166449 | 0.37541082 | 1.60268289 | 0.10900468 | 0.53262539 |
| MACF1 | 6891.41 | -0.1878492 | 0.28940265 | -0.6490928 | 0.51627842 | 0.85888182 |
| PABPC4 | 256.061004 | -0.1127228 | 0.23071039 | -0.4885902 | 0.62513182 | 0.89873262 |
| HPCAL4 | 113.567035 | 1.17677505 | 0.96501812 | 1.21943311 | 0.22267985 | 0.67568617 |
| PPIE | 206.569681 | 0.41106572 | 0.36943845 | 1.11267714 | 0.26584712 | 0.71641758 |
| TRIT1 | 609.76063 | -0.2832243 | 0.15337029 | -1.8466698 | 0.06479502 | 0.44041719 |
| CAP1 | 523.131065 | 0.05724623 | 0.18493753 | 0.30954359 | 7.57E-01 | 9.40E-01 |
| PPT1 | 150.100041 | 0.41308958 | 0.27442613 | 1.50528518 | 0.13225077 | 0.57615451 |
| RLF | 1413.89895 | -0.05433 | 0.24570645 | -0.2211175 | 0.82500092 | 0.96146415 |
| AL050341.2 | 40.64586 | 0.35962742 | 0.49013187 | 0.73373604 | 0.46310963 | 0.83531269 |
| ZMPSTE24 | 377.371469 | -0.311767 | 0.21976301 | -1.418651 | 0.15600078 | 0.60527706 |
| SMAP2 | 288.091322 | 0.09554883 | 0.20531124 | 0.46538526 | 0.64165559 | 0.90426796 |
| AL603839.3 | 248.153868 | 0.11907917 | 0.29384782 | 0.40524096 | 0.6853004 | 0.92053037 |
| EXO5 | 47.8581551 | -0.2869308 | 0.42871586 | -0.6692798 | 0.50331703 | 0.85525037 |
| ZNF684 | 296.357472 | 0.70189282 | 0.44791195 | 1.56703303 | 0.11710697 | 0.54756317 |
| NFYC | 1541.71995 | 0.23437776 | 0.24446768 | 0.95872699 | 3.38E-01 | 7.64E-01 |
| CITED4 | 113.263234 | 0.51823394 | 0.49550375 | 1.04587288 | 2.96E-01 | 7.37E-01 |
| CTPS1 | 159.910675 | -1.0714943 | 0.51460015 | -2.0821881 | 0.03732529 | 0.36263296 |
| SLFNL1-AS1 | 165.637802 | -0.2656088 | 0.51964187 | -0.5111381 | 0.60925434 | 0.8927935 |
| SLFNL1 | 61.4693503 | -0.5433203 | 0.63712752 | -0.8527655 | 0.39378938 | 0.79843808 |
| SCMH1 | 3249.4724 | 0.00582745 | 0.19562495 | 0.02978891 | 0.9762354 | 0.9941831 |
| AC093151.3 | 129.570802 | -1.5197031 | 0.92363424 | -1.6453516 | 9.99E-02 | 0.51665771 |
| HIVEP3 | 1799.45701 | -0.7503374 | 0.40173235 | -1.8677545 | 0.0617963 | 0.43264436 |
| FOXJ3 | 1965.47437 | -0.1159764 | 0.18567134 | -0.6246326 | 0.53221221 | 0.86397197 |
| RIMKLA | 68.1980323 | 1.85660822 | 0.8300725 | 2.236682 | 0.02530713 | 0.31080279 |
| ZMYND12 | 61.8580868 | 0.136999 | 0.58055647 | 0.23597876 | 0.81344915 | 0.9582375 |
| PPCS | 264.733632 | 0.50883401 | 0.22028316 | 2.30990885 | 0.0208932 | 0.2892543 |
| CCDC30 | 1831.66613 | 0.1488375 | 0.21361361 | 0.69676037 | 0.48595277 | 0.8447301 |
| PPIH | 166.349962 | -0.4214941 | 0.24601561 | -1.713282 | 0.08666067 | 0.49478465 |
| YBX1 | 786.566913 | 0.25115923 | 0.32188744 | 0.78027038 | 0.43523174 | 0.81919857 |
| CLDN19 | 138.57044 | 1.67116042 | 0.84101255 | 1.98708143 | 0.04691337 | 0.39245013 |
| P3H1 | 115.074492 | -0.452034 | 0.28808913 | -1.569077 | 0.11663 | 0.54715005 |

| | | | | | | |
|---|---|---|---|---|---|---|
| SVBP | 85.949621 | -0.0480133 | 0.43496198 | -0.1103851 | 0.91210395 | 0.98125148 |
| ERMAP | 259.587132 | -0.5569665 | 0.30531762 | -1.8242198 | 0.06811884 | 0.45260654 |
| ZNF691 | 55.8950109 | 0.15678128 | 0.35718524 | 0.4389355 | 0.66070827 | 0.91099119 |
| SLC2A1 | 595.553961 | 0.30130018 | 0.84403018 | 0.35697797 | 0.7211083 | 0.93044898 |
| SLC2A1-AS1 | 135.347479 | 0.87077224 | 0.85588316 | 1.01739616 | 0.30896501 | 0.74399658 |
| FAM183A | 55.0331405 | 0.82450766 | 0.49238649 | 1.67451319 | 0.09402979 | 0.5098684 |
| EBNA1BP2 | 215.645273 | 0.11708191 | 0.36959686 | 0.31678275 | 0.75140845 | 0.93794692 |
| CFAP57 | 66.0838377 | 0.13495446 | 0.42907268 | 0.31452587 | 0.75312167 | 0.93844712 |
| TMEM125 | 77.670145 | 0.67683628 | 0.45880339 | 1.47522075 | 0.1401532 | 0.58492375 |
| AL139289.2 | 38.3582357 | -0.1928695 | 0.70804416 | -0.2723976 | 0.78531633 | 0.94978829 |
| ELOVL1 | 125.121457 | 0.43114814 | 0.4113648 | 1.04809197 | 0.29459624 | 0.73556877 |
| MED8 | 110.820312 | 0.64135751 | 0.33037992 | 1.94127268 | 0.05222521 | 0.40866415 |
| SZT2 | 479.52192 | -0.3814495 | 0.20468177 | -1.8636225 | 0.06237474 | 0.43419765 |
| HYI | 85.2995125 | -0.0730258 | 0.28589901 | -0.2554253 | 0.79839463 | 0.95297658 |
| PTPRF | 750.834091 | 0.63311327 | 0.36244996 | 1.74676051 | 0.08067889 | 0.48456361 |
| KDM4A-AS1 | 57.9941908 | -0.0959301 | 0.36008058 | -0.2664129 | 0.78992123 | 0.95123253 |
| AL357079.1 | 43.6825209 | 0.43139546 | 0.48214957 | 0.89473368 | 0.37092947 | 0.78104287 |
| IPO13 | 99.5415928 | -0.0822136 | 0.36179398 | -0.2272387 | 0.82023816 | 0.96016713 |
| DPH2 | 43.3632541 | -0.0513429 | 0.50942016 | -0.100787 | 0.91971954 | 0.98215577 |
| ATP6V0B | 817.33821 | 0.77205635 | 0.36611628 | 2.10877361 | 0.03496413 | 0.35358029 |
| B4GALT2 | 104.682629 | -0.0207574 | 0.40640433 | -0.0510757 | 0.9592652 | 0.99117358 |
| CCDC24 | 328.745474 | -0.0017339 | 0.32036376 | -0.0054123 | 0.99568163 | 0.99882912 |
| SLC6A9 | 41.6618301 | -1.3018303 | 0.38341712 | -3.395337 | 0.00068544 | 0.04572727 |
| DMAP1 | 146.257644 | 0.13341694 | 0.34672522 | 0.38479157 | 0.70039185 | 0.92367966 |
| ERI3 | 1050.78884 | 0.1197949 | 0.23846336 | 0.50236189 | 0.61541298 | 0.8949877 |
| RNF220 | 163.875247 | -0.064862 | 0.18285765 | -0.3547131 | 0.72280451 | 0.93155672 |
| TMEM53 | 214.871729 | 0.12591416 | 0.34751002 | 0.36233248 | 0.71710359 | 0.9290703 |
| ARMH1 | 206.162581 | 0.15834128 | 0.3655983 | 0.43310181 | 0.66494081 | 0.91225625 |
| RPS8 | 1018.05107 | 0.51643195 | 0.40137228 | 1.28666572 | 0.19821083 | 0.65053266 |
| PLK3 | 64.0714165 | -0.1999526 | 0.52922592 | -0.3778209 | 7.06E-01 | 9.26E-01 |
| EIF2B3 | 612.061328 | -0.1965827 | 0.24589707 | -0.7994511 | 0.42402892 | 0.81421689 |
| HECTD3 | 100.134029 | -0.5161594 | 0.29545167 | -1.7470181 | 0.08063419 | 0.48456361 |
| UROD | 99.5355243 | 0.15538728 | 0.36806324 | 0.42217548 | 0.67289693 | 0.91608132 |
| ZSWIM5 | 1053.72968 | 0.19973898 | 0.35178953 | 0.56777979 | 0.57018451 | 0.87817509 |
| MUTYH | 106.247517 | -0.2971631 | 0.25086493 | -1.1845541 | 0.2361938 | 0.68856409 |
| TOE1 | 46.7112277 | -0.2215148 | 0.35027191 | -0.6324081 | 0.52712025 | 0.86151701 |
| TESK2 | 362.260032 | -0.1414132 | 0.26865509 | -0.5263745 | 0.59862803 | 0.88883692 |
| MMACHC | 57.4897024 | 0.35166383 | 0.39172045 | 0.89774181 | 0.36932321 | 0.78095617 |
| PRDX1 | 1283.6066 | 0.72990935 | 0.49522167 | 1.47390431 | 1.41E-01 | 0.58566162 |
| AKR1A1 | 199.466052 | -0.1931442 | 0.30043944 | -0.6428723 | 0.52030695 | 0.85935232 |
| NASP | 489.15187 | -0.0174607 | 0.20818279 | -0.0838718 | 0.9331584 | 0.98419748 |
| GPBP1L1 | 2250.43619 | -0.0494543 | 0.15773062 | -0.3135367 | 0.75387297 | 0.93874663 |
| TMEM69 | 144.477299 | 0.01814803 | 0.22564008 | 0.08042912 | 0.93589597 | 0.98613991 |
| IPP | 469.405829 | -0.2236357 | 0.20519177 | -1.0898862 | 0.27576328 | 0.72233844 |
| MAST2 | 510.587674 | -0.4129616 | 0.26027775 | -1.5866187 | 0.11259902 | 0.53737225 |
| PIK3R3 | 42.4609737 | -0.3125066 | 0.60255171 | -0.5186386 | 0.60401276 | 0.89001883 |
| TSPAN1 | 685.914916 | 0.55602271 | 0.46730117 | 1.18985945 | 2.34E-01 | 6.85E-01 |
| POMGNT1 | 389.994998 | 0.06633308 | 0.22674026 | 0.29255094 | 0.76986542 | 0.94386043 |
| LURAP1 | 413.998005 | 0.0856796 | 0.39090855 | 0.21918068 | 0.82650931 | 0.96214368 |
| LRRC41 | 418.229409 | -0.0656295 | 0.19989651 | -0.3283174 | 0.74267167 | 0.93598004 |
| UQCRH | 921.837596 | 0.25072642 | 0.22999138 | 1.09015571 | 0.27564456 | 0.72233844 |
| NSUN4 | 310.359835 | -0.0964134 | 0.14906922 | -0.6467693 | 0.51778128 | 0.85893097 |
| FAAH | 139.867653 | 0.03835138 | 0.28364606 | 0.13520859 | 0.89244695 | 0.97771356 |
| MKNK1 | 393.014333 | 0.08742982 | 0.15307191 | 0.57116825 | 0.5678856 | 0.87712536 |

| | | | | | | |
|---|---|---|---|---|---|---|
| ATPAF1 | 452.447953 | 0.28732475 | 0.15706389 | 1.82934947 | 0.06734727 | 0.45135397 |
| EFCAB14-AS | 44.9008026 | 0.68274318 | 0.62700527 | 1.08889544 | 0.2762 | 0.72241028 |
| EFCAB14 | 599.328906 | 0.00433611 | 0.1831901 | 0.02366998 | 0.98111585 | 0.99518016 |
| CYP4A11 | 52.8046355 | 2.13281654 | 0.87999925 | 2.42365724 | 0.0153651 | 0.25212125 |
| PDZK1IP1 | 480.097845 | 2.08138687 | 0.60856551 | 3.42015255 | 0.00062586 | 0.04487218 |
| CMPK1 | 624.353703 | 0.42197992 | 0.23437241 | 1.80046755 | 0.07178684 | 0.46757022 |
| TRABD2B | 2927.59469 | 1.00806227 | 0.68738129 | 1.46652563 | 0.14250515 | 0.58824095 |
| AL109659.1 | 54.8526834 | 2.59829225 | 0.83998035 | 3.09327741 | 0.00197959 | 0.0861633 |
| SLC5A9 | 54.9100849 | 0.80343559 | 0.41208521 | 1.94968314 | 0.0512139 | 0.40556617 |
| SPATA6 | 1585.63784 | -0.0599498 | 0.14427188 | -0.4155332 | 0.67775159 | 0.91753826 |
| AGBL4 | 4091.24997 | 1.40394021 | 0.37217045 | 3.77230437 | 0.00016175 | 0.0205434 |
| AC099788.1 | 68.1739648 | 2.05753627 | 0.62774535 | 3.27766071 | 0.00104671 | 0.05784941 |
| FAF1 | 5534.42809 | -0.1698094 | 0.14372921 | -1.1814536 | 0.23742259 | 0.68973903 |
| RNF11 | 630.458937 | 0.07371614 | 0.20982372 | 0.35132413 | 0.72534519 | 0.9326472 |
| TTC39A | 607.108874 | -0.425013 | 0.49719455 | -0.8548223 | 3.93E-01 | 7.98E-01 |
| TTC39A-AS1 | 73.588066 | -0.7871968 | 0.58876493 | -1.3370306 | 1.81E-01 | 6.36E-01 |
| EPS15 | 2359.58624 | -0.1281472 | 0.1627408 | -0.7874311 | 0.43102956 | 0.81706845 |
| OSBPL9 | 2553.23085 | -0.0344939 | 0.16901519 | -0.2040878 | 8.38E-01 | 9.66E-01 |
| NRDC | 1845.84377 | 0.01188575 | 0.16552334 | 0.07180708 | 0.94275544 | 0.98772472 |
| TXNDC12 | 331.633071 | -0.097449 | 0.22318809 | -0.4366226 | 0.6623851 | 0.91110616 |
| BTF3L4 | 556.316134 | -0.1037615 | 0.16051527 | -0.6464274 | 0.51800261 | 0.85893754 |
| ZFYVE9 | 1369.41248 | -0.2695814 | 0.16265573 | -1.6573744 | 0.09744381 | 0.51355946 |
| CC2D1B | 231.882612 | -0.4528493 | 0.23717765 | -1.9093255 | 0.05622011 | 0.41843207 |
| PRPF38A | 240.031742 | 0.11756682 | 0.17766137 | 0.66174663 | 0.50813361 | 0.85643442 |
| TUT4 | 2383.51919 | 0.09425144 | 0.17913864 | 0.52613682 | 0.59879315 | 0.88883692 |
| ZYG11B | 890.048259 | -0.2505005 | 0.21487307 | -1.1658071 | 0.24369246 | 0.6948422 |
| ECHDC2 | 1923.17832 | 0.62569994 | 0.27400799 | 2.28350978 | 0.02240036 | 0.29677665 |
| SCP2 | 1134.72779 | 0.26799456 | 0.17612018 | 1.52165737 | 0.12809496 | 0.56762585 |
| CPT2 | 184.16729 | 0.79584298 | 0.18651888 | 4.26682275 | 1.98E-05 | 0.0072549 |
| C1orf123 | 110.897143 | 0.39725957 | 0.30189281 | 1.31589611 | 0.18820891 | 0.64267818 |
| MAGOH | 226.31239 | 0.21802987 | 0.20054112 | 1.08720784 | 0.27694496 | 0.7230876 |
| LRP8 | 120.89843 | -1.6486413 | 0.99129536 | -1.6631182 | 0.09628878 | 0.51206987 |
| NDC1 | 302.203523 | -0.4209119 | 0.27179931 | -1.5486129 | 0.12147481 | 0.5564493 |
| YIPF1 | 233.043565 | -0.0697379 | 0.2544583 | -0.2740641 | 0.78403534 | 0.94978829 |
| HSPB11 | 324.768917 | 0.04007069 | 0.17322577 | 0.2313206 | 0.81706575 | 0.95925886 |
| LRRC42 | 154.412472 | 0.18972907 | 0.238706 | 0.79482318 | 0.42671636 | 0.81466629 |
| TMEM59 | 1180.45671 | 0.56598222 | 0.25440938 | 2.22469083 | 0.026102 | 0.31286872 |
| TCEANC2 | 220.156661 | -0.2238818 | 0.23346396 | -0.9589565 | 0.33758067 | 0.76409325 |
| MRPL37 | 125.841839 | 0.38778213 | 0.24420915 | 1.58790989 | 0.11230672 | 0.53670746 |
| SSBP3 | 2874.86718 | -0.0484601 | 0.38032661 | -0.1274171 | 0.89861028 | 0.9788023 |
| ACOT11 | 1982.18196 | 0.03310899 | 0.34011135 | 0.0973475 | 9.22E-01 | 9.82E-01 |
| FAM151A | 74.2800167 | 0.06395471 | 0.41008834 | 0.15595349 | 8.76E-01 | 9.76E-01 |
| MROH7-TTC4 | 150.941293 | 0.00581809 | 0.37407925 | 0.0155531 | 9.88E-01 | 9.97E-01 |
| TTC22 | 107.891492 | -0.1978172 | 0.38092967 | -0.5193012 | 0.60355074 | 0.89001883 |
| DHCR24 | 481.846744 | 0.11816683 | 0.48660867 | 0.2428375 | 0.80813128 | 0.95610623 |
| TMEM61 | 196.445422 | 0.22834785 | 0.45817382 | 0.49838694 | 0.61821134 | 0.89599505 |
| BSND | 137.379103 | 1.12438393 | 0.56202927 | 2.00057894 | 0.04543779 | 0.38897453 |
| USP24 | 3618.82586 | -0.5350565 | 0.31689431 | -1.6884383 | 0.09132712 | 0.50411197 |
| AC119674.2 | 91.4601307 | -0.4185413 | 0.57696386 | -0.7254203 | 0.46819423 | 0.83825637 |
| PLPP3 | 1444.46871 | -0.7059598 | 0.43325629 | -1.6294276 | 0.10322254 | 0.52337282 |
| PRKAA2 | 2992.57675 | -0.2623396 | 0.25963778 | -1.010406 | 3.12E-01 | 0.74687836 |
| DAB1 | 1922.32285 | 0.30075149 | 0.85500679 | 0.35175333 | 0.72502326 | 0.93241479 |
| TACSTD2 | 802.220979 | 0.39411429 | 0.6474619 | 0.60870654 | 0.54271897 | 0.86590899 |
| MYSM1 | 1372.88235 | -0.2313457 | 0.28464028 | -0.812765 | 0.41635281 | 0.80966301 |

| | | | | | | |
|---|---|---|---|---|---|---|
| AL136985.3 | 218.685717 | 0.10179228 | 0.64405823 | 0.15804825 | 0.87441878 | 0.97564728 |
| JUN | 479.891362 | 0.33964518 | 0.67200136 | 0.50542335 | 0.61326152 | 0.89441126 |
| LINC01135 | 440.053757 | 0.22945342 | 0.32722604 | 0.70120771 | 0.4831734 | 0.84427756 |
| LINC01358 | 79.9317675 | 0.3318836 | 0.46067047 | 0.72043602 | 0.47125658 | 0.83921324 |
| FGGY | 1674.81564 | -0.1799661 | 0.33671561 | -0.5344751 | 0.59301286 | 0.88740306 |
| HOOK1 | 953.818972 | 0.08773766 | 0.28031038 | 0.31300182 | 0.75427928 | 0.93901195 |
| NFIA | 3380.30261 | -0.4347827 | 0.43735992 | -0.9941072 | 0.32017066 | 0.75104498 |
| TM2D1 | 946.637354 | -0.1892577 | 0.15055099 | -1.2571002 | 0.20871736 | 0.6614379 |
| PATJ | 12520.3222 | 0.00775428 | 0.18550222 | 0.04180155 | 0.9666569 | 0.99248831 |
| USP1 | 207.355981 | 0.10043633 | 0.18315805 | 0.54835879 | 0.58344557 | 0.88379959 |
| DOCK7 | 1581.76343 | 0.01083283 | 0.1917604 | 0.0564915 | 0.95495027 | 0.99049376 |
| ANGPTL3 | 37.4582025 | 0.87740561 | 0.70641936 | 1.24204639 | 2.14E-01 | 6.68E-01 |
| ATG4C | 496.538716 | -0.1195658 | 0.17598586 | -0.6794056 | 4.97E-01 | 8.51E-01 |
| ALG6 | 246.849106 | -0.5432903 | 0.26526979 | -2.0480671 | 0.04055343 | 0.37344844 |
| ITGB3BP | 899.123245 | 0.59033016 | 0.44652604 | 1.32205091 | 0.1861512 | 0.64033103 |
| EFCAB7 | 463.724306 | 0.09782634 | 0.25380926 | 0.38543251 | 0.699917 | 0.92367966 |
| PGM1 | 1186.6382 | 0.17120845 | 0.24816223 | 0.68990533 | 0.49025372 | 0.84614483 |
| ROR1 | 549.263638 | -1.2264845 | 0.55849281 | -2.1960613 | 0.02808756 | 0.3228969 |
| UBE2U | 45.3716905 | -1.5219165 | 1.2618898 | -1.2060614 | 0.22779383 | 0.6795232 |
| CACHD1 | 129.277021 | -0.0017615 | 0.60761612 | -0.0028991 | 0.99768689 | 0.99957874 |
| RAVER2 | 785.589875 | -0.0959292 | 0.22674873 | -0.4230637 | 0.67224877 | 0.91576543 |
| JAK1 | 1996.26398 | -0.0454557 | 0.21114164 | -0.2152855 | 0.82954471 | 0.96295422 |
| AK4 | 528.923707 | 1.13647475 | 0.53836435 | 2.110977 | 0.03477429 | 0.35358029 |
| DNAJC6 | 902.423259 | -0.1532207 | 0.46532534 | -0.3292764 | 0.74194675 | 0.93598004 |
| AL139294.1 | 108.865929 | 0.28196183 | 0.82900312 | 0.34012155 | 0.73376499 | 0.93552616 |
| LEPR | 1214.65749 | -0.3974255 | 0.38511093 | -1.0319768 | 0.30208299 | 0.73905643 |
| SGIP1 | 7160.34328 | -0.4630536 | 0.40560434 | -1.1416386 | 0.25360426 | 0.70431689 |
| AL139147.1 | 119.98758 | -0.2642625 | 0.70089081 | -0.377038 | 0.70614538 | 0.92556842 |
| WDR78 | 912.905945 | 0.28019731 | 0.22975872 | 1.21952854 | 0.22264365 | 0.67568617 |
| MIER1 | 1366.18999 | 0.01496649 | 0.18755909 | 0.07979614 | 0.9363994 | 0.98621411 |
| SLC35D1 | 549.600983 | 0.19642858 | 0.24148382 | 0.81342338 | 0.41597536 | 0.80966301 |
| IL12RB2 | 43.0492465 | -0.069545 | 0.72734683 | -0.0956146 | 0.92382663 | 0.98215577 |
| SERBP1 | 849.96493 | 0.22276837 | 0.17053148 | 1.30631812 | 0.1914444 | 0.64571429 |
| GADD45A | 505.743169 | 0.44984443 | 0.30901612 | 1.45573127 | 0.14546692 | 0.59195088 |
| GNG12 | 1709.08539 | 0.06473557 | 0.22983615 | 0.28165967 | 7.78E-01 | 9.47E-01 |
| GNG12-AS1 | 1462.75825 | -0.1073484 | 0.31730697 | -0.338311 | 0.73512886 | 0.93552616 |
| DIRAS3 | 78.1323009 | 1.19317999 | 0.53457516 | 2.23201538 | 0.02561395 | 0.31211455 |
| WLS | 2788.16622 | 0.18639699 | 0.22447682 | 0.83036188 | 0.40633421 | 0.80499101 |
| LRRC7 | 71.7152312 | -0.1811233 | 0.42876636 | -0.4224289 | 0.67271198 | 0.91608132 |
| LRRC40 | 480.682429 | -0.1829021 | 0.18146985 | -1.0078925 | 0.31350605 | 0.74749123 |
| SRSF11 | 2647.92934 | 0.0241929 | 0.16086243 | 0.15039496 | 0.88045301 | 0.97664137 |
| ANKRD13C | 1220.6955 | -0.1246239 | 0.19263531 | -0.6469423 | 0.51766932 | 0.85893097 |
| HHLA3 | 136.170752 | 0.30432511 | 0.33661335 | 0.90407914 | 0.36595344 | 0.78095617 |
| CTH | 130.568416 | 0.36994448 | 0.37639121 | 0.98287227 | 0.32567031 | 0.75550719 |
| LINC01788 | 63.0728475 | 1.42499247 | 0.5843576 | 2.4385624 | 0.01474581 | 0.24869534 |
| PTGER3 | 9083.73995 | 0.73518046 | 0.48846371 | 1.50508716 | 1.32E-01 | 0.5761862 |
| AL031429.1 | 52.3414179 | 0.36490032 | 0.89944642 | 0.40569433 | 0.68496721 | 0.92053037 |
| ZRANB2-AS1 | 220.143667 | 0.49144239 | 0.52709847 | 0.93235405 | 0.35115358 | 0.77100859 |
| ZRANB2 | 2300.68059 | 0.09585856 | 0.15510968 | 0.61800503 | 0.53657202 | 0.86466169 |
| NEGR1 | 590.580643 | 0.13407154 | 0.40703716 | 0.32938403 | 0.74186544 | 0.93598004 |
| LRRIQ3 | 209.277921 | 0.070529 | 0.51498561 | 0.13695334 | 0.89106768 | 0.97729111 |
| CRYZ | 986.140907 | 0.07139897 | 0.34064867 | 0.2095971 | 0.83398214 | 0.96510927 |
| TYW3 | 287.974277 | 0.1249172 | 0.30976628 | 0.40326276 | 0.68675493 | 0.9208231 |
| SLC44A5 | 134.055136 | 1.37248414 | 0.682117 | 2.01209491 | 0.04420993 | 0.38531062 |

| | | | | | | |
|---|---|---|---|---|---|---|
| ACADM | 878.532269 | 0.04252339 | 0.23613922 | 0.18007762 | 0.85709164 | 0.97127317 |
| RABGGTB | 272.535086 | -0.115214 | 0.3688573 | -0.3123538 | 0.75477165 | 0.93922829 |
| MSH4 | 68.8646436 | 0.42115897 | 0.63002268 | 0.66848224 | 0.50382582 | 0.85525037 |
| ST6GALNAC3 | 79.9032462 | -0.6438521 | 0.45221397 | -1.4237775 | 0.15451089 | 0.60380955 |
| ST6GALNAC5 | 354.763385 | 0.41984223 | 0.70838149 | 0.59267815 | 5.53E-01 | 0.87104567 |
| PIGK | 567.541696 | -0.3770305 | 0.23872495 | -1.5793511 | 0.11425555 | 0.54132052 |
| AC118549.1 | 2805.2223 | -0.2372874 | 0.1049202 | -2.2615987 | 0.02372221 | 0.30187535 |
| USP33 | 1700.46409 | -0.161794 | 0.10834006 | -1.4933906 | 0.13533497 | 0.57708028 |
| MIGA1 | 1021.92829 | -0.0703883 | 0.11227968 | -0.6269013 | 0.53072395 | 0.86396107 |
| NEXN | 199.961237 | 0.567405 | 0.29760738 | 1.90655552 | 0.05657817 | 0.41843207 |
| FUBP1 | 1410.79795 | 0.00458447 | 0.17980022 | 0.02549757 | 0.97965809 | 0.99483846 |
| GIPC2 | 209.601141 | 1.0621421 | 0.31672954 | 3.35346717 | 0.00079806 | 0.04913638 |
| PTGFR | 480.070319 | -0.5160189 | 0.49175524 | -1.0493408 | 0.29402128 | 0.73527014 |
| IFI44L | 998.163173 | 0.98776752 | 1.72159841 | 0.57375025 | 0.56613682 | 0.87642772 |
| IFI44 | 365.190711 | 0.21032908 | 1.28617495 | 0.1635307 | 0.87010061 | 0.97463098 |
| AL136234.1 | 160.178948 | 2.54341099 | 0.90108906 | 2.82259667 | 0.00476365 | 0.1366507 |
| LINC01781 | 792.590153 | 4.25034868 | 1.1323421 | 3.75359062 | 0.00017432 | 0.02140101 |
| ADGRL2 | 2009.68588 | 0.68220785 | 0.2821422 | 2.41795745 | 0.0156079 | 0.25227487 |
| LINC01362 | 218.912976 | 0.84116941 | 0.66142306 | 1.27175699 | 0.20345948 | 0.65644753 |
| LINC01725 | 3016.27557 | 0.64207235 | 0.3893017 | 1.64929242 | 0.09908774 | 0.51618562 |
| TTLL7 | 195.607215 | 1.28555039 | 0.37226232 | 3.45334542 | 0.00055368 | 0.04085784 |
| PRKACB | 1110.23941 | 0.18294992 | 0.22259224 | 0.82190612 | 0.41113033 | 0.80675684 |
| SAMD13 | 72.0467889 | 0.25065505 | 0.43423718 | 0.57723075 | 0.56378358 | 0.87488456 |
| RPF1 | 181.902537 | -0.1167483 | 0.21910058 | -0.5328524 | 0.59413577 | 0.88807733 |
| GNG5 | 509.236953 | 0.05839514 | 0.3128049 | 0.18668231 | 0.85190973 | 0.97076831 |
| SPATA1 | 459.812425 | -0.0553541 | 0.52329296 | -0.1057804 | 0.91575659 | 0.98191348 |
| CTBS | 657.022976 | -0.2686831 | 0.19261034 | -1.394957 | 0.16302881 | 0.61387934 |
| SSX2IP | 305.484392 | -0.4798595 | 0.30427833 | -1.5770413 | 0.11478601 | 0.54247171 |
| WDR63 | 65.0766205 | 0.89115563 | 0.57748918 | 1.54315554 | 0.12279304 | 0.55947129 |
| SYDE2 | 230.006008 | -0.1984593 | 0.36705214 | -0.5406843 | 5.89E-01 | 8.86E-01 |
| C1orf52 | 333.502847 | -0.2142318 | 0.2347494 | -0.9125977 | 0.36145417 | 0.77772231 |
| BCL10 | 182.325716 | 0.09358497 | 0.25387892 | 0.36862048 | 0.71241063 | 0.92748721 |
| AL078459.1 | 515.607629 | 0.21675125 | 0.41431183 | 0.52315969 | 0.60086313 | 0.88897861 |
| DDAH1 | 2684.67867 | 0.10157255 | 0.29534333 | 0.34391347 | 0.73091136 | 0.93507101 |
| CYR61 | 168.185072 | -0.1431136 | 0.98921735 | -0.1446736 | 0.88496861 | 0.97664137 |
| ZNHIT6 | 637.480168 | -0.2464609 | 0.19663427 | -1.2533974 | 0.21006112 | 0.66269342 |
| COL24A1 | 219.897517 | 0.15341091 | 0.53125622 | 0.2887701 | 0.77275731 | 0.94530018 |
| AL136382.1 | 154.235645 | 0.56347955 | 0.49528567 | 1.13768597 | 0.25525164 | 0.70624609 |
| ODF2L | 1394.36877 | -0.2149767 | 0.16387767 | -1.3118117 | 0.18958366 | 0.64441858 |
| SH3GLB1 | 844.711263 | -0.1335403 | 0.19198715 | -0.695569 | 0.48669876 | 0.84474442 |
| SELENOF | 974.636328 | -0.0453377 | 0.15867536 | -0.2857263 | 0.77508778 | 0.94657308 |
| HS2ST1 | 1164.62134 | -0.2092395 | 0.21944922 | -0.9534759 | 0.34034901 | 0.76478309 |
| LMO4 | 517.211174 | -0.1107977 | 0.22855767 | -0.4847692 | 0.6278401 | 0.89919293 |
| PKN2-AS1 | 193.93303 | 0.24834319 | 0.28858859 | 0.86054405 | 0.38948921 | 0.79677297 |
| PKN2 | 2926.84258 | -0.1468143 | 0.12559893 | -1.168914 | 0.24243828 | 0.69337349 |
| GTF2B | 431.776053 | -0.0796888 | 0.24677123 | -0.3229259 | 0.74675133 | 0.93709275 |
| KYAT3 | 744.12141 | 0.01165746 | 0.20929395 | 0.05569898 | 0.95558161 | 0.99065996 |
| GBP3 | 170.326843 | 0.36037922 | 0.53997148 | 0.66740418 | 0.504514 | 0.85547181 |
| GBP2 | 596.799235 | -0.1526754 | 0.53539886 | -0.2851619 | 7.76E-01 | 9.47E-01 |
| LRRC8B | 599.84598 | 0.07852001 | 0.23183435 | 0.33869016 | 0.73484316 | 0.93552616 |
| LRRC8C-DT | 75.0248616 | 1.40729393 | 0.50956568 | 2.76175177 | 0.00574922 | 0.1552994 |
| LRRC8D | 71.9910534 | 0.5140489 | 0.51968291 | 0.98915875 | 0.32258548 | 0.7532337 |
| ZNF326 | 775.328385 | -0.0404747 | 0.13433595 | -0.3012943 | 0.76319006 | 0.94249976 |
| ZNF644 | 2460.9698 | -0.1244143 | 0.18977522 | -0.6555876 | 0.51208952 | 0.85762435 |

| | | | | | | |
|---|---|---|---|---|---|---|
| HFM1 | 403.58366 | -0.0155573 | 0.26057345 | -0.0597041 | 0.95239127 | 0.98938795 |
| CDC7 | 112.218627 | -1.7432563 | 0.53633587 | -3.2503072 | 0.0011528 | 0.06094516 |
| TGFBR3 | 454.819805 | -0.561315 | 0.36880081 | -1.5220005 | 0.12800895 | 0.56762585 |
| BTBD8 | 371.649503 | -0.3877037 | 0.40783417 | -0.9506405 | 0.34178691 | 0.76545664 |
| GLMN | 469.782863 | -0.2390411 | 0.19323444 | -1.2370523 | 0.21606768 | 0.66944638 |
| RPAP2 | 1558.7208 | -0.0862325 | 0.19461282 | -0.4430975 | 0.65769518 | 0.91037018 |
| EVI5 | 2686.7985 | -0.3301553 | 0.15830795 | -2.0855255 | 0.03702163 | 0.3605598 |
| RPL5 | 817.828164 | 0.54612621 | 0.27322046 | 1.99884816 | 0.04562478 | 0.38904776 |
| FAM69A | 324.696261 | 0.35615871 | 0.33961466 | 1.0487142 | 0.29430968 | 0.73527014 |
| MTF2 | 1402.52987 | -0.131529 | 0.17683934 | -0.7437769 | 0.45701147 | 0.83252855 |
| TMED5 | 459.759847 | 0.17109331 | 0.3006552 | 0.5690682 | 0.56930986 | 0.87789095 |
| CCDC18 | 215.436677 | -0.3645 | 0.41478163 | -0.8787756 | 0.37952295 | 0.78762967 |
| DR1 | 257.333035 | 0.23497271 | 0.2777302 | 0.84604667 | 0.39752671 | 0.80069929 |
| FNBP1L | 3798.99395 | 0.25405829 | 0.23736572 | 1.07032423 | 0.28447339 | 0.72738535 |
| BCAR3 | 1000.26394 | -0.7507174 | 0.32034488 | -2.3434661 | 0.0191055 | 0.27947345 |
| AL109613.1 | 69.8243779 | 0.28329052 | 0.4993136 | 0.56735991 | 0.57046968 | 0.87824406 |
| DNTTIP2 | 116.459892 | 0.14561017 | 0.25247064 | 0.576741 | 0.56411443 | 0.87508955 |
| GCLM | 537.439555 | -0.0698731 | 0.34724625 | -0.2012207 | 8.41E-01 | 0.96617873 |
| ABCA4 | 189.831078 | -2.2429454 | 0.74204123 | -3.0226695 | 0.00250556 | 0.09525007 |
| ARHGAP29 | 4365.16016 | -0.5890951 | 0.41297634 | -1.4264621 | 0.15373501 | 0.60285224 |
| ABCD3 | 1561.16069 | -0.0365012 | 0.22151944 | -0.1647765 | 0.86911993 | 0.9744678 |
| F3 | 86.4368372 | 0.75008445 | 0.78891395 | 0.95078107 | 0.34171553 | 0.76545664 |
| SLC44A3 | 1293.89064 | 0.46947853 | 0.36297895 | 1.29340427 | 0.19587128 | 0.64964978 |
| CNN3 | 1814.20682 | 0.17870886 | 0.30391752 | 0.58801764 | 0.55652045 | 0.87212955 |
| AC105942.1 | 110.225938 | -0.2583704 | 0.29222352 | -0.8841532 | 0.3766135 | 0.78527035 |
| ALG14 | 397.386055 | -0.4130248 | 0.31275143 | -1.3206169 | 0.18662913 | 0.6408069 |
| AC092802.1 | 99.8587127 | -0.0388437 | 0.44976801 | -0.0863639 | 0.93117713 | 0.9835214 |
| PTBP2 | 2495.41187 | -0.1865571 | 0.13489211 | -1.3830096 | 0.16666193 | 0.61925103 |
| DPYD | 3637.70106 | 0.35129017 | 0.37977467 | 0.9249963 | 3.55E-01 | 0.7726065 |
| DPYD-AS1 | 172.462035 | 0.64665008 | 0.52300858 | 1.23640435 | 0.2163083 | 0.66968853 |
| SNX7 | 431.108292 | 0.25147405 | 0.23027767 | 1.09204704 | 0.27481243 | 0.72164296 |
| FRRS1 | 224.938578 | -0.2530764 | 0.30860735 | -0.8200595 | 0.41218216 | 0.80695334 |
| AGL | 945.290826 | -0.0503916 | 0.13026916 | -0.3868268 | 6.99E-01 | 9.24E-01 |
| AC093019.2 | 51.9559107 | 0.13110509 | 0.62745239 | 0.20894827 | 0.83448862 | 0.96510927 |
| SASS6 | 102.675748 | -0.5594851 | 0.31897101 | -1.7540312 | 0.07942516 | 0.48169281 |
| TRMT13 | 622.839136 | -0.1395565 | 0.18018012 | -0.7745391 | 4.39E-01 | 0.82127804 |
| LRRC39 | 105.672218 | -0.5081132 | 0.31140442 | -1.6316825 | 0.10274639 | 0.52202889 |
| DBT | 679.583413 | -0.0468224 | 0.19047217 | -0.2458229 | 0.80581932 | 0.95522565 |
| RTCA | 405.95189 | 0.03841961 | 0.20153397 | 0.19063591 | 0.84881085 | 0.97017477 |
| CDC14A | 689.206765 | -0.121752 | 0.3036213 | -0.4009996 | 0.68842044 | 0.92136651 |
| VCAM1 | 107.789119 | 1.43686667 | 0.64022717 | 2.24430755 | 0.02481262 | 0.3089066 |
| EXTL2 | 375.995651 | -0.0156395 | 0.20612655 | -0.0758732 | 0.93952 | 0.98715467 |
| SLC30A7 | 891.586862 | -0.1183132 | 0.20789649 | -0.5690968 | 0.56929045 | 0.87789095 |
| DPH5 | 277.643663 | -0.0111898 | 0.18272054 | -0.0612401 | 9.51E-01 | 9.89E-01 |
| AC093157.1 | 356.429155 | 0.03429638 | 0.2100208 | 0.16329992 | 0.87028229 | 0.97463098 |
| RNPC3 | 1522.64709 | -0.1023565 | 0.20257711 | -0.5052716 | 0.61336811 | 0.89441126 |
| AMY2B | 381.283517 | -0.4399149 | 0.33695495 | -1.3055601 | 0.1917022 | 0.64596359 |
| PRMT6 | 44.5825544 | 0.32376088 | 0.52278748 | 0.61929731 | 0.53572051 | 0.8642718 |
| NTNG1 | 220.342709 | -1.2167957 | 0.66590648 | -1.8272771 | 6.77E-02 | 4.52E-01 |
| VAV3 | 5428.60293 | -0.109415 | 0.36900533 | -0.2965132 | 0.76683815 | 0.94312902 |
| SLC25A24 | 469.74919 | 0.13907332 | 0.24025955 | 0.57884618 | 0.56269297 | 0.87432201 |
| PRPF38B | 755.767693 | -0.2682868 | 0.23277958 | -1.1525359 | 0.24910094 | 0.69928042 |
| STXBP3 | 827.999042 | -0.1770075 | 0.20243326 | -0.8743995 | 0.38190071 | 0.78919376 |
| AL449266.1 | 169.457094 | -0.5139346 | 0.3205676 | -1.603202 | 0.10889006 | 0.53262539 |

| | | | | | | |
|---|---|---|---|---|---|---|
| GPSM2 | 203.247023 | -0.1465042 | 0.60052438 | -0.2439604 | 0.80726149 | 0.95582754 |
| WDR47 | 273.659175 | -0.2221341 | 0.25821508 | -0.8602678 | 0.38964142 | 0.79683752 |
| TAF13 | 160.04235 | -0.0823721 | 0.31954741 | -0.2577775 | 0.79657865 | 0.95278473 |
| TMEM167B | 153.273312 | 0.32038336 | 0.25320427 | 1.2653158 | 0.20575815 | 0.65813563 |
| SARS | 441.803866 | 0.39660668 | 0.2016787 | 1.96652738 | 0.04923772 | 0.39892272 |
| CELSR2 | 141.998855 | 0.15667888 | 0.47506044 | 0.3298083 | 0.74154482 | 0.93598004 |
| SORT1 | 1567.71272 | -0.2441989 | 0.3520046 | -0.6937379 | 0.48784659 | 0.8451268 |
| PSMA5 | 559.378197 | -0.4083272 | 0.28543422 | -1.4305473 | 0.15256001 | 0.60267503 |
| SYPL2 | 351.556585 | 0.12203309 | 0.50213023 | 0.24303076 | 0.80798157 | 0.95607172 |
| ATXN7L2 | 41.7794049 | -0.4697709 | 0.44296228 | -1.0605213 | 0.2889075 | 0.73176973 |
| CYB561D1 | 137.235975 | 0.02073509 | 0.32192564 | 0.06440955 | 0.94864413 | 0.98919519 |
| GNAI3 | 786.924849 | -0.0209552 | 0.20080179 | -0.1043576 | 0.91688557 | 0.98199511 |
| AMPD2 | 188.924438 | 0.06649454 | 0.43642371 | 0.15236234 | 0.87890116 | 0.9765717 |
| GSTM4 | 148.764522 | 0.30886217 | 0.41766473 | 0.73949785 | 0.45960474 | 0.83390278 |
| GSTM3 | 413.487641 | 0.02376895 | 0.5987867 | 0.03969519 | 0.96833614 | 0.99248831 |
| AHCYL1 | 1734.48853 | 0.90456564 | 0.19900478 | 4.54544668 | 5.48E-06 | 0.0032951 |
| STRIP1 | 262.719709 | -0.2067223 | 0.2308855 | -0.8953457 | 0.37060231 | 0.78104287 |
| KCNC4 | 516.823433 | 1.27624675 | 0.76575746 | 1.66664617 | 0.09558478 | 0.5117083 |
| RBM15-AS1 | 60.3621465 | 0.54724201 | 0.50095428 | 1.09239913 | 0.27465771 | 0.72164296 |
| RBM15 | 266.40979 | -0.0560333 | 0.22786682 | -0.2459037 | 0.80575681 | 0.95522565 |
| SLC16A4 | 277.875756 | 0.95897812 | 0.37104804 | 2.58451197 | 0.00975169 | 0.20084545 |
| AL355488.1 | 119.053144 | -0.2461464 | 0.27815339 | -0.8849303 | 0.37619421 | 0.78490077 |
| LAMTOR5 | 499.921959 | 0.71258197 | 0.33155639 | 2.14920293 | 0.03161832 | 0.33950635 |
| LRIF1 | 221.40039 | 0.11585289 | 0.28960594 | 0.40003629 | 0.68912979 | 0.92166785 |
| DRAM2 | 469.356248 | -0.0772146 | 0.17466845 | -0.4420636 | 0.65844318 | 0.91037018 |
| CEPT1 | 676.776802 | -0.1589143 | 0.19261578 | -0.8250327 | 0.40935301 | 0.80623405 |
| DENND2D | 234.199004 | 0.2035387 | 0.26262661 | 0.77501169 | 0.43833275 | 0.82127804 |
| WDR77 | 114.32306 | -0.0787973 | 0.30629675 | -0.2572582 | 0.79697948 | 0.95281098 |
| ATP5PB | 628.522523 | 0.64119506 | 0.27121629 | 2.36414657 | 0.01807166 | 0.26945723 |
| RAP1A | 1957.42082 | -0.0380174 | 0.23180057 | -0.1640089 | 0.8697241 | 0.97454113 |
| INKA2 | 131.498425 | -0.0919486 | 0.66469374 | -0.1383323 | 0.88997782 | 0.97729111 |
| DDX20 | 100.653974 | -0.1674485 | 0.33074098 | -0.5062829 | 0.61265808 | 0.89441126 |
| KCND3 | 472.821491 | 0.1282317 | 0.59312303 | 0.21619748 | 0.82883382 | 0.96288319 |
| CTTNBP2NL | 628.811695 | 0.00489545 | 0.19612942 | 0.02496032 | 9.80E-01 | 9.95E-01 |
| WNT2B | 186.930886 | 0.42851744 | 0.4884134 | 0.87736626 | 0.38028773 | 0.7880311 |
| ST7L | 764.480567 | -0.1528573 | 0.17017788 | -0.8982209 | 0.3690678 | 0.78095617 |
| CAPZA1 | 1131.20106 | -0.1940003 | 0.1704041 | -1.1384721 | 0.2549234 | 0.70578143 |
| MOV10 | 288.107297 | -0.0358141 | 0.33770549 | -0.1060513 | 0.91554164 | 0.98176629 |
| AL603832.1 | 61.4749675 | 0.1586251 | 0.47809189 | 0.33178789 | 0.74004943 | 0.93552616 |
| RHOC | 319.199721 | 0.22499373 | 0.42570119 | 0.52852501 | 5.97E-01 | 8.89E-01 |
| SLC16A1-AS1 | 74.5441627 | 0.27914544 | 0.48489459 | 0.57568273 | 0.56482965 | 0.87579905 |
| AL357055.3 | 88.5247151 | -0.011669 | 0.38680062 | -0.030168 | 9.76E-01 | 0.9941831 |
| LRIG2 | 663.622271 | -0.2991138 | 0.15550977 | -1.9234404 | 0.05442476 | 0.41578755 |
| MAGI3 | 4610.95269 | -0.4868401 | 0.30017961 | -1.6218293 | 0.10483989 | 0.52495457 |
| PHTF1 | 665.369642 | -0.1355431 | 0.18796015 | -0.7211267 | 0.47083155 | 0.83921324 |
| RSBN1 | 620.24851 | 0.07391444 | 0.19369174 | 0.38160863 | 0.70275169 | 0.92501157 |
| AP4B1 | 73.4259059 | 0.18251333 | 0.37447672 | 0.48738231 | 0.62598743 | 0.89873262 |
| HIPK1 | 464.341801 | 0.03065548 | 0.23835321 | 0.12861367 | 0.89766335 | 0.97823597 |
| TRIM33 | 1758.26872 | -0.3453799 | 0.200819 | -1.7198565 | 0.08545853 | 0.49346408 |
| BCAS2 | 205.236604 | 0.21779677 | 0.30194335 | 0.72131665 | 0.47071472 | 0.83921324 |
| DENND2C | 96.9750427 | -0.4788207 | 0.39489331 | -1.2125317 | 0.2253089 | 0.67798093 |
| NRAS | 136.033689 | 0.05653862 | 0.31260472 | 0.18086296 | 0.85647514 | 0.97118784 |
| CSDE1 | 1246.49651 | 0.12025112 | 0.13509131 | 0.89014698 | 0.37338697 | 0.78277446 |
| SIKE1 | 240.602747 | 0.17402687 | 0.2839921 | 0.61278772 | 0.54001671 | 0.86566791 |

| | | | | | | |
|---|---|---|---|---|---|---|
| VANGL1 | 104.200386 | -0.3433091 | 0.37226345 | -0.9222209 | 0.35641338 | 0.77368353 |
| SLC22A15 | 1014.99706 | 0.13807692 | 0.34437969 | 0.40094385 | 0.68846147 | 0.92136651 |
| MAB21L3 | 134.974815 | 2.80662094 | 1.01279914 | 2.77115258 | 0.00558582 | 0.15150546 |
| ATP1A1 | 15607.88 | 0.18967855 | 0.35782958 | 0.5300807 | 0.59605598 | 0.88859628 |
| LINC01762 | 7477.16215 | 1.41971411 | 0.64896526 | 2.18765808 | 0.02869452 | 0.32702837 |
| AL390066.1 | 69.5046911 | 0.10714598 | 0.60219229 | 0.17792652 | 0.85878068 | 0.97174503 |
| CD58 | 838.138823 | -0.1014951 | 0.18343551 | -0.5533011 | 5.80E-01 | 0.88258799 |
| IGSF3 | 1287.91085 | 0.30662947 | 0.3359597 | 0.91269719 | 0.36140182 | 0.77772231 |
| PTGFRN | 322.957413 | -0.0619102 | 0.33441199 | -0.1851315 | 0.8531259 | 0.97076831 |
| AL445231.1 | 41.5537699 | 0.27029605 | 0.45423451 | 0.59505837 | 0.55180445 | 0.87051638 |
| TTF2 | 250.917055 | -0.443192 | 0.28372905 | -1.5620256 | 0.11828196 | 0.54936233 |
| TRIM45 | 45.7130341 | -0.0723694 | 0.39545992 | -0.1830005 | 8.55E-01 | 0.97116182 |
| VTCN1 | 321.693416 | 1.99043865 | 0.70738465 | 2.8137996 | 0.00489597 | 0.13937698 |
| MAN1A2 | 2073.14117 | -0.0785221 | 0.16632677 | -0.4720951 | 0.63685892 | 0.90255504 |
| TENT5C | 87.7123276 | 0.87191061 | 0.57523826 | 1.51573821 | 0.12958556 | 0.57050365 |
| GDAP2 | 679.54482 | -0.2178196 | 0.19778514 | -1.101294 | 0.27076873 | 0.71965808 |
| WDR3 | 155.398913 | 0.02812064 | 0.29498815 | 0.09532804 | 0.92405427 | 0.98215577 |
| SPAG17 | 1193.808 | 0.36508124 | 0.32591498 | 1.12017325 | 0.26263994 | 0.7132328 |
| TBX15 | 109.384349 | 0.28741207 | 1.48679486 | 0.19330984 | 0.84671632 | 0.96883887 |
| WARS2 | 576.935982 | -0.2062754 | 0.13749171 | -1.5002753 | 0.1335431 | 0.57682553 |
| AL359915.2 | 1695.12979 | -0.3367767 | 0.345812 | -0.9738721 | 0.33012003 | 0.75885186 |
| AL359915.1 | 258.888721 | -0.2429023 | 0.89337743 | -0.2718922 | 7.86E-01 | 0.94978829 |
| ZNF697 | 217.819817 | -0.3053038 | 0.38786648 | -0.7871364 | 4.31E-01 | 8.17E-01 |
| PHGDH | 124.391402 | 0.92024379 | 0.50381856 | 1.82653811 | 0.06776925 | 0.45233147 |
| HMGCS2 | 86.5641802 | -0.4006826 | 1.1813609 | -0.3391704 | 0.73448138 | 0.93552616 |
| NOTCH2 | 556.402587 | 0.28476175 | 0.17269112 | 1.64896586 | 0.09915463 | 0.51618562 |
| SEC22B | 674.084533 | 0.03287953 | 0.18143307 | 0.18122127 | 0.8561939 | 0.97118784 |
| NBPF26 | 157.282079 | 0.09296841 | 0.63250418 | 0.14698465 | 0.88314414 | 0.97664137 |
| LINC00623 | 792.811501 | 1.23441786 | 0.57440767 | 2.14902748 | 0.03163222 | 0.33950635 |
| AC244453.3 | 53.0917526 | 0.88499542 | 0.45252618 | 1.95567784 | 0.05050312 | 0.40335713 |
| SRGAP2C | 120.832976 | -0.2223385 | 0.39059745 | -0.5692268 | 0.56920224 | 0.87789095 |
| AC239800.3 | 84.7762755 | 0.77047573 | 0.59795473 | 1.28851849 | 0.19756554 | 0.64964978 |
| NBPF15 | 232.113329 | -0.0125355 | 0.41980859 | -0.0298599 | 0.97617877 | 0.9941831 |
| FP700111.1 | 220.930666 | -0.6363336 | 0.36543007 | -1.7413279 | 0.08162612 | 0.48594425 |
| SRGAP2B | 143.764569 | -0.40375 | 0.36912235 | -1.0938108 | 2.74E-01 | 0.72135661 |
| GPR89A | 206.982976 | -0.0801798 | 0.27058309 | -0.2963222 | 0.76698402 | 0.94312902 |
| PDZK1 | 288.755536 | 0.70126584 | 0.49147387 | 1.42686292 | 0.15361941 | 0.60285224 |
| RNF115 | 1968.91764 | -0.4234841 | 0.15671466 | -2.702262 | 0.00688695 | 0.17059994 |
| POLR3C | 270.915421 | -0.485704 | 0.23826868 | -2.0384717 | 0.04150278 | 0.37579578 |
| PIAS3 | 112.968507 | -0.4720736 | 0.3281083 | -1.4387737 | 0.15021466 | 0.59663305 |
| ITGA10 | 100.864268 | -1.1396337 | 0.71235412 | -1.5998135 | 0.10963997 | 0.53361622 |
| PEX11B | 163.908895 | -0.3741767 | 0.28062886 | -1.3333508 | 0.1824167 | 0.63741052 |
| RBM8A | 268.141513 | 0.28442493 | 0.24424035 | 1.16452882 | 0.24420977 | 0.6948422 |
| LIX1L | 78.3071714 | -0.2483984 | 0.30274369 | -0.8204907 | 0.41193644 | 0.80679792 |
| POLR3GL | 46.236179 | 0.52905074 | 0.58000301 | 0.91215171 | 0.36168886 | 0.77772231 |
| TXNIP | 5191.7281 | 0.53949488 | 0.46373744 | 1.1633628 | 0.24468234 | 0.6948422 |
| NBPF12 | 433.93577 | -0.1391641 | 0.21139376 | -0.6583171 | 0.51033439 | 0.85762435 |
| PRKAB2 | 202.169722 | 0.09318062 | 0.17683123 | 0.52694661 | 0.59823066 | 0.88883692 |
| AC242426.2 | 571.703693 | -0.0654599 | 0.20158843 | -0.3247205 | 0.74539265 | 0.93675395 |
| FMO5 | 307.845487 | -0.1313577 | 0.41600837 | -0.3157574 | 0.75218666 | 0.93815321 |
| CHD1L | 532.258849 | 0.06788828 | 0.23990839 | 0.28297585 | 0.77719535 | 0.94694572 |
| LINC00624 | 119.234185 | 0.6331006 | 0.46649684 | 1.35713802 | 0.17473736 | 0.62932267 |
| BCL9 | 566.235133 | -0.0188466 | 0.27052568 | -0.0696665 | 0.94445913 | 0.98813503 |
| ACP6 | 600.31385 | -0.3539601 | 0.15894735 | -2.2269018 | 0.02595384 | 0.31222606 |

| | | | | | | |
|---|---|---|---|---|---|---|
| GPR89B | 266.14127 | -0.5330032 | 0.30823625 | -1.7292034 | 0.0837727 | 0.48884683 |
| NBPF11 | 350.914029 | -0.4161787 | 0.34403878 | -1.2096855 | 0.22639958 | 0.67880306 |
| AC239809.3 | 232.702694 | -0.9255843 | 0.30294859 | -3.0552521 | 0.00224871 | 0.09083568 |
| NBPF14 | 149.800103 | 0.18483925 | 0.29196265 | 0.63309211 | 0.52667349 | 0.86151701 |
| NUDT4B | 79.1586284 | -0.2270333 | 0.49076658 | -0.4626095 | 0.64364431 | 0.90478945 |
| PDE4DIP | 3157.38675 | 0.05692887 | 0.38053905 | 0.14960059 | 0.88107974 | 0.97664137 |
| NBPF9 | 158.965232 | -0.5657857 | 0.19861377 | -2.8486733 | 0.00439019 | 0.12886682 |
| AC245297.3 | 2808.98079 | 0.05286325 | 0.2377498 | 0.22234825 | 0.8240428 | 0.96141406 |
| NBPF19 | 283.740692 | 0.4467858 | 0.78998773 | 0.56556043 | 0.57169264 | 0.87899989 |
| BOLA1 | 57.1448981 | 0.0053993 | 0.37825115 | 0.01427437 | 0.98861109 | 0.99660871 |
| SF3B4 | 107.788109 | 0.35538416 | 0.36743824 | 0.96719427 | 0.33344693 | 0.76149731 |
| OTUD7B | 1128.70841 | -0.354045 | 0.24836134 | -1.4255239 | 0.1540058 | 0.60292313 |
| VPS45 | 1229.76803 | -0.0705277 | 0.12687339 | -0.5558906 | 0.57828566 | 0.88205258 |
| ANP32E | 296.200641 | 0.46110468 | 0.27492603 | 1.67719542 | 0.09350427 | 0.50887217 |
| APH1A | 221.183013 | 0.23675444 | 0.29067714 | 0.8144928 | 0.4153627 | 0.80966301 |
| C1orf54 | 116.493237 | 0.9258795 | 0.68762388 | 1.34649119 | 0.17814416 | 0.63204571 |
| MRPS21 | 523.57681 | 0.09251515 | 0.16568438 | 0.55838182 | 0.57658368 | 0.88147804 |
| PRPF3 | 679.932626 | -0.7626841 | 0.26714907 | -2.8549008 | 0.00430503 | 0.12803674 |
| RPRD2 | 2283.10896 | -0.283439 | 0.13172261 | -2.1517866 | 0.03141417 | 0.33928841 |
| TARS2 | 166.703103 | -0.6281993 | 0.31051542 | -2.0230856 | 0.04306433 | 0.38246672 |
| AL356356.1 | 42.3357877 | -0.9624679 | 0.54466332 | -1.7670878 | 0.07721352 | 0.47712313 |
| ADAMTSL4 | 62.1595237 | -0.230337 | 0.49233807 | -0.4678433 | 0.63989668 | 0.90330608 |
| MCL1 | 710.910253 | 0.13079317 | 0.40301955 | 0.32453307 | 0.74553449 | 0.93675395 |
| ENSA | 1538.91032 | -0.3508257 | 0.18174931 | -1.9302727 | 0.05357306 | 0.41262191 |
| GOLPH3L | 709.51099 | 0.07205868 | 0.1764388 | 0.4084061 | 0.68297555 | 0.9200778 |
| CTSS | 77.5260272 | 0.05515962 | 0.57284823 | 0.09629011 | 0.92329017 | 0.98215577 |
| CTSK | 49.5476181 | 0.35459827 | 0.37724235 | 0.9399747 | 0.34723054 | 0.76893725 |
| ARNT | 825.960966 | -0.0911388 | 0.18990919 | -0.4799073 | 0.63129328 | 0.90011746 |
| SETDB1 | 612.741303 | -0.3361716 | 0.1769155 | -1.9001819 | 0.05740925 | 0.42012124 |
| CERS2 | 333.609571 | -0.0351719 | 0.25746442 | -0.1366087 | 0.89134008 | 0.97729111 |
| MINDY1 | 203.208867 | 0.35002137 | 0.29382588 | 1.19125438 | 0.23355374 | 0.68493974 |
| PRUNE1 | 245.700097 | -0.1458356 | 0.22633413 | -0.6443375 | 0.5193566 | 0.85924115 |
| C1orf56 | 173.97059 | 0.03077856 | 0.64088276 | 0.04802526 | 0.96169611 | 0.99203779 |
| CDC42SE1 | 478.859907 | -0.1601534 | 0.26750597 | -0.5986909 | 0.54937905 | 0.86907161 |
| GABPB2 | 859.299426 | 0.17433182 | 0.34814619 | 0.50074315 | 0.6165519 | 0.89589954 |
| SEMA6C | 42.2886733 | -0.2620964 | 0.45177951 | -0.5801423 | 0.56181866 | 0.87403145 |
| SCNM1 | 167.079275 | -0.1591935 | 0.32597563 | -0.48836 | 0.62529488 | 0.89873262 |
| LYSMD1 | 62.6019042 | 0.21068778 | 0.27329328 | 0.77092192 | 0.44075321 | 0.82265742 |
| VPS72 | 151.973525 | -0.2101491 | 0.25209531 | -0.8336096 | 0.40450104 | 0.80437725 |
| PIP5K1A | 2301.08018 | 0.01858059 | 0.16944438 | 0.10965599 | 0.91268221 | 0.98135874 |
| PSMD4 | 384.343522 | 0.06136579 | 0.15739562 | 0.38988246 | 0.69662346 | 0.92322348 |
| ZNF687 | 66.6560627 | 0.41064662 | 0.35885436 | 1.14432668 | 0.25248818 | 0.70298696 |
| PI4KB | 551.283075 | 0.21408432 | 0.15992293 | 1.3386718 | 0.18067754 | 0.63503977 |
| RFX5 | 65.8077785 | -0.1171691 | 0.30178862 | -0.388249 | 0.69783179 | 0.92341984 |
| SELENBP1 | 215.055582 | -0.2731538 | 0.78425954 | -0.3482952 | 0.72761852 | 0.93420128 |
| PSMB4 | 205.391502 | 0.46363878 | 0.44382134 | 1.04465185 | 0.29618391 | 0.7366786 |
| POGZ | 3125.11413 | 0.04512127 | 0.17858289 | 0.25266287 | 8.01E-01 | 0.95370871 |
| CGN | 503.343769 | -0.070449 | 0.28402748 | -0.2480358 | 0.80410674 | 0.95450237 |
| TUFT1 | 506.900618 | -0.0709739 | 0.41960991 | -0.1691426 | 0.86568449 | 0.97379123 |
| SNX27 | 907.183746 | -0.0494424 | 0.1893452 | -0.2611232 | 0.7939975 | 0.95224951 |
| MRPL9 | 92.1029897 | -0.2011477 | 0.33055199 | -0.6085205 | 0.54284231 | 0.86590899 |
| TDRKH | 69.2108793 | 0.11299309 | 0.39502772 | 0.28603838 | 0.77484872 | 0.94636863 |
| RORC | 244.482893 | -0.0212231 | 0.51882103 | -0.0409064 | 0.96737054 | 0.99248831 |
| AL450992.2 | 70.0504036 | -0.0610185 | 0.41018654 | -0.1487578 | 0.88174472 | 0.97664137 |

| | | | | | | |
|---|---|---|---|---|---|---|
| THEM4 | 1026.60015 | 0.58969122 | 0.61554033 | 0.95800583 | 0.33805982 | 0.76436702 |
| S100A10 | 1562.53699 | -0.2859374 | 0.53330256 | -0.5361636 | 0.59184552 | 0.88736521 |
| S100A11 | 3061.8223 | 0.09456618 | 0.44414572 | 0.21291701 | 8.31E-01 | 9.64E-01 |
| FLG-AS1 | 1024.82447 | -0.1161821 | 0.40386325 | -0.2876768 | 0.77359416 | 0.94558571 |
| S100A9 | 93.5956255 | 0.66902814 | 0.83583987 | 0.80042621 | 4.23E-01 | 8.14E-01 |
| S100A6 | 5405.33713 | 0.09591143 | 0.62922268 | 0.15242844 | 8.79E-01 | 9.77E-01 |
| S100A4 | 153.178731 | 0.33479667 | 0.69743684 | 0.48003869 | 0.63119988 | 0.90011169 |
| S100A2 | 1317.04938 | -0.643961 | 1.22352144 | -0.5263177 | 0.59866745 | 0.88883692 |
| S100A16 | 318.843976 | 0.1062845 | 0.40843412 | 0.26022434 | 7.95E-01 | 9.53E-01 |
| S100A14 | 152.32635 | 0.31304227 | 0.42343591 | 0.73929079 | 4.60E-01 | 8.34E-01 |
| S100A13 | 675.298388 | 0.12825951 | 0.4052491 | 0.3164955 | 7.52E-01 | 9.38E-01 |
| S100A1 | 68.5407103 | 0.41612453 | 0.67137862 | 0.61980605 | 0.53538548 | 0.8642718 |
| AL162258.2 | 54.761086 | -1.5392565 | 0.43121339 | -3.5695936 | 0.00035754 | 0.03148458 |
| CHTOP | 506.202792 | -0.0201968 | 0.11906469 | -0.1696288 | 0.86530208 | 0.97379123 |
| SNAPIN | 75.636327 | 0.11062466 | 0.31275021 | 0.3537157 | 0.72355195 | 0.93188356 |
| ILF2 | 284.8882 | 0.42563788 | 0.28984075 | 1.46852324 | 0.14196215 | 0.5872148 |
| NPR1 | 46.0459572 | 0.47926924 | 0.73159026 | 0.65510609 | 0.51239945 | 0.85762435 |
| INTS3 | 781.702877 | -0.475273 | 0.25300639 | -1.8785018 | 0.06031255 | 0.42866735 |
| SLC27A3 | 59.192097 | -0.2840275 | 0.46079185 | -0.6163901 | 0.53763707 | 0.86489441 |
| GATAD2B | 2649.39717 | -0.2125643 | 0.23556064 | -0.902376 | 0.36685716 | 0.78095617 |
| DENND4B | 266.894166 | -0.2699032 | 0.24596416 | -1.0973274 | 0.27249832 | 0.72070318 |
| CRTC2 | 146.565208 | 0.02590533 | 0.33609855 | 0.07707658 | 0.93856262 | 0.9871386 |
| SLC39A1 | 239.424706 | 0.46271547 | 0.32989142 | 1.4026296 | 1.61E-01 | 0.61166537 |
| CREB3L4 | 72.011223 | -0.1337078 | 0.32620138 | -0.4098933 | 0.6818842 | 0.91964553 |
| JTB | 370.553379 | 0.42658166 | 0.41533697 | 1.02707365 | 0.30438578 | 0.74152784 |
| RAB13 | 362.315884 | 0.45749241 | 0.37820577 | 1.20963889 | 2.26E-01 | 0.67880306 |
| RPS27 | 347.451812 | 0.50947981 | 0.47533634 | 1.07183011 | 0.28379634 | 0.72690826 |
| NUP210L | 218.492842 | -1.6857198 | 1.38013406 | -1.2214174 | 0.22192802 | 0.67544503 |
| TPM3 | 487.044428 | 0.00697836 | 0.22992145 | 0.03035105 | 9.76E-01 | 9.94E-01 |
| C1orf43 | 580.427573 | -0.0294048 | 0.19756006 | -0.1488399 | 0.88167995 | 0.97664137 |
| UBAP2L | 736.027068 | -0.2387905 | 0.16120402 | -1.4812938 | 0.13852831 | 0.58206378 |
| HAX1 | 157.010241 | 0.41702145 | 0.25821141 | 1.61503881 | 0.10630229 | 0.52735777 |
| ATP8B2 | 109.503336 | 0.92597196 | 0.34453563 | 2.68759416 | 0.00719688 | 0.1729679 |
| IL6R | 227.827696 | -0.2387084 | 0.54231375 | -0.4401666 | 6.60E-01 | 9.11E-01 |
| UBE2Q1 | 174.031494 | -0.0514545 | 0.27362311 | -0.1880486 | 0.85083852 | 0.97056586 |
| ADAR | 973.831901 | -0.0960601 | 0.36472407 | -0.2633774 | 0.79225972 | 0.95212343 |
| KCNN3 | 635.163197 | -1.439056 | 0.78748727 | -1.8274022 | 0.06763931 | 0.45209626 |
| PMVK | 211.612032 | 0.57480393 | 0.2904938 | 1.97871325 | 0.0478483 | 0.39501762 |
| PBXIP1 | 242.933881 | 0.83907097 | 0.3385013 | 2.47878214 | 0.01318318 | 0.23563819 |
| PYGO2 | 85.5036035 | 0.58765539 | 0.29234158 | 2.01016693 | 0.04441352 | 0.38623091 |
| SHC1 | 51.6837033 | -0.7600912 | 0.33499716 | -2.2689482 | 0.02327148 | 0.30018842 |
| CKS1B | 46.9530346 | 0.43869946 | 0.47434258 | 0.92485784 | 0.35503986 | 0.7726065 |
| FLAD1 | 112.295699 | -0.3905875 | 0.2919347 | -1.3379277 | 0.18092001 | 0.63540877 |
| ZBTB7B | 137.899851 | 0.36190697 | 0.38317916 | 0.94448499 | 0.34492187 | 0.76724655 |
| ADAM15 | 65.7544908 | -0.1708276 | 0.52277142 | -0.3267731 | 0.74383953 | 0.93637438 |
| EFNA1 | 540.122766 | 0.39629895 | 0.44662498 | 0.88731927 | 3.75E-01 | 7.84E-01 |
| SLC50A1 | 137.89473 | -0.3235878 | 0.316102 | -1.0236815 | 0.30598574 | 0.74290739 |
| DPM3 | 180.986197 | 0.09120198 | 0.4418858 | 0.20639264 | 0.83648422 | 0.96511435 |
| MUC1 | 152.173723 | 0.75224771 | 0.67886403 | 1.10809775 | 0.2678196 | 0.71799256 |
| AC234582.1 | 98.3487824 | -0.3101836 | 0.40981072 | -0.7568949 | 0.44911285 | 0.8267002 |
| THBS3 | 131.54972 | -0.5410072 | 0.34106423 | -1.5862327 | 0.11268653 | 0.53755015 |
| MTX1 | 87.1157689 | 0.34982069 | 0.33535486 | 1.04313591 | 0.29688535 | 0.73673761 |
| GBA | 138.115055 | -0.0906382 | 0.27017987 | -0.3354734 | 0.73726802 | 0.93552616 |
| FAM189B | 101.079464 | -0.1482988 | 0.32226913 | -0.4601706 | 0.6453938 | 0.90537455 |

| | | | | | | |
|---|---|---|---|---|---|---|
| SCAMP3 | 263.603923 | -0.1845272 | 0.25159595 | -0.7334269 | 0.4632981 | 0.83531269 |
| CLK2 | 280.90995 | -0.1460354 | 0.20194364 | -0.7231493 | 0.46958815 | 0.83900851 |
| HCN3 | 49.5912513 | 0.17738908 | 0.29063291 | 0.61035443 | 0.54162705 | 0.86582694 |
| FDPS | 668.849637 | 0.12642396 | 0.28325324 | 0.44632837 | 0.65536006 | 0.90919768 |
| RUSC1 | 111.309646 | 0.12984978 | 0.36994539 | 0.35099714 | 0.72559049 | 0.93269023 |
| ASH1L | 4612.74488 | 0.08712044 | 0.12282828 | 0.70928649 | 0.47814671 | 0.84199814 |
| ASH1L-AS1 | 46.4519329 | 0.30542027 | 0.35659766 | 0.85648423 | 0.39172999 | 0.79765091 |
| MSTO1 | 78.9873094 | -0.2350231 | 0.34367943 | -0.6838439 | 0.49407373 | 0.84882719 |
| YY1AP1 | 819.449875 | -0.0056833 | 0.15861563 | -0.0358304 | 0.97141756 | 0.99327086 |
| DAP3 | 841.118634 | -0.1999152 | 0.19811966 | -1.0090632 | 0.31294435 | 0.74703371 |
| GON4L | 1117.59831 | -0.2065093 | 0.21898576 | -0.943026 | 0.34566762 | 0.76802752 |
| SYT11 | 63.6516074 | 0.84511001 | 0.45852244 | 1.84311593 | 0.06531209 | 0.44218726 |
| RIT1 | 315.996916 | 0.36669735 | 0.39818199 | 0.92092902 | 0.35708748 | 0.77448758 |
| KHDC4 | 1090.87532 | -0.1625634 | 0.24555204 | -0.6620323 | 0.50795055 | 0.85643442 |
| ARHGEF2 | 327.273044 | -0.0610212 | 0.26394902 | -0.2311856 | 0.81717062 | 0.95929672 |
| SSR2 | 347.248087 | 0.4438326 | 0.23975279 | 1.85120935 | 0.06413944 | 0.43894277 |
| UBQLN4 | 178.317711 | 0.26336188 | 0.23990012 | 1.09779801 | 0.2722927 | 0.72070318 |
| LAMTOR2 | 127.671542 | 0.27721338 | 0.27814254 | 0.99665943 | 0.31892985 | 0.75094569 |
| RAB25 | 173.287049 | 0.63627254 | 0.33186191 | 1.917281 | 0.05520224 | 0.41810002 |
| LMNA | 853.80552 | 0.56005596 | 0.33703612 | 1.661709 | 0.09657113 | 0.51229242 |
| SEMA4A | 289.461709 | 0.62387394 | 0.20812545 | 2.99758606 | 0.00272127 | 0.09842207 |
| SLC25A44 | 174.837959 | -0.1683976 | 0.2958324 | -0.5692333 | 0.56919785 | 0.87789095 |
| SMG5 | 499.332402 | -0.2171747 | 0.18487205 | -1.1747299 | 0.24010279 | 0.69159976 |
| TMEM79 | 47.5865233 | -0.2922594 | 0.40212578 | -0.726786 | 0.46735705 | 0.83762812 |
| GLMP | 93.5961981 | 0.44307751 | 0.31351624 | 1.41325216 | 0.15758159 | 0.6073933 |
| CCT3 | 507.108301 | 0.60628643 | 0.27784902 | 2.18207151 | 0.02910426 | 0.32942426 |
| MEF2D | 286.122836 | -0.333119 | 0.2794559 | -1.1920269 | 0.23325069 | 0.68484327 |
| NAXE | 126.074469 | 0.44533186 | 0.525913 | 0.84677857 | 0.39711855 | 0.80069929 |
| GPATCH4 | 96.9290506 | -0.1017272 | 0.34095473 | -0.2983598 | 0.76542854 | 0.94270299 |
| ISG20L2 | 130.663505 | -0.2386432 | 0.33815368 | -0.7057241 | 0.48035971 | 0.8433217 |
| RRNAD1 | 209.162853 | -0.4034595 | 0.202446 | -1.9929239 | 0.04626978 | 0.3904347 |
| MRPL24 | 102.148522 | 0.24549968 | 0.2690546 | 0.912453 | 0.3615303 | 0.77772231 |
| HDGF | 518.478158 | 0.12675134 | 0.24004717 | 0.52802679 | 5.97E-01 | 0.88883692 |
| PRCC | 403.356623 | -0.1821543 | 0.22849958 | -0.7971756 | 0.42534908 | 0.81466629 |
| ARHGEF11 | 1040.38754 | -0.1048852 | 0.1678368 | -0.624924 | 0.53202094 | 0.86397197 |
| ETV3 | 96.1599174 | 0.32128817 | 0.2595129 | 1.23804316 | 0.21570006 | 0.66885336 |
| KIRREL1 | 1073.03144 | -0.4046062 | 0.48390523 | -0.836127 | 0.40308348 | 0.80324472 |
| IFI16 | 209.043311 | -0.1822106 | 0.74453565 | -0.2447305 | 0.80666513 | 0.95566275 |
| DUSP23 | 726.411382 | 0.47551383 | 0.58398264 | 0.81426022 | 4.15E-01 | 0.80966301 |
| AL590560.2 | 52.7787673 | -0.515174 | 0.7801741 | -0.660332 | 0.50904077 | 0.8566594 |
| TAGLN2 | 383.456279 | 0.31348235 | 0.38196058 | 0.82071912 | 4.12E-01 | 8.07E-01 |
| KCNJ10 | 247.611949 | 1.65432191 | 0.32973393 | 5.01714185 | 5.24E-07 | 0.00076973 |
| IGSF8 | 86.1573787 | 0.33659491 | 0.41159998 | 0.81777194 | 0.41348742 | 0.80843359 |
| PEA15 | 116.056347 | 0.1157917 | 0.31968023 | 0.362211 | 7.17E-01 | 9.29E-01 |
| DCAF8 | 305.738425 | -0.1322835 | 0.30114042 | -0.439275 | 0.66046228 | 0.91099119 |
| AL139011.2 | 71.4934801 | 0.3723481 | 0.42788777 | 0.87020037 | 0.38419091 | 0.791697 |
| COPA | 1215.40675 | 0.06142212 | 0.17217091 | 0.35675084 | 0.72127834 | 0.93044898 |
| NCSTN | 190.317295 | 0.58751123 | 0.21854554 | 2.68827835 | 0.00718215 | 0.1729679 |
| VANGL2 | 50.7264729 | 1.29749915 | 0.62379211 | 2.08001854 | 0.03752383 | 0.36360429 |
| F11R | 258.972141 | 0.38051755 | 0.31799106 | 1.19662973 | 0.23145091 | 0.68363932 |
| AL591806.3 | 192.463531 | 0.45163864 | 0.36799786 | 1.22728608 | 0.21971509 | 0.67289974 |
| USF1 | 107.855056 | -0.9459538 | 0.39570651 | -2.390544 | 0.01682343 | 0.25960368 |
| NECTIN4 | 73.6529267 | -0.2075685 | 0.7373186 | -0.2815181 | 7.78E-01 | 0.94694572 |
| KLHDC9 | 57.8990969 | 0.48991836 | 0.43682779 | 1.1215366 | 0.26205952 | 0.712756 |

| | | | | | | |
|---|---|---|---|---|---|---|
| PFDN2 | 312.613195 | 0.40108411 | 0.27137067 | 1.4779936 | 0.13940949 | 0.58329426 |
| NIT1 | 344.077914 | 0.0879535 | 0.27576079 | 0.31894855 | 0.74976553 | 0.93775604 |
| DEDD | 164.29286 | 0.00878337 | 0.22085319 | 0.03977017 | 0.96827636 | 0.99248831 |
| UFC1 | 252.729262 | 0.43760236 | 0.36905249 | 1.18574558 | 0.23572279 | 0.68810212 |
| USP21 | 50.0938052 | -0.6738069 | 0.3138714 | -2.1467611 | 0.0318123 | 0.34024781 |
| PPOX | 104.820752 | 0.01883189 | 0.25334388 | 0.07433331 | 0.94074517 | 0.98715467 |
| B4GALT3 | 98.38792 | 0.31103694 | 0.35286833 | 0.88145326 | 0.37807254 | 0.78657429 |
| NDUFS2 | 448.270899 | 0.0891006 | 0.27468611 | 0.32437244 | 0.74565608 | 0.93675395 |
| TOMM40L | 44.271264 | -0.1503009 | 0.49510521 | -0.3035736 | 0.76145273 | 0.94202764 |
| SDHC | 739.017412 | 0.22765525 | 0.16242904 | 1.40156736 | 0.16104448 | 0.6121544 |
| AL592295.4 | 167.599747 | 0.00312569 | 0.36079372 | 0.00866337 | 0.99308772 | 0.99816851 |
| AL592295.3 | 59.1935588 | 0.23972798 | 0.35080032 | 0.68337447 | 0.49437025 | 0.84882719 |
| DUSP12 | 156.119911 | 0.05288174 | 0.32463292 | 0.16289703 | 0.87059951 | 0.9747202 |
| ATF6 | 1538.86211 | 0.00510589 | 0.23014427 | 0.02218561 | 0.9822999 | 0.99518326 |
| NOS1AP | 5071.18019 | -1.4274693 | 0.29574537 | -4.8266836 | 1.39E-06 | 0.0012225 |
| C1orf226 | 113.204485 | -0.6245176 | 0.44497341 | -1.4034943 | 0.1604695 | 0.61155269 |
| UHMK1 | 787.167637 | -0.150146 | 0.33899613 | -0.4429137 | 0.65782814 | 0.91037018 |
| UAP1 | 341.928731 | -0.9011965 | 0.47977387 | -1.8783777 | 0.06032952 | 0.42866735 |
| DDR2 | 123.943396 | -1.6657429 | 0.39644558 | -4.2016936 | 2.65E-05 | 0.00833191 |
| HSD17B7 | 319.823452 | -0.0727584 | 0.47992139 | -0.1516049 | 0.87949859 | 0.9765717 |
| RGS5 | 96.1267041 | -0.2915451 | 0.36951003 | -0.7890047 | 0.43010929 | 0.81675008 |
| NUF2 | 87.2677649 | 0.08089066 | 0.61158325 | 0.13226434 | 0.89477521 | 0.97788747 |
| PBX1 | 4883.00623 | 0.024665 | 0.20308263 | 0.12145304 | 9.03E-01 | 9.80E-01 |
| MGST3 | 1933.95075 | 0.44865085 | 0.21818301 | 2.05630518 | 0.03975311 | 0.37279391 |
| ALDH9A1 | 274.923475 | 0.84081904 | 0.28427256 | 2.95779175 | 0.00309851 | 0.1071796 |
| TMCO1 | 499.268522 | 0.40022521 | 0.17401946 | 2.29988761 | 0.02145459 | 0.29288643 |
| AL451074.2 | 40.7398102 | -0.4013739 | 0.5223705 | -0.7683702 | 0.44226729 | 0.82285053 |
| UCK2 | 250.612039 | -0.1286789 | 0.38242894 | -0.3364779 | 0.73651054 | 0.93552616 |
| FAM78B | 144.603566 | -0.2458989 | 0.4828008 | -0.5093174 | 0.61052976 | 0.89339478 |
| POGK | 258.137416 | -0.1186347 | 0.34532639 | -0.3435436 | 0.73118955 | 0.93515518 |
| TADA1 | 168.17113 | 0.07716588 | 0.19708582 | 0.39153443 | 0.69540224 | 0.92229578 |
| POU2F1 | 2238.21704 | -0.1143783 | 0.13385359 | -0.8545028 | 0.39282644 | 0.79842711 |
| CREG1 | 208.790052 | 0.87966921 | 0.32528836 | 2.70427513 | 0.00684536 | 0.17028313 |
| MPZL1 | 773.01944 | -0.3407931 | 0.17264081 | -1.974001 | 4.84E-02 | 0.39627811 |
| ADCY10 | 82.6083262 | -0.2261899 | 0.52092177 | -0.434211 | 6.64E-01 | 0.91206611 |
| MPC2 | 497.269568 | 0.83945072 | 0.29865573 | 2.81076384 | 0.0049424 | 0.14034467 |
| DCAF6 | 3174.69709 | -0.1052564 | 0.23974641 | -0.4390322 | 0.66063821 | 0.91099119 |
| TIPRL | 304.751773 | 0.02447867 | 0.25178442 | 0.09722074 | 0.9225511 | 0.98215577 |
| SFT2D2 | 452.702341 | 0.00114755 | 0.15940677 | 0.00719888 | 9.94E-01 | 9.98E-01 |
| TBX19 | 60.6491349 | -0.4357627 | 0.31256164 | -1.3941657 | 0.16326758 | 0.61387934 |
| DPT | 54.7628485 | 1.74970663 | 0.83787489 | 2.08826718 | 0.03677374 | 0.36038369 |
| AL031275.1 | 62.665804 | 1.58517079 | 0.79746675 | 1.98775785 | 0.04683848 | 0.39232052 |
| LINC00626 | 36.5615312 | 2.54907818 | 0.91338807 | 2.79079425 | 0.00525789 | 0.14621358 |
| LINC00970 | 2153.96578 | 1.0314513 | 0.80269027 | 1.28499291 | 0.19879476 | 0.65113421 |
| ATP1B1 | 10315.7315 | 1.73375154 | 0.49831776 | 3.4792088 | 0.0005029 | 0.03839746 |
| NME7 | 1933.31789 | 0.01827808 | 0.17189311 | 0.10633398 | 0.91531737 | 0.9817213 |
| BLZF1 | 485.226448 | 0.23286567 | 0.1842457 | 1.26388654 | 0.20627075 | 0.65831072 |
| CCDC181 | 101.637115 | 0.24164905 | 0.31929278 | 0.75682591 | 0.44915417 | 0.8267002 |
| SLC19A2 | 706.736972 | -0.0710691 | 0.48701098 | -0.1459292 | 0.88397731 | 0.97664137 |
| C1orf112 | 120.159636 | -0.6110462 | 0.40907906 | -1.4937118 | 0.13525097 | 0.57708028 |
| SCYL3 | 455.05676 | -0.1113852 | 0.17356343 | -0.6417549 | 0.52103235 | 0.85943874 |
| KIFAP3 | 1012.30335 | 0.01006126 | 0.14900004 | 0.0675252 | 0.9461636 | 0.98882138 |
| GORAB | 155.657037 | -0.5241894 | 0.29749255 | -1.7620254 | 0.078065 | 0.47980319 |
| FMO4 | 597.357708 | 0.97837729 | 0.39779036 | 2.45952989 | 0.01391191 | 0.24469033 |

| | | | | | | |
|---|---|---|---|---|---|---|
| PRRC2C | 2427.24826 | -0.1573935 | 0.13928984 | -1.1299711 | 0.2584884 | 0.70896455 |
| MYOCOS | 210.264896 | 1.10639617 | 0.64554692 | 1.71388963 | 0.086549 | 0.49478465 |
| VAMP4 | 306.90262 | -0.0642713 | 0.2773744 | -0.2317133 | 0.81676073 | 0.95909544 |
| METTL13 | 117.152577 | -0.0679924 | 0.32085101 | -0.2119126 | 0.83217523 | 0.96417211 |
| DNM3 | 3263.42242 | 0.16115082 | 0.25252526 | 0.63815723 | 0.52337134 | 0.86006369 |
| PIGC | 460.351005 | 0.21747763 | 0.28927442 | 0.75180389 | 0.452169 | 0.82885146 |
| SUCO | 862.693203 | -0.1115578 | 0.2420211 | -0.4609423 | 0.64484003 | 0.90507831 |
| AL645568.1 | 885.026638 | -0.1708865 | 0.25232812 | -0.6772391 | 4.98E-01 | 0.85092921 |
| PRDX6 | 998.527253 | 1.08752012 | 0.26413457 | 4.11729566 | 3.83E-05 | 0.00992861 |
| SLC9C2 | 55.1637482 | 0.75698146 | 0.58390066 | 1.29642166 | 0.19483025 | 0.64840333 |
| ANKRD45 | 64.021375 | 0.87155009 | 0.48993725 | 1.77890145 | 0.07525592 | 0.47201114 |
| KLHL20 | 758.804281 | -0.160601 | 0.20512884 | -0.7829277 | 0.43366958 | 0.81899321 |
| DARS2 | 158.444808 | -0.397865 | 0.28131014 | -1.4143288 | 0.15726537 | 0.60704801 |
| ZBTB37 | 457.345213 | -0.1019891 | 0.33077808 | -0.3083308 | 0.75783063 | 0.94056395 |
| RC3H1 | 1632.87892 | -0.3049871 | 0.18765662 | -1.6252405 | 0.10411133 | 0.52442862 |
| RABGAP1L | 6220.47015 | -0.0380621 | 0.17852868 | -0.2131986 | 0.83117207 | 0.96382687 |
| Z99127.1 | 48.5294556 | -0.6011975 | 0.45795164 | -1.312797 | 0.18925136 | 0.64432929 |
| CACYBP | 159.537513 | 0.22160869 | 0.27010662 | 0.82044894 | 0.41196023 | 0.80679792 |
| MRPS14 | 183.828735 | 0.36169219 | 0.22227461 | 1.62723124 | 0.10368799 | 0.52435479 |
| KIAA0040 | 357.992019 | -0.287085 | 0.34920399 | -0.8221127 | 0.41101276 | 0.80669651 |
| AL513329.1 | 68.7773878 | -0.9966678 | 0.6140158 | -1.6231958 | 0.10454756 | 0.52442862 |
| COP1 | 3073.33415 | -0.2974203 | 0.15900085 | -1.8705582 | 0.06140635 | 0.43236485 |
| AL359265.3 | 117.40291 | -1.0370417 | 0.65986192 | -1.5716042 | 0.11604238 | 0.54556096 |
| PAPPA2 | 4100.25713 | 2.74818387 | 0.82636648 | 3.32562362 | 0.00088221 | 0.05165902 |
| SEC16B | 125.44269 | -0.3648286 | 0.59533653 | -0.6128106 | 0.54000156 | 0.86566791 |
| RASAL2 | 3745.6303 | -0.3319505 | 0.27825972 | -1.1929521 | 0.23288814 | 0.68484327 |
| RALGPS2 | 2958.23561 | -0.103684 | 0.24852863 | -0.4171912 | 0.67653855 | 0.91658712 |
| ANGPTL1 | 285.365744 | 0.11128252 | 0.39761104 | 0.27987783 | 0.77957123 | 0.94758042 |
| FAM20B | 449.094782 | -0.0537138 | 0.15857632 | -0.3387253 | 0.73481666 | 0.93552616 |
| TOR3A | 105.089129 | -0.1702265 | 0.39282386 | -0.4333404 | 0.66476749 | 0.91220795 |
| ABL2 | 1747.99903 | -0.0702607 | 0.16346801 | -0.4298133 | 0.66733142 | 0.91360856 |
| SOAT1 | 447.417534 | 0.30298526 | 0.20587979 | 1.47166105 | 1.41E-01 | 0.5872148 |
| AXDND1 | 140.898082 | -0.6102192 | 0.66923873 | -0.911811 | 0.36186824 | 0.77784066 |
| TDRD5 | 156.147406 | 0.34667679 | 0.47313388 | 0.7327245 | 0.46372648 | 0.83531269 |
| TOR1AIP2 | 1280.58882 | 0.1913163 | 0.32996787 | 0.5798028 | 0.56204761 | 0.87403145 |
| TOR1AIP1 | 392.251413 | -0.1039002 | 0.25686546 | -0.4044926 | 0.68585055 | 0.92053037 |
| CEP350 | 1999.29093 | -0.0100552 | 0.12820894 | -0.0784279 | 0.93748765 | 0.98661414 |
| QSOX1 | 198.791186 | 0.17494397 | 0.21008139 | 0.83274375 | 4.05E-01 | 8.05E-01 |
| ACBD6 | 1827.84568 | 0.05520329 | 0.15327582 | 0.36015655 | 0.71873006 | 0.9298452 |
| XPR1 | 2913.5826 | -0.1328963 | 0.25339945 | -0.5244536 | 0.59996309 | 0.88883692 |
| KIAA1614 | 269.305667 | 0.49009892 | 0.4335369 | 1.13046645 | 0.25827973 | 0.70896455 |
| AL162431.2 | 53.050233 | -0.4820968 | 0.33812017 | -1.4258149 | 0.15392178 | 0.60292313 |
| STX6 | 542.30705 | -0.3296443 | 0.2452529 | -1.3440996 | 0.17891618 | 0.63284534 |
| MR1 | 202.774996 | 0.11208561 | 0.20050238 | 0.55902383 | 0.57614545 | 0.88123035 |
| CACNA1E | 190.945904 | -2.4349514 | 0.97279914 | -2.5030362 | 0.0123133 | 0.22749396 |
| GLUL | 711.340882 | 0.55461169 | 0.31943541 | 1.73622484 | 8.25E-02 | 4.86E-01 |
| RNASEL | 111.158645 | 0.44915188 | 0.44977563 | 0.9986132 | 0.3179821 | 0.75004029 |
| NPL | 99.0846899 | -0.6430992 | 0.32090324 | -2.0040283 | 0.04506703 | 0.38755881 |
| DHX9 | 1269.90208 | -0.0722822 | 0.27888151 | -0.259186 | 0.79549172 | 0.95278473 |
| LAMC1 | 1736.93458 | -0.0925704 | 0.25263954 | -0.3664129 | 7.14E-01 | 9.28E-01 |
| LAMC2 | 462.588297 | -0.3597281 | 0.71216062 | -0.5051221 | 6.13E-01 | 0.89441126 |
| SMG7 | 1629.95829 | -0.1454885 | 0.17589592 | -0.8271284 | 0.40816428 | 0.80588056 |
| ARPC5 | 439.847736 | 0.0463644 | 0.26465895 | 0.17518544 | 0.86093392 | 0.9720639 |
| RGL1 | 211.154239 | 0.47795287 | 0.35118801 | 1.36096008 | 0.1735263 | 0.6286116 |

| | | | | | | |
|---|---|---|---|---|---|---|
| TSEN15 | 494.299889 | -0.0080108 | 0.20080287 | -0.0398937 | 9.68E-01 | 0.99248831 |
| C1orf21 | 142.720677 | 0.32586837 | 0.52762975 | 0.61760802 | 0.53683376 | 0.86466169 |
| EDEM3 | 886.165502 | -0.0427222 | 0.265659 | -0.1608159 | 0.8722384 | 0.97548023 |
| FAM129A | 86.9895231 | 1.31288576 | 0.47740573 | 2.75004188 | 5.96E-03 | 1.58E-01 |
| RNF2 | 479.458259 | -0.0835943 | 0.19699381 | -0.4243498 | 0.67131073 | 0.91548043 |
| TRMT1L | 420.899641 | -0.058986 | 0.21178363 | -0.2785201 | 0.78061314 | 0.9480401 |
| SWT1 | 706.080266 | -0.5217235 | 0.15455773 | -3.3755895 | 0.00073658 | 0.04792832 |
| IVNS1ABP | 6950.45927 | -0.3470511 | 0.46053057 | -0.7535898 | 0.45109557 | 0.82814845 |
| LINC01350 | 172.547214 | -0.9784988 | 1.08494903 | -0.9018846 | 0.36711819 | 0.78095617 |
| AL133383.1 | 40.3175489 | -1.5344975 | 0.8765722 | -1.750566 | 0.0800207 | 0.48282462 |
| HMCN1 | 1691.1738 | -0.878088 | 0.78604317 | -1.117099 | 0.26395201 | 0.71445861 |
| TPR | 1347.90001 | -0.1422007 | 0.12573318 | -1.1309723 | 0.25806674 | 0.70896455 |
| ODR4 | 809.518579 | 0.07699939 | 0.15649886 | 0.49201248 | 0.62271052 | 0.89835991 |
| AL596220.1 | 241.333922 | 0.28052413 | 0.43894805 | 0.63908275 | 0.5227691 | 0.86006369 |
| PLA2G4A | 73.8696577 | 2.54258199 | 0.64335224 | 3.95208385 | 7.75E-05 | 0.01421317 |
| ERVMER61-1 | 1096.86582 | -0.4699601 | 0.31306963 | -1.5011362 | 0.13332034 | 0.57682553 |
| BRINP3 | 546.210796 | 1.12412168 | 0.75190735 | 1.49502685 | 0.13490744 | 0.57708028 |
| AL954650.1 | 45.9581213 | 1.49647569 | 1.02114972 | 1.46548118 | 1.43E-01 | 5.88E-01 |
| AL390957.1 | 152.965255 | -0.2338719 | 0.39458037 | -0.5927105 | 0.55337494 | 0.87104567 |
| AL136454.1 | 312.343416 | 0.46467297 | 0.47411952 | 0.98007558 | 0.32704881 | 0.75563893 |
| UCHL5 | 431.996835 | -0.0681937 | 0.16347955 | -0.417139 | 0.67657677 | 0.91658712 |
| TROVE2 | 701.236707 | 0.26922367 | 0.14440905 | 1.86431294 | 0.06227777 | 0.43419765 |
| GLRX2 | 101.539985 | 0.03491899 | 0.40709473 | 0.08577609 | 0.93164442 | 0.98357046 |
| CDC73 | 1560.95917 | -0.089355 | 0.19221358 | -0.4648733 | 0.64202216 | 0.90454089 |
| LINC01031 | 52.586666 | -1.8489895 | 0.68133465 | -2.7137758 | 0.00665212 | 0.16894224 |
| KCNT2 | 50.2480795 | 1.10818268 | 0.4835982 | 2.29153601 | 0.02193243 | 0.29509055 |
| ZBTB41 | 356.151282 | 0.13503553 | 0.16695789 | 0.80879992 | 0.41863024 | 0.81125469 |
| CRB1 | 226.841998 | 1.06297151 | 0.80797672 | 1.31559671 | 0.18830944 | 0.64273368 |
| DENND1B | 2980.18949 | -0.5167627 | 0.31833375 | -1.6233362 | 0.10451755 | 0.52442862 |
| NEK7 | 2375.21054 | -0.0120549 | 0.15776101 | -0.0764122 | 0.9390912 | 0.98715467 |
| LINC00862 | 167.210534 | 0.09769791 | 0.6741935 | 0.14491078 | 0.88478131 | 0.97664137 |
| AC104461.1 | 167.989857 | -0.3195939 | 0.60588547 | -0.5274823 | 0.59785867 | 0.88883692 |
| ZNF281 | 164.656346 | 0.34722218 | 0.40653102 | 0.85410994 | 0.39304408 | 0.79843808 |
| DDX59 | 310.830633 | -0.0655144 | 0.21899347 | -0.2991616 | 0.76481674 | 0.94256992 |
| CAMSAP2 | 1353.4334 | -0.267083 | 0.23037897 | -1.15932 | 0.24632576 | 0.6967274 |
| INAVA | 229.729277 | -0.1545605 | 0.32999435 | -0.4683733 | 6.40E-01 | 9.03E-01 |
| TMEM9 | 234.373344 | 0.96757661 | 0.36512829 | 2.64996337 | 0.00805005 | 0.1802126 |
| LAD1 | 113.192464 | 0.95073914 | 0.49922376 | 1.90443489 | 0.05685357 | 0.41872623 |
| CSRP1 | 155.722194 | 0.28492322 | 0.4188249 | 0.68029198 | 0.4963196 | 0.8506404 |
| AC096677.1 | 252.061434 | 0.50102885 | 0.45309247 | 1.10579821 | 0.26881387 | 0.71818262 |
| NAV1 | 147.547689 | -0.1504679 | 0.46204534 | -0.3256562 | 0.74468446 | 0.93675395 |
| IPO9-AS1 | 65.0671427 | 0.40398717 | 0.33624889 | 1.20145278 | 0.22957561 | 0.68191236 |
| IPO9 | 779.854508 | -0.2955273 | 0.21603205 | -1.3679789 | 0.17131868 | 0.62772493 |
| SHISA4 | 43.63191 | 0.40224607 | 0.49431183 | 0.81374964 | 0.41578839 | 0.80966301 |
| TIMM17A | 262.236123 | -0.0219324 | 0.31603216 | -0.0693993 | 0.94467175 | 0.98813503 |
| RNPEP | 331.549655 | -0.2900838 | 0.22395138 | -1.2952982 | 0.19521739 | 0.64871108 |
| ELF3 | 1752.72027 | -0.1195968 | 0.40511017 | -0.2952204 | 7.68E-01 | 9.43E-01 |
| ARL8A | 200.42531 | -0.1919152 | 0.23871326 | -0.8039571 | 0.42142175 | 0.81266383 |
| PPP1R12B | 1938.04621 | -0.1132079 | 0.1573033 | -0.7196789 | 0.47172273 | 0.83921324 |
| KDM5B | 1465.17584 | -0.2095354 | 0.21217611 | -0.9875541 | 0.32337105 | 0.75362713 |
| RABIF | 64.776672 | -0.3412722 | 0.44670764 | -0.7639721 | 0.44488384 | 0.8252311 |
| KLHL12 | 244.614449 | -0.301772 | 0.22346586 | -1.3504165 | 0.17688241 | 0.63025747 |
| ADIPOR1 | 172.064054 | 0.11615274 | 0.19737406 | 0.58849041 | 0.55620317 | 0.87212955 |
| CYB5R1 | 142.751505 | 0.88883926 | 0.29518345 | 3.01114195 | 0.00260267 | 0.09654879 |

| | | | | | | |
|---|---|---|---|---|---|---|
| TMEM183A | 138.503723 | -0.2858965 | 0.24163392 | -1.1831804 | 0.23673766 | 0.68878143 |
| PPFIA4 | 89.3048762 | 1.17483096 | 1.01575055 | 1.15661367 | 0.24743024 | 0.69761067 |
| BTG2 | 139.555641 | 0.62101814 | 0.6288458 | 0.98755234 | 0.32337194 | 0.75362713 |
| ATP2B4 | 596.645129 | 0.40896132 | 0.45056963 | 0.90765398 | 0.36406107 | 0.78015617 |
| SNRPE | 147.479358 | -0.0324589 | 0.33946724 | -0.0956171 | 0.9238247 | 0.98215577 |
| SOX13 | 257.837167 | -0.0127171 | 0.28556239 | -0.0445335 | 0.96447917 | 0.99226095 |
| ETNK2 | 104.14743 | 1.08435336 | 0.23420964 | 4.62984088 | 3.66E-06 | 0.00268544 |
| GOLT1A | 109.309051 | 1.03357245 | 0.57041319 | 1.81197151 | 0.0699906 | 0.45949593 |
| PLEKHA6 | 1576.45036 | -0.1129261 | 0.25521125 | -0.4424808 | 0.65814134 | 0.91037018 |
| AL606489.1 | 65.1838446 | -0.0283199 | 0.62068235 | -0.045627 | 0.96360755 | 0.99224805 |
| PPP1R15B | 217.60148 | 0.20784072 | 0.39793785 | 0.52229442 | 0.60146535 | 0.88920716 |
| PIK3C2B | 46.9271851 | -0.155752 | 0.53179689 | -0.2928787 | 0.76961488 | 0.94386043 |
| MDM4 | 1638.17994 | -0.2015423 | 0.20268609 | -0.9943566 | 0.32004927 | 0.75104498 |
| LRRN2 | 404.409068 | -0.7456181 | 0.75255487 | -0.9907824 | 0.32179184 | 0.75240254 |
| RBBP5 | 322.878633 | -0.0983922 | 0.18892021 | -0.5208137 | 0.60249659 | 0.88971776 |
| DSTYK | 767.430957 | -0.039786 | 0.20082932 | -0.1981085 | 0.84296015 | 0.96764236 |
| NUAK2 | 638.296671 | 0.22976853 | 0.58253866 | 0.39442624 | 0.69326638 | 0.92224682 |
| LEMD1 | 65.9616045 | -0.9183678 | 0.65078231 | -1.4111751 | 1.58E-01 | 6.08E-01 |
| CDK18 | 296.25464 | 0.3777282 | 0.57523894 | 0.65664574 | 0.51140873 | 0.85762435 |
| MFSD4A | 1505.78628 | 0.80766067 | 0.62774007 | 1.2866164 | 0.19822802 | 0.65053266 |
| ELK4 | 887.92919 | -0.0523687 | 0.37595989 | -0.1392933 | 0.88921835 | 0.97729111 |
| SLC45A3 | 58.8280265 | -0.6028969 | 0.5987243 | -1.0069692 | 0.31394958 | 0.74777437 |
| NUCKS1 | 1761.23255 | 0.05995846 | 0.160696 | 0.37311731 | 0.70906114 | 0.92577218 |
| RAB29 | 201.468227 | 0.07473055 | 0.33367292 | 0.22396348 | 0.82278572 | 0.96102012 |
| SLC41A1 | 115.223375 | 0.21950988 | 0.21620322 | 1.01529426 | 0.30996558 | 0.74455572 |
| RHEX | 609.308255 | -0.6643232 | 0.67770575 | -0.9802532 | 0.32696116 | 0.75556858 |
| SRGAP2 | 126.934283 | -0.9577144 | 0.52471045 | -1.8252246 | 0.06796714 | 0.45250883 |
| EIF2D | 181.005762 | 0.00513042 | 0.29244216 | 0.01754337 | 0.98600314 | 0.99618444 |
| DYRK3 | 170.492876 | 0.30092793 | 0.35101601 | 0.85730545 | 0.3912761 | 0.7973532 |
| MAPKAPK2 | 1133.42213 | -0.2121522 | 0.25714766 | -0.8250211 | 0.40935964 | 0.80623405 |
| PIGR | 594.338417 | 2.86879828 | 0.58353944 | 4.91620285 | 8.82E-07 | 0.00085043 |
| C1orf116 | 86.1619485 | 0.72689046 | 0.60955023 | 1.19250296 | 0.23306409 | 0.68484327 |
| PFKFB2 | 563.036701 | -0.0811476 | 0.30532422 | -0.2657752 | 0.79041238 | 0.95130361 |
| YOD1 | 142.701323 | -0.2853547 | 0.33700502 | -0.8467371 | 0.39714166 | 0.80069929 |
| CD55 | 314.816738 | 0.22042653 | 0.43008678 | 0.5125164 | 6.08E-01 | 0.89246894 |
| CD46 | 1617.42497 | -0.2609512 | 0.20289774 | -1.2861217 | 0.1984006 | 0.65077564 |
| MIR29B2CH( | 2568.69585 | 0.47685244 | 0.34455187 | 1.38397868 | 0.16636498 | 0.61901833 |
| PLXNA2 | 676.758503 | -0.628774 | 0.49719559 | -1.2646412 | 0.20599999 | 0.65816052 |
| LAMB3 | 239.39009 | -0.2484151 | 0.6326356 | -0.3926669 | 6.95E-01 | 9.22E-01 |
| AL031316.1 | 42.1210891 | 0.51193194 | 0.70613749 | 0.72497488 | 0.46846744 | 0.83825637 |
| G0S2 | 606.339449 | 0.92271185 | 0.58342391 | 1.58154618 | 0.11375321 | 0.54000194 |
| IRF6 | 136.786961 | 0.67640956 | 0.23143659 | 2.92265607 | 0.0034706 | 0.1138728 |
| UTP25 | 129.276981 | -0.4568214 | 0.34360884 | -1.329481 | 0.18368934 | 0.63834582 |
| SYT14 | 87.4678638 | 0.4287042 | 0.59640182 | 0.71881771 | 0.47225324 | 0.83921324 |
| SERTAD4-AS | 40.84922 | 0.46972626 | 0.85256835 | 0.55095438 | 0.58166495 | 0.88312785 |
| SERTAD4 | 40.4105513 | 0.94728624 | 0.86978294 | 1.08910648 | 0.27610693 | 0.72241028 |
| HHAT | 528.900805 | 0.35387419 | 0.51986587 | 0.68070287 | 0.49605952 | 0.85030498 |
| KCNH1 | 85.529346 | 0.96513652 | 0.56247514 | 1.7158741 | 0.08618509 | 0.49453468 |
| RCOR3 | 2113.52099 | -0.1284899 | 0.16310507 | -0.7877738 | 0.43082903 | 0.81706845 |
| TRAF5 | 298.0699 | 0.34361938 | 0.32692937 | 1.0510508 | 0.29323526 | 0.73527014 |
| LINC00467 | 442.560279 | -0.389018 | 0.31486472 | -1.2355082 | 0.21664142 | 0.66978787 |
| SLC30A1 | 118.473652 | 0.83290842 | 0.25234505 | 3.30067275 | 0.00096453 | 0.05563545 |
| LPGAT1 | 1525.89858 | -0.0030198 | 0.30202133 | -0.0099985 | 0.99202248 | 0.99776307 |
| INTS7 | 302.58193 | -0.4801243 | 0.35883753 | -1.3379992 | 0.18089669 | 0.63540877 |

| | | | | | | |
|---|---|---|---|---|---|---|
| PPP2R5A | 2491.76397 | -0.5012634 | 0.35414907 | -1.4154024 | 0.15695056 | 0.60654181 |
| AL360091.3 | 385.032117 | -0.424933 | 0.64425265 | -0.6595751 | 0.50952656 | 0.8568242 |
| TMEM206 | 107.274698 | -0.3045756 | 0.46632589 | -0.653139 | 0.51366666 | 0.85762435 |
| NENF | 243.43263 | 0.26517783 | 0.36403072 | 0.72844904 | 0.46633875 | 0.8367113 |
| ATF3 | 130.412374 | -0.1128322 | 0.87520619 | -0.1289207 | 0.89742039 | 0.97823597 |
| NSL1 | 540.583944 | -0.050908 | 0.17770096 | -0.2864813 | 0.77450953 | 0.94621715 |
| TATDN3 | 410.797948 | -0.0477314 | 0.34136882 | -0.1398235 | 0.88879943 | 0.97729111 |
| FLVCR1-DT | 56.6036944 | 0.26313562 | 0.39341163 | 0.66885572 | 0.50358752 | 0.85525037 |
| FLVCR1 | 320.855865 | -0.104647 | 0.16666602 | -0.6278842 | 0.53007978 | 0.86361777 |
| ANGEL2 | 443.030364 | -0.2506218 | 0.1552047 | -1.6147822 | 0.10635787 | 0.52735777 |
| RPS6KC1 | 1322.44092 | -0.4498331 | 0.14641583 | -3.0722986 | 0.00212417 | 0.08823327 |
| PROX1 | 1228.5397 | -0.5435001 | 0.59811139 | -0.9086938 | 3.64E-01 | 0.779589 |
| SMYD2 | 326.963304 | -0.0979266 | 0.3557031 | -0.2753043 | 0.78308244 | 0.94937658 |
| PTPN14 | 5119.94028 | -0.2747919 | 0.20962819 | -1.3108536 | 0.18990723 | 0.64452327 |
| KCTD3 | 1297.0181 | -0.5375025 | 0.2343831 | -2.2932649 | 0.02183276 | 0.29509055 |
| USH2A | 115.180688 | 0.25132236 | 0.42323926 | 0.59380684 | 5.53E-01 | 0.87104567 |
| ESRRG | 51368.916 | -0.4179547 | 0.21790651 | -1.918046 | 5.51E-02 | 4.18E-01 |
| GPATCH2 | 1802.89408 | -0.1768905 | 0.2716778 | -0.6511038 | 0.51497948 | 0.85869987 |
| SPATA17 | 438.167254 | 0.06497759 | 0.35007581 | 0.18561004 | 0.85275058 | 0.97076831 |
| RRP15 | 311.320165 | -0.1433666 | 0.35174411 | -0.4075878 | 0.6835763 | 0.9200778 |
| TGFB2 | 106.196315 | -0.6074789 | 0.90943383 | -0.6679749 | 0.50414963 | 0.85525037 |
| LYPLAL1-DT | 107.962876 | -1.0519333 | 0.44237121 | -2.3779425 | 1.74E-02 | 2.64E-01 |
| LYPLAL1 | 3025.35351 | -0.041798 | 0.34737848 | -0.1203242 | 0.90422636 | 0.97989015 |
| AC096642.1 | 72.0544289 | 0.63924014 | 0.65326792 | 0.97852677 | 0.32781386 | 0.75633196 |
| AL513283.1 | 72.5013315 | -0.9596321 | 1.05197457 | -0.9122198 | 0.361653 | 0.77772231 |
| ZC3H11B | 83.8858386 | 0.09689782 | 0.33005907 | 0.2935772 | 0.769081 | 0.94386043 |
| SLC30A10 | 48.7271522 | -0.0022989 | 0.9398537 | -0.002446 | 0.9980484 | 0.99969006 |
| EPRS | 610.476651 | 0.12412565 | 0.17633325 | 0.7039265 | 0.48147855 | 0.8437862 |
| BPNT1 | 195.382957 | 0.07809564 | 0.23264339 | 0.3356882 | 0.737106 | 0.93552616 |
| IARS2 | 454.629947 | -0.0242876 | 0.25166934 | -0.0965058 | 0.92311886 | 0.98215577 |
| RAB3GAP2 | 1812.92064 | -0.0324905 | 0.15675045 | -0.2072751 | 0.83579504 | 0.96511435 |
| MARK1 | 572.200769 | 0.40374937 | 0.51506397 | 0.78388199 | 0.43310936 | 0.81856368 |
| C1orf115 | 75.9897611 | 1.09616492 | 0.37507403 | 2.92252948 | 0.00347201 | 0.1138728 |
| 2-Mar | 679.800086 | 0.53877878 | 0.28632545 | 1.88170065 | 0.05987667 | 0.42682728 |
| 1-Mar | 145.689413 | -0.0059327 | 0.39070843 | -0.0151845 | 0.98788497 | 0.99653761 |
| AL513314.2 | 222.606368 | 0.01304453 | 0.34244143 | 0.03809273 | 0.96961375 | 0.99299658 |
| TAF1A | 92.4775163 | -0.3175468 | 0.31423885 | -1.0105269 | 0.31224294 | 0.74687836 |
| MIA3 | 534.896565 | -0.1140337 | 0.19369786 | -0.5887194 | 0.55604949 | 0.87212955 |
| AIDA | 224.268893 | 0.02268767 | 0.21257326 | 0.10672869 | 0.91500422 | 0.9816913 |
| BROX | 490.981282 | -0.0270984 | 0.18213847 | -0.1487793 | 0.88172775 | 0.97664137 |
| FAM177B | 325.540721 | 0.14289329 | 0.75987786 | 0.18804771 | 8.51E-01 | 0.97056586 |
| AC093152.1 | 413.407421 | -0.3058972 | 0.20114388 | -1.5207881 | 0.12831301 | 0.56762585 |
| TLR5 | 436.393009 | -0.1492491 | 0.31560272 | -0.4729019 | 0.63628313 | 0.90237131 |
| SUSD4 | 1602.60608 | -0.8270833 | 0.41974841 | -1.9704262 | 0.04878955 | 0.39762623 |
| CAPN8 | 96.9068487 | -0.2122516 | 0.55536529 | -0.3821838 | 7.02E-01 | 9.25E-01 |
| CAPN2 | 1514.02259 | -0.0631871 | 0.26849574 | -0.2353374 | 0.81394691 | 0.9582375 |
| TP53BP2 | 491.475302 | -0.1902602 | 0.19558691 | -0.9727654 | 0.33066993 | 0.75909265 |
| FBXO28 | 899.723241 | -0.0992622 | 0.1982671 | -0.5006491 | 0.61661813 | 0.89589954 |
| DEGS1 | 370.881312 | 0.78494801 | 0.3342246 | 2.34856442 | 0.01884594 | 0.27690323 |
| NVL | 718.557041 | -0.3168344 | 0.18261194 | -1.7350148 | 0.08273818 | 0.48594425 |
| CNIH4 | 587.768795 | -0.2121144 | 0.2668187 | -0.7949759 | 0.42662751 | 0.81466629 |
| WDR26 | 1621.60874 | -0.2353738 | 0.2191323 | -1.0741174 | 0.28277007 | 0.72604837 |
| CNIH3 | 690.414945 | -0.1222554 | 0.42865931 | -0.2852042 | 0.77548771 | 0.94680268 |
| DNAH14 | 3853.1588 | -0.1878634 | 0.2006788 | -0.9361397 | 0.34920126 | 0.76982416 |

| | | | | | | |
|---|---|---|---|---|---|---|
| LBR | 836.482168 | 0.00628227 | 0.23192347 | 0.02708769 | 0.97838979 | 0.99455895 |
| ENAH | 405.816243 | 0.01347095 | 0.33392965 | 0.04034069 | 0.96782152 | 0.99248831 |
| SRP9 | 511.594737 | 0.04717949 | 0.28573952 | 0.16511362 | 0.86885455 | 0.97442321 |
| EPHX1 | 210.092288 | -0.1596383 | 0.30047439 | -0.5312876 | 0.59521952 | 0.88859116 |
| AL591895.1 | 186.848475 | 0.08137478 | 0.27734057 | 0.29341102 | 7.69E-01 | 0.94386043 |
| TMEM63A | 553.061737 | -0.113065 | 0.28363443 | -0.3986293 | 6.90E-01 | 9.22E-01 |
| AL117348.2 | 49.4972381 | -0.7086942 | 0.540183 | -1.311952 | 0.18953634 | 0.64441858 |
| SDE2 | 98.0286482 | 0.03508402 | 0.24313992 | 0.1442956 | 0.88526705 | 0.97664137 |
| H3F3A | 225.848622 | -0.0536918 | 0.18991811 | -0.2827102 | 0.77739902 | 0.94694572 |
| ACBD3 | 881.087302 | 0.10016826 | 0.12474655 | 0.8029742 | 0.42198965 | 0.81266383 |
| LIN9 | 136.657844 | -0.3771271 | 0.2739212 | -1.3767724 | 0.16858264 | 0.62253512 |
| PARP1 | 476.642362 | 0.43711037 | 0.31172334 | 1.4022382 | 0.16084413 | 0.61166537 |
| STUM | 571.297783 | -0.5957676 | 0.50291346 | -1.1846324 | 0.23616284 | 0.68856409 |
| ITPKB | 1019.01057 | -0.1317472 | 0.20804672 | -0.6332578 | 0.52656529 | 0.86145663 |
| PSEN2 | 56.0580099 | 0.05015176 | 0.32299876 | 0.15526919 | 0.8766091 | 0.97631784 |
| COQ8A | 431.553906 | -0.4700696 | 0.28607768 | -1.6431538 | 0.10035111 | 0.51705755 |
| CDC42BPA | 5281.57155 | -0.1500442 | 0.31841249 | -0.4712259 | 0.63747942 | 0.90255504 |
| ZNF678 | 516.716094 | -0.3114252 | 0.1711157 | -1.8199688 | 0.06876376 | 0.45460483 |
| SNAP47 | 236.321811 | 0.06345699 | 0.17601573 | 0.36051886 | 0.71845916 | 0.9298398 |
| JMJD4 | 56.3291285 | -0.3488271 | 0.35206564 | -0.9908014 | 0.32178258 | 0.75240254 |
| ARF1 | 670.587393 | 0.32148025 | 0.19389453 | 1.65801613 | 9.73E-02 | 0.51355946 |
| C1orf35 | 108.629533 | -0.3592773 | 0.29919261 | -1.2008226 | 0.22982001 | 0.68217811 |
| MRPL55 | 113.612016 | 0.26546504 | 0.39837636 | 0.66636746 | 0.50517626 | 0.85582573 |
| GUK1 | 568.017043 | 0.61821801 | 0.30183891 | 2.04817203 | 0.04054315 | 0.37344844 |
| IBA57 | 60.9306466 | -0.0410611 | 0.32532175 | -0.1262168 | 0.89956032 | 0.97893329 |
| OBSCN-AS1 | 50.4786519 | 0.08416767 | 0.4208409 | 0.19999879 | 0.84148153 | 0.96661847 |
| OBSCN | 796.74138 | 0.00549798 | 0.2441999 | 0.02251427 | 0.98203773 | 0.99518016 |
| TRIM11 | 105.974701 | -0.087487 | 0.33877727 | -0.2582433 | 0.79621911 | 0.95278473 |
| RNF187 | 162.984459 | 0.63649566 | 0.31248412 | 2.03688963 | 0.0416611 | 0.37614595 |
| RAB4A | 340.64648 | 0.24921253 | 0.21101204 | 1.18103464 | 0.23758896 | 0.68973903 |
| CCSAP | 53.8497028 | -0.5653841 | 0.4425981 | -1.2774209 | 0.20145368 | 0.65525774 |
| NUP133 | 510.171105 | -0.1766677 | 0.14618009 | -1.2085621 | 0.22683111 | 0.67895131 |
| ABCB10 | 299.291218 | -0.1846791 | 0.20572716 | -0.8976892 | 0.36935127 | 0.78095617 |
| TAF5L | 157.563074 | -0.3519808 | 0.29679321 | -1.1859464 | 0.23564346 | 0.68802265 |
| URB2 | 75.0540392 | -0.2387167 | 0.40466154 | -0.5899169 | 0.55524639 | 0.87184917 |
| GALNT2 | 1158.24557 | -0.2046137 | 0.28985683 | -0.7059131 | 0.48024215 | 0.8433217 |
| PGBD5 | 113.32847 | 0.29522804 | 0.45904833 | 0.64313063 | 0.52013934 | 0.85935232 |
| COG2 | 347.745902 | 0.0373078 | 0.13772157 | 0.27089294 | 0.78647338 | 0.94978829 |
| C1orf198 | 120.103392 | 0.09807135 | 0.24169087 | 0.40577186 | 0.68491024 | 0.92053037 |
| TTC13 | 339.954021 | -0.5993729 | 0.24846686 | -2.4122851 | 0.01585288 | 0.25383866 |
| ARV1 | 213.359781 | 0.1621695 | 0.21305656 | 0.76115705 | 0.44656327 | 0.82571251 |
| FAM89A | 69.494501 | 0.33826026 | 0.35417696 | 0.95506004 | 3.40E-01 | 7.64E-01 |
| C1orf131 | 182.827984 | -0.1316633 | 0.258247 | -0.5098349 | 0.61016712 | 0.89339478 |
| GNPAT | 543.221695 | -0.1132159 | 0.14061971 | -0.805121 | 0.42074985 | 0.81260593 |
| EXOC8 | 71.8305045 | 0.62035928 | 0.32236186 | 1.92441898 | 0.05430209 | 0.41533078 |
| SPRTN | 146.320405 | -0.1841765 | 0.32777598 | -0.5618974 | 0.5741859 | 0.88010061 |
| EGLN1 | 1372.77252 | -0.210532 | 0.31452249 | -0.6693702 | 0.50325938 | 0.85525037 |
| TSNAX-DISC1 | 73.0949183 | -0.2153881 | 0.40166901 | -0.5362328 | 0.59179768 | 0.88736521 |
| DISC1 | 208.259763 | -0.2915167 | 0.36437315 | -0.8000498 | 4.24E-01 | 8.14E-01 |
| SIPA1L2 | 109.377112 | 1.27797863 | 0.65089006 | 1.96343241 | 0.04959595 | 0.39970282 |
| NTPCR | 321.70168 | -0.2192264 | 0.21252773 | -1.0315192 | 0.30229743 | 0.73905643 |
| PCNX2 | 2078.07472 | -0.119939 | 0.19138387 | -0.6266933 | 0.53086028 | 0.86396107 |
| MAP3K21 | 1163.59958 | 0.23350574 | 0.27807299 | 0.83972823 | 0.40106078 | 0.80243544 |
| KCNK1 | 984.684799 | 0.11407008 | 0.29604843 | 0.38530886 | 0.7000086 | 0.92367966 |

| | | | | | | |
|---|---|---|---|---|---|---|
| SLC35F3 | 120.765822 | -0.4276842 | 0.64372094 | -0.6643938 | 0.50643829 | 0.85626196 |
| AL355472.3 | 46.5194963 | 0.01741862 | 0.71178343 | 0.0244718 | 0.98047628 | 0.99518016 |
| AL355472.1 | 129.598525 | 0.23631909 | 0.48767838 | 0.48457979 | 0.62797445 | 0.89920892 |
| COA6 | 129.943675 | 0.15183783 | 0.34338078 | 0.442185 | 0.65835534 | 0.91037018 |
| TARBP1 | 588.966942 | -0.3249766 | 0.32995519 | -0.9849115 | 0.32466757 | 0.75478872 |
| IRF2BP2 | 738.001904 | 0.31225224 | 0.46734258 | 0.6681442 | 0.50404155 | 0.85525037 |
| TOMM20 | 487.834759 | 0.54145294 | 0.26863859 | 2.01554417 | 0.04384766 | 0.38458418 |
| ARID4B | 3707.64261 | -0.0147473 | 0.17737181 | -0.0831435 | 0.93373741 | 0.98457232 |
| TBCE | 67.8224122 | 0.15840352 | 0.43639241 | 0.36298413 | 0.71661674 | 0.92897478 |
| B3GALNT2 | 646.443395 | -0.062207 | 0.19249437 | -0.3231629 | 0.74657184 | 0.93695653 |
| LYST | 1310.88726 | 0.17605245 | 0.21315494 | 0.82593649 | 0.40884012 | 0.80602525 |
| GPR137B | 216.676833 | 0.65468765 | 0.49300579 | 1.32795125 | 0.18419421 | 0.63875594 |
| ERO1B | 529.054627 | 0.33801678 | 0.20950279 | 1.61342377 | 0.10665247 | 0.52775297 |
| EDARADD | 68.9287105 | 0.19953417 | 0.47561201 | 0.41953139 | 0.67482782 | 0.91636338 |
| LGALS8 | 414.525495 | -0.1706673 | 0.24773719 | -0.6889048 | 0.49088316 | 0.84635515 |
| AL359921.1 | 74.4105698 | -0.511726 | 0.3329049 | -1.5371536 | 0.1242557 | 0.56105491 |
| HEATR1 | 468.262439 | -0.3504875 | 0.16687067 | -2.1003539 | 0.03569772 | 0.35661333 |
| ACTN2 | 60.2067908 | -0.5542443 | 0.63533974 | -0.872359 | 0.38301255 | 0.79050201 |
| MTR | 4012.92635 | 0.1663328 | 0.28296951 | 0.58781172 | 0.55665867 | 0.87212955 |
| RYR2 | 119.84253 | -0.7764095 | 0.73294897 | -1.0592954 | 0.28946528 | 0.73204984 |
| LINC01139 | 66.8437152 | 0.44751514 | 0.50918958 | 0.87887726 | 0.37946783 | 0.78762967 |
| CHRM3 | 316.222984 | 0.78319656 | 0.60213381 | 1.30070185 | 0.19336053 | 0.64718686 |
| RGS7 | 510.000329 | 0.03290295 | 0.61551933 | 0.05345559 | 0.95736892 | 0.99065996 |
| FH | 195.133655 | 0.30846516 | 0.28708229 | 1.07448341 | 0.28260608 | 0.72604837 |
| OPN3 | 172.015857 | -0.5935106 | 0.33059248 | -1.7952938 | 0.0726069 | 0.4692634 |
| CEP170 | 744.024761 | -0.3797843 | 0.22270525 | -1.7053225 | 0.0881343 | 0.49729428 |
| SDCCAG8 | 2991.02154 | -0.0956338 | 0.16153585 | -0.5920283 | 0.55383161 | 0.87121136 |
| AKT3 | 4459.82836 | 0.01151096 | 0.30952881 | 0.03718864 | 0.9703346 | 0.99309664 |
| ZBTB18 | 206.839173 | -0.1693661 | 0.28406884 | -0.596215 | 0.55103158 | 0.87018163 |
| AL645465.1 | 234.736275 | -0.4900657 | 0.44115405 | -1.110872 | 0.26662346 | 0.71641758 |
| ADSS | 1021.34462 | -0.8460668 | 0.27392611 | -3.0886678 | 0.00201056 | 0.08650653 |
| CATSPERE | 832.452189 | -0.7780049 | 0.28940734 | -2.6882695 | 0.00718234 | 0.1729679 |
| DESI2 | 690.876749 | -0.1578394 | 0.22867552 | -0.6902329 | 0.49004772 | 0.84614483 |
| COX20 | 645.973055 | -0.0341723 | 0.29303388 | -0.1166154 | 0.90716485 | 0.98047754 |
| HNRNPU | 2849.63575 | 0.05525269 | 0.20880656 | 0.26461182 | 0.79130851 | 0.95168844 |
| EFCAB2 | 1628.15843 | -0.4519769 | 0.32643317 | -1.3845926 | 0.16617708 | 0.61901667 |
| KIF26B | 708.636771 | -1.7602254 | 0.58272768 | -3.0206654 | 0.0025222 | 0.09525007 |
| SMYD3 | 6033.79455 | -0.0600057 | 0.20705095 | -0.2898113 | 0.77196062 | 0.94478317 |
| TFB2M | 157.631849 | -0.1603165 | 0.24895037 | -0.6439695 | 0.51959519 | 0.85924779 |
| CNST | 701.984822 | -0.0889882 | 0.17943947 | -0.4959231 | 0.6199487 | 0.89702074 |
| SCCPDH | 261.712284 | -0.2597484 | 0.34938134 | -0.7434523 | 0.45720788 | 0.83255882 |
| AHCTF1 | 1054.87453 | -0.2346983 | 0.23204886 | -1.0114176 | 0.31181662 | 0.74629204 |
| ZNF124 | 470.35764 | -0.3534997 | 0.2418301 | -1.4617689 | 0.14380454 | 0.58881406 |
| AL390728.6 | 190.053209 | -0.1889617 | 0.30392946 | -0.6217287 | 0.53412031 | 0.86397197 |
| ZNF496 | 224.543939 | -0.5337299 | 0.37107707 | -1.4383262 | 0.15034153 | 0.59671313 |
| LYPD8 | 90.3496398 | 0.42203062 | 0.32280197 | 1.30739793 | 0.1910776 | 0.64567512 |
| SH3BP5L | 126.479749 | 0.23742886 | 0.31248658 | 0.75980498 | 0.44737116 | 0.82578615 |
| ZNF672 | 145.298355 | 0.26917106 | 0.22305775 | 1.2067326 | 0.22753514 | 0.67921637 |
| ZNF692 | 172.072766 | -0.2703946 | 0.249609 | -1.0832728 | 0.27868737 | 0.7244595 |
| PGBD2 | 70.4498805 | -0.1015833 | 0.43477526 | -0.2336456 | 0.81526013 | 0.95872901 |
| FAM110C | 1478.07195 | 0.69430131 | 0.55393532 | 1.25339779 | 2.10E-01 | 0.66269342 |
| SH3YL1 | 1146.43688 | 0.14681627 | 0.22254465 | 0.65971601 | 0.50943609 | 0.8568242 |
| ACP1 | 375.415263 | 0.11303619 | 0.20376699 | 0.55473261 | 0.57907757 | 0.8821758 |
| ALKAL2 | 61.4618231 | -0.5898493 | 0.91139009 | -0.6471974 | 0.51750419 | 0.85893097 |

| | | | | | | |
|---|---|---|---|---|---|---|
| TMEM18 | 85.5846927 | 0.12071124 | 0.30361083 | 0.3975854 | 0.69093582 | 0.92206635 |
| SNTG2 | 131.199138 | 0.59661604 | 0.86436532 | 0.69023597 | 0.4900458 | 0.84614483 |
| EIPR1 | 646.479396 | -0.1810938 | 0.29552571 | -0.6127851 | 0.54001844 | 0.86566791 |
| TRAPPC12 | 1129.54527 | -0.1182489 | 0.21582636 | -0.547889 | 0.58376811 | 0.88408541 |
| ADI1 | 449.088873 | 0.861879 | 0.31342521 | 2.74987132 | 0.00596187 | 0.15829975 |
| AC108488.2 | 108.726761 | -0.2526964 | 0.30576005 | -0.8264533 | 0.408547 | 0.80588056 |
| RNASEH1 | 45.6747662 | -0.3916045 | 0.40914116 | -0.957138 | 0.33849763 | 0.76449297 |
| RNASEH1-AS | 48.7240375 | 0.51113973 | 0.42275309 | 1.20907389 | 0.22663446 | 0.67880306 |
| RPS7 | 904.976035 | 0.28161113 | 0.45861595 | 0.61404566 | 0.53918515 | 0.86543657 |
| RNF144A | 401.752467 | -0.5709596 | 0.3830052 | -1.4907358 | 0.13603086 | 0.57783867 |
| ID2 | 599.930547 | 0.47220621 | 0.5258959 | 0.89790815 | 0.36923452 | 0.78095617 |
| KIDINS220 | 2009.82646 | -0.2274138 | 0.22540259 | -1.0089228 | 0.31301168 | 0.74703371 |
| MBOAT2 | 1101.43424 | 0.4076291 | 0.44310037 | 0.91994755 | 0.35760017 | 0.77484888 |
| ASAP2 | 3532.92708 | 0.44657992 | 0.40449136 | 1.10405305 | 0.26957012 | 0.71818309 |
| ITGB1BP1 | 949.559526 | 0.01458667 | 0.3538776 | 0.04121953 | 0.96712088 | 0.99248831 |
| CPSF3 | 294.546567 | -0.398072 | 0.16564412 | -2.4031764 | 0.01625334 | 0.25558383 |
| IAH1 | 354.429358 | 0.02285899 | 0.2543584 | 0.0898692 | 0.92839116 | 0.98314146 |
| ADAM17 | 1099.63808 | -0.3084637 | 0.19160071 | -1.60993 | 0.10741315 | 0.52901576 |
| AC080162.1 | 56.9508358 | -0.1804406 | 0.45847317 | -0.3935684 | 0.6938997 | 0.92229578 |
| YWHAQ | 1093.83175 | 0.18958436 | 0.15467598 | 1.22568714 | 0.22031643 | 0.67302974 |
| TAF1B | 574.633947 | -0.1088925 | 0.25986638 | -0.4190325 | 0.6751924 | 0.91649321 |
| GRHL1 | 165.210193 | -0.9315253 | 0.53480984 | -1.741788 | 0.08154555 | 0.48594425 |
| AC104794.2 | 44.5677695 | -0.8237552 | 0.48269757 | -1.7065658 | 0.08790278 | 0.49714378 |
| KLF11 | 261.085976 | 0.08663686 | 0.54495362 | 0.15898024 | 0.87368444 | 0.97564728 |
| AC104794.4 | 78.1150891 | 1.22172083 | 0.68281071 | 1.78925259 | 0.07357415 | 0.47028843 |
| CYS1 | 2692.24884 | 0.17460185 | 0.32658043 | 0.53463659 | 0.59290118 | 0.88740306 |
| HPCAL1 | 858.256092 | -0.3579629 | 0.36213996 | -0.9884655 | 0.3229247 | 0.7532337 |
| ODC1 | 151.565695 | 0.26652525 | 0.41512691 | 0.64203317 | 0.52085165 | 0.85943874 |
| NOL10 | 941.13219 | -0.3525379 | 0.23386605 | -1.5074351 | 0.13169916 | 0.57539433 |
| PDIA6 | 615.844075 | 0.85102989 | 0.24576381 | 3.46279585 | 0.00053459 | 0.04005562 |
| PQLC3 | 96.3598317 | -0.0343591 | 0.39833085 | -0.0862576 | 0.93126163 | 0.98353201 |
| ROCK2 | 1535.67911 | -0.0698757 | 0.17256667 | -0.4049202 | 0.6855362 | 0.92053037 |
| E2F6 | 178.902874 | 0.11296408 | 0.23395623 | 0.48284281 | 0.62920735 | 0.89996751 |
| LPIN1 | 807.861385 | -0.0409262 | 0.2523153 | -0.1622027 | 0.87114623 | 0.97500174 |
| MIR3681HG | 437.15413 | -1.7916673 | 0.65534555 | -2.7339276 | 0.00625838 | 0.16248472 |
| TRIB2 | 639.991228 | -0.8858093 | 0.38108244 | -2.324456 | 0.02010107 | 0.28397329 |
| LINC00276 | 136.94444 | 0.85895858 | 0.51873862 | 1.6558601 | 0.09775015 | 0.51355946 |
| FAM84A | 78.5811785 | 0.9126261 | 0.55310913 | 1.64999282 | 0.0989444 | 0.51618562 |
| NBAS | 2305.37984 | -0.2203465 | 0.16947444 | -1.3001757 | 0.19354077 | 0.64718686 |
| DDX1 | 640.858011 | 0.18509671 | 0.15196311 | 1.21803713 | 0.22320986 | 0.67624946 |
| SMC6 | 433.744399 | -0.2187302 | 0.33340467 | -0.6560501 | 0.51179191 | 0.85762435 |
| GEN1 | 158.535055 | -1.0811397 | 0.56329836 | -1.9193021 | 0.05494611 | 0.41687717 |
| KCNS3 | 78.5280166 | 1.41599179 | 0.48725797 | 2.90604132 | 0.00366033 | 0.11790188 |
| RDH14 | 118.33937 | 0.45515249 | 0.45937879 | 0.99079998 | 0.32178326 | 0.75240254 |
| LINC01376 | 889.726863 | -0.4881709 | 0.39728597 | -1.2287645 | 0.21916011 | 0.67275992 |
| TTC32 | 194.699911 | -0.2505286 | 0.18975114 | -1.3203007 | 0.18673464 | 0.64100257 |
| WDR35 | 774.902504 | 0.11500564 | 0.1813629 | 0.63411887 | 0.52600325 | 0.86119802 |
| LAPTM4A | 916.106331 | 0.74229955 | 0.32894565 | 2.25660239 | 0.02403293 | 0.30407182 |
| AC098828.3 | 70.2724599 | 3.23728188 | 1.3238847 | 2.4452899 | 0.01447358 | 0.24762526 |
| SDC1 | 771.733743 | 0.66380239 | 0.59732293 | 1.11129569 | 0.26644109 | 0.71641758 |
| PUM2 | 3427.74702 | -0.2460958 | 0.13889856 | -1.7717666 | 0.07643331 | 0.47578117 |
| RHOB | 337.934852 | 0.65788165 | 0.55356803 | 1.18843867 | 0.23466063 | 0.68606292 |
| HS1BP3 | 121.444525 | 0.42832604 | 0.35188497 | 1.21723313 | 0.22351553 | 0.67624946 |
| GDF7 | 84.2117636 | 0.8415162 | 0.56363666 | 1.49301184 | 0.13543409 | 0.57708028 |

| | | | | | | |
|---|---|---|---|---|---|---|
| LDAH | 1102.99975 | -0.0233094 | 0.23543297 | -0.0990064 | 0.92113316 | 0.98215577 |
| KLHL29 | 410.391288 | -0.7229379 | 0.43741892 | -1.6527359 | 9.84E-02 | 5.15E-01 |
| ATAD2B | 1912.02138 | -0.2103622 | 0.21118866 | -0.9960866 | 0.31920805 | 0.75094569 |
| UBXN2A | 826.127513 | 0.07386344 | 0.20987467 | 0.35194071 | 0.72488273 | 0.93241479 |
| WDCP | 42.8865156 | 0.4634145 | 0.43184748 | 1.07309761 | 0.28322732 | 0.72647048 |
| SF3B6 | 279.463476 | 0.34674699 | 0.36446427 | 0.95138814 | 0.34140738 | 0.76543778 |
| FAM228B | 207.851402 | -0.1636471 | 0.37281208 | -0.4389534 | 0.66069532 | 0.91099119 |
| TP53I3 | 306.00285 | -0.3685295 | 0.4437065 | -0.8305705 | 0.40621629 | 0.80499101 |
| ITSN2 | 2041.00136 | 0.12138395 | 0.1916586 | 0.63333421 | 0.52651542 | 0.86145663 |
| NCOA1 | 5039.20284 | 0.03227724 | 0.14741803 | 0.21895045 | 0.82668864 | 0.96214368 |
| PTRHD1 | 46.2238805 | 0.39715003 | 0.4624734 | 0.85875215 | 0.39047727 | 0.79701353 |
| CENPO | 54.8910876 | -0.5320349 | 0.36688145 | -1.4501548 | 0.14701536 | 0.59373375 |
| ADCY3 | 89.8917218 | -0.1110401 | 0.51765633 | -0.2145054 | 0.83015294 | 0.96332164 |
| DNAJC27 | 398.980492 | 0.06813899 | 0.2530162 | 0.26930684 | 0.78769357 | 0.95006812 |
| DNAJC27-AS | 265.980969 | -0.073404 | 0.40081236 | -0.1831381 | 8.55E-01 | 9.71E-01 |
| EFR3B | 98.9188656 | -1.0086676 | 0.44830253 | -2.2499707 | 0.0244508 | 0.3067148 |
| DNMT3A | 1460.33595 | -0.2087924 | 0.26235234 | -0.7958474 | 0.42612071 | 0.81466629 |
| DTNB | 1865.37256 | -0.5066474 | 0.16888487 | -2.9999573 | 0.00270017 | 0.09823475 |
| ASXL2 | 1438.37758 | -0.3005175 | 0.18578216 | -1.6175798 | 0.10575318 | 0.52694363 |
| KIF3C | 72.6840375 | -0.7322019 | 0.37697705 | -1.9422984 | 0.05210098 | 0.40866415 |
| RAB10 | 2419.30399 | -0.1837976 | 0.17453445 | -1.0530735 | 0.29230729 | 0.73404695 |
| HADHA | 1042.29297 | 0.02626928 | 0.29407061 | 0.08932983 | 0.92881979 | 0.98323294 |
| HADHB | 1527.03767 | 0.20426854 | 0.29637709 | 0.68921838 | 0.49068585 | 0.84614483 |
| ADGRF3 | 136.123785 | 0.44809547 | 0.43117836 | 1.03923462 | 0.29869563 | 0.73673761 |
| SELENOI | 742.998979 | -0.0422303 | 0.29174523 | -0.1447507 | 0.88490768 | 0.97664137 |
| AC013472.2 | 49.3307668 | 0.48879932 | 0.6360864 | 0.768448 | 0.44222107 | 0.82285053 |
| MAPRE3 | 246.62617 | 0.51259519 | 0.19840192 | 2.58362011 | 0.00977694 | 0.20084545 |
| TMEM214 | 138.842855 | 0.2537998 | 0.22206186 | 1.14292386 | 0.2530702 | 0.70374828 |
| AGBL5 | 275.928951 | 0.30321775 | 0.29795788 | 1.01765305 | 0.30884287 | 0.74399658 |
| OST4 | 302.354887 | 0.5081225 | 0.54714159 | 0.92868558 | 0.35305206 | 0.77216708 |
| KHK | 55.0062474 | 1.34632981 | 0.53111075 | 2.53493235 | 0.0112469 | 0.21687644 |
| PREB | 79.8379981 | 0.23288875 | 0.36090114 | 0.6452979 | 0.51873415 | 0.85916831 |
| SLC5A6 | 56.3333888 | -0.2586455 | 0.38844839 | -0.6658426 | 0.50551168 | 0.85595485 |
| ATRAID | 228.219285 | 0.68028661 | 0.34025416 | 1.99934842 | 0.04557067 | 0.38904776 |
| CAD | 125.772569 | -0.6322664 | 0.29844494 | -2.1185363 | 0.03412968 | 0.35153234 |
| MPV17 | 173.494499 | -0.0743606 | 0.31286837 | -0.2376737 | 0.81213422 | 0.95806331 |
| GTF3C2 | 526.557343 | -0.2375473 | 0.21218936 | -1.1195063 | 0.26292419 | 0.71328 |
| AC074117.1 | 76.084172 | -0.2835642 | 0.26404252 | -1.0739337 | 0.28285238 | 0.72604837 |
| EIF2B4 | 139.396491 | -0.2331492 | 0.20625521 | -1.1303918 | 0.25831118 | 0.70896455 |
| SNX17 | 135.745501 | 0.66268705 | 0.3511104 | 1.88740366 | 0.05910605 | 0.42419956 |
| ZNF513 | 77.1271043 | -0.0959826 | 0.40237886 | -0.238538 | 0.81146387 | 0.95761871 |
| PPM1G | 616.275303 | 0.28033632 | 0.26562239 | 1.05539414 | 0.2912451 | 0.73333141 |
| NRBP1 | 360.876968 | -0.308789 | 0.25695322 | -1.2017324 | 0.22946721 | 0.6817947 |
| KRTCAP3 | 100.808385 | 0.29290828 | 0.31442579 | 0.9315657 | 0.35156102 | 0.77126216 |
| IFT172 | 584.685227 | 0.14954924 | 0.26923182 | 0.55546643 | 0.57857566 | 0.8821758 |
| FNDC4 | 97.9634526 | 0.53932591 | 0.64020219 | 0.8424306 | 4.00E-01 | 8.02E-01 |
| ZNF512 | 510.483415 | 0.1372325 | 0.26791337 | 0.51222712 | 0.60849206 | 0.89266677 |
| GPN1 | 116.79109 | -0.2269751 | 0.21887728 | -1.0369969 | 0.29973732 | 0.73673761 |
| SUPT7L | 220.694175 | 0.21119313 | 0.23097325 | 0.91436184 | 0.36052674 | 0.77730375 |
| SLC4A1AP | 280.554261 | -0.1303072 | 0.1865335 | -0.6985725 | 0.48481921 | 0.8447301 |
| MRPL33 | 1213.72749 | -0.4566785 | 0.31823499 | -1.4350354 | 0.15127701 | 0.59916584 |
| RBKS | 998.01072 | 0.66823371 | 0.23367796 | 2.85963522 | 0.00424129 | 0.12646307 |
| BABAM2 | 5696.92351 | -0.0651655 | 0.29749849 | -0.2190449 | 0.82661504 | 0.96214368 |
| FOSL2 | 1207.29391 | 0.06462096 | 0.42446134 | 0.15224228 | 0.87899584 | 0.9765717 |

| | | | | | | |
|---|---|---|---|---|---|---|
| AC092164.1 | 80.5401165 | 0.1704186 | 0.36537103 | 0.46642614 | 0.64091051 | 0.90398151 |
| PPP1CB | 1294.39984 | -0.2347907 | 0.28552436 | -0.8223141 | 0.41089817 | 0.80666106 |
| SPDYA | 133.422481 | 0.53297395 | 0.52166634 | 1.02167595 | 0.3069343 | 0.74341546 |
| TRMT61B | 373.698256 | -0.1331502 | 0.1780619 | -0.7477748 | 0.45459603 | 0.83121393 |
| WDR43 | 587.541973 | -0.3684074 | 0.1653943 | -2.2274491 | 0.02591728 | 0.31222606 |
| CLIP4 | 688.118699 | 0.8880625 | 0.2392443 | 3.71194845 | 0.00020567 | 0.0232196 |
| ALK | 104.676827 | -0.4291566 | 0.5578536 | -0.7692997 | 0.44171544 | 0.82285053 |
| AC016907.2 | 45.1315889 | 0.47314235 | 0.62911005 | 0.75208201 | 0.45200174 | 0.82885146 |
| YPEL5 | 493.518841 | 0.14295124 | 0.29136697 | 0.49062265 | 0.62369336 | 0.89873262 |
| LBH | 135.975413 | 0.88409876 | 0.47117277 | 1.87637914 | 6.06E-02 | 4.30E-01 |
| LINC01936 | 95.9874838 | 1.45479099 | 0.74001528 | 1.96589318 | 0.04931095 | 0.39892272 |
| LCLAT1 | 1305.43485 | 0.31086161 | 0.19791254 | 1.57070197 | 0.11625189 | 0.54607796 |
| CAPN13 | 386.330341 | 0.72948477 | 0.48806878 | 1.4946352 | 0.13500968 | 0.57708028 |
| GALNT14 | 4385.16549 | 0.46125679 | 0.33087591 | 1.39404768 | 0.16330321 | 0.61387934 |
| MEMO1 | 102.624539 | 0.00719544 | 0.32289863 | 0.0222839 | 0.98222149 | 0.99518016 |
| DPY30 | 406.360819 | 0.01405761 | 0.24143306 | 0.0582257 | 0.95356885 | 0.98960488 |
| SPAST | 615.803623 | -0.3540471 | 0.20267522 | -1.746869 | 0.08066007 | 0.48456361 |
| SLC30A6 | 660.994561 | -0.2869119 | 0.16487811 | -1.7401455 | 0.08183347 | 0.48594425 |
| YIPF4 | 938.932856 | -0.115243 | 0.1810388 | -0.6365654 | 0.52440797 | 0.86006369 |
| BIRC6 | 5815.22295 | -0.3688181 | 0.13533468 | -2.72523 | 0.00642567 | 0.1657748 |
| TTC27 | 692.696125 | -0.3073395 | 0.21097847 | -1.4567339 | 0.14518984 | 0.59119271 |
| LINC00486 | 1086.26809 | -0.3209874 | 0.21348943 | -1.5035284 | 0.13270284 | 0.57682553 |
| LTBP1 | 420.550889 | -2.1566878 | 0.84078782 | -2.5650798 | 0.01031521 | 0.20627111 |
| RASGRP3 | 336.910264 | -0.9598976 | 0.30531126 | -3.1439968 | 0.00166657 | 0.07697119 |
| FAM98A | 196.717595 | 0.13181311 | 0.27357449 | 0.481818 | 0.62993524 | 0.90005881 |
| LINC01317 | 89.4604664 | 0.46538136 | 0.4574559 | 1.01732509 | 0.3089988 | 0.74399658 |
| AC068599.1 | 312.934384 | 0.18946448 | 0.60107034 | 0.31521183 | 0.75260082 | 0.93819406 |
| LINC01320 | 15940.276 | 0.56147419 | 0.32842251 | 1.70960934 | 8.73E-02 | 0.49566631 |
| AC073218.2 | 167.841249 | 0.18447384 | 0.77903478 | 0.23679795 | 0.81281354 | 0.95809779 |
| AC073218.1 | 141.458191 | 1.34813519 | 0.82992598 | 1.62440413 | 0.10428959 | 0.52442862 |
| AC012593.1 | 7575.71645 | 0.62692315 | 0.64086852 | 0.97823989 | 0.32795569 | 0.75641116 |
| CRIM1 | 3544.40026 | 0.23479181 | 0.33348595 | 0.70405309 | 0.48139971 | 0.8437862 |
| FEZ2 | 1207.94622 | 0.1598212 | 0.24814686 | 0.64405895 | 0.51953721 | 0.85924779 |
| STRN | 2160.6401 | -0.3051579 | 0.25791281 | -1.1831824 | 0.23673685 | 0.68878143 |
| HEATR5B | 939.017445 | -0.3165923 | 0.23018033 | -1.37541 | 0.16900437 | 0.62304738 |
| GPATCH11 | 90.0040376 | -0.059581 | 0.37731625 | -0.1579074 | 0.87452978 | 0.97564728 |
| EIF2AK2 | 904.1361 | 0.02050271 | 0.48365753 | 0.04239097 | 0.96618703 | 0.99248831 |
| CEBPZOS | 588.466495 | 0.17648387 | 0.19143814 | 0.92188456 | 0.35658879 | 0.77393713 |
| CEBPZ | 573.813262 | 0.3099848 | 0.19869575 | 1.56009782 | 0.11873677 | 0.55026203 |
| NDUFAF7 | 386.955189 | -0.0948031 | 0.20226202 | -0.4687144 | 0.63927378 | 0.90330608 |
| PRKD3 | 692.949419 | -0.1257819 | 0.22328818 | -0.5633166 | 0.57321934 | 0.87970887 |
| CDC42EP3 | 241.163342 | -0.3751915 | 0.38862232 | -0.9654399 | 0.33432452 | 0.76231532 |
| RMDN2 | 467.191226 | -0.0233653 | 0.34245008 | -0.0682299 | 0.94560265 | 0.98861299 |
| RMDN2-AS1 | 83.0599031 | 0.57608872 | 0.4711553 | 1.22271515 | 0.2214373 | 0.67442133 |
| CYP1B1 | 235.642701 | 0.37294368 | 0.79657742 | 0.46818259 | 0.63965402 | 0.90330608 |
| CYP1B1-AS1 | 279.150888 | 0.14107102 | 0.56333181 | 0.25042261 | 0.80226054 | 0.95443209 |
| ATL2 | 1077.65559 | 0.20457488 | 0.22561617 | 0.90673857 | 0.36454506 | 0.78068673 |
| AC011247.1 | 97.5603963 | 1.42011373 | 0.47850267 | 2.96782821 | 0.00299912 | 0.10480255 |
| HNRNPLL | 608.329909 | -0.0980465 | 0.25321488 | -0.3872069 | 0.69860304 | 0.92356678 |
| GALM | 326.72857 | -0.433469 | 0.26521393 | -1.6344126 | 0.10217224 | 0.52073581 |
| SRSF7 | 1009.24016 | 0.20505618 | 0.32168847 | 0.63743715 | 0.52384014 | 0.86006369 |
| GEMIN6 | 59.777109 | 0.04420002 | 0.34196648 | 0.1292525 | 0.89715786 | 0.97823597 |
| DHX57 | 569.539738 | -0.2870932 | 0.26286734 | -1.0921602 | 0.2747627 | 0.72164296 |
| MORN2 | 119.142662 | 0.60115261 | 0.37659548 | 1.59628208 | 0.11042583 | 0.53372881 |

| | | | | | | |
|---|---|---|---|---|---|---|
| ARHGEF33 | 61.3854465 | 0.23707925 | 0.410553 | 0.57746321 | 0.56362659 | 0.87487519 |
| SOS1 | 3365.04331 | -0.0448607 | 0.12504079 | -0.3587682 | 0.71976852 | 0.93001079 |
| CDKL4 | 51.1470687 | 0.01833378 | 0.6007374 | 0.03051879 | 0.97565331 | 0.9941831 |
| MAP4K3 | 3731.97896 | -0.082685 | 0.26054143 | -0.3173584 | 0.75097168 | 0.93775604 |
| AC007388.1 | 1891.92522 | 0.25389145 | 0.39728967 | 0.63905879 | 0.52278469 | 0.86006369 |
| TMEM178A | 90.2571533 | 0.33148838 | 0.599681 | 0.55277452 | 0.58041782 | 0.88258799 |
| THUMPD2 | 618.650853 | -0.1119325 | 0.20596522 | -0.5434536 | 0.5868175 | 0.88547501 |
| SLC8A1 | 158.541596 | 0.09483289 | 0.61940631 | 0.15310288 | 0.87831714 | 0.9765717 |
| EML4 | 1774.08154 | -0.228391 | 0.23175198 | -0.9854974 | 0.32437979 | 0.75473814 |
| COX7A2L | 608.531276 | -0.2120594 | 0.25265642 | -0.8393192 | 0.4012902 | 0.80259332 |
| KCNG3 | 78.2716324 | -0.0943215 | 0.7827777 | -0.1204959 | 0.90409035 | 0.97988471 |
| MTA3 | 1339.85819 | 0.05005101 | 0.16408459 | 0.30503178 | 0.76034195 | 0.94180015 |
| ZFP36L2 | 3244.98737 | 0.52872333 | 0.43937709 | 1.20334752 | 2.29E-01 | 6.81E-01 |
| THADA | 2547.32765 | -0.278563 | 0.22319103 | -1.2480921 | 0.21199735 | 0.66552984 |
| PLEKHH2 | 2634.09936 | -0.1453685 | 0.32071707 | -0.4532609 | 0.65036086 | 0.90825486 |
| DYNC2LI1 | 1176.81642 | 0.05122047 | 0.2888639 | 0.17731697 | 0.85925943 | 0.97174503 |
| LRPPRC | 1375.25732 | -0.3083145 | 0.2003731 | -1.5387018 | 0.12387711 | 0.56105491 |
| PPM1B | 1547.67527 | -0.1467056 | 0.16231064 | -0.903857 | 0.36607123 | 0.78095617 |
| SLC3A1 | 1494.62363 | 2.01862682 | 0.60040487 | 3.36210936 | 0.00077349 | 0.04906731 |
| PREPL | 1881.53408 | 0.21126007 | 0.26119592 | 0.80881841 | 0.41861961 | 0.81125469 |
| CAMKMT | 6331.70316 | -0.3481113 | 0.17744816 | -1.9617632 | 0.04979006 | 0.40017381 |
| SRBD1 | 1081.29405 | -0.1441229 | 0.14213988 | -1.0139514 | 3.11E-01 | 7.45E-01 |
| PRKCE | 5662.4027 | -0.3500099 | 0.20887769 | -1.675669 | 0.09380304 | 0.50926605 |
| EPAS1 | 936.886475 | 1.02036977 | 0.47702279 | 2.13903777 | 3.24E-02 | 0.3438118 |
| ATP6V1E2 | 71.0326568 | -0.0343542 | 0.40743698 | -0.0843179 | 0.93280368 | 0.9839805 |
| RHOQ | 644.289303 | 0.09523148 | 0.30618589 | 0.31102505 | 0.75578158 | 0.93957997 |
| AC018682.1 | 153.333166 | 0.05056218 | 0.60448652 | 0.08364484 | 0.93333881 | 0.98423059 |
| PIGF | 400.984899 | -0.1535031 | 0.20463485 | -0.7501318 | 0.4531753 | 0.82991556 |
| CRIPT | 165.358072 | -0.2325267 | 0.41475849 | -0.5606317 | 0.57504865 | 0.88057242 |
| SOCS5 | 722.81159 | -0.245339 | 0.25921977 | -0.9464518 | 0.34391821 | 0.76684937 |
| MCFD2 | 420.673868 | 0.24221389 | 0.23608311 | 1.02596871 | 0.30490633 | 0.74226092 |
| TTC7A | 827.270679 | 0.5838946 | 0.43073713 | 1.35557064 | 0.17523582 | 0.62944785 |
| AC106869.1 | 175.746114 | -0.1693243 | 0.52305575 | -0.3237213 | 0.74614905 | 0.93678195 |
| EPCAM | 1036.78267 | 0.17440018 | 0.33349138 | 0.52295259 | 0.60100724 | 0.8890585 |
| MSH2 | 732.742345 | -0.2487825 | 0.25853597 | -0.9622742 | 0.33591191 | 0.7637525 |
| MSH6 | 280.347064 | 0.08186757 | 0.19591395 | 0.41787514 | 0.67603841 | 0.91650575 |
| FBXO11 | 2690.72738 | -0.0292265 | 0.17634779 | -0.165732 | 0.86836783 | 0.97403792 |
| FOXN2 | 338.141965 | -0.1684171 | 0.21383156 | -0.7876158 | 0.43092147 | 0.81706845 |
| PPP1R21 | 1310.52985 | -0.2663348 | 0.23621693 | -1.1275009 | 0.25953076 | 0.70921184 |
| STON1-GTF2 | 105.214501 | -0.5557312 | 0.80950679 | -0.6865059 | 0.49239416 | 0.84734855 |
| AC009975.1 | 129.333909 | -0.7072104 | 0.93880336 | -0.7533104 | 0.45126338 | 0.82814845 |
| NRXN1 | 63.5846463 | -0.4116682 | 0.59243868 | -0.6948706 | 0.48713642 | 0.84475225 |
| AC007402.1 | 54.4413993 | -0.0799744 | 0.36885964 | -0.2168152 | 0.82835236 | 0.96288319 |
| ASB3 | 105.7976 | 0.41225615 | 0.23472631 | 1.75632696 | 0.0790326 | 0.48162624 |
| GPR75-ASB3 | 645.016369 | 0.38169052 | 0.40071436 | 0.95252518 | 0.3408307 | 0.76513132 |
| ERLEC1 | 322.569465 | 0.22111077 | 0.16635135 | 1.32917931 | 0.18378882 | 0.63843513 |
| PSME4 | 2316.89048 | -0.1380518 | 0.27752277 | -0.497443 | 0.61887667 | 0.89608494 |
| ACYP2 | 2756.16952 | -0.180656 | 0.26184111 | -0.6899451 | 0.49022872 | 0.84614483 |
| C2orf73 | 64.4248523 | 1.27005804 | 0.56117119 | 2.26322744 | 0.02362167 | 0.30185158 |
| SPTBN1 | 2475.24798 | -0.3236994 | 0.26090418 | -1.2406832 | 0.21472282 | 0.66830201 |
| AC092839.1 | 81.4439405 | -0.9328865 | 0.63724477 | -1.4639375 | 0.14321103 | 0.5883902 |
| AC093110.1 | 78.1541517 | -0.1404734 | 0.63260933 | -0.2220539 | 0.82427191 | 0.96146415 |
| EML6 | 647.19672 | -0.48824 | 0.27687751 | -1.7633789 | 0.07783661 | 0.47916634 |
| RTN4 | 3331.5258 | -0.1589322 | 0.19540707 | -0.8133388 | 0.41602384 | 0.80966301 |

| | | | | | | |
|---|---|---|---|---|---|---|
| CLHC1 | 626.228766 | -0.1723455 | 0.27213809 | -0.6333018 | 0.52653657 | 0.86145663 |
| RPS27A | 1444.02193 | 0.6033509 | 0.49110772 | 1.22855103 | 0.21924019 | 0.67278782 |
| MTIF2 | 380.297469 | -0.108666 | 0.19643125 | -0.5532012 | 0.58012564 | 0.88258799 |
| AC012358.3 | 340.87649 | -0.0448192 | 0.44090944 | -0.1016517 | 0.91903317 | 0.98215577 |
| CCDC88A | 398.383743 | 0.33968056 | 0.29360243 | 1.15694057 | 0.24729664 | 0.69761067 |
| CFAP36 | 650.780955 | -0.0089537 | 0.16813215 | -0.0532542 | 0.95752936 | 0.99065996 |
| PPP4R3B | 1400.48472 | -0.1328248 | 0.17437927 | -0.7617004 | 0.44623884 | 0.82571115 |
| PNPT1 | 551.825727 | -0.6091394 | 0.46870427 | -1.2996242 | 0.19372979 | 0.64718686 |
| EFEMP1 | 826.154361 | 1.3976921 | 0.55222604 | 2.53101445 | 0.01137332 | 0.21832654 |
| CCDC85A | 112.787036 | 0.32491379 | 0.49123062 | 0.6614282 | 0.50833775 | 0.85646191 |
| VRK2 | 1136.01588 | 0.00476485 | 0.16133063 | 0.02953471 | 0.97643813 | 0.9941831 |
| FANCL | 816.513922 | -0.0020447 | 0.18283004 | -0.0111835 | 0.99107702 | 0.99749592 |
| LINC01122 | 170.905433 | -1.2425809 | 0.53766757 | -2.311058 | 0.02082965 | 0.28870816 |
| BCL11A | 145.406763 | -1.2901989 | 0.6446947 | -2.0012557 | 0.04536484 | 0.3886245 |
| PAPOLG | 370.944487 | -0.0645313 | 0.20184479 | -0.3197074 | 0.74919013 | 0.93748128 |
| LINC01185 | 37.4792873 | -0.0037441 | 0.59088397 | -0.0063365 | 0.99494424 | 0.998438 |
| REL | 450.318907 | 0.18546092 | 0.2342164 | 0.79183576 | 0.42845645 | 0.81511869 |
| PUS10 | 379.617426 | 0.17536095 | 0.30550744 | 0.57399896 | 0.5659685 | 0.87642772 |
| PEX13 | 434.562566 | 0.06634318 | 0.24288054 | 0.27315149 | 0.78473677 | 0.94978829 |
| KIAA1841 | 248.440808 | -0.5851161 | 0.25651876 | -2.2809874 | 0.02254919 | 0.2981504 |
| C2orf74 | 202.576289 | -0.4581713 | 0.32768749 | -1.3981959 | 0.16205424 | 0.6135209 |
| USP34 | 6012.86991 | -0.2795874 | 0.15899143 | -1.7585063 | 0.07866139 | 0.48060389 |
| AC016727.1 | 162.973747 | -0.3283924 | 0.30685399 | -1.0701911 | 0.28453328 | 0.72738535 |
| XPO1 | 2591.13533 | -0.2951563 | 0.23038239 | -1.2811582 | 0.20013811 | 0.65306925 |
| FAM161A | 238.14562 | 0.28640805 | 0.22728939 | 1.26010301 | 0.2076322 | 0.6607067 |
| CCT4 | 494.040934 | 0.21352264 | 0.20518781 | 1.04062049 | 0.29805172 | 0.73673761 |
| COMMD1 | 2249.79029 | -0.2262053 | 0.18151955 | -1.2461759 | 0.21269984 | 0.66656043 |
| AC018462.1 | 158.526301 | -0.1382642 | 0.4490076 | -0.3079329 | 0.75813342 | 0.94056395 |
| B3GNT2 | 467.311202 | 0.67486338 | 0.28957895 | 2.33049875 | 0.01977981 | 0.28268153 |
| TMEM17 | 38.5518549 | -0.2934448 | 0.40480554 | -0.7249031 | 0.46851148 | 0.83825637 |
| EHBP1 | 3314.49274 | -0.3777625 | 0.16919527 | -2.2327012 | 0.02556865 | 0.31211455 |
| AC092567.1 | 67.7156707 | -0.3004006 | 0.52472526 | -0.5724912 | 0.56698926 | 0.87646121 |
| WDPCP | 2410.59417 | -0.2907786 | 0.23894121 | -1.2169464 | 0.22362462 | 0.67624946 |
| MDH1 | 649.605406 | 0.81400522 | 0.22655113 | 3.59303088 | 0.00032685 | 0.03094754 |
| UGP2 | 1383.62938 | 0.42511364 | 0.18196067 | 2.33629409 | 0.01947592 | 0.28049217 |
| VPS54 | 1645.47389 | 0.04278289 | 0.15908344 | 0.26893362 | 0.78798076 | 0.95010843 |
| PELI1 | 1241.60238 | -0.4955864 | 0.26147741 | -1.8953316 | 0.05804848 | 0.42087381 |
| AC012368.1 | 135.713747 | 0.68771377 | 0.81229334 | 0.84663229 | 0.39720011 | 0.80069929 |
| AC008074.1 | 217.087845 | 0.12613771 | 0.45923449 | 0.27466949 | 0.78357017 | 0.94955765 |
| AFTPH | 1758.18199 | 0.0392667 | 0.19152851 | 0.20501752 | 0.83755844 | 0.96527001 |
| SERTAD2 | 320.981431 | 0.42287293 | 0.25151213 | 1.68132219 | 0.09270035 | 0.50702841 |
| CEP68 | 315.623804 | 0.06803704 | 0.28890508 | 0.23549964 | 0.81382096 | 0.9582375 |
| RAB1A | 2128.42628 | -0.1235478 | 0.2363492 | -0.5227341 | 0.60115932 | 0.8890585 |
| ACTR2 | 1265.91262 | 0.04694971 | 0.18246937 | 0.25730185 | 7.97E-01 | 0.95281098 |
| SPRED2 | 1609.99334 | 0.10040291 | 0.19990791 | 0.50224583 | 0.61549461 | 0.8949877 |
| MEIS1 | 205.630452 | -1.1081374 | 0.39548799 | -2.8019497 | 0.00507948 | 0.14305936 |
| ETAA1 | 250.660648 | -0.1346534 | 0.19271851 | -0.6987049 | 0.48473646 | 0.8447301 |
| AC007422.2 | 83.9695069 | -1.5106428 | 0.50318562 | -3.0021581 | 2.68E-03 | 9.81E-02 |
| C1D | 325.46038 | -0.010304 | 0.16962406 | -0.0607459 | 0.95156157 | 0.98938795 |
| WDR92 | 76.886332 | -0.0593883 | 0.32137842 | -0.1847925 | 0.85339182 | 0.97076831 |
| PNO1 | 111.746719 | -0.1942532 | 0.32301672 | -0.6013719 | 0.54759232 | 0.86859163 |
| APLF | 512.961058 | 0.14496341 | 0.27397218 | 0.52911726 | 0.59672411 | 0.88883692 |
| ARHGAP25 | 108.327498 | 0.00831543 | 0.54732305 | 0.0151929 | 0.98787828 | 0.99653761 |
| ANTXR1 | 162.669306 | 0.7939993 | 0.60415225 | 1.3142371 | 0.18876643 | 0.64379441 |

| | | | | | | |
|---|---|---|---|---|---|---|
| GFPT1 | 1193.18731 | -0.0344332 | 0.26226082 | -0.1312935 | 0.8955431 | 0.97788747 |
| NFU1 | 423.871042 | -0.12084 | 0.18132021 | -0.6664451 | 0.50512667 | 0.85582573 |
| AAK1 | 4076.89243 | -0.0036214 | 0.18679731 | -0.0193867 | 0.98453259 | 0.995279 |
| ANXA4 | 3951.14543 | 0.52900901 | 0.51338854 | 1.03042621 | 0.30280998 | 0.73961113 |
| GMCL1 | 745.234164 | -0.2950526 | 0.23324405 | -1.2649953 | 0.20587302 | 0.65813563 |
| MXD1 | 277.415933 | 0.21952693 | 0.33494172 | 0.6554183 | 0.51219847 | 0.85762435 |
| PCBP1 | 457.884098 | 0.58548505 | 0.35121295 | 1.6670372 | 0.09550701 | 0.5117083 |
| C2orf42 | 243.29956 | -0.2197302 | 0.21356463 | -1.02887 | 0.30354077 | 0.7409078 |
| PCYOX1 | 313.945552 | 1.18562353 | 0.46565295 | 2.54615275 | 0.01089175 | 0.21282418 |
| SNRPG | 271.039307 | -0.0469378 | 0.30917574 | -0.1518158 | 0.87933223 | 0.9765717 |
| FAM136A | 217.320296 | 0.18287577 | 0.31301339 | 0.58424264 | 0.55905708 | 0.87295536 |
| TGFA | 411.194568 | 0.30102441 | 0.35326884 | 0.8521114 | 0.39415227 | 0.79852106 |
| ATP6V1B1 | 38.0440243 | 6.81568304 | 1.60524647 | 4.24587948 | 2.18E-05 | 0.00756868 |
| AC007040.2 | 190.836073 | 2.34862481 | 0.82384015 | 2.85082587 | 0.00436058 | 0.12885673 |
| NAGK | 202.091883 | -0.2053885 | 0.24573622 | -0.835809 | 0.40326236 | 0.80324472 |
| MCEE | 478.279936 | -0.0278488 | 0.21650418 | -0.1286292 | 0.89765108 | 0.97823597 |
| MPHOSPH10 | 259.461643 | -0.020522 | 0.24994178 | -0.0821071 | 0.93456157 | 0.98522769 |
| PAIP2B | 185.602773 | 0.02302462 | 0.31676508 | 0.07268674 | 0.9420554 | 0.98772472 |
| ZNF638 | 4853.12927 | 0.02121303 | 0.17748623 | 0.11951932 | 0.90486394 | 0.97992575 |
| DYSF | 47.4679753 | -1.3779072 | 0.62670885 | -2.19864 | 0.02790353 | 0.32224037 |
| EXOC6B | 4264.3743 | -0.3731482 | 0.20114672 | -1.8551044 | 0.06358131 | 0.43779712 |
| SPR | 60.2881983 | 0.70765677 | 0.55525598 | 1.27446943 | 0.2024971 | 0.65552334 |
| SFXN5 | 671.37183 | -0.4302852 | 0.23252708 | -1.8504737 | 0.06424531 | 0.43924239 |
| RAB11FIP5 | 272.628503 | -0.4499587 | 0.40597641 | -1.1083369 | 0.26771633 | 0.71787758 |
| SMYD5 | 71.1659177 | 0.05754365 | 0.33975217 | 0.1693695 | 0.86550602 | 0.97379123 |
| PRADC1 | 60.2530527 | 0.49315022 | 0.47009711 | 1.04903903 | 0.29416015 | 0.73527014 |
| CCT7 | 352.353086 | 0.6278041 | 0.31809213 | 1.97365495 | 0.04842099 | 0.39627811 |
| ALMS1 | 1059.873 | -0.331823 | 0.22742482 | -1.4590446 | 0.14455282 | 0.59035409 |
| NAT8 | 113.909642 | 3.61468452 | 1.14128714 | 3.1671999 | 0.00153914 | 0.07298272 |
| TPRKB | 109.127458 | 0.20272915 | 0.22512487 | 0.90051868 | 0.36784429 | 0.78095617 |
| DUSP11 | 204.46656 | 0.18090856 | 0.21070668 | 0.85858008 | 0.39057223 | 0.79701353 |
| STAMBP | 581.781482 | -0.4526258 | 0.19353371 | -2.338744 | 0.01934869 | 0.28045661 |
| DGUOK | 502.500926 | -0.1988099 | 0.18330867 | -1.0845635 | 0.27811502 | 0.72372365 |
| DGUOK-AS1 | 80.0378661 | -0.4190839 | 0.30017311 | -1.3961407 | 0.16267213 | 0.61387934 |
| TET3 | 584.980251 | -0.5073998 | 0.17274183 | -2.9373303 | 0.00331051 | 0.11183777 |
| BOLA3 | 101.723307 | 0.46965882 | 0.42256219 | 1.11145492 | 0.26637258 | 0.71641758 |
| MOB1A | 549.529104 | 0.30653549 | 0.23573135 | 1.30035948 | 0.19347779 | 0.64718686 |
| MTHFD2 | 52.8426511 | 0.27686163 | 0.72877198 | 0.37990159 | 0.70401847 | 0.92514314 |
| AC006030.1 | 190.433249 | -0.4842523 | 0.31361492 | -1.5440985 | 0.12256446 | 0.55930194 |
| DCTN1 | 121.337153 | -0.2060236 | 0.24704026 | -0.8339676 | 0.40429924 | 0.80437725 |
| WDR54 | 67.3214883 | 0.50090998 | 0.55348313 | 0.90501399 | 0.36545797 | 0.78095617 |
| RTKN | 333.717906 | -0.3096913 | 0.27743707 | -1.1162579 | 0.26431178 | 0.71488591 |
| MOGS | 275.790739 | 0.06551576 | 0.25996183 | 0.25202068 | 0.80102508 | 0.95370871 |
| CCDC142 | 122.499672 | -0.6245446 | 0.43669464 | -1.4301632 | 0.15267018 | 0.60267503 |
| TTC31 | 134.43045 | -0.5519821 | 0.33060795 | -1.6695973 | 0.09499907 | 0.51085035 |
| AUP1 | 191.037094 | 0.0323965 | 0.39969252 | 0.08105355 | 0.93539937 | 0.98586223 |
| HTRA2 | 73.9743662 | 0.26158027 | 0.41046775 | 0.63727362 | 0.52394663 | 0.86006369 |
| M1AP | 102.849458 | -0.7779898 | 0.71730457 | -1.0846018 | 0.27809806 | 0.72372365 |
| SEMA4F | 59.6839931 | 0.14188004 | 0.27879135 | 0.50891121 | 0.61081446 | 0.89339478 |
| POLE4 | 131.589063 | 0.22374582 | 0.3891587 | 0.57494751 | 0.5653268 | 0.8761426 |
| TACR1 | 109.314825 | 0.67476562 | 0.52287542 | 1.29049024 | 0.1968805 | 0.64964978 |
| AC007681.1 | 140.690904 | 0.248837 | 0.58432918 | 0.42585072 | 0.67021662 | 0.91474389 |
| EVA1A | 590.954603 | -0.3835218 | 0.39384612 | -0.9737858 | 0.33016289 | 0.75885186 |
| AC007099.1 | 157.975998 | -0.4241922 | 0.6619539 | -0.6408183 | 0.52164074 | 0.85957492 |

| | | | | | | |
|---|---|---|---|---|---|---|
| MRPL19 | 673.387031 | 0.1741917 | 0.31947404 | 0.54524524 | 0.58558486 | 0.88490909 |
| GCFC2 | 1216.87859 | 0.06102936 | 0.27850649 | 0.21913084 | 0.82654813 | 0.96214368 |
| CTNNA2 | 117.656145 | -0.7788264 | 0.47655659 | -1.634279 | 0.10220029 | 0.52073581 |
| SUCLG1 | 872.058045 | 0.46419274 | 0.23317875 | 1.99071628 | 0.04651209 | 0.39097111 |
| DNAH6 | 602.187282 | 0.14872187 | 0.45329476 | 0.32809086 | 0.74284297 | 0.93598004 |
| TMSB10 | 1630.20748 | 0.17823435 | 0.40307507 | 0.44218649 | 6.58E-01 | 9.10E-01 |
| KCMF1 | 1283.22853 | -0.2308623 | 0.19575139 | -1.1793647 | 0.23825299 | 0.68982886 |
| TCF7L1 | 184.713135 | -0.1937215 | 0.6369668 | -0.3041313 | 0.76102786 | 0.94202764 |
| TGOLN2 | 855.077544 | 0.30158756 | 0.20080405 | 1.50189977 | 0.13312299 | 0.57682553 |
| RETSAT | 133.556602 | 0.14741924 | 0.276631 | 0.53290934 | 0.59409634 | 0.88807733 |
| ELMOD3 | 535.072471 | -0.1406056 | 0.19277857 | -0.7293633 | 0.46577948 | 0.8367113 |
| CAPG | 176.061657 | 0.12023694 | 0.37777592 | 0.31827582 | 0.75027572 | 0.93775604 |
| MAT2A | 751.563105 | 0.02194388 | 0.33036039 | 0.06642406 | 0.94704022 | 0.98905425 |
| GGCX | 228.158159 | 0.06074171 | 0.21475516 | 0.28284168 | 0.7772982 | 0.94694572 |
| VAMP8 | 641.862229 | 0.69151048 | 0.33584383 | 2.0590239 | 0.03949195 | 0.37234059 |
| VAMP5 | 84.8370568 | -0.0912585 | 0.28475906 | -0.320476 | 0.7486075 | 0.93746253 |
| RNF181 | 85.118286 | 0.52849014 | 0.44383549 | 1.19073427 | 0.23375792 | 0.68493974 |
| TMEM150A | 75.597784 | -0.0472752 | 0.39976274 | -0.1182582 | 0.90586308 | 0.97997915 |
| USP39 | 594.454498 | -0.0757167 | 0.24994189 | -0.3029372 | 0.76193773 | 0.94210219 |
| C2orf68 | 273.030729 | -0.0322696 | 0.26638366 | -0.1211394 | 0.90358063 | 0.97975674 |
| ST3GAL5 | 178.835071 | 0.37968 | 0.57807774 | 0.65679748 | 0.51131115 | 0.85762435 |
| POLR1A | 383.255611 | -0.1727795 | 0.20676376 | -0.8356371 | 0.40335908 | 0.80324472 |
| PTCD3 | 831.718338 | -0.0551203 | 0.15042799 | -0.3664229 | 0.71404953 | 0.92760608 |
| IMMT | 1239.60036 | 0.00390996 | 0.19856818 | 0.01969078 | 0.98429004 | 0.99518552 |
| MRPL35 | 157.970518 | 0.32618582 | 0.2417727 | 1.34914249 | 0.1772912 | 0.63071358 |
| KDM3A | 1217.41742 | 0.00832885 | 0.36904607 | 0.02256859 | 0.9819944 | 0.99518016 |
| CHMP3 | 197.397825 | 0.32846151 | 0.27736341 | 1.18422794 | 2.36E-01 | 6.89E-01 |
| RNF103-CHM | 177.788716 | -0.9094575 | 0.88720185 | -1.0250852 | 3.05E-01 | 0.74229685 |
| AC015971.1 | 67.5668881 | 0.24061569 | 0.71581272 | 0.33614336 | 0.73676277 | 0.93552616 |
| RMND5A | 684.100747 | -0.2722892 | 0.20980167 | -1.2978409 | 0.19434202 | 0.64775769 |
| PLGLB1 | 239.621325 | 0.29242523 | 0.36744054 | 0.79584369 | 0.4261229 | 0.81466629 |
| CYTOR | 454.684932 | -0.0359327 | 0.68934647 | -0.0521257 | 0.95842851 | 0.99071816 |
| KRCC1 | 772.293495 | -0.037996 | 0.21003127 | -0.1809063 | 0.85644114 | 0.97118784 |
| THNSL2 | 170.888441 | 0.29018013 | 0.78881772 | 0.36786716 | 0.71297229 | 0.92760608 |
| EIF2AK3 | 707.092006 | -0.0767754 | 0.27836851 | -0.2758049 | 0.78269791 | 0.94919728 |
| RPIA | 262.828193 | 0.03989689 | 0.20375963 | 0.1958037 | 0.84476381 | 0.96853443 |
| MAL | 2516.61421 | 0.26090916 | 0.59807003 | 0.43625185 | 0.66265401 | 0.91110616 |
| MRPS5 | 919.995444 | -0.0562302 | 0.18686508 | -0.3009133 | 0.76348062 | 0.94255335 |
| ZNF514 | 100.467298 | 0.20015081 | 0.2596807 | 0.77075733 | 0.44085079 | 0.82265742 |
| ZNF2 | 83.8455088 | -0.4454235 | 0.33784011 | -1.3184448 | 0.18735478 | 0.64196349 |
| AC092835.1 | 156.079335 | -0.485925 | 0.22901376 | -2.1218157 | 0.03385322 | 0.35107893 |
| PROM2 | 517.213576 | -0.4897802 | 0.42665652 | -1.1479497 | 0.25098933 | 0.70200988 |
| KCNIP3 | 268.19754 | -0.1651262 | 0.37364201 | -0.4419369 | 0.65853486 | 0.91037018 |
| FAHD2A | 138.453664 | -0.0946683 | 0.313134 | -0.3023253 | 0.7624041 | 0.94212205 |
| LINC00342 | 41.3791141 | -0.0866376 | 0.52695252 | -0.1644125 | 0.86940644 | 0.97454113 |
| ANKRD36C | 456.409164 | -0.2211599 | 0.39602448 | -0.5584502 | 0.57653702 | 0.88147804 |
| STARD7 | 530.06249 | 0.10043856 | 0.18565236 | 0.54100342 | 0.58850522 | 0.88571413 |
| TMEM127 | 332.497147 | 0.17543708 | 0.23725188 | 0.73945495 | 0.45963078 | 0.83390278 |
| CIAO1 | 394.759563 | -0.1871454 | 0.25468078 | -0.7348232 | 0.46244716 | 0.83531269 |
| SNRNP200 | 617.409947 | -0.4876747 | 0.26725258 | -1.8247708 | 0.06803562 | 0.45250883 |
| NEURL3 | 61.0700189 | -1.1568435 | 0.75824235 | -1.5256909 | 0.12708686 | 0.56740575 |
| KANSL3 | 965.505354 | -0.4861964 | 0.20043367 | -2.4257223 | 0.01527796 | 0.25131572 |
| FER1L5 | 170.945029 | -0.502793 | 0.35319522 | -1.4235556 | 0.15457515 | 0.60380955 |
| LMAN2L | 316.460707 | 0.05840754 | 0.16494341 | 0.35410654 | 0.72325904 | 0.93177886 |

| | | | | | | |
|---|---|---|---|---|---|---|
| CNNM4 | 243.491896 | -0.5727533 | 0.32486233 | -1.7630647 | 0.07788958 | 0.47916634 |
| CNNM3 | 103.409858 | 0.02169002 | 0.31770241 | 0.06827149 | 0.94556952 | 0.98861299 |
| ANKRD23 | 236.447241 | -0.1048649 | 0.25784016 | -0.4067049 | 0.68422471 | 0.9200778 |
| SEMA4C | 80.0710736 | -0.1699457 | 0.38827167 | -0.4376979 | 0.66160533 | 0.91110616 |
| FAHD2B | 57.392295 | 2.00675018 | 0.51310469 | 3.91099557 | 9.19E-05 | 0.01517656 |
| ANKRD36 | 695.808507 | -0.2268256 | 0.31713742 | -0.7152282 | 0.47446805 | 0.84106098 |
| AC092683.1 | 527.579214 | 0.62389182 | 0.53621958 | 1.16350063 | 0.24462644 | 0.6948422 |
| ANKRD36B | 243.023854 | -0.7525511 | 0.59005649 | -1.2753883 | 0.20217184 | 0.65552334 |
| COX5B | 779.241748 | 0.66657318 | 0.40271875 | 1.65518289 | 0.0978874 | 0.51355946 |
| ACTR1B | 239.139635 | 0.0523159 | 0.28939586 | 0.18077626 | 0.8565432 | 0.97118784 |
| C2orf92 | 111.34717 | -0.7746023 | 0.38139531 | -2.0309696 | 0.04225808 | 0.37920308 |
| TMEM131 | 2380.15681 | -0.3173365 | 0.19759839 | -1.6059669 | 0.10828117 | 0.53091537 |
| INPP4A | 566.923519 | -0.3121587 | 0.15907328 | -1.9623578 | 0.04972085 | 0.40017381 |
| COA5 | 184.387162 | 0.16043412 | 0.21028234 | 0.76294623 | 0.44549546 | 0.82571115 |
| UNC50 | 236.980562 | 0.43078411 | 0.21667413 | 1.98816589 | 0.04679335 | 0.39219119 |
| MGAT4A | 1883.51738 | 0.28188574 | 0.19329807 | 1.45829572 | 0.14475905 | 0.59044137 |
| KIAA1211L | 406.476584 | 0.05737371 | 0.60813977 | 0.09434297 | 0.92483672 | 0.9822216 |
| TSGA10 | 1684.12321 | -0.0198151 | 0.18154842 | -0.1091451 | 0.91308744 | 0.98136469 |
| AC079447.1 | 817.720827 | -0.2189477 | 0.33510624 | -0.653368 | 0.51351905 | 0.85762435 |
| MITD1 | 370.085438 | 0.20002547 | 0.18107398 | 1.10466158 | 0.26930625 | 0.71818262 |
| LYG1 | 106.248134 | -0.3453655 | 0.52665219 | -0.6557753 | 0.51196873 | 0.85762435 |
| TXNDC9 | 302.342382 | -0.2494804 | 0.39730788 | -0.6279273 | 0.53005159 | 0.86361777 |
| EIF5B | 978.033896 | -0.0884992 | 0.1891245 | -0.4679416 | 0.63982634 | 0.90330608 |
| REV1 | 2051.16513 | -0.2380759 | 0.22304955 | -1.0673677 | 0.28580584 | 0.72866423 |
| AFF3 | 94.0679297 | -0.3010248 | 0.52892087 | -0.5691301 | 0.56926782 | 0.87789095 |
| LONRF2 | 548.078577 | 0.31349046 | 0.19140148 | 1.63786855 | 0.10144912 | 0.51919466 |
| PDCL3 | 46.4454675 | -0.0741608 | 0.46013916 | -0.1611703 | 0.87195925 | 0.97535872 |
| NPAS2 | 1166.76594 | 1.34723875 | 0.4513392 | 2.98498058 | 2.84E-03 | 1.01E-01 |
| RPL31 | 630.700245 | 0.27309467 | 0.34818163 | 0.78434541 | 0.43283746 | 0.81816686 |
| TBC1D8 | 2838.66977 | -0.7268238 | 0.24296181 | -2.9915147 | 0.00277597 | 0.09937076 |
| CNOT11 | 569.094242 | -0.2511871 | 0.31981933 | -0.7854032 | 0.43221718 | 0.81773248 |
| RNF149 | 2116.67503 | -0.6209595 | 0.30175618 | -2.0578188 | 0.03960753 | 0.37259071 |
| MAP4K4 | 1746.82472 | -0.9075083 | 0.37501256 | -2.4199411 | 1.55E-02 | 0.25214021 |
| LINC01127 | 48.9107879 | -1.6677212 | 0.67125605 | -2.4844785 | 0.01297413 | 0.23379993 |
| IL1R2 | 207.981754 | -1.8576413 | 0.60749115 | -3.0578904 | 0.00222901 | 0.09083568 |
| IL1R1 | 395.43812 | -0.5752935 | 0.30985182 | -1.8566728 | 0.06335772 | 0.43748783 |
| IL1RL2 | 318.922616 | -0.3022339 | 0.36012229 | -0.8392536 | 4.01E-01 | 0.80259332 |
| SLC9A2 | 44.8678566 | 0.35761539 | 0.57271199 | 0.62442449 | 0.53234885 | 0.86397197 |
| MFSD9 | 117.258775 | -0.0253822 | 0.32775189 | -0.0774434 | 0.93827084 | 0.98696247 |
| TMEM182 | 194.826443 | -0.0481398 | 0.3634547 | -0.1324505 | 0.89462794 | 0.97788747 |
| LINC01965 | 945.643768 | -0.8746234 | 0.69125784 | -1.2652637 | 0.20577682 | 0.65813563 |
| POU3F3 | 1177.86096 | -0.1908823 | 0.30941747 | -0.6169087 | 0.53729497 | 0.86487317 |
| AC010884.1 | 51.6864625 | 0.03970067 | 0.45799007 | 0.08668457 | 9.31E-01 | 0.98333082 |
| MRPS9 | 715.934004 | 0.10569289 | 0.16512367 | 0.64008323 | 5.22E-01 | 0.85993307 |
| TGFBRAP1 | 282.141752 | -0.3133596 | 0.23216971 | -1.3497005 | 0.17711206 | 0.63058577 |
| AC012360.3 | 93.8221766 | -0.4158105 | 0.30754403 | -1.3520356 | 0.17636392 | 0.63008941 |
| C2orf49 | 180.820549 | 0.1000589 | 0.21112014 | 0.47394295 | 0.63554059 | 0.90199374 |
| FHL2 | 786.576348 | -2.0992475 | 1.19166444 | -1.7616096 | 0.07813528 | 0.47981821 |
| AC108058.1 | 48.2470614 | -1.3169571 | 0.96008339 | -1.3717112 | 0.17015337 | 0.62606012 |
| NCK2 | 681.612746 | -0.2510346 | 0.20000404 | -1.2551476 | 0.20942517 | 0.66216422 |
| UXS1 | 1256.6741 | -0.3933529 | 0.37459682 | -1.0500701 | 2.94E-01 | 0.73527014 |
| AC018878.1 | 60.5115997 | -0.4270739 | 0.65707082 | -0.6499663 | 0.51571399 | 0.85869987 |
| LINC01594 | 431.261707 | -0.8910335 | 0.44190645 | -2.0163396 | 0.04376448 | 0.38453712 |
| SULT1C3 | 50.9560858 | -0.2669784 | 0.85703011 | -0.3115157 | 0.75540859 | 0.93953492 |

| | | | | | | |
|---|---|---|---|---|---|---|
| SULT1C2 | 2655.04576 | 0.81838641 | 0.50001291 | 1.63673054 | 0.10168678 | 0.5195058 |
| SULT1C4 | 168.865279 | 0.46652718 | 0.41378189 | 1.12747123 | 0.2595433 | 0.70921184 |
| GCC2 | 2871.95665 | -0.0861998 | 0.16100817 | -0.5353753 | 5.92E-01 | 8.87E-01 |
| GCC2-AS1 | 322.001044 | 0.42240343 | 0.40532634 | 1.04213171 | 0.29735062 | 0.73673761 |
| LIMS1 | 3125.5573 | -0.0246423 | 0.26321906 | -0.093619 | 0.92541185 | 0.9822216 |
| RANBP2 | 1178.70529 | 0.14864799 | 0.13070476 | 1.13728067 | 0.25542097 | 0.70643369 |
| CCDC138 | 314.258949 | -0.219945 | 0.28139399 | -0.7816264 | 0.43443418 | 0.81919857 |
| SH3RF3 | 199.623135 | 1.28343314 | 1.02867913 | 1.24765158 | 0.21215868 | 0.66552984 |
| 10-Sep | 1806.61153 | -0.5165169 | 0.3323479 | -1.5541453 | 0.12014977 | 0.55298199 |
| SOWAHC | 162.764638 | -0.1038426 | 0.51444997 | -0.2018517 | 0.84003266 | 0.96617873 |
| NPHP1 | 492.157153 | 0.16409206 | 0.26414643 | 0.62121629 | 0.53445733 | 0.86419602 |
| SMIM37 | 200.043025 | 0.44772668 | 0.35108844 | 1.27525328 | 0.20221961 | 0.65552334 |
| LINC01106 | 91.3945861 | -0.2167102 | 0.49916168 | -0.4341483 | 0.66418078 | 0.91206611 |
| ACOXL | 90.0867844 | 0.84086881 | 0.59530315 | 1.41250522 | 0.15780125 | 0.60779483 |
| MIR4435-2H | 2487.72131 | 0.65067444 | 0.5722798 | 1.13698657 | 0.2555439 | 0.70646282 |
| BCL2L11 | 644.033351 | 0.09010881 | 0.38509031 | 0.23399396 | 0.81498967 | 0.95872901 |
| AC017002.5 | 126.345236 | 0.28634966 | 0.57897589 | 0.4945796 | 0.62089691 | 0.89750791 |
| ANAPC1 | 853.822814 | -0.2686794 | 0.1606769 | -1.672172 | 0.09449041 | 0.51017802 |
| MERTK | 608.682852 | 0.3377707 | 0.36428116 | 0.92722528 | 0.35380958 | 0.77243419 |
| TMEM87B | 772.560521 | -0.4375984 | 0.16578843 | -2.6394987 | 8.30E-03 | 0.18347203 |
| ZC3H8 | 510.312133 | -0.3894616 | 0.26002268 | -1.4977986 | 0.13418559 | 0.57708028 |
| ZC3H6 | 966.351149 | -0.1195884 | 0.40873634 | -0.2925808 | 0.76984257 | 0.94386043 |
| RGPD8 | 82.8378189 | -0.1187748 | 0.39109312 | -0.3036996 | 0.76135679 | 0.94202764 |
| TTL | 366.54129 | -0.2598784 | 0.25420544 | -1.0223163 | 0.30663123 | 0.7430365 |
| POLR1B | 184.725289 | -0.3645929 | 0.29005763 | -1.256967 | 0.20876558 | 0.6614379 |
| CHCHD5 | 104.158467 | 0.58018311 | 0.27976842 | 2.07379775 | 0.0380981 | 0.36599111 |
| SLC20A1 | 850.911887 | -0.6097863 | 0.4923586 | -1.2385003 | 0.21553062 | 0.66885336 |
| PSD4 | 285.466859 | -0.4102061 | 0.26244658 | -1.5630082 | 0.11805067 | 0.54888439 |
| PAX8 | 14721.0745 | -0.6300462 | 0.27844751 | -2.2627107 | 0.02365353 | 0.30185158 |
| CBWD2 | 382.598068 | -0.1385633 | 0.2657239 | -0.5214557 | 0.60204932 | 0.88964714 |
| RABL2A | 276.793913 | -0.0926954 | 0.3280313 | -0.2825808 | 0.77749817 | 0.94694572 |
| AC017074.1 | 211.791138 | 0.25551338 | 0.46503799 | 0.54944626 | 0.58269923 | 0.88347959 |
| SLC35F5 | 3188.31073 | 0.01254045 | 0.18907232 | 0.0663262 | 0.94711813 | 0.98905425 |
| ACTR3 | 1658.39319 | -0.0436959 | 0.18368939 | -0.2378794 | 0.8119746 | 0.95806331 |
| LINC01191 | 54.4317876 | -0.1517437 | 0.43131912 | -0.3518131 | 7.25E-01 | 9.32E-01 |
| DPP10 | 555.673623 | -2.0148743 | 1.26934287 | -1.5873365 | 0.11243645 | 0.53693893 |
| DDX18 | 403.575792 | 0.08416694 | 0.17876636 | 0.47082092 | 6.38E-01 | 0.90273101 |
| AC009404.1 | 52.9185205 | -0.5278681 | 0.50400388 | -1.0473492 | 0.29493853 | 0.73575884 |
| CCDC93 | 957.586732 | -0.3111029 | 0.2099245 | -1.4819753 | 0.13834688 | 0.58206378 |
| INSIG2 | 518.298763 | -0.2138447 | 0.47334382 | -0.4517745 | 0.65143141 | 0.90825486 |
| STEAP3 | 114.669159 | 0.48531559 | 0.44365472 | 1.0939038 | 0.27399717 | 0.72135661 |
| C2orf76 | 195.455236 | -0.0600869 | 0.23125436 | -0.2598302 | 0.79499473 | 0.95272759 |
| DBI | 798.16106 | 0.32028815 | 0.45674002 | 0.70124826 | 0.48314809 | 0.84427756 |
| TMEM37 | 803.52828 | 0.86308327 | 0.40542934 | 2.12881308 | 0.03326973 | 0.34850106 |
| SCTR | 1743.45364 | -0.5964437 | 0.59375031 | -1.0045362 | 0.31512021 | 0.74785672 |
| AC013275.1 | 105.146522 | -0.5031082 | 0.72443802 | -0.6944807 | 0.48738081 | 0.84485737 |
| CFAP221 | 3028.02858 | -0.1470419 | 0.33449587 | -0.4395925 | 0.66023226 | 0.91099119 |
| TMEM177 | 76.1850267 | 0.57115465 | 0.52895044 | 1.07978859 | 0.28023633 | 0.72513763 |
| PTPN4 | 5499.42154 | -0.1987099 | 0.25933959 | -0.7662149 | 4.44E-01 | 8.24E-01 |
| EPB41L5 | 3507.79673 | -0.2535773 | 0.24556573 | -1.032625 | 0.30177945 | 0.73900718 |
| AC012363.1 | 44.6108512 | -0.3241243 | 0.38705062 | -0.837421 | 0.40235598 | 0.80310019 |
| RALB | 307.35199 | -0.4363346 | 0.20401591 | -2.1387285 | 0.03245767 | 0.3438118 |
| TFCP2L1 | 5671.24779 | 0.08748562 | 0.50709365 | 0.1725236 | 0.86302591 | 0.97319807 |
| CLASP1 | 6223.78925 | -0.0678873 | 0.27244326 | -0.2491795 | 0.80322195 | 0.95446295 |

| | | | | | | |
|---|---|---|---|---|---|---|
| NIFK-AS1 | 272.737162 | -0.3487585 | 0.37241306 | -0.936483 | 0.34902457 | 0.76982416 |
| NIFK | 111.879332 | -0.0109136 | 0.26944563 | -0.0405037 | 0.96769154 | 0.99248831 |
| TSN | 287.169545 | -0.0306549 | 0.22873318 | -0.1340205 | 0.89338637 | 0.97788747 |
| CNTNAP5 | 47.7413664 | -0.4219194 | 0.41489926 | -1.0169201 | 0.30919142 | 0.74420449 |
| LINC01889 | 94.2862993 | -0.5473082 | 0.62227474 | -0.8795283 | 3.79E-01 | 7.88E-01 |
| GYPC | 204.907417 | 0.40733082 | 0.2620735 | 1.5542618 | 0.12012198 | 0.55298199 |
| BIN1 | 148.373099 | 0.98045668 | 0.49462005 | 1.98224206 | 0.04745215 | 0.39411174 |
| ERCC3 | 374.969555 | -0.2987797 | 0.24323245 | -1.2283711 | 0.21930768 | 0.67278782 |
| MAP3K2 | 1861.76567 | -0.1396147 | 0.18588599 | -0.7510773 | 0.45260615 | 0.82942212 |
| IWS1 | 1245.4679 | -0.0748907 | 0.15387694 | -0.486692 | 0.62647665 | 0.89873262 |
| AC010976.1 | 159.98965 | 0.33778535 | 0.58239912 | 0.57998946 | 0.56192173 | 0.87403145 |
| WDR33 | 1744.02663 | -0.0019717 | 0.22909151 | -0.0086066 | 0.99313302 | 0.99816851 |
| SFT2D3 | 51.1895236 | 0.18765556 | 0.3734283 | 0.50252098 | 0.6153011 | 0.8949877 |
| POLR2D | 144.192775 | 0.20349056 | 0.25852163 | 0.78713166 | 0.4312048 | 0.81706845 |
| AMMECR1L | 210.410426 | -0.2333387 | 0.17689764 | -1.3190605 | 0.18714887 | 0.64195489 |
| SAP130 | 555.197269 | -0.3083828 | 0.20788077 | -1.4834602 | 0.13795218 | 0.58161836 |
| UGGT1 | 612.961741 | -0.0162158 | 0.16552121 | -0.097968 | 0.92195769 | 0.98215577 |
| HS6ST1 | 546.457025 | 0.19485366 | 0.40514795 | 0.48094447 | 0.63055596 | 0.90005881 |
| SMPD4 | 430.227588 | -0.2993398 | 0.19447448 | -1.539224 | 0.12374963 | 0.56102027 |
| MZT2B | 389.690781 | 0.50293661 | 0.33048925 | 1.52179417 | 0.12806066 | 0.56762585 |
| CCDC115 | 96.6589412 | 0.51700303 | 0.29020725 | 1.78149592 | 0.07483147 | 0.47190267 |
| IMP4 | 109.360475 | 0.16141382 | 0.33186039 | 0.48639074 | 0.62669015 | 0.89875738 |
| PTPN18 | 351.045228 | -0.0025997 | 0.2507729 | -0.0103669 | 0.99172855 | 0.99772135 |
| FAM168B | 969.840037 | -0.2025267 | 0.149624 | -1.3535709 | 0.17587333 | 0.62957039 |
| PLEKHB2 | 750.583773 | 0.1768553 | 0.17054315 | 1.03701202 | 0.29973025 | 0.73673761 |
| MZT2A | 269.131539 | 0.30889195 | 0.25918177 | 1.19179659 | 0.23334101 | 0.68484327 |
| CCDC74A | 42.3034563 | -0.0848575 | 0.42491732 | -0.1997035 | 0.84171251 | 0.96679831 |
| ANKRD30BL | 92.3892701 | 0.05952808 | 0.33810394 | 0.17606444 | 0.86024331 | 0.97200968 |
| GPR39 | 1099.38008 | -0.4642384 | 0.36665414 | -1.2661478 | 0.20546016 | 0.65813563 |
| LYPD1 | 50.1199544 | -1.6735656 | 1.014697 | -1.6493254 | 0.09908098 | 0.51618562 |
| NCKAP5 | 912.137965 | -0.0323709 | 0.45695996 | -0.0708397 | 0.9435253 | 0.98779628 |
| AC010890.1 | 39.4840974 | -0.2633261 | 0.79092638 | -0.3329337 | 0.7391843 | 0.93552616 |
| MGAT5 | 5915.0995 | -0.1208319 | 0.35443042 | -0.3409185 | 7.33E-01 | 0.93552616 |
| TMEM163 | 602.076237 | 0.59199741 | 0.72565818 | 0.81580753 | 0.41461023 | 0.8094275 |
| CCNT2-AS1 | 452.133989 | -0.6050567 | 0.41812451 | -1.447073 | 0.14787647 | 0.59424557 |
| ACMSD | 555.332094 | 0.88172152 | 0.73886621 | 1.19334394 | 0.2327347 | 0.68474693 |
| CCNT2 | 1405.48145 | -0.20779 | 0.18471797 | -1.1249041 | 0.26062966 | 0.71055875 |
| RAB3GAP1 | 2014.33896 | -0.0534681 | 0.18270627 | -0.2926452 | 0.76979339 | 0.94386043 |
| ZRANB3 | 955.286151 | -0.3536906 | 0.20112906 | -1.7585257 | 0.0786581 | 0.48060389 |
| R3HDM1 | 2039.8282 | -0.0118219 | 0.15550958 | -0.0760202 | 0.93940304 | 0.98715467 |
| UBXN4 | 1817.19791 | -0.1422837 | 0.17506112 | -0.812766 | 0.41635222 | 0.80966301 |
| MCM6 | 61.1273252 | -0.3403648 | 0.4616048 | -0.737351 | 0.46090892 | 0.83444983 |
| DARS | 1543.74385 | -0.006412 | 0.25625548 | -0.025022 | 0.98003744 | 0.99500147 |
| DARS-AS1 | 112.948498 | 0.07247538 | 0.72314523 | 0.10022245 | 0.92016773 | 0.98215577 |
| THSD7B | 223.488429 | 1.8202233 | 0.85088042 | 2.13922339 | 0.03241758 | 0.3438118 |
| HNMT | 997.044776 | -0.1060177 | 0.32701675 | -0.3241965 | 0.74578926 | 0.93675395 |
| AC114763.1 | 355.687143 | -0.310397 | 0.31184355 | -0.9953611 | 0.31956065 | 0.75094569 |
| SPOPL | 763.686012 | -0.2605913 | 0.16126605 | -1.6159092 | 0.10611395 | 0.52735777 |
| LRP1B | 8648.92436 | -0.1392558 | 0.5401118 | -0.2578276 | 0.79653993 | 0.95278473 |
| ARHGAP15 | 43.8275822 | 0.02192471 | 0.37307943 | 0.05876686 | 0.95313781 | 0.98949465 |
| GTDC1 | 738.757004 | 0.09508784 | 0.3017535 | 0.31511762 | 0.75267235 | 0.93819469 |
| TEX41 | 412.140227 | 1.28092072 | 0.6031889 | 2.12358139 | 3.37E-02 | 3.51E-01 |
| ACVR2A | 1047.50327 | -0.0662468 | 0.27357877 | -0.242149 | 0.80866472 | 0.95610623 |
| ORC4 | 1527.35072 | 0.02732964 | 0.20112826 | 0.13588165 | 0.89191484 | 0.97762038 |

| | | | | | | |
|---|---|---|---|---|---|---|
| MBD5 | 7239.73393 | -0.1002181 | 0.16445613 | -0.609391 | 0.54226532 | 0.86582694 |
| EPC2 | 1839.5209 | -0.0013796 | 0.25825361 | -0.0053422 | 0.99573759 | 0.99882912 |
| AC105402.3 | 61.8296273 | 0.38496927 | 1.15417228 | 0.33354576 | 0.73872236 | 0.93552616 |
| LYPD6B | 571.407032 | 1.03706254 | 0.38222764 | 2.71320655 | 0.00666356 | 0.16894224 |
| LYPD6 | 326.748867 | 0.17349791 | 0.45911592 | 0.37789566 | 7.06E-01 | 0.92556842 |
| MMADHC | 328.168573 | 0.32451392 | 0.32638239 | 0.99427522 | 0.32008889 | 0.75104498 |
| AC007364.1 | 185.432468 | -0.3736228 | 0.4357035 | -0.8575161 | 0.39115973 | 0.79735014 |
| RND3 | 121.159981 | -0.5894667 | 0.56755999 | -1.0385981 | 0.2989917 | 0.73673761 |
| RBM43 | 84.0178559 | 0.58445918 | 0.45891707 | 1.27356164 | 0.20281881 | 0.65552334 |
| NMI | 219.038834 | 0.1878524 | 0.42109505 | 0.44610452 | 0.65552175 | 0.90919768 |
| RIF1 | 1219.15229 | -0.2841253 | 0.19417478 | -1.4632449 | 0.14340037 | 0.58849865 |
| NEB | 327.015322 | 0.66992281 | 0.56382027 | 1.18818503 | 0.23476052 | 0.68620308 |
| ARL5A | 475.181157 | 0.27470981 | 0.37140046 | 0.7396593 | 0.45950675 | 0.83390278 |
| CACNB4 | 1276.21302 | -0.2549815 | 0.6455526 | -0.3949818 | 0.6928563 | 0.92224682 |
| STAM2 | 1297.76083 | -0.3596678 | 0.18894998 | -1.9035081 | 0.05697428 | 0.41872623 |
| FMNL2 | 3668.9885 | -0.0646183 | 0.28210845 | -0.2290549 | 0.81882623 | 0.96004577 |
| PRPF40A | 1042.86686 | -0.2301009 | 0.17693683 | -1.3004691 | 0.19344024 | 0.64718686 |
| ARL6IP6 | 590.363959 | -0.3375267 | 0.24613392 | -1.3713131 | 0.17027736 | 0.62634187 |
| GALNT13 | 536.784055 | 0.60573902 | 0.31324586 | 1.9337495 | 0.05314393 | 0.41099428 |
| AC009227.1 | 58.1811825 | 1.19821991 | 0.63390874 | 1.89020883 | 0.05873004 | 0.42299076 |
| KCNJ3 | 2020.02147 | -0.8738881 | 0.53494422 | -1.633606 | 0.1023416 | 0.52073581 |
| LINC01876 | 953.991396 | 0.42330003 | 0.72237714 | 0.58598204 | 0.55788758 | 0.87291366 |
| NR4A2 | 97.3102617 | -0.2052925 | 0.42498018 | -0.4830636 | 0.62905058 | 0.89984071 |
| GPD2 | 983.073581 | -0.2250033 | 0.23539189 | -0.955867 | 0.33913944 | 0.76449297 |
| CYTIP | 190.734917 | 2.61888871 | 0.69040214 | 3.79328009 | 0.00014867 | 0.02008985 |
| ACVR1C | 55.281011 | 1.45333911 | 0.48146089 | 3.01860265 | 0.00253943 | 0.09529369 |
| AC019186.1 | 62.2875716 | 0.5456028 | 0.42667772 | 1.27872343 | 0.20099447 | 0.6543242 |
| ACVR1 | 1522.37916 | -0.1105839 | 0.23225629 | -0.4761288 | 0.63398262 | 0.90136076 |
| UPP2 | 1099.11408 | 1.29497027 | 0.42362216 | 3.05689928 | 0.00223639 | 0.09083568 |
| CCDC148-AS1 | 147.704581 | 0.41838953 | 0.62054855 | 0.67422531 | 0.50016808 | 0.85270008 |
| CCDC148 | 5303.09064 | 0.45904955 | 0.31685272 | 1.44877896 | 0.14739932 | 0.59373375 |
| PKP4 | 15092.4726 | -0.1325954 | 0.27327278 | -0.4852126 | 0.62752552 | 0.89919293 |
| PKP4-AS1 | 417.095776 | 0.1003044 | 0.57219938 | 0.17529624 | 0.86084687 | 0.9720639 |
| TANC1 | 1975.50773 | -0.3363886 | 0.2324474 | -1.44716 | 0.14785213 | 0.59424557 |
| WDSUB1 | 855.305546 | -0.4119542 | 0.25119104 | -1.6400034 | 0.10100446 | 0.51819907 |
| BAZ2B | 5700.21768 | -0.1928923 | 0.18414748 | -1.0474882 | 0.29487446 | 0.73573797 |
| AC009961.1 | 216.2598 | 0.23572114 | 0.19498617 | 1.20891211 | 0.22669661 | 0.67880306 |
| 7-Mar | 1734.70745 | -0.1794935 | 0.13034593 | -1.3770547 | 0.16849533 | 0.62253512 |
| PLA2R1 | 199.082584 | 0.27247026 | 0.40734649 | 0.66889066 | 0.50356523 | 0.85525037 |
| ITGB6 | 14645.9371 | -0.3372348 | 0.77714919 | -0.4339383 | 0.66433326 | 0.91218067 |
| LINC02478 | 80.7009166 | 0.63327327 | 0.60981417 | 1.03846926 | 0.29905164 | 0.73673761 |
| RBMS1 | 5514.03512 | -0.5052292 | 0.29564732 | -1.7088916 | 0.08747103 | 0.49588191 |
| TANK | 1787.51021 | 0.02518012 | 0.26477438 | 0.09510028 | 0.92423518 | 0.98215577 |
| PSMD14 | 638.254431 | -0.2409677 | 0.25653372 | -0.9393216 | 0.34756566 | 0.76900643 |
| DPP4 | 349.147659 | 0.13399497 | 0.59145114 | 0.2265529 | 0.82077142 | 0.96045446 |
| IFIH1 | 247.996332 | 0.41414345 | 0.6934616 | 0.59721179 | 0.55036599 | 0.86959143 |
| GCA | 420.638405 | 0.10501203 | 0.23375707 | 0.44923574 | 0.65326161 | 0.908787 |
| FIGN | 72.4005533 | 1.04902764 | 0.54317131 | 1.93130163 | 0.05344576 | 0.4122367 |
| GRB14 | 2414.55402 | -0.0182989 | 0.37633142 | -0.0486244 | 0.96121864 | 0.99188706 |
| COBLL1 | 18887.582 | -0.6151311 | 0.30550733 | -2.0134743 | 0.04406476 | 0.3852094 |
| AC019197.1 | 7835.06212 | 1.79126877 | 0.46702986 | 3.83544806 | 0.00012534 | 0.01819294 |
| SLC38A11 | 703.894562 | -0.289578 | 0.65006494 | -0.4454602 | 0.65598725 | 0.90950195 |
| SCN3A | 693.923574 | -0.6581288 | 0.43416635 | -1.5158447 | 0.12955862 | 0.57050365 |
| SCN2A | 1191.72138 | 1.96669468 | 0.50724232 | 3.87722912 | 0.00010565 | 0.01624726 |

| | | | | | | |
|---|---|---|---|---|---|---|
| CSRNP3 | 1322.34234 | -0.3498972 | 0.28443576 | -1.2301449 | 2.19E-01 | 6.72E-01 |
| GALNT3 | 490.091043 | 0.02733287 | 0.25484374 | 0.10725346 | 9.15E-01 | 0.98156931 |
| AC009495.1 | 119.964838 | -0.5141831 | 0.47540215 | -1.081575 | 0.27944142 | 0.72489035 |
| TTC21B | 950.220452 | -0.1860796 | 0.19604794 | -0.9491536 | 0.3425425 | 0.76598 |
| AC010127.1 | 95.0234327 | 1.64197526 | 0.70273801 | 2.3365397 | 0.01946313 | 0.28049217 |
| SCN1A | 57.7378051 | 0.82290912 | 0.86799982 | 0.94805219 | 3.43E-01 | 7.66E-01 |
| SCN9A | 156.325254 | 0.99788301 | 0.79989749 | 1.24751361 | 0.21220923 | 0.66552984 |
| SCN7A | 161.922698 | 3.46487004 | 0.91419094 | 3.7900945 | 0.00015059 | 0.02008985 |
| STK39 | 5129.73029 | 0.95168312 | 0.52967567 | 1.79672802 | 0.07237881 | 0.46865279 |
| CERS6 | 1192.9435 | 0.44741641 | 0.35250412 | 1.26925159 | 0.20435135 | 0.65739819 |
| NOSTRIN | 155.598758 | 3.14181517 | 0.85249445 | 3.68543766 | 0.00022831 | 0.02471923 |
| LRP2 | 293.978224 | 2.98556914 | 0.69793229 | 4.27773463 | 1.89E-05 | 0.0071255 |
| FASTKD1 | 835.624827 | 0.17132267 | 0.22920152 | 0.74747615 | 4.55E-01 | 8.31E-01 |
| PPIG | 1128.50318 | 0.02779368 | 0.13547776 | 0.20515305 | 0.83745255 | 0.96527001 |
| CCDC173 | 84.1209485 | 0.04672684 | 0.45159124 | 0.10347154 | 0.91758873 | 0.98199511 |
| KLHL23 | 412.020359 | 0.06049409 | 0.18209289 | 0.33221557 | 0.73972649 | 0.93552616 |
| SSB | 352.512648 | 0.52369093 | 0.26582626 | 1.97004966 | 0.04883268 | 0.39762623 |
| METTL5 | 224.452425 | -0.2907289 | 0.16157555 | -1.7993375 | 0.07196531 | 0.46774933 |
| UBR3 | 3078.08318 | -0.0542231 | 0.20282509 | -0.2673392 | 0.78920799 | 0.95080703 |
| MYO3B | 3892.83594 | -0.160776 | 0.40683561 | -0.3951867 | 0.69270513 | 0.92224682 |
| AC012594.1 | 81.881344 | -0.1581537 | 0.65744314 | -0.2405588 | 0.80989708 | 0.95662439 |
| AC007277.1 | 84.677229 | -0.1540594 | 0.43166699 | -0.3568942 | 0.72117104 | 0.93044898 |
| AC007405.1 | 244.126472 | 0.21347133 | 0.46036764 | 0.46369751 | 0.64286448 | 0.90478945 |
| AC007405.3 | 73.0430593 | 0.33271466 | 0.41342719 | 0.80477209 | 0.42095121 | 0.81260593 |
| ERICH2 | 48.580392 | -0.1831952 | 0.43288143 | -0.4231994 | 0.6721498 | 0.91576543 |
| GAD1 | 701.898206 | 0.21870813 | 0.45590543 | 0.47972259 | 0.63142466 | 0.90011746 |
| GORASP2 | 540.918914 | 0.04403188 | 0.16302331 | 0.27009559 | 0.78708671 | 0.94978829 |
| TLK1 | 2407.32347 | -0.0733814 | 0.11944758 | -0.6143394 | 0.53899106 | 0.86542329 |
| METTL8 | 428.350992 | -0.4739462 | 0.32174113 | -1.4730668 | 0.14073304 | 0.58629918 |
| DCAF17 | 353.812284 | -0.1900781 | 0.26707993 | -0.7116898 | 0.47665686 | 0.84150767 |
| CYBRD1 | 168.200714 | 0.31863196 | 0.63340457 | 0.50304652 | 0.61493157 | 0.8949759 |
| DYNC1I2 | 1031.14826 | 0.04472489 | 0.09624886 | 0.46467966 | 0.64216088 | 0.90454089 |
| SLC25A12 | 2050.68758 | 0.13441671 | 0.1929744 | 0.69655206 | 0.48608316 | 0.8447301 |
| HAT1 | 462.985393 | -0.1150194 | 0.22468086 | -0.5119235 | 0.60870456 | 0.89273938 |
| METAP1D | 185.574352 | -0.6466783 | 0.40318005 | -1.6039443 | 0.10872633 | 0.53211043 |
| ITGA6 | 1103.57229 | 0.1997758 | 0.38422141 | 0.51994969 | 0.60309864 | 0.88978937 |
| AC078883.1 | 153.471899 | 0.29848994 | 0.6189257 | 0.48227104 | 0.62961342 | 0.90005881 |
| PDK1 | 829.945907 | -0.0198957 | 0.66669225 | -0.0298424 | 0.97619277 | 0.9941831 |
| RAPGEF4 | 158.791734 | -0.2351187 | 0.55901286 | -0.4205962 | 0.67404999 | 0.91636338 |
| MAP3K20 | 150.291309 | -0.3579509 | 0.47295546 | -0.7568385 | 4.49E-01 | 8.27E-01 |
| SP3 | 1137.94302 | -0.2319143 | 0.1804862 | -1.2849419 | 0.1988126 | 0.65113421 |
| OLA1 | 1967.69048 | -0.2722216 | 0.23404276 | -1.1631277 | 0.24477771 | 0.6948422 |
| CIR1 | 696.664515 | -0.0794124 | 0.14407635 | -0.5511828 | 0.58150837 | 0.88299162 |
| SCRN3 | 288.543816 | -0.0113511 | 0.30659435 | -0.0370233 | 0.97046645 | 0.99309664 |
| GPR155 | 509.500154 | 0.00399803 | 0.35471773 | 0.01127101 | 0.99100723 | 0.99749592 |
| AC010894.3 | 615.327336 | 0.5913202 | 0.38974613 | 1.51719326 | 1.29E-01 | 5.70E-01 |
| WIPF1 | 559.975977 | -0.322731 | 0.41501215 | -0.7776423 | 0.43677994 | 0.82013459 |
| CHN1 | 112.627107 | -0.3297591 | 0.32495041 | -1.0147983 | 0.31020197 | 0.74459794 |
| ATF2 | 1109.17247 | 0.07820093 | 0.1669436 | 0.46842722 | 0.63947911 | 0.90330608 |
| ATP5MC3 | 813.34898 | 1.0701442 | 0.4825925 | 2.21749032 | 0.0265896 | 0.31471512 |
| LNPK | 477.246856 | -0.0983034 | 0.31124428 | -0.3158401 | 0.75212387 | 0.93815321 |
| HOXD10 | 696.521055 | 0.06952045 | 0.54569197 | 0.12739871 | 0.89862484 | 0.9788023 |
| HOXD-AS2 | 180.799842 | 0.2752131 | 0.49122465 | 0.56025914 | 0.57530269 | 0.88065515 |
| HOXD9 | 253.738737 | -0.061722 | 0.61420677 | -0.1004906 | 0.91995481 | 0.98215577 |

| | | | | | | |
|---|---|---|---|---|---|---|
| HOXD8 | 255.765079 | 1.03424922 | 0.4460518 | 2.31867513 | 0.02041266 | 0.28653645 |
| HOXD3 | 874.699943 | 0.41069284 | 0.40664045 | 1.00996554 | 0.3125118 | 0.74693024 |
| HAGLR | 235.813876 | 0.16458835 | 0.24173424 | 0.68086486 | 0.495957 | 0.85023959 |
| HOXD1 | 50.1817132 | 0.54934875 | 0.48010552 | 1.14422502 | 0.25253033 | 0.70298696 |
| MTX2 | 2385.36569 | 0.17563023 | 0.20051138 | 0.87591154 | 0.38107812 | 0.78860425 |
| LINC01116 | 212.179544 | 0.45016379 | 0.49371204 | 0.91179421 | 0.36187706 | 0.77784066 |
| LINC01117 | 222.951076 | -0.5038797 | 0.42277163 | -1.1918484 | 0.23332068 | 0.68484327 |
| AC092162.2 | 49.4739172 | 0.12263932 | 0.54532092 | 0.22489385 | 0.82206185 | 0.96076933 |
| AC079305.3 | 104.725397 | 0.12929867 | 0.38237173 | 0.33814914 | 0.7352508 | 0.93552616 |
| HNRNPA3 | 1639.61865 | 0.15085922 | 0.15274969 | 0.98762376 | 0.32333694 | 0.75362713 |
| NFE2L2 | 3329.41848 | 0.24351467 | 0.29938476 | 0.81338366 | 0.41599813 | 0.80966301 |
| AGPS | 688.372537 | 0.05199429 | 0.1520442 | 0.34196825 | 0.73237479 | 0.93552616 |
| TTC30B | 41.3561907 | 1.03641929 | 0.56460327 | 1.83565939 | 0.06640805 | 0.4470866 |
| TTC30A | 70.7775294 | 0.29312614 | 0.31173391 | 0.9403088 | 0.34705919 | 0.76893725 |
| RBM45 | 268.901483 | -0.0292829 | 0.19897809 | -0.1471666 | 8.83E-01 | 0.97664137 |
| OSBPL6 | 50.2479131 | -0.3195322 | 0.43913014 | -0.7276481 | 0.46682904 | 0.83702251 |
| AC009948.1 | 174.189884 | -0.040211 | 0.33190526 | -0.1211522 | 0.90357051 | 0.97975674 |
| PRKRA | 654.175904 | 0.0099043 | 0.20259055 | 0.04888825 | 0.96100835 | 0.99179305 |
| PJVK | 164.097707 | -0.2622604 | 0.2731982 | -0.9599639 | 0.3370734 | 0.76409325 |
| PLEKHA3 | 315.070476 | 0.0724941 | 0.21468726 | 0.33767303 | 0.7356096 | 0.93552616 |
| TTN-AS1 | 1617.37887 | 0.43112509 | 0.23291356 | 1.85100902 | 0.06416826 | 0.43894277 |
| TTN | 169.982995 | 1.14449502 | 0.54091972 | 2.11583156 | 0.03435914 | 0.35209451 |
| SESTD1 | 3597.43797 | 0.03170939 | 0.42279315 | 0.07499976 | 0.9402149 | 0.98715467 |
| ZNF385B | 1530.96043 | 0.51713534 | 0.37246712 | 1.38840534 | 1.65E-01 | 6.17E-01 |
| CWC22 | 528.49683 | 0.08765273 | 0.22767358 | 0.38499298 | 0.70024261 | 0.92367966 |
| AC009478.1 | 920.3509 | 0.17499372 | 0.29494254 | 0.59331461 | 0.55297062 | 0.87104567 |
| SCHLAP1 | 3269.29313 | 0.01777315 | 0.44050276 | 0.04034742 | 0.96781615 | 0.99248831 |
| UBE2E3 | 2793.65633 | -0.1649651 | 0.23124623 | -0.7133741 | 0.47561428 | 0.84118672 |
| LINC01934 | 53.6519887 | 0.08118764 | 0.51824747 | 0.15665805 | 0.87551435 | 0.97614152 |
| CERKL | 403.874362 | -0.3348843 | 0.4029602 | -0.8310606 | 0.40593942 | 0.80499101 |
| SSFA2 | 1423.41215 | 0.80999514 | 0.36835306 | 2.19896403 | 0.02788048 | 0.32224037 |
| PPP1R1C | 101.35558 | 0.66401102 | 0.51088973 | 1.29971495 | 0.19369869 | 0.64718686 |
| PDE1A | 22859.8803 | -0.7072802 | 0.37989 | -1.8618027 | 0.0626309 | 0.43495877 |
| DNAJC10 | 592.697125 | -0.145739 | 0.20816697 | -0.7001063 | 0.48386092 | 0.8445691 |
| NCKAP1 | 2707.09288 | -0.1356798 | 0.21554265 | -0.6294802 | 0.52903476 | 0.86282506 |
| DUSP19 | 42.2807844 | -0.1243257 | 0.42670257 | -0.2913639 | 0.77077305 | 0.9443559 |
| NUP35 | 303.85299 | -0.3159446 | 0.22336226 | -1.4144942 | 0.15721686 | 0.60703814 |
| AC007966.1 | 66.7828097 | 0.70739539 | 0.33896917 | 2.08690182 | 0.03689701 | 0.36038369 |
| FSIP2 | 241.565475 | 0.11428172 | 0.64433061 | 0.17736503 | 0.85922167 | 0.97174503 |
| FSIP2-AS1 | 193.070497 | 0.65977919 | 0.33583491 | 1.96459384 | 0.04946126 | 0.39910437 |
| LINC01473 | 412.655545 | 0.28341019 | 0.28997668 | 0.97735512 | 0.32839337 | 0.75675996 |
| AC017071.1 | 55.1347814 | 0.3765884 | 0.37614386 | 1.00118185 | 0.3167389 | 0.74844439 |
| ZC3H15 | 544.813203 | 0.0514238 | 0.17950166 | 0.28648094 | 0.77450979 | 0.94621715 |
| ITGAV | 2113.06852 | 0.16714842 | 0.41575637 | 0.40203454 | 0.68765861 | 0.92103859 |
| AC017101.1 | 265.792166 | 0.43258845 | 0.85044806 | 0.50865946 | 0.61099094 | 0.89348926 |
| FAM171B | 123.970893 | 2.26062651 | 0.57512288 | 3.93068437 | 8.47E-05 | 0.01511702 |
| AC007319.1 | 2512.07907 | 0.63087476 | 0.66740912 | 0.94525943 | 0.34452644 | 0.76716956 |
| CALCRL | 278.920491 | 2.05887305 | 0.64740543 | 3.18019118 | 0.00147178 | 0.07095157 |
| TFPI | 6167.12535 | -0.1022031 | 0.60927342 | -0.1677459 | 8.67E-01 | 9.74E-01 |
| LINC01090 | 175.227099 | 0.75584868 | 0.58594085 | 1.28997438 | 0.19705955 | 0.64964978 |
| GULP1 | 3451.10326 | -0.338791 | 0.57608448 | -0.5880925 | 0.55647018 | 0.87212955 |
| COL5A2 | 237.806803 | -0.489247 | 0.48219202 | -1.0146311 | 0.31028172 | 0.74459794 |
| WDR75 | 344.840982 | -0.3355784 | 0.26416862 | -1.2703188 | 0.2039711 | 0.65712971 |
| SLC40A1 | 231.338294 | 0.21766205 | 0.43292216 | 0.50277409 | 0.61512311 | 0.8949877 |

| | | | | | | |
|---|---|---|---|---|---|---|
| ASNSD1 | 169.936859 | 0.20310311 | 0.31318813 | 0.64850194 | 0.51666036 | 0.8588826 |
| ANKAR | 883.334277 | -0.1756897 | 0.28298168 | -0.6208517 | 0.53469719 | 0.86419602 |
| OSGEPL1 | 112.522059 | -0.1548642 | 0.26966011 | -0.574294 | 0.56576886 | 0.87631812 |
| ORMDL1 | 564.041598 | 0.19370734 | 0.27045589 | 0.71622527 | 0.47385226 | 0.84075535 |
| PMS1 | 587.780421 | -0.006663 | 0.1978698 | -0.0336736 | 0.97313742 | 0.99327086 |
| C2orf88 | 310.14457 | 0.28396546 | 0.35774714 | 0.79376026 | 0.42733501 | 0.81466629 |
| HIBCH | 1938.97887 | 0.26139752 | 0.2583107 | 1.01195003 | 0.31156195 | 0.74607653 |
| INPP1 | 225.681636 | 0.24601472 | 0.29859915 | 0.82389624 | 0.40999852 | 0.80650342 |
| MFSD6 | 490.72969 | -0.1333512 | 0.26824198 | -0.4971304 | 6.19E-01 | 0.89618124 |
| NEMP2 | 95.3060649 | -0.7415024 | 0.32679351 | -2.2690242 | 0.02326686 | 0.30018842 |
| AC108047.1 | 106.593743 | -0.2404488 | 0.33708357 | -0.7133209 | 0.47564718 | 0.84118672 |
| NAB1 | 365.577767 | 0.06022464 | 0.39680189 | 0.15177509 | 0.87936433 | 0.9765717 |
| GLS | 5469.69674 | -0.5827301 | 0.35389824 | -1.6466036 | 0.09963956 | 0.51665771 |
| STAT1 | 1097.79631 | 0.46919286 | 0.69187555 | 0.67814632 | 0.49767892 | 0.85092921 |
| MYO1B | 461.227478 | 0.19682584 | 0.46707711 | 0.42139903 | 0.67346373 | 0.91636338 |
| NABP1 | 357.282047 | 1.69244724 | 0.67767461 | 2.49743347 | 0.01250959 | 0.22899031 |
| SLC39A10 | 745.278605 | 1.033477 | 0.2759895 | 3.74462432 | 0.00018066 | 0.021499 |
| DNAH7 | 2001.76686 | -0.1611649 | 0.16706782 | -0.9646674 | 0.33471144 | 0.76242642 |
| STK17B | 205.905972 | -0.0267204 | 0.23259512 | -0.1148794 | 0.90854068 | 0.98047754 |
| GTF3C3 | 443.812282 | -0.1544703 | 0.18321512 | -0.8431092 | 0.39916738 | 0.80174499 |
| PGAP1 | 483.808654 | 0.17651807 | 0.2046079 | 0.86271387 | 0.3882948 | 0.79634939 |
| ANKRD44 | 915.640405 | -0.5351348 | 0.28774282 | -1.8597677 | 0.0629184 | 0.43603835 |
| SF3B1 | 1337.38681 | 0.16461938 | 0.19196936 | 0.85752941 | 0.39115237 | 0.79735014 |
| COQ10B | 304.80543 | 0.01329373 | 0.2598742 | 0.05115449 | 0.95920241 | 0.99117358 |
| HSPD1 | 895.716081 | 0.2922904 | 0.28713812 | 1.01794358 | 0.30870477 | 0.74399658 |
| MOB4 | 125.698811 | 0.52454296 | 0.323181 | 1.6230625 | 0.10457605 | 0.52442862 |
| AC011997.1 | 132.32273 | -0.2335953 | 0.39190958 | -0.5960439 | 0.55114593 | 0.87018163 |
| PLCL1 | 18795.9735 | 0.31568418 | 0.41743447 | 0.75624848 | 4.50E-01 | 8.27E-01 |
| SATB2 | 483.020819 | -0.8035551 | 0.46236333 | -1.73793 | 0.08222314 | 0.48594425 |
| FTCDNL1 | 294.134345 | 0.66244214 | 0.56150539 | 1.17976096 | 0.2380953 | 0.68982886 |
| C2orf69 | 152.977195 | 0.33390049 | 0.3333473 | 1.00165949 | 0.31650808 | 0.74818634 |
| TYW5 | 392.813815 | -0.1701279 | 0.16338973 | -1.0412398 | 0.29776428 | 0.73673761 |
| MAIP1 | 173.827702 | -0.2758751 | 0.26592687 | -1.0374095 | 0.29954505 | 0.73673761 |
| SPATS2L | 1456.52927 | -0.0041224 | 0.61784111 | -0.0066723 | 0.99467631 | 0.998438 |
| KCTD18 | 179.238021 | -0.4379603 | 0.18757837 | -2.3348123 | 0.01955323 | 0.2807376 |
| SGO2 | 86.5236507 | -0.3813988 | 0.44235201 | -0.8622066 | 0.38857382 | 0.79634939 |
| AOX1 | 56.9606311 | 2.64352944 | 0.65900594 | 4.01138942 | 6.04E-05 | 0.01245824 |
| BZW1 | 238.712243 | -0.1400172 | 0.28047578 | -0.499213 | 0.61762935 | 0.89599505 |
| CLK1 | 1160.27198 | -0.1599695 | 0.299 | -0.5350149 | 0.59263955 | 0.88740306 |
| PPIL3 | 303.453865 | -0.4496525 | 0.33525234 | -1.3412359 | 0.17984387 | 0.63399761 |
| NIF3L1 | 147.157369 | -0.202969 | 0.27502068 | -0.7380136 | 0.46050621 | 0.83429248 |
| ORC2 | 1364.58223 | -0.4062786 | 0.22804821 | -1.7815471 | 0.07482313 | 0.47190267 |
| AC005037.1 | 316.873549 | -0.1221676 | 0.34260126 | -0.3565883 | 0.72140002 | 0.93044898 |
| FAM126B | 1201.24508 | 0.0040473 | 0.27627215 | 0.01464967 | 0.98831167 | 0.99653761 |
| NDUFB3 | 366.566221 | 0.66103032 | 0.46798711 | 1.41249684 | 0.15780372 | 0.60779483 |
| CFLAR | 3800.1795 | 0.1278311 | 0.21080159 | 0.60640482 | 0.54424597 | 0.86654229 |
| CFLAR-AS1 | 390.958322 | 0.0939391 | 0.61906186 | 0.15174429 | 0.87938863 | 0.9765717 |
| CASP10 | 540.514767 | 0.33025076 | 0.32062854 | 1.03001049 | 0.30300508 | 0.73995085 |
| CASP8 | 505.297177 | -0.0179297 | 0.29648828 | -0.0604736 | 0.95177844 | 0.98938795 |
| ALS2CR12 | 274.830425 | 0.45566321 | 0.43169138 | 1.05553 | 0.291183 | 0.73331481 |
| TRAK2 | 1112.46975 | 0.34674586 | 0.22478351 | 1.54257697 | 0.12293345 | 0.55947129 |
| STRADB | 612.249725 | 0.50822626 | 0.37364939 | 1.36016883 | 0.1737765 | 0.6286116 |
| C2CD6 | 100.158652 | 0.04249171 | 0.44528594 | 0.09542568 | 0.92397672 | 0.98215577 |
| TMEM237 | 150.33901 | -0.13744 | 0.21317869 | -0.6447174 | 0.51911035 | 0.85918126 |

| | | | | | | |
|---|---|---|---|---|---|---|
| ALS2 | 731.596882 | -0.0167544 | 0.17937438 | -0.0934049 | 0.92558193 | 0.9822216 |
| SUMO1 | 508.645097 | 0.08147526 | 0.22173594 | 0.36744275 | 0.71328879 | 0.92760608 |
| NOP58 | 753.285045 | -0.2251769 | 0.16707448 | -1.3477637 | 0.17773439 | 0.63160978 |
| BMPR2 | 2851.76435 | -0.2301756 | 0.1767511 | -1.3022582 | 0.19282812 | 0.64695623 |
| FAM117B | 1211.3143 | -0.5861555 | 0.30529972 | -1.9199346 | 0.05486616 | 0.41684503 |
| ICA1L | 432.529796 | -0.8470375 | 0.40167018 | -2.1087886 | 0.03496283 | 0.35358029 |
| WDR12 | 482.216354 | -0.0886664 | 0.26611766 | -0.3331851 | 0.73899457 | 0.93552616 |
| CARF | 776.320359 | -0.3779418 | 0.25373964 | -1.4894866 | 0.13635928 | 0.57787125 |
| NBEAL1 | 4969.50607 | -0.0207178 | 0.16856063 | -0.1229102 | 0.90217821 | 0.97941893 |
| CYP20A1 | 998.742271 | -0.2497415 | 0.23063022 | -1.0828653 | 0.27886822 | 0.72454177 |
| ABI2 | 1755.27937 | -0.0724082 | 0.16102388 | -0.4496739 | 0.65294559 | 0.908787 |
| RAPH1 | 895.655897 | -0.1195598 | 0.24884689 | -0.4804551 | 6.31E-01 | 9.00E-01 |
| PARD3B | 7681.73734 | -0.1699775 | 0.17076064 | -0.9954139 | 0.31953502 | 0.75094569 |
| AC007679.1 | 81.3965173 | 0.31399294 | 0.70778857 | 0.44362533 | 0.65731349 | 0.91026278 |
| INO80D | 2260.38872 | -0.1915541 | 0.54381819 | -0.3522393 | 0.72465882 | 0.93241479 |
| NDUFS1 | 1477.85755 | 0.17125445 | 0.14285744 | 1.19877861 | 0.23061404 | 0.68284709 |
| EEF1B2 | 327.472101 | 0.31629859 | 0.41214441 | 0.76744604 | 0.44281636 | 0.82301411 |
| ZDBF2 | 330.53068 | 0.01593266 | 0.28146867 | 0.05660546 | 0.95485948 | 0.99049376 |
| MDH1B | 86.8558016 | 0.57231919 | 0.40905787 | 1.39911547 | 0.16177836 | 0.61346689 |
| FASTKD2 | 478.527268 | -0.3412819 | 0.20904727 | -1.6325584 | 0.10256191 | 0.52165586 |
| CPO | 152.152822 | -0.2295402 | 0.4785948 | -0.4796127 | 0.63150279 | 0.90011794 |
| KLF7 | 922.671581 | -0.3822532 | 0.50304729 | -0.7598754 | 0.44732909 | 0.82578615 |
| AC009226.1 | 71.047477 | -1.4093789 | 0.59690467 | -2.3611457 | 0.01821857 | 0.2711921 |
| CREB1 | 1190.22325 | -0.0901414 | 0.14597178 | -0.6175264 | 0.53688755 | 0.86466169 |
| METTL21A | 304.680074 | -0.0783234 | 0.34373522 | -0.2278598 | 0.81975526 | 0.96014363 |
| CCNYL1 | 433.250353 | -0.6638905 | 0.40887683 | -1.623693 | 0.10444134 | 0.52442862 |
| FZD5 | 76.0849191 | 0.54436312 | 0.36915155 | 1.4746332 | 1.40E-01 | 0.58521327 |
| PLEKHM3 | 831.604394 | -0.1221989 | 0.19349732 | -0.6315276 | 0.52769564 | 0.86181403 |
| IDH1 | 351.020327 | -0.4671162 | 0.35080992 | -1.3315364 | 0.18301258 | 0.63767162 |
| PIKFYVE | 1168.16883 | -0.2741381 | 0.23231584 | -1.1800231 | 0.23799103 | 0.68982886 |
| PTH2R | 12015.8884 | 0.09141238 | 0.20459076 | 0.446806 | 6.55E-01 | 9.09E-01 |
| MAP2 | 510.736192 | -0.6002684 | 0.44844839 | -1.3385452 | 0.18071879 | 0.63503977 |
| RPE | 430.13783 | -0.2330213 | 0.21483718 | -1.0846415 | 0.27808046 | 0.72372365 |
| KANSL1L | 5280.92418 | 0.33845707 | 0.22547911 | 1.50105736 | 0.13334073 | 0.57682553 |
| AC007038.2 | 51.2589457 | -0.008822 | 0.38402555 | -0.0229725 | 0.98167218 | 0.99518016 |
| AC007038.1 | 682.281589 | -0.1049913 | 0.27497593 | -0.3818199 | 0.70259493 | 0.92501157 |
| ACADL | 338.431315 | 0.19120036 | 0.46298264 | 0.41297523 | 0.67962476 | 0.9182972 |
| LANCL1-AS1 | 169.771449 | 0.33176507 | 0.57894169 | 0.57305439 | 0.56660787 | 0.87646121 |
| LANCL1 | 867.890405 | -0.0998636 | 0.17117895 | -0.583387 | 0.55963282 | 0.87300289 |
| CPS1 | 120.982828 | 0.95786567 | 0.30365422 | 3.15446191 | 0.00160794 | 0.07531676 |
| ERBB4 | 70863.8385 | 1.04238333 | 0.41445082 | 2.51509536 | 0.01190002 | 0.22355405 |
| IKZF2 | 462.8565 | 0.19368494 | 0.23502661 | 0.82409792 | 0.40988392 | 0.80641944 |
| SPAG16 | 3913.94631 | 0.35258764 | 0.2909817 | 1.2117176 | 0.2256205 | 0.67798093 |
| AC068051.1 | 535.394202 | 0.69961599 | 0.43069609 | 1.62438436 | 0.1042938 | 0.52442862 |
| VWC2L | 81.6526099 | 1.07869784 | 0.49171662 | 2.19373883 | 0.02825419 | 0.32424814 |
| BARD1 | 258.620994 | -0.0438134 | 0.21743576 | -0.2015006 | 0.84030714 | 0.96617873 |
| AC016708.1 | 270.833151 | 0.0246067 | 0.3811413 | 0.06456058 | 0.94852387 | 0.98919519 |
| ATIC | 253.839047 | -0.2851102 | 0.2120378 | -1.3446195 | 0.17874816 | 0.63284534 |
| FN1 | 94.1941693 | -0.2268151 | 0.33599203 | -0.6750611 | 0.49963698 | 0.85244523 |
| MREG | 231.47851 | -0.2157902 | 0.28390503 | -0.7600789 | 0.44720745 | 0.82578615 |
| PECR | 278.77441 | -0.2134511 | 0.37003085 | -0.5768468 | 0.56404292 | 0.87508139 |
| TMEM169 | 51.8436667 | -0.3514226 | 0.66497739 | -0.528473 | 0.59717105 | 0.88883692 |
| XRCC5 | 2127.40514 | -0.2570213 | 0.18734729 | -1.3718977 | 0.1700953 | 0.62602085 |
| SMARCAL1 | 335.338955 | -0.4174096 | 0.28030279 | -1.4891384 | 0.13645094 | 0.57787125 |

| | | | | | | |
|---|---|---|---|---|---|---|
| RPL37A | 827.542498 | 0.18317805 | 0.36470422 | 0.50226469 | 0.61548135 | 0.8949877 |
| IGFBP2 | 287.262076 | 0.81212905 | 0.61226189 | 1.32644064 | 0.18469379 | 0.63998432 |
| IGFBP5 | 226.959411 | -1.1265603 | 0.35574673 | -3.166748 | 0.00154154 | 0.07298272 |
| DIRC3 | 767.564739 | -0.0197103 | 0.65490916 | -0.0300962 | 0.97599035 | 0.9941831 |
| TNS1 | 2621.12334 | 0.56695054 | 0.41650428 | 1.36121181 | 0.17344676 | 0.6286116 |
| ARPC2 | 913.579314 | -0.2025527 | 0.15904174 | -1.2735818 | 2.03E-01 | 6.56E-01 |
| AAMP | 226.803134 | 0.15867352 | 0.3269858 | 0.4852612 | 0.62749108 | 0.89919293 |
| PNKD | 531.697962 | 0.49476466 | 0.29348966 | 1.68579931 | 0.09183446 | 0.50476839 |
| TMBIM1 | 429.586356 | 0.37772077 | 0.19343108 | 1.95274085 | 5.09E-02 | 4.05E-01 |
| CATIP-AS2 | 70.9129144 | 0.92458693 | 0.6896955 | 1.34057267 | 0.18005924 | 0.63424065 |
| CATIP | 69.0006865 | 1.31241073 | 0.70700737 | 1.85629003 | 0.06341223 | 0.4376239 |
| CTDSP1 | 368.790045 | 0.11514758 | 0.17856495 | 0.64484986 | 0.51902449 | 0.85918126 |
| USP37 | 864.289068 | -0.283086 | 0.27877653 | -1.0154586 | 0.30988726 | 0.74450725 |
| CNOT9 | 422.295899 | -0.1761537 | 0.22209843 | -0.7931332 | 0.4277002 | 0.81466629 |
| PLCD4 | 238.055964 | 1.9775333 | 0.84865419 | 2.33019919 | 0.01979563 | 0.28268153 |
| ZNF142 | 176.520305 | 0.10218599 | 0.25123662 | 0.40673207 | 0.68420478 | 0.9200778 |
| BCS1L | 80.0689388 | -0.0858411 | 0.32709809 | -0.2624324 | 0.7929881 | 0.95224951 |
| RNF25 | 88.1200594 | -0.3647319 | 0.27359353 | -1.333116 | 0.18249375 | 0.63741052 |
| STK36 | 267.323077 | -0.4576391 | 0.19671889 | -2.3263608 | 0.01999931 | 0.2836927 |
| TTLL4 | 167.026809 | -0.1078992 | 0.27907594 | -0.3866303 | 0.69902992 | 0.92367966 |
| CYP27A1 | 169.818856 | 0.41445202 | 0.28729066 | 1.44262265 | 0.1491268 | 0.59583057 |
| CNPPD1 | 230.48432 | 0.31334062 | 0.21633063 | 1.44843388 | 0.14749575 | 0.59373375 |
| RETREG2 | 238.428334 | 0.06617802 | 0.32972876 | 0.20070441 | 0.84092971 | 0.96627616 |
| ZFAND2B | 147.626673 | -0.3837122 | 0.32905293 | -1.1661109 | 0.24356959 | 0.6948422 |
| ANKZF1 | 402.184134 | -0.1814883 | 0.38499568 | -0.4714034 | 0.63735268 | 0.90255504 |
| GLB1L | 49.4048784 | -0.5785855 | 0.40236342 | -1.4379674 | 0.15044331 | 0.59693772 |
| STK16 | 87.2047778 | -0.3437006 | 0.33001442 | -1.0414713 | 0.29765685 | 0.73673761 |
| TUBA4A | 215.83833 | 0.68684967 | 0.40198918 | 1.70862727 | 0.08752001 | 0.49594674 |
| DNAJB2 | 242.625749 | 0.09102538 | 0.32295143 | 0.28185471 | 0.77805492 | 0.94694572 |
| DNPEP | 180.58741 | -0.0861896 | 0.341493 | -0.2523905 | 0.80073928 | 0.95370871 |
| GMPPA | 104.868626 | 0.46245771 | 0.42155144 | 1.09703745 | 0.27262502 | 0.72070318 |
| CHPF | 107.956243 | 0.96634728 | 0.46537413 | 2.07649549 | 0.03784815 | 0.36437085 |
| OBSL1 | 688.688701 | -0.0782728 | 0.26081107 | -0.3001131 | 0.76409092 | 0.94256396 |
| INHA | 65.9871193 | 0.36705625 | 0.44379077 | 0.82709303 | 0.40818433 | 0.80588056 |
| STK11IP | 152.135387 | -0.8623365 | 0.22667726 | -3.8042477 | 0.00014224 | 0.01998715 |
| SLC4A3 | 108.740778 | 0.28128802 | 0.57539969 | 0.48885674 | 0.62494312 | 0.89873262 |
| AC019211.1 | 63.7990333 | -0.1040331 | 0.82919493 | -0.1254628 | 9.00E-01 | 9.79E-01 |
| EPHA4 | 786.668082 | -0.3731507 | 0.56308153 | -0.6626939 | 0.5075266 | 0.85632606 |
| SGPP2 | 2824.45998 | 0.31079195 | 0.3457806 | 0.89881258 | 3.69E-01 | 0.78095617 |
| FARSB | 392.419751 | -0.5575943 | 0.25631042 | -2.1754648 | 0.0295953 | 0.33073125 |
| MOGAT1 | 66.5774613 | -0.1997638 | 0.65300368 | -0.3059152 | 0.75966919 | 0.94133292 |
| ACSL3 | 1499.29374 | 0.19330826 | 0.30789449 | 0.62783927 | 0.53010923 | 0.86361777 |
| AP1S3 | 1863.97231 | -0.5831499 | 0.44868256 | -1.2996937 | 0.19370596 | 0.64718686 |
| WDFY1 | 802.454347 | -0.3128587 | 0.22287387 | -1.4037475 | 0.16039405 | 0.61151309 |
| MRPL44 | 111.172765 | 0.15586379 | 0.24002042 | 0.64937724 | 0.51609457 | 0.85888182 |
| SERPINE2 | 191.892828 | -1.9576899 | 0.73053685 | -2.6797962 | 0.0073667 | 0.17305714 |
| CUL3 | 2379.79198 | -0.1850525 | 0.16755938 | -1.1043998 | 0.26941976 | 0.71818262 |
| DOCK10 | 80.5747278 | -0.5304864 | 0.58996848 | -0.8991775 | 0.36855812 | 0.78095617 |
| AC062015.1 | 49.5534593 | 0.62673149 | 0.85510253 | 0.73293139 | 0.46360028 | 0.83531269 |
| IRS1 | 1285.04518 | -0.2071131 | 0.40787172 | -0.5077899 | 0.6116007 | 0.8941127 |
| AC010735.2 | 38.7732375 | -0.08812 | 0.37355741 | -0.2358942 | 0.81351476 | 0.9582375 |
| RHBDD1 | 1387.84266 | -0.175869 | 0.22624647 | -0.7773334 | 0.43696207 | 0.82013459 |
| COL4A4 | 4236.71464 | -0.1633911 | 0.29667279 | -0.550745 | 0.58180846 | 0.88324422 |
| COL4A3 | 7150.33117 | 0.1857831 | 0.38890617 | 0.47770674 | 0.63285895 | 0.90074133 |

| | | | | | | |
|---|---|---|---|---|---|---|
| AC097662.1 | 1038.69225 | -0.1514709 | 0.60933946 | -0.2485821 | 0.80368405 | 0.95450237 |
| MFF | 1269.67558 | 0.2965558 | 0.1951868 | 1.51934347 | 0.12867606 | 0.56864574 |
| AGFG1 | 2199.80587 | -0.0716631 | 0.20775355 | -0.3449427 | 0.73013742 | 0.93453345 |
| SLC19A3 | 335.073878 | 0.23512963 | 0.50524512 | 0.46537734 | 0.64166126 | 0.90426796 |
| DNER | 644.230676 | -0.2068121 | 0.44325842 | -0.4665724 | 0.64080584 | 0.9039304 |
| TRIP12 | 3038.70504 | -0.1165752 | 0.1467713 | -0.7942641 | 0.42704172 | 0.81466629 |
| FBXO36 | 382.828114 | -0.1242333 | 0.21111527 | -0.5884619 | 0.55622229 | 0.87212955 |
| AC009950.1 | 75.0343491 | -0.5099197 | 0.65332065 | -0.7805045 | 0.43509398 | 0.81919857 |
| SP110 | 544.639604 | -0.4547563 | 0.54020367 | -0.8418238 | 0.3998866 | 0.80215701 |
| SP140L | 271.613204 | -0.376314 | 0.44579365 | -0.8441438 | 0.39858904 | 0.80148305 |
| SP100 | 1543.56487 | -0.2362711 | 0.38733976 | -0.6099841 | 0.54187231 | 0.86582694 |
| AC010149.1 | 54.8639922 | -0.2210172 | 0.38464941 | -0.574594 | 0.5655659 | 0.87620925 |
| CAB39 | 1613.52629 | -0.1818548 | 0.1790287 | -1.0157856 | 0.30973148 | 0.74450725 |
| ITM2C | 955.822516 | 1.58936103 | 0.69640278 | 2.28224395 | 0.02247494 | 0.29746644 |
| PSMD1 | 1082.83162 | -0.2088611 | 0.24341181 | -0.8580565 | 0.39086129 | 0.79735014 |
| ARMC9 | 1358.04692 | -0.042267 | 0.19565784 | -0.2160252 | 0.82896807 | 0.96288319 |
| NCL | 600.916786 | 0.50230144 | 0.3173438 | 1.58283049 | 0.11346011 | 0.53989968 |
| PTMA | 2384.64246 | 0.32264619 | 0.3503986 | 0.92079762 | 3.57E-01 | 0.77448758 |
| PDE6D | 459.231805 | -0.2929462 | 0.33075876 | -0.8856793 | 0.37579037 | 0.78490077 |
| COPS7B | 453.239765 | -0.0980408 | 0.33023407 | -0.2968829 | 0.76655592 | 0.94312902 |
| DIS3L2 | 1954.54253 | -0.2390736 | 0.16770093 | -1.4255947 | 0.15398536 | 0.60292313 |
| AC019130.1 | 142.190012 | -0.1075049 | 0.55366654 | -0.1941691 | 0.8460435 | 0.96883382 |
| TIGD1 | 76.4348797 | 0.26404976 | 0.2516065 | 1.04945527 | 0.29396863 | 0.73527014 |
| EIF4E2 | 505.795267 | 0.29262503 | 0.20518935 | 1.42612193 | 0.15383315 | 0.60285224 |
| EFHD1 | 3424.41128 | 0.78789843 | 0.33009188 | 2.38690643 | 0.01699082 | 0.26127087 |
| GIGYF2 | 3230.70026 | -0.1370101 | 0.14616566 | -0.9373618 | 0.34857249 | 0.76982416 |
| KCNJ13 | 42.0571908 | 0.27923524 | 0.71159359 | 0.39240831 | 0.69475655 | 0.92229578 |
| SNORC | 72.0995684 | 0.42298465 | 0.46763624 | 0.9045164 | 0.36572164 | 0.78095617 |
| ATG16L1 | 213.02532 | -0.422151 | 0.27621632 | -1.5283348 | 0.12642943 | 0.56629581 |
| DGKD | 298.465043 | -0.4951289 | 0.3527352 | -1.4036845 | 0.16041281 | 0.61151309 |
| USP40 | 1439.58357 | -0.1555408 | 0.30130732 | -0.5162197 | 0.60570096 | 0.89081188 |
| ARL4C | 151.35314 | 0.05349534 | 0.85918558 | 0.06226285 | 9.50E-01 | 0.98929186 |
| SH3BP4 | 1282.94819 | -0.4553709 | 0.3188427 | -1.4281993 | 0.1532345 | 0.60285224 |
| AGAP1 | 7685.63712 | -0.5725173 | 0.32300957 | -1.7724468 | 0.07632042 | 0.47530241 |
| IQCA1 | 288.841946 | -1.5628544 | 0.34690895 | -4.5050854 | 6.63E-06 | 0.00349342 |
| AC105760.2 | 144.98203 | -0.6664932 | 0.34169853 | -1.95053 | 0.05111298 | 0.40556617 |
| COPS8 | 223.229381 | -0.2503873 | 0.24070636 | -1.0402189 | 0.29823822 | 0.73673761 |
| MLPH | 95.6201178 | -1.1590424 | 0.53656096 | -2.1601318 | 0.03076247 | 0.33646028 |
| RAB17 | 851.771581 | 0.07539442 | 0.46214636 | 0.16313971 | 0.87040843 | 0.9746715 |
| LRRFIP1 | 4314.63659 | -0.3867635 | 0.13024661 | -2.9694704 | 2.98E-03 | 1.05E-01 |
| RBM44 | 52.4872257 | -1.7970092 | 0.66877096 | -2.6870323 | 0.007209 | 0.1729679 |
| UBE2F-SCLY | 61.9361084 | -0.5783591 | 0.42461256 | -1.3620867 | 0.17317051 | 0.6286116 |
| ERFE | 74.6336826 | -1.2674859 | 0.56827086 | -2.2304257 | 2.57E-02 | 3.12E-01 |
| ILKAP | 773.210934 | -0.3004004 | 0.19909528 | -1.5088275 | 0.13134288 | 0.57506954 |
| PER2 | 312.678466 | -0.8043502 | 0.57025006 | -1.410522 | 0.15838561 | 0.60804051 |
| TRAF3IP1 | 554.997478 | -0.0381452 | 0.20601063 | -0.1851615 | 0.85310233 | 0.97076831 |
| ASB1 | 123.432583 | -0.0262283 | 0.28057528 | -0.0934805 | 0.92552181 | 0.9822216 |
| HDAC4 | 1648.30917 | -0.3808114 | 0.25964439 | -1.4666652 | 0.14246714 | 0.58824095 |
| NDUFA10 | 1143.05207 | -0.2835365 | 0.26259102 | -1.0797646 | 0.280247 | 0.72513763 |
| COPS9 | 250.856224 | 0.81704572 | 0.49304707 | 1.65713532 | 0.09749212 | 0.51355946 |
| AC124861.1 | 134.255902 | 1.40960333 | 0.59253462 | 2.37893834 | 0.01736258 | 0.26421926 |
| GPC1 | 253.142136 | -0.0100212 | 0.49717198 | -0.0201563 | 9.84E-01 | 9.95E-01 |
| ANKMY1 | 536.661745 | -0.7720671 | 0.27685975 | -2.7886577 | 0.0052927 | 0.14687236 |
| RNPEPL1 | 193.714819 | -0.0194163 | 0.36846218 | -0.0526956 | 9.58E-01 | 0.99069768 |

| | | | | | | |
|---|---|---|---|---|---|---|
| CAPN10-DT | 48.498142 | -0.1313648 | 0.46579513 | -0.2820227 | 0.77792609 | 0.94694572 |
| CAPN10 | 177.350251 | 0.08197745 | 0.25003406 | 0.32786512 | 0.74301364 | 0.93598059 |
| GPR35 | 38.9755581 | -0.6509364 | 0.43488801 | -1.4967908 | 0.1344477 | 0.57708028 |
| SNED1 | 1971.13462 | -0.6851871 | 0.50699833 | -1.3514583 | 0.17654865 | 0.63008941 |
| AC005237.1 | 362.710924 | -0.3363873 | 0.53701473 | -0.6264024 | 0.53105106 | 0.86397197 |
| MTERF4 | 603.077189 | -0.2673985 | 0.23156048 | -1.1547675 | 0.24818563 | 0.69838949 |
| PASK | 115.373351 | -0.4223654 | 0.39299168 | -1.0747439 | 0.28248942 | 0.72604837 |
| PPP1R7 | 309.541339 | 0.20277967 | 0.22745782 | 0.8915045 | 0.37265858 | 0.78269743 |
| HDLBP | 1624.12465 | -0.0046511 | 0.15696695 | -0.0296312 | 0.97636119 | 0.9941831 |
| 2-Sep | 2112.5711 | 0.10850342 | 0.16686532 | 0.65024549 | 0.51553366 | 0.85869987 |
| FARP2 | 1102.84685 | -0.0211421 | 0.16952083 | -0.1247165 | 0.90074797 | 0.97914987 |
| STK25 | 308.293852 | -0.1707858 | 0.19549776 | -0.8735945 | 0.38233911 | 0.78960558 |
| BOK | 95.9968059 | 0.21904066 | 0.34562911 | 0.63374484 | 0.52624736 | 0.86136324 |
| THAP4 | 541.74802 | -0.1519889 | 0.20042119 | -0.7583472 | 0.44824315 | 0.82623672 |
| ATG4B | 940.489204 | -0.4802537 | 0.20527928 | -2.3395139 | 0.01930885 | 0.28045661 |
| DTYMK | 85.8332277 | 0.59016549 | 0.28635704 | 2.0609428 | 0.0393085 | 0.37220498 |
| ING5 | 480.118914 | -0.2033537 | 0.23246063 | -0.8747877 | 0.38168941 | 0.78919376 |
| D2HGDH | 592.031399 | 0.05161688 | 0.35059952 | 0.1472246 | 0.88295474 | 0.97664137 |
| CHL1 | 1505.47555 | 0.84347108 | 0.43866434 | 1.92281661 | 0.05450309 | 0.41614524 |
| LINC01266 | 1502.99165 | 0.05619605 | 0.26603861 | 0.2112327 | 0.8327057 | 0.96441995 |
| CNTN6 | 106.563361 | -0.0008371 | 0.40298231 | -0.0020774 | 0.9983425 | 0.99972915 |
| CNTN4 | 2013.52582 | -1.139917 | 0.6174626 | -1.8461312 | 0.06487315 | 0.44041719 |
| TRNT1 | 603.49844 | -0.3924192 | 0.19635057 | -1.9985641 | 0.04565554 | 0.38904776 |
| CRBN | 647.643181 | 0.07510818 | 0.18990344 | 0.3955072 | 0.69246861 | 0.92224682 |
| AC024060.1 | 68.6227281 | 0.26742842 | 0.30438827 | 0.87857664 | 0.37963086 | 0.78762967 |
| AC034195.1 | 1069.09432 | 0.58800982 | 0.40914324 | 1.43717348 | 0.15066871 | 0.59747312 |
| SUMF1 | 4434.82446 | -0.3695969 | 0.34006502 | -1.0868419 | 0.2771067 | 0.72309412 |
| LRRN1 | 104.583855 | 0.42329427 | 0.37322521 | 1.13415242 | 0.25673062 | 0.70735159 |
| SETMAR | 176.087828 | 0.40825804 | 0.29495217 | 1.38414996 | 0.16631254 | 0.61901833 |
| ITPR1 | 4777.55664 | 0.69931457 | 0.23002962 | 3.04010665 | 0.00236494 | 0.09297185 |
| EGOT | 194.99765 | 1.02405387 | 0.63704031 | 1.60751816 | 0.10794075 | 0.53063658 |
| AC018816.1 | 230.142419 | 0.09720989 | 0.22957194 | 0.42343975 | 0.67197445 | 0.91574062 |
| BHLHE40-AS | 126.589413 | -0.876488 | 0.48752846 | -1.7978192 | 0.07220567 | 0.4679905 |
| BHLHE40 | 1530.55252 | 0.02217476 | 0.68011542 | 0.03260441 | 0.97399006 | 0.99346986 |
| ARL8B | 1168.07586 | -0.2778462 | 0.19459736 | -1.4278006 | 0.15334926 | 0.60285224 |
| AC026202.2 | 287.266777 | -0.0077458 | 0.41707152 | -0.018572 | 0.98518255 | 0.99572827 |
| EDEM1 | 274.548002 | 0.23446545 | 0.28143038 | 0.83312061 | 0.40477671 | 0.80437725 |
| AC087857.1 | 109.825898 | -0.5717823 | 0.85856876 | -0.6659714 | 0.50542935 | 0.85595485 |
| AC026167.1 | 168.44185 | -0.841953 | 0.84037763 | -1.0018746 | 0.31640415 | 0.74818634 |
| AC069277.1 | 279.634703 | 4.30181614 | 0.97655261 | 4.40510436 | 1.06E-05 | 0.00465542 |
| GRM7 | 216.009943 | 2.37695862 | 0.78277512 | 3.03657918 | 0.00239279 | 0.09378753 |
| LMCD1-AS1 | 1114.82175 | -0.2566179 | 0.31899384 | -0.8044603 | 0.4211312 | 0.81266383 |
| LMCD1 | 110.394492 | -0.9313569 | 0.79140887 | -1.176834 | 0.23926178 | 0.69029619 |
| RAD18 | 251.599599 | -0.5855378 | 0.30422044 | -1.9247156 | 0.05426495 | 0.41533078 |
| SRGAP3 | 1846.31703 | -0.7993352 | 0.4501267 | -1.7758005 | 0.0757658 | 0.47385913 |
| THUMPD3-A: | 1087.24096 | -0.19456 | 0.22798296 | -0.8533969 | 0.39343922 | 0.79843808 |
| THUMPD3 | 250.836004 | 0.0336346 | 0.25836543 | 0.13018227 | 0.89642222 | 0.97809691 |
| SETD5 | 3039.23771 | -0.036855 | 0.1227137 | -0.300333 | 0.76392314 | 0.94256396 |
| LHFPL4 | 46.7683147 | 0.69995127 | 0.8702327 | 0.80432655 | 0.42120841 | 0.81266383 |
| MTMR14 | 420.389758 | -0.1316089 | 0.22522221 | -0.5843512 | 0.55898405 | 0.87295536 |
| BRPF1 | 61.5960754 | -0.3222345 | 0.34231346 | -0.9413433 | 0.34652896 | 0.76864066 |
| OGG1 | 222.894712 | -0.206313 | 0.24007398 | -0.8593726 | 0.39013497 | 0.79701353 |
| CAMK1 | 116.32076 | 0.58361903 | 0.32638307 | 1.78814126 | 0.07375322 | 0.47028843 |
| TADA3 | 233.350911 | 0.06650853 | 0.19672752 | 0.33807437 | 0.73530714 | 0.93552616 |

| | | | | | | |
|---|---|---|---|---|---|---|
| TTLL3 | 84.9729226 | -0.308346 | 0.32876976 | -0.9378783 | 0.34830694 | 0.76974844 |
| RPUSD3 | 127.766492 | -0.1142528 | 0.30607041 | -0.3732892 | 0.70893321 | 0.92577218 |
| JAGN1 | 100.040337 | 0.56997525 | 0.46612056 | 1.2228065 | 0.22140279 | 0.67442133 |
| IL17RE | 164.660496 | 0.42832251 | 0.49280053 | 0.86915999 | 0.38475963 | 0.79237448 |
| IL17RC | 146.115072 | 0.50271512 | 0.30208446 | 1.66415416 | 0.09608163 | 0.5117083 |
| CRELD1 | 260.154178 | 0.63546291 | 0.23464386 | 2.70820171 | 0.00676489 | 0.16923754 |
| EMC3 | 487.86416 | 0.12059911 | 0.18671469 | 0.64590046 | 0.51834382 | 0.85916831 |
| FANCD2 | 75.2501999 | -0.5108798 | 0.36747327 | -1.3902504 | 0.16445287 | 0.61625736 |
| BRK1 | 305.076072 | 0.23630697 | 0.25742641 | 0.91795931 | 0.35864017 | 0.77558579 |
| VHL | 308.413469 | 0.14151252 | 0.24949777 | 0.56718954 | 0.57058541 | 0.87831986 |
| IRAK2 | 241.663338 | 0.55029349 | 0.55661881 | 0.98863617 | 3.23E-01 | 7.53E-01 |
| GHRL | 45.4248998 | -0.278652 | 0.41253186 | -0.6754679 | 0.49937854 | 0.85212391 |
| SEC13 | 407.642999 | 0.0479107 | 0.25738256 | 0.18614589 | 0.85233036 | 0.97076831 |
| HRH1 | 229.578237 | 1.42182134 | 0.73067789 | 1.94589348 | 5.17E-02 | 4.08E-01 |
| ATG7 | 2689.29042 | -0.3209258 | 0.26641622 | -1.204603 | 0.22835659 | 0.68012676 |
| VGLL4 | 2384.43209 | 0.08257962 | 0.18878948 | 0.43741641 | 0.66180939 | 0.91110616 |
| TAMM41 | 707.116835 | -0.3450138 | 0.33955009 | -1.016091 | 0.30958607 | 0.74450725 |
| PPARG | 307.90322 | -0.5382318 | 0.50490713 | -1.0660015 | 0.28642295 | 0.72934245 |
| TSEN2 | 679.18596 | -0.4288163 | 0.31535532 | -1.3597876 | 0.17389714 | 0.6286116 |
| MKRN2OS | 116.290261 | -0.2032416 | 0.45222758 | -0.4494233 | 0.65312631 | 0.908787 |
| MKRN2 | 188.861992 | -0.1915146 | 0.2096189 | -0.9136321 | 0.36091019 | 0.77730375 |
| RAF1 | 1484.23826 | -0.2430376 | 0.13046337 | -1.8628799 | 0.06247916 | 0.43419765 |
| CAND2 | 54.7070695 | 0.96266358 | 0.4146403 | 2.3216836 | 0.02024998 | 0.28534319 |
| RPL32 | 1066.19788 | 0.37587192 | 0.44971771 | 0.83579522 | 0.40327011 | 0.80324472 |
| IQSEC1 | 282.657867 | 0.43647346 | 0.2609532 | 1.67261203 | 0.0944037 | 0.51017802 |
| NUP210 | 517.467026 | 0.07589381 | 0.23842002 | 0.31831976 | 0.7502424 | 0.93775604 |
| HDAC11 | 181.085804 | 0.14170141 | 0.34278625 | 0.41338125 | 0.67932731 | 0.9182972 |
| CHCHD4 | 46.1410444 | 0.47709523 | 0.38537082 | 1.23801596 | 0.21571014 | 0.66885336 |
| TMEM43 | 128.730259 | -0.3829942 | 0.33995947 | -1.1265877 | 0.25991683 | 0.70934741 |
| XPC | 544.509025 | 0.268789 | 0.1523444 | 1.76435108 | 0.07767289 | 0.47898281 |
| LSM3 | 222.729881 | 0.46352203 | 0.31516373 | 1.47073405 | 0.14136305 | 0.5872148 |
| SLC6A6 | 320.383351 | 1.20613624 | 0.73581757 | 1.63917835 | 1.01E-01 | 0.51819907 |
| CCDC174 | 364.898083 | -0.0899238 | 0.20070652 | -0.448036 | 0.65412719 | 0.90900153 |
| C3orf20 | 66.7670914 | -3.1762673 | 0.61442878 | -5.1694638 | 2.35E-07 | 0.00051684 |
| FGD5-AS1 | 505.821279 | 0.18864047 | 0.31966976 | 0.59011045 | 0.5551166 | 0.87184917 |
| NR2C2 | 1100.76047 | -0.2326815 | 0.24120986 | -0.9646435 | 0.33472338 | 0.76242642 |
| MRPS25 | 843.819052 | -0.1638105 | 0.22208238 | -0.7376114 | 0.46075063 | 0.83439198 |
| RBSN | 343.16306 | -0.139519 | 0.21352577 | -0.6534059 | 0.51349463 | 0.85762435 |
| CAPN7 | 1378.40458 | -0.1988795 | 0.18897759 | -1.0523973 | 0.29261729 | 0.73454614 |
| SH3BP5-AS1 | 93.7563685 | -0.2887537 | 0.3005832 | -0.9606448 | 0.33673079 | 0.76409325 |
| SH3BP5 | 91.2110502 | 0.16672463 | 0.43888141 | 0.37988539 | 0.7040305 | 0.92514314 |
| METTL6 | 349.281655 | -0.1850631 | 0.28321306 | -0.6534414 | 0.51347174 | 0.85762435 |
| EAF1 | 132.467326 | -0.0539517 | 0.23180476 | -0.2327465 | 0.81595827 | 0.95879418 |
| COLQ | 219.529337 | -0.7623693 | 0.41355393 | -1.843458 | 0.06526217 | 0.44218726 |
| HACL1 | 511.099883 | -0.1813339 | 0.16223633 | -1.1177143 | 0.26368903 | 0.71432905 |
| BTD | 151.172738 | 0.28997806 | 0.25318624 | 1.14531523 | 0.2520786 | 0.70276117 |
| ANKRD28 | 3553.76879 | -0.2643258 | 0.20704804 | -1.2766397 | 0.20172946 | 0.65534788 |
| GALNT15 | 47.1002864 | -0.2837013 | 0.57256866 | -0.4954888 | 0.62025514 | 0.8972371 |
| DPH3 | 80.8624994 | 0.26012967 | 0.5231368 | 0.49724981 | 0.6190129 | 0.89615755 |
| OXNAD1 | 1172.28835 | -0.1657506 | 0.23655499 | -0.7006852 | 0.48349949 | 0.84454356 |
| RFTN1 | 111.010641 | 0.14083146 | 0.51786438 | 0.27194661 | 0.78566307 | 0.94978829 |
| PLCL2 | 704.061267 | 0.32679998 | 0.32225136 | 1.01411514 | 0.31052781 | 0.7446917 |
| TBC1D5 | 7098.9762 | -0.3315686 | 0.16257555 | -2.0394737 | 0.04140278 | 0.37577267 |
| KCNH8 | 105.510087 | -0.2724685 | 0.60731482 | -0.4486446 | 6.54E-01 | 9.09E-01 |

| | | | | | | |
|---|---|---|---|---|---|---|
| EFHB | 42.9782734 | -0.2103112 | 0.75989315 | -0.2767642 | 0.78196116 | 0.94882207 |
| RAB5A | 1109.9975 | -0.0540267 | 0.14483326 | -0.3730272 | 0.70912821 | 0.92577218 |
| KAT2B | 1017.1274 | 0.08275625 | 0.27894 | 0.29668119 | 7.67E-01 | 0.94312902 |
| ZNF385D | 2049.72553 | -1.6602047 | 0.60617112 | -2.7388384 | 0.00616567 | 0.16191312 |
| UBE2E2 | 3258.33315 | 0.90492642 | 0.2589578 | 3.49449375 | 0.00047496 | 0.03734387 |
| UBE2E1 | 2391.1681 | 0.00022782 | 0.19019989 | 0.00119777 | 0.99904432 | 0.99985365 |
| NKIRAS1 | 185.146426 | -0.0144776 | 0.26975285 | -0.05367 | 0.95719805 | 0.99065996 |
| RPL15 | 1708.79708 | 0.92984872 | 0.47671398 | 1.95053797 | 0.05111203 | 0.40556617 |
| NR1D2 | 481.299774 | -0.125645 | 0.47611711 | -0.2638952 | 0.79186064 | 0.95183249 |
| THRB | 5496.76239 | -0.0264228 | 0.25722547 | -0.1027223 | 9.18E-01 | 9.82E-01 |
| AC092422.1 | 69.2928822 | 0.79577653 | 0.69254612 | 1.14905926 | 0.25053155 | 0.70126536 |
| RARB | 778.47997 | -0.4711492 | 0.28016192 | -1.681703 | 0.09262644 | 0.50683623 |
| TOP2B | 1640.30707 | -0.2576414 | 0.20510611 | -1.2561372 | 0.20906624 | 0.66160901 |
| NGLY1 | 1171.52654 | -0.4712827 | 0.25054875 | -1.8810018 | 0.05997167 | 0.42727389 |
| OXSM | 135.104741 | 0.04492224 | 0.36572906 | 0.12282929 | 0.90224228 | 0.97941893 |
| NEK10 | 2862.13875 | -1.7787955 | 1.43260353 | -1.2416523 | 0.21436487 | 0.66827012 |
| SLC4A7 | 7179.52871 | -1.3467663 | 0.8085864 | -1.6655812 | 0.09579686 | 0.5117083 |
| CMC1 | 1757.05001 | -0.0655734 | 0.21439868 | -0.3058478 | 0.75972053 | 0.94133292 |
| AZI2 | 610.935704 | 0.15665828 | 0.27904034 | 0.56141806 | 0.57451258 | 0.88026176 |
| AC098650.1 | 63.2053576 | 0.73451217 | 0.32207424 | 2.2805679 | 0.02257403 | 0.29818035 |
| RBMS3 | 71.3414783 | -0.607474 | 0.59523604 | -1.0205599 | 0.30746299 | 0.74341546 |
| TGFBR2 | 1728.7962 | -0.4565082 | 0.29985795 | -1.522415 | 0.12790513 | 0.56762585 |
| GADL1 | 127.966945 | -0.5875772 | 0.98125057 | -0.5988044 | 5.49E-01 | 8.69E-01 |
| STT3B | 1363.29734 | -0.0128741 | 0.25966258 | -0.0495803 | 0.96045685 | 0.99153894 |
| OSBPL10 | 3172.15694 | -0.4958863 | 0.34403889 | -1.4413669 | 0.14948107 | 0.59625836 |
| OSBPL10-AS | 66.3445226 | -0.1009244 | 0.36338358 | -0.2777352 | 0.78121564 | 0.9482703 |
| ZNF860 | 54.3146341 | -0.3006982 | 0.534823 | -0.5622387 | 0.57395337 | 0.88002205 |
| GPD1L | 307.141049 | -0.0894066 | 0.33892149 | -0.2637974 | 0.79193603 | 0.95183649 |
| CMTM8 | 361.631905 | 0.30342305 | 0.43636493 | 0.69534243 | 0.48684073 | 0.84474442 |
| CMTM7 | 341.930332 | -0.0596076 | 0.52313509 | -0.113943 | 9.09E-01 | 9.81E-01 |
| CMTM6 | 439.307655 | 0.31947097 | 0.30050427 | 1.06311624 | 2.88E-01 | 7.31E-01 |
| DYNC1LI1 | 461.863239 | 0.02216397 | 0.19673833 | 0.11265711 | 0.91030241 | 0.98069436 |
| CNOT10 | 866.92194 | -0.2348247 | 0.14706493 | -1.5967414 | 0.11032335 | 0.53372881 |
| GLB1 | 771.485191 | 0.00858463 | 0.14333681 | 0.05989134 | 0.95224218 | 0.98938795 |
| CRTAP | 273.631964 | 0.25979552 | 0.28512875 | 0.91115163 | 0.36221549 | 0.77809471 |
| FBXL2 | 1271.22834 | 0.21007789 | 0.25579393 | 0.82127787 | 0.41148801 | 0.80675684 |
| UBP1 | 615.706071 | -0.3297895 | 0.22536399 | -1.4633638 | 0.14336787 | 0.58849865 |
| CLASP2 | 2895.99225 | 0.02394399 | 0.25547258 | 0.09372429 | 0.92532818 | 0.9822216 |
| PDCD6IP | 1634.15325 | -0.038984 | 0.13235335 | -0.2945449 | 0.76834159 | 0.9434809 |
| LINC01811 | 1031.6298 | 4.27002773 | 0.92834751 | 4.59960054 | 4.23E-06 | 0.00294284 |
| STAC | 314.882496 | 1.69557498 | 0.48504085 | 3.49573642 | 0.00047276 | 0.03734387 |
| TRANK1 | 193.006867 | 1.59727028 | 0.81868135 | 1.95102805 | 0.05105371 | 0.40556617 |
| EPM2AIP1 | 485.828377 | 0.63784198 | 0.2115 | 3.01580135 | 0.00256301 | 0.09563507 |
| MLH1 | 328.165049 | -0.4719643 | 0.17264183 | -2.7337772 | 0.00626124 | 0.16248472 |
| LRRFIP2 | 2524.73524 | 0.23674745 | 0.2734688 | 0.86572017 | 0.38664364 | 0.79523542 |
| GOLGA4 | 4444.56376 | -0.1972645 | 0.18573926 | -1.0620504 | 0.28821279 | 0.73099132 |
| ITGA9 | 179.862873 | -0.5310879 | 0.56295438 | -0.9433942 | 0.34547929 | 0.7678828 |
| CTDSPL | 4533.05479 | -0.2738037 | 0.21785676 | -1.2568062 | 0.20882383 | 0.6614379 |
| PLCD1 | 165.974866 | 0.58558836 | 0.4449963 | 1.31593983 | 0.18819424 | 0.64267818 |
| ACAA1 | 393.580145 | 0.98824606 | 0.27268218 | 3.62416816 | 0.00028989 | 0.02900906 |
| MYD88 | 50.6347442 | 0.3399443 | 0.40682326 | 0.83560683 | 0.40337612 | 0.80324472 |
| OXSR1 | 1367.73238 | 0.01298856 | 0.2062921 | 0.062962 | 0.94979677 | 0.98919519 |
| XYLB | 148.198546 | 0.05086325 | 0.52864389 | 0.09621458 | 0.92335015 | 0.98215577 |
| ACVR2B | 319.784505 | -0.1039606 | 0.27885283 | -0.3728153 | 0.70928591 | 0.92578632 |

| | | | | | | |
|---|---|---|---|---|---|---|
| EXOG | 342.078593 | -0.3013703 | 0.2361493 | -1.2761854 | 0.20188998 | 0.65552334 |
| SCN11A | 91.162745 | -0.3806485 | 0.35648412 | -1.0677853 | 0.28561735 | 0.72846488 |
| WDR48 | 718.524477 | 0.01304007 | 0.19258859 | 0.06770947 | 0.94601691 | 0.98877579 |
| GORASP1 | 204.175516 | 0.02386996 | 0.20580147 | 0.11598538 | 0.90766413 | 0.98047754 |
| TTC21A | 189.201776 | 0.30524625 | 0.35780897 | 0.85309837 | 0.39360476 | 0.79843808 |
| CSRNP1 | 114.9114 | 0.07153554 | 0.54681041 | 0.1308233 | 0.89591509 | 0.97788747 |
| SLC25A38 | 144.026508 | -0.1953205 | 0.30068827 | -0.6495781 | 0.51596477 | 0.85883493 |
| RPSA | 836.109144 | 0.61723861 | 0.36025247 | 1.71335012 | 0.08664815 | 0.49478465 |
| MYRIP | 324.625608 | 0.59964379 | 0.45860942 | 1.3075261 | 0.1910341 | 0.64567512 |
| EIF1B | 288.196585 | 0.21297197 | 0.48692515 | 0.43738133 | 0.66183482 | 0.91110616 |
| ENTPD3-AS1 | 411.019334 | -0.5515747 | 0.45196702 | -1.220387 | 0.22231822 | 0.67568617 |
| RPL14 | 1368.26679 | 0.53333622 | 0.44185603 | 1.20703617 | 0.22741821 | 0.67921637 |
| ZNF619 | 42.4119288 | -0.215285 | 0.40617814 | -0.530026 | 0.59609391 | 0.88859628 |
| ZNF620 | 81.937948 | 0.04321509 | 0.33039262 | 0.13079921 | 0.89593415 | 0.97788747 |
| ZNF621 | 531.232772 | -0.3846874 | 0.25220209 | -1.525314 | 0.12718081 | 0.56740575 |
| CTNNB1 | 1397.23693 | -0.1737172 | 0.13818909 | -1.2570975 | 0.20871833 | 0.6614379 |
| ULK4 | 1621.92943 | -0.5508521 | 0.2675184 | -2.0591187 | 0.03948287 | 0.37234059 |
| TRAK1 | 1360.28527 | -0.1145789 | 0.22286657 | -0.5141144 | 0.60717201 | 0.89174148 |
| SEC22C | 252.584004 | -0.3559048 | 0.23846251 | -1.4924977 | 0.13556871 | 0.57709645 |
| SS18L2 | 137.953341 | 0.1304395 | 0.3639127 | 0.35843624 | 0.72001687 | 0.93001079 |
| NKTR | 2901.90552 | -0.1499705 | 0.12963285 | -1.1568864 | 0.24731879 | 0.69761067 |
| AC006059.1 | 409.589983 | 0.00990021 | 0.50427054 | 0.01963273 | 0.98433635 | 0.99518552 |
| CCDC13 | 97.9916945 | -0.2305279 | 0.52153587 | -0.4420173 | 0.65847666 | 0.91037018 |
| AC099329.3 | 131.536791 | 0.13031658 | 0.43695434 | 0.29823844 | 0.76552118 | 0.94270299 |
| AC092042.3 | 72.9901724 | -0.4391089 | 0.49906552 | -0.8798622 | 0.37893399 | 0.78749827 |
| POMGNT2 | 79.2803815 | 0.20619407 | 0.25772577 | 0.8000522 | 0.42368055 | 0.81402399 |
| SNRK | 1525.57703 | -0.751197 | 0.33146029 | -2.2663257 | 0.02343145 | 0.30107587 |
| AC104184.1 | 35.1684728 | 1.14638458 | 0.52907784 | 2.16675976 | 0.03025318 | 0.3341256 |
| ANO10 | 1963.90399 | 0.03287111 | 0.2807948 | 0.11706452 | 0.90680893 | 0.9803357 |
| ABHD5 | 362.80456 | -0.261859 | 0.40259246 | -0.650432 | 0.5154132 | 0.85869987 |
| TCAIM | 2910.52816 | 0.15746001 | 0.2694356 | 0.58440683 | 0.55894663 | 0.87295536 |
| ZNF445 | 422.953137 | -0.3073239 | 0.18874935 | -1.6282117 | 0.10348 | 0.52383403 |
| ZNF852 | 113.686014 | -0.251305 | 0.25425359 | -0.988403 | 0.32295533 | 0.7532337 |
| ZKSCAN7 | 114.857615 | 0.00287084 | 0.25417521 | 0.01129473 | 0.9909883 | 0.99749592 |
| AC099669.1 | 769.088941 | -0.0339186 | 0.29177359 | -0.1162497 | 0.90745463 | 0.98047754 |
| ZNF197 | 409.880274 | -0.0757335 | 0.27164253 | -0.2787985 | 0.78039948 | 0.9480401 |
| ZNF35 | 91.7757603 | -0.3115966 | 0.28695005 | -1.0858914 | 0.27752703 | 0.72344438 |
| ZNF502 | 70.0237116 | 0.08513059 | 0.42267128 | 0.20141086 | 0.84037733 | 0.96617873 |
| KIAA1143 | 149.722501 | -0.0456934 | 0.30396778 | -0.150323 | 0.88050978 | 0.97664137 |
| ZDHHC3 | 483.045575 | -0.2336521 | 0.15304349 | -1.5267038 | 0.12683469 | 0.56735411 |
| EXOSC7 | 296.02437 | -0.3871818 | 0.24699339 | -1.5675795 | 0.1169793 | 0.54756317 |
| CDCP1 | 429.511771 | 0.92127209 | 0.5075511 | 1.81513169 | 6.95E-02 | 4.58E-01 |
| LARS2 | 753.29581 | -0.2854179 | 0.20106397 | -1.4195377 | 0.1557423 | 0.60505885 |
| LIMD1 | 549.00398 | -0.3774899 | 0.21956547 | -1.7192592 | 0.08556719 | 0.49356204 |
| SACM1L | 947.055111 | -0.1676917 | 0.18218638 | -0.9204405 | 0.35734259 | 0.77455501 |
| SLC6A20 | 224.957137 | 0.94632565 | 0.52771803 | 1.79324107 | 7.29E-02 | 4.70E-01 |
| LZTFL1 | 271.343363 | -0.196425 | 0.19848471 | -0.9896228 | 0.32235853 | 0.75310114 |
| FYCO1 | 370.651104 | -0.1299608 | 0.17662528 | -0.7357994 | 0.46185277 | 0.83531269 |
| LTF | 1199.36616 | 1.19500549 | 0.84250123 | 1.41840207 | 1.56E-01 | 6.05E-01 |
| LRRC2 | 689.689149 | 0.56917472 | 0.47240139 | 1.20485402 | 0.22825966 | 0.67999141 |
| TDGF1 | 80.5088823 | -0.296391 | 0.81823999 | -0.3622299 | 0.71718021 | 0.9290703 |
| AC104304.1 | 95.4937774 | 0.473391 | 0.5752006 | 0.82300158 | 0.4105071 | 0.80655687 |
| ALS2CL | 398.355755 | -0.5467181 | 0.20545132 | -2.6610591 | 7.79E-03 | 1.77E-01 |
| MYL3 | 86.4547131 | 2.55093629 | 0.70880007 | 3.5989504 | 0.0003195 | 0.03094754 |

| | | | | | | |
|---|---|---|---|---|---|---|
| PTH1R | 2119.66941 | 2.50526718 | 0.77649524 | 3.22637802 | 0.00125368 | 0.0645665 |
| CCDC12 | 791.509075 | -0.1042622 | 0.27410278 | -0.3803764 | 0.70366604 | 0.92513032 |
| NBEAL2 | 274.338103 | -0.2425064 | 0.38914582 | -0.6231761 | 0.53316879 | 0.86397197 |
| SETD2 | 3389.04519 | -0.0894842 | 0.13697428 | -0.6532918 | 0.51356818 | 0.85762435 |
| KIF9-AS1 | 1190.87388 | 0.25773348 | 0.28607043 | 0.90094413 | 0.36761802 | 0.78095617 |
| KIF9 | 163.36553 | -0.0495282 | 0.26191355 | -0.1891012 | 0.85001353 | 0.97041641 |
| KLHL18 | 413.946021 | -0.1336645 | 0.19218409 | -0.6955025 | 0.48674045 | 0.84474442 |
| AC099778.1 | 38.057616 | 0.4721001 | 0.47028947 | 1.00385004 | 0.3154509 | 0.74785672 |
| PTPN23 | 223.062416 | -0.2365015 | 0.24549138 | -0.96338 | 0.33535687 | 0.76322001 |
| SCAP | 646.095684 | -0.0866446 | 0.19378662 | -0.4471136 | 0.65479307 | 0.90902447 |
| ELP6 | 270.922122 | -0.027479 | 0.34159606 | -0.0804428 | 0.93588505 | 0.98613991 |
| SMARCC1 | 2291.03778 | -0.3380955 | 0.21595675 | -1.5655703 | 0.11744926 | 0.54780623 |
| DHX30 | 513.177822 | -0.0420801 | 0.16595362 | -0.2535652 | 0.7998315 | 0.95370871 |
| MAP4 | 3099.81676 | 0.06009399 | 0.23427118 | 0.25651466 | 0.79755346 | 0.95281098 |
| ZNF589 | 184.467856 | -0.3960775 | 0.24776604 | -1.5985949 | 0.10991064 | 0.53372881 |
| NME6 | 144.776569 | 0.10413319 | 0.27225927 | 0.38247803 | 0.70210682 | 0.9248705 |
| PLXNB1 | 1085.05553 | -0.3805193 | 0.22535826 | -1.6885084 | 0.09131367 | 0.50411197 |
| TMA7 | 433.014015 | 0.15127121 | 0.42718236 | 0.3541139 | 0.72325353 | 0.93177886 |
| ATRIP | 52.4467692 | 0.1535986 | 0.37461746 | 0.41001453 | 0.68179529 | 0.91961952 |
| SHISA5 | 508.330888 | 0.04862056 | 0.49835706 | 0.0975617 | 9.22E-01 | 0.98215577 |
| PFKFB4 | 349.04908 | -0.1231991 | 0.84158783 | -0.1463889 | 0.88361437 | 0.97664137 |
| COL7A1 | 393.202615 | -2.7647632 | 0.84557805 | -3.2696724 | 0.00107672 | 0.05901413 |
| UQCRC1 | 426.909073 | 0.36807329 | 0.26161375 | 1.40693404 | 0.15944695 | 0.60923771 |
| SLC26A6 | 56.6420225 | -0.2208597 | 0.36955595 | -0.5976354 | 0.55008323 | 0.86956073 |
| NCKIPSD | 113.155884 | 0.35193672 | 0.36348302 | 0.96823429 | 0.33292737 | 0.76111014 |
| IP6K2 | 809.600403 | -0.2946786 | 0.11387859 | -2.5876554 | 0.00966316 | 0.19975064 |
| PRKAR2A | 980.317068 | 0.15391774 | 0.20311613 | 0.75778196 | 0.44858152 | 0.82630667 |
| SLC25A20 | 181.813471 | 0.4811873 | 0.63858716 | 0.7535186 | 0.45113834 | 0.82814845 |
| ARIH2 | 1194.58634 | -0.2515875 | 0.15621475 | -1.6105232 | 0.10728369 | 0.52901576 |
| P4HTM | 309.030825 | 0.52639697 | 0.27218461 | 1.93397035 | 0.05311677 | 0.41099428 |
| WDR6 | 566.717396 | -0.4499686 | 0.33007996 | -1.3632108 | 0.17281609 | 0.6286116 |
| DALRD3 | 155.655154 | -0.2793973 | 0.298886 | -0.9347956 | 0.34989364 | 0.77041926 |
| NDUFAF3 | 138.705218 | 0.74212301 | 0.50451374 | 1.47096689 | 0.14130007 | 0.5872148 |
| IMPDH2 | 94.0689173 | 0.42489256 | 0.40718176 | 1.04349605 | 0.29671861 | 0.73672107 |
| QRICH1 | 795.03484 | -0.1787593 | 0.18075562 | -0.9889558 | 0.32268479 | 0.7532337 |
| QARS | 213.395635 | -0.0190671 | 0.29729455 | -0.0641353 | 0.94886252 | 0.98919519 |
| USP19 | 131.603709 | -0.0624159 | 0.218993 | -0.285013 | 0.77563417 | 0.94680268 |
| LAMB2 | 323.621352 | 0.39758969 | 0.32382134 | 1.2278057 | 0.21951991 | 0.67278782 |
| KLHDC8B | 59.9747357 | 0.72414279 | 0.39779529 | 1.82039057 | 0.06869955 | 0.45448131 |
| C3orf62 | 95.8584593 | -0.0566168 | 0.28164511 | -0.2010219 | 0.84068144 | 0.96617873 |
| USP4 | 744.230469 | -0.2444491 | 0.17481073 | -1.3983645 | 0.16200364 | 0.6135052 |
| RHOA | 1171.78441 | 0.33431987 | 0.19072923 | 1.75285068 | 7.96E-02 | 0.48176967 |
| TCTA | 160.74896 | 0.61477316 | 0.33655716 | 1.82665304 | 0.06775195 | 0.45233147 |
| DAG1 | 610.639618 | 0.45833222 | 0.23599817 | 1.94210071 | 0.05212491 | 0.40866415 |
| BSN | 40.7949794 | -0.310345 | 0.53074356 | -0.5847362 | 0.55872508 | 0.87295536 |
| APEH | 171.182568 | 0.82938363 | 0.42531822 | 1.95003083 | 0.05117244 | 0.40556617 |
| MST1 | 249.052582 | -0.8094083 | 0.35682242 | -2.2683786 | 0.02330614 | 0.30034225 |
| RNF123 | 181.581761 | -0.2273531 | 0.19799259 | -1.148291 | 0.25084846 | 0.70200367 |
| GMPPB | 50.6268659 | 0.11874276 | 0.42012762 | 0.28263498 | 0.77745667 | 0.94694572 |
| IP6K1 | 1323.76812 | -0.0480235 | 0.25231643 | -0.1903303 | 0.84905029 | 0.97017477 |
| MON1A | 68.1256057 | -0.1147208 | 0.34961231 | -0.3281372 | 0.74280795 | 0.93598004 |
| RBM6 | 4210.07524 | -0.1526903 | 0.18085338 | -0.8442766 | 0.39851486 | 0.80148305 |
| RBM5 | 1385.48209 | -0.0618667 | 0.13770061 | -0.4492841 | 0.65322676 | 0.908787 |
| SEMA3F | 87.5045522 | 0.65710931 | 0.59649184 | 1.1016233 | 0.27062547 | 0.71965808 |

| | | | | | | |
|---|---|---|---|---|---|---|
| GNAI2 | 334.497415 | 0.26583291 | 0.28206377 | 0.94245676 | 3.46E-01 | 7.68E-01 |
| SEMA3B | 510.556502 | 0.41390721 | 0.42631868 | 0.97088688 | 3.32E-01 | 0.76018142 |
| IFRD2 | 109.221437 | -0.1927512 | 0.31265447 | -0.6164992 | 0.53756511 | 0.86489441 |
| HYAL1 | 147.206547 | 0.63818528 | 0.53442454 | 1.19415415 | 0.23241767 | 0.68450837 |
| HYAL2 | 74.2312808 | 0.14134496 | 0.32057095 | 0.44091632 | 0.65927358 | 0.91099119 |
| TUSC2 | 111.888533 | 0.46138734 | 0.38790659 | 1.18942899 | 0.2342709 | 0.68546368 |
| RASSF1 | 112.545547 | -0.7579044 | 0.25268772 | -2.9993716 | 0.00270537 | 0.09823475 |
| NPRL2 | 59.7317876 | 0.47972752 | 0.34389011 | 1.3950024 | 0.16301512 | 0.61387934 |
| TMEM115 | 75.5671185 | 0.86236095 | 0.32738358 | 2.63409961 | 0.00843607 | 0.18510315 |
| C3orf18 | 45.2864203 | 0.9733868 | 0.45944063 | 2.11863459 | 0.03412136 | 0.35153234 |
| HEMK1 | 165.905152 | -0.2170957 | 0.28795634 | -0.7539187 | 0.45089802 | 0.82813014 |
| CISH | 216.108995 | 0.90509879 | 0.44169944 | 2.04912824 | 0.04044958 | 0.37344844 |
| MAPKAPK3 | 293.574899 | -0.2554741 | 0.36468578 | -0.7005322 | 0.48359501 | 0.84454356 |
| DOCK3 | 1030.37159 | 0.86016174 | 0.48485132 | 1.77407321 | 0.07605104 | 0.4745161 |
| MANF | 223.342701 | 1.30176459 | 0.34915805 | 3.72829607 | 0.00019278 | 0.02233698 |
| RBM15B | 75.2236207 | 0.11451719 | 0.29407493 | 0.38941499 | 0.69696918 | 0.92322348 |
| DCAF1 | 620.679608 | -0.4281335 | 0.22493667 | -1.9033511 | 0.05699475 | 0.41872623 |
| RAD54L2 | 713.348904 | -0.0684307 | 0.22583271 | -0.3030151 | 0.76187837 | 0.94210219 |
| TEX264 | 295.129496 | 0.25906111 | 0.31449514 | 0.82373645 | 0.41008933 | 0.80655687 |
| RRP9 | 39.1564446 | -0.9175228 | 0.51300174 | -1.7885374 | 0.07368934 | 0.47028843 |
| PARP3 | 58.8001493 | -0.0200308 | 0.46291215 | -0.0432713 | 0.96548524 | 0.99248831 |
| PCBP4 | 56.4920281 | 0.50049402 | 0.43824479 | 1.14204215 | 0.2534365 | 0.7041739 |
| ABHD14A | 42.4433359 | 0.26149295 | 0.45977396 | 0.56874242 | 0.56953096 | 0.87807251 |
| RPL29 | 1411.7906 | 0.77626112 | 0.47130712 | 1.64703881 | 0.09955007 | 0.51665771 |
| POC1A | 45.7254135 | -0.484599 | 0.52301775 | -0.9265441 | 0.35416327 | 0.77243419 |
| ALAS1 | 108.840292 | 0.24423321 | 0.27076036 | 0.90202721 | 0.36704242 | 0.78095617 |
| WDR82 | 471.010035 | 0.15766037 | 0.16961578 | 0.92951477 | 0.35262237 | 0.77210174 |
| GLYCTK | 47.1347532 | -0.3565391 | 0.3779377 | -0.9433807 | 0.34548622 | 0.7678828 |
| DNAH1 | 117.251553 | -0.7676801 | 0.28684348 | -2.6763031 | 0.00744393 | 0.17360516 |
| BAP1 | 150.78645 | 0.11348131 | 0.34162777 | 0.33217822 | 0.73975469 | 0.93552616 |
| PHF7 | 152.956721 | -0.3529485 | 0.41564992 | -0.8491485 | 0.39579864 | 0.79952657 |
| NISCH | 628.780895 | 0.27407125 | 0.21479836 | 1.27594665 | 0.20197438 | 0.65552334 |
| NT5DC2 | 61.7047093 | -0.0707832 | 0.55430656 | -0.1276969 | 0.89838888 | 0.97877011 |
| SMIM4 | 402.875097 | 0.06070555 | 0.31366181 | 0.19353821 | 0.84653748 | 0.96883382 |
| PBRM1 | 1711.99504 | -0.5051853 | 0.17085504 | -2.9568067 | 0.00310843 | 0.10720429 |
| GNL3 | 254.029486 | -0.200524 | 0.28848445 | -0.6950948 | 0.4869959 | 0.84474442 |
| GLT8D1 | 180.063332 | 0.15855343 | 0.2147175 | 0.73842807 | 0.46025436 | 0.83429248 |
| SPCS1 | 464.574888 | 0.86775519 | 0.39833768 | 2.17844115 | 0.02937321 | 0.33012415 |
| NEK4 | 445.790474 | -0.11014 | 0.1496978 | -0.735749 | 0.46188347 | 0.83531269 |
| SFMBT1 | 375.432253 | -0.0491041 | 0.17384576 | -0.2824579 | 0.77759243 | 0.94694572 |
| AC096887.1 | 76.2422774 | 0.05879117 | 0.51265584 | 0.11467961 | 0.90869907 | 0.98047754 |
| RFT1 | 112.624406 | -0.2364054 | 0.28880082 | -0.8185758 | 0.4130285 | 0.80777221 |
| PRKCD | 129.675669 | 0.38327175 | 0.2817292 | 1.36042605 | 0.17369514 | 0.6286116 |
| TKT | 629.192145 | 0.43447505 | 0.62377006 | 0.69653078 | 0.48609649 | 0.8447301 |
| DCP1A | 1065.16234 | -0.1071602 | 0.16557314 | -0.6472074 | 0.51749774 | 0.85893097 |
| CACNA1D | 2577.2151 | -0.0030162 | 0.4413968 | -0.0068334 | 0.9945478 | 0.998438 |
| CHDH | 269.503945 | -0.6715115 | 0.62356431 | -1.0768921 | 0.2815285 | 0.72604837 |
| IL17RB | 102.76796 | 0.07923082 | 0.71706563 | 0.11049312 | 0.9120183 | 0.98125148 |
| ACTR8 | 128.344241 | -0.290992 | 0.25646708 | -1.1346175 | 0.25653563 | 0.70735159 |
| SELENOK | 210.481939 | 0.83442842 | 0.41099569 | 2.03026077 | 0.04233004 | 0.37959096 |
| CACNA2D3 | 17909.8562 | -0.5905896 | 0.2850758 | -2.0716934 | 0.03829405 | 0.36662698 |
| CACNA2D3-A | 128.221047 | -0.7345046 | 0.46444706 | -1.5814603 | 0.11377284 | 0.54000194 |
| LRTM1 | 293.133274 | -0.7718366 | 0.46888502 | -1.6461106 | 0.099741 | 0.51665771 |
| ERC2 | 1308.68353 | -0.5329999 | 0.34962763 | -1.5244788 | 0.12738915 | 0.56740575 |

| | | | | | | |
|---|---|---|---|---|---|---|
| CCDC66 | 1417.85679 | -0.0166364 | 0.23139802 | -0.0718953 | 0.94268525 | 0.98772472 |
| FAM208A | 1452.40932 | -0.0685184 | 0.23281675 | -0.2943019 | 0.76852724 | 0.94362115 |
| ARHGEF3 | 5010.76077 | -0.1980043 | 0.366374 | -0.540443 | 0.58889153 | 0.88571413 |
| ARHGEF3-AS | 52.6634199 | -0.0132871 | 0.7971415 | -0.0166684 | 0.98670112 | 0.99643235 |
| SPATA12 | 132.327211 | -0.192752 | 0.60502605 | -0.3185846 | 0.75004156 | 0.93775604 |
| IL17RD | 253.810984 | 0.53586003 | 0.45856965 | 1.16854666 | 0.24258633 | 0.69357637 |
| HESX1 | 70.2356529 | -0.1913346 | 0.50154444 | -0.3814909 | 0.70283902 | 0.92501157 |
| APPL1 | 806.621492 | 0.20802816 | 0.11132415 | 1.86867059 | 0.06166866 | 0.43259761 |
| ASB14 | 134.013579 | 0.53624944 | 0.46443592 | 1.15462524 | 0.24824392 | 0.69838949 |
| DNAH12 | 79.1845606 | 0.10588615 | 0.43634504 | 0.24266611 | 0.80826407 | 0.95610623 |
| PDE12 | 135.987357 | 0.1810629 | 0.25609553 | 0.7070131 | 0.47955834 | 0.8430245 |
| ARF4 | 660.951304 | -0.1187267 | 0.31045347 | -0.3824298 | 0.70214258 | 0.9248705 |
| DENND6A | 955.352556 | -0.1238981 | 0.15679477 | -0.7901926 | 0.4294153 | 0.81613622 |
| DENND6A-AS | 217.186159 | 0.0090372 | 0.421982 | 0.02141609 | 0.98291374 | 0.99518552 |
| SLMAP | 3213.83866 | -0.2322718 | 0.1720168 | -1.3502855 | 0.17692442 | 0.63025747 |
| FLNB | 2938.43033 | 0.26678409 | 0.3639339 | 0.73305643 | 0.46352401 | 0.83531269 |
| ABHD6 | 196.259233 | -0.827231 | 0.44463786 | -1.8604602 | 0.06282045 | 0.43558808 |
| RPP14 | 107.358836 | 0.39232704 | 0.33453893 | 1.17273958 | 0.24090025 | 0.69178991 |
| PXK | 908.800548 | -0.3698004 | 0.29818131 | -1.2401864 | 0.21490647 | 0.66855854 |
| PDHB | 250.407664 | 0.39143165 | 0.29426377 | 1.33020675 | 0.18345016 | 0.63834582 |
| KCTD6 | 211.466021 | 0.52365543 | 0.24624192 | 2.12658931 | 0.03345422 | 0.34965509 |
| ACOX2 | 57.4314044 | 0.80598229 | 0.41203554 | 1.95609898 | 0.0504535 | 0.40335713 |
| FAM107A | 164.864523 | 2.15338818 | 0.84357784 | 2.55268462 | 0.01068962 | 0.21011792 |
| C3orf67 | 200.383076 | 0.67869478 | 0.49221252 | 1.37886532 | 0.16793628 | 0.62233912 |
| C3orf67-AS1 | 52.4018098 | 0.5151941 | 0.58382913 | 0.88243986 | 0.37753899 | 0.78633121 |
| FHIT | 17971.034 | 0.19605404 | 0.18116647 | 1.08217621 | 0.27917424 | 0.72482665 |
| PTPRG | 20125.5208 | 0.06907435 | 0.35569391 | 0.19419605 | 0.84602237 | 0.96883382 |
| C3orf14 | 43.2677855 | 0.04620323 | 0.44507202 | 0.10381067 | 0.91731959 | 0.98199511 |
| THOC7 | 339.388571 | 0.17306179 | 0.19630201 | 0.88160987 | 0.37798781 | 0.7865219 |
| ATXN7 | 288.61505 | 0.09826548 | 0.18849611 | 0.52131303 | 0.60214873 | 0.88968485 |
| PSMD6-AS2 | 134.709936 | 0.06693077 | 0.28496807 | 0.23487114 | 0.81430875 | 0.95832184 |
| PSMD6 | 197.627551 | 0.11778202 | 0.29749164 | 0.39591707 | 0.69216621 | 0.92224682 |
| PSMD6-AS1 | 49.1943944 | -0.0763673 | 0.43148247 | -0.1769883 | 0.85951758 | 0.97184883 |
| PRICKLE2 | 3223.32092 | 0.74931364 | 0.39178552 | 1.91256083 | 0.0558043 | 0.41843207 |
| ADAMTS9 | 3723.90908 | -0.9455883 | 0.52076855 | -1.8157554 | 0.06940789 | 0.45787876 |
| ADAMTS9-AS | 2222.14455 | 0.25886349 | 0.51602808 | 0.50164613 | 0.61591646 | 0.89540398 |
| ADAMTS9-AS | 13061.9409 | 1.06279241 | 0.45648911 | 2.32818784 | 0.01990213 | 0.28297874 |
| MAGI1 | 20414.0492 | -0.172951 | 0.28283314 | -0.6114947 | 0.54087213 | 0.86582694 |
| MAGI1-AS1 | 385.04892 | -0.0804908 | 0.7540669 | -0.1067422 | 0.91499351 | 0.9816913 |
| SLC25A26 | 1041.67046 | -0.3752829 | 0.2096614 | -1.7899475 | 0.07346235 | 0.4701377 |
| LRIG1 | 256.284031 | 0.07547277 | 0.34088017 | 0.22140556 | 0.82477666 | 0.96146415 |
| SUCLG2 | 2867.05222 | -0.1044568 | 0.19694635 | -0.5303818 | 0.59584725 | 0.88859628 |
| SUCLG2-AS1 | 1899.29113 | 0.44932549 | 0.28555352 | 1.57352462 | 0.11559739 | 0.54435863 |
| FAM19A1 | 754.84331 | 0.54631899 | 0.56561529 | 0.96588442 | 0.33410202 | 0.76220269 |
| EOGT | 135.720687 | -0.6395057 | 0.26761315 | -2.3896645 | 0.01686377 | 0.25961947 |
| AC109587.1 | 130.960696 | 0.27412468 | 0.75241268 | 0.36432757 | 0.71561342 | 0.92883553 |
| TMF1 | 973.059337 | -0.0291999 | 0.13494034 | -0.2163912 | 0.82868285 | 0.96288319 |
| UBA3 | 690.117772 | -0.2015816 | 0.14465177 | -1.3935649 | 0.16344902 | 0.61416689 |
| ARL6IP5 | 397.889035 | 0.29100024 | 0.24116246 | 1.20665646 | 2.28E-01 | 0.67921637 |
| FRMD4B | 1513.71724 | 0.07329341 | 0.26878045 | 0.27268878 | 0.78509246 | 0.94978829 |
| MITF | 4097.23022 | 0.22396446 | 0.19943936 | 1.1229702 | 2.61E-01 | 7.12E-01 |
| FOXP1 | 8997.18307 | 0.19666442 | 0.30704335 | 0.64051028 | 0.52184091 | 0.85969027 |
| AC097634.3 | 48.7435631 | 0.59984048 | 0.56925983 | 1.05372004 | 0.29201111 | 0.73384286 |
| EIF4E3 | 340.82623 | 0.67425667 | 0.33298358 | 2.02489467 | 0.04287818 | 0.38144955 |

| | | | | | | |
|---|---|---|---|---|---|---|
| RYBP | 1205.94785 | 0.3679581 | 0.26972147 | 1.36421509 | 0.17249988 | 0.6286116 |
| SHQ1 | 472.582841 | -0.2160259 | 0.29421962 | -0.7342335 | 0.46280643 | 0.83531269 |
| GXYLT2 | 153.788252 | 0.57731587 | 0.36561344 | 1.57903351 | 1.14E-01 | 0.54147129 |
| PPP4R2 | 1078.01442 | 0.09735465 | 0.20946642 | 0.46477449 | 0.64209296 | 0.90454089 |
| PDZRN3 | 1383.59338 | 1.65277121 | 0.57366392 | 2.88107926 | 0.00396316 | 0.12259807 |
| CNTN3 | 505.704276 | 0.36019253 | 0.3167531 | 1.13713972 | 0.25547988 | 0.70643369 |
| LINC02018 | 69.9704538 | -0.4666944 | 0.60839078 | -0.7670964 | 0.44302418 | 0.82310928 |
| ZNF717 | 669.465843 | -0.0102967 | 0.29093245 | -0.0353919 | 0.97176721 | 0.99327086 |
| ROBO2 | 492.077228 | 1.01554159 | 0.72046881 | 1.40955663 | 0.15867064 | 0.60804051 |
| ROBO1 | 185.005291 | 0.57932664 | 0.45099432 | 1.28455419 | 0.19894812 | 0.65113421 |
| LINC02027 | 128.115046 | 2.80633868 | 0.71602071 | 3.91935406 | 8.88E-05 | 0.01515778 |
| GBE1 | 4875.501 | -0.3884483 | 0.41811192 | -0.9290533 | 0.35286148 | 0.77216708 |
| CADM2 | 111.648809 | -0.6172644 | 0.69021364 | -0.8943092 | 0.37115649 | 0.78104287 |
| CHMP2B | 410.768897 | -0.0424649 | 0.1945263 | -0.2182991 | 0.82719611 | 0.96234221 |
| AC117513.1 | 72.5209484 | -4.3313955 | 1.09953647 | -3.9392923 | 8.17E-05 | 0.01478726 |
| CGGBP1 | 992.836697 | -0.0386157 | 0.29151262 | -0.1324666 | 0.89461526 | 0.97788747 |
| ZNF654 | 1320.84957 | 0.13771015 | 0.20449294 | 0.67342255 | 0.50067851 | 0.85323989 |
| C3orf38 | 241.352195 | 0.11128588 | 0.2118047 | 0.52541741 | 0.59929305 | 0.88883692 |
| PROS1 | 212.679813 | 0.30574944 | 0.34010412 | 0.89898777 | 3.69E-01 | 7.81E-01 |
| ARL13B | 573.796486 | -0.1554411 | 0.23457015 | -0.6626635 | 0.50754612 | 0.85632606 |
| STX19 | 47.4373782 | 0.5013821 | 0.63002825 | 0.79580891 | 0.42614311 | 0.81466629 |
| DHFR2 | 91.4749955 | 0.18059581 | 0.27441165 | 0.65812006 | 0.51046098 | 0.85762435 |
| NSUN3 | 607.004602 | -0.566837 | 0.16558996 | -3.4231364 | 0.00061903 | 0.04480846 |
| MTRNR2L12 | 888.673909 | 0.22121863 | 0.58758665 | 0.37648683 | 0.70655501 | 0.92556842 |
| EPHA6 | 263.52881 | 0.66982985 | 0.72618369 | 0.92239727 | 0.3563214 | 0.77368353 |
| ARL6 | 381.709276 | 0.00777183 | 0.23181807 | 0.03352557 | 0.97325547 | 0.99327086 |
| CRYBG3 | 1602.25414 | -0.0421243 | 0.318903 | -0.1320913 | 0.89491204 | 0.97788747 |
| AC110491.2 | 49.9527044 | 0.40958187 | 0.82166019 | 0.49848085 | 0.61814516 | 0.89599505 |
| RIOX2 | 520.841738 | 0.14672584 | 0.30137917 | 0.48684798 | 0.62636606 | 0.89873262 |
| CLDND1 | 280.945502 | -0.2739029 | 0.27205476 | -1.0067934 | 0.31403404 | 0.74777437 |
| CPOX | 106.845461 | 0.00573341 | 0.36174452 | 0.01584933 | 0.9873546 | 0.99653761 |
| ST3GAL6-AS | 92.7408013 | -0.5601809 | 0.32629439 | -1.7167961 | 0.08601643 | 0.49445601 |
| ST3GAL6 | 1588.7113 | -0.3216343 | 0.28220855 | -1.1397042 | 0.25440957 | 0.70509116 |
| DCBLD2 | 846.14391 | -0.2581137 | 0.22582535 | -1.1429793 | 0.25304719 | 0.70374828 |
| AC091212.1 | 45.9612224 | -1.3528854 | 0.49517746 | -2.7321222 | 0.00629278 | 0.16298298 |
| COL8A1 | 749.443338 | 0.48573389 | 0.57412699 | 0.84603912 | 0.39753092 | 0.80069929 |
| CMSS1 | 1465.8166 | -0.2091483 | 0.20105721 | -1.0402425 | 0.29822724 | 0.73673761 |
| FILIP1L | 457.173438 | 0.16704766 | 0.4847906 | 0.34457693 | 0.73041245 | 0.93467574 |
| TBC1D23 | 1003.83326 | 0.00616115 | 0.16927328 | 0.03639766 | 0.97096528 | 0.99317943 |
| NIT2 | 196.098407 | -0.2853673 | 0.22034634 | -1.2950854 | 0.19529079 | 0.64879175 |
| TOMM70 | 463.219057 | 0.0333622 | 0.18952149 | 0.17603388 | 0.86026733 | 0.97200968 |
| LNP1 | 69.8505722 | -0.4314445 | 0.44412786 | -0.971442 | 0.33132822 | 0.76000762 |
| TFG | 1190.01932 | 0.12821497 | 0.20682729 | 0.61991324 | 0.53531491 | 0.8642718 |
| ABI3BP | 314.385677 | 0.7057432 | 0.48822227 | 1.4455367 | 0.14830719 | 0.59438683 |
| IMPG2 | 65.5307891 | 0.35501622 | 0.50947187 | 0.69683184 | 0.48590804 | 0.8447301 |
| SENP7 | 1619.0823 | -0.1857188 | 0.18858385 | -0.9848074 | 0.32471869 | 0.75478872 |
| TRMT10C | 171.726429 | 0.60479761 | 0.35865897 | 1.68627485 | 0.09174287 | 0.50476839 |
| PCNP | 930.116715 | -0.0566629 | 0.14220461 | -0.3984606 | 0.69029073 | 0.92183073 |
| ZBTB11 | 398.975165 | -0.2219008 | 0.21922043 | -1.0122268 | 0.3114296 | 0.74590637 |
| RPL24 | 618.257488 | 0.37499089 | 0.35348421 | 1.06084196 | 0.28876173 | 0.73176973 |
| CEP97 | 265.016119 | -0.0042832 | 0.19586419 | -0.021868 | 0.98255327 | 0.99518552 |
| NXPE3 | 99.7642338 | 0.11471677 | 0.28390596 | 0.40406607 | 0.68616413 | 0.92053037 |
| NFKBIZ | 1219.43317 | 0.20747138 | 0.45205807 | 0.4589485 | 6.46E-01 | 9.06E-01 |
| ZPLD1 | 256.31923 | -1.2781249 | 0.85251227 | -1.4992452 | 1.34E-01 | 0.57704758 |

| | | | | | | |
|---|---|---|---|---|---|---|
| ALCAM | 1377.08878 | 0.5904432 | 0.4388617 | 1.34539696 | 0.17849708 | 0.63278795 |
| CBLB | 2452.82777 | 0.05186782 | 0.13023182 | 0.39827304 | 0.69042893 | 0.92183073 |
| LINC00882 | 452.512685 | 0.35321672 | 0.40102782 | 0.88077858 | 0.37843768 | 0.7869621 |
| DUBR | 98.6995175 | 0.55389413 | 0.34660076 | 1.59807532 | 0.11002622 | 0.53372881 |
| AC063944.1 | 105.874813 | 1.13301345 | 0.37805779 | 2.99693192 | 0.00272712 | 0.09842207 |
| BBX | 4284.96289 | 0.42905206 | 0.20403658 | 2.10281935 | 0.03548156 | 0.35586635 |
| CD47 | 1091.39873 | 0.40580645 | 0.30447869 | 1.332791 | 0.1826004 | 0.63741052 |
| IFT57 | 526.573974 | 0.38655728 | 0.20417318 | 1.89328139 | 0.05832046 | 0.42147872 |
| CIP2A | 67.529113 | -0.7355958 | 0.467309 | -1.57411 | 0.11546201 | 0.54427374 |
| DZIP3 | 730.081699 | 0.54601921 | 0.45620134 | 1.19688207 | 0.23135252 | 0.68357541 |
| AC078980.1 | 2068.1277 | -3.7005567 | 1.36538817 | -2.7102598 | 0.00672305 | 0.16906706 |
| NECTIN3-AS1 | 91.5551479 | -1.1254108 | 0.59557895 | -1.8896081 | 0.0588104 | 0.42333873 |
| NECTIN3 | 484.887399 | -0.1477418 | 0.28604123 | -0.5165053 | 0.60550151 | 0.89081188 |
| PHLDB2 | 4217.09368 | 0.14857576 | 0.15340396 | 0.96852628 | 3.33E-01 | 7.61E-01 |
| ABHD10 | 186.579507 | 0.72538861 | 0.20723488 | 3.5003211 | 0.0004647 | 0.03734387 |
| C3orf52 | 166.149866 | 0.45662503 | 0.57543905 | 0.79352457 | 0.42747226 | 0.81466629 |
| SLC9C1 | 114.717251 | 1.00660456 | 0.41992382 | 2.39711234 | 0.01652486 | 0.25721056 |
| ATG3 | 316.892915 | -0.1569129 | 0.21331882 | -0.7355795 | 0.46198667 | 0.83531269 |
| SLC35A5 | 195.481507 | -0.1671491 | 0.23824647 | -0.7015804 | 0.48294087 | 0.8441831 |
| GTPBP8 | 200.656148 | -0.0803933 | 0.32729695 | -0.2456279 | 0.8059703 | 0.95525039 |
| NEPRO | 535.985355 | 0.15926741 | 0.19235088 | 0.82800457 | 0.40766791 | 0.80588056 |
| AC078785.1 | 141.519497 | -0.2443492 | 0.25944448 | -0.9418171 | 0.34628631 | 0.76864066 |
| CFAP44 | 794.41609 | -0.0849428 | 0.23181157 | -0.3664304 | 0.71404392 | 0.92760608 |
| CFAP44-AS1 | 51.3749774 | 0.08004961 | 0.60759709 | 0.13174786 | 0.89518373 | 0.97788747 |
| SPICE1 | 586.903433 | -0.3513361 | 0.21884846 | -1.6053852 | 0.10840906 | 0.53134516 |
| SIDT1 | 187.46228 | 1.64631158 | 1.05785482 | 1.55627365 | 0.11964303 | 0.55232605 |
| USF3 | 559.685156 | 0.2112765 | 0.16320647 | 1.29453507 | 0.19548066 | 0.64893292 |
| NAA50 | 1346.03666 | -0.0137627 | 0.25946985 | -0.0530418 | 0.95769863 | 0.99069768 |
| ATP6V1A | 2658.40095 | 0.90840026 | 0.27587244 | 3.29282717 | 0.00099185 | 0.0559889 |
| GRAMD1C | 2781.60777 | 0.26224627 | 0.42693433 | 0.61425435 | 0.53904727 | 0.86542329 |
| ZDHHC23 | 213.522443 | -0.1834259 | 0.25069179 | -0.7316791 | 0.46436445 | 0.83586498 |
| CCDC191 | 705.93737 | -0.0745584 | 0.3573135 | -0.2086638 | 0.83471067 | 0.96511435 |
| QTRT2 | 356.666993 | -0.5941124 | 0.31154821 | -1.9069678 | 0.05652476 | 0.41843207 |
| ZBTB20 | 26385.9683 | -0.0770859 | 0.14136953 | -0.5452792 | 0.58556151 | 0.88490909 |
| ZBTB20-AS1 | 130.135247 | 0.20034008 | 0.68787211 | 0.29124611 | 0.7708631 | 0.9443559 |
| ZBTB20-AS5 | 624.911204 | 0.10728397 | 0.6961413 | 0.15411235 | 0.87752115 | 0.97655597 |
| ZBTB20-AS2 | 113.643474 | 0.05871471 | 0.62718365 | 0.09361646 | 0.92541384 | 0.9822216 |
| LSAMP | 676.370468 | -0.9307348 | 0.57080948 | -1.6305525 | 0.10298477 | 0.52280011 |
| IGSF11 | 72.6541452 | 0.03870986 | 0.51454986 | 0.07523054 | 0.94003129 | 0.98715467 |
| B4GALT4 | 178.017192 | 0.17970033 | 0.23727692 | 0.75734432 | 0.4488436 | 0.82665577 |
| ARHGAP31 | 48.1253815 | -0.544372 | 0.5028129 | -1.0826533 | 0.27896235 | 0.72464379 |
| TMEM39A | 358.184995 | -0.0235396 | 0.28495311 | -0.0826086 | 0.9341628 | 0.98494224 |
| POGLUT1 | 207.112445 | -0.1480054 | 0.30321639 | -0.488118 | 0.62546628 | 0.89873262 |
| TIMMDC1 | 485.812087 | 0.01418241 | 0.16199056 | 0.08755084 | 0.93023368 | 0.98331361 |
| PLA1A | 48.7985195 | 0.60662445 | 0.69903057 | 0.86780818 | 0.38549935 | 0.79365039 |
| COX17 | 227.726144 | 0.03001054 | 0.321842 | 0.09324618 | 0.92570798 | 0.9822216 |
| MAATS1 | 534.118243 | 0.44029082 | 0.28986778 | 1.51893678 | 0.12877841 | 0.56871749 |
| GSK3B | 2742.32587 | -0.1395386 | 0.1873624 | -0.7447522 | 0.45642156 | 0.83182139 |
| AC092910.3 | 144.521441 | -0.0757661 | 0.27218751 | -0.2783599 | 0.7807361 | 0.9480401 |
| LRRC58 | 336.047243 | 0.12299925 | 0.345082 | 0.35643483 | 0.72151495 | 0.93044898 |
| FSTL1 | 164.696499 | 0.24472251 | 0.46099468 | 0.53085755 | 0.5955175 | 0.88859628 |
| NDUFB4 | 386.579031 | 0.30364482 | 0.27252471 | 1.1141919 | 0.26519687 | 0.71512755 |
| HGD | 712.150891 | 1.19436158 | 0.55357192 | 2.15755451 | 0.03096248 | 0.33790537 |
| RABL3 | 317.731667 | -0.3108102 | 0.19285132 | -1.6116571 | 0.10703657 | 0.52854061 |

| | | | | | | |
|---|---|---|---|---|---|---|
| GTF2E1 | 118.92288 | -0.1035899 | 0.23740155 | -0.4363491 | 0.66258348 | 0.91110616 |
| GOLGB1 | 2402.98654 | -0.115982 | 0.18570907 | -0.6245359 | 0.53227567 | 0.86397197 |
| IQCB1 | 1135.66065 | -0.1865267 | 0.19438473 | -0.9595748 | 0.33726927 | 0.76409325 |
| EAF2 | 248.341023 | 0.42711051 | 0.30272819 | 1.41087126 | 0.15828258 | 0.60804051 |
| SLC15A2 | 503.594205 | 0.80398309 | 0.49403678 | 1.62737498 | 0.10365748 | 0.52435479 |
| ILDR1 | 328.112274 | 0.25337976 | 0.2664796 | 0.9508411 | 0.34168505 | 0.76545664 |
| CASR | 8411.38639 | -0.2033709 | 0.35125343 | -0.5789862 | 0.56259852 | 0.87432201 |
| CCDC58 | 204.513035 | -0.3035926 | 0.36278838 | -0.8368312 | 0.40268744 | 0.80324472 |
| FAM162A | 948.53431 | 0.02244034 | 0.47145531 | 0.04759803 | 0.96203661 | 0.99223192 |
| WDR5B | 40.596762 | 0.03694725 | 0.42972595 | 0.08597863 | 0.9314834 | 0.98357046 |
| AC083798.2 | 509.10807 | 0.12060791 | 0.31837265 | 0.37882623 | 0.7048169 | 0.92528913 |
| KPNA1 | 1087.85596 | -0.2694037 | 0.271959 | -0.9906041 | 0.32187894 | 0.75240254 |
| PARP9 | 667.913822 | 0.06914389 | 0.59144178 | 0.11690735 | 0.90693347 | 0.9803357 |
| DTX3L | 251.112187 | 0.50928117 | 0.5312814 | 0.95859027 | 0.3377652 | 0.76409325 |
| PARP14 | 1688.29393 | -0.1907734 | 0.6402736 | -0.297956 | 0.76573675 | 0.94273621 |
| HSPBAP1 | 929.92343 | 0.17987336 | 0.39597145 | 0.4542584 | 0.64964285 | 0.90824857 |
| DIRC2 | 1369.66938 | 0.16079975 | 0.40752047 | 0.39458079 | 0.6931523 | 0.92224682 |
| LINC02035 | 112.481326 | 0.42127172 | 0.56847136 | 0.7410606 | 0.4586567 | 0.833342 |
| SEMA5B | 90.2189851 | -0.8713966 | 1.17039331 | -0.7445331 | 0.456554 | 0.83192465 |
| PDIA5 | 515.161285 | -0.6753761 | 0.33810485 | -1.9975345 | 0.04576715 | 0.38904776 |
| SEC22A | 1251.71224 | -0.0250638 | 0.2184849 | -0.1147165 | 0.90866985 | 0.98047754 |
| ADCY5 | 1320.96721 | -1.0872567 | 0.47245696 | -2.3012821 | 0.02137569 | 0.29288643 |
| HACD2 | 775.652521 | -0.5311318 | 0.32880646 | -1.6153326 | 0.10623869 | 0.52735777 |
| MYLK-AS1 | 229.70661 | -0.5088611 | 0.28354843 | -1.7946179 | 0.0727146 | 0.4692634 |
| MYLK | 1695.81302 | -0.9798335 | 0.41901749 | -2.3384071 | 0.01936614 | 0.28045661 |
| CCDC14 | 1204.50981 | -0.2379721 | 0.16112025 | -1.4769844 | 0.13967981 | 0.58424021 |
| KALRN | 413.591776 | -2.1194106 | 0.62130975 | -3.411198 | 6.47E-04 | 0.04534035 |
| UMPS | 235.347024 | -0.0527411 | 0.21255658 | -0.2481275 | 0.80403579 | 0.95450237 |
| ITGB5 | 1560.06482 | 0.23586159 | 0.32087169 | 0.73506513 | 0.46229983 | 0.83531269 |
| ITGB5-AS1 | 97.4772688 | 0.36549602 | 0.38172987 | 0.95747294 | 0.3383286 | 0.76445133 |
| HEG1 | 63.0885995 | -0.2683982 | 0.48443154 | -0.5540476 | 0.57954624 | 0.88224343 |
| SLC12A8 | 666.033429 | -0.082013 | 0.35719247 | -0.2296046 | 0.81839904 | 0.95985564 |
| ZNF148 | 3518.67657 | -0.0768139 | 0.13352012 | -0.575298 | 0.56508981 | 0.87588257 |
| SNX4 | 1109.9452 | -0.1789351 | 0.13676306 | -1.3083587 | 0.19075166 | 0.64567512 |
| OSBPL11 | 416.012934 | -0.4966181 | 0.24988449 | -1.9873906 | 0.04687913 | 0.39241217 |
| AC092902.3 | 42.293619 | 0.23673083 | 0.74835429 | 0.31633524 | 0.75174807 | 0.93810488 |
| AC092903.2 | 39.7143794 | -0.5068293 | 0.41010687 | -1.2358468 | 0.21651552 | 0.66978787 |
| SLC41A3 | 185.660829 | -0.1607186 | 0.27671387 | -0.5808116 | 0.56136745 | 0.87401021 |
| AC079848.1 | 49.3401322 | -0.6503772 | 0.38110254 | -1.7065676 | 0.08790246 | 0.49714378 |
| ALDH1L1 | 976.627631 | 1.00387578 | 0.47599486 | 2.10900552 | 0.03494411 | 0.35358029 |
| ALDH1L1-AS: | 57.6910791 | 0.86329626 | 0.49588547 | 1.74091866 | 0.08169784 | 0.48594425 |
| KLF15 | 138.329335 | 0.88839484 | 0.45427292 | 1.95564121 | 0.05050743 | 0.40335713 |
| ZXDC | 1678.46745 | -0.2754825 | 0.22432572 | -1.228047 | 0.21942933 | 0.67278782 |
| TXNRD3NB | 124.790833 | -0.1110513 | 0.33287434 | -0.3336132 | 0.73867146 | 0.93552616 |
| TXNRD3 | 211.407424 | 0.07846561 | 0.25934036 | 0.30255842 | 0.76222641 | 0.94212205 |
| CHCHD6 | 412.831733 | -0.3120404 | 0.24164818 | -1.2913003 | 0.19659956 | 0.64964978 |
| PLXNA1 | 110.45147 | 0.04707682 | 0.33834096 | 0.13914019 | 0.88933937 | 0.97729111 |
| TPRA1 | 147.374782 | -0.0982896 | 0.25506311 | -0.3853541 | 0.69997505 | 0.92367966 |
| PODXL2 | 46.8433562 | 0.31334096 | 0.57140563 | 0.5483687 | 0.58343876 | 0.88379959 |
| ABTB1 | 155.393782 | 0.10230823 | 0.36191254 | 0.28268771 | 0.77741624 | 0.94694572 |
| MGLL | 4473.65281 | 0.26442601 | 0.37323148 | 0.70847724 | 0.47864894 | 0.84265945 |
| KBTBD12 | 362.937252 | -0.1803314 | 0.36634698 | -0.492242 | 0.62254831 | 0.89835355 |
| SEC61A1 | 563.03973 | 0.3316041 | 0.22628972 | 1.4653962 | 0.14281285 | 0.58824095 |
| RUVBL1 | 393.745612 | 0.16269676 | 0.20062469 | 0.81095084 | 0.4173939 | 0.81017256 |

| | | | | | | |
|---|---|---|---|---|---|---|
| EEFSEC | 675.201809 | -0.0757976 | 0.1781091 | -0.4255684 | 0.67042234 | 0.91492559 |
| RPN1 | 374.705269 | 0.40815614 | 0.23142157 | 1.76369102 | 0.07778401 | 0.47916634 |
| RAB7A | 1813.16776 | -0.2252001 | 0.13925552 | -1.6171718 | 0.1058412 | 0.52717062 |
| ACAD9 | 378.246051 | -0.0627691 | 0.17467809 | -0.3593417 | 0.71933951 | 0.93001079 |
| KIAA1257 | 64.3872157 | -0.6875122 | 0.39094244 | -1.7586021 | 0.07864512 | 0.48060389 |
| CNBP | 822.101772 | 0.53675939 | 0.46329405 | 1.15857173 | 0.2466308 | 0.69684411 |
| COPG1 | 331.830511 | -0.0197798 | 0.22743494 | -0.0869689 | 0.93069626 | 0.98331361 |
| HMCES | 211.603286 | 0.03288052 | 0.24069389 | 0.1366072 | 0.89134129 | 0.97729111 |
| H1FX | 264.569845 | 0.24854239 | 0.59644567 | 0.41670583 | 0.67689359 | 0.91675253 |
| MBD4 | 467.582572 | -0.026897 | 0.13375883 | -0.2010856 | 0.84063164 | 0.96617873 |
| IFT122 | 636.036257 | -0.1602148 | 0.19016566 | -0.8425012 | 0.39950746 | 0.80174499 |
| TMCC1 | 4514.70741 | 0.01520525 | 0.18986363 | 0.08008509 | 0.93616958 | 0.98621411 |
| LINC02021 | 49.7559358 | -0.0322712 | 0.56116971 | -0.057507 | 0.95414135 | 0.99004345 |
| COL6A6 | 45.9096049 | 0.85619731 | 0.72463588 | 1.18155523 | 0.23738223 | 0.68973903 |
| PIK3R4 | 511.878815 | -0.0964342 | 0.23437521 | -0.4114523 | 0.68074088 | 0.91867756 |
| ATP2C1 | 1885.19592 | 0.27560954 | 0.25646315 | 1.07465551 | 0.28252899 | 0.72604837 |
| AC055733.2 | 92.6052879 | 1.01344877 | 0.71595755 | 1.41551516 | 0.15691751 | 0.60654181 |
| ASTE1 | 80.3102378 | -0.1602164 | 0.353567 | -0.4531429 | 0.65044585 | 0.90825486 |
| NEK11 | 1085.05308 | -0.104829 | 0.34525812 | -0.303625 | 0.76141358 | 0.94202764 |
| NUDT16 | 130.278773 | 0.84190184 | 0.36995634 | 2.27567888 | 0.02286524 | 0.2983102 |
| MRPL3 | 461.359164 | -0.0146761 | 0.19292721 | -0.0760705 | 0.93936301 | 0.98715467 |
| CPNE4 | 121.334077 | 0.16441442 | 0.47632861 | 0.34517015 | 0.72996646 | 0.93453345 |
| ACPP | 1808.87395 | 0.33478167 | 0.4324751 | 0.77410623 | 0.43886797 | 0.82127804 |
| DNAJC13 | 985.911858 | -0.0245218 | 0.20647874 | -0.1187621 | 0.90546387 | 0.9799593 |
| NPHP3-ACAD | 75.6010595 | 0.15234568 | 0.46662923 | 0.32648123 | 0.74406029 | 0.93656303 |
| ACKR4 | 44.3037664 | 0.28338668 | 0.72321676 | 0.39184197 | 0.69517498 | 0.92229578 |
| UBA5 | 681.538352 | 0.36596957 | 0.20844738 | 1.75569282 | 0.07914087 | 0.48169281 |
| TMEM108 | 264.172009 | -1.3493676 | 0.66559271 | -2.0273172 | 0.04262998 | 0.38021566 |
| CDV3 | 426.840994 | 0.16970987 | 0.22463204 | 0.7555016 | 4.50E-01 | 8.27E-01 |
| TOPBP1 | 370.708318 | -0.4967557 | 0.22038036 | -2.2540832 | 0.02419093 | 0.30500338 |
| TF | 58.2116722 | -0.1509033 | 0.50387919 | -0.2994831 | 0.76457148 | 0.94256992 |
| SRPRB | 167.076087 | 0.53705837 | 0.3016134 | 1.78061842 | 0.07497481 | 0.47190267 |
| RAB6B | 114.349841 | 0.17224705 | 0.63795077 | 0.27000053 | 0.78715985 | 0.94978829 |
| SLCO2A1 | 121.98715 | 1.66853637 | 0.55633041 | 2.9991824 | 0.00270705 | 0.09823475 |
| RYK | 1663.11335 | -0.1824378 | 0.19770146 | -0.9227942 | 0.35611445 | 0.77367035 |
| AMOTL2 | 287.439112 | 0.74517332 | 0.46574793 | 1.59994983 | 0.10960971 | 0.53361622 |
| ANAPC13 | 126.026641 | -0.133514 | 0.32171817 | -0.4150029 | 0.67813976 | 0.91787561 |
| CEP63 | 693.398757 | -0.2319511 | 0.18960363 | -1.2233474 | 0.2211985 | 0.67400484 |
| EPHB1 | 319.869443 | -1.3442544 | 0.51370346 | -2.6167907 | 0.00887607 | 0.19125454 |
| PPP2R3A | 9811.2073 | -0.0379562 | 0.21679099 | -0.1750822 | 0.86101504 | 0.9720639 |
| MSL2 | 722.093693 | -0.0999031 | 0.23710166 | -0.4213515 | 0.67349843 | 0.91636338 |
| PCCB | 317.961354 | 0.12360832 | 0.23916913 | 0.51682388 | 0.60527911 | 0.89062401 |
| STAG1 | 5213.7948 | -0.3510709 | 0.20750505 | -1.6918668 | 0.09067136 | 0.50267341 |
| SLC35G2 | 239.937498 | 0.56668773 | 0.38681831 | 1.46499717 | 0.14292169 | 0.58829934 |
| NCK1-DT | 113.346585 | 0.26648913 | 0.29899181 | 0.89129242 | 0.37277232 | 0.78269743 |
| NCK1 | 602.681295 | 0.14252076 | 0.23439089 | 0.60804736 | 0.54315607 | 0.86612953 |
| IL20RB | 57.98475 | 0.19204129 | 0.61650815 | 0.31149838 | 0.75542177 | 0.93953492 |
| CLDN18 | 54.4518466 | 0.32611812 | 0.82264954 | 0.39642412 | 0.69179217 | 0.92224682 |
| DZIP1L | 440.810362 | 0.09763166 | 0.30375769 | 0.32141296 | 0.74789746 | 0.93746253 |
| DBR1 | 118.043951 | -0.2748796 | 0.23606691 | -1.1644139 | 0.24425632 | 0.6948422 |
| ARMC8 | 1513.1402 | -0.4104689 | 0.18950623 | -2.1659918 | 0.03031181 | 0.33449351 |
| NME9 | 190.988814 | -0.062551 | 0.39242274 | -0.1593971 | 0.87335604 | 0.97564728 |
| MRAS | 549.832502 | 0.19514116 | 0.25128467 | 0.77657409 | 0.43741009 | 0.82013459 |
| ESYT3 | 87.2935494 | 0.35041483 | 0.36541503 | 0.95895025 | 0.33758381 | 0.76409325 |

| | | | | | | |
|---|---|---|---|---|---|---|
| CEP70 | 2090.55988 | 0.24922029 | 0.32307192 | 0.77140808 | 0.44046508 | 0.82253723 |
| FAIM | 230.36797 | 0.25686894 | 0.27851978 | 0.92226463 | 0.35639056 | 0.77368353 |
| PIK3CB | 2078.16181 | 0.1297911 | 0.17684162 | 0.73393975 | 0.46298546 | 0.83531269 |
| MRPS22 | 127.480746 | 0.12546244 | 0.25658416 | 0.48897189 | 0.6248616 | 0.89873262 |
| COPB2 | 700.97136 | 0.06616647 | 0.17842029 | 0.37084609 | 0.71075217 | 0.92669287 |
| AC097103.2 | 964.226481 | -0.5017801 | 0.42640795 | -1.1767606 | 0.23929107 | 0.69029619 |
| NMNAT3 | 428.832929 | 0.36387619 | 0.29328589 | 1.24068768 | 0.21472115 | 0.66830201 |
| CLSTN2 | 80.9376164 | -0.1395499 | 0.46435338 | -0.3005252 | 0.76377658 | 0.94255335 |
| SLC25A36 | 1962.21929 | 0.2270628 | 0.31081018 | 0.73055137 | 0.46505322 | 0.8363817 |
| PXYLP1 | 118.743868 | 0.32185904 | 0.27873264 | 1.15472315 | 0.24820381 | 0.69838949 |
| AC117383.1 | 102.282978 | 0.66963736 | 0.68166412 | 0.98235678 | 3.26E-01 | 0.75550719 |
| ZBTB38 | 1259.08356 | 0.03355086 | 0.31150391 | 0.10770605 | 9.14E-01 | 0.98145965 |
| RASA2 | 2030.983 | -0.4698556 | 0.16768023 | -2.8020927 | 0.00507723 | 0.14305936 |
| RNF7 | 320.446039 | 0.42814747 | 0.27451293 | 1.5596623 | 0.11883971 | 0.55040451 |
| ATP1B3 | 1130.83193 | 0.07951804 | 0.29429357 | 0.27019971 | 0.78700661 | 0.94978829 |
| TFDP2 | 3045.62092 | 0.03247902 | 0.23229141 | 0.13982015 | 0.88880209 | 0.97729111 |
| GK5 | 861.959793 | -0.2998812 | 0.23582734 | -1.2716133 | 0.20351056 | 0.65645201 |
| XRN1 | 2256.48774 | -0.1491243 | 0.21416788 | -0.6962964 | 0.48624321 | 0.8447301 |
| ATR | 1787.13843 | -0.5167022 | 0.17514553 | -2.9501307 | 0.00317639 | 0.10846567 |
| PLS1 | 404.034246 | 0.01215329 | 0.20538575 | 0.05917299 | 0.95281432 | 0.98949465 |
| TRPC1 | 597.905124 | 0.76863226 | 0.37710843 | 2.03822614 | 0.04152732 | 0.37579578 |
| PCOLCE2 | 55.7557442 | 0.45628292 | 0.37360874 | 1.22128545 | 0.22197796 | 0.67544503 |
| U2SURP | 1490.40674 | -0.2149987 | 0.15800946 | -1.3606695 | 0.17361815 | 0.6286116 |
| SLC9A9 | 166.577537 | 0.34996568 | 0.64717558 | 0.54075848 | 0.58867407 | 0.88571413 |
| C3orf58 | 285.685478 | 0.03941527 | 0.36382234 | 0.10833659 | 0.9137287 | 0.98145965 |
| AC107021.1 | 463.007263 | -0.4773474 | 0.81252312 | -0.5874878 | 0.55687615 | 0.87221664 |
| LNCSRLR | 125.81761 | -0.5363354 | 0.57062745 | -0.9399047 | 0.34726643 | 0.76893725 |
| PLOD2 | 3239.64106 | -0.4813799 | 0.46638496 | -1.0321513 | 0.30200123 | 0.73905643 |
| PLSCR4 | 327.159122 | 0.85991164 | 0.48770441 | 1.76318201 | 0.0778698 | 0.47916634 |
| PLSCR2 | 446.077345 | 0.41335293 | 0.35053823 | 1.179195 | 0.23832054 | 0.68982886 |
| PLSCR1 | 2251.61474 | 0.16604494 | 0.39812961 | 0.41706252 | 0.67663268 | 0.91658712 |
| GYG1 | 140.864065 | -0.08688 | 0.26278839 | -0.3306083 | 7.41E-01 | 0.93598004 |
| HLTF | 499.311337 | 0.47445802 | 0.24502807 | 1.93634148 | 0.05282589 | 0.41066631 |
| HPS3 | 460.401792 | 0.37472576 | 0.17138774 | 2.18642106 | 0.02878481 | 0.32721051 |
| CP | 602.910048 | 0.15500802 | 1.23327227 | 0.1256884 | 9.00E-01 | 0.97898519 |
| TM4SF1 | 169.800538 | -0.0900715 | 0.74327827 | -0.1211813 | 0.9035474 | 0.97975674 |
| WWTR1 | 2716.64912 | -0.0814164 | 0.24127334 | -0.3374448 | 0.73578164 | 0.93552616 |
| COMMD2 | 283.24511 | 0.12988356 | 0.32251028 | 0.40272687 | 0.68714917 | 0.9208231 |
| ANKUB1 | 687.700728 | -0.2252879 | 0.55589589 | -0.40527 | 0.68527905 | 0.92053037 |
| RNF13 | 1613.07377 | -0.1689601 | 0.13010261 | -1.2986678 | 0.19405795 | 0.64749489 |
| PFN2 | 41.9253733 | 0.30205766 | 0.38432344 | 0.78594648 | 0.43189884 | 0.81761357 |
| TSC22D2 | 2716.53473 | 0.62969912 | 0.35453462 | 1.77612872 | 0.0757117 | 0.47374509 |
| SERP1 | 944.177693 | 0.46967855 | 0.26167605 | 1.7948855 | 0.07267195 | 0.4692634 |
| EIF2A | 477.803091 | -0.0213728 | 0.19951221 | -0.1071251 | 0.91468973 | 0.98156931 |
| SELENOT | 508.550785 | 0.33194227 | 0.23243428 | 1.42811235 | 0.15325952 | 0.60285224 |
| ERICH6 | 60.2755061 | -1.6616428 | 0.51720973 | -3.212706 | 0.00131491 | 0.06604038 |
| ERICH6-AS1 | 91.3632368 | -0.1570441 | 0.29706569 | -0.5286512 | 0.59704741 | 0.88883692 |
| SIAH2 | 196.467273 | 0.13425608 | 0.28302216 | 0.47436597 | 0.63523895 | 0.90199374 |
| MED12L | 241.23055 | -0.4159659 | 0.62989825 | -0.66037 | 0.50901641 | 0.8566594 |
| SUCNR1 | 468.272619 | 2.63038614 | 0.77764564 | 3.38249969 | 0.00071829 | 0.04697 |
| MBNL1 | 5162.56033 | 0.17772039 | 0.20271648 | 0.87669434 | 0.38065268 | 0.78826518 |
| RAP2B | 92.4412262 | 0.66811356 | 0.40439582 | 1.65212776 | 9.85E-02 | 5.15E-01 |
| ARHGEF26 | 146.512815 | 0.72464759 | 0.39078711 | 1.85432827 | 0.06369221 | 0.43779712 |
| DHX36 | 1336.43899 | -0.0152691 | 0.16674312 | -0.0915725 | 0.92703769 | 0.98252755 |

| | | | | | | |
|---|---|---|---|---|---|---|
| MME | 110.156936 | 0.28670374 | 0.55870863 | 0.5131543 | 0.60784339 | 0.89203142 |
| PLCH1 | 566.712137 | 0.75961864 | 0.79444139 | 0.956167 | 0.33898787 | 0.76449297 |
| GMPS | 683.988888 | -0.0335984 | 0.21043256 | -0.1596634 | 0.87314622 | 0.97564728 |
| KCNAB1 | 116.151182 | 0.47563308 | 0.62395199 | 0.76229115 | 0.44588626 | 0.82571115 |
| SSR3 | 410.475793 | 0.47744034 | 0.19794977 | 2.41192676 | 0.01586847 | 0.25383866 |
| TIPARP | 233.110073 | 0.1480698 | 0.36547564 | 0.40514272 | 0.6853726 | 0.92053037 |
| LINC00886 | 719.320059 | 0.73930163 | 0.41917683 | 1.76369871 | 0.07778272 | 0.47916634 |
| LEKR1 | 495.977899 | -0.1756922 | 0.4453623 | -0.3944928 | 0.69321725 | 0.92224682 |
| CCNL1 | 1648.542 | -0.1471959 | 0.43881364 | -0.3354406 | 0.73729273 | 0.93552616 |
| VEPH1 | 984.762549 | 0.39900426 | 0.45459428 | 0.87771511 | 3.80E-01 | 7.88E-01 |
| RSRC1 | 1769.6878 | -0.5217569 | 0.14950672 | -3.4898557 | 0.00048328 | 0.03777316 |
| MLF1 | 229.793881 | 0.63687877 | 0.29935764 | 2.12748462 | 0.03337984 | 0.3493774 |
| GFM1 | 387.709785 | -0.1211682 | 0.32539169 | -0.3723764 | 0.70961262 | 0.92593827 |
| LXN | 125.048478 | 0.00871534 | 0.44283162 | 0.01968092 | 0.98429791 | 0.99518552 |
| RARRES1 | 118.931692 | 0.45895002 | 0.55898585 | 0.8210405 | 0.4116232 | 0.80679792 |
| MFSD1 | 241.732224 | -0.0168602 | 0.19424534 | -0.0867987 | 0.93083151 | 0.98331361 |
| IQCJ-SCHIP1 | 941.652797 | -0.0941363 | 0.35514551 | -0.265064 | 0.79096014 | 0.95144272 |
| AC079594.2 | 60.6185101 | 0.44369625 | 0.8714531 | 0.5091453 | 0.61065038 | 0.89339478 |
| SMC4 | 235.164553 | -0.4170613 | 0.28373968 | -1.4698731 | 0.14159613 | 0.5872148 |
| KPNA4 | 1242.47238 | -0.2749221 | 0.26205357 | -1.0491063 | 0.29412919 | 0.73527014 |
| PPM1L | 3422.4354 | 0.27440843 | 0.42363139 | 0.64775283 | 0.51714483 | 0.85893097 |
| B3GALNT1 | 219.640797 | 0.3000619 | 0.29135536 | 1.0298829 | 0.30306498 | 0.73996031 |
| NMD3 | 1525.00732 | 0.18248592 | 0.29405697 | 0.62058015 | 0.5348759 | 0.8642718 |
| LINC01322 | 140.056125 | -0.1267689 | 0.4122846 | -0.3074791 | 0.75847874 | 0.94090399 |
| PDCD10 | 663.590612 | -0.0413928 | 0.20395956 | -0.2029461 | 0.83917712 | 0.966178 |
| SERPINI1 | 124.374553 | 0.00265952 | 0.3843405 | 0.00691969 | 0.99447893 | 0.998438 |
| GOLIM4 | 592.149679 | 0.34092471 | 0.30346843 | 1.12342726 | 0.26125608 | 0.71167902 |
| MECOM | 47927.8807 | 0.04277306 | 0.3046606 | 0.14039578 | 0.88834729 | 0.97729111 |
| AC074033.1 | 231.814313 | 0.26252916 | 0.60362319 | 0.43492226 | 0.66361886 | 0.91177281 |
| MYNN | 219.401486 | -0.2511097 | 0.26757062 | -0.9384802 | 0.3479977 | 0.76932244 |
| LRRC34 | 80.1492532 | 0.30879142 | 0.42484034 | 0.72684111 | 0.4673233 | 0.83762812 |
| SEC62 | 965.74377 | 0.10476749 | 0.20372141 | 0.51426842 | 0.60706434 | 0.89174148 |
| GPR160 | 236.583128 | 0.42347111 | 0.43188258 | 0.98052371 | 0.32682767 | 0.75556858 |
| PHC3 | 2399.76965 | -0.1416214 | 0.16996894 | -0.8332192 | 0.40472114 | 0.80437725 |
| PRKCI | 1819.77902 | -0.5097952 | 0.21261811 | -2.3977036 | 0.01649821 | 0.25721056 |
| SKIL | 769.089733 | 0.42447192 | 0.33387659 | 1.27134376 | 0.20360638 | 0.65660075 |
| RPL22L1 | 100.608054 | -0.061329 | 0.44136504 | -0.1389529 | 0.88948735 | 0.97729111 |
| TNIK | 437.276736 | 0.92320643 | 0.63347019 | 1.45737945 | 0.14501167 | 0.59091136 |
| PLD1 | 453.008445 | 0.44569894 | 0.25845577 | 1.7244689 | 0.08462323 | 0.49155155 |
| FNDC3B | 10286.0786 | 0.31546191 | 0.22144017 | 1.42459207 | 1.54E-01 | 0.60379867 |
| TNFSF10 | 2119.57228 | -0.7106286 | 0.42891774 | -1.6567945 | 9.76E-02 | 0.51355946 |
| LINC02068 | 75.4005717 | -0.1240728 | 0.640072 | -0.1938419 | 0.84629968 | 0.96883382 |
| NCEH1 | 1895.17795 | -0.8879509 | 0.3876688 | -2.2904883 | 0.02199302 | 0.29509055 |
| ECT2 | 269.454141 | -0.2449642 | 0.41092687 | -0.5961261 | 0.55109097 | 0.87018163 |
| NLGN1 | 3491.42944 | 2.73875829 | 0.64270112 | 4.26132491 | 2.03E-05 | 0.0072549 |
| NAALADL2 | 10571.8593 | -0.7328491 | 0.36574145 | -2.0037354 | 0.04509841 | 0.38757642 |
| AC092920.1 | 76.3836785 | -1.3270586 | 0.55748501 | -2.3804381 | 0.01729207 | 0.26361342 |
| TBL1XR1 | 5512.40486 | -0.6776478 | 0.30333271 | -2.2340083 | 0.02548253 | 0.31163914 |
| LINC00501 | 55.8303759 | 0.13362483 | 0.68484054 | 0.19511817 | 0.84530044 | 0.96859165 |
| LINC00578 | 128.909519 | 0.17925048 | 0.75140097 | 0.23855503 | 0.81145064 | 0.95761871 |
| KCNMB2-AS | 487.476713 | 0.32412672 | 0.58454921 | 0.55449004 | 0.57924352 | 0.8821758 |
| ZMAT3 | 414.889613 | 0.44115165 | 0.39752564 | 1.1097439 | 0.2671094 | 0.71712359 |
| AC076966.1 | 39.6413806 | -1.0199483 | 0.7449139 | -1.3692163 | 0.17093166 | 0.62735101 |
| PIK3CA | 1288.58188 | -0.270897 | 0.23802306 | -1.1381126 | 0.25507348 | 0.70604895 |

| | | | | | | |
|---|---|---|---|---|---|---|
| KCNMB3 | 237.878429 | -0.5994195 | 0.28327276 | -2.1160507 | 0.0343405 | 0.35209451 |
| ZNF639 | 173.472519 | 0.13511233 | 0.27399413 | 0.49312127 | 0.62192691 | 0.89830864 |
| MFN1 | 1135.42644 | -0.1980704 | 0.19730046 | -1.0039025 | 0.31542559 | 0.74785672 |
| GNB4 | 215.95424 | 1.09088325 | 0.39033781 | 2.79471582 | 0.00519454 | 0.14506262 |
| ACTL6A | 236.773411 | -0.2388348 | 0.30095209 | -0.7935975 | 0.42742977 | 0.81466629 |
| MRPL47 | 182.684884 | -0.0311832 | 0.2746325 | -0.113545 | 0.90959846 | 0.98069436 |
| NDUFB5 | 461.347275 | 0.53473591 | 0.23930664 | 2.23452177 | 0.02544876 | 0.31163914 |
| USP13 | 727.658605 | 0.25167407 | 0.30312044 | 0.83027747 | 4.06E-01 | 0.80499101 |
| PEX5L | 125.621467 | -0.249519 | 0.41878489 | -0.5958167 | 0.55129767 | 0.87023075 |
| AC125618.1 | 57.6881289 | -1.2250494 | 0.61734021 | -1.9843992 | 0.04721135 | 0.39394488 |
| TTC14 | 1693.34224 | 0.07070776 | 0.16204125 | 0.43635653 | 6.63E-01 | 9.11E-01 |
| CCDC39 | 51.3214943 | 0.16054146 | 0.76618757 | 0.20953284 | 0.8340323 | 0.96510927 |
| FXR1 | 1402.58505 | 0.28883965 | 0.2157627 | 1.33869129 | 0.18067119 | 0.63503977 |
| DNAJC19 | 383.806888 | 0.41338988 | 0.30792569 | 1.34249884 | 0.17943431 | 0.63322142 |
| ATP11B | 1357.30114 | 0.03492852 | 0.17133489 | 0.2038611 | 0.83846204 | 0.9657521 |
| DCUN1D1 | 855.443669 | 0.0362669 | 0.1491488 | 0.24315921 | 8.08E-01 | 0.95603962 |
| MCCC1 | 1511.0446 | -0.256801 | 0.22072781 | -1.1634283 | 0.24465578 | 0.6948422 |
| MCF2L2 | 156.965185 | 0.11129201 | 0.53699082 | 0.20725124 | 0.83581365 | 0.96511435 |
| B3GNT5 | 184.333534 | 0.43999826 | 0.41185786 | 1.06832552 | 0.2853737 | 0.72844615 |
| KLHL24 | 2202.67719 | -0.2512243 | 0.19954714 | -1.258972 | 0.20804044 | 0.66104323 |
| YEATS2 | 1135.91291 | -0.1923489 | 0.32363621 | -0.5943366 | 0.55228698 | 0.87095972 |
| ABCC5 | 2261.73657 | 0.01010814 | 0.18798475 | 0.05377106 | 0.95711757 | 0.99065996 |
| EIF2B5 | 212.75918 | -0.0459724 | 0.2682582 | -0.1713737 | 0.86392995 | 0.97349667 |
| DVL3 | 442.340285 | -0.3487027 | 0.42741274 | -0.8158453 | 0.41458862 | 0.8094275 |
| AP2M1 | 518.53897 | 0.4712642 | 0.38828102 | 1.2137194 | 0.2248549 | 0.67750405 |
| ABCF3 | 104.57326 | -0.0996691 | 0.3043059 | -0.3275294 | 0.74326748 | 0.93611803 |
| ALG3 | 84.8514524 | -0.2785688 | 0.40824275 | -0.6823607 | 0.49501089 | 0.84899096 |
| PSMD2 | 277.722303 | 0.22487656 | 0.3373713 | 0.6665551 | 0.50505635 | 0.85582573 |
| EIF4G1 | 550.85567 | -3.98E-05 | 0.25815474 | -0.0001542 | 0.99987697 | 0.99988492 |
| FAM131A | 129.669498 | 0.19704645 | 0.30213438 | 0.65218148 | 0.5142841 | 0.85777992 |
| CLCN2 | 76.4219381 | -0.3318542 | 0.3754427 | -0.8839009 | 0.37674969 | 0.78530639 |
| POLR2H | 414.312502 | 0.1472171 | 0.27287967 | 0.53949455 | 0.58954566 | 0.88621335 |
| EPHB3 | 69.6842434 | 0.40534316 | 0.66314497 | 0.61124366 | 0.54103828 | 0.86582694 |
| MAGEF1 | 179.563504 | 0.47607931 | 0.34265671 | 1.3893769 | 0.16471817 | 0.61673853 |
| VPS8 | 1453.7469 | -0.4080201 | 0.21701331 | -1.8801616 | 0.06008605 | 0.42785804 |
| EHHADH | 384.965703 | -0.1235463 | 0.28287037 | -0.4367593 | 0.66228593 | 0.91110616 |
| MAP3K13 | 1931.69266 | 0.41074368 | 0.23994027 | 1.71185808 | 0.08692283 | 0.49480766 |
| TMEM41A | 256.571024 | -0.0083063 | 0.24408367 | -0.0340304 | 0.97285287 | 0.99327086 |
| LIPH | 890.657927 | -1.1483147 | 0.40590292 | -2.8290378 | 0.00466882 | 0.13494621 |
| SENP2 | 684.545523 | -0.1905418 | 0.18477287 | -1.0312218 | 0.30243682 | 0.73905643 |
| IGF2BP2 | 1720.17966 | 0.73351568 | 0.48118152 | 1.52440536 | 0.12740748 | 0.56740575 |
| TRA2B | 959.566795 | 0.15983236 | 0.17935724 | 0.89113968 | 0.37285424 | 0.78270813 |
| ETV5 | 332.455719 | 1.03167182 | 0.53550367 | 1.92654482 | 0.05403638 | 0.41468669 |
| CRYGS | 76.1246195 | 0.11710151 | 0.29864758 | 0.39210602 | 0.69497988 | 0.92229578 |
| TBCCD1 | 141.321951 | -0.1077404 | 0.27816036 | -0.3873321 | 0.69851037 | 0.92356678 |
| DNAJB11 | 133.62322 | 0.41951295 | 0.3188235 | 1.31581566 | 0.18823592 | 0.64267818 |
| AC068631.1 | 1187.76516 | 0.95968603 | 0.33773089 | 2.84157022 | 0.0044892 | 0.13032483 |
| AC068631.2 | 72.9564662 | 1.15477166 | 0.53441578 | 2.1608113 | 0.03070992 | 0.33646028 |
| HRG | 61.0662896 | 1.78206825 | 0.47583048 | 3.74517464 | 0.00018027 | 0.021499 |
| KNG1 | 2181.03806 | 0.46260743 | 0.64831669 | 0.71355163 | 0.47550448 | 0.84118672 |
| EIF4A2 | 1052.88371 | -0.0139015 | 0.32284834 | -0.0430588 | 0.96565469 | 0.99248831 |
| RFC4 | 47.3446639 | -0.2589686 | 0.3413582 | -0.7586419 | 0.44806677 | 0.82614657 |
| ST6GAL1 | 956.389544 | 0.99086025 | 0.58782659 | 1.68563359 | 0.09186639 | 0.50476839 |
| AC072022.1 | 152.413446 | -0.4267082 | 0.95187874 | -0.44828 | 0.65395113 | 0.90898037 |

| | | | | | | |
|---|---|---|---|---|---|---|
| BCL6 | 1803.98613 | -0.6193646 | 0.49507624 | -1.251049 | 2.11E-01 | 6.64E-01 |
| LPP | 9219.96667 | -0.25463 | 0.2256023 | -1.1286674 | 0.25903816 | 0.70921184 |
| TPRG1 | 370.681771 | -1.5128106 | 0.35937631 | -4.2095446 | 2.56E-05 | 0.00824389 |
| P3H2 | 4752.31337 | 0.30148354 | 0.46027305 | 0.6550102 | 0.51246119 | 0.85762435 |
| P3H2-AS1 | 133.883505 | 0.32036644 | 0.64307464 | 0.49817924 | 0.61835771 | 0.89599505 |
| CLDN1 | 207.334861 | 0.83705113 | 0.51527338 | 1.62447966 | 1.04E-01 | 5.24E-01 |
| CLDN16 | 1251.80332 | 1.90281432 | 0.52398646 | 3.63141886 | 0.00028187 | 0.02863854 |
| TMEM207 | 336.974143 | 3.27718864 | 0.61873962 | 5.29655533 | 1.18E-07 | 0.00031175 |
| IL1RAP | 170.595489 | 0.3217434 | 0.6121519 | 0.52559405 | 0.59917029 | 0.88883692 |
| CCDC50 | 1807.21159 | -0.1176179 | 0.29683896 | -0.3962349 | 0.69193176 | 0.92224682 |
| FGF12 | 56.124146 | 0.17096831 | 0.41656277 | 0.41042628 | 0.68149327 | 0.91936732 |
| MB21D2 | 112.589625 | 1.00394187 | 0.52776232 | 1.90226135 | 0.057137 | 0.41929032 |
| OPA1 | 1374.15672 | -0.1401147 | 0.17434459 | -0.8036651 | 0.4215904 | 0.81266383 |
| OPA1-AS1 | 72.6423358 | 0.10548777 | 0.49738748 | 0.21208368 | 0.83204175 | 0.96417211 |
| HES1 | 245.039948 | 0.28559376 | 0.64816588 | 0.44061832 | 0.65948934 | 0.91099119 |
| ATP13A3 | 1044.61499 | -0.1617099 | 0.27296389 | -0.5924224 | 0.55356778 | 0.87121134 |
| LINC00884 | 74.0034125 | 0.30186837 | 0.46959203 | 0.64283112 | 0.52033369 | 0.85935232 |
| TMEM44-AS | 118.331399 | -0.2923423 | 0.5181978 | -0.5641519 | 0.57265073 | 0.87946834 |
| TMEM44 | 53.5568475 | -0.7988238 | 0.43530607 | -1.8350854 | 0.06649304 | 0.4472029 |
| LSG1 | 366.916386 | -0.0699366 | 0.30117445 | -0.2322131 | 0.81637252 | 0.9589293 |
| FAM43A | 238.378286 | -2.0695951 | 1.01437091 | -2.0402745 | 0.04132299 | 0.37577267 |
| LINC01968 | 38.1649636 | -2.2538136 | 0.93999963 | -2.397675 | 0.0164995 | 0.25721056 |
| XXYLT1 | 336.897386 | -0.5355258 | 0.28433148 | -1.8834558 | 0.05963862 | 0.42650032 |
| ACAP2 | 2031.2278 | -0.2552192 | 0.16740966 | -1.5245188 | 0.12737915 | 0.56740575 |
| PPP1R2 | 472.812119 | -0.0712894 | 0.16189315 | -0.4403488 | 0.65968454 | 0.91099119 |
| MUC20-OT1 | 5542.76703 | -0.1201417 | 0.47592562 | -0.2524379 | 0.80070259 | 0.95370871 |
| TNK2 | 131.225244 | 0.04612152 | 0.28364587 | 0.16260247 | 0.87083144 | 0.97473202 |
| TFRC | 667.305641 | 0.13411981 | 0.33984009 | 0.39465565 | 0.69309704 | 0.92224682 |
| LINC00885 | 89.8963877 | 3.23738789 | 1.19603667 | 2.70676306 | 0.00679427 | 0.16951739 |
| SLC51A | 46.3227013 | 0.01382719 | 0.48089667 | 0.02875292 | 0.97706165 | 0.99433432 |
| PCYT1A | 267.97326 | -0.0219887 | 0.22164268 | -0.099208 | 0.92097315 | 0.98215577 |
| UBXN7 | 770.1342 | 0.00895726 | 0.22082237 | 0.0405632 | 0.96764412 | 0.99248831 |
| RNF168 | 598.154746 | -0.5248518 | 0.23061376 | -2.275891 | 0.02285254 | 0.2983102 |
| WDR53 | 110.016362 | -0.1315769 | 0.31216489 | -0.421498 | 0.67339145 | 0.91636338 |
| FBXO45 | 58.3595167 | -0.0102309 | 0.30139911 | -0.0339446 | 0.97292136 | 0.99327086 |
| PIGX | 173.447549 | 0.07324633 | 0.21575813 | 0.33948354 | 0.73424549 | 0.93552616 |
| PAK2 | 952.245536 | -0.2040934 | 0.1462452 | -1.3955565 | 1.63E-01 | 6.14E-01 |
| SENP5 | 1562.19967 | -0.3365586 | 0.2563195 | -1.3130431 | 0.18916842 | 0.64432929 |
| NCBP2 | 248.628894 | 0.01292956 | 0.28330526 | 0.04563827 | 0.96359857 | 0.99224805 |
| NCBP2-AS2 | 107.68335 | 0.71189304 | 0.54835567 | 1.29823229 | 0.19420753 | 0.64760078 |
| MELTF | 272.376863 | -3.8367702 | 1.02412816 | -3.746377 | 0.00017941 | 0.021499 |
| DLG1 | 3504.04681 | -0.2296094 | 0.16139351 | -1.4226683 | 0.15483234 | 0.60380955 |
| BDH1 | 658.674856 | 0.24918991 | 0.56935458 | 0.43767086 | 0.6616249 | 0.91110616 |
| AC024560.1 | 54.4265126 | -1.4825108 | 2.08116463 | -0.7123467 | 0.47625009 | 0.84141268 |
| RUBCN | 413.728135 | -0.1023443 | 0.1822559 | -0.561542 | 0.57442814 | 0.88026176 |
| FYTTD1 | 508.183644 | -0.1466548 | 0.2834402 | -0.5174099 | 0.60487006 | 0.89044135 |
| LRCH3 | 1672.63361 | -0.1969202 | 0.13301429 | -1.4804437 | 0.13875488 | 0.58243213 |
| IQCG | 296.628441 | -0.3111807 | 0.20867302 | -1.4912358 | 0.13589961 | 0.57776571 |
| RPL35A | 686.128525 | 0.48655655 | 0.43593953 | 1.11611019 | 0.26437497 | 0.71488591 |
| LMLN | 474.340934 | -0.0069911 | 0.26016738 | -0.0268714 | 0.97856234 | 0.99455895 |
| ZNF595 | 395.947088 | 0.084422 | 0.19167919 | 0.44043383 | 0.65962293 | 0.91099119 |
| ZNF718 | 943.210186 | -0.2589126 | 0.17631302 | -1.4684825 | 0.1419732 | 0.5872148 |
| ZNF141 | 763.04768 | 0.04350532 | 0.15259754 | 0.28509841 | 0.77556875 | 0.94680268 |
| AC092574.2 | 140.932043 | 0.20800746 | 0.55501642 | 0.37477712 | 0.70782623 | 0.92577218 |

| | | | | | | |
|---|---|---|---|---|---|---|
| ZNF721 | 2139.92785 | -0.118061 | 0.19625151 | -0.6015802 | 0.54745358 | 0.86859163 |
| PIGG | 440.791409 | -0.2249568 | 0.21668858 | -1.038157 | 0.29919696 | 0.73673761 |
| ATP5ME | 258.775558 | 0.15424073 | 0.29707152 | 0.51920404 | 0.60361846 | 0.89001883 |
| SLC49A3 | 55.7437155 | -0.2693427 | 0.39774406 | -0.6771758 | 0.49829439 | 0.85092921 |
| PCGF3 | 676.324 | -0.3956594 | 0.23387893 | -1.6917275 | 0.09069793 | 0.50267341 |
| AC139887.2 | 1394.32134 | 0.82464053 | 0.34047625 | 2.42202074 | 0.01543447 | 0.25214021 |
| GAK | 840.996315 | -0.3684826 | 0.27576205 | -1.3362338 | 0.18147286 | 0.63616641 |
| TMEM175 | 155.474038 | 0.21232305 | 0.20565091 | 1.03244399 | 0.30186418 | 0.73905643 |
| DGKQ | 78.0968674 | -0.3816229 | 0.49365521 | -0.7730555 | 0.43948951 | 0.82177104 |
| IDUA | 135.625291 | -0.1113453 | 0.34749303 | -0.3204246 | 0.74864645 | 0.93746253 |
| FGFRL1 | 124.115639 | -0.4624747 | 0.31028773 | -1.4904706 | 0.13610053 | 0.57783867 |
| RNF212 | 525.8986 | -1.5423917 | 0.66685158 | -2.312946 | 0.02072561 | 0.28817321 |
| AC092535.3 | 1619.64124 | -0.7890992 | 0.54050208 | -1.4599373 | 0.14430731 | 0.58995828 |
| AC092535.1 | 48.9055662 | -0.7136012 | 0.79442755 | -0.8982584 | 0.36904782 | 0.78095617 |
| SPON2 | 134.185465 | -0.6334419 | 0.61273842 | -1.0337885 | 0.30123507 | 0.73820989 |
| CTBP1-AS | 179.959062 | 0.01818164 | 0.52386837 | 0.03470651 | 0.97231377 | 0.99327086 |
| CTBP1 | 445.144339 | -0.0323203 | 0.23076398 | -0.1400578 | 0.8886143 | 0.97729111 |
| CTBP1-DT | 141.182582 | -0.3004283 | 0.27604248 | -1.0883408 | 0.27644469 | 0.72245583 |
| MAEA | 665.770897 | 0.11281762 | 0.16811137 | 0.67108859 | 0.50216409 | 0.85510965 |
| UVSSA | 500.359479 | -0.6725083 | 0.20698896 | -3.2490056 | 0.00115809 | 0.06094516 |
| FAM53A | 63.461393 | 0.0679983 | 0.40322747 | 0.16863508 | 0.86608369 | 0.97379123 |
| SLBP | 418.637221 | -0.085984 | 0.32518456 | -0.264416 | 0.79145935 | 0.95178079 |
| TMEM129 | 164.767748 | 0.50187683 | 0.33327912 | 1.50587541 | 0.13209916 | 0.57606398 |
| FGFR3 | 227.498416 | 1.00476434 | 0.71290631 | 1.40939185 | 1.59E-01 | 0.60804051 |
| LETM1 | 356.523761 | -0.0654682 | 0.21863495 | -0.2994405 | 0.76460397 | 0.94256992 |
| NSD2 | 1379.45194 | -0.1082322 | 0.18490682 | -0.5853338 | 0.55832333 | 0.87291524 |
| NELFA | 636.488778 | -0.0516264 | 0.23156261 | -0.2229479 | 0.82357608 | 0.96135914 |
| C4orf48 | 108.112842 | 0.08711077 | 0.7299708 | 0.11933459 | 0.90501028 | 0.97992575 |
| NAT8L | 187.548347 | 0.80510049 | 0.71293675 | 1.12927337 | 0.25878253 | 0.70918225 |
| POLN | 262.178645 | -0.5962182 | 0.37533825 | -1.5884823 | 0.11217733 | 0.53638156 |
| MXD4 | 526.78885 | 0.211769 | 0.28408148 | 0.7454516 | 0.45599876 | 0.83148642 |
| ZFYVE28 | 933.830007 | -0.2394071 | 0.28970277 | -0.8263886 | 0.4085837 | 0.80588056 |
| AL158068.2 | 66.0967673 | -0.3051361 | 0.56244265 | -0.5425194 | 0.58746071 | 0.88571413 |
| RNF4 | 582.654893 | -0.4657383 | 0.21439304 | -2.1723575 | 0.02982871 | 0.33193548 |
| FAM193A | 1408.18683 | -0.2277144 | 0.17997897 | -1.2652277 | 0.20578973 | 0.65813563 |
| TNIP2 | 162.652349 | 0.23575056 | 0.24891235 | 0.94712278 | 3.44E-01 | 7.67E-01 |
| SH3BP2 | 244.412833 | -0.3054228 | 0.40301498 | -0.7578449 | 4.49E-01 | 8.26E-01 |
| ADD1 | 1331.56456 | -0.1303466 | 0.17889851 | -0.7286064 | 0.46624243 | 0.8367113 |
| MFSD10 | 249.084557 | 0.23327482 | 0.41443552 | 0.56287363 | 0.57352094 | 0.87986505 |
| NOP14-AS1 | 92.4284116 | -0.1502585 | 0.31982595 | -0.4698135 | 0.63848829 | 0.90307917 |
| NOP14 | 167.993304 | -0.1709887 | 0.22863112 | -0.7478801 | 0.45453248 | 0.83121393 |
| GRK4 | 275.773914 | -0.1753098 | 0.35236787 | -0.4975193 | 0.61882287 | 0.89608494 |
| HTT | 1091.30679 | -0.3083387 | 0.17225337 | -1.79003 | 0.07344908 | 0.4701377 |
| RGS12 | 503.384243 | -0.8400034 | 0.27771646 | -3.02468 | 0.00248896 | 0.09525007 |
| LRPAP1 | 422.334125 | 0.7931099 | 0.30889881 | 2.56753954 | 0.01024231 | 0.2062358 |
| TMEM128 | 164.623859 | -0.0714334 | 0.23520667 | -0.3037047 | 0.76135291 | 0.94202764 |
| ZBTB49 | 119.471386 | -0.2334161 | 0.28963967 | -0.8058843 | 0.42030957 | 0.8125045 |
| STX18 | 1785.88796 | 0.03864386 | 0.19828061 | 0.1948948 | 0.84547531 | 0.96859352 |
| STX18-AS1 | 841.369159 | -0.4930326 | 0.40388086 | -1.2207378 | 0.2221853 | 0.67560904 |
| STK32B | 3157.75803 | -0.3941153 | 0.48519646 | -0.8122799 | 0.41663104 | 0.80966301 |
| EVC2 | 755.492725 | -0.5954398 | 0.26745196 | -2.226343 | 0.02599122 | 0.31239124 |
| EVC | 1574.5666 | -1.0972072 | 0.41623772 | -2.636011 | 0.0083887 | 0.18436998 |
| CRMP1 | 496.66327 | -1.3910911 | 0.68201607 | -2.0396749 | 0.04138271 | 0.37577267 |
| WFS1 | 71.6939994 | 0.07911893 | 0.29709232 | 0.26631094 | 0.78999975 | 0.95123267 |

| | | | | | | |
|---|---|---|---|---|---|---|
| MAN2B2 | 160.225707 | 0.44162175 | 0.42373241 | 1.04221849 | 0.29731039 | 0.73673761 |
| MRFAP1 | 327.779801 | 0.84656731 | 0.40556507 | 2.08737729 | 0.03685404 | 0.36038369 |
| LINC02482 | 94.5493448 | -0.4561872 | 0.46948091 | -0.9716843 | 0.33120763 | 0.7599308 |
| AC093323.1 | 64.1575086 | -0.1024893 | 0.32303125 | -0.3172737 | 0.75103594 | 0.93775604 |
| MRFAP1L1 | 143.032018 | 0.49144095 | 0.30572718 | 1.60744933 | 0.10795584 | 0.53063658 |
| BLOC1S4 | 88.2132456 | 0.53124787 | 0.40504499 | 1.31157743 | 0.18966275 | 0.64447541 |
| KIAA0232 | 1935.6934 | -0.0340421 | 0.18252193 | -0.1865096 | 0.85204518 | 0.97076831 |
| TBC1D14 | 1732.88525 | 0.40938367 | 0.29523917 | 1.38661708 | 0.16555856 | 0.61845672 |
| TADA2B | 118.927953 | 0.44415711 | 0.31654939 | 1.40312104 | 0.16058075 | 0.61166537 |
| GRPEL1 | 196.953991 | 0.24966076 | 0.36058233 | 0.69238213 | 0.48869739 | 0.84569681 |
| SORCS2 | 880.512007 | -0.8472165 | 0.72218443 | -1.1731304 | 0.24074353 | 0.69178991 |
| AFAP1 | 257.289322 | -0.3293929 | 0.30905665 | -1.0658011 | 0.28651359 | 0.72934245 |
| ABLIM2 | 81.0090124 | 0.34479875 | 0.39836126 | 0.86554288 | 0.38674089 | 0.79523542 |
| SH3TC1 | 244.997399 | 1.63843613 | 0.63409639 | 2.58389126 | 0.00976926 | 0.20084545 |
| ACOX3 | 308.237751 | -0.2252979 | 0.22299759 | -1.0103152 | 0.31234431 | 0.74687836 |
| TRMT44 | 262.600955 | -0.1969259 | 0.21864083 | -0.9006824 | 0.36775722 | 0.78095617 |
| SLC2A9 | 269.267505 | -0.186983 | 0.61724209 | -0.302933 | 0.76194093 | 0.94210219 |
| WDR1 | 718.821001 | 0.27072439 | 0.2912501 | 0.92952548 | 0.35261683 | 0.77210174 |
| ZNF518B | 98.1221138 | 0.0645768 | 0.38253196 | 0.16881412 | 0.86594285 | 0.97379123 |
| CLNK | 41.5891405 | -0.2704142 | 0.4072424 | -0.6640129 | 0.50668202 | 0.85626196 |
| HS3ST1 | 562.219729 | -0.2874493 | 0.60436147 | -0.4756249 | 6.34E-01 | 9.01E-01 |
| AC005699.1 | 1003.74798 | -0.0199102 | 0.59753532 | -0.0333205 | 0.97341898 | 0.99327086 |
| RAB28 | 1993.6472 | -0.1367486 | 0.18437837 | -0.7416735 | 0.45828516 | 0.8331421 |
| BOD1L1 | 1358.11589 | -0.4172053 | 0.30361611 | -1.374121 | 0.16940414 | 0.62417273 |
| LINC01182 | 1718.45095 | 0.10718053 | 0.58988152 | 0.1816984 | 0.85581942 | 0.97118784 |
| AC073848.1 | 40.3251354 | 0.43835582 | 0.59603148 | 0.73545749 | 0.46206092 | 0.83531269 |
| AC092546.1 | 252.802757 | 1.00987999 | 0.74601516 | 1.35369902 | 0.17583242 | 0.62957039 |
| LINC00504 | 147.020394 | -1.0781655 | 0.66735892 | -1.6155706 | 0.10618718 | 0.52735777 |
| CPEB2-DT | 234.393469 | -1.1219065 | 0.41833645 | -2.6818283 | 0.0073221 | 0.17305714 |
| CPEB2 | 4404.05896 | 0.10111088 | 0.2974812 | 0.33988997 | 0.73393939 | 0.93552616 |
| AC098829.1 | 1142.60755 | -0.4396114 | 0.45831588 | -0.9591887 | 0.33746368 | 0.76409325 |
| C1QTNF7 | 213.850312 | -0.1650138 | 0.89849047 | -0.1836568 | 0.85428273 | 0.97085267 |
| CC2D2A | 717.065715 | -0.0105458 | 0.33246209 | -0.0317202 | 0.97469522 | 0.99402649 |
| FBXL5 | 1310.64315 | -0.0779577 | 0.17047368 | -0.4573006 | 0.64745503 | 0.90662922 |
| FAM200B | 364.154138 | 0.19046034 | 0.22326004 | 0.85308746 | 0.3936108 | 0.79843808 |
| PROM1 | 2052.06394 | -0.9823183 | 0.41681454 | -2.3567276 | 0.01843677 | 0.27301711 |
| TAPT1 | 1139.42918 | -0.216877 | 0.19726925 | -1.0993957 | 0.27159549 | 0.72040597 |
| TAPT1-AS1 | 2924.43854 | -0.042448 | 0.22144267 | -0.1916883 | 8.48E-01 | 9.70E-01 |
| AC097515.1 | 663.944843 | -0.2418372 | 0.5342023 | -0.4527071 | 0.65075968 | 0.90825486 |
| LDB2 | 217.212702 | -0.2685183 | 0.53431982 | -0.5025424 | 0.61528606 | 0.8949877 |
| QDPR | 103.300234 | 0.13935041 | 0.39051858 | 0.35683426 | 0.72121589 | 0.93044898 |
| LAP3 | 320.070048 | 0.69840723 | 0.5981543 | 1.16760379 | 0.24296662 | 0.69436307 |
| AC006160.1 | 80.1922238 | -0.2728593 | 0.5740101 | -0.4753562 | 0.63453308 | 0.90153248 |
| MED28 | 388.607449 | -0.1707449 | 0.14356095 | -1.189355 | 0.23429999 | 0.68546368 |
| FAM184B | 81.7541166 | 0.66227594 | 0.45711534 | 1.44881582 | 0.14738902 | 0.59373375 |
| DCAF16 | 414.439888 | 0.04199234 | 0.21538129 | 0.19496744 | 0.84541843 | 0.96859352 |
| LCORL | 1648.32599 | -0.2322443 | 0.23646242 | -0.9821616 | 0.32602025 | 0.75550719 |
| AC024230.1 | 142.643356 | 0.75987375 | 0.55797611 | 1.36183921 | 0.17324863 | 0.6286116 |
| SLIT2 | 813.633017 | 0.46646489 | 0.64778251 | 0.7200949 | 0.47146656 | 0.83921324 |
| PACRGL | 674.398225 | -0.2800384 | 0.22847255 | -1.2256982 | 0.22031226 | 0.67302974 |
| KCNIP4 | 84786.6426 | 0.78524939 | 0.53441196 | 1.46937092 | 0.14173221 | 0.5872148 |
| AC110296.1 | 97.2564257 | 0.83045358 | 0.82620644 | 1.00514053 | 0.31482919 | 0.74785672 |
| ADGRA3 | 1640.06756 | -0.4330094 | 0.28470554 | -1.5209026 | 0.12828428 | 0.56762585 |
| AC093607.1 | 78.3725903 | 0.39507171 | 0.38627167 | 1.02278201 | 0.30641093 | 0.74295884 |

| | | | | | | |
|---|---|---|---|---|---|---|
| PPARGC1A | 7234.8801 | 0.10040966 | 0.42872465 | 0.23420548 | 0.81482546 | 0.95872901 |
| DHX15 | 1849.41665 | -0.3594696 | 0.23091081 | -1.5567467 | 0.11953064 | 0.55224912 |
| SOD3 | 760.357734 | 1.37269745 | 0.95362515 | 1.43945181 | 1.50E-01 | 0.59663305 |
| CCDC149 | 494.364225 | -0.3747425 | 0.29875128 | -1.2543629 | 0.20971013 | 0.66237713 |
| SEPSECS | 420.000228 | -0.3866084 | 0.20950821 | -1.845314 | 0.06499188 | 0.44092331 |
| AC104662.2 | 137.302129 | -0.2428635 | 0.3522221 | -0.6895182 | 0.49049725 | 0.84614483 |
| PI4K2B | 60.9774377 | 0.21286321 | 0.3831776 | 0.55552102 | 0.57853833 | 0.8821758 |
| ZCCHC4 | 237.319795 | -0.4777252 | 0.2504667 | -1.9073401 | 0.05647656 | 0.41843207 |
| ANAPC4 | 1165.40272 | -0.210482 | 0.4222844 | -0.4984366 | 0.61817631 | 0.89599505 |
| SLC34A2 | 867.013449 | 0.4456796 | 0.72714676 | 0.61291561 | 5.40E-01 | 8.66E-01 |
| SEL1L3 | 424.622915 | -1.6523377 | 0.44314778 | -3.7286382 | 1.93E-04 | 2.23E-02 |
| SMIM20 | 169.323191 | -0.6763174 | 0.33497579 | -2.0190038 | 0.04348682 | 0.38387647 |
| RBPJ | 1733.21993 | -0.058014 | 0.19623509 | -0.2956352 | 0.76750866 | 0.94316227 |
| TBC1D19 | 1214.65808 | 0.22064507 | 0.21618106 | 1.02064939 | 0.30742058 | 0.74341546 |
| STIM2 | 836.244873 | 0.55020591 | 0.33085345 | 1.6629898 | 0.09631448 | 0.51206987 |
| PCDH7 | 305.135545 | -0.0992334 | 0.85830489 | -0.1156156 | 0.90795721 | 0.98047754 |
| AC016687.3 | 108.288137 | 0.20902792 | 0.92266105 | 0.22654898 | 8.21E-01 | 9.60E-01 |
| LINC02484 | 364.30876 | -0.7309106 | 0.84548657 | -0.8644852 | 3.87E-01 | 7.96E-01 |
| ARAP2 | 1572.03062 | 0.08621922 | 0.2407461 | 0.35813342 | 0.72024347 | 0.93001079 |
| C4orf19 | 4196.1255 | -0.0601512 | 0.3107439 | -0.1935717 | 0.84651125 | 0.96883382 |
| RELL1 | 611.814784 | -0.2160004 | 0.23136322 | -0.9335989 | 0.35051085 | 0.77044053 |
| PGM2 | 203.817665 | -0.0719047 | 0.36538246 | -0.1967931 | 0.84398947 | 0.96831902 |
| TBC1D1 | 4084.50813 | 0.22755697 | 0.31235974 | 0.72850926 | 0.4663019 | 0.8367113 |
| KLF3 | 886.23673 | -0.6672636 | 0.21642368 | -3.0831356 | 2.05E-03 | 8.76E-02 |
| TLR1 | 66.8337184 | 0.26390424 | 0.48102623 | 0.54862754 | 0.58326108 | 0.88379959 |
| FAM114A1 | 336.924415 | -0.7868623 | 0.35618093 | -2.2091645 | 0.0271632 | 0.31836623 |
| KLHL5 | 1177.55155 | 0.36041804 | 0.22875867 | 1.57553831 | 0.11513224 | 0.54371891 |
| WDR19 | 921.992469 | -0.3638734 | 0.18854711 | -1.9298806 | 0.05362163 | 0.41275533 |
| RFC1 | 732.165861 | -0.0610342 | 0.19689513 | -0.3099831 | 0.75657382 | 0.93989203 |
| RPL9 | 129.441633 | 0.29828063 | 0.3350884 | 0.89015505 | 0.37338264 | 0.78277446 |
| LIAS | 344.944435 | -0.0448167 | 0.24609741 | -0.1821095 | 0.8554968 | 0.97118784 |
| UGDH | 358.450329 | -0.1085024 | 0.36783216 | -0.294978 | 0.76801071 | 0.94316227 |
| UGDH-AS1 | 607.831294 | 0.16995322 | 0.3064676 | 0.55455526 | 0.57919889 | 0.8821758 |
| SMIM14 | 1217.49915 | -0.095549 | 0.12062908 | -0.792089 | 0.4283088 | 0.81497133 |
| AC108471.2 | 50.3413154 | -0.5656184 | 0.43136082 | -1.311242 | 0.18977603 | 0.64450954 |
| UBE2K | 2774.08612 | -0.1906566 | 0.18292169 | -1.0422852 | 0.29727948 | 0.73673761 |
| PDS5A | 2243.66177 | -0.2622316 | 0.16434035 | -1.5956614 | 0.1105644 | 0.53393773 |
| N4BP2 | 644.275038 | 0.16824264 | 0.14533446 | 1.15762384 | 0.24701758 | 0.69751478 |
| RBM47 | 9608.24714 | -0.5663953 | 0.22761114 | -2.4884341 | 0.0128307 | 0.23224886 |
| NSUN7 | 763.624172 | -0.1405323 | 0.24093124 | -0.583288 | 0.55969947 | 0.87300289 |
| APBB2 | 5283.88109 | -0.5078955 | 0.21119918 | -2.4048177 | 0.01618054 | 0.25504618 |
| AC131953.1 | 41.2019148 | -0.4853644 | 0.45861383 | -1.0583291 | 0.28990544 | 0.73204984 |
| UCHL1 | 769.442112 | 1.16537589 | 0.75142458 | 1.5508887 | 0.12092836 | 0.55498372 |
| LIMCH1 | 3208.03834 | 0.06800282 | 0.37532751 | 0.18118262 | 0.85622423 | 0.97118784 |
| TMEM33 | 415.95255 | -0.1090787 | 0.32786658 | -0.3326923 | 0.73936653 | 0.93552616 |
| SLC30A9 | 1617.42375 | -0.0548689 | 0.12279629 | -0.4468283 | 0.65499906 | 0.90902447 |
| AC024022.1 | 824.242917 | 1.55372587 | 0.80753678 | 1.92403109 | 0.05435069 | 0.41546193 |
| SHISA3 | 142.663278 | 0.06344852 | 0.66347396 | 0.09563077 | 0.92381383 | 0.98215577 |
| ATP8A1 | 1224.74778 | -0.5218495 | 0.30301182 | -1.7222085 | 0.08503176 | 0.49231347 |
| KCTD8 | 895.066074 | 1.37335095 | 0.55994025 | 2.4526741 | 0.01417987 | 0.24646911 |
| YIPF7 | 47.5558133 | 0.05384203 | 0.40909432 | 0.13161275 | 0.8952906 | 0.97788747 |
| GUF1 | 364.61025 | -0.1656596 | 0.24479587 | -0.6767255 | 0.49858015 | 0.85098142 |
| GNPDA2 | 297.545949 | 0.18761484 | 0.30082363 | 0.62367055 | 0.53284397 | 0.86397197 |
| GABRA2 | 174.340408 | 2.09054664 | 0.69193419 | 3.02130849 | 0.00251685 | 0.09525007 |

| | | | | | | |
|---|---|---|---|---|---|---|
| GABRB1 | 489.547506 | 1.40183331 | 0.55728273 | 2.51547954 | 0.01188706 | 0.22355405 |
| COMMD8 | 94.4197505 | 0.29736838 | 0.37320418 | 0.7967981 | 0.4255683 | 0.81466629 |
| ATP10D | 933.689157 | 0.09261383 | 0.26580675 | 0.34842543 | 0.72752071 | 0.93420128 |
| CORIN | 85.8177676 | -0.4422967 | 0.52487751 | -0.8426665 | 0.39941499 | 0.80174499 |
| AC107068.2 | 191.047857 | 0.30970508 | 0.62918318 | 0.49223357 | 0.62255423 | 0.89835355 |
| NFXL1 | 604.208781 | 0.09950682 | 0.21235531 | 0.46858647 | 0.63936525 | 0.90330608 |
| NIPAL1 | 330.103496 | 0.29719496 | 0.30619766 | 0.9705984 | 0.3317483 | 0.76035039 |
| CNGA1 | 117.700456 | 0.65357133 | 0.60556692 | 1.07927184 | 0.28046656 | 0.72527072 |
| TXK | 442.721409 | 0.19469782 | 0.44928078 | 0.43335443 | 0.6647573 | 0.91220795 |
| TEC | 511.658721 | -0.2100685 | 0.25516783 | -0.823256 | 0.41036241 | 0.80655687 |
| SLAIN2 | 947.01826 | -0.0994431 | 0.16591476 | -0.5993623 | 0.54893128 | 0.86904066 |
| FRYL | 2920.51012 | -0.1939262 | 0.13631328 | -1.4226506 | 0.15483748 | 0.60380955 |
| OCIAD1 | 1484.78148 | 0.0687573 | 0.14060546 | 0.48900871 | 0.62483553 | 0.89873262 |
| OCIAD2 | 260.615999 | -0.226253 | 0.24443683 | -0.9256095 | 0.35464896 | 0.77251948 |
| DCUN1D4 | 1209.5704 | 0.15065627 | 0.16347048 | 0.9216115 | 0.35673126 | 0.77411555 |
| SGCB | 158.236475 | 0.32276441 | 0.26216432 | 1.23115307 | 0.21826562 | 0.67173126 |
| SPATA18 | 44.3282912 | 1.29509773 | 0.56965609 | 2.273473 | 0.02299769 | 0.2983102 |
| USP46 | 320.768291 | -0.2293414 | 0.23505991 | -0.9756721 | 0.32922697 | 0.75815187 |
| SCFD2 | 1529.96559 | -0.5190066 | 0.34824903 | -1.4903318 | 0.13613702 | 0.57783867 |
| AC058822.1 | 5971.79728 | -0.2985585 | 0.72165482 | -0.4137138 | 0.67908375 | 0.91822574 |
| LNX1 | 4596.82291 | 0.35119216 | 0.40564011 | 0.86577276 | 0.38661479 | 0.79523542 |
| CHIC2 | 1137.10353 | -0.2721033 | 0.18177762 | -1.4969019 | 0.13441879 | 0.57708028 |
| SRD5A3 | 69.0996195 | 0.09909852 | 0.58017459 | 0.17080809 | 0.86437467 | 0.97361004 |
| TMEM165 | 1682.81097 | 0.01400475 | 0.18840179 | 0.07433448 | 0.94074424 | 0.98715467 |
| CLOCK | 1148.28397 | -0.0201502 | 0.22463668 | -0.0897012 | 0.92852465 | 0.98314146 |
| EXOC1 | 1384.90085 | -0.1994301 | 0.22099678 | -0.9024119 | 0.3668381 | 0.78095617 |
| CEP135 | 412.55957 | -0.4274994 | 0.2342175 | -1.8252239 | 0.06796724 | 0.45250883 |
| AASDH | 719.958839 | -0.0418667 | 0.25224039 | -0.1659794 | 0.86817312 | 0.97402505 |
| PPAT | 254.105352 | -0.7717523 | 0.36581234 | -2.1096946 | 0.03488467 | 0.35358029 |
| PAICS | 366.818921 | -0.2056594 | 0.30079328 | -0.6837233 | 0.49414992 | 0.84882719 |
| SRP72 | 541.658766 | 0.17663914 | 0.17370241 | 1.01690665 | 0.30919784 | 0.74420449 |
| REST | 959.536439 | 0.01292415 | 0.21947291 | 0.05888723 | 0.95304193 | 0.98949465 |
| NOA1 | 139.080093 | -0.1767694 | 0.26389063 | -0.6698585 | 0.50294802 | 0.85525037 |
| POLR2B | 731.914832 | -0.3150731 | 0.18545057 | -1.6989598 | 0.08932675 | 0.49927552 |
| IGFBP7 | 9523.13073 | 0.26076601 | 0.44731587 | 0.5829572 | 0.55992211 | 0.87300289 |
| ADGRL3 | 423.675526 | -1.9804871 | 0.58179418 | -3.4041026 | 0.00066382 | 0.04534035 |
| LINC02232 | 52.7586048 | 1.26428552 | 0.90267587 | 1.40059745 | 0.16133448 | 0.61272778 |
| AC116049.2 | 73.8055687 | 0.51530738 | 0.41235075 | 1.24968216 | 0.21141567 | 0.66484951 |
| AC104806.2 | 217.461801 | 0.64446458 | 0.38892209 | 1.65705317 | 0.09750873 | 0.51355946 |
| CENPC | 883.755397 | -0.2504585 | 0.17277445 | -1.449627 | 0.14716258 | 0.59373375 |
| UBA6 | 1281.50363 | -0.2760447 | 0.25770444 | -1.0711677 | 0.28409403 | 0.72738535 |
| UBA6-AS1 | 3176.6753 | -0.4676803 | 0.42726562 | -1.0945891 | 0.27369671 | 0.72135661 |
| TMPRSS11D | 64.29443 | 0.54991218 | 0.81129156 | 0.67782313 | 0.49788384 | 0.85092921 |
| YTHDC1 | 1977.81536 | -0.1207342 | 0.21524663 | -0.5609108 | 0.57485836 | 0.88046555 |
| UGT2B7 | 4183.51499 | 0.08638011 | 0.68012215 | 0.12700675 | 0.89893505 | 0.97889101 |
| AC111000.4 | 135.305084 | -0.8379303 | 0.83549391 | -1.0029161 | 0.31590133 | 0.74785672 |
| ENAM | 171.143529 | 0.94340545 | 0.46027848 | 2.04964056 | 0.04039952 | 0.37344844 |
| UTP3 | 125.37083 | 0.58909034 | 0.30894305 | 1.90679268 | 0.05654744 | 0.41843207 |
| RUFY3 | 3139.09083 | -0.5735176 | 0.28497528 | -2.012517 | 0.04416547 | 0.38531062 |
| GRSF1 | 477.399503 | -0.082306 | 0.18816226 | -0.4374204 | 0.66180649 | 0.91110616 |
| MOB1B | 1196.14346 | 0.16211711 | 0.27480363 | 0.589938 | 0.55523222 | 0.87184917 |
| SLC4A4 | 438.240646 | 0.81640844 | 0.45205777 | 1.80598252 | 0.07092107 | 0.46359457 |
| ADAMTS3 | 156.11889 | -0.4637337 | 0.72544703 | -0.6392385 | 0.52266777 | 0.86006369 |
| COX18 | 144.743155 | -0.4172026 | 0.22483886 | -1.8555627 | 0.06351592 | 0.43779712 |

| | | | | | | |
|---|---|---|---|---|---|---|
| ANKRD17 | 6561.16528 | -0.0177556 | 0.23827998 | -0.0745159 | 0.94059989 | 0.98715467 |
| AC053527.1 | 1202.23314 | 0.51558177 | 0.47199464 | 1.09234666 | 0.27468076 | 0.72164296 |
| RASSF6 | 490.467698 | 1.17793438 | 0.43955112 | 2.67985756 | 0.00736535 | 0.17305714 |
| MTHFD2L | 765.830752 | 0.78243673 | 0.33658785 | 2.32461373 | 0.02009262 | 0.28397329 |
| BTC | 224.599534 | -0.0765676 | 0.49492265 | -0.1547062 | 0.87705295 | 0.97639142 |
| RCHY1 | 578.193146 | 0.2239855 | 0.28246268 | 0.79297378 | 0.4277931 | 0.81466629 |
| THAP6 | 423.720553 | -0.1235314 | 0.20460041 | -0.6037691 | 0.54599716 | 0.86794271 |
| CDKL2 | 431.307263 | 0.1468145 | 0.46286536 | 0.3171862 | 0.75110232 | 0.93775604 |
| G3BP2 | 624.382 | 0.20581335 | 0.18626175 | 1.10496846 | 0.26917326 | 0.71818262 |
| USO1 | 1641.03517 | -0.1141181 | 0.19574567 | -0.5829915 | 0.55989905 | 0.87300289 |
| NAAA | 411.419623 | 1.08201371 | 0.42522532 | 2.54456555 | 0.01094138 | 0.21301305 |
| SDAD1 | 418.028228 | -0.1111524 | 0.21994891 | -0.5053555 | 0.61330914 | 0.89441126 |
| NUP54 | 745.365326 | -0.2337297 | 0.22046399 | -1.0601718 | 0.28906644 | 0.73189164 |
| SCARB2 | 827.847381 | 0.22428432 | 0.25047162 | 0.89544802 | 0.37054764 | 0.78104287 |
| FAM47E-STB | 59.246863 | -0.2956863 | 0.38546528 | -0.7670894 | 0.44302837 | 0.82310928 |
| SHROOM3 | 5477.98426 | 0.67590047 | 0.30963241 | 2.18291252 | 0.02904225 | 0.32900439 |
| AC107072.1 | 79.5365681 | 0.45277835 | 0.74182628 | 0.61035631 | 0.54162581 | 0.86582694 |
| 11-Sep | 1249.90663 | -0.4859452 | 0.52573739 | -0.9243116 | 0.35532411 | 0.77296991 |
| CCNI | 1732.11227 | 0.53932336 | 0.25955389 | 2.07788591 | 0.03771987 | 0.36396365 |
| CCNG2 | 656.627662 | 0.44082766 | 0.27991976 | 1.57483583 | 0.11529434 | 0.54409539 |
| CNOT6L | 1678.39224 | -0.2032857 | 0.33596009 | -0.6050888 | 0.54511999 | 0.86698116 |
| MRPL1 | 822.6463 | -0.0646121 | 0.21094946 | -0.3062918 | 0.7593825 | 0.94133292 |
| FRAS1 | 4070.37287 | 0.1076182 | 0.59108019 | 0.1820704 | 0.85552748 | 0.97118784 |
| ANXA3 | 272.746474 | 0.90052678 | 0.70427579 | 1.27865644 | 0.20101807 | 0.6543242 |
| BMP2K | 887.939831 | -0.4770791 | 0.26535557 | -1.797886 | 0.07219507 | 0.4679905 |
| PAQR3 | 166.157245 | -0.2248151 | 0.28855645 | -0.7791026 | 0.43591925 | 0.8196523 |
| LINC01088 | 109.615636 | -0.7857039 | 0.42415858 | -1.8523824 | 0.06397093 | 0.43872009 |
| ANTXR2 | 54.1260916 | -1.6577655 | 0.68514339 | -2.4195891 | 0.01553805 | 0.25214021 |
| CFAP299 | 283.859151 | -0.8633367 | 0.76975149 | -1.1215784 | 0.26204174 | 0.712756 |
| PRKG2 | 61.9175253 | -0.3418342 | 0.73132133 | -0.46742 | 0.6401994 | 0.90346088 |
| RASGEF1B | 3721.22569 | -0.3964556 | 0.6182155 | -0.6412903 | 0.52133408 | 0.85943874 |
| HNRNPD | 1756.81356 | 0.00023233 | 0.11849617 | 0.00196061 | 0.99843566 | 0.99972915 |
| AC124016.1 | 78.1867545 | -0.1396199 | 0.41390453 | -0.337324 | 7.36E-01 | 0.93552616 |
| HNRNPDL | 1363.73688 | -0.1619949 | 0.22841051 | -0.709227 | 0.47818365 | 0.84199814 |
| ENOPH1 | 264.912893 | -0.0539885 | 0.25431919 | -0.2122865 | 0.83188356 | 0.96414407 |
| TMEM150C | 369.017747 | 0.13815272 | 0.2987206 | 0.46248139 | 0.64373614 | 0.90478945 |
| SCD5 | 1672.04611 | 0.38640685 | 0.28996614 | 1.33259302 | 0.1826654 | 0.63741052 |
| SEC31A | 2349.95022 | -0.0915263 | 0.15208873 | -0.6017954 | 0.54731032 | 0.86859163 |
| THAP9 | 112.41173 | 0.07251756 | 0.32834656 | 0.22085677 | 0.82520396 | 0.96146415 |
| LIN54 | 372.631814 | -0.3109919 | 0.15452574 | -2.0125574 | 0.04416121 | 0.38531062 |
| COPS4 | 412.360181 | -0.1166574 | 0.18707513 | -0.623586 | 0.53289948 | 0.86397197 |
| AC114781.2 | 61.7937105 | -0.5561975 | 0.47323035 | -1.1753209 | 0.23986634 | 0.69118554 |
| COQ2 | 94.8903914 | 0.12815571 | 0.40235104 | 0.31851717 | 0.75009267 | 0.93775604 |
| HPSE | 85.244545 | 0.31503591 | 0.69857132 | 0.45097171 | 0.65200994 | 0.90838511 |
| HELQ | 562.170432 | -0.1057133 | 0.205463 | -0.5145128 | 0.60689353 | 0.89174148 |
| MRPS18C | 207.476237 | 0.07042492 | 0.25007299 | 0.28161745 | 0.77823686 | 0.94694572 |
| ABRAXAS1 | 225.407105 | -0.2225691 | 0.28135401 | -0.7910644 | 0.42890644 | 0.8155211 |
| GPAT3 | 2349.33552 | -1.2779214 | 0.37570842 | -3.4013647 | 6.71E-04 | 0.04534035 |
| CDS1 | 658.934755 | 0.08929047 | 0.24036645 | 0.37147643 | 0.71028271 | 0.92619854 |
| WDFY3 | 5034.78904 | -0.2607284 | 0.22648627 | -1.1511885 | 0.24965469 | 0.70032501 |
| WDFY3-AS1 | 61.502138 | -0.0968109 | 0.73500596 | -0.1317144 | 0.89521018 | 0.97788747 |
| ARHGAP24 | 12617.9106 | 0.3281183 | 0.27729518 | 1.18328169 | 0.23669753 | 0.68878143 |
| MAPK10 | 7040.81042 | 0.0676364 | 0.21758811 | 0.31084605 | 0.75591766 | 0.93957997 |
| AC104827.1 | 150.455922 | -0.2289802 | 0.5621161 | -0.4073539 | 0.68374806 | 0.9200778 |

| | | | | | | |
|---|---|---|---|---|---|---|
| PTPN13 | 5101.62269 | -0.0220026 | 0.2378638 | -0.092501 | 9.26E-01 | 9.82E-01 |
| AC093827.5 | 261.554839 | -0.1175643 | 0.38628672 | -0.3043446 | 0.7608654 | 0.94202764 |
| AFF1 | 7420.79631 | -0.1494905 | 0.21471184 | -0.6962378 | 0.48627989 | 0.8447301 |
| KLHL8 | 713.140033 | -0.1356975 | 0.3602406 | -0.3766858 | 0.70640714 | 0.92556842 |
| HSD17B11 | 172.507697 | -0.0726153 | 0.43098325 | -0.1684876 | 0.86619969 | 0.97383877 |
| NUDT9 | 397.733727 | -0.0877947 | 0.20417634 | -0.4299944 | 0.66719971 | 0.91355251 |
| SPARCL1 | 107.15341 | 0.03850488 | 0.52917833 | 0.07276353 | 0.94199429 | 0.98772472 |
| AC093895.1 | 275.484106 | 0.68190283 | 0.8975981 | 0.75969728 | 0.44743556 | 0.82578961 |
| SPP1 | 54676.147 | 0.20240689 | 0.66934106 | 0.30239724 | 7.62E-01 | 9.42E-01 |
| PKD2 | 3592.76213 | -0.3010295 | 0.23959669 | -1.2564011 | 0.20897059 | 0.66146477 |
| PPM1K | 1167.41799 | 0.51957809 | 0.41134153 | 1.26313064 | 2.07E-01 | 0.65883032 |
| AC107067.1 | 55.2499898 | -0.3966122 | 0.71439111 | -0.5551752 | 5.79E-01 | 0.8821758 |
| HERC6 | 371.903951 | -0.344868 | 0.91550389 | -0.3766974 | 0.70639848 | 0.92556842 |
| HERC5 | 258.197091 | 0.06501656 | 0.66166295 | 0.09826236 | 0.92172397 | 0.98215577 |
| PYURF | 136.075894 | 0.74890835 | 0.30999947 | 2.41583752 | 0.01569906 | 0.25227487 |
| AC098582.1 | 108.095474 | 0.0740338 | 0.53642484 | 0.13801338 | 0.89022984 | 0.97729111 |
| HERC3 | 1237.053 | -0.1805136 | 0.23520713 | -0.7674664 | 0.44280425 | 0.82301411 |
| FAM13A-AS1 | 460.310998 | -0.3414982 | 0.27584121 | -1.2380246 | 0.21570694 | 0.66885336 |
| FAM13A | 3773.62403 | -0.0234563 | 0.38909485 | -0.0602844 | 0.95192915 | 0.98938795 |
| GPRIN3 | 756.949425 | 0.79028784 | 0.52720778 | 1.4990064 | 0.13387197 | 0.57708028 |
| CCSER1 | 31289.809 | 0.69413584 | 0.3316456 | 2.09300482 | 0.03634872 | 0.35881276 |
| AC004054.1 | 71.5973106 | 0.74727733 | 0.53736627 | 1.3906294 | 0.16433784 | 0.61616193 |
| GRID2 | 46.967277 | -0.2894715 | 0.45065839 | -0.6423302 | 0.5206588 | 0.85943874 |
| SMARCAD1 | 1280.83055 | -0.185603 | 0.14788396 | -1.255058 | 0.2094577 | 0.66216422 |
| PDLIM5 | 2261.26467 | 0.16894495 | 0.18197414 | 0.92840087 | 0.35319967 | 0.77216708 |
| BMPR1B | 99.605337 | -0.2050925 | 0.4529568 | -0.4527861 | 0.6507028 | 0.90825486 |
| UNC5C | 111.087227 | 1.05561308 | 0.8444545 | 1.25005323 | 0.2112801 | 0.66484951 |
| STPG2 | 5690.76324 | -0.5755638 | 0.29801817 | -1.9313043 | 5.34E-02 | 4.12E-01 |
| STPG2-AS1 | 160.379436 | 0.24384015 | 0.60780427 | 0.40118202 | 0.68828611 | 0.92131853 |
| RAP1GDS1 | 1412.92796 | 0.00881657 | 0.1912655 | 0.04609597 | 0.96323376 | 0.99224805 |
| TSPAN5 | 102.855626 | 0.25773297 | 0.4683226 | 0.55033211 | 0.5820916 | 0.88336948 |
| EIF4E | 671.891028 | -0.036432 | 0.26401418 | -0.1379928 | 0.89024613 | 0.97729111 |
| METAP1 | 353.039473 | -0.0006953 | 0.23055455 | -0.0030157 | 0.99759385 | 0.99956134 |
| ADH5 | 442.860021 | 0.32225931 | 0.2151422 | 1.49788978 | 0.13416189 | 0.57708028 |
| AP002026.1 | 213.453931 | -0.823109 | 0.6084573 | -1.3527803 | 0.17612583 | 0.62957039 |
| TRMT10A | 118.478439 | -0.0857701 | 0.41476629 | -0.2067913 | 0.83617285 | 0.96511435 |
| MTTP | 40.0766263 | -0.4887776 | 0.62919314 | -0.7768324 | 0.43725766 | 0.82013459 |
| LAMTOR3 | 205.607004 | 0.21816169 | 0.24575208 | 0.88773083 | 0.37468556 | 0.78426576 |
| DNAJB14 | 1203.99389 | 0.03067019 | 0.15335306 | 0.19999721 | 0.84148276 | 0.96661847 |
| H2AFZ | 315.673904 | 0.34083001 | 0.60953209 | 0.55916663 | 0.576048 | 0.88123035 |
| AC097460.1 | 90.5426062 | -0.5079136 | 0.29350454 | -1.7305138 | 0.08353852 | 0.4882568 |
| DDIT4L | 187.181202 | -1.1431831 | 0.83829639 | -1.363698 | 0.17266263 | 0.6286116 |
| EMCN | 81.2554905 | -0.8899587 | 0.54138068 | -1.6438686 | 0.10020335 | 0.51692417 |
| PPP3CA | 3225.56813 | -0.1626085 | 0.30067057 | -0.5408193 | 0.58863213 | 0.88571413 |
| BANK1 | 164.455467 | -0.2309948 | 0.61842601 | -0.3735205 | 0.70876107 | 0.92577218 |
| AP002075.1 | 148.474417 | 0.32908476 | 0.68182483 | 0.48265294 | 0.62934218 | 0.90005881 |
| SLC39A8 | 163.834499 | 0.17368358 | 0.51436796 | 0.33766408 | 0.73561635 | 0.93552616 |
| NFKB1 | 1293.67622 | 0.0100627 | 0.18484305 | 0.05443914 | 0.9565853 | 0.99065996 |
| MANBA | 652.687779 | -0.0404805 | 0.21471389 | -0.1885321 | 0.85045955 | 0.97056586 |
| UBE2D3 | 2825.75949 | 0.07714493 | 0.1581497 | 0.48779685 | 0.62569374 | 0.89873262 |
| AC018797.2 | 114.425289 | -0.3516387 | 0.33482449 | -1.050218 | 0.29361788 | 0.73527014 |
| CISD2 | 259.595356 | 0.57305821 | 0.24518745 | 2.33722484 | 0.0194275 | 0.28045661 |
| SLC9B1 | 178.427539 | -0.0624855 | 0.52619584 | -0.1187495 | 0.90547379 | 0.9799593 |
| SLC9B2 | 89.6850209 | 0.62078723 | 0.29459458 | 2.10725947 | 0.03509509 | 0.35387107 |

| | | | | | | |
|---|---|---|---|---|---|---|
| BDH2 | 137.655137 | -0.2555994 | 0.25077873 | -1.0192228 | 0.30809719 | 0.74372365 |
| AC004053.1 | 41.8205427 | 0.47492642 | 0.57712695 | 0.82291499 | 0.41055634 | 0.80655687 |
| AC096577.1 | 44.0296466 | 1.78552932 | 0.7497121 | 2.38161998 | 0.01723667 | 0.26351764 |
| TET2 | 1939.3741 | -0.0584788 | 0.1633022 | -0.3581016 | 0.72026727 | 0.93001079 |
| PPA2 | 1984.24714 | 0.20462352 | 0.20874266 | 0.98026691 | 0.32695438 | 0.75556858 |
| ARHGEF38 | 5184.5471 | -0.1339368 | 0.33253493 | -0.4027752 | 0.68711361 | 0.9208231 |
| INTS12 | 764.010026 | -0.3268986 | 0.29543842 | -1.1064864 | 0.26851603 | 0.71818262 |
| GSTCD | 781.525183 | -0.5801569 | 0.2110025 | -2.7495262 | 0.00596815 | 0.15829975 |
| NPNT | 1296.61935 | 0.58801865 | 0.74871522 | 0.78537024 | 0.43223652 | 0.81773248 |
| AC109361.1 | 81.1654925 | 0.303995 | 0.43835879 | 0.69348444 | 0.48800559 | 0.845164 |
| TBCK | 3518.53992 | 0.09201788 | 0.13866732 | 0.66358737 | 0.50695444 | 0.85626196 |
| AIMP1 | 428.952896 | 0.16148911 | 0.17325288 | 0.93210054 | 0.35128457 | 0.77103986 |
| PAPSS1 | 619.451181 | -0.0663283 | 0.14681153 | -0.4517924 | 0.65141855 | 0.90825486 |
| SGMS2 | 992.174188 | 0.12418441 | 0.33192281 | 0.37413641 | 0.70830283 | 0.92577218 |
| AC096564.1 | 213.672124 | 0.52850121 | 0.49471668 | 1.06829067 | 0.28538942 | 0.72844615 |
| CYP2U1 | 44.6114121 | 1.03623086 | 0.47960251 | 2.1606035 | 0.03072598 | 0.33646028 |
| HADH | 334.127765 | 0.58690663 | 0.24266579 | 2.41858003 | 0.01558122 | 0.25227487 |
| LEF1 | 81.9101901 | -0.9969182 | 0.95247236 | -1.0466636 | 0.29525475 | 0.7364086 |
| RPL34-AS1 | 556.097533 | 0.1873373 | 0.40481049 | 0.46277777 | 0.64352367 | 0.90478945 |
| RPL34 | 697.778797 | 0.4922603 | 0.42943578 | 1.14629549 | 0.25167291 | 0.70276117 |
| OSTC | 317.220984 | 0.33308862 | 0.29898458 | 1.11406623 | 0.26525077 | 0.71512755 |
| COL25A1 | 134.079666 | 0.77842206 | 0.37167389 | 2.09436841 | 0.03622717 | 0.35817721 |
| SEC24B-AS1 | 137.77534 | 0.65644548 | 0.52304931 | 1.25503556 | 0.20946585 | 0.66216422 |
| SEC24B | 1821.60803 | -0.0428734 | 0.16403754 | -0.2613631 | 0.7938125 | 0.95224951 |
| MCUB | 286.407989 | -1.0761804 | 0.35163063 | -3.0605422 | 2.21E-03 | 9.04E-02 |
| CASP6 | 325.278556 | 0.33070325 | 0.17034093 | 1.9414198 | 0.05220738 | 0.40866415 |
| PLA2G12A | 479.782615 | 0.80085076 | 0.18695985 | 4.28354419 | 1.84E-05 | 0.0071255 |
| AC126283.2 | 164.989414 | 0.83308316 | 0.50299468 | 1.65624647 | 0.09767192 | 0.51355946 |
| CFI | 45.9874647 | 0.1541425 | 0.67786233 | 0.227395 | 0.82011661 | 0.96016713 |
| GAR1 | 158.316636 | -0.5347413 | 0.28096307 | -1.9032441 | 0.0570087 | 0.41872623 |
| RRH | 44.2083311 | -0.2603295 | 0.60826836 | -0.4279846 | 0.6686623 | 0.91395501 |
| LRIT3 | 37.1232975 | -0.5113852 | 0.81059286 | -0.6308781 | 0.52812026 | 0.86215467 |
| EGF | 16888.5984 | 0.16069643 | 0.583845 | 0.27523818 | 7.83E-01 | 9.49E-01 |
| ELOVL6 | 1227.77996 | -0.2805956 | 0.20711302 | -1.3547946 | 0.17548302 | 0.62957039 |
| ENPEP | 175.521701 | 0.23642082 | 0.58508211 | 0.40408144 | 0.68615283 | 0.92053037 |
| FAM241A | 195.8995 | 0.40113856 | 0.36257712 | 1.10635377 | 0.26857343 | 0.71818262 |
| AP1AR | 311.879289 | -0.5280491 | 0.27643433 | -1.9102152 | 0.05610551 | 0.41843207 |
| TIFA | 62.4384409 | -0.0409345 | 0.4588547 | -0.0892103 | 0.92891481 | 0.98323312 |
| ALPK1 | 1849.16775 | -0.0893735 | 0.28174467 | -0.3172144 | 0.75108091 | 0.93775604 |
| ZGRF1 | 100.038384 | -1.3266371 | 0.4311644 | -3.0768707 | 0.00209186 | 0.0878448 |
| LARP7 | 708.663518 | 0.01148792 | 0.16430133 | 0.06991983 | 0.94425747 | 0.98813503 |
| ANK2 | 4566.99999 | -1.1244438 | 0.61001997 | -1.8432902 | 0.06528665 | 0.44218726 |
| CAMK2D | 1960.17573 | 0.18251161 | 0.24749804 | 0.73742647 | 0.46086304 | 0.83444983 |
| ARSJ | 464.569654 | 1.29859456 | 0.48925848 | 2.65420963 | 0.00794944 | 0.17918803 |
| UGT8 | 3380.05827 | 0.19724453 | 0.27545092 | 0.71607867 | 4.74E-01 | 8.41E-01 |
| TRAM1L1 | 40.8126464 | 0.16975438 | 0.6244863 | 0.27183043 | 0.78575241 | 0.94978829 |
| LINC01378 | 61.1669907 | -1.1086525 | 0.64116552 | -1.7291206 | 8.38E-02 | 4.89E-01 |
| NDST3 | 129.850682 | -0.7365036 | 0.57339785 | -1.2844548 | 0.19898288 | 0.65113421 |
| SNHG8 | 199.852788 | 0.11311747 | 0.54433358 | 0.20780909 | 0.83537803 | 0.96511435 |
| METTL14 | 407.192712 | -0.0163602 | 0.22163042 | -0.0738173 | 0.94115578 | 0.98732467 |
| SEC24D | 609.471941 | 0.06751542 | 0.3326456 | 0.20296501 | 0.83916238 | 0.966178 |
| USP53 | 4454.39558 | -0.3159879 | 0.37345949 | -0.8461102 | 0.39749129 | 0.80069929 |
| C4orf3 | 408.667928 | 0.72663638 | 0.42713847 | 1.70117289 | 0.08891053 | 0.49869115 |
| PDE5A | 893.364747 | 0.2420998 | 0.33060632 | 0.73229028 | 0.46399141 | 0.83545018 |

| | | | | | | |
|---|---|---|---|---|---|---|
| AC108866.1 | 739.681898 | -0.7959677 | 0.55190785 | -1.4422112 | 0.14924279 | 0.5959335 |
| PRDM5 | 576.637975 | -0.3395713 | 0.26757774 | -1.2690568 | 0.20442081 | 0.65746152 |
| ANXA5 | 875.571969 | 0.52578218 | 0.33063007 | 1.59024309 | 0.11178002 | 0.53607455 |
| EXOSC9 | 130.438328 | -0.2300206 | 0.26734348 | -0.8603937 | 0.38957208 | 0.79681908 |
| BBS7 | 231.894702 | -0.0512644 | 0.32173524 | -0.1593372 | 0.87340323 | 0.97564728 |
| KIAA1109 | 3245.55412 | -0.0898832 | 0.17837823 | -0.5038914 | 0.61433771 | 0.89488165 |
| BBS12 | 74.1725618 | 0.55532695 | 0.38908643 | 1.42725859 | 0.15350537 | 0.60285224 |
| FGF2 | 450.180298 | -0.0119422 | 0.37117109 | -0.0321743 | 0.97433307 | 0.993743 |
| NUDT6 | 160.963145 | -0.0563621 | 0.29714541 | -0.1896787 | 0.84956093 | 0.97032861 |
| SPATA5 | 1611.33006 | -0.433934 | 0.18355151 | -2.3640992 | 0.01807397 | 0.26945723 |
| SPRY1 | 234.277914 | 1.16191499 | 0.39954921 | 2.90806477 | 0.00363673 | 0.11745127 |
| LINC01091 | 607.745133 | 1.88846588 | 0.72109315 | 2.61889313 | 0.00882156 | 0.19102288 |
| ANKRD50 | 171.608127 | -0.3627018 | 0.43677048 | -0.8304173 | 0.40630288 | 0.80499101 |
| FAT4 | 454.499295 | -1.1944218 | 0.49654035 | -2.4054879 | 1.62E-02 | 0.25504618 |
| INTU | 719.423037 | -0.0478414 | 0.32184329 | -0.1486481 | 0.88183133 | 0.97664137 |
| HSPA4L | 686.721292 | 1.24477121 | 0.38128452 | 3.26467804 | 0.00109589 | 0.05957018 |
| MFSD8 | 580.713054 | -0.4650101 | 0.18687817 | -2.4883063 | 0.01283531 | 0.23224886 |
| ABHD18 | 1500.28259 | -0.3351112 | 0.20931378 | -1.6009994 | 0.10937704 | 0.53319732 |
| LARP1B | 1586.90081 | 0.22817907 | 0.23566724 | 0.96822569 | 0.33293167 | 0.76111014 |
| PGRMC2 | 498.147084 | 0.38757656 | 0.19721769 | 1.96522215 | 0.04938853 | 0.39895974 |
| AC108062.1 | 632.428256 | 0.04127602 | 0.28382192 | 0.14542928 | 0.88437195 | 0.97664137 |
| AC078850.2 | 132.742435 | 2.29397647 | 0.72993182 | 3.14272704 | 0.00167382 | 0.0770051 |
| JADE1 | 1156.41112 | 0.89758039 | 0.29716606 | 3.02046733 | 0.00252385 | 0.09525007 |
| SCLT1 | 1372.15099 | -0.0997182 | 0.28900756 | -0.3450365 | 0.73006692 | 0.93453345 |
| C4orf33 | 189.065963 | 0.37281799 | 0.33709619 | 1.10596916 | 0.26873987 | 0.71818262 |
| AC096711.2 | 624.467886 | -0.0383713 | 0.34356966 | -0.1116842 | 0.91107378 | 0.98111638 |
| AC018680.1 | 68.1049491 | -0.3513787 | 0.67122596 | -0.5234879 | 0.60063474 | 0.88883692 |
| AC109927.1 | 173.250699 | -0.4390724 | 0.42814157 | -1.025531 | 0.30511272 | 0.74229685 |
| NOCT | 112.980182 | 0.05864758 | 0.39398001 | 0.14885928 | 0.88166467 | 0.97664137 |
| ELF2 | 3912.3093 | 0.08433273 | 0.24111929 | 0.34975524 | 0.7265224 | 0.93343394 |
| NDUFC1 | 409.447917 | 0.40850261 | 0.31896051 | 1.280731 | 0.20028817 | 0.65322946 |
| NAA15 | 1213.38567 | -0.1885612 | 0.24360158 | -0.7740557 | 0.43889785 | 0.82127804 |
| AC097376.2 | 1494.64513 | -0.0826345 | 0.28835408 | -0.2865732 | 0.77443915 | 0.94621715 |
| RAB33B | 92.0199424 | 0.18947715 | 0.28573304 | 0.66312652 | 0.50724953 | 0.85626313 |
| SETD7 | 287.763742 | -0.1707372 | 0.35176546 | -0.4853722 | 0.62741232 | 0.89919293 |
| MGST2 | 376.57506 | 0.03156733 | 0.24083689 | 0.13107347 | 0.89571719 | 0.97788747 |
| MAML3 | 2767.24972 | -0.4029526 | 0.36246089 | -1.1117133 | 0.26626146 | 0.71641758 |
| SCOC | 701.431733 | 0.29092622 | 0.17283756 | 1.68323492 | 0.09232962 | 0.50584073 |
| SCOC-AS1 | 146.640919 | 0.49773192 | 0.46543146 | 1.06939897 | 0.28488993 | 0.72787448 |
| ELMOD2 | 344.817982 | -0.100876 | 0.17472622 | -0.5773378 | 0.56371131 | 0.87487519 |
| TBC1D9 | 1064.85322 | -0.4119038 | 0.37554312 | -1.0968217 | 0.27271934 | 0.72070318 |
| RNF150 | 4828.75911 | -0.1029356 | 0.35872858 | -0.2869456 | 0.774154 | 0.94613251 |
| ZNF330 | 272.930841 | 0.04357376 | 0.22323413 | 0.19519308 | 0.84524179 | 0.96859165 |
| LINC02432 | 478.783481 | -0.000868 | 0.53013427 | -0.0016374 | 9.99E-01 | 0.99980325 |
| IL15 | 689.766791 | -0.3163503 | 0.30852194 | -1.0253739 | 0.30518681 | 0.74229685 |
| INPP4B | 3555.78298 | 0.24062683 | 0.67221245 | 0.35796247 | 0.7203714 | 0.93005432 |
| AC139720.1 | 99.4635453 | 0.15056497 | 0.76403674 | 0.19706509 | 0.8437766 | 0.96824299 |
| AC104596.1 | 626.778894 | -0.4434144 | 0.71263685 | -0.6222165 | 0.53379953 | 0.86397197 |
| USP38 | 331.481723 | 0.09746932 | 0.26013304 | 0.37469027 | 0.70789083 | 0.92577218 |
| GAB1 | 2633.50267 | -0.1437555 | 0.1880999 | -0.7642511 | 0.44471764 | 0.8252311 |
| SMARCA5 | 781.232984 | 0.25328608 | 0.16201743 | 1.56332615 | 0.1179759 | 0.54888439 |
| HHIP | 48.0945856 | 1.23101318 | 0.64467344 | 1.90951433 | 0.05619577 | 0.41843207 |
| ANAPC10 | 1125.85607 | -0.0787554 | 0.16395242 | -0.4803552 | 0.63097487 | 0.90005881 |
| ABCE1 | 345.877819 | -0.0460686 | 0.23568066 | -0.1954705 | 0.84502467 | 0.96858129 |

| | | | | | | |
|---|---|---|---|---|---|---|
| OTUD4 | 969.911514 | -0.0662966 | 0.11669644 | -0.5681113 | 0.5699594 | 0.87807251 |
| SMAD1 | 821.602907 | -0.1839915 | 0.13601012 | -1.3527777 | 1.76E-01 | 0.62957039 |
| MMAA | 224.725072 | -0.021235 | 0.23482741 | -0.0904279 | 0.92794718 | 0.98297055 |
| ZNF827 | 1751.96608 | 0.14132298 | 0.39380326 | 0.35886695 | 0.71969462 | 0.93001079 |
| LSM6 | 202.005267 | -0.1089292 | 0.24807634 | -0.4390953 | 0.66059248 | 0.91099119 |
| SLC10A7 | 1115.28435 | 0.3630393 | 0.22455873 | 1.61667864 | 0.10594767 | 0.52735777 |
| TMEM184C | 492.672677 | -0.1941258 | 0.31748824 | -0.6114424 | 0.54090676 | 0.86582694 |
| PRMT9 | 521.634706 | -0.0738667 | 0.23017395 | -0.320917 | 0.74827327 | 0.93746253 |
| ARHGAP10 | 542.728143 | -1.2485588 | 0.46212371 | -2.7017847 | 0.00689684 | 0.17059994 |
| NR3C2 | 10353.0557 | 0.37498211 | 0.36856497 | 1.01741116 | 0.30895788 | 0.74399658 |
| AC069272.1 | 108.986285 | 0.25426 | 0.40467123 | 0.62831252 | 0.52979923 | 0.86343221 |
| DCLK2 | 764.326769 | -0.8623569 | 0.37938794 | -2.2730215 | 0.02302489 | 0.2983102 |
| LRBA | 14033.9613 | -0.0708827 | 0.18488595 | -0.3833861 | 7.01E-01 | 0.92448967 |
| RPS3A | 493.28327 | 0.45973789 | 0.37883053 | 1.21357137 | 0.22491144 | 0.67750405 |
| SH3D19 | 2253.20596 | -0.4353588 | 0.24984513 | -1.7425146 | 0.08141844 | 0.48594425 |
| PRSS48 | 93.9418493 | -0.0772117 | 0.67541981 | -0.1143166 | 0.90898678 | 0.98054769 |
| FAM160A1 | 3652.73151 | 0.04662278 | 0.26889177 | 0.17338863 | 0.86234596 | 0.97281816 |
| AC092611.2 | 53.4562136 | 0.43045437 | 0.32003572 | 1.34501978 | 0.17861885 | 0.63284534 |
| GATB | 274.442974 | 0.03822402 | 0.30687036 | 0.12456081 | 0.90087125 | 0.97914987 |
| FBXW7 | 2946.06449 | -0.0916331 | 0.24437802 | -0.3749646 | 0.70768682 | 0.92577218 |
| TMEM154 | 65.2745507 | -0.0807008 | 0.52157682 | -0.1547246 | 8.77E-01 | 9.76E-01 |
| TIGD4 | 75.802465 | 0.46468592 | 0.30357769 | 1.53069852 | 0.12584392 | 0.56519704 |
| ARFIP1 | 1740.6441 | -0.0202081 | 0.17892663 | -0.1129407 | 0.91007757 | 0.98069436 |
| FHDC1 | 146.139395 | 0.16757412 | 0.39946493 | 0.41949647 | 0.67485334 | 0.91636338 |
| TRIM2 | 5660.60203 | -0.1926152 | 0.20872599 | -0.9228139 | 0.35610421 | 0.77367035 |
| MND1 | 135.26363 | 0.00569054 | 0.8307972 | 0.00684949 | 0.99453494 | 0.998438 |
| TMEM131L | 446.085307 | -0.1355958 | 0.2504995 | -0.5413017 | 0.58829966 | 0.88571413 |
| TLR2 | 1073.2145 | 1.35084302 | 0.67889993 | 1.98975276 | 0.04661818 | 0.39097111 |
| PLRG1 | 294.292486 | 0.09195423 | 0.25093503 | 0.36644636 | 0.71403203 | 0.92760608 |
| FGB | 91.5020639 | 2.61075547 | 1.20871939 | 2.15993512 | 3.08E-02 | 0.33646028 |
| MAP9 | 404.62446 | 0.19512615 | 0.20761631 | 0.93984019 | 0.34729954 | 0.76893725 |
| GUCY1A1 | 2108.58759 | 0.43601245 | 0.26148907 | 1.66742132 | 0.09543066 | 0.51158423 |
| GUCY1B1 | 167.464472 | 1.5771971 | 0.41628901 | 3.78870702 | 0.00015143 | 0.02008985 |
| TDO2 | 62.5642475 | 0.82640063 | 0.68009682 | 1.21512203 | 2.24E-01 | 0.67681479 |
| CTSO | 323.064317 | -0.313394 | 0.33373505 | -0.9390504 | 0.34770488 | 0.76900643 |
| PDGFC | 1606.40375 | 0.17233005 | 0.2024056 | 0.85140951 | 0.39454191 | 0.79876591 |
| GLRB | 257.135865 | 0.12674291 | 0.32302405 | 0.3923637 | 0.69478951 | 0.92229578 |
| TMEM144 | 105.534099 | -0.4072608 | 0.52163671 | -0.7807365 | 0.43495748 | 0.81919857 |
| ETFDH | 1049.82165 | 0.17246583 | 0.31750134 | 0.54319718 | 5.87E-01 | 0.8855551 |
| PPID | 303.137804 | 0.11565191 | 0.27944013 | 0.41387009 | 0.67896925 | 0.91822574 |
| FNIP2 | 4998.10169 | -1.2436682 | 0.37377176 | -3.3273468 | 0.00087677 | 0.05165902 |
| C4orf45 | 236.004936 | -0.3590083 | 0.78218832 | -0.4589793 | 0.64624901 | 0.90582727 |
| RAPGEF2 | 3183.03222 | -0.0334663 | 0.21793686 | -0.1535596 | 0.87795699 | 0.9765717 |
| FSTL5 | 43.6178804 | 0.67805446 | 0.50188036 | 1.35102807 | 0.17668644 | 0.63008941 |
| NAF1 | 376.027205 | -0.1472925 | 0.2291152 | -0.6428753 | 0.52030503 | 0.85935232 |
| NPY1R | 435.717738 | -1.2301062 | 0.63392618 | -1.9404566 | 0.05232423 | 0.40896493 |
| TMA16 | 245.690991 | -0.3267185 | 0.25033658 | -1.3051168 | 0.19185308 | 0.64596359 |
| 1-Mar | 92.2475214 | -0.2697433 | 0.33553447 | -0.8039213 | 0.42144243 | 0.81266383 |
| TMEM192 | 477.489987 | -0.4679112 | 0.35944569 | -1.3017578 | 0.19299918 | 0.64699503 |
| KLHL2 | 2295.68659 | -0.5656561 | 0.48531226 | -1.1655509 | 2.44E-01 | 0.6948422 |
| MSMO1 | 444.559365 | 0.46452206 | 0.57731582 | 0.80462382 | 0.42103679 | 0.81260593 |
| CPE | 103.949873 | -2.5773349 | 0.89549765 | -2.8781034 | 0.00400074 | 0.12328719 |
| DDX60 | 577.606263 | 0.24293528 | 1.177172 | 0.20637195 | 0.83650038 | 0.96511435 |
| DDX60L | 452.416461 | -0.1938763 | 0.98763729 | -0.1963032 | 0.84437289 | 0.96850656 |

| | | | | | | |
|---|---|---|---|---|---|---|
| PALLD | 267.934559 | -0.0570507 | 0.3552659 | -0.1605859 | 0.87241956 | 0.97554234 |
| CBR4 | 2677.53076 | -0.3234435 | 0.15857313 | -2.0397119 | 0.04137903 | 0.37577267 |
| SH3RF1 | 3197.13442 | 0.02579697 | 0.2694466 | 0.09574058 | 0.92372661 | 0.98215577 |
| AC096741.1 | 85.4255029 | 0.24162102 | 0.48903535 | 0.4940768 | 0.62125194 | 0.8978246 |
| NEK1 | 1795.55156 | 0.09381546 | 0.13989804 | 0.67059883 | 5.02E-01 | 8.55E-01 |
| CLCN3 | 1564.92255 | 0.43305878 | 0.19883983 | 2.17792777 | 0.02941141 | 0.33012415 |
| HPF1 | 243.405593 | 0.12177673 | 0.2467914 | 0.49343992 | 0.62170178 | 0.8982666 |
| LINC02275 | 128.245721 | -0.3730906 | 0.57369946 | -0.6503242 | 5.15E-01 | 8.59E-01 |
| MFAP3L | 805.96929 | -0.3510017 | 0.3961101 | -0.8861216 | 0.37555198 | 0.78490077 |
| LINC01612 | 71.610653 | -0.4006047 | 0.41646246 | -0.9619228 | 0.33608836 | 0.7637525 |
| LINC02512 | 69.3505531 | 0.88567834 | 0.49711675 | 1.78163044 | 0.07480952 | 0.47190267 |
| GALNTL6 | 49.7084456 | 0.04457611 | 0.46200827 | 0.09648335 | 0.9231367 | 0.98215577 |
| AC105285.1 | 59.8382571 | 0.23482077 | 0.42817299 | 0.548425 | 0.58340012 | 0.88379959 |
| GALNT7 | 1138.15189 | 0.51632307 | 0.27209328 | 1.89759586 | 0.05774934 | 0.42087381 |
| HMGB2 | 231.646216 | 0.68940078 | 0.249795 | 2.75986618 | 0.0057825 | 0.15587979 |
| SAP30 | 74.9002006 | -0.2120774 | 0.52087401 | -0.4071567 | 0.68389288 | 0.9200778 |
| FBXO8 | 227.680372 | 0.29721283 | 0.23311607 | 1.27495643 | 0.20232466 | 0.65552334 |
| CEP44 | 506.342516 | -0.0741073 | 0.19741655 | -0.3753853 | 0.70737396 | 0.92576069 |
| GPM6A | 51.1562253 | -0.6395049 | 0.38733561 | -1.6510357 | 0.09873129 | 0.51547101 |
| WDR17 | 400.116094 | 0.15863936 | 0.39690537 | 0.39969064 | 0.68938438 | 0.92166785 |
| AC019163.1 | 50.8870231 | 0.38992132 | 0.46022012 | 0.84724963 | 0.39685599 | 0.80068288 |
| SPCS3 | 553.058631 | 0.17792379 | 0.27268822 | 0.65248067 | 0.51409114 | 0.85762435 |
| AGA | 87.3924594 | 0.1990098 | 0.358986 | 0.55436648 | 0.57932806 | 0.8821758 |
| AC078881.1 | 602.660641 | -0.3397452 | 0.47020569 | -0.7225459 | 0.46995889 | 0.83921324 |
| TENM3 | 332.671186 | 0.61417398 | 0.39025385 | 1.57378069 | 0.11553816 | 0.54427374 |
| DCTD | 232.304315 | -0.2642733 | 0.20212591 | -1.3074689 | 0.1910535 | 0.64567512 |
| WWC2 | 4767.2367 | 0.09438104 | 0.17754567 | 0.53158739 | 0.5950118 | 0.88858235 |
| CDKN2AIP | 117.069409 | 0.19602234 | 0.31633829 | 0.6196605 | 0.53548133 | 0.8642718 |
| AC107214.1 | 96.8436203 | 0.06434032 | 0.41348347 | 0.15560554 | 0.87634396 | 0.97614152 |
| ING2 | 77.0158195 | 0.72939364 | 0.39266588 | 1.85754271 | 0.06323398 | 0.43730767 |
| RWDD4 | 211.989076 | 0.15876135 | 0.17520881 | 0.9061265 | 0.3648689 | 0.78094673 |
| TRAPPC11 | 727.156881 | -0.1179815 | 0.23324755 | -0.5058208 | 0.61298248 | 0.89441126 |
| STOX2 | 3636.40305 | 0.30872192 | 0.44003976 | 0.70157733 | 0.48294279 | 0.8441831 |
| IRF2 | 719.108951 | 0.0295805 | 0.22265566 | 0.13285314 | 0.89430953 | 0.97788747 |
| LINC02427 | 141.588431 | -0.7475691 | 0.61340173 | -1.2187268 | 0.22294791 | 0.67594033 |
| CASP3 | 276.370486 | -0.769618 | 0.34255214 | -2.2467179 | 0.02465806 | 0.30812502 |
| PRIMPOL | 355.033569 | -0.3203018 | 0.30894616 | -1.0367561 | 0.29984953 | 0.73673761 |
| ACSL1 | 2316.24252 | -0.2847478 | 0.41883485 | -0.679857 | 0.496595 | 0.85078123 |
| AC084871.2 | 44.1698781 | -0.3581691 | 0.67504015 | -0.5305893 | 0.59570341 | 0.88859628 |
| LINC02437 | 210.357683 | -0.0919758 | 0.76631154 | -0.1200241 | 0.90446408 | 0.97989015 |
| LINC02436 | 46.9697997 | 0.24067138 | 0.74980711 | 0.32097773 | 0.74822727 | 0.93746253 |
| SLC25A4 | 581.393086 | 1.11117224 | 0.38626412 | 2.87671614 | 0.00401837 | 0.12328719 |
| CFAP97 | 479.288086 | 0.17094529 | 0.24407127 | 0.70039085 | 0.48368325 | 0.84454356 |
| SNX25 | 1616.4384 | -0.4710802 | 0.35471643 | -1.3280473 | 0.18416247 | 0.63875594 |
| LRP2BP | 241.443312 | -0.1549389 | 0.45359651 | -0.3415786 | 0.73266804 | 0.93552616 |
| AC112722.1 | 244.882419 | -0.2836549 | 0.4645141 | -0.6106485 | 0.54143228 | 0.86582694 |
| ANKRD37 | 294.346764 | 0.12005558 | 0.87050056 | 0.13791557 | 0.89030714 | 0.97729111 |
| UFSP2 | 501.182366 | 0.1931892 | 0.20678167 | 0.93426657 | 0.35016641 | 0.77044053 |
| C4orf47 | 751.169204 | 0.80456005 | 0.64283885 | 1.25157347 | 0.21072533 | 0.66399591 |
| CCDC110 | 57.7437938 | 0.18839615 | 0.44329903 | 0.42498661 | 0.67084644 | 0.91516099 |
| SORBS2 | 2604.54304 | 0.80169377 | 0.47962482 | 1.67150183 | 0.0946226 | 0.51017802 |
| TLR3 | 97.1195307 | 0.29688039 | 0.33325585 | 0.89084826 | 0.37301058 | 0.78277446 |
| FAM149A | 481.494316 | -0.4928705 | 0.30750375 | -1.6028114 | 0.10897629 | 0.53262539 |
| CYP4V2 | 448.922394 | -0.3707179 | 0.26767536 | -1.3849534 | 0.1660667 | 0.61901667 |

| | | | | | | |
|---|---|---|---|---|---|---|
| KLKB1 | 477.542282 | -0.3762981 | 0.41188652 | -0.9135965 | 0.3609289 | 0.77730375 |
| F11 | 582.498466 | 0.28203582 | 0.59657282 | 0.47276009 | 0.63638435 | 0.90241555 |
| F11-AS1 | 105.280306 | 2.15800787 | 0.69454279 | 3.10709132 | 0.00188938 | 0.08431359 |
| AC018709.1 | 56.2874988 | 2.74665105 | 0.72796172 | 3.77307073 | 0.00016125 | 0.0205434 |
| FAT1 | 1129.08764 | 0.040931 | 0.24630363 | 0.16618105 | 0.86801447 | 0.97402505 |
| LINC01060 | 97.6145182 | 0.28470369 | 0.77512166 | 0.36730194 | 0.71339381 | 0.92760608 |
| FRG1-DT | 652.523408 | -1.0390367 | 0.54413945 | -1.9095044 | 0.05619706 | 0.41843207 |
| FRG1 | 192.51464 | 0.2936077 | 0.25087551 | 1.17033225 | 0.24186729 | 0.6933214 |
| PLEKHG4B | 203.197946 | -4.0861158 | 1.10984695 | -3.6816931 | 0.00023169 | 0.02488127 |
| CCDC127 | 89.5408453 | -0.3176947 | 0.32489896 | -0.9778263 | 0.32816025 | 0.75648668 |
| SDHA | 909.745351 | 0.15285464 | 0.13938252 | 1.09665575 | 0.27279191 | 0.72070318 |
| PDCD6 | 493.347308 | -0.4015298 | 0.2511328 | -1.5988743 | 0.10984853 | 0.53372881 |
| AHRR | 48.5634292 | -1.2854281 | 0.67771779 | -1.8967011 | 0.05786739 | 0.42087381 |
| EXOC3-AS1 | 47.0865768 | -0.1516003 | 0.37408621 | -0.405255 | 0.68529009 | 0.92053037 |
| EXOC3 | 723.905421 | -0.5085192 | 0.12435478 | -4.0892619 | 4.33E-05 | 0.01039303 |
| SLC9A3 | 1542.485 | 0.55370748 | 0.5066584 | 1.09286154 | 0.27445459 | 0.72164296 |
| CEP72 | 65.8480937 | -0.3157961 | 0.3487766 | -0.9054394 | 0.36523263 | 0.78095617 |
| TPPP | 81.0603319 | 2.48734354 | 0.70314271 | 3.53746616 | 0.00040399 | 0.03420673 |
| ZDHHC11B | 212.149211 | 2.44712838 | 0.76093775 | 3.2159377 | 0.00130019 | 0.06597468 |
| ZDHHC11 | 64.5346311 | 0.35654285 | 0.62720947 | 0.56845897 | 0.56972336 | 0.87807251 |
| BRD9 | 243.431667 | -0.2190777 | 0.24442233 | -0.8963082 | 0.37008817 | 0.78099673 |
| SLC12A7 | 254.041763 | 0.25622224 | 0.32546166 | 0.78725782 | 0.43113095 | 0.81706845 |
| CLPTM1L | 258.680209 | 0.24997736 | 0.28522912 | 0.87640898 | 0.38080774 | 0.78829172 |
| LPCAT1 | 417.446582 | -0.4178171 | 0.37952492 | -1.1008951 | 0.27094231 | 0.71977937 |
| MRPL36 | 296.861032 | 0.17539039 | 0.72883548 | 0.24064469 | 0.80983051 | 0.95662439 |
| NDUFS6 | 664.035067 | -0.021945 | 0.34207325 | -0.0641529 | 0.94884851 | 0.98919519 |
| AC126768.2 | 71.1817803 | 0.06001118 | 0.53165298 | 0.1128766 | 0.91012839 | 0.98069436 |
| IRX2 | 1256.87285 | 0.58465697 | 0.43525877 | 1.34323996 | 0.17919429 | 0.63288164 |
| C5orf38 | 214.573373 | 0.73401448 | 0.42534776 | 1.72568086 | 0.08440484 | 0.4907146 |
| IRX1 | 652.128319 | 0.76971784 | 0.37398324 | 2.05816134 | 0.03957465 | 0.37258841 |
| AC106799.2 | 47.3751845 | 0.29782884 | 0.47160246 | 0.63152521 | 0.52769717 | 0.86181403 |
| AC022424.1 | 89.6472261 | -0.779411 | 0.69814744 | -1.1163989 | 0.26425142 | 0.71488591 |
| ADAMTS16 | 679.181735 | -0.5317325 | 0.61464093 | -0.8651108 | 0.38697797 | 0.79542282 |
| AC022424.2 | 48.6157403 | 0.64460373 | 0.71004901 | 0.90782992 | 0.36396809 | 0.78015617 |
| ICE1 | 1119.19596 | -0.0317621 | 0.18569099 | -0.171048 | 0.86418603 | 0.97356361 |
| LINC02145 | 66.7726909 | -0.7827804 | 0.48990678 | -1.597815 | 0.11008417 | 0.53372881 |
| MED10 | 232.124361 | -0.0078187 | 0.20472818 | -0.0381908 | 0.96953556 | 0.99299658 |
| NSUN2 | 455.133921 | -0.2315514 | 0.24087273 | -0.961302 | 0.33640034 | 0.7637525 |
| SRD5A1 | 177.210595 | -0.5071659 | 0.3523473 | -1.4393919 | 0.15003951 | 0.59663305 |
| TENT4A | 726.420487 | -0.413206 | 0.16555897 | -2.4958237 | 0.0125665 | 0.22916272 |
| ADCY2 | 37.8271402 | -0.1834434 | 0.447764 | -0.4096878 | 0.682035 | 0.91969963 |
| MTRR | 423.10318 | -0.1876755 | 0.23659054 | -0.7932504 | 0.42763193 | 0.81466629 |
| FASTKD3 | 80.1302942 | -0.0386712 | 0.27843524 | -0.1388876 | 0.88953897 | 0.97729111 |
| MIR4458HG | 86.2548718 | 0.58651558 | 0.46295668 | 1.26689084 | 0.20519432 | 0.65813563 |
| SEMA5A | 338.708177 | -0.0313607 | 0.71216563 | -0.0440357 | 0.96487593 | 0.99237298 |
| SNHG18 | 74.3472684 | 0.99646066 | 0.37770017 | 2.63823196 | 0.00833396 | 0.18347203 |
| FAM173B | 354.501118 | -0.2346527 | 0.22948859 | -1.0225029 | 0.30654294 | 0.74295884 |
| CCT5 | 386.770589 | 0.51412847 | 0.28729618 | 1.7895416 | 0.07352763 | 0.47028843 |
| CMBL | 388.44469 | 1.0010837 | 0.43803521 | 2.2853955 | 0.02228965 | 0.29649945 |
| 6-Mar | 2780.92443 | -0.1405252 | 0.18079726 | -0.7772527 | 0.43700967 | 0.82013459 |
| ANKRD33B | 261.371839 | 1.75990754 | 0.52695715 | 3.33975452 | 0.00083852 | 0.05047093 |
| DAP | 1316.25011 | -0.4376179 | 0.29790609 | -1.4689793 | 0.14183841 | 0.5872148 |
| DNAH5 | 3219.98365 | 0.6392627 | 0.8023346 | 0.79675325 | 4.26E-01 | 0.81466629 |
| AC016576.1 | 183.014467 | 1.00078835 | 0.73745797 | 1.3570785 | 1.75E-01 | 0.62932267 |

| | | | | | | |
|---|---|---|---|---|---|---|
| TRIO | 7392.8602 | -0.5195556 | 0.22696679 | -2.289126 | 2.21E-02 | 0.29509055 |
| OTULINL | 98.0026585 | -0.5158523 | 0.5725183 | -0.9010233 | 0.36757594 | 0.78095617 |
| OTULIN | 407.423978 | -0.0433839 | 0.20005496 | -0.2168598 | 0.82831764 | 0.96288319 |
| ANKH | 778.783702 | -0.2986878 | 0.37661539 | -0.7930844 | 0.42772864 | 0.81466629 |
| FBXL7 | 524.010402 | -2.2824769 | 0.56296805 | -4.054363 | 5.03E-05 | 0.01125476 |
| ZNF622 | 82.0278519 | 0.14662855 | 0.30781922 | 0.47634631 | 0.63382768 | 0.90136076 |
| RETREG1 | 2130.2054 | 0.51986767 | 0.6818957 | 0.7623859 | 4.46E-01 | 8.26E-01 |
| MYO10 | 4223.14953 | -0.0598997 | 0.33416776 | -0.1792505 | 0.85774105 | 0.97127317 |
| CDH18 | 181.872394 | 0.39141606 | 0.86452587 | 0.45275228 | 6.51E-01 | 9.08E-01 |
| LINC02241 | 504.405144 | -0.0055096 | 0.28506138 | -0.0193279 | 0.9845795 | 0.995279 |
| AC093274.1 | 73.6773215 | 0.45296327 | 0.43455102 | 1.04237075 | 0.29723982 | 0.73673761 |
| CDH12 | 235.74593 | 0.51292127 | 0.46954403 | 1.09238161 | 0.2746654 | 0.72164296 |
| C5orf17 | 536.201623 | -1.3061155 | 0.52797105 | -2.4738393 | 0.01336698 | 0.23795747 |
| CDH9 | 104.286562 | 0.86670759 | 0.7453012 | 1.16289574 | 0.24487181 | 0.6948422 |
| CDH6 | 159.603781 | 1.86978485 | 0.78951286 | 2.36827662 | 0.01787117 | 0.26845799 |
| DROSHA | 800.282072 | -0.3235862 | 0.18068908 | -1.7908454 | 0.07331811 | 0.4701377 |
| C5orf22 | 227.355398 | -0.1150362 | 0.21178969 | -0.5431624 | 0.58701798 | 0.8855551 |
| PDZD2 | 800.487013 | 0.27828822 | 0.30366305 | 0.91643755 | 3.59E-01 | 0.77654715 |
| AC025178.1 | 88.981059 | -0.1409802 | 0.62595298 | -0.2252249 | 0.82180436 | 0.96066244 |
| GOLPH3 | 1936.13114 | -0.186068 | 0.34901067 | -0.5331299 | 0.59394366 | 0.88807733 |
| MTMR12 | 1011.01713 | -0.1295822 | 0.13798795 | -0.9390835 | 0.34768787 | 0.76900643 |
| ZFR | 2400.41535 | 0.00490137 | 0.14634737 | 0.03349132 | 0.97328279 | 0.99327086 |
| SUB1 | 824.950622 | 0.65129994 | 0.25065146 | 2.5984287 | 0.00936515 | 0.19669868 |
| NPR3 | 372.874658 | 2.58042005 | 0.71869094 | 3.59044467 | 0.00033011 | 0.03094754 |
| AC010343.3 | 69.8240855 | -0.2613976 | 0.57992073 | -0.450747 | 0.65217187 | 0.9085149 |
| TARS | 310.846642 | 0.06549734 | 0.27602451 | 0.23728813 | 0.81243327 | 0.95806331 |
| SLC45A2 | 59.6695576 | -0.2488241 | 0.60815626 | -0.409145 | 0.68243322 | 0.91969963 |
| C1QTNF3-AN | 1866.12237 | -0.1854975 | 0.29437672 | -0.6301366 | 0.52860524 | 0.86245226 |
| RAI14 | 1466.13858 | -0.4202463 | 0.23179358 | -1.8130194 | 0.06982882 | 0.45914671 |
| TTC23L | 139.583885 | 0.12845725 | 0.54684955 | 0.23490419 | 0.8142831 | 0.95832184 |
| RAD1 | 192.033258 | -0.0535523 | 0.34172571 | -0.1567112 | 0.87547243 | 0.97614152 |
| BRIX1 | 164.360191 | 0.03902514 | 0.24417281 | 0.15982589 | 0.87301823 | 0.97564728 |
| DNAJC21 | 508.005225 | 0.26088499 | 0.15171908 | 1.71952657 | 0.08551853 | 0.49349686 |
| PRLR | 333.33774 | 1.56469656 | 0.59094553 | 2.64778476 | 0.00810211 | 0.18047346 |
| SPEF2 | 789.339448 | 0.35865022 | 0.4820341 | 0.74403495 | 0.45685533 | 0.8323589 |
| AC137810.1 | 44.4090452 | 1.19064077 | 0.68218468 | 1.74533498 | 0.08092658 | 0.48522885 |
| LMBRD2 | 427.479226 | -0.0488753 | 0.12387113 | -0.3945661 | 0.69316315 | 0.92224682 |
| SKP2 | 132.403366 | -0.0326377 | 0.24132184 | -0.1352454 | 8.92E-01 | 0.97771356 |
| NADK2 | 808.3382 | -0.6748306 | 0.41280751 | -1.6347342 | 0.10210477 | 0.52073581 |
| RANBP3L | 4491.23661 | 0.44426688 | 0.69606583 | 0.63825411 | 0.52330828 | 0.86006369 |
| NIPBL | 4999.10051 | -0.0891851 | 0.14498349 | -0.6151395 | 0.53846261 | 0.86527404 |
| CPLANE1 | 1609.55776 | -0.2324548 | 0.16514615 | -1.4075703 | 0.15925834 | 0.60886928 |
| NUP155 | 331.093393 | -0.3589651 | 0.24511509 | -1.4644758 | 0.143064 | 0.5883902 |
| WDR70 | 2743.86444 | -0.2758101 | 0.15884959 | -1.736297 | 0.08251133 | 0.48594425 |
| LIFR | 2788.00855 | 0.00620723 | 0.27778206 | 0.0223457 | 9.82E-01 | 0.99518016 |
| LIFR-AS1 | 142.600018 | -0.214764 | 0.39314123 | -0.546277 | 0.58487557 | 0.8846469 |
| OSMR-AS1 | 1946.73568 | 0.46517325 | 0.37135888 | 1.25262456 | 2.10E-01 | 6.63E-01 |
| OSMR | 2251.08 | -0.1105363 | 0.45310825 | -0.2439511 | 8.07E-01 | 9.56E-01 |
| RICTOR | 2955.27398 | -0.2745528 | 0.16986182 | -1.61633 | 0.10602299 | 0.52735777 |
| FYB1 | 409.82635 | 1.64823118 | 0.61376862 | 2.68542757 | 0.00724371 | 0.17302374 |
| C9 | 69.3055786 | 0.25027694 | 0.52421405 | 0.47743271 | 0.63305402 | 0.90078752 |
| DAB2 | 129.16265 | 0.15465147 | 0.77708964 | 0.19901367 | 8.42E-01 | 0.96716572 |
| TTC33 | 522.779072 | -0.0763307 | 0.24587776 | -0.3104416 | 0.75622514 | 0.93964676 |
| PRKAA1 | 834.32389 | -0.3452728 | 0.18137541 | -1.9036361 | 0.05695759 | 0.41872623 |

| | | | | | | |
|---|---|---|---|---|---|---|
| RPL37 | 622.047799 | 0.29856129 | 0.27835131 | 1.07260603 | 0.28344792 | 0.72668316 |
| CARD6 | 64.4418307 | -2.0985517 | 0.8028076 | -2.6140157 | 0.00894849 | 0.19219616 |
| OXCT1 | 1182.11115 | -0.666788 | 0.33231883 | -2.0064706 | 4.48E-02 | 0.38683751 |
| C5orf51 | 184.771215 | 0.14620865 | 0.23803365 | 0.61423524 | 0.53905989 | 0.86542329 |
| FBXO4 | 146.178056 | -0.3269372 | 0.34197251 | -0.9560335 | 0.33905532 | 0.76449297 |
| GHR | 797.334846 | 0.58061876 | 0.42184117 | 1.3763919 | 1.69E-01 | 0.6226216 |
| SELENOP | 369.942459 | 0.29308052 | 0.41884231 | 0.69973952 | 0.48408999 | 0.8445691 |
| ZNF131 | 1184.91867 | -0.1744947 | 0.20067397 | -0.8695432 | 0.38455008 | 0.79217288 |
| NIM1K | 69.9354946 | -0.0186339 | 0.59125386 | -0.031516 | 0.97485804 | 0.99403355 |
| HMGCS1 | 712.653759 | 0.17565422 | 0.44603026 | 0.39381683 | 0.69371628 | 0.92229578 |
| CCL28 | 215.894177 | 0.42159224 | 0.80035204 | 0.5267585 | 0.59836131 | 0.88883692 |
| TMEM267 | 583.283257 | 0.14457182 | 0.21428728 | 0.67466358 | 0.49988953 | 0.85244523 |
| C5orf34 | 76.5181528 | 0.19717659 | 0.33779068 | 0.58372417 | 0.5594059 | 0.87295536 |
| PAIP1 | 435.39747 | -0.165353 | 0.2006519 | -0.8240788 | 0.40989478 | 0.80641944 |
| NNT-AS1 | 798.611682 | 0.29409104 | 0.21173998 | 1.38892539 | 0.16485544 | 0.61673853 |
| NNT | 2218.20414 | -0.1026387 | 0.24506323 | -0.4188256 | 6.75E-01 | 0.91649321 |
| MRPS30-DT | 380.565651 | -0.294043 | 0.29775578 | -0.9875309 | 0.32338243 | 0.75362713 |
| MRPS30 | 216.102801 | 0.35422345 | 0.20007817 | 1.77042527 | 0.07665632 | 0.47627158 |
| AC093297.2 | 69.3821593 | 0.31995159 | 0.28221688 | 1.13370822 | 0.25691696 | 0.70735159 |
| PARP8 | 1201.99955 | 0.15860269 | 0.35676076 | 0.44456316 | 0.65663548 | 0.90974386 |
| ITGA1 | 604.566037 | 0.12550061 | 0.2249826 | 0.55782363 | 0.57696482 | 0.88150331 |
| ITGA2 | 3129.93654 | -1.2095437 | 0.51420693 | -2.352251 | 0.01866018 | 0.27519285 |
| MOCS2 | 242.333219 | 0.92494699 | 0.28629735 | 3.23072144 | 0.00123478 | 0.06446734 |
| NDUFS4 | 1442.90868 | -0.0023437 | 0.18903545 | -0.0123983 | 9.90E-01 | 0.997268 |
| ARL15 | 7280.76812 | -0.3057727 | 0.38189181 | -0.8006788 | 0.42331762 | 0.8138418 |
| SNX18 | 284.314184 | 0.86261929 | 0.24683561 | 3.49471175 | 0.00047457 | 0.03734387 |
| AC026704.1 | 151.921249 | 0.2253657 | 0.52720781 | 0.42747033 | 0.66903679 | 0.91421919 |
| DHX29 | 664.416621 | -0.0903198 | 0.18460534 | -0.4892587 | 0.62465854 | 0.89873262 |
| MTREX | 1238.94343 | -0.037892 | 0.13134184 | -0.2884988 | 0.77296491 | 0.94534972 |
| PLPP1 | 824.14631 | -0.3560657 | 0.27159649 | -1.31101 | 0.18985439 | 0.64450954 |
| SLC38A9 | 808.046379 | -0.4492878 | 0.20036928 | -2.2422988 | 0.02494207 | 0.3093519 |
| IL6ST | 1415.41217 | 0.32154962 | 0.29057398 | 1.10660155 | 0.26846624 | 0.71818262 |
| AC008892.1 | 238.704955 | 0.88419512 | 0.41453103 | 2.13300102 | 0.03292464 | 0.34598377 |
| LINC01948 | 390.829311 | -0.3322821 | 0.5615687 | -0.5917033 | 0.55404928 | 0.8712425 |
| MAP3K1 | 2559.68458 | -0.2330017 | 0.23516703 | -0.9907924 | 0.32178697 | 0.75240254 |
| AC008937.2 | 116.814784 | 0.06462918 | 0.62499695 | 0.10340719 | 0.91763981 | 0.98199511 |
| SETD9 | 64.4359281 | 0.06504405 | 0.42862243 | 0.15175138 | 0.87938303 | 0.9765717 |
| MIER3 | 257.361832 | 0.08867498 | 0.27579513 | 0.32152482 | 0.7478127 | 0.93746253 |
| GPBP1 | 2578.58769 | 0.02732782 | 0.22196539 | 0.12311748 | 0.90201407 | 0.97941893 |
| PDE4D | 16278.1017 | -0.021083 | 0.51670778 | -0.0408026 | 0.96745327 | 0.99248831 |
| ELOVL7 | 1130.79774 | 0.04749517 | 0.36150284 | 0.13138258 | 0.89547267 | 0.97788747 |
| ERCC8 | 602.586087 | -0.1259459 | 0.21176814 | -0.594735 | 0.55202063 | 0.87064364 |
| NDUFAF2 | 946.714592 | 0.04877052 | 0.26419678 | 0.1845992 | 0.85354341 | 0.97076831 |
| SMIM15 | 135.109143 | 0.29153347 | 0.25049578 | 1.16382591 | 0.24449457 | 0.6948422 |
| SMIM15-AS1 | 138.203735 | 0.14581741 | 0.37561701 | 0.38820769 | 0.69786234 | 0.92341984 |
| ZSWIM6 | 4441.06647 | -0.3268061 | 0.28082086 | -1.163753 | 0.24452412 | 0.6948422 |
| KIF2A | 775.506419 | -0.1196458 | 0.18670153 | -0.6408399 | 0.52162669 | 0.85957492 |
| DIMT1 | 126.389679 | -0.1727373 | 0.30308312 | -0.5699337 | 0.56872268 | 0.87770276 |
| IPO11 | 507.33608 | -0.3182157 | 0.1688309 | -1.8848192 | 0.05945425 | 0.42578317 |
| RNF180 | 1220.21924 | 0.19767969 | 0.16861273 | 1.17238887 | 0.24104096 | 0.69178991 |
| SREK1IP1 | 448.997106 | -0.0083578 | 0.15042207 | -0.0555623 | 0.95569047 | 0.99065996 |
| CWC27 | 1217.23133 | -0.1574205 | 0.13097497 | -1.2019132 | 0.22939714 | 0.6817947 |
| PPWD1 | 567.813805 | 0.03982547 | 0.21204063 | 0.18781998 | 0.85101777 | 0.97056586 |
| TRIM23 | 423.627403 | 0.03375616 | 0.24252002 | 0.13918919 | 0.88930065 | 0.97729111 |

| | | | | | | |
|---|---|---|---|---|---|---|
| TRAPPC13 | 551.678091 | -0.1753905 | 0.20815278 | -0.8426046 | 0.39944963 | 0.80174499 |
| SGTB | 70.2629214 | 0.38605325 | 0.41345665 | 0.93372121 | 0.35044772 | 0.77044053 |
| NLN | 311.44988 | -0.6577759 | 0.32761561 | -2.0077672 | 0.04466803 | 0.38664485 |
| ERBIN | 4431.5847 | -0.0963603 | 0.16961483 | -0.5681124 | 0.56995862 | 0.87807251 |
| SREK1 | 1675.54235 | -0.3489509 | 0.20382305 | -1.7120284 | 0.08689144 | 0.49480766 |
| MAST4 | 16002.1959 | -0.8074329 | 0.30938557 | -2.6097948 | 0.00905965 | 0.19332628 |
| AC112206.2 | 125.63018 | 0.28243736 | 0.65404885 | 0.43182915 | 0.66586559 | 0.91305741 |
| AC106798.1 | 118.307675 | 0.07818605 | 0.74980877 | 0.10427465 | 0.91695139 | 0.98199511 |
| PIK3R1 | 1355.98694 | 0.03419202 | 0.32450216 | 0.1053676 | 0.91608412 | 0.98195017 |
| AC093523.1 | 179.518567 | -0.3775306 | 0.64330621 | -0.5868599 | 5.57E-01 | 0.87240458 |
| LINC02198 | 153.475196 | -0.9896671 | 0.62219972 | -1.5905939 | 0.111701 | 0.5359457 |
| SLC30A5 | 667.955803 | -0.2745627 | 0.284052 | -0.9665932 | 0.33374745 | 0.7617885 |
| AC010273.1 | 44.9741493 | -0.0381887 | 0.50174085 | -0.0761123 | 0.93932974 | 0.98715467 |
| MRPS36 | 358.921466 | 0.30141978 | 0.22235509 | 1.3555785 | 0.17523332 | 0.62944785 |
| CDK7 | 362.230781 | -0.1961286 | 0.18443129 | -1.0634238 | 0.28758983 | 0.73053346 |
| CCDC125 | 766.951586 | -0.0590091 | 0.40098642 | -0.1471599 | 0.88300581 | 0.97664137 |
| AK6 | 245.380092 | 0.15616787 | 0.26248717 | 0.59495427 | 0.55187404 | 0.87051638 |
| RAD17 | 586.27131 | -0.1909495 | 0.14604063 | -1.3075095 | 0.19103973 | 0.64567512 |
| MARVELD2 | 613.716671 | 0.17350304 | 0.22367487 | 0.77569303 | 4.38E-01 | 0.82074642 |
| AC145146.1 | 90.1288728 | 0.29569275 | 0.56263455 | 0.52555029 | 0.5992007 | 0.88883692 |
| OCLN | 1717.23203 | 0.10257795 | 0.30558088 | 0.33568184 | 0.7371108 | 0.93552616 |
| GTF2H2C | 696.728168 | -0.3264646 | 0.34894022 | -0.9355888 | 0.34948492 | 0.77016121 |
| AC146944.4 | 37.2677992 | -0.8755139 | 0.65323927 | -1.3402652 | 0.18015913 | 0.63442332 |
| NAIP | 2080.32957 | 0.14260227 | 0.31483524 | 0.45294252 | 6.51E-01 | 0.90825486 |
| GTF2H2 | 657.85909 | -0.4570201 | 0.55630846 | -0.8215228 | 0.41134857 | 0.80675684 |
| LINC02197 | 96.4271479 | -0.9687521 | 1.67387615 | -0.5787478 | 0.56275937 | 0.87432234 |
| BDP1 | 1703.40809 | -0.1315931 | 0.16437958 | -0.800544 | 0.42339571 | 0.8138418 |
| MCCC2 | 798.638106 | -0.0084886 | 0.11114862 | -0.0763713 | 0.93912373 | 0.98715467 |
| MRPS27 | 705.911032 | -0.0505099 | 0.15558853 | -0.3246377 | 0.74545533 | 0.93675395 |
| PTCD2 | 163.954058 | -0.3393338 | 0.26666973 | -1.2724869 | 0.20320016 | 0.65625206 |
| AC063979.2 | 78.3692571 | 1.43832745 | 0.70860978 | 2.02978775 | 0.04237812 | 0.3797643 |
| AC035140.1 | 41.1919218 | 0.32764814 | 0.45646932 | 0.71778787 | 0.47288809 | 0.83985854 |
| TNPO1 | 2021.45798 | -0.0722771 | 0.16048155 | -0.4503761 | 0.65243925 | 0.90868303 |
| FCHO2 | 2469.10528 | -0.07261 | 0.2176284 | -0.333642 | 7.39E-01 | 9.36E-01 |
| TMEM171 | 459.441864 | 0.31394246 | 0.60477694 | 0.51910455 | 0.60368784 | 0.89001883 |
| AC116345.1 | 293.297774 | 0.10877917 | 0.91419679 | 0.11898879 | 0.90528423 | 0.97992575 |
| BTF3 | 453.606709 | 0.54069477 | 0.30555769 | 1.76953416 | 0.07680477 | 0.47651607 |
| ANKRA2 | 245.531281 | 0.07963205 | 0.23713285 | 0.33581196 | 0.73701267 | 0.93552616 |
| UTP15 | 67.2412374 | -0.4439887 | 0.32176071 | -1.3798725 | 0.16762592 | 0.62191735 |
| ARHGEF28 | 5554.77275 | -0.3609163 | 0.1883975 | -1.9157172 | 0.05540111 | 0.41841807 |
| HEXB | 176.474818 | -0.2070082 | 0.24344699 | -0.8503215 | 0.39514639 | 0.79901033 |
| GFM2 | 396.243485 | 0.07388948 | 0.22088904 | 0.33450948 | 0.73799514 | 0.93552616 |
| NSA2 | 242.536735 | 0.0815298 | 0.29589511 | 0.27553615 | 0.78290436 | 0.94928378 |
| FAM169A | 383.114611 | 0.90430471 | 0.46514279 | 1.94414432 | 0.05187805 | 0.40837733 |
| AC116337.3 | 47.8252303 | -0.0748053 | 0.52619948 | -0.1421614 | 0.88695249 | 0.97729111 |
| ANKRD31 | 88.4563019 | -0.2333791 | 0.63355497 | -0.3683644 | 0.71260152 | 0.92754765 |
| HMGCR | 437.267552 | 0.15276705 | 0.48845314 | 0.31275682 | 0.75446542 | 0.93901195 |
| COL4A3BP | 1602.99541 | -0.0825213 | 0.21297354 | -0.387472 | 0.69840682 | 0.92356678 |
| POLK | 1387.41073 | 0.09517505 | 0.17336261 | 0.54899413 | 0.58300948 | 0.88379959 |
| ANKDD1B | 374.993341 | 0.62918363 | 0.51752091 | 1.21576466 | 0.22407458 | 0.67678006 |
| POC5 | 287.860273 | -0.31955 | 0.21573137 | -1.4812403 | 0.13854255 | 0.58206378 |
| IQGAP2 | 73.6087852 | 0.7812117 | 0.47298132 | 1.65167559 | 0.0986007 | 0.51540034 |
| F2RL1 | 129.75889 | 0.67455044 | 0.32921124 | 2.04898972 | 0.04046312 | 0.37344844 |
| AC008581.2 | 121.518109 | 0.25868345 | 0.47522347 | 0.54434065 | 0.58620708 | 0.88530153 |

| | | | | | | |
|---|---|---|---|---|---|---|
| ZBED3 | 249.487666 | 0.46871191 | 0.29390016 | 1.5947998 | 0.11075701 | 0.53393773 |
| AC022414.1 | 152.200001 | 0.97329069 | 0.37060565 | 2.62621653 | 8.63E-03 | 0.18819528 |
| PDE8B | 42.558914 | 1.88614901 | 0.61950899 | 3.04458696 | 0.00233 | 0.09242336 |
| WDR41 | 851.477584 | -0.0295824 | 0.19981965 | -0.1480454 | 0.88230695 | 0.97664137 |
| TBCA | 1682.68724 | -0.1393413 | 0.22460335 | -0.6203885 | 0.53500205 | 0.8642718 |
| AP3B1 | 3560.0006 | 0.18405484 | 0.27862197 | 0.66058983 | 0.50887539 | 0.85659934 |
| SCAMP1-AS1 | 56.5993234 | -0.053843 | 0.50013843 | -0.1076561 | 0.91426846 | 0.98145965 |
| SCAMP1 | 769.305788 | -0.004228 | 0.17398472 | -0.0243008 | 0.98061267 | 0.99518016 |
| LHFPL2 | 119.207709 | -0.3645948 | 0.38610808 | -0.9442817 | 0.3450257 | 0.76724655 |
| ARSB | 228.511399 | -0.6275126 | 0.30961679 | -2.0267395 | 0.04268906 | 0.38022913 |
| DMGDH | 1797.64484 | 0.03749896 | 0.32489979 | 0.11541699 | 9.08E-01 | 9.80E-01 |
| BHMT2 | 1132.65197 | 0.73830409 | 0.43489464 | 1.69766195 | 0.08957158 | 0.50007863 |
| BHMT | 184.266503 | 0.23550141 | 0.45770119 | 0.51453091 | 6.07E-01 | 0.89174148 |
| JMY | 999.809024 | -0.0870168 | 0.25199037 | -0.3453179 | 0.72985542 | 0.93453345 |
| HOMER1 | 1562.94046 | -0.4204172 | 0.50997163 | -0.8243933 | 4.10E-01 | 0.80641067 |
| TENT2 | 3510.92515 | -0.2676803 | 0.17608983 | -1.5201347 | 0.12847711 | 0.5681467 |
| CMYA5 | 915.003831 | 0.42948908 | 0.40324408 | 1.06508467 | 0.28683761 | 0.72949981 |
| MTX3 | 223.392059 | 0.00179911 | 0.21484201 | 0.00837409 | 0.99331852 | 0.99821837 |
| THBS4 | 366.016579 | -0.270312 | 0.49145438 | -0.5500247 | 0.58230244 | 0.88338315 |
| SERINC5 | 930.380403 | -0.8097998 | 0.25590427 | -3.1644638 | 0.00155369 | 0.07326389 |
| AC010260.1 | 84.4428489 | -0.2403477 | 0.64375702 | -0.3733515 | 0.70888681 | 0.92577218 |
| ZFYVE16 | 1495.77873 | -0.3816166 | 0.21849027 | -1.7466067 | 0.08070558 | 0.48456361 |
| AC008771.1 | 102.342903 | 0.01433223 | 0.34651478 | 0.0413611 | 0.96700803 | 0.99248831 |
| FAM151B | 370.142498 | -0.1365277 | 0.46562328 | -0.2932149 | 0.76935793 | 0.94386043 |
| DHFR | 1506.82598 | -0.0056793 | 0.35735068 | -0.0158929 | 0.98731983 | 0.99653761 |
| MSH3 | 1488.53611 | -0.3983879 | 0.23833738 | -1.6715291 | 0.09461721 | 0.51017802 |
| RASGRF2 | 363.176346 | 1.10446648 | 0.53128844 | 2.07884528 | 0.03763157 | 0.36389125 |
| CKMT2-AS1 | 700.539864 | -0.3028176 | 0.30691875 | -0.9866378 | 0.32382023 | 0.75402436 |
| CKMT2 | 241.544706 | 1.22506715 | 0.38300824 | 3.19853998 | 0.00138125 | 0.06807829 |
| ZCCHC9 | 190.378796 | 0.00046862 | 0.2400696 | 0.00195204 | 0.9984425 | 0.99972915 |
| SSBP2 | 5010.22956 | -0.1322487 | 0.21936734 | -0.6028642 | 0.54659903 | 0.86820907 |
| ATG10 | 1756.76338 | -0.112835 | 0.24973928 | -0.451811 | 0.65140515 | 0.90825486 |
| RPS23 | 1309.15406 | 0.37613217 | 0.54529514 | 0.68977723 | 0.49033429 | 0.84614483 |
| ATP6AP1L | 746.749074 | -0.5405907 | 0.39582892 | -1.3657179 | 0.17202752 | 0.6286116 |
| TMEM167A | 444.682457 | 0.06251326 | 0.23377581 | 0.26740686 | 0.78915592 | 0.95080703 |
| XRCC4 | 1036.04053 | -0.4817827 | 0.25152071 | -1.9154792 | 0.05543142 | 0.41841807 |
| VCAN | 173.162978 | 1.48016555 | 0.74100709 | 1.99750525 | 4.58E-02 | 3.89E-01 |
| EDIL3 | 162.331182 | 0.06854433 | 0.76478282 | 0.08962588 | 0.92858452 | 0.98314146 |
| COX7C | 861.499113 | 0.70218148 | 0.4330775 | 1.62137605 | 0.10493701 | 0.52524176 |
| RASA1 | 2117.47841 | -0.1508456 | 0.16791195 | -0.8983614 | 0.36899293 | 0.78095617 |
| CCNH | 3193.91136 | 0.57186495 | 0.190405 | 3.00341352 | 0.00266969 | 0.09795553 |
| TMEM161B | 936.166996 | -0.0495868 | 0.2381816 | -0.2081891 | 0.83508133 | 0.96511435 |
| TMEM161B-A | 3414.51685 | -0.3879464 | 0.26764637 | -1.4494737 | 0.14720534 | 0.59373375 |
| MEF2C | 1249.47182 | 0.8141708 | 0.48709832 | 1.67147119 | 0.09462865 | 0.51017802 |
| MEF2C-AS1 | 529.624211 | 1.62852944 | 0.57746437 | 2.82013838 | 0.00480029 | 0.13724478 |
| CETN3 | 181.509046 | 0.32750757 | 0.29062965 | 1.12688975 | 2.60E-01 | 0.70921184 |
| MBLAC2 | 75.3671437 | -0.2119266 | 0.36527248 | -0.5801876 | 0.56178812 | 0.87403145 |
| POLR3G | 67.1065632 | -0.1410415 | 0.29643704 | -0.475789 | 0.63422472 | 0.90136076 |
| LYSMD3 | 295.108505 | 0.05360908 | 0.29413127 | 0.18226244 | 0.85537677 | 0.97118784 |
| ADGRV1 | 1984.99139 | -0.0982222 | 0.32472278 | -0.3024801 | 0.76228614 | 0.94212205 |
| LUCAT1 | 103.115713 | -1.2013988 | 1.11199054 | -1.0804038 | 2.80E-01 | 7.25E-01 |
| ARRDC3 | 2896.41996 | 0.21041552 | 0.40447599 | 0.52021759 | 0.60291193 | 0.88971776 |
| ARRDC3-AS1 | 159.720783 | -0.2314792 | 0.38498257 | -0.6012719 | 0.54765891 | 0.86859163 |
| NR2F1-AS1 | 972.322219 | -0.042116 | 0.64491201 | -0.0653051 | 0.94793111 | 0.98919519 |

| | | | | | | |
|---|---|---|---|---|---|---|
| NR2F1 | 98.80792 | -0.1733183 | 0.68797704 | -0.2519245 | 0.80109939 | 0.95370871 |
| FAM172A | 4355.26984 | -0.1769502 | 0.1541328 | -1.1480375 | 0.25095309 | 0.70200988 |
| KIAA0825 | 1479.68279 | 0.04611627 | 0.34799983 | 0.1325181 | 0.89457451 | 0.97788747 |
| SLF1 | 240.302556 | -0.7509279 | 0.38596333 | -1.9455939 | 0.05170354 | 0.40795233 |
| MCTP1 | 94.4062715 | -0.0299057 | 0.34845876 | -0.0858228 | 0.93160724 | 0.98357046 |
| TTC37 | 1321.30947 | -0.1768779 | 0.19138498 | -0.9241996 | 0.35538241 | 0.77296991 |
| ARSK | 220.333109 | 0.20267596 | 0.31285807 | 0.64782078 | 0.51710087 | 0.85893097 |
| RFESD | 41.8834828 | -0.5633273 | 0.43237816 | -1.3028578 | 0.1926233 | 0.64676186 |
| RHOBTB3 | 4662.50544 | 0.40482073 | 0.30537235 | 1.32566271 | 0.18495144 | 0.64011182 |
| GLRX | 953.113095 | 0.921087 | 0.44794185 | 2.05626469 | 0.03975701 | 0.37279391 |
| ELL2 | 1681.07504 | -0.589726 | 0.40165699 | -1.4682329 | 0.14204097 | 0.58723604 |
| AC104123.1 | 1021.50372 | -0.2243893 | 0.40648867 | -0.5520186 | 0.58093562 | 0.88283233 |
| CAST | 2407.35403 | -0.228117 | 0.15734154 | -1.4498204 | 0.14710862 | 0.59373375 |
| ERAP1 | 511.548514 | 0.04825555 | 0.34083186 | 0.14158169 | 0.88741043 | 0.97729111 |
| AC009126.1 | 276.614375 | 0.21828292 | 0.54113288 | 0.40338137 | 0.68666769 | 0.9208231 |
| ERAP2 | 223.51256 | 0.60306356 | 0.63483875 | 0.94994762 | 0.34213887 | 0.76585533 |
| LNPEP | 2689.42807 | 0.14166498 | 0.30450945 | 0.46522359 | 0.64177134 | 0.90432661 |
| LIX1-AS1 | 940.639415 | -0.1247796 | 0.2704238 | -0.4614226 | 0.64449545 | 0.90507553 |
| LIX1 | 63.9687634 | 1.63909415 | 0.92010228 | 1.78142603 | 0.07484288 | 0.47190267 |
| RIOK2 | 179.970835 | 0.16397247 | 0.26214748 | 0.62549704 | 0.53164489 | 0.86397197 |
| AC008883.1 | 55.383284 | -0.3446754 | 0.56647959 | -0.6084516 | 0.54288801 | 0.86590899 |
| LINC01340 | 92.2520225 | -0.4288341 | 0.57681523 | -0.7434514 | 0.45720841 | 0.83255882 |
| RGMB | 82.2691389 | 0.14228842 | 0.2895359 | 0.49143619 | 0.62311797 | 0.89865327 |
| CHD1 | 1792.27906 | -0.1106933 | 0.25204029 | -0.4391888 | 0.66052477 | 0.91099119 |
| LINC02062 | 629.065259 | 0.56308898 | 0.29911823 | 1.88249636 | 0.05976865 | 0.42668737 |
| AC021086.1 | 162.43419 | 0.03784211 | 0.54395988 | 0.06956784 | 0.94453764 | 0.98813503 |
| FAM174A | 275.308087 | 0.32216111 | 0.27680027 | 1.16387571 | 0.24447439 | 0.6948422 |
| SLCO4C1 | 1434.18991 | 0.04215193 | 0.4086134 | 0.10315846 | 9.18E-01 | 9.82E-01 |
| PAM | 4442.26924 | 0.09537752 | 0.2616376 | 0.36454058 | 0.71545438 | 0.92880162 |
| GIN1 | 261.071701 | -0.2580223 | 0.20978402 | -1.2299427 | 0.21871855 | 0.67202914 |
| PPIP5K2 | 1327.37493 | -0.1869013 | 0.16358828 | -1.1425101 | 0.25324204 | 0.70407789 |
| C5orf30 | 85.5366615 | -0.1481864 | 0.3998297 | -0.3706239 | 7.11E-01 | 0.92681719 |
| NUDT12 | 220.71173 | -0.1066421 | 0.20837338 | -0.5117839 | 0.60880228 | 0.89273938 |
| LINC01950 | 351.980005 | -0.3745078 | 0.55508771 | -0.6746822 | 0.49987767 | 0.85244523 |
| EFNA5 | 23426.0176 | -0.3668263 | 0.33853076 | -1.0835834 | 0.27854956 | 0.7242837 |
| FBXL17 | 6468.15408 | -0.1739402 | 0.15731922 | -1.1056515 | 0.26887739 | 0.71818262 |
| LINC01023 | 44.4431269 | -0.9625133 | 0.59538709 | -1.6166177 | 0.10596084 | 0.52735777 |
| FER | 3547.16876 | -0.3272083 | 0.16132101 | -2.0283056 | 0.04252906 | 0.38021566 |
| AC008467.1 | 171.519265 | -0.7384187 | 0.49247408 | -1.4994061 | 0.13376831 | 0.57704758 |
| PJA2 | 1346.92893 | 0.25700062 | 0.16706465 | 1.53833031 | 0.12396787 | 0.56105491 |
| MAN2A1 | 2252.3812 | -0.1609834 | 0.25920559 | -0.6210645 | 0.53455717 | 0.86419602 |
| TMEM232 | 2094.27355 | 0.25659095 | 0.441448 | 0.58124841 | 0.56107304 | 0.87394741 |
| SLC25A46 | 705.619394 | -0.2879528 | 0.21610778 | -1.3324499 | 0.1827124 | 0.63741052 |
| WDR36 | 394.578495 | 0.00169171 | 0.2155244 | 0.00784925 | 0.99373727 | 0.9983477 |
| STARD4 | 159.230269 | 0.35631674 | 0.5082655 | 0.70104451 | 0.48327524 | 0.84427756 |
| STARD4-AS1 | 137.844875 | -0.9319568 | 0.36737672 | -2.5367879 | 0.01118747 | 0.21604575 |
| NREP | 560.987794 | -1.5290413 | 0.42502534 | -3.5975298 | 0.00032125 | 0.03094754 |
| EPB41L4A | 3532.35329 | -0.5039524 | 0.27047138 | -1.8632374 | 0.06242887 | 0.43419765 |
| EPB41L4A-AS | 81.0538979 | 0.09672665 | 0.36061337 | 0.26822813 | 0.78852372 | 0.95048754 |
| AC010261.2 | 75.6628045 | -0.6749225 | 0.64775549 | -1.0419402 | 0.29743939 | 0.73673761 |
| AC104126.1 | 83.5837154 | -0.3472119 | 0.61339388 | -0.5660505 | 0.57135944 | 0.8787896 |
| APC | 919.471066 | -0.3631636 | 0.21909841 | -1.6575364 | 0.09741108 | 0.51355946 |
| SRP19 | 399.543817 | 0.08501251 | 0.28415016 | 0.29918162 | 0.76480147 | 0.94256992 |
| REEP5 | 1114.53053 | 0.16833998 | 0.23114137 | 0.72829878 | 0.46643071 | 0.83676263 |

| | | | | | | |
|---|---|---|---|---|---|---|
| DCP2 | 1050.26057 | -0.3089941 | 0.26966121 | -1.1458603 | 0.25185295 | 0.70276117 |
| MCC | 724.279498 | 0.37083622 | 0.3004728 | 1.23417567 | 0.21713745 | 0.66978787 |
| YTHDC2 | 1236.52342 | 0.17401197 | 0.19259602 | 0.90350759 | 0.36625656 | 0.78095617 |
| KCNN2 | 44.86399 | 0.05573332 | 0.52434743 | 0.10629082 | 0.91535162 | 0.9817213 |
| PGGT1B | 438.611836 | -0.0159368 | 0.14340277 | -0.1111332 | 0.91151072 | 0.98125148 |
| CCDC112 | 161.470476 | -0.0457651 | 0.39019589 | -0.1172876 | 0.90663216 | 0.98032942 |
| FEM1C | 263.59251 | -0.2236286 | 0.26475513 | -0.844662 | 0.39829956 | 0.80148305 |
| TMED7-TICA | 115.03023 | 0.71613723 | 0.26819465 | 2.67021447 | 0.00758028 | 0.17504883 |
| AC010226.1 | 61.80929 | 1.13314584 | 0.45095548 | 2.51276652 | 0.01197886 | 0.22380303 |
| ATG12 | 438.412572 | -0.1038321 | 0.19575847 | -0.5304092 | 0.59582828 | 0.88859628 |
| AP3S1 | 593.083436 | 0.17245377 | 0.19593104 | 0.8801758 | 0.37876408 | 0.78739294 |
| COMMD10 | 2867.42952 | -0.2971784 | 0.14305311 | -2.0773994 | 0.03776471 | 0.36411243 |
| SEMA6A | 259.353367 | 0.7682517 | 0.36726454 | 2.09182109 | 0.03645452 | 0.35908113 |
| SEMA6A-AS1 | 377.656923 | -0.5256368 | 0.20288263 | -2.5908419 | 0.00957415 | 0.19884419 |
| LINC02147 | 51.2375021 | 0.75215483 | 0.69014364 | 1.08985259 | 0.27577809 | 0.72233844 |
| DTWD2 | 914.376154 | -0.1765788 | 0.22604878 | -0.7811537 | 4.35E-01 | 0.81919857 |
| AC008629.1 | 564.313082 | 0.9891398 | 0.49902322 | 1.98215185 | 0.04746225 | 0.39411174 |
| DMXL1 | 2475.79533 | -0.0138056 | 0.14532706 | -0.0949969 | 0.92431727 | 0.98215577 |
| TNFAIP8 | 1395.75637 | -0.1650449 | 0.43398469 | -0.3803012 | 7.04E-01 | 0.92513032 |
| HSD17B4 | 1166.5311 | 0.27653355 | 0.24642707 | 1.12217196 | 0.26178933 | 0.7126907 |
| SRFBP1 | 438.296328 | -0.3901617 | 0.25482812 | -1.5310778 | 0.12575016 | 0.56516974 |
| AC119150.1 | 81.686067 | 0.1345485 | 0.46035193 | 0.29227312 | 0.77007781 | 0.94387751 |
| AC022101.1 | 181.240631 | 0.1946438 | 0.82328108 | 0.23642448 | 0.8131033 | 0.95816239 |
| SNX2 | 981.435363 | 0.03958318 | 0.26224104 | 0.15094198 | 0.88002149 | 0.97664137 |
| SNX24 | 1998.39435 | 0.02863766 | 0.26918091 | 0.10638814 | 0.9152744 | 0.9817213 |
| PPIC | 183.430355 | 0.29443631 | 0.31874116 | 0.92374738 | 0.35561786 | 0.77310012 |
| AC106786.2 | 45.2216951 | 0.20053667 | 0.43428729 | 0.46176039 | 0.64425316 | 0.90507553 |
| PRDM6 | 135.460381 | -0.2197247 | 0.39381883 | -0.5579334 | 0.57688989 | 0.88150331 |
| CEP120 | 636.767453 | -0.1178297 | 0.21924115 | -0.5374434 | 0.59096137 | 0.8867913 |
| CSNK1G3 | 934.767243 | -0.0846317 | 0.28042337 | -0.3017998 | 0.76280468 | 0.94225911 |
| ZNF608 | 1892.34743 | 0.03152309 | 0.3121776 | 0.10097807 | 0.91956787 | 0.98215577 |
| AC112196.1 | 71.8347307 | 0.45881155 | 0.40411761 | 1.13534163 | 0.25623221 | 0.70709713 |
| LINC02240 | 71.5184599 | -0.3111311 | 0.48867206 | -0.6366868 | 0.52432885 | 0.86006369 |
| AC010235.1 | 116.478727 | -0.3952108 | 0.50814066 | -0.7777586 | 0.43671132 | 0.82013459 |
| GRAMD2B | 5226.09763 | -0.1017652 | 0.24010737 | -0.4238322 | 0.67168819 | 0.91574062 |
| AC093535.1 | 274.569319 | 0.16362541 | 0.74067812 | 0.22091298 | 0.82516019 | 0.96146415 |
| ALDH7A1 | 481.96942 | -0.17413 | 0.36275888 | -0.4800158 | 0.63121619 | 0.90011169 |
| PHAX | 197.038705 | 0.06409715 | 0.20043484 | 0.31979047 | 0.74912718 | 0.93748128 |
| 3-Mar | 177.1312 | -0.4948826 | 0.46322524 | -1.0683412 | 0.28536662 | 0.72844615 |
| C5orf63 | 273.669325 | 0.01679087 | 0.2535387 | 0.06622607 | 0.94719784 | 0.98905425 |
| PRRC1 | 362.397483 | 0.06601151 | 0.19469595 | 0.33904921 | 0.73457266 | 0.93552616 |
| CCDC192 | 49.7398328 | 0.55077691 | 0.54945827 | 1.00239988 | 0.3161505 | 0.74785672 |
| LINC01184 | 1213.88228 | -0.1632157 | 0.22763778 | -0.7169974 | 0.47337572 | 0.84043277 |
| SLC12A2 | 423.294973 | -0.1463782 | 0.39914787 | -0.3667268 | 0.71382285 | 0.92760608 |
| SLC27A6 | 142.966992 | 0.00790896 | 1.38243089 | 0.00572105 | 0.99543529 | 0.99868637 |
| ISOC1 | 192.261813 | 0.00410956 | 0.26025269 | 0.01579064 | 0.98740141 | 0.99653761 |
| CHSY3 | 146.050038 | 0.00734033 | 0.49997005 | 0.01468154 | 0.98828625 | 0.99653761 |
| HINT1 | 1340.06924 | 0.52078634 | 0.42083332 | 1.23751211 | 0.21589702 | 0.66911866 |
| LYRM7 | 304.007983 | 0.35030119 | 0.2026587 | 1.72852777 | 0.08389365 | 0.48924997 |
| CDC42SE2 | 4532.12266 | 0.08509575 | 0.43970071 | 0.19353107 | 8.47E-01 | 9.69E-01 |
| RAPGEF6 | 45.4825629 | 0.57283755 | 0.35965886 | 1.59272468 | 0.11122197 | 0.53461826 |
| P4HA2 | 146.709335 | 0.70668609 | 0.51035384 | 1.3846983 | 0.16614474 | 0.61901667 |
| PDLIM4 | 97.4633856 | 1.27650142 | 0.49795305 | 2.56349755 | 0.01036234 | 0.20676162 |
| SLC22A4 | 158.432243 | 0.11794875 | 0.41402233 | 0.284885 | 0.77573225 | 0.9468349 |

| | | | | | | |
|---|---|---|---|---|---|---|
| SLC22A5 | 363.462442 | 0.20533285 | 0.25888751 | 0.79313538 | 0.42769895 | 0.81466629 |
| AC116366.3 | 814.453102 | 0.41469943 | 0.7351199 | 0.56412489 | 0.57266914 | 0.87946834 |
| IRF1 | 154.352692 | 0.61831793 | 0.45459665 | 1.36014625 | 1.74E-01 | 0.6286116 |
| TH2LCRR | 110.19507 | 0.36895599 | 0.57912877 | 0.63708801 | 0.52406752 | 0.86006369 |
| AC004039.1 | 90.3400448 | -0.0120892 | 0.70341462 | -0.0171864 | 0.9862879 | 0.99624198 |
| KIF3A | 568.849688 | 0.21825123 | 0.18429884 | 1.18422469 | 0.23632413 | 0.68864007 |
| 8-Sep | 256.596843 | 0.13722748 | 0.32686376 | 0.41983082 | 0.67460905 | 0.91636338 |
| UQCRQ | 329.481501 | 0.71929436 | 0.44766348 | 1.60677472 | 0.1081038 | 0.53063658 |
| AFF4 | 2547.31444 | -0.1804963 | 0.15568807 | -1.1593455 | 0.24631539 | 0.6967274 |
| ZCCHC10 | 372.036889 | -0.4452601 | 0.21340963 | -2.0864106 | 0.03694144 | 0.36038369 |
| HSPA4 | 569.387641 | -0.0015523 | 0.21078605 | -0.0073644 | 0.99412412 | 0.998438 |
| FSTL4 | 213.016053 | 0.48091741 | 0.70045557 | 0.68657804 | 0.49234867 | 0.84734855 |
| C5orf15 | 383.395607 | 0.04594533 | 0.22515305 | 0.20406267 | 0.83830452 | 0.96566728 |
| VDAC1 | 721.529168 | 0.67840635 | 0.26212022 | 2.58814964 | 0.00964931 | 0.19975064 |
| TCF7 | 142.252371 | -0.0064681 | 0.30491854 | -0.0212126 | 0.98307603 | 0.99518552 |
| SKP1 | 939.365494 | 0.78275348 | 0.34445048 | 2.27247029 | 0.02305812 | 0.2983102 |
| CDKL3 | 539.532355 | -0.1941125 | 0.22031721 | -0.8810592 | 0.37828577 | 0.78689398 |
| UBE2B | 1242.85932 | -0.0929928 | 0.17367808 | -0.5354322 | 0.59235102 | 0.88740306 |
| CDKN2AIPNL | 73.2414928 | 0.24161203 | 0.32689221 | 0.73911836 | 0.45983513 | 0.83390278 |
| JADE2 | 406.257039 | 0.21460751 | 0.34074238 | 0.62982337 | 0.52881016 | 0.86267177 |
| SAR1B | 522.996193 | 0.29143089 | 0.25758208 | 1.13140982 | 0.25788264 | 0.70885276 |
| SEC24A | 674.780534 | -0.1055533 | 0.25628471 | -0.4118594 | 0.68044246 | 0.91867756 |
| CAMLG | 230.344759 | 0.32030157 | 0.18211299 | 1.75880682 | 0.07861033 | 0.48060389 |
| DDX46 | 1087.41449 | -0.2523139 | 0.17908519 | -1.4089045 | 0.1588634 | 0.60823961 |
| C5orf24 | 294.289128 | 0.35170373 | 0.32989218 | 1.06611722 | 0.28637065 | 0.72934245 |
| TXNDC15 | 320.006729 | -0.2315658 | 0.33436478 | -0.6925543 | 0.4885893 | 0.84562056 |
| PCBD2 | 1332.84579 | -0.6349394 | 0.24453642 | -2.5965023 | 0.00941783 | 0.19714749 |
| C5orf66 | 85.0183996 | 0.05483126 | 0.50602885 | 0.10835599 | 0.91371331 | 0.98145965 |
| H2AFY | 800.959316 | 0.0936566 | 0.17881634 | 0.5237586 | 0.60044646 | 0.88883692 |
| CXCL14 | 508.616415 | 1.44043074 | 0.56346054 | 2.55640039 | 0.01057613 | 0.20870812 |
| SLC25A48 | 248.450726 | 0.3230937 | 0.5023888 | 0.64311486 | 5.20E-01 | 8.59E-01 |
| SMAD5 | 770.557283 | 0.04009692 | 0.21955712 | 0.18262639 | 0.85509117 | 0.97118784 |
| SPOCK1 | 256.821825 | -2.6043867 | 0.65570983 | -3.9718585 | 7.13E-05 | 0.01385276 |
| KLHL3 | 248.279682 | 0.71668823 | 0.65196491 | 1.09927423 | 0.27164847 | 0.72040597 |
| HNRNPA0 | 254.394671 | 0.34552159 | 0.29775439 | 1.16042484 | 0.24587588 | 0.69605111 |
| AC106791.1 | 234.13711 | 0.25192852 | 0.44801861 | 0.56231709 | 0.5739 | 0.88002205 |
| PKD2L2 | 88.9800835 | -0.0634154 | 0.6605251 | -0.0960075 | 0.92351459 | 0.98215577 |
| FAM13B | 1824.05973 | -0.1670771 | 0.1476804 | -1.1313424 | 0.257911 | 0.70885276 |
| NME5 | 553.04523 | 0.27570447 | 0.44240271 | 0.62319797 | 0.53315444 | 0.86397197 |
| BRD8 | 605.90334 | -0.1068606 | 0.25083916 | -0.4260124 | 0.67009883 | 0.914701 |
| CDC23 | 205.622469 | 0.20618818 | 0.19517311 | 1.05643743 | 0.29076841 | 0.73296946 |
| FAM53C | 218.671048 | 0.16694641 | 0.39413985 | 0.42357151 | 0.67187834 | 0.91574062 |
| KDM3B | 1192.73696 | -0.1792669 | 0.14037429 | -1.277064 | 0.20157967 | 0.65534477 |
| EGR1 | 320.353726 | -0.1278482 | 0.82683892 | -0.1546228 | 0.87711867 | 0.97639142 |
| ETF1 | 681.053705 | -0.0953645 | 0.41163815 | -0.2316706 | 0.81679386 | 0.95909544 |
| HSPA9 | 1074.94175 | 0.13954424 | 0.18151432 | 0.76877812 | 0.44202504 | 0.82285053 |
| CTNNA1 | 4264.85979 | -0.2838153 | 0.17087725 | -1.6609308 | 0.09672734 | 0.51270925 |
| SIL1 | 2360.14791 | -0.1958813 | 0.19431354 | -1.0080681 | 0.31342175 | 0.74749123 |
| AC011405.1 | 95.7186862 | -0.2032002 | 0.64414661 | -0.3154565 | 0.75241508 | 0.93818313 |
| MATR3 | 52.2288104 | -0.9272186 | 0.410637 | -2.2580006 | 0.02394562 | 0.30325761 |
| PAIP2 | 884.678569 | -0.0396947 | 0.28894399 | -0.1373784 | 0.89073168 | 0.97729111 |
| SPATA24 | 44.5041427 | 0.44396647 | 0.41063286 | 1.08117618 | 0.27961875 | 0.72492327 |
| DNAJC18 | 90.0249737 | 0.17071232 | 0.29176264 | 0.58510685 | 0.55847587 | 0.87295536 |
| UBE2D2 | 4218.35491 | 0.62069223 | 0.53660806 | 1.15669569 | 0.24739671 | 0.69761067 |

| | | | | | | |
|---|---|---|---|---|---|---|
| CXXC5 | 747.114111 | 0.36910516 | 0.48973281 | 0.7536868 | 0.45103731 | 0.82814845 |
| AC008667.1 | 71.1566198 | 0.15350924 | 0.29128851 | 0.52700068 | 0.59819312 | 0.88883692 |
| NRG2 | 302.188799 | 1.42781001 | 0.60772741 | 2.34942508 | 0.01880243 | 0.2768799 |
| PURA | 957.323796 | 0.17947754 | 0.17048118 | 1.0527704 | 0.29244623 | 0.73425628 |
| IGIP | 135.010697 | 0.52810406 | 0.27749778 | 1.90309289 | 0.05702843 | 0.41872623 |
| CYSTM1 | 2303.84694 | 0.22862259 | 0.33050406 | 0.69173912 | 0.48910117 | 0.84606305 |
| PFDN1 | 532.621864 | -0.1324994 | 0.22123251 | -0.5989147 | 0.54922974 | 0.86904356 |
| SLC4A9 | 39.1132995 | 1.12505647 | 0.74255716 | 1.51511094 | 0.12974431 | 0.57050365 |
| AC008438.1 | 176.635883 | 0.04665725 | 0.42518225 | 0.1097347 | 0.91261978 | 0.98135874 |
| ANKHD1-EIF4 | 178.19086 | -0.0129099 | 0.30615118 | -0.0421684 | 0.96636448 | 0.99248831 |
| SRA1 | 231.280581 | 0.29171111 | 0.30554973 | 0.95470912 | 0.33972482 | 0.76449297 |
| APBB3 | 109.524766 | -0.1233109 | 0.41352763 | -0.2981925 | 0.76555621 | 0.94270299 |
| SLC35A4 | 132.027619 | 0.70280635 | 0.37555344 | 1.87138839 | 0.06129127 | 0.43236485 |
| NDUFA2 | 277.763454 | 0.85629064 | 0.38564654 | 2.22040277 | 0.02639144 | 0.31402818 |
| IK | 190.346475 | 0.10737896 | 0.25671967 | 0.4182732 | 0.67574738 | 0.91649321 |
| WDR55 | 147.832376 | -0.4271039 | 0.18628339 | -2.2927644 | 0.02186157 | 0.29509055 |
| HARS | 344.466623 | -0.1188762 | 0.15424299 | -0.770707 | 0.4408806 | 0.82265742 |
| HARS2 | 92.2842795 | -0.4818648 | 0.33376435 | -1.4437277 | 0.14881558 | 0.59502131 |
| ZMAT2 | 148.979776 | 0.36997585 | 0.28856781 | 1.28211062 | 0.19980384 | 0.65230079 |
| AC005609.5 | 106.453638 | -0.4273895 | 0.56193037 | -0.7605738 | 0.4469117 | 0.82571251 |
| AC244517.11 | 46.9020016 | 0.07541638 | 0.59683979 | 0.1263595 | 0.89944736 | 0.97889101 |
| AC244517.10 | 40.1240946 | -0.4017042 | 0.57499418 | -0.6986231 | 0.48478762 | 0.8447301 |
| TAF7 | 365.447952 | 0.41198121 | 0.2815775 | 1.46311839 | 0.14343499 | 0.58849865 |
| DIAPH1 | 2979.91395 | 0.13057499 | 0.28015196 | 0.4660863 | 0.64115373 | 0.90404588 |
| AC008781.2 | 115.485073 | 0.22312887 | 0.57266124 | 0.38963502 | 0.69680645 | 0.92322348 |
| AC008781.1 | 64.7743215 | -0.1221181 | 0.43927559 | -0.2779989 | 0.78101324 | 0.94811174 |
| HDAC3 | 371.35391 | -0.0366976 | 0.23725735 | -0.1546742 | 0.87707819 | 0.97639142 |
| FCHSD1 | 63.3686195 | -0.4076168 | 0.3380272 | -1.2058697 | 0.22786772 | 0.67959015 |
| PCDH1 | 305.636369 | 1.77323259 | 0.46053545 | 3.85037157 | 0.00011794 | 0.01730948 |
| DELE1 | 349.732075 | -0.0669164 | 0.2503032 | -0.2673412 | 0.78920648 | 0.95080703 |
| RNF14 | 342.925013 | 0.07398787 | 0.20793734 | 0.3558181 | 0.7219768 | 0.93079284 |
| GNPDA1 | 199.521451 | 0.04355034 | 0.31663333 | 0.13754186 | 0.89060251 | 0.97729111 |
| NDFIP1 | 1515.42111 | 0.04809619 | 0.18778746 | 0.25612038 | 0.79785789 | 0.95288536 |
| SPRY4 | 105.146554 | 0.12376714 | 0.46552996 | 0.26586289 | 7.90E-01 | 0.95130361 |
| SPRY4-AS1 | 236.578948 | -0.5453733 | 0.42773422 | -1.2750284 | 2.02E-01 | 0.65552334 |
| ARHGAP26 | 1863.86746 | -0.1107002 | 0.43851203 | -0.2524451 | 8.01E-01 | 0.95370871 |
| NR3C1 | 1894.97688 | -0.0258959 | 0.20602743 | -0.1256916 | 0.89997602 | 0.97898519 |
| YIPF5 | 229.408218 | -0.0138463 | 0.22761665 | -0.0608317 | 0.95149324 | 0.98938795 |
| KCTD16 | 2734.75325 | -1.0927 | 0.47528339 | -2.2990494 | 2.15E-02 | 2.93E-01 |
| PRELID2 | 1221.26426 | 0.50499121 | 0.45697639 | 1.10507066 | 0.26912897 | 0.71818262 |
| PLAC8L1 | 42.0463624 | -0.450466 | 0.43791581 | -1.028659 | 0.30363997 | 0.7409078 |
| AC091887.1 | 48.9679579 | -0.0686117 | 0.65930753 | -0.1040662 | 0.91711678 | 0.98199511 |
| LARS | 890.069105 | 0.03194787 | 0.16486957 | 0.19377663 | 0.84635079 | 0.96883382 |
| AC091959.3 | 54.8602678 | -0.1652966 | 0.33230983 | -0.4974171 | 0.61889492 | 0.89608494 |
| TCERG1 | 1173.32504 | -0.3293075 | 0.18655776 | -1.7651772 | 0.07753399 | 0.47834956 |
| PPP2R2B | 1454.26601 | -1.2074754 | 0.44545912 | -2.7106312 | 0.00671553 | 0.16906706 |
| STK32A-AS1 | 58.1557258 | 2.02679454 | 0.72123027 | 2.81019064 | 0.00495122 | 0.14034467 |
| STK32A | 787.593046 | 1.55198327 | 0.80133998 | 1.9367351 | 0.05277773 | 0.41066414 |
| AC091948.1 | 551.278569 | 0.46930115 | 0.30063671 | 1.5610241 | 1.19E-01 | 0.54998912 |
| FBXO38 | 676.074493 | -0.4099551 | 0.20471045 | -2.0026094 | 0.04521923 | 0.38785764 |
| GRPEL2 | 44.5739566 | 0.41469186 | 0.34721075 | 1.19435202 | 0.2323403 | 0.68450837 |
| PCYOX1L | 36.3478233 | 0.19243998 | 0.61525538 | 0.31278066 | 0.75444731 | 0.93901195 |
| CSNK1A1 | 3793.73313 | 0.02760177 | 0.15510415 | 0.17795634 | 0.85875727 | 0.97174503 |
| ARHGEF37 | 284.970542 | 0.49310038 | 0.47066991 | 1.04765648 | 0.29479691 | 0.73568342 |

| | | | | | | |
|---|---|---|---|---|---|---|
| SLC26A2 | 347.98549 | -0.915662 | 0.811225 | -1.1287399 | 0.25900756 | 0.70921184 |
| HMGXB3 | 469.396773 | -0.5587828 | 0.26379062 | -2.1182815 | 0.03415124 | 0.35153234 |
| TCOF1 | 148.700308 | -0.5298624 | 0.30591983 | -1.7320302 | 0.08326818 | 0.48775582 |
| CD74 | 1454.7124 | 0.57103359 | 0.49185752 | 1.16097358 | 0.24565264 | 0.69581163 |
| RPS14 | 1115.18195 | 0.63767968 | 0.34769713 | 1.83400903 | 0.06665265 | 0.4478204 |
| NDST1 | 382.343702 | -0.1041233 | 0.24334742 | -0.4278791 | 0.66873913 | 0.91395501 |
| SYNPO | 105.482684 | 0.59451526 | 0.38567598 | 1.54148896 | 0.12319783 | 0.56015422 |
| RBM22 | 165.039712 | -0.0169118 | 0.26627837 | -0.0635116 | 0.94935912 | 0.98919519 |
| DCTN4 | 764.800333 | -0.2085632 | 0.11720704 | -1.7794429 | 0.07516718 | 0.47190267 |
| SMIM3 | 98.2383191 | -0.6996435 | 0.49269156 | -1.4200436 | 0.155595 | 0.6050204 |
| ZNF300 | 92.2050127 | -1.1778943 | 0.32245754 | -3.6528663 | 0.00025933 | 0.02718635 |
| GPX3 | 4458.56018 | 0.89956906 | 0.70717886 | 1.27205309 | 0.20335426 | 0.65642873 |
| TNIP1 | 1027.66743 | 0.17624726 | 0.36467926 | 0.48329391 | 0.62888706 | 0.89970424 |
| ANXA6 | 196.726275 | 1.28567639 | 0.49531657 | 2.59566602 | 0.00944078 | 0.19731525 |
| GM2A | 41.1449678 | 0.85304535 | 0.49430376 | 1.72575127 | 0.08439217 | 0.4907146 |
| SLC36A1 | 105.31331 | -1.0295648 | 0.37501507 | -2.7453959 | 0.00604379 | 0.15966493 |
| SPARC | 72.4556977 | 0.00064724 | 0.43461203 | 0.00148924 | 0.99881176 | 0.99980325 |
| ATOX1 | 319.264948 | 0.2687022 | 0.21100971 | 1.27341155 | 0.20287204 | 0.65552334 |
| G3BP1 | 1248.91997 | 0.11228534 | 0.32390523 | 0.3466611 | 0.72884594 | 0.93453345 |
| LINC01933 | 349.103244 | 1.09017502 | 0.61436168 | 1.77448407 | 0.07598311 | 0.47431992 |
| LINC01470 | 133.779246 | -0.4263017 | 0.75713234 | -0.5630478 | 0.57340234 | 0.87985952 |
| FAM114A2 | 222.049422 | -0.1268833 | 0.1869923 | -0.6785484 | 0.49742403 | 0.85092921 |
| MFAP3 | 328.826439 | -0.0681827 | 0.27445883 | -0.248426 | 0.80380483 | 0.95450237 |
| GALNT10 | 1206.46813 | 0.74265933 | 0.54512461 | 1.36236618 | 0.17308235 | 0.6286116 |
| SAP30L-AS1 | 329.467599 | 0.39116934 | 0.28751045 | 1.36053956 | 0.17365924 | 0.6286116 |
| SAP30L | 151.918074 | -0.0470107 | 0.23973394 | -0.1960954 | 0.84453546 | 0.96853443 |
| LARP1 | 1770.89633 | -0.3505265 | 0.23582241 | -1.4864001 | 0.13717332 | 0.5794443 |
| FAXDC2 | 714.574148 | -0.4587582 | 0.46305333 | -0.9907243 | 0.32182022 | 0.75240254 |
| CNOT8 | 362.404753 | -0.4320096 | 0.21975744 | -1.9658474 | 0.04931624 | 0.39892272 |
| GEMIN5 | 227.101982 | -0.3863642 | 0.24193522 | -1.5969737 | 0.11027156 | 0.53372881 |
| MRPL22 | 229.103555 | 0.156455 | 0.2270028 | 0.68922056 | 0.49068448 | 0.84614483 |
| SGCD | 169.88218 | -0.4553133 | 0.78701915 | -0.5785289 | 5.63E-01 | 0.87436903 |
| TIMD4 | 114.009378 | 0.73368787 | 0.87095583 | 0.84239389 | 4.00E-01 | 8.02E-01 |
| HAVCR1 | 580.684223 | 1.25712826 | 0.87238698 | 1.44102136 | 1.50E-01 | 5.96E-01 |
| HAVCR2 | 204.100842 | -0.1190532 | 0.40736452 | -0.2922523 | 0.77009372 | 0.94387751 |
| AC009185.1 | 42.1954434 | -0.2555123 | 0.74065262 | -0.3449826 | 0.73010746 | 0.93453345 |
| CYFIP2 | 4251.82817 | 0.12194419 | 0.38798515 | 0.31430118 | 7.53E-01 | 9.38E-01 |
| AC008676.1 | 72.2784591 | 0.26187712 | 0.74816467 | 0.35002604 | 0.72631915 | 0.93335438 |
| THG1L | 45.7244323 | -0.7411551 | 0.393775 | -1.882179 | 0.05981171 | 0.42668737 |
| LSM11 | 43.4308915 | -0.7051785 | 0.31776916 | -2.2191534 | 0.02647629 | 0.31421858 |
| CLINT1 | 3445.3167 | -0.1803852 | 0.29615627 | -0.6090879 | 0.5424662 | 0.8658519 |
| RNF145 | 1090.99519 | 0.41895862 | 0.24529215 | 1.70799845 | 0.08763663 | 0.49639462 |
| UBLCP1 | 295.503558 | -0.1740362 | 0.27390497 | -0.6353889 | 0.52517477 | 0.8606763 |
| AC008691.1 | 84.6935364 | 0.10717441 | 0.62926682 | 0.17031633 | 0.86476137 | 0.97361767 |
| TTC1 | 694.448849 | -0.2421436 | 0.22550097 | -1.073803 | 0.28291097 | 0.72604837 |
| PWWP2A | 832.682871 | 0.17478771 | 0.16784581 | 1.04135875 | 0.29770908 | 0.73673761 |
| CCNJL | 471.704591 | -0.0529498 | 0.31791192 | -0.1665549 | 0.86772025 | 0.97402505 |
| ZBED8 | 70.4566744 | 0.24836687 | 0.28819617 | 0.86179796 | 0.38879871 | 0.79634939 |
| SLU7 | 215.362285 | -0.0810783 | 0.19020007 | -0.4262791 | 0.66990446 | 0.91465659 |
| GABRB2 | 177.829054 | 1.7724451 | 1.00542439 | 1.76288253 | 0.0779203 | 0.47916634 |
| AC113414.1 | 1326.39122 | 1.32452657 | 0.45186384 | 2.93125151 | 0.00337599 | 0.11318146 |
| CCNG1 | 281.84525 | 0.93481657 | 0.28873366 | 3.23764323 | 0.00120521 | 0.06317331 |
| NUDCD2 | 137.510908 | 0.68782829 | 0.29907398 | 2.29985999 | 0.02145615 | 0.29288643 |
| MAT2B | 332.030756 | 0.38922367 | 0.25041645 | 1.55430551 | 0.12011156 | 0.55298199 |

| | | | | | | |
|---|---|---|---|---|---|---|
| AC008415.1 | 205.489367 | -0.4196671 | 0.62507335 | -0.6713885 | 5.02E-01 | 8.55E-01 |
| TENM2 | 81.2958792 | 0.50645073 | 0.40355843 | 1.2549626 | 0.20949234 | 0.66216422 |
| WWC1 | 6635.38695 | -0.2950083 | 0.37799699 | -0.7804516 | 0.43512513 | 0.81919857 |
| RARS | 606.565034 | -0.2383132 | 0.25252874 | -0.9437074 | 0.34531921 | 0.76776999 |
| PANK3 | 540.120724 | -0.1018915 | 0.30525248 | -0.3337941 | 0.73853491 | 0.93552616 |
| SLIT3 | 173.395729 | -0.5162516 | 0.8082656 | -0.6387153 | 5.23E-01 | 0.86006369 |
| SPDL1 | 65.1091659 | -0.0954285 | 0.34305907 | -0.2781692 | 0.78088245 | 0.9480401 |
| KCNIP1 | 84.5699013 | 1.7970598 | 0.79076207 | 2.27256703 | 0.02305228 | 0.2983102 |
| GABRP | 71.5390562 | 0.77316205 | 0.82821084 | 0.93353288 | 3.51E-01 | 0.77044053 |
| RANBP17 | 641.076604 | 0.02798855 | 0.43967841 | 0.06365686 | 0.94924346 | 0.98919519 |
| NPM1 | 612.859806 | 0.31382527 | 0.22671705 | 1.38421557 | 0.16629246 | 0.61901833 |
| FBXW11 | 3510.70147 | -0.4882323 | 0.32090608 | -1.521418 | 0.12815498 | 0.56762585 |
| STK10 | 215.708702 | -0.6156391 | 0.33957087 | -1.8129915 | 0.06983312 | 0.45914671 |
| UBTD2 | 599.488311 | -0.2893103 | 0.20810124 | -1.3902381 | 0.1644566 | 0.61625736 |
| SH3PXD2B | 121.506208 | 0.57475318 | 0.51806971 | 1.10941283 | 0.26725213 | 0.71736098 |
| DUSP1 | 865.593433 | 0.37481294 | 0.68906877 | 0.54394127 | 0.58648189 | 0.88538429 |
| ERGIC1 | 1261.81293 | 0.27030803 | 0.31819348 | 0.84950837 | 0.39559847 | 0.79924445 |
| RPL26L1 | 72.7759324 | -0.0166595 | 0.36163448 | -0.0460672 | 0.96325666 | 0.99224805 |
| ATP6V0E1 | 2313.91974 | 0.99701221 | 0.26074839 | 3.82365621 | 0.00013149 | 0.01887842 |
| CREBRF | 2092.19037 | 0.24304407 | 0.29203455 | 0.83224425 | 0.40527109 | 0.80481948 |
| BNIP1 | 110.558848 | 0.180263 | 0.29123754 | 0.61895525 | 0.53594584 | 0.8642718 |
| BOD1 | 164.373555 | -0.0287297 | 0.27339464 | -0.1050849 | 0.91630841 | 0.98195017 |
| CPEB4 | 6988.76159 | -0.3328376 | 0.37327317 | -0.8916731 | 0.3725682 | 0.78269743 |
| SFXN1 | 793.987863 | 0.10233665 | 0.24357705 | 0.42014078 | 0.67438261 | 0.91636338 |
| THOC3 | 276.858567 | -0.5887329 | 0.32076935 | -1.8353776 | 0.06644976 | 0.4471395 |
| SIMC1 | 704.836429 | 0.00660841 | 0.18030966 | 0.03665032 | 0.97076382 | 0.99317943 |
| KIAA1191 | 346.6725 | 0.17854485 | 0.21122005 | 0.84530259 | 0.39794191 | 0.80116061 |
| ARL10 | 206.933805 | 0.14165671 | 0.33046327 | 0.42866099 | 0.66816995 | 0.91383898 |
| CLTB | 271.537742 | 0.39898006 | 0.29349707 | 1.35940047 | 0.17401972 | 0.6286158 |
| FAF2 | 846.527934 | -0.2436533 | 0.17962532 | -1.3564527 | 0.17495517 | 0.62944785 |
| RNF44 | 210.661556 | 0.11579617 | 0.31585528 | 0.36661147 | 0.71390885 | 0.92760608 |
| CDHR2 | 74.756913 | -0.833444 | 0.61185955 | -1.3621493 | 0.17315078 | 0.6286116 |
| TSPAN17 | 142.33039 | 0.21582591 | 0.39878299 | 0.54121143 | 0.58836186 | 0.88571413 |
| UIMC1 | 1121.8719 | -0.1583414 | 0.16324084 | -0.9699865 | 0.33205323 | 0.76037513 |
| ZNF346 | 616.635309 | -0.0353181 | 0.20861102 | -0.1693014 | 0.8655596 | 0.97379123 |
| FGFR4 | 68.9247924 | 0.80349839 | 0.55689948 | 1.44280686 | 0.14907489 | 0.59580338 |
| NSD1 | 2824.54663 | 0.11831754 | 0.19610456 | 0.60333906 | 0.54628315 | 0.86802047 |
| PRELID1 | 211.417383 | 0.64404253 | 0.42206858 | 1.52591914 | 0.12703 | 0.56740575 |
| LMAN2 | 361.608401 | 0.51862233 | 0.24595292 | 2.10862444 | 0.03497701 | 0.35358029 |
| RGS14 | 74.1308847 | 0.64583282 | 0.31732385 | 2.03524827 | 0.0418259 | 0.37686104 |
| GRK6 | 59.4732732 | 0.19529462 | 0.34098835 | 0.57273106 | 0.5668268 | 0.87646121 |
| DBN1 | 238.846201 | -0.711101 | 0.36087098 | -1.970513 | 0.0487796 | 0.39762623 |
| PDLIM7 | 158.790166 | -2.0307466 | 0.65281637 | -3.110747 | 0.00186615 | 0.08355912 |
| DDX41 | 96.6618441 | 0.12554386 | 0.42241257 | 0.29720673 | 0.76630868 | 0.94300088 |
| FAM193B | 687.313644 | -0.2747374 | 0.18864286 | -1.4563893 | 0.14528502 | 0.59139288 |
| TMED9 | 292.296818 | 0.57971682 | 0.32681101 | 1.77385953 | 0.07608639 | 0.4745161 |
| B4GALT7 | 97.0960419 | 0.1698541 | 0.4923127 | 0.34501263 | 0.73008488 | 0.93453345 |
| AC106795.3 | 97.4763487 | -0.3825129 | 0.33526819 | -1.1409163 | 0.25390475 | 0.70431689 |
| N4BP3 | 51.4441321 | 0.56724884 | 0.54604574 | 1.03883027 | 0.29888368 | 0.73673761 |
| RMND5B | 492.114242 | -0.4130663 | 0.30454415 | -1.356343 | 0.17499006 | 0.62944785 |
| NHP2 | 156.079928 | 0.54225606 | 0.43964829 | 1.23338603 | 0.21743177 | 0.67025818 |
| HNRNPAB | 219.944829 | -0.0151351 | 0.29719799 | -0.0509261 | 9.59E-01 | 9.91E-01 |
| PHYKPL | 1193.81077 | 0.45659449 | 0.30866254 | 1.47926759 | 0.13906881 | 0.58243213 |
| COL23A1 | 181.414172 | -0.6550813 | 0.60162269 | -1.0888573 | 0.2762168 | 0.72241028 |

| | | | | | | |
|---|---|---|---|---|---|---|
| CLK4 | 656.271332 | -0.0994835 | 0.22455391 | -0.4430272 | 6.58E-01 | 9.10E-01 |
| ZNF354A | 251.871378 | -0.3126534 | 0.36197173 | -0.863751 | 0.3877247 | 0.79612242 |
| ZNF354B | 222.651753 | -0.1870971 | 0.26932868 | -0.6946794 | 0.48725622 | 0.84475225 |
| ZFP2 | 179.806125 | 0.42579517 | 0.25174882 | 1.69134921 | 0.09077013 | 0.50267341 |
| ZNF454 | 86.0408296 | 0.24323995 | 0.2965806 | 0.82014787 | 0.41213182 | 0.80695334 |
| AC104117.3 | 57.74046 | 0.25943769 | 0.37246766 | 0.69653749 | 0.48609229 | 0.8447301 |
| ZNF879 | 40.0800292 | -0.459536 | 0.34716139 | -1.3236955 | 0.18560418 | 0.64030106 |
| ZNF354C | 156.957603 | -0.1499857 | 0.25018021 | -0.5995108 | 0.54883232 | 0.86904066 |
| RUFY1 | 862.176229 | -0.2481288 | 0.17050113 | -1.4552914 | 0.14558859 | 0.59208119 |
| AC136604.2 | 102.167419 | -0.3995891 | 0.34160333 | -1.1697459 | 0.24210323 | 0.69337349 |
| HNRNPH1 | 2851.14635 | -0.16872 | 0.28126357 | -0.5998642 | 0.54859673 | 0.8690392 |
| CANX | 1371.07481 | 0.90482608 | 0.20154501 | 4.48944913 | 7.14E-06 | 0.00349342 |
| MAML1 | 426.566182 | -0.1345746 | 0.22934923 | -0.5867674 | 0.55735996 | 0.87240458 |
| MGAT4B | 316.203982 | 0.02483873 | 0.33022707 | 0.07521711 | 9.40E-01 | 0.98715467 |
| SQSTM1 | 2005.35318 | 0.08145579 | 0.37106494 | 0.21951895 | 0.82624582 | 0.96214368 |
| MRNIP | 433.291246 | -0.0045916 | 0.21563241 | -0.0212936 | 0.98301143 | 0.99518552 |
| TBC1D9B | 491.108502 | -0.009036 | 0.3323438 | -0.0271886 | 0.9783093 | 0.99455895 |
| RNF130 | 1610.51655 | -0.2248035 | 0.29032107 | -0.7743272 | 0.4387373 | 0.82127804 |
| MAPK9 | 506.890503 | -0.1355977 | 0.15428063 | -0.8789027 | 0.37945405 | 0.78762967 |
| AC008610.1 | 52.6223934 | 0.58244718 | 0.45152336 | 1.28996024 | 0.19706446 | 0.64964978 |
| GFPT2 | 340.366021 | 0.56828803 | 0.29273412 | 1.94131121 | 5.22E-02 | 4.09E-01 |
| CNOT6 | 700.488978 | -0.1361273 | 0.15150711 | -0.898488 | 0.36892545 | 0.78095617 |
| MGAT1 | 304.941581 | 0.04097417 | 0.19857977 | 0.20633606 | 0.83652841 | 0.96511435 |
| LINC00847 | 44.1876423 | 0.27659521 | 0.40847484 | 0.67714136 | 0.49831627 | 0.85092921 |
| ZFP62 | 148.433941 | -0.0811649 | 0.26653103 | -0.3045234 | 0.76072917 | 0.94202764 |
| BTNL9 | 95.0298605 | 0.36643117 | 0.56006482 | 0.65426565 | 5.13E-01 | 8.58E-01 |
| TRIM41 | 187.575359 | -0.2128162 | 0.28371408 | -0.7501082 | 0.45318956 | 0.82991556 |
| RACK1 | 858.0182 | 0.43510918 | 0.38515184 | 1.12970817 | 0.25859922 | 0.70897406 |
| TRIM52 | 267.078854 | 0.07882669 | 0.22173305 | 0.35550269 | 0.72221303 | 0.93088524 |
| AC138035.1 | 112.720613 | -0.3964266 | 0.32794381 | -1.2088247 | 0.22673019 | 0.67880306 |
| DUSP22 | 367.647286 | -0.1234121 | 0.16413447 | -0.751896 | 4.52E-01 | 8.29E-01 |
| EXOC2 | 1085.82527 | -0.4180475 | 0.19559202 | -2.1373444 | 0.03256999 | 0.34456902 |
| FOXQ1 | 178.028967 | 0.69007141 | 0.56567112 | 1.2199163 | 0.22249661 | 0.67568617 |
| FOXC1 | 294.198995 | 0.15500984 | 0.54663707 | 0.28356995 | 0.77673997 | 0.94694572 |
| GMDS | 2941.48296 | 0.0644035 | 0.19836314 | 0.32467476 | 0.74542724 | 0.93675395 |
| GMDS-DT | 5167.46369 | 0.37410624 | 0.35850291 | 1.04352357 | 0.29670587 | 0.73672107 |
| LINC01600 | 83.1603824 | 1.0810198 | 0.5277821 | 2.04823125 | 0.04053735 | 0.37344844 |
| MYLK4 | 72.4730802 | -0.1514677 | 0.48200979 | -0.3142419 | 0.75333735 | 0.93844712 |
| WRNIP1 | 420.187209 | -0.215435 | 0.18806591 | -1.1455295 | 0.25198988 | 0.70276117 |
| SERPINB1 | 318.136361 | 0.7842545 | 0.24446301 | 3.20807024 | 1.34E-03 | 6.69E-02 |
| SERPINB6 | 585.645952 | 0.33882669 | 0.2446545 | 1.38491913 | 0.1660772 | 0.61901667 |
| NQO2 | 366.405272 | 0.38185988 | 0.38384801 | 0.99482053 | 0.31982356 | 0.75094569 |
| RIPK1 | 530.145696 | -0.1557688 | 0.28340329 | -0.5496365 | 0.58256875 | 0.88338315 |
| BPHL | 147.764976 | 0.01969203 | 0.31631732 | 0.06225404 | 0.95036053 | 0.98929186 |
| TUBB2A | 79.8520433 | 1.13414252 | 0.46021238 | 2.46438941 | 0.01372469 | 0.24270686 |
| LINC02525 | 81.9472391 | 0.91142345 | 0.98906252 | 0.92150236 | 0.35678821 | 0.77411555 |
| TUBB2B | 119.329139 | 1.42571133 | 0.56100435 | 2.5413552 | 0.01104237 | 0.21418301 |
| PSMG4 | 456.027378 | 0.48046119 | 0.43247648 | 1.11095334 | 0.26658844 | 0.71641758 |
| SLC22A23 | 190.697411 | 0.83095245 | 0.42993964 | 1.93271885 | 0.05327084 | 0.41173466 |
| PXDC1 | 67.8115154 | 0.66747731 | 0.41110516 | 1.62361694 | 0.10445759 | 0.52442862 |
| PRPF4B | 1424.19601 | -0.1661104 | 0.2018643 | -0.8228817 | 0.41057524 | 0.80655687 |
| FAM217A | 105.493893 | 0.35636884 | 0.55224386 | 0.64531065 | 0.51872589 | 0.85916831 |
| ECI2 | 630.550483 | -0.1593054 | 0.24115051 | -0.6606055 | 0.50886536 | 0.85659934 |
| CDYL | 1581.75826 | -0.3767177 | 0.2170382 | -1.7357207 | 0.08261323 | 0.48594425 |

| | | | | | | |
|---|---|---|---|---|---|---|
| LYRM4 | 677.204436 | -0.3689707 | 0.26797887 | -1.3768649 | 0.16855401 | 0.62253512 |
| FARS2 | 2852.28347 | -0.1056704 | 0.17266805 | -0.611986 | 0.54054702 | 0.86579684 |
| AL022097.1 | 62.5499335 | 0.26559887 | 0.58061092 | 0.45744725 | 0.64734961 | 0.90657771 |
| LY86-AS1 | 168.771756 | -0.0813529 | 0.53536565 | -0.1519577 | 0.8792203 | 0.9765717 |
| RREB1 | 1842.67542 | -0.1941634 | 0.30779553 | -0.6308193 | 0.52815865 | 0.86215467 |
| SSR1 | 491.818836 | 0.43915524 | 0.2733974 | 1.60628903 | 0.10821042 | 0.53091537 |
| AL139095.4 | 99.7383962 | 0.60386369 | 0.57057576 | 1.05834094 | 0.28990003 | 0.73204984 |
| RIOK1 | 148.709964 | -0.1977825 | 0.30424738 | -0.6500714 | 0.51564612 | 0.85869987 |
| DSP | 610.537604 | 0.50570819 | 0.24759704 | 2.04246461 | 4.11E-02 | 0.37523297 |
| SNRNP48 | 254.356735 | 0.09798824 | 0.21044468 | 0.4656247 | 0.64148416 | 0.90421132 |
| BMP6 | 2663.44946 | -1.7829928 | 0.54709546 | -3.259016 | 0.00111799 | 0.05996788 |
| BLOC1S5-TXI | 246.970069 | -0.5211696 | 0.42738091 | -1.2194499 | 0.22267347 | 0.67568617 |
| SLC35B3 | 384.122164 | -0.0692036 | 0.23026664 | -0.300537 | 0.76376757 | 0.94255335 |
| TFAP2A | 143.316729 | 0.07557963 | 0.34446472 | 0.21941182 | 0.82632926 | 0.96214368 |
| TFAP2A-AS1 | 44.9704888 | -0.5112538 | 0.48530811 | -1.0534623 | 0.29212916 | 0.73384286 |
| GCNT2 | 3601.64116 | -0.4823517 | 0.42885418 | -1.1247452 | 0.26069701 | 0.71059571 |
| C6orf52 | 50.7506241 | -0.4540934 | 0.48911502 | -0.9283981 | 0.35320111 | 0.77216708 |
| PAK1IP1 | 85.0897253 | -0.2091981 | 0.31342307 | -0.6674623 | 0.50447689 | 0.85547181 |
| TMEM14C | 296.728963 | 0.28260317 | 0.24109225 | 1.1721786 | 0.24112536 | 0.69178991 |
| AL024498.2 | 70.4724902 | -0.5233589 | 0.40249667 | -1.3002812 | 0.1935046 | 0.64718686 |
| MAK | 568.683445 | 0.13294925 | 0.48520275 | 0.27400763 | 0.78407876 | 0.94978829 |
| SMIM13 | 132.858735 | 0.01502076 | 0.31999964 | 0.04693992 | 0.96256111 | 0.99224805 |
| AL139807.1 | 761.396971 | 0.25697547 | 0.78542415 | 0.32718051 | 0.74353136 | 0.93633036 |
| NEDD9 | 2216.11085 | 0.01984881 | 0.47215646 | 0.04203862 | 0.96646791 | 0.99248831 |
| TMEM170B | 234.185125 | -0.0199272 | 0.30078445 | -0.0662509 | 0.94717805 | 0.98905425 |
| AL157373.2 | 104.629802 | -0.5784072 | 0.7782422 | -0.7432226 | 0.45734694 | 0.83255882 |
| HIVEP1 | 1790.49909 | 0.21481914 | 0.36698984 | 0.58535447 | 0.55830939 | 0.87291524 |
| EDN1 | 71.7064967 | 0.97803022 | 0.58617505 | 1.66849513 | 0.09521748 | 0.51085035 |
| PHACTR1 | 6461.09862 | 0.53177754 | 0.46000798 | 1.15601807 | 0.24767377 | 0.69800327 |
| TBC1D7 | 197.821071 | -0.3823956 | 0.21460122 | -1.781889 | 0.07476733 | 0.47190267 |
| GFOD1 | 411.950352 | -2.5824407 | 0.82122568 | -3.1446176 | 0.00166304 | 0.07697119 |
| SIRT5 | 512.369347 | -0.4149719 | 0.2832162 | -1.4652124 | 0.14286297 | 0.58824095 |
| NOL7 | 333.540026 | -0.0911754 | 0.31126377 | -0.2929202 | 0.76958315 | 0.94386043 |
| RANBP9 | 1415.82109 | -0.38028 | 0.25436207 | -1.4950343 | 0.1349055 | 0.57708028 |
| MCUR1 | 303.119347 | -0.0534782 | 0.24109917 | -0.2218101 | 0.82446171 | 0.96146415 |
| CD83 | 253.381117 | 0.47134923 | 0.73346616 | 0.64263256 | 0.52046255 | 0.85943874 |
| JARID2 | 1861.29665 | -0.3188168 | 0.30186876 | -1.0561438 | 0.29090253 | 0.73313067 |
| DTNBP1 | 1153.93214 | -0.5177045 | 0.23206881 | -2.2308231 | 0.02569285 | 0.31211455 |
| MYLIP | 406.85398 | 0.1025381 | 0.27429587 | 0.37382298 | 0.70853602 | 0.92577218 |
| GMPR | 1008.8466 | 0.27929086 | 0.37446701 | 0.74583571 | 0.45576667 | 0.83130183 |
| ATXN1 | 5170.95447 | 0.3472652 | 0.34619183 | 1.00310049 | 0.31581238 | 0.74785672 |
| CAP2 | 430.025245 | 1.26446011 | 0.45876675 | 2.75621566 | 0.00584744 | 0.15669584 |
| FAM8A1 | 181.474685 | 0.28516468 | 0.46453775 | 0.6138676 | 0.53930282 | 0.86546604 |
| NUP153 | 1109.52624 | -0.3099904 | 0.21231427 | -1.4600544 | 0.14427513 | 0.58995828 |
| KIF13A | 1291.54205 | -0.1345928 | 0.32085327 | -0.419484 | 0.67486245 | 0.91636338 |
| TPMT | 261.237923 | 0.26921134 | 0.21806994 | 1.23451834 | 0.21700982 | 0.66978787 |
| KDM1B | 228.843233 | -0.3889595 | 0.24474197 | -1.5892636 | 0.1120009 | 0.53621666 |
| DEK | 727.864158 | -0.3149093 | 0.19743813 | -1.5949772 | 0.11071733 | 0.53393773 |
| RNF144B | 2474.49112 | -0.196178 | 0.35238735 | -0.5567112 | 0.57772474 | 0.88190988 |
| MIR548A1HC | 86.3620463 | 0.85566215 | 0.70844391 | 1.20780508 | 2.27E-01 | 0.67921637 |
| AL022068.1 | 804.724136 | -0.4879884 | 0.81599852 | -0.5980261 | 0.5498225 | 0.86956073 |
| ID4 | 145.036328 | 0.19945205 | 0.65947395 | 0.30244113 | 0.7623158 | 0.94212205 |
| MBOAT1 | 249.463122 | 1.81625712 | 0.6898508 | 2.63282599 | 0.00846777 | 0.18546054 |
| E2F3 | 424.434408 | 0.43172104 | 0.31545128 | 1.36858231 | 1.71E-01 | 0.62772493 |

| | | | | | | |
|---|---|---|---|---|---|---|
| CDKAL1 | 4070.42403 | -0.3830167 | 0.17659928 | -2.1688466 | 0.03009433 | 0.33337391 |
| AL513188.1 | 95.047178 | 0.03515692 | 0.64187168 | 0.0547725 | 0.95631971 | 0.99065996 |
| AL512380.2 | 98.0501942 | -0.2253531 | 0.40793993 | -0.5524174 | 0.58066244 | 0.88272186 |
| SOX4 | 644.048636 | -0.5142432 | 0.49299237 | -1.0431059 | 0.29689926 | 0.73673761 |
| CASC15 | 1028.82826 | 0.23352344 | 0.39574788 | 0.59008133 | 0.55513612 | 0.87184917 |
| NBAT1 | 83.5519434 | 0.4087612 | 0.62012051 | 0.65916414 | 0.50979037 | 0.85704735 |
| DCDC2 | 7343.70497 | 0.52529232 | 0.41342095 | 1.27059919 | 0.20387127 | 0.65712971 |
| MRS2 | 544.29884 | -0.1013836 | 0.2364817 | -0.4287165 | 0.66812953 | 0.91383898 |
| GPLD1 | 65.616051 | -0.0495531 | 0.34997239 | -0.1415916 | 0.88740264 | 0.97729111 |
| ALDH5A1 | 142.035246 | 0.32986892 | 0.35444815 | 0.93065495 | 0.35203208 | 0.77151834 |
| TDP2 | 305.594069 | 0.41353268 | 0.21756376 | 1.90074251 | 0.05733575 | 0.42004874 |
| ACOT13 | 211.575519 | 0.31821119 | 0.26536178 | 1.19915985 | 0.2304658 | 0.68271422 |
| C6orf62 | 879.710998 | -0.1644778 | 0.23800367 | -0.6910725 | 0.48952 | 0.84614483 |
| GMNN | 101.071496 | -0.1729515 | 0.36792371 | -0.4700743 | 0.63830193 | 0.90303979 |
| RIPOR2 | 77.8554695 | 0.87043804 | 0.60844541 | 1.43059349 | 0.15254675 | 0.60267503 |
| CARMIL1 | 4008.76271 | -0.1701202 | 0.24654602 | -0.6900139 | 0.49018543 | 0.84614483 |
| SLC17A1 | 98.2894198 | 2.0547384 | 0.86757148 | 2.36837936 | 0.01786621 | 0.26845799 |
| TRIM38 | 689.848871 | 0.30848478 | 0.3570506 | 0.86398057 | 0.38759857 | 0.79598717 |
| U91328.2 | 76.7099417 | 0.86221419 | 0.46999511 | 1.83451737 | 0.06657723 | 0.4475413 |
| HIST1H1C | 160.463309 | 0.3833911 | 0.53088905 | 0.72216802 | 0.47019118 | 0.83921324 |
| HFE | 66.2832447 | 0.83300729 | 0.4467527 | 1.86458255 | 0.06223994 | 0.43419765 |
| HIST1H4C | 299.99102 | 1.13631257 | 0.76072086 | 1.49373131 | 1.35E-01 | 0.57708028 |
| HIST1H2AC | 303.294089 | 0.36625291 | 0.70074295 | 0.52266371 | 0.6012083 | 0.8890585 |
| HIST1H1E | 86.1126358 | -0.0065054 | 0.78428123 | -0.0082947 | 0.99338183 | 0.99821837 |
| HIST1H2BD | 73.1545286 | 1.04756573 | 0.80596173 | 1.29977104 | 0.19367945 | 0.64718686 |
| BTN3A2 | 83.6654082 | 1.93739834 | 0.81709168 | 2.37109052 | 0.01773569 | 0.26773793 |
| BTN2A2 | 48.8179749 | 0.84263953 | 0.50492901 | 1.66882772 | 0.09515153 | 0.51085035 |
| BTN3A1 | 90.1876312 | 1.54379238 | 0.7977335 | 1.93522321 | 0.05296291 | 0.41073851 |
| BTN2A1 | 175.925498 | -0.06022 | 0.28721509 | -0.2096686 | 0.83392633 | 0.96510927 |
| HCG11 | 52.5767508 | 0.74142443 | 0.52285277 | 1.41803673 | 0.15618004 | 0.60551282 |
| HMGN4 | 125.582317 | 0.37237694 | 0.34916432 | 1.06648051 | 0.28620649 | 0.72934245 |
| ABT1 | 64.6982289 | 0.50733244 | 0.38617752 | 1.31372857 | 0.18893756 | 0.64421174 |
| ZNF322 | 163.457113 | -0.004647 | 0.28412662 | -0.0163554 | 0.98695084 | 0.99653761 |
| AL513548.1 | 44.5745814 | 1.18025575 | 0.65613159 | 1.79880952 | 0.07204882 | 0.46774933 |
| LINC00240 | 377.87305 | 0.06814451 | 0.36167472 | 0.18841381 | 0.85055227 | 0.97056586 |
| ZNF391 | 58.5129673 | 0.31184528 | 0.28394405 | 1.09826316 | 0.27208959 | 0.72070318 |
| ZNF184 | 52.835649 | 0.09693167 | 0.28589996 | 0.33904053 | 0.73457919 | 0.93552616 |
| HIST1H2BN | 45.6386326 | -0.467144 | 0.54207492 | -0.8617701 | 0.38881403 | 0.79634939 |
| AL121944.1 | 102.099576 | -0.0593153 | 0.42979902 | -0.138007 | 0.89023485 | 0.97729111 |
| ZNF165 | 75.1260973 | -0.3287241 | 0.56164086 | -0.5852923 | 0.55835119 | 0.87291524 |
| ZSCAN16-AS | 188.807875 | 0.4431309 | 0.26429211 | 1.67667092 | 0.09360685 | 0.50887217 |
| ZSCAN16 | 44.6556491 | -0.2775138 | 0.4253117 | -0.6524952 | 0.51408176 | 0.85762435 |
| ZKSCAN8 | 291.963725 | -0.248757 | 0.18158289 | -1.3699362 | 0.17070681 | 0.62687414 |
| ZSCAN9 | 143.26355 | -0.2359004 | 0.22242738 | -1.0605728 | 0.28888409 | 0.73176973 |
| PGBD1 | 96.8588985 | -0.3526285 | 0.30475762 | -1.1570786 | 0.24724026 | 0.69761067 |
| ZSCAN31 | 50.4064216 | 1.85049344 | 0.5541951 | 3.33906495 | 0.00084061 | 0.05047093 |
| ZKSCAN3 | 70.1455119 | 0.17397557 | 0.36225319 | 0.48025959 | 0.63104282 | 0.90005881 |
| ZSCAN12 | 97.3937017 | -0.0165273 | 0.25488951 | -0.0648409 | 0.94830068 | 0.98919519 |
| ZSCAN23 | 119.256268 | 0.26131216 | 0.28381142 | 0.92072463 | 0.35719421 | 0.77448758 |
| ZBED9 | 42.6208681 | -0.3385196 | 0.42406019 | -0.798282 | 0.42470685 | 0.81466629 |
| TRIM27 | 441.252951 | -0.07822 | 0.1956863 | -0.3997213 | 0.68936181 | 0.92166785 |
| ZNF311 | 36.9617634 | -0.163825 | 0.56634803 | -0.2892656 | 0.77237811 | 0.94509888 |
| GABBR1 | 246.700201 | -0.1245064 | 0.61010998 | -0.2040721 | 0.83829718 | 0.96566728 |
| HLA-F | 144.31607 | 1.35382189 | 0.71297427 | 1.89883696 | 0.05758592 | 0.42087381 |

| | | | | | | |
|---|---|---|---|---|---|---|
| HLA-A | 2344.20027 | 1.27743853 | 0.45845214 | 2.78641634 | 0.00532944 | 0.14749423 |
| ZNRD1 | 75.2802305 | -0.1522641 | 0.45746283 | -0.3328447 | 0.73925148 | 0.93552616 |
| PPP1R11 | 132.470102 | 0.34128447 | 0.29077743 | 1.17369656 | 0.24051659 | 0.69178991 |
| TRIM26 | 662.120874 | -0.0055508 | 0.26087898 | -0.0212775 | 0.98302432 | 0.99518552 |
| HCG17 | 1415.52183 | -0.2263019 | 0.32757206 | -0.6908463 | 0.48966216 | 0.84614483 |
| HLA-E | 916.826265 | 0.6178899 | 0.46392967 | 1.33186113 | 1.83E-01 | 0.63763606 |
| GNL1 | 393.869448 | -0.3289899 | 0.2361931 | -1.3928852 | 0.16365451 | 0.61447199 |
| PRR3 | 164.524962 | -0.0953266 | 0.30876067 | -0.3087395 | 0.75751973 | 0.94056395 |
| ABCF1 | 327.902422 | 0.18228472 | 0.19200661 | 0.94936693 | 0.34243403 | 0.76591028 |
| PPP1R10 | 508.78375 | -0.4291536 | 0.29110233 | -1.4742361 | 0.14041804 | 0.58547408 |
| MRPS18B | 286.237047 | 0.15394769 | 0.21383142 | 0.71994884 | 0.4715565 | 0.83921324 |
| ATAT1 | 195.953697 | -0.2838529 | 0.20343793 | -1.3952802 | 0.16293137 | 0.61387934 |
| C6orf136 | 236.707435 | -0.3191851 | 0.24215259 | -1.3181157 | 0.18746492 | 0.64200436 |
| DHX16 | 230.218739 | -0.1585848 | 0.2457277 | -0.6453678 | 0.51868885 | 0.85916831 |
| PPP1R18 | 61.6217543 | -0.2649792 | 0.33564838 | -0.7894547 | 4.30E-01 | 8.17E-01 |
| MDC1 | 283.501998 | 0.03458599 | 0.27759337 | 0.12459229 | 0.90084633 | 0.97914987 |
| MDC1-AS1 | 51.9350807 | 0.09687219 | 0.44082017 | 0.21975444 | 0.8260624 | 0.96214368 |
| TUBB | 412.516839 | 0.31409067 | 0.52907872 | 0.59365584 | 0.55274233 | 0.87104567 |
| FLOT1 | 718.087528 | 0.10144336 | 0.25974016 | 0.39055709 | 0.69612465 | 0.92274064 |
| IER3 | 1939.72672 | 0.33312593 | 0.53472176 | 0.62298928 | 5.33E-01 | 8.64E-01 |
| DDR1 | 739.726412 | 0.51111312 | 0.33798783 | 1.51222344 | 1.30E-01 | 0.57223551 |
| VARS2 | 114.177204 | -0.2914592 | 0.41383624 | -0.7042862 | 0.48125454 | 0.8437862 |
| PSORS1C1 | 106.071133 | 1.09760275 | 1.05490092 | 1.04047948 | 0.29811719 | 0.73673761 |
| CCHCR1 | 161.199148 | 0.19191483 | 0.33022381 | 0.58116594 | 0.56112862 | 0.87394741 |
| POU5F1 | 898.520508 | 0.39820979 | 0.46768916 | 0.85144115 | 0.39452434 | 0.79876591 |
| HCG27 | 64.4064534 | 0.44193433 | 0.46663135 | 0.94707381 | 0.34360117 | 0.76666011 |
| HLA-C | 1233.35966 | 1.29661551 | 0.68413444 | 1.89526419 | 0.05805741 | 0.42087381 |
| HLA-B | 1751.92813 | 1.46680811 | 0.61954379 | 2.36756163 | 0.01790574 | 0.26845799 |
| MICA | 270.677404 | 0.187925 | 0.40159588 | 0.46794552 | 0.63982355 | 0.90330608 |
| NFKBIL1 | 121.837098 | 0.06932137 | 0.20659954 | 0.335535 | 0.73722155 | 0.93552616 |
| PRRC2A | 407.492146 | 0.04304784 | 0.25932455 | 0.16599985 | 0.86815707 | 0.97402505 |
| BAG6 | 253.199153 | -0.24256 | 0.2318994 | -1.0459707 | 0.29557455 | 0.73661928 |
| C6orf47 | 74.6034143 | 0.18256001 | 0.34118211 | 0.53508083 | 0.59259397 | 0.88740306 |
| GPANK1 | 90.1455113 | -0.0711922 | 0.30035924 | -0.2370236 | 0.81263845 | 0.95806331 |
| DDAH2 | 189.973719 | 0.84997196 | 0.44600179 | 1.90575909 | 0.05668147 | 0.41866502 |
| CLIC1 | 994.533192 | 1.03559006 | 0.38646633 | 2.67963853 | 7.37E-03 | 1.73E-01 |
| VARS | 128.349462 | 0.07240203 | 0.43226495 | 0.16749456 | 0.86698093 | 0.97402505 |
| LSM2 | 158.97854 | 0.05747033 | 0.25257343 | 0.22753909 | 0.82000458 | 0.96016713 |
| HSPA1A | 392.356573 | 0.01362207 | 0.50489621 | 0.02697995 | 0.97847573 | 0.99455895 |
| HSPA1B | 184.853451 | 0.15285616 | 0.63219718 | 0.24178557 | 0.80894632 | 0.95610623 |
| C6orf48 | 160.093941 | 0.29609083 | 0.35634494 | 0.83091072 | 0.40602407 | 0.80499101 |
| NEU1 | 143.610409 | 0.59155324 | 0.41745891 | 1.41703346 | 0.15647315 | 0.60611548 |
| SLC44A4 | 72.1305355 | 1.11836243 | 0.41739779 | 2.67936836 | 7.38E-03 | 1.73E-01 |
| EHMT2 | 189.030829 | -0.2525267 | 0.31745951 | -0.7954611 | 0.42634531 | 0.81466629 |
| C2 | 92.7256048 | -0.7071005 | 0.35693524 | -1.9810331 | 0.04758756 | 0.39447539 |
| NELFE | 120.6069 | -0.1851001 | 0.23656207 | -0.7824591 | 0.43394481 | 0.81919857 |
| SKIV2L | 101.965102 | -0.3014814 | 0.25538932 | -1.1804777 | 0.23781028 | 0.68982886 |
| DXO | 54.6512608 | -0.011565 | 0.42010224 | -0.027529 | 0.97803784 | 0.99455895 |
| STK19 | 175.377298 | 0.00824298 | 0.20902173 | 0.03943601 | 0.96854277 | 0.99250254 |
| ATF6B | 394.335743 | -0.3474653 | 0.26575433 | -1.3074681 | 0.1910538 | 0.64567512 |
| AGPAT1 | 118.608374 | 0.26566884 | 0.29771038 | 0.89237348 | 0.37219278 | 0.78256595 |
| RNF5 | 141.478568 | 0.52027152 | 0.35630264 | 1.46019551 | 0.14423635 | 0.58995828 |
| AGER | 41.011567 | -0.1806049 | 0.50399615 | -0.3583457 | 0.72008462 | 0.93001079 |
| PBX2 | 224.871307 | -0.1641579 | 0.27218736 | -0.6031064 | 0.54643793 | 0.86805757 |

| | | | | | | |
|---|---|---|---|---|---|---|
| AL662796.1 | 50.0904332 | -1.0513639 | 0.74466976 | -1.4118525 | 0.1579934 | 0.60779483 |
| HLA-DRA | 405.917 | 0.45483926 | 0.70740437 | 0.64296926 | 0.52024405 | 0.85935232 |
| HLA-DRB5 | 57.7847007 | 2.29095481 | 1.00129594 | 2.2879897 | 0.02213812 | 0.2955431 |
| HLA-DRB1 | 231.969391 | 1.12736533 | 0.96767533 | 1.16502436 | 0.24400913 | 0.6948422 |
| HLA-DQB1 | 72.2907463 | 1.33708513 | 0.93720135 | 1.42667863 | 0.15367255 | 0.60285224 |
| PSMB8 | 223.870075 | 1.02463035 | 0.46144267 | 2.22049328 | 0.0263853 | 0.31402818 |
| PSMB9 | 105.817217 | 1.64537221 | 0.71546421 | 2.29972678 | 0.0214637 | 0.29288643 |
| TAP1 | 151.378097 | 1.09974532 | 0.62936476 | 1.74738942 | 0.08056981 | 0.48456361 |
| HLA-DMB | 59.0277281 | 0.61709332 | 0.73106247 | 0.84410477 | 0.39861086 | 0.80148305 |
| HLA-DMA | 100.303069 | -0.088697 | 0.62507127 | -0.1418989 | 0.88715983 | 0.97729111 |
| BRD2 | 607.006071 | -0.0347422 | 0.2309271 | -0.1504467 | 0.88041217 | 0.97664137 |
| HLA-DPA1 | 66.5946219 | 0.78999791 | 0.86293046 | 0.9154827 | 0.35993828 | 0.77686679 |
| HLA-DPB1 | 47.5209389 | 0.50097061 | 0.79948123 | 0.62661959 | 0.53090862 | 0.86396107 |
| RXRB | 149.90174 | 0.1629695 | 0.26617323 | 0.61226854 | 0.5403601 | 0.86569031 |
| SLC39A7 | 181.756953 | 0.35636353 | 0.33674172 | 1.0582696 | 0.28993255 | 0.73204984 |
| HSD17B8 | 41.7114339 | 0.71716443 | 0.4174583 | 1.71793069 | 0.08580926 | 0.49387995 |
| RING1 | 242.941883 | -0.1361788 | 0.26910786 | -0.506038 | 0.61282995 | 0.89441126 |
| HCG25 | 76.4229434 | -0.2098876 | 0.37594206 | -0.5582978 | 0.57664106 | 0.88147804 |
| VPS52 | 301.49093 | -0.285709 | 0.15273086 | -1.8706696 | 0.06139089 | 0.43236485 |
| RPS18 | 650.442136 | 0.46693889 | 0.37500615 | 1.24514997 | 0.21307663 | 0.66710813 |
| B3GALT4 | 54.2954575 | 0.28748923 | 0.37111066 | 0.77467251 | 0.4385332 | 0.82127804 |
| WDR46 | 118.264578 | 0.40712328 | 0.35731202 | 1.1394055 | 0.25453406 | 0.70509116 |
| PFDN6 | 114.460493 | 0.2381809 | 0.35853278 | 0.66432111 | 0.50648481 | 0.85626196 |
| RGL2 | 283.31348 | -0.1745021 | 0.29559274 | -0.5903464 | 0.55495847 | 0.87184917 |
| TAPBP | 663.577241 | 0.53021591 | 0.34154928 | 1.55238482 | 1.21E-01 | 0.55433742 |
| DAXX | 96.9028239 | 0.11167646 | 0.30151828 | 0.37038039 | 0.71109908 | 0.92683165 |
| PHF1 | 270.549479 | -0.044034 | 0.23245948 | -0.1894264 | 0.84975863 | 0.9703866 |
| CUTA | 249.74824 | 0.67122291 | 0.463616 | 1.44779926 | 0.1476732 | 0.59397543 |
| SYNGAP1 | 192.777083 | -0.2796395 | 0.21566611 | -1.2966315 | 0.19475799 | 0.64840333 |
| SYNGAP1-AS | 84.0574335 | 0.37476482 | 0.35932292 | 1.04297501 | 0.29695987 | 0.73673761 |
| ITPR3 | 840.116702 | -0.2504296 | 0.36307817 | -0.6897401 | 4.90E-01 | 8.46E-01 |
| UQCC2 | 237.212007 | -0.0864543 | 0.20437513 | -0.4230177 | 6.72E-01 | 0.91576543 |
| LEMD2 | 272.539761 | -0.2463025 | 0.24761669 | -0.9946927 | 0.31988574 | 0.75094569 |
| HMGA1 | 161.569238 | -1.0518489 | 0.59389015 | -1.7711169 | 7.65E-02 | 0.4759127 |
| SMIM29 | 59.9634966 | -0.2561982 | 0.51179906 | -0.5005835 | 0.61666429 | 0.89589954 |
| NUDT3 | 265.109518 | -0.0262409 | 0.3308584 | -0.0793117 | 0.93678471 | 0.98630295 |
| C6orf106 | 924.703498 | -0.0467 | 0.14330066 | -0.3258884 | 0.74450879 | 0.93668127 |
| SNRPC | 186.812609 | 0.24677495 | 0.26649022 | 0.92601881 | 0.35443621 | 0.77243419 |
| UHRF1BP1 | 626.002958 | -0.1112333 | 0.16154911 | -0.6885418 | 0.49111167 | 0.84654758 |
| TAF11 | 374.329165 | -0.0661165 | 0.25592139 | -0.2583469 | 0.7961392 | 0.95278473 |
| ANKS1A | 1946.90893 | -0.1007612 | 0.15827866 | -0.6366066 | 0.52438113 | 0.86006369 |
| SCUBE3 | 54.8852481 | 0.93399235 | 0.56109872 | 1.66457757 | 0.09599706 | 0.5117083 |
| ZNF76 | 495.06533 | -0.1223022 | 0.24814934 | -0.4928573 | 0.62211339 | 0.89835355 |
| PPARD | 461.387174 | -0.0480533 | 0.1929222 | -0.249081 | 0.8032981 | 0.95446295 |
| RPL10A | 781.095913 | 0.68030844 | 0.44375857 | 1.53305982 | 0.12526112 | 0.56354703 |
| TEAD3 | 328.322863 | -0.6406005 | 0.2627483 | -2.4380767 | 0.01476564 | 0.24869534 |
| FKBP5 | 5601.25984 | -0.1223246 | 0.83196547 | -0.1470308 | 0.88310768 | 0.97664137 |
| SRPK1 | 927.934983 | -0.3927979 | 0.32492619 | -1.2088836 | 0.22670756 | 0.67880306 |
| SLC26A8 | 70.5281198 | 0.09290625 | 0.50956855 | 0.18232336 | 8.55E-01 | 0.97118784 |
| MAPK14 | 1978.26574 | -0.34275 | 0.38567062 | -0.8887116 | 0.37415809 | 0.78369843 |
| MAPK13 | 127.569133 | 0.34451331 | 0.2267917 | 1.5190737 | 1.29E-01 | 5.69E-01 |
| Z84485.1 | 67.3054082 | 0.09138338 | 0.36667079 | 0.24922459 | 0.80318706 | 0.95446295 |
| BRPF3 | 194.143463 | 0.09063318 | 0.32418346 | 0.27957374 | 0.77980456 | 0.9477748 |
| KCTD20 | 637.30458 | 0.06176916 | 0.17150709 | 0.36015513 | 0.71873113 | 0.9298452 |

| | | | | | | |
|---|---|---|---|---|---|---|
| STK38 | 1112.31626 | 0.00666536 | 0.18160628 | 0.03670226 | 0.97072241 | 0.99317943 |
| SRSF3 | 839.257493 | 0.04048066 | 0.21585182 | 0.1875391 | 0.85123797 | 0.97057139 |
| CDKN1A | 221.47363 | 0.55374587 | 0.56802436 | 0.9748629 | 3.30E-01 | 7.59E-01 |
| PPIL1 | 87.144088 | -0.3349383 | 0.4127446 | -0.8114904 | 0.41708412 | 0.81007111 |
| C6orf89 | 715.225296 | 0.01655074 | 0.14609375 | 0.1132885 | 0.90980183 | 0.98069436 |
| MTCH1 | 662.22275 | -0.3747831 | 0.23019793 | -1.6280905 | 0.1035057 | 0.52383403 |
| PIM1 | 118.322124 | -0.541515 | 0.23916891 | -2.264153 | 0.0235647 | 0.30185158 |
| TMEM217 | 108.377147 | -0.6616737 | 0.32228885 | -2.0530457 | 0.04006815 | 0.37342745 |
| TBC1D22B | 591.496306 | -0.0096 | 0.19318643 | -0.0496929 | 0.96036715 | 0.99153894 |
| RNF8 | 729.972878 | -0.712224 | 0.30734397 | -2.3173515 | 0.02048459 | 0.28691085 |
| CMTR1 | 252.853659 | -0.2619485 | 0.30885225 | -0.8481353 | 0.3963626 | 0.80029863 |
| ZFAND3 | 9941.2268 | -0.2527134 | 0.1834192 | -1.3777914 | 0.16826769 | 0.62253512 |
| BTBD9 | 5982.38957 | -0.3077762 | 0.21274986 | -1.4466574 | 0.1479929 | 0.59435642 |
| GLO1 | 464.303669 | 0.37597667 | 0.23309999 | 1.61294157 | 0.1067572 | 0.52778894 |
| SAYSD1 | 107.455858 | -0.3524956 | 0.28712931 | -1.2276545 | 0.21957668 | 0.67278782 |
| KCNK5 | 129.787981 | 1.62826971 | 0.7224052 | 2.25395625 | 0.02419892 | 0.30500338 |
| KIF6 | 216.361542 | 1.00952652 | 0.70632105 | 1.42927429 | 0.15292541 | 0.60285224 |
| DAAM2 | 45.7888442 | 1.74576824 | 0.48658193 | 3.58781972 | 0.00033345 | 0.03094754 |
| MOCS1 | 132.43409 | 1.12773263 | 0.35516976 | 3.17519329 | 0.00149737 | 0.07192263 |
| UNC5CL | 159.655113 | 1.52661364 | 0.64188535 | 2.3783276 | 0.01739137 | 0.26432489 |
| OARD1 | 452.5919 | -0.2772939 | 0.29274366 | -0.9472244 | 0.34352444 | 0.76666011 |
| NFYA | 225.328098 | 0.27756343 | 0.18545612 | 1.49665282 | 0.13448362 | 0.57708028 |
| FOXP4 | 128.608597 | 0.06776112 | 0.28807131 | 0.23522341 | 0.81403534 | 0.9582562 |
| TFEB | 701.398312 | 1.15826839 | 0.59561597 | 1.94465638 | 0.05181635 | 0.40820525 |
| FRS3 | 49.2478905 | 0.11417776 | 0.33891381 | 0.33689322 | 0.7361974 | 0.93552616 |
| USP49 | 462.934527 | 0.18518228 | 0.18249265 | 1.01473829 | 0.3102306 | 0.74459794 |
| MED20 | 83.8807573 | 0.13195676 | 0.25038447 | 0.52701656 | 0.59818209 | 0.88883692 |
| CCND3 | 2846.82741 | -0.423036 | 0.36182494 | -1.1691733 | 0.24233383 | 0.69337349 |
| TAF8 | 390.418181 | -0.2731352 | 0.19254456 | -1.4185555 | 0.15602864 | 0.60527706 |
| C6orf132 | 160.478224 | -0.1622493 | 0.47706124 | -0.3401017 | 7.34E-01 | 0.93552616 |
| MRPS10 | 140.585045 | 0.07022171 | 0.20789207 | 0.33777963 | 0.73552926 | 0.93552616 |
| TRERF1 | 327.003274 | -0.1595475 | 0.59106354 | -0.2699328 | 0.78721192 | 0.94978829 |
| UBR2 | 2830.58055 | -0.2188579 | 0.16093236 | -1.3599373 | 0.17384977 | 0.6286116 |
| TBCC | 52.3324047 | 0.91901564 | 0.49573641 | 1.85383929 | 0.06376216 | 0.43779712 |
| BICRAL | 1463.88031 | -0.3073753 | 0.28557159 | -1.0763513 | 0.28177019 | 0.72604837 |
| RPL7L1 | 503.479719 | 0.23609762 | 0.23085302 | 1.02271836 | 0.30644103 | 0.74295884 |
| C6orf226 | 62.9491508 | 0.07281138 | 0.45702262 | 0.15931681 | 0.87341928 | 0.97564728 |
| CNPY3 | 128.033014 | -0.2035434 | 0.36720285 | -0.5543078 | 0.57936824 | 0.8821758 |
| AL035587.1 | 66.9798008 | -0.3385598 | 0.5522943 | -0.6130062 | 0.53987221 | 0.86566791 |
| PEX6 | 278.83182 | 0.40746791 | 0.27791555 | 1.46615727 | 0.14260545 | 0.58824095 |
| PPP2R5D | 163.191748 | -0.4244918 | 0.22219911 | -1.9104117 | 0.05608023 | 0.41843207 |
| MEA1 | 74.1095141 | 0.04037215 | 0.42169651 | 0.09573746 | 0.92372909 | 0.98215577 |
| KLHDC3 | 256.501715 | -0.0174079 | 0.22215129 | -0.0783607 | 0.93754112 | 0.98661414 |
| RRP36 | 285.338482 | -0.0326201 | 0.19234419 | -0.1695926 | 0.86533057 | 0.97379123 |
| CUL7 | 170.471621 | -0.2389411 | 0.24514133 | -0.9747075 | 0.32970538 | 0.75860764 |
| KLC4 | 314.57059 | -0.2065674 | 0.21610004 | -0.9558878 | 0.33912895 | 0.76449297 |
| MRPL2 | 177.004148 | 0.27894739 | 0.23589297 | 1.18251676 | 0.23700072 | 0.68924318 |
| PTK7 | 839.089358 | 0.17870057 | 0.3647724 | 0.48989607 | 0.62420744 | 0.89873262 |
| SRF | 82.2676452 | 0.00831535 | 0.52845213 | 0.01573529 | 0.98744557 | 0.99653761 |
| CUL9 | 269.977784 | -0.3368416 | 0.18080889 | -1.8629702 | 0.06246646 | 0.43419765 |
| DNPH1 | 194.371286 | 1.18168171 | 0.43607786 | 2.70979523 | 0.00673248 | 0.16906706 |
| ZNF318 | 215.488813 | -0.2023745 | 0.20456039 | -0.9893142 | 0.32250941 | 0.7532337 |
| ABCC10 | 220.426137 | -0.5015757 | 0.21192763 | -2.3667309 | 0.01794598 | 0.26845799 |
| TJAP1 | 261.633676 | -0.1088913 | 0.19478744 | -0.5590265 | 0.57614366 | 0.88123035 |

| | | | | | | |
|---|---|---|---|---|---|---|
| POLR1C | 390.645365 | -0.1450007 | 0.26890398 | -0.5392286 | 0.58972914 | 0.88621335 |
| YIPF3 | 283.656399 | 0.43465613 | 0.33121115 | 1.31232336 | 0.18941105 | 0.64432929 |
| XPO5 | 772.528324 | 0.00415472 | 0.28091196 | 0.01479012 | 0.98819962 | 0.99653761 |
| POLH | 428.405484 | 0.08591585 | 0.29122921 | 0.29501109 | 0.76798541 | 0.94316227 |
| GTPBP2 | 339.031622 | -0.3604355 | 0.31097554 | -1.1590478 | 0.24643672 | 0.69677689 |
| MAD2L1BP | 54.7127773 | 0.65857284 | 0.31581568 | 2.08530764 | 0.03704138 | 0.3605598 |
| MRPS18A | 123.402538 | 0.4822823 | 0.32593352 | 1.47969529 | 0.13895458 | 0.58243213 |
| VEGFA | 1601.80441 | -0.2104839 | 0.52449668 | -0.4013064 | 0.68819455 | 0.92128933 |
| AL109615.3 | 58.5536568 | 0.06367403 | 0.72256231 | 0.08812255 | 0.92977928 | 0.98331361 |
| MRPL14 | 208.478359 | 0.54100396 | 0.31969133 | 1.69226974 | 0.09059455 | 0.50258861 |
| TMEM63B | 269.690723 | -0.2807386 | 0.24892189 | -1.1278182 | 0.25939673 | 0.70921184 |
| SLC29A1 | 284.61892 | 0.22176338 | 0.4997432 | 0.44375467 | 0.65721996 | 0.91026278 |
| HSP90AB1 | 1396.69838 | 0.84701923 | 0.40525639 | 2.09008235 | 0.0366104 | 0.3595441 |
| SLC35B2 | 88.8702751 | 0.39841087 | 0.41795845 | 0.95323079 | 0.34047315 | 0.76484861 |
| NFKBIE | 50.4506746 | 0.92463359 | 0.43787202 | 2.11165263 | 0.03471626 | 0.35358029 |
| AARS2 | 88.0166441 | -0.2092291 | 0.3965339 | -0.5276449 | 0.59774582 | 0.88883692 |
| CDC5L | 429.951957 | 0.02453634 | 0.20622938 | 0.11897595 | 0.90529441 | 0.97992575 |
| SUPT3H | 3418.1856 | -0.1491373 | 0.21854498 | -0.6824102 | 0.49497961 | 0.84899096 |
| RUNX2 | 157.605568 | -0.1398382 | 0.50514111 | -0.2768299 | 0.78191072 | 0.94882207 |
| CLIC5 | 90.3077343 | 0.95552928 | 0.474702 | 2.0129034 | 0.04412479 | 0.38531062 |
| AL035701.1 | 63.5105623 | 0.09328633 | 0.56652756 | 0.16466335 | 0.86920896 | 0.97448491 |
| ENPP4 | 279.460305 | 0.82005026 | 0.28547358 | 2.87259603 | 0.00407114 | 0.12448086 |
| ENPP5 | 204.60112 | 0.01139517 | 0.43038902 | 0.02647643 | 0.97887733 | 0.99455895 |
| RCAN2 | 526.359637 | -0.0822511 | 0.42608125 | -0.1930408 | 0.846927 | 0.96891207 |
| CYP39A1 | 230.297506 | 0.31715047 | 0.32740753 | 0.96867187 | 0.33270893 | 0.7610305 |
| SLC25A27 | 1127.4652 | 0.11051678 | 0.39409556 | 0.28043143 | 0.77914653 | 0.94732571 |
| AL591242.1 | 65.5540238 | 0.89771135 | 0.94020948 | 0.95479929 | 0.33967921 | 0.76449297 |
| TDRD6 | 115.039346 | 0.03771561 | 0.59980303 | 0.06287999 | 0.94986207 | 0.98919519 |
| ADGRF5 | 49.9245831 | -1.0637656 | 0.5293469 | -2.0095812 | 0.04447553 | 0.38623091 |
| ADGRF1 | 1504.38355 | -0.5185438 | 0.35015653 | -1.4808914 | 0.13863551 | 0.58226913 |
| TNFRSF21 | 475.862089 | -0.4684133 | 0.3639363 | -1.2870751 | 0.19806812 | 0.65049274 |
| AL355353.1 | 58.1699076 | 0.44962871 | 0.51844697 | 0.86726075 | 0.38579916 | 0.79402011 |
| CD2AP | 3157.39084 | -0.3532834 | 0.22355333 | -1.5803094 | 0.11403603 | 0.54047431 |
| PTCHD4 | 186.049225 | 2.08877632 | 0.53750413 | 3.88606564 | 0.00010188 | 0.01602094 |
| MUT | 363.889597 | 0.23141002 | 0.19824 | 1.16732252 | 0.24308015 | 0.6944117 |
| CENPQ | 59.727941 | -0.6375827 | 0.33528405 | -1.9016196 | 0.05722091 | 0.41944006 |
| TFAP2B | 1428.61284 | -0.4069562 | 0.48517397 | -0.8387841 | 0.40159046 | 0.80278824 |
| AL355997.1 | 114.617985 | 0.09683696 | 0.71668043 | 0.13511874 | 0.89251799 | 0.97771356 |
| PKHD1 | 21867.5805 | -0.4257899 | 0.24774816 | -1.7186398 | 0.08567997 | 0.49356597 |
| MCM3 | 191.156494 | -0.4859215 | 0.27878121 | -1.7430209 | 0.08132997 | 0.48594425 |
| EFHC1 | 717.171257 | -0.1264837 | 0.15685909 | -0.8063522 | 0.42003979 | 0.8125045 |
| TRAM2 | 212.435954 | -0.0655925 | 0.38975817 | -0.1682902 | 0.86635497 | 0.97386931 |
| TRAM2-AS1 | 45.5938659 | -0.3240263 | 0.3305106 | -0.9803809 | 0.32689814 | 0.75556858 |
| TMEM14A | 107.63683 | 0.00191529 | 0.29803398 | 0.00642643 | 0.99487249 | 0.998438 |
| GSTA4 | 300.9713 | 0.44722824 | 0.36989821 | 1.20905759 | 0.22664072 | 0.67880306 |
| ICK | 273.530096 | -0.1425562 | 0.21465459 | -0.6641192 | 0.50661402 | 0.85626196 |
| FBXO9 | 770.395007 | 0.00811182 | 0.16001018 | 0.05069565 | 0.95956804 | 0.991232 |
| ELOVL5 | 298.999396 | 0.58944325 | 0.48805049 | 1.20775056 | 0.22714322 | 0.67921637 |
| GCLC | 506.361566 | 0.02264426 | 0.52435709 | 0.0431848 | 0.96555422 | 0.99248831 |
| AL513211.1 | 39.9603005 | -0.2079418 | 0.40228734 | -0.5168986 | 0.60522697 | 0.89062401 |
| LRRC1 | 521.93911 | 0.07493375 | 0.17475784 | 0.42878617 | 0.66807884 | 0.91383898 |
| TINAG | 975.424887 | 1.97013109 | 0.69135901 | 2.84964985 | 0.00437674 | 0.12886682 |
| FAM83B | 1803.78616 | -0.2468596 | 0.24763729 | -0.9968597 | 0.31883262 | 0.75094569 |
| HCRTR2 | 103.77357 | 0.67486639 | 0.61108543 | 1.10437323 | 2.69E-01 | 7.18E-01 |

| | | | | | | |
|---|---|---|---|---|---|---|
| DST | 4208.89732 | 0.21019807 | 0.36343105 | 0.57837124 | 5.63E-01 | 0.87440863 |
| KIAA1586 | 103.649972 | 0.0960027 | 0.31376547 | 0.30596962 | 0.75962778 | 0.94133292 |
| ZNF451 | 1268.66062 | -0.0483597 | 0.13324525 | -0.3629373 | 0.71665176 | 0.92897478 |
| ZNF451-AS1 | 188.987121 | 0.1897408 | 0.63323849 | 0.29963561 | 0.76445512 | 0.94256396 |
| BAG2 | 62.2488825 | 0.44462879 | 0.43640771 | 1.01883807 | 0.30827985 | 0.74375973 |
| RAB23 | 99.812207 | 0.12408807 | 0.45927554 | 0.27018219 | 0.78702009 | 0.94978829 |
| PRIM2 | 991.979672 | -0.7474134 | 0.22539571 | -3.3160055 | 0.00091314 | 0.05313512 |
| AL021368.2 | 165.628754 | -0.1079873 | 0.29663954 | -0.3640354 | 0.7158316 | 0.92890921 |
| AL445250.1 | 76.9263899 | -0.3558283 | 0.63827426 | -0.5574849 | 0.57719618 | 0.88151051 |
| PTP4A1 | 839.740491 | 0.44032531 | 0.70530749 | 0.62430261 | 0.53242887 | 0.86397197 |
| PHF3 | 2236.94699 | 0.10461795 | 0.19124538 | 0.54703516 | 0.58435458 | 0.88416261 |
| EYS | 246.190066 | 0.71625284 | 0.56624518 | 1.26491645 | 0.20590128 | 0.65813563 |
| LMBRD1 | 1726.57321 | 0.16603898 | 0.15228653 | 1.0903064 | 0.2755782 | 0.72233844 |
| COL19A1 | 447.945161 | -0.2174467 | 0.34129369 | -0.6371248 | 0.52404355 | 0.86006369 |
| FAM135A | 4544.6158 | 0.04011095 | 0.29182095 | 0.13745055 | 8.91E-01 | 0.97729111 |
| SDHAF4 | 160.817203 | 0.00010616 | 0.2121788 | 0.00050032 | 1.00E+00 | 0.99987432 |
| SMAP1 | 1974.97926 | -0.1602292 | 0.19217766 | -0.8337558 | 0.40441863 | 0.80437725 |
| OGFRL1 | 272.190896 | -0.5235141 | 0.44909362 | -1.1657126 | 0.24373065 | 0.6948422 |
| AL136164.2 | 2060.20589 | 0.37693046 | 0.41996142 | 0.89753592 | 0.36943301 | 0.78095617 |
| LINC00472 | 917.114702 | -0.0691332 | 0.36864611 | -0.1875326 | 0.85124306 | 0.97057139 |
| RIMS1 | 126.604468 | 3.09318659 | 0.67751688 | 4.56547534 | 4.98E-06 | 0.00329144 |
| KHDC1 | 71.0283248 | -0.2033922 | 0.38545659 | -0.5276655 | 0.59773149 | 0.88883692 |
| AC019205.1 | 189.960598 | -0.2489591 | 0.2148695 | -1.1586526 | 0.24659784 | 0.69684411 |
| OOEP | 666.556024 | 0.33648643 | 0.43115423 | 0.78043171 | 0.43513681 | 0.81919857 |
| MTO1 | 414.442066 | 0.05745219 | 0.27435698 | 0.20940671 | 0.83413075 | 0.96510927 |
| EEF1A1 | 3472.3364 | 0.92077186 | 0.46454777 | 1.9820822 | 0.04747004 | 0.39411174 |
| AL121972.1 | 48.4534306 | -0.1500199 | 0.52440257 | -0.2860777 | 0.77481861 | 0.94636863 |
| SLC17A5 | 283.693229 | -0.1436376 | 0.33270437 | -0.4317274 | 0.66593952 | 0.91305741 |
| AL357507.1 | 55.7176335 | -0.2771755 | 0.38487908 | -0.7201626 | 0.47142488 | 0.83921324 |
| COX7A2 | 491.18205 | 0.39408002 | 0.38331738 | 1.02807761 | 0.30391332 | 0.74120958 |
| TMEM30A | 543.699784 | 0.41464446 | 0.26178188 | 1.5839311 | 0.1132094 | 0.53945994 |
| AL080250.1 | 101.044491 | 0.20079826 | 0.38406369 | 0.52282542 | 0.60109575 | 0.8890585 |
| FILIP1 | 86.787868 | 0.07089681 | 0.49036034 | 0.14458104 | 0.88504167 | 0.97664137 |
| SENP6 | 4050.453 | -0.108453 | 0.14235093 | -0.7618709 | 0.44613709 | 0.82571115 |
| MYO6 | 7032.49763 | -0.5563962 | 0.32520508 | -1.7109088 | 0.08709796 | 0.4952548 |
| MEI4 | 252.944934 | 0.1914474 | 0.41079693 | 0.46603902 | 0.64118757 | 0.90404588 |
| IRAK1BP1 | 1011.63046 | 0.5262321 | 0.21893817 | 2.4035649 | 0.01623608 | 0.25558383 |
| PHIP | 2693.93393 | -0.2214984 | 0.21137822 | -1.047877 | 0.29469528 | 0.73556877 |
| HMGN3 | 990.036025 | 0.25268854 | 0.21119151 | 1.19649004 | 0.23150538 | 0.68364735 |
| LCA5 | 139.289704 | 0.4447827 | 0.31618083 | 1.40673518 | 0.15950593 | 0.60928682 |
| SH3BGRL2 | 149.269979 | 0.14636051 | 0.32240459 | 0.45396535 | 0.64985377 | 0.90825486 |
| BCKDHB | 2443.20254 | 0.35106995 | 0.33637962 | 1.04367188 | 0.29663722 | 0.73672107 |
| AL359715.1 | 115.598024 | 0.47370462 | 0.52542062 | 0.90157219 | 0.36728417 | 0.78095617 |
| TENT5A | 260.344589 | -0.5321928 | 0.71067545 | -0.7488549 | 0.45394469 | 0.83060749 |
| IBTK | 1556.46054 | -0.5047374 | 0.21156209 | -2.3857651 | 0.01704363 | 0.26177833 |
| TPBG | 183.658145 | 0.38525836 | 0.48592642 | 0.79283271 | 4.28E-01 | 8.15E-01 |
| UBE3D | 1228.32072 | -0.0964242 | 0.21985123 | -0.4385884 | 0.66095983 | 0.91110616 |
| DOPEY1 | 1733.76661 | -0.0428045 | 0.17265538 | -0.2479189 | 0.80419717 | 0.95450237 |
| PGM3 | 410.384146 | 0.35990532 | 0.26543035 | 1.35593127 | 0.17512104 | 0.62944785 |
| RWDD2A | 72.4378977 | 0.78332224 | 0.3354534 | 2.33511496 | 0.01953742 | 0.2807376 |
| ME1 | 280.957488 | 1.44011436 | 0.37032795 | 3.88875417 | 0.00010076 | 0.01602094 |
| CYB5R4 | 519.055403 | -0.1798985 | 0.30338153 | -0.5929779 | 0.55319593 | 0.87104567 |
| MRAP2 | 205.414197 | -0.201096 | 0.60468482 | -0.3325634 | 0.73946391 | 0.93552616 |
| CEP162 | 951.766273 | -0.2450648 | 0.13924479 | -1.759957 | 0.0784151 | 0.48060389 |

| | | | | | | |
|---|---|---|---|---|---|---|
| NT5E | 55.6719273 | 1.49911059 | 0.57840794 | 2.59178772 | 9.55E-03 | 1.99E-01 |
| SNX14 | 1534.00185 | 0.01325358 | 0.12660902 | 0.10468114 | 0.91662883 | 0.98197488 |
| AL589666.1 | 84.5397058 | 0.0152748 | 0.39727342 | 0.03844909 | 0.96932962 | 0.99299658 |
| SYNCRIP | 535.587485 | -0.3190288 | 0.17268023 | -1.8475122 | 0.06467294 | 0.44001919 |
| ZNF292 | 5250.59933 | -0.2511896 | 0.29937381 | -0.8390501 | 0.40144121 | 0.80270011 |
| SMIM8 | 956.047829 | -0.2067241 | 0.2670354 | -0.7741451 | 0.43884501 | 0.82127804 |
| SLC35A1 | 68.8973998 | 0.47811581 | 0.44698998 | 1.06963427 | 0.28478396 | 0.72774451 |
| RARS2 | 1015.65454 | -0.131893 | 0.18665917 | -0.706598 | 0.47981636 | 0.84323863 |
| ORC3 | 742.9493 | -0.2922548 | 0.3825415 | -0.7639818 | 0.44487807 | 0.8252311 |
| AKIRIN2 | 331.284348 | 0.11435086 | 0.21582833 | 0.52982323 | 0.59623449 | 0.88859628 |
| RNGTT | 1733.75005 | -0.3664834 | 0.21084269 | -1.738184 | 0.08217841 | 0.48594425 |
| PNRC1 | 630.221363 | 0.55139799 | 0.40546944 | 1.35990024 | 0.17386149 | 0.6286116 |
| SRSF12 | 97.3328414 | 0.43548016 | 0.4435841 | 0.98173079 | 0.32623249 | 0.75554697 |
| PM20D2 | 427.928075 | 0.46636335 | 0.20369637 | 2.28950248 | 0.02205017 | 0.29509055 |
| UBE2J1 | 544.860089 | 0.23241578 | 0.21325364 | 1.0898561 | 0.27577654 | 0.72233844 |
| RRAGD | 686.692341 | 0.59839861 | 0.41923599 | 1.42735504 | 0.15347758 | 0.60285224 |
| ANKRD6 | 1734.42014 | -0.1506026 | 0.19507039 | -0.7720421 | 0.44008949 | 0.82232701 |
| LYRM2 | 413.879072 | 0.20166841 | 0.31221631 | 0.64592527 | 0.51832775 | 0.85916831 |
| MDN1 | 989.896388 | -0.2553072 | 0.1114053 | -2.2916973 | 0.02192312 | 0.29509055 |
| CASP8AP2 | 568.12191 | -0.1984038 | 0.18525928 | -1.0709521 | 0.28419098 | 0.72738535 |
| BACH2 | 1847.76413 | 0.3977361 | 0.48972627 | 0.81216003 | 0.41669981 | 0.80967753 |
| MAP3K7 | 1070.97371 | 0.0192889 | 0.2184864 | 0.08828421 | 0.92965079 | 0.98331361 |
| EPHA7 | 50.9050683 | 1.96640728 | 0.45072467 | 4.36276823 | 1.28E-05 | 0.00530122 |
| MANEA | 175.471497 | 0.5343929 | 0.24591331 | 2.17309462 | 0.0297732 | 0.33159709 |
| UFL1-AS1 | 372.147498 | 0.56681971 | 0.57530577 | 0.98524948 | 0.32450154 | 0.75477036 |
| UFL1 | 691.35388 | -0.0137401 | 0.2044832 | -0.0671943 | 0.94642704 | 0.98885769 |
| NDUFAF4 | 174.028821 | 0.40154567 | 0.36634443 | 1.09608783 | 0.27304034 | 0.72070318 |
| KLHL32 | 614.01196 | 0.41129716 | 0.40839339 | 1.00711023 | 0.3138818 | 0.74777437 |
| MMS22L | 996.239343 | 0.82970961 | 0.49354381 | 1.68112657 | 0.09273833 | 0.50702841 |
| AL589740.1 | 309.072756 | -0.0131753 | 0.56986844 | -0.0231198 | 0.9815547 | 0.99518016 |
| FBXL4 | 2166.82424 | -0.312104 | 0.20777566 | -1.5021201 | 0.13306609 | 0.57682553 |
| FAXC | 167.666599 | -0.8321811 | 0.43872378 | -1.8968225 | 0.05785137 | 0.42087381 |
| COQ3 | 122.804873 | 0.21538666 | 0.28405882 | 0.75824667 | 4.48E-01 | 8.26E-01 |
| PNISR | 3905.20646 | 0.14512876 | 0.15794821 | 0.91883762 | 3.58E-01 | 7.75E-01 |
| AL513550.1 | 282.861869 | 0.33617282 | 0.3421759 | 0.98245618 | 0.32587516 | 0.75550719 |
| USP45 | 770.883923 | 0.18703531 | 0.29825334 | 0.62710214 | 0.53059228 | 0.86396107 |
| TSTD3 | 131.054916 | 0.11031377 | 0.27280621 | 0.40436678 | 0.68594302 | 0.92053037 |
| CCNC | 1096.54887 | -0.0389311 | 0.23349143 | -0.1667345 | 0.86757896 | 0.97402505 |
| SIM1 | 3599.36576 | -0.4518041 | 0.51098758 | -0.8841782 | 0.37660002 | 0.78527035 |
| ASCC3 | 2428.00062 | 0.08125202 | 0.21377203 | 0.38008723 | 0.70388066 | 0.92513032 |
| GRIK2 | 551.258183 | -0.0574006 | 0.64765837 | -0.0886279 | 0.92937764 | 0.98331361 |
| HACE1 | 941.137569 | -0.0637769 | 0.12030757 | -0.5301157 | 5.96E-01 | 8.89E-01 |
| PREP | 481.531892 | -0.1288215 | 0.15972892 | -0.8065007 | 0.41995422 | 0.8125045 |
| ATG5 | 1262.26178 | -0.1590211 | 0.2178487 | -0.7299612 | 0.46541392 | 0.83652911 |
| CRYBG1 | 942.35229 | -1.0658671 | 0.41641344 | -2.5596365 | 0.01047817 | 0.20778283 |
| RTN4IP1 | 364.386332 | -0.4942125 | 0.36011214 | -1.372385 | 0.16994362 | 0.62581135 |
| QRSL1 | 619.428312 | -0.4193438 | 0.31113545 | -1.3477853 | 0.17772745 | 0.63160978 |
| LINC02532 | 11016.7901 | -0.0208559 | 0.37684344 | -0.0553436 | 0.95586471 | 0.99065996 |
| CD24 | 5013.94885 | 0.48729291 | 0.24957939 | 1.95245652 | 5.09E-02 | 0.40489586 |
| C6orf203 | 252.505756 | -0.170902 | 0.20811238 | -0.8212004 | 0.41153211 | 0.80675684 |
| BEND3 | 56.8623142 | -0.6216274 | 0.40402664 | -1.5385803 | 0.12390679 | 0.56105491 |
| PDSS2 | 2205.35012 | -0.3234232 | 0.2004719 | -1.6133094 | 0.1066773 | 0.52775297 |
| SOBP | 151.990838 | 1.62668966 | 0.69824504 | 2.3296831 | 0.01982291 | 0.28276541 |
| SEC63 | 1092.45754 | -0.002816 | 0.20875356 | -0.0134897 | 0.98923708 | 0.99670729 |

| | | | | | | |
|---|---|---|---|---|---|---|
| OSTM1 | 388.634158 | -0.0322062 | 0.31014022 | -0.1038439 | 0.9172932 | 0.98199511 |
| SNX3 | 765.581752 | 0.14374783 | 0.12154227 | 1.18269825 | 0.23692876 | 0.68918564 |
| AFG1L | 1022.78427 | -0.3826681 | 0.22384594 | -1.7095154 | 8.74E-02 | 0.49566631 |
| FOXO3 | 5197.82915 | 0.1061757 | 0.31221469 | 0.34007272 | 0.73380176 | 0.93552616 |
| LINC00222 | 45.7219749 | -0.0643108 | 0.57687343 | -0.1114817 | 0.91123436 | 0.98112934 |
| ARMC2 | 691.968622 | 0.558997 | 0.23063014 | 2.42378119 | 0.01535986 | 0.25212125 |
| SESN1 | 1233.71445 | 0.22732888 | 0.2350793 | 0.96703065 | 0.33352871 | 0.76155242 |
| CEP57L1 | 288.961152 | -0.4625703 | 0.16252108 | -2.8462173 | 0.0044242 | 0.12957705 |
| CD164 | 692.542893 | 0.30960704 | 0.1887555 | 1.64025442 | 0.10095228 | 0.51819907 |
| PPIL6 | 280.321985 | 0.42882266 | 0.31126907 | 1.37765908 | 0.16830857 | 0.62253512 |
| SMPD2 | 58.6562694 | 0.31142366 | 0.32663831 | 0.9534205 | 0.34037706 | 0.76478309 |
| MICAL1 | 115.207165 | 0.08342958 | 0.39009964 | 0.21386737 | 0.83065048 | 0.9635604 |
| ZBTB24 | 165.693558 | -0.0852637 | 0.21048264 | -0.4050868 | 0.68541374 | 0.92053037 |
| AK9 | 780.302355 | -0.0996206 | 0.24943593 | -0.3993835 | 0.68961062 | 0.92168526 |
| FIG4 | 675.830703 | -0.3144693 | 0.17805479 | -1.7661379 | 0.07737271 | 0.47763291 |
| WASF1 | 444.872336 | -0.2539776 | 0.3489969 | -0.7277358 | 0.46677531 | 0.83702251 |
| CDC40 | 401.176877 | -0.1618736 | 0.16268273 | -0.9950264 | 0.31972341 | 0.75094569 |
| METTL24 | 80.3547867 | 0.54403455 | 0.72373974 | 0.75169915 | 0.452232 | 0.82885146 |
| CDK19 | 1609.5822 | -0.2146859 | 0.25242138 | -0.850506 | 0.39504382 | 0.7989793 |
| AMD1 | 573.074609 | 0.10327662 | 0.29743981 | 0.34721855 | 0.72842713 | 0.93453345 |
| GTF3C6 | 280.336637 | 0.53028997 | 0.4137387 | 1.2817026 | 0.19994699 | 0.65260683 |
| RPF2 | 238.259705 | -0.2064395 | 0.30970189 | -0.6665747 | 0.50504382 | 0.85582573 |
| SLC16A10 | 411.055848 | -0.7142924 | 0.44878322 | -1.5916201 | 0.1114701 | 0.53522667 |
| MFSD4B | 268.554519 | 0.01090798 | 0.26139608 | 0.04172971 | 0.96671417 | 0.99248831 |
| AL080317.2 | 57.3409546 | -0.2153774 | 0.33734162 | -0.6384548 | 0.52317764 | 0.86006369 |
| REV3L | 2699.25829 | -0.0313396 | 0.19113687 | -0.163964 | 0.86975945 | 0.97454113 |
| AL080317.3 | 105.009103 | -0.5685427 | 0.22566649 | -2.5193933 | 0.01175573 | 0.22271223 |
| TRAF3IP2-AS | 1246.94917 | -0.0774731 | 0.24341376 | -0.3182776 | 0.75027437 | 0.93775604 |
| TRAF3IP2 | 236.402721 | -0.14924 | 0.24336823 | -0.6132272 | 0.53972611 | 0.86566791 |
| FYN | 82.881749 | 1.0764452 | 0.45981845 | 2.34102221 | 0.01923102 | 0.28037812 |
| TUBE1 | 417.484819 | 0.35781956 | 0.19973425 | 1.79147821 | 0.07321659 | 0.4701377 |
| FAM229B | 161.142557 | 0.4458678 | 0.27333556 | 1.63121035 | 0.10284594 | 0.52229606 |
| LAMA4 | 286.960945 | 2.50438136 | 0.9337275 | 2.68213302 | 0.00731544 | 0.17305714 |
| MARCKS | 156.9321 | 0.53007715 | 0.35054511 | 1.51215103 | 0.13049545 | 0.57223551 |
| HDAC2 | 480.072146 | -0.0203612 | 0.23150191 | -0.0879526 | 0.92991436 | 0.98331361 |
| HDAC2-AS2 | 73.6511766 | -0.1454534 | 0.44971319 | -0.323436 | 0.7463651 | 0.9368751 |
| FRK | 1946.97877 | 0.40855534 | 0.24430104 | 1.67234388 | 0.09445653 | 0.51017802 |
| NT5DC1 | 1989.11633 | 0.30319706 | 0.24783163 | 1.22339935 | 0.2211789 | 0.67400484 |
| COL10A1 | 99.0469709 | 0.85711475 | 0.72406784 | 1.18374924 | 0.23651234 | 0.6887327 |
| TSPYL4 | 138.830843 | 0.68809575 | 0.24123241 | 2.85241834 | 0.0043388 | 0.12850035 |
| DSE | 1299.36187 | -0.0248356 | 0.22212312 | -0.1118099 | 9.11E-01 | 9.81E-01 |
| TSPYL1 | 291.49272 | 1.09504918 | 0.39306414 | 2.78593 | 0.00533744 | 0.14749423 |
| RWDD1 | 237.380879 | 0.03054419 | 0.2704504 | 0.11293822 | 0.91007954 | 0.98069436 |
| ZUP1 | 250.885336 | -0.2968451 | 0.18608468 | -1.5952149 | 0.11066419 | 0.53393773 |
| KPNA5 | 761.199018 | 0.14965417 | 0.16159282 | 0.92611892 | 0.35438418 | 0.77243419 |
| DCBLD1 | 300.929851 | -0.5512692 | 0.29249934 | -1.8846854 | 0.05947233 | 0.42578317 |
| GOPC | 501.790875 | 0.07584948 | 0.22634773 | 0.33510157 | 0.73754847 | 0.93552616 |
| NUS1 | 475.856446 | 0.12819252 | 0.21386482 | 0.59940909 | 0.54890012 | 0.86904066 |
| SLC35F1 | 114.191484 | 0.11640313 | 0.44361508 | 0.26239669 | 0.79301562 | 0.95224951 |
| CEP85L | 752.159411 | -0.2208116 | 0.27558825 | -0.8012372 | 0.42299434 | 0.81379363 |
| MCM9 | 1246.09308 | -0.1260248 | 0.24496341 | -0.5144637 | 0.60692784 | 0.89174148 |
| ASF1A | 90.0983956 | 0.55420483 | 0.32425153 | 1.70918187 | 0.08741727 | 0.4957899 |
| AL365275.1 | 491.124205 | -0.211658 | 0.51672757 | -0.4096123 | 0.68209038 | 0.91969963 |
| FAM184A | 1732.42012 | 0.56220982 | 0.33367712 | 1.68489174 | 0.09200946 | 0.50534427 |

| | | | | | | |
|---|---|---|---|---|---|---|
| MAN1A1 | 2922.55492 | 0.1864612 | 0.26645867 | 0.69977529 | 0.48406765 | 0.8445691 |
| TBC1D32 | 1482.23764 | -0.3518657 | 0.26951657 | -1.3055439 | 0.19170771 | 0.64596359 |
| GJA1 | 70.1001106 | 1.34271606 | 0.67208059 | 1.99784977 | 0.04573295 | 0.38904776 |
| HSF2 | 218.149342 | -0.2931317 | 0.24160198 | -1.2132837 | 0.22502139 | 0.67768069 |
| SERINC1 | 808.76686 | 0.34669897 | 0.20462018 | 1.69435374 | 0.09019809 | 0.50138792 |
| PKIB | 260.820957 | 0.26626432 | 0.42817457 | 0.62185926 | 0.53403442 | 0.86397197 |
| SMPDL3A | 108.072447 | 0.5396266 | 0.30021992 | 1.79743771 | 0.07226616 | 0.46815286 |
| NKAIN2 | 526.830502 | -0.8362639 | 0.59744503 | -1.3997336 | 0.16159312 | 0.61311788 |
| RNF217-AS1 | 98.8729657 | -0.9360332 | 0.49941593 | -1.8742559 | 0.06089517 | 0.4312945 |
| RNF217 | 1288.19169 | -0.6956749 | 0.38159179 | -1.8230867 | 0.06829026 | 0.45323644 |
| TPD52L1 | 933.027657 | -0.0587451 | 0.35823062 | -0.1639868 | 0.86974152 | 0.97454113 |
| HDDC2 | 354.604008 | 0.19056941 | 0.2343873 | 0.81305348 | 0.4161874 | 0.80966301 |
| AL365259.1 | 74.3696162 | -0.0256338 | 0.78045598 | -0.0328447 | 0.97379843 | 0.99345929 |
| NCOA7 | 4171.98304 | 0.64854186 | 0.42194087 | 1.53704442 | 0.12428243 | 0.56105491 |
| NCOA7-AS1 | 63.3131173 | 0.62722128 | 0.85554664 | 0.73312341 | 0.46348316 | 0.83531269 |
| HINT3 | 260.58486 | 0.22350235 | 0.17700989 | 1.26265463 | 0.20671332 | 0.65921688 |
| TRMT11 | 745.456637 | -0.3700141 | 0.25059441 | -1.4765458 | 0.13979743 | 0.58431708 |
| AC020559.1 | 265.722441 | 0.07764718 | 0.45421087 | 0.17094964 | 0.86426336 | 0.97356769 |
| RNF146 | 413.596017 | 0.00836985 | 0.2866716 | 0.02919664 | 0.97670776 | 0.99418779 |
| ECHDC1 | 1057.48004 | -0.2003835 | 0.2059796 | -0.9728319 | 0.33063685 | 0.75909265 |
| PTPRK | 4596.12016 | -0.4809476 | 0.203787 | -2.3600502 | 0.01827246 | 0.2711921 |
| LAMA2 | 269.773635 | 0.25823876 | 0.50703228 | 0.50931424 | 6.11E-01 | 0.89339478 |
| AL356124.1 | 337.402523 | 0.20755958 | 0.44948087 | 0.46177622 | 0.64424181 | 0.90507553 |
| ARHGAP18 | 2326.69946 | -0.0944635 | 0.23906819 | -0.3951321 | 6.93E-01 | 0.92224682 |
| L3MBTL3 | 428.366819 | -0.2326592 | 0.33819682 | -0.6879401 | 0.49149048 | 0.84697949 |
| EPB41L2 | 620.584871 | -0.1199121 | 0.34161405 | -0.3510164 | 7.26E-01 | 0.93269023 |
| AKAP7 | 516.672902 | -0.0104826 | 0.37617979 | -0.027866 | 0.97776902 | 0.99455895 |
| MED23 | 724.902118 | -0.3802155 | 0.20508166 | -1.8539711 | 0.0637433 | 0.43779712 |
| ENPP3 | 47.3834399 | 0.31581684 | 0.43947292 | 0.71862639 | 0.47237115 | 0.83921324 |
| ENPP1 | 81.0577903 | 0.65872528 | 0.49538762 | 1.3297169 | 0.18361157 | 0.63834582 |
| CTGF | 128.143721 | -0.6085256 | 0.7328023 | -0.8304089 | 0.40630763 | 0.80499101 |
| AL133346.1 | 94.3602183 | 1.53250774 | 0.71429868 | 2.1454719 | 0.03191514 | 0.34024781 |
| STX7 | 1103.30245 | -0.0494873 | 0.21920606 | -0.2257568 | 0.82139057 | 0.96066244 |
| SLC18B1 | 578.256757 | 0.14148679 | 0.28580792 | 0.49504153 | 0.62057081 | 0.89731054 |
| RPS12 | 747.843324 | 0.83545505 | 0.42758651 | 1.95388544 | 0.05071477 | 0.40427963 |
| TBPL1 | 305.011148 | -0.2334287 | 0.28624113 | -0.8154967 | 0.41478803 | 0.80965493 |
| SLC2A12 | 213.198625 | 0.80562744 | 0.34882379 | 2.30955413 | 0.02091285 | 0.2892543 |
| SGK1 | 454.906778 | 0.23111104 | 0.56849306 | 0.40653275 | 6.84E-01 | 9.20E-01 |
| ALDH8A1 | 85.6803372 | -0.5041326 | 0.42774894 | -1.1785713 | 0.23856892 | 0.68985482 |
| HBS1L | 662.100582 | -0.119125 | 0.21299423 | -0.5592876 | 0.57596549 | 0.88123035 |
| AHI1 | 3289.2781 | -0.1710565 | 0.26720488 | -0.6401699 | 0.52206218 | 0.85993307 |
| LINC00271 | 1587.54212 | 0.14443156 | 0.44599059 | 0.3238444 | 0.74605584 | 0.93675395 |
| PDE7B | 7114.95948 | -0.3178399 | 0.34014384 | -0.934428 | 0.35008318 | 0.77044053 |
| AL138828.1 | 927.749098 | -0.3235226 | 0.65319119 | -0.4952954 | 0.62039165 | 0.89726851 |
| BCLAF1 | 1421.10996 | -0.0509984 | 0.19696658 | -0.2589189 | 0.7956978 | 0.95278473 |
| MAP7 | 2923.26935 | -0.1978102 | 0.17881268 | -1.1062424 | 0.26862163 | 0.71818262 |
| MAP3K5 | 1644.92635 | 1.34555188 | 0.6247299 | 2.15381381 | 0.03125478 | 0.3383722 |
| PEX7 | 409.470345 | 0.10238535 | 0.21679215 | 0.47227423 | 0.63673106 | 0.90255504 |
| IFNGR1 | 293.579474 | 0.34786643 | 0.21314432 | 1.63206994 | 0.10266475 | 0.52197795 |
| TNFAIP3 | 84.6470503 | 0.6628261 | 0.58443715 | 1.13412725 | 2.57E-01 | 0.70735159 |
| PERP | 353.663266 | 0.42702145 | 0.33509259 | 1.27433867 | 2.03E-01 | 6.56E-01 |
| ARFGEF3 | 1314.50297 | -0.7988291 | 0.35992142 | -2.2194542 | 0.02645584 | 0.31421858 |
| HEBP2 | 517.782956 | -0.0967679 | 0.24730141 | -0.3912954 | 6.96E-01 | 0.92229578 |
| NHSL1 | 1316.79501 | -0.916159 | 0.32012896 | -2.8618436 | 0.00421185 | 0.12615484 |

| | | | | | | |
|---|---|---|---|---|---|---|
| CCDC28A | 234.68135 | 0.01557481 | 0.21790229 | 0.07147613 | 0.94301882 | 0.98772472 |
| REPS1 | 1329.9678 | -0.3264671 | 0.18812172 | -1.7354036 | 0.08266934 | 0.48594425 |
| ABRACL | 59.5737722 | 0.51496112 | 0.58192809 | 0.88492226 | 0.37619856 | 0.78490077 |
| HECA | 227.46019 | -0.0710392 | 0.37101178 | -0.1914743 | 0.84815404 | 0.96972793 |
| AL031772.1 | 86.9656721 | 0.14977175 | 0.47773298 | 0.31350515 | 0.75389691 | 0.93874663 |
| CITED2 | 262.546287 | 0.34627519 | 0.30396408 | 1.13919771 | 0.2546207 | 0.70509116 |
| VTA1 | 558.776711 | -0.2137333 | 0.21164106 | -1.0098856 | 0.31255009 | 0.74693024 |
| HIVEP2 | 2876.85993 | 0.40759791 | 0.42527934 | 0.95842397 | 0.33784901 | 0.76415199 |
| LINC01277 | 91.146014 | -0.6287668 | 0.57357018 | -1.0962334 | 0.27297667 | 0.72070318 |
| AIG1 | 1955.77166 | 0.11457569 | 0.1569408 | 0.73005673 | 0.46535551 | 0.83652911 |
| ADAT2 | 186.10141 | -0.3319298 | 0.29858473 | -1.1116771 | 0.26627701 | 0.71641758 |
| PEX3 | 695.860685 | -0.0927124 | 0.24007315 | -0.386184 | 0.69936037 | 0.92367966 |
| FUCA2 | 226.206415 | -0.0496659 | 0.21445296 | -0.2315933 | 0.81685394 | 0.95909544 |
| PHACTR2 | 1195.19465 | 0.23539473 | 0.28143097 | 0.83642084 | 0.40291819 | 0.80324472 |
| PLAGL1 | 887.793155 | 0.49268149 | 0.21292251 | 2.3139004 | 0.02067319 | 0.28805076 |
| SF3B5 | 292.93057 | 1.01832284 | 0.45723995 | 2.22710818 | 0.02594004 | 0.31222606 |
| UTRN | 4862.12056 | -0.3278926 | 0.23219124 | -1.4121662 | 0.15790104 | 0.60779483 |
| AL024474.2 | 107.336593 | -0.0195746 | 0.49535773 | -0.039516 | 0.96847899 | 0.99250254 |
| EPM2A | 793.512311 | -0.6360149 | 0.34283592 | -1.8551583 | 0.06357363 | 0.43779712 |
| FBXO30 | 189.557528 | 0.06365185 | 0.39382172 | 0.16162604 | 0.87160035 | 0.97526209 |
| AL356599.1 | 598.790628 | 0.16704761 | 0.3407298 | 0.49026416 | 0.62394698 | 0.89873262 |
| SHPRH | 2400.0171 | -0.0692178 | 0.11904667 | -0.5814345 | 0.56094767 | 0.87394741 |
| GRM1 | 364.048104 | -0.0922629 | 0.50006763 | -0.1845009 | 8.54E-01 | 9.71E-01 |
| RAB32 | 143.31008 | 0.14967528 | 0.43742123 | 0.34217653 | 0.73221804 | 0.93552616 |
| ADGB | 74.4884942 | -0.4584161 | 0.63182455 | -0.7255433 | 4.68E-01 | 0.83825637 |
| STXBP5-AS1 | 205.890043 | -0.8111918 | 0.38740643 | -2.0939038 | 0.03626855 | 0.35831812 |
| STXBP5 | 1574.08698 | -0.0889362 | 0.18726711 | -0.4749163 | 0.63484662 | 0.90178395 |
| SAMD5 | 1115.79415 | -0.4158603 | 0.31334017 | -1.3271848 | 0.18444755 | 0.63929879 |
| AL033504.1 | 701.280042 | -0.7646499 | 0.43608488 | -1.7534428 | 0.07952602 | 0.48176967 |
| AL513164.1 | 53.0444104 | -0.3708877 | 0.76939072 | -0.4820538 | 0.62976774 | 0.90005881 |
| SASH1 | 7176.80114 | -0.5267347 | 0.35103112 | -1.5005356 | 0.13347571 | 0.57682553 |
| UST | 993.678985 | 0.13400588 | 0.31745191 | 0.42212969 | 0.67293036 | 0.91608132 |
| TAB2 | 1664.71772 | -0.0992525 | 0.1635527 | -0.6068531 | 0.54394838 | 0.86639505 |
| PPIL4 | 642.214521 | 0.08328547 | 0.19077836 | 0.43655617 | 0.66243326 | 0.91110616 |
| GINM1 | 716.496347 | 0.1339114 | 0.18509275 | 0.72348268 | 0.46938339 | 0.83881856 |
| AL078581.3 | 61.6749054 | 0.032179 | 0.56951531 | 0.05650243 | 0.95494156 | 0.99049376 |
| KATNA1 | 359.344028 | -0.0377969 | 0.27475188 | -0.1375675 | 0.89058221 | 0.97729111 |
| LATS1 | 1049.50596 | -0.1166524 | 0.14805086 | -0.7879212 | 0.43074282 | 0.81706845 |
| NUP43 | 358.068631 | -0.2251999 | 0.23679279 | -0.9510422 | 0.34158294 | 0.76545664 |
| PCMT1 | 587.570022 | -0.1253514 | 0.22528332 | -0.5564167 | 0.577926 | 0.88191134 |
| AL355312.2 | 45.7449525 | -0.3553454 | 0.38605847 | -0.9204445 | 0.35734051 | 0.77455501 |
| LRP11 | 244.732111 | -0.3404208 | 0.25161769 | -1.3529289 | 0.17607835 | 0.62957039 |
| PPP1R14C | 174.319397 | -0.4979331 | 0.60992746 | -0.8163809 | 4.14E-01 | 8.09E-01 |
| PLEKHG1 | 1557.7673 | 0.32136906 | 0.51803582 | 0.62036069 | 0.53502034 | 0.8642718 |
| MTHFD1L | 1112.886 | -0.3997611 | 0.43093868 | -0.9276519 | 3.54E-01 | 7.72E-01 |
| AL138733.1 | 56.7414174 | -0.2521243 | 0.32431038 | -0.7774165 | 0.43691305 | 0.82013459 |
| AKAP12 | 253.581148 | -0.0176068 | 0.65353128 | -0.0269411 | 0.97850674 | 0.99455895 |
| ZBTB2 | 136.34512 | -0.3032476 | 0.32111023 | -0.9443724 | 0.34497936 | 0.76724655 |
| RMND1 | 431.428269 | -0.4425767 | 0.26661429 | -1.6599888 | 0.09691671 | 0.51327103 |
| ARMT1 | 224.060533 | 0.16932963 | 0.30806217 | 0.54966058 | 0.5825522 | 0.88338315 |
| ESR1 | 226.817537 | 1.67740102 | 0.69446297 | 2.41539302 | 0.01571824 | 0.25227487 |
| SYNE1 | 6847.69627 | 0.7320314 | 0.38394944 | 1.9065828 | 0.05657463 | 0.41843207 |
| AL080276.2 | 82.2394206 | 0.20644059 | 0.36205459 | 0.57019187 | 0.56854757 | 0.87763759 |
| MTRF1L | 209.965765 | -0.0275532 | 0.25479118 | -0.1081401 | 0.91388453 | 0.98145965 |

| | | | | | | |
|---|---|---|---|---|---|---|
| RGS17 | 49.9684191 | -0.0720223 | 0.7274064 | -0.0990124 | 0.92112842 | 0.98215577 |
| OPRM1 | 173.788183 | 0.6195476 | 0.53087193 | 1.16703779 | 0.24319511 | 0.69452742 |
| IPCEF1 | 1665.57359 | 0.56941972 | 0.66082017 | 0.86168635 | 0.38886014 | 0.79634939 |
| CNKSR3 | 2878.90827 | -0.0315418 | 0.27488418 | -0.1147457 | 0.90864669 | 0.98047754 |
| SCAF8 | 2667.43076 | -0.1956253 | 0.21188556 | -0.9232591 | 0.35587222 | 0.77352578 |
| TIAM2 | 1548.04839 | 0.31897326 | 0.13195904 | 2.41721411 | 0.01563981 | 0.25227487 |
| TFB1M | 837.665318 | -0.3500743 | 0.24214164 | -1.4457419 | 0.1482496 | 0.59438683 |
| CLDN20 | 57.9612537 | -0.343643 | 0.43301786 | -0.7936001 | 0.42742827 | 0.81466629 |
| ARID1B | 9049.34524 | -0.0972411 | 0.17235502 | -0.5641908 | 0.57262426 | 0.87946834 |
| TMEM242 | 310.776978 | -0.0022342 | 0.19142118 | -0.0116717 | 0.99068753 | 0.99740789 |
| ZDHHC14 | 4810.67987 | -0.4382325 | 0.29333856 | -1.4939478 | 0.13518925 | 0.57708028 |
| SNX9 | 2291.91608 | -0.3881997 | 0.27295498 | -1.4222115 | 0.15496486 | 0.60399259 |
| SYNJ2 | 429.427073 | -0.8075573 | 0.46663302 | -1.7306047 | 8.35E-02 | 0.4882568 |
| SERAC1 | 222.306107 | -0.4214384 | 0.25032886 | -1.6835389 | 0.09227081 | 0.5058297 |
| GTF2H5 | 238.126888 | 0.10180071 | 0.36300141 | 0.28044164 | 0.7791387 | 0.94732571 |
| TULP4 | 1839.91724 | -0.4268531 | 0.16381691 | -2.6056716 | 0.00916943 | 0.19441258 |
| TMEM181 | 554.987337 | -0.2535991 | 0.21277159 | -1.1918843 | 0.2333066 | 0.68484327 |
| DYNLT1 | 604.385458 | 0.35051015 | 0.23537731 | 1.4891416 | 1.36E-01 | 0.57787125 |
| SYTL3 | 497.885521 | 1.2494534 | 0.65516595 | 1.90707928 | 0.05651032 | 0.41843207 |
| EZR | 1682.8765 | 0.21328702 | 0.32535009 | 0.65556157 | 0.51210626 | 0.85762435 |
| C6orf99 | 44.7640122 | -0.0324275 | 0.5487711 | -0.0590912 | 0.95287948 | 0.98949465 |
| RSPH3 | 548.997878 | 0.19176165 | 0.21273431 | 0.90141381 | 0.36736834 | 0.78095617 |
| AL035530.2 | 533.559101 | 0.36313644 | 0.4792202 | 0.7577653 | 0.4485915 | 0.82630667 |
| SOD2 | 131.243612 | -0.5474361 | 0.62647135 | -0.8738407 | 0.38220503 | 0.78957558 |
| SOD2.1 | 253.550839 | -0.1067076 | 0.56453663 | -0.189018 | 0.85007872 | 0.97041641 |
| WTAP | 1598.00067 | -0.2189512 | 0.30651882 | -0.7143158 | 0.47503192 | 0.84106098 |
| ACAT2 | 143.664835 | 0.16298757 | 0.43678716 | 0.37315101 | 0.70903606 | 0.92577218 |
| TCP1 | 349.172045 | 0.27760567 | 0.25823142 | 1.07502669 | 0.28236278 | 0.72604837 |
| MRPL18 | 194.944588 | 0.33469356 | 0.28265889 | 1.18408999 | 0.23637744 | 0.68864351 |
| IGF2R | 2550.15724 | -0.1146413 | 0.23369981 | -0.4905496 | 0.62374504 | 0.89873262 |
| SLC22A1 | 149.682129 | -0.2244594 | 0.54999743 | -0.4081098 | 0.68319303 | 0.9200778 |
| SLC22A2 | 464.240151 | -0.0138394 | 0.68165725 | -0.0203025 | 0.98380203 | 0.99518552 |
| SLC22A3 | 679.99071 | -0.197737 | 0.59388492 | -0.332955 | 0.73916822 | 0.93552616 |
| MAP3K4 | 1073.29369 | -0.2364416 | 0.15509146 | -1.5245303 | 0.1273763 | 0.56740575 |
| AGPAT4 | 1366.04118 | -0.0569009 | 0.36575519 | -0.1555711 | 8.76E-01 | 0.97614152 |
| PRKN | 8073.70409 | -0.2380722 | 0.2895369 | -0.8222518 | 0.41093363 | 0.80666106 |
| PACRG | 1697.26104 | -0.455503 | 0.23948709 | -1.9019939 | 0.05717196 | 0.41931393 |
| CAHM | 47.8448759 | -0.0695067 | 0.37722427 | -0.1842584 | 0.85381075 | 0.97077171 |
| QKI | 5162.54236 | -0.1024958 | 0.19521297 | -0.5250461 | 0.59955118 | 0.88883692 |
| PDE10A | 2232.66195 | 0.73920434 | 0.51995827 | 1.4216609 | 0.15512472 | 0.60416746 |
| SFT2D1 | 466.523265 | -0.0304213 | 0.17620481 | -0.1726475 | 0.86292849 | 0.97319807 |
| MPC1 | 2022.3764 | 0.40731167 | 0.2672244 | 1.52423086 | 0.12745105 | 0.56740848 |
| AL022069.1 | 89.5547913 | 0.01186001 | 0.50465496 | 0.02350123 | 0.98125046 | 0.99518016 |
| RPS6KA2 | 1287.82737 | -0.2654951 | 0.43788785 | -0.6063085 | 0.54430992 | 0.86654229 |
| AL159163.1 | 169.098856 | -0.6617845 | 0.60395186 | -1.095757 | 0.27318514 | 0.72070318 |
| Z94721.1 | 233.91486 | 0.08210681 | 0.51504617 | 0.15941641 | 0.87334081 | 0.97564728 |
| FGFR1OP | 999.098263 | -0.4324181 | 0.17264285 | -2.5046977 | 0.01225561 | 0.22749396 |
| LINC01558 | 47.647918 | 0.74713292 | 0.71109134 | 1.05068488 | 0.29340334 | 0.73527014 |
| AFDN | 3777.70255 | -0.0938349 | 0.21458242 | -0.4372909 | 0.66190039 | 0.91110616 |
| WDR27 | 1211.82694 | 0.00026281 | 0.23541183 | 0.00111638 | 0.99910926 | 0.99985365 |
| C6orf120 | 69.2052196 | 0.67191078 | 0.33992506 | 1.97664384 | 0.0480819 | 0.39546317 |
| PHF10 | 510.328832 | 0.13749214 | 0.25846333 | 0.53195995 | 0.59475374 | 0.8885412 |
| TCTE3 | 83.8773704 | -0.0201037 | 0.33138213 | -0.0606663 | 0.95162497 | 0.98938795 |
| ERMARD | 328.403905 | -0.6960886 | 0.33469533 | -2.0797677 | 0.03754684 | 0.36360429 |

| | | | | | | |
|---|---|---|---|---|---|---|
| DLL1 | 61.90343 | -0.4276252 | 0.50320034 | -0.849811 | 0.39543015 | 0.79924445 |
| FAM120B | 1679.03019 | -0.2241768 | 0.24094143 | -0.9304201 | 0.35215359 | 0.77151834 |
| PSMB1 | 553.041684 | 0.47004623 | 0.31163178 | 1.50833857 | 0.1314679 | 0.57519227 |
| TBP | 255.921724 | -0.4491425 | 0.19457595 | -2.3083147 | 0.02098164 | 0.28959923 |
| PDCD2 | 208.72061 | 0.31286018 | 0.29391605 | 1.06445424 | 0.28712298 | 0.72972899 |
| AC093627.5 | 55.5760518 | -0.7308296 | 0.4291129 | -1.7031174 | 0.0885461 | 0.49770444 |
| FAM20C | 654.383997 | -1.2543465 | 0.51534057 | -2.4340146 | 1.49E-02 | 2.50E-01 |
| PDGFA | 316.256808 | -0.6967964 | 0.27922058 | -2.4955053 | 0.01257779 | 0.22916272 |
| PRKAR1B | 624.974508 | -0.1881148 | 0.31959672 | -0.5886005 | 0.55612932 | 0.87212955 |
| DNAAF5 | 419.390631 | -0.3825916 | 0.18868521 | -2.0276711 | 0.04259383 | 0.38021566 |
| SUN1 | 1180.17665 | -0.2334682 | 0.17936311 | -1.3016513 | 0.19303561 | 0.64699503 |
| AC073957.3 | 61.398798 | 0.6454882 | 0.62212657 | 1.03755126 | 0.29947902 | 0.73673761 |
| ADAP1 | 83.509212 | 1.94808959 | 0.60390994 | 3.22579485 | 0.00125623 | 0.0645665 |
| COX19 | 320.456014 | 0.19361875 | 0.21854661 | 0.88593802 | 0.37565093 | 0.78490077 |
| C7orf50 | 665.590239 | 0.18307153 | 0.26389728 | 0.69372267 | 0.48785615 | 0.8451268 |
| ZFAND2A | 47.1071981 | -0.2121328 | 0.38671721 | -0.5485475 | 0.583316 | 0.88379959 |
| MICALL2 | 156.576986 | -0.4211974 | 0.46863951 | -0.8987663 | 0.36877717 | 0.78095617 |
| INTS1 | 324.750138 | -0.1953782 | 0.34154011 | -0.5720505 | 0.56728777 | 0.87671747 |
| MAFK | 215.179519 | -0.024008 | 0.24825776 | -0.0967059 | 0.92295995 | 0.98215577 |
| TMEM184A | 83.288927 | 0.13454379 | 0.47027062 | 0.28609866 | 0.77480256 | 0.94636863 |
| PSMG3 | 78.6333143 | 0.11004439 | 0.39279144 | 0.28015986 | 0.77935486 | 0.9474918 |
| PSMG3-AS1 | 125.465747 | -0.1529076 | 0.27548339 | -0.5550521 | 0.57885899 | 0.8821758 |
| ELFN1 | 127.566915 | 0.26347806 | 0.431459 | 0.61066765 | 0.54141963 | 0.86582694 |
| MAD1L1 | 698.528276 | -0.3547253 | 0.20381739 | -1.7404077 | 0.08178746 | 0.48594425 |
| MRM2 | 87.0811155 | -0.0875941 | 0.28441329 | -0.3079817 | 0.75809626 | 0.94056395 |
| SNX8 | 179.596739 | -0.7747242 | 0.41045804 | -1.8874627 | 0.05909812 | 0.42419956 |
| EIF3B | 528.883097 | -0.2886905 | 0.2949904 | -0.9786438 | 0.32775601 | 0.75633196 |
| CHST12 | 217.856471 | -0.1102286 | 0.28637552 | -0.3849093 | 0.70030459 | 0.92367966 |
| BRAT1 | 184.766121 | -0.2494934 | 0.33331401 | -0.7485236 | 0.45414439 | 0.83085778 |
| IQCE | 374.212115 | -0.2166489 | 0.23579393 | -0.9188062 | 0.35819694 | 0.77529959 |
| TTYH3 | 192.408666 | -0.3710021 | 0.2804028 | -1.3231041 | 0.18580075 | 0.64030106 |
| AMZ1 | 70.3846155 | 0.07017959 | 0.517596 | 0.13558758 | 0.89214732 | 0.97771293 |
| GNA12 | 460.108404 | -0.1216122 | 0.20624667 | -0.5896444 | 0.55542909 | 0.87184917 |
| SDK1 | 2424.2462 | 0.41262461 | 0.31184738 | 1.32316203 | 1.86E-01 | 0.64030106 |
| FOXK1 | 1134.57882 | -0.2969549 | 0.24468295 | -1.2136314 | 0.2248885 | 0.67750405 |
| AP5Z1 | 199.681648 | -0.3477262 | 0.32975402 | -1.0545017 | 0.29165329 | 0.73362416 |
| RBAK-RBAKI | 635.028894 | -0.3651173 | 0.25666621 | -1.4225374 | 0.15487029 | 0.60380955 |
| WIPI2 | 611.798665 | -0.2083143 | 0.2428647 | -0.8577381 | 0.3910371 | 0.79735014 |
| SLC29A4 | 43.8626948 | 0.09234516 | 0.54941788 | 0.16807819 | 0.86652177 | 0.97386931 |
| TNRC18 | 2145.97745 | -0.1272734 | 0.33565781 | -0.3791761 | 0.7045571 | 0.92528913 |
| AC092171.4 | 44.3101347 | 0.6361136 | 0.57027585 | 1.11544895 | 0.26465808 | 0.71489091 |
| FBXL18 | 247.551968 | -0.4477859 | 0.27970364 | -1.6009299 | 0.10939244 | 0.53319732 |
| ACTB | 2797.50851 | 0.55826772 | 0.52378376 | 1.06583624 | 0.28649767 | 0.72934245 |
| RNF216 | 1418.79929 | -0.3743232 | 0.18518826 | -2.0213116 | 0.04324751 | 0.38262319 |
| CCZ1 | 212.992408 | -0.3605139 | 0.36128592 | -0.9978632 | 0.31834571 | 0.75062987 |
| PMS2 | 367.529886 | -0.0614557 | 0.26329101 | -0.2334137 | 0.81544019 | 0.95872901 |
| AIMP2 | 229.431413 | -0.0038722 | 0.25857087 | -0.0149752 | 0.98805196 | 0.99653761 |
| EIF2AK1 | 548.70747 | 0.08833842 | 0.1331547 | 0.66342695 | 0.50705715 | 0.85626196 |
| USP42 | 548.598868 | -0.2753125 | 0.17325184 | -1.5890887 | 0.11204036 | 0.53621666 |
| CYTH3 | 638.269082 | 0.08848234 | 0.18422703 | 0.48028969 | 0.63102142 | 0.90005881 |
| FAM220A | 61.6090877 | 0.3701015 | 0.37718587 | 0.98121781 | 0.32648534 | 0.75554697 |
| RAC1 | 1339.62425 | 0.19645649 | 0.16807631 | 1.16885296 | 2.42E-01 | 6.93E-01 |
| DAGLB | 184.451786 | -0.4789138 | 0.25095855 | -1.9083381 | 0.05634753 | 0.41843207 |
| GRID2IP | 79.9888703 | -0.0982142 | 0.42767455 | -0.229647 | 0.81836609 | 0.95985564 |

| | | | | | | |
|---|---|---|---|---|---|---|
| ZDHHC4 | 125.786209 | 0.10668688 | 0.20383664 | 0.52339405 | 0.60070007 | 0.88883692 |
| C7orf26 | 164.90223 | -0.6581643 | 0.28689036 | -2.2941318 | 0.02178293 | 0.29480605 |
| ZNF316 | 181.120751 | -0.19594 | 0.26134061 | -0.7497497 | 0.45340547 | 0.83008079 |
| AC073343.2 | 204.950957 | -0.0303856 | 0.46356748 | -0.0655474 | 0.94773818 | 0.98919519 |
| ZNF12 | 307.52434 | 0.16034053 | 0.20492901 | 0.78241989 | 0.43396785 | 0.81919857 |
| CCZ1B | 299.904525 | -0.3244359 | 0.26601471 | -1.2196164 | 0.22261032 | 0.67568617 |
| C1GALT1 | 762.76025 | 0.35784768 | 0.3201148 | 1.11787293 | 0.26362127 | 0.71429196 |
| COL28A1 | 1612.1055 | 0.04226641 | 0.3163922 | 0.13358864 | 0.89372786 | 0.97788747 |
| AC004982.1 | 106.841545 | 0.50351022 | 0.31714326 | 1.58764283 | 0.11236713 | 0.53680195 |
| AC004982.2 | 300.795148 | 0.27380721 | 0.36232134 | 0.7557027 | 0.44982746 | 0.82719907 |
| MIOS | 811.357876 | -0.2331268 | 0.39455232 | -0.5908641 | 0.55461151 | 0.87160779 |
| RPA3 | 358.268047 | 0.32812928 | 0.51104525 | 0.64207479 | 0.52082462 | 0.85943874 |
| UMAD1 | 3178.8905 | 0.07626156 | 0.18203407 | 0.41894112 | 0.67525916 | 0.91649321 |
| GLCCI1 | 2024.79616 | 0.27754202 | 0.15595732 | 1.77960248 | 0.07514104 | 0.47190267 |
| ICA1 | 1794.31511 | 0.27777046 | 0.38045986 | 0.73009138 | 0.46533433 | 0.83652911 |
| AC004949.1 | 105.991303 | -0.6635721 | 1.03497849 | -0.6411458 | 0.52142797 | 0.85943874 |
| NDUFA4 | 978.662951 | 0.80606766 | 0.4753331 | 1.69579533 | 0.08992465 | 0.50134767 |
| PHF14 | 1994.44041 | -0.3915356 | 0.1799604 | -2.1756766 | 0.02957945 | 0.33073125 |
| THSD7A | 7583.58765 | -1.5327136 | 0.44276123 | -3.4617159 | 0.00053674 | 0.04005562 |
| TMEM106B | 960.426121 | -0.3668758 | 0.28470322 | -1.2886253 | 0.1975284 | 0.64964978 |
| SCIN | 2160.90017 | -0.236132 | 0.22709691 | -1.039785 | 0.2984398 | 0.73673761 |
| AC011287.1 | 340.515571 | 0.2985325 | 0.48732873 | 0.61258957 | 0.54014775 | 0.86569031 |
| ETV1 | 361.057793 | 0.84472292 | 0.58192661 | 1.45159699 | 0.1466137 | 0.59373375 |
| ISPD | 1236.43117 | -0.0368675 | 0.24546948 | -0.1501919 | 8.81E-01 | 0.97664137 |
| SOSTDC1 | 41.2108536 | 1.57133157 | 0.98362735 | 1.59748665 | 0.11015727 | 0.53372881 |
| ANKMY2 | 460.126837 | 0.32700642 | 0.24606585 | 1.32893866 | 0.18386821 | 0.63843513 |
| BZW2 | 957.673104 | 0.09554008 | 0.28657719 | 0.3333834 | 0.73884489 | 0.93552616 |
| AGR2 | 117.70722 | 4.56811674 | 0.82095785 | 5.56437427 | 2.63E-08 | 0.00016602 |
| AHR | 135.591891 | 0.96340593 | 0.59767487 | 1.6119231 | 0.10697867 | 0.52845225 |
| AC073332.1 | 218.896509 | -0.0923025 | 0.53898189 | -0.1712535 | 0.86402447 | 0.97349667 |
| AC019117.1 | 226.290789 | 2.54862567 | 0.78925991 | 3.22913358 | 1.24E-03 | 0.0645665 |
| SNX13 | 2039.48372 | 0.04721014 | 0.19767371 | 0.23882864 | 0.81123846 | 0.9575238 |
| HDAC9 | 1948.29378 | 0.22921199 | 0.71308502 | 0.32143711 | 0.74787917 | 0.93746253 |
| TWISTNB | 111.442561 | 0.32760888 | 0.2897205 | 1.13077562 | 0.25814954 | 0.70896455 |
| MACC1 | 1188.38257 | 0.30713483 | 0.51410355 | 0.59741822 | 0.55022819 | 0.86959143 |
| ITGB8 | 2490.68917 | 0.44324266 | 0.41177602 | 1.07641689 | 0.28174085 | 0.72604837 |
| SP4 | 519.489231 | -0.0280063 | 0.24387436 | -0.114839 | 0.90857272 | 0.98047754 |
| DNAH11 | 87.7885205 | -0.6526633 | 0.74776105 | -0.8728233 | 0.38275938 | 0.79022641 |
| CDCA7L | 105.636173 | -0.5554088 | 0.34336039 | -1.617568 | 0.10575574 | 0.52694363 |
| RAPGEF5 | 679.100098 | 0.50241199 | 0.66150572 | 0.75949758 | 4.48E-01 | 8.26E-01 |
| STEAP1B | 644.588376 | 0.54834998 | 0.46036452 | 1.19112127 | 0.23360598 | 0.68493974 |
| TOMM7 | 373.975523 | 0.52771421 | 0.39807192 | 1.32567552 | 0.1849472 | 0.64011182 |
| FAM126A | 89.7391309 | -0.3346559 | 0.38790763 | -0.8627206 | 0.38829113 | 0.79634939 |
| KLHL7 | 513.216255 | -0.1517026 | 0.246708 | -0.6149076 | 0.53861571 | 0.86541478 |
| NUPL2 | 582.750113 | 0.00377581 | 0.27609626 | 0.01367569 | 0.98908872 | 0.99670729 |
| AC005082.1 | 135.409876 | 0.54076822 | 0.45573999 | 1.1865718 | 0.23539657 | 0.68745373 |
| GPNMB | 201.730332 | 0.65086592 | 0.68175852 | 0.95468688 | 0.33973607 | 0.76449297 |
| MALSU1 | 308.870903 | 0.22407078 | 0.18022949 | 1.24325255 | 0.21377478 | 0.66800831 |
| TRA2A | 3074.43393 | 0.09735912 | 0.18699077 | 0.52066268 | 0.60260177 | 0.88971776 |
| CCDC126 | 260.668743 | -0.0948768 | 0.30795976 | -0.3080819 | 0.75802001 | 0.94056395 |
| FAM221A | 260.782099 | -0.488076 | 0.21107951 | -2.3122852 | 0.02076197 | 0.28837528 |
| STK31 | 59.7513894 | -1.1989119 | 0.55067301 | -2.1771757 | 0.02946746 | 0.3304208 |
| MPP6 | 1623.08143 | -0.3959223 | 0.28443775 | -1.3919473 | 0.16393837 | 0.61531922 |
| GSDME | 270.156389 | -0.0020531 | 0.32968195 | -0.0062276 | 0.99503116 | 0.998438 |

| | | | | | | |
|---|---|---|---|---|---|---|
| OSBPL3 | 6782.15885 | -0.6116532 | 0.30027274 | -2.0369919 | 0.04165085 | 0.37614595 |
| CYCS | 415.140273 | 0.64216217 | 0.50192558 | 1.27939719 | 0.20075722 | 0.6539586 |
| C7orf31 | 102.298597 | -0.3836131 | 0.33450436 | -1.1468105 | 0.25145996 | 0.70276117 |
| NFE2L3 | 250.744799 | -0.278668 | 0.46161669 | -0.6036783 | 0.54605754 | 0.86794271 |
| AC004520.1 | 45.5886602 | -0.7691236 | 0.61037242 | -1.2600891 | 0.20763724 | 0.6607067 |
| HNRNPA2B1 | 4353.82998 | -0.1233675 | 0.13518784 | -0.9125637 | 0.36147208 | 0.77772231 |
| CBX3 | 774.011008 | -0.0965467 | 0.13643294 | -0.7076492 | 4.79E-01 | 8.43E-01 |
| SNX10 | 2818.0173 | 0.14216728 | 0.37647773 | 0.37762467 | 0.70570944 | 0.92556842 |
| AC004540.2 | 47.3535304 | 1.42317943 | 0.44642881 | 3.1879202 | 0.001433 | 0.06933519 |
| AC004540.1 | 464.999279 | 1.11691774 | 0.52657807 | 2.12108671 | 0.03391451 | 0.35107893 |
| SKAP2 | 2699.07487 | 0.13192118 | 0.23992229 | 0.54984962 | 0.58242252 | 0.88338315 |
| HOTAIRM1 | 517.706023 | -0.1685167 | 0.43002086 | -0.3918803 | 0.69514663 | 0.92229578 |
| HOXA3 | 737.207125 | 0.1896158 | 0.33980299 | 0.55801687 | 0.57683287 | 0.88150331 |
| HOXA-AS2 | 320.345083 | -0.3448075 | 0.40873881 | -0.8435889 | 0.39889919 | 0.8016217 |
| HOXA-AS3 | 113.892201 | 0.49237374 | 0.47531109 | 1.03589785 | 0.3002498 | 0.73680045 |
| HOXA7 | 108.657955 | 0.17239348 | 0.59746896 | 0.28853963 | 0.7729337 | 0.94534972 |
| HOXA9 | 233.595694 | 0.01483767 | 0.55659488 | 0.02665794 | 0.97873256 | 0.99455895 |
| HOXA10-AS | 73.892168 | -0.2502146 | 0.61332033 | -0.4079673 | 0.68329768 | 0.9200778 |
| HIBADH | 3562.85164 | 0.0143626 | 0.29727502 | 0.04831419 | 0.96146585 | 0.99191712 |
| TAX1BP1 | 2880.07943 | 0.1377281 | 0.15472987 | 0.89011971 | 0.37340161 | 0.78277446 |
| JAZF1 | 1327.68296 | 0.1891622 | 0.45808459 | 0.41294162 | 0.67964938 | 0.9182972 |
| CREB5 | 474.767239 | -1.1015228 | 0.65982288 | -1.6694219 | 0.09503379 | 0.51085035 |
| AC005162.3 | 373.804399 | 0.12039222 | 0.50930587 | 0.23638491 | 0.81313401 | 0.95816239 |
| CPVL | 1130.32392 | -0.7776055 | 0.44532885 | -1.7461378 | 0.08078701 | 0.48483219 |
| AC007255.1 | 187.220959 | -0.3706976 | 0.37259396 | -0.9949104 | 0.31977984 | 0.75094569 |
| PRR15 | 78.1037485 | 0.18501404 | 0.38017523 | 0.48665464 | 6.27E-01 | 8.99E-01 |
| WIPF3 | 127.905692 | 1.57735749 | 0.56355916 | 2.79892084 | 0.00512737 | 0.14350465 |
| SCRN1 | 733.228702 | -0.0189652 | 0.19080559 | -0.0993956 | 0.92082418 | 0.98215577 |
| AC007285.1 | 112.2504 | 0.11556556 | 0.35673678 | 0.32395191 | 0.74597444 | 0.93675395 |
| FKBP14 | 194.796887 | 0.28262613 | 0.27518245 | 1.02705001 | 0.30439692 | 0.74152784 |
| PLEKHA8 | 590.238273 | -0.1532832 | 0.19952956 | -0.7682232 | 0.44235458 | 0.82285053 |
| MTURN | 1388.3835 | 0.08199989 | 0.28596662 | 0.28674638 | 0.77430652 | 0.94621715 |
| ZNRF2 | 907.235413 | -0.2816794 | 0.22610417 | -1.2457946 | 0.21283982 | 0.66684088 |
| AC006978.2 | 816.134882 | 0.08458967 | 0.26246147 | 0.32229368 | 0.74723022 | 0.93735494 |
| NOD1 | 144.08592 | 0.16972255 | 0.53073982 | 0.31978484 | 0.74913144 | 0.93748128 |
| GGCT | 58.816901 | 0.64778422 | 0.48988109 | 1.32232952 | 0.18605845 | 0.64033103 |
| AC005154.6 | 426.500276 | 0.06481058 | 0.26816113 | 0.24168519 | 0.8090241 | 0.95610623 |
| GARS | 366.524503 | -0.0699446 | 0.22265245 | -0.3141426 | 0.75341277 | 0.93844712 |
| AQP1 | 178.894346 | 1.0278045 | 0.69109464 | 1.48721238 | 0.13695873 | 0.57909342 |
| PDE1C | 53.2448092 | -0.0684156 | 0.54681612 | -0.1251163 | 0.90043148 | 0.97914987 |
| LSM5 | 171.926538 | 0.16643595 | 0.31214154 | 0.53320669 | 0.59389051 | 0.88807733 |
| AVL9 | 2004.26777 | -0.1449295 | 0.16038134 | -0.9036559 | 0.36617787 | 0.78095617 |
| KBTBD2 | 568.916946 | -0.1519304 | 0.23444499 | -0.6480428 | 0.51695724 | 0.85893097 |
| FKBP9 | 349.970977 | -0.097967 | 0.27540472 | -0.3557201 | 0.72205016 | 0.93079284 |
| NT5C3A | 745.852509 | -0.3161932 | 0.28537653 | -1.1079859 | 0.26786792 | 0.71799256 |
| RP9 | 146.258599 | 0.23999483 | 0.24456523 | 0.98131217 | 0.32643882 | 0.75554697 |
| BBS9 | 2382.86616 | 0.10268057 | 0.15282627 | 0.67187776 | 0.50166152 | 0.85452654 |
| BMPER | 133.963386 | 1.30313464 | 0.42323788 | 3.07896502 | 0.00207721 | 0.0878448 |
| NPSR1 | 109.129265 | 3.48320284 | 0.60438854 | 5.76318475 | 8.25E-09 | 0.00010903 |
| DPY19L1 | 1149.51274 | -0.2351345 | 0.30113098 | -0.7808379 | 0.43489781 | 0.81919857 |
| HERPUD2 | 753.14215 | -0.1013964 | 0.18959202 | -0.5348136 | 0.59277875 | 0.88740306 |
| AC018647.2 | 48.4600867 | -0.0047808 | 0.46687523 | -0.0102401 | 0.99182974 | 0.99772135 |
| SEPT7-AS1 | 229.014209 | 0.83162918 | 0.28753275 | 2.89229377 | 0.0038244 | 0.12084489 |
| 7-Sep | 1602.57766 | 0.04791078 | 0.18331496 | 0.2613577 | 0.79381668 | 0.95224951 |

| | | | | | | |
|---|---|---|---|---|---|---|
| EEPD1 | 379.391722 | -0.4080236 | 0.63999854 | -0.6375383 | 0.52377429 | 0.86006369 |
| KIAA0895 | 451.305441 | 0.13976024 | 0.34545729 | 0.40456589 | 0.68579663 | 0.92053037 |
| ELMO1 | 335.660377 | 1.93755473 | 0.94522072 | 2.0498437 | 0.04037969 | 0.37344844 |
| EPDR1 | 266.379556 | 0.32745772 | 0.39454213 | 0.82996896 | 0.40655633 | 0.80499101 |
| SFRP4 | 100.577569 | 0.2479519 | 0.73963007 | 0.33523772 | 0.73744577 | 0.93552616 |
| STARD3NL | 336.521115 | 0.13963665 | 0.15382707 | 0.90775083 | 0.36400988 | 0.78015617 |
| AMPH | 279.873104 | -1.5327488 | 0.78935514 | -1.9417734 | 5.22E-02 | 4.09E-01 |
| VPS41 | 1952.50646 | -0.1680218 | 0.18654038 | -0.900726 | 0.36773404 | 0.78095617 |
| YAE1D1 | 218.362245 | 0.53209029 | 0.21451401 | 2.48044535 | 0.01312184 | 0.23485956 |
| RALA | 793.535906 | -0.2971335 | 0.20213319 | -1.4699887 | 0.14156482 | 0.5872148 |
| LINC00265 | 175.804636 | -0.0138805 | 0.34648722 | -0.0400607 | 0.96804476 | 0.99248831 |
| CDK13 | 3423.52709 | -0.1715438 | 0.17722495 | -0.9679438 | 0.33307243 | 0.76124971 |
| MPLKIP | 309.830629 | 0.40947684 | 0.20388777 | 2.00834424 | 0.04460672 | 0.38636735 |
| SUGCT | 1242.24818 | 0.0649732 | 0.19486548 | 0.33342593 | 7.39E-01 | 0.93552616 |
| MRPL32 | 226.475163 | 0.11350719 | 0.30392744 | 0.37346806 | 0.70880012 | 0.92577218 |
| AC005537.1 | 83.5506327 | -2.1172988 | 0.71208421 | -2.9733826 | 0.00294537 | 0.10386419 |
| STK17A | 836.559988 | 0.09831317 | 0.29076758 | 0.33811601 | 0.73527576 | 0.93552616 |
| COA1 | 2151.0298 | -0.1993072 | 0.16083547 | -1.2391994 | 0.21527165 | 0.66885336 |
| BLVRA | 231.858395 | 0.39444273 | 0.2256457 | 1.74806223 | 0.08045325 | 0.4844196 |
| URGCP | 429.286743 | 0.1955429 | 0.23182924 | 0.84347815 | 0.39896108 | 0.80162411 |
| UBE2D4 | 124.830816 | -0.1376301 | 0.30773651 | -0.4472335 | 0.65470648 | 0.90902447 |
| SPDYE1 | 129.59931 | 0.15959489 | 0.32028148 | 0.49829572 | 0.61827562 | 0.89599505 |
| AC004951.1 | 120.753649 | -0.2421027 | 0.42768652 | -0.5660752 | 0.57134269 | 0.8787896 |
| DBNL | 298.810187 | 0.06289538 | 0.20083908 | 0.31316305 | 0.75415679 | 0.93898171 |
| POLM | 293.01239 | -0.2399532 | 0.30208634 | -0.7943198 | 0.4270093 | 0.81466629 |
| POLD2 | 142.992737 | 0.22140093 | 0.25676562 | 0.8622686 | 0.38853973 | 0.79634939 |
| YKT6 | 134.982743 | -0.1351032 | 0.27607616 | -0.4893695 | 0.62458015 | 0.89873262 |
| CAMK2B | 51.8768924 | 1.00778811 | 0.74975335 | 1.34415953 | 0.1788968 | 0.63284534 |
| NUDCD3 | 1096.04092 | -0.3170542 | 0.26159713 | -1.211994 | 0.22551466 | 0.67798093 |
| DDX56 | 189.71093 | -0.0051591 | 0.23387863 | -0.022059 | 0.98240086 | 0.99518552 |
| TMED4 | 371.01355 | 0.2684255 | 0.2367553 | 1.13376764 | 0.25689203 | 0.70735159 |
| OGDH | 2223.6061 | 0.02976307 | 0.13365836 | 0.22268018 | 8.24E-01 | 0.96141406 |
| ZMIZ2 | 557.007782 | -0.0200973 | 0.2885278 | -0.0696548 | 0.94446845 | 0.98813503 |
| PPIA | 1351.51426 | 0.39819982 | 0.39533428 | 1.00724841 | 0.31381542 | 0.74777437 |
| H2AFV | 556.991293 | 0.0786739 | 0.26028822 | 0.30225686 | 0.76245626 | 0.94212205 |
| PURB | 462.884683 | 0.37745936 | 0.17662939 | 2.13701336 | 0.0325969 | 0.34456902 |
| SNHG15 | 59.4298117 | -0.4777641 | 0.46499059 | -1.0274705 | 0.30419898 | 0.74152784 |
| CCM2 | 582.175183 | -0.2014212 | 0.15658024 | -1.2863766 | 0.19831166 | 0.65064548 |
| TBRG4 | 137.872807 | -0.0602352 | 0.27957173 | -0.2154553 | 0.82941238 | 0.96290106 |
| ADCY1 | 48.0075482 | -0.2048085 | 0.78350968 | -0.2613988 | 0.79378502 | 0.95224951 |
| IGFBP3 | 328.391599 | -0.0069915 | 0.59391312 | -0.0117719 | 0.99060759 | 0.99740343 |
| AC011294.1 | 84.2281257 | 3.82112732 | 1.30510637 | 2.92782826 | 0.00341339 | 0.11385709 |
| AC004870.4 | 755.042551 | 2.64742647 | 0.84093507 | 3.14819366 | 0.00164283 | 0.07640886 |
| AC004870.3 | 47.4851753 | 0.98248336 | 0.70871076 | 1.38629666 | 0.16565634 | 0.61864703 |
| TNS3 | 6356.96111 | -0.8247165 | 0.32603441 | -2.5295381 | 0.01142128 | 0.21832654 |
| HUS1 | 285.372412 | 0.21919644 | 0.23439232 | 0.93516902 | 0.3497012 | 0.77025232 |
| UPP1 | 1101.76682 | 0.33638922 | 0.82173022 | 0.40936698 | 0.68227037 | 0.91969963 |
| ZPBP | 144.575934 | 0.18323093 | 0.55135227 | 0.33233005 | 0.73964005 | 0.93552616 |
| FIGNL1 | 51.6136834 | 0.22322982 | 0.44817519 | 0.49808606 | 0.61842338 | 0.89599505 |
| GRB10 | 681.040825 | -1.2646702 | 0.38767883 | -3.2621595 | 0.00110567 | 0.05961135 |
| COBL | 2388.73922 | -0.7614708 | 0.43870447 | -1.7357261 | 0.08261228 | 0.48594425 |
| SEC61G | 229.064498 | 0.21788866 | 0.44296504 | 0.49188682 | 0.62279936 | 0.89838994 |
| EGFR | 1390.07039 | 0.58889569 | 0.26070986 | 2.25881635 | 2.39E-02 | 0.3030643 |
| LANCL2 | 387.073102 | -0.4167117 | 0.35373029 | -1.1780493 | 0.23877695 | 0.6901542 |

| | | | | | | |
|---|---|---|---|---|---|---|
| VOPP1 | 2158.5191 | 0.29617149 | 0.28560829 | 1.03698493 | 3.00E-01 | 7.37E-01 |
| AC092647.5 | 42.4422705 | 0.14135321 | 0.36004053 | 0.39260361 | 0.69461227 | 0.92229578 |
| NIPSNAP2 | 1135.32165 | 0.01894968 | 0.20536924 | 0.09227129 | 0.9264825 | 0.98238964 |
| PSPH | 235.418543 | -0.5556103 | 0.4563403 | -1.217535 | 0.22340073 | 0.67624946 |
| CCT6A | 476.863449 | 0.03866718 | 0.28744851 | 0.13451864 | 0.89299247 | 0.97782787 |
| SUMF2 | 652.124717 | 0.10980702 | 0.18060546 | 0.60799388 | 0.54319154 | 0.86612953 |
| PHKG1 | 163.774563 | 0.07875096 | 0.39246184 | 0.20065889 | 0.8409653 | 0.96627616 |
| CHCHD2 | 801.900849 | 0.67513918 | 0.37881012 | 1.78226276 | 0.07470639 | 0.47190267 |
| NUPR2 | 56.9702397 | 1.53418017 | 0.42765366 | 3.58743606 | 0.00033395 | 0.03094754 |
| ZNF727 | 145.040234 | 0.20289522 | 0.40126039 | 0.50564479 | 0.61310604 | 0.89441126 |
| ZNF736 | 865.073053 | 0.42035679 | 0.20005337 | 2.10122326 | 0.03562137 | 0.35661333 |
| ZNF680 | 905.027826 | 0.30680491 | 0.21986952 | 1.39539541 | 0.16289664 | 0.61387934 |
| ZNF107 | 308.381175 | -0.3679949 | 0.30792052 | -1.1950969 | 0.23204917 | 0.68448804 |
| ZNF138 | 680.851897 | 0.11084269 | 0.22159432 | 0.50020548 | 0.6169304 | 0.89599505 |
| ZNF273 | 101.386306 | -0.1768741 | 0.26768643 | -0.6607512 | 0.50877186 | 0.85659934 |
| ZNF117 | 325.38803 | -0.1961918 | 0.19556703 | -1.0031947 | 0.31576694 | 0.74785672 |
| ZNF92 | 114.400957 | 0.14194308 | 0.25262812 | 0.56186572 | 0.57420751 | 0.88010061 |
| VKORC1L1 | 881.54906 | -0.2370353 | 0.19326653 | -1.2264682 | 0.22002253 | 0.67301289 |
| GUSB | 222.90169 | 0.17884401 | 0.19496823 | 0.9172982 | 0.3589864 | 0.77608042 |
| ASL | 159.70188 | -0.3103526 | 0.38943227 | -0.7969361 | 0.42548814 | 0.81466629 |
| AC068533.4 | 186.837028 | 0.0660524 | 0.42392969 | 0.15580979 | 0.87618295 | 0.97614152 |
| CRCP | 139.85714 | -0.1326157 | 0.22707891 | -0.5840073 | 0.55921542 | 0.87295536 |
| AC068533.3 | 183.250851 | -0.0465075 | 0.51576503 | -0.0901719 | 0.92815064 | 0.98299726 |
| TPST1 | 7403.12251 | -0.4784114 | 0.30688065 | -1.5589493 | 0.11900838 | 0.55079946 |
| AC027644.4 | 863.440789 | -0.0857392 | 0.3258347 | -0.2631371 | 0.79244492 | 0.95215633 |
| AC027644.3 | 128.909468 | -0.1935148 | 0.46264598 | -0.4182783 | 0.67574365 | 0.91649321 |
| AC073335.2 | 52.8668335 | -0.3549993 | 0.40339736 | -0.8800238 | 0.37884639 | 0.78744013 |
| TMEM248 | 533.121913 | -0.0474718 | 0.18891076 | -0.2512921 | 0.80158827 | 0.95390307 |
| SBDS | 297.46211 | 0.7215949 | 0.29507681 | 2.44544767 | 0.01446724 | 0.24762526 |
| TYW1 | 1358.22943 | -0.2545909 | 0.16587438 | -1.5348416 | 0.12482275 | 0.56195762 |
| AC006480.2 | 186.042086 | 0.07714543 | 0.33453493 | 0.23060501 | 0.81762168 | 0.95957039 |
| LINC01372 | 112.08989 | -0.3698247 | 0.43890624 | -0.8426051 | 0.39944932 | 0.80174499 |
| AUTS2 | 3677.03197 | -1.3811559 | 0.37347669 | -3.6981047 | 0.00021722 | 0.02390998 |
| GALNT17 | 289.92556 | 1.10952536 | 1.23429811 | 0.89891198 | 0.36869955 | 0.78095617 |
| CALN1 | 136.219616 | 0.21995266 | 0.37144175 | 0.59215922 | 0.55374397 | 0.87121136 |
| TYW1B | 1002.7919 | -0.1045363 | 0.19206235 | -0.5442831 | 0.58624669 | 0.88530153 |
| POM121 | 1015.09495 | -0.1854602 | 0.20052036 | -0.9248947 | 0.35502067 | 0.7726065 |
| NSUN5 | 206.842771 | -0.0849057 | 0.31042274 | -0.2735165 | 0.78445622 | 0.94978829 |
| BAZ1B | 922.044274 | -0.3095311 | 0.16313253 | -1.8974213 | 0.05777236 | 0.42087381 |
| BCL7B | 174.173672 | -0.1325864 | 0.20433827 | -0.6488574 | 0.51643052 | 0.8588826 |
| TBL2 | 114.436088 | -0.0632363 | 0.24414817 | -0.259008 | 0.7956291 | 0.95278473 |
| MLXIPL | 249.138077 | 1.77416504 | 0.47123629 | 3.76491597 | 0.00016661 | 0.02095892 |
| VPS37D | 56.6955195 | 1.21341862 | 0.59015207 | 2.05611177 | 0.03977174 | 0.37279391 |
| DNAJC30 | 70.3531724 | 0.50152976 | 0.305914 | 1.6394469 | 0.10112022 | 0.51819907 |
| BUD23 | 303.128632 | 0.14873219 | 0.24616917 | 0.6041869 | 0.54571939 | 0.86764654 |
| ABHD11 | 243.699623 | -0.1711594 | 0.40044437 | -0.4274236 | 0.66907086 | 0.91421919 |
| CLDN3 | 393.726873 | 1.0550681 | 0.52806943 | 1.9979723 | 4.57E-02 | 3.89E-01 |
| CLDN4 | 622.540721 | 0.2691728 | 0.49908535 | 0.5393322 | 5.90E-01 | 0.88621335 |
| METTL27 | 77.3692139 | 0.14980213 | 0.40762629 | 0.36749871 | 0.71324705 | 0.92760608 |
| LIMK1 | 121.198385 | -0.209559 | 0.43800051 | -0.4784447 | 0.63233375 | 0.90044163 |
| EIF4H | 497.500875 | 0.11660415 | 0.17225482 | 0.67692819 | 0.49845152 | 0.85092921 |
| RFC2 | 101.037628 | -0.2574042 | 0.35499949 | -0.7250834 | 0.46840085 | 0.83825637 |
| CLIP2 | 768.019195 | -0.2311873 | 0.37475267 | -0.6169064 | 0.53729648 | 0.86487317 |
| GTF2IRD1 | 3697.1005 | -0.1904993 | 0.35639463 | -0.5345179 | 0.59298328 | 0.88740306 |

| | | | | | | |
|---|---|---|---|---|---|---|
| GTF2I | 4260.80318 | -0.2489704 | 0.19267469 | -1.2921799 | 0.19629486 | 0.64964978 |
| GTF2IRD2 | 500.157274 | -0.5793602 | 0.37234593 | -1.5559728 | 0.11971455 | 0.55232605 |
| CASTOR2 | 556.606893 | -0.2829716 | 0.42635745 | -0.6636957 | 0.50688508 | 0.85626196 |
| RCC1L | 370.807305 | -0.4157579 | 0.23247656 | -1.7883864 | 0.07371368 | 0.47028843 |
| GTF2IRD2B | 738.678568 | -0.5369696 | 0.35795161 | -1.5001179 | 0.13358387 | 0.57682553 |
| AC211486.5 | 171.428464 | 0.23646963 | 0.61039125 | 0.38740664 | 0.69845519 | 0.92356678 |
| TRIM73 | 348.121856 | 0.29702852 | 0.44094822 | 0.67361315 | 0.50055729 | 0.85314338 |
| POM121C | 895.289516 | -0.1674592 | 0.22357645 | -0.7490019 | 0.45385607 | 0.83060749 |
| SPDYE5 | 244.517577 | 0.04088434 | 0.37410722 | 0.10928508 | 0.91297638 | 0.98135874 |
| HIP1 | 8619.42008 | -0.0486344 | 0.37827875 | -0.1285678 | 0.89769968 | 0.97823597 |
| RHBDD2 | 195.389941 | 0.14379328 | 0.2798264 | 0.51386605 | 0.60734565 | 0.89174148 |
| POR | 412.748687 | -0.2506923 | 0.28002859 | -0.8952381 | 0.37065984 | 0.78104287 |
| TMEM120A | 118.918857 | -0.1168691 | 0.26742998 | -0.4370081 | 0.66210545 | 0.91110616 |
| STYXL1 | 558.851072 | -0.2285683 | 0.30109464 | -0.7591246 | 0.44777805 | 0.8260638 |
| MDH2 | 289.135998 | 0.67846932 | 0.29016984 | 2.33817998 | 0.01937791 | 0.28045661 |
| HSPB1 | 717.175781 | 0.65872297 | 0.49291661 | 1.33637813 | 0.18142571 | 0.63616641 |
| YWHAG | 470.025117 | 0.13680642 | 0.20075556 | 0.68145768 | 0.49558194 | 0.84970688 |
| SSC4D | 73.03033 | 0.62243442 | 0.722619 | 0.86135906 | 0.38904032 | 0.79659488 |
| ZP3 | 51.9140707 | -0.4485488 | 0.46621186 | -0.9621137 | 0.3359925 | 0.7637525 |
| DTX2 | 336.082275 | -0.3040069 | 0.22658338 | -1.3416997 | 0.17969338 | 0.63379703 |
| SPDYE16 | 76.2568784 | 0.05372595 | 0.41832855 | 0.12843004 | 0.89780866 | 0.97823597 |
| POMZP3 | 238.449219 | 0.13657183 | 0.41275711 | 0.33087698 | 0.74073741 | 0.93594801 |
| CCDC146 | 1229.70418 | 0.21723368 | 0.30487216 | 0.71254021 | 0.47613032 | 0.84141268 |
| FGL2 | 232.256271 | 3.03583963 | 0.66930087 | 4.53583698 | 5.74E-06 | 0.0032951 |
| GSAP | 1447.47545 | 0.16214124 | 0.29992463 | 0.54060661 | 0.58877876 | 0.88571413 |
| PTPN12 | 2202.72022 | -0.3310358 | 0.29680478 | -1.1153318 | 2.65E-01 | 0.71489091 |
| APTR | 428.061581 | 0.33069138 | 0.22788169 | 1.45115379 | 0.14673704 | 0.59373375 |
| RSBN1L | 1118.69764 | -0.0987547 | 0.15672144 | -0.6301288 | 0.5286103 | 0.86245226 |
| TMEM60 | 79.5054642 | 0.37231469 | 0.38662093 | 0.96299673 | 0.33554917 | 0.76339458 |
| PHTF2 | 813.014657 | -0.1783155 | 0.18381554 | -0.9700786 | 0.33200732 | 0.76037513 |
| MAGI2 | 4687.75902 | 0.58781429 | 0.42447704 | 1.38479644 | 0.16611472 | 0.61901667 |
| GNAI1 | 4065.80511 | -0.336845 | 0.3645565 | -0.9239857 | 0.35549378 | 0.77308485 |
| CD36 | 161.728334 | -1.0380072 | 0.52632287 | -1.9721871 | 0.04858826 | 0.39695534 |
| SEMA3C | 583.179285 | -0.9622583 | 0.7351616 | -1.3089072 | 0.19056578 | 0.64559718 |
| CACNA2D1 | 891.862426 | 0.98139625 | 0.55856928 | 1.75698215 | 0.07892085 | 0.48150833 |
| PCLO | 3686.35036 | -0.4007373 | 0.527337 | -0.7599265 | 0.44729855 | 0.82578615 |
| KIAA1324L | 247.120476 | 0.41442886 | 0.36443435 | 1.13718385 | 0.25546144 | 0.70643369 |
| DMTF1 | 1882.26237 | 0.16926444 | 0.25941864 | 0.65247601 | 0.51409415 | 0.85762435 |
| TMEM243 | 530.495032 | -0.1241509 | 0.21924134 | -0.5662749 | 5.71E-01 | 0.8787896 |
| TP53TG1 | 90.3902906 | 0.3863399 | 0.28020522 | 1.37877483 | 0.16796419 | 0.62233912 |
| CROT | 404.787088 | -0.2503382 | 0.22826076 | -1.0967203 | 0.2727637 | 0.72070318 |
| ABCB1 | 135.282498 | 1.69978942 | 0.50547715 | 3.36274239 | 0.00077172 | 0.04906731 |
| RUNDC3B | 86.3359995 | 0.9163529 | 0.44024483 | 2.08146205 | 0.03739163 | 0.36263296 |
| SLC25A40 | 496.337309 | 0.07146483 | 0.17709805 | 0.40353256 | 6.87E-01 | 9.21E-01 |
| DBF4 | 142.909188 | -0.3468995 | 0.2967125 | -1.1691434 | 0.24234587 | 0.69337349 |
| ADAM22 | 462.625026 | -0.4451217 | 0.44401598 | -1.0024904 | 0.31610682 | 0.74785672 |
| SRI | 409.513019 | 0.12154439 | 0.28623567 | 0.42463047 | 0.67110608 | 0.91529584 |
| AC003991.1 | 80.7424843 | -0.949157 | 0.40298648 | -2.3553074 | 0.01850739 | 0.27375602 |
| ZNF804B | 64.4928367 | -0.3432034 | 0.78654357 | -0.4363438 | 6.63E-01 | 0.91110616 |
| STEAP2 | 347.254998 | 0.31042709 | 0.30991813 | 1.00164224 | 0.31651641 | 0.74818634 |
| CFAP69 | 816.963776 | 0.11084431 | 0.25285005 | 0.43837964 | 0.6611111 | 0.91110616 |
| AC002064.1 | 52.9991356 | -0.4600981 | 0.51880865 | -0.8868359 | 0.37516726 | 0.78460803 |
| GTPBP10 | 465.960413 | -0.3719478 | 0.24113651 | -1.5424781 | 0.12295747 | 0.55947129 |
| CLDN12 | 474.741852 | -0.0148675 | 0.21504127 | -0.0691379 | 0.94487988 | 0.98827447 |

| | | | | | | |
|---|---|---|---|---|---|---|
| CDK14 | 2469.04946 | 0.15010635 | 0.29486388 | 0.50906999 | 0.61070316 | 0.89339478 |
| AC079760.1 | 341.034946 | -0.6768856 | 0.70017488 | -0.966738 | 0.33367503 | 0.76175485 |
| MTERF1 | 1437.74823 | -0.5640844 | 0.41672953 | -1.3535984 | 0.17586455 | 0.62957039 |
| AKAP9 | 4502.88906 | -0.1094405 | 0.17261929 | -0.6339994 | 0.52608122 | 0.86119802 |
| KRIT1 | 1515.4212 | 0.03378812 | 0.19847688 | 0.17023708 | 0.86482369 | 0.97361767 |
| ANKIB1 | 2800.78366 | -0.1536534 | 0.16502522 | -0.9310904 | 0.35180679 | 0.77151834 |
| GATAD1 | 447.665383 | -0.0805395 | 0.34262924 | -0.2350632 | 0.81415967 | 0.95831716 |
| AC007566.1 | 407.864012 | -0.4586939 | 0.36245398 | -1.2655231 | 0.20568388 | 0.65813563 |
| PEX1 | 599.865831 | -0.0539065 | 0.20423816 | -0.2639397 | 0.79182642 | 0.95183249 |
| RBM48 | 163.996411 | 0.03174508 | 0.30722196 | 0.10332947 | 0.91770149 | 0.98199511 |
| FAM133B | 699.020992 | -0.1422513 | 0.15447246 | -0.9208848 | 0.35711056 | 0.77448758 |
| CDK6 | 1619.17002 | 0.29228986 | 0.3675542 | 0.79522928 | 0.42648014 | 0.81466629 |
| SAMD9 | 140.793899 | 0.48563101 | 0.42190929 | 1.15103178 | 0.24971916 | 0.70032501 |
| SAMD9L | 307.509086 | 1.23659064 | 0.81360053 | 1.51989901 | 0.12853636 | 0.56821847 |
| VPS50 | 1297.01672 | -0.1705205 | 0.15033466 | -1.1342728 | 0.25668013 | 0.70735159 |
| CALCR | 2479.44112 | 1.71368158 | 0.41845304 | 4.09527811 | 4.22E-05 | 0.01031432 |
| GNGT1 | 1872.03806 | 0.93765491 | 0.30355755 | 3.08888681 | 0.00200908 | 0.08650653 |
| TFPI2 | 76.3021157 | -0.4170774 | 0.80913166 | -0.515463 | 0.60622953 | 0.89125807 |
| GNG11 | 65.4015811 | 0.47933655 | 0.4574186 | 1.04791661 | 0.29467703 | 0.73556877 |
| BET1 | 414.024 | 0.16043084 | 0.347687 | 0.46142316 | 0.64449504 | 0.90507553 |
| CASD1 | 833.943587 | -0.0220741 | 0.1675931 | -0.1317123 | 0.89521184 | 0.97788747 |
| SGCE | 1459.37692 | 0.31517844 | 0.26106107 | 1.20729775 | 0.2273175 | 0.67921637 |
| PEG10 | 90.0582861 | 1.74386523 | 0.51004768 | 3.41902396 | 0.00062846 | 0.04487218 |
| PPP1R9A | 3857.41847 | -0.1589392 | 0.24810276 | -0.6406184 | 0.52177063 | 0.8596817 |
| AC002429.2 | 326.743205 | -0.1216712 | 0.53843265 | -0.2259729 | 0.82122246 | 0.96066244 |
| PON2 | 1629.2063 | -0.6334692 | 0.42150912 | -1.5028601 | 0.13287512 | 0.57682553 |
| PDK4 | 3190.92568 | 0.86927504 | 1.31343368 | 0.66183398 | 0.50807762 | 0.85643442 |
| AC002451.1 | 66.8501048 | 0.60947357 | 0.72913233 | 0.83588884 | 0.40321744 | 0.80324472 |
| SLC25A13 | 2088.96205 | -0.2793482 | 0.15330503 | -1.8221725 | 0.06842881 | 0.45352545 |
| SEM1 | 968.295143 | -0.0852034 | 0.22717794 | -0.3750512 | 0.70762236 | 0.92577218 |
| SDHAF3 | 372.796151 | -0.8742305 | 0.45767239 | -1.9101666 | 0.05611177 | 0.41843207 |
| ASNS | 154.405505 | 0.3430836 | 0.3814601 | 0.89939576 | 0.3684419 | 0.78095617 |
| LMTK2 | 910.08115 | -0.2543404 | 0.25434915 | -0.9999655 | 0.31732719 | 0.74916441 |
| TECPR1 | 166.507573 | -0.3769271 | 0.45891889 | -0.8213371 | 0.41145429 | 0.80675684 |
| BRI3 | 1015.73837 | -0.09666 | 0.28642721 | -0.3374678 | 0.7357643 | 0.93552616 |
| BAIAP2L1 | 1811.32942 | 0.51008979 | 0.31606516 | 1.6138754 | 0.10655445 | 0.5275404 |
| TRRAP | 1466.93779 | -0.1937188 | 0.13736939 | -1.4102035 | 0.15847961 | 0.60804051 |
| SMURF1 | 2078.05595 | -0.2677623 | 0.20656828 | -1.2962414 | 0.19489235 | 0.6484466 |
| AC004834.1 | 68.6535111 | -0.6001008 | 0.38673373 | -1.5517158 | 0.12073025 | 0.55468726 |
| ARPC1B | 183.347993 | 0.28822727 | 0.47324275 | 0.6090474 | 5.42E-01 | 8.66E-01 |
| PDAP1 | 206.850732 | 0.20947768 | 0.28423442 | 0.73698915 | 0.46112895 | 0.83473377 |
| BUD31 | 168.288298 | 0.13290251 | 0.2352508 | 0.56493966 | 0.57211481 | 0.87909049 |
| CPSF4 | 147.01832 | 0.13500707 | 0.27047318 | 0.49915139 | 0.61767273 | 0.89599505 |
| ZNF789 | 378.732846 | 0.05830724 | 0.21417258 | 0.2722442 | 0.78543426 | 0.94978829 |
| ZNF394 | 247.357284 | 0.0535967 | 0.22172303 | 0.24172816 | 0.8089908 | 0.95610623 |
| ZKSCAN5 | 199.233246 | -0.1732595 | 0.2664158 | -0.6503351 | 0.51547579 | 0.85869987 |
| FAM200A | 169.480133 | 0.22397088 | 0.27080278 | 0.82706269 | 0.40820153 | 0.80588056 |
| ZSCAN25 | 151.003886 | -0.0492158 | 0.23572451 | -0.2087851 | 0.83461603 | 0.96510927 |
| CYP3A5 | 77.5099432 | 1.18015197 | 0.69911305 | 1.68807028 | 0.09139774 | 0.50429103 |
| TRIM4 | 454.277328 | -0.0356622 | 0.21599305 | -0.1651082 | 0.86885885 | 0.97442321 |
| ZKSCAN1 | 1316.43175 | 0.15029331 | 0.12455 | 1.20669057 | 0.22755133 | 0.67921637 |
| ZSCAN21 | 123.767651 | -0.3559035 | 0.27801163 | -1.2801748 | 0.20048365 | 0.65355097 |
| ZNF3 | 266.514183 | -0.5872147 | 0.32607374 | -1.8008648 | 0.07172419 | 0.4673926 |
| COPS6 | 274.499077 | 0.37673514 | 0.22744813 | 1.65635629 | 0.09764969 | 0.51355946 |

| | | | | | | |
|---|---|---|---|---|---|---|
| MCM7 | 143.872188 | -0.3606818 | 0.27730803 | -1.300654 | 0.19337691 | 0.64718686 |
| AP4M1 | 228.283911 | -0.4530993 | 0.20588055 | -2.2007874 | 0.02775108 | 0.32154734 |
| TAF6 | 117.578046 | -0.0609363 | 0.33988677 | -0.1792841 | 0.85771463 | 0.97127317 |
| AC073842.2 | 35.219415 | -0.47306 | 0.57509669 | -0.8225748 | 0.41074983 | 0.80665991 |
| LAMTOR4 | 433.879217 | 0.46733573 | 0.30639858 | 1.52525426 | 0.12719569 | 0.56740575 |
| C7orf43 | 63.3048231 | -0.4404397 | 0.48204511 | -0.9136899 | 0.36087983 | 0.77730375 |
| STAG3 | 161.165276 | -0.0740823 | 0.49914152 | -0.1484194 | 0.88201177 | 0.97664137 |
| CASTOR3 | 974.673502 | -0.1654723 | 0.35083538 | -0.4716523 | 6.37E-01 | 0.90255504 |
| SPDYE3 | 380.435205 | 0.15748743 | 0.34009487 | 0.46306912 | 0.64331482 | 0.90478945 |
| PILRB | 980.76789 | -0.225789 | 0.35898702 | -0.6289615 | 0.52937428 | 0.86316564 |
| PILRA | 263.105025 | -0.0261392 | 0.43901922 | -0.0595399 | 0.95252208 | 0.98938795 |
| ZCWPW1 | 132.167764 | -1.0720778 | 0.49279186 | -2.1755186 | 0.02959128 | 0.33073125 |
| MEPCE | 118.170037 | 0.08255655 | 0.30454446 | 0.27108208 | 0.7863279 | 0.94978829 |
| PPP1R35 | 178.15005 | -0.0975791 | 0.36134094 | -0.2700471 | 0.78712399 | 0.94978829 |
| TSC22D4 | 505.674765 | 0.19858617 | 0.30329471 | 0.65476306 | 0.51262032 | 0.85762435 |
| AGFG2 | 358.575765 | -1.062926 | 0.2894282 | -3.6725033 | 0.00024019 | 0.02558562 |
| SAP25 | 76.3764383 | 0.21265524 | 0.50613868 | 0.42015211 | 0.67437433 | 0.91636338 |
| LRCH4 | 159.235434 | -0.09018 | 0.34563995 | -0.2609075 | 0.79416386 | 0.95234775 |
| PCOLCE-AS1 | 49.9692374 | -0.0345043 | 0.68716285 | -0.0502126 | 0.95995294 | 0.99140097 |
| PCOLCE | 107.708401 | -0.5262146 | 0.3095834 | -1.6997506 | 0.08917785 | 0.49902981 |
| MOSPD3 | 53.8182418 | -0.2623498 | 0.38870685 | -0.6749296 | 0.4997205 | 0.85244523 |
| GNB2 | 548.18476 | 0.20558877 | 0.39146293 | 0.52518068 | 5.99E-01 | 0.88883692 |
| GIGYF1 | 602.493135 | -0.2967018 | 0.24246468 | -1.2236908 | 0.22106889 | 0.67392082 |
| POP7 | 62.8399525 | 0.18247778 | 0.56136482 | 0.32506094 | 0.74513495 | 0.93675395 |
| EPHB4 | 178.008397 | -0.172571 | 0.36405139 | -0.4740293 | 0.63547903 | 0.90199374 |
| SLC12A9 | 434.385097 | 0.01797048 | 0.43868993 | 0.04096396 | 0.96732463 | 0.99248831 |
| TRIP6 | 216.471881 | -0.1947362 | 0.28108333 | -0.6928061 | 0.48843127 | 0.84562056 |
| SRRT | 275.588119 | -0.4270008 | 0.21014801 | -2.0319048 | 0.04216329 | 0.37912522 |
| TRIM56 | 856.354045 | 0.27668124 | 0.49760127 | 0.55603001 | 0.57819034 | 0.88205258 |
| AP1S1 | 205.977598 | 1.04030554 | 0.41094675 | 2.53148498 | 0.01135807 | 0.21832654 |
| PLOD3 | 165.387218 | -0.0273986 | 0.26141349 | -0.1048094 | 0.91652702 | 0.98197488 |
| ZNHIT1 | 292.361443 | 0.25759086 | 0.24244096 | 1.06248902 | 0.28801374 | 0.73075337 |
| CLDN15 | 73.7403896 | -0.2652326 | 0.31951118 | -0.8301198 | 0.40647106 | 0.80499101 |
| FIS1 | 475.235459 | 0.49289889 | 0.28295833 | 1.74194867 | 0.08151743 | 0.48594425 |
| IFT22 | 132.377364 | 0.23776327 | 0.27738806 | 0.85715033 | 0.39136181 | 0.7973532 |
| COL26A1 | 126.623505 | -0.0737933 | 0.6127565 | -0.1204284 | 0.90414376 | 0.97988471 |
| CUX1 | 5220.11396 | 0.0495742 | 0.2324485 | 0.21326962 | 0.83111667 | 0.96382687 |
| SH2B2 | 84.5698568 | -0.2607985 | 0.44675235 | -0.5837652 | 5.59E-01 | 8.73E-01 |
| PRKRIP1 | 1584.69618 | -0.2594077 | 0.23160264 | -1.1200548 | 0.2626904 | 0.7132328 |
| ORAI2 | 234.548506 | 0.29647614 | 0.43330346 | 0.68422288 | 0.49383444 | 0.84869361 |
| ALKBH4 | 92.5180953 | 0.28655101 | 0.31946579 | 0.89696932 | 0.36973529 | 0.78095617 |
| LRWD1 | 41.4273172 | 0.45829739 | 0.46148527 | 0.99309213 | 0.32066506 | 0.75144499 |
| POLR2J | 137.825657 | 0.52498452 | 0.34009488 | 1.54364135 | 0.12267524 | 0.55947129 |
| AC093668.2 | 78.2413277 | -0.3740051 | 0.59161711 | -0.6321742 | 0.52727304 | 0.86151701 |
| SPDYE2 | 208.20004 | 0.5895042 | 0.59266817 | 0.99466149 | 0.31990093 | 0.75094569 |
| AC105052.1 | 83.631027 | -0.498991 | 0.54477119 | -0.9159644 | 0.35968559 | 0.77670213 |
| AC105052.3 | 492.972939 | 0.55127367 | 0.52468143 | 1.05068264 | 0.29340437 | 0.73527014 |
| FAM185A | 932.923031 | 0.10105195 | 0.180998 | 0.55830424 | 0.57663665 | 0.88147804 |
| FBXL13 | 690.14367 | -0.4705348 | 0.25324162 | -1.8580469 | 0.06316235 | 0.43727017 |
| LRRC17 | 114.963283 | -0.0228861 | 0.29964523 | -0.0763772 | 0.93911901 | 0.98715467 |
| ARMC10 | 452.57596 | -0.1894116 | 0.23314981 | -0.8124031 | 0.41656035 | 0.80966301 |
| NAPEPLD | 1056.01835 | -0.0872465 | 0.41195269 | -0.2117876 | 0.83227273 | 0.96417211 |
| PMPCB | 465.107425 | 0.09070971 | 0.23291511 | 0.38945396 | 0.69694036 | 0.92322348 |
| DNAJC2 | 413.142836 | -0.0810688 | 0.22613654 | -0.3584948 | 0.71997309 | 0.93001079 |

| | | | | | | |
|---|---|---|---|---|---|---|
| PSMC2 | 316.666256 | 0.12011433 | 0.1813508 | 0.66233138 | 0.50775887 | 0.85643442 |
| SLC26A5 | 121.031568 | 0.74095461 | 0.45745363 | 1.6197371 | 0.10528876 | 0.52613699 |
| ORC5 | 566.066305 | -0.3760918 | 0.15560993 | -2.4168884 | 0.01565381 | 0.25227487 |
| LHFPL3 | 572.211638 | 0.55096078 | 0.34383035 | 1.60242041 | 0.10906267 | 0.53262539 |
| LINC01004 | 1761.27676 | -0.1889759 | 0.23370285 | -0.8086163 | 0.41873591 | 0.81125469 |
| KMT2E-AS1 | 117.933192 | -0.0514396 | 0.31780234 | -0.1618605 | 0.87141574 | 0.97518512 |
| KMT2E | 3314.09994 | 0.17585693 | 0.18296506 | 0.96115035 | 0.33647658 | 0.76379431 |
| SRPK2 | 4810.373 | -0.0973977 | 0.18193339 | -0.5353482 | 0.59240912 | 0.88740306 |
| PUS7 | 271.256576 | -0.3988429 | 0.31308132 | -1.2739273 | 0.20268916 | 0.65552334 |
| RINT1 | 263.557492 | -0.1085511 | 0.29088825 | -0.3731713 | 0.70902098 | 0.92577218 |
| EFCAB10 | 77.679984 | -0.2150151 | 0.49886954 | -0.4310046 | 0.66646504 | 0.91318863 |
| ATXN7L1 | 1592.06456 | -0.3148018 | 0.25901609 | -1.2153757 | 0.22422285 | 0.67681479 |
| CDHR3 | 398.44581 | 1.21918377 | 0.60690509 | 2.00885411 | 0.04455261 | 0.38623091 |
| SYPL1 | 539.902436 | 0.33441365 | 0.302837 | 1.10426945 | 0.26947627 | 0.71818262 |
| NAMPT | 2947.96909 | 0.85157109 | 0.57312706 | 1.48583299 | 0.1373233 | 0.57989242 |
| AC005050.1 | 63.0272016 | 0.31561879 | 0.64775109 | 0.4872532 | 6.26E-01 | 8.99E-01 |
| CCDC71L | 167.493069 | 0.86815498 | 0.74404244 | 1.16680842 | 0.24328774 | 0.69452742 |
| HBP1 | 987.202306 | 0.02518995 | 0.22589391 | 0.11151228 | 0.91121013 | 0.98112934 |
| COG5 | 2616.2276 | -0.1751249 | 0.23838462 | -0.7346316 | 0.46256391 | 0.83531269 |
| DUS4L | 128.914395 | -0.3149957 | 0.22308451 | -1.4120017 | 0.15794945 | 0.60779483 |
| BCAP29 | 489.649924 | 0.26303683 | 0.26810842 | 0.98108381 | 0.32655142 | 0.75554697 |
| CBLL1 | 199.736484 | -0.1304386 | 0.30452846 | -0.4283298 | 0.66841106 | 0.91395501 |
| SLC26A3 | 2881.76756 | -0.2777679 | 0.6976147 | -0.3981681 | 0.6905063 | 0.92183073 |
| DLD | 674.249533 | 0.26601337 | 0.25961236 | 1.02465603 | 0.30552552 | 0.74251103 |
| LAMB1 | 5123.07176 | -1.1400745 | 0.40733114 | -2.7988886 | 0.00512788 | 0.14350465 |
| LAMB4 | 141.92334 | 0.50849448 | 0.45576639 | 1.11569106 | 0.2645544 | 0.71488591 |
| NRCAM | 2125.12146 | -0.9691734 | 0.47561462 | -2.0377283 | 0.04157711 | 0.37590145 |
| PNPLA8 | 1298.49443 | 0.29641665 | 0.37632075 | 0.78767023 | 0.43088962 | 0.81706845 |
| DNAJB9 | 158.422638 | 1.23262358 | 0.37627566 | 3.27585256 | 0.00105344 | 0.05797845 |
| IMMP2L | 25545.2378 | 0.27515186 | 0.18979076 | 1.44976422 | 0.14712428 | 0.59373375 |
| LRRN3 | 170.86298 | 0.68037261 | 0.71517143 | 0.95134199 | 0.3414308 | 0.76543778 |
| DOCK4 | 2728.12895 | -0.7413943 | 0.32588077 | -2.2750478 | 0.02290306 | 0.2983102 |
| ZNF277 | 1545.90283 | -0.0418874 | 0.27318399 | -0.1533305 | 0.87813763 | 0.9765717 |
| AC004112.1 | 87.7288428 | -0.1908879 | 0.47315652 | -0.403435 | 0.68662824 | 0.9208231 |
| IFRD1 | 1336.24176 | -0.1132087 | 0.50709677 | -0.2232488 | 0.82334191 | 0.96133701 |
| LSMEM1 | 156.390906 | 0.12615779 | 0.78839678 | 0.16001815 | 0.87286678 | 0.97564728 |
| TMEM168 | 753.304585 | 0.19037744 | 0.22667424 | 0.83987244 | 0.40097991 | 0.80243544 |
| BMT2 | 1036.5416 | -0.1839282 | 0.16921166 | -1.0869714 | 0.27704946 | 0.7230876 |
| SMIM30 | 119.299085 | 0.12554107 | 0.44739837 | 0.28060242 | 0.77901536 | 0.94732571 |
| FOXP2 | 6965.77193 | -1.1547926 | 0.64869835 | -1.7801689 | 0.07504833 | 0.47190267 |
| MDFIC | 814.541483 | 0.46519108 | 0.38151635 | 1.21932147 | 0.2227222 | 0.67568617 |
| TES | 523.306777 | 0.25506614 | 0.26633206 | 0.95769972 | 0.3382142 | 0.76444215 |
| AC002066.1 | 59.7102881 | 0.60224428 | 0.59298714 | 1.01561103 | 0.30981465 | 0.74450725 |
| CAV2 | 108.1447 | 1.40202084 | 0.32324048 | 4.33739259 | 1.44E-05 | 0.00577125 |
| MET | 2327.30067 | -0.9243555 | 0.50163488 | -1.8426858 | 0.0653749 | 0.44218998 |
| CAPZA2 | 1349.03644 | 0.12025471 | 0.21745581 | 0.55300754 | 0.58025824 | 0.88258799 |
| ST7 | 1955.22332 | -0.0417514 | 0.22223173 | -0.1878731 | 0.85097616 | 0.97056586 |
| ST7-AS2 | 126.950548 | 0.13670753 | 0.6536195 | 0.20915461 | 0.83432754 | 0.96510927 |
| CFTR | 57.3689886 | 0.47406436 | 0.47191516 | 1.00455419 | 0.31511156 | 0.74785672 |
| CTTNBP2 | 828.151989 | -0.4801037 | 0.36349555 | -1.3207967 | 0.18656918 | 0.6408069 |
| LSM8 | 493.540697 | -0.2510106 | 0.15588793 | -1.6101989 | 0.10735445 | 0.52901576 |
| ANKRD7 | 401.654168 | -0.3076031 | 0.31856649 | -0.9655852 | 0.33425178 | 0.76231532 |
| TSPAN12 | 1135.30676 | 0.56310366 | 0.39073724 | 1.44113127 | 0.1495476 | 0.59625836 |
| ING3 | 328.388984 | -0.0109619 | 0.27446906 | -0.0399386 | 0.96814206 | 0.99248831 |

| | | | | | | |
|---|---|---|---|---|---|---|
| FAM3C | 1017.72738 | 0.1271784 | 0.28461876 | 0.44683772 | 0.65499224 | 0.90902447 |
| AASS | 642.449167 | 0.70339819 | 0.32853464 | 2.1410168 | 0.03227268 | 0.34295243 |
| CADPS2 | 6101.65263 | -0.1619663 | 0.39987418 | -0.4050432 | 0.68544574 | 0.92053037 |
| AC004594.1 | 377.877944 | -0.1600001 | 0.72177086 | -0.2216771 | 0.82456527 | 0.96146415 |
| SLC13A1 | 68.3572108 | 3.99184366 | 0.81266602 | 4.91203467 | 9.01E-07 | 0.00085043 |
| IQUB | 307.944251 | 0.26617753 | 0.33045278 | 0.80549339 | 0.42053502 | 0.81260593 |
| NDUFA5 | 704.484782 | 0.44698093 | 0.25686048 | 1.74017007 | 0.08182916 | 0.48594425 |
| WASL | 1168.71297 | 0.01427385 | 0.22130158 | 0.06449955 | 0.94857247 | 0.98919519 |
| AC006333.2 | 52.9191271 | -0.0942694 | 0.46743921 | -0.201672 | 0.84017315 | 0.96617873 |
| POT1 | 635.808821 | -0.2504715 | 0.1949186 | -1.2850055 | 0.19879036 | 0.65113421 |
| POT1-AS1 | 342.560906 | 0.16385872 | 0.38137465 | 0.4296529 | 0.66744815 | 0.91360856 |
| GRM8 | 639.626703 | -0.379566 | 0.62699414 | -0.6053741 | 0.54493046 | 0.86680954 |
| ZNF800 | 818.30068 | 0.13951504 | 0.28094406 | 0.49659366 | 0.61947563 | 0.8966309 |
| GCC1 | 51.0763242 | 0.30449787 | 0.33786598 | 0.90123864 | 0.36746145 | 0.78095617 |
| ARF5 | 159.195199 | 0.43505913 | 0.4695843 | 0.92647718 | 0.35419805 | 0.77243419 |
| SND1 | 2765.15998 | 0.16378887 | 0.23550055 | 0.69549251 | 0.4867467 | 0.84474442 |
| RBM28 | 443.719341 | -0.4856416 | 0.16122429 | -3.0122112 | 0.00259352 | 0.09650091 |
| HILPDA | 472.522694 | 0.05982801 | 1.28915184 | 0.04640882 | 0.96298441 | 0.99224805 |
| METTL2B | 368.067407 | -0.1779449 | 0.22836011 | -0.7792294 | 0.43584459 | 0.8196523 |
| AC090114.2 | 111.350873 | -0.4177504 | 0.34989867 | -1.1939182 | 0.23250997 | 0.68462421 |
| AC018638.7 | 60.9357213 | 0.295427 | 0.30902114 | 0.95600901 | 0.33906768 | 0.76449297 |
| FAM71F2 | 164.808487 | 0.6068967 | 0.283158 | 2.1433147 | 0.03208784 | 0.34153767 |
| FAM71F1 | 44.0750314 | 0.06956716 | 0.67061155 | 0.1037369 | 0.91737815 | 0.98199511 |
| CALU | 578.264552 | -0.077246 | 0.22491248 | -0.3434493 | 0.73126045 | 0.93515532 |
| KCP | 1269.95072 | -0.5854334 | 0.66132739 | -0.88524 | 0.37602722 | 0.78490077 |
| ATP6V1F | 368.761903 | 0.90380095 | 0.39563004 | 2.28445985 | 0.02234452 | 0.29654015 |
| IRF5 | 55.7685745 | -0.4837617 | 0.51920128 | -0.9317421 | 0.35146985 | 0.77119024 |
| TNPO3 | 1291.78874 | -0.346079 | 0.16407544 | -2.1092677 | 0.03492148 | 0.35358029 |
| TSPAN33 | 509.091508 | 0.04846787 | 0.55666724 | 0.08706795 | 0.9306175 | 0.98331361 |
| SMO | 411.639839 | -0.1717743 | 0.31473766 | -0.5457698 | 5.85E-01 | 8.85E-01 |
| AHCYL2 | 2114.70614 | -0.1531568 | 0.14847693 | -1.0315195 | 0.30229728 | 0.73905643 |
| AC078846.1 | 93.9111772 | -0.227782 | 0.28241103 | -0.8065619 | 0.41991893 | 0.8125045 |
| NRF1 | 1697.71199 | -0.1108273 | 0.17271947 | -0.6416609 | 0.52109341 | 0.85943874 |
| UBE2H | 6443.81367 | -0.4686977 | 0.27363491 | -1.7128578 | 0.0867387 | 0.49478465 |
| AC073320.1 | 48.7183062 | -0.4126888 | 0.3543082 | -1.1647735 | 0.24411069 | 0.6948422 |
| ZC3HC1 | 359.873242 | -0.3320248 | 0.22767757 | -1.4583113 | 0.14475476 | 0.59044137 |
| KLHDC10 | 1942.28206 | 0.05710096 | 0.18725458 | 0.30493758 | 0.76041369 | 0.9418007 |
| TMEM209 | 388.719311 | 0.0419363 | 0.2267368 | 0.18495586 | 0.85326366 | 0.97076831 |
| AC087071.2 | 103.986895 | 0.22729182 | 0.51840175 | 0.43844725 | 0.6610621 | 0.91110616 |
| CEP41 | 159.094178 | -0.2523239 | 0.2731263 | -0.9238361 | 0.35557168 | 0.77310012 |
| MEST | 267.50429 | 2.35437366 | 0.63378339 | 3.7147923 | 0.00020337 | 0.02315796 |
| COPG2 | 599.010693 | -0.3744823 | 0.21820342 | -1.7162072 | 0.08612413 | 0.49453468 |
| AC016831.7 | 15047.5189 | -0.2764192 | 0.42634662 | -0.6483438 | 0.51676258 | 0.85893003 |
| LINC00513 | 1983.64782 | -0.5361625 | 0.72979909 | -0.7346714 | 0.46253965 | 0.83531269 |
| AC058791.1 | 406.448947 | -0.6489923 | 0.78043188 | -0.831581 | 0.40564549 | 0.80499101 |
| MKLN1 | 10904.0717 | -0.4497234 | 0.37321802 | -1.2049883 | 0.22820781 | 0.67999141 |
| PODXL | 142.166059 | 0.22732432 | 0.46963624 | 0.48404339 | 0.62835508 | 0.89962522 |
| PLXNA4 | 418.770925 | -1.2190339 | 0.57660717 | -2.1141498 | 0.03450248 | 0.35291306 |
| CHCHD3 | 3553.2331 | 0.09013452 | 0.1810486 | 0.49784708 | 0.61859183 | 0.89604387 |
| EXOC4 | 7612.95075 | -0.222769 | 0.11455881 | -1.9445822 | 0.05182529 | 0.40820525 |
| LRGUK | 457.042638 | 0.01494266 | 0.31454014 | 0.04750637 | 0.96210965 | 0.99223192 |
| SLC35B4 | 537.101924 | 0.15004597 | 0.22126877 | 0.67811635 | 4.98E-01 | 8.51E-01 |
| AKR1B1 | 621.124786 | -0.5675454 | 0.90521699 | -0.6269717 | 0.5306778 | 0.86396107 |
| BPGM | 149.615158 | 0.59371562 | 0.28346367 | 2.09450342 | 0.03621516 | 0.35817721 |

| | | | | | | |
|---|---|---|---|---|---|---|
| CALD1 | 241.916679 | -0.0020136 | 0.4717581 | -0.0042682 | 0.99659447 | 0.99931803 |
| AGBL3 | 404.262357 | -0.0434472 | 0.30714376 | -0.1414557 | 0.88750992 | 0.97729111 |
| CYREN | 411.00815 | -0.007285 | 0.25933458 | -0.0280911 | 0.97758952 | 0.99455895 |
| TMEM140 | 343.22637 | 0.28982619 | 0.49142021 | 0.58977263 | 0.5553431 | 0.87184917 |
| WDR91 | 1285.23064 | 0.30055742 | 0.18397413 | 1.63369397 | 0.10232312 | 0.52073581 |
| AC009542.1 | 222.863727 | 1.10281732 | 0.61452917 | 1.79457277 | 0.07272179 | 0.4692634 |
| CNOT4 | 2042.59052 | -0.0945151 | 0.14546138 | -0.6497609 | 0.51584665 | 0.85881251 |
| NUP205 | 436.727495 | -0.3654047 | 0.21463507 | -1.7024463 | 0.08867174 | 0.49819865 |
| STMP1 | 804.223689 | 0.26624581 | 0.29843786 | 0.89213146 | 0.37232247 | 0.78260237 |
| SLC13A4 | 164.069556 | 0.91437727 | 0.53323559 | 1.71477163 | 0.08638711 | 0.49478465 |
| MTPN | 494.55935 | 0.09696098 | 0.15608005 | 0.62122596 | 0.53445096 | 0.86419602 |
| DGKI | 137.368779 | -0.0318468 | 0.55239885 | -0.0576519 | 0.95402593 | 0.99000145 |
| CREB3L2 | 1846.70991 | 0.2976003 | 0.34899712 | 0.85272995 | 0.39380908 | 0.79843808 |
| TRIM24 | 1080.562 | -0.4575632 | 0.2181534 | -2.097438 | 0.03595482 | 0.35661333 |
| SVOPL | 677.253948 | -1.3819136 | 0.61463684 | -2.2483416 | 0.02455441 | 0.30772226 |
| ATP6V0A4 | 2639.2944 | -0.2051774 | 0.42198924 | -0.4862147 | 0.62681494 | 0.89878404 |
| TMEM213 | 1518.6696 | 0.4078875 | 0.34310958 | 1.18879658 | 0.23451972 | 0.68580275 |
| KIAA1549 | 336.179959 | -0.2919722 | 0.3621255 | -0.8062736 | 0.42008514 | 0.8125045 |
| ZC3HAV1L | 74.3313151 | 0.07126601 | 0.51041311 | 0.13962418 | 0.88895693 | 0.97729111 |
| ZC3HAV1 | 467.148751 | 0.06639349 | 0.37631665 | 0.17642984 | 0.85995626 | 0.97200968 |
| TTC26 | 304.310018 | 0.11344068 | 0.21456623 | 0.52869773 | 0.59701515 | 0.88883692 |
| UBN2 | 1338.77777 | -0.0128582 | 0.1745608 | -0.07366 | 0.94128091 | 0.98732467 |
| LUC7L2 | 323.266107 | 0.53639225 | 0.234048 | 2.29180442 | 0.02191693 | 0.29509055 |
| HIPK2 | 4938.28114 | -0.0890149 | 0.17616809 | -0.5052839 | 0.61335948 | 0.89441126 |
| TBXAS1 | 359.590134 | -0.2875286 | 0.40253315 | -0.7142979 | 0.475043 | 0.84106098 |
| PARP12 | 455.860384 | -0.3273271 | 0.66463144 | -0.4924942 | 0.62237003 | 0.89835355 |
| KDM7A | 2268.31907 | -1.1319451 | 0.27906454 | -4.0562124 | 4.99E-05 | 0.01125476 |
| SLC37A3 | 676.80277 | -0.454589 | 0.14987118 | -3.0331979 | 0.00241977 | 0.094008 |
| RAB19 | 54.0672626 | -0.1055505 | 0.34948443 | -0.3020176 | 0.76263864 | 0.9422053 |
| MKRN1 | 801.092307 | 0.00993268 | 0.21818999 | 0.04552308 | 0.96369038 | 0.99224805 |
| DENND2A | 3348.51712 | -0.5391729 | 0.31812877 | -1.6948258 | 0.09010846 | 0.50138792 |
| ADCK2 | 47.3899106 | 0.21667543 | 0.46129526 | 0.46971095 | 0.63856154 | 0.90307917 |
| NDUFB2 | 625.039624 | 0.64639245 | 0.33210044 | 1.94637639 | 0.05160954 | 0.40772157 |
| BRAF | 5047.99884 | 0.07495661 | 0.19596179 | 0.38250627 | 0.70208588 | 0.9248705 |
| MRPS33 | 194.9018 | 0.34742562 | 0.2870801 | 1.21020447 | 0.22620044 | 0.67875549 |
| TMEM178B | 884.951416 | 0.29463847 | 0.5856782 | 0.50307228 | 0.61491345 | 0.8949759 |
| AC073878.1 | 57.6404416 | -0.6763425 | 0.59137856 | -1.143671 | 0.25276009 | 0.7033301 |
| AGK | 539.408657 | -0.2146785 | 0.20107929 | -1.0676313 | 0.28568687 | 0.72850152 |
| AC004918.1 | 179.91052 | -0.0801242 | 0.44504174 | -0.1800375 | 0.85712311 | 0.97127317 |
| KIAA1147 | 523.71931 | -0.1715939 | 0.33365533 | -0.5142849 | 0.60705283 | 0.89174148 |
| WEE2-AS1 | 126.847596 | -1.3036761 | 0.38801927 | -3.3598231 | 0.00077992 | 0.04906731 |
| SSBP1 | 729.477811 | -0.2007738 | 0.21382921 | -0.9389449 | 0.34775903 | 0.76900643 |
| MGAM | 234.878174 | 1.38217714 | 0.81123159 | 1.70380094 | 0.08841828 | 0.49769182 |
| GSTK1 | 544.070949 | 0.30708043 | 0.26250046 | 1.16982815 | 0.24207013 | 0.69337349 |
| AC073342.1 | 127.03043 | -0.1447731 | 0.51317497 | -0.2821126 | 0.77785714 | 0.94694572 |
| TMEM139 | 88.0690862 | 0.65779634 | 0.64445967 | 1.02069434 | 0.30739928 | 0.74341546 |
| CASP2 | 157.427562 | -0.4036484 | 0.2375005 | -1.6995686 | 0.08921211 | 0.49902981 |
| ZYX | 138.319867 | -0.19493 | 0.49642224 | -0.3926697 | 0.69456344 | 0.92229578 |
| EPHA1 | 70.4925629 | -0.0002038 | 0.39030652 | -0.0005221 | 0.99958346 | 0.99987432 |
| TCAF2 | 230.68152 | 0.54724001 | 0.88577074 | 0.61781225 | 0.53669911 | 0.86466169 |
| TCAF1 | 703.500522 | 0.15387956 | 0.1915699 | 0.80325543 | 0.42182712 | 0.81266383 |
| AC004889.1 | 455.014462 | -0.3446345 | 0.37520461 | -0.9185243 | 0.35834445 | 0.77529959 |
| AC074386.1 | 939.007572 | -0.3092253 | 0.33652889 | -0.9188669 | 0.35816519 | 0.77529959 |
| ARHGEF5 | 189.747545 | -0.1610468 | 0.36307081 | -0.4435685 | 0.65735458 | 0.91026278 |

| | | | | | | |
|---|---|---|---|---|---|---|
| TPK1 | 543.721591 | -0.5213594 | 0.39027894 | -1.3358636 | 0.18159387 | 0.63635309 |
| CNTNAP2 | 156.907886 | -0.1941891 | 0.37621748 | -0.5161617 | 0.60574144 | 0.89081188 |
| CUL1 | 1516.16819 | -0.1869663 | 0.1314315 | -1.4225379 | 0.15487015 | 0.60380955 |
| EZH2 | 246.288933 | -0.776395 | 0.40427936 | -1.9204419 | 0.05480211 | 0.41684503 |
| PDIA4 | 208.547841 | 0.84854599 | 0.33157049 | 2.5591722 | 0.01049218 | 0.20778283 |
| ZNF786 | 180.185608 | -0.0939129 | 0.39733926 | -0.2363543 | 0.81315772 | 0.95816239 |
| ZNF425 | 151.330279 | -0.5796556 | 0.34445422 | -1.6828233 | 0.09240931 | 0.50585766 |
| ZNF398 | 424.060496 | -0.0413555 | 0.26260834 | -0.1574798 | 0.87486669 | 0.97585831 |
| ZNF282 | 249.286207 | -0.1826301 | 0.21536991 | -0.8479833 | 0.39644726 | 0.80030376 |
| ZNF212 | 172.79853 | -0.3714272 | 0.26152447 | -1.4202387 | 0.15553819 | 0.6049776 |
| ZNF783 | 226.864574 | -0.0351829 | 0.23982619 | -0.1467018 | 0.88336739 | 0.97664137 |
| ZNF777 | 126.423318 | -0.0009309 | 0.28544487 | -0.0032614 | 0.99739779 | 0.99948643 |
| ZNF746 | 184.882853 | 0.0612662 | 0.21729176 | 0.28195364 | 0.77797906 | 0.94694572 |
| KRBA1 | 77.4617202 | 0.29359594 | 0.3074616 | 0.95490278 | 0.33962686 | 0.76449297 |
| ZNF467 | 120.605914 | -0.305444 | 0.53777216 | -0.5679802 | 0.57004839 | 0.87810719 |
| SSPO | 43.5566515 | -1.6128084 | 0.49500552 | -3.2581625 | 0.00112136 | 0.05996788 |
| ZNF862 | 651.14448 | 0.37328619 | 0.24128838 | 1.54705415 | 0.12185019 | 0.5572644 |
| ATP6V0E2-A: | 49.7939125 | -0.3520227 | 0.52887746 | -0.6656035 | 0.50566454 | 0.85599422 |
| ATP6V0E2 | 79.3646572 | 0.75414031 | 0.61968857 | 1.21696663 | 0.22361691 | 0.67624946 |
| AC092666.1 | 135.311956 | 0.29780341 | 0.80827637 | 0.36844255 | 0.71254327 | 0.92754765 |
| ACTR3C | 2648.35978 | 0.19857679 | 0.26563072 | 0.74756711 | 0.45472131 | 0.83121393 |
| LRRC61 | 132.971818 | 0.2054792 | 0.42229467 | 0.48657778 | 0.62655757 | 0.89873262 |
| RARRES2 | 162.171838 | -1.1359113 | 0.76606452 | -1.482788 | 0.13813074 | 0.58206378 |
| AC073111.5 | 55.9204807 | 0.75030744 | 0.93717722 | 0.80060358 | 0.42336118 | 0.8138418 |
| TMEM176B | 581.935331 | 1.26243384 | 0.40722014 | 3.10012627 | 0.00193438 | 0.08545567 |
| TMEM176A | 264.092476 | 1.47383727 | 0.41487521 | 3.55248331 | 0.00038161 | 0.03316269 |
| AOC1 | 110.568798 | 2.43624598 | 0.65451176 | 3.72223407 | 0.00019747 | 0.02268133 |
| ABCB8 | 188.795903 | -0.3089842 | 0.25813948 | -1.1969662 | 0.23131974 | 0.68357541 |
| SLC4A2 | 291.474351 | -0.3216 | 0.29105503 | -1.1049459 | 0.26918305 | 0.71818262 |
| FASTK | 218.772577 | -0.3151931 | 0.36921483 | -0.8536848 | 0.39327963 | 0.79843808 |
| TMUB1 | 177.690928 | 0.27204693 | 0.36492456 | 0.74548813 | 0.45597668 | 0.83148642 |
| AGAP3 | 263.538408 | 0.16609181 | 0.27333218 | 0.60765554 | 0.54341596 | 0.86622038 |
| CHPF2 | 96.0974553 | 0.02207732 | 0.35506737 | 0.06217784 | 0.95042121 | 0.98929186 |
| SMARCD3 | 329.387368 | 1.11465304 | 0.38867203 | 2.8678499 | 0.00413272 | 0.12549204 |
| AC021097.1 | 53.6374731 | 0.17255701 | 0.3588068 | 0.48091902 | 0.63057406 | 0.90005881 |
| NUB1 | 483.424425 | -0.089542 | 0.32716746 | -0.2736887 | 0.78432384 | 0.94978829 |
| RHEB | 1165.13472 | -0.0410702 | 0.19119417 | -0.2148087 | 0.82991644 | 0.9631318 |
| PRKAG2 | 1694.57084 | 0.58334649 | 0.3775037 | 1.54527358 | 0.1222801 | 0.5585055 |
| GALNT11 | 1220.04156 | 0.04310954 | 0.14607858 | 0.29511196 | 0.76790835 | 0.94316227 |
| KMT2C | 5385.95873 | -0.1565133 | 0.14814031 | -1.0565207 | 0.29073039 | 0.73296946 |
| ACTR3B | 744.660868 | -0.2041952 | 0.16540257 | -1.2345345 | 0.21700381 | 0.66978787 |
| DPP6 | 1994.61632 | -0.1278496 | 0.45279405 | -0.2823572 | 0.77766965 | 0.94694572 |
| PAXIP1-AS2 | 227.631617 | 0.17860083 | 0.41209791 | 0.43339418 | 0.66472843 | 0.91220795 |
| PAXIP1 | 453.754495 | 0.05589263 | 0.21522782 | 0.25969056 | 0.79510248 | 0.95272759 |
| PAXIP1-AS1 | 138.583289 | 0.04595804 | 0.34402742 | 0.13358832 | 0.89372812 | 0.97788747 |
| INSIG1 | 228.068052 | -0.7341319 | 0.7474883 | -0.9821316 | 0.32603504 | 0.75550719 |
| AC009403.1 | 315.215296 | 0.16337363 | 0.32677143 | 0.49996302 | 0.61710112 | 0.89599505 |
| RBM33 | 2313.52325 | -0.1197606 | 0.19718835 | -0.6073409 | 0.54362468 | 0.86633953 |
| SHH | 77.865437 | -0.6389419 | 0.41931427 | -1.5237781 | 1.28E-01 | 0.56762585 |
| LINC01006 | 640.600355 | -0.8235141 | 0.43077202 | -1.9117167 | 0.05591254 | 0.41843207 |
| RNF32 | 64.6004688 | -0.1468581 | 0.48631521 | -0.3019813 | 0.76266629 | 0.9422053 |
| LMBR1 | 2056.72862 | -0.1686174 | 0.23421896 | -0.7199137 | 0.47157812 | 0.83921324 |
| NOM1 | 198.593366 | -0.5389799 | 0.29170441 | -1.8476921 | 0.06464689 | 0.44001919 |
| UBE3C | 2404.08182 | -0.3563748 | 0.19546823 | -1.8231856 | 0.06827529 | 0.45323644 |

| | | | | | | |
|---|---|---|---|---|---|---|
| DNAJB6 | 1224.05692 | -0.0453917 | 0.2274138 | -0.1995997 | 0.84179363 | 0.96680741 |
| PTPRN2 | 53.413058 | 0.66118833 | 0.44260574 | 1.49385395 | 0.1352138 | 0.57708028 |
| NCAPG2 | 98.0015228 | -0.3730658 | 0.37210443 | -1.0025835 | 0.31606184 | 0.74785672 |
| ESYT2 | 2048.7388 | -0.2158152 | 0.27934565 | -0.7725739 | 0.43977458 | 0.82210338 |
| WDR60 | 1998.06428 | -0.0347998 | 0.18722788 | -0.1858689 | 0.85254755 | 0.97076831 |
| PLCXD1 | 59.5263419 | -0.2551876 | 0.39629717 | -0.64393 | 0.51962081 | 0.85924779 |
| GTPBP6 | 308.759529 | -0.1065722 | 0.29612343 | -0.359891 | 0.71892863 | 0.92991855 |
| LINC00685 | 188.302824 | -0.1380158 | 0.57530548 | -0.2399001 | 0.8104077 | 0.95679972 |
| PPP2R3B | 134.969888 | -0.8577785 | 0.40845199 | -2.1000718 | 0.03572253 | 0.35661333 |
| IL3RA | 218.844066 | 0.33496395 | 0.5009171 | 0.66870137 | 0.50368599 | 0.85525037 |
| SLC25A6 | 953.280806 | 0.78154007 | 0.41167791 | 1.89842605 | 0.05763998 | 0.42087381 |
| LINC00106 | 414.749037 | -0.5047215 | 0.39112952 | -1.2904203 | 0.19690477 | 0.64964978 |
| ASMTL-AS1 | 202.279013 | -0.2767206 | 0.33189921 | -0.8337488 | 0.40442254 | 0.80437725 |
| ASMTL | 334.523879 | -0.3255958 | 0.23784683 | -1.3689308 | 0.17102092 | 0.62750424 |
| AKAP17A | 325.489241 | 0.06725238 | 0.24638598 | 0.27295537 | 0.78488752 | 0.94978829 |
| DHRSX | 1605.15013 | -0.3824829 | 0.24907038 | -1.5356418 | 0.12462626 | 0.56139092 |
| CD99 | 306.834929 | 0.0686201 | 0.40025376 | 0.1714415 | 0.86387663 | 0.97349667 |
| GYG2 | 49.7113346 | 0.12586123 | 0.50519605 | 0.24913345 | 0.80325756 | 0.95446295 |
| ARSD | 652.685655 | 0.3061564 | 0.43826443 | 0.69856546 | 0.48482364 | 0.8447301 |
| ARSE | 60.1586635 | 1.10846068 | 0.55597779 | 1.99371396 | 0.04618332 | 0.3904347 |
| MXRA5 | 47.2011905 | -0.4063513 | 0.80781029 | -0.5030282 | 0.61494445 | 0.8949759 |
| PRKX | 2688.40377 | -0.8106398 | 0.3699245 | -2.1913656 | 0.02842535 | 0.32580744 |
| PRKX-AS1 | 204.820647 | -1.1364451 | 0.65281296 | -1.7408434 | 0.08171104 | 0.48594425 |
| BX890604.1 | 203.268172 | -0.3812736 | 0.4228036 | -0.9017748 | 0.36717653 | 0.78095617 |
| FAM239A | 58.0904187 | -2.9518005 | 1.40514124 | -2.1007144 | 0.03566604 | 0.35661333 |
| NLGN4X | 224.363667 | 3.24877812 | 0.90805266 | 3.57774199 | 0.00034658 | 0.03115997 |
| AC097347.1 | 58.8974533 | -0.1469909 | 0.62098657 | -0.2367055 | 0.8128853 | 0.95809779 |
| PUDP | 404.559377 | -0.0610669 | 0.34172553 | -0.1787017 | 0.85817196 | 0.97143431 |
| STS | 126.265953 | -0.2990922 | 0.536392 | -0.5576001 | 0.57711752 | 0.88150331 |
| PNPLA4 | 104.100305 | 0.1875465 | 0.27580655 | 0.67999291 | 0.49650895 | 0.85074416 |
| AC078960.1 | 58.394651 | -0.4756888 | 0.50279186 | -0.9460948 | 0.34410022 | 0.76686685 |
| TBL1X | 3631.0499 | -0.0692642 | 0.37536803 | -0.1845235 | 0.85360283 | 0.97076831 |
| GPR143 | 93.9668973 | 0.71788018 | 0.59222246 | 1.21217993 | 0.22544351 | 0.67798093 |
| SHROOM2 | 382.983138 | 0.24866067 | 0.32310271 | 0.76960257 | 0.44153568 | 0.82285053 |
| WWC3 | 356.155492 | -0.1185689 | 0.322952 | -0.3671408 | 0.71351398 | 0.92760608 |
| CLCN4 | 71.3089607 | -0.3910496 | 0.48636094 | -0.8040317 | 0.42137867 | 0.81266383 |
| MID1 | 683.089457 | 1.16999228 | 0.42543095 | 2.75013437 | 0.00595708 | 0.15829975 |
| AC073529.1 | 236.834141 | 0.04644521 | 0.48008347 | 0.09674403 | 0.92292967 | 0.98215577 |
| HCCS | 59.4174964 | 0.50024013 | 0.33394749 | 1.49796044 | 0.13414353 | 0.57708028 |
| ARHGAP6 | 19651.9323 | 0.23701786 | 0.30704328 | 0.77193634 | 0.44015213 | 0.82232701 |
| MSL3 | 98.394113 | -0.3007537 | 0.35875517 | -0.8383257 | 0.40184778 | 0.80278824 |
| PRPS2 | 150.914347 | -0.1260904 | 0.39341055 | -0.3205059 | 0.74858484 | 0.93746253 |
| TMSB4X | 857.377309 | -0.3457453 | 0.52934469 | -0.6531572 | 0.51365494 | 0.85762435 |
| TCEANC | 38.3389062 | 0.30171105 | 0.4900104 | 0.61572377 | 0.53807685 | 0.86496983 |
| RAB9A | 429.970217 | -0.0231709 | 0.31725571 | -0.0730355 | 0.94177786 | 0.98768906 |
| TRAPPC2 | 219.109694 | -0.058429 | 0.2698107 | -0.2165556 | 0.82855472 | 0.96288319 |
| OFD1 | 1110.03162 | -0.0353458 | 0.23883247 | -0.147994 | 0.88234753 | 0.97664137 |
| GPM6B | 183.611191 | 0.33011819 | 0.46713674 | 0.70668429 | 0.4797627 | 0.84323863 |
| GEMIN8 | 311.40054 | 0.03487297 | 0.22800062 | 0.15295121 | 0.87843675 | 0.9765717 |
| MOSPD2 | 231.649567 | -0.1160825 | 0.25653215 | -0.4525067 | 0.65090398 | 0.90825486 |
| PIGA | 123.082595 | 0.09372443 | 0.34495674 | 0.27169908 | 0.78585341 | 0.94978829 |
| PIR | 175.299648 | 1.06786883 | 0.50045394 | 2.13380043 | 0.03285912 | 0.34596681 |
| BMX | 45.0163287 | 0.01705834 | 0.39460576 | 0.04322881 | 0.96551914 | 0.99248831 |
| CLTRN | 154.58266 | 0.50105665 | 0.71776963 | 0.69807446 | 0.48513063 | 0.8447301 |

| | | | | | | |
|---|---|---|---|---|---|---|
| CA5B | 608.205725 | -0.4299786 | 0.2168943 | -1.9824337 | 0.04743072 | 0.39411174 |
| ZRSR2 | 253.43186 | 0.02298875 | 0.32647824 | 0.07041433 | 0.94386388 | 0.98792866 |
| AP1S2 | 386.17475 | 1.03136394 | 0.25663253 | 4.0188356 | 5.85E-05 | 0.01226266 |
| CTPS2 | 571.548817 | -0.2935004 | 0.17430321 | -1.6838495 | 0.09221076 | 0.5058189 |
| SYAP1 | 519.2376 | 0.11950632 | 0.14456669 | 0.82665187 | 4.08E-01 | 0.80588056 |
| TXLNG | 362.424644 | -0.0394424 | 0.26301611 | -0.149962 | 0.88079462 | 0.97664137 |
| RBBP7 | 418.364895 | 0.25713264 | 0.17357514 | 1.48139091 | 0.13850244 | 0.58206378 |
| REPS2 | 305.603554 | -0.2511641 | 0.3702482 | -0.6783668 | 0.49753913 | 0.85092921 |
| NHS | 8926.37083 | -0.0192017 | 0.48054257 | -0.0399583 | 0.96812639 | 0.99248831 |
| NHS-AS1 | 47.7209454 | 0.28066183 | 0.51513273 | 0.544834 | 0.58586769 | 0.88523522 |
| SCML1 | 572.416724 | 0.65918734 | 0.32371885 | 2.03629581 | 0.04172066 | 0.37639126 |
| RAI2 | 230.325335 | 0.5767511 | 0.23048011 | 2.5023899 | 1.23E-02 | 0.22749396 |
| SCML2 | 66.208424 | 0.3655274 | 0.54836688 | 0.66657453 | 0.50504394 | 0.85582573 |
| CDKL5 | 1017.1716 | -0.3395844 | 0.25754093 | -1.3185646 | 0.18731473 | 0.64196349 |
| PHKA2 | 361.324899 | -0.2725671 | 0.17973053 | -1.5165321 | 0.12938485 | 0.57039829 |
| ADGRG2 | 61.0814945 | -0.0308094 | 0.58707928 | -0.0524791 | 0.95814694 | 0.99069768 |
| PDHA1 | 740.325722 | 0.78082185 | 0.31068244 | 2.51324742 | 0.01196254 | 0.22380303 |
| MAP3K15 | 735.471667 | 0.41083952 | 0.36966199 | 1.11139239 | 0.26639949 | 0.71641758 |
| SH3KBP1 | 163.129529 | 0.02312372 | 0.3252018 | 0.07110575 | 0.94331359 | 0.98772472 |
| BCLAF3 | 780.44802 | -0.2490971 | 0.25570396 | -0.9741622 | 0.32997603 | 0.75885186 |
| EIF1AX | 605.557426 | 0.47476987 | 0.25460115 | 1.86475933 | 0.06221515 | 0.43419765 |
| RPS6KA3 | 1040.16225 | -0.2936821 | 0.21740977 | -1.3508232 | 0.17675208 | 0.63015338 |
| MBTPS2 | 452.909104 | -0.4298064 | 0.19901186 | -2.1597025 | 0.03079571 | 0.33646028 |
| SMS | 1400.25108 | -0.6567498 | 0.18861691 | -3.481924 | 0.00049782 | 0.03839746 |
| PHEX | 116.160481 | -3.2645159 | 0.65379347 | -4.9931913 | 5.94E-07 | 0.00078448 |
| PTCHD1-AS | 396.302828 | -0.6047009 | 0.68818982 | -0.8786834 | 0.37957298 | 0.78762967 |
| PRDX4 | 116.376748 | 0.39008728 | 0.37370872 | 1.04382707 | 0.29656541 | 0.73672107 |
| ACOT9 | 293.177088 | -0.6027597 | 0.33927216 | -1.776626 | 0.07562979 | 0.47345685 |
| SAT1 | 7840.22703 | 0.10830928 | 0.42817718 | 0.25295436 | 8.00E-01 | 9.54E-01 |
| APOO | 2057.46872 | 0.36130598 | 0.35430332 | 1.01976457 | 0.30784013 | 0.74370953 |
| KLHL15 | 120.945569 | 0.39765708 | 0.43950504 | 0.90478389 | 0.36557989 | 0.78095617 |
| EIF2S3 | 530.983404 | -0.2368784 | 0.20689259 | -1.1449342 | 0.25223641 | 0.70276117 |
| ZFX | 890.347139 | -0.197272 | 0.33381399 | -0.5909638 | 0.55454465 | 0.87160642 |
| PDK3 | 586.264566 | -0.4939608 | 0.44272704 | -1.1157231 | 0.26454066 | 0.71488591 |
| POLA1 | 748.713466 | -0.5846893 | 0.24846348 | -2.3532202 | 0.01861161 | 0.27498967 |
| IL1RAPL1 | 78.2552679 | -0.2220872 | 0.41089164 | -0.5405007 | 0.58885181 | 0.88571413 |
| GK | 452.87037 | -0.3848853 | 0.18567217 | -2.0729296 | 0.03817883 | 0.36650015 |
| GK-AS1 | 44.735881 | -0.0536078 | 0.52345114 | -0.1024121 | 0.91842955 | 0.98215155 |
| TAB3 | 723.809938 | 0.29490901 | 0.17251162 | 1.70950227 | 0.08735795 | 0.49566631 |
| DMD | 727.619062 | -0.2168432 | 0.27262927 | -0.7953777 | 0.42639382 | 0.81466629 |
| TMEM47 | 422.920925 | 1.2323783 | 0.40258773 | 3.0611422 | 0.00220494 | 0.09035145 |
| CFAP47 | 197.299555 | 0.30452598 | 0.58778332 | 0.51809225 | 0.60439391 | 0.89013087 |
| AF241726.2 | 661.893924 | 0.2208926 | 0.41699111 | 0.52972978 | 0.5962993 | 0.88859628 |
| DYNLT3 | 247.600909 | -0.4493148 | 0.53238372 | -0.8439679 | 0.39868733 | 0.80148305 |
| RPGR | 469.502927 | -0.5917798 | 0.23896992 | -2.4763776 | 0.01327231 | 0.23659106 |
| MID1IP1 | 125.323303 | 0.59442296 | 0.34896422 | 1.70339228 | 0.08849468 | 0.49769182 |
| BCOR | 793.51138 | 0.12852463 | 0.34573921 | 0.37173864 | 0.71008745 | 0.92610043 |
| ATP6AP2 | 403.840227 | 0.56028559 | 0.25342347 | 2.21086705 | 0.02704504 | 0.3178271 |
| CXorf38 | 255.137898 | -0.3135832 | 0.16788211 | -1.8678777 | 0.06177913 | 0.43264436 |
| MED14 | 1204.19317 | -0.5026572 | 0.20090042 | -2.5020216 | 0.01234864 | 0.22749396 |
| USP9X | 2072.12209 | -0.1744334 | 0.21594238 | -0.8077776 | 0.41921862 | 0.8119441 |
| DDX3X | 955.930203 | 0.03476279 | 0.3446681 | 0.10085873 | 0.9196626 | 0.98215577 |
| CASK | 1477.46947 | 0.03952519 | 0.19478209 | 0.20292003 | 0.83919753 | 0.966178 |
| Z93403.1 | 147.200906 | 0.17276866 | 0.66492787 | 0.25983067 | 0.79499439 | 0.95272759 |

| | | | | | | |
|---|---|---|---|---|---|---|
| MAOA | 2222.82356 | 0.1665979 | 0.4543945 | 0.36663714 | 0.7138897 | 0.92760608 |
| MAOB | 611.295844 | -0.0997719 | 0.38148363 | -0.2615366 | 0.7936787 | 0.95224951 |
| EFHC2 | 408.491221 | 0.64212935 | 0.48720228 | 1.31799334 | 0.18750588 | 0.64200436 |
| FUNDC1 | 94.5532053 | 0.36675627 | 0.2954079 | 1.24152492 | 0.2144119 | 0.66827012 |
| KDM6A | 2955.42299 | -0.2038339 | 0.43563345 | -0.4679024 | 0.63985442 | 0.90330608 |
| MIR222HG | 724.947341 | -0.6051684 | 0.54292055 | -1.1146538 | 0.26499881 | 0.71512755 |
| KRBOX4 | 449.471331 | 0.07762461 | 0.17026061 | 0.45591641 | 0.64845009 | 0.90754156 |
| ZNF674 | 132.455139 | 0.19162356 | 0.19780883 | 0.96873108 | 0.33267938 | 0.7610305 |
| SLC9A7 | 552.796221 | -0.4862791 | 0.28647327 | -1.6974675 | 0.08960831 | 0.50007863 |
| RP2 | 138.498987 | -0.3118129 | 0.28046006 | -1.1117906 | 0.26622819 | 0.71641758 |
| JADE3 | 547.174044 | -0.2885904 | 0.25417544 | -1.1353984 | 0.25620843 | 0.70709713 |
| RGN | 383.735114 | 1.65880371 | 0.72615454 | 2.28436731 | 0.02234995 | 0.29654015 |
| NDUFB11 | 349.961497 | 0.96203997 | 0.51093071 | 1.88291673 | 0.05971165 | 0.42657179 |
| RBM10 | 465.875936 | 0.12463906 | 0.18788273 | 0.66338756 | 0.50708237 | 0.85626196 |
| UBA1 | 832.018398 | 0.15276891 | 0.22608122 | 0.67572578 | 0.49921476 | 0.8519545 |
| CDK16 | 255.37914 | 0.08571731 | 0.25993472 | 0.32976475 | 0.74157772 | 0.93598004 |
| USP11 | 303.350608 | 0.18776725 | 0.23477819 | 0.79976444 | 0.42384729 | 0.81413339 |
| ZNF41 | 255.002776 | -0.040103 | 0.18392658 | -0.2180382 | 0.82739935 | 0.96249389 |
| ARAF | 201.304803 | 0.07161715 | 0.32214363 | 0.22231434 | 0.82406919 | 0.96141406 |
| TIMP1 | 540.086293 | 0.09324584 | 0.48603238 | 0.19185109 | 8.48E-01 | 9.70E-01 |
| ELK1 | 105.796903 | 0.29981247 | 0.20454793 | 1.4657321 | 0.14272129 | 0.58824095 |
| UXT | 141.789219 | -0.1459924 | 0.24279971 | -0.6012875 | 0.54764854 | 0.86859163 |
| ZNF81 | 553.617058 | -0.1137164 | 0.20787388 | -0.547045 | 0.58434784 | 0.88416261 |
| ZNF182 | 113.021245 | -0.011749 | 0.21863628 | -0.0537377 | 0.95714414 | 0.99065996 |
| ZNF630 | 206.914691 | 0.28812109 | 0.33109596 | 0.87020421 | 0.38418882 | 0.791697 |
| FTSJ1 | 58.6742354 | -0.0650335 | 0.32342709 | -0.2010763 | 0.8406389 | 0.96617873 |
| PORCN | 37.8248813 | 0.10586974 | 0.42614219 | 0.2484376 | 0.80379585 | 0.95450237 |
| EBP | 169.620477 | 0.24726854 | 0.40113742 | 0.61641854 | 0.53761832 | 0.86489441 |
| TBC1D25 | 172.613869 | -0.2105363 | 0.25046802 | -0.8405716 | 0.40058797 | 0.80243544 |
| RBM3 | 368.445186 | 0.1211965 | 0.28786909 | 0.42101256 | 0.67374591 | 0.91636338 |
| WDR13 | 327.537295 | -0.0386768 | 0.27663661 | -0.1398107 | 0.88880958 | 0.97729111 |
| AC231533.1 | 47.5623124 | -0.7759756 | 0.41432017 | -1.8728889 | 0.06108373 | 0.43176766 |
| HDAC6 | 245.587094 | -0.2747131 | 0.26022921 | -1.0556582 | 0.29112442 | 0.73330711 |
| PCSK1N | 772.519498 | 0.93021971 | 0.55911932 | 1.66372307 | 0.09616778 | 0.51179704 |
| TIMM17B | 104.333269 | 0.39540465 | 0.32990672 | 1.19853468 | 0.23070893 | 0.68297496 |
| PQBP1 | 74.3426484 | -0.0701988 | 0.47772258 | -0.1469447 | 0.8831757 | 0.97664137 |
| SLC35A2 | 62.1438878 | 0.17807222 | 0.32377215 | 0.54999238 | 0.5823246 | 0.88338315 |
| OTUD5 | 491.039942 | -0.1757897 | 0.20901448 | -0.841041 | 0.40032498 | 0.80238662 |
| GRIPAP1 | 432.782379 | -0.4021315 | 0.22042001 | -1.8243875 | 0.06809351 | 0.45260654 |
| TFE3 | 238.958323 | 0.0267018 | 0.29566484 | 0.09031105 | 0.92804004 | 0.98297055 |
| CCDC120 | 147.597883 | -0.6031196 | 0.27398126 | -2.2013168 | 0.0277136 | 0.32139504 |
| WDR45 | 326.570395 | -0.2133341 | 0.22345497 | -0.9547072 | 0.33972576 | 0.76449297 |
| GPKOW | 63.9514137 | 0.08914247 | 0.25818947 | 0.34525988 | 0.729899 | 0.93453345 |
| MAGIX | 78.851389 | 0.32202434 | 0.3355003 | 0.95983324 | 0.33713915 | 0.76409325 |
| PLP2 | 141.616677 | 0.56619798 | 0.34646714 | 1.63420398 | 1.02E-01 | 5.21E-01 |
| PRICKLE3 | 53.0266643 | -0.271174 | 0.34202619 | -0.7928456 | 0.42786779 | 0.81466629 |
| SYP | 68.2964092 | -0.027723 | 0.53233956 | -0.0520777 | 0.95846675 | 0.99071816 |
| CCDC22 | 78.7736389 | -0.0384888 | 0.35760786 | -0.1076286 | 0.91429033 | 0.98145965 |
| PPP1R3F | 84.3998901 | -0.1656521 | 0.27567065 | -0.6009058 | 0.5479027 | 0.86859163 |
| CLCN5 | 5924.59299 | 0.95297899 | 0.3899975 | 2.44355153 | 0.01454349 | 0.24762526 |
| CCNB3 | 55.3548498 | -1.166181 | 0.48939279 | -2.3829142 | 0.0171762 | 0.26289734 |
| SHROOM4 | 1387.69809 | -0.4037043 | 0.31518869 | -1.2808337 | 0.2002521 | 0.65322946 |
| AL158055.1 | 74.6122823 | 1.80418942 | 0.62934013 | 2.86679545 | 0.00414651 | 0.12562211 |
| GSPT2 | 49.3844258 | 1.25498163 | 0.37717775 | 3.32729496 | 0.00087693 | 0.05165902 |

| | | | | | | |
|---|---|---|---|---|---|---|
| MAGED1 | 2100.01626 | 0.30919723 | 0.20775724 | 1.48826215 | 0.13668177 | 0.57847791 |
| FAM156A | 151.258678 | -0.2066523 | 0.23331855 | -0.8857087 | 0.37577451 | 0.78490077 |
| GPR173 | 67.5473102 | 0.21038344 | 0.64459176 | 0.32638245 | 0.74413501 | 0.93656783 |
| TSPYL2 | 175.224294 | -0.5662028 | 0.44747193 | -1.265337 | 0.20575055 | 0.65813563 |
| KANTR | 1052.15956 | 0.12123449 | 0.28540785 | 0.4247763 | 0.67099976 | 0.91524533 |
| KDM5C | 419.965581 | -0.3637101 | 0.29632494 | -1.2274028 | 0.21967123 | 0.67289974 |
| IQSEC2 | 155.947626 | 0.24518147 | 0.43206919 | 0.56745882 | 0.5704025 | 0.87824299 |
| SMC1A | 421.724597 | -0.3438283 | 0.26072426 | -1.3187429 | 0.18725507 | 0.64195489 |
| HSD17B10 | 153.682375 | 1.13756237 | 0.45382977 | 2.50658381 | 0.01219042 | 0.22679323 |
| HUWE1 | 1907.63918 | -0.1249666 | 0.14097666 | -0.8864348 | 0.37538328 | 0.78468709 |
| PHF8 | 1112.99299 | -0.5256602 | 0.41833184 | -1.2565627 | 0.20891202 | 0.6614379 |
| FAM120C | 414.407507 | -0.1817902 | 0.15361115 | -1.1834441 | 0.2366332 | 0.68878143 |
| WNK3 | 369.103622 | 0.32622679 | 0.2466269 | 1.32275427 | 0.18591711 | 0.64033103 |
| TSR2 | 93.4391895 | 0.66279712 | 0.28002894 | 2.36688798 | 0.01793836 | 0.26845799 |
| FGD1 | 131.863373 | -0.3618577 | 0.23459332 | -1.5424894 | 0.12295471 | 0.55947129 |
| GNL3L | 157.432274 | -0.1831395 | 0.30890446 | -0.5928678 | 0.55326962 | 0.87104567 |
| MAGED2 | 289.292349 | 0.91551407 | 0.33927685 | 2.69842777 | 0.00696679 | 0.17136735 |
| TRO | 40.3329378 | 0.43836865 | 0.68279992 | 0.64201626 | 0.52086262 | 0.85943874 |
| APEX2 | 55.1889937 | 0.31050483 | 0.42339604 | 0.73336734 | 0.46333441 | 0.83531269 |
| FAM104B | 96.7056751 | 0.21875114 | 0.3930988 | 0.55647878 | 0.5778836 | 0.88191134 |
| MAGEH1 | 46.8866375 | 0.61405415 | 0.58760917 | 1.04500438 | 0.29602094 | 0.7366786 |
| USP51 | 46.282563 | 0.64917013 | 0.59834249 | 1.0849474 | 0.27794495 | 0.72372365 |
| RRAGB | 730.241737 | 0.03527549 | 0.1658471 | 0.21269889 | 0.83156183 | 0.9639242 |
| AL445472.1 | 750.244137 | -0.1832502 | 0.21739222 | -0.8429472 | 0.39925796 | 0.80174499 |
| KLF8 | 106.305381 | 1.26503839 | 0.30465353 | 4.1523838 | 3.29E-05 | 0.00869231 |
| UBQLN2 | 207.064492 | 0.40202096 | 0.25237933 | 1.59292348 | 0.11117736 | 0.53461826 |
| AL354793.1 | 45.4095951 | -0.4286637 | 0.44689597 | -0.9592024 | 0.33745681 | 0.76409325 |
| NBDY | 946.960942 | 0.08548322 | 0.25537413 | 0.33473718 | 0.73782335 | 0.93552616 |
| SPIN3 | 276.614797 | -0.1072606 | 0.24048489 | -0.4460181 | 0.65558414 | 0.90919768 |
| FAAH2 | 2384.26541 | 0.2340606 | 0.3455107 | 0.67743373 | 0.49813081 | 0.85092921 |
| ZXDB | 47.9026787 | 0.45250884 | 0.35107311 | 1.28893049 | 0.19742225 | 0.64964978 |
| ARHGEF9 | 643.226955 | -0.2383971 | 0.29260313 | -0.8147457 | 0.4152179 | 0.80966301 |
| AMER1 | 40.4740629 | 0.28381019 | 0.47520886 | 0.59723253 | 0.55035214 | 0.86959143 |
| MTMR8 | 331.80803 | 0.1557617 | 0.20701324 | 0.75242387 | 0.4517962 | 0.82874267 |
| ZC4H2 | 761.825031 | 0.22276099 | 0.21501457 | 1.03602741 | 0.30018935 | 0.73680045 |
| ZC3H12B | 306.926426 | 0.17512801 | 0.21800877 | 0.80330717 | 0.42179721 | 0.81266383 |
| LAS1L | 132.855396 | -0.0181307 | 0.32744145 | -0.0553707 | 0.95584316 | 0.99065996 |
| MSN | 1361.95631 | 0.23517602 | 0.31880254 | 0.73768552 | 4.61E-01 | 0.83439198 |
| OPHN1 | 1309.74823 | -0.2799165 | 0.19950563 | -1.4030504 | 0.16060181 | 0.61166537 |
| YIPF6 | 450.543442 | 0.01808292 | 0.15999888 | 0.11301902 | 0.91001547 | 0.98069436 |
| STARD8 | 236.295795 | 0.02153695 | 0.36799704 | 0.0585248 | 0.95333061 | 0.98950524 |
| PJA1 | 42.6477988 | 0.30089467 | 0.45764766 | 0.65748107 | 0.51087164 | 0.85762435 |
| EDA | 1094.1319 | -0.5403751 | 0.28266643 | -1.911706 | 0.05591391 | 0.41843207 |
| IGBP1 | 416.756236 | 0.03166847 | 0.21848681 | 0.14494452 | 0.88475468 | 0.97664137 |
| PDZD11 | 48.3708223 | 0.35969004 | 0.3808953 | 0.94432785 | 0.34500214 | 0.76724655 |
| DLG3 | 831.610488 | -0.0060431 | 0.24537725 | -0.024628 | 0.98035171 | 0.99518016 |
| SNX12 | 114.636085 | 0.18487221 | 0.27682244 | 0.6678368 | 0.50423777 | 0.85525037 |
| MED12 | 122.332763 | -0.4133071 | 0.21531064 | -1.9195852 | 0.05491032 | 0.41684503 |
| ZMYM3 | 139.613798 | 0.03068587 | 0.37385097 | 0.08208049 | 0.93458271 | 0.98522769 |
| NONO | 461.352417 | -0.0620441 | 0.26136374 | -0.2373859 | 0.81235746 | 0.95806331 |
| TAF1 | 1066.97791 | -0.4255744 | 0.23875932 | -1.7824408 | 0.07467737 | 0.47190267 |
| OGT | 1680.11529 | -0.2944128 | 0.36078039 | -0.8160443 | 0.41447483 | 0.80940242 |
| GCNA | 230.494125 | 0.06133605 | 0.39955705 | 0.1535101 | 0.87799602 | 0.9765717 |
| NHSL2 | 122.452665 | 0.35202121 | 0.43130232 | 0.81618203 | 0.41439603 | 0.80936821 |

| | | | | | | |
|---|---|---|---|---|---|---|
| PIN4 | 367.68746 | -0.3817811 | 0.31365348 | -1.2172066 | 0.22352564 | 0.67624946 |
| RPS4X | 1160.40735 | 1.06673974 | 0.42987168 | 2.4815306 | 0.01308195 | 0.23485956 |
| HDAC8 | 2952.92779 | -0.0750703 | 0.22885168 | -0.3280306 | 0.74288854 | 0.93598004 |
| PHKA1 | 212.921149 | 0.69308792 | 0.223262 | 3.10437035 | 0.00190684 | 0.08480644 |
| NAP1L2 | 45.2536729 | 1.53976871 | 0.55396777 | 2.77952762 | 0.0054438 | 0.149384 |
| CHIC1 | 994.595162 | -0.0732883 | 0.24562676 | -0.2983728 | 0.76541866 | 0.94270299 |
| TSIX | 100.418882 | -0.5003394 | 2.79612094 | -0.1789406 | 0.85798438 | 0.97143431 |
| XIST | 3707.09931 | 0.17161108 | 4.22869357 | 0.04058253 | 0.96762871 | 0.99248831 |
| JPX | 2612.09954 | 0.02509905 | 0.37409227 | 0.06709322 | 0.94650749 | 0.98885769 |
| FTX | 17205.9332 | 0.12467511 | 0.24269782 | 0.51370513 | 0.60745817 | 0.89174427 |
| SLC16A2 | 316.322947 | -0.2090432 | 0.32590718 | -0.6414196 | 0.52125013 | 0.85943874 |
| RLIM | 372.346629 | 0.31756906 | 0.20309293 | 1.56366377 | 0.11789655 | 0.54888439 |
| NEXMIF | 1202.85644 | 0.44101374 | 0.26705739 | 1.65138189 | 0.09866062 | 0.51542775 |
| ABCB7 | 1004.53343 | -0.0036932 | 0.20511316 | -0.0180055 | 0.98563444 | 0.99594729 |
| UPRT | 217.489297 | 0.11723757 | 0.18277576 | 0.64142845 | 0.52124436 | 0.85943874 |
| ZDHHC15 | 262.999776 | 0.00954158 | 0.28632126 | 0.03332473 | 0.97341564 | 0.99327086 |
| PBDC1 | 90.7942957 | 0.27041436 | 0.39617356 | 0.68256538 | 0.49488152 | 0.84899096 |
| ATRX | 3764.42049 | -0.0388499 | 0.19257951 | -0.2017342 | 0.84012449 | 0.96617873 |
| MAGT1 | 580.676459 | 0.52568505 | 0.21393405 | 2.45722943 | 0.01400132 | 0.24537335 |
| COX7B | 783.226211 | 0.71940959 | 0.41000403 | 1.75464029 | 0.07932086 | 0.48169281 |
| PGK1 | 1603.04953 | 0.43393916 | 0.50636617 | 0.85696712 | 0.39146306 | 0.7973532 |
| TAF9B | 61.3013359 | -0.2427567 | 0.41463084 | -0.5854768 | 0.55822716 | 0.87291524 |
| TENT5D | 48.1068536 | -0.3544152 | 0.45496445 | -0.7789954 | 0.43598242 | 0.8196544 |
| BRWD3 | 1273.78031 | -0.1616036 | 0.17148096 | -0.9423998 | 0.34598796 | 0.76822239 |
| HMGN5 | 177.88125 | 0.13878178 | 0.2049113 | 0.67727733 | 0.49823001 | 0.85092921 |
| SH3BGRL | 876.522256 | 0.3406426 | 0.22821407 | 1.49264502 | 0.13553013 | 0.57709645 |
| RPS6KA6 | 1850.06498 | 0.16746719 | 0.31486866 | 0.53186362 | 0.59482045 | 0.8885412 |
| HDX | 444.053255 | -0.1307154 | 0.30452887 | -0.4292382 | 0.66774988 | 0.9137375 |
| APOOL | 646.607143 | 0.0660128 | 0.13937229 | 0.47364364 | 0.63575405 | 0.90207488 |
| SATL1 | 110.643307 | 0.69004214 | 0.35192868 | 1.96074424 | 4.99E-02 | 4.00E-01 |
| ZNF711 | 338.499855 | 0.43422291 | 0.23650402 | 1.83600649 | 6.64E-02 | 4.47E-01 |
| POF1B | 170.600976 | -0.2832268 | 0.59196528 | -0.4784517 | 0.63232874 | 0.90044163 |
| CHM | 1109.96749 | -0.0722795 | 0.212887 | -0.3395203 | 0.73421783 | 0.93552616 |
| PCDH11X | 82.8706711 | 0.60508429 | 0.41818881 | 1.44691649 | 0.14792031 | 0.59424557 |
| DIAPH2 | 1173.74656 | -0.5325012 | 0.45227135 | -1.1773932 | 0.23903861 | 0.69029619 |
| DIAPH2-AS1 | 117.120463 | -0.3603227 | 0.74257354 | -0.4852351 | 0.62750961 | 0.89919293 |
| PCDH19 | 43.8586654 | 0.19833688 | 0.52377927 | 0.37866501 | 0.70493664 | 0.92532128 |
| TSPAN6 | 180.01124 | 0.3638461 | 0.28163695 | 1.29189758 | 0.19639262 | 0.64964978 |
| SRPX2 | 164.894615 | 0.34870921 | 0.5137323 | 0.67877611 | 0.49727974 | 0.85092921 |
| SYTL4 | 980.408138 | 0.05677343 | 0.29883019 | 0.18998558 | 8.49E-01 | 9.70E-01 |
| CSTF2 | 46.8117967 | -0.6557856 | 0.41041487 | -1.5978603 | 0.11007407 | 0.53372881 |
| TRMT2B | 440.866422 | 0.00435312 | 0.28114405 | 0.0154836 | 0.98764637 | 0.99653761 |
| GLA | 59.8887649 | -0.7572678 | 0.38738372 | -1.9548261 | 0.0506036 | 0.40363702 |
| HNRNPH2 | 173.295676 | 0.7474365 | 0.37149326 | 2.01197864 | 0.04422219 | 0.38531062 |
| ARMCX4 | 1636.27533 | -0.1220087 | 0.41531806 | -0.2937717 | 0.76893235 | 0.94376765 |
| ARMCX1 | 69.5181143 | 0.18519999 | 0.33255307 | 0.55690356 | 0.57759332 | 0.88181117 |
| ARMCX6 | 95.4197732 | 0.1077222 | 0.29267676 | 0.36805861 | 0.71282953 | 0.92760608 |
| ARMCX3 | 341.834206 | 0.20698759 | 0.20318459 | 1.01871694 | 0.30833737 | 0.74375973 |
| ARMCX2 | 165.662317 | 0.14283151 | 0.31739834 | 0.4500071 | 0.65270532 | 0.90868303 |
| ZMAT1 | 1418.09302 | 0.12236188 | 0.37165093 | 0.32923873 | 0.74197525 | 0.93598004 |
| TCEAL2 | 53.6257854 | 2.38156992 | 0.60998544 | 3.90430619 | 9.45E-05 | 0.01540986 |
| BEX5 | 46.709883 | 1.06037802 | 0.52590693 | 2.01628456 | 0.04377023 | 0.38453712 |
| ARMCX5 | 85.9270178 | 0.5429908 | 0.26467425 | 2.05154372 | 0.04021403 | 0.37344844 |
| AL035427.1 | 112.049558 | 0.96830621 | 0.76016317 | 1.27381364 | 0.20272947 | 0.65552334 |

| | | | | | | |
|---|---|---|---|---|---|---|
| GPRASP1 | 82.6413437 | 0.87912175 | 0.39270096 | 2.23865447 | 0.02517841 | 0.31053369 |
| BHLHB9 | 160.809138 | 0.22294428 | 0.2015088 | 1.10637495 | 0.26856426 | 0.71818262 |
| LINC00630 | 612.302787 | -0.257734 | 0.21832929 | -1.180483 | 0.23780815 | 0.68982886 |
| Z68871.1 | 72.6675728 | -0.3199206 | 0.31273912 | -1.0229632 | 0.30632527 | 0.74295884 |
| BEX4 | 307.346366 | 0.40484613 | 0.24316352 | 1.66491309 | 0.0959301 | 0.5117083 |
| TCEAL8 | 122.695843 | 0.38411547 | 0.29847759 | 1.28691561 | 0.1981237 | 0.65051354 |
| BEX2 | 254.613763 | 0.91540871 | 0.35688086 | 2.56502605 | 0.01031681 | 0.20627111 |
| TCEAL9 | 331.839346 | 0.37178988 | 0.43569391 | 0.85332815 | 0.39347735 | 0.79843808 |
| BEX3 | 638.977535 | 0.36238113 | 0.33837705 | 1.07093887 | 0.28419692 | 0.72738535 |
| RAB40A | 51.0901855 | 0.74163381 | 0.3906987 | 1.89822441 | 0.05766653 | 0.42087381 |
| TCEAL4 | 385.961139 | 0.30028875 | 0.22394733 | 1.34089007 | 1.80E-01 | 0.63404661 |
| TCEAL3 | 204.700419 | 0.36825747 | 0.3420046 | 1.07676175 | 0.28158672 | 0.72604837 |
| TCEAL1 | 58.7564796 | -0.3412247 | 0.35937355 | -0.9494988 | 0.342367 | 0.76591028 |
| MORF4L2 | 936.374398 | 0.29249306 | 0.22963856 | 1.27371056 | 0.20276601 | 0.65552334 |
| MORF4L2-AS | 44.5979394 | -0.0404711 | 0.44109799 | -0.0917507 | 0.92689609 | 0.98252755 |
| SLC25A53 | 508.192834 | 0.27890158 | 0.19807438 | 1.40806485 | 0.15911187 | 0.60883797 |
| FAM199X | 675.915064 | 0.21686292 | 0.13861943 | 1.56444823 | 0.11771234 | 0.54883951 |
| IL1RAPL2 | 110.719988 | 0.79842549 | 0.48867279 | 1.63386524 | 0.10228715 | 0.52073581 |
| NRK | 2243.64241 | 0.51859964 | 0.29652125 | 1.74894595 | 8.03E-02 | 0.48389028 |
| CXorf57 | 212.726627 | 0.68124591 | 0.52118633 | 1.30710625 | 0.19117663 | 0.64571429 |
| RNF128 | 253.203103 | -1.7101315 | 0.67826669 | -2.5213261 | 0.01169134 | 0.22188357 |
| TBC1D8B | 448.027809 | 0.67237689 | 0.31885419 | 2.10872841 | 0.03496803 | 0.35358029 |
| MORC4 | 395.352668 | 0.76754424 | 0.29394757 | 2.61116039 | 0.00902356 | 0.19318015 |
| RBM41 | 861.825183 | 0.09708697 | 0.18423163 | 0.52698319 | 0.59820526 | 0.88883692 |
| NUP62CL | 194.037169 | 0.64562395 | 0.45594298 | 1.41601905 | 0.15676993 | 0.60654181 |
| PRPS1 | 71.5621844 | 0.46176635 | 0.36103307 | 1.279014 | 0.20089212 | 0.6542367 |
| TSC22D3 | 346.391236 | 0.86016643 | 0.52525022 | 1.63763175 | 0.10149854 | 0.51924638 |
| MID2 | 180.796551 | -0.1212271 | 0.31998039 | -0.378858 | 0.70479331 | 0.92528913 |
| PSMD10 | 57.278707 | 0.61305068 | 0.39532659 | 1.55074486 | 0.12096284 | 0.55498372 |
| ATG4A | 113.203554 | -0.0198278 | 0.25623539 | -0.077381 | 0.93832044 | 0.98696247 |
| COL4A6 | 159.533489 | -1.8176664 | 0.59648624 | -3.0472898 | 0.00230915 | 0.09214973 |
| COL4A5 | 1913.22092 | -0.9059127 | 0.4396754 | -2.0604126 | 0.03935912 | 0.37234059 |
| ACSL4 | 9408.76758 | 1.63422814 | 0.58362285 | 2.80014419 | 0.00510798 | 0.14350465 |
| TMEM164 | 1650.18257 | -0.0514749 | 0.30858802 | -0.1668079 | 0.86752121 | 0.97402505 |
| AMMECR1 | 1124.13766 | -0.2660842 | 0.20949683 | -1.2701107 | 0.20404521 | 0.65712971 |
| CAPN6 | 43.6947583 | -0.3190161 | 0.82175785 | -0.3882118 | 0.6978593 | 0.92341984 |
| ALG13 | 759.742383 | 0.18287664 | 0.27630983 | 0.66185354 | 0.50806509 | 0.85643442 |
| ALG13-AS1 | 68.0309573 | 0.64140664 | 0.5971258 | 1.07415663 | 0.28275249 | 0.72604837 |
| AMOT | 81.4600865 | 0.31115835 | 0.56842935 | 0.54740022 | 0.5841038 | 0.88416261 |
| LRCH2 | 128.013857 | 1.05025366 | 0.50804918 | 2.06722835 | 0.03871264 | 0.36916996 |
| PLS3-AS1 | 196.906872 | 1.91557481 | 0.4281812 | 4.47374802 | 7.69E-06 | 0.00350085 |
| PLS3 | 1228.58646 | 0.36910167 | 0.21895245 | 1.68576178 | 0.09184169 | 0.50476839 |
| DANT2 | 5262.89091 | -0.0793674 | 0.26452173 | -0.3000411 | 0.76414583 | 0.94256396 |
| KLHL13 | 6035.14239 | 0.28346331 | 0.29520755 | 0.96021698 | 0.33694602 | 0.76409325 |
| WDR44 | 706.937299 | -0.6118183 | 0.17109915 | -3.5758117 | 0.00034914 | 0.03115997 |
| DOCK11 | 469.956547 | -3.9735146 | 1.00256685 | -3.9633413 | 7.39E-05 | 0.01392432 |
| IL13RA1 | 605.816597 | -0.4265289 | 0.42816641 | -0.9961756 | 0.31916485 | 0.75094569 |
| LONRF3 | 247.902676 | -1.4457506 | 0.41044725 | -3.5223786 | 0.00042769 | 0.03553079 |
| PGRMC1 | 430.476827 | 0.6697513 | 0.38782856 | 1.72692618 | 0.08418092 | 0.49023622 |
| SLC25A43 | 471.100637 | 0.12692675 | 0.44742335 | 0.28368378 | 0.77665273 | 0.94694572 |
| SLC25A5 | 494.596833 | 0.87122999 | 0.52967421 | 1.64484123 | 0.10000256 | 0.51678168 |
| CXorf56 | 120.246723 | -0.4981316 | 0.2723659 | -1.8289058 | 0.06741373 | 0.45155573 |
| UBE2A | 301.039219 | 0.07013362 | 0.30576434 | 0.22937149 | 0.81858019 | 0.95992771 |
| NKRF | 170.698214 | 0.388478 | 0.21493946 | 1.80738339 | 0.07070252 | 0.4627897 |

| | | | | | | |
|---|---|---|---|---|---|---|
| 6-Sep | 1041.87631 | 0.39488953 | 0.34834109 | 1.13362892 | 0.25695024 | 0.70735159 |
| RPL39 | 47.5968822 | -0.3015876 | 0.51353869 | -0.5872733 | 0.55702014 | 0.87227967 |
| UPF3B | 675.477928 | -0.9815932 | 0.28077549 | -3.4960076 | 0.00047228 | 0.03734387 |
| NDUFA1 | 452.603341 | 0.58217439 | 0.36900027 | 1.57770725 | 0.11463288 | 0.54194192 |
| NKAP | 174.98007 | -0.1454148 | 0.2237146 | -0.6500013 | 0.51569139 | 0.85869987 |
| RHOXF1-AS1 | 250.607441 | 0.30780062 | 0.47604404 | 0.64658013 | 0.51790372 | 0.85893097 |
| ZBTB33 | 58.0251099 | 0.39659835 | 0.36914697 | 1.07436436 | 0.28265941 | 0.72604837 |
| LAMP2 | 517.837338 | 0.22108839 | 0.1540301 | 1.43535833 | 0.15118501 | 0.59898106 |
| CUL4B | 307.380375 | -0.3987202 | 0.16398397 | -2.4314581 | 0.01503819 | 0.25049106 |
| MCTS1 | 368.077264 | -0.1526742 | 0.21576837 | -0.7075837 | 0.4792038 | 0.84296219 |
| C1GALT1C1 | 88.3748447 | 0.29662581 | 0.30746019 | 0.96476169 | 0.33466419 | 0.76242642 |
| GRIA3 | 68.8021832 | -0.457609 | 0.64223044 | -0.7125308 | 0.47613612 | 0.84141268 |
| THOC2 | 1686.08901 | -0.3553028 | 0.14483063 | -2.4532296 | 0.01415799 | 0.24646911 |
| XIAP | 1230.83464 | 0.0878596 | 0.17332609 | 0.50690347 | 0.61222255 | 0.89440588 |
| STAG2 | 2062.36342 | -0.0239102 | 0.22308861 | -0.1071781 | 0.9146477 | 0.98156931 |
| TENM1 | 637.351103 | 1.17134328 | 0.5353055 | 2.1881772 | 0.0286567 | 0.32702837 |
| AL442647.1 | 107.278387 | 0.89743855 | 0.52827448 | 1.69881109 | 0.08935478 | 0.49927552 |
| SMARCA1 | 879.987878 | 0.19205154 | 0.21419999 | 0.89659919 | 0.36993283 | 0.78095617 |
| OCRL | 261.716084 | -0.3438692 | 0.2150424 | -1.5990764 | 0.10980363 | 0.53372881 |
| ZDHHC9 | 174.88478 | -0.3876769 | 0.31373046 | -1.2357006 | 2.17E-01 | 6.70E-01 |
| AL034405.1 | 52.9937009 | -0.1742052 | 0.44214771 | -0.3939978 | 0.69358269 | 0.92229578 |
| UTP14A | 124.997153 | -0.3570515 | 0.33234204 | -1.0743496 | 0.28266602 | 0.72604837 |
| BCORL1 | 151.697999 | 0.39523889 | 0.36380048 | 1.08641664 | 0.2772947 | 0.72344178 |
| ELF4 | 121.354826 | 0.33336342 | 0.29632732 | 1.12498376 | 0.26059592 | 0.71055875 |
| AIFM1 | 287.500262 | 0.52364859 | 0.20632365 | 2.53799602 | 0.01114893 | 0.21567475 |
| ZNF280C | 185.709938 | -0.120854 | 0.20127794 | -0.6004334 | 0.54821745 | 0.86869054 |
| SLC25A14 | 157.967821 | -0.0746591 | 0.20783914 | -0.3592159 | 0.71943355 | 0.93001079 |
| RBMX2 | 97.5983937 | 0.2838141 | 0.24667052 | 1.15057973 | 0.24990518 | 0.7004026 |
| ENOX2 | 718.362648 | -0.5820049 | 0.24325141 | -2.3926067 | 0.01672916 | 0.25905688 |
| STK26 | 111.395757 | 0.68267498 | 0.47990169 | 1.4225309 | 0.15487219 | 0.60380955 |
| RAP2C | 748.198449 | 0.76756651 | 0.38881422 | 1.97412151 | 4.84E-02 | 3.96E-01 |
| RAP2C-AS1 | 1396.25168 | -0.3244615 | 0.31798132 | -1.0203791 | 0.30754871 | 0.74348662 |
| MBNL3 | 710.930531 | 0.03193708 | 0.32000657 | 0.09980133 | 0.92050205 | 0.98215577 |
| HS6ST2 | 4185.52505 | -0.2605932 | 0.21518052 | -1.2110447 | 0.22587828 | 0.67809183 |
| GPC4 | 950.169463 | -0.540694 | 0.31593925 | -1.711386 | 0.08700988 | 0.49496705 |
| GPC3 | 1522.50275 | -0.53514 | 0.50723864 | -1.0550064 | 0.2914224 | 0.73349819 |
| CCDC160 | 143.246338 | 0.81940627 | 0.34611565 | 2.36743492 | 0.01791187 | 0.26845799 |
| PHF6 | 434.551107 | 0.15304278 | 0.22251983 | 0.68777143 | 0.49159673 | 0.84704258 |
| HPRT1 | 1005.39196 | 0.61150188 | 0.26401205 | 2.31618932 | 0.02054793 | 0.28691085 |
| MIR503HG | 369.251479 | 0.63616623 | 0.5849003 | 1.087649 | 0.27675009 | 0.72273466 |
| FAM122B | 199.450482 | -0.053508 | 0.25432589 | -0.2103914 | 0.83336219 | 0.96496852 |
| FAM122C | 164.317287 | 0.10565483 | 0.30036209 | 0.35175821 | 0.7250196 | 0.93241479 |
| MOSPD1 | 103.870027 | -0.035881 | 0.26488302 | -0.1354597 | 0.89224845 | 0.97771356 |
| RTL8C | 257.992366 | 0.41630734 | 0.45884508 | 0.9072939 | 0.3642514 | 0.78031086 |
| RTL8A | 149.67991 | 0.60765583 | 0.30975555 | 1.96172702 | 4.98E-02 | 0.40017381 |
| ZNF75D | 711.044971 | -0.4215633 | 0.31165169 | -1.3526746 | 0.1761596 | 0.62957039 |
| ZNF449 | 124.615104 | 0.11904802 | 0.26010914 | 0.45768488 | 0.64717885 | 0.9065308 |
| INTS6L | 245.67964 | -0.1915431 | 0.2651699 | -0.7223412 | 0.47008473 | 0.83921324 |
| MMGT1 | 102.314288 | 0.3643529 | 0.31067764 | 1.17276832 | 0.24088873 | 0.69178991 |
| SLC9A6 | 205.921139 | -0.139104 | 0.25220525 | -0.5515507 | 0.58125624 | 0.88290096 |
| MAP7D3 | 51.9968144 | 0.73140142 | 0.38587798 | 1.89542154 | 0.05803658 | 0.42087381 |
| HTATSF1 | 186.418709 | 0.40829673 | 0.26118669 | 1.5632371 | 0.11799684 | 0.54888439 |
| ARHGEF6 | 48.4339527 | -0.1528496 | 0.38159355 | -0.400556 | 0.68874705 | 0.92151057 |
| AL683813.1 | 268.429664 | -0.2384465 | 0.47268754 | -0.5044485 | 0.61394627 | 0.89477643 |

| | | | | | | |
|---|---|---|---|---|---|---|
| RBMX | 1376.6609 | -0.0349451 | 0.13575661 | -0.2574096 | 7.97E-01 | 0.95281098 |
| FGF13 | 3505.65567 | 1.14638047 | 0.72448435 | 1.58233985 | 0.11357201 | 0.53989968 |
| ATP11C | 538.681337 | -0.0565919 | 0.2204542 | -0.2567057 | 0.79740594 | 0.95281098 |
| LDOC1 | 58.8166202 | 0.97278779 | 0.49262772 | 1.97469152 | 0.04830317 | 0.39627811 |
| SLITRK4 | 179.551057 | 0.70915171 | 0.46879036 | 1.51272672 | 0.13034909 | 0.57221041 |
| AL589669.1 | 61.3071363 | 1.80051162 | 0.58657859 | 3.06951474 | 0.00214407 | 0.08878056 |
| FMR1 | 934.347707 | 0.17756914 | 0.24706268 | 0.71872101 | 0.47231284 | 0.83921324 |
| IDS | 319.886374 | 0.25706069 | 0.25049211 | 1.02622273 | 0.30478661 | 0.74210624 |
| LINC00893 | 66.6922756 | -0.5379413 | 0.36476088 | -1.4747779 | 0.14027226 | 0.58521327 |
| TMEM185A | 244.834538 | 0.16036694 | 0.24759231 | 0.64770563 | 0.51717536 | 0.85893097 |
| LINC00894 | 1295.46923 | 0.02322324 | 0.4079675 | 0.05692424 | 0.95460554 | 0.99036951 |
| MAMLD1 | 79.981322 | -1.125346 | 0.46771627 | -2.4060442 | 0.01612631 | 0.25504618 |
| MTM1 | 630.289889 | -0.1811496 | 0.17447867 | -1.0382334 | 0.2991614 | 0.73673761 |
| MTMR1 | 579.954938 | -0.0529944 | 0.1549944 | -0.3419118 | 0.73241726 | 0.93552616 |
| CD99L2 | 407.901613 | -0.23043 | 0.20557856 | -1.1208853 | 0.2623367 | 0.71300523 |
| VMA21 | 152.577788 | 0.32152638 | 0.28659156 | 1.12189757 | 0.26190599 | 0.71271452 |
| CETN2 | 88.6391476 | 1.05823071 | 0.48583863 | 2.17815268 | 0.02939467 | 0.33012415 |
| NSDHL | 72.1261765 | 0.18454116 | 0.36515829 | 0.50537305 | 0.61329684 | 0.89441126 |
| ZNF185 | 103.669902 | 0.7969749 | 0.51063684 | 1.56074696 | 0.11858347 | 0.54998912 |
| ZNF275 | 79.3398308 | -0.5461373 | 0.37348827 | -1.4622608 | 0.14366974 | 0.58876618 |
| BGN | 46.12554 | -0.8672569 | 0.90033452 | -0.9632607 | 0.33541669 | 0.76322465 |
| CCNQ | 70.5446769 | 0.25382894 | 0.41220512 | 0.61578309 | 0.53803769 | 0.86496983 |
| DUSP9 | 529.544623 | 1.72771741 | 0.64679332 | 2.67120476 | 0.00755795 | 0.17483887 |
| SLC6A8 | 705.540557 | 0.27269278 | 0.4475012 | 0.60936771 | 0.54228074 | 0.86582694 |
| BCAP31 | 371.810612 | 0.5168419 | 0.37063651 | 1.39447111 | 0.16317539 | 0.61387934 |
| ABCD1 | 42.8193506 | -0.0409818 | 0.45852404 | -0.0893777 | 0.92878171 | 0.98323294 |
| IDH3G | 150.691182 | 0.21510834 | 0.35299368 | 0.60938298 | 0.54227062 | 0.86582694 |
| SSR4 | 219.914842 | 0.78898308 | 0.47750054 | 1.65231873 | 0.09846959 | 0.51525577 |
| ARHGAP4 | 106.103488 | -0.5405839 | 0.49481508 | -1.0924968 | 0.27461481 | 0.72164296 |
| HCFC1 | 149.419552 | 0.16579686 | 0.27909421 | 0.5940534 | 0.5524764 | 0.87096727 |
| TMEM187 | 54.3548931 | 0.4580085 | 0.4020444 | 1.1391988 | 0.25462024 | 0.70509116 |
| IRAK1 | 139.907935 | 0.15475693 | 0.49771654 | 0.31093386 | 0.7558509 | 0.93957997 |
| MECP2 | 1108.39932 | -0.2535372 | 0.17587889 | -1.441544 | 0.14943106 | 0.59625836 |
| FLNA | 450.593728 | 0.22440711 | 0.71522141 | 0.31375894 | 0.75370413 | 0.93868357 |
| EMD | 222.635277 | 0.6234861 | 0.37168646 | 1.67745173 | 0.09345418 | 0.50887217 |
| RPL10 | 1660.12011 | 0.82711559 | 0.4967792 | 1.66495618 | 0.0959215 | 0.5117083 |
| DNASE1L1 | 114.447264 | -0.5984784 | 0.31378546 | -1.9072853 | 0.05648366 | 0.41843207 |
| TAZ | 296.79571 | -0.4781071 | 0.25826447 | -1.8512305 | 0.0641364 | 0.43894277 |
| AC244090.1 | 42.1715153 | -0.0033571 | 0.39280979 | -0.0085464 | 0.99318107 | 0.99816851 |
| ATP6AP1 | 259.5389 | 0.79082621 | 0.39414048 | 2.00645774 | 0.04480743 | 0.38683751 |
| GDI1 | 231.927837 | -0.1272778 | 0.33065601 | -0.3849252 | 0.70029281 | 0.92367966 |
| FAM50A | 137.512276 | 0.01418335 | 0.3550221 | 0.03995062 | 0.9681325 | 0.99248831 |
| PLXNA3 | 111.690914 | -0.3108185 | 0.4241494 | -0.7328043 | 4.64E-01 | 0.83531269 |
| UBL4A | 61.7222794 | 0.3564904 | 0.36522181 | 0.97609287 | 0.32901844 | 0.75793593 |
| SLC10A3 | 45.7848288 | 0.43853374 | 0.48161646 | 0.91054557 | 0.36253486 | 0.77849113 |
| FAM3A | 169.472896 | -0.0213036 | 0.22844549 | -0.0932545 | 0.9257014 | 0.9822216 |
| G6PD | 96.3054115 | 0.59762552 | 0.60743094 | 0.98385756 | 0.32518555 | 0.7552048 |
| IKBKG | 134.339964 | -0.4259826 | 0.25911889 | -1.6439659 | 0.10018325 | 0.51692417 |
| DKC1 | 140.264111 | -0.0589443 | 0.27351579 | -0.2155061 | 0.82937276 | 0.96290106 |
| MPP1 | 83.8728813 | 0.53466126 | 0.36803327 | 1.45275251 | 0.14629248 | 0.593482 |
| F8 | 219.806426 | -0.1032616 | 0.28364174 | -0.3640564 | 0.71581587 | 0.92890921 |
| FUNDC2 | 290.680662 | 0.21477479 | 0.3051901 | 0.70374101 | 0.48159407 | 0.8437862 |
| MTCP1 | 119.871205 | 0.35829806 | 0.71834145 | 0.49878517 | 0.61793074 | 0.89599505 |
| BRCC3 | 431.509922 | -0.0160906 | 0.21625463 | -0.074406 | 0.94068737 | 0.98715467 |

| | | | | | | |
|---|---|---|---|---|---|---|
| VBP1 | 140.314334 | 0.06052104 | 0.25916953 | 0.23351912 | 0.81535833 | 0.95872901 |
| CLIC2 | 53.3973388 | 0.51649439 | 0.52772724 | 0.97871466 | 0.32772099 | 0.75633196 |
| TMLHE-AS1 | 368.129351 | 0.16225264 | 0.59700401 | 0.27177815 | 0.78579261 | 0.94978829 |
| TMLHE | 797.026241 | -0.0231568 | 0.23169348 | -0.099946 | 0.9203872 | 0.98215577 |
| VAMP7 | 348.009274 | 0.17211332 | 0.23181263 | 0.74246741 | 0.45780419 | 0.83282548 |
| ZNF596 | 211.149854 | -0.2287125 | 0.23852364 | -0.958867 | 0.33762574 | 0.76409325 |
| FBXO25 | 667.027234 | -0.36264 | 0.22709467 | -1.5968672 | 0.1102953 | 0.53372881 |
| TDRP | 1030.51597 | -0.4935289 | 0.28321232 | -1.7426112 | 0.08140156 | 0.48594425 |
| ERICH1 | 2433.58611 | -0.6103452 | 0.49830059 | -1.2248534 | 0.22063044 | 0.67345736 |
| DLGAP2 | 108.910792 | -1.9552169 | 0.7301615 | -2.6777869 | 0.00741104 | 0.17326084 |
| AC100810.1 | 58.5136362 | 0.05321324 | 0.56009178 | 0.09500807 | 0.92430842 | 0.98215577 |
| AC019257.8 | 49.2015391 | -1.078462 | 0.48702771 | -2.2143749 | 0.026803 | 0.31695684 |
| KBTBD11 | 336.412518 | 0.18523797 | 0.39867951 | 0.46462878 | 0.64219732 | 0.90454089 |
| MYOM2 | 49.2950852 | -0.8598044 | 0.50520885 | -1.701879 | 0.08877804 | 0.49850673 |
| AC246817.2 | 276.818407 | 0.25274088 | 0.49690901 | 0.50862608 | 0.61101435 | 0.89348926 |
| AC246817.1 | 611.227362 | -0.0433742 | 0.35111197 | -0.1235337 | 0.90168446 | 0.97941893 |
| CSMD1 | 467.39957 | 0.22318154 | 0.94112978 | 0.23714215 | 0.81254651 | 0.95806331 |
| MCPH1 | 1718.7373 | -0.094809 | 0.21725152 | -0.4364021 | 0.66254505 | 0.91110616 |
| ANGPT2 | 301.722583 | 0.42059553 | 0.53479182 | 0.7864659 | 0.43159459 | 0.81733805 |
| MCPH1-AS1 | 416.875239 | -0.069402 | 0.23766331 | -0.2920181 | 0.77027277 | 0.94400938 |
| AGPAT5 | 711.427617 | -0.5323055 | 0.48308773 | -1.1018817 | 0.27051311 | 0.71954751 |
| AF233439.1 | 1280.82535 | 1.47824978 | 0.86984235 | 1.69944563 | 0.08923525 | 0.49902981 |
| DEFB1 | 2895.09626 | 1.52964816 | 0.44961241 | 3.40214841 | 0.00066858 | 0.04534035 |
| FAM66B | 59.5971812 | 0.01869656 | 0.52283624 | 0.03575989 | 0.97147382 | 0.99327086 |
| FAM85B | 1022.96736 | 0.13189173 | 0.43142158 | 0.30571426 | 0.75982222 | 0.94133292 |
| PRAG1 | 629.97825 | 1.99054437 | 0.7913998 | 2.51521969 | 0.01189582 | 0.22355405 |
| MFHAS1 | 5403.10283 | 0.25436367 | 0.3131095 | 0.81237927 | 0.41657404 | 0.80966301 |
| ERI1 | 480.565207 | -0.0489863 | 0.36570777 | -0.1339493 | 0.89344269 | 0.97788747 |
| PPP1R3B | 210.143768 | 0.20056094 | 0.41149134 | 0.48740015 | 0.62597479 | 0.89873262 |
| AC022784.3 | 105.632134 | 0.75388976 | 0.60128831 | 1.25379081 | 0.20991805 | 0.66255855 |
| AC022784.1 | 38.0053268 | 0.92978899 | 0.66528887 | 1.39757185 | 1.62E-01 | 6.14E-01 |
| TNKS | 3113.57406 | -0.2696778 | 0.19553745 | -1.3791617 | 0.16784492 | 0.62227466 |
| MSRA | 1514.39342 | -0.4838087 | 0.22752518 | -2.1263965 | 0.03347026 | 0.34965509 |
| PINX1 | 46.1447649 | -0.3664903 | 0.49086405 | -0.7466227 | 0.45529133 | 0.83121393 |
| XKR6 | 2863.285 | 0.15045965 | 0.33867848 | 0.44425511 | 0.65685815 | 0.90986151 |
| MTMR9 | 469.572583 | -0.0775314 | 0.19917473 | -0.3892631 | 0.69708151 | 0.92322348 |
| FAM167A | 119.647276 | 1.23980998 | 0.68414581 | 1.81220138 | 0.06995509 | 0.45949593 |
| NEIL2 | 76.8912512 | 0.00854633 | 0.3928142 | 0.02175669 | 0.98264205 | 0.99518552 |
| FDFT1 | 866.218002 | -0.0273039 | 0.38221887 | -0.0714354 | 0.94305126 | 0.98772472 |
| CTSB | 564.974882 | 0.19094784 | 0.35217632 | 0.54219387 | 5.88E-01 | 0.88571413 |
| FAM66D | 437.416194 | -0.2113184 | 0.69102804 | -0.3058029 | 0.75975469 | 0.94133292 |
| AC145124.1 | 120.112717 | 0.3994364 | 0.61055076 | 0.65422307 | 0.5129681 | 0.85762435 |
| FAM66A | 51.913566 | 1.02063733 | 0.99270956 | 1.02813287 | 0.30388733 | 0.74120958 |
| LONRF1 | 598.273237 | -0.6329679 | 0.3579942 | -1.7680953 | 0.07704497 | 0.47689411 |
| TRMT9B | 2849.59551 | 0.05865425 | 0.29045523 | 0.20193904 | 0.83996438 | 0.96617873 |
| DLC1 | 508.241775 | -1.0322654 | 0.73037339 | -1.4133393 | 0.15755599 | 0.6073933 |
| SGCZ | 41.3803128 | -0.0876208 | 0.47564146 | -0.184216 | 0.85384403 | 0.97077171 |
| TUSC3 | 1482.18994 | -0.3297027 | 0.29951377 | -1.1007932 | 0.27098666 | 0.71977937 |
| MSR1 | 1264.90501 | 0.96551455 | 0.79917826 | 1.20813415 | 0.22699567 | 0.67921637 |
| AC011586.1 | 91.1757377 | 1.60742971 | 0.56109402 | 2.86481348 | 0.00417255 | 0.12579326 |
| AC068992.1 | 617.272899 | 0.87717967 | 0.81824077 | 1.07203124 | 0.283706 | 0.72690826 |
| MICU3 | 1121.08837 | 0.14934123 | 0.23939121 | 0.62383755 | 0.53273428 | 0.86397197 |
| AC079193.2 | 59.4292641 | 0.21554879 | 0.4232862 | 0.50922707 | 0.61059307 | 0.89339478 |
| ZDHHC2 | 1290.61446 | -0.0729807 | 0.2668791 | -0.2734598 | 0.78449976 | 0.94978829 |

| | | | | | | |
|---|---|---|---|---|---|---|
| CNOT7 | 385.38491 | -0.1132892 | 0.20446461 | -0.5540773 | 0.57952593 | 0.88224343 |
| VPS37A | 586.874595 | -0.3018996 | 0.23275153 | -1.2970897 | 0.19460031 | 0.64814994 |
| MTMR7 | 256.343947 | 0.0860758 | 0.59092536 | 0.14566273 | 0.88418764 | 0.97664137 |
| SLC7A2 | 379.228208 | -0.3578497 | 0.65383654 | -0.5473076 | 0.58416742 | 0.88416261 |
| MTUS1 | 2032.87809 | -0.3179043 | 0.28088244 | -1.1318056 | 0.25771617 | 0.70875972 |
| AC027117.2 | 349.379307 | -0.1223358 | 0.47831631 | -0.2557635 | 0.79813348 | 0.95294186 |
| PCM1 | 3337.84653 | -0.0747667 | 0.12251124 | -0.610284 | 0.54167369 | 0.86582694 |
| ASAH1 | 479.171716 | 0.07555225 | 0.33162342 | 0.22782543 | 0.81978196 | 0.96014363 |
| NAT1 | 39.8876893 | -0.103156 | 0.4030795 | -0.2559199 | 0.79801272 | 0.95294186 |
| PSD3 | 594.18778 | -0.2482312 | 0.40607515 | -0.6112937 | 5.41E-01 | 0.86582694 |
| SH2D4A | 740.642601 | -0.474822 | 0.40988262 | -1.1584341 | 0.24668695 | 0.6968537 |
| CSGALNACT1 | 4171.09214 | -0.4848478 | 0.76842884 | -0.63096 | 0.52806669 | 0.86215467 |
| INTS10 | 1533.84438 | -0.2729467 | 0.23633737 | -1.1549027 | 0.24813028 | 0.69838949 |
| LPL | 242.185239 | -1.269051 | 0.71037683 | -1.7864476 | 7.40E-02 | 0.47070029 |
| ATP6V1B2 | 381.716454 | -0.1385966 | 0.22351363 | -0.6200814 | 0.5352042 | 0.8642718 |
| GFRA2 | 46.5248023 | 1.5567325 | 0.85308627 | 1.82482424 | 0.06802756 | 0.45250883 |
| XPO7 | 931.636816 | -0.2687922 | 0.12878427 | -2.0871508 | 0.03687451 | 0.36038369 |
| NPM2 | 39.9272504 | -0.0795769 | 0.59386177 | -0.133999 | 0.89340338 | 0.97788747 |
| DMTN | 445.392106 | 0.44884291 | 0.34224331 | 1.31147314 | 0.18969796 | 0.64447541 |
| FAM160B2 | 268.205518 | -0.0801689 | 0.23885513 | -0.3356381 | 0.73714379 | 0.93552616 |
| BMP1 | 81.8630821 | 0.10842992 | 0.44998959 | 0.24096095 | 0.80958538 | 0.95656232 |
| POLR3D | 68.0462579 | -0.0889102 | 0.33509022 | -0.265332 | 0.79075369 | 0.95144272 |
| SLC39A14 | 627.33079 | 0.47004343 | 0.52469982 | 0.89583302 | 0.37034195 | 0.78104287 |
| PPP3CC | 1229.32836 | -0.3048916 | 0.20612499 | -1.479159 | 0.13909782 | 0.58243213 |
| SORBS3 | 98.6172515 | 0.72169548 | 0.36851072 | 1.95841108 | 0.0501818 | 0.40148476 |
| CCAR2 | 190.598748 | -0.2841673 | 0.20342415 | -1.3969204 | 0.16243751 | 0.61374059 |
| AC037459.3 | 45.6116727 | -0.0730682 | 0.44577925 | -0.1639112 | 0.86980104 | 0.97454113 |
| BIN3 | 614.761816 | -0.3042441 | 0.24636456 | -1.2349345 | 0.21685489 | 0.66978787 |
| PEBP4 | 71.8150046 | 0.1524495 | 0.55141992 | 0.27646716 | 0.78218928 | 0.94884178 |
| RHOBTB2 | 80.8651706 | 0.23939263 | 0.52181872 | 0.4587659 | 0.64640229 | 0.90582727 |
| TNFRSF10B | 724.71483 | 0.14114169 | 0.29176383 | 0.48375321 | 0.62856102 | 0.89962754 |
| TNFRSF10D | 457.919462 | -0.0885564 | 0.38758648 | -0.2284816 | 8.19E-01 | 9.60E-01 |
| TNFRSF10A | 74.9150973 | 0.34366724 | 0.42229148 | 0.81381522 | 4.16E-01 | 8.10E-01 |
| CHMP7 | 131.709268 | -0.1585357 | 0.23682157 | -0.6694309 | 0.50322065 | 0.85525037 |
| R3HCC1 | 80.1831939 | 0.20490284 | 0.33946217 | 0.60361025 | 0.5461028 | 0.86794271 |
| LOXL2 | 71.1510224 | -0.0481873 | 0.48262988 | -0.0998431 | 0.92046886 | 0.98215577 |
| ENTPD4 | 420.920812 | -0.2110314 | 0.22499173 | -0.9379516 | 0.34826929 | 0.76974844 |
| SLC25A37 | 678.453488 | 0.69739296 | 0.34554121 | 2.01826278 | 0.0435639 | 0.38413589 |
| AC120193.1 | 68.5750584 | 0.48839468 | 0.68387619 | 0.71415657 | 0.47513037 | 0.84106098 |
| DOCK5 | 1343.87327 | 0.04246354 | 0.42599631 | 0.09968055 | 9.21E-01 | 0.98215577 |
| KCTD9 | 546.141018 | -0.0025669 | 0.21762196 | -0.0117952 | 0.99058898 | 0.99740343 |
| PPP2R2A | 1701.02303 | -0.1603327 | 0.1989094 | -0.8060588 | 0.42020894 | 0.8125045 |
| BNIP3L | 1627.48975 | 0.50962793 | 0.59036176 | 0.86324685 | 0.38800176 | 0.79631996 |
| DPYSL2 | 275.567564 | -0.5019928 | 0.60697361 | -0.8270423 | 0.40821309 | 0.80588056 |
| TRIM35 | 127.709309 | -0.0712627 | 0.34624994 | -0.205813 | 0.83693701 | 0.9652581 |
| PTK2B | 387.313748 | 0.12895984 | 0.47411832 | 0.27199928 | 0.78562257 | 0.94978829 |
| EPHX2 | 710.169287 | 0.54139556 | 0.45678297 | 1.18523585 | 0.23592421 | 0.68832244 |
| CLU | 1188.07582 | 1.14770592 | 0.56422056 | 2.03414411 | 4.19E-02 | 3.78E-01 |
| CCDC25 | 241.633126 | 0.26794968 | 0.22861144 | 1.17207467 | 0.24116708 | 0.69178991 |
| ELP3 | 759.913049 | -0.1491643 | 0.16300606 | -0.9150845 | 0.36014729 | 0.77706395 |
| ZNF395 | 272.305072 | 0.16753349 | 0.50708424 | 0.33038591 | 0.74110838 | 0.93598004 |
| FBXO16 | 431.890032 | -0.0418346 | 0.3298659 | -0.1268231 | 0.89908043 | 0.97889101 |
| FZD3 | 258.597424 | -0.3596884 | 0.34449517 | -1.0441029 | 0.29643776 | 0.73672107 |
| EXTL3 | 295.942091 | -0.4042484 | 0.23726094 | -1.7038136 | 0.08841592 | 0.49769182 |

| | | | | | | |
|---|---|---|---|---|---|---|
| INTS9 | 642.709774 | -0.192892 | 0.17226932 | -1.119712 | 0.26283653 | 0.71328 |
| HMBOX1 | 2061.74462 | -0.3419864 | 0.19844592 | -1.7233229 | 0.08483014 | 0.49198622 |
| KIF13B | 3177.09275 | -0.3984005 | 0.17272879 | -2.3065089 | 0.02108221 | 0.29038053 |
| AC108449.1 | 165.272504 | -0.2422187 | 0.52603741 | -0.4604591 | 0.64518673 | 0.90518019 |
| SARAF | 745.813575 | 0.63002703 | 0.26330665 | 2.3927502 | 0.01672262 | 0.25905688 |
| LEPROTL1 | 264.926051 | 0.39503473 | 0.25167542 | 1.56961983 | 0.11650358 | 0.54687129 |
| DCTN6 | 152.729304 | 0.0250606 | 0.28353395 | 0.08838659 | 0.92956942 | 0.98331361 |
| RBPMS-AS1 | 135.658094 | 0.06489101 | 0.55064065 | 0.11784638 | 0.90618938 | 0.98009133 |
| RBPMS | 13308.2277 | -0.3262228 | 0.23035814 | -1.4161549 | 1.57E-01 | 6.07E-01 |
| GTF2E2 | 583.596024 | -0.3869449 | 0.17106327 | -2.2619987 | 0.02369749 | 0.30185158 |
| GSR | 427.455655 | -0.3725028 | 0.23771158 | -1.567037 | 0.11710604 | 0.54756317 |
| UBXN8 | 77.9010464 | -0.383655 | 0.24841563 | -1.5444077 | 0.1224896 | 0.55926898 |
| PPP2CB | 1097.78935 | -0.1909024 | 0.19973528 | -0.9557769 | 0.33918496 | 0.76449297 |
| WRN | 1135.3906 | -0.5643845 | 0.17537356 | -3.2181844 | 0.00129005 | 0.06579248 |
| NRG1 | 1237.11717 | 0.38518127 | 0.28227407 | 1.36456485 | 0.17238986 | 0.6286116 |
| FUT10 | 583.748698 | 0.22413792 | 0.25034789 | 0.89530579 | 0.37062364 | 0.78104287 |
| TTI2 | 227.132591 | -0.2778009 | 0.35763726 | -0.7767672 | 0.43729611 | 0.82013459 |
| MAK16 | 90.0165552 | -0.5616004 | 0.38874996 | -1.4446313 | 0.14856147 | 0.59478974 |
| RNF122 | 167.085412 | -0.10194 | 0.41412679 | -0.2461565 | 0.80556111 | 0.95522565 |
| UNC5D | 106.175664 | 0.08888222 | 0.35572919 | 0.24985921 | 0.80269623 | 0.95446295 |
| ERLIN2 | 264.66435 | -0.226724 | 0.22368669 | -1.0135785 | 0.31078393 | 0.74495902 |
| PLPBP | 239.589719 | 0.2455155 | 0.16673744 | 1.47246772 | 0.14089462 | 0.58672037 |
| RAB11FIP1 | 611.539678 | 0.296755 | 0.2831722 | 1.04796656 | 0.29465401 | 0.73556877 |
| EIF4EBP1 | 96.3709677 | -0.3824247 | 0.3599701 | -1.062379 | 0.28806365 | 0.73075337 |
| ASH2L | 429.276609 | -0.4318287 | 0.22108196 | -1.9532517 | 0.05078978 | 0.40463338 |
| LSM1 | 222.846242 | 0.18353453 | 0.28468455 | 0.64469437 | 0.51912527 | 0.85918126 |
| BAG4 | 516.3164 | 0.08054902 | 0.20516103 | 0.39261362 | 0.69460488 | 0.92229578 |
| DDHD2 | 676.851523 | -0.0303078 | 0.17853305 | -0.1697601 | 0.86519878 | 0.97379123 |
| PLPP5 | 414.38673 | 0.23528282 | 0.40432146 | 0.58192019 | 0.56062044 | 0.873671 |
| NSD3 | 2411.53423 | -0.0484924 | 0.1383838 | -0.3504195 | 0.72602387 | 0.9330657 |
| LETM2 | 145.14087 | -0.0125514 | 0.3563469 | -0.0352224 | 0.97190237 | 0.99327086 |
| FGFR1 | 610.651936 | -0.236249 | 0.44000829 | -0.5369195 | 0.59132321 | 0.8868841 |
| AC069120.1 | 98.2888076 | -1.4113269 | 0.7949132 | -1.7754478 | 0.07582398 | 0.47399856 |
| TACC1 | 5376.78773 | -0.7007155 | 0.50350399 | -1.3916781 | 0.16401991 | 0.61531922 |
| PLEKHA2 | 763.423308 | -0.2659375 | 0.39723297 | -0.6694748 | 0.50319262 | 0.85525037 |
| TM2D2 | 78.043784 | 1.01657902 | 0.35040836 | 2.90112662 | 0.00371824 | 0.11865695 |
| ADAM9 | 823.760052 | -0.1047482 | 0.27162053 | -0.3856417 | 0.69976207 | 0.92367966 |
| ADAM32 | 93.9413191 | 0.30540749 | 0.35792308 | 0.85327686 | 3.94E-01 | 0.79843808 |
| TCIM | 4788.39262 | 0.34032617 | 0.58711403 | 0.5796594 | 0.56214433 | 0.87403145 |
| AC105999.2 | 233.506323 | -2.37975 | 1.47338716 | -1.6151559 | 0.10627694 | 0.52735777 |
| ZMAT4 | 218.485633 | -3.3940413 | 0.72896435 | -4.6559771 | 3.22E-06 | 0.00250542 |
| SFRP1 | 1737.71023 | 1.53882378 | 0.45449131 | 3.38581562 | 0.00070967 | 0.046637 |
| GOLGA7 | 229.007002 | 0.02298125 | 0.23757848 | 0.09673118 | 0.92293988 | 0.98215577 |
| AC009630.1 | 68.8189723 | -0.6990167 | 0.42913561 | -1.6288946 | 0.10333533 | 0.52337282 |
| GPAT4 | 280.373937 | -0.2895334 | 0.22857492 | -1.2666894 | 0.20526638 | 0.65813563 |
| KAT6A | 2268.58326 | -0.0280734 | 0.15965381 | -0.1758391 | 0.86042032 | 0.97200968 |
| AP3M2 | 188.998921 | 0.13267855 | 0.33841171 | 0.39206253 | 0.69501201 | 0.92229578 |
| IKBKB | 839.385472 | -0.3830615 | 0.18269759 | -2.0966971 | 0.0360204 | 0.35693431 |
| POLB | 222.979138 | -0.50045 | 0.18416878 | -2.7173445 | 0.00658081 | 0.1680054 |
| VDAC3 | 582.293006 | 0.07468815 | 0.28167977 | 0.2651527 | 0.79089183 | 0.95144272 |
| SLC20A2 | 4015.07362 | -0.5392483 | 0.32389369 | -1.664893 | 0.0959341 | 0.5117083 |
| SMIM19 | 490.157569 | -0.3280762 | 0.26471027 | -1.2393785 | 0.21520537 | 0.66885336 |
| THAP1 | 44.4658813 | 0.88384026 | 0.3568209 | 2.47698568 | 0.01324972 | 0.23650753 |
| RNF170 | 730.988589 | -0.1576457 | 0.26986205 | -0.5841715 | 0.55910496 | 0.87295536 |

| | | | | | | |
|---|---|---|---|---|---|---|
| HOOK3 | 1151.61587 | -0.1311402 | 0.12171766 | -1.0774135 | 0.28129558 | 0.72604837 |
| FNTA | 314.513374 | -0.0990244 | 0.2734401 | -0.3621429 | 0.71724525 | 0.9290703 |
| POMK | 80.199291 | 0.00249872 | 0.26998409 | 0.00925505 | 0.99261565 | 0.99805588 |
| HGSNAT | 1169.70517 | -0.3775684 | 0.18002011 | -2.097368 | 0.03596101 | 0.35661333 |
| SPIDR | 7075.00129 | -0.4671898 | 0.31750061 | -1.4714613 | 0.14116642 | 0.5872148 |
| CEBPD | 1944.34012 | 0.34421179 | 0.52089471 | 0.66080876 | 5.09E-01 | 8.57E-01 |
| PRKDC | 1361.38088 | -0.3484282 | 0.17706226 | -1.9678285 | 0.04908777 | 0.39852511 |
| MCM4 | 159.884589 | -0.3336443 | 0.36401954 | -0.9165559 | 0.35937538 | 0.77654006 |
| UBE2V2 | 534.427517 | -0.0615406 | 0.25138893 | -0.2448023 | 0.80660954 | 0.95566275 |
| SNTG1 | 48.7567846 | 0.28735244 | 0.30772813 | 0.93378669 | 0.35041393 | 0.77044053 |
| PXDNL | 1581.57656 | 0.23573844 | 0.57765913 | 0.40809264 | 6.83E-01 | 0.9200778 |
| PCMTD1 | 3199.50387 | 0.16198603 | 0.15007428 | 1.07937237 | 0.28042176 | 0.72527072 |
| AC064807.1 | 250.154769 | -0.1918287 | 0.36620748 | -0.5238253 | 0.60040004 | 0.88883692 |
| RB1CC1 | 2066.48276 | -0.3063139 | 0.21515126 | -1.4237142 | 0.15452921 | 0.60380955 |
| ATP6V1H | 2047.51895 | 0.20083766 | 0.23536456 | 0.85330458 | 0.39349042 | 0.79843808 |
| TCEA1 | 614.411649 | -0.304884 | 0.16744704 | -1.8207788 | 0.06864049 | 0.45447261 |
| LYPLA1 | 1123.54733 | -0.1180302 | 0.18041281 | -0.6542232 | 0.51296802 | 0.85762435 |
| MRPL15 | 299.59954 | 0.29283861 | 0.26760402 | 1.09429826 | 0.27382419 | 0.72135661 |
| RP1 | 2645.11131 | -0.0786995 | 0.56027857 | -0.140465 | 0.88829264 | 0.97729111 |
| XKR4 | 243.326469 | 2.08341645 | 0.53738772 | 3.87693349 | 1.06E-04 | 1.62E-02 |
| TMEM68 | 402.910095 | -0.0255702 | 0.21576063 | -0.1185117 | 0.90566222 | 0.97997915 |
| TGS1 | 461.970385 | 0.1167203 | 0.17591213 | 0.66351479 | 0.50700091 | 0.85626196 |
| LYN | 610.844988 | 0.19960048 | 0.3320038 | 0.60119938 | 5.48E-01 | 8.69E-01 |
| RPS20 | 650.5988 | 0.38782659 | 0.41726269 | 0.92945428 | 0.35265371 | 0.77210174 |
| PLAG1 | 434.771581 | 1.1281273 | 0.38599271 | 2.92266477 | 0.0034705 | 0.1138728 |
| CHCHD7 | 149.710169 | 0.44121954 | 0.26375151 | 1.67286067 | 0.09435473 | 0.51017802 |
| IMPAD1 | 378.56715 | 0.23585229 | 0.2087173 | 1.13000832 | 0.25847272 | 0.70896455 |
| LINC01606 | 5043.68312 | 1.53014765 | 0.4493725 | 3.40507632 | 0.00066146 | 0.04534035 |
| FAM110B | 1022.56408 | 0.03992212 | 0.26784365 | 0.1490501 | 0.8815141 | 0.97664137 |
| AC027698.1 | 83.4293554 | 0.54184243 | 0.60220602 | 0.89976257 | 0.36824662 | 0.78095617 |
| UBXN2B | 559.673911 | -0.2043166 | 0.15965376 | -1.2797479 | 0.20063382 | 0.65387914 |
| SDCBP | 744.08208 | 0.18688708 | 0.38866003 | 0.48084975 | 0.63062329 | 0.90005881 |
| NSMAF | 736.838898 | -0.2273757 | 0.16882002 | -1.3468529 | 0.17802761 | 0.63197168 |
| TOX | 2015.58418 | -0.3920978 | 0.41473383 | -0.9454202 | 0.34444437 | 0.76716956 |
| CA8 | 939.870708 | 0.21324798 | 0.70845963 | 0.3010023 | 7.63E-01 | 9.43E-01 |
| LINC01301 | 356.516534 | 0.8477801 | 0.46263137 | 1.83251755 | 0.06687435 | 0.4488533 |
| RAB2A | 2074.38152 | 0.01750721 | 0.18811164 | 0.09306821 | 0.92584937 | 0.98229272 |
| CHD7 | 1856.09174 | 0.04539072 | 0.21118651 | 0.21493192 | 0.8298204 | 0.96310497 |
| CLVS1 | 68.0129455 | 0.09611227 | 0.26285779 | 0.36564361 | 0.71463103 | 0.92808586 |
| ASPH | 1917.10999 | 0.02232342 | 0.178537 | 0.12503528 | 0.90049562 | 0.97914987 |
| NKAIN3 | 63.5624675 | -0.3194623 | 0.4490197 | -0.7114662 | 0.47679539 | 0.8415432 |
| YTHDF3 | 1056.94696 | 0.14702977 | 0.18889435 | 0.77837043 | 0.43635066 | 0.81999657 |
| AC022639.1 | 40.6454369 | -0.078696 | 0.40184311 | -0.1958375 | 0.84473732 | 0.96853443 |
| CYP7B1 | 1549.90405 | -0.2097511 | 0.26371793 | -0.7953616 | 0.4264032 | 0.81466629 |
| ARMC1 | 334.286764 | -0.0516164 | 0.19000704 | -0.2716555 | 0.78588696 | 0.94978829 |
| MTFR1 | 613.350341 | -0.2014307 | 0.34819058 | -0.5785071 | 0.56292181 | 0.87436903 |
| PDE7A | 1079.0252 | 0.01228583 | 0.27754176 | 0.04426661 | 0.96469189 | 0.99226095 |
| TRIM55 | 240.794984 | 0.6113416 | 0.48563199 | 1.25885778 | 0.20808171 | 0.66104323 |
| RRS1-AS1 | 97.4610238 | 0.69159214 | 0.41454598 | 1.66831227 | 0.09525375 | 0.51085035 |
| ADHFE1 | 652.048841 | 0.70769845 | 0.53894326 | 1.31312235 | 0.18914172 | 0.64432929 |
| VXN | 167.174428 | -0.5171536 | 0.41036752 | -1.2602205 | 2.08E-01 | 6.61E-01 |
| VCPIP1 | 320.18143 | 0.02983788 | 0.17500759 | 0.17049475 | 0.86462106 | 0.97361767 |
| SGK3 | 788.882423 | 0.0331028 | 0.26675324 | 0.12409523 | 0.90123987 | 0.97931857 |
| MCMDC2 | 113.066201 | 0.89395009 | 0.36943713 | 2.41976242 | 0.01553065 | 0.25214021 |

| | | | | | | |
|---|---|---|---|---|---|---|
| PPP1R42 | 191.124733 | 0.34307147 | 0.38907809 | 0.8817548 | 0.37790942 | 0.7865219 |
| COPS5 | 101.023011 | 0.10506482 | 0.39055765 | 0.26901231 | 0.78792021 | 0.95010843 |
| CSPP1 | 1658.09409 | -0.0081464 | 0.21336899 | -0.03818 | 0.9695442 | 0.99299658 |
| ARFGEF1 | 3963.24998 | 0.06054487 | 0.1844091 | 0.32831824 | 0.74267106 | 0.93598004 |
| C8orf34 | 84.8705317 | -0.2037766 | 0.33124937 | -0.6151759 | 0.53843857 | 0.86527404 |
| SULF1 | 72.9089168 | 0.39536565 | 0.60904363 | 0.64915816 | 0.51623615 | 0.85888182 |
| NCOA2 | 8331.62049 | -0.0842309 | 0.18918171 | -0.4452381 | 0.65614767 | 0.90950195 |
| TRAM1 | 898.100846 | 0.17898034 | 0.30901538 | 0.57919558 | 0.56245721 | 0.87426422 |
| LACTB2-AS1 | 122.162831 | 0.06274698 | 0.79434868 | 0.07899174 | 0.9370392 | 0.9864789 |
| LACTB2 | 674.616438 | -0.2719758 | 0.45533958 | -0.5973032 | 0.55030495 | 0.86959143 |
| XKR9 | 199.135821 | -0.7941133 | 0.56093112 | -1.4157055 | 0.15686176 | 0.60654181 |
| TERF1 | 491.769202 | 0.03283305 | 0.16549685 | 0.19839079 | 0.84273932 | 0.96747294 |
| AC022893.1 | 77.6912602 | -0.2961001 | 0.39118933 | -0.7569228 | 0.44909609 | 0.8267002 |
| SBSPON | 126.465523 | 0.33650092 | 0.46781089 | 0.71930973 | 0.4719501 | 0.83921324 |
| RPL7 | 1016.12977 | 0.73612937 | 0.51604782 | 1.42647513 | 0.15373124 | 0.60285224 |
| RDH10 | 368.444143 | -1.003132 | 0.40169621 | -2.4972405 | 0.0125164 | 0.22899031 |
| RDH10-AS1 | 145.009614 | -0.7434182 | 0.67064515 | -1.108512 | 0.26764074 | 0.71784829 |
| STAU2 | 1962.70019 | -0.053661 | 0.22324679 | -0.2403661 | 0.81004648 | 0.9567153 |
| AC027018.1 | 323.5202 | -0.0423872 | 0.59663001 | -0.0710444 | 0.9433624 | 0.98772472 |
| ELOC | 345.422236 | -0.0807386 | 0.22063092 | -0.3659441 | 0.7144068 | 0.92788588 |
| TMEM70 | 94.2396458 | 0.38029037 | 0.28388055 | 1.33961403 | 0.18037086 | 0.63479125 |
| GDAP1 | 1022.58147 | -0.7107622 | 0.47519543 | -1.4957261 | 0.13472503 | 0.57708028 |
| MIR2052HG | 274.60997 | -0.5237291 | 0.71877146 | -0.7286448 | 0.46621897 | 0.8367113 |
| CRISPLD1 | 47.3285161 | 0.17370211 | 0.57793177 | 0.30055817 | 0.76375144 | 0.94255335 |
| CASC9 | 77.8936919 | -0.1229798 | 0.89984705 | -0.1366674 | 0.89129367 | 0.97729111 |
| HNF4G | 126.269176 | -0.3044202 | 0.6774716 | -0.4493476 | 0.65318095 | 0.908787 |
| PEX2 | 388.047129 | 0.21222775 | 0.20905436 | 1.01517971 | 0.31002017 | 0.74455572 |
| PKIA | 73.2879634 | -0.2699455 | 0.539021 | -0.500807 | 0.61650694 | 0.89589954 |
| ZC2HC1A | 536.571086 | -0.2527409 | 0.32773513 | -0.7711742 | 0.44060367 | 0.82260549 |
| IL7 | 528.034851 | 1.04376373 | 0.28656835 | 3.64228547 | 0.00027023 | 0.02788628 |
| AC083837.1 | 59.4953843 | 1.47409876 | 0.79209042 | 1.86102335 | 0.06274088 | 0.43526485 |
| LINC01607 | 37.7817409 | -1.5363661 | 0.70379517 | -2.1829734 | 0.02903777 | 0.32900439 |
| TPD52 | 1343.93162 | -0.2612082 | 0.3651679 | -0.7153099 | 0.47441756 | 0.84106098 |
| ZBTB10 | 737.587093 | 0.55877248 | 0.39441869 | 1.4166988 | 0.15657101 | 0.60631676 |
| ZNF704 | 2781.79156 | -0.1406647 | 0.28517425 | -0.4932587 | 0.6218298 | 0.8982666 |
| PAG1 | 171.41973 | 1.44889049 | 0.69476147 | 2.08545026 | 0.03702845 | 0.3605598 |
| IMPA1 | 393.988445 | -0.0154762 | 0.19634852 | -0.0788199 | 0.93717589 | 0.98649645 |
| ZFAND1 | 477.780292 | -0.0703133 | 0.22557999 | -0.3117003 | 0.75526833 | 0.93953492 |
| CHMP4C | 212.222489 | 0.26922301 | 0.30083038 | 0.8949329 | 0.37082296 | 0.78104287 |
| SNX16 | 354.481372 | -0.3387124 | 0.23363605 | -1.4497439 | 0.14712996 | 0.59373375 |
| RALYL | 2442.94828 | 1.40001636 | 0.6829805 | 2.04986286 | 0.04037782 | 0.37344844 |
| LRRCC1 | 174.235519 | -0.5437263 | 0.36613922 | -1.4850262 | 0.13753689 | 0.58042326 |
| E2F5 | 279.589653 | -0.2135018 | 0.30493471 | -0.7001558 | 0.48383002 | 0.8445691 |
| C8orf59 | 282.806904 | -0.1739756 | 0.27371506 | -0.6356083 | 0.52503175 | 0.8606763 |
| CA13 | 467.584655 | 0.01786902 | 0.18671707 | 0.09570104 | 0.92375802 | 0.98215577 |
| CA1 | 151.358614 | 0.30907413 | 0.4918332 | 0.6284125 | 0.52973375 | 0.86343221 |
| CA3 | 588.290622 | 0.14901636 | 0.27585828 | 0.54019173 | 0.58906482 | 0.88581024 |
| CA2 | 383.359158 | -0.0166236 | 0.49347454 | -0.0336867 | 0.97312695 | 0.99327086 |
| ATP6V0D2 | 239.916548 | 1.34696139 | 0.82118962 | 1.64025623 | 0.1009519 | 0.51819907 |
| WWP1 | 2825.577 | 0.04366732 | 0.16615842 | 0.26280532 | 0.79270063 | 0.95224951 |
| RMDN1 | 1013.62697 | -0.1116018 | 0.15481081 | -0.7208913 | 4.71E-01 | 8.39E-01 |
| CPNE3 | 782.047071 | -0.0043001 | 0.20609545 | -0.0208644 | 0.98335385 | 0.99518552 |
| CNGB3 | 115.484204 | 0.26948348 | 0.46227592 | 0.58294941 | 0.55992736 | 0.87300289 |
| CNBD1 | 47.1206024 | -0.2698141 | 0.38342416 | -0.7036961 | 0.48162205 | 0.8437862 |

| | | | | | | |
|---|---|---|---|---|---|---|
| AF117829.1 | 2419.71853 | 0.48306846 | 0.26505023 | 1.82255444 | 0.0683709 | 0.4533691 |
| RIPK2 | 186.830585 | 0.07690533 | 0.35998777 | 0.21363316 | 0.83083314 | 0.96368764 |
| OSGIN2 | 278.277247 | 0.26451327 | 0.19527069 | 1.35459789 | 0.17554571 | 0.62957039 |
| NBN | 524.455048 | 0.01251364 | 0.25926253 | 0.04826627 | 0.96150404 | 0.99191712 |
| DECR1 | 1441.7702 | 0.17957039 | 0.19321983 | 0.92935797 | 0.3527036 | 0.77210174 |
| CALB1 | 50.3849536 | 1.81006081 | 0.62148304 | 2.91248623 | 0.00358564 | 0.11665695 |
| LINC00534 | 381.986243 | 1.11425751 | 0.66208273 | 1.68295813 | 0.0923832 | 0.50585766 |
| TMEM64 | 158.322674 | -0.0534155 | 0.61530018 | -0.0868121 | 0.93082085 | 0.98331361 |
| NECAB1 | 64.0801347 | 0.03321282 | 0.83757248 | 0.03965366 | 0.96836924 | 0.99248831 |
| C8orf88 | 66.3825359 | 0.32830402 | 0.73307583 | 0.44784456 | 0.65426538 | 0.90902447 |
| PIP4P2 | 425.991423 | -0.2652705 | 0.35926641 | -0.7383671 | 0.46029138 | 0.83429248 |
| OTUD6B-AS1 | 230.708132 | 0.44494379 | 0.36015864 | 1.23541055 | 0.21667775 | 0.66978787 |
| OTUD6B | 256.128343 | -0.1302072 | 0.18974027 | -0.6862391 | 0.49256232 | 0.84738939 |
| LRRC69 | 445.502167 | -0.0576579 | 0.40942248 | -0.1408274 | 0.8880063 | 0.97729111 |
| SLC26A7 | 216.891441 | 1.05365831 | 0.58726128 | 1.79419 | 0.07278285 | 0.46942805 |
| AC117834.1 | 132.1194 | 0.06501365 | 0.61796388 | 0.10520623 | 0.91621217 | 0.98195017 |
| TRIQK | 591.071189 | -0.2279792 | 0.14409985 | -1.5820916 | 0.11362867 | 0.53989968 |
| LINC00535 | 78.4737727 | -0.7828749 | 0.36114701 | -2.1677459 | 0.03017803 | 0.33385391 |
| FAM92A | 466.5592 | 0.31146647 | 0.35479013 | 0.87788935 | 0.38000377 | 0.7880311 |
| AC120053.1 | 63.6985639 | 0.18817972 | 0.33398877 | 0.56343126 | 0.57314125 | 0.87969124 |
| RBM12B | 520.219462 | 0.18946957 | 0.18235203 | 1.03903187 | 0.29878991 | 0.73673761 |
| TMEM67 | 803.973894 | 0.09961569 | 0.15894648 | 0.62672473 | 0.53083969 | 0.86396107 |
| PDP1 | 107.716503 | 0.06788182 | 0.26591465 | 0.25527673 | 0.79850937 | 0.95297658 |
| GEM | 69.2496929 | -0.0084892 | 0.46541646 | -0.01824 | 0.98544742 | 0.9958515 |
| RAD54B | 50.4298126 | -0.4143798 | 0.42885496 | -0.9662469 | 0.33392064 | 0.76205213 |
| VIRMA | 606.822775 | -0.1669828 | 0.14630256 | -1.1413524 | 0.2537233 | 0.70431689 |
| AC023632.2 | 97.0445041 | -0.6208481 | 0.33197151 | -1.8701847 | 0.06145817 | 0.43249565 |
| AC108860.2 | 56.161689 | 0.21509004 | 0.38062472 | 0.5650974 | 0.57200752 | 0.87907473 |
| ESRP1 | 1040.13389 | 0.2036407 | 0.29657647 | 0.6866381 | 0.49231081 | 0.84734855 |
| DPY19L4 | 660.46626 | -0.3887768 | 0.17270597 | -2.2510908 | 0.02437979 | 0.3064059 |
| INTS8 | 608.528943 | -0.1491299 | 0.15586473 | -0.9567907 | 0.33867291 | 0.76449297 |
| NDUFAF6 | 671.005104 | -0.1569827 | 0.19663362 | -0.7983513 | 0.42466665 | 0.81466629 |
| TP53INP1 | 317.859271 | 0.11385708 | 0.38808629 | 0.29338083 | 0.76923107 | 0.94386043 |
| PLEKHF2 | 182.69912 | 0.20250549 | 0.27119787 | 0.74670754 | 0.45524012 | 0.83121393 |
| C8orf37-AS1 | 466.647525 | 0.65884621 | 0.4143858 | 1.58993434 | 0.11184961 | 0.53607455 |
| C8orf37 | 60.4161863 | -0.0142401 | 0.34429179 | -0.0413605 | 0.96700852 | 0.99248831 |
| UQCRB | 805.016884 | 0.57341519 | 0.57327057 | 1.00025228 | 0.31718843 | 0.74897068 |
| MTERF3 | 202.927776 | -0.095899 | 0.22072117 | -0.4344802 | 0.66393979 | 0.91197775 |
| PTDSS1 | 176.258011 | -0.6870586 | 0.27511775 | -2.4973258 | 0.01251339 | 0.22899031 |
| SDC2 | 1217.4133 | 0.15337697 | 0.41980805 | 0.36535023 | 0.71484999 | 0.92827895 |
| CPQ | 1765.8757 | 0.86727338 | 0.30438601 | 2.84925511 | 0.00438217 | 0.12886682 |
| TSPYL5 | 100.782155 | 0.56892651 | 0.35112197 | 1.62031018 | 0.10516566 | 0.52578851 |
| MTDH | 2235.45848 | 0.09379402 | 0.1171793 | 0.80043168 | 0.42346073 | 0.8138418 |
| LAPTM4B | 925.448573 | -0.0318271 | 0.31309349 | -0.1016535 | 9.19E-01 | 0.98215577 |
| MATN2 | 45.5611811 | 0.14044631 | 0.61686186 | 0.22767871 | 0.81989603 | 0.96016713 |
| RPL30 | 761.833406 | 0.35323586 | 0.36731322 | 0.96167479 | 0.33621299 | 0.7637525 |
| RIDA | 199.372168 | 1.03309126 | 0.37960907 | 2.72146095 | 0.00649941 | 0.16670031 |
| POP1 | 138.272107 | -0.0284328 | 0.31329946 | -0.0907527 | 0.9276891 | 0.98297055 |
| NIPAL2 | 310.933762 | -0.690194 | 0.25614576 | -2.6945362 | 0.00704867 | 0.17262485 |
| AP003438.1 | 45.4751093 | -0.1049188 | 0.42810796 | -0.2450756 | 0.80639789 | 0.95565312 |
| STK3 | 4106.79434 | -0.2669681 | 0.20047411 | -1.3316837 | 0.18296413 | 0.63767103 |
| AC104986.2 | 394.226707 | 0.23835487 | 0.27002455 | 0.88271554 | 0.37738998 | 0.78614482 |
| VPS13B | 4855.52424 | -0.1796733 | 0.13368356 | -1.3440193 | 0.17894213 | 0.63284534 |
| COX6C | 705.455934 | 0.38366553 | 0.30212156 | 1.26990451 | 0.20411865 | 0.65712971 |

| | | | | | | |
|---|---|---|---|---|---|---|
| POLR2K | 159.950375 | 0.26499584 | 0.46391092 | 0.57122138 | 0.56784959 | 0.87712536 |
| SPAG1 | 530.999084 | 0.2772338 | 0.30124654 | 0.92028874 | 0.35742189 | 0.77459979 |
| RNF19A | 2400.65009 | -0.2174989 | 0.375248 | -0.5796137 | 5.62E-01 | 0.87403145 |
| AP000424.1 | 127.980485 | 1.05738843 | 0.97907121 | 1.07999134 | 2.80E-01 | 7.25E-01 |
| ANKRD46 | 339.68232 | 0.07751299 | 0.19470506 | 0.39810466 | 0.69055304 | 0.92183073 |
| PABPC1 | 1152.72843 | -0.0974794 | 0.28146885 | -0.3463241 | 0.72909918 | 0.93453345 |
| YWHAZ | 1816.43926 | 0.02823285 | 0.20735547 | 0.13615677 | 0.89169735 | 0.9774631 |
| AP003469.4 | 43.1383958 | -0.0387192 | 0.40308135 | -0.096058 | 0.92347448 | 0.98215577 |
| ZNF706 | 563.548632 | 0.29751327 | 0.19808945 | 1.50191378 | 0.13311938 | 0.57682553 |
| AP001207.3 | 58.4687709 | 0.35605391 | 0.50661607 | 0.70280816 | 0.48217531 | 0.84389635 |
| GRHL2 | 1995.45421 | 0.00069806 | 0.280103 | 0.00249215 | 0.99801155 | 0.99969006 |
| NCALD | 86.1423059 | -0.6557444 | 0.48620777 | -1.3486918 | 0.17743599 | 0.63105868 |
| RRM2B | 399.12668 | 0.31237679 | 0.48117924 | 0.64919008 | 0.51621553 | 0.85888182 |
| UBR5-AS1 | 106.885766 | 0.1286285 | 0.60852752 | 0.21137663 | 0.83259339 | 0.96438209 |
| UBR5 | 3356.41035 | -0.1228503 | 0.21681696 | -0.5666086 | 0.57098015 | 0.87863805 |
| KLF10 | 180.811675 | 0.5671997 | 0.60643381 | 0.93530355 | 0.34963188 | 0.77025232 |
| AZIN1 | 615.382026 | 0.47640136 | 0.24153154 | 1.9724188 | 0.04856182 | 0.39695534 |
| AZIN1-AS1 | 2078.06139 | 0.28979289 | 0.31983776 | 0.90606214 | 0.36490296 | 0.78094673 |
| AP003354.2 | 593.599425 | 0.49716479 | 0.4531389 | 1.0971576 | 0.27257251 | 0.72070318 |
| ATP6V1C1 | 843.832028 | -0.1667483 | 0.22550525 | -0.7394431 | 0.45963799 | 0.83390278 |
| BAALC-AS1 | 550.183957 | -0.2465476 | 0.33199819 | -0.7426174 | 0.45771336 | 0.83282548 |
| FZD6 | 1023.66184 | -0.1920233 | 0.24800578 | -0.7742693 | 0.43877155 | 0.82127804 |
| SLC25A32 | 336.731418 | 0.12818799 | 0.22384744 | 0.57265781 | 0.56687641 | 0.87646121 |
| DCAF13 | 197.072032 | -0.23362 | 0.29915715 | -0.7809273 | 0.43484526 | 0.81919857 |
| RIMS2 | 1184.62679 | -1.8742383 | 0.75267945 | -2.4900883 | 0.01277114 | 0.23204118 |
| DPYS | 76.7094895 | 1.74606426 | 0.71191039 | 2.45264612 | 0.01418098 | 0.24646911 |
| LRP12 | 290.44804 | -0.103007 | 0.28883809 | -0.3566253 | 0.72137232 | 0.93044898 |
| ZFPM2 | 85.7913253 | 0.74628542 | 0.47791767 | 1.56153553 | 0.11839745 | 0.54970541 |
| AC027031.2 | 380.383274 | -0.967589 | 0.43875732 | -2.2052942 | 0.02743345 | 0.31979365 |
| OXR1 | 18792.1481 | -0.8746834 | 0.3795746 | -2.3043782 | 0.02120142 | 0.29111184 |
| AC090579.1 | 1597.24432 | -0.2311887 | 0.50699726 | -0.455996 | 0.64839286 | 0.90754156 |
| ANGPT1 | 47.3998209 | 0.39439196 | 0.66691242 | 0.59136995 | 0.55427257 | 0.8714657 |
| EIF3E | 938.018536 | 0.24040363 | 0.145383 | 1.65358832 | 0.09821119 | 0.51479031 |
| EMC2 | 781.996197 | 0.21664646 | 0.24642172 | 0.8791695 | 0.37930938 | 0.78762967 |
| NUDCD1 | 363.723412 | -0.1463436 | 0.28351196 | -0.5161813 | 0.60572782 | 0.89081188 |
| ENY2 | 456.68916 | 0.10648201 | 0.29581201 | 0.35996514 | 0.7188732 | 0.92991855 |
| EBAG9 | 282.701015 | -0.2477205 | 0.21525797 | -1.1508074 | 0.24981147 | 0.70032501 |
| SYBU | 203.87946 | -0.2888934 | 0.90015793 | -0.3209364 | 0.74825861 | 0.93746253 |
| TRPS1 | 7059.12187 | -0.5971188 | 0.36398662 | -1.6404965 | 0.10090197 | 0.51819907 |
| EIF3H | 1166.13463 | 0.02128029 | 0.17950618 | 0.11854905 | 0.90563263 | 0.97997915 |
| UTP23 | 210.848768 | -0.2407197 | 0.29976951 | -0.8030158 | 0.4219656 | 0.81266383 |
| RAD21 | 680.626396 | 0.21396713 | 0.24099347 | 0.88785449 | 0.37461903 | 0.78426576 |
| MED30 | 160.930379 | 0.10204674 | 0.32373846 | 0.31521351 | 0.75259955 | 0.93819406 |
| EXT1 | 6098.38909 | -0.018202 | 0.2585556 | -0.070399 | 0.94387612 | 0.98792866 |
| SAMD12 | 7765.15821 | -0.057696 | 0.13298922 | -0.4338397 | 6.64E-01 | 9.12E-01 |
| AC023590.1 | 756.793608 | 0.26673217 | 0.54666116 | 0.48792961 | 0.6255997 | 0.89873262 |
| SAMD12-AS1 | 826.903446 | -0.7105583 | 0.46708738 | -1.5212536 | 1.28E-01 | 0.56762585 |
| TNFRSF11B | 596.912685 | 1.12740949 | 0.68953343 | 1.63503238 | 0.10204226 | 0.52073581 |
| COLEC10 | 311.690676 | 0.09735462 | 0.6649778 | 0.14640281 | 0.88360341 | 0.97664137 |
| MAL2 | 953.205486 | -0.1220644 | 0.32401068 | -0.3767295 | 0.70637461 | 0.92556842 |
| MAL2-AS1 | 113.888572 | 0.11974434 | 0.80972034 | 0.14788358 | 0.88243465 | 0.97664137 |
| ENPP2 | 93.2283369 | -0.5975881 | 0.58677906 | -1.018421 | 0.30847792 | 0.7438271 |
| TAF2 | 1027.37074 | -0.2872126 | 0.19960058 | -1.4389367 | 0.15016847 | 0.59663305 |
| DEPTOR | 11464.5596 | -0.1090158 | 0.41688853 | -0.2614987 | 0.79370793 | 0.95224951 |

| | | | | | | |
|---|---|---|---|---|---|---|
| AC091563.1 | 556.582515 | 0.46601981 | 0.46976675 | 0.99202383 | 0.3211859 | 0.75235761 |
| COL14A1 | 853.535314 | -0.0114206 | 0.61496833 | -0.018571 | 0.98518331 | 0.99572827 |
| MRPL13 | 303.859524 | 0.1290101 | 0.26667507 | 0.48377263 | 0.62854724 | 0.89962754 |
| MTBP | 178.938061 | -0.6176937 | 0.38441107 | -1.6068571 | 0.10808572 | 0.53063658 |
| SNTB1 | 1195.51087 | 0.7327945 | 0.39168102 | 1.87089612 | 6.14E-02 | 0.43236485 |
| ZHX2 | 3106.39215 | 0.19722823 | 0.24447211 | 0.80675144 | 0.4198097 | 0.8125045 |
| AC104316.1 | 175.41104 | -0.276847 | 0.42335254 | -0.6539395 | 0.51315076 | 0.85762435 |
| DERL1 | 393.849546 | -0.0672828 | 0.22512203 | -0.2988727 | 0.76503719 | 0.9426657 |
| TBC1D31 | 215.378622 | -0.5174893 | 0.1892861 | -2.7339 | 0.00625891 | 0.16248472 |
| C8orf76 | 82.4610276 | 0.01851214 | 0.36259976 | 0.05105393 | 0.95928254 | 0.99117358 |
| ATAD2 | 127.686121 | -0.0320765 | 0.30376462 | -0.1055965 | 0.91590249 | 0.98191348 |
| WDYHV1 | 219.865725 | -0.1574693 | 0.33941415 | -0.4639444 | 0.64268758 | 0.90477318 |
| FBXO32 | 414.043798 | -0.7192572 | 0.33516772 | -2.1459621 | 0.031876 | 0.34024781 |
| FAM91A1 | 776.460239 | -0.1215118 | 0.22974705 | -0.5288937 | 0.59687918 | 0.88883692 |
| TMEM65 | 1422.51223 | 0.10833509 | 0.20594506 | 0.52603877 | 0.59886127 | 0.88883692 |
| TRMT12 | 104.21839 | 0.11044613 | 0.23960885 | 0.46094344 | 0.64483918 | 0.90507831 |
| RNF139-AS1 | 58.753482 | 0.16997163 | 0.39822622 | 0.4268218 | 0.66950913 | 0.91443968 |
| RNF139 | 131.086351 | 0.17583618 | 0.31425971 | 0.55952505 | 0.57580344 | 0.88111534 |
| TATDN1 | 681.824399 | -0.0646545 | 0.17631325 | -0.3667026 | 0.71384088 | 0.92760608 |
| NDUFB9 | 435.743707 | 0.34097069 | 0.29963969 | 1.13793567 | 0.25514735 | 0.70610544 |
| MTSS1 | 1895.83245 | 0.27129383 | 0.4006355 | 0.67715874 | 0.49830524 | 0.85092921 |
| SQLE | 289.76934 | 0.38884648 | 0.49019877 | 0.79324245 | 0.42763658 | 0.81466629 |
| WASHC5 | 525.435679 | -0.2066938 | 0.20149453 | -1.0258035 | 0.30498424 | 0.74229685 |
| NSMCE2 | 3904.11933 | -0.2663613 | 0.19470354 | -1.3680351 | 0.1713011 | 0.62772493 |
| TRIB1 | 189.857525 | 0.35532276 | 0.30993521 | 1.14644204 | 2.52E-01 | 0.70276117 |
| FAM84B | 260.394804 | -0.1155598 | 0.38834751 | -0.297568 | 0.76603288 | 0.9428901 |
| PCAT1 | 3077.10487 | -0.5912093 | 0.52075763 | -1.1352869 | 0.25625513 | 0.70709713 |
| CASC8 | 46.2748919 | 1.07011278 | 0.60784733 | 1.76049598 | 0.07832375 | 0.48060389 |
| PVT1 | 126.709885 | 1.05424449 | 0.51092716 | 2.0633949 | 3.91E-02 | 0.37079261 |
| CCDC26 | 56.5136458 | -0.1307027 | 0.47437695 | -0.2755249 | 0.78291298 | 0.94928378 |
| FAM49B | 3168.37382 | -0.256953 | 0.16016129 | -1.6043389 | 0.10863937 | 0.53207912 |
| ASAP1 | 2161.8776 | -0.5895608 | 0.24215427 | -2.4346496 | 0.01490622 | 0.24967336 |
| EFR3A | 1582.16531 | -0.0171758 | 0.31549631 | -0.0544407 | 0.95658405 | 0.99065996 |
| KCNQ3 | 490.662145 | -0.947318 | 0.66553474 | -1.4233938 | 0.15462203 | 0.60380955 |
| LRRC6 | 736.672749 | 0.15205629 | 0.17851898 | 0.85176542 | 0.3943443 | 0.79876591 |
| PHF20L1 | 1003.81036 | 0.04339825 | 0.20467761 | 0.21203221 | 0.83208191 | 0.96417211 |
| TG | 109.527407 | 1.81379705 | 0.61044169 | 2.97128631 | 0.00296555 | 0.10418078 |
| NDRG1 | 2860.45813 | 0.69328481 | 0.83032409 | 0.83495687 | 0.40374199 | 0.80377211 |
| ST3GAL1 | 358.856164 | -0.4606388 | 0.39213321 | -1.1746999 | 0.24011481 | 0.69159976 |
| ZFAT | 517.831739 | -0.3556508 | 0.24026948 | -1.4802162 | 0.13881556 | 0.58243213 |
| AC083843.3 | 531.280921 | 0.12253576 | 0.39374173 | 0.31120845 | 0.75564216 | 0.93957997 |
| KHDRBS3 | 1363.76692 | -0.6583065 | 0.32168211 | -2.0464505 | 0.04071207 | 0.37344844 |
| LINC02055 | 147.489354 | 1.92987204 | 0.92217763 | 2.09273352 | 0.03637295 | 0.35881276 |
| TRAPPC9 | 4696.33401 | -0.1357123 | 0.21267646 | -0.6381162 | 0.52339804 | 0.86006369 |
| CHRAC1 | 86.674866 | 0.2740145 | 0.32883999 | 0.83327609 | 0.40468904 | 0.80437725 |
| AGO2 | 1183.30675 | -0.0518255 | 0.19417122 | -0.2669064 | 0.78954121 | 0.95090647 |
| PTK2 | 8080.56831 | -0.4548144 | 0.17060178 | -2.6659415 | 0.0076773 | 0.17544898 |
| DENND3 | 392.838426 | 0.34335983 | 0.55902778 | 0.61420889 | 0.5390773 | 0.86542329 |
| SLC45A4 | 919.3537 | 0.61376804 | 0.36033402 | 1.70333082 | 0.08850618 | 0.49769182 |
| TSNARE1 | 483.240584 | -0.219266 | 0.24005372 | -0.913404 | 0.36103008 | 0.77731806 |
| JRK | 159.197057 | -0.0822779 | 0.32851668 | -0.2504527 | 0.80223727 | 0.95443209 |
| THEM6 | 58.3356398 | 0.38337718 | 0.45638596 | 0.84002842 | 0.40089245 | 0.80243544 |
| LY6E | 699.088255 | 2.18640826 | 0.59320632 | 3.68574676 | 2.28E-04 | 0.02471923 |
| GLI4 | 96.3512514 | -0.0552957 | 0.33708989 | -0.1640384 | 0.86970094 | 0.97454113 |

| | | | | | | |
|---|---|---|---|---|---|---|
| TOP1MT | 120.66793 | 0.01193159 | 0.27986926 | 0.04263274 | 0.9659943 | 0.99248831 |
| RHPN1 | 261.458401 | -0.0309985 | 0.41399977 | -0.0748758 | 0.94031357 | 0.98715467 |
| ZC3H3 | 330.412099 | -0.0636383 | 0.34987131 | -0.1818907 | 0.85566848 | 0.97118784 |
| GSDMD | 204.078611 | 0.82588331 | 0.41098207 | 2.0095361 | 0.04448031 | 0.38623091 |
| NAPRT | 112.687425 | 0.59188259 | 0.44880365 | 1.31880076 | 0.18723573 | 0.64195489 |
| EEF1D | 915.941355 | 0.18189603 | 0.28119082 | 0.64687755 | 0.5177112 | 0.85893097 |
| TIGD5 | 40.100707 | 0.52147849 | 0.52272649 | 0.99761252 | 0.31846729 | 0.75064854 |
| TSTA3 | 106.88974 | 0.37101507 | 0.56117587 | 0.66113866 | 0.50852339 | 0.85659934 |
| ZNF623 | 165.309164 | 0.02384356 | 0.21551315 | 0.11063623 | 0.91190481 | 0.98125148 |
| ZNF707 | 69.8249144 | -0.2796703 | 0.3085362 | -0.9064425 | 0.36470166 | 0.78076892 |
| FAM83H | 163.000228 | 0.29544054 | 0.32445083 | 0.91058649 | 0.36251329 | 0.77849113 |
| IQANK1 | 186.479607 | -0.266036 | 0.44293137 | -0.6006258 | 0.54808927 | 0.86859163 |
| SCRIB | 158.694541 | 0.13119704 | 0.38108171 | 0.34427537 | 0.73063921 | 0.93481338 |
| PUF60 | 191.888727 | 0.44035697 | 0.3937251 | 1.11843764 | 0.26338013 | 0.71407802 |
| NRBP2 | 218.619071 | 0.37334633 | 0.40027689 | 0.93272018 | 0.35096446 | 0.77084962 |
| PLEC | 438.260873 | 0.03884945 | 0.34312086 | 0.11322381 | 0.90985312 | 0.98069436 |
| PARP10 | 226.297149 | 0.53864644 | 0.50445102 | 1.06778741 | 0.28561642 | 0.72846488 |
| GRINA | 274.357687 | 0.55459811 | 0.44026508 | 1.25969135 | 0.20778073 | 0.6607067 |
| SPATC1 | 332.227549 | -0.3485008 | 0.77249809 | -0.4511348 | 0.65189238 | 0.90838511 |
| OPLAH | 146.242784 | 0.94657591 | 0.46272038 | 2.04567587 | 0.04078827 | 0.37362849 |
| EXOSC4 | 64.7808882 | 0.61460337 | 0.45827288 | 1.34112969 | 0.17987835 | 0.63399761 |
| GPAA1 | 249.169922 | 0.59014526 | 0.47967315 | 1.23030703 | 0.21858215 | 0.67202914 |
| CYC1 | 356.670874 | 0.77193443 | 0.43632931 | 1.76915556 | 0.07686792 | 0.47651607 |
| SHARPIN | 270.145861 | -0.1522255 | 0.25542832 | -0.5959619 | 0.55120067 | 0.87018163 |
| MAF1 | 212.808972 | 0.52861027 | 0.37767743 | 1.39963426 | 0.16162287 | 0.61311788 |
| MROH1 | 1056.95729 | -0.4365984 | 0.27387929 | -1.5941269 | 0.11090761 | 0.53427373 |
| SCX | 85.3642146 | -0.5420696 | 0.38331299 | -1.4141696 | 0.1573121 | 0.607051 |
| HSF1 | 525.723022 | 0.02229955 | 0.18222405 | 0.12237437 | 0.90260253 | 0.97958071 |
| DGAT1 | 123.75316 | 0.23947921 | 0.29086436 | 0.82333638 | 0.41031673 | 0.80655687 |
| SLC52A2 | 76.2619903 | 0.24362151 | 0.49432709 | 0.49283463 | 0.62212944 | 0.89835355 |
| ADCK5 | 60.6110608 | -0.5783572 | 0.33581874 | -1.7222302 | 0.08502783 | 0.49231347 |
| CPSF1 | 141.638641 | -0.2322016 | 0.32278123 | -0.7193776 | 0.47190827 | 0.83921324 |
| SLC39A4 | 53.2219704 | 0.44450878 | 0.4580331 | 0.97047305 | 3.32E-01 | 0.76035039 |
| VPS28 | 433.376841 | 0.43203611 | 0.36682569 | 1.1777695 | 0.23888851 | 0.69019287 |
| CYHR1 | 335.767143 | 0.53392513 | 0.36157014 | 1.4766848 | 0.13976015 | 0.58431708 |
| KIFC2 | 166.181857 | 0.10703829 | 0.45075015 | 0.23746702 | 0.81229451 | 0.95806331 |
| PPP1R16A | 369.866805 | -0.058462 | 0.271864 | -0.2150413 | 0.82973512 | 0.96309061 |
| GPT | 39.336184 | -0.2371949 | 0.68017433 | -0.3487267 | 0.72729453 | 0.93415339 |
| MFSD3 | 61.636849 | 0.57421538 | 0.42206938 | 1.36047628 | 0.17367925 | 0.6286116 |
| LRRC14 | 66.828552 | 0.35821841 | 0.42755048 | 0.83783886 | 0.40212121 | 0.80296585 |
| ARHGAP39 | 200.737205 | -0.1118593 | 0.27421821 | -0.4079209 | 0.68333173 | 0.9200778 |
| ZNF251 | 798.731387 | -0.0205646 | 0.18591124 | -0.1106154 | 0.91192133 | 0.98125148 |
| ZNF34 | 57.1929075 | 0.13445945 | 0.44973896 | 0.29897221 | 0.76496125 | 0.94266005 |
| RPL8 | 1387.20047 | 0.47329152 | 0.52637622 | 0.89915065 | 0.36857242 | 0.78095617 |
| ZNF517 | 41.3067828 | 0.26181329 | 0.41285241 | 0.63415711 | 0.5259783 | 0.86119802 |
| ZNF7 | 219.766982 | -0.3616002 | 0.22245765 | -1.625479 | 0.10406054 | 0.52442862 |
| COMMD5 | 76.6026171 | 0.07826994 | 0.30611598 | 0.2556872 | 0.79819238 | 0.95294186 |
| ZNF250 | 178.539457 | -0.3020174 | 0.23630258 | -1.278096 | 0.20121557 | 0.65480574 |
| ZNF16 | 158.49174 | -0.3717958 | 0.35260165 | -1.0544358 | 0.29168346 | 0.73362416 |
| C8orf33 | 116.531651 | 0.39885198 | 0.29447536 | 1.3544494 | 0.17559305 | 0.62957039 |
| WASHC1 | 1049.12929 | -0.3993637 | 0.18053603 | -2.2120996 | 0.02695978 | 0.31767329 |
| CBWD1 | 241.833161 | 0.04225928 | 0.27904759 | 0.15144113 | 0.87962774 | 0.976633 |
| DOCK8 | 61.9920242 | 0.26354955 | 0.37410555 | 0.70447913 | 0.48113445 | 0.8437862 |
| AL161725.1 | 64.2126711 | -0.8996502 | 0.7508722 | -1.1981402 | 0.23086245 | 0.68321398 |

| | | | | | | |
|---|---|---|---|---|---|---|
| KANK1 | 5742.86387 | -0.3060824 | 0.34979902 | -0.8750237 | 3.82E-01 | 0.78910908 |
| SMARCA2 | 2196.26789 | -0.1796262 | 0.21705485 | -0.8275613 | 0.40791896 | 0.80588056 |
| VLDLR-AS1 | 1151.04873 | -0.6484867 | 0.59712003 | -1.0860241 | 0.27746835 | 0.72344438 |
| VLDLR | 505.128541 | -0.2061743 | 0.4359076 | -0.472977 | 0.63622955 | 0.90237131 |
| PUM3 | 1786.67477 | -0.097055 | 0.19699626 | -0.4926743 | 0.62224271 | 0.89835355 |
| RFX3 | 2822.82669 | -0.0407448 | 0.30246471 | -0.1347094 | 0.89284164 | 0.97782787 |
| RFX3-AS1 | 2325.39135 | -0.5063409 | 0.3458422 | -1.4640808 | 0.14317188 | 0.5883902 |
| GLIS3 | 10436.9176 | 0.30227032 | 0.18578893 | 1.62695547 | 0.10374655 | 0.52441026 |
| SLC1A1 | 647.125301 | 0.68832373 | 0.40978329 | 1.67972619 | 0.0930106 | 0.50795488 |
| SPATA6L | 1938.67391 | 0.1407914 | 0.29043723 | 0.48475673 | 6.28E-01 | 8.99E-01 |
| CDC37L1 | 687.678099 | 0.18174424 | 0.34600102 | 0.52527082 | 0.59939494 | 0.88883692 |
| AK3 | 5374.98805 | 0.4818314 | 0.40146425 | 1.20018505 | 0.23006748 | 0.68234886 |
| RCL1 | 1023.90955 | 0.17359869 | 0.2802217 | 0.61950479 | 0.53558387 | 0.8642718 |
| JAK2 | 1179.17374 | -0.0391758 | 0.27039218 | -0.1448851 | 0.88480158 | 0.97664137 |
| PLGRKT | 581.500877 | -0.2977355 | 0.20479593 | -1.4538156 | 0.14599743 | 0.59264906 |
| CD274 | 82.484131 | -0.6418438 | 0.58514964 | -1.0968884 | 0.27269019 | 0.72070318 |
| RIC1 | 2074.53064 | -0.4339817 | 0.27921114 | -1.5543136 | 0.12010962 | 0.55298199 |
| ERMP1 | 2136.50204 | -0.7627481 | 0.35256848 | -2.1634042 | 0.0305101 | 0.33556032 |
| KIAA2026 | 2899.35559 | -0.4250896 | 0.30605235 | -1.388944 | 0.16484979 | 0.61673853 |
| MLANA | 43.7616708 | -0.1933254 | 0.79041332 | -0.2445877 | 0.80677566 | 0.95566275 |
| RANBP6 | 216.72961 | 0.38378383 | 0.41170404 | 0.9321838 | 0.35124155 | 0.77103986 |
| UHRF2 | 1147.60167 | -0.1958898 | 0.18020234 | -1.0870549 | 0.27701255 | 0.7230876 |
| AL133480.1 | 131.873695 | 0.17139915 | 0.53602971 | 0.31975681 | 0.74915269 | 0.93748128 |
| GLDC | 735.87666 | 1.56152138 | 0.48053095 | 3.24957503 | 0.00115578 | 0.06094516 |
| AL162411.1 | 250.926821 | 1.40955642 | 0.57697454 | 2.4430132 | 0.0145652 | 0.24762526 |
| KDM4C | 4424.37949 | -0.1426324 | 0.21701754 | -0.6572393 | 0.51102709 | 0.85762435 |
| DMAC1 | 197.725531 | 0.6747689 | 0.28501116 | 2.36751752 | 0.01790787 | 0.26845799 |
| PTPRD | 285.31955 | 0.56500927 | 0.45037198 | 1.25453912 | 0.20964612 | 0.66233331 |
| TYRP1 | 114.376839 | -0.4197715 | 0.48220971 | -0.8705164 | 0.38401827 | 0.791697 |
| LURAP1L-AS | 74.890405 | 0.07418365 | 0.71996142 | 0.10303837 | 0.91793251 | 0.98201754 |
| LURAP1L | 327.037928 | -1.1555487 | 0.51922134 | -2.2255417 | 0.0260449 | 0.31246781 |
| MPDZ | 1702.43031 | -0.0411775 | 0.22752682 | -0.1809787 | 0.85638427 | 0.97118784 |
| NFIB | 3244.67992 | -0.2988349 | 0.27046385 | -1.1048979 | 0.26920382 | 0.71818262 |
| ZDHHC21 | 2408.03322 | -0.3887512 | 0.25269906 | -1.5383959 | 0.12395183 | 0.56105491 |
| FREM1 | 1986.35412 | -0.737244 | 0.51353417 | -1.435628 | 1.51E-01 | 5.99E-01 |
| TTC39B | 1060.85908 | -0.4880438 | 0.31089517 | -1.5698018 | 0.11646124 | 0.54686687 |
| SNAPC3 | 721.023343 | -0.1520271 | 0.2050291 | -0.7414904 | 0.45839618 | 0.8331421 |
| PSIP1 | 641.950089 | 0.18373838 | 0.22679641 | 0.81014678 | 0.41785582 | 0.81062751 |
| CCDC171 | 1578.23224 | -0.3372082 | 0.30491416 | -1.105912 | 0.26876462 | 0.71818262 |
| BNC2 | 2211.00232 | -2.3076538 | 0.45425453 | -5.0800899 | 3.77E-07 | 0.00071188 |
| CNTLN | 1018.06645 | -0.1448856 | 0.15791733 | -0.9174774 | 0.35889251 | 0.77600444 |
| SH3GL2 | 457.113114 | 0.23926604 | 0.39192276 | 0.61049285 | 0.54153538 | 0.86582694 |
| ADAMTSL1 | 89.1848451 | -0.1924043 | 0.3618734 | -0.5316896 | 0.59494102 | 0.88857711 |
| SAXO1 | 67.4829467 | -1.2864495 | 0.4734885 | -2.7169605 | 0.00658845 | 0.1680054 |
| RRAGA | 162.28843 | 0.92586702 | 0.45101702 | 2.05284277 | 0.04008783 | 0.37342745 |
| HAUS6 | 192.333193 | -0.2807789 | 0.21897973 | -1.2822141 | 0.19976755 | 0.65230079 |
| PLIN2 | 381.502542 | 0.31048618 | 0.64045562 | 0.48478953 | 0.62782565 | 0.89919293 |
| DENND4C | 2937.16621 | 0.14881731 | 0.26178477 | 0.56847198 | 0.56971453 | 0.87807251 |
| RPS6 | 1425.47088 | 0.48538065 | 0.50326432 | 0.96446465 | 0.33481302 | 0.76242642 |
| ACER2 | 104.738992 | 0.48947015 | 0.32307673 | 1.51502754 | 1.30E-01 | 5.71E-01 |
| MLLT3 | 3119.26048 | 0.11274656 | 0.26775974 | 0.42107362 | 0.67370132 | 0.91636338 |
| FOCAD | 2233.62144 | 0.01621409 | 0.16718417 | 0.0969834 | 0.92273957 | 0.98215577 |
| HACD4 | 40.2841798 | -0.4081437 | 0.59074127 | -0.6909009 | 0.48962781 | 0.84614483 |
| KLHL9 | 200.407055 | 0.60563875 | 0.27710318 | 2.18560738 | 0.02884434 | 0.3275529 |

| | | | | | | |
|---|---|---|---|---|---|---|
| LINC01239 | 249.696249 | 0.65626747 | 0.45832598 | 1.43187928 | 0.15217837 | 0.60219416 |
| AL391117.1 | 109.272028 | 0.82709774 | 0.53713729 | 1.53982559 | 0.12360287 | 0.56102027 |
| TUSC1 | 62.1123463 | 0.2753557 | 0.50234995 | 0.54813522 | 0.58359906 | 0.88393074 |
| CAAP1 | 1449.66161 | 0.02727012 | 0.24208537 | 0.11264669 | 0.91031067 | 0.98069436 |
| PLAA | 459.295056 | -0.180741 | 0.24204064 | -0.7467381 | 0.45522165 | 0.83121393 |
| IFT74 | 1412.40759 | 0.0855184 | 0.17113527 | 0.49971228 | 0.61727769 | 0.89599505 |
| LRRC19 | 154.322376 | 0.89971489 | 0.43953435 | 2.04697287 | 0.04066075 | 0.37344844 |
| TEK | 43.2664663 | 0.78695711 | 0.74016573 | 1.06321744 | 0.28768335 | 0.7306066 |
| MOB3B | 1975.52902 | 0.64411587 | 0.33727202 | 1.90978149 | 0.05616135 | 0.41843207 |
| C9orf72 | 428.615376 | 0.14891019 | 0.29473214 | 0.50523908 | 0.61339092 | 0.89441126 |
| LINGO2 | 464.835593 | 0.38208731 | 0.66629121 | 0.57345393 | 0.56633738 | 0.87642772 |
| ACO1 | 388.090465 | 0.21723584 | 0.19430812 | 1.11799669 | 0.26356841 | 0.71429196 |
| DDX58 | 796.691531 | -0.3624376 | 0.74663024 | -0.4854312 | 0.62737053 | 0.89919293 |
| TOPORS | 271.095465 | 0.19522201 | 0.24501404 | 0.79677886 | 0.42557947 | 0.81466629 |
| SMIM27 | 71.8296749 | 0.48947306 | 0.37976323 | 1.28889008 | 0.1974363 | 0.64964978 |
| NDUFB6 | 407.585951 | 0.03134233 | 0.26339571 | 0.11899334 | 0.90528063 | 0.97992575 |
| APTX | 699.869912 | -0.1149132 | 0.25343433 | -0.4534241 | 0.65024342 | 0.90825486 |
| DNAJA1 | 486.24619 | 0.57823114 | 0.29878019 | 1.93530616 | 0.05295274 | 0.41073851 |
| SMU1 | 410.909175 | 0.06290907 | 0.20428834 | 0.30794252 | 0.75812607 | 0.94056395 |
| B4GALT1 | 1068.32251 | 0.06002417 | 0.22906818 | 0.26203628 | 0.79329347 | 0.95224951 |
| BAG1 | 334.952368 | 0.46012221 | 0.34522011 | 1.33283724 | 0.18258522 | 0.63741052 |
| CHMP5 | 311.216577 | 0.2106882 | 0.20575695 | 1.02396637 | 0.30585116 | 0.74290739 |
| NFX1 | 1361.96836 | -0.2344288 | 0.24073364 | -0.9738098 | 0.33015099 | 0.75885186 |
| AQP7 | 51.5434645 | -0.048711 | 0.79613914 | -0.0611841 | 0.9512126 | 0.98938795 |
| AQP3 | 137.242759 | -0.805699 | 0.67606195 | -1.1917532 | 0.23335802 | 0.68484327 |
| NOL6 | 45.1965907 | -0.1810394 | 0.41740032 | -0.4337308 | 0.66448392 | 0.91219788 |
| UBE2R2 | 2025.70436 | -0.3404083 | 0.2689538 | -1.2656758 | 0.20562919 | 0.65813563 |
| UBAP2 | 1450.82425 | -0.3180897 | 0.15103601 | -2.106052 | 0.03519984 | 0.3546565 |
| DCAF12 | 283.369888 | -0.2743101 | 0.27408833 | -1.0008092 | 0.31691905 | 0.74873613 |
| UBAP1 | 2163.70218 | -0.2168308 | 0.22524473 | -0.9626453 | 0.33572553 | 0.76363358 |
| KIF24 | 118.305254 | -0.1501226 | 0.42205738 | -0.3556925 | 0.72207088 | 0.93079284 |
| NUDT2 | 155.640518 | 0.31935057 | 0.43144635 | 0.74018605 | 0.45918711 | 0.83384693 |
| C9orf24 | 58.9628519 | 0.64744079 | 0.43095083 | 1.5023542 | 0.13300566 | 0.57682553 |
| FAM219A | 403.402786 | -0.0283932 | 0.26773264 | -0.1060506 | 0.91554221 | 0.98176629 |
| DNAI1 | 189.406938 | 0.83071182 | 0.46357789 | 1.79195739 | 0.07313979 | 0.47001711 |
| RPP25L | 68.1289483 | 0.3623327 | 0.4731747 | 0.76574827 | 0.4438261 | 0.82410854 |
| DCTN3 | 132.865055 | 1.16944363 | 0.39972218 | 2.92564108 | 0.00343747 | 0.1138728 |
| SIGMAR1 | 50.1143072 | 0.28770094 | 0.41201286 | 0.69828147 | 0.48500119 | 0.8447301 |
| IL11RA | 56.315191 | -0.1587155 | 0.55173066 | -0.2876685 | 0.7736005 | 0.94558571 |
| AL162231.2 | 61.4263006 | 0.81686672 | 0.30699733 | 2.66082681 | 0.0077949 | 0.17721667 |
| VCP | 610.713988 | 0.15176192 | 0.23588605 | 0.64336961 | 0.5199843 | 0.85935232 |
| PIGO | 88.0674096 | 0.04589552 | 0.29012597 | 0.15819171 | 0.87430573 | 0.97564728 |
| STOML2 | 206.115365 | 0.62884521 | 0.39743804 | 1.58224717 | 0.11359315 | 0.53989968 |
| FAM214B | 101.740718 | -0.0003023 | 0.36283147 | -0.0008333 | 0.99933516 | 0.99987432 |
| UNC13B | 1542.38746 | 0.0507013 | 0.22901761 | 0.22138604 | 0.82479186 | 0.96146415 |
| RUSC2 | 982.942346 | -0.2688041 | 0.3370481 | -0.7975243 | 0.42514658 | 0.81466629 |
| TESK1 | 94.533498 | -0.3117895 | 0.34669224 | -0.8993263 | 0.36847886 | 0.78095617 |
| CD72 | 129.245842 | -0.2772223 | 0.41947881 | -0.6608732 | 0.50869364 | 0.85659934 |
| CCDC107 | 199.280554 | 0.43025265 | 0.43320785 | 0.99317833 | 3.21E-01 | 0.75144499 |
| TLN1 | 443.321519 | 0.26039406 | 0.19933259 | 1.30632958 | 0.1914405 | 0.64571429 |
| CREB3 | 103.530214 | 0.78478034 | 0.36955061 | 2.1236072 | 0.033703 | 0.35097935 |
| GBA2 | 514.232579 | -0.1882975 | 0.31061058 | -0.6062174 | 0.54437043 | 0.86654229 |
| RGP1 | 204.926254 | 0.08292099 | 0.19864556 | 0.41743188 | 0.67636254 | 0.91658712 |
| AL133410.1 | 244.001247 | -0.1866359 | 0.26548746 | -0.7029931 | 0.48206003 | 0.84389635 |

| | | | | | | |
|---|---|---|---|---|---|---|
| HINT2 | 101.025035 | 0.29303594 | 0.43425985 | 0.674794 | 0.49980666 | 0.85244523 |
| TMEM8B | 207.339376 | -0.1685691 | 0.34615514 | -0.4869757 | 0.62627558 | 0.89873262 |
| RECK | 389.90321 | 0.19180939 | 0.26352166 | 0.72786955 | 0.46669344 | 0.83700661 |
| CLTA | 1206.25573 | -0.2669816 | 0.21499565 | -1.2417999 | 0.21431042 | 0.66827012 |
| GNE | 907.226882 | -0.1617796 | 0.29335291 | -0.5514844 | 0.58130163 | 0.88290096 |
| RNF38 | 1993.3065 | -0.1165208 | 0.15224861 | -0.7653327 | 0.44407344 | 0.82437446 |
| EBLN3P | 498.690627 | 0.09967639 | 0.21988559 | 0.45331023 | 0.65032537 | 0.90825486 |
| ZCCHC7 | 4631.29125 | -0.2717179 | 0.15446774 | -1.7590591 | 0.07856747 | 0.48060389 |
| GRHPR | 242.58258 | 0.36961602 | 0.26720681 | 1.38325824 | 0.1665857 | 0.61925103 |
| ZBTB5 | 278.28553 | 0.10861572 | 0.18108444 | 0.59980701 | 0.54863486 | 0.8690392 |
| AL138752.2 | 563.052547 | -0.056366 | 0.22861837 | -0.2465506 | 0.80525601 | 0.95522565 |
| TRMT10B | 545.093302 | 0.1689154 | 0.23437205 | 0.72071477 | 0.47108502 | 0.83921324 |
| DCAF10 | 1416.31515 | 0.00022575 | 0.16299715 | 0.00138497 | 0.99889495 | 0.99980325 |
| FAM95C | 253.435986 | -0.5229501 | 0.29832342 | -1.7529637 | 0.07960823 | 0.48176967 |
| ANKRD18A | 240.701672 | -0.4784695 | 0.30538057 | -1.5667973 | 0.11716207 | 0.54757066 |
| CNTNAP3 | 201.741462 | -1.6588229 | 0.53560862 | -3.0970802 | 0.00195437 | 0.08583557 |
| GLIDR | 374.310314 | -0.1047086 | 0.46580811 | -0.2247892 | 0.82214329 | 0.9607795 |
| FAM95B1 | 48.5246577 | 0.17053388 | 0.9158499 | 0.18620287 | 0.85228567 | 0.97076831 |
| BMS1P14 | 307.135094 | -0.3016626 | 0.587884 | -0.5131328 | 0.60785844 | 0.89203142 |
| CBWD6 | 138.786555 | 0.42599175 | 0.44305283 | 0.961492 | 0.33630484 | 0.7637525 |
| AL591926.2 | 70.4507945 | 0.39520288 | 0.62027631 | 0.63714004 | 0.52403363 | 0.86006369 |
| CNTNAP3B | 281.055604 | 1.98031675 | 0.63971439 | 3.09562638 | 0.00196398 | 0.08590121 |
| FAM27C | 107.579003 | 0.31089941 | 0.45893288 | 0.67743983 | 0.49812693 | 0.85092921 |
| AL391987.2 | 100.604597 | 0.04331841 | 0.33584714 | 0.12898252 | 0.89737148 | 0.97823597 |
| AL512625.3 | 139.924688 | 0.36359426 | 0.49641603 | 0.73243859 | 0.46390091 | 0.83540111 |
| AL512625.1 | 40.1814196 | -0.0610578 | 0.68761618 | -0.0887963 | 0.92924382 | 0.98331361 |
| CBWD5 | 456.306006 | -0.5076973 | 0.33063386 | -1.5355273 | 0.12465437 | 0.56139092 |
| ZNF658 | 198.441933 | 0.21250749 | 0.38335535 | 0.55433553 | 0.57934924 | 0.8821758 |
| CBWD3 | 95.0629521 | 0.02661271 | 0.36939292 | 0.07204445 | 0.94256653 | 0.98772472 |
| PGM5 | 385.903575 | -1.7070884 | 1.29567612 | -1.3175271 | 0.18766199 | 0.64222928 |
| AL161457.2 | 142.370325 | -0.4154641 | 0.2874175 | -1.4455072 | 0.14831547 | 0.59438683 |
| AL353616.2 | 493.774316 | -0.4541069 | 0.93168778 | -0.4874025 | 0.62597313 | 0.89873262 |
| PIP5K1B | 728.062524 | 0.40003814 | 0.50979305 | 0.78470692 | 0.43262542 | 0.8178831 |
| FAM189A2 | 184.756652 | 1.22186307 | 0.71152817 | 1.71723779 | 0.08593573 | 0.49439246 |
| APBA1 | 83.2293377 | 0.62794628 | 0.38063364 | 1.64973932 | 0.09899626 | 0.51618562 |
| PTAR1 | 1441.71472 | 0.15810427 | 0.17383326 | 0.90951677 | 3.63E-01 | 0.77918595 |
| C9orf135 | 83.7422287 | 0.85745704 | 0.92092791 | 0.93107943 | 0.35181248 | 0.77151834 |
| MAMDC2-AS | 950.794016 | 0.08191764 | 0.29059558 | 0.28189568 | 0.7780235 | 0.94694572 |
| MAMDC2 | 848.070309 | 0.64213121 | 0.48482415 | 1.32446208 | 0.18534963 | 0.64030106 |
| SMC5-AS1 | 1175.13733 | 0.09328167 | 0.23659055 | 0.3942747 | 0.69337824 | 0.92224682 |
| SMC5 | 1912.74832 | -0.0820492 | 0.20171285 | -0.4067623 | 0.68418261 | 0.9200778 |
| KLF9 | 806.430764 | 0.14104367 | 0.47397294 | 0.29757748 | 0.76602566 | 0.9428901 |
| TRPM3 | 291.121161 | 0.90670692 | 0.37521263 | 2.41651492 | 0.01566988 | 0.25227487 |
| CEMIP2 | 1484.91121 | 0.01429915 | 0.42585283 | 0.03357769 | 9.73E-01 | 0.99327086 |
| ABHD17B | 358.890666 | -0.2373928 | 0.31195341 | -0.760988 | 0.44666422 | 0.82571251 |
| C9orf85 | 659.476442 | -0.090677 | 0.19385188 | -0.4677646 | 0.63995293 | 0.90330608 |
| GDA | 138.704273 | 1.8519408 | 0.48894471 | 3.78762827 | 0.00015209 | 0.02008985 |
| LINC01504 | 135.1045 | 0.14061562 | 1.55708719 | 0.09030684 | 0.92804339 | 0.98297055 |
| ZFAND5 | 1839.00497 | 0.21459924 | 0.46418401 | 0.46231502 | 0.64385543 | 0.90485013 |
| TMC1 | 49.3025596 | -0.8666161 | 0.43536418 | -1.9905544 | 0.0465299 | 0.39097111 |
| ALDH1A1 | 1061.34527 | 0.65229006 | 0.26657172 | 2.44695899 | 0.01440672 | 0.24762526 |
| ANXA1 | 109.89004 | 1.92859465 | 0.84606407 | 2.27949009 | 0.02263795 | 0.2983102 |
| C9orf41-AS1 | 86.4594195 | 0.12783905 | 0.50079279 | 0.25527335 | 0.79851199 | 0.95297658 |
| CARNMT1 | 692.810533 | 0.11990643 | 0.14531495 | 0.82514864 | 4.09E-01 | 0.80623405 |

| | | | | | | |
|---|---|---|---|---|---|---|
| NMRK1 | 471.624766 | -0.0056291 | 0.27683581 | -0.0203336 | 0.98377727 | 0.99518552 |
| OSTF1 | 306.845491 | -0.3150074 | 0.29197821 | -1.0788731 | 0.28064431 | 0.72530438 |
| PCSK5 | 421.494318 | -0.5294331 | 0.55283626 | -0.9576671 | 3.38E-01 | 7.64E-01 |
| RFK | 155.694782 | 1.01266124 | 0.29228734 | 3.46460857 | 0.000531 | 0.04005562 |
| GCNT1 | 127.690049 | 0.39380324 | 0.49122932 | 0.80166885 | 4.23E-01 | 0.8136431 |
| PRUNE2 | 630.837776 | 1.07899602 | 0.47104832 | 2.29062703 | 0.02198499 | 0.29509055 |
| PCA3 | 83.1872729 | 1.25335714 | 0.67170777 | 1.86592623 | 0.06205169 | 0.43390193 |
| VPS13A | 1869.44185 | -0.1807707 | 0.17426468 | -1.0373344 | 0.29958005 | 0.73673761 |
| GNA14 | 60.263281 | -0.5137542 | 0.48609287 | -1.0569055 | 0.29055471 | 0.73285033 |
| GNAQ | 4907.12012 | -0.2318562 | 0.17694326 | -1.3103421 | 1.90E-01 | 0.64494432 |
| CEP78 | 201.47884 | -0.3782992 | 0.40124783 | -0.9428068 | 0.34577976 | 0.76814746 |
| PSAT1 | 75.0615589 | 0.79662299 | 0.41179266 | 1.93452451 | 5.30E-02 | 4.11E-01 |
| TLE4 | 800.652032 | -0.2653738 | 0.28434936 | -0.9332667 | 0.35068226 | 0.77061421 |
| AL161782.1 | 59.2974565 | 0.36117247 | 0.60130471 | 0.600648 | 0.54807446 | 0.86859163 |
| TLE1 | 958.165211 | -0.0289376 | 0.38664362 | -0.074843 | 0.9403396 | 0.98715467 |
| AL591368.1 | 94.898428 | 0.41512254 | 0.80133619 | 0.51803793 | 0.6044318 | 0.89013087 |
| RASEF | 348.030971 | 0.37605174 | 0.39091301 | 0.96198318 | 0.33605806 | 0.7637525 |
| AL137847.1 | 81.7745074 | -0.0026456 | 0.56302677 | -0.0046988 | 0.99625091 | 0.99914308 |
| FRMD3 | 719.640702 | -0.2542069 | 0.22226743 | -1.1436984 | 0.25274875 | 0.7033301 |
| AL137847.2 | 57.997891 | -0.1240494 | 0.59314011 | -0.2091401 | 0.83433884 | 0.96510927 |
| IDNK | 176.757679 | -0.3272735 | 0.25862371 | -1.2654426 | 0.20571272 | 0.65813563 |
| UBQLN1 | 1029.65073 | 0.08310833 | 0.21336132 | 0.38951919 | 0.69689212 | 0.92322348 |
| GKAP1 | 662.932589 | -0.0289602 | 0.28133148 | -0.1029399 | 0.91801066 | 0.98202161 |
| AL354733.2 | 48.9871648 | -0.5424007 | 0.45957042 | -1.1802342 | 0.23790709 | 0.68982886 |
| KIF27 | 826.751351 | -0.4427006 | 0.1344157 | -3.2935188 | 0.00098942 | 0.0559889 |
| C9orf64 | 105.10441 | 0.13077043 | 0.27749385 | 0.47125524 | 0.63745847 | 0.90255504 |
| HNRNPK | 1304.9811 | 0.20637394 | 0.29371907 | 0.70262357 | 0.48229037 | 0.84389635 |
| AL354733.3 | 113.190238 | 0.27729121 | 0.60208987 | 0.46054787 | 0.64512302 | 0.90518019 |
| RMI1 | 212.991357 | -0.361685 | 0.24280466 | -1.489613 | 0.13632603 | 0.57787125 |
| NTRK2 | 1742.79458 | 0.03808286 | 0.30752283 | 0.12383751 | 0.90144393 | 0.97941893 |
| AGTPBP1 | 1101.9606 | -0.2862409 | 0.24904949 | -1.1493334 | 0.25041855 | 0.70109763 |
| NAA35 | 1220.13333 | -0.241137 | 0.13298816 | -1.8132217 | 0.06979763 | 0.45914671 |
| GOLM1 | 1493.93579 | -1.0345664 | 0.25490118 | -4.0586963 | 4.93E-05 | 0.01125476 |
| C9orf153 | 160.669858 | 0.31962898 | 0.43475911 | 0.73518638 | 4.62E-01 | 0.83531269 |
| ISCA1 | 338.451693 | 0.26960915 | 0.31629721 | 0.85239183 | 0.39399666 | 0.79845072 |
| TUT7 | 1029.91175 | -0.364824 | 0.20822751 | -1.7520449 | 0.07976609 | 0.48221063 |
| AL513318.2 | 192.642533 | 0.01904673 | 0.38166077 | 0.04990487 | 0.9601982 | 0.99153894 |
| DAPK1 | 7442.71946 | -0.3389224 | 0.35832954 | -0.9458399 | 0.34423023 | 0.76702717 |
| CTSL | 407.164323 | -0.3796796 | 0.41925199 | -0.9056118 | 0.36514136 | 0.78095617 |
| CDK20 | 58.7925362 | 0.04170763 | 0.36188754 | 0.11525026 | 0.90824676 | 0.98047754 |
| SPIN1 | 2962.32465 | -0.0802643 | 0.20210852 | -0.3971347 | 0.69126811 | 0.92206635 |
| SECISBP2 | 1427.49355 | -0.4149651 | 0.23593173 | -1.7588354 | 0.07860546 | 0.48060389 |
| SEMA4D | 2095.79782 | -0.2688112 | 0.29718413 | -0.9045275 | 0.36571578 | 0.78095617 |
| GADD45G | 45.5295189 | -0.0439077 | 0.69329548 | -0.0633319 | 0.94950222 | 0.98919519 |
| LINC01508 | 62.9540252 | 2.3762268 | 0.74341998 | 3.19634509 | 0.00139181 | 0.06809021 |
| SYK | 417.968751 | -0.0742774 | 0.3258907 | -0.2279213 | 0.81970741 | 0.96014363 |
| AL158071.1 | 265.481274 | -0.7157997 | 0.85897008 | -0.8333232 | 0.40466248 | 0.80437725 |
| AL158071.4 | 798.433841 | -0.5849751 | 0.54450488 | -1.0743249 | 0.28267711 | 0.72604837 |
| AL158071.5 | 66.9195218 | -0.4854536 | 0.48158466 | -1.0080338 | 0.31343821 | 0.74749123 |
| AUH | 3764.0865 | -0.2142836 | 0.27909497 | -0.7677802 | 0.44261778 | 0.82299244 |
| NFIL3 | 368.89816 | 0.10859071 | 0.62549581 | 0.17360742 | 0.862174 | 0.97270724 |
| SPTLC1 | 758.353957 | -0.1588754 | 0.15955188 | -0.9957604 | 0.3193666 | 0.75094569 |
| IARS | 618.260288 | -0.1687458 | 0.21588841 | -0.7816346 | 0.43442937 | 0.81919857 |
| NOL8 | 293.725531 | -0.1485491 | 0.1577836 | -0.9414737 | 0.34646216 | 0.76864066 |

| | | | | | | |
|---|---|---|---|---|---|---|
| CENPP | 617.536236 | -1.2565783 | 0.54320811 | -2.3132539 | 0.02070868 | 0.28817321 |
| IPPK | 240.256913 | -0.1808381 | 0.20343598 | -0.8889189 | 0.37404668 | 0.78362928 |
| BICD2 | 244.977614 | 0.24583648 | 0.28732667 | 0.85559925 | 0.39221948 | 0.79790962 |
| ZNF484 | 156.401735 | -0.3160992 | 0.25698071 | -1.2300502 | 0.21867832 | 0.67202914 |
| CARD19 | 137.481063 | 0.04796956 | 0.24790114 | 0.19350279 | 0.84656522 | 0.96883382 |
| NINJ1 | 246.664068 | 0.59744393 | 0.35884648 | 1.66490121 | 0.09593247 | 0.5117083 |
| WNK2 | 74.9854757 | 0.37311066 | 0.39734349 | 0.9390129 | 0.34772412 | 0.76900643 |
| FAM120AOS | 165.307288 | 0.02780531 | 0.22657916 | 0.12271785 | 0.90233053 | 0.97941893 |
| FAM120A | 1386.58472 | 0.14356268 | 0.14929059 | 0.96163249 | 0.33623424 | 0.7637525 |
| PHF2 | 372.13701 | -0.1020694 | 0.18863322 | -0.5410997 | 0.58843888 | 0.88571413 |
| PTPDC1 | 594.495731 | 0.0054796 | 0.27541626 | 0.01989572 | 0.98412656 | 0.99518552 |
| AL158152.2 | 739.472972 | -0.0515263 | 0.22715454 | -0.2268336 | 0.82055316 | 0.96036564 |
| MIRLET7DHG | 57.5572738 | 0.55579848 | 0.83111535 | 0.66873807 | 0.50366258 | 0.85525037 |
| ZNF169 | 292.508049 | -0.1786712 | 0.22075211 | -0.8093747 | 0.41829962 | 0.81099658 |
| MFSD14B | 658.851177 | -0.2962384 | 0.23513296 | -1.2598761 | 0.20771404 | 0.6607067 |
| FBP1 | 83.004497 | 1.16875158 | 0.47529166 | 2.45901974 | 0.0139317 | 0.24471246 |
| C9orf3 | 7143.50054 | -0.507407 | 0.2085794 | -2.4326802 | 0.01498753 | 0.25020868 |
| AL353768.1 | 87.8753264 | 0.09123302 | 0.51595643 | 0.1768231 | 0.85964734 | 0.97184883 |
| FANCC | 1617.2909 | -0.342193 | 0.21472315 | -1.5936473 | 0.11101506 | 0.53440159 |
| PTCH1 | 403.503364 | -0.0855991 | 0.30770336 | -0.2781871 | 0.78086873 | 0.9480401 |
| LINC00476 | 1320.63137 | -0.1142021 | 0.38601243 | -0.2958509 | 0.76734396 | 0.94316227 |
| ERCC6L2 | 1314.92535 | -0.2685318 | 0.15145717 | -1.7729885 | 0.07623061 | 0.47519121 |
| AL160269.1 | 50.0156766 | 0.01645981 | 0.33404962 | 0.04927354 | 0.96070131 | 0.99155365 |
| SLC35D2 | 284.676943 | 0.03113232 | 0.22246808 | 0.13994064 | 0.88870689 | 0.97729111 |
| ZNF367 | 69.7206355 | -0.5740242 | 0.48832165 | -1.1755043 | 0.23979303 | 0.69112505 |
| HABP4 | 523.026186 | 0.05270297 | 0.25324125 | 0.20811369 | 0.8351402 | 0.96511435 |
| CDC14B | 1179.30337 | 0.86715275 | 0.32858741 | 2.63903219 | 0.00831431 | 0.18347203 |
| AAED1 | 82.5016073 | 0.03785798 | 0.34584906 | 0.10946389 | 0.91283456 | 0.98135874 |
| AL589843.1 | 218.933053 | 0.50142421 | 0.65709943 | 0.76308728 | 0.44541135 | 0.82571115 |
| ZNF510 | 176.226382 | -0.539757 | 0.17706848 | -3.0482952 | 0.00230144 | 0.09212026 |
| ZNF782 | 679.370211 | -0.3092725 | 0.25056132 | -1.2343185 | 0.21708424 | 0.66978787 |
| MFSD14C | 1556.8323 | -0.2852133 | 0.21169184 | -1.3473044 | 0.17788223 | 0.63179521 |
| AL590705.1 | 103.439621 | -0.0817639 | 0.44136857 | -0.1852507 | 0.85303237 | 0.97076831 |
| CCDC180 | 185.488125 | -0.0778704 | 0.4439994 | -0.1753839 | 0.86077798 | 0.9720639 |
| TDRD7 | 369.45315 | -0.121664 | 0.2991153 | -0.4067463 | 0.68419432 | 0.9200778 |
| TMOD1 | 80.6270208 | -3.1153252 | 0.79564535 | -3.9154697 | 9.02E-05 | 0.01515778 |
| TSTD2 | 455.310636 | -0.249469 | 0.21339837 | -1.1690296 | 0.24239171 | 0.69337349 |
| NCBP1 | 431.157915 | -0.1695033 | 0.30638696 | -0.5532327 | 0.58010409 | 0.88258799 |
| XPA | 789.012209 | -0.0005869 | 0.25678949 | -0.0022855 | 9.98E-01 | 1.00E+00 |
| TRMO | 128.237284 | -0.3270403 | 0.26782764 | -1.2210848 | 0.22205391 | 0.67548814 |
| ANP32B | 463.439151 | -0.0310653 | 0.20336422 | -0.1527571 | 0.87858984 | 0.9765717 |
| NANS | 136.268358 | 0.27156915 | 0.32872803 | 0.82612107 | 0.40873542 | 0.80602525 |
| TRIM14 | 156.599644 | 0.76967842 | 0.65579072 | 1.17366469 | 0.24052936 | 0.69178991 |
| TBC1D2 | 126.517289 | -0.3692977 | 0.38465055 | -0.9600863 | 0.33701178 | 0.76409325 |
| ANKS6 | 515.146693 | -0.3685104 | 0.35275165 | -1.0446737 | 0.2961738 | 0.7366786 |
| GALNT12 | 90.7132056 | -1.1118704 | 0.59239179 | -1.8769174 | 0.06052942 | 0.42985653 |
| TGFBR1 | 610.609923 | -0.5333847 | 0.42261297 | -1.2621116 | 0.20690862 | 0.65951781 |
| ALG2 | 166.898085 | 0.66742297 | 0.2785816 | 2.39578987 | 0.01658459 | 0.2577246 |
| SEC61B | 426.551134 | 0.52389397 | 0.24306743 | 2.15534422 | 0.03113491 | 0.3383722 |
| AL359710.1 | 689.990606 | 0.46239241 | 0.38549051 | 1.19949105 | 0.23033706 | 0.68263905 |
| STX17 | 668.389795 | 0.10183782 | 0.20389046 | 0.49947321 | 0.61744606 | 0.89599505 |
| ERP44 | 966.225071 | 0.02817444 | 0.18915895 | 0.14894586 | 0.88159635 | 0.97664137 |
| INVS | 2207.71579 | 0.03209291 | 0.17139816 | 0.18724184 | 0.85147102 | 0.9707215 |
| TEX10 | 383.783094 | -0.2744105 | 0.2850849 | -0.9625571 | 0.33576984 | 0.76363358 |

| | | | | | | |
|---|---|---|---|---|---|---|
| MSANTD3 | 72.8315459 | -0.554652 | 0.41859349 | -1.3250372 | 0.18515881 | 0.64030106 |
| MSANTD3-TI | 50.9955011 | -0.4198042 | 0.46772754 | -0.89754 | 0.36943084 | 0.78095617 |
| PLPPR1 | 1510.83788 | -0.2314808 | 0.42976726 | -0.5386189 | 5.90E-01 | 8.86E-01 |
| MRPL50 | 136.504982 | 0.47976802 | 0.36634541 | 1.30960565 | 0.19032928 | 0.64549154 |
| ZNF189 | 271.153584 | -0.2392991 | 0.19633142 | -1.2188528 | 0.22290007 | 0.67594033 |
| ALDOB | 359.404896 | 1.79315679 | 0.9652738 | 1.85766649 | 0.06321639 | 0.43730767 |
| TMEM246 | 63.235942 | 0.9402429 | 0.55610353 | 1.69076952 | 0.09088083 | 0.50282313 |
| RNF20 | 230.848022 | 0.11182304 | 0.20567306 | 0.54369316 | 0.58665264 | 0.88547501 |
| SMC2 | 161.237216 | -0.4854524 | 0.290113 | -1.6733217 | 0.09426398 | 0.51017802 |
| NIPSNAP3A | 186.914926 | -0.1653736 | 0.28184133 | -0.5867612 | 0.55736409 | 0.87240458 |
| NIPSNAP3B | 105.673124 | 0.13207131 | 0.35498793 | 0.37204452 | 0.7098597 | 0.92607195 |
| ABCA1 | 586.481808 | 0.29439356 | 0.39509595 | 0.74511916 | 0.45619968 | 0.83173798 |
| SLC44A1 | 1094.67528 | -0.4006199 | 0.19645023 | -2.0392947 | 0.04142062 | 0.37577267 |
| FSD1L | 625.850358 | 0.13068516 | 0.19441696 | 0.67219012 | 0.50146267 | 0.85435578 |
| FKTN | 646.685377 | -0.0523501 | 0.24032985 | -0.217826 | 0.82756469 | 0.96260145 |
| TMEM38B | 643.890141 | -0.0174706 | 0.24715591 | -0.0706866 | 9.44E-01 | 0.9878456 |
| LINC01505 | 1056.37684 | 0.00205948 | 0.44038996 | 0.00467648 | 0.99626872 | 0.99914308 |
| ZNF462 | 1207.1416 | -0.0182901 | 0.33513582 | -0.0545751 | 0.95647699 | 0.99065996 |
| AL807761.4 | 47.9805868 | 0.32388177 | 0.69893861 | 0.46339088 | 0.64308422 | 0.90478945 |
| RAD23B | 1238.61168 | -0.1646294 | 0.16833065 | -0.978012 | 0.32806837 | 0.75648668 |
| ELP1 | 460.799836 | -0.0196691 | 0.15411807 | -0.1276233 | 0.89844709 | 0.97877011 |
| FAM206A | 145.338781 | 0.36241089 | 0.20268695 | 1.78803269 | 0.07377073 | 0.47028843 |
| CTNNAL1 | 872.453265 | -0.4693409 | 0.36035864 | -1.3024273 | 0.19277035 | 0.64695623 |
| TMEM245 | 1951.47853 | -0.2002011 | 0.20630945 | -0.9703923 | 0.33185101 | 0.76035039 |
| EPB41L4B | 494.463496 | 0.19533704 | 0.31547055 | 0.61919264 | 0.53578946 | 0.8642718 |
| PTPN3 | 3275.75017 | -0.2219889 | 0.26082327 | -0.8511084 | 0.39470915 | 0.7988451 |
| PALM2 | 718.994354 | -0.906524 | 0.33380673 | -2.7157152 | 0.00661328 | 0.16831367 |
| PALM2-AKAF | 201.246502 | -0.3680401 | 0.43527476 | -0.8455352 | 0.39781207 | 0.80102129 |
| TXN | 407.916919 | 0.07446284 | 0.2750755 | 0.27069963 | 0.78662206 | 0.94978829 |
| LPAR1 | 596.79281 | -0.4179467 | 0.25299401 | -1.6520025 | 0.09853404 | 0.51525577 |
| ECPAS | 1683.40421 | -0.0929424 | 0.18966886 | -0.4900245 | 0.62411659 | 0.89873262 |
| ZNF483 | 679.019509 | -0.1349831 | 0.26094597 | -0.5172839 | 0.60495799 | 0.89044908 |
| PTGR1 | 975.466385 | 0.18299592 | 0.20577378 | 0.88930629 | 0.37383849 | 0.78331736 |
| C9orf84 | 349.122706 | 0.63679292 | 0.60382939 | 1.05459079 | 0.29161252 | 0.73362416 |
| UGCG | 2554.47838 | 0.24960014 | 0.41510715 | 0.60129088 | 5.48E-01 | 0.86859163 |
| SUSD1 | 488.922791 | -0.6841961 | 0.31885614 | -2.145783 | 0.0318903 | 0.34024781 |
| PTBP3 | 1612.48156 | -0.6007398 | 0.25578915 | -2.3485743 | 1.88E-02 | 0.27690323 |
| HSDL2 | 1343.44945 | -0.022631 | 0.32392313 | -0.0698654 | 0.94430081 | 0.98813503 |
| C9orf147 | 309.249262 | 0.03344306 | 0.25542618 | 0.13093044 | 0.89583033 | 0.97788747 |
| KIAA1958 | 4008.12715 | -0.0760102 | 0.17409634 | -0.4365984 | 0.66240265 | 0.91110616 |
| INIP | 405.542858 | 0.05691821 | 0.20030229 | 0.28416153 | 0.7762866 | 0.94694572 |
| SNX30 | 498.958905 | -0.2808461 | 0.18350711 | -1.5304369 | 0.12590862 | 0.56519704 |
| ZNF883 | 50.6753866 | -0.2674358 | 0.51367254 | -0.5206347 | 0.60262126 | 0.88971776 |
| ZFP37 | 65.1082134 | -0.064401 | 0.44447832 | -0.1448913 | 0.8847967 | 0.97664137 |
| FKBP15 | 382.553787 | -0.2363137 | 0.15561687 | -1.5185611 | 0.12887301 | 0.56894506 |
| SLC31A1 | 265.978973 | 0.05278768 | 0.23391339 | 0.2256719 | 0.82145662 | 0.96066244 |
| CDC26 | 174.653441 | 0.5118854 | 0.36319238 | 1.40940564 | 0.15871526 | 0.60804051 |
| PRPF4 | 189.00501 | -0.594748 | 0.40421589 | -1.4713621 | 0.14119321 | 0.5872148 |
| WDR31 | 189.227914 | 0.30802785 | 0.34433956 | 0.89454681 | 0.3710294 | 0.78104287 |
| BSPRY | 250.776893 | 0.43391604 | 0.25876419 | 1.67687822 | 0.0935663 | 0.50887217 |
| HDHD3 | 333.134021 | -0.5372155 | 0.37194333 | -1.4443479 | 0.14864115 | 0.59478974 |
| ALAD | 132.436389 | 0.14978198 | 0.3442364 | 0.43511373 | 0.66347988 | 0.91176714 |
| POLE3 | 60.1020314 | -0.2180201 | 0.36607832 | -0.5955559 | 0.55147197 | 0.87039028 |
| RGS3 | 1130.02403 | -0.2393805 | 0.43176895 | -0.5544181 | 0.57929276 | 0.8821758 |

| | | | | | | |
|---|---|---|---|---|---|---|
| ZNF618 | 2562.09376 | 0.20083398 | 0.28885601 | 0.69527367 | 0.48688381 | 0.84474442 |
| KIF12 | 1078.67997 | 0.37192285 | 0.45875815 | 0.81071659 | 0.41752844 | 0.81021496 |
| COL27A1 | 1067.06002 | 0.06310726 | 0.46196145 | 0.13660721 | 0.89134128 | 0.97729111 |
| AKNA | 79.6629161 | 0.13468008 | 0.36946385 | 0.36452845 | 0.71546344 | 0.92880162 |
| WHRN | 501.965017 | -0.0559737 | 0.3028858 | -0.1848014 | 0.85338479 | 0.97076831 |
| ATP6V1G1 | 523.985672 | 0.59949306 | 0.27621593 | 2.1703783 | 0.0299782 | 0.33256306 |
| TMEM268 | 118.243478 | 0.92410642 | 0.4559518 | 2.02676338 | 0.04268662 | 0.38022913 |
| TNFSF15 | 52.3008217 | -0.9810366 | 0.54750282 | -1.7918385 | 7.32E-02 | 0.47001711 |
| TNC | 128.77585 | 0.07467581 | 1.00656943 | 0.07418843 | 0.94086045 | 0.98719721 |
| PAPPA | 638.484414 | 0.40569221 | 0.36938016 | 1.09830535 | 0.27207118 | 0.72070318 |
| AL137024.1 | 136.176557 | 0.56173854 | 0.51043184 | 1.10051625 | 0.27110725 | 0.7199549 |
| ASTN2 | 4916.2306 | -0.2590402 | 0.44689357 | -0.5796464 | 0.56215312 | 0.87403145 |
| ASTN2-AS1 | 143.324282 | -0.3278901 | 0.80094907 | -0.409377 | 0.68226303 | 0.91969963 |
| AL133284.1 | 40.3442013 | -0.4194556 | 0.92836314 | -0.4518228 | 0.65139667 | 0.90825486 |
| TRIM32 | 75.499051 | 0.43888547 | 0.39747681 | 1.10417882 | 0.26951558 | 0.71818262 |
| CDK5RAP2 | 676.540614 | -0.2894327 | 0.21211044 | -1.3645379 | 0.17239833 | 0.6286116 |
| MEGF9 | 781.790366 | 0.18473698 | 0.25208537 | 0.73283498 | 0.46365908 | 0.83531269 |
| FBXW2 | 501.243091 | -0.1941598 | 0.17740188 | -1.0944633 | 0.27375185 | 0.72135661 |
| PSMD5 | 440.16694 | -0.053175 | 0.25639616 | -0.2073938 | 0.83570234 | 0.96511435 |
| TRAF1 | 57.6091925 | 0.16038169 | 0.45311096 | 0.35395678 | 0.72337128 | 0.93183257 |
| C5 | 119.234723 | -0.0962783 | 0.3753824 | -0.2564806 | 0.79757976 | 0.95281098 |
| CNTRL | 359.350451 | -0.3739935 | 0.29036269 | -1.288022 | 0.1977383 | 0.64989432 |
| RAB14 | 706.835336 | 0.23061987 | 0.19666704 | 1.1726412 | 0.24093972 | 0.69178991 |
| GSN | 1359.23147 | 0.16880397 | 0.44068117 | 0.38305238 | 0.70168093 | 0.92463122 |
| STOM | 521.360858 | 0.87577674 | 0.4118957 | 2.12620996 | 3.35E-02 | 3.50E-01 |
| AL359644.1 | 97.9171103 | 0.94019034 | 0.59415263 | 1.58240542 | 0.11355705 | 0.53989968 |
| DAB2IP | 1692.1447 | 0.64515261 | 0.45401483 | 1.42099459 | 0.15531833 | 0.60447845 |
| TTLL11 | 313.613537 | 0.43422592 | 0.18032372 | 2.40803549 | 0.01603862 | 0.25504618 |
| NDUFA8 | 266.740704 | 0.11532804 | 0.23312399 | 0.49470686 | 0.62080706 | 0.89747624 |
| RBM18 | 157.289452 | 0.07404355 | 0.31653454 | 0.23391933 | 0.8150476 | 0.95872901 |
| MRRF | 865.493004 | -0.1361433 | 0.19538041 | -0.6968114 | 0.48592084 | 0.8447301 |
| PDCL | 156.416963 | 0.23497423 | 0.22407946 | 1.04862015 | 0.29435298 | 0.73527014 |
| RC3H2 | 906.383462 | -0.0555636 | 0.13433363 | -0.4136238 | 0.67914961 | 0.91822574 |
| ZBTB6 | 53.835585 | 0.03792012 | 0.29920892 | 0.1267346 | 0.89915046 | 0.97889101 |
| ZBTB26 | 144.813019 | -0.0159251 | 0.28447894 | -0.0559799 | 0.95535782 | 0.99065996 |
| RABGAP1 | 2339.03031 | -0.2172692 | 0.14530411 | -1.4952718 | 0.13484351 | 0.57708028 |
| STRBP | 2252.74676 | -0.0508921 | 0.20702828 | -0.2458217 | 0.80582023 | 0.95522565 |
| DENND1A | 3268.60788 | -0.0282068 | 0.18799247 | -0.150042 | 0.88073148 | 0.97664137 |
| AL390774.2 | 58.8352796 | 0.23043746 | 0.76687525 | 0.30048885 | 0.7638043 | 0.94255335 |
| NEK6 | 273.374495 | 1.21246956 | 0.71691658 | 1.69122823 | 9.08E-02 | 0.50267341 |
| PSMB7 | 647.963572 | -0.2069948 | 0.18922241 | -1.0939235 | 0.27398853 | 0.72135661 |
| NR6A1 | 4191.84742 | -0.5488553 | 0.22402439 | -2.44998 | 0.01428642 | 0.24700164 |
| MIR181A2HG | 564.308471 | -0.6970379 | 0.41626166 | -1.6745186 | 0.09402872 | 0.5098684 |
| RPL35 | 594.014849 | 0.23370867 | 0.46808025 | 0.49929187 | 0.61757378 | 0.89599505 |
| ARPC5L | 266.164306 | -0.2132438 | 0.34473969 | -0.6185647 | 0.53620315 | 0.8642718 |
| GOLGA1 | 501.690332 | -0.3118264 | 0.32489718 | -0.9597695 | 0.33717124 | 0.76409325 |
| SCAI | 859.9686 | -0.5275225 | 0.26260997 | -2.0087681 | 0.04456173 | 0.38623091 |
| PPP6C | 689.225332 | -0.1881913 | 0.18916899 | -0.9948316 | 0.3198182 | 0.75094569 |
| RABEPK | 209.894981 | -0.6588742 | 0.42381528 | -1.5546258 | 0.12003521 | 0.55298199 |
| HSPA5 | 677.852273 | 1.47814862 | 0.44139723 | 3.34879454 | 0.00081164 | 0.04963402 |
| GAPVD1 | 1191.53523 | -0.1860407 | 0.12686878 | -1.4664026 | 0.14253863 | 0.58824095 |
| MAPKAP1 | 2069.31496 | -0.1744671 | 0.18631652 | -0.9364016 | 0.34906644 | 0.76982416 |
| PBX3 | 1656.26902 | 1.05026557 | 0.38979065 | 2.69443501 | 0.00705081 | 0.17262485 |
| MVB12B | 376.003062 | 0.05466587 | 0.21200506 | 0.25785173 | 0.79652134 | 0.95278473 |

| | | | | | | |
|---|---|---|---|---|---|---|
| ZBTB43 | 649.404382 | 0.04139352 | 0.24053027 | 0.17209275 | 0.86336461 | 0.97319807 |
| ZBTB34 | 151.385731 | 0.04715324 | 0.27396522 | 0.17211397 | 0.86334793 | 0.97319807 |
| RALGPS1 | 2456.8131 | 0.13913204 | 0.2465656 | 0.56428003 | 0.57256357 | 0.87946834 |
| ANGPTL2 | 58.725755 | -0.2876085 | 0.60548327 | -0.4750066 | 0.63478227 | 0.90178395 |
| GARNL3 | 333.895076 | -0.3056608 | 0.35492768 | -0.8611917 | 0.38913245 | 0.79663535 |
| SLC2A8 | 58.154175 | -0.0022218 | 0.3621634 | -0.0061347 | 0.99510527 | 0.998438 |
| ZNF79 | 68.824888 | 0.07250552 | 0.35878218 | 0.20208784 | 0.83984805 | 0.96617873 |
| RPL12 | 287.294268 | 0.04926985 | 0.3891077 | 0.12662267 | 0.89923906 | 0.97889101 |
| LRSAM1 | 591.361365 | 0.00818931 | 0.27537762 | 0.02973847 | 0.97627563 | 0.9941831 |
| FAM129B | 791.979627 | -0.5286126 | 0.36421922 | -1.4513583 | 1.47E-01 | 5.94E-01 |
| STXBP1 | 448.302614 | 0.1390222 | 0.3643181 | 0.38159563 | 0.70276132 | 0.92501157 |
| PTRH1 | 109.432763 | 0.43091713 | 0.26329443 | 1.63663593 | 0.10170656 | 0.5195058 |
| CDK9 | 215.325733 | -0.032654 | 0.25919884 | -0.1259803 | 0.8997475 | 0.97897567 |
| FPGS | 316.040908 | -0.5486485 | 0.33071231 | -1.6589901 | 0.09711779 | 0.51343843 |
| ENG | 109.961057 | -0.3210886 | 0.21835774 | -1.4704704 | 0.14143441 | 0.5872148 |
| AL162586.1 | 90.1903189 | -0.812364 | 0.47149508 | -1.7229532 | 0.08489698 | 0.49205978 |
| ST6GALNAC4 | 47.4885633 | 1.01610711 | 0.47163625 | 2.15442961 | 0.0312065 | 0.3383722 |
| DPM2 | 143.527247 | -0.1187648 | 0.3907148 | -0.303968 | 0.76115225 | 0.94202764 |
| FAM102A | 338.608902 | -0.3706781 | 0.56223553 | -0.6592933 | 0.50970744 | 0.85701701 |
| SLC25A25 | 369.194883 | 0.63169817 | 0.41918058 | 1.50698338 | 0.13181492 | 0.57539433 |
| SLC25A25-AS | 202.651106 | 0.58194265 | 0.34426953 | 1.6903693 | 0.09095732 | 0.50282313 |
| PTGES2 | 266.74749 | 0.14640389 | 0.2849409 | 0.51380439 | 0.60738876 | 0.89174148 |
| LCN2 | 200.557956 | 0.93904722 | 0.7405048 | 1.26811766 | 2.05E-01 | 6.58E-01 |
| C9orf16 | 259.497901 | 0.33617096 | 0.41426563 | 0.81148648 | 0.41708635 | 0.81007111 |
| CIZ1 | 247.79584 | -0.2401052 | 0.33748631 | -0.7114517 | 0.4768044 | 0.8415432 |
| DNM1 | 240.844049 | -0.0283377 | 0.43110715 | -0.0657324 | 0.94759087 | 0.98919519 |
| GOLGA2 | 591.768909 | -0.0878399 | 0.2215834 | -0.3964193 | 0.69179572 | 0.92224682 |
| SWI5 | 111.149426 | 0.04748584 | 0.28811908 | 0.16481323 | 0.86909099 | 0.9744678 |
| TRUB2 | 138.315414 | -0.0290819 | 0.25274619 | -0.1150637 | 0.90839464 | 0.98047754 |
| COQ4 | 706.577494 | 0.23659681 | 0.24242914 | 0.97594213 | 0.32909314 | 0.75797582 |
| SLC27A4 | 286.990117 | -0.0141401 | 0.31210733 | -0.0453052 | 0.96386404 | 0.99224805 |
| URM1 | 210.124736 | -0.1768309 | 0.25474931 | -0.6941369 | 0.48759636 | 0.8451201 |
| CERCAM | 53.9156486 | -0.0112301 | 0.48443038 | -0.0231821 | 0.98150505 | 0.99518016 |
| ODF2 | 336.459982 | -0.245484 | 0.25822361 | -0.9506645 | 0.34177474 | 0.76545664 |
| GLE1 | 362.550163 | -0.2969036 | 0.26752891 | -1.1098 | 0.26708521 | 0.71712359 |
| AL356481.1 | 49.6021543 | -0.644143 | 0.54110896 | -1.1904127 | 0.23388424 | 0.68500598 |
| SPTAN1 | 1142.05893 | -0.5710206 | 0.37518825 | -1.5219576 | 0.1280197 | 0.56762585 |
| WDR34 | 312.441273 | 0.2846062 | 0.31661731 | 0.89889652 | 0.36870778 | 0.78095617 |
| SET | 602.089102 | 0.3446445 | 0.2872788 | 1.19968652 | 0.23026111 | 0.6825671 |
| PKN3 | 48.9158795 | -0.3808553 | 0.3447277 | -1.1048003 | 2.69E-01 | 0.71818262 |
| ZDHHC12 | 98.7753541 | 0.50639915 | 0.47507482 | 1.06593558 | 2.86E-01 | 0.72934245 |
| ZER1 | 1126.11216 | 0.01664264 | 0.29056477 | 0.05727687 | 0.95432464 | 0.99015586 |
| TBC1D13 | 200.956318 | -0.2117366 | 0.2249526 | -0.94125 | 0.34657674 | 0.76864066 |
| ENDOG | 121.39909 | 0.56321672 | 0.38870671 | 1.44895033 | 0.14735145 | 0.59373375 |
| SPOUT1 | 77.5418377 | 0.01631173 | 0.36816329 | 0.04430569 | 0.96466074 | 0.99226095 |
| KYAT1 | 268.690841 | 0.12082477 | 0.2273588 | 0.53142775 | 5.95E-01 | 8.89E-01 |
| LRRC8A | 124.322884 | 0.05715099 | 0.22252733 | 0.25682685 | 0.79731244 | 0.95281098 |
| PHYHD1 | 242.968285 | -0.2279475 | 0.28812145 | -0.7911507 | 0.42885604 | 0.8155211 |
| NUP188 | 186.841342 | -0.5130748 | 0.32287476 | -1.5890831 | 0.11204164 | 0.53621666 |
| SH3GLB2 | 350.675105 | -0.1169933 | 0.29941939 | -0.3907338 | 0.69599402 | 0.92266008 |
| MIGA2 | 298.156565 | -0.0936011 | 0.19036308 | -0.4916977 | 6.23E-01 | 8.98E-01 |
| CRAT | 84.7949004 | 0.39773074 | 0.28409169 | 1.40000837 | 0.16151081 | 0.61304492 |
| PTPA | 440.530846 | -0.1836553 | 0.34274574 | -0.5358355 | 0.59207222 | 0.88740306 |
| IER5L | 393.36938 | -0.1085271 | 0.43190068 | -0.2512779 | 8.02E-01 | 9.54E-01 |

| | | | | | | |
|---|---|---|---|---|---|---|
| LINC01503 | 979.803931 | 0.35625342 | 0.70182009 | 0.5076136 | 0.61172435 | 0.89413156 |
| NTMT1 | 212.836671 | 0.10739233 | 0.2370566 | 0.45302398 | 0.65053147 | 0.90825486 |
| ASB6 | 123.415693 | 0.23224535 | 0.38341664 | 0.6057258 | 0.54469685 | 0.86654229 |
| TOR1B | 178.358015 | 0.14386628 | 0.31787522 | 0.45258729 | 0.65084595 | 0.90825486 |
| TOR1A | 82.9140256 | 0.33315399 | 0.31988195 | 1.04149045 | 0.29764798 | 0.73673761 |
| C9orf78 | 200.713361 | 0.32680972 | 0.27539462 | 1.18669613 | 0.2353475 | 0.68745373 |
| USP20 | 305.193895 | -0.2203348 | 0.22427675 | -0.9824236 | 0.32589118 | 0.75550719 |
| FNBP1 | 1397.83473 | -0.2594677 | 0.24098509 | -1.0766959 | 0.28161615 | 0.72604837 |
| GPR107 | 778.287535 | -0.20335 | 0.19624287 | -1.0362162 | 0.30010128 | 0.73680045 |
| NCS1 | 307.302454 | -0.2985799 | 0.39138249 | -0.7628852 | 0.44553189 | 0.82571115 |
| ASS1 | 208.191515 | 0.350243 | 0.52301776 | 0.66965795 | 0.50307587 | 0.85525037 |
| FUBP3 | 791.372067 | -0.2862356 | 0.18633676 | -1.5361198 | 0.12450901 | 0.56111892 |
| EXOSC2 | 93.1305638 | -0.2340542 | 0.29364248 | -0.7970719 | 0.42540927 | 0.81466629 |
| ABL1 | 2065.34979 | -0.1063067 | 0.13724187 | -0.7745937 | 0.43857976 | 0.82127804 |
| AIF1L | 1200.98659 | 0.64701998 | 0.39265343 | 1.64781439 | 0.09939076 | 0.51665771 |
| NUP214 | 1804.78082 | -0.2931176 | 0.15489407 | -1.8923746 | 0.05844109 | 0.42147872 |
| AL157938.2 | 498.858573 | -0.0767781 | 0.583962 | -0.1314779 | 0.89539727 | 0.97788747 |
| AL157938.3 | 133.473063 | -0.4947945 | 0.4023475 | -1.229769 | 0.21878364 | 0.6720728 |
| PLPP7 | 79.2801277 | -0.3590057 | 0.33327853 | -1.0771943 | 0.28139348 | 0.72604837 |
| PRRC2B | 2062.41301 | -0.1918601 | 0.19827597 | -0.9676419 | 0.33322324 | 0.76124971 |
| POMT1 | 197.533425 | -0.3054141 | 0.23325372 | -1.3093642 | 0.19041101 | 0.64556957 |
| UCK1 | 95.8625202 | 0.09229252 | 0.28620414 | 0.32247094 | 0.74709595 | 0.93734711 |
| RAPGEF1 | 1691.37415 | -0.2271676 | 0.3160635 | -0.7187405 | 0.47230083 | 0.83921324 |
| MED27 | 1156.0967 | -0.3034324 | 0.11994449 | -2.5297739 | 0.0114136 | 0.21832654 |
| SETX | 1724.1758 | -0.1078705 | 0.19535979 | -0.5521634 | 0.58083639 | 0.88283233 |
| TTF1 | 345.108544 | -0.2706754 | 0.2449275 | -1.1051245 | 0.26910565 | 0.71818262 |
| CFAP77 | 45.3506784 | -0.3880775 | 0.54883346 | -0.7070953 | 0.47950728 | 0.8430245 |
| DDX31 | 443.217685 | -0.3894303 | 0.22189152 | -1.7550481 | 0.07925108 | 0.48169281 |
| GTF3C4 | 163.215418 | 0.20394568 | 0.23040712 | 0.88515353 | 0.37607383 | 0.78490077 |
| AK8 | 212.0951 | -0.3079089 | 0.4069944 | -0.7565434 | 0.44932346 | 0.82672287 |
| SPACA9 | 82.8081878 | -0.5697384 | 0.39951897 | -1.4260609 | 0.15385078 | 0.60285224 |
| TSC1 | 911.247036 | -0.3726244 | 0.18762399 | -1.9860166 | 0.04703148 | 0.39269202 |
| GTF3C5 | 211.506497 | -0.4192383 | 0.2808973 | -1.4924968 | 0.13556895 | 0.57709645 |
| RALGDS | 105.32469 | 0.07710523 | 0.33062564 | 0.23321006 | 0.81559829 | 0.95872901 |
| ABO | 587.155274 | -0.327276 | 0.51677744 | -0.6333017 | 0.52653662 | 0.86145663 |
| SURF6 | 121.018159 | 0.03717172 | 0.22329221 | 0.16647118 | 0.86778616 | 0.97402505 |
| MED22 | 93.3271035 | -0.013543 | 0.29544993 | -0.0458385 | 0.96343901 | 0.99224805 |
| RPL7A | 706.78801 | 0.59974104 | 0.48152767 | 1.24549654 | 0.21294929 | 0.66696598 |
| SURF1 | 141.874078 | 0.24052913 | 0.29315215 | 0.82049246 | 0.41193542 | 0.80679792 |
| SURF2 | 70.6662192 | 0.48572119 | 0.40097309 | 1.21135606 | 0.22575897 | 0.67804689 |
| SURF4 | 510.534962 | 0.3897558 | 0.25203096 | 1.54646 | 0.12199351 | 0.55758211 |
| REXO4 | 160.347606 | -0.0579373 | 0.26179545 | -0.2213075 | 0.82485302 | 0.96146415 |
| ADAMTS13 | 80.4097663 | -0.3767426 | 0.433312 | -0.8694487 | 0.38460176 | 0.79217288 |
| CACFD1 | 115.586834 | 0.2879708 | 0.39278672 | 0.733148 | 0.46346817 | 0.83531269 |
| ADAMTSL2 | 93.1632259 | -0.1323181 | 0.86946316 | -0.1521837 | 0.87904202 | 0.9765717 |
| VAV2 | 1000.55643 | -0.1229699 | 0.2872191 | -0.4281398 | 0.66854936 | 0.91395501 |
| BRD3OS | 89.4192295 | 0.37924322 | 0.35264747 | 1.07541738 | 0.2821879 | 0.72604837 |
| BRD3 | 308.786092 | -0.001956 | 0.18421053 | -0.0106183 | 0.99152799 | 0.99772135 |
| WDR5 | 215.990668 | -0.2560226 | 0.22537832 | -1.1359681 | 0.25596993 | 0.70709713 |
| RXRA | 323.997046 | 0.27718518 | 0.37719373 | 0.73486157 | 0.4624238 | 0.83531269 |
| COL5A1 | 74.4913034 | -1.0462345 | 0.92816473 | -1.1272078 | 0.25965463 | 0.70921184 |
| PPP1R26-AS1 | 334.687843 | -0.1157264 | 0.50643861 | -0.2285103 | 0.81924958 | 0.96006637 |
| PPP1R26 | 135.973609 | 0.29612229 | 0.25065708 | 1.1813841 | 0.23745017 | 0.68973903 |
| MRPS2 | 100.346502 | 0.49207108 | 0.42295356 | 1.16341633 | 0.24466063 | 0.6948422 |

| | | | | | | |
|---|---|---|---|---|---|---|
| CAMSAP1 | 431.858286 | -0.2952688 | 0.22385589 | -1.319013 | 0.18716475 | 0.64195489 |
| AL355574.1 | 43.9884616 | -0.1233129 | 0.47749871 | -0.2582476 | 0.79621583 | 0.95278473 |
| UBAC1 | 421.949588 | -0.0128199 | 0.24290776 | -0.0527768 | 0.95790973 | 0.99069768 |
| NACC2 | 421.646397 | -0.0674583 | 0.26824054 | -0.2514843 | 0.80143972 | 0.95388513 |
| TMEM250 | 64.1605887 | 0.08434769 | 0.45204986 | 0.18658935 | 0.85198262 | 0.97076831 |
| QSOX2 | 119.812377 | -0.4522691 | 0.29185783 | -1.5496214 | 0.12123242 | 0.55582251 |
| SNAPC4 | 188.765492 | -0.2590848 | 0.22609316 | -1.1459207 | 0.25182795 | 0.70276117 |
| ENTR1 | 189.976121 | -0.6315066 | 0.29413787 | -2.1469747 | 0.0317953 | 0.34024781 |
| PMPCA | 192.872152 | 0.14591084 | 0.27295773 | 0.53455473 | 0.59295779 | 0.88740306 |
| INPP5E | 115.041726 | -0.5363409 | 0.32617573 | -1.6443311 | 0.10010782 | 0.51692417 |
| SEC16A | 482.833545 | -0.071522 | 0.24679007 | -0.289809 | 0.77196234 | 0.94478317 |
| NOTCH1 | 350.804951 | -0.2102376 | 0.33128049 | -0.6346211 | 0.52567556 | 0.86119802 |
| EGFL7 | 69.6233357 | 0.30602222 | 0.45763052 | 0.66871026 | 0.50368032 | 0.85525037 |
| AGPAT2 | 362.813265 | 0.93945425 | 0.43897146 | 2.14012604 | 0.03234458 | 0.34344018 |
| FAM69B | 90.301619 | 1.1585285 | 0.60186106 | 1.92491021 | 0.05424059 | 0.41533078 |
| SNHG7 | 289.495545 | 1.0294657 | 0.35946792 | 2.86385976 | 0.00418513 | 0.12579326 |
| RABL6 | 686.195671 | -0.2135773 | 0.25731458 | -0.830024 | 0.4065252 | 0.80499101 |
| AJM1 | 82.3534623 | 0.87478065 | 0.6662279 | 1.31303516 | 0.1891711 | 0.64432929 |
| PHPT1 | 419.535158 | 0.99737381 | 0.40764292 | 2.44668498 | 0.01441768 | 0.24762526 |
| MAMDC4 | 45.3173922 | 0.15760596 | 0.62114491 | 0.25373461 | 0.7997006 | 0.95370871 |
| EDF1 | 632.295925 | 0.18370376 | 0.3688219 | 0.49808258 | 0.61842584 | 0.89599505 |
| TRAF2 | 111.716533 | 0.13224382 | 0.30072468 | 0.43975046 | 0.66011785 | 0.91099119 |
| FBXW5 | 324.958703 | 0.17096441 | 0.37844576 | 0.45175405 | 0.65144617 | 0.90825486 |
| LCN12 | 111.795543 | 0.72176663 | 0.67648345 | 1.06693907 | 0.28599935 | 0.72901687 |
| PAXX | 149.118402 | -0.0638854 | 0.51895999 | -0.1231027 | 0.90202576 | 0.97941893 |
| ABCA2 | 171.584657 | 0.8437852 | 0.52253304 | 1.61479781 | 0.10635449 | 0.52735777 |
| UAP1L1 | 87.4455719 | 0.80431268 | 0.6022642 | 1.33548146 | 0.18171882 | 0.63635309 |
| MAN1B1 | 249.73428 | -0.1337814 | 0.32333419 | -0.4137559 | 0.67905288 | 0.91822574 |
| DPP7 | 219.391113 | 0.07127813 | 0.38687366 | 0.18424137 | 0.8538241 | 0.97077171 |
| ANAPC2 | 45.5760375 | -0.3286342 | 0.38745697 | -0.8481825 | 0.39633635 | 0.80029863 |
| SSNA1 | 203.808639 | 0.42436634 | 0.36269982 | 1.17002084 | 0.24199258 | 0.69337349 |
| TPRN | 193.349802 | 0.3118984 | 0.30052665 | 1.03783939 | 0.29934483 | 0.73673761 |
| TMEM203 | 87.8639214 | 0.7921329 | 0.40431511 | 1.9591969 | 0.05008973 | 0.40103187 |
| NDOR1 | 93.772072 | -0.5687611 | 0.34973659 | -1.6262556 | 0.1038953 | 0.52442862 |
| BX255925.3 | 75.1616761 | -0.3643545 | 0.49652486 | -0.7338092 | 0.46306505 | 0.83531269 |
| RNF208 | 44.5806961 | 0.25624008 | 0.52823775 | 0.48508475 | 0.62761623 | 0.89919293 |
| TUBB4B | 498.959208 | 1.19093863 | 0.3715929 | 3.20495525 | 0.00135083 | 0.06712951 |
| NELFB | 323.761511 | -0.238056 | 0.24330527 | -0.9784252 | 0.32786409 | 0.75633196 |
| EXD3 | 597.799223 | -0.2395142 | 0.23319023 | -1.0271194 | 0.30436424 | 0.74152784 |
| NOXA1 | 179.484699 | 0.17996905 | 0.42627898 | 0.42218608 | 0.6728892 | 0.91608132 |
| NSMF | 109.898667 | 0.18944918 | 0.33013636 | 0.5738513 | 0.56606843 | 0.87642772 |
| PNPLA7 | 381.473993 | -0.0407831 | 0.46983426 | -0.0868032 | 0.93082793 | 0.98331361 |
| MRPL41 | 458.512583 | 0.53302036 | 0.41094168 | 1.29707056 | 0.19460691 | 0.64814994 |
| DPH7 | 208.20713 | -0.3041358 | 0.19784858 | -1.537215 | 0.12424066 | 0.56105491 |
| ZMYND19 | 56.6265509 | -0.7046549 | 0.42337157 | -1.6643888 | 0.09603476 | 0.5117083 |
| ARRDC1 | 296.298395 | -0.3874188 | 0.27913942 | -1.3879043 | 0.1651662 | 0.61734023 |
| ARRDC1-AS1 | 90.8082235 | 0.20011161 | 0.25108089 | 0.79700056 | 0.42545071 | 0.81466629 |
| EHMT1 | 2322.28735 | -0.3007807 | 0.18093596 | -1.6623601 | 0.09644058 | 0.512217 |
| LINC01001 | 87.8067737 | -0.7809368 | 0.49620712 | -1.5738122 | 0.11553088 | 0.54427374 |
| AC069287.3 | 103.46682 | -0.0869861 | 0.60532647 | -0.1437011 | 0.88573651 | 0.97664137 |
| BET1L | 141.657266 | -0.056421 | 0.22341109 | -0.2525435 | 0.80062102 | 0.95370871 |
| RIC8A | 127.988102 | -0.0592039 | 0.23886638 | -0.2478538 | 8.04E-01 | 0.95450237 |
| SIRT3 | 241.47198 | -0.2090642 | 0.2534121 | -0.824997 | 0.40937329 | 0.80623405 |
| PSMD13 | 365.116117 | -0.0309525 | 0.30517132 | -0.1014267 | 0.91921171 | 0.98215577 |

| | | | | | | |
|---|---|---|---|---|---|---|
| PGGHG | 158.88241 | -0.2787627 | 0.54314017 | -0.5132427 | 0.60778156 | 0.89203142 |
| IFITM2 | 1607.78534 | 1.20471049 | 0.45891639 | 2.62511978 | 8.66E-03 | 1.88E-01 |
| IFITM3 | 2461.40073 | 0.7160334 | 0.39649223 | 1.80592039 | 7.09E-02 | 4.64E-01 |
| SIGIRR | 243.919499 | 0.07367843 | 0.35130106 | 0.20973016 | 0.83387828 | 0.96510927 |
| ANO9 | 87.7729712 | -0.2761828 | 0.37912869 | -0.7284671 | 0.4663277 | 0.8367113 |
| PTDSS2 | 258.634783 | -0.0703245 | 0.16987546 | -0.4139767 | 0.67889117 | 0.91822574 |
| RNH1 | 384.781103 | 0.45422392 | 0.27581811 | 1.64682416 | 0.09959419 | 0.51665771 |
| HRAS | 170.081957 | 0.08530529 | 0.37148378 | 0.22963396 | 0.81837621 | 0.95985564 |
| LMNTD2 | 54.3922797 | -0.2088939 | 0.38284985 | -0.5456287 | 0.58532121 | 0.88490909 |
| AP006284.1 | 91.00676 | 0.39359347 | 0.57624247 | 0.68303448 | 0.49458506 | 0.84882719 |
| RASSF7 | 624.420171 | 0.25990777 | 0.35525849 | 0.73160184 | 0.46441162 | 0.83586498 |
| MIR210HG | 200.17505 | 0.59008558 | 0.63529974 | 0.92883019 | 0.3529771 | 0.77216708 |
| PHRF1 | 314.621 | -0.4783638 | 0.24011141 | -1.9922579 | 0.04634277 | 0.3904347 |
| IRF7 | 131.931566 | -0.0207655 | 0.59951708 | -0.034637 | 0.97236919 | 0.99327086 |
| DEAF1 | 783.363796 | -0.124425 | 0.22199465 | -0.5604866 | 0.57514755 | 0.88062177 |
| AC131934.1 | 169.234683 | -0.6733492 | 0.46136814 | -1.4594619 | 0.14443803 | 0.59012743 |
| EPS8L2 | 1297.68465 | 0.03466089 | 0.34313486 | 0.10101246 | 0.91954057 | 0.98215577 |
| AP006621.4 | 250.897568 | 0.66977906 | 0.60785652 | 1.10187032 | 0.27051805 | 0.71954751 |
| TALDO1 | 583.790981 | 0.49007668 | 0.43310478 | 1.131543 | 0.25782662 | 0.70885276 |
| GATD1 | 176.956485 | -0.0787761 | 0.23269242 | -0.3385415 | 0.73495512 | 0.93552616 |
| AP006621.3 | 229.321563 | 0.18759011 | 0.31209196 | 0.60107321 | 0.54779123 | 0.86859163 |
| PIDD1 | 114.298114 | -0.5685744 | 0.28349193 | -2.0056105 | 0.04489782 | 0.38711185 |
| RPLP2 | 690.402731 | 0.03579045 | 0.37088479 | 0.09650018 | 0.92312333 | 0.98215577 |
| PNPLA2 | 460.370752 | -0.3050562 | 0.24604347 | -1.2398468 | 0.21503206 | 0.66865355 |
| CRACR2B | 96.4864934 | -0.1214293 | 0.53881471 | -0.2253638 | 0.82169624 | 0.96066244 |
| CD151 | 568.566045 | 0.33105163 | 0.44445197 | 0.74485355 | 0.45636026 | 0.83182139 |
| POLR2L | 459.406372 | 0.94085162 | 0.51820525 | 1.81559649 | 0.06943229 | 0.45787876 |
| TSPAN4 | 147.739225 | 0.58382548 | 0.34962343 | 1.66986945 | 0.0949452 | 0.51085035 |
| CHID1 | 232.604209 | 0.20534061 | 0.27190029 | 0.75520556 | 0.45012566 | 0.82728674 |
| AP2A2 | 1599.02139 | 0.00679429 | 0.29665022 | 0.02290338 | 0.98172735 | 0.99518016 |
| MUC6 | 59.7768063 | -0.2099974 | 0.38564067 | -0.5445416 | 0.58606885 | 0.88530153 |
| TOLLIP | 272.65764 | 0.13264244 | 0.20968304 | 0.63258548 | 0.52700437 | 0.86151701 |
| MOB2 | 270.334486 | -0.084793 | 0.21973464 | -0.385888 | 0.69957962 | 0.92367966 |
| MRPL23 | 81.5022156 | 0.53516427 | 0.47347447 | 1.13029173 | 0.25835332 | 0.70896455 |
| CD81-AS1 | 98.5541512 | -0.5446803 | 0.41824494 | -1.3022998 | 0.19281392 | 0.64695623 |
| CD81 | 1466.00317 | 0.94291308 | 0.36040649 | 2.61624891 | 8.89E-03 | 0.19125454 |
| TSSC4 | 104.831049 | 0.07398667 | 0.3740601 | 0.19779353 | 0.84320661 | 0.96775707 |
| KCNQ1 | 2918.38457 | 0.11044247 | 0.3406117 | 0.32424744 | 0.74575071 | 0.93675395 |
| KCNQ1OT1 | 2828.9102 | 0.43504982 | 0.19953182 | 2.18035303 | 0.0292313 | 0.33001392 |
| CDKN1C | 255.271757 | 1.02038171 | 0.65297645 | 1.56266235 | 0.11813203 | 0.54905209 |
| SLC22A18 | 146.881833 | 0.03368444 | 0.74577018 | 0.04516733 | 0.96397394 | 0.99224805 |
| NAP1L4 | 1859.3702 | 0.03611052 | 0.18461884 | 0.19559499 | 0.84492718 | 0.96858129 |
| CARS | 496.670458 | -0.3908479 | 0.28260488 | -1.3830189 | 0.16665908 | 0.61925103 |
| OSBPL5 | 405.82813 | -0.3881246 | 0.30097197 | -1.2895704 | 0.19719985 | 0.64964978 |
| ZNF195 | 377.166313 | -0.5111829 | 0.26536371 | -1.9263483 | 0.0540609 | 0.41468669 |
| AC127526.1 | 188.742072 | -0.9320097 | 0.47796834 | -1.9499403 | 0.05118324 | 0.40556617 |
| NUP98 | 1194.58668 | -0.0947448 | 0.30829965 | -0.3073141 | 0.75860428 | 0.94097135 |
| PGAP2 | 198.515975 | 0.10316296 | 0.56081105 | 0.18395315 | 0.85405021 | 0.97085267 |
| RHOG | 70.4977895 | 0.17573836 | 0.29857971 | 0.58858106 | 0.55614234 | 0.87212955 |
| STIM1 | 542.376057 | 0.58539003 | 0.34949056 | 1.67498092 | 9.39E-02 | 5.10E-01 |
| RRM1 | 191.725397 | 0.01612418 | 0.31873237 | 0.05058846 | 0.95965346 | 0.991232 |
| TRIM21 | 52.8497043 | 0.248484 | 0.57198093 | 0.43442706 | 0.66397835 | 0.91197775 |
| TRIM68 | 58.5963154 | 0.38862117 | 0.38135041 | 1.01906583 | 0.30817172 | 0.74375973 |
| HBB | 63.4470885 | -1.8985527 | 1.04561804 | -1.815723 | 0.06941288 | 0.45787876 |

| | | | | | | |
|---|---|---|---|---|---|---|
| AC104389.4 | 476.074085 | 0.54378608 | 0.52056696 | 1.04460352 | 0.29620625 | 0.7366786 |
| TRIM5 | 153.81823 | 0.99867515 | 0.44010367 | 2.26918159 | 0.02325729 | 0.30018842 |
| TRIM22 | 407.828396 | 0.8718108 | 0.85224639 | 1.02295629 | 0.30632852 | 0.74295884 |
| FAM160A2 | 308.056687 | -0.1001819 | 0.2274254 | -0.4405044 | 0.6595718 | 0.91099119 |
| CAVIN3 | 75.4404535 | 0.93963163 | 0.6368857 | 1.47535362 | 0.1401175 | 0.58492375 |
| SMPD1 | 118.106204 | 0.72931348 | 0.44927442 | 1.62331406 | 0.10452229 | 0.52442862 |
| APBB1 | 219.849216 | 0.25153395 | 0.23980715 | 1.04890097 | 0.2942237 | 0.73527014 |
| TRIM3 | 44.7159299 | -0.0761793 | 0.50985926 | -0.1494125 | 0.88122817 | 0.97664137 |
| ARFIP2 | 149.758888 | -0.1474962 | 0.25084588 | -0.5879954 | 0.55653539 | 0.87212955 |
| AC084337.2 | 259.676494 | -0.5216199 | 0.37866201 | -1.3775342 | 0.16834715 | 0.62253512 |
| DNHD1 | 469.961625 | -0.7442969 | 0.28995158 | -2.5669696 | 0.01025916 | 0.20626067 |
| RRP8 | 103.665051 | 0.08660799 | 0.28061757 | 0.30863353 | 0.75760031 | 0.94056395 |
| ILK | 390.070755 | 0.1652604 | 0.1971162 | 0.83839077 | 4.02E-01 | 0.80278824 |
| TAF10 | 170.09997 | -0.1270752 | 0.37266953 | -0.3409862 | 0.73311393 | 0.93552616 |
| TPP1 | 226.121187 | 0.04959647 | 0.29769261 | 0.16660295 | 0.86768247 | 0.97402505 |
| MRPL17 | 141.046391 | 0.55893044 | 0.41908918 | 1.33367899 | 0.18230908 | 0.63741052 |
| AC087280.2 | 217.126597 | 0.21350974 | 0.61803104 | 0.34546767 | 0.72974281 | 0.93453345 |
| ZNF215 | 95.7446259 | 0.16300478 | 0.43757094 | 0.37252196 | 0.70950426 | 0.92588834 |
| ZNF214 | 139.544404 | -0.1302562 | 0.22471106 | -0.579661 | 0.56214324 | 0.87403145 |
| NLRP14 | 37.4365818 | -0.3782544 | 0.59265524 | -0.6382368 | 0.52331952 | 0.86006369 |
| SYT9 | 73.0672255 | 0.9848909 | 0.49450314 | 1.99167776 | 0.04640643 | 0.3904347 |
| AC107884.1 | 229.749906 | -0.1780628 | 0.32951306 | -0.5403817 | 0.58893385 | 0.88571413 |
| PPFIBP2 | 250.078892 | -0.3844997 | 0.28705943 | -1.3394427 | 0.18042658 | 0.63479125 |
| OVCH2 | 71.8742186 | 3.27992352 | 0.98502269 | 3.3297949 | 0.0008691 | 0.05165902 |
| AC044810.3 | 542.802373 | 2.73297734 | 0.94196946 | 2.90134388 | 0.00371566 | 0.11865695 |
| EIF3F | 381.169188 | 0.35036936 | 0.26974059 | 1.29891225 | 0.19397405 | 0.64749489 |
| TUB | 104.910931 | 0.49757704 | 0.52145143 | 0.9542155 | 0.33997457 | 0.76465348 |
| RIC3 | 1063.29925 | 0.01451495 | 0.28280483 | 0.05132498 | 0.95906657 | 0.99117358 |
| LMO1 | 139.32373 | 0.08091156 | 0.46721912 | 0.1731769 | 0.86251238 | 0.97283972 |
| STK33 | 1695.33204 | 0.00254714 | 0.35909797 | 0.00709317 | 0.99434052 | 0.998438 |
| TRIM66 | 927.460402 | -0.585573 | 0.47723559 | -1.2270104 | 0.21981867 | 0.67299198 |
| RPL27A | 844.040259 | 0.32156812 | 0.36622967 | 0.87805043 | 0.37991635 | 0.7880311 |
| ST5 | 2516.57113 | -1.0375993 | 0.30854559 | -3.3628719 | 7.71E-04 | 4.91E-02 |
| AC026894.1 | 248.776649 | -0.7461017 | 0.60347901 | -1.2363341 | 0.21633439 | 0.66968853 |
| AKIP1 | 123.05058 | -0.3161646 | 0.28292715 | -1.117477 | 0.26379045 | 0.71445725 |
| TMEM9B | 238.750596 | 0.21676896 | 0.21368475 | 1.01443344 | 0.31037597 | 0.74459794 |
| TMEM9B-AS | 253.97087 | 0.61073659 | 0.54396364 | 1.12275259 | 0.26154259 | 0.71216574 |
| SCUBE2 | 59.283484 | 0.33417576 | 0.49839184 | 0.67050809 | 0.50253395 | 0.85525037 |
| DENND5A | 1214.18103 | -0.1533375 | 0.30946318 | -0.4954952 | 0.62025061 | 0.8972371 |
| TMEM41B | 995.47386 | -0.3831901 | 0.23645456 | -1.6205656 | 0.10511083 | 0.52571334 |
| IPO7 | 911.082252 | -0.1046881 | 0.23296785 | -0.4493671 | 0.65316683 | 0.908787 |
| AC132192.2 | 48.5296776 | 0.37254592 | 0.58517425 | 0.63664099 | 0.52435872 | 0.86006369 |
| ZNF143 | 321.559172 | -0.0014876 | 0.36773584 | -0.0040452 | 0.99677243 | 0.99942061 |
| WEE1 | 440.695709 | -0.7195369 | 0.41802529 | -1.721276 | 0.08520075 | 0.49270382 |
| SWAP70 | 1087.00993 | -0.4360656 | 0.21289557 | -2.0482605 | 4.05E-02 | 0.37344844 |
| SBF2-AS1 | 401.816349 | -0.1541549 | 0.3369211 | -0.4575402 | 0.64728281 | 0.90657771 |
| SBF2 | 5992.97782 | 0.16368664 | 0.19765413 | 0.82814682 | 0.40758736 | 0.80588056 |
| AC011092.2 | 265.174361 | 0.06111425 | 0.51778713 | 0.11802968 | 0.90604414 | 0.98007461 |
| AC100763.1 | 77.9323171 | 0.57195788 | 0.65631195 | 0.87147259 | 0.38349616 | 0.79137648 |
| ADM | 339.97079 | 0.81097704 | 1.31691191 | 0.61581723 | 0.53801516 | 0.86496983 |
| AMPD3 | 495.522973 | -0.086816 | 0.58489516 | -0.14843 | 0.88200344 | 0.97664137 |
| MTRNR2L8 | 367.049984 | 0.80118767 | 0.74055645 | 1.08187252 | 0.27930918 | 0.724832 |
| RNF141 | 189.233141 | 0.27453609 | 0.18237119 | 1.50536987 | 0.13222901 | 0.57615451 |
| MRVI1-AS1 | 56.527658 | -0.2032964 | 0.60734887 | -0.3347276 | 0.73783058 | 0.93552616 |

| | | | | | | |
|---|---|---|---|---|---|---|
| CTR9 | 453.490101 | 0.21971914 | 0.21488238 | 1.02250889 | 0.30654011 | 0.74295884 |
| EIF4G2 | 1079.15885 | 0.08566118 | 0.22907262 | 0.3739477 | 0.70844323 | 0.92577218 |
| ZBED5 | 692.518491 | -0.0528135 | 0.17966933 | -0.2939484 | 0.76879731 | 0.94376765 |
| ZBED5-AS1 | 242.468157 | 0.1227402 | 0.35557071 | 0.3451921 | 0.72994996 | 0.93453345 |
| GALNT18 | 3618.057 | 0.11739457 | 0.50803848 | 0.23107416 | 0.81725719 | 0.9593122 |
| USP47 | 2413.57006 | 0.0241363 | 0.17932796 | 0.13459307 | 0.89293362 | 0.97782787 |
| MICAL2 | 87.7992498 | -0.2551171 | 0.36832765 | -0.6926362 | 4.89E-01 | 8.46E-01 |
| PARVA | 1214.0404 | 0.01453086 | 0.26991781 | 0.05383438 | 0.95706712 | 0.99065996 |
| TEAD1 | 4330.68053 | -0.6152848 | 0.2299324 | -2.6759377 | 0.00745205 | 0.17360516 |
| LINC00958 | 628.520025 | 0.46594547 | 0.28310315 | 1.64585055 | 0.09979453 | 0.51665771 |
| AC084859.1 | 63.0595108 | 0.66601368 | 0.56332263 | 1.18229527 | 0.23708856 | 0.68934687 |
| RASSF10 | 49.2890049 | 0.10758969 | 0.63759812 | 0.16874217 | 0.86599944 | 0.97379123 |
| ARNTL | 333.365894 | 0.0474837 | 0.33429757 | 0.14204023 | 0.88704823 | 0.97729111 |
| BTBD10 | 717.82153 | -0.323981 | 0.20716067 | -1.5639118 | 0.11783829 | 0.54888439 |
| FAR1 | 1157.7042 | 0.15427427 | 0.18833193 | 0.81916149 | 0.41269428 | 0.80735763 |
| SPON1 | 197.744541 | 0.87079183 | 0.53716941 | 1.62107488 | 0.10500158 | 0.52536586 |
| SPON1-AS1 | 44.087768 | -0.9410111 | 0.54416144 | -1.7292867 | 0.0837578 | 0.48884683 |
| RRAS2 | 957.367297 | -0.4320881 | 0.26687401 | -1.6190716 | 0.10543187 | 0.52640329 |
| COPB1 | 812.167475 | 0.08210386 | 0.190758 | 0.43040849 | 0.66689852 | 0.91331423 |
| PSMA1 | 46.5670608 | -0.5885088 | 0.50062693 | -1.1755437 | 0.23977728 | 0.69112505 |
| PDE3B | 69.2810153 | -0.5520173 | 0.40165986 | -1.3743402 | 0.16933608 | 0.62409608 |
| CYP2R1 | 176.228701 | -0.2129304 | 0.27604813 | -0.7713523 | 0.44049813 | 0.82253723 |
| SOX6 | 7661.43115 | 0.18658138 | 0.23902735 | 0.78058592 | 0.43504608 | 0.81919857 |
| C11orf58 | 1094.57956 | 0.05595067 | 0.1504227 | 0.37195628 | 0.7099254 | 0.92607195 |
| PLEKHA7 | 6310.98247 | -0.0624546 | 0.2877127 | -0.2170729 | 0.82815154 | 0.96288319 |
| RPS13 | 652.593024 | 0.3894131 | 0.47697708 | 0.81641888 | 0.4142606 | 0.80926578 |
| PIK3C2A | 2429.0507 | -0.072147 | 0.08587504 | -0.8401394 | 0.40083023 | 0.80243544 |
| NUCB2 | 391.947904 | 0.79781314 | 0.23769262 | 3.356491 | 0.00078938 | 0.04911302 |
| NCR3LG1 | 64.4415566 | 0.86840674 | 0.42155715 | 2.05999767 | 0.03939876 | 0.37234059 |
| KCNC1 | 134.697007 | -0.03075 | 0.51946174 | -0.0591959 | 0.95279609 | 0.98949465 |
| SERGEF | 1245.78757 | -0.2855142 | 0.27662125 | -1.0321485 | 0.30200255 | 0.73905643 |
| SAAL1 | 143.756196 | -0.4455638 | 0.27900169 | -1.596993 | 0.11026726 | 0.53372881 |
| SAA1 | 210.404974 | 4.0335213 | 0.81771217 | 4.93269082 | 8.11E-07 | 8.50E-04 |
| HPS5 | 289.922181 | -0.3838677 | 0.17832308 | -2.1526529 | 0.03134597 | 0.3388289 |
| GTF2H1 | 497.469762 | -0.258375 | 0.20129056 | -1.2835924 | 0.19928462 | 0.65150038 |
| LDHA | 2345.53088 | 0.3662927 | 0.42540041 | 0.86105395 | 3.89E-01 | 7.97E-01 |
| TSG101 | 852.852524 | -0.3292824 | 0.16767317 | -1.9638348 | 0.04954926 | 0.39957029 |
| UEVLD | 617.771099 | -0.1964036 | 0.1733207 | -1.13318 | 0.25713868 | 0.70746612 |
| SPTY2D1OS | 267.271132 | 0.02264136 | 0.20352448 | 0.11124635 | 0.911421 | 0.9812503 |
| SPTY2D1 | 210.124276 | 0.03775949 | 0.2333611 | 0.16180714 | 0.87145773 | 0.97518512 |
| TMEM86A | 48.1978563 | -0.3460165 | 0.46866818 | -0.7382975 | 0.46033369 | 0.83429248 |
| ZDHHC13 | 715.944408 | -0.1283194 | 0.20091931 | -0.6386613 | 0.52304332 | 0.86006369 |
| NAV2 | 3687.32446 | 0.11389965 | 0.41786401 | 0.27257588 | 0.78517926 | 0.94978829 |
| HTATIP2 | 152.939504 | 0.58435096 | 0.39041898 | 1.49672784 | 1.34E-01 | 5.77E-01 |
| PRMT3 | 776.494107 | -0.3574297 | 0.21480148 | -1.6640003 | 0.09611237 | 0.5117083 |
| NELL1 | 2920.727 | -0.5905414 | 0.63595331 | -0.9285924 | 3.53E-01 | 0.77216708 |
| ANO5 | 595.294289 | 0.01994539 | 0.2194634 | 0.09088255 | 0.92758592 | 0.98297055 |
| FANCF | 59.2691475 | 0.81002416 | 0.38732633 | 2.0913222 | 0.03649919 | 0.35908236 |
| GAS2 | 350.852552 | -0.398753 | 0.33388016 | -1.1942997 | 2.32E-01 | 0.68450837 |
| SVIP | 436.925627 | 0.50983942 | 0.19807621 | 2.57395589 | 0.01005431 | 0.20400517 |
| AC006299.1 | 162.708781 | -0.3936848 | 0.50350295 | -0.7818917 | 0.43427821 | 0.81919857 |
| MUC15 | 228.875653 | 0.61544769 | 0.50528481 | 1.21802137 | 0.22321585 | 0.67624946 |
| SLC5A12 | 94.0633557 | 2.44806167 | 0.91763875 | 2.66778366 | 0.00763534 | 0.17527709 |
| FIBIN | 115.629745 | 2.57804984 | 0.76890833 | 3.3528702 | 0.00079978 | 0.04913638 |

| | | | | | | |
|---|---|---|---|---|---|---|
| BBOX1 | 2166.26685 | 1.94957319 | 0.74373789 | 2.62131754 | 0.00875906 | 0.18998107 |
| BBOX1-AS1 | 406.53675 | 1.57675878 | 0.58219297 | 2.70830957 | 0.00676269 | 0.16923754 |
| CCDC34 | 134.248833 | 0.86133608 | 0.39414431 | 2.18533177 | 0.02886453 | 0.3275529 |
| LGR4 | 9093.09289 | -0.2550746 | 0.28746058 | -0.8873376 | 0.37489714 | 0.7844364 |
| LIN7C | 262.573653 | 0.19038904 | 0.33818278 | 0.56297674 | 0.57345072 | 0.87985952 |
| BDNF-AS | 1196.47845 | -0.1281619 | 0.21157651 | -0.6057475 | 0.54468241 | 0.86654229 |
| METTL15 | 2795.11005 | -0.031616 | 0.23931089 | -0.1321127 | 0.89489516 | 0.97788747 |
| AC090833.1 | 212.505637 | -0.7614256 | 0.42747596 | -1.7812127 | 0.07487771 | 0.47190267 |
| ARL14EP | 304.344596 | 0.41674051 | 0.20134801 | 2.06975236 | 0.03847554 | 0.36774487 |
| MPPED2 | 1956.17556 | -0.3469898 | 0.61032091 | -0.5685367 | 0.56967059 | 0.87807251 |
| AL136088.1 | 51.0268061 | 0.59206859 | 0.71411625 | 0.82909273 | 4.07E-01 | 0.80550547 |
| DCDC1 | 1122.24822 | 0.3389648 | 0.29375824 | 1.15389036 | 0.24854512 | 0.69881492 |
| DNAJC24 | 466.466609 | -0.4318919 | 0.25814478 | -1.6730606 | 0.09431537 | 0.51017802 |
| IMMP1L | 1292.44143 | 0.13769827 | 0.21017455 | 0.65516148 | 0.51236379 | 0.85762435 |
| ELP4 | 1899.7824 | 0.16024268 | 0.12981277 | 1.23441385 | 0.21704873 | 0.66978787 |
| AC131571.1 | 248.048846 | 0.44103496 | 0.60833772 | 0.72498375 | 0.468462 | 0.83825637 |
| RCN1 | 193.603663 | 0.52881663 | 0.36153441 | 1.4627007 | 0.1435493 | 0.58849865 |
| EIF3M | 432.323575 | 0.01837135 | 0.24911441 | 0.07374663 | 0.94121199 | 0.98732467 |
| CCDC73 | 81.2649126 | -0.4076819 | 0.51064013 | -0.7983742 | 0.42465337 | 0.81466629 |
| PRRG4 | 575.887707 | 0.56270134 | 0.38401463 | 1.46531225 | 0.14283575 | 0.58824095 |
| QSER1 | 812.941372 | -0.137409 | 0.12619995 | -1.0888201 | 0.27623323 | 0.72241028 |
| TCP11L1 | 381.029325 | -0.2581349 | 0.18225821 | -1.4163143 | 0.1566835 | 0.60654181 |
| CSTF3 | 778.482648 | -0.0963878 | 0.21209769 | -0.4544501 | 0.6495049 | 0.90815181 |
| HIPK3 | 2360.51826 | -0.2733356 | 0.2449465 | -1.1158991 | 0.26446534 | 0.71488591 |
| KIAA1549L | 78.8434304 | -0.2320738 | 0.48691574 | -0.47662 | 0.6336327 | 0.90136076 |
| AL049629.1 | 238.877552 | -0.1091215 | 0.35574968 | -0.3067367 | 0.75904379 | 0.94133292 |
| CD59 | 185.740704 | -0.0482991 | 0.28077122 | -0.1720229 | 0.8634195 | 0.97319807 |
| FBXO3 | 553.521086 | -0.0478169 | 0.2101949 | -0.2274885 | 0.8200439 | 0.96016713 |
| LMO2 | 51.1772651 | -0.8352594 | 0.484116 | -1.7253291 | 0.08446818 | 0.49086675 |
| CAPRIN1 | 1040.15681 | -0.0670413 | 0.16171703 | -0.4145591 | 0.67846468 | 0.91803319 |
| NAT10 | 340.50875 | -0.2727587 | 0.27155752 | -1.0044231 | 0.31517471 | 0.74785672 |
| ABTB2 | 2684.73431 | -0.2101579 | 0.34089043 | -0.6164968 | 0.53756665 | 0.86489441 |
| CAT | 1027.14979 | 0.65625472 | 0.54102375 | 1.21298691 | 0.22513482 | 0.67771328 |
| EHF | 1162.09971 | -0.1010196 | 0.23110489 | -0.437116 | 0.66202724 | 0.91110616 |
| APIP | 394.860755 | -1.0874391 | 0.31443674 | -3.4583717 | 0.00054345 | 0.04032835 |
| PDHX | 962.341728 | -0.1352791 | 0.20606148 | -0.6564989 | 0.5115032 | 0.85762435 |
| CD44 | 1112.54527 | -1.7080964 | 0.85999175 | -1.9861777 | 4.70E-02 | 3.93E-01 |
| TRIM44 | 2108.14618 | -0.1386982 | 0.15845607 | -0.8753102 | 0.38140514 | 0.78898747 |
| LDLRAD3 | 106.251381 | -0.0500318 | 0.62668447 | -0.0798357 | 0.9363679 | 0.98621411 |
| COMMD9 | 176.947266 | 0.02497912 | 0.28179643 | 0.08864242 | 0.92936609 | 0.98331361 |
| PRR5L | 215.448122 | 0.1329814 | 0.52554105 | 0.25303713 | 0.80023952 | 0.95370871 |
| TRAF6 | 330.382003 | 0.01647121 | 0.1730981 | 0.09515536 | 0.92419143 | 0.98215577 |
| RAG1 | 86.3335979 | -0.2031778 | 0.31257904 | -0.6500044 | 0.51568938 | 0.85869987 |
| C11orf74 | 327.998881 | -0.0268018 | 0.26824831 | -0.099914 | 9.20E-01 | 9.82E-01 |
| LRRC4C | 73.0805596 | -0.3541106 | 0.37136148 | -0.9535469 | 0.34031305 | 0.76478309 |
| API5 | 688.90654 | -0.1642526 | 0.17541898 | -0.9363447 | 0.34909572 | 0.76982416 |
| TTC17 | 3510.17387 | -0.0467247 | 0.18067632 | -0.2586102 | 0.79593604 | 0.95278473 |
| AC068205.1 | 96.1101845 | 0.39582323 | 0.23112816 | 1.71257032 | 8.68E-02 | 4.95E-01 |
| HSD17B12 | 2691.06421 | 0.13993667 | 0.29094047 | 0.48098043 | 0.63053041 | 0.90005881 |
| AC087521.2 | 67.4931772 | 1.00064651 | 0.49342702 | 2.02795241 | 0.0425651 | 0.38021566 |
| ALKBH3 | 274.799996 | -0.1901246 | 0.24953334 | -0.7619206 | 0.44610737 | 0.82571115 |
| ACCS | 138.008848 | 0.65760393 | 0.5521576 | 1.19097144 | 0.2336648 | 0.68493974 |
| EXT2 | 832.400147 | -0.5218886 | 0.34124415 | -1.5293701 | 0.12617273 | 0.5657215 |
| CD82 | 196.658723 | 0.04683344 | 0.46593608 | 0.10051474 | 0.91993568 | 0.98215577 |

| | | | | | | |
|---|---|---|---|---|---|---|
| PRDM11 | 221.365918 | 0.02278675 | 0.28652775 | 0.07952719 | 0.93661331 | 0.98630295 |
| SYT13 | 136.299951 | 0.44433402 | 0.41602797 | 1.06803882 | 0.285503 | 0.72846488 |
| CRY2 | 368.289789 | 0.19747297 | 0.41240854 | 0.47882851 | 0.63206063 | 0.90042718 |
| MAPK8IP1 | 79.7147982 | 0.73744893 | 0.53551195 | 1.37709147 | 0.16848397 | 0.62253512 |
| PEX16 | 103.137496 | -0.1768985 | 0.28692762 | -0.6165267 | 0.53754698 | 0.86489441 |
| LARGE2 | 258.033828 | 0.85483565 | 0.56021263 | 1.52591285 | 0.12703157 | 0.56740575 |
| PHF21A | 3246.21578 | -0.1952006 | 0.17816411 | -1.0956223 | 0.27324412 | 0.72070318 |
| DGKZ | 189.349844 | -0.2442201 | 0.34300125 | -0.7120093 | 0.47645903 | 0.84141268 |
| MDK | 75.4057476 | 0.49793518 | 0.81318488 | 0.61232715 | 0.54032133 | 0.86569031 |
| AMBRA1 | 2759.10419 | 0.19573233 | 0.19419757 | 1.00790304 | 0.31350101 | 0.74749123 |
| HARBI1 | 47.914902 | -0.1484093 | 0.45538193 | -0.3259008 | 0.74449943 | 0.93668127 |
| ATG13 | 457.910934 | -0.1437668 | 0.16240106 | -0.8852576 | 0.37601774 | 0.78490077 |
| ARHGAP1 | 376.83495 | -0.1243319 | 0.26884271 | -0.4624708 | 0.64374376 | 0.90478945 |
| CKAP5 | 657.679965 | -0.0858705 | 0.16825924 | -0.5103462 | 0.60980891 | 0.89330886 |
| LRP4 | 521.342139 | 0.2349655 | 0.48055067 | 0.48895052 | 0.62487673 | 0.89873262 |
| C11orf49 | 2579.84768 | -0.1267991 | 0.15125347 | -0.8383219 | 0.40184994 | 0.80278824 |
| ARFGAP2 | 312.068559 | -0.0395296 | 0.26616198 | -0.1485172 | 0.88193463 | 0.97664137 |
| PACSIN3 | 49.6383076 | 0.35122098 | 0.43544787 | 0.80657412 | 0.41991189 | 0.8125045 |
| DDB2 | 172.612066 | -0.1505382 | 0.43450917 | -0.3464557 | 0.72900024 | 0.93453345 |
| ACP2 | 93.4250499 | 0.35339677 | 0.34609723 | 1.02109098 | 0.30721134 | 0.74341546 |
| NR1H3 | 170.690051 | -0.3206409 | 0.29642921 | -1.0816777 | 0.27939576 | 0.72489035 |
| MADD | 399.304401 | -0.6410425 | 0.16328837 | -3.9258305 | 8.64E-05 | 0.01511702 |
| SPI1 | 51.4498453 | -0.0747319 | 0.78482178 | -0.0952214 | 0.92413893 | 0.98215577 |
| AC090559.1 | 40.5365162 | -0.2396412 | 0.57643848 | -0.4157273 | 0.67760954 | 0.91753403 |
| SLC39A13 | 270.416214 | 0.00999076 | 0.39264326 | 0.02544489 | 0.97970011 | 0.99483846 |
| PSMC3 | 117.235555 | 0.48161688 | 0.36621862 | 1.31510758 | 0.18847375 | 0.64312833 |
| CELF1 | 2816.93074 | -0.2080341 | 0.16781535 | -1.2396609 | 0.21510085 | 0.66869079 |
| NDUFS3 | 241.775397 | 0.5923444 | 0.35720459 | 1.65827769 | 0.09726143 | 0.51355946 |
| KBTBD4 | 66.7200079 | 0.30566697 | 0.27893887 | 1.09582067 | 0.27315726 | 0.72070318 |
| MTCH2 | 616.765538 | -0.2266219 | 0.25609218 | -0.8849231 | 0.37619808 | 0.78490077 |
| AGBL2 | 82.6734091 | -1.0408496 | 0.35146621 | -2.96145 | 0.00306194 | 0.10671555 |
| FNBP4 | 1902.31581 | -0.1577887 | 0.16070637 | -0.9818446 | 0.32617642 | 0.75554697 |
| NUP160 | 1055.14402 | -0.4886682 | 0.17807999 | -2.7440938 | 0.00606782 | 0.15969766 |
| PTPRJ | 4221.20595 | -0.9555522 | 0.25322439 | -3.7735392 | 0.00016095 | 0.0205434 |
| FOLH1 | 55.8792807 | -0.0806928 | 0.60452753 | -0.1334807 | 8.94E-01 | 9.78E-01 |
| TNKS1BP1 | 438.917263 | -0.0900581 | 0.16659165 | -0.5405918 | 5.89E-01 | 8.86E-01 |
| SSRP1 | 185.682007 | 0.32492906 | 0.29857599 | 1.08826251 | 0.27647924 | 0.72245583 |
| SLC43A1 | 99.7271575 | -1.2998971 | 0.58087838 | -2.2378128 | 2.52E-02 | 0.31063019 |
| TIMM10 | 106.433959 | 0.76331407 | 0.40954881 | 1.86379266 | 0.06235083 | 0.43419765 |
| UBE2L6 | 130.240929 | 0.61902603 | 0.6966019 | 0.88863672 | 0.37419835 | 0.78369843 |
| SERPING1 | 1728.2134 | 0.20597559 | 0.42735654 | 0.48197598 | 6.30E-01 | 9.00E-01 |
| ZDHHC5 | 557.345659 | -0.2647676 | 0.20608983 | -1.2847195 | 0.19889032 | 0.65113421 |
| MED19 | 88.799546 | -0.1005338 | 0.31415815 | -0.3200102 | 0.7489606 | 0.93748128 |
| CTNND1 | 1608.25276 | -0.1608207 | 0.17980677 | -0.8944084 | 0.3711034 | 0.78104287 |
| LPXN | 509.232959 | -0.2673085 | 0.23424108 | -1.1411681 | 2.54E-01 | 0.70431689 |
| GLYAT | 61.2460331 | 1.87390005 | 0.87841887 | 2.13326478 | 0.03290301 | 0.34598377 |
| GLYATL1 | 99.1586944 | 1.50281743 | 0.42011257 | 3.57717798 | 0.00034732 | 0.03115997 |
| FAM111A-D1 | 67.1942888 | 0.16246151 | 0.39071243 | 0.41580841 | 0.67755021 | 0.91753403 |
| FAM111A | 431.223669 | 0.13484535 | 0.20214475 | 0.66707324 | 0.50472535 | 0.85561052 |
| DTX4 | 82.0526621 | 0.3072371 | 0.90272086 | 0.34034563 | 0.73359626 | 0.93552616 |
| OSBP | 1030.63401 | -0.3706636 | 0.28412953 | -1.3045585 | 0.19204323 | 0.64596359 |
| AP000442.2 | 57.013272 | 0.29553837 | 0.56317426 | 0.52477251 | 0.59974135 | 0.88883692 |
| PATL1 | 576.517897 | -0.0599424 | 0.25821106 | -0.2321451 | 0.8164253 | 0.9589293 |
| STX3 | 778.879825 | 0.06743712 | 0.31037182 | 0.21727848 | 0.82799132 | 0.96284333 |

| | | | | | | |
|---|---|---|---|---|---|---|
| MRPL16 | 169.94894 | 0.17065921 | 0.30393837 | 0.56149281 | 0.57446163 | 0.88026176 |
| PRPF19 | 189.368804 | 0.27014566 | 0.37475706 | 0.72085543 | 0.47099847 | 0.83921324 |
| TMEM109 | 122.971972 | 0.94814373 | 0.3933776 | 2.41026363 | 0.015941 | 0.25461261 |
| TMEM132A | 46.8255455 | -0.0586486 | 0.49198963 | -0.1192071 | 0.90511131 | 0.97992575 |
| VPS37C | 285.167476 | 0.07360318 | 0.32060357 | 0.22957692 | 0.81842054 | 0.95985564 |
| DDB1 | 731.769447 | 0.18464657 | 0.25363796 | 0.72799262 | 0.4666181 | 0.83698513 |
| TKFC | 128.112425 | -0.0001543 | 0.3548056 | -0.0004347 | 0.99965312 | 0.99987432 |
| CYB561A3 | 98.9699331 | -0.2096095 | 0.25780225 | -0.8130631 | 0.41618186 | 0.80966301 |
| TMEM138 | 119.843716 | -0.2631495 | 0.29073588 | -0.9051153 | 0.36540432 | 0.78095617 |
| TMEM216 | 74.6524018 | -0.2190792 | 0.38875923 | -0.5635344 | 0.57307102 | 0.87968566 |
| CPSF7 | 658.217662 | 0.01366508 | 0.11102946 | 0.12307614 | 0.90204681 | 0.97941893 |
| AP003108.2 | 590.744327 | -0.2594428 | 0.41157521 | -0.6303655 | 0.52845549 | 0.86241272 |
| SYT7 | 307.385823 | 0.67168841 | 0.53843419 | 1.2474847 | 0.21221983 | 0.66552984 |
| DAGLA | 65.299267 | 0.07294241 | 0.54597106 | 0.13360123 | 0.89371791 | 0.97788747 |
| MYRF | 99.7945365 | 1.59562903 | 0.44328874 | 3.59952533 | 0.0003188 | 0.03094754 |
| TMEM258 | 190.531413 | 0.13100404 | 0.36068243 | 0.3632116 | 0.71644682 | 0.9289438 |
| FADS2 | 102.006538 | -0.6017801 | 0.33526426 | -1.7949426 | 0.07266285 | 0.4692634 |
| FADS1 | 109.740935 | -0.8496553 | 0.44906042 | -1.8920734 | 0.0584812 | 0.42147872 |
| FADS3 | 136.637924 | 0.06688166 | 0.38013149 | 0.17594349 | 8.60E-01 | 0.97200968 |
| FTH1 | 5963.00085 | 0.7556654 | 0.45279016 | 1.66890864 | 9.51E-02 | 5.11E-01 |
| INCENP | 68.7644822 | -0.7155837 | 0.35921131 | -1.9920969 | 0.04636043 | 0.3904347 |
| ASRGL1 | 756.240186 | 0.27308253 | 0.26544826 | 1.0287599 | 0.30359252 | 0.7409078 |
| AHNAK | 1103.71727 | 0.20530092 | 0.29241288 | 0.70209259 | 4.83E-01 | 8.44E-01 |
| MTA2 | 54.8074979 | -0.0936029 | 0.33588113 | -0.2786786 | 0.78049149 | 0.9480401 |
| EML3 | 148.812567 | -0.4150883 | 0.43175656 | -0.9613944 | 0.33635391 | 0.7637525 |
| B3GAT3 | 231.928455 | 0.16501129 | 0.29418077 | 0.56091801 | 0.57485343 | 0.88046555 |
| GANAB | 552.604122 | 0.26578672 | 0.18170928 | 1.46270304 | 0.14354865 | 0.58849865 |
| INTS5 | 53.0175106 | 1.11909142 | 0.45431083 | 2.46327257 | 0.01376752 | 0.24292128 |
| CSKMT | 58.7015274 | 0.22944458 | 0.26383707 | 0.86964497 | 0.38449446 | 0.79217288 |
| UQCC3 | 153.842852 | 0.39246553 | 0.33869299 | 1.15876486 | 0.24655205 | 0.69684411 |
| UBXN1 | 207.298538 | 0.44111022 | 0.31304397 | 1.40909987 | 0.15880564 | 0.60819475 |
| TTC9C | 302.060829 | -0.1758357 | 0.25644641 | -0.6856627 | 0.49292581 | 0.84779389 |
| ZBTB3 | 38.7140394 | -0.1403851 | 0.36127113 | -0.3885865 | 0.69758206 | 0.92341984 |
| POLR2G | 88.2594691 | 0.34641326 | 0.3312321 | 1.0458324 | 0.29563842 | 0.73661928 |
| TAF6L | 154.042725 | -0.3559244 | 0.32028907 | -1.1112598 | 0.26645655 | 0.71641758 |
| TMEM223 | 169.503865 | 0.20309217 | 0.24552975 | 0.82715911 | 0.40814688 | 0.80588056 |
| TMEM179B | 297.973749 | 0.25406723 | 0.28523672 | 0.89072415 | 0.37307718 | 0.78277446 |
| NXF1 | 312.250869 | 0.04780914 | 0.43613253 | 0.10962066 | 0.91271022 | 0.98135874 |
| STX5 | 616.802189 | -0.1181925 | 0.22616311 | -0.5225985 | 0.60125366 | 0.8890585 |
| WDR74 | 73.6881809 | -0.1468166 | 0.27826531 | -0.5276138 | 0.59776741 | 0.88883692 |
| SLC3A2 | 354.853371 | -0.045343 | 0.35303849 | -0.1284365 | 0.89780358 | 0.97823597 |
| SLC22A6 | 81.7987307 | 1.8681019 | 0.95068339 | 1.96500951 | 0.04941314 | 0.39895974 |
| RARRES3 | 257.054116 | 1.54084032 | 0.61168631 | 2.5190041 | 0.01176873 | 0.22271223 |
| PLA2G16 | 134.813123 | 0.98822027 | 0.48486625 | 2.03812963 | 0.04153697 | 0.37579578 |
| ATL3 | 573.586474 | 0.20468113 | 0.16903013 | 1.21091511 | 2.26E-01 | 0.67809183 |
| RTN3 | 1233.43059 | 0.03145926 | 0.3250878 | 0.09677158 | 0.92290779 | 0.98215577 |
| C11orf95 | 48.7697243 | 0.41093955 | 0.63367774 | 0.64849928 | 0.51666208 | 0.8588826 |
| SPINDOC | 89.0950689 | -0.3820419 | 0.29629315 | -1.2894051 | 0.19725728 | 0.64964978 |
| MARK2 | 933.673086 | -0.1260909 | 0.20987205 | -0.6007989 | 0.54797395 | 0.86859163 |
| NAA40 | 358.182655 | 0.09819365 | 0.25038946 | 0.39216367 | 0.69493728 | 0.92229578 |
| OTUB1 | 289.885273 | 0.05029239 | 0.21845399 | 0.23021958 | 0.81792115 | 0.9597513 |
| MACROD1 | 518.828532 | 0.58219839 | 0.42547583 | 1.36834658 | 0.17120362 | 0.62772493 |
| AP000721.2 | 178.012857 | 0.58168206 | 0.68324743 | 0.85134905 | 0.39457548 | 0.79876591 |
| FLRT1 | 55.9730058 | 1.08226959 | 0.61100083 | 1.77130625 | 0.07650979 | 0.4759127 |

| | | | | | | |
|---|---|---|---|---|---|---|
| STIP1 | 277.507616 | 0.16329523 | 0.2249004 | 0.72607801 | 0.46779095 | 0.83806465 |
| TRPT1 | 105.18451 | -0.367123 | 0.33223064 | -1.1050246 | 0.26914891 | 0.71818262 |
| NUDT22 | 212.338656 | 0.03312115 | 0.19895417 | 0.16647626 | 0.86778216 | 0.97402505 |
| AP001453.1 | 100.828565 | 0.07835779 | 0.38143386 | 0.20542955 | 0.83723654 | 0.96527001 |
| DNAJC4 | 154.006853 | 0.35650528 | 0.42719322 | 0.83452936 | 0.40398275 | 0.80413022 |
| VEGFB | 421.612263 | -0.0351815 | 0.30965509 | -0.113615 | 0.90954298 | 0.98069436 |
| FKBP2 | 365.262304 | 0.81639 | 0.34926842 | 2.33742865 | 0.01941691 | 0.28045661 |
| PPP1R14B | 165.270495 | 0.14727792 | 0.43858621 | 0.33580153 | 0.73702054 | 0.93552616 |
| PLCB3 | 61.5288988 | -0.0964086 | 0.37100968 | -0.2598547 | 0.79497589 | 0.95272759 |
| BAD | 174.392544 | 0.42654262 | 0.29856805 | 1.42862781 | 0.15311124 | 0.60285224 |
| GPR137 | 208.081621 | 0.01849256 | 0.28557464 | 0.0647556 | 0.94836859 | 0.98919519 |
| ESRRA | 616.919387 | -0.0555819 | 0.29926946 | -0.1857252 | 0.85266026 | 0.97076831 |
| TRMT112 | 226.662286 | 0.56942055 | 0.46994624 | 1.2116717 | 0.22563808 | 0.67798093 |
| PRDX5 | 367.978128 | 0.53370599 | 0.4396933 | 1.21381423 | 0.22481867 | 0.67750405 |
| RPS6KA4 | 76.6526135 | -0.2221034 | 0.45515423 | -0.487974 | 0.62556825 | 0.89873262 |
| SLC22A12 | 49.6141485 | 1.19934757 | 0.80524007 | 1.48942858 | 0.13637455 | 0.57787125 |
| SF1 | 1851.51854 | -0.2488721 | 0.20047468 | -1.2414143 | 0.21445274 | 0.66827012 |
| MAP4K2 | 110.660397 | 0.24416311 | 0.29339964 | 0.83218612 | 0.4053039 | 0.80481948 |
| MEN1 | 98.6118766 | -0.1664705 | 0.34658024 | -0.4803232 | 0.63099761 | 0.90005881 |
| CDC42BPG | 71.1297988 | -0.1959393 | 0.30207942 | -0.6486351 | 0.51657427 | 0.8588826 |
| EHD1 | 148.333514 | -0.0654738 | 0.36074837 | -0.1814945 | 0.85597949 | 0.97118784 |
| ATG2A | 225.431014 | -0.2416255 | 0.25698568 | -0.9402295 | 0.34709984 | 0.76893725 |
| PPP2R5B | 112.569527 | -0.0447089 | 0.27286507 | -0.1638499 | 0.86984934 | 0.97454113 |
| ZFPL1 | 94.1237927 | 0.05405841 | 0.37590571 | 0.14380844 | 0.88565174 | 0.97664137 |
| VPS51 | 555.164938 | 0.00016893 | 0.22772848 | 0.0007418 | 0.99940813 | 0.99987432 |
| TM7SF2 | 104.459588 | 0.09406845 | 0.47292748 | 0.19890671 | 0.84233572 | 0.96717773 |
| FAU | 772.820219 | 0.60071262 | 0.42097959 | 1.42694002 | 0.15359718 | 0.60285224 |
| SYVN1 | 150.557353 | 0.28197739 | 0.33089236 | 0.85217255 | 0.39411833 | 0.79852106 |
| MRPL49 | 96.4987993 | 0.76409019 | 0.37474087 | 2.03898279 | 0.04145175 | 0.37579578 |
| AP003068.2 | 49.7052194 | 0.26005575 | 0.49041935 | 0.5302722 | 0.59592322 | 0.88859628 |
| CAPN1 | 504.829983 | 0.11083867 | 0.29813593 | 0.37177226 | 0.71006242 | 0.92610043 |
| POLA2 | 87.7429678 | -1.1240018 | 0.39522307 | -2.8439682 | 0.00445555 | 0.12989304 |
| DPF2 | 298.76189 | -0.1470403 | 0.20270253 | -0.7253995 | 0.46820699 | 0.83825637 |
| SLC25A45 | 97.928657 | -0.1430377 | 0.70700599 | -0.2023147 | 0.83967069 | 0.96617873 |
| FRMD8 | 277.978553 | -0.4218237 | 0.29432548 | -1.4331879 | 0.15180414 | 0.60089326 |
| NEAT1 | 78404.0362 | -0.1908969 | 0.45511759 | -0.4194453 | 6.75E-01 | 9.16E-01 |
| MALAT1 | 1039802.28 | -0.1980843 | 0.44788414 | -0.4422668 | 0.65829618 | 0.91037018 |
| SCYL1 | 250.390332 | 0.00738088 | 0.31219102 | 0.02364218 | 0.98113803 | 0.99518016 |
| LTBP3 | 623.845876 | -0.6310864 | 0.37476512 | -1.6839519 | 0.09219097 | 0.5058189 |
| SSSCA1 | 87.6425503 | 0.74999404 | 0.49980278 | 1.50057999 | 0.13346423 | 0.57682553 |
| FAM89B | 81.6510387 | 0.26314137 | 0.45898851 | 0.57330709 | 0.56643678 | 0.87642772 |
| EHBP1L1 | 247.027044 | -0.9557895 | 0.43963542 | -2.1740504 | 2.97E-02 | 3.31E-01 |
| MAP3K11 | 148.410162 | 0.0975374 | 0.34847094 | 0.27990111 | 0.77955337 | 0.94758042 |
| PCNX3 | 143.426179 | -0.0481371 | 0.36894045 | -0.1304738 | 0.89619157 | 0.97800682 |
| SIPA1 | 94.0187647 | 0.00953412 | 0.36234343 | 0.02631239 | 9.79E-01 | 9.95E-01 |
| RELA | 204.201016 | 0.20548355 | 0.29918639 | 0.68680783 | 0.49220384 | 0.84734855 |
| KAT5 | 242.960323 | -0.1743172 | 0.27976857 | -0.6230763 | 0.53323442 | 0.86397197 |
| RNASEH2C | 158.693685 | 0.5377582 | 0.28817024 | 1.86611288 | 0.06202557 | 0.43390193 |
| AP5B1 | 71.4827027 | 0.73554143 | 0.39151967 | 1.87868321 | 0.06028776 | 0.42866735 |
| CFL1 | 675.615446 | 0.44725003 | 0.36045045 | 1.24080865 | 2.15E-01 | 0.66830201 |
| MUS81 | 152.429035 | -0.2209652 | 0.24237918 | -0.9116507 | 0.36195262 | 0.77784066 |
| EFEMP2 | 74.3918142 | 0.49258533 | 0.37046995 | 1.3296229 | 0.18364255 | 0.63834582 |
| FIBP | 61.5396273 | 0.37750396 | 0.48604554 | 0.77668434 | 0.43734503 | 0.82013459 |
| CCDC85B | 68.9982851 | 0.66811734 | 0.41350698 | 1.61573411 | 0.10615181 | 0.52735777 |

| | | | | | | |
|---|---|---|---|---|---|---|
| C11orf68 | 56.2082127 | 0.92514702 | 0.44223651 | 2.09197339 | 0.0364409 | 0.35908113 |
| DRAP1 | 131.808212 | 0.04685831 | 0.42293354 | 0.11079355 | 0.91178006 | 0.98125148 |
| SART1 | 158.371149 | -0.2847573 | 0.3166383 | -0.8993142 | 0.36848531 | 0.78095617 |
| EIF1AD | 70.110196 | 0.06176912 | 0.2905098 | 0.2126232 | 0.83162087 | 0.9639242 |
| BANF1 | 192.42414 | 0.34014858 | 0.4371978 | 0.77801987 | 0.4365573 | 0.82013459 |
| SF3B2 | 426.521205 | 0.02017525 | 0.21137321 | 0.09544848 | 0.92395861 | 0.98215577 |
| PACS1 | 3684.92721 | -0.029868 | 0.20468951 | -0.1459186 | 8.84E-01 | 0.97664137 |
| KLC2 | 131.659069 | 0.1059293 | 0.34694274 | 0.30532212 | 0.76012083 | 0.94161456 |
| RAB1B | 241.265509 | 0.56630036 | 0.33861395 | 1.67240705 | 0.09444408 | 0.51017802 |
| YIF1A | 100.343021 | 0.20894161 | 0.44245507 | 0.47223237 | 0.63676094 | 0.90255504 |
| BRMS1 | 102.626988 | 0.11156627 | 0.37413759 | 0.29819584 | 0.7655537 | 0.94270299 |
| B4GAT1 | 54.3933571 | 1.22108445 | 0.47431655 | 2.57440828 | 0.01004117 | 0.20400517 |
| AP001107.9 | 46.9421468 | -0.0564126 | 0.41135455 | -0.1371386 | 0.89092128 | 0.97729111 |
| SLC29A2 | 238.595042 | -0.216509 | 0.32825632 | -0.659573 | 0.50952791 | 0.8568242 |
| MRPL11 | 89.6622149 | 0.75754066 | 0.34491762 | 2.19629444 | 0.02807088 | 0.3228969 |
| DPP3 | 56.142303 | -0.5842594 | 0.48636222 | -1.2012844 | 0.2296409 | 0.68195292 |
| ZDHHC24 | 143.069204 | 0.427046 | 0.27173873 | 1.57153158 | 0.11605923 | 0.54556096 |
| CTSF | 162.093809 | 0.90315234 | 0.45156075 | 2.00006828 | 0.04549289 | 0.38904776 |
| CCS | 292.984865 | 0.39818677 | 0.20101391 | 1.98089157 | 0.04760343 | 0.39447539 |
| RBM14-RBM | 368.636432 | 0.09338729 | 0.29472856 | 0.31685865 | 0.75135086 | 0.93794692 |
| RBM4 | 220.694188 | -0.6786211 | 0.31993228 | -2.1211399 | 0.03391003 | 0.35107893 |
| RBM4B | 293.238862 | -0.0210009 | 0.23198272 | -0.0905277 | 0.92786791 | 0.98297055 |
| AP001157.1 | 67.8768717 | -0.6454847 | 0.4309678 | -1.4977561 | 0.13419663 | 0.57708028 |
| SPTBN2 | 142.636343 | 0.91591367 | 0.50915785 | 1.7988796 | 0.07203773 | 0.46774933 |
| C11orf80 | 1032.51635 | 0.3746235 | 0.24835812 | 1.50840045 | 1.31E-01 | 0.57519227 |
| RCE1 | 63.9722085 | 0.21493821 | 0.38324897 | 0.5608318 | 0.57491221 | 0.88046555 |
| PC | 118.855238 | 0.07126017 | 0.29178827 | 0.24421875 | 0.80706141 | 0.95575346 |
| RHOD | 130.173861 | -0.1429335 | 0.36267999 | -0.3941037 | 0.69350445 | 0.92229578 |
| KDM2A | 3584.28063 | -0.25933 | 0.16281774 | -1.5927624 | 0.11121351 | 0.53461826 |
| GRK2 | 262.08251 | -0.2259583 | 0.3046293 | -0.7417484 | 0.45823978 | 0.8331421 |
| ANKRD13D | 380.633723 | -0.3714632 | 0.29849846 | -1.2444392 | 0.21333795 | 0.66755797 |
| SSH3 | 140.188829 | -0.4846286 | 0.27874699 | -1.7385968 | 0.08210571 | 0.48594425 |
| RAD9A | 1368.27165 | 0.28160316 | 0.16970508 | 1.65936791 | 0.09704168 | 0.51334542 |
| CLCF1 | 67.5325737 | -0.4173336 | 0.603802 | -0.6911763 | 4.89E-01 | 0.84614483 |
| PPP1CA | 305.640494 | 0.55712716 | 0.44772657 | 1.24434688 | 0.21337192 | 0.66755797 |
| RPS6KB2 | 142.652485 | 0.17225243 | 0.41481454 | 0.41525168 | 0.67795767 | 0.91772319 |
| CORO1B | 171.430058 | 0.38958352 | 0.37254574 | 1.04573338 | 0.29568415 | 0.73661928 |
| TMEM134 | 230.19089 | -0.003812 | 0.26026783 | -0.0146465 | 0.98831423 | 0.99653761 |
| AIP | 98.1959397 | 0.52131602 | 0.4041294 | 1.28997302 | 0.19706003 | 0.64964978 |
| PITPNM1 | 157.403234 | 0.08738509 | 0.4600696 | 0.18993885 | 0.84935705 | 0.970224 |
| CDK2AP2 | 214.069847 | 0.82451273 | 0.47104418 | 1.75039363 | 0.08005041 | 0.48282462 |
| GSTP1 | 1980.77843 | 0.7471 | 0.54820637 | 1.36280794 | 0.17294305 | 0.6286116 |
| NDUFV1 | 656.264749 | 0.27514308 | 0.30747582 | 0.89484461 | 0.37087016 | 0.78104287 |
| NUDT8 | 71.8284934 | 0.63994411 | 0.4256705 | 1.50337904 | 0.13274133 | 0.57682553 |
| AP003716.1 | 44.5713072 | 0.21692542 | 0.65178606 | 0.33281691 | 0.73927249 | 0.93552616 |
| UNC93B1 | 144.932859 | -0.5465226 | 0.53396193 | -1.0235235 | 0.30606041 | 0.74290739 |
| NDUFS8 | 473.705864 | 0.52963876 | 0.32414888 | 1.6339367 | 0.10227214 | 0.52073581 |
| TCIRG1 | 60.1177543 | 0.45359495 | 0.64555985 | 0.70263811 | 0.48228131 | 0.84389635 |
| CHKA | 1420.10125 | -0.1666768 | 0.17907535 | -0.9307635 | 3.52E-01 | 7.72E-01 |
| KMT5B | 1441.93716 | -0.0401817 | 0.17228251 | -0.2332313 | 0.81558184 | 0.95872901 |
| C11orf24 | 44.5294574 | -0.1911208 | 0.38833333 | -0.4921566 | 0.62260865 | 0.89835355 |
| LRP5 | 539.509788 | 0.05499495 | 0.35612169 | 0.15442742 | 0.87727274 | 0.97639835 |
| PPP6R3 | 3835.78081 | -0.0964201 | 0.17324648 | -0.5565486 | 0.57783587 | 0.88191134 |
| AP003096.1 | 57.6431939 | 0.24063743 | 0.73055583 | 0.32938951 | 0.7418613 | 0.93598004 |

| | | | | | | |
|---|---|---|---|---|---|---|
| TESMIN | 99.1102762 | -0.0643217 | 0.50859863 | -0.1264685 | 0.89936108 | 0.97889101 |
| CPT1A | 1319.5998 | 0.23443815 | 0.57526334 | 0.40753188 | 0.68361738 | 0.9200778 |
| MRPL21 | 116.738354 | 0.31456337 | 0.2502062 | 1.25721654 | 0.20867524 | 0.6614379 |
| IGHMBP2 | 183.560827 | -0.1141322 | 0.34912113 | -0.326913 | 0.74373367 | 0.93633036 |
| TPCN2 | 171.827321 | -0.1335136 | 0.24827845 | -0.5377574 | 0.59074451 | 0.88672234 |
| CCND1 | 275.260106 | -0.0711528 | 0.70221125 | -0.1013267 | 0.91929109 | 0.98215577 |
| ORAOV1 | 234.370065 | -0.4058637 | 0.29791556 | -1.3623447 | 0.17308912 | 0.6286116 |
| FADD | 47.7717824 | 0.30005129 | 0.33673448 | 0.89106197 | 3.73E-01 | 0.78270813 |
| PPFIA1 | 1419.84185 | -0.4451592 | 0.20222232 | -2.2013355 | 0.02771228 | 0.32139504 |
| AP002336.2 | 562.912959 | -0.1449775 | 0.48348133 | -0.2998617 | 0.76428267 | 0.94256396 |
| AP000487.1 | 136.182799 | 0.11490136 | 0.28722573 | 0.40003855 | 0.68912812 | 0.92166785 |
| CTTN | 1489.36459 | -0.2699083 | 0.21205377 | -1.2728298 | 2.03E-01 | 6.56E-01 |
| SHANK2 | 4619.56821 | 0.53494989 | 0.33170575 | 1.61272421 | 0.10680444 | 0.52778894 |
| SHANK2-AS1 | 49.2893989 | 0.35137082 | 0.75093336 | 0.46791212 | 0.63984743 | 0.90330608 |
| DHCR7 | 54.4982852 | 0.40821996 | 0.60864351 | 0.67070452 | 0.50240878 | 0.85525037 |
| NADSYN1 | 309.931802 | -0.0912919 | 0.31502791 | -0.2897898 | 0.77197704 | 0.94478317 |
| FAM86C1 | 41.1800113 | -0.2424693 | 0.41996756 | -0.5773524 | 0.56370141 | 0.87487519 |
| AP002495.2 | 915.969631 | -0.1618733 | 0.15509426 | -1.0437094 | 0.29661986 | 0.73672107 |
| RNF121 | 857.407008 | -0.3469541 | 0.17971489 | -1.9305806 | 0.05353493 | 0.41256882 |
| IL18BP | 68.9505531 | -0.5470413 | 0.37601188 | -1.454851 | 0.14571051 | 0.59218144 |
| NUMA1 | 1012.37534 | -0.0332237 | 0.24686875 | -0.1345806 | 0.89294351 | 0.97782787 |
| LAMTOR1 | 362.782263 | 0.31227333 | 0.21220575 | 1.47155925 | 0.14113994 | 0.5872148 |
| ANAPC15 | 75.6706464 | 0.02397155 | 0.3869872 | 0.06194403 | 0.95060741 | 0.9893457 |
| FOLR3 | 67.8302012 | 1.6918571 | 0.54855568 | 3.08420307 | 0.00204098 | 0.08753031 |
| FOLR1 | 85.8740651 | 2.03323393 | 0.56723857 | 3.5844423 | 0.0003378 | 0.03098604 |
| INPPL1 | 281.394743 | -0.0731187 | 0.27474577 | -0.2661322 | 0.7901374 | 0.95123267 |
| CLPB | 197.128327 | -0.1465594 | 0.22972553 | -0.6379763 | 0.5234891 | 0.86006369 |
| ARAP1 | 307.060344 | 0.05284773 | 0.29158155 | 0.18124511 | 0.85617519 | 0.97118784 |
| ATG16L2 | 217.690194 | 0.08405301 | 0.32566465 | 0.25809682 | 0.79633218 | 0.95278473 |
| FCHSD2 | 2546.22362 | -0.6440485 | 0.20699224 | -3.111462 | 0.00186163 | 0.08355912 |
| P2RY6 | 350.704962 | 1.64718662 | 0.91561216 | 1.79900038 | 0.07201862 | 0.46774933 |
| ARHGEF17 | 501.324474 | 0.1218439 | 0.28541151 | 0.42690606 | 0.66944775 | 0.91443968 |
| FAM168A | 1525.27391 | -0.0040955 | 0.13713919 | -0.0298635 | 0.97617594 | 0.9941831 |
| RAB6A | 1860.90397 | -0.4131122 | 0.15411917 | -2.6804727 | 0.00735183 | 0.17305714 |
| MRPL48 | 379.999545 | -0.3264007 | 0.1708719 | -1.9102071 | 0.05610656 | 0.41843207 |
| COA4 | 221.381223 | 0.12786649 | 0.27732722 | 0.46106722 | 0.64475038 | 0.90507831 |
| PAAF1 | 376.900641 | -0.2645617 | 0.36078545 | -0.7332938 | 0.46337927 | 0.83531269 |
| UCP2 | 46.7731839 | -0.5887743 | 0.65820774 | -0.8945114 | 0.37104832 | 0.78104287 |
| C2CD3 | 844.840772 | -0.2328184 | 0.13349549 | -1.7440171 | 0.08115611 | 0.48572317 |
| PPME1 | 997.098161 | 0.26746438 | 0.30330946 | 0.8818201 | 0.3778741 | 0.7865219 |
| P4HA3 | 182.197514 | 0.04017742 | 0.53035611 | 0.07575555 | 0.93961358 | 0.98715467 |
| PGM2L1 | 478.982507 | -0.9596786 | 0.43214349 | -2.2207406 | 2.64E-02 | 0.31402818 |
| KCNE3 | 54.6894492 | 1.1935583 | 0.56732097 | 2.10385014 | 0.03539152 | 0.35577368 |
| POLD3 | 471.811818 | -0.3071158 | 0.3211776 | -0.956218 | 0.33896208 | 0.76449297 |
| RNF169 | 1128.43015 | -0.3182134 | 0.22539672 | -1.4117927 | 0.15801101 | 0.60779483 |
| XRRA1 | 1149.31049 | -0.4103639 | 0.49011179 | -0.8372862 | 0.40243169 | 0.80310019 |
| SPCS2 | 749.916835 | 0.40936417 | 0.23281605 | 1.75831592 | 0.07869377 | 0.48060389 |
| NEU3 | 199.247171 | 0.02517547 | 0.24801634 | 0.10150728 | 0.91914778 | 0.98215577 |
| ARRB1 | 183.297602 | -0.5459081 | 0.3486086 | -1.5659629 | 0.1173573 | 0.54757066 |
| RPS3 | 1241.69737 | 0.65922461 | 0.38999997 | 1.69031965 | 0.09096682 | 0.50282313 |
| SERPINH1 | 120.534624 | 0.17806471 | 0.28444544 | 0.62600657 | 0.53131063 | 0.86397197 |
| MAP6 | 102.751729 | 0.03428112 | 0.50469894 | 0.0679239 | 0.94584621 | 0.98873715 |
| UVRAG | 1638.61237 | -0.2160087 | 0.17422441 | -1.2398301 | 0.21503825 | 0.66865355 |
| THAP12 | 206.033637 | -0.1422568 | 0.26774398 | -0.5313165 | 0.59519948 | 0.88859116 |

| | | | | | | |
|---|---|---|---|---|---|---|
| GVQW3 | 104.189487 | -0.0666291 | 0.45752409 | -0.1456297 | 0.8842137 | 0.97664137 |
| AP002360.1 | 104.953667 | 0.40799766 | 0.43623535 | 0.93526958 | 0.34964939 | 0.77025232 |
| EMSY | 1120.11697 | -0.3040989 | 0.23868406 | -1.2740646 | 0.20264051 | 0.65552334 |
| ACER3 | 1491.07535 | 0.84183386 | 0.42750244 | 1.96919075 | 0.04893119 | 0.39768952 |
| CAPN5 | 147.450386 | 1.92905216 | 0.54865743 | 3.51595014 | 0.00043818 | 0.03595008 |
| PAK1 | 881.35009 | 0.00079 | 0.33455734 | 0.00236134 | 0.99811593 | 0.99969006 |
| CLNS1A | 642.867893 | 0.03744027 | 0.24029718 | 0.15580819 | 0.87618422 | 0.97614152 |
| AQP11 | 50.6916784 | 0.42507512 | 0.38227474 | 1.11196238 | 0.26615433 | 0.71641758 |
| RSF1 | 2265.66716 | -0.1527373 | 0.14131108 | -1.0808589 | 0.27975988 | 0.72500457 |
| AAMDC | 694.188145 | -0.0894425 | 0.18382888 | -0.4865532 | 0.62657496 | 0.89873262 |
| AP002812.2 | 297.473409 | -0.2168096 | 0.32580795 | -0.6654523 | 0.50576127 | 0.85604826 |
| INTS4 | 597.158571 | -0.1965672 | 0.24762826 | -0.7937997 | 0.42731206 | 0.81466629 |
| ALG8 | 305.729875 | -0.3709691 | 0.27893933 | -1.3299276 | 0.18354213 | 0.63834582 |
| KCTD21-AS1 | 59.0264566 | -0.6783349 | 0.80654786 | -0.8410349 | 0.40032839 | 0.80238662 |
| KCTD21 | 52.4384004 | 0.18292357 | 0.39091211 | 0.4679404 | 0.63982721 | 0.90330608 |
| GAB2 | 1293.04935 | -0.4756867 | 0.31706792 | -1.5002675 | 1.34E-01 | 0.57682553 |
| AP003086.2 | 496.267725 | -0.2767907 | 0.69441412 | -0.398596 | 0.69019092 | 0.92183073 |
| NARS2 | 801.431336 | -0.3581157 | 0.24966265 | -1.4343984 | 0.15145862 | 0.5997053 |
| TENM4 | 3488.10339 | -0.6724167 | 0.39796882 | -1.6896215 | 0.09110039 | 0.50328109 |
| MIR4300HG | 47.0450843 | -0.7905896 | 0.64520479 | -1.2253313 | 0.22045044 | 0.67327858 |
| PRCP | 505.116459 | 0.72578656 | 0.22869623 | 3.17358346 | 0.0015057 | 0.07206065 |
| RAB30 | 289.802804 | -0.2364402 | 0.29614568 | -0.7983915 | 0.42464335 | 0.81466629 |
| RAB30-AS1 | 994.946058 | -0.0871603 | 0.26020312 | -0.3349701 | 0.73764764 | 0.93552616 |
| AP001767.3 | 97.9768755 | 0.20312826 | 0.64640893 | 0.31424112 | 0.75333791 | 0.93844712 |
| PCF11 | 708.801101 | 0.08398887 | 0.24716156 | 0.33981362 | 0.73399689 | 0.93552616 |
| PCF11-AS1 | 75.4902579 | 0.10490669 | 0.39736647 | 0.26400488 | 0.79177616 | 0.95183249 |
| ANKRD42 | 414.190441 | 0.18689965 | 0.26617742 | 0.70216191 | 0.4825782 | 0.84407748 |
| CCDC90B | 317.650056 | -0.1140018 | 0.21417167 | -0.5322917 | 0.59452398 | 0.88845653 |
| AP000446.1 | 369.631216 | 0.13895009 | 0.35092082 | 0.39595853 | 0.69213562 | 0.92224682 |
| DLG2 | 2499.69121 | -0.4130099 | 0.23841602 | -1.7323078 | 0.08321877 | 0.48768268 |
| TMEM126B | 205.671657 | -0.1302737 | 0.23052523 | -0.5651169 | 0.57199425 | 0.87907473 |
| TMEM126A | 87.8749809 | 0.41055418 | 0.35312222 | 1.16264047 | 0.2449754 | 0.6948422 |
| CREBZF | 835.963618 | 0.02384973 | 0.17840231 | 0.1336851 | 0.89365159 | 0.97788747 |
| SYTL2 | 1851.54727 | -0.3985648 | 0.47701719 | -0.8355355 | 0.40341627 | 0.80324472 |
| PICALM | 4336.15958 | -0.1872232 | 0.15215608 | -1.2304681 | 0.21852188 | 0.67202914 |
| EED | 479.695975 | -0.0277438 | 0.26001496 | -0.1067008 | 0.91502637 | 0.9816913 |
| HIKESHI | 463.452899 | 0.18845278 | 0.17602208 | 1.07062009 | 0.28434029 | 0.72738535 |
| CCDC81 | 51.5691152 | -0.272098 | 0.38123453 | -0.7137286 | 0.475395 | 0.84118672 |
| AP001831.1 | 379.726593 | 0.49333253 | 0.40989106 | 1.20356987 | 0.22875587 | 0.68065126 |
| ME3 | 1326.42618 | 0.29876479 | 0.3199819 | 0.93369275 | 0.3504624 | 0.77044053 |
| AP003059.1 | 148.672544 | 0.82583208 | 0.48393997 | 1.70647628 | 0.08791944 | 0.49714378 |
| AP003059.2 | 153.240621 | 0.98239612 | 0.58045968 | 1.69244505 | 0.09056115 | 0.50258861 |
| PRSS23 | 387.937387 | 0.8408543 | 0.40392566 | 2.0817056 | 0.03736937 | 0.36263296 |
| AP001528.2 | 319.258514 | 1.17350158 | 0.69708413 | 1.68344325 | 0.09228932 | 0.5058297 |
| FZD4 | 168.060379 | 0.07077573 | 0.34420108 | 0.2056232 | 0.83708526 | 0.96527001 |
| AP001528.1 | 172.890557 | -0.2500452 | 0.42636199 | -0.5864623 | 0.55756489 | 0.87261547 |
| TMEM135 | 1551.60048 | 0.34219785 | 0.20668045 | 1.65568566 | 0.09778549 | 0.51355946 |
| RAB38 | 146.695901 | 0.28899003 | 0.46344153 | 0.62357386 | 0.53290748 | 0.86397197 |
| CTSC | 506.024933 | 0.67726953 | 0.36357334 | 1.86281403 | 0.06248843 | 0.43419765 |
| GRM5 | 128.375008 | 0.82990386 | 0.44236691 | 1.8760532 | 0.06064798 | 0.43023583 |
| NOX4 | 906.68353 | -0.0402979 | 0.5196601 | -0.0775467 | 0.93818863 | 0.98696247 |
| CHORDC1 | 452.455619 | 0.29949496 | 0.41068329 | 0.72926016 | 0.46584254 | 0.8367113 |
| SLC36A4 | 1168.51693 | 0.00319767 | 0.25951312 | 0.0123218 | 0.99016887 | 0.997268 |
| DEUP1 | 72.146123 | 0.33957753 | 0.47675171 | 0.71227332 | 0.47629554 | 0.84141268 |

| | | | | | | |
|---|---|---|---|---|---|---|
| SMCO4 | 344.223752 | 0.42992244 | 0.35892546 | 1.19780426 | 0.23099323 | 0.68321398 |
| CEP295 | 459.318738 | -0.3943848 | 0.2515763 | -1.5676548 | 0.11696172 | 0.54756317 |
| TAF1D | 1491.44067 | 0.07504006 | 0.20885832 | 0.35928689 | 0.71938048 | 0.93001079 |
| AP001273.2 | 56.1970161 | -0.2066647 | 0.33472441 | -0.6174176 | 0.53695932 | 0.86466169 |
| PANX1 | 119.08214 | 0.18413213 | 0.28168655 | 0.65367738 | 0.51331968 | 0.85762435 |
| MRE11 | 552.775307 | -0.4675827 | 0.22679169 | -2.0617278 | 0.03923366 | 0.37176281 |
| ANKRD49 | 131.255441 | -0.3446411 | 0.35072647 | -0.9826493 | 0.32578008 | 0.75550719 |
| PIWIL4 | 101.663421 | 0.91059589 | 0.37842787 | 2.40626008 | 0.01611679 | 0.25504618 |
| AMOTL1 | 287.828386 | 1.89279633 | 0.50439017 | 3.75264318 | 0.00017498 | 0.02140101 |
| CWC15 | 249.649173 | -0.084805 | 0.17528946 | -0.4837999 | 0.62852787 | 0.89962754 |
| KDM4D | 46.9383625 | -0.8932285 | 0.49834657 | -1.7923842 | 0.07307144 | 0.47001711 |
| SRSF8 | 232.890483 | 0.36476198 | 0.33839129 | 1.07792957 | 0.28106519 | 0.72582688 |
| ENDOD1 | 478.975697 | 0.39311933 | 0.44342544 | 0.88655115 | 0.37532061 | 0.78468026 |
| AP000787.1 | 345.511826 | -0.0545025 | 0.41713859 | -0.130658 | 0.89604589 | 0.97792862 |
| SESN3 | 594.87022 | 0.04864625 | 0.36721838 | 0.13247227 | 0.89461076 | 0.97788747 |
| FAM76B | 290.069543 | -0.2258377 | 0.24390899 | -0.9259098 | 0.35449285 | 0.77243419 |
| CEP57 | 795.408186 | 0.28057482 | 0.15639944 | 1.79396302 | 0.07281907 | 0.46943247 |
| MTMR2 | 621.730323 | -0.0606815 | 0.18061118 | -0.3359784 | 0.73688716 | 0.93552616 |
| MAML2 | 14811.2758 | 0.01543922 | 0.24635156 | 0.06267148 | 0.95002811 | 0.98919519 |
| CCDC82 | 482.340639 | -0.3198913 | 0.21404487 | -1.4945056 | 0.13504353 | 0.57708028 |
| JRKL | 136.488336 | 0.49326699 | 0.28787392 | 1.71348273 | 0.08662377 | 0.49478465 |
| CNTN5 | 168.781632 | 0.47853103 | 0.54964575 | 0.87061716 | 0.38396322 | 0.791697 |
| ARHGAP42 | 109.402566 | 1.05526116 | 0.32959893 | 3.20165222 | 0.00136642 | 0.06759934 |
| TRPC6 | 135.949211 | 0.3585484 | 0.47875518 | 0.74891806 | 0.45390659 | 0.83060749 |
| CEP126 | 435.108674 | 0.24601464 | 0.29423421 | 0.8361184 | 0.4030883 | 0.80324472 |
| CFAP300 | 61.5311987 | 2.53662309 | 0.72790741 | 3.4848156 | 0.00049248 | 0.03826541 |
| YAP1 | 4108.63794 | -0.0527217 | 0.2207178 | -0.2388646 | 0.81121055 | 0.9575238 |
| BIRC3 | 554.191611 | 1.28875104 | 0.55524618 | 2.32104442 | 2.03E-02 | 2.85E-01 |
| BIRC2 | 543.499138 | 0.02530967 | 0.25397661 | 0.09965356 | 0.92061937 | 0.98215577 |
| TMEM123 | 1140.50365 | 0.26821564 | 0.27311085 | 0.9820761 | 0.32606236 | 0.75550719 |
| MMP7 | 621.028705 | 1.01036944 | 0.78194881 | 1.29211711 | 1.96E-01 | 6.50E-01 |
| DCUN1D5 | 346.396108 | 0.1593686 | 0.23336835 | 0.68290582 | 0.49466635 | 0.84882719 |
| DYNC2H1 | 2202.18679 | 0.18697654 | 0.20054216 | 0.93235525 | 0.35115296 | 0.77100859 |
| PDGFD | 462.190349 | -0.5965563 | 0.4062585 | -1.4684156 | 0.14199138 | 0.5872148 |
| CASP4 | 623.990657 | 0.49109376 | 0.36543165 | 1.34387307 | 1.79E-01 | 6.33E-01 |
| MSANTD4 | 166.017363 | 0.16731931 | 0.20864631 | 0.80192797 | 0.42259462 | 0.8134731 |
| KBTBD3 | 209.983763 | 0.5005896 | 0.25112706 | 1.99337178 | 0.04622075 | 0.3904347 |
| AASDHPPT | 322.022104 | 0.33032464 | 0.27965505 | 1.18118603 | 0.23752883 | 0.68973903 |
| GUCY1A2 | 56.6373169 | 0.15087417 | 0.44987923 | 0.33536594 | 0.73734906 | 0.93552616 |
| AP000766.1 | 74.6973884 | -0.4657969 | 0.38175649 | -1.2201413 | 0.2224113 | 0.67568617 |
| CWF19L2 | 925.320656 | -0.2965366 | 0.20730436 | -1.4304407 | 0.15259056 | 0.60267503 |
| ALKBH8 | 224.652908 | -0.226744 | 0.22504953 | -1.0075295 | 0.31368039 | 0.74777193 |
| SLC35F2 | 299.187422 | 0.81851977 | 0.35745585 | 2.28984855 | 0.0220301 | 0.29509055 |
| CUL5 | 1731.86264 | -0.2241843 | 0.11760032 | -1.9063236 | 0.05660823 | 0.41843207 |
| AP002433.1 | 42.2075869 | 0.32579383 | 0.62586133 | 0.52055273 | 0.60267839 | 0.88971776 |
| ACAT1 | 743.551512 | 0.33686594 | 0.28968761 | 1.16285935 | 0.24488657 | 0.6948422 |
| NPAT | 522.087439 | -0.142435 | 0.17431943 | -0.817092 | 0.41387585 | 0.80874327 |
| ATM | 1624.31426 | -0.3961475 | 0.22781747 | -1.7388812 | 0.08205565 | 0.48594425 |
| C11orf65 | 646.428474 | -0.0785017 | 0.40756641 | -0.1926108 | 0.84726382 | 0.96905304 |
| KDELC2 | 111.302816 | -0.6254256 | 0.34251294 | -1.825991 | 0.06785161 | 0.45242399 |
| EXPH5 | 852.31908 | -0.5909629 | 0.41900098 | -1.4104094 | 0.15841884 | 0.60804051 |
| DDX10 | 1041.78861 | -0.3670824 | 0.24869354 | -1.476043 | 0.13993235 | 0.58437128 |
| AP003049.2 | 38.1055624 | -0.047819 | 0.59940592 | -0.0797773 | 0.93641437 | 0.98621411 |
| RDX | 1492.0998 | 0.12110679 | 0.23370165 | 0.51821109 | 0.604311 | 0.89013087 |

| | | | | | | |
|---|---|---|---|---|---|---|
| ZC3H12C | 718.457125 | 0.3501259 | 0.31416782 | 1.11445498 | 0.26508405 | 0.71512755 |
| FDX1 | 463.888334 | 0.02655167 | 0.20877425 | 0.12717888 | 0.89879882 | 0.97889101 |
| ARHGAP20 | 42.8138592 | 0.69743529 | 0.41617666 | 1.67581545 | 0.09377435 | 0.50926605 |
| LINC02550 | 399.374672 | -3.6251517 | 1.69849169 | -2.1343359 | 3.28E-02 | 0.34593557 |
| COLCA1 | 192.074174 | -0.9886557 | 0.59563895 | -1.6598238 | 0.0969499 | 0.51327103 |
| COLCA2 | 100.534341 | -1.582809 | 0.50845697 | -3.1129654 | 0.00185218 | 0.08349969 |
| POU2AF1 | 60.040102 | -0.3221785 | 0.9723746 | -0.3313316 | 0.74039401 | 0.93570489 |
| SIK2 | 2996.68787 | 0.11977839 | 0.32437033 | 0.36926432 | 0.71193072 | 0.92748721 |
| PPP2R1B | 724.836653 | -0.3585612 | 0.32912128 | -1.08945 | 0.27595548 | 0.72241028 |
| C11orf1 | 84.3768411 | 0.24312506 | 0.37522006 | 0.64795325 | 0.51701519 | 0.85893097 |
| CRYAB | 2394.05325 | 1.23240649 | 0.60184812 | 2.04770347 | 4.06E-02 | 0.37344844 |
| DIXDC1 | 587.830249 | 0.31392019 | 0.33726234 | 0.93078933 | 0.35196255 | 0.77151834 |
| DLAT | 382.922579 | 0.02089431 | 0.24756675 | 0.08439871 | 0.93273943 | 0.9839805 |
| PIH1D2 | 86.6147809 | 0.1348407 | 0.32519863 | 0.41464105 | 0.67840469 | 0.91803319 |
| NKAPD1 | 156.718904 | -0.1049605 | 0.2673168 | -0.3926445 | 0.69458205 | 0.92229578 |
| TIMM8B | 191.769576 | 0.3830219 | 0.38570008 | 0.9930563 | 0.32068252 | 0.75144499 |
| AP002884.3 | 279.418864 | -1.2092208 | 0.49659887 | -2.4350052 | 0.01489158 | 0.24967336 |
| IL18 | 70.212472 | 0.03332247 | 0.60381285 | 0.05518676 | 0.95598968 | 0.99065996 |
| BCO2 | 106.391827 | -0.8574277 | 0.6425307 | -1.334454 | 0.18205511 | 0.63706109 |
| PTS | 690.502088 | -0.4479935 | 0.20893705 | -2.1441554 | 0.03202043 | 0.3410951 |
| AP002884.1 | 412.441584 | -0.285368 | 0.43370619 | -0.6579754 | 0.51055395 | 0.85762435 |
| AP003063.1 | 60.4890603 | 0.73839092 | 0.62151955 | 1.18804134 | 0.23481712 | 0.68621668 |
| TTC12 | 488.140773 | -0.017776 | 0.22944985 | -0.0774721 | 0.93824802 | 0.98696247 |
| TMPRSS5 | 41.0320475 | -1.2284445 | 0.6639388 | -1.8502376 | 0.06427931 | 0.43924752 |
| ZW10 | 264.330816 | 0.00403421 | 0.26561659 | 0.0151881 | 0.98788211 | 0.99653761 |
| USP28 | 418.290201 | -0.0291591 | 0.24185852 | -0.1205626 | 0.90403752 | 0.97988471 |
| ZBTB16 | 2710.17512 | 0.22615333 | 0.53870612 | 0.41980836 | 6.75E-01 | 0.91636338 |
| AP002518.2 | 607.196582 | 1.62444416 | 1.21733941 | 1.33442173 | 1.82E-01 | 0.63706109 |
| NNMT | 2624.16997 | 1.8545622 | 1.08794396 | 1.70464865 | 8.83E-02 | 4.97E-01 |
| C11orf71 | 69.8164698 | 0.40702338 | 0.36100336 | 1.12747808 | 0.2595404 | 0.70921184 |
| AP002373.1 | 53.7931008 | -1.0671789 | 0.61226376 | -1.7430052 | 0.08133271 | 0.48594425 |
| REXO2 | 542.979208 | 0.1824507 | 0.46573634 | 0.39174676 | 0.69524534 | 0.92229578 |
| CADM1 | 6608.9211 | -0.8262439 | 0.378715 | -2.1817037 | 0.02913141 | 0.32944929 |
| AP000462.1 | 299.609896 | -0.5970775 | 0.63099553 | -0.9462468 | 0.34402274 | 0.76685327 |
| BUD13 | 125.987436 | -0.4490911 | 0.25920786 | -1.7325519 | 0.08317535 | 0.48764454 |
| ZPR1 | 74.5116819 | -0.1195825 | 0.36889068 | -0.3241678 | 0.74581099 | 0.93675395 |
| APOA1-AS | 60.3841872 | 0.38045859 | 0.6611224 | 0.57547375 | 0.56497094 | 0.87588257 |
| SIK3 | 8061.50504 | 0.02483953 | 0.29225831 | 0.0849917 | 0.93226799 | 0.98365108 |
| PAFAH1B2 | 800.030937 | -0.1798121 | 0.23390475 | -0.7687409 | 0.44204716 | 0.82285053 |
| SIDT2 | 180.745764 | 0.19339145 | 0.2375956 | 0.81395213 | 0.41567238 | 0.80966301 |
| TAGLN | 86.774375 | 0.0987235 | 0.38486549 | 0.25651429 | 0.79755375 | 0.95281098 |
| PCSK7 | 688.683506 | -0.0841657 | 0.27450016 | -0.3066143 | 0.75913698 | 0.94133292 |
| RNF214 | 484.295934 | -0.3772101 | 0.20867448 | -1.8076484 | 0.07066124 | 0.4627897 |
| BACE1 | 340.844839 | 0.3174642 | 0.25982523 | 1.22183745 | 0.2217691 | 0.67512055 |
| BACE1-AS | 39.3907773 | -0.5476421 | 0.40079466 | -1.3663907 | 0.17181635 | 0.6286116 |
| CEP164 | 492.481006 | -0.0730779 | 0.18921976 | -0.3862065 | 0.69934375 | 0.92367966 |
| AP000892.1 | 71.8820333 | 0.79035678 | 0.51645119 | 1.53036103 | 0.12592739 | 0.56519704 |
| DSCAML1 | 3597.57653 | -1.086971 | 0.63809687 | -1.7034577 | 0.08848244 | 0.49769182 |
| FXYD2 | 971.078912 | 1.14759942 | 0.51254837 | 2.23900706 | 0.02515546 | 0.31053369 |
| AP000757.2 | 89.7311406 | 0.37581687 | 0.48005558 | 0.78286116 | 0.43370865 | 0.81899321 |
| FXYD6-FXYD2 | 241.048463 | 2.81248845 | 0.87962738 | 3.19736345 | 0.0013869 | 0.06809021 |
| AP000757.1 | 281.490951 | 1.43644762 | 0.54993924 | 2.6120115 | 0.00900112 | 0.19301271 |
| SMIM35 | 43.4213389 | -0.657229 | 0.43408178 | -1.5140671 | 0.13000882 | 0.57109626 |
| TMPRSS4 | 844.664922 | -0.4165131 | 0.84440887 | -0.49326 | 0.62182892 | 0.8982666 |

| | | | | | | |
|---|---|---|---|---|---|---|
| MPZL3 | 246.447141 | -0.1091401 | 0.26545388 | -0.4111454 | 0.6809659 | 0.91887257 |
| MPZL2 | 83.3531525 | 0.01622805 | 0.45383461 | 0.03575763 | 0.97147562 | 0.99327086 |
| UBE4A | 616.687824 | -0.1018122 | 0.15877303 | -0.6412439 | 0.52136421 | 0.85943874 |
| ATP5MG | 924.066057 | 0.53963797 | 0.4085962 | 1.32071217 | 0.18659736 | 0.6408069 |
| KMT2A | 2275.12157 | 0.0702249 | 0.08884367 | 0.79043223 | 4.29E-01 | 8.16E-01 |
| AP001267.3 | 139.508592 | 0.20688647 | 0.39671851 | 0.52149437 | 0.60202243 | 0.88964714 |
| TMEM25 | 99.4659444 | 0.12578403 | 0.26153516 | 0.480945 | 0.63055559 | 0.90005881 |
| IFT46 | 196.798167 | 0.05134985 | 0.24121994 | 0.21287563 | 0.83142397 | 0.96386512 |
| ARCN1 | 383.824821 | 0.12399501 | 0.20873397 | 0.5940337 | 0.55248957 | 0.87096727 |
| PHLDB1 | 648.605161 | 0.01246085 | 0.49253083 | 0.02529963 | 0.97981597 | 0.99487963 |
| AP002954.1 | 123.060868 | 0.38406817 | 0.36458458 | 1.05344052 | 0.29213914 | 0.73384286 |
| DDX6 | 1170.19369 | 0.05549727 | 0.13197825 | 0.42050312 | 0.67411795 | 0.91636338 |
| BCL9L | 179.2146 | -0.0251829 | 0.29439745 | -0.0855404 | 0.93183182 | 0.98357046 |
| CCDC84 | 892.432921 | -0.2915032 | 0.20708035 | -1.4076817 | 0.15922535 | 0.60886928 |
| AP003392.1 | 58.9645851 | 0.2611887 | 0.48809303 | 0.53512073 | 0.59256638 | 0.88740306 |
| RPS25 | 853.018954 | 0.30576879 | 0.44473445 | 0.68753116 | 0.49174807 | 0.84704258 |
| TRAPPC4 | 216.515451 | 0.84681003 | 0.28027132 | 3.02139375 | 0.00251614 | 0.09525007 |
| SLC37A4 | 106.083022 | 0.60142989 | 0.46312245 | 1.29864119 | 0.1940671 | 0.64749489 |
| HYOU1 | 131.949982 | 0.67715063 | 0.39105483 | 1.73160024 | 0.08334476 | 0.48777181 |
| VPS11 | 147.437247 | -0.1026009 | 0.24740999 | -0.4146999 | 0.67836161 | 0.91803319 |
| HMBS | 54.8349053 | -0.2152837 | 0.46369714 | -0.4642766 | 0.64244958 | 0.90460682 |
| H2AFX | 51.4332457 | 0.21577597 | 0.47022036 | 0.45888267 | 0.64631842 | 0.90582727 |
| DPAGT1 | 68.0585375 | 0.14707691 | 0.42317024 | 0.34755968 | 0.72817089 | 0.93453345 |
| C2CD2L | 74.5835585 | -0.1245292 | 0.35156408 | -0.3542147 | 0.72317796 | 0.93177886 |
| HINFP | 177.099743 | -0.2586673 | 0.24602079 | -1.0514043 | 0.29307292 | 0.73527014 |
| NLRX1 | 141.921524 | -0.1057037 | 0.40427033 | -0.2614678 | 0.79373177 | 0.95224951 |
| CBL | 611.010031 | -0.3512132 | 0.29843692 | -1.1768423 | 0.23925846 | 0.69029619 |
| USP2 | 405.934953 | 0.67457325 | 0.7577765 | 0.89020081 | 0.37335807 | 0.78277446 |
| USP2-AS1 | 393.315748 | 0.24531839 | 0.57439092 | 0.42709309 | 0.66931153 | 0.91435887 |
| THY1 | 155.835197 | 0.7186469 | 1.03015212 | 0.6976124 | 0.48541962 | 0.8447301 |
| NECTIN1 | 66.4886886 | -0.0383532 | 0.50751975 | -0.0755699 | 0.93976129 | 0.98715467 |
| OAF | 276.120605 | 0.14437658 | 0.45481156 | 0.31744264 | 0.75090775 | 0.93775604 |
| TMEM136 | 84.0948529 | 0.53294373 | 0.39680443 | 1.34308917 | 0.1792431 | 0.63288483 |
| ARHGEF12 | 4672.68222 | -0.1737879 | 0.22574808 | -0.7698313 | 0.44139998 | 0.82285053 |
| TECTA | 55.4066972 | -0.7414956 | 0.39290067 | -1.8872343 | 0.05912881 | 0.42419956 |
| SC5D | 148.917799 | -0.1323467 | 0.41888526 | -0.3159496 | 0.75204073 | 0.93815321 |
| SORL1 | 1617.07168 | -0.2531862 | 0.54514709 | -0.4644364 | 0.64233511 | 0.90454205 |
| UBASH3B | 363.974248 | -0.4087653 | 0.46790787 | -0.8736021 | 3.82E-01 | 7.90E-01 |
| JHY | 312.97139 | 0.7219148 | 0.38616564 | 1.86944341 | 0.06156115 | 0.43249565 |
| HSPA8 | 573.888994 | 1.45880436 | 0.42817714 | 3.4070113 | 0.00065678 | 0.04534035 |
| CLMP | 46.0427737 | -0.8197415 | 0.4083695 | -2.0073525 | 0.04471215 | 0.38677325 |
| GRAMD1B | 346.903558 | -1.7610811 | 0.73309288 | -2.4022619 | 0.01629404 | 0.25591911 |
| ZNF202 | 100.539491 | -0.3602723 | 0.29176766 | -1.2347917 | 0.21690803 | 0.66978787 |
| VWA5A | 306.275154 | 0.09335969 | 0.30017493 | 0.31101759 | 7.56E-01 | 0.93957997 |
| TBRG1 | 681.61638 | 0.01086305 | 0.21505831 | 0.05051213 | 0.95971429 | 0.991232 |
| SIAE | 498.996983 | 0.250406 | 0.33428831 | 0.74907195 | 0.45381384 | 0.83060749 |
| SPA17 | 92.0828779 | -0.1051107 | 0.29532841 | -0.3559113 | 0.72190697 | 0.93079284 |
| ESAM | 173.652484 | 0.06649512 | 0.53588862 | 0.12408384 | 0.90124889 | 0.97931857 |
| AP000866.2 | 69.2264894 | -0.1339204 | 0.49968708 | -0.2680085 | 0.78869276 | 0.95053309 |
| MSANTD2 | 1070.36417 | -0.0719776 | 0.24920036 | -0.2888341 | 7.73E-01 | 0.94530018 |
| AP000866.1 | 67.2867813 | 0.063875 | 0.35916018 | 0.17784544 | 8.59E-01 | 9.72E-01 |
| ROBO3 | 46.3163553 | -0.243174 | 0.46955362 | -0.5178834 | 0.60453964 | 0.89013087 |
| CCDC15 | 55.947144 | -1.4832016 | 0.53369954 | -2.7790947 | 0.00545106 | 0.149384 |
| TMEM218 | 105.711668 | -0.00316 | 0.38783731 | -0.0081478 | 0.99349909 | 0.99826027 |

| | | | | | | |
|---|---|---|---|---|---|---|
| PKNOX2 | 391.97594 | 0.17870795 | 0.39883422 | 0.44807576 | 0.65409852 | 0.90900153 |
| EI24 | 273.522638 | 0.35894569 | 0.3289798 | 1.09108732 | 0.27523446 | 0.72220342 |
| STT3A-AS1 | 80.5686712 | -0.0682173 | 0.41024683 | -0.1662837 | 0.86793372 | 0.97402505 |
| STT3A | 271.066529 | 0.02722464 | 0.2299536 | 0.11839189 | 0.90575715 | 0.97997915 |
| HYLS1 | 59.7318683 | -0.2525557 | 0.45786015 | -0.5516001 | 0.58122237 | 0.88290096 |
| PUS3 | 61.4516486 | 0.3560722 | 0.41470469 | 0.85861627 | 0.39055226 | 0.79701353 |
| DDX25 | 59.1485698 | -0.5494932 | 0.52034824 | -1.0560104 | 0.29096345 | 0.73313067 |
| CDON | 63.7350072 | -0.187137 | 0.48380142 | -0.3868053 | 0.69890034 | 0.92367966 |
| RPUSD4 | 138.9017 | -0.3647643 | 0.31104026 | -1.1727236 | 0.24090664 | 0.69178991 |
| FAM118B | 258.046134 | -0.2567732 | 0.22051353 | -1.164433 | 0.24424858 | 0.6948422 |
| SRPRA | 224.282932 | 0.23781853 | 0.20971689 | 1.13399794 | 0.25679541 | 0.70735159 |
| FOXRED1 | 256.74048 | -0.1916559 | 0.32191085 | -0.5953696 | 0.55159642 | 0.87039028 |
| TIRAP | 118.351698 | 0.18114987 | 0.28984087 | 0.62499769 | 0.53197257 | 0.86397197 |
| AP001318.2 | 80.8286052 | -0.4457212 | 0.43382862 | -1.0274132 | 0.30422595 | 0.74152784 |
| DCPS | 431.907285 | -0.3912522 | 0.41098872 | -0.9519779 | 0.34110818 | 0.76543778 |
| GSEC | 46.3102666 | -0.6319131 | 0.55184618 | -1.1450892 | 0.25217221 | 0.70276117 |
| ST3GAL4 | 703.205432 | -1.60331 | 0.488094 | -3.2848385 | 0.00102041 | 0.0571126 |
| ETS1 | 290.984382 | -0.6203361 | 0.58059996 | -1.0684398 | 2.85E-01 | 0.72844615 |
| KCNJ1 | 1964.89306 | 1.3725853 | 0.59957138 | 2.28927754 | 0.02206323 | 0.29509055 |
| KCNJ5 | 122.874369 | -1.9411826 | 0.66738231 | -2.9086516 | 0.00362991 | 0.11745127 |
| ARHGAP32 | 2076.71195 | -0.4140247 | 0.19888848 | -2.0816926 | 0.03737056 | 0.36263296 |
| BARX2 | 141.078935 | -2.9616195 | 0.97965227 | -3.0231334 | 0.00250172 | 0.09525007 |
| TMEM45B | 979.808789 | -0.1773872 | 0.31250952 | -0.5676216 | 0.57029192 | 0.87817509 |
| NFRKB | 342.400685 | -0.0498296 | 0.1933029 | -0.25778 | 0.79657673 | 0.95278473 |
| PRDM10 | 499.823641 | -0.638605 | 0.30121834 | -2.1200736 | 0.03399984 | 0.35153234 |
| APLP2 | 1578.14772 | 0.53605566 | 0.27141247 | 1.97505905 | 0.04826145 | 0.39627811 |
| ST14 | 615.602158 | 0.01894627 | 0.26604962 | 0.07121328 | 0.94322801 | 0.98772472 |
| ZBTB44 | 1730.22185 | 0.10655986 | 0.32086299 | 0.33210393 | 0.73981078 | 0.93552616 |
| AP002986.1 | 297.96575 | 0.59475857 | 0.54593036 | 1.08944037 | 0.27595974 | 0.72241028 |
| ADAMTS15 | 288.241269 | 0.46147081 | 0.44647207 | 1.03359391 | 3.01E-01 | 0.73820989 |
| AP003486.1 | 45.7649608 | 0.04728454 | 0.37797981 | 0.12509806 | 0.90044592 | 0.97914987 |
| SNX19 | 727.938642 | -0.8034213 | 0.51032491 | -1.574333 | 0.11541048 | 0.54427374 |
| NTM | 199.42978 | -1.472811 | 0.74334152 | -1.9813383 | 0.04755335 | 0.39447539 |
| OPCML | 353.224097 | -1.9938139 | 0.63547601 | -3.1375125 | 0.00170388 | 0.07760881 |
| IGSF9B | 54.3972112 | 0.43827093 | 0.78601002 | 0.55758949 | 0.57712473 | 0.88150331 |
| JAM3 | 168.081277 | 0.2282684 | 0.45613751 | 0.50043769 | 0.61676692 | 0.8959501 |
| NCAPD3 | 206.70807 | -0.4748746 | 0.36705283 | -1.29375 | 0.19575179 | 0.64964978 |
| VPS26B | 232.601418 | 0.04914857 | 0.21930074 | 0.22411493 | 0.82266788 | 0.96096746 |
| THYN1 | 66.86654 | 0.30514798 | 0.35406487 | 0.86184201 | 0.38877446 | 0.79634939 |
| ACAD8 | 143.571946 | 0.31065944 | 0.30979914 | 1.00277696 | 0.31596849 | 0.74785672 |
| TUBB8 | 67.0680637 | 0.02598525 | 0.93205718 | 0.02787946 | 0.97775829 | 0.99455895 |
| ZMYND11 | 1954.79902 | -0.0382158 | 0.11071322 | -0.345178 | 7.30E-01 | 0.93453345 |
| DIP2C | 4876.10868 | 0.02391359 | 0.25302152 | 0.09451209 | 0.92470238 | 0.9822216 |
| PRR26 | 101.460486 | 0.20083206 | 0.51328658 | 0.39126692 | 0.69559995 | 0.92229578 |
| LARP4B | 2805.62516 | -0.2231173 | 0.15600822 | -1.4301638 | 0.15267003 | 0.60267503 |
| GTPBP4 | 250.941664 | -0.353151 | 0.19956828 | -1.7695749 | 0.07679798 | 0.47651607 |
| IDI1 | 296.892943 | 0.3730049 | 0.51351688 | 0.7263732 | 0.46761002 | 0.83785414 |
| WDR37 | 1068.78131 | -0.1343624 | 0.20116293 | -0.6679282 | 0.50417941 | 0.85525037 |
| ADARB2 | 236.69507 | -0.688868 | 1.08193058 | -0.6367026 | 0.52431856 | 0.86006369 |
| PFKP | 1413.3648 | 0.36547089 | 0.62587938 | 0.58393182 | 0.55926618 | 0.87295536 |
| PITRM1 | 561.940723 | -0.571791 | 0.2320237 | -2.4643645 | 0.01372564 | 0.24270686 |
| PITRM1-AS1 | 88.3037198 | -0.2243988 | 0.5350776 | -0.4193761 | 0.67494128 | 0.91636338 |
| KLF6 | 1261.04754 | -0.4038485 | 0.61824773 | -0.6532147 | 0.51361784 | 0.85762435 |
| AKR1C1 | 100.735191 | -0.3290917 | 0.71138295 | -0.4626083 | 0.64364516 | 0.90478945 |

| | | | | | | |
|---|---|---|---|---|---|---|
| AKR1C3 | 65.9688177 | 1.1821903 | 0.5300613 | 2.23028978 | 0.02572821 | 0.31211455 |
| NET1 | 137.224216 | 0.09810408 | 0.36961598 | 0.26542163 | 0.79068468 | 0.95144272 |
| ASB13 | 197.198555 | -0.1305824 | 0.47078642 | -0.2773708 | 0.78149541 | 0.94852273 |
| FAM208B | 1639.94631 | -0.2095467 | 0.18114237 | -1.1568069 | 0.24735128 | 0.69761067 |
| AL365356.1 | 112.80676 | 0.3097685 | 0.52345862 | 0.59177266 | 0.55400283 | 0.8712425 |
| GDI2 | 956.718099 | -0.0057023 | 0.1744583 | -0.032686 | 0.97392498 | 0.99346986 |
| ANKRD16 | 158.788727 | -0.2656246 | 0.29849145 | -0.8898901 | 0.37352488 | 0.78290863 |
| FBH1 | 741.696469 | -0.0067356 | 0.21290411 | -0.031637 | 0.97476154 | 0.99402649 |
| RBM17 | 678.047537 | -0.5819116 | 0.1897343 | -3.0669819 | 0.00216232 | 0.08925651 |
| PFKFB3 | 1748.7283 | -2.0839126 | 0.47458221 | -4.3910467 | 1.13E-05 | 0.00480664 |
| AL137145.2 | 324.15738 | -2.8202405 | 0.81901122 | -3.4434699 | 0.0005743 | 0.04214409 |
| PRKCQ | 2931.81556 | -0.6581483 | 0.28610326 | -2.3003873 | 0.02142629 | 0.29288643 |
| SFMBT2 | 1119.76265 | 0.19628017 | 0.30472424 | 0.6441239 | 0.51949509 | 0.85924779 |
| KIN | 301.427239 | -0.2878384 | 0.25012548 | -1.1507758 | 0.24982447 | 0.70032501 |
| ATP5F1C | 1214.8335 | 0.13018972 | 0.25190302 | 0.51682475 | 0.6052785 | 0.89062401 |
| TAF3 | 1184.24189 | -0.1732378 | 0.22539674 | -0.7685907 | 0.4421363 | 0.82285053 |
| CELF2 | 218.710722 | 0.6003309 | 0.61728631 | 0.97253234 | 0.33078578 | 0.75922666 |
| USP6NL | 1064.26077 | -0.1894981 | 0.1870969 | -1.0128338 | 0.31113956 | 0.74548204 |
| ECHDC3 | 188.540408 | 1.32082143 | 0.24009363 | 5.50127652 | 3.77E-08 | 0.00016602 |
| PROSER2 | 696.382253 | -0.6190311 | 0.45171981 | -1.3703874 | 0.170566 | 0.62661733 |
| PROSER2-AS | 51.6741833 | -0.7799331 | 0.67562446 | -1.1543885 | 0.24834094 | 0.69838949 |
| UPF2 | 1119.29251 | -0.2197926 | 0.19350487 | -1.1358503 | 0.25601924 | 0.70709713 |
| DHTKD1 | 486.263307 | -0.1102678 | 0.30033554 | -0.3671486 | 0.71350817 | 0.92760608 |
| SEC61A2 | 259.397765 | -0.2492499 | 0.27418408 | -0.9090604 | 0.36331822 | 0.77944947 |
| NUDT5 | 544.224063 | -0.1172797 | 0.21244091 | -0.5520582 | 0.58090847 | 0.88283233 |
| CDC123 | 459.968069 | -0.1623506 | 0.1880489 | -0.8633427 | 0.38794906 | 0.79631996 |
| CAMK1D | 214.502372 | 0.3940751 | 0.64928086 | 0.60694089 | 0.54389014 | 0.86639505 |
| OPTN | 367.649821 | 0.57510928 | 0.43984172 | 1.30753691 | 0.19103043 | 0.64567512 |
| PHYH | 182.348144 | 0.39909373 | 0.323412 | 1.23401027 | 0.21719908 | 0.66978787 |
| SEPHS1 | 296.008151 | -0.261277 | 0.2803801 | -0.931867 | 0.35140528 | 0.77117666 |
| BEND7 | 884.541786 | -0.3356222 | 0.25474426 | -1.317487 | 0.18767545 | 0.64222928 |
| PRPF18 | 155.519841 | -0.2621502 | 0.32425393 | -0.8084721 | 0.41881889 | 0.81128886 |
| AL157392.5 | 317.164011 | -0.1809306 | 0.20664337 | -0.8755696 | 0.38126408 | 0.78886549 |
| FRMD4A | 2279.76238 | -0.0689917 | 0.40070477 | -0.1721759 | 0.86329928 | 0.97319807 |
| FAM107B | 1191.3333 | -0.167169 | 0.33422513 | -0.5001688 | 0.6169562 | 0.89599505 |
| CDNF | 62.6159841 | -0.5595219 | 0.52023248 | -1.0755229 | 0.28214069 | 0.72604837 |
| HSPA14.1 | 133.647709 | 0.38928789 | 0.30336919 | 1.28321497 | 0.19941677 | 0.65150038 |
| SUV39H2 | 63.3832967 | -0.1548309 | 0.47012284 | -0.3293415 | 0.7418976 | 0.93598004 |
| DCLRE1C | 353.205121 | -0.5843418 | 0.21049409 | -2.7760486 | 0.0055024 | 0.15016771 |
| RPP38 | 499.978483 | 0.42784621 | 0.29918842 | 1.43002263 | 0.15271053 | 0.60267503 |
| NMT2 | 291.718585 | 0.33603594 | 0.26059655 | 1.28948729 | 0.19722873 | 0.64964978 |
| FAM171A1 | 2668.62907 | 0.02042035 | 0.29399168 | 0.06945893 | 0.94462432 | 0.98813503 |
| MINDY3 | 793.679097 | -0.1823454 | 0.20323772 | -0.8972023 | 0.36961098 | 0.78095617 |
| PTER | 713.151092 | -0.3810093 | 0.15641349 | -2.4359107 | 0.01485435 | 0.24963249 |
| RSU1 | 1014.05859 | -0.5708798 | 0.30154382 | -1.8931902 | 0.05833257 | 0.42147872 |
| CUBN | 378.555184 | 1.03743945 | 0.54129256 | 1.91659657 | 0.0552892 | 0.41827898 |
| TRDMT1 | 206.865201 | -0.0655946 | 0.30342406 | -0.2161812 | 0.82884647 | 0.96288319 |
| VIM | 249.787939 | -0.0156048 | 0.43567535 | -0.0358175 | 9.71E-01 | 0.99327086 |
| ST8SIA6 | 164.198898 | -1.2275878 | 1.14290912 | -1.0740905 | 0.28278212 | 0.72604837 |
| HACD1 | 109.306554 | 0.40404197 | 0.39457469 | 1.02399362 | 0.30583829 | 0.74290739 |
| STAM-AS1 | 40.0597025 | 0.69261281 | 0.4654662 | 1.48799807 | 0.1367514 | 0.57858721 |
| STAM | 433.204983 | -0.2137671 | 0.22581423 | -0.9466504 | 0.34381694 | 0.76680914 |
| CACNB2 | 2833.09278 | 0.49411757 | 0.49213339 | 1.00403179 | 3.15E-01 | 0.74785672 |
| AL450384.2 | 370.06572 | -0.1233979 | 0.36434216 | -0.3386868 | 0.73484568 | 0.93552616 |

| | | | | | | |
|---|---|---|---|---|---|---|
| NSUN6 | 859.524792 | -0.0930469 | 0.24113082 | -0.3858773 | 0.69958751 | 0.92367966 |
| ARL5B | 360.659648 | -0.175678 | 0.42921111 | -0.4093044 | 0.68231627 | 0.91969963 |
| MALRD1 | 1313.27754 | 0.71659837 | 0.3500886 | 2.04690575 | 4.07E-02 | 0.37344844 |
| PLXDC2 | 3491.13115 | 0.30339565 | 0.18891089 | 1.60602524 | 0.10826836 | 0.53091537 |
| NEBL | 5453.88131 | -0.3758913 | 0.37005731 | -1.015765 | 0.30974129 | 0.74450725 |
| MLLT10 | 3223.86407 | 0.03774966 | 0.16529574 | 0.22837649 | 0.81935356 | 0.96006637 |
| DNAJC1 | 2213.15091 | 0.14128538 | 0.21156475 | 0.66781153 | 0.50425391 | 0.85525037 |
| BMI1 | 129.574371 | 0.21698133 | 0.36007182 | 0.60260569 | 0.54677103 | 0.86837785 |
| PIP4K2A | 1134.55506 | -0.230113 | 0.32495043 | -0.708148 | 0.47885334 | 0.84279465 |
| MSRB2 | 218.125296 | 0.18943951 | 0.24272891 | 0.78045714 | 0.43512184 | 0.81919857 |
| C10orf67 | 79.4163648 | 0.40676025 | 0.6122354 | 0.66438538 | 0.50644369 | 0.85626196 |
| OTUD1 | 90.0617972 | 0.45906874 | 0.2949468 | 1.55644587 | 0.1196021 | 0.55232605 |
| KIAA1217 | 8556.73959 | -0.0838233 | 0.40068216 | -0.2092014 | 0.83429099 | 0.96510927 |
| ARHGAP21 | 1487.9273 | -0.339374 | 0.17929061 | -1.8928709 | 0.05837504 | 0.42147872 |
| PRTFDC1 | 95.1241414 | -0.1160873 | 0.40260119 | -0.2883431 | 0.77308408 | 0.94534972 |
| ENKUR | 52.4872955 | 0.29823922 | 0.68908367 | 0.43280552 | 0.66515606 | 0.91245679 |
| THNSL1 | 73.3292575 | -0.267959 | 0.47329352 | -0.5661581 | 0.57128631 | 0.8787896 |
| PDSS1 | 215.7753 | -0.1903714 | 0.37551289 | -0.5069636 | 0.61218038 | 0.89440588 |
| ABI1 | 1827.72234 | -0.0275951 | 0.16179357 | -0.1705575 | 0.86457169 | 0.97361767 |
| ANKRD26 | 1572.43314 | -0.2328829 | 0.14870271 | -1.5660971 | 1.17E-01 | 0.54757066 |
| YME1L1 | 1013.65213 | -0.0335693 | 0.1623742 | -0.2067404 | 0.83621261 | 0.96511435 |
| MASTL | 225.954479 | -0.6682942 | 0.32319484 | -2.0677751 | 0.03866118 | 0.36916996 |
| ACBD5 | 570.547364 | -0.246293 | 0.14321335 | -1.7197632 | 0.0854755 | 0.49346408 |
| RAB18 | 888.802243 | -0.0929851 | 0.10503655 | -0.8852643 | 0.37601412 | 0.78490077 |
| MPP7 | 1666.40065 | -0.5765271 | 0.39408782 | -1.4629408 | 0.14348358 | 0.58849865 |
| AL358234.1 | 220.452121 | -1.4548236 | 0.85078202 | -1.709984 | 0.08726883 | 0.49566631 |
| WAC-AS1 | 422.482614 | -0.270227 | 0.31637887 | -0.8541246 | 0.39303595 | 0.79843808 |
| WAC | 3625.64164 | -0.1748125 | 0.184437 | -0.9478171 | 0.34322258 | 0.76641095 |
| BAMBI | 42.4665239 | 0.02140775 | 0.60502857 | 0.03538305 | 0.9717743 | 0.99327086 |
| SVIL | 3590.60075 | 0.21077134 | 0.38297087 | 0.55035867 | 0.58207339 | 0.88336948 |
| MTPAP | 265.739331 | -0.1457292 | 0.18461925 | -0.7893498 | 0.4299076 | 0.81671932 |
| MAP3K8 | 231.968102 | 0.59138492 | 0.41308212 | 1.43164007 | 0.15224686 | 0.60228473 |
| ZNF438 | 897.455491 | -0.3451486 | 0.22576031 | -1.5288278 | 0.12630714 | 0.56613199 |
| ZEB1-AS1 | 76.9498486 | -0.9499093 | 0.41335076 | -2.2980709 | 0.02155776 | 0.29326096 |
| ZEB1 | 1376.03321 | -1.9413317 | 0.53915287 | -3.6007072 | 3.17E-04 | 0.03094754 |
| ARHGAP12 | 1828.52228 | -0.3009689 | 0.16719894 | -1.8000644 | 0.07185047 | 0.46774933 |
| KIF5B | 966.950668 | 0.10674082 | 0.19852889 | 0.53765887 | 0.59081258 | 0.88672234 |
| EPC1 | 1950.92755 | -0.2191546 | 0.1574691 | -1.3917307 | 0.16400396 | 0.61531922 |
| AL391839.1 | 64.6444785 | 0.1866712 | 0.53851226 | 0.34664243 | 0.72885996 | 0.93453345 |
| CCDC7 | 1797.96673 | -0.4257137 | 0.35331175 | -1.2049236 | 0.22823281 | 0.67999141 |
| ITGB1 | 1570.07404 | -0.1479241 | 0.27602652 | -0.5359053 | 0.59202402 | 0.88740306 |
| NRP1 | 4698.62108 | -0.761463 | 0.52129954 | -1.4607015 | 1.44E-01 | 5.90E-01 |
| PARD3 | 9733.78973 | -0.1729591 | 0.1654328 | -1.0454945 | 0.29579449 | 0.73664205 |
| CUL2 | 745.749985 | -0.2366255 | 0.168145 | -1.4072707 | 0.15934714 | 0.60903253 |
| CREM | 1462.62915 | -0.1589843 | 0.3025305 | -0.525515 | 0.59922523 | 0.88883692 |
| CCNY | 3461.87766 | -0.2746029 | 0.24979317 | -1.099321 | 0.27162807 | 0.72040597 |
| AL117336.3 | 460.419689 | 0.10538389 | 0.28650369 | 0.36782733 | 0.71300199 | 0.92760608 |
| FZD8 | 91.5787439 | 1.14258574 | 0.39066673 | 2.92470702 | 0.00344781 | 0.1138728 |
| ZNF248 | 1121.34797 | -0.0198244 | 0.26269627 | -0.0754653 | 0.93984454 | 0.98715467 |
| ZNF25 | 212.435509 | 0.09963804 | 0.2420294 | 0.41167745 | 0.68057586 | 0.91867756 |
| ZNF33A | 2284.84073 | -0.219958 | 0.211491 | -1.0400346 | 0.29832382 | 0.73673761 |
| AL117339.5 | 155.35304 | 0.03078176 | 0.35023639 | 0.08788852 | 0.92996528 | 0.98331361 |
| ZNF33B | 1390.57892 | -0.1545006 | 0.16496017 | -0.9365931 | 0.34896789 | 0.76982416 |
| BMS1 | 518.950004 | -0.428981 | 0.17672701 | -2.4273655 | 0.01520893 | 0.25080486 |

| | | | | | | |
|---|---|---|---|---|---|---|
| CSGALNACT2 | 363.53747 | 0.14293377 | 0.20288433 | 0.70450864 | 0.48111607 | 0.8437862 |
| FXYD4 | 61.1868011 | -0.6852987 | 0.80557939 | -0.8506905 | 0.39494131 | 0.7989793 |
| HNRNPF | 668.515594 | -0.1197899 | 0.20640176 | -0.5803724 | 5.62E-01 | 8.74E-01 |
| ZNF487 | 186.500059 | -0.1321543 | 0.25828923 | -0.5116526 | 0.6088942 | 0.89273938 |
| ZNF239 | 72.4480763 | 0.10895671 | 0.33416252 | 0.32605904 | 0.74437969 | 0.93668127 |
| ZNF32 | 232.665197 | 0.04747073 | 0.26584895 | 0.1785628 | 0.85828101 | 0.97143431 |
| CXCL12 | 459.335747 | 0.36334058 | 0.67644776 | 0.53713029 | 5.91E-01 | 8.87E-01 |
| TMEM72-AS | 1116.67032 | 0.28775927 | 0.5860905 | 0.49098094 | 6.23E-01 | 0.89873262 |
| TMEM72 | 1609.94746 | 1.40059021 | 0.63518981 | 2.20499476 | 0.02745446 | 0.31979365 |
| AL353801.1 | 89.6307145 | -0.049391 | 0.68938721 | -0.0716448 | 0.94288456 | 0.98772472 |
| RASSF4 | 250.01186 | 0.24060385 | 0.29859707 | 0.80578102 | 4.20E-01 | 0.8125045 |
| DEPP1 | 585.36729 | 0.30156473 | 0.79332683 | 0.38012672 | 0.70385135 | 0.92513032 |
| ZNF22 | 158.723195 | 0.74949334 | 0.28257804 | 2.65234107 | 0.00799358 | 0.17977198 |
| 8-Mar | 912.703407 | 0.05636204 | 0.19592333 | 0.28767398 | 0.77359631 | 0.94558571 |
| ZFAND4 | 348.871827 | 0.07947786 | 0.21396692 | 0.3714493 | 0.71030291 | 0.92619854 |
| WASHC2C | 350.195847 | -0.2071997 | 0.24846874 | -0.8339067 | 0.40433358 | 0.80437725 |
| AGAP4 | 411.989678 | -0.1204827 | 0.25477445 | -0.4728996 | 0.63628484 | 0.90237131 |
| TIMM23 | 403.856178 | -0.227349 | 0.22315508 | -1.0187937 | 0.30830094 | 0.74375973 |
| NCOA4 | 414.777307 | 0.70708293 | 0.24498992 | 2.88617156 | 0.0038996 | 0.12135113 |
| AL356056.2 | 71.6099826 | -0.3011372 | 0.36824935 | -0.8177534 | 0.41349801 | 0.80843359 |
| PTPN20 | 398.611289 | 1.41318586 | 0.46646482 | 3.0295658 | 0.00244906 | 0.0945894 |
| AGAP9 | 397.48674 | 0.6206504 | 0.30339435 | 2.04568874 | 0.040787 | 0.37362849 |
| FRMPD2 | 56.9902419 | 1.27466975 | 0.81083361 | 1.57204848 | 0.11593931 | 0.54556096 |
| MAPK8 | 3560.83889 | -0.5013477 | 0.19465883 | -2.57552 | 0.01000895 | 0.20371062 |
| ARHGAP22 | 244.153953 | -1.4114332 | 0.52227096 | -2.7024922 | 6.88E-03 | 1.71E-01 |
| AL138760.1 | 88.0823999 | 0.4435801 | 0.70521466 | 0.6290001 | 0.52934899 | 0.86316564 |
| ERCC6 | 643.857567 | -0.017959 | 0.15542931 | -0.1155443 | 0.90801371 | 0.98047754 |
| OGDHL | 599.044636 | 2.17647801 | 0.5618873 | 3.87351348 | 0.00010728 | 0.01628768 |
| PARG | 1285.52624 | -0.1791012 | 0.17442301 | -1.026821 | 0.30450478 | 0.74155671 |
| TIMM23B | 1024.82973 | -0.4665314 | 0.34360955 | -1.3577371 | 0.17454711 | 0.62908398 |
| LINC00843 | 277.782481 | 9.97E-05 | 0.23493676 | 0.0004244 | 0.99966138 | 0.99987432 |
| AGAP6 | 428.831343 | -0.1630987 | 0.18098271 | -0.9011841 | 0.36749042 | 0.78095617 |
| WASHC2A | 345.224453 | -0.5208685 | 0.29986985 | -1.7369818 | 0.08239039 | 0.48594425 |
| SGMS1 | 5566.60956 | 0.00566471 | 0.35904334 | 0.01577724 | 0.98741211 | 0.99653761 |
| SGMS1-AS1 | 140.315428 | -0.8744279 | 0.63933796 | -1.3677083 | 0.17140339 | 0.62786116 |
| ASAH2B | 100.228673 | -0.7390845 | 0.39990361 | -1.8481566 | 0.06457969 | 0.44001919 |
| PRKG1 | 4735.33513 | 0.38556911 | 0.50165098 | 0.76860032 | 0.44213061 | 0.82285053 |
| CSTF2T | 70.5936861 | 0.8390159 | 0.37342641 | 2.24680389 | 0.02465256 | 0.30812502 |
| PCDH15 | 73.8971022 | -0.0239908 | 0.51902732 | -0.0462226 | 0.9631328 | 0.99224805 |
| IPMK | 541.953692 | -0.0607147 | 0.42077304 | -0.1442932 | 0.88526892 | 0.97664137 |
| CISD1 | 849.708389 | 0.22014902 | 0.25125458 | 0.87619904 | 0.38092184 | 0.78840435 |
| UBE2D1 | 343.282148 | -0.5064516 | 0.32529409 | -1.5569037 | 0.11949335 | 0.55224912 |
| TFAM | 195.35968 | 0.09011059 | 0.19686605 | 0.45772539 | 0.64714975 | 0.9065308 |
| BICC1 | 22779.7227 | 0.10739669 | 0.28176824 | 0.38115259 | 0.70309003 | 0.92501157 |
| PHYHIPL | 937.618177 | -0.9391841 | 0.52789192 | -1.7791219 | 0.07521978 | 0.47200856 |
| SLC16A9 | 101.471737 | 1.21233172 | 0.76312454 | 1.58864204 | 0.11214123 | 0.53638156 |
| CCDC6 | 977.875022 | -0.2049504 | 0.19322928 | -1.060659 | 0.2888449 | 0.73176973 |
| ANK3 | 13636.3987 | -0.3232193 | 0.24411722 | -1.3240333 | 0.18549199 | 0.64030106 |
| AL592430.1 | 95.7811503 | -0.1212288 | 0.75156575 | -0.1613017 | 0.87185581 | 0.97535872 |
| RHOBTB1 | 151.562017 | -0.3730552 | 0.26774148 | -1.393341 | 0.16351671 | 0.61416689 |
| ARID5B | 5109.54879 | 0.18397148 | 0.40125438 | 0.4584909 | 0.6465998 | 0.90600793 |
| ADO | 49.8715206 | 0.42650056 | 0.3949968 | 1.07975699 | 0.28025041 | 0.72513763 |
| NRBF2 | 165.387535 | 0.51839616 | 0.29943469 | 1.73124952 | 0.08340727 | 0.48792145 |
| JMJD1C | 7762.96213 | -0.1970713 | 0.13196584 | -1.493351 | 0.13534534 | 0.57708028 |

| | | | | | | |
|---|---|---|---|---|---|---|
| REEP3 | 1381.74876 | -0.321202 | 0.27462445 | -1.1696044 | 0.2421602 | 0.69337349 |
| AC022387.1 | 425.743644 | -0.9424232 | 0.68119163 | -1.3834921 | 0.16651404 | 0.61922407 |
| LINC01515 | 858.039334 | -0.4506116 | 0.37498214 | -1.2016881 | 0.22948439 | 0.6817947 |
| CTNNA3 | 262.470845 | 0.12887168 | 0.44805695 | 0.28762344 | 7.74E-01 | 0.94558571 |
| SIRT1 | 493.160141 | -0.2875895 | 0.22757158 | -1.263732 | 0.20632624 | 0.65831072 |
| HERC4 | 3389.4308 | -0.2154575 | 0.13383033 | -1.6099304 | 0.10741305 | 0.52901576 |
| PBLD | 682.283236 | 0.50456921 | 0.37336601 | 1.35140638 | 0.17656529 | 0.63008941 |
| HNRNPH3 | 1132.10667 | -0.0475288 | 0.19515767 | -0.2435407 | 0.80758654 | 0.95594682 |
| RUFY2 | 932.190621 | -0.1056411 | 0.16841874 | -0.6272529 | 0.53049346 | 0.86396107 |
| DNA2 | 173.538024 | -0.2406646 | 0.42396561 | -0.5676513 | 0.57027175 | 0.87817509 |
| SLC25A16 | 1294.27738 | 0.06843495 | 0.21170535 | 0.32325563 | 0.74650166 | 0.93695653 |
| TET1 | 292.960866 | -0.2375228 | 0.42121007 | -0.5639057 | 5.73E-01 | 0.87946834 |
| CCAR1 | 1839.14478 | -0.3419157 | 0.19669176 | -1.7383327 | 0.08215221 | 0.48594425 |
| STOX1 | 226.541542 | -0.0147182 | 0.43297309 | -0.0339934 | 0.97288242 | 0.99327086 |
| DDX50 | 654.740148 | -0.1087181 | 0.16327417 | -0.665862 | 0.50549929 | 0.85595485 |
| DDX21 | 439.658361 | -0.1593984 | 0.2821777 | -0.5648866 | 5.72E-01 | 0.87909049 |
| KIF1BP | 376.021564 | 0.21644037 | 0.20482227 | 1.05672285 | 0.29063809 | 0.73292069 |
| SRGN | 42.9029771 | -0.7021672 | 0.65671756 | -1.0692073 | 0.28497629 | 0.72795431 |
| VPS26A | 598.928755 | -0.2332191 | 0.15259483 | -1.5283554 | 0.12642431 | 0.56629581 |
| SUPV3L1 | 339.361116 | -0.3355489 | 0.19061826 | -1.760319 | 0.07835373 | 0.48060389 |
| HKDC1 | 974.512349 | 0.53053102 | 0.69905972 | 0.75892088 | 0.44789989 | 0.8260638 |
| HK1 | 1477.39119 | -0.1239915 | 0.22765802 | -0.5446392 | 0.58600169 | 0.88530153 |
| TSPAN15 | 302.687676 | -0.0143527 | 0.38392846 | -0.0373839 | 0.97017893 | 0.99309664 |
| H2AFY2 | 495.773423 | 0.08015925 | 0.21733101 | 0.36883484 | 0.71225084 | 0.92748721 |
| AIFM2 | 141.62442 | 0.38309845 | 0.29790452 | 1.28597728 | 0.19845099 | 0.65077933 |
| TYSND1 | 114.853814 | 0.5302051 | 0.40147259 | 1.3206508 | 0.18661782 | 0.6408069 |
| SAR1A | 420.288287 | 0.10337832 | 0.20070223 | 0.51508303 | 0.606495 | 0.8914201 |
| PPA1 | 576.180189 | 0.42372254 | 0.34205832 | 1.23874355 | 2.15E-01 | 0.66885336 |
| EIF4EBP2 | 415.215689 | 0.24662406 | 0.23172761 | 1.06428431 | 0.28719992 | 0.72972899 |
| SGPL1 | 777.581617 | -0.0248807 | 0.19091034 | -0.1303264 | 0.8963082 | 0.97805329 |
| PCBD1 | 259.642513 | 1.0210002 | 0.39346886 | 2.59486917 | 0.00946269 | 0.19737334 |
| UNC5B | 85.7232425 | 1.91702273 | 0.81685891 | 2.34682232 | 0.01893428 | 0.27727597 |
| SLC29A3 | 92.5184874 | -0.4568629 | 0.62482523 | -0.7311851 | 0.46466608 | 0.83609511 |
| PSAP | 1416.9469 | 0.37278973 | 0.25001603 | 1.49106331 | 0.13594487 | 0.57776571 |
| CHST3 | 162.657012 | 0.61499898 | 0.75705881 | 0.81235298 | 0.41658912 | 0.80966301 |
| ASCC1 | 769.480657 | -0.5234044 | 0.1854174 | -2.8228437 | 0.00475998 | 0.1366507 |
| ANAPC16 | 1892.56546 | 0.36892973 | 0.21631545 | 1.70551725 | 0.088098 | 0.49729428 |
| DDIT4 | 358.352098 | 0.02862998 | 0.78273694 | 0.03657675 | 0.97082248 | 0.99317943 |
| DNAJB12 | 250.713199 | -0.2576836 | 0.27526197 | -0.9361395 | 0.34920139 | 0.76982416 |
| MICU1 | 1614.71317 | -0.0959744 | 0.12025016 | -0.7981226 | 0.42479937 | 0.81466629 |
| MCU | 1521.45003 | -0.3445449 | 0.26073854 | -1.3214191 | 0.18636167 | 0.6408069 |
| P4HA1 | 2399.64733 | 0.36452898 | 0.61810028 | 0.58975703 | 0.55535356 | 0.87184917 |
| NUDT13 | 137.637421 | 0.13593552 | 0.37910967 | 0.35856515 | 0.71992042 | 0.93001079 |
| ECD | 416.663425 | -0.2780626 | 0.2092763 | -1.3286868 | 0.18395134 | 0.63843513 |
| FAM149B1 | 708.87219 | -0.0598636 | 0.17603283 | -0.3400706 | 0.73380339 | 0.93552616 |
| DNAJC9 | 517.673313 | -0.1548832 | 0.25313346 | -0.6118636 | 0.54062799 | 0.86580445 |
| DNAJC9-AS1 | 51.5217759 | 0.43962139 | 0.40938494 | 1.07385825 | 0.28288622 | 0.72604837 |
| MRPS16 | 260.423791 | 0.2446912 | 0.24565104 | 0.99609267 | 0.31920512 | 0.75094569 |
| CFAP70 | 1178.7507 | 0.61873172 | 0.42900627 | 1.442244 | 0.14923355 | 0.5959335 |
| ANXA7 | 1051.07092 | 0.02032 | 0.12491179 | 0.16267479 | 0.8707745 | 0.97473202 |
| MSS51 | 62.081433 | 0.13658763 | 0.34105784 | 0.40048231 | 0.68880131 | 0.92151057 |
| PPP3CB | 1306.53791 | -0.2287793 | 0.18091778 | -1.2645483 | 0.20603329 | 0.65816052 |
| USP54 | 1054.98398 | -0.1781748 | 0.13132631 | -1.3567334 | 0.17486593 | 0.62944785 |
| AC073389.2 | 86.7275374 | -0.2922737 | 0.45409554 | -0.6436391 | 0.51980946 | 0.85935232 |

| | | | | | | |
|---|---|---|---|---|---|---|
| AGAP5 | 198.813435 | -0.0339102 | 0.17924198 | -0.1891865 | 0.84994667 | 0.97041641 |
| AC022400.6 | 75.6615926 | 0.01141479 | 0.26058443 | 0.04380457 | 0.96506019 | 0.99248521 |
| SEC24C | 354.911484 | -0.1077561 | 0.17973225 | -0.5995366 | 0.54881509 | 0.86904066 |
| FUT11 | 229.711548 | 0.12939653 | 0.53739766 | 0.24078357 | 0.80972286 | 0.95658969 |
| CHCHD1 | 177.564827 | 0.63934703 | 0.41461355 | 1.5420312 | 0.12306601 | 0.55977237 |
| ZSWIM8 | 641.539441 | -0.2319381 | 0.21281154 | -1.0898756 | 0.27576795 | 0.72233844 |
| CAMK2G | 468.170046 | -0.2893138 | 0.36026973 | -0.8030478 | 0.42194711 | 0.81266383 |
| PLAU | 589.36597 | -1.263388 | 0.68027283 | -1.8571784 | 0.06328578 | 0.4374369 |
| VCL | 2242.74862 | -0.3657234 | 0.51210286 | -0.7141601 | 0.47512819 | 0.84106098 |
| AP3M1 | 269.036857 | -0.1421778 | 0.18105078 | -0.7852925 | 0.43228211 | 0.81773248 |
| ADK | 6583.12695 | -0.436121 | 0.17094398 | -2.5512512 | 0.01073369 | 0.21067065 |
| AC022540.1 | 67.8477493 | -0.1050571 | 0.91826975 | -0.1144077 | 0.90891461 | 0.98054769 |
| KAT6B | 2410.31587 | -0.3357568 | 0.25489136 | -1.3172547 | 0.18775326 | 0.64232914 |
| SAMD8 | 520.393433 | -0.0190071 | 0.19391182 | -0.0980195 | 0.92191683 | 0.98215577 |
| VDAC2 | 515.224949 | 0.52537109 | 0.23295618 | 2.25523568 | 0.02411854 | 0.30486295 |
| COMTD1 | 204.059158 | 0.74600911 | 0.45943034 | 1.6237698 | 0.10442495 | 0.52442862 |
| AC010997.3 | 52.9024201 | -0.7481153 | 0.54932029 | -1.3618927 | 0.17323175 | 0.6286116 |
| LRMDA | 3084.00398 | -0.2289139 | 0.43157111 | -0.5304198 | 0.59582092 | 0.88859628 |
| KCNMA1 | 535.11553 | -0.1319315 | 0.84625564 | -0.1559003 | 0.87611164 | 0.97614152 |
| DLG5 | 926.230526 | -0.5580467 | 0.2640451 | -2.1134522 | 0.03456208 | 0.35307852 |
| POLR3A | 450.772674 | -0.3817569 | 0.1842081 | -2.0724219 | 0.03822611 | 0.36662698 |
| RPS24 | 1547.14354 | 0.37352535 | 0.46453969 | 0.80407629 | 0.42135291 | 0.81266383 |
| ZMIZ1-AS1 | 102.19648 | 0.32540222 | 0.59971328 | 0.54259632 | 0.58740778 | 0.88571413 |
| ZMIZ1 | 2312.38781 | 0.21900905 | 0.48888913 | 0.44797284 | 0.6541728 | 0.90900153 |
| PPIF | 186.657945 | 0.44170733 | 0.29308211 | 1.50711122 | 0.13178215 | 0.57539433 |
| ZCCHC24 | 50.2733412 | 0.588032 | 0.28835052 | 2.03929575 | 0.04142052 | 0.37577267 |
| NUTM2B-AS | 2009.42445 | -0.0932909 | 0.32110475 | -0.2905311 | 0.77140998 | 0.9446143 |
| AL135925.1 | 64.1340202 | -0.0209997 | 0.37880802 | -0.0554362 | 0.95579094 | 0.99065996 |
| SFTPD | 49.1420126 | -0.5067509 | 0.43971427 | -1.1524549 | 0.2491342 | 0.69928042 |
| TMEM254-A: | 97.411712 | -1.6992685 | 0.55810129 | -3.0447314 | 0.00232888 | 0.09242336 |
| TMEM254 | 180.631419 | 0.14362514 | 0.28201186 | 0.50928759 | 0.61055065 | 0.89339478 |
| ANXA11 | 2055.82321 | 0.16238084 | 0.34420355 | 0.47175818 | 0.63709941 | 0.90255504 |
| FAM213A | 692.075378 | 0.27284442 | 0.13370417 | 2.04065754 | 0.04128488 | 0.37577267 |
| TSPAN14 | 1106.88765 | -0.0993429 | 0.21835025 | -0.4549703 | 6.49E-01 | 0.90788153 |
| NRG3 | 1998.65598 | 1.16640084 | 0.54085894 | 2.15657125 | 0.03103909 | 0.33800731 |
| GHITM | 488.705588 | 0.51721993 | 0.31748826 | 1.62909939 | 0.10329198 | 0.52337282 |
| CCSER2 | 1867.32903 | -0.1231703 | 0.22156853 | -0.5559016 | 0.5782781 | 0.88205258 |
| WAPL | 1481.44973 | -0.1391529 | 0.17915259 | -0.7767281 | 0.4373192 | 0.82013459 |
| AL731569.1 | 40.11894 | -0.1106141 | 0.45243655 | -0.2444854 | 0.80685492 | 0.95566275 |
| BMPR1A | 2624.1828 | -0.1956154 | 0.15740922 | -1.2427187 | 0.21397152 | 0.66816781 |
| MMRN2 | 78.3668401 | -0.1514294 | 0.71089253 | -0.2130131 | 0.83131674 | 0.96386512 |
| ADIRF | 302.451028 | 0.71502459 | 0.79208233 | 0.90271499 | 0.36667718 | 0.78095617 |
| GLUD1 | 1386.89074 | -0.1031229 | 0.24681531 | -0.4178142 | 0.67608298 | 0.91650575 |
| SHLD2 | 2487.05892 | -0.1391012 | 0.22849755 | -0.6087647 | 0.54268043 | 0.86590899 |
| NUTM2A-AS | 2237.1203 | -0.4265062 | 0.20664899 | -2.0639161 | 0.03902567 | 0.37058958 |
| LINC00864 | 100.005021 | -2.0540207 | 0.98310044 | -2.0893294 | 0.03667808 | 0.3599411 |
| MINPP1 | 196.819305 | 0.23192824 | 0.30497577 | 0.76048088 | 0.44696719 | 0.82571251 |
| PAPSS2 | 747.17648 | 0.75952004 | 0.59861307 | 1.26879962 | 0.20451254 | 0.65759643 |
| ATAD1 | 1022.58795 | -0.0027009 | 0.25935875 | -0.0104139 | 0.99169106 | 0.99772135 |
| PTEN | 2744.49588 | -0.0072563 | 0.21315749 | -0.0340421 | 0.97284357 | 0.99327086 |
| RNLS | 941.486034 | -0.0248249 | 0.28237703 | -0.087914 | 0.92994503 | 0.98331361 |
| STAMBPL1 | 67.1437103 | 1.56019826 | 0.71342312 | 2.18691857 | 0.02874847 | 0.32721051 |
| ACTA2 | 79.5614586 | 1.464588 | 0.38662145 | 3.78817054 | 0.00015176 | 0.02008985 |
| FAS | 142.867214 | 0.90969666 | 0.50547634 | 1.79968198 | 7.19E-02 | 0.46774933 |

| | | | | | | |
|---|---|---|---|---|---|---|
| LIPA | 1190.95702 | 0.68277858 | 0.46777644 | 1.45962584 | 0.14439293 | 0.5901257 |
| IFIT2 | 252.688547 | 0.64927978 | 0.98163797 | 0.66142489 | 0.50833987 | 0.85646191 |
| IFIT3 | 559.549116 | 0.48730211 | 0.85968025 | 0.56684111 | 0.57082214 | 0.87858187 |
| IFIT1 | 555.915683 | 1.79351772 | 1.24056539 | 1.44572607 | 0.14825405 | 0.59438683 |
| IFIT5 | 113.174608 | 0.78798551 | 0.47608746 | 1.65512764 | 0.09789861 | 0.51355946 |
| SLC16A12 | 4018.03959 | 0.82194352 | 0.53962352 | 1.52317957 | 0.12771379 | 0.56762585 |
| SLC16A12-AS | 167.148054 | 0.65715449 | 0.69070824 | 0.95142123 | 0.34139059 | 0.76543778 |
| PANK1 | 648.580965 | 0.65559783 | 0.36319053 | 1.80510719 | 7.11E-02 | 0.46413489 |
| AL157400.3 | 80.1528392 | 0.39922961 | 0.44108975 | 0.90509835 | 0.36541329 | 0.78095617 |
| KIF20B | 134.847238 | -0.294098 | 0.370456 | -0.793881 | 0.42726469 | 0.81466629 |
| LINC00865 | 40.1851422 | 1.1018278 | 0.74122665 | 1.48649242 | 0.13714892 | 0.5794443 |
| RPP30 | 208.351688 | 0.02457367 | 0.25259319 | 0.09728555 | 0.92249963 | 0.98215577 |
| PCGF5 | 2203.93777 | -0.1965424 | 0.26111413 | -0.752707 | 0.45162598 | 0.8285455 |
| HECTD2 | 952.646233 | -0.3657407 | 0.21691561 | -1.6860967 | 0.09177718 | 0.50476839 |
| PPP1R3C | 58.7055616 | -0.0110248 | 1.13694494 | -0.0096968 | 0.99226315 | 0.99785337 |
| TNKS2 | 1556.6984 | -0.265197 | 0.16275449 | -1.6294298 | 0.10322206 | 0.52337282 |
| BTAF1 | 1387.95628 | -0.1401265 | 0.29558724 | -0.4740613 | 0.63545618 | 0.90199374 |
| CPEB3 | 4932.68465 | 0.40347586 | 0.40623418 | 0.99321003 | 0.32060761 | 0.75144499 |
| 5-Mar | 822.988722 | -0.1551433 | 0.25303935 | -0.6131193 | 0.53979746 | 0.86566791 |
| IDE | 670.600879 | 0.11416582 | 0.19625298 | 0.58172783 | 0.56075002 | 0.87376985 |
| EXOC6 | 947.974912 | -0.4082928 | 0.18723287 | -2.1806686 | 0.02920794 | 0.33001392 |
| MYOF | 2844.15441 | -0.1983789 | 0.36504668 | -0.5434343 | 5.87E-01 | 8.85E-01 |
| FRA10AC1 | 604.492402 | 0.03803613 | 0.21203154 | 0.17938904 | 0.85763224 | 0.97127317 |
| SLC35G1 | 73.3818737 | 1.07674365 | 0.47568288 | 2.26357454 | 0.0236003 | 0.30185158 |
| PLCE1 | 74.328958 | 1.3572925 | 0.33935956 | 3.99957056 | 6.35E-05 | 0.01289555 |
| NOC3L | 201.812668 | -0.1647076 | 0.25561376 | -0.6443611 | 0.51934132 | 0.85924115 |
| TBC1D12 | 494.403968 | -0.2933434 | 0.20950982 | -1.4001413 | 0.161471 | 0.61304492 |
| HELLS | 168.172085 | -0.4000786 | 0.32311165 | -1.2382055 | 0.21563988 | 0.66885336 |
| AL157834.1 | 57.2261499 | -0.0008735 | 0.59027118 | -0.0014798 | 0.99881933 | 0.99980325 |
| PDLIM1 | 89.2779538 | 2.25768413 | 0.67684607 | 3.33559464 | 0.00085117 | 0.05087388 |
| SORBS1 | 2664.34268 | -0.6041718 | 0.32892269 | -1.8368201 | 0.06623646 | 0.44661426 |
| ALDH18A1 | 590.360159 | -0.2029324 | 0.30205936 | -0.6718294 | 0.50169228 | 0.85452654 |
| TCTN3 | 480.5932 | 0.00307482 | 0.20898043 | 0.01471343 | 0.9882608 | 0.99653761 |
| ENTPD1 | 65.7263564 | 0.13754012 | 0.67037125 | 0.20517008 | 0.83743924 | 0.96527001 |
| ENTPD1-AS1 | 1493.75718 | 0.022966 | 0.36398052 | 0.06309678 | 0.94968944 | 0.98919519 |
| AL365273.2 | 97.6045204 | 0.05729623 | 0.51941948 | 0.1103082 | 0.91216495 | 0.98125148 |
| CC2D2B | 113.640606 | 0.45202285 | 0.58272836 | 0.7757008 | 0.43792567 | 0.82074642 |
| CCNJ | 79.3631248 | -0.4426248 | 0.28601107 | -1.5475793 | 0.12172362 | 0.55692668 |
| ZNF518A | 2123.60612 | 0.00741652 | 0.21812067 | 0.03400192 | 0.97287562 | 0.99327086 |
| BLNK | 2614.89664 | 0.20195187 | 0.42023168 | 0.4805727 | 0.63082022 | 0.90005881 |
| TM9SF3 | 1763.18226 | 0.07688784 | 0.15108014 | 0.50892092 | 0.61080766 | 0.89339478 |
| PIK3AP1 | 84.4074429 | 0.55875004 | 0.49422093 | 1.13056734 | 0.25823724 | 0.70896455 |
| LCOR | 3410.92659 | -0.2601313 | 0.22555266 | -1.1533061 | 0.24878478 | 0.6991991 |
| FRAT2 | 52.1114302 | 0.80360807 | 0.42443233 | 1.89337147 | 0.05830849 | 0.42147872 |
| RRP12 | 210.540833 | -0.3987393 | 0.2423211 | -1.6454998 | 0.09986678 | 0.51665771 |
| PGAM1 | 634.879891 | 0.6434076 | 0.3181384 | 2.02241413 | 0.04313359 | 0.38262319 |
| EXOSC1 | 307.908374 | -0.1793728 | 0.19088439 | -0.9396934 | 0.34737485 | 0.76897511 |
| ZDHHC16 | 230.978612 | -0.1108074 | 0.25831546 | -0.4289614 | 0.66795129 | 0.91383898 |
| MMS19 | 762.057035 | -0.3989373 | 0.15975151 | -2.4972366 | 0.01251654 | 0.22899031 |
| UBTD1 | 683.335955 | -0.5907394 | 0.33794885 | -1.7480143 | 0.08046155 | 0.4844196 |
| ANKRD2 | 105.406308 | 0.73850608 | 1.06959742 | 0.69045238 | 0.48990975 | 0.84614483 |
| AL355315.1 | 174.413996 | 0.82510534 | 0.41608017 | 1.98304415 | 0.0473625 | 0.39411174 |
| MORN4 | 328.569412 | 0.1593108 | 0.28830014 | 0.55258663 | 0.5805465 | 0.88264718 |
| PI4K2A | 75.5309858 | 0.50565 | 0.33646889 | 1.5028135 | 0.13288714 | 0.57682553 |

| | | | | | | |
|---|---|---|---|---|---|---|
| AVPI1 | 46.9949757 | -0.1262076 | 0.43260325 | -0.2917398 | 0.77048559 | 0.94409501 |
| ZFYVE27 | 313.202271 | -0.1411634 | 0.2171288 | -0.6501367 | 0.51560395 | 0.85869987 |
| R3HCC1L | 1189.76321 | -0.2507496 | 0.1710556 | -1.4658952 | 0.14267683 | 0.58824095 |
| PYROXD2 | 66.0746084 | -0.34552 | 0.42918646 | -0.805058 | 0.42078619 | 0.81260593 |
| HPS1 | 236.824451 | 0.1244139 | 0.25380529 | 0.49019427 | 0.62399643 | 0.89873262 |
| HPSE2 | 44.6657914 | 0.17426723 | 0.44277479 | 0.39357984 | 0.69389127 | 0.92229578 |
| GOT1 | 1689.44266 | 0.33958999 | 0.33641298 | 1.0094438 | 0.31276184 | 0.74693024 |
| AL391684.1 | 112.808142 | 0.06300688 | 0.49761705 | 0.12661721 | 8.99E-01 | 0.97889101 |
| SLC25A28 | 310.593511 | 0.44537429 | 0.30283986 | 1.47065941 | 0.14138325 | 0.5872148 |
| ENTPD7 | 140.043673 | -0.1201659 | 0.45955677 | -0.2614821 | 0.79372074 | 0.95224951 |
| CUTC | 365.809662 | -0.1235095 | 0.24292581 | -0.5084246 | 0.6111556 | 0.8935969 |
| COX15 | 312.89931 | -0.1889383 | 0.26888845 | -0.7026643 | 0.48226498 | 0.84389635 |
| ABCC2 | 117.388609 | -0.9578487 | 0.55161168 | -1.7364547 | 0.08248347 | 0.48594425 |
| DNMBP | 924.395416 | -0.6204709 | 0.24414152 | -2.5414395 | 0.0110397 | 0.21418301 |
| ERLIN1 | 300.017214 | 0.16664256 | 0.16999808 | 0.98026144 | 0.32695708 | 0.75556858 |
| CHUK | 325.844162 | -0.3616478 | 0.33947554 | -1.0653133 | 0.28673416 | 0.72949981 |
| CWF19L1 | 355.792898 | -0.2820639 | 0.24431471 | -1.1545106 | 0.24829091 | 0.69838949 |
| BLOC1S2 | 156.761885 | -0.0170411 | 0.2399499 | -0.0710196 | 0.94338217 | 0.98772472 |
| SCD | 62.4467376 | -0.4743461 | 0.62798702 | -0.7553438 | 4.50E-01 | 8.27E-01 |
| OLMALINC | 124.736963 | 0.63387896 | 0.62130497 | 1.02023803 | 0.30761558 | 0.74348936 |
| SEC31B | 173.956297 | 0.6809995 | 0.26365193 | 2.58294905 | 0.00979598 | 0.20092404 |
| HIF1AN | 306.547305 | -0.1356907 | 0.15676936 | -0.8655433 | 0.38674067 | 0.79523542 |
| PAX2 | 3186.64625 | -0.5089254 | 0.3359398 | -1.5149302 | 0.12979008 | 0.57050365 |
| SLF2 | 754.215908 | -0.2940863 | 0.252477 | -1.1648044 | 0.24409819 | 0.6948422 |
| MRPL43 | 184.554581 | 0.58599504 | 0.32728306 | 1.79048386 | 0.07337616 | 0.4701377 |
| SEMA4G | 68.0706012 | 0.52188262 | 0.44251115 | 1.17936603 | 0.23825245 | 0.68982886 |
| LZTS2 | 255.253651 | -0.3710652 | 0.22242043 | -1.6683056 | 0.09525508 | 0.51085035 |
| SFXN3 | 58.5149263 | -0.0034733 | 0.50212444 | -0.0069172 | 0.99448094 | 0.998438 |
| BTRC | 1817.11797 | -0.0373374 | 0.16732177 | -0.2231472 | 0.82342092 | 0.96133701 |
| DPCD | 176.515647 | 0.41468452 | 0.32136246 | 1.29039502 | 0.19691354 | 0.64964978 |
| POLL | 234.513166 | 0.27172558 | 0.29816215 | 0.91133492 | 0.36211893 | 0.77801382 |
| FBXW4 | 1337.01905 | -0.0257086 | 0.17189251 | -0.1495622 | 0.88111 | 0.97664137 |
| OGA | 2646.89401 | -0.1694673 | 0.2195609 | -0.7718463 | 0.44020549 | 0.82232701 |
| KCNIP2-AS1 | 41.3157338 | 0.40109831 | 0.72021076 | 0.55691797 | 0.57758348 | 0.88181117 |
| KCNIP2 | 42.2316951 | -0.0115986 | 0.44164794 | -0.026262 | 0.97904838 | 0.99455895 |
| ARMH3 | 1653.33748 | -0.2014004 | 0.16877198 | -1.1933285 | 0.23274075 | 0.68474693 |
| LDB1 | 239.060448 | 0.40123622 | 0.34403272 | 1.16627342 | 0.24350392 | 0.6948422 |
| PPRC1 | 163.482819 | -0.4989314 | 0.32415813 | -1.5391606 | 0.12376509 | 0.56102027 |
| NOLC1 | 146.176066 | 0.02329413 | 0.23298245 | 0.09998235 | 0.92035834 | 0.98215577 |
| GBF1 | 2446.76624 | -0.2750512 | 0.19515117 | -1.4094263 | 0.15870915 | 0.60804051 |
| NFKB2 | 166.235328 | 0.02420851 | 0.35929211 | 0.06737836 | 9.46E-01 | 9.89E-01 |
| FBXL15 | 178.912356 | 0.6674096 | 0.37104598 | 1.79872481 | 0.07206222 | 0.46774933 |
| CUEDC2 | 179.42548 | 0.20252125 | 0.24205163 | 0.83668614 | 0.40276901 | 0.80324472 |
| MFSD13A | 39.7647284 | -0.2786335 | 0.35886171 | -0.7764371 | 0.43749094 | 0.8201558 |
| ACTR1A | 236.549659 | -0.1194283 | 0.28103687 | -0.4249562 | 0.67086864 | 0.91516099 |
| SUFU | 746.692818 | 0.17835717 | 0.28220932 | 0.6320031 | 0.52738485 | 0.86151701 |
| TRIM8 | 491.685114 | 0.48006784 | 0.34360649 | 1.39714428 | 0.16237019 | 0.61373097 |
| ARL3 | 494.28708 | 0.11687937 | 0.23125121 | 0.50542166 | 0.61326271 | 0.89441126 |
| SFXN2 | 269.683906 | -0.2872761 | 0.30413978 | -0.9445528 | 0.34488726 | 0.76724655 |
| WBP1L | 893.292103 | 0.05394358 | 0.27982627 | 0.19277524 | 0.847135 | 0.96905304 |
| BORCS7-ASN | 53.4895181 | 0.17933925 | 0.47424342 | 0.37815864 | 0.70531275 | 0.92553905 |
| CNNM2 | 1705.72486 | 0.30804688 | 0.17943423 | 1.71676759 | 8.60E-02 | 4.94E-01 |
| NT5C2 | 2841.3357 | -0.137852 | 0.22882654 | -0.6024302 | 0.5468878 | 0.86845888 |
| PCGF6 | 299.356013 | -0.3198327 | 0.27411896 | -1.166766 | 0.24330489 | 0.69452742 |

| | | | | | | |
|---|---|---|---|---|---|---|
| TAF5 | 63.1982737 | -0.1911226 | 0.50878532 | -0.3756449 | 0.70718093 | 0.92573766 |
| ATP5MD | 385.759513 | 0.01818516 | 0.34193044 | 0.05318381 | 0.95758546 | 0.99065996 |
| PDCD11 | 225.040167 | -0.3901964 | 0.24973344 | -1.5624516 | 0.11818164 | 0.54908945 |
| SH3PXD2A | 356.764334 | -0.4321825 | 0.25219904 | -1.7136565 | 0.08659183 | 0.49478465 |
| STN1 | 324.757637 | -1.0131616 | 0.24862539 | -4.0750529 | 4.60E-05 | 0.01085117 |
| SLK | 1004.08727 | -0.2908475 | 0.19445003 | -1.4957444 | 0.13472028 | 0.57708028 |
| CFAP43 | 452.004092 | 0.09323872 | 0.34724353 | 0.26851103 | 0.78830599 | 0.95035277 |
| GSTO1 | 371.457264 | 0.30852943 | 0.24660135 | 1.2511263 | 0.2108884 | 0.66428174 |
| GSTO2 | 977.743855 | 0.27537732 | 0.28931197 | 0.95183521 | 0.34118056 | 0.76543778 |
| ITPRIP | 67.4717078 | 0.08167777 | 0.53803607 | 0.15180723 | 0.87933898 | 0.9765717 |
| SORCS3 | 804.908425 | 2.03414369 | 0.73778023 | 2.75711331 | 0.00583141 | 0.15669584 |
| SORCS1 | 61.1943648 | 0.04858583 | 0.5460841 | 0.08897133 | 0.9291047 | 0.98329813 |
| LINC01435 | 137.067182 | 1.00431329 | 0.7779877 | 1.29091153 | 0.19673436 | 0.64964978 |
| XPNPEP1 | 469.912225 | -0.2335057 | 0.27585165 | -0.8464902 | 0.39727932 | 0.80069929 |
| ADD3 | 2122.2009 | -0.0679701 | 0.18095182 | -0.3756252 | 0.70719558 | 0.92573766 |
| MXI1 | 1081.98807 | 0.07136456 | 0.3296503 | 0.21648567 | 0.8286092 | 0.96288319 |
| SMNDC1 | 240.750738 | 0.01661562 | 0.24824375 | 0.0669327 | 0.94663528 | 0.98885769 |
| DUSP5 | 176.350617 | 0.70586745 | 0.78759547 | 0.89623096 | 0.37012942 | 0.78099673 |
| SMC3 | 550.226228 | -0.0430718 | 0.21917436 | -0.1965185 | 0.84420434 | 0.96848143 |
| RBM20 | 553.274585 | 1.11164609 | 0.88385192 | 1.25772889 | 0.20848982 | 0.6614379 |
| PDCD4 | 752.723339 | 0.17508534 | 0.24007769 | 0.72928619 | 0.46582662 | 0.8367113 |
| BBIP1 | 441.072283 | 0.06783292 | 0.1366937 | 0.49624029 | 0.61972489 | 0.8968934 |
| SHOC2 | 1310.93049 | -0.0718609 | 0.2194152 | -0.3275108 | 0.74328154 | 0.93611803 |
| GPAM | 120.736536 | 0.09686216 | 0.26663644 | 0.36327426 | 0.71640002 | 0.9289438 |
| ACSL5 | 52.1485826 | 0.40924428 | 0.52244299 | 0.78332811 | 0.43343446 | 0.81882662 |
| ZDHHC6 | 302.198725 | -0.2845837 | 0.22452096 | -1.2675152 | 0.20497114 | 0.65813563 |
| VTI1A | 4113.02425 | -0.2664831 | 0.18322637 | -1.4543929 | 0.1458374 | 0.59218144 |
| TCF7L2 | 4116.86221 | 0.02258883 | 0.22354286 | 0.10104921 | 0.9195114 | 0.98215577 |
| AL158212.2 | 47.1706524 | 0.56863842 | 0.48687402 | 1.16793748 | 0.24283199 | 0.69412848 |
| CASP7 | 278.832069 | -0.5277788 | 0.21724547 | -2.4294123 | 0.01512332 | 0.25080486 |
| DCLRE1A | 52.1451186 | -0.3514416 | 0.34995124 | -1.0042587 | 0.31525392 | 0.74785672 |
| NHLRC2 | 821.875221 | -0.0541424 | 0.17045963 | -0.317626 | 0.75076863 | 0.93775604 |
| CCDC186 | 1071.92839 | -0.2647669 | 0.24209549 | -1.0936466 | 0.27411 | 0.72140248 |
| TDRD1 | 52.4682068 | -0.0739869 | 0.8406598 | -0.0880105 | 0.92986834 | 0.98331361 |
| VWA2 | 59.319986 | -2.091246 | 0.79445034 | -2.6323181 | 0.00848044 | 0.18546054 |
| AFAP1L2 | 282.479882 | -1.4349299 | 0.88887972 | -1.6143128 | 0.1064596 | 0.52743811 |
| ABLIM1 | 6744.33254 | 0.86435276 | 0.3696958 | 2.3380108 | 0.01938669 | 0.28045661 |
| FAM160B1 | 598.285125 | -0.2460254 | 0.19170126 | -1.283379 | 0.19935931 | 0.65150038 |
| TRUB1 | 269.956735 | 0.00801962 | 0.17888741 | 0.04483056 | 0.96424236 | 0.99224805 |
| ATRNL1 | 705.045437 | -0.125312 | 0.74531403 | -0.1681331 | 0.86647858 | 0.97386931 |
| HSPA12A | 1204.35343 | 0.03416025 | 0.34073162 | 0.10025558 | 0.92014142 | 0.98215577 |
| ENO4 | 160.603132 | 0.04675346 | 0.42278906 | 0.11058341 | 0.9119467 | 0.98125148 |
| SHTN1 | 1318.11331 | -0.4201963 | 0.15411102 | -2.7265821 | 0.0063994 | 0.1654202 |
| PDZD8 | 984.67552 | -0.3627997 | 0.23253775 | -1.5601754 | 0.11871843 | 0.55026203 |
| AC005871.2 | 78.5483087 | 0.67435658 | 0.61175547 | 1.10233028 | 0.27031811 | 0.71945032 |
| EMX2OS | 1455.24414 | 0.12085732 | 0.27523985 | 0.43909818 | 0.6605904 | 0.91099119 |
| EMX2 | 497.549375 | -0.2341063 | 0.28485844 | -0.8218338 | 0.41117152 | 0.80675684 |
| RAB11FIP2 | 982.046616 | 0.03928903 | 0.15705663 | 0.25015834 | 0.8024649 | 0.95446295 |
| AC022395.1 | 84.8729498 | 0.46299002 | 0.57899449 | 0.79964496 | 0.42391653 | 0.81413339 |
| CASC2 | 1188.0661 | 0.16019606 | 0.45460923 | 0.3523819 | 0.72455187 | 0.93241479 |
| FAM204A | 673.530107 | -0.1798771 | 0.13885023 | -1.2954754 | 0.19515627 | 0.64871108 |
| CACUL1 | 1183.39343 | -0.3920355 | 0.14461651 | -2.7108627 | 0.00671084 | 0.16906706 |
| EIF3A | 1302.04083 | -0.0801075 | 0.17137206 | -0.467448 | 0.64017936 | 0.90346088 |
| FAM45A | 291.707303 | -0.389419 | 0.22707114 | -1.7149647 | 0.0863517 | 0.49478465 |

| | | | | | | |
|---|---|---|---|---|---|---|
| SFXN4 | 376.410749 | -0.079832 | 0.38153446 | -0.2092394 | 0.83426136 | 0.96510927 |
| PRDX3 | 435.612331 | 0.87205925 | 0.43541165 | 2.00283856 | 0.04519462 | 0.38785764 |
| GRK5 | 114.260771 | -0.7433678 | 0.41800693 | -1.7783623 | 0.07534437 | 0.47234163 |
| RGS10 | 61.5846429 | 1.03500592 | 0.58442256 | 1.770989 | 0.07656253 | 0.4759127 |
| TIAL1 | 796.108157 | -0.1891632 | 0.16177909 | -1.1692682 | 0.24229559 | 0.69337349 |
| BAG3 | 283.604434 | 0.12354835 | 0.2603389 | 0.47456739 | 0.63509536 | 0.90199374 |
| INPP5F | 669.405073 | -0.0837241 | 0.17558944 | -0.4768177 | 0.63349195 | 0.90131357 |
| MCMBP | 476.442056 | -0.1796519 | 0.19998525 | -0.8983256 | 0.36901199 | 0.78095617 |
| SEC23IP | 664.12071 | -0.2592482 | 0.19486552 | -1.3303954 | 0.18338804 | 0.63834582 |
| WDR11-AS1 | 137.102549 | -0.7616066 | 0.3641938 | -2.0912125 | 0.03650902 | 0.35908236 |
| WDR11 | 1055.78713 | -0.1377563 | 0.14717286 | -0.9360173 | 0.34926428 | 0.76982416 |
| FGFR2 | 1932.95385 | 0.0438394 | 0.40526095 | 0.10817574 | 0.91385629 | 0.98145965 |
| ATE1 | 1676.23239 | -0.3149388 | 0.23275974 | -1.3530637 | 0.17603529 | 0.62957039 |
| NSMCE4A | 169.570749 | -0.2878558 | 0.26303515 | -1.0943626 | 0.27379598 | 0.72135661 |
| TACC2 | 1011.14765 | 0.74991928 | 0.4546091 | 1.64959143 | 0.09902653 | 0.51618562 |
| PLEKHA1 | 5035.29153 | -0.2231965 | 0.27200749 | -0.8205527 | 4.12E-01 | 8.07E-01 |
| HTRA1 | 93.0824893 | 1.15910699 | 0.4511638 | 2.56914889 | 0.01019486 | 0.20559384 |
| FAM24B | 55.8934201 | -0.365549 | 0.29259524 | -1.2493335 | 0.21154313 | 0.66484951 |
| C10orf88 | 204.916618 | 0.07479907 | 0.23955072 | 0.31224733 | 0.75485257 | 0.93922829 |
| PSTK | 199.949865 | 0.1144136 | 0.2346245 | 0.4876456 | 0.62580089 | 0.89873262 |
| IKZF5 | 251.088446 | 0.23511966 | 0.22938857 | 1.02498416 | 0.30537066 | 0.74229685 |
| ACADSB | 475.035763 | 0.39049728 | 0.17644474 | 2.21314214 | 0.02688785 | 0.31710856 |
| BUB3 | 312.762645 | 0.14327983 | 0.26610076 | 0.53844202 | 0.59027192 | 0.88631239 |
| CPXM2 | 541.343206 | -0.3038317 | 0.78328914 | -0.3878921 | 0.69809587 | 0.92356678 |
| CHST15 | 56.1674119 | -0.3364782 | 0.66935255 | -0.502692 | 6.15E-01 | 8.95E-01 |
| OAT | 212.90627 | 0.58938237 | 0.24556009 | 2.40015537 | 0.01638811 | 0.256786 |
| LHPP | 277.857855 | 0.12025293 | 0.43899888 | 0.27392538 | 0.78414197 | 0.94978829 |
| ABRAXAS2 | 280.795439 | -0.2978222 | 0.17815169 | -1.6717337 | 0.09457684 | 0.51017802 |
| ZRANB1 | 620.931754 | -0.1029089 | 0.2599683 | -0.3958515 | 0.69221455 | 0.92224682 |
| CTBP2 | 4495.33617 | -0.028693 | 0.29155452 | -0.0984139 | 0.92160361 | 0.98215577 |
| EDRF1 | 460.914686 | -0.18427 | 0.22885214 | -0.8051922 | 0.42070876 | 0.81260593 |
| UROS | 231.579478 | 0.02977704 | 0.29776835 | 0.10000068 | 0.92034378 | 0.98215577 |
| BCCIP | 472.481915 | -0.1499592 | 0.15233271 | -0.9844192 | 0.32490943 | 0.75478872 |
| DHX32 | 1233.0692 | -0.2648268 | 0.2522365 | -1.0499146 | 0.29375736 | 0.73527014 |
| FANK1 | 660.590758 | -0.0279813 | 0.29278054 | -0.095571 | 0.92386126 | 0.98215577 |
| ADAM12 | 161.329566 | 1.2705465 | 0.56415335 | 2.25212967 | 2.43E-02 | 3.06E-01 |
| DOCK1 | 8170.94607 | -0.3642751 | 0.15006294 | -2.4274819 | 0.01520405 | 0.25080486 |
| AL359094.1 | 119.506841 | -0.3437186 | 0.66359448 | -0.5179649 | 0.60448277 | 0.89013087 |
| FAM196A | 350.24224 | -0.3392832 | 0.65013355 | -0.5218669 | 0.60176302 | 0.88951296 |
| MGMT | 1204.89379 | -0.1274856 | 0.24480789 | -0.5207577 | 0.60253558 | 0.88971776 |
| AL157832.3 | 165.507792 | -1.0775998 | 0.3643391 | -2.9576838 | 0.0030996 | 0.1071796 |
| C10orf143 | 167.956073 | -0.7097345 | 0.41439098 | -1.7127172 | 0.08676459 | 0.49478465 |
| GLRX3 | 592.567038 | -0.3199548 | 0.1841429 | -1.7375352 | 0.08229275 | 0.48594425 |
| PPP2R2D | 1132.91693 | -0.2535986 | 0.17347532 | -1.4618715 | 0.14377643 | 0.58881406 |
| BNIP3 | 1052.95805 | 0.54513354 | 0.6495055 | 0.8393055 | 0.40129789 | 0.80259332 |
| STK32C | 252.207576 | -0.6414175 | 0.26025345 | -2.4645879 | 0.01371709 | 0.24270686 |
| LRRC27 | 204.247361 | -0.409355 | 0.41813454 | -0.979003 | 0.32757848 | 0.75620136 |
| PWWP2B | 90.4384895 | -0.192767 | 0.28793127 | -0.6694897 | 0.50318313 | 0.85525037 |
| INPP5A | 843.081329 | 0.00850296 | 0.32618185 | 0.02606815 | 0.97920298 | 0.99463083 |
| CFAP46 | 75.4987259 | 0.50273802 | 0.50075997 | 1.0039501 | 0.31540267 | 0.74785672 |
| KNDC1 | 94.6172429 | 1.51990658 | 0.52263544 | 2.90815829 | 0.00363564 | 0.11745127 |
| ADAM8 | 43.7289009 | -0.8282302 | 0.45325058 | -1.8273119 | 0.06765289 | 0.45209626 |
| TUBGCP2 | 747.595486 | -0.4176881 | 0.17865695 | -2.3379336 | 0.01939069 | 0.28045661 |
| ECHS1 | 359.56551 | 1.17714362 | 0.3586448 | 3.28219908 | 0.00103001 | 0.05732067 |

| | | | | | | |
|---|---|---|---|---|---|---|
| PAOX | 61.5131344 | -0.054064 | 0.24863633 | -0.2174419 | 8.28E-01 | 0.96278001 |
| SCART1 | 145.246506 | -1.0381629 | 0.35732912 | -2.9053408 | 0.00366853 | 0.11790188 |
| SLC6A12 | 489.234256 | -0.5327653 | 0.58635721 | -0.9086019 | 0.36356032 | 0.779589 |
| SLC6A13 | 61.3434858 | 1.54347522 | 0.55636437 | 2.77421651 | 0.00553348 | 0.15039463 |
| KDM5A | 1574.39567 | 0.02573306 | 0.13803466 | 0.18642466 | 0.85211176 | 0.97076831 |
| CCDC77 | 126.285054 | -0.6555316 | 0.29181725 | -2.2463772 | 0.02467986 | 0.30812502 |
| B4GALNT3 | 730.31664 | -0.552136 | 0.31522721 | -1.7515493 | 0.07985134 | 0.48228454 |
| WNK1 | 4596.51604 | 0.83130402 | 0.2138273 | 3.88773558 | 0.00010118 | 0.01602094 |
| RAD52 | 498.379901 | -0.2454582 | 0.19416988 | -1.2641416 | 0.2061792 | 0.65830819 |
| ERC1 | 4005.72562 | -0.148467 | 0.12251601 | -1.2118174 | 0.2255823 | 0.67798093 |
| WNT5B | 139.731024 | 0.16708476 | 0.44090628 | 0.37895755 | 0.70471939 | 0.92528913 |
| FBXL14 | 145.050368 | 0.04438228 | 0.22750317 | 0.19508422 | 0.84532702 | 0.96859165 |
| ADIPOR2 | 1236.5347 | -0.0282642 | 0.33679204 | -0.0839218 | 0.93311859 | 0.98419748 |
| DCP1B | 535.375946 | 0.32454245 | 0.16095281 | 2.01638261 | 0.04375998 | 0.38453712 |
| CACNA1C | 717.826539 | -0.7436385 | 0.86713954 | -0.8575765 | 0.39112638 | 0.79735014 |
| FKBP4 | 237.904104 | 0.89703016 | 0.45320301 | 1.97931202 | 0.04778089 | 0.39501762 |
| ITFG2-AS1 | 83.7070997 | -0.3693252 | 0.3750957 | -0.9846159 | 0.32481277 | 0.75478872 |
| ITFG2 | 910.013997 | -0.4918698 | 0.19969625 | -2.4630896 | 0.01377455 | 0.24292128 |
| TULP3 | 1429.20691 | 0.07899195 | 0.22162296 | 0.35642493 | 0.72152236 | 0.93044898 |
| TEAD4 | 584.036594 | -0.4902037 | 0.42583492 | -1.1511591 | 0.24966679 | 0.70032501 |
| TSPAN9 | 4068.44352 | 0.29237724 | 0.41658888 | 0.7018364 | 0.48278119 | 0.8441831 |
| LINC02417 | 61.8148951 | -1.298187 | 1.09869197 | -1.181575 | 0.23737437 | 0.68973903 |
| PRMT8 | 146.065339 | -1.3621295 | 0.96646475 | -1.4093939 | 0.15871872 | 0.60804051 |
| PARP11 | 292.269871 | -0.7080456 | 0.2301756 | -3.0761104 | 0.0020972 | 0.0878448 |
| CCND2 | 48.72158 | 0.16801502 | 1.06249273 | 0.15813286 | 0.87435211 | 0.97564728 |
| TIGAR | 180.16532 | -0.2884503 | 0.44614867 | -0.646534 | 0.51793362 | 0.85893097 |
| C12orf4 | 349.814209 | -0.1474041 | 0.18349541 | -0.8033121 | 0.42179437 | 0.81266383 |
| AKAP3 | 44.6026082 | -0.5791435 | 0.5846762 | -0.9905371 | 0.32191165 | 0.75240254 |
| AC005833.2 | 65.2971636 | 0.00348488 | 0.33951838 | 0.01026418 | 0.99181052 | 0.99772135 |
| GALNT8 | 50.1958888 | -0.5847732 | 0.43141696 | -1.3554709 | 0.17526756 | 0.62944785 |
| ANO2 | 53.4987558 | 0.11788758 | 0.58810853 | 0.20045209 | 0.84112702 | 0.96637791 |
| VWF | 44.8872081 | 0.65247665 | 0.44847458 | 1.45487989 | 0.14570252 | 0.59218144 |
| CD9 | 994.993848 | 0.82595647 | 0.40622847 | 2.03323139 | 0.04202915 | 0.37817647 |
| PLEKHG6 | 42.2282007 | 0.89482685 | 0.42560292 | 2.10249227 | 0.03551018 | 0.35588309 |
| TNFRSF1A | 501.635658 | -0.1385622 | 0.43515493 | -0.3184204 | 0.75016607 | 0.93775604 |
| SCNN1A | 2735.57698 | 0.07261876 | 0.35272253 | 0.2058807 | 0.8368841 | 0.9652581 |
| LTBR | 282.260566 | -0.1638133 | 0.25163105 | -0.651006 | 0.51504264 | 0.85869987 |
| TAPBPL | 139.436606 | 0.15872374 | 0.42092738 | 0.37708106 | 0.70611337 | 0.92556842 |
| MRPL51 | 253.780192 | 0.55930905 | 0.47365107 | 1.18084615 | 0.23766385 | 0.68980482 |
| NCAPD2 | 216.02849 | -0.4711681 | 0.30033063 | -1.5688314 | 0.11668724 | 0.54715005 |
| GAPDH | 2272.90492 | 0.78664596 | 0.51526159 | 1.52669243 | 0.12683751 | 0.56735411 |
| CHD4 | 313.446609 | -0.0460039 | 0.22267055 | -0.2066008 | 0.83632166 | 0.96511435 |
| AC006064.2 | 140.976316 | -0.2106453 | 0.47827688 | -0.4404254 | 0.65962905 | 0.91099119 |
| ING4 | 274.881844 | -0.1493195 | 0.20775043 | -0.7187447 | 0.47229825 | 0.83921324 |
| ZNF384 | 262.439445 | -0.2972291 | 0.21783991 | -1.3644381 | 0.17242972 | 0.6286116 |
| COPS7A | 200.479013 | 0.16305807 | 0.23861027 | 0.68336567 | 0.49437581 | 0.84882719 |
| MLF2 | 345.552249 | 0.49496429 | 0.40151575 | 1.23273943 | 0.217673 | 0.67068873 |
| PTMS | 511.922283 | -0.1951713 | 0.58353937 | -0.3344612 | 0.73803155 | 0.93552616 |
| P3H3 | 50.7170868 | 0.55944963 | 0.47542039 | 1.17674724 | 0.23929641 | 0.69029619 |
| USP5 | 159.377478 | 0.14186389 | 0.41154026 | 0.34471447 | 0.73030904 | 0.93466254 |
| TPI1 | 828.351336 | 0.78540136 | 0.42049608 | 1.86779712 | 0.06179036 | 0.43264436 |
| SPSB2 | 56.668704 | -0.1089569 | 0.36352701 | -0.2997217 | 0.76438943 | 0.94256396 |
| LRRC23 | 280.908117 | -0.2679846 | 0.29235262 | -0.9166487 | 0.35932675 | 0.77654006 |
| ENO2 | 137.727865 | 0.40815489 | 0.64371974 | 0.63405681 | 5.26E-01 | 0.86119802 |

| | | | | | | |
|---|---|---|---|---|---|---|
| ATN1 | 606.99151 | -0.5646089 | 0.32111427 | -1.7582802 | 0.07869984 | 0.48060389 |
| C12orf57 | 197.166055 | 0.72905745 | 0.4685381 | 1.55602597 | 0.11970192 | 0.55232605 |
| PTPN6 | 231.962095 | -0.2576368 | 0.41266164 | -0.6243294 | 0.5324113 | 0.86397197 |
| PHB2 | 322.09736 | 0.48968478 | 0.3380727 | 1.44846001 | 0.14748844 | 0.59373375 |
| EMG1.1 | 107.132232 | -0.2134285 | 0.35463289 | -0.6018294 | 0.54728771 | 0.86859163 |
| LPCAT3 | 988.929714 | 0.37596619 | 0.24349697 | 1.54402818 | 0.12258151 | 0.55930194 |
| C1S | 463.994264 | 0.05710727 | 0.38212703 | 0.14944579 | 0.88120188 | 0.97664137 |
| C1R | 252.372735 | 0.81466385 | 0.5542056 | 1.46996683 | 1.42E-01 | 5.87E-01 |
| C1RL | 884.923836 | 0.0771747 | 0.33055397 | 0.23347081 | 8.15E-01 | 9.59E-01 |
| C1RL-AS1 | 97.7125155 | -0.6693845 | 0.44557509 | -1.5022934 | 0.13302135 | 0.57682553 |
| RBP5 | 139.442483 | 0.57405923 | 0.29245725 | 1.96288253 | 0.04965982 | 0.39997355 |
| CLSTN3 | 308.834224 | 0.16161994 | 0.3375408 | 0.47881603 | 0.63206951 | 0.90042718 |
| PEX5 | 156.396882 | -0.2011377 | 0.23406474 | -0.8593252 | 0.39016113 | 0.79701353 |
| CD163L1 | 65.9805737 | 0.41399258 | 0.38300122 | 1.08091712 | 0.27973398 | 0.72500457 |
| FOXJ2 | 141.484351 | 0.15464566 | 0.29354524 | 0.52682055 | 0.59831821 | 0.88883692 |
| NECAP1 | 182.224815 | -0.2136897 | 0.22703079 | -0.9412365 | 0.34658368 | 0.76864066 |
| FAM66C | 198.803613 | 0.58238086 | 0.4326626 | 1.34603929 | 0.17828985 | 0.63222298 |
| LINC00937 | 212.660707 | 0.33248939 | 0.27758439 | 1.19779569 | 0.23099657 | 0.68321398 |
| RIMKLB | 1602.19931 | 0.70351743 | 0.55747297 | 1.26197585 | 0.20695748 | 0.65951781 |
| A2ML1-AS1 | 253.150596 | 0.78834055 | 0.69846631 | 1.12867369 | 0.25903551 | 0.70921184 |
| AC006581.2 | 103.687649 | -0.5574798 | 0.36060294 | -1.5459659 | 0.12211281 | 0.55793432 |
| PHC1 | 357.390998 | -0.2765677 | 0.18978039 | -1.4573038 | 0.14503252 | 0.59091136 |
| M6PR | 361.589219 | 0.18691046 | 0.25321204 | 0.73815787 | 0.46041852 | 0.83429248 |
| KLRG1 | 98.6489584 | -0.6961903 | 0.39916455 | -1.7441187 | 0.0811384 | 0.48572317 |
| A2M | 170.46574 | 0.14099534 | 0.643235 | 0.21919725 | 0.82649639 | 0.96214368 |
| CLEC2D | 449.043793 | -0.3062164 | 0.33694785 | -0.9087947 | 0.3634585 | 0.779589 |
| KLRF1 | 75.6508255 | 1.26388636 | 0.70705659 | 1.78753211 | 0.07385153 | 0.47056873 |
| TMEM52B | 684.610192 | 4.03542976 | 0.86415173 | 4.66981621 | 3.01E-06 | 0.00248882 |
| GABARAPL1 | 1166.08907 | 0.35770706 | 0.42053315 | 0.85060371 | 0.39498953 | 0.7989793 |
| KLRD1 | 579.617698 | 0.88771933 | 0.59532883 | 1.49114116 | 0.13592444 | 0.57776571 |
| AC022075.1 | 54.5345406 | -0.537132 | 0.28814446 | -1.8641068 | 0.0623067 | 0.43419765 |
| EIF2S3B | 70.8451214 | -0.5354433 | 0.3199021 | -1.6737724 | 0.09417534 | 0.51017802 |
| MAGOHB | 152.319993 | -0.0121608 | 0.32603748 | -0.0372989 | 0.97024672 | 0.99309664 |
| YBX3 | 2018.90374 | 0.01562372 | 0.36946986 | 0.04228685 | 9.66E-01 | 0.99248831 |
| PRH1 | 139.300245 | 1.07158867 | 0.47130687 | 2.27365381 | 0.02298681 | 0.2983102 |
| SMIM10L1 | 120.885363 | 0.47210758 | 0.27866021 | 1.69420523 | 0.09022629 | 0.50138792 |
| AC244131.2 | 337.070627 | -0.0549371 | 0.61628585 | -0.0891423 | 0.92896884 | 0.98323312 |
| PRB2 | 57.9146849 | 0.70906715 | 0.82460089 | 0.85989132 | 0.38984895 | 0.79698174 |
| LINC01252 | 146.19877 | 0.37357736 | 0.77596131 | 0.48143813 | 0.63020514 | 0.90005881 |
| ETV6 | 3606.60611 | -0.2869804 | 0.29346061 | -0.9779179 | 3.28E-01 | 7.56E-01 |
| BCL2L14 | 703.350756 | 0.56086502 | 0.47097253 | 1.19086567 | 0.23370632 | 0.68493974 |
| LRP6 | 2304.30104 | 0.14472818 | 0.18118026 | 0.79880765 | 4.24E-01 | 8.15E-01 |
| MANSC1 | 124.530955 | 0.35389643 | 0.21809973 | 1.62263581 | 0.10466728 | 0.52468695 |
| BORCS5 | 588.935846 | -0.3234428 | 0.24415591 | -1.3247386 | 0.18525787 | 0.64030106 |
| DUSP16 | 833.319118 | 0.11441793 | 0.39472264 | 0.28986917 | 0.77191633 | 0.94478317 |
| AC007619.1 | 52.0066792 | -0.4643976 | 0.46206798 | -1.0050417 | 0.31487679 | 0.74785672 |
| CREBL2 | 345.303084 | 0.09308002 | 0.19256965 | 0.48335767 | 0.6288418 | 0.89970424 |
| CDKN1B | 189.731644 | 0.5186864 | 0.34399207 | 1.50784404 | 0.13159445 | 0.57519227 |
| APOLD1 | 305.651201 | 0.51841837 | 0.5762044 | 0.89971261 | 0.36827321 | 0.78095617 |
| GPRC5A | 132.992315 | 0.1718294 | 0.96778026 | 0.17755002 | 0.85907638 | 0.97174503 |
| GPRC5D-AS1 | 414.994207 | -0.495099 | 0.38880008 | -1.2734025 | 0.20287527 | 0.65552334 |
| HEBP1 | 159.141224 | 0.0502952 | 0.3727063 | 0.13494595 | 0.89265461 | 0.97778214 |
| AC023790.2 | 91.142559 | -0.3881425 | 0.42779905 | -0.907301 | 0.36424763 | 0.78031086 |
| FAM234B | 114.801665 | -0.0197409 | 0.37106873 | -0.0532 | 0.95757255 | 0.99065996 |

| | | | | | | |
|---|---|---|---|---|---|---|
| EMP1 | 1053.77714 | -0.6741555 | 0.34376797 | -1.961077 | 0.04987004 | 0.40017381 |
| ATF7IP | 2449.35244 | -0.0990978 | 0.28822955 | -0.3438155 | 0.73098506 | 0.93507473 |
| PLBD1 | 133.468481 | -0.1821663 | 0.29669037 | -0.6139945 | 0.53921894 | 0.86543657 |
| PLBD1-AS1 | 119.692505 | -0.1674228 | 0.51950223 | -0.3222753 | 0.74724411 | 0.93735494 |
| AC010168.1 | 230.27998 | 0.05640567 | 0.5374377 | 0.10495296 | 0.91641314 | 0.98197488 |
| H2AFJ | 298.558907 | 0.78931492 | 0.40692756 | 1.93969393 | 0.0524169 | 0.4090401 |
| WBP11 | 321.917346 | 0.24463756 | 0.2185421 | 1.11940704 | 0.26296653 | 0.71328 |
| C12orf60 | 306.336645 | -0.4313828 | 0.3975034 | -1.0852306 | 0.27781955 | 0.72372365 |
| MGP | 127.523759 | -0.5716737 | 0.53701126 | -1.064547 | 0.28708095 | 0.72972899 |
| ERP27 | 55.0747579 | 1.5574818 | 0.70939165 | 2.19551754 | 0.02812649 | 0.32306336 |
| RERG | 792.526143 | -0.7554433 | 0.52312499 | -1.4440971 | 0.14871166 | 0.59489166 |
| PTPRO | 51.8172095 | -0.3759056 | 0.53951263 | -0.6967503 | 0.4859591 | 0.8447301 |
| EPS8 | 710.211692 | 0.03297847 | 0.43863561 | 0.07518421 | 0.94006815 | 0.98715467 |
| STRAP | 489.982972 | 0.28798949 | 0.27217479 | 1.05810496 | 0.29000759 | 0.73204984 |
| DERA | 1289.64619 | -0.4292591 | 0.1768214 | -2.4276423 | 0.01519732 | 0.25080486 |
| MGST1 | 583.309863 | 0.4382826 | 0.69310376 | 0.63234775 | 0.52715967 | 0.86151701 |
| LMO3 | 177.060789 | 0.63074043 | 0.59840819 | 1.0540304 | 0.291869 | 0.73368744 |
| PIK3C2G | 64.5649344 | 0.63350967 | 0.46211693 | 1.37088609 | 0.17041047 | 0.6265257 |
| PLEKHA5 | 9481.82787 | -0.1491148 | 0.2386762 | -0.6247577 | 0.53213009 | 0.86397197 |
| AEBP2 | 3269.6958 | 0.51654198 | 0.28779868 | 1.79480315 | 0.07268507 | 0.4692634 |
| AC024901.1 | 541.569731 | 0.69025777 | 0.64809656 | 1.06505389 | 0.28685154 | 0.72949981 |
| LINC02398 | 270.54487 | 1.64921494 | 0.72382976 | 2.27845694 | 0.02269937 | 0.2983102 |
| PDE3A | 451.049749 | 1.45049974 | 0.88035779 | 1.64762526 | 0.09942959 | 0.51665771 |
| PYROXD1 | 376.363423 | -0.0760671 | 0.15153595 | -0.5019741 | 0.61568577 | 0.89516713 |
| RECQL | 288.23495 | -0.4960085 | 0.18509417 | -2.6797632 | 0.00736743 | 0.17305714 |
| GOLT1B | 277.298461 | -0.1903721 | 0.18692015 | -1.0184674 | 0.30845587 | 0.7438271 |
| SPX | 93.3631681 | 0.89674408 | 0.71245574 | 1.25866637 | 0.20815086 | 0.66105261 |
| LDHB | 3187.10101 | 0.85844697 | 0.46665546 | 1.8395734 | 0.06583089 | 0.44463112 |
| AC010197.1 | 453.215327 | 0.38291941 | 0.44781056 | 0.8550924 | 0.3925 | 0.7982344 |
| AC010185.1 | 60.4447803 | 0.77367795 | 0.51677955 | 1.49711409 | 0.13436358 | 0.57708028 |
| ABCC9 | 228.476408 | 1.30322761 | 0.60193835 | 2.16505164 | 0.03038373 | 0.33500726 |
| AC008250.1 | 419.647345 | 0.90747628 | 0.37309026 | 2.43232373 | 0.01500229 | 0.25020868 |
| CMAS | 265.589096 | 0.0272035 | 0.22150466 | 0.12281233 | 0.90225571 | 0.97941893 |
| ST8SIA1 | 50.184627 | 0.14401562 | 0.48624507 | 0.29617909 | 0.76709332 | 0.94316227 |
| C2CD5 | 1843.13499 | -0.1995157 | 0.21294977 | -0.9369146 | 0.34880248 | 0.76982416 |
| ETNK1 | 3577.63361 | 0.0880134 | 0.32443715 | 0.27128029 | 0.78617547 | 0.94978829 |
| AC084816.1 | 213.937016 | -0.6604596 | 0.49699806 | -1.3288978 | 0.18388169 | 0.63843513 |
| SOX5 | 260.606304 | 1.20404967 | 0.5186649 | 2.32144042 | 0.02026309 | 0.28534319 |
| CASC1 | 541.77525 | 0.40921159 | 0.42361342 | 0.96600243 | 0.33404297 | 0.76219961 |
| ETFRF1 | 428.380303 | 0.3512597 | 0.21373042 | 1.64347074 | 0.10028557 | 0.51704612 |
| KRAS | 810.087553 | -0.0362009 | 0.23809283 | -0.1520455 | 0.87915104 | 0.9765717 |
| LMNTD1 | 58.0243092 | -1.8908677 | 0.92675542 | -2.0403093 | 0.04131953 | 0.37577267 |
| RASSF8-AS1 | 988.408846 | -0.9950744 | 0.38017114 | -2.6174381 | 0.00885926 | 0.19125454 |
| RASSF8 | 3625.66123 | -0.6778121 | 0.5071537 | -1.3365023 | 0.18138514 | 0.63616641 |
| BHLHE41 | 68.3039343 | -1.4275508 | 0.70711966 | -2.0188249 | 0.04350542 | 0.38387647 |
| SSPN | 94.6589534 | -0.2978463 | 0.44844242 | -0.6641797 | 0.50657533 | 0.85626196 |
| AC022509.2 | 59.9991429 | -0.8381112 | 0.41817588 | -2.0042074 | 0.04504785 | 0.38755881 |
| AC024145.1 | 177.501514 | -0.2605787 | 0.67107109 | -0.3883027 | 0.69779203 | 0.92341984 |
| ITPR2 | 7494.05068 | -0.6201705 | 0.1554159 | -3.9903931 | 6.60E-05 | 0.01300473 |
| AC023051.1 | 193.928995 | -0.1658201 | 0.64984537 | -0.2551686 | 0.79859287 | 0.95298701 |
| INTS13 | 185.822575 | -0.4604501 | 0.23982364 | -1.9199529 | 0.05486385 | 0.41684503 |
| FGFR1OP2 | 648.220675 | 0.20478853 | 0.20993611 | 0.97548027 | 0.32932208 | 0.75817321 |
| TM7SF3 | 541.102801 | -0.114285 | 0.28704524 | -0.3981426 | 0.69052504 | 0.92183073 |
| MED21 | 220.465283 | -0.0903687 | 0.29827629 | -0.3029699 | 0.7619128 | 0.94210219 |

| | | | | | | |
|---|---|---|---|---|---|---|
| STK38L | 707.917724 | -0.0254106 | 0.35058061 | -0.0724815 | 0.94221877 | 0.98772472 |
| ARNTL2 | 75.8293059 | 0.85506333 | 0.68882515 | 1.24133581 | 2.14E-01 | 0.66827012 |
| PPFIBP1 | 1724.35893 | -0.3987397 | 0.30379702 | -1.3125202 | 0.18934466 | 0.64432929 |
| MRPS35 | 429.845509 | 0.00029456 | 0.17337773 | 0.00169894 | 0.99864444 | 0.99980325 |
| KLHL42 | 491.436552 | 0.20977143 | 0.19564734 | 1.07219158 | 0.28363399 | 0.72690826 |
| CCDC91 | 3294.46417 | -0.2796678 | 0.15812909 | -1.7686046 | 0.07695989 | 0.47681197 |
| AC022364.1 | 84.239593 | 0.269152 | 0.28712737 | 0.93739581 | 0.348555 | 0.76982416 |
| FAR2 | 138.560043 | -0.3952601 | 0.47050071 | -0.8400839 | 0.40086134 | 0.80243544 |
| AC009318.1 | 205.280709 | -1.3606304 | 0.50514947 | -2.6935204 | 0.00707018 | 0.17262485 |
| ERGIC2 | 602.239387 | -0.2469165 | 0.17013547 | -1.4512935 | 0.14669816 | 0.59373375 |
| TMTC1 | 86.6400381 | -1.557137 | 0.63476367 | -2.4530972 | 0.0141632 | 0.24646911 |
| IPO8 | 757.730125 | 0.06229679 | 0.16467482 | 0.37830183 | 0.70520639 | 0.92549142 |
| CAPRIN2 | 503.426736 | 0.09966323 | 0.18427131 | 0.5408505 | 0.58861063 | 0.88571413 |
| AC010198.1 | 160.332587 | 0.18827843 | 0.51740447 | 0.36389021 | 0.71593999 | 0.92890921 |
| DDX11-AS1 | 50.7344581 | -0.4969825 | 0.43856025 | -1.1332136 | 0.25712455 | 0.70746612 |
| DDX11 | 145.396316 | -1.280011 | 0.47684898 | -2.6843112 | 0.00726794 | 0.17305714 |
| SINHCAF | 1241.06546 | 0.02713086 | 0.19863988 | 0.13658316 | 0.89136029 | 0.97729111 |
| DENND5B | 1445.11832 | -0.1170307 | 0.21374054 | -0.5475362 | 0.58401038 | 0.88416261 |
| DENND5B-AS | 69.5619271 | 0.4247265 | 0.47217154 | 0.89951736 | 0.36837715 | 0.78095617 |
| ETFBKMT | 103.818622 | 0.01959156 | 0.25148895 | 0.07790226 | 0.9379058 | 0.98691928 |
| AMN1 | 451.522638 | 0.15042532 | 0.24213172 | 0.6212541 | 0.53443245 | 0.86419602 |
| KIAA1551 | 516.447604 | 0.0232973 | 0.17968777 | 0.12965433 | 0.89683992 | 0.97823597 |
| BICD1 | 2085.86793 | 0.82063832 | 0.52100549 | 1.57510492 | 1.15E-01 | 5.44E-01 |
| FGD4 | 11626.6108 | -0.061272 | 0.20204558 | -0.3032583 | 0.76169301 | 0.94210219 |
| DNM1L | 1559.17378 | -0.0970345 | 0.16892263 | -0.5744316 | 0.56567579 | 0.87627671 |
| YARS2 | 454.90734 | -0.0884737 | 0.17489213 | -0.5058756 | 0.61294396 | 0.89441126 |
| PKP2 | 142.257885 | -0.1328091 | 0.29409534 | -0.4515852 | 0.65156787 | 0.90825486 |
| SYT10 | 79.0346718 | 1.09472449 | 0.61331354 | 1.7849345 | 0.07427194 | 0.47120945 |
| ALG10 | 42.6076518 | -0.4600314 | 0.39525968 | -1.1638714 | 0.24447612 | 0.6948422 |
| ALG10B | 155.119753 | -0.1673254 | 0.31939325 | -0.5238852 | 0.6003584 | 0.88883692 |
| CPNE8 | 2050.62365 | -0.3695437 | 0.29569064 | -1.2497646 | 0.21138553 | 0.66484951 |
| KIF21A | 702.351186 | 0.15830942 | 0.39404929 | 0.4017503 | 0.6878678 | 0.92113197 |
| C12orf40 | 205.591604 | 0.09080095 | 0.55808904 | 0.16269976 | 0.87075484 | 0.97473202 |
| SLC2A13 | 2491.52819 | -0.2225999 | 0.26486852 | -0.8404165 | 0.40067488 | 0.80243544 |
| LINC02471 | 106.095068 | 2.17043101 | 0.95754432 | 2.26666375 | 0.02341077 | 0.30107587 |
| AC079630.1 | 125.6449 | 1.17477813 | 0.45788897 | 2.56563971 | 0.01029857 | 0.20627111 |
| LRRK2 | 3576.64906 | 0.34541336 | 0.35789276 | 0.96513088 | 0.33447926 | 0.76240493 |
| CNTN1 | 634.207962 | 2.01917134 | 0.60173405 | 3.35558764 | 0.00079197 | 0.04911302 |
| PDZRN4 | 87.0845857 | 0.36266137 | 0.71469535 | 0.5074349 | 6.12E-01 | 8.94E-01 |
| AC090630.1 | 79.2033288 | 1.00694073 | 0.74326679 | 1.35475006 | 0.1754972 | 0.62957039 |
| GXYLT1 | 537.461515 | 0.12546797 | 0.2141912 | 0.58577559 | 0.55802633 | 0.87291524 |
| YAF2 | 1132.4728 | -0.2277426 | 0.24538189 | -0.928115 | 0.35334791 | 0.77219738 |
| PPHLN1 | 1597.67487 | 0.02577891 | 0.13330947 | 0.19337645 | 0.84666416 | 0.96883887 |
| ZCRB1 | 249.980653 | -0.0601231 | 0.22947746 | -0.2619999 | 0.79332151 | 0.95224951 |
| PRICKLE1 | 365.330097 | 0.80501941 | 0.6735338 | 1.19521753 | 2.32E-01 | 6.84E-01 |
| PUS7L | 384.390455 | 0.25721498 | 0.21149215 | 1.21619161 | 0.22391194 | 0.67665358 |
| IRAK4 | 478.66463 | 0.10394957 | 0.22919765 | 0.45353679 | 0.65016226 | 0.90825486 |
| TWF1 | 194.16418 | 0.24843921 | 0.30964222 | 0.80234282 | 0.42235468 | 0.81312972 |
| TMEM117 | 2419.30097 | -0.3689399 | 0.189651 | -1.9453622 | 0.05173141 | 0.40795233 |
| ANO6 | 2197.56001 | 0.08652765 | 0.23473714 | 0.36861507 | 0.71241466 | 0.92748721 |
| AC008124.1 | 209.344631 | 0.54728468 | 0.29562741 | 1.85126498 | 0.06413144 | 0.43894277 |
| ARID2 | 3092.48551 | -0.1136456 | 0.17782516 | -0.6390861 | 0.52276693 | 0.86006369 |
| SCAF11 | 2763.64145 | 0.02611387 | 0.13558428 | 0.19260247 | 0.84727031 | 0.96905304 |
| SLC38A1 | 1224.88881 | 0.1410722 | 0.34184469 | 0.41267922 | 6.80E-01 | 9.18E-01 |

| | | | | | | |
|---|---|---|---|---|---|---|
| SLC38A2 | 1324.0007 | -0.443547 | 0.47553044 | -0.9327416 | 0.35095339 | 0.77084962 |
| AC008014.1 | 4034.56389 | -0.2326982 | 0.31383923 | -0.7414568 | 0.45841655 | 0.8331421 |
| SLC38A4 | 96.0721815 | 0.19997793 | 0.70622924 | 0.28316292 | 0.77705195 | 0.94694572 |
| PCED1B | 321.594419 | -1.3377151 | 0.5593024 | -2.3917564 | 0.01676797 | 0.25935369 |
| RPAP3 | 304.130388 | -0.113246 | 0.23536607 | -0.4811484 | 0.63041105 | 0.90005881 |
| AC004241.1 | 61.3300144 | 0.10632774 | 0.36677353 | 0.28990025 | 0.77189255 | 0.94478317 |
| RAPGEF3 | 1186.95181 | -0.1539285 | 0.50700832 | -0.3036016 | 7.61E-01 | 0.94202764 |
| SLC48A1 | 421.422593 | 0.18866912 | 0.26551768 | 0.71057082 | 4.77E-01 | 8.42E-01 |
| HDAC7 | 1189.61056 | -0.5434998 | 0.39788059 | -1.3659873 | 1.72E-01 | 0.6286116 |
| VDR | 290.761991 | 1.2082847 | 0.67022923 | 1.80279319 | 0.0714207 | 0.46587458 |
| TMEM106C | 162.321138 | 0.56566065 | 0.32230335 | 1.75505668 | 0.07924961 | 0.48169281 |
| SENP1 | 400.797442 | -0.2083666 | 0.1713183 | -1.2162539 | 0.2238882 | 0.67665358 |
| PFKM | 202.529593 | 0.32938042 | 0.2840743 | 1.15948685 | 0.24625779 | 0.6967274 |
| ASB8 | 198.711628 | 0.26358209 | 0.20223144 | 1.30336852 | 0.19244898 | 0.64671221 |
| ZNF641 | 189.354508 | -0.0764095 | 0.29262022 | -0.2611218 | 0.79399857 | 0.95224951 |
| AC090115.1 | 152.361905 | 0.2025476 | 0.41627528 | 0.4865713 | 0.62656216 | 0.89873262 |
| KANSL2 | 289.639566 | -0.1477107 | 0.28006589 | -0.5274143 | 0.59790594 | 0.88883692 |
| CCNT1 | 460.373735 | -0.2058886 | 0.14561318 | -1.4139424 | 0.15737883 | 0.60713112 |
| ADCY6 | 533.493076 | 0.17331142 | 0.27782341 | 0.62381864 | 0.5327467 | 0.86397197 |
| DDX23 | 237.049442 | -0.2257648 | 0.24765778 | -0.9115998 | 0.36197945 | 0.77784066 |
| AC073610.3 | 67.5182293 | -0.170161 | 0.44681962 | -0.380827 | 0.70333161 | 0.92513032 |
| CCDC65 | 229.795369 | -0.1960236 | 0.43384683 | -0.4518267 | 0.65139383 | 0.90825486 |
| PRKAG1 | 328.076841 | -0.2859059 | 0.3015588 | -0.9480932 | 0.34308199 | 0.76641095 |
| KMT2D | 344.760832 | -0.4440164 | 0.23756765 | -1.8690103 | 0.06162138 | 0.43249565 |
| LMBR1L | 329.747355 | 0.13364413 | 0.27825843 | 0.48028779 | 0.63102277 | 0.90005881 |
| AC011603.2 | 161.73724 | -0.0969473 | 0.34096385 | -0.2843331 | 0.77615514 | 0.94694572 |
| TUBA1B | 447.421675 | 0.66959255 | 0.59706091 | 1.12148114 | 0.26208312 | 0.712756 |
| TUBA1A | 187.683335 | 0.49600728 | 0.51918569 | 0.95535622 | 0.33939758 | 0.76449297 |
| TUBA1C | 377.3182 | 0.38033211 | 0.33918069 | 1.12132595 | 0.26214914 | 0.71278881 |
| AC010173.1 | 68.7128603 | 0.13878093 | 0.43244384 | 0.32092243 | 0.74826918 | 0.93746253 |
| SPATS2 | 1000.9256 | 0.16275607 | 0.19341652 | 0.84147967 | 0.40007927 | 0.80238662 |
| KCNH3 | 41.9386099 | 0.12672659 | 0.80345113 | 0.15772782 | 0.87467128 | 0.97572274 |
| MCRS1 | 145.816189 | -0.1645948 | 0.28470964 | -0.5781146 | 0.56318672 | 0.87447201 |
| PRPF40B | 893.851107 | -0.0686843 | 0.20193645 | -0.3401281 | 0.73376009 | 0.93552616 |
| FMNL3 | 55.370702 | 1.29383794 | 0.47277419 | 2.73669326 | 0.00620601 | 0.16248472 |
| TMBIM6 | 1604.8832 | 0.40858933 | 0.33722162 | 1.21163447 | 0.22565234 | 0.67798093 |
| NCKAP5L | 239.109794 | -0.5027088 | 0.27609555 | -1.8207785 | 0.06864054 | 0.45447261 |
| BCDIN3D | 87.9150925 | -0.0435307 | 0.40205208 | -0.1082714 | 0.91378042 | 0.98145965 |
| AQP2 | 239.840419 | -1.5656929 | 0.5910294 | -2.6490948 | 0.00807077 | 0.1802126 |
| RACGAP1 | 43.2232607 | -0.6778639 | 0.49207636 | -1.3775583 | 0.1683397 | 0.62253512 |
| SMARCD1 | 232.06023 | -0.186494 | 0.26426275 | -0.7057143 | 0.48036585 | 0.8433217 |
| AC074032.1 | 1081.09068 | -0.0947554 | 0.25265579 | -0.3750375 | 0.70763256 | 0.92577218 |
| CERS5 | 709.502541 | 0.02590037 | 0.19323005 | 0.13403902 | 0.89337172 | 0.97788747 |
| LIMA1 | 976.890118 | -0.1388782 | 0.23235989 | -0.5976857 | 0.55004965 | 0.86956073 |
| LARP4 | 1497.14239 | -0.1064054 | 0.19153809 | -0.5555315 | 0.57853117 | 0.8821758 |
| DIP2B | 2418.32527 | -0.2282961 | 0.23976416 | -0.9521693 | 0.34101112 | 0.76540627 |
| ATF1 | 693.296584 | -0.3401131 | 0.23622267 | -1.4397989 | 0.1499243 | 0.59663305 |
| METTL7A | 446.085891 | 0.5509619 | 0.57077013 | 0.96529562 | 0.33439677 | 0.76234844 |
| SLC11A2 | 919.84309 | 0.18920309 | 0.20247875 | 0.93443434 | 0.35007989 | 0.77044053 |
| LETMD1 | 932.682179 | 0.18611039 | 0.23951518 | 0.77702964 | 0.43714128 | 0.82013459 |
| CSRNP2 | 308.177056 | -0.228478 | 0.26616046 | -0.8584221 | 0.39065944 | 0.79706836 |
| TFCP2 | 1900.04074 | 0.1999049 | 0.22668065 | 0.88187898 | 0.37784225 | 0.7865219 |
| POU6F1 | 363.506673 | -0.483482 | 0.27148877 | -1.7808544 | 0.07493624 | 0.47190267 |
| DAZAP2 | 910.963162 | 0.0742703 | 0.27364164 | 0.27141446 | 0.78607228 | 0.94978829 |

| | | | | | | |
|---|---|---|---|---|---|---|
| SMAGP | 255.726057 | -1.1106918 | 0.4962601 | -2.2381244 | 0.02521295 | 0.31063019 |
| GALNT6 | 148.34049 | 0.3812086 | 0.51298076 | 0.74312457 | 0.45740627 | 0.83255882 |
| SLC4A8 | 743.885984 | -1.333726 | 0.53262771 | -2.5040492 | 0.0122781 | 0.22749396 |
| SCN8A | 46.1664769 | -1.2510781 | 0.48487113 | -2.5802279 | 0.00987351 | 0.20220034 |
| AC068987.3 | 46.3512585 | -1.101416 | 0.53897263 | -2.0435472 | 0.0409983 | 0.37451348 |
| FIGNL2 | 239.139999 | -1.1460761 | 0.39189285 | -2.9244629 | 0.00345051 | 0.1138728 |
| ACVR1B | 2513.74072 | -0.3117579 | 0.27209039 | -1.1457879 | 0.25188293 | 0.70276117 |
| GRASP | 48.0386192 | -1.1926989 | 0.56168838 | -2.1234174 | 0.03371889 | 0.35097935 |
| NR4A1 | 182.923269 | 0.86030524 | 0.86404026 | 0.99567727 | 0.31940698 | 0.75094569 |
| ATG101 | 123.461998 | 0.31564868 | 0.31840478 | 0.99134402 | 0.32151762 | 0.75240254 |
| AC078864.2 | 70.7720275 | -0.672109 | 1.00558994 | -0.6683728 | 0.50389563 | 0.85525037 |
| AC078864.1 | 131.954254 | -0.4895353 | 0.98019716 | -0.4994253 | 0.61747981 | 0.89599505 |
| KRT7 | 1174.66954 | 1.1030719 | 0.67242656 | 1.64043476 | 0.1009148 | 0.51819907 |
| KRT8 | 705.396598 | 0.33977249 | 0.35570959 | 0.9551963 | 3.39E-01 | 0.76449297 |
| KRT18 | 768.287777 | 0.7558144 | 0.45877244 | 1.64747123 | 9.95E-02 | 5.17E-01 |
| EIF4B | 913.831035 | -0.0500377 | 0.1312316 | -0.3812933 | 0.70298564 | 0.92501157 |
| AC068888.1 | 472.792678 | -0.1435632 | 0.27346732 | -0.5249739 | 0.59960132 | 0.88883692 |
| TNS2 | 504.017586 | 0.0302337 | 0.37068592 | 0.08156149 | 0.93499542 | 0.98558411 |
| SPRYD3 | 78.0776244 | 0.54488195 | 0.403226 | 1.3513066 | 0.17659724 | 0.63008941 |
| IGFBP6 | 79.5813602 | 2.30207032 | 0.64410539 | 3.57405849 | 3.51E-04 | 0.03115997 |
| CSAD | 887.486653 | -0.0340291 | 0.19570518 | -0.1738796 | 0.86196006 | 0.97264823 |
| ZNF740 | 144.620525 | -0.0049557 | 0.21425209 | -0.0231301 | 0.98154648 | 0.99518016 |
| RARG | 156.183212 | -0.6618229 | 0.43771092 | -1.5120091 | 0.13053154 | 0.57223551 |
| MFSD5 | 52.4183565 | 0.4603507 | 0.51354933 | 0.89640989 | 0.37003389 | 0.78099673 |
| PFDN5 | 624.285029 | 0.27323406 | 0.37152922 | 0.73543087 | 0.46207713 | 0.83531269 |
| C12orf10 | 87.5284178 | 0.39801514 | 0.36285085 | 1.09691112 | 0.27268025 | 0.72070318 |
| AAAS | 141.990888 | -0.242302 | 0.27040595 | -0.8960675 | 0.37021674 | 0.78104287 |
| SP1 | 833.847032 | -0.1105315 | 0.20499768 | -0.539184 | 5.90E-01 | 0.88621335 |
| PRR13 | 543.094672 | 0.84449639 | 0.22497326 | 3.75376345 | 0.0001742 | 0.02140101 |
| PCBP2 | 1764.49055 | -0.1806867 | 0.2467837 | -0.7321661 | 0.46406721 | 0.83547278 |
| MAP3K12 | 165.028136 | 0.46312772 | 0.38987479 | 1.18788834 | 0.2348774 | 0.68624102 |
| TARBP2 | 55.7940574 | -0.1012178 | 0.41794344 | -0.2421806 | 0.80864025 | 0.95610623 |
| ATP5MC2 | 871.060475 | -0.0085722 | 0.23578374 | -0.036356 | 0.9709985 | 0.99317943 |
| CALCOCO1 | 489.69665 | -0.0415773 | 0.26167514 | -0.1588891 | 0.87375625 | 0.97564728 |
| HOXC4 | 395.998186 | -0.1305273 | 0.36119333 | -0.3613781 | 0.71781685 | 0.92945725 |
| AC023794.1 | 213.122366 | -0.5922582 | 0.33517564 | -1.7670083 | 0.07722683 | 0.47712313 |
| SMUG1 | 225.394689 | 0.14810862 | 0.22938792 | 0.64566882 | 0.51849386 | 0.85916831 |
| LINC02381 | 730.118922 | 0.31919743 | 0.2133103 | 1.49639951 | 0.13454958 | 0.57708028 |
| CBX5 | 804.049898 | 0.21021527 | 0.1694842 | 1.24032368 | 0.2148557 | 0.66855804 |
| AC078778.1 | 80.2402295 | 0.38436991 | 0.3904458 | 0.98443859 | 0.32489991 | 0.75478872 |
| HNRNPA1 | 552.820864 | 0.14514883 | 0.30331425 | 0.47854274 | 0.63226396 | 0.90044163 |
| COPZ1 | 441.411585 | -0.064015 | 0.16952456 | -0.3776147 | 0.70571687 | 0.92556842 |
| PPP1R1A | 1004.36797 | 0.97710971 | 0.50313251 | 1.9420524 | 0.05213076 | 0.40866415 |
| AC079310.1 | 173.779492 | 0.78702252 | 1.72794666 | 0.45546691 | 0.64877337 | 0.90754509 |
| CD63 | 812.286479 | 0.15300896 | 0.38453008 | 0.39791155 | 6.91E-01 | 0.92192759 |
| GDF11 | 55.8218866 | -0.2493743 | 0.55798274 | -0.4469212 | 0.65493199 | 0.90902447 |
| ORMDL2 | 118.852261 | -0.0157514 | 0.43170751 | -0.0364862 | 0.97089468 | 0.99317943 |
| PYM1 | 127.748014 | -0.4091221 | 0.19793268 | -2.0669758 | 0.03873643 | 0.36916996 |
| DGKA | 65.3604719 | 0.50216874 | 0.38566904 | 1.30207169 | 0.19289188 | 0.64699503 |
| CDK2 | 61.4796089 | -0.1270429 | 0.2784496 | -0.4562508 | 0.64820964 | 0.90739732 |
| RAB5B | 474.306454 | 0.06642226 | 0.20142014 | 0.3297697 | 0.74157399 | 0.93598004 |
| SUOX | 161.19456 | 0.6049686 | 0.32063182 | 1.88680152 | 0.05918703 | 0.42420045 |
| IKZF4 | 176.034171 | 0.24075395 | 0.25703106 | 0.93667261 | 0.34892699 | 0.76982416 |
| RPS26 | 184.277742 | 0.20831843 | 0.39988709 | 0.52094313 | 0.60240639 | 0.88971776 |

| | | | | | | |
|---|---|---|---|---|---|---|
| ERBB3 | 289.669514 | -0.3133681 | 0.31498299 | -0.9948731 | 0.31979799 | 0.75094569 |
| PA2G4 | 184.705334 | 0.33021046 | 0.36742888 | 0.8987058 | 0.36880939 | 0.78095617 |
| RPL41 | 2352.01135 | 0.70291427 | 0.40524697 | 1.73453309 | 0.08282354 | 0.48601338 |
| ESYT1 | 185.458256 | -0.3871615 | 0.2535215 | -1.5271346 | 0.12672755 | 0.56735041 |
| AC034102.5 | 45.5920353 | 0.01564265 | 0.55684262 | 0.02809168 | 0.97758903 | 0.99455895 |
| MYL6B | 143.269724 | -0.0748237 | 0.33225681 | -0.2251984 | 0.82182494 | 0.96066244 |
| MYL6 | 1321.4919 | 0.14374126 | 0.3988858 | 0.36035691 | 0.71858025 | 0.9298398 |
| SMARCC2 | 957.629878 | -0.3301714 | 0.18849442 | -1.7516242 | 0.07983845 | 0.48228454 |
| RNF41 | 207.053417 | 0.16417487 | 0.27040522 | 0.60714385 | 0.54375545 | 0.86639505 |
| NABP2 | 68.7478802 | 0.19054293 | 0.29540527 | 0.6450221 | 0.51891286 | 0.85918126 |
| ANKRD52 | 272.622887 | -0.0734922 | 0.15703607 | -0.4679957 | 0.6397877 | 0.90330608 |
| COQ10A | 123.472306 | 0.67674919 | 0.30933706 | 2.18774041 | 0.02868852 | 0.32702837 |
| CS | 406.063147 | -0.03255 | 0.24529466 | -0.1326976 | 0.89443254 | 0.97788747 |
| AC073896.1 | 105.17047 | 0.10712378 | 0.34723357 | 0.3085064 | 0.75769703 | 0.94056395 |
| AC073896.2 | 146.904806 | 0.17157639 | 0.33067002 | 0.51887497 | 0.60384794 | 0.89001883 |
| PAN2 | 397.428728 | -0.1805731 | 0.20282588 | -0.8902864 | 3.73E-01 | 0.78277446 |
| STAT2 | 602.961562 | -0.3993797 | 0.32852414 | -1.2156783 | 0.2241075 | 0.67678006 |
| TIMELESS | 83.5479954 | -0.659568 | 0.31597968 | -2.0873746 | 0.03685428 | 0.36038369 |
| SPRYD4 | 82.9395293 | 0.30810206 | 0.300515 | 1.02524685 | 0.30524673 | 0.74229685 |
| GLS2 | 69.7048283 | 0.23476159 | 0.32126567 | 0.73073973 | 0.46493814 | 0.83635679 |
| RBMS2 | 824.774027 | -0.1210592 | 0.18708791 | -0.647071 | 0.51758597 | 0.85893097 |
| BAZ2A | 686.265051 | -0.0355931 | 0.20073435 | -0.1773146 | 0.85926126 | 0.97174503 |
| ATP5F1B | 1212.8345 | 1.13838005 | 0.36250265 | 3.14033586 | 0.00168754 | 0.07713062 |
| PTGES3 | 1387.3772 | 0.13615362 | 0.1504663 | 0.90487783 | 0.36553012 | 0.78095617 |
| NACA | 589.307388 | 0.55374321 | 0.35776796 | 1.54777194 | 0.12167721 | 0.55692668 |
| PRIM1 | 52.0080213 | -0.2743611 | 0.33189354 | -0.826654 | 0.40843319 | 0.80588056 |
| NEMP1 | 145.251263 | -0.5453817 | 0.25080191 | -2.1745516 | 0.02966373 | 0.33121574 |
| NAB2 | 95.2636522 | -0.5033041 | 0.68519941 | -0.7345367 | 0.46262167 | 0.83531269 |
| STAT6 | 791.016568 | -0.2687345 | 0.21923692 | -1.225772 | 2.20E-01 | 6.73E-01 |
| LRP1 | 96.9161886 | 0.49511014 | 0.3471211 | 1.42633261 | 0.15377235 | 0.60285224 |
| SHMT2 | 65.0284477 | 1.00250685 | 0.53773731 | 1.86430591 | 0.06227876 | 0.43419765 |
| R3HDM2 | 2715.19171 | -0.1354498 | 0.15375702 | -0.880934 | 0.37835356 | 0.78691107 |
| MARS | 270.325171 | -0.5242343 | 0.23464108 | -2.2341964 | 0.02547015 | 0.31163914 |
| DDIT3 | 349.633122 | 0.1929145 | 0.7056396 | 0.27338956 | 0.78455378 | 0.94978829 |
| MBD6 | 271.692758 | 0.02459296 | 0.26513635 | 0.09275591 | 0.92609748 | 0.98234748 |
| DCTN2 | 493.782684 | -0.372721 | 0.15348659 | -2.4283619 | 0.0151672 | 0.25080486 |
| PIP4K2C | 312.714338 | 0.59931346 | 0.36506331 | 1.64166993 | 0.10065843 | 0.51815945 |
| DTX3 | 155.591487 | -0.0911507 | 0.3757772 | -0.2425658 | 0.80834176 | 0.95610623 |
| OS9 | 482.682293 | 0.18447073 | 0.24156245 | 0.7636565 | 0.44507197 | 0.82546414 |
| AC025165.2 | 62.6258679 | -0.1391418 | 0.55265569 | -0.2517694 | 0.80121929 | 0.95370871 |
| TSPAN31 | 163.755125 | 0.06929633 | 0.281786 | 0.2459183 | 0.80574548 | 0.95522565 |
| CDK4 | 127.89779 | -0.1697016 | 0.2269643 | -0.7477016 | 0.45464016 | 0.83121393 |
| 9-Mar | 185.981281 | 0.24873288 | 0.25152128 | 0.98891385 | 0.32270529 | 0.7532337 |
| TSFM | 242.692937 | 0.65066396 | 0.31683357 | 2.05364591 | 0.04000998 | 0.37342745 |
| AVIL | 723.514706 | 0.41307716 | 0.42583966 | 0.97002981 | 0.33203163 | 0.76037513 |
| CTDSP2 | 668.745749 | 0.09594344 | 0.19241326 | 0.49863218 | 6.18E-01 | 0.89599505 |
| AC084033.3 | 43.9689028 | -0.0548324 | 0.3929955 | -0.1395242 | 0.88903595 | 0.97729111 |
| ATP23 | 54.0229352 | 0.10740407 | 0.397031 | 0.2705181 | 0.7867617 | 0.94978829 |
| LRIG3 | 461.426991 | -0.2646727 | 0.33713029 | -0.7850756 | 0.43240927 | 0.81773248 |
| AC068305.2 | 92.755536 | -0.5744333 | 0.54128232 | -1.0612453 | 0.28857842 | 0.73163767 |
| SLC16A7 | 4725.03118 | -0.8780456 | 0.40092815 | -2.1900324 | 0.02852189 | 0.32600107 |
| FAM19A2 | 815.96968 | -0.0751833 | 0.49124859 | -0.1530453 | 0.87836259 | 0.9765717 |
| USP15 | 2070.48242 | -0.2136925 | 0.2200064 | -0.9713012 | 0.33139832 | 0.76000762 |
| MON2 | 3219.92224 | -0.1735929 | 0.16747686 | -1.0365187 | 0.29996024 | 0.73673761 |

| | | | | | | |
|---|---|---|---|---|---|---|
| AC048341.1 | 1120.59062 | -0.94964 | 0.4137482 | -2.2952124 | 0.02172095 | 0.29426879 |
| PPM1H | 2629.81194 | -0.5097858 | 0.22109029 | -2.3057811 | 0.02112287 | 0.29063746 |
| DPY19L2 | 671.904921 | 0.87800033 | 0.46004772 | 1.90849838 | 0.05632683 | 0.41843207 |
| RXYLT1 | 218.75083 | -0.1791292 | 0.35521542 | -0.5042834 | 0.61406226 | 0.89477643 |
| SRGAP1 | 2138.98595 | -0.8894234 | 0.38393617 | -2.3165918 | 0.02052598 | 0.28691085 |
| AC020611.2 | 106.37042 | -0.3418822 | 0.57283204 | -0.596828 | 0.55062224 | 0.86989226 |
| C12orf66 | 87.4866834 | -0.4001632 | 0.25165688 | -1.5901141 | 0.11180908 | 0.53607455 |
| C12orf56 | 44.6171269 | -0.2986197 | 0.42666373 | -0.6998948 | 0.483993 | 0.8445691 |
| XPOT | 251.533433 | -0.3509514 | 0.23305164 | -1.5058953 | 0.13209404 | 0.57606398 |
| TBK1 | 916.80224 | -0.2910124 | 0.22508032 | -1.292927 | 0.1960363 | 0.64964978 |
| RASSF3 | 174.216873 | -0.1425028 | 0.24518764 | -0.5811991 | 0.56110628 | 0.87394741 |
| GNS | 849.045618 | 0.08822288 | 0.23436969 | 0.37642617 | 0.70660009 | 0.92556842 |
| AC025262.1 | 40.6205564 | 0.74505863 | 0.51749505 | 1.43974058 | 0.14994081 | 0.59663305 |
| TBC1D30 | 582.476712 | 0.07531968 | 0.25337627 | 0.29726413 | 0.76626486 | 0.94300088 |
| LINC02389 | 309.206997 | -0.2239582 | 0.36164766 | -0.6192717 | 0.53573736 | 0.8642718 |
| LINC02231 | 41.5597454 | -0.132208 | 0.54036943 | -0.2446622 | 0.80671803 | 0.95566275 |
| LEMD3 | 1043.52476 | 0.03632743 | 0.23092518 | 0.15731254 | 0.87499854 | 0.97592296 |
| MSRB3 | 71.7675468 | 1.20167292 | 0.47497103 | 2.52999202 | 0.01140651 | 0.21832654 |
| HMGA2 | 394.811637 | -2.2674686 | 0.78491777 | -2.8887976 | 0.00386718 | 0.12104636 |
| AC090673.1 | 143.499826 | -0.1025577 | 0.61490798 | -0.1667855 | 0.86753886 | 0.97402505 |
| LLPH | 90.8982263 | 0.31218983 | 0.35697923 | 0.8745322 | 0.38182849 | 0.78919376 |
| HELB | 491.484233 | 0.33096276 | 0.29397202 | 1.12583081 | 0.26023714 | 0.70992821 |
| GRIP1 | 4984.28122 | -0.4953464 | 0.49687009 | -0.9969334 | 3.19E-01 | 0.75094569 |
| CAND1 | 1036.63494 | -0.0423631 | 0.16621677 | -0.2548663 | 0.79882634 | 0.95300733 |
| DYRK2 | 138.227337 | 0.1871902 | 0.23316153 | 0.80283485 | 0.4220702 | 0.81270047 |
| MDM1 | 167.041867 | -0.2790997 | 0.31313483 | -0.8913085 | 0.37276371 | 0.78269743 |
| RAP1B | 1700.90194 | -0.1654804 | 0.18453401 | -0.8967476 | 0.3698536 | 0.78095617 |
| NUP107 | 528.099527 | -0.524799 | 0.22568107 | -2.325401 | 0.02005053 | 0.28386651 |
| SLC35E3 | 505.030945 | -0.6489403 | 0.36404413 | -1.7825868 | 0.07465359 | 0.47190267 |
| MDM2 | 1244.22657 | 0.35359818 | 0.29998385 | 1.17872404 | 0.23850808 | 0.68982992 |
| CPM | 3705.29496 | -0.3639239 | 0.40527241 | -0.8979736 | 0.36919963 | 0.78095617 |
| CPSF6 | 1511.69618 | 0.00345227 | 0.20195991 | 0.01709382 | 0.98636177 | 0.99624198 |
| YEATS4 | 127.586337 | -0.0282135 | 0.21274865 | -0.1326143 | 0.8944984 | 0.97788747 |
| FRS2 | 1420.7087 | -0.0909101 | 0.1745464 | -0.5208362 | 0.60248085 | 0.88971776 |
| CCT2 | 396.40864 | 0.19848882 | 0.31187848 | 0.63642998 | 0.5244962 | 0.86009564 |
| RAB3IP | 1378.09638 | 0.05543052 | 0.23329762 | 0.23759576 | 0.81219464 | 0.95806331 |
| AC025263.2 | 54.1948847 | 0.84653927 | 0.70530548 | 1.20024485 | 0.23004426 | 0.68234886 |
| MYRFL | 618.191064 | 0.50430466 | 0.638802 | 0.7894538 | 0.42984682 | 0.81671932 |
| AC025159.1 | 967.711793 | 0.38333361 | 0.61282015 | 0.62552383 | 0.53162731 | 0.86397197 |
| CNOT2 | 1992.67407 | -0.1192526 | 0.21574773 | -0.552741 | 0.58044075 | 0.88258799 |
| TSPAN8 | 883.459666 | -0.3517057 | 0.32527755 | -1.081248 | 0.27958682 | 0.72492327 |
| LGR5 | 190.279988 | -5.9274244 | 1.17734624 | -5.0345635 | 4.79E-07 | 0.00076973 |
| ZFC3H1 | 1416.1555 | -0.1694855 | 0.17410857 | -0.9734473 | 0.33033106 | 0.75894644 |
| THAP2 | 77.3346538 | 0.54559687 | 0.45879894 | 1.18918513 | 0.23436682 | 0.68550739 |
| TMEM19 | 83.1940973 | 0.12711213 | 0.38698184 | 0.32847053 | 0.74255592 | 0.93598004 |
| RAB21 | 1174.39604 | 0.12420768 | 0.24804232 | 0.50075196 | 0.6165457 | 0.89589954 |
| TBC1D15 | 1357.17144 | -0.0521731 | 0.29700607 | -0.1756635 | 0.86055833 | 0.97204678 |
| TRHDE | 3268.00988 | 0.33975942 | 0.30115248 | 1.12819732 | 0.25923659 | 0.70921184 |
| ATXN7L3B | 314.23231 | 0.47379852 | 0.27790252 | 1.70490902 | 0.08821139 | 0.49730444 |
| CAPS2 | 1316.13548 | 0.39355141 | 0.24612202 | 1.59900934 | 1.10E-01 | 0.53372881 |
| GLIPR1L1 | 124.398638 | 0.47970244 | 0.64713547 | 0.74127051 | 0.45852944 | 0.83322539 |
| GLIPR1L2 | 122.158773 | 0.18139028 | 0.74829106 | 0.24240605 | 0.80846555 | 0.95610623 |
| KRR1 | 343.901496 | -0.1715811 | 0.26008138 | -0.6597209 | 0.50943295 | 0.8568242 |
| NAP1L1 | 1163.46466 | 0.28995315 | 0.18501414 | 1.56719452 | 0.11706923 | 0.54756317 |

| | | | | | | |
|---|---|---|---|---|---|---|
| BBS10 | 59.3796015 | 0.03964242 | 0.30302823 | 0.1308209 | 0.89591699 | 0.97788747 |
| OSBPL8 | 1683.29927 | -0.3348743 | 0.19841456 | -1.6877508 | 0.09145908 | 0.50441876 |
| AC107032.2 | 182.500157 | -1.2003379 | 0.5667196 | -2.1180455 | 0.03417121 | 0.35153234 |
| ZDHHC17 | 1321.3381 | -0.3712988 | 0.21429055 | -1.7326884 | 0.08315106 | 0.48764454 |
| CSRP2 | 351.434418 | 0.56626333 | 0.57621425 | 0.98273051 | 0.32574009 | 0.75550719 |
| NAV3 | 57.8179449 | -0.2403311 | 0.43663752 | -0.5504134 | 0.58203587 | 0.88336948 |
| SYT1 | 442.0408 | -0.7169754 | 0.51070701 | -1.4038879 | 0.16035223 | 0.61151309 |
| AC090709.1 | 682.118026 | -0.3008576 | 0.55737029 | -0.5397805 | 5.89E-01 | 8.86E-01 |
| AC027288.3 | 451.310694 | 0.24959671 | 0.61280725 | 0.40730052 | 0.68378728 | 0.9200778 |
| PAWR | 3463.23383 | -0.1899734 | 0.26992156 | -0.7038097 | 0.48155126 | 0.8437862 |
| AC073569.2 | 283.919197 | -0.167844 | 0.433523 | -0.3871627 | 0.69863573 | 0.92356678 |
| PPP1R12A | 2368.62141 | 0.01560272 | 0.14328645 | 0.10889178 | 0.91328833 | 0.98145965 |
| LIN7A | 375.352874 | 1.4791467 | 0.41183583 | 3.59159304 | 0.00032866 | 0.03094754 |
| ACSS3 | 2276.67006 | 0.40216018 | 0.26792891 | 1.5009958 | 0.13335665 | 0.57682553 |
| AC078955.1 | 50.99925 | 1.51664572 | 0.87061045 | 1.74204861 | 0.08149994 | 0.48594425 |
| PPFIA2 | 1146.89966 | 0.41783114 | 0.48658412 | 0.85870278 | 0.39050451 | 0.79701353 |
| CCDC59 | 191.572484 | 0.03257555 | 0.23456276 | 0.13887776 | 0.88954674 | 0.97729111 |
| METTL25 | 1070.04639 | 0.05615691 | 0.21986992 | 0.25540969 | 0.79840668 | 0.95297658 |
| TMTC2 | 3130.68096 | -0.1056328 | 0.20078139 | -0.5261087 | 0.59881271 | 0.88883692 |
| LRRIQ1 | 527.735199 | -0.1175314 | 0.42926087 | -0.2737995 | 0.78423869 | 0.94978829 |
| RASSF9 | 123.652183 | -0.0542978 | 0.62550164 | -0.0868069 | 0.93082502 | 0.98331361 |
| MGAT4C | 89.038883 | -0.4529053 | 0.7241847 | -0.6254003 | 0.53170835 | 0.86397197 |
| C12orf29 | 117.191479 | 0.32436798 | 0.40693273 | 0.79710466 | 0.42539025 | 0.81466629 |
| CEP290 | 1121.3905 | -0.3222885 | 0.16867101 | -1.9107519 | 0.05603647 | 0.41843207 |
| TMTC3 | 778.012996 | 0.21124591 | 0.16545084 | 1.2767896 | 0.20167654 | 0.65534788 |
| KITLG | 946.768412 | 0.22200796 | 0.46888585 | 0.47347977 | 0.63587093 | 0.90207488 |
| AC024941.2 | 58.5496545 | 0.63246341 | 0.67651246 | 0.93488804 | 0.349846 | 0.77041926 |
| AC024909.1 | 69.5091016 | 0.62595763 | 0.62414303 | 1.00290735 | 0.31590557 | 0.74785672 |
| DUSP6 | 235.069336 | 0.60387508 | 0.39698958 | 1.52113585 | 0.12822575 | 0.56762585 |
| POC1B | 741.096417 | -0.1586311 | 0.14852819 | -1.06802 | 0.28551149 | 0.72846488 |
| POC1B-AS1 | 337.959209 | -0.6157208 | 0.23432199 | -2.6276699 | 0.00859719 | 0.18770295 |
| AC025034.1 | 166.963857 | -0.7643655 | 0.30396388 | -2.5146589 | 0.01191476 | 0.22355405 |
| ATP2B1 | 1274.77644 | 0.258353 | 0.28119427 | 0.9187705 | 0.35821563 | 0.77529959 |
| ATP2B1-AS1 | 36.9497496 | -0.0635316 | 0.49028493 | -0.1295809 | 0.896898 | 0.97823597 |
| BTG1 | 1059.88392 | -0.0282505 | 0.35382623 | -0.079843 | 9.36E-01 | 0.98621411 |
| AC025164.1 | 165.877717 | -1.2944601 | 0.45774837 | -2.8278859 | 0.00468565 | 0.13513704 |
| LINC02391 | 118.558671 | 0.89708126 | 0.53716842 | 1.67001862 | 0.09491568 | 0.51085035 |
| PLEKHG7 | 212.844507 | -0.2121745 | 0.73740611 | -0.2877309 | 0.77355272 | 0.94558571 |
| EEA1 | 2350.74997 | -0.3706064 | 0.16625304 | -2.2291704 | 0.02580257 | 0.31222606 |
| LINC02413 | 72.9434008 | 0.13439003 | 0.57660283 | 0.23307209 | 8.16E-01 | 9.59E-01 |
| AC138123.1 | 3384.23489 | -0.465275 | 0.40025643 | -1.1624423 | 2.45E-01 | 6.95E-01 |
| AC138123.2 | 129.79482 | 0.07896264 | 0.54887186 | 0.14386352 | 8.86E-01 | 0.97664137 |
| AC124947.1 | 756.305342 | -1.0791662 | 0.35458364 | -3.0434744 | 0.00233863 | 0.09248804 |
| NUDT4 | 2991.86162 | -0.1854209 | 0.49502087 | -0.3745719 | 0.70797888 | 0.92577218 |
| UBE2N | 631.673952 | -0.0211199 | 0.24712439 | -0.0854625 | 0.93189373 | 0.98357046 |
| MRPL42 | 579.086661 | -0.0633546 | 0.16868029 | -0.3755901 | 0.70722164 | 0.92573766 |
| SOCS2 | 272.729445 | -0.8502249 | 0.69032952 | -1.2316218 | 0.21809041 | 0.67150495 |
| CRADD | 5188.0218 | -0.2604314 | 0.3468828 | -0.7507764 | 0.45278721 | 0.82963882 |
| AC012085.2 | 493.807075 | -0.1831231 | 0.81569134 | -0.2245005 | 0.8223679 | 0.96083745 |
| AC012464.1 | 47.342055 | 0.24932089 | 0.75671666 | 0.32947721 | 0.74179502 | 0.93598004 |
| PLXNC1 | 72.6225486 | 0.1280564 | 0.29715412 | 0.4309427 | 0.66651003 | 0.91318863 |
| CEP83 | 701.915443 | -0.2358601 | 0.16337628 | -1.4436617 | 0.14883416 | 0.59502131 |
| TMCC3 | 118.065004 | 0.36781675 | 0.6593862 | 0.55781687 | 0.57696944 | 0.88150331 |
| NDUFA12 | 432.104924 | 0.23081844 | 0.3283819 | 0.70289634 | 0.48212035 | 0.84389635 |

| | | | | | | |
|---|---|---|---|---|---|---|
| NR2C1 | 850.86253 | -0.4176852 | 0.17011522 | -2.4553076 | 0.01407641 | 0.24627187 |
| FGD6 | 498.397923 | -0.528832 | 0.27130769 | -1.9491967 | 0.05127194 | 0.40578257 |
| VEZT | 1677.39294 | 0.01037691 | 0.16073341 | 0.06455978 | 0.94852451 | 0.98919519 |
| METAP2 | 733.98003 | 0.08585684 | 0.11289484 | 0.76050279 | 0.4469541 | 0.82571251 |
| USP44 | 221.806514 | -0.1738516 | 0.42116106 | -0.4127912 | 0.67975959 | 0.9182972 |
| NTN4 | 3046.46605 | -0.4304296 | 0.43716475 | -0.9845936 | 0.32482376 | 0.75478872 |
| AC090001.1 | 75.7380032 | -1.480875 | 0.66190129 | -2.2373048 | 0.02526642 | 0.31074878 |
| SNRPF | 166.988974 | -0.2621803 | 0.27890187 | -0.940045 | 0.34719446 | 0.76893725 |
| LTA4H | 371.09778 | 0.25018815 | 0.19305997 | 1.29590901 | 0.19500683 | 0.64850083 |
| ELK3 | 393.880735 | -0.8021241 | 0.44836994 | -1.7889782 | 7.36E-02 | 4.70E-01 |
| CDK17 | 922.902442 | -0.1579413 | 0.17038634 | -0.9269596 | 0.35394749 | 0.77243419 |
| CFAP54 | 220.831119 | 0.99916087 | 0.68519752 | 1.45820853 | 0.14478307 | 0.59044137 |
| NEDD1 | 336.193997 | -0.1264832 | 0.19691203 | -0.6423336 | 0.52065661 | 0.85943874 |
| TMPO | 647.663135 | -0.14244 | 0.33783736 | -0.421623 | 0.67330021 | 0.91636338 |
| SLC25A3 | 1058.61616 | 0.90411504 | 0.35736875 | 2.52992194 | 0.01140879 | 0.21832654 |
| IKBIP | 126.470748 | 0.13576783 | 0.43869137 | 0.3094837 | 0.75695361 | 0.94015987 |
| APAF1 | 375.612339 | -0.0205262 | 0.21811657 | -0.0941066 | 0.92502447 | 0.9822216 |
| ANKS1B | 322.841421 | 0.98774121 | 0.49890246 | 1.9798283 | 0.04772283 | 0.39501762 |
| UHRF1BP1L | 625.628704 | -0.0938695 | 0.22022785 | -0.426238 | 0.66993446 | 0.91465659 |
| ACTR6 | 312.915199 | 0.19931602 | 0.20790861 | 0.95867129 | 0.33772436 | 0.76409325 |
| DEPDC4 | 124.411148 | -0.1600874 | 0.30438017 | -0.5259456 | 0.59892603 | 0.88883692 |
| SCYL2 | 1121.96508 | -0.2569564 | 0.22522244 | -1.1409005 | 0.25391133 | 0.70431689 |
| NR1H4 | 2069.9682 | 0.21659132 | 0.39170273 | 0.55294819 | 0.58029889 | 0.88258799 |
| GAS2L3 | 156.28882 | -0.9356779 | 0.48022431 | -1.9484184 | 5.14E-02 | 0.40627499 |
| SLC5A8 | 92.4478315 | -0.6896515 | 1.05326492 | -0.6547749 | 5.13E-01 | 0.85762435 |
| UTP20 | 407.491478 | -0.4747108 | 0.2157549 | -2.2002318 | 0.02779045 | 0.32172135 |
| ARL1 | 310.133179 | 0.3814505 | 0.20267464 | 1.88208304 | 0.05982474 | 0.42668737 |
| CHPT1 | 704.422847 | 0.81985216 | 0.33752553 | 2.42900787 | 0.0151402 | 0.25080486 |
| SYCP3 | 43.95146 | 1.72189138 | 0.58169676 | 2.96011856 | 0.00307521 | 0.1068958 |
| GNPTAB | 734.213467 | 0.14848773 | 0.30618603 | 0.48495918 | 0.6277053 | 0.89919293 |
| DRAM1 | 303.073554 | 0.23585954 | 0.41213086 | 0.57229285 | 5.67E-01 | 8.77E-01 |
| WASHC3 | 612.435536 | -0.3910391 | 0.18220818 | -2.1461115 | 0.03186408 | 0.34024781 |
| NUP37 | 200.065639 | 0.07742263 | 0.23406229 | 0.33077788 | 0.74081227 | 0.93595306 |
| PARPBP | 58.9239828 | -0.622508 | 0.57733824 | -1.0782379 | 0.2809276 | 0.72575254 |
| NT5DC3 | 131.945676 | -0.6779146 | 0.35156008 | -1.9283037 | 0.05381736 | 0.41353897 |
| HSP90B1 | 1223.70442 | 0.9044153 | 0.34698528 | 2.60649474 | 0.00914742 | 0.19425773 |
| C12orf73 | 141.753131 | 0.19653792 | 0.2968595 | 0.66205705 | 0.50793466 | 0.85643442 |
| TDG | 472.22714 | -0.1070647 | 0.17117214 | -0.6254797 | 0.53165624 | 0.86397197 |
| HCFC2 | 336.706951 | -0.2627235 | 0.19515426 | -1.3462352 | 0.17822668 | 0.63216868 |
| NFYB | 263.856134 | -0.2766065 | 0.25097428 | -1.1021308 | 0.27040479 | 0.71953603 |
| TXNRD1 | 2190.84698 | 0.11936651 | 0.25757125 | 0.46343105 | 0.64305543 | 0.90478945 |
| CHST11 | 273.405935 | -0.2461259 | 0.59802652 | -0.4115635 | 6.81E-01 | 9.19E-01 |
| SLC41A2 | 814.278643 | 0.0180474 | 0.33564982 | 0.05376855 | 0.95711957 | 0.99065996 |
| C12orf45 | 173.921008 | -0.5557489 | 0.20639412 | -2.6926588 | 0.00708848 | 0.1727522 |
| WASHC4 | 1647.39364 | -0.1468732 | 0.21504259 | -0.6829957 | 0.49460956 | 0.84882719 |
| APPL2 | 851.910325 | -0.254664 | 0.20479896 | -1.2434828 | 0.21368996 | 0.66800831 |
| C12orf75 | 446.987671 | 0.77657541 | 0.48890691 | 1.58839116 | 0.11219792 | 0.53638156 |
| NUAK1 | 383.724836 | -0.2667271 | 0.3831322 | -0.696175 | 0.48631922 | 0.8447301 |
| CKAP4 | 146.926609 | 0.58695479 | 0.32839958 | 1.78731895 | 0.07388595 | 0.47056873 |
| TCP11L2 | 647.844679 | -0.073432 | 0.26086592 | -0.2814932 | 0.77833217 | 0.94694572 |
| POLR3B | 459.107823 | -0.3093787 | 0.1865933 | -1.6580377 | 9.73E-02 | 0.51355946 |
| AC079385.1 | 49.8260646 | -0.2396388 | 0.35767742 | -0.6699857 | 0.50286688 | 0.85525037 |
| RIC8B | 1134.90946 | 0.05643636 | 0.19335381 | 0.29188131 | 0.77037737 | 0.94404998 |
| AC007541.1 | 46.2672131 | 0.74587143 | 0.56199119 | 1.32719416 | 0.18444446 | 0.63929879 |

| | | | | | | |
|---|---|---|---|---|---|---|
| TMEM263 | 771.161005 | 0.15037216 | 0.21352686 | 0.70423064 | 0.48128915 | 0.8437862 |
| MTERF2 | 214.185673 | 0.46595066 | 0.30715114 | 1.51700777 | 0.12926472 | 0.57010275 |
| CRY1 | 862.21031 | -0.3930056 | 0.63084634 | -0.6229814 | 0.53329673 | 0.86397197 |
| BTBD11 | 1556.43044 | -0.5719098 | 0.53128468 | -1.0764659 | 0.28171895 | 0.72604837 |
| PWP1 | 309.939364 | -0.0803148 | 0.17631131 | -0.4555284 | 0.64872916 | 0.90754509 |
| PRDM4 | 380.356514 | -0.177228 | 0.23489539 | -0.7544974 | 0.4505506 | 0.82783737 |
| FICD | 41.6707862 | 0.10198887 | 0.4880653 | 0.20896562 | 0.83447508 | 0.96510927 |
| SART3 | 350.596307 | 0.12816379 | 0.20466346 | 0.62621723 | 0.53117247 | 0.86397197 |
| ISCU | 607.307938 | 0.85596484 | 0.38708051 | 2.21133542 | 0.02701262 | 0.3178271 |
| CORO1C | 368.041541 | -0.0343859 | 0.34162318 | -0.1006545 | 9.20E-01 | 9.82E-01 |
| SSH1 | 1230.79924 | -0.0479503 | 0.32652522 | -0.1468501 | 0.88325034 | 0.97664137 |
| USP30 | 210.394531 | 0.12414291 | 0.26468038 | 0.46902951 | 0.63904854 | 0.90330608 |
| ALKBH2 | 51.3939906 | -0.5429942 | 0.3946292 | -1.3759604 | 0.16883389 | 0.62265166 |
| UNG | 154.259022 | -0.1096934 | 0.34374882 | -0.3191093 | 0.74964367 | 0.93775604 |
| ACACB | 3947.00808 | 0.15977277 | 0.50866452 | 0.31410244 | 0.75344323 | 0.93844712 |
| KCTD10 | 187.880445 | 0.42442155 | 0.22385667 | 1.89595218 | 0.05796637 | 0.42087381 |
| UBE3B | 440.119898 | -0.0277083 | 0.27877071 | -0.0993947 | 0.92082486 | 0.98215577 |
| MMAB | 418.641479 | -0.3308858 | 0.24704898 | -1.3393528 | 0.18045584 | 0.63479125 |
| MVK | 388.322529 | -0.4038149 | 0.33236445 | -1.2149761 | 0.2243752 | 0.67681479 |
| FAM222A | 252.06765 | -0.1140961 | 0.38120212 | -0.2993061 | 0.76470651 | 0.94256992 |
| FAM222A-AS | 104.544592 | -0.5026286 | 0.40729691 | -1.2340594 | 0.21718079 | 0.66978787 |
| TRPV4 | 152.695358 | -0.9211567 | 0.46531205 | -1.9796537 | 0.04774245 | 0.39501762 |
| GLTP | 621.113672 | -0.0573517 | 0.30202711 | -0.1898893 | 0.84939589 | 0.970224 |
| TCHP | 464.213007 | -0.2272707 | 0.30986879 | -0.7334416 | 0.46328912 | 0.83531269 |
| GIT2 | 757.291494 | -0.2546009 | 0.18303945 | -1.3909616 | 0.16423706 | 0.61595892 |
| ANKRD13A | 1093.11411 | 0.26303067 | 0.30629673 | 0.85874465 | 0.39048141 | 0.79701353 |
| C12orf76 | 203.368637 | 0.19447645 | 0.26251662 | 0.74081575 | 0.45880516 | 0.83340407 |
| IFT81 | 764.956638 | -0.2149875 | 0.19802256 | -1.0856716 | 0.27762431 | 0.72344438 |
| ATP2A2 | 2426.13029 | -0.1256322 | 0.39831325 | -0.3154105 | 0.75245 | 0.93818313 |
| ANAPC7 | 464.546313 | -0.5774828 | 0.18380473 | -3.141828 | 0.00167897 | 0.0770051 |
| ARPC3 | 954.920416 | 0.17525244 | 0.25291003 | 0.69294379 | 4.88E-01 | 8.46E-01 |
| GPN3 | 116.005601 | 0.29688534 | 0.28115928 | 1.0559329 | 0.29099887 | 0.73313067 |
| FAM216A | 68.0027341 | -0.1630384 | 0.49133933 | -0.3318245 | 0.74002179 | 0.93552616 |
| VPS29 | 597.721235 | 0.07954005 | 0.20955551 | 0.37956552 | 0.70426796 | 0.92528913 |
| PPTC7 | 787.423049 | 0.06283088 | 0.24805867 | 0.2532904 | 0.80004382 | 0.95370871 |
| TCTN1 | 417.521524 | 0.39672588 | 0.24153569 | 1.64251455 | 0.10048341 | 0.5174602 |
| HVCN1 | 125.990931 | 0.49723117 | 0.6860582 | 0.72476528 | 0.46859604 | 0.83825637 |
| PPP1CC | 302.212862 | 0.16774509 | 0.27112215 | 0.6187067 | 0.53610959 | 0.8642718 |
| SH2B3 | 141.297444 | 0.73014408 | 0.33971634 | 2.14927573 | 3.16E-02 | 0.33950635 |
| ATXN2 | 3392.91125 | -0.3495238 | 0.15730241 | -2.2219865 | 0.02628422 | 0.31391341 |
| BRAP | 400.935971 | -0.1326939 | 0.19601934 | -0.6769429 | 0.49844216 | 0.85092921 |
| ACAD10 | 509.961843 | -0.1985227 | 0.28776018 | -0.6898895 | 0.49026369 | 0.84614483 |
| ALDH2 | 564.588812 | 0.4265584 | 0.31222242 | 1.36620041 | 0.17187607 | 0.6286116 |
| MAPKAPK5-A | 112.531006 | 0.06730509 | 0.39309753 | 0.17121729 | 0.86405291 | 0.97349667 |
| MAPKAPK5 | 447.595082 | -0.2225708 | 0.1641556 | -1.3558523 | 0.17514616 | 0.62944785 |
| TMEM116 | 1267.24957 | -0.0999268 | 0.37756898 | -0.2646585 | 0.79127254 | 0.95168844 |
| ERP29 | 441.882182 | 0.93224724 | 0.31750961 | 2.93612288 | 0.00332343 | 0.1119876 |
| NAA25 | 816.571119 | -0.4875717 | 0.19586219 | -2.4893611 | 0.01279729 | 0.23219703 |
| TRAFD1 | 253.185632 | 0.03507586 | 0.15498573 | 0.22631673 | 0.82095509 | 0.96058072 |
| HECTD4 | 2815.35181 | -0.1359862 | 0.14320372 | -0.9495999 | 0.3423156 | 0.76591028 |
| RPL6 | 1039.86086 | 0.59576826 | 0.39712755 | 1.50019372 | 0.13356423 | 0.57682553 |
| PTPN11 | 792.693524 | 0.07450652 | 0.22386306 | 0.33282185 | 0.73926877 | 0.93552616 |
| DTX1 | 246.96701 | -0.4224469 | 0.40276092 | -1.0488776 | 2.94E-01 | 0.73527014 |
| RASAL1 | 601.349254 | -1.199942 | 0.42614396 | -2.8158138 | 4.87E-03 | 0.13880537 |

| | | | | | | |
|---|---|---|---|---|---|---|
| DDX54 | 184.695653 | -0.0968472 | 0.35113303 | -0.2758134 | 0.78269137 | 0.94919728 |
| RITA1 | 58.3789133 | 0.01623323 | 0.34602372 | 0.04691363 | 0.96258207 | 0.99224805 |
| IQCD | 99.6132477 | -0.3970154 | 0.53180439 | -0.746544 | 0.45533885 | 0.83121393 |
| TPCN1 | 1707.28093 | -0.4452148 | 0.28587461 | -1.5573778 | 0.11938082 | 0.55194302 |
| SLC8B1 | 531.275206 | -0.3512928 | 0.34179925 | -1.0277752 | 0.30405557 | 0.7414196 |
| PLBD2 | 159.392389 | -0.0084057 | 0.20916935 | -0.040186 | 0.96794481 | 0.99248831 |
| SDSL | 130.359374 | 0.66061863 | 0.46528648 | 1.41981049 | 0.15566286 | 0.60505885 |
| RBM19 | 395.333246 | -0.4033541 | 0.26881334 | -1.5004988 | 0.13348525 | 0.57682553 |
| MED13L | 7553.72104 | 0.21096455 | 0.21118267 | 0.99896718 | 0.31781059 | 0.74990356 |
| MAP1LC3B2 | 47.5598679 | 0.69192318 | 0.41021387 | 1.68673767 | 0.09165381 | 0.50476839 |
| C12orf49 | 386.791267 | 0.25792006 | 0.36296443 | 0.71059321 | 0.47733635 | 0.84174109 |
| RNFT2 | 86.4503126 | -0.4481425 | 0.38225007 | -1.1723804 | 0.24104435 | 0.69178991 |
| FBXW8 | 915.008105 | -0.2645105 | 0.21289981 | -1.2424176 | 0.21408252 | 0.66827012 |
| AC127164.1 | 71.8796562 | -0.1931095 | 0.61912051 | -0.3119094 | 0.75510934 | 0.93945929 |
| TESC | 813.210035 | 0.34034905 | 0.47350327 | 0.71878923 | 0.47227079 | 0.83921324 |
| FBXO21 | 1074.95835 | -0.3879301 | 0.27420887 | -1.4147249 | 0.15714916 | 0.60703814 |
| NOS1 | 752.485918 | 2.2847549 | 0.85637758 | 2.66792936 | 0.00763203 | 0.17527709 |
| KSR2 | 585.42299 | -0.1027113 | 0.31660808 | -0.3244114 | 0.74562662 | 0.93675395 |
| RFC5 | 86.5895769 | -0.4695143 | 0.36980635 | -1.2696221 | 0.20421929 | 0.65729352 |
| WSB2 | 330.860854 | -0.2331355 | 0.23301811 | -1.0005037 | 0.31706679 | 0.74881731 |
| VSIG10 | 740.224038 | -0.4643872 | 0.39168505 | -1.1856138 | 0.23577487 | 0.68810212 |
| PEBP1 | 2362.93858 | 0.96735222 | 0.4724856 | 2.04736868 | 0.0406219 | 0.37344844 |
| TAOK3 | 3892.02729 | 0.23939598 | 0.177438 | 1.34918097 | 0.17727885 | 0.63071358 |
| SUDS3 | 543.982525 | -0.2216059 | 0.17781233 | -1.246291 | 0.21265759 | 0.66656043 |
| HSPB8 | 60.5456686 | 1.43907283 | 0.42836767 | 3.35943379 | 7.81E-04 | 4.91E-02 |
| CCDC60 | 237.237657 | 0.50327741 | 0.54546203 | 0.92266259 | 0.35618307 | 0.77368353 |
| AC002070.1 | 1467.64447 | 0.26600474 | 0.39691625 | 0.6701785 | 0.50274401 | 0.85525037 |
| TMEM233 | 97.2152602 | 0.60768565 | 0.41913304 | 1.44986341 | 1.47E-01 | 0.59373375 |
| PRKAB1 | 952.841393 | -0.1304584 | 0.21256191 | -0.613743 | 0.53938518 | 0.86549306 |
| CIT | 1979.98217 | -0.1271434 | 0.24457332 | -0.519858 | 6.03E-01 | 8.90E-01 |
| BICDL1 | 7748.22127 | -0.9635118 | 0.32347942 | -2.9785877 | 0.0028958 | 0.10254865 |
| RAB35 | 240.703407 | -0.3832449 | 0.19561762 | -1.9591534 | 0.05009483 | 0.40103187 |
| GCN1 | 388.72835 | -0.3110171 | 0.16366723 | -1.9003015 | 0.05739357 | 0.42012124 |
| RPLP0 | 1429.54557 | 0.597046 | 0.49886116 | 1.19681795 | 0.23137752 | 0.68357541 |
| PXN-AS1 | 85.2779205 | 0.12288018 | 0.36125544 | 0.34014761 | 0.73374537 | 0.93552616 |
| PXN | 864.422279 | -0.128106 | 0.19722602 | -0.6495388 | 0.51599016 | 0.85883493 |
| SIRT4 | 79.176607 | 0.22656814 | 0.43245717 | 0.52390886 | 0.60034193 | 0.88883692 |
| GATC | 156.219137 | 0.23168836 | 0.24969088 | 0.92790076 | 0.35345905 | 0.77219738 |
| SRSF9 | 468.267765 | 0.22640329 | 0.29277291 | 0.77330683 | 0.43934081 | 0.82164134 |
| DYNLL1 | 683.774872 | 1.6107762 | 0.41046082 | 3.92431166 | 8.70E-05 | 0.01511702 |
| COQ5 | 248.600866 | -0.1320111 | 0.30633998 | -0.4309301 | 0.66651916 | 0.91318863 |
| RNF10 | 1696.98093 | 0.04626794 | 0.20281926 | 0.228124 | 0.81954984 | 0.96012717 |
| POP5 | 162.593669 | 0.74782112 | 0.32232191 | 2.32010637 | 0.02033512 | 0.28575175 |
| AC125616.1 | 51.0498034 | 0.24268733 | 0.76516145 | 0.3171714 | 0.75111355 | 0.93775604 |
| CABP1 | 425.913988 | 0.35502877 | 0.68441481 | 0.51873333 | 0.60394671 | 0.89001883 |
| MLEC | 672.398941 | 0.84180104 | 0.3116949 | 2.70072121 | 0.00691893 | 0.17082646 |
| UNC119B | 170.284291 | 0.43627415 | 0.26969867 | 1.61763557 | 0.10574116 | 0.52694363 |
| ACADS | 146.969929 | 0.61716208 | 0.25747776 | 2.39695301 | 0.01653204 | 0.25721056 |
| SPPL3 | 2212.83527 | -0.1793654 | 0.17913697 | -1.0012754 | 0.31669368 | 0.74844439 |
| C12orf43 | 78.1471019 | -0.171567 | 0.32902132 | -0.5214464 | 0.60205586 | 0.88964714 |
| P2RX4 | 150.076443 | 0.10650621 | 0.59157371 | 0.18003878 | 0.85712213 | 0.97127317 |
| CAMKK2 | 191.59975 | -0.0667553 | 0.23953985 | -0.2786814 | 0.7804893 | 0.9480401 |
| ANAPC5 | 981.080514 | -0.1870803 | 0.17672742 | -1.0585811 | 0.2897906 | 0.73204984 |
| RNF34 | 468.118698 | -0.4070108 | 0.27509648 | -1.4795202 | 0.13900133 | 0.58243213 |

| | | | | | | |
|---|---|---|---|---|---|---|
| KDM2B | 797.101574 | -0.4211189 | 0.32418732 | -1.2989986 | 0.19394439 | 0.64749489 |
| ORAI1 | 81.7450406 | 0.34959107 | 0.54165458 | 0.64541331 | 5.19E-01 | 0.85916831 |
| MORN3 | 59.3333664 | -0.4485052 | 0.39498804 | -1.1354906 | 0.25616982 | 0.70709713 |
| TMEM120B | 833.404645 | 0.96596982 | 0.43364965 | 2.22753511 | 0.02591153 | 0.31222606 |
| RHOF | 141.641798 | 0.54848267 | 0.28948122 | 1.89470901 | 0.05813096 | 0.42087381 |
| SETD1B | 310.027006 | 0.0059671 | 0.25791099 | 0.02313629 | 0.98154155 | 0.99518016 |
| HPD | 48.1381422 | 0.61067115 | 0.62318383 | 0.97992136 | 0.32712494 | 0.75568263 |
| WDR66 | 424.744551 | -0.1608762 | 0.3797686 | -0.4236165 | 0.67184553 | 0.91574062 |
| AC069503.1 | 39.1115884 | -0.887272 | 0.68437037 | -1.2964792 | 0.19481044 | 0.64840333 |
| BCL7A | 806.170846 | -0.3035104 | 0.35542747 | -0.8539307 | 0.39314337 | 0.79843808 |
| MLXIP | 1295.57628 | -0.535196 | 0.20990219 | -2.5497401 | 1.08E-02 | 2.11E-01 |
| CLIP1 | 1870.44584 | -0.3817346 | 0.26703746 | -1.4295171 | 0.15285565 | 0.60285224 |
| ZCCHC8 | 675.425151 | -0.1388424 | 0.17808671 | -0.7796337 | 0.43560654 | 0.81953094 |
| RSRC2 | 893.868094 | -0.1427945 | 0.17419736 | -0.8197283 | 0.41237104 | 0.80701706 |
| KNTC1 | 147.488602 | -0.7282677 | 0.29811484 | -2.4429098 | 0.01456938 | 0.24762526 |
| DENR | 936.589197 | -0.1296064 | 0.21486564 | -0.6031973 | 0.54637745 | 0.86805757 |
| CCDC62 | 52.9872905 | -0.1724887 | 0.44006105 | -0.3919655 | 0.69508371 | 0.92229578 |
| HIP1R | 750.40745 | 0.28160481 | 0.33902161 | 0.8306397 | 0.4061772 | 0.80499101 |
| VPS37B | 1377.45496 | -0.3984749 | 0.21662105 | -1.8395022 | 0.06584136 | 0.44463112 |
| ABCB9 | 184.476093 | -0.0779629 | 0.36106455 | -0.2159251 | 0.82904615 | 0.96288319 |
| OGFOD2 | 55.0993025 | -0.0594275 | 0.3760527 | -0.1580298 | 0.87443328 | 0.97564728 |
| ARL6IP4 | 524.183578 | 0.45464279 | 0.27406367 | 1.65889477 | 0.097137 | 0.51343843 |
| PITPNM2 | 2175.01868 | -0.3531088 | 0.3565071 | -0.9904677 | 0.32194558 | 0.75240254 |
| MPHOSPH9 | 867.67821 | -0.0797856 | 0.23907952 | -0.3337197 | 0.73859106 | 0.93552616 |
| C12orf65 | 309.333182 | 0.10070027 | 0.26157401 | 0.38497811 | 0.70025364 | 0.92367966 |
| AC068768.1 | 61.8790096 | -0.6882776 | 0.42040552 | -1.6371756 | 0.10159379 | 0.5195058 |
| SBNO1 | 949.638897 | -0.2385632 | 0.15256303 | -1.5637026 | 0.11788742 | 0.54888439 |
| KMT5A | 223.4143 | 0.43557521 | 0.19707741 | 2.21017323 | 0.02709314 | 0.3178271 |
| RILPL2 | 255.812823 | 0.13789868 | 0.22393038 | 0.6158105 | 0.5380196 | 0.86496983 |
| SNRNP35 | 364.984127 | 0.13198303 | 0.20169565 | 0.65436725 | 0.51287523 | 0.85762435 |
| RILPL1 | 978.695077 | 0.01761399 | 0.2365558 | 0.0744602 | 0.94064421 | 0.98715467 |
| AC055713.1 | 90.9005696 | -0.1392101 | 0.33389453 | -0.4169282 | 0.67673089 | 0.91662617 |
| TMED2 | 310.97637 | 0.30224121 | 0.2578734 | 1.17205268 | 0.24117591 | 0.69178991 |
| DDX55 | 214.657139 | -0.4854906 | 0.29039885 | -1.6718062 | 0.09456254 | 0.51017802 |
| EIF2B1 | 267.619736 | 0.01954757 | 0.22474236 | 0.08697769 | 0.93068925 | 0.98331361 |
| GTF2H3 | 258.996486 | -0.0825774 | 0.20016341 | -0.4125501 | 0.67993627 | 0.91833111 |
| TCTN2 | 228.736843 | -0.2139974 | 0.21060882 | -1.0160894 | 0.30958683 | 0.74450725 |
| ATP6V0A2 | 274.4175 | -0.6533424 | 0.30851808 | -2.1176795 | 0.03420222 | 0.3515775 |
| CCDC92 | 790.346895 | 0.07228842 | 0.14180354 | 0.5097787 | 0.61020651 | 0.89339478 |
| RFLNA | 167.386882 | -0.4128083 | 0.57788398 | -0.7143445 | 0.47501417 | 0.84106098 |
| NCOR2 | 1216.86211 | 0.04025199 | 0.37378694 | 0.10768699 | 0.91424399 | 0.98145965 |
| SCARB1 | 63.5993352 | -0.2061936 | 0.42892677 | -0.4807197 | 0.63071569 | 0.90005881 |
| UBC | 2190.87201 | 0.46836879 | 0.36328288 | 1.28926744 | 1.97E-01 | 6.50E-01 |
| DHX37 | 123.143621 | -0.5108746 | 0.33010018 | -1.5476351 | 0.12171019 | 0.55692668 |
| BRI3BP | 217.415597 | -0.1538734 | 0.35226507 | -0.4368114 | 0.6622481 | 0.91110616 |
| AACS | 530.737764 | -0.234052 | 0.44657959 | -0.5240991 | 0.60020962 | 0.88883692 |
| LINC02405 | 42.9169371 | 0.45617299 | 0.66856044 | 0.68232125 | 0.49503584 | 0.84899096 |
| TMEM132C | 150.419689 | 0.5502853 | 0.69411472 | 0.79278724 | 0.42790179 | 0.81466629 |
| SLC15A4 | 379.670297 | -0.0585114 | 0.28140233 | -0.2079278 | 0.83528536 | 0.96511435 |
| TMEM132D | 45.2650628 | -0.2408247 | 0.32862935 | -0.7328154 | 0.46367102 | 0.83531269 |
| RIMBP2 | 1404.23931 | 1.0856591 | 0.61070396 | 1.77771746 | 0.07545027 | 0.47278113 |
| STX2 | 344.420108 | -0.4551263 | 0.44872282 | -1.0142704 | 0.31045372 | 0.74464922 |
| RAN | 352.484089 | 0.78942459 | 0.48666672 | 1.62210515 | 0.10478084 | 0.52486729 |
| SFSWAP | 1043.04436 | -0.372098 | 0.19470939 | -1.9110428 | 0.05599908 | 0.41843207 |

| | | | | | | |
|---|---|---|---|---|---|---|
| ULK1 | 744.642103 | -0.1115255 | 0.27465468 | -0.4060572 | 0.68470057 | 0.92053026 |
| PUS1 | 155.360807 | -0.1384631 | 0.23358789 | -0.5927665 | 0.55333741 | 0.87104567 |
| EP400 | 1008.68591 | -0.2435072 | 0.20137153 | -1.2092435 | 0.2265693 | 0.67880306 |
| DDX51 | 112.371881 | -0.2500368 | 0.3522377 | -0.7098523 | 0.47779571 | 0.84190259 |
| NOC4L | 41.2335667 | -0.1019949 | 0.5683953 | -0.1794436 | 0.85758936 | 0.97127317 |
| GALNT9 | 209.179081 | -1.0025177 | 0.52338314 | -1.9154566 | 0.0554343 | 0.41841807 |
| AC148477.2 | 45.6481679 | 0.16153487 | 0.76795245 | 0.21034489 | 0.83339851 | 0.96496852 |
| FBRSL1 | 546.95735 | -0.3083467 | 0.24684265 | -1.2491628 | 0.21160553 | 0.66486617 |
| POLE | 136.067671 | -0.7117877 | 0.38396288 | -1.8537931 | 0.06376877 | 0.43779712 |
| AC135586.2 | 37.3334693 | 0.0675918 | 0.40625267 | 0.16637872 | 0.86785892 | 0.97402505 |
| PGAM5 | 54.1218091 | 0.04352617 | 0.37026081 | 0.11755543 | 0.90641992 | 0.98018017 |
| ANKLE2 | 731.085784 | -0.5441675 | 0.26219604 | -2.0754223 | 0.03794741 | 0.36480886 |
| GOLGA3 | 587.582662 | -0.3421765 | 0.22024282 | -1.5536328 | 0.12027203 | 0.55335186 |
| CHFR | 738.837824 | -0.4277551 | 0.1796974 | -2.3804188 | 0.01729297 | 0.26361342 |
| ZNF26 | 548.239153 | -0.3168164 | 0.23191587 | -1.3660832 | 0.17191285 | 0.6286116 |
| ZNF84 | 611.650754 | -0.2168879 | 0.17155767 | -1.2642275 | 0.20614837 | 0.65830819 |
| ZNF140 | 333.755726 | -0.1118682 | 0.20959179 | -0.5337433 | 0.59351915 | 0.88785893 |
| ZNF891 | 283.303928 | 0.12588816 | 0.18964675 | 0.66380344 | 0.50681612 | 0.85626196 |
| ZNF268 | 106.01996 | 0.22332443 | 0.35879861 | 0.62242278 | 0.5336639 | 0.86397197 |
| TPTE2 | 87.957782 | -0.3177165 | 0.44707107 | -0.7106623 | 0.47729355 | 0.84174109 |
| MPHOSPH8 | 1383.95057 | 0.37524303 | 0.26586853 | 1.41138566 | 0.15813093 | 0.60804051 |
| PSPC1 | 2009.74688 | 0.03881927 | 0.27961282 | 0.13883224 | 0.88958272 | 0.97729111 |
| ZMYM5 | 818.377416 | 0.10202996 | 0.14655872 | 0.69617119 | 0.48632163 | 0.8447301 |
| AL355001.1 | 67.7904418 | 0.21001267 | 0.51376208 | 0.40877417 | 0.68270539 | 0.91981391 |
| ZMYM2 | 4292.591 | -0.1414746 | 0.22310015 | -0.6341306 | 0.52599562 | 0.86119802 |
| CRYL1 | 1943.5245 | 0.09144099 | 0.31125508 | 0.29378153 | 0.76892485 | 0.94376765 |
| IFT88 | 2325.61779 | -0.3011156 | 0.19559755 | -1.5394652 | 0.12369076 | 0.56102027 |
| EEF1AKMT1 | 217.057948 | -0.0854718 | 0.25671097 | -0.3329495 | 0.73917241 | 0.93552616 |
| XPO4 | 1189.21494 | -0.0804109 | 0.15600613 | -0.515434 | 0.60624981 | 0.89125807 |
| LATS2 | 808.008319 | 0.14948947 | 0.28092455 | 0.53213389 | 0.59463327 | 0.88851932 |
| LATS2-AS1 | 51.6570372 | -0.1964411 | 0.60470322 | -0.3248553 | 0.74529058 | 0.93675395 |
| SAP18 | 452.366069 | 0.59849429 | 0.33685913 | 1.77669011 | 0.07561924 | 0.47345685 |
| MRPL57 | 305.754198 | 0.42273907 | 0.30431403 | 1.38915408 | 0.1647859 | 0.61673853 |
| ZDHHC20 | 1244.45729 | -0.23363 | 0.14680727 | -1.5914066 | 0.11151811 | 0.53526261 |
| MICU2 | 2277.22152 | -0.2008662 | 0.1885778 | -1.0651635 | 0.28680197 | 0.72949981 |
| FGF9 | 1205.74082 | 0.28936453 | 0.48902771 | 0.59171397 | 5.54E-01 | 0.8712425 |
| AL136962.1 | 464.233568 | 0.82443454 | 1.17858198 | 0.69951396 | 0.48423089 | 0.84458506 |
| LINC00621 | 437.232914 | -0.2703928 | 0.49175474 | -0.549853 | 0.58242019 | 0.88338315 |
| AL157931.1 | 189.900169 | -0.4009168 | 0.55640913 | -0.7205431 | 4.71E-01 | 8.39E-01 |
| SACS | 126.90408 | -0.7098445 | 0.44916545 | -1.5803631 | 0.11402374 | 0.54047431 |
| TNFRSF19 | 215.652817 | 0.09359729 | 0.56301412 | 0.16624324 | 0.86796554 | 0.97402505 |
| MIPEP | 1127.68409 | -0.346043 | 0.24929081 | -1.3881097 | 0.16510365 | 0.6172811 |
| PARP4 | 1031.92953 | -0.6792929 | 0.44184449 | -1.5374026 | 0.12419474 | 0.56105491 |
| CENPJ | 85.808644 | -0.831726 | 0.36567915 | -2.2744694 | 0.02293778 | 0.2983102 |
| MTMR6 | 338.382213 | -0.009469 | 0.15314412 | -0.0618307 | 0.95069763 | 0.9893457 |
| NUP58 | 954.789446 | -0.3959135 | 0.27077731 | -1.4621369 | 0.1437037 | 0.58876618 |
| RNF6 | 144.586076 | 0.10715984 | 0.25531296 | 0.41971956 | 0.67469034 | 0.91636338 |
| CDK8 | 1640.81693 | -0.1468974 | 0.19100933 | -0.769059 | 0.44185828 | 0.82285053 |
| WASF3 | 370.329536 | -0.1043746 | 0.19815972 | -0.5267194 | 0.5983885 | 0.88883692 |
| USP12 | 1212.7421 | 0.07749042 | 0.26342334 | 0.29416689 | 0.76863038 | 0.94366007 |
| LINC02340 | 420.298062 | -0.3431931 | 0.48196987 | -0.7120634 | 0.47642552 | 0.84141268 |
| RPL21 | 508.026803 | 0.62426852 | 0.39346886 | 1.58657668 | 0.11260855 | 0.53737225 |
| GTF3A | 199.611675 | -0.0840405 | 0.23492295 | -0.3577363 | 0.72054064 | 0.93009101 |
| MTIF3 | 492.912212 | -0.0887325 | 0.16258278 | -0.5457682 | 0.58522532 | 0.88487193 |

| | | | | | | |
|---|---|---|---|---|---|---|
| LNX2 | 637.568916 | 0.17566916 | 0.26387739 | 0.66572266 | 0.50558839 | 0.85597501 |
| POLR1D | 477.528151 | -0.1400706 | 0.16132359 | -0.8682587 | 0.38525274 | 0.79326632 |
| PAN3 | 4766.29292 | -0.2617505 | 0.15189531 | -1.7232294 | 0.08484704 | 0.49198622 |
| FLT1 | 67.4176943 | 0.10172455 | 0.42424254 | 0.23977923 | 0.81050141 | 0.95682484 |
| POMP | 320.009727 | 0.06544551 | 0.23657146 | 0.2766416 | 0.78205532 | 0.94882207 |
| SLC46A3 | 194.67833 | 1.09826804 | 0.27304706 | 4.02226645 | 5.76E-05 | 0.01226266 |
| MTUS2 | 51.4051174 | 1.10140341 | 0.64996561 | 1.69455643 | 0.0901596 | 0.50138792 |
| SLC7A1 | 381.354705 | 0.49879791 | 0.29931964 | 1.66643899 | 9.56E-02 | 5.12E-01 |
| UBL3 | 1157.59999 | -0.0317447 | 0.17772458 | -0.1786171 | 0.85823836 | 0.97143431 |
| KATNAL1 | 399.113961 | -0.4176679 | 0.31553226 | -1.3236932 | 0.18560496 | 0.64030106 |
| HMGB1 | 2185.78762 | 0.14357284 | 0.27430959 | 0.52339708 | 0.60069796 | 0.88883692 |
| USPL1 | 524.819555 | -0.3984766 | 0.27121326 | -1.4692371 | 0.1417685 | 0.5872148 |
| HSPH1 | 534.105647 | 0.18244478 | 0.43729408 | 0.41721303 | 0.6765226 | 0.91658712 |
| B3GLCT | 662.996534 | -0.3261754 | 0.29618071 | -1.1012717 | 0.27077845 | 0.71965808 |
| FRY | 2092.3387 | -1.6165537 | 0.7327958 | -2.2060084 | 0.02738341 | 0.31979365 |
| FRY-AS1 | 39.4491821 | -1.5539333 | 1.23660388 | -1.2566136 | 0.20889358 | 0.6614379 |
| N4BP2L1 | 771.982651 | 0.57725823 | 0.33245713 | 1.73633886 | 0.08250394 | 0.48594425 |
| N4BP2L2 | 8650.78926 | 0.03723646 | 0.19780074 | 0.18825238 | 0.85067881 | 0.97056586 |
| PDS5B | 2106.31216 | -0.1679071 | 0.21563 | -0.7786814 | 0.43616743 | 0.81976886 |
| KL | 184.64491 | 0.19883655 | 0.30350891 | 0.65512591 | 5.12E-01 | 8.58E-01 |
| STARD13 | 3704.51943 | -0.2308187 | 0.24431121 | -0.9447733 | 3.45E-01 | 0.76724655 |
| AL139383.1 | 1487.20299 | 0.20736996 | 0.3735403 | 0.55514748 | 0.57879379 | 0.8821758 |
| RFC3 | 301.999632 | 0.100877 | 0.40863174 | 0.24686532 | 0.80501246 | 0.95521106 |
| LINC02343 | 2342.72494 | 0.33444815 | 0.47434358 | 0.70507574 | 0.4807631 | 0.8437862 |
| LINC00457 | 136.357497 | 0.12259288 | 0.81207501 | 0.15096251 | 0.88000529 | 0.97664137 |
| NBEA | 2744.78501 | -0.1390744 | 0.33801782 | -0.411441 | 0.68074919 | 0.91867756 |
| DCLK1 | 65.3353945 | -1.3231152 | 0.67171399 | -1.9697598 | 0.0488659 | 0.39762623 |
| SPART | 899.552111 | 0.63279266 | 0.21043229 | 3.00710822 | 0.00263746 | 0.09727693 |
| SPART-AS1 | 77.1062308 | 1.005116 | 0.50829691 | 1.97741905 | 0.04799428 | 0.39511605 |
| RFXAP | 168.857358 | 0.05687112 | 0.27308764 | 0.20825227 | 0.835032 | 0.96511435 |
| SMAD9 | 172.690029 | 0.89233671 | 0.62486024 | 1.42805807 | 1.53E-01 | 6.03E-01 |
| ALG5 | 432.754433 | -0.1104733 | 0.14043292 | -0.7866627 | 0.43147932 | 0.81723693 |
| EXOSC8 | 236.927587 | -0.4452848 | 0.24966421 | -1.7835349 | 0.07449927 | 0.47190267 |
| SUPT20H | 1135.67424 | -0.0034 | 0.16972529 | -0.0200325 | 0.98401742 | 0.99518552 |
| LINC02334 | 125.274933 | 0.42892356 | 0.61540187 | 0.69698124 | 0.48581454 | 0.8447301 |
| LINC00571 | 422.746043 | -0.2042985 | 0.33990407 | -0.6010475 | 0.54780833 | 0.86859163 |
| UFM1 | 553.544876 | 0.18458907 | 0.20427017 | 0.90365164 | 0.36618015 | 0.78095617 |
| FREM2 | 1397.28842 | -0.107863 | 0.38748672 | -0.2783656 | 0.7807317 | 0.9480401 |
| PROSER1 | 400.472213 | -0.0904228 | 0.29999802 | -0.3014112 | 0.76310094 | 0.94247783 |
| NHLRC3 | 833.370491 | 0.29533663 | 0.32149275 | 0.91864167 | 0.35828303 | 0.77529959 |
| AL354809.1 | 57.978497 | -0.192431 | 0.37446903 | -0.5138769 | 0.60733803 | 0.89174148 |
| LHFPL6 | 58.5410372 | -0.3236944 | 0.65262678 | -0.495987 | 0.6199036 | 0.89702074 |
| COG6 | 1069.45484 | -0.1529673 | 0.23456567 | -0.6521301 | 0.51431728 | 0.85777992 |
| LINC00598 | 302.170852 | -0.0164093 | 0.3666583 | -0.0447536 | 0.96430375 | 0.99224805 |
| FOXO1 | 4280.27885 | 0.05219047 | 0.32870634 | 0.15877538 | 0.87384585 | 0.97564728 |
| MRPS31 | 915.774197 | 0.08052743 | 0.21742817 | 0.37036335 | 0.71111178 | 0.92683165 |
| SLC25A15 | 87.1822504 | -0.2778335 | 0.35010244 | -0.7935778 | 0.42744129 | 0.81466629 |
| ELF1 | 2985.75905 | -0.1373145 | 0.25661853 | -0.5350919 | 0.59258632 | 0.88740306 |
| WBP4 | 508.582428 | -0.1711903 | 0.16756646 | -1.0216264 | 0.30695776 | 0.74341546 |
| KBTBD6 | 52.8659317 | 1.24315748 | 0.45468324 | 2.73411768 | 0.00625477 | 0.16248472 |
| AL354696.2 | 212.098528 | -0.3205799 | 0.2680518 | -1.1959626 | 0.23171115 | 0.68394918 |
| KBTBD7 | 58.3738093 | 1.03201317 | 0.43485906 | 2.37321297 | 0.01763409 | 0.26681412 |
| MTRF1 | 290.126141 | -0.5225776 | 0.18960323 | -2.7561641 | 0.00584836 | 0.15669584 |
| NAA16 | 1185.20342 | -0.0771626 | 0.19179731 | -0.4023131 | 0.6874536 | 0.92094022 |

| | | | | | | |
|---|---|---|---|---|---|---|
| VWA8 | 4894.70441 | 0.22023199 | 0.22209103 | 0.99162937 | 0.32137836 | 0.75240254 |
| DGKH | 4028.65492 | 0.08393113 | 0.47688023 | 0.17600043 | 8.60E-01 | 9.72E-01 |
| AKAP11 | 891.881496 | -0.0051226 | 0.11138189 | -0.0459915 | 0.96331702 | 0.99224805 |
| LINC02341 | 72.4634516 | -0.5471475 | 0.54415762 | -1.0054945 | 0.31465879 | 0.74785672 |
| EPSTI1 | 625.33324 | 0.33033695 | 0.83451295 | 0.39584401 | 0.69222011 | 0.92224682 |
| DNAJC15 | 833.157291 | -0.0410636 | 0.29648974 | -0.1384992 | 0.88984593 | 0.97729111 |
| ENOX1 | 4013.33868 | 1.74975369 | 0.72637926 | 2.40887067 | 0.01600197 | 0.25496983 |
| CCDC122 | 693.615415 | 0.51370935 | 0.26584135 | 1.93239067 | 0.0533113 | 0.41180645 |
| LINC00284 | 66.977256 | -0.962573 | 0.97963335 | -0.982585 | 0.32581175 | 0.75550719 |
| SMIM2-AS1 | 188.423956 | 0.18110514 | 0.53982522 | 0.33548847 | 0.73725664 | 0.93552616 |
| SERP2 | 178.747381 | -0.5508351 | 0.31040812 | -1.7745512 | 0.07597203 | 0.47431992 |
| TSC22D1 | 4836.7496 | -0.0084699 | 0.33236942 | -0.0254833 | 0.97966945 | 0.99483846 |
| TSC22D1-AS1 | 47.7166911 | 0.30861365 | 0.53449152 | 0.57739672 | 0.56367149 | 0.87487519 |
| NUFIP1 | 158.588282 | -0.3704677 | 0.24479106 | -1.5134037 | 0.13017716 | 0.57164564 |
| AL359706.1 | 183.94988 | 0.01048927 | 0.52272035 | 0.0200667 | 0.98399017 | 0.99518552 |
| GPALPP1 | 481.065575 | -0.1261612 | 0.13617468 | -0.9264659 | 0.35420388 | 0.77243419 |
| GTF2F2 | 985.974595 | -0.3063027 | 0.22101944 | -1.3858633 | 0.16578864 | 0.61896611 |
| TPT1 | 2831.54807 | 0.55204251 | 0.48990661 | 1.12683212 | 0.25981347 | 0.70921184 |
| AL138963.1 | 57.5850399 | 0.23044044 | 0.45471128 | 0.50678409 | 0.61230632 | 0.89440588 |
| SLC25A30 | 548.337122 | 0.7446043 | 0.41552565 | 1.7919575 | 0.07313977 | 0.47001711 |
| COG3 | 707.717291 | 0.09875899 | 0.18526844 | 0.53305889 | 0.59399282 | 0.88807733 |
| LINC01055 | 115.639239 | 3.26664326 | 0.9231736 | 3.53849292 | 0.00040242 | 0.03420673 |
| SIAH3 | 198.345277 | -2.3741957 | 0.8254137 | -2.8763706 | 0.00402277 | 0.12328719 |
| ZC3H13 | 1298.53428 | -0.0615126 | 0.1186413 | -0.5184754 | 0.60412665 | 0.89001883 |
| CPB2-AS1 | 74.7320667 | 0.0996036 | 0.33419773 | 0.29803793 | 7.66E-01 | 0.94273621 |
| AL138686.1 | 139.356465 | 0.27208415 | 0.64933751 | 0.41901808 | 0.67520292 | 0.91649321 |
| LRCH1 | 3039.66663 | 0.3319446 | 0.20850615 | 1.59201346 | 0.11138168 | 0.53517349 |
| ESD | 447.905874 | 0.35320957 | 0.18897858 | 1.86904557 | 0.06161647 | 0.43249565 |
| SUCLA2 | 1320.65168 | -0.5616326 | 0.24866249 | -2.2586139 | 0.02390741 | 0.3030643 |
| NUDT15 | 58.5751227 | 0.07448085 | 0.32011189 | 0.23267129 | 0.81601667 | 0.95879418 |
| MED4 | 658.403477 | -0.1288312 | 0.21264417 | -0.6058536 | 0.54461198 | 0.86654229 |
| ITM2B | 3922.82138 | 0.58390409 | 0.30627241 | 1.90648612 | 0.05658716 | 0.41843207 |
| RB1 | 2455.39434 | -0.0928013 | 0.28383412 | -0.326956 | 0.74370119 | 0.93633036 |
| LPAR6 | 160.73702 | 0.27285161 | 0.56945921 | 0.47914163 | 0.63183787 | 0.90031785 |
| RCBTB2 | 424.169484 | -0.1211619 | 0.22093944 | -0.5483942 | 0.58342123 | 0.88379959 |
| FNDC3A | 3365.08671 | 0.18393937 | 0.34118306 | 0.53912222 | 0.58980252 | 0.88621335 |
| CDADC1 | 295.857124 | -0.3041966 | 0.22446721 | -1.3551937 | 0.17535585 | 0.62953084 |
| CAB39L | 521.68625 | -0.4253309 | 0.42698369 | -0.9961292 | 0.31918736 | 0.75094569 |
| SETDB2 | 1049.52568 | -0.3842149 | 0.30082523 | -1.2772029 | 0.20153063 | 0.65534477 |
| PHF11 | 564.405969 | -0.3313473 | 0.3315074 | -0.9995172 | 0.31754422 | 0.74948361 |
| RCBTB1 | 207.673831 | -0.615541 | 0.21934175 | -2.8063101 | 0.00501124 | 0.14174202 |
| EBPL | 214.4556 | -0.3829533 | 0.36810707 | -1.0403313 | 0.29818603 | 0.73673761 |
| KPNA3 | 930.893835 | -0.2408052 | 0.20061021 | -1.2003634 | 0.22999825 | 0.68234886 |
| SPRYD7 | 178.921191 | 0.23068739 | 0.27837192 | 0.82870209 | 0.407273 | 0.80561751 |
| DLEU2 | 1232.09777 | 0.00155751 | 0.48608428 | 0.00320421 | 0.99744342 | 0.99948643 |
| TRIM13 | 407.605486 | -0.1853035 | 0.20527885 | -0.9026915 | 0.36668963 | 0.78095617 |
| RNASEH2B | 450.73286 | -0.402194 | 0.23164762 | -1.7362321 | 0.0825228 | 0.48594425 |
| FAM124A | 152.07743 | -1.7030834 | 0.40734009 | -4.1809865 | 2.90E-05 | 0.00860682 |
| INTS6 | 2184.03882 | 0.07542187 | 0.41959457 | 0.17974939 | 0.85734931 | 0.97127317 |
| INTS6-AS1 | 303.596302 | 0.28288227 | 0.3327079 | 0.85024213 | 0.39519048 | 0.79901033 |
| WDFY2 | 2502.91299 | 0.07803119 | 0.29841043 | 0.2614895 | 7.94E-01 | 9.52E-01 |
| ATP7B | 145.378991 | -0.1656222 | 0.27041608 | -0.6124718 | 0.54022565 | 0.86569031 |
| ALG11 | 141.025232 | -0.0577106 | 0.24654245 | -0.2340799 | 0.81492299 | 0.95872901 |
| UTP14C | 162.910555 | 0.24918913 | 0.32641479 | 0.76341249 | 0.44521744 | 0.82561801 |

| | | | | | | |
|---|---|---|---|---|---|---|
| NEK5 | 118.949875 | 0.2902998 | 0.54707474 | 0.5306401 | 0.5956682 | 0.88859628 |
| NEK3 | 437.475785 | 0.02637702 | 0.42614805 | 0.06189638 | 0.95064535 | 0.9893457 |
| THSD1 | 59.8700564 | 0.29151478 | 0.4534868 | 0.6428297 | 0.52033461 | 0.85935232 |
| VPS36 | 872.455225 | -0.193987 | 0.12597942 | -1.5398306 | 0.12360165 | 0.56102027 |
| CKAP2 | 93.9138195 | 0.19950715 | 0.30071023 | 0.66345317 | 0.50704037 | 0.85626196 |
| HNRNPA1L2 | 38.7018358 | -0.0063075 | 0.45270354 | -0.013933 | 0.9888834 | 0.99670729 |
| SUGT1 | 616.651786 | 0.25378958 | 0.21524714 | 1.17906135 | 0.23837375 | 0.68982886 |
| OLFM4 | 336.745939 | 1.10222918 | 1.40012911 | 0.78723396 | 0.43114492 | 0.81706845 |
| DIAPH3 | 772.66819 | 0.45577965 | 0.63793158 | 0.71446479 | 0.47493982 | 0.84106098 |
| TDRD3 | 2488.05587 | 0.02711016 | 0.19568417 | 0.13854039 | 0.88981335 | 0.97729111 |
| PCDH9 | 9224.21489 | 1.35373038 | 0.32459543 | 4.17051586 | 3.04E-05 | 0.00860682 |
| PCDH9-AS2 | 201.888277 | 1.98917314 | 0.60870548 | 3.26787456 | 0.00108358 | 0.05914487 |
| DACH1 | 2248.78114 | 0.6011579 | 0.33653624 | 1.78630957 | 0.07404915 | 0.47070029 |
| MZT1 | 95.07591 | -0.1681995 | 0.3210902 | -0.5238387 | 0.60039074 | 0.88883692 |
| BORA | 97.2240209 | -0.3908756 | 0.34683669 | -1.1269732 | 0.25975382 | 0.70921184 |
| DIS3 | 421.080224 | -0.0613471 | 0.14924324 | -0.4110542 | 0.68103281 | 0.91887257 |
| PIBF1 | 2600.37141 | -0.1985144 | 0.14840793 | -1.3376267 | 0.18101815 | 0.63558446 |
| KLF5 | 475.697168 | -0.2239191 | 0.32999206 | -0.678559 | 4.97E-01 | 8.51E-01 |
| KLF12 | 4413.82942 | -0.2206559 | 0.26945612 | -0.8188937 | 0.41284707 | 0.80753695 |
| TBC1D4 | 6519.06486 | 0.15475416 | 0.22371324 | 0.69175234 | 4.89E-01 | 0.84606305 |
| COMMD6 | 401.564589 | 0.42356905 | 0.45192686 | 0.93725132 | 0.3486293 | 0.76982416 |
| UCHL3 | 504.242609 | -0.2568264 | 0.30369065 | -0.8456843 | 0.3977289 | 0.80097592 |
| LMO7-AS1 | 289.979661 | 0.08819085 | 0.5033643 | 0.17520283 | 0.86092026 | 0.9720639 |
| LMO7 | 5624.82234 | 0.10518167 | 0.40593615 | 0.25910891 | 0.79555121 | 0.95278473 |
| KCTD12 | 92.8766434 | 1.14709832 | 0.60196422 | 1.90559219 | 0.05670314 | 0.41866502 |
| AC001226.2 | 323.563842 | 0.24100455 | 0.45886677 | 0.52521683 | 0.59943247 | 0.88883692 |
| FBXL3 | 785.131444 | 0.26394923 | 0.26548564 | 0.99421285 | 0.32011926 | 0.75104498 |
| MYCBP2 | 2457.70697 | -0.1110172 | 0.15108603 | -0.7347945 | 0.46246467 | 0.83531269 |
| SLAIN1 | 333.643806 | 0.33045329 | 0.17798256 | 1.85666105 | 0.06335939 | 0.43748783 |
| RNF219 | 231.939885 | -0.2354233 | 0.19514768 | -1.2063854 | 0.22766892 | 0.67930398 |
| RBM26 | 2516.78113 | 0.00995132 | 0.08460051 | 0.1176272 | 9.06E-01 | 9.80E-01 |
| RBM26-AS1 | 90.9572187 | -0.5721795 | 0.43724681 | -1.3085962 | 0.19067115 | 0.64567512 |
| NDFIP2 | 1046.51964 | -0.1936822 | 0.32286492 | -0.5998861 | 0.54858212 | 0.8690392 |
| SPRY2 | 89.6081044 | -0.3863986 | 0.2961372 | -1.304796 | 0.19196231 | 0.64596359 |
| LINC00379 | 142.44302 | 1.66752597 | 0.79446021 | 2.09894208 | 3.58E-02 | 0.35661333 |
| GPC5 | 19699.5831 | 2.53223679 | 1.15176506 | 2.19857059 | 0.02790847 | 0.32224037 |
| GPC5-AS1 | 74.5760128 | 1.37339743 | 0.62383737 | 2.2015312 | 0.02769844 | 0.32139504 |
| GPC6 | 4177.1307 | -2.0261117 | 1.09674859 | -1.8473803 | 0.06469205 | 0.44001919 |
| TGDS | 274.804988 | -0.2377329 | 0.19377429 | -1.2268549 | 0.21987714 | 0.67299198 |
| GPR180 | 296.818265 | -0.5172834 | 0.28616299 | -1.8076529 | 0.07066055 | 0.4627897 |
| ABCC4 | 591.245015 | -0.0597819 | 0.47499319 | -0.1258585 | 0.89984394 | 0.97898519 |
| CLDN10 | 4339.36755 | -0.6826537 | 0.42041024 | -1.62378 | 0.10442278 | 0.52442862 |
| CLDN10-AS1 | 6061.76382 | -0.8517839 | 0.44935051 | -1.8955891 | 0.0580144 | 0.42087381 |
| DZIP1 | 2100.27829 | -0.5946425 | 0.3138667 | -1.8945704 | 0.05814935 | 0.42087381 |
| DNAJC3-DT | 224.366974 | 0.85024715 | 0.28281482 | 3.00637411 | 0.00264384 | 0.09727693 |
| DNAJC3 | 1266.72096 | 0.23402768 | 0.26738639 | 0.87524153 | 0.3814425 | 0.78898747 |
| UGGT2 | 2698.85835 | -0.4274448 | 0.23682909 | -1.8048664 | 0.0710956 | 0.46413489 |
| MBNL2 | 3807.20714 | 0.11302344 | 0.25393358 | 0.44509057 | 0.6562543 | 0.90950195 |
| RAP2A | 141.683349 | 0.23834277 | 0.31783398 | 0.74989709 | 0.45331669 | 0.83003329 |
| IPO5 | 519.76623 | 0.29710661 | 0.19108979 | 1.55480107 | 0.11999345 | 0.55298199 |
| FARP1 | 10572.457 | 0.02129698 | 0.28810614 | 0.07392062 | 0.94107355 | 0.98732467 |
| STK24 | 2571.52171 | -0.1631185 | 0.21001706 | -0.7766918 | 4.37E-01 | 8.20E-01 |
| SLC15A1 | 164.13156 | 0.76050206 | 0.76817007 | 0.99001782 | 3.22E-01 | 7.53E-01 |
| DOCK9 | 1621.67068 | -0.3863065 | 0.22307683 | -1.7317197 | 0.08332348 | 0.48777181 |

| | | | | | | |
|---|---|---|---|---|---|---|
| UBAC2 | 2270.03947 | -0.2206124 | 0.15461301 | -1.4268683 | 0.15361786 | 0.60285224 |
| TM9SF2 | 1120.44848 | -0.0602292 | 0.15544215 | -0.3874705 | 0.69840793 | 0.92356678 |
| CLYBL | 1947.97992 | -0.4145075 | 0.29734632 | -1.3940226 | 0.16331077 | 0.61387934 |
| AL355338.1 | 181.514658 | 0.71619489 | 0.31625331 | 2.26462421 | 0.02353575 | 0.30185158 |
| PCCA | 9502.18095 | 0.19790075 | 0.24397956 | 0.8111366 | 0.41728723 | 0.81017256 |
| PCCA-AS1 | 115.551996 | 0.82391491 | 0.50013843 | 1.64737374 | 0.09948125 | 0.51665771 |
| GGACT | 268.04119 | 0.07484001 | 0.35833304 | 0.20885602 | 0.83456064 | 0.96510927 |
| TMTC4 | 332.67897 | -0.6491217 | 0.32914622 | -1.9721376 | 0.0485939 | 0.39695534 |
| FGF14 | 4288.37788 | -1.1212931 | 0.47408158 | -2.3651902 | 0.01802081 | 0.26927252 |
| TPP2 | 1129.58088 | -0.0960213 | 0.1906634 | -0.5036167 | 0.61453076 | 0.89496546 |
| TEX30 | 143.286233 | 0.66729418 | 0.28950644 | 2.30493724 | 0.02117009 | 0.29098408 |
| KDELC1 | 57.103656 | 0.63264758 | 0.35112097 | 1.80179378 | 0.07157786 | 0.46666926 |
| ARGLU1 | 1694.08175 | 0.03938812 | 0.12331614 | 0.31940766 | 0.74941741 | 0.93758804 |
| FAM155A | 3886.78623 | 1.21631049 | 0.81210464 | 1.49772632 | 0.13420437 | 0.57708028 |
| LIG4 | 213.667536 | 0.38374976 | 0.38138137 | 1.00621004 | 0.31431454 | 0.74785672 |
| ABHD13 | 279.422588 | 0.23841604 | 0.33020496 | 0.7220244 | 0.47027948 | 0.83921324 |
| TNFSF13B | 368.797148 | 0.45436845 | 0.29872656 | 1.52101791 | 0.12825534 | 0.56762585 |
| MYO16 | 73.1928401 | 0.74314347 | 0.5312539 | 1.39884803 | 0.16185856 | 0.61346689 |
| IRS2 | 665.106675 | 0.15325355 | 0.40901943 | 0.37468526 | 0.70789456 | 0.92577218 |
| COL4A1 | 1690.22541 | -0.3183053 | 0.3909098 | -0.814268 | 0.41549143 | 0.80966301 |
| COL4A2 | 2650.09257 | -0.4550331 | 0.41428367 | -1.0983613 | 0.27204677 | 0.72070318 |
| RAB20 | 412.404564 | 0.48024291 | 0.4369479 | 1.09908507 | 0.27173096 | 0.72040597 |
| NAXD | 321.873693 | 0.15546283 | 0.19756735 | 0.78688524 | 0.43134905 | 0.81710736 |
| CARS2 | 645.982747 | -0.2465224 | 0.21325195 | -1.1560146 | 0.24767517 | 0.69800327 |
| ING1 | 73.8512999 | -0.0828182 | 0.39723462 | -0.2084869 | 0.83484884 | 0.96511435 |
| ANKRD10 | 3521.93076 | 0.0273237 | 0.24017104 | 0.11376768 | 0.90942196 | 0.98069436 |
| ARHGEF7 | 2217.81906 | 0.02889978 | 0.27907728 | 0.10355475 | 0.9175227 | 0.98199511 |
| TUBGCP3 | 734.552916 | -0.2299671 | 0.20090513 | -1.144655 | 0.25235211 | 0.70278705 |
| ATP11A | 2663.89115 | -0.0185768 | 0.37476801 | -0.0495687 | 0.96046611 | 0.99153894 |
| MCF2L | 1249.32589 | -0.6110345 | 0.25730332 | -2.3747634 | 0.0175602 | 0.26630624 |
| F10 | 48.9140346 | -0.1331071 | 0.66237449 | -0.2009544 | 0.84073422 | 0.96617873 |
| PCID2 | 439.098524 | 0.06284868 | 0.19906287 | 0.31572278 | 0.75221294 | 0.93815321 |
| CUL4A | 1270.19627 | -0.3838386 | 0.18183896 | -2.110871 | 0.0347834 | 0.35358029 |
| LAMP1 | 550.898313 | 0.52391827 | 0.22996995 | 2.2782032 | 0.02271447 | 0.2983102 |
| GRTP1 | 359.825725 | 0.09822385 | 0.26024672 | 0.3774259 | 0.70585712 | 0.92556842 |
| ADPRHL1 | 63.7943894 | -0.256659 | 0.3742239 | -0.6858433 | 0.49281191 | 0.84770837 |
| DCUN1D2 | 480.610941 | -0.2287572 | 0.19510567 | -1.1724783 | 0.24100508 | 0.69178991 |
| TMCO3 | 687.03551 | -0.2934704 | 0.26095695 | -1.1245932 | 0.26076144 | 0.71062469 |
| TFDP1 | 334.558783 | -0.4529323 | 0.22547237 | -2.0088151 | 0.04455675 | 0.38623091 |
| GAS6-AS1 | 60.0186958 | -0.2811111 | 0.71115848 | -0.3952861 | 0.69263174 | 0.92224682 |
| GAS6 | 570.845369 | -0.0876778 | 0.42457918 | -0.2065051 | 0.8363964 | 0.96511435 |
| RASA3 | 374.858524 | -1.2927495 | 0.55813521 | -2.3161942 | 0.02054767 | 0.28691085 |
| CDC16 | 737.205641 | -0.3771286 | 0.18657129 | -2.0213647 | 0.04324203 | 0.38262319 |
| UPF3A | 940.684368 | -0.0260783 | 0.18353337 | -0.1420903 | 0.88700872 | 0.97729111 |
| CHAMP1 | 151.062333 | 4.06E-05 | 0.28118353 | 0.00014423 | 0.99988492 | 0.99988492 |
| TTC5 | 209.908712 | -0.3096643 | 0.22963416 | -1.3485117 | 0.17749386 | 0.63109457 |
| CCNB1IP1 | 492.951645 | -0.237682 | 0.29486793 | -0.8060625 | 4.20E-01 | 8.13E-01 |
| PARP2 | 98.8707967 | -0.1006386 | 0.29995953 | -0.3355072 | 0.73724254 | 0.93552616 |
| TEP1 | 299.982266 | -0.4298493 | 0.24010532 | -1.7902531 | 7.34E-02 | 0.4701377 |
| OSGEP | 223.455515 | -0.4942524 | 0.33308583 | -1.483859 | 0.13784633 | 0.58135767 |
| APEX1 | 195.878731 | -0.1242385 | 0.45666235 | -0.2720578 | 0.7855776 | 0.94978829 |
| PIP4P1 | 101.916148 | -0.2179232 | 0.28972392 | -0.7521755 | 0.45194552 | 0.82885146 |
| PNP | 207.706819 | 0.24602408 | 0.41846435 | 0.58792123 | 0.55658516 | 0.87212955 |
| METTL17 | 232.45905 | -0.283371 | 0.23528349 | -1.2043813 | 0.22844224 | 0.68022848 |

| | | | | | | |
|---|---|---|---|---|---|---|
| NDRG2 | 385.071226 | 0.44798192 | 0.47008151 | 0.95298776 | 0.34059627 | 0.76494634 |
| ARHGEF40 | 167.071174 | -0.2956681 | 0.36160195 | -0.8176619 | 0.41355029 | 0.80843359 |
| ZNF219 | 99.3289263 | 0.15234318 | 0.38143852 | 0.39939119 | 0.68960499 | 0.92168526 |
| HNRNPC | 3955.55116 | -0.0654127 | 0.17538815 | -0.3729595 | 0.70917858 | 0.92577218 |
| SUPT16H | 553.684365 | 0.04163885 | 0.18677502 | 0.22293584 | 0.82358544 | 0.96135914 |
| CHD8 | 981.597913 | -0.1815543 | 0.15039695 | -1.2071676 | 0.2273676 | 0.67921637 |
| RAB2B | 382.494469 | -0.3560419 | 0.22338459 | -1.5938518 | 0.11096924 | 0.53437573 |
| TOX4 | 458.215134 | 0.01370867 | 0.13434198 | 0.10204308 | 0.91872248 | 0.98215577 |
| METTL3 | 214.101417 | -0.0671821 | 0.19831582 | -0.3387634 | 0.73478794 | 0.93552616 |
| DAD1 | 522.451339 | 0.26224747 | 0.27659908 | 0.94811401 | 0.34307141 | 0.76641095 |
| ABHD4 | 219.033347 | 0.61287647 | 0.20275682 | 3.02271698 | 0.00250516 | 0.09525007 |
| OXA1L | 220.675496 | 0.16026638 | 0.22704439 | 0.70588127 | 0.48026197 | 0.8433217 |
| MRPL52 | 179.445413 | 0.109052 | 0.30156018 | 0.36162599 | 0.71763154 | 0.92942397 |
| MMP14 | 149.378118 | 0.2009177 | 0.3002243 | 0.6692253 | 0.50335177 | 0.85525037 |
| LRP10 | 663.164924 | 0.25911964 | 0.29424954 | 0.88061185 | 3.79E-01 | 7.87E-01 |
| RBM23 | 714.466629 | -0.2085922 | 0.16005037 | -1.3032912 | 0.19247536 | 0.64671221 |
| PRMT5 | 127.611777 | 0.18176987 | 0.25207025 | 0.72110798 | 0.47084308 | 0.83921324 |
| AL132780.1 | 345.417922 | 1.30542904 | 0.50145992 | 2.60325699 | 0.00923427 | 0.19516077 |
| AL132780.3 | 42.3442842 | 0.33492833 | 0.55050665 | 0.60840015 | 0.54292211 | 0.86590899 |
| AL132780.2 | 69.2306232 | -0.548818 | 0.59169367 | -0.9275374 | 0.35364757 | 0.77224843 |
| C14orf93 | 217.661444 | 0.23584953 | 0.2609508 | 0.90380843 | 0.36609699 | 0.78095617 |
| PSMB5 | 326.462134 | 0.5463941 | 0.27794375 | 1.96584419 | 0.04931661 | 0.39892272 |
| ACIN1 | 1259.71965 | -0.317463 | 0.12473826 | -2.5450328 | 0.01092675 | 0.21301305 |
| C14orf119 | 153.085611 | 0.27081309 | 0.34373677 | 0.78785022 | 0.43078432 | 0.81706845 |
| SLC7A8 | 320.409475 | 0.67670605 | 0.28770116 | 2.35211447 | 0.01866703 | 0.27519285 |
| RNF212B | 77.4737185 | -0.0987804 | 0.51798935 | -0.1906997 | 0.84876086 | 0.97017477 |
| HOMEZ | 123.189229 | -0.0477071 | 0.32234726 | -0.1479992 | 0.88234342 | 0.97664137 |
| PPP1R3E | 106.627177 | -0.1400418 | 0.59062671 | -0.2371072 | 0.81257362 | 0.95806331 |
| BCL2L2 | 42.1097689 | -0.1548792 | 0.59648038 | -0.2596552 | 0.79512976 | 0.95272759 |
| BCL2L2-PABI | 268.322858 | 0.12455684 | 0.21328513 | 0.58399214 | 0.5592256 | 0.87295536 |
| PABPN1 | 569.073113 | -0.1401779 | 0.36978129 | -0.3790834 | 0.70462596 | 0.92528913 |
| NGDN | 209.825107 | -0.022138 | 0.27316697 | -0.081042 | 0.93540854 | 0.98586223 |
| ZFHX2 | 326.023584 | 0.44524402 | 0.37792422 | 1.17813042 | 0.23874462 | 0.6901542 |
| THTPA | 51.9912267 | 0.01917776 | 0.34489249 | 0.05560503 | 0.95565646 | 0.99065996 |
| AP1G2 | 220.831237 | -0.9962819 | 0.29538858 | -3.3727841 | 0.00074412 | 0.04818196 |
| AL135999.1 | 79.3545055 | -0.0895863 | 0.37962414 | -0.2359868 | 0.81344287 | 0.9582375 |
| DHRS4-AS1 | 251.902907 | -0.1822627 | 0.22084805 | -0.8252855 | 0.4092095 | 0.80623405 |
| DHRS4 | 92.8377811 | -0.2291749 | 0.30124035 | -0.760771 | 0.44679384 | 0.82571251 |
| DHRS4L2 | 173.649554 | 0.50806885 | 0.30614423 | 1.65957348 | 0.09700029 | 0.51333205 |
| NRL | 127.388963 | -0.8825989 | 0.34502666 | -2.5580601 | 0.01052579 | 0.20813646 |
| PCK2 | 64.1777289 | -0.074708 | 0.3667275 | -0.2037152 | 0.83857608 | 0.96579924 |
| DCAF11 | 72.9070143 | -0.2090026 | 0.38669041 | -0.5404907 | 0.58885864 | 0.88571413 |
| PSME1 | 283.772357 | 0.76004419 | 0.45309606 | 1.67744603 | 0.09345529 | 0.50887217 |
| EMC9 | 88.3723954 | 0.14714547 | 0.42247177 | 0.34829658 | 0.72761746 | 0.93420128 |
| PSME2 | 81.3805608 | 0.34375387 | 0.42445246 | 0.80987602 | 0.41801144 | 0.81067583 |
| REC8 | 68.1383085 | -0.1395584 | 0.38958076 | -0.3582271 | 0.72017339 | 0.93001079 |
| IPO4 | 45.4076698 | 0.31397365 | 0.49414558 | 0.63538693 | 0.52517609 | 0.8606763 |
| AL136295.1 | 236.329337 | -0.1073161 | 0.29049277 | -0.3694277 | 0.71180898 | 0.92748721 |
| GMPR2 | 136.024315 | 0.9209427 | 0.22842055 | 4.03178574 | 5.54E-05 | 0.01198655 |
| TINF2 | 79.8868059 | -0.0304847 | 0.32401839 | -0.0940833 | 0.92504297 | 0.9822216 |
| RABGGTA | 47.9662638 | -0.1812722 | 0.4232875 | -0.4282485 | 0.6684702 | 0.91395501 |
| DHRS1 | 137.508814 | 0.18921526 | 0.36126414 | 0.52375875 | 0.60044635 | 0.88883692 |
| LTB4R | 59.662209 | -1.6994135 | 0.49656088 | -3.4223668 | 0.00062078 | 0.04480846 |
| NFATC4 | 189.058814 | -0.7230454 | 0.51848352 | -1.3945388 | 0.16315495 | 0.61387934 |

| | | | | | | |
|---|---|---|---|---|---|---|
| NYNRIN | 97.3742596 | -0.5166525 | 0.64282664 | -0.8037198 | 0.42155884 | 0.81266383 |
| KHNYN | 191.025642 | -0.3670355 | 0.24222542 | -1.5152643 | 0.12970549 | 0.57050365 |
| SDR39U1 | 223.710837 | -0.1683914 | 0.29637455 | -0.568171 | 0.56991889 | 0.87807251 |
| NOVA1 | 1100.023 | -0.699148 | 0.31623189 | -2.2108713 | 0.02704475 | 0.3178271 |
| AL110292.1 | 159.60113 | -0.2609559 | 0.79082768 | -0.3299782 | 0.74141641 | 0.93598004 |
| LINC00645 | 1055.02258 | 0.25133145 | 0.30247945 | 0.8309042 | 4.06E-01 | 8.05E-01 |
| PRKD1 | 22141.5827 | 0.02237216 | 0.42554662 | 0.05257276 | 0.95807232 | 0.99069768 |
| AL133372.3 | 41.3575574 | 0.68360026 | 0.62607683 | 1.09187919 | 0.27488621 | 0.72164296 |
| G2E3-AS1 | 70.4254058 | -0.0671764 | 0.64835054 | -0.1036112 | 0.91747789 | 0.98199511 |
| G2E3 | 458.658855 | -0.371871 | 0.15172888 | -2.4508911 | 0.01425031 | 0.24670027 |
| SCFD1 | 1743.41715 | -0.2405359 | 0.17368991 | -1.3848584 | 0.16609578 | 0.61901667 |
| COCH | 129.156714 | -0.2107535 | 0.44021949 | -0.4787463 | 0.63211911 | 0.90042718 |
| STRN3 | 3609.71048 | -0.1604103 | 0.28961413 | -0.5538758 | 0.57966386 | 0.8823208 |
| AP4S1 | 267.289632 | 0.05785974 | 0.25911137 | 0.22330065 | 0.82330152 | 0.96133701 |
| HECTD1 | 3005.46106 | -0.2093469 | 0.2405291 | -0.8703599 | 0.38410376 | 0.791697 |
| AL136418.1 | 62.0123343 | -0.5407921 | 0.42977865 | -1.2583038 | 0.20828189 | 0.66118614 |
| HEATR5A | 488.242355 | -0.1980903 | 0.23222309 | -0.8530171 | 0.3936498 | 0.79843808 |
| AL139353.1 | 152.018859 | -0.2015089 | 0.27262843 | -0.7391337 | 0.45982584 | 0.83390278 |
| AL139353.2 | 52.1226321 | 0.16393288 | 0.51123567 | 0.32066011 | 0.74846798 | 0.93746253 |
| NUBPL | 1602.26541 | -0.267703 | 0.26909678 | -0.9948207 | 0.31982347 | 0.75094569 |
| ARHGAP5 | 2515.64517 | 0.32915839 | 0.10845206 | 3.035059 | 0.00240489 | 0.09386908 |
| AKAP6 | 1381.54121 | 0.34385224 | 0.36031755 | 0.95430333 | 0.33993012 | 0.76465348 |
| NPAS3 | 2109.01319 | 0.02179622 | 0.47083123 | 0.04629305 | 0.96307668 | 0.99224805 |
| EGLN3 | 389.21125 | -0.2001712 | 0.67541561 | -0.2963675 | 0.76694944 | 0.94312902 |
| EAPP | 524.189022 | -0.0542975 | 0.20145746 | -0.2695235 | 0.78752682 | 0.95004286 |
| SNX6 | 1167.6765 | 0.05841207 | 0.19932219 | 0.29305351 | 0.76948125 | 0.94386043 |
| CFL2 | 112.207405 | 0.64333982 | 0.37968763 | 1.69439235 | 0.09019075 | 0.50138792 |
| BAZ1A | 651.930596 | -0.3128598 | 0.28863179 | -1.0839409 | 2.78E-01 | 0.7242837 |
| SRP54 | 975.800243 | -0.161156 | 0.18327156 | -0.8793289 | 0.37922297 | 0.78762967 |
| FAM177A1 | 579.059426 | 0.2892916 | 0.18034165 | 1.60413083 | 0.10868522 | 0.53210639 |
| PPP2R3C | 398.206929 | -0.0881599 | 0.24400124 | -0.3613092 | 0.71786833 | 0.92945725 |
| NFKBIA | 475.068328 | 0.9336349 | 0.50780349 | 1.8385752 | 0.06597769 | 0.4450967 |
| RALGAPA1 | 3538.96534 | -0.1432164 | 0.1664171 | -0.8605872 | 0.38946543 | 0.79677297 |
| BRMS1L | 252.800028 | 0.36250175 | 0.34999975 | 1.03572001 | 0.30033278 | 0.73683056 |
| PTCSC3 | 41.732299 | -0.3514213 | 0.66840897 | -0.525758 | 5.99E-01 | 8.89E-01 |
| MBIP | 436.819247 | 0.32936152 | 0.19707133 | 1.67128075 | 0.09466624 | 0.51017802 |
| SLC25A21 | 216.524907 | -0.2696364 | 0.42619286 | -0.6326628 | 0.52695385 | 0.86151701 |
| MIPOL1 | 1518.54564 | 0.55677343 | 0.14681034 | 3.79246736 | 0.00014916 | 0.02008985 |
| SEC23A | 1023.13573 | 0.02118917 | 0.22631489 | 0.09362692 | 0.92540553 | 0.9822216 |
| GEMIN2 | 128.565884 | -0.1835537 | 0.21647696 | -0.8479132 | 0.39648632 | 0.80030376 |
| TRAPPC6B | 694.687618 | -0.1265303 | 0.21124555 | -0.5989726 | 0.54919115 | 0.86904356 |
| PNN | 1137.89834 | -0.2655627 | 0.17609218 | -1.5080892 | 0.1315317 | 0.57519227 |
| MIA2 | 2248.10404 | 0.10130725 | 0.22410535 | 0.45205191 | 0.65123159 | 0.90825486 |
| FBXO33 | 804.111577 | -0.0099528 | 0.21360381 | -0.0465946 | 0.96283634 | 0.99224805 |
| AL049875.1 | 380.086724 | -0.4587183 | 0.31910701 | -1.4375063 | 0.15057418 | 0.59727759 |
| LRFN5 | 259.282165 | 0.25760459 | 0.86917026 | 0.29637989 | 0.76693998 | 0.94312902 |
| C14orf28 | 71.343524 | -0.2734167 | 0.25657843 | -1.0656263 | 0.28659262 | 0.72940306 |
| KLHL28 | 746.076394 | 0.1330186 | 0.2701059 | 0.49246834 | 0.6223883 | 0.89835355 |
| TOGARAM1 | 1234.70351 | -0.2075711 | 0.15762481 | -1.3168679 | 0.18788289 | 0.64243984 |
| PRPF39 | 519.355219 | -0.2462107 | 0.24304158 | -1.0130394 | 0.31104133 | 0.74538189 |
| FKBP3 | 397.954018 | 0.27872998 | 0.23224749 | 1.20014206 | 0.23008417 | 0.68234886 |
| FANCM | 338.337678 | -0.751469 | 0.36194215 | -2.0762129 | 0.03787427 | 0.36437085 |
| MIS18BP1 | 428.056043 | -0.3539119 | 0.28899384 | -1.2246346 | 0.22071292 | 0.67345736 |
| LINC00871 | 173.218787 | 0.09347015 | 0.47688629 | 0.19600091 | 0.84460946 | 0.96853443 |

| | | | | | | |
|---|---|---|---|---|---|---|
| AL358335.2 | 112.43914 | 0.69519452 | 0.88965346 | 0.7814217 | 4.35E-01 | 0.81919857 |
| AL110505.1 | 35.3210152 | 1.29489422 | 1.19485416 | 1.08372575 | 0.27848642 | 0.7242837 |
| RPS29 | 757.359446 | 0.09858376 | 0.27936805 | 0.35288128 | 0.72417744 | 0.93223466 |
| LRR1 | 66.3868382 | -0.4431278 | 0.36206707 | -1.2238832 | 0.2209963 | 0.67392082 |
| RPL36AL | 702.112525 | 0.53318341 | 0.4133772 | 1.28982298 | 0.19711213 | 0.64964978 |
| MGAT2 | 48.8302026 | 0.40373885 | 0.47165064 | 0.85601251 | 0.39199086 | 0.79767768 |
| DNAAF2 | 172.001562 | 0.06672507 | 0.2866615 | 0.23276606 | 0.81594307 | 0.95879418 |
| POLE2 | 155.947792 | 0.12359499 | 0.37797727 | 0.32699053 | 0.74367504 | 0.93633036 |
| KLHDC1 | 597.995477 | 0.01048146 | 0.23255642 | 0.04507059 | 0.96405104 | 0.99224805 |
| KLHDC2 | 719.618266 | -0.2345048 | 0.24803832 | -0.9454379 | 0.34443539 | 0.76716956 |
| NEMF | 2096.29551 | -0.2022112 | 0.16774226 | -1.2054876 | 0.22801513 | 0.67987627 |
| ARF6 | 189.936572 | 0.52584906 | 0.31892096 | 1.64883817 | 0.09918079 | 0.51618562 |
| LINC01588 | 103.552918 | -0.0295274 | 0.40775745 | -0.0724142 | 0.94227228 | 0.98772472 |
| VCPKMT | 252.792522 | -0.0913446 | 0.22194768 | -0.411559 | 0.68066267 | 0.91867756 |
| SOS2 | 2185.80017 | 0.05494131 | 0.15702155 | 0.34989663 | 0.72641627 | 0.93338838 |
| L2HGDH | 775.782498 | -0.1759854 | 0.20855615 | -0.8438275 | 0.39876581 | 0.80148305 |
| ATP5S | 443.247945 | 0.13957373 | 0.19566155 | 0.71334266 | 0.47563374 | 0.84118672 |
| AL359397.2 | 73.3652937 | 0.16259736 | 0.39834156 | 0.40818577 | 0.6831373 | 0.9200778 |
| CDKL1 | 1836.00425 | -0.2311348 | 0.37928859 | -0.6093902 | 5.42E-01 | 0.86582694 |
| MAP4K5 | 1713.78793 | -0.0359813 | 0.10920844 | -0.3294737 | 0.74179769 | 0.93598004 |
| ATL1 | 178.437658 | 0.46272094 | 0.49872498 | 0.92780781 | 0.35350727 | 0.77219738 |
| SAV1 | 580.608444 | 0.13641802 | 0.31606714 | 0.43161091 | 0.66602423 | 0.91307878 |
| NIN | 602.248487 | -0.0382912 | 0.18682466 | -0.2049579 | 0.83760503 | 0.96527001 |
| PYGL | 241.960066 | 0.24367883 | 0.46971379 | 0.51878152 | 0.6039131 | 0.89001883 |
| AL358334.2 | 38.8902777 | 0.77213721 | 0.83386801 | 0.92597054 | 3.54E-01 | 0.77243419 |
| TRIM9 | 111.310922 | 0.34045383 | 0.68385881 | 0.49784229 | 0.61859521 | 0.89604387 |
| TMX1 | 285.695787 | -0.1752237 | 0.28172549 | -0.621966 | 0.53396421 | 0.86397197 |
| FRMD6 | 67.7037001 | -0.7954923 | 0.6901406 | -1.1526525 | 0.24905304 | 0.69928042 |
| GNG2 | 111.788461 | 1.93876606 | 0.79870744 | 2.42737949 | 0.01520834 | 0.25080486 |
| RTRAF | 311.247425 | 0.28226544 | 0.3440528 | 0.82041316 | 0.41198062 | 0.80679792 |
| TXNDC16 | 338.468583 | -0.0953546 | 0.25587972 | -0.3726539 | 0.70940607 | 0.92585167 |
| GPR137C | 92.3218454 | -0.527597 | 0.3661259 | -1.4410263 | 0.14957726 | 0.59625836 |
| ERO1A | 893.710524 | 0.30997006 | 0.68149962 | 0.45483526 | 0.64922776 | 0.90788153 |
| AL133453.1 | 83.494887 | 1.35125335 | 0.79825402 | 1.69276108 | 0.09050095 | 0.50249142 |
| PSMC6 | 586.087504 | 0.04931921 | 0.14869504 | 0.33168026 | 0.74013071 | 0.93553938 |
| STYX | 293.72253 | -0.0698338 | 0.20633209 | -0.3384533 | 0.73502162 | 0.93552616 |
| GNPNAT1 | 207.558632 | 0.0687644 | 0.26589886 | 0.25861111 | 0.79593531 | 0.95278473 |
| FERMT2 | 1585.50755 | -0.042634 | 0.34216562 | -0.1246006 | 0.90083978 | 0.97914987 |
| DDHD1 | 484.84909 | -0.0239392 | 0.21683735 | -0.1104017 | 0.91209084 | 0.98125148 |
| LINC02331 | 39.3492839 | -0.1792208 | 0.68855388 | -0.2602858 | 7.95E-01 | 0.95272008 |
| CNIH1 | 235.605726 | 0.1839129 | 0.27143859 | 0.67754883 | 0.4980578 | 0.85092921 |
| GMFB | 241.621972 | 0.08511367 | 0.21251495 | 0.40050675 | 0.68878331 | 0.92151057 |
| CGRRF1 | 192.79728 | 0.07909606 | 0.37094715 | 0.21322731 | 0.83114967 | 0.96382687 |
| SAMD4A | 4725.51627 | -0.1093643 | 0.44034678 | -0.2483596 | 0.8038562 | 0.95450237 |
| GCH1 | 130.361688 | -0.1425472 | 0.438165 | -0.3253277 | 0.74493307 | 0.93675395 |
| WDHD1 | 103.961685 | -1.1932223 | 0.41949855 | -2.8444015 | 0.00444949 | 0.12989304 |
| SOCS4 | 259.245818 | 0.32671921 | 0.25073526 | 1.30304455 | 0.19255955 | 0.64671221 |
| MAPK1IP1L | 1077.32016 | 0.19600456 | 0.16074486 | 1.219352 | 0.22271062 | 0.67568617 |
| LGALS3 | 634.34682 | 0.7540672 | 0.52151441 | 1.44591823 | 1.48E-01 | 5.94E-01 |
| FBXO34 | 2292.57404 | -0.4362136 | 0.20295866 | -2.149273 | 0.03161276 | 0.33950635 |
| AL158801.3 | 145.030525 | -0.4186741 | 0.49246973 | -0.8501519 | 0.39524063 | 0.79901033 |
| ATG14 | 556.549746 | -0.1789425 | 0.35871543 | -0.4988424 | 0.61789043 | 0.89599505 |
| KTN1-AS1 | 107.729647 | -0.5866357 | 0.35299957 | -1.6618595 | 0.09654095 | 0.51229242 |
| KTN1 | 2416.05654 | -0.1174037 | 0.18053523 | -0.6503089 | 0.51549268 | 0.85869987 |

| | | | | | | |
|---|---|---|---|---|---|---|
| PELI2 | 3222.59911 | -0.1673488 | 0.18618541 | -0.8988289 | 0.36874381 | 0.78095617 |
| LINC02284 | 71.4004865 | -0.5022543 | 0.55767868 | -0.9006159 | 0.36779259 | 0.78095617 |
| TMEM260 | 1145.00246 | -0.0060963 | 0.13497542 | -0.0451658 | 0.96397515 | 0.99224805 |
| EXOC5 | 748.931151 | 0.18563929 | 0.11419777 | 1.62559465 | 0.10403591 | 0.52442862 |
| AP5M1 | 730.502523 | 0.37811189 | 0.24934829 | 1.51640055 | 0.1294181 | 0.57039829 |
| NAA30 | 272.876174 | 0.17119276 | 0.20082291 | 0.85245637 | 0.39396085 | 0.79845072 |
| CCDC198 | 2689.4997 | 0.59865011 | 0.3787673 | 1.58052215 | 0.11398734 | 0.54047431 |
| ARMH4 | 1046.8436 | -0.3459444 | 0.24076616 | -1.4368479 | 0.1507612 | 0.59766048 |
| ACTR10 | 771.088393 | -0.1480117 | 0.16453338 | -0.8995848 | 0.36834123 | 0.78095617 |
| PSMA3 | 654.884188 | 0.15797376 | 0.26484578 | 0.59647451 | 0.55085828 | 0.87005704 |
| ARID4A | 1255.90778 | 0.08111193 | 0.17657937 | 0.4593511 | 0.64598206 | 0.90582727 |
| AL139021.2 | 56.8710823 | 0.24476521 | 0.51913528 | 0.47148638 | 0.63729345 | 0.90255504 |
| AL139021.1 | 167.416005 | -0.3114927 | 0.39857692 | -0.781512 | 0.4345014 | 0.81919857 |
| TIMM9 | 256.69421 | 0.13402181 | 0.20758709 | 0.64561726 | 0.51852726 | 0.85916831 |
| KIAA0586 | 821.859645 | -0.0434994 | 0.15463894 | -0.2812962 | 0.77848319 | 0.94704222 |
| DAAM1 | 1731.81469 | -0.0207361 | 0.25671551 | -0.0807745 | 0.93562127 | 0.98600777 |
| GPR135 | 231.316761 | -0.0309257 | 0.56682168 | -0.0545598 | 0.95648918 | 0.99065996 |
| L3HYPDH | 201.318218 | 0.12647113 | 0.36592724 | 0.34561824 | 0.72962964 | 0.93453345 |
| JKAMP | 429.605625 | 0.36745607 | 0.17881047 | 2.05500312 | 0.0398787 | 0.37279391 |
| CCDC175 | 43.2875696 | 0.38595808 | 0.89362961 | 0.43189939 | 0.66581454 | 0.91305741 |
| RTN1 | 85.2551062 | 0.74285055 | 0.42506085 | 1.74763345 | 0.08052752 | 0.48456361 |
| LRRC9 | 73.5156391 | 0.38553361 | 0.56794777 | 0.67881877 | 0.4972527 | 0.85092921 |
| AL157911.1 | 406.201399 | -0.145067 | 0.48367794 | -0.2999248 | 7.64E-01 | 0.94256396 |
| PCNX4 | 1660.1409 | -0.3783661 | 0.27912386 | -1.3555493 | 0.17524263 | 0.62944785 |
| DHRS7 | 193.653377 | 0.38775634 | 0.34108503 | 1.13683191 | 0.25560856 | 0.70649371 |
| PPM1A | 898.103682 | -0.2499157 | 0.28968274 | -0.8627222 | 0.38829022 | 0.79634939 |
| SIX4 | 121.771666 | 0.66375 | 0.36313287 | 1.82784336 | 0.06757306 | 0.45209626 |
| MNAT1 | 3310.93093 | -0.0414293 | 0.20050879 | -0.2066208 | 0.83630601 | 0.96511435 |
| TRMT5 | 103.140673 | 0.54828921 | 0.28702433 | 1.91025343 | 0.05610059 | 0.41843207 |
| SLC38A6 | 605.784069 | -0.2635059 | 0.12473856 | -2.1124652 | 0.03464657 | 0.35339604 |
| PRKCH | 447.168509 | 0.40627213 | 0.3087039 | 1.31605763 | 0.1881547 | 0.64267818 |
| TMEM30B | 151.299523 | 0.59141954 | 0.44363086 | 1.33313434 | 0.18248772 | 0.63741052 |
| AL359220.1 | 54.2547304 | 0.85287492 | 0.74619383 | 1.14296699 | 0.2530523 | 0.70374828 |
| HIF1A-AS1 | 46.4843392 | -0.7032567 | 0.41237929 | -1.7053637 | 0.08812661 | 0.49729428 |
| HIF1A | 13074.5635 | -0.2077916 | 0.40625521 | -0.5114805 | 6.09E-01 | 8.93E-01 |
| HIF1A-AS2 | 2515.76921 | -0.0140901 | 0.66427579 | -0.0212112 | 0.98307717 | 0.99518552 |
| SNAPC1 | 896.476226 | -0.0998829 | 0.45011709 | -0.2219043 | 0.82438838 | 0.96146415 |
| SYT16 | 300.675476 | 0.6764493 | 0.74047959 | 0.91352863 | 0.36096458 | 0.77730375 |
| PPP2R5E | 2222.33762 | -0.3340656 | 0.23595476 | -1.4158037 | 0.15683298 | 0.60654181 |
| WDR89 | 288.223515 | -0.148218 | 0.17294845 | -0.8570068 | 0.39144112 | 0.7973532 |
| SGPP1 | 551.347334 | -0.0355769 | 0.24832167 | -0.1432696 | 0.8860773 | 0.97681481 |
| SYNE2 | 16886.4949 | -0.3586549 | 0.22899751 | -1.5661958 | 0.11730279 | 0.54757066 |
| ESR2 | 1367.69822 | -0.2861086 | 0.6224588 | -0.4596427 | 0.64577272 | 0.9058099 |
| MTHFD1 | 286.198257 | -0.3324079 | 0.2071534 | -1.6046459 | 0.10857175 | 0.53194517 |
| AL122035.1 | 198.138381 | -0.5292272 | 0.40182036 | -1.317074 | 0.18781381 | 0.64236991 |
| ZBTB25 | 439.527743 | -0.2819067 | 0.31218581 | -0.9030093 | 0.36652094 | 0.78095617 |
| ZBTB1 | 1204.40067 | 0.05811613 | 0.23553393 | 0.24674208 | 0.80510784 | 0.95522565 |
| AL049869.3 | 176.776193 | -0.0107827 | 0.66660451 | -0.0161756 | 0.98709428 | 0.99653761 |
| HSPA2 | 286.386077 | 0.20489692 | 0.80518545 | 0.25447171 | 0.79913116 | 0.95319879 |
| PPP1R36 | 80.9693064 | -0.5192493 | 0.62988705 | -0.8243531 | 0.40973897 | 0.80641067 |
| PLEKHG3 | 759.538175 | -0.5984896 | 0.43960806 | -1.3614163 | 0.17338217 | 0.6286116 |
| SPTB | 55.7248553 | 0.09373181 | 0.40689386 | 0.23035936 | 0.81781254 | 0.95970911 |
| CHURC1-FNT | 307.048414 | -0.4031392 | 0.34671926 | -1.1627252 | 0.24494101 | 0.6948422 |
| RAB15 | 131.964844 | 0.46310985 | 0.3098343 | 1.49470169 | 0.13499231 | 0.57708028 |

| | | | | | | |
|---|---|---|---|---|---|---|
| FNTB | 44.2697438 | 0.39529349 | 0.3791159 | 1.0426719 | 0.29710028 | 0.73673761 |
| MAX | 736.420618 | -0.1065662 | 0.25355331 | -0.420291 | 0.67427291 | 0.91636338 |
| AL139022.1 | 47.8497897 | 0.54364444 | 0.69362615 | 0.78377155 | 0.43317417 | 0.81856905 |
| FUT8 | 758.059536 | 0.14095527 | 0.17753292 | 0.79396696 | 0.42721467 | 0.81466629 |
| AL359232.1 | 600.837779 | -0.2435696 | 0.54484396 | -0.4470448 | 0.65484274 | 0.90902447 |
| GPHN | 6339.78913 | 0.11518827 | 0.16947508 | 0.67967671 | 0.49670918 | 0.85081933 |
| MPP5 | 960.855412 | -0.1416863 | 0.17167563 | -0.8253141 | 0.4091933 | 0.80623405 |
| ATP6V1D | 630.802249 | -0.4543368 | 0.31740925 | -1.4313913 | 0.1523181 | 0.60238618 |
| EIF2S1 | 369.774678 | -0.1201146 | 0.19513882 | -0.615534 | 0.53820209 | 0.86506588 |
| PLEKHH1 | 364.394794 | -0.4312791 | 0.26992054 | -1.5978002 | 0.11008744 | 0.53372881 |
| PIGH | 286.575344 | 0.20686281 | 0.39265562 | 0.52683012 | 0.59831156 | 0.88883692 |
| ARG2 | 895.180813 | -0.3878708 | 1.13147892 | -0.3427999 | 0.73174902 | 0.93545724 |
| VTI1B | 212.035025 | 0.60341827 | 0.30028006 | 2.00951829 | 0.0444822 | 0.38623091 |
| RDH11 | 81.6747918 | -0.1179957 | 0.35846527 | -0.3291692 | 0.74202781 | 0.93598004 |
| ZFYVE26 | 354.738156 | -0.4148493 | 0.30534506 | -1.3586245 | 0.17426561 | 0.62892483 |
| RAD51B | 4178.96407 | -0.2624472 | 0.4102558 | -0.6397159 | 0.52235732 | 0.86006369 |
| ZFP36L1 | 2297.18055 | -0.1933952 | 0.31344256 | -0.6170036 | 0.53723235 | 0.86487317 |
| ACTN1 | 531.039331 | 0.03639086 | 0.3033169 | 0.11997635 | 9.05E-01 | 9.80E-01 |
| DCAF5 | 1910.81442 | 0.2155441 | 0.1495999 | 1.44080377 | 0.14964013 | 0.59625836 |
| AL359317.2 | 176.007854 | 0.11915496 | 0.53518028 | 0.22264452 | 0.82381219 | 0.96141406 |
| EXD2 | 408.349776 | 0.01535992 | 0.23722473 | 0.0647484 | 0.94837432 | 0.98919519 |
| AL359317.1 | 46.0212774 | 0.85029901 | 0.62443942 | 1.36169977 | 0.17329265 | 0.6286116 |
| GALNT16 | 471.32013 | 2.23377059 | 0.75134269 | 2.97303829 | 0.00294868 | 0.10386419 |
| ERH | 389.027224 | 0.36368147 | 0.262243 | 1.38681099 | 0.16549941 | 0.61841065 |
| SLC39A9 | 787.870116 | 0.01086906 | 0.16865718 | 0.06444468 | 0.94861615 | 0.98919519 |
| PLEKHD1 | 44.0754096 | -0.9478493 | 0.93889818 | -1.0095337 | 0.31271877 | 0.74693024 |
| SUSD6 | 1516.73459 | -0.1290035 | 0.22477292 | -0.5739281 | 0.56601646 | 0.87642772 |
| SRSF5 | 2029.17999 | 0.12603476 | 0.26828241 | 0.46978393 | 0.63850939 | 0.90307917 |
| AL357153.2 | 419.588986 | 0.02089071 | 0.45228955 | 0.04618879 | 0.96315978 | 0.99224805 |
| MED6 | 351.523066 | -0.3812083 | 0.25669306 | -1.4850744 | 0.13752412 | 0.58042326 |
| MAP3K9 | 481.785785 | -0.0205647 | 0.2184411 | -0.0941431 | 0.92499547 | 0.9822216 |
| AC004816.1 | 41.0047753 | -0.4980501 | 0.50379072 | -0.9886053 | 0.32285631 | 0.7532337 |
| PCNX1 | 2629.19877 | -0.1483015 | 0.24399534 | -0.6078046 | 0.5433171 | 0.86622038 |
| AC005476.2 | 40.2016091 | -0.7806459 | 0.51858149 | -1.5053486 | 0.13223449 | 0.57615451 |
| SIPA1L1 | 3729.28396 | -0.6368244 | 0.20434727 | -3.1163834 | 0.00183084 | 0.08282046 |
| RGS6 | 3070.80805 | -1.3806029 | 0.74708029 | -1.8479981 | 0.06460262 | 0.44001919 |
| DPF3 | 98.3522997 | 0.94286291 | 0.32961562 | 2.86049223 | 0.00422984 | 0.12640712 |
| DCAF4 | 115.379405 | -0.4384092 | 0.21987852 | -1.9938703 | 0.04616623 | 0.3904347 |
| ZFYVE1 | 436.044434 | -0.0777068 | 0.22670983 | -0.3427589 | 0.73177982 | 0.93545724 |
| RBM25 | 1481.1116 | -0.3064977 | 0.18010728 | -1.701751 | 0.08880205 | 0.49850673 |
| PSEN1 | 1120.88573 | -0.1356304 | 0.22026662 | -0.6157555 | 0.53805589 | 0.86496983 |
| PAPLN | 430.392929 | 1.14883054 | 0.79222898 | 1.45012436 | 0.14702384 | 0.59373375 |
| NUMB | 3341.12781 | 0.02374896 | 0.17161993 | 0.13838116 | 0.88993919 | 0.97729111 |
| HEATR4 | 153.140725 | -0.4104661 | 0.45254952 | -0.9070082 | 0.36440246 | 0.78050789 |
| AC005225.3 | 49.0189513 | -0.9646607 | 0.46939779 | -2.0551027 | 0.03986909 | 0.37279391 |
| ACOT2 | 50.1095288 | -0.1569488 | 0.63218273 | -0.2482649 | 0.80392949 | 0.95450237 |
| AC005225.2 | 60.228348 | 1.01436156 | 0.56808975 | 1.78556567 | 0.07416961 | 0.47078634 |
| DNAL1 | 385.548144 | 0.16744932 | 0.219989 | 0.76117132 | 0.44655474 | 0.82571251 |
| PNMA1 | 83.634893 | 0.81090252 | 0.53624907 | 1.51217514 | 0.13048931 | 0.57223551 |
| ELMSAN1 | 521.254654 | -0.3481037 | 0.2858042 | -1.2179795 | 0.22323177 | 0.67624946 |
| AC005520.2 | 72.5292414 | 1.27751477 | 0.94994639 | 1.34482828 | 0.1786807 | 0.63284534 |
| AC005480.1 | 313.314211 | -0.1029736 | 0.40875965 | -0.2519171 | 0.80110511 | 0.95370871 |
| COQ6 | 160.989011 | -0.166315 | 0.2154622 | -0.7718989 | 0.44017433 | 0.82232701 |
| ENTPD5 | 508.0951 | -0.3552687 | 0.26327336 | -1.3494291 | 0.17719919 | 0.63071358 |

| | | | | | | |
|---|---|---|---|---|---|---|
| BBOF1 | 428.125309 | 0.30349119 | 0.1701713 | 1.78344526 | 0.07451385 | 0.47190267 |
| ALDH6A1 | 371.066823 | 0.77326785 | 0.25257438 | 3.06154507 | 0.00220198 | 0.09035145 |
| LIN52 | 746.431295 | -0.2541556 | 0.23121301 | -1.0992272 | 0.27166897 | 0.72040597 |
| ABCD4 | 643.022049 | -0.0473551 | 0.19956351 | -0.2372933 | 0.81242925 | 0.95806331 |
| NPC2 | 341.193624 | 0.48047309 | 0.40006542 | 1.2009863 | 0.22975652 | 0.68214291 |
| ISCA2 | 123.363938 | 0.92026541 | 0.41199472 | 2.23368255 | 0.02550397 | 0.31163914 |
| LTBP2 | 287.986632 | 0.60675178 | 0.70730684 | 0.8578339 | 0.39098419 | 0.79735014 |
| AREL1 | 543.497873 | -0.1322803 | 0.21418947 | -0.6175857 | 0.53684851 | 0.86466169 |
| FCF1 | 244.790164 | -0.2945324 | 0.20206182 | -1.457635 | 0.14494117 | 0.59090366 |
| YLPM1 | 1282.80555 | 0.08762186 | 0.16154434 | 0.54240131 | 0.58754208 | 0.88571413 |
| DLST | 545.981795 | -0.0848359 | 0.22978034 | -0.3692044 | 0.71197541 | 0.92748721 |
| RPS6KL1 | 45.6967642 | -0.3999741 | 0.74762786 | -0.5349909 | 0.59265613 | 0.88740306 |
| EIF2B2 | 133.909404 | 0.1746205 | 0.28619289 | 0.61014968 | 0.54176266 | 0.86582694 |
| AL049780.1 | 40.9033503 | 0.06765692 | 0.42799795 | 0.15807767 | 0.8743956 | 0.97564728 |
| MLH3 | 897.923902 | -0.0165512 | 0.19292401 | -0.0857913 | 0.93163235 | 0.98357046 |
| ACYP1 | 579.772158 | -0.0560797 | 0.38853057 | -0.144338 | 0.8852336 | 0.97664137 |
| ZC2HC1C | 205.394295 | -0.2563582 | 0.61097615 | -0.4195879 | 0.67478657 | 0.91636338 |
| NEK9 | 724.943439 | 0.06997144 | 0.12888725 | 0.5428888 | 0.58720637 | 0.88563701 |
| AL049780.2 | 119.204251 | 0.42998009 | 0.4275793 | 1.00561483 | 0.31460089 | 0.74785672 |
| TMED10 | 817.562723 | 0.61521527 | 0.14672399 | 4.19301074 | 2.75E-05 | 0.00845611 |
| FOS | 233.436859 | 0.6159406 | 0.81170684 | 0.7588215 | 0.44795934 | 0.8260638 |
| JDP2 | 184.0059 | 0.50100299 | 0.37932445 | 1.32077694 | 0.18657575 | 0.6408069 |
| FLVCR2 | 58.013952 | -0.2869043 | 0.82417266 | -0.3481119 | 0.72775617 | 0.93420128 |
| TTLL5 | 1592.6864 | -0.5020848 | 0.15839564 | -3.1698144 | 0.00152536 | 0.07273836 |
| ERG28 | 123.358628 | 0.18406406 | 0.33747679 | 0.5454125 | 0.58546984 | 0.88490909 |
| AF107885.2 | 78.7320535 | -0.4978778 | 0.52669157 | -0.9452929 | 0.34450939 | 0.76716956 |
| IFT43 | 1017.04502 | -0.0863123 | 0.16621714 | -0.5192743 | 0.60356948 | 0.89001883 |
| GPATCH2L | 2276.22393 | -0.2503337 | 0.23382617 | -1.0705973 | 0.28435053 | 0.72738535 |
| AC016526.1 | 305.853362 | -0.0820016 | 0.77650104 | -0.105604 | 0.91589659 | 0.98191348 |
| ESRRB | 6085.30994 | -0.2365059 | 0.41288515 | -0.5728127 | 0.56677151 | 0.87646121 |
| AC008050.1 | 43.5093016 | -0.1119265 | 0.69458559 | -0.1611414 | 0.87198207 | 0.97535872 |
| VASH1-AS1 | 54.9312823 | -0.4544034 | 0.36557157 | -1.2429945 | 0.21386987 | 0.66800831 |
| ANGEL1 | 198.409887 | -0.298502 | 0.190483 | -1.5670797 | 0.11709607 | 0.54756317 |
| IRF2BPL | 346.64851 | 1.02014726 | 0.42134278 | 2.4211813 | 0.01547016 | 0.25214021 |
| AC007686.2 | 64.9516229 | 1.35131299 | 1.00762986 | 1.34108073 | 0.17989424 | 0.63399761 |
| POMT2 | 195.951478 | -0.5071704 | 0.23588364 | -2.1500871 | 0.03154833 | 0.33950635 |
| GSTZ1 | 87.029198 | 0.04223815 | 0.29960303 | 0.14098037 | 0.88788545 | 0.97729111 |
| TMED8 | 556.473567 | -0.1119761 | 0.14922207 | -0.7503994 | 0.45301421 | 0.82982454 |
| SAMD15 | 64.9365156 | 0.11015522 | 0.45254971 | 0.24341022 | 0.80768763 | 0.95598082 |
| NOXRED1 | 126.189665 | -0.6434425 | 0.34850415 | -1.8462978 | 0.06484897 | 0.44041719 |
| VIPAS39 | 265.399853 | -0.1336066 | 0.30541498 | -0.4374592 | 0.66177837 | 0.91110616 |
| AHSA1 | 204.674784 | 0.58398064 | 0.26449896 | 2.20787501 | 0.02725299 | 0.31913538 |
| SPTLC2 | 1206.40488 | -0.2017335 | 0.22301627 | -0.9045686 | 0.36569398 | 0.78095617 |
| ALKBH1 | 365.789904 | -0.3551055 | 0.24015347 | -1.4786606 | 0.13923104 | 0.58273218 |
| SLIRP | 412.76478 | 0.28911952 | 0.30192232 | 0.95759571 | 0.33826666 | 0.76444215 |
| SNW1 | 438.593529 | -0.2697241 | 0.15157451 | -1.7794818 | 0.07516081 | 0.47190267 |
| ADCK1 | 240.222159 | -0.863294 | 0.29916187 | -2.8857085 | 0.00390534 | 0.12135113 |
| NRXN3 | 996.208854 | -2.1816498 | 0.60533152 | -3.6040578 | 0.00031329 | 0.03094754 |
| CEP128 | 528.571893 | -0.5021375 | 0.40003548 | -1.2552323 | 0.20939443 | 0.66216422 |
| TSHR | 62.7970367 | 0.25746234 | 0.59975648 | 0.42927813 | 0.66772083 | 0.9137375 |
| AC007262.2 | 254.61471 | 0.1055861 | 0.55888021 | 0.18892439 | 0.85015207 | 0.97041641 |
| GTF2A1 | 472.812003 | 0.08727891 | 0.1793468 | 0.48664883 | 0.62650722 | 0.89873262 |
| STON2 | 1476.49051 | 0.29316075 | 0.22443005 | 1.30624554 | 0.19146907 | 0.64571429 |
| SEL1L | 747.390986 | 0.25341906 | 0.11731498 | 2.16015934 | 0.03076034 | 0.33646028 |

| | | | | | | |
|---|---|---|---|---|---|---|
| AL355838.1 | 60.5603153 | -1.3016662 | 0.7043209 | -1.8481153 | 0.06458567 | 0.44001919 |
| LINC02301 | 52.3918722 | 1.13663426 | 0.57484687 | 1.9772818 | 0.04800979 | 0.39511605 |
| GALC | 245.526806 | -0.0346064 | 0.28922323 | -0.1196529 | 0.90475813 | 0.97992575 |
| SPATA7 | 651.719922 | -0.2351271 | 0.21095025 | -1.1146091 | 0.26501797 | 0.71512755 |
| PTPN21 | 570.934283 | -0.5697947 | 0.23338201 | -2.441468 | 0.01462768 | 0.2480322 |
| AL162171.2 | 46.5140847 | -0.0836127 | 0.36030791 | -0.2320589 | 0.81649226 | 0.9589293 |
| ZC3H14 | 1069.641 | -0.1630691 | 0.24556503 | -0.6640568 | 0.50665397 | 0.85626196 |
| EML5 | 459.373773 | -0.8833224 | 0.43159416 | -2.0466505 | 0.04069242 | 0.37344844 |
| TTC8 | 586.714917 | -0.1364459 | 0.19546786 | -0.6980479 | 0.48514724 | 0.8447301 |
| FOXN3 | 2172.96693 | 0.02677379 | 0.18860773 | 0.14195487 | 0.88711566 | 0.97729111 |
| AL356805.1 | 66.3542623 | -0.2160405 | 0.54706534 | -0.394908 | 0.69291077 | 0.92224682 |
| EFCAB11 | 539.769615 | -0.5887741 | 0.51886337 | -1.1347383 | 0.256485 | 0.70735159 |
| TDP1 | 246.820993 | -0.1611097 | 0.26893968 | -0.5990552 | 0.5491361 | 0.86904356 |
| KCNK13 | 569.538788 | -0.6996909 | 0.47628708 | -1.4690529 | 1.42E-01 | 5.87E-01 |
| PSMC1 | 316.767855 | 0.25059787 | 0.24787333 | 1.01099166 | 0.31202042 | 0.74664452 |
| NRDE2 | 420.213644 | -0.1124225 | 0.26913761 | -0.4177139 | 0.67615629 | 0.91650575 |
| CALM1 | 669.980969 | 0.46980704 | 0.27743808 | 1.69337618 | 0.09038389 | 0.50205246 |
| TTC7B | 1355.91247 | -0.3676712 | 0.24522598 | -1.499316 | 0.13379167 | 0.57704758 |
| RPS6KA5 | 740.120489 | -0.1970628 | 0.19968532 | -0.9868668 | 3.24E-01 | 0.75398666 |
| DGLUCY | 1312.11286 | 0.05642236 | 0.41059199 | 0.1374171 | 0.89070112 | 0.97729111 |
| CCDC88C | 63.8060983 | -0.2702109 | 0.37571356 | -0.719194 | 0.47202141 | 0.83921324 |
| PPP4R3A | 1246.14138 | -0.2072996 | 0.17358561 | -1.194221 | 0.23239155 | 0.68450837 |
| CATSPERB | 217.356383 | 0.77150182 | 0.37124782 | 2.07813159 | 0.03769724 | 0.36396365 |
| TC2N | 293.724982 | 0.42806043 | 0.23622033 | 1.81212358 | 0.0699671 | 0.45949593 |
| FBLN5 | 50.4492182 | 0.40033191 | 0.55435237 | 0.72216146 | 0.47019521 | 0.83921324 |
| TRIP11 | 1481.23501 | 0.16833424 | 0.14420807 | 1.16730113 | 0.24308878 | 0.6944117 |
| ATXN3 | 1068.87343 | -0.2957595 | 0.2241755 | -1.3193211 | 0.1870618 | 0.64195489 |
| NDUFB1 | 448.015182 | -0.1998286 | 0.26010534 | -0.7682604 | 0.4423325 | 0.82285053 |
| CPSF2 | 490.058022 | -0.1062346 | 0.19784042 | -0.5369714 | 0.59128738 | 0.8868841 |
| RIN3 | 55.5869773 | 0.5656938 | 0.49193682 | 1.1499318 | 0.25017196 | 0.7008529 |
| LGMN | 255.47554 | 0.55285083 | 0.43209389 | 1.27946923 | 0.20073187 | 0.6539586 |
| GOLGA5 | 360.399716 | -0.191106 | 0.15975082 | -1.1962758 | 0.23158894 | 0.68374125 |
| ITPK1 | 781.537416 | 0.23956879 | 0.43011988 | 0.55698143 | 0.57754012 | 0.88181117 |
| MOAP1 | 176.938607 | 0.49776639 | 0.48762502 | 1.02079749 | 0.30735039 | 0.74341546 |
| AL110118.2 | 65.4110469 | -0.3842714 | 0.36750049 | -1.045635 | 0.29572958 | 0.73661928 |
| GON7 | 103.962525 | 0.19284189 | 0.25531073 | 0.75532231 | 0.45005561 | 0.82727311 |
| UBR7 | 122.861333 | 0.05940908 | 0.23786616 | 0.24975842 | 0.80277418 | 0.95446295 |
| BTBD7 | 3267.62636 | -0.1329389 | 0.14282883 | -0.9307568 | 0.3519794 | 0.77151834 |
| UNC79 | 175.891703 | 0.72482143 | 0.54238908 | 1.3363496 | 0.18143503 | 0.63616641 |
| DDX24 | 1100.82034 | 0.20996405 | 0.16069157 | 1.30662765 | 1.91E-01 | 6.46E-01 |
| IFI27L1 | 106.846285 | -0.3221571 | 0.29080376 | -1.1078162 | 0.26794119 | 0.71800072 |
| IFI27 | 219.266661 | -0.8442111 | 0.88258654 | -0.9565193 | 0.33880994 | 0.76449297 |
| IFI27L2 | 74.0881746 | -0.25162 | 0.5605356 | -0.4488921 | 0.65350946 | 0.90884465 |
| PPP4R4 | 91.0507926 | -3.1005447 | 0.89176436 | -3.4768654 | 0.00050731 | 0.038512 |
| SERPINA1 | 1214.05281 | -2.3015751 | 0.84876011 | -2.7116909 | 0.0066941 | 0.16906706 |
| DICER1 | 2553.26145 | -0.0751295 | 0.15266662 | -0.4921149 | 0.62263811 | 0.89835355 |
| DICER1-AS1 | 237.036368 | -0.2123239 | 0.36144442 | -0.5874316 | 0.55691385 | 0.87221664 |
| CLMN | 5653.37622 | 0.3442428 | 0.44489265 | 0.77376598 | 0.43906919 | 0.82136595 |
| SNHG10 | 55.487455 | 0.20429184 | 0.32997444 | 0.61911412 | 0.53584118 | 0.8642718 |
| GLRX5 | 255.679655 | 0.64383701 | 0.34728513 | 1.85391472 | 0.06375136 | 0.43779712 |
| C14orf132 | 133.798295 | -0.4799494 | 0.34750652 | -1.3811236 | 0.16724097 | 0.62087856 |
| AL355102.2 | 111.539652 | 0.02473945 | 0.31503114 | 0.07853019 | 0.93740632 | 0.98661414 |
| AL355102.4 | 81.0552881 | 0.52176283 | 0.58025067 | 0.89920245 | 0.36854484 | 0.78095617 |
| ATG2B | 1532.68602 | -0.1445908 | 0.20476359 | -0.7061353 | 0.480104 | 0.8433217 |

| | | | | | | |
|---|---|---|---|---|---|---|
| GSKIP | 258.600416 | 0.04377055 | 0.26527678 | 0.16499956 | 0.86894433 | 0.97443634 |
| AK7 | 311.814868 | 0.20702222 | 0.27144155 | 0.76267698 | 0.44565606 | 0.82571115 |
| PAPOLA | 1867.30649 | 0.08468924 | 0.23493371 | 0.36048146 | 0.71848712 | 0.9298398 |
| VRK1 | 200.932062 | -0.2918342 | 0.29102492 | -1.0027809 | 0.31596658 | 0.74785672 |
| LINC01550 | 418.18147 | 0.54522433 | 0.33917753 | 1.60748952 | 0.10794703 | 0.53063658 |
| BCL11B | 236.649877 | 0.91186416 | 0.48414402 | 1.88345642 | 0.05963854 | 0.42650032 |
| SETD3 | 1312.83243 | -0.1487469 | 0.17608125 | -0.8447628 | 0.39824329 | 0.80148305 |
| CCNK | 377.101895 | -0.2418218 | 0.18151929 | -1.3322098 | 0.18279126 | 0.63741052 |
| CCDC85C | 180.831481 | 0.46970087 | 0.23953827 | 1.96085936 | 0.04989543 | 0.40017381 |
| CYP46A1 | 109.85311 | 0.0299041 | 0.50239931 | 0.05952257 | 0.95253589 | 0.98938795 |
| EML1 | 570.390074 | 0.66395728 | 0.26692162 | 2.48746163 | 0.01286583 | 0.23248261 |
| EVL | 910.229459 | -0.4762641 | 0.33460203 | -1.4233748 | 0.15462752 | 0.60380955 |
| DEGS2 | 58.2289664 | 0.7123199 | 0.65006585 | 1.09576577 | 0.27318129 | 0.72070318 |
| AL133523.1 | 91.6972718 | -0.253046 | 0.25971455 | -0.9743234 | 0.329896 | 0.75885186 |
| YY1 | 1131.29338 | -0.0202588 | 0.17464843 | -0.1159977 | 0.90765439 | 0.98047754 |
| SLC25A29 | 1172.49628 | -0.0417976 | 0.34314202 | -0.1218085 | 0.90305068 | 0.97960781 |
| WARS | 447.030089 | -0.245755 | 0.44912678 | -0.547184 | 0.58425234 | 0.88416261 |
| AL157871.4 | 84.6607953 | -0.380865 | 0.42463835 | -0.8969161 | 0.36976366 | 0.78095617 |
| WDR25 | 424.050522 | -0.4015324 | 0.25892745 | -1.5507525 | 0.12096101 | 0.55498372 |
| PPP2R5C | 1706.93983 | 0.13513398 | 0.18445235 | 0.73262272 | 0.46378857 | 0.83531269 |
| DYNC1H1 | 1051.5289 | 0.10424771 | 0.29099037 | 0.3582514 | 0.72015518 | 0.93001079 |
| HSP90AA1 | 2781.39242 | 0.62171244 | 0.28525739 | 2.1794788 | 0.02929612 | 0.33012415 |
| WDR20 | 1511.14846 | -0.2162277 | 0.17813062 | -1.2138718 | 0.22479669 | 0.67750405 |
| MOK | 1348.8393 | -0.268492 | 0.27915932 | -0.9617877 | 0.33615625 | 0.7637525 |
| ZNF839 | 259.584178 | -0.3331341 | 0.29407718 | -1.1328119 | 0.25729325 | 0.70774396 |
| CINP | 135.384247 | 0.21747159 | 0.22635209 | 0.96076684 | 0.33666942 | 0.76409325 |
| TECPR2 | 710.908017 | -0.1849087 | 0.19106223 | -0.967793 | 0.33314776 | 0.76124971 |
| ANKRD9 | 255.385989 | 0.52712227 | 0.36636808 | 1.43877783 | 0.15021348 | 0.59663305 |
| RCOR1 | 1477.42099 | -0.0908957 | 0.23996181 | -0.3787923 | 0.70484209 | 0.92528913 |
| TRAF3 | 996.82927 | 0.32313503 | 0.28018319 | 1.15329913 | 2.49E-01 | 6.99E-01 |
| AMN | 100.365044 | 1.05879942 | 0.43298962 | 2.44532288 | 0.01447225 | 0.24762526 |
| CDC42BPB | 1458.20899 | -0.4430827 | 0.22888067 | -1.9358678 | 0.0528839 | 0.41066631 |
| AL117209.1 | 59.9792797 | -0.0752497 | 0.50925228 | -0.1477651 | 0.88252814 | 0.97664137 |
| TNFAIP2 | 323.503325 | -0.3550433 | 0.45349912 | -0.7828974 | 4.34E-01 | 8.19E-01 |
| AL161669.3 | 68.9682414 | 0.83468849 | 0.61984246 | 1.34661393 | 0.17810461 | 0.63204571 |
| EIF5 | 795.110593 | 0.03341533 | 0.31043913 | 0.10763892 | 0.91428212 | 0.98145965 |
| MARK3 | 3266.71628 | -0.1323971 | 0.19910518 | -0.6649606 | 0.50607569 | 0.85626196 |
| CKB | 1931.98168 | 1.48344456 | 0.45207196 | 3.28143454 | 0.0010328 | 0.05732067 |
| TRMT61A | 57.4677925 | 0.33159281 | 0.43525744 | 0.76183146 | 0.4461606 | 0.82571115 |
| AL139300.2 | 51.4535905 | 0.66146053 | 0.48557379 | 1.36222453 | 0.17312704 | 0.6286116 |
| BAG5 | 410.299334 | 0.79570414 | 0.2957868 | 2.69012727 | 0.00714248 | 0.17279872 |
| KLC1 | 765.398953 | -0.0819733 | 0.18253029 | -0.4490941 | 0.65336377 | 0.90883341 |
| AL049840.1 | 82.3218057 | 0.09109224 | 0.58110009 | 0.15675827 | 0.87543536 | 0.97614152 |
| ZFYVE21 | 229.061856 | 0.35415938 | 0.21475717 | 1.64911551 | 9.91E-02 | 0.51618562 |
| PPP1R13B | 2245.42526 | -0.288346 | 0.12834849 | -2.2465867 | 0.02466646 | 0.30812502 |
| ATP5MPL | 432.433954 | 0.50642782 | 0.26107063 | 1.93981154 | 0.0524026 | 0.4090401 |
| TDRD9 | 187.840063 | -0.1109528 | 0.78460634 | -0.141412 | 0.88754446 | 0.97729111 |
| AL590079.1 | 111.925669 | 0.91355954 | 0.84304433 | 1.08364354 | 0.27852288 | 0.7242837 |
| INF2 | 115.706951 | 0.41324408 | 0.39766313 | 1.0391813 | 2.99E-01 | 7.37E-01 |
| SIVA1 | 178.826196 | 0.25950121 | 0.26603998 | 0.97542187 | 0.32935104 | 0.75817321 |
| AKT1 | 288.945087 | -0.2459482 | 0.25895561 | -0.9497698 | 0.34222923 | 0.76591028 |
| AL583810.1 | 62.5465617 | 0.41304148 | 0.34143038 | 1.2097385 | 0.22637925 | 0.67880306 |
| CEP170B | 275.067929 | -0.0387767 | 0.29134563 | -0.1330952 | 0.89411809 | 0.97788747 |
| CLBA1 | 127.253241 | 0.09067906 | 0.2677243 | 0.33870313 | 0.73483338 | 0.93552616 |

| | | | | | | |
|---|---|---|---|---|---|---|
| CDCA4 | 46.3277531 | 0.34422805 | 0.41568284 | 0.82810262 | 0.40761239 | 0.80588056 |
| GPR132 | 505.930353 | 0.51432164 | 0.71464353 | 0.71968977 | 0.47171603 | 0.83921324 |
| JAG2 | 58.979112 | 0.81616406 | 0.47245704 | 1.72748841 | 0.08407998 | 0.48990407 |
| NUDT14 | 122.544344 | 1.09749684 | 0.45178373 | 2.42925268 | 0.01512998 | 0.25080486 |
| BRF1 | 364.438024 | -0.1539249 | 0.2474385 | -0.6220735 | 0.53389357 | 0.86397197 |
| BTBD6 | 91.5639183 | 0.42172267 | 0.35765155 | 1.17914396 | 0.23834086 | 0.68982886 |
| PACS2 | 684.061267 | -0.2416128 | 0.2553451 | -0.9462206 | 0.34403607 | 0.76685327 |
| TEX22 | 141.334435 | -0.3770613 | 0.35619793 | -1.0585723 | 0.28979459 | 0.73204984 |
| MTA1 | 401.502982 | -0.3641682 | 0.31878689 | -1.1423562 | 0.25330599 | 0.70408029 |
| CRIP2 | 328.540928 | 0.07303607 | 0.52305196 | 0.13963445 | 0.88894882 | 0.97729111 |
| AC087386.1 | 102.784795 | 0.05712703 | 0.48865939 | 0.11690563 | 0.90693484 | 0.9803357 |
| NIPA1 | 549.51209 | -0.0606888 | 0.1755262 | -0.3457532 | 0.72952817 | 0.93453345 |
| NIPA2 | 433.034707 | -0.1110051 | 0.18643033 | -0.5954241 | 0.55155999 | 0.87039028 |
| CYFIP1 | 931.760962 | -0.1331945 | 0.17963775 | -0.7414616 | 0.45841358 | 0.8331421 |
| TUBGCP5 | 428.708393 | -0.3710548 | 0.1766375 | -2.1006568 | 0.0356711 | 0.35661333 |
| SNRPN | 77.8203624 | -0.5521559 | 0.55943377 | -0.9869906 | 0.32364726 | 0.75397825 |
| AC124312.1 | 2557.36613 | 0.03933804 | 0.28205219 | 0.13947077 | 0.88907815 | 0.97729111 |
| AC124312.2 | 81.87391 | 0.15148232 | 1.03444365 | 0.14643845 | 0.88357528 | 0.97664137 |
| PWAR6 | 634.479322 | 0.15365794 | 0.34364827 | 0.44713724 | 0.65477597 | 0.90902447 |
| UBE3A | 1894.11035 | -0.089615 | 0.16404136 | -0.5462951 | 0.58486312 | 0.8846469 |
| LINC02250 | 59.3218341 | 0.81621263 | 0.63045999 | 1.29463033 | 0.19544779 | 0.64893292 |
| GABRB3 | 1084.60389 | 0.05401523 | 0.43482373 | 0.12422329 | 0.90113848 | 0.97931857 |
| OCA2 | 59.0709274 | 0.16531962 | 0.67176498 | 0.24609741 | 0.80560683 | 0.95522565 |
| HERC2 | 2233.97012 | -0.2981126 | 0.15436801 | -1.9311813 | 0.05346063 | 0.4122367 |
| GOLGA8M | 88.3308701 | -0.0338939 | 0.56757905 | -0.0597165 | 0.9523814 | 0.98938795 |
| FAM189A1 | 124.090169 | 1.0194148 | 0.38949888 | 2.61724708 | 0.00886422 | 0.19125454 |
| TJP1 | 1836.15768 | -0.1839197 | 0.20108089 | -0.9146552 | 0.36037268 | 0.77716943 |
| AC091057.2 | 50.9076893 | -0.4593558 | 0.43144382 | -1.0646944 | 0.28701425 | 0.72972899 |
| FAN1 | 548.414226 | -0.3036103 | 0.24730366 | -1.2276821 | 0.21956635 | 0.67278782 |
| MTMR10 | 1737.53477 | -0.2035054 | 0.27409384 | -0.7424661 | 0.45780497 | 0.83282548 |
| KLF13 | 361.163892 | 0.17719487 | 0.38422666 | 0.46117275 | 0.64467467 | 0.90507831 |
| OTUD7A | 1005.19451 | -0.5462229 | 0.2961241 | -1.8445742 | 0.0650995 | 0.44142675 |
| CHRNA7 | 55.7431976 | -0.2007849 | 0.67980103 | -0.2953583 | 0.76772015 | 0.94316227 |
| LINC02256 | 1645.40566 | 0.16201649 | 0.36329333 | 0.44596606 | 0.65562177 | 0.90919768 |
| AC123768.3 | 264.246402 | -0.2042783 | 0.31140235 | -0.6559947 | 0.51182754 | 0.85762435 |
| GOLGA8N | 199.455519 | 0.26403738 | 0.29618369 | 0.89146496 | 0.37267978 | 0.78269743 |
| FMN1 | 2698.15342 | -0.4537399 | 0.63857476 | -0.7105509 | 0.47736259 | 0.84174109 |
| RYR3 | 129.706194 | 1.24731439 | 0.46955479 | 2.65637665 | 0.00789853 | 0.17862547 |
| AC010809.1 | 55.4322451 | -0.5620636 | 0.40122958 | -1.4008529 | 0.16125806 | 0.61261368 |
| AVEN | 619.737511 | -0.2435725 | 0.29070107 | -0.8378796 | 0.40209835 | 0.80296585 |
| CHRM5 | 108.792823 | 0.00332457 | 0.50286219 | 0.0066113 | 0.99472498 | 0.998438 |
| EMC7 | 243.726571 | 0.41747624 | 0.24884747 | 1.67763904 | 0.09341759 | 0.50887217 |
| PGBD4 | 53.1854081 | -0.0813998 | 0.33455192 | -0.2433098 | 8.08E-01 | 0.9559872 |
| KATNBL1 | 1188.36174 | -0.2848474 | 0.1412924 | -2.0160135 | 0.04379856 | 0.38453712 |
| EMC4 | 139.464699 | -0.0001069 | 0.30786779 | -0.0003473 | 0.99972293 | 0.99987432 |
| SLC12A6 | 742.271857 | -0.5244921 | 0.26018103 | -2.0158737 | 0.04381319 | 0.38453712 |
| NOP10 | 257.345475 | 0.26286056 | 0.43330578 | 0.60663987 | 0.54408993 | 0.8664796 |
| LPCAT4 | 93.6396442 | -1.1251889 | 0.31888872 | -3.5284688 | 0.00041797 | 0.03514528 |
| GOLGA8A | 903.614568 | -0.1344206 | 0.26297671 | -0.5111502 | 0.60924591 | 0.8927935 |
| GOLGA8B | 970.246829 | 0.40162672 | 0.37120549 | 1.08195252 | 0.27927363 | 0.724832 |
| AQR | 991.451556 | -0.3138942 | 0.19177619 | -1.6367734 | 0.10167783 | 0.5195058 |
| ZNF770 | 457.965297 | 0.29710935 | 0.2856769 | 1.04001882 | 0.29833116 | 0.73673761 |
| AC021231.1 | 1317.86685 | 0.68772413 | 0.43578713 | 1.57811941 | 0.11453818 | 0.54179726 |
| DPH6 | 3869.0526 | 0.06286207 | 0.33029811 | 0.19031921 | 0.849059 | 0.97017477 |

| | | | | | | |
|---|---|---|---|---|---|---|
| DPH6-DT | 2224.11943 | -0.2968552 | 0.58571849 | -0.5068224 | 0.61227944 | 0.89440588 |
| AC021351.1 | 47.3920039 | -0.5904961 | 0.86424553 | -0.6832504 | 0.49444861 | 0.84882719 |
| C15orf41 | 783.365825 | -0.522799 | 0.22151985 | -2.3600551 | 0.01827222 | 0.2711921 |
| MEIS2 | 67.5108898 | -0.3740342 | 0.63290164 | -0.5909831 | 0.55453176 | 0.87160642 |
| SPRED1 | 1336.73347 | -0.23983 | 0.32346067 | -0.7414502 | 0.45842053 | 0.8331421 |
| FAM98B | 240.666641 | 0.08143603 | 0.21823083 | 0.37316466 | 0.70902589 | 0.92577218 |
| RASGRP1 | 114.719463 | 0.78988249 | 0.27227337 | 2.90106411 | 0.00371898 | 0.11865695 |
| C15orf53 | 231.800428 | 1.09407288 | 0.68002803 | 1.60886439 | 0.107646 | 0.52996496 |
| AC013652.1 | 506.173325 | 0.33784618 | 0.58429702 | 0.57820966 | 0.56312258 | 0.87447201 |
| THBS1 | 208.146469 | 0.2074311 | 0.59962658 | 0.3459338 | 0.72939247 | 0.93453345 |
| FSIP1 | 234.620267 | 0.20797383 | 0.17622559 | 1.18015683 | 0.23793785 | 0.68982886 |
| EIF2AK4 | 542.951024 | -0.178585 | 0.16852723 | -1.0596805 | 0.28928996 | 0.73204984 |
| SRP14 | 1287.85953 | 0.67442825 | 0.45527087 | 1.481378 | 0.13850587 | 0.58206378 |
| SRP14-AS1 | 207.682007 | 0.24977183 | 0.35179159 | 0.70999945 | 0.47770448 | 0.84190259 |
| BMF | 130.100194 | -0.35436 | 0.53110214 | -0.6672163 | 0.504634 | 0.85556546 |
| PAK6 | 41.6427893 | 1.17709429 | 0.61247889 | 1.92185282 | 0.05462428 | 0.41650243 |
| CCDC9B | 77.8156412 | -0.2509902 | 0.4692582 | -0.5348659 | 0.59274257 | 0.88740306 |
| KNSTRN | 45.9711976 | -0.2193904 | 0.46333246 | -0.4735053 | 0.63585272 | 0.90207488 |
| IVD | 455.984132 | 0.32606443 | 0.22856149 | 1.42659392 | 0.15369698 | 0.60285224 |
| BAHD1 | 231.597991 | 0.25242385 | 0.27867696 | 0.90579376 | 0.36504502 | 0.78095617 |
| AC091045.1 | 105.243762 | 0.08080341 | 0.7084944 | 0.11404946 | 0.90919858 | 0.98069436 |
| CCDC32 | 476.890505 | -0.2644123 | 0.2464875 | -1.072721 | 0.28339632 | 0.72668316 |
| RPUSD2 | 44.1675868 | 0.50431685 | 0.34810607 | 1.44874477 | 0.14740887 | 0.59373375 |
| KNL1 | 45.9296567 | -0.9231203 | 0.5768459 | -1.6002892 | 0.10953444 | 0.53361622 |
| RMDN3 | 231.590024 | -0.1278432 | 0.25019491 | -0.5109743 | 0.60936908 | 0.89286259 |
| GCHFR | 107.821078 | 1.04598655 | 0.50885926 | 2.05555178 | 3.98E-02 | 0.37279391 |
| DNAJC17 | 457.470698 | -0.2464343 | 0.24571547 | -1.0029255 | 0.31589678 | 0.74785672 |
| ZFYVE19 | 142.147107 | -0.5941443 | 0.28677506 | -2.0718129 | 0.03828289 | 0.36662698 |
| SPINT1-AS1 | 183.744214 | 0.64121715 | 0.22581378 | 2.83958383 | 0.00451724 | 0.13085144 |
| SPINT1 | 470.960304 | 0.10116826 | 0.27435512 | 0.36874931 | 7.12E-01 | 9.27E-01 |
| VPS18 | 79.1406342 | 0.13396874 | 0.33375254 | 0.4014014 | 0.68812462 | 0.92128908 |
| INO80 | 1079.21194 | -0.3523647 | 0.20986934 | -1.6789718 | 0.09315753 | 0.50826841 |
| AC020661.3 | 43.1374708 | 0.09815331 | 0.38882873 | 0.25243326 | 0.80070619 | 0.95370871 |
| INO80-AS1 | 46.4663324 | -0.4077927 | 0.56590658 | -0.7206008 | 0.47115515 | 0.83921324 |
| CHP1 | 879.149567 | 0.68192791 | 0.22793033 | 2.99182607 | 0.00277314 | 0.09937076 |
| OIP5 | 57.4558135 | 0.4450613 | 0.48681181 | 0.91423687 | 0.3605924 | 0.77730375 |
| NDUFAF1 | 158.285056 | 0.01614228 | 0.23966859 | 0.06735249 | 0.94630109 | 0.98882138 |
| RTF1 | 609.908858 | -0.1193825 | 0.22008045 | -0.5424495 | 0.58750889 | 0.88571413 |
| RPAP1 | 141.428634 | -0.0103482 | 0.27728308 | -0.0373198 | 0.97022999 | 0.99309664 |
| MGA | 1070.15984 | -0.0235824 | 0.21102112 | -0.1117536 | 0.91101878 | 0.98111638 |
| MAPKBP1 | 727.333992 | -0.4662428 | 0.29112164 | -1.6015394 | 0.1092575 | 0.53310139 |
| EHD4 | 236.075133 | 0.21310048 | 0.31832225 | 0.66944891 | 0.50320916 | 0.85525037 |
| PLA2G4F | 97.7409113 | 1.89854317 | 0.91325513 | 2.07887489 | 0.03762885 | 0.36389125 |
| VPS39 | 762.530268 | -0.1929073 | 0.16084397 | -1.1993444 | 0.23039405 | 0.68265478 |
| TMEM87A | 1062.36809 | -0.3563925 | 0.26954472 | -1.3222019 | 0.18610092 | 0.64033103 |
| GANC | 660.900751 | -0.2304668 | 0.35309646 | -0.6527021 | 0.51394835 | 0.85762435 |
| ZNF106 | 896.462191 | 0.23790485 | 0.17478948 | 1.36109368 | 0.17348409 | 0.6286116 |
| SNAP23 | 606.622498 | 0.0169394 | 0.21362309 | 0.07929574 | 0.9367974 | 0.98630295 |
| LRRC57 | 101.036402 | -0.0996545 | 0.22561657 | -0.4416983 | 0.65870751 | 0.91051355 |
| HAUS2 | 166.185448 | -0.1136815 | 0.25348652 | -0.4484716 | 0.65381287 | 0.90897911 |
| STARD9 | 45.9094929 | 0.16396364 | 0.34982922 | 0.46869623 | 6.39E-01 | 0.90330608 |
| CDAN1 | 82.6752686 | 0.23273416 | 0.32888851 | 0.70763847 | 0.4791698 | 0.84296219 |
| TTBK2 | 1258.63323 | 0.0029636 | 0.21649696 | 0.01368888 | 0.9890782 | 0.99670729 |
| UBR1 | 2064.69452 | -0.0403193 | 0.151477 | -0.2661744 | 0.79010491 | 0.95123267 |

| | | | | | | |
|---|---|---|---|---|---|---|
| AC068724.3 | 289.365363 | 0.05599539 | 0.38141405 | 0.14680998 | 0.88328201 | 0.97664137 |
| TMEM62 | 566.122678 | -0.4351011 | 0.25046131 | -1.7371987 | 0.0823521 | 0.48594425 |
| CCNDBP1 | 272.636998 | 0.50732187 | 0.19204189 | 2.64172498 | 0.0082485 | 0.18342499 |
| ADAL | 411.248852 | 0.07609717 | 0.32322542 | 0.23543065 | 0.8138745 | 0.9582375 |
| ZSCAN29 | 123.80451 | -0.1196467 | 0.29927302 | -0.3997911 | 0.68931035 | 0.92166785 |
| TUBGCP4 | 297.412961 | -0.2020102 | 0.2085734 | -0.9685331 | 0.33277821 | 0.7610305 |
| TP53BP1 | 1428.72597 | 0.18207883 | 0.1748371 | 1.04141983 | 0.29768074 | 0.73673761 |
| PPIP5K1 | 514.079594 | -0.008783 | 0.33177998 | -0.0264724 | 0.97888055 | 0.99455895 |
| AC011330.3 | 71.0301313 | -1.1308432 | 0.55772457 | -2.0276015 | 0.04260094 | 0.38021566 |
| CATSPER2 | 579.113085 | -0.1217011 | 0.49550959 | -0.245608 | 0.80598573 | 0.95525039 |
| PDIA3 | 812.633982 | 0.83802608 | 0.20450028 | 4.09792139 | 4.17E-05 | 0.01031432 |
| AC018512.1 | 45.7967386 | -0.6884283 | 0.39265084 | -1.7532837 | 0.07955333 | 0.48176967 |
| SERF2 | 2177.88609 | 0.53560654 | 0.48626125 | 1.10147897 | 0.27068825 | 0.71965808 |
| MFAP1 | 218.372336 | 0.2218342 | 0.23434815 | 0.94660103 | 0.34384212 | 0.76680914 |
| WDR76 | 53.999489 | -0.4174274 | 0.37728678 | -1.1063929 | 0.26855649 | 0.71818262 |
| FRMD5 | 82.9258467 | -0.0986888 | 0.29698125 | -0.3323065 | 0.73965781 | 0.93552616 |
| CASC4 | 2385.35377 | -0.3199067 | 0.17157534 | -1.8645259 | 0.06224788 | 0.43419765 |
| AC025043.1 | 115.744894 | 0.09025315 | 0.48334085 | 0.18672775 | 0.85187409 | 0.97076831 |
| CTDSPL2 | 1458.94243 | -0.3658099 | 0.17530137 | -2.0867487 | 0.03691086 | 0.36038369 |
| EIF3J-DT | 424.448967 | 0.33286641 | 0.29179235 | 1.1407647 | 0.25396785 | 0.70431689 |
| EIF3J | 561.254691 | -0.0240626 | 0.2034527 | -0.1182714 | 0.90585263 | 0.97997915 |
| SPG11 | 1862.15419 | -0.2839174 | 0.21370886 | -1.3285241 | 0.18400504 | 0.63843513 |
| B2M | 4055.12428 | 1.0339522 | 0.46422981 | 2.22724213 | 0.0259311 | 0.31222606 |
| TRIM69 | 130.104227 | 0.03494528 | 0.4897305 | 0.07135615 | 0.94311431 | 0.98772472 |
| SORD | 59.9941384 | 1.28136651 | 0.60455165 | 2.1195319 | 0.03404554 | 0.35153234 |
| DUOX1 | 168.992394 | 0.0832883 | 0.35047912 | 0.23764127 | 0.81215934 | 0.95806331 |
| SHF | 58.9501272 | -0.0977596 | 0.44931059 | -0.2175769 | 0.82775881 | 0.96274246 |
| AC051619.5 | 68.42027 | 0.50815222 | 0.62657476 | 0.81100014 | 0.41736559 | 0.81017256 |
| GATM | 208.180842 | 0.80602867 | 0.38551168 | 2.09080223 | 0.0365458 | 0.35917666 |
| SPATA5L1 | 174.03513 | -0.2386154 | 0.25253777 | -0.94487 | 0.34472523 | 0.76724655 |
| AC025580.2 | 79.2367119 | 1.32319055 | 0.69108394 | 1.91465967 | 0.05553592 | 0.41843207 |
| SLC30A4 | 111.494208 | 0.71600099 | 0.34122429 | 2.09832948 | 0.03587605 | 0.35661333 |
| SQOR | 133.34889 | 0.44909629 | 0.33030243 | 1.35965177 | 1.74E-01 | 6.29E-01 |
| SEMA6D | 1930.90753 | -0.132985 | 0.3854949 | -0.3449722 | 7.30E-01 | 9.35E-01 |
| AC092078.2 | 1190.2981 | 1.85409915 | 0.92958975 | 1.99453485 | 4.61E-02 | 0.3904347 |
| SLC24A5 | 91.349018 | 0.02675742 | 0.71689586 | 0.03732399 | 0.97022667 | 0.99309664 |
| MYEF2 | 1848.24336 | -0.4105821 | 0.2259891 | -1.8168225 | 0.0692443 | 0.45732396 |
| SLC12A1 | 33457.3288 | 0.61367632 | 0.56039162 | 1.09508474 | 2.73E-01 | 7.21E-01 |
| AC023355.1 | 100.422528 | 0.24362524 | 0.71002735 | 0.34312093 | 0.73150749 | 0.93529014 |
| DUT | 353.664548 | 0.16850508 | 0.27810119 | 0.60591285 | 0.54457263 | 0.86654229 |
| FBN1 | 139.800206 | -0.38914 | 0.36791435 | -1.0576918 | 0.29019597 | 0.73222514 |
| CEP152 | 619.40339 | -0.3866877 | 0.3172962 | -1.2186963 | 0.2229595 | 0.67594033 |
| EID1 | 470.52377 | 1.11733724 | 0.3684919 | 3.0321894 | 0.00242787 | 0.09404607 |
| SECISBP2L | 2100.99131 | 0.0174857 | 0.11819122 | 0.14794416 | 0.88238684 | 0.97664137 |
| COPS2 | 412.308101 | 0.15853119 | 0.18763003 | 0.84491378 | 0.39815898 | 0.80147546 |
| GALK2 | 1184.54563 | 0.26344817 | 0.2065229 | 1.27563662 | 0.202084 | 0.65552334 |
| FAM227B | 1070.20165 | 0.23491472 | 0.34565676 | 0.6796185 | 0.49674606 | 0.85081933 |
| FGF7 | 48.005022 | 1.32468506 | 0.50009927 | 2.64884421 | 0.00807676 | 0.1802126 |
| DTWD1 | 743.080921 | -0.1475667 | 0.23659308 | -0.6237153 | 5.33E-01 | 0.86397197 |
| SLC27A2 | 75.9175694 | 2.36700185 | 0.6391411 | 3.70341049 | 0.00021272 | 0.02361195 |
| GABPB1 | 525.585347 | -0.2117044 | 0.20928718 | -1.0115496 | 0.31175346 | 0.74627609 |
| GABPB1-AS1 | 1165.18995 | 0.32065477 | 0.44433765 | 0.72164663 | 0.47051176 | 0.83921324 |
| USP8 | 1433.84137 | -0.2106578 | 0.17037763 | -1.2364168 | 0.21630368 | 0.66968853 |
| USP50 | 263.030534 | -1.1839323 | 0.45543253 | -2.5995779 | 0.00933385 | 0.19663607 |

| | | | | | | |
|---|---|---|---|---|---|---|
| TRPM7 | 3518.87428 | -0.5472519 | 0.252347 | -2.1686482 | 0.0301094 | 0.33337391 |
| SPPL2A | 1919.24654 | -0.1467606 | 0.18728742 | -0.7836117 | 0.43326799 | 0.81862923 |
| AC021752.1 | 88.7251104 | 0.17710124 | 0.50066829 | 0.35372969 | 0.72354147 | 0.93188356 |
| AP4E1 | 627.143911 | -0.1767713 | 0.2664907 | -0.6633301 | 0.50711919 | 0.85626196 |
| DMXL2 | 710.030892 | -0.2123307 | 0.20590262 | -1.0312191 | 0.30243809 | 0.73905643 |
| LYSMD2 | 52.4445062 | 0.63378959 | 0.46017956 | 1.37726585 | 0.16843007 | 0.62253512 |
| TMOD2 | 117.706411 | 0.41070572 | 0.31502785 | 1.30371244 | 0.19233165 | 0.64660441 |
| TMOD3 | 1205.86726 | -0.2816091 | 0.13226687 | -2.1290976 | 0.03324618 | 0.34850106 |
| AC090971.2 | 76.2380314 | -0.1357197 | 0.47943302 | -0.2830837 | 0.77711266 | 0.94694572 |
| LEO1 | 154.446165 | -0.2113302 | 0.25560123 | -0.8267963 | 0.40835251 | 0.80588056 |
| MAPK6 | 665.362523 | -0.3738986 | 0.25084061 | -1.4905824 | 0.13607116 | 0.57783867 |
| GNB5 | 592.413729 | 0.30240375 | 0.21301362 | 1.41964515 | 1.56E-01 | 6.05E-01 |
| CERNA1 | 119.718973 | 0.03248363 | 0.57653797 | 0.05634256 | 0.95506891 | 0.99053904 |
| MYO5C | 1383.16027 | 0.11447269 | 0.26256287 | 0.43598202 | 0.66284977 | 0.91128046 |
| MYO5A | 955.085387 | 0.23977973 | 0.23054318 | 1.04006432 | 0.29831002 | 0.73673761 |
| ARPP19 | 647.14753 | 0.15739021 | 0.19201398 | 0.81968098 | 0.412398 | 0.80701706 |
| AC025917.1 | 67.9579236 | 0.37941974 | 0.75338623 | 0.50361917 | 0.61452902 | 0.89496546 |
| FAM214A | 1916.77405 | -0.0156169 | 0.2432218 | -0.0642084 | 0.94880432 | 0.98919519 |
| WDR72 | 3390.6106 | 1.65937414 | 0.45529747 | 3.6445934 | 0.00026781 | 0.02785486 |
| UNC13C | 691.537079 | -2.2435569 | 1.2788136 | -1.7544049 | 7.94E-02 | 0.48169281 |
| RSL24D1 | 229.926554 | 0.15170249 | 0.31727793 | 0.47813753 | 0.63255232 | 0.9005587 |
| RAB27A | 1022.24437 | 0.00438126 | 0.30651946 | 0.01429357 | 9.89E-01 | 0.99660871 |
| PIGB | 408.440045 | -0.1727253 | 0.20176033 | -0.8560916 | 0.39194714 | 0.79767768 |
| CCPG1 | 1177.7879 | 0.31615759 | 0.11617045 | 2.72149755 | 0.00649869 | 0.16670031 |
| C15orf65 | 76.01349 | 0.81611006 | 0.39867651 | 2.04704829 | 0.04065334 | 0.37344844 |
| DNAAF4 | 500.437209 | 0.20815382 | 0.30186573 | 0.68955763 | 0.49047242 | 0.84614483 |
| PYGO1 | 67.6461885 | 0.5305138 | 0.46761689 | 1.13450521 | 0.25658269 | 0.70735159 |
| PRTG | 205.460627 | 2.85023132 | 0.88310995 | 3.22749316 | 0.0012488 | 0.0645665 |
| NEDD4 | 273.372641 | 0.65210333 | 0.3433283 | 1.89935795 | 5.75E-02 | 4.21E-01 |
| RFX7 | 1450.40422 | -0.0239731 | 0.15733223 | -0.1523723 | 0.87889329 | 0.9765717 |
| TEX9 | 992.194908 | 0.11883226 | 0.2127887 | 0.55845195 | 0.57653581 | 0.88147804 |
| MNS1 | 176.644387 | 0.61845256 | 0.37163504 | 1.66413952 | 0.09608455 | 0.5117083 |
| AC090518.1 | 93.4590516 | 1.01203752 | 0.50470941 | 2.00518851 | 0.0449429 | 0.38724771 |
| ZNF280D | 803.046732 | 0.29459426 | 0.2110807 | 1.39564755 | 0.16282066 | 0.61387934 |
| AC090517.4 | 484.203946 | -0.5634549 | 0.30421501 | -1.85216 | 0.06400285 | 0.43872009 |
| TCF12 | 6997.18492 | -0.2291189 | 0.20825633 | -1.1001772 | 0.27125493 | 0.7200754 |
| CGNL1 | 11725.1648 | 0.44314651 | 0.39896798 | 1.11073201 | 0.26668372 | 0.71641758 |
| POLR2M | 125.221748 | -0.6333635 | 0.30355187 | -2.0865083 | 0.03693261 | 0.36038369 |
| ALDH1A2 | 4184.30232 | -0.1972064 | 0.63624339 | -0.3099544 | 0.75659565 | 0.93989203 |
| LIPC | 662.948973 | -1.5304215 | 0.43655402 | -3.5056865 | 0.00045543 | 0.03713452 |
| ADAM10 | 3487.55584 | -0.6676567 | 0.23657104 | -2.822225 | 0.00476917 | 0.1366507 |
| MINDY2 | 1772.86279 | -0.4135184 | 0.20360068 | -2.0310265 | 0.0422523 | 0.37920308 |
| RNF111 | 2012.36768 | -0.1400189 | 0.20363981 | -0.6875812 | 0.49171653 | 0.84704258 |
| SLTM | 2386.43273 | -0.2902309 | 0.15096177 | -1.9225455 | 0.05453715 | 0.41616478 |
| MYO1E | 4319.23873 | -0.8692749 | 0.34887841 | -2.4916272 | 0.01271594 | 0.23135662 |
| FAM81A | 222.867346 | -0.074107 | 0.46612758 | -0.1589843 | 0.87368121 | 0.97564728 |
| GCNT3 | 87.4437492 | 1.5063943 | 0.72870102 | 2.06723232 | 3.87E-02 | 3.69E-01 |
| GTF2A2 | 351.368551 | 0.25663672 | 0.27626162 | 0.92896265 | 0.35290844 | 0.77216708 |
| BNIP2 | 628.009675 | -0.1765054 | 0.11441512 | -1.5426759 | 0.12290943 | 0.55947129 |
| ANXA2 | 971.545496 | 0.18590753 | 0.56547137 | 0.32876559 | 7.42E-01 | 9.36E-01 |
| ICE2 | 994.444126 | -0.0420269 | 0.18998549 | -0.2212114 | 0.82492786 | 0.96146415 |
| RORA-AS1 | 685.176397 | 0.28513987 | 0.37079053 | 0.76900526 | 0.44189019 | 0.82285053 |
| RORA | 11124.5279 | 0.31470791 | 0.32123799 | 0.97967215 | 0.32724798 | 0.75583469 |
| AC012404.1 | 328.401398 | 0.76937224 | 0.38016244 | 2.02379866 | 0.04299087 | 0.3821443 |

| | | | | | | |
|---|---|---|---|---|---|---|
| AC009554.1 | 529.219699 | -0.3337633 | 0.75802206 | -0.4403082 | 0.65971393 | 0.91099119 |
| VPS13C | 3882.84046 | -0.4618123 | 0.34838035 | -1.325598 | 0.18497288 | 0.64011182 |
| C2CD4A | 280.818788 | 0.43690988 | 0.97821315 | 0.44664078 | 6.55E-01 | 9.09E-01 |
| TLN2 | 2051.6531 | -0.2895365 | 0.24601804 | -1.1768914 | 0.23923886 | 0.69029619 |
| TPM1 | 1064.58516 | 0.3017791 | 0.45824877 | 0.65854864 | 0.51018565 | 0.85760273 |
| LACTB | 275.231509 | 0.06295322 | 0.2291853 | 0.27468262 | 0.78356008 | 0.94955765 |
| RPS27L | 505.525909 | 0.63332509 | 0.32018777 | 1.97798028 | 0.04793093 | 0.39511605 |
| RAB8B | 463.739951 | -0.1190817 | 0.2029041 | -0.5868868 | 0.55727976 | 0.87240458 |
| APH1B | 332.537419 | 0.3443423 | 0.41324348 | 0.83326736 | 0.40469396 | 0.80437725 |
| CA12 | 7892.31211 | 1.02299449 | 0.69142793 | 1.47953886 | 0.13899635 | 0.58243213 |
| LINC02568 | 381.529714 | 0.3334167 | 0.7964335 | 0.41863721 | 0.67548129 | 0.91649321 |
| USP3 | 2136.9024 | -0.241833 | 0.16684857 | -1.4494162 | 0.14722138 | 0.59373375 |
| USP3-AS1 | 448.262577 | -0.0109541 | 0.37299109 | -0.0293682 | 0.97657096 | 0.9941831 |
| HERC1 | 2436.88098 | -0.2265241 | 0.2040429 | -1.1101787 | 0.26692202 | 0.71691195 |
| AC073167.1 | 108.962901 | -0.0252315 | 0.61077885 | -0.0413103 | 0.96704851 | 0.99248831 |
| DAPK2 | 55.3761907 | -2.3795218 | 0.58808074 | -4.0462501 | 5.20E-05 | 0.01145763 |
| FAM96A | 390.895264 | 0.00970773 | 0.23368274 | 0.04154235 | 0.96686353 | 0.99248831 |
| SNX1 | 977.452948 | 0.04654667 | 0.12233344 | 0.38049017 | 0.70358159 | 0.92513032 |
| PPIB | 678.857363 | 0.82339327 | 0.34025982 | 2.41989566 | 0.01552496 | 0.25214021 |
| CSNK1G1 | 254.17175 | 0.27669263 | 0.29198649 | 0.94762134 | 0.34332226 | 0.76642618 |
| TRIP4 | 629.838098 | -0.3291098 | 0.20674679 | -1.5918496 | 0.11141851 | 0.53517349 |
| ZNF609 | 3199.21902 | -0.1788902 | 0.16515219 | -1.0831838 | 0.27872687 | 0.7244595 |
| OAZ2 | 274.97543 | 0.47755508 | 0.2813855 | 1.69715595 | 0.08966718 | 0.50017474 |
| AC100830.1 | 62.8179816 | -0.3300805 | 0.61656366 | -0.5353551 | 0.59240434 | 0.88740306 |
| RBPMS2 | 74.3049511 | 0.33374913 | 0.49923576 | 0.66852008 | 0.50380167 | 0.85525037 |
| ANKDD1A | 64.9864981 | 0.05955765 | 0.49313421 | 0.12077372 | 0.90387026 | 0.97984491 |
| AC103691.1 | 41.4346449 | -0.5780639 | 0.51978308 | -1.1121253 | 0.26608429 | 0.71641758 |
| SPG21 | 415.694845 | -0.5099334 | 0.26356265 | -1.9347712 | 0.05301838 | 0.41073851 |
| MTFMT | 365.829578 | -0.2121627 | 0.20467852 | -1.0365655 | 0.29993842 | 0.73673761 |
| UBAP1L | 186.692217 | -0.0694732 | 0.41485696 | -0.1674631 | 0.8670057 | 0.97402505 |
| PDCD7 | 279.639193 | 0.12779664 | 0.28038585 | 0.45578849 | 0.64854208 | 0.90754509 |
| CLPX | 634.944346 | -0.0575748 | 0.17847607 | -0.3225911 | 0.74700496 | 0.93732199 |
| PARP16 | 238.862408 | -0.4416841 | 0.35086491 | -1.2588438 | 0.20808674 | 0.66104323 |
| DPP8 | 912.028568 | -0.2358902 | 0.17662338 | -1.3355548 | 0.18169484 | 0.63635309 |
| HACD3 | 597.418411 | 0.26410557 | 0.20059548 | 1.31660775 | 0.18797014 | 0.64257183 |
| INTS14 | 255.430268 | -0.0680824 | 0.19493926 | -0.3492493 | 0.72690216 | 0.93380972 |
| SLC24A1 | 232.032276 | -0.6372887 | 0.29262071 | -2.177866 | 0.02941601 | 0.33012415 |
| DENND4A | 3339.2538 | 0.11121004 | 0.33437218 | 0.33259357 | 0.7394411 | 0.93552616 |
| RAB11A | 1869.2386 | 0.18047212 | 0.47092651 | 0.38322778 | 0.70155088 | 0.92455209 |
| MEGF11 | 1356.21021 | 0.49617639 | 0.78657419 | 0.63080685 | 5.28E-01 | 8.62E-01 |
| DIS3L | 300.845331 | -0.1637589 | 0.23953118 | -0.6836641 | 0.49418728 | 0.84882719 |
| TIPIN | 305.352127 | 0.18306817 | 0.36569029 | 0.50060986 | 0.61664572 | 0.89589954 |
| MAP2K1 | 1120.2646 | -0.0926761 | 0.3079008 | -0.3009933 | 0.7634196 | 0.94255335 |
| SNAPC5 | 119.982411 | 0.47936833 | 0.26065806 | 1.83906966 | 0.06590494 | 0.4448331 |
| RPL4 | 773.477105 | 0.64429086 | 0.46716965 | 1.37913683 | 0.16785257 | 0.62227466 |
| ZWILCH | 108.930849 | -0.8549343 | 0.27024513 | -3.1635511 | 0.00155857 | 0.07326389 |
| SMAD6 | 305.008387 | -0.727504 | 0.55998079 | -1.2991588 | 0.19388943 | 0.64749489 |
| AC087482.1 | 67.0058853 | 0.6131288 | 0.93255119 | 0.65747468 | 0.51087575 | 0.85762435 |
| SMAD3 | 3344.31541 | -1.0893436 | 0.44433902 | -2.4516048 | 0.01422208 | 0.24653468 |
| AAGAB | 873.068276 | -0.3603816 | 0.18110026 | -1.9899565 | 0.04659573 | 0.39097111 |
| IQCH | 1043.3635 | -0.2633522 | 0.27674185 | -0.9516169 | 0.34129128 | 0.76543778 |
| IQCH-AS1 | 691.244397 | -0.1648885 | 0.26489824 | -0.6224599 | 0.53363954 | 0.86397197 |
| C15orf61 | 76.0119784 | 0.11943714 | 0.38775605 | 0.30802133 | 0.7580661 | 0.94056395 |
| MAP2K5 | 1873.18244 | -0.3382794 | 0.15966279 | -2.1187113 | 0.03411487 | 0.35153234 |

| | | | | | | |
|---|---|---|---|---|---|---|
| PIAS1 | 2445.71569 | -0.163458 | 0.15796465 | -1.0347761 | 0.3007735 | 0.73750086 |
| FEM1B | 247.055054 | 0.61940705 | 0.2404355 | 2.57618801 | 0.00998963 | 0.20363123 |
| CORO2B | 53.0851575 | -2.1937904 | 0.52525511 | -4.1766188 | 2.96E-05 | 0.00860682 |
| ANP32A | 520.154336 | 0.06997211 | 0.26986386 | 0.25928669 | 0.79541405 | 0.95278473 |
| SPESP1 | 192.67815 | 0.45899762 | 0.26454138 | 1.73506926 | 0.08272854 | 0.48594425 |
| NOX5 | 58.8282059 | 0.56045001 | 0.47636206 | 1.17652108 | 0.23938671 | 0.69040591 |
| GLCE | 782.643812 | -0.2406953 | 0.24739399 | -0.9729231 | 0.33059154 | 0.75909265 |
| AC026992.2 | 159.265677 | -0.0784628 | 0.50437736 | -0.1555638 | 0.87637689 | 0.97614152 |
| AC026992.1 | 119.856199 | -0.4268332 | 0.73185474 | -0.5832212 | 0.55974438 | 0.87300289 |
| PAQR5 | 3535.86869 | -0.7238513 | 0.25088648 | -2.8851747 | 0.00391196 | 0.12135113 |
| AC027237.3 | 116.216579 | -0.4595469 | 0.67295512 | -0.682879 | 0.4946833 | 0.84882719 |
| KIF23 | 518.382818 | -1.2264072 | 0.69918518 | -1.7540521 | 0.07942159 | 0.48169281 |
| RPLP1 | 1686.09696 | 0.48543477 | 0.46056763 | 1.05399238 | 0.2918864 | 0.73368744 |
| DRAIC | 60.4259364 | 0.09424468 | 0.52479992 | 0.17958213 | 8.57E-01 | 0.97127317 |
| TLE3 | 325.572396 | 0.41501022 | 0.28356902 | 1.46352452 | 0.14332391 | 0.58849865 |
| UACA | 1679.10257 | -0.0682257 | 0.19770337 | -0.3450913 | 0.73002572 | 0.93453345 |
| AC009269.2 | 47.261654 | 0.16800135 | 0.65131789 | 0.25794063 | 0.79645272 | 0.95278473 |
| LARP6 | 141.27679 | 0.36477377 | 0.45031162 | 0.81004743 | 0.41791292 | 0.81062751 |
| LRRC49 | 506.863722 | 0.26355782 | 0.27079619 | 0.97327003 | 0.33041911 | 0.75894644 |
| THSD4 | 2816.29811 | 0.79524806 | 0.56844846 | 1.39898005 | 0.16181896 | 0.61346689 |
| THSD4-AS1 | 423.507706 | 0.39107724 | 0.20395122 | 1.91750381 | 0.05517396 | 0.41810002 |
| MYO9A | 10171.5166 | 0.04890835 | 0.19084329 | 0.25627491 | 0.79773857 | 0.95288536 |
| AC022872.1 | 330.175568 | 0.09420804 | 0.5682002 | 0.16580079 | 0.86831372 | 0.97403792 |
| SENP8 | 259.200889 | -0.2150281 | 0.28794076 | -0.7467788 | 0.45519711 | 0.83121393 |
| AC020779.2 | 43.3511737 | 0.05640526 | 0.54348065 | 0.10378521 | 0.9173398 | 0.98199511 |
| GRAMD2A | 176.740332 | 0.29419501 | 0.35587983 | 0.82666952 | 0.4084244 | 0.80588056 |
| PKM | 3701.03428 | -0.0096241 | 0.35984826 | -0.0267449 | 9.79E-01 | 0.99455895 |
| PARP6 | 534.849646 | -0.4003366 | 0.21884271 | -1.829335 | 0.06734943 | 0.45135397 |
| AC009690.3 | 45.5586221 | -1.0927007 | 0.75173286 | -1.4535758 | 0.14606396 | 0.59273698 |
| AC009690.2 | 610.6491 | -0.9122605 | 0.73502906 | -1.2411217 | 0.2145608 | 0.66827012 |
| AC009690.1 | 2004.9591 | -0.6055129 | 0.49643659 | -1.2197186 | 0.22257157 | 0.67568617 |
| TMEM202-A: | 117.415191 | -0.5281536 | 0.48899045 | -1.0800899 | 0.28010215 | 0.72513763 |
| ARIH1 | 3613.49603 | -0.2544049 | 0.19966415 | -1.2741642 | 0.20260523 | 0.65552334 |
| BBS4 | 757.216588 | -0.5209171 | 0.21244398 | -2.4520212 | 0.01420563 | 0.24653468 |
| ADPGK | 432.345058 | -0.2061969 | 0.27599135 | -0.7471137 | 0.45499492 | 0.83121393 |
| NEO1 | 1211.10903 | -0.362595 | 0.19001778 | -1.9082162 | 0.05636328 | 0.41843207 |
| NPTN | 1257.75134 | -0.1557441 | 0.15907567 | -0.9790567 | 0.32755198 | 0.75620136 |
| CD276 | 92.6080545 | -0.2694147 | 0.28608988 | -0.9417136 | 0.34633929 | 0.76864066 |
| LOXL1-AS1 | 40.7972721 | 0.35619254 | 0.44770289 | 0.79560028 | 0.42626441 | 0.81466629 |
| LOXL1 | 78.1547666 | 0.25728739 | 0.34860516 | 0.73804815 | 0.46048519 | 0.83429248 |
| PML | 219.957827 | 0.13626804 | 0.40434253 | 0.33701139 | 0.73610831 | 0.93552616 |
| UBL7 | 64.7090067 | 0.08555048 | 0.26952653 | 0.31741022 | 0.75093235 | 0.93775604 |
| UBL7-AS1 | 63.0423219 | -0.4351867 | 0.47625504 | -0.9137683 | 0.36083862 | 0.77730375 |
| AC012435.2 | 89.3799412 | -0.5204424 | 0.31443336 | -1.6551755 | 0.0978889 | 0.51355946 |
| ARID3B | 189.097156 | -0.2742897 | 0.25076288 | -1.0938209 | 0.27403355 | 0.72135661 |
| CLK3 | 281.128356 | 0.02532313 | 0.19050477 | 0.13292649 | 0.89425152 | 0.97788747 |
| EDC3 | 376.809837 | -0.3242488 | 0.16394014 | -1.9778487 | 0.04794578 | 0.39511605 |
| CSK | 93.6291913 | -0.4542517 | 0.27619688 | -1.6446662 | 0.10003867 | 0.51678168 |
| ULK3 | 187.143403 | -0.6173653 | 0.28374735 | -2.1757569 | 0.02957344 | 0.33073125 |
| SCAMP2 | 315.51804 | -0.1448969 | 0.18812857 | -0.7702012 | 0.44118056 | 0.82285053 |
| MPI | 76.3703717 | 0.67117191 | 0.33319238 | 2.01436755 | 0.04397096 | 0.38494416 |
| FAM219B | 162.463801 | -0.045369 | 0.23594359 | -0.1922874 | 0.8475171 | 0.96925137 |
| COX5A | 504.6355 | 0.4366609 | 0.2331009 | 1.87326987 | 0.06103113 | 0.43176766 |
| SCAMP5 | 172.620665 | -0.1012683 | 0.39539428 | -0.2561197 | 0.79785844 | 0.95288536 |

| | | | | | | |
|---|---|---|---|---|---|---|
| PPCDC | 478.45582 | -0.6362792 | 0.33987168 | -1.872116 | 0.06119056 | 0.43222783 |
| COMMD4 | 207.380634 | -0.1187396 | 0.29461385 | -0.4030345 | 0.68692282 | 0.9208231 |
| NEIL1 | 258.941003 | -0.6378222 | 0.26919504 | -2.3693681 | 0.01781851 | 0.26845799 |
| MAN2C1 | 424.970023 | -0.3812352 | 0.29655037 | -1.2855666 | 0.19859437 | 0.65108786 |
| SIN3A | 788.347334 | -0.1495173 | 0.19544922 | -0.7649931 | 0.44427568 | 0.82456618 |
| PTPN9 | 897.102209 | -0.0710642 | 0.20956522 | -0.3391031 | 7.35E-01 | 0.93552616 |
| SNUPN | 157.837218 | 0.03093232 | 0.31563826 | 0.09799928 | 0.92193286 | 0.98215577 |
| IMP3 | 107.572812 | 0.63277463 | 0.39910131 | 1.58549876 | 0.11285306 | 0.5381502 |
| SNX33 | 77.1770859 | 0.43599353 | 0.26664671 | 1.63509809 | 0.10202849 | 0.52073581 |
| UBE2Q2 | 404.680888 | -0.2274501 | 0.21468841 | -1.0594429 | 2.89E-01 | 7.32E-01 |
| FBXO22 | 615.381388 | -0.089206 | 0.16960564 | -0.5259614 | 0.59891505 | 0.88883692 |
| NRG4 | 101.733492 | -0.370887 | 0.59895817 | -0.6192202 | 0.53577133 | 0.8642718 |
| TMEM266 | 209.155504 | -0.2639705 | 0.83150121 | -0.3174625 | 0.75089269 | 0.93775604 |
| ETFA | 1229.36879 | 0.04923049 | 0.23843661 | 0.20647203 | 0.83642221 | 0.96511435 |
| SCAPER | 2786.45164 | 0.47252263 | 0.16470862 | 2.86883978 | 0.0041198 | 0.12538824 |
| RCN2 | 568.515181 | 0.27534634 | 0.20092629 | 1.37038486 | 0.1705668 | 0.62661733 |
| TSPAN3 | 599.261175 | 0.09862299 | 0.200406 | 0.49211594 | 0.62263738 | 0.89835355 |
| PEAK1 | 1365.71673 | -0.015359 | 0.23551813 | -0.0652135 | 0.94800401 | 0.98919519 |
| HMG20A | 913.998053 | 0.11675017 | 0.11773917 | 0.99160013 | 0.32139263 | 0.75240254 |
| LINGO1 | 4882.46187 | -0.3101001 | 0.6107597 | -0.5077285 | 0.61164376 | 0.8941127 |
| TBC1D2B | 201.081367 | -0.1389406 | 0.25838258 | -0.537732 | 0.5907621 | 0.88672234 |
| IDH3A | 540.372831 | -0.0891181 | 0.27803479 | -0.3205287 | 0.74856761 | 0.93746253 |
| DNAJA4 | 85.3560532 | 0.00946333 | 0.31186153 | 0.03034466 | 0.97579218 | 0.9941831 |
| WDR61 | 358.080848 | 0.16458043 | 0.18896767 | 0.8709449 | 0.38378424 | 0.791697 |
| IREB2 | 1002.74455 | -0.2529419 | 0.1228922 | -2.0582421 | 0.0395669 | 0.37258841 |
| HYKK | 167.454064 | 0.37191892 | 0.35887115 | 1.03635781 | 0.30003525 | 0.73678482 |
| PSMA4 | 332.952264 | 0.32332312 | 0.29645907 | 1.09061638 | 0.27544172 | 0.72233844 |
| MORF4L1 | 1121.65877 | 0.13895467 | 0.15062124 | 0.92254369 | 0.35624505 | 0.77368353 |
| CTSH | 372.704169 | 0.06720361 | 0.3822754 | 0.17579894 | 0.8604519 | 0.97200968 |
| TMED3 | 292.863945 | -0.4187075 | 0.32844346 | -1.2748237 | 0.20237164 | 0.65552334 |
| MTHFS | 71.4728221 | 0.21079607 | 0.29651139 | 0.71092066 | 0.4771334 | 0.84174109 |
| ZFAND6 | 2319.11432 | -0.2085503 | 0.20749206 | -1.0051004 | 0.31484853 | 0.74785672 |
| FAH | 89.9807952 | 0.46997288 | 0.32177016 | 1.46058568 | 0.14412918 | 0.58977767 |
| CTXND1 | 108.78154 | -1.0987107 | 0.81309487 | -1.35127 | 0.17660896 | 0.63008941 |
| LINC00927 | 41.3623807 | -0.6912611 | 0.65326765 | -1.0581591 | 0.2899829 | 0.73204984 |
| AC016705.2 | 1445.42948 | -0.4664263 | 0.603122 | -0.7733532 | 0.43931339 | 0.82164134 |
| ARNT2 | 3162.6338 | 0.02079475 | 0.33028832 | 0.0629594 | 0.94979884 | 0.98919519 |
| ABHD17C | 993.157384 | 0.00227428 | 0.33746243 | 0.00673937 | 0.9946228 | 0.998438 |
| CEMIP | 144.25806 | 0.63973356 | 0.62681472 | 1.0206103 | 0.30743911 | 0.74341546 |
| AC027808.1 | 242.488725 | -0.018274 | 0.35000965 | -0.0522099 | 0.95836142 | 0.99071816 |
| MESD | 426.021804 | 0.50671534 | 0.15338778 | 3.30349228 | 0.00095489 | 0.05532057 |
| TLNRD1 | 60.2388978 | 0.36230779 | 0.46971156 | 0.771341 | 0.44050484 | 0.82253723 |
| CFAP161 | 70.4367497 | -0.6617187 | 0.60754258 | -1.0891725 | 0.27607781 | 0.72241028 |
| AC104041.1 | 441.851097 | -0.2698464 | 0.81303191 | -0.3319013 | 0.73996376 | 0.93552616 |
| EFL1 | 509.059462 | -0.2640851 | 0.25279834 | -1.0446473 | 0.29618603 | 0.7366786 |
| SAXO2 | 123.228485 | 1.09019354 | 0.46855796 | 2.32669942 | 0.01998127 | 0.2836927 |
| GOLGA6L10 | 39.7318262 | -0.2851498 | 0.82651699 | -0.3450018 | 0.73009303 | 0.93453345 |
| GOLGA6L9 | 70.389284 | -0.1636675 | 0.30390715 | -0.5385445 | 0.59020121 | 0.88631239 |
| RPS17 | 77.5903755 | 0.34883543 | 0.58818519 | 0.59307075 | 0.5531338 | 0.87104567 |
| AP3B2 | 108.275239 | 0.13927911 | 0.67013492 | 0.20783742 | 0.83535591 | 0.96511435 |
| AC105339.3 | 80.4348403 | -0.2657269 | 0.39210843 | -0.6776872 | 0.49797006 | 0.85092921 |
| SNHG21 | 73.6852946 | 0.16106646 | 0.33323441 | 0.48334282 | 0.62885234 | 0.89970424 |
| WHAMM | 570.726605 | -0.3066707 | 0.2145087 | -1.4296421 | 0.15281975 | 0.60285224 |
| HOMER2 | 438.585568 | -0.2342503 | 0.30000972 | -0.7808089 | 0.43491489 | 0.81919857 |

| | | | | | | |
|---|---|---|---|---|---|---|
| AC022558.2 | 141.572498 | 0.08507659 | 0.55246795 | 0.15399372 | 0.87761469 | 0.97655597 |
| C15orf40 | 285.455816 | 0.22893465 | 0.23742203 | 0.96425191 | 0.33491964 | 0.76248769 |
| BTBD1 | 643.255004 | -0.0309343 | 0.19789922 | -0.1563136 | 0.8757858 | 0.97614152 |
| HDGFL3 | 617.627076 | 0.14205579 | 0.19890532 | 0.71418799 | 0.47511094 | 0.84106098 |
| AC103876.1 | 94.8708716 | 0.21558012 | 0.46139566 | 0.46723483 | 0.64033188 | 0.9035513 |
| ADAMTSL3 | 644.745388 | 0.21047244 | 0.81870787 | 0.25707881 | 0.79711794 | 0.95281098 |
| UBE2Q2L | 49.4712712 | 0.61521123 | 0.74007799 | 0.83127891 | 0.40581608 | 0.80499101 |
| GOLGA6L4 | 58.0614171 | -0.5736452 | 0.42162978 | -1.3605425 | 0.17365832 | 0.6286116 |
| AC048382.5 | 116.654897 | -0.1547862 | 0.3937117 | -0.3931461 | 0.69421155 | 0.92229578 |
| ZSCAN2 | 98.196604 | -0.3255424 | 0.29801627 | -1.0923644 | 0.27467296 | 0.72164296 |
| WDR73 | 672.172603 | 0.08125221 | 0.23269621 | 0.34917722 | 0.72695627 | 0.93380972 |
| SEC11A | 1095.66846 | 0.01982244 | 0.17225538 | 0.11507589 | 0.90838497 | 0.98047754 |
| ZNF592 | 756.83261 | -0.2234276 | 0.17291533 | -1.2921212 | 0.19631516 | 0.64964978 |
| ALPK3 | 1039.91483 | -0.0173207 | 0.42923553 | -0.0403524 | 0.96781219 | 0.99248831 |
| SLC28A1 | 53.9192858 | 0.70563437 | 0.67122992 | 1.05125583 | 0.29314111 | 0.73527014 |
| PDE8A | 2521.62692 | -0.3625053 | 0.25470318 | -1.4232461 | 0.15466484 | 0.60380955 |
| AKAP13 | 5269.38703 | -0.488579 | 0.1935338 | -2.5245152 | 0.0115858 | 0.22051414 |
| AC087286.2 | 180.97693 | -0.10119 | 0.57431497 | -0.1761925 | 0.86014271 | 0.97200968 |
| KLHL25 | 91.9945457 | 0.12975964 | 0.29351643 | 0.44208645 | 0.65842665 | 0.91037018 |
| MRPL46 | 149.475652 | 0.54526265 | 0.2695471 | 2.02288452 | 0.04308505 | 0.38246672 |
| MRPS11 | 242.158706 | 0.12950932 | 0.28225408 | 0.4588395 | 0.64634943 | 0.90582727 |
| DET1 | 286.843323 | -0.0175112 | 0.50440994 | -0.0347161 | 0.97230609 | 0.99327086 |
| ABHD2 | 688.484106 | 0.0073192 | 0.35005886 | 0.02090847 | 0.98331867 | 0.99518552 |
| FANCI | 47.1504342 | -1.2205542 | 0.46790209 | -2.6085676 | 0.00909221 | 0.1936857 |
| POLG | 203.73429 | -0.3133718 | 0.22430427 | -1.3970834 | 0.1623885 | 0.61373097 |
| PEX11A | 106.563952 | 0.41364656 | 0.28740887 | 1.43922684 | 0.15008626 | 0.59663305 |
| WDR93 | 63.1405996 | 0.75331761 | 0.58054252 | 1.29760972 | 0.19442148 | 0.64785907 |
| ANPEP | 111.017306 | 0.47958794 | 0.73489959 | 0.65258975 | 0.5140208 | 0.85762435 |
| AP3S2 | 50.4326254 | 0.50240777 | 0.41468271 | 1.2115474 | 0.22568568 | 0.67798093 |
| ZNF710 | 562.173996 | -0.1948831 | 0.30204675 | -0.6452083 | 0.51879222 | 0.85916831 |
| ZNF710-AS1 | 347.403821 | 0.07765115 | 0.35944876 | 0.21602842 | 0.82896559 | 0.96288319 |
| IDH2 | 2753.49942 | 0.15131789 | 0.32078021 | 0.47171828 | 0.63712789 | 0.90255504 |
| IDH2-DT | 525.408033 | -0.4557648 | 0.63407125 | -0.7187911 | 0.47226966 | 0.83921324 |
| SEMA4B | 2089.51362 | -0.6765953 | 0.43666608 | -1.549457 | 1.21E-01 | 5.56E-01 |
| CIB1 | 772.169221 | 0.4833231 | 0.4868686 | 0.99271775 | 3.21E-01 | 0.75169828 |
| AC091167.6 | 106.618013 | -0.0823021 | 0.40504226 | -0.2031938 | 0.83898357 | 0.966178 |
| ZNF774 | 69.3525115 | 0.03565399 | 0.30225606 | 0.11795954 | 0.90609971 | 0.98007461 |
| IQGAP1 | 1844.18164 | 0.07864872 | 0.22619885 | 0.34769727 | 7.28E-01 | 9.35E-01 |
| CRTC3 | 1241.12016 | 0.00648011 | 0.24635729 | 0.02630372 | 0.97901509 | 0.99455895 |
| CRTC3-AS1 | 215.82379 | -1.5572608 | 0.43538754 | -3.5767233 | 0.00034793 | 0.03115997 |
| FURIN | 193.268183 | -0.1289669 | 0.44305222 | -0.2910874 | 0.77098448 | 0.9443559 |
| MAN2A2 | 185.572028 | -0.3703741 | 0.23298069 | -1.5897202 | 0.11189788 | 0.53611141 |
| HDDC3 | 79.6671671 | -0.0731149 | 0.32021553 | -0.2283303 | 0.8193895 | 0.96006637 |
| UNC45A | 277.933468 | 0.04402794 | 0.2268727 | 0.19406451 | 0.84612537 | 0.96883382 |
| RCCD1 | 94.3199471 | -0.0098398 | 0.26552316 | -0.0370582 | 0.97043859 | 0.99309664 |
| AC104035.1 | 700.511957 | -1.0536652 | 1.17377536 | -0.8976719 | 0.36936049 | 0.78095617 |
| SLCO3A1 | 2796.17444 | 0.22592907 | 0.28631477 | 0.78909332 | 0.43005746 | 0.81675008 |
| AC116903.2 | 60.210779 | 0.04175159 | 0.58504874 | 0.0713643 | 0.94310782 | 0.98772472 |
| FAM174B | 568.834307 | -1.1997782 | 0.31083582 | -3.8598455 | 0.00011346 | 0.01683906 |
| AC013394.1 | 1643.21469 | -0.3321393 | 0.14881811 | -2.2318471 | 0.02562507 | 0.31211455 |
| CHD2 | 2345.71202 | 0.05766853 | 0.32600185 | 0.17689633 | 0.85958982 | 0.97184883 |
| MCTP2 | 255.344738 | 2.33895056 | 0.62722136 | 3.72906712 | 0.00019219 | 0.02233698 |
| LINC01197 | 276.000891 | -0.9807972 | 0.91605562 | -1.0706743 | 0.28431592 | 0.72738535 |
| AC012409.2 | 986.635611 | 0.10472596 | 0.4741299 | 0.22088031 | 0.82518563 | 0.96146415 |

| | | | | | | |
|---|---|---|---|---|---|---|
| AC024337.2 | 62.8927247 | -0.1751023 | 0.40261015 | -0.4349178 | 6.64E-01 | 0.91177281 |
| NR2F2-AS1 | 6802.41482 | -0.0888659 | 0.29802931 | -0.2981784 | 7.66E-01 | 0.94270299 |
| NR2F2 | 1174.50733 | -0.2918052 | 0.27674705 | -1.0544114 | 0.29169461 | 0.73362416 |
| AC087477.2 | 262.72894 | -1.4168392 | 0.36196202 | -3.9143311 | 9.07E-05 | 0.01515778 |
| LINC00923 | 84.2189171 | 0.06471258 | 0.85589772 | 0.07560784 | 0.9397311 | 0.98715467 |
| ARRDC4 | 472.165188 | -0.1382507 | 0.3844341 | -0.3596212 | 0.71913042 | 0.93001079 |
| FAM169B | 208.02445 | 0.55928146 | 0.96475767 | 0.57971186 | 5.62E-01 | 0.87403145 |
| IGF1R | 9809.4182 | -0.122823 | 0.2791122 | -0.4400488 | 0.65990176 | 0.91099119 |
| AC036108.1 | 138.415627 | 0.27752481 | 0.61139567 | 0.45392015 | 0.6498863 | 0.90825486 |
| TTC23 | 636.680406 | -0.3270558 | 0.15879816 | -2.0595693 | 0.03943973 | 0.37234059 |
| LRRC28 | 1050.75993 | -0.1879719 | 0.16022753 | -1.1731562 | 2.41E-01 | 6.92E-01 |
| MEF2A | 4462.14988 | 0.03656847 | 0.26013629 | 0.14057428 | 0.88820627 | 0.97729111 |
| LYSMD4 | 119.661061 | 0.48400531 | 0.25210867 | 1.91982811 | 0.05487962 | 0.41684503 |
| AC090825.1 | 204.620203 | 1.48032034 | 0.43907731 | 3.37143438 | 0.00074778 | 0.04818247 |
| ADAMTS17 | 1298.00015 | 0.5951432 | 0.58921547 | 1.01006039 | 0.31246636 | 0.74693024 |
| AC027020.2 | 73.377735 | -0.1759581 | 0.5395857 | -0.3260985 | 0.74434985 | 0.93668127 |
| LINS1 | 319.862436 | -0.0373252 | 0.25972908 | -0.1437081 | 8.86E-01 | 0.97664137 |
| ASB7 | 413.26083 | -0.056458 | 0.19773491 | -0.2855239 | 0.77524279 | 0.94667486 |
| AC087762.1 | 48.806775 | 0.52919512 | 0.50069334 | 1.05692461 | 0.29054599 | 0.73285033 |
| CHSY1 | 806.08526 | 0.4812638 | 0.26951202 | 1.78568581 | 0.07415014 | 0.47078634 |
| SELENOS | 370.525576 | 0.64816276 | 0.28933573 | 2.24017534 | 0.02507954 | 0.31053369 |
| SNRPA1 | 339.272691 | -0.1997053 | 0.18320227 | -1.0900809 | 0.27567752 | 0.72233844 |
| AC023024.1 | 49.224642 | -0.0918655 | 0.36369685 | -0.2525882 | 0.80058647 | 0.95370871 |
| PCSK6 | 1195.8407 | -0.3004922 | 0.45268086 | -0.6638058 | 0.50681461 | 0.85626196 |
| TM2D3 | 373.022891 | 0.06756469 | 0.22709929 | 0.29751167 | 0.7660759 | 0.9428901 |
| TARSL2 | 628.242003 | -0.0301281 | 0.24949583 | -0.1207558 | 0.90388449 | 0.97984491 |
| POLR3K | 41.1045766 | 0.25128938 | 0.4204203 | 0.59770991 | 0.55003351 | 0.86956073 |
| SNRNP25 | 71.3101916 | 0.66920175 | 0.51874727 | 1.29003428 | 0.19703876 | 0.64964978 |
| RHBDF1 | 56.9919095 | -0.2669427 | 0.49682173 | -0.5373007 | 0.59105993 | 0.8867913 |
| MPG | 208.660535 | 0.23558845 | 0.23037997 | 1.02260822 | 3.06E-01 | 7.43E-01 |
| NPRL3 | 651.108065 | -0.1030313 | 0.27929042 | -0.3689038 | 0.71219942 | 0.92748721 |
| LUC7L | 1110.51391 | -0.1580578 | 0.17888867 | -0.8835537 | 0.37693716 | 0.78557321 |
| FAM234A | 269.047181 | -0.0230786 | 0.38384486 | -0.0601249 | 0.95205619 | 0.98938795 |
| AXIN1 | 285.581159 | -0.3989136 | 0.21565971 | -1.8497366 | 0.06435152 | 0.43948973 |
| MRPL28 | 170.527574 | 0.73879641 | 0.48044496 | 1.53773369 | 0.12411374 | 0.56105491 |
| TMEM8A | 596.757372 | 0.31072178 | 0.3443005 | 0.90247264 | 0.36680585 | 0.78095617 |
| NME4 | 136.040961 | 0.45610359 | 0.5542635 | 0.82290029 | 0.4105647 | 0.80655687 |
| DECR2 | 79.9331498 | 0.12521786 | 0.36671897 | 0.34145456 | 0.73276141 | 0.93552616 |
| RAB11FIP3 | 1845.06024 | 0.32287116 | 0.33155762 | 0.97380105 | 0.33015533 | 0.75885186 |
| CAPN15 | 275.534141 | -0.6025779 | 0.27765013 | -2.1702777 | 0.02998581 | 0.33256306 |
| PIGQ | 279.486957 | 0.11936711 | 0.37219144 | 0.32071427 | 0.74842693 | 0.93746253 |
| RAB40C | 670.533921 | 0.17170904 | 0.33955372 | 0.50569036 | 0.61307404 | 0.89441126 |
| WFIKKN1 | 39.5783757 | 0.27849232 | 0.76686365 | 0.36315755 | 0.7164872 | 0.9289438 |
| METTL26 | 101.483274 | 0.36099893 | 0.49221803 | 0.73341267 | 0.46330678 | 0.83531269 |
| MCRIP2 | 347.152495 | 0.17264422 | 0.31313287 | 0.55134493 | 5.81E-01 | 0.88292437 |
| WDR90 | 149.622973 | -0.4845482 | 0.25729653 | -1.8832288 | 0.05966936 | 0.42650032 |
| RHOT2 | 336.418425 | -0.0182759 | 0.29154648 | -0.0626862 | 0.95001636 | 0.98919519 |
| STUB1 | 293.678466 | 0.48179176 | 0.34091256 | 1.41324146 | 0.15758474 | 0.6073933 |
| JMJD8 | 148.332657 | 0.35815306 | 0.47194777 | 0.75888283 | 0.44792265 | 0.8260638 |
| WDR24 | 46.3627079 | -0.401846 | 0.39767527 | -1.0104877 | 0.31226169 | 0.74687836 |
| METRN | 224.151211 | 0.29893796 | 0.31266256 | 0.95610412 | 0.33901963 | 0.76449297 |
| FAM173A | 180.958129 | 0.19572172 | 0.45576499 | 0.42943562 | 0.66760624 | 0.91373027 |
| HAGHL | 84.7352903 | -0.2321908 | 0.60741412 | -0.3822612 | 0.70226764 | 0.92492636 |
| NARFL | 181.65487 | -0.5098663 | 0.24989153 | -2.0403505 | 0.04131543 | 0.37577267 |

| | | | | | | |
|---|---|---|---|---|---|---|
| PRR25 | 45.8578202 | -0.2860087 | 0.46069364 | -0.6208219 | 0.53471679 | 0.86419602 |
| LMF1 | 861.104778 | 0.10537479 | 0.59630395 | 0.17671321 | 0.85973366 | 0.97186324 |
| CACNA1H | 482.21616 | -0.9343105 | 0.42413937 | -2.2028385 | 0.02760614 | 0.32099424 |
| UBE2I | 697.931808 | -0.5338554 | 0.20620347 | -2.5889737 | 0.00962625 | 0.19961236 |
| BAIAP3 | 119.94984 | -0.0212887 | 0.39079371 | -0.0544756 | 0.95655621 | 0.99065996 |
| TSR3 | 125.377361 | 0.53669423 | 0.38637767 | 1.3890405 | 0.16482043 | 0.61673853 |
| GNPTG | 367.513 | -0.1561296 | 0.21422932 | -0.7287964 | 0.46612623 | 0.8367113 |
| UNKL | 377.831733 | -0.2124415 | 0.30543529 | -0.695537 | 0.48671882 | 0.84474442 |
| C16orf91 | 45.5719543 | 0.26340756 | 0.47026744 | 0.5601229 | 0.57539561 | 0.88069533 |
| CCDC154 | 61.6811252 | 0.11882899 | 0.42299662 | 0.28092184 | 0.77877035 | 0.94725685 |
| CLCN7 | 477.876489 | -0.4792673 | 0.27320155 | -1.7542628 | 0.07938549 | 0.48169281 |
| TELO2 | 129.488423 | -0.3198099 | 0.28505928 | -1.1219067 | 0.26190212 | 0.71271452 |
| IFT140 | 479.071193 | -0.2561757 | 0.3011582 | -0.8506348 | 0.39497222 | 0.7989793 |
| AL133297.2 | 55.7622751 | 0.1131154 | 0.43890165 | 0.2577238 | 0.79662007 | 0.95278473 |
| CRAMP1 | 107.792902 | -0.0776629 | 0.37193919 | -0.2088054 | 0.83460012 | 0.96510927 |
| JPT2 | 773.474933 | -0.090766 | 0.31991048 | -0.2837232 | 7.77E-01 | 0.94694572 |
| MAPK8IP3 | 1278.39712 | -0.1858252 | 0.28708606 | -0.6472805 | 0.51745042 | 0.85893097 |
| NME3 | 398.037593 | 0.49587079 | 0.35299971 | 1.40473426 | 0.16010032 | 0.61085068 |
| MRPS34 | 189.656682 | 0.12201879 | 0.35821375 | 0.34063122 | 0.73338122 | 0.93552616 |
| EME2 | 116.178856 | -0.7024082 | 0.28950391 | -2.4262477 | 0.01525585 | 0.25126505 |
| SPSB3 | 504.95678 | -0.2195217 | 0.23025077 | -0.9534027 | 0.34038609 | 0.76478309 |
| NUBP2 | 114.722352 | -0.0392697 | 0.29737892 | -0.1320526 | 0.89494269 | 0.97788747 |
| HAGH | 627.482414 | -0.1663579 | 0.19788784 | -0.8406677 | 0.40053414 | 0.80243544 |
| FAHD1 | 151.207215 | 0.42997415 | 0.49033253 | 0.87690318 | 0.38053923 | 0.78824206 |
| MSRB1 | 138.52177 | 0.11003202 | 0.50068674 | 0.2197622 | 8.26E-01 | 0.96214368 |
| NDUFB10 | 305.954442 | 0.72854508 | 0.42516011 | 1.71357815 | 0.08660623 | 0.49478465 |
| RPS2 | 867.329272 | 0.4237607 | 0.4196738 | 1.00973827 | 0.3126207 | 0.74693024 |
| SNHG9 | 88.6373778 | -0.011821 | 0.49551139 | -0.0238562 | 0.9809673 | 0.99518016 |
| TBL3 | 48.478632 | 0.11511193 | 0.40587539 | 0.28361397 | 0.77670623 | 0.94694572 |
| GFER | 48.3627866 | 0.17429827 | 0.47349232 | 0.36811215 | 0.71278961 | 0.92760608 |
| ZNF598 | 95.1707936 | -0.4083447 | 0.29255321 | -1.3957962 | 0.16277587 | 0.61387934 |
| SLC9A3R2 | 51.7093589 | 0.40562562 | 0.44747948 | 0.90646755 | 0.36468844 | 0.78076892 |
| TSC2 | 677.824764 | -0.5065105 | 0.32241541 | -1.5709874 | 0.11618558 | 0.54596061 |
| PKD1 | 428.873176 | -0.1436273 | 0.23218992 | -0.618577 | 0.53619505 | 0.8642718 |
| SNHG19 | 155.687598 | 0.38689176 | 0.33098 | 1.1689279 | 0.24243268 | 0.69337349 |
| TRAF7 | 124.437117 | 0.20577141 | 0.35914996 | 0.5729401 | 0.56668526 | 0.87646121 |
| MLST8 | 71.3398836 | -0.0979218 | 0.45337869 | -0.2159823 | 0.82900158 | 0.96288319 |
| BRICD5 | 59.7326861 | -0.6224044 | 0.48473344 | -1.2840138 | 0.19913713 | 0.65141218 |
| PGP | 409.05456 | 0.106537 | 0.24757121 | 0.43032871 | 0.66695655 | 0.91331423 |
| E4F1 | 188.210151 | -0.5757308 | 0.30140065 | -1.9101844 | 0.05610947 | 0.41843207 |
| ECI1 | 669.696615 | 0.25193447 | 0.26777194 | 0.94085463 | 0.34677936 | 0.76875481 |
| RNPS1 | 473.469488 | -0.0970211 | 0.18008154 | -0.5387622 | 0.59005096 | 0.88631239 |
| ABCA3 | 217.411975 | 0.13391818 | 0.20396369 | 0.65657853 | 0.51145196 | 0.85762435 |
| CCNF | 54.5879648 | 0.16601252 | 0.44200026 | 0.37559372 | 0.70721896 | 0.92573766 |
| TBC1D24 | 103.723245 | 0.01258303 | 0.68063066 | 0.01848732 | 0.9852501 | 0.99572827 |
| AMDHD2 | 75.6535185 | -0.0208957 | 0.46610314 | -0.0448307 | 0.96424227 | 0.99224805 |
| PDPK1 | 1034.43317 | -0.0436757 | 0.16523496 | -0.2643251 | 0.79152941 | 0.95178079 |
| AC093525.4 | 101.482409 | 0.03370796 | 0.39489101 | 0.08536015 | 0.93197508 | 0.98357772 |
| KCTD5 | 107.785927 | -0.2688319 | 0.34238252 | -0.7851801 | 0.43234801 | 0.81773248 |
| SRRM2-AS1 | 38.1883626 | -0.9013742 | 0.52575598 | -1.7144345 | 0.08644895 | 0.49478465 |
| SRRM2 | 3256.56835 | -0.2463095 | 0.20095696 | -1.2256829 | 0.22031801 | 0.67302974 |
| ELOB | 440.291019 | 0.68865761 | 0.49698362 | 1.38567466 | 0.16584627 | 0.61900632 |
| PRSS22 | 152.912823 | -0.525178 | 0.55391244 | -0.9481246 | 0.343066 | 0.76641095 |
| FLYWCH2 | 127.771414 | 0.5308155 | 0.31731862 | 1.6728155 | 0.09436362 | 0.51017802 |

| | | | | | | |
|---|---|---|---|---|---|---|
| FLYWCH1 | 339.325734 | 0.16558883 | 0.33890465 | 0.48860007 | 0.62512486 | 0.89873262 |
| TNFRSF12A | 505.745518 | -0.5351536 | 0.65788542 | -0.813445 | 0.41596297 | 0.80966301 |
| HCFC1R1 | 165.936058 | -0.3450285 | 0.49631275 | -0.6951836 | 0.48694027 | 0.84474442 |
| THOC6 | 51.8335218 | -1.0771583 | 0.46252313 | -2.3288744 | 0.01986572 | 0.28276543 |
| BICDL2 | 190.280323 | -0.3033974 | 0.50495775 | -0.6008372 | 0.54794842 | 0.86859163 |
| MMP25-AS1 | 96.6201906 | -0.9515132 | 0.28431106 | -3.3467331 | 0.0008177 | 0.04977413 |
| IL32 | 113.071696 | 0.68728456 | 0.50565475 | 1.35919729 | 0.17408408 | 0.6286158 |
| ZNF213-AS1 | 218.821788 | -0.461339 | 0.18340224 | -2.5154493 | 0.01188808 | 0.22355405 |
| ZNF205 | 82.0470988 | -0.1921404 | 0.35292165 | -0.544428 | 0.586147 | 0.88530153 |
| ZNF213 | 384.684409 | 0.03332793 | 0.33362981 | 0.09989494 | 0.92042773 | 0.98215577 |
| ZNF200 | 73.3468372 | -0.0035561 | 0.35775011 | -0.0099401 | 0.99206906 | 0.99776307 |
| ZNF263 | 192.382028 | 0.10218379 | 0.30030723 | 0.34026416 | 0.7336576 | 0.93552616 |
| TIGD7 | 154.565783 | 0.58152856 | 0.30350187 | 1.91606254 | 0.05535714 | 0.41841807 |
| ZNF75A | 554.538115 | -0.4443244 | 0.30836573 | -1.4409009 | 0.14961269 | 0.59625836 |
| AC025283.3 | 196.93246 | 0.39931051 | 0.50500859 | 0.79070043 | 0.42911882 | 0.81580751 |
| ZSCAN32 | 120.622008 | 0.15329525 | 0.26776137 | 0.57250695 | 0.56697858 | 0.87646121 |
| NAA60 | 156.075546 | 0.21319856 | 0.26687917 | 0.79885798 | 0.42437276 | 0.81457794 |
| CLUAP1 | 484.899405 | 0.11130976 | 0.25574478 | 0.43523766 | 0.66338993 | 0.91173838 |
| SLX4 | 75.3185673 | -0.498924 | 0.37735939 | -1.3221456 | 0.18611969 | 0.64033103 |
| DNASE1 | 1733.12347 | -0.2372835 | 0.38044806 | -0.6236949 | 0.53282795 | 0.86397197 |
| TRAP1 | 519.177814 | -0.1766032 | 0.15423609 | -1.1450185 | 0.2522015 | 0.70276117 |
| CREBBP | 2180.67551 | -0.1217846 | 0.19269384 | -0.632011 | 0.52737967 | 0.86151701 |
| ADCY9 | 698.186221 | 0.22135424 | 0.34412646 | 0.64323517 | 0.52007151 | 0.85935232 |
| TFAP4 | 47.4905659 | -0.5498817 | 0.58902146 | -0.9335513 | 0.3505354 | 0.77044053 |
| GLIS2 | 289.041913 | 0.0611054 | 0.54350024 | 0.11242938 | 0.91048296 | 0.98079999 |
| GLIS2-AS1 | 40.2421224 | 0.45225022 | 0.51594793 | 0.87654238 | 0.38073525 | 0.78826518 |
| PAM16 | 51.3242197 | -0.1533785 | 0.52765692 | -0.2906785 | 0.77129722 | 0.9446143 |
| VASN | 133.49764 | 1.6646087 | 0.56999497 | 2.92039191 | 0.00349591 | 0.11430083 |
| DNAJA3 | 375.206802 | -0.0870531 | 0.18802401 | -0.4629891 | 0.64337219 | 0.90478945 |
| NMRAL1 | 288.982515 | 0.08999255 | 0.23751436 | 0.37889307 | 0.70476727 | 0.92528913 |
| HMOX2 | 313.514064 | -0.0564022 | 0.27338004 | -0.2063143 | 0.8365454 | 0.96511435 |
| CDIP1 | 242.988377 | 0.58525696 | 0.24330043 | 2.40549081 | 0.01615076 | 0.25504618 |
| UBALD1 | 122.055207 | -0.1352144 | 0.30275783 | -0.4466092 | 0.65515728 | 0.90903073 |
| MGRN1 | 707.616389 | -0.1719594 | 0.19870166 | -0.8654149 | 0.38681108 | 0.79523542 |
| NUDT16L1 | 138.989077 | 0.56880802 | 0.37720448 | 1.50795672 | 0.1315656 | 0.57519227 |
| ANKS3 | 422.809397 | -0.358127 | 0.15966288 | -2.2430197 | 0.02489554 | 0.3093519 |
| ZNF500 | 201.504236 | -0.2683554 | 0.29515688 | -0.9091956 | 0.36324686 | 0.77942297 |
| ROGDI | 88.5082276 | 0.49410272 | 0.60079884 | 0.82240957 | 0.41084383 | 0.80666106 |
| GLYR1 | 1151.29794 | -0.113751 | 0.15740889 | -0.7226467 | 0.46989697 | 0.83921324 |
| UBN1 | 507.306051 | -0.1806902 | 0.23515157 | -0.768399 | 0.44225019 | 0.82285053 |
| PPL | 402.089519 | -0.4808497 | 0.29789565 | -1.6141548 | 0.10649386 | 0.52743811 |
| NAGPA | 107.581473 | -0.024352 | 0.36378363 | -0.0669409 | 0.94662876 | 0.98885769 |
| ALG1 | 166.92408 | 0.13164443 | 0.2925082 | 0.45005382 | 6.53E-01 | 0.90868303 |
| C16orf89 | 157.185208 | 3.22928658 | 0.7765339 | 4.15859061 | 3.20E-05 | 0.00863214 |
| EEF2KMT | 43.0382419 | 0.14382667 | 0.56251578 | 0.25568469 | 0.79819432 | 0.95294186 |
| AC026458.2 | 73.9981157 | 0.18507973 | 0.40849957 | 0.45307203 | 0.65049688 | 0.90825486 |
| RBFOX1 | 8820.73171 | 0.96897208 | 0.53080459 | 1.82547795 | 0.06792894 | 0.45250883 |
| METTL22 | 191.340564 | -0.2954048 | 0.34097001 | -0.866366 | 0.38628946 | 0.79490535 |
| ABAT | 233.288539 | -0.098216 | 0.3255021 | -0.301737 | 0.76285252 | 0.94225911 |
| TMEM186 | 55.7111304 | 0.14438149 | 0.36358263 | 0.39710777 | 0.69128799 | 0.92206635 |
| PMM2 | 788.211999 | -0.1509295 | 0.21398294 | -0.7053342 | 0.48060228 | 0.84362465 |
| AC022167.2 | 67.6263212 | -0.7381843 | 0.35658299 | -2.0701614 | 0.03843723 | 0.36764477 |
| CARHSP1 | 496.764535 | -0.0793044 | 0.3472721 | -0.2283638 | 0.8193634 | 0.96006637 |
| USP7 | 592.53472 | 0.07764983 | 0.1856138 | 0.41834081 | 0.67569796 | 0.91649321 |

| | | | | | | |
|---|---|---|---|---|---|---|
| C16orf72 | 607.013792 | 0.01886806 | 0.18511127 | 0.10192821 | 0.91881366 | 0.98215577 |
| AC087190.1 | 92.5931768 | 0.06064618 | 0.63973462 | 0.09479897 | 0.9244745 | 0.98215577 |
| ATF7IP2 | 1067.49118 | 0.52761931 | 0.29024004 | 1.81787225 | 0.06908366 | 0.45649128 |
| NUBP1 | 171.323445 | -0.3387028 | 0.22941418 | -1.4763813 | 0.13984157 | 0.58431708 |
| CIITA | 290.296024 | 1.12779852 | 0.53158707 | 2.12156875 | 0.03387397 | 0.35107893 |
| DEXI | 152.634929 | 0.545002 | 0.24125782 | 2.25900242 | 0.02388323 | 0.3030643 |
| CLEC16A | 3266.20681 | -0.2988435 | 0.21368297 | -1.3985369 | 0.16195191 | 0.61348517 |
| AC007014.1 | 210.34245 | -0.1408505 | 0.58045183 | -0.2426567 | 0.80827137 | 0.95610623 |
| AC007220.1 | 225.441804 | -0.727612 | 0.36930634 | -1.9702127 | 0.048814 | 0.39762623 |
| RMI2 | 58.7693638 | -0.55342 | 0.57186323 | -0.9677489 | 0.33316978 | 0.76124971 |
| SOCS1 | 322.079868 | -0.1743867 | 0.54479248 | -0.3200975 | 7.49E-01 | 0.93748128 |
| AC099489.1 | 237.155348 | -0.722583 | 0.3585795 | -2.0151263 | 0.04389142 | 0.38471248 |
| LITAF | 1512.96648 | 0.07621283 | 0.28282318 | 0.26947164 | 0.78756676 | 0.95004286 |
| SNN | 53.1732257 | 0.1590483 | 0.438643 | 0.36259168 | 0.71690992 | 0.9290703 |
| TXNDC11 | 1121.9402 | 0.09125209 | 0.18977467 | 0.4808444 | 0.63062709 | 0.90005881 |
| ZC3H7A | 1091.25192 | 0.13449675 | 0.14517515 | 0.92644472 | 0.35421491 | 0.77243419 |
| RSL1D1 | 197.447065 | 0.07667604 | 0.24076193 | 0.31847245 | 0.75012659 | 0.93775604 |
| AC007216.4 | 579.881028 | -0.012316 | 0.30321845 | -0.0406174 | 0.96760088 | 0.99248831 |
| GSPT1 | 1310.87104 | -0.1795912 | 0.18538578 | -0.9687432 | 0.33267332 | 0.7610305 |
| NPIPB2 | 128.726701 | 1.49002467 | 1.0682588 | 1.39481619 | 0.16307127 | 0.61387934 |
| SNX29 | 7638.09221 | -0.0919563 | 0.29114984 | -0.3158384 | 0.75212516 | 0.93815321 |
| AC010333.2 | 86.0514585 | -0.4768718 | 0.4566109 | -1.0443723 | 0.29631319 | 0.73668377 |
| CPPED1 | 1413.42484 | -0.5158519 | 0.33781142 | -1.5270411 | 0.12675078 | 0.56735041 |
| AC109597.1 | 95.4445987 | -0.3225399 | 0.57843782 | -0.5576052 | 0.57711401 | 0.88150331 |
| SHISA9 | 1233.39979 | -0.2551776 | 0.68252625 | -0.3738722 | 0.70849939 | 0.92577218 |
| U91319.1 | 720.981678 | 0.06927896 | 0.57723967 | 0.12001768 | 0.90446914 | 0.97989015 |
| ERCC4 | 224.380147 | -0.2211128 | 0.18856342 | -1.1726178 | 0.2409491 | 0.69178991 |
| MKL2 | 2744.8196 | -0.3658645 | 0.21052238 | -1.7378888 | 0.08223041 | 0.48594425 |
| AC130650.1 | 60.4509211 | -0.5955788 | 0.60832871 | -0.9790411 | 0.32755965 | 0.75620136 |
| MIR193BHG | 72.561095 | 0.20402541 | 0.44056712 | 0.46309723 | 0.64329467 | 0.90478945 |
| PARN | 1179.79844 | -0.4071915 | 0.17077235 | -2.3844112 | 0.01710648 | 0.26213393 |
| BFAR | 304.599042 | 0.06517791 | 0.24754185 | 0.26330056 | 0.79231893 | 0.95212343 |
| NOMO1 | 228.076426 | -0.4156334 | 0.31037303 | -1.3391414 | 0.18052465 | 0.63486424 |
| NPIPA1 | 99.887537 | -0.7828137 | 0.36542092 | -2.1422246 | 0.03217542 | 0.3421941 |
| PDXDC1 | 2386.32897 | -0.2742107 | 0.17595614 | -1.5584036 | 0.11913761 | 0.55120445 |
| NTAN1 | 153.692496 | -0.482076 | 0.29937221 | -1.6102896 | 0.10733465 | 0.52901576 |
| RRN3 | 343.430397 | -0.2579321 | 0.29056507 | -0.8876915 | 0.37470673 | 0.78426576 |
| C16orf45 | 833.789296 | -0.105346 | 0.28248651 | -0.372924 | 0.70920499 | 0.92577218 |
| MARF1 | 650.333628 | -0.4920372 | 0.21609028 | -2.2769981 | 0.02278634 | 0.2983102 |
| NDE1 | 232.617783 | -0.2061502 | 0.20515988 | -1.0048272 | 0.31498005 | 0.74785672 |
| MYH11 | 59.5485382 | 0.11313698 | 0.41787734 | 0.27074207 | 0.78658942 | 0.94978829 |
| FOPNL | 268.258949 | -0.1140394 | 0.19803886 | -0.5758433 | 0.5647211 | 0.87579905 |
| ABCC1 | 721.939366 | -0.1107148 | 0.26979631 | -0.4103643 | 0.68153871 | 0.91936732 |
| ABCC6 | 159.375691 | -0.6330911 | 0.71987589 | -0.8794448 | 0.37916013 | 0.78762967 |
| NOMO3 | 124.226198 | -0.8142411 | 0.38684154 | -2.1048441 | 0.03530488 | 0.355173 |
| NOMO2 | 142.410841 | -0.2573088 | 0.24523409 | -1.0492375 | 0.2940688 | 0.73527014 |
| RPS15A | 203.739139 | 0.61996042 | 0.46413554 | 1.33573142 | 0.18163708 | 0.63635309 |
| AC138811.1 | 53.869723 | 0.03411945 | 0.70668724 | 0.04828084 | 0.96149243 | 0.99191712 |
| SMG1 | 2802.12616 | -0.2914758 | 0.15720044 | -1.8541664 | 0.06371535 | 0.43779712 |
| COQ7 | 184.221745 | -0.2353913 | 0.21188374 | -1.1109456 | 0.26659175 | 0.71641758 |
| SYT17 | 300.410364 | 0.38514173 | 0.64827576 | 0.59410171 | 0.55244409 | 0.87096727 |
| GDE1 | 158.718788 | 0.43398478 | 0.2455562 | 1.76735417 | 0.07716893 | 0.47712313 |
| CCP110 | 332.515794 | 0.14257053 | 0.20602857 | 0.691994 | 0.4889411 | 0.84600773 |
| VPS35L | 1077.67854 | -0.2883068 | 0.26793241 | -1.076043 | 0.28190803 | 0.72604837 |

| | | | | | | |
|---|---|---|---|---|---|---|
| KNOP1 | 165.819179 | -0.2249416 | 0.46028737 | -0.4886982 | 0.62505537 | 0.89873262 |
| IQCK | 991.275788 | -0.1257661 | 0.30844371 | -0.4077439 | 0.68346168 | 0.9200778 |
| GPRC5B | 215.775552 | -0.0557373 | 0.5238513 | -0.1063991 | 9.15E-01 | 0.9817213 |
| GP2 | 1668.92653 | 2.60687177 | 1.11037674 | 2.3477363 | 0.01888789 | 0.27721124 |
| UMOD | 16978.8089 | 1.21734037 | 0.66779953 | 1.8229129 | 0.06831658 | 0.45323644 |
| ACSM2A | 117.110458 | 2.72884667 | 0.87967784 | 3.10209777 | 0.00192154 | 0.08517344 |
| ACSM2B | 130.030619 | 2.63089026 | 0.85543573 | 3.07549729 | 0.00210152 | 0.0878448 |
| ACSM3 | 468.50894 | 1.16738228 | 0.25966141 | 4.49578663 | 6.93E-06 | 0.00349342 |
| THUMPD1 | 481.520663 | 0.32549266 | 0.17172343 | 1.89544697 | 0.05803321 | 0.42087381 |
| ERI2 | 305.892264 | -0.2631715 | 0.36282289 | -0.7253444 | 0.4682408 | 0.83825637 |
| REXO5 | 51.0979486 | -0.3999541 | 0.37793926 | -1.0582496 | 0.28994167 | 0.73204984 |
| LYRM1 | 210.296112 | 0.24387075 | 0.30625959 | 0.79628771 | 0.42586483 | 0.81466629 |
| TMEM159 | 140.555062 | 0.3612368 | 0.41302929 | 0.87460334 | 0.38178977 | 0.78919376 |
| CRYM | 54.5136578 | 1.23330863 | 0.61568777 | 2.00313974 | 0.04516229 | 0.38785764 |
| NPIPB3 | 78.320936 | -0.5077768 | 0.52361082 | -0.96976 | 0.33216612 | 0.76041288 |
| METTL9 | 944.877848 | 0.20288564 | 0.27691776 | 0.73265667 | 0.46376786 | 0.83531269 |
| IGSF6 | 56.1146004 | 0.28007653 | 0.69710971 | 0.40176823 | 0.6878546 | 0.92113197 |
| NPIPB4 | 274.016431 | -0.9034715 | 0.29631178 | -3.0490571 | 0.00229561 | 0.09212026 |
| UQCRC2 | 1271.66323 | 0.21723908 | 0.18713122 | 1.16089174 | 2.46E-01 | 0.69581163 |
| MOSMO | 948.268747 | -0.0726189 | 0.2350065 | -0.3090079 | 0.7573155 | 0.94052092 |
| EEF2K | 200.107966 | -0.4420948 | 0.28099409 | -1.5733242 | 0.11564376 | 0.54438291 |
| POLR3E | 293.123141 | -0.3903694 | 0.15982212 | -2.4425245 | 0.01458494 | 0.24762526 |
| CDR2 | 2128.02317 | -0.5420241 | 0.25146155 | -2.1554949 | 0.03112313 | 0.3383722 |
| AC092338.2 | 57.0626173 | -0.8376621 | 0.75740181 | -1.1059679 | 0.26874043 | 0.71818262 |
| NPIPB5 | 742.572946 | -0.1118846 | 0.21289546 | -0.5255377 | 0.59920947 | 0.88883692 |
| USP31 | 540.781285 | -0.1732938 | 0.28143574 | -0.6157492 | 0.53806009 | 0.86496983 |
| COG7 | 846.662678 | -0.2330061 | 0.19169631 | -1.2154962 | 0.22417689 | 0.67681479 |
| GGA2 | 1014.58874 | -0.298614 | 0.32463542 | -0.9198443 | 0.35765415 | 0.77484888 |
| EARS2 | 159.480022 | -0.0135305 | 0.27345456 | -0.0494798 | 0.96053693 | 0.99153894 |
| UBFD1 | 206.099477 | -0.0134579 | 0.20710859 | -0.0649799 | 0.94818999 | 0.98919519 |
| NDUFAB1 | 299.95422 | 0.40310779 | 0.20371899 | 1.97874435 | 0.0478448 | 0.39501762 |
| PALB2 | 172.034659 | -0.3818682 | 0.35586568 | -1.0730684 | 0.28324044 | 0.72647048 |
| DCTN5 | 405.284786 | 0.29299838 | 0.22290922 | 1.31442913 | 0.18870183 | 0.64374031 |
| RBBP6 | 861.681334 | 0.34757439 | 0.17775654 | 1.95533947 | 0.05054302 | 0.40339739 |
| TNRC6A | 5747.5994 | -0.0517426 | 0.22208211 | -0.2329884 | 0.81577041 | 0.95876069 |
| AC008731.1 | 182.711884 | 0.02732843 | 0.50423072 | 0.05419825 | 0.95677721 | 0.99065996 |
| SLC5A11 | 89.009356 | 0.10731638 | 0.68710471 | 0.15618635 | 0.87588613 | 0.97614152 |
| ARHGAP17 | 757.178623 | -0.2433379 | 0.17165408 | -1.4176065 | 0.15630568 | 0.60564438 |
| LCMT1 | 536.976674 | -0.0670436 | 0.19692913 | -0.3404453 | 0.73352118 | 0.93552616 |
| AC008741.2 | 101.101577 | -0.1999131 | 0.39229334 | -0.5096011 | 0.61033093 | 0.89339478 |
| HS3ST4 | 52.3971539 | -0.1291992 | 0.42190387 | -0.306229 | 0.75943031 | 0.94133292 |
| AC009035.1 | 35.1006855 | 0.90641634 | 0.83834885 | 1.08119232 | 2.80E-01 | 7.25E-01 |
| NSMCE1 | 278.965872 | 0.33740626 | 0.2331293 | 1.44729237 | 0.14781506 | 0.59424557 |
| AC106739.1 | 87.9436822 | -0.0368532 | 0.5880451 | -0.0626706 | 0.95002879 | 0.98919519 |
| IL4R | 281.543055 | 1.0978267 | 0.54217958 | 2.02483962 | 4.29E-02 | 3.81E-01 |
| GTF3C1 | 455.757628 | -0.1176746 | 0.13699061 | -0.8589973 | 0.39034199 | 0.79701353 |
| KIAA0556 | 871.578387 | -0.1474856 | 0.20165588 | -0.7313729 | 0.46455139 | 0.83600264 |
| XPO6 | 794.893675 | -0.1894665 | 0.21432145 | -0.8840294 | 0.37668033 | 0.78528575 |
| SBK1 | 52.0921022 | -0.9436197 | 0.46164201 | -2.0440507 | 0.04094854 | 0.37443162 |
| NUPR1 | 207.941904 | 0.57885356 | 0.67413376 | 0.85866277 | 0.39052659 | 0.79701353 |
| SGF29 | 177.222506 | -0.5123626 | 0.31660803 | -1.6182866 | 0.10560084 | 0.52694363 |
| SULT1A2 | 49.5027466 | 1.6055092 | 0.69908372 | 2.29659073 | 0.02164213 | 0.29380356 |
| SULT1A1 | 385.477784 | 0.47230245 | 0.45039924 | 1.04863066 | 0.29434814 | 0.73527014 |
| ATXN2L | 447.181333 | -0.2870388 | 0.22669047 | -1.2662146 | 0.20543628 | 0.65813563 |

| | | | | | | |
|---|---|---|---|---|---|---|
| TUFM | 486.911464 | 0.8548852 | 0.44158751 | 1.93593608 | 0.05287553 | 0.41066631 |
| SH2B1 | 724.106251 | 0.07268114 | 0.20571321 | 0.35331294 | 0.72385385 | 0.93201977 |
| AC133550.2 | 119.511259 | -0.6702517 | 0.3709983 | -1.8066167 | 0.07082207 | 0.4633426 |
| ATP2A1 | 103.574283 | -0.6945861 | 0.32055556 | -2.1668196 | 0.03024861 | 0.3341256 |
| RABEP2 | 493.200812 | -0.6521378 | 0.29045068 | -2.2452617 | 0.02475134 | 0.30843438 |
| NFATC2IP | 913.608985 | -0.3286558 | 0.32426936 | -1.0135271 | 0.31080848 | 0.74495902 |
| AC109460.2 | 75.0103915 | -0.0878508 | 0.4267885 | -0.2058416 | 0.83691465 | 0.9652581 |
| SPNS1 | 188.832457 | -0.2523032 | 0.28200965 | -0.8946615 | 0.37096809 | 0.78104287 |
| LAT | 77.6276889 | -0.4312776 | 0.32581153 | -1.3237027 | 0.18560179 | 0.64030106 |
| NPIPB11 | 63.3191476 | -0.4235982 | 0.42576467 | -0.9949115 | 0.31977929 | 0.75094569 |
| QPRT | 85.3359867 | 1.17734703 | 0.55769382 | 2.1110993 | 0.03476378 | 0.35358029 |
| KIF22 | 158.744333 | -0.5934438 | 0.25454113 | -2.3314259 | 0.01973091 | 0.2823953 |
| MAZ | 165.027354 | 0.06152871 | 0.42351557 | 0.14528088 | 0.88448911 | 0.97664137 |
| AC009133.1 | 211.218484 | -0.089876 | 0.40961292 | -0.2194168 | 8.26E-01 | 9.62E-01 |
| MVP | 863.713417 | 0.2092905 | 0.48474994 | 0.43174942 | 6.66E-01 | 9.13E-01 |
| CDIPT | 221.677962 | 0.39744528 | 0.27251573 | 1.4584306 | 0.14472189 | 0.59044137 |
| ASPHD1 | 89.3971429 | -0.0970211 | 0.38472698 | -0.2521817 | 0.80090062 | 0.95370871 |
| KCTD13 | 332.263586 | -0.4306839 | 0.24054598 | -1.790443 | 0.07338272 | 0.4701377 |
| TMEM219 | 341.443064 | 0.61791176 | 0.31672303 | 1.95095306 | 0.05106263 | 0.40556617 |
| TAOK2 | 152.821186 | -0.1427533 | 0.33976941 | -0.4201477 | 0.67437758 | 0.91636338 |
| HIRIP3 | 41.9881465 | 0.92656021 | 0.38855973 | 2.38460173 | 0.01709762 | 0.26213393 |
| INO80E | 244.899098 | -0.033717 | 0.22069286 | -0.1527781 | 0.87857323 | 0.9765717 |
| DOC2A | 64.8317724 | -0.1418369 | 0.868676 | -0.1632794 | 0.87029847 | 0.97463098 |
| PPP4C | 263.39549 | 0.02764782 | 0.22655222 | 0.12203731 | 0.90286947 | 0.97960781 |
| YPEL3 | 508.051448 | -0.2112961 | 0.38591633 | -0.5475179 | 0.58402296 | 0.88416261 |
| MAPK3 | 106.051645 | 0.56763763 | 0.37999711 | 1.49379461 | 0.13522931 | 0.57708028 |
| NPIPB13 | 219.769758 | -0.8536304 | 0.44847541 | -1.9034051 | 0.05698771 | 0.41872623 |
| CD2BP2 | 169.618138 | 0.30726141 | 0.35052628 | 0.87657167 | 0.38071933 | 0.78826518 |
| TBC1D10B | 147.224666 | -0.3320182 | 0.27812441 | -1.1937758 | 0.23256567 | 0.68463559 |
| ZNF48 | 69.860575 | -0.4873353 | 0.396881 | -1.2279128 | 0.21947971 | 0.67278782 |
| 1-Sep | 58.3838071 | -0.428505 | 0.33605657 | -1.2750979 | 0.2022746 | 0.65552334 |
| ZNF771 | 132.222267 | 0.11017131 | 0.40932686 | 0.26915241 | 0.7878124 | 0.95007888 |
| DCTPP1 | 80.7757761 | 0.47087787 | 0.43282251 | 1.08792373 | 0.27662878 | 0.72258811 |
| SEPHS2 | 427.922609 | 1.07979026 | 0.33693524 | 3.2047412 | 0.00135184 | 0.06712951 |
| ZNF768 | 114.55019 | 0.47717481 | 0.28583774 | 1.6693905 | 0.09504002 | 0.51085035 |
| AC002310.4 | 50.6171327 | -0.4387402 | 0.34677705 | -1.2651938 | 0.20580187 | 0.65813563 |
| ZNF688 | 81.1094301 | -0.3241635 | 0.31273091 | -1.0365573 | 0.2999422 | 0.73673761 |
| AC002310.2 | 41.4266537 | 0.2991826 | 0.55104741 | 0.54293441 | 0.58717496 | 0.88563701 |
| ZNF785 | 236.949146 | -0.0182608 | 0.20203554 | -0.0903842 | 0.92798188 | 0.98297055 |
| PRR14 | 138.70225 | -0.2428405 | 0.32914953 | -0.7377816 | 0.46064718 | 0.83439198 |
| FBRS | 241.803575 | -0.0243896 | 0.23191663 | -0.1051654 | 0.91624458 | 0.98195017 |
| SRCAP | 51.7859764 | -0.4110072 | 0.33505826 | -1.2266739 | 0.21994518 | 0.67299198 |
| PHKG2 | 192.572335 | 0.07941629 | 0.27894883 | 0.28469842 | 0.7758752 | 0.94692188 |
| CCDC189 | 56.3209737 | -0.2228557 | 0.31165726 | -0.7150667 | 0.4745678 | 0.84106098 |
| RNF40 | 297.798239 | 0.24331154 | 0.273496 | 0.88963474 | 0.37366205 | 0.78307187 |
| ZNF629 | 87.1157358 | 0.27131167 | 0.347756 | 0.78017825 | 0.43528596 | 0.81919857 |
| BCL7C | 798.533698 | 0.015435 | 0.32656275 | 0.04726504 | 0.96230199 | 0.99224805 |
| MIR762HG | 172.417871 | -0.4694452 | 0.3600303 | -1.3039048 | 0.19226604 | 0.64654839 |
| CTF1 | 42.4893948 | -0.0029184 | 0.33917551 | -0.0086043 | 0.99313487 | 0.99816851 |
| FBXL19 | 222.659601 | -0.3559795 | 0.41173399 | -0.8645862 | 0.38726597 | 0.79554171 |
| ORAI3 | 74.0332472 | 0.38596788 | 0.31337535 | 1.23164722 | 0.21808089 | 0.67150495 |
| SETD1A | 173.400056 | -0.5377933 | 0.24251445 | -2.2175723 | 0.02658401 | 0.31471512 |
| HSD3B7 | 52.8494694 | 0.19020196 | 0.59518877 | 0.31956577 | 0.74929753 | 0.93752686 |
| STX4 | 153.581351 | -0.2251664 | 0.19905273 | -1.1311898 | 0.25797521 | 0.70888174 |

| | | | | | | |
|---|---|---|---|---|---|---|
| AC135050.3 | 67.4927224 | -0.5080884 | 0.342925 | -1.4816312 | 0.13843846 | 0.58206378 |
| ZNF668 | 49.9037428 | -0.3433049 | 0.39667521 | -0.8654558 | 0.38678868 | 0.79523542 |
| ZNF646 | 47.6577786 | -0.023074 | 0.42537567 | -0.0542439 | 0.95674084 | 0.99065996 |
| BCKDK | 154.307735 | 0.3795249 | 0.37803635 | 1.00393757 | 0.3154087 | 0.74785672 |
| KAT8 | 365.936769 | -0.5371374 | 0.21619012 | -2.4845602 | 0.01297116 | 0.23379993 |
| PRSS8 | 368.68579 | 0.29043362 | 0.46704633 | 0.62185184 | 0.5340393 | 0.86397197 |
| FUS | 1781.1325 | -0.2155966 | 0.13340443 | -1.6161125 | 0.10606998 | 0.52735777 |
| AC026471.4 | 165.867985 | -0.4257423 | 0.37160525 | -1.1456844 | 0.25192578 | 0.70276117 |
| AC026471.1 | 46.6377263 | -0.6160234 | 0.44077054 | -1.3976056 | 0.16223154 | 0.61370284 |
| ARMC5 | 42.6227377 | -0.0979327 | 0.42822692 | -0.2286935 | 0.81910711 | 0.96006637 |
| C16orf58 | 243.554436 | -0.206305 | 0.21913562 | -0.9414489 | 0.34647484 | 0.76864066 |
| ZNF720 | 528.094216 | 0.11013311 | 0.14360275 | 0.76692896 | 0.44312375 | 0.82312215 |
| ZNF267 | 145.594314 | -0.0515946 | 0.25015198 | -0.2062531 | 0.83659318 | 0.96511435 |
| VPS35 | 444.743769 | 0.12525521 | 0.20003173 | 0.62617669 | 0.53119905 | 0.86397197 |
| C16orf87 | 171.528903 | -0.1742701 | 0.26575944 | -0.655744 | 0.51198888 | 0.85762435 |
| GPT2 | 104.702465 | -0.306298 | 0.43835798 | -0.6987395 | 0.48471484 | 0.8447301 |
| DNAJA2 | 269.376689 | 0.29225963 | 0.21899559 | 1.33454572 | 0.18202507 | 0.63706109 |
| NETO2 | 80.0163421 | -0.2223048 | 0.44871767 | -0.4954224 | 0.62030201 | 0.8972371 |
| ITFG1 | 4123.71241 | -0.397731 | 0.1432454 | -2.7765712 | 0.00549356 | 0.15016771 |
| AC007533.1 | 49.7405621 | -0.1529404 | 0.64765178 | -0.2361461 | 0.81331933 | 0.9582375 |
| PHKB | 2593.40995 | 0.08620104 | 0.15918558 | 0.54151285 | 0.58815414 | 0.88571413 |
| LINC02133 | 120.615832 | -0.7597736 | 0.67251515 | -1.1297495 | 0.2585818 | 0.70897406 |
| LONP2 | 1861.89617 | -0.0274566 | 0.15374253 | -0.1785885 | 0.85826083 | 0.97143431 |
| SIAH1 | 395.300105 | -0.4429378 | 0.23223416 | -1.9072896 | 0.05648309 | 0.41843207 |
| N4BP1 | 817.392523 | -0.028525 | 0.21969514 | -0.1298391 | 0.89669373 | 0.97823153 |
| ZNF423 | 87.8198853 | 0.13478831 | 0.72457704 | 0.18602343 | 0.85242638 | 0.97076831 |
| CNEP1R1 | 120.922581 | 0.09336485 | 0.29141677 | 0.32038256 | 0.74867834 | 0.93746253 |
| HEATR3 | 172.115622 | -0.1630145 | 0.34542504 | -0.4719244 | 0.63698076 | 0.90255504 |
| TENT4B | 699.996276 | 0.18380328 | 0.25222121 | 0.72873838 | 0.46616171 | 0.8367113 |
| BRD7 | 559.49568 | -0.1875155 | 0.20315931 | -0.9229972 | 0.35600866 | 0.77367035 |
| CYLD | 488.180825 | 0.0849463 | 0.16017117 | 0.53034699 | 0.59587137 | 0.88859628 |
| SALL1 | 851.559498 | -0.0845526 | 0.46874882 | -0.1803794 | 0.85685469 | 0.97127317 |
| AC087564.1 | 2909.80008 | -0.7666698 | 0.36449238 | -2.1033906 | 0.03543164 | 0.35586635 |
| AC007344.1 | 406.707978 | -2.8144619 | 0.62182288 | -4.5261472 | 6.01E-06 | 0.00330603 |
| HNRNPA1P4 | 61.4129295 | 0.48204624 | 0.38777846 | 1.243097 | 0.21383209 | 0.66800831 |
| LINC01571 | 44.202502 | -0.8894806 | 0.69304782 | -1.2834333 | 0.19934032 | 0.65150038 |
| TOX3 | 942.579763 | 0.10602276 | 0.31846562 | 0.33291743 | 0.73919661 | 0.93552616 |
| AC007906.2 | 225.204453 | 0.54250748 | 0.63427851 | 0.85531431 | 0.39237717 | 0.79810749 |
| CHD9 | 6269.59058 | 0.30090078 | 0.14720261 | 2.04412673 | 0.04094103 | 0.37443162 |
| RBL2 | 1423.26213 | -0.1917608 | 0.14609278 | -1.3125959 | 0.18931915 | 0.64432929 |
| AKTIP | 221.777874 | 0.03385557 | 0.2322563 | 0.14576816 | 0.88410441 | 0.97664137 |
| RPGRIP1L | 272.744341 | -0.2112869 | 0.30314558 | -0.6969817 | 0.48581422 | 0.8447301 |
| FTO | 3280.87681 | 0.1110087 | 0.16220691 | 0.68436484 | 0.49374481 | 0.84865 |
| IRX3 | 70.3037707 | 0.53462454 | 0.48773124 | 1.09614578 | 2.73E-01 | 0.72070318 |
| CRNDE | 289.719476 | -0.4361114 | 0.42750179 | -1.0201393 | 0.30766242 | 0.74348936 |
| IRX5 | 60.7868498 | -0.5340092 | 0.35508957 | -1.5038718 | 0.13261439 | 0.57682553 |
| AMFR | 1513.76415 | -0.1950283 | 0.50534443 | -0.3859314 | 0.69954748 | 0.92367966 |
| AC092140.1 | 96.568423 | -0.1071618 | 0.84783192 | -0.1263951 | 0.89941919 | 0.97889101 |
| NUDT21 | 611.482187 | 0.23372673 | 0.26347172 | 0.88710365 | 0.3750231 | 0.78444261 |
| OGFOD1 | 273.319159 | 0.21942871 | 0.27672321 | 0.79295376 | 0.42780477 | 0.81466629 |
| BBS2 | 616.756658 | -0.4704074 | 0.12828901 | -3.6667786 | 0.00024563 | 0.02595571 |
| MT2A | 3555.52351 | 0.30623725 | 0.57582725 | 0.53182138 | 5.95E-01 | 8.89E-01 |
| MT1E | 907.322968 | -0.1011382 | 0.56381168 | -0.179383 | 0.85763696 | 0.97127317 |
| MT1M | 95.4149411 | -0.5308412 | 0.77327956 | -0.6864803 | 0.49241029 | 0.84734855 |

| | | | | | | |
|---|---|---|---|---|---|---|
| MT1F | 134.984883 | 0.6055411 | 0.56219868 | 1.07709449 | 0.28143805 | 0.72604837 |
| MT1G | 1271.11317 | -0.0775485 | 0.74765239 | -0.1037226 | 0.91738947 | 0.98199511 |
| MT1H | 403.585295 | -0.2587709 | 0.86250255 | -0.3000234 | 0.76415933 | 0.94256396 |
| MT1X | 504.490651 | 0.05071693 | 0.64375491 | 0.07878298 | 0.93720524 | 0.98649645 |
| NUP93 | 443.223207 | -0.6674689 | 0.1964464 | -3.3977152 | 0.00067951 | 0.04556174 |
| SLC12A3 | 91.0073739 | 0.86437373 | 0.65240503 | 1.32490353 | 1.85E-01 | 0.64030106 |
| HERPUD1 | 907.706634 | 0.16740402 | 0.3976256 | 0.42100917 | 0.67374839 | 0.91636338 |
| NLRC5 | 224.302087 | -1.0437215 | 0.62263466 | -1.6762985 | 0.09367974 | 0.50901512 |
| FAM192A | 642.012674 | -0.2049759 | 0.14681555 | -1.3961457 | 0.16267062 | 0.61387934 |
| RSPRY1 | 820.148893 | -0.2168602 | 0.126158 | -1.7189574 | 0.08562212 | 0.49356597 |
| ARL2BP | 287.02372 | 0.60543532 | 0.25018525 | 2.41994809 | 0.01552272 | 0.25214021 |
| PLLP | 94.7843975 | 2.32697384 | 0.6957958 | 3.34433441 | 0.0008248 | 0.04997622 |
| CX3CL1 | 63.900228 | 0.79940309 | 0.616813 | 1.29602179 | 0.19496798 | 0.64850083 |
| CIAPIN1 | 332.106409 | -0.2157992 | 0.25400562 | -0.8495843 | 0.39555627 | 0.79924445 |
| COQ9 | 649.555684 | 0.20337956 | 0.18374274 | 1.10687127 | 0.26834959 | 0.71818262 |
| POLR2C | 164.639105 | 0.29081197 | 0.23427634 | 1.24132029 | 0.21448745 | 0.66827012 |
| DOK4 | 92.6809767 | 0.97560664 | 0.50768093 | 1.92169249 | 0.05464446 | 0.41650243 |
| CCDC102A | 45.6788199 | 0.91697478 | 0.3834956 | 2.39109598 | 0.01679816 | 0.25951685 |
| ADGRG1 | 1430.04544 | 1.19502024 | 0.6473366 | 1.84605698 | 0.06488393 | 0.44041719 |
| KATNB1 | 49.1718347 | -0.2943789 | 0.37907998 | -0.7765615 | 0.43741753 | 0.82013459 |
| KIFC3 | 3063.52007 | 0.03052492 | 0.34955108 | 0.08732606 | 9.30E-01 | 0.98331361 |
| AC092118.1 | 76.5675539 | 0.51369138 | 0.51092322 | 1.00541795 | 0.31469564 | 0.74785672 |
| USB1 | 144.71379 | -0.0591212 | 0.35415505 | -0.1669358 | 0.86742056 | 0.97402505 |
| MMP15 | 63.7813152 | 0.6672075 | 0.55158173 | 1.20962583 | 0.2264225 | 0.67880306 |
| CFAP20 | 97.4885935 | 0.07716448 | 0.26717171 | 0.28881981 | 0.77271927 | 0.94530018 |
| CSNK2A2 | 428.250298 | -0.0967863 | 0.18450385 | -0.5245763 | 0.5998778 | 0.88883692 |
| CCDC113 | 82.2173378 | -0.135983 | 0.36553981 | -0.3720059 | 0.70988848 | 0.92607195 |
| SETD6 | 147.466933 | -0.0351062 | 0.33518941 | -0.1047354 | 0.91658575 | 0.98197488 |
| CNOT1 | 1504.32567 | 0.02700516 | 0.21337821 | 0.12656008 | 0.8992886 | 0.97889101 |
| AC009118.3 | 42.1143356 | -0.369079 | 0.46156088 | -0.7996324 | 0.4239238 | 0.81413339 |
| SLC38A7 | 157.975282 | -0.239701 | 0.30494936 | -0.7860353 | 0.43184679 | 0.81761357 |
| GOT2 | 424.001584 | 0.11836487 | 0.1886026 | 0.62758878 | 0.53027336 | 0.86377862 |
| AC092378.1 | 301.749015 | -0.0678058 | 0.41980413 | -0.1615178 | 0.87168558 | 0.97527484 |
| AC138305.1 | 5287.932 | 0.58916366 | 0.34122055 | 1.72663594 | 0.08423307 | 0.49023622 |
| BEAN1 | 72.9552336 | -0.2720397 | 0.33197107 | -0.8194682 | 0.41251936 | 0.80713497 |
| AC010542.3 | 156.302403 | -0.7535716 | 0.59023881 | -1.2767233 | 0.20169996 | 0.65534788 |
| TK2 | 200.576713 | -0.5357205 | 0.28360365 | -1.8889761 | 0.05889503 | 0.42348634 |
| CMTM2 | 71.8683614 | 0.45164695 | 0.69103407 | 0.65358132 | 0.51338158 | 0.85762435 |
| CMTM4 | 2088.6512 | 0.26654203 | 0.18392679 | 1.4491746 | 0.14728883 | 0.59373375 |
| DYNC1LI2 | 1338.18771 | -0.1004166 | 0.18541546 | -0.541576 | 0.58811064 | 0.88571413 |
| AC018557.1 | 113.147188 | 0.90374232 | 0.36429977 | 2.48076555 | 0.01311006 | 0.23485956 |
| NAE1 | 413.879659 | 0.04551328 | 0.17407411 | 0.26145919 | 0.79373842 | 0.95224951 |
| PDP2 | 322.359389 | 0.05333129 | 0.35421477 | 0.15056202 | 0.88032123 | 0.97664137 |
| CDH16 | 1988.44177 | 1.05372561 | 0.35175675 | 2.9956088 | 0.00273898 | 0.09858074 |
| RRAD | 826.673724 | 1.12001756 | 0.56046114 | 1.99838577 | 4.57E-02 | 3.89E-01 |
| FAM96B | 337.964298 | 0.33399036 | 0.32760532 | 1.01949001 | 0.30797039 | 0.74372365 |
| CES2 | 216.116277 | 0.36429907 | 0.33366177 | 1.09182142 | 0.2749116 | 0.72164296 |
| CBFB | 1519.19401 | 0.19582809 | 0.20533991 | 0.95367766 | 0.34024683 | 0.76478309 |
| C16orf70 | 264.487053 | 0.16237371 | 0.31479938 | 0.51580062 | 0.60599367 | 0.89107986 |
| TRADD | 131.972867 | 0.71075321 | 0.31587886 | 2.25008162 | 0.02444376 | 0.3067148 |
| FBXL8 | 42.213452 | -0.2492296 | 0.45821748 | -0.5439112 | 0.58650258 | 0.88538429 |
| HSF4 | 94.9420129 | -0.124296 | 0.49765064 | -0.2497655 | 0.80276871 | 0.95446295 |
| NOL3 | 246.408855 | 0.93736402 | 0.47690878 | 1.96549962 | 0.04935644 | 0.39895974 |
| KIAA0895L | 228.372568 | -0.6313399 | 0.24551844 | -2.571456 | 0.01012719 | 0.20516875 |

| | | | | | | |
|---|---|---|---|---|---|---|
| E2F4 | 220.083743 | -0.2177575 | 0.37475781 | -0.5810618 | 0.56119878 | 0.87395363 |
| ELMO3 | 157.20639 | -0.2031014 | 0.28316832 | -0.7172463 | 0.47322216 | 0.84027309 |
| LRRC29 | 167.892028 | -0.4731529 | 0.44734298 | -1.057696 | 0.29019408 | 0.73222514 |
| TMEM208 | 236.503999 | 0.47635941 | 0.42606003 | 1.118057 | 0.26354265 | 0.71429196 |
| FHOD1 | 107.296446 | -0.1734108 | 0.33127796 | -0.5234602 | 0.60065401 | 0.88883692 |
| SLC9A5 | 39.7023977 | -0.4821087 | 0.38271329 | -1.2597126 | 0.20777307 | 0.6607067 |
| LRRC36 | 118.612646 | 1.41310142 | 0.6178587 | 2.2870948 | 0.02219029 | 0.29577351 |
| ZDHHC1 | 300.91336 | 0.53359341 | 0.23360236 | 2.28419529 | 0.02236006 | 0.29654015 |
| HSD11B2 | 615.349596 | 3.22408114 | 0.91412981 | 3.52694015 | 0.00042039 | 0.03514528 |
| ATP6V0D1 | 446.918234 | 0.40155521 | 0.28110822 | 1.42847196 | 0.15315606 | 0.60285224 |
| AC009061.2 | 446.430067 | 0.19287848 | 0.44737613 | 0.4311327 | 0.66637188 | 0.91318863 |
| AC027682.6 | 66.6864184 | 1.82029383 | 0.61916757 | 2.939905 | 0.00328313 | 0.11119705 |
| RIPOR1 | 403.71634 | 0.22626064 | 0.3487655 | 0.6487472 | 0.51650179 | 0.8588826 |
| AC027682.4 | 161.304863 | 0.68478516 | 0.35778115 | 1.91397774 | 0.055623 | 0.41843207 |
| CTCF | 792.589861 | -0.3290493 | 0.19863704 | -1.6565354 | 0.09761345 | 0.51355946 |
| ENKD1 | 177.550636 | 0.09191851 | 0.36507114 | 0.25178245 | 0.80120922 | 0.95370871 |
| C16orf86 | 39.9914723 | 0.21288406 | 0.59217777 | 0.35949351 | 0.71922593 | 0.93001079 |
| GFOD2 | 273.235113 | 0.03508592 | 0.28834473 | 0.12168046 | 0.90315209 | 0.97961373 |
| RANBP10 | 918.834304 | -0.0416948 | 0.22901538 | -0.1820612 | 0.8555347 | 0.97118784 |
| TSNAXIP1 | 72.1391995 | 0.2283773 | 0.27143583 | 0.84136755 | 0.40014206 | 0.80238662 |
| CENPT | 104.574734 | -0.0045206 | 0.28279817 | -0.0159852 | 0.98724616 | 0.99653761 |
| NUTF2 | 147.175137 | 0.02311456 | 0.2714908 | 0.08513939 | 0.93215058 | 0.98360577 |
| PSKH1 | 250.12684 | 0.15225933 | 0.21855737 | 0.6966561 | 0.48601804 | 0.8447301 |
| LCAT | 88.8449643 | -0.4757179 | 0.43045539 | -1.1051503 | 0.26909447 | 0.71818262 |
| SLC12A4 | 89.160959 | 0.19014373 | 0.44689444 | 0.42547796 | 0.67048828 | 0.91492559 |
| DUS2 | 154.223524 | -0.4067182 | 0.21921199 | -1.8553647 | 0.06354417 | 0.43779712 |
| NFATC3 | 1460.03223 | -0.2510727 | 0.11033541 | -2.2755411 | 0.02287349 | 0.2983102 |
| AC020978.5 | 86.2229216 | -0.0528376 | 0.44082657 | -0.1198602 | 0.90459387 | 0.97989015 |
| ESRP2 | 127.737889 | -0.0477013 | 0.43664271 | -0.1092457 | 0.91300761 | 0.98135874 |
| PLA2G15 | 181.723002 | 0.32326087 | 0.3223914 | 1.00269693 | 0.31600711 | 0.74785672 |
| SLC7A6 | 613.344867 | 0.18397734 | 0.31298414 | 0.58781682 | 0.55665526 | 0.87212955 |
| SLC7A6OS | 330.285695 | -0.4200912 | 0.30351406 | -1.3840914 | 0.16633048 | 0.61901833 |
| PRMT7 | 371.618564 | -0.1275688 | 0.41380954 | -0.3082791 | 0.75786995 | 0.94056395 |
| ZFP90 | 435.702307 | 0.02480954 | 0.24191133 | 0.10255632 | 0.91831511 | 0.98210868 |
| CDH3 | 593.688225 | 1.38308743 | 0.64418512 | 2.14703413 | 0.03179056 | 0.34024781 |
| CDH1 | 979.081345 | 0.52260771 | 0.40046457 | 1.3050036 | 0.19189162 | 0.64596359 |
| TANGO6 | 859.760495 | -0.1867641 | 0.19106092 | -0.9775106 | 0.32831641 | 0.75671461 |
| CHTF8 | 335.151585 | 0.20454191 | 0.24141585 | 0.84725966 | 0.39685041 | 0.80068288 |
| UTP4 | 203.389588 | -0.162291 | 0.25660346 | -0.6324585 | 0.52708732 | 0.86151701 |
| SNTB2 | 519.62333 | -1.1229656 | 0.36531016 | -3.0740059 | 0.00211205 | 0.08800664 |
| VPS4A | 212.125068 | 0.22641168 | 0.20223726 | 1.11953496 | 0.26291199 | 0.71328 |
| COG8 | 114.815526 | 0.44838168 | 0.25492799 | 1.75885623 | 0.07860193 | 0.48060389 |
| NIP7 | 53.4775208 | 0.07608045 | 0.45087594 | 0.16873921 | 0.86600178 | 0.97379123 |
| TERF2 | 414.813761 | -0.3531109 | 0.23207568 | -1.5215335 | 0.128126 | 0.56762585 |
| CYB5B | 865.951335 | -0.2887745 | 0.16058877 | -1.7982233 | 0.07214163 | 0.4679905 |
| NFAT5 | 6935.33582 | -0.1317298 | 0.15410209 | -0.8548213 | 0.39265009 | 0.79829383 |
| NQO1 | 417.29665 | 1.73652955 | 0.76739891 | 2.26287729 | 0.02364325 | 0.30185158 |
| AC092115.3 | 64.9255422 | 0.11952533 | 0.37569339 | 0.31814595 | 0.75037423 | 0.93775604 |
| NOB1 | 92.8562083 | 0.10443378 | 0.27670531 | 0.37741877 | 0.70586242 | 0.92556842 |
| WWP2 | 2030.3039 | -0.1118943 | 0.24602186 | -0.4548143 | 0.64924286 | 0.90788153 |
| PDPR | 689.216041 | -0.6854647 | 0.29057824 | -2.3589678 | 0.01832585 | 0.27167913 |
| AC009060.1 | 306.758151 | 0.09452685 | 0.45991075 | 0.20553303 | 0.83715569 | 0.96527001 |
| EXOSC6 | 176.988062 | -0.1164983 | 0.28478168 | -0.4090795 | 0.68248132 | 0.91969963 |
| AARS | 404.785389 | 0.25296787 | 0.20192756 | 1.25276547 | 0.21029107 | 0.66326045 |

| | | | | | | |
|---|---|---|---|---|---|---|
| AC012184.2 | 73.1930526 | -0.5553357 | 0.31833727 | -1.7444884 | 0.08107396 | 0.48572317 |
| DDX19A | 87.7202849 | 0.36974734 | 0.29486196 | 1.2539676 | 0.20985378 | 0.66255855 |
| ST3GAL2 | 364.323532 | 0.20156842 | 0.18699633 | 1.07792712 | 0.28106628 | 0.72582688 |
| FUK | 227.720355 | 0.04618337 | 0.22526907 | 0.20501425 | 0.83756099 | 0.96527001 |
| COG4 | 508.208034 | -0.146561 | 0.29879492 | -0.4905069 | 0.62377521 | 0.89873262 |
| SF3B3 | 639.861979 | -0.0251588 | 0.12440654 | -0.2022307 | 0.83973634 | 0.96617873 |
| MTSS1L | 212.140433 | -0.0071499 | 0.36169353 | -0.0197677 | 0.98422866 | 0.99518552 |
| VAC14 | 426.553639 | -0.3655784 | 0.36034689 | -1.0145181 | 0.31033562 | 0.74459794 |
| HYDIN | 1198.29201 | 0.12834832 | 0.33298145 | 0.38545188 | 0.69990265 | 0.92367966 |
| CMTR2 | 355.17522 | 0.06199608 | 0.33044473 | 0.18761407 | 0.8511792 | 0.97057139 |
| ZNF19 | 435.506266 | -0.1184242 | 0.31063803 | -0.3812289 | 0.70303338 | 0.92501157 |
| MARVELD3 | 166.039721 | -0.6550253 | 0.49511042 | -1.3229884 | 0.18583924 | 0.64030106 |
| PHLPP2 | 343.277391 | 0.03264532 | 0.33041279 | 0.09880162 | 0.92129578 | 0.98215577 |
| AP1G1 | 1512.38779 | 0.00530193 | 0.22919045 | 0.02313331 | 0.98154393 | 0.99518016 |
| IST1 | 1012.44987 | -0.1602815 | 0.13972803 | -1.1470962 | 0.25134188 | 0.70276117 |
| ZNF821 | 389.976568 | -0.077566 | 0.31237521 | -0.2483104 | 0.80389424 | 0.95450237 |
| DHODH | 116.045546 | -1.2326462 | 0.49345137 | -2.4980094 | 0.01248929 | 0.22899031 |
| TXNL4B | 207.585727 | -0.0398997 | 0.17996613 | -0.2217066 | 0.8245423 | 0.96146415 |
| DHX38 | 205.803199 | -0.2000068 | 0.22613486 | -0.884458 | 0.37644903 | 0.78517531 |
| LINC01572 | 665.239227 | 0.02063785 | 0.41808749 | 0.04936252 | 0.9606304 | 0.99155365 |
| AC004158.1 | 46.361624 | 0.41250906 | 0.61211951 | 0.67390281 | 0.5003731 | 0.85293952 |
| AC004943.2 | 436.685452 | -0.0538022 | 0.20288892 | -0.2651808 | 0.79087018 | 0.95144272 |
| ZFHX3 | 3190.01979 | -0.6660319 | 0.24758479 | -2.6901162 | 0.00714271 | 0.17279872 |
| AC138627.1 | 515.71923 | -0.3784426 | 0.60608323 | -0.6244071 | 0.53236029 | 0.86397197 |
| PSMD7 | 342.479359 | 0.22244926 | 0.2282247 | 0.97469404 | 0.32971205 | 0.75860764 |
| NPIPB15 | 165.870934 | 0.13629139 | 1.00477078 | 0.13564426 | 0.89210251 | 0.97771293 |
| AC009053.3 | 208.215814 | 0.12634517 | 0.66478203 | 0.19005503 | 0.84926601 | 0.970224 |
| GLG1 | 3040.66577 | -0.1008948 | 0.15911673 | -0.6340928 | 0.52602023 | 0.86119802 |
| RFWD3 | 451.504853 | -0.1394337 | 0.20358438 | -0.684894 | 0.49341079 | 0.84818625 |
| MLKL | 74.667336 | 0.30200261 | 0.66071022 | 0.45708785 | 0.6476079 | 0.90674717 |
| FA2H | 570.906714 | 1.21591693 | 0.75074447 | 1.61961489 | 0.10531503 | 0.52613699 |
| WDR59 | 1267.2463 | -0.1565087 | 0.19756217 | -0.7921996 | 0.42824428 | 0.81497133 |
| ZNRF1 | 660.426869 | -0.4938273 | 0.32059981 | -1.5403232 | 0.1234816 | 0.56089013 |
| LDHD | 43.421992 | 1.01205376 | 0.46977929 | 2.15431753 | 0.03121528 | 0.3383722 |
| ZFP1 | 95.5275349 | -0.3761521 | 0.29919472 | -1.2572152 | 0.20867573 | 0.6614379 |
| BCAR1 | 737.996839 | -0.1243283 | 0.31304261 | -0.397161 | 6.91E-01 | 0.92206635 |
| CFDP1 | 2357.86919 | -0.6176103 | 0.28202465 | -2.1899159 | 0.02853034 | 0.32600107 |
| AC009163.4 | 45.6939588 | 0.15642672 | 0.34639983 | 0.45157853 | 0.65157264 | 0.90825486 |
| TMEM231 | 78.109525 | -0.3997345 | 0.48142529 | -0.8303147 | 0.40636087 | 0.80499101 |
| GABARAPL2 | 417.809424 | 0.72330265 | 0.35672939 | 2.0275948 | 0.04260162 | 0.38021566 |
| ADAT1 | 199.083281 | -0.3956916 | 0.25640397 | -1.5432352 | 0.12277373 | 0.55947129 |
| KARS | 266.441992 | 0.08517388 | 0.22103858 | 0.38533488 | 0.69998932 | 0.92367966 |
| TERF2IP | 1069.99121 | 0.25084173 | 0.31664799 | 0.79217849 | 0.42825661 | 0.81497133 |
| MON1B | 84.9328481 | -0.5052383 | 0.39253858 | -1.2871047 | 0.19805779 | 0.65049274 |
| NUDT7 | 164.043784 | -0.2356481 | 0.41886772 | -0.5625836 | 5.74E-01 | 0.88002205 |
| VAT1L | 808.758706 | -3.0243846 | 0.88645966 | -3.4117566 | 0.00064546 | 0.04534035 |
| WWOX | 18088.1508 | -0.4352392 | 0.24824767 | -1.753246 | 0.07955979 | 0.48176967 |
| AC046158.1 | 128.30765 | 0.29348964 | 0.65380652 | 0.44889371 | 0.65350833 | 0.90884465 |
| MAF | 95.5282406 | 0.86945298 | 0.59685278 | 1.45672939 | 0.14519109 | 0.59119271 |
| AC105411.1 | 106.47572 | -0.7042854 | 0.52430038 | -1.3432861 | 0.17917934 | 0.63288164 |
| DYNLRB2 | 95.2729087 | -0.719933 | 0.56218581 | -1.2805961 | 0.20033557 | 0.65322946 |
| CDYL2 | 1013.55467 | -0.3424091 | 0.59720642 | -0.5733513 | 0.56640685 | 0.87642772 |
| CMC2 | 706.454617 | 0.12617199 | 0.32089794 | 0.39318418 | 0.69418346 | 0.92229578 |
| ATMIN | 178.42705 | 0.36263655 | 0.26525133 | 1.3671432 | 0.17158042 | 0.62833541 |

| | | | | | | |
|---|---|---|---|---|---|---|
| GAN | 473.265818 | 0.01848105 | 0.18916794 | 0.09769652 | 0.92217328 | 0.98215577 |
| CMIP | 1873.56519 | -0.5007731 | 0.22622369 | -2.2136192 | 0.02685498 | 0.31710856 |
| PLCG2 | 1315.08117 | -0.0405709 | 0.5435613 | -0.074639 | 0.94050191 | 0.98715467 |
| SDR42E1 | 59.1953562 | 0.87246464 | 0.40456239 | 2.15656391 | 0.03103966 | 0.33800731 |
| AC092142.1 | 135.141122 | -0.0469789 | 0.39303761 | -0.1195278 | 0.90485719 | 0.97992575 |
| MPHOSPH6 | 228.386273 | -0.1476993 | 0.30997608 | -0.4764861 | 0.63372814 | 0.90136076 |
| CDH13 | 137.774979 | -0.8069863 | 0.59458194 | -1.357233 | 1.75E-01 | 0.62932267 |
| HSBP1 | 470.367831 | 0.30352988 | 0.38193252 | 0.79472123 | 0.42677568 | 0.81466629 |
| MLYCD | 100.303948 | -0.1467085 | 0.40492484 | -0.3623104 | 0.71712005 | 0.9290703 |
| AC009119.2 | 108.693506 | -0.6181424 | 0.58420734 | -1.0580873 | 0.29001566 | 0.73204984 |
| MBTPS1 | 885.924605 | -0.0849679 | 0.14626816 | -0.5809049 | 0.56130453 | 0.87401021 |
| HSDL1 | 179.479309 | -0.4212698 | 0.32409229 | -1.2998451 | 0.19365406 | 0.64718686 |
| DNAAF1 | 57.1330172 | 0.74905982 | 0.73998945 | 1.01225742 | 0.31141499 | 0.74590637 |
| TAF1C | 285.218232 | -0.2904452 | 0.25766884 | -1.1272036 | 0.25965642 | 0.70921184 |
| ATP2C2 | 51.5044093 | -0.3951874 | 0.5379556 | -0.7346097 | 0.46257725 | 0.83531269 |
| ATP2C2-AS1 | 39.3329607 | -0.3894764 | 0.53422191 | -0.7290535 | 0.46596893 | 0.8367113 |
| TLDC1 | 408.701236 | -0.5315254 | 0.39861498 | -1.3334305 | 0.18239055 | 0.63741052 |
| COTL1 | 172.892941 | -0.8136151 | 0.52032642 | -1.5636629 | 0.11789676 | 0.54888439 |
| KLHL36 | 445.651238 | 0.19096228 | 0.21726454 | 0.87893903 | 0.37943433 | 0.78762967 |
| USP10 | 996.378378 | -0.2904536 | 0.14706239 | -1.9750362 | 0.04826403 | 0.39627811 |
| ZDHHC7 | 976.242313 | -0.005819 | 0.29482568 | -0.0197372 | 0.98425304 | 0.99518552 |
| KIAA0513 | 322.36792 | 0.31317249 | 0.40060861 | 0.78174178 | 0.43436634 | 0.81919857 |
| GSE1 | 1492.61069 | 0.05214183 | 0.39780696 | 0.13107318 | 0.89571741 | 0.97788747 |
| EMC8 | 171.927794 | -0.2244603 | 0.28005272 | -0.8014929 | 0.42284638 | 0.81372055 |
| COX4I1 | 1245.56282 | 0.8668555 | 0.43984616 | 1.97081522 | 0.04874502 | 0.39762623 |
| IRF8 | 52.5816468 | 0.80392285 | 0.4684883 | 1.71599344 | 0.08616324 | 0.49453468 |
| MTHFSD | 175.66265 | -0.310265 | 0.2247899 | -1.3802444 | 0.1675114 | 0.62170779 |
| C16orf95 | 62.0795881 | -0.4046843 | 0.31994704 | -1.2648479 | 0.20592585 | 0.65813563 |
| FBXO31 | 213.884007 | -0.3315211 | 0.24495555 | -1.353393 | 0.17593011 | 0.62957039 |
| MAP1LC3B | 464.868584 | 0.24264722 | 0.41279908 | 0.58780949 | 0.55666017 | 0.87212955 |
| ZCCHC14 | 1391.98732 | -0.8065286 | 0.26201737 | -3.0781494 | 0.0020829 | 0.0878448 |
| KLHDC4 | 517.482089 | -0.3215583 | 0.32594461 | -0.9865428 | 0.32386682 | 0.75402436 |
| SLC7A5 | 425.478491 | 0.38499948 | 0.76304395 | 0.50455742 | 0.61386973 | 0.89477643 |
| BANP | 729.719083 | -0.4653339 | 0.29688399 | -1.5673931 | 0.11702284 | 0.54756317 |
| AC138512.1 | 72.5586644 | 0.9143022 | 0.75760154 | 1.20683783 | 0.22749461 | 0.67921637 |
| ZFPM1 | 290.753227 | 0.18713096 | 0.24860873 | 0.75271274 | 0.45162255 | 0.8285455 |
| AC116552.1 | 58.6260796 | -0.1031379 | 0.40440761 | -0.2550346 | 0.79869636 | 0.95300733 |
| ZC3H18 | 398.012859 | -0.1765489 | 0.2438158 | -0.7241079 | 0.46899951 | 0.83870782 |
| CYBA | 649.778902 | 1.43581536 | 0.59669445 | 2.40628239 | 0.0161158 | 0.25504618 |
| MVD | 212.262051 | 0.27272841 | 0.39081353 | 0.69784793 | 0.4852723 | 0.8447301 |
| SNAI3-AS1 | 78.7799626 | -0.0710161 | 0.35324202 | -0.2010408 | 0.84066664 | 0.96617873 |
| RNF166 | 63.0564679 | -0.5870013 | 0.32975994 | -1.7800868 | 0.07506176 | 0.47190267 |
| PIEZO1 | 141.300027 | -0.8926179 | 0.39806834 | -2.2423734 | 0.02493725 | 0.3093519 |
| APRT | 257.313085 | 0.69737097 | 0.45734764 | 1.52481591 | 0.12730502 | 0.56740575 |
| GALNS | 271.616789 | -0.1607425 | 0.31939768 | -0.5032676 | 0.61477612 | 0.8949759 |
| TRAPPC2L | 87.5146498 | 0.45616995 | 0.3889988 | 1.17267701 | 0.24092535 | 0.69178991 |
| ACSF3 | 324.369128 | 0.00131386 | 0.28888199 | 0.00454808 | 0.99637117 | 0.99916997 |
| AC009113.1 | 43.6960125 | 0.01607493 | 0.4612342 | 0.03485199 | 0.97219777 | 0.99327086 |
| ZNF778 | 114.685676 | -0.3100573 | 0.24601647 | -1.260311 | 0.2075572 | 0.6607067 |
| ANKRD11 | 3229.32112 | -0.0846995 | 0.20467055 | -0.4138333 | 0.67899621 | 0.91822574 |
| SPG7 | 1448.37689 | -0.6076788 | 0.287145 | -2.1162787 | 0.03432111 | 0.35209451 |
| AC092123.1 | 45.2039542 | -0.1822524 | 0.46005256 | -0.3961556 | 0.69199021 | 0.92224682 |
| RPL13 | 2155.47629 | 0.57775378 | 0.42635537 | 1.3550991 | 0.17538599 | 0.62953084 |
| CHMP1A | 250.106955 | 0.11159629 | 0.29016207 | 0.38459987 | 0.70053389 | 0.92376482 |

| | | | | | | |
|---|---|---|---|---|---|---|
| SPATA33 | 110.447673 | 0.34513671 | 0.28968692 | 1.1914128 | 0.23349157 | 0.68492492 |
| CDK10 | 391.771004 | -0.2109139 | 0.22786045 | -0.9256274 | 0.35463965 | 0.77251948 |
| SPATA2L | 73.4905952 | 0.22800609 | 0.29261849 | 0.77919235 | 0.43586641 | 0.8196523 |
| VPS9D1 | 418.404823 | -0.4763983 | 0.26974829 | -1.7660845 | 0.07738167 | 0.47763291 |
| ZNF276 | 241.785783 | -0.5892913 | 0.24025653 | -2.4527587 | 0.01417654 | 0.24646911 |
| FANCA | 62.261132 | -1.2865569 | 0.40299268 | -3.1925069 | 0.00141044 | 0.06874704 |
| SPIRE2 | 104.911041 | 0.17358654 | 0.360349 | 0.48171784 | 0.6300064 | 0.90005881 |
| TCF25 | 2141.69331 | -0.53191 | 0.3403152 | -1.5629923 | 0.1180544 | 0.54888439 |
| MC1R | 88.8200278 | -0.6740891 | 0.47105927 | -1.4310069 | 0.15242823 | 0.60264129 |
| DEF8 | 178.541557 | -0.1425561 | 0.25349501 | -0.5623627 | 0.5738689 | 0.88002205 |
| DBNDD1 | 56.0469467 | -0.0041651 | 0.5244048 | -0.0079424 | 0.99366292 | 0.9983477 |
| GAS8 | 458.029747 | -0.066457 | 0.32859528 | -0.2022457 | 0.83972468 | 0.96617873 |
| FAM157C | 461.35776 | -0.1060824 | 0.37611112 | -0.2820507 | 0.7779046 | 0.94694572 |
| AC133919.2 | 69.9933056 | -0.1326452 | 0.7480706 | -0.1773165 | 0.85925981 | 0.97174503 |
| LINC02193 | 123.604843 | -0.1512084 | 0.3360009 | -0.4500239 | 0.65269319 | 0.90868303 |
| AC240565.1 | 47.9446334 | -0.3857039 | 0.49122624 | -0.7851858 | 0.43234464 | 0.81773248 |
| RPH3AL | 701.508936 | 0.00152901 | 0.42035605 | 0.00363741 | 0.99709777 | 0.99948643 |
| AC129507.2 | 151.62977 | 0.63887713 | 0.58492784 | 1.09223237 | 0.27473098 | 0.72164296 |
| C17orf97 | 79.2631884 | 0.35811469 | 0.50123055 | 0.71447099 | 0.47493599 | 0.84106098 |
| VPS53 | 1728.16166 | -0.3706948 | 0.18575887 | -1.9955697 | 0.04598079 | 0.39025761 |
| AC015853.3 | 81.5604078 | -0.3506543 | 0.48923691 | -0.7167372 | 0.47353628 | 0.84060485 |
| FAM57A | 96.0430339 | -0.1179835 | 0.50090715 | -0.2355396 | 8.14E-01 | 9.58E-01 |
| GLOD4 | 223.711395 | 0.22386296 | 0.21594506 | 1.03666628 | 0.29989141 | 0.73673761 |
| MRM3 | 57.5976065 | 0.42047008 | 0.30075746 | 1.39803708 | 0.16210193 | 0.61352562 |
| NXN | 318.89521 | 0.1384776 | 0.49301476 | 0.28087922 | 0.77880304 | 0.94725685 |
| TIMM22 | 84.601068 | 0.03613899 | 0.32937596 | 0.10971959 | 0.91263177 | 0.98135874 |
| ABR | 399.84804 | -0.3411887 | 0.41990324 | -0.8125413 | 0.41648111 | 0.80966301 |
| YWHAE | 1858.3813 | -0.001598 | 0.11071817 | -0.0144328 | 0.98848472 | 0.99660871 |
| CRK | 820.488738 | -0.0519014 | 0.18357826 | -0.2827206 | 0.777391 | 0.94694572 |
| MYO1C | 617.137492 | 0.37101111 | 0.25790512 | 1.43855661 | 0.15027619 | 0.59663305 |
| INPP5K | 158.361219 | -0.368135 | 0.29877286 | -1.2321569 | 0.21789048 | 0.67120226 |
| PITPNA-AS1 | 44.3426944 | 0.27997968 | 0.42811796 | 0.6539779 | 0.51312604 | 0.85762435 |
| PITPNA | 1038.07534 | -0.0066792 | 0.19010784 | -0.0351336 | 0.97197324 | 0.99327086 |
| SLC43A2 | 1347.10124 | 0.57596799 | 0.44595546 | 1.29153701 | 0.19651753 | 0.64964978 |
| SCARF1 | 122.712622 | 0.42704438 | 0.42076611 | 1.01492104 | 0.31014347 | 0.74459794 |
| RILP | 63.2808825 | 0.2027532 | 0.45456597 | 0.44603692 | 0.65557058 | 0.90919768 |
| PRPF8 | 830.811991 | -0.0610685 | 0.20638502 | -0.2958962 | 0.76730937 | 0.94316227 |
| MIR22HG | 222.295155 | -0.1590197 | 0.59058598 | -0.2692575 | 0.78773155 | 0.95006812 |
| WDR81 | 131.068343 | -0.0752519 | 0.5322492 | -0.1413848 | 0.88756597 | 0.97729111 |
| SERPINF2 | 74.5383325 | 2.23930658 | 0.53150966 | 4.21310609 | 2.52E-05 | 0.00824389 |
| SMYD4 | 209.333991 | -0.3522632 | 0.31473763 | -1.1192281 | 0.26304283 | 0.71330996 |
| RPA1 | 332.896426 | 0.10078448 | 0.25047449 | 0.40237424 | 0.68740863 | 0.92094022 |
| RTN4RL1 | 185.048013 | 2.28254969 | 0.67281597 | 3.39253195 | 0.0006925 | 0.04573607 |
| DPH1 | 104.654089 | -0.1812887 | 0.35917122 | -0.5047418 | 0.61374018 | 0.89470191 |
| SMG6 | 2606.23809 | -0.385755 | 0.17712008 | -2.1779295 | 0.02941128 | 0.33012415 |
| AC090617.4 | 70.3568663 | -0.3783269 | 0.44321706 | -0.8535928 | 0.39333065 | 0.79843808 |
| SRR | 144.694525 | -0.2404006 | 0.30216505 | -0.7955938 | 0.42626817 | 0.81466629 |
| TSR1 | 180.938219 | 0.06291344 | 0.28000541 | 0.22468653 | 0.82222315 | 0.96078782 |
| SGSM2 | 394.053382 | 0.13425759 | 0.20438263 | 0.65689332 | 0.51124951 | 0.85762435 |
| MNT | 185.629467 | -0.1291557 | 0.29905749 | -0.4318758 | 6.66E-01 | 0.91305741 |
| METTL16 | 839.220266 | -0.3762216 | 0.25387348 | -1.4819256 | 0.13836009 | 0.58206378 |
| PAFAH1B1 | 2738.96918 | -0.2405914 | 0.11166256 | -2.1546287 | 0.0311909 | 0.3383722 |
| CLUH | 583.58213 | 0.17824481 | 0.3121226 | 0.57107305 | 0.56795013 | 0.87712536 |
| RAP1GAP2 | 1723.58339 | -0.2892064 | 0.21512725 | -1.3443504 | 0.17883512 | 0.63284534 |

| | | | | | | |
|---|---|---|---|---|---|---|
| TRPV1 | 87.0932739 | -0.6331318 | 0.36046451 | -1.756433 | 0.0790145 | 0.48162624 |
| CTNS | 209.22497 | -0.3173002 | 0.41741325 | -0.7601585 | 0.44715984 | 0.82578615 |
| TAX1BP3 | 119.512403 | 0.2234226 | 0.42174396 | 0.52975887 | 5.96E-01 | 0.88859628 |
| EMC6 | 146.964152 | 0.68563854 | 0.5704504 | 1.20192491 | 0.22939262 | 0.6817947 |
| ITGAE | 171.347594 | 0.00906933 | 0.31087407 | 0.02917366 | 0.97672609 | 0.99418779 |
| NCBP3 | 719.628762 | -0.142873 | 0.19812222 | -0.7211355 | 0.47082615 | 0.83921324 |
| CAMKK1 | 67.7105593 | 0.19900332 | 0.48640449 | 0.40913133 | 0.68244329 | 0.91969963 |
| ZZEF1 | 1113.37539 | -0.3801031 | 0.23791131 | -1.5976673 | 0.11011704 | 0.53372881 |
| CYB5D2 | 404.258137 | -0.3645214 | 0.31430238 | -1.1597792 | 0.24613873 | 0.69664593 |
| ANKFY1 | 811.624906 | -0.1442807 | 0.21949026 | -0.6573444 | 0.5109595 | 0.85762435 |
| UBE2G1 | 1152.8202 | -0.1781198 | 0.20301099 | -0.8773901 | 0.38027477 | 0.7880311 |
| SPNS2 | 668.820744 | -0.5894889 | 0.70097565 | -0.8409548 | 0.40037325 | 0.80238662 |
| MYBBP1A | 113.299409 | -0.323782 | 0.31267682 | -1.0355164 | 0.3004278 | 0.7369268 |
| GGT6 | 129.648902 | 1.57564645 | 0.56615007 | 2.78308976 | 0.00538439 | 0.14848104 |
| SMTNL2 | 618.771416 | 0.18354854 | 0.3266678 | 0.56188133 | 5.74E-01 | 8.80E-01 |
| PELP1 | 197.699181 | -0.0832218 | 0.35142677 | -0.2368113 | 0.81280317 | 0.95809779 |
| AC091153.2 | 58.549444 | -0.0454243 | 0.42299481 | -0.1073873 | 0.91448171 | 0.98156931 |
| ARRB2 | 46.5520185 | 0.29639373 | 0.47258374 | 0.62717717 | 0.5305431 | 0.86396107 |
| MED11 | 93.6266461 | 0.41914344 | 0.31921098 | 1.31306085 | 0.18916244 | 0.64432929 |
| CXCL16 | 72.6953844 | 0.80552058 | 0.53338734 | 1.51019815 | 1.31E-01 | 0.57389214 |
| PSMB6 | 211.449275 | 0.93450105 | 0.47102167 | 1.98398736 | 0.04725725 | 0.39407891 |
| PLD2 | 67.9934863 | -0.1204289 | 0.45282283 | -0.2659514 | 0.79027662 | 0.95130361 |
| MINK1 | 679.453448 | -0.3311856 | 0.23452837 | -1.4121346 | 0.15791032 | 0.60779483 |
| SLC25A11 | 160.588414 | 0.93468203 | 0.3682893 | 2.53790168 | 0.01115193 | 0.21567475 |
| RNF167 | 167.8895 | 0.11508025 | 0.26156868 | 0.43996191 | 0.6599647 | 0.91099119 |
| PFN1 | 657.471165 | 0.34651853 | 0.37453918 | 0.92518633 | 3.55E-01 | 0.7726065 |
| SPAG7 | 253.999842 | -0.3736268 | 0.31037665 | -1.2037852 | 0.22867262 | 0.68060764 |
| CAMTA2 | 226.097456 | -0.1014542 | 0.21584312 | -0.470037 | 0.6383286 | 0.90303979 |
| KIF1C | 285.526687 | 0.14645894 | 0.3427887 | 0.4272572 | 0.66919201 | 0.91429015 |
| ZFP3 | 109.777036 | 0.14985472 | 0.2500513 | 0.59929591 | 0.54897558 | 0.86904066 |
| ZNF232 | 87.9910482 | -0.2897759 | 0.84341015 | -0.3435765 | 0.73116479 | 0.93515518 |
| USP6 | 61.4627421 | -0.1378845 | 0.7567505 | -0.1822061 | 0.855421 | 0.97118784 |
| ZNF594 | 261.451246 | 0.19305027 | 0.2062505 | 0.93599902 | 0.34927369 | 0.76982416 |
| AC087500.1 | 551.982357 | 0.19232153 | 0.30930665 | 0.62178271 | 0.53408476 | 0.86397197 |
| SCIMP | 107.177031 | 0.46461824 | 0.35119834 | 1.32295111 | 0.18585164 | 0.64030106 |
| RABEP1 | 1463.64104 | -0.2448763 | 0.23672353 | -1.0344399 | 0.30093057 | 0.73774906 |
| NUP88 | 1333.18964 | -0.3442884 | 0.21817762 | -1.5780189 | 0.11456126 | 0.54179726 |
| RPAIN | 593.967964 | 0.27717315 | 0.23757426 | 1.16668003 | 0.24333961 | 0.69452742 |
| AC004148.2 | 67.0112409 | 0.24969472 | 0.6244748 | 0.39984756 | 0.6892688 | 0.92166785 |
| C1QBP | 183.406293 | 0.72446921 | 0.38045879 | 1.9041989 | 0.05688429 | 0.41872623 |
| DHX33 | 97.1625058 | -0.2550769 | 0.39766117 | -0.6414429 | 0.52123497 | 0.85943874 |
| DERL2 | 251.797324 | 0.2960078 | 0.20985669 | 1.41052351 | 0.15838516 | 0.60804051 |
| MIS12 | 173.730712 | 0.43445528 | 0.22666061 | 1.91676569 | 0.05526771 | 0.41827898 |
| AC007846.2 | 43.7536743 | 0.1251152 | 0.700897 | 0.17850726 | 0.85832462 | 0.97143431 |
| WSCD1 | 218.69755 | 0.33605213 | 0.46479938 | 0.72300469 | 0.469677 | 0.83905376 |
| PITPNM3 | 95.8368523 | 0.10990429 | 0.59832981 | 0.18368513 | 0.85426047 | 0.97085267 |
| KIAA0753 | 613.060339 | -0.0357918 | 0.20593025 | -0.1738056 | 0.86201825 | 0.97264823 |
| TXNDC17 | 125.466462 | 0.50514643 | 0.35631196 | 1.41770831 | 0.15627594 | 0.60564438 |
| MED31 | 154.571092 | 0.55013769 | 0.24311394 | 2.26288005 | 0.02364308 | 0.30185158 |
| XAF1 | 461.208151 | 1.58268022 | 1.59276465 | 0.9936686 | 0.32038423 | 0.75141253 |
| ALOX12-AS1 | 298.427425 | 0.89635174 | 0.36767571 | 2.43788672 | 0.0147734 | 0.24869534 |
| SLC16A13 | 39.6762157 | -0.3154689 | 0.38023876 | -0.8296599 | 0.40673109 | 0.80511179 |
| DLG4 | 56.1122044 | 0.35875261 | 0.81715738 | 0.43902512 | 0.66064334 | 0.91099119 |
| ACADVL | 1586.75515 | 0.30408256 | 0.52710274 | 0.57689429 | 0.56401087 | 0.87508139 |

| | | | | | | |
|---|---|---|---|---|---|---|
| DVL2 | 137.992834 | -0.2105422 | 0.20310011 | -1.0366424 | 0.29990254 | 0.73673761 |
| PHF23 | 87.1063896 | 0.47627075 | 0.35250668 | 1.35109708 | 0.17666434 | 0.63008941 |
| AC003688.1 | 45.2820363 | -0.0027212 | 0.41942692 | -0.0064879 | 0.99482347 | 0.998438 |
| ELP5 | 154.630235 | 0.27580339 | 0.31271801 | 0.88195556 | 0.37780084 | 0.7865219 |
| CLDN7 | 41.4181151 | 0.17193599 | 0.47282579 | 0.36363497 | 7.16E-01 | 0.92890921 |
| EIF5A | 204.754054 | 0.01765947 | 0.45958508 | 0.03842482 | 0.96934897 | 0.99299658 |
| GPS2 | 171.368925 | 0.26400657 | 0.36993083 | 0.71366468 | 0.47543455 | 0.84118672 |
| TNK1 | 41.6500372 | -0.2259155 | 0.31360863 | -0.7203739 | 0.4712948 | 0.83921324 |
| NLGN2 | 77.1060022 | -0.1261074 | 0.48915324 | -0.2578076 | 0.79655541 | 0.95278473 |
| FGF11 | 90.197922 | 1.34332454 | 0.72166082 | 1.86143476 | 0.06268281 | 0.43509048 |
| CHRNB1 | 59.7519887 | 0.34190029 | 0.47459582 | 0.72040307 | 0.47127686 | 0.83921324 |
| ZBTB4 | 573.193865 | 0.42464165 | 0.36213147 | 1.17261737 | 0.24094928 | 0.69178991 |
| POLR2A | 953.969517 | -0.0903602 | 0.25716187 | -0.3513747 | 0.72530727 | 0.9326472 |
| EIF4A1 | 275.062291 | 0.18825453 | 0.4701442 | 0.40041869 | 0.68884816 | 0.92151057 |
| CD68 | 67.8259405 | -0.1659088 | 0.6076265 | -0.273044 | 7.85E-01 | 9.50E-01 |
| MPDU1 | 207.968269 | -0.1057536 | 0.26506967 | -0.3989653 | 0.68991877 | 0.92176114 |
| FXR2 | 192.307354 | -0.0053794 | 0.29998714 | -0.0179321 | 0.98569301 | 0.99594729 |
| SAT2 | 446.312009 | 0.7273787 | 0.34652429 | 2.09906991 | 0.03581074 | 0.35661333 |
| TP53 | 265.921482 | 0.177146 | 0.3838444 | 0.46150472 | 0.64443654 | 0.90507553 |
| WRAP53 | 52.4658475 | -0.4849235 | 0.45823911 | -1.0582324 | 0.28994952 | 0.73204984 |
| KDM6B | 230.333341 | -0.3604297 | 0.29904903 | -1.2052528 | 0.22810572 | 0.67999141 |
| NAA38 | 652.009117 | 0.16103814 | 0.24412051 | 0.65966655 | 0.50946784 | 0.8568242 |
| CYB5D1 | 89.5813991 | 0.63025788 | 0.29747728 | 2.11867573 | 0.03411788 | 0.35153234 |
| CHD3 | 295.18443 | 0.14436482 | 0.28949159 | 0.498684 | 0.61800202 | 0.89599505 |
| KCNAB3 | 42.3122475 | -0.7022838 | 0.43832769 | -1.6021891 | 0.10911381 | 0.53262539 |
| TRAPPC1 | 84.144675 | 1.21569064 | 0.55341603 | 2.19670297 | 0.02804167 | 0.3228969 |
| CNTROB | 147.649412 | -0.5429401 | 0.36130618 | -1.5027148 | 0.1329126 | 0.57682553 |
| PER1 | 323.699696 | -0.096835 | 0.46947532 | -0.2062621 | 0.83658618 | 0.96511435 |
| TMEM107 | 108.017572 | 0.19789391 | 0.28356811 | 0.69787082 | 0.48525798 | 0.8447301 |
| CTC1 | 243.194977 | -0.1761723 | 0.18881388 | -0.9330475 | 0.35079545 | 0.77073471 |
| PFAS | 73.9108409 | -0.5110678 | 0.38338716 | -1.3330333 | 0.18252087 | 0.63741052 |
| SLC25A35 | 75.3837043 | -0.5067087 | 0.42769899 | -1.1847319 | 0.23612345 | 0.68856409 |
| RANGRF | 80.9826122 | 0.48322134 | 0.43099994 | 1.12116336 | 0.26221833 | 0.7128302 |
| AC135178.3 | 64.1880937 | 0.29430551 | 0.45518386 | 0.64656402 | 0.51791415 | 0.85893097 |
| NDEL1 | 738.495287 | -0.5186478 | 0.28146771 | -1.8426547 | 0.06537944 | 0.44218998 |
| MYH10 | 1314.13298 | 0.37235148 | 0.30329526 | 1.22768642 | 0.2195647 | 0.67278782 |
| STX8 | 2063.57062 | -0.2896336 | 0.2235853 | -1.2954054 | 0.19518042 | 0.64871108 |
| AC005695.2 | 121.858008 | -0.3989226 | 0.46244607 | -0.862636 | 0.38833763 | 0.79634939 |
| AC118755.2 | 53.322034 | 0.02130987 | 0.46403127 | 0.04592336 | 0.96337134 | 0.99224805 |
| USP43 | 436.508027 | -0.7897076 | 0.30825519 | -2.5618631 | 1.04E-02 | 0.20711146 |
| GAS7 | 82.434237 | 0.08803556 | 0.5986796 | 0.14704955 | 0.88309291 | 0.97664137 |
| MYH3 | 55.0168661 | -1.118873 | 0.39350806 | -2.8433295 | 0.00446449 | 0.12989304 |
| SCO1 | 246.638739 | -0.1232 | 0.23504286 | -0.5241598 | 0.60016738 | 0.88883692 |
| ADPRM | 72.7224584 | -0.0675102 | 0.31336279 | -0.2154378 | 0.82942601 | 0.96290106 |
| TMEM220 | 301.20143 | 0.22461473 | 0.31519211 | 0.71262802 | 0.47607597 | 0.84141268 |
| TMEM220-A! | 203.985312 | -0.2557666 | 0.55246728 | -0.4629533 | 0.64339781 | 0.90478945 |
| SHISA6 | 573.753228 | -0.9160275 | 0.63614866 | -1.4399582 | 0.14987922 | 0.59663305 |
| DNAH9 | 97.0731235 | 0.41454483 | 0.28659254 | 1.44646064 | 0.14804804 | 0.59438683 |
| ZNF18 | 88.2994039 | -0.3168073 | 0.30709228 | -1.0316354 | 0.30224294 | 0.73905643 |
| MAP2K4 | 2282.24453 | -0.1815832 | 0.14344484 | -1.265875 | 0.20555785 | 0.65813563 |
| ARHGAP44 | 880.910134 | 0.21569816 | 0.15805399 | 1.36471192 | 0.17234361 | 0.6286116 |
| ELAC2 | 206.573862 | -0.0570772 | 0.25510452 | -0.2237403 | 0.82295941 | 0.96113801 |
| COX10 | 777.466715 | -0.0007842 | 0.21712328 | -0.0036119 | 9.97E-01 | 0.99948643 |
| AC005224.1 | 36.2023027 | -0.2756055 | 0.55136032 | -0.4998647 | 0.61717037 | 0.89599505 |

| | | | | | | |
|---|---|---|---|---|---|---|
| PMP22 | 43.8388021 | 0.53542331 | 0.64367136 | 0.83182714 | 0.40550652 | 0.80485885 |
| TRIM16 | 206.538055 | -0.6386797 | 0.24048067 | -2.6558465 | 0.00791096 | 0.17862547 |
| ZSWIM7 | 505.634739 | -0.0732006 | 0.19023932 | -0.3847814 | 0.70039938 | 0.92367966 |
| TTC19 | 1315.34124 | 0.09390631 | 0.26973573 | 0.34814192 | 0.7277336 | 0.93420128 |
| NCOR1 | 3669.62156 | -0.0880701 | 0.15088717 | -0.583682 | 0.55943426 | 0.87295536 |
| PIGL | 1078.93353 | -0.3075807 | 0.28252783 | -1.0886739 | 0.27629773 | 0.72241028 |
| CENPV | 88.8673563 | -0.0918964 | 0.51206451 | -0.1794625 | 0.85757453 | 0.97127317 |
| UBB | 1041.39444 | 1.04809324 | 0.45652402 | 2.29581179 | 0.02168664 | 0.29410564 |
| LRRC75A | 126.162517 | 0.60457836 | 0.34106927 | 1.77259701 | 0.07629551 | 0.47530241 |
| ZNF287 | 92.7844984 | -0.2467529 | 0.26874894 | -0.9181539 | 0.35853828 | 0.77549242 |
| ZNF624 | 77.5498632 | -0.3047483 | 0.3610182 | -0.8441356 | 0.39859362 | 0.80148305 |
| AC098850.3 | 479.408826 | 0.31937821 | 0.68412673 | 0.46684071 | 0.64061385 | 0.9037561 |
| MPRIP | 1803.0888 | -0.4869935 | 0.18297214 | -2.6615717 | 0.00777768 | 0.17721667 |
| MPRIP-AS1 | 45.7413049 | -0.367404 | 0.55586647 | -0.6609572 | 0.50863975 | 0.85659934 |
| AC055811.1 | 47.5057102 | -0.3592184 | 0.59442694 | -0.6043105 | 0.54563724 | 0.86762035 |
| COPS3 | 218.006694 | -0.0585078 | 0.21818977 | -0.2681511 | 0.788583 | 0.95048754 |
| NT5M | 101.802433 | -0.6335762 | 0.37135154 | -1.7061359 | 0.08798279 | 0.4972891 |
| MED9 | 481.349857 | 0.22433617 | 0.31083179 | 0.72172852 | 0.4704614 | 0.83921324 |
| RASD1 | 2067.38401 | 0.82724536 | 0.64004711 | 1.29247575 | 1.96E-01 | 6.50E-01 |
| PEMT | 176.874643 | -0.3370199 | 0.4788286 | -0.7038424 | 0.48153095 | 0.8437862 |
| RAI1 | 932.33271 | -0.1401102 | 0.39222637 | -0.3572178 | 0.72092878 | 0.93041019 |
| SMCR5 | 107.613709 | -0.1064316 | 0.30121948 | -0.3533356 | 0.72383685 | 0.93201977 |
| SREBF1 | 541.845291 | -0.0822803 | 0.54089396 | -0.152119 | 0.87909304 | 0.9765717 |
| TOM1L2 | 2486.83922 | 0.01945437 | 0.21156774 | 0.09195337 | 9.27E-01 | 9.82E-01 |
| DRC3 | 175.513031 | 0.91501221 | 0.40956339 | 2.2341162 | 0.02547543 | 0.31163914 |
| ATPAF2 | 192.02053 | -0.0402537 | 0.2143045 | -0.1878341 | 0.85100667 | 0.97056586 |
| GID4 | 146.076766 | -0.1533678 | 0.26890308 | -0.5703461 | 0.56844296 | 0.87757867 |
| DRG2 | 136.58573 | -0.0818336 | 0.2940411 | -0.2783066 | 0.780777 | 0.9480401 |
| ALKBH5 | 436.465667 | -0.1493337 | 0.23262565 | -0.6419486 | 0.52090654 | 0.85943874 |
| FLII | 321.254694 | 0.03375734 | 0.26780468 | 0.12605208 | 0.8996907 | 0.97897567 |
| MIEF2 | 109.851107 | 0.28981029 | 0.26877163 | 1.07827708 | 0.28091012 | 0.72575254 |
| TOP3A | 309.992641 | -0.3458785 | 0.20451758 | -1.6911921 | 0.09080012 | 0.50267341 |
| SMCR8 | 116.088631 | 0.33041718 | 0.22874903 | 1.44445282 | 0.14861164 | 0.59478974 |
| SHMT1 | 552.066163 | 0.04319361 | 0.308388 | 0.14006255 | 0.88861057 | 0.97729111 |
| ZNF286B | 87.9047313 | 0.29992999 | 0.33506817 | 0.89513126 | 0.37071692 | 0.78104287 |
| TRIM16L | 86.7250569 | -0.5449938 | 0.43589274 | -1.2502934 | 0.21119239 | 0.66483325 |
| TVP23B | 292.193634 | -0.0703349 | 0.22659365 | -0.3104011 | 0.75625594 | 0.93964676 |
| PRPSAP2 | 761.658038 | -0.2474304 | 0.13982451 | -1.7695782 | 0.07679744 | 0.47651607 |
| SLC5A10 | 94.1359406 | -0.115844 | 0.36311168 | -0.3190314 | 0.74970272 | 0.93775604 |
| FAM83G | 63.4095304 | -0.2861048 | 0.36071909 | -0.7931512 | 4.28E-01 | 8.15E-01 |
| EPN2 | 1583.11101 | -0.4213539 | 0.25602826 | -1.6457318 | 0.09981899 | 0.51665771 |
| EPN2-AS1 | 82.3651685 | -0.3816604 | 0.49696632 | -0.7679804 | 0.4424988 | 0.82299244 |
| B9D1 | 333.307331 | -0.0998048 | 0.25648899 | -0.3891194 | 0.69718784 | 0.92322348 |
| MAPK7 | 55.9761494 | -0.1621303 | 0.42241028 | -0.3838218 | 0.70111056 | 0.92424844 |
| AC004448.2 | 88.8492097 | 0.17892862 | 0.40782036 | 0.4387437 | 0.66084726 | 0.91108771 |
| SLC47A1 | 332.100744 | 0.40775684 | 0.41882871 | 0.9735647 | 3.30E-01 | 0.75894644 |
| ALDH3A2 | 439.50035 | -0.1067379 | 0.35783697 | -0.2982863 | 0.76548467 | 0.94270299 |
| SLC47A2 | 64.2968092 | 0.99891733 | 0.57585464 | 1.73466923 | 0.08279941 | 0.48601338 |
| ULK2 | 583.150921 | 0.05875217 | 0.27740757 | 0.21179008 | 0.83227081 | 0.96417211 |
| AKAP10 | 1495.86678 | -0.2466696 | 0.16677077 | -1.4790938 | 0.13911525 | 0.58243213 |
| SPECC1 | 155.602066 | -0.2351464 | 0.30727194 | -0.7652714 | 4.44E-01 | 0.82437446 |
| CCDC144NL-/ | 186.423287 | 0.51994317 | 0.44147022 | 1.17775365 | 0.23889483 | 0.69019287 |
| USP22 | 1061.21733 | -0.0930051 | 0.13087562 | -0.7106373 | 0.47730904 | 0.84174109 |
| DHRS7B | 932.037754 | -0.0940234 | 0.21484683 | -0.43763 | 0.66165455 | 0.91110616 |

| | | | | | | |
|---|---|---|---|---|---|---|
| TMEM11 | 111.416341 | 0.28261079 | 0.28587365 | 0.98858634 | 0.32286557 | 0.7532337 |
| AC087294.1 | 165.948036 | -0.127438 | 0.28906772 | -0.4408588 | 0.65931523 | 0.91099119 |
| NATD1 | 61.4644046 | 0.34428543 | 0.4485907 | 0.76748232 | 0.4427948 | 0.82301411 |
| MAP2K3 | 320.515048 | -0.0976119 | 0.36072485 | -0.2705994 | 0.78669915 | 0.94978829 |
| KCNJ12 | 66.8186596 | 0.11480228 | 1.01089775 | 0.11356468 | 0.90958289 | 0.98069436 |
| C17orf51 | 54.456754 | 0.87394788 | 0.45407811 | 1.92466419 | 0.05427138 | 0.41533078 |
| WSB1 | 5360.54835 | 0.4106786 | 0.55026856 | 0.74632393 | 0.45547176 | 0.83121393 |
| KSR1 | 908.698797 | 0.50381123 | 0.29392858 | 1.71406006 | 0.0865177 | 0.49478465 |
| AC005697.1 | 104.916437 | -0.3405354 | 0.8054188 | -0.4228054 | 0.67243725 | 0.915882 |
| NLK | 2788.63633 | -0.364385 | 0.25886397 | -1.4076312 | 0.15924031 | 0.60886928 |
| IFT20 | 170.912517 | 0.12168946 | 0.23799603 | 0.51130878 | 0.60913486 | 0.8927935 |
| TNFAIP1 | 275.692204 | 0.1605188 | 0.20167982 | 0.79590908 | 0.42608488 | 0.81466629 |
| POLDIP2 | 229.163588 | 0.36895221 | 0.2669687 | 1.38200551 | 0.16697001 | 0.62022127 |
| TMEM199 | 93.9495736 | -0.1390371 | 0.27567657 | -0.5043485 | 0.61401652 | 0.89477643 |
| SARM1 | 239.281374 | 0.18879866 | 0.40998062 | 0.4605063 | 0.64515285 | 0.90518019 |
| AC005726.1 | 187.867448 | -0.4119439 | 0.27844427 | -1.4794484 | 0.13902051 | 0.58243213 |
| KIAA0100 | 499.324676 | -0.2030101 | 0.23226838 | -0.8740323 | 0.38210068 | 0.78948349 |
| SDF2 | 221.341947 | 0.21724354 | 0.24063799 | 0.90278155 | 0.36664185 | 0.78095617 |
| SUPT6H | 723.340089 | -0.3442865 | 0.21806887 | -1.5787972 | 0.11438258 | 0.54153385 |
| PROCA1 | 60.4756169 | -0.2879285 | 0.41896904 | -0.6872309 | 0.49193721 | 0.84719669 |
| RAB34 | 258.372329 | 0.71103562 | 0.44376767 | 1.60226999 | 0.10909592 | 0.53262539 |
| RPL23A | 821.404391 | 0.65634663 | 0.33860521 | 1.93838315 | 0.05257649 | 0.40972443 |
| TLCD1 | 67.8601587 | 0.62552684 | 0.57044963 | 1.09655051 | 0.27283793 | 0.72070318 |
| NEK8 | 314.15623 | 0.2915527 | 0.65967087 | 0.44196691 | 0.65851315 | 0.91037018 |
| TRAF4 | 493.04879 | 0.08282835 | 0.4513771 | 0.18350145 | 0.85440458 | 0.97090761 |
| FAM222B | 1826.68123 | -0.2461455 | 0.22515045 | -1.0932489 | 0.27428453 | 0.72164296 |
| ERAL1 | 94.8474419 | 0.17198449 | 0.26969152 | 0.6377082 | 0.52366364 | 0.86006369 |
| FLOT2 | 802.862988 | 0.54012704 | 0.28483256 | 1.89629673 | 5.79E-02 | 0.42087381 |
| DHRS13 | 144.655565 | 0.63938344 | 0.48767173 | 1.3110939 | 0.18982604 | 0.64450954 |
| PHF12 | 606.8947 | -0.1261916 | 0.2343014 | -0.5385866 | 0.59017217 | 0.88631239 |
| PIPOX | 63.5670084 | -0.6630617 | 0.40876837 | -1.6220963 | 0.10478273 | 0.52486729 |
| MYO18A | 420.768915 | -0.5744437 | 0.25220312 | -2.2777026 | 0.0227443 | 0.2983102 |
| AC024619.3 | 75.7817036 | -0.0147251 | 0.54097892 | -0.0272194 | 0.97828474 | 0.99455895 |
| NUFIP2 | 1368.14273 | -0.0030929 | 0.18298737 | -0.016902 | 0.98651478 | 0.99632034 |
| TAOK1 | 2467.87069 | -0.0538797 | 0.16786054 | -0.320979 | 0.74822628 | 0.93746253 |
| ABHD15-AS1 | 248.767518 | -0.0659581 | 0.38740008 | -0.1702582 | 0.86480706 | 0.97361767 |
| ABHD15 | 43.0921386 | 0.11326317 | 0.33714996 | 0.33594301 | 0.73691384 | 0.93552616 |
| GIT1 | 160.285899 | -0.2542728 | 0.30683137 | -0.8287054 | 0.40727111 | 0.80561751 |
| SSH2 | 4010.93117 | -0.2980256 | 0.20187933 | -1.4762562 | 0.13987513 | 0.58431708 |
| AC023389.1 | 164.365842 | -0.156108 | 0.3350165 | -0.465971 | 0.64123628 | 0.90404588 |
| EFCAB5 | 146.584126 | -1.1251191 | 0.53019549 | -2.1220835 | 0.03383073 | 0.35107893 |
| NSRP1 | 1741.41118 | -0.2395152 | 0.15280596 | -1.5674465 | 0.11701036 | 0.54756317 |
| AC104984.3 | 85.2606573 | 0.05392134 | 0.5671438 | 0.09507525 | 0.92425506 | 0.98215577 |
| BLMH | 317.106197 | 0.06234468 | 0.25063362 | 0.24874826 | 0.80355552 | 0.95450237 |
| CPD | 3076.45342 | 0.21391242 | 0.50536252 | 0.4232851 | 6.72E-01 | 0.91576543 |
| GOSR1 | 939.26293 | -0.1352715 | 0.19701219 | -0.6866148 | 0.49232549 | 0.84734855 |
| AC127024.2 | 56.055201 | 0.0844532 | 0.56356334 | 0.14985574 | 0.88087843 | 0.97664137 |
| CRLF3 | 761.807347 | 0.08506004 | 0.27962566 | 0.30419253 | 0.76098122 | 0.94202764 |
| ATAD5 | 184.813549 | -0.288758 | 0.39528292 | -0.7305098 | 0.46507862 | 0.8363817 |
| AC130324.2 | 85.7873791 | -0.8585052 | 0.4577525 | -1.875479 | 0.06072685 | 0.43056413 |
| TEFM | 385.972778 | 0.2063413 | 0.26389592 | 0.7819041 | 0.43427094 | 0.81919857 |
| ADAP2 | 84.9051489 | 0.68131836 | 0.5222075 | 1.30468894 | 0.19199879 | 0.64596359 |
| RNF135 | 175.565885 | 0.27926775 | 0.24010342 | 1.16311444 | 0.24478307 | 0.6948422 |
| NF1 | 5387.73222 | -0.175739 | 0.16240171 | -1.0821255 | 0.27919676 | 0.72482665 |

| | | | | | | |
|---|---|---|---|---|---|---|
| OMG | 142.51393 | -0.2952383 | 0.64823514 | -0.4554495 | 0.64878591 | 0.90754509 |
| RAB11FIP4 | 292.91984 | -0.1398747 | 0.35755282 | -0.3912 | 0.6956494 | 0.92229578 |
| COPRS | 126.849123 | -0.0164995 | 0.50294162 | -0.032806 | 0.97382928 | 0.99345929 |
| UTP6 | 336.767478 | -0.463113 | 0.20782847 | -2.2283425 | 0.02585768 | 0.31222606 |
| AC004253.1 | 122.949654 | 0.17334496 | 0.45443905 | 0.3814482 | 0.7028707 | 0.92501157 |
| SUZ12 | 1136.29819 | -0.1009035 | 0.17648305 | -0.5717464 | 0.56749381 | 0.87678876 |
| LRRC37B | 391.120139 | -0.2468777 | 0.23649907 | -1.0438843 | 0.29653891 | 0.73672107 |
| RHOT1 | 1061.18422 | 0.19575916 | 0.14588476 | 1.34187537 | 0.17963641 | 0.63376531 |
| C17orf75 | 180.006014 | -0.2353831 | 0.26291309 | -0.8952887 | 0.37063277 | 0.78104287 |
| ZNF207 | 1919.69422 | 0.25490339 | 0.15111913 | 1.68677119 | 0.09164736 | 0.50476839 |
| PSMD11 | 542.479598 | -0.3107475 | 0.29594824 | -1.0500063 | 0.29371523 | 0.73527014 |
| MYO1D | 2301.49413 | 0.30317542 | 0.24961879 | 1.21455368 | 2.25E-01 | 6.77E-01 |
| AC079336.4 | 50.3315721 | 0.11647797 | 0.48154518 | 0.24188379 | 0.80887021 | 0.95610623 |
| ASIC2 | 818.810002 | -0.4608626 | 0.64516329 | -0.7143348 | 0.47502017 | 0.84106098 |
| CCL2 | 165.4027 | -1.3074534 | 1.02798096 | -1.2718654 | 0.20342095 | 0.65644753 |
| CCT6B | 140.722111 | -1.5545228 | 0.47308636 | -3.2859176 | 0.00101651 | 0.0571126 |
| ZNF830 | 80.6482299 | 0.23819057 | 0.36471992 | 0.65307804 | 0.51370597 | 0.85762435 |
| LIG3 | 162.748798 | -0.1543375 | 0.21845083 | -0.706509 | 0.47987166 | 0.84323863 |
| RFFL | 92.5542185 | 0.16997231 | 0.31573056 | 0.5383461 | 0.59033813 | 0.88631239 |
| SLFN5 | 242.595565 | 0.33227376 | 0.54489683 | 0.60979205 | 5.42E-01 | 8.66E-01 |
| SLFN12 | 54.3460318 | -0.5685128 | 0.34076872 | -1.6683245 | 0.09525134 | 0.51085035 |
| SLFN13 | 76.6340515 | -1.6542342 | 0.79853103 | -2.0715967 | 0.03830307 | 0.36662698 |
| LINC02001 | 164.387589 | -0.0166838 | 0.27994284 | -0.0595972 | 0.95247643 | 0.98938795 |
| AP2B1 | 1422.80804 | 0.19309389 | 0.1824372 | 1.05841291 | 0.28986723 | 0.73204984 |
| TAF15 | 2175.22522 | -0.1275686 | 0.20943206 | -0.6091168 | 0.54244701 | 0.8658519 |
| AC015849.3 | 154.909748 | -0.2446204 | 0.53711348 | -0.4554352 | 0.64879615 | 0.90754509 |
| ZNHIT3 | 158.634065 | 0.32977025 | 0.25620461 | 1.28713626 | 0.1980468 | 0.65049274 |
| MYO19 | 902.303238 | -0.1539971 | 0.35229084 | -0.4371307 | 0.66201655 | 0.91110616 |
| GGNBP2 | 998.056633 | -0.1950773 | 0.18300486 | -1.0659677 | 0.28643824 | 0.72934245 |
| DHRS11 | 80.3720652 | 0.95174243 | 0.51834973 | 1.83610094 | 0.06634273 | 0.4469687 |
| MRM1 | 58.699701 | -0.6226599 | 0.38483914 | -1.6179745 | 0.1056681 | 0.52694363 |
| AATF | 796.315482 | -0.0989096 | 0.22428581 | -0.4409978 | 0.65921459 | 0.91099119 |
| ACACA | 2042.63596 | -0.6004597 | 0.30224247 | -1.9866821 | 0.04695764 | 0.39257181 |
| TADA2A | 390.666336 | -0.3992277 | 0.19415143 | -2.0562697 | 0.03975653 | 0.37279391 |
| DUSP14 | 43.0337507 | 0.76432526 | 0.37621658 | 2.03160974 | 0.04219318 | 0.37913585 |
| SYNRG | 1254.46474 | -0.137246 | 0.15471435 | -0.8870931 | 0.37502877 | 0.78444261 |
| DDX52 | 1025.76501 | -1.047865 | 0.59676915 | -1.7558967 | 0.07910605 | 0.48169281 |
| AC243585.2 | 61.8812331 | -0.7152559 | 0.88824972 | -0.8052419 | 0.42068012 | 0.81260593 |
| AC243571.2 | 100.80852 | -2.0046764 | 0.99362001 | -2.0175483 | 0.04363832 | 0.38443959 |
| HNF1B | 2664.38254 | -0.5176082 | 0.25736794 | -2.0111605 | 0.0443085 | 0.38580819 |
| MRPL45 | 317.173118 | -0.2088181 | 0.16143463 | -1.2935148 | 0.19583307 | 0.64964978 |
| SOCS7 | 321.243782 | -0.267659 | 0.24544861 | -1.0904892 | 0.27549773 | 0.72233844 |
| ARHGAP23 | 45.0914318 | -0.4377594 | 0.42094925 | -1.0399339 | 0.29837063 | 0.73673761 |
| MLLT6 | 1650.31069 | -0.0628555 | 0.24871791 | -0.2527182 | 0.80048601 | 0.95370871 |
| CISD3 | 138.20438 | 0.4638978 | 0.37619269 | 1.2331388 | 0.21752398 | 0.67038598 |
| PCGF2 | 246.124032 | -0.2434564 | 0.25100964 | -0.9699086 | 0.33209207 | 0.76037513 |
| PSMB3 | 293.350216 | 0.1075855 | 0.34385578 | 0.31287973 | 0.75437204 | 0.93901195 |
| PIP4K2B | 382.302195 | -0.2651012 | 0.24633999 | -1.0761596 | 0.28185587 | 0.72604837 |
| CWC25 | 276.693437 | -0.266564 | 0.18673253 | -1.4275177 | 0.15343073 | 0.60285224 |
| RPL23 | 621.988302 | 0.68018369 | 0.39181206 | 1.73599476 | 0.08256476 | 0.48594425 |
| LASP1 | 383.267403 | -0.2575801 | 0.22126025 | -1.16415 | 2.44E-01 | 6.95E-01 |
| RPL19 | 1157.27046 | 0.65096466 | 0.47866873 | 1.35994816 | 0.17384633 | 0.6286116 |
| STAC2 | 116.430715 | 2.44820309 | 0.81326065 | 3.0103548 | 2.61E-03 | 0.09654879 |
| FBXL20 | 2278.40182 | -0.2244832 | 0.20449767 | -1.0977298 | 0.27232248 | 0.72070318 |

| | | | | | | |
|---|---|---|---|---|---|---|
| MED1 | 384.931839 | -0.0974442 | 0.15395027 | -0.6329591 | 0.52676038 | 0.86151701 |
| CDK12 | 1274.24794 | -0.1145193 | 0.18096777 | -0.6328161 | 0.52685375 | 0.86151701 |
| PPP1R1B | 166.935224 | 3.2222256 | 0.90904172 | 3.54463994 | 0.00039315 | 0.03394193 |
| STARD3 | 268.1807 | -0.0784311 | 0.27593168 | -0.2842412 | 0.77622555 | 0.94694572 |
| PGAP3 | 180.079413 | 0.36811473 | 0.36588388 | 1.00609717 | 0.31436883 | 0.74785672 |
| ERBB2 | 1140.47945 | 0.43851493 | 0.35681242 | 1.22897888 | 0.21907972 | 0.67266947 |
| MIEN1 | 165.518544 | 0.21293455 | 0.473565 | 0.44964165 | 0.65296885 | 0.908787 |
| GRB7 | 1136.90895 | -0.1777856 | 0.23812851 | -0.7465952 | 4.55E-01 | 8.31E-01 |
| GSDMB | 54.2548826 | -0.6007573 | 0.38698145 | -1.5524188 | 0.12056204 | 0.55433742 |
| ORMDL3 | 71.5834064 | 0.6286686 | 0.45947015 | 1.36824688 | 0.17123481 | 0.62772493 |
| AC090844.2 | 82.7171715 | 0.27916474 | 0.50418704 | 0.55369281 | 0.5797891 | 0.88240975 |
| PSMD3 | 227.815984 | 0.07107376 | 0.28679141 | 0.24782388 | 0.80427068 | 0.95450237 |
| MED24 | 337.717143 | -0.0683366 | 0.21091825 | -0.3239956 | 0.74594138 | 0.93675395 |
| THRA | 238.484815 | -0.0201482 | 0.32941399 | -0.0611639 | 0.95122871 | 0.98938795 |
| NR1D1 | 57.9124834 | -0.1800828 | 0.65876799 | -0.273363 | 0.78457418 | 0.94978829 |
| MSL1 | 496.661037 | -0.2303211 | 0.26909356 | -0.8559145 | 0.39204509 | 0.79767768 |
| CASC3 | 489.225317 | -0.0593161 | 0.13219978 | -0.4486851 | 0.65365887 | 0.90890163 |
| RAPGEFL1 | 69.8689284 | -0.7694752 | 0.32087678 | -2.3980394 | 0.01648309 | 0.25721056 |
| WIPF2 | 1214.41823 | -0.0328324 | 0.23209339 | -0.1414619 | 0.8875051 | 0.97729111 |
| RARA | 165.175276 | 0.20130101 | 0.56304966 | 0.35751911 | 0.72070322 | 0.93020997 |
| IGFBP4 | 194.583988 | 0.5606378 | 0.31987429 | 1.75268163 | 0.07965667 | 0.48176967 |
| KRT10 | 247.196136 | 0.20050678 | 0.53241004 | 0.37660218 | 0.70646927 | 0.92556842 |
| TMEM99 | 81.9422615 | -0.0521683 | 0.3243627 | -0.1608332 | 0.87222474 | 0.97548023 |
| KRT19 | 393.098847 | 1.22394675 | 0.61397234 | 1.9934884 | 4.62E-02 | 3.90E-01 |
| EIF1 | 1535.70173 | 0.56537041 | 0.36791949 | 1.53666883 | 1.24E-01 | 0.56111892 |
| HAP1 | 139.320211 | 0.34051331 | 0.56543316 | 0.60221673 | 5.47E-01 | 8.69E-01 |
| JUP | 1506.70108 | -0.0481384 | 0.31265515 | -0.1539665 | 0.87763615 | 0.97655597 |
| P3H4 | 133.612043 | -0.1172428 | 0.32764499 | -0.357835 | 0.72046681 | 0.93008661 |
| FKBP10 | 49.6644255 | -1.8055056 | 0.63053008 | -2.8634725 | 0.00419025 | 0.12579326 |
| NT5C3B | 231.563482 | 0.28201853 | 0.33729151 | 0.83612696 | 0.40308348 | 0.80324472 |
| ACLY | 453.378026 | 0.0809671 | 0.28645367 | 0.28265341 | 0.77744254 | 0.94694572 |
| TTC25 | 147.145257 | -0.3269345 | 0.40282168 | -0.811611 | 0.41701486 | 0.81007111 |
| CNP | 292.071901 | -0.0087347 | 0.51013922 | -0.0171221 | 0.9863392 | 0.99624198 |
| DNAJC7 | 912.260872 | -0.0822529 | 0.12431438 | -0.6616526 | 0.50819388 | 0.85643442 |
| NKIRAS2 | 113.699485 | -0.1621483 | 0.23446684 | -0.6915619 | 0.4892125 | 0.84614482 |
| DHX58 | 69.4875207 | 0.49683621 | 0.66855276 | 0.74315184 | 0.45738976 | 0.83255882 |
| KAT2A | 207.909628 | -0.030514 | 0.46903986 | -0.0650564 | 0.94812912 | 0.98919519 |
| GHDC | 57.8810941 | 0.50228352 | 0.47372816 | 1.06027793 | 0.28901817 | 0.73189164 |
| STAT5B | 1304.13736 | 0.13842487 | 0.21469932 | 0.64473827 | 0.51909681 | 0.85918126 |
| STAT5A | 202.310167 | 0.26777638 | 0.35110105 | 0.76267609 | 0.4456566 | 0.82571115 |
| STAT3 | 5065.11645 | -0.0756042 | 0.28300719 | -0.2671457 | 7.89E-01 | 0.95089981 |
| CAVIN1 | 103.751683 | 1.13134149 | 0.67977606 | 1.66428556 | 0.09605538 | 0.5117083 |
| ATP6V0A1 | 676.240861 | 0.13767328 | 0.27166977 | 0.50676703 | 0.61231829 | 0.89440588 |
| NAGLU | 242.010075 | -0.3293081 | 0.40494511 | -0.8132167 | 0.41609381 | 0.80966301 |
| COASY | 224.304636 | 0.10178421 | 0.39591705 | 0.25708469 | 0.79711339 | 0.95281098 |
| MLX | 262.701663 | -0.5308011 | 0.24713139 | -2.1478496 | 0.0317257 | 0.34015001 |
| RETREG3 | 611.885496 | -0.0676453 | 0.2056097 | -0.3289987 | 0.74215669 | 0.93598004 |
| TUBG1 | 57.6260413 | 0.29055685 | 0.54909464 | 0.52915624 | 0.59669707 | 0.88883692 |
| TUBG2 | 218.090684 | -0.4557111 | 0.21123743 | -2.1573406 | 0.03097914 | 0.33790537 |
| PLEKHH3 | 93.7542988 | 0.28035393 | 0.4078025 | 0.68747477 | 0.4917836 | 0.84704258 |
| AC100793.2 | 65.212676 | 0.48300517 | 0.69329076 | 0.69668485 | 0.48600004 | 0.8447301 |
| EZH1 | 723.222397 | -0.3464496 | 0.20656505 | -1.6771936 | 0.09350463 | 0.50887217 |
| VPS25 | 222.232931 | -0.1322774 | 0.27065948 | -0.4887225 | 6.25E-01 | 0.89873262 |
| WNK4 | 1462.51123 | -0.7631063 | 0.54183545 | -1.4083729 | 0.15902069 | 0.60866541 |

| | | | | | | |
|---|---|---|---|---|---|---|
| COA3 | 401.521951 | 0.58364358 | 0.43772763 | 1.33334872 | 0.18241739 | 0.63741052 |
| CNTD1 | 79.1849846 | -0.2341825 | 0.50587183 | -0.4629285 | 0.64341563 | 0.90478945 |
| BECN1 | 477.24622 | -0.0837153 | 0.21806379 | -0.3839028 | 0.70105052 | 0.92424844 |
| PSME3 | 214.85665 | -0.1307203 | 0.20991419 | -0.6227321 | 0.5334606 | 0.86397197 |
| LINC00671 | 179.356499 | -0.3270226 | 0.82343246 | -0.3971456 | 6.91E-01 | 0.92206635 |
| RUNDC1 | 230.927322 | -0.0178887 | 0.23715818 | -0.0754292 | 0.93987319 | 0.98715467 |
| RPL27 | 420.476013 | 0.46528337 | 0.40769168 | 1.14126284 | 0.25376056 | 0.70431689 |
| IFI35 | 149.915954 | 0.46604613 | 0.65713014 | 0.70921436 | 0.47819146 | 0.84199814 |
| VAT1 | 92.8169591 | 0.34954413 | 0.36271209 | 0.96369584 | 3.35E-01 | 7.63E-01 |
| BRCA1 | 268.075754 | 0.12352516 | 0.4371774 | 0.28255156 | 0.77752061 | 0.94694572 |
| NBR1 | 838.509836 | 0.05237753 | 0.16665256 | 0.31429176 | 0.75329945 | 0.93844712 |
| TMEM106A | 76.6849028 | -0.2814003 | 0.44383835 | -0.6340152 | 0.5260709 | 0.86119802 |
| LINC00854 | 611.892569 | -1.8088524 | 0.724846 | -2.4954989 | 0.01257801 | 0.22916272 |
| ARL4D | 145.319371 | 1.64457214 | 0.46954616 | 3.5024717 | 0.00046096 | 0.03734387 |
| DHX8 | 363.31763 | -0.2333468 | 0.14958426 | -1.5599687 | 0.11876728 | 0.55026203 |
| ETV4 | 136.026067 | 0.93178479 | 0.85797235 | 1.08603125 | 0.27746517 | 0.72344438 |
| AC068675.1 | 64.7813815 | 0.37780233 | 0.57382676 | 0.65839092 | 0.51028697 | 0.85762435 |
| DUSP3 | 287.444507 | 0.53033147 | 0.23303268 | 2.27578157 | 0.02285909 | 0.2983102 |
| MPP3 | 55.4247076 | -0.3868945 | 0.39219521 | -0.9864846 | 0.32389539 | 0.75402436 |
| TMEM101 | 155.449645 | -0.2039169 | 0.55665291 | -0.3663269 | 0.71412115 | 0.92760608 |
| LSM12 | 344.316914 | -0.390713 | 0.17451857 | -2.238805 | 0.02516861 | 0.31053369 |
| G6PC3 | 115.906722 | 0.61056521 | 0.41009302 | 1.48884565 | 0.13652802 | 0.57801237 |
| HDAC5 | 469.579281 | 0.27910614 | 0.30353841 | 0.91950847 | 0.35782968 | 0.77510204 |
| ASB16-AS1 | 106.849797 | -0.3219979 | 0.3164741 | -1.0174543 | 0.30893737 | 0.74399658 |
| TMUB2 | 241.708601 | 0.14080728 | 0.25732898 | 0.5471878 | 0.58424972 | 0.88416261 |
| ATXN7L3 | 122.065897 | -0.3379111 | 0.33146278 | -1.019454 | 0.30798748 | 0.74372365 |
| AC004596.1 | 55.8132618 | -0.1040927 | 0.33329911 | -0.3123101 | 0.75480483 | 0.93922829 |
| UBTF | 362.07191 | 0.04544317 | 0.31526239 | 0.14414398 | 0.88538678 | 0.97664137 |
| AC003102.1 | 54.0158435 | -0.6539828 | 0.47530971 | -1.3759087 | 0.16884989 | 0.62265166 |
| SLC25A39 | 300.698866 | 0.04679273 | 0.34572424 | 0.13534699 | 0.89233754 | 0.97771356 |
| GRN | 388.904952 | 0.65227865 | 0.33180783 | 1.96583261 | 4.93E-02 | 3.99E-01 |
| GPATCH8 | 3057.13705 | -0.025427 | 0.17052625 | -0.1491087 | 0.88146784 | 0.97664137 |
| CCDC43 | 91.4323197 | -0.2699395 | 0.28512287 | -0.946748 | 0.34376719 | 0.76680914 |
| DBF4B | 151.49533 | -0.4879631 | 0.45190549 | -1.0797903 | 0.28023559 | 0.72513763 |
| EFTUD2 | 393.273191 | -0.4380711 | 0.25167163 | -1.7406455 | 0.08174574 | 0.48594425 |
| DCAKD | 596.287585 | -0.0135676 | 0.28731802 | -0.0472216 | 0.96233659 | 0.99224805 |
| NMT1 | 626.993266 | -0.1065104 | 0.1413747 | -0.7533907 | 0.45121518 | 0.82814845 |
| PLCD3 | 67.0416128 | 0.28880155 | 0.5066685 | 0.570001 | 0.56867702 | 0.87770276 |
| ACBD4 | 142.663079 | -0.0891165 | 0.27706155 | -0.3216489 | 0.74771872 | 0.93746253 |
| AC142472.1 | 44.5009327 | -0.3845935 | 0.49506263 | -0.7768583 | 0.43724235 | 0.82013459 |
| HEXIM1 | 155.923409 | 0.30147092 | 0.52080055 | 0.57886061 | 0.56268323 | 0.87432201 |
| AC138150.1 | 623.132701 | -0.1723491 | 0.41588782 | -0.4144124 | 0.67857208 | 0.91808446 |
| HEXIM2 | 49.3861328 | -0.1629858 | 0.3647454 | -0.446848 | 0.65498481 | 0.90902447 |
| MAP3K14 | 663.85862 | -0.0489612 | 0.33082671 | -0.1479964 | 0.88234557 | 0.97664137 |
| ARHGAP27 | 128.736423 | -0.1031411 | 0.31075686 | -0.3319029 | 0.73996259 | 0.93552616 |
| PLEKHM1 | 400.343758 | -0.287747 | 0.25506812 | -1.1281182 | 0.25927001 | 0.70921184 |
| LINC02210-C | 297.937984 | 0.08176669 | 0.39999607 | 0.20441873 | 0.83802629 | 0.96566728 |
| KANSL1 | 6688.35147 | -0.0633733 | 0.2998241 | -0.2113682 | 0.8326 | 0.96438209 |
| KANSL1-AS1 | 47.8523278 | 0.52014512 | 0.53444674 | 0.97324034 | 0.33043387 | 0.75894644 |
| ARL17B | 1419.17039 | 0.61845263 | 1.20321343 | 0.51400077 | 0.60725145 | 0.89174148 |
| LRRC37A | 76.932987 | 0.1384891 | 0.35734367 | 0.38755157 | 0.69834791 | 0.92356678 |
| LRRC37A2 | 388.331887 | 0.09550608 | 0.32924898 | 0.29007251 | 0.77176076 | 0.94478317 |
| ARL17A | 466.570779 | -0.5139496 | 0.48971552 | -1.049486 | 0.29395449 | 0.73527014 |
| NSF | 1689.7212 | -0.0290918 | 0.34450439 | -0.0844454 | 0.93270233 | 0.9839805 |

| | | | | | | |
|---|---|---|---|---|---|---|
| AC005670.2 | 288.25159 | 0.12717325 | 0.37761745 | 0.33677801 | 0.73628425 | 0.93552616 |
| CDC27 | 1599.52816 | -0.2189222 | 0.21227567 | -1.0313109 | 0.30239506 | 0.73905643 |
| AC068234.1 | 708.781355 | -0.2744449 | 0.84725053 | -0.3239241 | 7.46E-01 | 9.37E-01 |
| AC068234.2 | 129.849942 | 0.32293304 | 0.61397233 | 0.52597328 | 5.99E-01 | 8.89E-01 |
| EFCAB13 | 781.490988 | 0.11279685 | 0.44455552 | 0.2537295 | 8.00E-01 | 9.54E-01 |
| AC040934.1 | 134.298705 | -0.4060172 | 0.36515351 | -1.1119083 | 0.26617756 | 0.71641758 |
| NPEPPS | 3142.8486 | -0.3462065 | 0.17965073 | -1.9271086 | 0.0539661 | 0.41444084 |
| AC025682.1 | 415.19643 | 0.09798364 | 0.30526101 | 0.32098313 | 0.74822317 | 0.93746253 |
| KPNB1 | 1188.54932 | -0.022732 | 0.19514032 | -0.1164904 | 0.90726388 | 0.98047754 |
| TBKBP1 | 85.1366076 | -0.5414555 | 0.37454959 | -1.4456175 | 0.1482845 | 0.59438683 |
| OSBPL7 | 70.4196486 | -0.8135061 | 0.44825998 | -1.8148088 | 0.0695533 | 0.4580333 |
| MRPL10 | 73.7987927 | 0.43439911 | 0.3481597 | 1.24770072 | 0.21214068 | 0.66552984 |
| SCRN2 | 139.924013 | 0.1623224 | 0.33563062 | 0.48363406 | 0.6286456 | 0.8996511 |
| SP2 | 180.218756 | -0.3869714 | 0.25367827 | -1.5254415 | 0.12714902 | 0.56740575 |
| SP2-AS1 | 128.798158 | -0.3872039 | 0.28505024 | -1.3583708 | 0.17434607 | 0.62892483 |
| PNPO | 136.659699 | 0.79201704 | 0.28548602 | 2.77427613 | 0.00553247 | 0.15039463 |
| PRR15L | 177.220594 | 2.04646594 | 0.62865801 | 3.25529288 | 0.00113275 | 0.0603329 |
| CDK5RAP3 | 397.508692 | 0.02188584 | 0.23736245 | 0.09220432 | 0.92653571 | 0.98238964 |
| NFE2L1 | 662.221643 | 0.53502937 | 0.23595155 | 2.26753918 | 0.02335731 | 0.30070827 |
| CBX1 | 409.562946 | -0.0645719 | 0.27806448 | -0.2322193 | 0.81636772 | 0.9589293 |
| SNX11 | 119.828119 | -0.0631524 | 0.21901278 | -0.2883501 | 0.77307877 | 0.94534972 |
| SKAP1 | 4952.90701 | 0.26431438 | 0.21736712 | 1.21598142 | 0.223992 | 0.67674069 |
| SKAP1-AS1 | 103.053322 | 0.14041486 | 0.53524863 | 0.26233576 | 0.79306259 | 0.95224951 |
| HOXB2 | 240.939742 | -0.1942509 | 0.44700906 | -0.4345569 | 0.66388405 | 0.91197775 |
| HOXB-AS1 | 216.380948 | -0.3851924 | 0.3481279 | -1.1064682 | 0.26852393 | 0.71818262 |
| HOXB3 | 2656.15095 | 0.18246134 | 0.27824006 | 0.65576946 | 0.51197247 | 0.85762435 |
| HOXB-AS3 | 599.801943 | 0.19224257 | 0.3945236 | 0.48727773 | 0.62606152 | 0.89873262 |
| HOXB7 | 396.514278 | 0.11426666 | 0.29654814 | 0.38532244 | 7.00E-01 | 9.24E-01 |
| AC103702.2 | 60.8758681 | 1.24822981 | 0.64433577 | 1.937235 | 0.05271662 | 0.41057422 |
| LINC02086 | 128.473745 | 0.8868395 | 0.72852367 | 1.21731047 | 0.22348611 | 0.67624946 |
| TTLL6 | 42.4536363 | 2.30689084 | 0.83468546 | 2.7637846 | 0.00571352 | 0.15465152 |
| CALCOCO2 | 459.920983 | 0.16042183 | 0.1956725 | 0.81984861 | 0.41230242 | 0.80701706 |
| ATP5MC1 | 399.578563 | 0.79891637 | 0.43234049 | 1.84788699 | 0.06461869 | 0.44001919 |
| UBE2Z | 745.102531 | 0.00201298 | 0.21016312 | 0.00957819 | 0.99235783 | 0.99787261 |
| SNF8 | 325.583061 | -0.1257777 | 0.16461599 | -0.7640673 | 0.44482713 | 0.8252311 |
| AC091133.1 | 64.8927211 | -0.29799 | 0.3115191 | -0.9565705 | 0.33878407 | 0.76449297 |
| B4GALNT2 | 144.26396 | -0.629578 | 0.70557295 | -0.8922933 | 0.37223574 | 0.78256595 |
| ZNF652 | 1258.68395 | -0.1322863 | 0.15189911 | -0.8708828 | 0.38381813 | 0.791697 |
| AC091180.5 | 68.4105593 | 0.28178417 | 0.59803123 | 0.47118637 | 0.63750765 | 0.90255504 |
| PHB | 381.950457 | 1.05826172 | 0.41057572 | 2.57750685 | 0.00995159 | 0.20316934 |
| SPOP | 982.116413 | -0.0437221 | 0.19776237 | -0.2210842 | 0.82502691 | 0.96146415 |
| SLC35B1 | 117.208297 | 0.02842004 | 0.32665472 | 0.08700332 | 0.93066888 | 0.98331361 |
| FAM117A | 180.669411 | -0.4407334 | 0.33133294 | -1.3301829 | 0.18345801 | 0.63834582 |
| KAT7 | 398.982996 | -0.1322521 | 0.18196523 | -0.7267985 | 0.46734941 | 0.83762812 |
| TAC4 | 79.620686 | -0.060482 | 0.49664591 | -0.121781 | 0.90307247 | 0.97960781 |
| ITGA3 | 2580.37267 | -0.5903782 | 0.52697163 | -1.1203225 | 2.63E-01 | 7.13E-01 |
| PDK2 | 189.735912 | 0.72209745 | 0.31729391 | 2.27579993 | 0.02285799 | 0.2983102 |
| PPP1R9B | 96.0225322 | -0.4729685 | 0.36824149 | -1.2843976 | 0.19900286 | 0.65113421 |
| COL1A1 | 55.8124936 | -1.064168 | 0.47968782 | -2.2184595 | 2.65E-02 | 0.31449657 |
| XYLT2 | 171.846775 | 0.04940226 | 0.34216731 | 0.14438042 | 0.88520008 | 0.97664137 |
| MRPL27 | 265.883545 | 0.35779242 | 0.28274959 | 1.26540386 | 0.20572659 | 0.65813563 |
| LRRC59 | 117.437694 | 0.35699359 | 0.34355282 | 1.03912286 | 0.2987476 | 0.73673761 |
| ACSF2 | 251.073585 | 1.26615852 | 0.32726714 | 3.86888373 | 0.00010933 | 0.0164114 |
| RSAD1 | 164.188009 | 0.00515299 | 0.24778761 | 0.02079599 | 0.9834084 | 0.99518552 |

| | | | | | | |
|---|---|---|---|---|---|---|
| SPATA20 | 120.601911 | 0.30829332 | 0.45711131 | 0.67443818 | 0.50003278 | 0.85257945 |
| ABCC3 | 916.859695 | 0.75019867 | 0.7370386 | 1.01785533 | 3.09E-01 | 7.44E-01 |
| ANKRD40 | 214.263905 | 0.00800584 | 0.26057988 | 0.03072318 | 0.97549031 | 0.9941831 |
| AC005921.2 | 59.546552 | 0.24194648 | 0.50651398 | 0.47766989 | 0.63288518 | 0.90074133 |
| LUC7L3 | 2008.88312 | -0.2855165 | 0.11131912 | -2.5648466 | 1.03E-02 | 0.20627111 |
| TOB1 | 672.592574 | 0.67664505 | 0.41311877 | 1.63789472 | 0.10144366 | 0.51919466 |
| SPAG9 | 2757.78914 | -0.0504811 | 0.26000688 | -0.194153 | 0.84605606 | 0.96883382 |
| AC005920.2 | 261.099934 | 0.27416697 | 0.61146579 | 0.44837663 | 0.6538814 | 0.90897911 |
| MBTD1 | 1547.469 | -0.3069571 | 0.18225905 | -1.6841805 | 0.09214679 | 0.5058189 |
| UTP18 | 293.516951 | -0.122037 | 0.17382557 | -0.7020657 | 0.48263817 | 0.84407748 |
| CA10 | 3011.07034 | 0.09734839 | 0.3597162 | 0.27062553 | 0.78667906 | 0.94978829 |
| TOM1L1 | 2118.38615 | 0.26049375 | 0.1971495 | 1.32130059 | 0.18640116 | 0.6408069 |
| COX11 | 493.997354 | 0.45050233 | 0.21462803 | 2.09899112 | 0.03581768 | 0.35661333 |
| STXBP4 | 3266.48907 | 0.18815511 | 0.3883097 | 0.48454908 | 0.62799624 | 0.89920892 |
| HLF | 531.257251 | 0.14284834 | 0.60579951 | 0.23580134 | 0.81358683 | 0.9582375 |
| MMD | 106.88957 | 0.63321464 | 0.33779768 | 1.87453816 | 0.06085629 | 0.43125038 |
| PCTP | 131.103801 | -0.1750025 | 0.3664821 | -0.47752 | 0.63299191 | 0.90078752 |
| C17orf67 | 77.2201668 | 0.44901263 | 0.42253952 | 1.06265238 | 0.28793963 | 0.7307234 |
| DGKE | 244.297442 | 0.28094525 | 0.22434782 | 1.25227536 | 0.21046954 | 0.66334818 |
| TRIM25 | 207.625439 | 0.47522929 | 0.48455535 | 0.98075336 | 0.32671438 | 0.75556858 |
| COIL | 189.472051 | -0.0077079 | 0.20236687 | -0.0380888 | 0.96961692 | 0.99299658 |
| SCPEP1 | 254.536502 | 0.47096364 | 0.22118395 | 2.12928493 | 0.03323069 | 0.34850106 |
| AC007114.1 | 50.5565363 | -0.2270435 | 0.4767768 | -0.4762049 | 0.63392838 | 0.90136076 |
| AKAP1 | 547.945128 | 0.27023298 | 0.27085338 | 0.99770945 | 0.31842027 | 0.75064854 |
| AC007114.2 | 83.1731226 | -0.2600615 | 0.54835689 | -0.4742559 | 0.63531745 | 0.90199374 |
| MSI2 | 14816.6134 | 0.25455383 | 0.36682329 | 0.69394131 | 0.48771902 | 0.8451268 |
| AC015845.2 | 63.1349244 | -0.9089614 | 0.40182009 | -2.2621105 | 0.02369058 | 0.30185158 |
| MRPS23 | 195.253281 | 0.15269318 | 0.26623567 | 0.57352639 | 0.56628833 | 0.87642772 |
| CUEDC1 | 1177.87305 | -0.1764099 | 0.2971348 | -0.5937031 | 0.55271068 | 0.87104567 |
| VEZF1 | 607.569323 | 0.27386255 | 0.20106785 | 1.36204047 | 0.17318511 | 0.6286116 |
| AC015813.2 | 372.112929 | 0.04128307 | 0.24879207 | 0.16593402 | 8.68E-01 | 0.97402505 |
| SRSF1 | 175.558285 | -0.1564992 | 0.28690193 | -0.5454797 | 0.58542366 | 0.88490909 |
| DYNLL2 | 264.417668 | 0.71328777 | 0.27905685 | 2.55606611 | 0.0105863 | 0.20870812 |
| MKS1 | 155.994799 | -0.3842609 | 0.21892633 | -1.7552066 | 0.07922397 | 0.48169281 |
| TSPOAP1 | 370.074352 | 0.32990316 | 0.35195443 | 0.93734625 | 0.34858048 | 0.76982416 |
| TSPOAP1-AS | 151.501733 | 0.07754062 | 0.4965928 | 0.15614528 | 0.87591851 | 0.97614152 |
| SUPT4H1 | 121.746906 | 0.13651712 | 0.25118552 | 0.54349122 | 0.58679164 | 0.88547501 |
| RNF43 | 813.227476 | -0.0813016 | 0.3306104 | -0.2459135 | 0.80574919 | 0.95522565 |
| MTMR4 | 295.58628 | 0.28107525 | 0.37281427 | 0.75392836 | 0.45089224 | 0.82813014 |
| SEPT4-AS1 | 166.931166 | 0.01230083 | 0.52359435 | 0.02349306 | 0.98125698 | 0.99518016 |
| TEX14 | 89.4096636 | -0.329575 | 0.43580793 | -0.756239 | 0.44950592 | 0.82672287 |
| RAD51C | 219.350055 | 0.22031441 | 0.23088849 | 0.95420266 | 0.33998107 | 0.76465348 |
| PPM1E | 1769.82217 | 0.46568463 | 0.58138733 | 0.80098862 | 4.23E-01 | 0.8138418 |
| TRIM37 | 589.613862 | -0.4616385 | 0.25489178 | -1.8111155 | 0.07012297 | 0.4601363 |
| AC099850.1 | 46.6714291 | 0.4768535 | 0.45653872 | 1.04449737 | 0.29625533 | 0.7366786 |
| SKA2 | 306.468807 | -0.0431128 | 0.23349547 | -0.1846408 | 0.85351076 | 0.97076831 |
| GDPD1 | 158.207832 | -0.3870685 | 0.28806335 | -1.3436924 | 0.17904789 | 0.63287224 |
| YPEL2 | 1334.75144 | -0.1259286 | 0.27498331 | -0.4579499 | 6.47E-01 | 9.06E-01 |
| DHX40 | 577.810048 | 0.00479437 | 0.20363333 | 0.02354412 | 0.98121625 | 0.99518016 |
| CLTC | 1980.76933 | 0.08796713 | 0.1926909 | 0.45651939 | 0.64801653 | 0.90722315 |
| PTRH2 | 334.646066 | 0.07576807 | 0.303849 | 0.24936092 | 0.80308161 | 0.95446295 |
| VMP1 | 5593.71542 | -0.6726136 | 0.37063365 | -1.8147668 | 0.06955975 | 0.4580333 |
| TUBD1 | 384.75419 | 0.0583869 | 0.3145853 | 0.18559959 | 0.85275878 | 0.97076831 |
| RPS6KB1 | 579.952734 | -0.1942967 | 0.22783137 | -0.8528091 | 0.39376516 | 0.79843808 |

| | | | | | | |
|---|---|---|---|---|---|---|
| RNFT1 | 144.711203 | 0.05576984 | 0.42230382 | 0.13206095 | 0.89493608 | 0.97788747 |
| RNFT1-DT | 62.5282277 | 0.01123434 | 0.42662903 | 0.02633281 | 0.97899188 | 0.99455895 |
| HEATR6 | 153.023745 | 0.10142701 | 0.24301131 | 0.41737566 | 0.67640365 | 0.91658712 |
| AC025048.4 | 50.9204398 | 0.11881927 | 0.6460524 | 0.18391583 | 0.85407948 | 0.97085267 |
| CA4 | 354.364208 | 2.46147615 | 0.78075446 | 3.15268918 | 0.00161774 | 0.07550785 |
| USP32 | 2139.49311 | -0.245945 | 0.20531082 | -1.1979156 | 0.23094987 | 0.68321398 |
| APPBP2 | 1413.32967 | 0.04955947 | 0.15958403 | 0.31055408 | 0.75613964 | 0.93964676 |
| AC011921.1 | 191.692924 | -0.0701821 | 0.37494987 | -0.1871772 | 0.85152169 | 0.9707215 |
| PPM1D | 562.694933 | -0.1264555 | 0.33164375 | -0.3812993 | 0.70298121 | 0.92501157 |
| BCAS3 | 5899.96147 | -0.337765 | 0.15702373 | -2.1510443 | 0.03147271 | 0.33950635 |
| AC005884.1 | 44.5036693 | -0.3687443 | 0.5136718 | -0.7178597 | 0.47284379 | 0.83985854 |
| AC005746.2 | 70.8241401 | 0.24511233 | 0.55367129 | 0.4427037 | 0.65798007 | 0.91037018 |
| NACA2 | 71.0178653 | 0.75367886 | 0.59706647 | 1.26230311 | 0.20683974 | 0.6594608 |
| INTS2 | 317.223355 | -0.5377982 | 0.2220409 | -2.4220681 | 0.01543246 | 0.25214021 |
| MED13 | 3168.57485 | -0.1721889 | 0.24471823 | -0.7036212 | 0.48166871 | 0.8437862 |
| EFCAB3 | 62.8724757 | -0.4053926 | 0.51495161 | -0.7872441 | 0.43113898 | 0.81706845 |
| METTL2A | 159.209399 | -0.1821924 | 0.27351748 | -0.6661087 | 0.50534163 | 0.85595485 |
| TLK2 | 1648.63156 | -0.257659 | 0.21620354 | -1.1917429 | 0.23336207 | 0.68484327 |
| 10-Mar | 57.6675658 | 0.53595874 | 0.44768229 | 1.19718548 | 0.23123427 | 0.68357541 |
| TANC2 | 5970.80781 | 0.29359135 | 0.25620744 | 1.14591267 | 2.52E-01 | 0.70276117 |
| AC006270.1 | 58.6835451 | 0.8927865 | 0.59528695 | 1.49975824 | 0.13367704 | 0.57703921 |
| AC015923.1 | 468.298917 | 0.61991231 | 0.59474972 | 1.04230787 | 0.29726897 | 0.73673761 |
| CYB561 | 175.090239 | 0.74986951 | 0.28586813 | 2.62313085 | 0.00871258 | 0.18928367 |
| DCAF7 | 1000.39627 | 0.09153564 | 0.19403172 | 0.47175608 | 0.63710091 | 0.90255504 |
| TACO1 | 76.4802654 | 0.64538349 | 0.33459622 | 1.92884277 | 0.05375038 | 0.41326474 |
| MAP3K3 | 1567.82027 | 0.41000573 | 0.36345695 | 1.12807233 | 0.25928937 | 0.70921184 |
| CCDC47 | 297.256671 | -0.0154528 | 0.25478331 | -0.0606506 | 0.95163751 | 0.98938795 |
| DDX42 | 786.647054 | -0.2532642 | 0.13186223 | -1.9206731 | 0.05477294 | 0.41684503 |
| FTSJ3 | 148.587354 | 0.26878725 | 0.27195792 | 0.98834132 | 0.32298551 | 0.7532337 |
| PSMC5 | 168.282572 | 0.49697127 | 0.37707228 | 1.31797351 | 0.18751252 | 0.64200436 |
| SMARCD2 | 206.580514 | 0.15151257 | 0.29213959 | 0.51863073 | 0.60401827 | 0.89001883 |
| ERN1 | 572.843617 | 0.44365203 | 0.40963847 | 1.08303312 | 0.27879372 | 0.72449071 |
| SNHG25 | 226.832611 | 0.06709066 | 0.39670022 | 0.16912183 | 0.86570082 | 0.97379123 |
| TEX2 | 1016.80504 | 0.34331635 | 0.36036684 | 0.95268573 | 0.34074932 | 0.76507867 |
| PECAM1 | 53.3800779 | -0.5896394 | 0.30399848 | -1.9396129 | 0.05242675 | 0.4090401 |
| POLG2 | 277.233317 | -0.5600111 | 0.19382005 | -2.8893352 | 0.00386057 | 0.12104636 |
| DDX5 | 3673.84447 | 0.03169355 | 0.26598785 | 0.11915411 | 0.90515326 | 0.97992575 |
| CEP95 | 1273.8714 | -0.0869215 | 0.23042098 | -0.377229 | 0.70600343 | 0.92556842 |
| SMURF2 | 1977.70051 | -0.1262599 | 0.12742452 | -0.9908602 | 0.32175387 | 0.75240254 |
| LRRC37A3 | 1665.14246 | -0.0853979 | 0.34089288 | -0.2505125 | 0.80219106 | 0.95443209 |
| AC103810.2 | 57.3524491 | -0.8050076 | 0.66163092 | -1.2167019 | 0.22371765 | 0.67637593 |
| GNA13 | 801.089783 | -0.2731478 | 0.35478014 | -0.7699073 | 0.44135487 | 0.82285053 |
| AC004805.1 | 62.7046121 | 1.09454284 | 0.44900063 | 2.43773121 | 0.01477976 | 0.24869534 |
| CEP112 | 1962.93759 | 0.07401444 | 0.2221193 | 0.33321934 | 0.73896872 | 0.93552616 |
| PRKCA | 2252.55417 | 0.26227742 | 0.46738578 | 0.56115833 | 0.57468961 | 0.88043088 |
| PRKCA-AS1 | 90.6018087 | 0.58072382 | 0.41416576 | 1.40215314 | 0.16086952 | 0.61166537 |
| HELZ | 2664.98926 | -0.0049755 | 0.17330361 | -0.02871 | 0.97709588 | 0.99433432 |
| PSMD12 | 481.015487 | -0.2394385 | 0.15635183 | -1.5314086 | 0.12566843 | 0.56499465 |
| PITPNC1 | 4123.10325 | -1.2767375 | 0.47437411 | -2.6914149 | 0.00711496 | 0.17279872 |
| NOL11 | 301.098148 | -0.2932472 | 0.23087474 | -1.2701573 | 0.20402861 | 0.65712971 |
| BPTF | 3480.30159 | -0.360216 | 0.1968222 | -1.8301593 | 0.06722612 | 0.45098515 |
| KPNA2 | 67.1369487 | 0.23903463 | 0.54040414 | 0.44232568 | 0.65825355 | 0.91037018 |
| AC005332.7 | 123.814994 | 0.17272864 | 0.19346423 | 0.89281949 | 0.37195385 | 0.78247147 |
| AC005332.6 | 43.5846687 | -0.3387818 | 0.39873606 | -0.8496392 | 0.39552569 | 0.79924445 |

| | | | | | | |
|---|---|---|---|---|---|---|
| AMZ2 | 97.6284968 | 0.41960436 | 0.27308763 | 1.53651909 | 0.12441112 | 0.56111892 |
| ARSG | 322.530521 | -0.5583361 | 0.52090071 | -1.0718667 | 0.28377991 | 0.72690826 |
| WIPI1 | 372.14792 | -0.5926076 | 0.33855165 | -1.7504202 | 0.08004584 | 0.48282462 |
| PRKAR1A | 1124.91586 | 0.38650633 | 0.29091682 | 1.32858023 | 0.1839865 | 0.63843513 |
| FAM20A | 239.228095 | 0.11471271 | 0.326643 | 0.35118682 | 7.25E-01 | 9.33E-01 |
| LINC01482 | 277.033214 | 1.93885674 | 0.67052183 | 2.89156392 | 0.0038333 | 0.12084489 |
| ABCA9 | 139.552531 | 0.74566189 | 0.53830936 | 1.38519214 | 0.16599372 | 0.61901667 |
| ABCA6 | 160.805637 | 0.72591345 | 0.35938916 | 2.01985353 | 0.04339858 | 0.38352475 |
| ABCA10 | 592.544845 | 0.72230163 | 0.25158438 | 2.87101142 | 0.00409161 | 0.12481765 |
| ABCA5 | 1748.09049 | 0.51552076 | 0.26110064 | 1.97441398 | 0.04833469 | 0.39627811 |
| MAP2K6 | 396.246071 | 1.44354868 | 0.65460679 | 2.20521495 | 0.02743901 | 0.31979365 |
| AC005208.1 | 2549.53565 | 0.69512617 | 0.36667391 | 1.8957612 | 0.05799163 | 0.42087381 |
| KCNJ16 | 8368.17125 | 0.1974314 | 0.31977702 | 0.61740333 | 0.53696872 | 0.86466169 |
| SOX9 | 277.590223 | 1.68832448 | 0.58435702 | 2.88920032 | 3.86E-03 | 0.12104636 |
| LINC00511 | 615.306919 | 1.79358247 | 0.67053766 | 2.67484227 | 7.48E-03 | 1.74E-01 |
| SLC39A11 | 1322.21137 | -0.7679849 | 0.29967065 | -2.5627632 | 0.01038428 | 0.2068869 |
| COG1 | 193.551787 | -0.2461998 | 0.25024825 | -0.9838223 | 0.32520288 | 0.7552048 |
| FAM104A | 258.67984 | 0.0921392 | 0.17552689 | 0.52492924 | 0.59963238 | 0.88883692 |
| C17orf80 | 164.615337 | 0.01673626 | 0.27859026 | 0.06007481 | 0.95209605 | 0.98938795 |
| CDC42EP4 | 130.619421 | 0.12309668 | 0.25150389 | 0.48944246 | 6.25E-01 | 8.99E-01 |
| RPL38 | 340.791877 | 0.44945125 | 0.36539227 | 1.23005133 | 0.21867788 | 0.67202914 |
| GPRC5C | 852.432902 | 0.01541916 | 0.39181163 | 0.03935351 | 9.69E-01 | 0.99250254 |
| SLC9A3R1 | 135.080974 | 0.31583417 | 0.43948851 | 0.71864032 | 0.47236256 | 0.83921324 |
| NAT9 | 157.0348 | -0.2188116 | 0.31422676 | -0.6963492 | 0.48621014 | 0.8447301 |
| TMEM104 | 114.020926 | -0.2506367 | 0.37857326 | -0.6620561 | 0.50793529 | 0.85643442 |
| HID1 | 84.0719958 | 0.2605539 | 0.2717119 | 0.95893443 | 0.33759178 | 0.76409325 |
| CDR2L | 126.777742 | 0.08898649 | 0.41017782 | 0.21694614 | 0.82825032 | 0.96288319 |
| MRPL58 | 100.29249 | 0.4991426 | 0.22987101 | 2.17140301 | 0.02990072 | 0.33245676 |
| KCTD2 | 251.372822 | 0.14985991 | 0.28121309 | 0.53290517 | 0.59409922 | 0.88807733 |
| ATP5PD | 310.740367 | 0.46386635 | 0.35083737 | 1.32216915 | 0.18611183 | 0.64033103 |
| SLC16A5 | 1160.96731 | 0.49238303 | 0.51617185 | 0.95391299 | 0.34012769 | 0.76478309 |
| ARMC7 | 284.156602 | 0.40930868 | 0.51494237 | 0.79486309 | 0.42669315 | 0.81466629 |
| NT5C | 339.640567 | 0.0205579 | 0.32331014 | 0.06358568 | 0.94930013 | 0.98919519 |
| JPT1 | 234.731474 | 0.90170146 | 0.33568667 | 2.68614021 | 0.00722827 | 0.1729679 |
| SUMO2 | 237.341234 | -0.4857984 | 0.19076336 | -2.546602 | 0.01087774 | 0.21282418 |
| NUP85 | 282.21864 | -0.2399542 | 0.22873528 | -1.0490478 | 0.29415614 | 0.73527014 |
| GGA3 | 348.132967 | -0.1601205 | 0.1889237 | -0.8475403 | 0.39669402 | 0.80060066 |
| MRPS7 | 196.650444 | 0.67256021 | 0.28496623 | 2.36014006 | 0.01826804 | 0.2711921 |
| MIF4GD | 47.3381514 | -0.5803183 | 0.36873183 | -1.573822 | 0.1155286 | 0.54427374 |
| SLC25A19 | 57.5285569 | -0.1632283 | 0.34318933 | -0.4756216 | 6.34E-01 | 0.90136076 |
| GRB2 | 935.641624 | -0.1468777 | 0.20034578 | -0.7331209 | 0.46348471 | 0.83531269 |
| AC011933.4 | 156.247037 | 0.00583193 | 0.51824849 | 0.01125315 | 0.99102148 | 0.99749592 |
| TMEM94 | 311.986073 | -0.1441302 | 0.19309914 | -0.7464051 | 0.45542272 | 0.83121393 |
| CASKIN2 | 147.353403 | 0.06262978 | 0.40383005 | 0.15508945 | 0.8767508 | 0.9763537 |
| TSEN54 | 245.252313 | -0.1662413 | 0.29409361 | -0.5652665 | 0.57189249 | 0.87907473 |
| LLGL2 | 3073.35177 | 0.51720936 | 0.38691302 | 1.33675871 | 0.18130141 | 0.63616641 |
| MYO15B | 464.743574 | -0.3839609 | 0.4439173 | -0.8649379 | 0.38707285 | 0.79542282 |
| RECQL5 | 262.165108 | -0.5945937 | 0.2415953 | -2.4611145 | 0.01385062 | 0.24393706 |
| SMIM5 | 87.7322003 | 0.43791896 | 0.57190358 | 0.76572166 | 0.44384194 | 0.82410854 |
| SAP30BP | 441.820252 | -0.1862883 | 0.24911083 | -0.7478128 | 0.45457306 | 0.83121393 |
| ITGB4 | 91.7337382 | 1.27679526 | 0.47753546 | 2.67371821 | 0.00750155 | 0.17414396 |
| GALK1 | 58.9321205 | 0.51648068 | 0.50670838 | 1.01928585 | 0.30806728 | 0.74372365 |
| H3F3B | 1624.2767 | 0.42420311 | 0.34106491 | 1.24376065 | 2.14E-01 | 0.66791655 |
| UNK | 873.458519 | -0.2544652 | 0.26738883 | -0.9516673 | 0.34126574 | 0.76543778 |

| | | | | | | |
|---|---|---|---|---|---|---|
| UNC13D | 52.2546328 | -0.2475062 | 0.45761817 | -0.5408575 | 0.58860578 | 0.88571413 |
| WBP2 | 506.111771 | 0.36258657 | 0.27787699 | 1.30484559 | 1.92E-01 | 6.46E-01 |
| TRIM47 | 137.450874 | 0.63381533 | 0.53745169 | 1.17929732 | 2.38E-01 | 6.90E-01 |
| TRIM65 | 95.323189 | 0.20831502 | 0.31665318 | 0.65786492 | 0.51062493 | 0.85762435 |
| FBF1 | 103.43038 | -0.0934437 | 0.29388407 | -0.317961 | 0.75051448 | 0.93775604 |
| ACOX1 | 572.275044 | 0.25073313 | 0.17961578 | 1.39594153 | 0.1627321 | 0.61387934 |
| TEN1 | 106.287124 | 0.21725616 | 0.23261443 | 0.93397541 | 0.35031658 | 0.77044053 |
| CDK3 | 94.1775003 | -0.3448536 | 0.48234376 | -0.714954 | 0.47463748 | 0.84106098 |
| EVPL | 247.614239 | -0.4198499 | 0.53720985 | -0.781538 | 0.43448616 | 0.81919857 |
| SRP68 | 294.499673 | 0.11599811 | 0.1852749 | 0.62608646 | 0.53125823 | 0.86397197 |
| EXOC7 | 797.851106 | 0.05882673 | 0.29075505 | 0.20232403 | 0.83966342 | 0.96617873 |
| RNF157 | 111.67072 | 0.17235094 | 0.40459022 | 0.42598888 | 0.67011595 | 0.914701 |
| UBALD2 | 306.70734 | 0.07346278 | 0.29113496 | 0.25233239 | 8.01E-01 | 0.95370871 |
| QRICH2 | 158.075433 | -0.4399114 | 0.47973392 | -0.9169904 | 0.35914766 | 0.776302 |
| PRPSAP1 | 578.487271 | -0.3731849 | 0.31026815 | -1.202782 | 0.22906071 | 0.68114879 |
| UBE2O | 677.090949 | -0.0645201 | 0.23481094 | -0.2747745 | 0.78348946 | 0.94955765 |
| RHBDF2 | 191.617936 | -0.5524115 | 0.31425899 | -1.7578225 | 0.07877771 | 0.4808571 |
| AC015802.4 | 75.0768559 | 0.32539417 | 0.48072994 | 0.6768752 | 0.49848514 | 0.85092921 |
| ST6GALNAC2 | 119.609982 | -0.0441427 | 0.61672755 | -0.0715757 | 0.94293956 | 0.98772472 |
| AC005837.1 | 35.2356279 | -0.219991 | 0.78850817 | -0.2789964 | 0.78024756 | 0.9480401 |
| MXRA7 | 339.914248 | -0.1699892 | 0.48211988 | -0.3525869 | 0.72439816 | 0.93233706 |
| JMJD6 | 214.037177 | 0.0552535 | 0.34595951 | 0.15971089 | 0.87310882 | 0.97564728 |
| METTL23 | 117.889537 | -0.2337266 | 0.37703086 | -0.6199136 | 0.53531467 | 0.8642718 |
| AC005837.2 | 139.779578 | -0.1905184 | 0.74728446 | -0.2549476 | 0.79876354 | 0.95300733 |
| SRSF2 | 959.260489 | -0.0394323 | 0.32148645 | -0.1226562 | 0.90237931 | 0.97941893 |
| MFSD11 | 470.964957 | -0.3457994 | 0.19353052 | -1.7867953 | 0.07397058 | 0.47070029 |
| SEC14L1 | 2651.09858 | 0.34231832 | 0.41244802 | 0.82996717 | 0.40655735 | 0.80499101 |
| 9-Sep | 2228.96195 | 0.02537519 | 0.41736844 | 0.06079805 | 0.95152005 | 0.98938795 |
| TNRC6C | 1117.56711 | -0.1904154 | 0.24824527 | -0.7670454 | 0.44305451 | 0.82310928 |
| TMC6 | 100.281404 | 0.83366564 | 0.39551834 | 2.10778 | 3.51E-02 | 3.54E-01 |
| SYNGR2 | 240.666767 | 0.41589746 | 0.45779302 | 0.90848362 | 0.36362276 | 0.77959635 |
| AFMID | 736.610429 | -0.8224765 | 0.36867222 | -2.2309152 | 0.02568675 | 0.31211455 |
| SOCS3 | 246.992854 | 0.6147189 | 0.60883234 | 1.0096686 | 3.13E-01 | 7.47E-01 |
| AC061992.1 | 50.7971284 | -0.1453951 | 0.86462812 | -0.1681591 | 8.66E-01 | 9.74E-01 |
| PGS1 | 429.505799 | -0.3925671 | 0.23566 | -1.6658197 | 0.09574934 | 0.5117083 |
| CYTH1 | 1038.19891 | -0.2130933 | 0.2641437 | -0.8067325 | 0.41982063 | 0.8125045 |
| USP36 | 528.541104 | -0.476006 | 0.19672207 | -2.4196876 | 0.01553384 | 0.25214021 |
| TIMP2 | 315.472849 | -0.1471148 | 0.3459141 | -0.425293 | 0.67062313 | 0.91501507 |
| LGALS3BP | 405.13426 | 1.27330124 | 0.59111805 | 2.15405575 | 0.0312358 | 0.3383722 |
| CANT1 | 122.791812 | 0.43813452 | 0.27062106 | 1.61899641 | 0.10544804 | 0.52640329 |
| ENGASE | 164.073658 | 0.07644915 | 0.25848193 | 0.29576206 | 0.7674118 | 0.94316227 |
| CBX8 | 45.0911669 | -1.2959739 | 0.62826005 | -2.0627985 | 0.03913177 | 0.37106359 |
| CBX4 | 167.510739 | -0.5331215 | 0.51912316 | -1.0269653 | 0.30443678 | 0.74152784 |
| TBC1D16 | 1687.90423 | -0.5842278 | 0.41431702 | -1.4100985 | 0.1585106 | 0.60804051 |
| CCDC40 | 441.456562 | -0.3204325 | 0.37155655 | -0.8624056 | 0.38846435 | 0.79634939 |
| GAA | 244.910735 | 0.10489687 | 0.3498868 | 0.2998023 | 0.76432796 | 0.94256396 |
| EIF4A3 | 163.502718 | 0.09569932 | 0.30223287 | 0.31664101 | 0.75151601 | 0.93799254 |
| AC087741.1 | 157.234853 | 0.20005222 | 0.46440795 | 0.43076829 | 0.66663686 | 0.91321713 |
| SGSH | 96.6675841 | 0.10777659 | 0.32763145 | 0.32895678 | 0.74218836 | 0.93598004 |
| SLC26A11 | 309.329242 | 0.07067996 | 0.20956222 | 0.33727432 | 0.73591012 | 0.93552616 |
| RNF213 | 3809.50191 | -0.249948 | 0.50948291 | -0.4905915 | 0.62371537 | 0.89873262 |
| AC124319.2 | 1980.11681 | 0.3877889 | 0.34615281 | 1.12028238 | 0.26259345 | 0.7132328 |
| ENDOV | 243.036802 | -0.3787567 | 0.28470285 | -1.330358 | 0.18340036 | 0.63834582 |
| RPTOR | 1279.90006 | -0.2882847 | 0.24188874 | -1.191807 | 0.23333694 | 0.68484327 |

| | | | | | | |
|---|---|---|---|---|---|---|
| CHMP6 | 85.3649991 | 0.34887956 | 0.36059996 | 0.9674975 | 0.33329539 | 0.76128287 |
| BAIAP2-DT | 59.0451244 | 1.28997194 | 0.56166007 | 2.29671293 | 0.02163515 | 0.29380356 |
| BAIAP2 | 145.550525 | 0.39978702 | 0.26534386 | 1.50667524 | 0.13189392 | 0.575549 |
| CEP131 | 71.9493221 | 0.01393203 | 0.28675281 | 0.04858551 | 0.96124962 | 0.99188706 |
| TEPSIN | 162.413305 | -0.2327518 | 0.35523812 | -0.6551995 | 0.51233933 | 0.85762435 |
| NDUFAF8 | 204.901567 | 0.27705021 | 0.41472733 | 0.66802978 | 0.50411458 | 0.85525037 |
| SLC38A10 | 596.631595 | 0.36722951 | 0.19542029 | 1.87917796 | 0.0602202 | 0.42858222 |
| AC110285.6 | 68.1940651 | -0.0683823 | 0.51838722 | -0.1319136 | 0.89505262 | 0.97788747 |
| AC110285.2 | 89.4538389 | 0.11437951 | 0.63139714 | 0.18115303 | 0.85624746 | 0.97118784 |
| BAHCC1 | 176.288498 | -0.0376467 | 0.42899508 | -0.0877556 | 0.93007095 | 0.98331361 |
| ACTG1 | 2192.28282 | 0.34262671 | 0.56558731 | 0.60578925 | 0.54465471 | 0.86654229 |
| AC139149.1 | 45.5030741 | -0.0264776 | 0.48712148 | -0.0543553 | 0.95665213 | 0.99065996 |
| FAAP100 | 102.533303 | -0.5470289 | 0.32226556 | -1.6974475 | 0.08961209 | 0.50007863 |
| NPLOC4 | 961.575432 | -0.2254625 | 0.2006132 | -1.1238667 | 0.26106957 | 0.71131764 |
| TSPAN10 | 49.6284123 | -0.3513939 | 0.41948242 | -0.8376845 | 0.40220792 | 0.8030176 |
| OXLD1 | 108.811718 | 0.23380603 | 0.29712843 | 0.7868854 | 0.43134896 | 0.81710736 |
| CCDC137 | 192.21501 | -0.2626475 | 0.28870449 | -0.9097453 | 0.36295686 | 0.77905381 |
| ARL16 | 222.563262 | 0.41556889 | 0.27034516 | 1.53717898 | 0.12424949 | 0.56105491 |
| HGS | 425.69348 | -0.5827296 | 0.20862451 | -2.7931983 | 0.00521897 | 0.14543748 |
| AC139530.1 | 38.2950303 | 0.28666332 | 0.38377681 | 0.7469532 | 0.45509181 | 0.83121393 |
| SLC25A10 | 41.6814542 | 0.70544791 | 0.45501912 | 1.5503698 | 0.12105278 | 0.55520354 |
| GCGR | 404.823385 | 0.13094447 | 0.52973779 | 0.24718733 | 8.05E-01 | 0.95500115 |
| MCRIP1 | 233.677222 | -0.0775778 | 0.32324853 | -0.2399942 | 0.81033472 | 0.95679908 |
| P4HB | 835.441259 | 0.52001827 | 0.25176113 | 2.06552246 | 0.03887359 | 0.36967695 |
| ARHGDIA | 412.675783 | 0.45746662 | 0.44053256 | 1.03843997 | 0.29906527 | 0.73673761 |
| ALYREF | 45.2912018 | 0.37810202 | 0.3856731 | 0.98036918 | 0.32690392 | 0.75556858 |
| ANAPC11 | 282.646232 | 0.56811953 | 0.36211739 | 1.56888221 | 0.11667539 | 0.54715005 |
| PCYT2 | 93.1716959 | 1.0684321 | 0.5206741 | 2.05201699 | 0.04016801 | 0.37344844 |
| SIRT7 | 210.60637 | -0.1400662 | 0.28677373 | -0.4884206 | 0.62525196 | 0.89873262 |
| MAFG | 132.013281 | -0.1889099 | 0.38682021 | -0.4883661 | 6.25E-01 | 0.89873262 |
| ASPSCR1 | 316.750447 | -0.6163136 | 0.32328251 | -1.9064242 | 0.05659519 | 0.41843207 |
| CENPX | 102.829378 | 0.06967406 | 0.39350721 | 0.17705916 | 0.85946192 | 0.97184883 |
| LRRC45 | 38.9797384 | -0.111354 | 0.52875549 | -0.2105964 | 0.83320223 | 0.96491042 |
| DCXR | 170.485798 | 0.37249225 | 0.33593295 | 1.10882913 | 0.2675039 | 0.71774508 |
| RFNG | 171.723807 | 0.10537863 | 0.27123377 | 0.38851588 | 0.6976343 | 0.92341984 |
| GPS1 | 184.086108 | 0.06835329 | 0.43282884 | 0.15792222 | 0.87451809 | 0.97564728 |
| DUS1L | 199.487173 | -0.1293192 | 0.33467435 | -0.386403 | 0.69919823 | 0.92367966 |
| FASN | 92.0084915 | 0.25558615 | 0.49965 | 0.51153038 | 0.60897972 | 0.89273938 |
| CCDC57 | 1236.74661 | -0.1682573 | 0.22315553 | -0.7539912 | 0.45085453 | 0.82813014 |
| SLC16A3 | 268.604944 | 0.53449359 | 0.40829328 | 1.30909231 | 0.19050309 | 0.64559718 |
| CSNK1D | 695.886206 | -0.2743221 | 0.23921072 | -1.1467802 | 0.25147246 | 0.70276117 |
| AC132872.1 | 63.5985592 | -0.1486674 | 0.37266262 | -0.3989329 | 0.68994265 | 0.92176114 |
| SECTM1 | 239.45143 | -0.2253225 | 0.97098393 | -0.2320558 | 0.81649465 | 0.9589293 |
| OGFOD3 | 230.029362 | -0.0326671 | 0.33602035 | -0.0972176 | 0.92255357 | 0.98215577 |
| HEXDC | 495.38968 | -0.6027408 | 0.20625332 | -2.9223327 | 0.0034742 | 0.1138728 |
| CYBC1 | 231.472489 | 0.22388908 | 0.31777665 | 0.70454856 | 0.48109122 | 0.8437862 |
| NARF | 439.082986 | -0.2552191 | 0.1899144 | -1.343864 | 0.17899237 | 0.63284534 |
| FOXK2 | 1316.76274 | -0.4528094 | 0.19653462 | -2.3039678 | 0.02122445 | 0.29112539 |
| WDR45B | 823.982964 | -0.7324704 | 0.34673745 | -2.112464 | 0.03464667 | 0.35339604 |
| RAB40B | 617.812742 | 0.24859206 | 0.42671296 | 0.58257443 | 0.56017982 | 0.87329343 |
| FN3KRP | 152.726633 | -0.5103937 | 0.19132502 | -2.6676786 | 0.00763773 | 0.17527709 |
| FN3K | 386.39203 | 0.12692361 | 0.66381679 | 0.19120278 | 0.84836673 | 0.96988715 |
| TBCD | 571.657016 | -0.3832748 | 0.28223845 | -1.3579821 | 0.17446935 | 0.62897534 |
| B3GNTL1 | 79.9889928 | -0.3408622 | 0.48772515 | -0.6988817 | 0.48462596 | 0.8447301 |

| | | | | | | |
|---|---|---|---|---|---|---|
| METRNL | 167.196182 | 0.08633699 | 0.291024 | 0.29666621 | 0.76672135 | 0.94312902 |
| USP14 | 873.608623 | -0.1964341 | 0.15596084 | -1.2595091 | 0.2078465 | 0.66075678 |
| THOC1 | 852.018622 | -0.3780508 | 0.19981321 | -1.892021 | 0.05848819 | 0.42147872 |
| CLUL1 | 43.5046551 | 0.67447324 | 0.45167721 | 1.49326383 | 0.13536814 | 0.57708028 |
| TYMS | 56.1483394 | 0.69468103 | 0.55667015 | 1.24792217 | 0.21205956 | 0.66552984 |
| ENOSF1 | 2871.62286 | 0.25673702 | 0.45007513 | 0.57043147 | 0.56838509 | 0.87757867 |
| YES1 | 1059.85039 | -0.348761 | 0.20801311 | -1.6766297 | 0.09361491 | 0.50887217 |
| LINC00470 | 107.912308 | 1.19236585 | 1.05640274 | 1.12870386 | 0.25902278 | 0.70921184 |
| AP005230.1 | 50.3183338 | -0.6771486 | 0.61370285 | -1.1033818 | 0.26986139 | 0.71881411 |
| METTL4 | 342.470134 | -0.0541515 | 0.28610066 | -0.1892743 | 0.84987785 | 0.97041641 |
| SMCHD1 | 2202.02491 | -0.2278107 | 0.21437426 | -1.0626774 | 0.2879283 | 0.7307234 |
| AP001011.1 | 376.31982 | 0.21937295 | 0.60689283 | 0.36146901 | 0.71774887 | 0.92945725 |
| LPIN2 | 927.651312 | -0.5818826 | 0.39189294 | -1.4847998 | 0.13759686 | 0.58049087 |
| AP000919.2 | 87.2054443 | -0.5703195 | 0.66339204 | -0.8597021 | 0.38995329 | 0.79698174 |
| MYOM1 | 193.414983 | -0.533785 | 0.51410286 | -1.0382844 | 0.29913768 | 0.73673761 |
| AP005329.2 | 47.218921 | -0.6173988 | 0.67669891 | -0.9123686 | 0.36157471 | 0.77772231 |
| MYL12A | 646.134641 | 0.40619139 | 0.40143758 | 1.01184198 | 0.31161361 | 0.74607653 |
| MYL12B | 1310.08155 | 0.58964207 | 0.38352403 | 1.5374319 | 0.12418758 | 0.56105491 |
| TGIF1 | 357.369665 | -0.8696613 | 0.26361476 | -3.2989854 | 0.00097035 | 0.05572759 |
| DLGAP1 | 348.87187 | 0.94923032 | 0.57435527 | 1.65268845 | 0.09839429 | 0.51525577 |
| DLGAP1-AS1 | 132.124283 | -0.385565 | 0.33817739 | -1.1401266 | 0.25423357 | 0.70475787 |
| DLGAP1-AS2 | 60.0017235 | -0.5710771 | 0.44203643 | -1.2919231 | 0.19638377 | 0.64964978 |
| LINC00667 | 210.726671 | -0.2860392 | 0.31664595 | -0.9033408 | 0.36634503 | 0.78095617 |
| ZBTB14 | 42.2953374 | -0.3540111 | 0.44490123 | -0.7957072 | 0.42620223 | 0.81466629 |
| EPB41L3 | 1102.21706 | 0.801032 | 0.4614665 | 1.73584001 | 0.08259213 | 0.48594425 |
| L3MBTL4 | 578.832343 | 1.04887611 | 0.39260134 | 2.67160607 | 0.00754892 | 0.17483887 |
| LAMA1 | 2507.34339 | -0.8017392 | 0.41348017 | -1.9390027 | 0.05250101 | 0.40937769 |
| PTPRM | 8444.64438 | 0.07957048 | 0.23188846 | 0.34314119 | 0.73149225 | 0.93529014 |
| RAB12 | 727.228163 | -0.3260713 | 0.16781188 | -1.9430761 | 0.05200696 | 0.40866415 |
| MTCL1 | 370.977776 | 0.51163969 | 0.56582171 | 0.90424188 | 0.36586716 | 0.78095617 |
| NDUFV2 | 786.926981 | -0.117792 | 0.22114726 | -0.5326404 | 0.59428252 | 0.88819618 |
| AP005899.1 | 62.0059729 | -0.3577713 | 0.41745853 | -0.8570224 | 0.39143251 | 0.7973532 |
| NDUFV2-AS1 | 124.337677 | 0.32265253 | 0.36128355 | 0.89307286 | 0.37181815 | 0.78231061 |
| ANKRD12 | 4075.63798 | -0.1158389 | 0.21004725 | -0.5514895 | 0.58129815 | 0.88290096 |
| TWSG1 | 372.225992 | -0.1421234 | 0.19834537 | -0.7165449 | 0.47365495 | 0.84064485 |
| RALBP1 | 674.466177 | 0.26435365 | 0.13250076 | 1.99511041 | 0.04603084 | 0.39025761 |
| PPP4R1 | 1089.52513 | -0.3263693 | 0.25298814 | -1.2900578 | 0.19703058 | 0.64964978 |
| RAB31 | 330.788208 | -1.461074 | 0.35058083 | -4.1675809 | 3.08E-05 | 0.00860682 |
| VAPA | 2236.22851 | -0.1624517 | 0.34492748 | -0.4709735 | 0.63765964 | 0.90267348 |
| NAPG | 774.800092 | -0.433581 | 0.25800004 | -1.6805461 | 0.09285111 | 0.50743497 |
| GNAL | 230.020253 | 0.50384112 | 0.26552583 | 1.89752207 | 5.78E-02 | 4.21E-01 |
| MPPE1 | 677.895007 | -0.6016553 | 0.21633326 | -2.7811503 | 0.00541667 | 0.14905986 |
| IMPA2 | 1606.21106 | -0.0446772 | 0.41213244 | -0.1084049 | 0.91367454 | 0.98145965 |
| TUBB6 | 218.257806 | -0.9857848 | 0.39902129 | -2.4705068 | 1.35E-02 | 0.23970866 |
| AFG3L2 | 1083.83187 | -0.1177152 | 0.16921541 | -0.6956529 | 0.48664622 | 0.84474442 |
| SPIRE1 | 2734.12494 | 0.33999435 | 0.32110424 | 1.05882859 | 0.28967785 | 0.73204984 |
| PSMG2 | 617.950177 | 0.51008522 | 0.25228801 | 2.02183696 | 0.0431932 | 0.38262319 |
| CEP76 | 185.250386 | 0.12109965 | 0.30950978 | 0.39126276 | 0.69560302 | 0.92229578 |
| PTPN2 | 1515.10797 | -0.1980461 | 0.2830643 | -0.6996507 | 0.48414545 | 0.8445691 |
| SEH1L | 298.480267 | -0.3330148 | 0.2274455 | -1.4641523 | 0.14315234 | 0.5883902 |
| CEP192 | 1623.92784 | -0.5017556 | 0.22041483 | -2.2764148 | 0.02282119 | 0.2983102 |
| LDLRAD4 | 3791.23316 | -0.1701341 | 0.43057417 | -0.3951332 | 0.69274462 | 0.92224682 |
| AP005131.1 | 67.1211858 | -1.204602 | 0.92063731 | -1.3084436 | 0.19072291 | 0.64567512 |
| FAM210A | 504.274307 | -0.2383871 | 0.40806185 | -0.5841935 | 0.55909015 | 0.87295536 |

| | | | | | | |
|---|---|---|---|---|---|---|
| RNMT | 911.635913 | -0.23739 | 0.34902404 | -0.6801537 | 0.49640717 | 0.85068012 |
| ZNF519 | 88.8476307 | -0.5271257 | 0.54925805 | -0.9597049 | 0.33720374 | 0.76409325 |
| ANKRD30B | 59.3315475 | -0.0723359 | 0.64204267 | -0.1126652 | 0.91029598 | 0.98069436 |
| ROCK1 | 2030.76975 | -0.1326086 | 0.13883867 | -0.9551275 | 0.33951323 | 0.76449297 |
| GREB1L | 137.604427 | -0.5157708 | 0.40537919 | -1.272317 | 0.20326053 | 0.65628656 |
| ESCO1 | 939.524926 | 0.04775813 | 0.21194825 | 0.2253292 | 0.82172319 | 0.96066244 |
| SNRPD1 | 203.450376 | -0.0256197 | 0.29374949 | -0.0872163 | 0.9304996 | 0.98331361 |
| ABHD3 | 701.644977 | -0.1589281 | 0.24301545 | -0.6539836 | 0.51312238 | 0.85762435 |
| MIB1 | 2090.21813 | -0.272776 | 0.22270366 | -1.2248385 | 0.22063607 | 0.67345736 |
| AC090912.1 | 226.206426 | 0.2905698 | 0.42183649 | 0.68882093 | 0.49093596 | 0.84635515 |
| RBBP8 | 3213.18196 | 0.29041137 | 0.2884778 | 1.00670265 | 0.31407769 | 0.74777437 |
| CABLES1 | 798.055701 | -0.8750991 | 0.35615158 | -2.4570973 | 1.40E-02 | 0.24537335 |
| TMEM241 | 430.728831 | -0.3609044 | 0.22809004 | -1.5822891 | 1.14E-01 | 0.53989968 |
| RIOK3 | 1145.51647 | 0.04485143 | 0.28302977 | 0.15846894 | 0.87408729 | 0.97564728 |
| RMC1 | 324.422455 | -0.2990442 | 0.22604044 | -1.3229675 | 0.1858462 | 0.64030106 |
| NPC1 | 1243.94798 | -0.734423 | 0.30372361 | -2.4180637 | 0.01560334 | 0.25227487 |
| LAMA3 | 142.704375 | 0.02398258 | 0.79957211 | 0.02999427 | 0.97607162 | 0.9941831 |
| TTC39C | 549.815484 | 0.02395615 | 0.38169065 | 0.06276327 | 0.94995501 | 0.98919519 |
| OSBPL1A | 810.583792 | 0.21465133 | 0.3232871 | 0.66396504 | 0.50671269 | 0.85626196 |
| IMPACT | 172.671391 | 0.27540921 | 0.24368906 | 1.1301665 | 0.25840607 | 0.70896455 |
| SS18 | 1693.82943 | -0.0585481 | 0.26085742 | -0.2244449 | 0.8224111 | 0.96083745 |
| TAF4B | 347.584518 | -0.0791493 | 0.14077637 | -0.5622343 | 5.74E-01 | 8.80E-01 |
| KCTD1 | 4740.78716 | 0.41691702 | 0.24014394 | 1.73611305 | 0.08254385 | 0.48594425 |
| AQP4-AS1 | 540.483365 | 0.69681265 | 0.25938895 | 2.68636213 | 0.00722348 | 0.1729679 |
| AQP4 | 66.2334911 | 1.28814084 | 0.98948655 | 1.30182754 | 0.19297535 | 0.64699503 |
| CHST9 | 887.165174 | 0.8828728 | 0.42489405 | 2.07786578 | 0.03772172 | 0.36396365 |
| CDH2 | 61.3545031 | 0.50992543 | 0.54312569 | 0.93887187 | 0.34779653 | 0.76900643 |
| DSC2 | 180.515234 | 0.41616985 | 0.33380206 | 1.24675638 | 0.21248684 | 0.66620904 |
| DSG2 | 530.131344 | 0.98851303 | 0.31920091 | 3.09683646 | 0.00195598 | 0.08583557 |
| B4GALT6 | 565.982561 | -0.7531666 | 0.3798365 | -1.9828704 | 0.0473819 | 0.39411174 |
| TRAPPC8 | 1464.38105 | -0.1752926 | 0.23586023 | -0.7432054 | 0.4573573 | 0.83255882 |
| RNF138 | 529.151428 | -0.1722233 | 0.2097094 | -0.8212473 | 0.41150545 | 0.80675684 |
| GAREM1 | 4032.47848 | 0.02902442 | 0.26881935 | 0.10796997 | 0.91401951 | 0.98145965 |
| AC025887.2 | 151.176414 | 0.5627284 | 0.45339375 | 1.24114726 | 0.21455135 | 0.66827012 |
| KLHL14 | 628.618421 | 0.16596686 | 0.54154118 | 0.30647136 | 0.75924578 | 0.94133292 |
| CCDC178 | 2880.5269 | 2.44996282 | 0.57874082 | 4.23326421 | 2.30E-05 | 0.00780088 |
| ASXL3 | 1222.63467 | 0.19707014 | 0.30087446 | 0.65499125 | 0.51247339 | 0.85762435 |
| DTNA | 444.125084 | 1.54067876 | 0.59018428 | 2.61050459 | 0.00904088 | 0.19323775 |
| MAPRE2 | 378.74909 | 0.25119115 | 0.35722618 | 0.70317118 | 0.48194908 | 0.84389635 |
| ZNF397 | 555.50555 | -0.0427853 | 0.20704157 | -0.2066509 | 0.83628248 | 0.96511435 |
| ZSCAN30 | 432.884829 | 0.04991301 | 0.164118 | 0.30412879 | 0.76102977 | 0.94202764 |
| AC011815.2 | 47.9219054 | 0.03237189 | 0.59186163 | 0.05469504 | 0.95638142 | 0.99065996 |
| ZNF24 | 455.127954 | 0.14167489 | 0.19846405 | 0.71385669 | 0.4753158 | 0.84118672 |
| ZNF396 | 69.8001524 | 0.10845414 | 0.29830343 | 0.36356987 | 0.71617923 | 0.92890921 |
| GALNT1 | 632.885245 | -0.1955573 | 0.2162855 | -0.9041628 | 0.36590908 | 0.78095617 |
| C18orf21 | 58.5855885 | -0.0051323 | 0.37829046 | -0.013567 | 0.98917544 | 0.99670729 |
| RPRD1A | 1386.62481 | -0.1452627 | 0.18327366 | -0.7926002 | 0.42801078 | 0.81475637 |
| SLC39A6 | 212.63357 | 0.14952436 | 0.24621128 | 0.607301 | 0.54365117 | 0.86633953 |
| ELP2 | 839.936817 | -0.1478962 | 0.13824718 | -1.0697956 | 0.28471135 | 0.72769972 |
| MOCOS | 162.910263 | -0.4987599 | 0.5601517 | -0.8904015 | 3.73E-01 | 7.83E-01 |
| FHOD3 | 4349.48612 | -0.0772687 | 0.36100353 | -0.2140386 | 0.83051694 | 0.9635604 |
| TPGS2 | 500.815014 | -0.0486503 | 0.20268375 | -0.2400305 | 8.10E-01 | 0.95679908 |
| KIAA1328 | 2474.04741 | -0.5113119 | 0.22023789 | -2.3216348 | 0.02025261 | 0.28534319 |
| AC016493.1 | 95.1980603 | -0.112671 | 0.78042757 | -0.1443708 | 0.88520765 | 0.97664137 |

| | | | | | | |
|---|---|---|---|---|---|---|
| CELF4 | 60.7230313 | -0.1979013 | 0.6100934 | -0.3243787 | 0.74565132 | 0.93675395 |
| AC015961.2 | 44.8001385 | -0.4526001 | 0.35814819 | -1.2637229 | 0.2063295 | 0.65831072 |
| AC016205.1 | 195.581374 | 0.8205201 | 0.44363706 | 1.84953012 | 0.0643813 | 0.43948973 |
| AC021504.1 | 94.7384771 | -0.1484205 | 0.95269641 | -0.1557899 | 0.87619865 | 0.97614152 |
| PIK3C3 | 1847.52618 | -0.2942534 | 0.1436834 | -2.0479292 | 0.04056694 | 0.37344844 |
| LINC00907 | 376.738598 | -0.5206562 | 0.56918348 | -0.9147423 | 0.36032692 | 0.77716943 |
| SETBP1 | 7432.25649 | 0.16516995 | 0.23934008 | 0.69010567 | 0.49012774 | 0.84614483 |
| SLC14A2 | 187.821958 | 0.62931177 | 0.52568099 | 1.19713624 | 0.23125346 | 0.68357541 |
| EPG5 | 935.44854 | -0.2570267 | 0.10508313 | -2.4459365 | 0.01444765 | 0.24762526 |
| PSTPIP2 | 311.437514 | 0.62511891 | 0.41047704 | 1.52290833 | 0.12778164 | 0.56762585 |
| ATP5F1A | 1024.40522 | 0.87554895 | 0.32823615 | 2.66743605 | 0.00764324 | 0.17527709 |
| HAUS1 | 151.031458 | -0.8907433 | 0.28809957 | -3.0917896 | 0.00198954 | 0.0861633 |
| C18orf25 | 691.809516 | 0.05932714 | 0.2235966 | 0.26533114 | 0.79075438 | 0.95144272 |
| RNF165 | 246.699653 | 0.51707914 | 0.36441115 | 1.41894435 | 1.56E-01 | 0.60519671 |
| ST8SIA5 | 396.533033 | 0.32897929 | 0.4230488 | 0.77763909 | 0.43678181 | 0.82013459 |
| PIAS2 | 2759.30838 | -0.1318965 | 0.31507519 | -0.4186191 | 0.67549453 | 0.91649321 |
| KATNAL2 | 1639.98011 | -0.035545 | 0.35971804 | -0.0988134 | 0.92128641 | 0.98215577 |
| AC012254.1 | 129.253662 | -1.419719 | 0.46785423 | -3.0345327 | 0.00240909 | 0.09386908 |
| HDHD2 | 72.2557816 | 0.88112687 | 0.33369481 | 2.64051714 | 0.00827796 | 0.18347203 |
| SMAD2 | 2261.51951 | -0.2271871 | 0.18184934 | -1.249315 | 0.21154989 | 0.66484951 |
| ZBTB7C | 152.466615 | -0.3734862 | 0.90655375 | -0.4119847 | 0.68035067 | 0.91867756 |
| CTIF | 1261.99798 | 0.03794919 | 0.27302561 | 0.13899497 | 0.88945412 | 0.97729111 |
| SMAD7 | 307.579493 | -0.6241909 | 0.43003871 | -1.451476 | 0.14664736 | 0.59373375 |
| DYM | 3444.43045 | -0.0122535 | 0.14297058 | -0.0857067 | 0.93169956 | 0.98357046 |
| C18orf32 | 49.0552897 | 0.08221054 | 0.29155143 | 0.2819761 | 0.77796184 | 0.94694572 |
| LIPG | 472.787542 | -0.0228224 | 0.66299517 | -0.0344232 | 9.73E-01 | 0.99327086 |
| ACAA2 | 423.624469 | 0.41037758 | 0.39824416 | 1.03046728 | 0.3027907 | 0.73961113 |
| SNHG22 | 362.981024 | 0.19740558 | 0.30902747 | 0.63879624 | 0.5229555 | 0.86006369 |
| MYO5B | 1249.73833 | -0.1844962 | 0.28263514 | -0.6527716 | 0.51390355 | 0.85762435 |
| CFAP53 | 118.989454 | 0.31710518 | 0.32455051 | 0.97705959 | 0.32853965 | 0.75696499 |
| MBD1 | 281.434729 | -0.0323001 | 0.18004683 | -0.1793986 | 0.85762476 | 0.97127317 |
| CXXC1 | 178.558595 | -0.2193671 | 0.28346925 | -0.7738655 | 0.43901035 | 0.82136595 |
| MAPK4 | 337.62477 | 1.27011405 | 0.72988819 | 1.7401488 | 0.08183289 | 0.48594425 |
| MRO | 41.9240117 | 3.09105775 | 1.05490841 | 2.93016695 | 0.0033878 | 0.11328973 |
| ME2 | 487.767294 | 0.09881846 | 0.23178784 | 0.42633152 | 0.6698663 | 0.91465659 |
| AC091551.1 | 230.202602 | -0.2419177 | 0.40772045 | -0.593342 | 0.55295228 | 0.87104567 |
| SMAD4 | 670.240165 | -0.08895 | 0.27405431 | -0.3245708 | 0.74550596 | 0.93675395 |
| MEX3C | 157.551352 | 0.12298858 | 0.19037652 | 0.64602811 | 0.51826115 | 0.85916831 |
| DCC | 78.6961588 | 0.19540224 | 0.41192692 | 0.47436141 | 0.63524221 | 0.90199374 |
| AC103949.1 | 92.950734 | -0.7754462 | 0.77085487 | -1.0059561 | 0.31443669 | 0.74785672 |
| MBD2 | 1525.98783 | -0.2435662 | 0.17700341 | -1.3760539 | 1.69E-01 | 6.23E-01 |
| POLI | 572.276781 | 0.12907025 | 0.22231449 | 0.58057508 | 0.56152687 | 0.87403145 |
| TCF4 | 366.884848 | -1.2382071 | 0.55942881 | -2.2133417 | 0.0268741 | 0.31710856 |
| AC009271.1 | 65.6734038 | -0.390208 | 0.79575432 | -0.4903624 | 0.62387751 | 0.89873262 |
| TXNL1 | 562.045762 | -0.1691814 | 0.17243057 | -0.9811564 | 0.32651561 | 0.75554697 |
| WDR7 | 1895.32349 | -0.1997982 | 0.15934669 | -1.2538582 | 0.20989355 | 0.66255855 |
| FECH | 540.065372 | 0.42226204 | 0.35071005 | 1.20402035 | 2.29E-01 | 6.80E-01 |
| NARS | 698.697779 | 0.31521048 | 0.25066008 | 1.25752168 | 0.2085648 | 0.6614379 |
| AC027097.2 | 527.444805 | -0.7607591 | 0.25491035 | -2.984418 | 0.00284118 | 0.10088491 |
| ATP8B1 | 1490.30356 | -0.4628352 | 0.21950707 | -2.108521 | 0.03498595 | 0.35358029 |
| NEDD4L | 7701.71628 | -0.1941838 | 0.27055346 | -0.7177278 | 4.73E-01 | 0.83985854 |
| MALT1 | 694.708053 | -0.392125 | 0.17646936 | -2.2220569 | 0.02627946 | 0.31391341 |
| AC104365.1 | 120.60686 | -0.4421344 | 0.68225946 | -0.6480444 | 0.51695622 | 0.85893097 |
| ZNF532 | 2013.54117 | -0.0371565 | 0.36214755 | -0.1026005 | 0.91828001 | 0.98210868 |

| | | | | | | |
|---|---|---|---|---|---|---|
| SEC11C | 203.442167 | 0.05538576 | 0.37080975 | 0.14936435 | 0.88126614 | 0.97664137 |
| LMAN1 | 702.181475 | 0.07435322 | 0.23872851 | 0.31145514 | 0.75545464 | 0.93953492 |
| CDH20 | 99.3094568 | 1.12738375 | 0.65179549 | 1.7296587 | 0.08369127 | 0.48884683 |
| AC105094.2 | 116.480708 | 0.48174229 | 0.65970196 | 0.73024232 | 0.46524208 | 0.83652911 |
| RNF152 | 2058.78316 | 0.24244011 | 0.64383265 | 0.37655765 | 0.70650236 | 0.92556842 |
| PIGN | 1442.66438 | -0.1209421 | 0.13205813 | -0.9158247 | 0.35975884 | 0.77671671 |
| KIAA1468 | 1517.76908 | -0.1537631 | 0.15943574 | -0.9644208 | 0.33483501 | 0.76242642 |
| ZCCHC2 | 750.522493 | 0.11310742 | 0.20693741 | 0.5465779 | 0.58466876 | 0.88453666 |
| PHLPP1 | 1501.69716 | -0.4291157 | 0.1261942 | -3.4004392 | 0.00067278 | 0.04534035 |
| BCL2 | 3254.3508 | 0.55264484 | 0.23544473 | 2.34723805 | 0.01891317 | 0.27727416 |
| KDSR | 616.01125 | 0.01671811 | 0.22350766 | 0.07479882 | 9.40E-01 | 9.87E-01 |
| AC036176.1 | 57.4139762 | 0.11732205 | 0.40553233 | 0.28930381 | 0.7723489 | 0.94509888 |
| VPS4B | 675.88967 | -0.0399939 | 0.15317056 | -0.2611072 | 0.79400985 | 0.95224951 |
| SERPINB8 | 82.1081443 | -0.4688832 | 0.54363534 | -0.8624958 | 3.88E-01 | 7.96E-01 |
| TMX3 | 726.240092 | -0.5162743 | 0.23784232 | -2.1706577 | 0.02995705 | 0.33256306 |
| CCDC102B | 277.643011 | 0.55996259 | 0.53868189 | 1.03950512 | 0.29856988 | 0.73673761 |
| DOK6 | 85.2766772 | -0.1432016 | 0.3425691 | -0.4180226 | 0.67593057 | 0.91650575 |
| CD226 | 183.540107 | -1.8170701 | 0.66219311 | -2.7440185 | 0.00606921 | 0.15969766 |
| RTTN | 914.510869 | -0.8244264 | 0.30291482 | -2.7216441 | 0.00649581 | 0.16670031 |
| SOCS6 | 377.195414 | -0.0469484 | 0.28257122 | -0.166147 | 0.86804125 | 0.97402505 |
| FBXO15 | 133.88376 | 0.03041201 | 0.45560425 | 0.06675094 | 0.94677998 | 0.98893063 |
| TIMM21 | 149.484083 | -0.0741186 | 0.2445215 | -0.3031167 | 0.76180089 | 0.94210219 |
| CYB5A | 1329.56114 | 0.87630805 | 0.27183177 | 3.22371458 | 0.00126539 | 0.06478526 |
| CNDP2 | 1543.54137 | 0.65053044 | 0.28196457 | 2.30713539 | 2.10E-02 | 0.29020191 |
| LINC00909 | 159.522313 | 0.13464469 | 0.33422799 | 0.40285281 | 0.68705651 | 0.9208231 |
| ZNF407 | 2698.7301 | -0.4116411 | 0.16385717 | -2.5121945 | 0.0119983 | 0.22384955 |
| ZADH2 | 389.296599 | 0.46945318 | 0.18910712 | 2.48247224 | 0.01304742 | 0.23480029 |
| TSHZ1 | 879.363628 | -0.3049003 | 0.26523522 | -1.1495469 | 0.25033052 | 0.70099977 |
| ZNF516 | 997.771935 | 0.06376228 | 0.29195197 | 0.21839991 | 0.82711754 | 0.96233555 |
| ZNF236-DT | 99.3266298 | 0.03084216 | 0.30881817 | 0.09987158 | 0.92044628 | 0.98215577 |
| ZNF236 | 659.930437 | -0.6243062 | 0.21872633 | -2.85428 | 0.00431345 | 0.12803674 |
| MBP | 1338.98066 | -0.2308053 | 0.33841095 | -0.6820268 | 0.495222 | 0.84919997 |
| ATP9B | 2284.95186 | -0.1592694 | 0.25590875 | -0.6223678 | 0.53370005 | 0.86397197 |
| NFATC1 | 191.507799 | 0.34123848 | 0.41999869 | 0.8124751 | 0.41651907 | 0.80966301 |
| AC068473.1 | 42.832726 | -0.3091114 | 0.57617438 | -0.5364893 | 0.59162042 | 0.88722913 |
| CTDP1 | 236.11813 | -0.4686244 | 0.25203356 | -1.859373 | 0.06297428 | 0.43619679 |
| PQLC1 | 306.979662 | 0.3354821 | 0.17341179 | 1.934598 | 0.05303965 | 0.41073851 |
| HSBP1L1 | 392.818803 | 0.20766503 | 0.26627245 | 0.77989679 | 0.43545163 | 0.8193562 |
| TXNL4A | 480.120652 | -0.0754024 | 0.17405702 | -0.4332051 | 0.66486576 | 0.91224803 |
| ADNP2 | 567.656206 | -0.0355422 | 0.24910429 | -0.14268 | 0.88654293 | 0.97716502 |
| PARD6G-AS1 | 106.905187 | 0.8702041 | 0.28850735 | 3.01622857 | 0.0025594 | 0.09563507 |
| PARD6G | 58.8126893 | 0.22053422 | 0.41773406 | 0.5279297 | 0.59754813 | 0.88883692 |
| C20orf96 | 143.233683 | -0.0544963 | 0.3428997 | -0.1589277 | 0.87372581 | 0.97564728 |
| ZCCHC3 | 104.985969 | 0.17144779 | 0.32946157 | 0.52038781 | 0.6027933 | 0.88971776 |
| NRSN2-AS1 | 750.83376 | 0.03710368 | 0.37232433 | 0.0996542 | 0.92061886 | 0.98215577 |
| SOX12 | 128.590211 | 0.34554124 | 0.36163613 | 0.95549426 | 0.33932781 | 0.76449297 |
| NRSN2 | 137.727307 | -0.227217 | 0.34420553 | -0.66012 | 0.5091768 | 0.85677915 |
| RBCK1 | 459.134858 | 0.0929087 | 0.24542128 | 0.37856823 | 0.70500852 | 0.92532368 |
| TBC1D20 | 277.299173 | -0.1504439 | 0.21360717 | -0.7043019 | 0.48124478 | 0.8437862 |
| CSNK2A1 | 1563.73373 | -0.0320637 | 0.14222459 | -0.2254443 | 0.82163367 | 0.96066244 |
| RSPO4 | 101.028042 | -1.0918253 | 0.6378449 | -1.7117411 | 0.0869444 | 0.49480766 |
| PSMF1 | 651.159979 | -0.1020262 | 0.23539257 | -0.4334299 | 0.66470247 | 0.91220795 |
| AL136531.2 | 341.571063 | -0.3012538 | 0.33419602 | -0.9014285 | 0.36736051 | 0.78095617 |
| SDCBP2-AS1 | 517.178964 | -0.3438772 | 0.18013885 | -1.9089564 | 0.05626772 | 0.41843207 |

| | | | | | | |
|---|---|---|---|---|---|---|
| NSFL1C | 260.430654 | 0.04656621 | 0.30034802 | 0.15504084 | 0.87678912 | 0.9763537 |
| SIRPA | 161.031591 | -0.1468853 | 0.41584813 | -0.3532185 | 0.72392463 | 0.93201977 |
| STK35 | 332.73591 | -0.1861048 | 0.22425623 | -0.8298756 | 0.40660913 | 0.80499101 |
| SNRPB | 251.182304 | 0.32399772 | 0.27103269 | 1.19541934 | 2.32E-01 | 0.68442225 |
| AL049650.1 | 75.2639601 | 0.18612053 | 0.29217031 | 0.63702751 | 0.52410693 | 0.86006369 |
| NOP56 | 186.189556 | -0.3091164 | 0.44405056 | -0.6961289 | 0.48634813 | 0.8447301 |
| IDH3B | 165.754476 | 0.25617826 | 0.36572532 | 0.7004663 | 0.48363614 | 0.84454356 |
| EBF4 | 222.312473 | -0.200542 | 0.44468779 | -0.4509726 | 0.6520093 | 0.90838511 |
| PCED1A | 192.310773 | -0.2180548 | 0.24138598 | -0.903345 | 0.36634281 | 0.78095617 |
| VPS16 | 76.190027 | -0.5490737 | 0.36544068 | -1.5024975 | 0.13296868 | 0.57682553 |
| PTPRA | 1794.87321 | -0.1506211 | 0.17892529 | -0.84181 | 0.39989431 | 0.80215701 |
| MRPS26 | 230.343499 | 0.31370545 | 0.33044969 | 0.94932894 | 0.34245334 | 0.76591028 |
| UBOX5-AS1 | 179.300496 | 0.06224472 | 0.41893505 | 0.14857846 | 0.88188627 | 0.97664137 |
| UBOX5 | 199.437473 | -0.3566495 | 0.29902399 | -1.1927118 | 0.23298225 | 0.68484327 |
| LZTS3 | 230.677856 | -0.1084279 | 0.40385337 | -0.2684833 | 0.7883273 | 0.95035277 |
| DDRGK1 | 235.909633 | -0.0277294 | 0.21430129 | -0.1293946 | 8.97E-01 | 9.78E-01 |
| ITPA | 415.850788 | -0.3873791 | 0.26187052 | -1.4792773 | 0.13906621 | 0.58243213 |
| C20orf194 | 4610.63663 | 0.12854906 | 0.27116697 | 0.47405869 | 0.63545806 | 0.90199374 |
| ATRN | 1971.0916 | -0.2895067 | 0.14013424 | -2.0659243 | 3.88E-02 | 0.36967695 |
| CENPB | 143.639317 | 0.07063642 | 0.38612806 | 0.18293522 | 8.55E-01 | 0.97116182 |
| CDC25B | 166.939468 | -0.6054313 | 0.35402987 | -1.7101137 | 0.08724485 | 0.49566631 |
| AP5S1 | 63.4231137 | -0.2712115 | 0.42526333 | -0.6377496 | 0.52363668 | 0.86006369 |
| MAVS | 840.448713 | -0.0110117 | 0.20674123 | -0.0532632 | 0.9575222 | 0.99065996 |
| AL353194.1 | 53.4011007 | 0.51649156 | 0.51194463 | 1.00888167 | 0.31303139 | 0.74703371 |
| PANK2 | 719.156923 | -0.4048974 | 0.29345594 | -1.3797552 | 0.16766202 | 0.62191735 |
| RNF24 | 2883.32381 | -0.414507 | 0.44148031 | -0.9389026 | 0.34778077 | 0.76900643 |
| SMOX | 73.986843 | -0.7471553 | 0.45247385 | -1.6512674 | 9.87E-02 | 5.15E-01 |
| PRNP | 183.264152 | 0.51845394 | 0.3529111 | 1.46907804 | 0.14181162 | 0.5872148 |
| SLC23A2 | 539.771327 | -0.0969141 | 0.25024965 | -0.3872697 | 0.69855655 | 0.92356678 |
| TMEM230 | 443.574464 | 0.13693134 | 0.18495951 | 0.74033145 | 0.45909891 | 0.83380138 |
| PCNA | 121.418037 | 0.95649623 | 0.38720533 | 2.47025582 | 0.01350165 | 0.23970866 |
| CDS2 | 575.150966 | -0.0518807 | 0.13634278 | -0.3805166 | 0.70356196 | 0.92513032 |
| GPCPD1 | 1455.6764 | 0.26772607 | 0.31271467 | 0.85613533 | 0.39192293 | 0.79767768 |
| SHLD1 | 591.293409 | -0.6314281 | 0.17823836 | -3.5426052 | 0.0003962 | 0.03398276 |
| TRMT6 | 97.1221972 | -0.4597777 | 0.39236544 | -1.1718099 | 0.24127337 | 0.69191922 |
| MCM8 | 225.501268 | -1.092401 | 0.30457967 | -3.5865853 | 0.00033504 | 0.03094754 |
| CRLS1 | 945.92075 | -0.535018 | 0.33172399 | -1.6128409 | 0.10677908 | 0.52778894 |
| TMX4 | 691.868465 | -0.173977 | 0.40570949 | -0.4288216 | 0.66805307 | 0.91383898 |
| PLCB1 | 33273.0819 | -0.2993706 | 0.41351611 | -0.7239636 | 0.46908811 | 0.83870782 |
| PLCB4 | 445.98564 | 1.11087769 | 0.52007859 | 2.13598042 | 0.03268101 | 0.34507071 |
| ANKEF1 | 81.2963185 | -0.1471731 | 0.30261544 | -0.4863369 | 0.62672831 | 0.89875738 |
| SNAP25-AS1 | 49.4240955 | 1.21482111 | 0.49625713 | 2.44796706 | 0.01436648 | 0.24762526 |
| SNAP25 | 65.5007758 | 2.28650285 | 0.55717289 | 4.10375824 | 4.06E-05 | 0.01031432 |
| MKKS | 171.831018 | 0.51940507 | 0.19550832 | 2.65669037 | 0.00789119 | 0.17862547 |
| SLX4IP | 1699.32113 | -0.2577146 | 0.18264481 | -1.4110154 | 0.15824006 | 0.60804051 |
| JAG1 | 442.418134 | 0.61090104 | 0.38011833 | 1.60713389 | 0.108025 | 0.53063658 |
| AL050403.2 | 381.416004 | -0.2570787 | 0.64014236 | -0.4015961 | 0.68798128 | 0.92119054 |
| BTBD3 | 1259.24901 | -0.3949867 | 0.19912166 | -1.9836448 | 0.04729545 | 0.39411174 |
| SPTLC3 | 2395.35274 | 0.39825407 | 0.36877945 | 1.07992478 | 0.28017568 | 0.72513763 |
| ISM1 | 112.45129 | 1.04688785 | 0.47133323 | 2.22112038 | 0.02634281 | 0.31402818 |
| TASP1 | 2008.43888 | -0.1609088 | 0.18340356 | -0.8773481 | 0.38029759 | 0.7880311 |
| ESF1 | 592.377565 | -0.1445582 | 0.20999454 | -0.6883902 | 0.49120707 | 0.84660154 |
| NDUFAF5 | 530.534401 | -0.4483436 | 0.28343612 | -1.5818155 | 0.1136917 | 0.54000194 |
| MACROD2 | 7480.92954 | 0.29249598 | 0.28051309 | 1.04271775 | 0.29707904 | 0.73673761 |

| | | | | | | |
|---|---|---|---|---|---|---|
| FLRT3 | 728.390509 | 0.00363301 | 0.59304154 | 0.00612606 | 0.99511214 | 0.998438 |
| KIF16B | 2946.18291 | 0.48416591 | 0.37584548 | 1.28820467 | 0.19767473 | 0.64984705 |
| SNRPB2 | 233.369484 | 0.22548963 | 0.19520438 | 1.15514632 | 0.24803051 | 0.69838949 |
| BFSP1 | 52.9703916 | -0.0570689 | 0.68175353 | -0.083709 | 0.93328778 | 0.98423059 |
| DSTN | 2126.01835 | 0.5902363 | 0.33943722 | 1.73886734 | 0.0820581 | 0.48594425 |
| RRBP1 | 1513.337 | 0.1534501 | 0.27215174 | 0.56384024 | 0.57286287 | 0.87946834 |
| SNX5 | 464.782173 | 0.10510814 | 0.24660328 | 0.4262236 | 0.66994492 | 0.91465659 |
| OVOL2 | 682.687152 | -0.0198953 | 0.28900685 | -0.0688402 | 0.94511682 | 0.98836577 |
| MGME1 | 705.093969 | 0.1021991 | 0.20045879 | 0.50982598 | 0.61017339 | 0.89339478 |
| KAT14 | 138.804099 | -0.2618658 | 0.28244259 | -0.9271468 | 0.35385029 | 0.77243419 |
| ZNF133 | 535.536724 | 0.02953733 | 0.19397434 | 0.15227441 | 0.87897051 | 0.9765717 |
| DZANK1 | 349.268423 | -0.6290548 | 0.29210585 | -2.1535166 | 0.0312781 | 0.3383722 |
| POLR3F | 140.126736 | -0.9083112 | 0.26604221 | -3.414162 | 0.00063979 | 0.04534035 |
| RBBP9 | 86.6028592 | 0.54621512 | 0.2760425 | 1.97873555 | 0.04784579 | 0.39501762 |
| SEC23B | 605.893421 | 0.29797842 | 0.16646647 | 1.7900206 | 0.0734506 | 0.4701377 |
| DTD1 | 103.928458 | 0.49394746 | 0.33671837 | 1.4669454 | 0.14239091 | 0.58824095 |
| AL035563.1 | 38.823512 | -0.0320695 | 0.44639741 | -0.0718406 | 0.94272873 | 0.98772472 |
| SLC24A3 | 109.520154 | 0.87130329 | 0.60169511 | 1.44808104 | 0.14759439 | 0.59383925 |
| AL121761.1 | 402.080166 | 0.3483516 | 0.32268042 | 1.07955604 | 0.28033992 | 0.7252272 |
| RIN2 | 1048.91992 | 0.21212035 | 0.36034606 | 0.58865735 | 0.55609116 | 0.87212955 |
| NAA20 | 481.599586 | 0.21578237 | 0.24064081 | 0.89669899 | 0.36987956 | 0.78095617 |
| CRNKL1 | 169.513778 | 0.09432002 | 0.20245376 | 0.46588426 | 0.64129835 | 0.90404588 |
| CFAP61 | 78.6336084 | 0.02141638 | 0.32470481 | 0.06595647 | 0.94741248 | 0.98912199 |
| RALGAPA2 | 6178.40774 | -0.0481468 | 0.28539281 | -0.1687037 | 8.66E-01 | 0.97379123 |
| KIZ | 1527.85504 | 0.33298035 | 0.20675317 | 1.61052116 | 0.10728413 | 0.52901576 |
| KIZ-AS1 | 327.875031 | 0.56140824 | 0.60466319 | 0.92846439 | 0.35316673 | 0.77216708 |
| XRN2 | 1070.81648 | -0.1664381 | 0.2337671 | -0.7119824 | 0.47647565 | 0.84141268 |
| NXT1 | 44.4166461 | 0.30925704 | 0.39676558 | 0.77944524 | 0.43571748 | 0.81962293 |
| GZF1 | 94.81612 | 0.55054538 | 0.36772367 | 1.49717142 | 0.13434866 | 0.57708028 |
| NAPB | 738.65964 | 0.23392081 | 0.248199 | 0.94247281 | 3.46E-01 | 0.76822239 |
| CST3 | 1267.42423 | 1.06751276 | 0.44299151 | 2.40978154 | 0.01596208 | 0.25464137 |
| SYNDIG1 | 762.381418 | 0.30546926 | 0.43042144 | 0.70969804 | 0.4778914 | 0.84190259 |
| AL035661.1 | 49.0896774 | 1.05248435 | 0.64839856 | 1.62320588 | 0.1045454 | 0.52442862 |
| APMAP | 284.684454 | 0.00086592 | 0.25582501 | 0.0033848 | 0.99729933 | 0.99948643 |
| ACSS1 | 1354.19199 | 0.74082281 | 0.35307523 | 2.09820101 | 0.03588739 | 0.35661333 |
| ENTPD6 | 454.316482 | 0.02889641 | 0.31259108 | 0.09244158 | 0.9263472 | 0.98234748 |
| PYGB | 808.895852 | -0.393598 | 0.5996104 | -0.656423 | 0.51155202 | 0.85762435 |
| ABHD12 | 804.588244 | -0.3235682 | 0.22469146 | -1.4400554 | 0.14985172 | 0.59663305 |
| GINS1 | 91.3387306 | -0.5190044 | 0.3972475 | -1.3065014 | 0.19138211 | 0.64571429 |
| NINL | 2757.89729 | 0.08080523 | 0.27265974 | 0.29635923 | 0.76695576 | 0.94312902 |
| NANP | 58.8790154 | -0.2743407 | 0.31371425 | -0.8744923 | 0.3818502 | 0.78919376 |
| ZNF337-AS1 | 404.024796 | -0.5580107 | 0.23819785 | -2.3426353 | 0.01914809 | 0.27978663 |
| ZNF337 | 401.511857 | 0.13483531 | 0.23005656 | 0.58609634 | 0.55781077 | 0.87289687 |
| FAM182B | 313.048285 | 0.23750285 | 0.27504321 | 0.86351104 | 0.38785655 | 0.79626938 |
| HM13 | 832.016391 | 0.30649867 | 0.23784466 | 1.28865061 | 0.19751958 | 0.64964978 |
| ID1 | 221.843907 | 0.17557007 | 0.76010394 | 0.23098166 | 0.81732905 | 0.9593122 |
| BCL2L1 | 1622.69118 | -0.2640016 | 0.44925341 | -0.5876452 | 0.55677046 | 0.87219888 |
| AL117381.1 | 352.616675 | 0.06074616 | 0.32044735 | 0.18956674 | 0.84964865 | 0.97034489 |
| DUSP15 | 184.896047 | 1.30382544 | 0.63163638 | 2.06420258 | 0.03899851 | 0.37058958 |
| TTLL9 | 178.411159 | 0.56212199 | 0.3891579 | 1.44445735 | 0.14861037 | 0.59478974 |
| PDRG1 | 77.9224426 | 0.38785343 | 0.27729412 | 1.39870774 | 0.16190064 | 0.61346689 |
| TM9SF4 | 717.997986 | -0.1910768 | 0.17793596 | -1.0738517 | 0.28288916 | 0.72604837 |
| PLAGL2 | 115.231502 | 0.06323913 | 0.23697668 | 0.26685801 | 0.78957848 | 0.95090647 |
| POFUT1 | 224.796056 | -0.0724625 | 0.25935982 | -0.2793897 | 0.77994581 | 0.9477748 |

| | | | | | | |
|---|---|---|---|---|---|---|
| KIF3B | 742.590714 | 0.36919082 | 0.20880735 | 1.76809305 | 0.07704535 | 0.47689411 |
| ASXL1 | 2299.59863 | -0.3049252 | 0.2169008 | -1.4058278 | 0.15977526 | 0.60996285 |
| NOL4L | 183.386493 | 0.17766626 | 0.29201751 | 0.60840961 | 0.54291584 | 0.86590899 |
| FO393400.1 | 63.8699881 | -0.3292738 | 0.35989673 | -0.914912 | 0.36023781 | 0.77713233 |
| DNMT3B | 73.7105233 | 0.21101011 | 0.34964397 | 0.60349991 | 0.54617617 | 0.86795489 |
| MAPRE1 | 223.934756 | 0.25330403 | 0.2671787 | 0.9480697 | 0.34309397 | 0.76641095 |
| AL035071.2 | 80.1204828 | 0.03622431 | 0.60642426 | 0.05973426 | 0.95236728 | 0.98938795 |
| AL035071.1 | 60.376151 | -0.0099613 | 0.44031878 | -0.0226228 | 0.98195112 | 0.99518016 |
| CDK5RAP1 | 390.669873 | -0.2953784 | 0.15489752 | -1.9069277 | 0.05652995 | 0.41843207 |
| SNTA1 | 128.436911 | 0.71000856 | 0.36448459 | 1.94797966 | 0.0514174 | 0.40644668 |
| CBFA2T2 | 2595.91261 | -0.0738236 | 0.17250455 | -0.4279517 | 0.66868627 | 0.91395501 |
| NECAB3 | 90.6558531 | 0.32559755 | 0.38917452 | 0.83663637 | 0.402797 | 0.80324472 |
| PXMP4 | 77.5886648 | 0.7344847 | 0.48451993 | 1.51590194 | 0.12954415 | 0.57050365 |
| ZNF341 | 110.646055 | -0.5406737 | 0.36911938 | -1.4647665 | 0.14298463 | 0.58837508 |
| CHMP4B | 492.819304 | -0.1151458 | 0.18324225 | -0.6283801 | 5.30E-01 | 8.63E-01 |
| RALY | 903.618452 | -0.3136593 | 0.23544376 | -1.3322049 | 0.18279287 | 0.63741052 |
| EIF2S2 | 600.398577 | 0.31380171 | 0.25522888 | 1.22949141 | 0.21888762 | 0.67223588 |
| ASIP | 89.7647752 | -0.2618561 | 0.69807157 | -0.3751136 | 0.70757599 | 0.92577218 |
| AHCY | 144.0282 | 0.14094667 | 0.2883985 | 0.48872192 | 0.62503858 | 0.89873262 |
| ITCH | 2448.96043 | -0.268873 | 0.1349885 | -1.9918211 | 0.04639069 | 0.3904347 |
| DYNLRB1 | 642.572843 | 0.37894625 | 0.22996097 | 1.64787201 | 0.09937894 | 0.51665771 |
| MAP1LC3A | 195.644074 | 0.81473896 | 0.65208012 | 1.24944609 | 0.21150196 | 0.66484951 |
| PIGU | 557.162887 | -0.4441744 | 0.30334434 | -1.4642579 | 0.14312349 | 0.5883902 |
| NCOA6 | 1467.68349 | -0.2355521 | 0.12577377 | -1.8728238 | 0.06109272 | 0.43176766 |
| TP53INP2 | 187.237371 | 0.28039157 | 0.44475654 | 0.63043834 | 0.52840783 | 0.86241272 |
| GGT7 | 596.907108 | 0.85403275 | 0.2657156 | 3.21408589 | 0.00130861 | 0.06597468 |
| ACSS2 | 668.948652 | 0.17402457 | 0.23664207 | 0.7353915 | 0.4621011 | 0.83531269 |
| GSS | 182.645644 | 0.05790571 | 0.40150025 | 0.14422334 | 0.88532411 | 0.97664137 |
| MYH7B | 45.7860861 | 0.40937182 | 0.43194133 | 0.94774867 | 0.34325742 | 0.76641095 |
| TRPC4AP | 1458.47706 | -0.2051795 | 0.17918891 | -1.1450459 | 0.25219013 | 0.70276117 |
| EDEM2 | 294.649891 | -0.0766224 | 0.28204482 | -0.2716673 | 0.78587783 | 0.94978829 |
| PROCR | 187.918312 | -0.4892711 | 0.60018737 | -0.8151973 | 0.41495938 | 0.80965493 |
| MMP24OS | 514.627571 | -0.3591203 | 0.2058771 | -1.7443432 | 0.08109926 | 0.48572317 |
| MMP24 | 438.530089 | 0.26821079 | 0.35678344 | 0.75174674 | 0.45220337 | 0.82885146 |
| UQCC1 | 897.686273 | -0.3391332 | 0.13930761 | -2.43442 | 0.01491568 | 0.24967336 |
| CEP250 | 201.990363 | -0.2748258 | 0.34907786 | -0.7872908 | 0.43111168 | 0.81706845 |
| ERGIC3 | 365.844156 | -0.0064512 | 0.31547209 | -0.0204492 | 0.98368503 | 0.99518552 |
| CPNE1 | 359.480747 | -0.1939741 | 0.29614105 | -0.6550059 | 0.51246398 | 0.85762435 |
| NFS1 | 111.975493 | -0.0439961 | 0.28101752 | -0.15656 | 0.87559164 | 0.97614152 |
| ROMO1 | 208.925614 | 0.20522844 | 0.4586789 | 0.4474338 | 0.65456188 | 0.90902447 |
| RBM39 | 3538.87384 | -0.0100292 | 0.16384708 | -0.0612105 | 0.95119157 | 0.98938795 |
| PHF20 | 1322.21312 | -0.0508195 | 0.15711744 | -0.3234492 | 0.7463551 | 0.9368751 |
| SCAND1 | 378.783899 | 0.42472301 | 0.38764697 | 1.09564382 | 2.73E-01 | 0.72070318 |
| CNBD2 | 93.7434755 | -0.0907261 | 0.34670733 | -0.2616792 | 0.79356882 | 0.95224951 |
| NORAD | 875.706297 | 0.60668377 | 0.251154 | 2.41558473 | 0.01570997 | 0.25227487 |
| EPB41L1 | 3652.25085 | 0.03254892 | 0.60750219 | 0.05357827 | 0.95727117 | 0.99065996 |
| AAR2 | 181.46143 | -0.0084501 | 0.19518528 | -0.0432928 | 0.96546812 | 0.99248831 |
| DLGAP4 | 847.128485 | -0.4412604 | 0.15218107 | -2.8995746 | 0.00373669 | 0.11893493 |
| DLGAP4-AS1 | 423.344697 | -0.6997165 | 0.27831828 | -2.5140873 | 0.01193409 | 0.22359911 |
| MYL9 | 101.124018 | 0.89593737 | 0.3666242 | 2.44374857 | 0.01453555 | 0.24762526 |
| RAB5IF | 122.662666 | 0.0279294 | 0.53106945 | 0.05259086 | 0.9580579 | 0.99069768 |
| NDRG3 | 343.645795 | -0.1121258 | 0.17940862 | -0.6249744 | 0.53198783 | 0.86397197 |
| DSN1 | 95.7713611 | -0.3845028 | 0.32555148 | -1.1810814 | 0.23757037 | 0.68973903 |
| SOGA1 | 639.46228 | -0.4875582 | 0.33172823 | -1.4697518 | 0.14162898 | 0.5872148 |

| | | | | | | |
|---|---|---|---|---|---|---|
| SAMHD1 | 133.509613 | -0.50759 | 0.67076551 | -0.7567325 | 0.44921017 | 0.8267002 |
| RBL1 | 298.292017 | -0.2730258 | 0.25081828 | -1.0885404 | 0.27635663 | 0.72242128 |
| MROH8 | 146.689118 | -0.2228144 | 0.28825394 | -0.7729794 | 0.43953459 | 0.82177104 |
| RPN2 | 552.9283 | 0.1679372 | 0.18445193 | 0.91046594 | 0.36257684 | 0.77849113 |
| MANBAL | 159.07575 | 0.16439216 | 0.21684881 | 0.75809574 | 0.44839367 | 0.82628794 |
| SRC | 235.869966 | 0.29056495 | 0.22271133 | 1.30467069 | 0.19200501 | 0.64596359 |
| BLCAP | 619.682051 | 0.02472365 | 0.25005204 | 0.09887402 | 0.9212383 | 0.98215577 |
| CTNNBL1 | 555.120133 | -0.2257558 | 0.15797694 | -1.4290428 | 0.15299193 | 0.60285224 |
| TTI1 | 357.868316 | 0.12394151 | 0.19860658 | 0.62405541 | 0.53259119 | 0.86397197 |
| RPRD1B | 648.795628 | -0.4698423 | 0.15898891 | -2.9551894 | 0.00312477 | 0.10748727 |
| TGM2 | 607.989359 | 0.30946415 | 0.64534824 | 0.47953048 | 6.32E-01 | 9.00E-01 |
| RALGAPB | 1281.95638 | -0.1773525 | 0.17302816 | -1.0249919 | 0.30536699 | 0.74229685 |
| ARHGAP40 | 68.6664279 | 2.05745548 | 0.97309411 | 2.11434378 | 3.45E-02 | 0.35291306 |
| ACTR5 | 90.7983906 | -0.4961105 | 0.30550216 | -1.6239183 | 0.10439325 | 0.52442862 |
| DHX35 | 317.110301 | -0.4182445 | 0.24393542 | -1.7145708 | 0.08642395 | 0.49478465 |
| MAFB | 52.8667765 | 0.30127345 | 0.6086306 | 0.49500214 | 0.62059861 | 0.89731054 |
| TOP1 | 1773.43653 | -0.0057844 | 0.25285916 | -0.0228762 | 0.98174905 | 0.99518016 |
| PLCG1-AS1 | 73.193033 | -0.5023512 | 0.39151629 | -1.2830915 | 0.19946003 | 0.65150038 |
| PLCG1 | 480.415002 | -0.5116541 | 0.23350709 | -2.1911713 | 0.0284394 | 0.32580744 |
| ZHX3 | 4936.64678 | -0.420562 | 0.37101835 | -1.1335343 | 0.25698995 | 0.70735159 |
| LPIN3 | 517.163087 | 0.19656336 | 0.27939003 | 0.70354466 | 0.48171639 | 0.8437862 |
| CHD6 | 2578.37668 | -0.141921 | 0.13506477 | -1.0507625 | 0.29336769 | 0.73527014 |
| PTPRT | 48.6197802 | -0.3447611 | 0.54016199 | -0.6382551 | 0.52330766 | 0.86006369 |
| SRSF6 | 699.118703 | 0.20605237 | 0.29748834 | 0.69264016 | 4.89E-01 | 0.84562056 |
| L3MBTL1 | 279.188203 | 0.14528062 | 0.25325807 | 0.57364657 | 0.56620699 | 0.87642772 |
| Z98752.3 | 90.2697152 | -0.0134632 | 0.35764455 | -0.0376442 | 0.9699714 | 0.99309664 |
| SGK2 | 1045.25786 | 0.00535194 | 0.27170979 | 0.01969726 | 9.84E-01 | 0.99518552 |
| IFT52 | 389.208124 | -0.0300445 | 0.17626639 | -0.1704493 | 0.86465684 | 0.97361767 |
| OSER1 | 245.451569 | 0.10999902 | 0.30015216 | 0.36647751 | 0.71400879 | 0.92760608 |
| OSER1-DT | 66.7684328 | 0.16077708 | 0.38904716 | 0.41325859 | 0.67941717 | 0.9182972 |
| FITM2 | 77.9238417 | -0.1040473 | 0.35240147 | -0.2952522 | 0.76780119 | 0.94316227 |
| TTPAL | 123.376322 | -0.1605239 | 0.31855819 | -0.5039076 | 0.61432632 | 0.89488165 |
| SERINC3 | 511.273979 | 0.22089037 | 0.1902323 | 1.1611612 | 0.24557635 | 0.69579966 |
| PKIG | 717.222996 | 0.00558533 | 0.19002887 | 0.029392 | 0.97655195 | 0.9941831 |
| YWHAB | 885.503068 | 0.63516457 | 0.31694233 | 2.00403828 | 4.51E-02 | 0.38755881 |
| PABPC1L | 289.500557 | -0.2640891 | 0.22816014 | -1.1574724 | 0.24707939 | 0.69751478 |
| TOMM34 | 110.144619 | 0.22403176 | 0.29120043 | 0.76933872 | 0.44169225 | 0.82285053 |
| STK4 | 587.978276 | -0.2747959 | 0.23674563 | -1.1607222 | 0.24575488 | 0.69585772 |
| SLPI | 5909.96203 | 0.75189838 | 0.6841658 | 1.09900025 | 2.72E-01 | 7.20E-01 |
| SDC4 | 1787.06434 | 0.54656167 | 0.38637009 | 1.41460656 | 0.15718388 | 0.60703814 |
| SYS1-DBNDD | 104.308793 | -0.7334371 | 0.4107289 | -1.7856964 | 0.07414843 | 0.47078634 |
| TP53TG5 | 59.3464561 | -0.5476672 | 0.6796338 | -0.8058268 | 0.4203427 | 0.8125045 |
| PIGT | 177.30668 | 0.45107196 | 0.28928766 | 1.55925062 | 0.11893708 | 0.5506624 |
| WFDC2 | 9006.98496 | 2.15483396 | 0.64805051 | 3.32510185 | 8.84E-04 | 5.17E-02 |
| DNTTIP1 | 198.166545 | 0.10582256 | 0.27190186 | 0.38919393 | 0.69713269 | 0.92322348 |
| SNX21 | 200.776866 | 0.48022622 | 0.35268097 | 1.36164484 | 0.17331 | 0.6286116 |
| ACOT8 | 187.52416 | -0.1731192 | 0.23980621 | -0.721913 | 0.47034796 | 0.83921324 |
| ZSWIM3 | 82.7256827 | -0.4113102 | 0.3691741 | -1.114136 | 0.26522083 | 0.71512755 |
| CTSA | 304.232028 | 0.62781739 | 0.30668172 | 2.04713012 | 0.04064531 | 0.37344844 |
| PCIF1 | 72.9841044 | -0.3311695 | 0.26896766 | -1.2312616 | 0.21822505 | 0.67173126 |
| ZNF335 | 224.281919 | -0.2254151 | 0.24359167 | -0.9253808 | 0.35476785 | 0.77252365 |
| NCOA5 | 515.248154 | -0.1794135 | 0.28948128 | -0.6197759 | 0.53540534 | 0.8642718 |
| CD40 | 58.0588208 | 1.10278077 | 0.46331254 | 2.38020922 | 0.01730281 | 0.26361342 |
| SLC35C2 | 272.389344 | 0.12425134 | 0.25918733 | 0.47938817 | 0.63166251 | 0.90016507 |

| | | | | | | |
|---|---|---|---|---|---|---|
| AL133227.1 | 58.1033207 | -0.2718897 | 0.58654519 | -0.4635443 | 0.6429743 | 0.90478945 |
| ELMO2 | 242.120552 | -0.3799491 | 0.19290888 | -1.9695781 | 0.04888674 | 0.39762623 |
| ZNF334 | 183.341252 | -1.2413842 | 0.57751403 | -2.1495308 | 0.03159235 | 0.33950635 |
| SLC13A3 | 206.541704 | -0.1688726 | 0.57638309 | -0.2929868 | 0.76953224 | 0.94386043 |
| EYA2 | 1017.2734 | -5.5064578 | 1.21390177 | -4.5361642 | 5.73E-06 | 0.0032951 |
| ZMYND8 | 4737.98141 | -0.1913319 | 0.33601098 | -0.5694214 | 0.56907019 | 0.87789095 |
| NCOA3 | 2265.25458 | -0.1844574 | 0.1704586 | -1.0821242 | 0.27919736 | 0.72482665 |
| SULF2 | 674.964104 | 1.45858533 | 0.49508922 | 2.94610599 | 3.22E-03 | 0.10927208 |
| PREX1 | 137.054411 | -0.3251217 | 0.38717908 | -0.8397192 | 0.40106584 | 0.80243544 |
| ARFGEF2 | 2358.2825 | -0.018753 | 0.15274857 | -0.1227702 | 0.90228904 | 0.97941893 |
| CSE1L | 501.821691 | -0.4531492 | 0.21283211 | -2.1291393 | 0.03324274 | 0.34850106 |
| STAU1 | 1350.99493 | -0.0822976 | 0.20623884 | -0.3990404 | 0.68986341 | 0.92176114 |
| DDX27 | 225.136307 | 0.09359697 | 0.19093706 | 0.49019803 | 0.62399377 | 0.89873262 |
| ZNFX1 | 415.601512 | -0.1693287 | 0.37464672 | -0.451969 | 0.65129135 | 0.90825486 |
| ZFAS1 | 375.954375 | -0.1051015 | 0.15508411 | -0.6777063 | 0.49795795 | 0.85092921 |
| B4GALT5 | 2110.27879 | 0.18882452 | 0.22839355 | 0.82675066 | 0.4083784 | 0.80588056 |
| SLC9A8 | 792.23794 | -0.3943527 | 0.27179442 | -1.4509226 | 0.1468014 | 0.59373375 |
| SPATA2 | 52.3220149 | 0.02451675 | 0.35581339 | 0.06890338 | 0.94506653 | 0.98836577 |
| RNF114 | 624.209012 | 0.23442882 | 0.18996529 | 1.23406133 | 0.21718005 | 0.66978787 |
| CEBPB | 320.432274 | 0.96664246 | 0.60591408 | 1.59534575 | 1.11E-01 | 5.34E-01 |
| PTPN1 | 1632.4205 | 0.32485526 | 0.30617932 | 1.06099674 | 0.28869138 | 0.73176973 |
| RIPOR3 | 966.404631 | -2.7490333 | 0.83444213 | -3.2944565 | 0.00098612 | 0.0559889 |
| AL353653.1 | 51.3296125 | -3.6446914 | 1.04753393 | -3.4793063 | 0.00050271 | 0.03839746 |
| PARD6B | 1704.94396 | -0.2433992 | 0.31250818 | -0.7788571 | 0.43606391 | 0.81969093 |
| BCAS4 | 315.07123 | 0.11038722 | 0.31344297 | 0.3521764 | 0.72470598 | 0.93241479 |
| ADNP | 2198.79929 | -0.0058209 | 0.17032074 | -0.0341759 | 0.9727369 | 0.99327086 |
| ADNP-AS1 | 58.4774565 | 0.06712877 | 0.36805711 | 0.18238683 | 0.85527916 | 0.97118784 |
| DPM1 | 547.239706 | 0.07287235 | 0.1659867 | 0.43902522 | 0.66064326 | 0.91099119 |
| NFATC2 | 496.724956 | 0.59644536 | 0.39375315 | 1.51476977 | 0.12983072 | 0.57050365 |
| ATP9A | 885.661895 | -0.2655756 | 0.26462226 | -1.0036025 | 0.31557023 | 0.74785672 |
| ZFP64 | 443.643207 | -0.162711 | 0.28460319 | -0.5717117 | 0.56751729 | 0.87678876 |
| TSHZ2 | 85.6804322 | 0.60989793 | 0.52431717 | 1.16322327 | 0.24473893 | 0.6948422 |
| AL157838.1 | 62.5849369 | 0.36445534 | 0.78469396 | 0.46445539 | 0.64232151 | 0.90454205 |
| ZNF217 | 679.8475 | 0.0697601 | 0.26516892 | 0.26307798 | 7.92E-01 | 0.95215633 |
| BCAS1 | 109.825523 | 0.88255341 | 0.79110993 | 1.11558884 | 0.26459817 | 0.71488591 |
| CYP24A1 | 1108.31711 | -0.4430528 | 1.16542302 | -0.3801648 | 7.04E-01 | 9.25E-01 |
| PFDN4 | 153.242004 | -0.4410237 | 0.30317764 | -1.4546708 | 0.14576041 | 0.59218144 |
| DOK5 | 617.504162 | -0.4673167 | 0.48401052 | -0.9655094 | 0.33428973 | 0.76231532 |
| FAM210B | 113.221747 | 0.77598149 | 0.29870373 | 2.59782991 | 0.0093815 | 0.19669868 |
| CSTF1 | 114.793824 | -0.0043831 | 0.19518927 | -0.0224556 | 0.98208453 | 0.99518016 |
| RTF2 | 338.773823 | -0.1288323 | 0.2109287 | -0.6107859 | 0.5413413 | 0.86582694 |
| RAE1 | 210.036967 | 0.24839108 | 0.23130446 | 1.07387068 | 0.28288065 | 0.72604837 |
| RBM38 | 76.3666514 | 0.89620088 | 0.45710811 | 1.96058843 | 0.04992705 | 0.40017381 |
| PCK1 | 1785.25718 | 0.58210765 | 0.72704968 | 0.80064357 | 0.42333802 | 0.8138418 |
| PMEPA1 | 299.140137 | 0.42729899 | 0.93037189 | 0.45927763 | 0.64603481 | 0.90582727 |
| RAB22A | 627.098227 | -0.1929108 | 0.1884816 | -1.0234994 | 0.30607177 | 0.74290739 |
| VAPB | 827.228636 | -0.0885697 | 0.15025341 | -0.589469 | 0.55554672 | 0.87184917 |
| STX16-NPEPI | 99.97429 | -1.4503711 | 0.54558383 | -2.6583835 | 0.00785165 | 0.17820006 |
| NPEPL1 | 256.853518 | -0.1495661 | 0.34925082 | -0.4282485 | 0.66847023 | 0.91395501 |
| GNAS-AS1 | 49.235506 | 0.2844402 | 0.5817581 | 0.48893209 | 0.62488978 | 0.89873262 |
| GNAS | 3331.24294 | 0.49052631 | 0.23113938 | 2.12221006 | 0.0338201 | 0.35107893 |
| AL121917.1 | 64.7665035 | 0.40325686 | 0.4045059 | 0.99691219 | 0.31880713 | 0.75094569 |
| NELFCD | 407.446932 | -0.0735829 | 0.18260185 | -0.4029693 | 0.68697077 | 0.9208231 |
| CTSZ | 243.173759 | 0.84793637 | 0.34783377 | 2.43776324 | 0.01477845 | 0.24869534 |

| | | | | | | |
|---|---|---|---|---|---|---|
| ATP5F1E | 996.591923 | 0.26732156 | 0.52455309 | 0.50961775 | 0.61031929 | 0.89339478 |
| PRELID3B | 122.124877 | 0.25819948 | 0.35859246 | 0.72003599 | 0.47150284 | 0.83921324 |
| PHACTR3 | 490.384003 | 3.13331316 | 0.86330171 | 3.62945323 | 0.00028402 | 0.02863854 |
| SYCP2 | 202.749065 | -0.4217745 | 0.34081038 | -1.2375636 | 0.21587793 | 0.66911866 |
| FAM217B | 132.218792 | 0.20034414 | 0.34801797 | 0.57567184 | 0.56483701 | 0.87579905 |
| CDH26 | 56.9885364 | 0.34872078 | 0.57490086 | 0.60657551 | 0.54413266 | 0.8664796 |
| MIR646HG | 1665.61008 | 0.62727484 | 0.89646024 | 0.6997241 | 0.48409962 | 0.8445691 |
| CDH4 | 101.298222 | 0.00490909 | 0.56676849 | 0.00866154 | 0.99308918 | 0.99816851 |
| TAF4 | 534.756734 | -0.1713942 | 0.25300074 | -0.6774456 | 0.4981233 | 0.85092921 |
| LSM14B | 129.730167 | -0.0702166 | 0.28091901 | -0.2499531 | 0.80262359 | 0.95446295 |
| PSMA7 | 469.638206 | 0.57738638 | 0.2806155 | 2.05757126 | 0.03963131 | 0.37259071 |
| SS18L1 | 439.725541 | -0.2594385 | 0.18998636 | -1.3655636 | 0.17207598 | 0.6286116 |
| MTG2 | 505.691202 | 0.04195333 | 0.2201944 | 0.19052858 | 0.84889495 | 0.97017477 |
| OSBPL2 | 725.71633 | -0.3236092 | 0.24774616 | -1.3062126 | 0.19148027 | 0.64571429 |
| ADRM1 | 90.7779992 | -0.2197188 | 0.45630117 | -0.4815215 | 0.63014591 | 0.90005881 |
| AL354836.1 | 74.1724675 | 0.45498229 | 0.53375014 | 0.85242561 | 0.39397791 | 0.79845072 |
| LAMA5 | 516.534637 | -0.1066362 | 0.29592479 | -0.360349 | 0.71858617 | 0.9298398 |
| RPS21 | 102.375314 | -0.3079728 | 0.41285978 | -0.7459502 | 0.4556975 | 0.83130183 |
| CABLES2 | 50.1434642 | -0.2061265 | 0.39171644 | -0.5262134 | 0.59873992 | 0.88883692 |
| SLCO4A1 | 468.082153 | -0.0838711 | 0.46401399 | -0.1807513 | 8.57E-01 | 9.71E-01 |
| NTSR1 | 46.974436 | -0.5063491 | 0.72176711 | -0.7015409 | 0.48296553 | 0.8441831 |
| LINC00659 | 46.0565915 | -0.2077363 | 0.57647763 | -0.3603545 | 0.71858202 | 0.9298398 |
| MRGBP | 83.6892407 | -0.4695083 | 0.28633293 | -1.6397288 | 0.10106157 | 0.51819907 |
| OGFR | 335.038639 | -0.4716879 | 0.32406085 | -1.4555534 | 1.46E-01 | 0.59196873 |
| TCFL5 | 164.876766 | -0.0101419 | 0.23637987 | -0.0429053 | 0.96577705 | 0.99248831 |
| DIDO1 | 1025.52978 | -0.0514014 | 0.16798601 | -0.3059862 | 0.75961517 | 0.94133292 |
| GID8 | 270.184021 | -0.07289 | 0.17370649 | -0.4196159 | 0.67476606 | 0.91636338 |
| YTHDF1 | 343.421755 | -0.0851757 | 0.17986894 | -0.4735431 | 0.63582573 | 0.90207488 |
| NKAIN4 | 44.5303461 | 1.61505301 | 0.54729627 | 2.95096658 | 0.00316781 | 0.10846567 |
| ARFGAP1 | 275.135317 | -0.6220384 | 0.29315473 | -2.1218775 | 0.03384803 | 0.35107893 |
| PPDPF | 867.610691 | 0.79574827 | 0.46061281 | 1.72758606 | 0.08406246 | 0.48990407 |
| HELZ2 | 97.5501768 | 0.60719122 | 0.95237389 | 0.63755551 | 0.52376306 | 0.86006369 |
| GMEB2 | 196.334912 | -0.1559617 | 0.20472385 | -0.7618151 | 0.44617036 | 0.82571115 |
| MHENCR | 84.8021538 | -0.5008997 | 0.40219629 | -1.2454109 | 0.21298073 | 0.66696598 |
| STMN3 | 138.475327 | 0.59067625 | 0.7626731 | 0.77448156 | 0.43864607 | 0.82127804 |
| ARFRP1 | 177.276362 | 0.05576803 | 0.31842449 | 0.17513738 | 0.86097169 | 0.9720639 |
| SLC2A4RG | 250.251618 | 0.40772075 | 0.46161742 | 0.88324385 | 0.37710453 | 0.78579804 |
| ZBTB46 | 266.716935 | 1.10587184 | 0.4373759 | 2.52842428 | 0.01145758 | 0.21870401 |
| TPD52L2 | 428.342727 | -0.4001267 | 0.27552345 | -1.4522419 | 0.14643436 | 0.59373375 |
| DNAJC5 | 917.127411 | -0.0976413 | 0.21934551 | -0.4451485 | 0.65621245 | 0.90950195 |
| UCKL1 | 240.755685 | 0.16260241 | 0.30881947 | 0.526529 | 0.59852071 | 0.88883692 |
| ZNF512B | 165.091599 | -0.0551066 | 0.44174397 | -0.1247478 | 0.9007232 | 0.97914987 |
| SAMD10 | 41.3225133 | 0.45751319 | 0.52378968 | 0.87346736 | 0.38240839 | 0.7896252 |
| PRPF6 | 532.962446 | -0.1479386 | 0.25376172 | -0.5829825 | 0.55990507 | 0.87300289 |
| TCEA2 | 107.241982 | 0.28824669 | 0.3552929 | 0.81129313 | 0.41719735 | 0.81016758 |
| PCMTD2 | 923.370122 | 0.5275536 | 0.29632285 | 1.78033386 | 0.07502134 | 0.47190267 |
| LINC00266-1 | 48.7673364 | -0.3661753 | 0.33726194 | -1.0857296 | 0.27759865 | 0.72344438 |
| PLPP2 | 143.969395 | 0.74475651 | 0.58674142 | 1.2693096 | 2.04E-01 | 6.57E-01 |
| MIER2 | 154.515596 | -0.0924122 | 0.29682206 | -0.3113387 | 0.75554317 | 0.93955655 |
| SHC2 | 88.142538 | -1.5131653 | 0.56388416 | -2.6834683 | 0.00728629 | 0.17305714 |
| MADCAM1 | 53.6312463 | -0.6226392 | 0.37455038 | -1.6623643 | 0.09643975 | 0.512217 |
| TPGS1 | 176.064434 | 0.13478507 | 0.38946228 | 0.34607991 | 0.72928266 | 0.93453345 |
| CDC34 | 125.743023 | -0.2112998 | 0.27695831 | -0.7629299 | 0.44550519 | 0.82571115 |
| BSG | 781.879418 | 0.67421537 | 0.39049838 | 1.726551 | 0.08424833 | 0.49023622 |

| | | | | | | |
|---|---|---|---|---|---|---|
| POLRMT | 69.9808587 | -0.116571 | 0.3061739 | -0.3807346 | 0.70340016 | 0.92513032 |
| RNF126 | 183.755622 | -0.3201929 | 0.20798311 | -1.5395142 | 0.12367882 | 0.56102027 |
| FSTL3 | 131.800481 | -1.0830788 | 0.63700007 | -1.7002806 | 0.08907817 | 0.49895023 |
| PALM | 521.115271 | 0.54488266 | 0.45598357 | 1.19496116 | 0.2321022 | 0.68449163 |
| LINC01836 | 84.8163409 | 0.21216091 | 0.69142555 | 0.30684563 | 0.75896088 | 0.94132528 |
| PTBP1 | 592.203649 | -0.1177508 | 0.26886428 | -0.4379561 | 0.66141809 | 0.91110616 |
| MED16 | 178.93478 | 0.46553569 | 0.38128103 | 1.22097786 | 0.2220944 | 0.67548814 |
| R3HDM4 | 193.071415 | 0.34847409 | 0.36569223 | 0.95291631 | 0.34063248 | 0.76494634 |
| ARID3A | 733.900813 | 0.57787403 | 0.4495605 | 1.28541993 | 0.19864558 | 0.65109416 |
| WDR18 | 127.342783 | 0.71269366 | 0.51743214 | 1.37736644 | 0.16839898 | 0.62253512 |
| TMEM259 | 548.079566 | -0.0967598 | 0.34211134 | -0.2828312 | 0.77730626 | 0.94694572 |
| CNN2 | 173.669233 | 0.10364579 | 0.373907 | 0.27719671 | 0.78162907 | 0.94859779 |
| ABCA7 | 134.56848 | -0.7847292 | 0.39254798 | -1.9990656 | 0.04560126 | 0.38904776 |
| POLR2E | 215.428662 | 0.0769026 | 0.3144464 | 0.24456506 | 0.80679322 | 0.95566275 |
| GPX4 | 691.418205 | 0.29526733 | 0.41823233 | 0.70598878 | 0.48019512 | 0.8433217 |
| SBNO2 | 1091.57107 | -0.0177334 | 0.40023482 | -0.0443074 | 9.65E-01 | 9.92E-01 |
| STK11 | 412.320637 | -0.0022324 | 0.24878254 | -0.0089732 | 0.99284051 | 0.99816851 |
| ATP5F1D | 597.875606 | 0.80114325 | 0.41367863 | 1.93663196 | 0.05279035 | 0.41066414 |
| MIDN | 696.209946 | -0.0739017 | 0.42451112 | -0.1740866 | 8.62E-01 | 9.73E-01 |
| CIRBP | 1192.75233 | 0.18618895 | 0.29249248 | 0.63655981 | 0.52441161 | 0.86006369 |
| C19orf24 | 134.664358 | 0.24402624 | 0.40781826 | 0.59837005 | 0.54959304 | 0.86930601 |
| MUM1 | 564.50179 | 0.07267176 | 0.21019967 | 0.34572727 | 0.72954768 | 0.93453345 |
| NDUFS7 | 230.988536 | 0.45873982 | 0.33977867 | 1.35011367 | 0.17697952 | 0.63028378 |
| GAMT | 57.3376622 | 1.59507443 | 0.51078888 | 3.12276653 | 0.0017916 | 0.08132377 |
| DAZAP1 | 741.303633 | -0.0927411 | 0.22468125 | -0.4127673 | 0.67977713 | 0.9182972 |
| RPS15 | 1426.97057 | 0.55587264 | 0.44458872 | 1.25030758 | 0.21118721 | 0.66483325 |
| C19orf25 | 438.058912 | 0.01382346 | 0.27923635 | 0.04950453 | 0.96051723 | 0.99153894 |
| PCSK4 | 71.5074596 | -0.3755928 | 0.42598512 | -0.8817041 | 0.37793682 | 0.7865219 |
| MBD3 | 163.403868 | -0.1695642 | 0.40336007 | -0.4203792 | 0.67420847 | 0.91636338 |
| TCF3 | 257.578774 | -0.7193033 | 0.2647419 | -2.7169984 | 0.00658769 | 0.1680054 |
| REXO1 | 220.882526 | -0.267179 | 0.27201199 | -0.9822323 | 0.3259854 | 0.75550719 |
| KLF16 | 47.0902523 | 0.27351626 | 0.57189173 | 0.47826582 | 0.63246102 | 0.90052578 |
| AC012615.1 | 47.9718542 | 0.46886557 | 0.35912118 | 1.30559153 | 0.1916915 | 0.64596359 |
| ABHD17A | 224.676249 | -0.0174918 | 0.33486966 | -0.0522347 | 0.95834172 | 0.99071816 |
| SCAMP4 | 353.508609 | 0.25424948 | 0.31641737 | 0.80352566 | 0.42167097 | 0.81266383 |
| CSNK1G2 | 414.218239 | -0.0444566 | 0.22740619 | -0.1954943 | 8.45E-01 | 0.96858129 |
| BTBD2 | 463.287013 | -0.2551674 | 0.34489277 | -0.7398456 | 0.45939368 | 0.83390278 |
| MKNK2 | 184.099622 | 0.30765058 | 0.38225587 | 0.80482891 | 0.42091842 | 0.81260593 |
| MOB3A | 255.777611 | -0.0448026 | 0.38973969 | -0.1149552 | 0.9084806 | 0.98047754 |
| AP3D1 | 614.167874 | -0.1738093 | 0.23073795 | -0.7532759 | 0.45128417 | 0.82814845 |
| DOT1L | 481.558813 | -0.7295416 | 0.27513169 | -2.6516088 | 0.00801093 | 0.17977198 |
| PLEKHJ1 | 269.197651 | 0.36512778 | 0.32724017 | 1.1157792 | 0.26451666 | 0.71488591 |
| SF3A2 | 120.113107 | -0.464075 | 0.35437031 | -1.3095765 | 0.19033913 | 0.64549154 |
| LSM7 | 306.167549 | 0.38617937 | 0.36757451 | 1.05061523 | 0.29343534 | 0.73527014 |
| SPPL2B | 492.242197 | -0.0527804 | 0.21671962 | -0.2435422 | 0.80758543 | 0.95594682 |
| TMPRSS9 | 41.672413 | -0.6278077 | 0.45607426 | -1.3765471 | 0.16865231 | 0.62261832 |
| TIMM13 | 212.90643 | 0.61338671 | 0.43140891 | 1.42182205 | 0.15507792 | 0.60416746 |
| LMNB2 | 66.2726146 | -0.0976978 | 0.31026725 | -0.3148827 | 0.75285072 | 0.93832847 |
| GADD45B | 701.177508 | -0.1702526 | 0.42906053 | -0.3968031 | 6.92E-01 | 0.92224682 |
| GNG7 | 1310.5726 | 0.02392185 | 0.38016133 | 0.06292553 | 0.94982581 | 0.98919519 |
| SGTA | 230.712602 | 0.06037732 | 0.27974859 | 0.21582708 | 0.82912254 | 0.96288726 |
| THOP1 | 141.824899 | -0.2285942 | 0.33370322 | -0.6850225 | 0.49332973 | 0.84815728 |
| ZNF554 | 104.148402 | -0.2229302 | 0.37804744 | -0.5896883 | 0.55539963 | 0.87184917 |
| ZNF555 | 107.413822 | 0.14288318 | 0.35289017 | 0.40489419 | 0.68555529 | 0.92053037 |

| | | | | | | |
|---|---|---|---|---|---|---|
| ZNF57 | 71.6178268 | 0.22464953 | 0.26988763 | 0.83238174 | 0.4051935 | 0.80481948 |
| AC119403.1 | 48.7305819 | 0.20673388 | 0.29949967 | 0.69026413 | 0.4900281 | 0.84614483 |
| ZNF77 | 65.7781799 | -0.102351 | 0.28132812 | -0.3638138 | 0.71599707 | 0.92890921 |
| TLE2 | 236.370871 | 1.72334514 | 0.55936133 | 3.08091576 | 0.00206365 | 0.0878448 |
| AES | 1001.74033 | 0.48575272 | 0.42752822 | 1.13618868 | 0.25587761 | 0.7070894 |
| GNA11 | 296.756771 | 0.16268752 | 0.34897865 | 0.46618188 | 0.64108532 | 0.90404588 |
| NCLN | 208.201058 | -0.0721106 | 0.2863584 | -0.2518195 | 0.80118057 | 0.95370871 |
| NFIC | 2037.58185 | 0.03370204 | 0.26078696 | 0.12923209 | 0.89717401 | 0.97823597 |
| SMIM24 | 78.3133899 | 1.33443675 | 0.53169792 | 2.50976483 | 0.01208116 | 0.22507761 |
| DOHH | 46.1671298 | -0.1471341 | 0.32604111 | -0.4512746 | 0.65179167 | 0.90838511 |
| FZR1 | 182.261347 | -0.2145376 | 0.24419909 | -0.8785356 | 0.37965312 | 0.78762967 |
| MFSD12 | 281.335884 | 0.05179453 | 0.22976467 | 0.22542426 | 0.82164924 | 0.96066244 |
| C19orf71 | 70.3877927 | -0.0047974 | 0.48438379 | -0.0099041 | 0.99209783 | 0.99776307 |
| AC005786.3 | 115.49429 | -0.1135018 | 0.35354072 | -0.3210431 | 0.74817775 | 0.93746253 |
| HMG20B | 216.872138 | -0.0191289 | 0.28959917 | -0.0660532 | 0.9473355 | 0.9891198 |
| CACTIN | 66.2033013 | -0.2207873 | 0.37290591 | -0.5920725 | 0.55380204 | 0.87121136 |
| PIP5K1C | 211.236591 | -0.2037879 | 0.28789086 | -0.7078651 | 0.47902906 | 0.84296219 |
| TJP3 | 235.882823 | 0.00108979 | 0.33254275 | 0.00327713 | 0.99738524 | 0.99948643 |
| APBA3 | 81.8121964 | 0.16674667 | 0.33963466 | 0.49095894 | 0.62345549 | 0.89873262 |
| MRPL54 | 168.501203 | 0.29222811 | 0.40508183 | 0.72140514 | 0.47066029 | 0.83921324 |
| DAPK3 | 236.031666 | -0.1080249 | 0.24161495 | -0.4470954 | 0.65480619 | 0.90902447 |
| EEF2 | 1097.37519 | 0.90057954 | 0.41110558 | 2.19062835 | 0.0284787 | 0.32597498 |
| PIAS4 | 218.296698 | -0.2584161 | 0.31403542 | -0.8228884 | 0.41057144 | 0.80655687 |
| ZBTB7A | 548.686907 | -0.0540564 | 0.34179542 | -0.1581542 | 0.87433528 | 0.97564728 |
| MAP2K2 | 460.942635 | 0.24845243 | 0.22587828 | 1.09993942 | 0.27135852 | 0.72018779 |
| SIRT6 | 128.392033 | -0.0592032 | 0.28604844 | -0.2069693 | 0.83603386 | 0.96511435 |
| ANKRD24 | 128.815267 | -0.4197673 | 0.71289025 | -0.5888246 | 0.55597897 | 0.87212955 |
| YJU2 | 119.585139 | -0.3366177 | 0.3939424 | -0.8544846 | 0.39283653 | 0.79842711 |
| STAP2 | 82.9760162 | -0.3837028 | 0.32691876 | -1.1736946 | 0.24051738 | 0.69178991 |
| MPND | 41.5358611 | -0.1779317 | 0.32366053 | -0.5497479 | 0.5824923 | 0.88338315 |
| SH3GL1 | 366.983648 | -0.4733315 | 0.28400793 | -1.6666135 | 0.09559129 | 0.5117083 |
| CHAF1A | 55.434177 | -0.1588099 | 0.56841886 | -0.2793889 | 0.77994641 | 0.9477748 |
| UBXN6 | 259.208094 | 0.5252257 | 0.33952931 | 1.54692301 | 0.12188181 | 0.5572644 |
| HDGFL2 | 195.6566 | -0.0341586 | 0.2981997 | -0.1145495 | 0.90880224 | 0.98050877 |
| PLIN4 | 111.162512 | 0.47536095 | 0.31542124 | 1.50706704 | 0.13179347 | 0.57539433 |
| PLIN5 | 912.751432 | 0.68390158 | 0.25331426 | 2.69981473 | 0.00693781 | 0.17097301 |
| MYDGF | 282.310441 | 0.82950983 | 0.23589952 | 3.5163693 | 0.00043749 | 0.03595008 |
| DPP9 | 507.839501 | -0.2496909 | 0.22750815 | -1.0975032 | 0.2724215 | 0.72070318 |
| FEM1A | 52.4853316 | 0.04618885 | 0.33827473 | 0.13654243 | 0.89139248 | 0.97729111 |
| TICAM1 | 156.89975 | 0.46614309 | 0.29536692 | 1.5781831 | 0.11452355 | 0.54179726 |
| PLIN3 | 429.49948 | -0.1576389 | 0.29277892 | -0.538423 | 5.90E-01 | 8.86E-01 |
| KDM4B | 1157.12801 | -0.1252514 | 0.29099671 | -0.430422 | 0.66688869 | 0.91331423 |
| PTPRS | 667.670848 | -0.6159582 | 0.30136054 | -2.0439247 | 0.04096099 | 0.37443162 |
| TINCR | 88.0878207 | -0.1466288 | 0.36702067 | -0.3995109 | 0.68951677 | 0.92168526 |
| SAFB2 | 783.131798 | -0.1821936 | 0.15975818 | -1.1404333 | 0.25410582 | 0.7045516 |
| SAFB | 933.96109 | -0.1786415 | 0.15993408 | -1.1169696 | 0.26400731 | 0.71445861 |
| RPL36 | 830.245596 | 0.457965 | 0.48898836 | 0.93655604 | 0.34898697 | 0.76982416 |
| C19orf70 | 311.7152 | 0.50366732 | 0.30716157 | 1.63974721 | 0.10105774 | 0.51819907 |
| LONP1 | 431.603367 | -0.203758 | 0.24264603 | -0.8397335 | 0.40105785 | 0.80243544 |
| DUS3L | 53.4591227 | 0.09184748 | 0.37961493 | 0.24194908 | 0.80881962 | 0.95610623 |
| AC104532.2 | 122.483928 | 0.64760012 | 0.45564535 | 1.42128109 | 0.15523506 | 0.60433242 |
| CAPS | 172.162774 | 0.96457484 | 0.45870088 | 2.10284061 | 0.0354797 | 0.35586635 |
| RANBP3 | 325.840629 | -0.0341678 | 0.25748712 | -0.1326971 | 0.89443294 | 0.97788747 |
| AC011444.1 | 51.3427033 | 0.27031154 | 0.3561654 | 0.75894946 | 0.44788279 | 0.8260638 |

| | | | | | | |
|---|---|---|---|---|---|---|
| RFX2 | 617.685049 | 0.36660003 | 0.45111795 | 0.81264783 | 0.41642 | 0.80966301 |
| MLLT1 | 308.272261 | -0.0887889 | 0.25731449 | -0.3450599 | 0.7300493 | 0.93453345 |
| CLPP | 150.786722 | 0.74721522 | 0.2533627 | 2.94919186 | 0.00318606 | 0.10846567 |
| ALKBH7 | 132.214332 | 1.29177418 | 0.41959585 | 3.07861523 | 0.00207965 | 0.0878448 |
| GTF2F1 | 189.009023 | 0.20233139 | 0.28510083 | 0.70968364 | 0.47790034 | 0.84190259 |
| KHSRP | 287.837537 | -0.2780633 | 0.29049366 | -0.9572097 | 0.33846142 | 0.76449297 |
| SLC25A23 | 273.548905 | 0.43529064 | 0.35159159 | 1.23805761 | 0.2156947 | 0.66885336 |
| CRB3 | 69.3971843 | 0.49832268 | 0.34406575 | 1.4483356 | 0.14752322 | 0.59373375 |
| AC010503.4 | 91.4557457 | 0.4485864 | 0.3247924 | 1.3811481 | 0.16723343 | 0.62087856 |
| C3 | 382.579571 | 3.34425403 | 0.62110071 | 5.38439897 | 7.27E-08 | 2.40E-04 |
| GPR108 | 84.2961553 | 0.02258036 | 0.38638355 | 0.05844028 | 0.95339793 | 0.98950524 |
| TRIP10 | 334.542203 | -0.3130623 | 0.28291639 | -1.1065543 | 2.68E-01 | 0.71818262 |
| SH2D3A | 393.806052 | -0.1086626 | 0.31851527 | -0.3411535 | 7.33E-01 | 0.93552616 |
| ZNF557 | 125.992022 | -0.5086522 | 0.21774929 | -2.3359535 | 0.01949366 | 0.28049217 |
| INSR | 2481.32984 | 0.34687499 | 0.35220212 | 0.9848748 | 0.32468557 | 0.75478872 |
| AC119396.1 | 52.7014145 | -0.0993704 | 0.49326101 | -0.201456 | 0.84034206 | 0.96617873 |
| PEX11G | 46.0522486 | 0.16744783 | 0.43846402 | 0.3818964 | 0.70253821 | 0.92501157 |
| ZNF358 | 129.25693 | 0.43128937 | 0.43152593 | 0.9994518 | 0.31757588 | 0.74948361 |
| MCOLN1 | 110.451089 | -0.3562432 | 0.40351474 | -0.8828506 | 0.37731701 | 0.78611679 |
| PNPLA6 | 253.289503 | -0.2796194 | 0.3335155 | -0.8384001 | 0.40180605 | 0.80278824 |
| CAMSAP3 | 223.709006 | -0.012654 | 0.28155673 | -0.0449429 | 0.96415285 | 0.99224805 |
| XAB2 | 103.915201 | 0.35620923 | 0.36634261 | 0.97233907 | 0.33088189 | 0.75931531 |
| PCP2 | 73.0893778 | 0.19767712 | 0.56056823 | 0.35263704 | 0.72436057 | 0.93233706 |
| AC008763.1 | 53.421624 | 0.3822863 | 0.48495938 | 0.78828518 | 0.43052991 | 0.81706845 |
| STXBP2 | 129.268249 | 0.01437217 | 0.4217735 | 0.03407556 | 0.9728169 | 0.99327086 |
| EVI5L | 186.904955 | -0.1093658 | 0.33018546 | -0.3312255 | 0.74047416 | 0.93570489 |
| MAP2K7 | 395.238232 | -0.4278738 | 0.30806814 | -1.3888935 | 0.16486515 | 0.61673853 |
| TIMM44 | 223.731033 | 0.01048526 | 0.23563823 | 0.0444973 | 0.96450801 | 0.99226095 |
| ELAVL1 | 607.646281 | -0.1061321 | 0.19245482 | -0.551465 | 0.58131498 | 0.88290096 |
| AC008946.1 | 51.8447398 | 0.06065352 | 0.48553121 | 0.12492198 | 0.90058532 | 0.97914987 |
| FBN3 | 42.5707487 | -0.2849759 | 0.84656932 | -0.3366245 | 0.73640001 | 0.93552616 |
| CERS4 | 534.346053 | 0.38627074 | 0.3591869 | 1.07540317 | 0.28219426 | 0.72604837 |
| AC022146.2 | 170.6657 | 0.55079877 | 0.54962325 | 1.00213879 | 0.31627657 | 0.74802098 |
| CD320 | 95.3654408 | 0.44060302 | 0.42629668 | 1.0335596 | 0.30134209 | 0.73820989 |
| RPS28 | 365.283089 | 0.73141848 | 0.38238322 | 1.91278914 | 0.05577505 | 0.41843207 |
| KANK3 | 45.030563 | 0.20690968 | 0.60628399 | 0.34127519 | 0.73289643 | 0.93552616 |
| ANGPTL4 | 455.642664 | -1.1076339 | 1.28960421 | -0.8588945 | 0.39039874 | 0.79701353 |
| RAB11B-AS1 | 127.248806 | -0.4153695 | 0.5039048 | -0.8243016 | 0.40976822 | 0.80641067 |
| RAB11B | 320.935626 | 0.23777262 | 0.37619901 | 0.63203946 | 0.52736109 | 0.86151701 |
| 2-Mar | 181.600332 | -0.4200298 | 0.2443962 | -1.7186428 | 0.08567944 | 0.49356597 |
| HNRNPM | 1473.90627 | -0.0716602 | 0.15238725 | -0.4702507 | 0.63817594 | 0.90301725 |
| ZNF414 | 56.273374 | -0.5666679 | 0.3992968 | -1.4191646 | 0.15585104 | 0.60519671 |
| ZNF558 | 134.759729 | -0.2734529 | 0.23883341 | -1.1449525 | 0.25222883 | 0.70276117 |
| ZNF317 | 165.383885 | 0.02822172 | 0.18761239 | 0.15042566 | 0.8804288 | 0.97664137 |
| ZNF699 | 105.611452 | -0.2430538 | 0.34113686 | -0.7124818 | 0.47616647 | 0.84141268 |
| ZNF559-ZNF | 281.060516 | 0.002105 | 0.33801241 | 0.00622759 | 0.99503113 | 0.998438 |
| ZNF266 | 316.44181 | -0.0973709 | 0.23541982 | -0.4136054 | 0.67916311 | 0.91822574 |
| ZNF426 | 152.384098 | 0.1901164 | 0.2468488 | 0.77017349 | 0.44119699 | 0.82285053 |
| ZNF121 | 298.736387 | -0.1049343 | 0.20770141 | -0.5052171 | 0.61340637 | 0.89441126 |
| ZNF561 | 231.359018 | -0.0349132 | 0.28653717 | -0.1218452 | 0.90302164 | 0.97960781 |
| ZNF562 | 495.950433 | -0.377574 | 0.1718789 | -2.1967445 | 0.0280387 | 0.3228969 |
| ZNF846 | 588.814454 | 0.0763305 | 0.22480634 | 0.33953892 | 0.73420378 | 0.93552616 |
| FBXL12 | 177.224957 | -0.1689676 | 0.20024109 | -0.8438209 | 0.39876952 | 0.80148305 |
| UBL5 | 223.44407 | 0.67201962 | 0.46317644 | 1.45089336 | 0.14680956 | 0.59373375 |

| | | | | | | |
|---|---|---|---|---|---|---|
| PIN1 | 200.67079 | 0.34573977 | 0.27557641 | 1.25460582 | 0.20962189 | 0.66233331 |
| OLFM2 | 161.611365 | -2.0583243 | 0.57801369 | -3.5610304 | 0.0003694 | 0.03231414 |
| RDH8 | 292.414297 | -2.35312 | 1.33106993 | -1.767841 | 0.07708748 | 0.47693141 |
| C19orf66 | 422.983079 | -0.5068616 | 0.39964448 | -1.2682812 | 0.20469754 | 0.65803112 |
| EIF3G | 162.745262 | 0.24484068 | 0.33842921 | 0.72346202 | 0.46939608 | 0.83881856 |
| DNMT1 | 215.829337 | -0.8312422 | 0.24493299 | -3.3937534 | 0.00068942 | 0.04573607 |
| MRPL4 | 202.549658 | 0.35116722 | 0.29396229 | 1.19459957 | 0.23224352 | 0.68450837 |
| AC011511.2 | 67.4095271 | 0.58136013 | 0.61647936 | 0.94303259 | 0.34566423 | 0.76802752 |
| ICAM1 | 104.520654 | 0.32666066 | 0.55778014 | 0.58564413 | 5.58E-01 | 8.73E-01 |
| ZGLP1 | 37.9818508 | -0.9324696 | 0.63598795 | -1.466175 | 0.14260063 | 0.58824095 |
| RAVER1 | 75.9110829 | -0.5941973 | 0.34939026 | -1.7006693 | 0.0890051 | 0.49879864 |
| TYK2 | 365.437819 | -0.317809 | 0.3734269 | -0.8510608 | 0.39473556 | 0.7988451 |
| CDC37 | 484.143503 | -0.3653308 | 0.22898579 | -1.59543 | 0.11061612 | 0.53393773 |
| PDE4A | 148.411323 | -0.7483903 | 0.57905556 | -1.2924326 | 0.19620737 | 0.64964978 |
| KEAP1 | 152.042388 | -0.0057995 | 0.33636571 | -0.0172418 | 0.98624373 | 0.99624198 |
| ATG4D | 168.340666 | -0.0601823 | 0.22495603 | -0.2675293 | 0.78906163 | 0.95080703 |
| KRI1 | 181.559986 | -0.2395454 | 0.26853665 | -0.8920399 | 0.37237156 | 0.78260237 |
| CDKN2D | 44.0053053 | 0.52012995 | 0.41688535 | 1.24765705 | 0.21215667 | 0.66552984 |
| AP1M2 | 275.078796 | 0.53323632 | 0.32376091 | 1.64700649 | 0.09955671 | 0.51665771 |
| SLC44A2 | 416.140864 | 0.15884264 | 0.438356 | 0.3623599 | 0.7170831 | 0.9290703 |
| ILF3-DT | 263.588857 | 0.29631244 | 0.35148048 | 0.84304096 | 0.39920554 | 0.80174499 |
| ILF3 | 797.116073 | -0.1890755 | 0.25562483 | -0.7396602 | 0.45950618 | 0.83390278 |
| QTRT1 | 358.296035 | -0.3442955 | 0.3121876 | -1.102848 | 0.27009319 | 0.71914149 |
| DNM2 | 2811.7189 | 0.0213382 | 0.35428673 | 0.06022863 | 0.95197355 | 0.98938795 |
| TMED1 | 95.7313976 | 0.62713587 | 0.42098849 | 1.48967463 | 0.13630981 | 0.57787125 |
| CARM1 | 373.99099 | -0.4189124 | 0.3192817 | -1.3120463 | 0.1895045 | 0.64441858 |
| YIPF2 | 216.548734 | 0.17993726 | 0.23662271 | 0.76043951 | 0.44699191 | 0.82571251 |
| TIMM29 | 137.335181 | 0.55636529 | 0.26794847 | 2.07638911 | 0.03785798 | 0.36437085 |
| SMARCA4 | 1343.01484 | -0.1646506 | 0.41645103 | -0.3953661 | 0.69257271 | 0.92224682 |
| LDLR | 538.704832 | 0.11993538 | 0.61072706 | 0.1963813 | 0.84431173 | 0.96850656 |
| KANK2 | 378.717689 | -0.1762691 | 0.2948982 | -0.5977285 | 0.55002108 | 0.86956073 |
| DOCK6 | 640.333756 | 0.11288422 | 0.43503826 | 0.25948112 | 0.79526404 | 0.95278473 |
| AC011472.1 | 41.762712 | -0.4217642 | 0.45448125 | -0.9280123 | 0.35340119 | 0.77219738 |
| RAB3D | 202.463224 | 0.22432689 | 0.31595656 | 0.70999282 | 4.78E-01 | 0.84190259 |
| CCDC159 | 64.6503954 | 0.2286481 | 0.43673957 | 0.52353419 | 0.60060256 | 0.88883692 |
| PLPPR2 | 53.9574545 | -0.1783687 | 0.37486066 | -0.4758265 | 0.63419796 | 0.90136076 |
| SWSAP1 | 63.8349983 | -0.0069472 | 0.30437515 | -0.0228246 | 0.9817902 | 0.99518016 |
| EPOR | 252.264418 | 0.20599717 | 0.53489427 | 0.38511755 | 0.70015033 | 0.92367966 |
| RGL3 | 2354.5512 | 0.2894524 | 0.32693096 | 0.88536247 | 0.37596117 | 0.78490077 |
| CCDC151 | 45.5779376 | 0.02011469 | 0.46321501 | 0.04342409 | 0.96536347 | 0.99248831 |
| PRKCSH | 251.215923 | 0.23723902 | 0.27440053 | 0.86457201 | 0.38727374 | 0.79554171 |
| ECSIT | 310.810106 | -0.175505 | 0.29588696 | -0.5931487 | 0.55308162 | 0.87104567 |
| ELOF1 | 96.7905947 | 0.43713744 | 0.40660122 | 1.07510115 | 0.28232945 | 0.72604837 |
| ZNF627 | 378.198237 | -0.2603782 | 0.29031 | -0.8968973 | 0.36977374 | 0.78095617 |
| AC008543.1 | 182.027254 | 1.03729134 | 0.44810725 | 2.31482829 | 0.02062233 | 0.28764558 |
| ZNF823 | 345.14218 | -0.0993914 | 0.22032054 | -0.4511218 | 0.65190176 | 0.90838511 |
| ZNF441 | 265.987499 | -0.102995 | 0.30584034 | -0.3367606 | 0.73629735 | 0.93552616 |
| ZNF491 | 73.2897757 | -0.0229179 | 0.28670274 | -0.0799361 | 0.9362881 | 0.98621411 |
| ZNF440 | 325.640552 | -0.1101356 | 0.23885605 | -0.4610963 | 0.6447295 | 0.90507831 |
| ZNF439 | 547.980821 | 0.00673016 | 0.21582105 | 0.03118398 | 0.97512282 | 0.99408793 |
| ZNF69 | 819.818732 | -0.3521843 | 0.24647388 | -1.428891 | 0.15303558 | 0.60285224 |
| AC008770.2 | 121.19395 | -0.0056841 | 0.26237095 | -0.0216646 | 0.98271554 | 0.99518552 |
| ZNF763 | 57.482723 | 0.13841946 | 0.55450688 | 0.24962623 | 0.80287641 | 0.95446295 |
| ZNF433 | 106.211308 | 0.32639768 | 0.39191432 | 0.83282919 | 0.40494107 | 0.80455756 |

| | | | | | | |
|---|---|---|---|---|---|---|
| AC008770.1 | 39.7340958 | -0.1914064 | 0.41472736 | -0.4615234 | 0.64442314 | 0.90507553 |
| ZNF844 | 348.788496 | 0.41987098 | 0.26009076 | 1.61432485 | 0.10645698 | 0.52743811 |
| ZNF788P | 207.865481 | 0.52571249 | 0.28125648 | 1.86915692 | 0.06160099 | 0.43249565 |
| ZNF20 | 96.8950511 | 0.86420312 | 0.38857847 | 2.22401186 | 0.02614765 | 0.31313169 |
| ZNF136 | 185.491941 | -0.1387032 | 0.19386853 | -0.7154498 | 0.47433113 | 0.84106098 |
| ZNF44 | 2087.98228 | 0.13037742 | 0.21758091 | 0.59921348 | 0.54903053 | 0.86904066 |
| ZNF563 | 115.505564 | 0.43305309 | 0.39433407 | 1.09818838 | 0.27212224 | 0.72070318 |
| ZNF442 | 90.0041138 | 0.47477617 | 0.36569389 | 1.29828849 | 0.19418822 | 0.64760078 |
| AC008758.5 | 156.072099 | -1.2368838 | 0.49847975 | -2.481312 | 0.01308997 | 0.23485956 |
| AC008758.6 | 81.3665552 | -0.8766725 | 0.39146171 | -2.2394848 | 0.02512439 | 0.31053369 |
| AC008758.1 | 147.521944 | -1.0783671 | 0.36556493 | -2.9498647 | 0.00317913 | 0.10846567 |
| AC010422.6 | 43.529012 | -0.0315837 | 0.34977467 | -0.0902973 | 0.928051 | 0.98297055 |
| ZNF490 | 391.78003 | 0.05344387 | 0.27292246 | 0.19582071 | 0.8447505 | 0.96853443 |
| ZNF791 | 503.381653 | 0.06194409 | 0.14809516 | 0.41827221 | 0.67574811 | 0.91649321 |
| MAN2B1 | 124.219086 | -0.1851303 | 0.30307373 | -0.6108425 | 0.54130388 | 0.86582694 |
| WDR83 | 102.09712 | -0.014873 | 0.28290764 | -0.0525719 | 0.95807297 | 0.99069768 |
| DHPS | 156.120383 | 0.08294817 | 0.25559988 | 0.32452352 | 0.74554172 | 0.93675395 |
| FBXW9 | 56.8593935 | 0.25362595 | 0.43390504 | 0.5845195 | 0.55887085 | 0.87295536 |
| TNPO2 | 412.818254 | -0.2110994 | 0.22751959 | -0.9278295 | 0.35349603 | 0.77219738 |
| TRIR | 423.152057 | 0.25674101 | 0.22760597 | 1.12800651 | 0.25931717 | 0.70921184 |
| ASNA1 | 132.401251 | 0.78365701 | 0.33903283 | 2.31144875 | 0.02080808 | 0.28870816 |
| HOOK2 | 1239.79671 | -0.2010903 | 0.26955389 | -0.7460116 | 0.45566041 | 0.83130183 |
| JUNB | 441.241433 | 0.38347762 | 0.55287215 | 0.69360995 | 0.48792686 | 0.84513845 |
| DNASE2 | 62.1509931 | 0.97015313 | 0.41425168 | 2.34194135 | 0.01918373 | 0.27999766 |
| GCDH | 147.189056 | 0.09445893 | 0.36039635 | 0.26209737 | 0.79324638 | 0.95224951 |
| FARSA | 180.490145 | 0.24490716 | 0.29015149 | 0.84406652 | 0.39863224 | 0.80148305 |
| CALR | 650.734289 | 1.54664573 | 0.41708183 | 3.70825489 | 0.00020869 | 0.02336118 |
| AC092069.1 | 123.183178 | 1.12823312 | 0.60289438 | 1.87136115 | 0.06129504 | 0.43236485 |
| RAD23A | 286.279351 | -0.1679429 | 0.21392532 | -0.7850539 | 0.43242194 | 0.81773248 |
| GADD45GIP1 | 310.800177 | 0.84713944 | 0.45221455 | 1.8733131 | 0.06102516 | 0.43176766 |
| NFIX | 111.638848 | -0.1431482 | 0.35588713 | -0.4022291 | 0.68751545 | 0.92094022 |
| TRMT1 | 100.503115 | -0.3981073 | 0.38470135 | -1.0348476 | 0.30074009 | 0.73750086 |
| NACC1 | 151.877571 | 0.07830798 | 0.32505495 | 0.24090689 | 0.80962728 | 0.95656232 |
| STX10 | 119.075625 | -0.1935292 | 0.30155784 | -0.6417649 | 0.52102582 | 0.85943874 |
| IER2 | 357.536702 | 0.38638812 | 0.47998403 | 0.80500203 | 0.42081851 | 0.81260593 |
| CACNA1A | 63.4679673 | -0.0532852 | 0.37648926 | -0.1415319 | 0.88744976 | 0.97729111 |
| CCDC130 | 680.635971 | -0.0860262 | 0.25969972 | -0.3312524 | 0.74045385 | 0.93570489 |
| MRI1 | 370.0698 | 0.08033689 | 0.32976107 | 0.2436215 | 0.80752397 | 0.95594682 |
| AC008686.1 | 44.979373 | -0.2904705 | 0.7729598 | -0.3757899 | 0.70707311 | 0.92573766 |
| C19orf53 | 361.097141 | 0.46898077 | 0.49071954 | 0.9557002 | 0.33922372 | 0.76449297 |
| ZSWIM4 | 291.706396 | -0.7013012 | 0.28025184 | -2.5023965 | 1.23E-02 | 0.22749396 |
| AC020916.1 | 107.576097 | -0.3759987 | 0.57263647 | -0.6566098 | 0.51143184 | 0.85762435 |
| CC2D1A | 406.113962 | -0.3672916 | 0.22179823 | -1.6559716 | 0.09772758 | 0.51355946 |
| DCAF15 | 131.021546 | -0.2172851 | 0.41290927 | -0.5262297 | 0.59872865 | 0.88883692 |
| RFX1 | 335.151821 | -0.3673468 | 0.26934307 | -1.3638621 | 0.17261097 | 0.6286116 |
| PALM3 | 153.498534 | -1.2046244 | 0.56674018 | -2.1255321 | 0.03354224 | 0.34996799 |
| MISP3 | 83.5641175 | 0.21057468 | 0.58156062 | 0.36208551 | 0.71728813 | 0.9290703 |
| SAMD1 | 99.876217 | -0.0184376 | 0.54856256 | -0.0336107 | 0.97318756 | 0.99327086 |
| PRKACA | 265.205777 | 0.23801556 | 0.28862755 | 0.824646 | 0.40957259 | 0.80641067 |
| AC022098.1 | 103.331378 | -0.2049232 | 0.27662569 | -0.7407962 | 0.45881705 | 0.83340407 |
| ADGRL1 | 241.03816 | -0.6160931 | 0.3589767 | -1.7162482 | 0.08611663 | 0.49453468 |
| DDX39A | 89.9914767 | -0.6793284 | 0.26979204 | -2.5179704 | 0.01180332 | 0.22304735 |
| PKN1 | 191.112995 | -0.5980605 | 0.36714044 | -1.6289693 | 0.10331952 | 0.52337282 |
| GIPC1 | 762.512861 | -0.6964952 | 0.30420255 | -2.289577 | 0.02204585 | 0.29509055 |

| | | | | | | |
|---|---|---|---|---|---|---|
| DNAJB1 | 183.756387 | 0.2928764 | 0.33762545 | 0.86745949 | 0.3856903 | 0.79391977 |
| TECR | 277.631428 | -0.0267243 | 0.29176919 | -0.0915941 | 0.92702052 | 0.98252755 |
| NDUFB7 | 414.089535 | 0.45306375 | 0.38929515 | 1.1638053 | 0.24450293 | 0.6948422 |
| ADGRE3 | 53.6386167 | 0.31671168 | 0.34188362 | 0.92637278 | 0.35425229 | 0.77243419 |
| ZNF333 | 138.208933 | -0.5616507 | 0.43493806 | -1.2913349 | 0.19658758 | 0.64964978 |
| SYDE1 | 42.4634893 | 0.17890683 | 0.41514023 | 0.43095516 | 0.66650097 | 0.91318863 |
| ILVBL | 188.519325 | 0.46525418 | 0.30150061 | 1.54312847 | 0.12279961 | 0.55947129 |
| NOTCH3 | 234.093445 | -0.5418059 | 0.53693776 | -1.0090664 | 0.31294278 | 0.74703371 |
| BRD4 | 1987.99152 | -0.2004692 | 0.22161189 | -0.9045959 | 0.3656795 | 0.78095617 |
| AKAP8 | 623.858715 | -0.146741 | 0.23422625 | -0.6264927 | 0.53099185 | 0.86397197 |
| AKAP8L | 1209.84192 | -0.2065916 | 0.14785491 | -1.397259 | 0.1623357 | 0.61373097 |
| WIZ | 118.901149 | 0.19002238 | 0.32986078 | 0.57606843 | 0.56456893 | 0.87569175 |
| TPM4 | 146.619194 | 0.9153871 | 0.49740528 | 1.84032448 | 0.06572061 | 0.44427001 |
| RAB8A | 135.808103 | 0.07091849 | 0.27624636 | 0.25672188 | 0.79739348 | 0.95281098 |
| FAM32A | 117.876508 | 0.3663471 | 0.31532028 | 1.16182536 | 0.24530641 | 0.69518395 |
| AP1M1 | 195.25158 | -0.0956411 | 0.1932278 | -0.4949655 | 0.62062451 | 0.89731054 |
| KLF2 | 60.3857416 | 2.01799979 | 0.91872157 | 2.19653033 | 0.02805401 | 0.3228969 |
| EPS15L1 | 802.837163 | -0.2876335 | 0.19824174 | -1.4509229 | 0.14680134 | 0.59373375 |
| AC008764.1 | 55.2329087 | -0.1331133 | 0.38638158 | -0.3445125 | 0.73046087 | 0.93467574 |
| C19orf44 | 173.528185 | 0.14521447 | 0.26827327 | 0.5412931 | 0.58830557 | 0.88571413 |
| AC008764.8 | 47.5629608 | 0.32524654 | 0.37991252 | 0.85610902 | 0.39193748 | 0.79767768 |
| AC024075.3 | 41.6581957 | -0.0776539 | 0.50275776 | -0.154456 | 0.87725023 | 0.97639835 |
| TMEM38A | 236.72314 | -0.2180703 | 0.31773295 | -0.686332 | 0.49250377 | 0.84738939 |
| NWD1 | 93.6608077 | 0.69744179 | 0.45590404 | 1.52979954 | 1.26E-01 | 0.56562856 |
| SIN3B | 550.093922 | -0.2359008 | 0.13731611 | -1.7179399 | 0.08580758 | 0.49387995 |
| CPAMD8 | 68.3729989 | 1.56201626 | 0.78187884 | 1.99777277 | 0.0457413 | 0.38904776 |
| HAUS8 | 72.839034 | -0.4208564 | 0.32936806 | -1.2777695 | 0.20133073 | 0.6550191 |
| MYO9B | 704.704187 | -0.5103238 | 0.26966864 | -1.8924105 | 0.05843631 | 0.42147872 |
| USE1 | 102.632452 | -0.2865126 | 0.3127559 | -0.9160901 | 0.35961963 | 0.77668667 |
| OCEL1 | 83.5575951 | -0.1626891 | 0.34585896 | -0.4703916 | 0.63807528 | 0.90301725 |
| NR2F6 | 208.102617 | 0.52546302 | 0.3951802 | 1.32967952 | 0.18362389 | 0.63834582 |
| USHBP1 | 113.849357 | -0.0857386 | 0.30448525 | -0.2815855 | 0.77826135 | 0.94694572 |
| ABHD8 | 148.279661 | 0.02193545 | 0.25763798 | 0.0851406 | 0.93214961 | 0.98360577 |
| MRPL34 | 215.529009 | 0.46564657 | 0.47415911 | 0.98204709 | 0.32607665 | 0.75550719 |
| DDA1 | 145.289977 | 0.11738626 | 0.2376685 | 0.49390754 | 0.62137148 | 0.89789913 |
| ANO8 | 64.473951 | -0.2504875 | 0.30657056 | -0.8170631 | 0.41389238 | 0.80874327 |
| GTPBP3 | 92.4426899 | -0.6368934 | 0.2493699 | -2.5540108 | 0.01064899 | 0.20963126 |
| BST2 | 307.78168 | 2.02555824 | 0.69960076 | 2.89530596 | 0.00378789 | 0.12027465 |
| MVB12A | 212.847542 | 0.24076465 | 0.29130002 | 0.82651777 | 0.40851044 | 0.80588056 |
| SLC27A1 | 337.615975 | 0.12480221 | 0.34761787 | 0.35902127 | 0.71957918 | 0.93001079 |
| AC010618.3 | 262.037344 | -0.0421448 | 0.29080764 | -0.1449234 | 0.88477131 | 0.97664137 |
| PGLS | 276.468397 | 0.21365191 | 0.30035214 | 0.71133806 | 0.47687477 | 0.84155495 |
| COLGALT1 | 158.332296 | -0.6217133 | 0.23448599 | -2.6513876 | 0.00801618 | 0.17977198 |
| MAP1S | 71.3906036 | 0.07880337 | 0.29143208 | 0.27040047 | 0.78685218 | 0.94978829 |
| RPL18A | 93.4160866 | 0.40449345 | 0.52573971 | 0.76937968 | 0.44166795 | 0.82285053 |
| CCDC124 | 116.493225 | 0.11228734 | 0.34082325 | 0.32945916 | 0.74180866 | 0.93598004 |
| ARRDC2 | 136.708257 | -0.1183487 | 0.54090261 | -0.2187986 | 8.27E-01 | 9.62E-01 |
| MAST3 | 83.8032279 | 0.28927231 | 0.34415969 | 0.8405177 | 0.40061818 | 0.80243544 |
| IQCN | 91.4946402 | -0.2954073 | 0.57579873 | -0.5130392 | 0.6079239 | 0.89203142 |
| JUND | 663.555445 | -0.2058139 | 0.50583597 | -0.4068788 | 0.68409703 | 0.9200778 |
| LSM4 | 250.182434 | 0.54857168 | 0.33372684 | 1.64377452 | 0.10022279 | 0.51692417 |
| PGPEP1 | 377.36737 | -0.0553921 | 0.27809954 | -0.1991809 | 0.84212123 | 0.96709957 |
| GDF15 | 1038.76319 | 0.0188475 | 0.55532557 | 0.03393955 | 9.73E-01 | 9.93E-01 |
| SSBP4 | 295.338865 | -0.00062 | 0.26149377 | -0.0023709 | 0.99810827 | 0.99969006 |

| | | | | | | |
|---|---|---|---|---|---|---|
| ISYNA1 | 189.533533 | 0.33457202 | 0.57753498 | 0.57931041 | 0.56237974 | 0.87424668 |
| ELL | 713.433595 | 0.02952203 | 0.24191381 | 0.12203533 | 0.90287103 | 0.97960781 |
| FKBP8 | 302.76186 | 0.56400037 | 0.31880515 | 1.76910682 | 0.07687605 | 0.47651607 |
| KXD1 | 183.052198 | -0.0216917 | 0.21560136 | -0.1006102 | 0.91985989 | 0.98215577 |
| AC005253.1 | 46.3462034 | -0.0889173 | 0.61954095 | -0.1435213 | 0.88587846 | 0.97667712 |
| UBA52 | 488.23207 | 0.55195627 | 0.47622743 | 1.15901823 | 0.24644876 | 0.69677689 |
| CRLF1 | 54.1112068 | 0.34753108 | 0.38026329 | 0.91392225 | 0.3607577 | 0.77730375 |
| REX1BD | 170.08394 | 0.7587152 | 0.40960573 | 1.85230614 | 0.06398187 | 0.43872009 |
| KLHL26 | 79.0162381 | 0.34403733 | 0.50213258 | 0.68515238 | 0.49324777 | 0.84812676 |
| CRTC1 | 970.526413 | -0.1996027 | 0.27187175 | -0.7341796 | 0.46283931 | 0.83531269 |
| UPF1 | 415.801648 | 0.10964654 | 0.22357362 | 0.49042703 | 0.62383176 | 0.89873262 |
| COPE | 241.014496 | 0.24175587 | 0.27088068 | 0.89248104 | 0.37213515 | 0.78256595 |
| DDX49 | 128.099084 | -0.3877685 | 0.40795364 | -0.9505211 | 0.34184754 | 0.76546265 |
| SUGP2 | 1060.54859 | -0.1287211 | 0.18527539 | -0.6947558 | 0.48720836 | 0.84475225 |
| ARMC6 | 93.1400299 | -0.6276629 | 0.23299289 | -2.6939142 | 0.00706183 | 0.17262485 |
| SLC25A42 | 221.67288 | 0.78609611 | 0.37294536 | 2.10780501 | 0.03504786 | 0.3536866 |
| TMEM161A | 152.970061 | 0.2939407 | 0.25436943 | 1.15556615 | 0.24785866 | 0.69837138 |
| BORCS8-MEI | 85.7683967 | -0.4445322 | 0.40979839 | -1.0847582 | 0.27802877 | 0.72372365 |
| RFXANK | 96.2613306 | 0.18390386 | 0.26469255 | 0.6947829 | 0.48719137 | 0.84475225 |
| SUGP1 | 255.367746 | -0.3539622 | 0.19750492 | -1.7921691 | 0.07310588 | 0.47001711 |
| MAU2 | 747.58842 | -0.2856399 | 0.21567544 | -1.3243971 | 0.18537118 | 0.64030106 |
| GATAD2A | 1326.63148 | 0.00080606 | 0.22868239 | 0.00352479 | 0.99718763 | 0.99948643 |
| YJEFN3 | 58.3121445 | -0.7019794 | 0.47830662 | -1.4676348 | 0.14220346 | 0.58772386 |
| PBX4 | 412.022252 | 0.09098131 | 0.41540831 | 0.21901658 | 0.82663713 | 0.96214368 |
| ATP13A1 | 179.816515 | 0.24317163 | 0.40059249 | 0.60702994 | 0.54383105 | 0.86639505 |
| ZNF101 | 129.2743 | -0.1159623 | 0.30285517 | -0.3828969 | 0.70179621 | 0.92469089 |
| ZNF14 | 245.016799 | -0.1091029 | 0.31443 | -0.3469862 | 0.72860166 | 0.93453345 |
| AC011477.4 | 166.997244 | 0.058185 | 0.34763341 | 0.1673746 | 0.86707531 | 0.97402505 |
| ZNF253 | 432.296212 | 0.07979083 | 0.18525358 | 0.4307114 | 0.66667823 | 0.91321713 |
| ZNF93 | 137.381814 | -0.6023758 | 0.39081384 | -1.541337 | 0.12323478 | 0.56015422 |
| ZNF682 | 338.385954 | -0.2389705 | 0.45283127 | -0.5277252 | 0.59769009 | 0.88883692 |
| ZNF90 | 712.995929 | 0.44802787 | 0.38206632 | 1.17264426 | 0.24093849 | 0.69178991 |
| AC011447.3 | 487.935559 | -0.062229 | 0.33454069 | -0.1860132 | 0.85243438 | 0.97076831 |
| ZNF486 | 429.871656 | -0.3311377 | 0.5872553 | -0.5638735 | 0.57284021 | 0.87946834 |
| AC008554.1 | 534.771359 | 2.01812754 | 0.61854363 | 3.26270844 | 0.00110353 | 0.05961135 |
| ZNF737 | 416.326822 | 0.19263256 | 0.25517215 | 0.75491218 | 0.45030168 | 0.82749512 |
| AC010636.2 | 85.5235845 | -0.2728042 | 0.41930229 | -0.6506147 | 0.51529524 | 0.85869987 |
| ZNF66 | 138.818831 | -0.6825111 | 0.31547164 | -2.1634626 | 0.03050562 | 0.33556032 |
| ZNF85 | 169.504143 | 0.03643673 | 0.49707586 | 0.07330214 | 0.94156569 | 0.98754495 |
| ZNF430 | 428.320717 | -0.3382094 | 0.16148267 | -2.0944008 | 0.03622429 | 0.35817721 |
| ZNF714 | 244.219611 | -0.0848033 | 0.32537694 | -0.260631 | 0.79437709 | 0.95251697 |
| ZNF431 | 906.349541 | -0.0110894 | 0.17410042 | -0.0636951 | 0.94921297 | 0.98919519 |
| ZNF708 | 642.300497 | -0.0413771 | 0.22166379 | -0.1866662 | 0.85192233 | 0.97076831 |
| ZNF738 | 115.477064 | -0.533288 | 0.33449275 | -1.5943186 | 0.11086468 | 0.53426181 |
| ZNF493 | 382.323377 | 0.25681016 | 0.22312837 | 1.15095251 | 2.50E-01 | 0.70032501 |
| AC010615.4 | 537.567027 | 0.37031677 | 0.75380167 | 0.49126552 | 0.62323866 | 0.8987292 |
| ZNF429 | 678.939658 | 0.10469617 | 0.2487995 | 0.4208054 | 0.67389719 | 0.91636338 |
| AC123912.1 | 242.021582 | 0.23063522 | 0.67801686 | 0.34016149 | 0.73373492 | 0.93552616 |
| AC123912.4 | 62.2408196 | 0.22509472 | 0.51587925 | 0.43633219 | 0.66259573 | 0.91110616 |
| ZNF100 | 428.813238 | -0.1213136 | 0.49266241 | -0.2462408 | 0.80549584 | 0.95522565 |
| ZNF43 | 631.93004 | -0.3906763 | 0.28762296 | -1.358293 | 0.17437073 | 0.62892483 |
| ZNF208 | 661.758602 | 0.32587683 | 0.49175554 | 0.66268053 | 0.50753518 | 0.85632606 |
| ZNF257 | 134.268211 | -0.5099804 | 0.30774329 | -1.6571618 | 0.09748678 | 0.51355946 |
| ZNF676 | 157.612577 | 0.04076011 | 0.65542815 | 0.06218853 | 0.9504127 | 0.98929186 |

| | | | | | | |
|---|---|---|---|---|---|---|
| ZNF98 | 1343.79733 | -0.0247528 | 0.41442098 | -0.0597287 | 0.95237168 | 0.98938795 |
| ZNF99 | 63.5359701 | 0.85150734 | 0.75075829 | 1.13419639 | 0.25671218 | 0.70735159 |
| AC092329.3 | 138.487853 | -0.4108289 | 0.39720088 | -1.0343102 | 0.30099118 | 0.73776073 |
| ZNF91 | 1959.75554 | -0.0070778 | 0.17300313 | -0.0409113 | 0.96736664 | 0.99248831 |
| ZNF675 | 220.940897 | -0.1128782 | 0.23502405 | -0.4802834 | 0.63102587 | 0.90005881 |
| ZNF681 | 53.0775334 | 0.26988107 | 0.35587097 | 0.75836775 | 0.44823086 | 0.82623672 |
| AC139769.2 | 98.5941979 | 0.2849048 | 0.61700249 | 0.46175632 | 0.64425608 | 0.90507553 |
| ZNF726 | 138.064424 | 0.54223311 | 0.32166781 | 1.68569281 | 0.09185498 | 0.50476839 |
| AC011503.1 | 78.5933664 | 1.11166317 | 0.60645361 | 1.83305556 | 0.06679431 | 0.44854399 |
| ZNF254 | 1784.87747 | -0.1385092 | 0.19434622 | -0.7126931 | 0.47603567 | 0.84141268 |
| LINC00662 | 536.912121 | -0.2031347 | 0.22778081 | -0.891799 | 0.37250066 | 0.78269743 |
| AC006504.5 | 403.038001 | 0.63256901 | 0.45360669 | 1.39453193 | 0.16315704 | 0.61387934 |
| UQCRFS1 | 500.723859 | 0.73755444 | 0.43910893 | 1.6796617 | 0.09302316 | 0.50795488 |
| POP4 | 75.4704283 | 0.06012533 | 0.33144127 | 0.18140569 | 0.85604916 | 0.97118784 |
| C19orf12 | 346.449115 | 0.13084655 | 0.23376477 | 0.55973597 | 0.57565955 | 0.88099721 |
| URI1 | 2045.97879 | -0.2102954 | 0.10328593 | -2.0360507 | 0.04174526 | 0.37639126 |
| TSHZ3 | 80.11672 | -0.3863783 | 0.53409531 | -0.7234258 | 0.46941834 | 0.83881856 |
| ZNF507 | 505.971364 | -0.249241 | 0.1331943 | -1.871259 | 0.06130919 | 0.43236485 |
| DPY19L3 | 797.038928 | -0.234143 | 0.22384345 | -1.0460124 | 0.29555531 | 0.73661928 |
| PDCD5 | 177.474225 | -0.0033156 | 0.22489884 | -0.0147425 | 0.98823763 | 0.99653761 |
| ANKRD27 | 410.349352 | -0.2010248 | 0.19369301 | -1.0378528 | 0.29933858 | 0.73673761 |
| NUDT19 | 144.251242 | 0.16120361 | 0.20318794 | 0.79337193 | 4.28E-01 | 0.81466629 |
| CEP89 | 432.599732 | -0.140719 | 0.23585701 | -0.5966282 | 0.55075561 | 0.86999891 |
| RHPN2 | 1081.53523 | -0.4356363 | 0.25440507 | -1.7123728 | 8.68E-02 | 4.95E-01 |
| GPATCH1 | 170.625586 | -0.2141919 | 0.25580074 | -0.8373388 | 0.40240218 | 0.80310019 |
| WDR88 | 60.1864328 | -0.5729012 | 0.43196907 | -1.3262551 | 0.18475523 | 0.64002932 |
| LRP3 | 81.7021174 | 0.6379138 | 0.42496734 | 1.50108901 | 0.13333254 | 0.57682553 |
| CEBPG | 80.1730352 | -0.5282695 | 0.41472548 | -1.2737813 | 0.20274094 | 0.65552334 |
| PEPD | 350.706797 | -0.0059107 | 0.2790908 | -0.0211783 | 0.98310341 | 0.99518552 |
| KCTD15 | 157.48288 | -0.4161367 | 0.38119124 | -1.0916743 | 0.27497627 | 0.72166928 |
| LSM14A | 2487.22217 | -0.1914018 | 0.10514747 | -1.8203175 | 0.06871067 | 0.45448131 |
| KIAA0355 | 1295.15615 | -0.0167133 | 0.17125519 | -0.0975931 | 0.92225538 | 0.98215577 |
| AC010504.1 | 43.395341 | 0.53685144 | 0.48430959 | 1.10848814 | 0.26765106 | 0.71784829 |
| GPI | 2012.43435 | -0.358775 | 0.45937864 | -0.7810007 | 0.4348021 | 0.81919857 |
| UBA2 | 875.567108 | -0.2076693 | 0.18633818 | -1.1144752 | 0.26507537 | 0.71512755 |
| WTIP | 135.723142 | 0.81488919 | 0.3010973 | 2.70639823 | 0.00680174 | 0.16951739 |
| SCGB2B2 | 894.46819 | -0.3434066 | 0.35356445 | -0.9712702 | 0.33141372 | 0.76000762 |
| ZNF302 | 809.677892 | -0.0600689 | 0.22494154 | -0.2670422 | 0.78943668 | 0.95090647 |
| ZNF181 | 126.71304 | 0.2078062 | 0.22091019 | 0.94068181 | 0.34686795 | 0.76875481 |
| ZNF599 | 56.4523462 | 0.26993479 | 0.34868775 | 0.77414475 | 0.43884519 | 0.82127804 |
| ZNF30 | 94.6945959 | -0.2026016 | 0.45516355 | -0.4451183 | 0.65623429 | 0.90950195 |
| GRAMD1A | 422.233021 | -0.0485883 | 0.29089446 | -0.1670307 | 8.67E-01 | 0.97402505 |
| SCN1B | 49.0093034 | -0.2807654 | 0.47190641 | -0.5949599 | 0.55187025 | 0.87051638 |
| HPN | 1964.80456 | 0.48373284 | 0.41299225 | 1.17128794 | 0.24148306 | 0.69237024 |
| LSR | 337.273683 | -0.1101796 | 0.22067575 | -0.4992828 | 6.18E-01 | 8.96E-01 |
| USF2 | 293.032012 | -0.4481858 | 0.27560253 | -1.6262035 | 0.10390638 | 0.52442862 |
| DMKN | 451.051441 | 0.1479288 | 0.29721009 | 0.49772468 | 0.61867811 | 0.89606569 |
| TMEM147-A: | 55.1610749 | -0.0718879 | 0.3233413 | -0.2223283 | 0.82405833 | 0.96141406 |
| TMEM147 | 153.668355 | 0.50091605 | 0.49869301 | 1.00445773 | 0.31515804 | 0.74785672 |
| HAUS5 | 83.3732456 | -0.4492961 | 0.45785454 | -0.9813076 | 0.3264411 | 0.75554697 |
| RBM42 | 106.955875 | 0.27100542 | 0.31333016 | 0.86491965 | 0.38708289 | 0.79542282 |
| COX6B1 | 663.743709 | 0.56546603 | 0.34495986 | 1.63922269 | 0.10116689 | 0.51819907 |
| KMT2B | 297.28194 | -0.2941095 | 0.25008259 | -1.1760496 | 0.23957506 | 0.69079829 |
| PROSER3 | 179.410789 | -0.3640875 | 0.30034761 | -1.2122203 | 0.22542808 | 0.67798093 |

| | | | | | | |
|---|---|---|---|---|---|---|
| ARHGAP33 | 45.4523654 | 1.41867695 | 0.48347264 | 2.93434796 | 0.00334249 | 0.11234348 |
| NFKBID | 73.4675516 | -0.4475282 | 0.29942123 | -1.4946441 | 0.13500735 | 0.57708028 |
| SDHAF1 | 56.4511334 | 0.89261698 | 0.41304745 | 2.16105191 | 0.03069133 | 0.33646028 |
| SYNE4 | 89.6516337 | -0.0489226 | 0.37985575 | -0.1287925 | 0.89752183 | 0.97823597 |
| ALKBH6 | 60.2375244 | -0.3815721 | 0.28417855 | -1.3427196 | 0.17936278 | 0.63313816 |
| AC002116.2 | 53.8845981 | -0.4608054 | 0.45816619 | -1.0057605 | 0.31453081 | 0.74785672 |
| THAP8 | 51.6751634 | -0.2700951 | 0.43562657 | -0.6200151 | 0.53524783 | 0.8642718 |
| TBCB | 171.926025 | 0.00396709 | 0.33677665 | 0.01177959 | 0.99060146 | 0.99740343 |
| CAPNS1 | 525.442816 | 0.32807048 | 0.34873482 | 0.94074483 | 3.47E-01 | 0.76875481 |
| COX7A1 | 55.299423 | 0.80243299 | 0.68068783 | 1.17885607 | 0.23845549 | 0.68982886 |
| ZNF565 | 632.613831 | -0.0488108 | 0.30415093 | -0.1604821 | 0.87250128 | 0.97554234 |
| ZNF146 | 661.051829 | -0.1992198 | 0.12763967 | -1.5607988 | 0.11857123 | 0.54998912 |
| LINC00665 | 310.57484 | 0.25708049 | 0.36339634 | 0.7074383 | 0.47929414 | 0.84300883 |
| ZFP14 | 486.510908 | 0.03126389 | 0.25484417 | 0.12267847 | 0.90236172 | 0.97941893 |
| ZFP82 | 231.630968 | 0.48358902 | 0.28794638 | 1.67944122 | 0.09306609 | 0.50797931 |
| ZNF566 | 293.224532 | -0.361566 | 0.20763048 | -1.7413918 | 0.08161493 | 0.48594425 |
| ZNF260 | 241.738825 | -0.0703208 | 0.17380748 | -0.4045904 | 0.68577859 | 0.92053037 |
| ZNF529 | 447.229221 | 0.055859 | 0.24278984 | 0.23007141 | 0.81803628 | 0.95980114 |
| ZNF529-AS1 | 419.000363 | 0.18065486 | 0.21841612 | 0.82711324 | 0.40817288 | 0.80588056 |
| ZNF382 | 122.373997 | 0.38830356 | 0.23893681 | 1.62513078 | 0.1041347 | 0.52442862 |
| ZNF461 | 213.547501 | -0.1445183 | 0.28904877 | -0.499979 | 0.61708984 | 0.89599505 |
| ZNF567 | 246.244241 | 0.21214726 | 0.23318729 | 0.90977196 | 0.36294279 | 0.77905381 |
| ZNF850 | 78.4401243 | 0.02909445 | 0.37829849 | 0.07690872 | 0.93869616 | 0.98715467 |
| AC020928.1 | 498.439045 | 0.34369432 | 0.20274041 | 1.69524331 | 0.09002927 | 0.50134767 |
| ZNF790 | 416.670135 | 0.1825061 | 0.26031905 | 0.70108622 | 0.48324921 | 0.84427756 |
| ZNF345 | 658.658279 | -0.1294699 | 0.25283593 | -0.5120707 | 0.6086015 | 0.89272818 |
| ZNF829 | 62.0818187 | -0.29955 | 0.4184896 | -0.7157883 | 0.47412208 | 0.84096663 |
| ZNF568 | 371.529133 | -0.1657532 | 0.15163249 | -1.0931248 | 0.27433901 | 0.72164296 |
| ZNF420 | 721.239127 | 0.24995515 | 0.36801709 | 0.6791944 | 0.49701469 | 0.85092921 |
| AC010632.1 | 40.4461133 | 0.88445582 | 0.42818329 | 2.06560098 | 0.03886617 | 0.36967695 |
| AC012309.1 | 60.2157694 | 0.18316514 | 0.66224565 | 0.27658186 | 0.78210119 | 0.94882207 |
| ZNF383 | 240.009652 | -0.1953452 | 0.26230299 | -0.744731 | 0.45643436 | 0.83182139 |
| LINC01535 | 126.944302 | 0.34080132 | 0.33386267 | 1.02078292 | 0.3073573 | 0.74341546 |
| HKR1 | 787.785539 | 0.18908989 | 0.14734922 | 1.28327717 | 0.19939499 | 0.65150038 |
| ZNF527 | 92.9173978 | -0.0054689 | 0.27063213 | -0.0202077 | 9.84E-01 | 0.99518552 |
| ZNF569 | 226.822477 | 0.1702009 | 0.29834898 | 0.5704759 | 0.56835497 | 0.87757867 |
| ZNF570 | 83.1878087 | 0.29483037 | 0.32668803 | 0.90248293 | 0.36680039 | 0.78095617 |
| ZNF793 | 386.045236 | 0.08406938 | 0.24247824 | 0.34670898 | 0.72880997 | 0.93453345 |
| ZNF540 | 282.715528 | 0.73525194 | 0.34214796 | 2.14892978 | 0.03163997 | 0.33950635 |
| ZNF571 | 241.93316 | 0.11702413 | 0.18953099 | 0.61744061 | 0.53694415 | 0.86466169 |
| ZFP30 | 371.343977 | 0.13898425 | 0.23505084 | 0.59129441 | 0.55432317 | 0.8714657 |
| AC093227.1 | 52.6456326 | 0.57221624 | 0.34585597 | 1.65449287 | 0.09802741 | 0.51403099 |
| ZNF573 | 392.694559 | 0.32649339 | 0.26456063 | 1.23409666 | 0.21716689 | 0.66978787 |
| SIPA1L3 | 2812.49983 | 0.0533391 | 0.27032583 | 0.1973141 | 0.84358175 | 0.9681035 |
| SPINT2 | 231.311519 | -0.2691949 | 0.22303873 | -1.2069425 | 0.22745429 | 0.67921637 |
| AC011479.1 | 48.7825008 | 0.43836219 | 0.53927233 | 0.81287721 | 0.41628846 | 0.80966301 |
| YIF1B | 48.7084941 | 0.5084756 | 0.3758311 | 1.35293645 | 0.17607593 | 0.62957039 |
| CATSPERG | 203.06888 | -0.5749284 | 0.44289289 | -1.2981206 | 0.19424591 | 0.64760078 |
| PSMD8 | 363.280721 | 0.5332436 | 0.28525519 | 1.86935637 | 0.06157325 | 0.43249565 |
| SPRED3 | 143.308701 | -0.5813521 | 0.47739388 | -1.217762 | 2.23E-01 | 6.76E-01 |
| FAM98C | 81.3166985 | -0.1493262 | 0.35266456 | -0.4234228 | 0.67198682 | 0.91574062 |
| EIF3K | 624.57408 | 0.25282448 | 0.25657196 | 0.98539405 | 0.32443055 | 0.75473814 |
| ACTN4 | 2480.96174 | -0.0319573 | 0.32906479 | -0.0971155 | 0.92263465 | 0.98215577 |
| CAPN12 | 105.341652 | 0.84424679 | 0.50814797 | 1.66141921 | 0.09662928 | 0.5123951 |

| | | | | | | |
|---|---|---|---|---|---|---|
| ECH1 | 296.529004 | 1.08402675 | 0.5442677 | 1.99171612 | 0.04640221 | 0.3904347 |
| HNRNPL | 567.782688 | -0.3097909 | 0.20252792 | -1.5296208 | 0.12611061 | 0.565635 |
| SIRT2 | 324.527713 | 0.16768598 | 0.324101 | 0.51738802 | 0.60488532 | 0.89044135 |
| NFKBIB | 103.126076 | 0.00884233 | 0.25461956 | 0.0347276 | 0.97229696 | 0.99327086 |
| AC011455.2 | 40.9596163 | 0.51938216 | 0.50693228 | 1.02455925 | 0.3055712 | 0.74251103 |
| MRPS12 | 166.618148 | 0.70085725 | 0.40390763 | 1.73519191 | 0.08270682 | 0.48594425 |
| FBXO17 | 1221.12404 | 0.72322133 | 0.57464253 | 1.25855867 | 0.20818978 | 0.66105261 |
| FBXO27 | 78.0500834 | 0.01412963 | 0.3140209 | 0.04499583 | 0.96411063 | 0.99224805 |
| PAK4 | 684.805931 | -0.0623492 | 0.34415142 | -0.1811679 | 0.85623583 | 0.97118784 |
| AC011443.1 | 109.345543 | 0.00146151 | 0.38160593 | 0.0038299 | 0.99694419 | 0.99948643 |
| SAMD4B | 843.697497 | -0.0227639 | 0.24017807 | -0.0947795 | 0.92449 | 0.98215577 |
| PAF1 | 128.680771 | 0.66111654 | 0.38592596 | 1.71306573 | 0.08670045 | 0.49478465 |
| MED29 | 175.154609 | 0.10793831 | 0.19448662 | 0.55499093 | 0.57890087 | 0.8821758 |
| ZFP36 | 366.542641 | 0.03850704 | 0.59700506 | 0.06450036 | 0.94857182 | 0.98919519 |
| PLEKHG2 | 51.4299931 | -0.4640124 | 0.45406592 | -1.0219053 | 0.30682573 | 0.74337144 |
| RPS16 | 648.870663 | 0.39025225 | 0.46570637 | 0.83797919 | 0.40204239 | 0.80296585 |
| SUPT5H | 607.060419 | -0.1312187 | 0.17256661 | -0.7603947 | 0.44701871 | 0.82571251 |
| TIMM50 | 119.331423 | 0.19302069 | 0.20144628 | 0.95817449 | 0.33797478 | 0.76430557 |
| EID2 | 53.7929515 | 0.56120863 | 0.45766654 | 1.22623916 | 0.22010869 | 0.67301289 |
| DYRK1B | 113.953861 | -0.0918328 | 0.27981391 | -0.3281926 | 0.74276606 | 0.93598004 |
| FBL | 141.678375 | -0.3344918 | 0.27276126 | -1.2263172 | 0.22007933 | 0.67301289 |
| FCGBP | 439.082338 | -1.7356148 | 0.86990198 | -1.9951843 | 0.04602278 | 0.39025761 |
| PSMC4 | 204.513243 | 0.47133061 | 0.28886795 | 1.63164727 | 0.10275381 | 0.52202889 |
| ZNF546 | 187.884752 | -0.0560658 | 0.38502606 | -0.1456155 | 0.88422495 | 0.97664137 |
| ZNF780B | 1011.20877 | 0.30899346 | 0.20622297 | 1.49834648 | 0.13404325 | 0.57708028 |
| ZNF780A | 178.7449 | -0.202869 | 0.23677468 | -0.8568019 | 0.39155438 | 0.79741625 |
| MAP3K10 | 182.857359 | -0.2401764 | 0.30140412 | -0.7968583 | 0.42553332 | 0.81466629 |
| CNTD2 | 52.2284771 | -2.2649642 | 1.00587097 | -2.2517442 | 0.02433844 | 0.30617757 |
| AKT2 | 717.788687 | -0.291633 | 0.17096082 | -1.705847 | 0.08803657 | 0.49729428 |
| C19orf47 | 54.6536683 | -0.4434381 | 0.33441112 | -1.3260267 | 0.18483085 | 0.64011182 |
| PLD3 | 941.667524 | -0.2106634 | 0.24203528 | -0.8703829 | 0.3840912 | 0.791697 |
| SERTAD1 | 96.4449787 | 0.32100313 | 0.51585611 | 0.62227261 | 0.53376263 | 0.86397197 |
| SERTAD3 | 54.5203146 | 0.54021865 | 0.30338799 | 1.78061975 | 0.07497459 | 0.47190267 |
| BLVRB | 417.305576 | 0.42592452 | 0.32754341 | 1.30036053 | 0.19347743 | 0.64718686 |
| SPTBN4 | 150.769883 | 1.07640255 | 0.84733935 | 1.2703323 | 0.20396629 | 0.65712971 |
| SHKBP1 | 108.412339 | -0.0983544 | 0.31844832 | -0.3088551 | 0.75743175 | 0.94056395 |
| NUMBL | 48.910985 | -0.1088928 | 0.37469855 | -0.2906144 | 0.77134628 | 0.9446143 |
| COQ8B | 196.392101 | -0.0644181 | 0.29733165 | -0.216654 | 0.82847801 | 0.96288319 |
| ITPKC | 267.202084 | -0.0579474 | 0.2763331 | -0.2097012 | 8.34E-01 | 9.65E-01 |
| C19orf54 | 50.8991262 | -0.7087444 | 0.27276905 | -2.5983315 | 0.0093678 | 0.19669868 |
| SNRPA | 195.215908 | -0.1908907 | 0.24861756 | -0.7678087 | 0.44260083 | 0.82299244 |
| AC008537.4 | 49.0531364 | -0.1072179 | 0.50065415 | -0.2141556 | 0.83042575 | 0.96355356 |
| AC008537.1 | 231.199794 | -1.3745164 | 0.34707079 | -3.9603343 | 7.48E-05 | 0.01392432 |
| AXL | 93.8882226 | -0.2579776 | 0.84233036 | -0.3062665 | 0.75940172 | 0.94133292 |
| HNRNPUL1 | 504.542841 | -0.0991922 | 0.18992861 | -0.5222607 | 0.60148883 | 0.88920716 |
| TGFB1 | 186.905792 | 0.02890819 | 0.40166767 | 0.07197042 | 0.94262545 | 0.98772472 |
| CCDC97 | 105.017363 | -0.102117 | 0.27194706 | -0.3755032 | 0.70728628 | 0.92573766 |
| B9D2 | 46.4798637 | 0.15206835 | 0.38825249 | 0.39167386 | 0.69529921 | 0.92229578 |
| EXOSC5 | 40.8679111 | -0.4322094 | 0.54263859 | -0.7964958 | 0.42574391 | 0.81466629 |
| DMAC2 | 120.132406 | -0.1248303 | 0.27092335 | -0.4607588 | 0.64497163 | 0.90516684 |
| RPS19 | 1910.65501 | 0.75885732 | 0.33876695 | 2.24005711 | 0.02508722 | 0.31053369 |
| ARHGEF1 | 264.189215 | -0.3485049 | 0.26295413 | -1.3253447 | 0.18505685 | 0.64023465 |
| RABAC1 | 369.825642 | 0.37582635 | 0.52453491 | 0.71649445 | 0.47368609 | 0.84064485 |
| GRIK5 | 486.526267 | 0.15570785 | 0.27517055 | 0.56585944 | 0.57148934 | 0.8787896 |

| | | | | | | |
|---|---|---|---|---|---|---|
| POU2F2 | 48.9503145 | -0.7643408 | 0.57746237 | -1.3236201 | 0.18562923 | 0.64030106 |
| DEDD2 | 102.414572 | 0.2823965 | 0.26867728 | 1.05106209 | 0.29323008 | 0.73527014 |
| ZNF526 | 50.191189 | 0.48149049 | 0.45494329 | 1.05835276 | 0.28989465 | 0.73204984 |
| GSK3A | 96.1985776 | 0.39432319 | 0.35299412 | 1.11708147 | 0.26395949 | 0.71445861 |
| CIC | 211.603905 | -0.3333966 | 0.23424369 | -1.4232897 | 0.15465221 | 0.60380955 |
| PAFAH1B3 | 86.5690515 | -0.17665 | 0.501743 | -0.3520727 | 0.72478375 | 0.93241479 |
| MEGF8 | 252.388251 | -0.1226562 | 0.29694852 | -0.4130554 | 0.67956605 | 0.9182972 |
| LIPE-AS1 | 160.402284 | -0.2157115 | 0.48433452 | -0.4453772 | 0.65604721 | 0.90950195 |
| PHLDB3 | 107.70865 | 1.11375396 | 0.36893768 | 3.01881327 | 0.00253767 | 0.09529369 |
| ETHE1 | 243.758787 | 0.45159543 | 0.26238702 | 1.72110427 | 8.52E-02 | 0.49270382 |
| XRCC1 | 82.6416226 | -0.2552691 | 0.3161259 | -0.8074919 | 0.41938312 | 0.81214362 |
| PINLYP | 94.7058191 | -0.0686106 | 0.38263719 | -0.1793099 | 0.85769441 | 0.97127317 |
| ZNF428 | 213.944636 | 0.55591683 | 0.29180438 | 1.90510103 | 0.05676694 | 0.41872623 |
| CADM4 | 76.5512289 | 0.51578281 | 0.36717457 | 1.40473459 | 0.16010022 | 0.61085068 |
| PLAUR | 86.3907988 | -0.4193527 | 0.62671327 | -0.6691301 | 5.03E-01 | 0.85525037 |
| SMG9 | 133.934254 | -0.3710005 | 0.32360785 | -1.146451 | 0.25160859 | 0.70276117 |
| ZNF283 | 214.390556 | 0.21480707 | 0.18828596 | 1.14085548 | 0.25393007 | 0.70431689 |
| ZNF404 | 61.8540934 | 0.24160855 | 0.4731638 | 0.51062349 | 0.60961473 | 0.89312344 |
| AC006213.3 | 78.8085087 | 0.97476296 | 0.44102422 | 2.21022549 | 0.02708952 | 0.3178271 |
| ZNF45 | 132.876324 | -0.0553692 | 0.33907339 | -0.1632956 | 0.87028566 | 0.97463098 |
| ZNF155 | 248.812027 | 0.5521392 | 0.26952767 | 2.04854368 | 0.04050676 | 0.37344844 |
| ZNF230 | 57.2325334 | 0.10592851 | 0.30598971 | 0.34618323 | 0.72920501 | 0.93453345 |
| ZNF224 | 368.217666 | 0.00784401 | 0.19445237 | 0.04033898 | 0.96782288 | 0.99248831 |
| ZNF225 | 66.0202961 | 0.1124388 | 0.2792689 | 0.40261843 | 0.68722895 | 0.9208366 |
| ZNF234 | 162.536028 | 0.4294257 | 0.25453973 | 1.68706747 | 0.09159038 | 0.50476839 |
| ZNF226 | 267.76016 | 0.2715886 | 0.29712889 | 0.91404305 | 0.36069423 | 0.77730375 |
| ZNF227 | 179.038992 | -0.1090738 | 0.24235001 | -0.4500671 | 0.65266203 | 0.90868303 |
| ZNF235 | 633.928263 | 0.43945179 | 0.28862636 | 1.52256292 | 0.12786809 | 0.56762585 |
| ZNF233 | 49.5104449 | 1.09031504 | 0.48982562 | 2.2259249 | 0.02601921 | 0.31244343 |
| ZNF229 | 154.910367 | 0.06009787 | 0.34928777 | 0.17205832 | 0.86339168 | 0.97319807 |
| ZNF180 | 128.366766 | -0.392417 | 0.3030885 | -1.2947273 | 0.19541431 | 0.64893292 |
| AC243964.2 | 361.699873 | -0.0563847 | 0.39143907 | -0.1440447 | 0.88546516 | 0.97664137 |
| PVR | 133.314946 | -0.2570951 | 0.2809088 | -0.9152262 | 0.36007288 | 0.77703033 |
| CEACAM19 | 227.184618 | -0.516464 | 0.46816725 | -1.1031613 | 0.26995713 | 0.71892414 |
| BCL3 | 248.552106 | -0.0578414 | 0.62446764 | -0.0926252 | 9.26E-01 | 0.98234748 |
| CBLC | 144.142686 | 0.56281108 | 0.41353328 | 1.36098135 | 0.17351958 | 0.6286116 |
| BCAM | 1487.72286 | 1.04710728 | 0.46685091 | 2.24291581 | 0.02490224 | 0.3093519 |
| NECTIN2 | 637.791091 | 0.2043189 | 0.24361605 | 0.83869229 | 0.401642 | 0.80278824 |
| TOMM40 | 180.638227 | -0.2357232 | 0.33806145 | -0.6972793 | 0.48562804 | 0.8447301 |
| APOE | 504.385931 | 0.15542941 | 0.6906161 | 0.22505906 | 0.82193333 | 0.96070413 |
| CLPTM1 | 394.946671 | -0.1632186 | 0.21869187 | -0.7463403 | 0.45546185 | 0.83121393 |
| RELB | 339.414845 | 0.30255853 | 0.48626982 | 0.62220297 | 5.34E-01 | 8.64E-01 |
| CLASRP | 663.200646 | -0.3317299 | 0.21780318 | -1.5230719 | 0.12774072 | 0.56762585 |
| GEMIN7-AS1 | 224.195024 | -0.2208186 | 0.30505243 | -0.7238711 | 0.46914486 | 0.83870782 |
| MARK4 | 502.822057 | 0.07674 | 0.23335204 | 0.32885937 | 0.74226198 | 0.93598004 |
| TRAPPC6A | 192.132796 | -0.0764737 | 0.30701515 | -0.2490877 | 0.80329293 | 0.95446295 |
| KLC3 | 46.132574 | 0.6705639 | 0.431416 | 1.55433247 | 0.12010513 | 0.55298199 |
| ERCC2 | 111.084037 | 0.33925877 | 0.35748919 | 0.94900426 | 0.34261844 | 0.76602015 |
| PPP1R13L | 387.646167 | -0.1766187 | 0.33934848 | -0.5204641 | 0.60274014 | 0.88971776 |
| ERCC1 | 203.785418 | -0.4562654 | 0.20993195 | -2.1733966 | 0.02975048 | 0.33159709 |
| VASP | 285.662637 | -0.2630827 | 0.2525942 | -1.0415232 | 0.29763278 | 0.73673761 |
| OPA3 | 711.767718 | -0.2597513 | 0.32628959 | -0.7960759 | 0.4259879 | 0.81466629 |
| EML2 | 414.171609 | 1.19648037 | 0.29950491 | 3.99486062 | 6.47E-05 | 0.01295529 |
| SNRPD2 | 306.428182 | 0.1748003 | 0.37164544 | 0.47034157 | 0.638111 | 0.90301725 |

| | | | | | | |
|---|---|---|---|---|---|---|
| QPCTL | 101.706227 | -0.1975751 | 0.27859444 | -0.7091852 | 0.47820956 | 0.84199814 |
| FBXO46 | 269.724309 | -0.2261206 | 0.21814249 | -1.0365729 | 0.29993493 | 0.73673761 |
| SIX5 | 55.3464925 | 0.47282584 | 0.42564157 | 1.11085447 | 0.26663099 | 0.71641758 |
| DM1-AS | 69.3392814 | 0.18384722 | 0.49644639 | 0.37032643 | 0.71113928 | 0.92683165 |
| DMPK | 185.715669 | -0.2974022 | 0.36397664 | -0.8170915 | 0.41387614 | 0.80874327 |
| SYMPK | 740.117227 | -0.2659733 | 0.21963066 | -1.2110025 | 0.22589446 | 0.67809183 |
| MYPOP | 240.055845 | 0.13096962 | 0.33100429 | 0.39567347 | 0.69234593 | 0.92224682 |
| CCDC61 | 67.4886624 | 0.58849447 | 0.35342266 | 1.66512943 | 0.09588694 | 0.5117083 |
| HIF3A | 120.665894 | 1.71256971 | 0.7701912 | 2.22356438 | 0.02617777 | 0.31320845 |
| PPP5C | 513.846781 | 0.09174722 | 0.13117401 | 0.69943139 | 0.48428247 | 0.84458506 |
| AC007193.1 | 47.3447593 | 0.69407686 | 0.63032085 | 1.10114849 | 0.27083203 | 0.71965808 |
| CCDC8 | 122.924119 | 0.69366973 | 0.39371915 | 1.76183893 | 0.07809651 | 0.47980319 |
| PNMA8A | 51.3150756 | 1.98604356 | 0.59135658 | 3.35845348 | 0.0007838 | 0.04906731 |
| PPP5D1 | 294.578654 | -0.1309162 | 0.3931339 | -0.3330065 | 0.73912934 | 0.93552616 |
| CALM3 | 197.42787 | 0.90409085 | 0.35180884 | 2.56983549 | 0.01017468 | 0.20550057 |
| PRKD2 | 457.358082 | -0.3394572 | 0.35396336 | -0.9590178 | 0.33754978 | 0.76409325 |
| STRN4 | 164.015335 | -0.0485833 | 0.22009766 | -0.2207354 | 0.82529849 | 0.96148948 |
| FKRP | 183.96142 | 0.09039165 | 0.21880864 | 0.4131082 | 0.67952734 | 0.9182972 |
| SLC1A5 | 156.744252 | 1.01823694 | 0.50764473 | 2.00580619 | 0.04487693 | 0.38711185 |
| AP2S1 | 274.333872 | 0.44010688 | 0.2916924 | 1.50880475 | 0.13134869 | 0.57506954 |
| ARHGAP35 | 1108.133 | 0.15449799 | 0.23230339 | 0.66506991 | 0.50600578 | 0.85626196 |
| TMEM160 | 291.94962 | 1.10040982 | 0.47486951 | 2.31728886 | 0.020488 | 0.28691085 |
| ZC3H4 | 321.697963 | -0.3056949 | 0.20811325 | -1.4688872 | 0.14186339 | 0.5872148 |
| SAE1 | 290.375397 | -0.4135159 | 0.30422443 | -1.3592461 | 0.1740686 | 0.6286158 |
| BBC3 | 50.8438882 | 0.65013758 | 0.395281 | 1.64474787 | 1.00E-01 | 5.17E-01 |
| CCDC9 | 119.94049 | -0.011418 | 0.30699 | -0.0371935 | 0.97033075 | 0.99309664 |
| INAFM1 | 92.6387054 | 0.07346462 | 0.56948008 | 0.12900297 | 0.8973553 | 0.97823597 |
| C5AR1 | 424.536096 | 1.46637267 | 0.38769621 | 3.78227242 | 0.0001554 | 0.02032396 |
| DHX34 | 287.741746 | -0.4320217 | 0.26339366 | -1.640213 | 0.1009609 | 0.51819907 |
| KPTN | 41.6577885 | 0.07370536 | 0.44930463 | 0.16404318 | 0.86969715 | 0.97454113 |
| NAPA-AS1 | 112.896264 | 0.42059137 | 0.34951493 | 1.20335736 | 0.22883806 | 0.68065126 |
| NAPA | 535.996066 | 0.18192269 | 0.31653783 | 0.57472656 | 0.56547624 | 0.8761731 |
| AC010331.1 | 39.5710702 | -0.2785956 | 0.51520595 | -0.5407461 | 0.58868258 | 0.88571413 |
| BICRA-AS1 | 100.934888 | -0.1777151 | 0.54207454 | -0.3278426 | 0.7430307 | 0.93598059 |
| NOP53 | 133.028898 | 0.81316685 | 0.39051263 | 2.08230613 | 0.03731452 | 0.36263296 |
| SELENOW | 613.59498 | 0.58319907 | 0.29548497 | 1.97370129 | 0.04841572 | 0.39627811 |
| PLA2G4C | 85.9941502 | -0.6254473 | 0.67546166 | -0.9259553 | 0.35446919 | 0.77243419 |
| LIG1 | 122.266676 | -0.8134793 | 0.47667079 | -1.7065853 | 0.08789917 | 0.49714378 |
| ZSWIM9 | 158.252663 | -0.1074001 | 0.29079859 | -0.3693281 | 0.71188316 | 0.92748721 |
| CARD8 | 457.178062 | -0.1367309 | 0.16769584 | -0.8153507 | 0.41487161 | 0.80965493 |
| EMP3 | 60.9962591 | -0.6446366 | 0.42110709 | -1.5308139 | 0.12581539 | 0.56519704 |
| TMEM143 | 204.578144 | -0.6357471 | 0.39163358 | -1.6233213 | 0.10452073 | 0.52442862 |
| KDELR1 | 317.610401 | 0.34572906 | 0.23020502 | 1.5018311 | 0.13314073 | 0.57682553 |
| GRIN2D | 38.2884338 | -1.0418552 | 0.64042999 | -1.6268057 | 0.10377837 | 0.52441026 |
| GRWD1 | 59.1220418 | 0.56684857 | 0.46729496 | 1.21304234 | 0.22511364 | 0.67771328 |
| KCNJ14 | 71.0046246 | -0.4416674 | 0.39604883 | -1.1151842 | 0.2647715 | 0.71491553 |
| CYTH2 | 321.587028 | -0.6222402 | 0.25916276 | -2.4009631 | 0.01635198 | 0.25652417 |
| LMTK3 | 301.657603 | -1.0310792 | 0.3381199 | -3.0494484 | 0.00229262 | 0.09212026 |
| SULT2B1 | 103.12882 | -1.8809293 | 0.88976815 | -2.1139543 | 0.03451917 | 0.35291306 |
| RPL18 | 802.441215 | 0.72618684 | 0.44196643 | 1.64308146 | 0.10036608 | 0.51705755 |
| SPHK2 | 160.105554 | -0.4702264 | 0.32230793 | -1.4589354 | 0.1445829 | 0.59035409 |
| CA11 | 55.0546119 | -0.5131434 | 0.45796403 | -1.1204885 | 0.26250566 | 0.7132328 |
| MAMSTR | 51.8092745 | 0.48997353 | 0.4578142 | 1.07024537 | 0.28450888 | 0.72738535 |
| BCAT2 | 239.492226 | 0.16090086 | 0.41027636 | 0.39217677 | 0.6949276 | 0.92229578 |

| | | | | | | |
|---|---|---|---|---|---|---|
| HSD17B14 | 125.761926 | 0.62657984 | 0.55630978 | 1.12631463 | 0.26003237 | 0.70951612 |
| PLEKHA4 | 204.300645 | 0.11987835 | 0.41181885 | 0.29109486 | 7.71E-01 | 0.9443559 |
| PPP1R15A | 98.4495297 | -0.0763604 | 0.43004471 | -0.1775638 | 8.59E-01 | 0.97174503 |
| NUCB1 | 441.744173 | 0.38136967 | 0.28657945 | 1.33076418 | 0.18326662 | 0.63834582 |
| NUCB1-AS1 | 60.8421689 | -0.1873001 | 0.49769277 | -0.3763369 | 0.70666648 | 0.92556842 |
| BAX | 190.105551 | 0.20066534 | 0.36084893 | 0.55609239 | 0.57814769 | 0.88205258 |
| FTL | 6121.14695 | 0.83998465 | 0.61298775 | 1.37031228 | 0.17058944 | 0.62661733 |
| GYS1 | 508.409119 | 0.1648612 | 0.42472154 | 0.38816302 | 0.6978954 | 0.92341984 |
| RUVBL2 | 109.789232 | -0.1455792 | 0.26889011 | -0.5414077 | 0.5882266 | 0.88571413 |
| SNRNP70 | 1763.06072 | -0.038883 | 0.31630437 | -0.122929 | 0.90216334 | 0.97941893 |
| LIN7B | 52.056142 | 0.59137991 | 0.35929383 | 1.64595064 | 0.09977392 | 0.51665771 |
| PPFIA3 | 45.1897459 | 0.40899353 | 0.46619592 | 0.87729967 | 0.38032389 | 0.7880311 |
| TRPM4 | 464.419516 | 0.41787562 | 0.50231003 | 0.83190776 | 0.40546101 | 0.80485885 |
| SLC6A16 | 641.030324 | 0.04382327 | 0.42206618 | 0.10383032 | 0.917304 | 0.98199511 |
| AC011450.1 | 83.6919594 | 0.38835105 | 0.53577372 | 0.72484154 | 0.46854925 | 0.83825637 |
| TEAD2 | 141.726316 | -1.321941 | 0.36316351 | -3.6400712 | 0.00027256 | 0.02790915 |
| PIH1D1 | 236.122427 | 0.07066325 | 0.22393828 | 0.31554787 | 0.75234571 | 0.93818313 |
| RPL13A | 1062.54901 | 0.69042479 | 0.43249409 | 1.59637971 | 0.11040404 | 0.53372881 |
| RPS11 | 846.831224 | 0.51604866 | 0.44412528 | 1.16194389 | 0.24525826 | 0.69518395 |
| FCGRT | 280.370068 | 0.96491454 | 0.2432344 | 3.96701511 | 7.28E-05 | 0.01392432 |
| NOSIP | 329.505619 | -0.0450256 | 0.23188814 | -0.1941695 | 0.84604316 | 0.96883382 |
| PRRG2 | 119.062857 | -0.1414255 | 0.31893991 | -0.4434237 | 0.65745926 | 0.91031231 |
| PRR12 | 268.626808 | -0.2352943 | 0.34691847 | -0.678241 | 0.49761893 | 0.85092921 |
| RRAS | 80.3008421 | 1.65266626 | 0.51403876 | 3.21506155 | 0.00130417 | 0.06597468 |
| SCAF1 | 169.09276 | 0.40093062 | 0.26851516 | 1.49313961 | 0.13540065 | 0.57708028 |
| IRF3 | 469.830712 | -0.3465484 | 0.30063713 | -1.1527131 | 0.24902817 | 0.69928042 |
| BCL2L12 | 43.8034131 | -0.5379707 | 0.35583548 | -1.5118524 | 0.13057142 | 0.57223551 |
| PRMT1 | 316.966582 | 0.41339498 | 0.30433805 | 1.35834142 | 0.17435537 | 0.62892483 |
| AP2A1 | 358.793522 | -0.3232585 | 0.2619745 | -1.2339313 | 0.2172285 | 0.66978787 |
| FUZ | 86.0578673 | -0.0718896 | 0.3896131 | -0.1845154 | 0.85360912 | 0.97076831 |
| MED25 | 460.941994 | 0.04453916 | 0.29958562 | 0.14866921 | 0.88181466 | 0.97664137 |
| PTOV1-AS1 | 122.864274 | 0.05498061 | 0.33742957 | 0.16293953 | 0.87056605 | 0.9747202 |
| PTOV1 | 400.155632 | -0.0078634 | 0.25567938 | -0.0307547 | 0.97546514 | 0.9941831 |
| PNKP | 453.158656 | -0.3874453 | 0.33958096 | -1.1409513 | 0.2538902 | 0.70431689 |
| TBC1D17 | 315.794645 | -0.6277384 | 0.26172327 | -2.3984812 | 0.01646322 | 0.25721056 |
| AC011452.1 | 36.4182733 | 0.68232273 | 0.57572251 | 1.18515903 | 2.36E-01 | 6.88E-01 |
| ATF5 | 68.3832156 | 0.43683547 | 0.44412395 | 0.98358907 | 0.3253176 | 0.7553384 |
| VRK3 | 368.465506 | -0.1301464 | 0.27540337 | -0.4725665 | 0.63652252 | 0.9025146 |
| ZNF473 | 81.6098968 | -0.1352029 | 0.33925219 | -0.3985321 | 0.69023803 | 0.92183073 |
| MYH14 | 273.04036 | 1.37826105 | 0.50415271 | 2.73381663 | 0.00626049 | 0.16248472 |
| KCNC3 | 215.963226 | 1.31779339 | 0.57971549 | 2.27317262 | 0.02301578 | 0.2983102 |
| NR1H2 | 324.276136 | 0.43817382 | 0.28812538 | 1.52077481 | 0.12831636 | 0.56762585 |
| NAPSA | 171.833055 | 1.41271172 | 0.57714251 | 2.44776932 | 0.01437437 | 0.24762526 |
| EMC10 | 396.086443 | 0.67868419 | 0.28139975 | 2.41181522 | 0.01587332 | 0.25383866 |
| JOSD2 | 94.2155983 | 0.49421527 | 0.35421355 | 1.39524665 | 0.16294147 | 0.61387934 |
| CLEC11A | 113.430774 | 0.14676431 | 0.3640529 | 0.40314007 | 0.68684518 | 0.9208231 |
| C19orf48 | 73.2448704 | -0.5795015 | 0.46677315 | -1.2415057 | 0.214419 | 0.66827012 |
| CTU1 | 64.6740084 | 0.12218536 | 0.33902541 | 0.36040176 | 0.71854671 | 0.9298398 |
| ETFB | 366.682163 | 1.09610851 | 0.41542028 | 2.6385532 | 0.00832606 | 0.18347203 |
| ZNF175 | 194.837685 | -0.0520385 | 0.19078276 | -0.2727629 | 0.7850355 | 0.94978829 |
| SPACA6 | 244.379671 | -0.1331971 | 0.2994826 | -0.4447573 | 0.65649516 | 0.90974031 |
| ZNF577 | 481.081671 | 0.06559547 | 0.30669537 | 0.21387826 | 0.83064199 | 0.9635604 |
| ZNF649 | 65.3153118 | -0.1437755 | 0.33940008 | -0.4236165 | 0.67184553 | 0.91574062 |
| ZNF613 | 95.6413751 | 0.04021757 | 0.26387597 | 0.15241088 | 0.87886288 | 0.9765717 |

| | | | | | | |
|---|---|---|---|---|---|---|
| ZNF350-AS1 | 305.734667 | 1.17863237 | 0.68386725 | 1.72348121 | 0.08480154 | 0.49198622 |
| ZNF350 | 201.943502 | -0.1228881 | 0.40880715 | -0.3006016 | 0.76371833 | 0.94255335 |
| ZNF615 | 159.83709 | 0.16895857 | 0.25115329 | 0.67273086 | 0.50111853 | 0.8538796 |
| ZNF614 | 84.9441582 | 0.34200346 | 0.34241418 | 0.99880051 | 0.31789134 | 0.74996012 |
| ZNF432 | 359.92722 | 0.0472707 | 0.32835218 | 0.14396342 | 0.88552935 | 0.97664137 |
| ZNF616 | 105.892566 | -0.2833456 | 0.23771137 | -1.1919733 | 0.23327171 | 0.68484327 |
| AC011468.3 | 83.8916834 | 0.28810311 | 0.61538376 | 0.4681682 | 0.63966431 | 0.90330608 |
| ZNF836 | 90.2462561 | -0.4503995 | 0.27439185 | -1.6414462 | 0.10070482 | 0.51819632 |
| PPP2R1A | 739.330907 | 0.40959534 | 0.30270653 | 1.35311034 | 0.17602038 | 0.62957039 |
| ZNF766 | 324.617765 | -0.1455981 | 0.27259516 | -0.5341184 | 0.59325963 | 0.88761003 |
| AC010320.3 | 47.9013251 | -0.3568144 | 0.54422893 | -0.6556329 | 0.51206032 | 0.85762435 |
| ZNF480 | 266.334931 | -0.5004931 | 0.21896941 | -2.2856758 | 0.02227323 | 0.29649945 |
| AC010320.4 | 75.1083126 | -0.2175805 | 0.58290384 | -0.3732699 | 0.70894756 | 0.92577218 |
| ZNF610 | 112.30859 | 0.03354499 | 0.3825747 | 0.08768221 | 0.93012926 | 0.98331361 |
| ZNF880 | 94.4851367 | 0.1437853 | 0.33365136 | 0.43094476 | 0.66650854 | 0.91318863 |
| ZNF528 | 232.407078 | -0.301149 | 0.37139397 | -0.8108614 | 0.41744526 | 0.81017256 |
| ZNF578 | 58.6420114 | 0.37052764 | 0.49884398 | 0.74277261 | 0.45761936 | 0.83282548 |
| ZNF808 | 301.665461 | -0.2282713 | 0.26539087 | -0.8601325 | 0.389716 | 0.79686667 |
| ZNF701 | 414.93005 | 0.04374033 | 0.20003676 | 0.21866148 | 8.27E-01 | 0.96218322 |
| ZNF83 | 2381.39559 | -0.1795119 | 0.23723702 | -0.7566773 | 0.44924325 | 0.8267002 |
| ZNF611 | 1136.14096 | -0.2383059 | 0.3330899 | -0.7154403 | 0.47433701 | 0.84106098 |
| ZNF600 | 164.991938 | -0.1422283 | 0.3095672 | -0.4594424 | 0.6459165 | 0.90582727 |
| ZNF28 | 177.068329 | -0.1640972 | 0.20179422 | -0.8131907 | 0.4161087 | 0.80966301 |
| ZNF320 | 88.5287572 | 0.06800892 | 0.62137752 | 0.10944863 | 0.91284667 | 0.98135874 |
| AC010487.3 | 326.740568 | -0.2054173 | 0.54979872 | -0.3736227 | 0.70868503 | 0.92577218 |
| ZNF160 | 813.134816 | 0.07033726 | 0.32143924 | 0.21881977 | 0.82679045 | 0.96214368 |
| ZNF415 | 90.964292 | 0.16150784 | 0.27220102 | 0.59334034 | 0.55295341 | 0.87104567 |
| ZNF347 | 260.967727 | -0.0372567 | 0.23505768 | -0.1585003 | 0.87406255 | 0.97564728 |
| ZNF665 | 97.744299 | 0.36937106 | 0.44718509 | 0.82599145 | 0.40880894 | 0.80602525 |
| ZNF677 | 155.158487 | 0.3402646 | 0.33736466 | 1.00859587 | 0.31316849 | 0.74722589 |
| ZNF845 | 73.5917541 | -0.0047808 | 0.34243751 | -0.013961 | 0.98886109 | 0.99670729 |
| ZNF525 | 197.988006 | 0.2620134 | 0.24398222 | 1.07390368 | 0.28286586 | 0.72604837 |
| ZNF765 | 276.403442 | -0.2240108 | 0.27172581 | -0.8244002 | 0.40971221 | 0.80641067 |
| ZNF761 | 321.249696 | -0.2971591 | 0.36453393 | -0.8151754 | 0.4149719 | 0.80965493 |
| ZNF813 | 99.2280539 | -0.1233292 | 0.42137241 | -0.2926846 | 0.76976326 | 0.94386043 |
| ZNF331 | 352.597546 | 0.40965008 | 0.28816445 | 1.42158436 | 0.15514695 | 0.60416746 |
| MYADM | 57.3802863 | 0.28050656 | 0.84113088 | 0.33348741 | 0.73876639 | 0.93552616 |
| NDUFA3 | 142.076608 | 0.67650047 | 0.47631713 | 1.42027324 | 0.15552815 | 0.6049776 |
| TFPT | 69.4604225 | 0.04283143 | 0.35108284 | 0.12199808 | 0.90290054 | 0.97960781 |
| PRPF31 | 291.770418 | 0.07527012 | 0.23525353 | 0.31995322 | 0.7490038 | 0.93748128 |
| CNOT3 | 186.080663 | 0.13372096 | 0.29559463 | 0.45237955 | 0.65099558 | 0.90825486 |
| LENG1 | 59.113576 | -0.4874254 | 0.29494948 | -1.6525725 | 0.0984179 | 0.51525577 |
| TMC4 | 672.12296 | -0.210135 | 0.15599921 | -1.3470258 | 0.17797193 | 0.63194386 |
| MBOAT7 | 356.211925 | 0.12994231 | 0.30243353 | 0.42965578 | 0.66744606 | 0.91360856 |
| TSEN34 | 170.037602 | 0.5420122 | 0.36331253 | 1.49186211 | 0.1357353 | 0.57743238 |
| RPS9 | 1680.57774 | 0.55065888 | 0.38100607 | 1.44527587 | 0.14838041 | 0.59446675 |
| LENG8-AS1 | 209.146522 | 0.26821108 | 0.26075877 | 1.02857931 | 0.3036774 | 0.7409078 |
| LENG8 | 916.604469 | -0.1944877 | 0.34221892 | -0.5683136 | 0.56982204 | 0.87807251 |
| LENG9 | 38.9412411 | -0.0524319 | 0.69864936 | -0.0750475 | 0.94017688 | 0.98715467 |
| NLRP2 | 56.2759961 | -0.6596792 | 0.65930208 | -1.000572 | 0.31703379 | 0.74881731 |
| AC011476.3 | 168.227953 | 0.7934275 | 0.52977061 | 1.49768124 | 0.13421609 | 0.57708028 |
| GP6 | 52.2678281 | 1.06521752 | 0.85053311 | 1.25241158 | 0.21041993 | 0.66334818 |
| RDH13 | 384.218731 | -0.0465799 | 0.29176896 | -0.1596464 | 0.87315961 | 0.97564728 |
| EPS8L1 | 240.237297 | 0.57953968 | 0.56773918 | 1.02078508 | 0.30735627 | 0.74341546 |

| | | | | | | |
|---|---|---|---|---|---|---|
| PPP1R12C | 382.283024 | -0.2662784 | 0.22211407 | -1.1988361 | 0.23059167 | 0.68284709 |
| PPP6R1 | 285.111622 | -0.116652 | 0.2201787 | -0.5298061 | 0.59624637 | 0.88859628 |
| HSPBP1 | 148.104004 | 0.15045069 | 0.31194881 | 0.4822929 | 0.62959789 | 0.90005881 |
| KMT5C | 84.1099215 | 0.11145482 | 0.35265591 | 0.31604409 | 0.75196904 | 0.93815321 |
| TMEM238 | 148.614807 | 0.8594995 | 0.57154972 | 1.50380531 | 0.1326315 | 0.57682553 |
| RPL28 | 1260.05757 | 0.50619828 | 0.42420681 | 1.19328185 | 0.23275901 | 0.68474693 |
| UBE2S | 71.395261 | 0.56026501 | 0.39938647 | 1.4028142 | 0.16067225 | 0.61166537 |
| ISOC2 | 124.32576 | 0.81058729 | 0.42416561 | 1.91101604 | 0.05600252 | 0.41843207 |
| NAT14 | 97.6733641 | 0.85760084 | 0.4811895 | 1.78225178 | 0.07470818 | 0.47190267 |
| ZNF579 | 94.274459 | 0.22745772 | 0.44921263 | 0.50634757 | 0.61261267 | 0.89441126 |
| FIZ1 | 55.4514856 | -0.0321825 | 0.35880895 | -0.0896925 | 0.9285316 | 0.98314146 |
| ZNF524 | 76.4380383 | 0.33160376 | 0.43590124 | 0.7607314 | 0.44681752 | 0.82571251 |
| ZNF865 | 97.8110269 | -0.0343307 | 0.33273796 | -0.1031763 | 0.91782307 | 0.98199511 |
| CCDC106 | 89.9730213 | -0.0055306 | 0.41778449 | -0.0132379 | 0.989438 | 0.99683369 |
| U2AF2 | 265.625809 | -0.2452409 | 0.21757372 | -1.1271621 | 0.25967395 | 0.70921184 |
| AC008735.2 | 120.727947 | 0.21121507 | 0.57977522 | 0.3643051 | 0.71563019 | 0.92883553 |
| EPN1 | 767.198973 | -0.5864341 | 0.39903586 | -1.4696276 | 0.14166263 | 0.5872148 |
| ZNF787 | 172.807598 | -0.3628659 | 0.27087055 | -1.3396285 | 0.18036616 | 0.63479125 |
| ZNF444 | 311.945274 | -0.0064751 | 0.24919206 | -0.0259845 | 0.97926974 | 0.99463083 |
| ZSCAN5A | 972.668394 | -0.2343888 | 0.30798224 | -0.7610465 | 0.44662928 | 0.82571251 |
| EDDM13 | 140.951666 | 0.33808566 | 0.55030585 | 0.61435955 | 0.53897776 | 0.86542329 |
| ZNF582 | 97.346538 | 0.35182544 | 0.33964382 | 1.03586585 | 0.30026473 | 0.73680045 |
| ZNF583 | 100.779728 | -0.1319518 | 0.46791125 | -0.2820017 | 0.77794218 | 0.94694572 |
| ZNF667 | 249.75896 | 0.01337503 | 0.42542011 | 0.03143959 | 0.97491897 | 0.99403355 |
| ZNF667-AS1 | 216.412073 | 0.13684338 | 0.22279743 | 0.61420536 | 0.53907963 | 0.86542329 |
| ZNF471 | 228.459918 | 0.31438589 | 0.27593869 | 1.13933238 | 0.25456455 | 0.70509116 |
| ZFP28 | 79.0623975 | 0.38553963 | 0.31505184 | 1.22373398 | 0.2210526 | 0.67392082 |
| AC005498.3 | 155.907182 | 0.18361152 | 0.44182097 | 0.415579 | 0.6777181 | 0.91753826 |
| ZNF470 | 95.0189294 | 0.19732279 | 0.51845761 | 0.3805958 | 0.7035032 | 0.92513032 |
| ZNF71 | 65.2535487 | -0.0086323 | 0.3281099 | -0.0263092 | 0.97901068 | 0.99455895 |
| AC006115.3 | 71.55336 | -0.0530256 | 0.3078437 | -0.1722485 | 0.86324213 | 0.97319807 |
| ZIM2 | 40.3409476 | -0.8534734 | 0.47187336 | -1.8086915 | 0.07049894 | 0.46237363 |
| ZNF264 | 352.19307 | -0.0402789 | 0.18682745 | -0.2155943 | 0.82930398 | 0.96290106 |
| ZNF805 | 76.7052546 | -0.0420102 | 0.29243637 | -0.143656 | 0.88577209 | 0.97664137 |
| ZNF460-AS1 | 182.854845 | 0.10233259 | 0.36477791 | 0.28053395 | 0.77906788 | 0.94732571 |
| ZNF460 | 132.121238 | -0.0038367 | 0.30352076 | -0.0126407 | 0.98991444 | 0.99723763 |
| AC005261.1 | 422.036879 | 0.33727789 | 0.19954276 | 1.6902537 | 0.09097943 | 0.50282313 |
| ZNF304 | 70.9724085 | 0.1950923 | 0.38764872 | 0.50327085 | 0.61477386 | 0.8949759 |
| AC003002.2 | 112.461932 | 0.18675232 | 0.3261908 | 0.57252478 | 0.5669665 | 0.87646121 |
| ZNF749 | 45.1348376 | 0.38813723 | 0.37841599 | 1.0256893 | 0.30503806 | 0.74229685 |
| AC004076.1 | 249.306226 | -0.2540778 | 0.3579559 | -0.7098019 | 0.47782697 | 0.84190259 |
| ZNF773 | 47.8327172 | 0.21656245 | 0.52972224 | 0.40882265 | 0.68266981 | 0.91981391 |
| ZNF549 | 168.572046 | 0.40549091 | 0.23917501 | 1.69537323 | 0.09000464 | 0.50134767 |
| ZNF550 | 102.514614 | 0.37777403 | 0.29227185 | 1.29254332 | 0.19616905 | 0.64964978 |
| ZNF134 | 79.2394297 | 0.48812657 | 0.36047805 | 1.35410899 | 0.17570161 | 0.62957039 |
| ZNF211 | 86.8991947 | 0.50892838 | 0.23275923 | 2.18650143 | 0.02877894 | 0.32721051 |
| AC003006.1 | 144.321235 | -0.4397827 | 0.4326357 | -1.0165197 | 0.30938195 | 0.74450725 |
| ZNF154 | 97.0393499 | 0.75556457 | 0.55892291 | 1.35182251 | 0.1764321 | 0.63008941 |
| ZNF671 | 56.9416653 | 0.90189423 | 0.50316543 | 1.79244077 | 0.07306239 | 0.47001711 |
| ZNF776 | 92.5792362 | 0.28229986 | 0.28461904 | 0.99185165 | 0.3212699 | 0.75240254 |
| ZNF586 | 1329.01217 | -0.044248 | 0.23501863 | -0.1882746 | 0.85066138 | 0.97056586 |
| ZNF552 | 475.801261 | 0.43213013 | 0.27124736 | 1.59312197 | 0.11113283 | 0.53461826 |
| AC010522.1 | 71.8409597 | 0.20501423 | 0.2917682 | 0.70266131 | 0.48226685 | 0.84389635 |
| ZNF814 | 764.283053 | 0.15714716 | 0.24768419 | 0.63446583 | 0.52577686 | 0.86119802 |

| | | | | | | |
|---|---|---|---|---|---|---|
| ZNF587 | 393.703445 | 0.1671029 | 0.18298169 | 0.913222 | 0.3611258 | 0.77739743 |
| ZNF418 | 118.071218 | 0.15079425 | 0.3974812 | 0.37937454 | 0.70440975 | 0.92528913 |
| C19orf18 | 160.285818 | -0.5773132 | 0.50092408 | -1.1524965 | 0.24911713 | 0.69928042 |
| ZNF606 | 235.96824 | -0.081149 | 0.23976775 | -0.3384483 | 0.73502541 | 0.93552616 |
| ZSCAN1 | 50.4209741 | 0.58633425 | 0.64029352 | 0.91572729 | 0.35980995 | 0.77671671 |
| ZNF135 | 462.123751 | -0.0168864 | 0.40286162 | -0.0419161 | 0.96656559 | 0.99248831 |
| ZSCAN18 | 543.927072 | 0.57926351 | 0.30568364 | 1.89497717 | 0.05809543 | 0.42087381 |
| ZNF329 | 294.154451 | 0.19740755 | 0.30483857 | 0.64758063 | 0.51725623 | 0.85893097 |
| ZNF274 | 377.858849 | 0.08859943 | 0.26368434 | 0.33600565 | 0.7368666 | 0.93552616 |
| AC020915.3 | 186.705638 | 0.38010199 | 0.26787515 | 1.41895205 | 0.155913 | 0.60519671 |
| ERVK3-1 | 44.3603265 | 0.94745962 | 0.40607174 | 2.33323208 | 0.01963597 | 0.28161944 |
| AC010642.2 | 64.8599718 | 0.55825452 | 0.36601034 | 1.5252425 | 0.12719862 | 0.56740575 |
| ZNF497 | 72.3345158 | -0.1477556 | 0.3218618 | -0.4590655 | 0.64618714 | 0.90582727 |
| ZNF837 | 44.275689 | -0.3708496 | 0.33708642 | -1.1001617 | 0.27126166 | 0.7200754 |
| RPS5 | 523.417452 | 0.57436724 | 0.45347013 | 1.26660434 | 0.2052968 | 0.65813563 |
| ZNF584 | 112.620373 | 0.11435697 | 0.34465601 | 0.33180031 | 0.74004005 | 0.93552616 |
| ZNF132 | 43.440324 | 0.25895689 | 0.43034818 | 0.60173809 | 0.54734849 | 0.86859163 |
| ZNF324 | 94.9404741 | 0.04515875 | 0.28478673 | 0.1585704 | 0.87400735 | 0.97564728 |
| ZNF446 | 92.1417826 | -0.1968562 | 0.31330305 | -0.6283253 | 0.52979085 | 0.86343221 |
| SLC27A5 | 113.056861 | 0.01817574 | 0.23413564 | 0.07762908 | 0.93812311 | 0.98696247 |
| ZBTB45 | 54.5000239 | -0.433856 | 0.47204148 | -0.9191057 | 0.3580403 | 0.77529959 |
| TRIM28 | 282.976432 | 0.21146437 | 0.45093448 | 0.46894699 | 0.63910753 | 0.90330608 |
| CHMP2A | 222.809475 | 0.48563836 | 0.30423519 | 1.59625967 | 0.11043083 | 0.53372881 |
| UBE2M | 187.898457 | 0.43564792 | 0.3109269 | 1.4011265 | 0.16117625 | 0.61247902 |
| MZF1-AS1 | 211.882354 | -0.01558 | 0.54244842 | -0.0287216 | 0.97708664 | 0.99433432 |
| MZF1 | 637.895282 | -0.2604873 | 0.24728894 | -1.0533722 | 0.29217046 | 0.73384286 |
| RPS4Y1 | 483.867644 | 0.50579963 | 2.14590321 | 0.23570477 | 0.81366177 | 0.9582375 |
| ZFY | 331.24576 | -0.0601088 | 2.01199056 | -0.0298753 | 0.97616652 | 0.9941831 |
| ZFY-AS1 | 46.297898 | 0.32987067 | 2.05572877 | 0.1604641 | 0.8725155 | 0.97554234 |
| LINC00278 | 1415.34061 | -0.5082679 | 2.51001433 | -0.202496 | 0.83952898 | 0.96617873 |
| PCDH11Y | 47.9341188 | 0.08310977 | 0.49067481 | 0.16937851 | 0.86549893 | 0.97379123 |
| TBL1Y | 633.909327 | -0.5748165 | 2.10232021 | -0.27342 | 0.78453035 | 0.94978829 |
| TTTY15 | 329.050073 | 0.39940103 | 2.21759654 | 0.18010536 | 0.85706985 | 0.97127317 |
| USP9Y | 1359.75984 | 0.26717866 | 2.38952372 | 0.11181252 | 0.91097206 | 0.98111638 |
| DDX3Y | 315.535266 | 0.90231146 | 2.28549311 | 0.39479946 | 0.69299089 | 0.92224682 |
| UTY | 3264.18608 | 0.28141876 | 2.40635747 | 0.11694803 | 0.90690124 | 0.9803357 |
| NLGN4Y | 2311.08104 | 0.53040827 | 2.1264486 | 0.24943386 | 0.8030252 | 0.95446295 |
| TTTY14 | 2702.44213 | 0.09012341 | 1.33142576 | 0.0676894 | 0.94603289 | 0.98877579 |
| KDM5D | 374.914957 | 0.30557157 | 2.79137122 | 0.10947006 | 0.91282967 | 0.98135874 |
| TTTY10 | 199.272312 | -0.4803403 | 1.8181553 | -0.264191 | 0.79163273 | 0.95181838 |
| EIF1AY | 126.561631 | 0.49806678 | 2.56138429 | 0.19445219 | 0.84582183 | 0.96883382 |
| AP000547.3 | 126.489981 | -0.0983974 | 0.28497694 | -0.3452818 | 0.72988252 | 0.93453345 |
| IL17RA | 308.147959 | 0.02684535 | 0.23188467 | 0.11577027 | 0.90783462 | 0.98047754 |
| HDHD5 | 269.979335 | -0.2117071 | 0.39220581 | -0.5397858 | 0.58934477 | 0.88603498 |
| HDHD5-AS1 | 107.998102 | -0.1061707 | 0.61783052 | -0.1718443 | 0.86355993 | 0.9732733 |
| ADA2 | 119.318294 | 0.69468341 | 0.70368928 | 0.98720192 | 0.32354366 | 0.75386985 |
| CECR3 | 64.0327134 | 1.64301596 | 0.63923021 | 2.570304 | 0.01016093 | 0.20550057 |
| CECR2 | 1774.69206 | 0.655775 | 0.33436055 | 1.96128103 | 0.04984625 | 0.40017381 |
| ATP6V1E1 | 486.680515 | 0.14832488 | 0.17212181 | 0.86174365 | 3.89E-01 | 0.79634939 |
| BCL2L13 | 1028.48167 | -0.1550518 | 0.1648281 | -0.9406882 | 0.34686465 | 0.76875481 |
| MICAL3 | 2371.32027 | -0.0201926 | 0.37346593 | -0.0540682 | 0.95688085 | 0.99065996 |
| PEX26 | 371.122029 | 0.00669933 | 0.17669502 | 0.03791465 | 0.96975573 | 0.99306175 |
| USP18 | 201.942629 | 2.14169141 | 0.74730676 | 2.8658799 | 0.00415852 | 0.12569768 |
| AC023490.3 | 103.334616 | 2.24078952 | 0.94492335 | 2.37139818 | 0.01772093 | 0.26773793 |

| | | | | | | |
|---|---|---|---|---|---|---|
| AC007326.4 | 38.1508644 | 0.19780861 | 0.63663879 | 0.31070776 | 0.7560228 | 0.93962224 |
| DGCR2 | 648.300978 | 0.0112385 | 0.2413072 | 0.04657343 | 0.96285321 | 0.99224805 |
| ESS2 | 53.4938072 | -0.07829 | 0.3955715 | -0.1979163 | 0.84311055 | 0.9677309 |
| SLC25A1 | 210.513749 | 0.49918247 | 0.39306971 | 1.26995913 | 0.20409919 | 0.65712971 |
| CLTCL1 | 244.526237 | -0.4157676 | 0.31763686 | -1.3089399 | 0.19055472 | 0.64559718 |
| HIRA | 257.651482 | -0.4718016 | 0.24306781 | -1.9410286 | 0.05225481 | 0.40866415 |
| MRPL40 | 173.155475 | 0.93750662 | 0.46555309 | 2.01374806 | 0.04403599 | 0.3852094 |
| UFD1 | 237.549417 | -0.3352427 | 0.24719854 | -1.3561677 | 0.17504581 | 0.62944785 |
| GNB1L | 45.3591292 | -0.265144 | 0.38114286 | -0.6956552 | 0.48664476 | 0.84474442 |
| TXNRD2 | 298.673571 | -0.2815386 | 0.28343653 | -0.993304 | 0.32056185 | 0.75144499 |
| COMT | 781.571143 | 0.4048039 | 0.2634908 | 1.53631133 | 0.12446204 | 0.56111892 |
| ARVCF | 70.1992482 | -0.5163125 | 0.38399942 | -1.3445658 | 0.17876549 | 0.63284534 |
| TANGO2 | 338.291804 | -0.2813695 | 0.23314311 | -1.206853 | 0.22748877 | 0.67921637 |
| AC006547.1 | 73.6241095 | 0.19859215 | 0.50335899 | 0.39453383 | 0.69318695 | 0.92224682 |
| DGCR8 | 309.325212 | -0.4318241 | 0.34596048 | -1.2481892 | 0.21196179 | 0.66552984 |
| TRMT2A | 77.017455 | -0.0806562 | 0.34566233 | -0.233338 | 0.81549896 | 0.95872901 |
| RANBP1 | 125.954774 | 0.22237974 | 0.33891363 | 0.65615461 | 0.51172465 | 0.85762435 |
| ZDHHC8 | 166.937722 | -0.603408 | 0.48882779 | -1.2343979 | 0.21705468 | 0.66978787 |
| DGCR6L | 87.1374512 | 0.80286286 | 0.49938524 | 1.6077024 | 0.10790037 | 0.53063658 |
| ZNF74 | 46.6026512 | -0.5355282 | 0.44999733 | -1.1900698 | 0.23401897 | 0.68509675 |
| KLHL22 | 635.46121 | 0.20832118 | 0.21122142 | 0.98626921 | 0.32400103 | 0.75413737 |
| MED15 | 941.429145 | -0.3493073 | 0.25366705 | -1.3770306 | 0.1685028 | 0.62253512 |
| PI4KA | 1371.81878 | 0.0422913 | 0.16372455 | 0.25830761 | 0.79616952 | 0.95278473 |
| SNAP29 | 217.449036 | -0.0669437 | 0.3011068 | -0.2223254 | 0.82406058 | 0.96141406 |
| CRKL | 256.707013 | 0.12878381 | 0.27062755 | 0.47587101 | 0.63416628 | 0.90136076 |
| LZTR1 | 138.645013 | -0.0263501 | 0.28409596 | -0.0927507 | 0.92610165 | 0.98234748 |
| THAP7 | 77.7082727 | -0.0043712 | 0.31852496 | -0.0137231 | 0.98905087 | 0.99670729 |
| GGT2 | 3305.37979 | 1.50222964 | 0.88597079 | 1.69557469 | 9.00E-02 | 0.50134767 |
| HIC2 | 197.824231 | 0.48219934 | 0.28282376 | 1.70494636 | 0.08820443 | 0.49730444 |
| UBE2L3 | 634.971122 | 0.05531081 | 0.18287284 | 0.30245501 | 0.76230523 | 0.94212205 |
| YDJC | 68.5321893 | 0.32801956 | 0.3975038 | 0.82519855 | 0.40925888 | 0.80623405 |
| SDF2L1 | 122.298727 | 1.28425414 | 0.50475797 | 2.54429691 | 0.0109498 | 0.21301305 |
| PPIL2 | 716.103622 | -0.5875839 | 0.30683612 | -1.9149764 | 0.05549551 | 0.41843207 |
| YPEL1 | 255.121624 | -0.4675329 | 0.35877359 | -1.3031419 | 0.19252631 | 0.64671221 |
| MAPK1 | 1586.207 | -0.0859352 | 0.23407776 | -0.3671223 | 0.71352781 | 0.92760608 |
| PPM1F | 169.953163 | -0.2537631 | 0.40610853 | -0.6248652 | 0.53205956 | 0.86397197 |
| AC245452.1 | 71.8201885 | -0.2022392 | 0.3427146 | -0.5901096 | 0.55511715 | 0.87184917 |
| TOP3B | 188.663366 | -0.0391559 | 0.32410586 | -0.120812 | 0.90383991 | 0.97984491 |
| AC245060.5 | 328.062167 | -0.149241 | 0.4397154 | -0.3394037 | 0.73430566 | 0.93552616 |
| RSPH14 | 91.8135287 | 0.23521656 | 0.59657817 | 0.39427618 | 0.69337715 | 0.92224682 |
| RAB36 | 109.384503 | 0.7792553 | 0.52073751 | 1.4964455 | 0.1345376 | 0.57708028 |
| BCR | 2646.89386 | -0.3720049 | 0.36008265 | -1.0331096 | 0.3015526 | 0.73858859 |
| PCAT14 | 96.522281 | 0.57216475 | 0.45793951 | 1.24943302 | 0.21150674 | 0.66484951 |
| AP000345.2 | 35.9927282 | -1.7144986 | 0.65738581 | -2.6080553 | 0.00910582 | 0.1936857 |
| DRICH1 | 86.167955 | -1.6265465 | 0.47719748 | -3.4085395 | 0.00065312 | 0.04534035 |
| CHCHD10 | 854.52342 | 0.82269706 | 0.28599316 | 2.87663196 | 0.00401944 | 0.12328719 |
| SMARCB1 | 404.561205 | -0.2232451 | 0.27569586 | -0.8097515 | 0.41808301 | 0.81069561 |
| AP000350.4 | 44.0917619 | 0.34857165 | 0.44680396 | 0.78014449 | 0.43530583 | 0.81919857 |
| GSTT2B | 99.4936935 | -0.4514189 | 0.72501359 | -0.622635 | 0.53352438 | 0.86397197 |
| DDT | 437.412247 | 0.93818765 | 0.36388298 | 2.57826748 | 0.00992971 | 0.20303644 |
| CABIN1 | 1526.96357 | 0.10372461 | 0.28445809 | 0.36463935 | 0.71538064 | 0.92880162 |
| GUCD1 | 249.151184 | 0.14193058 | 0.19988725 | 0.71005321 | 4.78E-01 | 0.84190259 |
| SNRPD3 | 418.70745 | 0.15770444 | 0.22409742 | 0.70373159 | 0.48159994 | 0.8437862 |
| GGT1 | 281.609598 | 1.5981628 | 0.7167041 | 2.22987813 | 0.02575553 | 0.31211455 |

| | | | | | | |
|---|---|---|---|---|---|---|
| LRRC75B | 90.17143 | 1.22239726 | 0.63966658 | 1.91099129 | 0.0560057 | 0.41843207 |
| SGSM1 | 706.199274 | 0.89863306 | 0.55773687 | 1.6112133 | 0.10713324 | 0.52882025 |
| KIAA1671 | 1873.03091 | -0.3076798 | 0.19640975 | -1.5665199 | 0.11722695 | 0.54757066 |
| AL022323.4 | 145.561929 | -0.4986646 | 0.36621735 | -1.361663 | 0.17330427 | 0.6286116 |
| AL022323.1 | 43.683098 | 0.17017292 | 0.4818059 | 0.35319808 | 0.72393995 | 0.93201977 |
| LRP5L | 621.000347 | -0.2788708 | 0.54501866 | -0.511672 | 0.60888057 | 0.89273938 |
| GRK3 | 437.473658 | 0.35933267 | 0.29339167 | 1.22475418 | 0.22066785 | 0.67345736 |
| ASPHD2 | 76.0341874 | -0.0612086 | 0.56247319 | -0.1088204 | 0.91334492 | 0.98145965 |
| HPS4 | 445.410751 | -0.3694824 | 0.2808065 | -1.31579 | 0.18824452 | 0.64267818 |
| SRRD | 60.1164869 | 0.03394457 | 0.34878486 | 0.09732236 | 0.9224704 | 0.98215577 |
| TFIP11 | 174.04062 | 0.4036817 | 0.25311892 | 1.59483021 | 0.11075021 | 0.53393773 |
| TPST2 | 832.884024 | 1.13167143 | 0.43527875 | 2.5998775 | 0.0093257 | 0.19663607 |
| MIATNB | 433.074164 | -0.0192393 | 0.44703848 | -0.0430373 | 0.96567183 | 0.99248831 |
| PITPNB | 1223.81412 | -0.0424459 | 0.25120068 | -0.168972 | 0.86581868 | 0.97379123 |
| TTC28 | 2925.93334 | 0.26280317 | 0.22075432 | 1.19047805 | 0.23385855 | 0.68500598 |
| CHEK2 | 120.602836 | -0.4464877 | 0.42744915 | -1.04454 | 0.29623564 | 0.7366786 |
| HSCB | 415.220432 | -0.285309 | 0.24751576 | -1.1526904 | 0.24903747 | 0.69928042 |
| CCDC117 | 171.690759 | -0.0604027 | 0.34599014 | -0.1745793 | 0.86141024 | 0.97234387 |
| XBP1 | 327.903262 | 0.95689147 | 0.28079158 | 3.40783533 | 0.0006548 | 0.04534035 |
| Z93930.2 | 749.32726 | 0.34310673 | 0.47055498 | 0.72915332 | 0.46590788 | 0.8367113 |
| ZNRF3 | 2752.86521 | -0.489621 | 0.32723605 | -1.4962319 | 0.13459324 | 0.57708028 |
| ZNRF3-AS1 | 61.9965027 | -0.1719622 | 0.58145017 | -0.295747 | 0.76742327 | 0.94316227 |
| KREMEN1 | 465.837806 | 0.267813 | 0.37818478 | 0.70815381 | 0.47884975 | 0.84279465 |
| RHBDD3 | 140.479203 | -0.2387755 | 0.28216204 | -0.8462354 | 0.39742142 | 0.80069929 |
| EWSR1 | 1233.53132 | -0.2362158 | 0.14762541 | -1.6001029 | 0.10957576 | 0.53361622 |
| GAS2L1 | 74.2742634 | 0.29394275 | 0.40217191 | 0.73088832 | 0.46484737 | 0.8363074 |
| AP1B1 | 416.078557 | 0.17883767 | 0.17778236 | 1.00593593 | 0.31444639 | 0.74785672 |
| THOC5 | 170.843137 | -0.3442697 | 0.23151558 | -1.4870259 | 0.13700797 | 0.57911626 |
| NIPSNAP1 | 220.34969 | -0.0835087 | 0.34477414 | -0.2422127 | 0.80861536 | 0.95610623 |
| NF2 | 511.360996 | -0.0660797 | 0.22359828 | -0.2955287 | 0.76758999 | 0.94316227 |
| ZMAT5 | 99.2975674 | -0.0611222 | 0.24420481 | -0.2502907 | 0.80236255 | 0.95446295 |
| UQCR10 | 608.63943 | 0.76159214 | 0.49095616 | 1.55124268 | 0.12084354 | 0.55498372 |
| ASCC2 | 361.925523 | 0.14154821 | 0.24623344 | 0.57485372 | 0.56539023 | 0.8761426 |
| MTMR3 | 2070.77245 | 0.05878166 | 0.15631214 | 0.37605307 | 0.70687744 | 0.92573766 |
| LIF | 93.3232969 | 0.81653173 | 0.96475301 | 0.84636349 | 0.39735 | 0.80069929 |
| TBC1D10A | 219.917506 | -0.2118448 | 0.29152433 | -0.7266798 | 0.46742215 | 0.83763115 |
| SF3A1 | 176.181851 | 0.17491821 | 0.22658093 | 0.77198997 | 0.44012037 | 0.82232701 |
| CCDC157 | 66.3160085 | -0.3756066 | 0.27612625 | -1.3602711 | 0.17374416 | 0.6286116 |
| RNF215 | 256.755023 | -0.6517087 | 0.34496947 | -1.8891779 | 0.05886799 | 0.42348634 |
| SEC14L2 | 69.6487333 | -0.6192575 | 0.40619523 | -1.5245316 | 0.12737597 | 0.56740575 |
| SEC14L6 | 226.76367 | 0.8123339 | 0.51335026 | 1.58241645 | 0.11355453 | 0.53989968 |
| GAL3ST1 | 169.100907 | 0.02691663 | 0.37398826 | 0.07197186 | 9.43E-01 | 9.88E-01 |
| PES1 | 189.780145 | -0.2882516 | 0.27759164 | -1.0384017 | 0.29908307 | 0.73673761 |
| TCN2 | 128.389028 | 1.10593576 | 0.45250863 | 2.44401033 | 0.01452501 | 0.24762526 |
| SLC35E4 | 92.3325508 | 0.37392269 | 0.34462306 | 1.08501934 | 0.27791309 | 0.72372365 |
| DUSP18 | 235.434433 | 0.42417177 | 0.27681815 | 1.53231198 | 0.12544547 | 0.56418425 |
| OSBP2 | 790.963771 | -0.175709 | 0.22152756 | -0.79317 | 0.4276788 | 0.81466629 |
| MORC2-AS1 | 48.2766384 | -0.1579048 | 0.38791855 | -0.4070566 | 0.68396639 | 0.9200778 |
| MORC2 | 342.249206 | -0.1637863 | 0.2525663 | -0.6484885 | 0.51666903 | 0.8588826 |
| SMTN | 99.863834 | 0.05265375 | 0.61535359 | 0.08556666 | 0.93181091 | 0.98357046 |
| SELENOM | 236.845346 | 1.39518334 | 0.46710926 | 2.98684583 | 0.00281872 | 0.10062824 |
| INPP5J | 78.4072273 | 1.0692286 | 0.61251101 | 1.74564798 | 0.08087214 | 0.48512266 |
| RNF185 | 452.94792 | -0.0971845 | 0.18858998 | -0.5153219 | 0.6063281 | 0.89127397 |
| LIMK2 | 495.907592 | -0.2007314 | 0.25053399 | -0.8012143 | 4.23E-01 | 8.14E-01 |

| | | | | | | |
|---|---|---|---|---|---|---|
| PIK3IP1 | 168.131972 | 0.24849197 | 0.42677572 | 0.58225422 | 0.56039546 | 0.8735265 |
| PIK3IP1-AS1 | 107.033872 | -0.6221632 | 0.2880225 | -2.1601203 | 3.08E-02 | 3.36E-01 |
| PATZ1 | 271.662418 | 0.24942444 | 0.37190568 | 0.67066584 | 0.50243343 | 0.85525037 |
| DRG1 | 643.536922 | -0.1754722 | 0.20988145 | -0.8360538 | 0.40312463 | 0.80324472 |
| EIF4ENIF1 | 563.008435 | -0.1554838 | 0.14410468 | -1.0789642 | 0.2806037 | 0.72530438 |
| SFI1 | 585.121772 | -0.6109973 | 0.21178135 | -2.885038 | 0.00391366 | 0.12135113 |
| PISD | 1134.41375 | -0.007802 | 0.19908496 | -0.0391895 | 0.9687393 | 0.99255953 |
| PRR14L | 1042.15946 | -0.0834739 | 0.19157344 | -0.4357277 | 0.66303429 | 0.91138507 |
| YWHAH | 160.999203 | 0.67348423 | 0.49090393 | 1.37192674 | 1.70E-01 | 0.62602085 |
| RTCB | 478.093601 | 0.20856315 | 0.18169115 | 1.14789934 | 0.25101012 | 0.70200988 |
| FBXO7 | 534.812231 | 0.11659614 | 0.19083197 | 0.61098849 | 0.5412072 | 0.86582694 |
| SYN3 | 897.718364 | -0.9059538 | 0.90747062 | -0.9983285 | 3.18E-01 | 0.75023175 |
| TIMP3 | 689.494374 | 0.25251076 | 0.51784967 | 0.48761401 | 0.62582327 | 0.89873262 |
| LARGE1 | 1991.04067 | -0.5136749 | 0.2135743 | -2.4051342 | 0.01616653 | 0.25504618 |
| HMGXB4 | 351.027748 | -0.4272174 | 0.30463962 | -1.4023696 | 0.16080489 | 0.61166537 |
| TOM1 | 620.171151 | -0.0513315 | 0.27686039 | -0.1854056 | 0.85291094 | 0.97076831 |
| HMOX1 | 185.925782 | 1.0798056 | 0.98620635 | 1.09490838 | 2.74E-01 | 0.72123981 |
| MCM5 | 61.4892464 | -0.9095364 | 0.42965488 | -2.1168999 | 0.03426834 | 0.35198331 |
| APOL6 | 213.116344 | 1.11844291 | 0.65753148 | 1.70097241 | 0.08894817 | 0.49869115 |
| RBFOX2 | 3231.39361 | -0.0192094 | 0.16615849 | -0.1156089 | 0.90796254 | 0.98047754 |
| APOL4 | 154.452176 | 1.27623291 | 1.0687364 | 1.19415125 | 0.23241881 | 0.68450837 |
| APOL2 | 487.650705 | 0.08496379 | 0.61804387 | 0.1374721 | 0.89065764 | 0.97729111 |
| APOL1 | 227.500087 | 0.71824704 | 0.82498279 | 0.87062064 | 3.84E-01 | 0.791697 |
| MYH9 | 1740.10039 | -0.5100843 | 0.29642653 | -1.7207783 | 0.08529105 | 0.49283006 |
| TXN2 | 257.292643 | 0.79900548 | 0.24243222 | 3.29578917 | 0.00098146 | 0.0559889 |
| EIF3D | 366.506422 | 0.17161342 | 0.28159128 | 0.60944153 | 0.54223182 | 0.86582694 |
| IFT27 | 197.575654 | -0.2233107 | 0.31583805 | -0.7070418 | 0.47954051 | 0.8430245 |
| PVALB | 55.3888435 | 1.53628625 | 0.9238677 | 1.66288554 | 0.09633535 | 0.51206987 |
| TST | 211.824432 | 0.75627844 | 0.32438955 | 2.33138964 | 0.01973282 | 0.2823953 |
| MPST | 255.498156 | 0.50946053 | 0.27881613 | 1.82722761 | 0.06766555 | 0.45209626 |
| C1QTNF6 | 91.7184953 | 0.2751969 | 0.45132126 | 0.60975833 | 0.54202191 | 0.86582694 |
| CARD10 | 87.1085077 | 1.39605849 | 0.65331499 | 2.13688422 | 0.03260741 | 0.34456902 |
| CDC42EP1 | 52.385687 | 0.53146595 | 0.59544268 | 0.89255603 | 3.72E-01 | 7.83E-01 |
| GGA1 | 557.36506 | -0.216295 | 0.18331727 | -1.1798943 | 0.23804227 | 0.68982886 |
| LGALS1 | 56.0374501 | 0.25035526 | 0.46024532 | 0.54396046 | 5.86E-01 | 0.88538429 |
| TRIOBP | 661.409708 | -0.2088879 | 0.34526125 | -0.6050141 | 0.54516962 | 0.86698116 |
| H1F0 | 440.419022 | -0.2785392 | 0.7389657 | -0.3769312 | 0.70622472 | 0.92556842 |
| GCAT | 77.0122943 | 0.26439027 | 0.64860619 | 0.40762835 | 0.68354654 | 0.9200778 |
| ANKRD54 | 156.799597 | -0.3099304 | 0.18563491 | -1.6695695 | 0.09500457 | 0.51085035 |
| EIF3L | 558.589647 | -0.1215468 | 0.24193072 | -0.5024032 | 0.61538394 | 0.8949877 |
| MICALL1 | 115.203423 | -0.3416413 | 0.35606746 | -0.9594848 | 0.33731456 | 0.76409325 |
| POLR2F | 157.599017 | -0.6126946 | 0.26948961 | -2.2735371 | 0.02299383 | 0.2983102 |
| PICK1 | 219.65799 | -0.4159007 | 0.27961882 | -1.4873844 | 0.13691331 | 0.57908675 |
| AL031587.2 | 115.176788 | -0.0245767 | 0.44193146 | -0.055612 | 0.95565086 | 0.99065996 |
| BAIAP2L2 | 91.5059327 | 0.75893539 | 0.33797618 | 2.24552925 | 0.02473418 | 0.30843438 |
| PLA2G6 | 260.149099 | -0.1838706 | 0.25397606 | -0.7239681 | 0.46908532 | 0.83870782 |
| MAFF | 155.644004 | 0.29794341 | 0.66186635 | 0.45015645 | 0.65259763 | 0.90868303 |
| TMEM184B | 389.498314 | -0.4304456 | 0.18641251 | -2.3091025 | 0.0209379 | 0.28929778 |
| CSNK1E | 143.8392 | -0.2788068 | 0.4805756 | -0.5801518 | 0.56181228 | 0.87403145 |
| TPTEP2-CSNI | 1807.46029 | -0.2004098 | 0.26352438 | -0.7604982 | 0.44695684 | 0.82571251 |
| DDX17 | 7845.14199 | 0.1437571 | 0.23222025 | 0.61905498 | 0.53588014 | 0.8642718 |
| FAM227A | 300.51802 | -0.1383617 | 0.27127117 | -0.5100493 | 0.61001693 | 0.89339478 |
| CBY1 | 292.719148 | 0.11747085 | 0.201255 | 0.58369161 | 0.55942782 | 0.87295536 |
| AL021707.2 | 69.6981694 | -0.0428064 | 0.30880901 | -0.1386178 | 0.88975215 | 0.97729111 |

| | | | | | | |
|---|---|---|---|---|---|---|
| TOMM22 | 71.2653128 | 0.82695263 | 0.43787697 | 1.88855019 | 0.05895213 | 0.42366629 |
| JOSD1 | 292.37141 | -0.0509217 | 0.23014061 | -0.2212636 | 0.82488722 | 0.96146415 |
| GTPBP1 | 250.243791 | -0.0565283 | 0.22198213 | -0.2546523 | 0.79899164 | 0.95311845 |
| SUN2 | 237.214412 | 0.12040476 | 0.29746751 | 0.40476607 | 0.68564947 | 0.92053037 |
| AL021707.1 | 182.751335 | -0.3179847 | 0.3042452 | -1.0451591 | 0.29594943 | 0.7366786 |
| CBX6 | 402.067336 | 0.38762998 | 0.54804016 | 0.70730215 | 0.47937873 | 0.8430245 |
| CBX7 | 310.227043 | 0.30083143 | 0.28541146 | 1.05402717 | 0.29187048 | 0.73368744 |
| AL031846.2 | 111.371851 | 0.12989285 | 0.42707682 | 0.30414399 | 0.7610182 | 0.94202764 |
| PDGFB | 63.1915453 | -0.0196849 | 0.72165553 | -0.0272774 | 0.97823846 | 0.99455895 |
| AL031590.1 | 43.457381 | 0.30007119 | 0.72892008 | 0.41166542 | 0.68058468 | 0.91867756 |
| RPL3 | 814.185163 | 0.76499066 | 0.51363943 | 1.48935344 | 0.13639432 | 0.57787125 |
| SYNGR1 | 233.621045 | 0.81127558 | 0.4065592 | 1.99546728 | 0.04599194 | 0.39025761 |
| TAB1 | 240.526581 | -0.2128483 | 0.21918202 | -0.9711028 | 0.3314971 | 0.76006686 |
| MIEF1 | 125.046692 | 0.3283872 | 0.25254417 | 1.30031589 | 0.19349272 | 0.64718686 |
| ATF4 | 426.383676 | -0.2825877 | 0.2551059 | -1.1077271 | 0.26797968 | 0.71800072 |
| RPS19BP1 | 281.30407 | 0.50025054 | 0.40238166 | 1.24322403 | 0.21378529 | 0.66800831 |
| FAM83F | 229.585749 | 0.39562234 | 0.50163571 | 0.78866462 | 0.43030805 | 0.81700326 |
| TNRC6B | 7556.61751 | -0.1933388 | 0.17403457 | -1.1109219 | 0.26660196 | 0.71641758 |
| AL022238.2 | 74.098154 | -0.092797 | 0.69911917 | -0.1327342 | 0.89440363 | 0.97788747 |
| MKL1 | 2550.30403 | -0.2034634 | 0.23840368 | -0.8534408 | 0.39341488 | 0.79843808 |
| SLC25A17 | 588.504802 | -0.2210504 | 0.20046438 | -1.1026919 | 0.27016101 | 0.71917709 |
| ST13 | 760.244933 | 0.48405017 | 0.1673915 | 2.89172498 | 0.00383133 | 0.12084489 |
| XPNPEP3 | 831.340455 | -0.0263487 | 0.19100707 | -0.1379463 | 0.89028285 | 0.97729111 |
| RBX1 | 511.651233 | -0.291664 | 0.19517295 | -1.4943872 | 0.13507444 | 0.57708028 |
| EP300 | 1684.01138 | -0.1974852 | 0.19151247 | -1.0311871 | 0.30245308 | 0.73905643 |
| L3MBTL2 | 154.338311 | 0.10909804 | 0.30018791 | 0.36343248 | 0.71628184 | 0.9289438 |
| AL035681.1 | 60.0187324 | 0.07285644 | 0.43149209 | 0.16884767 | 0.86591646 | 0.97379123 |
| RANGAP1 | 266.866624 | -0.1962988 | 0.29615302 | -0.6628288 | 0.50744019 | 0.85632606 |
| ZC3H7B | 486.150745 | -0.1646241 | 0.24824823 | -0.6631429 | 0.50723903 | 0.85626313 |
| TEF | 177.673502 | -0.2523339 | 0.63195 | -0.3992941 | 0.68967652 | 0.92168526 |
| TOB2 | 337.198609 | -0.058825 | 0.28320392 | -0.2077126 | 0.83545338 | 0.96511435 |
| PHF5A | 89.2624831 | -0.0761501 | 0.29678239 | -0.2565855 | 0.79749874 | 0.95281098 |
| ACO2 | 1015.68088 | 0.10754441 | 0.18811716 | 0.57168848 | 0.56753304 | 0.87678876 |
| POLR3H | 140.463749 | -0.1706582 | 0.27935423 | -0.6109025 | 0.54126411 | 0.86582694 |
| CSDC2 | 206.127967 | 0.124434 | 0.62649411 | 0.19861958 | 0.84256034 | 0.96735154 |
| PMM1 | 212.077621 | 0.44995411 | 0.35524477 | 1.26660307 | 0.20529726 | 0.65813563 |
| DESI1 | 331.605135 | 0.14709462 | 0.23947678 | 0.61423331 | 0.53906116 | 0.86542329 |
| XRCC6 | 828.677386 | 0.27556396 | 0.19914207 | 1.3837556 | 0.1664333 | 0.61909814 |
| SNU13 | 597.642879 | 0.38514343 | 0.23967808 | 1.60691971 | 0.10807198 | 0.53063658 |
| C22orf46 | 96.6834927 | -0.2247678 | 0.45485639 | -0.4941512 | 0.62119939 | 0.8978246 |
| CCDC134 | 172.107061 | -0.8699651 | 0.44228281 | -1.9669883 | 0.04918456 | 0.39892272 |
| SREBF2 | 2919.72269 | -0.3307572 | 0.32928855 | -1.0044601 | 0.31515691 | 0.74785672 |
| WBP2NL | 403.620318 | 0.00673735 | 0.54690576 | 0.01231904 | 0.99017108 | 0.997268 |
| NAGA | 50.6707148 | 0.31129836 | 0.34568751 | 0.90051955 | 0.36784382 | 0.78095617 |
| SMDT1 | 271.256645 | 0.73715579 | 0.36497058 | 2.01976771 | 0.04340749 | 0.38352475 |
| NDUFA6 | 264.286911 | 0.18894333 | 0.36605707 | 0.51615811 | 0.60574399 | 0.89081188 |
| TCF20 | 1306.92202 | -0.1631748 | 0.21419725 | -0.7617967 | 0.44618132 | 0.82571115 |
| AL022316.1 | 51.7309093 | 0.20377786 | 0.8270181 | 0.24640072 | 8.05E-01 | 0.95522565 |
| RRP7A | 196.791606 | 0.1087383 | 0.44922476 | 0.24205768 | 0.80873547 | 0.95610623 |
| SERHL2 | 79.4704546 | 0.2786385 | 0.32843794 | 0.84837489 | 0.39622922 | 0.80027397 |
| POLDIP3 | 354.417848 | -0.35777 | 0.14936457 | -2.39528 | 0.01660768 | 0.25778007 |
| CYB5R3 | 641.44004 | 0.31870204 | 0.22724648 | 1.4024509 | 1.61E-01 | 6.12E-01 |
| A4GALT | 423.97491 | 1.36118437 | 0.59222515 | 2.29842378 | 2.15E-02 | 2.93E-01 |
| ARFGAP3 | 2058.15646 | -0.0617263 | 0.35522331 | -0.1737677 | 8.62E-01 | 0.97264823 |

| | | | | | | |
|---|---|---|---|---|---|---|
| PACSIN2 | 2049.33193 | -0.4731608 | 0.15562148 | -3.0404593 | 0.00236218 | 0.09297185 |
| TTLL1 | 104.497615 | -0.4067764 | 0.37904933 | -1.0731489 | 0.28320433 | 0.72647048 |
| MCAT | 106.894386 | 0.32789409 | 0.39911263 | 0.82155778 | 0.41132863 | 0.80675684 |
| TSPO | 408.461824 | 0.89886702 | 0.35898958 | 2.50388056 | 1.23E-02 | 2.27E-01 |
| TTLL12 | 84.119372 | 0.41368627 | 0.3834315 | 1.07890528 | 0.28062996 | 0.72530438 |
| EFCAB6 | 1289.37139 | -0.2165126 | 0.26816585 | -0.8073831 | 0.4194458 | 0.81214593 |
| SAMM50 | 168.585408 | -0.0173036 | 0.27471441 | -0.0629875 | 0.94977646 | 0.98919519 |
| PARVB | 65.2213306 | -1.5431287 | 0.67902982 | -2.2725492 | 0.02305336 | 0.2983102 |
| RTL6 | 53.3805942 | 0.82687646 | 0.41209106 | 2.0065382 | 0.04479886 | 0.38683751 |
| PHF21B | 1200.06176 | -0.0560524 | 0.43140881 | -0.1299287 | 0.89662287 | 0.97823153 |
| NUP50-DT | 690.594548 | 0.14564951 | 0.31434644 | 0.46334073 | 0.64312016 | 0.90478945 |
| NUP50 | 294.469966 | -0.0787872 | 0.18661817 | -0.4221838 | 0.67289083 | 0.91608132 |
| KIAA0930 | 164.752012 | -0.2067804 | 0.3309761 | -0.6247593 | 5.32E-01 | 8.64E-01 |
| FAM118A | 261.206799 | -0.5090767 | 0.40897439 | -1.2447642 | 0.21321845 | 0.66739395 |
| FBLN1 | 283.7204 | 0.57328177 | 1.29013524 | 0.44435789 | 6.57E-01 | 0.909854 |
| ATXN10 | 2542.40882 | -0.3969787 | 0.20354755 | -1.9502996 | 0.05114043 | 0.40556617 |
| MIRLET7BHG | 1515.31292 | -0.0018455 | 0.35095341 | -0.0052584 | 0.99580443 | 0.99882912 |
| PPARA | 2187.4203 | -0.2116598 | 0.25380342 | -0.8339519 | 0.40430811 | 0.80437725 |
| TTC38 | 385.280016 | 0.28677384 | 0.24775718 | 1.15747945 | 0.24707653 | 0.69751478 |
| TRMU | 402.916954 | -0.0557954 | 0.24549378 | -0.2272784 | 0.82020729 | 0.96016713 |
| CELSR1 | 726.561681 | -0.2308671 | 0.42081252 | -0.5486222 | 0.58326474 | 0.88379959 |
| GRAMD4 | 369.288901 | -0.137346 | 0.25972303 | -0.5288173 | 0.59693222 | 0.88883692 |
| CERK | 268.543783 | 0.34152218 | 0.39669781 | 0.86091268 | 0.38928614 | 0.79663535 |
| AL118516.1 | 87.1992642 | -0.3154324 | 0.55799732 | -0.5652938 | 0.57187395 | 0.87907473 |
| TBC1D22A | 3017.11744 | -0.3752675 | 0.33586715 | -1.1173092 | 0.26386215 | 0.71445861 |
| BRD1 | 642.918118 | -0.3391129 | 0.22875579 | -1.4824231 | 0.13822774 | 0.58206378 |
| ZBED4 | 365.87593 | -0.5030415 | 0.33715167 | -1.4920332 | 0.13569043 | 0.57742748 |
| ALG12 | 123.949646 | 0.24761164 | 0.29910575 | 0.82783979 | 0.40776124 | 0.80588056 |
| CRELD2 | 97.8955457 | 0.73708254 | 0.41212321 | 1.78850041 | 0.07369531 | 0.47028843 |
| PIM3 | 502.269609 | 0.06476148 | 0.40989967 | 0.15799348 | 0.87446193 | 0.97564728 |
| TRABD | 303.399867 | 0.43308066 | 0.2858076 | 1.51528742 | 0.12969963 | 0.57050365 |
| SELENOO | 552.442261 | -0.5057908 | 0.22919028 | -2.2068596 | 0.02732387 | 0.31968201 |
| TUBGCP6 | 180.480581 | -0.3860064 | 0.29508536 | -1.3081178 | 0.19083336 | 0.64567512 |
| HDAC10 | 43.9589174 | 0.12306236 | 0.53723803 | 0.22906488 | 0.81881849 | 0.96004577 |
| PLXNB2 | 819.748075 | 0.09871376 | 0.31045589 | 0.3179639 | 0.75051232 | 0.93775604 |
| DENND6B | 173.259639 | 0.41798811 | 0.44103032 | 0.9477537 | 0.34325486 | 0.76641095 |
| PPP6R2 | 2761.66171 | -0.2150121 | 0.17696098 | -1.2150254 | 0.22435641 | 0.67681479 |
| SBF1 | 126.515902 | -0.0132318 | 0.22593008 | -0.0585658 | 0.95329798 | 0.98950524 |
| MIOX | 108.406552 | -0.3416007 | 0.87203711 | -0.3917272 | 0.69525978 | 0.92229578 |
| LMF2 | 132.200362 | 0.08925519 | 0.42905507 | 0.20802735 | 0.83520762 | 0.96511435 |
| NCAPH2 | 72.0978214 | -0.0045216 | 0.37603925 | -0.0120242 | 0.99040632 | 0.99740343 |
| TYMP | 82.4399368 | 0.704191 | 0.43973918 | 1.60138334 | 0.10929204 | 0.53310139 |
| ARSA | 106.952718 | 0.00636728 | 0.27886695 | 0.02283267 | 0.98178375 | 0.99518016 |
| RABL2B | 310.604085 | -0.3941066 | 0.24655478 | -1.5984547 | 0.10994182 | 0.53372881 |
| CU633967.1 | 210.643805 | 0.60226583 | 0.74018449 | 0.81366988 | 0.41583409 | 0.80966301 |
| U2AF1L5 | 50.5895253 | -0.201875 | 1.1550795 | -0.1747715 | 0.86125914 | 0.97225639 |
| FP475955.1 | 47.3819069 | -0.5827279 | 0.92537691 | -0.6297195 | 0.52887811 | 0.86267608 |
| FP671120.1 | 3531.63061 | -0.6606604 | 0.41715773 | -1.5837184 | 0.11325782 | 0.53949604 |
| FP236383.1 | 2071.25225 | -0.8451492 | 0.49035451 | -1.7235473 | 0.08478961 | 0.49198622 |
| CR381653.1 | 399.172673 | -0.7196329 | 0.56403305 | -1.2758701 | 0.20200145 | 0.65552334 |
| AP001347.1 | 100.206076 | -0.6579902 | 0.51898803 | -1.267833 | 0.20485758 | 0.65806513 |
| RBM11 | 109.420129 | 0.26024853 | 0.38076327 | 0.68349169 | 0.4942962 | 0.84882719 |
| HSPA13 | 115.218574 | 0.65022273 | 0.3205836 | 2.02824703 | 0.04253504 | 0.38021566 |
| AF127577.4 | 127.127558 | 0.07309559 | 0.39772712 | 0.18378327 | 0.85418348 | 0.97085267 |

| | | | | | | |
|---|---|---|---|---|---|---|
| NRIP1 | 1386.37426 | -0.5051518 | 0.22673317 | -2.2279573 | 0.02588336 | 0.31222606 |
| AF127577.2 | 54.7229935 | -0.2335902 | 0.60146211 | -0.3883707 | 0.69774176 | 0.92341984 |
| USP25 | 3680.76022 | -0.1469572 | 0.23000685 | -0.6389254 | 0.52287144 | 0.86006369 |
| MIR99AHG | 4895.10665 | -0.3721069 | 0.34964409 | -1.0642447 | 2.87E-01 | 0.72972899 |
| CXADR | 511.662633 | 0.14167529 | 0.22220244 | 0.63759555 | 0.52373699 | 0.86006369 |
| BTG3 | 230.268005 | 0.46254757 | 0.25589452 | 1.80757121 | 0.07067326 | 0.4627897 |
| C21orf91 | 162.382489 | -0.1122984 | 0.33581072 | -0.3344098 | 0.73807038 | 0.93552616 |
| CHODL | 508.990441 | 1.16740745 | 0.53930294 | 2.16465991 | 0.03041374 | 0.33505845 |
| MRPL39 | 201.729276 | -0.0850282 | 0.24422731 | -0.3481517 | 0.72772626 | 0.93420128 |
| JAM2 | 112.059954 | -0.0815735 | 0.72146764 | -0.1130661 | 0.90997814 | 0.98069436 |
| ATP5PF | 714.133769 | 0.5759741 | 0.37949181 | 1.51775108 | 0.12907716 | 0.56965595 |
| GABPA | 338.754302 | -0.1265671 | 0.19346244 | -0.6542204 | 0.51296985 | 0.85762435 |
| APP | 8889.13847 | -0.0223554 | 0.23812486 | -0.093881 | 0.92520366 | 0.9822216 |
| AP001439.1 | 206.714325 | 0.05654825 | 0.47149506 | 0.11993392 | 0.90453549 | 0.97989015 |
| AP000229.1 | 47.5490811 | 0.95712135 | 0.60891133 | 1.57185669 | 0.11598379 | 0.54556096 |
| CYYR1 | 282.505045 | 0.80036977 | 0.46478453 | 1.72202324 | 0.08506531 | 0.49231347 |
| ADAMTS1 | 5363.47415 | 0.32902606 | 0.66027954 | 0.49831328 | 6.18E-01 | 8.96E-01 |
| ADAMTS5 | 121.488937 | 1.79270769 | 0.99895019 | 1.79459167 | 0.07271878 | 0.4692634 |
| LINC00113 | 196.743752 | 0.51469276 | 0.90346124 | 0.56968992 | 5.69E-01 | 0.87785537 |
| LINC01695 | 84.8819798 | 0.12571508 | 0.55752594 | 0.22548742 | 0.82160011 | 0.96066244 |
| AF165147.1 | 298.459267 | 0.47257699 | 0.62015202 | 0.76203411 | 0.44603965 | 0.82571115 |
| N6AMT1 | 114.902436 | 0.34137277 | 0.20335384 | 1.67871321 | 0.09320795 | 0.50833353 |
| LTN1 | 920.325796 | -0.2173982 | 0.16857807 | -1.2895996 | 0.19718972 | 0.64964978 |
| RWDD2B | 97.8174432 | 0.162027 | 0.32183129 | 0.50345322 | 0.61464566 | 0.8949759 |
| USP16 | 486.993065 | -0.1063761 | 0.1879645 | -0.5659373 | 0.57143639 | 0.8787896 |
| CCT8 | 335.807577 | 0.32995658 | 0.24435828 | 1.35029832 | 0.17692031 | 0.63025747 |
| MAP3K7CL | 397.805079 | 0.18959565 | 0.34300736 | 0.55274514 | 0.58043794 | 0.88258799 |
| BACH1 | 3551.26314 | -0.1582251 | 0.30427243 | -0.5200114 | 0.60305561 | 0.88978937 |
| GRIK1 | 210.072485 | 0.70071753 | 0.58874252 | 1.19019351 | 0.23397034 | 0.68509675 |
| CLDN8 | 37.1133839 | 1.05426078 | 0.68749095 | 1.53349041 | 0.12515507 | 0.56326176 |
| TIAM1 | 126.966046 | -1.4946429 | 0.33392835 | -4.4759389 | 7.61E-06 | 0.00350085 |
| SOD1 | 671.618395 | 0.91575579 | 0.34681291 | 2.64048932 | 0.00827864 | 0.18347203 |
| SCAF4 | 731.482481 | -0.1802067 | 0.17734673 | -1.0161264 | 0.3095692 | 0.74450725 |
| HUNK | 598.970523 | 0.36453597 | 0.76289053 | 0.47783523 | 0.63276748 | 0.90074133 |
| MIS18A | 44.1653132 | -0.2666887 | 0.31901001 | -0.8359887 | 0.40316129 | 0.80324472 |
| URB1 | 278.254909 | -0.5634872 | 0.24102163 | -2.3379112 | 0.01939186 | 0.28045661 |
| EVA1C | 498.124584 | 0.3880105 | 0.40058211 | 0.96861664 | 3.33E-01 | 0.7610305 |
| SYNJ1 | 854.008754 | 0.18148406 | 0.20468916 | 0.88663251 | 0.37527678 | 0.78468026 |
| PAXBP1-AS1 | 97.8537645 | -0.1003471 | 0.4495201 | -0.2232317 | 0.82335521 | 0.96133701 |
| PAXBP1 | 1410.48093 | -0.2542819 | 0.19677064 | -1.2922759 | 0.19626163 | 0.64964978 |
| C21orf62-AS | 346.336988 | 0.28346403 | 0.40160668 | 0.70582499 | 0.48029698 | 0.8433217 |
| C21orf62 | 281.215134 | 1.25560821 | 0.6468436 | 1.94113106 | 0.05224238 | 0.40866415 |
| AP000282.1 | 521.928675 | 0.11701609 | 0.83289829 | 0.14049266 | 0.88827075 | 0.97729111 |
| IFNAR1 | 636.69712 | 0.08262012 | 0.13000528 | 0.63551355 | 5.25E-01 | 8.61E-01 |
| IFNGR2 | 811.965246 | 0.40245832 | 0.28288141 | 1.42271039 | 0.15482013 | 0.60380955 |
| TMEM50B | 702.641874 | 0.17709776 | 0.17028611 | 1.0400012 | 0.29833934 | 0.73673761 |
| DNAJC28 | 41.945149 | -0.1651232 | 0.45003729 | -0.3669101 | 0.71368607 | 0.92760608 |
| GART | 493.000494 | -0.2824477 | 0.20620923 | -1.3697144 | 0.17077608 | 0.62695422 |
| SON | 2207.21604 | 0.02680931 | 0.28381093 | 0.09446186 | 0.92474228 | 0.9822216 |
| DONSON | 92.5618532 | 0.02849676 | 0.47359899 | 0.06017065 | 0.95201972 | 0.98938795 |
| AP000311.1 | 470.804431 | 0.03704179 | 0.46665775 | 0.07937678 | 0.93673294 | 0.98630295 |
| ITSN1 | 1609.87952 | -0.2383883 | 0.14399727 | -1.6555055 | 0.09782199 | 0.51355946 |
| LINC00649 | 136.851206 | -0.1120524 | 0.68538154 | -0.1634891 | 0.87013336 | 0.97463098 |
| MRPS6 | 5740.90708 | -1.1835308 | 0.50815125 | -2.3290916 | 0.01985421 | 0.28276543 |

| | | | | | | |
|---|---|---|---|---|---|---|
| LINC00310 | 881.564984 | -1.0632009 | 0.58216567 | -1.8262857 | 0.06780723 | 0.45235643 |
| AP000317.1 | 938.636934 | -0.9775205 | 0.40622491 | -2.406353 | 0.01611269 | 0.25504618 |
| KCNE2 | 48.4715642 | -0.7116998 | 0.67024449 | -1.0618511 | 0.28830329 | 0.73108047 |
| RCAN1 | 444.571748 | 0.71899842 | 0.6034865 | 1.19140764 | 0.23349359 | 0.68492492 |
| CLIC6 | 192.366066 | -0.4543191 | 0.637615 | -0.7125289 | 0.47613734 | 0.84141268 |
| LINC01426 | 49.1366249 | -0.1744204 | 0.92658244 | -0.1882406 | 0.85068809 | 0.97056586 |
| RUNX1 | 6036.74426 | -1.2891144 | 0.51109627 | -2.5222535 | 0.01166056 | 0.22161781 |
| SETD4 | 436.944391 | -0.651094 | 0.25180181 | -2.5857398 | 0.00971702 | 0.20055028 |
| CBR1 | 656.99856 | 0.2063135 | 0.38629635 | 0.53408091 | 0.59328556 | 0.88761003 |
| AP000688.2 | 171.351492 | -0.8504713 | 0.42157356 | -2.0173735 | 0.04365655 | 0.38443959 |
| CBR3 | 290.841819 | -0.0296422 | 0.48601983 | -0.0609898 | 0.95136734 | 0.98938795 |
| DOPEY2 | 271.758793 | -0.7398624 | 0.35140217 | -2.1054578 | 0.03525148 | 0.35490605 |
| MORC3 | 1020.98387 | -0.1372627 | 0.13692985 | -1.0024305 | 0.3161357 | 0.74785672 |
| CHAF1B | 48.183944 | -0.3652862 | 0.3627209 | -1.0070723 | 0.31390003 | 0.74777437 |
| AP000695.1 | 45.7823836 | -0.0959581 | 0.87730071 | -0.1093788 | 0.91290208 | 0.98135874 |
| AP000695.2 | 356.431044 | -0.0527888 | 0.48933837 | -0.1078778 | 0.91409261 | 0.98145965 |
| CLDN14 | 1700.88282 | 0.66610724 | 0.4252792 | 1.56628221 | 0.11728256 | 0.54757066 |
| AP000696.1 | 148.581442 | -0.4502097 | 0.32841682 | -1.3708485 | 0.1704222 | 0.6265257 |
| SIM2 | 4962.1858 | -0.5095462 | 0.43855771 | -1.1618681 | 0.24528903 | 0.69518395 |
| HLCS | 2198.92894 | -0.1054196 | 0.18004621 | -0.5855141 | 0.55820211 | 0.87291524 |
| PIGP | 154.650673 | 0.40019679 | 0.29790626 | 1.34336483 | 0.17915387 | 0.63288164 |
| TTC3 | 2863.29249 | 0.06889455 | 0.17049805 | 0.40407826 | 0.68615517 | 0.92053037 |
| VPS26C | 564.855776 | -0.0374555 | 0.16625542 | -0.2252888 | 0.8217546 | 0.96066244 |
| DYRK1A | 4048.04541 | -0.1426647 | 0.19574387 | -0.7288336 | 0.46610347 | 0.8367113 |
| KCNJ6 | 55.2760935 | 0.12280769 | 0.50246368 | 0.24441108 | 8.07E-01 | 0.95566275 |
| KCNJ15 | 178.860042 | -0.5548524 | 0.510874 | -1.0860846 | 0.27744155 | 0.72344438 |
| ETS2 | 575.994841 | -0.1604362 | 0.43481962 | -0.368972 | 7.12E-01 | 9.27E-01 |
| PSMG1 | 90.5126046 | 0.26297529 | 0.31935501 | 0.82345754 | 0.41024786 | 0.80655687 |
| BRWD1 | 3781.74776 | -0.1307578 | 0.18970929 | -0.6892534 | 0.49066384 | 0.84614483 |
| HMGN1 | 747.28889 | 0.13519966 | 0.1972544 | 0.68540757 | 0.49308677 | 0.84796031 |
| WRB | 254.779463 | -0.3759332 | 0.27548718 | -1.3646123 | 0.17237494 | 0.6286116 |
| LCA5L | 165.65142 | -0.1976796 | 0.24974424 | -0.7915283 | 0.42863578 | 0.81524117 |
| SH3BGR | 136.484569 | 0.26539101 | 0.32258993 | 0.82268844 | 4.11E-01 | 8.07E-01 |
| B3GALT5 | 43.5990218 | -0.0189554 | 0.66702304 | -0.0284179 | 0.97732883 | 0.99449476 |
| PCP4 | 466.503567 | -0.5244323 | 0.98392467 | -0.5330005 | 0.59403325 | 0.88807733 |
| DSCAM | 48.4163845 | 0.12050432 | 0.35662083 | 0.337906 | 0.73543403 | 0.93552616 |
| BACE2 | 4682.13474 | -0.4483912 | 0.58312828 | -0.768941 | 0.44192833 | 0.82285053 |
| PLAC4 | 51.4715497 | 0.24708656 | 1.07642312 | 0.22954408 | 0.81844606 | 0.95985564 |
| FAM3B | 181.24265 | 0.55512428 | 0.32651998 | 1.70012349 | 0.0891077 | 0.49895023 |
| MX1 | 856.321375 | 1.06588435 | 1.20475408 | 0.88473189 | 0.37630125 | 0.78499104 |
| TMPRSS2 | 1195.63427 | -0.6835543 | 0.38261584 | -1.7865291 | 0.07401363 | 0.47070029 |
| RIPK4 | 296.557098 | -0.3491939 | 0.33623303 | -1.0385474 | 0.2990153 | 0.73673761 |
| PRDM15 | 174.673423 | -0.4506413 | 0.2529219 | -1.781741 | 0.07479147 | 0.47190267 |
| C2CD2 | 212.092298 | 0.33999474 | 0.25906835 | 1.3123747 | 1.89E-01 | 6.44E-01 |
| ZBTB21 | 219.514783 | 0.43740487 | 0.37164715 | 1.17693589 | 0.2392211 | 0.69029619 |
| ABCG1 | 302.535203 | 0.62961736 | 0.57343465 | 1.0979758 | 0.27221506 | 0.72070318 |
| TMPRSS3 | 109.334267 | 1.45301422 | 0.52874555 | 2.74804058 | 0.00599526 | 0.15870015 |
| RSPH1 | 121.067515 | 0.79779824 | 0.48956142 | 1.62961828 | 0.1031822 | 0.52337282 |
| SLC37A1 | 415.130882 | -0.0338303 | 0.19530844 | -0.1732146 | 8.62E-01 | 9.73E-01 |
| LINC01671 | 205.389891 | 0.83720776 | 0.4716514 | 1.77505623 | 0.0758886 | 0.47417812 |
| PDE9A | 2267.74045 | -0.2471135 | 0.26002431 | -0.9503475 | 0.34193571 | 0.76553029 |
| AP001628.1 | 77.1398283 | -0.9600877 | 0.62496163 | -1.5362346 | 0.12448085 | 0.56111892 |
| WDR4 | 185.563936 | -0.7541235 | 0.31769583 | -2.3737279 | 0.01760952 | 0.26674791 |
| NDUFV3 | 670.296538 | -0.0612469 | 0.15730267 | -0.3893569 | 6.97E-01 | 0.92322348 |

| | | | | | | |
|---|---|---|---|---|---|---|
| PKNOX1 | 619.049368 | -0.1199296 | 0.26532802 | -0.452005 | 0.6512654 | 0.90825486 |
| U2AF1 | 62.7419433 | 0.49250333 | 1.16239938 | 0.42369545 | 0.67178793 | 0.91574062 |
| HSF2BP | 46.2477568 | -1.4713928 | 0.42843955 | -3.4343066 | 0.00059407 | 0.04335412 |
| RRP1B | 270.892402 | -0.2869697 | 0.23482409 | -1.2220625 | 0.22168401 | 0.67501708 |
| PDXK | 1562.92828 | 0.25480456 | 0.24710742 | 1.03114896 | 0.30247097 | 0.73905643 |
| CSTB | 774.76256 | 0.74345725 | 0.42766944 | 1.73839228 | 8.21E-02 | 4.86E-01 |
| RRP1 | 98.4439281 | -0.3188885 | 0.356311 | -0.8949724 | 0.37080186 | 0.78104287 |
| AGPAT3 | 1104.10865 | 0.12894174 | 0.22410345 | 0.57536702 | 0.5650431 | 0.87588257 |
| TRAPPC10 | 1067.49792 | -0.2284824 | 0.19598371 | -1.1658234 | 0.24368587 | 0.6948422 |
| PFKL | 1237.66492 | 0.15543093 | 0.30712848 | 0.50607788 | 0.61280198 | 0.89441126 |
| C21orf2 | 130.542328 | 0.31596977 | 0.37336013 | 0.84628684 | 0.39739275 | 0.80069929 |
| AP001062.1 | 47.886367 | 0.60808743 | 0.39268495 | 1.54853765 | 0.1214929 | 0.5564493 |
| UBE2G2 | 937.519713 | 0.16884667 | 0.16136577 | 1.04635987 | 0.29539491 | 0.7366191 |
| SUMO3 | 170.949406 | 0.1355445 | 0.38236396 | 0.35449079 | 7.23E-01 | 0.93168052 |
| PTTG1IP | 1660.02842 | 0.43614668 | 0.36146897 | 1.20659505 | 0.22758813 | 0.67921637 |
| FAM207A | 132.92206 | 0.20802305 | 0.24566812 | 0.84676456 | 0.39712637 | 0.80069929 |
| ADARB1 | 471.291029 | -0.1814667 | 0.38145959 | -0.4757166 | 0.63427626 | 0.90136076 |
| POFUT2 | 265.200872 | -0.4155035 | 0.18161905 | -2.287775 | 0.02215063 | 0.2955431 |
| BX322562.1 | 48.7020859 | -0.1944473 | 0.83172193 | -0.2337888 | 0.81514896 | 0.95872901 |
| COL18A1 | 1906.46476 | 0.3730002 | 0.41580852 | 0.89704801 | 0.3696933 | 0.78095617 |
| SLC19A1 | 56.2229573 | -0.6222677 | 0.41995268 | -1.4817567 | 0.13840504 | 0.58206378 |
| PCBP3 | 533.642412 | -4.7240802 | 0.95931054 | -4.9244535 | 8.46E-07 | 0.00085043 |
| COL6A1 | 467.606951 | 1.75978105 | 0.56907566 | 3.09234987 | 0.00198579 | 0.0861633 |
| FTCD | 71.8018202 | 0.34054935 | 0.51406848 | 0.66245911 | 5.08E-01 | 0.85643442 |
| LSS | 367.216702 | 0.2625295 | 0.41178855 | 0.63753473 | 0.5237766 | 0.86006369 |
| MCM3AP-AS | 151.289254 | 0.24809384 | 0.32804942 | 0.75626973 | 0.44948749 | 0.82672287 |
| MCM3AP | 709.402048 | -0.2173018 | 0.13605081 | -1.5972103 | 0.11021883 | 0.53372881 |
| YBEY | 160.980861 | -0.6270151 | 0.35702385 | -1.7562275 | 0.07904956 | 0.48162624 |
| C21orf58 | 50.6526293 | -0.5509392 | 0.43794574 | -1.2580078 | 0.20838894 | 0.66136702 |
| PCNT | 372.396488 | -0.4635526 | 0.27245897 | -1.7013667 | 0.08887415 | 0.49869115 |
| DIP2A | 1164.14807 | -0.3901449 | 0.17573388 | -2.2200894 | 0.0264127 | 0.31402818 |
| PRMT2 | 908.46061 | -0.0916338 | 0.22645723 | -0.4046408 | 0.68574159 | 0.92053037 |
| AC011043.1 | 180.500103 | 0.50009976 | 0.39640417 | 1.26159057 | 0.20709615 | 0.65980054 |
| AL592183.1 | 898.697904 | -0.2534995 | 0.5429517 | -0.4668914 | 0.64057758 | 0.9037561 |
| AL354822.1 | 81.7576291 | -0.6301744 | 0.68613835 | -0.9184364 | 0.35839044 | 0.77529959 |

**Distal convoluted tubules**

*This means that positive fold changes imply higher mean value in non-COVID AKI than in COVID AKI and vice versa.

| | baseMean | log2 fold cha | lfcSE | stat | pvalue | padj |
|---|---|---|---|---|---|---|
| AL669831.5 | 37.7512535 | -0.009993 | 0.32138012 | -0.0310939 | 0.97519464 | 0.99037778 |
| NOC2L | 22.2146633 | 0.56144893 | 0.55842997 | 1.00540616 | 0.31470132 | 0.57214023 |
| ISG15 | 83.7123448 | -2.1011923 | 1.35894905 | -1.5461892 | 0.12205889 | 0.35134155 |
| AGRN | 81.1115278 | -0.0582374 | 0.47838997 | -0.1217362 | 0.90310792 | 0.95661107 |
| C1orf159 | 32.4346201 | -0.0732515 | 0.36261023 | -0.2020116 | 0.83990768 | 0.92315337 |
| AL390719.2 | 23.2684139 | -0.8022305 | 0.61677482 | -1.3006862 | 0.1933659 | 0.4452101 |
| SDF4 | 41.711187 | -0.1434084 | 0.44770501 | -0.3203189 | 0.74872659 | 0.87556192 |
| UBE2J2 | 41.0119848 | -0.5115509 | 0.4079478 | -1.2539617 | 0.20985591 | 0.46495709 |
| ACAP3 | 17.6617193 | -1.1597136 | 0.48629584 | -2.3847904 | 0.01708886 | NA |
| INTS11 | 38.9613273 | 0.1839294 | 0.30262586 | 0.6077782 | 0.54333459 | 0.75189614 |
| AURKAIP1 | 31.3272465 | 0.57319772 | 0.44344997 | 1.29258712 | 0.1961539 | 0.44947798 |
| CCNL2 | 119.184147 | -0.5313554 | 0.20627167 | -2.5759982 | 0.00999512 | 0.09901212 |
| MRPL20 | 37.9688801 | -0.0154814 | 0.39047753 | -0.0396473 | 0.96837428 | 0.98741848 |
| ATAD3B | 34.263048 | -0.3492248 | 0.34086549 | -1.0245237 | 0.305588 | 0.56557634 |
| ATAD3A | 16.2446781 | 0.2719361 | 0.61776066 | 0.44019653 | 0.65979477 | NA |
| SSU72 | 42.685554 | 0.235898 | 0.41135295 | 0.5734686 | 0.56632746 | 0.77020309 |
| MIB2 | 20.3861767 | 0.63441573 | 0.4344674 | 1.46021479 | 0.14423105 | NA |
| CDK11B | 27.4942725 | -1.2206445 | 0.42219071 | -2.891216 | 0.00383754 | 0.06110382 |
| SLC35E2B | 26.8833805 | -0.1587503 | 0.42241343 | -0.3758173 | 0.70705269 | 0.8547469 |
| CDK11A | 18.457347 | -0.0631051 | 0.47440425 | -0.1330197 | 0.89417778 | NA |
| SLC35E2A | 32.2079983 | -0.0803404 | 0.35091305 | -0.2289467 | 0.81891034 | 0.913067 |
| NADK | 29.9025058 | 0.36382525 | 0.43195673 | 0.84227245 | 0.39963547 | 0.64766685 |
| GNB1 | 220.314499 | -0.5742413 | 0.3086094 | -1.8607383 | 0.06278114 | 0.24960868 |
| PRKCZ | 65.2804535 | -0.8612287 | 0.29791785 | -2.8908259 | 0.00384231 | 0.06110382 |
| FAAP20 | 26.0485945 | 0.18163673 | 0.38737906 | 0.46888628 | 0.63915092 | 0.81390792 |
| SKI | 85.7238378 | -0.3563532 | 0.58839655 | -0.6056345 | 0.54475751 | 0.75328235 |
| MORN1 | 28.8976537 | -0.1649048 | 0.36770726 | -0.4484677 | 0.6538157 | 0.82242514 |
| RER1 | 55.8600929 | 0.30822984 | 0.35851971 | 0.85972913 | 0.38993837 | 0.6386799 |
| PEX10 | 12.3028773 | 0.2601737 | 0.54607962 | 0.47643913 | 0.63376157 | NA |
| TNFRSF14 | 25.8165348 | -1.143542 | 0.74221422 | -1.5407169 | 0.12338571 | 0.35349809 |
| LINC00982 | 59.6924952 | -0.2417892 | 0.60674155 | -0.3985044 | 0.69025838 | 0.84627568 |
| PRDM16 | 280.54792 | -0.7426012 | 0.45098819 | -1.6466089 | 0.09963845 | 0.31793995 |
| ARHGEF16 | 31.1733735 | -0.5459684 | 0.42315794 | -1.2902237 | 0.19697299 | 0.45014879 |
| LRRC47 | 17.0226089 | -0.0380553 | 0.46485425 | -0.081865 | 0.93475407 | NA |
| CEP104 | 60.3455956 | -0.7099544 | 0.29679661 | -2.3920568 | 0.01675425 | 0.1285308 |
| DFFB | 11.6469611 | 1.46723958 | 0.699938 | 2.09624222 | 0.03606071 | NA |
| AJAP1 | 52.8823468 | -0.2736042 | 0.98066066 | -0.2789999 | 0.7802449 | 0.89076609 |
| NPHP4 | 42.7468475 | -0.1907614 | 0.41500634 | -0.459659 | 0.64576103 | 0.81777273 |
| RPL22 | 26.3328227 | 0.05308656 | 0.51715136 | 0.10265189 | 0.91823926 | 0.96421386 |
| RNF207 | 22.2928014 | 0.89010778 | 0.43854257 | 2.02969526 | 0.04238752 | 0.20555189 |
| ICMT | 15.3380692 | 1.2700808 | 0.50645548 | 2.50778369 | 0.0121491 | NA |
| NOL9 | 39.4444687 | -0.0042608 | 0.32443783 | -0.0131329 | 0.98952179 | 0.99583841 |
| ZBTB48 | 17.096483 | 0.57307137 | 0.48135941 | 1.19052699 | 0.23383933 | NA |
| KLHL21 | 33.4805719 | 0.13337646 | 0.40018044 | 0.33329081 | 0.73891478 | 0.87054913 |
| DNAJC11 | 126.268844 | -0.2347464 | 0.25742794 | -0.9118915 | 3.62E-01 | 0.61764768 |
| CAMTA1 | 426.101238 | -1.16579 | 0.42188941 | -2.7632598 | 0.00572272 | 0.0741545 |
| VAMP3 | 17.7993446 | -0.1717625 | 0.39884153 | -0.4306536 | 0.66672026 | NA |
| PER3 | 99.2047566 | -0.521993 | 0.53060906 | -0.9837619 | 0.32523262 | 0.58122021 |
| PARK7 | 52.9308574 | 0.57718938 | 0.37308317 | 1.54707967 | 0.12184404 | 0.35111702 |
| ERRFI1 | 121.731756 | -0.1917157 | 0.55401702 | -0.3460467 | 0.72930765 | 0.86533507 |
| RERE | 1351.91417 | -0.0950529 | 0.21689312 | -0.4382479 | 0.6612066 | 0.82766421 |
| ENO1 | 235.601653 | -0.3348227 | 0.37003378 | -0.9048437 | 0.36554822 | 0.62110919 |

| | | | | | | |
|---|---|---|---|---|---|---|
| GPR157 | 33.3378966 | 0.75669184 | 0.35303777 | 2.14337361 | 0.03208311 | 0.1766407 |
| H6PD | 34.6219096 | -0.124179 | 0.34938625 | -0.3554204 | 0.72227464 | 0.86153309 |
| SPSB1 | 45.8485804 | -1.8027438 | 0.44224143 | -4.0763792 | 4.57E-05 | 0.00423103 |
| SLC25A33 | 80.9286143 | -0.0069555 | 0.37433812 | -0.0185808 | 0.9851755 | 0.99408703 |
| PIK3CD | 21.4933439 | -0.5623957 | 0.66281059 | -0.8485014 | 0.39615878 | 0.64452212 |
| CLSTN1 | 104.001204 | 0.25376205 | 0.30294274 | 0.8376568 | 0.40222348 | 0.65012897 |
| CTNNBIP1 | 34.2452033 | -0.0059944 | 0.33928735 | -0.0176675 | 0.98590407 | 0.9942828 |
| LZIC | 23.0201226 | 0.04457853 | 0.44189258 | 0.10088091 | 0.919645 | 0.9650317 |
| NMNAT1 | 63.9459435 | 0.13114793 | 0.30584179 | 0.42880971 | 0.66806171 | 0.83285892 |
| UBE4B | 177.993772 | 0.05011651 | 0.23731374 | 0.2111825 | 0.83274487 | 0.91881268 |
| KIF1B | 203.631003 | 0.28716503 | 0.2127545 | 1.34974832 | 0.17709673 | 0.42569745 |
| PGD | 27.2746927 | -0.3616333 | 0.37448089 | -0.9656923 | 0.33419818 | 0.58942664 |
| DFFA | 36.8410371 | 0.25600299 | 0.37087654 | 0.69026472 | 0.49002773 | 0.7163358 |
| PEX14 | 128.786202 | -0.2982288 | 0.25629511 | -1.1636149 | 0.24458009 | 0.50200879 |
| CASZ1 | 218.424067 | 1.04785463 | 0.37758256 | 2.77516693 | 0.00551734 | 0.07254189 |
| EXOSC10 | 31.5505061 | -0.6634413 | 0.35572385 | -1.8650458 | 0.06217498 | 0.24899105 |
| MTOR | 86.0107608 | -0.3687096 | 0.27873824 | -1.3227809 | 0.18590824 | 0.43607799 |
| FBXO44 | 22.4467998 | 0.79788195 | 0.49898089 | 1.59902306 | 0.10981548 | 0.33253594 |
| MAD2L2 | 14.4256247 | 0.07447128 | 0.63234334 | 0.11777032 | 0.90624965 | NA |
| AGTRAP | 17.1388518 | -0.4351088 | 0.49770969 | -0.874222 | 0.38199734 | NA |
| MTHFR | 22.1808484 | -0.0907361 | 0.43657507 | -0.2078362 | 0.83535688 | 0.92037547 |
| CLCN6 | 31.6619692 | -0.7972481 | 0.46820747 | -1.7027667 | 0.08861173 | 0.29770601 |
| KIAA2013 | 16.3176484 | 0.4544611 | 0.51797868 | 0.87737415 | 0.38028345 | NA |
| PLOD1 | 13.7166917 | 0.14075098 | 0.54132424 | 0.26001233 | 0.79485426 | NA |
| MFN2 | 45.7761056 | 0.23901961 | 0.38818013 | 0.61574405 | 0.53806346 | 0.74878549 |
| VPS13D | 408.235957 | -0.1510209 | 0.20401828 | -0.7402324 | 0.45915902 | 0.69296999 |
| DHRS3 | 94.1616146 | -0.9650854 | 0.47603761 | -2.0273301 | 0.04262867 | 0.20610104 |
| PRDM2 | 105.661076 | -0.2676361 | 0.21214218 | -1.2615882 | 0.207097 | 0.46066753 |
| KAZN | 1471.97281 | -0.7156374 | 0.34207445 | -2.0920516 | 0.0364339 | 0.19061356 |
| TMEM51-AS | 33.3392523 | -0.8986462 | 0.6521854 | -1.3778999 | 0.16823421 | 0.41623231 |
| TMEM51 | 251.611706 | -0.8170722 | 0.51776322 | -1.5780809 | 0.11454703 | 0.3409391 |
| CASP9 | 24.5750924 | -0.7191565 | 0.38268376 | -1.879245 | 0.06021105 | 0.24444898 |
| DNAJC16 | 62.4305497 | -0.1085563 | 0.26525021 | -0.4092602 | 0.68234876 | 0.84194267 |
| DDI2 | 67.1823243 | -0.4455269 | 0.23114479 | -1.9274797 | 0.05391988 | 0.23287156 |
| PLEKHM2 | 60.0012419 | -1.3895226 | 0.33934906 | -4.0946705 | 4.23E-05 | 0.00401254 |
| AL450998.2 | 11.7912633 | 0.14733834 | 0.5601124 | 0.26305138 | 0.79251098 | NA |
| SPEN | 148.052062 | -0.7060812 | 0.32791369 | -2.1532533 | 0.03129878 | 0.17469667 |
| ZBTB17 | 21.4068263 | -0.5283173 | 0.38181688 | -1.3836928 | 0.16645255 | 0.41429299 |
| CLCNKA | 123.072369 | 1.14244547 | 0.63768215 | 1.79155943 | 7.32E-02 | 0.26946334 |
| CLCNKB | 184.920379 | 2.37306521 | 0.79105887 | 2.99985917 | 0.00270104 | 0.05029571 |
| FBXO42 | 151.238878 | -0.7217281 | 0.2888093 | -2.4989781 | 0.0124552 | 0.10951947 |
| SZRD1 | 39.6984238 | -0.2081862 | 0.32243017 | -0.6456784 | 0.51848766 | 0.73536015 |
| NECAP2 | 22.029023 | -0.0849718 | 0.35395832 | -0.2400615 | 0.81028254 | 0.90835199 |
| NBPF1 | 34.4114376 | 0.59666911 | 0.37171435 | 1.60518181 | 0.1084538 | 0.33024913 |
| CROCC | 37.7465235 | 0.57138837 | 0.4332797 | 1.31875178 | 0.18725211 | 0.43794576 |
| ATP13A2 | 14.5549482 | 0.56758041 | 0.50775119 | 1.11783176 | 0.26363885 | NA |
| SDHB | 76.7735569 | 0.04060058 | 0.24136647 | 0.16821133 | 0.86641702 | 0.93793856 |
| PADI2 | 12.9277227 | 1.91731942 | 1.08785718 | 1.76247347 | 0.07798934 | NA |
| RCC2 | 30.9798382 | -0.1534988 | 0.46750361 | -0.3283372 | 0.74265668 | 0.87154797 |
| ARHGEF10L | 234.870454 | -0.2184175 | 0.35325487 | -0.6182999 | 0.53637766 | 0.74845881 |
| UBR4 | 141.651676 | -0.5298698 | 0.28695898 | -1.8465 | 0.06481964 | 0.25312875 |
| AL035413.1 | 27.0527767 | -0.2477669 | 0.42174612 | -0.5874788 | 0.55688217 | 0.76323398 |
| EMC1 | 16.2127884 | 0.06419382 | 0.46744366 | 0.13732953 | 0.89077033 | NA |
| AKR7A2 | 14.038456 | 1.09445995 | 0.58660558 | 1.86575102 | 0.06207621 | NA |

| | | | | | | |
|---|---|---|---|---|---|---|
| CAPZB | 150.217445 | -0.508273 | 0.31549891 | -1.6110134 | 0.10717681 | 0.32918592 |
| TMCO4 | 99.910813 | 0.65866684 | 0.50909693 | 1.29379455 | 0.1957364 | 0.44885942 |
| OTUD3 | 39.1230079 | -0.3440441 | 0.41668407 | -0.8256713 | 0.40899056 | 0.65472852 |
| CAMK2N1 | 68.4821145 | 0.16200851 | 0.5167923 | 0.31348863 | 0.75390946 | 0.87868581 |
| MUL1 | 14.6938437 | -0.7808887 | 0.45889862 | -1.7016585 | 0.08881941 | NA |
| PINK1 | 26.8085725 | 0.33759666 | 0.39571302 | 0.85313506 | 0.39358441 | 0.64243226 |
| DDOST | 18.5390663 | 0.59698443 | 0.42806281 | 1.39461877 | 0.16313083 | NA |
| HP1BP3 | 229.421898 | -0.0461509 | 0.17988431 | -0.2565585 | 0.7975196 | 0.90121835 |
| EIF4G3 | 901.01262 | -0.8256877 | 0.4403136 | -1.8752265 | 0.06076156 | 0.24588946 |
| ECE1 | 79.5324165 | -0.3474558 | 0.49115913 | -0.70742 | 0.4793055 | 0.70695585 |
| NBPF3 | 16.7439629 | 1.52254071 | 0.60541035 | 2.51489046 | 0.01190694 | NA |
| RAP1GAP | 50.6853698 | 1.29540303 | 0.58055595 | 2.2313147 | 0.02566029 | 0.15662748 |
| USP48 | 136.156381 | -0.0832215 | 0.24499696 | -0.3396836 | 0.73409478 | 0.86800594 |
| HSPG2 | 64.5897302 | 0.23200757 | 0.39731228 | 0.58394261 | 0.55925892 | 0.76485713 |
| CDC42 | 107.717484 | -0.2007517 | 0.2582207 | -0.7774423 | 0.43689787 | 0.67734041 |
| ZBTB40 | 50.3977142 | 0.07311232 | 0.37525066 | 0.19483594 | 0.84552138 | 0.92531372 |
| EPHB2 | 43.7277707 | -2.0984166 | 0.57565894 | -3.6452427 | 0.00026714 | 0.01346273 |
| KDM1A | 81.1732075 | 0.33983979 | 0.3473309 | 0.97843235 | 0.32786054 | 0.58415337 |
| AL031428.1 | 21.6652599 | 0.66486307 | 0.50553422 | 1.31516927 | 0.18845302 | 0.43914317 |
| LUZP1 | 147.586827 | 0.19824532 | 0.26458277 | 0.74927525 | 4.54E-01 | 6.89E-01 |
| LINC01355 | 20.903928 | 0.51679068 | 0.41449949 | 1.24678242 | 0.21247729 | NA |
| HNRNPR | 114.461203 | -0.2716437 | 0.22983938 | -1.181885 | 0.23725135 | 0.49515284 |
| TCEA3 | 56.467264 | 2.10859854 | 0.54679179 | 3.85630982 | 0.00011511 | 0.00784083 |
| ASAP3 | 63.3049494 | 0.72627898 | 0.31413942 | 2.31196385 | 2.08E-02 | 0.14123712 |
| RPL11 | 88.7359097 | 0.66821446 | 0.32782518 | 2.03832562 | 0.04151738 | 0.20310896 |
| ELOA | 24.8713467 | -0.306227 | 0.4069453 | -0.7525017 | 0.45174941 | 0.68853766 |
| ELOA-AS1 | 24.8993049 | 0.36364474 | 0.6409462 | 0.5673561 | 0.57047227 | 0.77196291 |
| PITHD1 | 19.5640683 | 0.10039501 | 0.41497703 | 0.24192909 | 0.80883511 | NA |
| LYPLA2 | 18.3897874 | 0.06184885 | 0.59199691 | 0.10447496 | 0.91679244 | NA |
| GALE | 23.0275384 | -0.9668463 | 0.37131289 | -2.6038586 | 9.22E-03 | 0.09482608 |
| HMGCL | 32.9843768 | -0.1038869 | 0.35833687 | -0.2899142 | 0.77188189 | 0.88727224 |
| FUCA1 | 24.2563681 | 0.77247788 | 0.4131163 | 1.86987994 | 0.06150049 | 0.24796715 |
| PNRC2 | 34.9888794 | -0.894112 | 0.43074163 | -2.0757501 | 0.03791707 | 0.19457113 |
| SRSF10 | 141.095368 | -0.2579946 | 0.3024319 | -0.8530668 | 0.39362226 | 0.64243226 |
| IFNLR1 | 19.0011207 | 0.31496434 | 0.54140501 | 0.58175365 | 0.56073263 | NA |
| STPG1 | 14.5221871 | 0.37674913 | 0.56739255 | 0.66400083 | 0.50668978 | NA |
| NIPAL3 | 49.9663732 | 0.35652466 | 0.46911287 | 0.75999763 | 0.447256 | 0.68660592 |
| RCAN3 | 26.2240626 | -0.0723959 | 0.47624445 | -0.1520141 | 0.87917585 | 0.94360719 |
| NCMAP | 22.5721907 | -0.3856378 | 0.63741626 | -0.6050015 | 5.45E-01 | 7.53E-01 |
| SRRM1 | 92.1078946 | -0.6372931 | 0.22498497 | -2.832603 | 0.00461707 | 0.06581034 |
| CLIC4 | 61.9799372 | -0.2652654 | 0.72791654 | -0.3644173 | 0.71554639 | 0.85868129 |
| SYF2 | 24.1632212 | -0.4117666 | 0.42220355 | -0.9752799 | 0.32942146 | 0.58535385 |
| RSRP1 | 279.556805 | 0.63122292 | 0.32455674 | 1.94487695 | 0.05178979 | 0.22830814 |
| TMEM50A | 33.6930384 | -0.2139922 | 0.40653858 | -0.5263762 | 0.59862686 | 0.78956448 |
| MACO1 | 69.9311308 | -0.3068993 | 0.24121419 | -1.2723103 | 0.2032629 | 0.45705947 |
| LDLRAP1 | 15.7306427 | 1.13749291 | 0.49032468 | 2.31987693 | 0.02034754 | NA |
| MAN1C1 | 27.2694307 | 0.43433238 | 0.62682585 | 0.69290757 | 0.48836756 | 0.71517601 |
| MTFR1L | 26.3457302 | 1.75882264 | 0.64532042 | 2.72550285 | 0.00642036 | 0.07805175 |
| STMN1 | 15.7159396 | 0.34470762 | 0.52758417 | 0.65336991 | 0.51351784 | NA |
| PAFAH2 | 23.9734828 | 0.38371963 | 0.44386043 | 0.86450516 | 0.38731044 | 0.63701577 |
| CNKSR1 | 19.0479271 | 0.19327574 | 0.43144575 | 0.44797229 | 0.65417319 | NA |
| CEP85 | 22.5257108 | -0.5815377 | 0.53383955 | -1.0893492 | 0.27599994 | 0.53482093 |
| SH3BGRL3 | 22.5682654 | -0.566753 | 0.65796911 | -0.8613672 | 0.38903584 | 0.63805857 |
| DHDDS | 39.5185136 | 0.14702753 | 0.35688589 | 0.4119735 | 0.68035885 | 0.84089068 |

| | | | | | | |
|---|---|---|---|---|---|---|
| HMGN2 | 33.8601836 | -0.29095 | 0.42162019 | -0.6900761 | 0.49014631 | 0.7163358 |
| RPS6KA1 | 31.0707828 | 0.23617905 | 0.32510475 | 0.72647063 | 0.46755031 | 0.69742921 |
| ARID1A | 186.581077 | 0.51867819 | 0.20832669 | 2.48973468 | 0.01278385 | 0.11046442 |
| ZDHHC18 | 14.1473709 | 0.13001417 | 0.56699245 | 0.22930494 | 0.81863192 | NA |
| GPN2 | 18.8076684 | -0.8760929 | 0.50626376 | -1.7305069 | 0.08353975 | NA |
| NUDC | 47.3816428 | -0.662903 | 0.29986348 | -2.2106826 | 0.02705782 | 0.16195658 |
| TRNP1 | 58.1375021 | 0.50849952 | 0.48646229 | 1.04530101 | 2.96E-01 | 5.56E-01 |
| SLC9A1 | 36.9971572 | -0.527414 | 0.50489136 | -1.0446088 | 0.2962038 | 0.55582698 |
| WDTC1 | 60.0110137 | 0.08202251 | 0.26512735 | 0.30937022 | 0.75703992 | 0.87904624 |
| TMEM222 | 19.4470846 | -0.2191194 | 0.46622983 | -0.4699814 | 0.63836828 | NA |
| WASF2 | 129.346353 | 0.07785483 | 0.21653131 | 0.35955462 | 0.71918023 | 0.85995641 |
| AHDC1 | 58.572462 | 0.52147257 | 0.34136485 | 1.5276106 | 0.12660925 | 0.35827624 |
| IFI6 | 79.6207149 | 1.80465184 | 1.56704823 | 1.15162495 | 0.24947522 | 0.5058511 |
| FAM76A | 20.767325 | 0.27629094 | 0.56130355 | 0.49223088 | 0.62255614 | NA |
| STX12 | 84.0024666 | -0.3254208 | 0.39881711 | -0.8159649 | 0.41452019 | 0.6585267 |
| PPP1R8 | 20.5568664 | -0.3857698 | 0.41089321 | -0.9388565 | 0.34780444 | NA |
| RPA2 | 20.0112751 | 0.41913604 | 0.43333219 | 0.96723958 | 0.33342428 | NA |
| EYA3 | 254.445494 | 0.33055751 | 0.28730829 | 1.15053242 | 0.24992465 | 0.5063622 |
| DNAJC8 | 38.371278 | -0.3921656 | 0.38492866 | -1.0188006 | 0.30829764 | 0.5682392 |
| ATP5IF1 | 58.0276126 | 0.78178122 | 0.44549854 | 1.75484577 | 0.07928569 | 0.2808912 |
| PHACTR4 | 247.936219 | -0.0794758 | 0.21406828 | -0.3712636 | 0.71044121 | 0.8562897 |
| RCC1 | 27.1530613 | -0.1133964 | 0.34454346 | -0.3291207 | 0.74206447 | 0.87152705 |
| TRNAU1AP | 48.8421078 | -0.4176523 | 0.398521 | -1.0480058 | 0.29463595 | 0.55396961 |
| TAF12 | 33.8489906 | -0.8351657 | 0.37143042 | -2.2485119 | 0.02454357 | 0.15333754 |
| GMEB1 | 32.590517 | -1.0319898 | 0.36125216 | -2.8567022 | 0.00428067 | 0.06301187 |
| YTHDF2 | 75.5971025 | -0.2430781 | 0.30123058 | -0.8069501 | 0.4196952 | 0.6636918 |
| EPB41 | 195.187213 | 0.17428192 | 0.2007867 | 0.86799536 | 0.38539687 | 0.63547552 |
| SRSF4 | 272.243983 | -0.455023 | 0.26645918 | -1.7076651 | 0.0876985 | 0.29621428 |
| MECR | 26.4402998 | 0.87339462 | 0.46153308 | 1.89237705 | 0.05844076 | 0.24020328 |
| PUM1 | 358.505001 | -0.2802291 | 0.23931733 | -1.1709518 | 0.24161814 | 0.49885591 |
| SNRNP40 | 37.0275602 | 0.13885243 | 0.39519505 | 0.35135164 | 0.72532456 | 0.86314519 |
| ZCCHC17 | 82.2601577 | -0.3017007 | 0.27787244 | -1.0857526 | 0.27758846 | 0.53602567 |
| FABP3 | 18.4426252 | 0.49120167 | 0.58396161 | 0.84115404 | 0.40026164 | NA |
| SERINC2 | 92.7426203 | -0.1508421 | 0.30568139 | -0.4934617 | 0.62168637 | 0.80354659 |
| TINAGL1 | 44.9480644 | -0.6241378 | 0.59085163 | -1.0563358 | 0.29081481 | 0.55011098 |
| PEF1 | 21.7312176 | 0.28438966 | 0.42196288 | 0.67396841 | 0.5003314 | 0.72238806 |
| PTP4A2 | 34.4069963 | -0.8866852 | 0.40391947 | -2.1952029 | 0.02814904 | 0.16513963 |
| KHDRBS1 | 128.929642 | -0.6985285 | 0.25245073 | -2.7669893 | 0.00565766 | 0.07350161 |
| TMEM39B | 15.0949307 | -0.1583513 | 0.49245259 | -0.3215564 | 0.74778876 | NA |
| KPNA6 | 117.833799 | -0.6002296 | 0.31467989 | -1.9074291 | 0.05646505 | 0.23671649 |
| TXLNA | 36.2072299 | -0.1947904 | 0.39532609 | -0.4927335 | 0.62220092 | 0.80366942 |
| EIF3I | 16.8414226 | 0.17704653 | 0.45879277 | 0.38589651 | 0.69957331 | NA |
| HDAC1 | 48.0880916 | 0.00639305 | 0.34073815 | 0.01876236 | 0.98503068 | 0.99407122 |
| BSDC1 | 63.4424215 | 0.11884771 | 0.29466992 | 0.40332487 | 0.68670925 | 0.8446294 |
| ZBTB8OS | 113.911972 | -0.6141327 | 0.25975849 | -2.3642449 | 0.01806686 | 0.13331596 |
| RBBP4 | 92.7599476 | -0.2462212 | 0.33692362 | -0.7307923 | 0.46490606 | 0.69577716 |
| SYNC | 15.8788712 | 0.50642193 | 0.5062314 | 1.00037638 | 0.3171284 | NA |
| KIAA1522 | 91.6533006 | 0.00376371 | 0.29442919 | 0.01278307 | 0.98980086 | 0.99583841 |
| YARS | 50.3447139 | -0.9299831 | 0.52560348 | -1.7693624 | 0.07683341 | 0.27670206 |
| S100PBP | 101.779862 | -0.3300749 | 0.23671476 | -1.3943992 | 0.16319709 | 0.40987726 |
| RNF19B | 41.2857587 | -1.1028916 | 0.45219566 | -2.4389698 | 0.0147292 | 0.11971557 |
| AL020995.1 | 32.5567013 | -0.1856248 | 0.57506231 | -0.3227908 | 0.7468537 | 0.87469995 |
| AK2 | 112.457236 | -0.1379796 | 0.2879025 | -0.479258 | 0.63175512 | 0.80908805 |
| ZNF362 | 22.826344 | 1.44264138 | 0.66348009 | 2.1743552 | 0.02967847 | 0.1702177 |

| | | | | | | |
|---|---|---|---|---|---|---|
| AL513327.1 | 43.3456108 | 0.02562108 | 0.70030804 | 0.03658544 | 0.97081555 | 0.98827765 |
| PHC2 | 206.127224 | -1.0071775 | 0.29157016 | -3.4543228 | 0.00055168 | 0.02102106 |
| SMIM12 | 33.9251853 | -0.7469469 | 0.38991858 | -1.9156484 | 0.05540986 | 0.23522598 |
| ZMYM1 | 42.1784265 | 0.90881421 | 0.38567318 | 2.35643614 | 0.01845124 | 0.13411314 |
| SFPQ | 221.051841 | -0.3333939 | 0.26978382 | -1.2357818 | 0.21653968 | 0.47282437 |
| ZMYM4 | 324.997492 | -0.1468132 | 0.19985403 | -0.7346021 | 0.46258186 | 0.69418845 |
| KIAA0319L | 162.469001 | -0.1812167 | 0.24638992 | -0.7354875 | 0.46204266 | 0.69416206 |
| PSMB2 | 25.7651951 | 0.09925815 | 0.39818506 | 0.24927644 | 0.80314696 | 0.90408418 |
| AGO4 | 78.6596924 | 0.19088479 | 0.31453349 | 0.60688223 | 0.54392907 | 0.75258351 |
| AGO1 | 23.6530401 | 0.39825839 | 0.41074566 | 0.96959855 | 0.33224664 | 0.58794039 |
| AGO3 | 241.764855 | -0.4818485 | 0.19203147 | -2.5092162 | 0.01209994 | 0.1074201 |
| ADPRHL2 | 18.6947661 | -0.7031564 | 0.4566214 | -1.5399111 | 0.12358203 | NA |
| TRAPPC3 | 32.4700073 | -0.5940083 | 0.49461412 | -1.2009531 | 0.22976941 | 0.48827585 |
| MAP7D1 | 19.9639947 | -0.5172467 | 0.62595393 | -0.8263335 | 0.40861494 | NA |
| THRAP3 | 174.573584 | -0.3537594 | 0.27180509 | -1.3015185 | 0.19308102 | 0.44483671 |
| SH3D21 | 24.5424977 | -0.7264282 | 0.50763019 | -1.4310185 | 0.15242492 | 0.39409292 |
| EVA1B | 16.3134184 | 0.22268876 | 0.56732626 | 0.39252328 | 0.69467162 | NA |
| AL591845.1 | 23.6028297 | 0.11364912 | 0.5407549 | 0.21016753 | 0.83353692 | 0.919423 |
| STK40 | 50.8067846 | -0.7218187 | 0.4656092 | -1.5502673 | 0.12107736 | 0.35008733 |
| LINC01137 | 17.2249122 | 0.42361275 | 0.47144292 | 0.89854516 | 0.36889498 | NA |
| MEAF6 | 27.8321551 | 0.01688253 | 0.38451772 | 0.04390572 | 0.96497956 | 0.9869039 |
| SNIP1 | 14.1151572 | -0.2763774 | 0.46858621 | -0.5898111 | 0.55531728 | NA |
| GNL2 | 13.2480086 | -0.4917158 | 0.57301459 | -0.8581209 | 0.39082571 | NA |
| C1orf122 | 24.0102175 | 0.49240651 | 0.52351849 | 0.94057139 | 0.34692455 | 0.60386211 |
| MTF1 | 28.0474046 | 0.1664467 | 0.40479947 | 0.41118309 | 0.68093829 | 0.84114621 |
| INPP5B | 103.003987 | 0.42621384 | 0.37079818 | 1.14944965 | 0.25037062 | 0.50659901 |
| SF3A3 | 30.9061999 | -0.7056726 | 0.32409217 | -2.1773825 | 0.02945204 | 0.16942626 |
| RRAGC | 22.8907961 | -1.1779134 | 0.58075177 | -2.0282563 | 0.04253409 | 0.20590247 |
| AL139260.1 | 76.3005875 | -1.8097172 | 0.65249767 | -2.7735228 | 0.00554529 | 0.07254189 |
| AKIRIN1 | 105.522215 | 0.14498727 | 0.29788777 | 0.48671776 | 0.62645836 | 0.8053173 |
| NDUFS5 | 53.7495369 | 0.44287251 | 0.38313656 | 1.15591294 | 0.24771677 | 0.50401162 |
| MACF1 | 899.953329 | -0.8690923 | 0.2461689 | -3.5304715 | 0.00041482 | 0.01783725 |
| PABPC4 | 34.2764283 | -0.7779526 | 0.39199926 | -1.9845768 | 0.04719157 | 0.21733813 |
| PPIE | 22.7212737 | -0.0136307 | 0.4242801 | -0.0321265 | 0.97437114 | 0.9901947 |
| TRIT1 | 58.6961508 | -0.5837694 | 0.36820053 | -1.5854659 | 0.11286051 | 0.33755464 |
| CAP1 | 45.9651933 | -0.1754038 | 0.34432359 | -0.5094156 | 0.61046094 | 0.79720357 |
| PPT1 | 17.810095 | 0.39193476 | 0.49666178 | 0.78913815 | 0.43003126 | NA |
| RLF | 173.200669 | -0.4611283 | 0.18626376 | -2.4756737 | 0.01329851 | 0.11268704 |
| ZMPSTE24 | 37.0754174 | -0.290255 | 0.35637891 | -0.8144562 | 0.41538365 | 0.65906718 |
| SMAP2 | 29.0996384 | 0.05687897 | 0.38752697 | 0.14677422 | 0.88331023 | 0.94552272 |
| AL603839.3 | 26.0444723 | 0.95266668 | 0.45086327 | 2.11298355 | 0.03460218 | 0.18447938 |
| ZNF684 | 32.4484966 | 1.9877627 | 0.60240139 | 3.29973128 | 0.00096777 | 0.02829008 |
| NFYC | 172.958259 | 0.46745522 | 0.24290389 | 1.92444519 | 5.43E-02 | 0.23287156 |
| SCMH1 | 305.269698 | -0.2104934 | 0.17564481 | -1.1984039 | 0.23075982 | 0.48936597 |
| HIVEP3 | 160.75948 | -0.7734608 | 0.27574086 | -2.805028 | 5.03E-03 | 0.06878385 |
| FOXJ3 | 212.074421 | -0.8200343 | 0.26045192 | -3.1485056 | 0.00164108 | 0.03697992 |
| PPCS | 21.6311736 | 0.81955243 | 0.48409564 | 1.69295561 | 0.09046392 | 0.3003886 |
| CCDC30 | 138.848626 | 1.22396648 | 0.41754806 | 2.9313188 | 0.00337526 | 0.05734965 |
| PPIH | 15.8786324 | -0.0421433 | 0.58006387 | -0.0726529 | 0.94208235 | NA |
| YBX1 | 63.8764799 | 0.02573678 | 0.32282741 | 0.07972303 | 0.93645754 | 0.97405271 |
| ERMAP | 24.5834951 | -0.2612758 | 0.46591271 | -0.5607828 | 0.5749456 | 0.77388388 |
| SLC2A1 | 62.8819231 | -0.7712344 | 0.56760565 | -1.3587504 | 0.17422569 | 0.42295604 |
| SLC2A1-AS1 | 19.6843178 | 1.02172555 | 0.73038869 | 1.39887921 | 0.16184921 | NA |
| EBNA1BP2 | 17.3045671 | 0.41439521 | 0.52151448 | 0.79459962 | 0.42684644 | NA |

| | | | | | | |
|---|---|---|---|---|---|---|
| SZT2 | 49.2863376 | -0.4403502 | 0.33720556 | -1.3058807 | 0.19159313 | 0.44319545 |
| PTPRF | 95.6618066 | 0.66773015 | 0.43797182 | 1.52459615 | 0.12735986 | 0.35947519 |
| ATP6V0B | 84.3810146 | 0.34245334 | 0.4193674 | 0.81659503 | 0.41415989 | 0.65845716 |
| CCDC24 | 29.0877672 | 1.02458537 | 0.45010023 | 2.27634938 | 0.02282511 | 0.14701661 |
| DMAP1 | 13.7403295 | 0.49032213 | 0.50296388 | 0.9748655 | 0.32962698 | NA |
| ERI3 | 91.1584706 | 1.00822989 | 0.38286956 | 2.63335087 | 0.0084547 | 0.09005263 |
| RNF220 | 16.0747664 | 0.54578557 | 0.4755104 | 1.14778892 | 0.25105571 | NA |
| TMEM53 | 22.5171112 | 0.54864807 | 0.41066995 | 1.335983 | 0.18155483 | 0.4309058 |
| ARMH1 | 15.7150239 | 0.75406731 | 0.66438282 | 1.13498918 | 0.25637985 | NA |
| RPS8 | 78.709234 | 0.33830908 | 0.38065641 | 0.88875185 | 0.37413646 | 0.62575308 |
| EIF2B3 | 53.6830756 | -0.02872 | 0.27623064 | -0.1039711 | 0.91719227 | 0.96401363 |
| ZSWIM5 | 94.5670927 | -0.2417859 | 0.44277895 | -0.5460646 | 0.58502152 | 0.78080643 |
| TESK2 | 24.442997 | 0.38898791 | 0.36286647 | 1.07198635 | 0.28372616 | 0.54270385 |
| PRDX1 | 94.4405285 | 0.49255225 | 0.41258882 | 1.19380902 | 0.23255268 | 0.49082729 |
| AKR1A1 | 18.9964463 | -0.3151033 | 0.50232404 | -0.6272908 | 0.53046861 | NA |
| NASP | 51.5646612 | -0.1179377 | 0.30338038 | -0.3887452 | 0.69746466 | 0.85050468 |
| GPBP1L1 | 226.549744 | -0.2653957 | 0.17969177 | -1.4769496 | 0.13968914 | 0.37697679 |
| TMEM69 | 14.5962113 | -0.0097264 | 0.44434654 | -0.0218892 | 0.98253636 | NA |
| IPP | 40.8761234 | 0.09361339 | 0.30211137 | 0.30986385 | 0.7566645 | 0.87904624 |
| MAST2 | 69.3379514 | -0.5641493 | 0.47218029 | -1.1947752 | 0.23217488 | 0.49066933 |
| TSPAN1 | 66.8925765 | 0.28515718 | 0.35184422 | 0.8104643 | 4.18E-01 | 6.62E-01 |
| POMGNT1 | 39.4090349 | 0.4084615 | 0.38617222 | 1.05771852 | 0.29018379 | 0.54959267 |
| LURAP1 | 42.2662934 | 0.23983659 | 0.4894508 | 0.49001165 | 0.62412566 | 0.80467256 |
| LRRC41 | 42.1540822 | -0.6858308 | 0.32016082 | -2.1421447 | 0.03218184 | 0.17693114 |
| UQCRH | 81.769519 | 0.69615407 | 0.32703685 | 2.12867162 | 0.03328144 | 0.18078139 |
| NSUN4 | 32.4272862 | 0.11757832 | 0.3629776 | 0.3239272 | 0.74599315 | 0.87409184 |
| FAAH | 14.7748871 | 0.99702391 | 0.50330554 | 1.9809516 | 0.0475967 | NA |
| MKNK1 | 51.4029195 | 0.57076745 | 0.30566791 | 1.86727961 | 0.06186255 | 0.24877537 |
| ATPAF1 | 55.9727505 | 0.27161777 | 0.29900686 | 0.90839978 | 0.36366704 | 0.61996572 |
| EFCAB14 | 79.4475307 | 0.07968244 | 0.41017883 | 0.19426268 | 0.8459702 | 0.92531372 |
| PDZK1IP1 | 43.5141714 | 0.84342808 | 0.60654542 | 1.39054398 | 0.16436375 | 0.41120418 |
| CMPK1 | 55.8641546 | 0.1720585 | 0.34793685 | 0.49451071 | 0.62094555 | 0.80353114 |
| TRABD2B | 774.049077 | -0.2323134 | 0.32292455 | -0.7194044 | 0.47189177 | 0.70118744 |
| SPATA6 | 152.074991 | 0.81879269 | 0.42015879 | 1.94876961 | 0.05132294 | 0.2274224 |
| AGBL4 | 456.512184 | 2.11814349 | 0.59714457 | 3.5471201 | 0.00038947 | 0.01710271 |
| FAF1 | 625.60612 | -0.2178598 | 0.16802117 | -1.2966209 | 0.19476165 | 0.44735315 |
| RNF11 | 68.3627481 | -0.2608556 | 0.3203338 | -0.8143242 | 0.41545922 | 0.65906718 |
| TTC39A | 76.0285556 | -1.1543083 | 0.34104587 | -3.3846132 | 7.13E-04 | 2.39E-02 |
| EPS15 | 239.051589 | 0.07893743 | 0.23365421 | 0.3378387 | 0.73548475 | 0.86838246 |
| OSBPL9 | 215.78403 | 0.19765051 | 0.2524516 | 0.78292438 | 0.43367152 | 0.67472483 |
| NRDC | 185.35788 | -0.0609601 | 0.22181821 | -0.2748201 | 0.78345444 | 0.89323786 |
| TXNDC12 | 35.7175787 | 0.07712971 | 0.34592037 | 0.22296954 | 0.82355921 | 0.91484128 |
| BTF3L4 | 56.9323199 | -0.2845684 | 0.28631453 | -0.9939013 | 0.32027091 | 0.57724473 |
| ZFYVE9 | 162.434378 | -0.0156726 | 0.25257691 | -0.0620509 | 0.95052233 | 0.97993174 |
| CC2D1B | 25.6625455 | -1.0487883 | 0.39296524 | -2.6689085 | 0.00760982 | 0.08575808 |
| PRPF38A | 26.012217 | -0.5267123 | 0.38691139 | -1.3613254 | 0.17341088 | 0.42238719 |
| TUT4 | 300.599298 | -0.1277586 | 0.24411711 | -0.5233496 | 0.600731 | 0.79096587 |
| ZYG11B | 89.4066198 | -0.148469 | 0.24681688 | -0.601535 | 0.54748371 | 0.75546471 |
| ECHDC2 | 219.099153 | 0.81437072 | 0.42500786 | 1.91613097 | 0.05534843 | 0.23510863 |
| SCP2 | 124.514349 | 0.36025022 | 0.21211372 | 1.69838245 | 0.0894356 | 0.29807849 |
| CPT2 | 15.6374669 | 1.10283051 | 0.57732977 | 1.9102263 | 0.05610408 | NA |
| MAGOH | 20.9700687 | -0.4794974 | 0.45009569 | -1.0653231 | 0.28672972 | NA |
| NDC1 | 33.2595432 | -0.6751921 | 0.35149758 | -1.9209012 | 0.05474417 | 0.23396216 |
| YIPF1 | 27.6790789 | -0.1402698 | 0.34341059 | -0.408461 | 0.68293526 | 0.84226129 |

| | | | | | | |
|---|---|---|---|---|---|---|
| HSPB11 | 33.1194213 | 0.38502206 | 0.38056659 | 1.01170746 | 0.31167795 | 0.57088166 |
| LRRC42 | 15.5760827 | 0.15338295 | 0.4653241 | 0.32962607 | 0.74168252 | NA |
| TMEM59 | 111.502826 | 0.40480295 | 0.28449442 | 1.42288537 | 0.15476939 | 0.39755983 |
| TCEANC2 | 23.4569451 | 1.30953928 | 0.55257949 | 2.36986591 | 0.01779454 | 0.13271759 |
| MRPL37 | 13.309099 | 0.31198879 | 0.4732972 | 0.65918158 | 0.50977917 | NA |
| SSBP3 | 360.067094 | -0.4960067 | 0.16752659 | -2.9607642 | 0.00306877 | 0.05464622 |
| ACOT11 | 241.741919 | 0.14289973 | 0.30819282 | 0.4636699 | 6.43E-01 | 8.16E-01 |
| MROH7-TTC4 | 20.6486472 | -0.7884708 | 0.59601463 | -1.3229052 | 0.18586692 | NA |
| DHCR24 | 34.9753101 | -0.7354696 | 0.65018466 | -1.1311704 | 0.25798338 | 0.51619036 |
| BSND | 11.5751771 | 1.7051917 | 0.72518949 | 2.35137397 | 0.01870422 | NA |
| USP24 | 640.252642 | -0.5758075 | 0.26897806 | -2.1407229 | 0.03229639 | 0.17730764 |
| PLPP3 | 288.89634 | -1.8975068 | 0.50250922 | -3.7760637 | 0.00015933 | 0.00936133 |
| PRKAA2 | 272.233442 | -0.662105 | 0.28826352 | -2.2968741 | 0.02162596 | 0.14349647 |
| DAB1 | 216.574928 | 2.15258708 | 0.84260945 | 2.55466762 | 0.01062892 | 0.1014484 |
| TACSTD2 | 47.2706421 | -0.2322842 | 0.56429328 | -0.4116374 | 0.68060524 | 0.84094592 |
| MYSM1 | 134.561282 | -0.0348525 | 0.28463613 | -0.122446 | 0.90254583 | 0.95661107 |
| AL136985.3 | 21.4826754 | -1.581542 | 0.52012366 | -3.0407038 | 0.00236026 | 0.04622624 |
| JUN | 38.9548985 | -0.9848896 | 0.63083682 | -1.5612431 | 0.1184664 | 0.34566454 |
| LINC01135 | 34.3320659 | 1.69265326 | 0.60624336 | 2.79203597 | 0.00523775 | 0.07072805 |
| FGGY | 138.703501 | -0.1881553 | 0.42525346 | -0.4424544 | 0.6581604 | 0.82521413 |
| HOOK1 | 105.161139 | -0.0927851 | 0.28373313 | -0.3270155 | 0.74365619 | 0.87230965 |
| NFIA | 20.6767898 | -0.5149214 | 0.66278369 | -0.7769071 | 0.43721356 | NA |
| TM2D1 | 80.4136527 | -0.2428567 | 0.27588571 | -0.8802801 | 0.37870761 | 0.63024389 |
| PATJ | 1296.62041 | -0.2478912 | 0.15524209 | -1.5968039 | 0.11030944 | 0.33347479 |
| USP1 | 18.4852948 | -0.5758555 | 0.4330189 | -1.3298623 | 0.18356365 | NA |
| DOCK7 | 310.2562 | 0.47864381 | 0.34410578 | 1.39097869 | 0.16423189 | 0.41120418 |
| AC103923.1 | 16.8063484 | 0.3863457 | 0.64402878 | 0.59988888 | 5.49E-01 | NA |
| ATG4C | 55.6090137 | 0.96072756 | 0.36714203 | 2.61677359 | 0.00887652 | 0.09320951 |
| ALG6 | 19.1407255 | 0.24462828 | 0.46240045 | 0.52903989 | 0.59677778 | NA |
| ITGB3BP | 67.1696184 | 1.14221353 | 0.69075222 | 1.65357923 | 0.09821303 | 0.31550323 |
| EFCAB7 | 37.0944637 | 1.10081208 | 0.61973172 | 1.77627198 | 0.0756881 | 0.27489837 |
| PGM1 | 98.5219581 | 0.6341618 | 0.36051525 | 1.75904295 | 0.07857021 | 0.27941662 |
| CACHD1 | 308.261427 | 0.3415194 | 0.26356039 | 1.29579182 | 1.95E-01 | 4.48E-01 |
| RAVER2 | 88.7773546 | 0.66289198 | 0.29588572 | 2.24036491 | 0.02506724 | 0.15502387 |
| JAK1 | 165.106164 | -0.6800328 | 0.22147096 | -3.0705278 | 0.00213681 | 0.04362602 |
| AK4 | 31.7665314 | -0.6611032 | 0.80663327 | -0.8195833 | 0.41245369 | 0.65722052 |
| DNAJC6 | 23.7983621 | -1.2687813 | 0.49273126 | -2.5749964 | 0.01002411 | 0.09908219 |
| LEPR | 99.189432 | 0.03753215 | 0.48236066 | 0.0778093 | 0.93797975 | 0.97466007 |
| SGIP1 | 313.764654 | -0.2409602 | 0.58012617 | -0.4153582 | 0.67787968 | 0.83969138 |
| WDR78 | 60.1718359 | 1.23132568 | 0.51051897 | 2.41190972 | 0.01586921 | 0.12463807 |
| MIER1 | 133.514937 | -0.3764407 | 0.2306609 | -1.6320092 | 0.10267755 | 0.3217871 |
| SLC35D1 | 60.6983186 | 0.41871511 | 0.3407412 | 1.22883616 | 0.21913324 | 0.47565385 |
| SERBP1 | 83.5575657 | -0.3439298 | 0.31064711 | -1.1071399 | 0.26823343 | 0.52717484 |
| GADD45A | 22.2173585 | 0.33152426 | 0.3638448 | 0.91116943 | 3.62E-01 | 0.61802271 |
| GNG12 | 178.288916 | -0.4350622 | 0.24730817 | -1.7591905 | 0.07854516 | 0.27941662 |
| GNG12-AS1 | 94.4474938 | 0.27363789 | 0.43114951 | 0.63467054 | 0.52564331 | 0.74059611 |
| WLS | 203.520435 | 0.59906245 | 0.22169788 | 2.70215685 | 0.00688913 | 0.08095511 |
| LRRC40 | 41.3598275 | 0.28033145 | 0.35264223 | 0.79494578 | 0.42664504 | 0.66963436 |
| SRSF11 | 230.341021 | -0.0833829 | 0.22051697 | -0.3781245 | 0.70533812 | 0.8538829 |
| ANKRD13C | 142.517999 | -0.2311309 | 0.22918065 | -1.0085096 | 0.31320987 | 0.57102302 |
| PTGER3 | 928.528996 | 1.87150185 | 0.38883526 | 4.8130971 | 1.49E-06 | 0.00051083 |
| ZRANB2-AS1 | 16.9300226 | 1.19101905 | 0.58693077 | 2.02923257 | 0.04243461 | NA |
| ZRANB2 | 250.25174 | -0.1246069 | 0.19374056 | -0.6431638 | 0.52011785 | 0.73699321 |
| NEGR1 | 18.3033385 | 0.16577503 | 0.60481283 | 0.27409312 | 0.78401306 | NA |

| | | | | | | |
|---|---|---|---|---|---|---|
| LRRIQ3 | 18.124793 | 3.08977576 | 1.08211301 | 2.85531707 | 0.00429939 | NA |
| CRYZ | 93.051584 | 0.89571739 | 0.4149615 | 2.15855539 | 0.03088468 | 0.17358581 |
| TYW3 | 28.1890896 | 0.1556545 | 0.43228711 | 0.36007203 | 0.71879327 | 0.85995641 |
| ACADM | 69.1529173 | 0.34133146 | 0.27807541 | 1.22747805 | 0.21964297 | 0.47614454 |
| RABGGTB | 24.8226168 | -0.2286424 | 0.49207244 | -0.4646519 | 0.64218078 | 0.81594508 |
| PIGK | 45.7031387 | 0.87685648 | 0.32295123 | 2.71513588 | 0.00662486 | 0.07957541 |
| AC118549.1 | 255.869004 | -0.0302405 | 0.16139715 | -0.1873672 | 0.85137273 | 0.92884705 |
| USP33 | 196.734073 | 0.05195031 | 0.17357948 | 0.29928831 | 0.76472007 | 0.88317118 |
| MIGA1 | 116.612142 | 0.26040398 | 0.2224838 | 1.17044017 | 0.24182388 | 0.49914679 |
| NEXN | 51.4213964 | 0.08396771 | 0.46575414 | 0.18028334 | 0.85693014 | 0.93203598 |
| FUBP1 | 158.634635 | -0.1999524 | 0.2109942 | -0.9476677 | 0.34329866 | 0.60053858 |
| GIPC2 | 27.6179942 | 2.34194885 | 0.61972982 | 3.77898366 | 0.00015747 | 0.00932342 |
| PTGFR | 63.6773293 | 0.15588795 | 0.49349562 | 0.31588518 | 7.52E-01 | 0.87816052 |
| IFI44L | 90.2922598 | 1.39659342 | 2.00145981 | 0.6977874 | 0.48531016 | 0.71218919 |
| IFI44 | 28.3670274 | 0.33673664 | 1.58123113 | 0.21295852 | 0.83135932 | 0.9181376 |
| LINC01781 | 93.4603819 | 1.8893138 | 1.34990009 | 1.39959528 | 0.16163455 | 0.40750118 |
| ADGRL2 | 701.549089 | 0.89687404 | 0.58346061 | 1.53716298 | 0.1242534 | 0.35460505 |
| LINC01362 | 17.3264278 | 2.33529757 | 0.88632753 | 2.63480203 | 0.00841864 | NA |
| LINC01725 | 198.306171 | 1.38060451 | 0.47827731 | 2.88661928 | 0.00389405 | 0.06131052 |
| TTLL7 | 19.005611 | 0.50590087 | 0.45845416 | 1.10349281 | 0.26981321 | NA |
| PRKACB | 124.818769 | 0.93358067 | 0.39345765 | 2.37276026 | 0.01765572 | 0.13219462 |
| RPF1 | 15.8519345 | -0.194201 | 0.44854157 | -0.4329609 | 0.66504316 | NA |
| GNG5 | 37.0690804 | 0.16780819 | 0.37810365 | 0.44381531 | 0.65717612 | 0.82476514 |
| SPATA1 | 21.8100468 | 1.12163445 | 0.64275091 | 1.74505308 | 0.08097563 | 0.28382032 |
| CTBS | 41.30191 | -0.1438071 | 0.35872364 | -0.4008855 | 0.68850441 | 0.84543868 |
| SSX2IP | 16.2095454 | 0.26564708 | 0.47551387 | 0.55865265 | 0.5763988 | NA |
| SYDE2 | 20.1996731 | 0.4765498 | 0.48444554 | 0.9837015 | 0.3252623 | NA |
| C1orf52 | 32.5350852 | 0.26773491 | 0.33763411 | 0.79297352 | 0.42779325 | 0.67061602 |
| BCL10 | 21.1480926 | -0.5452307 | 0.38738771 | -1.4074548 | 0.15929257 | NA |
| AL078459.1 | 54.2524004 | 0.13593669 | 0.43425767 | 0.31303232 | 0.75425611 | 0.87872046 |
| DDAH1 | 321.551971 | 0.50009203 | 0.43354856 | 1.15348563 | 0.24871111 | 0.50492395 |
| CYR61 | 25.011754 | -1.6817254 | 0.92803889 | -1.8121282 | 0.0699664 | 0.26406366 |
| ZNHIT6 | 62.6633087 | -0.173113 | 0.24782438 | -0.698531 | 0.48484515 | 0.7118397 |
| ODF2L | 126.633107 | 0.21028726 | 0.26976293 | 0.77952613 | 0.43566985 | 0.676367 |
| SH3GLB1 | 71.8756243 | -0.2599281 | 0.25161907 | -1.0330221 | 0.30159356 | 0.56180195 |
| SELENOF | 93.9581402 | 0.00698122 | 0.23447017 | 0.02977446 | 0.97624693 | 0.99092346 |
| HS2ST1 | 118.856426 | -0.1203337 | 0.2243747 | -0.5363068 | 0.59174652 | 0.78528845 |
| PKN2-AS1 | 26.5893967 | 1.03672441 | 0.68212491 | 1.5198454 | 0.12854984 | 0.36084904 |
| PKN2 | 319.697363 | -0.4964213 | 0.32040065 | -1.5493767 | 0.12129119 | 0.35057391 |
| GTF2B | 41.0753907 | -0.6079788 | 0.38263864 | -1.5889112 | 0.11208044 | 0.33659117 |
| KYAT3 | 67.2571134 | -0.5711458 | 0.39319761 | -1.4525668 | 0.14634408 | 0.38641866 |
| GBP3 | 18.3873542 | -0.1063336 | 0.45947876 | -0.2314222 | 8.17E-01 | NA |
| GBP2 | 42.0721574 | 0.02616014 | 0.71931835 | 0.03636796 | 9.71E-01 | 9.88E-01 |
| LRRC8B | 81.3951445 | -0.2521502 | 0.29118204 | -0.8659538 | 0.38651552 | 0.6367744 |
| LRRC8C-DT | 11.7339324 | 1.14600808 | 0.84172728 | 1.36149571 | 0.17335709 | NA |
| ZNF326 | 64.9878728 | 0.07240664 | 0.31733175 | 0.22817332 | 0.8195115 | 0.913067 |
| ZNF644 | 288.242643 | -0.6212589 | 0.24705056 | -2.5147034 | 0.01191326 | 0.10687217 |
| HFM1 | 18.490416 | 0.43844032 | 0.69825291 | 0.62791048 | 0.53006259 | NA |
| TGFBR3 | 18.1703106 | -1.2291955 | 0.5809908 | -2.1156884 | 0.03437131 | NA |
| BTBD8 | 27.6064078 | 0.97845479 | 0.62628186 | 1.56232337 | 0.11821183 | 0.34566454 |
| GLMN | 44.3668177 | 0.28182652 | 0.37061694 | 0.76042536 | 0.44700037 | 0.68660184 |
| RPAP2 | 161.893115 | -0.0067281 | 0.19470276 | -0.0345555 | 0.97243419 | 0.98887911 |
| EVI5 | 296.014352 | -0.4106577 | 0.20664693 | -1.987243 | 0.04689547 | 0.21661307 |
| RPL5 | 76.7615627 | 0.38066858 | 0.27612968 | 1.37858624 | 0.16802236 | 0.41617965 |

| | | | | | | |
|---|---|---|---|---|---|---|
| FAM69A | 32.5437548 | -0.1138962 | 0.34488185 | -0.3302469 | 0.74121339 | 0.87152705 |
| MTF2 | 150.004753 | -0.3384587 | 0.2064782 | -1.6391983 | 0.10117198 | 0.32019766 |
| TMED5 | 44.2303444 | 0.01016978 | 0.40169593 | 0.02531711 | 0.97980202 | 0.99256859 |
| CCDC18 | 21.158835 | 0.4663454 | 0.58638931 | 0.79528292 | 0.42644895 | NA |
| DR1 | 22.7234163 | 0.05531524 | 0.39770161 | 0.1390873 | 0.88938116 | 0.94945448 |
| FNBP1L | 373.3459 | 0.07966205 | 0.20951786 | 0.38021605 | 0.70378505 | 0.85295217 |
| BCAR3 | 177.721566 | -0.4224746 | 0.44970211 | -0.9394544 | 0.34749751 | 0.60403983 |
| GCLM | 46.9301781 | -0.3147364 | 0.38121218 | -0.82562 | 4.09E-01 | 0.65472852 |
| ARHGAP29 | 424.11885 | -0.6358231 | 0.65175936 | -0.9755489 | 0.32928806 | 0.58534184 |
| ABCD3 | 166.616761 | 0.47005995 | 0.23361322 | 2.01212907 | 0.04420633 | 0.21042433 |
| SLC44A3 | 150.245493 | 1.07047278 | 0.43485835 | 2.46165854 | 0.01382963 | 0.11532965 |
| CNN3 | 154.587482 | -0.2908992 | 0.24967944 | -1.1650907 | 0.24398228 | 0.50145036 |
| ALG14 | 39.4431572 | 0.04337663 | 0.32495922 | 0.13348332 | 0.89381116 | 0.95193918 |
| PTBP2 | 229.933226 | -0.166836 | 0.20097677 | -0.8301259 | 0.40646758 | 0.65333469 |
| DPYD | 363.540689 | -0.2705944 | 0.37739071 | -0.717014 | 0.47336548 | 0.70242802 |
| SNX7 | 38.6858327 | 0.0698646 | 0.35939085 | 0.19439727 | 0.84586483 | 0.92531372 |
| PALMD | 12.109348 | 0.40458887 | 0.79976011 | 0.50588778 | 6.13E-01 | NA |
| FRRS1 | 16.6030406 | -0.7311015 | 0.5012053 | -1.4586867 | 0.14465136 | NA |
| AGL | 98.0018243 | 0.15389837 | 0.23261161 | 0.66161086 | 0.50822064 | 0.72894696 |
| TRMT13 | 66.4353835 | 0.35931557 | 0.30194465 | 1.19000475 | 0.23404453 | 0.49243138 |
| LRRC39 | 12.2469573 | 0.31655579 | 0.61393785 | 0.51561536 | 0.60612308 | NA |
| DBT | 75.7205573 | 0.56842887 | 0.26504683 | 2.14463563 | 0.03198199 | 0.17658923 |
| RTCA | 44.173198 | 0.40889394 | 0.33079863 | 1.23608113 | 0.21642841 | 0.47271551 |
| CDC14A | 68.780213 | 0.1053758 | 0.26348715 | 0.39992767 | 0.68920979 | 0.84569856 |
| EXTL2 | 29.3906204 | 0.00544118 | 0.3752501 | 0.01450015 | 0.98843096 | 0.99581595 |
| SLC30A7 | 101.241713 | -0.5428784 | 0.23483193 | -2.3117741 | 0.02079013 | 0.14123712 |
| DPH5 | 24.9515473 | 0.00407351 | 0.36974619 | 0.01101703 | 0.99120986 | 0.99583841 |
| AC093157.1 | 33.1666297 | 0.42725109 | 0.34754471 | 1.22934134 | 0.21894385 | 0.47565385 |
| RNPC3 | 428.153059 | -0.3618995 | 0.45911208 | -0.7882597 | 4.31E-01 | 6.72E-01 |
| AMY2B | 140.0477 | -0.1988725 | 0.58327458 | -0.3409587 | 7.33E-01 | 0.86776236 |
| NTNG1 | 59.8087709 | 0.86530682 | 0.95230372 | 0.90864585 | 3.64E-01 | 6.20E-01 |
| VAV3 | 652.343686 | 0.53316569 | 0.48768076 | 1.09326784 | 0.27427622 | 0.53337647 |
| SLC25A24 | 51.5538611 | -0.4170467 | 0.31127529 | -1.3398 | 0.18031036 | 0.42954159 |
| PRPF38B | 80.8430994 | -0.7248909 | 0.37049232 | -1.9565613 | 0.05039906 | 0.22527387 |
| STXBP3 | 77.729593 | -0.2282699 | 0.26493767 | -0.8615986 | 0.38890845 | 0.63798557 |
| AL449266.1 | 18.8567857 | 0.11047517 | 0.41659306 | 0.26518724 | 0.79086522 | NA |
| GPSM2 | 17.0846476 | -0.2332849 | 0.62148837 | -0.3753649 | 0.70738907 | NA |
| WDR47 | 23.2882667 | -0.379762 | 0.40132899 | -0.946261 | 0.3440155 | 0.60110949 |
| TAF13 | 14.7304028 | -0.9793983 | 0.56570042 | -1.731302 | 0.08339791 | NA |
| TMEM167B | 13.7965773 | 0.31433814 | 0.4387725 | 0.71640347 | 0.47374225 | NA |
| SARS | 40.4238405 | 0.008011 | 0.46130234 | 0.01736606 | 0.98614459 | 0.9942828 |
| SORT1 | 79.9963937 | 0.26921446 | 0.34536406 | 0.7795092 | 0.43567982 | 0.676367 |
| PSMA5 | 51.2482255 | -0.9729241 | 0.40620563 | -2.3951517 | 0.01661349 | 0.12787404 |
| GNAI3 | 79.6640313 | -0.4307908 | 0.2339218 | -1.8416019 | 0.06553341 | 0.25452134 |
| AMPD2 | 17.5919861 | -0.4564461 | 0.48300452 | -0.9450142 | 0.34465165 | NA |
| GSTM4 | 14.3132533 | 1.93367261 | 0.61373289 | 3.15067459 | 0.00162894 | NA |
| GSTM3 | 60.720757 | -0.1202664 | 0.62804101 | -0.1914946 | 0.84813814 | 0.92660264 |
| AHCYL1 | 199.55695 | -0.2695229 | 0.35577945 | -0.757556 | 0.44871681 | 0.68731807 |
| STRIP1 | 26.3241896 | 0.32505033 | 0.44855052 | 0.72466827 | 0.46865557 | 0.69853638 |
| SLC6A17 | 21.2253452 | -1.0256774 | 0.68105824 | -1.5060054 | 0.1320658 | 0.36551976 |
| KCNC4 | 69.0737876 | 0.89177323 | 0.51910996 | 1.71788889 | 0.08581688 | 0.29278925 |
| RBM15 | 29.4884844 | -0.1337378 | 0.40535902 | -0.3299244 | 0.74145705 | 0.87152705 |
| SLC16A4 | 32.5640898 | 0.98841441 | 0.48515868 | 2.03730129 | 0.04161986 | 0.20335702 |
| AL355488.1 | 14.0606863 | 0.79402949 | 0.495183 | 1.60350718 | 0.10882273 | NA |

| | | | | | | |
|---|---|---|---|---|---|---|
| LAMTOR5 | 41.9672692 | 0.6565446 | 0.33734708 | 1.94619914 | 0.05163082 | 0.22799911 |
| LRIF1 | 17.8784358 | -0.3663826 | 0.49488499 | -0.7403389 | 0.45909439 | NA |
| DRAM2 | 48.7829558 | 0.47472828 | 0.32589503 | 1.45669076 | 0.14520176 | 0.38485467 |
| CEPT1 | 66.527549 | -0.0583964 | 0.27470545 | -0.2125783 | 0.83165589 | 0.9181376 |
| DENND2D | 26.9771693 | -0.5184253 | 0.4458907 | -1.1626736 | 0.24496196 | 0.5022131 |
| ATP5PB | 63.1153393 | 0.12854938 | 0.41404216 | 0.31047413 | 0.75620043 | 0.87904624 |
| RAP1A | 145.695764 | -0.33511 | 0.26184175 | -1.2798188 | 0.20060888 | 0.45454417 |
| KCND3 | 21.923194 | 0.00396586 | 0.70343513 | 0.00563784 | 0.99550168 | 0.99796515 |
| CTTNBP2NL | 63.9797164 | 0.1922853 | 0.33705514 | 0.57048619 | 0.56834799 | 0.77103518 |
| WNT2B | 17.959338 | 1.16551063 | 0.58993073 | 1.97567369 | 0.04819175 | NA |
| ST7L | 76.2000197 | -0.1600981 | 0.27406627 | -0.5841583 | 0.55911378 | 0.76485713 |
| CAPZA1 | 127.219054 | -0.6121236 | 0.28416573 | -2.1541079 | 0.03123171 | 0.17457551 |
| MOV10 | 31.6087368 | 0.18379809 | 0.38403327 | 0.4785994 | 0.63222365 | 0.80928412 |
| RHOC | 32.9295933 | -0.7878932 | 0.51507354 | -1.5296713 | 0.12609812 | 0.35730275 |
| LRIG2 | 60.670102 | -0.0565197 | 0.28471125 | -0.1985157 | 0.84264158 | 0.92390487 |
| MAGI3 | 286.883152 | 0.46545472 | 0.22502498 | 2.06845799 | 0.03859698 | 0.1958098 |
| PHTF1 | 64.736194 | 0.21216851 | 0.269134 | 0.78833782 | 0.43049913 | 0.67197671 |
| RSBN1 | 63.2940421 | 0.23414959 | 0.30666694 | 0.76353059 | 0.44514702 | 0.68514266 |
| HIPK1 | 45.0647719 | 0.3898833 | 0.38670273 | 1.00822483 | 0.31334654 | 0.57102302 |
| TRIM33 | 222.361339 | -1.0060371 | 0.426181 | -2.3605864 | 0.01824607 | 0.13368407 |
| BCAS2 | 20.3209238 | -0.218904 | 0.60337174 | -0.3628012 | 0.71675341 | NA |
| CSDE1 | 130.112775 | -0.2033741 | 0.24472133 | -0.8310435 | 0.40594906 | 0.65318894 |
| SIKE1 | 25.3831678 | 0.10237057 | 0.37135567 | 0.27566718 | 0.78280371 | 0.89289273 |
| SLC22A15 | 149.960278 | 0.37761032 | 0.26747395 | 1.41176484 | 0.15801921 | 0.40275854 |
| ATP1A1 | 1298.34454 | 0.56999266 | 0.31572547 | 1.80534267 | 0.07102108 | 0.26639826 |
| LINC01762 | 852.294932 | 1.52491555 | 0.47191662 | 3.23132407 | 0.00123218 | 0.0311977 |
| CD58 | 79.4141141 | 0.40297423 | 0.22348772 | 1.80311578 | 7.14E-02 | 0.26675397 |
| IGSF3 | 144.707155 | 0.43085342 | 0.25755147 | 1.67288275 | 0.09435038 | 0.30830642 |
| PTGFRN | 44.1686804 | 0.99097805 | 0.38467565 | 2.57613928 | 0.00999104 | 0.09901212 |
| TTF2 | 32.0701747 | 0.56616425 | 0.41904658 | 1.35107712 | 0.17667073 | 0.42569745 |
| VTCN1 | 33.9698522 | 2.62701762 | 0.91009481 | 2.88653181 | 3.90E-03 | 0.06131052 |
| MAN1A2 | 199.295018 | 0.01991136 | 0.2041016 | 0.09755614 | 0.92228476 | 0.96656839 |
| GDAP2 | 73.362802 | -0.0094523 | 0.23837347 | -0.0396533 | 0.96836952 | 0.98741848 |
| WDR3 | 12.0956125 | 0.7024815 | 0.59151054 | 1.18760606 | 0.23498865 | NA |
| SPAG17 | 68.7176676 | 0.51094844 | 0.57875146 | 0.88284604 | 0.37731946 | 0.62875685 |
| WARS2 | 44.2990373 | -0.4138259 | 0.32713628 | -1.2649956 | 0.20587292 | 0.45902815 |
| AL359915.2 | 131.716554 | 0.20036913 | 0.44139735 | 0.45394276 | 0.64987003 | 0.81947559 |
| ZNF697 | 27.072743 | -0.5823129 | 0.48696377 | -1.1958034 | 0.2317733 | 0.49050291 |
| PHGDH | 22.1324514 | 0.84340436 | 0.64136616 | 1.31501225 | 0.18850578 | 0.43914317 |
| NOTCH2 | 52.1389875 | 0.42829783 | 0.29669563 | 1.44355962 | 0.14886289 | 0.38937375 |
| SEC22B | 68.1322032 | -0.1398901 | 0.27056377 | -0.5170319 | 0.60513386 | 0.79401898 |
| NBPF26 | 18.6395227 | -0.6247152 | 0.60557272 | -1.0316106 | 0.30225458 | NA |
| LINC00623 | 123.547925 | 1.20652487 | 0.50037637 | 2.41123468 | 0.01589861 | 0.1247389 |
| SRGAP2C | 22.2849223 | 0.0419452 | 0.42054949 | 0.09973904 | 0.9205515 | 0.96558802 |
| NBPF15 | 21.2478899 | 0.24640273 | 0.43990184 | 0.56013115 | 0.57538998 | 0.77388388 |
| FP700111.1 | 24.6775805 | -0.135572 | 0.42132668 | -0.321774 | 0.74762394 | 0.87507018 |
| SRGAP2B | 27.5598843 | -0.1512924 | 0.40234421 | -0.3760272 | 0.70689664 | 0.8547469 |
| GPR89A | 19.7206397 | 0.41314699 | 0.46370256 | 0.89097414 | 0.37294304 | NA |
| PDZK1 | 22.0099438 | 0.75033141 | 0.55795673 | 1.34478424 | 0.17869492 | 0.42767873 |
| RNF115 | 217.123942 | -0.1915589 | 0.18881841 | -1.0145139 | 0.3103376 | 0.56995192 |
| POLR3C | 26.2335028 | -0.5142084 | 0.35341606 | -1.4549662 | 0.14567863 | 0.38558748 |
| PEX11B | 12.9413919 | 1.12072513 | 0.51082211 | 2.19396364 | 0.02823802 | NA |
| RBM8A | 20.0036257 | -0.0965357 | 0.46251761 | -0.208718 | 0.83466841 | NA |
| TXNIP | 453.287708 | 1.57576903 | 0.37393381 | 4.21403197 | 2.51E-05 | 0.00292522 |

| | | | | | | |
|---|---|---|---|---|---|---|
| NBPF12 | 50.0263996 | -0.1477155 | 0.3232846 | -0.456921 | 0.64772783 | 0.81885075 |
| PRKAB2 | 23.1845511 | 0.96881745 | 0.51853657 | 1.8683686 | 0.06171071 | 0.24842434 |
| AC242426.2 | 43.457836 | 0.79230038 | 0.49023924 | 1.61615047 | 0.10606178 | 0.32732861 |
| FMO5 | 23.2031836 | 1.26541039 | 0.7306739 | 1.73184013 | 0.08330203 | 0.28855622 |
| CHD1L | 54.7777393 | 0.18335418 | 0.37724306 | 0.48603724 | 0.62694076 | 0.80546871 |
| BCL9 | 87.2898982 | 0.69025622 | 0.35051253 | 1.96927689 | 0.0489213 | 0.22162876 |
| ACP6 | 66.6061607 | 0.7619214 | 0.33279945 | 2.28943104 | 0.02205432 | 0.14508727 |
| GPR89B | 25.2257615 | -0.2319231 | 0.40542746 | -0.5720458 | 0.56729093 | 0.77077463 |
| NBPF11 | 39.5068263 | -0.3392692 | 0.34606661 | -0.9803582 | 0.32690935 | 0.58313355 |
| NBPF14 | 17.1377677 | -0.4596621 | 0.44130347 | -1.0416009 | 0.29759676 | NA |
| PDE4DIP | 215.280355 | 0.37780275 | 0.2655458 | 1.42274045 | 0.15481141 | 0.39755983 |
| NBPF9 | 18.2100706 | -0.2297128 | 0.42114751 | -0.545445 | 0.58544747 | NA |
| AC245297.3 | 334.496742 | 0.27383166 | 0.27566064 | 0.9933651 | 0.32053206 | 0.57737779 |
| NBPF19 | 27.1468337 | -0.5163335 | 0.90404777 | -0.5711352 | 0.56790798 | 0.77103518 |
| OTUD7B | 142.623103 | -0.8266379 | 0.25253783 | -3.2733231 | 0.00106291 | 0.02953506 |
| VPS45 | 121.24982 | 0.40578176 | 0.28248866 | 1.43645328 | 0.1508734 | 0.39152818 |
| ANP32E | 25.404478 | 0.54615133 | 0.41442456 | 1.31785464 | 0.18755231 | 0.43816783 |
| APH1A | 25.071575 | 0.14075244 | 0.51732886 | 0.27207536 | 0.78556407 | 0.89452315 |
| C1orf54 | 32.2317152 | 0.32800423 | 0.59428956 | 0.55192663 | 0.58099862 | 0.77786509 |
| MRPS21 | 54.9645868 | 0.03114945 | 0.25155342 | 0.12382836 | 0.90145117 | 0.95637074 |
| PRPF3 | 68.5309829 | -0.8288469 | 0.29529229 | -2.8068695 | 0.00500255 | 0.06866339 |
| RPRD2 | 238.138585 | -0.2624963 | 0.1578882 | -1.6625454 | 0.09640347 | 0.31164112 |
| TARS2 | 14.6724253 | -0.3316781 | 0.48917398 | -0.6780371 | 0.49774817 | NA |
| MCL1 | 51.927109 | -0.4847833 | 0.29458744 | -1.6456346 | 9.98E-02 | 3.18E-01 |
| ENSA | 187.46707 | -0.9637623 | 0.41537981 | -2.3201953 | 0.02033032 | 0.14033487 |
| GOLPH3L | 66.315524 | 0.14096407 | 0.27900299 | 0.50524215 | 0.61338877 | 0.79899363 |
| ARNT | 84.1505647 | -0.2234408 | 0.28977323 | -0.7710884 | 0.44065452 | 0.68065781 |
| SETDB1 | 63.3532821 | -0.1477937 | 0.26496657 | -0.5577823 | 0.57699308 | 0.77465825 |
| CERS2 | 29.2867935 | -0.1186148 | 0.41763275 | -0.2840169 | 0.77639741 | 0.88914312 |
| MINDY1 | 19.1603228 | 1.30233844 | 0.54440835 | 2.39220877 | 0.01674731 | NA |
| PRUNE1 | 25.5270863 | -0.1285573 | 0.44789226 | -0.2870272 | 0.77409147 | 0.88782329 |
| CDC42SE1 | 40.7873743 | -0.0853276 | 0.40780787 | -0.2092347 | 0.83426499 | 0.91992307 |
| GABPB2 | 75.0366046 | 0.30547464 | 0.45919654 | 0.66523725 | 0.50589876 | 0.72769627 |
| SCNM1 | 13.543945 | 0.99163552 | 0.5469419 | 1.81305458 | 0.0698234 | NA |
| VPS72 | 14.0222728 | -0.6329031 | 0.47150842 | -1.3422943 | 0.17950059 | NA |
| PIP5K1A | 251.552121 | -0.2982777 | 0.19038608 | -1.5666992 | 0.11718503 | 0.34465921 |
| PSMD4 | 41.3843764 | -0.1937622 | 0.29923372 | -0.6475281 | 0.51729023 | 0.73469647 |
| PI4KB | 51.3906621 | 0.7210776 | 0.3166063 | 2.27752137 | 0.02275511 | 0.14700765 |
| PSMB4 | 18.4841159 | 0.19570628 | 0.50629487 | 0.38654604 | 0.69909231 | NA |
| POGZ | 323.66991 | 0.03184173 | 0.21138657 | 0.15063269 | 0.88026547 | 0.94376451 |
| CGN | 52.9338179 | -0.9056789 | 0.35139843 | -2.5773562 | 0.00995593 | 0.0990062 |
| TUFT1 | 43.6945759 | -0.2889836 | 0.35216931 | -0.8205814 | 0.41188474 | 0.65707964 |
| SNX27 | 78.5807829 | 0.52165468 | 0.25097674 | 2.07849807 | 0.03766351 | 0.19425568 |
| RORC | 37.4963299 | 0.25822599 | 0.5040352 | 0.51231738 | 0.60842891 | 0.79566688 |
| THEM4 | 108.328526 | 1.08789403 | 0.57447862 | 1.89370674 | 0.05826395 | 0.24020328 |
| S100A10 | 103.16598 | -0.0190092 | 0.47233668 | -0.040245 | 0.9678978 | 0.98741848 |
| S100A11 | 211.570238 | -0.3979043 | 0.46860762 | -0.8491205 | 3.96E-01 | 6.44E-01 |
| FLG-AS1 | 83.8065696 | 0.8582819 | 0.42573187 | 2.01601518 | 0.04379839 | 0.20912915 |
| S100A6 | 302.943367 | -0.1273548 | 0.71052248 | -0.1792411 | 0.8577484 | 0.93263023 |
| S100A2 | 54.0410976 | -1.8608579 | 1.36960991 | -1.3586773 | 0.17424888 | 0.42295604 |
| S100A16 | 26.9682245 | -0.6936037 | 0.45300237 | -1.5311259 | 1.26E-01 | 0.35686116 |
| S100A13 | 33.1134929 | 0.10816131 | 0.44565818 | 0.24270016 | 0.80823768 | 0.90713122 |
| CHTOP | 42.7689189 | -0.3872125 | 0.3546425 | -1.0918389 | 0.27490392 | 0.5337282 |
| ILF2 | 23.179734 | -0.0106392 | 0.3721893 | -0.0285854 | 0.97719523 | 0.99136308 |

| | | | | | | |
|---|---|---|---|---|---|---|
| INTS3 | 56.9557886 | 0.25694865 | 0.30517422 | 0.84197365 | 0.39980271 | 0.64766685 |
| GATAD2B | 300.297691 | -0.525325 | 0.32536391 | -1.6145767 | 0.1064024 | 0.32732986 |
| DENND4B | 30.6064564 | -0.635184 | 0.41229635 | -1.5406007 | 0.12341401 | 0.35349809 |
| CRTC2 | 16.6943596 | -0.7299409 | 0.53256662 | -1.3706096 | 0.1704967 | NA |
| SLC39A1 | 28.8007929 | -0.4162079 | 0.60094209 | -0.6925924 | 0.48856538 | 0.71532961 |
| JTB | 31.4033548 | -0.0381537 | 0.43888136 | -0.0869339 | 0.93072408 | 0.97162394 |
| RAB13 | 28.256187 | 0.26350213 | 0.342739 | 0.76881279 | 4.42E-01 | 0.68190353 |
| RPS27 | 27.0399203 | 0.41344041 | 0.49915414 | 0.82828205 | 0.40751079 | 0.65373292 |
| TPM3 | 55.1357501 | -1.0373673 | 0.31444704 | -3.2990207 | 0.00097023 | 0.02829008 |
| C1orf43 | 65.3186915 | -0.2948201 | 0.3141416 | -0.9384943 | 0.34799044 | 0.60448713 |
| UBAP2L | 82.2435295 | -0.7529683 | 0.34691792 | -2.1704508 | 0.02997271 | 0.17114241 |
| HAX1 | 17.8572458 | 0.65565639 | 0.48392479 | 1.35487252 | 0.17545818 | NA |
| ATP8B2 | 14.3601226 | -0.3481962 | 0.5554825 | -0.6268355 | 0.53076706 | NA |
| IL6R | 26.574408 | -1.9907367 | 0.6886325 | -2.890855 | 0.00384195 | 0.06110382 |
| UBE2Q1 | 27.415611 | -1.3995769 | 0.55251079 | -2.5331214 | 0.01130518 | 0.1036885 |
| ADAR | 104.832623 | -0.5623436 | 0.30437954 | -1.8475077 | 0.06467359 | 0.25284322 |
| KCNN3 | 81.9430389 | 0.81176478 | 0.54825988 | 1.48062043 | 1.39E-01 | 0.37517394 |
| PMVK | 19.7739269 | 0.80393575 | 0.43523948 | 1.84711128 | 0.06473101 | NA |
| PBXIP1 | 28.9249775 | 1.35001922 | 0.50789846 | 2.65804946 | 0.00785944 | 0.08691678 |
| ZBTB7B | 12.0701818 | 0.67005312 | 0.56675988 | 1.1822522 | 0.23710565 | NA |
| EFNA1 | 41.0834853 | 0.02877076 | 0.38626329 | 0.07448485 | 9.41E-01 | 9.75E-01 |
| SLC50A1 | 15.4102118 | -0.6252579 | 0.43315552 | -1.4434952 | 0.14888104 | NA |
| DPM3 | 15.464502 | 0.41046305 | 0.52419172 | 0.78303995 | 0.43360365 | NA |
| SCAMP3 | 26.0687252 | -0.2412248 | 0.54052466 | -0.4462789 | 0.65539578 | 0.8237396 |
| CLK2 | 26.4155754 | 0.27395771 | 0.37361357 | 0.73326488 | 0.46339689 | 0.69473429 |
| FDPS | 60.1462172 | 0.39816238 | 0.3281649 | 1.21329972 | 0.22501525 | 0.48216659 |
| RUSC1 | 13.3206307 | 0.88892271 | 0.52964267 | 1.67834421 | 0.09327992 | NA |
| ASH1L | 537.183166 | -0.3091528 | 0.20025123 | -1.5438246 | 0.12263084 | 0.35251484 |
| YY1AP1 | 89.2003682 | -0.6028641 | 0.23802594 | -2.5327665 | 0.01131663 | 0.1036885 |
| DAP3 | 95.3282581 | -0.1157907 | 0.24713264 | -0.4685366 | 0.63940092 | 0.81400411 |
| GON4L | 116.58701 | -0.7070102 | 0.2364441 | -2.9901792 | 0.00278814 | 0.0514788 |
| RIT1 | 36.9876677 | -1.0886497 | 0.38022518 | -2.863171 | 0.00419424 | 0.06301187 |
| KHDC4 | 103.981975 | -0.6405284 | 0.23701091 | -2.7025272 | 0.00688145 | 0.08095511 |
| ARHGEF2 | 31.411436 | -0.7436365 | 0.43003841 | -1.7292328 | 0.08376745 | 0.28873024 |
| SSR2 | 32.5615431 | 0.22537769 | 0.4486505 | 0.5023458 | 0.6154243 | 0.80075958 |
| UBQLN4 | 15.250341 | 0.16052685 | 0.45078654 | 0.35610391 | 0.72176275 | NA |
| RAB25 | 14.0612749 | 0.88566323 | 0.60688226 | 1.45936584 | 0.14446444 | NA |
| LMNA | 84.8846923 | -0.2948467 | 0.39978491 | -0.7375134 | 0.46081018 | 0.69410104 |
| SLC25A44 | 17.8695893 | -0.3571125 | 0.42712133 | -0.8360915 | 0.40310341 | NA |
| SMG5 | 53.1591246 | -0.4105993 | 0.25576874 | -1.6053539 | 0.10841595 | 0.33024913 |
| CCT3 | 48.3883758 | 1.09571969 | 0.33286366 | 3.29179725 | 0.00099549 | 0.02848444 |
| RHBG | 58.0153479 | 0.95714061 | 0.80828487 | 1.18416247 | 0.23634875 | 0.49420711 |
| MEF2D | 43.2791159 | -1.0226953 | 0.49325273 | -2.0733698 | 0.03813788 | 0.19510866 |
| ISG20L2 | 12.8915062 | -0.891185 | 0.5339159 | -1.6691486 | 0.09508793 | NA |
| RRNAD1 | 18.5076641 | -0.1622932 | 0.42088824 | -0.3855968 | 0.69979527 | NA |
| HDGF | 42.5417415 | -0.2425741 | 0.29686759 | -0.8171121 | 0.41386437 | 0.65845053 |
| PRCC | 47.4488261 | -0.5045816 | 0.32846924 | -1.5361608 | 0.12449894 | 0.35478281 |
| ARHGEF11 | 107.504895 | 0.36487897 | 0.25316419 | 1.44127401 | 0.14950729 | 0.38995513 |
| KIRREL1 | 72.1151215 | -0.7216532 | 0.38101459 | -1.8940302 | 0.058221 | 0.24020328 |
| IFI16 | 30.7146053 | -0.7763469 | 0.69883009 | -1.1109237 | 0.26660119 | 0.52548766 |
| DUSP23 | 39.6714806 | -0.316425 | 0.49632935 | -0.6375304 | 0.52377943 | 0.73909151 |
| TAGLN2 | 39.0719832 | -0.2101816 | 0.53236418 | -0.394808 | 6.93E-01 | 0.84786241 |
| KCNJ10 | 48.3117258 | 2.51762881 | 0.64396881 | 3.90955082 | 9.25E-05 | 0.00652959 |
| IGSF8 | 14.6139774 | 0.17476315 | 0.6102897 | 0.28636097 | 0.77460166 | NA |

| | | | | | | |
|---|---|---|---|---|---|---|
| DCAF8 | 29.8195568 | -0.0374824 | 0.42618344 | -0.087949 | 0.9299172 | 0.97116166 |
| COPA | 125.442676 | -0.6055564 | 0.20605372 | -2.9388278 | 0.00329456 | 0.0569848 |
| NCSTN | 21.2387333 | 0.6739878 | 0.5312907 | 1.26858574 | 0.20458885 | 0.45773044 |
| F11R | 37.289245 | -0.5825659 | 0.58271745 | -0.99974 | 0.31743636 | 0.57495296 |
| AL591806.3 | 21.3078062 | 0.78824539 | 0.4627786 | 1.70328833 | 0.08851412 | 0.29750795 |
| PFDN2 | 39.7869036 | -0.650163 | 0.31979853 | -2.0330395 | 0.04204854 | 0.20441466 |
| NIT1 | 30.5006825 | 0.01369194 | 0.34633079 | 0.03953428 | 0.96846442 | 0.98741848 |
| DEDD | 15.8552199 | -0.4062469 | 0.43681416 | -0.9300222 | 0.35235959 | NA |
| UFC1 | 18.943181 | 0.2046514 | 0.46947312 | 0.43591718 | 0.66289682 | NA |
| NDUFS2 | 47.9833627 | -0.1155465 | 0.32481129 | -0.3557343 | 0.72203957 | 0.86153309 |
| SDHC | 79.2540425 | 0.41704192 | 0.24620064 | 1.69391078 | 0.09028224 | 0.30003934 |
| AL592295.4 | 17.3679961 | 0.54479371 | 0.58217611 | 0.93578851 | 0.34938209 | NA |
| DUSP12 | 15.867992 | -0.3353474 | 0.44841522 | -0.7478502 | 0.45455055 | NA |
| ATF6 | 176.291314 | -0.5163357 | 0.37999898 | -1.3587818 | 0.17421576 | 0.42295604 |
| NOS1AP | 823.521198 | -1.0113161 | 0.32105984 | -3.1499302 | 0.00163309 | 0.03697039 |
| C1orf226 | 15.4540475 | -0.3250812 | 0.50657037 | -0.6417296 | 0.52104875 | NA |
| UHMK1 | 70.3386403 | 0.0956269 | 0.26169924 | 0.36540762 | 0.71480715 | 0.85843895 |
| UAP1 | 35.4632392 | -0.7538991 | 0.47978734 | -1.5713194 | 0.11610849 | 0.34306603 |
| HSD17B7 | 38.4108443 | -1.169657 | 0.32913074 | -3.5537763 | 0.00037974 | 0.01697016 |
| PBX1 | 228.232244 | -0.0663689 | 0.40239678 | -0.164934 | 0.86899595 | 0.93875956 |
| MGST3 | 160.230978 | 0.44828037 | 0.28577951 | 1.56862323 | 0.11673576 | 0.34414008 |
| ALDH9A1 | 21.6332824 | 0.6302596 | 0.47552929 | 1.32538544 | 0.18504334 | 0.43476148 |
| TMCO1 | 45.4585565 | 0.47815435 | 0.30730979 | 1.55593599 | 0.11972331 | 0.3480474 |
| UCK2 | 31.1145556 | 0.01155749 | 0.38363642 | 0.03012616 | 0.97596644 | 0.99088558 |
| FAM78B | 11.9073003 | 0.98312156 | 0.56587449 | 1.73734913 | 0.08232556 | NA |
| POGK | 34.0423388 | -0.1111277 | 0.36792796 | -0.3020366 | 0.76262415 | 0.88247846 |
| TADA1 | 17.7015834 | 0.4368418 | 0.45451404 | 0.96111839 | 0.33649265 | NA |
| POU2F1 | 229.188417 | 0.31831494 | 0.28874843 | 1.10239538 | 0.27028982 | 0.52870255 |
| CREG1 | 22.8344125 | 0.75974168 | 0.4078243 | 1.8629142 | 0.06247434 | 0.24933194 |
| MPZL1 | 69.3682253 | -0.2995996 | 0.25859223 | -1.1585792 | 0.24662777 | 0.50296216 |
| MPC2 | 42.3497932 | 1.0821211 | 0.35092327 | 3.0836402 | 0.00204485 | 0.04242369 |
| DCAF6 | 294.800038 | 0.10406969 | 0.30099126 | 0.34575653 | 0.72952569 | 0.86533507 |
| TIPRL | 27.1442783 | 0.09859847 | 0.4052512 | 0.24330211 | 0.80777137 | 0.90698997 |
| SFT2D2 | 40.1211627 | -0.2463681 | 0.31282687 | -0.787554 | 0.43095763 | 0.67197671 |
| LINC00970 | 257.120814 | 1.78540475 | 0.57014329 | 3.13150181 | 0.00173915 | 0.03863922 |
| ATP1B1 | 1530.61446 | 0.97306581 | 0.51834725 | 1.87724696 | 0.06048425 | 0.24514087 |
| NME7 | 194.279901 | 0.16077468 | 0.20603643 | 0.78032161 | 0.4352016 | 0.67618228 |
| BLZF1 | 54.10012 | -0.4298712 | 0.33773745 | -1.272797 | 0.20309008 | 0.45693784 |
| SLC19A2 | 89.0544055 | -0.6898441 | 0.52381348 | -1.3169652 | 0.18785031 | 0.43841231 |
| SCYL3 | 38.357797 | 0.5312297 | 0.39674103 | 1.33898351 | 0.18057604 | 0.42977544 |
| KIFAP3 | 90.0482271 | 0.5187694 | 0.27975308 | 1.85438316 | 0.06368436 | 0.25118809 |
| GORAB | 15.0356812 | -0.1138918 | 0.60789465 | -0.1873544 | 0.85138274 | NA |
| FMO4 | 57.0420853 | 0.81078568 | 0.50207716 | 1.6148627 | 0.10634043 | 0.32732986 |
| PRRC2C | 288.245146 | -0.7587129 | 0.28027058 | -2.7070728 | 0.00678794 | 0.08022018 |
| VAMP4 | 25.1481497 | 0.45181798 | 0.37715088 | 1.19797674 | 0.23092607 | 0.48951748 |
| DNM3 | 264.489926 | -0.028883 | 0.49490413 | -0.0583608 | 0.95346128 | 0.98153155 |
| PIGC | 34.161491 | 0.68289088 | 0.41722809 | 1.63673275 | 0.10168632 | 0.32089434 |
| SUCO | 95.0998568 | -1.1826941 | 0.29408461 | -4.0216116 | 5.78E-05 | 0.00496891 |
| AL645568.1 | 68.6679647 | 1.43053682 | 0.38364335 | 3.7288196 | 0.00019238 | 0.01080832 |
| PRDX6 | 82.0624341 | 0.91280017 | 0.35627546 | 2.56206299 | 0.01040524 | 0.10033015 |
| KLHL20 | 77.9941195 | -0.4462662 | 0.29480891 | -1.5137473 | 0.13008994 | 0.36252797 |
| DARS2 | 16.7791371 | 0.68103145 | 0.530908 | 1.28276734 | 0.1995736 | NA |
| ZBTB37 | 41.3754015 | 0.48286107 | 0.31206644 | 1.54730218 | 0.1217904 | 0.35109389 |
| RC3H1 | 160.342884 | -0.3257675 | 0.23252958 | -1.4009723 | 0.16122236 | 0.40699525 |

| | | | | | | |
|---|---|---|---|---|---|---|
| RABGAP1L | 781.166627 | 0.30478728 | 0.1721512 | 1.77046272 | 0.07665009 | 0.27659435 |
| CACYBP | 13.4511336 | 1.16498369 | 0.61316302 | 1.89995752 | 0.05743869 | NA |
| MRPS14 | 14.548537 | 1.09576061 | 0.48016386 | 2.28205556 | 0.02248606 | NA |
| KIAA0040 | 23.2601056 | -0.1911633 | 0.43451346 | -0.4399479 | 0.65997488 | 0.82676279 |
| COP1 | 320.691801 | -0.2681458 | 0.21379949 | -1.2541928 | 0.20977192 | 0.46490483 |
| PAPPA2 | 396.318428 | 1.05410946 | 0.54931143 | 1.91896509 | 5.50E-02 | 2.34E-01 |
| RASAL2 | 326.6233 | -1.5613249 | 0.59869869 | -2.6078643 | 0.00911091 | 0.09427267 |
| RALGPS2 | 217.410139 | 0.45505864 | 0.32062415 | 1.41928995 | 0.1558145 | 0.39897013 |
| FAM20B | 43.2010865 | 0.26902104 | 0.29788407 | 0.90310653 | 0.36646936 | 0.62157013 |
| ABL2 | 174.936833 | -0.2345198 | 0.24782378 | -0.946317 | 0.34398694 | 0.60110949 |
| SOAT1 | 53.230779 | -0.5741894 | 0.40054175 | -1.433532 | 1.52E-01 | 0.39273741 |
| TOR1AIP2 | 101.077798 | 0.03385192 | 0.20579385 | 0.16449434 | 0.869342 | 0.9390016 |
| TOR1AIP1 | 46.0126206 | -1.2007119 | 0.36838744 | -3.2593724 | 0.00111659 | 0.03002592 |
| CEP350 | 220.854437 | -0.1984922 | 0.25168101 | -0.7886658 | 0.43030738 | 0.67197671 |
| QSOX1 | 31.556147 | -0.0022415 | 0.38016572 | -0.005896 | 0.99529569 | 0.99796515 |
| ACBD6 | 178.62684 | 0.15497558 | 0.30101329 | 0.51484631 | 0.60666042 | 0.79507326 |
| XPR1 | 194.374367 | -0.4632091 | 0.30802505 | -1.5038033 | 0.13263201 | 0.3666913 |
| KIAA1614 | 29.426976 | 0.70596499 | 0.56766914 | 1.24362051 | 0.21363926 | 0.46920405 |
| STX6 | 55.6152544 | 0.21352538 | 0.26521541 | 0.80510168 | 0.42076101 | 0.66405525 |
| MR1 | 18.4215301 | 0.48826728 | 0.45130159 | 1.08190906 | 0.27929294 | NA |
| GLUL | 54.0702166 | -0.0728088 | 0.33610838 | -0.2166229 | 8.29E-01 | 9.17E-01 |
| DHX9 | 127.050132 | -0.5161629 | 0.33003896 | -1.5639452 | 0.11783044 | 0.34551224 |
| LAMC1 | 160.181474 | -1.050234 | 0.47281286 | -2.2212468 | 0.02633425 | 0.15910105 |
| LAMC2 | 21.6551915 | -1.785946 | 0.84639587 | -2.1100599 | 3.49E-02 | 1.86E-01 |
| SMG7 | 154.189973 | -0.2839444 | 0.23891606 | -1.1884691 | 0.23464866 | 0.49243138 |
| ARPC5 | 34.2628045 | -0.1076344 | 0.36570639 | -0.2943192 | 0.76851403 | 0.88545674 |
| RGL1 | 15.688218 | -0.187665 | 0.55281635 | -0.3394708 | 0.73425505 | NA |
| TSEN15 | 48.8618962 | 0.86296816 | 0.33031178 | 2.61258673 | 0.00898599 | 0.09383967 |
| EDEM3 | 86.1480551 | 0.09726053 | 0.35719008 | 0.27229348 | 0.78539637 | 0.89452315 |
| RNF2 | 48.3457604 | -0.325936 | 0.29469381 | -1.1060157 | 0.26871971 | 0.52744122 |
| TRMT1L | 37.8928915 | 0.82304331 | 0.35549137 | 2.31522728 | 0.0206005 | 0.14099415 |
| SWT1 | 72.3240736 | -0.5846782 | 0.35761727 | -1.6349273 | 0.10206429 | 0.32100708 |
| IVNS1ABP | 699.180668 | -1.2356942 | 0.32604232 | -3.7899811 | 0.00015066 | 0.00913087 |
| HMCN1 | 210.959261 | 0.01894997 | 0.31040981 | 0.06104823 | 0.9513208 | 0.98040586 |
| TPR | 135.560969 | -0.3112013 | 0.22748422 | -1.3680126 | 0.17130813 | 0.41952233 |
| ODR4 | 83.1795258 | -0.0913498 | 0.27468327 | -0.332564 | 0.73946346 | 0.87054913 |
| AL596220.1 | 17.8200964 | 1.26913468 | 0.76891746 | 1.65054735 | 0.09883104 | NA |
| ERVMER61-1 | 126.157236 | 0.33957533 | 0.40869719 | 0.83087269 | 0.40604555 | 0.65318894 |
| BRINP3 | 54.0848082 | 0.13433895 | 0.52155199 | 0.25757538 | 0.79673463 | 0.90077357 |
| AL390957.1 | 22.4581093 | 0.96704364 | 0.51129532 | 1.89136025 | 0.05857627 | 0.24020328 |
| AL136454.1 | 19.0871578 | 0.38437139 | 0.51885436 | 0.74080787 | 0.45880994 | NA |
| UCHL5 | 42.6932814 | 1.17667888 | 0.36612637 | 3.21385993 | 0.00130964 | 0.03230854 |
| TROVE2 | 70.2604475 | 0.28906391 | 0.26002848 | 1.11166252 | 0.26628328 | 0.52522169 |
| CDC73 | 169.060196 | -0.3595916 | 0.21267358 | -1.6908146 | 0.09087223 | 0.30109493 |
| ZBTB41 | 36.7601316 | 0.26405998 | 0.36236906 | 0.72870454 | 0.46618241 | 0.69633341 |
| CRB1 | 18.3478104 | 3.78655873 | 1.09787649 | 3.44898427 | 0.0005627 | NA |
| DENND1B | 229.654862 | 0.60843955 | 0.18598119 | 3.27151125 | 1.07E-03 | 0.02960163 |
| NEK7 | 202.558409 | 0.06207151 | 0.19076663 | 0.32537931 | 0.74489401 | 0.87306978 |
| AC104461.1 | 24.1258647 | -1.2127228 | 0.56097884 | -2.1617978 | 0.03063377 | 0.17299202 |
| ZNF281 | 15.9362497 | 0.07927831 | 0.53622064 | 0.14784643 | 0.88246396 | NA |
| DDX59 | 32.017981 | 0.22520433 | 0.40889252 | 0.55076657 | 0.5817937 | 0.7783237 |
| CAMSAP2 | 154.903181 | -0.0035992 | 0.27731484 | -0.0129788 | 0.98964467 | 0.99583841 |
| INAVA | 26.806218 | -0.876418 | 0.43135565 | -2.0317759 | 0.04217635 | 0.20468559 |
| TMEM9 | 24.0203367 | 0.77743572 | 0.46481605 | 1.67256643 | 0.09441268 | 0.30830642 |

| | | | | | | |
|---|---|---|---|---|---|---|
| CSRP1 | 25.259063 | -0.2129495 | 0.67865684 | -0.3137808 | 0.7536875 | 0.87868581 |
| AC096677.1 | 38.9278991 | 0.64497658 | 0.37172199 | 1.73510471 | 0.08272226 | 0.28758502 |
| NAV1 | 40.2578257 | -0.1063196 | 0.43274454 | -0.2456868 | 0.80592469 | 0.90570921 |
| IPO9 | 81.2627569 | -0.659849 | 0.28630079 | -2.3047405 | 0.02118111 | 0.1427136 |
| TIMM17A | 27.5433453 | -0.0462076 | 0.44469058 | -0.1039097 | 0.91724104 | 0.96401363 |
| RNPEP | 27.8211298 | -0.2240479 | 0.35023625 | -0.639705 | 0.52236438 | 0.7380123 |
| ELF3 | 111.001087 | 0.17029638 | 0.42010837 | 0.40536299 | 6.85E-01 | 8.44E-01 |
| ARL8A | 23.9472521 | -0.3380232 | 0.42761813 | -0.7904791 | 0.42924803 | 0.6716654 |
| PPP1R12B | 218.429884 | 0.44770765 | 0.20420632 | 2.19242794 | 0.02834862 | 0.16567907 |
| KDM5B | 144.812846 | -0.4230954 | 0.19340333 | -2.1876326 | 0.02869637 | 0.16632227 |
| KLHL12 | 27.8713302 | -0.2518488 | 0.37493513 | -0.6717129 | 0.50176647 | 0.72337451 |
| ADIPOR1 | 15.8715559 | 0.44879794 | 0.46323903 | 0.96882583 | 0.33263209 | NA |
| CYB5R1 | 16.1566009 | 0.38735787 | 0.54037509 | 0.71683147 | 0.47347809 | NA |
| TMEM183A | 14.9470901 | -0.685784 | 0.51488426 | -1.3319187 | 0.1828869 | NA |
| BTG2 | 25.4070726 | 0.85909427 | 0.62080985 | 1.38382835 | 0.16641102 | 0.41429299 |
| ATP2B4 | 171.562905 | -0.0144047 | 0.2873718 | -0.0501258 | 0.96002214 | 0.98458234 |
| SNRPE | 12.5487514 | 0.07452511 | 0.48135915 | 0.15482225 | 0.87696145 | NA |
| SOX13 | 36.6962526 | -0.6235827 | 0.34031555 | -1.8323663 | 0.06689686 | 0.25822726 |
| GOLT1A | 18.7080876 | 0.38893385 | 0.54172776 | 0.71795076 | 0.47278765 | NA |
| PLEKHA6 | 126.40223 | -0.7425423 | 0.25954516 | -2.8609367 | 0.00422391 | 0.06301187 |
| PPP1R15B | 24.8520981 | -0.7664502 | 0.48970218 | -1.5651354 | 0.11755117 | 0.34520844 |
| MDM4 | 170.739358 | -0.252261 | 0.21072423 | -1.1971145 | 0.23126192 | 0.48982471 |
| LRRN2 | 52.1278076 | 1.27076285 | 0.48017741 | 2.64644446 | 0.00813429 | 0.08868215 |
| RBBP5 | 41.7318742 | -0.8125415 | 0.31830826 | -2.5526874 | 0.01068954 | 0.101577 |
| DSTYK | 77.744107 | 0.49371126 | 0.29454991 | 1.67615481 | 0.09370787 | 0.30744651 |
| NUAK2 | 55.1380385 | -0.1407834 | 0.6166789 | -0.2282929 | 0.81941853 | 0.913067 |
| CDK18 | 27.9294698 | -0.1006412 | 0.52588578 | -0.1913747 | 0.84823207 | 0.92660264 |
| MFSD4A | 92.9214632 | 1.60152795 | 0.51267317 | 3.12387705 | 0.00178485 | 0.03909384 |
| ELK4 | 60.8096879 | 0.14253716 | 0.36632771 | 0.3890974 | 0.6972041 | 0.85041636 |
| NUCKS1 | 170.767728 | -0.2306589 | 0.27303842 | -0.8447854 | 0.39823065 | 0.64642108 |
| RAB29 | 14.9345029 | 0.7788765 | 0.47340495 | 1.64526481 | 0.09991521 | NA |
| SRGAP2 | 44.5206266 | -0.5424055 | 0.2962356 | -1.8309938 | 0.06710147 | 0.25862794 |
| EIF2D | 21.2897551 | 0.17276623 | 0.43921618 | 0.39335123 | 0.69406008 | 0.84823443 |
| DYRK3 | 22.8239888 | -1.8110786 | 0.61604139 | -2.9398651 | 0.00328355 | 0.0569223 |
| MAPKAPK2 | 127.51982 | -0.7347166 | 0.35826042 | -2.0507893 | 0.04028747 | 0.19980197 |
| PIGR | 44.3014489 | 2.51478729 | 0.81811 | 3.07389871 | 0.00211281 | 0.04325083 |
| PFKFB2 | 97.8683008 | -1.0432767 | 0.47338489 | -2.2038657 | 0.02753379 | 0.16347426 |
| CD55 | 36.843581 | -1.3630759 | 0.59231888 | -2.3012536 | 0.0213773 | 0.14320374 |
| CD46 | 146.351317 | 0.57133182 | 0.25657844 | 2.22673353 | 0.02596509 | 0.157613 |
| MIR29B2CH( | 246.269776 | 0.71816543 | 0.35610332 | 2.01673334 | 0.04372335 | 0.20890044 |
| PLXNA2 | 14.4824694 | -0.2825918 | 0.58781201 | -0.4807521 | 0.63069272 | NA |
| LAMB3 | 57.4873008 | -3.1749896 | 1.28693294 | -2.467098 | 1.36E-02 | 0.11413904 |
| G0S2 | 93.0450224 | -1.0851694 | 0.661881 | -1.6395235 | 0.10110429 | 0.32011506 |
| IRF6 | 18.0909935 | 0.02014557 | 0.51158622 | 0.03937864 | 0.96858851 | NA |
| UTP25 | 13.0521574 | -0.5898895 | 0.51787373 | -1.1390605 | 0.25467791 | NA |
| SYT14 | 20.3030299 | -1.3713774 | 0.77130718 | -1.7779912 | 0.0754053 | NA |
| HHAT | 81.9112777 | 2.02899473 | 0.55963749 | 3.62555184 | 0.00028835 | 0.01395844 |
| KCNH1 | 16.0692997 | 1.21267631 | 0.78370822 | 1.54735688 | 1.22E-01 | NA |
| RCOR3 | 174.471839 | 0.51475037 | 0.21664737 | 2.37598255 | 0.01750229 | 0.1316147 |
| TRAF5 | 33.5992945 | -0.7093186 | 0.46447191 | -1.5271508 | 0.12672352 | 0.35846782 |
| LINC00467 | 29.9735374 | 0.71477956 | 0.51614049 | 1.38485464 | 0.16609692 | 0.41400518 |
| LPGAT1 | 129.679233 | -0.7787894 | 0.29396475 | -2.6492613 | 0.00806679 | 0.08827138 |
| INTS7 | 32.6103507 | -0.1383673 | 0.41573201 | -0.3328281 | 0.73926406 | 0.87054913 |
| PPP2R5A | 188.917381 | -0.2903134 | 0.29111856 | -0.9972344 | 0.31865076 | 0.57587641 |

| | | | | | | |
|---|---|---|---|---|---|---|
| AL360091.3 | 25.6535627 | 0.80390321 | 0.68756068 | 1.16921056 | 0.24231881 | 0.49963243 |
| NENF | 29.5133846 | 1.07197246 | 0.60352054 | 1.7761988 | 0.07570016 | 0.27489837 |
| NSL1 | 61.7562788 | -0.0279528 | 0.28274485 | -0.0988623 | 0.92124761 | 0.96605489 |
| TATDN3 | 45.7870445 | 0.02434974 | 0.3098305 | 0.07859053 | 0.93735832 | 0.97445259 |
| FLVCR1 | 37.4836292 | -0.2372198 | 0.37465357 | -0.633171 | 0.52662198 | 0.74154041 |
| ANGEL2 | 51.0742012 | 0.04505375 | 0.34255297 | 0.13152345 | 0.89536124 | 0.95279904 |
| RPS6KC1 | 158.089036 | -0.5879971 | 0.19101781 | -3.078232 | 0.00208233 | 0.04285475 |
| PROX1 | 57.3184084 | 0.71997946 | 0.52582569 | 1.36923599 | 0.17092552 | 0.41898315 |
| SMYD2 | 38.6046405 | 0.10368177 | 0.40604604 | 0.25534487 | 0.79845675 | 0.90179334 |
| PTPN14 | 529.810489 | -0.56312 | 0.42578212 | -1.3225544 | 0.18598362 | 0.43607799 |
| KCTD3 | 95.4517717 | -0.342455 | 0.24176014 | -1.4165074 | 0.156627 | 0.40065072 |
| ESRRG | 5317.15071 | 0.12030721 | 0.28302955 | 0.42506945 | 0.67078605 | 0.83476802 |
| GPATCH2 | 176.263884 | 0.23334765 | 0.20163344 | 1.15728643 | 0.24715535 | 0.50340163 |
| SPATA17 | 28.3687775 | 1.61939758 | 0.83678593 | 1.93525909 | 0.05295851 | 0.2312091 |
| RRP15 | 24.3416274 | 0.09182757 | 0.40985478 | 0.22404905 | 0.82271914 | 0.91475524 |
| LYPLAL1 | 221.713407 | 0.29896854 | 0.34870875 | 0.85735887 | 0.39124658 | 0.63984683 |
| EPRS | 58.4520277 | 0.1564996 | 0.32517639 | 0.48127603 | 0.63032033 | 0.80805722 |
| BPNT1 | 16.3660979 | 0.43945309 | 0.48960963 | 0.8975581 | 0.36942119 | NA |
| IARS2 | 50.9004773 | 0.37298519 | 0.29975878 | 1.24428448 | 0.21339488 | 0.46890289 |
| RAB3GAP2 | 203.068353 | -0.3442623 | 0.2037873 | -1.6893215 | 0.09115783 | 0.30165168 |
| MARK1 | 161.708683 | -0.9584091 | 0.57065692 | -1.6794839 | 0.09305777 | 0.30571879 |
| 2-Mar | 71.8236333 | 1.06123937 | 0.31208378 | 3.40049506 | 0.00067264 | 0.02351966 |
| 1-Mar | 13.1371093 | 1.03048859 | 0.55821077 | 1.84605646 | 0.06488401 | NA |
| AL513314.2 | 19.2034698 | 0.45108331 | 0.45501171 | 0.99136637 | 0.32150671 | NA |
| MIA3 | 45.1708839 | 0.19653825 | 0.30902799 | 0.63598851 | 0.52478391 | 0.73968378 |
| AIDA | 18.7414452 | -0.1483264 | 0.3838597 | -0.3864079 | 0.69919457 | NA |
| BROX | 43.8287312 | 0.35134062 | 0.27821159 | 1.26285401 | 0.20664165 | 0.460087 |
| FAM177B | 32.1579362 | -1.3149684 | 0.85484801 | -1.5382482 | 0.12398793 | 0.35457643 |
| AC093152.1 | 35.8156783 | 0.24189588 | 0.39268613 | 0.61600312 | 0.53789246 | 0.74878549 |
| TLR5 | 26.547939 | 0.92957729 | 0.37402358 | 2.4853441 | 0.01294263 | 0.11079049 |
| SUSD4 | 109.581939 | 0.65457447 | 0.33512731 | 1.95321133 | 0.05079456 | 0.22622461 |
| CAPN2 | 149.860457 | -0.5100522 | 0.27901988 | -1.8280139 | 0.06754746 | 0.25898011 |
| TP53BP2 | 69.7010667 | -1.5810826 | 0.39679733 | -3.9846101 | 6.76E-05 | 0.00530264 |
| FBXO28 | 110.156777 | -0.7525731 | 0.27627893 | -2.7239613 | 0.00645041 | 0.07818705 |
| DEGS1 | 46.4045904 | -0.3637689 | 0.49323707 | -0.7375134 | 0.46081022 | 0.69410104 |
| NVL | 80.5166083 | -0.4399144 | 0.25445778 | -1.7288305 | 0.08383943 | 0.28873024 |
| CNIH4 | 65.9108348 | -0.1775504 | 0.27731434 | -0.6402498 | 0.52201022 | 0.7380123 |
| WDR26 | 164.481996 | -0.3476423 | 0.23984164 | -1.4494659 | 0.14720751 | 0.38715156 |
| CNIH3 | 49.4903327 | -0.0604983 | 0.68043716 | -0.0889109 | 0.92915272 | 0.97064176 |
| DNAH14 | 374.791787 | 0.25594367 | 0.23481076 | 1.08999977 | 0.27571325 | 0.53465985 |
| LBR | 50.2700249 | 0.11772467 | 0.33953095 | 0.34672736 | 0.72879616 | 0.86526979 |
| ENAH | 75.6507191 | -0.2313744 | 0.36676401 | -0.6308535 | 0.52813629 | 0.74274895 |
| SRP9 | 48.0023295 | -0.0254384 | 0.31838593 | -0.0798981 | 0.93631833 | 0.97405271 |
| EPHX1 | 19.186864 | 0.55737089 | 0.4710123 | 1.18334678 | 0.23667174 | NA |
| AL591895.1 | 18.9046352 | 0.3554836 | 0.45538597 | 0.78062045 | 0.43502576 | NA |
| TMEM63A | 61.698654 | 0.35359804 | 0.29377235 | 1.20364642 | 0.22872627 | 0.48701178 |
| H3F3A | 22.27977 | 0.01877718 | 0.38687551 | 0.04853545 | 0.96128951 | 0.98496344 |
| ACBD3 | 87.2327726 | -0.4993934 | 0.26112581 | -1.9124629 | 0.05581685 | 0.23575965 |
| LIN9 | 15.5384638 | 0.78194666 | 0.48190748 | 1.62260745 | 0.10467335 | NA |
| PARP1 | 46.8671505 | -0.3366006 | 0.33296905 | -1.0109064 | 0.31206121 | 0.57101636 |
| STUM | 45.2770436 | 0.74785118 | 0.59725559 | 1.25214598 | 0.21051668 | 0.46548315 |
| ITPKB | 292.795765 | -0.3803496 | 0.48342564 | -0.78678 | 0.43141069 | 0.67234892 |
| COQ8A | 37.8880812 | 0.1522565 | 0.35996264 | 0.42297863 | 0.67231086 | 0.83624371 |
| CDC42BPA | 374.012199 | 0.09455282 | 0.27780521 | 0.34035654 | 0.73358804 | 0.86776236 |

| | | | | | | |
|---|---|---|---|---|---|---|
| ZNF678 | 53.0525023 | -0.0981741 | 0.27704241 | -0.3543648 | 0.72306549 | 0.86192273 |
| SNAP47 | 26.1512346 | 0.36480065 | 0.40977694 | 0.89024203 | 0.37333594 | 0.62575308 |
| ARF1 | 72.9900005 | -0.5252933 | 0.4135669 | -1.2701532 | 0.20403005 | 0.45744809 |
| GUK1 | 57.7865469 | 0.59982256 | 0.36718564 | 1.63356761 | 0.10234967 | 0.32120262 |
| OBSCN | 133.510115 | 0.55426613 | 0.28189096 | 1.96624302 | 0.04927055 | 0.22242545 |
| TRIM11 | 12.6088143 | -0.3327167 | 0.50523258 | -0.6585416 | 0.51019018 | NA |
| RNF187 | 15.8462815 | -0.1368979 | 0.62813059 | -0.2179449 | 0.82747204 | NA |
| RAB4A | 26.9637735 | -0.0727915 | 0.428894 | -0.1697191 | 0.86523109 | 0.93738047 |
| NUP133 | 48.5106777 | 0.12226383 | 0.28289198 | 0.43219262 | 0.66560142 | 0.83059892 |
| ABCB10 | 23.5709533 | 1.1837753 | 0.47846846 | 2.47409266 | 0.0133575 | 0.11285699 |
| TAF5L | 17.1759512 | -0.7056331 | 0.58214073 | -1.212135 | 0.22546072 | NA |
| GALNT2 | 105.677671 | -0.9363706 | 0.31554198 | -2.9674992 | 0.00300233 | 0.05399285 |
| PGBD5 | 24.5001905 | -0.1654749 | 0.46098409 | -0.3589601 | 0.71962497 | 0.85995641 |
| COG2 | 36.1319413 | 0.00859187 | 0.31923942 | 0.02691357 | 0.97852867 | 0.9916702 |
| TTC13 | 25.0783532 | 0.5414551 | 0.40754068 | 1.32859155 | 0.18398276 | 0.43412487 |
| ARV1 | 17.7883574 | 1.12736728 | 0.50640658 | 2.2262098 | 0.02600013 | NA |
| C1orf131 | 19.3943063 | 0.72723161 | 0.51630392 | 1.40853397 | 0.15897302 | NA |
| GNPAT | 54.5766201 | 0.06935257 | 0.27783943 | 0.24961383 | 0.802886 | 0.90408418 |
| SPRTN | 13.532094 | 0.05835267 | 0.49607263 | 0.11762929 | 0.9063614 | NA |
| EGLN1 | 142.438201 | -0.0120337 | 0.28383757 | -0.0423965 | 0.9661826 | 0.98723631 |
| DISC1 | 20.9823098 | 0.60630625 | 0.67409166 | 0.89944185 | 0.36841735 | NA |
| NTPCR | 28.7398464 | 0.16390569 | 0.42745132 | 0.38344878 | 0.70138704 | 0.85190818 |
| PCNX2 | 184.434658 | -0.0559276 | 0.25054743 | -0.2232218 | 0.82336291 | 0.91475524 |
| MAP3K21 | 89.8505785 | 0.79573436 | 0.28934433 | 2.7501294 | 0.00595717 | 0.07594487 |
| KCNK1 | 40.4549629 | 0.43427107 | 0.41403151 | 1.04888411 | 2.94E-01 | 0.55371627 |
| AL355472.1 | 11.4112237 | -0.5734172 | 0.83571491 | -0.6861397 | 0.49262501 | NA |
| TARBP1 | 63.6755579 | 0.99873959 | 0.42667547 | 2.34074763 | 0.01924517 | 0.13639344 |
| IRF2BP2 | 75.7398002 | 0.59990005 | 0.41563541 | 1.4433324 | 0.14892686 | 0.38937375 |
| TOMM20 | 48.3109607 | 0.46087263 | 0.31983156 | 1.44098549 | 0.14958878 | 0.39003552 |
| ARID4B | 397.274835 | -0.5327206 | 0.19512428 | -2.7301606 | 0.00633035 | 0.07758708 |
| B3GALNT2 | 71.7296518 | 0.45791454 | 0.28010761 | 1.63478076 | 0.10209502 | 0.32100708 |
| LYST | 144.531971 | -0.4028985 | 0.21728315 | -1.8542554 | 0.06370263 | 0.25118809 |
| GPR137B | 19.8124903 | -0.1082151 | 0.46490571 | -0.2327678 | 0.8159417 | NA |
| ERO1B | 44.9352426 | -0.3596827 | 0.37251346 | -0.9655563 | 0.33426624 | 0.58942664 |
| LGALS8 | 40.2300885 | -0.7306521 | 0.30548108 | -2.3918079 | 0.01676561 | 0.1285308 |
| HEATR1 | 46.4361412 | -0.0758413 | 0.36852836 | -0.2057949 | 0.83695111 | 0.92119416 |
| MTR | 388.694462 | 0.37257442 | 0.28106919 | 1.32556123 | 0.18498508 | 0.43476148 |
| CHRM3 | 158.917575 | 0.47954266 | 0.35292719 | 1.35875803 | 0.17422328 | 0.42295604 |
| RGS7 | 31.3984904 | -0.4040977 | 0.54185627 | -0.7457654 | 0.45580914 | 0.6908246 |
| FH | 17.7804858 | 0.51425002 | 0.55757774 | 0.92229295 | 0.35637579 | NA |
| CEP170 | 79.4800167 | 0.17887439 | 0.25659706 | 0.69710225 | 0.48573881 | 0.71268235 |
| SDCCAG8 | 285.111223 | 0.58169621 | 0.28577794 | 2.03548327 | 0.04180227 | 0.20402795 |
| AKT3 | 316.845597 | -0.0371325 | 0.30410712 | -0.1221032 | 0.90281726 | 0.95661107 |
| ZBTB18 | 15.0625272 | -0.3148735 | 0.5386162 | -0.5845972 | 0.55881858 | NA |
| ADSS | 60.2953563 | -0.3907979 | 0.29818778 | -1.3105764 | 0.1900009 | 0.44127941 |
| CATSPERE | 45.3034965 | -0.5330187 | 0.49801901 | -1.0702778 | 0.28449428 | 0.54316116 |
| DESI2 | 85.4908816 | -0.0929546 | 0.35615781 | -0.2609926 | 0.79409819 | 0.89884908 |
| COX20 | 65.8641727 | 0.0343409 | 0.32030494 | 0.10721314 | 0.91461989 | 0.96388328 |
| HNRNPU | 308.853218 | -0.2128516 | 0.19796325 | -1.0752076 | 0.28228177 | 0.54101664 |
| EFCAB2 | 182.437877 | -0.5865982 | 0.48196171 | -1.2171053 | 0.22356414 | 0.48054488 |
| KIF26B | 49.4590927 | -0.2399338 | 0.52519394 | -0.456848 | 0.6477803 | 0.81885075 |
| SMYD3 | 502.893829 | 0.35628327 | 0.39924056 | 0.89240251 | 0.37217723 | 0.62492323 |
| TFB2M | 15.1489658 | -0.817653 | 0.48083717 | -1.700478 | 0.08904105 | NA |
| CNST | 90.6016762 | -0.9270098 | 0.28241016 | -3.2824946 | 0.00102893 | 0.02922388 |

| | | | | | | |
|---|---|---|---|---|---|---|
| SCCPDH | 20.5160811 | -0.0802628 | 0.48846 | -0.1643181 | 0.86948073 | NA |
| AHCTF1 | 108.557571 | -0.6941447 | 0.30896105 | -2.2467061 | 0.02465881 | 0.15380786 |
| ZNF124 | 52.9267208 | -0.3888973 | 0.33830523 | -1.1495457 | 0.25033104 | 0.50659901 |
| AL390728.6 | 15.413382 | 0.15896872 | 0.48763026 | 0.32600257 | 0.74442241 | NA |
| ZNF496 | 26.9876047 | -0.9377825 | 0.36760146 | -2.5510848 | 0.01073882 | 0.10191948 |
| SH3BP5L | 10.4617998 | -0.0479303 | 0.566747 | -0.0845709 | 0.9326025 | NA |
| ZNF672 | 14.2420436 | -0.7231326 | 0.49798687 | -1.4521119 | 0.1464705 | NA |
| ZNF692 | 18.846058 | 0.768422 | 0.45014016 | 1.70707275 | 0.08780853 | NA |
| FAM110C | 110.956442 | -1.2839828 | 0.6002598 | -2.1390451 | 3.24E-02 | 1.78E-01 |
| SH3YL1 | 102.646344 | 0.84396317 | 0.31015172 | 2.7211301 | 0.00650592 | 0.07861229 |
| ACP1 | 31.7326943 | -0.1962908 | 0.32191176 | -0.6097659 | 0.54201692 | 0.7506125 |
| EIPR1 | 80.703525 | -0.0862146 | 0.34845089 | -0.2474225 | 0.80458126 | 0.90446356 |
| TRAPPC12 | 113.799008 | -0.1062882 | 0.26081265 | -0.4075268 | 0.68362109 | 0.8424322 |
| ADI1 | 36.7148848 | 0.6874647 | 0.3923372 | 1.75222919 | 0.07973441 | 0.28190893 |
| RPS7 | 71.2785662 | 0.15128404 | 0.39869801 | 0.37944518 | 0.7043573 | 0.85317002 |
| RNF144A | 18.5904901 | 0.28743201 | 0.45411796 | 0.6329457 | 0.5267691 | NA |
| ID2 | 46.4960316 | -0.1134699 | 0.43609644 | -0.2601946 | 0.79471369 | 0.89901694 |
| KIDINS220 | 162.500018 | 0.05695647 | 0.22000019 | 0.25889283 | 0.79571793 | 0.8998778 |
| MBOAT2 | 83.6849269 | 0.40590137 | 0.46985985 | 0.86387752 | 0.38765518 | 0.63701577 |
| ASAP2 | 497.858576 | -0.0330269 | 0.21827679 | -0.1513075 | 0.87973315 | 0.94376451 |
| ITGB1BP1 | 67.5115624 | -0.4775887 | 0.24803825 | -1.9254639 | 0.05417134 | 0.23287156 |
| CPSF3 | 28.3410563 | -0.6599223 | 0.39903034 | -1.6538148 | 0.09816515 | 0.31548106 |
| IAH1 | 26.9736634 | 0.16304626 | 0.3510935 | 0.46439555 | 0.64236438 | 0.81595394 |
| ADAM17 | 118.568004 | -0.834541 | 0.28946661 | -2.8830304 | 0.00393869 | 0.06143969 |
| YWHAQ | 80.1242535 | -0.4561013 | 0.28971015 | -1.5743368 | 0.11540961 | 0.34185848 |
| TAF1B | 99.7109872 | -0.176013 | 0.32797395 | -0.5366678 | 0.59149713 | 0.78517152 |
| GRHL1 | 15.4373904 | 0.68416053 | 0.66880514 | 1.02295945 | 0.30632702 | NA |
| KLF11 | 23.4745558 | 0.49780007 | 0.5408422 | 0.92041648 | 0.35735516 | 0.61328042 |
| CYS1 | 261.241435 | 1.74708187 | 0.60670391 | 2.87962848 | 0.00398144 | 0.06175835 |
| HPCAL1 | 109.225604 | -0.4466674 | 0.28843596 | -1.5485845 | 0.12148164 | 0.35059775 |
| NOL10 | 117.565082 | -0.6334785 | 0.21765325 | -2.910494 | 0.00360858 | 0.05973168 |
| PDIA6 | 59.1723857 | 0.01438606 | 0.55690688 | 0.02583207 | 0.97939128 | 0.9922831 |
| ROCK2 | 183.346374 | -0.50857 | 0.34565623 | -1.4713173 | 0.14120533 | 0.3792245 |
| E2F6 | 20.9177405 | -0.2837546 | 0.49085839 | -0.5780783 | 0.56321128 | NA |
| LPIN1 | 70.5469307 | -0.905764 | 0.38808269 | -2.3339458 | 0.01959856 | 0.13704876 |
| TRIB2 | 58.2625597 | 0.33848511 | 0.45088921 | 0.75070572 | 4.53E-01 | 6.89E-01 |
| NBAS | 287.61242 | -0.3480503 | 0.22868031 | -1.521995 | 0.12801035 | 0.36035989 |
| DDX1 | 67.0587646 | 0.0970905 | 0.28750857 | 0.33769603 | 0.73559227 | 0.86838246 |
| SMC6 | 53.6372313 | -0.2800775 | 0.35778999 | -0.7827986 | 0.43374542 | 0.67472483 |
| GEN1 | 17.508111 | -1.2412517 | 0.50614853 | -2.4523468 | 0.01419278 | NA |
| LINC01376 | 65.5331033 | 0.25923686 | 0.53191226 | 0.48736772 | 0.62599776 | 0.8053173 |
| TTC32 | 16.8573681 | -0.5264339 | 0.44657484 | -1.1788257 | 0.23846759 | NA |
| WDR35 | 59.925694 | -0.0070096 | 0.36607452 | -0.019148 | 0.98472303 | 0.99407122 |
| LAPTM4A | 70.5218852 | 0.48690117 | 0.25314054 | 1.92344211 | 0.05442455 | 0.23311395 |
| SDC1 | 25.3529966 | -0.3816049 | 0.74300859 | -0.5135942 | 0.60753571 | 0.79552231 |
| PUM2 | 330.975842 | -0.5364459 | 0.17188353 | -3.1209852 | 0.00180247 | 0.03909384 |
| RHOB | 39.7139504 | -0.4245375 | 0.69206759 | -0.6134336 | 0.53958967 | 0.74929145 |
| LDAH | 97.7248176 | 0.5714809 | 0.46011007 | 1.24205257 | 0.21421717 | 0.4699161 |
| KLHL29 | 115.176011 | -2.2405613 | 1.15255325 | -1.943998 | 5.19E-02 | 0.22838257 |
| ATAD2B | 197.921588 | -0.300926 | 0.23085709 | -1.3035166 | 0.19239847 | 0.44417846 |
| UBXN2A | 87.0061875 | 0.27997263 | 0.29014736 | 0.96493256 | 0.33457859 | 0.58984229 |
| SF3B6 | 21.3751057 | 0.47520238 | 0.41807321 | 1.13664873 | 0.25568516 | 0.5136319 |
| FAM228B | 13.9243003 | -0.0176834 | 0.57842698 | -0.0305715 | 0.97561126 | NA |
| TP53I3 | 18.2998328 | 1.25242251 | 0.60774446 | 2.06077159 | 0.03932484 | NA |

| | | | | | | |
|---|---|---|---|---|---|---|
| ITSN2 | 214.283233 | -0.3893352 | 0.26697267 | -1.4583336 | 0.14474861 | 0.38431531 |
| NCOA1 | 510.845546 | 0.07335923 | 0.18599 | 0.39442568 | 0.69326679 | 0.84791601 |
| DNAJC27 | 40.8973 | -0.0042677 | 0.30133201 | -0.0141627 | 0.9887002 | 0.99581595 |
| DNAJC27-AS | 18.8923076 | 0.45519006 | 0.73695302 | 0.61766496 | 0.53679622 | NA |
| DNMT3A | 122.026564 | 0.28389349 | 0.2822777 | 1.00572411 | 0.3145483 | 0.57208844 |
| DTNB | 168.905733 | -0.3961013 | 0.24874255 | -1.5924147 | 0.11129155 | 0.33461909 |
| ASXL2 | 171.771616 | -0.4217952 | 0.27851937 | -1.5144195 | 0.12991947 | 0.36231527 |
| RAB10 | 294.752945 | -0.6124783 | 0.28531948 | -2.1466402 | 0.03182194 | 0.17583164 |
| HADHA | 88.2680613 | 0.26578571 | 0.26422644 | 1.00590126 | 0.31446307 | 0.5720686 |
| HADHB | 122.259577 | 0.68801845 | 0.23490574 | 2.92891283 | 0.0034015 | 0.05757539 |
| SELENOI | 72.9658728 | 0.01538684 | 0.35302677 | 0.04358546 | 0.96523484 | 0.9869039 |
| MAPRE3 | 32.2576232 | 1.17553691 | 0.46811049 | 2.51123815 | 0.01203085 | 0.10726863 |
| TMEM214 | 13.5865995 | 0.26984444 | 0.45978888 | 0.58688771 | 0.55727913 | NA |
| AGBL5 | 25.2785392 | -0.0661502 | 0.58271432 | -0.1135209 | 0.90961762 | 0.96026976 |
| OST4 | 25.9172644 | 0.50145711 | 0.5121695 | 0.97908428 | 0.32753834 | 0.58371442 |
| ATRAID | 20.8790262 | 0.75732041 | 0.4187043 | 1.80872375 | 0.07049393 | NA |
| MPV17 | 14.5683551 | -0.8565301 | 0.55155089 | -1.5529484 | 0.12043545 | NA |
| GTF3C2 | 53.881943 | -0.20423 | 0.27904186 | -0.7318975 | 0.46423115 | 0.69530788 |
| EIF2B4 | 16.22778 | -0.3221374 | 0.44447298 | -0.7247626 | 0.4685977 | NA |
| PPM1G | 50.4445052 | 0.17443395 | 0.36722501 | 0.47500565 | 0.63478295 | 0.81099192 |
| NRBP1 | 47.8133629 | -1.0184045 | 0.41665597 | -2.4442335 | 0.01451603 | 0.11873638 |
| IFT172 | 54.0167285 | 0.97322067 | 0.5105033 | 1.90639446 | 0.05659905 | 0.23714908 |
| ZNF512 | 50.5906205 | -0.4898345 | 0.36620552 | -1.3375944 | 0.18102869 | 0.43032052 |
| SUPT7L | 20.8116477 | 0.26785278 | 0.3818662 | 0.70143098 | 0.48303409 | NA |
| SLC4A1AP | 31.2943301 | -0.5637255 | 0.34926952 | -1.6140129 | 0.10652463 | 0.32738142 |
| MRPL33 | 94.5955897 | -0.4405313 | 0.28486354 | -1.5464642 | 0.12199249 | 0.35128178 |
| RBKS | 110.044526 | -0.006657 | 0.31031236 | -0.0214527 | 0.98288452 | 0.99382437 |
| BABAM2 | 635.164353 | -0.2850456 | 0.25748573 | -1.1070344 | 0.26827905 | 0.52717484 |
| FOSL2 | 182.407858 | -0.798058 | 0.52676459 | -1.5150183 | 0.12976777 | 0.3622089 |
| PPP1CB | 125.153728 | -1.1706364 | 0.26325323 | -4.4468073 | 8.72E-06 | 0.00146459 |
| TRMT61B | 41.9863963 | -0.1012994 | 0.33531342 | -0.3021037 | 0.76257304 | 0.88247846 |
| WDR43 | 68.6556414 | -0.8153491 | 0.30879137 | -2.6404528 | 0.00827953 | 0.08912945 |
| CLIP4 | 117.174801 | -0.3510053 | 0.33069114 | -1.0614293 | 0.28849486 | 0.54774172 |
| YPEL5 | 62.6411902 | -0.1713569 | 0.30678013 | -0.5585659 | 0.57645805 | 0.77421002 |
| LCLAT1 | 103.364446 | 0.4111129 | 0.27475166 | 1.49630727 | 0.1345736 | 0.36839609 |
| CAPN13 | 47.0780169 | -2.814184 | 1.30238373 | -2.1607948 | 0.0307112 | 0.17330211 |
| GALNT14 | 224.822612 | -0.5231071 | 0.39452294 | -1.325923 | 0.18486519 | 0.43476148 |
| DPY30 | 35.8036893 | 0.60155867 | 0.31223224 | 1.92663856 | 0.05402469 | 0.23287156 |
| SPAST | 64.1777552 | 0.16481486 | 0.26243602 | 0.62801919 | 0.52999137 | 0.74381523 |
| SLC30A6 | 74.9702746 | -0.0800724 | 0.35085705 | -0.2282196 | 0.81947555 | 0.913067 |
| YIPF4 | 94.674674 | -0.1640122 | 0.28757167 | -0.570335 | 0.56845049 | 0.77103518 |
| BIRC6 | 575.593722 | -0.4455596 | 0.16555558 | -2.6912992 | 0.00711743 | 0.08238028 |
| TTC27 | 67.0312761 | 0.15925356 | 0.3234258 | 0.49239598 | 0.62243944 | 0.80371018 |
| LINC00486 | 106.361877 | -0.1961416 | 0.30966699 | -0.6333951 | 0.52647563 | 0.74149734 |
| RASGRP3 | 42.0911511 | -1.4629278 | 0.44430572 | -3.2926152 | 0.0009926 | 0.02848444 |
| FAM98A | 15.3612184 | 1.06473293 | 0.60033051 | 1.77357791 | 0.07613299 | NA |
| AC068599.1 | 23.56259 | 0.41104947 | 0.71515946 | 0.57476617 | 0.56544945 | 0.76965382 |
| LINC01320 | 360.856037 | 0.55837309 | 0.62895389 | 0.88778065 | 3.75E-01 | 0.62575308 |
| AC012593.1 | 106.950232 | 1.65156968 | 0.93792256 | 1.76088064 | 0.07825861 | 0.27886875 |
| CRIM1 | 227.958937 | -0.3121429 | 0.31408242 | -0.9938247 | 0.3203082 | 0.57724473 |
| FEZ2 | 108.700522 | 0.37324909 | 0.24566869 | 1.51931889 | 0.12868225 | 0.36108905 |
| STRN | 157.471121 | -0.3178498 | 0.25790611 | -1.2324244 | 0.21779059 | 0.47407641 |
| HEATR5B | 91.8151745 | 0.05405939 | 0.27573975 | 0.19605221 | 0.8445693 | 0.92496441 |
| EIF2AK2 | 88.9555167 | 0.26815502 | 0.56082367 | 0.47814497 | 0.63254702 | 0.80953494 |

| | | | | | | |
|---|---|---|---|---|---|---|
| CEBPZOS | 54.2626253 | 0.19490524 | 0.31322827 | 0.62224666 | 0.53377969 | 0.74672888 |
| CEBPZ | 56.8692144 | -0.1312326 | 0.39068223 | -0.3359062 | 0.73694163 | 0.86917557 |
| NDUFAF7 | 43.1692403 | -0.0448723 | 0.30650576 | -0.1463996 | 0.88360593 | 0.94565001 |
| PRKD3 | 52.2089981 | 0.37238685 | 0.27718123 | 1.34347786 | 0.17911729 | 0.42802774 |
| RMDN2 | 33.2775593 | 0.62594064 | 0.46613584 | 1.34282881 | 0.17932741 | 0.42839327 |
| ATL2 | 186.530603 | -0.2773305 | 0.32684375 | -0.848511 | 0.39615347 | 0.64452212 |
| AC011247.1 | 16.457146 | -0.5432114 | 0.90069787 | -0.6031005 | 0.54644181 | NA |
| HNRNPLL | 62.235253 | -0.2781241 | 0.28451993 | -0.9775205 | 0.32831153 | 0.58467456 |
| GALM | 31.5465575 | 0.52714637 | 0.46478588 | 1.13417036 | 0.25672309 | 0.51485087 |
| SRSF7 | 94.2416827 | 0.49927922 | 0.36930322 | 1.35194926 | 0.17639155 | 0.42560681 |
| DHX57 | 58.3282552 | 0.09046463 | 0.26555244 | 0.34066579 | 0.73335519 | 0.86776236 |
| SOS1 | 339.939219 | 0.36847261 | 0.1978031 | 1.86282527 | 0.06248685 | 0.24933194 |
| MAP4K3 | 568.630913 | -0.4008218 | 0.19850766 | -2.0191753 | 0.04346901 | 0.20859161 |
| AC007388.1 | 306.40082 | -0.1464043 | 0.20474202 | -0.7150673 | 0.47456746 | 0.70288437 |
| TMEM178A | 47.9990164 | 0.77450513 | 0.62221475 | 1.24475533 | 2.13E-01 | 0.46890289 |
| THUMPD2 | 77.7635883 | 0.11168157 | 0.22136532 | 0.50451249 | 0.61390128 | 0.79925544 |
| SLC8A1-AS1 | 79.8350602 | 1.1271782 | 0.60043661 | 1.87726427 | 0.06048188 | 0.24514087 |
| SLC8A1 | 2520.77743 | 0.73224484 | 0.57624767 | 1.27071202 | 2.04E-01 | 0.45736371 |
| EML4 | 217.552826 | -0.7092775 | 0.25255307 | -2.8084295 | 0.00497838 | 0.06860371 |
| COX7A2L | 64.0535784 | -0.4292281 | 0.51007868 | -0.8414939 | 0.40007132 | 0.64773852 |
| MTA3 | 137.530117 | 0.24261866 | 0.30794803 | 0.78785588 | 0.43078101 | 0.67197671 |
| ZFP36L2 | 178.274468 | 0.34417988 | 0.38383427 | 0.89668877 | 3.70E-01 | 0.62368335 |
| THADA | 237.230293 | -0.1667803 | 0.23053615 | -0.7234454 | 0.46940627 | 0.69910103 |
| PLEKHH2 | 132.561462 | 0.71109556 | 0.40454969 | 1.75774589 | 0.07879076 | 0.27998728 |
| DYNC2LI1 | 95.5430541 | 0.89091715 | 0.5112829 | 1.74251311 | 0.0814187 | 0.28447225 |
| LRPPRC | 184.37996 | 0.05880637 | 0.28185576 | 0.20863993 | 0.83472933 | 0.91994726 |
| PPM1B | 220.101046 | 0.07495875 | 0.19385682 | 0.38667069 | 0.69900002 | 0.85102865 |
| SLC3A1 | 190.41764 | 0.70708544 | 0.40342347 | 1.75271268 | 0.07965134 | 0.28174465 |
| PREPL | 208.959967 | 0.35469338 | 0.19906148 | 1.78182825 | 0.07477724 | 0.2729068 |
| CAMKMT | 619.383899 | 0.02576539 | 0.24808587 | 0.10385674 | 0.91728303 | 0.96401363 |
| SRBD1 | 114.678085 | -0.1248789 | 0.32762052 | -0.3811692 | 0.70307767 | 0.85275589 |
| PRKCE | 545.191565 | -0.7949563 | 0.22742181 | -3.495515 | 0.00047315 | 0.01957967 |
| EPAS1 | 129.106628 | -0.3501153 | 0.5349508 | -0.6544813 | 5.13E-01 | 7.31E-01 |
| RHOQ | 67.5495474 | -0.9827635 | 0.50848448 | -1.9327305 | 0.0532694 | 0.23189906 |
| PIGF | 32.0325373 | 0.27285716 | 0.38896928 | 0.70148768 | 0.48299872 | 0.71069416 |
| CRIPT | 14.3190632 | 1.22057169 | 0.50927848 | 2.39666851 | 0.01654488 | NA |
| SOCS5 | 75.0572803 | -0.3775053 | 0.32744314 | -1.1528882 | 0.24895626 | 0.50506493 |
| MCFD2 | 45.3831474 | 0.43021239 | 0.33600852 | 1.28036155 | 0.20041801 | 0.45451309 |
| TTC7A | 87.9033775 | 0.16461338 | 0.3348073 | 0.49166604 | 0.62295545 | 0.80410667 |
| AC106869.1 | 13.4202433 | -0.5855723 | 0.69423089 | -0.8434836 | 0.39895806 | NA |
| EPCAM | 108.909279 | -0.0656896 | 0.32115367 | -0.2045426 | 0.83792947 | 0.92175835 |
| MSH2 | 98.4724786 | 0.93397236 | 0.43490867 | 2.14751376 | 0.0317524 | 0.17557345 |
| MSH6 | 27.5043812 | 0.2137759 | 0.39633186 | 0.5393861 | 0.58962047 | 0.78422477 |
| FBXO11 | 279.622546 | -0.5237062 | 0.28714373 | -1.823847 | 0.06817521 | 0.25987613 |
| FOXN2 | 41.0020926 | -2.179794 | 0.34633942 | -6.2938086 | 3.10E-10 | 1.19E-06 |
| PPP1R21 | 113.703241 | -0.2237495 | 0.22608877 | -0.9896535 | 0.32234352 | 0.57901472 |
| GPR75-ASB3 | 53.1672352 | 1.08572063 | 0.34371173 | 3.15881168 | 0.00158414 | 0.03639735 |
| ERLEC1 | 31.558542 | 0.29006108 | 0.32880343 | 0.8821717 | 0.37768396 | 0.62906718 |
| PSME4 | 193.933214 | -0.4466062 | 0.28517886 | -1.5660566 | 0.11733537 | 0.34472743 |
| ACYP2 | 266.84996 | -0.574139 | 0.38271835 | -1.5001606 | 0.13357282 | 0.36762309 |
| SPTBN1 | 686.964855 | -1.1995006 | 0.25661169 | -4.6743801 | 2.95E-06 | 0.00076688 |
| EML6 | 52.34588 | 0.67489346 | 0.43468126 | 1.55261688 | 0.12051469 | 0.34903213 |
| RTN4 | 357.418533 | -0.558849 | 0.25509228 | -2.1907719 | 0.0284683 | 0.16587474 |
| CLHC1 | 46.1893544 | 0.51478748 | 0.34769638 | 1.4805661 | 0.13872223 | 0.37517394 |

| | | | | | | |
|---|---|---|---|---|---|---|
| RPS27A | 109.319878 | 0.8314903 | 0.46729259 | 1.77937832 | 0.07517776 | 0.27367272 |
| MTIF2 | 37.101935 | 0.27887495 | 0.35382413 | 0.78817391 | 0.43059498 | 0.67197671 |
| AC012358.3 | 27.6132348 | 1.67006005 | 0.67412576 | 2.47737167 | 0.0132354 | 0.11256671 |
| CCDC88A | 49.6087815 | 0.24967652 | 0.31459758 | 0.79363776 | 0.42740634 | 0.67024083 |
| CFAP36 | 67.2100094 | -0.6236617 | 0.31849173 | -1.9581722 | 0.05020981 | 0.22507781 |
| PPP4R3B | 150.293053 | -0.0651859 | 0.19044766 | -0.3422774 | 0.73214212 | 0.86750276 |
| PNPT1 | 66.9894428 | -0.9351865 | 0.41876445 | -2.2332042 | 0.02553548 | 0.15648615 |
| EFEMP1 | 109.78772 | 0.48604658 | 0.54540806 | 0.89116133 | 0.37284262 | 0.62543211 |
| VRK2 | 119.292227 | 0.29968019 | 0.27852347 | 1.07596026 | 0.28194503 | 0.54064048 |
| FANCL | 83.7342943 | 0.56623894 | 0.43844482 | 1.29147138 | 0.19654027 | 0.44992056 |
| PAPOLG | 40.0146214 | -0.3468441 | 0.29984082 | -1.1567607 | 0.24737015 | 0.50358016 |
| REL | 51.2646584 | -0.7673458 | 0.53089872 | -1.4453713 | 0.14835361 | 0.38865817 |
| PUS10 | 29.4104374 | 0.00822436 | 0.34563729 | 0.02379478 | 0.98101631 | 0.99304653 |
| PEX13 | 45.2105617 | -0.3122772 | 0.32219244 | -0.9692256 | 0.33243266 | 0.58794039 |
| KIAA1841 | 20.5048375 | -0.2507443 | 0.52788601 | -0.4749971 | 0.63478904 | NA |
| C2orf74 | 13.7813864 | 0.73896039 | 0.55256864 | 1.33731873 | 0.18111862 | NA |
| USP34 | 658.099202 | -0.481516 | 0.18314425 | -2.6291625 | 0.00855954 | 0.09049837 |
| AC016727.1 | 14.4216415 | -0.0223749 | 0.57416815 | -0.0389693 | 0.96891486 | NA |
| XPO1 | 257.297356 | -0.0458105 | 0.23535499 | -0.1946442 | 0.84567152 | 0.92531372 |
| FAM161A | 19.5590676 | 1.45410305 | 0.4584873 | 3.1715231 | 0.00151642 | NA |
| CCT4 | 50.2453929 | -0.1266045 | 0.36321997 | -0.3485615 | 0.72741854 | 0.86461401 |
| COMMD1 | 260.194108 | -0.310495 | 0.20530854 | -1.5123338 | 0.13044897 | 0.36313409 |
| AC018462.1 | 17.755405 | -0.4357054 | 0.58287616 | -0.7475094 | 0.45475611 | NA |
| B3GNT2 | 60.4629844 | 0.25887342 | 0.43078022 | 0.60094083 | 0.5478794 | 0.7554786 |
| EHBP1 | 305.394179 | -0.2806463 | 0.24612073 | -1.1402789 | 0.25417013 | 0.51173097 |
| WDPCP | 195.619361 | 0.40189066 | 0.39133076 | 1.0269846 | 0.30442771 | 0.56489395 |
| MDH1 | 67.5448194 | 0.83277138 | 0.25677186 | 3.24323452 | 0.00118181 | 0.03062755 |
| UGP2 | 109.193564 | -0.1583684 | 0.25171959 | -0.629146 | 0.52925345 | 0.74350499 |
| VPS54 | 155.683092 | 0.24836505 | 0.23534761 | 1.05531154 | 0.29128286 | 0.55045524 |
| PELI1 | 54.7895794 | -0.9592244 | 0.39153479 | -2.4499085 | 0.01428925 | 0.11775628 |
| AC008074.1 | 29.7410336 | -1.3805459 | 0.56152301 | -2.458574 | 0.013949 | 0.11594544 |
| AFTPH | 158.162259 | -0.4141832 | 0.23059054 | -1.7961847 | 0.07246515 | 0.26831077 |
| SERTAD2 | 33.4512709 | -0.5689391 | 0.37532406 | -1.515861 | 0.1295545 | 0.36208459 |
| CEP68 | 23.5744481 | -0.3758024 | 0.36362829 | -1.0334796 | 0.30137952 | 0.56180195 |
| RAB1A | 222.729481 | -0.8263651 | 0.33344204 | -2.4782872 | 0.01320148 | 0.11240246 |
| ACTR2 | 122.085465 | -0.2450935 | 0.26070062 | -0.940134 | 0.34714881 | 0.6039793 |
| SPRED2 | 189.915697 | -0.1586554 | 0.23880134 | -0.6643823 | 0.50644565 | 0.7281071 |
| LINC01828 | 85.6055555 | -1.0535109 | 1.39921705 | -0.7529289 | 4.51E-01 | 6.89E-01 |
| ETAA1 | 27.4542193 | 0.04543497 | 0.38639561 | 0.11758666 | 0.90639517 | 0.9580376 |
| C1D | 34.7319982 | 0.29544256 | 0.36535908 | 0.80863616 | 0.41872446 | 0.66260735 |
| APLF | 59.5421833 | 0.53905474 | 0.31718436 | 1.69949973 | 0.08922507 | 0.29794449 |
| ANTXR1 | 23.455802 | 0.30772618 | 0.42008054 | 0.73254092 | 0.46383848 | 0.69499022 |
| GFPT1 | 134.559694 | 0.21403514 | 0.26848876 | 0.79718473 | 4.25E-01 | 6.68E-01 |
| NFU1 | 48.2522135 | 0.03600006 | 0.33700782 | 0.10682262 | 0.9149297 | 0.96388328 |
| AAK1 | 335.206277 | 0.01841697 | 0.19773661 | 0.09313888 | 0.92579323 | 0.96902431 |
| ANXA4 | 523.946852 | -0.8962376 | 0.57379163 | -1.5619566 | 0.11829822 | 0.34566454 |
| GMCL1 | 97.3815696 | -1.0086933 | 0.45447622 | -2.2194633 | 0.02645522 | 0.15970654 |
| MXD1 | 48.807262 | -0.333466 | 0.37194597 | -0.8965441 | 0.36996224 | 0.62368335 |
| PCBP1 | 37.8389715 | -0.1550013 | 0.45660797 | -0.3394626 | 0.73426127 | 0.86800937 |
| C2orf42 | 20.7847329 | -0.2567822 | 0.52108893 | -0.49278 | 0.62216807 | NA |
| PCYOX1 | 38.5384037 | 2.31308911 | 0.57337505 | 4.03416425 | 5.48E-05 | 0.00479287 |
| SNRPG | 25.7702644 | -0.1959914 | 0.36525317 | -0.5365905 | 0.59155051 | 0.78517152 |
| FAM136A | 18.8952339 | 0.25423659 | 0.45413684 | 0.55982376 | 0.57559966 | NA |
| TGFA | 60.3092789 | -0.1803697 | 0.53642923 | -0.3362414 | 0.7366888 | 0.86914373 |

| | | | | | | |
|---|---|---|---|---|---|---|
| AC007040.2 | 46.020087 | 2.47526622 | 0.8938004 | 2.76937247 | 0.00561644 | 0.07314675 |
| NAGK | 18.2043713 | -0.7374943 | 0.46643077 | -1.5811441 | 0.11384508 | NA |
| MCEE | 41.1656065 | -0.039689 | 0.43209933 | -0.0918516 | 0.92681596 | 0.9695235 |
| MPHOSPH10 | 24.911671 | -0.7112487 | 0.39074869 | -1.8202203 | 0.06872546 | 0.2608382 |
| PAIP2B | 27.4225263 | -1.0816777 | 0.51936783 | -2.0826813 | 0.03728028 | 0.19296996 |
| ZNF638 | 554.807594 | -0.1190992 | 0.17142145 | -0.6947744 | 0.48719668 | 0.7138688 |
| EXOC6B | 612.19228 | -0.6909081 | 0.43892947 | -1.5740754 | 0.11547002 | 0.34185848 |
| EMX1 | 175.722901 | 0.4901406 | 0.49052433 | 0.99921773 | 0.31768923 | 0.57515952 |
| SFXN5 | 198.339587 | 0.20149207 | 0.28186068 | 0.71486405 | 0.47469305 | 0.70288437 |
| RAB11FIP5 | 49.3147979 | -0.1409255 | 0.42881673 | -0.328638 | 0.74242931 | 0.87152705 |
| CCT7 | 27.9087949 | 0.91018525 | 0.39919294 | 2.28006348 | 0.02260392 | 0.14632666 |
| ALMS1 | 101.624111 | -0.2575187 | 0.20474064 | -1.2577803 | 0.20847124 | 0.46255495 |
| DUSP11 | 20.6844694 | 0.02842488 | 0.44749056 | 0.06352063 | 0.94935193 | NA |
| STAMBP | 58.5065002 | -0.3788096 | 0.26293731 | -1.4406841 | 0.14967394 | 0.39012541 |
| DGUOK | 50.9009049 | -0.5400276 | 0.28988711 | -1.8628893 | 0.06247785 | 0.24933194 |
| TET3 | 75.2178982 | -0.9866099 | 0.5327816 | -1.8518093 | 0.06405321 | 0.25205396 |
| MOB1A | 53.0680137 | -0.0277416 | 0.35684125 | -0.0777422 | 0.93803312 | 0.97466007 |
| AC006030.1 | 22.7211738 | -0.1239871 | 0.47365725 | -0.2617654 | 0.79350231 | 0.89875118 |
| DCTN1 | 13.58214 | 0.19945031 | 0.52192805 | 0.38214139 | 0.7023565 | NA |
| RTKN | 34.1812674 | -0.1303405 | 0.46263376 | -0.2817359 | 0.77814603 | 0.88986706 |
| MOGS | 24.0635326 | 0.09805348 | 0.48173878 | 0.20354076 | 0.83871239 | 0.92222418 |
| CCDC142 | 10.3674596 | -0.5307396 | 0.70675188 | -0.7509561 | 0.45267907 | NA |
| TTC31 | 12.4390195 | 0.01157357 | 0.49485655 | 0.02338772 | 0.981341 | NA |
| AUP1 | 18.7955676 | -0.3095562 | 0.44892325 | -0.6895525 | 0.49047563 | NA |
| POLE4 | 17.5916605 | 0.80897408 | 0.46475664 | 1.74064016 | 0.08174667 | NA |
| TACR1 | 17.6064129 | 2.47755776 | 0.92695323 | 2.67279694 | 0.00752218 | NA |
| AC007681.1 | 26.4530146 | 1.4860361 | 0.70124372 | 2.11914353 | 0.03407834 | 0.18312899 |
| EVA1A | 65.1944139 | -0.4674048 | 0.41371114 | -1.1297853 | 0.25856669 | 0.51706621 |
| MRPL19 | 67.8437012 | 0.99489787 | 0.36951166 | 2.69246677 | 0.00709256 | 0.08221602 |
| GCFC2 | 122.663824 | 0.38456026 | 0.2772236 | 1.38718441 | 0.16538554 | 0.41243204 |
| SUCLG1 | 98.3687327 | 0.66743303 | 0.33890854 | 1.9693603 | 0.04891173 | 0.22162876 |
| DNAH6 | 33.5792308 | 2.70969446 | 0.92796984 | 2.92002427 | 0.00350004 | 0.05869303 |
| TMSB10 | 124.912461 | -0.1172929 | 0.44932203 | -0.2610441 | 0.79405851 | 0.89884908 |
| KCMF1 | 121.608931 | -0.2147322 | 0.28792088 | -0.7458027 | 0.45578661 | 0.6908246 |
| TGOLN2 | 89.9522237 | 0.19712876 | 0.39686822 | 0.49671088 | 0.61939295 | 0.8027391 |
| RETSAT | 13.0474722 | -0.039311 | 0.49675344 | -0.0791358 | 0.93692464 | NA |
| ELMOD3 | 45.9645843 | 0.19609474 | 0.30355409 | 0.64599604 | 0.51828192 | 0.73536015 |
| CAPG | 17.5346603 | -0.0308678 | 0.62577513 | -0.0493272 | 0.96065851 | NA |
| MAT2A | 116.549758 | -1.8272689 | 0.43961682 | -4.1565037 | 3.23E-05 | 0.00345462 |
| GGCX | 25.547051 | -0.6883599 | 0.47193787 | -1.4585816 | 0.1446803 | 0.38426647 |
| VAMP8 | 44.1259599 | 0.32223928 | 0.35112484 | 0.91773422 | 0.35875803 | 0.61513935 |
| USP39 | 55.0859367 | -0.0004712 | 0.39126534 | -0.0012042 | 0.99903918 | 0.99968858 |
| C2orf68 | 27.7967533 | -0.1483706 | 0.37631898 | -0.3942681 | 0.69338308 | 0.84791601 |
| POLR1A | 37.8406144 | 0.08636727 | 0.31842002 | 0.27123695 | 0.7862088 | 0.89452315 |
| PTCD3 | 90.8512629 | 0.06121633 | 0.23869825 | 0.25645906 | 0.79759639 | 0.90121835 |
| IMMT | 110.774468 | -0.1123876 | 0.26786786 | -0.4195637 | 0.67480419 | 0.83760166 |
| MRPL35 | 15.0860518 | 0.28504965 | 0.48250886 | 0.59076564 | 0.55467746 | NA |
| KDM3A | 92.3581886 | 0.09127933 | 0.35080405 | 0.26020032 | 0.79470925 | 0.89901694 |
| CHMP3 | 15.6203002 | 0.77419836 | 0.49317513 | 1.56982441 | 0.11645597 | NA |
| RMND5A | 54.1840969 | 0.05536701 | 0.3862351 | 0.14335054 | 0.88601334 | 0.94717288 |
| PLGLB1 | 19.5376351 | 0.5114048 | 0.51762624 | 0.98798082 | 0.32316204 | NA |
| CYTOR | 60.2756752 | -2.4389248 | 0.36504889 | -6.6810909 | 2.37E-11 | 1.83E-07 |
| KRCC1 | 62.9011044 | 0.02112766 | 0.38772898 | 0.05449079 | 0.95654415 | 0.98284879 |
| THNSL2 | 12.641161 | 0.70731622 | 0.8985772 | 0.78715131 | 0.43119329 | NA |

| | | | | | | |
|---|---|---|---|---|---|---|
| EIF2AK3 | 97.0291306 | -1.3107477 | 0.30306772 | -4.3249335 | 1.53E-05 | 0.00207296 |
| RPIA | 33.5683722 | 0.81619379 | 0.35793067 | 2.28031251 | 0.02258916 | 0.14632666 |
| MAL | 211.477447 | 1.63004884 | 0.61546265 | 2.64849353 | 0.00808514 | 0.08827138 |
| MRPS5 | 93.8587938 | 0.10881665 | 0.28635728 | 0.38000309 | 0.70394312 | 0.85295217 |
| AC092835.1 | 16.0770937 | 0.35646327 | 0.46427562 | 0.76778373 | 0.44261568 | NA |
| PROM2 | 54.1186737 | 1.34955364 | 0.55400042 | 2.43601553 | 0.01485005 | 0.12019013 |
| KCNIP3 | 23.2055788 | 0.36060806 | 0.58797812 | 0.61330184 | 0.53967678 | 0.74929145 |
| FAHD2A | 13.7286988 | 1.09391707 | 0.56182546 | 1.94707635 | 0.05152558 | NA |
| ANKRD36C | 57.8500629 | 0.9255737 | 0.46121494 | 2.00681641 | 0.04476922 | 0.21170763 |
| STARD7 | 54.0990179 | 0.14871128 | 0.33810718 | 0.43983472 | 0.66005682 | 0.82676279 |
| TMEM127 | 30.7684809 | 0.4069021 | 0.40387654 | 1.00749132 | 0.31369873 | 0.57113617 |
| CIAO1 | 38.868185 | -0.1686773 | 0.34582413 | -0.4877544 | 0.62572379 | 0.8053173 |
| SNRNP200 | 63.2205412 | -0.6244303 | 0.39101571 | -1.5969443 | 0.11027812 | 0.33347479 |
| KANSL3 | 111.609732 | -0.7236251 | 0.21641025 | -3.3437654 | 0.0008265 | 0.02565549 |
| FER1L5 | 12.9392506 | 0.1203931 | 0.53993557 | 0.22297679 | 0.82355357 | NA |
| LMAN2L | 29.2826033 | 0.61903866 | 0.35052404 | 1.76603767 | 0.07738953 | 0.27731248 |
| CNNM4 | 24.0666954 | -0.397823 | 0.35859529 | -1.1093926 | 0.26726086 | 0.52637204 |
| ANKRD23 | 16.8194493 | 1.20433955 | 0.47979702 | 2.51010217 | 0.01206962 | NA |
| FAHD2B | 11.105899 | 0.91534044 | 0.57882143 | 1.58138659 | 0.11378967 | NA |
| ANKRD36 | 77.9785738 | 0.38686155 | 0.4109947 | 0.94128111 | 0.34656082 | 0.60363852 |
| AC092683.1 | 112.899741 | 0.45302883 | 0.42750426 | 1.05970599 | 0.28927838 | 0.54868794 |
| ANKRD36B | 33.3363549 | 0.42747383 | 0.37365508 | 1.14403324 | 0.25260985 | 0.5096561 |
| COX5B | 75.4501664 | 0.65702752 | 0.3093939 | 2.12359563 | 0.03370397 | 0.18191827 |
| ACTR1B | 29.5238345 | -0.3784357 | 0.42558924 | -0.8892042 | 0.37389336 | 0.62575308 |
| C2orf92 | 46.6655527 | -2.3026032 | 0.78863534 | -2.9197312 | 0.00350333 | 0.05869303 |
| TMEM131 | 242.633801 | -0.4331465 | 0.21585869 | -2.0066204 | 0.04479009 | 0.21170763 |
| INPP4A | 60.154591 | -0.4187786 | 0.27056783 | -1.5477767 | 0.12167607 | 0.35089572 |
| COA5 | 18.6825609 | 0.72356201 | 0.42345453 | 1.70871242 | 0.08750423 | NA |
| UNC50 | 25.3325275 | 0.51747608 | 0.46471868 | 1.11352545 | 0.26548282 | 0.524719 |
| MGAT4A | 65.4915426 | 0.85578647 | 0.48074433 | 1.78012804 | 0.07505501 | 0.27353144 |
| KIAA1211L | 19.6222064 | 0.62611247 | 0.76517117 | 0.81826458 | 0.41320613 | NA |
| TSGA10 | 139.899515 | 0.68568204 | 0.34595208 | 1.98201453 | 0.04747761 | 0.21778021 |
| AC079447.1 | 60.3701263 | -0.0655587 | 0.38166468 | -0.1717704 | 0.86361801 | 0.93643831 |
| MITD1 | 38.3972627 | 0.33158585 | 0.30625964 | 1.08269522 | 0.27894373 | 0.53717915 |
| TXNDC9 | 26.647117 | 0.16337483 | 0.46070636 | 0.35461813 | 0.72287569 | 0.86192273 |
| EIF5B | 94.8596536 | -0.6453326 | 0.27268183 | -2.3666139 | 0.01795165 | 0.13297634 |
| REV1 | 238.53983 | 0.01597013 | 0.19372411 | 0.08243749 | 0.93429882 | 0.97327579 |
| LONRF2 | 30.8354838 | 0.98632728 | 0.59150214 | 1.66749571 | 0.09541588 | 0.31001098 |
| NPAS2 | 308.798996 | -1.1345659 | 0.55055594 | -2.0607641 | 0.03932555 | 0.19770658 |
| RPL31 | 58.1485589 | 0.19073444 | 0.37119204 | 0.51384303 | 0.60736174 | 0.79545062 |
| TBC1D8 | 223.512074 | -0.4147235 | 0.30612148 | -1.3547679 | 0.17549152 | 0.4243664 |
| CNOT11 | 73.2545972 | -1.343754 | 0.4174142 | -3.2192339 | 0.00128534 | 0.03201692 |
| RNF149 | 255.957172 | -0.8422547 | 0.21139872 | -3.9841997 | 6.77E-05 | 0.00530264 |
| MAP4K4 | 133.854426 | -0.9041388 | 0.38478355 | -2.3497334 | 0.01878686 | 0.13549695 |
| IL1R2 | 19.3147076 | -3.3103057 | 0.65645875 | -5.0426713 | 4.59E-07 | NA |
| IL1R1 | 88.9919611 | -1.5471945 | 0.38483652 | -4.0203942 | 5.81E-05 | 0.00496891 |
| IL1RL2 | 48.0183767 | -1.5233713 | 0.74362869 | -2.0485644 | 0.04050473 | 0.20046817 |
| SLC9A2 | 27.3325089 | 1.32589461 | 0.55931743 | 2.37055837 | 0.01776124 | 0.13259773 |
| MFSD9 | 13.7507671 | 0.0215455 | 0.45490705 | 0.04736241 | 0.96222438 | NA |
| TMEM182 | 31.0801568 | 0.49691546 | 0.40462406 | 1.22809174 | 0.21941253 | 0.4758575 |
| POU3F3 | 82.8774869 | 0.14136074 | 0.33610256 | 0.42058811 | 0.67405588 | 0.83715334 |
| MRPS9 | 80.1909614 | 0.61070989 | 0.35670148 | 1.71210362 | 0.08687757 | 0.29483981 |
| TGFBRAP1 | 25.4796018 | -0.5206291 | 0.37452552 | -1.390103 | 0.16449759 | 0.41121726 |
| C2orf49 | 18.2606268 | -0.5586156 | 0.45508322 | -1.227502 | 0.21963396 | NA |

| | | | | | | |
|---|---|---|---|---|---|---|
| NCK2 | 93.4110242 | -0.0784955 | 0.29038307 | -0.2703171 | 0.78691633 | 0.89489881 |
| UXS1 | 127.830278 | -0.4740592 | 0.2618896 | -1.8101489 | 7.03E-02 | 0.26462278 |
| LINC01594 | 36.810031 | 0.07783232 | 0.53508378 | 0.1454582 | 0.88434912 | 0.94605075 |
| SULT1C2 | 348.042013 | -0.4478484 | 0.42846014 | -1.0452511 | 0.29590695 | 0.55551117 |
| GCC2 | 257.865562 | 0.05130801 | 0.24352687 | 0.21068727 | 0.83313131 | 0.91910731 |
| GCC2-AS1 | 25.2641834 | 1.64029332 | 0.7086239 | 2.31475869 | 0.02062614 | 0.14099415 |
| LIMS1 | 364.594987 | -0.1771448 | 0.28458132 | -0.6224751 | 0.53362953 | 0.74672888 |
| RANBP2 | 139.170435 | -0.0595345 | 0.26215719 | -0.2270945 | 0.82035024 | 0.91338577 |
| CCDC138 | 37.3067358 | -0.1599028 | 0.48070588 | -0.3326416 | 0.73940485 | 0.87054913 |
| SH3RF3 | 30.7393783 | -1.1502608 | 0.90720521 | -1.2679169 | 0.20482764 | 0.45803555 |
| 10-Sep | 242.248636 | -1.5067599 | 0.46627033 | -3.2315157 | 1.23E-03 | 3.12E-02 |
| NPHP1 | 44.2925663 | 1.41899221 | 0.63479269 | 2.23536319 | 0.02539351 | 0.15636311 |
| SMIM37 | 21.4607343 | 1.29675209 | 0.61371045 | 2.11297051 | 0.03460329 | 0.18447938 |
| ACOXL | 25.1984755 | 1.30166511 | 0.68959591 | 1.88757661 | 0.05908281 | 0.24138026 |
| MIR4435-2H | 338.066897 | -0.788185 | 0.34316136 | -2.2968349 | 2.16E-02 | 1.43E-01 |
| BCL2L11 | 54.0165895 | -0.5384983 | 0.40598431 | -1.3264018 | 0.18470666 | 0.43476148 |
| ANAPC1 | 87.2150914 | 0.07760544 | 0.27799797 | 0.2791583 | 0.78012334 | 0.89075944 |
| MERTK | 72.7003925 | 0.12829282 | 0.31660362 | 0.40521591 | 0.68531881 | 0.84355496 |
| TMEM87B | 76.0525188 | -0.3333343 | 0.22397357 | -1.4882753 | 0.13667831 | 0.37226218 |
| ZC3H8 | 63.6323561 | -0.1328727 | 0.30886849 | -0.4301919 | 0.66705607 | 0.83187469 |
| ZC3H6 | 75.5189545 | 1.35359509 | 0.5122192 | 2.64260907 | 0.008227 | 0.08906215 |
| TTL | 36.1897253 | 0.11677464 | 0.3970911 | 0.29407518 | 0.76870045 | 0.88546654 |
| POLR1B | 17.0587562 | -0.2110059 | 0.496045 | -0.4253764 | 0.67056228 | NA |
| SLC20A1 | 191.503379 | -1.0201406 | 0.50038648 | -2.0387054 | 0.04147944 | 0.20309619 |
| PSD4 | 36.74901 | -0.4297699 | 0.36003571 | -1.1936868 | 0.23260052 | 0.49082729 |
| PAX8 | 801.145814 | -0.6091556 | 0.39618361 | -1.5375589 | 0.1241565 | 0.35457643 |
| CBWD2 | 39.7761452 | -0.2746476 | 0.33167712 | -0.8280571 | 0.40763818 | 0.65378997 |
| RABL2A | 27.5185619 | 0.08014696 | 0.41093947 | 0.19503349 | 0.84536673 | 0.92531372 |
| AC017074.1 | 27.3321485 | -0.1910105 | 0.52830644 | -0.3615525 | 0.71768649 | 0.8593704 |
| SLC35F5 | 418.221912 | -0.8188146 | 0.16817357 | -4.8688659 | 1.12E-06 | 0.00045469 |
| ACTR3 | 194.271036 | -0.4649999 | 0.20361916 | -2.2836748 | 0.02239065 | 0.14604538 |
| DDX18 | 40.0130037 | -0.4286813 | 0.33275131 | -1.2882933 | 0.19764389 | 0.45090075 |
| CCDC93 | 128.418606 | -0.727061 | 0.31499004 | -2.308203 | 0.02098785 | 0.14220374 |
| INSIG2 | 65.5003079 | -0.4004813 | 0.54876889 | -0.7297813 | 0.46552386 | 0.69602508 |
| C2orf76 | 22.9428425 | -0.41748 | 0.41026706 | -1.017581 | 0.30887712 | 0.56849049 |
| DBI | 67.4426915 | 0.90619209 | 0.47128327 | 1.9228183 | 0.05450287 | 0.23331959 |
| TMEM37 | 63.8394924 | 1.09994263 | 0.60820434 | 1.80850836 | 0.07052742 | 0.26532235 |
| SCTR | 41.1427231 | 2.53967303 | 0.89257812 | 2.84532297 | 0.00443664 | 0.0642158 |
| AC013275.1 | 17.3078698 | 1.5996307 | 0.82721926 | 1.93374452 | 0.05314455 | NA |
| CFAP221 | 192.595356 | 1.17021921 | 0.54111848 | 2.16259333 | 0.03057247 | 0.17296809 |
| PTPN4 | 437.419848 | -0.1586295 | 0.40973494 | -0.3871515 | 0.69864402 | 0.85102865 |
| EPB41L5 | 340.830706 | -0.5010881 | 0.29505632 | -1.6982795 | 0.08945502 | 0.29807849 |
| RALB | 29.3505231 | -0.1302816 | 0.33807577 | -0.3853622 | 0.6999691 | 0.85153503 |
| TFCP2L1 | 602.053013 | -0.1237262 | 0.47520227 | -0.2603654 | 0.79458192 | 0.89901694 |
| CLASP1 | 868.346513 | -0.1272703 | 0.32337699 | -0.3935664 | 0.69390118 | 0.8481749 |
| NIFK-AS1 | 31.7866234 | -0.1775506 | 0.501923 | -0.3537407 | 0.72353319 | 0.86194629 |
| TSN | 26.4934635 | -0.2865357 | 0.37004795 | -0.7743204 | 0.43874132 | 0.67892881 |
| ERCC3 | 39.6814787 | -0.6878626 | 0.33766485 | -2.0371163 | 0.04163839 | 0.20335702 |
| MAP3K2 | 182.232842 | -0.3874738 | 0.21753249 | -1.7812229 | 0.07487604 | 0.27312881 |
| IWS1 | 173.942634 | -0.5658311 | 0.2414439 | -2.3435305 | 0.0191022 | 0.1359475 |
| AC010976.1 | 17.540281 | 0.4118455 | 0.70265235 | 0.58612983 | 0.55778827 | NA |
| WDR33 | 213.268339 | -0.3358993 | 0.2755929 | -1.2188243 | 0.22291089 | 0.48006298 |
| POLR2D | 13.5538381 | 0.26187762 | 0.45069472 | 0.58105323 | 0.56120458 | NA |
| AMMECR1L | 27.131059 | -0.2509396 | 0.40043522 | -0.626667 | 0.53087751 | 0.74456344 |

| | | | | | | |
|---|---|---|---|---|---|---|
| SAP130 | 68.2959393 | -0.8871207 | 0.33357889 | -2.6594029 | 0.00782793 | 0.08671693 |
| UGGT1 | 62.6940589 | -0.0217116 | 0.34479599 | -0.0629693 | 0.94979094 | 0.97993174 |
| HS6ST1 | 81.6152106 | 0.62116939 | 0.38788166 | 1.60144046 | 1.09E-01 | 3.32E-01 |
| SMPD4 | 42.7543663 | -0.5083296 | 0.33079115 | -1.5367087 | 0.12436465 | 0.35460505 |
| MZT2B | 33.6972984 | 0.54274933 | 0.43471489 | 1.24851793 | 0.21184145 | 0.46653608 |
| PTPN18 | 23.1700443 | 0.45320537 | 0.51560408 | 0.87897941 | 0.37941244 | 0.63087871 |
| FAM168B | 108.478588 | -0.7080847 | 0.31780617 | -2.2280396 | 0.02587787 | 0.15729431 |
| PLEKHB2 | 98.4482557 | -0.2670895 | 0.33796708 | -0.7902824 | 0.42936284 | 0.67170849 |
| MZT2A | 24.6673422 | 1.11281462 | 0.43934229 | 2.53291031 | 0.01131199 | 0.1036885 |
| ANKRD30BL | 14.2256884 | 0.45344149 | 0.56912049 | 0.79674074 | 0.42560162 | NA |
| GPR39 | 132.398188 | -0.1661248 | 0.34434238 | -0.4824407 | 0.62949288 | 0.80726536 |
| NCKAP5 | 210.053006 | -1.1869217 | 0.55831534 | -2.1258984 | 0.03351172 | 0.1813922 |
| MGAT5 | 181.349278 | -0.0462887 | 0.24135519 | -0.1917866 | 0.84790938 | 0.92651313 |
| TMEM163 | 39.5769283 | -0.804526 | 1.30254529 | -0.6176568 | 0.53680159 | 0.7486432 |
| CCNT2-AS1 | 44.2368052 | -1.1962621 | 0.59302818 | -2.0172094 | 0.04367366 | 0.20890044 |
| ACMSD | 28.6552482 | -0.2812612 | 0.91375412 | -0.3078084 | 0.7582281 | 0.87985552 |
| CCNT2 | 137.581072 | -0.328588 | 0.26596477 | -1.2354568 | 0.21666054 | 0.47286334 |
| RAB3GAP1 | 192.189411 | -0.0297473 | 0.21817827 | -0.1363438 | 0.89154948 | 0.95026032 |
| ZRANB3 | 85.8593507 | 0.15908655 | 0.33037197 | 0.48153767 | 0.63013441 | 0.80795345 |
| R3HDM1 | 197.80239 | 0.16238442 | 0.199507 | 0.81392845 | 0.41568594 | 0.65929007 |
| UBXN4 | 180.762578 | -0.7109244 | 0.25635712 | -2.7731797 | 0.00555115 | 0.07254189 |
| DARS | 167.095878 | -0.1930583 | 0.215006 | -0.8979206 | 0.36922789 | 0.6236443 |
| THSD7B | 18.9248008 | 0.01943145 | 0.71468025 | 0.02718901 | 0.97830898 | NA |
| HNMT | 82.3525356 | 1.07133905 | 0.46758994 | 2.29119353 | 0.02195222 | 0.14478686 |
| AC114763.1 | 27.0336946 | 0.94516865 | 0.55200729 | 1.71223944 | 0.08685255 | 0.29483981 |
| SPOPL | 81.9732324 | 0.13469673 | 0.23800817 | 0.56593325 | 0.57143916 | 0.77245539 |
| LRP1B | 738.258554 | 0.9556206 | 0.59927353 | 1.59463174 | 0.11079461 | 0.33416384 |
| GTDC1 | 88.1795164 | 0.02908877 | 0.31399224 | 0.09264169 | 0.92618822 | 0.96925503 |
| TEX41 | 530.448126 | 0.27963715 | 0.47941707 | 0.58328575 | 5.60E-01 | 7.65E-01 |
| ACVR2A | 102.113981 | -0.090671 | 0.40953522 | -0.2213997 | 0.8247812 | 0.91537186 |
| ORC4 | 150.329214 | 0.14078418 | 0.21347314 | 0.65949363 | 5.10E-01 | 0.72990216 |
| MBD5 | 740.029229 | 0.17335371 | 0.29648157 | 0.58470315 | 0.55874734 | 0.76483696 |
| EPC2 | 179.913317 | -0.3268827 | 0.30277012 | -1.0796399 | 0.28030257 | 0.53856437 |
| LYPD6 | 23.4392392 | -0.7110232 | 0.7706718 | -0.9226018 | 0.35621475 | 0.61227891 |
| MMADHC | 28.6848264 | -0.0203697 | 0.41423698 | -0.049174 | 0.96078062 | 0.98496344 |
| AC007364.1 | 30.5583775 | -1.4534718 | 0.70200499 | -2.0704579 | 0.03840948 | 0.19552762 |
| RND3 | 34.6366426 | -2.5872552 | 0.73735557 | -3.5088298 | 0.00045008 | 0.01893054 |
| NMI | 16.1534194 | 0.37354255 | 0.67963249 | 0.54962432 | 0.58257707 | NA |
| RIF1 | 129.014007 | -0.6832381 | 0.2520042 | -2.7112169 | 0.00670368 | 0.07974991 |
| NEB | 19.2424563 | 0.19374101 | 0.58645533 | 0.33035937 | 0.74112844 | NA |
| ARL5A | 49.4578341 | 0.1835621 | 0.32851803 | 0.55875807 | 0.57632684 | 0.77421002 |
| CACNB4 | 544.840118 | -0.1261094 | 0.31772507 | -0.3969137 | 6.91E-01 | 8.47E-01 |
| STAM2 | 142.506585 | -0.1428851 | 0.26964561 | -0.5298995 | 0.59618163 | 0.78913289 |
| FMNL2 | 492.903384 | -0.8597019 | 0.33372184 | -2.5761031 | 9.99E-03 | 9.90E-02 |
| PRPF40A | 111.687851 | -0.1388357 | 0.22525047 | -0.6163612 | 0.53765613 | 0.74878549 |
| ARL6IP6 | 50.4384559 | -0.1610811 | 0.43695389 | -0.3686456 | 0.71239189 | 0.857224 |
| GALNT13 | 31.6910432 | 1.92508552 | 0.7403869 | 2.6001075 | 0.00931946 | 0.09528654 |
| KCNJ3 | 128.098378 | 0.3574937 | 0.46318608 | 0.77181441 | 0.44022435 | 0.68012984 |
| LINC01876 | 126.744187 | 1.40513436 | 1.18192735 | 1.18885003 | 0.23449869 | 0.49243138 |
| GPD2 | 93.4565747 | 0.07457189 | 0.29425994 | 0.25342183 | 0.79994226 | 0.90280874 |
| CYTIP | 19.8007943 | 2.37248974 | 0.74350637 | 3.19094742 | 0.00141807 | NA |
| ACVR1C | 13.6470038 | 1.02897977 | 0.59465898 | 1.73036949 | 0.08356428 | NA |
| ACVR1 | 164.720786 | -0.4245121 | 0.18591047 | -2.2834224 | 0.0224055 | 0.14604538 |
| UPP2 | 52.581794 | 1.85885127 | 0.66752494 | 2.78469185 | 0.00535786 | 0.07122532 |

| | | | | | | |
|---|---|---|---|---|---|---|
| CCDC148 | 356.230189 | 0.84710287 | 0.39518535 | 2.14355839 | 3.21E-02 | 0.1766407 |
| PKP4 | 1126.89727 | -0.2262163 | 0.23884824 | -0.947113 | 3.44E-01 | 6.01E-01 |
| PKP4-AS1 | 27.0548377 | 0.51803018 | 0.69797276 | 0.74219254 | 0.45797069 | 0.69231672 |
| TANC1 | 375.13787 | -0.3507009 | 0.17535958 | -1.9998957 | 0.04551153 | 0.21335013 |
| WDSUB1 | 80.9043536 | 0.04235277 | 0.27124938 | 0.15613959 | 0.87592299 | 0.94201191 |
| BAZ2B | 599.612295 | -0.0914682 | 0.15134178 | -0.6043817 | 0.5455899 | 0.75352691 |
| AC009961.1 | 29.6777454 | 0.35915358 | 0.56127194 | 0.63989227 | 0.52224264 | 0.7380123 |
| 7-Mar | 211.060316 | -0.5684653 | 0.29404941 | -1.9332304 | 0.05320783 | 0.23177172 |
| PLA2R1 | 44.5702926 | 0.6008202 | 0.32401678 | 1.85428728 | 6.37E-02 | 0.25118809 |
| ITGB6 | 361.338407 | -0.4275395 | 0.55777904 | -0.7665033 | 4.43E-01 | 6.83E-01 |
| RBMS1 | 505.858948 | -0.9986269 | 0.29889319 | -3.3410829 | 0.00083452 | 0.0256933 |
| TANK | 149.616955 | -0.4000582 | 0.26643833 | -1.501504 | 0.13322524 | 0.36753932 |
| PSMD14 | 66.7899174 | -0.4696525 | 0.34036388 | -1.3798541 | 0.16763158 | 0.41581059 |
| IFIH1 | 29.4184633 | -0.0763688 | 0.65025174 | -0.117445 | 0.90650745 | 0.9580376 |
| GCA | 48.4484624 | -0.2610853 | 0.35116521 | -0.7434829 | 0.45718936 | 0.69176066 |
| KCNH7 | 137.974986 | 0.86951938 | 0.78362111 | 1.10961709 | 0.26716406 | 0.52632757 |
| FIGN | 19.5318788 | 1.06790016 | 0.66914362 | 1.59592071 | 0.1105065 | NA |
| GRB14 | 488.895987 | 0.52580168 | 0.22493123 | 2.33761083 | 1.94E-02 | 1.37E-01 |
| COBLL1 | 2465.0763 | -0.0116636 | 0.24619063 | -0.0473762 | 0.96221336 | 0.98535004 |
| AC019197.1 | 5568.98734 | 1.44264449 | 0.43795826 | 3.29402276 | 0.00098765 | 0.02847155 |
| SLC38A11 | 56.3053887 | 1.18494319 | 0.70376955 | 1.68370908 | 0.09223791 | 0.30444046 |
| SCN3A | 186.697013 | 2.34394591 | 0.69259671 | 3.38428681 | 0.00071363 | 0.02388192 |
| SCN2A | 903.324381 | 1.70295334 | 0.44978482 | 3.78615117 | 0.000153 | 0.00920024 |
| CSRNP3 | 73.7154769 | 0.96144043 | 0.64144275 | 1.49887176 | 0.1339069 | 0.36774274 |
| GALNT3 | 36.7560261 | -0.2768187 | 0.32015098 | -0.8646504 | 0.38723071 | 0.63701577 |
| TTC21B | 65.7658702 | 0.40925978 | 0.35482397 | 1.15341639 | 0.24873952 | 0.50492395 |
| SCN7A | 58.4179221 | 0.7977917 | 0.59168192 | 1.34834558 | 0.17754726 | 0.42593802 |
| STK39 | 1008.88829 | -0.915812 | 0.23206668 | -3.9463314 | 7.94E-05 | 0.00593026 |
| CERS6 | 104.038574 | -0.0079525 | 0.35397912 | -0.0224659 | 0.98207629 | 0.99382437 |
| LRP2 | 43.8727376 | 1.77803567 | 0.70159099 | 2.5342909 | 0.01126752 | 0.1036885 |
| FASTKD1 | 100.850115 | 0.40457101 | 0.26936263 | 1.50195669 | 1.33E-01 | 3.67E-01 |
| PPIG | 134.986821 | -0.4333354 | 0.21087874 | -2.0549032 | 0.03988836 | 0.19882936 |
| KLHL23 | 38.5528939 | 0.37511503 | 0.32273045 | 1.1623168 | 0.2451068 | 0.5022131 |
| SSB | 33.9824561 | 0.31382275 | 0.38271669 | 0.81998711 | 0.41222346 | 0.65722052 |
| METTL5 | 23.6780447 | 0.1706366 | 0.44151409 | 0.38648053 | 0.69914082 | 0.85106546 |
| UBR3 | 357.211125 | 0.24709126 | 0.22005042 | 1.12288475 | 0.26148645 | 0.52104231 |
| MYO3B | 260.241087 | 0.78853392 | 0.53913866 | 1.46258093 | 1.44E-01 | 3.82E-01 |
| AC007405.1 | 19.8145319 | 1.39605417 | 0.7010214 | 1.99145728 | 0.04643064 | NA |
| GAD1 | 126.809207 | -0.6543644 | 0.54694632 | -1.1963961 | 2.32E-01 | 4.90E-01 |
| GORASP2 | 53.7824997 | -0.0781743 | 0.28879247 | -0.2706936 | 0.78662673 | 0.89473415 |
| TLK1 | 311.755697 | -0.4710029 | 0.24732638 | -1.9043778 | 0.05686099 | 0.23747101 |
| METTL8 | 54.0244084 | -0.1521723 | 0.30201477 | -0.5038571 | 0.61436176 | 0.79971969 |
| DCAF17 | 30.7255858 | -0.013581 | 0.36571278 | -0.0371357 | 0.97037679 | 0.98796166 |
| CYBRD1 | 14.4646465 | 1.00915105 | 0.55961222 | 1.80330418 | 0.07134046 | NA |
| DYNC1I2 | 110.847978 | -0.6103629 | 0.24492078 | -2.4920829 | 0.01269964 | 0.11020195 |
| SLC25A12 | 227.68763 | 0.85989356 | 0.34937677 | 2.46122132 | 0.01384649 | 0.11532965 |
| HAT1 | 50.5073866 | -0.4275132 | 0.36072123 | -1.1851622 | 0.23595333 | 0.49381285 |
| METAP1D | 16.7276599 | 1.62023498 | 0.69481002 | 2.33191079 | 0.01970538 | NA |
| ITGA6 | 176.760772 | -0.2539729 | 0.29306832 | -0.8665997 | 0.38616135 | 0.63638899 |
| PDK1 | 69.2567931 | -0.4667245 | 0.64066552 | -0.7284995 | 0.46630786 | 0.69638565 |
| RAPGEF4 | 18.1080027 | -0.4024965 | 0.63669296 | -0.6321674 | 0.52727749 | NA |
| MAP3K20 | 25.0057674 | -0.6830433 | 0.51082263 | -1.3371438 | 1.81E-01 | 0.4304342 |
| SP3 | 106.510905 | -0.2661399 | 0.27179609 | -0.9791895 | 0.32748633 | 0.58371442 |
| OLA1 | 199.863048 | -0.4156715 | 0.27733235 | -1.498821 | 0.13392009 | 0.36774274 |

| | | | | | | |
|---|---|---|---|---|---|---|
| CIR1 | 58.8367166 | 0.13752078 | 0.30584688 | 0.4496393 | 0.65297055 | 0.82176493 |
| SCRN3 | 22.9382466 | 1.72683784 | 0.54422351 | 3.17303059 | 0.00150857 | 0.03507986 |
| GPR155 | 35.6513896 | 1.22555795 | 0.42836656 | 2.86100287 | 0.00422303 | 0.06301187 |
| AC010894.3 | 35.880725 | 1.03910224 | 0.47235899 | 2.19981469 | 0.02782005 | 0.16446305 |
| ATF2 | 112.914559 | -0.0726642 | 0.28520612 | -0.2547779 | 0.79889463 | 0.90203341 |
| ATP5MC3 | 71.6776585 | 1.32790492 | 0.55458513 | 2.39441135 | 0.01664707 | 0.12787672 |
| LNPK | 44.5442267 | 0.71226233 | 0.43609001 | 1.63329203 | 0.10240759 | 0.32120262 |
| HOXD10 | 79.3100555 | 1.36946402 | 0.44046277 | 3.10914816 | 0.00187628 | 0.04011582 |
| HOXD-AS2 | 27.5170611 | 1.36956714 | 0.62096624 | 2.20554204 | 0.02741608 | 0.16307695 |
| HOXD9 | 38.9894542 | 1.41218468 | 0.55725236 | 2.53419237 | 0.01127068 | 0.1036885 |
| HOXD8 | 73.7893078 | 2.03107975 | 0.57955025 | 3.5045792 | 0.00045733 | 0.01913079 |
| HOXD3 | 148.650867 | 0.83612181 | 0.40928882 | 2.04286503 | 0.0410658 | 0.20235817 |
| HAGLR | 40.0337841 | -0.162748 | 0.42742443 | -0.3807645 | 0.703378 | 0.8528536 |
| MTX2 | 293.278012 | 0.19635149 | 0.22422194 | 0.87570152 | 0.38119232 | 0.63258566 |
| LINC01116 | 34.6781958 | 0.41652576 | 0.64499722 | 0.64577916 | 0.51842238 | 0.73536015 |
| LINC01117 | 28.3163397 | 0.08671955 | 0.55904681 | 0.15512038 | 0.87672642 | 0.94225833 |
| AC079305.3 | 16.0472595 | -0.7204529 | 0.66403673 | -1.0849594 | 0.27793964 | NA |
| HNRNPA3 | 187.740266 | -0.512684 | 0.19056885 | -2.6902824 | 0.00713916 | 0.08250765 |
| NFE2L2 | 855.563434 | 0.66893349 | 0.47674776 | 1.40311826 | 0.16058158 | 0.40604349 |
| AGPS | 77.7314271 | 0.21663114 | 0.29271994 | 0.74006281 | 0.45926189 | 0.69297787 |
| RBM45 | 27.2200894 | 0.62266853 | 0.39604098 | 1.57223258 | 0.11589662 | 0.34268368 |
| AC009948.1 | 15.3587874 | 0.09706678 | 0.65425799 | 0.14836163 | 0.88205738 | NA |
| PRKRA | 67.9213807 | -0.3059409 | 0.235562 | -1.2987702 | 0.1940228 | 0.44613935 |
| PJVK | 13.3683673 | -0.0760249 | 0.61334882 | -0.1239505 | 0.90135445 | NA |
| PLEKHA3 | 31.5448141 | 0.27959622 | 0.3557727 | 0.78588441 | 0.43193521 | 0.67275181 |
| TTN-AS1 | 196.823397 | 0.70399307 | 0.31991168 | 2.20058573 | 0.02776536 | 0.16439232 |
| TTN | 22.357554 | 1.28616544 | 0.64109109 | 2.00621326 | 0.0448335 | 0.21170763 |
| SESTD1 | 462.821482 | -0.3002611 | 0.45687537 | -0.6572058 | 5.11E-01 | 7.31E-01 |
| ZNF385B | 65.4685528 | 0.74692615 | 0.52262269 | 1.42918814 | 0.15295017 | 0.3947713 |
| CWC22 | 64.4578299 | -0.5633386 | 0.33143293 | -1.6997063 | 0.08918619 | 0.29794449 |
| SCHLAP1 | 63.525505 | 0.44031901 | 0.3727466 | 1.18128243 | 2.37E-01 | 4.95E-01 |
| UBE2E3 | 210.498346 | 0.40781609 | 0.20274706 | 2.01145258 | 0.04427767 | 0.21063366 |
| CERKL | 35.5305264 | 0.97867405 | 0.63229281 | 1.54781776 | 0.12166618 | 0.35089572 |
| SSFA2 | 58.8680268 | -0.0601043 | 0.28960128 | -0.2075414 | 0.83558705 | 0.92049715 |
| PPP1R1C | 41.0960437 | -1.9938995 | 0.70534572 | -2.8268401 | 0.00470098 | 0.06664966 |
| PDE1A | 68.0275687 | 0.46285204 | 0.51292629 | 0.90237534 | 0.36685751 | 0.62174959 |
| DNAJC10 | 61.0565652 | 0.08880668 | 0.30082488 | 0.29521056 | 0.76783304 | 0.88499714 |
| NCKAP1 | 264.787436 | -0.292761 | 0.20910223 | -1.4000854 | 0.16148774 | 0.40727295 |
| NUP35 | 30.941209 | -0.4249239 | 0.41926295 | -1.0135022 | 0.31082034 | 0.57002243 |
| FSIP2 | 19.3964613 | 0.1971997 | 0.68353172 | 0.28850117 | 0.77296314 | NA |
| FSIP2-AS1 | 17.2543741 | 0.74213683 | 0.58058931 | 1.27824749 | 0.20116218 | NA |
| LINC01473 | 37.465212 | 1.15552797 | 0.66779922 | 1.73035239 | 0.08356733 | 0.28861463 |
| ZC3H15 | 54.0146449 | -0.3185021 | 0.28752017 | -1.1077556 | 0.26796738 | 0.52710067 |
| ITGAV | 278.764888 | -0.7673151 | 0.35956071 | -2.1340348 | 0.03283994 | 0.17901486 |
| AC017101.1 | 24.8004122 | 0.38992851 | 0.67472692 | 0.57790567 | 0.56332781 | 0.76782967 |
| AC007319.1 | 66.8037041 | 1.37768071 | 0.76462961 | 1.80176217 | 0.07158283 | 0.26694431 |
| TFPI | 59.7220507 | -0.4027306 | 0.88763212 | -0.4537135 | 0.65003508 | 0.81947559 |
| GULP1 | 473.775747 | -1.3999197 | 0.65596679 | -2.134132 | 0.03283197 | 0.17901486 |
| COL5A2 | 23.928691 | 0.59949593 | 0.71995077 | 0.83269018 | 0.40501949 | 0.65245604 |
| WDR75 | 35.1060361 | 0.02253721 | 0.43656582 | 0.05162385 | 0.95882841 | 0.98401364 |
| SLC40A1 | 15.4484055 | 1.55173024 | 0.60376608 | 2.57008515 | 0.01016735 | NA |
| ASNSD1 | 16.3189836 | 1.28176814 | 0.49540184 | 2.58733019 | 0.00967229 | NA |
| ANKAR | 75.9220455 | 0.72717009 | 0.42595285 | 1.70716101 | 0.08779213 | 0.29624552 |
| ORMDL1 | 58.0269244 | 0.67298682 | 0.3514923 | 1.91465598 | 0.05553639 | 0.235266 |

| | | | | | | |
|---|---|---|---|---|---|---|
| PMS1 | 56.1354753 | 0.36409114 | 0.30721731 | 1.18512574 | 0.23596774 | 0.49381285 |
| C2orf88 | 29.3566176 | 1.08712625 | 0.46880245 | 2.3189432 | 0.02039812 | 0.14043319 |
| HIBCH | 200.668312 | 0.92904854 | 0.35661082 | 2.6052169 | 0.00918161 | 0.09473306 |
| INPP1 | 24.4522979 | -0.7537565 | 0.43061643 | -1.7504127 | 0.08004713 | 0.28223671 |
| MFSD6 | 45.9933566 | 0.1984841 | 0.27938884 | 0.71042242 | 0.47744223 | 0.70521451 |
| NAB1 | 35.7652188 | -0.2421881 | 0.54706715 | -0.4427028 | 0.65798074 | 0.82521413 |
| GLS | 907.19636 | -1.612686 | 0.31142758 | -5.1783661 | 2.24E-07 | 0.00014357 |
| STAT1 | 127.270431 | 0.3273292 | 0.79010105 | 0.41428777 | 0.67866337 | 0.84008877 |
| MYO1B | 292.332629 | 0.32571562 | 0.3668857 | 0.88778499 | 0.37465642 | 0.62575308 |
| NABP1 | 127.426579 | -0.7742506 | 0.47790375 | -1.6200974 | 1.05E-01 | 0.32614246 |
| SLC39A10 | 107.007052 | 2.14939656 | 0.53704555 | 4.0022612 | 6.27E-05 | 0.00519258 |
| DNAH7 | 159.792849 | -0.0245419 | 0.31529757 | -0.0778372 | 0.93795757 | 0.97466007 |
| STK17B | 16.3350305 | -0.3221025 | 0.53326866 | -0.6040155 | 0.54583334 | NA |
| GTF3C3 | 44.3412433 | -0.0890596 | 0.33788041 | -0.2635833 | 0.79210104 | 0.89843821 |
| PGAP1 | 28.8719122 | -0.0982131 | 0.36206668 | -0.2712569 | 0.78619345 | 0.89452315 |
| ANKRD44 | 47.3242883 | -0.3955181 | 0.35507498 | -1.1139002 | 0.265322 | 0.5247131 |
| SF3B1 | 157.495222 | 0.0168201 | 0.22658916 | 0.07423169 | 0.94082603 | 0.97515997 |
| COQ10B | 41.6486961 | -1.0131938 | 0.45209625 | -2.241102 | 0.02501947 | 0.15492746 |
| HSPD1 | 87.3418601 | 0.10966017 | 0.3695555 | 0.29673533 | 0.76666858 | 0.88444965 |
| PLCL1 | 1482.40687 | -0.1046584 | 0.23958623 | -0.4368298 | 6.62E-01 | 8.28E-01 |
| FTCDNL1 | 38.8149348 | 1.33898376 | 0.744231 | 1.79915076 | 0.07199484 | 0.26757327 |
| C2orf69 | 15.6384765 | 0.50476946 | 0.52972912 | 0.95288222 | 0.34064975 | NA |
| TYW5 | 43.9274766 | -0.4828619 | 0.29267279 | -1.6498354 | 0.0989766 | 0.31649984 |
| MAIP1 | 20.4108015 | -0.1311174 | 0.43471284 | -0.3016184 | 0.76294299 | NA |
| SPATS2L | 161.729066 | -0.5789038 | 0.54028229 | -1.071484 | 0.28395185 | 0.54273452 |
| KCTD18 | 15.361941 | -0.0926997 | 0.50514941 | -0.1835094 | 0.85439835 | NA |
| BZW1 | 30.0205229 | -1.151995 | 0.53199089 | -2.1654412 | 0.03035391 | 0.17229651 |
| CLK1 | 115.506853 | -0.6807247 | 0.25815519 | -2.6368817 | 0.0083672 | 0.08952533 |
| PPIL3 | 27.4925872 | 0.03269462 | 0.53603406 | 0.06099355 | 0.95136435 | 0.98040586 |
| NIF3L1 | 13.3359263 | 0.19549407 | 0.50449365 | 0.38750552 | 0.698382 | NA |
| ORC2 | 127.940482 | -0.4521143 | 0.23199463 | -1.9488137 | 0.05131768 | 0.2274224 |
| AC005037.1 | 26.5767814 | -0.3513423 | 0.48316855 | -0.7271629 | 0.46712615 | 0.69693157 |
| FAM126B | 106.552665 | -0.2659296 | 0.27688403 | -0.9604369 | 0.33683539 | 0.59259931 |
| NDUFB3 | 30.2079807 | 0.71919707 | 0.46108489 | 1.55979319 | 0.11880876 | 0.34612832 |
| CFLAR | 252.812797 | 0.16105177 | 0.2067418 | 0.77899954 | 0.43597999 | 0.6764237 |
| CFLAR-AS1 | 19.2110909 | 1.11399733 | 0.60554304 | 1.83966664 | 0.06581719 | NA |
| CASP10 | 18.0088068 | -0.4012182 | 0.42134971 | -0.9522213 | 3.41E-01 | NA |
| CASP8 | 28.8874741 | -0.2356051 | 0.34275966 | -0.6873769 | 4.92E-01 | 0.71672078 |
| ALS2CR12 | 21.2416971 | -0.5087342 | 0.47661796 | -1.0673837 | 0.28579861 | 0.54477264 |
| TRAK2 | 139.607674 | -0.9206001 | 0.38912561 | -2.3658174 | 0.01799031 | 0.13301773 |
| STRADB | 58.9611364 | -0.197493 | 0.35515929 | -0.5560689 | 0.57816376 | 0.77555358 |
| TMEM237 | 14.3863312 | -0.0679566 | 0.61719885 | -0.1101048 | 0.91232624 | NA |
| ALS2 | 78.627669 | -0.2949095 | 0.32569484 | -0.9054781 | 0.36521216 | 0.62097479 |
| SUMO1 | 47.4856293 | -0.2467641 | 0.2723186 | -0.9061597 | 0.36485133 | 0.62061009 |
| NOP58 | 86.4305367 | -0.9938906 | 0.24758966 | -4.0142653 | 5.96E-05 | 0.00498893 |
| BMPR2 | 271.437811 | -0.1378988 | 0.17123168 | -0.8053344 | 0.42062676 | 0.66399954 |
| FAM117B | 79.7333466 | 0.50234971 | 0.29533121 | 1.70097062 | 0.0889485 | 0.29779758 |
| ICA1L | 21.8343348 | -0.150658 | 0.63966202 | -0.2355276 | 0.81379928 | 0.90970797 |
| WDR12 | 43.3629492 | 0.4054947 | 0.38094732 | 1.06443772 | 0.28713045 | 0.5463983 |
| CARF | 64.3865364 | -0.2841844 | 0.60021324 | -0.4734725 | 0.63587613 | 0.81188871 |
| NBEAL1 | 433.401004 | 0.51201867 | 0.26578232 | 1.92645874 | 0.05404712 | 0.23287156 |
| CYP20A1 | 101.812478 | -0.7035966 | 0.29525583 | -2.3830067 | 0.01717188 | 0.13009053 |
| ABI2 | 179.371392 | -0.2003361 | 0.220923 | -0.906814 | 0.36450516 | 0.62015831 |
| RAPH1 | 101.403212 | -0.4754272 | 0.29665963 | -1.6026016 | 1.09E-01 | 0.33102453 |

| | | | | | | |
|---|---|---|---|---|---|---|
| PARD3B | 892.408906 | 0.51952984 | 0.230754 | 2.25144456 | 0.02435739 | 0.15259218 |
| INO80D | 250.112453 | -0.7493402 | 0.5194034 | -1.4426941 | 0.14910667 | 0.38943809 |
| NDUFS1 | 162.689432 | 0.1018621 | 0.25782643 | 0.39508011 | 0.69278377 | 0.84775145 |
| EEF1B2 | 23.2638206 | 0.56584984 | 0.42111452 | 1.34369588 | 0.17904675 | 0.42802774 |
| ZDBF2 | 36.9079702 | 0.31951958 | 0.39114206 | 0.81688883 | 0.41399196 | 0.65845716 |
| FASTKD2 | 48.8133316 | -0.4327864 | 0.28400131 | -1.5238887 | 0.12753651 | 0.35967181 |
| CPO | 11.9936406 | 0.5625587 | 0.51851806 | 1.08493559 | 0.27795018 | NA |
| KLF7 | 41.2021548 | -0.399421 | 0.52695864 | -0.7579741 | 0.44846648 | 0.68731807 |
| CREB1 | 108.832411 | -0.1153715 | 0.19993765 | -0.5770375 | 0.56391411 | 0.76835668 |
| METTL21A | 22.6358237 | 0.84752369 | 0.41534277 | 2.04054038 | 0.04129653 | 0.20301425 |
| CCNYL1 | 40.8659445 | -0.7763109 | 0.51907674 | -1.4955609 | 0.13476811 | 0.36875584 |
| PLEKHM3 | 84.551507 | 0.39113162 | 0.27765866 | 1.4086779 | 0.15893043 | 0.40385855 |
| IDH1 | 28.4286066 | 1.22669959 | 0.40532749 | 3.02644068 | 0.00247451 | 0.04726137 |
| PIKFYVE | 118.04745 | -0.0857881 | 0.26010535 | -0.3298206 | 0.74153553 | 0.87152705 |
| PTH2R | 1270.49578 | -1.2192458 | 0.3571148 | -3.4141566 | 0.0006398 | 0.02310022 |
| RPE | 43.8695991 | -0.4960783 | 0.28469285 | -1.7425037 | 0.08142034 | 0.28447225 |
| KANSL1L | 450.219119 | 0.2449974 | 0.36224944 | 0.67632239 | 0.49883598 | 0.72176422 |
| AC007038.1 | 49.4613486 | 0.5544285 | 0.29908793 | 1.85373076 | 0.06377769 | 0.25122665 |
| LANCL1 | 64.7905502 | 0.68837413 | 0.31522534 | 2.18375248 | 0.02898044 | 0.16771613 |
| ERBB4 | 6360.59089 | 0.5033836 | 0.35762798 | 1.4075621 | 0.15926078 | 0.40403105 |
| IKZF2 | 38.2833835 | 0.80376541 | 0.57386349 | 1.40062126 | 0.16132736 | 0.40712678 |
| SPAG16 | 278.373437 | 1.30245103 | 0.51521584 | 2.52797163 | 0.01147236 | 0.10425357 |
| AC068051.1 | 73.6609159 | 1.8705594 | 0.75163595 | 2.48865078 | 0.01282288 | 0.11046442 |
| BARD1 | 26.1186955 | 0.16296952 | 0.44114188 | 0.36942655 | 0.71180981 | 0.857224 |
| AC016708.1 | 17.6672158 | 0.59051303 | 0.70691358 | 0.83533978 | 0.40352643 | NA |
| ATIC | 23.2129928 | -0.448291 | 0.38135072 | -1.1755346 | 0.23978089 | 0.49733052 |
| MREG | 16.0475106 | 0.1533637 | 0.48204188 | 0.3181543 | 0.7503679 | NA |
| PECR | 24.7906555 | 2.10826227 | 0.47188398 | 4.4677555 | 7.90E-06 | 0.00144859 |
| XRCC5 | 203.833805 | -0.4943386 | 0.27432643 | -1.8020088 | 0.07154403 | 0.26692893 |
| SMARCAL1 | 32.4157517 | 0.90447591 | 0.44143949 | 2.04892387 | 0.04046956 | 0.20044672 |
| RPL37A | 76.7586595 | -0.0289403 | 0.44955163 | -0.064376 | 0.94867084 | 0.97959746 |
| IGFBP5 | 21.9936077 | -1.4162379 | 0.69677045 | -2.0325745 | 0.04209552 | 0.20442224 |
| DIRC3 | 66.5558095 | 0.49843244 | 0.8053112 | 0.61893146 | 0.53596151 | 0.74808305 |
| TNS1 | 201.178877 | 0.66613608 | 0.41392427 | 1.60931875 | 0.10754666 | 0.32954311 |
| ARPC2 | 116.484866 | -0.9119651 | 0.33634257 | -2.7114173 | 0.00669963 | 0.07974991 |
| AAMP | 19.3386484 | -0.1198422 | 0.53145274 | -0.2254993 | 0.82159088 | NA |
| PNKD | 56.2240462 | 1.09143659 | 0.35600811 | 3.06576328 | 0.00217115 | 0.04403601 |
| TMBIM1 | 54.6690583 | 0.05175785 | 0.29666412 | 0.17446616 | 8.61E-01 | 9.36E-01 |
| CTDSP1 | 38.6767757 | -0.1645665 | 0.35055116 | -0.4694507 | 0.63874748 | 0.8138255 |
| USP37 | 99.565933 | -0.5935788 | 0.26295972 | -2.2572996 | 0.02398936 | 0.15134929 |
| CNOT9 | 47.9190363 | -0.6394267 | 0.27226774 | -2.3485215 | 0.01884811 | 0.1355543 |
| PLCD4 | 57.6491411 | 0.65365316 | 0.4071142 | 1.60557694 | 0.1083669 | 0.33024913 |
| ZNF142 | 20.2106873 | -0.0370072 | 0.40091656 | -0.0923066 | 0.92645448 | NA |
| STK36 | 18.1628447 | 0.96518914 | 0.67492286 | 1.43007327 | 0.15269599 | NA |
| TTLL4 | 24.1897637 | -0.6503376 | 0.53078386 | -1.22524 | 0.2204848 | 0.47710754 |
| CNPPD1 | 22.4261974 | 0.00782906 | 0.50466975 | 0.01551324 | 0.98762272 | 0.99527537 |
| RETREG2 | 22.5052439 | 0.28829354 | 0.50954445 | 0.56578682 | 0.57153871 | 0.77245539 |
| ZFAND2B | 12.5626206 | -0.346916 | 0.56578663 | -0.6131569 | 0.53977262 | NA |
| ANKZF1 | 33.6636584 | 0.18641464 | 0.53489096 | 0.34850961 | 0.7274575 | 0.86461401 |
| TUBA4A | 21.9125943 | -0.0477045 | 0.54597641 | -0.0873746 | 0.93037373 | 0.97138994 |
| DNAJB2 | 26.0196148 | -0.310148 | 0.40982572 | -0.7567802 | 0.44918153 | 0.68734597 |
| DNPEP | 18.553168 | -0.1467385 | 0.44040015 | -0.3331935 | 0.73898819 | NA |
| OBSL1 | 36.1611971 | 1.65203612 | 0.57392391 | 2.87849326 | 0.0039958 | 0.06175835 |
| SGPP2 | 249.816357 | -0.3165105 | 0.41657565 | -0.7597913 | 0.44737936 | 0.68660592 |

| | | | | | | |
|---|---|---|---|---|---|---|
| FARSB | 46.1715505 | -0.2700577 | 0.32957767 | -0.8194052 | 0.41255526 | 0.65722052 |
| MOGAT1 | 24.0326638 | -3.2728378 | 1.0139386 | -3.2278462 | 0.00124726 | 0.03140706 |
| ACSL3 | 208.143997 | -0.8950545 | 0.45024148 | -1.9879432 | 0.04681797 | 0.21661307 |
| AP1S3 | 49.5152517 | -1.0438408 | 0.50476693 | -2.0679659 | 0.03864323 | 0.1958098 |
| WDFY1 | 90.4812663 | -0.9548999 | 0.3034852 | -3.1464464 | 0.00165268 | 0.03708643 |
| CUL3 | 237.931096 | -0.5026798 | 0.21495437 | -2.338542 | 0.01935915 | 0.1366689 |
| IRS1 | 47.1272478 | 0.05971435 | 0.49929096 | 0.1195983 | 0.90480137 | 0.95673178 |
| RHBDD1 | 124.455705 | -0.0782448 | 0.35018768 | -0.2234369 | 0.82319552 | 0.91475524 |
| COL4A4 | 581.197847 | 0.39027791 | 0.36083141 | 1.08160735 | 0.27942704 | 0.53755309 |
| COL4A3 | 1070.71572 | 0.72145565 | 0.5257793 | 1.37216442 | 0.17001226 | 0.41834538 |
| AC097662.1 | 124.4742 | 1.19383819 | 0.60069348 | 1.98743325 | 0.04687441 | 0.21661307 |
| MFF | 149.378788 | 0.3962769 | 0.29053182 | 1.36397072 | 0.17257678 | 0.42128877 |
| AGFG1 | 302.965499 | -1.0330058 | 0.34671966 | -2.9793688 | 0.00288843 | 0.05257034 |
| SLC19A3 | 222.274787 | -0.5088125 | 0.71575853 | -0.7108717 | 0.47716375 | 0.7051944 |
| PID1 | 98.1185331 | -0.9791603 | 0.652082 | -1.5015908 | 1.33E-01 | 0.36753932 |
| DNER | 254.942589 | -0.8411475 | 0.56950083 | -1.4769909 | 0.13967807 | 0.37697679 |
| TRIP12 | 310.833833 | -0.4227214 | 0.18335267 | -2.3055102 | 0.02113802 | 0.1427136 |
| FBXO36 | 33.3633775 | 0.15155529 | 0.40355707 | 0.37554859 | 0.70725252 | 0.8547469 |
| SP110 | 40.5656955 | -0.2365454 | 0.66335843 | -0.3565876 | 0.72140055 | 0.86114805 |
| SP140L | 16.2287004 | -0.7783273 | 0.6500804 | -1.1972786 | 0.23119799 | NA |
| SP100 | 152.281009 | -0.4315748 | 0.49173647 | -0.8776547 | 0.38013116 | 0.63170913 |
| CAB39 | 223.165142 | -0.7361275 | 0.31538856 | -2.3340337 | 0.01959396 | 0.13704876 |
| ITM2C | 55.7211301 | 1.54351092 | 0.65801687 | 2.34570113 | 0.01899133 | 0.13585154 |
| PSMD1 | 118.801702 | -0.3126727 | 0.296109 | -1.0559379 | 0.29099659 | 0.55031959 |
| ARMC9 | 103.497871 | -0.074376 | 0.23169513 | -0.3210082 | 0.74820416 | 0.87522263 |
| NCL | 53.3576628 | -0.0353533 | 0.41709613 | -0.0847606 | 0.93245175 | 0.97276784 |
| PTMA | 175.34039 | -0.0791687 | 0.4029917 | -0.1964524 | 0.84425609 | 0.92488459 |
| PDE6D | 31.3223637 | 0.29944314 | 0.37518204 | 0.79812759 | 0.42479645 | 0.66754967 |
| COPS7B | 37.183521 | 0.55641539 | 0.34405525 | 1.61722684 | 0.10582932 | 0.32713587 |
| DIS3L2 | 202.819602 | -0.0977896 | 0.19203293 | -0.5092336 | 0.61058853 | 0.79723493 |
| EIF4E2 | 48.7773313 | -0.3987842 | 0.28528006 | -1.3978692 | 0.16215234 | 0.40813819 |
| EFHD1 | 182.724545 | 1.57859699 | 0.3608687 | 4.37443587 | 1.22E-05 | 0.00187417 |
| GIGYF2 | 305.193499 | -0.6791686 | 0.19321034 | -3.5151771 | 0.00043946 | 0.01858533 |
| ATG16L1 | 25.187954 | -0.8586858 | 0.35669069 | -2.4073682 | 0.01606796 | 0.12530404 |
| DGKD | 36.4889687 | -0.7723691 | 0.48718723 | -1.5853639 | 0.11288367 | 0.33755464 |
| USP40 | 151.982993 | 0.02451815 | 0.31834394 | 0.07701781 | 0.93860938 | 0.97481036 |
| AC114812.2 | 27.1248739 | 1.74302391 | 0.90545919 | 1.92501653 | 0.05422729 | 0.23287156 |
| SH3BP4 | 183.061393 | -1.1969269 | 0.46024725 | -2.6006171 | 0.00930562 | 0.09528654 |
| AGAP1 | 965.425204 | -0.9041479 | 0.38796186 | -2.3305071 | 0.01977937 | 0.13777536 |
| COPS8 | 19.8664527 | 0.23684536 | 0.42465062 | 0.55774172 | 0.57702077 | NA |
| RAB17 | 101.820407 | -0.9479372 | 0.40205438 | -2.3577337 | 0.01838688 | 0.13391856 |
| LRRFIP1 | 544.056348 | -0.8153422 | 0.19363785 | -4.2106553 | 2.55E-05 | 0.00292522 |
| ILKAP | 75.3761515 | -0.7772366 | 0.26202243 | -2.9662978 | 0.00301408 | 0.05407788 |
| PER2 | 23.3293878 | -0.9599124 | 0.52891741 | -1.8148625 | 0.06954504 | 0.26273569 |
| TRAF3IP1 | 48.8594309 | 0.0900341 | 0.34253862 | 0.26284364 | 0.7926711 | 0.8986875 |
| ASB1 | 14.3221453 | 0.07141321 | 0.52239139 | 0.13670442 | 0.89126444 | NA |
| HDAC4 | 193.891908 | -0.8188909 | 0.3498904 | -2.3404211 | 0.01926201 | 0.13639344 |
| NDUFA10 | 126.694906 | -0.2019178 | 0.25234254 | -0.8001736 | 0.42361022 | 0.66695091 |
| COPS9 | 17.323694 | -0.1379834 | 0.68908997 | -0.20024 | 0.84129291 | NA |
| GPC1 | 27.1471111 | -1.2659244 | 0.61371603 | -2.0627201 | 3.91E-02 | 0.19721202 |
| ANKMY1 | 48.7892357 | -0.5334188 | 0.36488269 | -1.4618913 | 0.14377099 | 0.38252489 |
| RNPEPL1 | 17.2012704 | -0.2734681 | 0.55820798 | -0.4899037 | 0.62420206 | NA |
| CAPN10 | 17.6645614 | -0.7436925 | 0.53952729 | -1.3784151 | 0.16807518 | NA |
| SNED1 | 78.9197648 | -1.9016048 | 0.62383412 | -3.048254 | 2.30E-03 | 0.04566132 |

| | | | | | | |
|---|---|---|---|---|---|---|
| MTERF4 | 55.087631 | -0.2814382 | 0.42449844 | -0.6629898 | 0.50733707 | 0.72850275 |
| PPP1R7 | 28.6811966 | -0.1310379 | 0.35284671 | -0.3713733 | 0.71035948 | 0.8562897 |
| HDLBP | 173.006029 | -0.5065339 | 0.30698065 | -1.6500516 | 0.09893238 | 0.31649984 |
| 2-Sep | 212.62788 | -0.016729 | 0.19430804 | -0.0860952 | 0.93139076 | 0.97192444 |
| FARP2 | 106.651232 | -0.3568423 | 0.30739188 | -1.1608711 | 0.24569433 | 0.50278177 |
| STK25 | 27.3165364 | 0.19364144 | 0.37188515 | 0.52070225 | 0.6025742 | 0.79255189 |
| THAP4 | 46.0153702 | 0.22629241 | 0.34580052 | 0.6544016 | 0.51285311 | 0.73124162 |
| ATG4B | 106.573868 | -0.4233399 | 0.28813796 | -1.4692266 | 0.14177134 | 0.37968477 |
| ING5 | 49.649783 | 0.34271618 | 0.28831743 | 1.18867661 | 0.23456695 | 0.49243138 |
| D2HGDH | 51.7020525 | 1.08944801 | 0.46252136 | 2.35545449 | 0.01850006 | 0.13433489 |
| LINC01266 | 164.836125 | 0.75810671 | 0.44613822 | 1.69926421 | 0.08926941 | 0.29794449 |
| CNTN4 | 43.2663459 | -0.3797287 | 0.44696436 | -0.8495726 | 0.39556275 | 0.64402157 |
| TRNT1 | 49.700002 | -0.2468426 | 0.30651779 | -0.8053124 | 0.42063944 | 0.66399954 |
| CRBN | 59.8474579 | 0.33703577 | 0.30934728 | 1.08950618 | 0.27593073 | 0.53482093 |
| AC034195.1 | 108.326433 | 1.78020918 | 0.55615087 | 3.20094647 | 0.00136977 | 0.03328762 |
| SUMF1 | 461.414311 | 0.22221924 | 0.32915787 | 0.67511448 | 0.49960303 | 0.72201362 |
| SETMAR | 15.7412263 | 1.35084047 | 0.55149165 | 2.44943048 | 0.01430823 | NA |
| ITPR1 | 961.800612 | 0.11380919 | 0.29714661 | 0.38300686 | 0.70171468 | 0.85190818 |
| EGOT | 18.2162894 | -0.7406669 | 0.45917855 | -1.613026 | 0.10673886 | NA |
| AC018816.1 | 24.5259123 | -0.0124771 | 0.47561791 | -0.0262335 | 0.97907112 | 0.99208931 |
| BHLHE40 | 143.108847 | -0.4977536 | 0.55075769 | -0.9037614 | 0.36612191 | 0.62157013 |
| ARL8B | 150.911575 | -1.3837847 | 0.43745151 | -3.1632871 | 0.00155998 | 0.03616626 |
| AC026202.2 | 21.7359478 | -0.5436315 | 0.53140536 | -1.0230072 | 0.30630446 | 0.56595793 |
| EDEM1 | 30.6652754 | -0.3685613 | 0.4150511 | -0.8879901 | 0.37454606 | 0.62575308 |
| AC069277.1 | 53.9847139 | 1.1454037 | 1.51698558 | 0.75505246 | 0.4502175 | 0.68770076 |
| LMCD1-AS1 | 76.6286542 | 1.51123093 | 0.67717638 | 2.23166515 | 0.0256371 | 0.15662748 |
| RAD18 | 24.137243 | -0.1465244 | 0.52429408 | -0.2794699 | 0.77988427 | 0.89074516 |
| SRGAP3 | 110.630798 | 0.7054778 | 0.45163742 | 1.56204464 | 0.11827747 | 0.34566454 |
| THUMPD3-A: | 88.6335218 | -0.0489157 | 0.28979752 | -0.1687926 | 0.86595976 | 0.93778686 |
| THUMPD3 | 22.2145721 | -0.0382131 | 0.41055063 | -0.0930777 | 0.92584186 | 0.96902431 |
| SETD5 | 291.437307 | -0.2079219 | 0.15500302 | -1.3414057 | 0.17978875 | 0.42896282 |
| MTMR14 | 39.5010605 | -0.4631446 | 0.30464559 | -1.5202733 | 0.12844231 | 0.36084904 |
| OGG1 | 25.8970885 | 0.25776116 | 0.49283906 | 0.52301284 | 0.60096532 | 0.79111169 |
| TADA3 | 24.0082601 | -0.0018039 | 0.41432288 | -0.0043539 | 0.99652608 | 0.99847191 |
| IL17RE | 17.7699395 | 1.77643476 | 0.65697783 | 2.70394931 | 0.00685207 | NA |
| IL17RC | 11.1957266 | 2.33877142 | 0.71267591 | 3.28167597 | 0.00103192 | NA |
| CRELD1 | 24.118379 | -0.4061351 | 0.53311929 | -0.7618091 | 0.44617398 | 0.686017 |
| EMC3 | 47.1021296 | 0.39707006 | 0.37359633 | 1.06283181 | 0.28785823 | 0.54734308 |
| BRK1 | 28.0268638 | -0.1508598 | 0.37739096 | -0.3997442 | 0.68934494 | 0.84569856 |
| VHL | 29.2081332 | 0.23844611 | 0.36432281 | 0.65449131 | 0.51279532 | 0.73124162 |
| SEC13 | 37.5294287 | -0.2808178 | 0.33738362 | -0.8323398 | 0.40521717 | 0.65250136 |
| ATG7 | 292.70553 | -0.4323693 | 0.2453488 | -1.7622639 | 0.07802472 | 0.27868041 |
| VGLL4 | 267.232102 | 0.56628329 | 0.21333947 | 2.65437657 | 0.00794551 | 0.08725829 |
| TAMM41 | 78.2993242 | 0.52179687 | 0.31703036 | 1.64588928 | 0.09978656 | 0.31793995 |
| SYN2 | 20.5547847 | -0.1724583 | 0.83016232 | -0.2077405 | 0.83543163 | NA |
| PPARG | 23.1387743 | -0.8746843 | 0.71450177 | -1.2241878 | 0.2208814 | 0.47769714 |
| TSEN2 | 62.4484078 | 0.00679769 | 0.41712315 | 0.0162966 | 0.98699777 | 0.99501268 |
| MKRN2 | 15.3323601 | 0.35146139 | 0.47906688 | 0.73363742 | 0.46316975 | NA |
| RAF1 | 159.815038 | -0.5440685 | 0.24232746 | -2.2451788 | 0.02475666 | 0.15424817 |
| RPL32 | 75.3156747 | 0.24135334 | 0.43091075 | 0.56010053 | 0.57541087 | 0.77388388 |
| IQSEC1 | 62.7365657 | 0.7533207 | 0.43900032 | 1.71599121 | 0.08616365 | 0.29319258 |
| NUP210 | 46.7161777 | 0.53799952 | 0.35045334 | 1.53515308 | 0.12474623 | 0.35504829 |
| HDAC11 | 12.7172981 | 0.99173628 | 0.53258747 | 1.86210967 | 0.06258763 | NA |
| XPC | 52.304227 | -0.3769903 | 0.33435907 | -1.1275015 | 0.25953053 | 0.51818585 |

| | | | | | | |
|---|---|---|---|---|---|---|
| LSM3 | 17.2716517 | 0.27826744 | 0.50562712 | 0.55034121 | 0.58208536 | NA |
| SLC6A6 | 28.552309 | -0.7535874 | 0.65723464 | -1.1466033 | 2.52E-01 | 0.50804157 |
| CCDC174 | 40.6658553 | -0.6875975 | 0.34785682 | -1.9766681 | 0.04807916 | 0.21962333 |
| FGD5-AS1 | 42.9594583 | 0.26175403 | 0.38529746 | 0.67935569 | 0.49691252 | 0.72101702 |
| NR2C2 | 115.866954 | -0.4333777 | 0.30250044 | -1.4326516 | 0.15195742 | 0.39301622 |
| MRPS25 | 89.3134351 | -0.1004066 | 0.28003296 | -0.3585528 | 0.71992964 | 0.85995641 |
| RBSN | 35.758004 | 0.40250073 | 0.38278766 | 1.05149873 | 0.29302959 | 0.55234367 |
| CAPN7 | 149.536424 | -0.3466487 | 0.24759311 | -1.4000742 | 0.16149111 | 0.40727295 |
| METTL6 | 39.8170254 | -0.812348 | 0.38837777 | -2.0916439 | 0.03647039 | 0.19061356 |
| COLQ | 17.8688454 | -0.1239761 | 0.73411184 | -0.1688791 | 0.86589177 | NA |
| HACL1 | 50.7474681 | -0.59546 | 0.35107258 | -1.6961165 | 0.08986382 | 0.29929978 |
| ANKRD28 | 400.122083 | -0.1664843 | 0.15983789 | -1.0415823 | 0.29760536 | 0.5574759 |
| OXNAD1 | 145.382073 | -0.1241201 | 0.36386016 | -0.3411202 | 0.73301306 | 0.86776236 |
| PLCL2 | 68.2182987 | 0.18682989 | 0.50689222 | 0.36857912 | 7.12E-01 | 0.857224 |
| TBC1D5 | 568.364702 | 0.96917146 | 0.2605317 | 3.71997513 | 0.00019924 | 0.01103287 |
| RAB5A | 122.291855 | -0.5725396 | 0.24014877 | -2.3841039 | 0.01712077 | 0.12988597 |
| KAT2B | 87.1843231 | 0.86018777 | 0.42387467 | 2.02934462 | 0.0424232 | 0.20555189 |
| ZNF385D | 961.81339 | -0.9394019 | 0.51301261 | -1.8311478 | 0.06707848 | 0.25862794 |
| UBE2E2 | 529.914577 | 0.37001822 | 0.36480844 | 1.01428084 | 0.31044876 | 0.57001845 |
| UBE2E1 | 227.23672 | -0.011822 | 0.22777665 | -0.0519015 | 0.95860715 | 0.98401364 |
| NKIRAS1 | 17.2502838 | 0.39776874 | 0.48810283 | 0.81492816 | 0.41511344 | NA |
| RPL15 | 120.012011 | 1.26535671 | 0.3628397 | 3.48737112 | 0.00048779 | 0.01986534 |
| NR1D2 | 62.6208826 | -0.1715696 | 0.58908355 | -0.2912483 | 0.77086141 | 0.88662885 |
| THRB | 479.520451 | -0.3661151 | 0.46492526 | -0.7874709 | 0.43100628 | 0.67197671 |
| RARB | 25.7111743 | 0.3101357 | 0.49581734 | 0.62550393 | 0.53164037 | 0.74523961 |
| TOP2B | 158.363337 | 0.17203835 | 0.23827871 | 0.72200473 | 0.47029157 | 0.70002596 |
| NGLY1 | 128.841389 | -0.3942549 | 0.23861297 | -1.652278 | 0.0984779 | 0.31609025 |
| SLC4A7 | 45.9750554 | -1.0012769 | 0.99599802 | -1.0053001 | 0.31475236 | 0.57214023 |
| CMC1 | 181.684564 | 0.26712084 | 0.26198737 | 1.01959433 | 0.3079209 | 0.56808896 |
| AZI2 | 55.9141324 | -0.232262 | 0.32300294 | -0.7190709 | 0.47209723 | 0.701222 |
| TGFBR2 | 86.9927086 | 0.31072619 | 0.36056189 | 0.86178321 | 0.38880683 | 0.63795484 |
| STT3B | 137.767952 | -0.4782754 | 0.32458371 | -1.4735041 | 0.14061517 | 0.37865124 |
| OSBPL10 | 173.773732 | -0.4597176 | 0.24958504 | -1.8419279 | 0.0654857 | 0.25452134 |
| GPD1L | 30.3621085 | 0.73514285 | 0.48516957 | 1.51522868 | 0.1297145 | 0.3622089 |
| CMTM8 | 122.208597 | 0.87600551 | 0.34172761 | 2.56346129 | 0.01036343 | 0.10015449 |
| CMTM7 | 73.8743533 | 0.35224851 | 0.46078851 | 0.76444725 | 4.45E-01 | 6.85E-01 |
| CMTM6 | 51.3779428 | 0.25323059 | 0.38780107 | 0.65299096 | 0.51376211 | 0.73176309 |
| DYNC1LI1 | 54.4049411 | -0.672508 | 0.29536925 | -2.2768382 | 0.02279589 | 0.14700765 |
| CNOT10 | 96.1607169 | -0.5016861 | 0.22508086 | -2.2289152 | 0.02581955 | 0.15729431 |
| TRIM71 | 40.6717894 | -1.1982526 | 0.61403099 | -1.9514529 | 0.05100319 | 0.22661979 |
| GLB1 | 77.7947463 | 0.57056875 | 0.30714439 | 1.85765645 | 0.06321782 | 0.25056001 |
| CRTAP | 36.1957423 | 0.47268169 | 0.35878594 | 1.31744763 | 0.18768862 | 0.43816783 |
| FBXL2 | 122.737796 | 0.11977621 | 0.38308224 | 0.31266447 | 0.75453559 | 0.87875026 |
| UBP1 | 65.3280919 | -0.5211383 | 0.27819852 | -1.8732605 | 0.06103242 | 0.24672613 |
| CLASP2 | 325.652949 | 0.53155541 | 0.2474025 | 2.14854498 | 0.03167049 | 0.17539916 |
| PDCD6IP | 166.108409 | -0.1937558 | 0.18898369 | -1.0252517 | 0.30524443 | 0.56557634 |
| STAC | 142.276923 | -0.283365 | 0.44789564 | -0.6326586 | 0.52695658 | 0.74176753 |
| TRANK1 | 18.2538531 | -0.7321812 | 1.04436593 | -0.7010772 | 0.48325483 | NA |
| EPM2AIP1 | 48.3968599 | 0.18691986 | 0.31024313 | 0.60249478 | 0.54684484 | 0.75478822 |
| MLH1 | 31.1856351 | -0.5105173 | 0.35206255 | -1.4500756 | 0.14703743 | 0.38705441 |
| LRRFIP2 | 296.221593 | -0.3553491 | 0.25683441 | -1.3835729 | 0.16648926 | 0.41429299 |
| GOLGA4 | 523.963912 | -0.7202808 | 0.32463547 | -2.2187372 | 0.02650461 | 0.15987928 |
| ITGA9 | 56.903164 | -1.7318799 | 0.45771459 | -3.7837551 | 0.00015448 | 0.0092173 |
| CTDSPL | 525.221493 | 0.52321355 | 0.32303662 | 1.61967255 | 0.10530264 | 0.32629404 |

| | | | | | | |
|---|---|---|---|---|---|---|
| PLCD1 | 18.0796763 | 1.50285359 | 0.52728533 | 2.85017148 | 0.00436957 | NA |
| ACAA1 | 37.1676948 | 0.76643141 | 0.39810662 | 1.92519134 | 0.05420543 | 0.23287156 |
| OXSR1 | 155.287177 | -0.1889188 | 0.2901567 | -0.6510923 | 0.5149869 | 0.73279894 |
| XYLB | 16.4026073 | 0.10149498 | 0.62506029 | 0.16237631 | 8.71E-01 | NA |
| ACVR2B | 29.627672 | 0.3090055 | 0.40876477 | 0.75594945 | 0.44967951 | 0.68770076 |
| EXOG | 24.0460268 | 0.66246166 | 0.66038846 | 1.00313936 | 0.31579362 | 0.57299942 |
| WDR48 | 81.8574124 | -0.0932241 | 0.34721918 | -0.2684878 | 0.78832389 | 0.89534145 |
| GORASP1 | 17.7383667 | 0.53431185 | 0.41973041 | 1.27298818 | 0.20302224 | NA |
| TTC21A | 13.4444848 | -0.0255584 | 0.75678491 | -0.0337723 | 0.97305871 | NA |
| RPSA | 62.0063669 | 0.49969075 | 0.43169769 | 1.15750156 | 0.2470675 | 0.5033559 |
| MYRIP | 23.9298777 | 1.03248316 | 0.74236779 | 1.39079735 | 0.16428689 | 0.41120418 |
| EIF1B | 20.3199465 | 0.07848122 | 0.48599701 | 0.16148498 | 0.87171144 | NA |
| ENTPD3-AS1 | 28.723764 | 1.13402201 | 0.65142052 | 1.74084479 | 0.08171079 | 0.28509879 |
| RPL14 | 93.4730163 | 0.43406583 | 0.48412126 | 0.89660558 | 0.36992942 | 0.62368335 |
| ZNF621 | 49.2732165 | -0.1852195 | 0.33258089 | -0.5569158 | 0.57758499 | 0.77491227 |
| CTNNB1 | 122.671738 | -0.2505605 | 0.21129304 | -1.1858434 | 0.23568414 | 0.49360921 |
| ULK4 | 168.741872 | 0.29681612 | 0.31769959 | 0.93426661 | 0.35016638 | 0.60717068 |
| TRAK1 | 240.037084 | 0.44848754 | 0.19644261 | 2.2830461 | 0.02242765 | 0.14604538 |
| SEC22C | 23.7866013 | -0.4760869 | 0.43937604 | -1.0835524 | 0.27856332 | 0.53717915 |
| NKTR | 329.481762 | -0.1736458 | 0.17439518 | -0.9957031 | 0.31939444 | 0.57666635 |
| AC006059.1 | 31.1625437 | 1.16371326 | 0.47766354 | 2.43626144 | 0.01483995 | 0.12019013 |
| SNRK | 206.887498 | -0.645356 | 0.43819189 | -1.4727702 | 0.140813 | 0.37896421 |
| ANO10 | 254.279199 | -0.2113154 | 0.34068333 | -0.6202694 | 0.53508046 | 0.74775242 |
| ABHD5 | 34.8791694 | -0.9724572 | 0.65180095 | -1.4919542 | 0.13571115 | 0.37028313 |
| TCAIM | 253.674626 | 0.66523984 | 0.40669779 | 1.63571046 | 0.1019002 | 0.32089434 |
| ZNF445 | 35.8652173 | 0.21989197 | 0.41198596 | 0.53373656 | 0.59352381 | 0.78655148 |
| AC099669.1 | 90.8987426 | 0.34604124 | 0.29486401 | 1.17356216 | 0.24057044 | 0.49822614 |
| ZNF197 | 41.3044761 | 0.01824167 | 0.30035763 | 0.06073316 | 0.95157172 | 0.98048829 |
| KIAA1143 | 11.32944 | -0.5172194 | 0.50071847 | -1.0329546 | 0.30162514 | NA |
| ZDHHC3 | 77.3411511 | 0.23441087 | 0.24893748 | 0.94164555 | 0.34637414 | 0.60358654 |
| EXOSC7 | 38.2336162 | -0.3677 | 0.36397539 | -1.0102331 | 0.31238361 | 0.57101636 |
| CDCP1 | 131.195727 | -0.6965634 | 0.39250937 | -1.7746415 | 0.07595709 | 0.27525506 |
| LARS2 | 79.327246 | 0.10692999 | 0.28447863 | 0.37588059 | 0.70700567 | 0.8547469 |
| LIMD1 | 45.7973469 | 0.46265744 | 0.39112468 | 1.18288992 | 0.23685278 | 0.49485772 |
| SACM1L | 92.1238153 | 0.09532196 | 0.27306187 | 0.34908558 | 0.72702506 | 0.86450052 |
| LZTFL1 | 23.2842038 | 0.00029699 | 0.43468371 | 0.00068324 | 0.99945486 | 0.99980616 |
| FYCO1 | 36.4994912 | 0.42011752 | 0.36746796 | 1.14327661 | 0.25292376 | 0.51005615 |
| LTF | 46.1363324 | 0.57601466 | 1.34885643 | 0.42703927 | 0.66935073 | 0.83376893 |
| ALS2CL | 22.4889966 | -1.6625423 | 0.44654178 | -3.7231507 | 1.97E-04 | 0.01097391 |
| PTH1R | 236.844389 | 0.6539523 | 0.4845828 | 1.34951612 | 0.17717125 | 0.42569745 |
| CCDC12 | 84.5191874 | -0.0300207 | 0.28880417 | -0.1039484 | 0.91721032 | 0.96401363 |
| NBEAL2 | 42.8359298 | -0.7203021 | 0.44087656 | -1.6337953 | 0.10230183 | 0.32120262 |
| SETD2 | 361.229794 | -0.3175517 | 0.18458728 | -1.7203334 | 0.08537185 | 0.29152934 |
| KIF9-AS1 | 132.053787 | 0.74277086 | 0.22744583 | 3.26570448 | 0.00109192 | 0.02988759 |
| KLHL18 | 47.8044753 | -0.639112 | 0.36001422 | -1.7752409 | 0.07585812 | 0.27514133 |
| PTPN23 | 23.8697223 | -0.4793729 | 0.54031913 | -0.8872033 | 0.37496948 | 0.62603002 |
| SCAP | 76.0545079 | 0.19228592 | 0.29428684 | 0.65339627 | 0.51350084 | 0.73176309 |
| ELP6 | 23.697795 | 0.05511094 | 0.39339598 | 0.14009026 | 0.88858868 | 0.94900334 |
| SMARCC1 | 276.973798 | -0.7904525 | 0.22846945 | -3.4597734 | 0.00054063 | 0.02091071 |
| DHX30 | 54.5231672 | 0.12910734 | 0.33614304 | 0.38408451 | 0.70091581 | 0.85187889 |
| MAP4 | 334.578499 | 0.05220755 | 0.25786757 | 0.20245877 | 0.83955809 | 0.92302222 |
| ZNF589 | 28.2162269 | 0.77929915 | 0.43755757 | 1.78102085 | 0.07490904 | 0.27312881 |
| NME6 | 15.3747928 | 0.72131659 | 0.5343978 | 1.34977464 | 0.17708828 | NA |
| PLXNB1 | 130.851674 | 0.51824237 | 0.39653915 | 1.30691351 | 0.19124209 | 0.44283704 |

| | | | | | | |
|---|---|---|---|---|---|---|
| TMA7 | 42.8037608 | 0.61793876 | 0.50901205 | 1.21399633 | 0.22474913 | 0.48213323 |
| SHISA5 | 68.138324 | 0.22826783 | 0.41632239 | 0.54829584 | 5.83E-01 | 7.79E-01 |
| PFKFB4 | 51.889432 | -1.9966384 | 0.57244347 | -3.4879224 | 0.00048679 | 0.01986534 |
| COL7A1 | 27.8624963 | -1.5082649 | 0.72194086 | -2.0891806 | 3.67E-02 | 0.19119673 |
| UQCRC1 | 44.4602978 | 0.10194397 | 0.40710486 | 0.25041207 | 0.80226869 | 0.90378853 |
| NCKIPSD | 14.447431 | 0.21540402 | 0.61464763 | 0.35045123 | 0.72600008 | NA |
| IP6K2 | 83.1184864 | -0.5747606 | 0.27878055 | -2.0616956 | 0.03923673 | 0.19738894 |
| PRKAR2A | 120.508842 | -0.2102428 | 0.30551276 | -0.6881638 | 0.49134962 | 0.71672078 |
| SLC25A20 | 18.3556951 | 0.73701339 | 0.6118989 | 1.20446922 | 0.22840827 | NA |
| ARIH2 | 123.829213 | -0.4316781 | 0.26844568 | -1.6080651 | 0.10782093 | 0.32997921 |
| P4HTM | 27.7452052 | 0.37792305 | 0.48633754 | 0.77707974 | 0.43711173 | 0.67749677 |
| WDR6 | 55.4827171 | -0.4291264 | 0.38626443 | -1.1109652 | 0.26658331 | 0.52548766 |
| DALRD3 | 14.668395 | 0.38319133 | 0.51087068 | 0.75007501 | 0.45320953 | NA |
| QRICH1 | 92.4187624 | -0.6587396 | 0.25396932 | -2.5937764 | 0.00949282 | 0.09644302 |
| QARS | 19.7505873 | -0.055409 | 0.47291237 | -0.1171654 | 0.90672898 | NA |
| LAMB2 | 29.7049669 | 0.31528002 | 0.46238657 | 0.68185376 | 0.49533142 | 0.71935207 |
| USP4 | 79.8380873 | -0.3943968 | 0.27068559 | -1.4570292 | 0.14510833 | 0.38473951 |
| RHOA | 108.345043 | -0.1015527 | 0.34774811 | -0.2920294 | 0.77026416 | 0.88651526 |
| TCTA | 15.2689043 | 0.60044162 | 0.55900542 | 1.07412487 | 0.28276672 | NA |
| DAG1 | 76.1983536 | 0.62621642 | 0.3621155 | 1.72932784 | 0.08375044 | 0.28873024 |
| APEH | 14.3360563 | 0.40506768 | 0.51606136 | 0.78492154 | 0.43249957 | NA |
| MST1 | 20.1137053 | 0.57463395 | 0.52699603 | 1.09039521 | 0.27553909 | NA |
| RNF123 | 16.2712881 | -0.0591639 | 0.52539941 | -0.1126075 | 0.91034173 | NA |
| IP6K1 | 133.773653 | -0.1574519 | 0.28593532 | -0.5506556 | 0.5818698 | 0.7783237 |
| RBM6 | 453.849863 | -0.2026443 | 0.22014644 | -0.9204977 | 0.35731276 | 0.61328042 |
| RBM5 | 141.047907 | -0.3690975 | 0.22427739 | -1.6457189 | 0.09982165 | 0.31793995 |
| GNAI2 | 29.4720762 | -0.0678971 | 0.41645069 | -0.1630376 | 0.87048883 | 0.9398207 |
| SEMA3B | 63.6693204 | -1.0249301 | 0.32512093 | -3.152458 | 0.00161902 | 0.03677143 |
| HYAL1 | 16.3141593 | 0.67406109 | 0.60061196 | 1.1222905 | 0.26173894 | NA |
| HEMK1 | 18.7041384 | 1.18251234 | 0.47123568 | 2.50938628 | 0.01209411 | NA |
| CISH | 23.1254851 | 0.82552412 | 0.4661265 | 1.77103025 | 0.07655567 | 0.27638321 |
| MAPKAPK3 | 26.8657388 | -0.1643518 | 0.4707532 | -0.3491253 | 0.72699526 | 0.86450052 |
| DOCK3 | 103.286423 | -0.5321562 | 0.53828436 | -0.9886154 | 0.32285135 | 0.57912936 |
| MANF | 20.1262945 | -0.5590095 | 0.69336954 | -0.8062216 | 0.42011511 | NA |
| DCAF1 | 71.1275367 | -1.0173332 | 0.25696593 | -3.9590197 | 7.53E-05 | 0.00573526 |
| RAD54L2 | 104.963975 | 0.14284268 | 0.29725626 | 0.48053716 | 0.63084549 | 0.80819203 |
| TEX264 | 27.4052851 | 0.25925437 | 0.41194985 | 0.62933479 | 0.5291299 | 0.74346711 |
| RPL29 | 101.54116 | 0.75071068 | 0.42328271 | 1.7735444 | 0.07613854 | 0.27578275 |
| ALAS1 | 16.0426421 | 0.10491392 | 0.4385739 | 0.23921605 | 0.81093806 | NA |
| WDR82 | 53.0269827 | -0.2930438 | 0.37891374 | -0.7733787 | 0.43929832 | 0.67938098 |
| DNAH1 | 12.8182411 | -0.7366951 | 0.56727066 | -1.2986659 | 0.19405861 | NA |
| BAP1 | 13.4380991 | -0.2251302 | 0.54189521 | -0.4154498 | 0.67781269 | NA |
| PHF7 | 16.789385 | 0.4729963 | 0.54335396 | 0.87051229 | 0.3840205 | NA |
| NISCH | 75.1387159 | 0.18163812 | 0.24088628 | 0.75404094 | 0.45082464 | 0.6884212 |
| SMIM4 | 38.5390397 | 0.38079723 | 0.4640749 | 0.82055122 | 0.41190195 | 0.65707964 |
| PBRM1 | 188.894848 | -0.5270767 | 0.30196737 | -1.7454755 | 0.08090213 | 0.28382032 |
| GNL3 | 22.6838716 | -0.4944543 | 0.4023534 | -1.2289055 | 0.21910722 | 0.47565385 |
| GLT8D1 | 17.0243701 | 0.06689658 | 0.46906515 | 0.14261681 | 0.88659282 | NA |
| SPCS1 | 32.674874 | 0.86299944 | 0.3924472 | 2.19902048 | 0.02787647 | 0.16467012 |
| NEK4 | 47.9678308 | 0.15235488 | 0.32055862 | 0.47527931 | 0.63458791 | 0.81099192 |
| SFMBT1 | 39.879568 | 0.01417704 | 0.40417586 | 0.03507641 | 0.97201881 | 0.98878882 |
| RFT1 | 11.3979607 | 0.30712191 | 0.59092222 | 0.51973322 | 0.60324953 | NA |
| TKT | 55.5639279 | 0.4420031 | 0.45610729 | 0.96907703 | 0.33250676 | 0.58794039 |
| DCP1A | 138.362708 | -0.6144341 | 0.19484096 | -3.1535162 | 0.00161316 | 0.03677143 |

| | | | | | | |
|---|---|---|---|---|---|---|
| CACNA1D | 894.428844 | 0.28589619 | 0.4068211 | 0.70275654 | 0.48220748 | 0.7098013 |
| CHDH | 35.5750252 | 0.60675146 | 0.43965495 | 1.38006284 | 0.1675673 | 0.41578513 |
| SELENOK | 16.4615133 | 0.2949546 | 0.49096161 | 0.60076916 | 0.54799375 | NA |
| CACNA2D3 | 1265.81226 | -0.7625102 | 0.29716048 | -2.5659878 | 0.01028824 | 0.09994074 |
| ERC2 | 96.7676789 | 0.62628368 | 0.38393146 | 1.63123826 | 0.10284006 | 0.32202997 |
| CCDC66 | 142.456529 | 0.05930133 | 0.26826924 | 0.22105156 | 0.82505228 | 0.91544291 |
| FAM208A | 153.437138 | -0.2096399 | 0.25355344 | -0.8268077 | 0.40834605 | 0.65425469 |
| ARHGEF3 | 964.730308 | -0.5778698 | 0.31069218 | -1.8599432 | 6.29E-02 | 0.24991833 |
| SPATA12 | 17.863779 | 0.52065585 | 0.54662473 | 0.95249231 | 3.41E-01 | NA |
| IL17RD | 15.9330749 | 0.3008078 | 0.56959578 | 0.5281075 | 0.59742473 | NA |
| APPL1 | 71.5229676 | 0.44483366 | 0.27674324 | 1.60738761 | 0.10796937 | 0.32999829 |
| ARF4 | 66.4741195 | -0.771347 | 0.35983627 | -2.1436054 | 0.03206452 | 0.1766407 |
| DENND6A | 91.6449552 | 0.01267135 | 0.26686379 | 0.04748248 | 0.96212869 | 0.98535004 |
| DENND6A-A! | 17.5210228 | 0.85654811 | 0.57280006 | 1.49537015 | 0.13481787 | NA |
| SLMAP | 341.1461 | -0.4263687 | 0.25840306 | -1.6500143 | 0.09894002 | 0.31649984 |
| FLNB | 301.088495 | -0.523502 | 0.45837197 | -1.1420899 | 0.25341667 | 0.51069785 |
| ABHD6 | 13.4091996 | 0.470018 | 0.5281921 | 0.88986185 | 0.37354007 | NA |
| PXK | 83.9797997 | 0.50808626 | 0.33619555 | 1.51128193 | 0.13071663 | 0.36322236 |
| PDHB | 23.3869591 | 0.27324603 | 0.42070022 | 0.64950294 | 0.51601335 | 0.73374372 |
| KCTD6 | 26.6871258 | 0.89975253 | 0.46621155 | 1.92992328 | 0.05361635 | 0.2324986 |
| FAM107A | 19.6924593 | 2.20450156 | 0.66962356 | 3.29215056 | 0.00099424 | NA |
| FAM3D-AS1 | 13.4577058 | -0.3950074 | 0.66751742 | -0.5917559 | 0.55401404 | NA |
| FHIT | 2499.2198 | -0.1970291 | 0.16471164 | -1.1962066 | 0.23161595 | 0.49030472 |
| PTPRG | 2422.37625 | -0.2996642 | 0.26404006 | -1.1349195 | 0.25640907 | 0.51462336 |
| THOC7 | 28.0694464 | 0.26218642 | 0.36585617 | 0.71663795 | 0.47359752 | 0.7026171 |
| ATXN7 | 32.2139951 | -0.1201466 | 0.32904951 | -0.3651323 | 0.71501262 | 0.85843895 |
| PSMD6-AS2 | 16.4574137 | 0.562228 | 0.44262535 | 1.27021194 | 0.20400915 | NA |
| PSMD6 | 22.0695468 | 0.54146475 | 0.47849866 | 1.13159095 | 0.25780645 | 0.51619036 |
| PRICKLE2 | 393.895157 | 1.4929255 | 0.34809181 | 4.28888426 | 1.80E-05 | 0.00230362 |
| ADAMTS9 | 677.358883 | -1.3538013 | 0.31316367 | -4.3229834 | 1.54E-05 | 0.00207296 |
| ADAMTS9-A! | 254.014042 | 0.47445569 | 0.59892067 | 0.79218454 | 0.42825308 | 0.67072527 |
| ADAMTS9-A! | 1572.95676 | 0.50259143 | 0.35667265 | 1.4091112 | 0.15880229 | 0.40367819 |
| MAGI1 | 1946.76498 | -0.4300115 | 0.18800233 | -2.2872668 | 0.02218026 | 0.14554257 |
| MAGI1-AS1 | 26.68443 | 0.49035417 | 0.77843835 | 0.62992037 | 0.52874669 | 0.74333575 |
| SLC25A26 | 112.35712 | -0.9687475 | 0.37637652 | -2.5738787 | 0.01005655 | 0.09911049 |
| LRIG1 | 26.6305814 | 0.86893098 | 0.46074274 | 1.88593528 | 0.05930368 | 0.2418974 |
| SUCLG2 | 292.547443 | 0.16066075 | 0.22814847 | 0.70419383 | 0.48131207 | 0.7087866 |
| SUCLG2-AS1 | 186.819331 | 2.33515045 | 0.4998638 | 4.67157342 | 2.99E-06 | 0.00076688 |
| FAM19A1 | 72.8553284 | 1.93487194 | 0.63234241 | 3.05984844 | 0.00221449 | 0.04462024 |
| TMF1 | 95.5699852 | -0.1874857 | 0.22976762 | -0.8159795 | 0.41451188 | 0.6585267 |
| UBA3 | 65.0414076 | -0.0259199 | 0.26596855 | -0.0974548 | 0.92236519 | 0.96656839 |
| ARL6IP5 | 38.817216 | 0.45203661 | 0.30125177 | 1.50052768 | 0.13347777 | 0.36762309 |
| FRMD4B | 130.336263 | 0.27162427 | 0.26021342 | 1.0438519 | 0.29655392 | 0.55618311 |
| MITF | 701.64415 | 0.49862627 | 0.38937051 | 1.28059586 | 2.00E-01 | 0.4544844 |
| FOXP1 | 820.096987 | 0.58265812 | 0.33865479 | 1.72050753 | 0.08534023 | 0.29152934 |
| EIF4E3 | 39.7965843 | 0.59237384 | 0.33408298 | 1.77313387 | 0.07620653 | 0.27589917 |
| RYBP | 184.251586 | -0.7613335 | 0.23020408 | -3.3072113 | 0.0009423 | 0.02822127 |
| SHQ1 | 54.3321866 | 0.0668831 | 0.30910815 | 0.21637443 | 0.8286959 | 0.91688285 |
| PPP4R2 | 119.175085 | -0.145227 | 0.2733664 | -0.5312541 | 0.59524271 | 0.78829717 |
| PDZRN3 | 311.026322 | 0.62991148 | 0.51848168 | 1.2149156 | 0.22439828 | 0.48173901 |
| CNTN3 | 20.5201748 | 1.38552172 | 0.86382757 | 1.6039332 | 0.10872878 | NA |
| ZNF717 | 68.3389754 | 0.42291006 | 0.37172303 | 1.13770208 | 0.2552449 | 0.51322362 |
| ROBO1 | 55.8172752 | 1.62610011 | 0.57177513 | 2.84395038 | 0.0044558 | 0.06422525 |
| GBE1 | 779.741066 | -1.0401868 | 0.38923721 | -2.6723724 | 0.0075317 | 0.08512703 |

| | | | | | | |
|---|---|---|---|---|---|---|
| CHMP2B | 37.4826572 | 0.11686785 | 0.36395546 | 0.32110482 | 0.74813096 | 0.87522263 |
| CGGBP1 | 82.1638449 | 0.21795177 | 0.35513181 | 0.61372078 | 0.53939986 | 0.74929145 |
| ZNF654 | 135.298389 | 0.0764999 | 0.22525399 | 0.33961617 | 0.7341456 | 0.86800594 |
| C3orf38 | 26.7419369 | -0.1672033 | 0.4034825 | -0.4144003 | 0.67858094 | 0.84008877 |
| ARL13B | 45.4500212 | -0.5059888 | 0.3025241 | -1.6725568 | 0.09441457 | 0.30830642 |
| NSUN3 | 48.706733 | -0.1313351 | 0.31375756 | -0.418588 | 0.67551727 | 0.83808131 |
| MTRNR2L12 | 97.3161765 | -0.2181335 | 0.58157685 | -0.3750726 | 0.7076065 | 0.8547469 |
| ARL6 | 30.3615466 | 0.25313291 | 0.58897849 | 0.42978294 | 0.66735354 | 0.83211084 |
| CRYBG3 | 232.067441 | -0.3560599 | 0.3288816 | -1.0826385 | 2.79E-01 | 0.53717915 |
| RIOX2 | 58.8259212 | 0.8933246 | 0.43924478 | 2.03377397 | 0.04197439 | 0.20441466 |
| CLDND1 | 31.9622872 | -0.4955079 | 0.35127645 | -1.4105924 | 0.15836483 | 0.40308667 |
| ST3GAL6 | 125.961614 | -1.3745375 | 0.65826864 | -2.0881103 | 0.03678789 | 0.19145123 |
| DCBLD2 | 73.8784008 | -0.2418411 | 0.32417966 | -0.7460096 | 0.4556616 | 0.6908246 |
| COL8A1 | 45.325338 | 1.40383526 | 0.75700872 | 1.85445059 | 0.06367472 | 0.25118809 |
| CMSS1 | 153.988208 | -0.1064474 | 0.30031139 | -0.3544567 | 0.72299665 | 0.86192273 |
| FILIP1L | 43.5076164 | 1.68501133 | 0.56923987 | 2.96010771 | 0.00307532 | 0.05464622 |
| TBC1D23 | 109.111738 | -0.2062136 | 0.20324016 | -1.0146301 | 0.31028219 | 0.56995192 |
| NIT2 | 13.2370996 | 0.90431408 | 0.64229972 | 1.4079316 | 0.15915133 | NA |
| TOMM70 | 48.7891369 | 0.448616 | 0.32698405 | 1.37198131 | 0.17006926 | 0.41835189 |
| TFG | 103.438893 | -0.4003107 | 0.26022541 | -1.5383228 | 0.1239697 | 0.35457643 |
| SENP7 | 146.692687 | 0.07985135 | 0.28023841 | 0.28494078 | 0.77568951 | 0.88880003 |
| TRMT10C | 14.0423553 | -0.178496 | 0.5173273 | -0.345035 | 0.73006806 | NA |
| PCNP | 100.558022 | -0.4632652 | 0.27249349 | -1.7000965 | 0.08911278 | 0.29794449 |
| ZBTB11 | 42.9166905 | -0.4459112 | 0.36539602 | -1.2203506 | 0.22233201 | 0.4793528 |
| RPL24 | 47.6172973 | 0.35535151 | 0.37709744 | 0.94233339 | 0.34602198 | 0.60338268 |
| CEP97 | 25.1754101 | 0.47517166 | 0.39983902 | 1.18840743 | 0.23467293 | 0.49243138 |
| NFKBIZ | 151.692356 | -1.1200321 | 0.48077995 | -2.3296149 | 1.98E-02 | 1.38E-01 |
| ALCAM | 61.2396944 | 0.38291886 | 0.56541969 | 0.67722944 | 0.49826039 | 0.72149172 |
| CBLB | 471.759775 | 0.19492272 | 0.24726667 | 0.78830972 | 4.31E-01 | 6.72E-01 |
| LINC00882 | 56.5062812 | 1.96627606 | 0.62799686 | 3.13102849 | 0.00174195 | 0.03863922 |
| BBX | 451.64842 | 0.06275607 | 0.29245666 | 0.21458247 | 0.83009286 | 0.91772835 |
| CD47 | 123.194428 | 0.29219275 | 0.42366544 | 0.68967804 | 0.49039668 | 0.7163358 |
| IFT57 | 52.6077997 | 0.49819218 | 0.37520492 | 1.32778689 | 0.18424852 | 0.43435248 |
| DZIP3 | 62.375567 | 1.3406001 | 0.53281479 | 2.51607148 | 0.01186711 | 0.10677891 |
| AC078980.1 | 474.372592 | 1.24234883 | 0.54087765 | 2.29691286 | 2.16E-02 | 1.43E-01 |
| NECTIN3 | 46.0050955 | 0.19727589 | 0.30962564 | 0.63714327 | 0.52403153 | 0.73927248 |
| PHLDB2 | 468.382576 | -0.1375833 | 0.22300531 | -0.6169506 | 0.53726733 | 0.74878549 |
| ABHD10 | 18.5909184 | 1.3560884 | 0.5171439 | 2.62226512 | 0.00873475 | NA |
| C3orf52 | 131.50721 | -1.7822175 | 0.570796 | -3.1223371 | 0.00179421 | 0.03909384 |
| AC128688.2 | 23.3803269 | -1.4691838 | 0.67141705 | -2.1881836 | 0.02865623 | 0.16622949 |
| SLC9C1 | 68.6538985 | 0.06226886 | 0.82192865 | 0.07575945 | 0.93961048 | 0.97491804 |
| AC112487.1 | 27.5093077 | -0.5650763 | 0.61206992 | -0.9232217 | 0.35589166 | 0.61199689 |
| ATG3 | 29.8731884 | -0.0685715 | 0.38037959 | -0.1802713 | 0.85693958 | 0.93203598 |
| SLC35A5 | 18.6220875 | -0.3599393 | 0.49678453 | -0.7245381 | 0.46873546 | NA |
| GTPBP8 | 22.3491299 | -0.7691341 | 0.4492066 | -1.7122058 | 0.08685875 | 0.29483981 |
| NEPRO | 49.5754332 | -0.4294167 | 0.28958456 | -1.4828715 | 0.13810854 | 0.37473724 |
| CFAP44 | 60.8306677 | 0.95779636 | 0.45678726 | 2.0968106 | 0.03601034 | 0.18932488 |
| SPICE1 | 54.6007693 | 0.19755742 | 0.36771579 | 0.53725574 | 0.59109097 | 0.78517152 |
| SIDT1 | 37.7297027 | 1.4569516 | 0.72834173 | 2.00036816 | 0.04546052 | 0.21335013 |
| USF3 | 60.3886158 | 0.28629444 | 0.26067756 | 1.09827037 | 0.27208644 | 0.53072716 |
| NAA50 | 126.288533 | -0.1029135 | 0.28957181 | -0.3553987 | 0.7222909 | 0.86153309 |
| ATP6V1A | 299.996399 | 0.11163832 | 0.24702002 | 0.45194038 | 0.65131194 | 0.820214 |
| GRAMD1C | 221.708182 | -0.1096641 | 0.57414259 | -0.1910049 | 0.84852173 | 0.92678754 |
| ZDHHC23 | 23.7012413 | 0.27255304 | 0.42059207 | 0.6480223 | 0.51697052 | 0.73456196 |

| | | | | | | |
|---|---|---|---|---|---|---|
| CCDC191 | 61.8670873 | 1.08418488 | 0.65902497 | 1.64513473 | 0.09994203 | 0.31813639 |
| QTRT2 | 37.9648685 | -0.4131227 | 0.43264253 | -0.9548823 | 0.33963722 | 0.59535132 |
| ZBTB20 | 1832.68118 | 0.14155022 | 0.20219324 | 0.70007396 | 0.48388111 | 0.71144851 |
| ZBTB20-AS5 | 34.0118187 | 0.67932576 | 0.68475261 | 0.99207473 | 0.32116107 | 0.57791615 |
| LSAMP | 119.226161 | 2.59340768 | 1.179618 | 2.19851485 | 0.02791244 | 0.16475616 |
| B4GALT4 | 20.6295813 | 0.39582819 | 0.41193797 | 0.96089271 | 0.33660612 | NA |
| TMEM39A | 32.695499 | -0.3747521 | 0.35294534 | -1.0617851 | 0.28833324 | 0.54770507 |
| POGLUT1 | 21.7371437 | 0.33458825 | 0.42285599 | 0.79125816 | 0.42879336 | 0.67122686 |
| TIMMDC1 | 46.3743864 | 0.72169273 | 0.29110869 | 2.47911781 | 0.01317078 | 0.11234472 |
| COX17 | 20.2941898 | 0.1239977 | 0.44814903 | 0.27668854 | 0.78201927 | NA |
| MAATS1 | 46.1308363 | 0.3618873 | 0.35334378 | 1.02417907 | 0.3057507 | 0.56557634 |
| GSK3B | 345.337948 | -0.2264844 | 0.24841598 | -0.9117143 | 0.36191915 | 0.61767 |
| AC092910.3 | 11.521932 | -0.082526 | 0.56019489 | -0.1473167 | 0.88288208 | NA |
| LRRC58 | 32.6412834 | 0.1172989 | 0.41124603 | 0.28522805 | 0.77546943 | 0.88874154 |
| FSTL1 | 20.528956 | -0.7069065 | 0.62961875 | -1.1227533 | 0.26154228 | NA |
| NDUFB4 | 36.9477993 | 0.2615266 | 0.39173377 | 0.66761309 | 0.5043806 | 0.72605526 |
| HGD | 66.233213 | 2.11106991 | 0.64484578 | 3.27375935 | 0.00106127 | 0.02953506 |
| RABL3 | 30.2600673 | -0.1470398 | 0.38550363 | -0.3814225 | 0.70288976 | 0.85275589 |
| GOLGB1 | 232.077925 | -0.4613792 | 0.21442049 | -2.1517495 | 0.03141709 | 0.17484986 |
| IQCB1 | 105.600734 | 0.01102188 | 0.2912479 | 0.03784364 | 0.96981235 | 0.98788821 |
| EAF2 | 14.593732 | 1.06756143 | 0.51593227 | 2.06918911 | 0.03852834 | NA |
| SLC15A2 | 16.4671696 | 0.89394977 | 0.6496405 | 1.37606842 | 0.16880045 | NA |
| ILDR1 | 23.1881296 | 0.27380065 | 0.54804449 | 0.49959566 | 0.61735982 | 0.80205143 |
| CASR | 290.791906 | 0.05874954 | 0.35399212 | 0.16596285 | 8.68E-01 | 9.39E-01 |
| CCDC58 | 18.1564852 | -0.3134001 | 0.50851187 | -0.6163083 | 0.53769106 | NA |
| FAM162A | 82.8952241 | 0.18740021 | 0.46199297 | 0.40563432 | 0.6850113 | 0.84355496 |
| AC083798.2 | 35.4624274 | 1.07219857 | 0.52472053 | 2.04337073 | 0.04101575 | 0.20224103 |
| KPNA1 | 116.195953 | -0.7137167 | 0.32547762 | -2.1928287 | 0.02831972 | 0.16563595 |
| PARP9 | 53.4561485 | -0.1406138 | 0.75008802 | -0.187463 | 0.85129761 | 0.92884705 |
| DTX3L | 19.7639558 | 0.42515526 | 0.59369202 | 0.7161209 | 0.4739167 | NA |
| PARP14 | 171.222444 | 0.17429264 | 0.6935828 | 0.25129319 | 0.80158744 | 0.90378853 |
| HSPBAP1 | 81.7385387 | -0.6379512 | 0.52698142 | -1.2105762 | 0.22605787 | 0.48386192 |
| DIRC2 | 145.704983 | -0.4973799 | 0.4514131 | -1.1018287 | 0.27053615 | 0.52890952 |
| PDIA5 | 26.7089749 | 0.49051595 | 0.41309055 | 1.18742962 | 0.2350582 | 0.49257908 |
| SEC22A | 118.675699 | -0.040409 | 0.23951651 | -0.1687106 | 0.8660243 | 0.93778686 |
| ADCY5 | 85.5264704 | -0.0220173 | 0.51245166 | -0.0429647 | 0.96572966 | 0.98714756 |
| HACD2 | 66.6587843 | 0.58996405 | 0.32313419 | 1.82575557 | 0.06788709 | 0.25944733 |
| MYLK-AS1 | 17.8321589 | 0.15331718 | 0.49439053 | 0.31011351 | 0.75647464 | NA |
| MYLK | 49.9839096 | 0.14484351 | 0.47735498 | 0.30342935 | 0.76156268 | 0.88239469 |
| CCDC14 | 83.2118985 | 0.57276086 | 0.30110985 | 1.90216581 | 0.05714948 | 0.23764428 |
| KALRN | 74.3679783 | -0.8303456 | 0.78796566 | -1.053784 | 0.29198183 | 0.55137 |
| UMPS | 24.5025714 | 0.08176981 | 0.36377528 | 0.22478109 | 0.82214958 | 0.91475524 |
| ITGB5 | 165.034017 | 0.72500939 | 0.21483967 | 3.37465333 | 0.00073909 | 0.02425576 |
| SLC12A8 | 32.6498802 | 0.37635206 | 0.55498096 | 0.67813509 | 0.49768605 | 0.72141045 |
| ZNF148 | 315.908544 | -0.3069369 | 0.16780675 | -1.8291091 | 0.06738327 | 0.25898011 |
| SNX4 | 134.663694 | -0.0263711 | 0.24931147 | -0.1057756 | 0.91576038 | 0.96388328 |
| OSBPL11 | 34.1711433 | -0.3150022 | 0.36697143 | -0.8583836 | 0.3906807 | 0.63925794 |
| SLC41A3 | 15.2389213 | 0.76919256 | 0.53332498 | 1.44225864 | 0.14922942 | NA |
| ALDH1L1 | 133.769438 | 2.06367507 | 0.55977002 | 3.68664811 | 0.00022723 | 0.01206185 |
| ZXDC | 162.274449 | -0.1586376 | 0.29110434 | -0.5449511 | 0.58578714 | 0.78146492 |
| TXNRD3NB | 14.4724811 | -0.2819689 | 0.62189131 | -0.4534054 | 0.65025684 | NA |
| TXNRD3 | 23.9861129 | 0.12159517 | 0.45377115 | 0.26796585 | 0.7887256 | 0.89544661 |
| CHCHD6 | 46.0959137 | 0.67796982 | 0.35479817 | 1.91086053 | 0.05602251 | 0.23590316 |
| TPRA1 | 20.4733813 | -1.1754344 | 0.60807446 | -1.9330435 | 0.05323084 | NA |

| | | | | | | |
|---|---|---|---|---|---|---|
| MGLL | 330.157328 | -1.1903969 | 0.43594919 | -2.7305864 | 6.32E-03 | 7.76E-02 |
| SEC61A1 | 53.4611867 | -0.3852764 | 0.38138328 | -1.010208 | 0.31239565 | 0.57101636 |
| RUVBL1 | 32.5510947 | 0.12887527 | 0.35477182 | 0.36326243 | 0.71640886 | 0.8589283 |
| EEFSEC | 69.3291835 | -0.1088882 | 0.23914625 | -0.4553206 | 0.6488786 | 0.81913126 |
| RPN1 | 40.6768553 | -0.0816776 | 0.34609749 | -0.235996 | 0.81343575 | 0.90970797 |
| RAB7A | 219.424925 | -0.6778109 | 0.33256165 | -2.0381512 | 0.04153482 | 0.20310896 |
| ACAD9 | 38.3546557 | -0.0523489 | 0.34155834 | -0.1532648 | 0.87818943 | 0.94339485 |
| CNBP | 76.2021175 | 0.02802425 | 0.40321481 | 0.06950203 | 0.94459002 | 0.9774035 |
| COPG1 | 33.9191864 | -0.4603532 | 0.31772465 | -1.4489062 | 0.14736379 | 0.38722988 |
| HMCES | 24.2175334 | -0.1592893 | 0.42122456 | -0.3781576 | 0.70531353 | 0.8538829 |
| MBD4 | 30.3234291 | 0.05304582 | 0.33801719 | 0.15693232 | 0.87529819 | 0.94169674 |
| IFT122 | 31.3698473 | 1.11085092 | 0.51458423 | 2.15873488 | 0.03087074 | 0.17358581 |
| TMCC1 | 493.807251 | -0.3576623 | 0.22096823 | -1.6186142 | 0.1055303 | 0.32665298 |
| PIK3R4 | 54.5578007 | -0.3274544 | 0.33412088 | -0.9800478 | 0.32706252 | 0.58316036 |
| ATP2C1 | 183.752022 | 0.38395113 | 0.19921754 | 1.92729582 | 0.05394277 | 0.23287156 |
| NEK11 | 112.614487 | 0.61726094 | 0.43848538 | 1.40771157 | 0.15921649 | 0.40403105 |
| NUDT16 | 15.3631401 | 1.07542873 | 0.52672293 | 2.04173518 | 0.04117781 | NA |
| MRPL3 | 47.5505076 | 0.20084527 | 0.29607227 | 0.67836567 | 0.49753987 | 0.72133441 |
| ACPP | 51.1240663 | -0.9511719 | 0.70503829 | -1.3491067 | 0.17730269 | 0.42576289 |
| DNAJC13 | 112.343296 | -0.1403827 | 0.2702843 | -0.5193889 | 0.60348957 | 0.79301836 |
| UBA5 | 79.7962464 | 0.48584053 | 0.31512475 | 1.54174031 | 0.12313671 | 0.35325504 |
| TMEM108 | 62.8644922 | 0.12477028 | 0.53360499 | 0.23382518 | 0.8151207 | 0.91046836 |
| CDV3 | 64.0222781 | -0.9667945 | 0.3700848 | -2.6123593 | 0.00899197 | 0.09383967 |
| TOPBP1 | 35.2712535 | -0.1865275 | 0.36045451 | -0.5174786 | 0.60482212 | 0.794016 |
| SRPRB | 12.8085692 | 0.26835993 | 0.52600593 | 0.51018423 | 0.6099224 | NA |
| RYK | 165.035168 | -0.4198319 | 0.19354101 | -2.1692142 | 0.03006643 | 0.17145264 |
| AMOTL2 | 33.9537217 | 0.07495816 | 0.5691065 | 0.13171201 | 0.89521209 | 0.95277205 |
| CEP63 | 56.5169771 | 0.14309617 | 0.34142084 | 0.41911962 | 0.67512871 | 0.83786935 |
| EPHB1 | 59.8594935 | -1.8661612 | 0.51346013 | -3.6344813 | 0.00027854 | 0.01365558 |
| PPP2R3A | 934.099257 | -0.083234 | 0.22645179 | -0.3675573 | 0.71320338 | 0.8577966 |
| MSL2 | 72.1958641 | -0.108125 | 0.30560518 | -0.3538062 | 0.72348413 | 0.86194629 |
| PCCB | 38.7354509 | 0.4676564 | 0.39304303 | 1.18983512 | 0.2341112 | 0.49243138 |
| STAG1 | 575.904219 | -0.5515252 | 0.28504569 | -1.9348661 | 0.05300673 | 0.23122687 |
| SLC35G2 | 24.7000844 | 0.31169071 | 0.63487884 | 0.49094518 | 0.62346522 | 0.8044129 |
| NCK1-DT | 15.5522665 | 1.00453055 | 0.53377974 | 1.8819196 | 0.05984693 | NA |
| NCK1 | 73.5314774 | -0.0345263 | 0.284162 | -0.1215021 | 0.90329339 | 0.95661107 |
| DZIP1L | 28.3324386 | -0.219817 | 0.48336991 | -0.4547594 | 0.64928237 | 0.81913234 |
| ARMC8 | 133.606636 | 0.04634351 | 0.26256897 | 0.17650035 | 0.85990087 | 0.93444261 |
| MRAS | 57.6108819 | -1.4644593 | 0.60080236 | -2.437506 | 0.01478897 | 0.12007458 |
| CEP70 | 161.120148 | 0.93097193 | 0.37431041 | 2.48716545 | 0.01287655 | 0.11061474 |
| FAIM | 13.7407983 | 0.65654307 | 0.61822731 | 1.06197682 | 0.28824621 | NA |
| PIK3CB | 171.648509 | 0.34158486 | 0.26684795 | 1.28007303 | 0.20051945 | 0.45454417 |
| COPB2 | 68.8200371 | -0.1978619 | 0.26870287 | -0.7363593 | 0.46151205 | 0.69416206 |
| AC097103.2 | 86.415223 | 0.66938813 | 0.55030945 | 1.21638495 | 0.22383832 | 0.48098367 |
| NMNAT3 | 74.6487782 | 0.69815378 | 0.32495178 | 2.14848426 | 0.03167531 | 0.17539916 |
| SLC25A36 | 233.43992 | -0.1249508 | 0.34106738 | -0.3663521 | 0.71410235 | 0.85788134 |
| SPSB4 | 14.5503211 | -0.5146726 | 0.78070717 | -0.659239 | 5.10E-01 | NA |
| PXYLP1 | 13.2776836 | 0.79088794 | 0.58389054 | 1.35451406 | 0.17557243 | NA |
| ZBTB38 | 190.130116 | -0.9206788 | 0.34657459 | -2.6565098 | 0.00789542 | 0.0871894 |
| RASA2 | 237.524423 | -0.7889524 | 0.20613853 | -3.8272921 | 0.00012956 | 0.00831024 |
| RNF7 | 27.903556 | 0.73472132 | 0.37597018 | 1.95420104 | 0.05067745 | 0.22586236 |
| ATP1B3 | 189.134596 | -0.7685351 | 0.46217367 | -1.6628707 | 0.09633832 | 0.31156136 |
| TFDP2 | 361.880395 | 0.23445971 | 0.3406134 | 0.68834552 | 0.49123523 | 0.71672078 |
| GK5 | 82.1616419 | 0.55394758 | 0.28879743 | 1.91811815 | 0.05509603 | 0.23466461 |

| | | | | | | |
|---|---|---|---|---|---|---|
| XRN1 | 258.706389 | -0.5061363 | 0.3148344 | -1.607627 | 0.10791689 | 0.32999829 |
| ATR | 181.935536 | -0.6899681 | 0.29794314 | -2.3157712 | 0.02057077 | 0.14099127 |
| PLS1 | 41.9168053 | 0.9316007 | 0.36664391 | 2.54088685 | 0.01105717 | 0.10282338 |
| TRPC1 | 83.1336728 | 0.35463575 | 0.41254681 | 0.85962548 | 0.38999552 | 0.6386799 |
| U2SURP | 147.726211 | -0.1719969 | 0.18032518 | -0.953815 | 0.34017732 | 0.59602659 |
| AC107021.1 | 21.0425131 | -0.2199809 | 0.65193858 | -0.3374257 | 0.73579599 | NA |
| PLOD2 | 217.535092 | -0.9136929 | 0.34657599 | -2.6363422 | 0.00838052 | 0.08952533 |
| PLSCR4 | 27.0271885 | -0.3584886 | 0.39233443 | -0.9137323 | 0.36085752 | 0.61731435 |
| PLSCR2 | 46.2795457 | 0.37749331 | 0.36722545 | 1.02796065 | 0.30396834 | 0.56472225 |
| PLSCR1 | 170.342316 | -0.2544987 | 0.41977256 | -0.6062775 | 0.54433048 | 0.75286822 |
| HLTF | 43.3656623 | 0.42065736 | 0.40445055 | 1.04007118 | 0.29830683 | 0.55811077 |
| HPS3 | 64.3835781 | -0.4694586 | 0.32155617 | -1.4599584 | 0.1443015 | 0.38365755 |
| WWTR1 | 667.117544 | -0.6745673 | 0.3578357 | -1.8851313 | 0.05941211 | 0.24195504 |
| COMMD2 | 26.3715921 | 0.54422758 | 0.4379592 | 1.24264448 | 0.21399887 | 0.4698087 |
| ANKUB1 | 55.3544446 | 0.16992284 | 0.57222215 | 0.29695258 | 0.76650271 | 0.88444965 |
| RNF13 | 140.25467 | 0.27812109 | 0.24924475 | 1.11585539 | 0.26448404 | 0.5237724 |
| AC117386.2 | 93.1916642 | 1.57990663 | 0.57757581 | 2.73540995 | 0.00623026 | 0.07721978 |
| TSC22D2 | 686.675393 | -0.0989075 | 0.2017857 | -0.4901611 | 6.24E-01 | 8.05E-01 |
| SERP1 | 89.0192506 | 0.55865201 | 0.34259628 | 1.63064238 | 0.1029658 | 0.32216576 |
| EIF2A | 45.4387331 | -0.1112375 | 0.30486641 | -0.3648729 | 0.71520626 | 0.85853753 |
| SELENOT | 48.9608155 | 0.13740485 | 0.40562917 | 0.33874498 | 0.73480185 | 0.86814805 |
| SIAH2 | 23.9481306 | 0.9342083 | 0.58874281 | 1.58678507 | 0.11256133 | 0.33724584 |
| MED12L | 95.6823181 | -0.150458 | 0.43704006 | -0.344266 | 0.73064627 | 0.86626377 |
| MBNL1 | 602.359702 | 0.26847204 | 0.24147252 | 1.11181194 | 0.26621902 | 0.52522169 |
| DHX36 | 138.82119 | -0.6687299 | 0.21213585 | -3.1523665 | 0.00161953 | 0.03677143 |
| PLCH1 | 34.853532 | 2.09681678 | 0.70833248 | 2.96021548 | 0.00307424 | 0.05464622 |
| GMPS | 59.3747941 | 0.19110853 | 0.29171142 | 0.6551287 | 0.51238489 | 0.73101511 |
| SSR3 | 39.0647096 | -0.1886798 | 0.39498801 | -0.4776849 | 0.63287447 | 0.80971323 |
| TIPARP | 28.2970883 | -1.078966 | 0.37307715 | -2.8920722 | 0.0038271 | 0.06110382 |
| LINC00886 | 59.9392434 | 1.85739687 | 0.73050791 | 2.54261022 | 0.01100279 | 0.10267384 |
| LEKR1 | 32.6411028 | 0.58859768 | 0.54046021 | 1.08906756 | 0.27612409 | 0.53482093 |
| CCNL1 | 212.235656 | -0.7166329 | 0.3624491 | -1.977196 | 0.04801948 | 0.21948095 |
| VEPH1 | 62.3301223 | 1.40183283 | 0.59575647 | 2.35302999 | 0.01862113 | 0.13495937 |
| RSRC1 | 148.243907 | 0.05561975 | 0.2440637 | 0.22789031 | 0.81973152 | 0.91308622 |
| MLF1 | 31.2930707 | 1.32411222 | 0.50757915 | 2.60868126 | 0.00908919 | 0.09427267 |
| GFM1 | 39.8644732 | -0.8010891 | 0.31887067 | -2.5122696 | 0.01199574 | 0.10726863 |
| MFSD1 | 24.3339219 | -0.2579567 | 0.49346932 | -0.5227411 | 0.6011544 | 0.79122528 |
| IQCJ-SCHIP1 | 173.568286 | -1.250102 | 0.68417384 | -1.8271701 | 0.06767419 | 0.25898011 |
| SMC4 | 19.1065008 | -0.9291394 | 0.4702304 | -1.9759238 | 0.04816341 | NA |
| KPNA4 | 153.757108 | -0.8303271 | 0.39478995 | -2.1032124 | 0.03544721 | 0.18721309 |
| PPM1L | 223.056284 | 1.03088031 | 0.3887509 | 2.651776 | 0.00800696 | 0.08779146 |
| B3GALNT1 | 15.7670773 | 1.75611634 | 0.55281998 | 3.17665137 | 0.00148986 | NA |
| NMD3 | 109.292459 | 0.83111438 | 0.40877787 | 2.03316875 | 0.04203548 | 0.20441466 |
| LINC01322 | 14.6415771 | 0.59471913 | 0.59914385 | 0.99261493 | 0.32089765 | NA |
| PDCD10 | 69.2684231 | -0.1300062 | 0.24678696 | -0.5267951 | 0.59833587 | 0.7894568 |
| SERPINI1 | 13.98431 | 1.05982541 | 0.53973855 | 1.96359036 | 0.04957762 | NA |
| GOLIM4 | 55.8010519 | 0.75858866 | 0.40904714 | 1.85452624 | 0.06366391 | 0.25118809 |
| MECOM | 6209.8852 | 0.52691219 | 0.24806807 | 2.12406291 | 0.03366488 | 0.18191827 |
| AC074033.1 | 25.4218703 | 1.82318802 | 0.78585837 | 2.31999568 | 0.02034111 | 0.14033487 |
| MYNN | 22.6217743 | 0.03139826 | 0.5047798 | 0.0622019 | 0.95040205 | 0.97993174 |
| SEC62 | 97.6948389 | -0.163494 | 0.27882324 | -0.5863714 | 0.55762596 | 0.76343775 |
| GPR160 | 61.5679913 | 0.35016094 | 0.36615258 | 0.95632521 | 0.33890796 | 0.59462212 |
| PHC3 | 261.151918 | -0.2048287 | 0.17594542 | -1.1641603 | 0.24435906 | 0.5016889 |
| PRKCI | 136.997272 | -0.4627752 | 0.25119158 | -1.8423199 | 0.06542837 | 0.25452134 |

| | | | | | | |
|---|---|---|---|---|---|---|
| SKIL | 60.6592064 | -0.4088507 | 0.40420981 | -1.0114814 | 0.31178609 | 0.57088166 |
| PLD1 | 79.8796027 | -0.2879771 | 0.32968546 | -0.8734906 | 0.38239575 | 0.63351272 |
| FNDC3B | 1804.69679 | -0.4811675 | 0.2482568 | -1.9381848 | 0.05260068 | 0.23030004 |
| TNFSF10 | 155.201674 | -1.4753726 | 0.51295881 | -2.876201 | 0.00402493 | 0.06202123 |
| NCEH1 | 154.118336 | -1.1370776 | 0.42739283 | -2.6604977 | 0.00780253 | 0.08671693 |
| ECT2 | 35.1499791 | -0.562215 | 0.52231363 | -1.0763934 | 0.28175134 | 0.5405384 |
| NLGN1 | 385.233277 | 0.68923883 | 0.745911 | 0.92402288 | 0.3554744 | 0.61168935 |
| NAALADL2 | 1338.57864 | 0.41820611 | 0.44834385 | 0.93277987 | 0.35093364 | 0.60767969 |
| TBL1XR1 | 615.787598 | -1.0848423 | 0.28555718 | -3.7990372 | 0.00014526 | 0.00894448 |
| ZMAT3 | 32.6011965 | 1.2649717 | 0.43647594 | 2.89814758 | 0.00375374 | 0.0609547 |
| PIK3CA | 116.62417 | -0.0453364 | 0.21592558 | -0.2099631 | 0.83369643 | 0.91946718 |
| ZNF639 | 17.6577951 | -0.1319482 | 0.47134384 | -0.2799405 | 0.77952318 | NA |
| MFN1 | 100.714691 | -0.5064191 | 0.2764613 | -1.8317902 | 0.06698268 | 0.25842892 |
| GNB4 | 43.6518356 | 0.46742365 | 0.3693369 | 1.26557526 | 0.20566519 | 0.45902815 |
| ACTL6A | 22.5740353 | -0.1573735 | 0.42544323 | -0.3699049 | 0.71145336 | 0.857224 |
| MRPL47 | 18.5102926 | -1.035937 | 0.47040145 | -2.2022403 | 0.02764834 | NA |
| NDUFB5 | 39.8551661 | 0.87727852 | 0.38654023 | 2.26956588 | 0.02323394 | 0.1486547 |
| TTC14 | 155.7793 | 0.3035883 | 0.2141206 | 1.41783785 | 0.15623811 | 0.39978881 |
| FXR1 | 159.967837 | 0.14531094 | 0.1960019 | 0.74137516 | 0.458466 | 0.6928053 |
| DNAJC19 | 23.8548647 | 0.88105926 | 0.44704153 | 1.97086669 | 0.04873913 | 0.22149053 |
| ATP11B | 144.477351 | -0.0880499 | 0.26419504 | -0.333276 | 0.73892596 | 0.87054913 |
| DCUN1D1 | 113.757556 | -0.0960881 | 0.250035 | -0.3842987 | 7.01E-01 | 8.52E-01 |
| MCCC1 | 137.350225 | -0.2291208 | 0.27236233 | -0.8412354 | 0.40021607 | 0.64782669 |
| KLHL24 | 186.200471 | -0.3474575 | 0.30017656 | -1.1575105 | 0.24706387 | 0.5033559 |
| YEATS2 | 91.2835089 | -0.0392399 | 0.4717717 | -0.0831756 | 0.93371188 | 0.9730757 |
| ABCC5 | 313.323897 | 0.28498854 | 0.50164677 | 0.56810599 | 0.569963 | 0.77196291 |
| EIF2B5 | 22.6585278 | 0.3696517 | 0.4344784 | 0.8507942 | 0.39488368 | 0.64334162 |
| DVL3 | 46.2147204 | -0.4894662 | 0.477683 | -1.0246673 | 0.30552019 | 0.56557634 |
| AP2M1 | 45.121933 | 0.4752996 | 0.50187508 | 0.94704761 | 0.34361452 | 0.60081803 |
| PSMD2 | 25.4371935 | -0.0520816 | 0.48097821 | -0.1082828 | 0.9137714 | 0.96346555 |
| EIF4G1 | 52.1722766 | -0.252457 | 0.36814991 | -0.6857452 | 0.49287379 | 0.71695332 |
| FAM131A | 15.9048569 | 1.31503769 | 0.49972934 | 2.63149985 | 0.00850089 | NA |
| POLR2H | 47.0720672 | -0.1844593 | 0.37446949 | -0.4925884 | 0.62230346 | 0.80366942 |
| VPS8 | 177.951341 | -0.4511834 | 0.17668327 | -2.5536282 | 0.0106607 | 0.10155374 |
| EHHADH | 27.9419659 | 0.54301694 | 0.5462542 | 0.99407371 | 0.32018698 | 0.57724473 |
| MAP3K13 | 241.250193 | 0.03496604 | 0.23228264 | 0.15053228 | 0.88034468 | 0.94376451 |
| TMEM41A | 34.7788851 | -0.4257911 | 0.43422685 | -0.980573 | 0.32680335 | 0.58313355 |
| LIPH | 15.4641373 | -0.436417 | 0.71289463 | -0.612176 | 0.54042134 | NA |
| SENP2 | 84.3477566 | -0.4967242 | 0.30905147 | -1.607254 | 0.10799866 | 0.32999829 |
| IGF2BP2 | 396.599662 | -0.3028564 | 0.29242551 | -1.0356704 | 0.30035595 | 0.56044599 |
| TRA2B | 100.684958 | -0.1734622 | 0.23031185 | -0.7531624 | 0.45135235 | 0.68853766 |
| ETV5 | 33.3226154 | 0.72750822 | 0.66851815 | 1.08824004 | 2.76E-01 | 5.35E-01 |
| TBCCD1 | 14.6050051 | 0.27419338 | 0.50819668 | 0.53954186 | 0.58951302 | NA |
| AC068631.1 | 104.894237 | 1.71784861 | 0.49431242 | 3.47522849 | 0.00051042 | 0.02046193 |
| KNG1 | 347.082851 | 0.6678752 | 0.61872461 | 1.07943855 | 0.28039227 | 0.53860228 |
| EIF4A2 | 105.926741 | -0.0329396 | 0.3751299 | -0.0878085 | 0.93002892 | 0.97116166 |
| ST6GAL1 | 239.954184 | 0.71122904 | 0.41473113 | 1.71491599 | 8.64E-02 | 0.29360326 |
| BCL6 | 167.635758 | -1.4916219 | 0.60254965 | -2.4755169 | 1.33E-02 | 1.13E-01 |
| LPP | 1455.52481 | -0.6909806 | 0.28824083 | -2.3972336 | 0.01651939 | 0.12745979 |
| TPRG1 | 26.7347309 | -1.6173968 | 0.66870988 | -2.4186823 | 0.01557684 | 0.12347403 |
| P3H2 | 130.742294 | 0.13233681 | 0.54761794 | 0.24165901 | 0.8090444 | 0.90756938 |
| CCDC50 | 176.824597 | -0.2489233 | 0.31728789 | -0.7845345 | 0.43272652 | 0.67354824 |
| MB21D2 | 51.1740949 | -1.1622983 | 0.43972576 | -2.6432344 | 0.00821182 | 0.08906215 |
| OPA1 | 153.308409 | -0.250458 | 0.20165454 | -1.2420152 | 0.21423095 | 0.4699161 |

| | | | | | | |
|---|---|---|---|---|---|---|
| HES1 | 27.3140677 | 0.02544136 | 0.55495645 | 0.04584389 | 0.96343468 | 0.98600419 |
| ATP13A3 | 145.461857 | -1.1192482 | 0.37156079 | -3.0122884 | 0.00259286 | 0.04879525 |
| LSG1 | 33.0095653 | -0.2587037 | 0.41097986 | -0.6294803 | 0.52903467 | 0.74346711 |
| XXYLT1 | 26.2717753 | 0.61909516 | 0.40779107 | 1.51816751 | 0.12897218 | 0.3611257 |
| ACAP2 | 237.781628 | -0.242944 | 0.21660465 | -1.121601 | 0.26203214 | 0.52137312 |
| PPP1R2 | 45.2506968 | -0.2066762 | 0.31478215 | -0.656569 | 0.51145809 | 0.73064085 |
| MUC20-OT1 | 576.669295 | 0.16427658 | 0.4260493 | 0.38558115 | 0.69980689 | 0.85147228 |
| TNK2 | 17.8560371 | 0.54541938 | 0.51395764 | 1.06121466 | 0.28859236 | NA |
| TFRC | 103.166828 | -0.6179303 | 0.39836719 | -1.5511577 | 0.1208639 | 0.34986439 |
| PCYT1A | 26.9584708 | 0.23915008 | 0.36255624 | 0.65962203 | 0.50949642 | 0.72990216 |
| UBXN7 | 73.6393249 | 0.06648985 | 0.26808678 | 0.24801612 | 0.80412193 | 0.90408418 |
| RNF168 | 58.8798194 | -0.3777447 | 0.34879672 | -1.0829938 | 0.27881118 | 0.53717915 |
| PIGX | 20.154882 | 0.05795841 | 0.4450846 | 0.13021885 | 0.89639328 | NA |
| PAK2 | 111.583405 | -0.8539312 | 0.25740778 | -3.3174257 | 0.00090851 | 0.02742277 |
| SENP5 | 177.7802 | -0.7153278 | 0.3094961 | -2.3112658 | 0.02081818 | 0.14130292 |
| NCBP2 | 24.7690927 | 0.23510705 | 0.4391216 | 0.53540306 | 0.59237117 | 0.7855446 |
| DLG1 | 375.024608 | -0.2951877 | 0.20366165 | -1.4494024 | 0.14722523 | 0.38715156 |
| BDH1 | 66.9018568 | 0.55795154 | 0.43602741 | 1.27962493 | 0.20067708 | 0.45456488 |
| RUBCN | 41.8326517 | -0.6125469 | 0.3241917 | -1.8894589 | 0.05883037 | 0.2411168 |
| FYTTD1 | 41.9655931 | -0.4267932 | 0.35397069 | -1.2057303 | 0.22792151 | 0.48609361 |
| LRCH3 | 184.639392 | -0.6197273 | 0.26265486 | -2.3594739 | 0.01830087 | 0.13368407 |
| IQCG | 25.4175644 | 0.91538808 | 0.41778315 | 2.19106035 | 0.02844743 | 0.16587474 |
| RPL35A | 55.5344132 | 0.1926265 | 0.3904783 | 0.49330911 | 0.62179419 | 0.80355102 |
| LMLN | 42.1625066 | 0.21937833 | 0.46189715 | 0.4749506 | 0.63482218 | 0.81099192 |
| ZNF595 | 45.7594705 | -0.2310133 | 0.28482917 | -0.8110591 | 0.41733173 | 0.66149141 |
| ZNF718 | 112.514717 | 0.04791421 | 0.23851462 | 0.20088583 | 0.84078785 | 0.92355925 |
| ZNF141 | 94.0705612 | -0.2077403 | 0.2711467 | -0.7661547 | 0.44358428 | 0.68340036 |
| ZNF721 | 251.084183 | -0.3448205 | 0.19924963 | -1.7305954 | 0.08352396 | 0.28861463 |
| PIGG | 44.3014366 | -0.4780568 | 0.30548315 | -1.5649204 | 0.11760158 | 0.34520992 |
| ATP5ME | 25.581679 | 0.4122407 | 0.49088094 | 0.83979773 | 0.40102181 | 0.64859526 |
| PCGF3 | 69.7170963 | -0.2005208 | 0.24874225 | -0.8061387 | 0.42016288 | 0.66385094 |
| AC139887.2 | 124.035481 | 0.11894196 | 0.37390542 | 0.31810708 | 0.75040371 | 0.87712337 |
| GAK | 106.612442 | -0.9290458 | 0.41224086 | -2.253648 | 0.02421832 | 0.15204601 |
| TMEM175 | 12.12438 | 1.0771914 | 0.59757885 | 1.80259292 | 0.07145217 | NA |
| IDUA | 14.1456227 | -0.5696896 | 0.53823449 | -1.0584413 | 0.28985429 | NA |
| RNF212 | 24.2539525 | -0.5487529 | 0.55818751 | -0.9830977 | 0.32555936 | 0.58154857 |
| AC092535.3 | 87.9955971 | 0.15742769 | 0.41064896 | 0.38336317 | 0.70145051 | 0.85190818 |
| CTBP1-AS | 14.6544328 | 0.60919959 | 0.57114093 | 1.06663619 | 0.28613615 | NA |
| CTBP1 | 47.4310659 | -0.4468422 | 0.35681561 | -1.2523056 | 0.21045852 | 0.46548315 |
| CTBP1-DT | 12.1894585 | 0.57206811 | 0.59460931 | 0.96209073 | 0.33600403 | NA |
| MAEA | 76.1911993 | -0.5411143 | 0.33680427 | -1.6066134 | 0.1081392 | 0.33009771 |
| UVSSA | 42.4032348 | -0.2843455 | 0.32029653 | -0.8877571 | 0.3746714 | 0.62575308 |
| SLBP | 44.7307561 | -0.0773187 | 0.51891969 | -0.1489993 | 0.88155418 | 0.94443055 |
| TMEM129 | 16.5529456 | 0.30276322 | 0.5256607 | 0.575967 | 0.56463748 | NA |
| LETM1 | 33.541305 | -0.1091541 | 0.46057422 | -0.2369957 | 0.81266009 | 0.90970797 |
| NSD2 | 155.128583 | -0.3183177 | 0.30265195 | -1.0517616 | 0.29290896 | 0.55234367 |
| NELFA | 56.7831015 | 0.76607675 | 0.44765959 | 1.71129304 | 0.08702703 | 0.29500251 |
| NAT8L | 19.6266428 | 2.22530993 | 0.8753145 | 2.54229757 | 0.01101264 | NA |
| POLN | 16.361608 | 0.07375713 | 0.42359525 | 0.17412171 | 0.86176981 | NA |
| MXD4 | 25.7112459 | 1.17698192 | 0.50674016 | 2.32265375 | 0.02019776 | 0.13990198 |
| ZFYVE28 | 56.9584266 | -0.1432561 | 0.26688079 | -0.5367792 | 0.59142014 | 0.78517152 |
| RNF4 | 61.3593442 | -0.6973598 | 0.35484779 | -1.9652366 | 0.04938686 | 0.22281986 |
| FAM193A | 154.891262 | -0.4289691 | 0.3117339 | -1.3760745 | 0.16879856 | 0.41629954 |
| TNIP2 | 17.8435044 | -0.9856158 | 0.45595503 | -2.1616513 | 0.03064507 | NA |

| | | | | | | |
|---|---|---|---|---|---|---|
| ADD1 | 123.044754 | -0.3423446 | 0.25079184 | -1.3650547 | 0.17223585 | 0.42085694 |
| MFSD10 | 28.0546681 | -0.046033 | 0.55974088 | -0.0822398 | 0.93445604 | 0.97327579 |
| NOP14 | 19.7581963 | -0.7567833 | 0.42261776 | -1.790704 | 0.07334081 | NA |
| GRK4 | 28.3426798 | 0.1186594 | 0.59049357 | 0.20094952 | 0.84073805 | 0.92355925 |
| HTT | 133.555544 | 0.00012621 | 0.22198413 | 0.00056854 | 0.99954637 | 0.99980616 |
| RGS12 | 66.6913195 | -1.0174231 | 0.45535169 | -2.2343677 | 0.02545889 | 0.15648615 |
| LRPAP1 | 42.4286858 | 0.8564013 | 0.42749396 | 2.00330621 | 0.04514443 | 0.21296142 |
| TMEM128 | 18.3131191 | -0.3647163 | 0.42864525 | -0.8508581 | 0.39484816 | NA |
| ZBTB49 | 13.3906623 | -0.0768783 | 0.52945165 | -0.1452036 | 0.88455012 | NA |
| STX18 | 213.891102 | 0.10652573 | 0.26454314 | 0.40267811 | 0.68718504 | 0.84493023 |
| STX18-AS1 | 92.6693569 | 0.50423569 | 0.37763021 | 1.33526313 | 0.18179024 | 0.43122793 |
| STK32B | 239.28782 | 0.63542035 | 0.43172291 | 1.47182447 | 0.14106828 | 0.37904789 |
| EVC2 | 58.7222614 | -0.4869229 | 0.29698325 | -1.6395634 | 0.10109597 | 0.32011506 |
| EVC | 75.7344234 | -0.8824712 | 0.29527646 | -2.9886269 | 0.00280234 | 0.0514788 |
| CRMP1 | 19.5261743 | 0.52775952 | 0.71353022 | 0.73964565 | 0.45951504 | NA |
| MAN2B2 | 17.9879761 | 0.61349845 | 0.52259541 | 1.17394535 | 0.24041692 | NA |
| MRFAP1 | 23.8978264 | 0.31302552 | 0.45213833 | 0.69232246 | 0.48873485 | 0.71544163 |
| KIAA0232 | 198.29514 | -0.1026881 | 0.26269674 | -0.3908998 | 0.69587133 | 0.84977338 |
| TBC1D14 | 178.330326 | -0.7663946 | 0.30130232 | -2.5436068 | 0.01097145 | 0.10267384 |
| TADA2B | 12.7128954 | 0.66523669 | 0.68282332 | 0.97424424 | 0.32993528 | NA |
| GRPEL1 | 18.6054404 | -0.9170313 | 0.52163519 | -1.7579935 | 0.07874861 | NA |
| SORCS2 | 26.2233204 | -1.8246001 | 0.9149497 | -1.9942081 | 0.04612932 | 0.21505596 |
| SH3TC1 | 43.5748981 | 0.07952727 | 0.33737038 | 0.23572689 | 0.8136446 | 0.90970797 |
| ACOX3 | 34.0978533 | -0.2882717 | 0.39176955 | -0.7358195 | 0.46184053 | 0.69416206 |
| TRMT44 | 26.0251785 | 0.15962219 | 0.35958065 | 0.44391205 | 0.65710617 | 0.82476514 |
| WDR1 | 71.727846 | -0.151411 | 0.34690489 | -0.4364625 | 0.6625012 | 0.8285925 |
| HS3ST1 | 66.9700656 | -0.9190249 | 0.40394344 | -2.2751327 | 2.29E-02 | 1.47E-01 |
| AC005699.1 | 54.5273886 | -0.7325515 | 0.72701705 | -1.0076125 | 0.31364051 | 0.57113617 |
| RAB28 | 231.031308 | -0.4912821 | 0.26866396 | -1.8286118 | 0.06745779 | 0.25898011 |
| BOD1L1 | 147.227209 | -0.8015453 | 0.37013705 | -2.1655365 | 0.03034663 | 0.17229651 |
| LINC01182 | 211.846352 | -0.917918 | 0.3740375 | -2.4540801 | 0.01412455 | 0.11677408 |
| AC092546.1 | 34.0963599 | 0.27380092 | 0.56296972 | 0.48635106 | 0.62671828 | 0.8053173 |
| CPEB2 | 450.022757 | -0.4493088 | 0.35341271 | -1.271343 | 0.20360666 | 0.45736371 |
| AC098829.1 | 139.705908 | -0.4244489 | 0.56333711 | -0.7534545 | 0.45117685 | 0.68853766 |
| CC2D2A | 67.2313135 | 1.28847948 | 0.5205103 | 2.47541593 | 0.01330811 | 0.11268704 |
| FBXL5 | 163.118649 | -0.0968074 | 0.19929713 | -0.4857442 | 0.62714853 | 0.80560118 |
| FAM200B | 38.0859179 | 0.41982168 | 0.38803321 | 1.08192203 | 0.27928718 | 0.53741835 |
| PROM1 | 49.1218785 | 0.72124005 | 0.96070436 | 0.75074089 | 0.45280861 | 0.68922439 |
| TAPT1 | 94.5658937 | 0.35308556 | 0.26253334 | 1.34491699 | 0.17865204 | 0.42767873 |
| TAPT1-AS1 | 242.988447 | 1.41543471 | 0.51407033 | 2.75338729 | 0.00589821 | 0.07567727 |
| AC097515.1 | 52.6428974 | 0.9066227 | 0.68091088 | 1.3314851 | 0.18302944 | 0.43276095 |
| LDB2 | 16.0917473 | -1.036505 | 0.60657902 | -1.7087715 | 0.08749328 | NA |
| QDPR | 13.7182892 | -1.0312039 | 0.68679875 | -1.5014645 | 0.13323547 | NA |
| LAP3 | 31.9097378 | 0.42778833 | 0.60643658 | 0.70541315 | 0.48055317 | 0.70817877 |
| MED28 | 50.7310541 | -0.5629104 | 0.27811663 | -2.0240085 | 0.04296927 | 0.20709737 |
| DCAF16 | 42.9717214 | -0.0351126 | 0.32374804 | -0.1084566 | 0.91363354 | 0.96345217 |
| LCORL | 204.030216 | -0.1957758 | 0.25159306 | -0.7781448 | 0.43648365 | 0.67706866 |
| SLIT2 | 61.7748262 | -0.5554598 | 0.58010674 | -0.957513 | 0.33830837 | 0.59410438 |
| PACRGL | 60.7823387 | 0.52676194 | 0.34316083 | 1.53502935 | 0.12477662 | 0.35504829 |
| KCNIP4 | 6748.14251 | 0.66423416 | 0.45250737 | 1.46789687 | 0.14213224 | 0.38005954 |
| ADGRA3 | 115.458815 | 0.41854735 | 0.40740134 | 1.02735878 | 0.30425155 | 0.56489395 |
| PPARGC1A | 802.494464 | 9.97E-05 | 0.34681601 | 0.00028762 | 0.99977052 | 0.99990042 |
| DHX15 | 195.128794 | -0.4301886 | 0.27524127 | -1.562951 | 0.11806413 | 0.34552837 |
| CCDC149 | 26.9785273 | -0.2650033 | 0.39361751 | -0.6732509 | 0.50078769 | 0.72277571 |

| | | | | | | |
|---|---|---|---|---|---|---|
| SEPSECS | 26.7687318 | 0.18715894 | 0.38470854 | 0.48649542 | 0.62661595 | 0.8053173 |
| ZCCHC4 | 21.2073322 | 0.60192997 | 0.4581866 | 1.31372235 | 0.18893966 | NA |
| ANAPC4 | 129.376653 | 0.06890705 | 0.45180413 | 0.15251532 | 0.8787805 | 0.94350307 |
| SEL1L3 | 39.590668 | -1.8698302 | 0.65717505 | -2.8452544 | 4.44E-03 | 6.42E-02 |
| SMIM20 | 15.5017811 | -0.9292044 | 0.75685004 | -1.2277259 | 0.21954987 | NA |
| RBPJ | 180.649319 | -0.157963 | 0.21485039 | -0.7352233 | 0.46220353 | 0.69416206 |
| TBC1D19 | 100.545759 | 0.83624646 | 0.46839181 | 1.78535672 | 0.07420347 | 0.27171463 |
| STIM2 | 70.3914629 | 0.49406035 | 0.47652264 | 1.03680352 | 0.29982743 | 0.559867 |
| ARAP2 | 82.0054939 | -0.5223709 | 0.48319764 | -1.0810709 | 0.27966559 | 0.53787757 |
| C4orf19 | 572.996788 | -0.1876649 | 0.23355545 | -0.8035131 | 0.42167825 | 0.66509374 |
| RELL1 | 163.594331 | -0.9614215 | 0.33517293 | -2.8684343 | 0.00412509 | 0.06262486 |
| PGM2 | 22.6834698 | 0.53748572 | 0.39030964 | 1.37707519 | 0.168489 | 0.41629954 |
| TBC1D1 | 406.776196 | -0.1751248 | 0.26225706 | -0.6677599 | 0.50428685 | 0.72605526 |
| KLF3 | 72.4221673 | -0.6846421 | 0.29598853 | -2.3130698 | 0.0207188 | 0.14118898 |
| FAM114A1 | 15.9393766 | -0.3179348 | 0.58029558 | -0.5478843 | 0.58377137 | NA |
| KLHL5 | 154.386008 | -0.100812 | 0.40279165 | -0.2502832 | 0.80236834 | 0.90378853 |
| WDR19 | 81.9868845 | -0.1561979 | 0.23516717 | -0.6641993 | 0.50656276 | 0.7281071 |
| RFC1 | 65.8702033 | 0.20883152 | 0.27870028 | 0.74930501 | 0.45367339 | 0.68944958 |
| LIAS | 27.8875252 | 1.08634118 | 0.46506634 | 2.33588434 | 0.01949727 | 0.13692563 |
| UGDH | 43.1717425 | -0.6612055 | 0.40161922 | -1.6463493 | 0.09969187 | 0.31793995 |
| UGDH-AS1 | 44.5351534 | 0.63238588 | 0.43555094 | 1.45192174 | 0.14652337 | 0.38645 |
| SMIM14 | 104.518468 | 0.05690934 | 0.35192922 | 0.16170677 | 0.87153677 | 0.9400893 |
| UBE2K | 373.594389 | -0.0458705 | 0.25206673 | -0.1819778 | 0.85560015 | 0.93162353 |
| PDS5A | 235.600117 | -0.2665696 | 0.19966359 | -1.3350935 | 0.18184574 | 0.43122793 |
| N4BP2 | 59.7260903 | -0.1141571 | 0.30420863 | -0.3752593 | 0.70746763 | 0.8547469 |
| RBM47 | 882.966167 | -0.9418361 | 0.38777925 | -2.4287945 | 0.01514911 | 0.12124045 |
| NSUN7 | 51.4940885 | 0.61208549 | 0.4424429 | 1.38342255 | 0.16653534 | 0.41429299 |
| APBB2 | 1433.83321 | -0.4172339 | 0.24304549 | -1.7166906 | 8.60E-02 | 2.93E-01 |
| UCHL1 | 45.5654963 | -0.1809022 | 0.61063145 | -0.2962544 | 0.76703584 | 0.88460817 |
| LIMCH1 | 971.391199 | 0.82902757 | 0.39355198 | 2.10652621 | 0.03515867 | 0.18663191 |
| TMEM33 | 49.9539933 | -0.0887617 | 0.4905687 | -0.1809364 | 0.85641753 | 0.93201099 |
| SLC30A9 | 214.362146 | -0.3547195 | 0.19895197 | -1.7829403 | 0.07459603 | 0.27250386 |
| AC024022.1 | 93.7889908 | 1.79733584 | 0.62223062 | 2.88853647 | 0.00387039 | 0.06118306 |
| SHISA3 | 40.3835262 | -0.1412856 | 0.73066732 | -0.1933652 | 0.84667299 | 0.92563742 |
| ATP8A1 | 189.865746 | 0.29492876 | 0.37935375 | 0.77745051 | 0.43689302 | 0.67734041 |
| KCTD8 | 262.681032 | 0.59901162 | 0.40879488 | 1.46531095 | 0.1428361 | 0.38081381 |
| GUF1 | 36.0862188 | -0.1308923 | 0.4375932 | -0.2991188 | 0.76484943 | 0.88317118 |
| GNPDA2 | 30.5191031 | 0.64830193 | 0.44478094 | 1.45757578 | 0.14495751 | 0.38459378 |
| GABRA2 | 51.6945199 | 0.21357153 | 0.68885412 | 0.31003884 | 0.75653142 | 0.87904624 |
| ATP10D | 79.7727522 | 0.20280645 | 0.34172736 | 0.59347444 | 0.55286369 | 0.759349 |
| AC107068.2 | 15.3171809 | 1.93859571 | 0.76437226 | 2.53619316 | 0.01120649 | NA |
| NFXL1 | 57.8609141 | 0.88089405 | 0.30954506 | 2.84577004 | 0.00443042 | 0.0642158 |
| NIPAL1 | 28.3022429 | 0.65741536 | 0.46913153 | 1.40134551 | 0.16111078 | 0.406847 |
| TXK | 64.73365 | -0.9740935 | 0.39156175 | -2.4877135 | 0.01285673 | 0.11056784 |
| TEC | 74.0992811 | -0.7139217 | 0.27891867 | -2.5596054 | 0.01047911 | 0.1006115 |
| SLAIN2 | 97.6399804 | -0.1651732 | 0.23946335 | -0.6897641 | 0.49034255 | 0.7163358 |
| FRYL | 330.027057 | -0.104249 | 0.18883801 | -0.5520549 | 0.58091074 | 0.77786509 |
| OCIAD1 | 164.071481 | 0.17443498 | 0.20395027 | 0.85528193 | 0.39239509 | 0.64124523 |
| OCIAD2 | 23.9575629 | -0.6617445 | 0.49091813 | -1.3479732 | 0.17766701 | 0.42601339 |
| CWH43 | 28.355788 | 1.1408462 | 0.68726814 | 1.65997249 | 0.09691999 | 0.3125608 |
| DCUN1D4 | 123.613614 | 0.02423538 | 0.25560115 | 0.09481717 | 0.92446004 | 0.96808157 |
| SGCB | 13.1219279 | 0.76513741 | 0.4820271 | 1.58733278 | 0.1124373 | NA |
| USP46 | 30.2944708 | 0.66766643 | 0.38612393 | 1.72915062 | 0.08378214 | 0.28873024 |
| SCFD2 | 127.838115 | -0.253453 | 0.32964063 | -0.7688768 | 0.44196647 | 0.68190353 |

| | | | | | | |
|---|---|---|---|---|---|---|
| AC058822.1 | 80.3724698 | 0.34502047 | 0.57484421 | 0.60019821 | 0.54837414 | 0.7554786 |
| LNX1 | 227.408167 | -0.2014957 | 0.20764908 | -0.9703665 | 0.33186385 | 0.58761353 |
| CHIC2 | 108.312114 | -0.4224277 | 0.21599278 | -1.9557494 | 0.05049469 | 0.22545112 |
| TMEM165 | 176.12529 | -0.5069995 | 0.25907105 | -1.9569902 | 0.05034862 | 0.22527387 |
| CLOCK | 99.8378676 | 0.43333196 | 0.26292588 | 1.64811453 | 0.09932917 | 0.31710354 |
| EXOC1 | 134.981436 | 0.1093772 | 0.25079413 | 0.43612346 | 0.66274715 | 0.8285925 |
| CEP135 | 31.3017906 | -0.0168278 | 0.44138724 | -0.0381248 | 0.96958817 | 0.98781207 |
| AASDH | 54.865546 | 0.61904321 | 0.36856041 | 1.67962484 | 0.09303033 | 0.30571879 |
| PPAT | 17.1767006 | -0.3500529 | 0.47492555 | -0.737069 | 0.4610804 | NA |
| PAICS | 31.2463457 | 0.1568388 | 0.32174399 | 0.48746458 | 0.62592914 | 0.8053173 |
| SRP72 | 52.3576967 | -0.2341115 | 0.25011924 | -0.9359995 | 0.34927346 | 0.60589538 |
| REST | 100.973256 | 0.15958556 | 0.30860576 | 0.5171179 | 0.60507386 | 0.79401898 |
| NOA1 | 12.6750892 | -0.0670972 | 0.51048991 | -0.1314368 | 0.89542978 | NA |
| POLR2B | 69.7044095 | -0.4530374 | 0.24425026 | -1.8548083 | 0.06362361 | 0.25118809 |
| IGFBP7 | 1511.7823 | 0.23080631 | 0.37414108 | 0.61689646 | 0.53730302 | 0.74878549 |
| ADGRL3 | 52.9958814 | -0.7609334 | 0.87194662 | -0.8726835 | 0.38283562 | 0.63396854 |
| AC104806.2 | 18.8315846 | 1.24341944 | 0.57265032 | 2.17134154 | 0.02990536 | NA |
| CENPC | 88.0677739 | 0.35470877 | 0.30135063 | 1.17706333 | 0.23917023 | 0.49686728 |
| UBA6 | 140.198736 | -0.6498731 | 0.21035059 | -3.0894761 | 0.0020051 | 0.04196595 |
| UBA6-AS1 | 353.577307 | 0.1339365 | 0.49765984 | 0.26913262 | 0.78782763 | 0.89506208 |
| YTHDC1 | 196.906302 | -0.2350941 | 0.21390154 | -1.0990763 | 0.2717348 | 0.5301756 |
| UGT2B7 | 114.719141 | -0.5557732 | 0.6637483 | -0.8373252 | 0.40240979 | 0.65015704 |
| RUFY3 | 312.316921 | -0.8198055 | 0.42750005 | -1.9176734 | 0.05515243 | 0.23466461 |
| GRSF1 | 47.0931119 | -0.5756394 | 0.29947115 | -1.9221866 | 0.05458228 | 0.23352755 |
| MOB1B | 135.431313 | 0.03087997 | 0.44219408 | 0.06983352 | 0.94432617 | 0.97734013 |
| SLC4A4 | 22.1639343 | 0.08659763 | 0.50571942 | 0.17123651 | 0.8640378 | 0.93668999 |
| ADAMTS3 | 23.3254025 | -1.963185 | 0.64802765 | -3.0294772 | 0.00244977 | 0.04713978 |
| COX18 | 12.009547 | 0.45756266 | 0.58169179 | 0.7866067 | 0.43151214 | NA |
| ANKRD17 | 739.279953 | -0.2070477 | 0.24232738 | -0.8544132 | 0.39287609 | 0.64175877 |
| AC053527.1 | 123.541618 | 0.95675464 | 0.53795481 | 1.77850372 | 0.07532116 | 0.2739825 |
| RASSF6 | 88.2506505 | -0.9810152 | 0.26972836 | -3.6370487 | 2.76E-04 | 1.36E-02 |
| MTHFD2L | 63.3942461 | 0.24891427 | 0.29543073 | 0.84254698 | 0.39948185 | 0.64760148 |
| BTC | 13.1606713 | 1.4160239 | 0.8804916 | 1.60821966 | 0.10778708 | NA |
| RCHY1 | 61.0410851 | 0.1285936 | 0.35878383 | 0.35841525 | 0.72003258 | 0.85995641 |
| THAP6 | 36.7580261 | -0.0082285 | 0.39515271 | -0.0208235 | 0.98338647 | 0.99382437 |
| G3BP2 | 55.7989423 | -0.5879142 | 0.32834004 | -1.7905652 | 0.0733631 | 0.26979255 |
| USO1 | 143.5293 | 0.03358691 | 0.268763 | 0.12496851 | 0.90054848 | 0.95621962 |
| NAAA | 29.3107231 | 1.58094633 | 0.5730164 | 2.75898966 | 0.00579804 | 0.07462791 |
| SDAD1 | 47.7495475 | -0.5048562 | 0.2967298 | -1.7014005 | 0.08886781 | 0.29778649 |
| NUP54 | 77.5965618 | -0.4509333 | 0.28818222 | -1.5647507 | 0.11764137 | 0.34520992 |
| SCARB2 | 89.0347655 | -0.5253714 | 0.37014674 | -1.4193597 | 0.15579417 | 0.39897013 |
| SHROOM3 | 816.59023 | -0.6681687 | 0.49035805 | -1.3626138 | 0.17300424 | 0.42179716 |
| 11-Sep | 186.638842 | -0.009829 | 0.45556552 | -0.0215753 | 0.9827867 | 0.99382437 |
| CCNI | 180.0398 | 0.01791721 | 0.32443889 | 0.05522521 | 0.95595904 | 0.98250992 |
| CCNG2 | 63.1123239 | 0.37919967 | 0.32109022 | 1.18097547 | 0.23761247 | 0.49536923 |
| CNOT6L | 309.38086 | 0.39494432 | 0.31053261 | 1.2718288 | 2.03E-01 | 0.4573105 |
| MRPL1 | 111.092745 | 0.58019653 | 0.38724859 | 1.49825342 | 0.13406742 | 0.36801602 |
| FRAS1 | 501.215672 | 1.4871905 | 0.64068768 | 2.32124098 | 0.02027384 | 0.14020463 |
| ANXA3 | 45.1079074 | 0.23881945 | 0.33764455 | 0.70731025 | 0.47937369 | 0.70695585 |
| BMP2K | 91.8791201 | -0.351773 | 0.27194256 | -1.2935563 | 0.19581872 | 0.44885942 |
| PAQR3 | 14.9613982 | 0.46968274 | 0.46537355 | 1.00925965 | 0.31285013 | NA |
| RASGEF1B | 573.311084 | -1.3210294 | 0.61621652 | -2.1437747 | 0.03205094 | 0.1766407 |
| HNRNPD | 165.934535 | -0.2958699 | 0.19824134 | -1.4924731 | 0.13557517 | 0.3700433 |
| HNRNPDL | 135.710039 | -0.2119603 | 0.23624737 | -0.8971965 | 0.36961408 | 0.62368335 |

| | | | | | | |
|---|---|---|---|---|---|---|
| ENOPH1 | 31.4499298 | 0.19437272 | 0.3476529 | 0.55909996 | 0.5760935 | 0.77421002 |
| TMEM150C | 39.9752422 | 0.79762287 | 0.51495824 | 1.54890785 | 0.12140387 | 0.35059775 |
| SCD5 | 117.591787 | 0.25638148 | 0.40036912 | 0.64036278 | 0.52193677 | 0.7380123 |
| SEC31A | 207.867393 | -0.3279003 | 0.20160125 | -1.6264795 | 0.10384769 | 0.32339043 |
| LIN54 | 36.6135763 | 0.14879235 | 0.36981971 | 0.40233754 | 0.68743563 | 0.84510334 |
| COPS4 | 47.907262 | -0.2712495 | 0.29546784 | -0.9180339 | 0.35860111 | 0.6150073 |
| HELQ | 51.2259041 | -0.2946148 | 0.33836883 | -0.8706913 | 0.38392272 | 0.63451053 |
| MRPS18C | 20.3600418 | 0.11636745 | 0.37684631 | 0.30879287 | 0.75747909 | NA |
| ABRAXAS1 | 18.3607669 | 1.150539 | 0.66770401 | 1.72312728 | 0.08486551 | NA |
| GPAT3 | 254.729792 | -1.1659473 | 0.53604309 | -2.1750999 | 2.96E-02 | 1.70E-01 |
| CDS1 | 60.2447378 | 0.47274928 | 0.32792317 | 1.44164646 | 0.14940213 | 0.38981296 |
| WDFY3 | 405.305833 | 2.12E-02 | 0.15478502 | 1.37E-01 | 0.89083307 | 0.94981883 |
| ARHGAP24 | 973.22839 | -0.8347488 | 0.15876188 | -5.257867 | 1.46E-07 | 0.00010198 |
| MAPK10 | 590.544998 | 1.42045687 | 0.52169341 | 2.72278094 | 0.0064735 | 0.07834356 |
| PTPN13 | 387.390133 | 0.63737267 | 0.26883668 | 2.37085459 | 0.01774701 | 0.13259773 |
| AC093827.5 | 20.4079594 | 0.954376 | 0.47514724 | 2.00859 | 0.04458063 | NA |
| AFF1 | 697.987571 | -0.3967683 | 0.23746412 | -1.6708559 | 0.09475015 | 0.30876032 |
| KLHL8 | 53.501468 | 0.92992227 | 0.4234677 | 2.19596976 | 0.02809411 | 0.16508057 |
| HSD17B11 | 18.6720974 | 0.12305832 | 0.44181294 | 0.27853037 | 0.78060525 | NA |
| NUDT9 | 52.812601 | -0.2756187 | 0.34902038 | -0.7896924 | 0.42970744 | 0.67197443 |
| SPARCL1 | 23.2672015 | 0.0073742 | 0.71588031 | 0.01030088 | 0.99178123 | 0.99605169 |
| AC093895.1 | 74.979436 | 0.31960744 | 0.79777403 | 0.40062401 | 0.68869696 | 0.84543868 |
| SPP1 | 6572.93451 | 0.09861848 | 0.58777917 | 0.16778152 | 0.86675516 | 0.93793856 |
| PKD2 | 283.487177 | -0.3011551 | 0.20928745 | -1.4389546 | 0.15016339 | 0.39073955 |
| PPM1K | 82.5915255 | 0.61548446 | 0.46756566 | 1.31635943 | 0.18805343 | 0.43860181 |
| HERC6 | 33.2656291 | -0.0801728 | 0.97942116 | -0.0818573 | 0.93476018 | 0.97332915 |
| HERC5 | 43.2207239 | -0.9385674 | 0.55162957 | -1.7014451 | 0.08885944 | 0.29778649 |
| PYURF | 14.5236331 | 1.13515667 | 0.49380037 | 2.29881697 | 0.02151533 | NA |
| HERC3 | 139.325464 | -0.4087158 | 0.33177962 | -1.2318894 | 0.21799041 | 0.47437721 |
| FAM13A-AS1 | 55.2730549 | -0.3099972 | 0.30262912 | -1.0243468 | 0.30567152 | 0.56557634 |
| FAM13A | 211.89028 | 0.37744983 | 0.57182809 | 0.66007571 | 0.50920524 | 0.72986085 |
| GPRIN3 | 328.651038 | -0.278401 | 0.5128317 | -0.5428701 | 0.58721925 | 0.78270395 |
| CCSER1 | 3464.41782 | 0.46993065 | 0.35128245 | 1.33775728 | 0.18097557 | 0.43032052 |
| SMARCAD1 | 114.700051 | -0.0714008 | 0.26450725 | -0.2699389 | 0.78720725 | 0.89499767 |
| PDLIM5 | 281.952347 | -0.4491965 | 0.20356981 | -2.206597 | 0.02734223 | 0.16284037 |
| BMPR1B | 37.1037445 | -0.0541889 | 0.421273 | -0.1286314 | 8.98E-01 | 0.9539772 |
| UNC5C | 298.09196 | 0.31403036 | 0.55704731 | 0.56374092 | 0.57293047 | 0.7733858 |
| STPG2 | 318.01196 | 1.94790301 | 0.51898141 | 3.75331944 | 0.00017451 | 0.0098764 |
| RAP1GDS1 | 158.032981 | 0.32863859 | 0.24000564 | 1.3692953 | 0.17090699 | 0.41898315 |
| TSPAN5 | 53.8008623 | -0.3377007 | 0.59463371 | -0.5679139 | 5.70E-01 | 0.77196291 |
| EIF4E | 89.0780396 | -0.6860888 | 0.32386454 | -2.1184438 | 0.0341375 | 0.18312899 |
| METAP1 | 56.5605304 | 0.50919652 | 0.30416519 | 1.67407888 | 9.41E-02 | 0.30825698 |
| ADH5 | 36.7966645 | 0.97460163 | 0.34051498 | 2.86214027 | 0.00420791 | 0.06301187 |
| LAMTOR3 | 21.7962232 | -0.4079822 | 0.369505 | -1.1041318 | 0.26953596 | 0.52764898 |
| DNAJB14 | 121.488259 | 0.18532817 | 0.20212413 | 0.91690273 | 0.35919361 | 0.61561194 |
| H2AFZ | 25.5810725 | 0.16990659 | 0.48598599 | 0.34961211 | 0.72662982 | 0.86429759 |
| PPP3CA | 463.105103 | 0.190532 | 0.32862516 | 0.57978518 | 0.56205949 | 0.76693811 |
| BANK1 | 30.3258389 | 0.03319735 | 0.84195513 | 0.03942888 | 0.96854845 | 0.98741848 |
| AP002075.1 | 16.0600645 | 0.68883637 | 0.85488998 | 0.80576026 | 0.4203811 | NA |
| NFKB1 | 152.540958 | -0.3325665 | 0.27322038 | -1.2172097 | 0.22352446 | 0.48054488 |
| MANBA | 81.0589081 | -0.0456653 | 0.3082999 | -0.1481198 | 0.88224821 | 0.94485383 |
| UBE2D3 | 292.695718 | -0.3642906 | 0.22068953 | -1.6506926 | 0.09880136 | 0.31647454 |
| CISD2 | 25.7615473 | 0.62593176 | 0.41647753 | 1.50291843 | 0.13286008 | 0.36692646 |
| SLC9B1 | 13.1837827 | 0.22487734 | 0.75255801 | 0.29881728 | 0.76507946 | NA |

| | | | | | | |
|---|---|---|---|---|---|---|
| BDH2 | 14.183372 | 0.46710057 | 0.50428464 | 0.92626372 | 0.35430895 | NA |
| TET2 | 190.315001 | 0.04347596 | 0.21671338 | 0.20061503 | 0.8409996 | 0.92355925 |
| PPA2 | 205.05693 | 0.15031258 | 0.28605857 | 0.52546083 | 0.59926287 | 0.78981615 |
| ARHGEF38 | 392.815995 | 0.39845926 | 0.39255906 | 1.01503009 | 0.31009148 | 0.56995192 |
| INTS12 | 73.124678 | 0.01601956 | 0.37506525 | 0.04271138 | 0.96593161 | 0.98722289 |
| GSTCD | 70.405825 | 0.3984473 | 0.3297034 | 1.20850226 | 0.22685413 | 0.48461808 |
| NPNT | 183.568076 | 0.6760918 | 0.52796345 | 1.28056554 | 0.20034631 | 0.4544844 |
| TBCK | 370.992367 | 0.33471744 | 0.26335136 | 1.27099188 | 0.20373154 | 0.45736371 |
| AIMP1 | 37.8500751 | 0.02502104 | 0.34312045 | 0.07292204 | 0.94186815 | 0.97576674 |
| PAPSS1 | 71.8663622 | 0.32293463 | 0.25447286 | 1.26903368 | 0.20442905 | 0.45773044 |
| SGMS2 | 142.135334 | -1.1170338 | 0.3293159 | -3.3919825 | 0.00069389 | 0.02384313 |
| AC096564.1 | 21.3651447 | 0.03716679 | 0.73225031 | 0.05075695 | 0.95951919 | 0.98419766 |
| HADH | 27.0652501 | 0.88117985 | 0.42776006 | 2.05998627 | 0.03939985 | 0.1978217 |
| RPL34-AS1 | 56.9910684 | 1.62900117 | 0.51531552 | 3.16117235 | 0.00157135 | 0.03632048 |
| RPL34 | 53.2543178 | 0.52043446 | 0.34701712 | 1.49973715 | 0.1336825 | 0.36762309 |
| OSTC | 31.9534753 | -0.5064125 | 0.34194482 | -1.4809773 | 0.13861263 | 0.37517394 |
| SEC24B-AS1 | 13.1354985 | 0.67539568 | 0.77077609 | 0.87625407 | 0.38089193 | NA |
| SEC24B | 195.535491 | -0.1362748 | 0.20332266 | -0.6702392 | 0.50270531 | 0.72445662 |
| MCUB | 27.9318265 | -1.1731614 | 0.62283385 | -1.8835865 | 0.05962093 | 0.24254878 |
| CASP6 | 30.3191764 | 0.62215842 | 0.41810251 | 1.48805233 | 0.13673709 | 0.37229056 |
| PLA2G12A | 52.4025298 | 0.19622216 | 0.50423554 | 0.38914782 | 0.6971668 | 0.85041636 |
| GAR1 | 18.9549867 | -0.1840305 | 0.41962643 | -0.438558 | 0.66098183 | NA |
| EGF | 2416.16125 | 0.27620692 | 0.53492963 | 0.51634252 | 6.06E-01 | 7.94E-01 |
| ELOVL6 | 69.8933381 | -0.9393845 | 0.34025028 | -2.7608632 | 0.00576488 | 0.07451815 |
| ENPEP | 18.5514044 | 0.29313777 | 0.6555569 | 0.4471584 | 0.6547607 | NA |
| FAM241A | 22.6068097 | -0.3421641 | 0.49818969 | -0.6868149 | 0.4921994 | 0.71672078 |
| AP1AR | 40.5613955 | -1.0746864 | 0.58757712 | -1.8290134 | 0.0673976 | 0.25898011 |
| ALPK1 | 170.987692 | 0.01677903 | 0.30370087 | 0.05524855 | 0.95594045 | 0.98250992 |
| LARP7 | 61.7278837 | -0.2437448 | 0.26691953 | -0.9131772 | 0.36114935 | 0.61731435 |
| ANK2 | 171.279084 | 0.07450008 | 0.47427207 | 0.157083 | 0.87517944 | 0.94169674 |
| CAMK2D | 155.111831 | -0.2912697 | 0.38973417 | -0.7473548 | 0.4548494 | 0.69080028 |
| ARSJ | 63.9607352 | 0.44021671 | 0.3738284 | 1.17759034 | 0.23895996 | 0.49681334 |
| UGT8 | 118.141447 | 0.55564991 | 0.29779435 | 1.86588466 | 0.0620575 | 0.24884382 |
| SNHG8 | 16.3417093 | 0.07278497 | 0.64301121 | 0.11319394 | 0.9098768 | NA |
| METTL14 | 46.1089557 | -0.0780342 | 0.3119222 | -0.250172 | 0.80245431 | 0.90378853 |
| SEC24D | 47.0588094 | -0.1257474 | 0.37690989 | -0.3336271 | 0.73866096 | 0.87054913 |
| USP53 | 457.611271 | -1.0512063 | 0.31331077 | -3.3551554 | 0.0007932 | 0.02533316 |
| C4orf3 | 31.7226122 | -0.0627165 | 0.49009847 | -0.1279671 | 0.89817501 | 0.95434195 |
| PDE5A | 90.3937151 | 0.38129894 | 0.26745378 | 1.42566292 | 1.54E-01 | 0.39634572 |
| AC108866.1 | 51.1970463 | 0.50762216 | 0.41503736 | 1.22307582 | 0.22130106 | 0.47833593 |
| PRDM5 | 43.4469911 | 1.25748616 | 0.43804709 | 2.87066432 | 0.0040961 | 0.06243109 |
| ANXA5 | 80.8230322 | -0.6434541 | 0.41999404 | -1.5320554 | 0.12550877 | 0.35647269 |
| EXOSC9 | 13.04722 | 0.10197736 | 0.50273713 | 0.20284431 | 0.83925673 | NA |
| BBS7 | 18.3170457 | -0.2446463 | 0.43633776 | -0.5606811 | 0.57501495 | NA |
| KIAA1109 | 298.613762 | 0.32741361 | 0.22246658 | 1.47174289 | 0.14109032 | 0.37904789 |
| FGF2 | 36.0502815 | -2.0875022 | 0.82540425 | -2.5290665 | 0.01143664 | 0.10405175 |
| NUDT6 | 11.0175327 | 0.31872383 | 0.61215968 | 0.52065472 | 0.60260732 | NA |
| SPATA5 | 157.542337 | -0.0359729 | 0.27602978 | -0.1303226 | 0.89631117 | 0.9533797 |
| ANKRD50 | 32.4899543 | 0.50649725 | 0.46313083 | 1.09363752 | 0.27411399 | 0.53327316 |
| FAT4 | 24.6982184 | 0.57736341 | 0.56648029 | 1.01921182 | 0.30810241 | 0.56815148 |
| INTU | 65.2560342 | 1.09420721 | 0.54817005 | 1.99610907 | 0.04592205 | 0.21446595 |
| HSPA4L | 106.15236 | 0.62617376 | 0.38187911 | 1.63971725 | 0.10106397 | 0.32011506 |
| MFSD8 | 53.4564252 | -0.078872 | 0.27381152 | -0.2880521 | 0.77330682 | 0.88771702 |
| ABHD18 | 155.601616 | -0.1649699 | 0.26915051 | -0.6129279 | 0.53992401 | 0.74933197 |

| | | | | | | |
|---|---|---|---|---|---|---|
| LARP1B | 141.648098 | -0.4626092 | 0.28949859 | -1.5979668 | 0.11005036 | 0.3329629 |
| PGRMC2 | 58.5112169 | -0.5372838 | 0.4093771 | -1.3124422 | 0.18937099 | 0.44049215 |
| AC108062.1 | 64.4710981 | 0.19919283 | 0.43560267 | 0.45728102 | 0.64746907 | 0.81885075 |
| AC078850.2 | 13.9421302 | 2.40919836 | 1.19988601 | 2.00785604 | 0.04465859 | NA |
| JADE1 | 121.92143 | 1.4597334 | 0.31587641 | 4.62121684 | 3.81E-06 | 0.00086364 |
| SCLT1 | 113.36584 | 0.18550936 | 0.39730738 | 0.46691648 | 0.64055963 | 0.81489142 |
| C4orf33 | 15.9774807 | 0.97368817 | 0.70096247 | 1.38907317 | 0.1648105 | NA |
| AC096711.2 | 73.9947162 | -0.2314173 | 0.43791338 | -0.5284546 | 0.59718386 | 0.7894568 |
| AC109927.1 | 23.3868121 | -0.255017 | 0.52403925 | -0.4866373 | 0.62651541 | 0.8053173 |
| NOCT | 20.2141531 | -0.6589777 | 0.41368778 | -1.5929348 | 0.11117482 | NA |
| ELF2 | 430.436733 | -0.0467735 | 0.19874384 | -0.2353458 | 0.81394034 | 0.90970797 |
| NDUFC1 | 32.7285223 | 0.15453967 | 0.45626232 | 0.33870795 | 0.73482975 | 0.86814805 |
| NAA15 | 125.456193 | -0.369611 | 0.28053657 | -1.3175145 | 0.18766623 | 0.43816783 |
| AC097376.2 | 183.140797 | -0.6357112 | 0.37269349 | -1.7057211 | 0.08806001 | 0.29675763 |
| SETD7 | 18.9830975 | -0.2301855 | 0.50378363 | -0.4569134 | 0.6477333 | NA |
| MGST2 | 29.3156412 | 1.75851449 | 0.48247352 | 3.64478969 | 0.00026761 | 0.01346273 |
| MAML3 | 285.159296 | 0.54148179 | 0.33112614 | 1.63527346 | 0.10199173 | 0.32094455 |
| SCOC | 61.4570356 | 0.41598457 | 0.26072442 | 1.59549525 | 0.11060153 | 0.3339741 |
| ELMOD2 | 41.126201 | -0.0572492 | 0.36050246 | -0.1588039 | 0.87382339 | 0.94120049 |
| TBC1D9 | 524.545935 | -0.1739435 | 0.42406259 | -0.4101835 | 0.68167136 | 0.84151154 |
| RNF150 | 421.220855 | 0.69351631 | 0.3347889 | 2.07150332 | 0.03831178 | 0.19533626 |
| ZNF330 | 27.6002587 | -0.2639571 | 0.44157428 | -0.5977638 | 0.54999755 | 0.75662755 |
| IL15 | 23.9849514 | 0.18357654 | 0.59012675 | 0.31107985 | 0.75573992 | 0.87904624 |
| INPP4B | 300.105636 | 0.01703019 | 0.33379986 | 0.05101915 | 0.95931026 | 0.9841145 |
| AC104596.1 | 37.6023281 | 2.7989647 | 1.02302597 | 2.73596642 | 0.00621974 | 0.07721978 |
| USP38 | 33.8546627 | -0.7363801 | 0.37743307 | -1.9510217 | 0.05105447 | 0.22662412 |
| GAB1 | 405.523452 | -0.9285589 | 0.24659152 | -3.7655752 | 0.00016617 | 0.00964706 |
| SMARCA5 | 79.808525 | -0.2532149 | 0.2877982 | -0.8798349 | 0.37894878 | 0.63024389 |
| ANAPC10 | 118.153432 | 0.06886183 | 0.28823373 | 0.23890969 | 0.81117561 | 0.90882368 |
| ABCE1 | 37.7732101 | -0.1020329 | 0.41408817 | -0.2464039 | 0.80536962 | 0.90521758 |
| OTUD4 | 96.7350925 | 0.10813122 | 0.2416711 | 0.44743133 | 0.65456366 | 0.82296251 |
| SMAD1 | 86.1101185 | 0.77447808 | 0.24815972 | 3.12088556 | 0.00180308 | 0.03909384 |
| MMAA | 19.7819343 | -0.1264466 | 0.43644664 | -0.2897183 | 0.77203176 | NA |
| ZNF827 | 158.932274 | 0.59947025 | 0.25677949 | 2.33457211 | 0.01956578 | 0.13704876 |
| LSM6 | 17.3208013 | 0.33847734 | 0.4835622 | 0.6999665 | 0.48394823 | NA |
| SLC10A7 | 96.9754153 | 0.42562549 | 0.26372305 | 1.61391085 | 0.10654676 | 0.32738142 |
| TMEM184C | 43.5749818 | 0.05454159 | 0.37390683 | 0.14586947 | 0.88402444 | 0.94596637 |
| PRMT9 | 44.351475 | 0.37436306 | 0.28673108 | 1.3056243 | 0.19168035 | 0.44319545 |
| ARHGAP10 | 26.2200971 | -1.2965946 | 0.59964171 | -2.1622823 | 0.03059642 | 0.17296809 |
| NR3C2 | 1325.04746 | 0.8246164 | 0.25945583 | 3.17825343 | 0.00148165 | 0.03500548 |
| DCLK2 | 85.6270487 | -0.5361941 | 0.57989555 | -0.9246391 | 0.35515364 | 0.61168935 |
| LRBA | 1591.25331 | 0.052352 | 0.1306667 | 0.40065296 | 6.89E-01 | 8.45E-01 |
| RPS3A | 38.5347259 | 0.67852219 | 0.39539207 | 1.71607435 | 0.08614844 | 0.29319258 |
| SH3D19 | 296.645635 | -1.0110118 | 0.39860973 | -2.5363449 | 0.01120163 | 0.1036885 |
| FAM160A1 | 415.984527 | -0.7170923 | 0.325933 | -2.2001217 | 0.02779827 | 0.16446061 |
| GATB | 31.2334005 | 0.31155612 | 0.44480259 | 0.70043685 | 0.48365453 | 0.71144239 |
| FBXW7 | 308.022935 | -0.3317394 | 0.30008097 | -1.1054996 | 0.26894316 | 0.52756125 |
| ARFIP1 | 176.855299 | -0.0325627 | 0.21110305 | -0.1542504 | 0.87741232 | 0.9428232 |
| FHDC1 | 18.9938831 | -1.2569261 | 0.61592064 | -2.0407273 | 0.04127794 | NA |
| TRIM2 | 420.956641 | 0.05123116 | 0.20620409 | 0.24844881 | 0.80378717 | 0.90408418 |
| TMEM131L | 30.0730233 | -0.1619253 | 0.39712335 | -0.4077456 | 0.68346046 | 0.8424322 |
| PLRG1 | 26.514761 | -0.6224855 | 0.42428108 | -1.4671535 | 0.1423343 | 0.38005954 |
| MAP9 | 27.220781 | 0.49227274 | 0.4329264 | 1.13708182 | 0.25550408 | 0.51347648 |
| GUCY1A1 | 48.5862963 | 0.30741668 | 0.53890392 | 0.57044804 | 0.56837385 | 0.77103518 |

| | | | | | | |
|---|---|---|---|---|---|---|
| CTSO | 13.858839 | -0.1829196 | 0.59128945 | -0.3093571 | 0.75704993 | NA |
| PDGFC | 88.3846463 | -0.191787 | 0.65061149 | -0.2947796 | 0.76816226 | 0.88524403 |
| ETFDH | 87.1166011 | 0.58114282 | 0.32374434 | 1.79506711 | 7.26E-02 | 0.26848318 |
| PPID | 33.7726638 | 0.00050706 | 0.33239705 | 0.00152546 | 0.99878286 | 0.99956204 |
| FNIP2 | 888.893715 | -1.9031561 | 0.51336328 | -3.7072306 | 0.00020954 | 0.01135785 |
| C4orf45 | 30.1117908 | -0.6120894 | 0.68297791 | -0.8962068 | 0.37014233 | 0.62368335 |
| RAPGEF2 | 378.975308 | -0.027901 | 0.27502789 | -0.101448 | 0.9191948 | 0.9647335 |
| NAF1 | 48.1725923 | -0.4872487 | 0.36725929 | -1.3267159 | 0.18460269 | 0.43476148 |
| NPY1R | 69.054956 | 0.29935502 | 0.65423877 | 0.45756233 | 0.64726691 | 0.81885075 |
| TMA16 | 23.9581413 | 0.65968188 | 0.36856264 | 1.78987725 | 0.07347365 | 0.26995071 |
| TMEM192 | 42.1296133 | -0.4829663 | 0.45276556 | -1.0667028 | 0.28610604 | 0.54522362 |
| KLHL2 | 136.541507 | -0.6951758 | 0.41542557 | -1.6734063 | 9.42E-02 | 0.30830642 |
| MSMO1 | 36.4326905 | 0.53063654 | 0.34221838 | 1.55057872 | 0.12100268 | 0.35000285 |
| DDX60 | 52.5818969 | 0.93561268 | 1.0255193 | 0.91233064 | 0.36159469 | 0.61739005 |
| DDX60L | 43.3480366 | 0.98705083 | 1.49211155 | 0.66151276 | 0.50828353 | 0.72894696 |
| PALLD | 163.717519 | 0.68372415 | 0.37519546 | 1.82231457 | 0.06840727 | 0.26001518 |
| CBR4 | 392.656185 | 0.9064377 | 0.32120839 | 2.8219615 | 0.00477309 | 0.06667601 |
| SH3RF1 | 616.26788 | 0.68298159 | 0.2971297 | 2.29859755 | 0.0215278 | 0.14349647 |
| NEK1 | 183.681511 | 0.16594703 | 0.20005395 | 0.82951137 | 0.40681511 | 0.65367876 |
| CLCN3 | 128.319624 | 0.35236079 | 0.26657171 | 1.32182363 | 0.18622689 | 0.43607799 |
| HPF1 | 20.4402172 | 0.68812693 | 0.40555142 | 1.69676863 | 0.08974041 | NA |
| MFAP3L | 36.3403799 | -0.0518211 | 0.32938626 | -0.1573263 | 0.87498769 | 0.94169674 |
| LINC02512 | 12.5061403 | 1.65195967 | 1.02410491 | 1.6130766 | 0.10672787 | NA |
| GALNT7 | 91.3871799 | 0.81920247 | 0.27048292 | 3.02866615 | 0.00245636 | 0.04714863 |
| HMGB2 | 18.6379921 | 0.19803748 | 0.51938549 | 0.3812919 | 0.70298666 | NA |
| FBXO8 | 23.6241144 | 0.27320529 | 0.42195212 | 0.64747936 | 0.51732175 | 0.73469647 |
| CEP44 | 48.4703348 | 0.50854604 | 0.35837276 | 1.41904213 | 0.15588673 | 0.39902235 |
| WDR17 | 28.7668248 | 0.60952974 | 0.51027276 | 1.19451748 | 0.23227561 | 0.49066933 |
| SPCS3 | 51.5833883 | -0.0679099 | 0.39554598 | -0.1716865 | 0.86368398 | 0.93643831 |
| AC078881.1 | 51.0075955 | 0.23395569 | 0.34548237 | 0.6771856 | 0.4982882 | 0.72149172 |
| LINC01099 | 50.3241111 | 2.23138404 | 0.86030431 | 2.59371481 | 0.00949452 | 0.09644302 |
| DCTD | 14.8743066 | 0.68829858 | 0.53865563 | 1.2778082 | 0.20131706 | NA |
| WWC2 | 241.179483 | -0.3037661 | 0.23579459 | -1.2882656 | 0.19765352 | 0.45090075 |
| RWDD4 | 27.0825469 | 0.20978235 | 0.38848385 | 0.54000274 | 0.58919514 | 0.78397234 |
| TRAPPC11 | 78.7779102 | -0.1402588 | 0.23346796 | -0.6007627 | 0.54799807 | 0.7554786 |
| STOX2 | 192.144996 | 0.89820606 | 0.67508443 | 1.33050922 | 0.18335055 | 0.4332917 |
| IRF2 | 69.6965586 | -0.0706908 | 0.25024466 | -0.2824869 | 0.77757017 | 0.8896927 |
| LINC02427 | 12.0745946 | -0.5238106 | 0.85304202 | -0.6140502 | 0.53918213 | NA |
| CASP3 | 29.4428482 | -0.913935 | 0.36284528 | -2.5188007 | 0.01177553 | 0.10625585 |
| PRIMPOL | 30.5860172 | 1.09902036 | 0.57605639 | 1.90783467 | 0.05641259 | 0.23662544 |
| ACSL1 | 210.956029 | -0.4419736 | 0.44436236 | -0.9946243 | 0.31991903 | 0.57721912 |
| LINC02437 | 19.6979232 | -0.126162 | 0.94993179 | -0.1328116 | 0.89434235 | NA |
| SLC25A4 | 47.2725762 | 1.77077083 | 0.43251926 | 4.09408551 | 4.24E-05 | 0.00401254 |
| CFAP97 | 65.1842209 | 0.18525024 | 0.39351296 | 0.47076021 | 0.63781199 | 0.81305711 |
| SNX25 | 189.629536 | -0.5902628 | 0.27065935 | -2.1808328 | 0.02919579 | 0.16858212 |
| LRP2BP | 22.8125874 | 0.89160264 | 0.48410266 | 1.84176356 | 0.06550974 | 0.25452134 |
| AC112722.1 | 25.6184554 | -0.4061104 | 0.36697929 | -1.1066303 | 0.26845378 | 0.52726441 |
| ANKRD37 | 25.5663585 | -1.1030587 | 0.89892935 | -1.2270805 | 0.21979233 | 0.47614454 |
| UFSP2 | 51.126939 | 0.40524656 | 0.34012037 | 1.19147984 | 0.23346527 | 0.49233972 |
| C4orf47 | 72.7863994 | -0.0615835 | 0.5751255 | -0.1070784 | 0.91472677 | 0.96388328 |
| SORBS2 | 265.611162 | 1.87428669 | 0.57636064 | 3.25193388 | 0.00114623 | 0.03021158 |
| FAM149A | 15.7477535 | -0.0208744 | 0.58133306 | -0.0359079 | 0.97135581 | NA |
| CYP4V2 | 25.6317388 | 0.49511984 | 0.42380748 | 1.16826595 | 0.24269951 | 0.50001555 |
| KLKB1 | 14.2805854 | 0.66928223 | 0.52022248 | 1.28653079 | 0.19825788 | NA |

| | | | | | | |
|---|---|---|---|---|---|---|
| F11 | 30.0607927 | 1.24847285 | 0.59175093 | 2.10979447 | 3.49E-02 | 0.18551559 |
| FAT1 | 146.814777 | -0.3652557 | 0.30438807 | -1.1999673 | 0.23015204 | 0.48881906 |
| FRG1-DT | 39.7914018 | 1.12576416 | 0.65216008 | 1.72620834 | 0.08430994 | 0.28931012 |
| FRG1 | 17.696095 | 0.04722644 | 0.44938048 | 0.10509233 | 0.91630255 | NA |
| SDHA | 99.625257 | -0.0709715 | 0.32368182 | -0.2192633 | 0.82644496 | 0.9158 |
| PDCD6 | 61.9090135 | -0.8872916 | 0.3174786 | -2.7948075 | 0.00519306 | 0.07024781 |
| EXOC3 | 56.2718539 | -0.7354335 | 0.33398467 | -2.2019978 | 0.02766547 | 0.16405324 |
| ZDHHC11B | 38.4740899 | 0.92596125 | 0.48716929 | 1.90069707 | 0.0573417 | 0.23805776 |
| BRD9 | 30.5727385 | -0.507505 | 0.3652233 | -1.3895746 | 0.1646581 | 0.41135131 |
| SLC12A7 | 51.211024 | -0.8202643 | 0.58117702 | -1.4113845 | 0.15813126 | 0.40275854 |
| CLPTM1L | 27.6078348 | -0.4437654 | 0.43738231 | -1.014594 | 0.31029942 | 0.56995192 |
| LPCAT1 | 45.619478 | -1.4261973 | 0.50526102 | -2.8226941 | 0.0047622 | 0.06664966 |
| NDUFS6 | 39.8680037 | -0.3215533 | 0.49426543 | -0.6505681 | 0.51532534 | 0.73303625 |
| AC022424.1 | 39.5309026 | 0.75976236 | 0.7548748 | 1.00647467 | 0.31418729 | 0.57172406 |
| ADAMTS16 | 301.72658 | 0.68720268 | 0.45884494 | 1.49767955 | 0.13421653 | 0.36808465 |
| ICE1 | 104.814421 | -0.256121 | 0.21349988 | -1.1996308 | 0.23028277 | 0.48896178 |
| MED10 | 23.8148526 | -0.1370133 | 0.4025791 | -0.3403389 | 0.73360136 | 0.86776236 |
| NSUN2 | 50.7991666 | -0.65408 | 0.36145373 | -1.8095817 | 0.07036069 | 0.26482455 |
| SRD5A1 | 33.5420106 | 0.08018189 | 0.56038574 | 0.14308338 | 0.88622433 | 0.94726686 |
| TENT4A | 75.36676 | -0.3639977 | 0.26430327 | -1.3771973 | 0.16845127 | 0.41629954 |
| MTRR | 37.7564208 | -0.5092344 | 0.34034069 | -1.496249 | 0.13458878 | 0.36839609 |
| SEMA5A | 80.7009897 | -1.9746704 | 0.62899308 | -3.1394151 | 0.00169285 | 0.03787762 |
| FAM173B | 40.8853684 | 0.63069919 | 0.40466928 | 1.55855463 | 0.11910183 | 0.34671965 |
| CCT5 | 32.8269227 | 0.29825991 | 0.43408518 | 0.68709996 | 0.49201974 | 0.71672078 |
| CMBL | 39.9632828 | 0.29220458 | 0.61021372 | 0.47885613 | 0.63204098 | 0.80927518 |
| 6-Mar | 311.461068 | -0.530291 | 0.17800318 | -2.9791095 | 0.00289087 | 0.05257034 |
| ANKRD33B | 50.8799878 | -0.0419509 | 0.67603813 | -0.0620541 | 0.95051978 | 0.97993174 |
| DAP | 78.4203769 | 0.19206906 | 0.28230731 | 0.68035452 | 0.49628001 | 0.72045779 |
| TRIO | 630.90415 | -0.4663788 | 0.29583055 | -1.5765066 | 0.1149091 | 0.34128941 |
| OTULIN | 39.6506672 | -0.1419315 | 0.30178234 | -0.4703107 | 0.63813306 | 0.81325069 |
| ANKH | 42.0793675 | 0.17322782 | 0.31182689 | 0.5555256 | 0.57853521 | 0.77576204 |
| RETREG1 | 320.995686 | -0.6678677 | 0.60021848 | -1.1127077 | 0.26583398 | 0.52505111 |
| MYO10 | 430.434672 | -0.3360682 | 0.40132259 | -0.8374016 | 0.40236684 | 0.65015704 |
| LINC02241 | 45.3512051 | 0.42633424 | 0.34405806 | 1.23913458 | 0.21529566 | 0.47115856 |
| CDH12 | 17.5126851 | 1.59574837 | 0.7152092 | 2.23116311 | 0.02567033 | NA |
| C5orf17 | 81.8169563 | 0.44446091 | 0.71191824 | 0.62431454 | 0.53242104 | 0.74604856 |
| CDH6 | 12.7987287 | 0.05706455 | 0.92238484 | 0.06186632 | 0.95066929 | NA |
| DROSHA | 79.3631243 | 0.38563161 | 0.27015022 | 1.42747101 | 0.15344418 | 0.39526768 |
| C5orf22 | 26.209926 | 0.05304275 | 0.34168314 | 0.15523958 | 0.87663244 | 0.94225833 |
| PDZD2 | 48.2184837 | 0.69509801 | 0.47127852 | 1.47491979 | 0.14023411 | 0.37806723 |
| GOLPH3 | 133.931463 | -0.0932495 | 0.25699815 | -0.3628411 | 0.71672361 | 0.8589283 |
| MTMR12 | 90.5694512 | 0.71058169 | 0.29227596 | 2.43120129 | 0.01504885 | 0.12124045 |
| ZFR | 248.821404 | -0.1346302 | 0.20481523 | -0.6573253 | 0.5109718 | 0.73060499 |
| SUB1 | 77.6652085 | -0.1048221 | 0.26075815 | -0.4019899 | 0.68769144 | 0.84528282 |
| NPR3 | 33.7844629 | 1.40184626 | 0.5391681 | 2.60001705 | 0.00932191 | 0.09528654 |
| TARS | 37.3541095 | -0.6890729 | 0.33476762 | -2.058362 | 0.0395554 | 0.19795704 |
| C1QTNF3-AN | 161.810543 | 0.54992424 | 0.25145159 | 2.1869985 | 0.02874263 | 0.16646505 |
| RAI14 | 155.034178 | -1.0257713 | 0.2959663 | -3.4658381 | 0.00052858 | 0.02087746 |
| RAD1 | 15.5370911 | -0.1266622 | 0.49237846 | -0.2572456 | 0.79698921 | NA |
| BRIX1 | 17.0936966 | 0.4546231 | 0.49228876 | 0.92348867 | 0.3557526 | NA |
| DNAJC21 | 55.5451006 | 0.14751522 | 0.26964715 | 0.54706761 | 0.58433229 | 0.78002178 |
| PRLR | 46.6447772 | 0.01326168 | 0.38077698 | 0.03482795 | 9.72E-01 | 0.98878882 |
| SPEF2 | 64.4851059 | 1.23239463 | 0.75337498 | 1.63583164 | 0.10187483 | 0.32089434 |
| LMBRD2 | 39.9599044 | 0.61545698 | 0.3176246 | 1.93768676 | 0.05266145 | 0.23043501 |

| | | | | | | |
|---|---|---|---|---|---|---|
| SKP2 | 14.0326264 | 0.64356269 | 0.59116223 | 1.08863972 | 0.27631279 | NA |
| NADK2 | 55.3042857 | 0.4460627 | 0.38024459 | 1.17309414 | 0.24075806 | 0.49828308 |
| RANBP3L | 159.746119 | 1.75315043 | 0.43232635 | 4.05515516 | 5.01E-05 | 0.00448404 |
| NIPBL | 539.966587 | -0.7095527 | 0.22359081 | -3.1734431 | 0.00150642 | 0.03507986 |
| CPLANE1 | 165.278049 | 0.17784453 | 0.26058449 | 0.68248317 | 0.49493348 | 0.719004 |
| NUP155 | 35.8134314 | -0.1431808 | 0.3399726 | -0.4211538 | 0.67364278 | 0.83710502 |
| WDR70 | 288.068571 | -0.1175367 | 0.16811276 | -0.6991539 | 0.48445588 | 0.71174975 |
| LIFR | 228.44605 | -0.1332119 | 0.20322553 | -0.6554882 | 5.12E-01 | 7.31E-01 |
| OSMR-AS1 | 188.834406 | 0.38038874 | 0.37068489 | 1.02617815 | 3.05E-01 | 0.56546258 |
| OSMR | 222.852402 | -0.406701 | 0.29191185 | -1.3932323 | 1.64E-01 | 4.10E-01 |
| RICTOR | 274.461466 | -0.3463319 | 0.20448082 | -1.6937134 | 0.09031976 | 0.30003934 |
| TTC33 | 43.5749531 | 0.81461068 | 0.35771998 | 2.27723003 | 0.02277249 | 0.14700765 |
| PRKAA1 | 66.8250721 | 0.116835 | 0.29220101 | 0.39984461 | 0.68927097 | 0.84569856 |
| RPL37 | 52.8941833 | 0.0628526 | 0.4024851 | 0.15616132 | 0.87590587 | 0.94201191 |
| OXCT1 | 71.2301957 | -0.8380746 | 0.54582667 | -1.5354225 | 0.12468008 | 0.35504829 |
| C5orf51 | 15.6865337 | 0.29196067 | 0.49770707 | 0.58661145 | 0.55746469 | NA |
| GHR | 198.355804 | 0.18054242 | 0.49876124 | 0.36198165 | 7.17E-01 | 0.85911998 |
| SELENOP | 42.9582425 | 1.10834337 | 0.34034281 | 3.25655002 | 0.00112775 | 0.03009201 |
| ZNF131 | 121.963963 | -0.2137462 | 0.29482898 | -0.7249838 | 0.46846196 | 0.69838306 |
| HMGCS1 | 63.4352998 | -0.1721884 | 0.37938306 | -0.4538643 | 0.64992647 | 0.81947559 |
| TMEM267 | 44.3468799 | 0.26968242 | 0.33106281 | 0.81459595 | 0.41530363 | 0.65906718 |
| PAIP1 | 51.6431643 | -0.2622041 | 0.31634596 | -0.8288523 | 0.40718798 | 0.65367876 |
| NNT-AS1 | 89.9479689 | 0.15132164 | 0.28088931 | 0.53872338 | 0.59007774 | 0.78469736 |
| NNT | 278.635232 | -0.1603255 | 0.22780646 | -0.7037795 | 4.82E-01 | 7.09E-01 |
| MRPS30-DT | 39.861086 | 0.28165031 | 0.4986077 | 0.56487358 | 0.57215976 | 0.77272682 |
| MRPS30 | 24.0691271 | 0.13976672 | 0.38948245 | 0.35885242 | 0.7197055 | 0.85995641 |
| PARP8 | 113.976088 | -0.1364437 | 0.31326336 | -0.4355559 | 0.66315898 | 0.8287603 |
| ITGA1 | 56.7218675 | -1.2889627 | 0.39515636 | -3.2619055 | 0.00110666 | 0.02988759 |
| ITGA2 | 798.783465 | -2.5230532 | 0.51497512 | -4.8993691 | 9.61E-07 | 0.00043531 |
| MOCS2 | 23.3312026 | 0.8920925 | 0.39205982 | 2.27539894 | 0.02288201 | 0.1471166 |
| NDUFS4 | 151.60266 | 0.05093295 | 0.23892021 | 0.21317974 | 0.83118677 | 0.9181376 |
| ARL15 | 439.048143 | 0.66140993 | 0.5704035 | 1.15954746 | 0.2462331 | 0.50296216 |
| SNX18 | 24.5830587 | 0.89755806 | 0.50179138 | 1.78870761 | 0.07366192 | 0.27037471 |
| DHX29 | 62.0148953 | 0.16133122 | 0.27616974 | 0.58417415 | 0.55910315 | 0.76485713 |
| MTREX | 126.454414 | -0.0186082 | 0.21168126 | -0.0879065 | 0.92995099 | 0.97116166 |
| PLPP1 | 102.088353 | -0.1116985 | 0.29985999 | -0.372502 | 0.70951909 | 0.85607462 |
| SLC38A9 | 66.0378339 | 0.24473621 | 0.36691361 | 0.66701317 | 0.50476372 | 0.72633508 |
| IL6ST | 145.202742 | -0.497635 | 0.28417694 | -1.751145 | 0.07992095 | 0.28223671 |
| AC008892.1 | 46.6486915 | -0.2064837 | 0.4668586 | -0.4422831 | 6.58E-01 | 0.82521413 |
| LINC01948 | 15.0304605 | 0.80981457 | 0.57903143 | 1.39856754 | 0.16194271 | NA |
| MAP3K1 | 197.47001 | -0.0239234 | 0.28704348 | -0.0833441 | 0.93357794 | 0.9730757 |
| MIER3 | 28.5538899 | -0.5888507 | 0.36314658 | -1.6215233 | 0.10490545 | 0.32571894 |
| GPBP1 | 251.186796 | -0.3841941 | 0.19745349 | -1.9457447 | 0.05168542 | 0.22810932 |
| PDE4D | 4568.64577 | 0.08046591 | 0.25833394 | 0.31148022 | 0.75543558 | 0.87904624 |
| DEPDC1B | 159.219719 | -1.2208437 | 0.61839793 | -1.9742041 | 0.04835855 | 0.22050695 |
| ELOVL7 | 294.995628 | -0.6888094 | 0.29476105 | -2.33684 | 0.0194475 | 0.13682581 |
| ERCC8 | 61.6831459 | 0.34048556 | 0.3268246 | 1.04179906 | 0.29750485 | 0.55742328 |
| NDUFAF2 | 117.883166 | -0.2442452 | 0.34111384 | -0.7160226 | 0.47397737 | 0.7026171 |
| SMIM15-AS1 | 11.5465515 | 0.97312214 | 0.65290654 | 1.49044632 | 0.13610692 | NA |
| ZSWIM6 | 627.933031 | -0.9185966 | 0.30626179 | -2.9993836 | 0.00270526 | 0.05029571 |
| KIF2A | 71.8725649 | -0.4393517 | 0.30259923 | -1.4519261 | 0.14652215 | 0.38645 |
| IPO11 | 53.3900378 | -0.3540476 | 0.32554015 | -1.0875698 | 0.27678509 | 0.53501126 |
| RNF180 | 138.447653 | 0.08207475 | 0.24864595 | 0.33008682 | 0.74133436 | 0.87152705 |
| SREK1IP1 | 52.0559714 | -0.0780466 | 0.26397885 | -0.2956547 | 0.76749377 | 0.88473859 |

| | | | | | | |
|---|---|---|---|---|---|---|
| CWC27 | 117.217859 | 0.53636357 | 0.27475935 | 1.95212128 | 0.05092381 | 0.2264359 |
| PPWD1 | 56.7526841 | 0.36009166 | 0.28174282 | 1.27808639 | 0.20121897 | 0.45528981 |
| TRIM23 | 39.3641583 | 0.32818099 | 0.36170341 | 0.90732069 | 0.36423723 | 0.62003028 |
| TRAPPC13 | 59.3261709 | 0.00094734 | 0.27987877 | 0.00338483 | 0.9972993 | 0.99868755 |
| NLN | 32.3237433 | -0.1894941 | 0.37109173 | -0.5106395 | 0.60960351 | 0.79648926 |
| ERBIN | 467.72294 | -0.3043486 | 0.15212266 | -2.0006789 | 0.045427 | 0.21335013 |
| SREK1 | 150.085099 | -0.2075694 | 0.29697523 | -0.6989453 | 0.48458621 | 0.71180536 |
| MAST4 | 1325.86291 | -1.2913653 | 0.37926499 | -3.4049157 | 0.00066184 | 0.02347567 |
| PIK3R1 | 141.702336 | 0.8313535 | 0.39240035 | 2.11863599 | 0.03412124 | 0.18312899 |
| AC093523.1 | 16.7958793 | -0.3623381 | 0.71971403 | -0.5034473 | 6.15E-01 | NA |
| SLC30A5 | 64.6427388 | -0.6431466 | 0.34481317 | -1.8652031 | 0.06215294 | 0.24899105 |
| MRPS36 | 35.6273122 | 0.57930063 | 0.32138244 | 1.80252734 | 0.07146248 | 0.26675397 |
| CDK7 | 36.6774765 | 0.14585198 | 0.39199553 | 0.37207561 | 0.70983656 | 0.85607462 |
| CCDC125 | 72.4130939 | 0.65803663 | 0.40014785 | 1.64448374 | 0.10007632 | 0.31841979 |
| AK6 | 23.125924 | -0.526066 | 0.38523583 | -1.3655688 | 0.17207436 | 0.42066785 |
| RAD17 | 49.3040656 | -0.1303218 | 0.28048624 | -0.4646283 | 0.64219764 | 0.81594508 |
| MARVELD2 | 45.1328318 | 0.0390418 | 0.36814414 | 0.10605031 | 0.91554243 | 0.96388328 |
| OCLN | 118.020243 | 0.00831166 | 0.25824421 | 0.03218527 | 0.9743243 | 0.9901947 |
| GTF2H2C | 34.3067603 | 0.20903786 | 0.44590922 | 0.46879018 | 0.63921962 | 0.81390792 |
| NAIP | 149.072342 | 0.90266704 | 0.38992843 | 2.31495569 | 0.02061536 | 0.14099415 |
| GTF2H2 | 31.5225295 | 0.38462188 | 0.58043676 | 0.66264218 | 0.50755975 | 0.72858773 |
| BDP1 | 182.052724 | -0.320463 | 0.21431412 | -1.4952959 | 0.13483722 | 0.36881383 |
| MCCC2 | 70.3298924 | -0.0218086 | 0.27414422 | -0.0795514 | 0.93659401 | 0.97405271 |
| MRPS27 | 63.7483578 | 0.38008034 | 0.31107811 | 1.22181641 | 0.22177706 | 0.47896129 |
| PTCD2 | 18.991911 | 0.92777239 | 0.59034105 | 1.57158712 | 0.11604634 | NA |
| TNPO1 | 204.443898 | -0.1395326 | 0.19796383 | -0.7048389 | 0.48091049 | 0.70856969 |
| FCHO2 | 186.633564 | 0.00978577 | 0.25950665 | 0.03770912 | 0.9699196 | 0.98788821 |
| BTF3 | 33.6465487 | 0.41636068 | 0.33414745 | 1.24603879 | 0.21275016 | 0.46813549 |
| ANKRA2 | 20.5824936 | 0.39881456 | 0.37943545 | 1.05107354 | 0.29322482 | NA |
| ARHGEF28 | 471.85136 | -0.1732439 | 0.32286576 | -0.536582 | 0.5915564 | 0.78517152 |
| HEXB | 20.0426972 | -0.7011221 | 0.42026401 | -1.6682896 | 0.09525826 | NA |
| GFM2 | 40.9691899 | -0.0444375 | 0.31855812 | -0.1394958 | 0.88905839 | 0.94924156 |
| NSA2 | 24.487941 | -0.0579728 | 0.38158555 | -0.151926 | 0.87924526 | 0.94360719 |
| FAM169A | 58.0369766 | 0.90221957 | 0.38030262 | 2.37237272 | 0.01767425 | 0.13220477 |
| HMGCR | 56.9288433 | -0.2217727 | 0.42053624 | -0.527357 | 0.59794567 | 0.7894568 |
| COL4A3BP | 172.29221 | -0.5105178 | 0.2959706 | -1.7248937 | 0.08454662 | 0.28931012 |
| POLK | 130.724119 | 0.34657909 | 0.3034405 | 1.1421649 | 0.25338548 | 0.51069785 |
| ANKDD1B | 20.5154246 | 1.44054107 | 0.9027185 | 1.59578105 | 0.11053769 | NA |
| POC5 | 23.4914034 | -0.4934083 | 0.49356875 | -0.9996749 | 0.31746786 | 0.57495296 |
| IQGAP2 | 15.4224472 | 0.79915403 | 0.64658833 | 1.23595493 | 0.21647532 | NA |
| ZBED3 | 24.0518995 | 0.87109547 | 0.38464249 | 2.26468861 | 0.0235318 | 0.14961264 |
| WDR41 | 84.9138965 | 0.23257763 | 0.23235756 | 1.00094714 | 0.31685236 | 0.57464954 |
| TBCA | 163.605642 | -0.2883907 | 0.27069854 | -1.0653573 | 0.28671428 | 0.5461123 |
| AP3B1 | 340.682829 | 0.35698692 | 0.18845313 | 1.89430083 | 0.05818509 | 0.24020328 |
| SCAMP1 | 82.0637926 | 0.40595826 | 0.26941285 | 1.50682592 | 0.13185528 | 0.36533122 |
| LHFPL2 | 20.3440501 | -0.2978707 | 0.42831202 | -0.6954525 | 0.48677175 | NA |
| ARSB | 20.8125637 | -1.567982 | 0.47919924 | -3.2720879 | 0.00106756 | NA |
| DMGDH | 105.002199 | 1.22477247 | 0.4750285 | 2.57831364 | 0.00992838 | 0.09885999 |
| BHMT2 | 40.693288 | 1.05552082 | 0.54288758 | 1.94427143 | 0.05186273 | 0.22838257 |
| BHMT | 12.1159062 | 2.59026836 | 0.9290327 | 2.78813476 | 0.00530125 | NA |
| JMY | 82.7856182 | -0.2828683 | 0.27995675 | -1.0104002 | 0.31230361 | 0.57101636 |
| HOMER1 | 98.9118188 | -1.7861249 | 0.75675638 | -2.3602377 | 0.01826323 | 0.13368407 |
| TENT2 | 369.953034 | -0.4497762 | 0.20773928 | -2.1650994 | 0.03038007 | 0.17231791 |
| CMYA5 | 100.599374 | -0.0958532 | 0.32832168 | -0.2919491 | 0.77032553 | 0.88651526 |

| | | | | | | |
|---|---|---|---|---|---|---|
| MTX3 | 21.7191346 | 0.81937217 | 0.48759367 | 1.68044056 | 0.09287163 | 0.30571879 |
| THBS4 | 33.6104539 | -0.0939567 | 0.57432509 | -0.1635949 | 0.87005006 | 0.93963453 |
| SERINC5 | 70.2121384 | -0.2302683 | 0.33421317 | -0.6889863 | 0.49083191 | 0.7166095 |
| ZFYVE16 | 151.523873 | -0.5468138 | 0.19353871 | -2.8253457 | 0.00472296 | 0.06664966 |
| FAM151B | 29.6926577 | -0.1564454 | 0.55923751 | -0.2797476 | 0.77967116 | 0.89074516 |
| DHFR | 174.071921 | 0.10872081 | 0.48928053 | 0.22220547 | 0.82415394 | 0.91510572 |
| MSH3 | 130.106632 | 0.51549378 | 0.27908198 | 1.84710523 | 0.06473189 | 0.2529144 |
| RASGRF2 | 155.076907 | -1.18362 | 0.48384862 | -2.4462609 | 0.01443465 | 0.11844724 |
| CKMT2-AS1 | 67.6802369 | -0.2705838 | 0.41237444 | -0.6561605 | 0.51172089 | 0.73088062 |
| CKMT2 | 69.7361189 | 0.76975917 | 0.27649526 | 2.78398684 | 0.00536952 | 0.07124746 |
| ZCCHC9 | 23.7697085 | 0.11758012 | 0.47409343 | 0.24801045 | 0.80412632 | 0.90408418 |
| SSBP2 | 198.215696 | 0.84305818 | 0.33256654 | 2.53500599 | 0.01124454 | 0.1036885 |
| ATG10 | 119.0637 | 0.98091383 | 0.39088827 | 2.50944811 | 0.012092 | 0.1074201 |
| RPS23 | 82.8211069 | 0.59502458 | 0.38066737 | 1.56310899 | 0.11802696 | 0.34552837 |
| ATP6AP1L | 78.1882586 | 0.50446941 | 0.62866748 | 0.80244235 | 0.42229712 | 0.665797 |
| TMEM167A | 37.8676986 | 0.07039443 | 0.36581105 | 0.19243385 | 0.84740238 | 0.92622211 |
| XRCC4 | 88.8946796 | -0.1885066 | 0.36164743 | -0.5212442 | 0.60219665 | 0.79219067 |
| COX7C | 70.7402121 | 0.94438838 | 0.3837615 | 2.46087316 | 0.01385994 | 0.11532965 |
| RASA1 | 221.849326 | 0.2875438 | 0.20132457 | 1.42825986 | 0.15321708 | 0.39507934 |
| CCNH | 297.030131 | 0.22105717 | 0.34031787 | 0.64956087 | 0.51597591 | 0.73374372 |
| TMEM161B | 102.2279 | -0.3426244 | 0.28231046 | -1.2136441 | 0.22488366 | 0.48216659 |
| TMEM161B-. | 326.830374 | 0.09884056 | 0.24656548 | 0.40086941 | 0.68851628 | 0.84543868 |
| MEF2C | 72.8041257 | 0.22455051 | 0.63350773 | 0.35445583 | 0.7229973 | 0.86192273 |
| MEF2C-AS1 | 24.1463012 | 2.1171271 | 0.87171226 | 2.42869946 | 0.01515309 | 0.12124045 |
| LYSMD3 | 20.83187 | 0.13434843 | 0.38500021 | 0.34895678 | 0.72712176 | NA |
| ADGRV1 | 82.059493 | 0.33119428 | 0.40482086 | 0.81812554 | 0.41328551 | 0.65805928 |
| ARRDC3 | 262.786719 | -0.2030199 | 0.49450688 | -0.4105502 | 0.68140241 | 0.84151154 |
| ARRDC3-AS1 | 18.6521379 | -0.8238152 | 0.53310836 | -1.5453053 | 0.12227244 | NA |
| NR2F1-AS1 | 111.332744 | -0.9798024 | 0.52344987 | -1.8718171 | 0.06123192 | 0.24714319 |
| FAM172A | 391.444758 | 0.39721621 | 0.28838829 | 1.37736596 | 0.16839913 | 0.41629954 |
| KIAA0825 | 106.194959 | 1.87094927 | 0.60562645 | 3.08927933 | 0.00200643 | 0.04196595 |
| SLF1 | 22.7984553 | -0.3204281 | 0.39283424 | -0.8156827 | 0.41468166 | 0.65864728 |
| TTC37 | 139.273923 | -0.2932555 | 0.26482585 | -1.1073522 | 0.26814168 | 0.52717407 |
| ARSK | 19.0930023 | 1.18407803 | 0.54282786 | 2.18131404 | 0.0291602 | NA |
| RHOBTB3 | 501.508691 | 1.09062869 | 0.30529666 | 3.57235711 | 0.00035378 | 0.01620871 |
| GLRX | 63.8310198 | 0.93678023 | 0.65612725 | 1.42774168 | 0.15336623 | 0.39519914 |
| ELL2 | 105.948373 | -1.6901104 | 0.5418956 | -3.1188857 | 0.00181536 | 0.03924959 |
| AC104123.1 | 38.0129065 | 0.14940492 | 0.59352941 | 0.25172286 | 0.80125528 | 0.90376054 |
| CAST | 209.784447 | -0.5524443 | 0.25346667 | -2.1795541 | 0.02929053 | 0.16874939 |
| ERAP1 | 40.4191794 | 0.33920355 | 0.47380077 | 0.71592022 | 0.47404061 | 0.7026171 |
| AC009126.1 | 17.1836986 | 2.3598844 | 0.70810957 | 3.33265428 | 0.00086022 | NA |
| ERAP2 | 18.0404784 | 0.26355715 | 0.65214143 | 0.40414109 | 0.68610897 | NA |
| LNPEP | 238.329304 | 0.15319693 | 0.37582788 | 0.40762524 | 0.68354883 | 0.8424322 |
| LIX1-AS1 | 54.1129645 | 0.86756061 | 0.30982897 | 2.80012747 | 0.00510824 | 0.06958964 |
| RIOK2 | 12.8265034 | 0.88609564 | 0.50640691 | 1.74977004 | 0.080158 | NA |
| CHD1 | 177.647843 | -0.4416034 | 0.2958862 | -1.4924773 | 0.13557406 | 0.3700433 |
| LINC02062 | 63.4289708 | 0.00683448 | 0.60498237 | 0.01129699 | 0.9909865 | 0.99583841 |
| AC021086.1 | 12.2432786 | 0.67422236 | 0.72150547 | 0.93446604 | 0.35006355 | NA |
| FAM174A | 27.0715776 | 0.253757 | 0.38162614 | 0.66493611 | 0.50609136 | 0.72783729 |
| SLCO4C1 | 29.9853731 | -0.1006745 | 0.41214995 | -0.2442666 | 0.80702437 | 0.90641566 |
| PAM | 402.483766 | -0.5811341 | 0.26877673 | -2.1621442 | 0.03060706 | 0.17296809 |
| GIN1 | 24.4339358 | 0.45823542 | 0.47081427 | 0.97328278 | 0.33041278 | 0.5865284 |
| PPIP5K2 | 124.361653 | 0.3102229 | 0.25125288 | 1.23470388 | 0.21694073 | 0.47304841 |
| NUDT12 | 19.2413617 | 1.22658885 | 0.52210007 | 2.34933668 | 0.01880689 | NA |

| | | | | | | |
|---|---|---|---|---|---|---|
| LINC01950 | 27.5842341 | 0.85766194 | 0.83302504 | 1.02957523 | 0.30320945 | 0.56372056 |
| EFNA5 | 2474.06132 | 0.07750877 | 0.36198095 | 0.21412391 | 0.83045043 | 0.91785999 |
| FBXL17 | 723.769517 | 0.75980645 | 0.1389698 | 5.46742134 | 4.57E-08 | 7.03E-05 |
| FER | 333.545508 | -0.2838289 | 0.16891375 | -1.6803186 | 0.09289535 | 0.30571879 |
| AC008467.1 | 21.7475524 | -1.0242209 | 0.77275093 | -1.3254217 | 0.18503131 | 0.43476148 |
| PJA2 | 135.588821 | -0.1674579 | 0.20898242 | -0.8013012 | 0.42295727 | 0.66656472 |
| MAN2A1 | 246.612418 | -0.6044942 | 0.36417188 | -1.6599147 | 0.09693161 | 0.3125608 |
| TMEM232 | 144.349445 | 1.88394547 | 0.71709886 | 2.62717678 | 0.00860966 | 0.09090336 |
| SLC25A46 | 63.7570755 | 0.00751551 | 0.27524796 | 0.02730452 | 0.97821685 | 0.99161532 |
| WDR36 | 39.5831605 | 0.00054813 | 0.32675963 | 0.00167747 | 0.99866157 | 0.99956204 |
| STARD4 | 15.4729021 | -1.0847073 | 0.60882328 | -1.7816456 | 0.07480704 | NA |
| NREP | 53.9467432 | -1.0382521 | 0.42792181 | -2.426266 | 0.01525508 | 0.12155112 |
| EPB41L4A | 250.313608 | 0.23936476 | 0.26638961 | 0.89855143 | 0.36889164 | 0.62348681 |
| APC | 106.226795 | -0.1573994 | 0.28071838 | -0.5607023 | 0.57500052 | 0.77388388 |
| SRP19 | 31.8936315 | -0.3190987 | 0.43754032 | -0.7293013 | 0.46581738 | 0.69622439 |
| REEP5 | 100.6197 | 0.14969308 | 0.26248821 | 0.57028497 | 0.56848443 | 0.77103518 |
| DCP2 | 116.853294 | -0.9229854 | 0.29945351 | -3.0822326 | 0.00205454 | 0.04251024 |
| MCC | 138.458763 | -0.3565492 | 0.31017058 | -1.1495261 | 0.25033912 | 0.50659901 |
| YTHDC2 | 130.253348 | 0.43240786 | 0.2744567 | 1.57550487 | 0.11513996 | 0.34138376 |
| PGGT1B | 43.0168608 | 0.53015242 | 0.2938666 | 1.80405808 | 0.0712222 | 0.2665798 |
| CCDC112 | 15.5084316 | 0.222223 | 0.54678577 | 0.40641694 | 0.68443627 | NA |
| FEM1C | 34.1771645 | -1.1471096 | 0.48626666 | -2.3590134 | 0.01832359 | 0.13368407 |
| ATG12 | 49.0239302 | 0.10050325 | 0.30482529 | 0.32970772 | 0.74162082 | 0.87152705 |
| AP3S1 | 97.032533 | -0.2172497 | 0.26099223 | -0.832399 | 0.40518377 | 0.65250136 |
| COMMD10 | 312.032356 | -0.3285868 | 0.21858476 | -1.5032467 | 0.13277545 | 0.36682435 |
| SEMA6A | 118.01747 | -1.0309935 | 0.65426638 | -1.5758008 | 1.15E-01 | 3.41E-01 |
| SEMA6A-AS1 | 37.9090126 | 0.28516486 | 0.34412195 | 0.82867385 | 0.40728899 | 0.65367876 |
| DTWD2 | 89.4279365 | 0.29105774 | 0.39526892 | 0.7363537 | 0.46151547 | 0.69416206 |
| AC008629.1 | 35.4337262 | 1.48552016 | 0.72339995 | 2.05352539 | 0.04002165 | 0.19912517 |
| DMXL1 | 252.928535 | -0.0697282 | 0.22544766 | -0.3092879 | 0.75710254 | 0.87904624 |
| TNFAIP8 | 73.4265749 | -1.4313068 | 0.54981303 | -2.6032609 | 9.23E-03 | 9.48E-02 |
| HSD17B4 | 94.2860765 | 0.87146171 | 0.35317129 | 2.4675327 | 0.01360478 | 0.11413904 |
| SRFBP1 | 44.8185724 | -0.5980405 | 0.33989027 | -1.7595105 | 0.07849084 | 0.27941662 |
| SNX2 | 95.1169574 | -0.081331 | 0.28200663 | -0.2884011 | 0.77303969 | 0.88767514 |
| SNX24 | 189.003616 | 0.46576294 | 0.29741716 | 1.56602577 | 0.11734258 | 0.34472743 |
| PPIC | 18.2536397 | 0.7590288 | 0.43459758 | 1.74650949 | 0.08072246 | NA |
| CEP120 | 48.9290732 | -0.117166 | 0.3057464 | -0.3832129 | 0.70156189 | 0.85190818 |
| CSNK1G3 | 101.770369 | 0.49305781 | 0.38194826 | 1.2909021 | 0.19673763 | 0.45014552 |
| ZNF608 | 199.060193 | 0.45963733 | 0.41350861 | 1.11155444 | 0.26632977 | 0.52522169 |
| GRAMD2B | 294.189174 | 0.24508882 | 0.27693488 | 0.88500525 | 0.3761538 | 0.6276297 |
| ALDH7A1 | 37.1343774 | 0.80062179 | 0.49571668 | 1.61507938 | 0.10629351 | 0.32732986 |
| PHAX | 18.8748237 | -0.4283118 | 0.41886958 | -1.0225421 | 0.30652439 | NA |
| C5orf63 | 21.5270307 | 1.44330164 | 0.53320339 | 2.70685008 | 0.00679249 | 0.08022018 |
| PRRC1 | 37.3373934 | 0.08055279 | 0.3338624 | 0.24127543 | 0.80934166 | 0.90756938 |
| LINC01184 | 138.704184 | 0.69847778 | 0.29504226 | 2.36738211 | 0.01791443 | 0.13296754 |
| SLC12A2 | 124.165179 | 0.11763273 | 0.37579869 | 0.3130206 | 7.54E-01 | 8.79E-01 |
| SLC27A6 | 17.0991845 | -1.464555 | 1.76009994 | -0.8320863 | 0.40536024 | NA |
| ISOC1 | 20.8690485 | 0.76913446 | 0.51696002 | 1.48780261 | 0.13680296 | NA |
| HINT1 | 93.2527851 | 0.50220493 | 0.28987816 | 1.73246899 | 0.08319009 | 0.2884298 |
| LYRM7 | 28.0351374 | 0.81046471 | 0.41462109 | 1.95471175 | 0.0506171 | 0.22572413 |
| CDC42SE2 | 334.514846 | -0.4970026 | 0.41033401 | -1.2112148 | 2.26E-01 | 0.48347239 |
| SLC22A5 | 32.2976877 | 0.5522709 | 0.43649211 | 1.26524831 | 0.20578233 | 0.45902815 |
| AC116366.3 | 103.753918 | 0.08420124 | 0.56977567 | 0.14777964 | 0.88251668 | 0.94500986 |
| IRF1 | 23.4109332 | -1.2490096 | 0.76910118 | -1.6239861 | 0.10437878 | 0.32447637 |

| | | | | | | |
|---|---|---|---|---|---|---|
| KIF3A | 61.8033198 | 0.54464049 | 0.31232605 | 1.74382025 | 0.08119044 | 0.284185 |
| 8-Sep | 20.9090081 | 1.13924308 | 0.47916095 | 2.37757912 | 0.0174267 | NA |
| UQCRQ | 35.4477673 | 0.40093097 | 0.45749997 | 0.8763519 | 0.38083876 | 0.63229421 |
| AFF4 | 319.690317 | -0.7155324 | 0.30479408 | -2.3475928 | 0.01889517 | 0.1355543 |
| ZCCHC10 | 40.3016861 | -0.0654923 | 0.38653371 | -0.1694348 | 0.86545467 | 0.9374338 |
| HSPA4 | 61.2140445 | -0.1901127 | 0.33189554 | -0.5728089 | 0.56677408 | 0.77048041 |
| FSTL4 | 130.556976 | -1.5300561 | 0.6833057 | -2.239197 | 0.0251431 | 0.15519363 |
| C5orf15 | 54.7704987 | -0.1285434 | 0.3082852 | -0.4169625 | 0.67670584 | 0.83887358 |
| VDAC1 | 78.4923322 | 0.2867336 | 0.2934299 | 0.9771792 | 0.32848044 | 0.58471646 |
| TCF7 | 20.936284 | -0.1625554 | 0.43311718 | -0.3753151 | 7.07E-01 | NA |
| SKP1 | 78.757094 | 0.65235819 | 0.36933855 | 1.76628786 | 0.07734757 | 0.27731248 |
| CDKL3 | 42.9898745 | -0.1105996 | 0.30793605 | -0.3591642 | 0.71947223 | 0.85995641 |
| UBE2B | 149.62844 | -0.646853 | 0.3540491 | -1.8270148 | 0.06769754 | 0.25898011 |
| JADE2 | 31.0443557 | 0.02530127 | 0.51644706 | 0.04899102 | 0.96092645 | 0.98496344 |
| SAR1B | 50.7751289 | -0.4812628 | 0.4279317 | -1.1246252 | 0.2607479 | 0.52007685 |
| SEC24A | 105.142075 | -1.5279254 | 0.4171598 | -3.662686 | 0.00024958 | 0.01289505 |
| CAMLG | 22.6649686 | -0.3118691 | 0.3960494 | -0.7874501 | 0.43101845 | 0.67197671 |
| DDX46 | 118.893308 | -0.4598694 | 0.23684265 | -1.9416663 | 0.05217751 | 0.22883777 |
| C5orf24 | 23.6718014 | 0.58648167 | 0.40444331 | 1.45009612 | 0.14703172 | 0.38705441 |
| TXNDC15 | 32.9899801 | 0.05803586 | 0.56987166 | 0.10184023 | 0.9188835 | 0.9647335 |
| PCBD2 | 114.076262 | -0.287637 | 0.29336967 | -0.9804593 | 0.32685944 | 0.58313355 |
| H2AFY | 88.8209323 | -0.7287537 | 0.28125878 | -2.5910435 | 0.00956854 | 0.09652563 |
| CXCL14 | 35.2378175 | 1.82234814 | 0.665632 | 2.73777122 | 0.00618571 | 0.07721978 |
| SMAD5 | 82.3149352 | 0.07946104 | 0.24204271 | 0.32829349 | 0.74268977 | 0.87154797 |
| SPOCK1 | 180.773744 | -2.532451 | 0.85590377 | -2.9588035 | 0.00308836 | 0.05464622 |
| KLHL3 | 386.294043 | 1.16937846 | 0.41114176 | 2.84422207 | 4.45E-03 | 6.42E-02 |
| HNRNPA0 | 28.9998617 | -0.1699153 | 0.49797742 | -0.3412108 | 0.73294487 | 0.86776236 |
| AC106791.1 | 15.7386529 | 1.37031943 | 0.79213885 | 1.72989802 | 0.0836485 | NA |
| FAM13B | 227.222954 | -0.4743501 | 0.20425076 | -2.3223908 | 0.0202119 | 0.13990198 |
| NME5 | 44.9902019 | 1.03863743 | 0.61269791 | 1.69518682 | 0.09003999 | 0.29957137 |
| BRD8 | 53.4189222 | 0.68055412 | 0.30332474 | 2.24364854 | 0.02485502 | 0.15453079 |
| CDC23 | 21.683299 | -0.0256084 | 0.41701657 | -0.0614086 | 0.95103383 | 0.98032776 |
| FAM53C | 19.2071517 | 0.06424197 | 0.43021102 | 0.14932665 | 0.88129589 | NA |
| KDM3B | 124.070847 | -0.4853993 | 0.2398207 | -2.0240092 | 0.04296921 | 0.20709737 |
| ETF1 | 69.4920746 | -0.7097413 | 0.49136709 | -1.4444217 | 0.1486204 | 0.38909225 |
| HSPA9 | 145.057695 | -0.791232 | 0.44054095 | -1.7960465 | 0.07248712 | 0.26831077 |
| CTNNA1 | 371.426148 | -0.4738585 | 0.2097442 | -2.2592209 | 0.02386965 | 0.15095838 |
| SIL1 | 195.839937 | 0.05049122 | 0.25321365 | 0.19940164 | 0.84194858 | 0.92367136 |
| PAIP2 | 77.6625701 | -0.0668214 | 0.28284474 | -0.2362475 | 0.81324064 | 0.90970797 |
| UBE2D2 | 330.770314 | -0.6535739 | 0.43183339 | -1.5134862 | 0.13015619 | 0.36258133 |
| CXXC5 | 74.8804916 | 0.10668635 | 0.4190713 | 0.25457803 | 0.79904903 | 0.90206517 |
| NRG2 | 73.9384033 | -0.2966915 | 0.42433281 | -0.6991952 | 0.48443004 | 0.71174975 |
| PURA | 87.3850511 | 0.70868214 | 0.28956003 | 2.44744461 | 0.01438732 | 0.11818488 |
| CYSTM1 | 217.850831 | -0.2207903 | 0.32769931 | -0.673759 | 0.50046457 | 0.72244483 |
| PFDN1 | 56.5593786 | 0.18018628 | 0.29578652 | 0.60917677 | 0.54240728 | 0.75101796 |
| AC008438.1 | 14.7394923 | 0.1318195 | 0.66312706 | 0.19878468 | 0.84243118 | NA |
| ANKHD1-EIF4 | 15.8479676 | 0.57634135 | 0.50918886 | 1.13188129 | 0.25768434 | NA |
| SRA1 | 17.170158 | 0.09643876 | 0.43144233 | 0.22352642 | 0.82312583 | NA |
| NDUFA2 | 22.9138741 | 0.50362893 | 0.49519077 | 1.01704022 | 0.3091343 | 0.56869184 |
| IK | 18.3244546 | -0.2729754 | 0.44490246 | -0.6135623 | 0.53950462 | NA |
| WDR55 | 14.6465728 | -0.4360789 | 0.47691611 | -0.9143724 | 0.36052122 | NA |
| HARS | 35.2230157 | -0.053945 | 0.34753452 | -0.155222 | 0.8766463 | 0.94225833 |
| TAF7 | 25.5837692 | -0.3365933 | 0.45537113 | -0.7391626 | 0.45980824 | 0.69299864 |
| DIAPH1 | 321.926844 | -0.3428885 | 0.26049215 | -1.3163104 | 0.18806988 | 0.43860181 |

| | | | | | | |
|---|---|---|---|---|---|---|
| HDAC3 | 31.6219511 | 0.55779893 | 0.44090717 | 1.26511649 | 0.20582957 | 0.45902815 |
| PCDH1 | 39.5171656 | 0.27035529 | 0.40042977 | 0.67516283 | 0.49957232 | 0.72201362 |
| DELE1 | 33.6346864 | -0.2571831 | 0.37231488 | -0.6907678 | 0.4897115 | 0.7163358 |
| RNF14 | 39.4528629 | -0.3629736 | 0.30388098 | -1.1944598 | 0.23229818 | 0.49066933 |
| GNPDA1 | 22.1936552 | -0.9442911 | 0.56738987 | -1.6642721 | 0.09605807 | 0.31117801 |
| NDFIP1 | 187.687847 | -0.2534327 | 0.26566217 | -0.9539659 | 0.34010088 | 0.59602659 |
| SPRY4-AS1 | 20.6368205 | 1.55341833 | 0.6546275 | 2.37298056 | 1.76E-02 | NA |
| ARHGAP26 | 77.3875617 | -0.6593204 | 0.40936867 | -1.6105785 | 0.10727163 | 0.32921441 |
| YIPF5 | 15.082396 | 0.01791944 | 0.55555675 | 0.03225492 | 0.97426876 | NA |
| PRELID2 | 78.0254076 | 1.45976308 | 0.81121072 | 1.79948692 | 0.07194169 | 0.26750492 |
| LARS | 87.8813367 | 0.25857766 | 0.29676958 | 0.87130782 | 0.3835861 | 0.6343924 |
| TCERG1 | 123.29571 | -0.1820788 | 0.20609623 | -0.8834648 | 0.3769852 | 0.62858285 |
| STK32A | 72.5504082 | 1.94614509 | 0.59399509 | 3.27636559 | 0.00105152 | 0.02953506 |
| AC091948.1 | 51.8057109 | 1.36556359 | 0.55241472 | 2.47198985 | 0.01343633 | 0.11302673 |
| FBXO38 | 65.6227073 | -0.3725744 | 0.28965515 | -1.2862689 | 0.19834923 | 0.4514175 |
| CSNK1A1 | 436.902575 | -0.4995882 | 0.21122985 | -2.3651404 | 0.01802324 | 0.13313325 |
| ARHGEF37 | 20.2669003 | 1.3951844 | 0.60602268 | 2.30219831 | 0.02132399 | NA |
| SLC26A2 | 33.566518 | 0.59308493 | 0.78285031 | 0.75759686 | 0.44869236 | 0.68731807 |
| HMGXB3 | 46.1294518 | -0.485203 | 0.29873784 | -1.6241766 | 0.10433812 | 0.32447637 |
| TCOF1 | 13.507042 | -0.8931749 | 0.49306575 | -1.8114721 | 0.0700678 | NA |
| CD74 | 74.3838706 | 0.69043102 | 0.46961815 | 1.47019663 | 0.14150851 | 0.37950905 |
| RPS14 | 92.0663943 | 0.59370956 | 0.39928736 | 1.486923 | 0.13703515 | 0.37283829 |
| NDST1 | 30.1585219 | 0.16857993 | 0.37631233 | 0.44797876 | 0.65416853 | 0.82260009 |
| RBM22 | 13.5768916 | -0.3578874 | 0.4852235 | -0.7375723 | 0.46077442 | NA |
| DCTN4 | 71.6687215 | 0.25455033 | 0.26176979 | 0.97242058 | 0.33084136 | 0.58655523 |
| GPX3 | 302.562688 | -0.2934277 | 0.68700659 | -0.4271105 | 0.66929886 | 0.83376893 |
| TNIP1 | 92.6725902 | -0.6528831 | 0.35974004 | -1.8148748 | 0.06954315 | 0.26273569 |
| ANXA6 | 27.8225436 | 0.89489051 | 0.38641917 | 2.31585437 | 0.02056622 | 0.14099127 |
| ATOX1 | 26.1274314 | 0.36948931 | 0.37170453 | 0.99404037 | 0.32020321 | 0.57724473 |
| G3BP1 | 137.390245 | -0.1158158 | 0.30307299 | -0.3821384 | 0.7023587 | 0.85228676 |
| FAM114A2 | 22.0733479 | 0.28576083 | 0.39894486 | 0.71629153 | 0.47381136 | 0.7026171 |
| MFAP3 | 35.9393661 | 0.94288522 | 0.41779447 | 2.25681594 | 0.02401958 | 0.15136029 |
| GALNT10 | 109.640157 | 0.47164813 | 0.41949885 | 1.12431329 | 0.26088014 | 0.52020582 |
| SAP30L-AS1 | 29.8563983 | 0.55589762 | 0.39624303 | 1.40292086 | 0.16064044 | 0.40605894 |
| SAP30L | 12.9141287 | 0.53986757 | 0.49397355 | 1.09290785 | 0.27443426 | NA |
| LARP1 | 166.866521 | -0.0231489 | 0.20166567 | -0.1147887 | 0.90861263 | 0.95957056 |
| FAXDC2 | 40.1433449 | 1.62525156 | 0.5859904 | 2.77351224 | 0.00554548 | 0.07254189 |
| CNOT8 | 30.0228388 | -0.2412305 | 0.34119405 | -0.7070185 | 0.479555 | 0.70695585 |
| GEMIN5 | 21.8154069 | -0.2279181 | 0.38232055 | -0.596144 | 0.55107905 | 0.75768657 |
| MRPL22 | 21.8293968 | 0.38735383 | 0.40694386 | 0.95186062 | 0.34116767 | 0.59735386 |
| SGCD | 61.3635203 | -2.2819063 | 0.79783762 | -2.8601137 | 0.00423489 | 0.06301187 |
| HAVCR1 | 54.9238555 | -1.9919455 | 1.49556284 | -1.3319036 | 0.18289188 | 0.43261243 |
| HAVCR2 | 26.5031884 | -0.9227222 | 0.58559934 | -1.5756886 | 0.11509758 | 0.34138376 |
| CYFIP2 | 322.822221 | 0.63249932 | 0.26324766 | 2.40267786 | 1.63E-02 | 0.12641035 |
| CLINT1 | 345.173777 | 0.39340621 | 0.32706501 | 1.20283798 | 0.22903902 | 0.48726185 |
| RNF145 | 106.947003 | 0.61723751 | 0.24966382 | 2.47227455 | 1.34E-02 | 0.11302673 |
| UBLCP1 | 28.2430613 | 0.37276046 | 0.44143242 | 0.84443381 | 0.39842704 | 0.64642108 |
| TTC1 | 75.9589009 | 0.12702481 | 0.27804801 | 0.45684489 | 0.64778254 | 0.81885075 |
| PWWP2A | 116.07616 | 0.62542946 | 0.28775525 | 2.17347719 | 0.02974442 | 0.1702177 |
| CCNJL | 21.9405624 | 1.53973618 | 0.47857593 | 3.21732891 | 0.0012939 | 0.03212633 |
| SLU7 | 17.2906116 | -0.598727 | 0.49643995 | -1.2060411 | 0.22780166 | NA |
| AC113414.1 | 102.059173 | 1.84544847 | 0.63629351 | 2.90031003 | 0.00372794 | 0.06079223 |
| CCNG1 | 24.8348093 | 0.37221952 | 0.36008535 | 1.03369805 | 0.30127735 | 0.56175672 |
| MAT2B | 22.914712 | 0.42780195 | 0.4916583 | 0.87012046 | 0.38423458 | 0.63451053 |

| | | | | | | |
|---|---|---|---|---|---|---|
| WWC1 | 553.503257 | -1.1611545 | 0.32801063 | -3.5399905 | 0.00040014 | 0.01738515 |
| RARS | 54.0223256 | 0.03338655 | 0.2982764 | 0.1119316 | 0.91087764 | 0.96120444 |
| PANK3 | 65.3154523 | -0.5057228 | 0.31685966 | -1.5960465 | 0.11047841 | 0.33373324 |
| KCNIP1 | 30.7470077 | 2.07164255 | 1.11839254 | 1.85233939 | 0.0639771 | 0.25188326 |
| RANBP17 | 31.3794924 | -0.0352704 | 0.65510278 | -0.0538395 | 0.95706302 | 0.98311946 |
| NPM1 | 63.9740486 | -0.1887675 | 0.31173838 | -0.6055319 | 0.54482563 | 0.75328235 |
| FBXW11 | 296.44922 | -0.7715693 | 0.27055685 | -2.8517825 | 0.00434748 | 0.06373824 |
| STK10 | 19.5818622 | -1.2817252 | 0.43145757 | -2.9706866 | 0.00297135 | NA |
| UBTD2 | 62.0029124 | 0.05454925 | 0.31376633 | 0.17385311 | 0.8619809 | 0.93551424 |
| SH3PXD2B | 15.2985659 | -0.2759629 | 0.50058711 | -0.5512784 | 0.58144281 | NA |
| DUSP1 | 60.1835052 | -0.9375844 | 0.61758385 | -1.5181491 | 0.12897682 | 0.3611257 |
| ERGIC1 | 113.563558 | 0.32329313 | 0.28094317 | 1.1507421 | 0.24983835 | 0.5063622 |
| ATP6V0E1 | 256.530922 | -0.1405107 | 0.24235973 | -0.5797609 | 0.56207586 | 0.76693811 |
| CREBRF | 216.517575 | -0.7251708 | 0.25897297 | -2.8001796 | 0.00510742 | 0.06958964 |
| BOD1 | 18.9102458 | 0.84074344 | 0.5279326 | 1.5925204 | 0.11126782 | NA |
| LINC01485 | 18.3753254 | 0.86910641 | 1.2349112 | 0.70378049 | 4.82E-01 | NA |
| CPEB4 | 1238.20832 | -0.0681635 | 0.23746579 | -0.2870456 | 7.74E-01 | 8.88E-01 |
| SFXN1 | 127.353145 | -4.31E-07 | 0.25929949 | -1.66E-06 | 0.99999867 | 0.99999867 |
| THOC3 | 36.6318049 | -0.0958045 | 0.36757946 | -0.2606362 | 0.79437305 | 0.89901694 |
| SIMC1 | 52.2282608 | -0.036518 | 0.27446187 | -0.1330532 | 0.89415132 | 0.95216972 |
| KIAA1191 | 31.1078051 | 0.27931331 | 0.32274588 | 0.86542797 | 0.38680394 | 0.63697687 |
| ARL10 | 16.4905991 | 0.68029175 | 0.51404562 | 1.32340733 | 0.18569996 | NA |
| CLTB | 27.301204 | -0.8036636 | 0.41753384 | -1.9247868 | 0.05425603 | 0.23287156 |
| FAF2 | 92.6681925 | -0.3347345 | 0.36157784 | -0.9257605 | 0.35457045 | 0.61168935 |
| RNF44 | 24.6651184 | -0.2709359 | 0.44343226 | -0.6109972 | 0.54120145 | 0.74994873 |
| UIMC1 | 123.912266 | -0.4058484 | 0.23719377 | -1.7110415 | 0.08707345 | 0.29500251 |
| ZNF346 | 62.6918669 | -0.042742 | 0.28059417 | -0.1523266 | 0.87892934 | 0.94353126 |
| FGFR4 | 16.0143883 | 1.17591581 | 0.64079958 | 1.83507581 | 0.06649446 | NA |
| NSD1 | 321.006263 | -0.1100376 | 0.19912185 | -0.5526144 | 0.58052745 | 0.77770268 |
| PRELID1 | 19.3419408 | 0.63246666 | 0.52963506 | 1.19415557 | 0.23241712 | NA |
| LMAN2 | 46.1601782 | 0.36369662 | 0.40262447 | 0.90331473 | 0.36635888 | 0.62157013 |
| DBN1 | 27.4112817 | 0.18556308 | 0.50265928 | 0.36916274 | 0.71200643 | 0.857224 |
| PDLIM7 | 14.9320734 | -2.2556891 | 0.53032701 | -4.2533928 | 2.11E-05 | NA |
| FAM193B | 71.7427789 | -0.211512 | 0.27845962 | -0.7595785 | 0.44750655 | 0.68660592 |
| TMED9 | 25.5216387 | 0.77691018 | 0.48536127 | 1.60068433 | 0.10944685 | 0.33178907 |
| RMND5B | 42.7860814 | -0.648341 | 0.43481858 | -1.491061 | 0.13594547 | 0.37065968 |
| HNRNPAB | 20.410858 | -0.8470396 | 0.47584985 | -1.7800564 | 0.07506673 | NA |
| PHYKPL | 104.657849 | 0.40542753 | 0.27313109 | 1.48436976 | 0.13771085 | 0.37407407 |
| CLK4 | 57.9835729 | -0.0350273 | 0.28957577 | -0.1209607 | 9.04E-01 | 0.95671001 |
| ZNF354A | 17.0761277 | 0.51808889 | 0.49318296 | 1.0505004 | 0.29348811 | NA |
| ZNF354B | 20.311291 | -0.5894265 | 0.4460708 | -1.3213743 | 0.18637658 | NA |
| ZFP2 | 14.5498413 | 1.33430518 | 0.56004686 | 2.38248847 | 0.01719607 | NA |
| ZNF354C | 13.4453044 | 0.55125583 | 0.5839602 | 0.94399555 | 0.34517192 | NA |
| RUFY1 | 94.1169295 | -0.4339988 | 0.22647647 | -1.9163086 | 0.05532582 | 0.23510863 |
| HNRNPH1 | 259.457583 | -0.3468132 | 0.25076113 | -1.3830422 | 0.16665192 | 0.41444906 |
| CANX | 143.30932 | 0.17978834 | 0.30824566 | 0.58326316 | 0.55971616 | 0.76507463 |
| MAML1 | 47.5749987 | -0.2514904 | 0.33954516 | -0.7406686 | 0.45889438 | 0.6928053 |
| MGAT4B | 33.7037749 | -0.0004415 | 0.51615675 | -0.0008553 | 0.99931758 | 0.99980616 |
| SQSTM1 | 246.716962 | -1.3645789 | 0.42593138 | -3.2037529 | 0.00135649 | 0.03314569 |
| MRNIP | 41.3975185 | 0.20452678 | 0.39320854 | 0.52014838 | 0.60296016 | 0.79291384 |
| TBC1D9B | 42.8376655 | 0.53497272 | 0.44037717 | 1.21480575 | 0.22444019 | 0.48173901 |
| RNF130 | 293.685328 | -1.0858412 | 0.53242042 | -2.0394432 | 0.04140582 | 0.20301425 |
| MAPK9 | 53.5795421 | 0.48766735 | 0.31191626 | 1.56345599 | 0.11794538 | 0.34552837 |
| GFPT2 | 37.9306271 | 2.37132966 | 0.52816298 | 4.48976882 | 7.13E-06 | 0.00135324 |

| | | | | | | |
|---|---|---|---|---|---|---|
| CNOT6 | 85.3307278 | -0.4451862 | 0.26553053 | -1.6765912 | 0.09362244 | 0.3072972 |
| MGAT1 | 29.5930119 | -0.1875701 | 0.35914786 | -0.5222644 | 0.60148625 | 0.7915267 |
| ZFP62 | 15.5042269 | 0.23730484 | 0.47715726 | 0.49733045 | 0.61895604 | NA |
| TRIM41 | 18.3674366 | -0.1555727 | 0.44217125 | -0.3518381 | 0.72495966 | NA |
| RACK1 | 73.670098 | 0.22712653 | 0.3660871 | 0.62041664 | 0.53498352 | 0.74775242 |
| TRIM52 | 27.9751598 | 0.06409025 | 0.35880182 | 0.17862299 | 0.85823374 | 0.93289438 |
| AC138035.1 | 10.8890443 | -0.1269006 | 0.54569382 | -0.232549 | 0.81611163 | NA |
| DUSP22 | 41.8507179 | -0.1231561 | 0.29769731 | -0.4136957 | 0.67909699 | 0.84034192 |
| EXOC2 | 117.597705 | -0.2102193 | 0.23591534 | -0.8910794 | 0.37288657 | 0.62543211 |
| FOXQ1 | 28.7491632 | -1.4148598 | 0.76563801 | -1.8479487 | 0.06460976 | 0.25284322 |
| GMDS | 255.145656 | -0.4946227 | 0.24930509 | -1.9840058 | 0.0472552 | 0.21733813 |
| GMDS-DT | 471.650872 | 0.02893016 | 0.37310932 | 0.07753803 | 0.93819554 | 0.97466007 |
| WRNIP1 | 35.5070219 | 0.11047975 | 0.38804408 | 0.28470928 | 0.77586688 | 0.88880003 |
| SERPINB1 | 27.3351155 | -0.5129173 | 0.43970958 | -1.1664911 | 0.24341594 | 0.50068747 |
| SERPINB6 | 57.0169121 | 0.42678793 | 0.31383586 | 1.35990811 | 0.173859 | 0.42294335 |
| NQO2 | 49.2752453 | 0.09758595 | 0.37287746 | 0.26171052 | 0.79354463 | 0.89875118 |
| RIPK1 | 61.5610808 | -0.8025633 | 0.28594149 | -2.8067396 | 0.00500457 | 0.06866339 |
| BPHL | 17.8440515 | 0.19540568 | 0.5575357 | 0.35048102 | 0.72597773 | NA |
| PSMG4 | 73.6462552 | 0.35152972 | 0.44157826 | 0.79607569 | 0.42598804 | 0.66887596 |
| SLC22A23 | 186.51645 | -1.4183885 | 0.4936814 | -2.8730848 | 0.00406485 | 0.06207769 |
| PXDC1 | 18.7375095 | -1.4933538 | 0.44973276 | -3.320536 | 0.00089845 | NA |
| PRPF4B | 146.916545 | -0.6877586 | 0.24696902 | -2.7847971 | 0.00535612 | 0.07122532 |
| ECI2 | 50.5063845 | 0.07327283 | 0.35710586 | 0.20518516 | 0.83742746 | 0.9214695 |
| CDYL | 310.129068 | -1.5552943 | 0.48061005 | -3.2360836 | 0.00121182 | 0.03090537 |
| LYRM4 | 71.4396212 | 0.0117507 | 0.33385402 | 0.03519711 | 0.97192256 | 0.98878882 |
| FARS2 | 233.943809 | 0.78503893 | 0.3794646 | 2.06880677 | 0.03856422 | 0.19579738 |
| RREB1 | 186.464878 | 0.06167464 | 0.26422499 | 0.23341712 | 0.81543752 | 0.91060816 |
| SSR1 | 49.9147335 | 0.63541776 | 0.3799496 | 1.6723738 | 0.09445064 | 0.30830642 |
| RIOK1 | 16.5658914 | 0.15448698 | 0.54226717 | 0.28489089 | 0.77572773 | NA |
| DSP | 86.9438424 | -0.0561587 | 0.31933161 | -0.1758634 | 0.86040129 | 0.93485442 |
| SNRNP48 | 23.3038382 | 0.48157529 | 0.373887 | 1.28802365 | 0.19773773 | 0.45094881 |
| BLOC1S5-TXI | 19.0579367 | -0.22953 | 0.52748507 | -0.4351402 | 0.66346066 | NA |
| SLC35B3 | 33.4027479 | 0.32079896 | 0.36772523 | 0.87238769 | 0.38299691 | 0.63409921 |
| TFAP2A | 87.4850459 | 0.68534089 | 0.37737925 | 1.81605346 | 0.06936217 | 0.26260569 |
| GCNT2 | 469.149333 | -0.5788052 | 0.37087888 | -1.5606315 | 0.11861073 | 0.34581317 |
| TMEM14C | 22.0541546 | 0.94079672 | 0.42229371 | 2.22782554 | 0.02589215 | 0.15729431 |
| MAK | 45.9606763 | 0.52179416 | 0.59660047 | 0.87461238 | 0.38178485 | 0.63318207 |
| SMIM13 | 14.1539115 | 1.08960237 | 0.47454142 | 2.29611646 | 0.02166922 | NA |
| AL139807.1 | 35.6150856 | 0.29220585 | 0.61963958 | 0.4715739 | 0.63723096 | 0.81285494 |
| NEDD9 | 143.796908 | -0.6884322 | 0.3622859 | -1.9002455 | 0.05740091 | 0.23817084 |
| TMEM170B | 18.9033266 | 0.99402969 | 0.54733692 | 1.81612028 | 0.06935192 | NA |
| HIVEP1 | 145.618329 | -0.6481532 | 0.3869467 | -1.6750451 | 0.09392539 | 0.30789766 |
| PHACTR1 | 158.733338 | 0.19059132 | 0.57982267 | 0.32870623 | 0.74237775 | 0.87152705 |
| TBC1D7 | 19.0208643 | 0.83212495 | 0.51048044 | 1.63008194 | 0.10308418 | NA |
| SIRT5 | 50.9677002 | 0.79332609 | 0.35551944 | 2.23145631 | 0.02565092 | 0.15662748 |
| NOL7 | 24.6009755 | 0.21215241 | 0.40202276 | 0.52771245 | 0.59769893 | 0.7894568 |
| RANBP9 | 138.610461 | 0.37107734 | 0.25261509 | 1.46894365 | 0.14184807 | 0.37975813 |
| MCUR1 | 29.719455 | 0.91898084 | 0.40659626 | 2.26018026 | 0.02381007 | 0.15078654 |
| CD83 | 41.1336169 | -0.2746963 | 0.69707801 | -0.3940682 | 0.69353066 | 0.84791601 |
| JARID2 | 179.794771 | 0.31835739 | 0.29722807 | 1.07108792 | 0.2841299 | 0.54290897 |
| DTNBP1 | 118.00698 | -0.457543 | 0.27704657 | -1.651502 | 0.09863612 | 0.31646453 |
| MYLIP | 35.2538623 | -0.0905406 | 0.38401076 | -0.2357763 | 0.81360629 | 0.90970797 |
| GMPR | 48.1091601 | 2.3945075 | 0.80478237 | 2.97534784 | 0.00292656 | 0.05300181 |
| ATXN1 | 761.974914 | 0.57780619 | 0.28562041 | 2.02298634 | 0.04307456 | 0.20739942 |

| | | | | | | |
|---|---|---|---|---|---|---|
| CAP2 | 68.9582763 | 0.94005463 | 0.31626888 | 2.97232727 | 0.00295551 | 0.05340046 |
| NUP153 | 108.988983 | -0.2852951 | 0.31799833 | -0.8971593 | 0.36963395 | 0.62368335 |
| KIF13A | 134.221053 | 0.32205485 | 0.30428876 | 1.05838562 | 0.28987967 | 0.54928701 |
| TPMT | 22.0984984 | 0.34734892 | 0.39539366 | 0.87848885 | 0.37967849 | 0.63118473 |
| KDM1B | 16.9221108 | 0.55000296 | 0.47145847 | 1.16659895 | 0.24337237 | NA |
| DEK | 73.6720283 | -0.5674908 | 0.35377199 | -1.6041144 | 0.10868883 | 0.33040203 |
| RNF144B | 157.642253 | 0.12907588 | 0.35222832 | 0.36645514 | 7.14E-01 | 8.58E-01 |
| MBOAT1 | 56.4804895 | 0.82837893 | 0.4915416 | 1.68526717 | 0.09193704 | 0.30383829 |
| E2F3 | 58.1974256 | -0.0961434 | 0.30596642 | -0.3142286 | 0.7533474 | 0.87842978 |
| CDKAL1 | 440.975883 | -0.1578818 | 0.16179089 | -0.9758389 | 0.32914432 | 0.58534184 |
| SOX4 | 62.7788451 | -1.3929139 | 0.604572 | -2.303967 | 0.0212245 | 0.1427136 |
| CASC15 | 118.075861 | -0.9736322 | 0.80177139 | -1.2143513 | 0.22461359 | 0.48197681 |
| DCDC2 | 1820.37877 | 0.63269131 | 0.49357229 | 1.28186149 | 0.19989124 | 0.45385336 |
| MRS2 | 61.7656729 | -0.1623712 | 0.30877819 | -0.5258505 | 0.59899208 | 0.78963276 |
| ALDH5A1 | 15.7068919 | 1.19117764 | 0.55087007 | 2.16235679 | 0.03059069 | NA |
| TDP2 | 31.0257841 | 0.00699938 | 0.34616404 | 0.02021984 | 0.983868 | 0.99382437 |
| ACOT13 | 16.3475851 | 0.64543288 | 0.55577723 | 1.16131581 | 0.24551349 | NA |
| C6orf62 | 90.6283501 | -0.7439761 | 0.27784329 | -2.6776825 | 0.00741334 | 0.08440905 |
| CARMIL1 | 427.201889 | -0.1992634 | 0.50798549 | -0.3922619 | 0.69486468 | 0.84881343 |
| SLC17A1 | 12.9866414 | 1.17697602 | 1.06699275 | 1.10307781 | 0.26999338 | NA |
| TRIM38 | 73.5127083 | -0.1029132 | 0.31031195 | -0.3316444 | 0.74015779 | 0.87123329 |
| HIST1H4C | 25.2694234 | 0.93541998 | 0.86596815 | 1.08020137 | 0.28005252 | 0.53828072 |
| HIST1H2AC | 26.2654067 | -0.620201 | 0.62661154 | -0.9897696 | 0.32228676 | 0.57901472 |
| BTN2A1 | 13.6751346 | 0.31317709 | 0.64903208 | 0.48252945 | 0.62942988 | NA |
| ZNF322 | 13.8364507 | 0.32728633 | 0.50935794 | 0.64254682 | 0.5205182 | NA |
| LINC00240 | 25.1901116 | 0.73686145 | 0.54578118 | 1.35010419 | 0.17698257 | 0.42569745 |
| ZNF165 | 30.8302312 | -2.911458 | 0.96550525 | -3.0154761 | 0.00256576 | 0.04846819 |
| ZSCAN16-AS | 19.3420531 | 0.88701995 | 0.45671171 | 1.94218788 | 0.05211436 | NA |
| ZKSCAN8 | 26.2839672 | -0.5160678 | 0.36109946 | -1.4291569 | 0.15295914 | 0.3947713 |
| ZSCAN9 | 17.7102431 | -0.6278595 | 0.41905649 | -1.4982694 | 0.13406326 | NA |
| TRIM27 | 39.4075066 | 0.22569757 | 0.37940923 | 0.59486579 | 0.55193319 | 0.75820628 |
| HLA-A | 229.754179 | 0.5259106 | 0.41749315 | 1.25968679 | 0.20778237 | 0.46155871 |
| TRIM26 | 54.7043852 | 0.32900484 | 0.29909054 | 1.10001753 | 0.27132448 | 0.52964356 |
| HCG17 | 135.353673 | -0.3946943 | 0.33220757 | -1.1880955 | 0.23479579 | 0.49243138 |
| HLA-E | 87.8886151 | -0.2235167 | 0.39325167 | -0.5683808 | 5.70E-01 | 0.77196291 |
| GNL1 | 37.1852868 | -0.5533618 | 0.36320106 | -1.5235689 | 0.12761644 | 0.35967181 |
| PRR3 | 15.3056631 | 0.26525692 | 0.57919395 | 0.45797598 | 0.6469697 | NA |
| ABCF1 | 34.6688762 | -1.1523581 | 0.38539921 | -2.9900376 | 0.00278943 | 0.0514788 |
| PPP1R10 | 42.6901107 | -0.2731699 | 0.35195835 | -0.7761426 | 0.43766478 | 0.67793318 |
| MRPS18B | 25.1626388 | 0.06541956 | 0.36294655 | 0.1802457 | 0.85695968 | 0.93203598 |
| ATAT1 | 23.6666753 | -0.1930294 | 0.38936386 | -0.4957558 | 0.62006669 | 0.8030714 |
| C6orf136 | 28.6579254 | -0.5779156 | 0.36952798 | -1.5639293 | 0.11783417 | 0.34551224 |
| DHX16 | 23.6011558 | -0.0931163 | 0.3617911 | -0.257376 | 0.79688856 | 0.90081528 |
| MDC1 | 21.8930662 | -0.6172172 | 0.49316028 | -1.251555 | 0.21073206 | 0.4656925 |
| TUBB | 30.5077356 | 0.4994022 | 0.55003636 | 0.90794399 | 0.36390782 | 0.62003028 |
| FLOT1 | 75.8451062 | -0.492371 | 0.3489761 | -1.4109018 | 0.15827356 | 0.40298762 |
| IER3 | 196.749062 | -0.7754523 | 0.55123708 | -1.4067492 | 1.60E-01 | 4.04E-01 |
| DDR1 | 59.4295805 | 0.88713843 | 0.38629258 | 2.29654535 | 0.02164472 | 0.14349647 |
| CCHCR1 | 17.0624057 | 0.005855 | 0.66752466 | 0.0087712 | 0.99300168 | NA |
| POU5F1 | 32.6302507 | 0.55169996 | 0.54763075 | 1.00743056 | 0.31372791 | 0.57113617 |
| HLA-C | 98.9085281 | 0.74790727 | 0.62608174 | 1.19458405 | 0.23224958 | 0.49066933 |
| HLA-B | 156.779117 | 1.01629911 | 0.55681124 | 1.82521297 | 0.0679689 | 0.25963107 |
| MICA | 30.1303891 | 0.33829755 | 0.50642691 | 0.66800865 | 0.50412807 | 0.72596329 |
| NFKBIL1 | 12.4001365 | 0.4407495 | 0.52521919 | 0.83917249 | 0.40137252 | NA |

| | | | | | | |
|---|---|---|---|---|---|---|
| PRRC2A | 44.5285728 | -0.5143546 | 0.32975164 | -1.5598243 | 0.11880141 | 0.34612832 |
| BAG6 | 25.8976101 | -0.1855324 | 0.47537367 | -0.3902875 | 0.69632398 | 0.85012783 |
| DDAH2 | 22.4514858 | 0.56153959 | 0.55587328 | 1.01019354 | 0.31240257 | 0.57101636 |
| CLIC1 | 67.7920126 | 0.36797219 | 0.349527 | 1.05277187 | 2.92E-01 | 0.55197486 |
| LSM2 | 15.1417558 | 0.56226644 | 0.48992659 | 1.14765447 | 0.25111124 | NA |
| HSPA1A | 39.0442934 | 0.5043162 | 0.69070973 | 0.73014202 | 0.46530338 | 0.69589923 |
| HSPA1B | 26.525739 | 0.0760315 | 0.74789981 | 0.10166 | 0.91902655 | 0.9647335 |
| C6orf48 | 19.567791 | 0.01936248 | 0.54928695 | 0.03525021 | 0.97188022 | NA |
| NEU1 | 19.5243276 | 0.16620912 | 0.69878781 | 0.23785349 | 0.81199473 | NA |
| EHMT2 | 16.4776792 | 1.11455083 | 0.55453281 | 2.00989158 | 0.04444267 | NA |
| STK19 | 14.0593862 | -0.2875019 | 0.50212342 | -0.5725722 | 0.56693439 | NA |
| ATF6B | 40.6104173 | -0.3041876 | 0.3320564 | -0.9160722 | 0.35962901 | 0.6159629 |
| PBX2 | 22.7735502 | -0.1251866 | 0.52199093 | -0.2398252 | 0.81046579 | 0.9084251 |
| HLA-DRA | 20.3089761 | 0.21888388 | 0.61628836 | 0.35516471 | 0.7224662 | NA |
| TAP1 | 17.6977379 | 0.28539629 | 0.70006647 | 0.40767028 | 0.68351576 | NA |
| BRD2 | 72.7653613 | -0.4622264 | 0.40186252 | -1.1502103 | 0.25005728 | 0.5064976 |
| RXRB | 12.0315727 | 1.33799097 | 0.53487742 | 2.50149083 | 0.01236716 | NA |
| SLC39A7 | 17.890782 | 0.4280967 | 0.65080927 | 0.65779134 | 0.51067222 | NA |
| RING1 | 22.2286707 | -0.1540755 | 0.49953054 | -0.3084405 | 0.75774715 | 0.87943001 |
| VPS52 | 27.9502698 | 0.17482694 | 0.37281388 | 0.46893893 | 0.63911329 | 0.81390792 |
| RPS18 | 51.596628 | 0.44611016 | 0.45233169 | 0.98624563 | 0.32401259 | 0.58021133 |
| RGL2 | 19.2145492 | 0.24431332 | 0.50111026 | 0.48754404 | 0.62587284 | NA |
| TAPBP | 72.9302838 | -0.0706267 | 0.32795837 | -0.2153527 | 0.82949236 | 0.91745979 |
| PHF1 | 40.0829073 | -0.7202253 | 0.45665896 | -1.5771624 | 0.11475817 | 0.34109892 |
| CUTA | 25.3737586 | 0.54246108 | 0.46672389 | 1.16227408 | 0.24512415 | 0.5022131 |
| SYNGAP1 | 26.6533777 | -0.7944734 | 0.47833835 | -1.6609026 | 0.09673302 | 0.31231293 |
| ITPR3 | 59.3848237 | -0.8486112 | 0.54542523 | -1.5558709 | 1.20E-01 | 0.3480474 |
| UQCC2 | 16.810764 | -0.3296043 | 0.44644712 | -0.738283 | 0.46034248 | NA |
| LEMD2 | 29.6813202 | -0.8806615 | 0.374571 | -2.3511203 | 0.01871698 | 0.13535558 |
| NUDT3 | 26.4666855 | 0.35811667 | 0.56979946 | 0.62849597 | 0.52967908 | 0.74370906 |
| C6orf106 | 101.878277 | -0.303038 | 0.31570299 | -0.9598831 | 0.33711405 | 0.59276835 |
| SNRPC | 17.0433937 | -0.4057631 | 0.41621436 | -0.9748896 | 0.32961501 | NA |
| UHRF1BP1 | 68.4221213 | 0.19996917 | 0.32060092 | 0.62373235 | 0.53280338 | 0.74615054 |
| TAF11 | 41.3083759 | 0.03318494 | 0.35411874 | 0.09371134 | 0.92533846 | 0.9687609 |
| ANKS1A | 394.069267 | 0.04126301 | 0.36200421 | 0.11398489 | 0.90924977 | 0.9600131 |
| SCUBE3 | 34.6902262 | 4.05705834 | 1.03330544 | 3.92629146 | 8.63E-05 | 0.00614802 |
| ZNF76 | 53.5190442 | 0.22647218 | 0.3280091 | 0.69044481 | 0.4899145 | 0.7163358 |
| PPARD | 57.1668074 | -1.111801 | 0.51008304 | -2.1796471 | 0.02928363 | 0.16874939 |
| RPL10A | 55.2963766 | 1.0274117 | 0.46793211 | 2.19564266 | 0.02811753 | 0.16508057 |
| TEAD3 | 31.2613986 | -0.1090753 | 0.45541848 | -0.2395057 | 0.81071351 | 0.90857045 |
| SRPK1 | 74.3624855 | -0.2969107 | 0.32212195 | -0.9217339 | 0.35666739 | 0.61264648 |
| MAPK14 | 183.181802 | -0.8592338 | 0.27022885 | -3.1796523 | 0.00147452 | 0.03500548 |
| BRPF3 | 19.0985218 | -0.6050746 | 0.46109277 | -1.3122623 | 0.18943166 | NA |
| KCTD20 | 64.3809253 | 0.07283696 | 0.278233 | 0.26178404 | 0.79348795 | 0.89875118 |
| STK38 | 108.206926 | 0.27391493 | 0.22063456 | 1.24148696 | 0.21442591 | 0.46999219 |
| SRSF3 | 79.9847533 | -0.3292664 | 0.2772414 | -1.1876523 | 0.23497043 | 0.49257908 |
| CDKN1A | 22.4012992 | -1.1624634 | 0.64699124 | -1.7967221 | 7.24E-02 | 0.26822676 |
| CPNE5 | 22.6579414 | 0.2924974 | 0.84253648 | 0.34716289 | 0.72846895 | 0.86501473 |
| C6orf89 | 71.1195005 | -0.2014415 | 0.27180961 | -0.7411124 | 0.45862527 | 0.6928053 |
| MTCH1 | 75.510518 | -1.1250089 | 0.49414185 | -2.2766921 | 0.02280462 | 0.14700765 |
| TBC1D22B | 59.7802834 | -0.2796835 | 0.30249681 | -0.9245834 | 0.35518266 | 0.61168935 |
| RNF8 | 31.2265732 | 0.2435461 | 0.41969823 | 0.5802886 | 0.56172002 | 0.76672441 |
| CMTR1 | 22.5165241 | 0.06574799 | 0.60568474 | 0.10855151 | 0.91355822 | 0.96345217 |
| ZFAND3 | 1063.19797 | -0.6454104 | 0.30718328 | -2.1010596 | 0.03563573 | 0.18774007 |

| | | | | | | |
|---|---|---|---|---|---|---|
| BTBD9 | 717.267233 | -0.4877144 | 0.2089746 | -2.3338452 | 0.01960383 | 0.13704876 |
| GLO1 | 47.7873336 | 0.85422537 | 0.30562857 | 2.7949788 | 0.00519031 | 0.07024781 |
| KIF6 | 15.1254363 | 1.51959479 | 1.04433614 | 1.45508206 | 0.14564654 | NA |
| MOCS1 | 18.0456163 | 1.16382399 | 0.42888738 | 2.71358882 | 0.00665587 | NA |
| UNC5CL | 12.8678561 | 0.63653005 | 0.60025357 | 1.06043526 | 0.28894663 | NA |
| OARD1 | 38.9569975 | 0.60600258 | 0.39281621 | 1.54271277 | 0.12290048 | 0.35290359 |
| NFYA | 20.7631137 | 0.08071803 | 0.48844402 | 0.16525543 | 0.86874294 | NA |
| FOXP4 | 14.4715432 | 0.40369765 | 0.51084776 | 0.7902504 | 0.42938155 | NA |
| TFEB | 75.9538067 | 1.206513 | 0.42767701 | 2.82108456 | 0.00478616 | 0.06673743 |
| USP49 | 55.2504381 | -0.1592091 | 0.29346638 | -0.5425123 | 0.5874656 | 0.78270395 |
| CCND3 | 123.39036 | 0.57400731 | 0.47530031 | 1.2076729 | 0.2271731 | 0.48503506 |
| TAF8 | 44.2576096 | -0.172726 | 0.30982347 | -0.5574981 | 0.57718716 | 0.77478367 |
| C6orf132 | 17.2936694 | -0.0129194 | 0.47735249 | -0.0270647 | 0.97840814 | NA |
| MRPS10 | 12.0198473 | 0.36186617 | 0.57224808 | 0.63235891 | 0.52715238 | NA |
| TRERF1 | 37.565797 | 1.42362951 | 0.57430692 | 2.4788653 | 0.01318011 | 0.11234472 |
| UBR2 | 290.452881 | -0.1478338 | 0.16960741 | -0.8716232 | 0.38341394 | 0.6343472 |
| BICRAL | 123.348338 | 0.57560991 | 0.37119435 | 1.55069687 | 0.12097435 | 0.35000285 |
| RPL7L1 | 47.7520143 | -0.1251558 | 0.35724473 | -0.3503362 | 0.72608639 | 0.86378469 |
| PEX6 | 25.5363573 | 0.6188055 | 0.4535961 | 1.36422139 | 0.1724979 | 0.42128877 |
| PPP2R5D | 13.2924202 | 0.50612608 | 0.52394829 | 0.96598479 | 0.33405179 | NA |
| KLHDC3 | 24.587828 | -0.0933351 | 0.39382985 | -0.2369934 | 0.81266188 | 0.90970797 |
| RRP36 | 28.0036266 | -0.679449 | 0.37939166 | -1.7908907 | 0.07331084 | 0.26972923 |
| CUL7 | 15.6953854 | -0.3895357 | 0.50280633 | -0.7747231 | 0.43850331 | NA |
| KLC4 | 31.7438943 | -0.1169732 | 0.3542697 | -0.3301811 | 0.74126314 | 0.87152705 |
| MRPL2 | 17.580866 | 0.13999367 | 0.51155478 | 0.27366311 | 0.78434353 | NA |
| PTK7 | 72.7017086 | 0.49739538 | 0.44870762 | 1.10850665 | 0.26764307 | 0.52673196 |
| CUL9 | 27.8274124 | -0.9366477 | 0.41920294 | -2.2343538 | 0.02545981 | 0.15648615 |
| DNPH1 | 16.3453118 | 1.10846631 | 0.73161529 | 1.51509451 | 0.12974847 | NA |
| ZNF318 | 20.41919 | 1.12232332 | 0.47229363 | 2.37632534 | 0.01748604 | NA |
| ABCC10 | 19.5789694 | 0.08170932 | 0.48215124 | 0.16946823 | 0.86542836 | NA |
| TJAP1 | 28.1893526 | -0.5477972 | 0.34891139 | -1.570018 | 0.11641092 | 0.34369575 |
| POLR1C | 33.7583929 | 0.05857526 | 0.34916528 | 0.16775798 | 0.86677368 | 0.93793856 |
| YIPF3 | 22.1300365 | 0.09263783 | 0.47821213 | 0.19371702 | 0.84639746 | 0.92563742 |
| XPO5 | 72.9892274 | 0.13788674 | 0.28967789 | 0.47600024 | 0.63407421 | 0.81084386 |
| POLH | 37.9525294 | 1.02625722 | 0.38826207 | 2.6432075 | 0.00821247 | 0.08906215 |
| GTPBP2 | 35.5836441 | -0.7289702 | 0.43509279 | -1.6754363 | 0.09384866 | 0.30777722 |
| MRPS18A | 12.1345952 | 0.41296357 | 0.51577293 | 0.80066935 | 0.42332309 | NA |
| VEGFA | 244.752726 | -0.6361022 | 0.40415764 | -1.5738961 | 0.11551146 | 0.34185848 |
| MRPL14 | 19.3534576 | 0.76368921 | 0.48056057 | 1.58916329 | 0.11202353 | NA |
| TMEM63B | 33.299949 | -0.2788089 | 0.31948348 | -0.8726864 | 0.38283402 | 0.63396854 |
| SLC29A1 | 23.0613114 | 0.1886245 | 0.48631535 | 0.38786457 | 0.69811626 | 0.85089483 |
| HSP90AB1 | 130.599472 | 0.44007403 | 0.40409329 | 1.08904067 | 0.27613595 | 0.53482093 |
| CDC5L | 37.5110592 | -0.2946757 | 0.38880176 | -0.7579072 | 0.44850654 | 0.68731807 |
| SUPT3H | 391.105759 | -0.1220281 | 0.21723926 | -0.5617222 | 0.57430534 | 0.77388388 |
| RUNX2 | 19.591744 | 0.74048155 | 0.56024179 | 1.32171781 | 0.18626214 | NA |
| ENPP4 | 42.0102683 | 1.55320668 | 0.58888757 | 2.63752668 | 0.00835131 | 0.08952533 |
| ENPP5 | 17.0391521 | 1.36932355 | 0.5669448 | 2.41526785 | 0.01572364 | NA |
| RCAN2 | 21.2653253 | 0.31810623 | 0.49067167 | 0.64830772 | 5.17E-01 | 0.7344633 |
| CYP39A1 | 16.3943484 | 2.53939118 | 0.72007916 | 3.52654449 | 0.00042102 | NA |
| SLC25A27 | 106.267018 | 0.62082266 | 0.38169467 | 1.62649025 | 0.10384541 | 0.32339043 |
| ADGRF1 | 29.1205071 | 0.03645993 | 0.70312918 | 0.05185381 | 9.59E-01 | 0.98401364 |
| TNFRSF21 | 16.969048 | -0.7934545 | 0.68874101 | -1.152036 | 0.24930626 | NA |
| CD2AP | 534.937659 | -0.5067796 | 0.26492674 | -1.9129049 | 0.05576023 | 0.23568726 |
| PTCHD4 | 71.5252538 | 2.75671718 | 0.93693032 | 2.94228624 | 0.00325799 | 0.05673466 |

| | | | | | | |
|---|---|---|---|---|---|---|
| MUT | 39.7983385 | 0.22148891 | 0.39252236 | 0.5642708 | 0.57256985 | 0.77303458 |
| TFAP2B | 75.5266039 | 0.3227584 | 0.44788788 | 0.72062322 | 0.47114136 | 0.70047809 |
| PKHD1 | 2367.7482 | 1.07825369 | 0.42445275 | 2.54033859 | 0.01107452 | 0.10282338 |
| MCM3 | 13.4076776 | 0.17723449 | 0.48820845 | 0.36303035 | 0.71658221 | NA |
| EFHC1 | 58.7137398 | 0.5018194 | 0.36060288 | 1.3916123 | 0.16403983 | 0.41100735 |
| TRAM2 | 16.0258678 | 0.24285188 | 0.4530355 | 0.53605485 | 0.59192064 | NA |
| GSTA4 | 22.9810752 | 0.53723722 | 0.59216923 | 0.90723596 | 0.36428203 | 0.62003028 |
| ICK | 23.7068737 | 1.02875238 | 0.56029518 | 1.83609001 | 0.06634435 | 0.25673827 |
| FBXO9 | 94.5426469 | 0.08753011 | 0.31103767 | 0.28141321 | 0.77839348 | 0.88997247 |
| ELOVL5 | 41.8706221 | -0.3353709 | 0.59605434 | -0.5626516 | 0.57367214 | 0.77388388 |
| GCLC | 68.3647231 | 0.13209293 | 0.41145624 | 0.32103761 | 0.74818189 | 0.87522263 |
| LRRC1 | 63.1475069 | -0.120157 | 0.29903345 | -0.4018179 | 0.68781801 | 0.84530341 |
| TINAG | 61.7285932 | 0.98588323 | 0.61084855 | 1.61395689 | 0.10653678 | 0.32738142 |
| FAM83B | 187.253137 | -0.6053782 | 0.30105223 | -2.0108742 | 0.04433874 | 0.21075748 |
| HMGCLL1 | 23.5073463 | 1.18759036 | 0.71370266 | 1.66398477 | 0.09611547 | 0.31123298 |
| DST | 1007.29938 | -1.6402526 | 0.52102261 | -3.1481409 | 0.00164312 | 0.03697992 |
| ZNF451 | 121.736177 | -0.0679291 | 0.20499447 | -0.3313704 | 0.74036468 | 0.87134357 |
| PRIM2 | 78.432388 | -0.094644 | 0.40558751 | -0.2333503 | 0.81548942 | 0.91060816 |
| AL021368.2 | 15.8947701 | 0.90992747 | 0.56925798 | 1.59844481 | 0.10994402 | NA |
| PTP4A1 | 85.5434802 | -1.4716617 | 0.92524659 | -1.5905616 | 0.11170827 | 0.33573549 |
| PHF3 | 206.05694 | 0.34577665 | 0.17430085 | 1.9837921 | 0.04727902 | 0.21733813 |
| EYS | 22.9929324 | 0.85169639 | 0.58564177 | 1.45429584 | 0.1458643 | 0.38581359 |
| LMBRD1 | 191.089943 | -0.0883953 | 0.2036646 | -0.4340237 | 0.66427126 | 0.8292079 |
| COL19A1 | 45.3659191 | -0.4239195 | 0.37759362 | -1.1226871 | 0.26157042 | 0.52104231 |
| FAM135A | 443.98015 | -0.7045562 | 0.28491098 | -2.4728995 | 0.01340218 | 0.11302673 |
| SDHAF4 | 15.1657678 | 0.74175852 | 0.53787216 | 1.37906101 | 0.16787594 | NA |
| SMAP1 | 260.757387 | -0.5022359 | 0.32794147 | -1.5314803 | 0.12565073 | 0.35674425 |
| OGFRL1 | 36.7513078 | -1.2583924 | 0.72610886 | -1.7330629 | 0.08308448 | 0.28832338 |
| AL136164.2 | 202.26091 | 0.68062667 | 0.53413753 | 1.2742536 | 0.20257356 | 0.45644281 |
| LINC00472 | 130.756545 | 0.14604362 | 0.50488193 | 0.28926293 | 0.77238018 | 0.88744742 |
| AC019205.1 | 18.6119145 | 0.53331071 | 0.69665919 | 0.76552598 | 0.4439584 | NA |
| OOEP | 49.6366784 | 0.0969152 | 0.38859349 | 0.24939995 | 0.80305142 | 0.90408418 |
| MTO1 | 40.9073734 | -0.1248921 | 0.34651765 | -0.3604207 | 0.71853257 | 0.85995641 |
| EEF1A1 | 255.693877 | 0.83400413 | 0.43593041 | 1.91315886 | 0.05572772 | 0.23568726 |
| SLC17A5 | 27.1658569 | -0.4578154 | 0.38753612 | -1.181349 | 0.2374641 | 0.49524918 |
| COX7A2 | 39.0488336 | 5.23E-01 | 0.39572202 | 1.32229421 | 0.1860702 | 0.43607799 |
| TMEM30A | 57.178076 | -0.0649906 | 0.40823506 | -0.159199 | 0.8735121 | 0.94114315 |
| SENP6 | 435.138977 | -0.0536382 | 0.16116303 | -0.3328195 | 0.73927053 | 0.87054913 |
| MYO6 | 810.90087 | -0.6807357 | 0.29692576 | -2.2926124 | 0.02187033 | 0.14449437 |
| MEI4 | 25.0821514 | 0.97949311 | 0.60193834 | 1.62723163 | 0.10368791 | 0.32339043 |
| IRAK1BP1 | 87.4541206 | 1.00042772 | 0.52276321 | 1.91373016 | 0.05565464 | 0.23562915 |
| PHIP | 248.00213 | 0.030773 | 0.22051713 | 0.13954925 | 0.88901614 | 0.94924156 |
| HMGN3 | 107.307883 | 0.70227021 | 0.35387581 | 1.98451034 | 0.04719898 | 0.21733813 |
| LCA5 | 14.7592545 | 0.8870073 | 0.55583382 | 1.59581382 | 0.11053037 | NA |
| SH3BGRL2 | 18.6321814 | 0.25098715 | 0.443497 | 0.56592751 | 0.57144306 | NA |
| BCKDHB | 267.62214 | 0.77889909 | 0.32059041 | 2.42957701 | 0.01511645 | 0.12124045 |
| AL359715.1 | 13.893756 | 1.31168611 | 0.60350305 | 2.17345398 | 0.02974617 | NA |
| TENT5A | 20.8834031 | -0.3750494 | 0.68936164 | -0.5440532 | 0.58640485 | NA |
| IBTK | 196.650047 | -1.2174249 | 0.25761952 | -4.7256701 | 2.29E-06 | 0.00065384 |
| UBE3D | 123.799979 | 0.1125694 | 0.36386911 | 0.30936783 | 0.75704174 | 0.87904624 |
| DOPEY1 | 172.88387 | -0.1650024 | 0.22439616 | -0.7353173 | 0.46214628 | 0.69416206 |
| PGM3 | 42.9908264 | -0.3188477 | 0.39154932 | -0.8143233 | 0.41545978 | 0.65906718 |
| CYB5R4 | 54.092458 | -0.7778891 | 0.27671404 | -2.811166 | 0.00493623 | 0.06830815 |
| CEP162 | 78.0755344 | 0.52785326 | 0.37128008 | 1.4217118 | 0.15510993 | 0.39796039 |

| | | | | | | |
|---|---|---|---|---|---|---|
| SNX14 | 144.019287 | 0.1301994 | 0.26142803 | 0.49803151 | 0.61846183 | 0.80247821 |
| SYNCRIP | 60.4938926 | -0.8814703 | 0.26315102 | -3.3496747 | 0.00080907 | 0.02562706 |
| ZNF292 | 786.200905 | -1.3741737 | 0.36159292 | -3.8003336 | 0.0001445 | 0.00894448 |
| SMIM8 | 95.9758735 | 0.68482066 | 0.37124153 | 1.84467685 | 0.06508456 | 0.25390566 |
| RARS2 | 129.168242 | -0.2684928 | 0.21784158 | -1.2325139 | 0.21775717 | 0.47407641 |
| ORC3 | 64.2004246 | 0.08317694 | 0.49956604 | 0.16649839 | 0.86776475 | 0.93827484 |
| AKIRIN2 | 46.3655986 | -0.8875824 | 0.38101417 | -2.3295259 | 0.01983122 | 0.13786014 |
| RNGTT | 183.963321 | -0.2278974 | 0.21418852 | -1.064004 | 0.28732687 | 0.54660279 |
| PNRC1 | 51.0462495 | 0.09203796 | 0.39094404 | 0.2354249 | 0.81387896 | 0.90970797 |
| PM20D2 | 36.9065289 | 0.14124364 | 0.39896395 | 0.35402607 | 0.72331934 | 0.86194629 |
| UBE2J1 | 49.2988438 | -0.1122947 | 0.31247893 | -0.3593674 | 0.71932026 | 0.85995641 |
| RRAGD | 67.8942886 | -0.290122 | 0.42022934 | -0.6903896 | 0.4899492 | 0.7163358 |
| ANKRD6 | 107.631607 | 0.16770806 | 0.36969284 | 0.45364163 | 0.65008678 | 0.81947559 |
| LYRM2 | 26.4932903 | 0.72494986 | 0.41674078 | 1.73957026 | 0.08193451 | 0.28574984 |
| MDN1 | 109.733823 | -0.0716189 | 0.20608136 | -0.3475275 | 0.72819508 | 0.86482295 |
| CASP8AP2 | 47.7944728 | 0.25409804 | 0.28165716 | 0.90215368 | 0.36697524 | 0.62174959 |
| BACH2 | 345.461113 | -1.0484371 | 0.39206852 | -2.6741171 | 0.00749263 | 0.08493485 |
| MAP3K7 | 117.870994 | -0.0939823 | 0.22563003 | -0.4165329 | 0.67702008 | 0.8388857 |
| MANEA | 17.9050707 | 0.64123424 | 0.50128261 | 1.27918708 | 0.20083118 | NA |
| UFL1-AS1 | 27.0220541 | 1.0420206 | 0.69253144 | 1.50465459 | 0.1324129 | 0.36634871 |
| UFL1 | 69.0313411 | 0.06712448 | 0.31564449 | 0.21265849 | 0.83159335 | 0.9181376 |
| NDUFAF4 | 15.7747535 | 0.44199931 | 0.48538139 | 0.91062271 | 0.3624942 | NA |
| KLHL32 | 60.8862949 | 0.55126446 | 0.55116785 | 1.00017528 | 0.31722569 | 0.57492021 |
| MMS22L | 150.539384 | 0.72464281 | 0.81596975 | 0.88807558 | 0.3745001 | 0.62575308 |
| AL589740.1 | 24.3850129 | 2.63881891 | 0.89870398 | 2.93624926 | 0.00332207 | 0.05707588 |
| FBXL4 | 140.363131 | 0.20737644 | 0.2750465 | 0.75396867 | 0.45086804 | 0.6884212 |
| FAXC | 22.79032 | 0.18376938 | 0.45280699 | 0.40584484 | 0.68485661 | 0.84354958 |
| PNISR | 447.68255 | -0.2605386 | 0.15928585 | -1.6356669 | 0.10190932 | 0.32089434 |
| AL513550.1 | 28.8589224 | 0.82886261 | 0.44241819 | 1.8734822 | 0.06100183 | 0.24672613 |
| USP45 | 83.1673421 | 0.24576759 | 0.35695889 | 0.6885039 | 0.49113551 | 0.71672078 |
| TSTD3 | 16.5314638 | 0.36703793 | 0.51259417 | 0.71603999 | 0.47396665 | NA |
| CCNC | 116.608933 | -0.6378787 | 0.27808458 | -2.2938297 | 0.02180028 | 0.14415531 |
| SIM1 | 469.794529 | 0.00625816 | 0.30322467 | 0.0206387 | 0.98353387 | 0.99382437 |
| ASCC3 | 213.956704 | 0.28278337 | 0.21208427 | 1.33335379 | 0.18241573 | 0.43214954 |
| HACE1 | 91.0655129 | 0.44703947 | 0.30483501 | 1.46649648 | 0.14251308 | 0.38032372 |
| PREP | 43.4254194 | -0.0177886 | 0.30485954 | -0.0583501 | 0.95346973 | 0.98153155 |
| ATG5 | 129.677645 | 0.04044356 | 0.24096074 | 0.16784292 | 0.86670685 | 0.93793856 |
| CRYBG1 | 158.49442 | -0.7985152 | 0.54429069 | -1.4670749 | 0.14235568 | 0.38005954 |
| RTN4IP1 | 32.5627607 | -0.1139307 | 0.48630542 | -0.2342782 | 0.81476904 | 0.91033202 |
| QRSL1 | 57.0948305 | 0.05147789 | 0.30340056 | 0.16966972 | 0.86526989 | 0.93738047 |
| LINC02532 | 799.154309 | -0.4572085 | 0.58292073 | -0.7843407 | 0.43284021 | 0.67358899 |
| CD24 | 341.603868 | 0.56815279 | 0.43327838 | 1.31128811 | 0.18976044 | 0.44113142 |
| C6orf203 | 28.8111461 | 0.08779346 | 0.36830878 | 0.23836917 | 0.81159477 | 0.90902866 |
| PDSS2 | 223.925465 | 0.10359988 | 0.18239162 | 0.56800789 | 0.57002961 | 0.77196291 |
| SOBP | 74.8108765 | 0.88460498 | 0.46657704 | 1.89594622 | 0.05796716 | 0.23962041 |
| SEC63 | 97.9097147 | -0.206296 | 0.27907848 | -0.7392041 | 0.45978308 | 0.69299864 |
| OSTM1 | 41.3004431 | -0.7395066 | 0.56747953 | -1.3031424 | 0.19252615 | 0.44433997 |
| SNX3 | 72.7116284 | 0.23446807 | 0.24733873 | 0.94796343 | 0.34314807 | 0.60053858 |
| AFG1L | 108.33184 | 0.2725125 | 0.29430137 | 0.92596408 | 3.54E-01 | 0.61168935 |
| FOXO3 | 349.249835 | -0.1989366 | 0.26343531 | -0.7551629 | 0.45015125 | 0.68770076 |
| ARMC2 | 63.4019862 | 0.96504301 | 0.54713781 | 1.76380246 | 0.07776524 | 0.27801165 |
| SESN1 | 117.759422 | 0.14132187 | 0.30273184 | 0.46682197 | 0.64062726 | 0.81489142 |
| CEP57L1 | 23.6446804 | 0.26053402 | 0.42162172 | 0.61793311 | 0.53661944 | 0.74852479 |
| CD164 | 73.9101128 | -0.187054 | 0.32558577 | -0.5745153 | 0.56561913 | 0.76965382 |

| | | | | | | |
|---|---|---|---|---|---|---|
| PPIL6 | 21.0877599 | 1.17500794 | 0.47836626 | 2.45629349 | 0.01403785 | NA |
| ZBTB24 | 13.4852881 | 0.59836006 | 0.51526621 | 1.16126392 | 0.24553459 | NA |
| AK9 | 73.3789913 | 0.01186669 | 0.30107895 | 0.03941388 | 0.96856042 | 0.98741848 |
| FIG4 | 73.2770666 | 0.05081924 | 0.25912056 | 0.19612197 | 0.8445147 | 0.92496441 |
| WASF1 | 32.4764208 | -0.2191468 | 0.40270845 | -0.5441823 | 0.586316 | 0.78199173 |
| CDC40 | 38.795105 | 0.04921507 | 0.30297881 | 0.16243732 | 0.87096148 | 0.93986793 |
| METTL24 | 13.7530804 | 0.70871571 | 0.73159945 | 0.96872093 | 0.33268444 | NA |
| CDK19 | 246.533802 | -0.180358 | 0.22850491 | -0.7892959 | 0.42993908 | 0.67197671 |
| AMD1 | 114.704242 | -0.4655535 | 0.45015875 | -1.0341986 | 0.30104333 | 0.56145639 |
| GTF3C6 | 24.3723994 | -0.0088817 | 0.51127087 | -0.0173718 | 0.98613998 | 0.9942828 |
| RPF2 | 32.3597399 | -0.9198726 | 0.33731178 | -2.7270693 | 0.00638996 | 0.07793129 |
| SLC16A10 | 91.2535652 | -0.1488061 | 0.30111369 | -0.4941859 | 6.21E-01 | 0.80353114 |
| MFSD4B | 27.3968012 | 0.67806365 | 0.36628218 | 1.85120568 | 0.06413997 | 0.25213756 |
| REV3L | 270.661366 | 0.19563708 | 0.20072812 | 0.97463715 | 0.32974028 | 0.58560474 |
| TRAF3IP2-AS | 119.932111 | 0.09515097 | 0.35130371 | 0.27085102 | 0.78650562 | 0.8947286 |
| TRAF3IP2 | 22.8289144 | -0.7259123 | 0.42054453 | -1.7261248 | 0.08432497 | 0.28931012 |
| TUBE1 | 44.5079458 | 0.4005752 | 0.36591141 | 1.09473273 | 0.27363374 | 0.53266538 |
| MARCKS | 17.634207 | 0.45042881 | 0.44310957 | 1.01651788 | 0.30938284 | NA |
| HDAC2 | 49.4001044 | -0.0201408 | 0.29225541 | -0.068915 | 0.94505729 | 0.97757102 |
| FRK | 235.080606 | 0.48940829 | 0.2214464 | 2.21005309 | 0.02710148 | 0.16195658 |
| NT5DC1 | 249.681342 | 0.61611772 | 0.24029756 | 2.56397826 | 0.010348 | 0.10015449 |
| DSE | 94.366654 | 0.71902709 | 0.44191491 | 1.62707137 | 0.10372194 | 0.32339043 |
| TSPYL1 | 27.2919263 | 0.1165274 | 0.58885955 | 0.19788657 | 0.84313381 | 0.92431291 |
| RWDD1 | 19.7074958 | -0.060394 | 0.45344448 | -0.1331895 | 0.89404353 | NA |
| ZUP1 | 27.3597514 | 0.14976508 | 0.34429987 | 0.43498442 | 0.66357374 | 0.82887489 |
| KPNA5 | 79.9763316 | 0.31532061 | 0.37631636 | 0.83791364 | 0.40207921 | 0.65003228 |
| DCBLD1 | 18.8342994 | 0.75896741 | 0.61723803 | 1.22961869 | 0.21883993 | NA |
| GOPC | 44.4967024 | 0.4384517 | 0.40141143 | 1.0922751 | 0.2747122 | 0.5337282 |
| NUS1 | 52.6540945 | -0.3334514 | 0.2829607 | -1.1784371 | 0.23862238 | 0.49653323 |
| SLC35F1 | 21.7221721 | 0.06198832 | 0.73568287 | 0.08425956 | 0.93285006 | 0.97291964 |
| CEP85L | 78.6012262 | 0.17348147 | 0.26585547 | 0.65254052 | 0.51405254 | 0.73176668 |
| MCM9 | 114.112347 | 0.28582468 | 0.25578399 | 1.11744552 | 0.26380388 | 0.52345926 |
| AL365275.1 | 28.393717 | 0.74707729 | 0.62772235 | 1.19013969 | 0.23399149 | 0.49243138 |
| FAM184A | 119.028004 | 1.44338418 | 0.44072215 | 3.27504345 | 0.00105646 | 0.02953506 |
| MAN1A1 | 398.383982 | 0.28447119 | 0.26903924 | 1.05735946 | 0.29034756 | 0.54976758 |
| TBC1D32 | 110.501317 | 0.48003122 | 0.44559181 | 1.07728914 | 0.28135111 | 0.54003978 |
| HSF2 | 19.6037708 | -0.0800973 | 0.43876731 | -0.1825507 | 0.85515058 | NA |
| SERINC1 | 69.8373699 | -0.3712353 | 0.31575319 | -1.1757135 | 0.23970939 | 0.49733052 |
| PKIB | 26.402752 | 0.69270819 | 0.74943085 | 0.92431236 | 0.35532371 | 0.61168935 |
| RNF217 | 101.653771 | -0.8452116 | 0.52555029 | -1.6082411 | 0.10778238 | 0.32997921 |
| TPD52L1 | 195.533866 | 0.09012225 | 0.28793175 | 0.31299866 | 0.75428168 | 0.87872046 |
| HDDC2 | 40.4654339 | 1.00344461 | 0.366764 | 2.73594084 | 0.00622022 | 0.07721978 |
| NCOA7 | 117.091805 | 1.31456939 | 0.30087007 | 4.36922613 | 1.25E-05 | 0.0018818 |
| HINT3 | 25.6080394 | 0.90425342 | 0.40682276 | 2.22272085 | 0.02623463 | 0.15862368 |
| TRMT11 | 51.0943732 | -0.3253561 | 0.26638149 | -1.2213915 | 0.22193783 | 0.47914579 |
| AC020559.1 | 14.5697984 | 1.06353728 | 0.6346292 | 1.67584044 | 0.09376945 | NA |
| RNF146 | 40.1230453 | 0.3151009 | 0.31206375 | 1.00973247 | 0.31262348 | 0.57101636 |
| ECHDC1 | 99.7758135 | 0.25190926 | 0.26199705 | 0.96149658 | 0.33630255 | 0.59220332 |
| PTPRK | 427.844952 | 0.12612171 | 0.25678269 | 0.49116128 | 0.62331239 | 0.8044129 |
| AL356124.1 | 29.9834004 | 0.5728418 | 0.56512118 | 1.0136619 | 0.31074411 | 0.57001845 |
| ARHGAP18 | 199.574626 | 0.73595896 | 0.36015116 | 2.04347243 | 0.04100569 | 0.20224103 |
| L3MBTL3 | 32.8696566 | 0.05654336 | 0.46141181 | 0.12254424 | 0.90246802 | 0.95661107 |
| EPB41L2 | 42.3328376 | -0.3172292 | 0.36478476 | -0.869634 | 3.85E-01 | 0.63468687 |
| AKAP7 | 46.6307972 | 1.37629784 | 0.53625144 | 2.56651589 | 0.01027259 | 0.09994074 |

| | | | | | | |
|---|---|---|---|---|---|---|
| MED23 | 69.6202731 | -0.1399576 | 0.28246172 | -0.4954924 | 0.62025259 | 0.80317702 |
| CTGF | 20.0823036 | -1.052328 | 0.75459419 | -1.3945615 | 0.16314812 | NA |
| STX7 | 103.311274 | 0.32373431 | 0.25837528 | 1.25296161 | 0.21021968 | 0.46522739 |
| SLC18B1 | 47.9495737 | 0.71197604 | 0.46033494 | 1.54664784 | 0.12194819 | 0.35128178 |
| RPS12 | 51.3485874 | 0.67460842 | 0.45173522 | 1.49337132 | 0.13534002 | 0.36966363 |
| TBPL1 | 27.7354359 | -0.1183141 | 0.44007013 | -0.2688527 | 0.78804302 | 0.89515453 |
| SLC2A12 | 26.5761572 | 0.77416882 | 0.5345084 | 1.4483754 | 0.14751209 | 0.38722988 |
| SGK1 | 108.119903 | -1.5225922 | 0.68065433 | -2.2369536 | 0.02528937 | 0.15597139 |
| HBS1L | 70.0885505 | 0.28597749 | 0.27242839 | 1.04973454 | 0.29384018 | 0.55311516 |
| AHI1 | 444.565096 | -0.5566826 | 0.23446479 | -2.3742696 | 0.01758371 | 0.13204078 |
| LINC00271 | 144.539645 | 1.41308491 | 0.64301923 | 2.197578 | 0.02797919 | 0.16477111 |
| PDE7B | 469.730502 | -0.5702514 | 0.33711673 | -1.6915547 | 0.09073092 | 0.30098894 |
| AL138828.1 | 193.203183 | -2.9127249 | 0.94673691 | -3.0765938 | 0.0020938 | 0.04297603 |
| BCLAF1 | 152.521371 | -0.3508758 | 0.20095298 | -1.7460593 | 0.08080065 | 0.28378572 |
| MAP7 | 327.518135 | -0.5202123 | 0.23874985 | -2.1789012 | 0.02933901 | 0.16890227 |
| MAP3K5 | 118.091316 | 0.36013835 | 0.38792895 | 0.92836161 | 0.35322003 | 0.6104029 |
| PEX7 | 52.5251616 | 0.20281032 | 0.30920582 | 0.65590719 | 0.51188384 | 0.73097773 |
| IFNGR1 | 35.817907 | 0.41848528 | 0.33560185 | 1.24696953 | 0.21240867 | 0.46751774 |
| PERP | 32.5602251 | 0.57161445 | 0.47275284 | 1.20911901 | 0.22661713 | 0.48449513 |
| ARFGEF3 | 63.049921 | -0.8371363 | 0.36972157 | -2.2642344 | 0.0235597 | 0.14961965 |
| HEBP2 | 36.8465367 | 0.34186406 | 0.35899874 | 0.95227093 | 0.3409596 | 0.59712537 |
| NHSL1 | 106.065832 | -1.4556224 | 0.5216467 | -2.7904373 | 0.00526369 | 0.07082974 |
| CCDC28A | 20.4340198 | 1.03255683 | 0.47762955 | 2.16183613 | 0.03063081 | NA |
| REPS1 | 132.083805 | -0.2971663 | 0.19339172 | -1.5366033 | 0.1243905 | 0.35460505 |
| HECA | 19.9922702 | -0.1861908 | 0.52863736 | -0.3522089 | 0.72468163 | NA |
| CITED2 | 21.6302922 | -0.8839355 | 0.38714638 | -2.2832074 | 0.02241816 | 0.14604538 |
| VTA1 | 60.9393363 | -0.0177805 | 0.27394405 | -0.0649056 | 0.94824919 | 0.97942485 |
| HIVEP2 | 364.585414 | -0.7645922 | 0.46156235 | -1.6565308 | 0.09761437 | 0.31397319 |
| AIG1 | 227.544598 | -0.019544 | 0.25838343 | -0.0756394 | 0.93970599 | 0.97491804 |
| ADAT2 | 21.4459521 | 0.94640988 | 0.56125664 | 1.68623374 | 0.09175079 | 0.30348337 |
| PEX3 | 74.6343311 | -0.1745028 | 0.2671847 | -0.6531169 | 0.51368092 | 0.73176309 |
| FUCA2 | 22.7769134 | -0.0288963 | 0.36613413 | -0.0789226 | 0.93709418 | 0.97430959 |
| PHACTR2 | 230.023569 | 0.0940505 | 0.22412991 | 0.41962492 | 0.67475948 | 0.83760166 |
| PLAGL1 | 83.4077197 | 0.5688512 | 0.42029713 | 1.35345012 | 0.17591188 | 0.42498233 |
| SF3B5 | 21.6388011 | 0.78017596 | 0.56896104 | 1.37122915 | 0.17030353 | 0.41839333 |
| UTRN | 401.644648 | 0.21233957 | 0.38499459 | 0.55153911 | 0.58126416 | 0.77808527 |
| EPM2A | 85.9449308 | -0.6857025 | 0.37573415 | -1.8249673 | 0.06800597 | 0.25964381 |
| FBXO30 | 15.5614337 | 0.47329538 | 0.48634054 | 0.9731769 | 0.33046539 | NA |
| AL356599.1 | 45.9205448 | 0.72627419 | 0.63370791 | 1.14607089 | 0.25176582 | 0.50835297 |
| SHPRH | 211.746384 | 0.65602562 | 0.23981353 | 2.73556551 | 0.00622732 | 0.07721978 |
| STXBP5-AS1 | 30.2239598 | 0.19744723 | 0.65695649 | 0.30054842 | 0.76375887 | 0.88294563 |
| STXBP5 | 164.620977 | 0.04072053 | 0.21960178 | 0.18542895 | 0.8528926 | 0.92984623 |
| SAMD5 | 901.103969 | 0.40215092 | 0.31514022 | 1.27610154 | 2.02E-01 | 0.45588081 |
| AL033504.1 | 527.360001 | 0.20834847 | 0.35148128 | 0.59277258 | 0.55333336 | 0.75972295 |
| AL513164.1 | 25.5183355 | 1.08367741 | 0.53264496 | 2.03452112 | 0.04189908 | 0.20424144 |
| SASH1 | 369.32054 | -0.0036403 | 0.33074477 | -0.0110064 | 9.91E-01 | 9.96E-01 |
| UST | 132.218431 | 0.19202376 | 0.36257975 | 0.52960419 | 0.59638639 | 0.78913289 |
| TAB2 | 204.305223 | -0.4827343 | 0.31088945 | -1.5527523 | 0.12048233 | 0.34903213 |
| PPIL4 | 80.0313245 | -0.4030231 | 0.24662366 | -1.6341624 | 0.10222476 | 0.32120262 |
| GINM1 | 77.9447128 | -0.0750021 | 0.36729927 | -0.2041988 | 0.83819812 | 0.92179039 |
| KATNA1 | 37.2979449 | -0.3849015 | 0.32012984 | -1.2023294 | 0.22923595 | 0.48741135 |
| LATS1 | 103.054761 | -0.0134323 | 0.19885437 | -0.0675485 | 0.94614507 | 0.97812984 |
| NUP43 | 37.6180417 | 0.3223355 | 0.33138539 | 0.97269073 | 0.33070704 | 0.58655523 |
| PCMT1 | 59.9321159 | -0.0828466 | 0.37280619 | -0.2222243 | 0.82413928 | 0.91510572 |

| | | | | | | |
|---|---|---|---|---|---|---|
| LRP11 | 24.0375296 | 1.04542462 | 0.45206207 | 2.31256875 | 0.02074636 | 0.14118898 |
| PLEKHG1 | 57.3518221 | 0.64076837 | 0.50272136 | 1.27459946 | 2.02E-01 | 0.45643401 |
| MTHFD1L | 134.488189 | -0.1701237 | 0.60501041 | -0.2811914 | 7.79E-01 | 0.89003475 |
| RMND1 | 40.4988898 | 0.29142754 | 0.34858457 | 0.83603108 | 0.40313742 | 0.65053591 |
| ARMT1 | 20.4832347 | 0.74183838 | 0.46834346 | 1.58396227 | 0.1132023 | NA |
| SYNE1 | 389.52572 | 0.8867642 | 0.39306683 | 2.25601381 | 0.02406977 | 0.15136029 |
| MTRF1L | 20.4997235 | 0.35693712 | 0.43771107 | 0.81546286 | 0.41480742 | NA |
| IPCEF1 | 165.828783 | 0.84626681 | 0.46034567 | 1.83832902 | 0.06601394 | 0.25609634 |
| CNKSR3 | 293.068596 | 0.11943292 | 0.32665766 | 0.36562105 | 0.71464786 | 0.85840272 |
| SCAF8 | 277.561609 | -0.2149401 | 0.22021581 | -0.9760429 | 0.32904319 | 0.5853121 |
| TIAM2 | 161.362012 | 0.55409052 | 0.22655704 | 2.44569989 | 0.01445713 | 0.11850534 |
| TFB1M | 83.2335381 | -0.0101812 | 0.34986034 | -0.0291008 | 0.97678416 | 0.99130791 |
| ARID1B | 966.604319 | 0.09453993 | 0.14449195 | 0.65429204 | 0.51292367 | 0.73124162 |
| TMEM242 | 33.5798895 | 0.00257507 | 0.33027929 | 0.00779664 | 0.99377924 | 0.99727755 |
| ZDHHC14 | 609.431035 | -0.7863104 | 0.37529108 | -2.0952013 | 0.0361531 | 0.18968672 |
| SNX9 | 380.739152 | -1.4130261 | 0.27782119 | -5.0860989 | 3.66E-07 | 0.00020095 |
| SYNJ2 | 118.740482 | -1.9327488 | 0.47168464 | -4.0975445 | 4.18E-05 | 0.00401254 |
| SERAC1 | 23.4079921 | 0.07532912 | 0.41398154 | 0.18196251 | 0.85561215 | 0.93162353 |
| GTF2H5 | 18.4503204 | 1.10980026 | 0.57150368 | 1.94189522 | 0.05214979 | NA |
| TULP4 | 282.241889 | -0.6333719 | 0.43073316 | -1.4704508 | 0.14143969 | 0.37950905 |
| TMEM181 | 55.8211374 | -0.0158525 | 0.2756504 | -0.0575094 | 0.95413941 | 0.98153155 |
| DYNLT1 | 40.882471 | 0.66584807 | 0.35810562 | 1.85936226 | 0.06297581 | 0.249987 |
| SYTL3 | 41.583093 | -0.5323424 | 0.44395565 | -1.1990891 | 0.2304933 | 0.48927384 |
| EZR | 163.163942 | -0.1160564 | 0.35487175 | -0.3270377 | 0.7436394 | 0.87230965 |
| RSPH3 | 47.4106516 | 0.02416285 | 0.42265064 | 0.05716978 | 0.95440995 | 0.98153155 |
| AL035530.2 | 47.1092235 | 1.86570459 | 0.58759437 | 3.17515737 | 0.00149755 | 0.03503545 |
| SOD2.1 | 19.8543092 | 0.74558226 | 0.52730703 | 1.41394334 | 0.15737854 | NA |
| WTAP | 171.283376 | -1.0762746 | 0.29075378 | -3.7016701 | 0.00021418 | 0.01144848 |
| TCP1 | 32.0868828 | 0.1907907 | 0.33433591 | 0.57065574 | 0.56823303 | 0.77103518 |
| MRPL18 | 16.5862992 | 0.26497432 | 0.42369983 | 0.62538217 | 0.53172026 | NA |
| IGF2R | 285.055902 | -0.6590261 | 0.36591636 | -1.8010293 | 0.07169827 | 0.26724532 |
| SLC22A1 | 15.5318641 | -1.1271838 | 0.79352095 | -1.420484 | 0.15546682 | NA |
| SLC22A2 | 16.0810579 | -0.3570484 | 0.6309815 | -0.5658619 | 0.57148769 | NA |
| SLC22A3 | 85.725073 | -0.931526 | 0.73390714 | -1.2692696 | 0.20434494 | 0.45773044 |
| MAP3K4 | 106.612879 | 0.31007363 | 0.22862406 | 1.3562598 | 0.17501652 | 0.42343073 |
| AGPAT4 | 45.966017 | 0.34693574 | 0.31810554 | 1.09063092 | 0.27543532 | 0.53428066 |
| PRKN | 573.738096 | 1.04335407 | 0.39307872 | 2.65431328 | 0.007947 | 0.08725829 |
| PACRG | 81.2493133 | 0.68577534 | 0.5156383 | 1.32995422 | 0.18353335 | 0.4334631 |
| QKI | 587.363295 | 0.00833489 | 0.29119587 | 0.02862298 | 0.97716529 | 0.99136308 |
| PDE10A | 706.740629 | -0.173419 | 0.36950761 | -0.4693245 | 0.63883769 | 0.8138255 |
| SFT2D1 | 43.0420189 | -0.2585469 | 0.33808445 | -0.7647405 | 0.44442611 | 0.68456029 |
| MPC1 | 184.465335 | 0.93489416 | 0.29023317 | 3.22118306 | 0.00127663 | 0.03190321 |
| RPS6KA2 | 86.7018149 | -0.6173552 | 0.21279169 | -2.9012185 | 0.00371715 | 0.06079223 |
| Z94721.1 | 26.384406 | -0.1565669 | 0.45412507 | -0.3447661 | 0.73027019 | 0.86595126 |
| FGFR1OP | 92.3141164 | 0.37916208 | 0.37394256 | 1.01395808 | 0.31060275 | 0.57001845 |
| AFDN | 354.322181 | -0.1394381 | 0.22672389 | -0.6150129 | 0.53854622 | 0.74904053 |
| WDR27 | 123.078895 | 0.64274794 | 0.39487591 | 1.62772132 | 0.10358399 | 0.32331143 |
| PHF10 | 54.4177026 | -0.0938339 | 0.30671193 | -0.305935 | 0.75965411 | 0.88058098 |
| ERMARD | 22.653273 | -0.2080533 | 0.41728673 | -0.4985859 | 0.61807117 | 0.80237709 |
| FAM120B | 181.826481 | -0.826727 | 0.25330157 | -3.2638052 | 0.00109927 | 0.02988759 |
| PSMB1 | 48.4594059 | 0.06663375 | 0.30208633 | 0.2205785 | 0.82542064 | 0.9154963 |
| TBP | 24.4629968 | 0.01139565 | 0.40334794 | 0.02825265 | 0.97746064 | 0.99142241 |
| PDCD2 | 16.4910112 | 0.3712357 | 0.50055109 | 0.74165398 | 0.45829701 | NA |
| FAM20C | 104.08467 | 0.07636996 | 0.3445045 | 0.22168058 | 8.25E-01 | 9.15E-01 |

| | | | | | | |
|---|---|---|---|---|---|---|
| PDGFA | 27.6443492 | -0.3007993 | 0.39084167 | -0.7696193 | 0.44152574 | 0.68172992 |
| PRKAR1B | 84.7529734 | -0.6334065 | 0.38016957 | -1.6661158 | 0.09569035 | 0.31077157 |
| DNAAF5 | 38.2832131 | -0.1441688 | 0.36223933 | -0.3979933 | 0.69063511 | 0.84646791 |
| SUN1 | 95.8242274 | -0.4581925 | 0.26263237 | -1.7446155 | 0.08105181 | 0.28395803 |
| COX19 | 40.3388067 | -0.0252776 | 0.40179389 | -0.062912 | 0.94983661 | 0.97993174 |
| C7orf50 | 61.6679873 | -0.1280494 | 0.30456559 | -0.420433 | 0.67416916 | 0.83715334 |
| INTS1 | 32.8845317 | 0.21580678 | 0.48613095 | 0.44392725 | 0.65709518 | 0.82476514 |
| MAFK | 26.5393083 | -0.6119183 | 0.63288054 | -0.966878 | 0.33360501 | 0.58883681 |
| MAD1L1 | 48.8214245 | -0.2267591 | 0.37094829 | -0.6112957 | 0.54100384 | 0.74994873 |
| SNX8 | 17.5124791 | -1.0749677 | 0.52758016 | -2.0375439 | 0.04159557 | NA |
| EIF3B | 52.2633368 | -0.4202315 | 0.34509466 | -1.2177282 | 0.22332727 | 0.48054488 |
| CHST12 | 18.8115202 | 0.25995893 | 0.44829365 | 0.57988537 | 0.56199192 | NA |
| BRAT1 | 18.2979638 | -0.3443737 | 0.49657402 | -0.6934993 | 0.48799629 | NA |
| IQCE | 32.740213 | 0.06148611 | 0.38657915 | 0.15905181 | 0.87362805 | 0.94114315 |
| TTYH3 | 17.4720038 | -1.2312757 | 0.53702171 | -2.2927856 | 0.02186035 | NA |
| GNA12 | 44.3113525 | 0.12362327 | 0.31611835 | 0.3910664 | 0.69574815 | 0.84975778 |
| SDK1 | 83.2579705 | 0.60513007 | 0.42071769 | 1.43832809 | 0.15034098 | 0.39093735 |
| FOXK1 | 112.039387 | -0.4923244 | 0.23186545 | -2.1233195 | 0.03372708 | 0.18191827 |
| AP5Z1 | 26.4973558 | -1.465569 | 0.51451002 | -2.8484751 | 0.00439293 | 0.0642158 |
| RBAK-RBAKD | 71.9254667 | -0.2695252 | 0.27867435 | -0.9671689 | 0.33345963 | 0.58883681 |
| WIPI2 | 48.3755834 | -0.2551721 | 0.33195981 | -0.7686836 | 0.44208115 | 0.68190353 |
| TNRC18 | 239.235903 | 0.84372883 | 0.32558416 | 2.59143079 | 0.00955778 | 0.09652563 |
| FBXL18 | 34.1226222 | -0.430697 | 0.35196013 | -1.2237096 | 0.22106179 | 0.47795297 |
| ACTB | 239.26961 | -0.0845983 | 0.55769057 | -0.151694 | 0.8794283 | 0.94367205 |
| RNF216 | 176.889115 | -0.0251979 | 0.20715819 | -0.121636 | 0.9031873 | 0.95661107 |
| CCZ1 | 24.9686374 | -0.7613176 | 0.40804472 | -1.86577 | 0.06207355 | 0.24884382 |
| PMS2 | 41.8829038 | 0.40194546 | 0.3709107 | 1.08367179 | 0.27851035 | 0.53717915 |
| AIMP2 | 21.6728713 | -0.2150978 | 0.44748173 | -0.4806851 | 0.63074032 | 0.80819181 |
| EIF2AK1 | 54.5117711 | 0.03636427 | 0.28733913 | 0.12655523 | 0.89929244 | 0.95528285 |
| USP42 | 63.1986919 | -0.3170245 | 0.25537149 | -1.2414249 | 0.21444882 | 0.46999219 |
| CYTH3 | 73.3832785 | 0.39494459 | 0.3197315 | 1.2352383 | 0.21674183 | 0.47286334 |
| RAC1 | 111.799749 | -0.3663635 | 0.21616953 | -1.6947972 | 0.0901139 | 0.29961411 |
| DAGLB | 19.3007516 | -0.5189353 | 0.44222484 | -1.1734649 | 0.24060942 | NA |
| C7orf26 | 15.5888816 | -0.2316166 | 0.48569198 | -0.4768796 | 0.63344784 | NA |
| ZNF316 | 21.143852 | -0.719731 | 0.56556835 | -1.2725801 | 0.20316707 | NA |
| AC073343.2 | 17.472493 | -0.1410176 | 0.67797079 | -0.2079995 | 0.83522935 | NA |
| ZNF12 | 29.5335948 | -0.2030884 | 0.3791298 | -0.5356699 | 0.5921867 | 0.7855446 |
| CCZ1B | 30.8840789 | -0.7326768 | 0.41481962 | -1.7662539 | 0.07735327 | 0.27731248 |
| C1GALT1 | 71.3689689 | 0.14808328 | 0.37725318 | 0.39253024 | 0.69466648 | 0.84870601 |
| COL28A1 | 45.7310053 | -0.2525107 | 0.43735697 | -0.577356 | 0.56369901 | 0.76819959 |
| AC004982.2 | 11.2655782 | -0.1525644 | 0.68173823 | -0.2237873 | 0.82292278 | NA |
| MIOS | 55.762872 | 0.29279454 | 0.32089355 | 0.91243509 | 0.36153973 | 0.61739005 |
| RPA3 | 29.0670581 | 0.62614296 | 0.57563148 | 1.08774968 | 0.27670563 | 0.53501126 |
| UMAD1 | 340.828808 | 0.0277846 | 0.22608805 | 0.12289282 | 0.90219197 | 0.95661107 |
| GLCCI1 | 127.470808 | 0.37069981 | 0.39423874 | 0.94029268 | 0.34706745 | 0.60397426 |
| ICA1 | 473.676787 | 0.21701618 | 0.28076181 | 0.77295479 | 4.40E-01 | 6.80E-01 |
| NDUFA4 | 70.7672547 | 0.91990809 | 0.38875305 | 2.36630449 | 0.01796666 | 0.13297634 |
| PHF14 | 141.02055 | 0.19324971 | 0.2864543 | 0.67462668 | 0.49991298 | 0.72205483 |
| THSD7A | 598.817119 | 0.01299847 | 0.6174118 | 0.02105316 | 0.98320325 | 0.99382437 |
| TMEM106B | 69.2044734 | -0.0593627 | 0.29685989 | -0.1999688 | 0.84150496 | 0.92367136 |
| SCIN | 232.862331 | 0.35102232 | 0.30905106 | 1.13580689 | 0.25603739 | 0.51414553 |
| AC011287.1 | 23.1632471 | 0.41873937 | 0.51885749 | 0.8070412 | 4.20E-01 | 0.6636918 |
| ISPD | 61.9839594 | 0.11813042 | 0.33585866 | 0.35172659 | 0.72504331 | 0.86294393 |
| ANKMY2 | 41.0875983 | 0.53968456 | 0.35360926 | 1.52621728 | 0.12695576 | 0.35861434 |

| | | | | | | |
|---|---|---|---|---|---|---|
| BZW2 | 55.2214682 | 0.60422223 | 0.33422507 | 1.80783038 | 7.06E-02 | 2.66E-01 |
| AC073332.1 | 20.1738725 | 1.786684 | 0.88947393 | 2.00869743 | 0.04456923 | NA |
| SNX13 | 195.292148 | 0.10764817 | 0.22645644 | 0.47535931 | 0.63453089 | 0.81099192 |
| HDAC9 | 34.6786738 | -0.5073285 | 0.57053489 | -0.8892156 | 0.37388723 | 0.62575308 |
| MACC1 | 100.203029 | -1.1095579 | 0.57449492 | -1.9313623 | 0.05343826 | 0.23211867 |
| ITGB8 | 74.5476202 | 0.64145267 | 0.5574696 | 1.15065049 | 0.24987605 | 0.5063622 |
| SP4 | 61.5438567 | -0.2790886 | 0.3013589 | -0.9261004 | 0.35439383 | 0.61168935 |
| RAPGEF5 | 232.495536 | 1.05029456 | 0.58703025 | 1.789166 | 0.07358809 | 0.27023258 |
| STEAP1B | 54.3117506 | 0.36467204 | 0.5240167 | 0.69591683 | 0.48648091 | 0.71336323 |
| TOMM7 | 34.4068609 | 0.95128799 | 0.50975017 | 1.86618473 | 0.06201552 | 0.24884382 |
| KLHL7 | 48.653573 | 0.07106828 | 0.29493729 | 0.24096064 | 0.80958562 | 0.90770292 |
| NUPL2 | 56.5097889 | 0.42803862 | 0.29750507 | 1.43876073 | 0.15021833 | 0.39075041 |
| GPNMB | 28.9063134 | -0.9318783 | 0.75851931 | -1.2285493 | 0.21924085 | 0.47575326 |
| MALSU1 | 35.8260523 | -0.4297656 | 0.33092906 | -1.2986639 | 0.19405931 | 0.44613935 |
| TRA2A | 265.969264 | -0.0975673 | 0.26618004 | -0.3665464 | 0.71395739 | 0.85788134 |
| CCDC126 | 26.4107763 | -0.521738 | 0.47596147 | -1.0961769 | 0.27300135 | 0.53186453 |
| FAM221A | 16.3709792 | 0.06926773 | 0.51254212 | 0.13514544 | 0.89249688 | NA |
| MPP6 | 29.6389738 | 0.00540898 | 0.42763316 | 0.01264865 | 0.98990811 | 0.99583841 |
| GSDME | 44.9322803 | -0.2909268 | 0.53693567 | -0.5418281 | 0.58793695 | 0.78279721 |
| OSBPL3 | 591.96416 | -0.6597168 | 0.30684589 | -2.1499938 | 0.03155571 | 0.17516458 |
| CYCS | 37.6459731 | 0.10427326 | 0.51979065 | 0.20060626 | 0.84100646 | 0.92355925 |
| NFE2L3 | 20.2830189 | 0.01824064 | 0.60565972 | 0.03011697 | 0.97597377 | NA |
| HNRNPA2B1 | 468.694253 | -0.3549323 | 0.20441494 | -1.7363326 | 0.08250504 | 0.28695946 |
| CBX3 | 78.4930781 | -0.4048163 | 0.27161587 | -1.4904001 | 0.13611906 | 0.37100157 |
| SNX10 | 467.00234 | -0.1706262 | 0.22464839 | -0.7595255 | 0.44753825 | 0.68660592 |
| AC004540.1 | 54.3938141 | 2.54942924 | 0.68813683 | 3.70482896 | 0.00021153 | 0.01138582 |
| SKAP2 | 118.179256 | 0.47097086 | 0.37654899 | 1.25075589 | 0.21102355 | 0.46593468 |
| HOXA3 | 17.2366245 | 2.51844559 | 0.79900077 | 3.15199395 | 0.0016216 | NA |
| HOXA9 | 12.5479709 | 0.47504086 | 0.72747787 | 0.65299698 | 0.51375822 | NA |
| HIBADH | 277.322594 | 0.54876636 | 0.31682544 | 1.73207796 | 8.33E-02 | 0.2885411 |
| TAX1BP1 | 275.623057 | -0.0168568 | 0.16419751 | -0.1026618 | 0.91823136 | 0.96421386 |
| JAZF1 | 80.4518004 | 0.75868974 | 0.51044355 | 1.48633428 | 0.13719072 | 0.37299788 |
| CREB5 | 42.2082335 | -0.6090699 | 0.87587489 | -0.6953846 | 0.4868143 | 0.71361615 |
| AC005162.3 | 21.9805374 | 0.78588081 | 0.72727269 | 1.08058617 | 0.27988124 | 0.53815785 |
| CPVL | 55.2400936 | 0.67784661 | 0.49709996 | 1.36360222 | 0.17269279 | 0.42133629 |
| AC007255.1 | 22.2718329 | 1.47556818 | 0.73975905 | 1.99466054 | 0.04607992 | 0.21495583 |
| SCRN1 | 56.8963504 | -0.1174149 | 0.31554178 | -0.3721056 | 0.70981423 | 0.85607462 |
| FKBP14 | 19.137332 | 0.3624091 | 0.45728101 | 0.79253039 | 0.42805148 | NA |
| PLEKHA8 | 65.8399053 | -0.133627 | 0.30964432 | -0.43155 | 0.66606851 | 0.83091237 |
| MTURN | 122.028475 | 0.57307435 | 0.30747564 | 1.86380405 | 0.06234923 | 0.24933194 |
| ZNRF2 | 108.752614 | -0.440085 | 0.26331383 | -1.6713325 | 0.09465602 | 0.30858458 |
| AC006978.2 | 90.4668932 | 0.20287014 | 0.33930333 | 0.59790199 | 0.54990533 | 0.75662755 |
| NOD1 | 18.610678 | 0.82272187 | 0.48155143 | 1.70848183 | 0.08754697 | NA |
| AC005154.6 | 41.7756971 | 0.86736845 | 0.40654759 | 2.13349794 | 0.0328839 | 0.17912076 |
| GARS | 43.7248048 | -0.3533377 | 0.49735593 | -0.7104322 | 0.47743616 | 0.70521451 |
| AQP1 | 22.4175797 | -0.7403103 | 0.58974316 | -1.2553097 | 0.20936637 | 0.46413968 |
| PDE1C | 49.69614 | 0.23681708 | 0.72064851 | 0.32861662 | 0.74244548 | 0.87152705 |
| LSM5 | 14.7173126 | -0.6429549 | 0.46499946 | -1.3827004 | 0.16675674 | NA |
| AVL9 | 227.25216 | -0.4407735 | 0.20126311 | -2.1900361 | 0.02852162 | 0.16593415 |
| KBTBD2 | 53.612881 | -0.3523245 | 0.26771497 | -1.3160435 | 0.18815944 | 0.43860181 |
| FKBP9 | 33.9094894 | 1.01899797 | 0.43483129 | 2.34343293 | 0.0191072 | 0.1359475 |
| NT5C3A | 64.5477932 | -0.3638657 | 0.40987121 | -0.8877563 | 0.37467188 | 0.62575308 |
| RP9 | 16.4800978 | 0.60161687 | 0.44956064 | 1.33823297 | 0.18082051 | NA |
| BBS9 | 218.17093 | 1.26698389 | 0.35084901 | 3.61119413 | 0.00030479 | 0.01457125 |

| | | | | | | |
|---|---|---|---|---|---|---|
| DPY19L1 | 121.443894 | -0.5871632 | 0.27136918 | -2.1637064 | 0.03048689 | 0.17266931 |
| HERPUD2 | 74.3378106 | -0.158466 | 0.2630496 | -0.6024188 | 0.5468954 | 0.75478822 |
| SEPT7-AS1 | 23.1812756 | 1.15437323 | 0.60680161 | 1.90238986 | 0.05712021 | 0.23764428 |
| 7-Sep | 164.889954 | -0.2647076 | 0.18811513 | -1.4071573 | 0.15938074 | 0.40420216 |
| EEPD1 | 49.6678015 | -1.4788223 | 0.6201873 | -2.3844769 | 0.01710342 | 0.12988597 |
| KIAA0895 | 37.381835 | 0.33435747 | 0.37303338 | 0.89632049 | 0.37008162 | 0.62368335 |
| AOAH | 20.5587193 | -2.0582838 | 0.82609139 | -2.4915934 | 0.01271715 | NA |
| ELMO1 | 26.3882596 | 1.64456666 | 0.96264337 | 1.70838621 | 0.0875647 | 0.29612721 |
| EPDR1 | 21.8964625 | 2.29793214 | 0.47511487 | 4.83658218 | 1.32E-06 | 0.00048414 |
| STARD3NL | 30.6175302 | 0.23355159 | 0.39176588 | 0.59615091 | 0.55107441 | 0.75768657 |
| AMPH | 37.8599471 | -1.7552592 | 0.77586205 | -2.2623341 | 0.02367677 | 0.15011539 |
| VPS41 | 151.013231 | -0.2060019 | 0.23016109 | -0.8950336 | 0.37076915 | 0.62398581 |
| YAE1D1 | 17.8777942 | 0.40251676 | 0.40930515 | 0.98341484 | 0.32540331 | NA |
| RALA | 80.9046788 | -0.6723528 | 0.32433481 | -2.0730206 | 0.03817036 | 0.19510866 |
| LINC00265 | 14.209792 | -0.8530633 | 0.52008311 | -1.6402442 | 0.1009544 | NA |
| CDK13 | 351.322155 | -0.200043 | 0.22168629 | -0.9023699 | 0.36686041 | 0.62174959 |
| MPLKIP | 27.0300179 | 0.8089018 | 0.38902644 | 2.07929772 | 0.03758999 | 0.19425568 |
| SUGCT | 103.451977 | 0.67064358 | 0.34338839 | 1.95301763 | 0.05081751 | 0.22622461 |
| MRPL32 | 17.8217552 | 0.45431152 | 0.43950628 | 1.03368606 | 0.30128295 | NA |
| STK17A | 52.6329122 | -0.8326002 | 0.51368807 | -1.6208284 | 0.10505445 | 0.32605003 |
| COA1 | 204.338546 | -0.2828676 | 0.25279534 | -1.118959 | 0.26315762 | 0.52284475 |
| BLVRA | 20.8139014 | 0.43221555 | 0.42036654 | 1.02818735 | 0.30386171 | NA |
| URGCP | 39.1775808 | 0.64843901 | 0.44010183 | 1.47338404 | 0.14064751 | 0.37865124 |
| UBE2D4 | 11.0331242 | 0.38843803 | 0.54237508 | 0.71617971 | 0.47388039 | NA |
| AC004951.1 | 10.4734888 | -0.0466422 | 0.55725438 | -0.0837001 | 0.93329489 | NA |
| DBNL | 23.9760446 | -0.3579184 | 0.34518839 | -1.0368784 | 0.29979252 | 0.559867 |
| POLM | 21.6688377 | 0.43403299 | 0.57025264 | 0.76112404 | 0.44658298 | 0.68637165 |
| YKT6 | 14.4738666 | -0.3110367 | 0.6543612 | -0.4753287 | 0.63455267 | NA |
| CAMK2B | 10.6176023 | 1.13971573 | 0.85450333 | 1.33377565 | 1.82E-01 | NA |
| NUDCD3 | 114.157253 | -0.2279716 | 0.23173083 | -0.9837775 | 0.32522492 | 0.58122021 |
| DDX56 | 19.6130495 | -0.5996382 | 0.40950148 | -1.4643127 | 0.14310852 | NA |
| TMED4 | 33.7257278 | 0.19272069 | 0.38633124 | 0.49884833 | 0.61788624 | 0.80236132 |
| OGDH | 317.483623 | -0.2617508 | 0.30774033 | -0.8505574 | 0.39501527 | 0.64334162 |
| ZMIZ2 | 54.3234096 | -0.1115609 | 0.32992806 | -0.3381371 | 0.73525986 | 0.86838246 |
| PPIA | 109.508798 | 0.0633564 | 0.42701962 | 0.14836883 | 0.8820517 | 0.94477483 |
| H2AFV | 58.8910812 | -0.0284335 | 0.3589387 | -0.0792154 | 0.93686132 | 0.97419907 |
| PURB | 44.7068422 | -0.1115504 | 0.29646168 | -0.3762726 | 0.70671424 | 0.8547469 |
| CCM2 | 55.152093 | 0.28885189 | 0.3225514 | 0.89552204 | 0.37050809 | 0.62376026 |
| TBRG4 | 16.698891 | -0.4556335 | 0.5534887 | -0.8232028 | 0.41039267 | NA |
| IGFBP3 | 125.243396 | -1.3287616 | 0.87314113 | -1.5218177 | 0.12805476 | 0.36035989 |
| AC004870.4 | 34.9297203 | -0.4325007 | 0.93561202 | -0.462265 | 0.64389131 | 0.81701804 |
| TNS3 | 520.695403 | -0.5519614 | 0.38512322 | -1.4332073 | 0.1517986 | 0.39273741 |
| HUS1 | 31.9294133 | -0.8606957 | 0.34099272 | -2.5240881 | 0.01159988 | 0.10516409 |
| UPP1 | 146.105197 | -0.5686065 | 0.69001024 | -0.8240552 | 4.10E-01 | 0.65566569 |
| GRB10 | 65.6006304 | -1.1100154 | 0.68726703 | -1.6151151 | 0.10628577 | 0.32732986 |
| COBL | 197.353118 | -0.945164 | 0.43752408 | -2.1602559 | 0.03075287 | 0.17341013 |
| SEC61G | 19.1841357 | 0.14446841 | 0.41158986 | 0.35100088 | 0.72558768 | NA |
| EGFR | 50.0912251 | 1.03182248 | 0.49441988 | 2.08693566 | 0.03689395 | 0.19174392 |
| LANCL2 | 39.1067828 | -0.2878234 | 0.32226297 | -0.8931321 | 0.37178641 | 0.62455659 |
| VOPP1 | 166.938235 | 0.20188892 | 0.22451808 | 0.89921006 | 0.36854078 | 0.62316748 |
| NIPSNAP2 | 112.292348 | 0.11261867 | 0.26447973 | 0.4258121 | 0.67024477 | 0.83436422 |
| PSPH | 16.528292 | 1.21475377 | 0.57465236 | 2.1138933 | 0.03452438 | NA |
| CCT6A | 40.5256768 | -0.5533053 | 0.37583077 | -1.4722192 | 0.14096171 | 0.37904789 |
| SUMF2 | 43.8232383 | 0.45231601 | 0.31432887 | 1.43898973 | 0.15015343 | 0.39073955 |

| | | | | | | |
|---|---|---|---|---|---|---|
| CHCHD2 | 65.2872608 | 0.47643612 | 0.44018312 | 1.08235891 | 0.27909308 | 0.53717915 |
| ZNF727 | 15.476178 | -0.1048269 | 0.6490699 | -0.1615032 | 0.87169711 | NA |
| ZNF736 | 79.5235421 | 0.19066216 | 0.34076539 | 0.55951151 | 0.57581268 | 0.77401855 |
| ZNF680 | 94.1389153 | 0.84596894 | 0.29426777 | 2.87482706 | 0.00404249 | 0.06207769 |
| ZNF107 | 22.6984182 | 0.86142611 | 0.49338311 | 1.74595784 | 0.08081828 | 0.28378572 |
| ZNF138 | 62.8143733 | 0.73033047 | 0.34431545 | 2.12110863 | 0.03391266 | 0.1825355 |
| ZNF117 | 26.4568537 | 0.40607413 | 0.49743219 | 0.81634067 | 0.41430532 | 0.65845716 |
| VKORC1L1 | 86.4248244 | 0.08893355 | 0.29554227 | 0.30091652 | 0.76347816 | 0.88288632 |
| GUSB | 21.6155124 | 0.48625155 | 0.38608014 | 1.25945758 | 0.2078651 | 0.46160926 |
| ASL | 17.9415844 | -0.0592206 | 0.54998665 | -0.1076764 | 0.91425237 | NA |
| AC068533.4 | 22.5932259 | 0.06035707 | 0.42608492 | 0.14165502 | 0.88735251 | 0.94807777 |
| CRCP | 12.4235945 | -0.3570195 | 0.5432243 | -0.657223 | 0.51103753 | NA |
| TPST1 | 475.691894 | 0.19219185 | 0.25629717 | 0.74987896 | 0.45332761 | 0.68943529 |
| AC027644.4 | 89.0882658 | -0.1508124 | 0.32961352 | -0.4575431 | 0.64728074 | 0.81885075 |
| TMEM248 | 55.6775051 | -0.0858353 | 0.35457162 | -0.2420816 | 0.80871694 | 0.90752213 |
| SBDS | 24.8865046 | -0.1988593 | 0.39373764 | -0.5050555 | 0.61351988 | 0.7990292 |
| TYW1 | 147.284094 | 0.09200242 | 0.23759936 | 0.38721659 | 0.69859587 | 0.85102865 |
| AC006480.2 | 19.4981344 | 0.71332872 | 0.50747011 | 1.40565661 | 0.15982612 | NA |
| AUTS2 | 219.086541 | -0.262824 | 0.22819965 | -1.1517283 | 2.49E-01 | 0.5058511 |
| GALNT17 | 215.152097 | 1.11974629 | 0.91252106 | 1.22709089 | 0.21978843 | 0.47614454 |
| CALN1 | 19.0940518 | 0.69451504 | 0.57944522 | 1.19858619 | 0.23068889 | NA |
| TYW1B | 113.103505 | 0.20817998 | 0.28439204 | 0.7320176 | 0.46415783 | 0.69530788 |
| POM121 | 111.053153 | -0.381945 | 0.24586963 | -1.5534452 | 0.12031682 | 0.34903213 |
| NSUN5 | 23.2986025 | -0.0784254 | 0.4622715 | -0.1696522 | 0.86528365 | 0.93738047 |
| TRIM50 | 74.0475331 | -0.2654093 | 1.55771487 | -0.1703837 | 0.86470837 | 0.93715296 |
| BAZ1B | 96.4148206 | -0.5725701 | 0.32774827 | -1.7469814 | 0.08064056 | 0.28378572 |
| BCL7B | 17.2585864 | -0.1286211 | 0.52708862 | -0.2440217 | 0.80721401 | NA |
| MLXIPL | 31.0278622 | 0.59950813 | 0.52611043 | 1.13951007 | 0.25449047 | 0.51197417 |
| BUD23 | 32.5136334 | -0.7931717 | 0.39487249 | -2.0086781 | 0.04457129 | 0.21133145 |
| ABHD11 | 22.4806918 | -1.415962 | 0.41957245 | -3.3747736 | 0.00073876 | 0.02425576 |
| CLDN3 | 22.4319019 | 0.1514947 | 0.48926922 | 0.30963464 | 7.57E-01 | 8.79E-01 |
| CLDN4 | 52.9811925 | -0.7890814 | 0.54712853 | -1.442223 | 0.14923947 | 0.3896527 |
| LIMK1 | 15.4200957 | -0.8611448 | 0.51343612 | -1.6772189 | 0.09349968 | NA |
| EIF4H | 45.8108101 | -0.3638708 | 0.32552673 | -1.1177908 | 0.26365636 | 0.52333673 |
| CLIP2 | 85.8602375 | -0.9383145 | 0.46013612 | -2.0392107 | 0.041429 | 0.20301425 |
| GTF2IRD1 | 740.612562 | -1.3844957 | 0.34940584 | -3.9624286 | 7.42E-05 | 0.0057105 |
| GTF2I | 357.607284 | 0.37706758 | 0.2119257 | 1.77924426 | 0.07519972 | 0.27367272 |
| GTF2IRD2 | 41.4198162 | -0.4636306 | 0.45314492 | -1.0231398 | 0.30624176 | 0.56595793 |
| CASTOR2 | 56.3946624 | -0.4418126 | 0.43379891 | -1.0184734 | 0.30845306 | 0.56825351 |
| RCC1L | 38.9043489 | -0.5288379 | 0.37716338 | -1.4021453 | 0.16087186 | 0.406377 |
| GTF2IRD2B | 72.4479721 | -0.9478437 | 0.34553349 | -2.7431311 | 0.00608564 | 0.07661007 |
| TRIM73 | 40.873538 | 0.25000718 | 0.44605064 | 0.56049057 | 0.57514487 | 0.77388388 |
| POM121C | 96.1390949 | -0.5701276 | 0.22217687 | -2.566098 | 0.01028498 | 0.09994074 |
| SPDYE5 | 21.2659968 | -0.3734409 | 0.73028217 | -0.5113653 | 0.60909532 | 0.79623076 |
| HIP1 | 449.110129 | 0.6769445 | 0.30228342 | 2.23943644 | 0.02512753 | 0.15519363 |
| RHBDD2 | 20.7065535 | -0.4695332 | 0.48355858 | -0.9709956 | 0.33155049 | NA |
| POR | 66.0549753 | -0.8949606 | 0.70724509 | -1.2654179 | 0.20572158 | 0.45902815 |
| STYXL1 | 49.6625375 | 0.06151744 | 0.36853347 | 0.16692496 | 0.86742909 | 0.93827484 |
| MDH2 | 24.8293173 | 0.7278887 | 0.48497883 | 1.50086696 | 0.13338998 | 0.36762309 |
| HSPB1 | 72.7916326 | 0.10231283 | 0.52243353 | 0.19583894 | 0.84473623 | 0.92501562 |
| YWHAG | 59.1886154 | -0.7437277 | 0.32603432 | -2.281133 | 0.02254058 | 0.14632666 |
| SSC4D | 35.1607076 | 1.77263519 | 0.53732616 | 3.2989929 | 0.00097032 | 0.02829008 |
| DTX2 | 35.4429528 | 0.16065693 | 0.41192637 | 0.39001371 | 0.69652641 | 0.85016869 |
| POMZP3 | 23.9840918 | 0.01056213 | 0.43367414 | 0.024355 | 0.98056944 | 0.99299791 |

| | | | | | | |
|---|---|---|---|---|---|---|
| CCDC146 | 104.263866 | 0.8554107 | 0.3173796 | 2.69522902 | 0.00703403 | 0.08190757 |
| GSAP | 49.9763988 | 1.38605964 | 0.60472654 | 2.29204367 | 0.02190312 | 0.14458691 |
| PTPN12 | 176.109087 | -0.4500698 | 0.27672508 | -1.626415 | 0.10386139 | 0.32339043 |
| APTR | 29.9260601 | 0.41818851 | 0.44399857 | 0.94186904 | 0.34625969 | 0.60358654 |
| RSBN1L | 91.9760387 | -0.1171568 | 0.22742054 | -0.5151548 | 0.60644482 | 0.79492605 |
| PHTF2 | 88.9606767 | 0.03656665 | 0.24181636 | 0.1512166 | 0.87980486 | 0.94376451 |
| MAGI2 | 1012.25843 | -0.425699 | 0.40825243 | -1.0427348 | 0.29707116 | 0.55674621 |
| GNAI1 | 237.059319 | -1.2755538 | 0.55502559 | -2.2981891 | 0.02155103 | 0.14349647 |
| SEMA3C | 77.7095519 | -2.5034083 | 0.68349751 | -3.6626444 | 0.00024962 | 0.01289505 |
| CACNA2D1 | 101.686607 | 0.20675942 | 0.26950194 | 0.76719087 | 4.43E-01 | 0.68272427 |
| PCLO | 213.560098 | 0.48843657 | 0.52794967 | 0.92515744 | 0.35488402 | 0.61168935 |
| KIAA1324L | 28.9899483 | 0.1296576 | 0.71676805 | 0.18089199 | 0.85645235 | 0.93201099 |
| DMTF1 | 155.737028 | -0.1528494 | 0.32932941 | -0.4641231 | 0.64255954 | 0.81595394 |
| TMEM243 | 34.832759 | 0.59547099 | 0.30936324 | 1.9248279 | 0.05425089 | 0.23287156 |
| CROT | 32.6531783 | 1.19217761 | 0.43351938 | 2.74999839 | 0.00595956 | 0.07594487 |
| SLC25A40 | 45.0528652 | 1.22604483 | 0.3437734 | 3.56643305 | 0.00036187 | 0.01648129 |
| DBF4 | 15.5946551 | -0.3548036 | 0.42385646 | -0.8370842 | 0.40254522 | NA |
| ADAM22 | 63.4338249 | 1.43852599 | 0.42853887 | 3.35681569 | 0.00078846 | 0.02532098 |
| SRI | 37.7051809 | 0.61210982 | 0.46151489 | 1.3263057 | 0.18473846 | 0.43476148 |
| STEAP2 | 33.9386462 | 0.73162154 | 0.48408262 | 1.51135675 | 0.13069758 | 0.36322236 |
| CFAP69 | 62.6741983 | 0.43603928 | 0.42792025 | 1.01897323 | 0.30821568 | 0.5682242 |
| GTPBP10 | 47.6397969 | 0.02106007 | 0.29559525 | 0.0712463 | 0.94320174 | 0.97657032 |
| CLDN12 | 55.4575233 | -0.1381966 | 0.28343896 | -0.487571 | 0.62585372 | 0.8053173 |
| CDK14 | 675.38874 | -0.0473706 | 0.21839364 | -0.2169046 | 8.28E-01 | 9.17E-01 |
| AC079760.1 | 39.0365111 | 1.10701531 | 0.61493491 | 1.80021543 | 0.07182663 | 0.26733538 |
| MTERF1 | 165.716852 | 0.41481682 | 0.32181337 | 1.28899808 | 0.19739875 | 0.45058666 |
| AKAP9 | 376.176242 | -0.0489272 | 0.19119712 | -0.2558991 | 0.79802873 | 0.90153736 |
| KRIT1 | 149.51168 | -0.2362157 | 0.20935325 | -1.1283118 | 0.25918826 | 0.5179055 |
| ANKIB1 | 284.258052 | -0.1949251 | 0.19144792 | -1.0181623 | 0.30860081 | 0.56838967 |
| GATAD1 | 39.0328757 | 0.145074 | 0.38284254 | 0.37893909 | 0.70473309 | 0.85341891 |
| AC007566.1 | 33.3584274 | -0.1451828 | 0.4112953 | -0.3529893 | 0.72409646 | 0.86235038 |
| PEX1 | 55.0644191 | 0.34750716 | 0.38246807 | 0.90859131 | 0.3635659 | 0.6199306 |
| RBM48 | 15.9953244 | 0.8349328 | 0.45052098 | 1.85326064 | 0.06384501 | NA |
| FAM133B | 67.3526722 | -0.1099843 | 0.295165 | -0.3726199 | 0.70943138 | 0.85607462 |
| CDK6 | 210.430018 | 0.44908176 | 0.44028866 | 1.01997122 | 0.30774211 | 0.56799861 |
| SAMD9L | 23.5571864 | 1.53780474 | 1.11337124 | 1.38121471 | 0.16721295 | 0.41517358 |
| VPS50 | 151.442876 | 0.03511107 | 0.20344206 | 0.17258513 | 0.86297754 | 0.93629034 |
| CALCR | 238.054564 | 1.83238549 | 0.88171436 | 2.0782076 | 0.03769024 | 0.19425568 |
| GNGT1 | 152.669321 | 1.13700481 | 0.70248695 | 1.61854224 | 0.10554579 | 0.32665298 |
| BET1 | 37.0311122 | 0.47769003 | 0.42973777 | 1.11158495 | 0.26631665 | 0.52522169 |
| CASD1 | 80.8940665 | 0.24342346 | 0.26957914 | 0.90297589 | 0.36653869 | 0.62157013 |
| SGCE | 111.144451 | 0.99864272 | 0.41782485 | 2.39009888 | 0.01684384 | 0.129002 |
| PPP1R9A | 323.28495 | 0.62587557 | 0.26912406 | 2.32560241 | 0.02003977 | 0.13896047 |
| AC002429.2 | 23.1319822 | 2.10488017 | 0.75782972 | 2.77751073 | 0.0054777 | 0.07231884 |
| PON2 | 68.6644176 | 0.50296495 | 0.44151822 | 1.13917143 | 0.25463166 | 0.51212435 |
| PDK4 | 120.183149 | -0.0585138 | 0.79208731 | -0.073873 | 0.94111147 | 0.97532449 |
| SLC25A13 | 226.89928 | -0.4439855 | 0.19592862 | -2.2660574 | 0.02344787 | 0.14954518 |
| SEM1 | 86.8730989 | 0.45498554 | 0.23578765 | 1.92964109 | 0.05365132 | 0.23251928 |
| SDHAF3 | 18.3454426 | 0.26440513 | 0.49376943 | 0.535483 | 0.59231591 | NA |
| LMTK2 | 91.6550514 | -0.2705985 | 0.27052617 | -1.0002675 | 0.31718105 | 0.57492021 |
| TECPR1 | 15.3810028 | 0.36375235 | 0.5584256 | 0.65138911 | 0.51479534 | NA |
| BRI3 | 102.072192 | -0.2960973 | 0.34028087 | -0.8701556 | 0.38421537 | 0.63451053 |
| BAIAP2L1 | 180.097418 | 0.20395466 | 0.42238572 | 0.48286354 | 0.62919263 | 0.80701478 |
| TRRAP | 140.382363 | -0.2460261 | 0.22612298 | -1.0880191 | 0.27658668 | 0.53501126 |

| | | | | | | |
|---|---|---|---|---|---|---|
| SMURF1 | 253.607539 | -0.5674146 | 0.31341963 | -1.8103989 | 0.07023395 | 0.26460632 |
| PDAP1 | 23.7161939 | -0.7112933 | 0.38206615 | -1.8617021 | 0.06264509 | 0.24960868 |
| BUD31 | 13.6648278 | 0.15068787 | 0.55645297 | 0.27080073 | 0.7865443 | NA |
| CPSF4 | 14.6808446 | 0.11016093 | 0.51625109 | 0.21338635 | 0.83102563 | NA |
| ZNF789 | 38.7084033 | 0.17942889 | 0.40287435 | 0.44537183 | 0.65605107 | 0.82442858 |
| ZNF394 | 28.0897234 | -1.0008523 | 0.40071557 | -2.4976626 | 0.01250151 | 0.10951947 |
| ZKSCAN5 | 21.6049609 | -0.1194657 | 0.42031785 | -0.284227 | 0.77623645 | 0.88909107 |
| FAM200A | 15.7716115 | -0.8480229 | 0.58145042 | -1.4584611 | 0.14471348 | NA |
| ZSCAN25 | 14.7196981 | 0.15278752 | 0.48908629 | 0.31239378 | 0.75474128 | NA |
| TRIM4 | 44.9371375 | -0.2169205 | 0.35357275 | -0.6135103 | 0.53953895 | 0.74929145 |
| ZKSCAN1 | 132.310708 | -0.1016673 | 0.25120612 | -0.4047165 | 0.68568589 | 0.84376087 |
| ZNF3 | 27.3411628 | -0.8912594 | 0.51495501 | -1.7307519 | 0.08349603 | 0.28861463 |
| COPS6 | 24.4754241 | 0.36205989 | 0.3659593 | 0.9893447 | 0.32249452 | 0.57912936 |
| AP4M1 | 19.6881835 | 0.04661906 | 0.56846905 | 0.08200809 | 0.93464028 | NA |
| LAMTOR4 | 37.856223 | 0.32492289 | 0.36090406 | 0.90030267 | 0.3679592 | 0.62245758 |
| STAG3 | 15.1600263 | 0.42731454 | 0.57752374 | 0.73990818 | 0.45935571 | NA |
| CASTOR3 | 92.6634972 | 0.10260071 | 0.32286383 | 0.31778323 | 0.75064937 | 0.8772395 |
| SPDYE3 | 36.1224981 | 0.88951525 | 0.57857268 | 1.53743046 | 0.12418793 | 0.35457643 |
| PILRB | 86.4086214 | -0.596132 | 0.41835464 | -1.4249441 | 0.15417337 | 0.39652407 |
| PILRA | 37.441334 | -0.7226772 | 0.52037264 | -1.3887686 | 0.16490313 | 0.4116962 |
| ZCWPW1 | 12.5210497 | -1.188615 | 0.63618976 | -1.868334 | 0.06171553 | NA |
| PPP1R35 | 17.2555071 | 0.10693441 | 0.49845744 | 0.21453067 | 0.83013325 | NA |
| TSC22D4 | 46.4447932 | 0.66083686 | 0.32788499 | 2.01545321 | 0.04385718 | 0.20924217 |
| AGFG2 | 19.3277951 | 0.09027013 | 0.54228764 | 0.16646172 | 0.86779361 | NA |
| LRCH4 | 12.9457549 | -0.6175999 | 0.60589698 | -1.0193151 | 0.3080534 | NA |
| GNB2 | 44.1089928 | 0.26017744 | 0.42282561 | 0.61533037 | 0.53833656 | 0.74901962 |
| GIGYF1 | 54.3534731 | -0.2675885 | 0.35917955 | -0.7449993 | 0.45627216 | 0.69129178 |
| EPHB4 | 17.0351579 | 0.30216855 | 0.48096672 | 0.62825251 | 0.52983854 | NA |
| SLC12A9 | 32.4337155 | 1.06502408 | 0.48159803 | 2.21143779 | 0.02700554 | 0.16195658 |
| TRIP6 | 21.8792818 | -0.5033974 | 0.54186865 | -0.9290026 | 0.35288775 | 0.61004394 |
| SRRT | 23.544145 | -1.0302627 | 0.39003314 | -2.6414747 | 0.0082546 | 0.08912945 |
| TRIM56 | 75.4124769 | -0.0945633 | 0.42539908 | -0.2222931 | 0.82408576 | 0.91510572 |
| AP1S1 | 22.3829174 | 0.56667166 | 0.50365618 | 1.12511607 | 0.26053985 | 0.51979659 |
| PLOD3 | 20.5829719 | 0.34132297 | 0.44384549 | 0.76901305 | 0.44188556 | NA |
| ZNHIT1 | 22.0173731 | 1.07141496 | 0.430424 | 2.4892082 | 0.0128028 | 0.11046442 |
| FIS1 | 38.8070159 | 0.19648538 | 0.39118935 | 0.50227692 | 0.61547274 | 0.80075958 |
| COL26A1 | 14.0646232 | 0.9729702 | 0.75275868 | 1.29253933 | 0.19617043 | NA |
| CUX1 | 633.144182 | -0.5300592 | 0.20958011 | -2.5291484 | 0.01143397 | 0.10405175 |
| PRKRIP1 | 171.517725 | -0.4150711 | 0.19947065 | -2.0808632 | 0.03744643 | 0.1936997 |
| ORAI2 | 24.1714046 | -0.5766759 | 0.42128661 | -1.3688446 | 0.17104786 | 0.41901826 |
| AC093668.2 | 10.3228161 | -0.197788 | 0.76385959 | -0.2589325 | 0.79568736 | NA |
| SPDYE2 | 17.1754184 | 1.23132906 | 0.66477747 | 1.85224246 | 0.06399101 | NA |
| AC105052.1 | 11.3006823 | -0.9815367 | 0.66112113 | -1.4846549 | 0.13763527 | NA |
| AC105052.3 | 44.4271495 | 0.82023068 | 0.56280045 | 1.45740942 | 0.1450034 | 0.38459378 |
| FAM185A | 79.4812386 | 1.15581674 | 0.39993459 | 2.89001441 | 0.00385224 | 0.06113547 |
| FBXL13 | 56.185454 | -0.2289428 | 0.40624773 | -0.5635547 | 0.5730572 | 0.77342122 |
| ARMC10 | 43.6082214 | -0.1916516 | 0.34655577 | -0.5530181 | 0.58025105 | 0.77754044 |
| NAPEPLD | 73.2921427 | 1.03285359 | 0.35060999 | 2.94587612 | 0.00322041 | 0.0564636 |
| PMPCB | 42.7247092 | 0.33050563 | 0.29079019 | 1.13657767 | 0.25571488 | 0.5136319 |
| DNAJC2 | 41.5940656 | -0.4927593 | 0.32713698 | -1.5062782 | 0.13199578 | 0.36545739 |
| PSMC2 | 26.5793636 | 0.06366121 | 0.35123873 | 0.18124768 | 0.85617317 | 0.93197071 |
| ORC5 | 52.9391816 | 0.52255978 | 0.28152429 | 1.85618005 | 0.0634279 | 0.25113402 |
| LHFPL3 | 112.983766 | -0.5561033 | 0.50708491 | -1.0966669 | 0.27278702 | 0.53182413 |
| LINC01004 | 159.853207 | -0.0298832 | 0.22678928 | -0.1317662 | 0.8951692 | 0.95277205 |

| | | | | | | |
|---|---|---|---|---|---|---|
| KMT2E-AS1 | 12.1504779 | -1.2249209 | 0.59897604 | -2.0450249 | 0.0408524 | NA |
| KMT2E | 346.557115 | -0.0534869 | 0.18666642 | -0.2865375 | 0.77446646 | 0.88786322 |
| SRPK2 | 540.574176 | -0.0944973 | 0.25011052 | -0.3778221 | 0.70556278 | 0.85402056 |
| PUS7 | 27.7766089 | -0.2187728 | 0.38959664 | -0.5615367 | 0.57443173 | 0.77388388 |
| RINT1 | 28.5628994 | -0.048696 | 0.4605639 | -0.1057313 | 0.91579556 | 0.96388328 |
| ATXN7L1 | 172.473367 | -0.5544129 | 0.45354903 | -1.2223881 | 0.22156089 | 0.47873331 |
| CDHR3 | 35.4873701 | -0.4331983 | 0.68727079 | -0.6303168 | 0.52848735 | 0.74310689 |
| SYPL1 | 53.3165073 | -0.3980558 | 0.339231 | -1.1734064 | 0.24063288 | 0.49822614 |
| NAMPT | 166.632074 | -0.5041313 | 0.49079985 | -1.0271626 | 0.30434389 | 0.56489395 |
| HBP1 | 89.8217955 | -0.0981834 | 0.27254792 | -0.3602426 | 0.71866575 | 0.85995641 |
| COG5 | 296.117442 | -0.2266068 | 0.25986974 | -0.8720014 | 0.38320764 | 0.63431165 |
| DUS4L | 14.119805 | -0.5430394 | 0.52479145 | -1.0347718 | 0.30077549 | NA |
| BCAP29 | 46.579514 | 0.17210364 | 0.28620671 | 0.60132636 | 0.54762264 | 0.7554786 |
| CBLL1 | 21.1242943 | -0.4474984 | 0.43234364 | -1.0350526 | 0.30064437 | NA |
| SLC26A3 | 304.21319 | -0.8483105 | 0.7252995 | -1.1696003 | 0.24216185 | 0.49957647 |
| DLD | 65.3387027 | 0.08775815 | 0.28133205 | 0.31193799 | 0.75508765 | 0.87904624 |
| LAMB1 | 564.527963 | -1.2042429 | 0.32760399 | -3.6759104 | 0.000237 | 0.01240959 |
| PNPLA8 | 94.1025133 | -0.5352735 | 0.3025 | -1.7694993 | 0.07681059 | 0.27670206 |
| IMMP2L | 2419.07997 | -0.301102 | 0.37901982 | -0.7944228 | 0.42694935 | 0.66997536 |
| DOCK4 | 268.658569 | -0.5830823 | 0.4106212 | -1.4200005 | 0.15560753 | 0.39879246 |
| ZNF277 | 135.393739 | 0.09310657 | 0.28218333 | 0.32995063 | 0.74143726 | 0.87152705 |
| IFRD1 | 124.173136 | -0.6167132 | 0.52922926 | -1.1653044 | 0.24389581 | 0.50140654 |
| LSMEM1 | 17.1609581 | -0.8066713 | 0.66020245 | -1.2218545 | 0.22176265 | NA |
| TMEM168 | 86.0442487 | 0.2115608 | 0.28753628 | 0.73577079 | 0.4618702 | 0.69416206 |
| BMT2 | 106.028458 | -0.007706 | 0.226847 | -0.0339702 | 0.97290095 | 0.98922306 |
| MDFIC | 40.4600627 | 1.41210768 | 0.52507018 | 2.68936943 | 0.00715871 | 0.08259723 |
| TES | 53.3452401 | -0.6053552 | 0.33775276 | -1.7923029 | 0.07308446 | 0.26939652 |
| MET | 217.781262 | -2.2180609 | 0.52993516 | -4.1855326 | 2.84E-05 | 0.00322027 |
| CAPZA2 | 147.34441 | -0.8362279 | 0.36309405 | -2.3030615 | 0.02127538 | 0.14276951 |
| ST7 | 132.57755 | -0.2428619 | 0.25626027 | -0.9477157 | 0.34327418 | 0.60053858 |
| LSM8 | 37.7227913 | -0.3374979 | 0.40764588 | -0.8279192 | 0.40771623 | 0.65378997 |
| ANKRD7 | 54.0578388 | 1.61146958 | 0.57060394 | 2.82414732 | 0.00474066 | 0.06664966 |
| TSPAN12 | 86.4513048 | 0.92034188 | 0.60404171 | 1.52363962 | 0.12759876 | 0.35967181 |
| ING3 | 27.364329 | 0.02900624 | 0.4539375 | 0.06389918 | 0.9490505 | 0.97977013 |
| FAM3C | 110.303118 | -0.6860383 | 0.37749851 | -1.8173272 | 6.92E-02 | 0.26199732 |
| AASS | 45.9865936 | -0.2523885 | 0.37995222 | -0.6642638 | 0.50652149 | 0.7281071 |
| CADPS2 | 684.59032 | 0.89499566 | 0.34557892 | 2.58984451 | 0.00960193 | 0.09673566 |
| AC004594.1 | 33.9681843 | 1.63848273 | 0.82906174 | 1.97630966 | 0.04811972 | 0.21967821 |
| IQUB | 23.1203792 | 1.88964009 | 0.67792928 | 2.78737052 | 0.00531377 | 0.07113057 |
| NDUFA5 | 69.2423163 | 0.42544924 | 0.3013466 | 1.41182692 | 0.15800093 | 0.40275854 |
| WASL | 108.018247 | -0.2942155 | 0.25369704 | -1.1597121 | 0.24616603 | 0.50296216 |
| POT1 | 61.4186687 | 0.30261515 | 0.25540262 | 1.18485529 | 0.23607467 | 0.49390235 |
| POT1-AS1 | 30.7694825 | 0.65126441 | 0.6078177 | 1.07147982 | 0.28395374 | 0.54273452 |
| ZNF800 | 79.8592107 | 0.20925785 | 0.2893053 | 0.7233115 | 0.46948853 | 0.69910103 |
| ARF5 | 14.2628768 | -0.1704596 | 0.60670509 | -0.2809595 | 0.77874143 | NA |
| SND1 | 248.269878 | -0.398174 | 0.21796973 | -1.8267401 | 0.06773886 | 0.25900943 |
| RBM28 | 46.4397568 | 0.193298 | 0.29332163 | 0.65899673 | 0.50989787 | 0.72990216 |
| HILPDA | 38.0214772 | -1.1430233 | 1.53970601 | -0.7423646 | 0.45786644 | 0.69231672 |
| METTL2B | 37.39398 | -0.094771 | 0.38148554 | -0.2484261 | 0.80380474 | 0.90408418 |
| FAM71F2 | 14.1617526 | -0.6248386 | 0.50573915 | -1.2354958 | 0.21664603 | NA |
| CALU | 51.7910367 | -0.3090771 | 0.32198558 | -0.9599098 | 0.33710061 | 0.59276835 |
| KCP | 23.5415052 | -1.6485903 | 0.64317202 | -2.5632184 | 1.04E-02 | 1.00E-01 |
| ATP6V1F | 29.2715827 | 0.80057751 | 0.42866601 | 1.86760203 | 0.06181756 | 0.2487244 |
| TNPO3 | 136.835894 | -0.5899037 | 0.25655172 | -2.2993559 | 0.02148474 | 0.14349647 |

| | | | | | | |
|---|---|---|---|---|---|---|
| TSPAN33 | 29.2743755 | 0.88048782 | 0.56154154 | 1.56798342 | 0.116885 | 0.34414008 |
| SMO | 27.4894492 | 1.10319061 | 0.47062828 | 2.34408059 | 1.91E-02 | 0.1359475 |
| AHCYL2 | 309.361601 | 0.13825267 | 0.21674178 | 0.63786814 | 0.52355952 | 0.73904282 |
| STRIP2 | 39.6303375 | -3.5416451 | 0.85964527 | -4.1198913 | 3.79E-05 | 0.00383215 |
| NRF1 | 168.510181 | -0.1900171 | 0.22086169 | -0.8603443 | 0.38959929 | 0.63857447 |
| UBE2H | 864.37545 | -0.6664765 | 0.31880149 | -2.090569 | 0.03656671 | 0.19068699 |
| ZC3HC1 | 32.3253196 | -0.1747336 | 0.37813 | -0.4620992 | 0.64401019 | 0.81703418 |
| KLHDC10 | 175.381909 | 0.21747078 | 0.25180313 | 0.863654 | 0.38777799 | 0.63702067 |
| TMEM209 | 31.5331655 | 0.02029309 | 0.39542293 | 0.05131997 | 0.95907055 | 0.98404745 |
| COPG2 | 55.400575 | 0.28306407 | 0.39989807 | 0.70784056 | 0.47904428 | 0.70690257 |
| AC016831.7 | 971.684588 | -0.8205985 | 0.37386681 | -2.1948954 | 0.0281711 | 0.16514315 |
| LINC00513 | 112.42501 | -0.946649 | 0.67151264 | -1.4097263 | 0.15862052 | 0.40347064 |
| AC058791.1 | 18.8151438 | -0.3277247 | 0.82133988 | -0.3990122 | 0.6898842 | NA |
| MKLN1 | 829.313703 | -0.6115461 | 0.31608757 | -1.9347363 | 0.05302266 | 0.23122687 |
| PODXL | 15.7942022 | -0.9670479 | 0.49429606 | -1.9564143 | 0.05041637 | NA |
| CHCHD3 | 401.773164 | 0.09616591 | 0.23344214 | 0.41194752 | 0.68037789 | 0.84089068 |
| EXOC4 | 737.787243 | 0.20403299 | 0.28029781 | 0.72791506 | 0.46666559 | 0.69672484 |
| LRGUK | 45.0973511 | 0.97254926 | 0.47831199 | 2.03329475 | 0.04202275 | 0.20441466 |
| SLC35B4 | 111.494103 | 0.55683813 | 0.34572225 | 1.6106517 | 1.07E-01 | 0.32921441 |
| AKR1B1 | 77.9889668 | -0.9939439 | 0.8908585 | -1.1157146 | 0.2645443 | 0.5237724 |
| BPGM | 20.7080471 | 0.28675289 | 0.44090084 | 0.65037956 | 0.51544708 | NA |
| CALD1 | 36.9254711 | -0.4374114 | 0.45806958 | -0.9549016 | 0.33962747 | 0.59535132 |
| AGBL3 | 22.2957833 | 0.94723396 | 0.49210716 | 1.92485306 | 0.05424774 | 0.23287156 |
| CYREN | 29.203022 | 0.2710444 | 0.41278806 | 0.6566188 | 0.51142605 | 0.73064085 |
| TMEM140 | 37.8025861 | -1.4746526 | 0.47145406 | -3.1278818 | 0.00176071 | 0.03887902 |
| WDR91 | 135.136706 | 0.31602467 | 0.42531429 | 0.74303797 | 0.45745869 | 0.69203214 |
| AC009542.1 | 15.5136364 | 0.65300596 | 0.80247442 | 0.81374054 | 0.41579361 | NA |
| CNOT4 | 234.904824 | -0.3808256 | 0.24157741 | -1.5764125 | 0.11493077 | 0.34128941 |
| NUP205 | 56.2724148 | -0.5615513 | 0.36525873 | -1.537407 | 0.12419367 | 0.35457643 |
| STMP1 | 60.4971359 | 0.83840467 | 0.36863694 | 2.27433709 | 0.02294573 | 0.14730047 |
| MTPN | 55.9206105 | -0.07954 | 0.3148647 | -0.2526165 | 0.80056455 | 0.90333444 |
| CREB3L2 | 77.0494179 | 1.09330153 | 0.30417159 | 3.59435783 | 0.00032519 | 0.01498807 |
| TRIM24 | 95.967477 | -0.3434594 | 0.26227708 | -1.309529 | 0.19035521 | 0.44158049 |
| SVOPL | 63.0901198 | 0.35686185 | 0.6480369 | 0.55068137 | 0.58185212 | 0.7783237 |
| ATP6V0A4 | 447.262718 | 0.75237783 | 0.33966233 | 2.21507587 | 0.02675486 | 0.16063349 |
| TMEM213 | 285.076944 | 1.12257006 | 0.44620198 | 2.51583389 | 0.01187511 | 0.10677891 |
| KIAA1549 | 33.4963694 | 0.45004464 | 0.40866998 | 1.10124225 | 0.27079124 | 0.52912064 |
| ZC3HAV1 | 55.0203356 | 0.03570922 | 0.48029837 | 0.07434799 | 0.9407335 | 0.97515997 |
| TTC26 | 22.4568918 | -0.1090868 | 0.57193071 | -0.1907343 | 0.84873376 | 0.9268876 |
| UBN2 | 112.78377 | 0.14012035 | 0.20623979 | 0.67940504 | 0.49688126 | 0.72101702 |
| LUC7L2 | 34.8071376 | -0.3356392 | 0.39877331 | -0.8416791 | 0.39996757 | 0.6477068 |
| HIPK2 | 384.416159 | -0.2149436 | 0.25009217 | -0.8594576 | 3.90E-01 | 0.63869565 |
| TBXAS1 | 34.8747512 | -0.0058884 | 0.5228804 | -0.0112615 | 9.91E-01 | 9.96E-01 |
| PARP12 | 37.1263076 | 0.15626382 | 0.68613294 | 0.2277457 | 0.81984395 | 0.91308622 |
| KDM7A | 195.934108 | -1.9217524 | 0.32812334 | -5.856799 | 4.72E-09 | 1.21E-05 |
| SLC37A3 | 79.9152521 | -0.5737572 | 0.24966474 | -2.2981108 | 0.02155548 | 0.14349647 |
| MKRN1 | 69.6314781 | -0.466737 | 0.28887551 | -1.6157028 | 0.10615859 | 0.32732986 |
| DENND2A | 256.832709 | -1.116101 | 0.44511398 | -2.5074499 | 0.01216058 | 0.10783409 |
| NDUFB2 | 55.2678836 | 0.93838365 | 0.40037898 | 2.34373859 | 0.01909155 | 0.1359475 |
| BRAF | 525.814551 | -0.3644221 | 0.20268457 | -1.7979765 | 0.07218073 | 0.26800535 |
| MRPS33 | 16.5652858 | 1.02365315 | 0.47503786 | 2.15488752 | 0.03117064 | NA |
| TMEM178B | 80.046176 | 1.35228686 | 0.62428232 | 2.16614632 | 0.03030001 | 0.17229651 |
| AGK | 53.8789643 | 0.35567568 | 0.29934949 | 1.18816199 | 0.23476959 | 0.49243138 |
| AC004918.1 | 14.7618042 | 0.75375325 | 0.56485628 | 1.33441598 | 0.18206757 | NA |

| | | | | | | |
|---|---|---|---|---|---|---|
| KIAA1147 | 43.4738522 | 1.06917339 | 0.44833447 | 2.38476731 | 0.01708993 | 0.12988597 |
| SSBP1 | 69.0484313 | -0.2382722 | 0.28790564 | -0.8276052 | 0.40789413 | 0.65393899 |
| MGAM | 16.0946401 | 1.72526313 | 0.93466706 | 1.84585849 | 0.06491276 | NA |
| TRPV6 | 17.414514 | 0.761135 | 0.71584853 | 1.06326264 | 0.28766286 | NA |
| TRPV5 | 64.6570741 | 3.7026694 | 1.0062989 | 3.67949263 | 0.0002337 | 0.01232039 |
| GSTK1 | 49.8598418 | 1.08318295 | 0.38057835 | 2.8461497 | 0.00442514 | 0.0642158 |
| CASP2 | 14.6197286 | -0.4234757 | 0.44028764 | -0.9618159 | 0.33614209 | NA |
| TCAF2 | 17.8522292 | 0.50973175 | 0.84880999 | 0.60052515 | 0.5481563 | NA |
| TCAF1 | 63.0369126 | 1.31254304 | 0.36080221 | 3.63784641 | 0.00027493 | 0.01360691 |
| AC004889.1 | 30.4642987 | 0.30896865 | 0.47757008 | 0.64695982 | 0.51765795 | 0.73486042 |
| AC074386.1 | 67.9749684 | -0.2247008 | 0.30580049 | -0.7347953 | 0.46246416 | 0.69418845 |
| ARHGEF5 | 15.2960325 | -0.278867 | 0.63211008 | -0.4411684 | 0.65909106 | NA |
| TPK1 | 41.6756555 | -0.1358483 | 0.41301737 | -0.3289167 | 0.74221862 | 0.87152705 |
| CNTNAP2 | 15.9670664 | 0.07158197 | 0.59479018 | 0.12034827 | 0.90420727 | NA |
| CUL1 | 145.338382 | -0.2093744 | 0.23471493 | -0.892037 | 0.37237311 | 0.62511578 |
| EZH2 | 24.9575181 | 0.00708935 | 0.57021062 | 0.01243286 | 0.99008027 | 0.99583841 |
| PDIA4 | 21.4625378 | -0.0816251 | 0.72650481 | -0.1123531 | 0.91054346 | 0.96104245 |
| ZNF786 | 18.5243973 | 0.31756615 | 0.54941712 | 0.57800557 | 0.56326036 | NA |
| ZNF425 | 14.2798163 | -0.0488139 | 0.51460991 | -0.0948561 | 0.92442912 | NA |
| ZNF398 | 37.395391 | -0.1602221 | 0.34842763 | -0.4598431 | 0.64562882 | 0.81777273 |
| ZNF282 | 25.047608 | -0.7063024 | 0.39461296 | -1.789861 | 0.07347625 | 0.26995071 |
| ZNF212 | 17.573893 | -0.5869148 | 0.51351891 | -1.1429272 | 0.2530688 | NA |
| ZNF783 | 18.1500094 | 0.02065456 | 0.47228846 | 0.04373293 | 0.96511729 | NA |
| ZNF746 | 16.2044141 | 0.2533116 | 0.4851098 | 0.52217375 | 0.60154936 | NA |
| ZNF467 | 14.4039782 | -0.108335 | 0.5706075 | -0.189859 | 0.84941963 | NA |
| ZNF862 | 51.0502101 | 1.05557677 | 0.46736966 | 2.25854794 | 0.02391152 | 0.15098192 |
| ACTR3C | 342.889586 | 0.05073364 | 0.29472324 | 0.17213995 | 0.8633275 | 0.93643831 |
| LRRC61 | 17.4837919 | 0.45945945 | 0.47761555 | 0.96198596 | 0.33605665 | NA |
| RARRES2 | 16.1115404 | -0.7606767 | 0.62152752 | -1.2238826 | 0.22099653 | NA |
| TMEM176B | 32.1488996 | 1.0326841 | 0.51857201 | 1.99139961 | 0.04643697 | 0.21595574 |
| TMEM176A | 15.8257982 | 0.81803305 | 0.62758219 | 1.30346759 | 0.19241517 | NA |
| AOC1 | 12.4162088 | 0.60070311 | 0.85455287 | 0.70294434 | 0.48209043 | NA |
| ABCB8 | 15.315575 | -0.5692562 | 0.59720956 | -0.9531933 | 0.34049213 | NA |
| SLC4A2 | 28.3502639 | -0.3740991 | 0.55136358 | -0.6784979 | 0.49745603 | 0.72133441 |
| FASTK | 19.9511729 | 0.10605204 | 0.60637146 | 0.17489615 | 0.86116123 | NA |
| TMUB1 | 17.3410489 | 0.55688099 | 0.53948916 | 1.03223757 | 0.30196084 | NA |
| AGAP3 | 21.3117712 | 0.29839591 | 0.5488182 | 0.54370629 | 0.5866436 | 0.78229311 |
| SMARCD3 | 42.9948386 | 0.62281787 | 0.3976693 | 1.56617037 | 0.11730874 | 0.34472743 |
| NUB1 | 49.9176836 | -0.4704506 | 0.39028296 | -1.205409 | 0.22804547 | 0.48611194 |
| RHEB | 125.973139 | -0.9204777 | 0.32131317 | -2.864737 | 0.00417356 | 0.06301187 |
| PRKAG2 | 243.362977 | -0.7567115 | 0.35761604 | -2.1159887 | 0.03434577 | 0.18371048 |
| GALNT11 | 169.316276 | 0.02693658 | 0.47207083 | 0.05706046 | 0.95449703 | 0.98153155 |
| KMT2C | 521.00899 | -0.4210726 | 0.21454398 | -1.9626402 | 0.049688 | 0.22378497 |
| ACTR3B | 81.548352 | -0.2582657 | 0.25183788 | -1.0255238 | 0.30511611 | 0.56557634 |
| DPP6 | 221.892943 | -0.166712 | 0.27050446 | -0.6163003 | 0.53769635 | 0.74878549 |
| PAXIP1-AS2 | 19.2351722 | -0.1296173 | 0.54221793 | -0.2390501 | 0.8110667 | NA |
| PAXIP1 | 47.7043864 | -0.2745281 | 0.30426034 | -0.9022802 | 0.36690802 | 0.62174959 |
| INSIG1 | 33.2984627 | -1.9580146 | 0.76151535 | -2.5712083 | 0.01013443 | 0.09949585 |
| AC009403.1 | 26.1581326 | 0.06858684 | 0.44896088 | 0.15276797 | 0.87858126 | 0.94342075 |
| RBM33 | 223.917902 | -0.3531563 | 0.25358278 | -1.3926667 | 0.16372061 | 0.41042742 |
| LINC01006 | 33.8593484 | -0.2686195 | 0.50995234 | -0.526754 | 0.59836441 | 0.7894568 |
| LMBR1 | 216.301548 | -0.0903298 | 0.24811061 | -0.3640708 | 0.71580517 | 0.85868129 |
| NOM1 | 21.9279941 | -0.6299555 | 0.58106928 | -1.0841316 | 0.27830647 | 0.537008 |
| UBE3C | 289.747842 | -0.4869233 | 0.28883772 | -1.6858023 | 0.09183389 | 0.30362776 |

| | | | | | | |
|---|---|---|---|---|---|---|
| DNAJB6 | 123.939661 | -0.5997261 | 0.32219631 | -1.8613685 | 0.06269215 | 0.24960868 |
| ESYT2 | 160.304243 | -0.2700333 | 0.23745065 | -1.1372184 | 0.25544698 | 0.51347648 |
| WDR60 | 160.665149 | -0.1550997 | 0.18980092 | -0.8171703 | 0.4138311 | 0.65845053 |
| GTPBP6 | 33.0195925 | -0.8920871 | 0.41884508 | -2.1298737 | 0.03318205 | 0.18036879 |
| LINC00685 | 12.2603578 | -0.1794164 | 0.78907175 | -0.2273766 | 0.82013092 | NA |
| PPP2R3B | 12.7580034 | 0.47460044 | 0.49679369 | 0.95532702 | 0.33941235 | NA |
| IL3RA | 16.4188225 | 0.84606354 | 0.54047195 | 1.56541617 | 0.11748536 | NA |
| SLC25A6 | 80.6484801 | 0.76795549 | 0.51816118 | 1.48207841 | 0.13831943 | 0.37487489 |
| LINC00106 | 37.4392231 | -0.2244112 | 0.43001099 | -0.5218732 | 0.60175863 | 0.79174977 |
| ASMTL-AS1 | 24.341851 | -0.7768371 | 0.48301767 | -1.6082996 | 0.10776958 | 0.32997921 |
| ASMTL | 25.8466944 | 0.15696786 | 0.36971368 | 0.42456601 | 0.67115307 | 0.83508975 |
| AKAP17A | 36.6985494 | 0.08845917 | 0.30343419 | 0.2915267 | 0.77064853 | 0.88651647 |
| DHRSX | 204.078664 | -0.3473846 | 0.25855127 | -1.3435811 | 0.17908387 | 0.42802774 |
| CD99 | 36.8679032 | 0.78190698 | 0.39537089 | 1.97765442 | 0.04796771 | 0.21943015 |
| ARSD | 50.8840048 | 0.53562238 | 0.56989666 | 0.93985878 | 0.34729 | 0.60403983 |
| PRKX | 45.0032648 | 1.36978153 | 0.43878989 | 3.12172534 | 0.00179795 | 0.03909384 |
| PUDP | 32.0256188 | 0.45042397 | 0.61285668 | 0.73495808 | 0.46236502 | 0.69418845 |
| TBL1X | 431.389459 | -0.2291043 | 0.34327183 | -0.6674136 | 0.50450797 | 0.72610282 |
| SHROOM2 | 33.0535631 | 0.77060591 | 0.4202942 | 1.83349165 | 0.06672949 | 0.25794497 |
| WWC3 | 29.4498249 | 0.37926077 | 0.46070341 | 0.82322111 | 0.41038227 | 0.65612215 |
| MID1 | 340.079812 | -0.1140224 | 0.30365641 | -0.3754982 | 7.07E-01 | 8.55E-01 |
| AC073529.1 | 31.7422825 | 0.65146793 | 0.44989543 | 1.44804301 | 0.14760502 | 0.38722988 |
| ARHGAP6 | 2051.7259 | 0.25536468 | 0.35725634 | 0.71479397 | 4.75E-01 | 0.70288437 |
| PRPS2 | 33.6051576 | -0.3122071 | 0.43523629 | -0.7173279 | 0.47317182 | 0.70227603 |
| TMSB4X | 117.998277 | -0.7179863 | 0.51006394 | -1.4076398 | 0.15923776 | 0.40403105 |
| RAB9A | 40.7577549 | -0.4898943 | 0.37384496 | -1.3104211 | 0.19005343 | 0.44127941 |
| TRAPPC2 | 23.1809171 | -0.0237949 | 0.42854556 | -0.0555249 | 0.95572031 | 0.98250992 |
| OFD1 | 131.815194 | -0.6736552 | 0.26232001 | -2.5680665 | 0.01022675 | 0.09994074 |
| GEMIN8 | 35.7845633 | 0.7922576 | 0.36463687 | 2.17273039 | 0.02980062 | 0.17041258 |
| MOSPD2 | 21.2460019 | -0.0107863 | 0.44386893 | -0.0243006 | 0.98061285 | 0.99299791 |
| PIR | 23.914181 | 1.61503562 | 0.94670882 | 1.70594757 | 0.08801784 | 0.29675763 |
| CLTRN | 17.2855929 | -0.6114708 | 0.79824767 | -0.7660165 | 0.44366651 | NA |
| CA5B | 62.5386435 | -0.6989531 | 0.31897458 | -2.1912502 | 0.02843369 | 0.16587474 |
| ZRSR2 | 28.9308092 | -0.2963188 | 0.37424224 | -0.7917834 | 0.428487 | 0.67088373 |
| AP1S2 | 27.6685525 | -0.6281403 | 0.50924315 | -1.2334781 | 0.21739744 | 0.47375655 |
| CTPS2 | 50.4278636 | 0.59059755 | 0.3087323 | 1.91297622 | 0.0557511 | 0.23568726 |
| SYAP1 | 48.3471565 | 0.05268139 | 0.28673497 | 0.18372851 | 0.85422644 | 0.93090484 |
| TXLNG | 45.1875123 | -1.237236 | 0.39761882 | -3.1116132 | 0.00186068 | 0.03989322 |
| RBBP7 | 40.5800434 | -0.1091885 | 0.29772429 | -0.3667436 | 0.7138103 | 0.85788134 |
| REPS2 | 23.6727649 | 0.20620177 | 0.63988223 | 0.32224956 | 0.74726364 | 0.87478148 |
| NHS | 1027.26458 | -1.0378207 | 0.19188532 | -5.4085465 | 6.35E-08 | 7.50E-05 |
| SCML1 | 137.046716 | -0.1012006 | 0.32899981 | -0.3076007 | 0.75838622 | 0.87990635 |
| RAI2 | 29.1443095 | 0.70272376 | 0.44771739 | 1.56956993 | 0.1165152 | 0.34387174 |
| CDKL5 | 69.2943501 | 0.0414025 | 0.26400973 | 0.15682187 | 0.87538524 | 0.94169674 |
| PHKA2 | 35.5576786 | 0.17304592 | 0.33738466 | 0.51290393 | 0.60801853 | 0.7956339 |
| PDHA1 | 63.7242324 | 0.73539206 | 0.41675138 | 1.76458217 | 0.07763401 | 0.27780056 |
| MAP3K15 | 95.2136023 | 1.0185063 | 0.48856629 | 2.08468392 | 0.037098 | 0.19247901 |
| BCLAF3 | 89.5292567 | -0.3593313 | 0.29103191 | -1.2346801 | 0.21694957 | 0.47304841 |
| EIF1AX | 60.5044611 | 0.00260795 | 0.33120932 | 0.00787404 | 0.99371749 | 0.99727755 |
| RPS6KA3 | 157.868265 | -1.4738249 | 0.35931486 | -4.1017644 | 4.10E-05 | 0.00401254 |
| MBTPS2 | 40.117781 | -0.3761304 | 0.36951612 | -1.0179 | 0.30872549 | 0.56848327 |
| SMS | 193.661683 | -1.5370646 | 0.42086884 | -3.6521226 | 0.00026008 | 0.01334566 |
| PTCHD1-AS | 127.334277 | 1.35754934 | 0.82303566 | 1.64944169 | 0.09905718 | 0.31649984 |
| ACOT9 | 30.7399399 | -1.0483382 | 0.44076912 | -2.3784294 | 0.01738656 | 0.13115162 |

| | | | | | | |
|---|---|---|---|---|---|---|
| SAT1 | 1198.91712 | -0.1726302 | 0.28111391 | -0.6140933 | 0.53915369 | 0.74929145 |
| APOO | 199.949287 | 0.39906362 | 0.33170744 | 1.203059 | 0.22895349 | 0.48721455 |
| EIF2S3 | 58.8014317 | -0.1419638 | 0.37350574 | -0.3800845 | 0.70388268 | 0.85295217 |
| ZFX | 86.7773538 | -0.6036155 | 0.3873952 | -1.5581389 | 0.11920033 | 0.34687522 |
| PDK3 | 38.062795 | 0.04177216 | 0.57738425 | 0.07234725 | 0.94232557 | 0.97605704 |
| POLA1 | 82.3161599 | -0.340083 | 0.28453145 | -1.1952386 | 0.23199381 | 0.49066933 |
| GK | 36.4308077 | -1.0738693 | 0.4471604 | -2.4015305 | 0.01632665 | 0.12667965 |
| TAB3 | 79.6434207 | 0.51212327 | 0.36886471 | 1.38837698 | 0.16502228 | 0.41186008 |
| DMD | 37.796641 | 0.52117108 | 0.32593768 | 1.59898995 | 0.10982284 | 0.33253594 |
| TMEM47 | 40.812489 | 0.63332309 | 0.75110834 | 0.84318474 | 0.39912513 | 0.64729586 |
| CFAP47 | 37.9471202 | 0.62393724 | 0.59953135 | 1.04070827 | 0.29801097 | 0.5578284 |
| AF241726.2 | 47.1507814 | -0.359308 | 0.40147921 | -0.8949604 | 0.37080825 | 0.62398581 |
| DYNLT3 | 26.6153285 | -0.6530099 | 0.47619886 | -1.3712967 | 0.17028249 | 0.41839333 |
| RPGR | 54.2283762 | -0.5581617 | 0.28335705 | -1.9698175 | 0.04885929 | 0.22162876 |
| BCOR | 70.9215615 | -0.6955576 | 0.35868808 | -1.9391713 | 0.05248048 | 0.22990454 |
| ATP6AP2 | 40.052272 | 0.47757224 | 0.36288362 | 1.31604795 | 0.18815795 | 0.43860181 |
| CXorf38 | 21.0966221 | 0.64053352 | 0.38933033 | 1.64521865 | 0.09992473 | NA |
| MED14 | 106.781481 | -0.8221593 | 0.22148929 | -3.7119597 | 0.00020566 | 0.01130693 |
| USP9X | 230.848729 | -0.6864493 | 0.22672052 | -3.0277333 | 0.00246395 | 0.04717675 |
| DDX3X | 92.0751947 | -0.3603658 | 0.31858532 | -1.1311438 | 0.25799459 | 0.51619036 |
| CASK | 143.498597 | -0.2345729 | 0.21147031 | -1.1092475 | 0.26732341 | 0.52637204 |
| MAOA | 45.0487655 | 1.07044735 | 0.62063731 | 1.72475507 | 0.08457162 | 0.28931012 |
| MAOB | 32.9440193 | 1.54900227 | 0.6207415 | 2.49540632 | 0.0125813 | 0.10970808 |
| EFHC2 | 18.7363124 | 0.30605721 | 0.62856223 | 0.48691632 | 0.62631763 | NA |
| KDM6A | 294.24203 | -0.5442485 | 0.35807342 | -1.5199356 | 0.12852715 | 0.36084904 |
| MIR222HG | 111.385669 | -1.9637636 | 0.42449238 | -4.6261458 | 3.73E-06 | 0.00086364 |
| KRBOX4 | 40.4753337 | 0.17027127 | 0.41844209 | 0.40691718 | 0.68406882 | 0.8427141 |
| ZNF674 | 13.2395828 | 0.067171 | 0.59008295 | 0.11383316 | 0.90937006 | NA |
| SLC9A7 | 58.5568953 | -0.7215897 | 0.29374326 | -2.456532 | 0.01402853 | 0.11638862 |
| RP2 | 17.4534315 | -0.9707175 | 0.45023073 | -2.1560446 | 0.03108019 | NA |
| JADE3 | 75.1775368 | -0.4518405 | 0.35009638 | -1.2906173 | 0.19683641 | 0.45014879 |
| RGN | 36.0359936 | 2.81197761 | 0.73018634 | 3.85104112 | 0.00011762 | 0.00794119 |
| NDUFB11 | 28.3301736 | 1.01038773 | 0.72674275 | 1.39029626 | 0.16443893 | 0.41120418 |
| RBM10 | 48.4990152 | -0.4827683 | 0.33656187 | -1.4344118 | 0.15145477 | 0.39263974 |
| UBA1 | 82.3811627 | -0.0069221 | 0.35067659 | -0.0197391 | 0.98425147 | 0.99382437 |
| CDK16 | 35.0472717 | -0.368217 | 0.57903886 | -0.6359107 | 0.52483466 | 0.73968378 |
| USP11 | 26.6911175 | -0.5709702 | 0.520889 | -1.0961457 | 0.27301504 | 0.53186453 |
| ZNF41 | 26.1513371 | -0.2213648 | 0.39142078 | -0.5655419 | 0.57170524 | 0.7725448 |
| ARAF | 18.6980887 | -0.1541758 | 0.49971049 | -0.3085303 | 0.75767883 | NA |
| TIMP1 | 22.3931884 | -0.0756425 | 0.5485317 | -0.1378999 | 8.90E-01 | 9.50E-01 |
| UXT | 12.7404729 | -0.8234004 | 0.48072817 | -1.7128191 | 0.08674582 | NA |
| ZNF81 | 49.1953215 | 0.60368679 | 0.32293263 | 1.86938925 | 0.06156868 | 0.24806975 |
| ZNF630 | 26.9551485 | 1.24794201 | 0.54716276 | 2.28075099 | 0.02256319 | 0.14632666 |
| EBP | 15.1316892 | 0.81284505 | 0.6096072 | 1.3333915 | 0.18240336 | NA |
| TBC1D25 | 16.7898282 | 0.13725212 | 0.46723095 | 0.29375649 | 0.76894398 | NA |
| RBM3 | 29.7095035 | -0.0328582 | 0.41279914 | -0.0795986 | 0.93655648 | 0.97405271 |
| WDR13 | 23.8354531 | 0.53561756 | 0.44207946 | 1.21158662 | 0.22567066 | 0.48330191 |
| HDAC6 | 20.0026444 | 0.26934427 | 0.42583675 | 0.63250594 | 0.52705633 | NA |
| PCSK1N | 58.1598224 | 1.18766374 | 0.67752341 | 1.75294865 | 0.07961082 | 0.2817308 |
| OTUD5 | 48.4882468 | -0.6603333 | 0.31811988 | -2.0757375 | 0.03791824 | 0.19457113 |
| GRIPAP1 | 49.9873635 | -0.977978 | 0.29664862 | -3.2967557 | 0.00097809 | 0.0283787 |
| TFE3 | 22.245414 | -0.9372199 | 0.42529692 | -2.2036838 | 0.02754659 | 0.16347426 |
| CCDC120 | 14.5403147 | -0.7418217 | 0.49203791 | -1.5076516 | 0.13164373 | NA |
| WDR45 | 21.795421 | -0.1788417 | 0.43280633 | -0.413214 | 0.6794498 | 0.84052156 |

| | | | | | | |
|---|---|---|---|---|---|---|
| CLCN5 | 405.893414 | 0.8219707 | 0.34841512 | 2.35917053 | 0.01831584 | 0.13368407 |
| SHROOM4 | 184.011404 | -0.0059822 | 0.28460232 | -0.0210195 | 0.98323006 | 0.99382437 |
| AL158055.1 | 13.5454224 | 2.6646823 | 0.82824038 | 3.21728132 | 0.00129412 | NA |
| MAGED1 | 225.680907 | 0.17441877 | 0.19204197 | 0.90823258 | 3.64E-01 | 0.61997896 |
| FAM156A | 14.9460316 | 0.89089502 | 0.47693452 | 1.86796087 | 0.06176752 | NA |
| KANTR | 89.3295647 | 0.27870989 | 0.28646285 | 0.97293554 | 0.33058534 | 0.58655523 |
| KDM5C | 51.8300796 | -0.9378199 | 0.3773289 | -2.4854177 | 0.01293995 | 0.11079049 |
| IQSEC2 | 15.7449301 | 1.20129701 | 0.56784729 | 2.11552829 | 0.03438495 | NA |
| SMC1A | 46.8437389 | -0.8894039 | 0.44470348 | -1.9999932 | 0.045501 | 0.21335013 |
| HUWE1 | 209.680442 | -0.3439252 | 0.24175721 | -1.4226056 | 0.15485051 | 0.39755983 |
| PHF8 | 63.3615471 | -0.1630519 | 0.29776162 | -0.5475921 | 0.58397204 | 0.77981138 |
| FAM120C | 37.2512644 | 0.09881 | 0.32224699 | 0.30662815 | 0.75912643 | 0.88058098 |
| WNK3 | 40.551657 | 0.39355546 | 0.40605899 | 0.96920759 | 0.33244162 | 0.58794039 |
| FGD1 | 13.4841838 | 1.02753644 | 0.50215469 | 2.04625479 | 0.04073131 | NA |
| GNL3L | 15.8327785 | -0.2780033 | 0.46290938 | -0.6005566 | 0.54813535 | NA |
| MAGED2 | 24.917182 | 0.64949278 | 0.49860796 | 1.30261213 | 0.19270722 | 0.44446998 |
| PFKFB1 | 14.2216801 | 3.54087638 | 0.94244447 | 3.75711937 | 0.00017188 | NA |
| RRAGB | 72.5588883 | 0.00741835 | 0.26789051 | 0.02769173 | 0.97790802 | 0.99143283 |
| AL445472.1 | 80.2656197 | 0.81863107 | 0.32024832 | 2.55623847 | 0.01058106 | 0.10129652 |
| UBQLN2 | 17.5269787 | -0.1924529 | 0.50204299 | -0.3833395 | 0.70146803 | NA |
| NBDY | 101.218712 | -0.0372632 | 0.2863195 | -0.1301454 | 0.89645137 | 0.9533797 |
| SPIN3 | 29.142181 | 0.32860858 | 0.33806558 | 0.97202615 | 0.33103754 | 0.58655523 |
| FAAH2 | 372.816032 | 0.44710626 | 0.36735813 | 1.21708552 | 0.22357168 | 0.48054488 |
| ARHGEF9 | 79.9931442 | 0.12394912 | 0.32343931 | 0.3832222 | 0.70155502 | 0.85190818 |
| MTMR8 | 36.3440272 | 1.51146702 | 0.45341464 | 3.33352057 | 0.00085754 | 0.02623258 |
| ZC4H2 | 60.9251915 | 1.39360731 | 0.4276689 | 3.2586127 | 0.00111958 | 0.03002592 |
| ZC3H12B | 26.9310063 | 1.58589646 | 0.59009809 | 2.68751327 | 0.00719862 | 0.08282184 |
| LAS1L | 16.0265819 | -0.120057 | 0.57071924 | -0.2103609 | 0.83338602 | NA |
| MSN | 121.304018 | -0.4935182 | 0.28959858 | -1.7041458 | 0.08835385 | 0.29722885 |
| OPHN1 | 366.00045 | -0.1412998 | 0.1836793 | -0.7692747 | 4.42E-01 | 6.82E-01 |
| YIPF6 | 52.814372 | -0.507114 | 0.32074899 | -1.5810306 | 0.11387105 | 0.33971529 |
| STARD8 | 20.5485678 | -0.0600398 | 0.56739426 | -0.1058168 | 0.91572774 | NA |
| EDA | 94.0811958 | 0.42184353 | 0.43215366 | 0.97614245 | 0.32899388 | 0.5853121 |
| IGBP1 | 43.0984719 | 0.04367993 | 0.29313237 | 0.14901095 | 0.88154499 | 0.94443055 |
| DLG3 | 58.5704763 | 0.61105464 | 0.3795995 | 1.60973507 | 0.1074557 | 0.32951656 |
| ZMYM3 | 12.6201242 | 0.97092406 | 0.65807379 | 1.47540302 | 0.14010423 | NA |
| NONO | 43.07963 | -0.5499006 | 0.33508653 | -1.6410705 | 0.10078279 | 0.31985368 |
| TAF1 | 112.796958 | -0.6562638 | 0.32537227 | -2.016963 | 0.04369937 | 0.20890044 |
| OGT | 345.091014 | -0.2749264 | 0.38371842 | -0.7164794 | 0.47369535 | 0.7026171 |
| GCNA | 44.3458475 | 0.13097295 | 0.58043781 | 0.2256451 | 0.82147746 | 0.91434507 |
| PIN4 | 37.9758065 | 0.03579533 | 0.31215697 | 0.11467092 | 0.90870597 | 0.95957056 |
| RPS4X | 85.9016093 | 0.57727905 | 0.48864783 | 1.18138055 | 0.23745158 | 0.49524918 |
| HDAC8 | 314.661127 | 0.19391363 | 0.26804096 | 0.72344776 | 0.46940484 | 0.69910103 |
| PHKA1 | 28.7098909 | 0.6902323 | 0.47508581 | 1.45285817 | 0.14626314 | 0.38641495 |
| CHIC1 | 100.866407 | -0.585013 | 0.29737787 | -1.9672377 | 0.04915581 | 0.22229864 |
| XIST | 394.694639 | -1.7409717 | 4.09436147 | -0.425212 | 0.67068213 | 0.83476802 |
| JPX | 245.809756 | 0.168275 | 0.45730534 | 0.36797076 | 0.71289504 | 0.85763568 |
| FTX | 1729.81101 | 0.38392566 | 0.30855717 | 1.24426102 | 0.21340351 | 0.46890289 |
| SLC16A2 | 32.9424268 | 0.09764975 | 0.4176346 | 0.23381624 | 0.81512764 | 0.91046836 |
| RLIM | 39.1868268 | -0.2865889 | 0.35088797 | -0.8167532 | 0.41406951 | 0.65845716 |
| NEXMIF | 126.040367 | 0.39339614 | 0.33900841 | 1.1604318 | 0.24587305 | 0.50291917 |
| ABCB7 | 122.056069 | 0.11938585 | 0.22399328 | 0.53298853 | 0.59404152 | 0.78697721 |
| UPRT | 26.8261371 | 0.01329695 | 0.38053216 | 0.03494303 | 0.97212517 | 0.98878882 |
| ZDHHC15 | 26.5444064 | 0.19711378 | 0.43243604 | 0.45582182 | 0.64851811 | 0.81913126 |

| | | | | | | |
|---|---|---|---|---|---|---|
| ATRX | 417.967854 | -0.2217923 | 0.21538783 | -1.0297346 | 0.30313461 | 0.5637176 |
| MAGT1 | 70.9524992 | -0.4752489 | 0.3420868 | -1.3892641 | 0.16475246 | 0.41145351 |
| COX7B | 66.5165385 | 0.6511819 | 0.43971232 | 1.4809271 | 0.138626 | 0.37517394 |
| PGK1 | 155.041197 | -0.45149 | 0.53956126 | -0.8367724 | 0.4027205 | 0.65024957 |
| BRWD3 | 121.764792 | -0.3862039 | 0.38259357 | -1.0094365 | 0.31276534 | 0.57102302 |
| HMGN5 | 14.685922 | 0.06650972 | 0.50108712 | 0.13273084 | 0.89440625 | NA |
| SH3BGRL | 132.049276 | 1.30161196 | 0.36656374 | 3.55084751 | 0.00038399 | 0.01698617 |
| RPS6KA6 | 134.230014 | 0.78595966 | 0.28183466 | 2.78872605 | 0.00529158 | 0.07098247 |
| HDX | 56.5278311 | -0.5210683 | 0.29418758 | -1.771211 | 0.07652561 | 0.27638321 |
| APOOL | 72.5114195 | -0.03211 | 0.26764651 | -0.1199718 | 0.90450548 | 0.95671001 |
| ZNF711 | 19.4596736 | -0.6442855 | 0.46307465 | -1.3913211 | 0.16412808 | NA |
| CHM | 110.762064 | -0.1533692 | 0.2544335 | -0.6027869 | 0.54665049 | 0.75472086 |
| DIAPH2 | 127.420918 | 0.35549063 | 0.38837677 | 0.91532412 | 0.3600215 | 0.61634464 |
| TSPAN6 | 15.1317015 | 0.63248245 | 0.45725094 | 1.3832283 | 0.16659487 | NA |
| SYTL4 | 33.538906 | 0.45405419 | 0.51737871 | 0.87760508 | 0.38015807 | 0.63170913 |
| TRMT2B | 52.7021488 | -0.0092548 | 0.32831226 | -0.0281891 | 0.97751132 | 0.99142241 |
| ARMCX4 | 141.816246 | 0.26014215 | 0.49289235 | 0.52778695 | 0.59764722 | 0.7894568 |
| ARMCX3 | 33.6550878 | -0.1877512 | 0.37133565 | -0.5056106 | 0.61313005 | 0.7987918 |
| ZMAT1 | 136.489743 | 0.79235437 | 0.53341687 | 1.48543179 | 0.13742948 | 0.37351508 |
| GPRASP1 | 17.5141768 | 0.7158558 | 0.59807548 | 1.1969322 | 0.23133298 | NA |
| BHLHB9 | 22.3388143 | 0.47931766 | 0.46185007 | 1.03782091 | 0.29935344 | 0.55938903 |
| LINC00630 | 60.5523638 | 0.1075471 | 0.35648429 | 0.30168819 | 0.76288977 | 0.88247846 |
| BEX4 | 23.4294545 | 0.48539249 | 0.35425267 | 1.37018725 | 0.17062846 | 0.41879058 |
| BEX2 | 25.6243592 | -0.3729807 | 0.47063411 | -0.7925068 | 0.42806522 | 0.67072527 |
| TCEAL9 | 29.5960098 | -0.4302462 | 0.43117345 | -0.9978495 | 0.31835234 | 0.57587641 |
| BEX3 | 54.1673593 | 0.50539274 | 0.36842356 | 1.37177093 | 0.17013476 | 0.41837931 |
| TCEAL4 | 35.3595503 | -0.0253298 | 0.31251391 | -0.0810517 | 0.93540083 | 0.9738645 |
| TCEAL3 | 20.8520936 | -0.0409968 | 0.50751258 | -0.0807798 | 0.93561708 | NA |
| MORF4L2 | 92.2732529 | -0.2842019 | 0.31662902 | -0.8975862 | 0.36940617 | 0.62368335 |
| SLC25A53 | 52.9405204 | 0.50184217 | 0.28900481 | 1.73644917 | 0.08248444 | 0.28695946 |
| FAM199X | 66.9010538 | 0.09196108 | 0.35029856 | 0.26252201 | 0.79291902 | 0.89875118 |
| IL1RAPL2 | 216.644821 | 0.38303163 | 0.60241097 | 0.6358311 | 5.25E-01 | 7.40E-01 |
| NRK | 145.452182 | 0.64460017 | 0.32368257 | 1.99145777 | 4.64E-02 | 0.21595574 |
| RNF128 | 28.5329783 | -0.4094998 | 0.65633071 | -0.6239229 | 0.53267822 | 0.74615054 |
| TBC1D8B | 31.9718383 | 0.69910754 | 0.65651389 | 1.06487852 | 0.2869309 | 0.54625456 |
| MORC4 | 30.4363791 | 1.60277599 | 0.44428005 | 3.60758041 | 0.00030907 | 0.01457249 |
| RBM41 | 84.5505996 | 0.77622246 | 0.31186544 | 2.48896592 | 0.01281152 | 0.11046442 |
| NUP62CL | 20.5707225 | 0.51488647 | 0.41331737 | 1.2457412 | 0.21285942 | NA |
| MID2 | 23.4651769 | -0.0409278 | 0.56174148 | -0.0728587 | 0.94191852 | 0.97576674 |
| COL4A6 | 59.5840295 | 0.9351465 | 0.49441518 | 1.89141948 | 5.86E-02 | 2.40E-01 |
| COL4A5 | 294.665493 | 0.83751299 | 0.49166586 | 1.70341904 | 0.08848968 | 0.29750795 |
| ACSL4 | 1543.17495 | -0.2442221 | 0.67322698 | -0.3627634 | 0.7167816 | 0.8589283 |
| TMEM164 | 203.179526 | -0.2024165 | 0.20415796 | -0.9914699 | 0.3214562 | 0.57796038 |
| AMMECR1 | 60.7936062 | 0.28067456 | 0.37018407 | 0.75820269 | 0.44832965 | 0.68731807 |
| PAK3 | 13.3839888 | -0.4740792 | 0.53049911 | -0.8936474 | 0.37151057 | NA |
| ALG13 | 77.8097122 | -0.2124276 | 0.28093824 | -0.7561362 | 0.44956751 | 0.68770076 |
| AMOT | 29.0171464 | -0.2885576 | 0.67355042 | -0.4284128 | 0.66835062 | 0.83308417 |
| LRCH2 | 14.5084375 | 0.30022194 | 0.58864768 | 0.51001974 | 0.61003763 | NA |
| PLS3-AS1 | 34.4326807 | 0.65634794 | 0.45332423 | 1.44785541 | 0.14765749 | 0.38722988 |
| PLS3 | 139.019484 | 0.19426119 | 0.32630148 | 0.59534265 | 0.55161445 | 0.758039 |
| DANT2 | 687.439687 | -0.0026658 | 0.20836031 | -0.0127941 | 0.98979209 | 0.99583841 |
| KLHL13 | 652.913277 | -0.6407562 | 0.40744646 | -1.5726144 | 0.11580812 | 0.34257306 |
| WDR44 | 69.5783964 | -0.4663798 | 0.31097675 | -1.4997257 | 0.13368546 | 0.36762309 |
| IL13RA1 | 58.2930253 | -0.1101029 | 0.38929167 | -0.2828287 | 7.77E-01 | 8.90E-01 |

| | | | | | | |
|---|---|---|---|---|---|---|
| LONRF3 | 41.6621061 | -2.3394887 | 0.89844899 | -2.6039193 | 9.22E-03 | 0.09482608 |
| PGRMC1 | 38.0999101 | 0.50837647 | 0.51449759 | 0.98810272 | 0.32310235 | 0.57912936 |
| SLC25A43 | 37.7860889 | 0.60797197 | 0.54577669 | 1.11395737 | 0.26529747 | 0.5247131 |
| SLC25A5 | 38.732466 | 1.15188534 | 0.55543746 | 2.07383447 | 0.03809469 | 0.19510866 |
| UBE2A | 25.9906099 | 0.38533409 | 0.43135238 | 0.89331625 | 0.37168784 | 0.62455659 |
| NKRF | 16.0784182 | 0.67052115 | 0.50353734 | 1.3316215 | 0.1829846 | NA |
| 6-Sep | 149.294991 | 1.03684766 | 0.38258427 | 2.71011573 | 0.00672597 | 0.0798917 |
| UPF3B | 53.4905835 | -0.617273 | 0.46251842 | -1.3345913 | 0.18201016 | 0.43145433 |
| NDUFA1 | 38.692976 | 0.677518 | 0.39474124 | 1.71635979 | 0.08609621 | 0.29319258 |
| NKAP | 18.743173 | 0.66884456 | 0.42139631 | 1.5872103 | 0.11246502 | NA |
| RHOXF1-AS1 | 28.2810366 | 1.52310577 | 0.79321481 | 1.92016809 | 0.05483667 | 0.23407018 |
| LAMP2 | 63.4894889 | 0.15265475 | 0.29510558 | 0.51728857 | 0.60495473 | 0.79401898 |
| CUL4B | 33.1810017 | -0.2953227 | 0.33919788 | -0.8706501 | 0.38394523 | 0.63451053 |
| MCTS1 | 29.7600017 | 0.19859961 | 0.41417754 | 0.47950358 | 0.63158042 | 0.80899892 |
| THOC2 | 180.230091 | -0.3354434 | 0.21605985 | -1.5525485 | 0.12053104 | 0.34903213 |
| XIAP | 129.720354 | -0.4977695 | 0.22172571 | -2.2449789 | 0.02476949 | 0.15424817 |
| STAG2 | 208.178041 | -0.1752869 | 0.19999124 | -0.8764731 | 0.38077288 | 0.63229421 |
| TENM1 | 35.6714847 | 1.81154929 | 0.5953138 | 3.04301581 | 0.0023422 | 0.04598958 |
| SMARCA1 | 73.4314287 | 0.62992174 | 0.30366013 | 2.07443018 | 0.03803938 | 0.19506268 |
| OCRL | 17.9258777 | 0.79531713 | 0.51408109 | 1.54706552 | 0.12184745 | NA |
| ZDHHC9 | 25.1189042 | -0.700342 | 0.4121863 | -1.6990909 | 0.08930206 | 0.29794449 |
| UTP14A | 12.3611292 | -0.7629053 | 0.56995101 | -1.3385454 | 0.1807187 | NA |
| BCORL1 | 14.7999671 | 0.6923705 | 0.69163149 | 1.00106851 | 0.31679369 | NA |
| AIFM1 | 31.7494376 | 0.89266894 | 0.35510206 | 2.51383768 | 0.01194254 | 0.10701015 |
| ZNF280C | 13.571477 | 0.65015128 | 0.52266013 | 1.24392744 | 0.21352627 | NA |
| SLC25A14 | 14.2254267 | 0.19098281 | 0.59219556 | 0.32249956 | 0.74707427 | NA |
| ENOX2 | 63.6514789 | -0.2671253 | 0.29222153 | -0.9141193 | 0.36065415 | 0.61715317 |
| RAP2C | 71.8646272 | 0.7792523 | 0.37890741 | 2.05657708 | 3.97E-02 | 0.19847343 |
| RAP2C-AS1 | 104.05955 | 0.67197051 | 0.52081376 | 1.29023187 | 0.19697016 | 0.45014879 |
| MBNL3 | 56.7012516 | 0.75062199 | 0.38321411 | 1.9587535 | 0.05014166 | 0.22503812 |
| HS6ST2 | 409.394184 | -0.2805232 | 0.33555161 | -0.8360062 | 0.4031514 | 0.65053591 |
| PHF6 | 32.469776 | 0.49720995 | 0.34275729 | 1.45061816 | 0.14688622 | 0.3869457 |
| HPRT1 | 143.851879 | -0.0500506 | 0.2238788 | -0.223561 | 0.82309891 | 0.91475524 |
| MIR503HG | 38.0254421 | 1.37338848 | 0.65140188 | 2.1083582 | 0.03500002 | 0.18604636 |
| FAM122B | 18.436616 | 0.54019266 | 0.41245942 | 1.30968682 | 0.1903018 | NA |
| FAM122C | 14.1205007 | 0.9160202 | 0.63097469 | 1.45175428 | 0.14656994 | NA |
| RTL8C | 37.7757755 | -0.0887583 | 0.54468709 | -0.1629528 | 0.87055556 | 0.9398207 |
| ZNF75D | 62.7132481 | 0.18140694 | 0.27514956 | 0.65930304 | 0.50970119 | 0.72990216 |
| INTS6L | 27.3631542 | -0.5662599 | 0.43845166 | -1.2914991 | 0.19653067 | 0.44992056 |
| SLC9A6 | 19.9172468 | 0.09832311 | 0.41984322 | 0.23419007 | 0.81483743 | NA |
| HTATSF1 | 13.1298769 | 1.13123817 | 0.54527195 | 2.07463113 | 0.03802073 | NA |
| AL683813.1 | 27.283698 | -0.9215216 | 0.47425203 | -1.9431053 | 0.05200344 | 0.22846489 |
| RBMX | 135.310745 | -0.1674852 | 0.20399167 | -0.8210396 | 0.41162369 | 0.65707964 |
| FGF13 | 1199.89651 | -1.3287159 | 0.45836418 | -2.8988215 | 0.00374568 | 0.06095244 |
| ATP11C | 34.9668276 | -0.353139 | 0.44345358 | -0.7963381 | 0.42583552 | 0.66877291 |
| FMR1 | 79.2910603 | 0.04087863 | 0.30878766 | 0.13238428 | 0.89468034 | 0.95246952 |
| IDS | 38.0638176 | 0.49606349 | 0.42356537 | 1.17116158 | 0.24153384 | 0.49881565 |
| TMEM185A | 19.9146424 | 0.58254661 | 0.4904973 | 1.18766529 | 0.23496531 | NA |
| LINC00894 | 145.425846 | 0.46069433 | 0.40413004 | 1.13996557 | 0.25430065 | 0.51185985 |
| MAMLD1 | 28.5457899 | -2.4295462 | 0.70922333 | -3.4256433 | 0.00061335 | 0.02248056 |
| MTM1 | 63.7241605 | -0.3651062 | 0.24165648 | -1.5108478 | 0.13082722 | 0.36326737 |
| MTMR1 | 56.904207 | 0.09412528 | 0.27816146 | 0.33838363 | 0.73507411 | 0.86830347 |
| CD99L2 | 38.6495928 | -0.2583805 | 0.44894102 | -0.5755333 | 0.56493069 | 0.7693258 |
| VMA21 | 17.3412734 | 0.30361993 | 0.55931609 | 0.5428414 | 0.587239 | NA |

| | | | | | | |
|---|---|---|---|---|---|---|
| DUSP9 | 33.1011426 | 2.37981953 | 0.61875167 | 3.84616262 | 1.20E-04 | 0.00800536 |
| SLC6A8 | 66.1074895 | -0.429719 | 0.45434549 | -0.9457978 | 0.34425172 | 0.60138572 |
| BCAP31 | 32.4210123 | 0.32383582 | 0.45810995 | 0.70689542 | 0.47963147 | 0.70695585 |
| IDH3G | 16.3281995 | -0.1659684 | 0.61754066 | -0.2687571 | 0.78811663 | NA |
| SSR4 | 16.578311 | 0.2496146 | 0.50127067 | 0.4979637 | 0.61850963 | NA |
| ARHGAP4 | 12.8880095 | 0.60226488 | 0.59231592 | 1.0167967 | 0.30925015 | NA |
| HCFC1 | 13.4580513 | 0.6190427 | 0.57302368 | 1.08030911 | 0.28000455 | NA |
| IRAK1 | 14.8766444 | 0.19422751 | 0.58735485 | 0.33068172 | 0.74088491 | NA |
| MECP2 | 129.345537 | -0.7292797 | 0.26728624 | -2.7284596 | 0.00636309 | 0.07774078 |
| FLNA | 38.4371359 | -0.9453441 | 0.59057016 | -1.6007313 | 0.10943645 | 0.33178907 |
| EMD | 23.5504804 | -0.7540645 | 0.57353662 | -1.3147625 | 0.18858973 | 0.43920578 |
| RPL10 | 113.041162 | 0.58447695 | 0.43934225 | 1.33034541 | 0.18340449 | 0.4332917 |
| TAZ | 28.6907635 | 0.24499889 | 0.36959634 | 0.66288235 | 0.5074059 | 0.72850275 |
| ATP6AP1 | 25.5352514 | 0.61261587 | 0.5270957 | 1.1622479 | 0.24513478 | 0.5022131 |
| GDI1 | 22.487879 | -0.4223697 | 0.46158631 | -0.9150394 | 0.36017093 | 0.61646334 |
| FAM50A | 16.9009054 | -0.9374202 | 0.52040417 | -1.8013311 | 0.07165072 | NA |
| FAM3A | 15.0344952 | -0.0419785 | 0.44140664 | -0.0951016 | 0.92423417 | NA |
| DKC1 | 16.8405291 | -0.2422261 | 0.45124713 | -0.5367926 | 0.59141088 | NA |
| MPP1 | 17.4580028 | 0.88837531 | 0.50222473 | 1.76888005 | 0.07691389 | NA |
| F8 | 18.4998629 | -0.0360284 | 0.57608364 | -0.0625402 | 0.95013265 | NA |
| FUNDC2 | 24.8775286 | -0.2054646 | 0.47136033 | -0.4358971 | 0.66291142 | 0.8285925 |
| BRCC3 | 41.1774021 | -0.1338136 | 0.32356406 | -0.4135613 | 0.67919541 | 0.84034192 |
| VBP1 | 13.5773733 | -0.0216663 | 0.53556105 | -0.0404554 | 0.96773006 | NA |
| TMLHE-AS1 | 28.0019893 | 1.1482853 | 0.72299674 | 1.58823026 | 0.11223428 | 0.33671279 |
| TMLHE | 81.6237199 | 0.96394783 | 0.35724951 | 2.69824815 | 0.00697055 | 0.08149858 |
| VAMP7 | 39.3112849 | -0.1219534 | 0.37354545 | -0.3264755 | 0.74406464 | 0.87236337 |
| ZNF596 | 20.6212265 | 0.05952255 | 0.39285924 | 0.15151113 | 0.87957253 | NA |
| FBXO25 | 47.535352 | -0.7142886 | 0.37701683 | -1.8945803 | 0.05814803 | 0.24020328 |
| TDRP | 68.4990294 | -0.0937033 | 0.33741353 | -0.2777105 | 0.78123462 | 0.89136716 |
| ERICH1 | 238.246403 | -0.9884126 | 0.54466361 | -1.8147212 | 0.06956676 | 0.26273569 |
| AC246817.2 | 27.6728929 | 1.44073823 | 0.58066724 | 2.48117707 | 0.01309493 | 0.11186646 |
| AC246817.1 | 59.2008075 | 0.62545843 | 0.51214125 | 1.22126159 | 0.22198699 | 0.47914579 |
| CSMD1 | 50.2034951 | -0.1759564 | 0.95185687 | -0.184856 | 0.853342 | 0.93020441 |
| MCPH1 | 180.204332 | -0.1667826 | 0.27931789 | -0.5971068 | 0.55043611 | 0.75709555 |
| ANGPT2 | 23.5520075 | 1.2642671 | 0.61520432 | 2.05503614 | 0.03987551 | 0.19882936 |
| MCPH1-AS1 | 41.0008253 | -0.949707 | 0.38590698 | -2.4609738 | 0.01385605 | 0.11532965 |
| AGPAT5 | 56.4592581 | -1.6539205 | 0.30926519 | -5.3479039 | 8.90E-08 | 7.61E-05 |
| AF233439.1 | 228.790142 | 0.9008284 | 0.61862053 | 1.45618897 | 0.14534039 | 0.38495697 |
| DEFB1 | 479.144038 | 0.66246696 | 0.40989776 | 1.61617609 | 0.10605625 | 0.32732861 |
| FAM85B | 109.686212 | 0.0198217 | 0.495159 | 0.04003098 | 0.96806842 | 0.98741848 |
| MFHAS1 | 729.042821 | 0.86586068 | 0.41165389 | 2.10337058 | 0.03543339 | 0.18721309 |
| ERI1 | 53.8606491 | 0.1610336 | 0.28742447 | 0.56026405 | 5.75E-01 | 7.74E-01 |
| PPP1R3B | 19.8010448 | -0.3042172 | 0.43461666 | -0.6999666 | 0.48394816 | NA |
| TNKS | 326.176582 | -0.2770326 | 0.19437319 | -1.4252611 | 0.15408173 | 0.3965119 |
| MSRA | 141.019825 | -0.5480067 | 0.28345968 | -1.9332792 | 0.05320181 | 0.23177172 |
| XKR6 | 555.912651 | -0.5215826 | 0.1859177 | -2.8054492 | 0.00502465 | 0.06878385 |
| MTMR9 | 54.2140979 | 0.429568 | 0.28146366 | 1.5261935 | 0.12696168 | 0.35861434 |
| FDFT1 | 82.6783732 | 0.16982164 | 0.25359035 | 0.66966917 | 0.50306871 | 0.7248446 |
| CTSB | 62.3887043 | -0.3006837 | 0.48316678 | -0.6223186 | 0.53373237 | 0.74672888 |
| FAM66D | 54.1386313 | 1.05811517 | 0.8123189 | 1.30258593 | 0.19271616 | 0.44446998 |
| AC145124.1 | 12.6256398 | 0.7710977 | 0.70883402 | 1.08783958 | 0.27666593 | NA |
| LONRF1 | 77.0626127 | -0.4252624 | 0.43773375 | -0.9715092 | 0.33129476 | 0.5868759 |
| TRMT9B | 233.067435 | 0.88520841 | 0.46514639 | 1.90307487 | 0.05703078 | 0.23764428 |
| TUSC3 | 133.029719 | -0.1643925 | 0.31207334 | -0.5267751 | 0.59834977 | 0.7894568 |

| | | | | | | |
|---|---|---|---|---|---|---|
| MSR1 | 169.041248 | 0.9371679 | 0.56833673 | 1.64896593 | 0.09915461 | 0.31667762 |
| AC068992.1 | 70.9882196 | -0.1272406 | 0.64475208 | -0.1973481 | 0.84355515 | 0.92437984 |
| MICU3 | 104.485544 | 0.26377211 | 0.321186 | 0.8212441 | 0.41150724 | 0.65707964 |
| ZDHHC2 | 130.930153 | -0.0076942 | 0.25011301 | -0.0307629 | 0.97545865 | 0.9905152 |
| CNOT7 | 36.6738731 | -0.1794602 | 0.35548065 | -0.5048382 | 0.61367248 | 0.79909273 |
| VPS37A | 60.4986857 | -0.3439362 | 0.28285602 | -1.2159409 | 0.22400744 | 0.48121274 |
| MTMR7 | 25.5170024 | -1.356832 | 0.88665594 | -1.5302801 | 0.1259474 | 0.35705972 |
| MTUS1 | 247.987778 | -0.5012538 | 0.31040467 | -1.6148399 | 0.10634537 | 0.32732986 |
| AC027117.2 | 42.0773456 | -1.3047399 | 0.63324136 | -2.0604148 | 0.0393589 | 0.19774509 |
| PCM1 | 313.639015 | -0.2705959 | 0.18438787 | -1.4675366 | 0.14223014 | 0.38005954 |
| ASAH1 | 56.3961346 | -0.3022556 | 0.41087747 | -0.7356344 | 0.46195323 | 0.69416206 |
| PSD3 | 27.3442276 | -0.0343278 | 0.44693795 | -0.0768065 | 0.93877746 | 0.97481036 |
| SH2D4A | 29.0918653 | 0.66363389 | 0.58836731 | 1.12792448 | 0.25935182 | 0.51809783 |
| CSGALNACT1 | 727.946667 | -1.091077 | 0.52366689 | -2.0835325 | 0.03720271 | 0.19282779 |
| INTS10 | 148.192537 | -0.2898051 | 0.26336044 | -1.1004125 | 0.27115245 | 0.52944592 |
| LPL | 19.4867373 | -1.8054658 | 0.83206933 | -2.1698502 | 3.00E-02 | NA |
| ATP6V1B2 | 36.4586035 | -1.0885884 | 0.39803878 | -2.7348801 | 0.0062403 | 0.07721978 |
| XPO7 | 101.935794 | 0.00839308 | 0.26492859 | 0.03168055 | 0.97472681 | 0.99032688 |
| DMTN | 44.343571 | 1.16181526 | 0.45016966 | 2.58083865 | 0.00985606 | 0.09830513 |
| FAM160B2 | 27.0737194 | 0.38898986 | 0.40388253 | 0.96312623 | 0.33548418 | 0.59130335 |
| SLC39A14 | 72.6627308 | -1.0384879 | 0.62930112 | -1.6502241 | 0.0988971 | 0.31649984 |
| PPP3CC | 154.897974 | -0.2245054 | 0.2740301 | -0.8192728 | 0.41263078 | 0.65722052 |
| CCAR2 | 16.4346421 | -0.402961 | 0.45689889 | -0.881948 | 0.37780495 | NA |
| BIN3 | 75.1858244 | -0.5977315 | 0.2619587 | -2.2817777 | 0.02250247 | 0.14628503 |
| PEBP4 | 19.9051005 | -0.2609292 | 0.75148584 | -0.3472177 | 7.28E-01 | NA |
| RHOBTB2 | 11.9728939 | -0.7978435 | 0.63401001 | -1.2584084 | 0.20824408 | NA |
| TNFRSF10B | 97.8749155 | -0.8222958 | 0.4943039 | -1.6635431 | 0.09620377 | 0.3113879 |
| TNFRSF10D | 59.0053458 | -0.7862076 | 0.24102381 | -3.2619499 | 1.11E-03 | 2.99E-02 |
| TNFRSF10A | 16.8801162 | -1.37332 | 0.49860745 | -2.7543111 | 0.00588158 | NA |
| CHMP7 | 14.2597873 | 0.14501776 | 0.46801281 | 0.30985852 | 0.75666855 | NA |
| ENTPD4 | 47.1868572 | -0.6413675 | 0.43651464 | -1.4692921 | 0.14175357 | 0.37968477 |
| SLC25A37 | 111.969584 | -0.1987335 | 0.25879255 | -0.7679257 | 0.44253131 | 0.68246113 |
| DOCK5 | 84.8963922 | -0.8240593 | 0.47749314 | -1.7258035 | 0.08438278 | 0.28931012 |
| KCTD9 | 43.2606677 | 0.23569747 | 0.32747976 | 0.71973141 | 0.47169038 | 0.70102354 |
| PPP2R2A | 184.068172 | -0.4074757 | 0.26448889 | -1.5406155 | 0.1234104 | 0.35349809 |
| BNIP3L | 172.551462 | -0.5526233 | 0.53940042 | -1.0245141 | 0.30559251 | 0.56557634 |
| TRIM35 | 13.6969575 | -0.620274 | 0.45254319 | -1.3706404 | 0.1704871 | NA |
| PTK2B | 61.8957299 | 0.08947246 | 0.32583883 | 0.27459116 | 0.78363036 | 0.89330611 |
| EPHX2 | 90.8929099 | 1.67944443 | 0.57679956 | 2.91166038 | 0.00359513 | 0.05963737 |
| CLU | 95.1545029 | 0.73184045 | 0.47725136 | 1.53344863 | 1.25E-01 | 0.35575987 |
| CCDC25 | 24.7043785 | 0.66713713 | 0.39116425 | 1.70551663 | 0.08809811 | 0.29675763 |
| ELP3 | 113.812737 | -0.3634353 | 0.32640365 | -1.1134534 | 0.26551374 | 0.524719 |
| ZNF395 | 33.9152478 | -0.5999325 | 0.3569764 | -1.6805943 | 0.09284174 | 0.30571879 |
| FBXO16 | 37.2609514 | 0.88411625 | 0.58005496 | 1.52419392 | 0.12746028 | 0.35962674 |
| FZD3 | 27.5715238 | 1.05778101 | 0.37968532 | 2.78594128 | 0.00533725 | 0.07122532 |
| EXTL3 | 32.7233901 | -1.2127411 | 0.42805603 | -2.8331363 | 0.00460937 | 0.06581034 |
| INTS9 | 65.7519782 | -0.1721919 | 0.27082245 | -0.6358109 | 0.52489967 | 0.73968378 |
| HMBOX1 | 188.885117 | -0.1051724 | 0.21114032 | -0.4981161 | 0.61840219 | 0.80247821 |
| KIF13B | 379.833066 | -0.5605904 | 0.2420387 | -2.3161189 | 0.02055178 | 0.14099127 |
| AC108449.1 | 15.9066088 | -0.0476781 | 0.52775262 | -0.0903418 | 0.9280156 | NA |
| SARAF | 70.0550151 | 0.28454047 | 0.39479737 | 0.72072535 | 0.47107851 | 0.70047809 |
| LEPROTL1 | 25.3771351 | 0.10892761 | 0.38957865 | 0.27960365 | 0.77978161 | 0.89074516 |
| DCTN6 | 13.4655239 | -0.4330838 | 0.55009276 | -0.7872924 | 0.43111074 | NA |
| RBPMS | 1318.53496 | -0.4219752 | 0.25354682 | -1.664289 | 9.61E-02 | 3.11E-01 |

| | | | | | | |
|---|---|---|---|---|---|---|
| GTF2E2 | 65.6882525 | 0.05430096 | 0.27230336 | 0.19941347 | 0.84193933 | 0.92367136 |
| GSR | 78.5144892 | 0.77316563 | 0.35725587 | 2.16417894 | 0.03045062 | 0.17259088 |
| PPP2CB | 112.468079 | -0.3454643 | 0.2440701 | -1.4154307 | 0.15694227 | 0.40105731 |
| WRN | 103.611749 | 0.10186107 | 0.28118345 | 0.36225842 | 0.71715893 | 0.85900596 |
| NRG1 | 168.495184 | -0.9059332 | 0.64164141 | -1.4118995 | 0.15797956 | 0.40275854 |
| FUT10 | 58.2315772 | 0.73207458 | 0.37413541 | 1.95671024 | 0.05038155 | 0.22527387 |
| TTI2 | 21.725082 | -0.3706381 | 0.44695885 | -0.8292443 | 0.40696619 | 0.65367876 |
| RNF122 | 18.1446116 | -1.0376582 | 0.47861246 | -2.1680551 | 0.03015449 | NA |
| ERLIN2 | 25.9831178 | 0.20251032 | 0.41592551 | 0.48689083 | 0.6263357 | 0.8053173 |
| PLPBP | 26.9201361 | -0.1153025 | 0.36394916 | -0.3168094 | 0.7513882 | 0.8776233 |
| RAB11FIP1 | 100.083104 | -0.14414 | 0.25451984 | -0.5663213 | 0.57117535 | 0.77245539 |
| ASH2L | 42.6440939 | -0.0693725 | 0.32440345 | -0.2138465 | 0.83066679 | 0.91794754 |
| LSM1 | 18.2806304 | -0.3683209 | 0.42897176 | -0.8586133 | 0.39055388 | NA |
| BAG4 | 54.8093795 | -0.0909724 | 0.29369354 | -0.3097527 | 0.75674904 | 0.87904624 |
| DDHD2 | 53.0074768 | -0.1371263 | 0.28974461 | -0.473266 | 0.63602344 | 0.81188871 |
| PLPP5 | 36.0006955 | -0.1556967 | 0.38604805 | -0.403309 | 0.6867209 | 0.8446294 |
| NSD3 | 246.097345 | -0.0934752 | 0.20275972 | -0.4610147 | 0.64478808 | 0.81734747 |
| LETM2 | 16.3199395 | -0.7374313 | 0.63115647 | -1.1683811 | 0.24265306 | NA |
| FGFR1 | 118.202742 | -1.718575 | 0.39784129 | -4.3197503 | 1.56E-05 | 0.00207296 |
| TACC1 | 565.357878 | -0.7220533 | 0.48139576 | -1.4999163 | 0.13363609 | 0.36762309 |
| PLEKHA2 | 55.2135759 | -0.2525747 | 0.41645709 | -0.6064843 | 0.54419322 | 0.75281366 |
| ADAM9 | 99.3080576 | -0.8914369 | 0.37358739 | -2.3861537 | 0.01702564 | 0.12988597 |
| TCIM | 163.841641 | -0.4889972 | 0.51164992 | -0.9557262 | 0.33921059 | 0.59487445 |
| SFRP1 | 434.978887 | 0.02059484 | 0.51224262 | 0.04020525 | 0.96792949 | 0.98741848 |
| GOLGA7 | 20.9866468 | 0.17813432 | 0.48068637 | 0.37058326 | 0.71094795 | NA |
| GPAT4 | 28.7849582 | -0.7365079 | 0.3986602 | -1.8474578 | 0.06468082 | 0.25284322 |
| KAT6A | 261.426408 | -0.2395798 | 0.21151882 | -1.1326642 | 0.2573553 | 0.51571564 |
| AP3M2 | 19.6012436 | 0.11919627 | 0.43143318 | 0.2762798 | 0.78233317 | NA |
| IKBKB | 79.5765503 | -0.2010016 | 0.24408267 | -0.8234981 | 0.4102248 | 0.6560358 |
| POLB | 23.9895385 | -0.2046178 | 0.3782853 | -0.5409088 | 0.58857047 | 0.78344371 |
| VDAC3 | 59.8607144 | 0.71432906 | 0.41200991 | 1.7337667 | 0.08295949 | 0.28801947 |
| SLC20A2 | 225.546134 | -1.1927741 | 0.31682439 | -3.7647797 | 0.0001667 | 0.00964706 |
| SMIM19 | 33.1305844 | -0.0989215 | 0.395111 | -0.2503637 | 0.80230607 | 0.90378853 |
| RNF170 | 70.0115986 | 0.43634009 | 0.38842611 | 1.12335417 | 0.26128711 | 0.52088238 |
| HOOK3 | 119.936262 | -0.1017239 | 0.25704665 | -0.3957411 | 0.69229607 | 0.84755891 |
| FNTA | 32.108589 | -0.3936255 | 0.32876814 | -1.1972738 | 0.23119984 | 0.48982471 |
| HGSNAT | 105.928969 | 0.38452642 | 0.30267383 | 1.27043167 | 0.20393091 | 0.45736371 |
| SPIDR | 539.200684 | -0.094405 | 0.30578508 | -0.30873 | 0.75752694 | 0.87930702 |
| CEBPD | 112.602226 | -0.093553 | 0.47083336 | -0.1986967 | 0.8425 | 0.92390487 |
| PRKDC | 139.676235 | -0.2194292 | 0.19598782 | -1.1196063 | 0.26288155 | 0.52266675 |
| MCM4 | 15.4607954 | -0.2778251 | 0.57266715 | -0.4851424 | 0.62757531 | NA |
| UBE2V2 | 61.7497428 | -0.6653743 | 0.27549832 | -2.4151665 | 0.01572802 | 0.12390845 |
| PXDNL | 69.2608254 | -0.1135558 | 0.43433657 | -0.2614465 | 0.79374817 | 0.89884908 |
| PCMTD1 | 254.918011 | 0.48700812 | 0.17503049 | 2.78241885 | 0.00539554 | 0.07135642 |
| AC064807.1 | 18.9768053 | 1.26178268 | 0.5096923 | 2.47557729 | 0.0133021 | NA |
| RB1CC1 | 224.712124 | -1.0043306 | 0.36565336 | -2.746674 | 0.00602029 | 0.07621415 |
| ATP6V1H | 251.953239 | 0.1225871 | 0.23840737 | 0.51419175 | 0.60711794 | 0.79532486 |
| TCEA1 | 65.8140663 | -0.3647872 | 0.35982463 | -1.0137915 | 0.31068223 | 0.57001845 |
| LYPLA1 | 126.246995 | 0.07552427 | 0.26286919 | 0.28730742 | 0.77387694 | 0.88782329 |
| MRPL15 | 27.0002098 | 0.1402926 | 0.41098098 | 0.34136033 | 0.73283233 | 0.86776236 |
| XKR4 | 39.6751996 | -0.1744609 | 0.6928341 | -0.2518076 | 8.01E-01 | 9.04E-01 |
| TMEM68 | 34.874576 | 0.28943059 | 0.3678842 | 0.78674374 | 0.4314319 | 0.67234892 |
| TGS1 | 46.7567896 | -0.7779479 | 0.29781905 | -2.6121495 | 0.00899749 | 0.09383967 |
| LYN | 53.1439943 | 0.02069126 | 0.31340528 | 0.06602077 | 9.47E-01 | 9.79E-01 |

| | | | | | | |
|---|---|---|---|---|---|---|
| RPS20 | 48.5785228 | 0.29223426 | 0.34591605 | 0.84481265 | 0.39821545 | 0.64642108 |
| PLAG1 | 45.7531004 | 0.80515816 | 0.41780034 | 1.9271362 | 0.05396266 | 0.23287156 |
| CHCHD7 | 14.7075845 | 0.44122064 | 0.42553538 | 1.03686005 | 0.29980108 | NA |
| IMPAD1 | 38.3139062 | 0.26469721 | 0.35780597 | 0.73977862 | 0.45943433 | 0.69297787 |
| LINC01606 | 384.988174 | 0.62615479 | 0.55931911 | 1.11949471 | 0.26292915 | 0.52266675 |
| FAM110B | 86.6586034 | -0.748521 | 0.40462125 | -1.8499299 | 0.06432365 | 0.25247277 |
| UBXN2B | 55.031894 | -0.53091 | 0.2600906 | -2.0412501 | 0.04122598 | 0.20301425 |
| SDCBP | 61.5117663 | -0.4220575 | 0.38259341 | -1.1031489 | 0.26996251 | 0.52832481 |
| NSMAF | 67.2406674 | 0.33613832 | 0.25452942 | 1.32062659 | 0.1866259 | 0.4367466 |
| TOX | 196.351674 | 1.75171218 | 0.36499911 | 4.79922315 | 1.59E-06 | 0.00051083 |
| LINC01301 | 25.3772613 | 2.09233421 | 0.86709192 | 2.41304777 | 0.01581975 | 0.12437649 |
| RAB2A | 204.60494 | -0.3725255 | 0.2650722 | -1.4053738 | 0.15991016 | 0.40470667 |
| CHD7 | 182.991224 | 0.07617558 | 0.23979044 | 0.31767562 | 0.75073101 | 0.8772395 |
| ASPH | 129.087999 | 0.07150406 | 0.21493558 | 0.33267669 | 0.73937835 | 0.87054913 |
| YTHDF3 | 110.752498 | -0.2795602 | 0.34054761 | -0.8209138 | 0.41169538 | 0.65707964 |
| CYP7B1 | 102.181668 | 0.08732761 | 0.35984184 | 0.24268331 | 0.80825074 | 0.90713122 |
| ARMC1 | 34.9392581 | -0.1048749 | 0.33707672 | -0.3111308 | 0.75570119 | 0.87904624 |
| MTFR1 | 64.3445961 | -0.1893621 | 0.32147362 | -0.589044 | 0.55583176 | 0.76247318 |
| PDE7A | 153.006727 | 0.20339587 | 0.35120192 | 0.57914226 | 0.56249319 | 0.76709958 |
| ADHFE1 | 49.1244784 | 0.95827212 | 0.52447135 | 1.82712003 | 0.06768172 | 0.25898011 |
| VCPIP1 | 29.8517875 | 0.3632382 | 0.38587854 | 0.94132781 | 0.34653689 | 0.60363852 |
| SGK3 | 92.4041404 | 0.02866422 | 0.27325493 | 0.10489917 | 0.91645582 | 0.96401363 |
| TCF24 | 31.7162453 | 0.89711823 | 0.43036598 | 2.0845473 | 3.71E-02 | 1.92E-01 |
| PPP1R42 | 23.6129445 | 0.94462176 | 0.65120077 | 1.45058452 | 0.14689559 | 0.3869457 |
| CSPP1 | 187.319198 | 0.11913918 | 0.20681783 | 0.57605858 | 0.56457559 | 0.76912182 |
| ARFGEF1 | 501.08507 | -0.3627799 | 0.20836784 | -1.7410551 | 0.08167392 | 0.28509879 |
| SULF1 | 14.5846317 | 0.55515401 | 0.67712609 | 0.81986799 | 0.41229136 | NA |
| NCOA2 | 710.111919 | 0.03318998 | 0.32011578 | 0.10368119 | 0.91742235 | 0.96401363 |
| TRAM1 | 83.5804717 | -0.3718478 | 0.39202424 | -0.9485327 | 0.34285833 | 0.60017752 |
| LACTB2 | 47.5204615 | -0.3191098 | 0.37635638 | -0.8478926 | 3.96E-01 | 6.45E-01 |
| EYA1 | 34.8885213 | -0.1164845 | 0.62562456 | -0.1861891 | 0.85229644 | 0.92945957 |
| TERF1 | 56.4301302 | 0.24542513 | 0.30529782 | 0.80388759 | 0.4214619 | 0.66488875 |
| RPL7 | 86.5961648 | 0.53717499 | 0.54312961 | 0.98903647 | 0.3226453 | 0.57912936 |
| RDH10 | 79.3718503 | -0.8964166 | 0.50650828 | -1.7697965 | 0.07676105 | 0.27670206 |
| RDH10-AS1 | 23.6834502 | 0.3992404 | 0.67430198 | 0.59207953 | 0.55379734 | 0.76022438 |
| STAU2 | 202.01048 | 1.00781251 | 0.19158041 | 5.26051976 | 1.44E-07 | 0.00010198 |
| AC027018.1 | 27.4330261 | 0.74161517 | 0.5998759 | 1.23628099 | 0.21635413 | 0.47268742 |
| ELOC | 36.0143963 | -0.3229749 | 0.3450302 | -0.9360772 | 0.34923344 | 0.60589538 |
| GDAP1 | 63.0724241 | 0.34202059 | 0.49801128 | 0.68677278 | 0.49222593 | 0.71672078 |
| PEX2 | 40.6760062 | 0.41521623 | 0.33504057 | 1.23930136 | 0.21523391 | 0.47115856 |
| ZC2HC1A | 48.0542732 | 0.74795905 | 0.40401498 | 1.85131515 | 0.06412423 | 0.25213756 |
| IL7 | 23.7342682 | 1.3680368 | 0.62304283 | 2.19573475 | 2.81E-02 | 0.16508057 |
| TPD52 | 230.947974 | 0.14600836 | 0.28444258 | 0.51331401 | 6.08E-01 | 0.79552231 |
| ZBTB10 | 82.6017374 | -0.0397878 | 0.4742248 | -0.0839007 | 0.93313538 | 0.9730757 |
| ZNF704 | 402.886869 | -0.1817518 | 0.32179071 | -0.5648138 | 0.57220043 | 0.77272682 |
| IMPA1 | 40.2742692 | -0.1827888 | 0.37647792 | -0.4855233 | 0.62730516 | 0.80566792 |
| ZFAND1 | 50.0070561 | -0.1684275 | 0.35816446 | -0.4702518 | 0.63817516 | 0.81325069 |
| CHMP4C | 23.0962338 | -0.8091948 | 0.36769386 | -2.2007296 | 0.02775517 | 0.16439232 |
| SNX16 | 30.1114114 | -1.4748176 | 0.38505778 | -3.8301201 | 0.00012808 | 0.00828435 |
| RALYL | 30.9107531 | 0.58006823 | 0.62124891 | 0.93371307 | 3.50E-01 | 0.60737992 |
| LRRCC1 | 17.2488132 | -0.9041061 | 0.49931458 | -1.8106944 | 0.07018818 | NA |
| E2F5 | 33.572757 | 0.5811131 | 0.36835771 | 1.57757823 | 0.11466254 | 0.34103102 |
| C8orf59 | 26.385449 | 0.43352546 | 0.43128209 | 1.00520162 | 0.31479978 | 0.57214023 |
| CA13 | 41.043693 | 0.13580301 | 0.36746716 | 0.36956503 | 0.71170661 | 0.857224 |

| | | | | | | |
|---|---|---|---|---|---|---|
| CA3 | 55.8863837 | 0.3063138 | 0.3194229 | 0.95896004 | 0.33757888 | 0.59336483 |
| CA2 | 127.916493 | 1.64608122 | 0.37331722 | 4.40933637 | 1.04E-05 | 0.00166268 |
| ATP6V0D2 | 154.496717 | 0.32722842 | 0.65431086 | 0.50011154 | 0.61699654 | 0.80179341 |
| AC023194.3 | 29.9822539 | 1.2772669 | 0.87033585 | 1.46755636 | 0.14222477 | 0.38005954 |
| WWP1 | 359.581554 | 0.33059429 | 0.25959965 | 1.27347737 | 0.2028487 | 0.45686328 |
| RMDN1 | 91.2577183 | 0.29649577 | 0.29725958 | 0.99743048 | 0.3185556 | 0.57587641 |
| CPNE3 | 74.5962211 | 0.51087588 | 0.30016829 | 1.70196485 | 0.08876196 | 0.29778649 |
| AF117829.1 | 264.305301 | 0.66739741 | 0.40078358 | 1.66523141 | 0.0958666 | 0.31098513 |
| RIPK2 | 21.9963956 | -0.0426579 | 0.40119441 | -0.1063273 | 0.91532265 | 0.96388328 |
| OSGIN2 | 26.5896084 | 0.16532172 | 0.40490926 | 0.40829325 | 0.68305839 | 0.84227819 |
| NBN | 44.0665247 | 0.07667099 | 0.34814679 | 0.22022604 | 0.82569511 | 0.9154963 |
| DECR1 | 143.846405 | 0.62512401 | 0.32813064 | 1.90510709 | 0.05676615 | 0.23723351 |
| CALB1 | 191.954416 | 1.31848873 | 0.82233668 | 1.60334418 | 1.09E-01 | 0.33065719 |
| LINC00534 | 30.0037184 | 1.94570331 | 0.88014673 | 2.2106579 | 0.02705954 | 0.16195658 |
| PIP4P2 | 45.4385256 | 0.46537128 | 0.36615125 | 1.270981 | 0.20373541 | 0.45736371 |
| OTUD6B-AS1 | 20.7486642 | 0.30546133 | 0.5352727 | 0.57066487 | 0.56822683 | NA |
| OTUD6B | 23.3394708 | 0.34238956 | 0.41325174 | 0.82852538 | 0.40737303 | 0.65367876 |
| LRRC69 | 38.8837215 | 0.88868052 | 0.42764088 | 2.07810003 | 0.03770015 | 0.19425568 |
| TRIQK | 48.929062 | 1.03696766 | 0.3932604 | 2.63684739 | 0.00836805 | 0.08952533 |
| FAM92A | 34.8038101 | 1.60996577 | 0.48399399 | 3.32641689 | 0.0008797 | 0.02668352 |
| RBM12B | 45.7443932 | 0.57216286 | 0.30243116 | 1.89187803 | 0.05850723 | 0.24020328 |
| TMEM67 | 78.5169667 | -0.1245704 | 0.41793418 | -0.2980624 | 0.76565557 | 0.88394344 |
| PDP1 | 19.5253639 | -1.5874512 | 0.59056708 | -2.6880117 | 0.00718789 | NA |
| GEM | 34.9689009 | -2.0570962 | 0.90821842 | -2.2649796 | 0.02351393 | 0.14961264 |
| VIRMA | 60.2221626 | -0.2380831 | 0.26738226 | -0.8904223 | 0.37323918 | 0.62575082 |
| ESRP1 | 90.9923556 | 0.29454249 | 0.3455233 | 0.85245334 | 0.39396253 | 0.6425014 |
| DPY19L4 | 65.2277752 | 0.0923297 | 0.23266497 | 0.39683542 | 0.69148883 | 0.846991 |
| INTS8 | 67.3325995 | -0.4778899 | 0.24743436 | -1.9313805 | 0.05343602 | 0.23211867 |
| NDUFAF6 | 58.6519228 | 0.48516272 | 0.32722993 | 1.48263553 | 0.13817127 | 0.37473724 |
| TP53INP1 | 29.4510332 | -0.164273 | 0.45806396 | -0.3586245 | 0.71987603 | 0.85995641 |
| PLEKHF2 | 18.2733456 | 0.51157626 | 0.48368244 | 1.05766969 | 0.29020606 | NA |
| C8orf37-AS1 | 38.2213416 | 2.10501008 | 0.67339382 | 3.12597177 | 0.00177219 | 0.03897289 |
| UQCRB | 58.5490797 | 0.84748992 | 0.49356357 | 1.71708362 | 0.08596389 | 0.29308354 |
| MTERF3 | 24.2468119 | 0.31731134 | 0.39711383 | 0.79904379 | 0.42426502 | 0.66715549 |
| PTDSS1 | 16.9171054 | -0.2449631 | 0.53220962 | -0.4602757 | 0.64531836 | NA |
| SDC2 | 139.347992 | -0.0382048 | 0.40348695 | -0.0946865 | 9.25E-01 | 9.68E-01 |
| CPQ | 212.281274 | 1.91695739 | 0.45926444 | 4.17397305 | 2.99E-05 | 0.00329138 |
| MTDH | 237.562727 | -0.3544933 | 0.17962963 | -1.9734678 | 0.04844229 | 0.22062742 |
| LAPTM4B | 97.4462823 | -0.943983 | 0.43305716 | -2.1798116 | 0.02927143 | 0.16874939 |
| RPL30 | 61.7275646 | 0.30193119 | 0.35761124 | 0.84430005 | 0.39850176 | 0.64642108 |
| RIDA | 17.4273143 | 1.23589188 | 0.59159383 | 2.08908851 | 0.03669976 | NA |
| POP1 | 14.0623306 | 0.69599473 | 0.49027379 | 1.4196042 | 0.15572294 | NA |
| NIPAL2 | 24.0368593 | 0.30734361 | 0.52274005 | 0.58794732 | 0.55656765 | 0.76317239 |
| STK3 | 486.577665 | -0.9720155 | 0.36083128 | -2.6938228 | 0.00706377 | 0.08210773 |
| AC104986.2 | 39.9629698 | 0.82104337 | 0.32691229 | 2.51150968 | 0.0120216 | 0.10726863 |
| VPS13B | 518.196863 | -0.0800884 | 0.19678232 | -0.4069899 | 0.68401543 | 0.8427141 |
| COX6C | 79.7961837 | 0.33754291 | 0.34912096 | 0.96683657 | 0.33362574 | 0.58883681 |
| SPAG1 | 54.9906848 | -0.1168412 | 0.3688264 | -0.3167919 | 0.75140153 | 0.8776233 |
| RNF19A | 373.193784 | -0.9692747 | 0.45495822 | -2.1304698 | 0.03313285 | 0.18022864 |
| AP000424.1 | 40.4612104 | -0.9032913 | 0.66245013 | -1.3635611 | 1.73E-01 | 4.21E-01 |
| ANKRD46 | 35.6033255 | 0.52314628 | 0.32765777 | 1.5966241 | 0.11034952 | 0.33347479 |
| PABPC1 | 116.912528 | -0.2856485 | 0.34703854 | -0.8231032 | 0.41044929 | 0.65612215 |
| YWHAZ | 199.410576 | -0.2934949 | 0.32486141 | -0.9034464 | 0.36628902 | 0.62157013 |
| ZNF706 | 55.2706274 | 0.23735937 | 0.34360971 | 0.69078191 | 0.48970261 | 0.7163358 |

| | | | | | | |
|---|---|---|---|---|---|---|
| GRHL2 | 181.695639 | 0.83860659 | 0.29065376 | 2.88524254 | 0.00391112 | 0.06143655 |
| RRM2B | 27.5925499 | 1.01216423 | 0.43200985 | 2.34291935 | 0.01913352 | 0.1359475 |
| UBR5 | 304.814166 | -0.2544426 | 0.20332825 | -1.2513882 | 0.21079287 | 0.4656925 |
| AZIN1 | 55.3370433 | 0.18361151 | 0.30628357 | 0.59948207 | 0.54885146 | 0.75579697 |
| AZIN1-AS1 | 134.083507 | 0.41870269 | 0.44373061 | 0.94359658 | 0.34537584 | 0.60277826 |
| AP003354.2 | 31.2491251 | 1.04499984 | 0.61074942 | 1.7110124 | 0.08707882 | 0.29500251 |
| ATP6V1C1 | 89.7243347 | -0.7867455 | 0.2658473 | -2.9593886 | 0.0030825 | 0.05464622 |
| BAALC-AS1 | 64.5055765 | 0.34994188 | 0.49236272 | 0.71074 | 0.47724537 | 0.7051944 |
| FZD6 | 109.978632 | -0.2780139 | 0.25685392 | -1.0823813 | 0.27908313 | 0.53717915 |
| SLC25A32 | 38.9074226 | -0.3428163 | 0.31037088 | -1.1045375 | 0.26936005 | 0.52764898 |
| DCAF13 | 21.7030248 | 0.14633801 | 0.38957963 | 0.37563054 | 0.70719158 | 0.8547469 |
| LRP12 | 34.2085518 | 0.00424172 | 0.38723158 | 0.01095396 | 0.99126018 | 0.99583841 |
| ZFPM2 | 32.5821633 | 3.4224206 | 0.94562519 | 3.6192147 | 0.0002955 | 0.01421532 |
| AC027031.2 | 29.4826349 | -1.0758569 | 0.66024468 | -1.6294822 | 0.10321098 | 0.32267056 |
| OXR1 | 1085.00781 | -0.1704735 | 0.3608436 | -0.4724304 | 0.63661963 | 0.81220973 |
| AC090579.1 | 81.8874818 | 1.40640057 | 0.71361961 | 1.97079866 | 0.04874691 | 0.22149053 |
| ANGPT1 | 39.7392432 | 1.41909755 | 0.77625629 | 1.82813018 | 0.06753002 | 0.25898011 |
| EIF3E | 95.158494 | -0.1675724 | 0.24731026 | -0.6775796 | 0.49803827 | 0.72149172 |
| EMC2 | 82.1527703 | 0.45114415 | 0.31891993 | 1.41460006 | 0.15718579 | 0.40154631 |
| NUDCD1 | 38.3577554 | -0.2065851 | 0.31434209 | -0.6571985 | 0.5110533 | 0.73060499 |
| ENY2 | 46.52586 | -0.1078785 | 0.32390442 | -0.3330567 | 0.73909151 | 0.87054913 |
| EBAG9 | 27.0695017 | 0.93302612 | 0.38310012 | 2.43546288 | 0.01487275 | 0.12024744 |
| SYBU | 23.1050131 | 1.04021473 | 0.49886005 | 2.08518348 | 0.03705265 | 0.19243876 |
| TRPS1 | 584.487102 | 0.08497331 | 0.30329121 | 0.28017071 | 0.77934654 | 0.89066523 |
| EIF3H | 124.728606 | -0.2347065 | 0.23320249 | -1.0064495 | 0.3141994 | 0.57172406 |
| UTP23 | 21.2711526 | 0.38612225 | 0.50756572 | 0.76073352 | 0.44681625 | 0.68656074 |
| RAD21 | 62.035806 | 0.41011078 | 0.2636938 | 1.55525376 | 0.11988565 | 0.3483427 |
| MED30 | 14.2173265 | 0.41812899 | 0.72432703 | 0.57726548 | 0.56376013 | NA |
| EXT1 | 658.118682 | -0.6036767 | 0.35915129 | -1.6808424 | 0.09279353 | 0.30571879 |
| SAMD12 | 603.695487 | 0.47416846 | 0.21864453 | 2.16867291 | 0.03010753 | 0.17153044 |
| AC023590.1 | 46.6042917 | 1.40055844 | 0.67409446 | 2.0776887 | 0.03773804 | 0.19425568 |
| SAMD12-AS1 | 40.3089718 | 0.66864072 | 0.49385604 | 1.35391828 | 0.17576245 | 0.42480254 |
| TNFRSF11B | 26.8145474 | 0.31225993 | 0.69218284 | 0.45112348 | 0.65190056 | 0.82061848 |
| MAL2 | 83.8811581 | -0.4203948 | 0.33083226 | -1.2707188 | 0.2038287 | 0.45736371 |
| TAF2 | 107.596962 | -0.7345911 | 0.26483725 | -2.7737453 | 0.0055415 | 0.07254189 |
| DEPTOR | 968.79877 | -0.014246 | 0.31054597 | -0.0458741 | 9.63E-01 | 9.86E-01 |
| AC091563.1 | 50.0679633 | 0.27776225 | 0.45949595 | 0.60449336 | 5.46E-01 | 7.54E-01 |
| COL14A1 | 55.1320095 | 0.73761405 | 0.54661727 | 1.34941593 | 1.77E-01 | 0.42569745 |
| MRPL13 | 26.9235692 | 0.28919499 | 0.45925198 | 0.62970876 | 0.52888515 | 0.74339464 |
| MTBP | 15.6222399 | -0.2527425 | 0.61130779 | -0.4134456 | 0.67928017 | NA |
| SNTB1 | 131.185649 | 0.02362523 | 0.58598766 | 0.04031694 | 0.96784045 | 0.98741848 |
| ZHX2 | 288.748176 | -0.9478753 | 0.35865558 | -2.6428568 | 0.00822098 | 0.08906215 |
| DERL1 | 23.3857265 | 0.15279669 | 0.43927873 | 0.3478354 | 0.7279638 | 0.86468169 |
| TBC1D31 | 16.2360567 | 0.70685847 | 0.51425372 | 1.37453254 | 0.16927642 | NA |
| WDYHV1 | 23.5269397 | -0.3250831 | 0.49357616 | -0.6586279 | 0.51013471 | 0.73010541 |
| FBXO32 | 22.1011104 | -0.1955984 | 0.41548922 | -0.4707664 | 0.63780756 | 0.81305711 |
| FAM91A1 | 74.3026531 | -0.5899274 | 0.30890277 | -1.9097511 | 0.05616527 | 0.23625346 |
| TMEM65 | 106.20407 | 0.20627181 | 0.32977681 | 0.62548913 | 0.53165008 | 0.74523961 |
| RNF139 | 12.283805 | 0.05057589 | 0.58199794 | 0.08690046 | 0.93075064 | NA |
| TATDN1 | 71.0083379 | -0.2681559 | 0.309459 | -0.8665312 | 0.3861989 | 0.63638899 |
| NDUFB9 | 44.2142095 | 0.0381991 | 0.39004617 | 0.09793483 | 0.92198404 | 0.96643214 |
| MTSS1 | 381.521405 | -0.0429369 | 0.33015465 | -0.1300508 | 0.89652621 | 0.9533797 |
| SQLE | 37.1019229 | 0.1281571 | 0.57380189 | 0.22334729 | 0.82326523 | 0.91475524 |
| WASHC5 | 58.935968 | -0.4887449 | 0.27352289 | -1.7868517 | 0.07396146 | 0.2712155 |

| | | | | | | |
|---|---|---|---|---|---|---|
| NSMCE2 | 426.639517 | -0.7931184 | 0.27987539 | -2.8338269 | 0.00459942 | 0.06580253 |
| TRIB1 | 22.3326599 | 0.36619656 | 0.43661539 | 0.83871656 | 0.40162838 | 0.64943985 |
| FAM84B | 69.4747788 | -1.3725527 | 0.57023898 | -2.4069781 | 0.01608513 | 0.12531101 |
| PCAT1 | 399.361082 | -1.1499736 | 0.47953846 | -2.3980842 | 0.01648107 | 0.12745979 |
| FAM49B | 392.392763 | -0.2583979 | 0.26181333 | -0.9869548 | 0.3236648 | 0.57976448 |
| ASAP1 | 198.258132 | -0.6347588 | 0.29666023 | -2.139683 | 0.0323804 | 0.17754567 |
| EFR3A | 160.505206 | -0.5519399 | 0.40657571 | -1.3575329 | 0.17461196 | 0.42330339 |
| KCNQ3 | 37.6167147 | 0.90240104 | 0.91382163 | 0.98750239 | 0.32339641 | 0.57941858 |
| LRRC6 | 51.9703453 | 1.098775 | 0.37881415 | 2.90056487 | 0.00372491 | 0.06079223 |
| PHF20L1 | 105.823563 | -0.297125 | 0.25357021 | -1.1717661 | 0.24129098 | 0.49867735 |
| NDRG1 | 361.09699 | -0.7019435 | 0.54617437 | -1.2852004 | 0.19872226 | 0.45213279 |
| ST3GAL1 | 63.1703068 | -1.1054079 | 0.36287717 | -3.0462315 | 2.32E-03 | 0.04573387 |
| ZFAT | 61.0425225 | -0.5225188 | 0.35316629 | -1.4795262 | 0.13899974 | 0.37579241 |
| AC083843.3 | 69.9403094 | -0.489662 | 0.47673254 | -1.027121 | 0.30436349 | 0.56489395 |
| KHDRBS3 | 191.078877 | -0.3636314 | 0.49102175 | -0.7405606 | 0.45895989 | 0.6928053 |
| TRAPPC9 | 483.588864 | 0.11939605 | 0.14694183 | 0.8125396 | 0.41648208 | 0.66028065 |
| AGO2 | 122.513334 | -0.6364221 | 0.3085528 | -2.0626035 | 0.03915031 | 0.19721202 |
| PTK2 | 756.236709 | -0.5090032 | 0.17056783 | -2.9841686 | 0.0028435 | 0.05198675 |
| DENND3 | 14.8559717 | 0.22565552 | 0.52536556 | 0.42952097 | 0.66754414 | NA |
| SLC45A4 | 78.9506165 | 0.69571649 | 0.29838168 | 2.33163274 | 0.01972002 | 0.13748641 |
| TSNARE1 | 42.9732325 | -0.5612768 | 0.47582702 | -1.1795817 | 0.23816664 | 0.49612142 |
| JRK | 11.6796948 | -0.1358741 | 0.53733259 | -0.2528678 | 0.80037034 | NA |
| LY6E | 32.5546921 | 0.09938197 | 0.75864861 | 0.13099869 | 0.89577635 | 0.953109 |
| GLI4 | 10.5630083 | 0.97684533 | 0.65148521 | 1.49941289 | 0.13376655 | NA |
| TOP1MT | 16.3718117 | -0.5989936 | 0.47876684 | -1.2511175 | 0.2108916 | NA |
| RHPN1 | 22.6242463 | 0.4971806 | 0.55600557 | 0.89420075 | 0.37121449 | 0.6243964 |
| ZC3H3 | 31.7247832 | -0.1281933 | 0.54515541 | -0.23515 | 0.8140923 | 0.90970797 |
| GSDMD | 14.6882405 | 0.42734566 | 0.54261946 | 0.7875605 | 0.43095383 | NA |
| EEF1D | 85.6879655 | -0.5330976 | 0.39765797 | -1.3405932 | 0.18005257 | 0.42915841 |
| ZNF623 | 17.070944 | -0.2119513 | 0.42731799 | -0.4960037 | 0.61989178 | NA |
| FAM83H | 13.5852454 | 0.60850811 | 0.49917094 | 1.21903753 | 0.22282995 | NA |
| IQANK1 | 15.4369922 | 0.42184141 | 0.80558722 | 0.52364462 | 0.60052574 | NA |
| SCRIB | 15.1215729 | 1.08691806 | 0.50241669 | 2.16337968 | 0.03051199 | NA |
| PUF60 | 14.5707297 | -0.027234 | 0.55596066 | -0.0489856 | 0.9609308 | NA |
| NRBP2 | 14.4446788 | 1.3373464 | 0.49180325 | 2.71927116 | 0.00654259 | NA |
| PLEC | 33.0778189 | -0.2258782 | 0.3932106 | -0.574446 | 0.56566606 | 0.76965382 |
| PARP10 | 15.531019 | 0.6619643 | 0.77562248 | 0.85346198 | 0.39340316 | NA |
| GRINA | 22.7277676 | 0.71278612 | 0.59862201 | 1.19071152 | 0.23376685 | 0.49243138 |
| GPAA1 | 20.6832454 | 0.8102523 | 0.53218109 | 1.52251238 | 0.12788075 | NA |
| CYC1 | 26.8824506 | 0.50716411 | 0.54721972 | 0.92680159 | 0.35402956 | 0.61144431 |
| SHARPIN | 27.9519714 | 0.01636864 | 0.33721548 | 0.0485406 | 0.96128541 | 0.98496344 |
| MAF1 | 18.8786725 | -0.0821342 | 0.68449451 | -0.1199926 | 0.90448905 | NA |
| MROH1 | 130.977044 | -0.9026636 | 0.38444912 | -2.3479403 | 0.01887755 | 0.1355543 |
| HSF1 | 57.7556781 | -0.1524933 | 0.26917726 | -0.5665164 | 0.57104276 | 0.77245539 |
| DGAT1 | 13.0009767 | 0.75766031 | 0.54188567 | 1.39819218 | 0.16205536 | NA |
| CPSF1 | 11.9810026 | 0.99202915 | 0.65484784 | 1.51490024 | 0.12979767 | NA |
| VPS28 | 37.2510154 | 0.53846379 | 0.46674346 | 1.15366113 | 0.24863913 | 0.50492395 |
| CYHR1 | 34.043809 | 0.73503666 | 0.49122796 | 1.49632496 | 0.13456899 | 0.36839609 |
| KIFC2 | 18.109221 | 1.09771881 | 0.56428928 | 1.94531217 | 0.05173742 | NA |
| PPP1R16A | 30.4982023 | 0.31419197 | 0.42134207 | 0.74569333 | 0.45585269 | 0.6908246 |
| ARHGAP39 | 23.1773826 | 0.23171467 | 0.45067436 | 0.514151 | 0.60714643 | 0.79532486 |
| ZNF251 | 71.0577023 | 0.32484428 | 0.25854883 | 1.25641365 | 0.20896603 | 0.46338564 |
| RPL8 | 100.376273 | 0.25582414 | 0.47654969 | 0.53682573 | 0.591388 | 0.78517152 |
| ZNF7 | 24.3022713 | 0.16096395 | 0.37272288 | 0.4318596 | 0.66584346 | 0.83076627 |

| | | | | | | |
|---|---|---|---|---|---|---|
| ZNF250 | 17.1672308 | 0.52935704 | 0.46693945 | 1.13367385 | 0.25693138 | NA |
| ZNF16 | 18.1372874 | -0.4697071 | 0.48350185 | -0.9714691 | 0.33131476 | NA |
| WASHC1 | 101.792986 | 0.41647618 | 0.34574359 | 1.20458106 | 0.22836507 | 0.48663509 |
| CBWD1 | 29.7247747 | 0.20090352 | 0.39686808 | 0.50622241 | 0.61270052 | 0.79866179 |
| DOCK8 | 36.1010819 | 0.33974991 | 0.52053681 | 0.65269143 | 0.51395523 | 0.73176668 |
| KANK1 | 1277.48317 | 0.4057697 | 0.25554637 | 1.58785159 | 1.12E-01 | 3.37E-01 |
| SMARCA2 | 160.651439 | 0.69710193 | 0.27928698 | 2.4960058 | 0.01256005 | 0.10970808 |
| VLDLR-AS1 | 67.5490325 | -0.6519138 | 0.66151554 | -0.9854852 | 0.32438579 | 0.58025777 |
| VLDLR | 38.6038588 | 0.34053935 | 0.39392126 | 0.86448584 | 0.38732105 | 0.63701577 |
| PUM3 | 207.769363 | 0.20303675 | 0.27268298 | 0.744589 | 0.45652022 | 0.69129178 |
| RFX3 | 262.428783 | 0.53077335 | 0.28525104 | 1.86072364 | 0.06278321 | 0.24960868 |
| RFX3-AS1 | 211.94267 | 0.07782921 | 0.41838402 | 0.18602339 | 0.85242642 | 0.92946964 |
| GLIS3 | 1011.35865 | 0.33880841 | 0.40858138 | 0.82923114 | 0.40697363 | 0.65367876 |
| SLC1A1 | 75.5351805 | 0.91135762 | 0.42342623 | 2.15234098 | 0.03137051 | 0.17484345 |
| SPATA6L | 111.311756 | -0.1139569 | 0.37309784 | -0.3054343 | 0.7600354 | 0.8808903 |
| CDC37L1 | 67.0736936 | -0.8245798 | 0.33001191 | -2.4986365 | 0.01246721 | 0.10951947 |
| AK3 | 522.345192 | -0.2488358 | 0.43468418 | -0.572452 | 0.56701582 | 0.77060355 |
| RCL1 | 119.729135 | -0.7015067 | 0.28149102 | -2.4921103 | 0.01269866 | 0.11020195 |
| JAK2 | 130.148399 | -0.2383012 | 0.21292281 | -1.1191904 | 0.2630589 | 0.52278966 |
| PLGRKT | 55.5584721 | -0.4014804 | 0.2974388 | -1.3497916 | 0.17708283 | 0.42569745 |
| RIC1 | 221.714258 | -0.2551818 | 0.27574451 | -0.9254283 | 0.35474317 | 0.61168935 |
| ERMP1 | 185.372043 | 0.10415452 | 0.31662574 | 0.32895154 | 0.74219232 | 0.87152705 |
| KIAA2026 | 283.343424 | 0.03940943 | 0.22662622 | 0.17389616 | 0.86194707 | 0.93551424 |
| RANBP6 | 15.1526646 | 0.03874009 | 0.63989875 | 0.06054096 | 0.95172479 | NA |
| UHRF2 | 144.975375 | -0.2314605 | 0.20455732 | -1.1315192 | 0.25783664 | 0.51619036 |
| GLDC | 176.245745 | -0.5571202 | 0.45906061 | -1.2136093 | 0.22489695 | 0.48216659 |
| AL162411.1 | 27.513749 | -0.132809 | 0.38692619 | -0.3432411 | 0.73141711 | 0.86703081 |
| KDM4C | 452.913488 | -0.5204754 | 0.20078202 | -2.5922411 | 0.00953529 | 0.09644302 |
| DMAC1 | 16.8425629 | 0.462562 | 0.44394118 | 1.04194436 | 0.29743748 | NA |
| PTPRD | 35.6334634 | 0.78743314 | 0.53823066 | 1.46300313 | 0.14346652 | 0.38222977 |
| LURAP1L | 91.7926003 | -1.4934005 | 0.72758727 | -2.052538 | 4.01E-02 | 1.99E-01 |
| MPDZ | 141.70127 | 0.42306306 | 0.3067611 | 1.37912879 | 0.16785505 | 0.41609672 |
| NFIB | 257.933603 | 0.5431826 | 0.37014341 | 1.4674923 | 0.14224218 | 0.38005954 |
| ZDHHC21 | 251.671639 | 0.04912217 | 0.27655305 | 0.17762296 | 0.85901909 | 0.9336162 |
| FREM1 | 336.179646 | 2.48420626 | 0.61025702 | 4.07075412 | 4.69E-05 | 0.00424342 |
| TTC39B | 93.7785377 | -0.0475822 | 0.34742189 | -0.1369579 | 0.89106404 | 0.94993351 |
| SNAPC3 | 68.511401 | 0.04382116 | 0.23945074 | 0.18300701 | 0.85479252 | 0.93125803 |
| PSIP1 | 69.0641176 | 0.08274039 | 0.40196231 | 0.20584115 | 0.83691499 | 0.92119416 |
| CCDC171 | 153.372057 | 1.27709155 | 0.51260312 | 2.49138465 | 0.01272463 | 0.11029444 |
| BNC2 | 75.0682747 | -1.23552 | 0.44779069 | -2.7591463 | 5.80E-03 | 0.07462791 |
| CNTLN | 97.1768101 | -0.0486938 | 0.33191404 | -0.1467059 | 0.88336415 | 0.94552272 |
| SH3GL2 | 58.4699214 | -1.2810679 | 0.50587653 | -2.5323726 | 0.01132935 | 0.1036885 |
| HAUS6 | 19.0618484 | -0.0523178 | 0.44872417 | -0.1165923 | 0.90718317 | NA |
| PLIN2 | 47.1368139 | -0.341936 | 0.62499875 | -0.5470986 | 0.58431097 | 0.78002178 |
| DENND4C | 285.352858 | -0.2041465 | 0.16185343 | -1.2613045 | 0.20719916 | 0.46066753 |
| RPS6 | 107.229918 | 0.78854802 | 0.48145435 | 1.63784588 | 0.10145385 | 0.32069415 |
| MLLT3 | 480.650272 | 0.29565057 | 0.26576956 | 1.11243205 | 0.26595243 | 0.5251503 |
| FOCAD | 195.906279 | 0.99636078 | 0.36894084 | 2.7005977 | 0.0069215 | 0.08121157 |
| KLHL9 | 17.2250552 | 0.27734708 | 0.48375978 | 0.57331571 | 0.56643095 | NA |
| LINC01239 | 24.731759 | 1.42963562 | 0.72059403 | 1.98396817 | 0.04725939 | 0.21733813 |
| CAAP1 | 150.638087 | -0.0757781 | 0.30233523 | -0.2506425 | 0.8020905 | 0.90378853 |
| PLAA | 48.0214667 | -0.3105008 | 0.2878689 | -1.0786189 | 0.28075766 | 0.5390351 |
| IFT74 | 116.582651 | 0.2284847 | 0.28568991 | 0.79976467 | 0.42384715 | 0.6670468 |
| MOB3B | 122.11284 | -0.0578304 | 0.33823971 | -0.1709746 | 0.86424374 | 0.93678131 |

| | | | | | | |
|---|---|---|---|---|---|---|
| C9orf72 | 37.7425021 | -0.4402549 | 0.37000461 | -1.1898632 | 0.23410018 | 0.49243138 |
| LINGO2 | 104.427008 | 0.96810967 | 0.35697704 | 2.71196623 | 0.00668854 | 0.07974991 |
| ACO1 | 34.8182006 | 0.80534391 | 0.46640184 | 1.72671685 | 0.08421853 | 0.28931012 |
| DDX58 | 77.9071435 | 0.13237066 | 0.922151 | 0.14354553 | 0.88585935 | 0.9471398 |
| TOPORS | 25.3489115 | 0.14994449 | 0.45291048 | 0.33106871 | 0.74059258 | 0.87147854 |
| NDUFB6 | 39.3960921 | 0.58387925 | 0.33141795 | 1.76176107 | 0.07810967 | 0.27885444 |
| APTX | 66.9588009 | 0.53001978 | 0.35061017 | 1.51170682 | 0.13060846 | 0.36322236 |
| DNAJA1 | 37.553333 | 0.26794803 | 0.35699655 | 0.75056195 | 0.45291633 | 0.68922439 |
| SMU1 | 40.1617694 | -0.0331013 | 0.33661904 | -0.0983346 | 0.92166663 | 0.96635254 |
| B4GALT1 | 106.934303 | -0.1254594 | 0.31027636 | -0.4043473 | 0.68595733 | 0.84395997 |
| BAG1 | 28.4210982 | -0.0883238 | 0.52975455 | -0.166726 | 0.86758568 | 0.93827484 |
| CHMP5 | 25.8719865 | -0.124261 | 0.43033069 | -0.288757 | 0.77276737 | 0.88749485 |
| NFX1 | 149.612615 | -0.8240555 | 0.25573321 | -3.2223247 | 0.00127155 | 0.03187985 |
| UBE2R2 | 236.269396 | -0.585845 | 0.35801301 | -1.6363791 | 0.10176026 | 0.32089434 |
| UBAP2 | 148.798989 | -0.3995433 | 0.29453301 | -1.3565315 | 0.17493013 | 0.42340793 |
| DCAF12 | 29.4612312 | -0.2218693 | 0.39502715 | -0.5616558 | 0.57435053 | 0.77388388 |
| UBAP1 | 220.465425 | -0.6156762 | 0.28940664 | -2.1273742 | 0.03338901 | 0.18111006 |
| NUDT2 | 14.2115932 | 0.53931074 | 0.69754921 | 0.77315081 | 0.43943313 | NA |
| FAM219A | 40.2681439 | -0.0070755 | 0.31779216 | -0.0222645 | 0.98223696 | 0.99382437 |
| VCP | 58.0368 | -0.4171134 | 0.32827414 | -1.2706252 | 0.20386201 | 0.45736371 |
| STOML2 | 15.0201383 | 0.96876675 | 0.63780775 | 1.51890086 | 0.12878745 | NA |
| UNC13B | 242.136688 | 0.14659886 | 0.22301227 | 0.65735783 | 0.51095086 | 0.73060499 |
| RUSC2 | 157.35262 | -0.712599 | 0.33530548 | -2.1252231 | 0.033568 | 0.18156913 |
| CCDC107 | 16.4334564 | 0.01565282 | 0.42045749 | 0.03722808 | 0.97030315 | NA |
| TLN1 | 39.6590778 | -0.0126892 | 0.35590807 | -0.0356531 | 0.97155895 | 0.98877288 |
| GBA2 | 59.5702773 | -0.2136255 | 0.36340677 | -0.5878411 | 0.55663892 | 0.76317239 |
| RGP1 | 18.4099434 | 0.97027084 | 0.51821468 | 1.87233377 | 0.06116044 | NA |
| AL133410.1 | 20.8313861 | 0.0931975 | 0.46860647 | 0.19888222 | 0.84235487 | NA |
| TMEM8B | 18.898896 | 0.32147465 | 0.42817028 | 0.75081028 | 0.45276684 | NA |
| RECK | 39.9454352 | 0.1607741 | 0.42852746 | 0.37517807 | 0.70752804 | 0.8547469 |
| CLTA | 98.4900935 | -0.695126 | 0.32186137 | -2.1597064 | 0.03079541 | 0.17345817 |
| GNE | 78.4344416 | -1.7767089 | 0.48939367 | -3.6304288 | 0.00028295 | 0.013784 |
| RNF38 | 254.005465 | -0.7074686 | 0.36623764 | -1.93172 | 0.05339408 | 0.23211867 |
| EBLN3P | 41.1653421 | 0.63828608 | 0.39117148 | 1.6317296 | 0.10273646 | 0.32184067 |
| ZCCHC7 | 476.249845 | -0.3734475 | 0.16905259 | -2.2090611 | 0.02717039 | 0.16201805 |
| GRHPR | 22.6516576 | 0.89569629 | 0.42198447 | 2.12258118 | 0.03378896 | 0.1821244 |
| ZBTB5 | 25.6818751 | 0.15176177 | 0.38014899 | 0.39921657 | 0.68973364 | 0.84603662 |
| AL138752.2 | 63.3772897 | 0.06613407 | 0.32739745 | 0.20199933 | 0.83991725 | 0.92315337 |
| FRMPD1 | 14.6548024 | 2.63626268 | 0.89783884 | 2.9362315 | 0.00332226 | NA |
| TRMT10B | 53.5518403 | 0.65700507 | 0.47646556 | 1.37891408 | 0.16792125 | 0.4161268 |
| DCAF10 | 158.613315 | 0.10596166 | 0.21507553 | 0.49267184 | 0.62224448 | 0.80366942 |
| FAM95C | 18.3729303 | -0.0333052 | 0.49847479 | -0.0668143 | 0.94672954 | NA |
| ANKRD18A | 20.1306761 | -0.3072796 | 0.489217 | -0.6281049 | 0.52993523 | NA |
| GLIDR | 37.2682488 | -0.0731786 | 0.44997329 | -0.1626287 | 0.87081076 | 0.93986793 |
| BMS1P14 | 33.6165879 | -1.0508737 | 0.55385395 | -1.8973841 | 0.05777726 | 0.23909224 |
| CBWD6 | 16.7272429 | -0.2685526 | 0.66069638 | -0.406469 | 0.68439804 | NA |
| CNTNAP3B | 33.4513358 | 0.73126242 | 0.63721052 | 1.14759941 | 0.25113397 | 0.50787656 |
| AL512625.3 | 13.3341207 | 0.59526013 | 0.5595276 | 1.06386196 | 0.28739124 | NA |
| CBWD5 | 46.6607434 | -0.5999474 | 0.31772335 | -1.8882697 | 0.05898976 | 0.2412562 |
| ZNF658 | 17.0176424 | 1.22748242 | 0.62270893 | 1.97119771 | 0.04870127 | NA |
| PGM5 | 22.2207128 | 0.96483623 | 0.72088255 | 1.33840974 | 0.1807629 | 0.4300872 |
| AL161457.2 | 11.7835928 | 1.18645362 | 0.67054054 | 1.76939879 | 0.07682735 | NA |
| PIP5K1B | 207.7087 | -0.0687706 | 0.38153724 | -0.1802462 | 0.85695929 | 0.93203598 |
| FAM189A2 | 39.4623974 | 1.05739345 | 0.5258865 | 2.01068758 | 0.04435847 | 0.21075748 |

| | | | | | | |
|---|---|---|---|---|---|---|
| PTAR1 | 138.449389 | 0.0631581 | 0.18915361 | 0.33389846 | 0.73845618 | 0.87054913 |
| MAMDC2-AS | 69.9593944 | 1.87271461 | 0.54632049 | 3.42786818 | 0.00060834 | 0.02240382 |
| MAMDC2 | 55.9218231 | 0.87221861 | 0.51132992 | 1.70578444 | 0.08804822 | 0.29675763 |
| SMC5-AS1 | 86.5958339 | 1.05946865 | 0.31695905 | 3.34260418 | 0.00082996 | 0.02565549 |
| SMC5 | 189.54772 | -0.3882361 | 0.20051849 | -1.9361611 | 0.05284797 | 0.23105954 |
| KLF9 | 47.7891119 | -0.2594494 | 0.40356611 | -0.6428919 | 0.52029425 | 0.73710746 |
| TRPM3 | 108.204808 | 0.89511997 | 0.66766897 | 1.34066434 | 0.18002946 | 0.42915841 |
| CEMIP2 | 153.137459 | -0.6383131 | 0.41105213 | -1.5528762 | 1.20E-01 | 3.49E-01 |
| ABHD17B | 33.5622594 | -0.4233289 | 0.4396232 | -0.9629358 | 0.33557973 | 0.59131149 |
| C9orf85 | 68.7965277 | 0.02241469 | 0.26831658 | 0.08353824 | 0.93342357 | 0.9730757 |
| ZFAND5 | 136.038429 | -0.6078611 | 0.49557498 | -1.2265775 | 0.21998141 | 0.47642007 |
| ALDH1A1 | 40.8299046 | 1.39269049 | 0.46891867 | 2.97000433 | 0.00297796 | 0.05367991 |
| TRPM6 | 739.203867 | -1.0598417 | 0.56409154 | -1.878847 | 6.03E-02 | 2.45E-01 |
| CARNMT1 | 101.851015 | 0.1914665 | 0.22171487 | 0.86357085 | 0.38782368 | 0.63702067 |
| NMRK1 | 42.0926836 | 1.07430646 | 0.62919945 | 1.70741799 | 0.08774439 | 0.29621428 |
| OSTF1 | 28.627447 | -0.1915322 | 0.38289063 | -0.5002268 | 0.61691537 | 0.80179341 |
| PCSK5 | 46.2619561 | -0.9656897 | 0.82922022 | -1.1645757 | 0.24419078 | 0.5015083 |
| VPS13A | 183.951816 | -0.006572 | 0.23545266 | -0.0279123 | 0.97773207 | 0.99142598 |
| GNAQ | 553.414471 | -0.4479465 | 0.17940918 | -2.496787 | 0.01253242 | 0.10961596 |
| TLE4 | 26.6758513 | -0.0920694 | 0.56719509 | -0.1623241 | 0.87105063 | 0.93986793 |
| TLE1 | 59.352779 | -0.0347862 | 0.4490524 | -0.0774659 | 0.93825295 | 0.97466007 |
| AL591368.1 | 10.9978533 | 0.7242934 | 1.07036488 | 0.67667896 | 0.49860967 | NA |
| RASEF | 26.057963 | -0.3971695 | 0.39849368 | -0.9966771 | 0.31892128 | 0.57595896 |
| FRMD3 | 497.521507 | -0.5707569 | 0.42008082 | -1.3586835 | 1.74E-01 | 0.42295604 |
| IDNK | 16.3603913 | -0.0062994 | 0.43300281 | -0.0145482 | 0.98839265 | NA |
| UBQLN1 | 124.458021 | -0.2416179 | 0.2788095 | -0.8666058 | 0.38615804 | 0.63638899 |
| GKAP1 | 49.6164603 | 0.4586799 | 0.34743258 | 1.32019831 | 0.18676882 | 0.4369482 |
| KIF27 | 71.807873 | 0.59220331 | 0.31911398 | 1.85577365 | 0.06348583 | 0.25118809 |
| HNRNPK | 135.241297 | -0.2747409 | 0.27255404 | -1.0080235 | 0.3134432 | 0.57102302 |
| RMI1 | 13.83588 | 0.24449975 | 0.57330886 | 0.42647126 | 0.66976448 | NA |
| AGTPBP1 | 118.625486 | -0.0391111 | 0.22543911 | -0.1734886 | 0.86226738 | 0.93569322 |
| NAA35 | 132.939095 | -0.3214891 | 0.22562405 | -1.4248883 | 0.15418949 | 0.39652407 |
| GOLM1 | 93.4276225 | -0.5280826 | 0.38369099 | -1.3763226 | 0.16872179 | 0.41629954 |
| C9orf153 | 15.1285497 | 0.98146511 | 0.58307104 | 1.68326848 | 0.09232313 | NA |
| ISCA1 | 35.4802685 | 0.09284434 | 0.31956206 | 0.29053619 | 0.77140607 | 0.88685773 |
| TUT7 | 106.069718 | -0.3828041 | 0.22244118 | -1.7209228 | 0.08526483 | 0.29142247 |
| AL513318.2 | 18.6389734 | -0.2955398 | 0.56259783 | -0.5253127 | 0.59936582 | NA |
| DAPK1 | 445.013303 | -0.9187324 | 0.54182104 | -1.6956381 | 0.08995443 | 0.299472 |
| CTSL | 45.9805606 | -0.586584 | 0.56458283 | -1.0389688 | 0.29881923 | 0.55866203 |
| SPIN1 | 348.969865 | 0.12907451 | 0.24940352 | 0.51753281 | 0.60478427 | 0.794016 |
| SHC3 | 131.878791 | -1.9815585 | 0.94470572 | -2.0975405 | 0.03594575 | 0.18911447 |
| SECISBP2 | 150.858364 | -0.6691563 | 0.23374687 | -2.8627391 | 0.00419996 | 0.06301187 |
| SEMA4D | 136.98109 | -0.9421364 | 0.26623526 | -3.5387363 | 0.00040205 | 0.01738515 |
| SYK | 53.6230215 | 0.58042953 | 0.36590524 | 1.58628373 | 0.11267496 | 0.33745493 |
| AL158071.1 | 55.0501193 | -0.359247 | 0.80809948 | -0.4445579 | 0.65663925 | 0.82462919 |
| AL158071.4 | 268.442512 | -1.3604639 | 0.51035646 | -2.6657131 | 0.00768252 | 0.08614009 |
| AL158071.5 | 18.3996949 | -0.5554747 | 0.63232642 | -0.8784619 | 0.3796931 | NA |
| AUH | 406.818433 | 0.06257626 | 0.25936578 | 0.24126646 | 0.80934861 | 0.90756938 |
| NFIL3 | 54.9390961 | -1.0558711 | 0.71968576 | -1.4671279 | 0.14234126 | 0.38005954 |
| SPTLC1 | 81.1325862 | -0.3368215 | 0.25488958 | -1.3214408 | 0.18635443 | 0.43624393 |
| IARS | 88.6182618 | -1.2610502 | 0.56222168 | -2.2429768 | 0.02489831 | 0.1546391 |
| NOL8 | 29.4813299 | 0.03901279 | 0.36625192 | 0.106519 | 0.91517058 | 0.96388328 |
| CENPP | 49.6250336 | -0.7254449 | 0.70735582 | -1.0255728 | 0.30509301 | 0.56557634 |
| IPPK | 29.6028928 | -0.4260032 | 0.483666 | -0.8807796 | 0.37843711 | 0.62993738 |

| | | | | | | |
|---|---|---|---|---|---|---|
| BICD2 | 27.1385475 | 0.5178627 | 0.40080117 | 1.29206881 | 0.19633331 | 0.44975522 |
| ZNF484 | 18.460873 | -0.367738 | 0.44150381 | -0.8329216 | 0.40488896 | NA |
| NINJ1 | 21.5269754 | 1.96194389 | 0.57394023 | 3.41837664 | 0.00062996 | 0.02287166 |
| FAM120AOS | 12.4988779 | -0.0548699 | 0.52563645 | -0.1043875 | 0.91686182 | NA |
| FAM120A | 138.948091 | 0.39861923 | 0.27511218 | 1.4489334 | 0.14735618 | 0.38722988 |
| PHF2 | 42.6211652 | -0.0011522 | 0.35224376 | -0.0032711 | 0.99739005 | 0.99868755 |
| PTPDC1 | 58.8950345 | 0.19841785 | 0.33654644 | 0.58957049 | 0.55547864 | 0.76212461 |
| AL158152.2 | 74.571716 | -0.260698 | 0.27354074 | -0.95305 | 0.34056473 | 0.59656958 |
| ZNF169 | 29.9461133 | 0.0630282 | 0.41880123 | 0.15049669 | 0.88037276 | 0.94376451 |
| MFSD14B | 54.3739206 | -0.1620606 | 0.3319675 | -0.4881822 | 0.62542081 | 0.8053173 |
| C9orf3 | 648.396667 | -0.3234732 | 0.21690796 | -1.4912926 | 0.1358847 | 0.37062528 |
| FANCC | 144.616863 | 0.38176362 | 0.27370597 | 1.39479462 | 0.16307778 | 0.40979748 |
| PTCH1 | 48.6151827 | -0.6096973 | 0.3557523 | -1.7138252 | 0.08656083 | 0.29415396 |
| LINC00476 | 89.7625761 | 0.39544145 | 0.4674971 | 0.84586932 | 0.39762565 | 0.64588458 |
| ERCC6L2 | 133.14663 | -0.5430138 | 0.22343894 | -2.430256 | 0.01508816 | 0.12124045 |
| SLC35D2 | 27.998072 | 0.07353809 | 0.43969785 | 0.16724686 | 0.86717581 | 0.93824181 |
| HABP4 | 78.9053804 | -0.5120311 | 0.46915187 | -1.0913973 | 0.27509808 | 0.53376102 |
| CDC14B | 165.318706 | 1.08506966 | 0.42148971 | 2.57436809 | 0.01004234 | 0.09909727 |
| AL589843.1 | 25.2698981 | -0.4576493 | 0.49521546 | -0.9241419 | 0.35541245 | 0.61168935 |
| ZNF510 | 18.4477953 | 0.03586282 | 0.45545037 | 0.07874145 | 0.93723827 | NA |
| ZNF782 | 66.7838443 | -0.4181467 | 0.25803236 | -1.6205203 | 0.10512056 | 0.32611764 |
| MFSD14C | 156.579183 | -0.3719424 | 0.2129898 | -1.7462921 | 0.08076021 | 0.28378572 |
| CCDC180 | 14.7396055 | 0.85281063 | 0.68034068 | 1.25350527 | 0.21002188 | NA |
| TDRD7 | 32.2448445 | 0.46026154 | 0.46398179 | 0.99198191 | 0.32120635 | 0.57791615 |
| TSTD2 | 37.0620133 | -0.0008789 | 0.31683653 | -0.0027739 | 0.99778679 | 0.99882493 |
| NCBP1 | 40.3474645 | -0.3667677 | 0.37700806 | -0.9728377 | 0.33063397 | 0.58655523 |
| XPA | 54.52131 | 0.82440916 | 0.33941096 | 2.42894091 | 0.015143 | 0.12124045 |
| ANP32B | 42.8685014 | -0.0260974 | 0.38245498 | -0.0682366 | 0.94559727 | 0.97792536 |
| TRIM14 | 15.6441025 | -0.8919542 | 0.82819296 | -1.0769883 | 0.28148548 | NA |
| ANKS6 | 25.9064903 | 0.42598219 | 0.47607611 | 0.89477751 | 0.37090604 | 0.62401394 |
| TGFBR1 | 75.6972175 | -0.1649898 | 0.47757919 | -0.345471 | 0.7297403 | 0.86545625 |
| ALG2 | 16.4002255 | 0.42335008 | 0.51017262 | 0.82981732 | 0.40664208 | NA |
| SEC61B | 37.7459763 | -0.0898308 | 0.35876886 | -0.2503864 | 0.80228857 | 0.90378853 |
| AL359710.1 | 69.3232833 | 0.55389372 | 0.47340738 | 1.17001499 | 0.24199494 | 0.49936596 |
| STX17 | 62.825751 | -0.002763 | 0.37983814 | -0.0072743 | 0.99419602 | 0.99730572 |
| ERP44 | 117.302638 | -0.571164 | 0.29091332 | -1.9633479 | 0.04960576 | 0.22367637 |
| INVS | 219.875698 | 0.31355901 | 0.30157256 | 1.03974649 | 0.2984577 | 0.55825732 |
| TEX10 | 37.6072063 | -0.1708976 | 0.37059703 | -0.4611413 | 0.64469723 | 0.81734747 |
| PLPPR1 | 183.392239 | 1.11192095 | 0.46607091 | 2.38573344 | 1.70E-02 | 1.30E-01 |
| ZNF189 | 27.9350019 | -0.3762891 | 0.35165823 | -1.0700421 | 0.28460036 | 0.54316116 |
| ALDOB | 50.6100816 | 0.63578098 | 0.88549015 | 0.71799894 | 0.47275794 | 0.70193246 |
| RNF20 | 22.9009305 | 0.17482825 | 0.46950903 | 0.37236398 | 0.70962187 | 0.85607462 |
| SMC2 | 15.3664293 | 0.5996274 | 0.50300956 | 1.19207954 | 0.23323006 | NA |
| NIPSNAP3A | 19.4558246 | 0.76514806 | 0.43813465 | 1.74637652 | 0.08074555 | NA |
| ABCA1 | 60.8145535 | -0.3906171 | 0.37273811 | -1.0479667 | 0.29465397 | 0.55396961 |
| SLC44A1 | 153.358216 | 0.19290397 | 0.39060597 | 0.49385822 | 0.62140632 | 0.80353114 |
| FSD1L | 95.0372424 | 0.29613075 | 0.27956459 | 1.05925701 | 0.28948275 | 0.5488051 |
| FKTN | 67.7497135 | 0.48472443 | 0.37673282 | 1.28665304 | 0.19821525 | 0.45124601 |
| TMEM38B | 99.6824514 | -0.856091 | 0.57160134 | -1.4977065 | 0.13420952 | 0.36808465 |
| LINC01505 | 174.990495 | -1.2732633 | 0.43257217 | -2.9434703 | 0.00324555 | 0.05664625 |
| ZNF462 | 107.407815 | -0.4349652 | 0.32330284 | -1.34538 | 0.17850254 | 0.42765983 |
| RAD23B | 132.094574 | -0.476011 | 0.23803425 | -1.9997582 | 0.04552638 | 0.21335013 |
| ELP1 | 49.4822815 | 0.43807356 | 0.35094651 | 1.24826302 | 0.21193475 | 0.46660806 |
| FAM206A | 17.0248309 | -0.2703963 | 0.47224227 | -0.5725796 | 0.5669294 | NA |

| | | | | | | |
|---|---|---|---|---|---|---|
| CTNNAL1 | 110.760482 | -0.6871785 | 0.31613867 | -2.1736616 | 0.02973056 | 0.1702177 |
| TMEM245 | 243.918864 | 0.17933012 | 0.29350615 | 0.61099272 | 0.5412044 | 0.74994873 |
| EPB41L4B | 71.9226371 | 0.63423156 | 0.33125351 | 1.914641 | 0.0555383 | 0.235266 |
| PTPN3 | 478.276475 | -1.2733661 | 0.29865656 | -4.263647 | 2.01E-05 | 0.00245714 |
| PALM2 | 62.191866 | -0.5262019 | 0.25262389 | -2.0829461 | 3.73E-02 | 1.93E-01 |
| TXN | 40.2225125 | -0.0421061 | 0.39712303 | -0.1060279 | 0.9155602 | 0.96388328 |
| ECPAS | 187.498658 | -0.4126089 | 0.25554343 | -1.6146332 | 0.10639016 | 0.32732986 |
| ZNF483 | 96.7913982 | -0.8170931 | 0.32606227 | -2.5059419 | 0.01221257 | 0.10817047 |
| PTGR1 | 177.966644 | -1.0722434 | 0.31055827 | -3.4526319 | 0.00055515 | 0.02104906 |
| C9orf84 | 33.5682849 | 1.21040006 | 0.6571322 | 1.84194301 | 0.06548349 | 0.25452134 |
| UGCG | 438.151885 | -1.2445437 | 0.41369079 | -3.0083911 | 0.00262635 | 0.0493049 |
| SUSD1 | 140.974808 | -1.3005973 | 0.38130506 | -3.4109102 | 0.00064746 | 0.02317922 |
| PTBP3 | 298.36559 | -0.5004941 | 0.29275706 | -1.7095884 | 0.08734201 | 0.29567725 |
| HSDL2 | 132.275285 | 0.0273515 | 0.27308105 | 0.1001589 | 0.92021818 | 0.96536995 |
| C9orf147 | 30.2383873 | 0.34937675 | 0.45933187 | 0.76061943 | 0.44688441 | 0.68656074 |
| KIAA1958 | 500.984926 | 0.3530749 | 0.29344591 | 1.20320268 | 0.2288979 | 0.48721455 |
| INIP | 39.4638939 | 0.37563566 | 0.29984669 | 1.25275907 | 0.2102934 | 0.46525677 |
| SNX30 | 51.6164251 | -0.22301 | 0.33366773 | -0.6683594 | 0.5039042 | 0.72577669 |
| FKBP15 | 40.1436072 | 0.10317114 | 0.3199189 | 0.32249153 | 0.74708035 | 0.87469995 |
| SLC31A1 | 28.602101 | 0.09905687 | 0.36004564 | 0.27512308 | 0.78322168 | 0.89310478 |
| CDC26 | 14.0508247 | 0.86471919 | 0.54459768 | 1.58781284 | 0.11232867 | NA |
| PRPF4 | 19.5768317 | -0.5143786 | 0.51509734 | -0.9986047 | 0.31798621 | NA |
| WDR31 | 15.8792704 | 0.56283297 | 0.61994935 | 0.90786928 | 0.36394729 | NA |
| BSPRY | 29.0251982 | 0.18789667 | 0.33830913 | 0.55539937 | 0.57862153 | 0.77576204 |
| HDHD3 | 30.6611489 | -0.5351248 | 0.43527677 | -1.2293897 | 0.21892574 | 0.47565385 |
| RGS3 | 128.978952 | 0.06761945 | 0.29648153 | 0.22807307 | 0.81958943 | 0.913067 |
| ZNF618 | 198.413337 | 0.7950801 | 0.45842588 | 1.73437002 | 0.08285245 | 0.28790759 |
| KIF12 | 80.7656278 | 0.43382598 | 0.50982704 | 0.85092776 | 0.39480949 | 0.64334162 |
| COL27A1 | 93.0685236 | -0.0493959 | 0.57111913 | -0.0864897 | 0.93107717 | 0.97177475 |
| WHRN | 48.8764124 | -0.705523 | 0.40165887 | -1.7565229 | 0.07899916 | 0.28033956 |
| ATP6V1G1 | 52.091119 | -0.1435788 | 0.37315506 | -0.3847699 | 0.70040787 | 0.85179956 |
| TMEM268 | 14.4331455 | 0.4330355 | 0.66696023 | 0.64926735 | 0.51616558 | NA |
| PAPPA | 25.6638729 | 0.62736164 | 0.48638461 | 1.28984683 | 0.19710385 | 0.45031229 |
| ASTN2 | 105.213359 | -0.5028173 | 0.52476223 | -0.9581812 | 0.33797137 | 0.59378353 |
| CDK5RAP2 | 71.7060713 | 0.02282083 | 0.334786 | 0.06816543 | 0.94565395 | 0.97792536 |
| MEGF9 | 119.42214 | 0.77363312 | 0.28771152 | 2.68891947 | 0.00716837 | 0.08259723 |
| FBXW2 | 61.0480073 | -0.615229 | 0.25602437 | -2.4030094 | 0.01626077 | 0.12641035 |
| PSMD5 | 46.5671944 | 0.24147208 | 0.40242234 | 0.60004641 | 0.54847531 | 0.7554786 |
| CNTRL | 35.2857379 | 0.10613383 | 0.36890738 | 0.28769776 | 0.7735781 | 0.88776363 |
| RAB14 | 79.8067071 | 0.29690285 | 0.28648827 | 1.03635256 | 0.30003769 | 0.56012373 |
| GSN | 127.424275 | -0.1886381 | 0.41424703 | -0.4553757 | 0.64883894 | 0.81913126 |
| STOM | 87.9805774 | -1.6072212 | 0.42280316 | -3.8013463 | 0.00014391 | 0.00894448 |
| DAB2IP | 181.440983 | -0.6869509 | 0.34177046 | -2.0099774 | 0.04443358 | 0.21098411 |
| TTLL11 | 33.3056716 | -0.7818787 | 0.56772661 | -1.37721 | 0.16844734 | 0.41629954 |
| NDUFA8 | 27.3666247 | 0.9902624 | 0.38933684 | 2.5434593 | 0.01097608 | 0.10267384 |
| RBM18 | 17.5084454 | 0.22163175 | 0.52549063 | 0.42176157 | 0.67319906 | NA |
| MRRF | 98.0926367 | 0.27009494 | 0.22555467 | 1.19746995 | 0.23112344 | 0.48980096 |
| PDCL | 12.7980412 | 0.53182385 | 0.48981957 | 1.08575461 | 0.27758757 | NA |
| RC3H2 | 102.221415 | -0.5339469 | 0.23225513 | -2.2989671 | 0.02150681 | 0.14349647 |
| ZBTB26 | 18.1529088 | -0.0688129 | 0.43183286 | -0.1593507 | 0.87339256 | NA |
| RABGAP1 | 267.203307 | -0.3712016 | 0.31155167 | -1.1914607 | 0.23347278 | 0.49233972 |
| STRBP | 244.287204 | 0.54655974 | 0.18704321 | 2.92210418 | 0.00347675 | 0.05855703 |
| DENND1A | 298.33091 | 0.27069314 | 0.32153659 | 0.84187351 | 0.39985876 | 0.64766685 |
| NEK6 | 27.3100185 | 1.95179411 | 0.723733 | 2.69684275 | 0.00700003 | 0.08163522 |

| | | | | | | |
|---|---|---|---|---|---|---|
| PSMB7 | 66.6501331 | -0.3877865 | 0.33083883 | -1.1721313 | 0.24114435 | 0.49867735 |
| NR6A1 | 515.443862 | 0.17631487 | 0.20711011 | 0.85130984 | 0.39459726 | 0.6432568 |
| MIR181A2HG | 45.334718 | -0.0258262 | 0.44734896 | -0.0577316 | 0.9539624 | 0.98153155 |
| RPL35 | 51.2343725 | 0.01064018 | 0.5396616 | 0.0197164 | 0.98426961 | 0.99382437 |
| ARPC5L | 25.8006941 | -0.1680469 | 0.45822662 | -0.3667332 | 0.71381803 | 0.85788134 |
| GOLGA1 | 51.6891231 | -0.2176327 | 0.38656565 | -0.5629903 | 0.57344146 | 0.77380416 |
| SCAI | 66.3302099 | 1.13168181 | 0.32371004 | 3.49597377 | 0.00047234 | 0.01957967 |
| PPP6C | 70.5962795 | -0.3981495 | 0.28255688 | -1.4090952 | 0.15880701 | 0.40367819 |
| RABEPK | 20.534408 | 0.21381027 | 0.4892163 | 0.4370465 | 0.66207763 | NA |
| HSPA5 | 82.4411994 | -0.9978391 | 0.68156643 | -1.4640379 | 0.14318359 | 0.38160805 |
| GAPVD1 | 120.175034 | -0.4478474 | 0.26495309 | -1.6902894 | 0.0909726 | 0.30125735 |
| MAPKAP1 | 189.272332 | -0.389372 | 0.20394439 | -1.9092065 | 0.05623546 | 0.23639777 |
| PBX3 | 61.5164469 | 0.53676994 | 0.33062813 | 1.62348541 | 0.10448568 | 0.32467754 |
| MVB12B | 42.0290924 | 1.1376599 | 0.4431439 | 2.56724714 | 0.01025095 | 0.09994074 |
| ZBTB43 | 63.9212383 | -0.8132267 | 0.37789056 | -2.1520166 | 0.03139605 | 0.17484986 |
| ZBTB34 | 11.8693766 | 0.55846578 | 0.52344419 | 1.06690605 | 0.28601426 | NA |
| RALGPS1 | 221.921415 | 0.4455402 | 0.22612534 | 1.9703241 | 0.04880124 | 0.22160657 |
| GARNL3 | 24.1120708 | 0.6638876 | 0.41334645 | 1.60612872 | 0.10824563 | 0.33009771 |
| RPL12 | 22.559203 | -0.2108116 | 0.45299213 | -0.4653759 | 0.64166229 | 0.81566881 |
| LRSAM1 | 58.0300119 | 0.12325118 | 0.34020001 | 0.36229035 | 0.71713507 | 0.85900596 |
| FAM129B | 55.2234506 | -0.7455584 | 0.36995472 | -2.0152694 | 0.04387643 | 0.20924217 |
| STXBP1 | 37.9619209 | 0.62829564 | 0.35984433 | 1.74602065 | 0.08080736 | 0.28378572 |
| CDK9 | 18.8696534 | -0.1134111 | 0.45607386 | -0.2486683 | 0.80361736 | NA |
| FPGS | 24.1655205 | -0.6402228 | 0.41594376 | -1.5392052 | 0.12375421 | 0.35423434 |
| DPM2 | 26.208299 | -2.0140934 | 0.67132547 | -3.0001742 | 0.00269825 | 0.05029571 |
| FAM102A | 69.9019101 | -1.7589855 | 0.5261059 | -3.3434058 | 0.00082757 | 0.02565549 |
| SLC25A25 | 45.0245842 | -0.8475896 | 0.57588764 | -1.4717967 | 0.14107577 | 0.37904789 |
| SLC25A25-AS | 19.9521043 | 0.92565685 | 0.60032487 | 1.54192654 | 0.12309144 | NA |
| PTGES2 | 25.4543412 | 0.27955149 | 0.40921286 | 0.68314446 | 0.49451556 | 0.71857397 |
| C9orf16 | 20.3630755 | 0.45416945 | 0.41647071 | 1.09051955 | 0.27548435 | NA |
| CIZ1 | 25.0219009 | 0.18257817 | 0.4455195 | 0.4098096 | 0.68194562 | 0.84158016 |
| DNM1 | 17.9219257 | 0.8638581 | 0.68745027 | 1.25661176 | 0.20889425 | NA |
| GOLGA2 | 64.9230429 | -0.5281226 | 0.36224583 | -1.4579121 | 0.14486478 | 0.38449111 |
| TRUB2 | 12.7692108 | -0.3722709 | 0.51478923 | -0.7231521 | 0.46958646 | NA |
| COQ4 | 62.657346 | 0.5288448 | 0.33004246 | 1.60235381 | 0.10907739 | 0.33106021 |
| SLC27A4 | 27.251898 | 0.66653297 | 0.41004521 | 1.62551094 | 0.10405373 | 0.32385831 |
| URM1 | 19.237474 | 0.63247718 | 0.41984699 | 1.50644685 | 0.1319525 | NA |
| ODF2 | 39.6888447 | -0.5581482 | 0.32865858 | -1.6982616 | 0.0894584 | 0.29807849 |
| GLE1 | 38.0359716 | -0.3519012 | 0.33977884 | -1.0356773 | 0.30035273 | 0.56044599 |
| SPTAN1 | 99.402775 | 0.19217018 | 0.30604079 | 0.62792341 | 0.53005411 | 0.74381523 |
| WDR34 | 24.8801636 | 0.87193919 | 0.47112783 | 1.85074863 | 0.06420572 | 0.25217177 |
| SET | 54.4057356 | -0.1499378 | 0.40311125 | -0.3719513 | 0.7099291 | 0.85607462 |
| ZER1 | 116.511557 | -0.0174322 | 0.27066047 | -0.0644062 | 0.94864682 | 0.97959746 |
| TBC1D13 | 18.3551919 | -0.1328407 | 0.43925717 | -0.3024212 | 0.76233096 | NA |
| KYAT1 | 27.511961 | 0.28584694 | 0.39664303 | 0.72066548 | 0.47111536 | 0.70047809 |
| PHYHD1 | 24.881884 | 0.38640877 | 0.45685907 | 0.84579425 | 0.39766753 | 0.64588458 |
| NUP188 | 24.4681277 | -0.795042 | 0.39906953 | -1.9922393 | 0.04634481 | 0.21579917 |
| SH3GLB2 | 34.8302046 | -0.0577471 | 0.48333965 | -0.1194751 | 0.90489897 | 0.95673178 |
| MIGA2 | 32.1967279 | -0.1433171 | 0.39414612 | -0.3636142 | 0.71614614 | 0.85886208 |
| PTPA | 49.7975382 | -0.5852929 | 0.50754192 | -1.1531912 | 0.24883192 | 0.50494577 |
| IER5L | 27.9645211 | -0.6314314 | 0.49600291 | -1.2730397 | 2.03E-01 | 4.57E-01 |
| LINC01503 | 108.605252 | -0.6248769 | 0.48893199 | -1.2780445 | 2.01E-01 | 0.45528981 |
| NTMT1 | 24.7969138 | -0.3405625 | 0.4240951 | -0.8030333 | 0.42195549 | 0.66539467 |
| ASB6 | 15.4028368 | -0.3114108 | 0.55117934 | -0.56499 | 0.57208055 | NA |

| | | | | | | |
|---|---|---|---|---|---|---|
| TOR1B | 16.108762 | 0.59390694 | 0.48791143 | 1.21724334 | 0.22351165 | NA |
| C9orf78 | 17.3131959 | 0.09951039 | 0.4617694 | 0.21549802 | 0.82937906 | NA |
| USP20 | 29.713707 | -0.4536348 | 0.35342141 | -1.2835522 | 0.19929869 | 0.45290877 |
| FNBP1 | 96.1086671 | 0.00288939 | 0.2620842 | 0.01102465 | 0.99120378 | 0.99583841 |
| GPR107 | 89.4410447 | -0.2612353 | 0.23861857 | -1.0947819 | 0.27361221 | 0.53266538 |
| NCS1 | 19.3382689 | -0.6225111 | 0.4192762 | -1.484728 | 0.13761589 | NA |
| ASS1 | 15.4299904 | 0.19056826 | 0.46339184 | 0.41124648 | 0.68089181 | NA |
| FUBP3 | 84.7671414 | -0.3956219 | 0.32686783 | -1.2103422 | 0.2261476 | 0.48391941 |
| ABL1 | 197.59864 | -0.0786846 | 0.21547499 | -0.365168 | 0.71498603 | 0.85843895 |
| AIF1L | 129.812349 | 1.01952749 | 0.48252528 | 2.11289964 | 0.03460936 | 0.18447938 |
| NUP214 | 186.564994 | -0.3212649 | 0.22620052 | -1.4202662 | 0.15553021 | 0.39879246 |
| AL157938.2 | 41.1627518 | 0.81513349 | 0.650226 | 1.25361565 | 0.20998174 | 0.46499349 |
| PRRC2B | 200.442127 | -0.4203936 | 0.27285654 | -1.5407129 | 0.12338667 | 0.35349809 |
| POMT1 | 19.5606434 | 0.78340734 | 0.44029853 | 1.77926404 | 0.07519648 | NA |
| RAPGEF1 | 181.100513 | -0.9804262 | 0.4752431 | -2.0629993 | 0.0391127 | 0.19721202 |
| MED27 | 119.873105 | -0.6751517 | 0.24890314 | -2.7125077 | 0.00667762 | 0.07974991 |
| SETX | 179.850419 | -1.0837249 | 0.25217361 | -4.2975349 | 1.73E-05 | 0.00225311 |
| TTF1 | 31.7316043 | -0.3154079 | 0.42149635 | -0.7483051 | 0.45427612 | 0.690202 |
| DDX31 | 36.2585207 | -0.4613597 | 0.36130843 | -1.2769137 | 0.20163273 | 0.45567274 |
| GTF3C4 | 13.7218033 | -0.0582154 | 0.48162014 | -0.120874 | 0.90379082 | NA |
| AK8 | 15.0062988 | 0.9369784 | 0.82314953 | 1.13828457 | 0.25500168 | NA |
| TSC1 | 122.962601 | -0.4291131 | 0.23370629 | -1.8361215 | 0.06633969 | 0.25673827 |
| GTF3C5 | 25.8390614 | -0.642228 | 0.49617065 | -1.2943691 | 0.19553796 | 0.44873454 |
| RALGDS | 17.0285509 | -1.4362663 | 0.46879281 | -3.0637549 | 0.00218578 | NA |
| ABO | 49.7235243 | 1.38947083 | 0.62727793 | 2.21508004 | 0.02675457 | 0.16063349 |
| RPL7A | 54.5234461 | 0.3086009 | 0.47780159 | 0.64587667 | 0.51835923 | 0.73536015 |
| SURF1 | 12.9255765 | 0.87529595 | 0.53272668 | 1.64304884 | 0.10037283 | NA |
| SURF4 | 51.1876703 | -0.2300535 | 0.32233516 | -0.713709 | 0.47540716 | 0.70355873 |
| REXO4 | 15.002799 | 0.30657053 | 0.4795694 | 0.63926207 | 0.52265246 | NA |
| VAV2 | 143.060165 | -0.6744133 | 0.40139559 | -1.6801712 | 0.09292402 | 0.30571879 |
| BRD3 | 30.752463 | 0.31283397 | 0.41089167 | 0.76135387 | 0.44644573 | 0.68629774 |
| WDR5 | 22.5174657 | 0.06312943 | 0.42388188 | 0.14893164 | 0.88160757 | 0.94443055 |
| RXRA | 24.5866007 | 0.8357305 | 0.40315613 | 2.07296984 | 0.03817509 | 0.19510866 |
| PPP1R26-AS1 | 41.0896778 | 0.07584841 | 0.35811222 | 0.21180068 | 0.83226254 | 0.91853705 |
| PPP1R26 | 15.0660768 | 0.67412818 | 0.61878818 | 1.08943286 | 0.27596305 | NA |
| CAMSAP1 | 43.5114404 | -1.0207519 | 0.34926381 | -2.9225813 | 0.00347143 | 0.05855703 |
| UBAC1 | 38.9131906 | 0.64334846 | 0.36859552 | 1.74540497 | 0.0809144 | 0.28382032 |
| NACC2 | 40.4582421 | -0.3587004 | 0.39092786 | -0.9175615 | 0.35884846 | 0.61515737 |
| SNAPC4 | 19.2453278 | -1.0713821 | 0.50325957 | -2.1288857 | 0.03326372 | NA |
| ENTR1 | 20.8389557 | -0.6198399 | 0.40808325 | -1.5189055 | 0.12878628 | NA |
| PMPCA | 24.1087098 | -0.1645555 | 0.43314802 | -0.379906 | 0.70401522 | 0.85295217 |
| SEC16A | 53.0140477 | -0.5517519 | 0.42853452 | -1.287532 | 0.19790893 | 0.45094881 |
| NOTCH1 | 33.0539429 | 0.21702052 | 0.39995153 | 0.54261706 | 0.58739349 | 0.78270395 |
| AGPAT2 | 34.6603994 | 1.01773078 | 0.48194915 | 2.11169742 | 0.03471241 | 0.18490064 |
| SNHG7 | 27.1495571 | 0.87746224 | 0.36082174 | 2.43184415 | 0.01502217 | 0.12124045 |
| RABL6 | 64.6881318 | -0.094068 | 0.3834986 | -0.245289 | 0.80623267 | 0.9057619 |
| PHPT1 | 35.9072089 | 1.19126094 | 0.52409874 | 2.27297045 | 0.02302796 | 0.14770518 |
| EDF1 | 45.2480673 | -0.1777441 | 0.37126162 | -0.478757 | 0.63211152 | 0.80927518 |
| FBXW5 | 32.0286323 | 0.01427174 | 0.58482065 | 0.02440362 | 0.98053066 | 0.99299791 |
| ABCA2 | 18.5788815 | 2.23313582 | 0.71146308 | 3.13879366 | 0.00169645 | NA |
| UAP1L1 | 12.7941572 | 0.03619842 | 0.66796689 | 0.05419194 | 0.95678224 | NA |
| MAN1B1 | 24.6720454 | 0.61270552 | 0.3701795 | 1.65515789 | 0.09789248 | 0.31473617 |
| DPP7 | 22.9391929 | -0.5166512 | 0.56724088 | -0.9108144 | 0.36239315 | 0.61820481 |
| SSNA1 | 16.4567644 | 0.59161804 | 0.51834459 | 1.14136051 | 0.25371993 | NA |

| | | | | | | |
|---|---|---|---|---|---|---|
| TPRN | 15.0658266 | -0.6397439 | 0.50956578 | -1.2554688 | 0.20930862 | NA |
| TUBB4B | 33.2637598 | 0.19420067 | 0.49768455 | 0.39020836 | 0.69638248 | 0.85012783 |
| NELFB | 31.5383496 | 0.27467752 | 0.40699478 | 0.67489201 | 0.49974438 | 0.72203669 |
| EXD3 | 58.0634741 | 0.02408061 | 0.37092943 | 0.06491965 | 0.94823797 | 0.97942485 |
| NOXA1 | 16.5789057 | 1.17544697 | 0.54186973 | 2.16924271 | 0.03006426 | NA |
| PNPLA7 | 25.8269 | 0.63004162 | 0.61913497 | 1.01761595 | 0.3088605 | 0.56849049 |
| MRPL41 | 43.669084 | 0.55502025 | 0.56911825 | 0.97522833 | 0.32944701 | 0.58535385 |
| DPH7 | 27.726656 | -0.1434106 | 0.36938863 | -0.3882378 | 0.69784003 | 0.8508196 |
| ARRDC1 | 20.258175 | 0.50818996 | 0.47784254 | 1.06350925 | 0.28755107 | NA |
| EHMT1 | 227.141571 | -0.4246302 | 0.26925153 | -1.5770762 | 0.11477799 | 0.34109892 |
| BET1L | 13.110234 | -0.3290866 | 0.58608029 | -0.5615043 | 0.57445378 | NA |
| SIRT3 | 19.0945932 | 0.21227221 | 0.48187366 | 0.44051425 | 0.65956469 | NA |
| PSMD13 | 36.2888689 | -0.1515425 | 0.33595125 | -0.4510849 | 0.65192836 | 0.82061848 |
| IFITM2 | 104.864161 | -0.2877681 | 0.61812795 | -0.4655478 | 6.42E-01 | 0.81566881 |
| IFITM3 | 163.140897 | -0.3568647 | 0.3681995 | -0.9692157 | 3.32E-01 | 5.88E-01 |
| SIGIRR | 32.4861118 | -0.2726954 | 0.50208291 | -0.5431282 | 0.58704155 | 0.78268817 |
| PTDSS2 | 30.8874759 | -0.1915301 | 0.35769015 | -0.5354636 | 0.59232931 | 0.7855446 |
| RNH1 | 33.9452259 | 0.21632678 | 0.38752181 | 0.55823124 | 0.57668649 | 0.77438171 |
| RASSF7 | 49.9116539 | -0.1580334 | 0.45369201 | -0.3483276 | 0.72759419 | 0.86464296 |
| MIR210HG | 17.6474101 | -0.0249381 | 0.75476112 | -0.033041 | 0.97364187 | NA |
| PHRF1 | 32.985663 | -0.6886324 | 0.38609692 | -1.7835739 | 0.07449294 | 0.27225647 |
| DEAF1 | 87.9113058 | -0.4603989 | 0.28989617 | -1.5881511 | 0.11225218 | 0.33671279 |
| AC131934.1 | 17.2512641 | 0.23369057 | 0.46102875 | 0.50688937 | 0.61223244 | NA |
| EPS8L2 | 99.1112683 | 0.14618838 | 0.37103415 | 0.39400248 | 0.69357921 | 0.84791601 |
| AP006621.4 | 17.5531008 | 0.76375792 | 0.65540511 | 1.16532189 | 0.24388872 | NA |
| TALDO1 | 40.3945544 | 0.35256043 | 0.38039383 | 0.92683006 | 0.35401477 | 0.61144431 |
| GATD1 | 21.3663112 | -0.2882821 | 0.42855993 | -0.6726763 | 0.50115323 | 0.72289663 |
| AP006621.3 | 21.3651553 | 0.75348577 | 0.47277667 | 1.59374568 | 0.11099301 | 0.33449169 |
| PIDD1 | 10.9528188 | 0.56403551 | 0.59909559 | 0.94147832 | 0.34645979 | NA |
| RPLP2 | 59.3040436 | -0.0787125 | 0.39426505 | -0.1996437 | 0.84175922 | 0.92367136 |
| PNPLA2 | 25.4750788 | 0.1361868 | 0.37497796 | 0.36318615 | 0.71646583 | 0.8589283 |
| CD151 | 51.0543554 | 0.00956868 | 0.5049415 | 0.01895007 | 0.98488094 | 0.99407122 |
| POLR2L | 33.7095142 | 0.76197013 | 0.41656681 | 1.82916668 | 0.06737464 | 0.25898011 |
| TSPAN4 | 16.7763212 | -0.4904078 | 0.4742635 | -1.0340407 | 0.30111715 | NA |
| CHID1 | 16.9363243 | -0.2914185 | 0.45030651 | -0.6471559 | 0.51753102 | NA |
| AP2A2 | 213.387094 | 0.42311802 | 0.30956372 | 1.36682044 | 0.17168159 | 0.42016955 |
| TOLLIP | 24.8087093 | -0.6351929 | 0.41151221 | -1.5435578 | 0.1226955 | 0.35251484 |
| MOB2 | 26.8844042 | -0.5355428 | 0.41079741 | -1.3036665 | 0.19234732 | 0.44417846 |
| MRPL23 | 12.3062637 | -0.0473118 | 0.61296251 | -0.0771855 | 0.93847598 | NA |
| CD81 | 106.060809 | 0.90884379 | 0.5033604 | 1.80555281 | 0.07098822 | 0.26639826 |
| KCNQ1 | 273.010871 | -0.0066948 | 0.56558865 | -0.0118369 | 0.99055575 | 0.99583841 |
| KCNQ1OT1 | 215.548487 | 0.386036 | 0.2963876 | 1.30247013 | 1.93E-01 | 0.44446998 |
| NAP1L4 | 248.413208 | 0.12367196 | 0.16568686 | 0.74641983 | 0.45541385 | 0.6908246 |
| CARS | 45.8789544 | -0.6877205 | 0.48855145 | -1.4076726 | 0.15922805 | 0.40403105 |
| OSBPL5 | 20.8979406 | 0.98619384 | 0.51316885 | 1.92177259 | 0.05463438 | NA |
| ZNF195 | 31.3777746 | 0.31516908 | 0.35731513 | 0.88204795 | 0.37775088 | 0.62906718 |
| AC127526.1 | 16.3952112 | -0.7842616 | 0.52193133 | -1.5026145 | 0.13293849 | NA |
| NUP98 | 147.129207 | -0.6063048 | 0.36970225 | -1.6399812 | 0.10100907 | 0.32011506 |
| PGAP2 | 18.9140683 | 1.10511097 | 0.70072422 | 1.57709828 | 0.11477291 | NA |
| STIM1 | 44.9877195 | 0.28465151 | 0.52482647 | 0.54237262 | 5.88E-01 | 0.78270395 |
| RRM1 | 15.1900694 | 0.78600277 | 0.72849532 | 1.07894003 | 0.28061446 | NA |
| AC104389.4 | 39.0289332 | 2.09247683 | 0.73055944 | 2.86421162 | 0.00418049 | 0.06301187 |
| TRIM5 | 15.3830785 | 0.72106061 | 0.93884922 | 0.768026 | 0.44247174 | NA |
| TRIM22 | 27.437691 | 0.33440487 | 1.17414502 | 0.28480712 | 0.77579191 | 0.88880003 |

| | | | | | | |
|---|---|---|---|---|---|---|
| FAM160A2 | 27.9971609 | -0.222276 | 0.37592226 | -0.5912819 | 0.55433152 | 0.76082199 |
| SMPD1 | 13.0310287 | 0.14860991 | 0.67981027 | 0.21860498 | 0.82695778 | NA |
| APBB1 | 23.1298062 | -0.2890052 | 0.38371563 | -0.7531755 | 0.45134446 | 0.68853766 |
| ARFIP2 | 13.6560651 | -1.1031944 | 0.55099432 | -2.0021884 | 0.04526447 | NA |
| AC084337.2 | 34.7696652 | -0.8007627 | 0.5603532 | -1.429032 | 0.15299504 | 0.3947713 |
| DNHD1 | 47.4385789 | -0.4964198 | 0.4402252 | -1.1276497 | 0.25946789 | 0.51818585 |
| ILK | 35.9615879 | 0.26553823 | 0.37228729 | 0.7132616 | 0.47568389 | 0.70369766 |
| TPP1 | 28.6311946 | -0.6420209 | 0.38552089 | -1.6653337 | 0.0958462 | 0.31098513 |
| AC107884.1 | 29.3519109 | -0.0339976 | 0.48013695 | -0.0708082 | 0.94355039 | 0.97679991 |
| PPFIBP2 | 26.1192414 | 0.31158043 | 0.50702781 | 0.61452336 | 0.53886954 | 0.74929145 |
| EIF3F | 30.4146591 | 0.22921141 | 0.44924593 | 0.51021365 | 0.60990179 | 0.79660853 |
| TUB | 18.4374249 | -0.4187106 | 0.55332677 | -0.7567149 | 0.44922069 | NA |
| RIC3 | 99.1903999 | -0.3849164 | 0.4471031 | -0.860912 | 0.38928651 | 0.63833368 |
| STK33 | 164.80753 | 1.23119893 | 0.58543919 | 2.10303469 | 0.03546274 | 0.18721309 |
| TRIM66 | 70.9742545 | 0.15826151 | 0.54377544 | 0.29104203 | 0.77101917 | 0.88667781 |
| RPL27A | 66.9237513 | 0.30203607 | 0.36353811 | 0.83082368 | 0.40607325 | 0.65318894 |
| ST5 | 380.158896 | -0.5593643 | 0.41818178 | -1.3376104 | 1.81E-01 | 4.30E-01 |
| AC026894.1 | 31.4460778 | 0.50520916 | 0.48798738 | 1.03529143 | 3.01E-01 | 5.61E-01 |
| TMEM9B | 27.2895763 | 0.26509049 | 0.4516198 | 0.58697714 | 0.55721906 | 0.763236 |
| TMEM9B-AS | 24.2763808 | 0.93712837 | 0.68351214 | 1.37104861 | 0.1703598 | 0.41839803 |
| DENND5A | 129.35294 | -0.6735069 | 0.27724043 | -2.4293243 | 0.01512699 | 0.12124045 |
| TMEM41B | 129.042732 | -1.6537446 | 0.34538808 | -4.7880767 | 1.68E-06 | 0.00051843 |
| IPO7 | 105.309235 | -0.6582936 | 0.36448955 | -1.8060698 | 0.07090744 | 0.26623151 |
| ZNF143 | 33.7417242 | -0.5541255 | 0.41307682 | -1.3414588 | 0.17977154 | 0.42896282 |
| WEE1 | 19.3088186 | -1.9003551 | 0.52578092 | -3.6143478 | 0.0003011 | NA |
| SWAP70 | 72.1463408 | -0.2425341 | 0.2619003 | -0.9260549 | 3.54E-01 | 6.12E-01 |
| SBF2-AS1 | 55.8795176 | -1.5987313 | 0.57198883 | -2.7950393 | 0.00518934 | 0.07024781 |
| SBF2 | 718.931018 | 0.20590963 | 0.30085773 | 0.68440864 | 0.49371716 | 0.71768479 |
| AC011092.2 | 27.6509919 | -0.2544716 | 0.58036019 | -0.4384718 | 0.66104434 | 0.82759568 |
| ADM | 41.2731917 | -1.5830542 | 0.65519061 | -2.4161735 | 0.01568458 | 0.12386371 |
| AMPD3 | 73.1511127 | -1.0552992 | 0.45973482 | -2.295452 | 0.02170724 | 0.14366346 |
| MTRNR2L8 | 50.6403271 | 0.5584587 | 0.74878992 | 0.74581493 | 0.45577922 | 0.6908246 |
| RNF141 | 19.0710426 | 0.12380611 | 0.45810625 | 0.27025632 | 0.78696306 | NA |
| CTR9 | 59.9972663 | 0.08847609 | 0.29262266 | 0.30235555 | 0.76238103 | 0.88247846 |
| EIF4G2 | 102.807023 | -0.4015913 | 0.2701819 | -1.4863739 | 0.13718026 | 0.37299788 |
| ZBED5 | 64.6665959 | -0.2833149 | 0.30284313 | -0.935517 | 0.34952193 | 0.60618979 |
| ZBED5-AS1 | 14.2419243 | 1.80881469 | 0.6938816 | 2.60680594 | 0.00913911 | NA |
| GALNT18 | 232.109428 | -0.8768693 | 0.47837243 | -1.8330264 | 0.06679865 | 0.25797753 |
| USP47 | 264.744731 | -0.1318042 | 0.20315913 | -0.648773 | 0.51648511 | 0.73427889 |
| PARVA | 118.972063 | 0.02314946 | 0.21575536 | 0.10729494 | 0.914555 | 0.96388328 |
| TEAD1 | 487.901715 | -1.3825642 | 0.40953471 | -3.3759389 | 0.00073564 | 0.02425576 |
| LINC00958 | 68.3505839 | -0.9439708 | 0.33019883 | -2.8587949 | 0.00425254 | 0.06301187 |
| ARNTL | 33.2554854 | -0.02343 | 0.4101402 | -0.0571268 | 0.95444418 | 0.98153155 |
| BTBD10 | 77.2651232 | -0.5340662 | 0.28050249 | -1.9039626 | 0.05691506 | 0.23751276 |
| FAR1 | 114.650246 | 0.28168009 | 0.25454274 | 1.10661218 | 0.26846164 | 0.52726441 |
| RRAS2 | 92.5328376 | -0.972717 | 0.36350075 | -2.6759696 | 0.00745134 | 0.08459138 |
| COPB1 | 75.4600479 | 0.27930009 | 0.285764 | 0.97738023 | 0.32838094 | 0.58467456 |
| PSMA1 | 31.1283288 | 1.34482779 | 0.68560591 | 1.96151722 | 0.04981872 | 0.2242425 |
| PDE3B | 70.144161 | 0.84941607 | 0.68630957 | 1.23765733 | 0.21584315 | 0.47170491 |
| CYP2R1 | 15.1068618 | 0.589233 | 0.53926433 | 1.09266081 | 0.27454275 | NA |
| SOX6 | 772.54488 | 1.21105294 | 0.34962835 | 3.46382933 | 0.00053254 | 0.02088709 |
| C11orf58 | 107.611354 | -0.1888404 | 0.20097909 | -0.9396024 | 0.34742157 | 0.60403983 |
| PLEKHA7 | 512.443447 | -0.1410026 | 0.1905923 | -0.7398129 | 0.45941354 | 0.69297787 |
| RPS13 | 42.4592213 | 0.28757883 | 0.40825268 | 0.70441381 | 0.48117511 | 0.7087866 |

| | | | | | | |
|---|---|---|---|---|---|---|
| PIK3C2A | 327.386909 | -0.6649922 | 0.144805 | -4.5923292 | 4.38E-06 | 0.00096394 |
| NUCB2 | 40.2264111 | 0.48464173 | 0.4183431 | 1.15847908 | 0.24666858 | 0.50296216 |
| NCR3LG1 | 13.7865646 | -0.1181032 | 0.80389502 | -0.1469137 | 0.88320011 | NA |
| SERGEF | 156.136849 | -0.6317759 | 0.24128885 | -2.6183384 | 0.00883591 | 0.09290987 |
| SAAL1 | 13.1121756 | 0.36820246 | 0.55074694 | 0.66855109 | 0.50378188 | NA |
| HPS5 | 32.531982 | -0.965475 | 0.35258499 | -2.7382759 | 0.00617622 | 0.07721978 |
| GTF2H1 | 60.9653522 | -0.5640111 | 0.32883904 | -1.7151585 | 0.08631617 | 0.29358176 |
| LDHA | 134.896764 | 0.10836049 | 0.31871179 | 0.33999522 | 7.34E-01 | 8.68E-01 |
| TSG101 | 94.4002024 | -0.6334888 | 0.26199897 | -2.4179058 | 0.01561012 | 0.12348518 |
| UEVLD | 78.9865011 | 0.16779691 | 0.24817457 | 0.67612451 | 0.4989616 | 0.72176422 |
| SPTY2D1OS | 34.1685336 | 0.64452449 | 0.40431978 | 1.59409585 | 0.11091457 | 0.3343946 |
| SPTY2D1 | 23.8016502 | -0.2021547 | 0.38485438 | -0.5252758 | 0.59939146 | 0.78985038 |
| ZDHHC13 | 93.6707028 | -0.5148264 | 0.36840521 | -1.397446 | 0.1622795 | 0.40832472 |
| NAV2 | 502.276116 | -0.8453224 | 0.26549824 | -3.1839095 | 0.001453 | 0.03484042 |
| HTATIP2 | 16.0265121 | 0.43208534 | 0.58083731 | 0.7439008 | 0.45693649 | NA |
| PRMT3 | 68.6428297 | -0.1778736 | 0.26573026 | -0.6693765 | 0.50325535 | 0.72497781 |
| NELL1 | 235.669317 | 0.51361575 | 0.64725119 | 0.79353389 | 4.27E-01 | 6.70E-01 |
| ANO5 | 63.5974099 | 1.01555871 | 0.31131359 | 3.26217275 | 0.00110562 | 0.02988759 |
| GAS2 | 22.8955186 | 1.38006986 | 0.79376547 | 1.73863681 | 0.08209867 | 0.28580436 |
| SVIP | 42.9538622 | 0.46019211 | 0.37107521 | 1.24015858 | 0.21491674 | 0.47074963 |
| SLC5A12 | 12.0081829 | 1.77175888 | 1.05388048 | 1.68117629 | 0.09272868 | NA |
| BBOX1 | 26.8235872 | 1.25339913 | 0.82953892 | 1.5109588 | 0.13079895 | 0.36326737 |
| CCDC34 | 16.1757749 | 1.48456826 | 0.69210873 | 2.14499282 | 0.03195342 | NA |
| LGR4 | 1172.95804 | -0.6808049 | 0.21375696 | -3.1849482 | 0.0014478 | 0.03482409 |
| LIN7C | 26.6444149 | 0.10700191 | 0.42374425 | 0.2525153 | 0.80064279 | 0.90333444 |
| BDNF-AS | 105.938953 | 0.53301944 | 0.35309326 | 1.50957127 | 0.13115286 | 0.36377786 |
| METTL15 | 261.668333 | 0.69060901 | 0.3734589 | 1.84922362 | 0.06442553 | 0.25274378 |
| ARL14EP | 30.9142361 | 0.27653427 | 0.37050723 | 0.74636675 | 0.4554459 | 0.6908246 |
| MPPED2 | 121.353186 | 0.30725152 | 0.74193104 | 0.4141241 | 0.67878323 | 0.84010202 |
| DCDC1 | 73.2336927 | 1.0877304 | 0.720835 | 1.50898666 | 0.13130219 | 0.36392977 |
| DNAJC24 | 45.4075028 | -0.3140302 | 0.29377993 | -1.0689301 | 0.28510117 | 0.54384727 |
| IMMP1L | 127.217453 | -0.010427 | 0.27050111 | -0.0385471 | 0.96925145 | 0.98781207 |
| ELP4 | 182.056035 | 0.3083133 | 0.25794364 | 1.19527393 | 0.23198002 | 0.49066933 |
| AC131571.1 | 19.2093921 | 1.45320419 | 0.655451 | 2.21710575 | 0.02661587 | NA |
| EIF3M | 44.1577201 | -0.1458441 | 0.27550702 | -0.5293662 | 0.59655147 | 0.78921565 |
| PRRG4 | 37.0300597 | 0.57650542 | 0.42429113 | 1.35874965 | 0.17422593 | 0.42295604 |
| QSER1 | 76.8298587 | 0.99864164 | 0.28015236 | 3.56463764 | 0.00036436 | 0.0164969 |
| TCP11L1 | 33.7063751 | 0.57138501 | 0.44559925 | 1.28228447 | 0.19974287 | 0.45365031 |
| CSTF3 | 80.7365937 | 0.23188844 | 0.3311934 | 0.70016021 | 0.48382726 | 0.71144851 |
| HIPK3 | 211.282332 | -0.3407125 | 0.25179776 | -1.3531196 | 0.17601741 | 0.42508357 |
| CD59 | 17.0488051 | 0.36951939 | 0.43009533 | 0.85915695 | 0.39025393 | NA |
| FBXO3 | 50.9906881 | 0.52870263 | 0.27994033 | 1.88862614 | 0.05894194 | 0.2411888 |
| CAPRIN1 | 108.521261 | -0.1973472 | 0.27813995 | -0.7095249 | 0.47799879 | 0.70576571 |
| NAT10 | 41.6109908 | -0.5414749 | 0.32803072 | -1.6506835 | 0.09880321 | 0.31647454 |
| ABTB2 | 365.858053 | -0.8882825 | 0.37586497 | -2.3633022 | 0.01811289 | 0.13341143 |
| CAT | 59.2706136 | 1.60020996 | 0.39828507 | 4.01775027 | 5.88E-05 | 0.00496976 |
| EHF | 19.5082558 | -0.8684425 | 0.68003744 | -1.277051 | 2.02E-01 | NA |
| APIP | 38.9950139 | -0.8576814 | 0.35745581 | -2.3994055 | 0.01642172 | 0.12723517 |
| PDHX | 106.613822 | -0.233083 | 0.21490398 | -1.0845913 | 0.2781027 | 0.53674936 |
| CD44 | 79.5472176 | -2.5556757 | 0.7852138 | -3.2547513 | 1.13E-03 | 3.01E-02 |
| TRIM44 | 241.913009 | -0.6481748 | 0.25440262 | -2.5478306 | 0.01083951 | 0.10244908 |
| COMMD9 | 16.9995059 | -0.0003499 | 0.48122128 | -0.0007271 | 0.99941985 | NA |
| TRAF6 | 39.3480578 | -0.1115406 | 0.37579378 | -0.2968133 | 0.76660909 | 0.88444965 |
| C11orf74 | 19.5667086 | 1.80095023 | 0.60980794 | 2.95330726 | 0.00314389 | NA |

| | | | | | | |
|---|---|---|---|---|---|---|
| API5 | 68.6035205 | -0.3642813 | 0.24100202 | -1.511528 | 0.13065399 | 0.36322236 |
| TTC17 | 352.745638 | 0.0015555 | 0.1616641 | 0.00962182 | 0.99232302 | 0.99646579 |
| HSD17B12 | 307.409285 | -0.0097998 | 0.18129083 | -0.0540557 | 0.95689082 | 0.98307376 |
| ALKBH3 | 30.1738126 | 0.48503238 | 0.41868884 | 1.15845548 | 0.24667821 | 0.50296216 |
| EXT2 | 88.1882375 | -0.7568497 | 0.30624561 | -2.4713815 | 0.01345922 | 0.11309562 |
| PRDM11 | 21.7203348 | 0.16562314 | 0.43657975 | 0.37936516 | 0.70441672 | 0.85317002 |
| SYT13 | 18.0174643 | -0.1800501 | 0.68831148 | -0.2615823 | 0.7936435 | NA |
| CRY2 | 40.2887345 | 0.09979118 | 0.32173497 | 0.31016579 | 0.75643489 | 0.87904624 |
| LARGE2 | 25.7103304 | 1.41236662 | 0.5369162 | 2.63051592 | 8.53E-03 | 0.0903651 |
| PHF21A | 329.353111 | 0.06672204 | 0.26084691 | 0.25579003 | 0.79811298 | 0.90153736 |
| DGKZ | 24.7940484 | -0.0873496 | 0.3710273 | -0.2354264 | 0.81387778 | 0.90970797 |
| AMBRA1 | 258.537676 | 0.08198154 | 0.27154902 | 0.30190329 | 0.76272579 | 0.88247846 |
| ATG13 | 56.8008328 | -1.088011 | 0.2983566 | -3.6466799 | 0.00026565 | 0.01346273 |
| ARHGAP1 | 35.70229 | -0.2222562 | 0.38027404 | -0.5844631 | 0.55890877 | 0.76485713 |
| CKAP5 | 72.2358832 | -0.1645804 | 0.2445915 | -0.6728787 | 0.50102443 | 0.72289663 |
| LRP4 | 45.5269082 | 0.38211807 | 0.53639739 | 0.7123787 | 0.4762303 | 0.70423528 |
| C11orf49 | 235.758015 | 0.39366016 | 0.2450755 | 1.60628115 | 0.10821215 | 0.33009771 |
| ARFGAP2 | 34.0563581 | -0.7026815 | 0.33796802 | -2.079136 | 0.03760485 | 0.19425568 |
| DDB2 | 19.2936611 | 1.33097605 | 0.55810555 | 2.38481062 | 0.01708792 | NA |
| NR1H3 | 21.4632531 | -0.5220736 | 0.53397041 | -0.9777201 | 0.3282128 | 0.58464566 |
| MADD | 31.964264 | -0.7854759 | 0.41940542 | -1.872832 | 0.06109158 | 0.24683566 |
| SLC39A13 | 26.626737 | 0.06949388 | 0.51987309 | 0.1336747 | 0.89365981 | 0.95191019 |
| CELF1 | 315.379314 | -0.3661152 | 0.2698087 | -1.3569436 | 0.17479913 | 0.42340329 |
| NDUFS3 | 20.7506029 | 0.72737851 | 0.45477888 | 1.59941137 | 0.10972923 | NA |
| MTCH2 | 65.0663711 | -0.5774341 | 0.31251582 | -1.8476957 | 0.06464638 | 0.25284322 |
| FNBP4 | 203.417738 | -0.6252285 | 0.31211274 | -2.0032137 | 0.04515435 | 0.21296142 |
| NUP160 | 100.760015 | -0.8106238 | 0.28962957 | -2.7988296 | 0.00512882 | 0.0697465 |
| PTPRJ | 215.272521 | -0.5826598 | 0.38588774 | -1.5099204 | 0.13106375 | 0.36366174 |
| TNKS1BP1 | 27.8587818 | -0.9311277 | 0.33729607 | -2.7605651 | 0.00577015 | 0.07451815 |
| SSRP1 | 16.3455941 | 0.74669376 | 0.55785851 | 1.33850027 | 0.18073341 | NA |
| SLC43A1 | 23.5441224 | -0.4661934 | 0.76643843 | -0.6082594 | 0.54301545 | 0.7517248 |
| SERPING1 | 107.010082 | -0.2792676 | 0.30055232 | -0.9291812 | 3.53E-01 | 6.10E-01 |
| ZDHHC5 | 55.3504283 | -0.8360126 | 0.3659303 | -2.2846224 | 0.02233498 | 0.14604538 |
| CTNND1 | 161.214744 | -0.2791299 | 0.21271725 | -1.3122107 | 0.18944904 | 0.44054056 |
| LPXN | 41.4831751 | 0.74837621 | 0.43309605 | 1.72796821 | 0.08399392 | 0.28888262 |
| FAM111A | 36.8309693 | 0.805353 | 0.5754987 | 1.39940021 | 0.161693 | 0.40751508 |
| OSBP | 82.4950933 | -0.5018589 | 0.28080965 | -1.7871854 | 0.07390753 | 0.27114691 |
| PATL1 | 58.5841729 | -0.3581078 | 0.31214739 | -1.1472393 | 0.25128272 | 0.50802481 |
| STX3 | 110.651137 | -0.9814268 | 0.4241681 | -2.3137684 | 0.02068043 | 0.14112911 |
| MRPL16 | 16.5678135 | 1.52532521 | 0.60765494 | 2.51018319 | 0.01206685 | NA |
| PRPF19 | 20.7543151 | -0.7694817 | 0.48702004 | -1.5799794 | 0.11411158 | NA |
| VPS37C | 30.6375582 | -0.927425 | 0.39152245 | -2.3687659 | 0.01784754 | 0.13283085 |
| DDB1 | 65.0515973 | -0.2772119 | 0.39614561 | -0.6997727 | 0.48406924 | 0.71145331 |
| TKFC | 10.6055044 | 0.97683731 | 0.59855953 | 1.6319802 | 0.10268366 | NA |
| CPSF7 | 68.0361642 | -0.523861 | 0.27134716 | -1.9305934 | 0.05353336 | 0.2322696 |
| AP003108.2 | 57.4859471 | -0.4439842 | 0.3520059 | -1.2612976 | 0.20720164 | 0.46066753 |
| SYT7 | 22.9911444 | -0.6929188 | 0.73795229 | -0.938975 | 0.34774357 | 0.60433106 |
| MYRF | 21.3869004 | 0.31523266 | 0.43349523 | 0.72718829 | 0.46711062 | 0.69693157 |
| TMEM258 | 18.0826864 | 0.24444859 | 0.45065658 | 0.54242766 | 0.58752393 | NA |
| FTH1 | 437.844526 | 0.16907817 | 0.43427227 | 0.38933679 | 6.97E-01 | 0.85041636 |
| ASRGL1 | 47.5310175 | 0.6319403 | 0.50777133 | 1.2445372 | 0.21330192 | 0.46890289 |
| AHNAK | 137.576044 | 0.10438746 | 0.34075615 | 0.30634066 | 0.75934529 | 0.88058098 |
| B3GAT3 | 24.9679895 | 0.08253868 | 0.43029148 | 0.19182039 | 0.8478829 | 0.92651313 |
| GANAB | 62.3323577 | -0.2270926 | 0.35261678 | -0.644021 | 0.51956179 | 0.73634084 |

| | | | | | | |
|---|---|---|---|---|---|---|
| UBXN1 | 17.4164934 | -0.670746 | 0.47550147 | -1.4106075 | 0.15836037 | NA |
| TTC9C | 32.6197258 | 0.25612471 | 0.36768545 | 0.69658646 | 0.48606163 | 0.71289302 |
| TAF6L | 15.2103234 | -0.6904525 | 0.58249008 | -1.1853464 | 0.23588052 | NA |
| TMEM223 | 16.805073 | 1.0760672 | 0.49333961 | 2.18118954 | 0.0291694 | NA |
| TMEM179B | 25.6488816 | 0.95105869 | 0.39473263 | 2.40937438 | 0.0159799 | 0.12499722 |
| NXF1 | 24.5315221 | 0.02962087 | 0.4719126 | 0.06276771 | 0.94995148 | 0.97993174 |
| STX5 | 60.7540059 | -0.7085872 | 0.35489329 | -1.9966204 | 0.04586644 | 0.2143497 |
| SLC3A2 | 52.3862929 | -0.8672806 | 0.57254212 | -1.5147891 | 0.12982583 | 0.3622089 |
| RARRES3 | 20.8079492 | 1.56719037 | 0.80041321 | 1.95797666 | 0.05023275 | NA |
| ATL3 | 57.2124775 | 0.18931359 | 0.2777427 | 0.681615 | 0.49548243 | 0.71943562 |
| RTN3 | 145.762417 | 0.55449486 | 0.27961566 | 1.98306084 | 0.04736063 | 0.21737317 |
| MARK2 | 90.0120241 | -0.4101149 | 0.25803167 | -1.5893974 | 0.1119707 | 0.33639285 |
| NAA40 | 31.8975778 | 0.50304422 | 0.36652522 | 1.37246821 | 0.16991773 | 0.41832117 |
| OTUB1 | 24.9252366 | -0.3667156 | 0.40178534 | -0.9127152 | 0.36139237 | 0.61731848 |
| MACROD1 | 46.7112957 | 0.99266925 | 0.43997447 | 2.25619738 | 0.02405827 | 0.15136029 |
| STIP1 | 34.0161961 | -0.1446729 | 0.3766164 | -0.3841387 | 0.70087562 | 0.85187889 |
| NUDT22 | 16.5053084 | 0.41717954 | 0.45710182 | 0.91266218 | 0.36142024 | NA |
| DNAJC4 | 13.2935554 | 1.19418717 | 0.74757824 | 1.59740762 | 0.11017488 | NA |
| VEGFB | 41.846796 | -0.1628025 | 0.42499424 | -0.3830699 | 0.70166794 | 0.85190818 |
| FKBP2 | 28.2134539 | 0.33124146 | 0.48285756 | 0.68600243 | 0.49271158 | 0.7169 |
| PPP1R14B | 17.0652469 | 0.25123065 | 0.73266995 | 0.34289744 | 0.73167562 | NA |
| GPR137 | 19.2248265 | 0.0073135 | 0.60639369 | 0.01206065 | 0.99037723 | NA |
| ESRRA | 59.635548 | -0.00559 | 0.44663329 | -0.0125159 | 0.99001399 | 0.99583841 |
| TRMT112 | 20.1991598 | 0.0633128 | 0.56241953 | 0.11257219 | 0.91036973 | NA |
| PRDX5 | 28.793549 | 0.53580662 | 0.51894992 | 1.03248232 | 0.30184623 | 0.56201529 |
| SF1 | 170.625939 | -0.5025444 | 0.23776445 | -2.1136229 | 0.03454749 | 0.18447938 |
| ATG2A | 20.8852167 | -1.2009041 | 0.56332493 | -2.1318142 | 0.03302213 | NA |
| VPS51 | 43.4775457 | 0.03309538 | 0.32026875 | 0.10333629 | 0.91769608 | 0.96416963 |
| FAU | 61.0085393 | 0.17813355 | 0.40157859 | 0.44358329 | 0.65734389 | 0.8248412 |
| SYVN1 | 14.4613276 | 0.69283714 | 0.55885953 | 1.23973397 | 0.21507381 | NA |
| CAPN1 | 38.4322325 | 0.56934885 | 0.37623244 | 1.51329018 | 0.13020597 | 0.36258875 |
| DPF2 | 31.4969125 | 0.07840904 | 0.36952465 | 0.21218893 | 0.83195964 | 0.91834123 |
| FRMD8 | 26.846359 | -1.0518128 | 0.42267891 | -2.4884439 | 0.01283035 | 0.11046442 |
| NEAT1 | 11322.8536 | -0.441471 | 0.5105076 | -0.8647688 | 3.87E-01 | 0.63701577 |
| MALAT1 | 115875.25 | -0.3717227 | 0.37816649 | -0.9829606 | 0.32562685 | 0.58154857 |
| SCYL1 | 22.5938581 | -0.6616458 | 0.49819448 | -1.3280874 | 0.18414924 | 0.43435248 |
| LTBP3 | 46.8521586 | -0.1777587 | 0.3654307 | -0.4864362 | 0.6266579 | 0.8053173 |
| EHBP1L1 | 18.0200428 | -1.0952736 | 0.53020398 | -2.0657589 | 3.89E-02 | NA |
| MAP3K11 | 14.1726898 | -0.3672343 | 0.49157005 | -0.7470639 | 0.45502497 | NA |
| PCNX3 | 13.9811816 | -0.3722593 | 0.59699153 | -0.6235588 | 0.53291739 | NA |
| RELA | 18.0589799 | -0.3418292 | 0.4725565 | -0.7233616 | 0.46945777 | NA |
| KAT5 | 24.4037096 | -0.7368152 | 0.4218912 | -1.7464579 | 0.08073142 | 0.28378572 |
| CFL1 | 49.1473857 | 0.07201024 | 0.46179951 | 0.155934 | 0.87608505 | 0.94205457 |
| SART1 | 20.1335645 | -1.076475 | 0.41293162 | -2.6069086 | 0.00913637 | NA |
| BANF1 | 17.4884904 | 0.24650488 | 0.55629891 | 0.44311587 | 0.65768192 | NA |
| SF3B2 | 44.2094673 | -0.7870639 | 0.33499852 | -2.3494548 | 0.01880093 | 0.13549695 |
| PACS1 | 341.29079 | 0.04437866 | 0.20009883 | 0.2217837 | 0.82448227 | 0.91529534 |
| RAB1B | 20.4491074 | 0.0787669 | 0.473305 | 0.16641891 | 0.8678273 | NA |
| SLC29A2 | 18.6369246 | 0.33877464 | 0.70817753 | 0.4783753 | 0.63238311 | NA |
| ZDHHC24 | 13.6702739 | 0.87045136 | 0.48657897 | 1.78892105 | 0.07362753 | NA |
| CTSF | 14.7176468 | 1.857941 | 0.67306973 | 2.76039896 | 0.00577308 | NA |
| CCS | 23.5212401 | 0.33364517 | 0.48623708 | 0.68617797 | 0.49260089 | 0.71687447 |
| RBM14-RBM | 42.9060876 | -0.127281 | 0.40119024 | -0.3172585 | 0.75104747 | 0.87747607 |
| RBM4 | 22.2166427 | -0.1257741 | 0.46547988 | -0.2702031 | 0.78700404 | 0.89489881 |

| | | | | | | |
|---|---|---|---|---|---|---|
| RBM4B | 30.2179163 | 0.1305019 | 0.38318179 | 0.34057438 | 0.73342402 | 0.86776236 |
| SPTBN2 | 40.9589429 | 1.0581996 | 0.54757948 | 1.93250411 | 0.05329732 | 0.23189906 |
| C11orf80 | 173.354274 | 0.72826784 | 0.42850422 | 1.69955814 | 0.08921407 | 0.29794449 |
| RHOD | 12.7630504 | 0.43752694 | 0.53162085 | 0.82300561 | 0.4105048 | NA |
| KDM2A | 344.063699 | -0.2656851 | 0.17665926 | -1.5039409 | 0.13259659 | 0.3666913 |
| GRK2 | 18.3391467 | -0.2539634 | 0.48021717 | -0.5288512 | 0.5969087 | NA |
| ANKRD13D | 26.8930915 | 0.43817242 | 0.37516668 | 1.16794068 | 0.24283069 | 0.50005426 |
| RAD9A | 114.131735 | 1.10377628 | 0.381094 | 2.89633606 | 0.00377548 | 0.06101472 |
| PPP1CA | 23.981482 | 0.41721513 | 0.52898529 | 0.78870839 | 0.43028247 | 0.67197671 |
| RPS6KB2 | 14.5396221 | 0.06182907 | 0.61554971 | 0.10044529 | 0.91999082 | NA |
| TMEM134 | 22.3590303 | 0.47020705 | 0.4698485 | 1.00076313 | 0.31694134 | 0.5746755 |
| PITPNM1 | 17.9032449 | -0.3990814 | 0.69237852 | -0.5763919 | 0.56435029 | NA |
| CDK2AP2 | 18.0395825 | -0.1907328 | 0.56422352 | -0.3380448 | 0.73532946 | NA |
| GSTP1 | 111.106855 | 0.78253222 | 0.54042963 | 1.44798172 | 0.14762216 | 0.38722988 |
| NDUFV1 | 58.1383591 | 1.01013987 | 0.4332244 | 2.33167813 | 0.01971763 | 0.13748641 |
| NDUFS8 | 43.5507922 | 0.60560484 | 0.4099261 | 1.47735125 | 0.13958151 | 0.37696802 |
| CHKA | 150.62549 | -0.9700542 | 0.41475085 | -2.3388843 | 0.01934142 | 0.1366689 |
| KMT5B | 132.371231 | 0.05125549 | 0.2104059 | 0.24360292 | 0.80753837 | 0.90686064 |
| LRP5 | 36.2764106 | 0.78144217 | 0.41855497 | 1.8670001 | 0.06190157 | 0.24880231 |
| PPP6R3 | 443.30396 | -0.3156307 | 0.24585228 | -1.2838224 | 0.1992041 | 0.45282752 |
| CPT1A | 115.499258 | 1.16964875 | 0.42739183 | 2.73671294 | 0.00620564 | 0.07721978 |
| IGHMBP2 | 16.5787444 | -0.5538096 | 0.54856153 | -1.0095669 | 0.31270283 | NA |
| TPCN2 | 23.4557908 | -0.5039998 | 0.42389787 | -1.1889652 | 0.23445334 | 0.49243138 |
| CCND1 | 15.7552048 | -0.6618329 | 0.59913238 | -1.1046522 | 0.26931033 | NA |
| ORAOV1 | 25.0493168 | -0.7675391 | 0.44499245 | -1.7248362 | 0.08455699 | 0.28931012 |
| PPFIA1 | 164.329528 | -1.0143641 | 0.3338121 | -3.0387278 | 0.00237579 | 0.0462949 |
| AP002336.2 | 57.7901299 | -0.5949093 | 0.40685797 | -1.4622038 | 0.14368536 | 0.38252489 |
| AP000487.1 | 14.0014731 | 0.72909343 | 0.48615609 | 1.49971058 | 0.13368939 | NA |
| CTTN | 155.975128 | -0.6815455 | 0.30751715 | -2.2162846 | 0.02667201 | 0.16052735 |
| SHANK2 | 364.099019 | -0.0750372 | 0.47035541 | -0.1595329 | 0.87324901 | 0.94114315 |
| NADSYN1 | 32.1521214 | 0.05387374 | 0.37870909 | 0.14225627 | 0.88687759 | 0.9477019 |
| AP002495.2 | 58.2082248 | 0.1242987 | 0.41090689 | 0.30249845 | 0.76227212 | 0.88247846 |
| RNF121 | 70.6997073 | -0.2010178 | 0.27086485 | -0.7421333 | 0.45800659 | 0.69231672 |
| NUMA1 | 100.938244 | 0.1730892 | 0.2882948 | 0.60038962 | 0.54824661 | 0.7554786 |
| LAMTOR1 | 29.2250465 | -0.1223744 | 0.32473238 | -0.3768469 | 0.70628737 | 0.85458443 |
| INPPL1 | 24.6940455 | 0.89074373 | 0.38630279 | 2.30581752 | 0.02112083 | 0.1427136 |
| CLPB | 18.8945042 | 0.1488537 | 0.50061344 | 0.2973426 | 0.76620496 | NA |
| ARAP1 | 35.4468194 | 0.13318222 | 0.44090399 | 0.30206627 | 0.76260154 | 0.88247846 |
| ATG16L2 | 23.9329922 | -0.0911327 | 0.44035984 | -0.2069505 | 0.83604854 | 0.92074197 |
| FCHSD2 | 241.71986 | -0.017358 | 0.18938691 | -0.0916534 | 0.92697341 | 0.96954944 |
| P2RY6 | 49.8013782 | 2.34662595 | 0.79421969 | 2.9546308 | 0.00313043 | 0.05513719 |
| ARHGEF17 | 39.206409 | 0.43539962 | 0.42713092 | 1.01935868 | 0.30803271 | 0.56815148 |
| FAM168A | 151.174267 | 0.8484789 | 0.24119217 | 3.51785431 | 0.00043505 | 0.01850049 |
| RAB6A | 178.665329 | -0.4385248 | 0.21906406 | -2.0018106 | 0.04530511 | 0.21333202 |
| MRPL48 | 41.8264311 | -0.0710585 | 0.3172285 | -0.2239978 | 0.82275902 | 0.91475524 |
| COA4 | 19.2062487 | 0.27838392 | 0.42862477 | 0.64948164 | 0.51602711 | NA |
| PAAF1 | 36.580455 | 0.08116269 | 0.50252735 | 0.161509 | 0.87169252 | 0.9400893 |
| C2CD3 | 74.2775978 | 0.39057542 | 0.27090075 | 1.44176573 | 0.14936847 | 0.38981296 |
| PPME1 | 86.9042725 | 0.07952455 | 0.45345394 | 0.17537514 | 0.86078488 | 0.93500722 |
| PGM2L1 | 23.4344384 | -0.3990341 | 0.55624345 | -0.7173731 | 0.47314389 | 0.70227603 |
| POLD3 | 37.290974 | 0.15031773 | 0.40744879 | 0.36892422 | 0.71218421 | 0.857224 |
| RNF169 | 129.728189 | -0.6280336 | 0.33310202 | -1.885409 | 0.05937463 | 0.24193042 |
| XRRA1 | 122.037649 | -0.4402345 | 0.5696171 | -0.7728603 | 0.43960505 | 0.67958226 |
| SPCS2 | 68.4242757 | 0.1376245 | 0.29898347 | 0.46030806 | 0.64529512 | 0.81777273 |

| | | | | | | |
|---|---|---|---|---|---|---|
| NEU3 | 23.0460069 | -0.7703936 | 0.40834263 | -1.8866353 | 0.0592094 | 0.24164091 |
| RPS3 | 92.1631414 | 0.42041204 | 0.38232814 | 1.0996105 | 0.27150187 | 0.52985544 |
| UVRAG | 215.872254 | -0.7781773 | 0.32619164 | -2.3856446 | 0.01704922 | 0.12988597 |
| THAP12 | 20.1585181 | 0.04898302 | 0.39795798 | 0.12308592 | 0.90203906 | NA |
| AP002360.1 | 11.3858335 | 1.2384443 | 0.6338709 | 1.95378002 | 0.05072724 | NA |
| EMSY | 130.538907 | -0.4286016 | 0.31858597 | -1.3453249 | 0.17852034 | 0.42765983 |
| ACER3 | 130.603504 | 0.81421089 | 0.36027173 | 2.25999107 | 0.02382181 | 0.15078654 |
| PAK1 | 89.7467449 | 0.02474657 | 0.27423077 | 0.09023996 | 0.92809653 | 0.97006504 |
| CLNS1A | 58.1519613 | 0.41263712 | 0.3892137 | 1.06018139 | 0.28906209 | 0.54841284 |
| RSF1 | 206.804046 | 0.05610403 | 0.19296197 | 0.29075173 | 0.7712412 | 0.88680065 |
| AAMDC | 52.9524271 | 1.00959657 | 0.43532532 | 2.31917721 | 0.02038543 | 0.14043319 |
| AP002812.2 | 21.9300296 | 0.36283226 | 0.45780936 | 0.79254006 | 0.42804585 | 0.67072527 |
| INTS4 | 63.1858966 | -0.3959633 | 0.29729144 | -1.3319028 | 0.18289215 | 0.43261243 |
| ALG8 | 28.7435942 | 0.39011721 | 0.53494393 | 0.72926747 | 0.46583807 | 0.69622439 |
| GAB2 | 187.680806 | -1.1819646 | 0.47766074 | -2.4744856 | 0.01334282 | 0.11285681 |
| AP003086.2 | 43.7694211 | -0.0260596 | 0.63269911 | -0.041188 | 0.96714599 | 0.98741848 |
| NARS2 | 75.0069268 | 0.06540078 | 0.30601659 | 0.21371647 | 0.83076817 | 0.91794754 |
| TENM4 | 26.9376796 | -1.6174944 | 0.81777499 | -1.9779211 | 0.0479376 | 0.21943015 |
| PRCP | 62.908442 | 1.14089996 | 0.35237662 | 3.23772891 | 0.00120485 | 0.03090537 |
| RAB30 | 24.5017262 | -1.0255898 | 0.49488765 | -2.072369 | 0.03823104 | 0.19526497 |
| RAB30-AS1 | 97.3858882 | -0.033641 | 0.33524332 | -0.1003481 | 0.92006801 | 0.96534398 |
| PCF11 | 81.4235401 | -0.2542819 | 0.33335962 | -0.7627857 | 0.44559123 | 0.68538685 |
| ANKRD42 | 35.9904634 | -0.0321492 | 0.42181352 | -0.0762167 | 0.93924668 | 0.97483572 |
| CCDC90B | 30.9269614 | -0.0494358 | 0.41182627 | -0.1200404 | 0.90445118 | 0.95671001 |
| AP000446.1 | 34.4646399 | 0.68633117 | 0.4997109 | 1.37345647 | 0.16961049 | 0.41775741 |
| DLG2 | 162.28561 | 0.2102993 | 0.39051241 | 0.53852144 | 0.59021711 | 0.78474712 |
| TMEM126B | 20.8378486 | 0.23483731 | 0.40165355 | 0.5846763 | 0.55876539 | NA |
| CREBZF | 81.4355358 | 0.01723271 | 0.25884134 | 0.06657633 | 0.94691899 | 0.97857619 |
| SYTL2 | 185.217258 | -1.0091562 | 0.38324582 | -2.6331825 | 0.00845889 | 0.09005263 |
| PICALM | 490.635786 | -0.6568764 | 0.21911747 | -2.9978276 | 0.00271911 | 0.05043138 |
| EED | 52.0909815 | 0.51524533 | 0.31454793 | 1.63805032 | 0.1014112 | 0.32069104 |
| HIKESHI | 38.8233138 | 0.45167857 | 0.38271274 | 1.18020261 | 0.23791964 | 0.49574106 |
| AP001831.1 | 51.0681606 | 0.96375496 | 0.55027777 | 1.75139725 | 0.07987751 | 0.28223671 |
| ME3 | 303.000155 | -0.9008376 | 0.32363804 | -2.7834726 | 0.00537804 | 0.07124746 |
| AP003059.1 | 17.0547374 | 0.71364984 | 0.64582054 | 1.10502808 | 0.26914742 | NA |
| AP003059.2 | 16.1338139 | 0.37375413 | 0.68956044 | 0.54201793 | 0.58780615 | NA |
| PRSS23 | 129.617702 | 1.35522652 | 0.49391127 | 2.74386639 | 0.00607202 | 0.07661007 |
| AP001528.2 | 78.2899701 | 2.88591618 | 0.83271631 | 3.46566548 | 0.00052892 | 0.02087746 |
| FZD4 | 83.9991654 | 0.91084411 | 0.43044965 | 2.11602941 | 0.03434231 | 0.18371048 |
| AP001528.1 | 19.6817569 | 0.86181068 | 0.51033519 | 1.68871498 | 0.09127406 | NA |
| TMEM135 | 193.319155 | -0.021994 | 0.2880141 | -0.0763642 | 0.93912938 | 0.97483572 |
| RAB38 | 32.0555304 | -0.2697144 | 0.48925727 | -0.5512731 | 0.58144649 | 0.77819399 |
| CTSC | 49.1504222 | 0.67891286 | 0.51194996 | 1.32613129 | 0.18479621 | 0.43476148 |
| NOX4 | 18.949581 | -0.4971978 | 0.7496212 | -0.6632654 | 0.50716057 | NA |
| CHORDC1 | 44.2633245 | 0.14720489 | 0.45026236 | 0.32693137 | 0.74371979 | 0.87230965 |
| SLC36A4 | 124.128938 | -0.2591241 | 0.26787384 | -0.9673363 | 0.33337596 | 0.58883681 |
| SMCO4 | 55.7559787 | -0.9816475 | 0.75174293 | -1.3058287 | 0.1916108 | 0.44319545 |
| CEP295 | 58.6026177 | -0.2257029 | 0.43462366 | -0.5193065 | 0.60354704 | 0.79301836 |
| TAF1D | 134.276522 | -0.057728 | 0.30715544 | -0.1879439 | 0.85092063 | 0.92874559 |
| MRE11 | 43.666755 | -0.1868025 | 0.3394196 | -0.5503587 | 0.58207337 | 0.7783237 |
| AMOTL1 | 42.1807444 | 0.82085576 | 0.45206228 | 1.81580236 | 0.0694007 | 0.26260569 |
| CWC15 | 22.919875 | -0.0157601 | 0.41085927 | -0.0383588 | 0.96940161 | 0.98781207 |
| SRSF8 | 17.4754101 | 0.40070016 | 0.41587681 | 0.96350686 | 0.33529322 | NA |
| ENDOD1 | 33.8996495 | -0.2670374 | 0.44462929 | -0.6005843 | 0.54811688 | 0.7554786 |

| | | | | | | |
|---|---|---|---|---|---|---|
| AP000787.1 | 28.8366374 | -0.0650418 | 0.46947218 | -0.1385424 | 0.8898118 | 0.94965078 |
| SESN3 | 47.8443188 | 0.62338407 | 0.57793212 | 1.07864583 | 0.28074564 | 0.5390351 |
| FAM76B | 29.3101109 | -0.1502991 | 0.37483511 | -0.4009739 | 0.68843936 | 0.84543868 |
| CEP57 | 97.8880238 | 0.40506135 | 0.24265711 | 1.6692746 | 0.09506297 | 0.30939937 |
| MTMR2 | 68.7800598 | -0.0773179 | 0.28021863 | -0.2759199 | 0.78260959 | 0.89280362 |
| MAML2 | 2323.20742 | -0.2249992 | 0.25854746 | -0.8702434 | 3.84E-01 | 6.35E-01 |
| CCDC82 | 47.5320232 | -0.3139185 | 0.40664609 | -0.7719697 | 0.44013235 | 0.68012421 |
| ARHGAP42 | 21.3056531 | 0.13760271 | 0.452621 | 0.30401309 | 0.76111792 | 0.88201214 |
| CEP126 | 27.4276933 | 1.19876035 | 0.50392926 | 2.37882663 | 0.01736784 | 0.13115162 |
| YAP1 | 530.522762 | -0.2635964 | 0.21521037 | -1.2248315 | 0.22063871 | 0.4773064 |
| BIRC3 | 81.4538946 | -0.2387141 | 0.56510819 | -0.4224219 | 6.73E-01 | 0.83624371 |
| BIRC2 | 65.4218172 | -0.8850618 | 0.30800987 | -2.8734853 | 0.0040597 | 0.06207769 |
| TMEM123 | 118.016848 | -0.0194642 | 0.33352199 | -0.0583597 | 0.95346215 | 0.98153155 |
| DCUN1D5 | 48.0926636 | -0.0912581 | 0.30249974 | -0.3016798 | 0.76289615 | 0.88247846 |
| DYNC2H1 | 538.996116 | 0.84029817 | 0.42150732 | 1.99355535 | 0.04620067 | 0.2152582 |
| CASP4 | 69.9449728 | -0.5804085 | 0.33951566 | -1.7095192 | 8.74E-02 | 2.96E-01 |
| MSANTD4 | 20.9908768 | -0.3352194 | 0.43913734 | -0.7633589 | 0.44524938 | NA |
| KBTBD3 | 21.469079 | 0.17407023 | 0.44056326 | 0.39510837 | 0.69276291 | 0.84775145 |
| AASDHPPT | 30.769408 | 0.69036673 | 0.35254565 | 1.95823357 | 0.05020261 | 0.22507781 |
| CWF19L2 | 117.667237 | -0.2518725 | 0.22621582 | -1.1134167 | 0.26552949 | 0.524719 |
| ALKBH8 | 21.3007769 | 1.41513954 | 0.64616832 | 2.19004785 | 0.02852077 | 0.16593415 |
| SLC35F2 | 70.0569862 | -0.0866039 | 0.30978179 | -0.2795643 | 0.77981181 | 0.89074516 |
| CUL5 | 178.458998 | 0.01012769 | 0.19151051 | 0.05288319 | 0.95782498 | 0.98363961 |
| ACAT1 | 70.36254 | -0.0040361 | 0.30655257 | -0.0131662 | 0.98949517 | 0.99583841 |
| NPAT | 53.6770544 | -0.4589826 | 0.25796553 | -1.7792399 | 0.07520044 | 0.27367272 |
| ATM | 150.686858 | 0.0638091 | 0.22623745 | 0.28204479 | 0.77790917 | 0.88981526 |
| C11orf65 | 40.8378384 | 1.23020171 | 0.63800418 | 1.92820321 | 0.05382986 | 0.23287156 |
| KDELC2 | 41.9653637 | 0.99739255 | 0.52404033 | 1.90327442 | 5.70E-02 | 0.23764428 |
| EXPH5 | 207.27219 | -0.1502853 | 0.25078501 | -0.5992595 | 5.49E-01 | 0.75579697 |
| DDX10 | 98.2024863 | -0.1181493 | 0.282245 | -0.4186055 | 0.67550444 | 0.83808131 |
| RDX | 167.43821 | 0.21781156 | 0.29609592 | 0.73561147 | 0.46196718 | 0.69416206 |
| ZC3H12C | 92.3710786 | 0.16142663 | 0.37343153 | 0.43227907 | 0.6655386 | 0.83059892 |
| FDX1 | 94.5811858 | 0.08852224 | 0.38803516 | 0.22812943 | 0.81954562 | 0.913067 |
| COLCA1 | 20.7037682 | 1.62820083 | 0.75682918 | 2.15134521 | 0.03144896 | NA |
| SIK2 | 364.464225 | -0.3097469 | 0.38232105 | -0.8101748 | 0.41783974 | 0.66175153 |
| PPP2R1B | 80.1434992 | -0.7253715 | 0.38019941 | -1.9078711 | 0.05640788 | 0.23662544 |
| CRYAB | 162.703976 | 0.80520271 | 0.80147121 | 1.00465582 | 0.31506261 | 0.57225891 |
| DIXDC1 | 38.9870298 | 1.17406417 | 0.46135464 | 2.54481926 | 0.01093343 | 0.10262759 |
| DLAT | 39.4891402 | 0.17551932 | 0.30936162 | 0.5673597 | 0.57046982 | 0.77196291 |
| NKAPD1 | 17.0662281 | -0.1685795 | 0.45222855 | -0.3727749 | 0.70931596 | NA |
| AP002884.3 | 26.8973331 | -0.6692609 | 0.46543857 | -1.4379145 | 0.15045831 | 0.39108044 |
| PTS | 55.792089 | -0.6868627 | 0.29786563 | -2.3059483 | 0.02111352 | 0.1427136 |
| AP002884.1 | 33.2762538 | -0.5459726 | 0.44083491 | -1.2384967 | 0.21553193 | 0.47115856 |
| TTC12 | 47.7406626 | 0.5542427 | 0.52536569 | 1.05496553 | 0.29144109 | 0.55061906 |
| ZW10 | 25.1627421 | 0.48460564 | 0.37575629 | 1.2896807 | 0.19716154 | 0.45031229 |
| USP28 | 29.3427332 | 0.28410343 | 0.3616997 | 0.78546769 | 0.4321794 | 0.6728327 |
| NNMT | 82.3861472 | 0.63859866 | 1.39024477 | 0.45934261 | 6.46E-01 | 0.81779125 |
| REXO2 | 42.1286896 | 1.2852492 | 0.45305496 | 2.83685054 | 0.00455609 | 0.06530401 |
| CADM1 | 643.499113 | -0.4388751 | 0.30737076 | -1.4278361 | 0.15333903 | 0.39519914 |
| AP000462.1 | 19.1066638 | 0.28679212 | 0.63919743 | 0.44867533 | 0.65366588 | NA |
| BUD13 | 14.704144 | -0.1445064 | 0.56203278 | -0.2571138 | 0.79709093 | NA |
| SIK3 | 876.884733 | -0.3484343 | 0.35713488 | -0.9756377 | 0.32924401 | 0.58534184 |
| PAFAH1B2 | 71.9490655 | -0.3076262 | 0.25252983 | -1.2181776 | 0.22315649 | 0.48036053 |
| SIDT2 | 15.0311007 | 0.23262896 | 0.5007406 | 0.46456979 | 0.64223957 | NA |

| | | | | | | |
|---|---|---|---|---|---|---|
| PCSK7 | 68.1457609 | 0.0718935 | 0.30254648 | 0.23762795 | 0.81216967 | 0.90954022 |
| RNF214 | 50.4772521 | -0.1591847 | 0.38059628 | -0.4182508 | 0.67576378 | 0.83825202 |
| BACE1 | 33.6990935 | 0.95552888 | 0.46562527 | 2.05214138 | 0.04015592 | 0.19927798 |
| CEP164 | 41.4129603 | 0.77683685 | 0.44016441 | 1.76487883 | 0.07758413 | 0.27775119 |
| DSCAML1 | 344.390645 | 0.2558248 | 0.66599194 | 0.38412598 | 0.70088508 | 0.85187889 |
| FXYD2 | 73.8696694 | 1.2989352 | 0.63218563 | 2.05467371 | 0.03991053 | 0.19882936 |
| FXYD6-FXYD2 | 22.9921119 | 3.30069639 | 1.22303896 | 2.69876636 | 0.0069597 | 0.08149858 |
| AP000757.1 | 26.3037749 | 1.35591939 | 0.4623727 | 2.93252474 | 0.00336218 | 0.05733014 |
| TMPRSS4 | 43.5754327 | -0.9469508 | 0.70830049 | -1.3369337 | 1.81E-01 | 4.30E-01 |
| MPZL3 | 23.1194991 | -0.7545502 | 0.3819278 | -1.9756357 | 0.04819605 | 0.21989628 |
| UBE4A | 65.8817745 | -0.3703074 | 0.28362635 | -1.3056172 | 0.19168275 | 0.44319545 |
| ATP5MG | 78.5680637 | 0.35814414 | 0.33887107 | 1.05687432 | 0.29056895 | 0.54991621 |
| KMT2A | 206.198025 | -0.512721 | 0.16621305 | -3.0847216 | 0.00203743 | 0.042384 |
| IFT46 | 20.879554 | -0.1058623 | 0.45991793 | -0.2301766 | 0.81795456 | NA |
| ARCN1 | 41.1027898 | -0.3717125 | 0.32369404 | -1.1483453 | 0.25082604 | 0.50738714 |
| PHLDB1 | 63.6709125 | 0.62196797 | 0.53104591 | 1.17121319 | 0.2415131 | 0.49881565 |
| DDX6 | 125.467201 | -0.0943869 | 0.26304321 | -0.3588266 | 0.71972479 | 0.85995641 |
| BCL9L | 24.5419035 | -1.5049221 | 0.51325553 | -2.9321109 | 0.00336667 | 0.05733014 |
| CCDC84 | 109.060404 | -0.5024841 | 0.30625823 | -1.6407202 | 0.10085551 | 0.31985368 |
| RPS25 | 70.0997953 | -0.2308578 | 0.45550344 | -0.5068191 | 0.61228175 | 0.79863288 |
| TRAPPC4 | 13.8092391 | 1.30785526 | 0.54403449 | 2.40399329 | 0.01621707 | NA |
| VPS11 | 13.9721046 | -0.5863004 | 0.49703694 | -1.1795912 | 0.23816285 | NA |
| HINFP | 15.694854 | -0.3348761 | 0.50196497 | -0.6671304 | 0.50468887 | NA |
| CBL | 58.7150639 | -0.1664661 | 0.44686536 | -0.3725195 | 0.70950606 | 0.85607462 |
| USP2 | 28.2862124 | 0.93149305 | 0.66615019 | 1.39832288 | 1.62E-01 | 4.08E-01 |
| USP2-AS1 | 15.8362702 | 1.17986727 | 0.82721329 | 1.42631567 | 1.54E-01 | NA |
| OAF | 46.8281925 | -0.514863 | 0.38914027 | -1.3230781 | 1.86E-01 | 4.36E-01 |
| ARHGEF12 | 468.130192 | -0.2680443 | 0.22269614 | -1.2036325 | 0.22873166 | 0.48701178 |
| SC5D | 17.0407992 | 0.45289135 | 0.46429564 | 0.97543744 | 0.32934332 | NA |
| SORL1 | 202.839377 | -0.1855058 | 0.59165787 | -0.3135356 | 0.75387375 | 0.87868581 |
| UBASH3B | 27.394283 | -1.4236851 | 0.559119 | -2.5463007 | 1.09E-02 | 1.02E-01 |
| JHY | 15.7746187 | 0.85651463 | 0.63572728 | 1.34729884 | 0.17788401 | NA |
| HSPA8 | 53.1975376 | 1.90767546 | 0.42208863 | 4.51960878 | 6.20E-06 | 0.001256 |
| GRAMD1B | 498.352781 | -1.1107978 | 0.62499765 | -1.7772831 | 0.07552167 | 0.27458208 |
| VWA5A | 21.3450746 | 0.38336286 | 0.41608593 | 0.92135502 | 3.57E-01 | 0.6128044 |
| TBRG1 | 56.8986209 | -0.0145683 | 0.32435671 | -0.0449145 | 0.96417546 | 0.98634483 |
| SIAE | 34.3958844 | 0.91719735 | 0.40477025 | 2.26597029 | 0.0234532 | 0.14954518 |
| MSANTD2 | 72.1771724 | 0.56843134 | 0.32424624 | 1.75308538 | 0.07958735 | 0.2817308 |
| EI24 | 23.8327791 | 0.65004184 | 0.44780407 | 1.4516211 | 0.14660699 | 0.38645 |
| STT3A | 25.218035 | -0.7054991 | 0.39733354 | -1.7755841 | 0.07580149 | 0.27507972 |
| RPUSD4 | 15.1659925 | -0.6559654 | 0.47092208 | -1.3929382 | 0.16363849 | NA |
| FAM118B | 24.9239102 | -0.8183506 | 0.42799549 | -1.9120542 | 0.05586925 | 0.23575965 |
| SRPRA | 21.5512387 | -0.4548004 | 0.38761697 | -1.1733242 | 0.24066582 | 0.49822614 |
| FOXRED1 | 23.8771105 | -0.2042844 | 0.54720023 | -0.3733266 | 0.70890538 | 0.8559129 |
| TIRAP | 13.152846 | 0.25300972 | 0.51277573 | 0.49341203 | 0.62172148 | NA |
| DCPS | 28.1583185 | -0.10627 | 0.423883 | -0.250706 | 0.8020414 | 0.90378853 |
| ST3GAL4 | 39.772317 | 0.645932 | 0.71512618 | 0.903242 | 0.36639747 | 0.62157013 |
| ETS1 | 88.4552679 | -1.8667232 | 0.43634291 | -4.2781105 | 1.88E-05 | 0.00237833 |
| KCNJ1 | 200.98779 | 2.02442289 | 0.63249398 | 3.20069905 | 0.00137095 | 0.03328762 |
| ARHGAP32 | 192.447511 | -0.1084221 | 0.29168304 | -0.3717121 | 0.71010721 | 0.85615527 |
| BARX2 | 48.1377072 | 0.49447776 | 0.56420739 | 0.87641136 | 0.38080644 | 0.63229421 |
| TMEM45B | 44.0536206 | 0.18226223 | 0.49381094 | 0.36909314 | 0.7120583 | 0.857224 |
| NFRKB | 37.3642641 | 0.11364191 | 0.38381132 | 0.29608795 | 0.76716291 | 0.88462217 |
| PRDM10 | 54.8533865 | -0.1097708 | 0.30143823 | -0.3641568 | 0.71574094 | 0.85868129 |

| | | | | | | |
|---|---|---|---|---|---|---|
| APLP2 | 173.649878 | 0.56986324 | 0.37231001 | 1.5306149 | 0.1258646 | 0.35705972 |
| ST14 | 52.8714017 | -0.0672498 | 0.38446698 | -0.1749171 | 0.8611448 | 0.93526619 |
| ZBTB44 | 113.948609 | 0.70234452 | 0.25758925 | 2.72660644 | 0.00639893 | 0.07793129 |
| AP002986.1 | 17.5450804 | 0.71431425 | 0.76025967 | 0.93956615 | 0.34744015 | NA |
| ADAMTS15 | 19.3829575 | -1.9005026 | 0.78143424 | -2.4320698 | 0.01501281 | NA |
| SNX19 | 54.7802405 | -0.4683273 | 0.38370831 | -1.2205296 | 0.22226418 | 0.4793528 |
| NCAPD3 | 19.9880376 | -0.1318276 | 0.52014601 | -0.2534435 | 0.79992552 | NA |
| VPS26B | 20.7202856 | 1.13276692 | 0.42740936 | 2.65030913 | 0.00804182 | NA |
| ACAD8 | 15.1219145 | 0.55678242 | 0.49307295 | 1.12920903 | 0.25880967 | NA |
| ZMYND11 | 208.735421 | -0.2805929 | 0.27742048 | -1.0114355 | 0.31180804 | 0.57088166 |
| DIP2C | 705.007274 | -1.2843975 | 0.38922988 | -3.299843 | 9.67E-04 | 0.02829008 |
| LARP4B | 273.407414 | -0.3611433 | 0.17765347 | -2.0328527 | 0.04206741 | 0.20441466 |
| GTPBP4 | 28.5278821 | -0.8693115 | 0.46427113 | -1.8724221 | 0.06114823 | 0.2469349 |
| IDI1 | 28.11896 | -0.0115851 | 0.56048422 | -0.0206698 | 0.98350908 | 0.99382437 |
| WDR37 | 128.410558 | -0.5119157 | 0.27923052 | -1.8333085 | 0.06675671 | 0.25794497 |
| PFKP | 158.687223 | -0.2824095 | 0.38482781 | -0.7338593 | 0.46303447 | 0.69434917 |
| PITRM1 | 43.9439199 | -0.9796686 | 0.40510415 | -2.4183128 | 0.01559267 | 0.12347403 |
| KLF6 | 123.812466 | -1.2517607 | 0.46884948 | -2.6698562 | 0.00758837 | 0.08564181 |
| NET1 | 16.0858835 | -0.6347533 | 0.47014932 | -1.3501101 | 0.17698068 | NA |
| ASB13 | 13.2394633 | 0.71136456 | 0.53648376 | 1.32597594 | 0.18484767 | NA |
| FAM208B | 210.82363 | -0.7398014 | 0.35639406 | -2.075796 | 0.03791282 | 0.19457113 |
| GDI2 | 120.45017 | 0.23616248 | 0.30304774 | 0.77929133 | 0.43580812 | 0.67638148 |
| ANKRD16 | 18.3615084 | 0.69515072 | 0.44367969 | 1.56678508 | 0.11716494 | NA |
| FBH1 | 91.5377019 | -0.1523366 | 0.31175724 | -0.4886385 | 0.62509764 | 0.8053173 |
| RBM17 | 60.6568421 | -0.2828963 | 0.26550169 | -1.0655161 | 0.28664244 | 0.54611061 |
| PFKFB3 | 59.0963391 | -1.5134092 | 0.3936242 | -3.8448072 | 0.00012065 | 0.00800536 |
| PRKCQ | 155.501384 | -0.8922289 | 0.29449919 | -3.0296481 | 0.00244839 | 0.04713978 |
| SFMBT2 | 170.246019 | -1.2139045 | 0.6136822 | -1.9780671 | 0.04792114 | 0.21943015 |
| KIN | 31.2274884 | -0.3102923 | 0.35502127 | -0.8740104 | 0.38211257 | 0.6334526 |
| ATP5F1C | 133.373524 | 0.39446178 | 0.23795651 | 1.65770536 | 9.74E-02 | 0.31373395 |
| TAF3 | 161.499626 | -0.0997459 | 0.2585008 | -0.3858632 | 0.699598 | 0.8513527 |
| GATA3 | 144.597604 | 0.96508887 | 0.39955473 | 2.41541093 | 0.01571747 | 0.12390845 |
| GATA3-AS1 | 15.183886 | 0.99742155 | 0.71142167 | 1.40201177 | 0.16091173 | NA |
| USP6NL | 109.502568 | -0.1780946 | 0.22277267 | -0.7994455 | 0.42403211 | 0.66715549 |
| PROSER2 | 39.2389719 | 0.01903702 | 0.38982912 | 0.04883427 | 0.96105137 | 0.98496344 |
| UPF2 | 116.612778 | -0.4701649 | 0.27367749 | -1.7179523 | 0.08580531 | 0.29278925 |
| DHTKD1 | 45.2010078 | -0.0232134 | 0.32282141 | -0.071908 | 0.94267513 | 0.97628774 |
| SEC61A2 | 21.7499481 | -0.1877102 | 0.41875345 | -0.4482594 | 0.65396598 | 0.82247976 |
| NUDT5 | 45.8874648 | -0.4843306 | 0.30979433 | -1.5633939 | 0.11795997 | 0.34552837 |
| CDC123 | 44.4425706 | -0.7494848 | 0.3357562 | -2.2322293 | 0.02559981 | 0.15662748 |
| OPTN | 42.3618843 | -0.7702066 | 0.45857041 | -1.6795821 | 0.09303866 | 0.30571879 |
| PHYH | 22.2323364 | 0.14769759 | 0.45640146 | 0.32361331 | 0.74623081 | 0.87410417 |
| SEPHS1 | 26.7898712 | 0.39526265 | 0.38566025 | 1.02489861 | 0.30541103 | 0.56557634 |
| BEND7 | 81.0753473 | 0.24277415 | 0.2694807 | 0.90089626 | 0.36764348 | 0.6222756 |
| PRPF18 | 16.8313559 | -0.1824463 | 0.44842735 | -0.4068582 | 0.68411217 | NA |
| AL157392.5 | 28.1514064 | 0.19450778 | 0.356609 | 0.54543711 | 0.58545292 | 0.78124673 |
| FRMD4A | 361.158794 | -0.275467 | 0.41845433 | -0.6582965 | 0.5103476 | 0.7302743 |
| FAM107B | 208.200248 | -0.0388302 | 0.31197958 | -0.1244641 | 0.90094785 | 0.95623215 |
| HSPA14.1 | 12.695071 | -0.4926591 | 0.47590061 | -1.0352142 | 0.3005689 | NA |
| DCLRE1C | 34.0077282 | -0.0769645 | 0.3619346 | -0.2126476 | 0.83160185 | 0.9181376 |
| RPP38 | 34.8606541 | 0.36301116 | 0.36407743 | 0.99707132 | 0.31872989 | 0.57588356 |
| NMT2 | 33.816804 | 0.39742556 | 0.36717644 | 1.08238308 | 0.27908234 | 0.53717915 |
| FAM171A1 | 300.02126 | 1.04966939 | 0.37333613 | 2.81159337 | 0.00492968 | 0.06830815 |
| MINDY3 | 85.5556235 | -0.0740727 | 0.27861754 | -0.265858 | 0.7903486 | 0.89697923 |

| | | | | | | |
|---|---|---|---|---|---|---|
| PTER | 74.8229892 | -0.0448512 | 0.28140278 | -0.1593844 | 0.87336602 | 0.94114315 |
| RSU1 | 75.9642572 | -0.055789 | 0.2298438 | -0.2427258 | 0.8082178 | 0.90713122 |
| CUBN | 39.9427342 | 0.3714722 | 0.63226045 | 0.58753034 | 0.55684758 | 0.76323398 |
| TRDMT1 | 16.1110129 | 1.43132448 | 0.62440613 | 2.29229729 | 0.02188849 | NA |
| VIM | 27.7785233 | -0.5699603 | 0.48802266 | -1.1678972 | 0.24284823 | 0.50005426 |
| STAM | 58.6068576 | -0.8121089 | 0.27958236 | -2.9047216 | 0.0036758 | 0.06071379 |
| CACNB2 | 15.6162337 | 1.27981616 | 0.72520355 | 1.7647682 | 0.07760273 | NA |
| AL450384.2 | 28.3046556 | 0.22190217 | 0.51962777 | 0.42704063 | 0.66934974 | 0.83376893 |
| NSUN6 | 88.326766 | -0.1513377 | 0.37543523 | -0.4030994 | 0.68687507 | 0.84468404 |
| ARL5B | 61.2206664 | -2.2640521 | 0.46407847 | -4.8785976 | 1.07E-06 | 0.00045469 |
| MALRD1 | 89.7309312 | 1.25468016 | 0.71039255 | 1.76617865 | 0.07736588 | 0.27731248 |
| PLXDC2 | 343.312987 | 0.44574938 | 0.23758755 | 1.87614792 | 0.06063497 | 0.24550625 |
| NEBL | 335.708666 | 0.41690428 | 0.36521004 | 1.14154661 | 0.25364253 | 0.51093602 |
| MLLT10 | 349.975858 | -0.0541933 | 0.19152846 | -0.2829515 | 0.77721404 | 0.8896927 |
| DNAJC1 | 241.587282 | -0.1452275 | 0.17915988 | -0.8106027 | 0.41759387 | 0.66167578 |
| PIP4K2A | 87.6045367 | 0.41027923 | 0.25383141 | 1.61634538 | 0.10601966 | 0.32732861 |
| MSRB2 | 20.8342007 | 0.55634095 | 0.43603011 | 1.27592326 | 0.20198265 | NA |
| KIAA1217 | 684.964598 | 0.14432352 | 0.2965158 | 0.48673128 | 0.62644877 | 0.8053173 |
| ARHGAP21 | 160.225935 | -0.8884608 | 0.18980618 | -4.6808846 | 2.86E-06 | 0.00076688 |
| PRTFDC1 | 32.1162759 | -0.9082067 | 0.5810833 | -1.5629545 | 0.1180633 | 0.34552837 |
| PDSS1 | 28.750263 | -0.4186161 | 0.37896681 | -1.1046246 | 0.26932227 | 0.52764898 |
| ABI1 | 185.093575 | -0.5159334 | 0.21243138 | -2.428706 | 0.01515281 | 0.12124045 |
| ANKRD26 | 164.348548 | 0.49643821 | 0.27007997 | 1.83811561 | 0.06604537 | 0.25609634 |
| YME1L1 | 103.858618 | -0.3253393 | 0.23607278 | -1.3781314 | 0.16816271 | 0.4161892 |
| MASTL | 25.2256319 | -1.0827741 | 0.44611772 | -2.427104 | 0.01521989 | 0.12139638 |
| ACBD5 | 48.9106139 | 0.30513093 | 0.31347656 | 0.9733772 | 0.33036587 | 0.5865284 |
| RAB18 | 99.749595 | -0.9448163 | 0.23702204 | -3.9861959 | 6.71E-05 | 0.00530264 |
| MPP7 | 95.1507739 | 0.30169382 | 0.24222369 | 1.24551742 | 0.21294162 | 0.46842287 |
| WAC-AS1 | 28.2950696 | 0.29918227 | 0.43578636 | 0.68653426 | 0.49237627 | 0.71672078 |
| WAC | 367.513975 | -0.3723235 | 0.20339868 | -1.8305109 | 0.06717358 | 0.25864684 |
| SVIL | 571.218846 | -1.0715604 | 0.46264077 | -2.3161824 | 0.02054831 | 0.14099127 |
| MTPAP | 30.3303111 | -0.5782747 | 0.40346339 | -1.4332767 | 0.15177878 | 0.39273741 |
| MAP3K8 | 33.0974803 | -0.3890786 | 0.45530587 | -0.8545434 | 0.39280399 | 0.64175877 |
| ZNF438 | 80.624302 | 0.16197041 | 0.3558316 | 0.45518837 | 0.64897373 | 0.81913126 |
| ZEB1 | 107.139758 | -2.3289768 | 0.71643548 | -3.2507838 | 1.15E-03 | 0.03021158 |
| ARHGAP12 | 194.440872 | -0.1271599 | 0.18429295 | -0.689988 | 0.49020176 | 0.7163358 |
| KIF5B | 112.466149 | -0.3410133 | 0.29367994 | -1.1611732 | 0.24557148 | 0.50275206 |
| EPC1 | 193.296456 | -0.3083478 | 0.2102828 | -1.4663482 | 0.14255345 | 0.38032372 |
| CCDC7 | 142.702771 | 0.67747041 | 0.60373174 | 1.12213814 | 0.26180371 | 0.52123724 |
| ITGB1 | 177.39358 | -0.7632954 | 0.34033829 | -2.2427549 | 0.02491263 | 0.1546391 |
| NRP1 | 240.992134 | 0.44980229 | 0.44534739 | 1.0100032 | 3.12E-01 | 0.57101636 |
| PARD3 | 1291.36463 | -0.2509055 | 0.22465648 | -1.1168408 | 0.2640624 | 0.52361258 |
| CUL2 | 83.494934 | -0.3150655 | 0.23412597 | -1.3457095 | 0.17839622 | 0.42762867 |
| CREM | 196.711245 | -0.716062 | 0.45671956 | -1.5678373 | 0.1169191 | 0.34414008 |
| CCNY | 376.439699 | -0.3120207 | 0.19758321 | -1.5791862 | 0.11429336 | 0.34044736 |
| AL117336.3 | 55.6245642 | 0.26788792 | 0.36742319 | 0.72909909 | 0.46594105 | 0.69624311 |
| ZNF248 | 155.234021 | 0.2436097 | 0.52841093 | 0.46102321 | 0.64478195 | 0.81734747 |
| ZNF25 | 24.7507539 | 0.72727566 | 0.4629324 | 1.57101916 | 0.1161782 | 0.34314029 |
| ZNF33A | 239.712106 | -0.300676 | 0.23472533 | -1.2809695 | 0.20020439 | 0.45443032 |
| AL117339.5 | 28.2438764 | 0.44177429 | 0.38158298 | 1.15774108 | 0.24696972 | 0.5033559 |
| ZNF33B | 124.731457 | 0.09961194 | 0.28634272 | 0.34787662 | 0.72793284 | 0.86468169 |
| BMS1 | 54.2423746 | -0.4746426 | 0.26575637 | -1.7860063 | 0.07409824 | 0.27145842 |
| CSGALNACT2 | 40.9117951 | 0.14370686 | 0.28366637 | 0.50660521 | 0.61243185 | 0.79866179 |
| HNRNPF | 68.0019286 | -0.685352 | 0.38670075 | -1.7723059 | 0.07634379 | 0.27600666 |

| | | | | | | |
|---|---|---|---|---|---|---|
| ZNF487 | 16.1516006 | 0.21281916 | 0.47072416 | 0.45211013 | 0.65118965 | NA |
| ZNF32 | 23.7468754 | -0.1152906 | 0.3874362 | -0.2975732 | 0.76602896 | 0.8841093 |
| TMEM72-AS | 32.815253 | 0.02874532 | 0.63596215 | 0.04519974 | 0.9639481 | 0.98624332 |
| TMEM72 | 137.616233 | 0.95172388 | 0.40220113 | 2.36628842 | 0.01796744 | 0.13297634 |
| RASSF4 | 22.9091084 | 2.42592392 | 0.52463023 | 4.6240643 | 3.76E-06 | 0.00086364 |
| DEPP1 | 24.5534405 | 0.57508157 | 0.41279039 | 1.39315638 | 0.16357251 | 0.41042742 |
| 8-Mar | 105.911565 | -0.0856671 | 0.24409869 | -0.3509527 | 0.7256238 | 0.86336781 |
| ZFAND4 | 32.6045762 | 0.14884663 | 0.36135513 | 0.41191232 | 0.68040369 | 0.84089068 |
| WASHC2C | 41.7989013 | -0.7027042 | 0.55394293 | -1.2685497 | 0.2046017 | 0.45773044 |
| AGAP4 | 46.2351133 | 0.48692679 | 0.4884843 | 0.99681153 | 0.318856 | 0.57595896 |
| TIMM23 | 44.5752679 | -0.436306 | 0.33360488 | -1.3078525 | 0.19092335 | 0.44249835 |
| NCOA4 | 37.2297063 | 0.56390317 | 0.43332875 | 1.30132879 | 0.19314594 | 0.44483671 |
| PTPN20 | 48.5833611 | 1.86206492 | 0.38498715 | 4.83669364 | 1.32E-06 | 0.00048414 |
| AGAP9 | 39.7872299 | 1.41016239 | 0.60154993 | 2.34421502 | 0.01906718 | 0.1359475 |
| MAPK8 | 434.23717 | -0.8292513 | 0.24372716 | -3.4023753 | 0.00066803 | 0.02351966 |
| ARHGAP22 | 25.8906051 | -1.1037061 | 0.38503424 | -2.866514 | 4.15E-03 | 6.29E-02 |
| ERCC6 | 70.0951447 | -0.283688 | 0.2877652 | -0.9858315 | 0.32421579 | 0.58021133 |
| OGDHL | 99.1726958 | 2.13818706 | 0.51322634 | 4.16616783 | 3.10E-05 | 0.00335809 |
| PARG | 133.573613 | -0.1838996 | 0.27704894 | -0.6637802 | 0.50683101 | 0.72824214 |
| TIMM23B | 149.899143 | -0.8193067 | 0.40833823 | -2.0064413 | 0.04480919 | 0.21170763 |
| LINC00843 | 29.8491629 | 0.0946807 | 0.42786468 | 0.22128655 | 0.82486932 | 0.91537186 |
| AGAP6 | 48.9183997 | -0.204221 | 0.31889015 | -0.6404117 | 0.521905 | 0.7380123 |
| WASHC2A | 42.3767327 | -1.3964919 | 0.33670828 | -4.1474832 | 3.36E-05 | 0.00354431 |
| SGMS1 | 376.62873 | 0.20672339 | 0.20965112 | 0.98603522 | 0.32411583 | 0.58021133 |
| ASAH2B | 15.0559621 | -0.7554247 | 0.5392677 | -1.4008344 | 0.1612636 | NA |
| PRKG1 | 261.280009 | -0.8355754 | 0.61909809 | -1.3496656 | 0.17712326 | 0.42569745 |
| IPMK | 58.6181757 | -0.6047782 | 0.33525587 | -1.8039302 | 0.07124225 | 0.2665798 |
| CISD1 | 83.7609127 | 0.4950188 | 0.24762035 | 1.99910392 | 0.04559711 | 0.21335013 |
| UBE2D1 | 39.0631245 | -0.4578389 | 0.41760128 | -1.0963542 | 0.2729238 | 0.53186453 |
| TFAM | 20.911792 | 0.07327239 | 0.51328055 | 0.1427531 | 0.88648518 | NA |
| BICC1 | 2034.05542 | -0.3776454 | 0.46090156 | -0.8193624 | 0.41257966 | 0.65722052 |
| SLC16A9 | 14.4753494 | 1.41864488 | 1.02901274 | 1.37864657 | 0.16800375 | NA |
| CCDC6 | 97.1997246 | 0.06144498 | 0.23746529 | 0.25875353 | 0.79582541 | 0.8998778 |
| ANK3 | 1215.47249 | 0.63977755 | 0.30065134 | 2.12797175 | 0.03333943 | 0.18096866 |
| ARID5B | 880.636266 | -1.1407259 | 0.51898338 | -2.1980008 | 0.02794905 | 0.16477111 |
| NRBF2 | 22.1051817 | -0.6141968 | 0.54363501 | -1.1297962 | 0.25856212 | 0.51706621 |
| JMJD1C | 850.417105 | -0.5666319 | 0.21195029 | -2.6734191 | 0.00750824 | 0.08498664 |
| REEP3 | 152.807863 | -0.3038365 | 0.34251063 | -0.8870864 | 0.37503241 | 0.62603002 |
| AC022387.1 | 27.4184633 | 0.06550126 | 0.63589449 | 0.10300649 | 0.91795782 | 0.96421386 |
| LINC01515 | 33.2064166 | 2.12625299 | 1.01429017 | 2.09629656 | 0.03605589 | 0.18943495 |
| CTNNA3 | 30.2730351 | 0.91166414 | 0.70373401 | 1.29546693 | 0.1951592 | 0.44799892 |
| SIRT1 | 55.4316962 | -1.0379155 | 0.33898774 | -3.0618082 | 0.00220004 | 0.0444455 |
| HERC4 | 407.3922 | -0.9095771 | 0.24212841 | -3.7565897 | 0.00017224 | 0.00982049 |
| PBLD | 40.7719897 | 0.2908267 | 0.36855433 | 0.7891013 | 0.4300528 | 0.67197671 |
| HNRNPH3 | 133.661477 | -0.6653737 | 0.31177716 | -2.1341321 | 0.03283197 | 0.17901486 |
| RUFY2 | 100.007641 | 0.15734629 | 0.27552819 | 0.57107146 | 0.56795121 | 0.77103518 |
| DNA2 | 21.5532201 | -0.5614551 | 0.44094849 | -1.2732896 | 0.20291529 | 0.45686328 |
| SLC25A16 | 165.068704 | -0.3428461 | 0.33975074 | -1.0091106 | 0.31292158 | 0.57102302 |
| TET1 | 29.0997975 | 1.22113527 | 0.62533619 | 1.95276601 | 0.05084733 | 0.22622652 |
| CCAR1 | 220.042505 | -0.6678395 | 0.27547208 | -2.4243454 | 0.01533601 | 0.12206959 |
| STOX1 | 18.4230244 | 1.83195483 | 0.56195667 | 3.25995748 | 0.00111429 | NA |
| DDX50 | 70.174577 | -0.3703859 | 0.28371778 | -1.3054729 | 0.19173187 | 0.44319545 |
| DDX21 | 46.8779426 | -0.9083882 | 0.35480259 | -2.5602637 | 0.01045928 | 0.1006115 |
| KIF1BP | 33.1726267 | 0.02750208 | 0.34297083 | 0.08018782 | 0.93608788 | 0.97405271 |

| | | | | | | |
|---|---|---|---|---|---|---|
| VPS26A | 59.1179486 | -0.4987154 | 0.31035658 | -1.6069109 | 0.10807393 | 0.33009722 |
| SUPV3L1 | 40.619715 | -0.7619144 | 0.36437762 | -2.0910022 | 0.03652787 | 0.19061356 |
| HKDC1 | 179.035863 | -1.7990331 | 0.61027209 | -2.9479197 | 3.20E-03 | 5.62E-02 |
| HK1 | 163.492323 | -0.9069971 | 0.27460008 | -3.3029748 | 0.00095665 | 0.02829008 |
| H2AFY2 | 48.1563379 | 0.53146946 | 0.33895318 | 1.56797309 | 0.11688741 | 0.34414008 |
| AIFM2 | 16.8476279 | 0.07880183 | 0.43098184 | 0.18284258 | 0.85492153 | NA |
| TYSND1 | 14.2544952 | 1.29635193 | 0.51894795 | 2.49803842 | 0.01248827 | NA |
| SAR1A | 45.774189 | -0.5714707 | 0.31888558 | -1.792087 | 0.07311903 | 0.26939652 |
| PPA1 | 55.2811282 | -0.3940444 | 0.31313669 | -1.2583783 | 2.08E-01 | 0.4622639 |
| EIF4EBP2 | 41.7555725 | -0.0042045 | 0.38603698 | -0.0108914 | 0.9913101 | 0.99583841 |
| SGPL1 | 76.6970143 | 0.0946166 | 0.31198654 | 0.30327141 | 0.76168303 | 0.88240131 |
| PCBD1 | 23.6311425 | 0.58337626 | 0.41807505 | 1.39538645 | 0.16289934 | 0.40961653 |
| UNC5B | 34.1430404 | -0.6490528 | 0.53534978 | -1.2123901 | 0.22536308 | 0.48277751 |
| PSAP | 142.341462 | 0.25999913 | 0.45376372 | 0.57298351 | 0.56665586 | 0.77045578 |
| CHST3 | 33.110727 | -0.2493467 | 0.39520416 | -0.6309314 | 0.52808541 | 0.74274895 |
| ASCC1 | 64.4423779 | -0.3176781 | 0.27530656 | -1.1539068 | 0.24853838 | 0.50492395 |
| ANAPC16 | 171.533894 | 0.30805444 | 0.25971797 | 1.18611141 | 0.2355783 | 0.49353462 |
| DDIT4 | 19.1081226 | -0.4497594 | 0.68830598 | -0.6534294 | 0.51347947 | NA |
| DNAJB12 | 23.865866 | -0.435291 | 0.38535152 | -1.1295946 | 0.25864709 | 0.51709263 |
| MICU1 | 177.155545 | -0.1746309 | 0.26321195 | -0.6634613 | 0.50703516 | 0.72824214 |
| MCU | 155.440021 | -0.3760242 | 0.31497487 | -1.1938228 | 0.2325473 | 0.49082729 |
| P4HA1 | 187.791998 | -0.9882291 | 0.56113238 | -1.7611336 | 0.0782158 | 0.27886875 |
| ECD | 42.8969776 | -0.4004859 | 0.30161269 | -1.3278153 | 0.18423913 | 0.43435248 |
| FAM149B1 | 68.1539603 | 0.30361563 | 0.4474701 | 0.67851602 | 0.49744457 | 0.72133441 |
| DNAJC9 | 54.2662106 | -0.4133406 | 0.36194749 | -1.1419906 | 0.25345795 | 0.51069785 |
| MRPS16 | 24.8370138 | 0.01339143 | 0.36030914 | 0.03716651 | 0.97035224 | 0.98796166 |
| CFAP70 | 87.1373926 | 1.00213404 | 0.61665492 | 1.62511319 | 0.10413844 | 0.32399095 |
| ANXA7 | 99.4785797 | -0.3647425 | 0.26790894 | -1.3614422 | 0.17337399 | 0.42238719 |
| PPP3CB | 131.352037 | -0.0478167 | 0.21369949 | -0.2237569 | 0.8229465 | 0.91475524 |
| USP54 | 80.750521 | -0.3250949 | 0.27969199 | -1.1623318 | 0.24510073 | 0.5022131 |
| AGAP5 | 16.68439 | 0.54046595 | 0.49857195 | 1.08402798 | 0.27835239 | NA |
| SEC24C | 35.5639334 | -0.5263562 | 0.33045708 | -1.5928127 | 0.11120221 | 0.33460648 |
| FUT11 | 24.2851769 | -0.9359375 | 0.57018742 | -1.6414559 | 0.10070281 | 0.31976466 |
| ZSWIM8 | 56.9429073 | -0.8082768 | 0.33986842 | -2.3782052 | 0.01739714 | 0.13115162 |
| CAMK2G | 40.651622 | 0.99960292 | 0.36444332 | 2.74282135 | 0.00609138 | 0.07661007 |
| VCL | 177.969204 | -1.0276992 | 0.35975855 | -2.856636 | 0.00428157 | 0.06301187 |
| AP3M1 | 24.077903 | 0.267343 | 0.38802743 | 0.68897965 | 0.49083607 | 0.7166095 |
| ADK | 627.245172 | -0.1091375 | 0.11639563 | -0.9376421 | 0.34842835 | 0.60511124 |
| KAT6B | 242.390243 | -0.3290588 | 0.24098484 | -1.365475 | 0.17210382 | 0.42066785 |
| SAMD8 | 50.7910864 | -0.4387493 | 0.39812092 | -1.1020504 | 0.27043978 | 0.52885543 |
| VDAC2 | 48.1927431 | 0.14308882 | 0.30354816 | 0.47138753 | 0.63736402 | 0.81288995 |
| COMTD1 | 16.8886874 | 0.14167585 | 0.60906565 | 0.23261178 | 0.81606288 | NA |
| LRMDA | 1740.16663 | 0.40519206 | 0.35812412 | 1.13142912 | 0.25787452 | 0.51619036 |
| KCNMA1 | 75.0645676 | -3.2998209 | 1.2959853 | -2.546187 | 0.01089068 | 0.10244908 |
| DLG5 | 128.71518 | -0.5023832 | 0.42219591 | -1.1899291 | 0.23407425 | 0.49243138 |
| POLR3A | 45.7556165 | -0.4636975 | 0.35538152 | -1.304788 | 0.19196505 | 0.44344387 |
| RPS24 | 117.783366 | 0.23776023 | 0.41882637 | 0.56768209 | 0.57025086 | 0.77196291 |
| ZMIZ1-AS1 | 25.8535399 | -2.197369 | 0.71862833 | -3.0577267 | 0.00223023 | 0.04482003 |
| ZMIZ1 | 273.385023 | -0.4508358 | 0.3340819 | -1.3494768 | 0.17718386 | 0.42569745 |
| PPIF | 20.0116161 | 0.10555347 | 0.50931494 | 0.20724596 | 0.83581777 | NA |
| NUTM2B-AS | 210.148196 | 0.19789211 | 0.34948311 | 0.56624228 | 0.57122909 | 0.77245539 |
| TMEM254 | 18.5412442 | 0.47251079 | 0.57987334 | 0.81485171 | 0.41515719 | NA |
| ANXA11 | 220.448622 | -0.1439782 | 0.23294696 | -0.6180728 | 0.53652735 | 0.74852479 |
| FAM213A | 42.7586622 | 0.32214841 | 0.31748778 | 1.01467971 | 0.31025853 | 0.56995192 |

| | | | | | | |
|---|---|---|---|---|---|---|
| TSPAN14 | 54.9516728 | 0.65944858 | 0.33301739 | 1.98022263 | 0.04767852 | 0.2185715 |
| NRG3 | 116.972105 | -0.9667531 | 0.54963389 | -1.7589037 | 0.07859387 | 0.27941662 |
| GHITM | 46.2706368 | 0.42883561 | 0.42739547 | 1.00336957 | 0.31568258 | 0.57293299 |
| CCSER2 | 195.108066 | -0.1484967 | 0.25943746 | -0.5723795 | 0.5670649 | 0.77060355 |
| WAPL | 161.604325 | -0.5094149 | 0.20684289 | -2.4628107 | 0.01378527 | 0.11520651 |
| BMPR1A | 277.318673 | 0.31690823 | 0.284227 | 1.11498285 | 0.26485777 | 0.52419909 |
| GLUD1 | 115.479062 | 0.70109021 | 0.23867781 | 2.93739168 | 0.00330986 | 0.05707588 |
| SHLD2 | 196.323806 | 0.03270274 | 0.22592065 | 0.14475321 | 0.88490573 | 0.94638313 |
| NUTM2A-AS | 228.281409 | 0.08813525 | 0.26227688 | 0.33603895 | 0.73684149 | 0.86917557 |
| MINPP1 | 16.5290042 | 0.87597482 | 0.45972623 | 1.90542708 | 0.05672458 | NA |
| ATAD1 | 106.936765 | 0.26388979 | 0.32712368 | 0.80669731 | 0.4198409 | 0.6636918 |
| PTEN | 281.275944 | -0.3480205 | 0.19298779 | -1.803329 | 0.07133655 | 0.26675397 |
| RNLS | 89.7520411 | 0.47868099 | 0.41304738 | 1.15890091 | 0.24649658 | 0.50296216 |
| LIPA | 47.9891505 | -0.4126047 | 0.42706138 | -0.9661486 | 0.33396984 | 0.58917394 |
| IFIT2 | 23.9538407 | -2.0460374 | 0.98955556 | -2.0676326 | 0.03867458 | 0.1958098 |
| IFIT3 | 49.0434213 | -2.9323495 | 1.39922624 | -2.0956936 | 0.03610938 | 0.18958655 |
| IFIT1 | 37.4763867 | -0.569885 | 1.03660508 | -0.5497609 | 0.58248337 | 0.7783237 |
| SLC16A12 | 23.6925026 | 1.73831192 | 0.54391702 | 3.19591383 | 0.00139389 | 0.03373821 |
| PANK1 | 52.6187765 | 0.93015255 | 0.47792377 | 1.94623621 | 0.05162637 | 0.22799911 |
| RPP30 | 17.2418613 | 0.38378156 | 0.48848683 | 0.78565384 | 0.43207031 | NA |
| PCGF5 | 234.774313 | -0.8263934 | 0.25759042 | -3.2081682 | 0.00133583 | 0.03284955 |
| HECTD2 | 100.334978 | 0.37221584 | 0.30966629 | 1.20199017 | 0.22936734 | 0.48755603 |
| TNKS2 | 174.104567 | -0.5748967 | 0.21842229 | -2.6320422 | 0.00848733 | 0.09010619 |
| BTAF1 | 166.381016 | -0.8862968 | 0.34450389 | -2.5726756 | 0.01009157 | 0.0992926 |
| CPEB3 | 409.809976 | 0.98726066 | 0.22854558 | 4.31975384 | 1.56E-05 | 0.00207296 |
| 5-Mar | 83.2655258 | 0.03462756 | 0.3305738 | 0.10474988 | 0.91657428 | 0.96401363 |
| IDE | 67.6473093 | 0.72454058 | 0.29844203 | 2.42774309 | 0.0151931 | 0.1213084 |
| EXOC6 | 113.8439 | -0.5292344 | 0.23711283 | -2.231994 | 0.02561536 | 0.15662748 |
| MYOF | 262.229569 | -1.0665907 | 0.31375737 | -3.3994125 | 6.75E-04 | 0.02351966 |
| FRA10AC1 | 56.9407044 | 0.31293184 | 0.29510438 | 1.0604107 | 0.2889578 | 0.54835014 |
| NOC3L | 21.7993688 | -0.5416962 | 0.41226947 | -1.3139371 | 0.18886736 | 0.43958636 |
| TBC1D12 | 60.4342771 | -0.167548 | 0.29118851 | -0.5753935 | 0.56502517 | 0.7693258 |
| HELLS | 19.0982423 | 1.37412958 | 0.66866922 | 2.05502144 | 0.03987693 | NA |
| SORBS1 | 62.1029194 | -2.0317336 | 0.42326592 | -4.8001351 | 1.59E-06 | 0.00051083 |
| ALDH18A1 | 53.3808117 | 0.22206279 | 0.43031047 | 0.51605249 | 0.60581775 | 0.79431477 |
| TCTN3 | 57.1724147 | 0.05331806 | 0.30383075 | 0.17548605 | 0.86069773 | 0.93500722 |
| ENTPD1-AS1 | 131.187357 | 1.45186858 | 0.47908614 | 3.03049586 | 0.00244153 | 0.04713978 |
| ZNF518A | 237.577762 | 0.31283938 | 0.30277941 | 1.0332254 | 0.30149843 | 0.56180195 |
| BLNK | 478.039874 | 0.47520808 | 0.49382559 | 0.96229942 | 0.33589922 | 0.59162844 |
| TM9SF3 | 169.482756 | -0.2233182 | 0.23923774 | -0.933457 | 0.35058404 | 0.60737992 |
| LCOR | 317.131286 | -0.5118094 | 0.21770877 | -2.3508903 | 0.01872856 | 0.13535558 |
| RRP12 | 26.2264033 | -0.5585328 | 0.47447073 | -1.1771702 | 0.23912759 | 0.49686728 |
| PGAM1 | 53.6304878 | -0.0561829 | 0.45309385 | -0.1239984 | 0.90131653 | 0.95637074 |
| EXOSC1 | 28.6723134 | -0.2953955 | 0.39112411 | -0.7552475 | 0.45010048 | 0.68770076 |
| ZDHHC16 | 17.5223935 | 0.22856785 | 0.420458 | 0.54361638 | 0.58670549 | NA |
| MMS19 | 71.1773175 | -0.0008984 | 0.24706158 | -0.0036365 | 0.99709849 | 0.99868755 |
| UBTD1 | 35.5340921 | 1.08973844 | 0.34296554 | 3.17739918 | 0.00148602 | 0.03500548 |
| AL355315.1 | 13.3635749 | 1.42473179 | 0.68520087 | 2.0792907 | 0.03759064 | NA |
| MORN4 | 28.5590356 | 1.39362635 | 0.41317904 | 3.37293573 | 0.00074371 | 0.02425576 |
| ZFYVE27 | 35.0207925 | -0.200418 | 0.34420011 | -0.5822719 | 0.56038356 | 0.76571495 |
| R3HCC1L | 119.039024 | -0.2269369 | 0.18462571 | -1.2291727 | 0.21900704 | 0.47565385 |
| HPS1 | 18.6719506 | -0.4054358 | 0.44648233 | -0.9080669 | 0.36384289 | NA |
| GOT1 | 128.383569 | 0.10028476 | 0.29335706 | 0.34185222 | 0.73246211 | 0.86774832 |
| AL391684.1 | 11.4579655 | 1.14793384 | 0.77550366 | 1.48024297 | 0.13880842 | NA |

| | | | | | | |
|---|---|---|---|---|---|---|
| SLC25A28 | 28.8711928 | 0.32860259 | 0.53692973 | 0.61200297 | 0.54053579 | 0.74994017 |
| CUTC | 33.2684904 | 0.34142255 | 0.36470349 | 0.93616476 | 0.34918836 | 0.60589538 |
| COX15 | 33.1036216 | 0.46912508 | 0.36175474 | 1.29680426 | 0.19469854 | 0.4473417 |
| DNMBP | 85.5824083 | -1.2590491 | 0.36196637 | -3.4783593 | 0.00050449 | 0.02033028 |
| ERLIN1 | 27.6389965 | 0.71197768 | 0.40471791 | 1.7591949 | 0.07854441 | 0.27941662 |
| CHUK | 29.7481959 | -0.7490959 | 0.39245042 | -1.9087658 | 0.05629231 | 0.23642261 |
| CWF19L1 | 41.1906564 | -0.208981 | 0.45945981 | -0.4548408 | 0.64922381 | 0.81913234 |
| BLOC1S2 | 16.141538 | 0.1032575 | 0.48260918 | 0.21395677 | 0.83058077 | NA |
| SEC31B | 18.9355722 | 0.86651479 | 0.48034579 | 1.80393958 | 0.07124078 | NA |
| HIF1AN | 33.7728612 | -0.5417787 | 0.32578537 | -1.6629927 | 0.09631389 | 0.31156136 |
| PAX2 | 154.34314 | 0.28593018 | 0.45166848 | 0.63305321 | 0.5266989 | 0.74154041 |
| SLF2 | 80.7819701 | -0.2536274 | 0.28712268 | -0.8833416 | 0.37705171 | 0.62858285 |
| MRPL43 | 17.4841733 | 0.54046806 | 0.47994135 | 1.12611273 | 0.26011781 | NA |
| LZTS2 | 25.9642899 | -0.2105434 | 0.39307452 | -0.5356324 | 0.59221264 | 0.7855446 |
| BTRC | 192.704325 | 0.53259856 | 0.20754036 | 2.56624086 | 0.01028074 | 0.09994074 |
| DPCD | 15.0483672 | 0.63782736 | 0.5254716 | 1.21381891 | 0.22481689 | NA |
| POLL | 20.9349423 | 0.69487496 | 0.48083247 | 1.44514982 | 0.14841581 | NA |
| FBXW4 | 137.577511 | -0.0326275 | 0.23592411 | -0.1382968 | 0.89000588 | 0.94967256 |
| OGA | 292.540385 | -0.326849 | 0.18114759 | -1.8043246 | 0.07118044 | 0.2665798 |
| ARMH3 | 174.766961 | -0.1639003 | 0.20466904 | -0.8008066 | 0.42324362 | 0.66684898 |
| LDB1 | 32.2436563 | 1.17362712 | 0.45477002 | 2.5807047 | 0.00985989 | 0.09830513 |
| PPRC1 | 20.4173213 | -1.8226732 | 0.581147 | -3.1363376 | 0.00171072 | NA |
| GBF1 | 228.104856 | -0.1488496 | 0.26637247 | -0.5588025 | 0.57629649 | 0.77421002 |
| NFKB2 | 25.7753947 | -0.9203766 | 0.69378702 | -1.3265982 | 0.18464164 | 0.43476148 |
| FBXL15 | 16.5054209 | 1.17108898 | 0.64242458 | 1.82292056 | 0.06831542 | NA |
| CUEDC2 | 14.5992481 | 0.17397157 | 0.45179648 | 0.38506623 | 0.70018834 | NA |
| ACTR1A | 25.396079 | -0.288397 | 0.36714878 | -0.7855045 | 0.43215781 | 0.6728327 |
| SUFU | 77.2166577 | 0.93259138 | 0.36508918 | 2.55442078 | 0.01063646 | 0.1014484 |
| TRIM8 | 40.8974328 | 0.84566914 | 0.3964267 | 2.13322949 | 0.0329059 | 0.17912076 |
| ARL3 | 38.8840338 | 0.71142289 | 0.37732518 | 1.8854371 | 0.05937085 | 0.24193042 |
| SFXN2 | 18.1962958 | 0.40658394 | 0.517511 | 0.78565275 | 0.43207095 | NA |
| WBP1L | 88.2623753 | -0.8231951 | 0.2666544 | -3.0871237 | 0.00202103 | 0.04215692 |
| CNNM2 | 1090.24453 | 0.22901053 | 0.26488705 | 0.86455918 | 3.87E-01 | 6.37E-01 |
| NT5C2 | 311.050821 | -0.8581081 | 0.37246384 | -2.3038695 | 0.02122997 | 0.1427136 |
| PCGF6 | 28.6590012 | -0.8209218 | 0.39243309 | -2.0918772 | 0.0364495 | 0.19061356 |
| ATP5MD | 37.2539223 | 0.48942748 | 0.42235026 | 1.15881895 | 0.24652999 | 0.50296216 |
| PDCD11 | 23.1256959 | -0.366055 | 0.46398964 | -0.7889293 | 0.43015335 | 0.67197671 |
| SH3PXD2A | 135.329949 | 0.30697025 | 0.39557729 | 0.77600574 | 4.38E-01 | 0.67793318 |
| STN1 | 43.6321138 | -0.709658 | 0.34159296 | -2.0774959 | 0.03775581 | 0.19425568 |
| SLK | 116.66724 | -0.300165 | 0.35644208 | -0.8421142 | 0.39972401 | 0.64766685 |
| CFAP43 | 24.4561493 | 1.70522024 | 0.7928502 | 2.15074706 | 0.03149617 | 0.17516331 |
| GSTO1 | 38.0046567 | -0.9905082 | 0.4811151 | -2.058776 | 0.03951571 | 0.19795704 |
| GSTO2 | 101.77099 | 0.98101415 | 0.34079159 | 2.87863365 | 0.00399402 | 0.06175835 |
| SORCS3 | 32.2415075 | -1.2711209 | 0.72905553 | -1.7435173 | 8.12E-02 | 2.84E-01 |
| XPNPEP1 | 49.6463698 | -0.0870391 | 0.38068043 | -0.2286408 | 0.81914812 | 0.913067 |
| ADD3 | 248.73628 | 0.49046756 | 0.31807396 | 1.54199218 | 0.12307549 | 0.35321106 |
| MXI1 | 82.9418962 | 0.84841839 | 0.39113013 | 2.16914609 | 0.03007159 | 0.17145264 |
| SMNDC1 | 26.3750912 | -0.3383658 | 0.47579318 | -0.7111616 | 0.47698411 | 0.7051944 |
| SMC3 | 47.431159 | -0.3341417 | 0.35693743 | -0.9361352 | 0.34920361 | 0.60589538 |
| PDCD4 | 85.9807266 | -0.1059585 | 0.46636162 | -0.2272025 | 0.82026628 | 0.91338577 |
| BBIP1 | 44.4068805 | 0.37396725 | 0.35206182 | 1.06222041 | 0.28813564 | 0.54746482 |
| SHOC2 | 116.79919 | -0.1473633 | 0.29361596 | -0.5018913 | 0.61574398 | 0.80084174 |
| GPAM | 20.9873216 | 0.80580315 | 0.48070662 | 1.67628886 | 0.09368163 | NA |
| ZDHHC6 | 28.9255887 | 0.04061267 | 0.32585054 | 0.1246359 | 0.90081181 | 0.95621962 |

| | | | | | | |
|---|---|---|---|---|---|---|
| VTI1A | 443.245277 | -0.1133796 | 0.12615611 | -0.8987244 | 0.36879945 | 0.62346791 |
| TCF7L2 | 284.76113 | 0.07635723 | 0.2321617 | 0.32889677 | 0.74223372 | 0.87152705 |
| CASP7 | 27.584089 | -0.8968356 | 0.37454506 | -2.3944666 | 0.01664456 | 0.12787672 |
| NHLRC2 | 93.2448283 | 0.06505657 | 0.2474262 | 0.26293322 | 0.79260205 | 0.8986875 |
| CCDC186 | 69.7428332 | -0.4120484 | 0.29588863 | -1.3925794 | 0.16374702 | 0.41042742 |
| ABLIM1 | 857.584623 | 0.09224552 | 0.34735165 | 0.2655681 | 0.79057185 | 0.89710033 |
| FAM160B1 | 52.2373736 | 0.62345244 | 0.30910567 | 2.01695567 | 0.04370014 | 0.20890044 |
| TRUB1 | 28.2351057 | 0.43323915 | 0.41255552 | 1.0501354 | 0.29365587 | 0.55303871 |
| HSPA12A | 105.081151 | 0.4991426 | 0.49475174 | 1.00887488 | 0.31303465 | 0.57102302 |
| SHTN1 | 111.442431 | -0.2299418 | 0.27566998 | -0.8341199 | 0.40421342 | 0.65170312 |
| PDZD8 | 150.921652 | -0.8683043 | 0.35939087 | -2.4160443 | 0.01569015 | 0.12386371 |
| EMX2OS | 176.590803 | 0.73627255 | 0.4507897 | 1.63329498 | 0.10240697 | 0.32120262 |
| EMX2 | 51.4154563 | 1.02892213 | 0.5023182 | 2.0483473 | 0.04052598 | 0.20046817 |
| RAB11FIP2 | 102.216526 | 0.54143839 | 0.41330372 | 1.31002545 | 0.19018722 | 0.44142207 |
| CASC2 | 89.6857721 | 2.04345781 | 0.69997188 | 2.91934271 | 0.0035077 | 0.05869303 |
| FAM204A | 62.0970121 | -0.2161141 | 0.35038117 | -0.6167971 | 0.53736855 | 0.74878549 |
| CACUL1 | 129.218722 | -0.5343217 | 0.24976181 | -2.1393253 | 0.03240933 | 0.17754567 |
| EIF3A | 135.075534 | -0.4603023 | 0.34122083 | -1.3489866 | 0.17734128 | 0.42576289 |
| FAM45A | 24.8327073 | 0.43089598 | 0.47979015 | 0.89809259 | 0.36913619 | 0.6236443 |
| SFXN4 | 38.1544037 | 0.2565657 | 0.44589043 | 0.57540078 | 0.56502028 | 0.7693258 |
| PRDX3 | 34.0857763 | 0.97716172 | 0.41583976 | 2.34985159 | 0.0187809 | 0.13549695 |
| TIAL1 | 85.3222446 | 0.10498057 | 0.25313795 | 0.41471685 | 0.67834919 | 0.83997004 |
| BAG3 | 23.6493737 | 0.29404529 | 0.41983878 | 0.70037668 | 0.48369209 | 0.71144239 |
| INPP5F | 46.1708808 | 0.85064815 | 0.34230407 | 2.48506586 | 0.01295275 | 0.11079049 |
| MCMBP | 43.2900785 | 0.4652907 | 0.32212799 | 1.44442805 | 0.1486186 | 0.38909225 |
| SEC23IP | 80.0378808 | -0.8126408 | 0.31961794 | -2.5425382 | 0.01100506 | 0.10267384 |
| PLPP4 | 14.7027932 | -0.4196381 | 0.62712513 | -0.6691457 | 5.03E-01 | NA |
| WDR11-AS1 | 14.6733784 | 1.25992974 | 0.71734472 | 1.75637975 | 0.07902359 | NA |
| WDR11 | 113.644424 | 0.53245493 | 0.28143526 | 1.89192685 | 0.05850073 | 0.24020328 |
| FGFR2 | 152.293144 | -0.2886781 | 0.42001052 | -0.6873115 | 0.49188642 | 0.71672078 |
| ATE1 | 170.244291 | -0.1279154 | 0.25639207 | -0.4989055 | 0.61784599 | 0.80236132 |
| NSMCE4A | 19.8147548 | -1.2201848 | 0.46811953 | -2.6065667 | 0.0091455 | NA |
| TACC2 | 239.567642 | -0.1467262 | 0.73059544 | -0.200831 | 0.84083076 | 0.92355925 |
| PLEKHA1 | 560.279109 | -0.4371731 | 0.30039771 | -1.4553145 | 1.46E-01 | 3.85E-01 |
| HTRA1 | 17.8155055 | 2.04386201 | 0.60888308 | 3.3567397 | 0.00078867 | NA |
| C10orf88 | 23.8756393 | -0.1586461 | 0.47005279 | -0.3375069 | 0.73573481 | 0.86841755 |
| PSTK | 19.5446301 | 0.10781843 | 0.54507882 | 0.19780338 | 0.8431989 | NA |
| IKZF5 | 25.4304539 | -0.2470376 | 0.34671556 | -0.7125081 | 0.47615019 | 0.70423528 |
| ACADSB | 63.2270431 | 0.04003551 | 0.29582166 | 0.13533663 | 0.89234572 | 0.95077312 |
| BUB3 | 27.9923862 | 0.5989042 | 0.37322799 | 1.60466045 | 0.10856855 | 0.33029728 |
| CPXM2 | 495.176793 | -1.059886 | 0.52335225 | -2.0251867 | 0.04284819 | 0.20681879 |
| OAT | 17.0637578 | 0.21967813 | 0.54797462 | 0.40089108 | 0.68850032 | NA |
| LHPP | 31.6082045 | 0.93151155 | 0.51103931 | 1.82277866 | 0.06833692 | 0.25987613 |
| ABRAXAS2 | 30.0720857 | -0.1149784 | 0.38192811 | -0.3010472 | 0.76337853 | 0.88288632 |
| ZRANB1 | 73.2797382 | -0.3709617 | 0.32867264 | -1.1286662 | 0.25903865 | 0.51774098 |
| CTBP2 | 480.403571 | -0.7309789 | 0.30800128 | -2.3732983 | 0.01763002 | 0.13213074 |
| EDRF1 | 52.3838258 | -0.6724767 | 0.2706127 | -2.4850153 | 0.01295459 | 0.11079049 |
| UROS | 21.5862436 | 0.74380599 | 0.44546098 | 1.66974443 | 0.09496994 | 0.30934559 |
| BCCIP | 44.2161105 | -0.1911538 | 0.28344353 | -0.6743983 | 0.50005815 | 0.722129 |
| DHX32 | 134.476213 | -0.5563218 | 0.21773271 | -2.5550677 | 0.01061671 | 0.1014484 |
| FANK1 | 53.6211974 | 0.25953764 | 0.48251819 | 0.53788156 | 0.59065882 | 0.78506319 |
| DOCK1 | 681.432307 | -0.2272427 | 0.22437419 | -1.0127843 | 0.31116321 | 0.57037944 |
| FAM196A | 19.0283625 | 0.92234157 | 0.60667477 | 1.52032293 | 0.12842983 | NA |
| MGMT | 96.5134888 | 0.69265293 | 0.44598346 | 1.55309106 | 0.12040138 | 0.34903213 |

| | | | | | | |
|---|---|---|---|---|---|---|
| AL157832.3 | 13.7193165 | 1.04842325 | 0.56983347 | 1.83987657 | 0.06578636 | NA |
| C10orf143 | 15.8232623 | 0.11286827 | 0.6138419 | 0.18387189 | 0.85411395 | NA |
| GLRX3 | 77.2542369 | -0.9837828 | 0.39527571 | -2.4888521 | 0.01281563 | 0.11046442 |
| PPP2R2D | 136.208861 | -0.3751418 | 0.36672021 | -1.0229645 | 0.30632464 | 0.56595793 |
| BNIP3 | 74.6310502 | -0.2639358 | 0.68196512 | -0.3870224 | 0.69873964 | 0.85102865 |
| STK32C | 23.7847719 | -0.4828666 | 0.36022285 | -1.3404664 | 0.18009376 | 0.42915841 |
| LRRC27 | 22.1789375 | 1.27663892 | 0.61045874 | 2.09127797 | 0.03650315 | 0.19061356 |
| INPP5A | 95.5152459 | -0.4050994 | 0.38285944 | -1.0580892 | 0.2900148 | 0.5494078 |
| KNDC1 | 22.7329088 | 0.39098606 | 0.63730973 | 0.61349457 | 0.53954937 | 0.74929145 |
| TUBGCP2 | 61.9433866 | -0.5563204 | 0.29237633 | -1.9027546 | 0.05707258 | 0.23764428 |
| ECHS1 | 36.6849465 | 0.5406771 | 0.37708045 | 1.43385078 | 0.15161485 | 0.39273741 |
| SCART1 | 22.8200512 | -0.6746147 | 0.52871045 | -1.2759625 | 0.20196876 | 0.45588081 |
| SLC6A12 | 50.5499416 | -1.1285774 | 0.54686249 | -2.0637316 | 0.03904318 | 0.19705923 |
| KDM5A | 160.313196 | -0.0924365 | 0.18619828 | -0.4964414 | 0.61958302 | 0.80279998 |
| B4GALNT3 | 178.252741 | -0.6440684 | 0.3654213 | -1.7625365 | 0.07797869 | 0.27864531 |
| WNK1 | 1504.36638 | 2.02456229 | 0.49692379 | 4.07419071 | 4.62E-05 | 0.00423103 |
| RAD52 | 42.3917638 | 0.13065664 | 0.37730182 | 0.3462921 | 0.72912321 | 0.86533507 |
| ERC1 | 343.302817 | 0.03579853 | 0.13134274 | 0.27255808 | 0.78519294 | 0.89452315 |
| ADIPOR2 | 120.162512 | -0.2513008 | 0.39183191 | -0.6413484 | 0.52129633 | 0.73784808 |
| DCP1B | 49.3851011 | 0.44314747 | 0.3902802 | 1.1354598 | 0.25618271 | 0.51430317 |
| CACNA1C | 32.2948827 | 1.78377253 | 0.58447328 | 3.0519317 | 0.00227374 | 0.04522213 |
| FKBP4 | 22.6528531 | 0.79722103 | 0.49266785 | 1.6181714 | 0.10562566 | 0.32676878 |
| ITFG2 | 68.3893374 | 0.3259146 | 0.39858337 | 0.81768237 | 0.41353858 | 0.65832605 |
| TULP3 | 121.098976 | -0.4001999 | 0.27237177 | -1.469315 | 0.14174737 | 0.37968477 |
| TEAD4 | 56.8291044 | -0.7740807 | 0.6692165 | -1.156697 | 2.47E-01 | 0.50358016 |
| TSPAN9 | 473.822312 | 0.22656985 | 0.35137713 | 0.64480533 | 0.51905335 | 0.73602682 |
| PARP11 | 24.9508791 | -0.6033683 | 0.38147225 | -1.5816832 | 0.1137219 | 0.33940189 |
| TIGAR | 20.9187173 | -0.6393794 | 0.58266028 | -1.0973451 | 0.27249056 | NA |
| C12orf4 | 34.9293436 | 0.44531682 | 0.40468342 | 1.10040787 | 0.27115445 | 0.52944592 |
| CD9 | 70.0388171 | 0.52554257 | 0.41394781 | 1.26958655 | 0.20423194 | 0.45771763 |
| TNFRSF1A | 42.5921509 | -0.005894 | 0.33138008 | -0.0177862 | 9.86E-01 | 9.94E-01 |
| SCNN1A | 197.615105 | 0.88974291 | 0.26254523 | 3.38891294 | 0.0007017 | 0.02388192 |
| LTBR | 22.7621859 | -0.4698833 | 0.46602099 | -1.0082879 | 0.31331629 | 0.57102302 |
| TAPBPL | 11.8907027 | 0.05144108 | 0.5658607 | 0.09090767 | 0.92756595 | NA |
| MRPL51 | 18.9414483 | 0.58287424 | 0.50715315 | 1.14930616 | 0.25042976 | NA |
| NCAPD2 | 21.9363582 | -0.0824576 | 0.54670432 | -0.1508267 | 0.88011243 | 0.94376451 |
| GAPDH | 167.543287 | 0.56094052 | 0.49799311 | 1.12640216 | 0.25999533 | 0.51897927 |
| CHD4 | 33.1772899 | -0.3222386 | 0.34644794 | -0.9301213 | 0.35230829 | 0.60964858 |
| ING4 | 26.9827554 | -0.3394756 | 0.38834742 | -0.8741543 | 0.38203424 | 0.6334526 |
| ZNF384 | 23.3589165 | -0.1825979 | 0.44696101 | -0.4085322 | 0.68288303 | 0.84226129 |
| COPS7A | 18.519654 | 0.5679137 | 0.49413405 | 1.14931102 | 0.25042776 | NA |
| MLF2 | 31.3164955 | 0.12180213 | 0.38643175 | 0.315197 | 0.75261208 | 0.87837077 |
| PTMS | 47.7035631 | 0.2841712 | 0.59911948 | 0.47431474 | 0.63527548 | 0.81143634 |
| USP5 | 14.8196207 | 0.39636141 | 0.55570683 | 0.71325632 | 0.47568716 | NA |
| TPI1 | 66.9947133 | 0.50265523 | 0.50821712 | 0.98905608 | 0.32263571 | 0.57912936 |
| LRRC23 | 23.687885 | -0.4706812 | 0.44049334 | -1.068532 | 0.28528062 | 0.54405474 |
| ATN1 | 48.7428391 | -0.1932449 | 0.33362653 | -0.5792252 | 0.56243724 | 0.76709958 |
| PTPN6 | 20.9763916 | 0.25001238 | 0.47916378 | 0.52176811 | 0.6018318 | NA |
| PHB2 | 30.0547744 | 0.17777778 | 0.39978318 | 0.4446855 | 0.65654705 | 0.82462919 |
| LPCAT3 | 111.674944 | -0.1927645 | 0.21414729 | -0.9001493 | 0.36804079 | 0.62245879 |
| C1S | 38.2188803 | -0.7569955 | 0.3290197 | -2.3007605 | 0.02140517 | 0.14326576 |
| C1RL | 66.1154964 | 0.27165396 | 0.32297613 | 0.84109609 | 4.00E-01 | 0.64782669 |
| CLSTN3 | 26.3064277 | 0.55461944 | 0.60166087 | 0.92181405 | 0.35662558 | 0.61264648 |
| PEX5 | 17.3925099 | -0.3602807 | 0.45761653 | -0.7872982 | 0.43110731 | NA |

| | | | | | | |
|---|---|---|---|---|---|---|
| FOXJ2 | 15.0116379 | -0.4968605 | 0.4844753 | -1.0255641 | 0.30509707 | NA |
| NECAP1 | 20.2581426 | 0.13185008 | 0.44822748 | 0.29415884 | 0.76863653 | NA |
| FAM66C | 18.1447139 | 0.42763213 | 0.54866556 | 0.77940399 | 0.43574177 | NA |
| RIMKLB | 653.607188 | -1.7571554 | 1.08669317 | -1.6169748 | 0.10588373 | 0.32717264 |
| A2ML1-AS1 | 54.6689218 | -0.4938431 | 0.84146336 | -0.586886 | 0.55728028 | 0.763236 |
| PHC1 | 32.726422 | -0.0300405 | 0.38819751 | -0.0773845 | 0.93831767 | 0.97466007 |
| M6PR | 36.8992452 | -0.4971039 | 0.40272659 | -1.2343459 | 0.21707403 | 0.47318572 |
| A2M | 21.4140506 | 0.89829195 | 0.57599104 | 1.55955888 | 0.11886416 | 0.3461587 |
| CLEC2D | 40.3364908 | 0.14478713 | 0.44319985 | 0.32668587 | 0.74390549 | 0.87230965 |
| OLR1 | 63.9081348 | 2.0087569 | 0.74258728 | 2.70507852 | 0.00682882 | 0.08049228 |
| TMEM52B | 488.914749 | 2.42074084 | 0.53239201 | 4.5469143 | 5.44E-06 | 0.00116392 |
| GABARAPL1 | 183.2639 | 0.40228355 | 0.25422361 | 1.58240042 | 0.11355819 | 0.33904475 |
| KLRD1 | 152.196988 | 0.29908257 | 0.34689692 | 0.86216553 | 0.38859643 | 0.63787614 |
| YBX3 | 217.030262 | -0.4173636 | 0.29967197 | -1.3927348 | 1.64E-01 | 0.41042742 |
| PRH1 | 14.4455274 | 2.13684915 | 0.68424695 | 3.12292099 | 0.00179066 | NA |
| AC244131.2 | 35.7182553 | 0.47161065 | 0.60458344 | 0.78005883 | 0.43535625 | 0.67626555 |
| ETV6 | 351.091836 | -0.6976857 | 0.35848133 | -1.946226 | 0.05162759 | 0.22799911 |
| BCL2L14 | 49.5836612 | 1.38288133 | 0.51891196 | 2.66496328 | 0.00769967 | 0.08614009 |
| LRP6 | 231.318126 | 0.41279193 | 0.29393089 | 1.40438431 | 0.16020444 | 0.405223 |
| BORCS5 | 62.0065214 | 0.22900941 | 0.30899967 | 0.74113158 | 0.45861366 | 0.6928053 |
| DUSP16 | 91.671506 | -0.0904294 | 0.41850927 | -0.2160749 | 0.82892935 | 0.91696884 |
| CREBL2 | 31.807494 | 0.50004778 | 0.31681067 | 1.57838051 | 0.11447822 | 0.3408661 |
| CDKN1B | 16.3719226 | -0.1687098 | 0.47426171 | -0.3557314 | 0.7220417 | NA |
| APOLD1 | 14.0042572 | -0.5271687 | 0.70580259 | -0.7469067 | 0.45511986 | NA |
| GPRC5D-AS1 | 61.5386886 | -0.6513757 | 0.41722936 | -1.5611934 | 0.11847812 | 0.34566454 |
| HEBP1 | 22.9528958 | 0.38172982 | 0.41256975 | 0.92524919 | 0.3548363 | 0.61168935 |
| EMP1 | 38.9825218 | -1.9268973 | 0.37687459 | -5.1128343 | 3.17E-07 | 1.88E-04 |
| ATF7IP | 230.11895 | -0.2168933 | 0.24063597 | -0.9013338 | 0.36741087 | 0.62220175 |
| AC010168.1 | 16.5390459 | 1.09124961 | 0.71208289 | 1.53247553 | 0.12540513 | NA |
| H2AFJ | 24.0525495 | -0.0222047 | 0.46183132 | -0.0480796 | 0.96165277 | 0.98520449 |
| WBP11 | 28.1621055 | -0.0961704 | 0.33460992 | -0.2874105 | 0.773798 | 0.88782329 |
| C12orf60 | 39.2232692 | 0.39001584 | 0.43669254 | 0.89311315 | 0.37179658 | 0.62455659 |
| EPS8 | 96.4656315 | -1.0081526 | 0.52337346 | -1.9262586 | 0.05407209 | 0.23287156 |
| STRAP | 47.9840613 | -0.1554164 | 0.28783122 | -0.5399566 | 0.58922697 | 0.78397234 |
| DERA | 130.915758 | -0.5566273 | 0.2354677 | -2.3639222 | 0.01808261 | 0.13331596 |
| MGST1 | 28.367727 | 0.46291716 | 0.68568486 | 0.67511649 | 0.49960176 | 0.72201362 |
| PIK3C2G | 418.728206 | 3.51133548 | 0.65366517 | 5.3717647 | 7.80E-08 | 7.50E-05 |
| AC087242.1 | 15.8596232 | 3.12261041 | 0.92622847 | 3.37131767 | 0.0007481 | NA |
| PLEKHA5 | 1273.01487 | -0.5618252 | 0.26056217 | -2.156204 | 0.03106774 | 0.17412703 |
| AEBP2 | 399.099943 | 0.53862029 | 0.29139355 | 1.84842897 | 0.06454031 | 0.25284322 |
| AC024901.1 | 54.0405505 | 0.60501478 | 0.43991538 | 1.3752981 | 0.16903905 | 0.41675001 |
| LINC02398 | 24.7411811 | 1.6476929 | 0.66090673 | 2.49307933 | 0.01266405 | 0.1101415 |
| PYROXD1 | 32.8704694 | -0.6508167 | 0.32752424 | -1.9870794 | 0.0469136 | 0.21661307 |
| RECQL | 21.3505657 | -0.472866 | 0.38174937 | -1.2386818 | 0.21546335 | 0.47115856 |
| GOLT1B | 23.8414443 | -0.4728506 | 0.40766201 | -1.1599084 | 0.24608608 | 0.50296216 |
| LDHB | 203.642238 | 1.16293318 | 0.40176701 | 2.8945462 | 0.00379707 | 0.06101472 |
| AC010197.1 | 78.2333462 | 0.85809261 | 0.6957679 | 1.23330295 | 0.21746276 | 0.47376475 |
| ABCC9 | 34.7530909 | 2.26711252 | 0.95227612 | 2.38073021 | 0.01727836 | 0.13076848 |
| AC008250.1 | 77.4198078 | 1.39212112 | 0.79234239 | 1.75696913 | 0.07892307 | 0.28019874 |
| CMAS | 24.0594128 | -0.1000646 | 0.40191187 | -0.2489715 | 0.80338281 | 0.90408418 |
| C2CD5 | 134.765878 | -0.248386 | 0.28738356 | -0.8643014 | 0.38742235 | 0.63701577 |
| ETNK1 | 312.350805 | -0.158636 | 0.37245989 | -0.4259143 | 0.67017029 | 0.83436422 |
| SOX5 | 87.0192066 | 0.47348799 | 0.57150603 | 0.82849168 | 0.4073921 | 0.65367876 |
| CASC1 | 29.3622183 | 1.7576865 | 0.72992972 | 2.40802153 | 0.01603923 | 0.12520689 |

| | | | | | | |
|---|---|---|---|---|---|---|
| ETFRF1 | 37.7128607 | 0.41731369 | 0.31084725 | 1.34250404 | 0.17943262 | 0.4285116 |
| KRAS | 79.4298762 | 0.04780155 | 0.34748621 | 0.13756388 | 0.8905851 | 0.949686 |
| RASSF8-AS1 | 50.6811847 | -0.4672883 | 0.31131755 | -1.5010021 | 0.13335502 | 0.36762309 |
| RASSF8 | 307.081519 | -0.6945353 | 0.53442174 | -1.2996015 | 0.19373758 | 0.44566591 |
| ITPR2 | 778.105213 | -0.9526841 | 0.36129905 | -2.6368297 | 0.00836848 | 0.08952533 |
| INTS13 | 18.7163815 | 0.14092232 | 0.45390632 | 0.31046564 | 0.75620689 | NA |
| FGFR1OP2 | 59.4236402 | 0.13655818 | 0.30428066 | 0.4487902 | 0.65358301 | 0.82226683 |
| TM7SF3 | 54.5183391 | 1.19988462 | 0.42617664 | 2.81546311 | 0.0048707 | 0.06779343 |
| MED21 | 17.784639 | -0.0755301 | 0.44601801 | -0.1693431 | 0.86552674 | NA |
| STK38L | 78.8224013 | 0.46896719 | 0.29194509 | 1.60635408 | 0.10819614 | 0.33009771 |
| PPFIBP1 | 263.996269 | -0.254084 | 0.23276428 | -1.0915935 | 0.27501183 | 0.5337282 |
| MRPS35 | 42.1375043 | 0.24399596 | 0.34322742 | 0.71088712 | 0.47715419 | 0.7051944 |
| KLHL42 | 43.994065 | 0.7154682 | 0.3217791 | 2.22347631 | 0.0261837 | 0.15845355 |
| CCDC91 | 307.506669 | -0.4080924 | 0.2056681 | -1.984228 | 0.04723043 | 0.21733813 |
| AC009318.1 | 17.0995471 | -0.8509025 | 0.71305902 | -1.1933128 | 0.23274688 | NA |
| ERGIC2 | 68.915115 | 0.00156943 | 0.31363615 | 0.00500397 | 0.99600742 | 0.99816804 |
| TMTC1 | 210.915132 | -1.530281 | 0.626659 | -2.4419676 | 0.01460746 | 0.11902037 |
| IPO8 | 78.3431507 | 0.03445394 | 0.28771045 | 0.11975214 | 0.9046795 | 0.95673178 |
| CAPRIN2 | 55.9450559 | 0.12273502 | 0.26317111 | 0.46636965 | 0.64095093 | 0.81516843 |
| DDX11 | 16.2173305 | -0.2407851 | 0.5970246 | -0.4033084 | 0.68672133 | NA |
| SINHCAF | 114.058384 | -0.0955135 | 0.21111358 | -0.4524272 | 0.65096123 | 0.81992533 |
| DENND5B | 369.673201 | -0.5082155 | 0.27873846 | -1.8232702 | 0.06826247 | 0.25987613 |
| AMN1 | 84.6509514 | -0.8538343 | 0.26056377 | -3.2768727 | 0.00104964 | 0.02953506 |
| KIAA1551 | 126.502322 | -1.0835386 | 0.26290654 | -4.1213831 | 3.77E-05 | 0.00383215 |
| BICD1 | 379.914811 | -0.3892932 | 0.46864293 | -0.8306819 | 4.06E-01 | 0.65318894 |
| FGD4 | 1050.35302 | -0.1231243 | 0.16829965 | -0.7315778 | 0.46442631 | 0.69532957 |
| DNM1L | 161.996471 | -0.0127551 | 0.21087788 | -0.0604858 | 0.95176873 | 0.98050223 |
| YARS2 | 44.9403647 | 0.2933625 | 0.36412428 | 0.80566588 | 0.42043553 | 0.66394999 |
| PKP2 | 16.4918528 | -0.6005326 | 0.45441857 | -1.3215407 | 0.18632115 | NA |
| ALG10B | 12.3017208 | 0.21253784 | 0.51585033 | 0.41201454 | 0.68032876 | NA |
| CPNE8 | 300.548212 | -1.0644752 | 0.38935817 | -2.733923 | 0.00625847 | 0.07721978 |
| KIF21A | 70.6418045 | -0.628178 | 0.46908768 | -1.3391483 | 0.18052241 | 0.42977544 |
| SLC2A13 | 82.2213408 | 0.43214484 | 0.42794512 | 1.00981371 | 0.31258455 | 0.57101636 |
| LRRK2 | 123.466016 | 0.62435592 | 0.64621132 | 0.96617918 | 0.33395453 | 0.58917394 |
| GXYLT1 | 47.496273 | 0.4654824 | 0.30277312 | 1.53739673 | 0.12419619 | 0.35457643 |
| YAF2 | 126.466539 | -0.166291 | 0.31492159 | -0.5280394 | 0.597472 | 0.7894568 |
| PPHLN1 | 159.986276 | 0.03599414 | 0.22186441 | 0.16223484 | 0.87112093 | 0.93986793 |
| ZCRB1 | 21.0835512 | 0.87950745 | 0.51836323 | 1.69670108 | 0.08975319 | NA |
| PUS7L | 37.0564419 | 0.63687097 | 0.39715758 | 1.60357251 | 0.10880831 | 0.33063466 |
| IRAK4 | 36.3393857 | 0.65617637 | 0.38003719 | 1.72661093 | 0.08423756 | 0.28931012 |
| TWF1 | 17.132792 | -0.0766203 | 0.57314974 | -0.1336829 | 0.8936533 | NA |
| TMEM117 | 324.47712 | 0.76853152 | 0.30000377 | 2.56173951 | 0.01041494 | 0.10033015 |
| ANO6 | 192.720117 | -1.0627325 | 0.31662505 | -3.3564385 | 0.00078953 | 0.02532098 |
| AC008124.1 | 17.325304 | 0.69839766 | 0.45100829 | 1.54852511 | 0.12149592 | NA |
| ARID2 | 285.883028 | 0.17931587 | 0.23917818 | 0.7497167 | 0.45342535 | 0.68943529 |
| SCAF11 | 274.956654 | -0.298127 | 0.19019112 | -1.5675129 | 0.11699485 | 0.34423141 |
| SLC38A1 | 179.963968 | -1.0242477 | 0.57064614 | -1.7948912 | 0.07267104 | 0.26848318 |
| SLC38A2 | 157.543249 | -0.666119 | 0.28137214 | -2.3673951 | 0.0179138 | 0.13296754 |
| AC008014.1 | 473.836812 | 0.60330607 | 0.39340602 | 1.53354558 | 0.12514148 | 0.35575987 |
| PCED1B | 57.051132 | -1.1630335 | 0.68110101 | -1.7075786 | 0.08771456 | 0.29621428 |
| RPAP3 | 28.6050131 | -0.4180976 | 0.39484711 | -1.0588847 | 0.28965229 | 0.5489913 |
| RAPGEF3 | 45.4700421 | -0.3789552 | 0.61223622 | -0.6189689 | 5.36E-01 | 0.74808305 |
| SLC48A1 | 25.3834608 | -0.5358 | 0.37660627 | -1.4227059 | 1.55E-01 | 3.98E-01 |
| HDAC7 | 86.0755233 | -1.1599948 | 0.30235179 | -3.8365732 | 1.25E-04 | 8.14E-03 |

| | | | | | | |
|---|---|---|---|---|---|---|
| VDR | 152.15427 | -0.4657204 | 0.29228637 | -1.5933703 | 1.11E-01 | 3.34E-01 |
| TMEM106C | 15.9436104 | 1.36455479 | 0.52083346 | 2.61994458 | 0.00879441 | NA |
| SENP1 | 42.3132203 | -0.4072125 | 0.30456648 | -1.3370234 | 0.18121501 | 0.4304342 |
| PFKM | 20.0009876 | 1.02540593 | 0.48666141 | 2.10702126 | 0.03511574 | NA |
| ASB8 | 17.1923065 | 0.65381746 | 0.65734735 | 0.99463009 | 0.31991621 | NA |
| ZNF641 | 18.0528596 | -0.0491496 | 0.51813917 | -0.094858 | 0.92442764 | NA |
| AC090115.1 | 14.1553371 | 1.06889722 | 0.6714458 | 1.59193374 | 0.11139959 | NA |
| KANSL2 | 29.7236735 | -0.5828804 | 0.41051273 | -1.4198839 | 0.15564149 | 0.39879246 |
| CCNT1 | 46.6265207 | -0.3829739 | 0.3255192 | -1.1765018 | 0.23939441 | 0.49706468 |
| ADCY6 | 44.6332319 | 0.53265345 | 0.38901615 | 1.36923223 | 0.1709267 | 0.41898315 |
| DDX23 | 23.5633574 | -0.8853823 | 0.40308671 | -2.1965058 | 0.02805576 | 0.16508057 |
| CCDC65 | 20.9982487 | 0.57331906 | 0.52449428 | 1.09308925 | 0.27435462 | NA |
| PRKAG1 | 32.8121089 | -0.4636497 | 0.32271684 | -1.4367074 | 0.15080114 | 0.39147264 |
| KMT2D | 37.0083192 | -0.1982498 | 0.37880396 | -0.5233573 | 0.60072565 | 0.79096587 |
| LMBR1L | 36.8042045 | -0.1622465 | 0.32326403 | -0.5019009 | 0.61573721 | 0.80084174 |
| AC011603.2 | 16.2041129 | 0.50166697 | 0.47621329 | 1.05345015 | 0.29213473 | NA |
| TUBA1B | 43.3426503 | 0.10184664 | 0.54667227 | 0.18630293 | 0.85220721 | 0.92945957 |
| TUBA1A | 16.7692278 | -0.2874632 | 0.63617315 | -0.4518632 | 0.65136753 | NA |
| TUBA1C | 38.7261573 | -0.389118 | 0.35940792 | -1.0826639 | 0.27895764 | 0.53717915 |
| SPATS2 | 166.431387 | 0.40602755 | 0.32475364 | 1.25026325 | 0.2112034 | 0.46604977 |
| MCRS1 | 14.8598182 | -0.3999114 | 0.46282534 | -0.8640655 | 0.38755189 | NA |
| PRPF40B | 88.1992528 | 0.3414078 | 0.33771128 | 1.0109458 | 0.31204237 | 0.57101636 |
| TMBIM6 | 164.767833 | -0.0086267 | 0.42050541 | -0.0205151 | 0.98363243 | 0.99382437 |
| NCKAP5L | 27.3716217 | 0.21245696 | 0.43610573 | 0.48716846 | 0.62613895 | 0.8053173 |
| AQP2 | 28.6214269 | -0.4954607 | 0.60027721 | -0.8253866 | 0.40915216 | 0.65472852 |
| SMARCD1 | 22.2755119 | -0.7382316 | 0.42215325 | -1.7487289 | 0.0803379 | 0.28313224 |
| AC074032.1 | 101.16797 | 0.11875108 | 0.31497969 | 0.37701187 | 0.70616478 | 0.85458443 |
| CERS5 | 65.0998748 | -0.1384177 | 0.3179229 | -0.4353813 | 0.66328569 | 0.82878409 |
| LIMA1 | 111.84893 | -0.3792893 | 0.32195583 | -1.1780785 | 0.2387653 | 0.49669635 |
| LARP4 | 173.282124 | -0.2950945 | 0.26375822 | -1.118807 | 0.26322248 | 0.52284475 |
| DIP2B | 248.907703 | -0.6975909 | 0.38588745 | -1.8077575 | 0.07064425 | 0.26550233 |
| ATF1 | 76.0823017 | -0.660574 | 0.2703178 | -2.443694 | 0.01453775 | 0.11878669 |
| METTL7A | 26.3916587 | 1.73558644 | 0.52756973 | 3.28977639 | 0.00100267 | 0.02858353 |
| SLC11A2 | 121.064439 | -0.2344674 | 0.21707368 | -1.0801284 | 0.280085 | 0.53828072 |
| LETMD1 | 85.4306113 | 0.47415697 | 0.32554404 | 1.45650639 | 0.14525268 | 0.38485711 |
| CSRNP2 | 29.6182176 | -0.1823405 | 0.45955348 | -0.3967775 | 0.69153152 | 0.846991 |
| TFCP2 | 196.116163 | -0.1675227 | 0.23223745 | -0.7213424 | 0.47069887 | 0.70036133 |
| POU6F1 | 27.645433 | -0.6084516 | 0.41271662 | -1.47426 | 0.1404116 | 0.37835958 |
| DAZAP2 | 78.9196329 | -0.297836 | 0.27266951 | -1.0922968 | 0.27470268 | 0.5337282 |
| SMAGP | 18.3341148 | -0.7919626 | 0.55973301 | -1.4148935 | 0.15709973 | NA |
| ACVR1B | 308.046807 | -0.2460553 | 0.28419549 | -0.8657959 | 0.38660208 | 0.63678069 |
| NR4A1 | 37.7471506 | -2.2373366 | 1.08092486 | -2.0698354 | 0.03846776 | 0.19556562 |
| KRT8 | 56.8886926 | -0.0033727 | 0.32055059 | -0.0105216 | 9.92E-01 | 0.99600481 |
| KRT18 | 49.7127644 | 0.20057616 | 0.5074515 | 0.39526174 | 0.69264973 | 0.84775145 |
| EIF4B | 107.376612 | -0.537149 | 0.23174293 | -2.3178659 | 0.02045661 | 0.14071003 |
| AC068888.1 | 48.9702301 | -0.0048918 | 0.31593164 | -0.0154837 | 0.98764626 | 0.99527537 |
| TNS2 | 28.4483682 | 0.87529057 | 0.57562935 | 1.52058017 | 0.12836523 | 0.36084904 |
| CSAD | 93.7036543 | 0.38371514 | 0.33861274 | 1.13319759 | 0.25713128 | 0.51553516 |
| ZNF740 | 14.1006812 | 0.14268501 | 0.58729725 | 0.24295194 | 0.80804263 | NA |
| RARG | 13.1810038 | -0.0773176 | 0.60893481 | -0.1269719 | 0.89896263 | NA |
| PFDN5 | 45.7153529 | 0.08518873 | 0.38739114 | 0.21990365 | 0.82594619 | 0.91551092 |
| AAAS | 11.4988566 | 0.04838659 | 0.58607735 | 0.08256008 | 0.93420135 | NA |
| SP1 | 71.4768864 | -0.0511894 | 0.22697517 | -0.2255284 | 8.22E-01 | 0.91434507 |
| PRR13 | 51.6566127 | 0.63935078 | 0.35878741 | 1.78197663 | 0.07475304 | 0.2729068 |

| | | | | | | |
|---|---|---|---|---|---|---|
| PCBP2 | 179.526197 | -0.3101888 | 0.23088574 | -1.3434732 | 0.1791188 | 0.42802774 |
| MAP3K12 | 13.128957 | 0.42108855 | 0.48682885 | 0.86496219 | 0.38705954 | NA |
| ATP5MC2 | 78.4635391 | 0.26257077 | 0.34645964 | 0.75786827 | 0.44852985 | 0.68731807 |
| CALCOCO1 | 36.3946381 | 0.44728257 | 0.43047906 | 1.03903444 | 0.29878872 | 0.55866203 |
| SMUG1 | 17.9239112 | 0.38648668 | 0.56681216 | 0.68186025 | 0.49532732 | NA |
| LINC02381 | 18.5824309 | 0.71136164 | 0.55074646 | 1.2916318 | 0.19648469 | NA |
| CBX5 | 71.8007945 | 0.63910079 | 0.33465838 | 1.90971101 | 0.05617043 | 0.23625346 |
| HNRNPA1 | 48.5859713 | 0.0262443 | 0.34779464 | 0.07545919 | 0.93984937 | 0.97493539 |
| COPZ1 | 35.0762974 | -0.1429394 | 0.3467802 | -0.4121903 | 0.68019992 | 0.84089068 |
| PPP1R1A | 27.3447494 | 0.34271465 | 0.55227326 | 0.62055268 | 0.53489398 | 0.74775242 |
| CD63 | 77.6119762 | -0.4959142 | 0.4915208 | -1.0089384 | 0.31300416 | 0.57102302 |
| RAB5B | 45.0282235 | 0.04528899 | 0.38772551 | 0.11680685 | 0.90701312 | 0.95830885 |
| SUOX | 14.7288565 | 1.78695664 | 0.66842721 | 2.67337508 | 0.00750922 | NA |
| IKZF4 | 17.7936942 | -0.2485272 | 0.43881953 | -0.5663541 | 0.57115308 | NA |
| RPS26 | 15.4566362 | 0.35742078 | 0.46146221 | 0.77453966 | 0.43861173 | NA |
| ERBB3 | 27.2742658 | 0.80927672 | 0.42754242 | 1.89285712 | 0.05837687 | 0.24020328 |
| PA2G4 | 15.4589849 | -0.3379932 | 0.51291355 | -0.6589673 | 0.5099168 | NA |
| RPL41 | 166.344567 | 0.48763193 | 0.36104728 | 1.35060406 | 0.1768223 | 0.42569745 |
| ESYT1 | 20.9009337 | -0.7108495 | 0.48529737 | -1.4647709 | 0.14298342 | NA |
| MYL6B | 13.5273561 | 0.8477069 | 0.58128943 | 1.45832156 | 0.14475193 | NA |
| MYL6 | 98.2159828 | 0.21632484 | 0.44991489 | 0.4808128 | 0.63064955 | 0.80819181 |
| SMARCC2 | 82.4089626 | -0.1659081 | 0.30055813 | -0.5520001 | 0.58094826 | 0.77786509 |
| RNF41 | 19.4127313 | 0.63906272 | 0.41673904 | 1.53348415 | 0.12515661 | NA |
| ANKRD52 | 27.339878 | 0.54820663 | 0.37365515 | 1.46714591 | 0.14233637 | 0.38005954 |
| COQ10A | 13.9277132 | 0.55098358 | 0.58497625 | 0.94189051 | 0.3462487 | NA |
| CS | 43.415813 | 0.03959057 | 0.46963585 | 0.08430058 | 0.93281745 | 0.97291964 |
| AC073896.2 | 16.2633257 | 0.148133 | 0.58077333 | 0.25506164 | 0.79867549 | NA |
| PAN2 | 29.6779126 | 0.39153311 | 0.44807284 | 0.87381577 | 0.38221859 | 0.63349193 |
| STAT2 | 53.4848186 | -0.2488225 | 0.3882922 | -0.6408125 | 0.52164452 | 0.7380123 |
| RBMS2 | 83.677976 | -0.314217 | 0.35362407 | -0.8885623 | 0.37423836 | 0.62575308 |
| BAZ2A | 70.9332653 | -0.0217961 | 0.2401057 | -0.0907769 | 0.92766984 | 0.96988247 |
| ATP5F1B | 102.978686 | 0.98138001 | 0.36893604 | 2.66002748 | 0.00781343 | 0.08671693 |
| PTGES3 | 145.757426 | -0.0826632 | 0.3083176 | -0.2681107 | 0.78861414 | 0.89544661 |
| NACA | 47.5555094 | 0.64999742 | 0.36055105 | 1.80278884 | 0.07142138 | 0.26675397 |
| NEMP1 | 12.7951472 | -0.2067546 | 0.48539776 | -0.4259489 | 0.67014508 | NA |
| STAT6 | 61.3236041 | -0.3229167 | 0.31314012 | -1.0312211 | 0.30243717 | 0.56282855 |
| R3HDM2 | 294.938041 | -0.2896498 | 0.17987965 | -1.610242 | 0.10734503 | 0.32930838 |
| MARS | 35.5946169 | -0.8774201 | 0.50681027 | -1.7312594 | 0.08340551 | 0.28861463 |
| DDIT3 | 44.6463566 | -1.7689061 | 0.5125508 | -3.451182 | 0.00055814 | 0.02105873 |
| MBD6 | 23.5625915 | 0.1799064 | 0.51970792 | 0.34616829 | 0.72921625 | 0.86533507 |
| DCTN2 | 42.0247021 | -0.197484 | 0.31682693 | -0.6233182 | 0.53307546 | 0.74615054 |
| PIP4K2C | 33.7168744 | -0.124554 | 0.38962663 | -0.3196752 | 0.74921457 | 0.87588883 |
| DTX3 | 18.8317949 | -0.0821115 | 0.42795838 | -0.1918679 | 0.84784567 | NA |
| OS9 | 49.5817797 | 0.27011212 | 0.35689436 | 0.75684053 | 0.44914541 | 0.68734597 |
| 9-Mar | 15.5414089 | 0.34145645 | 0.44606781 | 0.76548102 | 0.44398516 | NA |
| TSFM | 17.8973844 | 0.64450378 | 0.48037929 | 1.34165604 | 0.17970755 | NA |
| AVIL | 50.5828347 | -0.0771681 | 0.32331249 | -0.2386797 | 8.11E-01 | 0.90889121 |
| CTDSP2 | 57.3102857 | 0.1721719 | 0.26799121 | 0.64245353 | 0.52057875 | 0.73723913 |
| LRIG3 | 16.9221685 | 0.89327036 | 0.52116399 | 1.71399096 | 0.08653039 | NA |
| SLC16A7 | 318.502641 | -0.2997501 | 0.51518489 | -0.5818301 | 0.5606811 | 0.76573742 |
| FAM19A2 | 59.8320569 | 1.19056389 | 0.66252597 | 1.79700713 | 0.07233449 | 0.26822676 |
| USP15 | 216.800187 | -0.3623493 | 0.19014767 | -1.9056206 | 0.05669945 | 0.23723351 |
| MON2 | 324.02956 | -0.2854137 | 0.15531107 | -1.8376906 | 0.06610801 | 0.25621016 |
| AC048341.1 | 97.9582864 | -1.3284361 | 0.29393931 | -4.519423 | 6.20E-06 | 0.001256 |

| | | | | | | |
|---|---|---|---|---|---|---|
| PPM1H | 377.772151 | -0.8157405 | 0.31185641 | -2.6157566 | 0.008903 | 0.09323317 |
| DPY19L2 | 231.094657 | 1.2483823 | 0.48483417 | 2.57486452 | 0.01002794 | 0.09908219 |
| RXYLT1 | 21.5113779 | -0.2201349 | 0.4260775 | -0.5166546 | 0.6053973 | 0.79422925 |
| SRGAP1 | 136.672285 | -1.5653878 | 0.45520522 | -3.4388616 | 0.00058417 | 0.02172137 |
| XPOT | 27.5010399 | -0.4437144 | 0.51691707 | -0.858386 | 0.39067933 | 0.63925794 |
| TBK1 | 94.442019 | -0.2883553 | 0.29808829 | -0.9673488 | 0.33336972 | 0.58883681 |
| RASSF3 | 22.6146749 | -1.0305463 | 0.48960633 | -2.1048468 | 0.03530465 | 0.18721309 |
| GNS | 79.0516063 | 0.01147889 | 0.39614356 | 0.02897659 | 0.97688326 | 0.99130791 |
| TBC1D30 | 51.3896422 | 0.27644837 | 0.30531603 | 0.90544991 | 0.36522709 | 0.62097479 |
| LINC02389 | 25.6380987 | 1.24766024 | 0.65137207 | 1.91543404 | 0.05543717 | 0.23522598 |
| LEMD3 | 108.222813 | -0.1317259 | 0.25713802 | -0.5122769 | 0.60845722 | 0.79566688 |
| AC090673.1 | 25.0714033 | -0.2580558 | 0.89273631 | -0.2890617 | 0.77253419 | 0.88749189 |
| HELB | 57.7579302 | -0.0189845 | 0.41538082 | -0.0457038 | 0.96354632 | 0.98600419 |
| GRIP1 | 695.990513 | -0.3354992 | 0.30722496 | -1.0920311 | 0.27481941 | 0.5337282 |
| CAND1 | 95.4462785 | -0.1595787 | 0.21198811 | -0.7527721 | 0.45158689 | 0.68853766 |
| MDM1 | 13.5376489 | 0.37264808 | 0.55487395 | 0.67159051 | 0.50184442 | NA |
| RAP1B | 204.987389 | -0.9502642 | 0.24130041 | -3.9380959 | 8.21E-05 | 0.00598111 |
| NUP107 | 46.8614218 | -0.4005974 | 0.39427438 | -1.0160372 | 0.30961167 | 0.5692979 |
| SLC35E3 | 39.9067132 | 0.47172998 | 0.37636504 | 1.25338417 | 0.21006593 | 0.46502084 |
| MDM2 | 94.6577219 | 0.38913674 | 0.25931319 | 1.50064386 | 0.1334477 | 0.36762309 |
| CPM | 490.17671 | -1.2159698 | 0.52837249 | -2.3013496 | 0.02137188 | 0.14320374 |
| CPSF6 | 169.007286 | -0.4025366 | 0.21420458 | -1.8792156 | 0.06021505 | 0.24444898 |
| FRS2 | 141.19581 | -0.4352791 | 0.19892575 | -2.1881488 | 0.02865877 | 0.16622949 |
| CCT2 | 38.9721403 | -0.6444789 | 0.49233704 | -1.3090198 | 0.19052764 | 0.44184732 |
| RAB3IP | 149.775113 | -0.6244201 | 0.36837418 | -1.6950701 | 0.09006212 | 0.29957137 |
| MYRFL | 59.4468551 | 0.32419371 | 0.50692834 | 0.63952571 | 0.52248099 | 0.7380123 |
| AC025159.1 | 45.1245016 | 1.86203867 | 0.6879338 | 2.70671202 | 0.00679532 | 0.08022018 |
| CNOT2 | 202.754437 | -0.5431979 | 0.21513889 | -2.5248708 | 0.01157408 | 0.10505392 |
| TSPAN8 | 15.7164074 | -0.2500704 | 0.57024112 | -0.4385345 | 0.66099888 | NA |
| ZFC3H1 | 175.446196 | -0.6169067 | 0.32587239 | -1.8930928 | 0.05834553 | 0.24020328 |
| RAB21 | 128.180604 | -0.4967402 | 0.25899371 | -1.9179625 | 0.05511577 | 0.23466461 |
| TBC1D15 | 131.120678 | -0.7077663 | 0.26557166 | -2.6650671 | 0.0076973 | 0.08614009 |
| TRHDE | 245.797518 | 1.56658668 | 0.35987171 | 4.35318095 | 1.34E-05 | 0.00194859 |
| ATXN7L3B | 24.9050404 | 0.03461893 | 0.37437423 | 0.09247145 | 0.92632347 | 0.96926479 |
| CAPS2 | 112.910722 | 1.51433456 | 0.56847305 | 2.66386343 | 0.00772489 | 0.08618711 |
| KRR1 | 36.8368046 | -0.4453178 | 0.30779017 | -1.4468227 | 0.14794659 | 0.38772383 |
| NAP1L1 | 116.773946 | 0.02177525 | 0.29677976 | 0.07337176 | 0.94151029 | 0.97560645 |
| OSBPL8 | 238.897085 | 0.09317818 | 0.20433223 | 0.45601314 | 0.64838053 | 0.81913126 |
| ZDHHC17 | 117.36989 | 0.11700893 | 0.22758708 | 0.51412817 | 0.60716238 | 0.79532486 |
| AC027288.3 | 56.3144772 | 0.09286418 | 0.59177589 | 0.15692457 | 0.87530429 | 0.94169674 |
| PAWR | 366.826679 | -0.1968513 | 0.21352693 | -0.9219038 | 0.35657876 | 0.61264648 |
| AC073569.2 | 26.0185976 | -0.4152788 | 0.54452435 | -0.7626451 | 0.4456751 | 0.68538685 |
| PPP1R12A | 278.746077 | 0.05236302 | 0.21900537 | 0.2390947 | 0.81103215 | 0.90879523 |
| LIN7A | 83.1923117 | 1.65263419 | 0.41972736 | 3.9373992 | 8.24E-05 | 0.00598111 |
| ACSS3 | 107.941776 | 1.67635474 | 0.49023184 | 3.41951419 | 0.00062733 | 0.02287166 |
| PPFIA2 | 119.287524 | 0.15134602 | 0.70523175 | 0.21460466 | 0.83007556 | 0.91772835 |
| CCDC59 | 24.7607053 | -0.2888509 | 0.42644187 | -0.6773512 | 0.49818314 | 0.72149172 |
| METTL25 | 107.653566 | 0.32451861 | 0.23544519 | 1.37831911 | 0.16810478 | 0.41617965 |
| TMTC2 | 542.890757 | 1.55141773 | 0.48742133 | 3.18290899 | 0.00145803 | 0.03485245 |
| LRRIQ1 | 35.1930968 | 1.6297648 | 0.89380393 | 1.82340303 | 0.06824237 | 0.25987613 |
| CEP290 | 94.2317543 | 0.2479867 | 0.28931745 | 0.85714394 | 0.39136534 | 0.63984683 |
| TMTC3 | 81.7747877 | 0.72416408 | 0.26306593 | 2.75278548 | 0.00590906 | 0.07567727 |
| KITLG | 386.692042 | 1.77700938 | 0.44620808 | 3.98246796 | 6.82E-05 | 0.00530264 |
| DUSP6 | 27.6985125 | -1.2163322 | 0.68523905 | -1.775048 | 7.59E-02 | 0.27514133 |

| | | | | | | |
|---|---|---|---|---|---|---|
| POC1B | 73.5557994 | -0.4841315 | 0.24350256 | -1.9881987 | 0.04678971 | 0.21661307 |
| POC1B-AS1 | 26.6066782 | -0.8890369 | 0.50149074 | -1.7727883 | 0.0762638 | 0.27590775 |
| ATP2B1 | 142.278644 | -0.1833982 | 0.28682868 | -0.6393997 | 0.52256295 | 0.7380123 |
| BTG1 | 74.3239853 | -0.6743437 | 0.2996582 | -2.2503764 | 2.44E-02 | 0.15276639 |
| LINC02391 | 23.6218216 | -1.1695665 | 0.63269919 | -1.8485349 | 0.064525 | 0.25284322 |
| EEA1 | 164.44061 | -0.3064379 | 0.37836355 | -0.8099034 | 0.41799572 | 0.66186239 |
| AC138123.1 | 148.730096 | 0.71244739 | 0.53449025 | 1.33294739 | 0.18254907 | 0.43233236 |
| AC124947.1 | 56.6964384 | 0.40074631 | 0.39141107 | 1.02385021 | 0.30590603 | 0.5657277 |
| NUDT4 | 215.707591 | 0.96714374 | 0.47834817 | 2.02184059 | 0.04319282 | 0.20765469 |
| UBE2N | 69.0969614 | -0.2698175 | 0.2521386 | -1.070116 | 0.28456711 | 0.54316116 |
| MRPL42 | 67.3305145 | -0.229876 | 0.29501728 | -0.7791949 | 0.4358649 | 0.67638148 |
| SOCS2 | 31.8311781 | 0.12294207 | 0.41895984 | 0.29344596 | 0.7691813 | 0.88575962 |
| CRADD | 490.804765 | 0.26622498 | 0.18982473 | 1.40247781 | 0.16077261 | 0.40625962 |
| AC012085.2 | 34.8477279 | 0.77052074 | 0.64768635 | 1.18965104 | 0.23418357 | 0.49243138 |
| CEP83 | 69.7857918 | 0.59515416 | 0.37098571 | 1.60425091 | 0.10865876 | 0.33040203 |
| TMCC3 | 34.1319465 | -0.289942 | 0.53509871 | -0.5418477 | 0.58792346 | 0.78279721 |
| NDUFA12 | 39.2718629 | 0.58687795 | 0.32186167 | 1.82338562 | 0.06824501 | 0.25987613 |
| NR2C1 | 75.1362391 | -0.199669 | 0.26368081 | -0.7572375 | 0.44890756 | 0.68733668 |
| FGD6 | 41.142781 | -0.7899537 | 0.38976151 | -2.0267618 | 0.04268678 | 0.20625243 |
| VEZT | 143.916095 | -0.2481635 | 0.20702931 | -1.198688 | 0.2306493 | 0.48933508 |
| METAP2 | 73.3913075 | -0.3048069 | 0.24113096 | -1.2640721 | 0.20620414 | 0.45924573 |
| USP44 | 16.0788888 | 1.58511682 | 0.57473204 | 2.75801019 | 0.00581544 | NA |
| NTN4 | 208.636723 | 0.42582062 | 0.47853932 | 0.88983414 | 0.37355495 | 0.62575308 |
| SNRPF | 16.6419405 | -0.4107388 | 0.48433883 | -0.8480403 | 0.39641554 | NA |
| LTA4H | 32.6152576 | -0.2988313 | 0.42272986 | -0.7069083 | 0.47962345 | 0.70695585 |
| ELK3 | 38.5791859 | -1.5730557 | 0.53934327 | -2.9166132 | 0.00353854 | 0.05908061 |
| CDK17 | 98.1655066 | -0.3137896 | 0.23383685 | -1.3419167 | 0.17962302 | 0.42883325 |
| CFAP54 | 16.6053167 | 0.87452953 | 0.9817441 | 0.89079173 | 0.37304091 | NA |
| NEDD1 | 34.2566925 | 0.48789724 | 0.33132601 | 1.47255944 | 0.14086987 | 0.37898475 |
| TMPO | 38.1141913 | 0.40626555 | 0.32495006 | 1.25023994 | 0.21121191 | 0.46604977 |
| SLC25A3 | 89.5838577 | 0.51476458 | 0.39794292 | 1.29356387 | 0.19581611 | 0.44885942 |
| APAF1 | 33.7664905 | 0.42687218 | 0.39731668 | 1.07438778 | 0.28264892 | 0.54138498 |
| ANKS1B | 71.7516011 | 0.5233922 | 0.70089099 | 0.74675264 | 0.45521289 | 0.6908246 |
| UHRF1BP1L | 56.7340245 | -0.613961 | 0.29908832 | -2.0527748 | 0.04009442 | 0.19926552 |
| ACTR6 | 29.8942019 | 0.40151522 | 0.36915054 | 1.08767341 | 0.27673931 | 0.53501126 |
| SCYL2 | 106.003424 | -0.4960924 | 0.27456174 | -1.806852 | 0.07078536 | 0.26590283 |
| NR1H4 | 156.01938 | -1.0066899 | 0.63747077 | -1.5791939 | 1.14E-01 | 3.40E-01 |
| UTP20 | 37.8871804 | 0.10737069 | 0.35720464 | 0.30058594 | 0.76373025 | 0.88294563 |
| ARL1 | 31.2793912 | 0.1042593 | 0.4504831 | 0.23143888 | 0.81697386 | 0.91186888 |
| CHPT1 | 75.885336 | 1.03525951 | 0.36351153 | 2.84794133 | 0.0044003 | 0.0642158 |
| GNPTAB | 79.7580848 | 0.09690222 | 0.2672104 | 0.3626439 | 0.71687091 | 0.8589283 |
| DRAM1 | 51.2090811 | -1.3616061 | 0.56240036 | -2.421062 | 0.01547524 | 0.12292353 |
| WASHC3 | 52.5498536 | 0.33900473 | 0.43751936 | 0.77483366 | 0.43843796 | 0.67873229 |
| NUP37 | 17.789232 | 0.63772144 | 0.5855264 | 1.08914207 | 0.27609124 | NA |
| NT5DC3 | 54.6722289 | -1.3514226 | 0.37453355 | -3.608282 | 0.00030823 | 0.01457249 |
| HSP90B1 | 149.004765 | -1.0198644 | 0.62154429 | -1.6408556 | 0.10082739 | 0.31985368 |
| C12orf73 | 12.0203131 | 0.29757731 | 0.57515443 | 0.51738679 | 0.60488618 | NA |
| TDG | 51.9423146 | -0.8355223 | 0.27371315 | -3.0525471 | 0.00226908 | 0.04522213 |
| HCFC2 | 26.6475765 | -0.1045497 | 0.38522719 | -0.2713975 | 0.7860853 | 0.89452315 |
| NFYB | 19.0561681 | 0.44035751 | 0.42901716 | 1.02643333 | 0.30468737 | NA |
| TXNRD1 | 177.195868 | -0.3492745 | 0.26916024 | -1.2976451 | 1.94E-01 | 4.47E-01 |
| SLC41A2 | 44.628827 | -0.3775142 | 0.29233979 | -1.2913543 | 1.97E-01 | 4.50E-01 |
| C12orf45 | 12.4661981 | -0.0518677 | 0.48715711 | -0.1064702 | 0.91520932 | NA |
| WASHC4 | 151.188533 | -0.3343993 | 0.24106903 | -1.3871515 | 0.16539557 | 0.41243204 |

| | | | | | | |
|---|---|---|---|---|---|---|
| APPL2 | 102.217962 | -0.3118888 | 0.2724342 | -1.1448226 | 0.25228264 | 0.50912938 |
| C12orf75 | 91.7688096 | 0.3328806 | 0.58328366 | 0.57070106 | 5.68E-01 | 0.77103518 |
| CKAP4 | 16.8657629 | 0.65867285 | 0.63456121 | 1.03799733 | 0.2992713 | NA |
| TCP11L2 | 60.8291539 | -0.4814085 | 0.36890184 | -1.3049771 | 0.19190063 | 0.44342814 |
| POLR3B | 37.6633768 | 0.23974453 | 0.35385763 | 0.67751693 | 0.49807803 | 0.72149172 |
| RIC8B | 96.909708 | 0.39589367 | 0.30326405 | 1.30544211 | 0.19174235 | 0.44319545 |
| TMEM263 | 57.2245774 | 0.59919252 | 0.34385357 | 1.7425805 | 0.08140692 | 0.28447225 |
| MTERF2 | 18.0008019 | 0.17948842 | 0.50193689 | 0.35759161 | 0.72064895 | NA |
| CRY1 | 77.3461227 | -0.848756 | 0.64571664 | -1.3144403 | 0.18869806 | 0.43932516 |
| BTBD11 | 310.752584 | -0.7117625 | 0.52468635 | -1.3565486 | 0.1749247 | 0.42340793 |
| PWP1 | 28.4172085 | -0.0193271 | 0.37330793 | -0.0517727 | 0.95870984 | 0.98401364 |
| PRDM4 | 57.1355474 | -0.3869293 | 0.32483335 | -1.1911626 | 0.23358976 | 0.49243138 |
| SART3 | 40.0796508 | -0.5706308 | 0.2984833 | -1.9117678 | 0.05590598 | 0.2357854 |
| ISCU | 54.1197182 | 0.93784277 | 0.32827452 | 2.85688565 | 0.0042782 | 0.06301187 |
| CORO1C | 60.677473 | -1.2879902 | 0.42588829 | -3.0242441 | 2.49E-03 | 0.04746918 |
| SSH1 | 170.648258 | -0.7127481 | 0.27099339 | -2.6301309 | 0.0085352 | 0.0903651 |
| USP30 | 23.4604893 | 0.24724092 | 0.39781616 | 0.62149541 | 0.53427371 | 0.74728417 |
| UNG | 14.492673 | 0.74912246 | 0.47151694 | 1.58874982 | 0.11211689 | NA |
| ACACB | 326.742062 | 1.11676475 | 0.27123824 | 4.11728364 | 3.83E-05 | 0.00383215 |
| KCTD10 | 22.2728875 | -0.1953516 | 0.3910679 | -0.4995336 | 0.6174035 | 0.80205143 |
| UBE3B | 41.1363561 | 0.03374102 | 0.31542679 | 0.10696941 | 0.91481324 | 0.96388328 |
| MMAB | 25.7660724 | -0.3941526 | 0.46761713 | -0.8428959 | 0.39928664 | 0.64742138 |
| MVK | 20.9451878 | 0.88712923 | 0.42242887 | 2.1000677 | 0.03572289 | NA |
| FAM222A | 36.2006031 | 0.93878435 | 0.4877458 | 1.92474102 | 0.05426177 | 0.23287156 |
| TRPV4 | 37.9818068 | 2.87086547 | 0.79640002 | 3.60480337 | 0.00031239 | 0.01457249 |
| GLTP | 34.3384352 | 0.38137543 | 0.34595382 | 1.10238826 | 0.27029291 | 0.52870255 |
| TCHP | 34.2755205 | 0.58541114 | 0.36709132 | 1.5947289 | 0.11077287 | 0.33416384 |
| GIT2 | 75.7464598 | 0.21114794 | 0.27059554 | 0.78030829 | 0.43520944 | 0.67618228 |
| ANKRD13A | 93.3946174 | 0.33039813 | 0.27325027 | 1.20914112 | 0.22660863 | 0.48449513 |
| C12orf76 | 15.939702 | 1.24968163 | 0.54647394 | 2.28680921 | 0.02220696 | NA |
| IFT81 | 58.9577465 | 0.20218939 | 0.32868756 | 0.61514159 | 0.53846122 | 0.74904053 |
| ATP2A2 | 185.182168 | -0.6602109 | 0.40559746 | -1.627749 | 0.10357812 | 0.32331143 |
| ANAPC7 | 41.2390114 | -0.0400971 | 0.38629747 | -0.1037985 | 0.91732922 | 0.96401363 |
| ARPC3 | 78.3313459 | -0.3443464 | 0.27062178 | -1.2724268 | 2.03E-01 | 0.45705947 |
| VPS29 | 48.926194 | 0.40599115 | 0.26215726 | 1.54865502 | 0.12146467 | 0.35059775 |
| PPTC7 | 98.2250195 | -0.2647937 | 0.25389057 | -1.0429443 | 0.29697411 | 0.55674621 |
| TCTN1 | 31.3326383 | 1.49243666 | 0.62484314 | 2.38849811 | 0.0169174 | 0.12943657 |
| PPP1CC | 27.5574587 | 0.4660549 | 0.36545067 | 1.27528812 | 0.20220728 | 0.45628538 |
| ATXN2 | 355.298915 | -0.3503892 | 0.2402155 | -1.4586452 | 0.14466278 | 0.38426647 |
| BRAP | 45.6666457 | -0.2175454 | 0.28928035 | -0.7520226 | 0.45203745 | 0.68880381 |
| ACAD10 | 56.7785155 | -0.2892169 | 0.35891771 | -0.8058029 | 0.42035648 | 0.66394999 |
| ALDH2 | 57.8141573 | 1.22825962 | 0.42952546 | 2.85957346 | 0.00424211 | 0.06301187 |
| MAPKAPK5 | 48.7413745 | -0.1866798 | 0.32775637 | -0.5695689 | 0.56897014 | 0.77142208 |
| TMEM116 | 132.794378 | 0.57032968 | 0.26045817 | 2.18971695 | 2.85E-02 | 1.66E-01 |
| ERP29 | 41.7941942 | 0.73326028 | 0.35294278 | 2.07756134 | 0.03774978 | 0.19425568 |
| NAA25 | 99.5303138 | -0.6138709 | 0.28746057 | -2.1354958 | 0.03272053 | 0.17874373 |
| TRAFD1 | 36.7538854 | -0.4072769 | 0.31628451 | -1.2876916 | 0.19785332 | 0.45094881 |
| HECTD4 | 249.80915 | 0.08258868 | 0.1765467 | 0.46780073 | 0.6399271 | 0.81426994 |
| RPL6 | 84.3685862 | 0.56035969 | 0.36656396 | 1.52868191 | 0.12634332 | 0.3577868 |
| PTPN11 | 83.1928642 | -0.6659703 | 0.26523476 | -2.510871 | 0.01204337 | 0.10726863 |
| RASAL1 | 23.0774362 | -1.1150159 | 0.61283733 | -1.8194321 | 6.88E-02 | 2.61E-01 |
| DDX54 | 17.8903555 | -0.2528497 | 0.49391904 | -0.5119253 | 0.60870326 | NA |
| TPCN1 | 90.5007586 | -0.4097761 | 0.30354278 | -1.3499781 | 0.177023 | 0.42569745 |
| SLC8B1 | 36.0737797 | 0.21141657 | 0.59928312 | 0.35278245 | 0.72425154 | 0.86240162 |

| | | | | | | |
|---|---|---|---|---|---|---|
| PLBD2 | 14.8289051 | -0.2004227 | 0.5114609 | -0.3918631 | 0.69515936 | NA |
| RBM19 | 44.6104952 | -0.9202077 | 0.28083671 | -3.2766644 | 0.00105041 | 0.02953506 |
| MED13L | 969.888755 | -0.2049635 | 0.23569225 | -0.8696233 | 0.38450629 | 0.63468687 |
| C12orf49 | 67.3771189 | -1.4937249 | 0.58211416 | -2.5660342 | 0.01028687 | 0.09994074 |
| FBXW8 | 93.495835 | -0.0447591 | 0.24553118 | -0.1822948 | 0.85535138 | 0.93160317 |
| TESC | 108.867569 | 1.13552573 | 0.41540511 | 2.73353819 | 6.27E-03 | 7.72E-02 |
| FBXO21 | 79.6472974 | -0.0311482 | 0.22422294 | -0.1389161 | 0.88951642 | 0.94946719 |
| NOS1 | 69.6412152 | 2.28001815 | 0.68187387 | 3.34375351 | 8.27E-04 | 0.02565549 |
| KSR2 | 101.156552 | 1.32300406 | 0.56210164 | 2.35367409 | 0.0185889 | 0.13485275 |
| WSB2 | 36.2078551 | -0.6888691 | 0.37690055 | -1.827721 | 0.06759143 | 0.25898011 |
| VSIG10 | 42.627931 | 1.39539131 | 0.54872215 | 2.54298338 | 0.01099105 | 0.10267384 |
| PEBP1 | 138.730621 | 0.86087426 | 0.366594 | 2.34830431 | 0.01885911 | 0.1355543 |
| TAOK3 | 359.161655 | 0.55014399 | 0.19086018 | 2.88244514 | 0.00394602 | 0.06143969 |
| SUDS3 | 51.7519991 | -0.1099331 | 0.28206847 | -0.3897392 | 0.69672939 | 0.85028161 |
| SRRM4 | 72.1047004 | 3.26269341 | 1.03270117 | 3.15937805 | 1.58E-03 | 3.64E-02 |
| CCDC60 | 23.0137592 | 2.70110902 | 0.83168748 | 3.24774519 | 0.00116323 | 0.03031862 |
| AC002070.1 | 167.855723 | 0.36767087 | 0.40255365 | 0.91334627 | 0.36106045 | 0.61731435 |
| PRKAB1 | 71.3068113 | 0.03782113 | 0.34779404 | 0.10874578 | 0.91340413 | 0.96345217 |
| CIT | 102.586328 | -0.1369183 | 0.38075684 | -0.3595951 | 0.71914998 | 0.85995641 |
| BICDL1 | 298.300256 | -0.4690841 | 0.3705125 | -1.2660411 | 2.05E-01 | 0.45886886 |
| RAB35 | 29.2644022 | -0.5278752 | 0.57087438 | -0.9246783 | 0.35513325 | 0.61168935 |
| GCN1 | 37.4244941 | -0.1509448 | 0.35708171 | -0.4227178 | 0.67250116 | 0.83624371 |
| RPLP0 | 107.920171 | 0.48328577 | 0.51807211 | 0.93285425 | 0.35089522 | 0.60767969 |
| PXN | 73.3667563 | -0.9088249 | 0.3014273 | -3.0150717 | 0.00256919 | 0.04846819 |
| GATC | 14.24202 | 0.12949834 | 0.52767207 | 0.24541443 | 0.80613555 | NA |
| SRSF9 | 42.4633678 | 0.12110367 | 0.28809796 | 0.42035589 | 0.67422548 | 0.83715334 |
| DYNLL1 | 53.9186215 | 1.66798952 | 0.52507788 | 3.17665163 | 0.00148986 | 0.03500548 |
| COQ5 | 34.0338414 | 1.34029304 | 0.64996326 | 2.06210586 | 0.03919766 | 0.19732137 |
| RNF10 | 201.92784 | -0.0181815 | 0.18721626 | -0.097115 | 0.92263507 | 0.96671959 |
| POP5 | 26.4140924 | 1.64661386 | 0.67130055 | 2.45287131 | 0.0141721 | 0.1170415 |
| MLEC | 77.502442 | 0.56605325 | 0.4103235 | 1.37952919 | 0.16773165 | 0.41592478 |
| UNC119B | 16.9322054 | 0.35188344 | 0.4901989 | 0.71783809 | 0.47285712 | NA |
| ACADS | 22.939205 | 1.06197761 | 0.45544366 | 2.33174307 | 0.01971421 | 0.13748641 |
| SPPL3 | 302.802515 | -0.4658656 | 0.21089471 | -2.2089961 | 0.02717491 | 0.16201805 |
| P2RX4 | 33.9668189 | -0.940915 | 0.47064989 | -1.9991824 | 0.04558862 | 0.21335013 |
| CAMKK2 | 18.1569162 | 0.9957929 | 0.43429704 | 2.29288437 | 0.02185466 | NA |
| ANAPC5 | 78.417765 | 0.24726956 | 0.32964795 | 0.7501019 | 0.45319334 | 0.68943192 |
| RNF34 | 53.395596 | -0.4810731 | 0.32949942 | -1.460012 | 0.14428678 | 0.38365755 |
| KDM2B | 65.9884683 | 0.16287327 | 0.30057938 | 0.54186441 | 0.58791192 | 0.78279721 |
| TMEM120B | 114.687529 | 0.46911237 | 0.429691 | 1.09174355 | 0.27494584 | 0.5337282 |
| RHOF | 17.9960718 | -0.4211186 | 0.676528 | -0.6224704 | 0.53363261 | NA |
| SETD1B | 23.4321382 | 0.30797555 | 0.37312224 | 0.82540121 | 0.40914385 | 0.65472852 |
| WDR66 | 35.0969189 | -0.7558331 | 0.52202587 | -1.4478845 | 0.14764937 | 0.38722988 |
| BCL7A | 68.7693027 | 0.74359306 | 0.37045575 | 2.00723855 | 0.04472427 | 0.21170763 |
| MLXIP | 87.0847126 | -0.5331203 | 0.43685144 | -1.2203699 | 0.22232469 | 0.4793528 |
| CLIP1 | 268.013488 | -0.9133504 | 0.44139503 | -2.0692357 | 0.03852397 | 0.19572213 |
| ZCCHC8 | 69.3106619 | -0.3323147 | 0.26378355 | -1.2598007 | 0.20774127 | 0.46155871 |
| RSRC2 | 91.677842 | -0.0960129 | 0.32917361 | -0.2916787 | 0.7705323 | 0.88651526 |
| KNTC1 | 14.183261 | 0.4019874 | 0.54799213 | 0.73356419 | 0.46321439 | NA |
| DENR | 95.6052695 | 0.0972511 | 0.3177898 | 0.30602336 | 0.75958686 | 0.88058098 |
| HIP1R | 89.4400585 | 0.63710677 | 0.38260588 | 1.66517765 | 0.09587732 | 0.31098513 |
| VPS37B | 142.653421 | -0.6583491 | 0.26113474 | -2.5211087 | 0.01169857 | 0.10593399 |
| ABCB9 | 22.6403975 | 1.23499825 | 0.52351528 | 2.35904911 | 0.01832183 | 0.13368407 |
| ARL6IP4 | 43.5203394 | 0.29601929 | 0.35914122 | 0.82424203 | 0.40980205 | 0.65563217 |

| | | | | | | |
|---|---|---|---|---|---|---|
| PITPNM2 | 209.024831 | 0.00112211 | 0.32863656 | 0.00341444 | 0.99727567 | 0.99868755 |
| MPHOSPH9 | 88.1998211 | 0.23179123 | 0.25939583 | 0.89358116 | 0.37154603 | 0.62454462 |
| C12orf65 | 31.0697138 | 0.01644393 | 0.38955387 | 0.04221222 | 0.96632952 | 0.98723631 |
| SBNO1 | 97.7212341 | -0.2564627 | 0.29046446 | -0.8829402 | 0.3772686 | 0.62875685 |
| KMT5A | 17.9583798 | 0.46537686 | 0.52740831 | 0.88238439 | 0.37756897 | NA |
| RILPL2 | 23.0456399 | 0.57646077 | 0.40833131 | 1.41174764 | 0.15802428 | 0.40275854 |
| SNRNP35 | 32.5155346 | 0.32081239 | 0.35340972 | 0.90776332 | 0.36400328 | 0.62003028 |
| RILPL1 | 85.9058326 | 0.5681341 | 0.29465273 | 1.92814808 | 0.05383671 | 0.23287156 |
| TMED2 | 31.9460765 | 0.0871476 | 0.37940625 | 0.22969468 | 0.81832903 | 0.913067 |
| DDX55 | 22.2826851 | -0.1133797 | 0.43409568 | -0.2611859 | 0.79394912 | 0.89884908 |
| EIF2B1 | 28.1141168 | -0.0310031 | 0.33937472 | -0.0913537 | 0.92721153 | 0.96966673 |
| GTF2H3 | 24.9535821 | -0.1879285 | 0.37989919 | -0.4946798 | 0.62082615 | 0.80353114 |
| TCTN2 | 19.8961116 | 0.91646019 | 0.4712429 | 1.94477243 | 0.05180237 | NA |
| ATP6V0A2 | 29.9251013 | -0.6225564 | 0.40474926 | -1.5381286 | 0.12401717 | 0.35457643 |
| CCDC92 | 82.8314947 | -0.166465 | 0.23087385 | -0.7210214 | 0.47089637 | 0.70047809 |
| RFLNA | 22.6642146 | 0.1429112 | 0.56095043 | 0.25476618 | 0.7989037 | 0.90203341 |
| NCOR2 | 138.20335 | -0.1198332 | 0.3233605 | -0.3705871 | 0.71094512 | 0.8567629 |
| UBC | 156.243483 | -0.6904744 | 0.29473759 | -2.3426752 | 1.91E-02 | 0.1359475 |
| BRI3BP | 20.4692837 | 1.06815774 | 0.50713538 | 2.10625757 | 0.03518198 | NA |
| AACS | 42.5066498 | -0.166153 | 0.33659654 | -0.4936264 | 6.22E-01 | 8.04E-01 |
| SLC15A4 | 41.7937023 | -0.4268838 | 0.32114082 | -1.329273 | 0.18375791 | 0.43386032 |
| RIMBP2 | 72.0655368 | -0.2873389 | 0.26397199 | -1.0885204 | 2.76E-01 | 5.35E-01 |
| STX2 | 17.0546366 | 0.22579765 | 0.57310229 | 0.39399189 | 0.69358703 | NA |
| RAN | 23.740978 | 0.09251529 | 0.53086622 | 0.17427232 | 0.86165145 | 0.93551424 |
| SFSWAP | 116.841218 | -0.5722475 | 0.29926706 | -1.9121633 | 0.05585525 | 0.23575965 |
| ULK1 | 58.5475988 | 0.26263293 | 0.35828162 | 0.7330349 | 0.46353715 | 0.69473502 |
| PUS1 | 15.2573236 | -0.6165263 | 0.55274069 | -1.1153989 | 0.26467953 | NA |
| EP400 | 105.1116 | -0.329998 | 0.26574895 | -1.2417661 | 0.21432288 | 0.46998382 |
| FBRSL1 | 62.230629 | -0.6649531 | 0.52600213 | -1.2641642 | 0.20617108 | 0.45924573 |
| ANKLE2 | 87.0635595 | -1.4294574 | 0.42373846 | -3.3734427 | 0.00074235 | 0.02425576 |
| GOLGA3 | 58.4785967 | -0.717975 | 0.43050322 | -1.6677575 | 0.09536388 | 0.30997288 |
| CHFR | 78.7871966 | -0.7030526 | 0.29133019 | -2.4132501 | 0.01581097 | 0.12437649 |
| ZNF26 | 56.7808601 | -0.6107203 | 0.24881256 | -2.4545397 | 0.01410651 | 0.11675032 |
| ZNF84 | 60.6732289 | 0.1511183 | 0.31612824 | 0.47802847 | 0.63262994 | 0.80953494 |
| ZNF140 | 34.4386211 | -0.2463067 | 0.37720015 | -0.6529868 | 0.51376481 | 0.73176309 |
| ZNF891 | 23.1702217 | 1.0043188 | 0.46582489 | 2.15600073 | 0.03108361 | 0.17412703 |
| MPHOSPH8 | 173.851194 | 0.00151992 | 0.22272466 | 0.00682423 | 0.99455509 | 0.99753591 |
| PSPC1 | 241.706352 | -0.6200462 | 0.30284976 | -2.0473724 | 0.04062153 | 0.20081179 |
| ZMYM5 | 83.3355418 | -0.2325355 | 0.29108453 | -0.7988589 | 0.42437226 | 0.66715549 |
| ZMYM2 | 448.36666 | -0.1911615 | 0.2569129 | -0.7440712 | 0.45683341 | 0.69135799 |
| CRYL1 | 123.065685 | 0.5700715 | 0.362625 | 1.57206896 | 0.11593456 | 0.34268368 |
| IFT88 | 197.27876 | 0.10236853 | 0.31627696 | 0.32366736 | 0.74618989 | 0.87410417 |
| EEF1AKMT1 | 19.783619 | 0.60768912 | 0.40767072 | 1.49063718 | 0.13605678 | NA |
| XPO4 | 147.075768 | -0.1231492 | 0.21133483 | -0.5827207 | 0.56008135 | 0.76543789 |
| LATS2 | 113.542395 | -0.1566152 | 0.33762103 | -0.4638787 | 0.64273467 | 0.81595394 |
| SAP18 | 37.2641813 | 0.86405514 | 0.41080249 | 2.10333471 | 0.03543652 | 0.18721309 |
| MRPL57 | 25.5033591 | 0.60492588 | 0.40944463 | 1.47743024 | 0.13956035 | 0.37696802 |
| ZDHHC20 | 130.07618 | -0.1798964 | 0.25547705 | -0.7041587 | 0.48133397 | 0.7087866 |
| MICU2 | 235.470607 | -0.523479 | 0.19262898 | -2.7175506 | 0.00657671 | 0.07921899 |
| FGF9 | 78.2287386 | 0.53132744 | 0.53409755 | 0.99481346 | 0.319827 | 0.57718837 |
| AL136962.1 | 53.3287484 | -0.0871694 | 0.72375332 | -0.1204408 | 0.90413399 | 0.95671001 |
| TNFRSF19 | 127.602739 | 0.51712549 | 0.38276621 | 1.35102179 | 1.77E-01 | 4.26E-01 |
| MIPEP | 119.012865 | 0.16530267 | 0.27734365 | 0.59602112 | 0.55116112 | 0.75768657 |
| PARP4 | 64.340481 | 0.42643545 | 0.4912729 | 0.86802151 | 0.38538256 | 0.63547552 |

| | | | | | | |
|---|---|---|---|---|---|---|
| MTMR6 | 31.6621517 | 0.02258313 | 0.39219657 | 0.05758114 | 0.95408227 | 0.98153155 |
| NUP58 | 91.2649787 | -0.6746812 | 0.34625945 | -1.9484847 | 0.05135699 | 0.22744234 |
| RNF6 | 16.8121592 | 0.21447424 | 0.60600855 | 0.3539129 | 0.72340416 | NA |
| CDK8 | 246.147228 | -0.0493474 | 0.22371009 | -0.2205863 | 0.8254146 | 0.9154963 |
| WASF3 | 43.48525 | 0.41841514 | 0.49451992 | 0.84610372 | 0.39749489 | 0.64588458 |
| USP12 | 122.441689 | 0.04890956 | 0.28115219 | 0.17396115 | 0.861896 | 0.93551424 |
| LINC02340 | 37.8113324 | 1.18990405 | 0.71819976 | 1.65678703 | 0.09756254 | 0.31393766 |
| RPL21 | 45.0506773 | 0.43537402 | 0.4616269 | 0.94312966 | 0.34561459 | 0.60294548 |
| GTF3A | 21.3589322 | 0.60458958 | 0.44320987 | 1.36411579 | 0.17253113 | 0.42128877 |
| MTIF3 | 45.7702243 | 0.05900002 | 0.32882945 | 0.17942437 | 0.8576045 | 0.93260551 |
| LNX2 | 61.862026 | -0.2736255 | 0.36118106 | -0.7575853 | 0.44869929 | 0.68731807 |
| POLR1D | 41.4808321 | -0.1475832 | 0.36919817 | -0.3997398 | 0.68934816 | 0.84569856 |
| PAN3 | 481.842321 | -0.2446969 | 0.14299132 | -1.7112707 | 0.08703115 | 0.29500251 |
| POMP | 32.4451351 | -0.5331993 | 0.38289337 | -1.3925529 | 0.16375505 | 0.41042742 |
| SLC7A1 | 64.728105 | -0.4293148 | 0.35967571 | -1.1936165 | 2.33E-01 | 0.49082729 |
| UBL3 | 131.068277 | -0.3493979 | 0.29406575 | -1.1881626 | 0.23476937 | 0.49243138 |
| KATNAL1 | 22.7307113 | 1.82303703 | 0.57395191 | 3.1762888 | 0.00149172 | 0.03500548 |
| HMGB1 | 185.959144 | 0.31123383 | 0.2906112 | 1.07096297 | 0.28418608 | 0.54290897 |
| USPL1 | 43.233723 | -0.1243784 | 0.33013779 | -0.3767469 | 0.70636172 | 0.85458443 |
| HSPH1 | 60.6797171 | -0.085557 | 0.42874002 | -0.1995546 | 0.84182895 | 0.92367136 |
| B3GLCT | 73.690248 | 0.59110506 | 0.31245542 | 1.89180606 | 5.85E-02 | 0.24020328 |
| FRY | 91.3315364 | 0.8467364 | 0.69081008 | 1.22571518 | 0.22030587 | 0.47685442 |
| N4BP2L1 | 52.5359768 | 0.62470252 | 0.43267792 | 1.44380495 | 0.14879385 | 0.38937375 |
| N4BP2L2 | 901.012053 | -0.1356857 | 0.21859775 | -0.6207096 | 0.5347907 | 0.74775242 |
| PDS5B | 260.108701 | 0.10732809 | 0.19518194 | 0.54988741 | 0.5823966 | 0.7783237 |
| KL | 258.534174 | 0.35018335 | 0.51010463 | 0.68649318 | 4.92E-01 | 7.17E-01 |
| STARD13 | 186.963527 | -0.8527887 | 0.37366835 | -2.2822076 | 0.02247709 | 0.14624356 |
| AL139383.1 | 54.9214688 | 1.86567935 | 0.5580503 | 3.34321004 | 0.00082815 | 0.02565549 |
| RFC3 | 29.0411232 | 1.50060113 | 0.7425644 | 2.02083636 | 0.04329671 | 0.20789442 |
| LINC02343 | 68.9682689 | 1.68871236 | 0.63327989 | 2.66661297 | 0.00766199 | 0.08614009 |
| NBEA | 409.239483 | -0.2806489 | 0.32684533 | -0.8586596 | 0.39052837 | 0.63925794 |
| SPART | 103.596071 | 0.04269839 | 0.2620969 | 0.16291069 | 0.87058875 | 0.9398207 |
| RFXAP | 14.1816478 | 1.61270576 | 0.57657064 | 2.79706534 | 0.00515691 | NA |
| ALG5 | 42.3698432 | 0.18036226 | 0.32548159 | 0.55413967 | 0.57948326 | 0.77674437 |
| EXOSC8 | 20.1096385 | -0.1919564 | 0.46680135 | -0.4112165 | 0.68091382 | NA |
| SUPT20H | 121.260526 | 0.04498103 | 0.28285218 | 0.15902663 | 0.87364789 | 0.94114315 |
| LINC00571 | 45.120887 | -0.798689 | 0.40733768 | -1.960754 | 0.04990772 | 0.2242948 |
| UFM1 | 55.3971612 | 0.53626791 | 0.31232855 | 1.71699932 | 0.0859793 | 0.29308354 |
| FREM2 | 233.702561 | 1.96546499 | 0.50693801 | 3.87713082 | 0.0001057 | 0.0073958 |
| PROSER1 | 42.317663 | 0.08637041 | 0.38613551 | 0.22367902 | 0.82300707 | 0.91475524 |
| NHLRC3 | 67.6851628 | 1.37705408 | 0.52805109 | 2.60780466 | 0.00911249 | 0.09427267 |
| COG6 | 106.960476 | 0.60486261 | 0.2614232 | 2.31372966 | 0.02068256 | 0.14112911 |
| LINC00598 | 27.608767 | -0.3447986 | 0.56342328 | -0.6119707 | 0.54055711 | 0.74994017 |
| FOXO1 | 518.174837 | -0.7872501 | 0.54225873 | -1.451798 | 0.14655778 | 0.38645 |
| MRPS31 | 82.9838132 | 0.20678075 | 0.2732061 | 0.75686723 | 0.44912941 | 0.68734597 |
| ELF1 | 255.964665 | -0.0889179 | 0.16770651 | -0.5301997 | 0.59597346 | 0.78899341 |
| WBP4 | 40.2755703 | -0.2004884 | 0.40947272 | -0.4896258 | 0.62439874 | 0.80488982 |
| AL354696.2 | 20.3956622 | 1.1502657 | 0.67280844 | 1.70964815 | 0.08733096 | NA |
| MTRF1 | 22.9075395 | 0.447219 | 0.41172112 | 1.08621827 | 0.27738244 | 0.53576226 |
| NAA16 | 75.5117607 | 0.3310446 | 0.28322774 | 1.16882832 | 0.24247281 | 0.49968225 |
| VWA8 | 409.368587 | 0.79737436 | 0.29447246 | 2.70780628 | 0.00677295 | 0.08022018 |
| DGKH | 201.621789 | -0.3533064 | 0.27026135 | -1.3072767 | 1.91E-01 | 4.43E-01 |
| AKAP11 | 74.0265703 | 0.32418941 | 0.23835183 | 1.36012975 | 0.17378887 | 0.42294335 |
| EPSTI1 | 31.8306005 | 0.87790227 | 1.50323174 | 0.58400993 | 0.55921363 | 0.76485713 |

| | | | | | | |
|---|---|---|---|---|---|---|
| DNAJC15 | 53.6364596 | 0.40868646 | 0.32016121 | 1.27650213 | 0.20177808 | 0.45571768 |
| ENOX1 | 18.9800146 | 0.65281924 | 0.79065485 | 0.82566906 | 0.40899185 | NA |
| CCDC122 | 43.13153 | 1.32072295 | 0.55110874 | 2.3964834 | 0.01655324 | 0.12753785 |
| TSC22D1 | 768.797562 | -0.0171126 | 0.21165176 | -0.0808525 | 9.36E-01 | 9.74E-01 |
| NUFIP1 | 16.403616 | -0.7046221 | 0.47957661 | -1.4692586 | 0.14176265 | NA |
| AL359706.1 | 17.5072399 | 0.93467314 | 0.80243154 | 1.16480111 | 0.24409951 | NA |
| GPALPP1 | 52.4457625 | 0.16132649 | 0.25558841 | 0.63119641 | 0.52791211 | 0.74270508 |
| GTF2F2 | 101.102974 | -0.3652821 | 0.29139014 | -1.2535842 | 0.20999316 | 0.46499349 |
| TPT1 | 204.083738 | 0.38896715 | 0.36144379 | 1.07614835 | 0.28186091 | 0.54061387 |
| SLC25A30 | 80.7938335 | 0.48154191 | 0.39812613 | 1.20952098 | 0.22646276 | 0.4844591 |
| COG3 | 74.590719 | -0.7319224 | 0.25217935 | -2.9023883 | 0.00370329 | 0.06079223 |
| LINC01055 | 170.711729 | 0.89192339 | 0.61932426 | 1.44015574 | 0.14982334 | 0.39038262 |
| ZC3H13 | 136.777202 | 0.0853365 | 0.18725057 | 0.45573428 | 0.64858107 | 0.81913126 |
| LRCH1 | 646.264764 | 0.14184613 | 0.26862749 | 0.52804027 | 5.97E-01 | 7.89E-01 |
| ESD | 42.13938 | 0.62017716 | 0.3647627 | 1.7002209 | 0.08908938 | 0.29794449 |
| SUCLA2 | 130.653468 | 0.05612208 | 0.26889481 | 0.20871389 | 0.8346716 | 0.91994726 |
| MED4 | 67.4551085 | -0.0686859 | 0.27652497 | -0.2483895 | 0.80383307 | 0.90408418 |
| ITM2B | 435.014874 | 1.19061164 | 0.34535188 | 3.44753197 | 0.00056573 | 0.02124122 |
| RB1 | 242.683678 | 0.24787957 | 0.29797117 | 0.83189112 | 0.4054704 | 0.65277258 |
| RCBTB2 | 27.1388902 | -0.4083922 | 0.45804973 | -0.8915892 | 0.37261317 | 0.62538237 |
| FNDC3A | 601.042546 | -0.7700378 | 0.58242532 | -1.3221227 | 0.18612728 | 0.43607799 |
| CDADC1 | 28.6760135 | 0.1269276 | 0.3559615 | 0.35657675 | 0.72140868 | 0.86114805 |
| CAB39L | 61.9972356 | -0.5420542 | 0.46626614 | -1.1625426 | 0.24501515 | 0.5022131 |
| SETDB2 | 116.824898 | -0.3025233 | 0.41220828 | -0.7339088 | 0.46300434 | 0.69434917 |
| PHF11 | 54.0981129 | -0.5036617 | 0.34076825 | -1.4780182 | 0.13940291 | 0.37675008 |
| RCBTB1 | 21.434423 | -0.4687961 | 0.36717663 | -1.2767591 | 0.20168731 | 0.45567274 |
| EBPL | 16.9856085 | 0.49508547 | 0.48330687 | 1.02437086 | 0.30566014 | NA |
| KPNA3 | 89.1560999 | -0.1919536 | 0.23386002 | -0.8208054 | 0.41175713 | 0.65707964 |
| SPRYD7 | 16.1876592 | 0.68986743 | 0.61964212 | 1.11333204 | 0.26556585 | NA |
| DLEU2 | 104.393045 | -0.3339063 | 0.47935307 | -0.6965769 | 0.48606763 | 0.71289302 |
| TRIM13 | 36.9494273 | -0.0988762 | 0.34318962 | -0.2881097 | 0.77326279 | 0.88771702 |
| RNASEH2B | 34.5315578 | 0.46898412 | 0.35474349 | 1.32203729 | 0.18615573 | 0.43607799 |
| INTS6 | 178.483353 | 0.24269074 | 0.37485057 | 0.64743331 | 0.51735154 | 0.73469647 |
| INTS6-AS1 | 19.3629278 | 0.53775326 | 0.50810062 | 1.05835978 | 0.28989145 | NA |
| WDFY2 | 424.782428 | -0.3553226 | 0.37730367 | -0.9417418 | 3.46E-01 | 0.60358654 |
| ATP7B | 14.3090787 | 1.02348113 | 0.53923892 | 1.89801049 | 0.0576947 | NA |
| ALG11 | 15.7362028 | -0.3560863 | 0.54187932 | -0.657132 | 0.51109603 | NA |
| UTP14C | 17.0445754 | -0.5185465 | 0.45592901 | -1.1373404 | 0.255396 | NA |
| NEK3 | 39.3565807 | 0.54742696 | 0.58323196 | 0.93860933 | 0.34793136 | 0.60448713 |
| VPS36 | 89.9359737 | 0.03778263 | 0.25970788 | 0.14548127 | 0.8843309 | 0.94605075 |
| SUGT1 | 59.6282156 | 0.18887418 | 0.30039354 | 0.62875579 | 0.52950894 | 0.74370906 |
| DIAPH3 | 110.542859 | -1.2887487 | 0.73610016 | -1.750779 | 0.07998399 | 0.28223671 |
| TDRD3 | 230.923442 | 0.85968575 | 0.19735219 | 4.35609932 | 1.32E-05 | 0.00194859 |
| PCDH9 | 90.1668986 | -0.2705326 | 0.56683108 | -0.4772721 | 0.63316841 | 0.80995467 |
| DACH1 | 1465.48041 | 0.92358286 | 0.52550465 | 1.75751604 | 0.07882989 | 0.27999708 |
| DIS3 | 41.9333475 | -0.5874162 | 0.3977653 | -1.4767909 | 0.13973169 | 0.37697679 |
| PIBF1 | 255.44541 | -0.147036 | 0.19032829 | -0.7725391 | 0.43979522 | 0.67973973 |
| KLF5 | 57.9504946 | -1.368296 | 0.51984114 | -2.6321425 | 0.00848483 | 0.09010619 |
| KLF12 | 380.280689 | 0.06867417 | 0.19703153 | 0.34854407 | 7.27E-01 | 8.65E-01 |
| TBC1D4 | 480.125774 | 1.22158579 | 0.40449845 | 3.02000114 | 0.00252774 | 0.04792117 |
| COMMD6 | 36.9615393 | 0.45440265 | 0.45773319 | 0.99272383 | 0.32084456 | 0.57769945 |
| UCHL3 | 55.1600212 | -0.6488337 | 0.30870813 | -2.1017708 | 0.03557336 | 0.18754755 |
| LMO7-AS1 | 21.2959314 | 1.31056308 | 0.50540391 | 2.59310041 | 0.0095115 | 0.09644302 |
| LMO7 | 366.113183 | 0.12936602 | 0.415932 | 0.31102686 | 0.75578021 | 0.87904624 |

| | | | | | | |
|---|---|---|---|---|---|---|
| AC001226.2 | 24.0208672 | 0.87237414 | 0.60866062 | 1.43326857 | 0.1517811 | 0.39273741 |
| FBXL3 | 65.0972806 | 0.16836075 | 0.33259983 | 0.50619614 | 0.61271896 | 0.79866179 |
| MYCBP2 | 240.202504 | -0.1652465 | 0.24133097 | -0.6847297 | 0.49351451 | 0.71752572 |
| SLAIN1 | 27.2695522 | -0.0616433 | 0.37037809 | -0.1664335 | 0.86781585 | 0.93827484 |
| RNF219 | 19.837755 | -0.8227169 | 0.40674342 | -2.0226926 | 0.04310485 | NA |
| RBM26 | 249.128073 | 0.59668491 | 0.21703479 | 2.74925929 | 0.00597301 | 0.07599053 |
| NDFIP2 | 140.495432 | -0.915207 | 0.31532722 | -2.9024041 | 0.00370311 | 0.06079223 |
| GPC5 | 1660.85762 | 1.84129527 | 1.35297073 | 1.36092764 | 0.17353656 | 0.4225596 |
| GPC6 | 58.1452264 | -0.6040106 | 0.98759022 | -0.6116005 | 0.54080213 | 0.74994873 |
| TGDS | 24.507683 | -0.2123921 | 0.39718606 | -0.534742 | 0.59282828 | 0.785911 |
| GPR180 | 20.9716401 | -0.0536291 | 0.47810399 | -0.1121704 | 0.91068832 | NA |
| ABCC4 | 26.3583692 | -0.0178829 | 0.46634781 | -0.0383467 | 9.69E-01 | 0.98781207 |
| CLDN10 | 31.8792812 | 0.6806863 | 0.66686853 | 1.02072038 | 0.30738694 | 0.56764809 |
| CLDN10-AS1 | 39.479504 | 1.02891725 | 0.83288513 | 1.23536514 | 0.21669464 | 0.47286334 |
| DZIP1 | 37.9395526 | 0.18346096 | 0.33377349 | 0.54965707 | 0.58255461 | 0.7783237 |
| DNAJC3 | 123.605648 | -0.5007654 | 0.4144228 | -1.2083443 | 0.22691485 | 0.48461808 |
| UGGT2 | 252.94102 | -0.2940526 | 0.2321362 | -1.2667244 | 0.20525384 | 0.45870784 |
| MBNL2 | 366.138415 | -1.2756782 | 0.43437641 | -2.9368037 | 0.00331614 | 0.05707588 |
| RAP2A | 13.6687363 | 1.32271571 | 0.53309434 | 2.48120384 | 0.01309395 | NA |
| IPO5 | 54.0082463 | -0.0380473 | 0.27013074 | -0.1408478 | 0.88799017 | 0.94862739 |
| FARP1 | 890.284239 | 0.07114699 | 0.24391514 | 0.29168745 | 0.7705256 | 0.88651526 |
| STK24 | 329.012535 | -0.6513834 | 0.29990648 | -2.1719552 | 2.99E-02 | 0.17061993 |
| DOCK9 | 132.365106 | -0.3204374 | 0.26507807 | -1.2088418 | 0.22672363 | 0.48449513 |
| UBAC2 | 276.732156 | -0.145656 | 0.17417935 | -0.8362415 | 0.40301906 | 0.65053591 |
| TM9SF2 | 118.161923 | -0.247932 | 0.19263253 | -1.2870723 | 0.1980691 | 0.45104671 |
| CLYBL | 151.638273 | 0.72464606 | 0.3360263 | 2.15651588 | 0.03104341 | 0.17412703 |
| AL355338.1 | 20.1198254 | -0.1813829 | 0.45820269 | -0.3958573 | 0.69221034 | NA |
| PCCA | 1349.87887 | -0.4406864 | 0.16036446 | -2.7480301 | 0.00599545 | 0.07605145 |
| GGACT | 19.9827271 | 0.69309052 | 0.49780195 | 1.39230173 | 0.16383105 | NA |
| TMTC4 | 23.5236283 | 0.77560271 | 0.43862159 | 1.76827297 | 0.07701528 | 0.27700309 |
| FGF14 | 108.831512 | 0.14486452 | 0.70439301 | 0.20565866 | 0.83705755 | 0.92119416 |
| TPP2 | 132.508563 | -0.1972487 | 0.24646248 | -0.8003192 | 4.24E-01 | 0.66695091 |
| ARGLU1 | 199.163896 | -0.03914 | 0.2279715 | -0.1716883 | 0.86368262 | 0.93643831 |
| FAM155A | 1354.59069 | -1.7127562 | 0.55614627 | -3.0796865 | 0.00207219 | 0.04276036 |
| LIG4 | 16.9728008 | 0.28730438 | 0.51750452 | 0.55517271 | 0.57877654 | NA |
| ABHD13 | 33.1227328 | 0.30593281 | 0.45639886 | 0.67031896 | 0.50265448 | 0.72445662 |
| TNFSF13B | 41.9393857 | 0.81587822 | 0.52035499 | 1.5679262 | 0.11689836 | 0.34414008 |
| IRS2 | 97.4408101 | -0.4890417 | 0.4421999 | -1.105929 | 0.26875724 | 0.52744122 |
| COL4A1 | 93.9863915 | -0.8336132 | 0.41428463 | -2.012175 | 0.04420149 | 0.21042433 |
| COL4A2 | 141.248673 | -1.0309998 | 0.28338365 | -3.6381766 | 0.00027458 | 0.01360691 |
| RAB20 | 36.6929369 | 0.21277179 | 0.38280233 | 0.55582679 | 0.57832927 | 0.77564042 |
| NAXD | 34.7370411 | -0.3121537 | 0.33682942 | -0.9267411 | 0.35406097 | 0.61144431 |
| CARS2 | 67.4039244 | 0.19969359 | 0.29124757 | 0.68564893 | 0.49293452 | 0.71695332 |
| ANKRD10 | 337.461536 | -0.5133581 | 0.28090182 | -1.8275358 | 0.06761925 | 0.25898011 |
| ARHGEF7 | 245.701917 | -0.5022307 | 0.22405839 | -2.2415171 | 0.0249926 | 0.15492746 |
| TUBGCP3 | 60.1260636 | 0.56973756 | 0.30231881 | 1.88455879 | 0.05948943 | 0.2421418 |
| ATP11A | 94.9240507 | -0.1175019 | 0.78385937 | -0.1499018 | 8.81E-01 | 0.94400471 |
| MCF2L | 69.9139486 | 0.23439613 | 0.43629658 | 0.53724035 | 0.5911016 | 0.78517152 |
| PCID2 | 47.4345498 | 0.72967174 | 0.28136224 | 2.59335344 | 0.0095045 | 0.09644302 |
| CUL4A | 135.876181 | -0.4460251 | 0.24998115 | -1.7842347 | 0.07438553 | 0.27199309 |
| LAMP1 | 70.6616885 | -0.2001945 | 0.39531816 | -0.5064136 | 0.61256631 | 0.79866179 |
| GRTP1 | 28.1605465 | 0.8349208 | 0.36235992 | 2.30412018 | 0.0212159 | 0.1427136 |
| DCUN1D2 | 52.1292155 | 0.06515839 | 0.32980291 | 0.19756766 | 0.84338334 | 0.92437726 |
| TMCO3 | 42.0692993 | 0.01991976 | 0.2957343 | 0.06735695 | 0.94629754 | 0.97812984 |

| | | | | | | |
|---|---|---|---|---|---|---|
| TFDP1 | 37.2943741 | 0.03870455 | 0.34470629 | 0.11228268 | 0.91059927 | 0.96104245 |
| GAS6 | 90.555162 | 0.28401177 | 0.4305555 | 0.65964033 | 5.09E-01 | 0.72990216 |
| RASA3 | 47.7923399 | -2.2045868 | 0.38978323 | -5.6559303 | 1.55E-08 | 2.98E-05 |
| CDC16 | 72.017017 | -0.2403189 | 0.27677914 | -0.8682696 | 0.38524674 | 0.63547552 |
| UPF3A | 83.312101 | 0.61384756 | 0.23746583 | 2.58499323 | 0.00973809 | 0.09772371 |
| CHAMP1 | 10.961059 | -0.1524238 | 0.6038236 | -0.252431 | 0.80070795 | NA |
| TTC5 | 24.5537017 | 0.6667482 | 0.46378001 | 1.43763893 | 0.15053653 | 0.39108044 |
| CCNB1IP1 | 72.2124293 | -0.3204905 | 0.39722179 | -0.8068302 | 4.20E-01 | 6.64E-01 |
| TEP1 | 30.3896314 | -0.8595754 | 0.37616602 | -2.2850958 | 0.02230721 | 0.14604538 |
| OSGEP | 22.8195747 | -0.523193 | 0.45125558 | -1.1594162 | 0.24628657 | 0.50296216 |
| APEX1 | 18.6374831 | -0.091298 | 0.63018775 | -0.1448744 | 0.88481008 | NA |
| PNP | 17.4819068 | 0.01580934 | 0.41563204 | 0.03803687 | 0.96965829 | NA |
| METTL17 | 23.3857681 | 0.08529192 | 0.41431569 | 0.20586216 | 0.83689858 | 0.92119416 |
| NDRG2 | 31.5764668 | 0.8396525 | 0.52466544 | 1.60035795 | 0.1095192 | 0.33187766 |
| ARHGEF40 | 16.510828 | -0.527277 | 0.54590036 | -0.965885 | 0.33410171 | NA |
| HNRNPC | 504.339633 | -0.6500652 | 0.2569366 | -2.5300608 | 0.01140427 | 0.1040032 |
| SUPT16H | 53.5817539 | -0.0672707 | 0.31096684 | -0.2163275 | 0.82873249 | 0.91688285 |
| CHD8 | 98.2464257 | -0.1870312 | 0.24614537 | -0.7598404 | 0.44735001 | 0.68660592 |
| RAB2B | 36.2383275 | -0.4293224 | 0.35617163 | -1.2053806 | 0.22805641 | 0.48611194 |
| TOX4 | 44.4063293 | -0.3539323 | 0.30854466 | -1.1471024 | 0.25133928 | 0.50802481 |
| METTL3 | 19.7586969 | 0.50579009 | 0.40941351 | 1.23540156 | 0.21668109 | NA |
| DAD1 | 44.1240996 | -0.0844848 | 0.31140875 | -0.2712987 | 0.78616129 | 0.89452315 |
| ABHD4 | 26.0767705 | 0.41313899 | 0.3578728 | 1.1544297 | 0.24832405 | 0.50492395 |
| OXA1L | 22.8103062 | -0.3489197 | 0.48512987 | -0.7192294 | 0.47199956 | 0.70121225 |
| MRPL52 | 16.4839761 | 1.03213805 | 0.47876435 | 2.15583729 | 0.03109637 | NA |
| MMP14 | 22.407098 | -0.9051175 | 0.5234864 | -1.7290181 | 0.08380585 | 0.28873024 |
| LRP10 | 54.3020181 | 0.06949532 | 0.43267498 | 0.16061785 | 8.72E-01 | 0.94058267 |
| RBM23 | 70.0958222 | -0.6704825 | 0.28108298 | -2.3853545 | 0.01706267 | 0.12988597 |
| AL132780.1 | 44.4800588 | 1.75591334 | 0.75689835 | 2.31987999 | 0.02034737 | 0.14033487 |
| C14orf93 | 15.9262546 | 0.43488954 | 0.47946721 | 0.90702667 | 0.36439269 | NA |
| PSMB5 | 27.2646771 | 0.38957969 | 0.4055507 | 0.96061896 | 0.33674379 | 0.59259931 |
| ACIN1 | 136.706917 | -0.571727 | 0.25470981 | -2.2446212 | 0.02479246 | 0.15426642 |
| C14orf119 | 13.3544093 | 0.12077831 | 0.46225911 | 0.26127838 | 0.79387784 | NA |
| SLC7A8 | 69.1939587 | -1.3428677 | 0.56579211 | -2.3734295 | 0.01762376 | 0.13213074 |
| HOMEZ | 14.417681 | -0.7588918 | 0.56037306 | -1.3542618 | 0.17565288 | NA |
| PPP1R3E | 10.6782724 | 1.77905696 | 0.82054393 | 2.16814347 | 0.03014777 | NA |
| BCL2L2-PABI | 28.0843446 | -0.3657516 | 0.3960455 | -0.9235089 | 0.35574206 | 0.61187634 |
| PABPN1 | 72.3735133 | -0.550553 | 0.47601331 | -1.1565915 | 0.24743928 | 0.50358016 |
| NGDN | 18.6219756 | -0.312899 | 0.41621035 | -0.751781 | 0.45218274 | NA |
| ZFHX2 | 32.7209512 | 0.64264031 | 0.5753803 | 1.11689661 | 0.26403853 | 0.52361258 |
| AP1G2 | 15.5403041 | -0.3073208 | 0.53075949 | -0.5790208 | 0.5625751 | NA |
| DHRS4-AS1 | 22.7258324 | 0.92520799 | 0.49775125 | 1.85877583 | 0.06305892 | 0.25018789 |
| DHRS4L2 | 13.0100643 | 1.49292393 | 0.68028902 | 2.19454363 | 0.02819635 | NA |
| PSME1 | 20.2900223 | 0.47303476 | 0.45333805 | 1.04344818 | 0.29674077 | NA |
| AL136295.1 | 18.3113941 | -0.7985638 | 0.45670168 | -1.7485458 | 0.08036956 | NA |
| DHRS1 | 15.3304819 | 0.19338564 | 0.46685387 | 0.41423162 | 0.67870448 | NA |
| KHNYN | 18.8943649 | -0.220216 | 0.44622701 | -0.4935067 | 0.62165459 | NA |
| SDR39U1 | 21.0363776 | -0.2514432 | 0.45154725 | -0.5568479 | 0.57763134 | NA |
| NOVA1 | 91.92041 | 0.56724057 | 0.26514864 | 2.13933047 | 0.03240891 | 0.17754567 |
| LINC00645 | 159.411788 | 1.53755452 | 0.53847567 | 2.85538345 | 0.00429849 | 0.06314022 |
| PRKD1 | 1168.33454 | -0.7130433 | 0.49193352 | -1.4494708 | 0.14720615 | 0.38715156 |
| G2E3 | 40.5406381 | 0.0212811 | 0.31318551 | 0.06795045 | 0.94582508 | 0.97797094 |
| SCFD1 | 170.592968 | -0.3598713 | 0.19752181 | -1.8219321 | 0.0684653 | 0.26010729 |
| STRN3 | 393.279771 | -0.8463544 | 0.30978343 | -2.7320843 | 0.0062935 | 0.07738194 |

| | | | | | | |
|---|---|---|---|---|---|---|
| AP4S1 | 24.2299008 | -0.7616534 | 0.44078308 | -1.7279551 | 0.08399627 | 0.28888262 |
| HECTD1 | 394.80346 | -1.5261868 | 0.44930718 | -3.3967558 | 0.0006819 | 0.0236422 |
| HEATR5A | 48.6323103 | 0.22202167 | 0.36309473 | 0.61147038 | 0.54088822 | 0.74994873 |
| AL139353.1 | 12.9413875 | 1.36566392 | 0.5764269 | 2.3691884 | 0.01782717 | NA |
| NUBPL | 161.627559 | 0.79397925 | 0.32184973 | 2.46692532 | 0.01362788 | 0.11413904 |
| ARHGAP5 | 420.137633 | -0.7086757 | 0.2040879 | -3.4724042 | 0.00051582 | 0.02057129 |
| AKAP6 | 116.657642 | 0.94669231 | 0.35514838 | 2.66562473 | 0.00768454 | 0.08614009 |
| NPAS3 | 206.344343 | 0.76833381 | 0.60166957 | 1.27700294 | 0.20160121 | 0.45567274 |
| EGLN3 | 27.1501602 | -1.4898091 | 0.54670815 | -2.7250537 | 0.0064291 | 0.07805175 |
| EAPP | 52.4430854 | -0.4331201 | 0.37265849 | -1.162244 | 0.24513637 | 0.5022131 |
| SNX6 | 141.03057 | -0.3351067 | 0.26100359 | -1.283916 | 0.19917136 | 0.45282752 |
| BAZ1A | 74.4358191 | -1.190405 | 0.38454095 | -3.0956521 | 0.00196381 | 0.04142115 |
| SRP54 | 105.959504 | -0.7574391 | 0.26102733 | -2.9017618 | 0.00371071 | 0.06079223 |
| FAM177A1 | 56.4518571 | 0.03073907 | 0.25433598 | 0.12086008 | 0.90380186 | 0.95671001 |
| PPP2R3C | 45.9099675 | -0.0552909 | 0.47231793 | -0.1170628 | 0.90681028 | 0.95822608 |
| NFKBIA | 38.6742752 | -0.5130477 | 0.49054557 | -1.0458716 | 0.29562032 | 0.55524392 |
| RALGAPA1 | 384.576789 | -0.1071312 | 0.2344039 | -0.4570368 | 0.64764459 | 0.81885075 |
| BRMS1L | 25.1336691 | 0.13589247 | 0.4436252 | 0.3063227 | 0.75935896 | 0.88058098 |
| MBIP | 44.314244 | -0.0046129 | 0.32426915 | -0.0142256 | 0.98864997 | 0.99581595 |
| SLC25A21 | 20.8225605 | -1.1581417 | 0.71134922 | -1.6280916 | 0.10350547 | NA |
| MIPOL1 | 170.208944 | 0.18673788 | 0.24812572 | 0.75259383 | 0.45169403 | 0.68853766 |
| SEC23A | 110.08005 | -0.1940364 | 0.25693775 | -0.7551882 | 0.45013608 | 0.68770076 |
| TRAPPC6B | 84.1989402 | -0.3097083 | 0.29996963 | -1.0324654 | 0.30185413 | 0.56201529 |
| PNN | 130.130905 | -0.8408387 | 0.30678383 | -2.740818 | 0.00612864 | 0.07682764 |
| MIA2 | 181.184544 | 0.0202824 | 0.26779436 | 0.07573871 | 0.93962698 | 0.97491804 |
| FBXO33 | 84.9887892 | -0.2412831 | 0.26151509 | -0.9226354 | 0.35619723 | 0.61227891 |
| AL049875.1 | 27.8822717 | -0.4157538 | 0.35012071 | -1.1874585 | 0.23504683 | 0.49257908 |
| KLHL28 | 68.7759207 | 0.26488585 | 0.27674369 | 0.95715227 | 0.33849041 | 0.59428847 |
| TOGARAM1 | 123.388257 | -0.0809778 | 0.27710041 | -0.2922329 | 0.77010859 | 0.88651526 |
| PRPF39 | 52.6359328 | -0.1236207 | 0.31078562 | -0.3977685 | 0.69080087 | 0.84653627 |
| FKBP3 | 32.1342799 | 0.5158821 | 0.33044089 | 1.56119328 | 0.11847815 | 0.34566454 |
| FANCM | 32.3327929 | -0.237395 | 0.38554884 | -0.6157326 | 0.53807101 | 0.74878549 |
| MIS18BP1 | 27.1410613 | -0.0802339 | 0.52504215 | -0.1528142 | 0.87854483 | 0.94342075 |
| AL358335.2 | 23.111112 | 1.3310456 | 0.8399149 | 1.58473864 | 0.11302573 | 0.33771702 |
| RPS29 | 63.6286526 | 0.3956234 | 0.33004305 | 1.1987024 | 0.23064369 | 0.48933508 |
| RPL36AL | 48.6626565 | 0.04658822 | 0.48080353 | 0.09689658 | 0.92280852 | 0.96676973 |
| DNAAF2 | 16.2933631 | 1.08820329 | 0.46353915 | 2.34759738 | 0.01889493 | NA |
| KLHDC1 | 44.4349237 | 0.83576058 | 0.51099708 | 1.63554864 | 0.10193409 | 0.32089434 |
| KLHDC2 | 45.3546953 | 0.50925592 | 0.38847805 | 1.31090011 | 0.18989151 | 0.44120467 |
| NEMF | 213.966247 | -0.3747789 | 0.19287078 | -1.9431607 | 0.05199675 | 0.22846489 |
| VCPKMT | 29.9349349 | -0.3460613 | 0.42243899 | -0.8191982 | 0.41267335 | 0.65722052 |
| SOS2 | 315.713194 | -0.6562373 | 0.28646884 | -2.2907806 | 0.02197611 | 0.14481901 |
| L2HGDH | 81.6108244 | 0.57013984 | 0.25793637 | 2.21038945 | 0.02707815 | 0.16195658 |
| ATP5S | 47.6286522 | 0.2877118 | 0.3218387 | 0.89396272 | 0.37134184 | 0.62447414 |
| CDKL1 | 76.5009905 | -0.3691358 | 0.38433418 | -0.9604552 | 0.33682617 | 0.59259931 |
| MAP4K5 | 151.598092 | -0.2310191 | 0.20838027 | -1.1086418 | 0.26758476 | 0.52673196 |
| SAV1 | 47.8085609 | -0.2914346 | 0.30797525 | -0.9462924 | 0.34399946 | 0.60110949 |
| NIN | 55.6763955 | 0.65598756 | 0.39725453 | 1.6513029 | 0.09867674 | 0.31646453 |
| PYGL | 16.6484901 | 0.82634587 | 0.68545928 | 1.20553605 | 0.22799643 | NA |
| TMX1 | 26.7389123 | 0.32862965 | 0.36598793 | 0.89792483 | 0.36922563 | 0.6236443 |
| FRMD6 | 18.4696583 | 0.68798522 | 0.42348138 | 1.62459378 | 1.04E-01 | NA |
| RTRAF | 25.5560499 | -0.2801302 | 0.37302205 | -0.7509748 | 0.4526678 | 0.68922439 |
| TXNDC16 | 28.0395831 | 0.17642004 | 0.37907887 | 0.46539138 | 0.64165121 | 0.81566881 |
| ERO1A | 173.014084 | -1.008346 | 0.61711463 | -1.6339687 | 0.10226541 | 0.32120262 |

| | | | | | | |
|---|---|---|---|---|---|---|
| PSMC6 | 68.0381155 | -0.4993146 | 0.288575 | -1.7302765 | 0.08358088 | 0.28861463 |
| STYX | 36.0504003 | 0.3583625 | 0.39237848 | 0.91330825 | 0.36108045 | 0.61731435 |
| GNPNAT1 | 25.9591582 | 0.80975149 | 0.48449471 | 1.67133197 | 0.09465612 | 0.30858458 |
| FERMT2 | 319.065092 | -0.1608402 | 0.28049972 | -0.5734058 | 0.56636995 | 0.77020309 |
| DDHD1 | 55.4826747 | 0.2490995 | 0.30972647 | 0.80425642 | 0.4212489 | 0.66468897 |
| CNIH1 | 23.2257254 | 0.67875301 | 0.38540309 | 1.76115095 | 0.07821286 | 0.27886875 |
| GMFB | 22.6226637 | 0.16826543 | 0.38676115 | 0.43506291 | 0.66351676 | 0.82887489 |
| CGRRF1 | 15.9717471 | 0.04934796 | 0.53070938 | 0.0929849 | 0.92591556 | NA |
| SAMD4A | 224.090663 | -1.1282743 | 0.70186414 | -1.6075395 | 0.10793607 | 0.32999829 |
| SOCS4 | 29.6128785 | -0.1025535 | 0.38574101 | -0.265861 | 0.79034624 | 0.89697923 |
| MAPK1IP1L | 126.955207 | -0.1621485 | 0.26452441 | -0.6129811 | 0.53988884 | 0.74933197 |
| LGALS3 | 69.2178553 | -0.0204501 | 0.38436227 | -0.0532053 | 9.58E-01 | 9.84E-01 |
| FBXO34 | 198.257341 | 0.06865038 | 0.24683704 | 0.27812025 | 0.78092005 | 0.89127248 |
| ATG14 | 57.2843605 | 0.46728479 | 0.37666739 | 1.24057671 | 0.21476216 | 0.47054493 |
| KTN1 | 268.053657 | -0.4904872 | 0.27716173 | -1.7696786 | 0.07678069 | 0.27670206 |
| PELI2 | 215.853002 | -0.2400987 | 0.28379122 | -0.8460401 | 0.39753039 | 0.64588458 |
| TMEM260 | 112.307666 | 0.2941398 | 0.29487498 | 0.99750682 | 0.31851856 | 0.57587641 |
| EXOC5 | 83.8049522 | -0.1116752 | 0.25566631 | -0.4368005 | 0.66225602 | 0.82847174 |
| AP5M1 | 73.7048793 | 0.4705906 | 0.30797186 | 1.52803115 | 0.1265048 | 0.35811235 |
| NAA30 | 30.3781819 | -0.2832176 | 0.34728363 | -0.8155225 | 0.41477328 | 0.65865688 |
| CCDC198 | 229.630925 | 0.46386533 | 0.56126575 | 0.82646291 | 0.40854155 | 0.6544317 |
| ARMH4 | 73.8982181 | -0.2781596 | 0.28148855 | -0.9881739 | 0.32306751 | 0.57912936 |
| ACTR10 | 68.8823291 | -0.2968474 | 0.23590979 | -1.2583091 | 0.20828001 | 0.4622639 |
| PSMA3 | 59.6831074 | -0.0375548 | 0.30702358 | -0.122319 | 0.90264641 | 0.95661107 |
| ARID4A | 113.87769 | 0.05386796 | 0.26021289 | 0.20701497 | 0.83599817 | 0.92074197 |
| TIMM9 | 25.5224338 | -0.8026582 | 0.39158635 | -2.0497604 | 0.04038782 | 0.20017066 |
| KIAA0586 | 76.8907034 | 0.46455679 | 0.27645483 | 1.68040758 | 0.09287804 | 0.30571879 |
| DAAM1 | 195.534142 | -0.2529275 | 0.26039319 | -0.9713292 | 0.33138438 | 0.58689959 |
| GPR135 | 20.8711305 | 0.58251303 | 0.47150054 | 1.23544509 | 0.2166649 | NA |
| L3HYPDH | 21.9619908 | 0.22596639 | 0.48986861 | 0.4612796 | 0.64459802 | 0.81734747 |
| JKAMP | 35.8404246 | 0.65735833 | 0.36583059 | 1.79689274 | 0.07235265 | 0.26822676 |
| AL157911.1 | 26.2357214 | 1.0859867 | 0.69795764 | 1.5559493 | 0.11972015 | 0.3480474 |
| PCNX4 | 163.535923 | -0.5203873 | 0.2737386 | -1.9010375 | 0.05729711 | 0.23800098 |
| DHRS7 | 17.0932425 | 0.59149158 | 0.45037617 | 1.31332786 | 0.18907249 | NA |
| PPM1A | 84.3619405 | -0.508085 | 0.42090122 | -1.2071359 | 0.22737983 | 0.48520724 |
| MNAT1 | 319.937771 | -0.0510943 | 0.20543298 | -0.248715 | 0.80358127 | 0.90408418 |
| SLC38A6 | 54.0024336 | -0.3543588 | 0.31751458 | -1.1160394 | 0.26440526 | 0.5237724 |
| PRKCH | 144.348461 | -1.8424598 | 0.76422981 | -2.4108714 | 0.01591446 | 0.1247389 |
| TMEM30B | 19.2752878 | -0.5356172 | 0.66908625 | -0.8005204 | 0.42340935 | NA |
| HIF1A | 1119.94709 | -0.8307868 | 0.33294308 | -2.4952818 | 0.01258571 | 0.10970808 |
| HIF1A-AS2 | 246.393482 | -0.7128897 | 0.63491949 | -1.1228033 | 2.62E-01 | 0.52104231 |
| SNAPC1 | 146.161724 | -1.1516461 | 0.66613067 | -1.7288591 | 0.08383432 | 0.28873024 |
| SYT16 | 33.6359805 | 0.05827267 | 0.76107944 | 0.07656582 | 9.39E-01 | 0.97481036 |
| PPP2R5E | 209.363236 | -0.2334315 | 0.23748594 | -0.9829277 | 0.32564302 | 0.58154857 |
| WDR89 | 36.6113037 | 0.21744826 | 0.42847067 | 0.5074986 | 0.61180502 | 0.79828161 |
| SGPP1 | 190.03465 | -0.1604351 | 0.49097198 | -0.3267705 | 0.74384149 | 0.87230965 |
| SYNE2 | 1056.65717 | -0.8175626 | 0.36032813 | -2.2689393 | 0.02327202 | 0.1487747 |
| ESR2 | 61.1664498 | -0.214168 | 0.6194052 | -0.345764 | 0.7295201 | 0.86533507 |
| MTHFD1 | 27.6567101 | 0.31459987 | 0.36908039 | 0.85238847 | 0.39399852 | 0.6425014 |
| AL122035.1 | 23.9622258 | -0.7130124 | 0.6083901 | -1.1719658 | 0.24121081 | 0.49867735 |
| ZBTB25 | 41.6403747 | -0.7167585 | 0.40457696 | -1.7716245 | 0.07645691 | 0.27628583 |
| ZBTB1 | 112.291847 | 0.13611754 | 0.27839952 | 0.48892878 | 0.62489212 | 0.8053173 |
| AL049869.3 | 13.2711593 | 1.21436333 | 0.74786519 | 1.62377304 | 0.10442426 | NA |
| HSPA2 | 23.7861938 | 0.64992637 | 0.63216451 | 1.02809691 | 0.30390425 | 0.56472225 |

| | | | | | | |
|---|---|---|---|---|---|---|
| PLEKHG3 | 69.6285249 | -0.5174451 | 0.35618252 | -1.4527525 | 0.14629247 | 0.38641495 |
| CHURC1-FNT | 45.5396333 | -0.2535522 | 0.37325377 | -0.6793024 | 0.49694625 | 0.72101702 |
| RAB15 | 44.1597918 | 0.45609909 | 0.45706742 | 0.99788142 | 0.31833686 | 0.57587641 |
| MAX | 74.056268 | -0.3414549 | 0.27052558 | -1.2621909 | 0.2068801 | 0.46048471 |
| FUT8 | 101.900233 | 0.80379831 | 0.30808993 | 2.60897303 | 0.00908144 | 0.09427267 |
| AL359232.1 | 58.9949246 | 0.61149752 | 0.6170035 | 0.99107626 | 0.32164834 | 0.57817079 |
| GPHN | 1017.0093 | 0.07287096 | 0.25748231 | 0.28301347 | 0.77716651 | 0.8896927 |
| MPP5 | 78.7902434 | -0.0177771 | 0.26423186 | -0.0672785 | 0.94636 | 0.97812984 |
| ATP6V1D | 56.7548594 | -0.9965196 | 0.48458781 | -2.0564273 | 0.03974135 | 0.19847343 |
| EIF2S1 | 35.8902246 | -0.6137083 | 0.33303486 | -1.8427751 | 0.06536186 | 0.25452134 |
| PLEKHH1 | 29.1968432 | 0.24264969 | 0.39164357 | 0.61956767 | 0.53554246 | 0.74789968 |
| PIGH | 20.0050665 | -0.1371803 | 0.43616299 | -0.3145161 | 0.75312912 | NA |
| ARG2 | 54.2670474 | -1.1712867 | 0.62400798 | -1.877038 | 0.06051288 | 0.24514087 |
| VTI1B | 23.1681498 | 0.35170825 | 0.50657437 | 0.69428749 | 0.48750193 | 0.71418012 |
| ZFYVE26 | 44.8147777 | -0.7362619 | 0.49614595 | -1.4839623 | 0.13781893 | 0.37417718 |
| RAD51B | 526.795033 | 0.04775825 | 0.37149326 | 0.12855751 | 0.89770779 | 0.9539772 |
| ZFP36L1 | 256.79031 | -0.711991 | 0.36871644 | -1.9309989 | 0.05348319 | 0.23218282 |
| ACTN1 | 60.3432222 | -0.8288599 | 0.42679625 | -1.9420506 | 0.05213097 | 0.22876401 |
| DCAF5 | 186.404415 | 0.02127026 | 0.22202579 | 0.09580087 | 0.92367873 | 0.9675497 |
| AL359317.2 | 14.4600473 | 0.18537953 | 0.60057964 | 0.30866769 | 0.75757432 | NA |
| EXD2 | 41.5366136 | 0.59593788 | 0.36641116 | 1.62641848 | 0.10386066 | 0.32339043 |
| GALNT16 | 33.4942048 | 2.89220311 | 0.83645164 | 3.45770512 | 0.0005448 | 0.02096653 |
| ERH | 32.3481353 | 0.0549869 | 0.33166059 | 0.16579268 | 0.86832011 | 0.93855636 |
| SLC39A9 | 83.948915 | -0.0680193 | 0.2585501 | -0.2630799 | 0.79248896 | 0.8986875 |
| PLEKHD1 | 36.8404481 | 0.97138009 | 0.83188445 | 1.16768632 | 0.24293332 | 0.50009568 |
| SUSD6 | 104.499427 | 0.18505758 | 0.20475584 | 0.90379632 | 0.36610341 | 0.62157013 |
| SRSF5 | 241.315316 | -0.0134769 | 0.2355711 | -0.0572095 | 0.95437828 | 0.98153155 |
| AL357153.2 | 51.2908013 | 0.32117775 | 0.40112603 | 0.80069038 | 0.42331091 | 0.66684898 |
| MED6 | 35.2606101 | -0.4004161 | 0.32199244 | -1.2435574 | 0.21366249 | 0.46920405 |
| MAP3K9 | 42.7032766 | -0.1734722 | 0.42822769 | -0.4050933 | 0.68540895 | 0.84355496 |
| PCNX1 | 269.45688 | -0.3999923 | 0.27888942 | -1.4342324 | 0.15150594 | 0.39264015 |
| SIPA1L1 | 448.68563 | -0.9061984 | 0.29091448 | -3.1149993 | 0.00183945 | 0.03965902 |
| RGS6 | 151.774344 | -3.2551392 | 1.15692639 | -2.8136096 | 0.00489887 | 0.06806244 |
| DPF3 | 14.4485513 | -0.5661452 | 0.58036521 | -0.9754982 | 0.32931319 | NA |
| ZFYVE1 | 40.3991439 | -0.5025154 | 0.37153429 | -1.3525411 | 0.17620229 | 0.42528349 |
| RBM25 | 178.518361 | -0.5101879 | 0.21171139 | -2.4098271 | 0.01596008 | 0.12496923 |
| PSEN1 | 123.360501 | -0.8441193 | 0.29223693 | -2.8884758 | 0.00387114 | 0.06118306 |
| PAPLN | 24.2741694 | 0.22939813 | 0.56104982 | 0.40887302 | 0.68263285 | 0.84215821 |
| NUMB | 307.017333 | 0.01455714 | 0.22894357 | 0.06358396 | 0.94930151 | 0.97985432 |
| HEATR4 | 14.2412936 | 0.88282871 | 0.46220508 | 1.91003681 | 0.05612847 | NA |
| DNAL1 | 31.6240699 | 0.62830181 | 0.40888782 | 1.53661168 | 0.12438843 | 0.35460505 |
| ELMSAN1 | 56.1650215 | -1.0670585 | 0.27918715 | -3.8220189 | 0.00013236 | 0.00841985 |
| AC005480.1 | 28.575123 | -0.0241263 | 0.51721001 | -0.046647 | 0.96279461 | 0.98571829 |
| COQ6 | 16.5005388 | -0.5066773 | 0.48906195 | -1.0360187 | 0.30019344 | NA |
| ENTPD5 | 45.1623467 | 0.70930298 | 0.40179145 | 1.76535112 | 0.07750477 | 0.2775962 |
| BBOF1 | 36.7401157 | 0.44096179 | 0.4966734 | 0.88783051 | 0.37463193 | 0.62575308 |
| ALDH6A1 | 41.8106073 | 0.67500808 | 0.46477054 | 1.45234697 | 0.14640515 | 0.38644735 |
| LIN52 | 79.8543826 | -0.5683903 | 0.28117482 | -2.0214838 | 0.0432297 | 0.20770226 |
| ABCD4 | 69.3316836 | 0.2510245 | 0.37140232 | 0.67588295 | 0.49911496 | 0.72185041 |
| NPC2 | 28.2899476 | 0.03516495 | 0.49927137 | 0.07043254 | 0.94384939 | 0.97697805 |
| LTBP2 | 34.2789294 | 2.15485103 | 0.747163 | 2.88404408 | 0.00392604 | 0.06143969 |
| AREL1 | 57.3883619 | -0.4026933 | 0.30674199 | -1.3128079 | 0.18924768 | 0.4403384 |
| FCF1 | 25.2361227 | -0.1276208 | 0.3767275 | -0.3387615 | 0.73478937 | 0.86814805 |
| YLPM1 | 134.805829 | 0.0486264 | 0.22070534 | 0.22032271 | 0.82561983 | 0.9154963 |

| | | | | | | |
|---|---|---|---|---|---|---|
| DLST | 48.2306448 | -0.1096754 | 0.30604954 | -0.3583583 | 0.72007523 | 0.85995641 |
| MLH3 | 88.4405873 | 0.07152888 | 0.29175189 | 0.24517025 | 0.80632461 | 0.9057619 |
| ACYP1 | 48.7962532 | 0.62873701 | 0.39696681 | 1.58385286 | 0.11322721 | 0.33818774 |
| ZC2HC1C | 14.6536206 | 0.34267886 | 0.53723621 | 0.63785511 | 0.523568 | NA |
| NEK9 | 72.815945 | -0.4051992 | 0.24416094 | -1.6595579 | 0.09700342 | 0.31266135 |
| TMED10 | 92.2033435 | 0.16343929 | 0.31672168 | 0.51603443 | 0.60583036 | 0.79431477 |
| JDP2 | 20.2700129 | 1.0597149 | 0.73802207 | 1.43588511 | 0.15103504 | NA |
| TTLL5 | 148.596004 | 0.21674213 | 0.24510862 | 0.88426971 | 0.37655064 | 0.62801956 |
| IFT43 | 107.947136 | 0.3045499 | 0.36389448 | 0.83691814 | 0.40263858 | 0.65024957 |
| GPATCH2L | 240.618161 | -0.5265796 | 0.24200754 | -2.175881 | 0.02956416 | 0.16994425 |
| AC016526.1 | 38.0871112 | 1.21274539 | 0.60616494 | 2.00068549 | 4.54E-02 | 0.21335013 |
| ESRRB | 397.533861 | -0.2133593 | 0.33555177 | -0.635846 | 0.52487678 | 0.73968378 |
| ANGEL1 | 19.2648046 | 0.40758694 | 0.47546371 | 0.85724091 | 0.39131176 | NA |
| IRF2BPL | 22.8109438 | 0.03707395 | 0.74600339 | 0.04969676 | 0.96036404 | 0.98480176 |
| POMT2 | 18.3486529 | 0.39788035 | 0.43154284 | 0.92199503 | 0.35653117 | NA |
| TMED8 | 53.4754777 | 0.15531585 | 0.31316186 | 0.49596029 | 0.61992243 | 0.80301968 |
| VIPAS39 | 23.8825302 | 0.02466957 | 0.54036306 | 0.0456537 | 0.96358627 | 0.98600419 |
| AHSA1 | 19.3402286 | 0.6135166 | 0.45539221 | 1.34722681 | 0.1779072 | NA |
| SPTLC2 | 100.168928 | -0.0728575 | 0.26831311 | -0.2715392 | 0.78597634 | 0.89452315 |
| ALKBH1 | 38.4172081 | -0.3785897 | 0.44474411 | -0.8512529 | 0.39462889 | 0.6432568 |
| SLIRP | 33.4379014 | 0.45205617 | 0.33297607 | 1.35762361 | 0.17458315 | 0.42330339 |
| SNW1 | 44.8802472 | -0.618472 | 0.30704132 | -2.0142957 | 0.04397849 | 0.20959906 |
| ADCK1 | 24.4763051 | -0.3854065 | 0.42951443 | -0.8973075 | 0.36955488 | 0.62368335 |
| NRXN3 | 36.1894833 | -0.4549271 | 0.64311499 | -0.7073807 | 0.47932996 | 0.70695585 |
| DIO2 | 38.063948 | -0.1006528 | 0.96302655 | -0.1045172 | 0.91675891 | 0.96401363 |
| CEP128 | 39.5775108 | 0.23414704 | 0.44518916 | 0.52594956 | 0.59892325 | 0.78963276 |
| AC007262.2 | 20.659485 | 1.73294147 | 0.79324866 | 2.18461317 | 0.02891722 | NA |
| GTF2A1 | 41.0840768 | 0.54982182 | 0.39528877 | 1.39093709 | 0.1642445 | 0.41120418 |
| STON2 | 96.2659738 | -0.0616979 | 0.50165253 | -0.1229894 | 0.90211553 | 0.95661107 |
| SEL1L | 80.1207598 | -0.513128 | 0.32603359 | -1.57385 | 0.11552214 | 0.34185848 |
| GALC | 23.5314592 | 0.19199151 | 0.40997598 | 0.46829942 | 0.63957048 | 0.81408534 |
| SPATA7 | 58.0096822 | 1.00711851 | 0.37094194 | 2.71503 | 0.00662698 | 0.07957541 |
| PTPN21 | 51.4227085 | 0.05795968 | 0.34813751 | 0.16648503 | 0.86777526 | 0.93827484 |
| ZC3H14 | 104.343907 | 0.39562275 | 0.26098431 | 1.51588711 | 0.1295479 | 0.36208459 |
| EML5 | 155.865793 | -3.1183502 | 1.20272326 | -2.5927412 | 0.00952144 | 0.09644302 |
| TTC8 | 46.1160256 | -0.1397824 | 0.46165519 | -0.3027853 | 0.76205346 | 0.88247846 |
| FOXN3 | 237.775629 | 0.13691631 | 0.30264003 | 0.45240648 | 0.65097618 | 0.81992533 |
| EFCAB11 | 46.5951488 | 0.80760017 | 0.59428709 | 1.35893945 | 0.17416578 | 0.42295604 |
| TDP1 | 25.4922666 | 0.38365315 | 0.42270504 | 0.90761433 | 0.36408202 | 0.62003028 |
| PSMC1 | 33.5175625 | -0.4981732 | 0.35424666 | -1.4062891 | 0.15963828 | 0.40445247 |
| NRDE2 | 48.0988447 | -0.0403834 | 0.32865287 | -0.1228754 | 0.90220574 | 0.95661107 |
| CALM1 | 66.2456519 | 0.38877827 | 0.40131632 | 0.96875769 | 0.3326661 | 0.58796066 |
| TTC7B | 170.110166 | -0.5509951 | 0.4516374 | -1.2199944 | 0.22246698 | 0.47950948 |
| RPS6KA5 | 73.4952936 | -1.345989 | 0.33322301 | -4.0393037 | 5.36E-05 | 0.00474295 |
| DGLUCY | 148.313609 | 0.23941776 | 0.44726798 | 0.53528929 | 0.59244983 | 0.7855446 |
| PPP4R3A | 137.253571 | -0.5639644 | 0.31426791 | -1.7945338 | 0.072728 | 0.26848318 |
| CATSPERB | 25.384052 | 0.00286727 | 0.38749314 | 0.00739953 | 0.99409608 | 0.99730572 |
| TC2N | 32.7843707 | 0.56173075 | 0.45572817 | 1.23260044 | 0.21772487 | 0.47407641 |
| TRIP11 | 152.385977 | 0.08290382 | 0.20207985 | 0.41025277 | 0.68162053 | 0.84151154 |
| ATXN3 | 96.5492316 | -0.6246722 | 0.33088273 | -1.8878961 | 0.0590399 | 0.24133304 |
| NDUFB1 | 47.2764279 | 0.24427518 | 0.39189426 | 0.62331911 | 0.53307484 | 0.74615054 |
| CPSF2 | 51.3140196 | -0.1049048 | 0.3217144 | -0.3260807 | 0.74436333 | 0.87258065 |
| LGMN | 29.6696957 | -0.008927 | 0.45195496 | -0.019752 | 0.98424121 | 0.99382437 |
| GOLGA5 | 36.0465305 | -0.1902616 | 0.39051975 | -0.4872009 | 0.62611593 | 0.8053173 |

| | | | | | | |
|---|---|---|---|---|---|---|
| ITPK1 | 60.0854572 | -0.0188245 | 0.39673109 | -0.0474489 | 0.96215544 | 0.98535004 |
| BTBD7 | 311.061934 | -0.1125788 | 0.21364328 | -0.5269476 | 5.98E-01 | 0.7894568 |
| UNC79 | 30.1053281 | 0.60913534 | 0.70854982 | 0.85969303 | 0.38995828 | 0.6386799 |
| DDX24 | 94.1275496 | -0.3428118 | 0.22572279 | -1.5187294 | 0.12883062 | 0.3611257 |
| IFI27 | 21.5932321 | -1.6291388 | 1.15860894 | -1.4061162 | 0.15968963 | 0.40445247 |
| SERPINA1 | 35.9161243 | -1.050499 | 0.91365784 | -1.1497729 | 0.25023743 | 0.50659901 |
| DICER1 | 218.62186 | 0.14641827 | 0.19617057 | 0.74638245 | 0.45543642 | 0.6908246 |
| DICER1-AS1 | 23.888174 | 0.37726551 | 0.50299523 | 0.75003795 | 0.45323185 | 0.68943192 |
| CLMN | 514.726716 | 0.87993523 | 0.36035788 | 2.44183709 | 0.01461274 | 0.11902037 |
| GLRX5 | 21.4309741 | 0.39921381 | 0.44114777 | 0.90494351 | 0.36549532 | 0.62110919 |
| ATG2B | 110.656359 | -0.2286655 | 0.26117419 | -0.8755288 | 0.38128624 | 0.63258566 |
| GSKIP | 19.4994106 | -0.4144182 | 0.38904134 | -1.0652291 | 0.28677225 | NA |
| AK7 | 18.1970871 | -0.6866137 | 0.58260149 | -1.1785306 | 0.23858515 | NA |
| PAPOLA | 195.14213 | -0.0821236 | 0.27769965 | -0.2957283 | 0.76743761 | 0.88473859 |
| VRK1 | 14.831508 | 0.43871018 | 0.62504694 | 0.70188357 | 0.48275177 | NA |
| LINC01550 | 21.3688911 | 0.65127844 | 0.60623105 | 1.0743073 | 0.28268497 | 0.54138498 |
| SETD3 | 102.736401 | 0.20481034 | 0.34079315 | 0.6009814 | 0.54785238 | 0.7554786 |
| CCNK | 38.6975005 | -0.2267978 | 0.30173228 | -0.7516525 | 0.45226005 | 0.68890671 |
| CCDC85C | 20.8444811 | -0.4526544 | 0.53122958 | -0.8520881 | 0.39416521 | NA |
| EVL | 107.052802 | -0.6045994 | 0.33713875 | -1.793325 | 0.07292097 | 0.26906649 |
| YY1 | 118.845457 | -0.6314365 | 0.30737606 | -2.0542801 | 0.03994859 | 0.19889024 |
| SLC25A29 | 144.530092 | -0.3483436 | 0.39380728 | -0.8845536 | 0.37639745 | 0.62790013 |
| WARS | 65.2834835 | -0.9345121 | 0.45531898 | -2.0524338 | 0.04012752 | 0.19926552 |
| WDR25 | 34.297149 | -0.5001292 | 0.37089549 | -1.348437 | 0.17751788 | 0.42593802 |
| PPP2R5C | 158.692781 | -0.1885756 | 0.24962482 | -0.7554361 | 0.44998735 | 0.68770076 |
| DYNC1H1 | 110.840958 | -0.5342644 | 0.32655285 | -1.6360732 | 0.10182426 | 0.32089434 |
| HSP90AA1 | 189.704788 | 0.75794445 | 0.26355142 | 2.87588832 | 0.00402892 | 0.06202123 |
| WDR20 | 140.273348 | -0.3110895 | 0.26658293 | -1.1669522 | 0.24322969 | 0.5004381 |
| MOK | 70.1393238 | 0.64184233 | 0.44342709 | 1.44745854 | 0.14776854 | 0.38738912 |
| ZNF839 | 22.1161096 | 0.82036923 | 0.5143284 | 1.59503001 | 0.11070552 | 0.33415703 |
| TECPR2 | 71.4039154 | -0.1447961 | 0.27947068 | -0.5181083 | 0.60438274 | 0.79370993 |
| ANKRD9 | 40.281903 | 0.62615565 | 0.53442174 | 1.17165078 | 0.24133729 | 0.49867735 |
| RCOR1 | 155.932998 | 0.094242 | 0.28131558 | 0.33500455 | 0.73762165 | 0.86984432 |
| TRAF3 | 98.3085324 | -0.6703776 | 0.36322322 | -1.8456354 | 0.06494517 | 0.25349034 |
| CDC42BPB | 164.26981 | -0.9625615 | 0.30130308 | -3.1946621 | 0.00139995 | 0.03377864 |
| EIF5 | 88.6151324 | -0.4496676 | 0.43981553 | -1.0224005 | 0.30659136 | 0.56631479 |
| MARK3 | 339.167015 | -0.3478322 | 0.25519095 | -1.3630271 | 0.17287396 | 0.42161308 |
| CKB | 119.358174 | 1.20924908 | 0.48141969 | 2.51183968 | 0.01201036 | 0.10726863 |
| BAG5 | 29.8293183 | 0.50379878 | 0.37814613 | 1.33228597 | 0.18276624 | 0.43258048 |
| KLC1 | 72.501509 | 0.35376145 | 0.38723285 | 0.9135626 | 0.36094673 | 0.61731435 |
| ZFYVE21 | 16.4879986 | 0.91452676 | 0.56161936 | 1.62837472 | 0.10344546 | NA |
| PPP1R13B | 100.969492 | -0.1068835 | 0.29100488 | -0.367291 | 7.13E-01 | 0.85784326 |
| ATP5MPL | 36.1383405 | 0.7053859 | 0.38000983 | 1.85623065 | 0.06342069 | 0.25113402 |
| AL590079.1 | 12.8980729 | 0.9605828 | 0.66610119 | 1.4420974 | 0.1492749 | NA |
| SIVA1 | 13.8623148 | 0.21495129 | 0.65646301 | 0.32743854 | 0.74333622 | NA |
| AKT1 | 25.1447787 | -0.3989366 | 0.40213599 | -0.9920441 | 0.32117601 | 0.57791615 |
| CEP170B | 23.1320362 | -0.2452448 | 0.51881592 | -0.4727011 | 0.63642646 | 0.81209789 |
| NUDT14 | 10.9185233 | 0.48898705 | 0.78105345 | 0.62606094 | 0.53127497 | NA |
| BRF1 | 46.0499584 | -0.1431617 | 0.29527032 | -0.4848497 | 0.62778295 | 0.80601257 |
| PACS2 | 64.5324439 | 0.17784815 | 0.29385681 | 0.60522045 | 0.54503253 | 0.75343308 |
| TEX22 | 12.6895592 | -0.3572595 | 0.60531969 | -0.5901997 | 0.5550568 | NA |
| MTA1 | 34.6658654 | 0.27048786 | 0.45137149 | 0.59925775 | 0.54900102 | 0.75579697 |
| CRIP2 | 15.6235833 | 0.6287967 | 0.58052392 | 1.08315381 | 0.27874016 | NA |
| AC087386.1 | 13.8103208 | 0.10706378 | 0.66859587 | 0.16013228 | 0.87277688 | NA |

| | | | | | | |
|---|---|---|---|---|---|---|
| NIPA1 | 62.7745752 | 0.32342026 | 0.29523649 | 1.09546169 | 0.27331442 | 0.53231303 |
| NIPA2 | 52.2362818 | -0.5989467 | 0.36608137 | -1.6361028 | 0.10181808 | 0.32089434 |
| CYFIP1 | 101.683845 | -0.6089606 | 0.2643951 | -2.3032219 | 0.02126636 | 0.14276951 |
| TUBGCP5 | 38.6408871 | -0.4524688 | 0.33420746 | -1.3538562 | 0.17578227 | 0.42480254 |
| SNRPN | 13.83805 | 0.75052739 | 0.88428595 | 0.84873834 | 0.39602691 | NA |
| AC124312.1 | 334.109024 | -0.1590304 | 0.25931691 | -0.6132665 | 0.53970011 | 0.74929145 |
| PWAR6 | 79.5507558 | 0.81812932 | 0.27898782 | 2.93249121 | 0.00336254 | 0.05733014 |
| UBE3A | 222.822718 | -0.3520781 | 0.24962939 | -1.4104031 | 0.15842069 | 0.40309554 |
| GABRB3 | 54.5404713 | 1.27168808 | 0.52343822 | 2.4294903 | 0.01512007 | 0.12124045 |
| HERC2 | 235.769782 | -0.1392555 | 0.15823057 | -0.8800797 | 0.37881614 | 0.63024389 |
| FAM189A1 | 121.246956 | 0.78034918 | 0.4678433 | 1.66797124 | 0.09532144 | 0.30996582 |
| TJP1 | 284.134006 | -0.577728 | 0.36257125 | -1.5934193 | 0.11106616 | 0.33449169 |
| FAN1 | 52.3823583 | 1.37607471 | 0.36295891 | 3.79126856 | 0.00014988 | 0.00913087 |
| MTMR10 | 197.035275 | 0.19408326 | 0.37360404 | 0.51948918 | 0.60341966 | 0.79301836 |
| KLF13 | 36.3409583 | 0.63677639 | 0.39432464 | 1.61485315 | 0.1063425 | 0.32732986 |
| OTUD7A | 91.9378977 | -0.4642798 | 0.2669941 | -1.7389142 | 0.08204985 | 0.28580436 |
| LINC02256 | 140.950872 | 0.9383538 | 0.37638074 | 2.49309729 | 0.01266341 | 0.1101415 |
| AC123768.3 | 17.7677619 | 1.50208568 | 0.6566311 | 2.28756402 | 0.02216292 | NA |
| GOLGA8N | 19.0251688 | 1.17179451 | 0.46782515 | 2.50477021 | 0.0122531 | NA |
| FMN1 | 553.504845 | 0.56768537 | 0.50580668 | 1.12233665 | 0.26171933 | 0.52120406 |
| AVEN | 66.2644208 | -0.698709 | 0.27058257 | -2.5822395 | 0.00981614 | 0.0981232 |
| EMC7 | 22.0807228 | 0.44908754 | 0.40624084 | 1.10547117 | 0.26895548 | 0.52756125 |
| KATNBL1 | 113.619519 | -0.1208311 | 0.23069897 | -0.5237607 | 0.60044497 | 0.79096587 |
| SLC12A6 | 52.8302205 | -0.4730121 | 0.45473708 | -1.0401881 | 0.29825253 | 0.55811077 |
| NOP10 | 23.0535108 | 0.27139278 | 0.5018097 | 0.54082809 | 5.89E-01 | 0.78344371 |
| LPCAT4 | 13.0469881 | -0.0255662 | 0.50667788 | -0.0504586 | 0.95975698 | NA |
| GOLGA8A | 80.3801164 | 0.6441353 | 0.47424993 | 1.35821907 | 0.17439418 | 0.42317528 |
| GOLGA8B | 84.9910266 | 0.98396703 | 0.51248869 | 1.91997804 | 0.05486067 | 0.23407018 |
| AQR | 101.770937 | -0.2134068 | 0.20800633 | -1.0259632 | 0.30490893 | 0.56551423 |
| ZNF770 | 51.5610833 | 0.79687687 | 0.38975812 | 2.04454206 | 0.04090003 | 0.20192913 |
| AC021231.1 | 94.8060298 | 2.58301792 | 0.59611003 | 4.33312269 | 1.47E-05 | 0.00207296 |
| DPH6 | 294.335352 | 1.61962519 | 0.47248287 | 3.42790246 | 0.00060826 | 0.02240382 |
| DPH6-DT | 186.808803 | 1.05826201 | 0.66750034 | 1.58541045 | 0.11287311 | 0.33755464 |
| C15orf41 | 86.9692992 | 0.97500382 | 0.30208917 | 3.2275365 | 0.00124861 | 0.03140706 |
| SPRED1 | 175.101696 | -0.1121132 | 0.28947308 | -0.3873009 | 0.69853348 | 0.85102865 |
| FAM98B | 31.4037241 | 0.08818271 | 0.40001243 | 0.22044992 | 0.82552077 | 0.9154963 |
| RASGRP1 | 19.9388001 | 0.9310383 | 0.45679921 | 2.03817841 | 0.04153209 | NA |
| C15orf53 | 110.519961 | 1.07341588 | 0.63063738 | 1.70211268 | 8.87E-02 | 0.29778649 |
| AC013652.1 | 46.1969898 | -0.0757424 | 0.45817961 | -0.1653115 | 0.86869881 | 0.93857029 |
| THBS1 | 17.9302988 | -0.7622033 | 0.56719267 | -1.3438173 | 0.17900747 | NA |
| FSIP1 | 15.5934302 | 0.02317538 | 0.6135073 | 0.03777523 | 0.96986689 | NA |
| EIF2AK4 | 48.9094932 | 0.65659479 | 0.29309121 | 2.24024045 | 0.02507532 | 0.15502387 |
| SRP14 | 93.2993596 | 0.80914434 | 0.42047099 | 1.92437616 | 0.05430745 | 0.23287156 |
| SRP14-AS1 | 17.6521964 | 0.1383982 | 0.49664645 | 0.27866544 | 0.78050159 | NA |
| IVD | 39.4150992 | 0.48938426 | 0.38572536 | 1.2687376 | 0.20453467 | 0.45773044 |
| BAHD1 | 20.332087 | 0.34025684 | 0.4822968 | 0.70549264 | 0.48050371 | NA |
| CCDC32 | 41.9782092 | -0.3503995 | 0.43469899 | -0.8060738 | 0.42020031 | 0.66385094 |
| RMDN3 | 24.0374597 | -0.049875 | 0.38148487 | -0.1307392 | 0.89598162 | 0.95319565 |
| DNAJC17 | 39.0954323 | 0.22984007 | 0.31416428 | 0.73159197 | 0.46441765 | 0.69532957 |
| SPINT1-AS1 | 13.8538961 | 0.4105507 | 0.49925774 | 0.82232216 | 0.41089357 | NA |
| SPINT1 | 47.0688393 | -0.1392516 | 0.44071601 | -0.3159667 | 0.75202779 | 0.87816052 |
| INO80 | 130.323674 | -0.5485531 | 0.25301259 | -2.168086 | 0.03015214 | 0.17165758 |
| CHP1 | 86.3099263 | 0.83392224 | 0.38500777 | 2.1659881 | 0.0303121 | 0.17229651 |
| NDUFAF1 | 13.5955685 | 0.66656359 | 0.54296115 | 1.22764511 | 0.21958022 | NA |

| | | | | | | |
|---|---|---|---|---|---|---|
| RTF1 | 64.9759981 | -0.4726539 | 0.282766 | -1.6715372 | 0.09461562 | 0.30858458 |
| RPAP1 | 15.2712999 | 0.45205831 | 0.55540384 | 0.81392724 | 0.41568664 | NA |
| MGA | 122.979994 | -0.7061749 | 0.27518742 | -2.56616 | 0.01028314 | 0.09994074 |
| MAPKBP1 | 96.0273279 | -1.6964299 | 0.35661948 | -4.7569749 | 1.97E-06 | 0.00058176 |
| EHD4 | 22.7377899 | -0.7040301 | 0.45100133 | -1.5610378 | 0.11851484 | 0.34566454 |
| PLA2G4F | 15.6604933 | 2.1606072 | 0.71291663 | 3.03065899 | 0.00244021 | NA |
| VPS39 | 78.2523601 | -0.0714226 | 0.25700762 | -0.2779008 | 0.78108847 | 0.89133254 |
| TMEM87A | 112.973398 | -0.3294756 | 0.2472407 | -1.3326107 | 0.18265959 | 0.43246104 |
| GANC | 63.6048803 | 0.10274706 | 0.38731638 | 0.26527941 | 0.79079422 | 0.89722039 |
| ZNF106 | 80.6215664 | 0.13670551 | 0.25999097 | 0.52580869 | 0.59902114 | 0.78963276 |
| SNAP23 | 57.3132058 | -0.1771901 | 0.26769291 | -0.6619155 | 0.50802536 | 0.72884831 |
| HAUS2 | 16.2666643 | 0.17283488 | 0.48333266 | 0.35758991 | 0.72065023 | NA |
| TTBK2 | 161.20442 | -0.6720959 | 0.27936743 | -2.4057775 | 0.01613809 | 0.12559644 |
| UBR1 | 184.155344 | -0.3307505 | 0.21677549 | -1.5257746 | 0.12706601 | 0.35877736 |
| AC068724.3 | 18.229566 | 0.92431541 | 0.57994039 | 1.59381105 | 0.11097836 | NA |
| TMEM62 | 47.2020218 | -0.1324006 | 0.3048963 | -0.4342481 | 0.66410827 | 0.8292079 |
| CCNDBP1 | 21.089565 | 0.30091231 | 0.42618631 | 0.70605814 | 0.48015198 | NA |
| ADAL | 39.5202038 | 0.91570262 | 0.43734769 | 2.09376348 | 0.03628106 | 0.19022839 |
| TUBGCP4 | 28.8413749 | -0.2908981 | 0.421616 | -0.6899599 | 0.49021941 | 0.7163358 |
| TP53BP1 | 170.875475 | -0.1000843 | 0.22946308 | -0.4361672 | 0.66271544 | 0.8285925 |
| PPIP5K1 | 47.5083557 | 0.08944601 | 0.35283897 | 0.25350377 | 0.79987895 | 0.90280874 |
| CATSPER2 | 61.4838478 | 0.593637 | 0.50297928 | 1.18024146 | 0.23790419 | 0.49574106 |
| PDIA3 | 80.5807774 | -0.0329866 | 0.33149511 | -0.0995084 | 0.92073461 | 0.96564849 |
| SERF2 | 190.015876 | -0.0204361 | 0.46769771 | -0.043695 | 0.96514749 | 0.9869039 |
| MFAP1 | 19.3604093 | -0.0630716 | 0.38747268 | -0.1627768 | 0.87069416 | NA |
| CASC4 | 201.52397 | -0.0468398 | 0.18868368 | -0.2482452 | 0.80394469 | 0.90408418 |
| CTDSPL2 | 148.928457 | -0.5400938 | 0.23528581 | -2.2954796 | 0.02170565 | 0.14366346 |
| EIF3J-DT | 37.0474646 | 0.39416021 | 0.51844983 | 0.76026684 | 0.4470951 | 0.68660592 |
| EIF3J | 64.2406565 | -0.4378059 | 0.30122284 | -1.4534287 | 0.14610477 | 0.38623836 |
| SPG11 | 198.874213 | -0.4821493 | 0.20623729 | -2.3378374 | 0.01939568 | 0.1366689 |
| B2M | 341.798377 | 0.49285514 | 0.31926696 | 1.54370857 | 0.12265895 | 0.35251484 |
| TRIM69 | 12.6332776 | 0.10365599 | 0.62613751 | 0.16554828 | 0.86851245 | NA |
| GATM | 17.6980868 | 0.38865529 | 0.57862671 | 0.67168569 | 0.50178381 | NA |
| SPATA5L1 | 22.4454919 | -0.2040741 | 0.46956068 | -0.4346065 | 0.66384807 | 0.82908302 |
| SEMA6D | 168.679076 | 0.26740914 | 0.45551068 | 0.58705349 | 5.57E-01 | 7.63E-01 |
| AC092078.2 | 151.233331 | 0.33893118 | 1.56005902 | 0.21725535 | 0.82800935 | 0.91685523 |
| MYEF2 | 125.635069 | 0.02400434 | 0.23673667 | 0.1013968 | 0.91923548 | 0.9647335 |
| SLC12A1 | 47.8973048 | 1.4675225 | 0.83016497 | 1.76774805 | 0.07710303 | 0.27718919 |
| DUT | 22.6835298 | 0.60270557 | 0.51406945 | 1.1724205 | 0.24102827 | 0.49867735 |
| CEP152 | 56.5989659 | 0.35577412 | 0.35408638 | 1.00476646 | 0.31500932 | 0.57225891 |
| EID1 | 35.4071247 | 1.25666676 | 0.49462467 | 2.54064717 | 0.01106475 | 0.10282338 |
| SECISBP2L | 220.198433 | -0.105298 | 0.23254702 | -0.4528031 | 0.65069052 | 0.81985999 |
| COPS2 | 41.2742045 | -0.0490342 | 0.30961068 | -0.1583738 | 0.87416225 | 0.94143373 |
| GALK2 | 131.015061 | 0.43881134 | 0.32032471 | 1.36989535 | 0.17071957 | 0.41888063 |
| FAM227B | 84.5469702 | 1.56483402 | 0.61296635 | 2.55288731 | 0.01068341 | 0.101577 |
| DTWD1 | 63.0088476 | 0.40584792 | 0.36065281 | 1.12531472 | 0.26045569 | 0.51976341 |
| GABPB1 | 59.7774149 | -0.8846131 | 0.33211009 | -2.6636142 | 0.00773062 | 0.08618711 |
| GABPB1-AS1 | 109.897692 | 0.23178467 | 0.39427626 | 0.58787376 | 0.55661703 | 0.76317239 |
| USP8 | 152.132921 | 0.16997548 | 0.19929023 | 0.85290426 | 0.3937124 | 0.64244315 |
| USP50 | 39.5092545 | -0.8149835 | 0.42854697 | -1.9017367 | 0.05720558 | 0.23774912 |
| TRPM7 | 1614.59516 | 0.03992288 | 0.27949414 | 0.14283978 | 8.86E-01 | 9.47E-01 |
| SPPL2A | 340.45207 | -0.5637667 | 0.35790817 | -1.5751713 | 0.11521692 | 0.34148041 |
| AP4E1 | 61.412448 | -0.2604537 | 0.29555092 | -0.8812481 | 0.37818353 | 0.62965144 |
| DMXL2 | 76.8486441 | 0.61957199 | 0.3329352 | 1.86093865 | 0.06275284 | 0.24960868 |

| | | | | | | |
|---|---|---|---|---|---|---|
| TMOD3 | 125.069813 | -0.1067307 | 0.20570995 | -0.5188406 | 0.60387187 | 0.79317437 |
| LEO1 | 17.0486103 | -0.0142817 | 0.49142439 | -0.0290618 | 0.9768153 | NA |
| MAPK6 | 87.7042225 | -0.7408569 | 0.37463433 | -1.9775468 | 0.04797985 | 0.21943015 |
| GNB5 | 62.7870054 | 0.2139054 | 0.28040152 | 0.76285391 | 0.44555052 | 0.68538685 |
| MYO5C | 163.176212 | 0.23375385 | 0.35591579 | 0.65676729 | 0.51133056 | 0.73064085 |
| MYO5A | 215.456168 | -0.010003 | 0.29826879 | -0.0335368 | 0.97324655 | 0.98944375 |
| ARPP19 | 75.0741209 | -0.3375098 | 0.30086301 | -1.1218057 | 0.26194505 | 0.52137312 |
| FAM214A | 218.829499 | -0.5132187 | 0.31974423 | -1.6050912 | 0.10847373 | 0.33024913 |
| WDR72 | 552.651238 | 1.07260957 | 0.46378136 | 2.31274834 | 0.02073648 | 0.14118898 |
| UNC13C | 85.3824855 | 0.0480992 | 0.72813992 | 0.06605764 | 9.47E-01 | 9.79E-01 |
| RSL24D1 | 20.0215043 | 0.09941379 | 0.40341335 | 0.24643159 | 0.80534816 | NA |
| RAB27A | 62.43166 | -0.1458352 | 0.32209241 | -0.4527743 | 0.65071127 | 0.81985999 |
| PIGB | 36.0966443 | 0.05652614 | 0.31104627 | 0.18172904 | 0.85579537 | 0.93169123 |
| CCPG1 | 108.497077 | 0.79918604 | 0.26246727 | 3.04489786 | 0.00232759 | 0.04581966 |
| DNAAF4 | 49.6484644 | 1.35571111 | 0.56556897 | 2.39707476 | 0.01652655 | 0.12745979 |
| PYGO1 | 16.9362581 | 0.42762374 | 0.54155162 | 0.78962693 | 0.42974568 | NA |
| PRTG | 79.436575 | 1.72785421 | 0.52312244 | 3.30296329 | 0.00095669 | 0.02829008 |
| NEDD4 | 74.4222124 | -0.969794 | 0.38310947 | -2.5313757 | 0.01136161 | 0.103737 |
| RFX7 | 182.383711 | 0.35653536 | 0.28002543 | 1.27322495 | 0.20293823 | 0.45686328 |
| TEX9 | 106.273971 | 0.96504225 | 0.43738076 | 2.2064122 | 0.02735515 | 0.16284037 |
| ZNF280D | 72.2817722 | 1.1648029 | 0.41951025 | 2.77657795 | 0.00549345 | 0.07240249 |
| AC090517.4 | 32.8421865 | 0.77297698 | 0.44239715 | 1.74724675 | 0.08059455 | 0.28377686 |
| TCF12 | 733.860242 | -0.2129639 | 0.18341339 | -1.1611144 | 0.24559539 | 0.50275206 |
| CGNL1 | 887.049462 | 0.10169893 | 0.37260219 | 0.27294238 | 0.78489751 | 0.89448566 |
| ALDH1A2 | 322.149254 | -0.924069 | 0.56215728 | -1.6437909 | 0.1002194 | 0.31860564 |
| LIPC | 21.0806324 | -0.4915474 | 0.52806 | -0.9308552 | 0.3519285 | NA |
| ADAM10 | 352.179843 | -0.9140493 | 0.30687458 | -2.9785761 | 0.00289591 | 0.05257034 |
| MINDY2 | 194.323918 | -0.4663147 | 0.18617246 | -2.5047459 | 0.01225394 | 0.10833956 |
| RNF111 | 214.680654 | -0.4206571 | 0.24861934 | -1.6919725 | 0.09065121 | 0.3008807 |
| SLTM | 285.639402 | -0.5499721 | 0.28063477 | -1.9597433 | 0.0500258 | 0.22464912 |
| MYO1E | 567.596075 | -1.8675736 | 0.41482182 | -4.5021103 | 6.73E-06 | 0.00132787 |
| FAM81A | 37.3551192 | -0.2913573 | 0.52072964 | -0.5595174 | 0.57580868 | 0.77401855 |
| GTF2A2 | 31.6308788 | -0.0712464 | 0.36658629 | -0.194351 | 0.84590104 | 0.92531372 |
| BNIP2 | 60.7440224 | -0.6616542 | 0.30310426 | -2.182926 | 0.02904126 | 0.16794185 |
| ANXA2 | 98.5656724 | -0.5259369 | 0.47538647 | -1.1063355 | 2.69E-01 | 5.27E-01 |
| ICE2 | 97.1326641 | 0.32188746 | 0.34147497 | 0.94263854 | 0.34586582 | 0.60324705 |
| RORA-AS1 | 60.4050233 | 1.67101569 | 0.55103214 | 3.03251947 | 0.00242521 | 0.04701984 |
| RORA | 1266.701 | 0.31367285 | 0.42727554 | 0.73412312 | 0.4628737 | 0.69434917 |
| AC012404.1 | 31.3558398 | 1.54516166 | 0.65572454 | 2.35641883 | 0.0184521 | 0.13411314 |
| AC009554.1 | 36.2455007 | 0.63190294 | 0.60194372 | 1.04977079 | 0.29382351 | 0.55311516 |
| VPS13C | 362.867333 | -0.4359936 | 0.31552126 | -1.3818199 | 0.167027 | 0.41508187 |
| C2CD4A | 32.1979316 | -1.0114037 | 0.92953047 | -1.0880802 | 2.77E-01 | 5.35E-01 |
| TLN2 | 158.339224 | 0.70605906 | 0.43511137 | 1.62270883 | 0.10465166 | 0.32506209 |
| TPM1 | 73.389916 | 0.68741424 | 0.41594363 | 1.65266204 | 0.09839966 | 0.31597089 |
| LACTB | 28.2687026 | -0.3550794 | 0.42608701 | -0.8333494 | 0.40464768 | 0.65220185 |
| RPS27L | 39.3713784 | 0.78671838 | 0.41804983 | 1.88187702 | 0.05985272 | 0.2433631 |
| RAB8B | 56.6188459 | -0.8958422 | 0.36575353 | -2.4493056 | 0.01431319 | 0.11781182 |
| APH1B | 26.4959861 | -0.1566322 | 0.46780763 | -0.3348217 | 0.73775956 | 0.86987367 |
| CA12 | 1030.24944 | 0.67017777 | 0.32858316 | 2.03959869 | 0.04139031 | 0.20301425 |
| USP3 | 245.491067 | -0.2956175 | 0.19505833 | -1.515534 | 0.12963723 | 0.36218431 |
| USP3-AS1 | 45.0974168 | 0.23446253 | 0.41513877 | 0.56478111 | 0.57222266 | 0.77272682 |
| HERC1 | 310.673322 | -0.8887518 | 0.49805465 | -1.7844463 | 0.07435117 | 0.27199309 |
| FAM96A | 42.844478 | 0.17457428 | 0.31085518 | 0.56159359 | 0.57439295 | 0.77388388 |
| SNX1 | 97.9841051 | -0.2514945 | 0.21326585 | -1.1792536 | 0.23829721 | 0.49612486 |

| | | | | | | |
|---|---|---|---|---|---|---|
| PPIB | 54.6351409 | 0.33768308 | 0.39546071 | 0.85389792 | 0.39316155 | 0.64195258 |
| CSNK1G1 | 27.7469362 | 0.52256865 | 0.34459288 | 1.5164813 | 0.1293977 | 0.36208459 |
| TRIP4 | 63.3423228 | -0.4634561 | 0.34254645 | -1.3529731 | 0.17606423 | 0.42508357 |
| ZNF609 | 276.648735 | -0.2162671 | 0.22306337 | -0.9695322 | 0.3322797 | 0.58794039 |
| OAZ2 | 27.1988874 | 0.91160314 | 0.43891241 | 2.07695912 | 0.03780533 | 0.19438052 |
| RBPMS2 | 14.9872166 | 0.83000607 | 0.60982002 | 1.36106726 | 0.17349243 | NA |
| SPG21 | 43.7042567 | -0.9442046 | 0.36419117 | -2.592607 | 0.00952515 | 0.09644302 |
| MTFMT | 39.1761839 | -0.1866471 | 0.30415747 | -0.6136528 | 0.53944476 | 0.74929145 |
| UBAP1L | 22.3022544 | 0.49108278 | 0.52012613 | 0.94416094 | 0.34508741 | 0.60270883 |
| PDCD7 | 26.4197374 | 0.12663299 | 0.40958294 | 0.30917545 | 0.75718807 | 0.87904624 |
| CLPX | 58.3138329 | -0.5112256 | 0.27434797 | -1.8634204 | 0.06240314 | 0.24933194 |
| PARP16 | 22.4069647 | 0.31903392 | 0.48302893 | 0.66048615 | 0.5089419 | 0.72967436 |
| DPP8 | 94.263105 | -0.1419246 | 0.31080685 | -0.4566328 | 0.64793502 | 0.81890901 |
| HACD3 | 78.8612033 | 0.6798121 | 0.31057594 | 2.18887559 | 0.02860588 | 0.1661732 |
| INTS14 | 24.2687645 | 0.04035262 | 0.3713304 | 0.10867039 | 0.91346393 | 0.96345217 |
| SLC24A1 | 18.4954418 | 0.98868535 | 0.56288333 | 1.75646586 | 0.0790089 | NA |
| DENND4A | 255.634926 | -0.1901378 | 0.22515571 | -0.8444725 | 0.39840543 | 0.64642108 |
| RAB11A | 125.169525 | 0.34694301 | 0.443472 | 0.78233352 | 0.43401859 | 0.67475797 |
| MEGF11 | 200.596825 | -0.732296 | 0.86377646 | -0.8477842 | 3.97E-01 | 6.45E-01 |
| DIS3L | 29.9309541 | 0.41305831 | 0.38155457 | 1.08256679 | 0.27900075 | 0.53717915 |
| TIPIN | 25.064967 | 0.44812314 | 0.53310017 | 0.84059839 | 0.40057296 | 0.64800548 |
| MAP2K1 | 131.583384 | -0.9680995 | 0.28144831 | -3.4397061 | 0.00058235 | 0.02172137 |
| RPL4 | 64.2216768 | 0.52129798 | 0.50604609 | 1.03013933 | 0.3029446 | 0.56350039 |
| SMAD6 | 17.424941 | 0.00063876 | 0.53166875 | 0.00120143 | 0.9990414 | NA |
| SMAD3 | 552.065264 | -2.045636 | 0.38078343 | -5.3721771 | 7.78E-08 | 7.50E-05 |
| AAGAB | 98.0498382 | -0.3867861 | 0.28310143 | -1.3662457 | 0.17186186 | 0.42045926 |
| IQCH | 89.5561482 | 0.60821314 | 0.5157532 | 1.17927167 | 0.23829002 | 0.49612486 |
| IQCH-AS1 | 53.6428747 | 0.62326453 | 0.51071186 | 1.2203839 | 0.22231938 | 0.4793528 |
| MAP2K5 | 175.306289 | 0.75956592 | 0.32422175 | 2.34273585 | 0.01914293 | 0.1359475 |
| PIAS1 | 293.974924 | -0.4771555 | 0.20855528 | -2.2879091 | 0.02214281 | 0.14542084 |
| FEM1B | 28.6836815 | -0.3250589 | 0.37356568 | -0.8701518 | 0.38421744 | 0.63451053 |
| ANP32A | 50.086906 | 0.71477121 | 0.37519773 | 1.90505206 | 0.0567733 | 0.23723351 |
| SPESP1 | 16.6477226 | 1.2816972 | 0.59100445 | 2.16867605 | 0.03010729 | NA |
| GLCE | 112.35046 | 0.41920921 | 0.3413114 | 1.2282309 | 0.2193603 | 0.4758575 |
| AC026992.2 | 15.4226488 | -0.6771947 | 0.63358591 | -1.0688286 | 0.28514693 | NA |
| PAQR5 | 384.686254 | -1.207635 | 0.3955103 | -3.0533591 | 0.00226295 | 0.04522213 |
| KIF23 | 41.3496372 | -1.402003 | 0.65948359 | -2.1259103 | 0.03351073 | 0.1813922 |
| RPLP1 | 126.120625 | 0.42815944 | 0.45369485 | 0.94371678 | 0.3453144 | 0.60277826 |
| TLE3 | 60.2775009 | -0.0316557 | 0.35017931 | -0.0903987 | 0.92797043 | 0.97006497 |
| UACA | 140.687448 | 0.2441651 | 0.28035692 | 0.87090803 | 0.38380437 | 0.63451053 |
| LARP6 | 19.7109372 | 0.28830967 | 0.54360979 | 0.53036143 | 0.59586136 | NA |
| LRRC49 | 56.6981712 | 0.14236064 | 0.34688816 | 0.41039349 | 0.68151732 | 0.84151154 |
| THSD4 | 25.0503403 | -0.4177719 | 0.50076531 | -0.8342668 | 0.40413064 | 0.65170312 |
| THSD4-AS1 | 28.695457 | 0.48899212 | 0.44752248 | 1.09266494 | 0.27454094 | 0.5337282 |
| MYO9A | 465.299699 | 0.32711604 | 0.25471508 | 1.28424288 | 0.19905698 | 0.45276052 |
| SENP8 | 19.6091265 | 0.07120389 | 0.42315915 | 0.16826739 | 0.86637293 | NA |
| PKM | 299.625541 | -0.336974 | 0.31022006 | -1.0862419 | 2.77E-01 | 0.53576226 |
| PARP6 | 51.7286339 | -0.8353981 | 0.42947203 | -1.9451747 | 0.05175395 | 0.22828091 |
| AC009690.2 | 58.5807499 | 0.0534665 | 0.58215004 | 0.09184316 | 0.92682265 | 0.9695235 |
| AC009690.1 | 238.816108 | -0.8140392 | 0.47747294 | -1.7048907 | 0.08821481 | 0.29702074 |
| ARIH1 | 473.653863 | -0.5276216 | 0.26978282 | -1.9557272 | 0.0504973 | 0.22545112 |
| BBS4 | 71.2345772 | -0.0509458 | 0.28484191 | -0.1788563 | 0.85805054 | 0.93282698 |
| ADPGK | 45.1192776 | -0.3309746 | 0.28247384 | -1.1716998 | 0.2413176 | 0.49867735 |
| NEO1 | 94.5558497 | 0.55007715 | 0.29722818 | 1.85068977 | 0.0642142 | 0.25217177 |

| | | | | | | |
|---|---|---|---|---|---|---|
| NPTN | 119.777203 | 0.19837499 | 0.2399111 | 0.82686875 | 0.40831146 | 0.65425469 |
| PML | 19.4263796 | 0.44502768 | 0.57461223 | 0.77448348 | 0.43864493 | NA |
| ARID3B | 35.5341086 | -1.4780135 | 0.35000457 | -4.2228404 | 2.41E-05 | 0.00285668 |
| CLK3 | 31.9907584 | -0.262057 | 0.33834604 | -0.7745236 | 0.4386212 | 0.67887942 |
| EDC3 | 41.3403975 | -0.0678933 | 0.31319724 | -0.2167748 | 0.82838382 | 0.91688285 |
| ULK3 | 21.1888346 | -0.008342 | 0.54211029 | -0.015388 | 0.98772267 | NA |
| SCAMP2 | 30.4567638 | -0.4603534 | 0.44789982 | -1.0278044 | 0.30404183 | 0.56472248 |
| FAM219B | 20.3893959 | 0.47385912 | 0.44669436 | 1.06081285 | 0.28877496 | NA |
| COX5A | 47.3596257 | 0.43684814 | 0.3237207 | 1.34946002 | 0.17718925 | 0.42569745 |
| SCAMP5 | 18.9353698 | 0.79125378 | 0.47692215 | 1.65908373 | 0.09709892 | NA |
| PPCDC | 55.4343581 | -0.4345696 | 0.37676431 | -1.1534257 | 0.24873571 | 0.50492395 |
| COMMD4 | 20.0051031 | -0.5045988 | 0.46673072 | -1.0811349 | 0.27963713 | NA |
| NEIL1 | 21.783478 | 1.03937136 | 0.51072879 | 2.03507495 | 0.04184333 | 0.20409894 |
| MAN2C1 | 49.6603103 | -0.4004792 | 0.35709249 | -1.1214998 | 0.26207517 | 0.52137312 |
| SIN3A | 77.8218175 | 0.00273453 | 0.23821867 | 0.01147909 | 0.99084121 | 0.99583841 |
| PTPN9 | 82.8229849 | 0.2964391 | 0.36761665 | 0.80638105 | 0.42002317 | 0.6638436 |
| SNUPN | 14.5600485 | 1.04655782 | 0.51216674 | 2.04339276 | 0.04101357 | NA |
| UBE2Q2 | 55.7408982 | -0.9075689 | 0.25539613 | -3.5535733 | 0.00038004 | 0.01697016 |
| FBXO22 | 76.3762419 | -0.4628014 | 0.26057112 | -1.7761038 | 0.0757158 | 0.27489837 |
| ETFA | 107.535291 | 0.17372728 | 0.21712385 | 0.80012984 | 0.42363557 | 0.66695091 |
| SCAPER | 294.387867 | 0.76925979 | 0.2377268 | 3.23589846 | 0.00121261 | 0.03090537 |
| RCN2 | 59.7478988 | 0.54351942 | 0.30808601 | 1.76418077 | 0.07770155 | 0.27791302 |
| TSPAN3 | 56.9718807 | -0.7202828 | 0.37861081 | -1.902436 | 0.05711418 | 0.23764428 |
| PEAK1 | 158.187507 | -0.1091268 | 0.30912132 | -0.3530225 | 0.7240716 | 0.86235038 |
| HMG20A | 99.5908912 | 0.16115751 | 0.24712271 | 0.65213557 | 0.51431372 | 0.73200309 |
| LINGO1 | 506.323609 | -0.8227382 | 0.6002437 | -1.3706737 | 0.17047671 | 0.41855159 |
| TBC1D2B | 28.9532872 | -0.8576526 | 0.55675513 | -1.5404485 | 0.12345107 | 0.35349809 |
| IDH3A | 73.7655061 | -0.1300581 | 0.48295376 | -0.2692971 | 0.78770105 | 0.89506208 |
| DNAJA4 | 22.696746 | -1.7761955 | 0.75923832 | -2.339444 | 0.01931247 | 0.13662506 |
| WDR61 | 38.0614803 | 0.68525902 | 0.3972058 | 1.72519894 | 0.08449163 | 0.28931012 |
| IREB2 | 94.2819541 | -0.4340426 | 0.25195411 | -1.7227049 | 0.08494191 | 0.29044775 |
| HYKK | 15.729243 | 1.60574756 | 0.62725732 | 2.55995032 | 0.01046871 | NA |
| PSMA4 | 29.8902761 | 0.20129619 | 0.41063918 | 0.4902021 | 0.62399089 | 0.80467102 |
| MORF4L1 | 116.336875 | -0.4305399 | 0.21508617 | -2.0017086 | 0.04531608 | 0.21333202 |
| CTSH | 32.5802244 | 0.25429914 | 0.43767911 | 0.5810173 | 0.56122879 | 0.76632571 |
| TMED3 | 27.5573042 | 0.25818743 | 0.37544315 | 0.68768714 | 0.49164982 | 0.71672078 |
| ZFAND6 | 222.663995 | -0.3722681 | 0.25231793 | -1.475393 | 0.14010692 | 0.37785668 |
| AC016705.2 | 153.985437 | -1.1100124 | 0.56776441 | -1.955058 | 0.05057622 | 0.22567257 |
| ARNT2 | 332.817563 | -0.0818687 | 0.35339314 | -0.2316647 | 0.81679845 | 0.91186888 |
| ABHD17C | 74.2544627 | -0.1722345 | 0.3464463 | -0.4971462 | 0.61908598 | 0.8027391 |
| AC027808.1 | 24.0458715 | 0.04017428 | 0.48450867 | 0.08291756 | 0.93391709 | 0.9731095 |
| MESD | 36.9361944 | 0.44989422 | 0.34011702 | 1.32276305 | 0.18591419 | 0.43607799 |
| EFL1 | 52.5209684 | -0.3501135 | 0.29016223 | -1.2066128 | 0.22758128 | 0.48550252 |
| WHAMM | 46.2609371 | -0.2205404 | 0.2910069 | -0.7578526 | 0.4485392 | 0.68731807 |
| HOMER2 | 38.445932 | -0.7752343 | 0.44156643 | -1.7556458 | 0.0791489 | 0.28074153 |
| C15orf40 | 21.9770897 | 0.63978082 | 0.39259765 | 1.6296094 | 0.10318408 | 0.32267056 |
| BTBD1 | 80.2408859 | -0.3548007 | 0.29193853 | -1.2153267 | 0.22424151 | 0.48158117 |
| HDGFL3 | 82.0134159 | -0.3301383 | 0.22266003 | -1.4827012 | 0.13815381 | 0.37473724 |
| ADAMTSL3 | 31.6441123 | 1.94364238 | 0.74557712 | 2.60689651 | 0.0091367 | 0.09439618 |
| AC048382.5 | 12.950298 | 1.07452517 | 0.59464976 | 1.80698831 | 0.0707641 | NA |
| WDR73 | 65.5561559 | -0.1692708 | 0.31991496 | -0.5291119 | 0.59672782 | 0.78931328 |
| SEC11A | 107.42719 | -0.1476882 | 0.28419164 | -0.5196783 | 0.60328781 | 0.79301836 |
| ZNF592 | 58.5743258 | 0.24842255 | 0.25208656 | 0.9854653 | 0.32439557 | 0.58025777 |
| ALPK3 | 156.377791 | -1.0809732 | 0.5300944 | -2.0392089 | 0.04142918 | 0.20301425 |

| | | | | | | |
|---|---|---|---|---|---|---|
| PDE8A | 473.553307 | 0.38479933 | 0.17898639 | 2.14988046 | 3.16E-02 | 0.17516458 |
| AKAP13 | 426.111828 | -1.1292741 | 0.33213153 | -3.4000811 | 0.00067366 | 0.02351966 |
| MRPS11 | 22.9108079 | 0.0435413 | 0.40576619 | 0.10730637 | 0.91454593 | 0.96388328 |
| DET1 | 22.486928 | 1.1721111 | 0.64412038 | 1.81970813 | 0.06880346 | 0.26100555 |
| ABHD2 | 61.8137481 | -0.5869736 | 0.52737068 | -1.1130191 | 0.26570024 | 0.52492165 |
| POLG | 19.7752843 | -0.2676603 | 0.40872257 | -0.6548703 | 0.51255126 | NA |
| RHCG | 58.309865 | 1.58294317 | 0.78411674 | 2.01875956 | 0.04351222 | 0.20866886 |
| ANPEP | 16.5558138 | -0.3909844 | 0.55398977 | -0.7057611 | 0.48033672 | NA |
| ZNF710 | 45.3906501 | -0.1437202 | 0.48977522 | -0.2934411 | 0.76918505 | 0.88575962 |
| ZNF710-AS1 | 33.873176 | 0.50020118 | 0.53170235 | 0.94075413 | 0.34683087 | 0.60383561 |
| IDH2 | 253.55186 | 0.76219971 | 0.37946679 | 2.00860716 | 0.04457881 | 0.21133145 |
| IDH2-DT | 43.88773 | -0.2724817 | 0.36484042 | -0.7468517 | 0.4551531 | 0.6908246 |
| SEMA4B | 223.46512 | -1.4189463 | 0.31879357 | -4.4509878 | 8.55E-06 | 0.00146459 |
| CIB1 | 70.7903891 | -0.5192808 | 0.58382408 | -0.8894474 | 0.37376267 | 0.62575308 |
| AC091167.6 | 15.0206179 | 0.17099861 | 0.70603361 | 0.24219613 | 0.80862819 | NA |
| ZNF774 | 11.7630252 | 1.03835106 | 0.691111 | 1.50243745 | 0.13298417 | NA |
| IQGAP1 | 208.01932 | -0.1355762 | 0.20693647 | -0.6551586 | 0.51236564 | 0.73101511 |
| CRTC3 | 109.243033 | 0.35531406 | 0.25866069 | 1.37366856 | 0.1695446 | 0.41775657 |
| CRTC3-AS1 | 19.1372404 | -0.4134545 | 0.45673689 | -0.9052357 | 0.36534054 | NA |
| FURIN | 19.2421841 | 0.17926943 | 0.581856 | 0.30809932 | 0.75800676 | NA |
| MAN2A2 | 17.1977771 | 0.71827727 | 0.44245704 | 1.62338308 | 0.10450754 | NA |
| UNC45A | 29.5575121 | -0.0368726 | 0.40593969 | -0.0908328 | 0.92762547 | 0.96988247 |
| AC104035.1 | 37.4968081 | -2.9476273 | 1.11277805 | -2.6488906 | 0.00807565 | 0.08827138 |
| FAM174B | 38.5436953 | -0.6121116 | 0.37845742 | -1.6173856 | 0.10579506 | 0.32713587 |
| AC013394.1 | 145.944595 | -0.0949916 | 0.23171085 | -0.4099575 | 0.68183715 | 0.84158016 |
| CHD2 | 255.158266 | -0.7058973 | 0.32964896 | -2.1413606 | 0.03224497 | 0.17715171 |
| MCTP2 | 21.7227093 | 0.28942357 | 0.46051251 | 0.62848146 | 0.52968859 | 0.74370906 |
| AC012409.2 | 62.0312605 | 1.64868054 | 0.67664377 | 2.43655616 | 0.01482787 | 0.12019013 |
| NR2F2-AS1 | 532.726084 | 0.87388332 | 0.45993436 | 1.90001746 | 0.05743083 | 0.23817084 |
| NR2F2 | 86.1641606 | 0.30360487 | 0.33689142 | 0.90119504 | 0.36748463 | 0.62220175 |
| AC087477.2 | 15.05013 | -0.0479767 | 0.63670314 | -0.0753518 | 0.93993478 | NA |
| ARRDC4 | 37.611622 | 0.44171701 | 0.47814691 | 0.92381025 | 0.35558512 | 0.61174311 |
| FAM169B | 20.9791803 | -0.5022581 | 0.66330081 | -0.7572102 | 4.49E-01 | NA |
| IGF1R | 817.556143 | -0.3582413 | 0.21183336 | -1.6911468 | 0.09080877 | 0.30101425 |
| TTC23 | 60.6782999 | 0.31543777 | 0.29843386 | 1.05697715 | 0.29052202 | 0.54991621 |
| LRRC28 | 178.076494 | 0.05092359 | 0.42308123 | 0.12036363 | 0.9041951 | 0.95671001 |
| MEF2A | 500.815694 | -0.0049769 | 0.18414704 | -0.027027 | 9.78E-01 | 0.9916702 |
| LYSMD4 | 12.5403633 | 0.65507182 | 0.55561233 | 1.17900879 | 0.23839468 | NA |
| AC090825.1 | 41.7881617 | 1.54023932 | 0.63015317 | 2.44423008 | 0.01451617 | 0.11873638 |
| ADAMTS17 | 561.266879 | -0.9198398 | 0.61846666 | -1.4872909 | 0.136938 | 0.37270573 |
| LINS1 | 28.1070752 | 0.17254073 | 0.34594561 | 0.49875105 | 0.61795477 | 0.80236132 |
| ASB7 | 37.8051085 | 0.13825935 | 0.33594522 | 0.41155326 | 0.68066689 | 0.84094592 |
| CHSY1 | 179.496594 | -0.0078956 | 0.3904256 | -0.0202231 | 9.84E-01 | 0.99382437 |
| SELENOS | 35.2815022 | -0.0487079 | 0.36437774 | -0.1336741 | 0.89366026 | 0.95191019 |
| SNRPA1 | 33.0088053 | -0.1237036 | 0.32031464 | -0.3861942 | 0.69935288 | 0.85118899 |
| PCSK6 | 64.8925938 | -0.6732392 | 0.60283555 | -1.1167874 | 0.26408523 | 0.52361258 |
| TM2D3 | 40.7288149 | -0.9002225 | 0.49385082 | -1.8228632 | 0.06832411 | 0.25987613 |
| TARSL2 | 76.0713407 | -1.0408519 | 0.33546376 | -3.1027253 | 0.00191748 | 0.04077019 |
| MPG | 14.2919328 | -0.1201701 | 0.428043 | -0.2807432 | 0.7789074 | NA |
| NPRL3 | 65.4664887 | 0.38074308 | 0.3125631 | 1.21813186 | 0.22317387 | 0.48036053 |
| LUC7L | 134.414311 | -0.2737704 | 0.27802438 | -0.9846993 | 0.32477182 | 0.58066636 |
| FAM234A | 26.0275401 | 0.3791749 | 0.52452008 | 0.72289874 | 0.4697421 | 0.69934331 |
| AXIN1 | 25.6774764 | -0.473158 | 0.44063962 | -1.0737981 | 0.28291318 | 0.54168724 |
| TMEM8A | 51.9759641 | -0.1728021 | 0.44686629 | -0.3866976 | 0.69898008 | 0.85102865 |

| | | | | | | |
|---|---|---|---|---|---|---|
| RAB11FIP3 | 152.664807 | 0.35611899 | 0.26046936 | 1.36722028 | 0.17155627 | 0.41999638 |
| CAPN15 | 23.0286785 | -0.4459406 | 0.37723543 | -1.1821282 | 0.23715485 | 0.49508567 |
| PIGQ | 35.3732883 | 1.01058158 | 0.4731773 | 2.13573554 | 0.03270098 | 0.17874373 |
| RAB40C | 62.6300468 | 0.41022309 | 0.28543312 | 1.43719513 | 0.15066256 | 0.39124483 |
| MCRIP2 | 41.0176912 | -0.0292818 | 0.50196268 | -0.0583346 | 0.9534821 | 0.98153155 |
| RHOT2 | 30.6932287 | -0.2393805 | 0.41914976 | -0.5711097 | 0.5679253 | 0.77103518 |
| STUB1 | 26.9465772 | -0.2040372 | 0.44224891 | -0.4613628 | 0.64453836 | 0.81734747 |
| METRN | 19.3184199 | 1.33148216 | 0.58234173 | 2.28642752 | 0.02222926 | NA |
| FAM173A | 18.8734804 | 0.43163789 | 0.7056588 | 0.61168073 | 0.54074902 | NA |
| NARFL | 18.5736323 | -0.1596193 | 0.4034241 | -0.3956613 | 0.6923549 | NA |
| LMF1 | 71.6442625 | -0.3424883 | 0.38581675 | -0.8876967 | 0.3747039 | 0.62575308 |
| CACNA1H | 28.1540604 | -1.0070369 | 0.75485853 | -1.3340737 | 0.1821797 | 0.43172326 |
| UBE2I | 88.4027684 | -1.0594039 | 0.47645772 | -2.2235004 | 0.02618208 | 0.15845355 |
| BAIAP3 | 21.5054449 | 0.12722918 | 0.40474304 | 0.31434557 | 0.75325859 | 0.87842978 |
| GNPTG | 40.1390919 | -1.2226076 | 0.34414328 | -3.5526123 | 0.00038143 | 0.01697016 |
| UNKL | 35.4233868 | 0.1254818 | 0.40466136 | 0.31009088 | 0.75649184 | 0.87904624 |
| CLCN7 | 51.1544083 | -1.0638695 | 0.51980204 | -2.0466821 | 0.04068931 | 0.20101774 |
| TELO2 | 11.2987079 | -0.7079555 | 0.63369517 | -1.1171862 | 0.2639147 | NA |
| IFT140 | 38.4585136 | 0.36907672 | 0.47177533 | 0.78231459 | 0.43402972 | 0.67475797 |
| JPT2 | 33.4998595 | 0.28602361 | 0.40018218 | 0.71473349 | 0.47477374 | 0.70288437 |
| MAPK8IP3 | 88.0940951 | 0.11806857 | 0.23566914 | 0.50099291 | 0.61637611 | 0.80152846 |
| NME3 | 45.3705901 | 0.81656292 | 0.54420998 | 1.50045562 | 0.13349642 | 0.36762309 |
| MRPS34 | 19.8338603 | -0.1827283 | 0.48787743 | -0.3745374 | 0.70800455 | NA |
| SPSB3 | 40.7991261 | 0.93734679 | 0.41155755 | 2.27755945 | 0.02275284 | 0.14700765 |
| HAGH | 59.5222098 | -0.1635775 | 0.32156283 | -0.5086953 | 0.6109658 | 0.79759223 |
| FAHD1 | 13.1116421 | 0.34203596 | 0.45537249 | 0.75111247 | 0.45258497 | NA |
| NDUFB10 | 26.9670259 | 1.08158545 | 0.6177708 | 1.7507876 | 0.0799825 | 0.28223671 |
| RPS2 | 66.6984113 | 0.69750373 | 0.41700799 | 1.67263874 | 0.09439844 | 0.30830642 |
| TSC2 | 54.0187146 | -0.1569949 | 0.46453255 | -0.3379632 | 0.73539093 | 0.86838246 |
| PKD1 | 34.7957207 | 0.07352949 | 0.33421581 | 0.22000603 | 0.82586646 | 0.91551092 |
| SNHG19 | 19.4679609 | -0.130509 | 0.63562274 | -0.2053247 | 0.83731847 | NA |
| PGP | 34.8505731 | 0.70166013 | 0.46111537 | 1.52165851 | 0.12809467 | 0.36035989 |
| E4F1 | 17.1482876 | 0.21398285 | 0.49947173 | 0.42841834 | 0.66834657 | NA |
| ECI1 | 67.7656002 | 0.31863052 | 0.30604092 | 1.04113699 | 0.29781197 | 0.55772719 |
| RNPS1 | 49.4305938 | -0.4067666 | 0.32841809 | -1.2385633 | 0.21550728 | 0.47115856 |
| ABCA3 | 20.0701046 | 0.15875642 | 0.51054591 | 0.31095425 | 0.7558354 | NA |
| TBC1D24 | 17.9665067 | -1.2739451 | 0.59937552 | -2.1254539 | 0.03354875 | NA |
| PDPK1 | 114.718938 | -0.4350087 | 0.4350552 | -0.9998931 | 0.31736224 | 0.57495296 |
| AC093525.4 | 11.8258082 | 0.33136977 | 0.58311518 | 0.56827499 | 0.56984826 | NA |
| KCTD5 | 13.6538582 | -0.1433984 | 0.49480234 | -0.2898094 | 0.77196204 | NA |
| SRRM2 | 330.932869 | -0.7521448 | 0.28894776 | -2.6030478 | 0.00923991 | 0.09482608 |
| ELOB | 38.046207 | 0.11467339 | 0.57496958 | 0.19944253 | 0.8419166 | 0.92367136 |
| PRSS22 | 14.6783317 | -1.7217383 | 0.63845922 | -2.6967084 | 7.00E-03 | NA |
| FLYWCH1 | 37.9921678 | 0.47470935 | 0.38725914 | 1.22581834 | 0.22026704 | 0.47685442 |
| TNFRSF12A | 48.6929781 | -1.8274184 | 0.56278796 | -3.2470816 | 0.00116595 | 0.03031862 |
| HCFC1R1 | 14.6117556 | -0.1996984 | 0.59209391 | -0.3372748 | 0.73590975 | NA |
| ZNF213-AS1 | 20.3145596 | -0.7260771 | 0.4504928 | -1.6117395 | 0.10701863 | NA |
| ZNF213 | 32.6225713 | 0.61079483 | 0.41540354 | 1.470365 | 0.14146293 | 0.37950905 |
| ZNF263 | 15.6535251 | 0.34892529 | 0.44148599 | 0.79034284 | 0.42932757 | NA |
| TIGD7 | 16.269517 | 0.05325943 | 0.45582695 | 0.11684134 | 0.90698579 | NA |
| ZNF75A | 56.6530433 | -0.8339961 | 0.49308278 | -1.6913916 | 0.09076203 | 0.30098894 |
| AC025283.3 | 19.3082977 | -0.2275542 | 0.58052635 | -0.3919792 | 0.6950736 | NA |
| ZSCAN32 | 12.8495522 | 0.37950106 | 0.50302672 | 0.7544352 | 0.45058794 | NA |
| NAA60 | 17.0666495 | 0.49282607 | 0.50793443 | 0.9702553 | 0.33191925 | NA |

| | | | | | | |
|---|---|---|---|---|---|---|
| CLUAP1 | 41.2140707 | 0.15705384 | 0.45816158 | 0.34279139 | 0.7317554 | 0.86717803 |
| DNASE1 | 145.62838 | -0.1815943 | 0.27611593 | -0.6576743 | 0.51074747 | 0.73060499 |
| TRAP1 | 52.5518692 | 0.28912423 | 0.32501122 | 0.88958231 | 0.37369021 | 0.62575308 |
| CREBBP | 238.564056 | -0.3114411 | 0.21706739 | -1.4347669 | 0.15135354 | 0.39250951 |
| ADCY9 | 69.2084525 | 0.23606726 | 0.39310948 | 0.60051277 | 0.54816455 | 0.7554786 |
| GLIS2 | 29.083788 | -0.5518223 | 0.46675104 | -1.1822626 | 0.23710153 | 0.49508567 |
| VASN | 17.2571668 | 1.18586102 | 0.5341618 | 2.22004086 | 0.02641599 | NA |
| DNAJA3 | 42.7954545 | -0.2287367 | 0.27863198 | -0.8209276 | 0.4116875 | 0.65707964 |
| NMRAL1 | 27.7836279 | 0.29176111 | 0.38521166 | 0.75740467 | 0.44880746 | 0.68732014 |
| HMOX2 | 32.370706 | -0.6532439 | 0.37669629 | -1.7341395 | 0.08289333 | 0.28791966 |
| CDIP1 | 28.5886663 | 0.44248271 | 0.42829362 | 1.03312935 | 0.30154337 | 0.56180195 |
| MGRN1 | 64.6781454 | -0.4394211 | 0.30739766 | -1.4294875 | 0.15286416 | 0.3947713 |
| ANKS3 | 39.5844044 | -0.1497622 | 0.29529047 | -0.5071692 | 0.61203613 | 0.79844781 |
| ZNF500 | 16.3387981 | 0.18840822 | 0.45195282 | 0.41687587 | 0.67676921 | NA |
| GLYR1 | 113.816802 | -0.4424032 | 0.31894177 | -1.387097 | 0.1654122 | 0.41243204 |
| UBN1 | 47.63437 | 0.01002448 | 0.32054483 | 0.03127325 | 0.97505162 | 0.99037778 |
| PPL | 22.4785772 | -0.2823762 | 0.44389916 | -0.6361269 | 0.52469371 | 0.73968378 |
| ALG1 | 15.4623861 | 0.79855019 | 0.60371833 | 1.3227198 | 0.18592858 | NA |
| RBFOX1 | 19.7606858 | -0.6593216 | 0.6105985 | -1.0797957 | 0.28023317 | NA |
| METTL22 | 22.0236353 | -0.3326201 | 0.41745238 | -0.7967858 | 0.42557542 | 0.66850082 |
| ABAT | 20.2276249 | 0.94030162 | 0.51083582 | 1.840712 | 0.06566377 | NA |
| PMM2 | 67.8513334 | -0.6238239 | 0.37549114 | -1.6613547 | 0.09664224 | 0.31216475 |
| CARHSP1 | 32.8734784 | -0.0377124 | 0.36155701 | -0.1043056 | 9.17E-01 | 0.96401363 |
| USP7 | 60.7132182 | -0.5242967 | 0.34277641 | -1.5295588 | 0.12612597 | 0.35730275 |
| C16orf72 | 66.5389939 | -0.6079595 | 0.32188561 | -1.8887439 | 0.05892616 | 0.2411888 |
| ATF7IP2 | 94.4122536 | 0.33635038 | 0.44509106 | 0.75568891 | 0.44983574 | 0.68770076 |
| NUBP1 | 17.4195842 | -0.0152729 | 0.50159695 | -0.0304485 | 0.97570936 | NA |
| CIITA | 15.5707295 | 0.03022953 | 0.47119687 | 0.06415478 | 0.94884698 | NA |
| DEXI | 15.6105799 | 0.260532 | 0.43554531 | 0.59817428 | 0.54972365 | NA |
| CLEC16A | 268.611028 | 0.10748435 | 0.25884891 | 0.41523973 | 0.67796641 | 0.83969138 |
| AC007220.1 | 24.2759125 | -0.3281825 | 0.49237163 | -0.666534 | 0.50506981 | 0.72663969 |
| SOCS1 | 17.9155118 | -0.4360237 | 0.76979213 | -0.5664175 | 0.57111002 | NA |
| AC099489.1 | 45.7816216 | -2.3684865 | 0.94566347 | -2.5045765 | 0.01225981 | 0.10833956 |
| LITAF | 133.043289 | 0.01972121 | 0.2457009 | 0.0802651 | 0.93602642 | 0.97405271 |
| TXNDC11 | 108.097374 | 0.2747261 | 0.24624867 | 1.11564498 | 0.26457413 | 0.5237724 |
| ZC3H7A | 106.055103 | -0.0180645 | 0.24290117 | -0.0743698 | 0.94071615 | 0.97515997 |
| RSL1D1 | 20.067296 | -0.0432973 | 0.4098804 | -0.1056339 | 0.91587285 | NA |
| AC007216.4 | 48.5340381 | 0.37328932 | 0.30371201 | 1.22908973 | 0.21903816 | 0.47565385 |
| GSPT1 | 142.334633 | -0.4377707 | 0.28748839 | -1.5227422 | 0.12782322 | 0.35999098 |
| SNX29 | 1211.78539 | -0.8666463 | 0.4420158 | -1.960668 | 0.04991776 | 0.2242948 |
| AC010333.2 | 26.1908524 | -2.4471707 | 0.76104367 | -3.2155457 | 0.00130197 | 0.03222265 |
| CPPED1 | 181.214229 | -1.7424169 | 0.45356359 | -3.8416155 | 0.00012223 | 0.00804088 |
| SHISA9 | 57.8593431 | -1.7556674 | 0.68616715 | -2.5586585 | 0.01050769 | 0.10071942 |
| U91319.1 | 64.8329819 | -2.2127102 | 0.66524713 | -3.3261477 | 0.00088055 | 0.02668352 |
| ERCC4 | 19.9664825 | -0.2727971 | 0.43111048 | -0.6327777 | 0.52687884 | NA |
| MKL2 | 248.950486 | 0.51069996 | 0.24339193 | 2.09826169 | 0.03588204 | 0.18890836 |
| PARN | 113.219526 | 0.15370453 | 0.24583133 | 0.62524385 | 0.53181103 | 0.74532947 |
| BFAR | 34.7665766 | -0.263576 | 0.30094455 | -0.8758293 | 0.38112285 | 0.63258566 |
| NOMO1 | 18.7280945 | -0.1819604 | 0.47515262 | -0.3829514 | 0.70175577 | NA |
| PDXDC1 | 219.537292 | -0.4349157 | 0.22111472 | -1.9669231 | 0.04919207 | 0.22233199 |
| RRN3 | 32.8732019 | -0.228553 | 0.35734017 | -0.639595 | 0.52243593 | 0.7380123 |
| C16orf45 | 134.815934 | -0.4890957 | 0.36239514 | -1.3496199 | 0.17713792 | 0.42569745 |
| MARF1 | 59.6427924 | -0.5261388 | 0.3269357 | -1.6093035 | 0.10754999 | 0.32954311 |
| NDE1 | 24.7942521 | -0.1176334 | 0.50825798 | -0.2314442 | 0.8169697 | 0.91186888 |

| | | | | | | |
|---|---|---|---|---|---|---|
| FOPNL | 26.5326086 | -0.1057509 | 0.38641733 | -0.2736701 | 0.78433819 | 0.8939806 |
| ABCC1 | 54.7410089 | -0.3450535 | 0.4734889 | -0.7287467 | 0.4661566 | 0.69633341 |
| ABCC6 | 14.2651014 | 1.50898349 | 0.84294057 | 1.79014222 | 0.07343105 | NA |
| RPS15A | 13.5547976 | 1.69464107 | 0.55232015 | 3.0682224 | 0.00215336 | NA |
| SMG1 | 284.567667 | -0.8537848 | 0.25615373 | -3.3330952 | 0.00085886 | 0.02623258 |
| COQ7 | 15.0974619 | -0.3655331 | 0.51863601 | -0.704797 | 0.48093656 | NA |
| SYT17 | 31.0126106 | 1.60599912 | 0.84536657 | 1.89976654 | 0.05746376 | 0.2381791 |
| GDE1 | 17.0889484 | -0.0344009 | 0.48822724 | -0.0704608 | 0.94382694 | NA |
| CCP110 | 26.3270451 | 0.54438351 | 0.43523052 | 1.2507935 | 0.21100983 | 0.46593468 |
| VPS35L | 122.536453 | -0.4318833 | 0.24095364 | -1.7923917 | 0.07307025 | 0.26939652 |
| KNOP1 | 16.5670272 | 0.08534318 | 0.52274892 | 0.16325846 | 0.87031494 | NA |
| IQCK | 147.729138 | 0.94098503 | 0.39911822 | 2.35765993 | 0.01839053 | 0.13391856 |
| GPRC5B | 100.044326 | 0.99011271 | 0.34192507 | 2.89570078 | 0.00378313 | 0.06101472 |
| UMOD | 117.374312 | -0.1757115 | 0.78405145 | -0.2241071 | 0.82267395 | 0.91475524 |
| ACSM2A | 12.4013065 | 2.17681024 | 1.09249396 | 1.99251467 | 0.04631462 | NA |
| ACSM2B | 14.8266194 | 2.5872515 | 1.01132246 | 2.55828541 | 0.01051897 | NA |
| ACSM3 | 35.4452725 | 0.5954059 | 0.42185104 | 1.41141268 | 0.15812297 | 0.40275854 |
| THUMPD1 | 44.9140541 | 0.04326719 | 0.32564358 | 0.13286671 | 0.89429879 | 0.95219502 |
| ERI2 | 21.3226884 | 0.10460472 | 0.50330416 | 0.20783599 | 0.83535703 | 0.92037547 |
| LYRM1 | 20.8238316 | 0.31529361 | 0.48793991 | 0.64617303 | 0.5181673 | NA |
| TMEM159 | 22.431618 | -0.5600086 | 0.38800022 | -1.4433203 | 0.14893027 | 0.38937375 |
| METTL9 | 94.0619604 | -0.563732 | 0.33694584 | -1.673064 | 0.09431469 | 0.30830642 |
| NPIPB4 | 37.1500582 | -1.4821258 | 0.57617397 | -2.5723582 | 0.01010083 | 0.0992926 |
| UQCRC2 | 126.293943 | 0.18896694 | 0.24361772 | 0.77566995 | 0.43794389 | 0.67810382 |
| MOSMO | 84.2555487 | 0.22582514 | 0.2748857 | 0.8215238 | 0.41134797 | 0.65707964 |
| EEF2K | 15.796869 | 0.54431014 | 0.44678072 | 1.2182937 | 0.22311238 | NA |
| POLR3E | 30.0835329 | -0.2427356 | 0.33116807 | -0.7329681 | 0.4635779 | 0.69473502 |
| CDR2 | 194.548534 | -0.9918976 | 0.36120988 | -2.7460423 | 0.0060319 | 0.07623568 |
| NPIPB5 | 90.6077454 | -0.854479 | 0.44138581 | -1.9359004 | 0.0528799 | 0.23105954 |
| USP31 | 28.6870933 | -0.4137948 | 0.39887371 | -1.0374081 | 0.29954572 | 0.55961248 |
| COG7 | 94.6438239 | -0.2891982 | 0.26168661 | -1.1051317 | 0.26910255 | 0.52764898 |
| GGA2 | 97.8677204 | -1.03304 | 0.41359205 | -2.497727 | 0.01249924 | 0.10951947 |
| EARS2 | 14.7732514 | -0.2321034 | 0.55891674 | -0.4152737 | 0.67794154 | NA |
| UBFD1 | 20.4036895 | -0.1043167 | 0.42833429 | -0.2435404 | 0.80758676 | NA |
| NDUFAB1 | 27.1918591 | 0.41272842 | 0.41862599 | 0.98591208 | 0.32417626 | 0.58021133 |
| PALB2 | 18.8222721 | -0.9825983 | 0.45780912 | -2.1463057 | 0.0318486 | NA |
| DCTN5 | 37.4945972 | 0.38841403 | 0.37081917 | 1.04744861 | 0.29489272 | 0.55409516 |
| RBBP6 | 89.7134004 | -0.1852005 | 0.27420554 | -0.6754076 | 0.49941682 | 0.72201362 |
| TNRC6A | 792.706601 | -1.405841 | 0.52390935 | -2.683367 | 0.0072885 | 0.0836059 |
| ARHGAP17 | 72.3276169 | -0.2185472 | 0.30870391 | -0.7079508 | 0.47897579 | 0.70690257 |
| LCMT1 | 57.1423208 | 0.05236357 | 0.33241786 | 0.15752334 | 0.87483241 | 0.94169674 |
| AC008741.2 | 12.1987887 | 0.05475583 | 0.57677341 | 0.09493473 | 0.92436667 | NA |
| NSMCE1 | 21.9448915 | 0.58536599 | 0.40796448 | 1.4348455 | 0.15133113 | 0.39250951 |
| IL4R | 32.6007415 | -0.0782531 | 0.65126594 | -0.1201554 | 9.04E-01 | 9.57E-01 |
| GTF3C1 | 51.6848003 | -0.541749 | 0.2856736 | -1.8963916 | 0.05790828 | 0.23950565 |
| KIAA0556 | 96.2490233 | -0.2227963 | 0.24099979 | -0.9244668 | 0.35524333 | 0.61168935 |
| XPO6 | 106.762571 | -0.8519347 | 0.32217216 | -2.644346 | 0.00818489 | 0.08906215 |
| SGF29 | 18.7741622 | -0.6696688 | 0.42781301 | -1.5653307 | 0.11750538 | NA |
| SULT1A1 | 39.0822495 | 1.39313938 | 0.61488476 | 2.26569182 | 0.02347026 | 0.14954518 |
| ATXN2L | 50.0445396 | -0.4320085 | 0.30970708 | -1.3948938 | 0.16304785 | 0.40979748 |
| TUFM | 44.5238627 | 0.33948366 | 0.54268974 | 0.62555753 | 0.5316052 | 0.74523961 |
| SH2B1 | 67.9764497 | 0.65585417 | 0.34636896 | 1.89351309 | 0.05828967 | 0.24020328 |
| RABEP2 | 57.5506973 | -1.0021643 | 0.3934898 | -2.5468622 | 0.01086964 | 0.10244908 |
| NFATC2IP | 90.4538671 | 0.01245088 | 0.30564278 | 0.04073671 | 0.96750579 | 0.98741848 |

| | | | | | | |
|---|---|---|---|---|---|---|
| SPNS1 | 22.6001623 | -0.4920046 | 0.38820852 | -1.2673719 | 0.20502235 | 0.45833779 |
| QPRT | 11.3906083 | 0.43858245 | 0.68067103 | 0.64433836 | 0.51935605 | NA |
| KIF22 | 12.1418349 | -0.4668872 | 0.5754873 | -0.8112901 | 0.41719908 | NA |
| MAZ | 17.3706144 | 0.08582297 | 0.5985728 | 0.14337933 | 0.88599061 | NA |
| AC009133.1 | 19.3968358 | -1.2673699 | 0.58799607 | -2.1554053 | 3.11E-02 | NA |
| MVP | 66.5768313 | -0.2782211 | 0.33573617 | -0.8286898 | 4.07E-01 | 6.54E-01 |
| CDIPT | 20.1097644 | 0.4291743 | 0.46525411 | 0.9224514 | 0.35629317 | NA |
| KCTD13 | 31.6087314 | 0.09741679 | 0.44403414 | 0.21939032 | 0.82634601 | 0.9158 |
| TMEM219 | 32.2389279 | 0.52112158 | 0.40883971 | 1.27463543 | 0.20243831 | 0.45643401 |
| TAOK2 | 19.2099337 | -0.3248293 | 0.56670497 | -0.5731894 | 0.56651648 | NA |
| INO80E | 23.4255609 | 0.30362186 | 0.53259717 | 0.57007787 | 0.56862488 | 0.7710898 |
| PPP4C | 25.5734521 | -0.1614318 | 0.47301292 | -0.3412842 | 0.73288966 | 0.86776236 |
| YPEL3 | 38.4629484 | -0.0074008 | 0.34819415 | -0.0212549 | 0.98304234 | 0.99382437 |
| NPIPB13 | 23.8463677 | -1.33645 | 0.63179581 | -2.1153195 | 0.03440272 | 0.18388733 |
| CD2BP2 | 17.7062937 | -0.6896676 | 0.53182799 | -1.2967869 | 0.19470452 | NA |
| TBC1D10B | 14.9514005 | 0.01761361 | 0.57515213 | 0.03062426 | 0.97556919 | NA |
| SEPHS2 | 31.8529212 | -0.0382763 | 0.70132527 | -0.0545771 | 0.95647542 | 0.98284879 |
| ZNF785 | 22.8117212 | -0.0859047 | 0.37556574 | -0.228734 | 0.81907567 | 0.913067 |
| FBRS | 23.7691772 | 0.01339586 | 0.3572327 | 0.03749899 | 0.97008715 | 0.98792813 |
| PHKG2 | 15.5308069 | -0.3710774 | 0.51510859 | -0.7203868 | 0.47128687 | NA |
| RNF40 | 30.2708298 | 0.11588059 | 0.43894623 | 0.26399724 | 0.79178204 | 0.89820875 |
| BCL7C | 62.5946416 | 0.78751887 | 0.38158405 | 2.06381496 | 0.03903526 | 0.19705923 |
| MIR762HG | 15.8895592 | 0.08331963 | 0.61320541 | 0.13587556 | 0.89191966 | NA |
| FBXL19 | 18.167589 | 0.18863785 | 0.51103273 | 0.36913067 | 0.71203033 | NA |
| SETD1A | 17.1789561 | -0.4206695 | 0.43810526 | -0.960202 | 0.33695356 | NA |
| STX4 | 15.0769986 | -0.9082587 | 0.43954219 | -2.0663743 | 0.03879315 | NA |
| KAT8 | 33.016297 | -0.8055987 | 0.42314699 | -1.9038271 | 0.05693271 | 0.23751276 |
| PRSS8 | 43.1833317 | -1.2590396 | 0.63081944 | -1.9958795 | 0.04594704 | 0.21446595 |
| FUS | 177.688741 | -0.511415 | 0.24504333 | -2.087039 | 0.03688461 | 0.19174392 |
| AC026471.4 | 18.2710772 | -0.7681677 | 0.63937685 | -1.2014319 | 0.22958372 | NA |
| C16orf58 | 26.6577235 | -0.3383084 | 0.39307409 | -0.8606733 | 0.389418 | 0.63841327 |
| ZNF720 | 64.1716529 | -0.1333946 | 0.26135889 | -0.5103886 | 0.60977921 | 0.7965836 |
| VPS35 | 49.8206707 | -0.0065214 | 0.30644929 | -0.0212804 | 0.98302198 | 0.99382437 |
| C16orf87 | 14.2825334 | 0.20100524 | 0.48063975 | 0.41820353 | 0.67579832 | NA |
| DNAJA2 | 25.3228049 | 0.66646623 | 0.36297013 | 1.8361462 | 0.06633604 | 0.25673827 |
| ITFG1 | 374.595759 | 0.00574716 | 0.18163771 | 0.03164076 | 0.97475853 | 0.99032688 |
| PHKB | 236.910188 | 0.24432649 | 0.24176856 | 1.01058006 | 0.31221746 | 0.57101636 |
| LONP2 | 204.017616 | 0.08072102 | 0.16320568 | 0.49459686 | 0.62088472 | 0.80353114 |
| SIAH1 | 39.7054732 | -0.2989829 | 0.44197516 | -0.6764698 | 0.4987424 | 0.72176422 |
| N4BP1 | 93.3939128 | -0.1759834 | 0.29106 | -0.6046293 | 0.54542537 | 0.75352691 |
| HEATR3 | 15.3910791 | 0.44228031 | 0.47713754 | 0.92694511 | 0.35395504 | NA |
| TENT4B | 58.5164915 | 0.75009785 | 0.30973297 | 2.42175656 | 0.01544569 | 0.1228156 |
| BRD7 | 51.2061566 | -0.1571992 | 0.26419232 | -0.5950182 | 0.55183132 | 0.75820166 |
| CYLD | 57.9499699 | -1.2342674 | 0.59933017 | -2.0594113 | 0.03945485 | 0.19795704 |
| SALL1 | 48.9930441 | -0.1820687 | 0.35478979 | -0.5131736 | 0.6078299 | 0.79552231 |
| AC087564.1 | 165.232589 | -1.0329273 | 0.47344945 | -2.1817055 | 0.02913128 | 0.16833591 |
| TOX3 | 501.576126 | -0.228312 | 0.32002955 | -0.713409 | 4.76E-01 | 7.04E-01 |
| CHD9 | 628.992967 | 0.26086352 | 0.22866831 | 1.14079437 | 0.25395551 | 0.51143263 |
| RBL2 | 120.800055 | 0.04674345 | 0.25494271 | 0.18334882 | 0.85452433 | 0.93109765 |
| AKTIP | 23.3851157 | 0.00771827 | 0.40840472 | 0.01889857 | 0.98492202 | 0.99407122 |
| RPGRIP1L | 21.666325 | 0.99478405 | 0.76436213 | 1.30145647 | 0.19310225 | 0.44483671 |
| FTO | 336.438779 | 0.23688098 | 0.31454686 | 0.75308645 | 0.45139797 | 0.68853766 |
| LPCAT2 | 62.4350876 | 0.72859163 | 0.5822057 | 1.25143335 | 0.21077642 | 0.4656925 |
| AMFR | 105.761296 | 0.47044682 | 0.48395356 | 0.97209084 | 0.33100536 | 0.58655523 |

| | | | | | | |
|---|---|---|---|---|---|---|
| NUDT21 | 60.7949197 | 0.2437224 | 0.32270016 | 0.75525961 | 0.45009323 | 0.68770076 |
| OGFOD1 | 28.7195334 | 0.14840917 | 0.41753341 | 0.35544263 | 0.72225801 | 0.86153309 |
| BBS2 | 57.3959583 | -0.1677806 | 0.28164899 | -0.595708 | 0.55137032 | 0.75783881 |
| MT2A | 210.649825 | -0.7566794 | 0.41474258 | -1.8244555 | 0.06808324 | 0.25980998 |
| MT1E | 67.6468538 | -1.1291109 | 0.58798736 | -1.920298 | 0.05482027 | 0.23407018 |
| MT1F | 15.9909302 | -0.6275044 | 0.59005987 | -1.0634589 | 0.28757392 | NA |
| MT1G | 160.327644 | -0.9985465 | 0.75411025 | -1.3241386 | 0.18545702 | 0.43546756 |
| MT1H | 52.7184693 | -1.5016817 | 0.91326383 | -1.6443022 | 0.1001138 | 0.31841979 |
| MT1X | 40.0404212 | -0.9725472 | 0.7678556 | -1.2665756 | 0.20530707 | 0.45870784 |
| NUP93 | 44.5312119 | -0.3306709 | 0.30813166 | -1.0731482 | 0.28320461 | 0.54207638 |
| SLC12A3 | 5322.16201 | 3.13349761 | 0.79799146 | 3.92673077 | 8.61E-05 | 0.00614802 |
| HERPUD1 | 76.3590596 | -0.8053436 | 0.58184747 | -1.3841147 | 0.16632334 | 0.41429299 |
| NLRC5 | 58.4836493 | -3.3618068 | 0.93251256 | -3.6051061 | 0.00031203 | 0.01457249 |
| FAM192A | 68.9074914 | -0.4720721 | 0.25598619 | -1.8441312 | 0.06516403 | 0.25408689 |
| RSPRY1 | 78.9193963 | -0.3291289 | 0.29592257 | -1.1122129 | 0.26604662 | 0.52520154 |
| ARL2BP | 27.1316546 | 0.83333481 | 0.38608924 | 2.15839948 | 0.03089679 | 0.17358581 |
| PLLP | 19.078401 | 0.72495373 | 0.71939285 | 1.00772995 | 0.31358412 | NA |
| CIAPIN1 | 41.4336883 | -0.3530669 | 0.35519989 | -0.9939948 | 0.32022538 | 0.57724473 |
| COQ9 | 66.0423089 | 0.24396512 | 0.27325701 | 0.8928046 | 0.37196182 | 0.62469783 |
| POLR2C | 15.3040884 | 0.34574951 | 0.51857026 | 0.6667361 | 0.50494072 | NA |
| ADGRG1 | 157.42236 | 0.1996531 | 0.42013222 | 0.47521493 | 0.63463379 | 0.81099192 |
| KIFC3 | 184.935546 | 0.2483175 | 0.24494395 | 1.01377276 | 3.11E-01 | 0.57001845 |
| CSNK2A2 | 50.503938 | -0.4879833 | 0.39636322 | -1.2311518 | 0.2182661 | 0.47484289 |
| SETD6 | 12.5922118 | -0.6444553 | 0.58863801 | -1.0948245 | 0.27359352 | NA |
| CNOT1 | 167.212577 | -0.1931439 | 0.26116438 | -0.7395491 | 0.45957366 | 0.69299864 |
| SLC38A7 | 20.118547 | -0.727502 | 0.45557476 | -1.5968883 | 0.11029061 | NA |
| GOT2 | 41.3450432 | 0.24619209 | 0.37571814 | 0.65525741 | 0.51230204 | 0.73101511 |
| AC092378.1 | 23.6560309 | 1.12513684 | 0.54312469 | 2.07159949 | 0.03830281 | 0.19533626 |
| AC138305.1 | 843.417789 | 0.32744163 | 0.25439923 | 1.28711719 | 0.19805344 | 0.45104671 |
| AC010542.3 | 27.1303867 | -0.0688526 | 0.65482932 | -0.1051459 | 0.91626003 | 0.96401363 |
| TK2 | 19.0698823 | -0.4773092 | 0.45482987 | -1.0494235 | 0.29398325 | NA |
| CMTM4 | 244.521205 | 0.41343014 | 0.24948565 | 1.65712994 | 9.75E-02 | 0.31384576 |
| DYNC1LI2 | 122.62575 | -0.3538471 | 0.36243251 | -0.9763118 | 0.32890997 | 0.5853121 |
| NAE1 | 41.3504981 | 0.02658636 | 0.35322262 | 0.07526799 | 0.94000149 | 0.97496179 |
| PDP2 | 31.308225 | -0.1873576 | 0.48439154 | -0.3867896 | 0.69891195 | 0.85102865 |
| CDH16 | 231.743953 | 0.56530376 | 0.50635642 | 1.11641471 | 0.26424466 | 0.5237724 |
| RRAD | 111.480928 | -0.8032298 | 0.62305728 | -1.2891749 | 0.19733729 | 0.45057998 |
| FAM96B | 29.1864725 | -0.2061099 | 0.42341997 | -0.486774 | 0.62641846 | 0.8053173 |
| CES2 | 30.0144537 | -0.2114899 | 0.44889071 | -0.4711388 | 0.63754159 | 0.81298171 |
| CBFB | 161.196718 | 0.09395926 | 0.25632239 | 0.36656671 | 0.71394224 | 0.85788134 |
| C16orf70 | 21.3973087 | 0.31949719 | 0.4823296 | 0.66240428 | 0.50771216 | 0.7286503 |
| NOL3 | 27.8955615 | 2.0503466 | 0.70823456 | 2.89501065 | 0.00379146 | 0.06101472 |
| KIAA0895L | 21.9909461 | 0.40841091 | 0.47704629 | 0.85612426 | 0.39192905 | 0.64061964 |
| E2F4 | 21.8995044 | -0.4909114 | 0.49133854 | -0.9991307 | 0.31773136 | 0.57515952 |
| LRRC29 | 16.5228413 | 0.02253919 | 0.52481827 | 0.04294665 | 0.96574406 | NA |
| TMEM208 | 16.8957957 | 0.79906407 | 0.48112916 | 1.66080989 | 0.09675163 | NA |
| FHOD1 | 10.8913618 | 0.73355671 | 0.58511784 | 1.25369056 | 0.2099545 | NA |
| LRRC36 | 15.9465646 | 1.33829026 | 0.84976303 | 1.57489819 | 0.11527995 | NA |
| ZDHHC1 | 25.4566023 | 0.67966962 | 0.39086926 | 1.7388669 | 0.08205818 | 0.28580436 |
| HSD11B2 | 217.057938 | 4.12506512 | 0.81720149 | 5.04779444 | 4.47E-07 | 0.00022934 |
| ATP6V0D1 | 50.6282328 | -0.3125869 | 0.49305502 | -0.6339797 | 0.52609405 | 0.74109552 |
| AC009061.2 | 41.6534622 | 1.10340986 | 0.54100105 | 2.03957064 | 0.04139311 | 0.20301425 |
| RIPOR1 | 43.2939336 | 0.35174898 | 0.39712623 | 0.88573595 | 0.37575983 | 0.62710829 |
| AC027682.4 | 15.9315046 | -0.3391467 | 0.51780132 | -0.6549746 | 0.51248414 | NA |

| | | | | | | |
|---|---|---|---|---|---|---|
| CTCF | 91.3465188 | -0.2758406 | 0.27461139 | -1.0044763 | 0.31514909 | 0.57225891 |
| GFOD2 | 22.7639418 | -0.3101881 | 0.37130921 | -0.8353902 | 0.40349804 | 0.65082239 |
| RANBP10 | 94.4374594 | -0.2660148 | 0.34271465 | -0.7761992 | 0.4376314 | 0.67793318 |
| PSKH1 | 26.7772676 | -0.040417 | 0.39350807 | -0.1027095 | 0.91819357 | 0.96421386 |
| DUS2 | 17.6889171 | -0.6271224 | 0.64838126 | -0.9672124 | 0.33343787 | NA |
| NFATC3 | 143.305035 | 0.41979742 | 0.18880507 | 2.22344356 | 0.02618591 | 0.15845355 |
| PLA2G15 | 20.9559788 | 0.64445499 | 0.45792401 | 1.40734046 | 0.15932646 | NA |
| SLC7A6 | 63.7938755 | 0.60191277 | 0.36948563 | 1.62905595 | 0.10330118 | 0.32282142 |
| SLC7A6OS | 34.2109345 | -0.2841451 | 0.43100609 | -0.6592601 | 0.50972878 | 0.72990216 |
| PRMT7 | 36.8306829 | 1.11930754 | 0.56892732 | 1.96739987 | 0.04913713 | 0.22229864 |
| ZFP90 | 43.3132372 | 1.11630318 | 0.43165723 | 2.58608708 | 0.00970724 | 0.09754129 |
| CDH3 | 59.6909509 | 1.11999914 | 0.57654267 | 1.94261274 | 5.21E-02 | 0.22859592 |
| CDH1 | 76.4520532 | -0.1797548 | 0.40208364 | -0.4470583 | 0.65483296 | 0.82316664 |
| TANGO6 | 106.187165 | -0.0716952 | 0.38171534 | -0.1878238 | 0.85101475 | 0.92874559 |
| CHTF8 | 32.0328833 | 0.41622678 | 0.32552031 | 1.27865071 | 0.20102008 | 0.45520788 |
| UTP4 | 24.0639077 | -0.4072705 | 0.34643077 | -1.1756186 | 0.2397473 | 0.49733052 |
| SNTB2 | 30.6662523 | -0.898559 | 0.39269247 | -2.2882001 | 0.02212587 | 0.14542084 |
| VPS4A | 19.2602061 | 0.11453479 | 0.43841532 | 0.26124724 | 0.79390185 | NA |
| TERF2 | 42.651635 | -0.1343533 | 0.33692925 | -0.3987581 | 0.69007143 | 0.84618126 |
| CYB5B | 85.7218769 | 0.17893356 | 0.28309798 | 0.63205523 | 0.52735079 | 0.74205101 |
| NFAT5 | 696.293025 | -0.1051132 | 0.18500945 | -0.5681503 | 0.56993288 | 0.77196291 |
| NQO1 | 28.6900251 | 0.73188752 | 0.66084197 | 1.10750762 | 0.26807452 | 0.52717407 |
| WWP2 | 203.287162 | -0.3195297 | 0.26204361 | -1.2193759 | 0.22270155 | 0.47974639 |
| PDPR | 72.2797541 | -0.4682487 | 0.34431036 | -1.3599611 | 0.17384222 | 0.42294335 |
| AC009060.1 | 25.7760293 | 0.38604019 | 0.51500305 | 0.74958817 | 0.45350278 | 0.68943529 |
| EXOSC6 | 20.8624345 | -0.2033657 | 0.46996754 | -0.4327229 | 0.66521612 | NA |
| AARS | 53.3282934 | -1.10051 | 0.56528016 | -1.9468399 | 0.05155393 | 0.22799911 |
| ST3GAL2 | 38.1709963 | 0.58656956 | 0.32606599 | 1.798929 | 0.07202991 | 0.26757444 |
| FUK | 20.0044058 | 0.92002984 | 0.47328944 | 1.94390529 | 0.05190687 | NA |
| COG4 | 49.36424 | 0.13848092 | 0.32649434 | 0.42414492 | 0.67146013 | 0.83533677 |
| SF3B3 | 64.9418462 | -0.0914275 | 0.27792038 | -0.32897 | 0.74217834 | 0.87152705 |
| MTSS1L | 55.6209869 | -0.9892771 | 0.50401253 | -1.9628026 | 0.04966911 | 0.22378497 |
| VAC14 | 51.4952157 | -0.6064813 | 0.57519935 | -1.0543845 | 0.29170693 | 0.55098608 |
| HYDIN | 94.2289399 | -0.3761249 | 0.55739213 | -0.6747941 | 0.49980661 | 0.72203669 |
| CMTR2 | 78.3422428 | -1.0234522 | 0.46185871 | -2.2159421 | 0.02669547 | 0.16052735 |
| ZNF19 | 28.9302736 | 0.68756824 | 0.43327045 | 1.58692624 | 0.11252935 | 0.33724584 |
| MARVELD3 | 16.147078 | -0.6755777 | 0.44421831 | -1.5208236 | 0.12830411 | NA |
| PHLPP2 | 38.4319937 | -0.2227928 | 0.45994344 | -0.4843918 | 0.62810783 | 0.80629519 |
| AP1G1 | 176.848583 | -0.5052296 | 0.30434189 | -1.6600726 | 0.09689986 | 0.3125608 |
| IST1 | 114.210038 | -0.4733094 | 0.38197357 | -1.2391155 | 0.21530273 | 0.47115856 |
| ZNF821 | 37.8900877 | 0.56508306 | 0.36358438 | 1.55420057 | 0.12013658 | 0.34880847 |
| TXNL4B | 17.8358838 | 0.65918284 | 0.47419286 | 1.39011549 | 0.16449381 | NA |
| DHX38 | 18.7814856 | -0.8524932 | 0.44589038 | -1.9118897 | 0.05589034 | NA |
| LINC01572 | 48.4487848 | 0.99717369 | 0.64378301 | 1.54892826 | 0.12139897 | 0.35059775 |
| AC004943.2 | 31.3680628 | 0.68260162 | 0.63804347 | 1.0698356 | 0.28469332 | 0.54320388 |
| ZFHX3 | 229.568859 | -0.3015157 | 0.18116785 | -1.664289 | 0.09605469 | 0.31117801 |
| AC138627.1 | 51.6075104 | -0.2531724 | 0.72723515 | -0.3481301 | 0.72774249 | 0.86468169 |
| PSMD7 | 35.2463144 | -0.4144388 | 0.3592465 | -1.1536335 | 0.24865045 | 0.50492395 |
| NPIPB15 | 18.4917233 | 0.0635019 | 0.97361349 | 0.06522291 | 0.94799652 | NA |
| GLG1 | 308.174309 | -0.1007057 | 0.18848425 | -0.5342923 | 0.59313933 | 0.78618794 |
| RFWD3 | 42.1254488 | -0.3422776 | 0.30709851 | -1.114553 | 0.26504202 | 0.52442889 |
| FA2H | 77.3321211 | -0.2211536 | 0.43134417 | -0.512708 | 0.60815557 | 0.79566688 |
| WDR59 | 132.114692 | -0.0573507 | 0.25045774 | -0.2289837 | 0.81888162 | 0.913067 |
| ZNRF1 | 81.1795209 | -0.8559102 | 0.39850338 | -2.1478117 | 0.03172872 | 0.1755686 |

| | | | | | | |
|---|---|---|---|---|---|---|
| BCAR1 | 79.1247066 | -0.8736449 | 0.36728423 | -2.3786618 | 0.01737561 | 0.13115162 |
| CFDP1 | 204.181101 | -0.4126341 | 0.21261284 | -1.9407768 | 0.05228536 | 0.22918019 |
| GABARAPL2 | 33.8624443 | 0.64141794 | 0.43307282 | 1.48108564 | 0.13858375 | 0.37517394 |
| ADAT1 | 16.3976908 | -0.1890301 | 0.51339891 | -0.3681935 | 0.71272897 | NA |
| KARS | 24.7793948 | -0.4092218 | 0.45112767 | -0.9071086 | 0.36434935 | 0.62003028 |
| TERF2IP | 118.880791 | -0.0068107 | 0.38397545 | -0.0177374 | 0.98584834 | 0.9942828 |
| WWOX | 1200.64208 | 0.52950753 | 0.36014522 | 1.47026118 | 0.14149103 | 0.37950905 |
| CDYL2 | 70.5204943 | -1.0909223 | 0.80442246 | -1.3561559 | 0.17504957 | 0.42343073 |
| CMC2 | 78.3706415 | -0.021361 | 0.33456879 | -0.0638463 | 0.94909264 | 0.97977013 |
| ATMIN | 15.0969308 | -0.0255335 | 0.49859002 | -0.0512114 | 0.95915704 | NA |
| GAN | 48.0853806 | -0.0296599 | 0.47581431 | -0.0623349 | 0.95029611 | 0.97993174 |
| CMIP | 281.055931 | -0.6971969 | 0.4226874 | -1.6494385 | 0.09905783 | 0.31649984 |
| PLCG2 | 248.23392 | -0.1744447 | 0.22674756 | -0.7693344 | 0.44169482 | 0.68177214 |
| MPHOSPH6 | 25.0485267 | 0.5012452 | 0.38928706 | 1.2875979 | 0.19788597 | 0.45094881 |
| HSBP1 | 30.2472588 | -0.0574816 | 0.47144122 | -0.1219274 | 0.90295649 | 0.95661107 |
| MBTPS1 | 77.2921087 | -0.0250044 | 0.27872291 | -0.0897105 | 0.92851731 | 0.97037308 |
| HSDL1 | 15.3830931 | -0.5139418 | 0.45972018 | -1.1179448 | 0.26359057 | NA |
| TAF1C | 18.8945396 | -0.0985543 | 0.52438978 | -0.1879409 | 0.850923 | NA |
| TLDC1 | 30.1894664 | 0.18790701 | 0.44485459 | 0.42240097 | 0.67273237 | 0.83624371 |
| KLHL36 | 45.3465124 | 0.01602026 | 0.30738143 | 0.05211849 | 0.95843428 | 0.98401364 |
| USP10 | 112.452066 | -0.7639533 | 0.23564113 | -3.2420204 | 0.00118686 | 0.03065512 |
| ZDHHC7 | 94.8443621 | -0.502361 | 0.41921976 | -1.1983237 | 0.23079102 | 0.48936597 |
| KIAA0513 | 37.1295663 | -1.0291982 | 0.44814076 | -2.2965958 | 0.02164184 | 0.14349647 |
| GSE1 | 162.510453 | -0.2622629 | 0.36170163 | -0.7250807 | 0.46840251 | 0.69838306 |
| EMC8 | 22.8434097 | -0.6005771 | 0.54383959 | -1.1043277 | 0.26945102 | 0.52764898 |
| COX4I1 | 102.123525 | 0.77903839 | 0.47724671 | 1.63235988 | 0.10260369 | 0.32168661 |
| MTHFSD | 15.3098427 | -0.5734561 | 0.48313052 | -1.1869591 | 0.23524377 | NA |
| FBXO31 | 22.4755707 | -0.486563 | 0.41621624 | -1.1690148 | 0.24239766 | 0.49966116 |
| MAP1LC3B | 38.5266726 | -0.3618863 | 0.46242435 | -0.7825848 | 0.43387094 | 0.67475797 |
| ZCCHC14 | 188.108847 | -1.188262 | 0.53263899 | -2.2308956 | 0.02568804 | 0.15667263 |
| KLHDC4 | 56.128539 | -0.2765794 | 0.54153694 | -0.5107305 | 0.60953978 | 0.79648926 |
| SLC7A5 | 49.2664828 | -1.0796795 | 0.64712608 | -1.6684221 | 9.52E-02 | 0.30980573 |
| BANP | 76.6135132 | -0.9520867 | 0.48548123 | -1.9611196 | 0.04986507 | 0.2242948 |
| ZFPM1 | 25.9406431 | -0.1281373 | 0.34746807 | -0.3687744 | 0.71229591 | 0.857224 |
| ZC3H18 | 42.1615325 | -0.7467711 | 0.31605506 | -2.362788 | 0.01813804 | 0.13346893 |
| CYBA | 46.8653218 | 0.64672474 | 0.690481 | 0.9366293 | 0.34894927 | 0.60587921 |
| MVD | 28.4772514 | -0.2424284 | 0.55844466 | -0.4341136 | 0.66420599 | 0.8292079 |
| PIEZO1 | 32.6243558 | -1.0560125 | 0.54944867 | -1.9219493 | 0.05461213 | 0.23352755 |
| APRT | 21.1681163 | 0.5569993 | 0.44142047 | 1.26183386 | 0.20700857 | NA |
| GALNS | 30.3261906 | -0.3312361 | 0.3564076 | -0.9293745 | 0.35269503 | 0.61004394 |
| ACSF3 | 34.5183864 | -0.0097707 | 0.35068694 | -0.0278617 | 0.97777246 | 0.99142598 |
| ZNF778 | 15.9470386 | -0.5790012 | 0.4932578 | -1.1738308 | 0.24046279 | NA |
| ANKRD11 | 370.13334 | -0.6404728 | 0.28404395 | -2.254837 | 0.02414356 | 0.15170042 |
| SPG7 | 150.031736 | -0.4120515 | 0.35469282 | -1.1617135 | 0.24535184 | 0.50252079 |
| RPL13 | 163.104471 | 0.40827389 | 0.39091025 | 1.04441849 | 0.29629181 | 0.55582698 |
| CHMP1A | 24.007659 | -0.1119273 | 0.49612011 | -0.2256052 | 0.82150849 | 0.91434507 |
| CDK10 | 38.8749835 | -0.4711686 | 0.37715174 | -1.2492812 | 0.21156223 | 0.46623687 |
| VPS9D1 | 38.6900545 | 0.2229145 | 0.40620613 | 0.54877187 | 0.58316201 | 0.779 |
| ZNF276 | 24.7264453 | 0.23574121 | 0.51804834 | 0.45505639 | 0.64906867 | 0.81913126 |
| SPIRE2 | 10.0881846 | 1.39955187 | 0.73982806 | 1.89172587 | 0.05852752 | NA |
| TCF25 | 275.814196 | -0.9053097 | 0.38769601 | -2.3351018 | 0.0195381 | 0.13704876 |
| DEF8 | 15.0855137 | -0.2757067 | 0.46386226 | -0.5943719 | 0.55226339 | NA |
| GAS8 | 49.8152118 | -0.3686906 | 0.350716 | -1.0512511 | 0.29314329 | 0.55234367 |
| FAM157C | 46.5613295 | -0.3776645 | 0.49420648 | -0.7641836 | 0.44475783 | 0.68479716 |

| | | | | | | |
|---|---|---|---|---|---|---|
| RPH3AL | 71.1486216 | -0.1056411 | 0.33048992 | -0.3196499 | 0.74923373 | 0.87588883 |
| VPS53 | 194.160801 | -0.7683859 | 0.23217315 | -3.3095381 | 0.0009345 | 0.02809707 |
| GLOD4 | 21.0801911 | 0.77479755 | 0.38838285 | 1.99493243 | 0.04605025 | NA |
| NXN | 27.1775231 | 1.06065323 | 0.60228616 | 1.76104534 | 0.07823073 | 0.27886875 |
| ABR | 22.0842042 | -0.2118712 | 0.47785893 | -0.4433761 | 0.65749371 | 0.82489472 |
| YWHAE | 206.372219 | -0.2762552 | 0.27897713 | -0.9902432 | 0.32205524 | 0.57876703 |
| CRK | 88.6865761 | -0.741511 | 0.35177436 | -2.1079165 | 0.03503821 | 0.18612085 |
| MYO1C | 62.7235421 | 0.11404019 | 0.29815713 | 0.38248352 | 0.70210275 | 0.85211052 |
| INPP5K | 14.2199213 | -0.3844084 | 0.48047467 | -0.8000597 | 0.42367621 | NA |
| PITPNA | 105.921947 | -0.2734194 | 0.24907148 | -1.0977546 | 0.27231168 | 0.53103192 |
| SLC43A2 | 94.2304958 | 1.17373743 | 0.41253356 | 2.84519263 | 0.00443846 | 0.0642158 |
| SCARF1 | 15.089714 | -0.0747378 | 0.6131283 | -0.1218958 | 0.90298153 | NA |
| PRPF8 | 82.3122824 | -0.4108146 | 0.32386733 | -1.2684656 | 0.20463173 | 0.45773044 |
| MIR22HG | 21.3922248 | -1.2233355 | 0.66429414 | -1.8415569 | 0.06553999 | 0.25452134 |
| WDR81 | 10.5284536 | -0.8375801 | 0.79958987 | -1.0475121 | 0.29486345 | NA |
| SMYD4 | 21.1115574 | 0.27875988 | 0.42069392 | 0.66261922 | 0.50757446 | NA |
| RPA1 | 32.38114 | 0.20559637 | 0.39286952 | 0.52331971 | 0.60075178 | 0.79096587 |
| SMG6 | 275.824333 | -0.373432 | 0.27335733 | -1.3660948 | 0.17190922 | 0.42045926 |
| TSR1 | 19.1476584 | -0.5935618 | 0.41963049 | -1.4144867 | 0.15721905 | NA |
| SGSM2 | 47.3752276 | 0.29157621 | 0.29512195 | 0.9879855 | 0.32315975 | 0.57912936 |
| MNT | 17.6710015 | -0.9674546 | 0.49908043 | -1.9384743 | 0.05256538 | NA |
| METTL16 | 82.0010784 | -0.1182815 | 0.23895983 | -0.4949849 | 0.62061081 | 0.8035057 |
| PAFAH1B1 | 311.212206 | -0.4755968 | 0.22117937 | -2.1502766 | 0.03153334 | 0.17516458 |
| CLUH | 65.0696245 | 0.08475647 | 0.42419459 | 0.19980563 | 0.84163259 | 0.92367136 |
| RAP1GAP2 | 180.704527 | 0.30968839 | 0.2575324 | 1.20252206 | 2.29E-01 | 0.48738732 |
| CTNS | 21.3412048 | -0.2327538 | 0.43847692 | -0.5308235 | 0.59554111 | 0.78855668 |
| ITGAE | 16.3535402 | 0.12390702 | 0.49251736 | 0.25157898 | 0.8013665 | NA |
| NCBP3 | 66.2514607 | -0.1775401 | 0.26154399 | -0.6788155 | 0.4972548 | 0.72132872 |
| ZZEF1 | 124.822995 | -0.5148534 | 0.35983205 | -1.4308159 | 0.15248298 | 0.39411063 |
| CYB5D2 | 36.6085326 | 0.30974468 | 0.41194184 | 0.75191362 | 0.45210301 | 0.68880381 |
| ANKFY1 | 107.790699 | -0.3147232 | 0.26425184 | -1.190997 | 0.23365475 | 0.49243138 |
| UBE2G1 | 149.385707 | -0.3443085 | 0.34031222 | -1.0117429 | 0.311661 | 0.57088166 |
| SPNS2 | 19.7428116 | -1.1000533 | 0.67756193 | -1.6235465 | 0.10447264 | NA |
| GGT6 | 14.915343 | 0.28050702 | 0.66682541 | 0.42066037 | 0.67400311 | NA |
| SMTNL2 | 34.5156133 | 0.33210942 | 0.4098054 | 0.81040762 | 0.41770594 | 0.66167578 |
| PELP1 | 23.8741826 | -0.2114585 | 0.37950729 | -0.5571923 | 0.57739604 | 0.77491227 |
| PSMB6 | 17.5644198 | 1.06935678 | 0.54649219 | 1.95676498 | 0.05037511 | NA |
| MINK1 | 70.9201837 | -0.3317874 | 0.32180845 | -1.0310089 | 0.30253663 | 0.56287755 |
| SLC25A11 | 15.8392331 | 0.55749416 | 0.57896958 | 0.96290752 | 0.33559394 | NA |
| RNF167 | 15.4567195 | 0.63008088 | 0.53729087 | 1.17269976 | 0.24091623 | NA |
| PFN1 | 49.3705301 | -0.4348292 | 0.44369387 | -0.9800207 | 0.32707591 | 0.58316036 |
| SPAG7 | 24.6496816 | -0.7008222 | 0.49017841 | -1.4297288 | 0.15279487 | 0.3947713 |
| CAMTA2 | 20.4570122 | 0.46190496 | 0.47356906 | 0.97536979 | 0.32937686 | NA |
| KIF1C | 27.5378685 | 0.07342954 | 0.46655259 | 0.15738748 | 0.87493948 | 0.94169674 |
| ZNF594 | 23.3285508 | -0.4180282 | 0.48493971 | -0.862021 | 0.38867599 | 0.63787614 |
| AC087500.1 | 49.9421471 | 0.8409881 | 0.48220556 | 1.74404479 | 0.08115128 | 0.28417716 |
| RABEP1 | 186.445862 | -0.6561955 | 0.3174961 | -2.0667828 | 0.03875462 | 0.1959884 |
| NUP88 | 124.791017 | -0.3366762 | 0.22144788 | -1.5203406 | 0.12842539 | 0.36084904 |
| RPAIN | 58.1645291 | 0.06635857 | 0.31131858 | 0.21315325 | 0.83120743 | 0.9181376 |
| DERL2 | 19.6988738 | 0.35509869 | 0.43264111 | 0.82076965 | 0.41177749 | NA |
| MIS12 | 15.9865879 | -0.433613 | 0.50115044 | -0.8652351 | 0.38690976 | NA |
| KIAA0753 | 56.2467098 | -0.1493212 | 0.42208522 | -0.3537703 | 0.72351105 | 0.86194629 |
| XAF1 | 51.5743608 | 1.84163125 | 2.24487667 | 0.82037079 | 0.41200476 | 0.65710747 |
| ALOX12-AS1 | 36.9587374 | 1.74120343 | 0.61265315 | 2.84207049 | 0.00448216 | 0.06436412 |

| | | | | | | |
|---|---|---|---|---|---|---|
| ACADVL | 130.958861 | -0.0417366 | 0.4828698 | -0.0864345 | 0.93112106 | 0.97177475 |
| DVL2 | 16.3696742 | -0.2752756 | 0.44128119 | -0.62381 | 0.53275239 | NA |
| ELP5 | 15.5932602 | 0.62100241 | 0.50243114 | 1.23599508 | 0.21646039 | NA |
| EIF5A | 21.5237744 | -0.7052056 | 0.6233064 | -1.1313948 | 0.25788898 | 0.51619036 |
| ZBTB4 | 59.486375 | 0.56401153 | 0.31475771 | 1.7918911 | 0.07315041 | 0.26939652 |
| POLR2A | 97.3733744 | -0.3056368 | 0.3889228 | -0.7858547 | 0.43195263 | 0.67275181 |
| EIF4A1 | 21.4500171 | -0.5273953 | 0.64855062 | -0.8131907 | 0.41610875 | 0.65982469 |
| MPDU1 | 23.8328904 | -0.0175843 | 0.39450371 | -0.0445732 | 0.96444749 | 0.986492 |
| FXR2 | 20.8788956 | 0.09793614 | 0.39178867 | 0.24997186 | 0.80260911 | NA |
| SAT2 | 50.1406613 | 0.1394729 | 0.40962534 | 0.34048894 | 0.73348835 | 0.86776236 |
| TP53 | 28.5600576 | 0.11526171 | 0.38383907 | 0.30028654 | 0.7639586 | 0.8830439 |
| KDM6B | 46.0312929 | -1.4238645 | 0.63755478 | -2.233321 | 0.02552778 | 0.15648615 |
| NAA38 | 50.4346661 | 0.32264437 | 0.32898887 | 0.98071514 | 0.32673323 | 0.58313355 |
| CHD3 | 24.5568775 | 0.29207143 | 0.465368 | 0.62761392 | 0.53025689 | 0.74396414 |
| CNTROB | 15.8029364 | -0.6023441 | 0.50998768 | -1.1810954 | 0.23756485 | NA |
| PER1 | 36.6442419 | -0.9483404 | 0.47225813 | -2.0080975 | 0.04463294 | 0.21140906 |
| CTC1 | 24.5000676 | -0.2293187 | 0.39403867 | -0.58197 | 0.56058691 | 0.76573742 |
| NDEL1 | 78.4377249 | 0.04365057 | 0.28989463 | 0.15057391 | 0.88031184 | 0.94376451 |
| MYH10 | 112.607106 | 0.26880891 | 0.36407261 | 0.73833872 | 0.46030864 | 0.69361699 |
| NTN1 | 26.1836557 | 0.3523263 | 0.63254616 | 0.55699698 | 0.57752949 | 0.77491227 |
| STX8 | 229.105421 | 0.01351218 | 0.22372541 | 0.06039628 | 0.95184002 | 0.98050223 |
| USP43 | 116.236082 | -1.1861526 | 0.58765872 | -2.0184378 | 0.04354569 | 0.20869935 |
| SCO1 | 30.0783959 | -0.3974604 | 0.41281252 | -0.9628109 | 0.33564244 | 0.59131149 |
| TMEM220 | 28.7648247 | -0.4906544 | 0.58088198 | -0.8446713 | 0.39829437 | 0.64642108 |
| TMEM220-A! | 21.8636058 | 0.22259373 | 0.45162729 | 0.49287041 | 0.62210415 | 0.80366942 |
| SHISA6 | 24.4850714 | 0.71709875 | 0.69070301 | 1.03821575 | 0.29916962 | 0.5591813 |
| MAP2K4 | 260.628753 | -0.5760317 | 0.24605439 | -2.3410745 | 0.01922833 | 0.13639344 |
| ARHGAP44 | 136.438195 | -0.7084916 | 0.358945 | -1.9738165 | 0.04840261 | 0.22057721 |
| ELAC2 | 17.3174463 | -0.0125233 | 0.42400661 | -0.0295357 | 0.97643732 | NA |
| COX10 | 81.0201816 | 0.09685111 | 0.26908676 | 0.35992522 | 0.71890306 | 0.85995641 |
| TRIM16 | 19.3258043 | 1.889657 | 0.58381722 | 3.23672707 | 0.00120909 | NA |
| ZSWIM7 | 48.1415171 | 0.40190177 | 0.38147418 | 1.05354907 | 0.29208942 | 0.55143788 |
| TTC19 | 129.597922 | 0.17893288 | 0.25737839 | 0.69521331 | 0.48692163 | 0.71361615 |
| NCOR1 | 348.755315 | 0.00150313 | 0.17160716 | 0.00875912 | 0.99301132 | 0.99689685 |
| PIGL | 110.174347 | -0.0894954 | 0.30898851 | -0.28964 | 0.77209163 | 0.88734195 |
| UBB | 79.4101596 | 1.11911428 | 0.4900228 | 2.28380045 | 0.02238326 | 0.14604538 |
| AC098850.3 | 63.7660256 | -0.1614194 | 0.6581196 | -0.2452736 | 0.80624457 | 0.9057619 |
| MPRIP | 182.675461 | -0.6390561 | 0.39439825 | -1.620332 | 0.10516097 | 0.32611764 |
| COPS3 | 21.8177997 | -0.7922461 | 0.5626011 | -1.4081843 | 0.15907649 | 0.40403105 |
| MED9 | 49.1850242 | -0.4190417 | 0.44977641 | -0.9316667 | 0.3515088 | 0.60853875 |
| RASD1 | 120.26871 | -0.1479565 | 0.74413879 | -0.1988292 | 0.84239638 | 0.92390487 |
| PEMT | 19.6319711 | 0.23929628 | 0.50432683 | 0.47448652 | 0.63515301 | NA |
| RAI1 | 101.520825 | -0.4156149 | 0.29358982 | -1.4156311 | 0.15688353 | 0.40104036 |
| SREBF1 | 57.0760969 | -1.2736576 | 0.44047218 | -2.8915734 | 0.00383318 | 0.06110382 |
| TOM1L2 | 327.707223 | -0.5381508 | 0.37802759 | -1.4235753 | 0.15456944 | 0.39736839 |
| DRC3 | 16.8335488 | 0.90684498 | 0.72301661 | 1.25425194 | 0.20975045 | NA |
| ATPAF2 | 19.2144156 | 0.39676743 | 0.47783472 | 0.83034448 | 0.40634405 | NA |
| GID4 | 13.125176 | 0.61352219 | 0.48533716 | 1.26411543 | 0.2061886 | NA |
| DRG2 | 12.5899791 | -0.5059764 | 0.50540628 | -1.001128 | 0.31676491 | NA |
| ALKBH5 | 42.7387087 | -0.4109749 | 0.34246739 | -1.2000409 | 0.23012346 | 0.48881906 |
| FLII | 25.9297042 | -0.5682966 | 0.47811756 | -1.1886126 | 0.23459214 | 0.49243138 |
| TOP3A | 34.2054861 | -0.7459176 | 0.3462149 | -2.154493 | 0.03120153 | 0.17453355 |
| SMCR8 | 14.1840903 | -0.6809278 | 0.49981802 | -1.3623515 | 0.17308699 | NA |
| SHMT1 | 30.5452187 | 0.80136783 | 0.44580239 | 1.79758532 | 0.07224275 | 0.26810628 |

| | | | | | | |
|---|---|---|---|---|---|---|
| TVP23B | 31.552963 | -0.6302602 | 0.33703515 | -1.8700134 | 0.06148196 | 0.24796715 |
| PRPSAP2 | 76.0317554 | -0.4131119 | 0.27251998 | -1.5158958 | 0.12954571 | 0.36208459 |
| EPN2 | 142.086659 | -0.1152334 | 0.25115353 | -0.4588164 | 0.64636599 | 0.81813501 |
| B9D1 | 23.1415538 | 0.13223436 | 0.5044423 | 0.26213971 | 0.79321373 | 0.89875118 |
| SLC47A1 | 35.216964 | 0.61072188 | 0.39114712 | 1.56136104 | 1.18E-01 | 3.46E-01 |
| ALDH3A2 | 46.9732634 | 0.30278843 | 0.31662273 | 0.95630665 | 0.33891733 | 0.59462212 |
| ULK2 | 66.4515997 | 0.76228259 | 0.44021039 | 1.73163245 | 0.08333902 | 0.28855622 |
| AKAP10 | 158.223567 | -0.486142 | 0.21361484 | -2.2757878 | 0.02285872 | 0.14710998 |
| SPECC1 | 21.5456434 | -0.7246284 | 0.40259437 | -1.7998969 | 0.07187692 | 0.26739325 |
| CCDC144NL-/ | 14.7602763 | 0.56610414 | 0.49168019 | 1.15136656 | 0.24958146 | NA |
| USP22 | 134.729683 | -0.1672781 | 0.30430297 | -0.5497091 | 0.58251892 | 0.7783237 |
| DHRS7B | 100.495148 | -0.4392887 | 0.20738298 | -2.1182488 | 0.034154 | 0.18312899 |
| AC087294.1 | 16.3249449 | 0.00775583 | 0.51055957 | 0.01519084 | 0.98787993 | NA |
| MAP2K3 | 35.8942308 | -1.3891191 | 0.48618046 | -2.8572089 | 0.00427385 | 0.06301187 |
| WSB1 | 925.858233 | -0.6212607 | 0.36375788 | -1.7078961 | 0.08765562 | 0.29621428 |
| KSR1 | 111.536747 | 0.04719597 | 0.30775322 | 0.15335654 | 0.87811712 | 0.94339485 |
| AC005697.1 | 13.8218987 | 1.81407862 | 1.23368016 | 1.47046105 | 0.14143693 | NA |
| NLK | 307.991537 | -0.6794202 | 0.44496436 | -1.5269092 | 0.12678358 | 0.35850596 |
| IFT20 | 13.7400138 | 0.01504726 | 0.49802826 | 0.03021367 | 0.97589664 | NA |
| TNFAIP1 | 29.5001222 | -0.4104421 | 0.38745845 | -1.059319 | 0.28945452 | 0.5488051 |
| POLDIP2 | 19.5029676 | 0.43374847 | 0.44001086 | 0.98576766 | 0.32424714 | NA |
| SARM1 | 22.5813991 | -0.7115316 | 0.55006696 | -1.2935363 | 0.19582564 | 0.44885942 |
| AC005726.1 | 22.6802623 | 0.20352773 | 0.42092478 | 0.48352517 | 0.62872289 | 0.80681562 |
| KIAA0100 | 56.2392517 | 0.00086204 | 0.29921865 | 0.00288097 | 0.99770133 | 0.99882493 |
| SDF2 | 21.9849916 | -0.4247631 | 0.4384661 | -0.9687479 | 0.332671 | 0.58796066 |
| SUPT6H | 86.0933989 | -0.8997508 | 0.26611279 | -3.3810882 | 0.00072199 | 0.02405708 |
| RAB34 | 20.6308382 | 0.46191452 | 0.42231021 | 1.09378015 | 0.27405141 | NA |
| RPL23A | 61.4283458 | 0.58921226 | 0.35549115 | 1.65745969 | 0.09742657 | 0.31376248 |
| NEK8 | 16.6143264 | 0.35336351 | 0.86685267 | 0.40763963 | 0.68353826 | NA |
| TRAF4 | 41.4358983 | 0.03090298 | 0.49140282 | 0.06288727 | 0.94985628 | 0.97993174 |
| FAM222B | 191.499776 | -0.2345463 | 0.24527798 | -0.9562468 | 0.33894758 | 0.59462212 |
| FLOT2 | 73.6897373 | 0.2420554 | 0.28394842 | 0.85246257 | 3.94E-01 | 0.6425014 |
| PHF12 | 60.563028 | -0.2703323 | 0.27810328 | -0.9720572 | 0.33102207 | 0.58655523 |
| MYO18A | 49.4260552 | -1.2424808 | 0.52444871 | -2.3691179 | 0.01783057 | 0.13283085 |
| NUFIP2 | 160.179225 | -0.3340667 | 0.31896276 | -1.0473532 | 0.29493671 | 0.55409516 |
| TAOK1 | 267.556534 | -0.505158 | 0.27140883 | -1.8612437 | 0.06270976 | 0.24960868 |
| ABHD15-AS1 | 16.9481969 | 0.54517133 | 0.49364023 | 1.10439 | 0.269424 | NA |
| GIT1 | 14.6603303 | 0.07160104 | 0.54118489 | 0.13230422 | 0.89474367 | NA |
| SSH2 | 427.917598 | -0.3478966 | 0.26559015 | -1.3099003 | 0.19022957 | 0.44142207 |
| AC023389.1 | 15.0875297 | 0.46335885 | 0.50194289 | 0.9231306 | 0.35593915 | NA |
| EFCAB5 | 15.0619249 | -0.889157 | 0.55853609 | -1.591942 | 1.11E-01 | NA |
| NSRP1 | 179.900765 | -0.1594067 | 0.18807718 | -0.8475602 | 0.39668297 | 0.64496595 |
| BLMH | 29.2554348 | 0.38991886 | 0.38102024 | 1.0233547 | 0.30614017 | 0.56595793 |
| CPD | 168.934469 | -0.5468494 | 0.42382676 | -1.2902663 | 0.19695821 | 0.45014879 |
| GOSR1 | 105.942033 | -0.8456189 | 0.21403666 | -3.9508136 | 7.79E-05 | 0.00587734 |
| CRLF3 | 80.1201375 | 0.37189068 | 0.40500221 | 0.91824357 | 0.35849136 | 0.6149561 |
| ATAD5 | 16.7850858 | 0.32185444 | 0.65533578 | 0.49112906 | 0.62333517 | NA |
| TEFM | 38.9908564 | 0.60774772 | 0.31155463 | 1.9506939 | 0.05109347 | 0.22666653 |
| RNF135 | 16.5091137 | 0.2982566 | 0.45229626 | 0.65942752 | 0.50962127 | NA |
| NF1 | 552.923278 | -0.1632269 | 0.16004943 | -1.0198531 | 0.30779813 | 0.56799861 |
| RAB11FIP4 | 30.3114663 | 0.78863454 | 0.48351452 | 1.63104624 | 0.10288056 | 0.32202997 |
| COPRS | 18.1810472 | 0.46228503 | 0.60350038 | 0.7660062 | 0.44367261 | NA |
| UTP6 | 37.000868 | -0.8803516 | 0.35899071 | -2.4522963 | 0.01419477 | 0.11710305 |
| SUZ12 | 130.440373 | -0.2050364 | 0.23007265 | -0.891181 | 0.37283206 | 0.62543211 |

| | | | | | | |
|---|---|---|---|---|---|---|
| LRRC37B | 37.8220063 | 0.52726936 | 0.35609429 | 1.48070155 | 0.13868612 | 0.37517394 |
| RHOT1 | 121.643858 | 0.50139873 | 0.25498712 | 1.96636884 | 0.04925602 | 0.22242545 |
| C17orf75 | 16.2534401 | 0.41012466 | 0.51483334 | 0.79661636 | 0.42567387 | NA |
| ZNF207 | 212.213067 | -0.5784049 | 0.17684694 | -3.2706525 | 0.001073 | 0.02960163 |
| PSMD11 | 56.8702048 | -0.7065723 | 0.41538914 | -1.7009889 | 0.08894507 | 0.29779758 |
| MYO1D | 216.663679 | -0.349255 | 0.20860401 | -1.6742489 | 0.09408169 | 0.30825698 |
| LIG3 | 16.9290531 | -0.3653846 | 0.44238155 | -0.8259489 | 0.40883306 | NA |
| SLFN5 | 19.448667 | -0.1442884 | 0.58904189 | -0.2449544 | 8.06E-01 | NA |
| LINC02001 | 13.6145585 | 0.33756717 | 0.53733835 | 0.62822087 | 0.52985926 | NA |
| AP2B1 | 163.380175 | 0.0728276 | 0.26072672 | 0.27932541 | 0.77999511 | 0.89074516 |
| TAF15 | 235.734039 | -0.2619475 | 0.18221207 | -1.437597 | 0.15054844 | 0.39108044 |
| ZNHIT3 | 14.8873105 | 0.19095473 | 0.52458621 | 0.3640102 | 0.71585039 | NA |
| MYO19 | 84.676608 | 0.37642073 | 0.26088986 | 1.4428339 | 0.14906727 | 0.38943809 |
| GGNBP2 | 106.106869 | -0.7311286 | 0.37341918 | -1.9579299 | 0.05023823 | 0.22507781 |
| LHX1-DT | 180.687292 | 0.22069658 | 0.44402869 | 0.49703226 | 0.6191663 | 0.8027391 |
| LHX1 | 215.205334 | 1.19121554 | 0.47369153 | 2.5147495 | 0.0119117 | 0.10687217 |
| AATF | 113.558381 | -0.2781023 | 0.22255364 | -1.2495967 | 0.21144692 | 0.46620079 |
| ACACA | 125.474898 | -0.1675567 | 0.25573781 | -0.6551895 | 0.51234572 | 0.73101511 |
| TADA2A | 42.5152014 | 0.27767351 | 0.33982962 | 0.81709627 | 0.41387341 | 0.65845053 |
| SYNRG | 143.237061 | -0.1652914 | 0.18965414 | -0.8715413 | 0.38345869 | 0.6343472 |
| DDX52 | 108.927498 | -1.4268897 | 0.48465486 | -2.9441357 | 0.00323858 | 0.05664625 |
| HNF1B | 291.25177 | -1.0291991 | 0.39343729 | -2.6159165 | 0.00889883 | 0.09323317 |
| MRPL45 | 32.2478684 | -0.1885585 | 0.3354085 | -0.5621757 | 0.57399629 | 0.77388388 |
| SOCS7 | 33.1106502 | 0.65415872 | 0.35779601 | 1.82830078 | 0.06750442 | 0.25898011 |
| MLLT6 | 157.114332 | -0.153884 | 0.29118116 | -0.528482 | 0.59716482 | 0.7894568 |
| PCGF2 | 28.0717946 | -0.3104249 | 0.44993773 | -0.6899286 | 0.49023906 | 0.7163358 |
| PSMB3 | 24.2033153 | -0.3445473 | 0.42025637 | -0.8198502 | 0.41230152 | 0.65722052 |
| PIP4K2B | 39.2186621 | -0.1812731 | 0.32388662 | -0.5596809 | 0.57569715 | 0.77401855 |
| CWC25 | 29.6617519 | -0.210122 | 0.34112581 | -0.6159664 | 0.5379167 | 0.74878549 |
| RPL23 | 52.2939219 | 0.32272901 | 0.40354969 | 0.79972558 | 0.42386981 | 0.6670468 |
| LASP1 | 35.075766 | -0.3838104 | 0.36803464 | -1.0428649 | 0.29701085 | 0.55674621 |
| RPL19 | 69.4136122 | 1.03644428 | 0.49537107 | 2.09225841 | 0.03641541 | 0.19061356 |
| FBXL20 | 201.160194 | 0.12694315 | 0.19452348 | 0.65258521 | 0.51402372 | 0.73176668 |
| MED1 | 41.4175829 | -0.234299 | 0.3308894 | -0.7080887 | 0.47889017 | 0.70690257 |
| CDK12 | 154.757382 | -0.3545542 | 0.22474837 | -1.5775609 | 0.11466653 | 0.34103102 |
| STARD3 | 30.6471728 | -0.2847196 | 0.33861809 | -0.8408281 | 0.40044425 | 0.64793345 |
| PGAP3 | 18.5393583 | 0.77883795 | 0.58316019 | 1.33554719 | 0.18169732 | NA |
| ERBB2 | 108.796944 | 1.24199277 | 0.47616263 | 2.60833732 | 0.00909832 | 0.09427267 |
| GRB7 | 97.3345597 | -0.095535 | 0.33823243 | -0.2824537 | 0.77759565 | 0.8896927 |
| PSMD3 | 23.9122692 | -0.4980638 | 0.48811544 | -1.0203812 | 0.30754771 | 0.56780876 |
| MED24 | 36.6377385 | -0.0232981 | 0.40414581 | -0.0576477 | 0.95402925 | 0.98153155 |
| THRA | 17.2329393 | 1.22016888 | 0.51954633 | 2.34852758 | 0.0188478 | NA |
| MSL1 | 48.1303122 | 0.14524138 | 0.30563391 | 0.47521357 | 0.63463476 | 0.81099192 |
| CASC3 | 50.4679033 | -0.3409651 | 0.29730541 | -1.1468513 | 0.25144309 | 0.50804157 |
| WIPF2 | 156.333307 | -0.7851398 | 0.33149566 | -2.3684769 | 0.0178615 | 0.13283085 |
| RARA | 25.6812686 | -1.9589151 | 0.44029621 | -4.4490847 | 8.62E-06 | 0.00146459 |
| IGFBP4 | 19.62078 | -0.7468735 | 0.43739811 | -1.7075371 | 0.08772227 | NA |
| KRT10 | 20.227872 | -0.1506475 | 0.547934 | -0.2749374 | 0.78336433 | NA |
| KRT19 | 70.8595357 | 1.68552927 | 0.80592453 | 2.09142321 | 0.03649014 | 0.19061356 |
| EIF1 | 132.68272 | -0.1798912 | 0.37190668 | -0.4836999 | 0.62859885 | 0.80679095 |
| JUP | 104.233377 | -0.3749641 | 0.35765012 | -1.0484104 | 0.29444958 | 0.55385592 |
| NT5C3B | 22.4574242 | 0.36881312 | 0.63391261 | 0.58180436 | 0.56069847 | 0.76573742 |
| ACLY | 33.5853636 | 0.09725189 | 0.36299662 | 0.26791403 | 0.78876549 | 0.89544661 |
| DNAJC7 | 86.4736713 | -0.2201383 | 0.25259443 | -0.8715089 | 0.38347636 | 0.6343472 |

| | | | | | | |
|---|---|---|---|---|---|---|
| KAT2A | 25.4513876 | 0.2050557 | 0.51480108 | 0.39832026 | 0.69039413 | 0.84630731 |
| STAT5B | 171.864814 | -0.6641219 | 0.30839057 | -2.1535093 | 0.03127868 | 0.17469667 |
| STAT5A | 22.5180092 | -0.9416848 | 0.46500798 | -2.0250938 | 0.04285773 | 0.20681879 |
| STAT3 | 552.679316 | -0.7162927 | 0.27352883 | -2.6187102 | 0.00882629 | 0.09290987 |
| ATP6V0A1 | 76.136943 | -0.6617318 | 0.29698657 | -2.228154 | 0.02587025 | 0.15729431 |
| NAGLU | 23.3469371 | -0.9151632 | 0.5979776 | -1.5304305 | 0.12591021 | 0.35705972 |
| COASY | 16.9333559 | 0.00479405 | 0.57571992 | 0.00832706 | 0.99335604 | NA |
| MLX | 16.1770928 | 0.40695954 | 0.45319123 | 0.89798637 | 0.36919282 | NA |
| RETREG3 | 57.4174818 | -0.2050713 | 0.29498282 | -0.6951975 | 0.48693153 | 0.71361615 |
| TUBG2 | 28.365489 | 0.138685 | 0.34582834 | 0.40102267 | 0.68840344 | 0.84543868 |
| EZH1 | 74.0423665 | -0.202047 | 0.31959841 | -0.6321903 | 0.52726253 | 0.74205101 |
| VPS25 | 18.5939731 | -0.5519482 | 0.41913119 | -1.3168864 | 0.1878767 | NA |
| WNK4 | 200.727067 | 0.35552085 | 0.6588197 | 0.539633 | 0.58945015 | 0.78413374 |
| COA3 | 32.6016401 | 0.98811185 | 0.39968855 | 2.47220453 | 0.01342827 | 0.11302673 |
| BECN1 | 55.332447 | -0.0720078 | 0.26725594 | -0.2694338 | 0.78759585 | 0.89506208 |
| PSME3 | 21.6299012 | -0.2801096 | 0.39198447 | -0.7145936 | 0.47486017 | 0.70288437 |
| LINC00671 | 12.9615169 | 0.42257042 | 0.99664579 | 0.42399258 | 6.72E-01 | NA |
| RUNDC1 | 27.9541007 | -1.2612416 | 0.42012533 | -3.0020603 | 0.00268159 | 0.05021947 |
| RPL27 | 33.4298098 | 0.43409852 | 0.34320342 | 1.26484324 | 0.20592753 | 0.45902815 |
| BRCA1 | 21.9427904 | 1.15553548 | 0.64345667 | 1.79582487 | 0.07252237 | 0.26831077 |
| NBR1 | 74.7700465 | -0.0202677 | 0.24643948 | -0.082242 | 0.93445429 | 0.97327579 |
| LINC00854 | 23.3537506 | -0.6580738 | 0.68492592 | -0.9607956 | 0.33665495 | 0.59259931 |
| DHX8 | 36.703509 | -0.7320125 | 0.30537196 | -2.3971175 | 0.01652462 | 0.12745979 |
| DUSP3 | 36.5008849 | -0.5074017 | 0.39564803 | -1.2824572 | 0.1996823 | 0.4536466 |
| PYY | 21.320565 | 1.47055224 | 0.93092949 | 1.57966018 | 0.11418471 | 0.34044736 |
| TMEM101 | 293.905931 | 2.08011099 | 0.78362555 | 2.6544706 | 0.0079433 | 0.08725829 |
| LSM12 | 41.5132894 | -0.5336525 | 0.31308438 | -1.7045006 | 0.08828761 | 0.29713586 |
| HDAC5 | 51.4249683 | 0.22280973 | 0.30362408 | 0.73383419 | 0.4630498 | 0.69434917 |
| ASB16-AS1 | 10.7353825 | -0.073431 | 0.55120282 | -0.1332195 | 0.89401976 | NA |
| TMUB2 | 24.6169412 | 0.13116659 | 0.36433316 | 0.36001827 | 0.71883347 | 0.85995641 |
| UBTF | 29.8731206 | 0.15866147 | 0.38064709 | 0.41682039 | 0.67680978 | 0.83887358 |
| SLC25A39 | 32.0296927 | -0.4781975 | 0.52713386 | -0.9071652 | 0.36431943 | 0.62003028 |
| GRN | 37.4863791 | -0.1701824 | 0.54138556 | -0.314346 | 0.75325828 | 0.87842978 |
| GPATCH8 | 323.421744 | -0.0771081 | 0.1932528 | -0.3990013 | 0.68989225 | 0.84609634 |
| DBF4B | 12.2139044 | 0.0947158 | 0.68179962 | 0.13892029 | 0.88951313 | NA |
| EFTUD2 | 44.0796976 | -0.4837869 | 0.34667682 | -1.3954982 | 0.16286565 | 0.40961653 |
| DCAKD | 55.9162395 | 0.09905445 | 0.3201129 | 0.30943599 | 0.7569899 | 0.87904624 |
| NMT1 | 69.7019008 | -0.1031949 | 0.28323651 | -0.3643417 | 0.71560287 | 0.85868129 |
| ACBD4 | 12.4537973 | 1.27215499 | 0.57340771 | 2.21858719 | 0.02651482 | NA |
| HEXIM1 | 18.3510583 | -0.3956307 | 0.5645045 | -0.7008459 | 0.48339917 | NA |
| AC138150.1 | 57.4529228 | 0.33390147 | 0.49680452 | 0.6720983 | 0.50152112 | 0.72315625 |
| MAP3K14 | 45.6270409 | -0.5808345 | 0.45444064 | -1.2781307 | 0.20120334 | 0.45528981 |
| PLEKHM1 | 42.5490993 | -1.2267555 | 0.45739697 | -2.6820368 | 0.00731754 | 0.08373567 |
| LINC02210-C | 36.8654608 | 1.10411767 | 0.50333501 | 2.19360394 | 0.02826389 | 0.16543513 |
| KANSL1 | 710.715127 | -0.2094132 | 0.30572285 | -0.6849773 | 0.49335822 | 0.71743401 |
| ARL17B | 154.891923 | 0.798772 | 1.14361219 | 0.69846404 | 0.48488704 | 0.7118397 |
| LRRC37A2 | 33.4150953 | 0.40941351 | 0.5157569 | 0.79381103 | 0.42730545 | 0.67024083 |
| ARL17A | 40.8630963 | -0.5839541 | 0.65208787 | -0.8955145 | 0.37051214 | 0.62376026 |
| NSF | 163.829545 | 0.07126363 | 0.35895402 | 0.19853137 | 0.84262934 | 0.92390487 |
| AC005670.2 | 32.8495761 | 0.09448912 | 0.4806068 | 0.19660379 | 0.84413761 | 0.92488459 |
| CDC27 | 164.416344 | -0.5141509 | 0.23883486 | -2.1527463 | 0.03133862 | 0.17479229 |
| AC068234.1 | 40.0827974 | -1.0811312 | 0.65827545 | -1.642369 | 1.01E-01 | 3.19E-01 |
| EFCAB13 | 63.7273399 | 0.16402549 | 0.38434545 | 0.4267658 | 0.66954992 | 0.83376893 |
| NPEPPS | 344.308641 | -0.8538078 | 0.22062559 | -3.8699401 | 0.00010886 | 0.00754875 |

| | | | | | | |
|---|---|---|---|---|---|---|
| AC025682.1 | 38.6989193 | 0.10666986 | 0.39234988 | 0.27187434 | 0.78571864 | 0.89452315 |
| KPNB1 | 134.000206 | -0.4332278 | 0.2293574 | -1.8888766 | 0.05890837 | 0.2411888 |
| TBKBP1 | 14.5195436 | -0.6554019 | 0.61551738 | -1.0647984 | 0.28696717 | NA |
| SCRN2 | 13.4914754 | 0.98163562 | 0.5544542 | 1.77045393 | 0.07665155 | NA |
| SP2 | 18.6755352 | -0.5751616 | 0.42789008 | -1.3441807 | 0.17888996 | NA |
| CDK5RAP3 | 33.5954125 | 0.27199581 | 0.34268747 | 0.79371391 | 0.427362 | 0.67024083 |
| NFE2L1 | 60.4168596 | 0.0950312 | 0.31678414 | 0.29998725 | 0.76418688 | 0.88317118 |
| CBX1 | 45.3434579 | -0.2971133 | 0.30721939 | -0.9671045 | 0.33349178 | 0.58883681 |
| SNX11 | 12.7803531 | 0.22369509 | 0.47200885 | 0.47392139 | 0.63555596 | NA |
| SKAP1 | 428.01927 | 0.35289049 | 0.34693293 | 1.01717209 | 0.30907157 | 0.56869184 |
| HOXB2 | 17.9596474 | 0.62799659 | 0.5265321 | 1.19270335 | 0.23298557 | NA |
| HOXB-AS1 | 15.3386329 | 1.20118531 | 0.4993003 | 2.4057372 | 0.01613987 | NA |
| HOXB3 | 229.034874 | 0.45625352 | 0.41788661 | 1.09181177 | 0.27491585 | 0.5337282 |
| HOXB-AS3 | 44.8406572 | 1.15023947 | 0.45716184 | 2.51604439 | 0.01186802 | 0.10677891 |
| HOXB7 | 27.2558474 | 1.11543093 | 0.38697879 | 2.88240841 | 0.00394648 | 0.06143969 |
| LINC02086 | 16.3696109 | 0.15586522 | 0.77789643 | 0.20036758 | 0.84119311 | NA |
| CALCOCO2 | 42.1568402 | 0.39971148 | 0.33642954 | 1.18809865 | 0.23479455 | 0.49243138 |
| ATP5MC1 | 33.8301964 | 1.28171048 | 0.47794812 | 2.68169375 | 0.00732505 | 0.08373567 |
| UBE2Z | 81.6502445 | -0.5297072 | 0.2637314 | -2.0085103 | 0.0445891 | 0.21133145 |
| SNF8 | 26.4841719 | -0.2597367 | 0.39218652 | -0.6622786 | 0.50779267 | 0.7286503 |
| ZNF652 | 122.159574 | -0.0934114 | 0.22373338 | -0.4175119 | 0.676304 | 0.83865183 |
| PHB | 32.1133009 | 0.79567422 | 0.45018916 | 1.76742198 | 0.07715758 | 0.2772558 |
| SPOP | 82.8845946 | 0.19610388 | 0.25225611 | 0.77739994 | 0.43692285 | 0.67734041 |
| KAT7 | 45.52253 | -0.3756086 | 0.31743659 | -1.1832555 | 0.2367079 | 0.4946893 |
| ITGA3 | 372.474866 | -1.6344086 | 0.48523763 | -3.3682643 | 7.56E-04 | 2.45E-02 |
| PDK2 | 13.557227 | 1.65979891 | 0.61185485 | 2.7127331 | 0.00667308 | NA |
| XYLT2 | 16.5884588 | 0.91075622 | 0.53201201 | 1.71190915 | 0.08691341 | NA |
| MRPL27 | 23.6182299 | 0.32114062 | 0.43647207 | 0.73576442 | 0.46187408 | 0.69416206 |
| ACSF2 | 23.5611136 | 1.57966374 | 0.57874431 | 2.72946741 | 0.00634367 | 0.07762678 |
| RSAD1 | 15.2029111 | 1.34706475 | 0.48011758 | 2.80569759 | 0.00502078 | NA |
| SPATA20 | 15.4104537 | 1.29756578 | 0.57465604 | 2.25798685 | 0.02394648 | NA |
| ABCC3 | 37.6793158 | 0.7941672 | 0.69261765 | 1.14661704 | 2.52E-01 | 0.50804157 |
| ANKRD40 | 16.1449112 | 0.29298379 | 0.45078646 | 0.6499392 | 0.51573149 | NA |
| LUC7L3 | 267.710894 | -0.7112123 | 0.21883344 | -3.2500166 | 0.00115398 | 0.03021158 |
| TOB1 | 50.9444821 | 1.18642959 | 0.47442494 | 2.5007741 | 0.01239222 | 0.10925878 |
| SPAG9 | 283.954075 | -1.1464216 | 0.42493023 | -2.6979054 | 0.00697773 | 0.08149858 |
| AC005920.2 | 21.4443706 | -0.6506371 | 0.61172042 | -1.0636184 | 0.2875016 | 0.54680004 |
| MBTD1 | 160.152628 | -0.2506238 | 0.23270124 | -1.0770197 | 0.28147146 | 0.54013608 |
| UTP18 | 36.2449327 | -0.7850081 | 0.31431726 | -2.4975024 | 0.01250716 | 0.10951947 |
| CA10 | 436.179175 | -0.7361647 | 0.54973043 | -1.3391377 | 0.18052585 | 0.42977544 |
| TOM1L1 | 226.606374 | 0.66436253 | 0.28140547 | 2.36087286 | 0.01823198 | 0.13368407 |
| COX11 | 50.0465694 | 0.68698191 | 0.36403478 | 1.88713261 | 0.05914249 | 0.24149589 |
| STXBP4 | 253.507488 | 0.7608943 | 0.30191529 | 2.52022443 | 0.011728 | 0.10607572 |
| HLF | 206.655837 | 0.10282725 | 0.53203821 | 0.19327043 | 0.84674718 | 0.92563742 |
| PCTP | 15.7182648 | 0.16630447 | 0.44334417 | 0.3751137 | 0.70757591 | NA |
| DGKE | 33.8728807 | -0.1073696 | 0.41777922 | -0.2570008 | 0.79717812 | 0.90101028 |
| TRIM25 | 45.1414997 | -1.9889575 | 0.67278298 | -2.9563137 | 0.0031134 | 0.05496298 |
| COIL | 19.6079349 | -0.06006 | 0.43401781 | -0.1383814 | 0.88993901 | NA |
| SCPEP1 | 56.3284279 | 0.77409537 | 0.45952972 | 1.6845382 | 0.0920777 | 0.30404208 |
| AKAP1 | 56.9975396 | 1.07126181 | 0.34566726 | 3.09911273 | 0.00194101 | 0.04115693 |
| MSI2 | 1834.07305 | 0.44185881 | 0.3144349 | 1.40524737 | 0.15994773 | 0.40470667 |
| MRPS23 | 17.7973671 | -0.4382988 | 0.4331707 | -1.0118385 | 0.3116153 | NA |
| CUEDC1 | 122.95385 | -0.6966322 | 0.35109155 | -1.9841895 | 0.04723472 | 0.21733813 |
| VEZF1 | 56.4835038 | -0.0099589 | 0.31900673 | -0.0312185 | 0.97509525 | 0.99037778 |

| | | | | | | |
|---|---|---|---|---|---|---|
| AC015813.2 | 43.3966104 | -0.3891993 | 0.33137172 | -1.1745097 | 0.24019094 | 0.4977947 |
| SRSF1 | 19.4182778 | -0.1842761 | 0.49255196 | -0.3741252 | 0.70831114 | NA |
| DYNLL2 | 25.3990113 | 0.60770658 | 0.45888046 | 1.32432436 | 0.18539534 | 0.43545559 |
| MKS1 | 14.414477 | 0.34877717 | 0.50228862 | 0.69437602 | 0.48744643 | NA |
| TSPOAP1 | 28.3913509 | 2.27549123 | 0.80050359 | 2.84257468 | 0.00447507 | 0.06436412 |
| TSPOAP1-AS | 12.4402972 | 1.13537733 | 0.78074382 | 1.45422519 | 0.14588388 | NA |
| RNF43 | 79.4654693 | 1.04033878 | 0.41554811 | 2.50353389 | 0.01229599 | 0.10853468 |
| MTMR4 | 27.0498149 | 0.52486733 | 0.41085817 | 1.27749029 | 0.20142921 | 0.45559818 |
| RAD51C | 16.6862005 | 0.69363474 | 0.5182955 | 1.33829974 | 0.18079875 | NA |
| TRIM37 | 53.2769807 | -0.8474966 | 0.31869045 | -2.6593096 | 0.0078301 | 0.08671693 |
| SKA2 | 29.5762536 | 0.18082775 | 0.42994471 | 0.42058373 | 0.67405908 | 0.83715334 |
| GDPD1 | 19.7754152 | 1.10479502 | 0.47147968 | 2.34325055 | 0.01911654 | NA |
| YPEL2 | 99.168177 | 0.04125655 | 0.37683152 | 0.10948275 | 0.91281961 | 0.96312166 |
| DHX40 | 63.2344209 | -0.3131074 | 0.30806287 | -1.0163751 | 0.30945078 | 0.56913803 |
| CLTC | 222.925992 | -0.0326934 | 0.30852488 | -0.1059669 | 0.91560857 | 0.96388328 |
| PTRH2 | 33.4992639 | -0.2780947 | 0.34375443 | -0.8089923 | 0.41851956 | 0.66255555 |
| VMP1 | 629.645525 | -1.6739855 | 0.36050006 | -4.6435095 | 3.43E-06 | 0.00085049 |
| TUBD1 | 47.3716393 | -0.2064954 | 0.40337808 | -0.5119153 | 0.60871027 | 0.79586257 |
| RPS6KB1 | 64.1610585 | -0.7331959 | 0.27021056 | -2.7134245 | 0.00665918 | 0.07974991 |
| RNFT1 | 15.9840544 | 0.05967056 | 0.80245128 | 0.07436035 | 0.94072366 | NA |
| HEATR6 | 16.5502211 | 0.59093087 | 0.49669068 | 1.18973617 | 0.2341501 | NA |
| USP32 | 237.68277 | -1.2199483 | 0.2747916 | -4.4395398 | 9.02E-06 | 0.00147637 |
| APPBP2 | 169.405482 | -0.3809782 | 0.20412683 | -1.8663798 | 0.06198825 | 0.24884382 |
| AC011921.1 | 14.5631764 | 1.14871119 | 0.62246462 | 1.84542408 | 0.06497587 | NA |
| PPM1D | 62.0483937 | 0.08840467 | 0.39341257 | 0.22471236 | 0.82220305 | 0.91475524 |
| BCAS3 | 534.690059 | 0.21404862 | 0.29309203 | 0.73031199 | 0.4651995 | 0.69589923 |
| TBX2 | 21.9288761 | 0.75611109 | 0.63880993 | 1.18362452 | 0.23656173 | 0.4945181 |
| INTS2 | 37.4547986 | -0.0713742 | 0.34828507 | -0.2049303 | 0.83762655 | 0.92155682 |
| MED13 | 355.029352 | -0.5697769 | 0.29119618 | -1.956677 | 0.05038546 | 0.22527387 |
| METTL2A | 14.8829424 | 0.39849249 | 0.51865963 | 0.76831213 | 0.44230177 | NA |
| TLK2 | 160.505584 | -0.3706775 | 0.25394068 | -1.4597011 | 0.14437225 | 0.38371312 |
| TANC2 | 249.050446 | 0.13128543 | 0.21498453 | 0.61067385 | 5.41E-01 | 7.50E-01 |
| CYB561 | 15.0840376 | 0.32427934 | 0.51148207 | 0.63399943 | 0.52608119 | NA |
| DCAF7 | 106.880058 | -0.1029163 | 0.23610644 | -0.4358893 | 0.66291706 | 0.8285925 |
| MAP3K3 | 114.632889 | -0.0011159 | 0.3377891 | -0.0033035 | 0.99736422 | 0.99868755 |
| CCDC47 | 28.2872587 | -0.3819843 | 0.33520895 | -1.139541 | 0.25447759 | 0.51197417 |
| DDX42 | 84.5929248 | -0.5706275 | 0.29243791 | -1.9512775 | 0.05102405 | 0.22661979 |
| FTSJ3 | 15.1581894 | -0.5587096 | 0.58515657 | -0.9548036 | 0.33967702 | NA |
| PSMC5 | 13.8676801 | 0.13796619 | 0.58165342 | 0.23719657 | 0.8125043 | NA |
| SMARCD2 | 17.0139047 | -0.0519643 | 0.53877828 | -0.0964483 | 0.92316451 | NA |
| ERN1 | 57.9460516 | -1.3659257 | 0.4839125 | -2.8226707 | 0.00476255 | 0.06664966 |
| SNHG25 | 26.02168 | -0.7325834 | 0.50888343 | -1.4395898 | 0.14998349 | 0.3905355 |
| TEX2 | 121.67521 | -1.0883601 | 0.47370344 | -2.2975558 | 0.02158709 | 0.14349647 |
| POLG2 | 28.7035927 | 0.24791709 | 0.34176504 | 0.72540215 | 0.46820535 | 0.698271 |
| DDX5 | 400.956596 | 0.03248167 | 0.25871363 | 0.12555069 | 0.9000876 | 0.95597823 |
| CEP95 | 150.995042 | -0.5615135 | 0.27272392 | -2.058908 | 0.03950305 | 0.19795704 |
| SMURF2 | 195.372185 | -0.0994448 | 0.20263712 | -0.4907529 | 0.62360122 | 0.8044129 |
| LRRC37A3 | 146.392049 | 0.3444171 | 0.3738574 | 0.92125261 | 0.35691856 | 0.6128044 |
| GNA13 | 86.8273365 | -0.6586157 | 0.31299676 | -2.1042252 | 0.03535881 | 0.18721309 |
| CEP112 | 144.754543 | 1.48872878 | 0.45941789 | 3.24046758 | 0.00119334 | 0.03071948 |
| PRKCA | 397.203441 | -0.0944006 | 0.25936695 | -0.3639654 | 0.71588383 | 0.85868129 |
| HELZ | 279.081632 | -0.6520932 | 0.20348878 | -3.2045661 | 0.00135266 | 0.03314569 |
| PSMD12 | 58.1284476 | -0.5547165 | 0.36620856 | -1.5147558 | 0.12983427 | 0.3622089 |
| PITPNC1 | 426.084259 | -1.7763747 | 0.50094647 | -3.5460369 | 0.00039107 | 0.01710271 |

| | | | | | | |
|---|---|---|---|---|---|---|
| NOL11 | 26.1707227 | -0.1033336 | 0.42821749 | -0.241311 | 0.80931412 | 0.90756938 |
| BPTF | 358.638068 | -0.0094615 | 0.23958032 | -0.039492 | 0.96849811 | 0.98741848 |
| AC005332.7 | 16.9267985 | -0.3946481 | 0.44666971 | -0.8835345 | 0.37694756 | NA |
| ARSG | 44.9910089 | 0.46388441 | 0.4986785 | 0.93022741 | 0.35225335 | 0.60964858 |
| WIPI1 | 43.8825243 | -1.3961185 | 0.63525917 | -2.1977148 | 0.02796944 | 0.16477111 |
| PRKAR1A | 146.794283 | -0.0627758 | 0.33116571 | -0.18956 | 0.84965392 | 0.92776085 |
| FAM20A | 81.3801172 | 0.10555719 | 0.56513742 | 0.18678146 | 8.52E-01 | 9.29E-01 |
| LINC01482 | 139.091682 | 1.67238153 | 0.79570753 | 2.10175406 | 0.03557483 | 0.18754755 |
| ABCA9 | 75.8222178 | 0.45127302 | 0.57124517 | 0.78998132 | 0.42953868 | 0.67184702 |
| ABCA6 | 68.199887 | 0.30052393 | 0.49191149 | 0.61093091 | 0.54124532 | 0.74994873 |
| ABCA10 | 234.90249 | -0.2072885 | 0.37414441 | -0.5540333 | 5.80E-01 | 0.77674437 |
| ABCA5 | 662.9753 | -0.4310731 | 0.32046985 | -1.3451285 | 1.79E-01 | 4.28E-01 |
| MAP2K6 | 106.873722 | 1.59596694 | 0.5683299 | 2.80816996 | 0.00498239 | 0.06860371 |
| AC005208.1 | 323.775493 | 2.05900645 | 0.61121786 | 3.36869485 | 0.00075525 | 0.02446321 |
| KCNJ16 | 1225.1092 | 1.09761527 | 0.39618186 | 2.77048343 | 0.00559731 | 0.07302124 |
| SOX9 | 51.7438069 | 2.08354448 | 0.61045307 | 3.41311165 | 6.42E-04 | 2.31E-02 |
| LINC00511 | 104.890347 | 1.07760768 | 0.9725014 | 1.10807827 | 2.68E-01 | 5.27E-01 |
| SLC39A11 | 134.355356 | -0.1907811 | 0.34016758 | -0.5608443 | 0.57490366 | 0.77388388 |
| COG1 | 18.3253014 | 0.03266831 | 0.43549255 | 0.07501462 | 0.94020308 | NA |
| FAM104A | 26.6276116 | 0.55654239 | 0.36248083 | 1.53537054 | 0.12469284 | 0.35504829 |
| C17orf80 | 17.0987025 | 0.83872939 | 0.44151684 | 1.89965434 | 0.0574785 | NA |
| SDK2 | 55.342124 | 0.43174677 | 0.57967273 | 0.74481125 | 0.45638583 | 0.69129178 |
| RPL38 | 26.5844283 | 0.17969135 | 0.48896605 | 0.3674925 | 0.71325169 | 0.8577966 |
| GPRC5C | 52.985125 | -0.0059314 | 0.33662494 | -0.0176203 | 0.98594174 | 0.9942828 |
| SLC9A3R1 | 15.4467199 | -0.2290146 | 0.47876237 | -0.4783471 | 0.6324032 | NA |
| NAT9 | 14.4345014 | -0.4819508 | 0.52069805 | -0.925586 | 0.35466115 | NA |
| KCTD2 | 26.7612077 | 0.38451457 | 0.38143495 | 1.00807376 | 0.31341905 | 0.57102302 |
| ATP5PD | 26.8995788 | 0.63266642 | 0.38492289 | 1.64361857 | 0.10025502 | 0.31860564 |
| SLC16A5 | 51.8940393 | -1.0167498 | 0.57545672 | -1.7668572 | 0.07725215 | 0.27731248 |
| ARMC7 | 18.144184 | 0.31303767 | 0.44781387 | 0.69903523 | 0.48453002 | NA |
| NT5C | 26.4041634 | 0.01149929 | 0.38353106 | 0.02998267 | 0.97608087 | 0.99088558 |
| JPT1 | 15.1022013 | -0.0096182 | 0.47563756 | -0.0202218 | 0.98386646 | NA |
| SUMO2 | 23.9617978 | -0.8690648 | 0.45480903 | -1.9108345 | 0.05602585 | 0.23590316 |
| NUP85 | 28.1327882 | -0.4893117 | 0.35418604 | -1.3815104 | 0.16712209 | 0.41508187 |
| GGA3 | 36.5974228 | -0.7278543 | 0.34367927 | -2.11783 | 0.03418947 | 0.18312899 |
| MRPS7 | 18.4538176 | 0.61037553 | 0.46660697 | 1.30811489 | 0.19083434 | NA |
| GRB2 | 100.560204 | -0.3222639 | 0.25560483 | -1.2607894 | 0.20738472 | 0.46094144 |
| TMEM94 | 32.5904459 | 0.12008229 | 0.3352743 | 0.35816133 | 0.72022259 | 0.85999895 |
| CASKIN2 | 13.3345498 | -0.0770188 | 0.59994734 | -0.128376 | 0.89785141 | NA |
| TSEN54 | 24.4811586 | 0.74555602 | 0.49899566 | 1.49411323 | 0.13514603 | 0.36934198 |
| LLGL2 | 359.697866 | 0.87850614 | 0.29391716 | 2.98895828 | 0.0027993 | 0.0514788 |
| MYO15B | 40.0416124 | 0.09687639 | 0.52637668 | 0.18404385 | 0.85397906 | 0.93076703 |
| RECQL5 | 21.4622836 | -0.1219453 | 0.43191671 | -0.2823353 | 0.77768643 | 0.8896927 |
| SAP30BP | 52.1369458 | -0.5703869 | 0.2710197 | -2.1045957 | 0.03532652 | 0.18721309 |
| H3F3B | 121.772088 | -0.2747516 | 0.34260708 | -0.8019436 | 0.42258557 | 0.66611533 |
| UNK | 72.6076335 | -0.0715575 | 0.32827907 | -0.2179778 | 0.82744644 | 0.91638205 |
| WBP2 | 61.0825084 | -0.7468876 | 0.37880521 | -1.9716932 | 0.04864465 | 0.22128714 |
| ACOX1 | 53.0001955 | 0.14385052 | 0.31003535 | 0.46398101 | 0.64266135 | 0.81595394 |
| EVPL | 12.6283881 | 2.0663657 | 0.71586893 | 2.886514 | 0.00389535 | NA |
| SRP68 | 28.1164165 | -0.3618828 | 0.35883496 | -1.0084938 | 0.31321748 | 0.57102302 |
| EXOC7 | 73.1437404 | 0.41239376 | 0.34780718 | 1.18569653 | 0.23574217 | 0.49360921 |
| RNF157 | 18.2171013 | 0.35382533 | 0.58453828 | 0.60530737 | 0.54497478 | NA |
| UBALD2 | 27.7390095 | 0.22528635 | 0.37255775 | 0.60470181 | 0.54537714 | 0.75352691 |
| PRPSAP1 | 38.8813705 | -0.2195205 | 0.41304974 | -0.5314627 | 0.59509816 | 0.78824136 |

| | | | | | | |
|---|---|---|---|---|---|---|
| UBE2O | 63.873829 | -0.2641967 | 0.35899481 | -0.7359345 | 0.46177055 | 0.69416206 |
| RHBDF2 | 18.7238793 | 0.40188336 | 0.46715686 | 0.86027499 | 0.38963748 | NA |
| MXRA7 | 25.378383 | -0.0524527 | 0.45740526 | -0.1146744 | 0.90870319 | 0.95957056 |
| JMJD6 | 22.0654827 | -0.145591 | 0.50584474 | -0.2878176 | 0.77348636 | 0.88776363 |
| SRSF2 | 84.4587595 | -0.0685772 | 0.34721849 | -0.1975046 | 0.8434327 | 0.92437726 |
| MFSD11 | 53.4700439 | -0.3253083 | 0.36526674 | -0.8906048 | 0.37314124 | 0.62572291 |
| SEC14L1 | 111.239083 | -0.078284 | 0.2872079 | -0.2725692 | 0.78518436 | 0.89452315 |
| 9-Sep | 242.575365 | -0.352894 | 0.24073128 | -1.465925 | 1.43E-01 | 3.80E-01 |
| TNRC6C | 120.114783 | -0.4093812 | 0.35808984 | -1.1432359 | 0.25294067 | 0.51005615 |
| SYNGR2 | 21.3585477 | 0.55369802 | 0.61777864 | 0.89627252 | 0.37010723 | 0.62368335 |
| AFMID | 71.2364045 | 1.10624206 | 0.42166125 | 2.62353264 | 8.70E-03 | 9.18E-02 |
| PGS1 | 42.7259033 | -0.7850635 | 0.34939544 | -2.2469198 | 0.02464515 | 0.15380786 |
| CYTH1 | 101.033254 | -0.2007313 | 0.29216978 | -0.6870365 | 0.49205972 | 0.71672078 |
| USP36 | 97.5185832 | -1.6564784 | 0.58623098 | -2.8256412 | 0.00471861 | 0.06664966 |
| TIMP2 | 32.3106344 | -0.4646332 | 0.60890065 | -0.7630689 | 0.4454223 | 0.68538685 |
| LGALS3BP | 31.5118732 | 1.29361783 | 0.71042217 | 1.82091422 | 0.0686199 | 0.26056603 |
| CANT1 | 10.8080984 | 0.7076986 | 0.58873499 | 1.20206649 | 0.22933777 | NA |
| ENGASE | 12.825683 | 0.05623613 | 0.60510426 | 0.09293626 | 0.9259542 | NA |
| TBC1D16 | 49.862539 | 0.45705806 | 0.33684397 | 1.3568836 | 0.1748182 | 0.42340329 |
| CCDC40 | 21.1425055 | 0.52818309 | 0.47607101 | 1.10946283 | 0.26723057 | NA |
| GAA | 20.3980315 | 0.60256173 | 0.4398043 | 1.37006784 | 0.17066573 | NA |
| EIF4A3 | 15.6976328 | -0.2131203 | 0.43216308 | -0.4931479 | 0.62190808 | NA |
| AC087741.1 | 15.2276444 | -0.6635501 | 0.64345484 | -1.0312303 | 0.30243282 | NA |
| SLC26A11 | 37.5425463 | -0.6063094 | 0.42113836 | -1.4396918 | 0.14995463 | 0.3905355 |
| RNF213 | 411.65506 | -0.188381 | 0.5191468 | -0.3628666 | 0.71670456 | 0.8589283 |
| AC124319.2 | 276.375018 | 0.04207884 | 0.34521359 | 0.12189218 | 0.90298441 | 0.95661107 |
| ENDOV | 28.3717818 | -0.1083844 | 0.3440992 | -0.31498 | 0.75277683 | 0.87842978 |
| RPTOR | 153.980767 | -0.3668656 | 0.26577093 | -1.3803825 | 0.1674689 | 0.41567497 |
| BAIAP2 | 14.9207084 | -1.0044369 | 0.44626266 | -2.2507752 | 0.02439978 | NA |
| TEPSIN | 13.7306589 | -0.2243146 | 0.56413694 | -0.3976244 | 0.69090705 | NA |
| NDUFAF8 | 17.5609012 | 1.10873344 | 0.55721745 | 1.98976799 | 0.0466165 | NA |
| SLC38A10 | 63.2416124 | 0.70423892 | 0.31420671 | 2.24132359 | 0.02500512 | 0.15492746 |
| BAHCC1 | 16.7051917 | 0.07865735 | 0.52060407 | 0.15108862 | 0.87990581 | NA |
| ACTG1 | 159.07964 | -0.2164348 | 0.57824161 | -0.3742982 | 0.70818248 | 0.85517423 |
| NPLOC4 | 116.10128 | -0.5366513 | 0.24888265 | -2.1562423 | 0.03106475 | 0.17412703 |
| CCDC137 | 19.1243901 | -0.9022611 | 0.53134597 | -1.6980672 | 0.08949508 | NA |
| ARL16 | 19.711302 | 0.52768355 | 0.45770451 | 1.15289132 | 0.24895499 | NA |
| HGS | 45.0765097 | -1.1078708 | 0.33531191 | -3.3040007 | 0.00095316 | 0.02829008 |
| GCGR | 54.1805303 | 0.95308199 | 0.56822006 | 1.67731142 | 0.0934816 | 0.30696582 |
| MCRIP1 | 21.9098855 | -0.3715619 | 0.44698632 | -0.8312601 | 0.40582671 | 0.65318894 |
| P4HB | 85.2551624 | -0.5540216 | 0.37582408 | -1.4741513 | 0.14044086 | 0.37835958 |
| ARHGDIA | 33.2642858 | -0.67389 | 0.5574768 | -1.2088216 | 0.22673138 | 0.48449513 |
| ANAPC11 | 22.86228 | 0.34790844 | 0.48582018 | 0.71612595 | 0.47391358 | 0.7026171 |
| SIRT7 | 16.456418 | -0.2154443 | 0.50652125 | -0.425341 | 0.67058811 | NA |
| MAFG | 17.5027308 | -0.3596702 | 0.76759428 | -0.4685681 | 0.63937838 | NA |
| ASPSCR1 | 25.2365383 | -0.8795234 | 0.50762465 | -1.7326256 | 0.08316224 | 0.2884298 |
| DCXR | 13.9312729 | 0.29174688 | 0.63016254 | 0.46297085 | 0.64338526 | NA |
| RFNG | 15.8186421 | 0.7557773 | 0.52145826 | 1.44935338 | 0.14723892 | NA |
| GPS1 | 15.8315438 | 0.67873332 | 0.6474701 | 1.04828521 | 0.29450722 | NA |
| DUS1L | 20.0437567 | -0.4256197 | 0.52860795 | -0.8051708 | 0.42072114 | NA |
| CCDC57 | 114.356595 | -0.0931929 | 0.28195843 | -0.33052 | 0.74100711 | 0.87152705 |
| SLC16A3 | 23.6367116 | -0.465479 | 0.4406514 | -1.056343 | 0.29081155 | 0.55011098 |
| CSNK1D | 87.8520637 | -0.8734682 | 0.47017888 | -1.8577359 | 0.06320652 | 0.25056001 |
| OGFOD3 | 22.1967363 | -0.1602195 | 0.49444027 | -0.3240422 | 0.74590612 | 0.87409184 |

| | | | | | | |
|---|---|---|---|---|---|---|
| HEXDC | 47.6733748 | -0.4792526 | 0.38131224 | -1.2568508 | 0.20880767 | 0.46316791 |
| CYBC1 | 16.7798416 | 0.14939092 | 0.52391993 | 0.28514075 | 0.77553631 | NA |
| NARF | 44.2542946 | -0.5963113 | 0.39486629 | -1.5101601 | 0.13100257 | 0.36362306 |
| FOXK2 | 134.105733 | -0.4249658 | 0.26972637 | -1.5755442 | 0.11513088 | 0.34138376 |
| WDR45B | 95.1013249 | -1.0825 | 0.36271501 | -2.9844367 | 0.00284101 | 0.05198675 |
| RAB40B | 59.4441319 | 0.53319488 | 0.30078343 | 1.77268699 | 0.07628059 | 0.27590775 |
| FN3KRP | 16.0418421 | -0.7307235 | 0.4952146 | -1.4755694 | 0.14005953 | NA |
| FN3K | 43.5778235 | 1.19160815 | 0.6129033 | 1.94420255 | 0.05187103 | 0.22838257 |
| TBCD | 57.1951813 | -0.2553622 | 0.33987608 | -0.7513392 | 0.45244853 | 0.68905745 |
| METRNL | 26.0787633 | -1.1515923 | 0.62574157 | -1.8403641 | 0.0657148 | 0.25507153 |
| USP14 | 101.934721 | -0.3190641 | 0.24927559 | -1.2799651 | 0.20055739 | 0.45454417 |
| THOC1 | 91.8029111 | -0.4941179 | 0.25525472 | -1.9357834 | 0.05289423 | 0.23105954 |
| ENOSF1 | 339.095206 | -0.1950934 | 0.36832342 | -0.5296796 | 0.59633413 | 0.78913289 |
| YES1 | 106.715694 | -0.8939473 | 0.2811464 | -3.1796506 | 0.00147453 | 0.03500548 |
| METTL4 | 33.7003736 | 0.22463404 | 0.3908486 | 0.57473415 | 0.56547111 | 0.76965382 |
| SMCHD1 | 216.966351 | -0.4119757 | 0.23835891 | -1.7283838 | 0.08391944 | 0.28887652 |
| AP001011.1 | 28.7160347 | 0.25286736 | 0.67760876 | 0.37317605 | 0.70901742 | 0.85591391 |
| LPIN2 | 99.0827078 | -0.5909727 | 0.38717281 | -1.5263796 | 0.12691535 | 0.35861434 |
| MYOM1 | 38.7513557 | 0.59535678 | 0.54424059 | 1.09392206 | 2.74E-01 | 5.33E-01 |
| MYL12A | 53.9922919 | -0.1718256 | 0.33958214 | -0.5059913 | 0.61286273 | 0.79871392 |
| MYL12B | 103.944079 | 0.20930087 | 0.32570905 | 0.64260072 | 0.52048322 | 0.73723913 |
| TGIF1 | 43.8036931 | -1.6730906 | 0.49399405 | -3.3868638 | 0.00070696 | 0.02388192 |
| DLGAP1 | 133.583735 | 1.34184344 | 0.81285992 | 1.65076837 | 9.88E-02 | 0.31647454 |
| DLGAP1-AS1 | 17.1977111 | -0.2563406 | 0.44349144 | -0.5780058 | 0.56326019 | NA |
| LINC00667 | 18.6641951 | -0.1235386 | 0.47458588 | -0.2603083 | 0.79462599 | NA |
| EPB41L3 | 225.650913 | 0.26636935 | 0.36135266 | 0.73714512 | 0.4610341 | 0.69416206 |
| L3MBTL4 | 232.197717 | 0.54116458 | 0.27427427 | 1.97307822 | 0.04848665 | 0.22069885 |
| LAMA1 | 254.980962 | -0.0849095 | 0.35916655 | -0.2364071 | 0.81311681 | 0.90970797 |
| PTPRM | 458.964043 | 0.042762 | 0.31466916 | 0.1358951 | 0.89190421 | 0.95043427 |
| RAB12 | 68.4162598 | 0.01113302 | 0.25714859 | 0.0432941 | 0.96546709 | 0.98701025 |
| MTCL1 | 39.6483347 | 0.10364918 | 0.34597092 | 0.29958927 | 0.76449047 | 0.88317118 |
| NDUFV2 | 91.7101791 | -0.5600215 | 0.28126218 | -1.9911014 | 0.04646974 | 0.21595574 |
| ANKRD12 | 455.063416 | -0.4408164 | 0.19890358 | -2.2162318 | 0.02667563 | 0.16052735 |
| TWSG1 | 59.6716674 | -0.3374551 | 0.36323489 | -0.9290272 | 0.35287499 | 0.61004394 |
| RALBP1 | 83.4524365 | 0.70482343 | 0.2495094 | 2.82483718 | 0.00473046 | 0.06664966 |
| PPP4R1 | 141.457044 | -0.3969026 | 0.31264929 | -1.269482 | 0.2042692 | 0.45771763 |
| RAB31 | 84.0582747 | -1.6847101 | 0.66867225 | -2.5194855 | 0.01175265 | 0.10617385 |
| VAPA | 243.54687 | -0.1947868 | 0.27750299 | -0.7019269 | 0.48272475 | 0.71042684 |
| NAPG | 60.0108911 | 0.00553863 | 0.27725571 | 0.01997662 | 0.98406202 | 0.99382437 |
| GNAL | 33.6092174 | 1.12598615 | 0.49846151 | 2.25892295 | 0.02388818 | 0.15095838 |
| MPPE1 | 47.6404365 | 0.48210865 | 0.41768741 | 1.15423315 | 0.24840459 | 0.50492395 |
| IMPA2 | 112.785183 | -0.015462 | 0.32717626 | -0.0472589 | 0.96230691 | 0.98535004 |
| AFG3L2 | 116.063878 | -0.4094387 | 0.29754368 | -1.3760626 | 0.16880224 | 0.41629954 |
| SPIRE1 | 491.025228 | -0.8548681 | 0.3452216 | -2.476288 | 0.01327564 | 0.11268704 |
| PSMG2 | 65.5698461 | -0.1947517 | 0.3470332 | -0.5611903 | 0.57466778 | 0.77388388 |
| CEP76 | 19.724143 | 0.4699266 | 0.52216721 | 0.89995424 | 0.3681446 | NA |
| PTPN2 | 171.229068 | -0.6662001 | 0.23115663 | -2.882029 | 0.00395123 | 0.06143969 |
| SEH1L | 30.3034553 | -0.515276 | 0.33932809 | -1.5185186 | 0.12888371 | 0.3611257 |
| CEP192 | 158.217401 | 0.21500467 | 0.31005952 | 0.69343031 | 0.48803955 | 0.71483166 |
| LDLRAD4 | 74.6956451 | -0.2568137 | 0.54877869 | -0.4679732 | 0.63980377 | 0.81424762 |
| FAM210A | 64.2489432 | -1.0611088 | 0.30381342 | -3.4926331 | 0.00047828 | 0.01968634 |
| RNMT | 100.962185 | -0.7865474 | 0.35302177 | -2.2280423 | 0.02587769 | 0.15729431 |
| ROCK1 | 215.819897 | -0.2646194 | 0.20972985 | -1.2617154 | 0.2070512 | 0.46066753 |
| ESCO1 | 95.0018386 | -0.2876941 | 0.23894622 | -1.204012 | 0.22858493 | 0.48696878 |

| | | | | | | |
|---|---|---|---|---|---|---|
| SNRPD1 | 16.7309531 | 0.40610801 | 0.41873049 | 0.96985536 | 0.3321186 | NA |
| ABHD3 | 101.126332 | -1.308707 | 0.87429983 | -1.496863 | 0.13442892 | 0.36835151 |
| MIB1 | 197.726178 | -0.7673775 | 0.43974 | -1.745071 | 0.08097252 | 0.28382032 |
| AC090912.1 | 24.6123446 | 0.6478779 | 0.55138478 | 1.17500142 | 0.23999415 | 0.49763873 |
| RBBP8 | 429.426732 | -0.3670559 | 0.34361517 | -1.0682179 | 0.28542224 | 0.54418999 |
| CABLES1 | 195.555523 | -0.4618695 | 0.39498306 | -1.1693401 | 0.24226665 | 0.49963243 |
| TMEM241 | 53.2422837 | 0.73141202 | 0.40987595 | 1.78447169 | 0.07434705 | 0.27199309 |
| RIOK3 | 110.630633 | -0.5535999 | 0.20671774 | -2.6780474 | 0.00740527 | 0.08440905 |
| RMC1 | 32.2520894 | -0.6283778 | 0.36075057 | -1.7418621 | 0.08153257 | 0.28473512 |
| NPC1 | 177.428907 | -1.9203843 | 0.57288716 | -3.3521161 | 0.00080196 | 0.02550709 |
| TTC39C | 66.9014614 | 0.11213249 | 0.40220793 | 0.27879234 | 0.78040419 | 0.89081582 |
| OSBPL1A | 79.0687865 | 0.56329251 | 0.31532977 | 1.78636007 | 0.07404098 | 0.27137781 |
| IMPACT | 15.1127236 | -0.5209609 | 0.58306228 | -0.893491 | 0.3715943 | NA |
| SS18 | 158.351567 | 0.00916542 | 0.22399391 | 0.04091817 | 0.96736113 | 0.98741848 |
| TAF4B | 61.8065471 | 0.61626233 | 0.32334328 | 1.90590736 | 0.05666223 | 0.23723351 |
| KCTD1 | 501.972353 | 1.49920511 | 0.35090951 | 4.27234108 | 1.93E-05 | 0.00240136 |
| AQP4-AS1 | 45.5980512 | 1.51247839 | 0.64300495 | 2.35220333 | 0.01866257 | 0.13513246 |
| CHST9 | 54.786834 | -0.0657274 | 0.66939995 | -0.0981886 | 0.92178256 | 0.96635254 |
| DSC2 | 16.4928483 | 0.47257563 | 0.44360212 | 1.06531417 | 0.28673378 | NA |
| DSG2 | 52.439812 | 0.98010353 | 0.35182339 | 2.78578276 | 0.00533986 | 0.07122532 |
| B4GALT6 | 29.4742691 | 0.30592077 | 0.46789653 | 0.65382142 | 0.51322687 | 0.73153837 |
| TRAPPC8 | 139.309247 | -0.1649161 | 0.26252737 | -0.6281862 | 0.52988196 | 0.74381523 |
| RNF138 | 54.2205175 | -0.6904954 | 0.34740761 | -1.9875655 | 0.04685977 | 0.21661307 |
| GAREM1 | 638.28719 | -0.5113331 | 0.46438119 | -1.1011063 | 0.27085038 | 0.52912064 |
| KLHL14 | 37.3212907 | 0.6194034 | 0.46395377 | 1.33505414 | 0.18185863 | 0.43122793 |
| CCDC178 | 294.946367 | 1.18182648 | 0.75652706 | 1.56217345 | 0.11824713 | 0.34566454 |
| ASXL3 | 125.785149 | 1.1563988 | 0.39037366 | 2.96228694 | 0.00305363 | 0.05464622 |
| DTNA | 50.5038888 | -0.3374717 | 0.55502592 | -0.6080287 | 0.54316844 | 0.75180139 |
| MAPRE2 | 43.7441009 | -0.9784997 | 0.48881755 | -2.0017688 | 0.0453096 | 0.21333202 |
| ZNF397 | 57.5437864 | 0.09805678 | 0.31616099 | 0.31014826 | 0.75644821 | 0.87904624 |
| ZSCAN30 | 43.1800249 | 0.51983745 | 0.3696277 | 1.40638121 | 0.15961096 | 0.40445247 |
| ZNF24 | 45.9784277 | -0.0302605 | 0.36406941 | -0.0831174 | 0.93375823 | 0.9730757 |
| GALNT1 | 68.5380714 | 0.40369204 | 0.26965809 | 1.49705146 | 0.13437987 | 0.36834824 |
| RPRD1A | 168.053368 | -0.2139795 | 0.23860919 | -0.896778 | 0.3698374 | 0.62368335 |
| SLC39A6 | 26.9240633 | 0.22268658 | 0.5214264 | 0.42707193 | 0.66932694 | 0.83376893 |
| ELP2 | 97.8953796 | -0.1391024 | 0.20384877 | -0.6823806 | 0.49499834 | 0.719004 |
| MOCOS | 26.0843193 | -0.2813047 | 0.62044834 | -0.4533894 | 0.65026834 | 0.81957023 |
| FHOD3 | 829.39836 | -0.1681398 | 0.39395532 | -0.4267992 | 0.6695256 | 0.83376893 |
| TPGS2 | 43.0733334 | 0.50968186 | 0.38953313 | 1.3084429 | 0.19072313 | 0.44216745 |
| KIAA1328 | 194.377851 | -0.0170028 | 0.34407144 | -0.0494166 | 0.96058733 | 0.98489952 |
| PIK3C3 | 164.081496 | -0.094108 | 0.22667155 | -0.4151733 | 0.67801506 | 0.83969138 |
| LINC00907 | 35.3407536 | 0.11271721 | 0.47869968 | 0.2354654 | 0.81384753 | 0.90970797 |
| SETBP1 | 597.932043 | 0.12406281 | 0.30620125 | 0.40516753 | 0.68535437 | 0.84355496 |
| SLC14A2 | 17.0227508 | 0.4923612 | 0.58235121 | 0.84547123 | 0.39784779 | NA |
| EPG5 | 103.981424 | -0.7460872 | 0.37529491 | -1.9880024 | 0.04681143 | 0.21661307 |
| PSTPIP2 | 28.9659132 | 0.6798526 | 0.55249738 | 1.23050827 | 0.21850683 | 0.47523229 |
| ATP5F1A | 98.898459 | 0.82871972 | 0.35123376 | 2.35945352 | 0.01830187 | 0.13368407 |
| C18orf25 | 74.7119064 | -0.1999802 | 0.32603113 | -0.6133777 | 0.53962666 | 0.74929145 |
| ST8SIA5 | 18.0139415 | 0.56605849 | 0.43109904 | 1.31305903 | 0.18916306 | NA |
| PIAS2 | 247.890538 | -0.2305203 | 0.29102187 | -0.7921064 | 0.42829865 | 0.67072527 |
| KATNAL2 | 112.415045 | 0.5331291 | 0.45392382 | 1.17449024 | 0.24019872 | 0.4977947 |
| SMAD2 | 268.162279 | -0.4090512 | 0.38503403 | -1.0623767 | 0.28806469 | 0.54746482 |
| CTIF | 100.879802 | 0.30411796 | 0.26199958 | 1.16075745 | 0.24574055 | 0.50278177 |
| SMAD7 | 38.781539 | -0.1262147 | 0.55256687 | -0.2284152 | 0.81932345 | 0.913067 |

| | | | | | | |
|---|---|---|---|---|---|---|
| DYM | 365.996004 | -0.4142501 | 0.20146238 | -2.0562157 | 0.03976173 | 0.19847343 |
| LIPG | 89.2066448 | -1.7294979 | 0.72547626 | -2.3839484 | 0.01712801 | 0.12988597 |
| ACAA2 | 28.4401492 | 1.11511989 | 0.44046587 | 2.53168285 | 0.01135166 | 0.103737 |
| SNHG22 | 30.1270905 | -0.0755684 | 0.34489814 | -0.2191035 | 0.82656944 | 0.9158061 |
| MYO5B | 135.580155 | -0.3903521 | 0.40135238 | -0.972592 | 0.33075612 | 0.58655523 |
| MBD1 | 28.9067498 | 0.14890045 | 0.34087283 | 0.43682111 | 0.66224109 | 0.82847174 |
| CXXC1 | 14.9286669 | 0.13860195 | 0.53240954 | 0.26032958 | 0.79460956 | NA |
| MAPK4 | 56.0761276 | 1.10335634 | 0.58333465 | 1.89146376 | 0.05856246 | 0.24020328 |
| ME2 | 39.1979868 | 0.06732895 | 0.35624123 | 0.18899821 | 0.85009422 | 0.92810996 |
| AC091551.1 | 18.7552744 | 0.9134412 | 0.71399961 | 1.2793301 | 0.20078084 | NA |
| SMAD4 | 65.7082873 | -0.1262028 | 0.27158023 | -0.4646981 | 0.64214769 | 0.81594508 |
| MEX3C | 17.4890973 | 0.55865295 | 0.42655131 | 1.30969694 | 0.19029838 | NA |
| MBD2 | 153.769052 | -1.1241801 | 0.25683968 | -4.3769719 | 1.20E-05 | 0.00187417 |
| POLI | 53.0294835 | 0.18413178 | 0.40053693 | 0.45971237 | 0.64572269 | 0.81777273 |
| TCF4 | 26.234686 | -0.1026392 | 0.50982432 | -0.2013227 | 8.40E-01 | 0.92355925 |
| TXNL1 | 64.6985517 | -0.4708605 | 0.3105158 | -1.5163818 | 0.12942284 | 0.36208459 |
| WDR7 | 187.994924 | -0.1345467 | 0.19869263 | -0.6771599 | 0.49830453 | 0.72149172 |
| FECH | 37.6701076 | 1.10226305 | 0.44872584 | 2.45642874 | 0.01403256 | 0.11638862 |
| NARS | 69.3813539 | 0.37089821 | 0.31185817 | 1.18931695 | 0.23431497 | 0.49243138 |
| AC027097.2 | 52.1058287 | -0.2356016 | 0.40666266 | -0.5793539 | 0.5623504 | 0.76709958 |
| ATP8B1 | 102.286387 | -0.0603293 | 0.32125319 | -0.1877937 | 0.85103841 | 0.92874559 |
| NEDD4L | 991.339804 | -1.3117959 | 0.38724313 | -3.3875253 | 0.00070526 | 0.02388192 |
| MALT1 | 54.8373534 | -0.1668068 | 0.37666735 | -0.4428492 | 0.65787482 | 0.82521413 |
| ZNF532 | 198.904882 | -0.6588922 | 0.28854346 | -2.2835109 | 0.02240029 | 0.14604538 |
| SEC11C | 13.0930776 | 0.42857371 | 0.52314004 | 0.81923324 | 0.41265335 | NA |
| LMAN1 | 63.8245675 | 0.46203542 | 0.33572863 | 1.37621694 | 0.16875448 | 0.41629954 |
| RNF152 | 125.005681 | 1.11911778 | 0.39576874 | 2.82770636 | 0.00468828 | 0.06664966 |
| PIGN | 136.848207 | 0.12367329 | 0.21858592 | 0.56578801 | 0.5715379 | 0.77245539 |
| KIAA1468 | 154.564794 | -0.1027417 | 0.19254417 | -0.5336005 | 0.59361797 | 0.78655148 |
| ZCCHC2 | 71.5747917 | 0.15180689 | 0.25294757 | 0.6001516 | 0.5484052 | 0.7554786 |
| PHLPP1 | 107.309737 | -0.3327633 | 0.24140706 | -1.3784324 | 0.16806982 | 0.41617965 |
| BCL2 | 130.721131 | 0.1071356 | 0.49925666 | 0.21459022 | 0.83008682 | 0.91772835 |
| KDSR | 63.7467229 | -0.8053182 | 0.34876211 | -2.3090758 | 0.02093937 | 0.14200031 |
| VPS4B | 67.0571196 | -0.2747896 | 0.32347003 | -0.8495055 | 0.3956001 | 0.64402157 |
| TMX3 | 76.3401515 | -0.5188658 | 0.27320093 | -1.8992093 | 0.05753696 | 0.23835412 |
| CCDC102B | 35.8905843 | 1.56601664 | 0.58354551 | 2.68362386 | 0.0072829 | 0.0836059 |
| RTTN | 58.3317101 | -0.1732732 | 0.39424063 | -0.4395113 | 0.66029111 | 0.82678716 |
| SOCS6 | 35.7565611 | 0.08776675 | 0.40422464 | 0.21712371 | 0.82811193 | 0.91685523 |
| TIMM21 | 13.9062255 | 0.44685663 | 0.51649484 | 0.86517153 | 0.38694465 | NA |
| CYB5A | 90.8127135 | 1.31008318 | 0.42716858 | 3.06689966 | 0.00216291 | 0.04403601 |
| CNDP2 | 86.1481809 | 0.55971711 | 0.32190235 | 1.73877921 | 0.08207361 | 0.28580436 |
| LINC00909 | 12.9688489 | -0.5471154 | 0.58597163 | -0.9336893 | 0.35046418 | NA |
| ZNF407 | 300.036067 | -0.2675906 | 0.29308434 | -0.9130156 | 0.36123431 | 0.61731848 |
| ZADH2 | 30.3690879 | 1.40704182 | 0.42062547 | 3.34511801 | 0.00082248 | 0.02565549 |
| TSHZ1 | 80.2176098 | 0.40860401 | 0.29522378 | 1.38404843 | 0.16634362 | 0.41429299 |
| ZNF516 | 190.935182 | -0.5168586 | 0.34687307 | -1.490051 | 0.13621084 | 0.37112029 |
| ZNF236 | 66.511629 | -0.1464779 | 0.30209152 | -0.4848793 | 0.62776196 | 0.80601257 |
| MBP | 188.223007 | -1.0556732 | 0.3838753 | -2.7500421 | 0.00595876 | 0.07594487 |
| LINC01896 | 73.0762692 | 0.98290752 | 0.41346637 | 2.3772369 | 0.01744288 | 0.13136777 |
| SALL3 | 113.318949 | 0.71895678 | 0.32526805 | 2.21035167 | 0.02708077 | 0.16195658 |
| ATP9B | 353.221873 | -0.0266069 | 0.21331219 | -0.1247323 | 0.90073548 | 0.95621962 |
| NFATC1 | 31.2576387 | 0.94120558 | 0.37660533 | 2.49918283 | 0.01244801 | 0.10951947 |
| CTDP1 | 30.2549134 | -0.547261 | 0.42030612 | -1.3020533 | 0.19289815 | 0.44466519 |
| PQLC1 | 48.7550487 | 0.23864134 | 0.38521724 | 0.61949808 | 0.53558829 | 0.74789968 |

| | | | | | | |
|---|---|---|---|---|---|---|
| HSBP1L1 | 58.5481748 | -0.2102234 | 0.28383361 | -0.740657 | 0.45890142 | 0.6928053 |
| TXNL4A | 59.5646939 | 0.01338445 | 0.3223868 | 0.04151675 | 0.96688394 | 0.98741848 |
| ADNP2 | 91.3323503 | -0.9582076 | 0.52240737 | -1.8342153 | 0.06662204 | 0.25768334 |
| PARD6G-AS1 | 16.9897849 | -0.9792342 | 0.67994247 | -1.4401722 | 0.14981869 | NA |
| C20orf96 | 13.2886312 | 1.80044412 | 0.71152348 | 2.53040718 | 0.01139302 | NA |
| NRSN2-AS1 | 87.7565417 | -0.3252337 | 0.57900706 | -0.5617094 | 0.57431404 | 0.77388388 |
| SOX12 | 13.9908767 | 1.50599196 | 0.55574473 | 2.70986278 | 0.00673111 | NA |
| RBCK1 | 45.4363105 | 0.31564632 | 0.36581518 | 0.86285735 | 0.3882159 | 0.63752887 |
| TBC1D20 | 29.2986658 | -0.4590234 | 0.34477529 | -1.3313697 | 0.18306738 | 0.43276095 |
| CSNK2A1 | 191.887165 | 0.0186986 | 0.228225 | 0.08193055 | 0.93470194 | 0.97332915 |
| PSMF1 | 72.4590541 | -0.5240213 | 0.23140005 | -2.2645687 | 0.02353916 | 0.14961264 |
| AL136531.2 | 41.2376986 | -1.2631218 | 0.43128666 | -2.9287292 | 0.00340351 | 0.05757539 |
| SDCBP2-AS1 | 48.0662817 | -0.1588003 | 0.3954526 | -0.4015661 | 0.6880034 | 0.84539626 |
| NSFL1C | 35.1508999 | -0.403405 | 0.29313324 | -1.3761832 | 0.16876493 | 0.41629954 |
| STK35 | 32.3280777 | -0.3524752 | 0.34249735 | -1.0291328 | 0.3034173 | 0.56397076 |
| SNRPB | 25.3787788 | -0.0053688 | 0.48414484 | -0.0110892 | 0.99115231 | 0.99583841 |
| NOP56 | 19.6898449 | -1.3627147 | 0.54767129 | -2.4881982 | 0.01283922 | NA |
| IDH3B | 15.3063305 | 0.1317648 | 0.60492793 | 0.21781901 | 0.82757013 | NA |
| EBF4 | 20.4602817 | -0.8604438 | 0.7924731 | -1.0857704 | 0.27758059 | NA |
| PCED1A | 22.525317 | -0.0488929 | 0.42321871 | -0.1155263 | 0.90802794 | 0.95924939 |
| PTPRA | 187.5914 | -0.0864678 | 0.19550182 | -0.4422865 | 0.65828193 | 0.82521413 |
| MRPS26 | 16.4559202 | 0.87772383 | 0.46508363 | 1.88723869 | 0.05912823 | NA |
| UBOX5-AS1 | 14.0824187 | 0.34667289 | 0.65428288 | 0.52985169 | 0.59621476 | NA |
| UBOX5 | 21.6279575 | -0.5035772 | 0.45120056 | -1.1160828 | 0.26438671 | 0.5237724 |
| LZTS3 | 19.2048881 | 0.442412 | 0.39733761 | 1.11344104 | 0.26551906 | NA |
| DDRGK1 | 21.6262797 | 0.06742415 | 0.45879876 | 0.146958 | 0.88316517 | 0.94552272 |
| ITPA | 35.6720842 | 0.00548574 | 0.46842035 | 0.01171115 | 0.99065607 | 0.99583841 |
| C20orf194 | 544.797231 | 0.62248022 | 0.24578135 | 2.5326585 | 0.01132012 | 0.1036885 |
| ATRN | 257.665954 | 0.20180757 | 0.21389924 | 0.94347026 | 3.45E-01 | 6.03E-01 |
| MAVS | 76.1874478 | 0.02818767 | 0.29606729 | 0.09520697 | 0.92415043 | 0.96791208 |
| PANK2 | 63.9644371 | -0.3574022 | 0.32365212 | -1.1042789 | 0.26947217 | 0.52764898 |
| RNF24 | 225.17304 | -0.3775175 | 0.40184479 | -0.939461 | 0.34749408 | 0.60403983 |
| PRNP | 19.066482 | -1.4364846 | 0.67713666 | -2.1214102 | 0.0338873 | NA |
| SLC23A2 | 45.7359261 | 0.41726822 | 0.3295642 | 1.26612119 | 0.2054697 | 0.45886886 |
| TMEM230 | 48.7913295 | 0.51138679 | 0.37669808 | 1.35755082 | 0.17460626 | 0.42330339 |
| CDS2 | 65.293708 | -0.6801037 | 0.35463091 | -1.917779 | 0.05513904 | 0.23466461 |
| GPCPD1 | 116.724815 | -0.046265 | 0.38224012 | -0.1210366 | 0.90366205 | 0.95671001 |
| SHLD1 | 41.4265103 | -0.1513002 | 0.32896278 | -0.4599311 | 0.64556565 | 0.81777273 |
| MCM8 | 15.466511 | 0.31245341 | 0.50766958 | 0.6154661 | 0.53824695 | NA |
| CRLS1 | 70.5072514 | -0.5290271 | 0.35988902 | -1.4699728 | 0.14156912 | 0.37953937 |
| TMX4 | 43.0265497 | 0.10639309 | 0.4522257 | 0.23526548 | 0.81400269 | 0.90970797 |
| PLCB1 | 113.4183 | 0.44364219 | 0.76040472 | 0.58342903 | 0.55960452 | 0.76507463 |
| PLCB4 | 264.092159 | 0.36673675 | 0.49245905 | 0.74470508 | 0.45645003 | 0.69129178 |
| MKKS | 16.4918961 | 0.00427786 | 0.47083996 | 0.0090856 | 0.99275084 | NA |
| SLX4IP | 152.777137 | 0.49085478 | 0.34716446 | 1.413897 | 0.15739215 | 0.40194007 |
| JAG1 | 101.877012 | 0.34946253 | 0.54090213 | 0.6460735 | 0.51823175 | 0.73536015 |
| AL050403.2 | 41.5406546 | -0.3284218 | 0.67187176 | -0.4888162 | 0.62497185 | 0.8053173 |
| BTBD3 | 145.940244 | -0.2938086 | 0.24929007 | -1.1785813 | 0.23856494 | 0.49653323 |
| SPTLC3 | 73.0636512 | 0.05812151 | 0.45933294 | 0.1265346 | 0.89930876 | 0.95528285 |
| TASP1 | 179.434022 | -0.1475584 | 0.23065032 | -0.6397492 | 0.52233567 | 0.7380123 |
| ESF1 | 66.144778 | -0.2433557 | 0.2921297 | -0.83304 | 0.40482219 | 0.65227474 |
| NDUFAF5 | 51.3462942 | -0.1107985 | 0.42239456 | -0.2623104 | 0.79308211 | 0.89875118 |
| MACROD2 | 613.066182 | 0.85780907 | 0.39440623 | 2.17493793 | 0.02963477 | 0.17009606 |
| FLRT3 | 19.1801859 | 0.21490095 | 0.59604086 | 0.36054734 | 0.71843786 | NA |

| | | | | | | |
|---|---|---|---|---|---|---|
| KIF16B | 187.738506 | 0.21420206 | 0.37739968 | 0.56757349 | 0.57032461 | 0.77196291 |
| SNRPB2 | 22.004021 | -0.0805844 | 0.38561869 | -0.2089742 | 0.83446835 | 0.91992307 |
| DSTN | 228.149506 | 0.78517514 | 0.33709203 | 2.32926051 | 0.01984527 | 0.13786014 |
| RRBP1 | 121.170987 | -0.3635599 | 0.36456838 | -0.9972336 | 0.31865112 | 0.57587641 |
| SNX5 | 36.728729 | 0.66851092 | 0.38635889 | 1.73028479 | 0.08357941 | 0.28861463 |
| OVOL2 | 67.1814134 | 0.99033544 | 0.31665178 | 3.12752208 | 0.00176287 | 0.03887902 |
| MGME1 | 64.7799279 | 0.22801531 | 0.33195848 | 0.686879 | 0.49215898 | 0.71672078 |
| KAT14 | 14.1394907 | -0.0680853 | 0.56777144 | -0.1199168 | 0.90454907 | NA |
| ZNF133 | 49.0423904 | 0.45412968 | 0.45615391 | 0.99556238 | 0.31946282 | 0.57666635 |
| DZANK1 | 27.1894169 | -0.0290235 | 0.42095933 | -0.068946 | 0.94503259 | 0.97757102 |
| POLR3F | 15.5317255 | -0.8193436 | 0.4813398 | -1.7022145 | 0.08871516 | NA |
| SEC23B | 72.9427092 | -0.1980992 | 0.25337116 | -0.7818537 | 0.43430058 | 0.67504272 |
| SLC24A3 | 189.642376 | -0.4478346 | 0.61253046 | -0.7311221 | 0.46470456 | 0.69561086 |
| AL121761.1 | 38.5543464 | 0.82627818 | 0.44276042 | 1.866197 | 0.06201381 | 0.24884382 |
| RIN2 | 175.639275 | 1.65669827 | 0.47851749 | 3.4621478 | 0.00053588 | 0.02088709 |
| NAA20 | 47.3970841 | 0.31792192 | 0.28016722 | 1.13475774 | 0.25647684 | 0.51462518 |
| CRNKL1 | 12.3059953 | -0.1947396 | 0.50632267 | -0.3846156 | 0.70052222 | NA |
| RALGAPA2 | 493.637865 | -0.0129156 | 0.30765593 | -0.0419805 | 9.67E-01 | 9.87E-01 |
| KIZ | 96.9843019 | 0.29158909 | 0.4338265 | 0.67213297 | 0.50149905 | 0.72315625 |
| KIZ-AS1 | 18.5776194 | 0.94360643 | 0.62004119 | 1.52184475 | 0.12804799 | NA |
| XRN2 | 115.245549 | -0.5879288 | 0.26321851 | -2.2336149 | 0.02550842 | 0.15648615 |
| NAPB | 67.7809343 | 0.12462805 | 0.34663509 | 0.35953673 | 0.7191936 | 0.85995641 |
| CST3 | 83.399389 | 1.03904516 | 0.56759798 | 1.83060052 | 0.06716019 | 0.25864684 |
| APMAP | 20.0599149 | 0.86034036 | 0.43705077 | 1.96851353 | 0.04900898 | NA |
| ACSS1 | 132.071635 | 1.06782222 | 0.27640063 | 3.86331325 | 0.00011186 | 0.00768734 |
| ENTPD6 | 36.9902168 | 0.29026526 | 0.39740761 | 0.73039685 | 0.46514765 | 0.69589923 |
| PYGB | 47.2902906 | -0.0555347 | 0.42988746 | -0.1291843 | 0.89721185 | 0.95374748 |
| ABHD12 | 66.7668648 | 0.02527459 | 0.24064178 | 0.10502994 | 0.91635205 | 0.96401363 |
| NINL | 264.522266 | -0.31966 | 0.24590994 | -1.2999069 | 0.1936329 | 0.44555827 |
| ZNF337-AS1 | 26.9371511 | -0.45103 | 0.5034227 | -0.8959271 | 0.37029172 | 0.6236924 |
| ZNF337 | 33.7811098 | 0.7438698 | 0.42391571 | 1.75475875 | 0.07930058 | 0.2808912 |
| FAM182B | 34.4454318 | 1.25219219 | 0.60831437 | 2.05846228 | 0.03954578 | 0.19795704 |
| HM13 | 76.5094791 | -0.14853 | 0.24088837 | -0.6165928 | 0.53750335 | 0.74878549 |
| ID1 | 17.2625609 | -0.0279659 | 0.85682506 | -0.0326389 | 0.97396252 | NA |
| BCL2L1 | 362.747405 | -1.3381898 | 0.31596191 | -4.2352884 | 2.28E-05 | 0.00274517 |
| AL117381.1 | 70.8363482 | -0.7799728 | 0.56676987 | -1.376172 | 0.16876838 | 0.41629954 |
| DUSP15 | 32.4444249 | 1.9055518 | 0.54735814 | 3.48136197 | 0.00049887 | 0.02020952 |
| TTLL9 | 16.2088315 | 1.02107933 | 0.63957496 | 1.59649672 | 0.11037794 | NA |
| TM9SF4 | 78.9327384 | -0.3828709 | 0.24401042 | -1.5690762 | 0.11663019 | 0.34407918 |
| PLAGL2 | 13.3095175 | -0.6237359 | 0.50036565 | -1.2465603 | 0.21255878 | NA |
| POFUT1 | 21.219467 | 0.42569175 | 0.42733563 | 0.99615319 | 0.31917572 | NA |
| KIF3B | 71.5323119 | 0.48834758 | 0.27091125 | 1.80261096 | 0.07144933 | 0.26675397 |
| ASXL1 | 224.039837 | -0.3118378 | 0.20739243 | -1.5036123 | 0.13268122 | 0.36669565 |
| NOL4L | 17.5005312 | 0.18082992 | 0.49239125 | 0.36724843 | 0.71343371 | NA |
| MAPRE1 | 25.6442302 | -0.0711126 | 0.42368029 | -0.167845 | 0.86670523 | 0.93793856 |
| CDK5RAP1 | 38.2629439 | 0.07099157 | 0.36502838 | 0.19448232 | 0.84579823 | 0.92531372 |
| SNTA1 | 17.0660396 | 0.75635338 | 0.56258657 | 1.34442131 | 0.17881219 | NA |
| CBFA2T2 | 326.922203 | -0.1000124 | 0.19688579 | -0.5079715 | 0.61147333 | 0.7979841 |
| NECAB3 | 11.5951455 | 1.11555117 | 0.61070438 | 1.82666312 | 0.06775044 | NA |
| CHMP4B | 55.3514683 | -0.7936971 | 0.45227912 | -1.7548833 | 0.07927926 | 0.2808912 |
| RALY | 112.31247 | -0.8238014 | 0.36291775 | -2.2699397 | 0.02321124 | 0.14863305 |
| EIF2S2 | 60.9581763 | -0.3942783 | 0.35274309 | -1.1177492 | 0.26367414 | 0.52333673 |
| ITCH | 274.43527 | -0.4576543 | 0.27161393 | -1.684944 | 0.09199937 | 0.3039138 |
| DYNLRB1 | 51.5020747 | 0.00412623 | 0.29791126 | 0.01385053 | 0.98894923 | 0.99583841 |

| | | | | | | |
|---|---|---|---|---|---|---|
| MAP1LC3A | 24.7290991 | 0.59331439 | 0.74264958 | 0.79891567 | 0.42433931 | 0.66715549 |
| PIGU | 56.1269162 | -0.3213764 | 0.33611013 | -0.9561639 | 0.33898946 | 0.59462212 |
| NCOA6 | 162.40728 | -0.1927116 | 0.26231267 | -0.7346638 | 0.46254428 | 0.69418845 |
| TP53INP2 | 21.2381889 | -0.2227107 | 0.66172499 | -0.3365608 | 0.73644798 | 0.86899281 |
| GGT7 | 67.6628883 | 1.11751766 | 0.44128557 | 2.53241377 | 0.01132802 | 0.1036885 |
| ACSS2 | 51.1217393 | 0.76815915 | 0.34385891 | 2.23393703 | 0.02548722 | 0.15648615 |
| GSS | 15.8845294 | 0.04208153 | 0.52609611 | 0.0799883 | 0.93624656 | NA |
| TRPC4AP | 155.501129 | -0.4853818 | 0.24643908 | -1.9695813 | 0.04888638 | 0.22162876 |
| EDEM2 | 38.9099572 | 0.2730217 | 0.40348974 | 0.6766509 | 0.49862748 | 0.72176422 |
| PROCR | 16.8080818 | 0.58309051 | 0.64958892 | 0.89763002 | 0.36938283 | NA |
| MMP24OS | 47.2274443 | -0.6170005 | 0.4095986 | -1.506354 | 0.13197632 | 0.36545739 |
| MMP24 | 43.4108326 | -0.0430712 | 0.36621332 | -0.1176124 | 0.90637477 | 0.9580376 |
| UQCC1 | 105.024537 | -0.3495795 | 0.20775533 | -1.68265 | 0.09244287 | 0.3049862 |
| CEP250 | 40.6379838 | -1.9956175 | 0.78742743 | -2.534351 | 0.01126558 | 0.1036885 |
| ERGIC3 | 35.4264476 | 0.31609417 | 0.36640835 | 0.86268276 | 0.38831191 | 0.63755051 |
| CPNE1 | 38.5956659 | -0.2155289 | 0.34550084 | -0.6238158 | 0.53274858 | 0.74615054 |
| NFS1 | 12.5484777 | 0.11916327 | 0.52879539 | 0.22534854 | 0.82170814 | NA |
| ROMO1 | 19.6356684 | -0.5504663 | 0.63298325 | -0.8696379 | 0.38449829 | NA |
| RBM39 | 366.884509 | -0.5532274 | 0.17001148 | -3.254059 | 0.00113769 | 0.03009201 |
| PHF20 | 158.370658 | -0.5671885 | 0.25855101 | -2.1937201 | 0.02825554 | 0.16543513 |
| SCAND1 | 27.4280401 | -0.1085782 | 0.36901626 | -0.2942369 | 0.76857691 | 0.88545674 |
| NORAD | 71.4765482 | 0.17624378 | 0.3415775 | 0.51597012 | 0.60587528 | 0.79431477 |
| EPB41L1 | 299.425229 | 0.01920265 | 0.37460099 | 0.05126162 | 0.95911705 | 0.98404745 |
| AAR2 | 18.4067044 | -0.3172792 | 0.47875173 | -0.6627218 | 0.50750873 | NA |
| DLGAP4 | 94.9030171 | -0.7572585 | 0.29255165 | -2.5884609 | 0.00964059 | 0.09699819 |
| DLGAP4-AS1 | 33.427585 | 0.4964212 | 0.47343077 | 1.04856132 | 0.29438006 | 0.55385592 |
| NDRG3 | 28.4400691 | 1.06795931 | 0.48167479 | 2.21717919 | 0.02661085 | 0.16039444 |
| SOGA1 | 37.9056443 | 0.96905554 | 0.36140503 | 2.68135598 | 0.00733245 | 0.08373567 |
| RBL1 | 28.3736684 | 0.41990902 | 0.44986033 | 0.93342089 | 0.3506027 | 0.60737992 |
| RPN2 | 56.9819044 | 0.15046668 | 0.30809656 | 0.48837506 | 0.6252842 | 0.8053173 |
| MANBAL | 14.3578953 | -0.6355322 | 0.45635552 | -1.3926251 | 0.16373319 | NA |
| SRC | 25.3316428 | 0.15771781 | 0.39970822 | 0.39458235 | 0.69315114 | 0.84791601 |
| BLCAP | 74.9215678 | -0.1463499 | 0.34804288 | -0.4204939 | 0.67412467 | 0.83715334 |
| CTNNBL1 | 56.1071402 | -0.1303201 | 0.27538622 | -0.4732265 | 0.63605157 | 0.81188871 |
| TTI1 | 33.5996409 | 0.37941119 | 0.42012698 | 0.90308694 | 0.36647976 | 0.62157013 |
| RPRD1B | 73.6761169 | -0.5111889 | 0.28337547 | -1.8039277 | 0.07124265 | 0.2665798 |
| TGM2 | 59.8127758 | -0.9594658 | 0.5866006 | -1.6356373 | 1.02E-01 | 3.21E-01 |
| RALGAPB | 127.470204 | -0.2320666 | 0.26462519 | -0.8769632 | 0.38050663 | 0.63215185 |
| DHX35 | 29.9345909 | -0.2490476 | 0.3893585 | -0.6396357 | 0.52240944 | 0.7380123 |
| TOP1 | 240.95597 | -1.0984848 | 0.26512087 | -4.1433358 | 3.42E-05 | 0.00356028 |
| PLCG1 | 32.6430156 | -0.1828331 | 0.35673109 | -0.5125237 | 0.60828457 | 0.79566688 |
| ZHX3 | 249.948301 | 0.21876627 | 0.47932366 | 0.45640616 | 0.64809794 | 0.81898044 |
| LPIN3 | 35.8141606 | 1.27545197 | 0.53684641 | 2.37582284 | 0.01750987 | 0.1316147 |
| CHD6 | 229.97403 | -0.0983293 | 0.21406475 | -0.4593438 | 0.64598733 | 0.81779125 |
| SRSF6 | 58.6829563 | -0.1900313 | 0.31124752 | -0.6105474 | 0.54149929 | 0.75003059 |
| L3MBTL1 | 28.2632956 | 0.15078917 | 0.39417353 | 0.38254514 | 0.70205706 | 0.85211052 |
| SGK2 | 93.6616376 | 1.2335474 | 0.37892542 | 3.25538309 | 0.0011324 | 0.03009201 |
| IFT52 | 35.8403648 | 0.12886653 | 0.33594651 | 0.38359242 | 0.70128056 | 0.85190818 |
| OSER1 | 23.7829012 | -0.2385723 | 0.43782432 | -0.5449041 | 0.58581948 | 0.78146492 |
| SERINC3 | 48.5390685 | -0.2548177 | 0.27797407 | -0.916696 | 0.35930196 | 0.61566055 |
| PKIG | 70.0277014 | 0.42789312 | 0.29441752 | 1.45335485 | 0.14612526 | 0.38623836 |
| YWHAB | 73.5309745 | 0.45029213 | 0.3829767 | 1.17576902 | 0.23968719 | 0.49733052 |
| PABPC1L | 27.4090279 | 0.55069184 | 0.39001133 | 1.41198934 | 0.1579531 | 0.40275854 |
| STK4 | 55.1220167 | -0.1367439 | 0.25980837 | -0.5263262 | 0.5986616 | 0.78956448 |

| | | | | | | |
|---|---|---|---|---|---|---|
| SLPI | 67.3003207 | -0.3763739 | 0.85475879 | -0.4403276 | 6.60E-01 | 8.27E-01 |
| SDC4 | 180.074615 | -0.300725 | 0.42968622 | -0.6998711 | 0.48400778 | 0.71145331 |
| PIGT | 14.5833667 | 0.61180581 | 0.60498729 | 1.01127053 | 0.31188697 | NA |
| WFDC2 | 496.574447 | 1.74488343 | 0.66885988 | 2.60874287 | 9.09E-03 | 9.43E-02 |
| DNTTIP1 | 15.6364976 | -0.2998412 | 0.43665826 | -0.6866723 | 0.49228923 | NA |
| SNX21 | 17.0952317 | 0.51456249 | 0.49534003 | 1.0388066 | 0.29889469 | NA |
| ACOT8 | 18.0140942 | 0.23586259 | 0.42988346 | 0.54866634 | 0.58323445 | NA |
| CTSA | 37.5578693 | -0.4548874 | 0.53695397 | -0.8471627 | 0.39690445 | 0.64518977 |
| ZNF335 | 21.6399922 | -0.6986445 | 0.48915045 | -1.4282814 | 0.15321087 | 0.39507934 |
| NCOA5 | 49.4367404 | -0.5944701 | 0.37196183 | -1.5982018 | 0.10999806 | 0.33293555 |
| SLC35C2 | 25.7649105 | -0.2879293 | 0.42797075 | -0.672778 | 0.50108852 | 0.72289663 |
| ELMO2 | 23.2929194 | 0.04060127 | 0.47487858 | 0.08549821 | 0.93186532 | 0.97228783 |
| SLC13A3 | 24.5482311 | -0.1357604 | 0.47349599 | -0.2867192 | 0.77432733 | 0.88786322 |
| EYA2 | 123.666474 | 0.45810909 | 0.74903157 | 0.61160184 | 0.54080122 | 0.74994873 |
| ZMYND8 | 367.813446 | -0.2767716 | 0.26054985 | -1.0622597 | 0.28811781 | 0.54746482 |
| NCOA3 | 233.006842 | -0.7716984 | 0.25522365 | -3.0236164 | 0.00249773 | 0.04746918 |
| SULF2 | 71.2722612 | 0.71379839 | 0.41950722 | 1.70151634 | 0.08884607 | 0.29778649 |
| ARFGEF2 | 285.958047 | -0.2760193 | 0.16614325 | -1.6613331 | 0.09664656 | 0.31216475 |
| CSE1L | 50.0008043 | 0.01701177 | 0.29325949 | 0.05800929 | 0.95374123 | 0.98153155 |
| STAU1 | 138.000801 | -0.20321 | 0.20490549 | -0.9917253 | 0.32133155 | 0.57793564 |
| DDX27 | 27.9336521 | -0.3343467 | 0.47087449 | -0.7100549 | 0.47767012 | 0.70541576 |
| ZNFX1 | 35.7796573 | -1.0610504 | 0.41614267 | -2.5497275 | 0.01078071 | 0.10206537 |
| ZFAS1 | 38.0795068 | -0.1175154 | 0.37255211 | -0.3154333 | 0.75243266 | 0.87837077 |
| B4GALT5 | 196.18307 | -0.8514473 | 0.26100289 | -3.2622142 | 0.00110546 | 0.02988759 |
| SLC9A8 | 74.5851167 | -1.1416137 | 0.37466481 | -3.0470267 | 0.00231117 | 0.04573031 |
| RNF114 | 71.5391502 | -0.2574782 | 0.24487117 | -1.0514844 | 0.29303615 | 0.55234367 |
| CEBPB | 33.6017932 | -0.7938006 | 0.72589542 | -1.0935468 | 2.74E-01 | 0.53327316 |
| PTPN1 | 187.206745 | -1.0467312 | 0.34485585 | -3.0352718 | 0.00240319 | 0.04671049 |
| PARD6B | 130.614126 | -0.0721526 | 0.31275522 | -0.2307 | 0.8175479 | 0.9123773 |
| BCAS4 | 35.0662804 | 0.69871151 | 0.37802196 | 1.84833576 | 0.06455379 | 0.25284322 |
| ADNP | 240.802163 | 0.02978367 | 0.19206293 | 0.15507244 | 0.87676421 | 0.94225833 |
| DPM1 | 59.8531333 | -0.666584 | 0.26551524 | -2.5105301 | 0.012055 | 0.10726863 |
| NFATC2 | 18.0533016 | 0.30486925 | 0.44506495 | 0.68499946 | 0.49334426 | NA |
| ATP9A | 77.9453924 | -0.080953 | 0.29751564 | -0.2720968 | 0.78554763 | 0.89452315 |
| ZFP64 | 42.5221898 | 0.47088318 | 0.40435182 | 1.16453829 | 0.24420594 | 0.5015083 |
| ZNF217 | 52.4462475 | -0.4585361 | 0.34592951 | -1.3255188 | 0.18499915 | 0.43476148 |
| DOK5 | 25.4970212 | -0.9174623 | 0.61380266 | -1.4947187 | 0.13498787 | 0.36909471 |
| RTF2 | 33.0521979 | -0.6305197 | 0.3513167 | -1.794733 | 0.07269625 | 0.26848318 |
| RAE1 | 20.3681537 | 0.2002543 | 0.44891347 | 0.44608664 | 0.65553467 | NA |
| PCK1 | 133.539541 | -0.7517412 | 1.64478837 | -0.4570443 | 0.64763917 | 0.81885075 |
| PMEPA1 | 26.5912528 | -2.1929577 | 0.89760149 | -2.4431307 | 0.01456046 | 0.11884612 |
| RAB22A | 65.309414 | -0.6594195 | 0.33150394 | -1.9891754 | 0.04668185 | 0.21658238 |
| VAPB | 88.5109091 | -0.1639655 | 0.21369791 | -0.7672769 | 0.44291692 | 0.68272427 |
| NPEPL1 | 18.3297707 | 0.77194834 | 0.62554995 | 1.23403149 | 0.21719117 | NA |
| GNAS | 242.2012 | -0.0013361 | 0.2599855 | -0.005139 | 0.99589968 | 0.99816804 |
| NELFCD | 38.2126511 | -0.0699713 | 0.33059032 | -0.2116556 | 0.83237572 | 0.91853705 |
| CTSZ | 17.6529634 | 0.31587579 | 0.41160521 | 0.76742417 | 0.44282936 | NA |
| ATP5F1E | 75.6725419 | -0.1298229 | 0.47120428 | -0.275513 | 0.78292214 | 0.89289549 |
| MIR646HG | 339.876404 | -1.005468 | 1.11534908 | -0.9014828 | 0.36733168 | 0.62220175 |
| TAF4 | 57.6563295 | 0.12265153 | 0.36440135 | 0.33658364 | 0.73643079 | 0.86899281 |
| PSMA7 | 40.0781329 | -0.1090446 | 0.36447352 | -0.299184 | 0.76479969 | 0.88317118 |
| SS18L1 | 55.2735589 | -0.4991496 | 0.36337934 | -1.3736323 | 0.16955587 | 0.41775657 |
| MTG2 | 42.8393987 | 0.04186629 | 0.34688869 | 0.12069084 | 0.90393592 | 0.95671001 |
| OSBPL2 | 71.9807388 | -0.5536576 | 0.29776661 | -1.8593677 | 0.06297504 | 0.249987 |

| | | | | | | |
|---|---|---|---|---|---|---|
| LAMA5 | 60.3410032 | -0.2165587 | 0.38762374 | -0.5586827 | 0.57637831 | 0.77421002 |
| SLCO4A1 | 60.460068 | -1.7055344 | 0.48348251 | -3.5276032 | 4.19E-04 | 1.79E-02 |
| OGFR | 29.6641389 | -0.531569 | 0.40005192 | -1.32875 | 0.18393047 | 0.43412487 |
| TCFL5 | 13.3368134 | 0.55652898 | 0.51964471 | 1.07097978 | 0.28417852 | NA |
| DIDO1 | 112.95365 | 0.01658655 | 0.23177143 | 0.07156426 | 0.94294868 | 0.97643966 |
| GID8 | 26.0144358 | -0.6017063 | 0.37966372 | -1.58484 | 0.11300269 | 0.33771702 |
| YTHDF1 | 35.2573656 | -0.561728 | 0.33048636 | -1.6997011 | 0.08918717 | 0.29794449 |
| ARFGAP1 | 31.4261148 | -0.6618068 | 0.41560163 | -1.5924067 | 0.11129334 | 0.33461909 |
| PPDPF | 76.2350912 | 0.66910048 | 0.53563517 | 1.24917204 | 0.21160216 | 0.46623687 |
| GMEB2 | 22.6740545 | -1.1365254 | 0.43720463 | -2.5995274 | 0.00933522 | 0.09529602 |
| ARFRP1 | 17.8382584 | -0.3043944 | 0.47723216 | -0.6378329 | 0.52358243 | NA |
| SLC2A4RG | 27.465767 | 1.1352466 | 0.49717906 | 2.28337576 | 0.02240824 | 0.14604538 |
| ZBTB46 | 30.5997623 | -0.0562113 | 0.43483541 | -0.1292704 | 0.89714371 | 0.95374748 |
| TPD52L2 | 43.4262944 | -0.8692355 | 0.33961618 | -2.5594644 | 0.01048336 | 0.1006115 |
| DNAJC5 | 110.728148 | -0.5995446 | 0.33299091 | -1.8004833 | 0.07178435 | 0.26730728 |
| UCKL1 | 25.4711937 | 1.14546927 | 0.42528609 | 2.6934087 | 0.00707255 | 0.08210773 |
| ZNF512B | 15.9278177 | 0.49638735 | 0.56205399 | 0.88316667 | 0.37714623 | NA |
| PRPF6 | 57.5055093 | -0.2611821 | 0.30059716 | -0.8688773 | 0.38491424 | 0.63508787 |
| PCMTD2 | 78.711501 | 1.02255885 | 0.43775283 | 2.33592744 | 0.01949502 | 0.13692563 |
| PLPP2 | 22.5865471 | 0.59701377 | 0.60414819 | 0.98819095 | 3.23E-01 | 5.79E-01 |
| MIER2 | 17.6937146 | -0.4951089 | 0.4425498 | -1.1187642 | 0.26324076 | NA |
| BSG | 71.9532883 | 0.77702092 | 0.3902841 | 1.99091106 | 0.04649067 | 0.21595574 |
| RNF126 | 21.4166223 | -0.4349005 | 0.36935375 | -1.1774632 | 0.23901066 | 0.49681334 |
| PALM | 58.4029715 | 1.03854137 | 0.51461889 | 2.0180786 | 0.04358308 | 0.20874856 |
| PTBP1 | 58.0947122 | -0.4575503 | 0.30103789 | -1.5199093 | 0.12853378 | 0.36084904 |
| MED16 | 14.1739956 | 0.63904895 | 0.55010652 | 1.16168218 | 0.24536459 | NA |
| R3HDM4 | 23.8034873 | -0.3510548 | 0.51381477 | -0.6832322 | 0.49446013 | 0.71857397 |
| ARID3A | 113.149793 | -0.4242247 | 0.52456961 | -0.80871 | 0.418682 | 0.66260735 |
| WDR18 | 13.8328339 | 0.43394784 | 0.64619762 | 0.67154045 | 0.5018763 | NA |
| TMEM259 | 69.9300022 | -0.1803673 | 0.38972686 | -0.4628044 | 0.64350458 | 0.81666196 |
| CNN2 | 15.0660429 | -0.6062738 | 0.51230928 | -1.1834136 | 0.23664528 | NA |
| POLR2E | 18.8658811 | 0.46052553 | 0.45306668 | 1.01646302 | 0.30940895 | NA |
| GPX4 | 65.6979418 | 0.18459822 | 0.42264948 | 0.43676434 | 0.66228226 | 0.82847174 |
| SBNO2 | 115.229712 | -0.4340805 | 0.31419738 | -1.3815535 | 1.67E-01 | 4.15E-01 |
| STK11 | 44.8244627 | 0.03098338 | 0.40425164 | 0.07664379 | 0.93890693 | 0.97481036 |
| ATP5F1D | 49.622304 | 0.96854253 | 0.45543936 | 2.1266114 | 0.03345238 | 0.18132606 |
| MIDN | 78.7179275 | -0.997733 | 0.47762902 | -2.0889287 | 3.67E-02 | 0.19119673 |
| CIRBP | 139.802482 | 0.24329973 | 0.36665735 | 0.66356158 | 0.50697096 | 0.72824214 |
| MUM1 | 49.3333017 | 0.89420542 | 0.41484675 | 2.15550781 | 0.03112212 | 0.17421596 |
| NDUFS7 | 20.6001787 | 0.48014152 | 0.4689348 | 1.02389823 | 0.30588334 | NA |
| DAZAP1 | 79.8043997 | -0.5424435 | 0.37298593 | -1.4543269 | 0.14585569 | 0.38581359 |
| RPS15 | 107.630724 | 0.27806814 | 0.48843504 | 0.56930423 | 0.56914969 | 0.77149801 |
| C19orf25 | 44.4575961 | -0.8893494 | 0.31108456 | -2.8588671 | 0.00425157 | 0.06301187 |
| MBD3 | 17.6614994 | -0.7925732 | 0.55681271 | -1.4234108 | 0.15461709 | NA |
| TCF3 | 20.7549879 | -0.1452234 | 0.4197066 | -0.3460117 | 0.72933393 | NA |
| REXO1 | 21.6227553 | -0.3196951 | 0.43075024 | -0.7421821 | 0.45797703 | 0.69231672 |
| ABHD17A | 20.0821718 | 0.11789186 | 0.49200582 | 0.23961476 | 0.81062892 | NA |
| SCAMP4 | 31.2469339 | 0.20597769 | 0.45357451 | 0.45412096 | 0.64974176 | 0.81947559 |
| CSNK1G2 | 38.5769706 | 0.00206106 | 0.33760874 | 0.00610489 | 0.99512903 | 0.99796515 |
| BTBD2 | 48.2146987 | -0.1414152 | 0.36022119 | -0.3925789 | 0.69463053 | 0.84870601 |
| MKNK2 | 21.9202417 | -0.5370812 | 0.49422795 | -1.0867075 | 0.27716608 | 0.53561319 |
| MOB3A | 25.1348202 | -0.2574031 | 0.43630141 | -0.5899663 | 0.55521327 | 0.76189633 |
| AP3D1 | 77.9134399 | -0.7571946 | 0.38172071 | -1.9836352 | 0.04729653 | 0.21733813 |
| DOT1L | 84.911558 | -1.5726474 | 0.46132811 | -3.4089564 | 0.00065212 | 0.02323777 |

| | | | | | | |
|---|---|---|---|---|---|---|
| PLEKHJ1 | 32.8317188 | 0.68847266 | 0.49817797 | 1.38198133 | 0.16697743 | 0.41508187 |
| SF3A2 | 12.0157151 | -1.0337214 | 0.55548514 | -1.8609343 | 0.06275345 | NA |
| LSM7 | 27.8384086 | 0.28689287 | 0.43675143 | 0.65687907 | 0.51125868 | 0.73064085 |
| SPPL2B | 45.8307188 | -0.0031214 | 0.37734534 | -0.008272 | 0.99340001 | 0.997157 |
| TIMM13 | 21.5386482 | 0.47302531 | 0.52799292 | 0.89589328 | 0.37030977 | 0.6236924 |
| GADD45B | 42.1011485 | -0.5596959 | 0.47688355 | -1.1736532 | 0.24053396 | 0.49822614 |
| GNG7 | 102.562404 | -0.2323056 | 0.44756623 | -0.5190419 | 0.60373154 | 0.79312539 |
| SGTA | 25.281313 | -0.0677884 | 0.48184497 | -0.1406852 | 0.88811867 | 0.948633 |
| ZNF554 | 9.94005284 | 0.93291058 | 0.64568825 | 1.44483127 | 0.14850529 | NA |
| ZNF555 | 13.3635579 | 0.11843185 | 0.60209021 | 0.19670118 | 0.84406139 | NA |
| AES | 82.5808137 | 0.35558878 | 0.33383971 | 1.06514823 | 0.28680886 | 0.54615729 |
| GNA11 | 22.6943957 | -0.4244779 | 0.41936212 | -1.0121991 | 0.31144288 | 0.57075616 |
| NCLN | 20.5949895 | 0.71715248 | 0.45498981 | 1.57619458 | 0.11498097 | NA |
| NFIC | 200.565878 | 0.04729474 | 0.29663108 | 0.1594396 | 0.87332254 | 0.94114315 |
| FZR1 | 21.2896644 | -1.0540478 | 0.44636559 | -2.3614002 | 0.01820607 | 0.13368407 |
| MFSD12 | 31.0603651 | -0.8165874 | 0.40841938 | -1.9993846 | 0.04556676 | 0.21335013 |
| HMG20B | 20.5761463 | -0.1794389 | 0.47436109 | -0.3782749 | 0.70522641 | NA |
| PIP5K1C | 21.3066719 | 0.36725486 | 0.46356633 | 0.79223799 | 0.42822192 | 0.67072527 |
| TJP3 | 38.4166408 | 0.13024327 | 0.37960038 | 0.34310626 | 0.73151852 | 0.86703081 |
| DAPK3 | 24.9615964 | -0.3867141 | 0.4981407 | -0.776315 | 0.43756301 | 0.67793318 |
| EEF2 | 98.7880845 | 0.87711164 | 0.41687663 | 2.10400767 | 0.03537778 | 0.18721309 |
| PIAS4 | 25.7972798 | -0.3096172 | 0.45790208 | -0.6761646 | 0.49893615 | 0.72176422 |
| ZBTB7A | 52.9717885 | -0.208538 | 0.36838165 | -0.5660923 | 0.57133107 | 0.77245539 |
| MAP2K2 | 43.5471616 | 0.24001315 | 0.3616444 | 0.66367169 | 0.50690046 | 0.72824214 |
| YJU2 | 11.4227808 | -0.7771397 | 0.65992597 | -1.1776164 | 0.23894956 | NA |
| SH3GL1 | 31.1376001 | -0.3672269 | 0.33260211 | -1.1041026 | 0.26954863 | 0.52764898 |
| UBXN6 | 23.5046935 | 0.49797858 | 0.47396256 | 1.05067071 | 0.29340985 | 0.55271063 |
| HDGFL2 | 15.8073864 | 0.47784372 | 0.48525177 | 0.98473359 | 0.32475495 | NA |
| PLIN5 | 56.2224145 | 1.05239939 | 0.3829811 | 2.74791467 | 0.00599756 | 0.07605145 |
| MYDGF | 22.2989392 | 0.09474701 | 0.46064581 | 0.205683 | 0.83703854 | 0.92119416 |
| DPP9 | 41.1685497 | -0.246279 | 0.31840626 | -0.773474 | 0.43924188 | 0.67938098 |
| PLIN3 | 31.5351365 | -0.6316268 | 0.40905532 | -1.5441109 | 1.23E-01 | 0.35251484 |
| KDM4B | 102.177781 | -0.6210773 | 0.24213068 | -2.56505 | 0.01031609 | 0.10000374 |
| PTPRS | 66.7245451 | 0.52947323 | 0.38101873 | 1.38962522 | 0.16464271 | 0.41135131 |
| SAFB2 | 98.1890641 | -0.48951 | 0.30866668 | -1.5858853 | 0.11276532 | 0.33755464 |
| SAFB | 115.886371 | -0.6673707 | 0.27188026 | -2.4546494 | 0.0141022 | 0.11675032 |
| RPL36 | 63.8235656 | 0.24288978 | 0.46400474 | 0.52346401 | 0.60065139 | 0.79096587 |
| C19orf70 | 29.4947551 | 0.8694735 | 0.52087517 | 1.66925504 | 0.09506685 | 0.30939937 |
| LONP1 | 58.3766371 | -1.2521187 | 0.40348379 | -3.1032689 | 0.00191396 | 0.04077019 |
| RANBP3 | 31.9294868 | -0.4787673 | 0.34336566 | -1.3943364 | 0.16321603 | 0.40987726 |
| RFX2 | 71.3609009 | -0.4891889 | 0.50968494 | -0.9597868 | 0.33716251 | 0.59276835 |
| MLLT1 | 30.0692811 | -0.3263547 | 0.38209277 | -0.8541242 | 0.39303618 | 0.64188404 |
| GTF2F1 | 16.3909962 | -0.4050881 | 0.45824414 | -0.8840005 | 0.37669594 | NA |
| KHSRP | 38.1362634 | -0.8908123 | 0.42054491 | -2.1182336 | 0.03415529 | 0.18312899 |
| SLC25A23 | 27.0590069 | 1.74417425 | 0.43757419 | 3.98600804 | 6.72E-05 | 0.00530264 |
| TRIP10 | 30.9947536 | -0.9362461 | 0.41498452 | -2.2560988 | 0.02406444 | 0.15136029 |
| SH2D3A | 47.5513099 | -1.4207689 | 0.28361941 | -5.0094206 | 5.46E-07 | 0.00026263 |
| INSR | 344.089279 | -0.3880881 | 0.31044175 | -1.2501156 | 0.21125733 | 0.46604977 |
| PNPLA6 | 26.6679203 | -0.2768421 | 0.42975222 | -0.6441899 | 0.51945227 | 0.7363212 |
| CAMSAP3 | 13.3592425 | 0.38440493 | 0.47187021 | 0.81464123 | 0.4152777 | NA |
| EVI5L | 14.4521174 | 0.41575967 | 0.48006763 | 0.86604397 | 0.38646605 | NA |
| MAP2K7 | 42.025498 | -1.0679605 | 0.43756317 | -2.4407001 | 0.01465882 | 0.1192695 |
| TIMM44 | 26.4679995 | -0.8028175 | 0.6720338 | -1.1946089 | 0.23223988 | 0.49066933 |
| ELAVL1 | 77.6673733 | -0.9958095 | 0.54006282 | -1.8438771 | 0.06520105 | 0.25410253 |

| | | | | | | |
|---|---|---|---|---|---|---|
| CERS4 | 62.898234 | -0.4569441 | 0.3880788 | -1.1774519 | 0.23901518 | 0.49681334 |
| AC022146.2 | 17.4758891 | 0.28761404 | 0.62470757 | 0.46039788 | 0.64523066 | NA |
| RPS28 | 27.7279494 | 0.24644964 | 0.38620585 | 0.63813027 | 0.52338889 | 0.73904282 |
| ANGPTL4 | 35.0691888 | -1.9910259 | 0.98425456 | -2.022877 | 4.31E-02 | 0.20739942 |
| RAB11B | 27.2444516 | -0.1732077 | 0.41545562 | -0.4169103 | 0.676744 | 0.83887358 |
| 2-Mar | 14.9603553 | -0.5376571 | 0.48137968 | -1.1169086 | 0.26403339 | NA |
| HNRNPM | 152.898197 | -0.4976938 | 0.24029568 | -2.0711726 | 0.03834267 | 0.19533626 |
| ZNF558 | 15.6081401 | -0.1550519 | 0.455659 | -0.3402807 | 0.73364518 | NA |
| ZNF317 | 20.849951 | -0.2893663 | 0.42901082 | -0.6744964 | 0.49999574 | NA |
| ZNF559-ZNF | 31.536053 | 0.68035062 | 0.47909621 | 1.42007098 | 0.15558702 | 0.39879246 |
| ZNF266 | 34.2592155 | -0.1752128 | 0.39035687 | -0.4488528 | 0.65353785 | 0.82226683 |
| ZNF426 | 17.1987217 | -0.3012073 | 0.54493647 | -0.5527384 | 0.58044258 | NA |
| ZNF121 | 30.6172781 | -0.0281763 | 0.3368069 | -0.0836572 | 0.93332901 | 0.9730757 |
| ZNF561 | 25.4344143 | -0.1167173 | 0.43357275 | -0.2691988 | 0.78777669 | 0.89506208 |
| ZNF562 | 58.2440476 | -0.8045428 | 0.29671308 | -2.7115178 | 0.00669759 | 0.07974991 |
| ZNF846 | 51.2794037 | 1.08243072 | 0.31271165 | 3.46143398 | 0.00053731 | 0.02088709 |
| FBXL12 | 20.3794044 | -0.0561134 | 0.4395659 | -0.1276563 | 0.89842098 | NA |
| UBL5 | 14.8772515 | 0.70592533 | 0.54034006 | 1.3064464 | 0.1914008 | NA |
| PIN1 | 17.9095908 | 0.43496882 | 0.41615133 | 1.0452179 | 0.29592227 | NA |
| C19orf66 | 30.2451041 | -0.2766069 | 0.55665029 | -0.4969133 | 0.6192502 | 0.8027391 |
| EIF3G | 13.5198537 | -0.0861217 | 0.56327396 | -0.1528948 | 0.87848125 | NA |
| DNMT1 | 19.9115257 | -1.0330455 | 0.49389027 | -2.0916498 | 0.03646986 | NA |
| MRPL4 | 22.9739897 | 0.87566869 | 0.42355468 | 2.06742773 | 0.03869387 | 0.1958098 |
| AC011511.2 | 15.872706 | 0.43187066 | 0.81027888 | 0.53299015 | 0.5940404 | NA |
| TYK2 | 40.736388 | 0.22997481 | 0.31036612 | 0.74097911 | 0.45870611 | 0.6928053 |
| CDC37 | 56.4710016 | -0.5310072 | 0.34422377 | -1.5426221 | 0.1229225 | 0.35290359 |
| KEAP1 | 13.9401602 | 0.20086298 | 0.54150033 | 0.37093788 | 0.7106838 | NA |
| ATG4D | 23.0580888 | -0.1578158 | 0.38245888 | -0.4126348 | 0.6798742 | 0.8407763 |
| KRI1 | 16.3684511 | 0.07977368 | 0.50836628 | 0.15692166 | 0.87530659 | NA |
| AP1M2 | 29.7011717 | 0.9015611 | 0.40799856 | 2.20971638 | 0.02712485 | 0.16197051 |
| SLC44A2 | 40.3856692 | 1.19522075 | 0.5186191 | 2.30462154 | 0.02118778 | 0.1427136 |
| ILF3-DT | 21.9483706 | 0.24149351 | 0.52512487 | 0.45987826 | 0.6456036 | 0.81777273 |
| ILF3 | 94.263629 | -0.6404651 | 0.32530773 | -1.9687976 | 0.04897634 | 0.22174758 |
| QTRT1 | 33.8013012 | 0.26402771 | 0.47106252 | 0.56049398 | 0.57514254 | 0.77388388 |
| DNM2 | 211.922763 | 0.17696248 | 0.27666382 | 0.63962999 | 0.52241318 | 0.7380123 |
| CARM1 | 32.6926726 | 0.39425801 | 0.39716101 | 0.99269062 | 0.32086075 | 0.57769945 |
| YIPF2 | 19.0494387 | -0.2894811 | 0.46624315 | -0.6208801 | 0.5346785 | NA |
| TIMM29 | 14.1333586 | 1.02620047 | 0.49977057 | 2.05334313 | 0.04003931 | NA |
| SMARCA4 | 145.157523 | -0.0673751 | 0.27123319 | -0.2484027 | 0.80382286 | 0.90408418 |
| LDLR | 58.3124341 | -1.3054669 | 0.64548073 | -2.0224723 | 0.04312758 | 0.20747061 |
| KANK2 | 28.0679119 | -0.3895276 | 0.52905377 | -0.7362722 | 0.46156505 | 0.69416206 |
| DOCK6 | 80.0862317 | -0.4440164 | 0.31221305 | -1.4221584 | 0.15498027 | 0.39776029 |
| EPOR | 28.8082178 | 1.31531192 | 0.49806074 | 2.64086647 | 0.00826943 | 0.08912945 |
| RGL3 | 272.030691 | 0.27286003 | 0.3469354 | 0.78648656 | 0.43158249 | 0.67244746 |
| PRKCSH | 25.5681258 | 0.39064346 | 0.47228573 | 0.82713375 | 0.40816126 | 0.65423098 |
| ECSIT | 29.886856 | 0.40042094 | 0.37745386 | 1.06084738 | 0.28875926 | 0.54810852 |
| ZNF627 | 37.6347799 | -0.0260675 | 0.37540997 | -0.0694375 | 0.94464137 | 0.9774035 |
| AC008543.1 | 25.6003724 | 0.5963735 | 0.42419039 | 1.40590996 | 0.15975086 | 0.40447447 |
| ZNF823 | 29.3041742 | 0.39749991 | 0.45990786 | 0.86430337 | 0.38742126 | 0.63701577 |
| ZNF441 | 27.8726928 | 0.15327779 | 0.45220702 | 0.33895492 | 0.73464369 | 0.86814805 |
| ZNF440 | 39.5544721 | -0.3768639 | 0.3422304 | -1.1011993 | 0.27080991 | 0.52912064 |
| ZNF439 | 53.8356081 | 0.15147217 | 0.3150498 | 0.48078803 | 0.63066716 | 0.80819181 |
| ZNF69 | 69.949927 | 0.54338328 | 0.36609002 | 1.48428868 | 0.13773235 | 0.37407407 |
| ZNF433 | 11.7849457 | 0.2359039 | 0.51226559 | 0.46051091 | 0.64514954 | NA |

| | | | | | | |
|---|---|---|---|---|---|---|
| ZNF844 | 35.5854321 | -0.3610348 | 0.38866284 | -0.9289152 | 0.35293305 | 0.61004394 |
| ZNF788P | 16.7720178 | -0.0810931 | 0.48702519 | -0.166507 | 0.86775798 | NA |
| ZNF136 | 18.1723553 | -0.5501243 | 0.44433571 | -1.2380826 | 0.21568543 | NA |
| ZNF44 | 241.299671 | -0.0616832 | 0.17502359 | -0.3524279 | 0.72451739 | 0.86245134 |
| ZNF563 | 14.9977533 | 1.2941949 | 0.59276039 | 2.18333568 | 0.0290111 | NA |
| AC008758.5 | 21.5066116 | -1.5543479 | 0.66211272 | -2.3475578 | 0.01889694 | 0.1355543 |
| AC008758.1 | 16.9812077 | -0.8957122 | 0.56163223 | -1.5948376 | 0.11074855 | NA |
| ZNF490 | 33.1886442 | 0.36681758 | 0.44183832 | 0.83020771 | 0.40642136 | 0.65333469 |
| ZNF791 | 56.9342986 | -0.9387841 | 0.27161952 | -3.456247 | 0.00054775 | 0.0209754 |
| DHPS | 13.2130799 | -0.3974093 | 0.51160062 | -0.7767961 | 0.4372791 | NA |
| TNPO2 | 42.4001615 | -0.4466321 | 0.33884608 | -1.3180973 | 0.18747109 | 0.43816783 |
| TRIR | 37.101596 | -0.2109373 | 0.34225831 | -0.6163102 | 0.53768978 | 0.74878549 |
| HOOK2 | 124.608627 | -0.3576247 | 0.36477931 | -0.9803866 | 0.32689533 | 0.58313355 |
| JUNB | 34.4704504 | -0.4618815 | 0.524952 | -0.8798546 | 0.37893806 | 0.63024389 |
| GCDH | 16.6228506 | -0.0710195 | 0.49645724 | -0.1430526 | 0.88624861 | NA |
| FARSA | 19.3335753 | -0.4611261 | 0.46436841 | -0.9930179 | 0.32070122 | NA |
| CALR | 56.4212051 | 0.36950005 | 0.4839502 | 0.76350841 | 0.44516025 | 0.68514266 |
| RAD23A | 28.8486635 | -0.2678999 | 0.34349728 | -0.7799187 | 0.43543875 | 0.67626555 |
| GADD45GIP1 | 19.0060073 | 0.43483332 | 0.50555051 | 0.86011843 | 0.38972376 | NA |
| NFIX | 18.3032397 | 0.22042872 | 0.50789319 | 0.43400604 | 0.66428405 | NA |
| NACC1 | 17.983004 | -0.0642153 | 0.45998784 | -0.1396021 | 0.88897438 | NA |
| IER2 | 40.5743312 | -1.0338562 | 0.47180985 | -2.191256 | 0.02843327 | 0.16587474 |
| CCDC130 | 70.1579583 | -0.0849503 | 0.27431031 | -0.3096868 | 0.75679912 | 0.87904624 |
| MRI1 | 26.6700659 | 0.19699256 | 0.46746169 | 0.42140898 | 0.67345646 | 0.83700862 |
| C19orf53 | 22.1774983 | 0.75443443 | 0.47491445 | 1.58856913 | 0.1121577 | 0.33669182 |
| ZSWIM4 | 26.0825379 | -1.1754291 | 0.38478538 | -3.0547656 | 0.00225236 | 0.04514701 |
| CC2D1A | 36.1845413 | 0.15165029 | 0.31806743 | 0.4767866 | 0.63351409 | 0.8102622 |
| DCAF15 | 12.8118774 | 0.13038719 | 0.49857028 | 0.26152219 | 0.79368984 | NA |
| RFX1 | 28.0571875 | 0.30718427 | 0.42424618 | 0.7240708 | 0.46902227 | 0.69884046 |
| PRKACA | 30.3445116 | 0.28926582 | 0.49325144 | 0.58644699 | 0.55757518 | 0.76343775 |
| ADGRL1 | 27.3768577 | 0.15009384 | 0.42579986 | 0.35249857 | 0.72446439 | 0.86245134 |
| PKN1 | 19.9228892 | 0.06172383 | 0.42844658 | 0.14406425 | 0.88544974 | NA |
| GIPC1 | 58.366406 | -0.2080342 | 0.28732413 | -0.72404 | 0.46904116 | 0.69884046 |
| DNAJB1 | 15.7238256 | -0.1453001 | 0.42474594 | -0.3420871 | 0.73228535 | NA |
| TECR | 22.5777665 | 0.76583736 | 0.42529347 | 1.80072682 | 0.07174595 | 0.26729359 |
| NDUFB7 | 35.1205052 | 0.62063267 | 0.40439409 | 1.53472239 | 0.12485203 | 0.3551316 |
| ZNF333 | 12.8190979 | -1.1776239 | 0.54965228 | -2.1424888 | 0.03215417 | NA |
| ILVBL | 17.7773956 | 0.3162106 | 0.48529514 | 0.65158412 | 0.5146695 | NA |
| NOTCH3 | 27.415883 | 0.09701901 | 0.60185149 | 0.16120091 | 0.87193517 | 0.94021925 |
| BRD4 | 220.148675 | -0.3466672 | 0.22758805 | -1.5232224 | 0.12770308 | 0.35978425 |
| AKAP8 | 57.9693978 | -0.4655671 | 0.31162351 | -1.4940052 | 0.13517427 | 0.36934198 |
| AKAP8L | 110.923449 | -0.3890591 | 0.32064597 | -1.2133603 | 0.2249921 | 0.48216659 |
| TPM4 | 20.8092157 | 0.14814997 | 0.3806372 | 0.38921569 | 0.6971166 | NA |
| RAB8A | 14.4164671 | -0.0372265 | 0.5248214 | -0.0709317 | 0.94345209 | NA |
| AP1M1 | 17.1321384 | -0.2309429 | 0.46773985 | -0.4937422 | 0.62148828 | NA |
| EPS15L1 | 86.1699053 | -0.5430493 | 0.31476698 | -1.7252424 | 0.0844838 | 0.28931012 |
| C19orf44 | 18.0076853 | 0.27820434 | 0.51067169 | 0.54478121 | 0.585904 | NA |
| TMEM38A | 32.8507715 | 0.6167956 | 0.43221964 | 1.42704204 | 0.15356778 | 0.39545372 |
| SIN3B | 67.9147418 | -0.6874348 | 0.24352044 | -2.8229039 | 0.00475908 | 0.06664966 |
| MYO9B | 84.3834296 | -1.1278312 | 0.36433512 | -3.0955873 | 0.00196424 | 0.04142115 |
| NR2F6 | 16.3574886 | 1.34954418 | 0.54414613 | 2.48011352 | 0.01313406 | NA |
| MRPL34 | 21.8235591 | 0.69849147 | 0.5691458 | 1.22726282 | 0.21972383 | 0.47614454 |
| BST2 | 22.3967082 | 1.7605399 | 1.06968906 | 1.64584267 | 0.09979615 | 0.31793995 |
| MVB12A | 17.3591765 | 0.29983973 | 0.44003305 | 0.68140274 | 0.49561669 | NA |

| | | | | | | |
|---|---|---|---|---|---|---|
| SLC27A1 | 38.8951529 | 1.51918146 | 0.38029934 | 3.99469915 | 6.48E-05 | 0.00530264 |
| AC010618.3 | 28.798323 | -0.5110983 | 0.41903204 | -1.2197118 | 0.22257416 | 0.4796062 |
| PGLS | 20.6266721 | 0.38481527 | 0.45331893 | 0.84888417 | 0.39594575 | NA |
| JUND | 89.5497812 | -1.1210492 | 0.63372294 | -1.7689894 | 0.07689564 | 0.27670206 |
| LSM4 | 22.094252 | 0.43871362 | 0.49084528 | 0.89379206 | 0.37143316 | 0.62449127 |
| PGPEP1 | 54.44801 | -0.276365 | 0.33166303 | -0.8332704 | 0.40469222 | 0.65220185 |
| GDF15 | 192.902623 | -1.1150478 | 0.61779429 | -1.8048853 | 7.11E-02 | 0.26653676 |
| SSBP4 | 23.7530074 | 0.49409501 | 0.40595747 | 1.21711028 | 0.22356226 | 0.48054488 |
| ISYNA1 | 24.3552787 | 0.28212022 | 0.69153991 | 0.40795942 | 0.68330347 | 0.8424322 |
| ELL | 58.848809 | -0.4819908 | 0.36569522 | -1.3180123 | 0.18749952 | 0.43816783 |
| FKBP8 | 27.7601721 | 0.9522041 | 0.50940898 | 1.86923305 | 0.0615904 | 0.24806975 |
| KXD1 | 20.1018255 | -0.3767646 | 0.49220744 | -0.765459 | 0.44399828 | NA |
| UBA52 | 37.2317112 | 0.3119136 | 0.43308332 | 0.72021614 | 0.47139193 | 0.70071527 |
| REX1BD | 12.6030995 | 2.0592838 | 0.75756303 | 2.71830027 | 0.00656183 | NA |
| CRTC1 | 86.7706587 | 0.14548027 | 0.34928478 | 0.41650905 | 0.67703754 | 0.8388857 |
| UPF1 | 39.7280182 | -0.6011765 | 0.31555976 | -1.9051115 | 0.05676557 | 0.23723351 |
| COPE | 22.6533352 | 0.07119169 | 0.47395326 | 0.15020824 | 0.88060032 | 0.94387699 |
| DDX49 | 12.5164606 | -0.1878739 | 0.59975066 | -0.3132533 | 0.75408822 | NA |
| SUGP2 | 110.836033 | -0.1270399 | 0.23640266 | -0.5373876 | 0.59099991 | 0.78517152 |
| SLC25A42 | 21.600332 | 1.47952485 | 0.47530639 | 3.11278128 | 0.00185333 | 0.03984666 |
| TMEM161A | 18.5818984 | 1.77210501 | 0.52392333 | 3.38237467 | 0.00071862 | NA |
| SUGP1 | 28.0611424 | -0.203234 | 0.39783409 | -0.5108512 | 0.60945525 | 0.79648926 |
| MAU2 | 71.2569459 | -0.073236 | 0.30336727 | -0.2414104 | 0.80923706 | 0.90756938 |
| GATAD2A | 155.305164 | -0.5910187 | 0.29590582 | -1.9973202 | 0.04579041 | 0.21412441 |
| ATP13A1 | 17.0328521 | -0.5108508 | 0.54640212 | -0.9349355 | 0.34982153 | NA |
| ZNF101 | 13.6363729 | 0.6649703 | 0.48210168 | 1.37931546 | 0.16779751 | NA |
| ZNF14 | 29.2794914 | -0.3251719 | 0.36097772 | -0.9008089 | 0.36768993 | 0.6222756 |
| AC011477.4 | 14.2834183 | -0.4450266 | 0.57115383 | -0.7791711 | 0.43587893 | NA |
| ZNF253 | 37.6631882 | 0.69953888 | 0.32979437 | 2.1211365 | 0.03391032 | 0.1825355 |
| ZNF682 | 16.5234479 | -0.4279074 | 0.49514844 | -0.8642002 | 0.38747794 | NA |
| ZNF90 | 41.7840696 | 0.72560145 | 0.47923501 | 1.51408272 | 0.13000486 | 0.3624221 |
| AC011447.3 | 18.2713901 | 0.80533752 | 0.49519307 | 1.62631015 | 0.10388369 | NA |
| ZNF737 | 35.8590636 | 0.69849183 | 0.38557669 | 1.81155098 | 0.0700556 | 0.26406366 |
| ZNF85 | 15.8063624 | 0.81951669 | 0.62269774 | 1.31607462 | 0.188149 | NA |
| ZNF430 | 34.4459108 | -0.8245465 | 0.40633487 | -2.029229 | 0.04243497 | 0.20555189 |
| ZNF714 | 22.1001773 | -0.3629628 | 0.49880473 | -0.7276652 | 0.46681858 | 0.6967428 |
| ZNF431 | 77.89282 | 0.12298822 | 0.42478053 | 0.28953356 | 0.7721731 | 0.88734195 |
| ZNF708 | 60.1284174 | -0.0965127 | 0.3375214 | -0.2859455 | 0.7749199 | 0.888244 |
| ZNF493 | 37.4493734 | 0.99964809 | 0.37857402 | 2.64056179 | 0.00827687 | 0.08912945 |
| AC010615.4 | 58.419703 | 0.51916043 | 0.80829598 | 0.64229001 | 0.5206849 | 0.7372538 |
| ZNF429 | 48.7395399 | 0.45537559 | 0.35667447 | 1.27672606 | 0.20169898 | 0.45567274 |
| AC123912.1 | 19.2164243 | 0.07795086 | 0.61995706 | 0.12573589 | 0.89994099 | NA |
| ZNF100 | 37.3653657 | 0.22809966 | 0.51883105 | 0.43964149 | 0.66019679 | 0.82678716 |
| ZNF43 | 57.707846 | -0.2939319 | 0.34022006 | -0.8639463 | 0.38761741 | 0.63701577 |
| ZNF208 | 72.1062048 | -0.1027997 | 0.59601267 | -0.1724791 | 0.86306093 | 0.93629034 |
| ZNF676 | 16.7887363 | 0.03473635 | 0.91141838 | 0.0381124 | 0.96959806 | NA |
| ZNF98 | 86.9415929 | 0.64167814 | 0.35294469 | 1.81806995 | 0.06905344 | 0.26169589 |
| ZNF91 | 263.489214 | -0.7955051 | 0.41623917 | -1.9111731 | 0.05598234 | 0.23590316 |
| ZNF675 | 17.9458281 | 0.34782453 | 0.52839428 | 0.65826703 | 0.51036656 | NA |
| ZNF254 | 154.918459 | -0.4582819 | 0.26180025 | -1.7505022 | 0.08003169 | 0.28223671 |
| LINC00662 | 58.8539851 | 0.21095112 | 0.28208383 | 0.7478313 | 0.45456192 | 0.69049992 |
| AC006504.5 | 46.5437557 | 1.37924887 | 0.4371747 | 3.15491465 | 0.00160545 | 0.03677143 |
| UQCRFS1 | 36.214405 | 0.94539582 | 0.35610419 | 2.65482922 | 0.00793486 | 0.08725829 |
| C19orf12 | 33.5659339 | 0.23797012 | 0.32695719 | 0.72783266 | 0.46671603 | 0.69672484 |

| | | | | | | |
|---|---|---|---|---|---|---|
| URI1 | 182.134945 | 0.05251705 | 0.20291836 | 0.25880878 | 0.79578278 | 0.8998778 |
| ZNF507 | 49.8084528 | -0.3125055 | 0.26974423 | -1.1585251 | 0.24664983 | 0.50296216 |
| DPY19L3 | 74.5638954 | -0.6880181 | 0.31649831 | -2.1738443 | 0.02971683 | 0.1702177 |
| PDCD5 | 19.051026 | 0.16794091 | 0.45821418 | 0.36651182 | 0.7139832 | NA |
| ANKRD27 | 38.0073141 | 0.17509079 | 0.33240519 | 0.52673905 | 0.59837482 | 0.7894568 |
| NUDT19 | 14.4673769 | 0.4418023 | 0.54323313 | 0.81328307 | 0.41605578 | NA |
| CEP89 | 34.7049094 | 0.39869843 | 0.42372105 | 0.94094553 | 0.34673278 | 0.6038014 |
| RHPN2 | 124.114875 | -0.7725258 | 0.32368031 | -2.3866937 | 1.70E-02 | 0.12988597 |
| GPATCH1 | 19.2411223 | -0.0750345 | 0.44787093 | -0.167536 | 0.86694829 | NA |
| PEPD | 33.2310912 | -0.4962363 | 0.46246227 | -1.0730309 | 0.28325727 | 0.54207638 |
| LSM14A | 241.25125 | -0.4884474 | 0.17515805 | -2.7886096 | 0.00529348 | 0.07098247 |
| KIAA0355 | 141.334199 | -0.6757512 | 0.23922926 | -2.8247015 | 0.00473247 | 0.06664966 |
| GPI | 203.99804 | -0.6274564 | 0.27620171 | -2.2717324 | 0.02310268 | 0.14806105 |
| UBA2 | 94.5374746 | -0.4382565 | 0.23514792 | -1.8637482 | 0.06235707 | 0.24933194 |
| WTIP | 22.4802471 | 0.87701034 | 0.5591721 | 1.56840863 | 0.1167858 | 0.34414008 |
| SCGB2B2 | 74.7953067 | 0.36946395 | 0.39984061 | 0.92402808 | 0.3554717 | 0.61168935 |
| ZNF302 | 78.2459683 | -0.29259 | 0.36622247 | -0.7989405 | 0.42432493 | 0.66715549 |
| ZNF181 | 13.2069918 | -0.0755956 | 0.49008763 | -0.1542491 | 0.87741333 | NA |
| GRAMD1A | 55.3790465 | -0.205515 | 0.3738448 | -0.5497336 | 5.83E-01 | 0.7783237 |
| HPN | 228.520086 | 1.29853331 | 0.34117199 | 3.8060959 | 0.00014118 | 0.00890692 |
| LSR | 32.3946939 | -0.3681881 | 0.38436715 | -0.9579072 | 0.33810955 | 0.59389074 |
| USF2 | 25.3576215 | 0.26430505 | 0.53742497 | 0.49179897 | 0.62286146 | 0.80410667 |
| DMKN | 47.8155775 | 0.15021517 | 0.50464773 | 0.29766341 | 0.76596007 | 0.8841093 |
| TMEM147 | 16.6036453 | 0.58427386 | 0.56038174 | 1.04263543 | 0.29711717 | NA |
| COX6B1 | 59.0946463 | 0.53948945 | 0.43191813 | 1.24905486 | 0.21164501 | 0.46623687 |
| KMT2B | 30.4206816 | -0.2021022 | 0.40937785 | -0.4936814 | 0.62153123 | 0.80353114 |
| PROSER3 | 17.6541703 | -0.8578374 | 0.416662 | -2.0588328 | 0.03951026 | NA |
| TBCB | 16.8571725 | -0.1555878 | 0.45063003 | -0.3452673 | 0.7298934 | NA |
| CAPNS1 | 47.5054356 | -0.4119124 | 0.42244168 | -0.9750752 | 0.32952296 | 0.58535385 |
| ZNF565 | 60.0902181 | 0.20852304 | 0.41230501 | 0.50574947 | 0.61303254 | 0.7987918 |
| ZNF146 | 77.737052 | -0.5056112 | 0.36150142 | -1.3986424 | 0.16192024 | 0.40782071 |
| LINC00665 | 29.3897702 | 0.11158233 | 0.57728937 | 0.19328666 | 0.84673447 | 0.92563742 |
| ZFP14 | 56.9224146 | 0.98085417 | 0.37963472 | 2.58367879 | 0.00977528 | 0.09787489 |
| ZFP82 | 31.0851098 | 0.1188704 | 0.37997562 | 0.3128369 | 0.75440458 | 0.87873063 |
| ZNF566 | 31.789058 | 0.46747853 | 0.44278052 | 1.05577934 | 0.29106904 | 0.55032139 |
| ZNF260 | 25.4027701 | 0.54163445 | 0.37539289 | 1.44284685 | 0.14906362 | 0.38943809 |
| ZNF529 | 46.9169663 | 0.18834046 | 0.32073863 | 0.58720855 | 0.55706365 | 0.763236 |
| ZNF529-AS1 | 37.1942038 | 0.18560012 | 0.38398324 | 0.48335473 | 0.62884389 | 0.80683637 |
| ZNF382 | 14.9733077 | 0.13264823 | 0.54249276 | 0.24451613 | 0.80683111 | NA |
| ZNF461 | 28.1272013 | -0.3990489 | 0.34188823 | -1.1671912 | 0.24313316 | 0.50037324 |
| ZNF567 | 25.7513504 | 0.23019702 | 0.42779115 | 0.53810609 | 0.59050381 | 0.78499271 |
| AC020928.1 | 48.3093532 | 0.28392717 | 0.32176234 | 0.88241271 | 0.37755367 | 0.62901095 |
| ZNF790 | 43.1914736 | 0.00959885 | 0.40407473 | 0.02375515 | 0.98104792 | 0.99304653 |
| ZNF345 | 71.9094794 | 0.12259548 | 0.40033612 | 0.30623137 | 0.75942849 | 0.88058098 |
| ZNF568 | 42.5760593 | -0.3786382 | 0.33807572 | -1.1199805 | 0.26272205 | 0.52252497 |
| ZNF420 | 74.3534174 | 0.09891684 | 0.34987758 | 0.28271844 | 0.77739268 | 0.8896927 |
| ZNF383 | 23.49809 | 0.31634818 | 0.48798287 | 0.64827722 | 0.51680567 | 0.7344633 |
| LINC01535 | 15.3426611 | 0.47738795 | 0.51065047 | 0.93486245 | 0.34985919 | NA |
| HKR1 | 79.8602023 | 0.34515741 | 0.33699611 | 1.02421778 | 0.30573242 | 0.56557634 |
| ZNF569 | 25.5256305 | 0.28630487 | 0.43437592 | 0.65911774 | 0.50982017 | 0.72990216 |
| ZNF793 | 39.3043509 | 0.23986344 | 0.36319489 | 0.66042625 | 0.50898033 | 0.72967436 |
| ZNF540 | 29.8291087 | 1.185816 | 0.46552594 | 2.54726083 | 0.01085722 | 0.10244908 |
| ZNF571 | 33.305068 | 0.4995544 | 0.45877192 | 1.08889491 | 0.27620023 | 0.53482093 |
| ZFP30 | 37.576269 | 0.46043234 | 0.4337619 | 1.06148636 | 0.28846893 | 0.54774172 |

| | | | | | | |
|---|---|---|---|---|---|---|
| ZNF573 | 50.337284 | 0.61543164 | 0.40719967 | 1.51137558 | 0.13069279 | 0.36322236 |
| SIPA1L3 | 253.284065 | 0.03570821 | 0.23330717 | 0.15305233 | 0.87835701 | 0.94342075 |
| SPINT2 | 23.6602488 | -0.0331626 | 0.37288006 | -0.0889364 | 0.92913248 | 0.97064176 |
| CATSPERG | 26.0777397 | -0.6104803 | 0.61763484 | -0.9884162 | 0.32294886 | 0.57912936 |
| PSMD8 | 33.7675559 | 0.27605951 | 0.46130667 | 0.59842948 | 0.5495534 | 0.75628688 |
| SPRED3 | 19.0852142 | -1.1499641 | 0.40736062 | -2.8229634 | 4.76E-03 | NA |
| EIF3K | 66.0806103 | -0.1503135 | 0.28462572 | -0.5281092 | 0.59742354 | 0.7894568 |
| ACTN4 | 291.827401 | -1.0569196 | 0.28498963 | -3.7086248 | 0.00020839 | 0.01135785 |
| ECH1 | 19.077063 | 1.30593425 | 0.56156468 | 2.32552776 | 0.02004376 | NA |
| HNRNPL | 70.1387156 | -0.6122842 | 0.34660897 | -1.7664984 | 0.07731228 | 0.27731248 |
| SIRT2 | 32.2153061 | 1.2505231 | 0.4840306 | 2.58356206 | 0.00977859 | 0.09787489 |
| MRPS12 | 18.7295694 | 0.38099893 | 0.48498171 | 0.78559443 | 0.43210513 | NA |
| FBXO17 | 139.572225 | 0.0066684 | 0.46271785 | 0.01441136 | 0.98850179 | 0.99581595 |
| PAK4 | 72.0650983 | 0.29029336 | 0.38998312 | 0.74437417 | 0.45665014 | 0.69135251 |
| SAMD4B | 86.5594988 | -0.0768678 | 0.25943539 | -0.2962888 | 0.76700957 | 0.88460817 |
| MED29 | 13.6765353 | 0.50396037 | 0.48057277 | 1.0486661 | 0.29433182 | NA |
| ZFP36 | 27.8159453 | -1.5657884 | 0.67892241 | -2.3062847 | 0.02109473 | 0.1427136 |
| RPS16 | 48.9988862 | 0.27606137 | 0.43304581 | 0.63748769 | 0.52380722 | 0.73909151 |
| SUPT5H | 64.0277673 | -0.5095488 | 0.25694948 | -1.9830701 | 0.0473596 | 0.21737317 |
| TIMM50 | 13.5719041 | -0.5117887 | 0.6103184 | -0.8385601 | 0.40171622 | NA |
| FBL | 11.6074793 | 0.71150587 | 0.52212411 | 1.36271407 | 0.17297264 | NA |
| PSMC4 | 18.521518 | -0.4720423 | 0.46788538 | -1.0088845 | 0.31303004 | NA |
| ZNF546 | 19.1204267 | -0.1459153 | 0.48831675 | -0.2988127 | 0.76508294 | NA |
| ZNF780B | 119.701257 | 0.26083777 | 0.33991408 | 0.76736384 | 0.44286522 | 0.68272427 |
| ZNF780A | 13.9546094 | 0.62343673 | 0.58109517 | 1.07286511 | 0.28333164 | NA |
| MAP3K10 | 14.1952651 | 0.56632832 | 0.5967616 | 0.94900261 | 0.34261928 | NA |
| AKT2 | 66.5614194 | 0.15413468 | 0.32595352 | 0.47287318 | 0.63630366 | 0.81207582 |
| PLD3 | 90.1761405 | 0.10846551 | 0.27883709 | 0.38899242 | 0.69728175 | 0.85041636 |
| BLVRB | 42.5023299 | -0.0027001 | 0.35821613 | -0.0075375 | 0.99398598 | 0.99730572 |
| SPTBN4 | 29.8822703 | -1.8231534 | 0.75685041 | -2.4088688 | 0.01600205 | 0.12504341 |
| COQ8B | 17.2240014 | -0.0840137 | 0.51130875 | -0.1643111 | 0.86948624 | NA |
| ITPKC | 31.586813 | -1.1287161 | 0.33328134 | -3.3866766 | 0.00070745 | 0.02388192 |
| SNRPA | 19.258146 | -0.5989083 | 0.45356275 | -1.3204529 | 0.18668385 | NA |
| AC008537.1 | 27.0243176 | -1.8777195 | 0.61256003 | -3.065364 | 0.00217405 | 0.04403601 |
| HNRNPUL1 | 54.0675958 | -0.5618086 | 0.28907689 | -1.9434571 | 0.05196095 | 0.22846489 |
| TGFB1 | 21.2221015 | 0.30366378 | 0.46920884 | 0.64718256 | 0.5175138 | 0.73479131 |
| DMAC2 | 11.7713637 | 1.47902686 | 0.56966801 | 2.59629615 | 0.00942348 | NA |
| RPS19 | 133.771834 | 0.73781541 | 0.39798473 | 1.8538787 | 0.06375652 | 0.25122665 |
| ARHGEF1 | 26.9419412 | -0.1520872 | 0.38341167 | -0.3966683 | 0.69161211 | 0.846991 |
| RABAC1 | 23.1138169 | 0.28700467 | 0.58484122 | 0.49073946 | 0.62361073 | 0.8044129 |
| GRIK5 | 54.4882508 | 0.29350567 | 0.45548764 | 0.64437679 | 0.51933113 | 0.73628508 |
| CIC | 20.5152528 | -0.2778796 | 0.42277801 | -0.6572708 | 0.51100683 | NA |
| MEGF8 | 29.1565584 | 0.73514681 | 0.38217723 | 1.92357564 | 0.0544078 | 0.23311395 |
| LIPE-AS1 | 20.7636028 | -0.1395921 | 0.5373599 | -0.2597739 | 0.79503819 | NA |
| PHLDB3 | 13.5659743 | 0.06611174 | 0.48558099 | 0.13614977 | 0.89170288 | NA |
| ETHE1 | 18.8510169 | 0.00983081 | 0.43200792 | 0.0227561 | 0.98184483 | NA |
| ZNF428 | 18.6449291 | 0.25255599 | 0.47262815 | 0.53436511 | 0.59308895 | NA |
| SMG9 | 13.8618578 | 0.09001712 | 0.47071819 | 0.19123357 | 0.8483426 | NA |
| ZNF283 | 26.248665 | 0.2535233 | 0.42000407 | 0.60362106 | 0.54609561 | 0.75409004 |
| ZNF45 | 12.9716334 | 0.57308399 | 0.53967959 | 1.06189672 | 0.28828257 | NA |
| ZNF155 | 26.7833568 | 0.85189115 | 0.49390561 | 1.72480555 | 0.08456252 | 0.28931012 |
| ZNF224 | 37.8377214 | -0.0395969 | 0.3194768 | -0.1239428 | 0.90136055 | 0.95637074 |
| ZNF234 | 18.4616804 | 0.72249259 | 0.53201278 | 1.35803615 | 0.17445221 | NA |
| ZNF226 | 28.8527458 | -0.0489948 | 0.3558634 | -0.1376787 | 0.89049437 | 0.949686 |

| | | | | | | |
|---|---|---|---|---|---|---|
| ZNF227 | 22.3894571 | 0.28015637 | 0.40107847 | 0.69850762 | 0.48485979 | 0.7118397 |
| ZNF235 | 87.5469089 | 0.38544953 | 0.45313005 | 0.85063775 | 0.39497061 | 0.64334162 |
| ZNF229 | 16.7371822 | -0.1075211 | 0.52827203 | -0.2035336 | 0.83871799 | NA |
| ZNF180 | 14.5089699 | -0.9340253 | 0.4919862 | -1.8984786 | 0.05763306 | NA |
| AC243964.2 | 31.6552417 | -0.9055552 | 0.41064713 | -2.2051906 | 0.02744072 | 0.16309747 |
| CEACAM19 | 27.6718503 | -1.6200839 | 0.56376234 | -2.8737002 | 4.06E-03 | 0.06207769 |
| BCL3 | 19.7450622 | -0.4342698 | 0.63512477 | -0.6837552 | 4.94E-01 | NA |
| CBLC | 18.9404415 | 0.5191714 | 0.47002487 | 1.10456157 | 0.26934961 | NA |
| BCAM | 111.855283 | 0.54466705 | 0.508168 | 1.07182477 | 0.28379875 | 0.54270782 |
| NECTIN2 | 78.4587433 | -0.2228105 | 0.35922891 | -0.6202465 | 0.53509548 | 0.74775242 |
| TOMM40 | 15.6252276 | -0.4280001 | 0.57422651 | -0.7453506 | 0.45605978 | NA |
| APOE | 66.5300739 | -0.7593183 | 0.67027955 | -1.1328382 | 0.25728221 | 0.51570343 |
| CLPTM1 | 38.6063868 | -0.2699199 | 0.37722284 | -0.715545 | 0.47427235 | 0.70282524 |
| RELB | 63.1330864 | -1.4147664 | 0.51135626 | -2.7666942 | 0.00566279 | 0.07350161 |
| CLASRP | 65.9059129 | -0.3060473 | 0.28994786 | -1.0555252 | 0.29118518 | 0.55040579 |
| GEMIN7-AS1 | 20.3868014 | 0.22673183 | 0.50306189 | 0.45070365 | 0.65220315 | NA |
| MARK4 | 58.1084218 | 0.33775919 | 0.32125923 | 1.05136024 | 0.29309317 | 0.55234367 |
| TRAPPC6A | 15.5199939 | -0.1192875 | 0.50820008 | -0.2347254 | 0.81442188 | NA |
| PPP1R13L | 38.6129214 | -0.6220001 | 0.32863945 | -1.8926519 | 5.84E-02 | 0.24020328 |
| ERCC1 | 24.6342111 | -2.0345393 | 0.45338516 | -4.4874413 | 7.21E-06 | 0.00135324 |
| VASP | 31.3692082 | -1.207793 | 0.53373829 | -2.2628937 | 0.02364224 | 0.15002007 |
| OPA3 | 64.0303766 | -0.488389 | 0.38309671 | -1.2748452 | 0.20236404 | 0.45643401 |
| EML2 | 43.4925941 | 0.81003043 | 0.33464126 | 2.42059339 | 0.0154952 | 0.1229552 |
| SNRPD2 | 24.8736224 | -0.2929955 | 0.45008796 | -0.6509739 | 0.51506331 | 0.73279894 |
| FBXO46 | 27.9580696 | -0.5093778 | 0.48941471 | -1.0407896 | 0.29797319 | 0.5578284 |
| DMPK | 16.3184792 | 0.02918186 | 0.43266216 | 0.06744722 | 0.94622568 | NA |
| SYMPK | 88.6416342 | -0.558652 | 0.29428976 | -1.8983059 | 0.0576558 | 0.23871796 |
| MYPOP | 26.2735026 | 0.73150952 | 0.4934699 | 1.48237923 | 0.13823942 | 0.37479 |
| PPP5C | 41.7997782 | 0.27419058 | 0.30815984 | 0.88976738 | 0.37359081 | 0.62575308 |
| PPP5D1 | 21.2594905 | 0.34453944 | 0.61301616 | 0.56203972 | 0.57408896 | 0.77388388 |
| CALM3 | 19.2916599 | 0.84379517 | 0.60121985 | 1.4034719 | 0.16047617 | NA |
| PRKD2 | 40.4677107 | -0.0752079 | 0.33587865 | -0.2239139 | 0.82282432 | 0.91475524 |
| STRN4 | 18.150745 | 0.44774228 | 0.43322146 | 1.03351824 | 0.30136144 | NA |
| FKRP | 17.7066168 | -0.6708652 | 0.53503543 | -1.2538706 | 0.20988903 | NA |
| SLC1A5 | 17.5574533 | -0.164363 | 0.57276498 | -0.2869641 | 0.7741398 | NA |
| AP2S1 | 20.9336006 | 0.49997267 | 0.5144397 | 0.9718781 | 0.3311112 | NA |
| ARHGAP35 | 108.255168 | 0.34387512 | 0.31551894 | 1.08987155 | 0.27576973 | 0.53465985 |
| TMEM160 | 22.0038462 | 0.31269981 | 0.63321275 | 0.49383056 | 0.62142585 | 0.80353114 |
| ZC3H4 | 29.4018437 | -0.3555727 | 0.39283693 | -0.9051406 | 0.36539088 | 0.62110919 |
| SAE1 | 30.7559158 | -0.302217 | 0.40871377 | -0.7394343 | 0.45964334 | 0.69299864 |
| CCDC9 | 14.2529974 | -0.3494677 | 0.51076489 | -0.6842046 | 0.493846 | NA |
| C5AR1 | 38.1670529 | 1.27577213 | 0.38706736 | 3.29599512 | 0.00098074 | 0.0283787 |
| DHX34 | 33.6413571 | -0.4689218 | 0.39657459 | -1.1824304 | 0.23703498 | 0.49508567 |
| NAPA-AS1 | 15.3809805 | 0.22998323 | 0.59091768 | 0.38919673 | 0.69713062 | NA |
| NAPA | 73.0635082 | -0.7852344 | 0.3428284 | -2.2904589 | 0.02199473 | 0.14481901 |
| NOP53 | 12.8806978 | 0.85392587 | 0.54785863 | 1.55866098 | 0.11907664 | NA |
| SELENOW | 60.5987746 | 1.00335431 | 0.36443947 | 2.75314394 | 0.00590259 | 0.07567727 |
| PLA2G4C | 38.6237312 | -3.6945237 | 0.88362894 | -4.1810805 | 2.90E-05 | 0.00323639 |
| ZSWIM9 | 15.2574601 | 0.12590494 | 0.50528925 | 0.24917399 | 0.8032262 | NA |
| CARD8 | 38.5771447 | 0.13768743 | 0.44258966 | 0.311095 | 0.7557284 | 0.87904624 |
| TMEM143 | 21.5793927 | 0.24133718 | 0.50926817 | 0.47389018 | 0.63557822 | 0.81168834 |
| KDELR1 | 27.759229 | 0.24146694 | 0.35095169 | 0.68803471 | 0.49143092 | 0.71672078 |
| CYTH2 | 29.1706671 | -0.566159 | 0.36450677 | -1.5532194 | 0.12037073 | 0.34903213 |
| LMTK3 | 16.2684772 | -0.3804873 | 0.43867072 | -0.8673642 | 0.3857425 | NA |

| | | | | | | |
|---|---|---|---|---|---|---|
| RPL18 | 61.1754226 | 0.35177561 | 0.47269338 | 0.74419406 | 0.45675908 | 0.69135799 |
| BCAT2 | 25.8705942 | -0.4484719 | 0.54505958 | -0.8227942 | 0.41062502 | 0.65626678 |
| HSD17B14 | 27.3300956 | -0.8223932 | 0.65798092 | -1.2498739 | 0.21134561 | 0.46611094 |
| NUCB1 | 48.921442 | 0.02403866 | 0.38708327 | 0.06210204 | 0.95048157 | 0.97993174 |
| BAX | 17.1435964 | -0.4585286 | 0.51856679 | -0.8842228 | 0.37657595 | NA |
| FTL | 629.435705 | 0.45534721 | 0.55118762 | 0.82612016 | 0.40873593 | 0.65460684 |
| GYS1 | 59.9210039 | -0.3849335 | 0.37572526 | -1.0245078 | 0.30559549 | 0.56557634 |
| SNRNP70 | 187.50541 | -0.1587507 | 0.35673857 | -0.4450057 | 0.65631564 | 0.82449183 |
| TRPM4 | 42.6126681 | 1.39091583 | 0.47296417 | 2.94084821 | 0.00327315 | 0.05687004 |
| SLC6A16 | 72.994305 | -0.1099101 | 0.297842 | -0.3690215 | 0.71211174 | 0.857224 |
| TEAD2 | 13.7388389 | -0.1128823 | 0.50595475 | -0.2231074 | 0.82345189 | NA |
| PIH1D1 | 20.5706028 | 0.34214146 | 0.46324324 | 0.73857841 | 0.46016304 | NA |
| RPL13A | 83.2717139 | 0.44489378 | 0.44863906 | 0.99165192 | 0.32136735 | 0.57793564 |
| RPS11 | 68.1814374 | 0.17209299 | 0.45901212 | 0.37492035 | 0.7077197 | 0.8547495 |
| FCGRT | 22.940994 | 1.55412915 | 0.41368691 | 3.7567762 | 0.00017212 | 0.00982049 |
| NOSIP | 32.9190129 | 0.2962908 | 0.41237531 | 0.71849792 | 0.47245033 | 0.70161107 |
| PRR12 | 24.9750748 | 0.52216009 | 0.52104576 | 1.00213863 | 0.31627664 | 0.57374058 |
| SCAF1 | 20.126751 | -0.5365803 | 0.43515877 | -1.2330679 | 0.21755042 | NA |
| IRF3 | 42.1061635 | -0.2436721 | 0.42450319 | -0.5740171 | 0.5659562 | 0.76991249 |
| PRMT1 | 32.6902068 | -0.2488904 | 0.44414501 | -0.560381 | 0.57521957 | 0.77388388 |
| AP2A1 | 26.0473418 | -0.8972022 | 0.43341461 | -2.0700784 | 0.03844501 | 0.19556562 |
| MED25 | 34.3189644 | 0.44324489 | 0.4212691 | 1.05216568 | 0.29272354 | 0.5522287 |
| PTOV1 | 37.5519947 | 0.15300714 | 0.30791541 | 0.49691292 | 0.61925046 | 0.8027391 |
| PNKP | 37.2812408 | -0.6384535 | 0.50503087 | -1.2641871 | 0.20616287 | 0.45924573 |
| TBC1D17 | 29.7995479 | -0.5633873 | 0.44210651 | -1.2743248 | 0.20254831 | 0.45644281 |
| VRK3 | 39.6957245 | -0.4155726 | 0.35315071 | -1.1767572 | 0.23929244 | 0.49698703 |
| MYH14 | 37.5865059 | 0.40193025 | 0.34658087 | 1.15970121 | 0.24617048 | 0.50296216 |
| NR1H2 | 23.5520033 | -0.1647539 | 0.37349371 | -0.4411155 | 0.65912938 | 0.82613888 |
| EMC10 | 45.876518 | -0.0418955 | 0.33578036 | -0.1247707 | 0.90070509 | 0.95621962 |
| ETFB | 30.2500187 | 1.24465617 | 0.53244384 | 2.337629 | 0.0194065 | 0.1366689 |
| ZNF175 | 19.6014354 | -0.2544384 | 0.41423286 | -0.6142401 | 0.53905667 | NA |
| SPACA6 | 28.5356647 | -0.427263 | 0.427789 | -0.9987704 | 0.31790593 | 0.57534022 |
| ZNF577 | 57.1134103 | -0.1328713 | 0.46000135 | -0.2888499 | 0.77269624 | 0.88749485 |
| ZNF350-AS1 | 45.8514242 | 0.12945346 | 0.66556988 | 0.19450018 | 0.84578425 | 0.92531372 |
| ZNF350 | 21.7345465 | -0.3635567 | 0.45695712 | -0.7956035 | 0.42626251 | 0.66917041 |
| ZNF615 | 18.9180145 | 0.27621534 | 0.51182079 | 0.53967199 | 0.58942326 | NA |
| ZNF432 | 38.8161338 | -0.8142895 | 0.37706204 | -2.1595637 | 0.03080646 | 0.17345817 |
| ZNF616 | 13.1230983 | 0.48507948 | 0.494107 | 0.98172964 | 0.32623306 | NA |
| PPP2R1A | 77.5801322 | -0.0574255 | 0.36340032 | -0.1580227 | 0.87443887 | 0.94159989 |
| ZNF766 | 37.0726354 | -0.4218275 | 0.36842207 | -1.1449572 | 0.25222688 | 0.50912938 |
| ZNF480 | 29.084487 | -0.9458903 | 0.42503779 | -2.2254264 | 0.02605263 | 0.15801977 |
| ZNF528 | 25.4888994 | -0.1295909 | 0.41181154 | -0.314685 | 0.75300086 | 0.87842978 |
| ZNF808 | 28.6466956 | 0.18164217 | 0.40804599 | 0.44515122 | 0.65621047 | 0.82449183 |
| ZNF701 | 43.625858 | -0.2279309 | 0.41251007 | -0.5525462 | 0.5805742 | 0.77770268 |
| ZNF83 | 357.934802 | -1.5193921 | 0.67435319 | -2.2531102 | 0.0242522 | 0.15213462 |
| ZNF611 | 151.007403 | -1.8320671 | 0.44770781 | -4.0921043 | 4.27E-05 | 0.00401254 |
| ZNF28 | 19.1197709 | -1.0451019 | 0.46766068 | -2.234744 | 0.02543416 | NA |
| AC010487.3 | 17.6972256 | 0.64355118 | 0.59841464 | 1.07542687 | 0.28218366 | NA |
| ZNF160 | 87.5930236 | 0.04973063 | 0.35987676 | 0.13818795 | 0.89009188 | 0.94967256 |
| ZNF347 | 29.9963864 | -0.8339325 | 0.36759686 | -2.2686063 | 0.02329228 | 0.14878065 |
| ZNF677 | 16.5172943 | 0.48323916 | 0.55748046 | 0.86682708 | 0.38603676 | NA |
| ZNF525 | 25.8179603 | -0.2171852 | 0.35247245 | -0.6161763 | 0.53777819 | 0.74878549 |
| ZNF765 | 32.167822 | 0.04907148 | 0.39862486 | 0.12310192 | 0.90202639 | 0.95661107 |
| ZNF761 | 36.9281785 | -0.5705975 | 0.3411144 | -1.6727451 | 0.09437748 | 0.30830642 |

| | | | | | | |
|---|---|---|---|---|---|---|
| ZNF331 | 43.8425991 | -0.0283773 | 0.31874416 | -0.0890285 | 0.92905928 | 0.97064176 |
| PRPF31 | 26.6540045 | 0.28460862 | 0.34665694 | 0.82100944 | 0.41164089 | 0.65707964 |
| CNOT3 | 22.9178583 | -1.1797578 | 0.42271745 | -2.7908898 | 0.00525634 | 0.07082974 |
| TMC4 | 53.3344131 | 0.6327583 | 0.33039231 | 1.91517262 | 0.05547049 | 0.23523768 |
| MBOAT7 | 45.0080382 | -0.4199026 | 0.42263336 | -0.9935387 | 0.32044748 | 0.57736055 |
| RPS9 | 120.617356 | 0.5282898 | 0.45355028 | 1.16478773 | 0.24410493 | 0.5015083 |
| LENG8-AS1 | 16.749606 | 0.26040809 | 0.45019904 | 0.5784288 | 0.56297465 | NA |
| LENG8 | 125.776523 | -1.0471898 | 0.39122733 | -2.6766787 | 0.00743559 | 0.08453729 |
| AC011476.3 | 18.501048 | 1.40970062 | 0.6750981 | 2.08814188 | 0.03678504 | NA |
| RDH13 | 41.1825758 | 0.05597618 | 0.38663287 | 0.14477864 | 0.88488565 | 0.94638313 |
| EPS8L1 | 16.4389858 | 0.59891254 | 0.70810763 | 0.84579309 | 0.39766818 | NA |
| PPP1R12C | 45.1751088 | -0.532581 | 0.40291223 | -1.3218288 | 0.18622518 | 0.43607799 |
| PPP6R1 | 31.2036454 | -0.609434 | 0.38586217 | -1.5794084 | 0.1142424 | 0.34044736 |
| HSPBP1 | 12.5815383 | -0.6152701 | 0.51798521 | -1.1878139 | 0.23490672 | NA |
| RPL28 | 91.1871711 | 0.46837125 | 0.44500646 | 1.05250437 | 0.29256821 | 0.55207097 |
| U2AF2 | 25.8868257 | -0.7656198 | 0.37055733 | -2.0661305 | 0.03881616 | 0.1961707 |
| EPN1 | 55.5016615 | -0.3097586 | 0.34836229 | -0.8891851 | 0.3739036 | 0.62575308 |
| ZNF787 | 21.8228162 | -0.5207422 | 0.48923629 | -1.0643981 | 0.28714839 | 0.5463983 |
| ZNF444 | 22.8372828 | 0.36204545 | 0.41898861 | 0.86409376 | 0.38753639 | 0.63701577 |
| ZSCAN5A | 96.9214591 | -0.3581035 | 0.26113609 | -1.3713288 | 0.17027247 | 0.41839333 |
| ZNF667 | 25.5532556 | 0.10111682 | 0.38613392 | 0.26186981 | 0.79342181 | 0.89875118 |
| ZNF667-AS1 | 22.511465 | 0.11866617 | 0.39675833 | 0.2990893 | 0.76487191 | 0.88317118 |
| ZNF471 | 26.7147235 | -0.0586677 | 0.42546137 | -0.137892 | 0.89032574 | 0.94967256 |
| ZNF264 | 36.6646116 | -0.4423847 | 0.3181819 | -1.3903516 | 0.16442213 | 0.41120418 |
| ZNF460-AS1 | 17.8226758 | -0.0340944 | 0.56341259 | -0.0605141 | 0.95174621 | NA |
| ZNF460 | 18.9798877 | -1.414185 | 0.66551767 | -2.1249399 | 0.03359163 | NA |
| AC005261.1 | 59.2660103 | -1.1052754 | 0.41499971 | -2.6633161 | 0.00773747 | 0.08618711 |
| AC003002.2 | 13.9541076 | 0.83682926 | 0.5206253 | 1.60735421 | 0.10797669 | NA |
| AC004076.1 | 28.096686 | -0.5586608 | 0.42402443 | -1.3175202 | 0.1876643 | 0.43816783 |
| ZNF549 | 19.9215511 | 0.025541 | 0.38062759 | 0.06710234 | 0.94650023 | NA |
| ZNF550 | 10.6525132 | 0.18386568 | 0.53694755 | 0.34242763 | 0.7320291 | NA |
| AC003006.1 | 14.2951392 | 0.44332317 | 0.50307654 | 0.8812241 | 0.37819654 | NA |
| ZNF154 | 14.9291656 | 0.0074811 | 0.54533636 | 0.01371832 | 0.98905471 | NA |
| ZNF586 | 147.797234 | -0.5152329 | 0.252672 | -2.0391373 | 0.04143632 | 0.20301425 |
| ZNF552 | 65.1611829 | 0.85317064 | 0.41810967 | 2.04054272 | 0.0412963 | 0.20301425 |
| ZNF814 | 80.1959571 | -0.476487 | 0.35528665 | -1.3411339 | 0.17987699 | 0.42904035 |
| ZNF587 | 38.8728163 | -0.3282306 | 0.31804561 | -1.0320235 | 0.30206109 | 0.56226462 |
| ZNF418 | 15.2252637 | 0.13875746 | 0.53537439 | 0.25917836 | 0.79549762 | NA |
| C19orf18 | 14.6896998 | -0.6045894 | 0.5382344 | -1.1232826 | 0.26131747 | NA |
| ZNF606 | 19.9874885 | 0.25224605 | 0.4564477 | 0.5526286 | 0.58051776 | NA |
| ZNF135 | 39.9096302 | 0.15649405 | 0.4394807 | 0.35608855 | 0.72177425 | 0.86145083 |
| ZSCAN18 | 66.0790309 | 0.51135314 | 0.36117514 | 1.4158038 | 0.15683296 | 0.40104036 |
| ZNF329 | 30.4407044 | 0.0966189 | 0.38540748 | 0.25069286 | 0.80205158 | 0.90378853 |
| ZNF274 | 31.621118 | -0.9790165 | 0.41730835 | -2.3460267 | 0.01897475 | 0.13585154 |
| AC020915.3 | 19.9361606 | -0.398209 | 0.40674794 | -0.9790067 | 0.32757665 | NA |
| RPS5 | 36.9503927 | 0.66045073 | 0.49142157 | 1.34395959 | 0.17896145 | 0.42802774 |
| TRIM28 | 27.5241967 | -0.4381379 | 0.52355131 | -0.8368576 | 0.40267261 | 0.65024957 |
| CHMP2A | 16.7201531 | 0.31055752 | 0.44930854 | 0.69118989 | 0.48944622 | NA |
| UBE2M | 15.7235784 | 0.40532911 | 0.50501763 | 0.80260388 | 0.42220373 | NA |
| MZF1-AS1 | 15.4231221 | 0.06984075 | 0.53985676 | 0.12936904 | 0.89706565 | NA |
| MZF1 | 64.8237459 | -0.5859565 | 0.23783324 | -2.4637286 | 0.01375002 | 0.11503685 |
| RPS4Y1 | 50.5808917 | 0.64456948 | 2.04915484 | 0.31455382 | 0.75310045 | 0.87842978 |
| ZFY | 30.60898 | -0.7365445 | 2.46175711 | -0.2991946 | 0.76479156 | 0.88317118 |
| LINC00278 | 97.5651367 | 0.57105184 | 2.73084741 | 0.20911159 | 0.83436113 | 0.91992307 |

| | | | | | | |
|---|---|---|---|---|---|---|
| TBL1Y | 59.8463455 | -0.109077 | 2.58318813 | -0.0422257 | 0.96631875 | 0.98723631 |
| TTTY15 | 35.5959816 | 0.08764563 | 2.4945616 | 0.03513468 | 0.97197235 | 0.98878882 |
| USP9Y | 161.435456 | 0.42353472 | 2.55833757 | 0.16555075 | 0.8685105 | 0.93857029 |
| DDX3Y | 36.2594854 | 0.08305336 | 2.4994768 | 0.0332283 | 0.97349253 | 0.98956313 |
| UTY | 352.431842 | 0.24169245 | 3.01453354 | 0.08017574 | 0.93609749 | 0.97405271 |
| NLGN4Y | 208.933368 | -0.0126197 | 2.57724765 | -0.0048966 | 0.99609311 | 0.99816804 |
| TTTY14 | 244.755253 | 0.18238796 | 1.41228491 | 0.12914389 | 0.8972438 | 0.95374748 |
| KDM5D | 41.6802319 | 0.54559242 | 2.49794135 | 0.21841683 | 0.82710436 | 0.91613502 |
| TTTY10 | 20.8252938 | 0.61520089 | 1.56864666 | 0.39218577 | 0.69492096 | NA |
| EIF1AY | 13.690566 | 0.37979473 | 1.91615066 | 0.19820714 | 0.842883 | NA |
| IL17RA | 29.9104065 | 0.21732243 | 0.31632915 | 0.68701361 | 0.49207415 | 0.71672078 |
| HDHD5 | 25.3945044 | -0.2123432 | 0.40301384 | -0.5268882 | 0.59827121 | 0.7894568 |
| ADA2 | 15.2768889 | 0.30299674 | 0.70698207 | 0.42857768 | 0.66823059 | NA |
| CECR2 | 236.15723 | 0.42726106 | 0.28500576 | 1.4991313 | 0.13383957 | 0.36774274 |
| ATP6V1E1 | 61.0839864 | -0.2129238 | 0.33186275 | -0.6416019 | 0.52113169 | 0.73775071 |
| BCL2L13 | 110.110527 | -0.1546383 | 0.24937611 | -0.6201009 | 0.53519139 | 0.74775242 |
| MICAL3 | 237.495155 | -0.9698418 | 0.38159115 | -2.541573 | 0.01103549 | 0.10282338 |
| PEX26 | 39.5698711 | 0.533748 | 0.3457417 | 1.54377676 | 0.12264242 | 0.35251484 |
| DGCR2 | 69.6463253 | -0.3050639 | 0.30236667 | -1.0089204 | 0.31301279 | 0.57102302 |
| SLC25A1 | 22.9707337 | 0.14988748 | 0.53446229 | 0.28044537 | 0.77913583 | 0.89055665 |
| CLTCL1 | 20.4261285 | 0.32807453 | 0.49852941 | 0.65808461 | 0.51048377 | NA |
| HIRA | 28.9673519 | -0.3740764 | 0.36930766 | -1.0129126 | 0.3111019 | 0.57037944 |
| UFD1 | 23.310977 | -0.0912125 | 0.42543792 | -0.2143967 | 0.83023775 | 0.91775671 |
| TXNRD2 | 29.309112 | -0.359437 | 0.39915126 | -0.9005032 | 0.36785253 | 0.62241391 |
| COMT | 66.0515253 | 0.68917175 | 0.39894064 | 1.72750449 | 0.0840771 | 0.28903146 |
| TANGO2 | 58.6748514 | 0.10335204 | 0.5058127 | 0.20432868 | 0.83809666 | 0.92179039 |
| DGCR8 | 34.7298154 | -0.9816491 | 0.4043096 | -2.4279639 | 0.01518385 | 0.1213084 |
| ZDHHC8 | 13.797065 | 0.7182394 | 0.62866847 | 1.14247721 | 0.25325571 | NA |
| KLHL22 | 66.1087754 | 0.15744728 | 0.28617776 | 0.55017303 | 0.5822007 | 0.7783237 |
| MED15 | 96.1797728 | -0.8734022 | 0.24248376 | -3.6018997 | 0.0003159 | 0.0146475 |
| PI4KA | 162.621135 | -0.5119682 | 0.28258476 | -1.8117333 | 0.07002742 | 0.26406366 |
| SNAP29 | 23.5183199 | 0.00980195 | 0.41112232 | 0.02384194 | 0.98097869 | 0.99304653 |
| CRKL | 33.4054316 | -0.1025458 | 0.35087291 | -0.2922591 | 0.77008854 | 0.88651526 |
| LZTR1 | 12.7332258 | 0.74567911 | 0.56622444 | 1.31693205 | 0.1878614 | NA |
| GGT2 | 132.960986 | 0.82382069 | 0.9123541 | 0.90296157 | 3.67E-01 | 0.62157013 |
| HIC2 | 34.0870396 | 0.26081775 | 0.4618259 | 0.5647534 | 0.57224151 | 0.77272682 |
| UBE2L3 | 67.3556327 | -0.2797045 | 0.34258753 | -0.8164468 | 0.41424466 | 0.65845716 |
| PPIL2 | 58.1548184 | -0.2185014 | 0.35042878 | -0.6235259 | 0.53293896 | 0.74615054 |
| YPEL1 | 15.8931212 | -0.6180853 | 0.61015654 | -1.0129946 | 0.31106273 | NA |
| MAPK1 | 153.952346 | -0.3293949 | 0.29667134 | -1.1103026 | 0.26686867 | 0.52588023 |
| PPM1F | 19.2343641 | 0.62142402 | 0.50904749 | 1.22075843 | 0.2221775 | NA |
| TOP3B | 16.8535056 | 0.20246646 | 0.58721742 | 0.34478959 | 0.73025256 | NA |
| AC245060.5 | 47.1764397 | 0.53039742 | 0.53640278 | 0.98880438 | 0.32275886 | 0.57912936 |
| BCR | 314.74246 | -1.0185251 | 0.40007223 | -2.5458532 | 0.0109011 | 0.10244908 |
| CHCHD10 | 76.5295986 | 0.80188563 | 0.40353398 | 1.98715761 | 0.04690493 | 0.21661307 |
| SMARCB1 | 41.2098643 | -0.18325 | 0.36791348 | -0.498079 | 0.61842839 | 0.80247821 |
| DDT | 36.4801425 | 0.8442735 | 0.49952619 | 1.69014863 | 0.09099952 | 0.30125735 |
| CABIN1 | 133.972661 | 0.17322118 | 0.33742418 | 0.51336327 | 0.60769723 | 0.79552231 |
| GUCD1 | 20.6537073 | -0.2450846 | 0.45985669 | -0.5329586 | 0.59406224 | NA |
| SNRPD3 | 34.6794333 | -0.0176357 | 0.31518726 | -0.0559531 | 0.95537919 | 0.98230746 |
| GGT1 | 16.4686987 | 1.06770534 | 0.66312282 | 1.61011703 | 0.10737231 | NA |
| SGSM1 | 143.912713 | 0.07380415 | 0.40434738 | 0.1825266 | 0.85516948 | 0.93153686 |
| KIAA1671 | 200.652314 | -0.5372424 | 0.26496997 | -2.0275596 | 0.04260521 | 0.20610104 |
| AL022323.4 | 14.6974308 | -0.2724675 | 0.5118667 | -0.5323016 | 0.59451713 | NA |

| | | | | | | |
|---|---|---|---|---|---|---|
| LRP5L | 51.2140506 | 0.22178242 | 0.37056343 | 0.59850056 | 0.54950598 | 0.75628688 |
| GRK3 | 45.6217459 | -0.6196271 | 0.4671249 | -1.3264699 | 0.18468411 | 0.43476148 |
| HPS4 | 55.004128 | -0.5813116 | 0.29214737 | -1.9897888 | 0.0466142 | 0.21639898 |
| TFIP11 | 17.8364348 | 0.02636064 | 0.53376546 | 0.04938618 | 0.96061154 | NA |
| TPST2 | 137.61269 | -0.545528 | 0.36148999 | -1.5091096 | 0.13127078 | 0.36392977 |
| MIATNB | 40.8419081 | -0.8424135 | 0.36191469 | -2.3276577 | 0.01993029 | 0.13832589 |
| PITPNB | 125.937788 | -0.3244336 | 0.24004949 | -1.3515281 | 0.17652632 | 0.42569745 |
| TTC28 | 289.75735 | -0.3616968 | 0.27817251 | -1.3002608 | 0.19351159 | 0.4454123 |
| HSCB | 39.5890253 | -0.2575039 | 0.36023889 | -0.7148142 | 0.47472384 | 0.70288437 |
| CCDC117 | 15.3911578 | 0.15150741 | 0.44728258 | 0.33872863 | 0.73481417 | NA |
| XBP1 | 28.4224275 | -0.0303336 | 0.40493085 | -0.0749105 | 0.94028596 | 0.97512544 |
| Z93930.2 | 66.8305469 | 0.97298548 | 0.51750156 | 1.88015952 | 0.06008634 | 0.24418404 |
| ZNRF3 | 274.034743 | 0.25842032 | 0.35093899 | 0.73636822 | 0.46150664 | 0.69416206 |
| KREMEN1 | 64.4763257 | 0.86344496 | 0.3385546 | 2.55038612 | 0.01076037 | 0.10199821 |
| RHBDD3 | 14.862589 | -0.5626682 | 0.49737164 | -1.1312831 | 0.25793594 | NA |
| EWSR1 | 123.053461 | -0.6434981 | 0.22083572 | -2.9139224 | 0.00356919 | 0.0593726 |
| AP1B1 | 37.6399594 | 0.23891362 | 0.34708192 | 0.68834936 | 0.4912328 | 0.71672078 |
| THOC5 | 16.9368277 | -0.4700347 | 0.46192978 | -1.0175457 | 0.30889388 | NA |
| NIPSNAP1 | 25.7663276 | 0.81673342 | 0.44803918 | 1.82290622 | 0.0683176 | 0.25987613 |
| NF2 | 60.1247621 | -0.3300824 | 0.39496723 | -0.835721 | 0.40331187 | 0.65065844 |
| UQCR10 | 54.049489 | 0.60631863 | 0.52568977 | 1.15337727 | 0.24875557 | 0.50492395 |
| ASCC2 | 33.972139 | -0.2744324 | 0.33061311 | -0.830071 | 0.40649862 | 0.65333469 |
| MTMR3 | 225.90467 | -0.2633616 | 0.21199495 | -1.2423011 | 0.21412548 | 0.4699161 |
| TBC1D10A | 19.400448 | -0.3968639 | 0.49713703 | -0.7982988 | 0.42469714 | NA |
| SF3A1 | 16.4747286 | 0.20401164 | 0.57293288 | 0.35608297 | 0.72177843 | NA |
| RNF215 | 23.4089896 | 0.14341498 | 0.37708043 | 0.38032995 | 0.7037005 | 0.85295217 |
| SEC14L6 | 22.0571235 | 0.79251802 | 0.58393677 | 1.35719836 | 0.17471819 | 0.42340329 |
| GAL3ST1 | 17.8376068 | 1.21129816 | 0.6545286 | 1.85064207 | 0.06422106 | NA |
| PES1 | 16.9109118 | 0.01098664 | 0.48004877 | 0.0228865 | 0.98174081 | NA |
| TCN2 | 13.5245773 | 0.33793489 | 0.64316176 | 0.52542752 | 0.59928603 | NA |
| DUSP18 | 19.3063845 | 0.86991446 | 0.56432409 | 1.54151571 | 0.12319132 | NA |
| OSBP2 | 105.086025 | 0.19239243 | 0.30163324 | 0.63783563 | 0.52358068 | 0.73904282 |
| MORC2 | 33.9250378 | 0.10545869 | 0.4211457 | 0.25040903 | 0.80227104 | 0.90378853 |
| SMTN | 23.8016186 | -0.4296313 | 0.4917605 | -0.8736597 | 0.38230362 | 0.63349644 |
| SELENOM | 35.0119387 | 0.67584163 | 0.66353725 | 1.01854362 | 0.30841968 | 0.56825351 |
| INPP5J | 47.173693 | 1.92560309 | 0.65607195 | 2.93504867 | 0.00333495 | 0.05716957 |
| RNF185 | 56.2586108 | -0.3146666 | 0.39906204 | -0.7885155 | 0.43039524 | 0.67197671 |
| LIMK2 | 47.8226896 | 0.58884932 | 0.35061792 | 1.67946159 | 9.31E-02 | 0.30571879 |
| PIK3IP1 | 18.0218409 | 0.74629475 | 0.597778 | 1.24844801 | 0.21186704 | NA |
| PATZ1 | 24.0304908 | 0.66488299 | 0.63656962 | 1.04447804 | 0.29626427 | 0.55582698 |
| DRG1 | 60.3693234 | 0.18879837 | 0.30457244 | 0.61988003 | 0.53533677 | 0.74781981 |
| EIF4ENIF1 | 52.6661168 | -0.1393029 | 0.28027295 | -0.4970259 | 0.61917081 | 0.8027391 |
| SFI1 | 50.4801276 | -0.6715031 | 0.29841104 | -2.2502623 | 0.0244323 | 0.15276639 |
| PISD | 151.799889 | -0.9920418 | 0.32075185 | -3.0928639 | 0.00198235 | 0.04168893 |
| PRR14L | 114.679388 | -0.3573766 | 0.21845905 | -1.6358974 | 0.10186107 | 0.32089434 |
| RTCB | 47.99919 | -0.1602321 | 0.4202559 | -0.3812728 | 0.70300085 | 0.85275589 |
| FBXO7 | 55.779903 | 0.06383226 | 0.29215671 | 0.21848636 | 0.82705019 | 0.91613502 |
| TIMP3 | 110.268544 | -0.138705 | 0.4321005 | -0.3210018 | 0.74820903 | 0.87522263 |
| LARGE1 | 319.550568 | -1.0774908 | 0.45920135 | -2.3464451 | 0.01895346 | 0.13583312 |
| HMGXB4 | 43.1296776 | -0.582142 | 0.42521151 | -1.3690646 | 0.17097909 | 0.41898315 |
| TOM1 | 60.8816429 | -0.8439827 | 0.40706638 | -2.0733293 | 0.03814164 | 0.19510866 |
| HMOX1 | 19.4092164 | -1.2430003 | 0.78402156 | -1.5854159 | 1.13E-01 | NA |
| APOL6 | 14.3666096 | 0.4019727 | 0.84864429 | 0.47366454 | 0.63573915 | NA |
| RBFOX2 | 368.972861 | 0.21192834 | 0.27396582 | 0.77355761 | 0.43919245 | 0.67938098 |

| | | | | | | |
|---|---|---|---|---|---|---|
| APOL4 | 17.4143547 | 0.28923504 | 1.59584716 | 0.18124232 | 0.85617738 | NA |
| APOL2 | 39.6933602 | 0.24882453 | 0.64113947 | 0.38809735 | 0.69794399 | 0.8508196 |
| MYH9 | 192.419002 | -0.9383896 | 0.32400992 | -2.8961755 | 0.00377741 | 0.06101472 |
| TXN2 | 23.0526996 | 1.17697841 | 0.5264357 | 2.23574958 | 0.02536818 | 0.15633216 |
| EIF3D | 34.3990636 | -0.1073715 | 0.37473072 | -0.2865298 | 0.77447235 | 0.88786322 |
| IFT27 | 15.0790177 | 0.60655515 | 0.62651239 | 0.96814549 | 0.33297171 | NA |
| PVALB | 14.0232436 | 1.68754478 | 0.86226921 | 1.95709734 | 0.05033602 | NA |
| TST | 14.972062 | 0.66601303 | 0.65136153 | 1.02249366 | 0.30654731 | NA |
| MPST | 24.1462165 | 0.08414624 | 0.50872355 | 0.16540661 | 0.86862395 | 0.93857029 |
| CARD10 | 14.4011426 | 0.8563323 | 0.50114639 | 1.70874682 | 0.08749786 | NA |
| GGA1 | 56.391753 | -0.1370828 | 0.26963646 | -0.5083985 | 0.61117393 | 0.79772863 |
| TRIOBP | 62.1435072 | 0.35715763 | 0.40798228 | 0.87542436 | 0.38134305 | 0.63258566 |
| H1F0 | 25.9962099 | -1.3948843 | 0.51032937 | -2.7333019 | 0.00627028 | 0.07721978 |
| ANKRD54 | 15.8250592 | -0.1829734 | 0.50330967 | -0.3635403 | 0.7162013 | NA |
| EIF3L | 66.380341 | -0.0923978 | 0.40276437 | -0.2294091 | 0.81855099 | 0.913067 |
| MICALL1 | 15.83986 | -0.6046975 | 0.68284651 | -0.885554 | 0.37585789 | NA |
| POLR2F | 25.2865319 | 0.12768773 | 0.54160897 | 0.23575631 | 0.81362177 | 0.90970797 |
| PICK1 | 26.6973796 | -0.6686856 | 0.3655925 | -1.8290462 | 0.06739268 | 0.25898011 |
| PLA2G6 | 26.5716038 | -0.3758748 | 0.48758941 | -0.7708839 | 0.44077577 | 0.68070849 |
| TMEM184B | 42.8836956 | -0.2238559 | 0.39878171 | -0.5613495 | 0.57455927 | 0.77388388 |
| CSNK1E | 15.8907707 | -0.5069647 | 0.55569433 | -0.9123085 | 0.36160634 | NA |
| TPTEP2-CSNI | 145.523826 | 0.65055176 | 0.30715839 | 2.11796835 | 0.03417775 | 0.18312899 |
| DDX17 | 841.181413 | 0.05439762 | 0.16964749 | 0.3206509 | 7.48E-01 | 0.87540066 |
| FAM227A | 24.278509 | 0.8095541 | 0.59466987 | 1.36135046 | 0.17340296 | 0.42238719 |
| CBY1 | 31.0494734 | 0.35294733 | 0.44618831 | 0.79102774 | 0.42892781 | 0.6713008 |
| JOSD1 | 34.501059 | -0.4902906 | 0.4061061 | -1.2072968 | 0.22731786 | 0.48520724 |
| GTPBP1 | 23.8699891 | -0.758883 | 0.36705778 | -2.0674756 | 0.03868936 | 0.1958098 |
| SUN2 | 19.2425773 | 0.31863615 | 0.50820712 | 0.62698089 | 0.53067176 | NA |
| AL021707.1 | 14.7798458 | -0.711452 | 0.52762566 | -1.3484029 | 0.17752883 | NA |
| CBX6 | 31.4482829 | 0.23350264 | 0.46643098 | 0.50061562 | 0.61664166 | 0.80172727 |
| CBX7 | 24.7512419 | 0.72596082 | 0.39984369 | 1.81561153 | 0.06942998 | 0.26260569 |
| RPL3 | 67.2696255 | 0.333675 | 0.57722505 | 0.57806742 | 0.5632186 | 0.76781678 |
| SYNGR1 | 22.0903792 | 1.57989248 | 0.54222453 | 2.91372371 | 0.00357146 | 0.0593726 |
| TAB1 | 25.3816545 | -0.4033594 | 0.43877072 | -0.9192943 | 0.35794168 | 0.61415005 |
| MIEF1 | 12.2498132 | 0.15603136 | 0.55298954 | 0.28215969 | 0.77782106 | NA |
| ATF4 | 46.8122672 | -1.1684696 | 0.56819857 | -2.0564459 | 0.03973955 | 0.19847343 |
| RPS19BP1 | 21.5317782 | 0.00720777 | 0.45488489 | 0.01584527 | 0.98735784 | 0.99524532 |
| FAM83F | 29.9012457 | -0.3842078 | 0.73884455 | -0.5200117 | 0.60305542 | 0.79291384 |
| TNRC6B | 719.073661 | -0.0749721 | 0.19690025 | -0.380762 | 0.70337984 | 0.8528536 |
| MKL1 | 296.754729 | -0.7228688 | 0.33818658 | -2.1374851 | 0.03255856 | 0.17811174 |
| SLC25A17 | 55.7663582 | 0.26105843 | 0.28230439 | 0.92474094 | 0.35510068 | 0.61168935 |
| ST13 | 81.1044513 | 0.34609471 | 0.25218748 | 1.37237069 | 0.16994807 | 0.41832117 |
| XPNPEP3 | 71.9183334 | 0.39833442 | 0.37214235 | 1.07038186 | 0.28444746 | 0.54316116 |
| RBX1 | 50.1988066 | -0.5614865 | 0.40132625 | -1.3990774 | 0.16178978 | 0.40762552 |
| EP300 | 176.876491 | -0.3115721 | 0.26300095 | -1.1846807 | 0.23614373 | 0.49391258 |
| L3MBTL2 | 15.3620946 | -0.4368233 | 0.45520397 | -0.959621 | 0.33724602 | NA |
| RANGAP1 | 29.4446391 | -0.5511084 | 0.4858681 | -1.1342757 | 0.25667893 | 0.51485087 |
| ZC3H7B | 47.8231674 | -0.2967997 | 0.31449651 | -0.9437296 | 0.34530782 | 0.60277826 |
| TEF | 15.3376673 | 0.35363334 | 0.69228154 | 0.51082301 | 0.609475 | NA |
| TOB2 | 35.0819269 | 0.3840514 | 0.36656705 | 1.04769756 | 0.29477797 | 0.55406741 |
| ACO2 | 133.21895 | 0.22543889 | 0.30579736 | 0.73721659 | 0.46099065 | 0.69416206 |
| CSDC2 | 26.2377091 | -0.3755204 | 0.84563807 | -0.4440675 | 0.65699379 | 0.82476514 |
| PMM1 | 20.0455684 | 1.53184557 | 0.55293282 | 2.77040087 | 0.00559873 | NA |
| DESI1 | 28.6248241 | 0.23065232 | 0.42520289 | 0.54245239 | 0.5875069 | 0.78270395 |

| | | | | | | |
|---|---|---|---|---|---|---|
| XRCC6 | 93.4696449 | -0.362463 | 0.27733534 | -1.3069484 | 0.19123023 | 0.44283704 |
| SNU13 | 50.5194469 | -0.1456102 | 0.32282448 | -0.4510508 | 0.65195297 | 0.82061848 |
| CCDC134 | 14.7207999 | -0.1086652 | 0.45321289 | -0.2397664 | 0.81051136 | NA |
| SREBF2 | 284.976663 | -0.5235175 | 0.16684486 | -3.1377504 | 0.0017025 | 0.03798297 |
| WBP2NL | 36.5507727 | 0.75615588 | 0.65258291 | 1.15871235 | 0.24657345 | 0.50296216 |
| SMDT1 | 20.2351274 | 0.69865506 | 0.40701876 | 1.71651808 | 0.08606726 | NA |
| NDUFA6 | 24.1541255 | 0.2132639 | 0.42966561 | 0.49634854 | 0.61964852 | 0.80279998 |
| TCF20 | 131.889791 | -0.4164656 | 0.28488497 | -1.4618728 | 0.14377608 | 0.38252489 |
| RRP7A | 21.6604223 | 0.25516631 | 0.52818377 | 0.48310137 | 0.62902376 | 0.80693265 |
| POLDIP3 | 31.253388 | -0.7032828 | 0.36846464 | -1.9086845 | 0.0563028 | 0.23642261 |
| CYB5R3 | 49.1448398 | 0.26912594 | 0.37303234 | 0.72145472 | 0.47062979 | 0.70036133 |
| A4GALT | 35.1979822 | 1.17107245 | 0.46231595 | 2.53305655 | 1.13E-02 | 1.04E-01 |
| ARFGAP3 | 153.563283 | -0.5168354 | 0.34040123 | -1.5183124 | 1.29E-01 | 3.61E-01 |
| PACSIN2 | 207.457338 | -0.6020464 | 0.31225934 | -1.9280334 | 0.05385098 | 0.23287156 |
| TSPO | 34.1595048 | 1.17957891 | 0.43023015 | 2.74173933 | 0.00611148 | 0.07673748 |
| EFCAB6 | 111.365061 | 0.41998318 | 0.48989981 | 0.85728382 | 0.39128805 | 0.63984683 |
| SAMM50 | 20.7143347 | 0.44154336 | 0.40517839 | 1.08975053 | 0.27582305 | NA |
| PHF21B | 203.750689 | -0.3949325 | 0.24397431 | -1.6187463 | 0.10550185 | 0.32665298 |
| NUP50-DT | 74.4792036 | 0.15350061 | 0.33725562 | 0.45514619 | 0.64900407 | 0.81913126 |
| NUP50 | 25.2131639 | -0.2977386 | 0.36845389 | -0.8080757 | 0.419047 | 0.66298145 |
| KIAA0930 | 25.1394811 | -1.8380463 | 0.67618103 | -2.7182755 | 6.56E-03 | 0.07916953 |
| FAM118A | 26.0949564 | -0.1787006 | 0.57187258 | -0.3124832 | 0.75467335 | 0.87877773 |
| ATXN10 | 184.722488 | -0.3168827 | 0.24943034 | -1.2704258 | 0.20393299 | 0.45736371 |
| MIRLET7BHG | 82.2766927 | 0.23152734 | 0.27011932 | 0.85712987 | 0.39137312 | 0.63984683 |
| PPARA | 205.27747 | 0.30571446 | 0.20126903 | 1.51893441 | 1.29E-01 | 3.61E-01 |
| TTC38 | 31.82672 | 0.80015872 | 0.43133972 | 1.85505458 | 0.06358843 | 0.25118809 |
| TRMU | 43.6832294 | 0.30817834 | 0.30702476 | 1.00375729 | 0.31549562 | 0.57272872 |
| CELSR1 | 46.7330846 | 0.22366176 | 0.54150575 | 0.41303672 | 0.67957971 | 0.84054717 |
| GRAMD4 | 39.9111048 | 0.55230405 | 0.51391208 | 1.07470533 | 0.28250668 | 0.5413129 |
| CERK | 28.0655255 | 0.39379283 | 0.39966065 | 0.98531798 | 0.3244679 | 0.58025777 |
| TBC1D22A | 302.741434 | -0.7963065 | 0.31036547 | -2.5657059 | 0.01029661 | 0.09994074 |
| BRD1 | 65.5210883 | -0.6602719 | 0.35452382 | -1.862419 | 0.06254405 | 0.24943086 |
| ZBED4 | 29.2384111 | -0.4559008 | 0.53012289 | -0.8599908 | 0.3897941 | 0.6386799 |
| PIM3 | 65.5057692 | -0.6464686 | 0.39445781 | -1.6388789 | 0.10123848 | 0.32027643 |
| TRABD | 29.6515187 | 0.96546404 | 0.40241219 | 2.39919181 | 0.01643131 | 0.12723517 |
| SELENOO | 61.1934466 | 0.11682115 | 0.30336955 | 0.38507871 | 0.7001791 | 0.8516559 |
| TUBGCP6 | 18.3284937 | 0.27559779 | 0.53781097 | 0.5124436 | 0.60834058 | NA |
| PLXNB2 | 67.1627946 | 0.16262921 | 0.3231667 | 0.50323627 | 0.61479816 | 0.80015243 |
| DENND6B | 13.8445539 | 0.8641113 | 0.58835262 | 1.46869628 | 0.14191519 | NA |
| PPP6R2 | 285.475134 | -0.3373653 | 0.16869642 | -1.999837 | 0.04551787 | 0.21335013 |
| MIOX | 17.0159684 | -1.4029097 | 0.79430822 | -1.7662031 | 0.07736178 | NA |
| RABL2B | 27.3544264 | -0.1812029 | 0.42018764 | -0.4312429 | 0.66629177 | 0.83105619 |
| CU633967.1 | 15.5671853 | 1.22004723 | 0.80520035 | 1.51520952 | 0.12971935 | NA |
| FP671120.1 | 482.139949 | -1.39596 | 0.48664584 | -2.8685337 | 0.00412379 | 0.06262486 |
| FP236383.1 | 268.096977 | -1.5397603 | 0.50655961 | -3.0396429 | 0.00236859 | 0.04627162 |
| CR381653.1 | 40.0082935 | -0.7327908 | 0.55901724 | -1.3108555 | 0.1899066 | 0.44120467 |
| NRIP1 | 134.929073 | -1.3573433 | 0.34462261 | -3.9386368 | 8.19E-05 | 0.00598111 |
| USP25 | 379.236414 | -0.2702949 | 0.28206048 | -0.9582869 | 0.33791813 | 0.59378353 |
| MIR99AHG | 121.680468 | 0.94807583 | 0.4157437 | 2.2804334 | 0.022582 | 0.14632666 |
| CXADR | 54.061292 | 0.15322925 | 0.36670722 | 0.41785175 | 0.67605551 | 0.83847878 |
| BTG3 | 28.2783535 | -0.2095658 | 0.39680847 | -0.5281284 | 0.59741025 | 0.7894568 |
| C21orf91 | 15.9761328 | 0.97319394 | 0.47077442 | 2.06721922 | 0.0387135 | NA |
| CHODL | 56.5202404 | 1.052148 | 0.98130425 | 1.07219347 | 0.28363314 | 0.54266078 |
| MRPL39 | 21.0431482 | -0.2096292 | 0.40117278 | -0.5225408 | 0.60129383 | NA |

| | | | | | | |
|---|---|---|---|---|---|---|
| ATP5PF | 55.8258423 | 0.65930452 | 0.38572115 | 1.70927763 | 0.08739953 | 0.29569856 |
| GABPA | 34.4836941 | -0.6651912 | 0.34711668 | -1.9163332 | 0.0553227 | 0.23510863 |
| APP | 916.818723 | -0.4607185 | 0.26354568 | -1.7481543 | 8.04E-02 | 0.28335284 |
| CYYR1 | 39.2811969 | 0.19163388 | 0.55362418 | 0.34614434 | 0.72923424 | 0.86533507 |
| ADAMTS1 | 350.633162 | -0.329238 | 0.44146877 | -0.7457787 | 4.56E-01 | 6.91E-01 |
| LINC00113 | 22.2697292 | -0.0424537 | 1.05587837 | -0.040207 | 0.9679281 | 0.98741848 |
| LINC01695 | 27.4457327 | -1.5113684 | 0.57333142 | -2.6361165 | 0.00838609 | 0.08952533 |
| AF165147.1 | 33.9275575 | 0.56143118 | 0.5589447 | 1.00444853 | 0.31516247 | 0.57225891 |
| N6AMT1 | 12.6540887 | 0.30379108 | 0.59883418 | 0.50730418 | 0.61194141 | NA |
| LTN1 | 91.9108814 | -0.1617679 | 0.25813049 | -0.6266904 | 0.53086221 | 0.74456344 |
| USP16 | 55.2542415 | -0.4515034 | 0.49287845 | -0.9160542 | 0.35963847 | 0.6159629 |
| CCT8 | 31.0306691 | 0.25164428 | 0.36492745 | 0.68957344 | 0.49046247 | 0.7163358 |
| MAP3K7CL | 37.0642287 | 0.32663522 | 0.5649594 | 0.57815698 | 0.56315814 | 0.76781678 |
| BACH1 | 354.530291 | -0.2399834 | 0.30053296 | -0.7985262 | 0.4245652 | 0.6673225 |
| GRIK1 | 23.2787032 | 1.48185358 | 0.77001245 | 1.92445406 | 0.05429769 | 0.23287156 |
| CLDN8 | 24.6518478 | 1.47186849 | 0.92401088 | 1.59291251 | 0.11117982 | 0.33460648 |
| SOD1 | 46.6688304 | 0.84340115 | 0.30006656 | 2.81071353 | 0.00494318 | 0.06830815 |
| SCAF4 | 77.6541952 | -0.3807864 | 0.28490345 | -1.3365455 | 0.18137105 | 0.43060239 |
| HUNK | 29.4689078 | 0.27299288 | 0.84643351 | 0.32252135 | 0.74705777 | 0.87469995 |
| URB1 | 24.4126782 | -0.9113885 | 0.40482266 | -2.2513277 | 0.02436479 | 0.15259218 |
| EVA1C | 28.0460873 | 0.7946389 | 0.4669488 | 1.70176882 | 0.08879871 | 0.29778649 |
| SYNJ1 | 61.6974883 | 0.38711682 | 0.31481318 | 1.22967157 | 0.21882012 | 0.47565385 |
| PAXBP1 | 112.216765 | -0.0011878 | 0.20505181 | -0.0057925 | 0.99537831 | 0.99796515 |
| C21orf62-AS | 30.5227517 | 1.14264355 | 0.73482206 | 1.55499353 | 0.11994761 | 0.34839123 |
| AP000282.1 | 40.9499541 | 2.17855114 | 0.64184858 | 3.39418238 | 0.00068834 | 0.02375848 |
| IFNAR1 | 55.0093853 | -0.083592 | 0.27619812 | -0.3026524 | 0.76215475 | 0.88247846 |
| IFNGR2 | 82.0790408 | 0.05457914 | 0.40055802 | 0.13625776 | 0.89161752 | 0.95026032 |
| TMEM50B | 69.9224719 | -0.4699185 | 0.42009098 | -1.1186113 | 0.263306 | 0.52287572 |
| GART | 47.601996 | -0.1542896 | 0.26574493 | -0.5805927 | 0.561515 | 0.76658052 |
| SON | 222.455043 | -0.1601901 | 0.28065213 | -0.5707781 | 0.56815008 | 0.77103518 |
| AP000311.1 | 53.3814314 | -0.1762481 | 0.62569387 | -0.2816842 | 0.77818568 | 0.88986706 |
| ITSN1 | 204.030097 | -0.4576022 | 0.25483185 | -1.7957024 | 0.07254186 | 0.26831077 |
| MRPS6 | 362.58063 | -0.7306834 | 0.44866763 | -1.6285627 | 0.10340562 | 0.32301666 |
| LINC00310 | 39.9033503 | -0.4941699 | 0.65638884 | -0.7528615 | 0.45153314 | 0.68853766 |
| AP000317.1 | 51.0923341 | -0.4082009 | 0.65960559 | -0.6188561 | 0.53601117 | 0.74808305 |
| RCAN1 | 56.5273386 | 0.65368013 | 0.60239086 | 1.08514284 | 0.27785839 | 0.53641236 |
| RUNX1 | 397.182499 | -2.0732773 | 0.84659209 | -2.4489684 | 0.0143266 | 0.11781182 |
| SETD4 | 31.2159273 | -0.1914873 | 0.38541475 | -0.4968345 | 0.61930577 | 0.8027391 |
| CBR1 | 38.6743113 | 0.52455916 | 0.42917148 | 1.22226006 | 0.2216093 | 0.47873331 |
| CBR3 | 31.0505259 | -0.5214009 | 0.55840258 | -0.9337366 | 0.35043977 | 0.60737992 |
| DOPEY2 | 34.6951686 | -0.4060287 | 0.51535479 | -0.7878626 | 0.43077709 | 0.67197671 |
| MORC3 | 98.1368282 | -0.4030497 | 0.26912952 | -1.4976051 | 0.13423589 | 0.36808465 |
| AP000695.2 | 22.1011347 | -1.5571868 | 0.83556686 | -1.8636293 | 0.06237379 | 0.24933194 |
| CLDN14 | 69.7358164 | -1.4838688 | 0.69266002 | -2.1422758 | 0.03217129 | 0.17693114 |
| HLCS | 175.124555 | 0.05637864 | 0.34889997 | 0.16158969 | 0.87162898 | 0.9400893 |
| PIGP | 11.9970286 | 1.7105036 | 0.75242703 | 2.27331493 | 0.02300721 | NA |
| TTC3 | 262.595786 | 0.01100651 | 0.28760952 | 0.03826895 | 0.96947325 | 0.98781207 |
| VPS26C | 55.3246362 | -0.0983146 | 0.31179633 | -0.3153167 | 0.7525212 | 0.87837077 |
| DYRK1A | 466.936452 | -0.6932652 | 0.33472777 | -2.0711315 | 0.03834651 | 0.19533626 |
| KCNJ15 | 20.8646921 | 0.0107657 | 0.54537882 | 0.01973986 | 0.98425089 | NA |
| ETS2 | 108.567958 | -1.0453374 | 0.49230499 | -2.1233532 | 3.37E-02 | 1.82E-01 |
| BRWD1 | 350.018917 | -0.0018299 | 0.20544233 | -0.008907 | 0.99289331 | 0.99689685 |
| HMGN1 | 72.210942 | 0.16544161 | 0.35659728 | 0.46394524 | 0.64268698 | 0.81595394 |
| WRB | 25.2227929 | 0.16049636 | 0.40506436 | 0.39622434 | 0.69193954 | 0.84725718 |

| | | | | | | |
|---|---|---|---|---|---|---|
| LCA5L | 13.8616098 | -0.1965014 | 0.53027117 | -0.3705677 | 0.71095954 | NA |
| SH3BGR | 14.7281174 | 1.09875594 | 0.60091117 | 1.82848312 | 0.06747708 | NA |
| PCP4 | 32.9065017 | -1.892485 | 1.06972367 | -1.7691345 | 0.07687144 | 0.27670206 |
| BACE2 | 949.636725 | -0.9706682 | 0.71995052 | -1.348243 | 0.17758024 | 0.42593802 |
| FAM3B | 17.3611168 | -0.3599977 | 0.50114323 | -0.718353 | 0.47253966 | NA |
| MX1 | 88.0205837 | -1.7698309 | 0.90872786 | -1.9475918 | 0.05146383 | 0.2277844 |
| TMPRSS2 | 266.867091 | -0.1945418 | 0.38871072 | -0.5004797 | 6.17E-01 | 8.02E-01 |
| RIPK4 | 26.4216403 | -0.2849681 | 0.39033141 | -0.7300669 | 0.46534927 | 0.69589923 |
| PRDM15 | 16.772608 | 0.08226458 | 0.59359337 | 0.13858743 | 0.88977618 | NA |
| ZBTB21 | 24.479817 | 0.29934077 | 0.42885238 | 0.69800423 | 0.48517455 | 0.71212596 |
| ABCG1 | 27.0871455 | -1.1319561 | 0.70530058 | -1.6049273 | 0.10850981 | 0.33024913 |
| SLC37A1 | 26.2888408 | 0.36700272 | 0.33229667 | 1.10444295 | 2.69E-01 | 5.28E-01 |
| PDE9A | 148.700352 | 0.32836421 | 0.35851518 | 0.91590045 | 0.35971911 | 0.61596396 |
| WDR4 | 22.9696446 | -0.5535903 | 0.51473713 | -1.0754817 | 0.28215914 | 0.54091629 |
| NDUFV3 | 70.2116312 | -0.149326 | 0.26234776 | -0.5691909 | 0.56922661 | 0.77149801 |
| PKNOX1 | 70.9883664 | 0.03769463 | 0.26193612 | 0.14390774 | 0.88557333 | 0.94696553 |
| RRP1B | 25.6895377 | -0.1586605 | 0.35985621 | -0.4408998 | 0.65928554 | 0.82620006 |
| PDXK | 114.986376 | -0.531902 | 0.24746286 | -2.1494216 | 0.03160099 | 0.17523981 |
| CSTB | 47.2500683 | 0.15852125 | 0.37515861 | 0.42254461 | 6.73E-01 | 8.36E-01 |
| AGPAT3 | 122.678912 | 0.22280484 | 0.2736072 | 0.81432372 | 0.41545952 | 0.65906718 |
| TRAPPC10 | 126.625257 | -0.2719776 | 0.31300032 | -0.8689373 | 0.38488145 | 0.63508787 |
| PFKL | 140.030924 | -0.4151381 | 0.33507167 | -1.2389532 | 0.21536283 | 0.47115856 |
| UBE2G2 | 101.48811 | -0.0758308 | 0.28178392 | -0.2691096 | 0.78784534 | 0.89506208 |
| SUMO3 | 18.9709634 | -0.1560467 | 0.60370093 | -0.2584834 | 0.79603385 | NA |
| PTTG1IP | 129.972437 | -0.6237628 | 0.34430331 | -1.8116666 | 0.07003773 | 0.26406366 |
| ADARB1 | 55.2690981 | -0.8190659 | 0.58029935 | -1.4114541 | 0.15811076 | 0.40275854 |
| POFUT2 | 25.4744495 | -0.5286529 | 0.36505762 | -1.4481355 | 0.14757917 | 0.38722988 |
| COL18A1 | 242.329294 | 0.50674433 | 0.33059019 | 1.53284746 | 0.12531344 | 0.35604931 |
| COL6A1 | 67.4815602 | 1.69444906 | 0.38112713 | 4.44588937 | 8.75E-06 | 0.00146459 |
| LSS | 38.2122981 | -0.5394791 | 0.46263428 | -1.1661027 | 0.24357294 | 0.50087656 |
| MCM3AP-AS | 15.3403345 | 0.22982752 | 0.59681996 | 0.38508685 | 0.70017307 | NA |
| MCM3AP | 76.8638619 | -0.3607896 | 0.24070827 | -1.4988666 | 0.13390825 | 0.36774274 |
| YBEY | 14.0790158 | -1.3405311 | 0.5391088 | -2.4865688 | 0.01289816 | NA |
| PCNT | 32.7093427 | -0.3271451 | 0.35841632 | -0.9127517 | 0.36137315 | 0.61731848 |
| DIP2A | 125.797517 | -0.1140279 | 0.23087567 | -0.4938932 | 0.62138161 | 0.80353114 |
| PRMT2 | 99.7750724 | 0.15479885 | 0.26090765 | 0.59330896 | 0.5529744 | 0.75936556 |
| AC011043.1 | 19.3678039 | 0.54606443 | 0.6176718 | 0.88406891 | 0.37665902 | NA |
| AL592183.1 | 68.4249855 | -0.0131654 | 0.59034262 | -0.0223014 | 0.98220757 | 0.99382437 |

**Connecting tubules**

*This means that positive fold changes imply higher mean value in non-COVID AKI than in COVID AKI and vice versa.

| | baseMean | log2 fold cha | lfcSE | stat | pvalue | padj |
|---|---|---|---|---|---|---|
| AL669831.5 | 44.0210642 | -0.0778137 | 0.30779887 | -0.252807 | 0.80041739 | 0.97019967 |
| SAMD11 | 20.726368 | 0.55963157 | 0.62930011 | 0.88929203 | 3.74E-01 | 0.85056774 |
| NOC2L | 29.2722437 | 0.55731389 | 0.30334224 | 1.83724456 | 0.0661738 | 0.59523726 |
| HES4 | 27.2135181 | -1.9723391 | 0.52105182 | -3.7853033 | 0.00015352 | 0.05491436 |
| ISG15 | 141.642756 | -5.5516342 | 1.71027421 | -3.2460492 | 0.00117019 | 0.14394871 |
| AGRN | 112.768069 | -0.1581009 | 0.45330533 | -0.3487736 | 0.72725927 | 0.95211709 |
| C1orf159 | 32.282756 | -0.3403972 | 0.31494674 | -1.0808087 | 0.27978221 | 0.80291849 |
| LINC01342 | 11.2489136 | 0.18558511 | 0.59684884 | 0.31094156 | 0.75584505 | NA |
| AL390719.2 | 15.3804117 | -0.5632234 | 0.45621667 | -1.2345524 | 0.21699712 | NA |
| SDF4 | 53.8263123 | -0.0934981 | 0.36863437 | -0.2536338 | 0.79977852 | 0.97019967 |
| UBE2J2 | 43.8341854 | -0.2219032 | 0.28158159 | -0.7880602 | 0.43066152 | 0.86560207 |
| ACAP3 | 16.1378878 | -0.1671772 | 0.46432835 | -0.3600409 | 0.71881652 | NA |
| INTS11 | 37.3101774 | 0.26116475 | 0.27910444 | 0.9357241 | 0.34941526 | 0.83687845 |
| CPTP | 11.9249236 | -0.0511035 | 0.44397807 | -0.1151038 | 0.90836288 | NA |
| DVL1 | 13.8374438 | -0.2931367 | 0.44966864 | -0.6518949 | 0.51446899 | NA |
| AURKAIP1 | 57.154339 | 0.15868354 | 0.42544891 | 0.37297908 | 0.70916401 | 0.94558664 |
| CCNL2 | 101.280223 | -0.3829765 | 0.23373058 | -1.6385383 | 0.10130945 | 0.65863153 |
| MRPL20 | 57.843837 | 0.23882911 | 0.29902393 | 0.79869566 | 0.4244669 | 0.86190547 |
| VWA1 | 22.2996216 | 1.05228898 | 0.71071874 | 1.48059833 | 0.13871364 | 0.69023184 |
| ATAD3C | 11.2319531 | 1.04325706 | 0.47010176 | 2.2192154 | 0.02647207 | NA |
| ATAD3B | 27.3281183 | -0.4099319 | 0.36855685 | -1.1122624 | 0.26602535 | 0.79666376 |
| ATAD3A | 15.6381034 | -0.2818453 | 0.52457051 | -0.5372878 | 0.59106884 | NA |
| SSU72 | 59.4391194 | -0.0036492 | 0.35221442 | -0.0103608 | 0.99173346 | 0.99814944 |
| MIB2 | 20.6421402 | 0.18736361 | 0.39582905 | 0.47334478 | 0.63596722 | 0.93251095 |
| CDK11B | 28.8836561 | -0.3299487 | 0.46103905 | -0.7156633 | 0.47419931 | 0.88403192 |
| SLC35E2B | 30.5154438 | -0.6313237 | 0.31644967 | -1.9950208 | 0.04604061 | 0.52984532 |
| CDK11A | 18.5704888 | -0.0889491 | 0.42193695 | -0.2108114 | 0.83303447 | 0.97600352 |
| SLC35E2A | 31.3236577 | -0.0704144 | 0.30383499 | -0.2317521 | 0.81673057 | 0.97288403 |
| NADK | 39.4955212 | -0.3101722 | 0.23992141 | -1.2928074 | 0.19607769 | 0.75350056 |
| GNB1 | 275.592998 | -0.6049793 | 0.27602665 | -2.1917425 | 0.02839811 | 0.46148943 |
| PRKCZ | 63.1323577 | -0.3204831 | 0.28453532 | -1.1263386 | 0.26002224 | 0.79535701 |
| FAAP20 | 32.3955021 | 0.56163649 | 0.29921976 | 1.87700335 | 0.06051763 | 0.57774794 |
| SKI | 74.9409621 | -0.0566035 | 0.32195623 | -0.175811 | 0.86044241 | 0.97718459 |
| MORN1 | 31.1670492 | -0.2605469 | 0.30386028 | -0.8574562 | 0.39119282 | 0.85605713 |
| RER1 | 71.3177887 | 0.09892569 | 0.24347081 | 0.40631438 | 0.68451162 | 0.94205398 |
| PEX10 | 19.6798616 | -0.2026973 | 0.36752775 | -0.5515157 | 0.58128023 | 0.92034873 |
| TNFRSF14-A | 13.8215883 | 0.08037956 | 0.50149128 | 0.16028106 | 0.87265968 | NA |
| TNFRSF14 | 33.2217397 | -0.4011208 | 0.43963735 | -0.9123902 | 0.36156333 | 0.84395756 |
| FAM213B | 11.6918607 | 0.29917537 | 0.57885686 | 0.51683826 | 0.60526907 | NA |
| LINC00982 | 108.111997 | 0.7702986 | 0.32211336 | 2.39138976 | 1.68E-02 | 3.97E-01 |
| PRDM16 | 321.223399 | -0.1371269 | 0.28872228 | -0.474944 | 0.63482691 | 0.93251095 |
| ARHGEF16 | 23.627701 | -0.2605018 | 0.35927759 | -0.725071 | 0.46840846 | 0.88170434 |
| TPRG1L | 13.2126936 | 0.59848649 | 0.47982799 | 1.24729382 | 0.21228978 | NA |
| WRAP73 | 11.1572594 | 0.32595042 | 0.44747998 | 0.72841342 | 0.46636055 | NA |
| LRRC47 | 17.9846091 | -0.183298 | 0.34442473 | -0.5321858 | 0.59459731 | 0.9229638 |
| CEP104 | 66.2682141 | -0.8908279 | 0.37694441 | -2.3632871 | 0.01811363 | 0.41119297 |
| DFFB | 13.476099 | 0.35519225 | 0.49578262 | 0.7164274 | 0.47372748 | NA |
| C1orf174 | 12.6580701 | -0.4249714 | 0.42183805 | -1.0074279 | 0.31372919 | NA |
| NPHP4 | 20.8977575 | 0.38413188 | 0.3989346 | 0.96289437 | 0.33560054 | 0.82974237 |
| RPL22 | 43.754267 | -0.072271 | 0.3731631 | -0.1936715 | 0.84643313 | 0.97612951 |
| RNF207 | 34.371161 | 0.12605002 | 0.37809906 | 0.3333783 | 0.73884874 | 0.9542764 |
| ICMT | 28.6273691 | 0.15739389 | 0.39924661 | 0.39422724 | 0.69341328 | 0.94394214 |

| | | | | | | |
|---|---|---|---|---|---|---|
| ESPN | 18.2956137 | 1.50182978 | 0.57744469 | 2.60082013 | 0.00930012 | 0.32071089 |
| NOL9 | 34.5607552 | 0.06319993 | 0.32589709 | 0.19392603 | 0.8462338 | 0.97612951 |
| ZBTB48 | 13.908182 | 0.11930267 | 0.40126103 | 0.29731936 | 0.76622271 | NA |
| KLHL21 | 24.62217 | -0.2881004 | 0.42270545 | -0.681563 | 0.49551533 | 0.89230917 |
| PHF13 | 13.1075208 | 0.66452752 | 0.45707711 | 1.45386304 | 0.14598428 | NA |
| THAP3 | 11.0317388 | 0.26693794 | 0.42935336 | 0.62172086 | 0.53412543 | NA |
| DNAJC11 | 96.5753594 | -0.1456025 | 0.29177999 | -0.4990145 | 0.61776913 | 0.92749951 |
| CAMTA1 | 371.935075 | -0.2410308 | 0.26006016 | -0.926827 | 0.35401635 | 0.83979358 |
| VAMP3 | 24.195392 | 0.39353702 | 0.30274681 | 1.29988825 | 0.19363927 | 0.75067401 |
| PER3 | 90.9762286 | -0.1646041 | 0.60281875 | -0.2730573 | 0.78480917 | 0.96738224 |
| PARK7 | 88.6123018 | 0.6015145 | 0.2931831 | 2.05166843 | 0.0402019 | 0.50338084 |
| ERRFI1 | 78.1494281 | -0.4667398 | 0.48497199 | -0.9624057 | 0.33584587 | 0.82974237 |
| RERE | 1128.23004 | -0.224311 | 0.18832051 | -1.1911129 | 0.23360927 | 0.77650198 |
| ENO1 | 362.380156 | -0.2374957 | 0.29049859 | -0.8175451 | 0.41361697 | 0.85867915 |
| GPR157 | 29.6188511 | -0.2010335 | 0.3573324 | -0.5625951 | 5.74E-01 | 0.91813957 |
| MIR34AHG | 16.8815704 | 1.37959301 | 0.61761443 | 2.23374479 | 0.02549987 | 0.44138123 |
| H6PD | 40.6060445 | 0.0296348 | 0.27748353 | 0.10679842 | 0.9149489 | 0.98691474 |
| SPSB1 | 42.8087218 | -1.4253024 | 0.38151022 | -3.7359481 | 1.87E-04 | 0.05491436 |
| SLC25A33 | 59.6447987 | -0.1473843 | 0.37455189 | -0.3934949 | 0.69395399 | 0.94395238 |
| PIK3CD | 14.301563 | -0.4251397 | 0.43171433 | -0.9847708 | 0.32473668 | NA |
| CLSTN1 | 110.345495 | 0.24249102 | 0.23565968 | 1.02898815 | 0.30348524 | 0.81568231 |
| CTNNBIP1 | 30.9609797 | -0.049212 | 0.31311861 | -0.1571672 | 0.87511307 | 0.97918624 |
| LZIC | 28.7808292 | -0.456233 | 0.39183103 | -1.1643615 | 0.24427755 | 0.78561483 |
| NMNAT1 | 49.3366757 | 0.17626278 | 0.32941376 | 0.53508018 | 0.59259441 | 0.92278087 |
| UBE4B | 165.808087 | -0.1664487 | 0.27406295 | -0.6073375 | 0.54362699 | 0.90965439 |
| KIF1B | 177.69987 | -0.1864918 | 0.17489936 | -1.0662808 | 0.28629672 | 0.80463985 |
| PGD | 37.4606466 | -0.3113087 | 0.25055194 | -1.2424916 | 0.21405524 | 0.7643572 |
| DFFA | 38.0876339 | 0.56499127 | 0.30443845 | 1.85584731 | 0.06347532 | 0.58688097 |
| PEX14 | 121.010175 | -0.1917931 | 0.24529462 | -0.7818886 | 0.43428003 | 0.86729771 |
| CASZ1 | 238.710087 | 0.20075539 | 0.25242379 | 0.7953109 | 0.42643267 | 0.86239557 |
| SRM | 13.6740638 | -0.5127276 | 0.5178292 | -0.9901481 | 0.32210174 | NA |
| EXOSC10 | 32.6390192 | -0.7091938 | 0.39649463 | -1.7886592 | 0.07366971 | 0.60985192 |
| MTOR | 71.519547 | -0.1338464 | 0.27331524 | -0.4897144 | 0.624336 | 0.92981013 |
| FBXO2 | 27.2785392 | 0.27836144 | 0.36565942 | 0.76125875 | 0.44650253 | 0.87231627 |
| FBXO44 | 23.5850546 | -0.0806418 | 0.39913455 | -0.2020417 | 0.83988411 | 0.97600352 |
| MAD2L2 | 9.08338931 | 0.69308119 | 0.63662276 | 1.08868428 | 0.27629314 | NA |
| AGTRAP | 30.3711459 | -0.6747606 | 0.5660371 | -1.1920784 | 0.23323049 | 0.77650198 |
| MTHFR | 27.1082671 | 0.36356233 | 0.3075637 | 1.18207165 | 0.23717727 | 0.77930556 |
| CLCN6 | 30.7387379 | -0.9311399 | 0.47563223 | -1.9576888 | 0.05026654 | 0.54473367 |
| KIAA2013 | 19.7849889 | 0.53865776 | 0.57374322 | 0.93884815 | 0.34780871 | 0.83687845 |
| PLOD1 | 14.8727839 | 0.54917643 | 0.46079886 | 1.19179207 | 0.23334279 | NA |
| MFN2 | 49.9077597 | -0.2383872 | 0.30388737 | -0.7844592 | 0.43277072 | 0.86729771 |
| VPS13D | 348.254052 | -0.1686458 | 0.23873388 | -0.7064177 | 0.4799284 | 0.88660781 |
| DHRS3 | 53.3409941 | -0.7967178 | 0.55171569 | -1.444073 | 0.14871843 | 0.70006785 |
| PRDM2 | 136.549902 | 0.06802595 | 0.23306448 | 0.29187608 | 0.77038137 | 0.96342652 |
| KAZN | 2972.47758 | -0.4858607 | 0.22931096 | -2.1187852 | 3.41E-02 | 0.4908133 |
| TMEM51-AS | 51.0659576 | -0.45646 | 0.37974737 | -1.2020095 | 2.29E-01 | 0.77185154 |
| TMEM51 | 342.652194 | -0.4294174 | 0.51361533 | -0.8360681 | 0.40311657 | 0.85782098 |
| EFHD2 | 12.8933645 | 0.59523627 | 0.3837599 | 1.55106427 | 0.12088628 | NA |
| CASP9 | 28.6560633 | -0.7435839 | 0.53430134 | -1.3916939 | 0.16401511 | 0.71368524 |
| DNAJC16 | 62.8842557 | -0.1081203 | 0.2151618 | -0.5025068 | 0.61531109 | 0.92662475 |
| DDI2 | 56.6269048 | -0.3916739 | 0.23112941 | -1.6946088 | 0.09014966 | 0.63862437 |
| PLEKHM2 | 57.6428297 | -0.6429011 | 0.25950071 | -2.477454 | 0.01323235 | 0.36723795 |
| FBLIM1 | 15.9263663 | 0.23785776 | 0.56406208 | 0.42168721 | 0.67325334 | NA |

| | | | | | | |
|---|---|---|---|---|---|---|
| UQCRHL | 13.3737285 | 0.43126045 | 0.52284497 | 0.82483426 | 0.40946569 | NA |
| SPEN | 126.82479 | -0.1103563 | 0.22236888 | -0.4962759 | 0.61969977 | 0.92798856 |
| ZBTB17 | 17.8150059 | -0.2769356 | 0.43412501 | -0.6379166 | 0.52352795 | 0.90487731 |
| CLCNKA | 103.580627 | 0.50191331 | 0.49111191 | 1.02199376 | 3.07E-01 | 8.16E-01 |
| CLCNKB | 132.156684 | 1.01278511 | 0.57656151 | 1.75659506 | 0.07898686 | 0.61367304 |
| EPHA2 | 20.8920497 | -0.9725015 | 0.62984917 | -1.5440228 | 0.12258281 | 0.66929384 |
| FBXO42 | 143.733772 | -0.4168025 | 0.2261633 | -1.8429274 | 0.06533961 | 0.59241679 |
| SZRD1 | 45.1059142 | -0.3013537 | 0.32811134 | -0.9184494 | 0.35838362 | 0.84204868 |
| NECAP2 | 24.9071802 | -0.2161533 | 0.28986474 | -0.7457041 | 0.45584621 | 0.87394768 |
| NBPF1 | 35.2554079 | 0.19824847 | 0.3046282 | 0.65078829 | 0.51518316 | 0.90174188 |
| CROCC | 39.0172438 | -0.1162791 | 0.35094426 | -0.3313321 | 0.74039364 | 0.9544376 |
| ATP13A2 | 18.6606133 | 0.14814939 | 0.47281502 | 0.31333478 | 0.75402633 | 0.95826617 |
| SDHB | 84.5009944 | 0.10025919 | 0.26411305 | 0.3796071 | 0.70423709 | 0.94476609 |
| RCC2 | 39.1533293 | -0.1305935 | 0.29817809 | -0.4379713 | 0.66140708 | 0.93729379 |
| ARHGEF10L | 370.310747 | 0.14853648 | 0.29032358 | 0.5116239 | 0.60891426 | 0.92573951 |
| KLHDC7A | 12.8435265 | 0.41866742 | 0.66196523 | 0.63246134 | 0.52708547 | NA |
| UBR4 | 143.013927 | -0.4798084 | 0.271467 | -1.7674649 | 0.0771504 | 0.61329066 |
| AL035413.1 | 31.0877525 | -0.2791645 | 0.32506657 | -0.8587919 | 0.39045535 | 0.85605713 |
| EMC1 | 19.705369 | 0.15313602 | 0.34980875 | 0.43777069 | 0.66155252 | 0.93729379 |
| MRTO4 | 11.6446446 | -0.1034363 | 0.44121941 | -0.2344327 | 0.81464905 | NA |
| AKR7A2 | 24.0303452 | 0.89517721 | 0.37303833 | 2.39969233 | 0.01640886 | 0.39748373 |
| CAPZB | 154.966016 | -0.103525 | 0.23918467 | -0.4328245 | 0.6651423 | 0.93860542 |
| NBL1 | 16.342794 | -0.2850403 | 0.45440581 | -0.6272814 | 0.53047482 | 0.9052447 |
| TMCO4 | 130.983853 | -0.0583737 | 0.27193408 | -0.2146611 | 0.83003152 | 0.97567084 |
| OTUD3 | 31.7006795 | -0.9690982 | 0.3785641 | -2.5599316 | 0.01046928 | 0.33556978 |
| VWA5B1 | 25.6904984 | 0.20921749 | 0.57130013 | 0.36621292 | 0.7142062 | 0.94841799 |
| CAMK2N1 | 177.684046 | 0.05395951 | 0.37041886 | 0.14567162 | 0.88418063 | 0.98154805 |
| MUL1 | 14.6407906 | -0.1764906 | 0.38563269 | -0.457665 | 0.64719313 | NA |
| PINK1 | 34.1955657 | 0.22175934 | 0.36420294 | 0.60888948 | 0.5425977 | 0.90945809 |
| DDOST | 29.4203343 | 0.44698354 | 0.35066076 | 1.27468934 | 0.20241922 | 0.76065252 |
| HP1BP3 | 203.023268 | -0.0838348 | 0.15314164 | -0.5474331 | 0.5840812 | 0.92071026 |
| EIF4G3 | 730.830471 | -0.7434697 | 0.39558879 | -1.8794003 | 0.06018985 | 0.57733412 |
| ECE1 | 38.4244836 | 0.15077217 | 0.39695674 | 0.37982014 | 0.70407893 | 0.94476609 |
| NBPF3 | 17.5801131 | 0.11859575 | 0.36744881 | 0.32275447 | 0.74688119 | 0.95563105 |
| RAP1GAP | 77.5270057 | 0.74739116 | 0.32748986 | 2.28218107 | 0.02247865 | 0.425412 |
| USP48 | 123.980607 | -0.1056409 | 0.23044403 | -0.4584231 | 0.6466485 | 0.93328692 |
| HSPG2 | 82.6950299 | 0.53825534 | 0.50010327 | 1.07628839 | 0.2817983 | 0.80425482 |
| CDC42 | 136.364376 | 0.00219903 | 0.20806725 | 0.01056886 | 0.99156742 | 0.99814944 |
| ZBTB40 | 44.6093357 | -0.2503839 | 0.31278177 | -0.8005068 | 0.42341725 | 0.86190547 |
| EPHB2 | 40.927607 | -0.4861713 | 0.34096089 | -1.4258859 | 0.15390127 | 0.70572801 |
| KDM1A | 88.8385344 | 0.17984023 | 0.22855207 | 0.78686764 | 0.43135936 | 0.86604857 |
| AL031428.1 | 19.3723305 | 0.94798432 | 0.4782512 | 1.98218908 | 0.04745808 | 0.53530393 |
| LUZP1 | 122.92579 | 0.10308246 | 0.22787819 | 0.45235772 | 0.6510113 | 0.93453685 |
| LINC01355 | 16.6447042 | 0.6559082 | 0.48148488 | 1.36226125 | 0.17311545 | 0.72721271 |
| HNRNPR | 128.027657 | -0.0357653 | 0.20972259 | -0.1705362 | 0.86458846 | 0.97788853 |
| TCEA3 | 68.4306432 | 0.69805185 | 0.41112303 | 1.69791472 | 0.08952386 | 0.63653714 |
| ASAP3 | 64.3657933 | 0.68924568 | 0.31234965 | 2.20664789 | 0.02733867 | 0.45247982 |
| RPL11 | 180.837291 | 0.481232 | 0.42066792 | 1.14397122 | 0.25263557 | 0.79056088 |
| ELOA | 21.9996572 | 0.01877757 | 0.33786497 | 0.05557716 | 0.95567866 | 0.99148072 |
| ELOA-AS1 | 23.3061766 | 0.64088317 | 0.53802766 | 1.19117143 | 0.23358629 | 0.77650198 |
| PITHD1 | 21.9097806 | 0.29744215 | 0.35472675 | 0.83851064 | 0.40174398 | 0.85778581 |
| LYPLA2 | 24.4306873 | -0.2583837 | 0.43729459 | -0.5908686 | 0.55460845 | 0.9137961 |
| GALE | 34.0361412 | -0.5576981 | 0.25489819 | -2.1879248 | 2.87E-02 | 4.61E-01 |
| HMGCL | 43.4375953 | -0.2886213 | 0.30359088 | -0.9506917 | 0.34176092 | 0.83294501 |

| | | | | | | |
|---|---|---|---|---|---|---|
| FUCA1 | 45.7865094 | 0.45504529 | 0.34858432 | 1.3054095 | 0.19175344 | 0.75034462 |
| PNRC2 | 48.0953422 | -0.0624677 | 0.25950981 | -0.2407143 | 0.80977657 | 0.97257655 |
| SRSF10 | 103.378962 | 0.131116 | 0.19572872 | 0.66988634 | 0.50293025 | 0.89564545 |
| IFNLR1 | 18.8288368 | 0.03653441 | 0.38982477 | 0.09372009 | 0.92533151 | 0.98925753 |
| STPG1 | 17.8001762 | 0.39830294 | 0.42464215 | 0.93797315 | 0.34825821 | 0.83687845 |
| NIPAL3 | 68.1325523 | 0.24019818 | 0.36802274 | 0.65267212 | 0.51396768 | 0.90156229 |
| RCAN3 | 33.0483053 | -0.6807215 | 0.35554432 | -1.9145898 | 5.55E-02 | 0.56684377 |
| NCMAP | 52.4179984 | 0.14892453 | 0.7638425 | 0.19496759 | 8.45E-01 | 9.76E-01 |
| SRRM1 | 101.839085 | 0.06302475 | 0.20630932 | 0.30548668 | 0.75999551 | 0.95913478 |
| CLIC4 | 40.1771811 | -0.6532123 | 0.48023814 | -1.3601842 | 0.17377165 | 0.72794035 |
| SYF2 | 35.9666303 | 0.16551524 | 0.34983984 | 0.47311718 | 0.63612958 | 0.93251095 |
| RSRP1 | 241.920978 | 0.54447046 | 0.26832207 | 2.02916764 | 0.04244122 | 0.51343649 |
| TMEM50A | 48.887853 | 0.22983412 | 0.23922894 | 0.96072875 | 3.37E-01 | 0.83037432 |
| MACO1 | 79.2569403 | -0.4363805 | 0.28904237 | -1.5097458 | 0.13110831 | 0.67492558 |
| LDLRAP1 | 14.3701954 | 0.10881668 | 0.46014466 | 0.23648363 | 0.81305741 | NA |
| MAN1C1 | 47.128605 | 0.32237306 | 0.46408782 | 0.69463805 | 0.48728216 | 0.88838332 |
| MTFR1L | 34.9870504 | 0.14683199 | 0.45391334 | 0.32348023 | 0.74633158 | 0.95563105 |
| PAQR7 | 24.3496438 | 0.25060018 | 0.44101519 | 0.56823481 | 0.56987554 | 0.91778542 |
| STMN1 | 21.2727114 | 0.52817442 | 0.52937413 | 0.99773371 | 0.3184085 | 0.82354084 |
| PAFAH2 | 23.8057706 | 0.20067514 | 0.42541628 | 0.47171476 | 0.6371304 | 0.93259128 |
| PDIK1L | 11.5218368 | 0.17529738 | 0.50279711 | 0.34864436 | 0.72735632 | NA |
| ZNF593 | 15.769816 | 0.23863899 | 0.47743771 | 0.49983272 | 0.61719287 | NA |
| CNKSR1 | 22.2431472 | -0.2617285 | 0.32295567 | -0.8104162 | 0.41770102 | 0.85950165 |
| CEP85 | 15.659103 | -0.0529197 | 0.46753959 | -0.1131877 | 0.90988174 | NA |
| SH3BGRL3 | 45.4686445 | -0.4673889 | 0.43786944 | -1.067416 | 2.86E-01 | 0.80431986 |
| UBXN11 | 10.5796904 | -0.4992738 | 0.43641537 | -1.1440336 | 0.2526097 | NA |
| DHDDS | 42.8529707 | -0.5334917 | 0.30916996 | -1.7255611 | 0.0844264 | 0.62840022 |
| HMGN2 | 64.6000193 | 0.06467138 | 0.34131864 | 0.1894751 | 0.84972047 | 0.97612951 |
| RPS6KA1 | 43.7961898 | 0.03435082 | 0.32515039 | 0.10564596 | 0.91586326 | 0.98691474 |
| ARID1A | 178.530833 | 0.09654784 | 0.21123687 | 0.45705962 | 0.64762819 | 0.93328692 |
| ZDHHC18 | 15.1092053 | 0.40406697 | 0.38112975 | 1.06018218 | 0.28906173 | NA |
| SFN | 19.2962862 | -0.7229357 | 0.61988122 | -1.1662488 | 2.44E-01 | 7.85E-01 |
| GPN2 | 20.5742474 | -0.2033271 | 0.36281528 | -0.5604149 | 0.57519649 | 0.91854298 |
| NUDC | 51.7318895 | -0.144267 | 0.31372479 | -0.459852 | 0.64562248 | 0.93328692 |
| NR0B2 | 34.9177932 | -0.6739236 | 0.74313088 | -0.9068707 | 3.64E-01 | 0.8468666 |
| KDF1 | 10.852572 | 0.5798398 | 0.42826621 | 1.35392379 | 0.17576069 | NA |
| TRNP1 | 97.0832575 | 0.45175223 | 0.5445589 | 0.82957458 | 4.07E-01 | 8.59E-01 |
| SLC9A1 | 45.3278239 | -0.0667675 | 0.31742296 | -0.2103425 | 0.83340039 | 0.97600352 |
| WDTC1 | 60.6655803 | -0.020429 | 0.25503222 | -0.0801036 | 0.93615483 | 0.99012269 |
| TMEM222 | 22.8907377 | -0.9010743 | 0.32347691 | -2.7855908 | 0.00534303 | 0.28183167 |
| WASF2 | 131.775803 | -0.027608 | 0.15822761 | -0.1744826 | 0.86148619 | 0.97767895 |
| AHDC1 | 48.3316168 | 0.72250526 | 0.43435889 | 1.66338314 | 0.09623577 | 0.65104726 |
| IFI6 | 127.248864 | -4.0391521 | 1.62200624 | -2.4902198 | 0.01276641 | 0.36544853 |
| FAM76A | 21.5899041 | 0.21437454 | 0.37585168 | 0.57037006 | 0.56842674 | 0.91698355 |
| STX12 | 105.890121 | 0.10278556 | 0.29397081 | 0.34964546 | 0.72660479 | 0.95209193 |
| PPP1R8 | 18.2798846 | -0.4383205 | 0.3807931 | -1.1510726 | 0.24970235 | 0.78799766 |
| RPA2 | 30.0219564 | 0.37440839 | 0.37795368 | 0.99061976 | 0.32187128 | 0.82511808 |
| EYA3 | 463.941907 | 0.2056854 | 0.20011853 | 1.02781784 | 3.04E-01 | 0.81568231 |
| PTAFR | 49.0822447 | 0.118889 | 0.3769584 | 0.31539025 | 0.75246537 | 0.95782508 |
| DNAJC8 | 41.0217909 | -0.0514829 | 0.28391847 | -0.18133 | 0.85610854 | 0.97634735 |
| ATP5IF1 | 121.740341 | 0.28364412 | 0.39783941 | 0.71296134 | 0.47586968 | 0.88495683 |
| SESN2 | 12.9389225 | 0.18326627 | 0.49860164 | 0.36756051 | 0.71320097 | NA |
| PHACTR4 | 208.361791 | -0.0237456 | 0.20010949 | -0.1186632 | 0.90554219 | 0.98531749 |
| RCC1 | 32.5759664 | 0.10354188 | 0.31895565 | 0.32462783 | 0.74546276 | 0.95563105 |

| | | | | | | |
|---|---|---|---|---|---|---|
| TRNAU1AP | 44.2589971 | -0.3791196 | 0.27999966 | -1.3540003 | 0.17573627 | 0.72841249 |
| SNHG12 | 13.289271 | -0.8424711 | 0.45522796 | -1.8506577 | 0.06421882 | NA |
| TAF12 | 27.2447012 | -0.1978049 | 0.36052955 | -0.5486509 | 0.58324506 | 0.92034873 |
| GMEB1 | 30.6590113 | -0.5685873 | 0.41562675 | -1.3680238 | 0.17130464 | 0.72463203 |
| YTHDF2 | 76.0818556 | -0.1499813 | 0.24342729 | -0.6161238 | 0.53781281 | 0.90763121 |
| EPB41 | 191.196762 | 0.15746116 | 0.19850686 | 0.79322778 | 0.42764513 | 0.86239557 |
| SRSF4 | 227.155288 | -0.2014894 | 0.27225584 | -0.7400737 | 0.45925525 | 0.87574928 |
| MECR | 27.7938165 | 0.33219639 | 0.40144663 | 0.82749826 | 0.40795471 | 0.85867915 |
| PTPRU | 17.4642555 | -0.0390731 | 0.46548366 | -0.0839408 | 0.93310352 | 0.98960607 |
| PUM1 | 394.233467 | -0.1804284 | 0.18738767 | -0.9628618 | 0.3356169 | 0.82974237 |
| SNRNP40 | 34.0839394 | -0.0186372 | 0.29404911 | -0.0633811 | 0.94946304 | 0.99105535 |
| ZCCHC17 | 63.2795774 | -0.2665489 | 0.28432317 | -0.9374856 | 0.34850883 | 0.83687845 |
| SERINC2 | 107.691887 | -0.1182062 | 0.34953511 | -0.3381811 | 0.73522674 | 0.95341665 |
| TINAGL1 | 41.9692584 | -0.6233005 | 0.46309309 | -1.3459507 | 0.17831842 | 0.72841249 |
| PEF1 | 30.9243017 | -0.1794159 | 0.39614407 | -0.4529056 | 0.65061674 | 0.93453685 |
| PTP4A2 | 59.4615971 | -0.5518298 | 0.3843482 | -1.435755 | 0.15107208 | 0.70572801 |
| KHDRBS1 | 148.666651 | -0.4481403 | 0.23378842 | -1.9168629 | 0.05525535 | 0.5645224 |
| TMEM39B | 11.1899399 | 0.32225056 | 0.50703624 | 0.63555724 | 0.52506504 | NA |
| KPNA6 | 124.591859 | -0.1783829 | 0.21555338 | -0.8275581 | 0.40792083 | 0.85867915 |
| TXLNA | 40.1461935 | -0.028079 | 0.27323227 | -0.1027662 | 0.91814852 | 0.98735642 |
| TMEM234 | 13.8513271 | -0.0425867 | 0.42130122 | -0.1010838 | 0.91948394 | NA |
| EIF3I | 28.7839193 | 0.55782836 | 0.49117953 | 1.13569139 | 0.25608575 | 0.7940072 |
| HDAC1 | 47.7545373 | 0.42488873 | 0.30314593 | 1.40159801 | 0.16103532 | 0.71101785 |
| BSDC1 | 59.1320721 | -0.1917319 | 0.30530073 | -0.62801 | 0.52999742 | 0.9052447 |
| ZBTB8OS | 90.5302037 | -0.1222616 | 0.22332064 | -0.5474713 | 0.58405502 | 0.92071026 |
| RBBP4 | 92.7274355 | -0.1099135 | 0.19014715 | -0.5780445 | 0.56323408 | 0.91624774 |
| SYNC | 20.4281386 | -0.0205274 | 0.50060468 | -0.0410052 | 0.96729178 | 0.99314878 |
| KIAA1522 | 112.602138 | 0.06157714 | 0.27739904 | 0.22198038 | 0.82432916 | 0.97518529 |
| YARS | 52.413763 | -0.7300976 | 0.49882087 | -1.4636469 | 0.14329044 | 0.69428142 |
| S100PBP | 85.875865 | 0.16187802 | 0.26201995 | 0.61780798 | 0.53670192 | 0.90763121 |
| TMEM54 | 13.4652685 | 0.51259218 | 0.6411081 | 0.79954094 | 0.42397682 | NA |
| RNF19B | 34.3904919 | -1.5027916 | 0.44703079 | -3.3617182 | 0.00077459 | 0.12591256 |
| AL020995.1 | 22.3207829 | 0.57319039 | 0.66554765 | 0.86123119 | 0.38911073 | 0.85605713 |
| AK2 | 108.319278 | -0.2389964 | 0.25030117 | -0.9548352 | 0.33966106 | 0.83071861 |
| TRIM62 | 12.3302778 | -0.025653 | 0.41741644 | -0.0614565 | 0.95099564 | NA |
| ZNF362 | 23.6835894 | 0.48871971 | 0.45089022 | 1.08389956 | 0.27840934 | 0.80210301 |
| AL513327.1 | 21.3749936 | -0.0799898 | 0.64636322 | -0.1237536 | 0.90151038 | 0.98531749 |
| PHC2 | 154.799123 | -0.9717715 | 0.22142689 | -4.3886787 | 1.14E-05 | 0.00954327 |
| SMIM12 | 32.5956803 | 0.11002155 | 0.48508816 | 0.22680733 | 0.82057356 | 0.97368535 |
| ZMYM1 | 38.96879 | -0.3274407 | 0.30795367 | -1.063279 | 0.28765546 | 0.80523851 |
| SFPQ | 203.835672 | -0.0932656 | 0.20625528 | -0.4521851 | 0.65113563 | 0.93453685 |
| ZMYM4 | 341.564126 | 0.08462057 | 0.1774443 | 0.47688528 | 0.63344382 | 0.93251095 |
| KIAA0319L | 186.759314 | -0.1173806 | 0.16771533 | -0.6998798 | 0.48400239 | 0.88838332 |
| PSMB2 | 40.3779764 | -0.0114564 | 0.35863592 | -0.0319443 | 0.97451644 | 0.99431961 |
| AGO4 | 70.2576393 | -0.3537015 | 0.19876919 | -1.7794586 | 0.0751646 | 0.61020315 |
| AGO1 | 26.9434803 | 0.09785347 | 0.29204344 | 0.33506479 | 0.73757621 | 0.9542764 |
| AGO3 | 221.446063 | -0.3381236 | 0.19228552 | -1.7584456 | 0.07867171 | 0.61367304 |
| ADPRHL2 | 21.2402506 | -0.2495637 | 0.47738668 | -0.5227706 | 0.60113393 | 0.92356492 |
| TRAPPC3 | 30.1909509 | -0.438871 | 0.27972709 | -1.5689255 | 0.11666529 | 0.6668162 |
| MAP7D1 | 20.3127871 | -0.1775288 | 0.42483091 | -0.417881 | 0.67603414 | 0.9402691 |
| THRAP3 | 159.587415 | -0.2501083 | 0.26244782 | -0.9529831 | 0.34059865 | 0.8319325 |
| SH3D21 | 12.2497538 | 0.1239714 | 0.54998427 | 0.22540898 | 0.82166112 | NA |
| EVA1B | 15.4696032 | 0.24311989 | 0.45208234 | 0.53777789 | 0.5907304 | NA |
| STK40 | 47.5393277 | -0.4726781 | 0.34127766 | -1.3850251 | 0.1660448 | 0.71756194 |

| | | | | | | |
|---|---|---|---|---|---|---|
| OSCP1 | 11.3593147 | -0.0520856 | 0.51317695 | -0.1014964 | 0.91915638 | NA |
| MRPS15 | 20.0474747 | -0.0266083 | 0.33803227 | -0.0787153 | 0.93725911 | 0.99012269 |
| LINC01137 | 19.3013827 | 0.76737446 | 0.48980131 | 1.56670562 | 0.11718352 | 0.6668162 |
| ZC3H12A | 25.787307 | 0.4053741 | 0.57629105 | 0.70341904 | 0.48179464 | 0.88708902 |
| MEAF6 | 35.9828081 | -0.1083264 | 0.31581308 | -0.3430081 | 0.7315924 | 0.95279138 |
| SNIP1 | 14.82084 | 0.04076529 | 0.43426825 | 0.09387122 | 0.92521146 | NA |
| DNALI1 | 12.9960135 | 0.31446051 | 0.47675785 | 0.65958118 | 0.50952263 | NA |
| GNL2 | 14.4732727 | -0.9872759 | 0.42769722 | -2.3083524 | 0.02097955 | NA |
| C1orf122 | 33.0824031 | 0.33681211 | 0.40111333 | 0.83969312 | 0.40108047 | 0.85778581 |
| MTF1 | 28.7234817 | -0.1917239 | 0.3596241 | -0.5331232 | 0.5939483 | 0.9229638 |
| INPP5B | 80.4334998 | 0.0696426 | 0.32407758 | 0.21489484 | 0.82984931 | 0.97567084 |
| SF3A3 | 34.8814675 | -0.2258247 | 0.2801585 | -0.8060604 | 0.42020802 | 0.86124817 |
| RRAGC | 27.2921114 | -0.8060856 | 0.3910288 | -2.0614483 | 0.03926029 | 0.50068767 |
| AL139260.1 | 42.9812536 | -0.708024 | 0.47371227 | -1.4946289 | 0.13501133 | 0.68329553 |
| AKIRIN1 | 78.5252921 | 0.24146713 | 0.33768093 | 0.7150748 | 0.47456282 | 0.88424821 |
| NDUFS5 | 113.314627 | -0.1678868 | 0.43746344 | -0.3837734 | 0.70114638 | 0.94440669 |
| MACF1 | 967.094779 | -0.3708505 | 0.21946838 | -1.6897672 | 0.09107251 | 0.64297142 |
| PABPC4 | 39.472452 | -0.0750693 | 0.29744335 | -0.2523818 | 0.80074593 | 0.97021033 |
| PPIE | 23.7640233 | 0.0855614 | 0.34255008 | 0.24977779 | 0.8027592 | 0.9710986 |
| TRIT1 | 56.3153891 | -0.3930988 | 0.2407556 | -1.632771 | 0.10251716 | 0.65946648 |
| CAP1 | 65.8545967 | 0.09765589 | 0.30407499 | 0.32115724 | 0.74809123 | 0.95563105 |
| PPT1 | 25.9407399 | -0.1624641 | 0.36532401 | -0.4447123 | 0.65652772 | 0.93570875 |
| RLF | 174.9686 | -0.2963812 | 0.28989525 | -1.0223735 | 0.30660415 | 0.81568231 |
| ZMPSTE24 | 53.9855693 | -0.3597139 | 0.25818246 | -1.3932546 | 0.16354283 | 0.71332671 |
| COL9A2 | 11.5400777 | 0.46115122 | 0.74084522 | 0.62246635 | 0.53363526 | NA |
| SMAP2 | 30.5792904 | 0.16309942 | 0.32499602 | 0.5018505 | 0.61577269 | 0.92662475 |
| AL603839.3 | 26.0751417 | 0.45465971 | 0.35292172 | 1.28827353 | 0.19765076 | 0.75559912 |
| ZNF684 | 45.1853898 | 0.94913891 | 0.34828153 | 2.72520601 | 0.00642614 | 0.29935381 |
| NFYC | 161.693192 | 0.4132198 | 0.24468729 | 1.6887669 | 9.13E-02 | 0.64297142 |
| CITED4 | 17.0073572 | 0.81377631 | 0.52379175 | 1.55362568 | 1.20E-01 | 0.6668162 |
| SLFNL1-AS1 | 16.7178439 | 0.6232997 | 0.42234236 | 1.47581621 | 0.13999324 | 0.6912882 |
| SCMH1 | 502.642061 | 0.01520897 | 0.14316441 | 0.10623428 | 0.91539647 | 0.98691474 |
| AC093151.3 | 16.4383335 | 0.124471 | 0.62357765 | 0.19960786 | 0.84178728 | 0.97600352 |
| HIVEP3 | 104.114557 | 0.16851907 | 0.27517619 | 0.61240425 | 0.54027033 | 0.90763121 |
| FOXJ3 | 203.649604 | -0.5371648 | 0.25385758 | -2.1160084 | 0.03434409 | 0.4908133 |
| PPCS | 26.2349538 | 0.60986622 | 0.39286383 | 1.55236032 | 0.12057603 | 0.6668162 |
| CCDC30 | 149.693308 | 0.30685482 | 0.31061863 | 0.98788286 | 0.32321002 | 0.82558302 |
| PPIH | 15.266307 | 0.11431538 | 0.43572099 | 0.26235913 | 0.79304458 | NA |
| YBX1 | 99.3558033 | -0.0534636 | 0.35760114 | -0.1495062 | 0.8811542 | 0.98130006 |
| ERMAP | 23.6086158 | 0.20904841 | 0.37352803 | 0.55965922 | 0.5757119 | 0.91854298 |
| SLC2A1 | 120.943809 | -1.654063 | 0.51779085 | -3.1944615 | 0.00140092 | 0.15940712 |
| SLC2A1-AS1 | 20.1180537 | -0.0406419 | 0.51574063 | -0.078803 | 0.93718928 | 0.99012269 |
| EBNA1BP2 | 18.9216276 | 0.28562135 | 0.41011886 | 0.69643555 | 0.48615611 | 0.88838332 |
| TMEM125 | 11.8038228 | 0.1152182 | 0.50804376 | 0.22678794 | 0.82058864 | NA |
| ELOVL1 | 17.1302198 | -0.0889948 | 0.44666842 | -0.1992414 | 0.84207395 | 0.97600352 |
| SZT2 | 42.8921361 | -0.2866677 | 0.29564938 | -0.9696205 | 0.3322357 | 0.82890589 |
| PTPRF | 145.630186 | 0.42099301 | 0.39487554 | 1.06614102 | 0.2863599 | 0.80463985 |
| IPO13 | 10.444541 | 0.36491619 | 0.5410102 | 0.67450888 | 0.49998784 | NA |
| ATP6V0B | 109.322116 | 0.24683088 | 0.33564128 | 0.73540083 | 0.46209542 | 0.87689276 |
| B4GALT2 | 14.4536453 | 0.29565982 | 0.58218009 | 0.50784942 | 0.61155895 | NA |
| CCDC24 | 26.6157233 | 0.01101943 | 0.33686418 | 0.03271178 | 0.97390443 | 0.99431961 |
| DMAP1 | 17.3958409 | 0.01587132 | 0.4311294 | 0.03681336 | 0.97063382 | 0.99403127 |
| ERI3 | 90.6675649 | 0.23951598 | 0.36902301 | 0.64905434 | 0.51630325 | 0.90225158 |
| RNF220 | 20.777935 | -0.2525939 | 0.35800418 | -0.7055613 | 0.48046098 | 0.88660781 |

| | | | | | | |
|---|---|---|---|---|---|---|
| TMEM53 | 17.8790116 | 0.42310845 | 0.42855684 | 0.98728666 | 0.32350213 | 0.82558302 |
| ARMH1 | 19.7422496 | 0.47807445 | 0.51446461 | 0.92926597 | 0.35275126 | 0.83872681 |
| RPS8 | 145.113195 | 0.36408503 | 0.26105097 | 1.39468944 | 0.16310951 | 0.71325312 |
| EIF2B3 | 59.1666292 | -0.2385179 | 0.29898186 | -0.797767 | 0.4250057 | 0.86207532 |
| HECTD3 | 12.5975089 | -0.0887858 | 0.49356222 | -0.1798877 | 0.85724073 | NA |
| ZSWIM5 | 93.9160183 | 0.1972772 | 0.34988599 | 0.56383283 | 0.57286791 | 0.91813957 |
| TESK2 | 30.2333691 | -0.0923542 | 0.35025767 | -0.263675 | 0.79203038 | 0.96931299 |
| PRDX1 | 172.090696 | 0.25768493 | 0.42436007 | 0.6072318 | 0.54369709 | 0.90965439 |
| AKR1A1 | 21.7192157 | -0.3136081 | 0.34387287 | -0.9119885 | 0.36177476 | 0.84398658 |
| NASP | 61.6790039 | -0.2658374 | 0.34949482 | -0.7606334 | 0.44687607 | 0.87231627 |
| GPBP1L1 | 208.762033 | 0.03844317 | 0.15794151 | 0.24340132 | 0.80769452 | 0.97241677 |
| TMEM69 | 12.3712023 | 0.22742766 | 0.47663272 | 0.47715495 | 0.63325179 | NA |
| IPP | 46.0859131 | 0.38313691 | 0.2778175 | 1.37909568 | 0.16786525 | 0.72008028 |
| MAST2 | 58.8371542 | -0.398686 | 0.37353133 | -1.067343 | 0.285817 | 0.80431986 |
| PIK3R3 | 12.2720481 | 0.28926471 | 0.59919057 | 0.48275912 | 0.62926678 | NA |
| TSPAN1 | 290.296938 | 0.34287287 | 0.46323723 | 0.74016693 | 4.59E-01 | 8.76E-01 |
| POMGNT1 | 53.8614751 | 0.25794172 | 0.31906584 | 0.80842787 | 0.41884431 | 0.86022698 |
| LURAP1 | 60.8927814 | 0.23043684 | 0.2509213 | 0.91836299 | 0.35842886 | 0.84204868 |
| LRRC41 | 45.1656527 | -0.0921559 | 0.25682891 | -0.3588223 | 0.71972804 | 0.94995363 |
| UQCRH | 131.514804 | 0.03732027 | 0.32543482 | 0.11467816 | 0.90870023 | 0.9854228 |
| NSUN4 | 31.441604 | -0.078979 | 0.32350758 | -0.2441335 | 0.80712744 | 0.97216031 |
| FAAH | 13.5223075 | -0.0414615 | 0.43394661 | -0.0955451 | 0.92388187 | NA |
| MKNK1 | 45.3070998 | -0.2015795 | 0.27918492 | -0.7220287 | 0.47027685 | 0.88260202 |
| ATPAF1 | 57.0128605 | 0.16219765 | 0.29995961 | 0.54073165 | 0.58869256 | 0.92187655 |
| EFCAB14 | 150.185979 | 0.08491302 | 0.3156378 | 0.26902043 | 0.78791396 | 0.96767512 |
| PDZK1IP1 | 75.5879573 | 0.68212633 | 0.5367898 | 1.27075129 | 0.20381714 | 0.76173533 |
| CMPK1 | 71.9941481 | 0.04856139 | 0.28559713 | 0.17003459 | 0.86498294 | 0.97788853 |
| TRABD2B | 936.711774 | -0.0485254 | 0.25199199 | -0.1925673 | 0.84729784 | 0.97612951 |
| SPATA6 | 164.314842 | 0.37714981 | 0.23792044 | 1.58519296 | 0.1129225 | 0.6656824 |
| AGBL4 | 778.119499 | 0.65405584 | 0.40379265 | 1.61978143 | 0.10527924 | 0.65946648 |
| FAF1 | 610.965659 | -0.1598611 | 0.16609276 | -0.9624809 | 0.33580808 | 0.82974237 |
| RNF11 | 80.4650149 | -0.202235 | 0.30369339 | -0.6659184 | 0.50546328 | 0.89723255 |
| TTC39A | 79.7684766 | -0.1039506 | 0.40792527 | -0.2548277 | 7.99E-01 | 9.70E-01 |
| EPS15 | 245.365856 | -0.3871745 | 0.20635481 | -1.876256 | 0.06062013 | 0.57774794 |
| OSBPL9 | 201.315741 | -0.2223096 | 0.16813907 | -1.3221769 | 0.18610924 | 0.73819275 |
| NRDC | 168.917197 | -0.0526999 | 0.18379573 | -0.286731 | 0.77431829 | 0.96435639 |
| TXNDC12 | 37.4235878 | -0.220329 | 0.29110205 | -0.7568788 | 0.44912249 | 0.87314185 |
| BTF3L4 | 63.9151708 | -0.0367951 | 0.2577684 | -0.1427449 | 0.88649163 | 0.98216235 |
| ZFYVE9 | 139.875399 | 0.06916575 | 0.19062594 | 0.36283491 | 0.71672822 | 0.94913834 |
| CC2D1B | 25.9593935 | -0.553831 | 0.36762455 | -1.5065126 | 0.13193565 | 0.6765887 |
| PRPF38A | 24.9750788 | -0.1165125 | 0.3172026 | -0.3673126 | 0.71338584 | 0.94829221 |
| TUT4 | 218.079423 | 0.01003613 | 0.23918537 | 0.04195965 | 0.96653086 | 0.99301184 |
| ZYG11B | 95.6476055 | -0.0384629 | 0.25269248 | -0.1522125 | 0.87901936 | 0.98061726 |
| ECHDC2 | 193.432647 | 0.45882695 | 0.32217117 | 1.42417135 | 0.15439688 | 0.70572801 |
| SCP2 | 142.116299 | 0.19193062 | 0.25829708 | 0.74306151 | 0.45744444 | 0.87420547 |
| CPT2 | 20.5207549 | 0.71575654 | 0.46830053 | 1.52841282 | 0.12641007 | 0.66929384 |
| C1orf123 | 10.8764507 | -0.1376828 | 0.42904392 | -0.3209061 | 0.74828158 | NA |
| MAGOH | 23.3444859 | 0.02308814 | 0.32313825 | 0.07144973 | 0.94303983 | 0.99058259 |
| LRP8 | 13.6634801 | -0.0590251 | 0.3939388 | -0.1498332 | 0.8808962 | NA |
| NDC1 | 34.8834814 | -0.4748993 | 0.32237719 | -1.473117 | 0.14071948 | 0.69227547 |
| YIPF1 | 25.9142551 | 0.10767574 | 0.33221708 | 0.3241126 | 0.74585279 | 0.95563105 |
| HSPB11 | 30.5295408 | 0.21365396 | 0.44253434 | 0.48279634 | 0.62924035 | 0.9315295 |
| LRRC42 | 18.5192928 | 0.15661921 | 0.36940158 | 0.42398088 | 0.67157976 | 0.93878847 |
| TMEM59 | 160.65356 | 0.51213764 | 0.2433253 | 2.10474471 | 0.03531354 | 0.49209205 |

| | | | | | | |
|---|---|---|---|---|---|---|
| TCEANC2 | 29.8284601 | 0.11930097 | 0.41099455 | 0.29027386 | 0.77160673 | 0.9637673 |
| MRPL37 | 16.1020805 | -0.0931837 | 0.39238561 | -0.23748 | 0.81228444 | NA |
| SSBP3 | 239.619672 | -0.3136053 | 0.26944717 | -1.1638843 | 0.24447091 | 0.78564467 |
| ACOT11 | 229.270454 | -0.3984716 | 0.33746342 | -1.1807846 | 0.23768831 | 0.77930556 |
| MROH7-TTC4 | 21.5679454 | -0.9377285 | 0.55052781 | -1.7033263 | 0.08850702 | 0.63466198 |
| TTC22 | 11.4042834 | 0.02686762 | 0.41761057 | 0.06433654 | 0.94870226 | NA |
| DHCR24 | 41.7955165 | -0.2030363 | 0.51356553 | -0.3953464 | 0.69258725 | 0.94374959 |
| USP24 | 380.826391 | -0.6399309 | 0.2437608 | -2.6252413 | 0.00865876 | 0.31119527 |
| PLPP3 | 167.206959 | -1.0028927 | 0.46187845 | -2.1713347 | 0.02990588 | 0.46381645 |
| PRKAA2 | 288.91245 | -0.2491193 | 0.26077381 | -0.9553079 | 0.33942203 | 0.83066816 |
| FYB2 | 315.595842 | 0.75614115 | 0.40648726 | 1.86018412 | 0.06285948 | 0.58508165 |
| DAB1 | 254.020468 | 1.3402864 | 1.0809505 | 1.23991468 | 0.21500695 | 0.7643572 |
| TACSTD2 | 171.089832 | 0.6077672 | 0.46772912 | 1.29939996 | 0.19380671 | 0.75067401 |
| MYSM1 | 121.79729 | -0.0326134 | 0.30787373 | -0.1059312 | 0.91563696 | 0.98691474 |
| AL136985.3 | 14.2060695 | -0.7077194 | 0.45366772 | -1.559995 | 0.11876107 | NA |
| JUN | 79.9237643 | -0.1043233 | 0.56652019 | -0.1841476 | 0.85389765 | 0.97612951 |
| LINC01135 | 38.1159131 | 0.83924272 | 0.56235811 | 1.49236351 | 0.13560388 | 0.6835006 |
| FGGY | 166.070114 | 0.18933328 | 0.2767796 | 0.68405793 | 0.49393859 | 0.89177369 |
| HOOK1 | 129.477062 | 0.0624745 | 0.2929679 | 0.21324691 | 0.83113438 | 0.97567084 |
| AC097066.1 | 15.0012111 | 2.49096225 | 0.80182769 | 3.10660542 | 0.00189249 | NA |
| NFIA | 137.427851 | 1.18267829 | 0.52736182 | 2.2426316 | 0.02492058 | 0.43948277 |
| TM2D1 | 84.3908124 | 0.05141923 | 0.25407877 | 0.20237515 | 0.83962346 | 0.97600352 |
| PATJ | 1210.34399 | -0.1367244 | 0.15972293 | -0.8560099 | 0.3919923 | 0.85605713 |
| USP1 | 18.8506389 | 0.18001211 | 0.35065493 | 0.51335968 | 0.60769974 | 0.92569157 |
| DOCK7 | 364.050106 | 0.01986481 | 0.20417973 | 0.09729078 | 0.92249548 | 0.98886909 |
| AC103923.1 | 14.7389028 | 0.61877214 | 0.44388592 | 1.3939891 | 0.1633209 | NA |
| ATG4C | 42.0603425 | 0.53472389 | 0.3468464 | 1.54167345 | 0.12315297 | 0.66929384 |
| ALG6 | 22.4702553 | 0.00017114 | 0.36012059 | 0.00047523 | 0.99962082 | 0.99973064 |
| ITGB3BP | 63.3736577 | 0.11736004 | 0.48615146 | 0.24140632 | 0.80924021 | 0.97257655 |
| EFCAB7 | 35.4970281 | -0.1824367 | 0.39322454 | -0.4639505 | 0.64268321 | 0.93328692 |
| PGM1 | 66.7556577 | -0.1216351 | 0.23873495 | -0.5094985 | 0.61040288 | 0.92606167 |
| ROR1 | 55.1422887 | -0.1537948 | 0.55640517 | -0.276408 | 0.78223474 | 0.96652357 |
| CACHD1 | 216.141259 | 0.40341026 | 0.17215927 | 2.3432386 | 0.01911715 | 0.41544117 |
| RAVER2 | 67.0021233 | 0.23147116 | 0.30082949 | 0.76944304 | 0.44163035 | 0.86925451 |
| JAK1 | 230.953277 | -0.3157918 | 0.30743078 | -1.0271963 | 0.30432803 | 0.81568231 |
| AK4 | 26.3804389 | -0.6923965 | 0.62093345 | -1.1150898 | 0.26481195 | 0.79666376 |
| DNAJC6 | 19.3523588 | -0.3932032 | 0.62386015 | -0.6302746 | 0.52851496 | 0.90487731 |
| LEPR | 88.5042095 | 0.27166732 | 0.33713276 | 0.80581702 | 0.42034836 | 0.86124817 |
| SGIP1 | 34.4692748 | -1.0733256 | 0.81667338 | -1.3142654 | 0.18875689 | 0.74318944 |
| WDR78 | 54.5874123 | 0.6915005 | 0.42198612 | 1.63868066 | 0.10127978 | 0.65863153 |
| MIER1 | 132.687782 | -0.1961678 | 0.27826673 | -0.7049633 | 0.48083307 | 0.88675516 |
| SLC35D1 | 43.1703574 | -0.1284563 | 0.25824359 | -0.497423 | 0.6188908 | 0.92766755 |
| SERBP1 | 120.93739 | 0.07490558 | 0.28802917 | 0.26006249 | 0.79481557 | 0.96974891 |
| GADD45A | 28.5796567 | -0.0192134 | 0.40137729 | -0.0478686 | 0.96182094 | 0.99256424 |
| GNG12 | 193.779423 | -0.1321651 | 0.16156296 | -0.818041 | 0.4133338 | 0.85867915 |
| GNG12-AS1 | 69.7047492 | 0.03667379 | 0.30258934 | 0.12119988 | 0.90353272 | 0.98531749 |
| WLS | 105.87396 | 0.36222426 | 0.24335551 | 1.48845718 | 0.13663037 | 0.68412884 |
| LRRC40 | 44.6975787 | -0.142192 | 0.32880317 | -0.4324534 | 0.66541194 | 0.93860542 |
| SRSF11 | 206.218189 | 0.16915944 | 0.23896985 | 0.70786939 | 0.47902637 | 0.88622003 |
| ANKRD13C | 133.700484 | -0.3514421 | 0.230061 | -1.5276038 | 0.12661094 | 0.66929384 |
| LINC01788 | 56.2908164 | -0.3004434 | 0.50445252 | -0.595583 | 5.51E-01 | 0.9128014 |
| PTGER3 | 634.303121 | 1.11573277 | 0.43939632 | 2.53924013 | 0.01110935 | 0.34397433 |
| ZRANB2-AS1 | 17.1027671 | 0.9111258 | 0.58567873 | 1.55567507 | 0.11978538 | 0.6668162 |
| ZRANB2 | 211.621966 | -0.1663075 | 0.20760381 | -0.8010813 | 0.42308456 | 0.86190547 |

| | | | | | | |
|---|---|---|---|---|---|---|
| NEGR1 | 15.4807554 | 0.14819593 | 0.52316782 | 0.28326653 | 0.77697253 | NA |
| LRRIQ3 | 13.9793412 | 0.77642024 | 0.92848884 | 0.83621925 | 0.40303157 | NA |
| CRYZ | 105.413829 | 0.30585125 | 0.33163833 | 0.92224335 | 0.35640166 | 0.84192356 |
| TYW3 | 30.9098255 | -0.1319915 | 0.36238074 | -0.3642343 | 0.71568305 | 0.94885853 |
| ACADM | 81.2051037 | 0.5966742 | 0.23217426 | 2.56994125 | 0.01017158 | 0.33306084 |
| RABGGTB | 23.0968603 | 0.20357306 | 0.34786785 | 0.58520227 | 0.55841172 | 0.91425998 |
| ST6GALNAC3 | 29.4666473 | 0.12182438 | 0.35518599 | 0.34298757 | 0.73160781 | 0.95279138 |
| PIGK | 46.5347042 | -0.0302456 | 0.2645255 | -0.1143391 | 0.90896901 | 0.9854228 |
| AC118549.1 | 241.92711 | -0.1198837 | 0.18582633 | -0.6451383 | 0.51883759 | 0.90368693 |
| USP33 | 177.295672 | -0.0167421 | 0.18608376 | -0.0899705 | 0.92831065 | 0.98960607 |
| MIGA1 | 104.843231 | 0.07885885 | 0.22557988 | 0.34958282 | 0.72665181 | 0.95209193 |
| NEXN | 25.8451268 | 0.78759022 | 0.52308228 | 1.50567176 | 0.13215146 | 0.6765887 |
| FUBP1 | 132.041651 | 0.02827918 | 0.21571577 | 0.13109465 | 0.89570043 | 0.98503595 |
| GIPC2 | 23.3107404 | 0.70743583 | 0.43370768 | 1.63113511 | 0.10286181 | 0.65946648 |
| PTGFR | 61.552397 | 0.17685315 | 0.43151735 | 0.40984018 | 0.68192319 | 0.94205398 |
| IFI44L | 102.662232 | -4.181569 | 1.64068481 | -2.5486729 | 0.01081337 | 0.33942942 |
| IFI44 | 40.2897682 | -2.8866587 | 1.26630128 | -2.2795986 | 0.0226315 | 0.42564995 |
| LINC01781 | 33.6738283 | 1.0205579 | 0.72804203 | 1.40178432 | 1.61E-01 | 0.71101785 |
| ADGRL2 | 438.166923 | -0.0042983 | 0.3132708 | -0.0137208 | 9.89E-01 | 9.98E-01 |
| LINC01362 | 19.0445223 | 0.74527664 | 0.75925452 | 0.98159 | 0.32630188 | 0.82567895 |
| LINC01725 | 298.6756 | 0.79059794 | 0.4326797 | 1.82721293 | 0.06766776 | 0.60095573 |
| PRKACB | 110.115088 | 0.35625242 | 0.3277548 | 1.08694799 | 0.2770598 | 0.80159799 |
| RPF1 | 15.2310482 | 0.16407479 | 0.38821226 | 0.42264197 | 0.6725565 | NA |
| GNG5 | 58.5974727 | 0.26117121 | 0.3030676 | 0.86175893 | 0.38882019 | 0.85605713 |
| SPATA1 | 22.8331178 | 0.45921414 | 0.64504538 | 0.71190982 | 0.47652062 | 0.88495683 |
| CTBS | 39.1165349 | 0.07349446 | 0.29186591 | 0.251809 | 0.8011887 | 0.97045451 |
| SSX2IP | 22.3350676 | -0.1761907 | 0.34885785 | -0.50505 | 0.61352372 | 0.92662475 |
| SYDE2 | 21.9479039 | 0.07790642 | 0.50007925 | 0.15578815 | 0.87620001 | 0.97925705 |
| C1orf52 | 35.7036809 | 0.11193586 | 0.36439368 | 0.30718386 | 0.75870343 | 0.95903525 |
| BCL10 | 31.6518623 | -0.1807001 | 0.35752241 | -0.5054232 | 0.61326161 | 0.92662475 |
| AL078459.1 | 50.1852191 | 0.44296406 | 0.4558518 | 0.97172822 | 0.33118577 | 0.82879734 |
| DDAH1 | 245.551024 | -0.0291768 | 0.32530194 | -0.0896915 | 0.9285324 | 0.98960607 |
| ZNHIT6 | 56.2143357 | -0.2610143 | 0.25944173 | -1.0060613 | 0.31438609 | 0.82040655 |
| ODF2L | 130.438896 | -0.5219975 | 0.24927527 | -2.0940606 | 0.03625458 | 0.49485984 |
| SH3GLB1 | 99.3683334 | -0.1176285 | 0.18760299 | -0.6270075 | 0.5306543 | 0.9052447 |
| SELENOF | 101.209272 | -0.0155018 | 0.18626698 | -0.0832235 | 0.93367383 | 0.98960607 |
| HS2ST1 | 109.230799 | -0.4156042 | 0.24159556 | -1.7202475 | 0.08538746 | 0.63062869 |
| LMO4 | 25.4958291 | 0.14799207 | 0.35275948 | 0.41952684 | 0.67483114 | 0.93936166 |
| PKN2-AS1 | 21.1239921 | 1.51010553 | 0.73808161 | 2.04598721 | 0.04075763 | 0.50616193 |
| PKN2 | 282.675235 | -0.1763967 | 0.16339117 | -1.0795973 | 0.28032153 | 0.80311628 |
| GTF2B | 35.4537906 | -0.3058045 | 0.28217875 | -1.0837261 | 0.27848626 | 0.80210301 |
| KYAT3 | 54.7922802 | -0.1240689 | 0.26265318 | -0.4723677 | 0.63666434 | 0.93251095 |
| GBP3 | 28.4330631 | -0.0042019 | 0.57801008 | -0.0072695 | 0.99419979 | 0.99921857 |
| GBP2 | 53.4408446 | -0.4242167 | 0.58911845 | -0.7200873 | 4.71E-01 | 8.83E-01 |
| LRRC8B | 73.8563175 | -0.1268033 | 0.23484192 | -0.5399516 | 0.58923042 | 0.92197072 |
| LRRC8D | 10.1812693 | 0.44761631 | 0.53667831 | 0.83404956 | 0.40425306 | NA |
| ZNF326 | 63.8664449 | -0.0392706 | 0.24970191 | -0.15727 | 0.87503204 | 0.97918624 |
| ZNF644 | 229.473223 | -0.1695588 | 0.24223409 | -0.6999792 | 0.48394027 | 0.88838332 |
| HFM1 | 17.2517985 | -0.1900876 | 0.69212779 | -0.2746424 | 0.78359102 | 0.96717296 |
| TGFBR3 | 29.042097 | -0.6131063 | 0.45984944 | -1.3332762 | 0.18244118 | 0.73398872 |
| BTBD8 | 36.2103222 | 0.49083283 | 0.5505522 | 0.89152822 | 0.37264586 | 0.8491102 |
| GLMN | 42.7159566 | -0.0619775 | 0.25417695 | -0.243836 | 0.80735786 | 0.97216031 |
| RPAP2 | 136.565783 | 0.00874906 | 0.26041961 | 0.03359601 | 0.97319931 | 0.99431961 |
| EVI5 | 272.768501 | -0.3043188 | 0.17814094 | -1.7083035 | 0.08758004 | 0.63466198 |

| | | | | | | |
|---|---|---|---|---|---|---|
| RPL5 | 106.338239 | 0.76010089 | 0.27130512 | 2.801646 | 0.00508426 | 0.28057141 |
| FAM69A | 28.81521 | 0.55307608 | 0.29853735 | 1.8526194 | 0.06393693 | 0.58781448 |
| MTF2 | 138.773931 | 0.06157996 | 0.22923398 | 0.26863364 | 0.78821162 | 0.96768123 |
| TMED5 | 61.2218494 | 0.12972728 | 0.24231294 | 0.53537081 | 0.59239347 | 0.92278087 |
| CCDC18 | 23.9740662 | -0.3388963 | 0.40117788 | -0.8447533 | 0.3982486 | 0.85758974 |
| DR1 | 29.3506577 | 0.27765976 | 0.33753841 | 0.82260195 | 0.41073439 | 0.85867915 |
| FNBP1L | 407.919571 | -0.0122194 | 0.2168925 | -0.0563387 | 0.95507197 | 0.99148072 |
| BCAR3 | 228.446245 | -0.2764224 | 0.35125051 | -0.7869666 | 0.43130141 | 0.86604857 |
| DNTTIP2 | 12.3005697 | -0.2015091 | 0.38139666 | -0.5283453 | 0.59725968 | NA |
| GCLM | 44.6942187 | -0.2113568 | 0.34917163 | -0.6053091 | 0.54497366 | 0.90999223 |
| ARHGAP29 | 651.605593 | -0.0734146 | 0.45290423 | -0.1620974 | 0.87122915 | 0.97860207 |
| ABCD3 | 202.249028 | 0.2876707 | 0.19615162 | 1.46657315 | 0.14249221 | 0.69328517 |
| SLC44A3 | 111.450624 | 0.38174207 | 0.39262149 | 0.97229031 | 0.33090614 | 0.82867637 |
| CNN3 | 172.250143 | -0.2239679 | 0.23396585 | -0.9572674 | 0.33843232 | 0.83055261 |
| AC105942.1 | 12.8912892 | -0.0673879 | 0.59677612 | -0.1129199 | 0.91009404 | NA |
| ALG14 | 32.5338247 | -0.3227294 | 0.30023218 | -1.0749326 | 0.2824049 | 0.80431986 |
| PTBP2 | 215.588273 | -0.3337884 | 0.18930864 | -1.763197 | 0.07786727 | 0.61367304 |
| DPYD | 201.937127 | -0.5178793 | 0.33112508 | -1.5639989 | 0.11781783 | 0.6668162 |
| SNX7 | 49.2750457 | -0.1852455 | 0.25466913 | -0.7273966 | 0.46698306 | 0.88090126 |
| FRRS1 | 17.6310654 | 0.12276418 | 0.48068468 | 0.25539441 | 0.79841849 | 0.97019967 |
| AGL | 107.624309 | -0.1754095 | 0.18563359 | -0.9449234 | 0.34469797 | 0.83545751 |
| TRMT13 | 54.9300363 | 0.10005081 | 0.240191 | 0.41654688 | 0.67700987 | 0.94034237 |
| DBT | 66.9936091 | 0.24729613 | 0.24082637 | 1.02686484 | 0.30448411 | 0.81568231 |
| RTCA | 36.3431385 | 0.13840654 | 0.32350063 | 0.42784009 | 0.66876754 | 0.93878847 |
| CDC14A | 60.3616461 | 0.18676409 | 0.4007231 | 0.46606769 | 0.64116705 | 0.93328692 |
| VCAM1 | 15.4008927 | 1.35508747 | 0.79006584 | 1.71515764 | 0.08631633 | NA |
| EXTL2 | 28.4638648 | -0.3175511 | 0.34309162 | -0.9255578 | 0.35467583 | 0.84012856 |
| SLC30A7 | 101.824741 | -0.2162619 | 0.25138727 | -0.8602738 | 0.38963811 | 0.85605713 |
| DPH5 | 26.3122242 | 0.21812192 | 0.303396 | 0.71893473 | 0.47218114 | 0.88277854 |
| AC093157.1 | 27.9455342 | 0.1363713 | 0.34370068 | 0.39677343 | 0.69153455 | 0.94321831 |
| RNPC3 | 168.648222 | -0.0360939 | 0.22093457 | -0.163369 | 0.87022794 | 0.97854266 |
| AMY2B | 45.024985 | 0.05360824 | 0.33307168 | 0.16095107 | 0.87213194 | 0.97866716 |
| NTNG1 | 25.5124086 | -1.6661435 | 0.86454471 | -1.9271918 | 5.40E-02 | 0.56390244 |
| VAV3 | 733.237085 | 0.39528311 | 0.25467628 | 1.5521002 | 0.12063825 | 0.6668162 |
| SLC25A24 | 42.5027741 | -0.1959329 | 0.27783591 | -0.7052108 | 0.48067909 | 0.88675516 |
| PRPF38B | 70.0070163 | -0.1555554 | 0.26687478 | -0.582878 | 0.55997546 | 0.91516276 |
| STXBP3 | 87.6661776 | -0.1957778 | 0.25792171 | -0.7590589 | 0.44781734 | 0.87291341 |
| AL449266.1 | 17.8271088 | -0.0077254 | 0.38552093 | -0.020039 | 0.98401229 | 0.99685229 |
| GPSM2 | 20.1280717 | 0.40380445 | 0.4233637 | 0.95380038 | 0.3401847 | 0.83155245 |
| WDR47 | 22.420537 | 0.3468559 | 0.4446859 | 0.78000201 | 0.43538969 | 0.86729771 |
| TAF13 | 15.4944186 | -0.1152658 | 0.4018674 | -0.2868254 | 0.77424599 | NA |
| TMEM167B | 21.1683683 | 0.18564483 | 0.37446406 | 0.4957614 | 0.62006276 | 0.92798856 |
| SARS | 49.9177795 | -0.1420905 | 0.29518921 | -0.4813541 | 0.63026484 | 0.9320416 |
| CELSR2 | 22.4386708 | 0.19650321 | 0.47639788 | 0.41247709 | 0.67998977 | 0.94195357 |
| SORT1 | 104.906124 | 0.3198067 | 0.22711282 | 1.40814023 | 0.15908955 | 0.71094363 |
| PSMA5 | 53.2097864 | -0.6399412 | 0.29233188 | -2.1890915 | 0.02859019 | 0.46148943 |
| SYPL2 | 15.863572 | -0.2971507 | 0.48696585 | -0.6102085 | 0.54172369 | NA |
| CYB561D1 | 9.73521464 | -0.2398564 | 0.49752496 | -0.4820993 | 0.62973539 | NA |
| GNAI3 | 83.9867179 | -0.3430961 | 0.24256448 | -1.4144533 | 0.15722884 | 0.7098097 |
| AMPD2 | 15.928367 | -0.535581 | 0.53224258 | -1.0062723 | 0.31428459 | NA |
| GSTM4 | 25.701981 | 0.43973223 | 0.30833882 | 1.42613318 | 0.1538299 | 0.70572801 |
| GSTM3 | 97.1679422 | -0.435375 | 0.53396182 | -0.8153673 | 0.4148621 | 0.85867915 |
| AHCYL1 | 184.27587 | 0.15075227 | 0.24307749 | 0.62018193 | 0.53513802 | 0.90763121 |
| STRIP1 | 26.9393625 | -0.3728971 | 0.39194278 | -0.951407 | 0.3413978 | 0.8325058 |

| | | | | | | |
|---|---|---|---|---|---|---|
| KCNC4 | 106.028717 | 0.11375155 | 0.43955531 | 0.25878779 | 0.79579898 | 0.96974891 |
| RBM15 | 28.6211954 | 0.75043185 | 0.31362768 | 2.39274751 | 0.01672274 | 0.39748373 |
| SLC16A4 | 98.7113561 | -0.4273805 | 0.41834542 | -1.0215971 | 0.30697162 | 0.81568231 |
| AL355488.1 | 12.5500126 | 0.58688458 | 0.48513604 | 1.20973198 | 0.22638175 | NA |
| LAMTOR5 | 89.3097184 | 0.06332418 | 0.35689153 | 0.17743255 | 0.85916864 | 0.97691173 |
| LRIF1 | 20.687793 | 0.21905949 | 0.3820342 | 0.57340282 | 0.56637198 | 0.91698355 |
| DRAM2 | 49.2300071 | 0.16385773 | 0.25458374 | 0.64362999 | 0.5198154 | 0.90368693 |
| CEPT1 | 58.9624409 | 0.04460282 | 0.26277488 | 0.16973776 | 0.86521638 | 0.97788853 |
| DENND2D | 33.6271539 | 0.12344447 | 0.2856752 | 0.43211477 | 0.665658 | 0.93860542 |
| WDR77 | 13.1379107 | 0.72998948 | 0.49021104 | 1.4891331 | 0.13645232 | NA |
| ATP5PB | 87.1696378 | 0.38390499 | 0.37572019 | 1.0217843 | 0.306883 | 0.81568231 |
| RAP1A | 156.313695 | -0.0132881 | 0.28653323 | -0.0463755 | 0.96301096 | 0.99256424 |
| KCND3 | 15.3198264 | 1.34439555 | 0.45825908 | 2.93370192 | 3.35E-03 | NA |
| CTTNBP2NL | 70.0159847 | -0.1325794 | 0.24763119 | -0.5353907 | 0.59237975 | 0.92278087 |
| ST7L | 68.9693551 | -0.4004978 | 0.24672671 | -1.6232448 | 0.10453708 | 0.65946648 |
| CAPZA1 | 109.190747 | -0.1961723 | 0.19784795 | -0.9915307 | 0.32142651 | 0.82490712 |
| MOV10 | 31.1394898 | -0.5008963 | 0.43297841 | -1.156862 | 0.24732876 | 0.78583441 |
| RHOC | 54.434503 | 0.14274769 | 0.41917281 | 0.34054616 | 0.73344527 | 0.95329896 |
| LRIG2 | 63.113711 | -0.2264643 | 0.27874727 | -0.8124358 | 0.41654162 | 0.85867915 |
| MAGI3 | 414.769584 | 0.01515388 | 0.28887877 | 0.05245756 | 0.95816411 | 0.99183169 |
| PHTF1 | 56.4366663 | -0.071406 | 0.3055344 | -0.2337085 | 0.81521129 | 0.97284589 |
| RSBN1 | 59.322297 | 0.24534439 | 0.27205145 | 0.90183083 | 0.36714674 | 0.84777627 |
| HIPK1 | 61.086583 | 0.18826502 | 0.29875423 | 0.63016687 | 0.52858541 | 0.90487731 |
| TRIM33 | 198.100796 | -0.4214943 | 0.25969996 | -1.6230048 | 0.10458837 | 0.65946648 |
| BCAS2 | 20.9085439 | 0.35700882 | 0.39649845 | 0.90040409 | 0.36790525 | 0.84777627 |
| DENND2C | 13.0286164 | 0.24220265 | 0.49603648 | 0.48827588 | 0.62535444 | NA |
| NRAS | 14.487058 | -0.3263327 | 0.43589779 | -0.7486449 | 0.45407126 | NA |
| CSDE1 | 164.430633 | 0.17757795 | 0.21714693 | 0.81777789 | 0.41348403 | 0.85867915 |
| SIKE1 | 29.1344214 | -0.0052232 | 0.30455639 | -0.0171503 | 0.98631672 | 0.99771543 |
| SLC22A15 | 123.698842 | -0.1748278 | 0.26682716 | -0.6552099 | 0.51233261 | 0.9014941 |
| ATP1A1 | 1384.36523 | 0.28145938 | 0.31517966 | 0.89301253 | 0.37185046 | 0.8491102 |
| LINC01762 | 371.333737 | 1.25211672 | 0.36296587 | 3.44968171 | 0.00056125 | 0.10017724 |
| CD58 | 64.0112697 | 0.03703901 | 0.29084644 | 0.12734901 | 8.99E-01 | 9.85E-01 |
| IGSF3 | 97.6136255 | 0.34568365 | 0.33736046 | 1.0246715 | 0.30551822 | 0.81568231 |
| PTGFRN | 23.9450447 | 0.07605987 | 0.35064524 | 0.21691403 | 0.82827534 | 0.97567084 |
| TTF2 | 33.0934531 | 0.18227816 | 0.33045143 | 0.55160349 | 0.58122005 | 0.92034873 |
| VTCN1 | 72.3188336 | 3.27502227 | 0.82053837 | 3.99130912 | 6.57E-05 | 0.03323079 |
| MAN1A2 | 189.049804 | -0.1494718 | 0.21488353 | -0.6955944 | 0.48668285 | 0.88838332 |
| GDAP2 | 68.6710669 | 0.24883729 | 0.27786998 | 0.895517 | 0.37051078 | 0.84852785 |
| WDR3 | 16.6722691 | 0.38492186 | 0.36617744 | 1.05118945 | 0.29317159 | 0.80935782 |
| SPAG17 | 83.4561697 | -0.1970459 | 0.37099198 | -0.5311324 | 0.59532703 | 0.9229638 |
| WARS2 | 48.8738217 | -0.351361 | 0.27401812 | -1.2822548 | 0.19975329 | 0.75605362 |
| AL359915.2 | 113.344677 | -0.2619726 | 0.3170313 | -0.8263303 | 0.40861674 | 0.85867915 |
| ZNF697 | 16.8841839 | -0.0602476 | 0.48326925 | -0.1246667 | 0.90078738 | 0.98531749 |
| PHGDH | 24.2848416 | -0.3907488 | 0.51179465 | -0.7634874 | 0.44517277 | 0.87129815 |
| HMGCS2 | 98.3040796 | 1.48426538 | 0.82838153 | 1.79176542 | 0.07317055 | 0.60952146 |
| NOTCH2 | 59.5295351 | -0.0100544 | 0.27567671 | -0.0364719 | 0.97090612 | 0.99403127 |
| SEC22B | 70.8481426 | -0.5401676 | 0.19403974 | -2.7837986 | 0.00537264 | 0.28183167 |
| NBPF26 | 14.1489209 | 0.07258507 | 0.52989595 | 0.13697985 | 0.89104672 | NA |
| LINC00623 | 102.665923 | 0.14650694 | 0.49904267 | 0.29357597 | 0.76908194 | 0.96289288 |
| SRGAP2C | 29.5237106 | -0.1533643 | 0.38449953 | -0.3988672 | 0.68999105 | 0.94252529 |
| AC239800.3 | 13.3621379 | -0.3724569 | 0.45279293 | -0.8225768 | 0.41074871 | NA |
| NBPF15 | 24.4678232 | 0.59924535 | 0.35194242 | 1.70268006 | 0.08862796 | 0.63466198 |
| FP700111.1 | 23.0119297 | -0.4668714 | 0.33264759 | -1.4035016 | 0.16046731 | 0.71101785 |

| | | | | | | |
|---|---|---|---|---|---|---|
| SRGAP2B | 24.1474105 | 0.16101485 | 0.38332181 | 0.42005138 | 0.67444792 | 0.9393359 |
| GPR89A | 17.9068463 | -0.1369546 | 0.39425948 | -0.3473718 | 0.72831201 | 0.95272318 |
| PDZK1 | 48.4017386 | 1.54486939 | 0.48800926 | 3.16565592 | 0.00154734 | 0.16768345 |
| RNF115 | 188.677096 | -0.1976164 | 0.20997213 | -0.9411552 | 0.34662533 | 0.83687845 |
| POLR3C | 20.7993817 | -0.0677278 | 0.39514774 | -0.1713988 | 0.8639102 | 0.97788853 |
| PEX11B | 17.2359745 | -0.2894588 | 0.39130384 | -0.739729 | 0.45946442 | 0.8759172 |
| RBM8A | 33.3174407 | -0.256303 | 0.31636495 | -0.8101499 | 0.41785403 | 0.85950165 |
| TXNIP | 801.450077 | 0.64359026 | 0.57232207 | 1.12452461 | 0.26079054 | 0.79574359 |
| NBPF12 | 40.0132301 | 0.21285876 | 0.29914711 | 0.71155212 | 0.47674217 | 0.88495683 |
| PRKAB2 | 29.2116298 | 0.06497236 | 0.38645181 | 0.1681254 | 0.86648463 | 0.97788853 |
| AC242426.2 | 52.3864204 | 0.1084696 | 0.33440465 | 0.32436631 | 0.74566072 | 0.95563105 |
| FMO5 | 34.871861 | 0.64610658 | 0.42864591 | 1.50732007 | 0.13172864 | 0.67632587 |
| CHD1L | 52.6772747 | 0.35412239 | 0.24984282 | 1.41738073 | 0.15637164 | 0.70818462 |
| BCL9 | 74.4243533 | 0.21945674 | 0.2265265 | 0.96879061 | 0.33264967 | 0.82928059 |
| ACP6 | 60.6350134 | 0.12525378 | 0.22611844 | 0.55392994 | 0.57962679 | 0.92027213 |
| GPR89B | 24.5110021 | 0.01146644 | 0.33124721 | 0.03461597 | 0.97238597 | 0.99431961 |
| NBPF11 | 39.6375216 | 0.10819284 | 0.30956039 | 0.34950481 | 0.72671036 | 0.95209193 |
| AC239809.3 | 22.8474568 | -0.5560752 | 0.38808696 | -1.4328624 | 0.15189715 | 0.70572801 |
| NBPF14 | 16.8911841 | -0.0063151 | 0.38936713 | -0.016219 | 0.98705969 | 0.99803728 |
| PDE4DIP | 156.066658 | 0.24600617 | 0.2742701 | 0.89694857 | 0.36974636 | 0.84852785 |
| NBPF9 | 17.3212184 | -0.6571544 | 0.3526084 | -1.8636946 | 0.0623646 | 0.58393911 |
| AC245297.3 | 212.385283 | -0.1781873 | 0.33598059 | -0.5303501 | 0.59586922 | 0.9229638 |
| NBPF19 | 27.7649264 | 0.18049244 | 0.69775848 | 0.25867466 | 0.79588627 | 0.96974891 |
| SF3B4 | 13.9973623 | 0.0994738 | 0.38528201 | 0.25818439 | 0.79626461 | NA |
| OTUD7B | 103.957312 | -0.5562853 | 0.2416708 | -2.3018311 | 0.0213447 | 0.4212027 |
| VPS45 | 101.35166 | -0.0384937 | 0.21425379 | -0.179664 | 0.85741639 | 0.97685375 |
| ANP32E | 43.3403272 | 0.33203555 | 0.37221283 | 0.89205833 | 0.37236167 | 0.8491102 |
| APH1A | 48.0532715 | 0.32175906 | 0.40809577 | 0.78844007 | 0.43043934 | 0.86534658 |
| C1orf54 | 32.6263572 | 0.48143753 | 0.47722626 | 1.00882448 | 0.31305882 | 0.81913609 |
| MRPS21 | 69.6923742 | -0.149307 | 0.23825476 | -0.6266696 | 0.53087583 | 0.90525279 |
| PRPF3 | 55.9930921 | -0.5423985 | 0.22479764 | -2.4128299 | 0.0158292 | 0.39477603 |
| RPRD2 | 213.453905 | -0.1841456 | 0.19689163 | -0.9352636 | 0.34965249 | 0.83687845 |
| TARS2 | 17.6018363 | -0.3976179 | 0.38668432 | -1.0282751 | 0.30382043 | 0.81568231 |
| MCL1 | 77.0721141 | -0.3384775 | 0.36292218 | -0.932645 | 0.35100331 | 0.83797093 |
| ENSA | 174.171753 | -0.0291052 | 0.32148921 | -0.0905324 | 0.92786418 | 0.98960607 |
| GOLPH3L | 64.4438631 | -0.1587603 | 0.25038219 | -0.6340717 | 0.52603405 | 0.90487731 |
| CTSS | 18.2002767 | -0.1802382 | 0.58867642 | -0.3061753 | 0.75947117 | 0.95903525 |
| ARNT | 80.73055 | -0.122561 | 0.28906001 | -0.4239984 | 0.67156696 | 0.93878847 |
| SETDB1 | 56.0062837 | -0.4673323 | 0.28901816 | -1.6169651 | 0.10588581 | 0.65946648 |
| CERS2 | 37.7033112 | 0.19740758 | 0.36633309 | 0.53887457 | 0.58997341 | 0.92229571 |
| ANXA9 | 10.736788 | 1.02413656 | 0.65521538 | 1.5630533 | 0.11804006 | NA |
| MINDY1 | 20.8769938 | 0.5433423 | 0.41167935 | 1.31981917 | 0.1868954 | 0.74034327 |
| PRUNE1 | 26.8365303 | -0.0191492 | 0.31905301 | -0.0600188 | 0.95214064 | 0.99141043 |
| C1orf56 | 16.0579933 | -0.4243455 | 0.66689495 | -0.6363003 | 0.52458068 | NA |
| CDC42SE1 | 44.2173235 | -0.1522094 | 0.33057207 | -0.4604425 | 0.64519865 | 0.93328692 |
| GABPB2 | 69.8791503 | -0.0720631 | 0.27438373 | -0.2626363 | 0.79283092 | 0.96947477 |
| SCNM1 | 13.4779249 | 0.21806986 | 0.41410742 | 0.52660215 | 0.5984699 | NA |
| VPS72 | 17.1416917 | -0.4017565 | 0.40215398 | -0.9990116 | 0.31778908 | 0.82351215 |
| PIP5K1A | 220.636187 | -0.1330433 | 0.21767697 | -0.6111961 | 0.54106975 | 0.90783815 |
| PSMD4 | 44.0023441 | 0.14182027 | 0.24318097 | 0.5831882 | 0.55976661 | 0.91515005 |
| PI4KB | 53.1721978 | 0.1591036 | 0.27818593 | 0.57193259 | 0.56736765 | 0.91698355 |
| SELENBP1 | 15.8576448 | 0.52676834 | 0.53638677 | 0.98206811 | 0.32606629 | NA |
| PSMB4 | 28.9021899 | -0.0715988 | 0.47383236 | -0.1511057 | 0.87989234 | 0.98093344 |
| POGZ | 318.778039 | 0.1326577 | 0.22159434 | 0.59865113 | 0.54940555 | 0.91221128 |

| | | | | | | |
|---|---|---|---|---|---|---|
| CGN | 40.1472103 | -0.001531 | 0.25962402 | -0.0058969 | 0.995295 | 0.99927028 |
| TUFT1 | 53.3294325 | -0.2677893 | 0.37223456 | -0.7194101 | 0.47188828 | 0.88267439 |
| SNX27 | 87.6905461 | -0.2330474 | 0.30174404 | -0.7723349 | 0.43991612 | 0.86895963 |
| MRPL9 | 14.0726695 | -0.4591612 | 0.41297625 | -1.1118345 | 0.26620931 | NA |
| RORC | 24.935893 | -0.4602031 | 0.56627795 | -0.8126805 | 0.41640126 | 0.85867915 |
| THEM4 | 92.7455751 | 0.99103331 | 0.45117567 | 2.1965575 | 0.02805206 | 0.45940027 |
| S100A10 | 206.331582 | -0.4156698 | 0.38092136 | -1.091222 | 0.2751752 | 0.7994854 |
| S100A11 | 586.161926 | -0.1045511 | 0.41401106 | -0.2525321 | 8.01E-01 | 9.70E-01 |
| FLG-AS1 | 115.190375 | 0.26065259 | 0.34249122 | 0.76104897 | 4.47E-01 | 0.87231627 |
| S100A6 | 795.213806 | -0.5599015 | 0.59561666 | -0.9400367 | 0.34719872 | 0.83687845 |
| S100A4 | 22.692235 | 0.15191025 | 0.81888107 | 0.18550953 | 0.8528294 | 0.97612951 |
| S100A2 | 245.095159 | -0.7900391 | 1.05478385 | -0.7490056 | 0.45385381 | 0.87378486 |
| S100A16 | 55.8574884 | -0.5288446 | 0.45233725 | -1.1691378 | 0.2423481 | 0.78464787 |
| S100A14 | 31.4601853 | -0.8564872 | 0.53360675 | -1.6050907 | 1.08E-01 | 0.65986731 |
| S100A13 | 59.4506743 | 0.43984546 | 0.38849165 | 1.13218769 | 0.25755553 | 0.79532468 |
| CHTOP | 40.3403621 | -0.0646979 | 0.27193631 | -0.2379156 | 0.81194657 | 0.97257655 |
| ILF2 | 31.2369422 | 0.46764436 | 0.35903243 | 1.30251287 | 0.19274112 | 0.75061775 |
| INTS3 | 62.3312813 | -0.0387245 | 0.26214007 | -0.1477245 | 0.88256023 | 0.98137082 |
| GATAD2B | 240.89454 | -0.2813292 | 0.26624539 | -1.0566537 | 0.29066966 | 0.80595978 |
| DENND4B | 27.8676609 | -1.0709351 | 0.28626828 | -3.741019 | 0.00018328 | 0.05491436 |
| CRTC2 | 16.7752708 | -0.4422419 | 0.40375484 | -1.0953227 | 0.27337528 | 0.79726027 |
| SLC39A1 | 40.352123 | -0.0182962 | 0.40423245 | -0.0452615 | 0.96389889 | 0.99257597 |
| JTB | 38.5621289 | -0.0499722 | 0.375325 | -0.1331437 | 0.89407971 | 0.98408497 |
| RAB13 | 53.7608351 | 0.55060024 | 0.27793496 | 1.98103988 | 0.0475868 | 0.53545446 |
| RPS27 | 45.763811 | 0.26644011 | 0.34731107 | 0.76715121 | 0.44299162 | 0.86991418 |
| TPM3 | 73.9449894 | -0.2112081 | 0.35110754 | -0.6015481 | 0.54747497 | 0.91139843 |
| C1orf43 | 60.3434321 | -0.1535618 | 0.26721109 | -0.5746835 | 0.56550539 | 0.91698355 |
| UBAP2L | 67.3132928 | -0.0601982 | 0.22328346 | -0.2696043 | 0.78746466 | 0.96767512 |
| HAX1 | 18.0390256 | 0.2979738 | 0.43841236 | 0.67966558 | 0.49671623 | 0.89306891 |
| IL6R | 22.7643072 | -0.6975004 | 0.39994975 | -1.7439701 | 0.08116431 | 0.61925304 |
| UBE2Q1 | 22.2019661 | 0.10670922 | 0.38635342 | 0.27619589 | 0.78239762 | 0.96652357 |
| ADAR | 115.716105 | -0.6286991 | 0.32661634 | -1.9248855 | 0.05424368 | 0.5645224 |
| KCNN3 | 19.9507267 | 1.04748683 | 0.63372496 | 1.65290448 | 0.09835031 | 0.65695163 |
| PMVK | 23.1981643 | 0.79845296 | 0.51046115 | 1.56417967 | 0.11777538 | 0.6668162 |
| PBXIP1 | 24.9847162 | 0.60162057 | 0.50089482 | 1.20109161 | 0.22971567 | 0.77219413 |
| PYGO2 | 10.5386471 | -0.061432 | 0.5131201 | -0.1197224 | 0.90470307 | NA |
| FLAD1 | 11.1952427 | -0.1187485 | 0.45456986 | -0.2612327 | 0.79391304 | NA |
| ZBTB7B | 15.0604574 | 0.01536291 | 0.4094623 | 0.03751972 | 0.97007062 | NA |
| EFNA1 | 53.4906059 | -0.5353536 | 0.33343923 | -1.6055506 | 1.08E-01 | 0.65986731 |
| SLC50A1 | 17.0412461 | -0.159918 | 0.43258051 | -0.3696837 | 0.71161819 | 0.94676706 |
| DPM3 | 31.6766847 | -0.1466296 | 0.53570565 | -0.2737129 | 0.78430523 | 0.96732016 |
| MUC1 | 18.7177182 | 0.25677378 | 0.41784042 | 0.61452595 | 0.53886783 | 0.90763121 |
| THBS3 | 13.5530985 | -0.476855 | 0.53751122 | -0.8871536 | 0.37499619 | NA |
| MTX1 | 11.4977161 | 0.22314351 | 0.48814029 | 0.45712988 | 0.64757769 | NA |
| GBA | 17.5132587 | 0.06205069 | 0.46094636 | 0.13461585 | 0.89291561 | 0.98377746 |
| FAM189B | 11.251277 | 0.09152485 | 0.46857167 | 0.19532733 | 0.8451367 | NA |
| SCAMP3 | 27.1325427 | 0.12330836 | 0.42024469 | 0.29342037 | 0.76920085 | 0.96289288 |
| CLK2 | 21.1814679 | -0.1273005 | 0.35416369 | -0.3594399 | 0.71926607 | 0.94959812 |
| FDPS | 70.9136059 | 0.01157873 | 0.38768016 | 0.02986671 | 0.97617336 | 0.99440179 |
| RUSC1 | 15.99733 | -0.0522106 | 0.42076176 | -0.1240859 | 0.90124724 | NA |
| ASH1L | 473.483612 | -0.0776295 | 0.18159119 | -0.4274959 | 0.66901819 | 0.93878847 |
| YY1AP1 | 78.748128 | -0.0155202 | 0.2429856 | -0.0638731 | 0.94907126 | 0.99105535 |
| DAP3 | 96.0441284 | -0.2164544 | 0.21316029 | -1.0154535 | 0.30988971 | 0.81568231 |
| GON4L | 106.851603 | -0.4533503 | 0.23106147 | -1.9620334 | 0.0497586 | 0.54357315 |

| | | | | | | |
|---|---|---|---|---|---|---|
| RIT1 | 33.4893328 | -0.6527622 | 0.38822741 | -1.6813913 | 0.09268693 | 0.64572229 |
| KHDC4 | 93.546956 | -0.5050287 | 0.22251218 | -2.2696678 | 0.02322775 | 0.43085291 |
| ARHGEF2 | 29.3178481 | -0.1838386 | 0.27277157 | -0.6739655 | 0.50033328 | 0.89469323 |
| SSR2 | 50.0183648 | 0.05249745 | 0.28476202 | 0.1843555 | 0.85373458 | 0.97612951 |
| UBQLN4 | 19.7808968 | 0.12316223 | 0.33310196 | 0.36974334 | 0.71157373 | 0.94676706 |
| LAMTOR2 | 18.1988155 | 0.50823324 | 0.56146758 | 0.90518715 | 0.36536625 | 0.84720508 |
| RAB25 | 35.3197204 | -0.0246335 | 0.48523036 | -0.0507666 | 0.95951147 | 0.99245244 |
| LMNA | 92.0430671 | -0.0154503 | 0.29043172 | -0.0531978 | 0.95757432 | 0.99170927 |
| SEMA4A | 23.4377374 | 0.21401502 | 0.34139565 | 0.6268827 | 0.53073613 | 0.9052447 |
| SLC25A44 | 20.5978871 | -0.2376849 | 0.32174611 | -0.7387342 | 0.4600684 | 0.87607759 |
| SMG5 | 53.0071777 | -0.397166 | 0.24224221 | -1.6395408 | 0.10110068 | 0.65863153 |
| GLMP | 12.6108451 | 0.22011663 | 0.58560998 | 0.37587582 | 0.70700921 | NA |
| CCT3 | 73.568022 | 0.46470905 | 0.27411396 | 1.69531333 | 0.090016 | 0.63817417 |
| RHBG | 103.615037 | 0.92108725 | 0.40205686 | 2.29093775 | 2.20E-02 | 4.21E-01 |
| MEF2D | 38.2152876 | -0.1716139 | 0.32494225 | -0.5281365 | 0.59740457 | 0.9229638 |
| NAXE | 20.4510508 | 0.07879117 | 0.52089239 | 0.15126189 | 0.87976913 | 0.98093344 |
| GPATCH4 | 11.0263215 | 0.37023922 | 0.47866112 | 0.77348923 | 0.4392329 | NA |
| ISG20L2 | 14.189264 | 0.11144245 | 0.43901391 | 0.2538472 | 0.7996136 | NA |
| RRNAD1 | 20.5267951 | 0.23625389 | 0.35839593 | 0.65919802 | 0.50976862 | 0.89963034 |
| MRPL24 | 12.5666332 | 0.38774509 | 0.54125347 | 0.71638356 | 0.47375454 | NA |
| HDGF | 58.7978148 | 0.13908008 | 0.31123667 | 0.44686276 | 0.65497415 | 0.93548446 |
| PRCC | 45.9274048 | -0.1243888 | 0.26484351 | -0.4696692 | 0.63859137 | 0.9331046 |
| ARHGEF11 | 119.275756 | 0.10190521 | 0.19552663 | 0.52118328 | 0.6022391 | 0.92407937 |
| ETV3 | 10.9467534 | 0.82138664 | 0.47078259 | 1.74472602 | 0.08103257 | NA |
| KIRREL1 | 30.8324795 | 0.3092824 | 0.52669403 | 0.58721456 | 0.55705961 | 0.9141017 |
| IFI16 | 17.0796431 | -1.775685 | 1.07920434 | -1.645365 | 0.09989456 | 0.65863153 |
| DUSP23 | 41.0577533 | -0.1169991 | 0.52786286 | -0.2216467 | 0.82458891 | 0.97518529 |
| TAGLN2 | 90.0170378 | -0.2622856 | 0.47780139 | -0.5489428 | 5.83E-01 | 0.92034873 |
| KCNJ10 | 14.7410225 | -0.0332031 | 0.63015067 | -0.0526907 | 9.58E-01 | NA |
| IGSF8 | 14.8891614 | 0.13509945 | 0.60137883 | 0.2246495 | 0.82225195 | NA |
| PEA15 | 15.2573021 | 0.05449535 | 0.48360252 | 0.11268625 | 0.91027931 | NA |
| DCAF8 | 33.1026341 | 0.20552602 | 0.35287843 | 0.58242728 | 0.56027891 | 0.91533003 |
| COPA | 133.534661 | -0.4460808 | 0.20524083 | -2.1734508 | 0.02974641 | 0.46381645 |
| NCSTN | 30.6069729 | 0.78003755 | 0.35168369 | 2.21800891 | 0.02655422 | 0.44763538 |
| VANGL2 | 9.67963375 | 1.03878865 | 0.6106274 | 1.7011825 | 0.08890872 | NA |
| F11R | 73.9075829 | -0.2046076 | 0.41928068 | -0.4879968 | 0.62555214 | 0.92995158 |
| AL591806.3 | 25.0018295 | 0.21898551 | 0.43914311 | 0.49866548 | 0.61801506 | 0.92749951 |
| NECTIN4 | 29.9535394 | -0.5680274 | 0.40457213 | -1.40402 | 0.16031288 | 0.71101785 |
| PFDN2 | 43.961515 | 0.06949198 | 0.27300019 | 0.25454921 | 0.79907129 | 0.97019967 |
| NIT1 | 32.946332 | 0.35779115 | 0.33135314 | 1.07978802 | 0.28023659 | 0.80311628 |
| DEDD | 15.8624534 | 0.15021749 | 0.4199084 | 0.35773872 | 0.72053885 | NA |
| UFC1 | 29.9844458 | 0.69016012 | 0.41774235 | 1.65211912 | 0.09851027 | 0.65695163 |
| B4GALT3 | 12.7977181 | 0.31335568 | 0.52500605 | 0.59686109 | 0.55060013 | NA |
| NDUFS2 | 50.6076919 | -0.1126392 | 0.32110514 | -0.3507859 | 0.72574894 | 0.95164932 |
| SDHC | 98.881769 | 0.44981831 | 0.19648683 | 2.28930518 | 0.02206163 | 0.42190544 |
| AL592295.4 | 16.5769713 | 0.69925348 | 0.51006281 | 1.37091641 | 0.17040102 | 0.72463203 |
| DUSP12 | 12.3290113 | -0.4953248 | 0.41970912 | -1.180162 | 0.23793577 | NA |
| ATF6 | 218.969874 | -0.1545114 | 0.3225295 | -0.4790612 | 0.63189508 | 0.93251095 |
| NOS1AP | 380.842686 | -0.9953041 | 0.34284492 | -2.9030739 | 0.00369519 | 0.24199539 |
| UHMK1 | 89.378358 | -0.2710016 | 0.28000358 | -0.9678503 | 0.33311912 | 0.82942652 |
| UAP1 | 21.2394947 | -0.5780936 | 0.42777187 | -1.3514064 | 0.17656528 | 0.72841249 |
| HSD17B7 | 29.5409094 | -0.1391778 | 0.43761498 | -0.3180371 | 0.75045681 | 0.95718206 |
| PBX1 | 192.76913 | 0.44604652 | 0.3498825 | 1.2748466 | 0.20236354 | 0.76065252 |
| MGST3 | 214.917945 | -0.0066743 | 0.19241054 | -0.0346879 | 0.97232865 | 0.99431961 |

| | | | | | | |
|---|---|---|---|---|---|---|
| ALDH9A1 | 38.6677809 | 0.71860346 | 0.34273188 | 2.0966928 | 0.03602078 | 0.493821 |
| TMCO1 | 74.7359473 | 0.3809702 | 0.28581502 | 1.33292574 | 0.18255617 | 0.73398872 |
| UCK2 | 31.8467479 | 0.27589659 | 0.30595969 | 0.90174163 | 0.36719413 | 0.84777627 |
| POGK | 38.686797 | 0.17584724 | 0.32569985 | 0.53990579 | 0.589262 | 0.92197072 |
| TADA1 | 20.0521339 | 0.40366547 | 0.38661489 | 1.04410224 | 0.29643809 | 0.81120086 |
| POU2F1 | 208.859892 | -0.1100145 | 0.19700519 | -0.5584346 | 0.57654765 | 0.91854298 |
| CREG1 | 33.8000805 | 0.46949716 | 0.40535273 | 1.15824349 | 0.24676468 | 0.78583441 |
| MPZL1 | 73.0754475 | -0.4798728 | 0.27545703 | -1.7420966 | 0.08149154 | 0.61925304 |
| MPC2 | 58.9865067 | 0.35793214 | 0.32108103 | 1.11477201 | 0.26494813 | 0.79666376 |
| DCAF6 | 258.830307 | -0.184433 | 0.22958313 | -0.8033388 | 0.42177896 | 0.86190547 |
| TIPRL | 31.7824228 | -0.425757 | 0.28917529 | -1.4723146 | 0.14093594 | 0.69227547 |
| SFT2D2 | 44.2083393 | 0.03546494 | 0.24638213 | 0.14394281 | 0.88554563 | 0.98210549 |
| LINC00970 | 154.163599 | 1.50234497 | 0.47847635 | 3.13985207 | 1.69E-03 | 0.17205462 |
| ATP1B1 | 2345.9703 | 0.52221095 | 0.32804867 | 1.59187032 | 1.11E-01 | 6.64E-01 |
| NME7 | 222.492952 | 0.15812646 | 0.1782805 | 0.88695319 | 0.3751041 | 0.85172677 |
| BLZF1 | 46.8876306 | -0.3664766 | 0.25527222 | -1.4356306 | 0.15110747 | 0.70572801 |
| CCDC181 | 15.030632 | -0.0621095 | 0.4693145 | -0.1323408 | 0.89471471 | NA |
| SLC19A2 | 86.3363159 | -0.2167268 | 0.42696189 | -0.5076022 | 0.61173235 | 0.92630153 |
| C1orf112 | 11.555576 | -0.3324107 | 0.4223498 | -0.7870507 | 0.43125221 | NA |
| SCYL3 | 40.1039295 | 0.46471958 | 0.28527244 | 1.62903779 | 0.10330502 | 0.65946648 |
| KIFAP3 | 87.6010749 | 0.08319599 | 0.24937531 | 0.3336176 | 0.73866813 | 0.9542764 |
| GORAB | 14.9453496 | -0.3866622 | 0.36688029 | -1.0539191 | 0.29191994 | NA |
| FMO4 | 58.7479107 | 0.68795765 | 0.3836346 | 1.79326277 | 0.07293092 | 0.60952146 |
| PRRC2C | 277.895099 | -0.2633473 | 0.20261059 | -1.2997705 | 0.19367964 | 0.75067401 |
| VAMP4 | 26.2652111 | 7.77E-06 | 0.36049479 | 2.16E-05 | 0.9999828 | 0.9999828 |
| METTL13 | 10.8323614 | 0.09515035 | 0.49491803 | 0.19225477 | 0.84754265 | NA |
| DNM3 | 64.8108377 | 0.39901479 | 0.32625688 | 1.22300804 | 0.22132666 | 0.76680432 |
| PIGC | 31.2600048 | 0.36703067 | 0.40343959 | 0.90975373 | 0.3629524 | 0.84543391 |
| SUCO | 85.2507613 | -0.5458463 | 0.2114303 | -2.5816846 | 0.00983194 | 0.32904468 |
| AL645568.1 | 42.6973998 | 0.2540349 | 0.40515135 | 0.62701235 | 0.53065114 | 0.9052447 |
| PRDX6 | 107.22173 | 0.82521467 | 0.3738126 | 2.20756246 | 0.02727479 | 0.45224481 |
| KLHL20 | 69.2356374 | 0.04832751 | 0.23178061 | 0.20850541 | 0.83483436 | 0.97600352 |
| DARS2 | 15.4299585 | 0.3499787 | 0.43884208 | 0.79750488 | 0.42515787 | NA |
| ZBTB37 | 37.6733668 | -0.0449763 | 0.30569007 | -0.1471304 | 0.88302912 | 0.98140267 |
| RC3H1 | 143.254236 | 0.0973274 | 0.23127236 | 0.42083456 | 0.6738759 | 0.93901638 |
| RABGAP1L | 748.587699 | -0.2769005 | 0.23812741 | -1.162825 | 0.24490049 | 0.78583441 |
| CACYBP | 17.4878335 | 0.4254889 | 0.39146237 | 1.08692158 | 0.27707147 | 0.80159799 |
| MRPS14 | 20.3235887 | 0.48612763 | 0.35510265 | 1.36897774 | 0.17100623 | 0.72463203 |
| KIAA0040 | 24.6253828 | 0.15565106 | 0.35215197 | 0.44199968 | 0.65848944 | 0.93630574 |
| COP1 | 315.223436 | -0.2773073 | 0.17356672 | -1.5976988 | 0.11011003 | 0.66283855 |
| PAPPA2 | 46.8282012 | 0.95273646 | 0.51437216 | 1.85223178 | 0.06399255 | 0.58781448 |
| RASAL2 | 322.871043 | -0.971144 | 0.47385426 | -2.0494572 | 0.04041743 | 0.50399977 |
| RALGPS2 | 270.866343 | 0.07970517 | 0.250449 | 0.3182491 | 0.75029599 | 0.95716475 |
| ANGPTL1 | 18.9712913 | 0.45239888 | 0.53969423 | 0.83825034 | 0.40189012 | 0.85778581 |
| FAM20B | 47.01696 | -0.378393 | 0.26920503 | -1.405594 | 0.15984473 | 0.71101785 |
| TOR3A | 14.0614628 | -0.3633502 | 0.5067215 | -0.7170609 | 0.47333654 | NA |
| ABL2 | 145.263786 | -0.095656 | 0.26739647 | -0.3577309 | 0.72054474 | 0.95048937 |
| SOAT1 | 66.9509384 | 0.44510173 | 0.28327004 | 1.57129828 | 1.16E-01 | 0.6668162 |
| TDRD5 | 14.8787311 | 0.48761309 | 0.60491892 | 0.80608007 | 0.42019668 | NA |
| TOR1AIP2 | 99.9220131 | -0.1092916 | 0.23996485 | -0.4554484 | 0.64878665 | 0.934183 |
| TOR1AIP1 | 46.1762001 | 0.16088607 | 0.31142316 | 0.51661563 | 0.60542451 | 0.92569157 |
| CEP350 | 205.050595 | -0.269217 | 0.21669746 | -1.2423634 | 0.2141025 | 0.7643572 |
| QSOX1 | 46.6792307 | 0.05458559 | 0.38503112 | 0.1417693 | 0.88726224 | 0.98229432 |
| ACBD6 | 199.508291 | -0.2465641 | 0.16890124 | -1.4598124 | 0.14434164 | 0.69512617 |

| | | | | | | |
|---|---|---|---|---|---|---|
| XPR1 | 184.764472 | -0.3936398 | 0.29599172 | -1.3299014 | 0.18355075 | 0.73573866 |
| KIAA1614 | 30.4787734 | 0.82009183 | 0.38956585 | 2.10514303 | 0.03527886 | 0.49209205 |
| STX6 | 46.3365381 | 0.12583068 | 0.27897684 | 0.45104345 | 0.65195823 | 0.93490482 |
| MR1 | 24.9555264 | 0.59963117 | 0.42570696 | 1.40855382 | 0.15896714 | 0.71094363 |
| GLUL | 70.0790574 | 0.60447815 | 0.23349483 | 2.58882886 | 9.63E-03 | 0.32468359 |
| RNASEL | 12.8611018 | -0.1769111 | 0.57359348 | -0.3084259 | 0.75775827 | NA |
| DHX9 | 122.942163 | -0.0887323 | 0.30014516 | -0.2956314 | 0.76751156 | 0.96283923 |
| LAMC1 | 177.23871 | -0.6992876 | 0.39250683 | -1.7815935 | 0.07481554 | 0.61020315 |
| LAMC2 | 30.0922993 | -0.3948152 | 0.63201741 | -0.6246903 | 5.32E-01 | 0.90612014 |
| SMG7 | 149.32787 | -0.2677728 | 0.18823364 | -1.4225556 | 0.15486502 | 0.70572801 |
| ARPC5 | 43.7292005 | 0.10006307 | 0.2832414 | 0.35327841 | 7.24E-01 | 0.95093722 |
| RGL1 | 27.3181186 | 0.2449311 | 0.42633518 | 0.57450362 | 0.56562705 | 0.91698355 |
| TSEN15 | 43.573538 | 0.02484489 | 0.29758468 | 0.08348846 | 0.93346315 | 0.98960607 |
| EDEM3 | 81.6768379 | 0.09147541 | 0.20821112 | 0.43933968 | 0.66041543 | 0.93728744 |
| RNF2 | 44.0319561 | -0.3290943 | 0.26496774 | -1.2420163 | 0.21423053 | 0.7643572 |
| TRMT1L | 37.1479506 | 0.26263578 | 0.44486303 | 0.59037448 | 0.55493961 | 0.9137961 |
| SWT1 | 57.9924958 | -0.2911522 | 0.31159538 | -0.9343918 | 0.35010182 | 0.83687845 |
| IVNS1ABP | 352.406585 | -1.1518974 | 0.53933337 | -2.13578 | 0.03269735 | 0.48384156 |
| HMCN1 | 54.1427891 | -0.1586607 | 0.43591503 | -0.3639717 | 7.16E-01 | 0.94885853 |
| TPR | 125.289697 | -0.2260323 | 0.18402891 | -1.2282436 | 0.21935552 | 0.76680432 |
| ODR4 | 72.2118698 | 0.11641505 | 0.21269235 | 0.54734007 | 0.58414512 | 0.92071026 |
| AL596220.1 | 10.581527 | 1.58097823 | 0.76198538 | 2.07481438 | 0.03800374 | NA |
| ERVMER61-1 | 116.633497 | 0.18112818 | 0.392611 | 0.46134261 | 6.45E-01 | 0.93328692 |
| BRINP3 | 46.0554177 | -0.2015883 | 0.6045936 | -0.3334278 | 0.73881137 | 0.9542764 |
| AL390957.1 | 21.1531967 | -0.022956 | 0.47464683 | -0.0483644 | 0.96142587 | 0.99256424 |
| AL136454.1 | 42.4679341 | 0.24617818 | 0.36632913 | 0.67201367 | 0.50157499 | 0.89492452 |
| UCHL5 | 42.0947393 | -0.1366922 | 0.26122694 | -0.5232701 | 0.60078631 | 0.92352528 |
| TROVE2 | 73.7228148 | 0.36619978 | 0.22485589 | 1.62859772 | 0.10339821 | 0.65946648 |
| GLRX2 | 10.5898277 | 0.12108441 | 0.4362602 | 0.27755089 | 0.78135713 | NA |
| CDC73 | 167.938465 | -0.3967688 | 0.25065891 | -1.5829034 | 0.11344349 | 0.6656824 |
| ZBTB41 | 41.7607541 | 0.28092896 | 0.27353154 | 1.02704415 | 0.30439968 | 0.81568231 |
| CRB1 | 20.369947 | 0.84421071 | 0.81023753 | 1.04192991 | 0.29744418 | 0.81214575 |
| DENND1B | 222.21419 | 0.09531325 | 0.22707657 | 0.41974059 | 0.67467497 | 0.93936166 |
| NEK7 | 202.092345 | 0.00127084 | 0.18594348 | 0.00683454 | 0.99454687 | 0.99926712 |
| ZNF281 | 14.7180473 | 0.19203366 | 0.378312 | 0.50760656 | 0.61172929 | NA |
| DDX59 | 23.1730899 | -0.3236836 | 0.32462569 | -0.9970979 | 0.31871698 | 0.82354084 |
| CAMSAP2 | 142.307951 | 0.07311915 | 0.2158748 | 0.33871091 | 0.73482752 | 0.95329896 |
| TMEM9 | 30.4606513 | 0.0811319 | 0.51955077 | 0.1561578 | 0.87590864 | 0.97918624 |
| LAD1 | 17.9546587 | 0.78967354 | 0.51440094 | 1.5351324 | 0.12475131 | 0.66929384 |
| CSRP1 | 40.9633483 | -0.1077875 | 0.53032125 | -0.2032495 | 0.83894004 | 0.97600352 |
| AC096677.1 | 40.3197461 | 0.11312453 | 0.34801441 | 0.32505702 | 0.74513792 | 0.95563105 |
| NAV1 | 46.5519657 | -0.0344639 | 0.40480039 | -0.085138 | 0.93215165 | 0.98960607 |
| IPO9-AS1 | 13.0569213 | 0.00671122 | 0.49760611 | 0.01348701 | 0.98923925 | NA |
| IPO9 | 72.8864114 | -0.2608078 | 0.24601231 | -1.0601411 | 0.28908042 | 0.80536924 |
| TIMM17A | 25.3615892 | 0.29751561 | 0.34695015 | 0.85751689 | 0.39115929 | 0.85605713 |
| RNPEP | 34.05096 | 0.07245654 | 0.33947855 | 0.2134348 | 0.83098784 | 0.97567084 |
| ELF3 | 239.923971 | -0.1017042 | 0.39509441 | -0.2574175 | 0.79685649 | 0.97010563 |
| ARL8A | 23.8641901 | -0.3371708 | 0.37305097 | -0.9038197 | 0.36609101 | 0.84741115 |
| PPP1R12B | 218.400433 | -0.068883 | 0.17819925 | -0.3865504 | 0.69908905 | 0.94395238 |
| KDM5B | 165.863052 | -0.3426113 | 0.21579705 | -1.5876552 | 0.11236434 | 0.6656824 |
| KLHL12 | 28.0389956 | -0.3810558 | 0.33632083 | -1.1330128 | 0.25720889 | 0.79530316 |
| ADIPOR1 | 19.537317 | 0.17107141 | 0.38621977 | 0.44293799 | 0.65781058 | 0.93593674 |
| CYB5R1 | 17.5683955 | 0.2480149 | 0.4918029 | 0.50429735 | 0.61405244 | 0.92662475 |
| TMEM183A | 13.4241849 | -0.3238151 | 0.44474206 | -0.7280965 | 0.46655451 | NA |

| | | | | | | |
|---|---|---|---|---|---|---|
| BTG2 | 148.692199 | 0.21740965 | 0.59666855 | 0.36437257 | 0.71557982 | 0.94885853 |
| ATP2B4 | 133.897615 | -0.0902951 | 0.27664341 | -0.3263954 | 0.74412525 | 0.95563105 |
| SNRPE | 17.4355255 | -0.0480246 | 0.37757373 | -0.1271928 | 0.89878784 | 0.98531749 |
| SOX13 | 74.5301858 | -0.1912542 | 0.30723906 | -0.6224931 | 0.53361767 | 0.90760867 |
| ETNK2 | 12.1447256 | 0.44292268 | 0.48848799 | 0.90672173 | 0.36455397 | NA |
| GOLT1A | 18.0171155 | -0.2746041 | 0.3948082 | -0.6955379 | 0.48671829 | 0.88838332 |
| PLEKHA6 | 129.645316 | -0.1539716 | 0.20286852 | -0.7589726 | 0.44786897 | 0.87291341 |
| PPP1R15B | 32.3396796 | -0.3203203 | 0.4377679 | -0.7317127 | 0.46434394 | 0.87877814 |
| PIK3C2B | 27.575702 | -0.6996457 | 0.63428549 | -1.1030454 | 0.27000747 | 0.79666376 |
| MDM4 | 162.205629 | -0.1645286 | 0.19928813 | -0.8255817 | 0.4090414 | 0.85867915 |
| LRRN2 | 101.245526 | 0.56950651 | 0.37130025 | 1.53381663 | 0.12507477 | 0.66929384 |
| RBBP5 | 38.4190826 | -0.2020174 | 0.25228539 | -0.8007496 | 0.42327662 | 0.86190547 |
| DSTYK | 75.3736272 | 0.26160082 | 0.25878312 | 1.01088826 | 0.31206991 | 0.81701824 |
| NUAK2 | 69.9671867 | 0.09003572 | 0.46758685 | 0.19255401 | 0.84730827 | 0.97612951 |
| CDK18 | 63.5830492 | -0.6452227 | 0.42108474 | -1.532287 | 0.12545163 | 0.66929384 |
| MFSD4A | 37.461215 | 1.31441446 | 0.41930378 | 3.13475462 | 0.00171998 | 0.17205462 |
| ELK4 | 66.7787414 | -0.3999627 | 0.30717195 | -1.3020807 | 0.1928888 | 0.75061775 |
| NUCKS1 | 233.814256 | -0.0702483 | 0.21802419 | -0.3222042 | 0.74729797 | 0.95563105 |
| RAB29 | 20.5928127 | 0.48060349 | 0.41341549 | 1.16251932 | 0.24502458 | 0.78583441 |
| SLC41A1 | 13.994409 | 0.5039099 | 0.43354495 | 1.16230138 | 0.24511307 | NA |
| RHEX | 61.662417 | -0.0923726 | 0.6613003 | -0.1396833 | 8.89E-01 | 0.98270788 |
| SRGAP2 | 46.5958759 | -0.1766867 | 0.34042724 | -0.5190145 | 0.60375061 | 0.92508698 |
| EIF2D | 19.5489113 | -0.0693135 | 0.45274206 | -0.153097 | 0.87832176 | 0.98061726 |
| MAPKAPK2 | 117.545757 | -0.4908184 | 0.25911144 | -1.8942369 | 0.05819358 | 0.57141408 |
| PIGR | 291.033091 | 1.6124664 | 0.59238057 | 2.72201095 | 0.0064886 | 0.29935381 |
| PFKFB2 | 68.3448572 | -0.5443863 | 0.4115492 | -1.3227732 | 0.18591081 | 0.73804265 |
| CD55 | 54.1376898 | -0.5931014 | 0.49138083 | -1.2070097 | 0.2274284 | 0.77185154 |
| CD46 | 199.833893 | 0.20035569 | 0.15368763 | 1.30365532 | 0.19235113 | 0.75061775 |
| MIR29B2CH( | 199.687061 | 0.65613535 | 0.36873573 | 1.77941899 | 0.0751711 | 0.61020315 |
| PLXNA2 | 20.7391177 | 0.97547369 | 0.59302735 | 1.64490507 | 0.09998939 | 0.65863153 |
| LAMB3 | 20.0796175 | -0.7155481 | 0.73893112 | -0.9683556 | 3.33E-01 | 8.29E-01 |
| G0S2 | 65.2702038 | -1.352201 | 1.12686464 | -1.1999676 | 0.23015193 | 0.77249016 |
| IRF6 | 21.9988987 | 0.65243267 | 0.41608474 | 1.56802837 | 0.11687451 | 0.6668162 |
| UTP25 | 10.6317451 | 0.22452395 | 0.48053634 | 0.46723615 | 0.64033093 | NA |
| HHAT | 65.1724219 | 0.60637844 | 0.42287979 | 1.43392628 | 0.1515933 | 0.70572801 |
| RCOR3 | 168.787772 | 0.05366369 | 0.19846498 | 0.27039375 | 0.78685735 | 0.96767512 |
| TRAF5 | 34.5878705 | 0.27954103 | 0.38667624 | 0.72293303 | 0.46972103 | 0.88207878 |
| LINC00467 | 38.317842 | 0.01547995 | 0.42508559 | 0.03641607 | 0.9709506 | 0.99403127 |
| SLC30A1 | 16.3719443 | 0.00174107 | 0.52161991 | 0.00333782 | 0.99733681 | 0.99953287 |
| LPGAT1 | 151.751766 | -0.3155278 | 0.21490765 | -1.4682017 | 1.42E-01 | 0.69328517 |
| INTS7 | 26.9392006 | -0.0683443 | 0.38731582 | -0.1764562 | 0.85993559 | 0.97691173 |
| PPP2R5A | 146.48406 | -0.1314761 | 0.24258683 | -0.5419754 | 0.58783548 | 0.92148551 |
| AL360091.3 | 19.1273601 | 1.01159641 | 0.56741166 | 1.78282628 | 0.07461459 | 0.61020315 |
| TMEM206 | 11.479141 | 0.32122157 | 0.51561603 | 0.622986 | 0.53329372 | NA |
| NENF | 35.5383011 | -0.1420826 | 0.44239616 | -0.321166 | 0.74808458 | 0.95563105 |
| ATF3 | 32.0723948 | -1.0120461 | 0.80412291 | -1.2585714 | 0.2081852 | 0.76231289 |
| NSL1 | 59.2301795 | -0.0171135 | 0.26694388 | -0.064109 | 0.94888342 | 0.99105535 |
| TATDN3 | 41.6477587 | -0.1854487 | 0.32980348 | -0.5623007 | 0.57391113 | 0.91813957 |
| FLVCR1 | 36.1889935 | -0.0986669 | 0.31077442 | -0.3174871 | 0.75087402 | 0.95740778 |
| ANGEL2 | 41.5673528 | -0.122593 | 0.25662728 | -0.4777084 | 0.63285777 | 0.93251095 |
| RPS6KC1 | 135.544464 | -0.5690754 | 0.20373887 | -2.7931608 | 0.00521957 | 0.28057141 |
| SMYD2 | 31.3553157 | -0.4841615 | 0.30052498 | -1.6110524 | 0.1071683 | 0.65986731 |
| PTPN14 | 334.350968 | -0.6888984 | 0.30851627 | -2.2329403 | 0.02555288 | 0.44138123 |
| KCTD3 | 79.8986505 | -0.1797266 | 0.22242358 | -0.8080377 | 0.41906885 | 0.86031916 |

| | | | | | | |
|---|---|---|---|---|---|---|
| ESRRG | 3810.07894 | 0.03825136 | 0.17766525 | 0.21530015 | 8.30E-01 | 9.76E-01 |
| GPATCH2 | 194.204549 | -0.0596517 | 0.17957851 | -0.3321761 | 0.73975626 | 0.9544376 |
| SPATA17 | 53.1011954 | -0.3750658 | 0.41883931 | -0.8954885 | 0.370526 | 0.84852785 |
| RRP15 | 20.2668407 | -0.0277052 | 0.3701661 | -0.0748452 | 0.94033787 | 0.99012269 |
| LYPLAL1 | 190.174977 | -0.040916 | 0.32379001 | -0.1263658 | 0.89944237 | 0.98531749 |
| EPRS | 64.1389466 | -0.4097824 | 0.2953394 | -1.3874966 | 0.1652904 | 0.71615349 |
| BPNT1 | 16.5482723 | 0.65302517 | 0.37973306 | 1.71969531 | 0.08548784 | 0.63062869 |
| IARS2 | 60.9123942 | -0.043577 | 0.23001091 | -0.1894563 | 0.84973524 | 0.97612951 |
| RAB3GAP2 | 182.650686 | -0.3990364 | 0.18173151 | -2.1957468 | 0.02811007 | 0.45940027 |
| MARK1 | 78.9484272 | -0.1082365 | 0.26134084 | -0.4141584 | 0.67875812 | 0.94144983 |
| C1orf115 | 12.0390967 | 0.56454207 | 0.5255649 | 1.07416243 | 0.28274989 | NA |
| 2-Mar | 81.5957947 | 0.51162188 | 0.29110884 | 1.75749346 | 0.07883374 | 0.61367304 |
| 1-Mar | 12.3201897 | 0.44007798 | 0.4810109 | 0.9149023 | 0.36024292 | NA |
| AL513314.2 | 18.643524 | -0.3956458 | 0.37763909 | -1.0476824 | 0.29478496 | 0.8101516 |
| MIA3 | 48.1658818 | 0.06437173 | 0.26292543 | 0.24482883 | 0.80658897 | 0.97216031 |
| AIDA | 20.6329879 | -0.283515 | 0.36344653 | -0.7800734 | 0.43534766 | 0.86729771 |
| BROX | 47.3847594 | 0.17573871 | 0.26152591 | 0.67197437 | 0.50160001 | 0.89492452 |
| FAM177B | 33.8525749 | -0.8494508 | 0.64027446 | -1.3266979 | 1.85E-01 | 0.73705812 |
| AC093152.1 | 44.6917259 | 0.01394011 | 0.30372794 | 0.04589671 | 0.96339258 | 0.99256424 |
| TLR5 | 17.1915635 | 1.06471197 | 0.47120922 | 2.2595313 | 0.02385036 | 0.43508975 |
| SUSD4 | 45.7757602 | 0.23529044 | 0.41126644 | 0.57211194 | 0.56724614 | 0.91698355 |
| CAPN2 | 145.983288 | -0.3748503 | 0.19813374 | -1.8919056 | 0.05850356 | 0.57141408 |
| TP53BP2 | 54.5254592 | -0.3596663 | 0.32757917 | -1.0979522 | 0.27222535 | 0.7968545 |
| FBXO28 | 85.3285008 | -0.2386253 | 0.2011512 | -1.186298 | 0.23550465 | 0.77848938 |
| DEGS1 | 20.7539711 | -1.0107603 | 0.43858119 | -2.3046139 | 0.0211882 | 0.4212027 |
| NVL | 76.1427569 | -0.46337 | 0.2617896 | -1.7700092 | 0.07672561 | 0.61188697 |
| CNIH4 | 49.5939658 | -0.283935 | 0.2413585 | -1.1764037 | 0.2394336 | 0.78098673 |
| WDR26 | 151.362427 | -0.2655458 | 0.23898211 | -1.1111533 | 0.26650235 | 0.79666376 |
| CNIH3 | 46.7476083 | 0.24607992 | 0.43470695 | 0.56608232 | 0.57133782 | 0.91791025 |
| DNAH14 | 253.818591 | -0.1032882 | 0.18410775 | -0.5610203 | 0.57478369 | 0.91824492 |
| LBR | 51.7203677 | 0.03194167 | 0.26407648 | 0.12095613 | 0.90372578 | 0.98531749 |
| ENAH | 77.4495285 | 0.19071327 | 0.26532904 | 0.71878024 | 0.47227634 | 0.88277854 |
| SRP9 | 71.649489 | -0.0045904 | 0.33054453 | -0.0138874 | 0.98891984 | 0.99814944 |
| EPHX1 | 23.9039771 | 0.07136141 | 0.36756629 | 0.19414568 | 0.84606181 | 0.97612951 |
| AL591895.1 | 25.7525554 | 0.16756049 | 0.30833208 | 0.54344164 | 0.58682576 | 0.92076087 |
| TMEM63A | 59.9312713 | -0.0577209 | 0.22747325 | -0.253748 | 0.79969029 | 0.97019967 |
| SDE2 | 11.6396422 | -0.0891796 | 0.42129578 | -0.2116794 | 0.83235717 | NA |
| H3F3A | 15.7358948 | 0.72527902 | 0.41500813 | 1.74762605 | 0.0805288 | NA |
| ACBD3 | 87.1295049 | -0.2236028 | 0.22762365 | -0.9823356 | 0.32593453 | 0.82567895 |
| LIN9 | 12.2191722 | 0.71831902 | 0.51598026 | 1.3921444 | 0.16387867 | NA |
| PARP1 | 46.9263805 | 0.40193704 | 0.24825414 | 1.61905471 | 0.10543549 | 0.65946648 |
| STUM | 21.1298195 | -0.6275178 | 0.82291943 | -0.7625507 | 0.44573142 | 0.8718292 |
| ITPKB | 106.544876 | 0.24493866 | 0.42405124 | 0.57761572 | 0.56352359 | 0.91624774 |
| COQ8A | 29.7271058 | 0.3327927 | 0.42257149 | 0.78754177 | 0.43096479 | 0.86602041 |
| CDC42BPA | 316.175993 | -0.2290459 | 0.18480086 | -1.2394199 | 0.21519002 | 0.7643572 |
| ZNF678 | 47.2105139 | -0.2939916 | 0.26456334 | -1.1112333 | 0.26646792 | 0.79666376 |
| SNAP47 | 25.3718425 | 0.74234472 | 0.34149312 | 2.17382044 | 0.02971862 | 0.46381645 |
| ARF1 | 107.25066 | -0.2759039 | 0.27662383 | -0.9973974 | 0.31857165 | 0.82354084 |
| C1orf35 | 14.8906315 | -0.0801711 | 0.39680915 | -0.2020395 | 0.83988585 | NA |
| MRPL55 | 15.642766 | 0.44624787 | 0.44259999 | 1.00824192 | 0.31333834 | NA |
| GUK1 | 108.514902 | -0.1097376 | 0.41146174 | -0.2667018 | 0.78969877 | 0.96868722 |
| OBSCN | 106.37047 | 0.29566668 | 0.21835939 | 1.3540369 | 0.17572461 | 0.72841249 |
| TRIM11 | 13.0226036 | -0.5426677 | 0.44060454 | -1.2316434 | 0.21808232 | NA |
| RNF187 | 33.3303346 | -0.3913521 | 0.50436388 | -0.7759321 | 0.4377891 | 0.86821458 |

| | | | | | | |
|---|---|---|---|---|---|---|
| RAB4A | 36.6133176 | 0.2636393 | 0.32120489 | 0.82078232 | 0.41177027 | 0.85867915 |
| NUP133 | 53.0712598 | -0.0978017 | 0.28234483 | -0.346391 | 0.72904885 | 0.95279138 |
| ABCB10 | 27.0676123 | 0.15378111 | 0.33916822 | 0.4534066 | 0.65025599 | 0.93453685 |
| TAF5L | 16.4418308 | -0.2845469 | 0.40983274 | -0.6943002 | 0.48749399 | 0.88838332 |
| GALNT2 | 96.9263335 | -0.3936042 | 0.28653137 | -1.3736863 | 0.1695391 | 0.72354169 |
| COG2 | 37.2600438 | 0.34187636 | 0.27995564 | 1.22118047 | 0.2220177 | 0.76680432 |
| C1orf198 | 14.6202919 | -0.77994 | 0.44690012 | -1.7452222 | 0.0809462 | NA |
| TTC13 | 31.2701101 | -0.0117558 | 0.32820553 | -0.0358183 | 0.97142724 | 0.99403127 |
| ARV1 | 25.920654 | -0.0893791 | 0.38445103 | -0.2324851 | 0.81616125 | 0.97288403 |
| C1orf131 | 17.5879802 | 0.17088535 | 0.38410081 | 0.44489715 | 0.65639408 | 0.93570875 |
| GNPAT | 49.3651771 | -0.0344126 | 0.24826823 | -0.1386107 | 0.88975781 | 0.9827634 |
| EGLN1 | 140.938652 | -0.5248715 | 0.25220212 | -2.0811543 | 0.03741979 | 0.49538341 |
| DISC1 | 25.6872618 | -0.1922178 | 0.41567984 | -0.4624179 | 0.64378167 | 0.93328692 |
| SIPA1L2 | 35.2309447 | 0.24722985 | 0.56012448 | 0.44138375 | 0.6589352 | 0.93664697 |
| NTPCR | 31.8167362 | 0.36992642 | 0.29359137 | 1.2600044 | 0.20766777 | 0.76231289 |
| PCNX2 | 175.186848 | 0.03399836 | 0.22925042 | 0.14830227 | 0.88210423 | 0.98137082 |
| MAP3K21 | 270.307037 | 0.68370748 | 0.35573214 | 1.92197275 | 0.05460919 | 0.5645224 |
| KCNK1 | 81.2445754 | -0.1033249 | 0.32043528 | -0.3224517 | 0.74711053 | 0.95563105 |
| AL355472.1 | 12.1191968 | 0.02668319 | 0.66360857 | 0.04020922 | 0.96792632 | NA |
| COA6 | 13.3323431 | 0.40950283 | 0.41866801 | 0.97810871 | 0.32802056 | NA |
| TARBP1 | 62.2371626 | 0.05900717 | 0.30726589 | 0.19203944 | 0.84771131 | 0.97612951 |
| IRF2BP2 | 75.2425858 | 0.25317766 | 0.34159248 | 0.74116873 | 0.45859114 | 0.87549589 |
| TOMM20 | 65.900281 | 0.59493758 | 0.29186076 | 2.03842953 | 0.041507 | 0.51036486 |
| ARID4B | 373.86977 | -0.2903611 | 0.19199247 | -1.5123567 | 0.13044313 | 0.67375513 |
| B3GALNT2 | 60.470619 | 0.36768164 | 0.29519029 | 1.24557499 | 0.21292047 | 0.7643572 |
| LYST | 140.877991 | -0.0341214 | 0.22705242 | -0.15028 | 0.88054373 | 0.98093344 |
| GPR137B | 38.0599331 | -0.5960742 | 0.32014621 | -1.8618812 | 0.06261984 | 0.5842021 |
| ERO1B | 36.7835202 | 0.45493725 | 0.43924068 | 1.0357357 | 0.30032546 | 0.81399689 |
| LGALS8 | 33.4399701 | -0.0504917 | 0.27508153 | -0.1835517 | 0.85436518 | 0.97612951 |
| HEATR1 | 43.8546006 | -0.2859174 | 0.29721094 | -0.9620015 | 0.33604885 | 0.82974237 |
| MTR | 513.130088 | 0.45077266 | 0.33951493 | 1.32769614 | 0.18427851 | 0.73705812 |
| CHRM3 | 279.006716 | 0.271893 | 0.43837882 | 0.62022386 | 0.53511042 | 0.90763121 |
| RGS7 | 57.0233165 | -0.9366403 | 0.51666152 | -1.8128702 | 6.99E-02 | 6.03E-01 |
| FH | 22.4617664 | 0.48657169 | 0.39719998 | 1.22500431 | 0.2205736 | 0.76680432 |
| OPN3 | 12.1245736 | 0.62007754 | 0.41804348 | 1.48328482 | 0.13799875 | NA |
| CEP170 | 74.9500376 | -0.4680599 | 0.26902093 | -1.7398643 | 0.08188285 | 0.61927158 |
| SDCCAG8 | 287.134088 | -0.1403196 | 0.22880041 | -0.6132838 | 0.53968867 | 0.90763121 |
| AKT3 | 427.399949 | -0.3835888 | 0.24711844 | -1.5522467 | 0.12060322 | 0.6668162 |
| ZBTB18 | 19.3448027 | 0.16233237 | 0.41913732 | 0.38730115 | 0.69853327 | 0.94395238 |
| AL645465.1 | 12.5269693 | 0.18535773 | 0.43030162 | 0.43076234 | 0.66664119 | NA |
| ADSS | 71.3576287 | -0.3029493 | 0.26819542 | -1.129584 | 0.25865156 | 0.79532468 |
| CATSPERE | 49.4607985 | -0.4599943 | 0.301965 | -1.5233365 | 0.12767455 | 0.66929384 |
| DESI2 | 70.0728706 | 0.03525526 | 0.27973464 | 0.12603109 | 0.89970731 | 0.98531749 |
| COX20 | 77.8840713 | -0.0012303 | 0.32735153 | -0.0037582 | 0.99700138 | 0.99941676 |
| HNRNPU | 311.939217 | -0.0190705 | 0.19435481 | -0.0981223 | 0.92183519 | 0.98846691 |
| EFCAB2 | 161.786449 | -0.1528994 | 0.43602368 | -0.3506675 | 0.72583777 | 0.95164932 |
| KIF26B | 195.296244 | 0.42960668 | 0.58934578 | 0.72895522 | 0.46602905 | 0.88032008 |
| SMYD3 | 649.420184 | -0.1006619 | 0.14394597 | -0.6993032 | 0.48436255 | 0.88838332 |
| TFB2M | 12.0214475 | -0.0848544 | 0.5006497 | -0.1694886 | 0.86541232 | NA |
| CNST | 72.1523359 | 0.01983479 | 0.24785228 | 0.08002665 | 0.93621606 | 0.99012269 |
| SCCPDH | 68.4104162 | -0.2953622 | 0.42705089 | -0.6916324 | 0.48916818 | 0.88930178 |
| AHCTF1 | 100.388565 | -0.354627 | 0.26232848 | -1.3518433 | 0.17642544 | 0.72841249 |
| ZNF124 | 58.0388698 | 0.28615502 | 0.37202559 | 0.76918101 | 0.44178586 | 0.86925451 |
| AL390728.6 | 19.6037915 | 0.12259447 | 0.35582849 | 0.34453249 | 0.73044587 | 0.95279138 |

| | | | | | | |
|---|---|---|---|---|---|---|
| ZNF496 | 26.1085489 | 0.42702662 | 0.32178822 | 1.32704245 | 0.18449464 | 0.73705812 |
| ZNF672 | 19.2798789 | 0.24365797 | 0.35327105 | 0.68971959 | 0.49037054 | 0.8897498 |
| ZNF692 | 18.0573561 | -0.3040637 | 0.41020122 | -0.741255 | 0.45853885 | 0.87549589 |
| FAM110C | 63.8041328 | -0.595655 | 0.39972854 | -1.4901487 | 1.36E-01 | 6.84E-01 |
| SH3YL1 | 108.141165 | 0.45255357 | 0.20434877 | 2.21461358 | 0.0267866 | 0.44988636 |
| ACP1 | 33.5926607 | 0.33881756 | 0.30246745 | 1.12017858 | 0.26263767 | 0.79666376 |
| EIPR1 | 118.013397 | 0.25873824 | 0.3017751 | 0.85738765 | 0.39123068 | 0.85605713 |
| TRAPPC12 | 123.825123 | 0.1246115 | 0.19357577 | 0.64373502 | 0.51974727 | 0.90368693 |
| ADI1 | 66.3753798 | 0.67031976 | 0.42227988 | 1.58738266 | 0.112426 | 0.6656824 |
| AC108488.2 | 12.0284673 | 0.05186156 | 0.46248828 | 0.11213595 | 0.91071561 | NA |
| RPS7 | 110.48542 | 0.17166659 | 0.40607132 | 0.42274986 | 0.67247777 | 0.93878847 |
| RSAD2 | 20.0979944 | -4.5153806 | 1.92836094 | -2.341564 | 0.01920313 | 0.41544117 |
| RNF144A | 50.9662761 | -0.0699529 | 0.35585916 | -0.1965746 | 0.84416042 | 0.97612951 |
| ID2 | 76.7813071 | 0.30575127 | 0.4404378 | 0.69419853 | 0.48755771 | 0.88838332 |
| KIDINS220 | 155.262383 | -0.1163806 | 0.20033901 | -0.5809183 | 0.56129549 | 0.91608386 |
| MBOAT2 | 139.310307 | 0.07409913 | 0.33876843 | 0.21873093 | 0.82685965 | 0.97567084 |
| ASAP2 | 620.240406 | 0.04771246 | 0.2363689 | 0.2018559 | 8.40E-01 | 9.76E-01 |
| ITGB1BP1 | 53.8621599 | 0.16514353 | 0.22159803 | 0.74523917 | 0.45612715 | 0.87394768 |
| CPSF3 | 25.0092584 | -0.2311824 | 0.33708361 | -0.685831 | 0.49281969 | 0.89092098 |
| IAH1 | 31.7022197 | -0.2947151 | 0.28489274 | -1.0344773 | 0.30091309 | 0.81437329 |
| ADAM17 | 109.20094 | -0.4079156 | 0.24722313 | -1.6499897 | 0.09894504 | 0.65744285 |
| YWHAQ | 89.6704692 | -0.0042413 | 0.27484911 | -0.0154314 | 0.98768801 | 0.99814944 |
| TAF1B | 105.540208 | -0.2609938 | 0.26192881 | -0.9964304 | 0.31904108 | 0.82354084 |
| GRHL1 | 25.1157706 | 0.42038803 | 0.45556473 | 0.92278441 | 0.35611957 | 0.84192356 |
| KLF11 | 24.6665781 | 0.02311664 | 0.49905565 | 0.04632077 | 0.96305458 | 0.99256424 |
| CYS1 | 243.217455 | 0.18987355 | 0.33170262 | 0.57242102 | 0.56703678 | 0.91698355 |
| HPCAL1 | 110.637979 | -0.2740315 | 0.26189014 | -1.0463605 | 0.29539464 | 0.81042115 |
| ODC1 | 21.4512105 | 0.77422948 | 0.44045729 | 1.75778559 | 0.078784 | 0.61367304 |
| NOL10 | 105.184226 | -0.21727 | 0.22032987 | -0.9861122 | 0.32407805 | 0.82567895 |
| PDIA6 | 90.7362863 | -0.0325203 | 0.42159151 | -0.077137 | 0.93851457 | 0.99012269 |
| PQLC3 | 16.347092 | 0.08991802 | 0.49015898 | 0.18344665 | 0.85444758 | 0.97612951 |
| ROCK2 | 251.156877 | -0.5608959 | 0.30950946 | -1.8122092 | 0.06995388 | 0.60302096 |
| E2F6 | 18.8359032 | -0.3578481 | 0.3438158 | -1.0408135 | 0.29796211 | 0.81214575 |
| LPIN1 | 77.8215319 | -0.1100813 | 0.25551221 | -0.430826 | 0.66659487 | 0.93860542 |
| MIR3681HG | 47.5034235 | 0.74613711 | 0.51309683 | 1.45418383 | 0.14589535 | 0.69679188 |
| TRIB2 | 53.2001436 | 0.16177002 | 0.27338139 | 0.59173749 | 0.55402639 | 0.9137961 |
| LINC00276 | 30.6528134 | 0.11108836 | 0.55164165 | 0.20137776 | 0.8404032 | 0.97600352 |
| FAM84A | 76.0636264 | 0.04779805 | 0.36348725 | 0.13149855 | 8.95E-01 | 9.85E-01 |
| NBAS | 240.598649 | -0.47921 | 0.16096091 | -2.9771826 | 0.00290911 | 0.21399685 |
| DDX1 | 68.5933153 | -0.0002836 | 0.21002921 | -0.0013505 | 0.99892246 | 0.99957534 |
| SMC6 | 61.3343242 | -0.117256 | 0.29141785 | -0.4023637 | 0.68741636 | 0.94205398 |
| GEN1 | 18.5381611 | -0.5363696 | 0.4119906 | -1.3018977 | 0.19295136 | 0.75061775 |
| KCNS3 | 13.870177 | 1.50624072 | 0.63118389 | 2.38637382 | 0.01701545 | NA |
| RDH14 | 13.5432833 | 0.51175125 | 0.44990877 | 1.1374556 | 0.25534787 | NA |
| LINC01376 | 69.1312886 | -0.2331532 | 0.42355149 | -0.5504719 | 0.58199573 | 0.92034873 |
| TTC32 | 18.2704383 | -0.5663485 | 0.36843193 | -1.5371862 | 0.12424771 | 0.66929384 |
| WDR35 | 61.471406 | -0.2276755 | 0.27284706 | -0.8344437 | 0.404031 | 0.85782098 |
| LAPTM4A | 136.012857 | 0.21943416 | 0.325879 | 0.67336083 | 0.50071776 | 0.89492452 |
| SDC1 | 39.8724457 | -0.1710664 | 0.55567684 | -0.3078523 | 0.75819474 | 0.95903525 |
| PUM2 | 306.990139 | -0.4238138 | 0.22744979 | -1.863329 | 0.06241599 | 0.58393911 |
| RHOB | 35.3264266 | -0.1369672 | 0.4434057 | -0.3088981 | 0.75739902 | 0.95903525 |
| HS1BP3 | 13.6075333 | -0.4086295 | 0.41532952 | -0.9838682 | 0.32518033 | NA |
| LDAH | 100.762373 | -0.0251107 | 0.20790886 | -0.1207775 | 0.90386729 | 0.98531749 |
| KLHL29 | 36.1412151 | -0.8283569 | 0.67423383 | -1.2285899 | 2.19E-01 | 7.67E-01 |

| | | | | | | |
|---|---|---|---|---|---|---|
| ATAD2B | 197.150578 | -0.1392752 | 0.24535796 | -0.5676409 | 0.57027883 | 0.91778542 |
| UBXN2A | 71.5525746 | 0.29555174 | 0.34438883 | 0.85819201 | 0.39078645 | 0.85605713 |
| SF3B6 | 35.433607 | 0.48350341 | 0.32382226 | 1.49311356 | 0.13540746 | 0.6835006 |
| FAM228B | 14.0806409 | -0.2155885 | 0.43058257 | -0.5006903 | 0.61658911 | NA |
| TP53I3 | 23.7145409 | 0.69469102 | 0.40619672 | 1.71023294 | 0.08722281 | 0.63466198 |
| ITSN2 | 230.322108 | -0.1260325 | 0.2266025 | -0.5561833 | 0.57808559 | 0.91878391 |
| NCOA1 | 459.204201 | 0.20651536 | 0.19539137 | 1.05693184 | 0.29054269 | 0.80585318 |
| DNAJC27 | 35.681907 | 0.15222488 | 0.28464457 | 0.53478934 | 0.59279553 | 0.92278087 |
| DNAJC27-AS | 21.3646432 | 0.13026124 | 0.66216887 | 0.19671906 | 0.84404739 | 0.97612951 |
| DNMT3A | 115.091503 | 0.07833465 | 0.2654331 | 0.29512015 | 0.7679021 | 0.96283923 |
| DTNB | 178.784769 | 0.48242525 | 0.29120769 | 1.65663638 | 0.09759301 | 0.65521765 |
| ASXL2 | 143.358683 | -0.3815127 | 0.20954295 | -1.8206899 | 0.06865401 | 0.60275677 |
| RAB10 | 358.551291 | -0.5403551 | 0.26608583 | -2.0307549 | 0.04227986 | 0.51343649 |
| HADHA | 103.301424 | 0.24231729 | 0.20990966 | 1.15438847 | 0.24834094 | 0.78713356 |
| HADHB | 127.47448 | 0.50319131 | 0.18802452 | 2.67620044 | 0.00744621 | 0.30654023 |
| SELENOI | 79.3724625 | -0.4906994 | 0.25553781 | -1.9202614 | 0.05482489 | 0.5645224 |
| MAPRE3 | 26.4429573 | 0.46608449 | 0.30450087 | 1.5306508 | 0.12585572 | 0.66929384 |
| TMEM214 | 15.6885671 | 0.08950882 | 0.38306599 | 0.23366423 | 0.81524566 | NA |
| AGBL5 | 25.4083913 | 0.20052996 | 0.36769197 | 0.54537487 | 0.58549572 | 0.92076087 |
| OST4 | 51.1560704 | -0.1238812 | 0.57941138 | -0.2138052 | 0.83069895 | 0.97567084 |
| ATRAID | 39.3879479 | 0.22566081 | 0.38234091 | 0.59020839 | 0.55505095 | 0.9137961 |
| CAD | 11.5303542 | -0.4730235 | 0.5422821 | -0.8722831 | 0.38305393 | NA |
| MPV17 | 17.3573598 | -0.3043973 | 0.38937758 | -0.7817535 | 0.43435945 | 0.86729771 |
| GTF3C2 | 53.2395438 | -0.0506333 | 0.24022301 | -0.2107761 | 0.83306196 | 0.97600352 |
| EIF2B4 | 13.2915636 | 0.25565458 | 0.45325796 | 0.5640377 | 0.57272847 | NA |
| SNX17 | 21.2336414 | -0.0622509 | 0.34331477 | -0.1813232 | 0.85611387 | 0.97634735 |
| PPM1G | 65.0575724 | 0.24976919 | 0.3036162 | 0.82264776 | 0.41070833 | 0.85867915 |
| NRBP1 | 46.8180186 | -0.4900238 | 0.26187931 | -1.8711819 | 0.06131987 | 0.58024409 |
| KRTCAP3 | 10.7325026 | 0.38798685 | 0.49477563 | 0.78416725 | 0.43294198 | NA |
| IFT172 | 55.6531626 | 0.05872112 | 0.35528416 | 0.16527932 | 0.86872414 | 0.9783491 |
| FNDC4 | 14.0572579 | 0.42506282 | 0.50522088 | 0.84134056 | 4.00E-01 | NA |
| ZNF512 | 51.0481035 | -0.3531579 | 0.27249046 | -1.2960378 | 0.19496248 | 0.7523287 |
| GPN1 | 12.3864037 | 0.2159305 | 0.41520948 | 0.52005194 | 0.60302737 | NA |
| SUPT7L | 20.1373261 | -0.1492081 | 0.40734875 | -0.3662908 | 0.71414813 | 0.94841799 |
| SLC4A1AP | 26.9860285 | 0.41720206 | 0.30722534 | 1.3579676 | 0.17447396 | 0.72841249 |
| MRPL33 | 162.120808 | -0.4065269 | 0.23589381 | -1.723347 | 0.08482579 | 0.62880224 |
| RBKS | 103.576652 | -0.0959299 | 0.3505794 | -0.2736324 | 0.78436711 | 0.96732016 |
| BABAM2 | 777.678848 | 0.01353165 | 0.1781759 | 0.07594543 | 0.93946251 | 0.99012269 |
| FOSL2 | 257.296198 | -1.057428 | 0.55076423 | -1.9199286 | 0.05486692 | 0.5645224 |
| PPP1CB | 144.175646 | -0.527856 | 0.49150519 | -1.0739582 | 0.28284141 | 0.80431986 |
| TRMT61B | 37.7288206 | -0.3834718 | 0.31581841 | -1.2142161 | 0.2246652 | 0.76931755 |
| WDR43 | 64.0904513 | -0.090819 | 0.34179869 | -0.2657089 | 0.79046341 | 0.96904219 |
| CLIP4 | 107.691147 | -0.321461 | 0.30951388 | -1.0385997 | 0.29899095 | 0.81301499 |
| ALK | 21.2974486 | -1.0972466 | 0.75005597 | -1.4628863 | 0.14349849 | 0.69460324 |
| YPEL5 | 61.1547217 | 0.36778286 | 0.25690893 | 1.43156901 | 0.1522672 | 0.70572801 |
| LCLAT1 | 88.9723628 | -0.2520834 | 0.19337297 | -1.3036123 | 0.19236582 | 0.75061775 |
| GALNT14 | 246.038603 | 0.31712135 | 0.39722388 | 0.79834412 | 0.42467082 | 0.861935 |
| DPY30 | 42.5861601 | 0.0984104 | 0.29545652 | 0.33307913 | 0.73907455 | 0.95443263 |
| SPAST | 73.4440482 | -0.5096158 | 0.26152136 | -1.9486585 | 0.05133622 | 0.55238015 |
| SLC30A6 | 62.2600565 | 0.09682136 | 0.22679215 | 0.4269167 | 0.66944 | 0.93878847 |
| YIPF4 | 85.2431026 | -0.0014077 | 0.25635683 | -0.0054911 | 0.99561873 | 0.99927028 |
| BIRC6 | 546.91703 | -0.2193908 | 0.15323071 | -1.4317679 | 0.15221025 | 0.70572801 |
| TTC27 | 67.7521261 | -0.4431205 | 0.24724725 | -1.7922163 | 0.07309833 | 0.60952146 |
| LINC00486 | 104.402646 | -0.0144789 | 0.22289288 | -0.0649588 | 0.94820678 | 0.99105535 |

| | | | | | | |
|---|---|---|---|---|---|---|
| LTBP1 | 72.5766274 | 1.10897406 | 0.71850604 | 1.5434443 | 0.12272301 | 0.66929384 |
| RASGRP3 | 21.9152547 | -1.0033567 | 0.43416356 | -2.3110108 | 0.02083226 | 0.4212027 |
| FAM98A | 19.5438261 | -0.6008529 | 0.39161149 | -1.5343087 | 0.12495373 | 0.66929384 |
| LINC01317 | 20.1756978 | 1.48110215 | 0.57399374 | 2.58034549 | 0.00987015 | 0.32911351 |
| AC068599.1 | 202.032053 | 2.06016842 | 0.57553182 | 3.57959083 | 3.44E-04 | 0.0763464 |
| LINC01320 | 321.907169 | 0.24258164 | 0.56344658 | 0.43053174 | 0.66680888 | 0.93860542 |
| AC012593.1 | 90.3099232 | 2.71734868 | 1.07886928 | 2.51870058 | 0.01177888 | 0.35687659 |
| CRIM1 | 245.245495 | 0.09518473 | 0.24617318 | 0.38665761 | 0.6990097 | 0.94395238 |
| FEZ2 | 94.8989819 | 0.10751663 | 0.23625689 | 0.45508358 | 0.64904911 | 0.93437275 |
| STRN | 156.142454 | -0.4632947 | 0.23465556 | -1.9743604 | 0.04834078 | 0.53861209 |
| HEATR5B | 85.8689418 | -0.2309531 | 0.27531689 | -0.8388627 | 0.40154637 | 0.85778581 |
| EIF2AK2 | 93.3113858 | -0.4410436 | 0.60314811 | -0.731236 | 0.46463504 | 0.87896358 |
| CEBPZOS | 52.2803751 | 0.4059811 | 0.33329053 | 1.21809972 | 0.22318608 | 0.76834258 |
| CEBPZ | 58.3715048 | -0.0287672 | 0.26864413 | -0.1070831 | 0.91472306 | 0.98691474 |
| NDUFAF7 | 40.5193625 | -0.0756311 | 0.35075785 | -0.2156221 | 0.82928232 | 0.97567084 |
| PRKD3 | 64.1302221 | 0.19354608 | 0.25093521 | 0.77129904 | 0.4405297 | 0.86895963 |
| CDC42EP3 | 20.6260788 | -0.2678263 | 0.34905139 | -0.7672975 | 0.44290465 | 0.86991418 |
| RMDN2 | 45.9446018 | 0.23207345 | 0.32217167 | 0.72034096 | 0.47131509 | 0.88267439 |
| CYP1B1-AS1 | 12.4805914 | 1.00856459 | 0.5568095 | 1.81132792 | 0.07009011 | NA |
| ATL2 | 125.191617 | 0.099574 | 0.28766154 | 0.34614985 | 0.7292301 | 0.95279138 |
| HNRNPLL | 60.2647189 | -0.2425754 | 0.30010405 | -0.8083045 | 0.41891532 | 0.86022698 |
| GALM | 47.3940769 | 0.55943023 | 0.38155135 | 1.46619906 | 0.14259406 | 0.69328517 |
| SRSF7 | 69.4965063 | 0.70002465 | 0.35324736 | 1.981684 | 0.04751462 | 0.53530393 |
| DHX57 | 52.638825 | -0.1723465 | 0.27340648 | -0.6303674 | 0.52845425 | 0.90487731 |
| MORN2 | 14.4558591 | 0.1498492 | 0.436197 | 0.34353561 | 0.73119556 | NA |
| SOS1 | 310.428281 | 0.21528814 | 0.18235307 | 1.18061156 | 0.23775707 | 0.77930556 |
| MAP4K3 | 743.583159 | -0.0330286 | 0.21158631 | -0.1560999 | 8.76E-01 | 9.79E-01 |
| AC007388.1 | 366.865125 | 0.47449838 | 0.25167402 | 1.88536897 | 0.05938004 | 0.57442774 |
| TMEM178A | 177.832083 | 1.20362181 | 0.47643996 | 2.52628223 | 1.15E-02 | 3.53E-01 |
| THUMPD2 | 67.7116551 | 0.13688622 | 0.25282335 | 0.54143031 | 0.58821102 | 0.92175048 |
| SLC8A1-AS1 | 291.003153 | 1.08698842 | 0.46561118 | 2.33454108 | 1.96E-02 | 0.41544117 |
| SLC8A1 | 9040.06803 | 0.72462205 | 0.39394446 | 1.83940155 | 6.59E-02 | 5.95E-01 |
| EML4 | 267.1023 | -0.34254 | 0.31088456 | -1.1018236 | 0.27053835 | 0.79666376 |
| COX7A2L | 74.7567238 | -0.2808018 | 0.32153324 | -0.8733212 | 0.38248806 | 0.8548566 |
| MTA3 | 133.164781 | -0.1686717 | 0.1844117 | -0.9146476 | 0.36037668 | 0.84288514 |
| ZFP36L2 | 296.109709 | -0.1474963 | 0.31675742 | -0.4656443 | 6.41E-01 | 0.93328692 |
| THADA | 250.117897 | -0.5190856 | 0.17345935 | -2.9925493 | 0.00276658 | 0.21163175 |
| PLEKHH2 | 122.69501 | 0.3165639 | 0.34210949 | 0.92532921 | 0.35479469 | 0.84019149 |
| DYNC2LI1 | 100.825373 | 0.00410744 | 0.30850101 | 0.01331419 | 0.98937713 | 0.99814944 |
| LRPPRC | 149.694582 | -0.0047354 | 0.17012381 | -0.0278353 | 0.97779351 | 0.99469237 |
| PPM1B | 176.969118 | 0.15885058 | 0.18388954 | 0.86383697 | 0.38767746 | 0.85605713 |
| SLC3A1 | 139.878976 | 1.37250492 | 0.6027403 | 2.27710825 | 0.02277976 | 0.42667526 |
| PREPL | 181.591886 | 0.11956588 | 0.28452675 | 0.42022719 | 0.6743195 | 0.93930074 |
| CAMKMT | 551.657766 | -0.1727365 | 0.19858714 | -0.8698271 | 0.38439488 | 0.85511218 |
| SRBD1 | 109.539087 | -0.4270277 | 0.21158145 | -2.0182663 | 0.04356354 | 0.51770086 |
| PRKCE | 531.701446 | -0.1677574 | 0.18809376 | -0.8918818 | 0.37245627 | 0.8491102 |
| EPAS1 | 59.2839505 | 0.04846752 | 0.34699146 | 0.13967928 | 0.8889134 | 0.98270788 |
| RHOQ | 77.5742943 | -0.4608682 | 0.33423783 | -1.378863 | 0.16793699 | 0.72008028 |
| PIGF | 35.1923183 | 0.14258871 | 0.31131089 | 0.45802674 | 0.64693323 | 0.93328692 |
| CRIPT | 12.8267516 | 0.52905042 | 0.46413923 | 1.13985284 | 0.25434762 | NA |
| SOCS5 | 52.35474 | 0.1387407 | 0.2306426 | 0.60153977 | 0.54748053 | 0.91139843 |
| MCFD2 | 62.3550158 | -0.3219362 | 0.30859847 | -1.0432202 | 0.29684631 | 0.81137495 |
| TTC7A | 110.161799 | 0.06685928 | 0.29146578 | 0.22938982 | 8.19E-01 | 0.97288403 |
| AC106869.1 | 18.7417621 | 0.25183162 | 0.55189971 | 0.45629961 | 0.64817455 | 0.93359698 |

| | | | | | | |
|---|---|---|---|---|---|---|
| EPCAM | 230.557689 | 0.45510748 | 0.30802984 | 1.47747853 | 0.13954741 | 0.69054855 |
| MSH2 | 83.2171301 | 0.59637064 | 0.28618829 | 2.0838401 | 0.03717471 | 0.49537143 |
| MSH6 | 23.2136244 | 0.53399138 | 0.33767995 | 1.58135353 | 0.11379723 | 0.6656824 |
| FBXO11 | 260.50664 | -0.1701454 | 0.23224849 | -0.7326007 | 0.46380199 | 0.87824803 |
| FOXN2 | 27.962218 | 0.03524301 | 0.30856016 | 0.11421762 | 0.90906529 | 0.9854228 |
| PPP1R21 | 141.77943 | -0.204765 | 0.18873104 | -1.0849565 | 0.27794092 | 0.80193223 |
| GPR75-ASB3 | 43.2643492 | 0.45366746 | 0.40556617 | 1.11860281 | 0.26330962 | 0.79666376 |
| ERLEC1 | 31.7609196 | -0.0084392 | 0.34651588 | -0.0243543 | 0.98056997 | 0.99592962 |
| PSME4 | 180.440479 | -0.327071 | 0.29910144 | -1.0935121 | 0.27416904 | 0.79838796 |
| ACYP2 | 230.235303 | -0.2771042 | 0.30924234 | -0.8960747 | 0.37021287 | 0.84852785 |
| SPTBN1 | 644.101118 | -0.9623659 | 0.23159489 | -4.1553847 | 3.25E-05 | 0.0211151 |
| AC093110.1 | 21.5107609 | -0.2483632 | 0.5780151 | -0.4296829 | 6.67E-01 | 0.93878847 |
| EML6 | 134.961939 | 0.3271036 | 0.30659051 | 1.06690713 | 0.28601377 | 0.80431986 |
| RTN4 | 639.481614 | -0.2573149 | 0.35767361 | -0.7194126 | 0.47188675 | 0.88267439 |
| CLHC1 | 42.7560696 | 0.38527264 | 0.28665579 | 1.34402533 | 0.1789402 | 0.72924345 |
| RPS27A | 218.094067 | 0.15059277 | 0.45006765 | 0.33460029 | 0.73792663 | 0.9542764 |
| MTIF2 | 36.1086423 | 0.57602729 | 0.32480084 | 1.77347843 | 0.07614946 | 0.61051335 |
| AC012358.3 | 31.3697761 | 0.43232564 | 0.50752031 | 0.85183909 | 0.3943034 | 0.85605713 |
| CCDC88A | 35.4548783 | -0.2971319 | 0.32046431 | -0.9271917 | 0.353827 | 0.83979358 |
| CFAP36 | 65.6885689 | -0.0224171 | 0.25813142 | -0.0868438 | 0.93079571 | 0.98960607 |
| PPP4R3B | 132.851327 | -0.0792207 | 0.19859608 | -0.3989039 | 0.68996403 | 0.94252529 |
| PNPT1 | 61.9978737 | -1.0627815 | 0.4397147 | -2.4169796 | 0.01564989 | 0.39416332 |
| EFEMP1 | 52.1463064 | 0.3619799 | 0.60761301 | 0.59574086 | 5.51E-01 | 0.9128014 |
| VRK2 | 105.951557 | 0.02931187 | 0.20831333 | 0.14071048 | 0.88809867 | 0.9825428 |
| FANCL | 73.0188472 | -0.2496112 | 0.30478667 | -0.8189701 | 0.41280348 | 0.85867915 |
| PAPOLG | 39.3462038 | -0.135412 | 0.36724937 | -0.3687195 | 0.71233683 | 0.94731953 |
| REL | 53.4179011 | 0.26707982 | 0.34148331 | 0.78211678 | 0.43414595 | 0.86729771 |
| PUS10 | 21.7931998 | 0.82222209 | 0.44867154 | 1.83257019 | 0.06686651 | 0.59652811 |
| PEX13 | 36.5693053 | 0.0796519 | 0.29527139 | 0.26975826 | 0.78734624 | 0.96767512 |
| KIAA1841 | 21.4274327 | -0.2492811 | 0.40136088 | -0.6210896 | 0.53454065 | 0.90763121 |
| C2orf74 | 16.9474806 | -0.0266574 | 0.41620344 | -0.0640489 | 0.9489313 | 0.99105535 |
| USP34 | 570.264875 | -0.3920751 | 0.19825173 | -1.9776631 | 0.04796673 | 0.53707392 |
| AC016727.1 | 15.0462023 | 0.00213812 | 0.43663496 | 0.00489681 | 0.99609292 | NA |
| XPO1 | 239.534253 | -0.2911069 | 0.33874317 | -0.8593736 | 0.39013443 | 0.85605713 |
| FAM161A | 18.2306257 | 0.1588144 | 0.35837818 | 0.4431475 | 0.65765905 | 0.93593674 |
| CCT4 | 55.3496458 | -0.0362575 | 0.26377153 | -0.1374579 | 0.89066885 | 0.98296447 |
| COMMD1 | 221.941964 | -0.2707509 | 0.25894049 | -1.0456106 | 0.29574085 | 0.81054924 |
| AC018462.1 | 14.299221 | -0.924707 | 0.4652204 | -1.9876751 | 0.04684763 | NA |
| B3GNT2 | 46.4697502 | 0.18937596 | 0.28008782 | 0.67613067 | 0.49895768 | 0.89378207 |
| EHBP1 | 267.461173 | -0.2541107 | 0.22412438 | -1.1337931 | 0.25688137 | 0.79456034 |
| WDPCP | 223.982796 | 0.09848346 | 0.26922909 | 0.365798 | 0.71451582 | 0.94841799 |
| MDH1 | 75.2668558 | 0.59786997 | 0.33150672 | 1.80349278 | 0.07131086 | 0.60665346 |
| UGP2 | 116.733871 | -0.0164004 | 0.19232602 | -0.0852742 | 0.9320434 | 0.98960607 |
| VPS54 | 156.595417 | -0.0141938 | 0.20359429 | -0.0697159 | 0.9444198 | 0.99069421 |
| PELI1 | 33.6905465 | 0.28111942 | 0.4246059 | 0.66207141 | 0.50792546 | 0.89884243 |
| AC008074.1 | 15.1267669 | 1.25912298 | 0.49368642 | 2.55045091 | 0.01075837 | NA |
| AFTPH | 130.037299 | -0.3240161 | 0.24412174 | -1.3272726 | 0.18441854 | 0.73705812 |
| SERTAD2 | 41.9251118 | 0.06723647 | 0.37024272 | 0.18160107 | 0.85589581 | 0.97634735 |
| CEP68 | 23.7623703 | -0.4097853 | 0.35566981 | -1.1521508 | 0.24925911 | 0.78763828 |
| RAB1A | 232.267763 | -0.4801003 | 0.26459324 | -1.8144843 | 0.06960319 | 0.60275677 |
| ACTR2 | 154.858448 | -0.0908772 | 0.24587291 | -0.3696104 | 0.7116728 | 0.94676706 |
| SPRED2 | 162.968967 | -0.1744694 | 0.27851798 | -0.6264204 | 0.53103921 | 0.90525279 |
| ETAA1 | 21.5116083 | 0.07156124 | 0.35106487 | 0.20384051 | 0.83847813 | 0.97600352 |
| C1D | 32.432234 | 0.25644431 | 0.28180851 | 0.90999491 | 0.36282519 | 0.84543391 |

| | | | | | | |
|---|---|---|---|---|---|---|
| PNO1 | 13.4563261 | -0.0596499 | 0.43561386 | -0.136933 | 0.89108374 | NA |
| APLF | 56.9786732 | 0.09264799 | 0.31016742 | 0.29870317 | 0.76516654 | 0.96205953 |
| ANTXR1 | 31.2482457 | 0.21431607 | 0.37153685 | 0.57683664 | 0.56404981 | 0.91650701 |
| GFPT1 | 124.996884 | -0.3820725 | 0.24349296 | -1.5691317 | 1.17E-01 | 0.6668162 |
| NFU1 | 45.2134332 | -0.026202 | 0.25180989 | -0.1040546 | 0.91712605 | 0.98710975 |
| AAK1 | 459.631141 | -0.0602262 | 0.31787375 | -0.1894658 | 0.84972773 | 0.97612951 |
| ANXA4 | 271.441341 | -0.2903335 | 0.29891899 | -0.9712783 | 0.33140972 | 0.82890589 |
| GMCL1 | 60.7167377 | -0.2913683 | 0.26883008 | -1.0838383 | 0.27843649 | 0.80210301 |
| MXD1 | 35.5340866 | -0.4110188 | 0.47637133 | -0.8628118 | 0.38824095 | 0.85605713 |
| PCBP1 | 62.4276573 | 0.0335623 | 0.41017804 | 0.08182372 | 0.93478689 | 0.99012269 |
| C2orf42 | 22.8315982 | -0.1391929 | 0.41110587 | -0.3385816 | 0.73492497 | 0.95329896 |
| PCYOX1 | 60.8607771 | 0.87082051 | 0.51052563 | 1.70573319 | 0.08805777 | 0.63466198 |
| SNRPG | 29.5760592 | -0.2937816 | 0.33670009 | -0.8725321 | 0.38291816 | 0.85510589 |
| FAM136A | 26.0841707 | 0.08236258 | 0.33190427 | 0.24815161 | 0.8040171 | 0.97175786 |
| TGFA | 72.7900376 | 0.51384831 | 0.36659009 | 1.40169723 | 0.16100568 | 0.71101785 |
| ATP6V1B1 | 9.37461384 | 0.8565567 | 0.98836282 | 0.86664197 | 0.38613821 | NA |
| AC007040.2 | 52.9381135 | 1.3441323 | 0.77306468 | 1.73870613 | 0.08208647 | 0.61959627 |
| NAGK | 22.9025744 | 0.31510363 | 0.38530541 | 0.81780224 | 0.41347012 | 0.85867915 |
| MCEE | 38.5236767 | 0.32671325 | 0.36619989 | 0.89217188 | 0.37230081 | 0.8491102 |
| MPHOSPH10 | 23.7891766 | -0.0512053 | 0.41363501 | -0.1237933 | 0.90147891 | 0.98531749 |
| PAIP2B | 15.5196629 | 0.27215493 | 0.48952038 | 0.55596241 | 0.57823655 | NA |
| ZNF638 | 476.467264 | -0.0012789 | 0.1783654 | -0.0071703 | 0.994279 | 0.99921857 |
| EXOC6B | 541.447551 | -0.6979663 | 0.30047432 | -2.3228816 | 0.02018551 | 0.41761076 |
| SPR | 19.1243192 | 1.00980699 | 0.47187091 | 2.1400069 | 0.03235421 | 0.48175799 |
| EMX1 | 243.838265 | 0.01000756 | 0.41640029 | 0.02403351 | 9.81E-01 | 0.99592962 |
| SFXN5 | 199.419217 | -0.2735218 | 0.25554177 | -1.0703605 | 0.28445709 | 0.80431986 |
| RAB11FIP5 | 41.2771443 | -0.347312 | 0.3313606 | -1.0481391 | 0.29457453 | 0.81012446 |
| SMYD5 | 9.71955169 | -0.2993217 | 0.50362084 | -0.5943395 | 0.55228507 | NA |
| CCT7 | 47.7693968 | 0.56700249 | 0.34874508 | 1.62583653 | 0.10398443 | 0.65946648 |
| ALMS1 | 97.1191265 | -0.0936738 | 0.20487026 | -0.4572348 | 0.64750226 | 0.93328692 |
| NAT8 | 10.2180889 | -0.4588558 | 1.07220004 | -0.4279573 | 0.6686822 | NA |
| DUSP11 | 17.2612289 | -0.0430079 | 0.44930074 | -0.0957219 | 0.92374144 | 0.98925753 |
| STAMBP | 54.4471501 | -0.4144619 | 0.27061864 | -1.5315349 | 0.12563724 | 0.66929384 |
| DGUOK | 50.5223951 | -0.5116303 | 0.35334877 | -1.4479469 | 0.1476319 | 0.69842926 |
| TET3 | 47.1080682 | -0.2246141 | 0.34418627 | -0.6525947 | 0.51401763 | 0.90156229 |
| BOLA3 | 16.6717642 | 0.06999462 | 0.62360058 | 0.11224272 | 0.91063096 | 0.98566868 |
| MOB1A | 78.8165967 | 0.32009403 | 0.24332891 | 1.31547881 | 0.18834903 | 0.74222565 |
| AC006030.1 | 24.4881636 | -0.3907557 | 0.44963746 | -0.8690461 | 0.38482192 | 0.85531718 |
| DCTN1 | 10.8972356 | 0.43501969 | 0.55840088 | 0.7790455 | 0.43595291 | NA |
| RTKN | 30.9032085 | -0.4310776 | 0.34781216 | -1.2393976 | 0.21519829 | 0.7643572 |
| MOGS | 23.4097925 | 0.22210033 | 0.30967371 | 0.71720756 | 0.47324604 | 0.88359968 |
| CCDC142 | 11.4097881 | 0.41149914 | 0.50337613 | 0.81747845 | 0.41365506 | NA |
| TTC31 | 13.2988958 | -0.6177358 | 0.44750263 | -1.3804071 | 0.16746133 | NA |
| AUP1 | 18.1924239 | -0.2773708 | 0.38159513 | -0.7268719 | 0.46730443 | 0.88090126 |
| POLE4 | 26.380703 | -0.1314293 | 0.35096365 | -0.3744813 | 0.70804629 | 0.94558664 |
| TACR1 | 24.6457913 | 0.28439562 | 0.59806919 | 0.47552294 | 0.63441429 | 0.93251095 |
| AC007681.1 | 23.0826734 | -0.0906261 | 0.57159967 | -0.1585482 | 0.87402481 | 0.97918624 |
| EVA1A | 52.2358154 | -0.7622344 | 0.39274559 | -1.9407841 | 0.05228447 | 0.55403388 |
| MRPL19 | 77.6107097 | 0.41303919 | 0.27622153 | 1.49531859 | 0.13483132 | 0.68328989 |
| GCFC2 | 98.5293185 | 0.57289983 | 0.3005203 | 1.90635982 | 0.05660354 | 0.56976762 |
| CTNNA2 | 11.9850856 | -0.2889845 | 0.62199424 | -0.4646097 | 0.64221099 | NA |
| SUCLG1 | 92.0533932 | 0.01110785 | 0.29089771 | 0.03818473 | 0.96954039 | 0.99398482 |
| DNAH6 | 36.1964534 | 0.15219852 | 0.4587442 | 0.33177209 | 0.74006136 | 0.9544376 |
| TMSB10 | 221.341937 | -0.6258188 | 0.4375847 | -1.4301662 | 1.53E-01 | 0.70572801 |

| | | | | | | |
|---|---|---|---|---|---|---|
| KCMF1 | 130.911593 | -0.072541 | 0.24801187 | -0.29249 | 0.76991199 | 0.96310415 |
| TCF7L1 | 57.1907491 | -0.2610908 | 0.68674117 | -0.380188 | 0.70380586 | 0.94476609 |
| TGOLN2 | 166.384003 | 0.58826969 | 0.29847644 | 1.97090826 | 0.04873437 | 0.53969461 |
| RETSAT | 16.9519465 | 0.64364007 | 0.39694478 | 1.62148515 | 0.10491363 | 0.65946648 |
| ELMOD3 | 49.2234664 | 0.31791919 | 0.25095414 | 1.26684178 | 0.20521187 | 0.76231289 |
| CAPG | 40.0943692 | 0.35504278 | 0.4317855 | 0.82226655 | 0.41092521 | 0.85867915 |
| MAT2A | 70.5577213 | -0.3785713 | 0.28129025 | -1.3458388 | 0.17835449 | 0.72841249 |
| GGCX | 25.4108431 | 0.18287497 | 0.3111419 | 0.58775423 | 0.55669727 | 0.9141017 |
| VAMP8 | 65.9888586 | 0.25950297 | 0.37176459 | 0.69803036 | 0.48515821 | 0.88838332 |
| RNF181 | 11.3402082 | -0.4679838 | 0.44495329 | -1.0517595 | 0.29290991 | NA |
| USP39 | 52.0091923 | -0.1395355 | 0.23844409 | -0.5851919 | 0.55841871 | 0.91425998 |
| C2orf68 | 25.0025795 | 0.19259247 | 0.36683349 | 0.52501333 | 0.59957392 | 0.92298075 |
| POLR1A | 36.712359 | -0.1308101 | 0.28926892 | -0.4522095 | 0.65111808 | 0.93453685 |
| PTCD3 | 83.7791071 | -0.0403934 | 0.21713693 | -0.1860274 | 0.85242328 | 0.97612951 |
| IMMT | 117.251542 | 0.12720924 | 0.22832179 | 0.55714894 | 0.57742567 | 0.91859435 |
| MRPL35 | 18.9547322 | 0.79906356 | 0.38320605 | 2.085206 | 0.03705061 | 0.49537143 |
| KDM3A | 83.0738349 | -0.2054155 | 0.30289122 | -0.6781825 | 0.49765602 | 0.89378207 |
| CHMP3 | 32.831445 | 0.0909344 | 0.47737032 | 0.19049026 | 0.84892497 | 0.97612951 |
| RMND5A | 55.922439 | 0.06151133 | 0.31110698 | 0.19771761 | 0.84326601 | 0.97612951 |
| PLGLB1 | 14.9108799 | 0.57672361 | 0.60566054 | 0.95222253 | 0.34098414 | NA |
| CYTOR | 56.0351348 | -0.4006574 | 0.57556496 | -0.6961115 | 0.48635901 | 0.88838332 |
| KRCC1 | 60.489218 | 0.35870105 | 0.24693219 | 1.45262978 | 0.14632657 | 0.69713616 |
| THNSL2 | 16.8352264 | 1.62076243 | 0.95805191 | 1.69172715 | 0.09069801 | 0.64151047 |
| EIF2AK3 | 108.91452 | -0.7037423 | 0.4968481 | -1.4164133 | 0.15665452 | 0.70821492 |
| RPIA | 28.6696487 | 0.210928 | 0.3233869 | 0.65224658 | 0.51424211 | 0.90156229 |
| MAL | 376.798015 | 0.36600598 | 0.51214253 | 0.71465647 | 0.47482134 | 0.88444826 |
| MRPS5 | 96.4372518 | -0.2493625 | 0.21073813 | -1.1832816 | 0.23669758 | 0.77864547 |
| ZNF514 | 10.4570182 | -0.3403124 | 0.52870479 | -0.643672 | 0.51978818 | NA |
| AC092835.1 | 16.2943087 | 0.7163542 | 0.43242705 | 1.65658972 | 0.09760245 | 0.65521765 |
| PROM2 | 60.0488684 | 0.36825258 | 0.51608741 | 0.71354691 | 0.4755074 | 0.88495683 |
| KCNIP3 | 13.296345 | 0.63282179 | 0.49753082 | 1.27192482 | 0.20339983 | NA |
| FAHD2A | 15.9745253 | 0.15192928 | 0.43962949 | 0.34558483 | 0.72965475 | NA |
| ANKRD36C | 37.1566667 | 0.32271327 | 0.46472637 | 0.69441567 | 0.48742157 | 0.88838332 |
| STARD7 | 67.4406983 | -0.3630743 | 0.23317654 | -1.5570791 | 0.11945171 | 0.6668162 |
| TMEM127 | 43.1801813 | -0.0106615 | 0.29655366 | -0.0359512 | 0.97132127 | 0.99403127 |
| CIAO1 | 35.908706 | 0.22642906 | 0.26619061 | 0.85062753 | 0.39497629 | 0.85605713 |
| SNRNP200 | 66.219293 | -0.1204439 | 0.27439901 | -0.438937 | 0.66070721 | 0.93729379 |
| NEURL3 | 17.2024334 | -0.000796 | 0.71966974 | -0.0011061 | 0.99911745 | 0.99957534 |
| KANSL3 | 105.644885 | -0.3051339 | 0.20351601 | -1.4993116 | 0.13379282 | 0.68154225 |
| FER1L5 | 18.8243733 | 0.07864699 | 0.33646044 | 0.23374809 | 0.81518055 | 0.97284589 |
| LMAN2L | 33.5283864 | -0.2925348 | 0.28093159 | -1.0413024 | 0.29773521 | 0.81214575 |
| CNNM4 | 24.9202371 | -0.4440632 | 0.31837796 | -1.3947675 | 0.16308595 | 0.71325312 |
| ANKRD23 | 18.5427434 | 0.54026752 | 0.42517841 | 1.27068427 | 0.20384099 | 0.76173533 |
| ANKRD36 | 57.1664912 | 0.19026208 | 0.34853026 | 0.54589832 | 0.58513583 | 0.92076087 |
| AC092683.1 | 55.9458706 | 0.5170037 | 0.43553907 | 1.18704321 | 0.23521058 | 0.77830675 |
| ANKRD36B | 16.4937536 | 0.16294212 | 0.66513533 | 0.24497589 | 0.8064751 | 0.97216031 |
| COX5B | 109.348112 | 0.33414292 | 0.43815162 | 0.76261938 | 0.44569042 | 0.8718292 |
| ACTR1B | 27.5362291 | 0.46352712 | 0.38557798 | 1.20216181 | 0.22930084 | 0.77185154 |
| TMEM131 | 265.556823 | -0.1594154 | 0.20652203 | -0.7719053 | 0.44017052 | 0.86895963 |
| INPP4A | 52.2105891 | -0.4306502 | 0.27035473 | -1.5929079 | 0.11118085 | 0.66409404 |
| COA5 | 27.2611976 | 0.33514277 | 0.31299832 | 1.07074941 | 0.28428212 | 0.80431986 |
| UNC50 | 30.0038069 | 0.0452231 | 0.2809051 | 0.16099067 | 0.87210075 | 0.97866716 |
| MGAT4A | 118.724115 | 0.20561243 | 0.52554976 | 0.39123304 | 0.69562499 | 0.94395238 |
| TSGA10 | 132.885619 | 0.40625213 | 0.33179641 | 1.22440182 | 0.22080068 | 0.76680432 |

| | | | | | | |
|---|---|---|---|---|---|---|
| AC079447.1 | 56.3130487 | 0.11407283 | 0.28342716 | 0.40247669 | 0.68733324 | 0.94205398 |
| MITD1 | 39.2703272 | 0.22128476 | 0.31329353 | 0.70631767 | 0.47999061 | 0.88660781 |
| TXNDC9 | 26.9050482 | -0.1518612 | 0.28113007 | -0.5401812 | 0.58907208 | 0.92197072 |
| EIF5B | 115.027715 | -0.3748301 | 0.22702444 | -1.6510562 | 0.0987271 | 0.65744285 |
| REV1 | 189.4834 | 0.06109712 | 0.21892931 | 0.27907236 | 0.78018929 | 0.96650724 |
| LONRF2 | 33.2093079 | 1.05166809 | 0.38818389 | 2.70920079 | 0.00674455 | 0.30102597 |
| NPAS2 | 282.213915 | -0.5063969 | 0.25820809 | -1.9611969 | 0.04985606 | 0.54357315 |
| RPL31 | 74.0388561 | 0.39061202 | 0.25187775 | 1.55080004 | 0.12094961 | 0.66727534 |
| TBC1D8 | 250.752093 | -0.3208894 | 0.25642829 | -1.2513806 | 0.21079564 | 0.76397692 |
| CNOT11 | 54.7936388 | -0.6011048 | 0.2651931 | -2.2666683 | 0.0234105 | 0.43088993 |
| RNF149 | 254.505173 | -0.8492286 | 0.18663187 | -4.550287 | 5.36E-06 | 0.00541859 |
| MAP4K4 | 121.832954 | -0.3252045 | 0.35995143 | -0.9034677 | 0.36627773 | 0.84741115 |
| IL1R1 | 86.5878648 | -0.7588299 | 0.36324911 | -2.0890069 | 0.0367071 | 0.49537143 |
| IL1RL2 | 12.8591235 | -1.353781 | 0.6677272 | -2.0274462 | 0.0426168 | NA |
| SLC9A4 | 90.3677744 | 0.17374796 | 0.62315524 | 0.27881971 | 7.80E-01 | 9.67E-01 |
| SLC9A2 | 85.6632318 | 0.63492062 | 0.40863992 | 1.55374106 | 0.12024619 | 0.6668162 |
| MFSD9 | 12.7197004 | -0.3431286 | 0.42018128 | -0.8166204 | 0.41414538 | NA |
| TMEM182 | 31.3107046 | -0.1444276 | 0.35586948 | -0.4058444 | 0.68485693 | 0.94205398 |
| POU3F3 | 126.814961 | -0.3706645 | 0.30409991 | -1.2188904 | 0.22288579 | 0.76834258 |
| MRPS9 | 67.5643101 | 0.23500486 | 0.23664357 | 0.99307521 | 0.3206733 | 0.82464962 |
| TGFBRAP1 | 24.9301689 | 0.46935314 | 0.30839032 | 1.52194512 | 0.12802283 | 0.66929384 |
| C2orf49 | 22.6122624 | -0.0450386 | 0.39834486 | -0.1130642 | 0.90997964 | 0.98556715 |
| NCK2 | 95.3834241 | 0.03803949 | 0.22815537 | 0.16672625 | 0.86758545 | 0.97805828 |
| UXS1 | 94.4288995 | -0.1704596 | 0.23983777 | -0.7107287 | 0.47725237 | 0.88512746 |
| LINC01594 | 29.2021002 | 0.1405847 | 0.63179304 | 0.22251702 | 0.82391143 | 0.97518529 |
| SULT1C2 | 310.105231 | -0.0198888 | 0.3190794 | -0.0623319 | 0.95029853 | 0.99105535 |
| SULT1C4 | 10.9058872 | 1.54868356 | 0.66528339 | 2.32785545 | 0.01991978 | NA |
| GCC2 | 255.701112 | 0.13478189 | 0.19656047 | 0.68570193 | 0.49290109 | 0.89092098 |
| GCC2-AS1 | 29.8888227 | 0.73438904 | 0.47412406 | 1.54893859 | 0.12139648 | 0.66771734 |
| LIMS1 | 290.374751 | -0.5721424 | 0.29494774 | -1.9398094 | 0.05240285 | 0.55403388 |
| RANBP2 | 222.563732 | -0.1554093 | 0.66850381 | -0.2324733 | 0.81617044 | 0.97288403 |
| CCDC138 | 46.7423985 | -0.4353378 | 0.36430629 | -1.1949775 | 0.23209582 | 0.77390779 |
| 10-Sep | 206.15182 | -1.2723804 | 0.42565366 | -2.9892386 | 0.00279674 | 0.21215575 |
| SOWAHC | 29.7076456 | -1.5311801 | 0.59471991 | -2.5746238 | 0.01003492 | 0.33306084 |
| NPHP1 | 47.9683976 | 0.57777726 | 0.40134152 | 1.43961496 | 0.14997637 | 0.70336675 |
| SMIM37 | 33.6073591 | 0.43856146 | 0.41775821 | 1.04979733 | 0.2938113 | 0.8097379 |
| LINC01106 | 11.6720704 | -1.1038345 | 0.4777294 | -2.3105853 | 0.02085577 | NA |
| ACOXL | 14.3229332 | 1.63487865 | 0.70117514 | 2.33162665 | 0.01972034 | NA |
| MIR4435-2H | 316.006461 | -0.3439572 | 0.33146202 | -1.0376973 | 0.29941099 | 0.81335071 |
| BCL2L11 | 29.8145085 | 0.13453673 | 0.3431986 | 0.39200839 | 0.69505201 | 0.94395238 |
| AC017002.5 | 23.0851891 | -0.7853667 | 0.88643793 | -0.8859805 | 0.37562802 | 0.85178063 |
| ANAPC1 | 88.7456367 | -0.0202283 | 0.25878644 | -0.0781658 | 0.93769615 | 0.99012269 |
| MERTK | 45.2177231 | 0.65557175 | 0.36579981 | 1.79215991 | 0.07310735 | 0.60952146 |
| TMEM87B | 79.2641761 | -0.252141 | 0.2129698 | -1.1839284 | 0.23644141 | 0.77864547 |
| ZC3H8 | 55.1991465 | -0.0440683 | 0.26262549 | -0.1677991 | 0.86674133 | 0.97788853 |
| ZC3H6 | 86.1863026 | 0.6309282 | 0.42101022 | 1.49860543 | 0.13397602 | 0.68205729 |
| TTL | 33.6884948 | 0.10918807 | 0.31889834 | 0.34239145 | 0.73205632 | 0.95279138 |
| POLR1B | 18.6987246 | -0.123668 | 0.35886708 | -0.3446066 | 0.73039015 | 0.95279138 |
| CHCHD5 | 13.3654876 | 0.25403684 | 0.54544032 | 0.46574636 | 0.64139707 | NA |
| SLC20A1 | 99.7023409 | -1.2202704 | 0.52382889 | -2.3295209 | 0.01983148 | 0.41544117 |
| PSD4 | 33.9861325 | -0.6185749 | 0.39212024 | -1.5775133 | 0.11467748 | 0.6668162 |
| PAX8 | 1363.01732 | -0.0658431 | 0.31679748 | -0.2078397 | 0.83535411 | 0.97600352 |
| CBWD2 | 32.8524381 | -0.4800248 | 0.30259077 | -1.5863829 | 0.11265248 | 0.6656824 |
| RABL2A | 24.9243782 | 0.06012 | 0.3609442 | 0.16656314 | 0.8677138 | 0.97805828 |

| | | | | | | |
|---|---|---|---|---|---|---|
| AC017074.1 | 23.3407015 | 0.32983777 | 0.474437 | 0.69521931 | 0.48691787 | 0.88838332 |
| SLC35F5 | 338.467823 | -0.3779518 | 0.14372963 | -2.6296025 | 0.00854848 | 0.31119527 |
| ACTR3 | 202.040851 | -0.1251856 | 0.23714022 | -0.5278972 | 0.59757072 | 0.9229638 |
| DDX18 | 45.2158815 | -0.0385066 | 0.24431146 | -0.1576127 | 0.87476201 | 0.97918624 |
| CCDC93 | 103.386952 | -0.5297897 | 0.20071644 | -2.6394931 | 0.00830301 | 0.30976355 |
| INSIG2 | 79.9753853 | -1.0431905 | 0.49245797 | -2.1183341 | 0.03414678 | 0.4908133 |
| C2orf76 | 20.7419868 | 0.31302646 | 0.34837645 | 0.89852933 | 0.36890341 | 0.84844057 |
| DBI | 116.877356 | 0.02884121 | 0.42561962 | 0.06776287 | 0.9459744 | 0.99083449 |
| TMEM37 | 39.3844968 | 0.35678034 | 0.55135559 | 0.64709662 | 0.51756941 | 0.90320951 |
| SCTR | 25.8846609 | 2.04116875 | 0.85482092 | 2.387832 | 0.01694809 | 0.39865229 |
| CFAP221 | 175.84298 | 0.74561479 | 0.3835167 | 1.94415208 | 0.05187712 | 0.55403388 |
| PTPN4 | 198.650888 | 0.05803896 | 0.19805068 | 0.29305103 | 7.69E-01 | 9.63E-01 |
| EPB41L5 | 325.106842 | -0.214211 | 0.17642019 | -1.2142089 | 0.22466795 | 0.76931755 |
| RALB | 28.9483522 | 0.06108048 | 0.3189357 | 0.19151345 | 0.84812334 | 0.97612951 |
| TFCP2L1 | 618.51039 | 0.18172996 | 0.29973465 | 0.60630282 | 5.44E-01 | 9.10E-01 |
| CLASP1 | 503.606172 | -0.3067427 | 0.25478551 | -1.2039253 | 0.22861844 | 0.77185154 |
| NIFK-AS1 | 24.5377057 | -0.1233652 | 0.47713964 | -0.2585515 | 0.79598133 | 0.96974891 |
| NIFK | 12.8664396 | -0.268351 | 0.50375847 | -0.5326977 | 0.59424285 | NA |
| TSN | 29.1685806 | 0.28101307 | 0.28507538 | 0.98575005 | 0.32425578 | 0.82567895 |
| BIN1 | 17.7650546 | 0.34056321 | 0.45772708 | 0.74403116 | 0.45685762 | 0.87405946 |
| ERCC3 | 35.6470793 | -0.1226231 | 0.28149092 | -0.4356202 | 0.66311234 | 0.93848128 |
| MAP3K2 | 206.467904 | -0.2232449 | 0.19597947 | -1.1391237 | 0.25465156 | 0.79251048 |
| IWS1 | 144.297526 | -0.4119912 | 0.22125775 | -1.8620419 | 0.06259718 | 0.5842021 |
| AC010976.1 | 15.5941322 | 1.02252477 | 0.73779806 | 1.38591416 | 0.16577312 | NA |
| WDR33 | 166.76282 | -0.1265091 | 0.21185526 | -0.5971489 | 0.55040795 | 0.91248567 |
| POLR2D | 14.5936637 | 0.1285882 | 0.37327041 | 0.34449075 | 0.73047725 | NA |
| AMMECR1L | 20.448653 | 0.15458633 | 0.34411552 | 0.44922801 | 0.65326719 | 0.93532021 |
| SAP130 | 54.8128938 | -0.4074783 | 0.2659124 | -1.532378 | 0.12542918 | 0.66929384 |
| UGGT1 | 58.7132676 | -0.1231363 | 0.30990478 | -0.3973358 | 0.69111982 | 0.94293839 |
| HS6ST1 | 226.182644 | 0.84498141 | 0.38203659 | 2.21178139 | 0.02698177 | 0.45084589 |
| SMPD4 | 39.0863832 | -0.0463534 | 0.32451412 | -0.1428393 | 0.88641711 | 0.98216235 |
| MZT2B | 50.4149554 | 0.12769156 | 0.37923685 | 0.33670663 | 0.73633807 | 0.95376039 |
| CCDC115 | 17.5231185 | 0.46931009 | 0.49533544 | 0.94745915 | 0.34340486 | 0.83449398 |
| IMP4 | 13.0173656 | 0.56007317 | 0.44426167 | 1.26068309 | 0.20742305 | NA |
| PTPN18 | 37.4057405 | 0.14529801 | 0.32102545 | 0.45260589 | 0.65083256 | 0.93453685 |
| FAM168B | 107.124666 | -0.2350594 | 0.26852592 | -0.8753694 | 0.38137297 | 0.85432452 |
| PLEKHB2 | 127.944673 | -0.1750514 | 0.23235212 | -0.7533885 | 0.45121649 | 0.87346041 |
| MZT2A | 34.4447664 | 0.14264623 | 0.35338202 | 0.40366012 | 0.68646267 | 0.94205398 |
| ANKRD30BL | 9.9390733 | -0.4740626 | 0.49739683 | -0.9530873 | 0.34054582 | NA |
| GPR39 | 119.141662 | -0.0191972 | 0.31292362 | -0.0613479 | 0.95108214 | 0.99105535 |
| NCKAP5 | 80.1230948 | 0.13438516 | 0.46175753 | 0.29102971 | 7.71E-01 | 0.96364125 |
| MGAT5 | 101.04974 | -0.2896451 | 0.22798721 | -1.2704447 | 0.20392627 | 0.76173533 |
| CCNT2-AS1 | 32.5286655 | -1.5413934 | 0.50801233 | -3.0341653 | 0.00241202 | 0.19505808 |
| ACMSD | 15.8789514 | -1.2917806 | 0.73163587 | -1.7656059 | 0.07746199 | NA |
| CCNT2 | 117.751174 | -0.3401791 | 0.244956 | -1.3887355 | 0.1649132 | 0.71485946 |
| RAB3GAP1 | 193.655048 | -0.1585869 | 0.17701655 | -0.8958875 | 0.37031283 | 0.84852785 |
| ZRANB3 | 98.2028317 | 0.04912074 | 0.26347581 | 0.18643357 | 0.85210477 | 0.97612951 |
| R3HDM1 | 172.987093 | 0.18676255 | 0.17584422 | 1.0620909 | 0.28819442 | 0.80523851 |
| UBXN4 | 166.01968 | -0.0817124 | 0.21305338 | -0.3835301 | 0.70132678 | 0.94440669 |
| DARS | 143.239204 | -0.3383214 | 0.20922621 | -1.6170126 | 0.10587557 | 0.65946648 |
| THSD7B | 31.4981711 | 0.88622718 | 0.8059672 | 1.09958219 | 0.27151421 | 0.79666376 |
| HNMT | 64.0615373 | 0.36022723 | 0.36938861 | 0.97519851 | 0.3294618 | 0.82831342 |
| AC114763.1 | 26.4377217 | -0.1105105 | 0.4809398 | -0.2297803 | 0.8182625 | 0.97288403 |
| SPOPL | 72.8002776 | -0.0913864 | 0.22331965 | -0.4092178 | 0.68237984 | 0.94205398 |

| | | | | | | |
|---|---|---|---|---|---|---|
| LRP1B | 1009.09209 | 0.09736035 | 0.49862674 | 0.19525697 | 0.84519178 | 0.97612951 |
| GTDC1 | 97.8487232 | -0.2097064 | 0.23353141 | -0.8979796 | 0.36919643 | 0.84848455 |
| TEX41 | 1104.98071 | -0.3409735 | 0.30231693 | -1.1278677 | 2.59E-01 | 7.95E-01 |
| ACVR2A | 93.2040628 | -0.0643454 | 0.35270573 | -0.1824336 | 0.85524249 | 0.97612951 |
| ORC4 | 144.626304 | -0.2019038 | 0.30999578 | -0.6513113 | 0.51484554 | 0.90156229 |
| MBD5 | 767.661352 | -0.1382386 | 0.1350357 | -1.0237186 | 0.30596822 | 0.81568231 |
| EPC2 | 173.072655 | -0.2206146 | 0.2709035 | -0.814366 | 0.41543531 | 0.85867915 |
| AC105402.3 | 15.0551055 | -2.0215972 | 1.57600849 | -1.2827324 | 0.19958583 | NA |
| LYPD6B | 58.6316751 | -0.2108302 | 0.58508553 | -0.3603408 | 0.71859231 | 0.94925929 |
| MMADHC | 24.4253757 | 0.46432345 | 0.30378664 | 1.52845248 | 0.12640023 | 0.66929384 |
| NMI | 21.184089 | -0.262611 | 0.74457669 | -0.3526984 | 0.72431456 | 0.95093722 |
| RIF1 | 127.452981 | -0.520525 | 0.25743984 | -2.0219287 | 0.04318372 | 0.51638379 |
| NEB | 11.5761134 | 0.40566404 | 0.45365445 | 0.89421374 | 0.37120754 | NA |
| ARL5A | 37.8845529 | 0.39306673 | 0.33413899 | 1.17635696 | 0.23945226 | 0.78098673 |
| CACNB4 | 310.214779 | -0.1588084 | 0.23185016 | -0.6849612 | 4.93E-01 | 8.91E-01 |
| STAM2 | 116.270596 | -0.3531507 | 0.22343622 | -1.5805439 | 0.11398237 | 0.66597015 |
| FMNL2 | 388.46487 | -0.5468112 | 0.24454154 | -2.2360668 | 2.53E-02 | 4.40E-01 |
| PRPF40A | 115.098546 | -0.1805026 | 0.17584363 | -1.0264947 | 0.30465848 | 0.81568231 |
| ARL6IP6 | 47.8295118 | -0.183124 | 0.25890258 | -0.7073087 | 0.47937465 | 0.88622003 |
| KCNJ3 | 317.405947 | 0.36915372 | 0.33257084 | 1.11000026 | 2.67E-01 | 7.97E-01 |
| LINC01876 | 94.3419194 | 0.14073561 | 1.1200982 | 0.12564578 | 0.90001232 | 0.98531749 |
| GPD2 | 106.515872 | -0.1586984 | 0.25691863 | -0.6176992 | 0.53677367 | 0.90763121 |
| ACVR1 | 110.215396 | -0.3467541 | 0.28391906 | -1.2213132 | 0.22196747 | 0.76680432 |
| UPP2 | 36.360216 | 1.66084283 | 0.62230914 | 2.66883886 | 0.0076114 | 0.30654023 |
| CCDC148 | 170.434364 | 0.05965924 | 0.4836397 | 0.12335471 | 0.90182622 | 0.98531749 |
| PKP4 | 632.784761 | -0.4672409 | 0.24353028 | -1.9186154 | 0.05503303 | 0.5645224 |
| PKP4-AS1 | 17.9913143 | 0.4816101 | 0.73670816 | 0.65373254 | 0.51328413 | 0.90156229 |
| TANC1 | 304.194833 | -0.2235644 | 0.17177269 | -1.3015128 | 0.19308299 | 0.75061775 |
| WDSUB1 | 88.7126946 | -0.2330464 | 0.28017823 | -0.8317791 | 0.40553365 | 0.8583057 |
| BAZ2B | 584.91728 | -0.026685 | 0.15011981 | -0.1777581 | 0.85891296 | 0.97691173 |
| AC009961.1 | 38.6045349 | 0.1302232 | 0.46625417 | 0.27929658 | 0.78001723 | 0.96650724 |
| 7-Mar | 199.766347 | -0.145488 | 0.26706358 | -0.5447693 | 0.58591217 | 0.92076087 |
| PLA2R1 | 34.6477371 | 0.33516888 | 0.27396848 | 1.22338483 | 0.22118438 | 0.76680432 |
| ITGB6 | 462.059327 | -0.3363466 | 0.65573829 | -0.5129282 | 0.60800158 | 0.92573951 |
| RBMS1 | 485.52909 | -0.6762142 | 0.25557046 | -2.6459011 | 0.00814736 | 0.30976355 |
| TANK | 159.497534 | -0.2356943 | 0.1621112 | -1.453905 | 0.14597264 | 0.69679548 |
| PSMD14 | 68.7959442 | -0.4873893 | 0.30354825 | -1.6056402 | 0.10835298 | 0.65986731 |
| DPP4 | 16.1913273 | 0.12604677 | 0.54595948 | 0.23087203 | 8.17E-01 | 9.73E-01 |
| IFIH1 | 35.1779711 | -0.8689665 | 0.83539559 | -1.0401856 | 0.29825368 | 0.81214575 |
| GCA | 40.7660516 | -0.1685972 | 0.39301711 | -0.4289819 | 0.66793642 | 0.93878847 |
| KCNH7 | 63.7422168 | 1.19439681 | 0.76523451 | 1.56082455 | 0.11856515 | 0.6668162 |
| FIGN | 19.7642315 | 1.10348416 | 0.6686408 | 1.65033925 | 0.09887357 | 0.65744285 |
| GRB14 | 214.70462 | 0.7086945 | 0.29648604 | 2.39031327 | 0.01683401 | 0.39748373 |
| COBLL1 | 3251.57532 | -0.3564896 | 0.26713713 | -1.3344816 | 1.82E-01 | 0.73398872 |
| AC019197.1 | 8368.61065 | 0.50124309 | 0.23828555 | 2.1035396 | 0.03541863 | 0.49209205 |
| SLC38A11 | 165.988872 | 0.70318763 | 0.49363479 | 1.42450987 | 0.15429893 | 0.70572801 |
| SCN3A | 296.61585 | 1.40368331 | 0.68207812 | 2.05795094 | 0.03959484 | 0.50156625 |
| SCN2A | 1187.37198 | 0.72263491 | 0.23210977 | 3.11333257 | 0.00184987 | 0.17725691 |
| CSRNP3 | 98.3516113 | 0.76481027 | 0.42348906 | 1.80597407 | 0.07092239 | 0.60572821 |
| GALNT3 | 53.0693665 | -0.1460897 | 0.33734705 | -0.4330546 | 0.66497512 | 0.93860542 |
| TTC21B | 60.1875865 | 0.38904342 | 0.31239191 | 1.24536969 | 0.21299589 | 0.7643572 |
| SCN7A | 36.1948615 | 0.75939267 | 0.5547562 | 1.36887641 | 1.71E-01 | 0.72463203 |
| STK39 | 401.145007 | -0.3831867 | 0.24852572 | -1.5418391 | 0.1231127 | 0.66929384 |
| CERS6 | 37.2734364 | 1.06086979 | 0.47286413 | 2.24349811 | 0.02486471 | 0.43948277 |

| | | | | | | |
|---|---|---|---|---|---|---|
| LRP2 | 42.2537188 | 1.68010171 | 0.47554645 | 3.53299183 | 0.00041089 | 0.07958059 |
| FASTKD1 | 96.0708973 | 0.06235588 | 0.27110169 | 0.23000919 | 8.18E-01 | 0.97288403 |
| PPIG | 134.076798 | -0.198826 | 0.2263905 | -0.8782435 | 0.37981162 | 0.85431805 |
| KLHL23 | 33.4717142 | -0.0136003 | 0.36688073 | -0.0370701 | 0.97042908 | 0.99401551 |
| SSB | 41.3832827 | -0.18799 | 0.41750829 | -0.4502666 | 0.65251823 | 0.93532021 |
| METTL5 | 23.9079186 | -0.4622224 | 0.36708002 | -1.2591871 | 0.20796274 | 0.76231289 |
| UBR3 | 281.252687 | -0.0479421 | 0.20142815 | -0.2380108 | 0.8118727 | 0.97257655 |
| MYO3B | 237.883714 | 0.86438843 | 0.42077054 | 2.05429882 | 0.03994678 | 0.50156625 |
| GORASP2 | 55.1613256 | 0.00226183 | 0.290155 | 0.00779526 | 0.99378034 | 0.99921857 |
| TLK1 | 362.997987 | -0.2020753 | 0.22100755 | -0.9143368 | 0.36053992 | 0.84305031 |
| METTL8 | 51.1746216 | -0.0472245 | 0.2918118 | -0.1618321 | 0.87143805 | 0.97860207 |
| DCAF17 | 32.5538393 | 0.06793472 | 0.35068126 | 0.19372212 | 0.84639347 | 0.97612951 |
| CYBRD1 | 29.9988528 | -0.1276836 | 0.51107728 | -0.2498322 | 0.80271712 | 0.9710986 |
| DYNC1I2 | 132.473079 | -0.28419 | 0.18173912 | -1.5637251 | 0.11788214 | 0.6668162 |
| SLC25A12 | 195.898409 | 0.39183537 | 0.25946783 | 1.51015009 | 0.13100513 | 0.67492558 |
| HAT1 | 50.7533934 | 0.0139936 | 0.35876976 | 0.03900439 | 0.96888689 | 0.99379133 |
| METAP1D | 24.7987198 | 0.04399573 | 0.37732978 | 0.11659757 | 0.90717897 | 0.9854228 |
| ITGA6 | 164.204904 | -0.1702878 | 0.25450505 | -0.6690938 | 0.50343563 | 0.89612329 |
| AC078883.1 | 13.8190844 | 0.21326085 | 0.40103635 | 0.53177435 | 5.95E-01 | NA |
| PDK1 | 60.5305126 | -0.3756091 | 0.42125886 | -0.8916349 | 3.73E-01 | 8.49E-01 |
| MAP3K20 | 25.5676455 | -0.6007484 | 0.49863358 | -1.2047894 | 0.22828463 | 0.77185154 |
| SP3 | 96.6682673 | 0.13222672 | 0.25008308 | 0.52873117 | 0.59699195 | 0.9229638 |
| OLA1 | 193.154899 | -0.261754 | 0.20424657 | -1.2815587 | 0.19999749 | 0.75605362 |
| CIR1 | 59.0850715 | 0.10988551 | 0.28498212 | 0.38558737 | 0.69980229 | 0.94419257 |
| SCRN3 | 23.9904774 | -0.0166401 | 0.46200689 | -0.036017 | 0.97126882 | 0.99403127 |
| GPR155 | 25.0274003 | 1.00666192 | 0.61617616 | 1.63372422 | 0.10231677 | 0.65946648 |
| AC010894.3 | 89.2341685 | 1.49693904 | 0.49355053 | 3.03300056 | 0.00242135 | 0.19505808 |
| WIPF1 | 27.0210194 | 0.61173877 | 0.53534804 | 1.14269359 | 0.25316583 | 0.7912221 |
| ATF2 | 104.330223 | -0.1697346 | 0.20863126 | -0.8135625 | 0.41589561 | 0.85867915 |
| ATP5MC3 | 128.342098 | 0.81868265 | 0.43832259 | 1.86776286 | 0.06179513 | 0.5835281 |
| LNPK | 52.9356567 | -0.5477013 | 0.29037434 | -1.8861903 | 0.05926932 | 0.57442774 |
| HOXD10 | 108.899741 | 0.3403321 | 0.3590663 | 0.94782523 | 0.34321843 | 0.83426365 |
| HOXD-AS2 | 26.7365921 | -0.1318442 | 0.45057151 | -0.2926154 | 0.76981615 | 0.96310415 |
| HOXD9 | 42.0947668 | 0.02298252 | 0.32321807 | 0.07110531 | 9.43E-01 | 0.99065376 |
| HOXD8 | 68.0273215 | 0.92253966 | 0.39066034 | 2.3614879 | 0.01820177 | 0.41186103 |
| HOXD3 | 126.835057 | 0.16538176 | 0.26729192 | 0.61873088 | 0.53609366 | 0.90763121 |
| HAGLR | 30.6745557 | -0.0619858 | 0.3289837 | -0.1884161 | 0.85055046 | 0.97612951 |
| MTX2 | 273.569673 | -0.0343375 | 0.2380725 | -0.1442311 | 0.88531801 | 0.98197268 |
| LINC01116 | 37.2091745 | 0.72598707 | 0.43566234 | 1.66639849 | 0.09563407 | 0.65011451 |
| LINC01117 | 22.6923641 | 0.20511121 | 0.47573132 | 0.43114927 | 0.66635984 | 0.93860542 |
| AC079305.3 | 14.8705038 | 0.93575829 | 0.78334486 | 1.19456747 | 0.23225606 | NA |
| HNRNPA3 | 186.120244 | -0.2223707 | 0.26167616 | -0.8497934 | 0.39543997 | 0.85605713 |
| NFE2L2 | 1618.84316 | 0.29882557 | 0.27940743 | 1.06949759 | 2.85E-01 | 8.04E-01 |
| AGPS | 79.0346271 | -0.1617827 | 0.26346088 | -0.6140674 | 0.5391708 | 0.90763121 |
| RBM45 | 28.6756866 | 0.39057677 | 0.30783447 | 1.26878829 | 0.20451658 | 0.76173533 |
| AC009948.1 | 14.5320298 | -0.418653 | 0.67447188 | -0.6207124 | 0.53478889 | NA |
| PRKRA | 58.5137898 | -0.0141012 | 0.32628693 | -0.0432171 | 0.96552848 | 0.99270395 |
| PJVK | 9.944113 | 0.20740355 | 0.60137669 | 0.34488126 | 0.73018364 | NA |
| PLEKHA3 | 39.2306989 | 0.48292547 | 0.26895542 | 1.79555951 | 0.0725646 | 0.60824637 |
| TTN-AS1 | 171.039488 | 0.04025017 | 0.27689696 | 0.14536155 | 0.88442542 | 0.98154805 |
| TTN | 13.7445698 | 1.66764823 | 0.88141622 | 1.89200992 | 0.05848966 | NA |
| SESTD1 | 219.094878 | -0.4615138 | 0.37069195 | -1.2450062 | 2.13E-01 | 0.7643572 |
| ZNF385B | 48.8309575 | 0.76747874 | 0.40420931 | 1.89871613 | 0.05760181 | 0.57118657 |
| CWC22 | 50.0286254 | 0.11356853 | 0.27462928 | 0.41353395 | 0.67921546 | 0.94176503 |

| | | | | | | |
|---|---|---|---|---|---|---|
| UBE2E3 | 171.900661 | 0.02070522 | 0.16659915 | 0.12428169 | 0.90109225 | 0.98531749 |
| CERKL | 18.2157129 | 0.25136352 | 0.48877252 | 0.51427507 | 0.60705969 | 0.92569157 |
| SSFA2 | 40.8924029 | 0.28443662 | 0.26911079 | 1.05694988 | 0.29053446 | 0.80585318 |
| PDE1A | 462.292012 | -0.4911555 | 0.42143298 | -1.1654414 | 0.24384036 | 0.78544896 |
| DNAJC10 | 71.3216758 | -0.2246376 | 0.25644072 | -0.8759824 | 0.38103961 | 0.85432452 |
| NCKAP1 | 272.952518 | -0.3186708 | 0.24945429 | -1.2774715 | 0.20143582 | 0.75928375 |
| NUP35 | 23.8859393 | -0.0699426 | 0.32153653 | -0.2175262 | 0.82779831 | 0.97567084 |
| FSIP2 | 15.6165421 | 0.88265158 | 0.79596361 | 1.10890946 | 0.26746924 | NA |
| FSIP2-AS1 | 12.3414185 | 0.07202745 | 0.55342405 | 0.13014876 | 0.89644874 | NA |
| LINC01473 | 34.8574719 | 0.42974252 | 0.50492841 | 0.85109594 | 0.39471606 | 0.85605713 |
| ZC3H15 | 67.8725354 | -0.1858205 | 0.26228032 | -0.7084805 | 0.47864691 | 0.88588465 |
| ITGAV | 332.014033 | 0.06665821 | 0.28502161 | 0.23387073 | 0.81508533 | 0.97284589 |
| AC017101.1 | 31.2384591 | 1.16968577 | 0.73313657 | 1.59545413 | 0.11061072 | 0.66311481 |
| AC007319.1 | 56.2072934 | 0.44072548 | 0.61104595 | 0.72126406 | 0.47074707 | 0.88263863 |
| TFPI | 32.1450133 | 0.11043442 | 0.66103146 | 0.16706378 | 0.86731986 | 0.97805828 |
| LINC01090 | 16.0382789 | 0.65010471 | 0.68377835 | 0.95075358 | 0.34172948 | NA |
| GULP1 | 350.050248 | -0.9543115 | 0.42136834 | -2.2647917 | 0.02352546 | 0.43088993 |
| COL5A2 | 16.2357226 | -0.8926636 | 0.58615787 | -1.5229064 | 0.12778213 | 0.66929384 |
| WDR75 | 32.718138 | -0.2108662 | 0.29221592 | -0.7216111 | 0.47053363 | 0.88260202 |
| SLC40A1 | 22.8055218 | 0.63747286 | 0.4942208 | 1.28985438 | 0.19710122 | 0.75418766 |
| ASNSD1 | 16.6606905 | -0.058101 | 0.39892001 | -0.1456458 | 0.884201 | 0.98154805 |
| ANKAR | 70.0164192 | 0.30863848 | 0.35355073 | 0.87296803 | 0.38268051 | 0.8548566 |
| OSGEPL1 | 10.6240005 | -0.114342 | 0.44190513 | -0.2587479 | 0.79582976 | NA |
| ORMDL1 | 51.4695875 | 0.34220697 | 0.34419418 | 0.99422649 | 0.32011262 | 0.82449332 |
| PMS1 | 61.8451482 | 0.20669006 | 0.25537974 | 0.80934399 | 0.41831731 | 0.86010695 |
| C2orf88 | 26.6303185 | 0.37472997 | 0.38179862 | 0.98148593 | 0.32635317 | 0.82567895 |
| HIBCH | 215.583583 | 0.21916198 | 0.27699795 | 0.79120435 | 0.42882475 | 0.86357236 |
| INPP1 | 43.1381403 | -0.2305754 | 0.35367011 | -0.6519506 | 0.51443305 | 0.90156229 |
| MFSD6 | 51.6865934 | 0.03852117 | 0.27682419 | 0.13915391 | 0.88932853 | 0.9827634 |
| NAB1 | 35.6013497 | -0.0547472 | 0.43718236 | -0.1252273 | 0.90034357 | 0.98531749 |
| GLS | 1368.86657 | -0.8624775 | 0.4674531 | -1.8450567 | 0.06502929 | 0.59078006 |
| STAT1 | 296.426661 | -1.1368423 | 0.78658027 | -1.4452973 | 0.1483744 | 0.69945736 |
| MYO1B | 473.075116 | -0.4984377 | 0.45997029 | -1.0836302 | 0.2785288 | 0.80210301 |
| NABP1 | 28.8515199 | -0.3566094 | 0.79459856 | -0.448792 | 0.65358174 | 0.93532021 |
| SLC39A10 | 83.0795826 | 1.0419393 | 0.37856979 | 2.75230437 | 0.00591775 | 0.2943676 |
| DNAH7 | 121.525348 | -0.0633852 | 0.30381608 | -0.2086301 | 0.83473702 | 0.97600352 |
| GTF3C3 | 37.4047182 | 0.15301687 | 0.31903363 | 0.47962616 | 0.63149324 | 0.93251095 |
| PGAP1 | 83.6037519 | -0.4604494 | 0.74454702 | -0.6184289 | 0.53629265 | 0.90763121 |
| ANKRD44 | 74.3168025 | 0.06345113 | 0.25588602 | 0.24796638 | 0.80416042 | 0.97175786 |
| SF3B1 | 137.892435 | 0.18546331 | 0.22050657 | 0.84107836 | 0.40030403 | 0.85778581 |
| COQ10B | 34.3752325 | -0.2348282 | 0.29730737 | -0.7898499 | 0.42961545 | 0.86426286 |
| HSPD1 | 109.949734 | -0.0462688 | 0.26158448 | -0.1768788 | 0.85960356 | 0.97691173 |
| MOB4 | 13.5084785 | 0.30456379 | 0.46795812 | 0.65083556 | 0.51515264 | NA |
| AC011997.1 | 12.8729548 | 0.15902284 | 0.59459719 | 0.26744635 | 0.78912552 | NA |
| PLCL1 | 504.02698 | 0.08539835 | 0.27989849 | 0.30510472 | 7.60E-01 | 9.59E-01 |
| FTCDNL1 | 19.6786217 | 1.15960416 | 0.62184543 | 1.86477878 | 0.06221242 | 0.58370968 |
| C2orf69 | 17.7512467 | 0.42908014 | 0.42945673 | 0.9991231 | 0.31773506 | 0.82351215 |
| TYW5 | 36.4802607 | -0.0992903 | 0.31768502 | -0.3125431 | 0.75462779 | 0.95832511 |
| MAIP1 | 18.2810225 | 0.3255241 | 0.43458819 | 0.74904037 | 0.45383288 | 0.87378486 |
| SPATS2L | 243.628132 | -0.5180819 | 0.41587045 | -1.2457772 | 0.21284622 | 0.7643572 |
| KCTD18 | 16.8437457 | -0.1140122 | 0.35379433 | -0.3222557 | 0.74725898 | 0.95563105 |
| BZW1 | 31.173011 | -0.637802 | 0.36137489 | -1.7649317 | 0.07757524 | 0.61352514 |
| CLK1 | 96.1493458 | -0.4309357 | 0.29179185 | -1.47686 | 0.13971316 | 0.69082503 |
| PPIL3 | 26.6530297 | 0.10374578 | 0.41861452 | 0.2478313 | 0.80426494 | 0.97175786 |

| | | | | | | |
|---|---|---|---|---|---|---|
| NIF3L1 | 14.6582719 | 0.28785002 | 0.44926024 | 0.64072 | 0.52170462 | NA |
| ORC2 | 117.928722 | -0.1389905 | 0.33711574 | -0.412293 | 0.68012466 | 0.94195357 |
| AC005037.1 | 23.8014683 | 0.25029644 | 0.53657316 | 0.46647216 | 0.64087757 | 0.93328692 |
| FAM126B | 94.1739108 | -0.067109 | 0.25832436 | -0.2597857 | 0.79502904 | 0.96974891 |
| NDUFB3 | 51.3905605 | 0.41415816 | 0.44271577 | 0.93549447 | 0.34953353 | 0.83687845 |
| CFLAR | 216.258596 | 0.21631403 | 0.28785063 | 0.75148015 | 0.45236374 | 0.87346041 |
| CFLAR-AS1 | 17.6475031 | 0.41842992 | 0.58700871 | 0.71281723 | 0.47595886 | 0.88495683 |
| CASP10 | 52.0865393 | -0.2025555 | 0.24756122 | -0.8182035 | 4.13E-01 | 0.85867915 |
| CASP8 | 32.0482396 | -0.3976039 | 0.40018161 | -0.9935587 | 0.32043776 | 0.82464962 |
| ALS2CR12 | 15.4405246 | 0.15355768 | 0.42795971 | 0.35881339 | 0.71973469 | NA |
| TRAK2 | 136.214554 | -0.3265285 | 0.21854529 | -1.4941001 | 0.13514946 | 0.68329553 |
| STRADB | 63.320696 | -0.051826 | 0.38251168 | -0.1354888 | 0.89222542 | 0.98340332 |
| TMEM237 | 14.1755265 | -0.6115052 | 0.4556515 | -1.3420458 | 0.17958114 | NA |
| ALS2 | 69.3316502 | -0.2098931 | 0.25353574 | -0.8278639 | 0.40774755 | 0.85867915 |
| SUMO1 | 62.607225 | -0.0078974 | 0.349163 | -0.0226182 | 0.98195485 | 0.99626066 |
| NOP58 | 72.0033771 | -0.3193474 | 0.27249328 | -1.1719461 | 0.24121869 | 0.78310048 |
| BMPR2 | 330.198823 | -0.2390059 | 0.18598268 | -1.2850975 | 0.19875821 | 0.75605362 |
| FAM117B | 137.693926 | -0.1944523 | 0.22852818 | -0.8508899 | 0.39483054 | 0.85605713 |
| ICA1L | 46.4429369 | 0.0194807 | 0.36965454 | 0.05269976 | 0.95797113 | 0.99183169 |
| WDR12 | 41.6233667 | 0.16435703 | 0.35003793 | 0.46954065 | 0.63868324 | 0.9331046 |
| CARF | 62.5899187 | 0.45226042 | 0.36231163 | 1.24826362 | 0.21193453 | 0.7643572 |
| NBEAL1 | 522.090446 | -0.1683057 | 0.24741004 | -0.6802704 | 0.49633326 | 0.89255664 |
| CYP20A1 | 103.120727 | -0.3657896 | 0.39512372 | -0.9257598 | 0.35457084 | 0.84009848 |
| ABI2 | 210.666552 | -0.152769 | 0.26679711 | -0.5726036 | 0.5669131 | 0.91698355 |
| RAPH1 | 224.501381 | -0.2995736 | 0.44297849 | -0.6762712 | 0.49886846 | 0.89378207 |
| PARD3B | 767.793225 | -0.00554 | 0.16627272 | -0.0333189 | 9.73E-01 | 9.94E-01 |
| INO80D | 236.587321 | -0.9298693 | 0.60549524 | -1.5357169 | 0.12460782 | 0.66929384 |
| NDUFS1 | 167.924235 | 0.07400456 | 0.25065617 | 0.29524334 | 0.767808 | 0.96283923 |
| EEF1B2 | 39.3336191 | 0.18603319 | 0.36696548 | 0.50695011 | 0.61218983 | 0.92644418 |
| ZDBF2 | 37.5735225 | -0.1386637 | 0.50360223 | -0.2753437 | 0.78305218 | 0.96678746 |
| FASTKD2 | 43.1406841 | -0.1022636 | 0.28679118 | -0.3565785 | 0.72140739 | 0.95090812 |
| KLF7 | 24.2311938 | 0.61758417 | 0.65115853 | 0.94843903 | 0.342906 | 0.83417245 |
| CREB1 | 103.575088 | -0.2530267 | 0.20347398 | -1.2435332 | 0.21367141 | 0.7643572 |
| METTL21A | 25.9033167 | 0.10002196 | 0.32058862 | 0.31199473 | 0.75504453 | 0.95832511 |
| CCNYL1 | 56.108153 | 0.15874911 | 0.38114312 | 0.41650787 | 0.67703841 | 0.94034237 |
| PLEKHM3 | 65.5416315 | 0.45733287 | 0.32881115 | 1.39086787 | 0.1642655 | 0.71397803 |
| IDH1 | 24.4534627 | 0.68086208 | 0.38897786 | 1.75038774 | 0.08005143 | 0.61650437 |
| PIKFYVE | 111.142586 | 0.15460333 | 0.24377512 | 0.63420471 | 0.52594723 | 0.90487731 |
| PTH2R | 1217.66906 | -0.5832624 | 0.22493032 | -2.5930802 | 0.00951206 | 0.32430062 |
| RPE | 54.1441598 | -0.2326508 | 0.27843814 | -0.8355564 | 0.40340453 | 0.85782098 |
| KANSL1L | 513.487067 | 0.31345976 | 0.25054309 | 1.25112118 | 0.21089027 | 0.76397692 |
| AC007038.1 | 53.9948045 | 0.24236971 | 0.32511809 | 0.74548209 | 0.45598034 | 0.87394768 |
| ACADL | 41.0239275 | 0.06885796 | 0.45266761 | 0.15211595 | 0.87909548 | 0.98061726 |
| LANCL1 | 65.3765597 | -0.0441427 | 0.28289416 | -0.1560398 | 0.87600168 | 0.97918624 |
| ERBB4 | 3062.64568 | 0.2755687 | 0.16792638 | 1.641009 | 0.10079555 | 0.65863153 |
| IKZF2 | 91.4161542 | 0.3556452 | 0.37857137 | 0.93944028 | 0.34750474 | 0.83687845 |
| SPAG16 | 371.850455 | 0.70796562 | 0.29737839 | 2.38068952 | 0.01728027 | 0.40063153 |
| AC068051.1 | 57.2885535 | 0.94980691 | 0.56424059 | 1.68333674 | 0.09230992 | 0.64572229 |
| BARD1 | 23.1927619 | 0.6779187 | 0.365712 | 1.85369552 | 0.06378273 | 0.58781448 |
| AC016708.1 | 24.9310011 | 0.01644154 | 0.43754347 | 0.03757692 | 0.97002501 | 0.99398482 |
| ATIC | 35.4277547 | -0.3567832 | 0.38998077 | -0.9148738 | 0.36025787 | 0.84282379 |
| MREG | 34.7933745 | -0.5533104 | 0.30541844 | -1.8116471 | 0.07004074 | 0.60319855 |
| PECR | 53.2903482 | 0.03627262 | 0.28276291 | 0.12827929 | 0.89792796 | 0.98531749 |
| XRCC5 | 229.208619 | -0.2473491 | 0.17976045 | -1.375993 | 0.1688238 | 0.72184269 |

| | | | | | | |
|---|---|---|---|---|---|---|
| SMARCAL1 | 35.592022 | 0.02444034 | 0.33811693 | 0.07228368 | 0.94237615 | 0.99058259 |
| RPL37A | 126.293091 | -0.2435216 | 0.28849066 | -0.8441229 | 0.39860073 | 0.85758979 |
| IGFBP2 | 18.0910373 | 0.49854009 | 0.64176724 | 0.77682384 | 0.43726271 | 0.86794646 |
| IGFBP5 | 35.3184837 | -0.67749 | 0.60716171 | -1.1158312 | 0.2644944 | 0.79666376 |
| DIRC3 | 54.0259697 | -0.4153433 | 0.62971442 | -0.659574 | 0.50952725 | 0.89963034 |
| TNS1 | 195.293633 | 0.59909832 | 0.29636009 | 2.02152159 | 0.0432258 | 0.51638379 |
| ARPC2 | 129.56022 | -0.4701758 | 0.2780052 | -1.6912482 | 0.0907894 | 0.64165834 |
| AAMP | 28.4445861 | -0.0477987 | 0.35716178 | -0.1338293 | 0.89353752 | 0.98377746 |
| PNKD | 78.125039 | 0.29705073 | 0.30906063 | 0.96114062 | 0.33648147 | 0.83037432 |
| TMBIM1 | 73.671036 | -0.3456165 | 0.23731005 | -1.4563922 | 0.14528424 | 0.69606446 |
| CATIP-AS2 | 14.2078462 | 0.51921592 | 0.57251053 | 0.9069107 | 0.36445402 | NA |
| CATIP | 16.9417577 | -0.1081154 | 0.53308452 | -0.202811 | 8.39E-01 | 0.97600352 |
| CTDSP1 | 51.6764028 | -0.1800825 | 0.25448436 | -0.707637 | 0.47917073 | 0.88622003 |
| USP37 | 91.9807515 | -0.6104242 | 0.27909738 | -2.1871368 | 0.02873254 | 0.46148943 |
| CNOT9 | 47.294268 | -0.3056351 | 0.22510645 | -1.3577357 | 0.17454757 | 0.72841249 |
| PLCD4 | 19.4454529 | 0.58666253 | 0.68378676 | 0.85796124 | 0.39091387 | 0.85605713 |
| ZNF142 | 17.2498492 | -0.4680081 | 0.41440003 | -1.1293632 | 0.25874465 | 0.79532468 |
| STK36 | 16.4581668 | -0.4088593 | 0.42468108 | -0.9627442 | 0.3356759 | 0.82974237 |
| TTLL4 | 18.0234306 | 0.01920027 | 0.35858896 | 0.05354396 | 0.95729851 | 0.99170927 |
| CYP27A1 | 16.6758089 | 0.09130035 | 0.45813612 | 0.19928651 | 0.84203863 | 0.97600352 |
| CNPPD1 | 27.504921 | 0.4732598 | 0.324451 | 1.45864799 | 0.14466202 | 0.69512617 |
| RETREG2 | 39.9066857 | 0.00105218 | 0.38217573 | 0.00275313 | 0.99780332 | 0.99956021 |
| ZFAND2B | 14.5619158 | -0.3827843 | 0.46507427 | -0.8230605 | 0.41047358 | NA |
| ANKZF1 | 28.4984296 | -0.3601603 | 0.29129022 | -1.2364311 | 0.21629837 | 0.76616336 |
| STK16 | 12.0092553 | 0.36076356 | 0.44095217 | 0.8181467 | 0.41327343 | NA |
| TUBA4A | 31.9546518 | -0.3993587 | 0.48971388 | -0.815494 | 0.41478961 | 0.85867915 |
| DNAJB2 | 31.7642668 | -0.4608845 | 0.39087846 | -1.1790993 | 0.23835863 | 0.78077674 |
| DNPEP | 21.425765 | -0.2123715 | 0.40292894 | -0.5270694 | 0.5981454 | 0.92296674 |
| CHPF | 21.5673687 | 0.11496429 | 0.5088281 | 0.22593935 | 0.8212486 | 0.97396757 |
| OBSL1 | 29.3748539 | 1.03605972 | 0.37083648 | 2.79384517 | 0.00520854 | 0.28057141 |
| SGPP2 | 273.862536 | -0.1353696 | 0.31970718 | -0.4234175 | 0.6719907 | 0.93878847 |
| FARSB | 40.8283385 | -0.5485802 | 0.32669778 | -1.6791671 | 0.09311949 | 0.64572229 |
| ACSL3 | 136.303388 | -0.2977908 | 0.28384841 | -1.0491193 | 0.29412323 | 0.81010703 |
| AP1S3 | 52.8109653 | -0.697677 | 0.49700872 | -1.4037521 | 0.1603927 | 0.71101785 |
| WDFY1 | 82.5279524 | -0.426741 | 0.19217242 | -2.2206153 | 0.02637703 | 0.44713242 |
| MRPL44 | 12.4508866 | -0.0339573 | 0.55385107 | -0.0613113 | 0.9511113 | NA |
| CUL3 | 247.151942 | -0.0428851 | 0.21717214 | -0.1974704 | 0.84345943 | 0.97612951 |
| IRS1 | 24.2943156 | 0.30907253 | 0.44562163 | 0.69357614 | 0.48794806 | 0.88866642 |
| RHBDD1 | 128.849506 | -0.4564038 | 0.2349944 | -1.9421902 | 0.05211408 | 0.55403388 |
| COL4A4 | 304.888211 | 0.08497673 | 0.33547401 | 0.25330346 | 0.80003372 | 0.97019967 |
| COL4A3 | 455.103627 | 0.16067572 | 0.4008978 | 0.40078973 | 0.68857494 | 0.94205398 |
| AC097662.1 | 66.7665494 | -0.0319614 | 0.57493971 | -0.0555909 | 0.95566773 | 0.99148072 |
| MFF | 132.795281 | 0.24994671 | 0.26483007 | 0.94380032 | 0.3452717 | 0.83564825 |
| AGFG1 | 249.478745 | -0.4493524 | 0.28804959 | -1.5599827 | 0.11876396 | 0.6668162 |
| SLC19A3 | 49.74776 | -0.7949762 | 0.85446409 | -0.9303799 | 0.35217442 | 0.83872681 |
| DNER | 25.5392159 | -0.2436187 | 0.67418491 | -0.361353 | 0.71783558 | 0.94925929 |
| TRIP12 | 287.03261 | -0.1505465 | 0.18766975 | -0.8021885 | 0.42244391 | 0.86190547 |
| FBXO36 | 34.4596826 | 0.65805128 | 0.28423517 | 2.31516485 | 0.02060391 | 0.4212027 |
| SP110 | 48.6605321 | -1.3829924 | 0.63577457 | -2.1752873 | 0.02960859 | 0.46381645 |
| SP140L | 39.2434059 | -0.6788757 | 0.43574088 | -1.5579802 | 0.11923794 | 0.6668162 |
| SP100 | 144.689372 | -0.7694088 | 0.43641882 | -1.7630055 | 0.07789956 | 0.61367304 |
| CAB39 | 236.186537 | -0.2120254 | 0.2829233 | -0.7494094 | 0.45361051 | 0.87378486 |
| ITM2C | 115.64945 | 0.93143913 | 0.46755871 | 1.99213299 | 0.04635647 | 0.53087707 |
| PSMD1 | 124.748247 | -0.1993737 | 0.26628728 | -0.7487165 | 0.45402811 | 0.87378486 |

| | | | | | | |
|---|---|---|---|---|---|---|
| ARMC9 | 131.177857 | 0.07740557 | 0.27113458 | 0.28548763 | 0.77527058 | 0.96469469 |
| NCL | 86.1238524 | 0.13233064 | 0.33047265 | 0.40042841 | 0.688841 | 0.94205398 |
| PTMA | 383.766537 | 0.10815457 | 0.26848878 | 0.40282715 | 0.68707539 | 0.94205398 |
| PDE6D | 30.7198761 | 0.1562847 | 0.29541672 | 0.52903132 | 0.59678372 | 0.9229638 |
| COPS7B | 29.9767395 | 0.02548414 | 0.27823121 | 0.09159339 | 0.9270211 | 0.98925753 |
| DIS3L2 | 206.206832 | -0.189027 | 0.18274117 | -1.0343974 | 0.30095043 | 0.81437329 |
| EIF4E2 | 48.6021377 | 0.05669754 | 0.27357602 | 0.20724602 | 0.83581773 | 0.97600352 |
| EFHD1 | 81.2675981 | 0.6492919 | 0.42190969 | 1.53893572 | 0.12381999 | 0.66929384 |
| GIGYF2 | 258.438 | -0.2432862 | 0.22358184 | -1.0881305 | 0.2765375 | 0.80159799 |
| ATG16L1 | 19.7746392 | -0.1884118 | 0.39698881 | -0.4746022 | 0.63507053 | 0.93251095 |
| DGKD | 38.5861048 | -0.3866788 | 0.34254384 | -1.1288446 | 0.25896338 | 0.79532468 |
| USP40 | 146.784518 | 0.11691741 | 0.30067413 | 0.3888509 | 0.69738644 | 0.94395238 |
| AC114812.2 | 56.9076918 | -0.0277731 | 0.85761935 | -0.0323839 | 0.97416588 | 0.99431961 |
| ARL4C | 72.7220582 | -0.0918527 | 0.71566868 | -0.1283453 | 0.89787573 | 0.98531749 |
| SH3BP4 | 120.830738 | -0.4942887 | 0.3574201 | -1.3829348 | 0.16668486 | 0.71877417 |
| AGAP1 | 747.26154 | -0.7091816 | 0.33604223 | -2.1103943 | 0.0348244 | 0.49148303 |
| IQCA1 | 36.3909154 | -0.2845473 | 0.35034151 | -0.8121998 | 0.41667698 | 0.85867915 |
| AC105760.2 | 13.758433 | -0.5599686 | 0.454806 | -1.2312251 | 0.21823868 | NA |
| COPS8 | 20.6405405 | 0.38326942 | 0.33917241 | 1.13001355 | 0.25847052 | 0.79532468 |
| RAB17 | 103.32881 | 0.28340101 | 0.33911395 | 0.83571027 | 0.40331792 | 0.85782098 |
| LRRFIP1 | 636.223696 | -0.5672822 | 0.36797532 | -1.5416311 | 0.12316325 | 0.66929384 |
| ILKAP | 74.6437506 | -0.1552343 | 0.21664183 | -0.716548 | 0.47365303 | 0.88366688 |
| PER2 | 26.4353057 | -0.4894636 | 0.40025287 | -1.2228858 | 0.22137282 | 0.76680432 |
| TRAF3IP1 | 51.1740204 | -0.1106412 | 0.27090798 | -0.4084086 | 0.68297372 | 0.94205398 |
| ASB1 | 14.6896149 | -0.2963106 | 0.44954751 | -0.6591308 | 0.50981181 | NA |
| HDAC4 | 362.386873 | -0.6362663 | 0.27133709 | -2.3449292 | 0.01903069 | 0.41544117 |
| NDUFA10 | 141.428652 | -0.0086154 | 0.20495146 | -0.0420363 | 0.96646974 | 0.99301184 |
| COPS9 | 28.0302447 | 0.04579632 | 0.44521123 | 0.10286425 | 0.91807071 | 0.98735642 |
| GPC1 | 31.42709 | -0.5357391 | 0.49998686 | -1.0715064 | 0.2839418 | 0.80431986 |
| ANKMY1 | 45.7550456 | 0.29566057 | 0.30424277 | 0.97179159 | 0.33115424 | 0.82879734 |
| RNPEPL1 | 30.4102068 | -0.4088504 | 0.35452329 | -1.15324 | 0.24881188 | 0.78729484 |
| CAPN10 | 12.4748117 | -0.3690068 | 0.47816932 | -0.7717074 | 0.44028772 | NA |
| SNED1 | 33.4897759 | -0.0515503 | 0.46583254 | -0.1106628 | 0.91188373 | 0.9859256 |
| MTERF4 | 43.1145515 | 0.40038101 | 0.36689975 | 1.09125453 | 0.2751609 | 0.7994854 |
| PASK | 11.5074769 | 0.59303255 | 0.46107125 | 1.28620588 | 0.19837122 | NA |
| PPP1R7 | 34.3245075 | 0.20952685 | 0.28381075 | 0.73826258 | 0.4603549 | 0.87607759 |
| HDLBP | 187.378266 | -0.2821429 | 0.24540089 | -1.1497222 | 0.25025829 | 0.78799766 |
| 2-Sep | 222.483654 | -0.0753312 | 0.15753918 | -0.4781741 | 0.63252631 | 0.93251095 |
| FARP2 | 89.2790385 | -0.0992719 | 0.21555351 | -0.4605442 | 0.64512567 | 0.93328692 |
| STK25 | 34.5165374 | -0.0805812 | 0.3118556 | -0.2583927 | 0.79610388 | 0.96974891 |
| BOK | 13.4253177 | 0.34679194 | 0.6533764 | 0.530769 | 0.59557886 | NA |
| THAP4 | 58.0376968 | 0.02881826 | 0.24597912 | 0.11715735 | 0.90673537 | 0.9854228 |
| ATG4B | 92.2658082 | -0.2582002 | 0.20736295 | -1.2451605 | 0.21307274 | 0.7643572 |
| DTYMK | 12.7665291 | 0.49891642 | 0.4316652 | 1.15579487 | 0.24776507 | NA |
| ING5 | 43.1636121 | -0.2525586 | 0.33924982 | -0.744462 | 0.456597 | 0.87396417 |
| D2HGDH | 62.6763063 | 0.34951725 | 0.3314401 | 1.05454126 | 0.29163518 | 0.80716177 |
| CHL1 | 45.6363423 | 1.16515094 | 0.69934119 | 1.66606937 | 0.0956996 | 0.65011451 |
| LINC01266 | 122.081171 | 1.20758831 | 0.46343864 | 2.60571348 | 0.00916831 | 0.32071089 |
| CNTN4 | 148.963443 | 0.76536827 | 0.40173807 | 1.90514251 | 0.05676155 | 0.56976762 |
| TRNT1 | 40.4221636 | -0.1594214 | 0.25760125 | -0.6188689 | 0.53600272 | 0.90763121 |
| CRBN | 50.5326606 | 0.07160584 | 0.25230587 | 0.2838057 | 0.77655929 | 0.9649632 |
| AC034195.1 | 97.5763444 | 0.28914292 | 0.39759758 | 0.72722504 | 0.46708811 | 0.88090126 |
| SUMF1 | 457.225842 | -0.1442312 | 0.30374541 | -0.4748423 | 0.63489935 | 0.93251095 |
| SETMAR | 16.1762888 | 0.59488554 | 0.5182135 | 1.14795454 | 0.25098733 | 0.78892875 |

| | | | | | | |
|---|---|---|---|---|---|---|
| ITPR1 | 657.351101 | 0.49122201 | 0.25797053 | 1.90417877 | 0.05688691 | 0.56976762 |
| EGOT | 35.3969743 | 0.42077886 | 0.56598952 | 0.74343931 | 0.45721575 | 0.87419345 |
| AC018816.1 | 24.5411129 | 0.18816915 | 0.36591151 | 0.51424769 | 0.60707883 | 0.92569157 |
| BHLHE40 | 325.795943 | -0.881371 | 0.40728256 | -2.1640284 | 0.03046217 | 0.46919989 |
| ARL8B | 120.942661 | -0.6701318 | 0.29081375 | -2.3043334 | 0.02120394 | 0.4212027 |
| AC026202.2 | 15.9487501 | 0.02831236 | 0.55673945 | 0.05085387 | 0.95944196 | NA |
| EDEM1 | 30.4844145 | 0.03626169 | 0.30902928 | 0.11734062 | 0.90659014 | 0.98539583 |
| AC087857.1 | 21.4423589 | -1.5582946 | 0.62714738 | -2.4847343 | 0.01296482 | 0.36664851 |
| AC026167.1 | 61.1456354 | -1.5498487 | 0.58233833 | -2.6614232 | 7.78E-03 | 3.07E-01 |
| AC069277.1 | 37.5674299 | 1.99027307 | 1.38735666 | 1.43457925 | 0.15140703 | 0.70572801 |
| LMCD1-AS1 | 109.787137 | 0.40984728 | 0.29027373 | 1.41193374 | 1.58E-01 | 0.71089007 |
| LMCD1 | 13.7719948 | 0.14676964 | 0.49914116 | 0.29404435 | 0.76872401 | NA |
| RAD18 | 22.394974 | -0.4626433 | 0.40198979 | -1.1508832 | 0.24978028 | 0.78799766 |
| SRGAP3 | 222.278844 | 0.32298132 | 0.27741616 | 1.16424839 | 0.24432336 | 0.78561483 |
| THUMPD3-A: | 89.7530875 | 0.03895303 | 0.26360369 | 0.14777118 | 0.88252336 | 0.98137082 |
| THUMPD3 | 28.1936724 | -0.0522849 | 0.28090354 | -0.1861311 | 0.85234198 | 0.97612951 |
| SETD5 | 278.595611 | -0.0459472 | 0.17332193 | -0.2650976 | 0.79093427 | 0.96909068 |
| MTMR14 | 42.7247791 | -0.124464 | 0.31320869 | -0.3973835 | 0.69108467 | 0.94293839 |
| OGG1 | 20.6550305 | -0.2160083 | 0.41778821 | -0.5170282 | 0.6051365 | 0.92569157 |
| CAMK1 | 11.8700217 | 0.28335623 | 0.43447093 | 0.65218686 | 0.51428063 | NA |
| TADA3 | 26.6127917 | 0.04728924 | 0.33247563 | 0.1422337 | 0.88689542 | 0.98216235 |
| RPUSD3 | 14.3615689 | -0.1832132 | 0.3961149 | -0.4625254 | 0.64370461 | NA |
| JAGN1 | 12.0006935 | 0.67392101 | 0.46828639 | 1.43912151 | 0.1501161 | NA |
| IL17RE | 16.6914508 | 0.84059308 | 0.55870414 | 1.50454064 | 0.13244222 | 0.67747802 |
| IL17RC | 14.4876217 | 0.83526333 | 0.43588235 | 1.91625864 | 0.05533218 | NA |
| CRELD1 | 27.3105016 | 0.10548974 | 0.38160724 | 0.27643537 | 0.78221369 | 0.96652357 |
| EMC3 | 44.9603747 | 0.28983896 | 0.3026453 | 0.95768532 | 0.33822146 | 0.83055261 |
| BRK1 | 46.2286612 | -0.1590206 | 0.32533396 | -0.488792 | 0.62498896 | 0.92981013 |
| VHL | 27.8354181 | -0.0214076 | 0.34654561 | -0.0617743 | 0.95074254 | 0.99105535 |
| IRAK2 | 18.903945 | 0.4110935 | 0.69807008 | 0.58890005 | 0.55592833 | 0.91398748 |
| SEC13 | 46.8322944 | 0.03680937 | 0.31052393 | 0.11853956 | 0.90564015 | 0.98531749 |
| ATG7 | 242.388207 | -0.09987 | 0.27518911 | -0.362914 | 0.7166691 | 0.94913834 |
| VGLL4 | 277.171111 | -0.0564188 | 0.17855099 | -0.3159814 | 0.75201663 | 0.95769549 |
| TAMM41 | 75.679626 | -0.2853921 | 0.29075426 | -0.9815579 | 0.3263177 | 0.82567895 |
| PPARG | 257.205194 | -0.9166403 | 0.37872644 | -2.4203231 | 0.01550672 | 0.39401695 |
| TSEN2 | 62.3054867 | 0.17056383 | 0.2945478 | 0.57907012 | 0.56254186 | 0.91624774 |
| MKRN2OS | 10.6011261 | 1.17308841 | 0.66376733 | 1.76731869 | 0.07717487 | NA |
| MKRN2 | 16.3497969 | 0.21942378 | 0.44137236 | 0.49713983 | 0.61909045 | 0.92766755 |
| RAF1 | 130.35775 | -0.2831816 | 0.1858423 | -1.5237736 | 0.12756528 | 0.66929384 |
| RPL32 | 156.979886 | 0.32434107 | 0.40109249 | 0.80864409 | 0.4187199 | 0.86022698 |
| IQSEC1 | 58.8585236 | 0.88249828 | 0.34498583 | 2.55807108 | 0.01052546 | 0.33618681 |
| NUP210 | 39.8706937 | 0.06335198 | 0.36746695 | 0.17240185 | 0.86312161 | 0.97788853 |
| HDAC11 | 20.2097536 | 0.86642759 | 0.48158107 | 1.79913133 | 0.07199791 | 0.60821941 |
| XPC | 47.3681438 | 0.06344859 | 0.29648127 | 0.21400538 | 0.83054286 | 0.97567084 |
| LSM3 | 32.3698999 | -0.3146389 | 0.28198857 | -1.115786 | 0.26451375 | 0.79666376 |
| SLC6A6 | 103.182542 | -0.19638 | 0.40524336 | -0.4845976 | 0.62796182 | 0.93061955 |
| CCDC174 | 31.737063 | 0.05947224 | 0.29069891 | 0.20458362 | 0.83789745 | 0.97600352 |
| FGD5-AS1 | 62.4077435 | 0.25086121 | 0.32390156 | 0.77449833 | 0.43863616 | 0.86879768 |
| NR2C2 | 107.88746 | -0.4858761 | 0.28959538 | -1.6777756 | 0.09339092 | 0.64572229 |
| MRPS25 | 108.344577 | -0.2381975 | 0.24229894 | -0.9830729 | 0.32557155 | 0.82567895 |
| RBSN | 39.2441694 | -0.1126586 | 0.31507533 | -0.3575608 | 0.72067204 | 0.95048937 |
| CAPN7 | 139.774795 | -0.0268911 | 0.27025808 | -0.0995016 | 0.92074004 | 0.98804116 |
| SH3BP5 | 40.5103708 | -1.4240287 | 0.81871904 | -1.7393375 | 0.08197542 | 0.61927158 |
| METTL6 | 31.0187689 | 0.09533765 | 0.36369905 | 0.26213335 | 0.79321864 | 0.96947761 |

| | | | | | | |
|---|---|---|---|---|---|---|
| EAF1 | 12.6852024 | 0.30437703 | 0.44193066 | 0.68874386 | 0.49098447 | NA |
| COLQ | 16.6917615 | -0.0895715 | 0.39825264 | -0.2249112 | 0.82204837 | 0.97448969 |
| HACL1 | 47.9688464 | 0.03488066 | 0.28689075 | 0.12158168 | 0.90323033 | 0.98531749 |
| BTD | 15.2782877 | 0.56295122 | 0.44533883 | 1.26409643 | 0.20619542 | NA |
| ANKRD28 | 360.265208 | -0.1684133 | 0.18862118 | -0.8928651 | 0.37192941 | 0.8491102 |
| OXNAD1 | 124.510872 | -0.1969971 | 0.29839128 | -0.6601973 | 0.5091272 | 0.89963034 |
| RFTN1 | 15.3112642 | 1.36805013 | 0.58270615 | 2.34775304 | 0.01888704 | NA |
| PLCL2 | 133.745727 | -0.5534081 | 0.52308851 | -1.0579626 | 0.29007251 | 0.80553084 |
| TBC1D5 | 539.157293 | 0.23946267 | 0.19676022 | 1.21702784 | 0.22359362 | 0.76834258 |
| RAB5A | 105.823208 | -0.2884925 | 0.20919234 | -1.3790778 | 0.16787078 | 0.72008028 |
| KAT2B | 85.6703724 | 0.36235573 | 0.40603557 | 0.89242362 | 0.37216592 | 0.8491102 |
| ZNF385D | 194.515263 | -0.5917921 | 0.78761555 | -0.7513717 | 0.45242896 | 0.87346041 |
| UBE2E2 | 365.257866 | -0.1757559 | 0.21718976 | -0.8092273 | 0.41838444 | 0.86010695 |
| UBE2E1 | 189.849087 | -0.2880029 | 0.26116585 | -1.1027587 | 0.27013197 | 0.79666376 |
| NKIRAS1 | 19.5615676 | 0.07985117 | 0.39834563 | 0.20045701 | 0.84112317 | 0.97600352 |
| RPL15 | 254.759228 | 0.79600894 | 0.41301931 | 1.92729232 | 0.05394321 | 0.56390244 |
| NR1D2 | 56.0254087 | 0.04219292 | 0.49582023 | 0.08509721 | 0.93218411 | 0.98960607 |
| THRB | 290.846725 | 0.30476217 | 0.27505718 | 1.10799568 | 0.26786368 | 0.79666376 |
| RARB | 46.1963566 | -0.300915 | 0.27882951 | -1.079208 | 0.28049502 | 0.80319164 |
| TOP2B | 166.276806 | -0.12714 | 0.22376034 | -0.568197 | 0.56990119 | 0.91778542 |
| NGLY1 | 114.230395 | -0.5803944 | 0.38446812 | -1.5096034 | 0.13114467 | 0.67492558 |
| OXSM | 13.7437455 | -0.5882215 | 0.44149252 | -1.3323476 | 0.18274598 | NA |
| NEK10 | 30.6393872 | 0.16126492 | 0.29919155 | 0.53900227 | 0.58988529 | 0.92229571 |
| SLC4A7 | 46.6485061 | -0.5720743 | 0.58217143 | -0.9826561 | 0.32577673 | 0.82567895 |
| CMC1 | 136.261788 | -0.0918272 | 0.26309153 | -0.3490314 | 0.72706575 | 0.95211709 |
| AZI2 | 61.4037238 | -0.2184907 | 0.35356639 | -0.6179624 | 0.53660013 | 0.90763121 |
| TGFBR2 | 78.605553 | -0.2163941 | 0.40305972 | -0.5368786 | 0.59135149 | 0.92254886 |
| STT3B | 130.880105 | 0.00011454 | 0.23727273 | 0.00048272 | 0.99961485 | 0.99973064 |
| OSBPL10 | 45.2258339 | -0.4123085 | 0.29486047 | -1.3983173 | 0.16201782 | 0.71179932 |
| GPD1L | 25.3361836 | 0.39227109 | 0.38458801 | 1.01997744 | 0.30773916 | 0.81568231 |
| CMTM8 | 66.9776428 | 0.55949365 | 0.25599307 | 2.18558122 | 0.02884626 | 0.46148943 |
| CMTM7 | 55.2407737 | 0.31503373 | 0.23444351 | 1.34375111 | 1.79E-01 | 7.29E-01 |
| CMTM6 | 63.4826933 | 0.13860577 | 0.31291886 | 0.44294476 | 0.65780569 | 0.93593674 |
| DYNC1LI1 | 36.8301338 | -0.1219995 | 0.30887329 | -0.3949823 | 0.69285593 | 0.94388919 |
| CNOT10 | 88.2838084 | -0.6370679 | 0.20189246 | -3.1554812 | 0.00160234 | 0.16785081 |
| GLB1 | 86.7357348 | 0.44582875 | 0.25145074 | 1.77302619 | 0.07622437 | 0.61051335 |
| CRTAP | 52.1785439 | 0.36300268 | 0.29873077 | 1.21514996 | 0.2243089 | 0.76907116 |
| FBXL2 | 135.119501 | -0.3969934 | 0.24604664 | -1.6134884 | 0.10663845 | 0.65986731 |
| UBP1 | 64.2607492 | -0.1354525 | 0.24422993 | -0.5546106 | 0.57916104 | 0.91976673 |
| CLASP2 | 268.529588 | -0.0407885 | 0.2000035 | -0.203939 | 0.83840113 | 0.97600352 |
| PDCD6IP | 157.245913 | 0.02662262 | 0.17880984 | 0.1488879 | 0.88164209 | 0.98137082 |
| STAC | 99.9372724 | -0.6103374 | 0.27644714 | -2.2077906 | 0.02725887 | 0.45224481 |
| TRANK1 | 24.4608666 | -1.7013997 | 1.23829286 | -1.3739881 | 0.16944537 | 0.72348087 |
| EPM2AIP1 | 50.8513154 | 0.30561526 | 0.30895282 | 0.98919716 | 0.32256669 | 0.82558302 |
| MLH1 | 31.6310252 | -0.4293454 | 0.26490408 | -1.620758 | 0.10506954 | 0.65946648 |
| LRRFIP2 | 233.25671 | -0.1908666 | 0.21181445 | -0.9011028 | 0.36753364 | 0.84777627 |
| GOLGA4 | 454.743001 | -0.3888982 | 0.25140382 | -1.5469066 | 0.12188577 | 0.66879213 |
| ITGA9 | 20.8715904 | -1.3512115 | 0.50740553 | -2.6629815 | 0.00774517 | 0.30654023 |
| CTDSPL | 289.44867 | -0.2854967 | 0.26756785 | -1.0670068 | 0.28596877 | 0.80431986 |
| PLCD1 | 15.0172084 | 0.88666653 | 0.39927567 | 2.22068758 | 0.02637213 | NA |
| ACAA1 | 40.9572304 | 0.64763435 | 0.31523014 | 2.05448106 | 0.03992915 | 0.50156625 |
| OXSR1 | 155.585404 | -0.447163 | 0.26101678 | -1.7131581 | 0.08668345 | 0.63466198 |
| ACVR2B | 25.4854711 | 0.20331611 | 0.37304963 | 0.54501087 | 0.58574604 | 0.92076087 |
| EXOG | 26.273192 | 0.1235277 | 0.33000061 | 0.37432568 | 0.70816203 | 0.94558664 |

| | | | | | | |
|---|---|---|---|---|---|---|
| WDR48 | 59.4867136 | 0.17312051 | 0.25557369 | 0.67738003 | 0.49816487 | 0.89378207 |
| GORASP1 | 17.1201952 | -0.0103989 | 0.37878758 | -0.027453 | 0.97809842 | 0.9946931 |
| TTC21A | 15.2789901 | 0.22741768 | 0.53325204 | 0.42647315 | 0.66976311 | NA |
| CSRNP1 | 14.3113424 | -0.4711266 | 0.47907078 | -0.9834175 | 0.32540199 | NA |
| SLC25A38 | 15.189836 | -0.178876 | 0.4392253 | -0.4072533 | 0.68382196 | NA |
| RPSA | 115.39192 | 0.28705177 | 0.44705427 | 0.64209603 | 0.52081083 | 0.90441454 |
| MYRIP | 13.4347598 | 1.07471192 | 0.67324609 | 1.59631366 | 0.11041878 | NA |
| EIF1B | 27.7270897 | -0.0196453 | 0.2957407 | -0.0664274 | 0.94703753 | 0.99083449 |
| ENTPD3-AS1 | 34.1535124 | 0.23710517 | 0.52445949 | 0.45209435 | 0.65120101 | 0.93453685 |
| RPL14 | 173.809619 | 0.21337254 | 0.40343445 | 0.52889022 | 0.59688161 | 0.9229638 |
| ZNF621 | 41.0328611 | -0.361862 | 0.28632112 | -1.2638327 | 0.20629008 | 0.76231289 |
| CTNNB1 | 121.024105 | -0.2155606 | 0.22211318 | -0.9704987 | 0.33179797 | 0.82890589 |
| ULK4 | 292.386453 | 0.16942308 | 0.1572627 | 1.07732527 | 0.28133498 | 0.80416687 |
| TRAK1 | 409.147773 | 0.18803613 | 0.22849279 | 0.82294122 | 0.41054142 | 0.85867915 |
| SEC22C | 24.6626175 | -0.5116898 | 0.31250371 | -1.6373878 | 0.10154947 | 0.65863153 |
| SS18L2 | 12.3987603 | -0.3708928 | 0.43672926 | -0.849251 | 0.39574164 | NA |
| NKTR | 283.969713 | -0.0826513 | 0.17287869 | -0.4780887 | 0.63258707 | 0.93251095 |
| AC006059.1 | 32.5761514 | 0.18336382 | 0.56550094 | 0.32425025 | 0.74574858 | 0.95563105 |
| AC099329.3 | 16.4666975 | -0.0136319 | 0.45410711 | -0.0300192 | 0.97605173 | 0.99440179 |
| SNRK | 126.31217 | -0.5077542 | 0.29796818 | -1.7040552 | 0.08837077 | 0.63466198 |
| ANO10 | 261.399331 | 0.12723455 | 0.2578571 | 0.49343046 | 0.62170846 | 0.92868594 |
| ABHD5 | 36.3431425 | -0.2395004 | 0.3752693 | -0.6382093 | 0.52333744 | 0.90487731 |
| TCAIM | 160.315035 | 0.45346259 | 0.29409525 | 1.54189018 | 0.12310028 | 0.66929384 |
| ZNF445 | 35.0306704 | 0.14906065 | 0.28875739 | 0.51621416 | 0.60570484 | 0.92569157 |
| ZNF852 | 9.89994681 | 0.3840734 | 0.45012109 | 0.85326685 | 0.39351134 | NA |
| AC099669.1 | 80.2322157 | -0.0280342 | 0.27942216 | -0.1003293 | 0.92008286 | 0.98790868 |
| ZNF197 | 34.6307311 | 0.18822806 | 0.33647353 | 0.55941417 | 0.57587909 | 0.91854298 |
| KIAA1143 | 16.8362962 | 0.32514148 | 0.37084538 | 0.87675752 | 0.38061836 | 0.85432452 |
| ZDHHC3 | 104.623875 | 0.41814487 | 0.25614976 | 1.63242345 | 0.10259031 | 0.65946648 |
| EXOSC7 | 34.1189649 | 0.32139608 | 0.31949904 | 1.00593754 | 0.31444562 | 0.82040655 |
| CDCP1 | 70.7588118 | 0.13054559 | 0.35553589 | 0.36717977 | 7.13E-01 | 9.48E-01 |
| LARS2 | 78.1389512 | 0.03640132 | 0.24890972 | 0.14624308 | 0.8837295 | 0.98154805 |
| LIMD1 | 60.8135556 | 0.13933578 | 0.27109703 | 0.51397013 | 0.60727288 | 0.92569157 |
| SACM1L | 77.7643846 | 0.26928874 | 0.26006651 | 1.03546105 | 0.30045365 | 0.81399689 |
| SLC6A20 | 17.4294605 | 1.63668913 | 0.49707129 | 3.29266477 | 9.92E-04 | 1.39E-01 |
| LZTFL1 | 19.9064654 | 0.08228703 | 0.34957681 | 0.23539041 | 0.81390573 | 0.97282508 |
| FYCO1 | 34.7471507 | -0.3267372 | 0.34532097 | -0.9461841 | 0.34405468 | 0.83495861 |
| LTF | 324.668173 | 0.46489828 | 1.23271601 | 0.37713332 | 7.06E-01 | 9.46E-01 |
| ALS2CL | 18.8726611 | -0.3738724 | 0.34927968 | -1.0704098 | 0.2844349 | 0.80431986 |
| PTH1R | 132.61025 | 0.94094706 | 0.44337941 | 2.12221639 | 0.03381957 | 0.4908133 |
| CCDC12 | 106.163679 | 0.07430429 | 0.21714062 | 0.34219434 | 0.73220464 | 0.95279138 |
| NBEAL2 | 53.9937161 | -0.7596016 | 0.41121255 | -1.8472239 | 0.0647147 | 0.58968763 |
| SETD2 | 327.640462 | -0.1974208 | 0.16842717 | -1.1721431 | 0.2411396 | 0.78310048 |
| KIF9-AS1 | 109.86031 | 0.58541701 | 0.25036074 | 2.33829402 | 0.019372 | 0.41544117 |
| KIF9 | 11.1315988 | 0.01300863 | 0.44238289 | 0.02940582 | 0.97654093 | NA |
| KLHL18 | 41.485204 | -0.3142006 | 0.31498034 | -0.9975246 | 0.31850994 | 0.82354084 |
| PTPN23 | 25.7786424 | -0.0424695 | 0.37036072 | -0.1146707 | 0.90870616 | 0.9854228 |
| SCAP | 60.2450193 | 0.39800314 | 0.25056837 | 1.58840135 | 0.11219561 | 0.6656824 |
| ELP6 | 21.2975108 | 0.50300093 | 0.45898981 | 1.09588693 | 0.27312826 | 0.79726027 |
| SMARCC1 | 275.190154 | -0.4418011 | 0.2333474 | -1.8933193 | 0.05831542 | 0.57141408 |
| DHX30 | 53.6975193 | -0.134602 | 0.23638365 | -0.5694217 | 0.56906999 | 0.91718202 |
| MAP4 | 530.282174 | -0.4583733 | 0.23145672 | -1.9803846 | 0.04766033 | 0.53561977 |
| ZNF589 | 19.2955347 | -0.1958955 | 0.33901545 | -0.5778365 | 0.56337452 | 0.91624774 |
| NME6 | 15.5120564 | 0.53864117 | 0.42132992 | 1.27843087 | 0.20109755 | NA |

| | | | | | | |
|---|---|---|---|---|---|---|
| PLXNB1 | 101.928165 | -0.1461341 | 0.30053585 | -0.4862451 | 0.62679342 | 0.93032781 |
| TMA7 | 64.8779665 | 0.53862021 | 0.40863651 | 1.31809127 | 0.1874731 | 0.74166345 |
| SHISA5 | 92.2681227 | -0.381752 | 0.39882729 | -0.9571863 | 0.33847321 | 0.83055261 |
| PFKFB4 | 40.4163759 | -1.4700754 | 0.58461083 | -2.5146222 | 0.011916 | 0.35710117 |
| UQCRC1 | 52.8289917 | 0.23245097 | 0.36114013 | 0.64365865 | 0.51979681 | 0.90368693 |
| NCKIPSD | 16.5291496 | -0.371753 | 0.36757405 | -1.011369 | 0.31183984 | 0.81701824 |
| IP6K2 | 79.4950132 | -0.3022966 | 0.22424524 | -1.3480625 | 0.17763827 | 0.72841249 |
| PRKAR2A | 124.806442 | -0.0979203 | 0.27670206 | -0.3538837 | 0.72342605 | 0.95093722 |
| SLC25A20 | 14.3806389 | 0.07974915 | 0.58627134 | 0.13602771 | 0.89179938 | NA |
| ARIH2 | 103.022549 | 0.08892302 | 0.24648059 | 0.36077088 | 0.71827073 | 0.94925929 |
| P4HTM | 34.2370627 | 0.46064473 | 0.34004074 | 1.35467514 | 0.17552108 | 0.72841249 |
| WDR6 | 65.9660942 | -0.2356501 | 0.34887217 | -0.6754627 | 0.49938186 | 0.89380123 |
| DALRD3 | 16.8351161 | 0.19598879 | 0.38309905 | 0.51158777 | 0.60893955 | 0.92573951 |
| NDUFAF3 | 21.7738019 | 0.65175976 | 0.48671902 | 1.33908832 | 0.18054192 | 0.73303885 |
| IMPDH2 | 12.9267618 | 0.21031711 | 0.42134745 | 0.49915364 | 0.61767115 | NA |
| QRICH1 | 71.584189 | -0.1566656 | 0.22703128 | -0.6900616 | 0.49015545 | 0.88970788 |
| QARS | 21.4909736 | 0.49734594 | 0.39150327 | 1.27034938 | 0.20396021 | 0.76173533 |
| USP19 | 12.5374909 | -0.1112034 | 0.46179858 | -0.2408049 | 0.80970634 | NA |
| LAMB2 | 35.8226414 | 0.24993833 | 0.292576 | 0.85426804 | 0.39295649 | 0.85605713 |
| USP4 | 88.5097606 | -0.3425888 | 0.24953173 | -1.3729268 | 0.1697751 | 0.72416949 |
| RHOA | 159.828136 | 0.18144078 | 0.32226431 | 0.56301854 | 0.57342226 | 0.91813957 |
| TCTA | 18.7186539 | 0.60228922 | 0.4493765 | 1.3402775 | 0.18015514 | 0.73244853 |
| DAG1 | 68.6352927 | 0.12747938 | 0.26873728 | 0.47436434 | 0.63524012 | 0.93251095 |
| APEH | 23.1003189 | -0.2253347 | 0.55636571 | -0.4050119 | 0.68546879 | 0.94205398 |
| MST1 | 17.7129146 | -0.3676075 | 0.41644603 | -0.8827253 | 0.37738469 | 0.85310518 |
| RNF123 | 16.5563764 | 0.1171432 | 0.39849341 | 0.29396522 | 0.76878448 | 0.96289288 |
| IP6K1 | 118.135543 | -0.3351821 | 0.29294882 | -1.144166 | 0.2525548 | 0.79056088 |
| RBM6 | 418.211879 | -0.3392089 | 0.16659661 | -2.0361094 | 0.04173937 | 0.51069022 |
| RBM5 | 122.047449 | -0.0787075 | 0.21313986 | -0.3692765 | 0.71192165 | 0.94690572 |
| SEMA3F-AS1 | 14.6094584 | -1.3839067 | 0.76490068 | -1.8092633 | 0.07041011 | NA |
| SEMA3F | 65.058006 | -0.8896433 | 0.42968666 | -2.0704467 | 0.03841053 | 0.49671741 |
| GNAI2 | 49.7055047 | -0.2563987 | 0.35457734 | -0.7231109 | 0.46961174 | 0.88207878 |
| SEMA3B | 81.5371508 | -1.2782041 | 0.45930205 | -2.782927 | 0.00538709 | 0.28183167 |
| IFRD2 | 14.4899962 | 0.01490581 | 0.38405792 | 0.03881135 | 0.9690408 | NA |
| HYAL1 | 20.5959371 | 1.10119683 | 0.47948856 | 2.29660711 | 0.02164119 | 0.4212027 |
| HYAL2 | 12.5891243 | 0.74723739 | 0.61251283 | 1.21995385 | 0.22248237 | NA |
| TUSC2 | 15.4348284 | 0.34352462 | 0.50297373 | 0.6829872 | 0.49461493 | NA |
| TMEM115 | 14.8476081 | 0.13198869 | 0.43234103 | 0.30528837 | 0.76014653 | NA |
| CACNA2D2 | 33.8860779 | 1.56080369 | 0.53554051 | 2.91444562 | 0.00356321 | 0.23862997 |
| HEMK1 | 16.4751608 | 0.12449978 | 0.38380583 | 0.32438221 | 0.74564869 | 0.95563105 |
| CISH | 39.5924965 | 0.28706037 | 0.37948734 | 0.75644256 | 0.4493839 | 0.87315723 |
| MAPKAPK3 | 37.5795916 | -0.2975162 | 0.42749367 | -0.6959546 | 0.48645726 | 0.88838332 |
| DOCK3 | 54.2013392 | -0.0559628 | 0.71373079 | -0.0784088 | 0.93750283 | 0.99012269 |
| MANF | 30.9351272 | 0.0472015 | 0.59081181 | 0.07989261 | 0.93632267 | 0.99012269 |
| RBM15B | 11.5798567 | -0.0523867 | 0.52397984 | -0.0999784 | 0.92036145 | NA |
| DCAF1 | 56.7650123 | -0.4041041 | 0.34949878 | -1.1562391 | 0.24758339 | 0.78593073 |
| RAD54L2 | 70.7005811 | -0.140787 | 0.29774054 | -0.4728514 | 0.63631922 | 0.93251095 |
| TEX264 | 38.4723644 | 0.65203915 | 0.43610252 | 1.49515107 | 0.13487502 | 0.68328989 |
| RPL29 | 217.310898 | 0.63452853 | 0.39416733 | 1.60979481 | 0.10744266 | 0.65986731 |
| ALAS1 | 35.5136977 | 1.07670614 | 0.50597006 | 2.12800365 | 0.03333678 | 0.49058916 |
| WDR82 | 62.9622494 | -0.0216407 | 0.21461563 | -0.1008348 | 0.9196816 | 0.98759722 |
| BAP1 | 19.7711138 | -0.1302904 | 0.3614601 | -0.3604559 | 0.71850625 | 0.94925929 |
| PHF7 | 18.7116072 | 0.22471216 | 0.52941592 | 0.42445297 | 0.6712355 | 0.93878847 |
| NISCH | 72.1095523 | -0.184223 | 0.2473954 | -0.74465 | 0.45648335 | 0.87396417 |

| | | | | | | |
|---|---|---|---|---|---|---|
| SMIM4 | 39.5012788 | 0.05191995 | 0.33421771 | 0.15534768 | 0.87654723 | 0.97948831 |
| PBRM1 | 173.765664 | -0.4483762 | 0.24023722 | -1.8663893 | 0.06198693 | 0.58370968 |
| GNL3 | 23.8404572 | -0.2088328 | 0.36261834 | -0.5759024 | 0.56468113 | 0.91676339 |
| GLT8D1 | 18.3820335 | 0.60792462 | 0.37879641 | 1.60488486 | 0.10851914 | 0.65986731 |
| SPCS1 | 68.6811629 | 0.86070194 | 0.33495774 | 2.56958371 | 0.01018208 | 0.33306084 |
| NEK4 | 45.5712416 | 0.26095282 | 0.2443838 | 1.06779917 | 0.28561111 | 0.80431986 |
| SFMBT1 | 37.6303001 | -0.2956659 | 0.29440864 | -1.0042704 | 0.31524828 | 0.82084199 |
| AC096887.1 | 10.5475884 | 0.10322123 | 0.7399867 | 0.13949066 | 0.88906244 | NA |
| PRKCD | 14.5075977 | 0.64632325 | 0.48207219 | 1.34071879 | 0.18001177 | NA |
| TKT | 93.2002652 | -0.1303756 | 0.37156989 | -0.3508777 | 0.72568007 | 0.95164932 |
| DCP1A | 110.054315 | -0.32765 | 0.17865458 | -1.8339862 | 0.06665604 | 0.59652811 |
| CACNA1D | 172.860661 | 1.12922643 | 0.48205944 | 2.34250456 | 0.0191548 | 0.41544117 |
| CHDH | 34.2100362 | 0.58601813 | 0.34941326 | 1.67714909 | 0.09351333 | 0.64572229 |
| ACTR8 | 13.0320172 | 0.1732272 | 0.49566998 | 0.34948092 | 0.72672829 | NA |
| SELENOK | 28.2020321 | 0.32238762 | 0.36154549 | 0.89169311 | 0.37255745 | 0.8491102 |
| CACNA2D3 | 188.358524 | -0.0550101 | 0.60807155 | -0.0904664 | 0.92791658 | 0.98960607 |
| ERC2 | 68.7361306 | 0.5073451 | 0.31443921 | 1.61349183 | 0.1066377 | 0.65986731 |
| CCDC66 | 108.000013 | 0.26111655 | 0.2429459 | 1.07479299 | 0.28246742 | 0.80431986 |
| FAM208A | 123.044129 | -0.1026143 | 0.26209808 | -0.3915109 | 0.69541963 | 0.94395238 |
| ARHGEF3 | 832.392286 | -0.1363591 | 0.24365302 | -0.5596446 | 0.5757219 | 0.91854298 |
| SPATA12 | 18.2528998 | 0.89302655 | 0.57434812 | 1.5548524 | 0.11998123 | 0.6668162 |
| APPL1 | 70.4230455 | 0.30055606 | 0.23180547 | 1.29658743 | 0.19477318 | 0.75191699 |
| PDE12 | 18.0658236 | 0.22886446 | 0.43711776 | 0.52357621 | 0.60057333 | 0.92352528 |
| ARF4 | 82.6007433 | -0.6871722 | 0.3123668 | -2.1998888 | 0.02781479 | 0.45703613 |
| DENND6A | 87.639841 | -0.1548436 | 0.24145306 | -0.641299 | 0.52132842 | 0.90451015 |
| DENND6A-A! | 19.0830323 | -0.4826936 | 0.52297845 | -0.9229704 | 0.35602263 | 0.84192356 |
| SLMAP | 292.653423 | -0.301743 | 0.18665596 | -1.6165731 | 0.10597047 | 0.65946648 |
| FLNB | 267.747504 | -0.3875212 | 0.27809531 | -1.3934835 | 0.16347364 | 0.71332671 |
| RPP14 | 10.1947894 | 0.52254632 | 0.50268073 | 1.03951929 | 0.29856329 | NA |
| PXK | 85.6162625 | 0.16289698 | 0.35172371 | 0.46313905 | 0.6432647 | 0.93328692 |
| PDHB | 26.0531422 | 0.46675911 | 0.28718573 | 1.62528656 | 0.10410151 | 0.65946648 |
| KCTD6 | 20.8768132 | 1.07916159 | 0.47830465 | 2.25622224 | 0.02405672 | 0.43623165 |
| FAM107A | 75.3821748 | 1.77330876 | 0.50001717 | 3.5464957 | 0.00039039 | 0.07897178 |
| FAM3D-AS1 | 18.7701069 | 1.82145956 | 0.76020426 | 2.39601335 | 0.01657449 | 0.39748373 |
| C3orf67 | 18.0453918 | -0.3608957 | 0.64009214 | -0.5638183 | 0.57287779 | 0.91813957 |
| FHIT | 1838.25288 | 0.51495153 | 0.28330826 | 1.81763686 | 0.06911965 | 0.60275677 |
| PTPRG | 1154.92131 | 0.1255417 | 0.2971186 | 0.42253061 | 0.67263776 | 0.93878847 |
| THOC7 | 34.7609372 | -0.1168912 | 0.34193951 | -0.3418476 | 0.73246557 | 0.95279138 |
| ATXN7 | 21.7442808 | -0.0082204 | 0.33885326 | -0.0242594 | 0.9806457 | 0.99592962 |
| PSMD6-AS2 | 15.9185353 | 0.53422621 | 0.51261453 | 1.04215971 | 0.29733764 | NA |
| PSMD6 | 19.6964917 | 0.33660156 | 0.38107225 | 0.88330116 | 0.37707358 | 0.85310518 |
| PRICKLE2 | 345.274815 | 0.01657068 | 0.25942065 | 0.06387571 | 0.9490692 | 0.99105535 |
| ADAMTS9 | 506.097547 | -0.4873049 | 0.3159781 | -1.5422111 | 0.12302229 | 0.66929384 |
| ADAMTS9-A! | 77.5880461 | 0.37293458 | 0.52559784 | 0.70954359 | 0.47798721 | 0.88545331 |
| ADAMTS9-A! | 1486.48657 | 0.6143816 | 0.34209763 | 1.79592474 | 0.07250649 | 0.60824637 |
| MAGI1 | 1595.51061 | -0.0436385 | 0.17365527 | -0.2512936 | 0.80158711 | 0.97045451 |
| MAGI1-AS1 | 24.1365913 | 0.4149616 | 0.74324421 | 0.55831125 | 0.57663186 | 0.91854298 |
| SLC25A26 | 90.4996416 | -0.4288053 | 0.24650461 | -1.7395426 | 0.08193937 | 0.61927158 |
| LRIG1 | 27.5304687 | 0.38916518 | 0.3530186 | 1.10239283 | 0.27029093 | 0.79666376 |
| SUCLG2 | 264.170914 | -0.0338796 | 0.16248366 | -0.2085108 | 0.83483017 | 0.97600352 |
| SUCLG2-AS1 | 145.604868 | 0.74102433 | 0.35755343 | 2.0724856 | 0.03822018 | 0.49671741 |
| FAM19A1 | 47.6007554 | 0.60355713 | 0.47436309 | 1.27235264 | 0.20324785 | 0.76169832 |
| EOGT | 13.695152 | -0.2868837 | 0.40884575 | -0.7016917 | 0.48287144 | NA |
| TMF1 | 88.5226904 | -0.1494888 | 0.22483655 | -0.6648778 | 0.50612865 | 0.89723255 |

| | | | | | | |
|---|---|---|---|---|---|---|
| UBA3 | 53.7118418 | -0.0717435 | 0.24627015 | -0.2913204 | 0.7708063 | 0.96364125 |
| ARL6IP5 | 54.2715197 | 0.73845213 | 0.28327759 | 2.60681451 | 0.00913889 | 0.32071089 |
| FRMD4B | 68.2874929 | 0.0687872 | 0.31380534 | 0.21920341 | 0.8264916 | 0.97567084 |
| MITF | 532.28882 | 0.11767142 | 0.27412707 | 0.42925865 | 6.68E-01 | 0.93878847 |
| FOXP1 | 722.361961 | -0.107325 | 0.18765167 | -0.5719376 | 0.56736428 | 0.91698355 |
| EIF4E3 | 36.2331235 | 0.30707934 | 0.3184368 | 0.96433371 | 0.33487865 | 0.82974237 |
| RYBP | 139.436547 | -0.3055167 | 0.25415965 | -1.2020662 | 0.22933787 | 0.77185154 |
| SHQ1 | 61.4772266 | -0.0862818 | 0.21044289 | -0.4100011 | 0.68180511 | 0.94205398 |
| GXYLT2 | 17.0953731 | 0.16068457 | 0.54686626 | 0.2938279 | 0.76888941 | 0.96289288 |
| PPP4R2 | 125.180417 | -0.1901122 | 0.24714473 | -0.7692343 | 0.44175423 | 0.86925451 |
| PDZRN3 | 253.101624 | 0.68862699 | 0.34007229 | 2.02494297 | 4.29E-02 | 0.51555473 |
| CNTN3 | 57.5272465 | 0.60860507 | 0.45016318 | 1.35196547 | 0.17638636 | 0.72841249 |
| ZNF717 | 77.0970018 | -0.0454887 | 0.30768596 | -0.1478413 | 0.88246804 | 0.98137082 |
| ROBO2 | 21.1754067 | -0.5378041 | 0.60039659 | -0.8957482 | 0.37038727 | 0.84852785 |
| ROBO1 | 92.7034167 | 0.74525227 | 0.33278283 | 2.2394553 | 0.02512631 | 0.4403385 |
| GBE1 | 549.006447 | -1.2770713 | 0.41809414 | -3.0545066 | 0.00225431 | 0.18826595 |
| VGLL3 | 14.3275898 | -0.8575562 | 0.48191083 | -1.7794915 | 0.07515922 | NA |
| CHMP2B | 54.5928943 | 0.0004288 | 0.2259964 | 0.00189737 | 0.99848612 | 0.99957534 |
| CGGBP1 | 87.2169721 | 0.23140643 | 0.29669472 | 0.77994792 | 0.43542153 | 0.86729771 |
| ZNF654 | 130.417811 | -0.1568618 | 0.22595968 | -0.6942025 | 0.4875552 | 0.88838332 |
| C3orf38 | 21.4212927 | 0.08181374 | 0.36995729 | 0.22114376 | 0.8249805 | 0.97518529 |
| PROS1 | 15.0931321 | 0.06743432 | 0.5809092 | 0.1160841 | 9.08E-01 | NA |
| ARL13B | 45.5065054 | -0.1234548 | 0.26132643 | -0.4724161 | 0.63662983 | 0.93251095 |
| NSUN3 | 39.461464 | -0.0517531 | 0.26974007 | -0.1918627 | 0.84784972 | 0.97612951 |
| MTRNR2L12 | 236.239612 | -0.6315163 | 0.69356916 | -0.9105312 | 0.36254244 | 0.84512775 |
| ARL6 | 31.077059 | -0.0951799 | 0.35931525 | -0.2648925 | 0.79109226 | 0.96909068 |
| CRYBG3 | 210.747909 | -0.52954 | 0.26173691 | -2.0231767 | 0.04305494 | 0.51637559 |
| RIOX2 | 52.452021 | 0.11301751 | 0.29236044 | 0.3865691 | 0.69907523 | 0.94395238 |
| CLDND1 | 26.8270455 | -0.1820889 | 0.34907796 | -0.5216281 | 0.60192929 | 0.92400714 |
| DCBLD2 | 59.5885512 | -0.2208905 | 0.253022 | -0.873009 | 0.38265819 | 0.8548566 |
| CMSS1 | 151.57897 | -0.0530358 | 0.2257832 | -0.234897 | 0.81428868 | 0.97282508 |
| FILIP1L | 44.9937359 | 0.83949652 | 0.48483986 | 1.73149236 | 0.08336399 | 0.62487197 |
| TBC1D23 | 77.8736049 | -0.2091213 | 0.27107904 | -0.7714403 | 0.44044601 | 0.86895963 |
| NIT2 | 16.7947296 | 0.10565081 | 0.40842813 | 0.25867663 | 0.79588476 | 0.96974891 |
| TOMM70 | 49.8922688 | 0.14765778 | 0.24902439 | 0.59294505 | 0.55321793 | 0.91322379 |
| TFG | 104.154347 | -0.2656731 | 0.24595616 | -1.0801645 | 0.28006895 | 0.80311628 |
| SENP7 | 132.32161 | -0.0384694 | 0.23148207 | -0.1661876 | 0.86800934 | 0.97810461 |
| TRMT10C | 15.8712611 | 0.69995868 | 0.56749221 | 1.2334243 | 0.2174175 | NA |
| PCNP | 86.9369852 | -0.2813334 | 0.22066986 | -1.2749062 | 0.20234243 | 0.76065252 |
| ZBTB11 | 36.2002582 | -0.353728 | 0.33385761 | -1.0595177 | 0.28936408 | 0.80536924 |
| RPL24 | 86.0812047 | 0.278606 | 0.33924258 | 0.82125895 | 0.41149878 | 0.85867915 |
| CEP97 | 20.9252049 | 0.43904349 | 0.41414337 | 1.0601244 | 0.28908801 | 0.80536924 |
| NFKBIZ | 277.728081 | -0.8740753 | 0.40213423 | -2.1735909 | 2.97E-02 | 0.46381645 |
| ZPLD1 | 249.622478 | -1.5392439 | 0.79088484 | -1.9462302 | 5.16E-02 | 5.53E-01 |
| ALCAM | 67.1043764 | 0.70331696 | 0.38746625 | 1.8151696 | 0.06949785 | 0.60275677 |
| CBLB | 305.298876 | 0.10789651 | 0.27164811 | 0.3971922 | 6.91E-01 | 9.43E-01 |
| LINC00882 | 54.5838066 | 0.1550526 | 0.52512291 | 0.29526915 | 0.76778828 | 0.96283923 |
| BBX | 426.043745 | 0.03235819 | 0.240884 | 0.13433102 | 0.89314083 | 0.98377746 |
| CD47 | 104.447399 | -0.2207489 | 0.41453313 | -0.5325242 | 0.59436298 | 0.9229638 |
| IFT57 | 51.2436369 | 0.44507667 | 0.27782551 | 1.60200069 | 0.10915546 | 0.6606663 |
| DZIP3 | 125.724777 | 1.13626193 | 0.47167604 | 2.40898804 | 0.01599682 | 0.39716901 |
| AC078980.1 | 33.1450764 | 0.86717055 | 0.74129127 | 1.16981081 | 2.42E-01 | 7.84E-01 |
| NECTIN3 | 29.0658441 | 0.3563305 | 0.46602916 | 0.76460989 | 0.44450388 | 0.87079259 |
| PHLDB2 | 417.909872 | -0.1227474 | 0.19102536 | -0.6425712 | 5.21E-01 | 0.90427107 |

| | | | | | | |
|---|---|---|---|---|---|---|
| ABHD10 | 20.8301761 | 0.3367181 | 0.43498394 | 0.77409318 | 0.43887569 | 0.86879768 |
| C3orf52 | 60.2451506 | 0.01727638 | 0.40000664 | 0.04319022 | 0.9655499 | 0.99270395 |
| SLC9C1 | 38.6986927 | 1.12584242 | 0.7483579 | 1.5044171 | 0.13247401 | 0.67747802 |
| ATG3 | 31.299976 | -0.1026643 | 0.33512921 | -0.3063424 | 0.75934392 | 0.95903525 |
| SLC35A5 | 17.8766734 | -0.2297721 | 0.49484868 | -0.464328 | 0.64241273 | 0.93328692 |
| GTPBP8 | 17.7651191 | -0.0769754 | 0.37479211 | -0.2053816 | 0.83727397 | 0.97600352 |
| NEPRO | 42.8555589 | -0.2350909 | 0.29473891 | -0.7976242 | 0.42508858 | 0.86207532 |
| CFAP44 | 64.6601936 | 0.15812107 | 0.26020651 | 0.60767531 | 0.54340284 | 0.90965439 |
| SPICE1 | 50.6198009 | -0.4814996 | 0.2544867 | -1.8920421 | 0.05848537 | 0.57141408 |
| SIDT1 | 40.9572309 | 0.04073369 | 0.78901086 | 0.05162627 | 0.95882648 | 0.99229276 |
| USF3 | 57.0108201 | 0.17224826 | 0.27281089 | 0.63138337 | 0.52778989 | 0.90487731 |
| NAA50 | 131.302383 | -0.1421449 | 0.2675563 | -0.531271 | 0.59523103 | 0.9229638 |
| ATP6V1A | 257.01579 | 0.25038964 | 0.28475929 | 0.87930278 | 0.37923713 | 0.85429312 |
| GRAMD1C | 147.8664 | -0.126062 | 0.33415611 | -0.3772548 | 0.70598426 | 0.94558664 |
| ZDHHC23 | 26.4571794 | -0.1832902 | 0.31844968 | -0.5755705 | 0.56490552 | 0.91680067 |
| CCDC191 | 58.4544249 | -0.1316967 | 0.42089599 | -0.3128961 | 0.75435959 | 0.95826617 |
| QTRT2 | 32.8006776 | -0.5516277 | 0.34963477 | -1.5777255 | 0.11462869 | 0.6668162 |
| ZBTB20 | 2839.63486 | 0.12844001 | 0.18978796 | 0.67675532 | 4.99E-01 | 0.89378207 |
| ZBTB20-AS5 | 51.2770931 | 0.75567818 | 0.62812079 | 1.20307781 | 0.22894621 | 0.77185154 |
| LSAMP | 381.757718 | -1.6714703 | 0.54037924 | -3.0931432 | 0.00198049 | 0.18210466 |
| AC092691.1 | 131.638514 | -2.2253081 | 0.55462448 | -4.012279 | 6.01E-05 | 0.03220072 |
| AC068633.1 | 52.884024 | -3.453351 | 0.73570175 | -4.6939551 | 2.68E-06 | 0.00350412 |
| IGSF11 | 23.1183576 | 0.52679618 | 0.41524159 | 1.26864986 | 0.20456597 | 0.76173533 |
| B4GALT4 | 17.2557188 | 0.33123972 | 0.37319385 | 0.88758086 | 0.37476625 | 0.85117195 |
| TMEM39A | 30.5412135 | -0.5199475 | 0.36633622 | -1.4193178 | 0.15580639 | 0.70690214 |
| POGLUT1 | 19.9225837 | 0.18056638 | 0.42251655 | 0.42735931 | 0.66911764 | 0.93878847 |
| TIMMDC1 | 46.8124908 | 0.46944096 | 0.37247406 | 1.26033197 | 0.20754963 | 0.76231289 |
| COX17 | 24.1209677 | 0.13632729 | 0.32579791 | 0.41844127 | 0.67562452 | 0.94011158 |
| MAATS1 | 43.8608344 | 0.25778476 | 0.35675488 | 0.72258231 | 0.46993654 | 0.88220919 |
| GSK3B | 353.735807 | -0.1653516 | 0.17241666 | -0.9590234 | 0.33754697 | 0.83055261 |
| LRRC58 | 49.9237915 | 0.23301262 | 0.29030416 | 0.80264995 | 0.42217709 | 0.86190547 |
| FSTL1 | 12.3499547 | 1.29451047 | 0.5887182 | 2.19886267 | 0.02788769 | NA |
| NDUFB4 | 44.5234439 | 0.35547085 | 0.29669428 | 1.19810482 | 0.23087621 | 0.77249016 |
| HGD | 53.3024501 | 0.23840915 | 0.53243777 | 0.44776904 | 0.65431989 | 0.93548446 |
| RABL3 | 34.3281974 | -0.336295 | 0.27396285 | -1.2275205 | 0.21962703 | 0.76680432 |
| GTF2E1 | 10.9160692 | 0.31937838 | 0.44740769 | 0.71384195 | 0.47532492 | NA |
| GOLGB1 | 215.182829 | -0.2459326 | 0.23735819 | -1.0361243 | 0.30014417 | 0.81388512 |
| IQCB1 | 90.0128404 | 0.08676194 | 0.26870198 | 0.32289284 | 0.7467764 | 0.95563105 |
| EAF2 | 14.0065638 | 0.75151689 | 0.42213756 | 1.78026539 | 0.07503254 | NA |
| SLC15A2 | 35.1997081 | 1.3988539 | 0.48397829 | 2.89032367 | 0.00384845 | 0.24454118 |
| ILDR1 | 17.9413199 | 0.25819611 | 0.42966326 | 0.60092667 | 0.54788883 | 0.91157238 |
| CCDC58 | 15.9415694 | -0.2451843 | 0.43612061 | -0.5621938 | 0.57398397 | NA |
| FAM162A | 104.843823 | -0.7407622 | 0.44268301 | -1.6733467 | 0.09425906 | 0.64660155 |
| AC083798.2 | 30.2348665 | 0.39366496 | 0.47728317 | 0.82480377 | 0.409483 | 0.85867915 |
| KPNA1 | 110.236858 | -0.2572969 | 0.29391073 | -0.8754253 | 0.38134256 | 0.85432452 |
| PARP9 | 76.5261279 | -0.8966452 | 0.6283573 | -1.4269671 | 0.15358937 | 0.70572801 |
| DTX3L | 29.0511744 | -0.17662 | 0.64427608 | -0.2741372 | 0.78397921 | 0.96727606 |
| PARP14 | 199.561462 | -1.407728 | 0.71866746 | -1.958803 | 0.05013586 | 0.54461422 |
| HSPBAP1 | 39.066594 | -0.1612193 | 0.37711773 | -0.427504 | 0.66901226 | 0.93878847 |
| DIRC2 | 107.691602 | -0.6657233 | 0.51169026 | -1.3010278 | 0.19324893 | 0.75067401 |
| PDIA5 | 23.4991663 | -0.6165674 | 0.37951138 | -1.6246347 | 0.10424042 | 0.65946648 |
| SEC22A | 94.7824694 | -0.1812471 | 0.21624855 | -0.8381426 | 0.4019506 | 0.85778581 |
| ADCY5 | 14.0905596 | 0.48810581 | 0.5219281 | 0.93519742 | 0.34968657 | NA |
| HACD2 | 66.6994158 | 0.09971985 | 0.26202165 | 0.38057867 | 0.70351591 | 0.94476609 |

| | | | | | | |
|---|---|---|---|---|---|---|
| MYLK-AS1 | 16.6293247 | 0.40461477 | 0.38218886 | 1.05867756 | 0.28974665 | 0.80536924 |
| MYLK | 23.9436526 | 1.0372383 | 0.65046319 | 1.59461491 | 0.11079838 | 0.66311481 |
| CCDC14 | 83.3701822 | -0.0660238 | 0.27966033 | -0.2360858 | 0.81336606 | 0.97282508 |
| KALRN | 150.87765 | 0.15264626 | 0.4371316 | 0.34919977 | 7.27E-01 | 9.52E-01 |
| UMPS | 19.8553871 | 0.54318174 | 0.34622054 | 1.56888944 | 0.1166737 | 0.6668162 |
| ITGB5 | 89.611579 | 0.09749822 | 0.24740343 | 0.39408599 | 0.69351755 | 0.94394293 |
| SLC12A8 | 27.4735478 | 0.5524248 | 0.47712958 | 1.15780873 | 0.2469421 | 0.78583441 |
| ZNF148 | 309.431 | -0.061833 | 0.18236752 | -0.3390571 | 0.73456673 | 0.95329896 |
| SNX4 | 118.921973 | -0.2789306 | 0.2369434 | -1.1772035 | 0.23911428 | 0.78098673 |
| OSBPL11 | 42.7820193 | 0.03225543 | 0.29242215 | 0.11030434 | 0.91216801 | 0.9859256 |
| SLC41A3 | 19.6329473 | 0.4374941 | 0.38930242 | 1.12378983 | 0.2611022 | 0.79589391 |
| ALDH1L1 | 139.250295 | -0.1326481 | 0.44871253 | -0.2956194 | 0.76752074 | 0.96283923 |
| ZXDC | 138.354379 | -0.1494359 | 0.21418955 | -0.6976807 | 0.4853769 | 0.88838332 |
| TXNRD3NB | 11.222533 | 0.33832691 | 0.47928335 | 0.70590166 | 0.4802493 | NA |
| TXNRD3 | 21.4461371 | 0.23704184 | 0.36772304 | 0.64462058 | 0.5191731 | 0.90368693 |
| CHCHD6 | 50.2596854 | -0.0292002 | 0.32606416 | -0.0895536 | 0.92864194 | 0.98960607 |
| PLXNA1 | 12.1739105 | -0.2379236 | 0.48881538 | -0.4867351 | 0.62644604 | NA |
| TPRA1 | 16.226889 | 0.24486733 | 0.49288113 | 0.49680809 | 0.61932439 | 0.92786536 |
| PODXL2 | 9.60630978 | 0.06113729 | 0.73618095 | 0.08304655 | 0.93381452 | NA |
| ABTB1 | 14.9206896 | 0.33110833 | 0.46713203 | 0.70881102 | 0.47844176 | NA |
| MGLL | 249.571969 | -0.386307 | 0.35811045 | -1.078737 | 2.81E-01 | 0.80354009 |
| SEC61A1 | 77.2464531 | -0.3605782 | 0.3289691 | -1.0960854 | 0.27304141 | 0.79726027 |
| RUVBL1 | 35.9661608 | 0.53405882 | 0.37628985 | 1.41927511 | 0.15581883 | 0.70690214 |
| EEFSEC | 75.7525993 | 0.04911673 | 0.20454606 | 0.24012555 | 0.81023293 | 0.97257655 |
| GATA2 | 72.7641699 | 0.12905188 | 0.43915642 | 0.29386312 | 0.7688625 | 0.96289288 |
| GATA2-AS1 | 23.7733295 | -0.5160371 | 0.4033444 | -1.2793958 | 0.2007577 | 0.75829766 |
| RPN1 | 54.5101218 | 0.22610143 | 0.30297055 | 0.74628187 | 0.45549716 | 0.8739446 |
| RAB7A | 197.117358 | -0.5622945 | 0.25726964 | -2.1856231 | 0.02884319 | 0.46148943 |
| ACAD9 | 41.2280661 | -0.3091516 | 0.2455575 | -1.2589784 | 0.20803815 | 0.76231289 |
| CNBP | 92.429742 | 0.31487225 | 0.43299136 | 0.72720217 | 0.46710212 | 0.88090126 |
| COPG1 | 35.1649162 | -0.4726205 | 0.43769133 | -1.0798032 | 0.28022984 | 0.80311628 |
| HMCES | 26.7610591 | 0.45732829 | 0.35942581 | 1.27238578 | 0.20323608 | 0.76169832 |
| H1FX | 33.4963833 | -0.7535063 | 0.57963079 | -1.2999764 | 0.19360907 | 0.75067401 |
| MBD4 | 33.4001758 | 0.43864902 | 0.25978314 | 1.68851998 | 0.09131146 | 0.64297142 |
| IFT122 | 36.9704035 | -0.0414662 | 0.35522998 | -0.1167305 | 0.90707359 | 0.9854228 |
| TMCC1 | 388.154116 | -0.0896906 | 0.15111259 | -0.5935349 | 0.55282323 | 0.91322379 |
| PIK3R4 | 45.2284442 | 0.08589646 | 0.31909834 | 0.26918492 | 0.78778738 | 0.96767512 |
| ATP2C1 | 181.769849 | -0.176635 | 0.2186352 | -0.8078981 | 0.41914923 | 0.86031916 |
| NEK11 | 125.391442 | -0.2813208 | 0.29794428 | -0.9442062 | 0.34506429 | 0.83562656 |
| NUDT16 | 20.7316346 | 0.42883753 | 0.4553758 | 0.94172226 | 0.34633486 | 0.83687845 |
| MRPL3 | 44.1400916 | 0.20276616 | 0.31964271 | 0.63435253 | 0.52585078 | 0.90487731 |
| DNAJC13 | 102.893826 | -0.4661313 | 0.20160125 | -2.312145 | 0.02076969 | 0.4212027 |
| UBA5 | 86.1083651 | 0.04458382 | 0.33811412 | 0.13186027 | 0.89509482 | 0.98473305 |
| CDV3 | 94.1630486 | -0.8200678 | 0.35853829 | -2.2872531 | 0.02218105 | 0.4232651 |
| TOPBP1 | 38.76503 | -0.3411923 | 0.27631688 | -1.2347863 | 0.21691004 | 0.76666749 |
| SRPRB | 22.3925731 | -0.3120886 | 0.37042087 | -0.8425245 | 0.39949444 | 0.85778581 |
| SLCO2A1 | 47.7677702 | -0.2168016 | 0.78572576 | -0.2759253 | 0.78260541 | 0.96652357 |
| RYK | 168.924215 | -0.2206843 | 0.18516632 | -1.1918165 | 0.23333321 | 0.77650198 |
| AMOTL2 | 29.2024137 | 0.17916896 | 0.46715962 | 0.38352836 | 0.70132805 | 0.94440669 |
| ANAPC13 | 19.478147 | -0.5833463 | 0.43771773 | -1.3326996 | 0.18263039 | 0.73398872 |
| CEP63 | 52.227256 | -0.0016155 | 0.26900444 | -0.0060054 | 0.9952084 | 0.99927028 |
| EPHB1 | 34.1503668 | -0.4971095 | 0.42729095 | -1.1633982 | 0.24466796 | 0.78583441 |
| PPP2R3A | 864.103323 | -0.0829375 | 0.1944412 | -0.4265427 | 0.66971247 | 0.93878847 |
| MSL2 | 54.6302219 | 0.24412586 | 0.29470559 | 0.828372 | 0.40745986 | 0.85867915 |

| | | | | | | |
|---|---|---|---|---|---|---|
| PCCB | 42.8930858 | 0.60225304 | 0.27930408 | 2.15626292 | 0.03106314 | 0.47444257 |
| STAG1 | 510.776116 | -0.4740419 | 0.32734639 | -1.4481354 | 0.14757918 | 0.69842926 |
| NCK1-DT | 12.9961566 | -0.1880166 | 0.41651276 | -0.4514065 | 0.65169661 | NA |
| NCK1 | 54.8342673 | 0.10849627 | 0.34789469 | 0.31186527 | 0.75514292 | 0.95832511 |
| DZIP1L | 21.1676342 | 0.62396976 | 0.3996988 | 1.56109991 | 0.11850018 | 0.6668162 |
| DBR1 | 13.4781632 | 0.20269506 | 0.5836047 | 0.34731567 | 0.72835418 | NA |
| ARMC8 | 122.30918 | 0.02519657 | 0.19622346 | 0.12840751 | 0.89782649 | 0.98531749 |
| NME9 | 19.0082243 | 0.9971733 | 0.53866324 | 1.85119982 | 0.06414081 | 0.58858247 |
| MRAS | 39.3063164 | -0.0368303 | 0.30862425 | -0.1193369 | 0.90500842 | 0.98531749 |
| CEP70 | 137.599997 | 0.14797886 | 0.28010169 | 0.52830404 | 0.59728833 | 0.9229638 |
| FAIM | 15.7343257 | -0.2605247 | 0.39810361 | -0.6544143 | 0.51284495 | NA |
| PIK3CB | 164.593533 | -0.0948449 | 0.20350188 | -0.4660639 | 0.64116974 | 0.93328692 |
| MRPS22 | 11.8642839 | 0.51883714 | 0.53752788 | 0.96522834 | 0.33443046 | NA |
| COPB2 | 74.5432759 | -0.2057696 | 0.21079696 | -0.9761505 | 0.32898988 | 0.82781307 |
| AC097103.2 | 77.1036996 | 0.05438754 | 0.42897306 | 0.12678543 | 0.89911022 | 0.98531749 |
| NMNAT3 | 45.2976276 | 0.52118023 | 0.34608129 | 1.50594747 | 0.13208066 | 0.6765887 |
| CLSTN2 | 31.4280056 | -0.7613359 | 0.76621558 | -0.9936315 | 0.3204023 | 0.82464962 |
| SLC25A36 | 165.565349 | -0.197556 | 0.24886168 | -0.7938384 | 0.42728951 | 0.86239557 |
| PXYLP1 | 20.9290597 | 0.55951767 | 0.36603667 | 1.52858367 | 0.12636769 | 0.66929384 |
| AC117383.1 | 19.1625344 | -0.1275584 | 0.48786924 | -0.2614603 | 0.79373757 | 0.96960502 |
| ZBTB38 | 220.73325 | -0.81488 | 0.29092512 | -2.8009955 | 0.00509452 | 0.28057141 |
| RASA2 | 203.416636 | -0.6545714 | 0.2329874 | -2.8094713 | 0.00496229 | 0.27957621 |
| RNF7 | 38.8620755 | 0.11630326 | 0.33870416 | 0.34337712 | 0.73131477 | 0.95279138 |
| ATP1B3 | 409.369715 | -0.506201 | 0.76681397 | -0.6601354 | 0.50916697 | 0.89963034 |
| TFDP2 | 337.258426 | 0.11491954 | 0.30715846 | 0.37413762 | 0.70830193 | 0.94558664 |
| GK5 | 101.916422 | 0.07625274 | 0.21072684 | 0.36185585 | 0.71745976 | 0.94925929 |
| XRN1 | 236.589222 | -0.6235981 | 0.33312473 | -1.8719657 | 0.06121135 | 0.58024409 |
| ATR | 155.11533 | -0.2566188 | 0.2969798 | -0.8640951 | 0.38753566 | 0.85605713 |
| PLS1 | 70.5600131 | 0.16505092 | 0.43411072 | 0.38020466 | 0.7037935 | 0.94476609 |
| TRPC1 | 77.5040138 | 0.78069929 | 0.30450014 | 2.56387165 | 0.01035118 | 0.33440378 |
| U2SURP | 137.247909 | -0.2275025 | 0.19001155 | -1.1973087 | 0.23118624 | 0.77260653 |
| SLC9A9 | 15.8071396 | -0.716764 | 0.70974057 | -1.0098958 | 0.31254523 | NA |
| C3orf58 | 21.1107675 | -0.3129561 | 0.42413662 | -0.7378662 | 0.46059579 | 0.87623897 |
| PLOD2 | 149.857195 | -0.9210063 | 0.31889498 | -2.8881179 | 0.00387555 | 0.24454118 |
| PLSCR4 | 37.5311135 | 0.52312106 | 0.42563672 | 1.22903179 | 0.21905989 | 0.76666749 |
| PLSCR2 | 46.153944 | 0.73232934 | 0.43348832 | 1.68938656 | 0.09114537 | 0.64297142 |
| PLSCR1 | 211.330994 | -0.2306484 | 0.43374493 | -0.5317606 | 0.59489183 | 0.9229638 |
| GYG1 | 14.2249255 | 0.06766798 | 0.46697822 | 0.14490608 | 0.88478503 | NA |
| HLTF | 36.9931838 | 0.40934511 | 0.39686441 | 1.03144827 | 0.30233065 | 0.81568231 |
| HPS3 | 43.6575204 | 0.05025793 | 0.33445077 | 0.15027004 | 0.88055157 | 0.98093344 |
| WWTR1 | 526.316266 | -0.5770512 | 0.42373076 | -1.3618345 | 0.17325012 | 0.72744273 |
| COMMD2 | 26.313973 | 0.19013899 | 0.30471608 | 0.62398738 | 0.53263588 | 0.90661637 |
| ANKUB1 | 55.8692991 | 0.47161295 | 0.49012306 | 0.96223376 | 0.3359322 | 0.82974237 |
| RNF13 | 138.918937 | -0.0052995 | 0.18241952 | -0.0290514 | 0.97682364 | 0.99441127 |
| AC117386.2 | 71.014078 | 2.28808799 | 0.68268362 | 3.35160813 | 0.00080344 | 0.12609798 |
| TSC22D2 | 272.69625 | 0.14225174 | 0.28485588 | 0.49938143 | 0.6175107 | 0.92749951 |
| SERP1 | 109.283935 | 0.33796825 | 0.32545548 | 1.03844695 | 0.29906202 | 0.81301499 |
| EIF2A | 42.8177597 | 0.01790039 | 0.32363029 | 0.05531124 | 0.95589051 | 0.99148072 |
| SELENOT | 55.3317523 | 0.40290922 | 0.32406374 | 1.24330237 | 0.21375643 | 0.7643572 |
| SIAH2 | 21.3040907 | -0.2349666 | 0.47411923 | -0.4955855 | 0.62018689 | 0.92798856 |
| MED12L | 30.0674963 | 0.71681787 | 0.44502235 | 1.61074576 | 0.10723515 | 0.65986731 |
| SUCNR1 | 41.1165405 | -0.7827928 | 0.70867521 | -1.1045861 | 0.26933898 | 0.79666376 |
| MBNL1 | 523.850606 | -0.0416607 | 0.23136553 | -0.1800644 | 0.85710199 | 0.97675334 |
| ARHGEF26 | 18.7768893 | 0.6324193 | 0.448599 | 1.40976528 | 0.158609 | 0.71089007 |

| | | | | | | |
|---|---|---|---|---|---|---|
| DHX36 | 141.112139 | -0.1696132 | 0.21711245 | -0.7812229 | 0.43467139 | 0.86729771 |
| PLCH1 | 82.6158947 | 0.1786449 | 0.59292514 | 0.3012942 | 0.76319016 | 0.96091323 |
| GMPS | 56.6417558 | 0.21348035 | 0.24181693 | 0.88281801 | 0.37733461 | 0.85310518 |
| SSR3 | 46.7587813 | -0.0325412 | 0.29975231 | -0.1085603 | 0.91355126 | 0.986403 |
| TIPARP | 21.5697587 | -0.0435078 | 0.37466047 | -0.1161259 | 0.9075528 | 0.9854228 |
| LINC00886 | 46.1883962 | 0.78052895 | 0.49877999 | 1.56487624 | 0.11761193 | 0.6668162 |
| LEKR1 | 26.759775 | -0.0517767 | 0.55841019 | -0.0927217 | 0.92612466 | 0.98925753 |
| CCNL1 | 159.578931 | -0.3559787 | 0.35786083 | -0.9947405 | 0.31986247 | 0.82449332 |
| VEPH1 | 152.539413 | 0.83137977 | 0.62501398 | 1.33017788 | 0.18345967 | 0.73569753 |
| RSRC1 | 170.258595 | -0.2324039 | 0.19903645 | -1.1676447 | 0.2429501 | 0.78495944 |
| MLF1 | 22.3909674 | 0.83147281 | 0.42916392 | 1.93742474 | 0.05269344 | 0.55581509 |
| GFM1 | 46.6704194 | -0.0954122 | 0.37642213 | -0.2534712 | 0.7999041 | 0.97019967 |
| LXN | 11.8551459 | 1.11057303 | 0.55873156 | 1.98766834 | 0.04684838 | NA |
| RARRES1 | 13.6168031 | -0.0327585 | 0.56242547 | -0.0582451 | 0.95355343 | NA |
| MFSD1 | 22.2777605 | -0.0283604 | 0.35272073 | -0.0804047 | 0.93591539 | 0.99012269 |
| IQCJ-SCHIP1 | 80.5216105 | -0.2684826 | 0.64626989 | -0.4154342 | 0.67782408 | 0.94093842 |
| SMC4 | 26.6015192 | -0.0637063 | 0.31135696 | -0.2046086 | 0.83787794 | 0.97600352 |
| KPNA4 | 130.089057 | -0.3842493 | 0.31153735 | -1.2333971 | 0.21742763 | 0.76666749 |
| PPM1L | 134.064595 | 0.76348425 | 0.31347194 | 2.43557443 | 0.01486817 | 0.389664 |
| B3GALNT1 | 13.7229709 | 0.88331858 | 0.45306937 | 1.94963206 | 0.05121999 | NA |
| NMD3 | 106.023168 | 0.4183281 | 0.37857052 | 1.10502029 | 0.2691508 | 0.79666376 |
| LINC01322 | 17.1553522 | -0.7073999 | 0.50107428 | -1.4117665 | 0.15801871 | 0.71089007 |
| PDCD10 | 72.8820516 | -0.141645 | 0.21892818 | -0.6469931 | 0.51763638 | 0.90320951 |
| GOLIM4 | 57.6323722 | -0.1328916 | 0.37671728 | -0.3527621 | 0.72426679 | 0.95093722 |
| MECOM | 5670.2243 | -0.3329088 | 0.25775569 | -1.2915671 | 0.19650709 | 0.75350056 |
| AC074033.1 | 22.3314022 | 0.43296106 | 0.6306102 | 0.68657478 | 0.49235073 | 0.89067343 |
| MYNN | 17.4389336 | 0.55173836 | 0.4188868 | 1.31715385 | 0.18778706 | 0.7419 |
| LRRC34 | 10.7964591 | 0.68284397 | 0.50265095 | 1.35848538 | 0.17430972 | NA |
| SEC62 | 138.800306 | -0.0282127 | 0.23282908 | -0.1211733 | 0.9035538 | 0.98531749 |
| GPR160 | 103.554389 | -0.2843519 | 0.23592597 | -1.2052589 | 0.22810336 | 0.77185154 |
| PHC3 | 235.137458 | -0.1153791 | 0.16380078 | -0.7043869 | 0.48119189 | 0.88689225 |
| PRKCI | 135.090378 | -0.3129766 | 0.29472348 | -1.0619331 | 0.28826606 | 0.80523851 |
| SKIL | 82.9897879 | -0.3530047 | 0.42535329 | -0.8299094 | 0.40659003 | 0.85867915 |
| TNIK | 67.6040464 | 0.83912425 | 0.80916623 | 1.03702332 | 0.29972499 | 0.81371804 |
| PLD1 | 60.6042507 | 0.04725521 | 0.3731146 | 0.12665067 | 0.89921689 | 0.98531749 |
| FNDC3B | 1261.9181 | -0.2699851 | 0.19498796 | -1.3846247 | 0.16616726 | 0.71756194 |
| TNFSF10 | 139.061181 | -0.8411434 | 0.57419579 | -1.4649069 | 1.43E-01 | 6.94E-01 |
| NCEH1 | 126.627718 | -0.2912322 | 0.36241917 | -0.8035784 | 0.42164049 | 0.86190547 |
| ECT2 | 27.636719 | -0.2307305 | 0.37592671 | -0.6137645 | 5.39E-01 | 0.90763121 |
| NLGN1 | 1083.19283 | 0.19238914 | 0.58217845 | 0.3304642 | 7.41E-01 | 9.55E-01 |
| NAALADL2 | 1641.35317 | 0.05829796 | 0.30359944 | 0.1920226 | 0.8477245 | 0.97612951 |
| TBL1XR1 | 656.374365 | -0.8792177 | 0.34054048 | -2.5818303 | 0.00982779 | 0.32904468 |
| ZMAT3 | 47.7935058 | 0.79299802 | 0.29776022 | 2.66321012 | 0.00773991 | 0.30654023 |
| PIK3CA | 110.277076 | -0.4794817 | 0.16846129 | -2.8462427 | 0.00442385 | 0.26149529 |
| KCNMB3 | 18.5386776 | -0.7378044 | 0.40816191 | -1.8076269 | 0.07066459 | 0.60513619 |
| ZNF639 | 15.7132018 | 0.2881561 | 0.38403267 | 0.75034268 | 0.45304834 | NA |
| MFN1 | 121.540429 | -0.1618118 | 0.19590845 | -0.8259562 | 0.40882895 | 0.85867915 |
| GNB4 | 17.9280516 | -0.0601043 | 0.54726949 | -0.1098258 | 0.91254753 | 0.9862187 |
| ACTL6A | 23.0747868 | 0.2011006 | 0.34868983 | 0.57673204 | 0.56412048 | 0.91650701 |
| MRPL47 | 23.9670555 | -0.4488099 | 0.37345058 | -1.2017919 | 0.22944417 | 0.77185154 |
| NDUFB5 | 58.2210799 | 0.773471 | 0.26456355 | 2.92357357 | 0.00346038 | 0.23507372 |
| USP13 | 14.9527938 | 0.75153537 | 0.4063955 | 1.84927089 | 0.06441871 | NA |
| TTC14 | 148.796194 | -0.0662486 | 0.20344685 | -0.325631 | 0.74470352 | 0.95563105 |
| FXR1 | 143.21555 | 0.01824691 | 0.27401255 | 0.06659151 | 0.94690691 | 0.99083449 |

| | | | | | | |
|---|---|---|---|---|---|---|
| DNAJC19 | 20.4214987 | -0.0388061 | 0.40406872 | -0.0960383 | 0.92349017 | 0.98923641 |
| ATP11B | 160.468139 | -0.0172688 | 0.26093931 | -0.0661795 | 0.94723493 | 0.99083449 |
| DCUN1D1 | 96.2071326 | -0.1723284 | 0.24559022 | -0.7016906 | 0.48287211 | 0.88781758 |
| MCCC1 | 125.283215 | -0.2611207 | 0.22743629 | -1.1481047 | 0.25092536 | 0.78892875 |
| KLHL24 | 159.667709 | -0.3287148 | 0.28395335 | -1.1576368 | 0.24701229 | 0.78583441 |
| YEATS2 | 83.968181 | -0.6353002 | 0.32715366 | -1.9419015 | 0.05214902 | 0.55403388 |
| ABCC5 | 245.592528 | -0.1212701 | 0.28603567 | -0.4239685 | 0.67158876 | 0.93878847 |
| EIF2B5 | 20.7574299 | -0.56821 | 0.33363699 | -1.7030786 | 0.08855337 | 0.63466198 |
| DVL3 | 32.0809547 | 0.20416195 | 0.35618407 | 0.57319226 | 0.56651453 | 0.91698355 |
| AP2M1 | 68.5232363 | 0.22697581 | 0.37858942 | 0.59953025 | 0.54881934 | 0.91208861 |
| PSMD2 | 29.6649961 | 0.16091709 | 0.31894266 | 0.50453297 | 0.6138869 | 0.92662475 |
| EIF4G1 | 69.947217 | -0.0535274 | 0.3417856 | -0.156611 | 0.8755514 | 0.97918624 |
| FAM131A | 14.3837138 | 0.36440611 | 0.57950072 | 0.6288277 | 0.52946186 | NA |
| POLR2H | 44.7320062 | -0.1719335 | 0.30096837 | -0.5712675 | 0.56781832 | 0.91698355 |
| MAGEF1 | 16.5671705 | 0.44555654 | 0.43413909 | 1.02629906 | 0.30475064 | 0.81568231 |
| VPS8 | 149.250323 | -0.4064319 | 0.23074224 | -1.7614109 | 0.07816888 | 0.61367304 |
| EHHADH | 41.2459326 | 0.6405547 | 0.3948596 | 1.62223407 | 0.10475324 | 0.65946648 |
| MAP3K13 | 189.695948 | 0.21583951 | 0.26262358 | 0.82185885 | 0.41115723 | 0.85867915 |
| TMEM41A | 27.3391302 | -0.1968913 | 0.33598345 | -0.5860148 | 0.5578656 | 0.91425998 |
| LIPH | 53.4456726 | -0.371887 | 0.40644181 | -0.9149822 | 0.36020096 | 0.84282379 |
| SENP2 | 68.683378 | -0.4530229 | 0.25597642 | -1.7697839 | 0.07676315 | 0.61188697 |
| IGF2BP2 | 273.335376 | -0.3445899 | 0.32135406 | -1.0723061 | 0.28358258 | 0.80431986 |
| TRA2B | 78.8243571 | 0.26860785 | 0.25397718 | 1.05760627 | 0.29023498 | 0.80559136 |
| TBCCD1 | 11.828223 | -0.1623535 | 0.4587969 | -0.353868 | 0.72343782 | NA |
| AC068631.2 | 12.957701 | 0.46985036 | 0.64365971 | 0.72996702 | 0.46541034 | NA |
| KNG1 | 72.6659686 | 0.04865808 | 0.72328559 | 0.06727367 | 9.46E-01 | 9.91E-01 |
| EIF4A2 | 108.307795 | 0.3353755 | 0.31264944 | 1.07268862 | 0.28341085 | 0.80431986 |
| ST6GAL1 | 436.18296 | 0.15100437 | 0.33911153 | 0.44529413 | 0.65610721 | 0.93570875 |
| BCL6 | 99.1692347 | -0.4396983 | 0.31302955 | -1.4046543 | 1.60E-01 | 7.11E-01 |
| LPP | 1112.30652 | -0.5084098 | 0.2611096 | -1.9471128 | 0.05152121 | 0.55289583 |
| TPRG1 | 16.3637995 | -0.4475637 | 0.43932286 | -1.0187581 | 0.30831783 | 0.81568231 |
| P3H2 | 241.64444 | -0.1236002 | 0.42528855 | -0.2906267 | 0.77133687 | 0.96366407 |
| CCDC50 | 125.211936 | -0.1175718 | 0.21228092 | -0.5538501 | 0.57968147 | 0.92027213 |
| FGF12 | 30.534891 | 0.24822976 | 0.78491933 | 0.31624875 | 0.75181371 | 0.95769549 |
| MB21D2 | 20.7551688 | 0.63428277 | 0.37106873 | 1.70934042 | 0.08738791 | 0.63466198 |
| OPA1 | 141.336371 | -0.1480078 | 0.1834845 | -0.8066503 | 0.41986796 | 0.86082389 |
| HES1 | 151.739217 | -0.4539334 | 0.44273251 | -1.0252995 | 0.30522188 | 0.81568231 |
| ATP13A3 | 181.033176 | -0.2404073 | 0.27843982 | -0.8634084 | 0.38791294 | 0.85605713 |
| LINC00884 | 14.7244605 | 0.32882595 | 0.47043156 | 0.69898786 | 0.48455962 | NA |
| LSG1 | 29.9719671 | -0.1580792 | 0.38070588 | -0.4152267 | 0.67797595 | 0.94093842 |
| XXYLT1 | 26.8500336 | -0.0872003 | 0.30296832 | -0.2878198 | 0.7734847 | 0.96435639 |
| ACAP2 | 213.68551 | -0.2718911 | 0.22601598 | -1.2029729 | 0.22898679 | 0.77185154 |
| PPP1R2 | 49.9149055 | -0.1557909 | 0.25467233 | -0.6117307 | 0.54071596 | 0.90783815 |
| MUC20-OT1 | 587.630919 | -0.1730195 | 0.41100358 | -0.4209684 | 0.67377818 | 0.93901638 |
| TNK2 | 12.9115169 | 0.48517088 | 0.43483742 | 1.11575237 | 0.26452815 | NA |
| TFRC | 87.4359852 | -0.2724559 | 0.31645696 | -0.8609572 | 0.3892616 | 0.85605713 |
| LINC00885 | 12.8542356 | 2.70174111 | 1.00104148 | 2.69893023 | 0.00695628 | NA |
| PCYT1A | 34.7103859 | -0.2271638 | 0.27618479 | -0.8225065 | 0.41078868 | 0.85867915 |
| UBXN7 | 66.9811907 | -0.1085262 | 0.27555606 | -0.3938443 | 0.69369599 | 0.94395238 |
| RNF168 | 60.8923653 | -0.3967001 | 0.24789288 | -1.6002883 | 0.10953465 | 0.66067258 |
| PIGX | 24.4422317 | 0.35916269 | 0.41587895 | 0.86362314 | 0.38779495 | 0.85605713 |
| PAK2 | 131.972974 | -0.0156884 | 0.1879277 | -0.0834811 | 0.93346899 | 0.98960607 |
| SENP5 | 147.499654 | -0.2567167 | 0.35181111 | -0.7297004 | 0.46557335 | 0.87981788 |
| NCBP2 | 27.9713538 | 0.22923092 | 0.35145886 | 0.65222688 | 0.51425482 | 0.90156229 |

| | | | | | | |
|---|---|---|---|---|---|---|
| NCBP2-AS2 | 12.9646304 | 0.93512205 | 0.51911026 | 1.80139389 | 0.07164082 | NA |
| DLG1 | 409.425497 | -0.2413752 | 0.21155326 | -1.1409665 | 0.25388387 | 0.7912221 |
| BDH1 | 59.7513117 | 0.1150209 | 0.36687403 | 0.31351606 | 0.75388862 | 0.95826617 |
| RUBCN | 43.3054413 | -0.3759379 | 0.40012232 | -0.9395574 | 0.34744465 | 0.83687845 |
| FYTTD1 | 48.3956862 | -0.0491444 | 0.34433702 | -0.1427219 | 0.88650982 | 0.98216235 |
| LRCH3 | 152.168338 | -0.4491678 | 0.21110231 | -2.1277255 | 0.03335985 | 0.49058916 |
| IQCG | 27.9575817 | 0.02995291 | 0.31045774 | 0.09647982 | 0.9231395 | 0.98915831 |
| RPL35A | 93.6299036 | 0.57802552 | 0.33174511 | 1.7423784 | 0.08144225 | 0.61925304 |
| LMLN | 39.9039412 | 0.59102844 | 0.35289263 | 1.67481092 | 0.09397133 | 0.64572229 |
| ZNF595 | 33.5054141 | 0.42967616 | 0.41243042 | 1.04181491 | 0.2974975 | 0.81214575 |
| ZNF718 | 102.460993 | -0.058988 | 0.19191626 | -0.3073634 | 0.75856679 | 0.95903525 |
| ZNF141 | 92.094288 | -0.0566834 | 0.36510449 | -0.1552527 | 0.87662213 | 0.97948831 |
| AC092574.2 | 11.8341439 | 1.06847 | 0.86077051 | 1.24129485 | 0.21449685 | NA |
| ZNF721 | 187.958829 | -0.0372754 | 0.21110102 | -0.176576 | 0.85984146 | 0.97691173 |
| PIGG | 42.4790576 | -0.1078584 | 0.25007041 | -0.4313122 | 0.66624135 | 0.93860542 |
| ATP5ME | 36.3929085 | -0.0942605 | 0.40548137 | -0.2324658 | 0.81617625 | 0.97288403 |
| PCGF3 | 69.3324761 | -0.1564978 | 0.20997893 | -0.7453023 | 0.45608902 | 0.87394768 |
| AC139887.2 | 113.980055 | -0.1681156 | 0.26337816 | -0.6383051 | 0.52327508 | 0.90487731 |
| GAK | 91.1963522 | -0.6126041 | 0.35196747 | -1.7405135 | 0.08176888 | 0.61925304 |
| TMEM175 | 13.5515095 | 0.24153126 | 0.50467993 | 0.47858304 | 0.63223529 | NA |
| IDUA | 14.6567622 | 0.14494715 | 0.43863954 | 0.33044706 | 0.74106219 | NA |
| FGFRL1 | 14.2370157 | 0.08482218 | 0.41052009 | 0.20662126 | 0.83630565 | NA |
| RNF212 | 30.7394992 | -0.5797532 | 0.4293602 | -1.3502724 | 1.77E-01 | 0.72841249 |
| AC092535.3 | 91.6075168 | 0.10500094 | 0.34994753 | 0.30004766 | 0.7641408 | 0.96130096 |
| CTBP1-AS | 18.4426284 | -0.2669852 | 0.529229 | -0.5044795 | 0.61392445 | 0.92662475 |
| CTBP1 | 52.5964334 | -0.2451723 | 0.25363123 | -0.9666489 | 0.33371959 | 0.82974237 |
| CTBP1-DT | 14.7458078 | 0.25688805 | 0.42826719 | 0.59983127 | 0.54861869 | NA |
| MAEA | 77.9423018 | -0.1277485 | 0.25128125 | -0.5083885 | 0.6111809 | 0.92619266 |
| UVSSA | 43.1657265 | -0.4762995 | 0.26505778 | -1.7969648 | 0.07234121 | 0.60824637 |
| SLBP | 57.7162878 | 0.20585352 | 0.36691622 | 0.56103686 | 0.57477241 | 0.91824492 |
| TMEM129 | 23.2504568 | 0.52416043 | 0.44738916 | 1.17159842 | 0.24135832 | 0.78323911 |
| LETM1 | 37.2562779 | 0.10322753 | 0.29716295 | 0.34737684 | 0.72830823 | 0.95272318 |
| NSD2 | 116.703996 | -0.2294273 | 0.18851507 | -1.2170236 | 0.22359522 | 0.76834258 |
| NELFA | 75.1516477 | 0.11665062 | 0.24966688 | 0.46722507 | 0.64033886 | 0.93328692 |
| C4orf48 | 20.1289612 | -0.6051526 | 0.78351942 | -0.7723518 | 0.43990609 | 0.86895963 |
| NAT8L | 24.575189 | -0.3276688 | 0.61348429 | -0.5341112 | 0.59326459 | 0.9229638 |
| POLN | 11.5865549 | -0.0577862 | 0.49254923 | -0.1173207 | 0.90660588 | NA |
| MXD4 | 45.0492025 | 0.54280191 | 0.32496799 | 1.67032424 | 0.09485523 | 0.64796091 |
| ZFYVE28 | 74.0706067 | -0.0504608 | 0.28442641 | -0.1774124 | 0.85918449 | 0.97691173 |
| RNF4 | 59.8451602 | -0.5338604 | 0.25327199 | -2.1078539 | 0.03504363 | 0.49209205 |
| FAM193A | 148.887329 | -0.5650178 | 0.24832825 | -2.275286 | 0.02288878 | 0.42783689 |
| TNIP2 | 20.7121109 | -0.0795365 | 0.34527842 | -0.2303547 | 0.81781613 | 0.97288403 |
| SH3BP2 | 74.7192094 | -0.5794619 | 0.35157779 | -1.6481754 | 9.93E-02 | 6.58E-01 |
| ADD1 | 146.763322 | -0.1287591 | 0.23255853 | -0.5536634 | 0.57980926 | 0.92031451 |
| MFSD10 | 35.3490314 | -0.6441687 | 0.57808412 | -1.1143166 | 0.2651434 | 0.79666376 |
| NOP14 | 18.1240813 | -0.6146434 | 0.35264799 | -1.7429375 | 0.08134454 | 0.61925304 |
| GRK4 | 26.6857373 | 0.39466093 | 0.56218864 | 0.70200801 | 0.48267417 | 0.88781228 |
| HTT | 143.990491 | -0.0723028 | 0.19821167 | -0.3647756 | 0.71527893 | 0.9487373 |
| RGS12 | 67.3678027 | -0.6485085 | 0.38217823 | -1.6968746 | 0.08972038 | 0.63707066 |
| LRPAP1 | 66.3161116 | 0.9500399 | 0.26676437 | 3.56134483 | 0.00036896 | 0.0763464 |
| TMEM128 | 16.3233737 | 0.78932757 | 0.3894142 | 2.02696144 | 0.04266636 | 0.51510854 |
| ZBTB49 | 12.4454823 | -0.8843655 | 0.41865897 | -2.1123767 | 0.03465415 | NA |
| STX18 | 170.664533 | -0.1472615 | 0.20043747 | -0.7347007 | 0.46252182 | 0.87713916 |
| STX18-AS1 | 105.249332 | 0.06926935 | 0.35399562 | 0.19567857 | 0.84486176 | 0.97612951 |

| | | | | | | |
|---|---|---|---|---|---|---|
| STK32B | 35.9722509 | 0.64562333 | 0.42017605 | 1.53655433 | 1.24E-01 | 6.69E-01 |
| EVC2 | 49.5216242 | -0.5038117 | 0.28022062 | -1.7979109 | 0.07219113 | 0.60821941 |
| EVC | 55.4882147 | 0.1062662 | 0.31638328 | 0.33587806 | 0.73696282 | 0.95387069 |
| CRMP1 | 18.1064141 | -0.2642726 | 0.68949417 | -0.3832848 | 0.7015086 | 0.94451009 |
| WFS1 | 10.2075451 | 1.08431122 | 0.70492642 | 1.53819063 | 0.12400201 | NA |
| MAN2B2 | 19.2251351 | 1.07547839 | 0.53851533 | 1.99711749 | 0.04581242 | 0.52835471 |
| MRFAP1 | 57.3584767 | 0.38858274 | 0.40207801 | 0.9664362 | 0.33382596 | 0.82974237 |
| LINC02482 | 11.3366322 | 0.3206327 | 0.54686114 | 0.58631465 | 0.55766409 | NA |
| MRFAP1L1 | 18.0244191 | 0.47185434 | 0.47031887 | 1.00326476 | 0.31573314 | 0.82094223 |
| KIAA0232 | 207.919463 | -0.1738174 | 0.21539862 | -0.8069569 | 0.41969132 | 0.86065558 |
| TBC1D14 | 126.721242 | -0.0821437 | 0.21863629 | -0.3757091 | 0.70713315 | 0.94558664 |
| TADA2B | 15.6283424 | 0.22435971 | 0.43814288 | 0.51206974 | 0.6086022 | NA |
| GRPEL1 | 19.7050063 | -0.2835683 | 0.40856175 | -0.6940647 | 0.48764161 | 0.88838332 |
| SORCS2 | 95.6352925 | 0.30849643 | 0.3650691 | 0.84503572 | 0.3980909 | 0.85758974 |
| AFAP1 | 22.9080758 | 0.04955451 | 0.55970498 | 0.08853684 | 0.92945001 | 0.98960607 |
| SH3TC1 | 86.2629885 | 0.22631293 | 0.28325199 | 0.79898088 | 0.4243015 | 0.86190547 |
| ACOX3 | 30.2523914 | 0.04930595 | 0.34853303 | 0.14146709 | 0.88750096 | 0.9823591 |
| TRMT44 | 25.2801901 | -0.2931993 | 0.34711945 | -0.8446639 | 0.39829854 | 0.85758974 |
| WDR1 | 86.8349803 | 0.04001965 | 0.32554903 | 0.12292972 | 0.90216275 | 0.98531749 |
| ZNF518B | 13.3359929 | -0.1900641 | 0.50965207 | -0.3729292 | 0.70920117 | NA |
| HS3ST1 | 66.5896351 | -0.0070327 | 0.36963963 | -0.0190258 | 9.85E-01 | 9.97E-01 |
| AC005699.1 | 44.5164786 | -0.6427609 | 0.39025693 | -1.6470198 | 0.09955398 | 0.65860455 |
| RAB28 | 198.324147 | -0.2135562 | 0.24865031 | -0.8588617 | 0.39041684 | 0.85605713 |
| BOD1L1 | 124.060675 | -0.420294 | 0.18730208 | -2.2439365 | 0.02483649 | 0.43948277 |
| LINC01182 | 75.7122918 | -1.5600033 | 0.43420017 | -3.5928205 | 0.00032712 | 0.0763464 |
| CPEB2 | 222.296934 | -1.1661942 | 0.40909123 | -2.8506948 | 0.00436238 | 0.26125499 |
| AC098829.1 | 69.8708111 | -0.494559 | 0.56789393 | -0.8708651 | 0.38382784 | 0.85511218 |
| CC2D2A | 41.3926004 | 0.6698777 | 0.3938575 | 1.70081235 | 0.08897823 | 0.63542189 |
| FBXL5 | 109.132271 | -0.2115468 | 0.2075463 | -1.0192751 | 0.3080724 | 0.81568231 |
| FAM200B | 37.8803972 | 0.34765616 | 0.34356741 | 1.01190087 | 0.31158545 | 0.81701824 |
| PROM1 | 126.852695 | 0.90020088 | 0.50991603 | 1.76539044 | 0.07749817 | 0.61344855 |
| TAPT1 | 88.6465192 | 0.34682371 | 0.25497073 | 1.36024915 | 0.1737511 | 0.72794035 |
| TAPT1-AS1 | 203.427462 | 0.65152797 | 0.40809234 | 1.59652094 | 0.11037253 | 0.66311481 |
| AC097515.1 | 34.3349153 | 0.39693506 | 0.77268455 | 0.51370906 | 0.60745542 | 0.92569157 |
| QDPR | 17.4361735 | 0.33475874 | 0.51228638 | 0.65346016 | 0.51345967 | 0.90156229 |
| LAP3 | 43.6348059 | -0.7769483 | 0.7046065 | -1.1026698 | 0.27017058 | 0.79666376 |
| MED28 | 46.6642974 | -0.3395806 | 0.26407899 | -1.2859056 | 0.19847602 | 0.75563666 |
| FAM184B | 11.790174 | 0.50150127 | 0.57996263 | 0.86471308 | 0.38719629 | NA |
| DCAF16 | 55.4512653 | -0.0624153 | 0.2890579 | -0.2159267 | 0.82904487 | 0.97567084 |
| LCORL | 267.527364 | 0.0255532 | 0.31256069 | 0.08175435 | 0.93484206 | 0.99012269 |
| AC024230.1 | 22.6170759 | -0.0238698 | 0.52114157 | -0.0458029 | 0.96346736 | 0.99256424 |
| SLIT2 | 370.423466 | 0.07843483 | 0.40586636 | 0.19325284 | 8.47E-01 | 0.97612951 |
| PACRGL | 56.9921955 | -0.4305088 | 0.25214221 | -1.7074046 | 0.08774688 | 0.63466198 |
| KCNIP4 | 7250.11125 | 0.70055152 | 0.30690752 | 2.28261436 | 0.02245309 | 0.425412 |
| ADGRA3 | 86.0131037 | 0.25521325 | 0.36645636 | 0.6964356 | 0.48615607 | 0.88838332 |
| PPARGC1A | 854.21757 | -0.2833192 | 0.31383848 | -0.9027549 | 0.36665599 | 0.84741115 |
| DHX15 | 145.740641 | -0.2787364 | 0.24903713 | -1.1192563 | 0.26303081 | 0.79666376 |
| CCDC149 | 28.1094242 | -0.5536704 | 0.34472786 | -1.6061086 | 0.10825005 | 0.65986731 |
| SEPSECS | 26.702386 | -0.4340344 | 0.34318935 | -1.2647081 | 0.20597598 | 0.76231289 |
| AC104662.2 | 9.73694889 | 0.362768 | 0.65050511 | 0.55767124 | 0.5770689 | NA |
| ZCCHC4 | 17.3800912 | 0.22724908 | 0.42351731 | 0.53657567 | 0.59156076 | 0.9227172 |
| ANAPC4 | 53.6167718 | -0.0899337 | 0.37288554 | -0.2411832 | 0.80941316 | 0.97257655 |
| SEL1L3 | 20.2029101 | -0.4295749 | 0.63788186 | -0.6734396 | 0.50066769 | 0.89492452 |
| SMIM20 | 20.2026103 | -0.4812692 | 0.42385616 | -1.1354541 | 0.25618512 | 0.7940072 |

| | | | | | | |
|---|---|---|---|---|---|---|
| RBPJ | 159.457328 | -0.076936 | 0.18912944 | -0.4067901 | 0.68416217 | 0.94205398 |
| TBC1D19 | 120.625732 | -0.2361125 | 0.24007399 | -0.9834989 | 0.32536194 | 0.82567895 |
| STIM2 | 363.454563 | 0.53171495 | 0.44106202 | 1.20553329 | 0.22799749 | 0.77185154 |
| PCDH7 | 449.339247 | 0.3207364 | 0.37137344 | 0.86364927 | 0.38778059 | 0.85605713 |
| ARAP2 | 105.462137 | -0.4997553 | 0.55573673 | -0.8992662 | 0.36851088 | 0.84818067 |
| C4orf19 | 628.135912 | 0.04633588 | 0.24802037 | 0.1868229 | 0.8517995 | 0.97612951 |
| RELL1 | 79.9435956 | -0.0560099 | 0.33027026 | -0.1695882 | 0.86533401 | 0.97788853 |
| PGM2 | 25.5559456 | -0.1592514 | 0.28952323 | -0.5500471 | 0.58228704 | 0.92034873 |
| TBC1D1 | 521.128608 | -0.2423599 | 0.18025359 | -1.3445497 | 0.17877069 | 0.72924345 |
| KLF3 | 76.0335484 | -0.5248173 | 0.26566114 | -1.9755141 | 4.82E-02 | 0.5380949 |
| FAM114A1 | 27.9208656 | -0.7357263 | 0.35312085 | -2.0834972 | 0.03720592 | 0.49537143 |
| KLHL5 | 138.029199 | 0.43095643 | 0.30561434 | 1.41013159 | 0.15850083 | 0.71089007 |
| WDR19 | 67.641403 | -0.3054801 | 0.29329277 | -1.0415533 | 0.29761883 | 0.81214575 |
| RFC1 | 69.7407089 | -0.0735149 | 0.23300753 | -0.3155044 | 0.75237871 | 0.95782508 |
| RPL9 | 14.8475065 | 0.05212057 | 0.42140548 | 0.1236827 | 0.90156651 | NA |
| LIAS | 30.7027323 | 0.75620868 | 0.43352988 | 1.74430576 | 0.08110579 | 0.61925304 |
| UGDH | 63.05471 | -0.7110447 | 0.43989556 | -1.6163945 | 0.10600906 | 0.65946648 |
| UGDH-AS1 | 53.3357037 | 0.12263728 | 0.3615109 | 0.33923536 | 0.73443243 | 0.95329896 |
| SMIM14 | 164.766986 | -0.1625489 | 0.21060735 | -0.77181 | 0.44022697 | 0.86895963 |
| UBE2K | 355.807292 | -0.4058184 | 0.2242079 | -1.8100091 | 0.07029437 | 0.60423953 |
| PDS5A | 228.549825 | -0.1187421 | 0.19327114 | -0.6143808 | 0.5389637 | 0.90763121 |
| N4BP2 | 48.4628175 | -0.2032821 | 0.25596727 | -0.7941722 | 0.42709519 | 0.86239557 |
| RBM47 | 821.12199 | -0.4476759 | 0.27712729 | -1.615416 | 0.10622064 | 0.65986731 |
| NSUN7 | 56.2250341 | 0.37449474 | 0.30794662 | 1.21610278 | 0.22394577 | 0.76840496 |
| APBB2 | 576.608103 | -0.3922218 | 0.21940301 | -1.7876775 | 7.38E-02 | 6.10E-01 |
| UCHL1 | 54.9054494 | -0.4275953 | 0.60240742 | -0.7098108 | 0.47782145 | 0.88545331 |
| LIMCH1 | 1088.98277 | 0.39751423 | 0.26066405 | 1.52500596 | 0.12725761 | 0.66929384 |
| TMEM33 | 54.9628211 | -0.2024729 | 0.34966026 | -0.579056 | 0.56255141 | 0.91624774 |
| SLC30A9 | 202.387567 | -0.2488712 | 0.15049677 | -1.6536644 | 0.09819572 | 0.65695163 |
| AC024022.1 | 21.6060954 | 2.71302947 | 0.79445775 | 3.41494493 | 0.00063795 | 0.10957077 |
| ATP8A1 | 113.102027 | -0.3563038 | 0.22738187 | -1.5669843 | 0.11711836 | 0.6668162 |
| KCTD8 | 328.714537 | 0.57176497 | 0.31597757 | 1.80951129 | 0.07037161 | 0.60433278 |
| GUF1 | 34.6286807 | -0.2286658 | 0.29326158 | -0.7797331 | 0.43554797 | 0.86729771 |
| GNPDA2 | 27.2395079 | 0.35721495 | 0.33390125 | 1.06982214 | 0.28469937 | 0.80431986 |
| GABRA2 | 78.8252166 | 0.27581261 | 0.56091053 | 0.491723 | 0.62291518 | 0.92911632 |
| ATP10D | 77.9762079 | 0.12260197 | 0.28468236 | 0.43066233 | 0.66671391 | 0.93860542 |
| AC107068.2 | 15.8734058 | 1.38203455 | 0.62593304 | 2.20795911 | 0.02724712 | NA |
| NFXL1 | 65.7878068 | -0.232967 | 0.36958438 | -0.6303487 | 0.52846649 | 0.90487731 |
| NIPAL1 | 58.7383167 | -0.590725 | 0.38014174 | -1.5539598 | 0.12019401 | 0.6668162 |
| CNGA1 | 40.4268555 | -0.0978946 | 0.40757253 | -0.2401895 | 0.81018338 | 0.97257655 |
| TXK | 33.3901628 | 0.14406161 | 0.3049103 | 0.47247212 | 6.37E-01 | 0.93251095 |
| TEC | 54.9213961 | -0.2695337 | 0.25546338 | -1.0550776 | 0.29138986 | 0.80688911 |
| SLAIN2 | 101.90412 | -0.0605604 | 0.21119061 | -0.2867571 | 0.77429828 | 0.96435639 |
| FRYL | 316.015757 | -0.1434579 | 0.16380068 | -0.8758077 | 0.3811346 | 0.85432452 |
| OCIAD1 | 164.741473 | 0.12146373 | 0.16975596 | 0.71551966 | 0.474288 | 0.88403192 |
| OCIAD2 | 32.7281116 | -0.0289249 | 0.27684217 | -0.1044816 | 0.91678715 | 0.98710975 |
| CWH43 | 65.073132 | 0.41972933 | 0.37971275 | 1.10538643 | 2.69E-01 | 7.97E-01 |
| DCUN1D4 | 123.549766 | -0.0323388 | 0.28007395 | -0.1154651 | 0.90807648 | 0.9854228 |
| SGCB | 13.8657574 | 0.93301552 | 0.40973701 | 2.27710822 | 0.02277976 | NA |
| USP46 | 38.3229307 | 0.06547248 | 0.31136654 | 0.21027463 | 0.83345334 | 0.97600352 |
| SCFD2 | 129.569618 | -0.2942762 | 0.28088481 | -1.0476757 | 0.29478805 | 0.8101516 |
| AC058822.1 | 56.4630089 | 0.52336877 | 0.57843925 | 0.90479471 | 0.36557416 | 0.84720508 |
| LNX1 | 136.260502 | -0.1962548 | 0.20524116 | -0.9562158 | 3.39E-01 | 0.83066816 |
| CHIC2 | 87.3078076 | -0.2556971 | 0.24594751 | -1.039641 | 0.29850671 | 0.81246117 |

| | | | | | | |
|---|---|---|---|---|---|---|
| TMEM165 | 163.127208 | -0.133407 | 0.26329309 | -0.5066864 | 0.61237489 | 0.92644418 |
| CLOCK | 112.393454 | 0.27300703 | 0.22688803 | 1.20326767 | 0.22887276 | 0.77185154 |
| EXOC1 | 120.056186 | -0.3716769 | 0.2287907 | -1.624528 | 0.10426317 | 0.65946648 |
| CEP135 | 31.6839008 | -0.5241246 | 0.35197714 | -1.489087 | 0.13646445 | 0.68406472 |
| AASDH | 51.312176 | 0.2742728 | 0.30263684 | 0.90627697 | 0.36478927 | 0.8468666 |
| PPAT | 12.9751541 | -0.1750701 | 0.43391395 | -0.4034674 | 0.68660441 | NA |
| PAICS | 23.1765935 | 0.13542756 | 0.33438582 | 0.40500391 | 0.68547464 | 0.94205398 |
| SRP72 | 54.0011749 | -0.0509321 | 0.27399968 | -0.1858839 | 0.8525358 | 0.97612951 |
| REST | 77.9827757 | 0.3871628 | 0.299499 | 1.29270149 | 0.19611432 | 0.75350056 |
| NOA1 | 14.2269117 | 0.43833952 | 0.40834173 | 1.07346248 | 0.28306366 | NA |
| POLR2B | 73.465096 | -0.429046 | 0.2402139 | -1.7860999 | 0.07408308 | 0.61020315 |
| IGFBP7 | 1054.422 | 0.04967935 | 0.18615209 | 0.26687507 | 0.78956535 | 0.96865409 |
| AC104806.2 | 19.8473428 | 1.20070419 | 0.48810011 | 2.45995476 | 0.01389545 | 0.37638271 |
| CENPC | 101.888002 | 0.02025643 | 0.34091832 | 0.05941727 | 0.95261976 | 0.99141043 |
| UBA6 | 138.059115 | -0.4486436 | 0.26774734 | -1.6756229 | 0.09381208 | 0.64572229 |
| UBA6-AS1 | 303.162813 | -0.0891319 | 0.36379207 | -0.2450077 | 0.80645044 | 0.97216031 |
| YTHDC1 | 164.612255 | -0.1964447 | 0.20748475 | -0.9467909 | 0.34374533 | 0.83465492 |
| UGT2B7 | 137.941513 | -0.3092153 | 0.68415088 | -0.4519695 | 0.65129098 | 0.93453685 |
| ENAM | 11.626195 | 0.20318166 | 0.44277133 | 0.45888623 | 0.64631587 | NA |
| UTP3 | 11.679838 | 0.26207835 | 0.47311311 | 0.55394439 | 0.57961691 | NA |
| RUFY3 | 416.345607 | -0.4793232 | 0.29762608 | -1.6104878 | 0.1072914 | 0.65986731 |
| GRSF1 | 54.9386818 | -0.0858709 | 0.28088411 | -0.3057163 | 0.75982066 | 0.95913478 |
| MOB1B | 250.53046 | -0.3257757 | 0.30406259 | -1.0714101 | 0.28398505 | 0.80431986 |
| SLC4A4 | 159.84277 | -0.1680984 | 0.35628752 | -0.4718055 | 0.63706565 | 0.93259128 |
| ADAMTS3 | 166.295189 | 0.28691453 | 0.61110193 | 0.46950356 | 0.63870974 | 0.9331046 |
| COX18 | 14.4292342 | 0.1901385 | 0.41407973 | 0.45918331 | 0.64610254 | NA |
| ANKRD17 | 645.989719 | -0.0372217 | 0.21791483 | -0.1708086 | 0.86437426 | 0.97788853 |
| AC053527.1 | 96.2174865 | 0.25909649 | 0.44339331 | 0.58434912 | 0.55898545 | 0.91469433 |
| RASSF6 | 63.8923179 | -0.2373758 | 0.33873507 | -0.7007711 | 0.48344585 | 0.88833419 |
| CXCL2 | 23.8156542 | 0.6519443 | 0.61370809 | 1.06230358 | 0.28809789 | 0.80523851 |
| MTHFD2L | 75.0571704 | 0.12669877 | 0.26784084 | 0.47303754 | 0.6361864 | 0.93251095 |
| BTC | 21.4280931 | -0.3795361 | 0.59524954 | -0.6376084 | 0.52372861 | 0.90487731 |
| PARM1 | 62.1955959 | 0.04461916 | 0.40239834 | 0.11088307 | 0.91170907 | 0.9858978 |
| RCHY1 | 57.3179309 | 0.05979879 | 0.26373687 | 0.22673655 | 0.8206286 | 0.97368535 |
| THAP6 | 35.7453635 | -0.0606581 | 0.32221356 | -0.1882543 | 0.85067728 | 0.97612951 |
| CDKL2 | 39.1205478 | -0.0703937 | 0.35147637 | -0.2002802 | 0.84126149 | 0.97600352 |
| G3BP2 | 73.9425938 | 0.12257297 | 0.25833313 | 0.47447639 | 0.63516023 | 0.93251095 |
| USO1 | 147.439282 | -0.2025052 | 0.26343845 | -0.7687001 | 0.44207137 | 0.86925451 |
| NAAA | 26.4540708 | 0.99091408 | 0.46766309 | 2.11886316 | 0.03410203 | 0.4908133 |
| SDAD1 | 39.3943939 | -0.2525341 | 0.27163593 | -0.9296785 | 0.35253755 | 0.83872681 |
| NUP54 | 60.88698 | 0.17490058 | 0.25144853 | 0.6955721 | 0.48669684 | 0.88838332 |
| SCARB2 | 142.66903 | -0.1857712 | 0.30051344 | -0.6181792 | 0.53645723 | 0.90763121 |
| SHROOM3 | 564.870521 | 0.09561256 | 0.33290075 | 0.28721039 | 0.77395123 | 0.96435639 |
| 11-Sep | 216.011727 | -0.2861136 | 0.31837928 | -0.8986565 | 0.36883567 | 0.84844057 |
| CCNI | 228.433534 | 0.08172907 | 0.28410464 | 0.28767241 | 0.77359751 | 0.96435639 |
| CCNG2 | 88.3904761 | -0.1976181 | 0.35384924 | -0.558481 | 0.57651598 | 0.91854298 |
| CNOT6L | 162.370771 | 0.11504957 | 0.25698584 | 0.44768838 | 0.65437811 | 0.93548446 |
| MRPL1 | 73.6695972 | -0.0611205 | 0.21660778 | -0.2821713 | 0.77781216 | 0.96518573 |
| FRAS1 | 1060.21366 | 0.27603164 | 0.40121996 | 0.68798082 | 4.91E-01 | 8.90E-01 |
| ANXA3 | 40.4738498 | 0.1341028 | 0.54872452 | 0.24439003 | 0.80692876 | 0.97216031 |
| BMP2K | 85.7544279 | -0.5330188 | 0.30707171 | -1.7358122 | 0.08259704 | 0.62228971 |
| PAQR3 | 15.7334755 | 0.42975614 | 0.45776956 | 0.93880455 | 0.3478311 | NA |
| RASGEF1B | 447.513177 | -1.1730046 | 0.67899804 | -1.7275523 | 0.08406852 | 0.62624856 |
| HNRNPD | 155.273363 | -0.3732832 | 0.17450996 | -2.1390368 | 0.03243269 | 0.48175799 |

| | | | | | | |
|---|---|---|---|---|---|---|
| HNRNPDL | 132.209041 | -0.0924403 | 0.19625423 | -0.4710234 | 0.63762405 | 0.93271601 |
| ENOPH1 | 28.4297679 | -0.007831 | 0.31640656 | -0.0247498 | 0.98025455 | 0.9957881 |
| TMEM150C | 34.5712541 | 0.0718043 | 0.34325443 | 0.20918682 | 0.8343024 | 0.97600352 |
| SCD5 | 122.346799 | 1.55486977 | 0.43660197 | 3.56129813 | 0.00036903 | 0.0763464 |
| SEC31A | 210.884867 | -0.4141193 | 0.25835919 | -1.6028819 | 0.10896074 | 0.6603659 |
| LIN54 | 36.5805633 | -0.4460752 | 0.28420734 | -1.5695414 | 0.11652183 | 0.6668162 |
| COPS4 | 34.9375917 | 0.1269885 | 0.295646 | 0.4295289 | 0.66753837 | 0.93878847 |
| HELQ | 41.1420799 | 0.32769523 | 0.26344448 | 1.24388723 | 0.21354107 | 0.7643572 |
| MRPS18C | 18.670173 | -0.0221716 | 0.43212425 | -0.0513085 | 0.95907969 | 0.99232808 |
| ABRAXAS1 | 19.8195281 | -0.3972532 | 0.44391016 | -0.8948954 | 0.37084299 | 0.84889051 |
| GPAT3 | 165.1488 | -0.7507745 | 0.69120924 | -1.0861755 | 2.77E-01 | 8.02E-01 |
| CDS1 | 80.4196131 | -0.212042 | 0.33106486 | -0.6404849 | 0.52185742 | 0.90451015 |
| WDFY3 | 505.86684 | 0.0303255 | 0.1288846 | 0.23529187 | 0.81398221 | 0.97282508 |
| ARHGAP24 | 581.773423 | -0.2988376 | 0.24327344 | -1.2284022 | 0.21929604 | 0.76680432 |
| MAPK10 | 938.522102 | 0.40434719 | 0.36094096 | 1.12025852 | 0.26260361 | 0.79666376 |
| AC104827.1 | 14.4490617 | 0.491339 | 0.69645985 | 0.70548072 | 0.48051112 | NA |
| PTPN13 | 784.647162 | 0.01534728 | 0.31659384 | 0.04847623 | 0.96133671 | 0.99256424 |
| AC093827.5 | 21.744871 | -0.3160141 | 0.46530406 | -0.6791561 | 0.49703899 | 0.89347273 |
| AFF1 | 950.148447 | -0.5199429 | 0.19210422 | -2.7065668 | 0.00679829 | 0.30102597 |
| KLHL8 | 88.2324118 | 0.37875259 | 0.28379992 | 1.33457612 | 0.18201512 | 0.73398872 |
| HSD17B11 | 18.6940776 | -0.0245605 | 0.40827439 | -0.0601568 | 0.95203074 | 0.99141043 |
| NUDT9 | 42.8797556 | -0.3341803 | 0.27477866 | -1.21618 | 0.22391636 | 0.76840496 |
| SPP1 | 7134.45954 | -0.0833624 | 0.54215002 | -0.1537627 | 8.78E-01 | 9.80E-01 |
| PKD2 | 230.959974 | -0.1683755 | 0.19335361 | -0.8708164 | 0.38385439 | 0.85511218 |
| PPM1K | 82.6918382 | -0.0589599 | 0.39134901 | -0.1506582 | 0.88024534 | 0.98093344 |
| HERC6 | 39.5302532 | -1.8639073 | 0.87820093 | -2.1224155 | 0.03380286 | 0.4908133 |
| HERC5 | 34.2623831 | -1.7557463 | 0.73005092 | -2.4049642 | 0.01617405 | 0.39748373 |
| PYURF | 21.9336665 | 0.62218641 | 0.34952787 | 1.78007667 | 0.07506341 | 0.61020315 |
| HERC3 | 157.776596 | -0.217284 | 0.32370768 | -0.6712352 | 0.50207069 | 0.89499455 |
| FAM13A-AS1 | 59.7577116 | -0.0943611 | 0.27768536 | -0.3398131 | 0.73399726 | 0.95329896 |
| FAM13A | 220.225536 | -0.1013291 | 0.39648274 | -0.2555701 | 0.79828279 | 0.97019967 |
| GPRIN3 | 165.035192 | -0.8211808 | 0.49929839 | -1.6446694 | 0.100038 | 0.65863153 |
| CCSER1 | 2895.01609 | 0.10462845 | 0.29604551 | 0.35342016 | 7.24E-01 | 9.51E-01 |
| SMARCAD1 | 111.058547 | -0.2987505 | 0.18618838 | -1.6045607 | 0.10859053 | 0.65986731 |
| PDLIM5 | 242.761155 | -0.2017825 | 0.22878337 | -0.8819807 | 0.37778724 | 0.85310518 |
| BMPR1B | 718.53482 | -0.0916864 | 0.19985616 | -0.4587617 | 6.46E-01 | 9.33E-01 |
| UNC5C | 44.7639551 | 0.78062633 | 0.38566271 | 2.02411669 | 0.04295815 | 0.51589447 |
| STPG2 | 246.613297 | 0.37631365 | 0.47331595 | 0.79505804 | 0.42657974 | 0.86239557 |
| RAP1GDS1 | 158.300886 | 0.025454 | 0.19316337 | 0.13177445 | 0.8951627 | 0.98473305 |
| TSPAN5 | 105.493685 | -1.1122663 | 0.44490597 | -2.5000031 | 0.01241922 | 0.36046953 |
| EIF4E | 85.0894258 | -0.2512786 | 0.27724304 | -0.9063479 | 0.36475175 | 0.8468666 |
| METAP1 | 54.9700766 | 0.34345688 | 0.24647281 | 1.39348789 | 0.1634723 | 0.71332671 |
| ADH5 | 48.9110873 | 0.31870108 | 0.35377135 | 0.9008674 | 0.36765883 | 0.84777627 |
| TRMT10A | 9.25974876 | 0.12438643 | 0.59635563 | 0.2085776 | 0.834778 | NA |
| LAMTOR3 | 29.7555264 | -0.4698028 | 0.36349781 | -1.29245 | 0.19620134 | 0.75350056 |
| DNAJB14 | 121.61174 | 0.17676117 | 0.20396178 | 0.86663867 | 0.38614002 | 0.85586379 |
| H2AFZ | 37.7622662 | 0.3492242 | 0.48531453 | 0.71958323 | 0.47178164 | 0.88267439 |
| PPP3CA | 299.74622 | -0.3584317 | 0.33010728 | -1.0858038 | 0.2775658 | 0.80176531 |
| BANK1 | 43.5240571 | -0.3605246 | 0.43989198 | -0.8195754 | 0.41245821 | 0.85867915 |
| AP002075.1 | 15.0995556 | 0.96526396 | 0.676669 | 1.42649354 | 0.15372593 | NA |
| SLC39A8 | 13.6787906 | 1.17747793 | 0.56925991 | 2.06843641 | 0.038599 | NA |
| NFKB1 | 132.963983 | 0.18872525 | 0.24223665 | 0.77909457 | 0.43592401 | 0.86729771 |
| MANBA | 83.0252593 | 0.19937236 | 0.24472403 | 0.81468239 | 0.41525413 | 0.85867915 |
| UBE2D3 | 297.195203 | -0.1225703 | 0.19308946 | -0.6347853 | 0.52556848 | 0.90487731 |

| | | | | | | |
|---|---|---|---|---|---|---|
| CISD2 | 26.2550839 | 0.523609 | 0.33310657 | 1.57189636 | 0.11597459 | 0.6668162 |
| SLC9B1 | 10.250159 | 0.84169949 | 0.66860293 | 1.2588929 | 0.20806902 | NA |
| SLC9B2 | 10.0214116 | 1.61362569 | 0.5331906 | 3.02635809 | 0.00247519 | NA |
| BDH2 | 10.7227911 | 0.45828845 | 0.55303644 | 0.82867678 | 0.40728733 | NA |
| TET2 | 193.667371 | -0.1659223 | 0.2692488 | -0.6162416 | 0.53773505 | 0.90763121 |
| PPA2 | 203.990864 | 0.13543853 | 0.24147166 | 0.56088787 | 0.57487398 | 0.91824492 |
| ARHGEF38 | 388.577299 | -0.0987088 | 0.33976547 | -0.2905203 | 0.77141822 | 0.96366407 |
| INTS12 | 50.1568008 | -0.0552122 | 0.31207214 | -0.1769212 | 0.85957032 | 0.97691173 |
| GSTCD | 69.1248982 | 5.20E-03 | 0.23337165 | 0.022289 | 0.98221743 | 0.99626066 |
| NPNT | 150.325307 | 0.09565502 | 0.36200172 | 0.26423913 | 0.79159566 | 0.96925674 |
| TBCK | 316.656663 | 0.2106381 | 0.22296772 | 0.94470222 | 0.34481092 | 0.83545751 |
| AIMP1 | 41.6344838 | 0.10591535 | 0.32296435 | 0.32794749 | 0.74295137 | 0.95563105 |
| PAPSS1 | 95.5264083 | 0.13782292 | 0.23085964 | 0.59699878 | 0.5505082 | 0.91248567 |
| SGMS2 | 114.864252 | -0.9950194 | 0.44774232 | -2.2223036 | 0.02626279 | 0.44686017 |
| AC096564.1 | 12.6523109 | 0.5362197 | 0.70965881 | 0.75560212 | 0.44988779 | NA |
| HADH | 30.0947838 | 0.54862769 | 0.34635573 | 1.58400064 | 0.11319357 | 0.6656824 |
| RPL34-AS1 | 51.3853173 | 0.96678639 | 0.52045798 | 1.85756859 | 0.0632303 | 0.58673337 |
| RPL34 | 104.150257 | 0.43168831 | 0.41094224 | 1.05048416 | 0.29349557 | 0.80935782 |
| OSTC | 37.4727955 | -0.1308794 | 0.36687075 | -0.3567452 | 0.7212826 | 0.95088329 |
| SEC24B-AS1 | 14.3197579 | 0.23704077 | 0.72461433 | 0.32712681 | 0.74357198 | NA |
| SEC24B | 196.52012 | -0.2651867 | 0.23377616 | -1.1343617 | 0.25664287 | 0.79437752 |
| MCUB | 16.140318 | -0.7005371 | 0.41291075 | -1.6965824 | 8.98E-02 | NA |
| CASP6 | 22.6254886 | 0.69707252 | 0.32555859 | 2.1411584 | 0.03226127 | 0.48175799 |
| PLA2G12A | 32.4440764 | 0.43878634 | 0.32763969 | 1.3392344 | 0.18049438 | 0.73303885 |
| AC126283.2 | 18.3251461 | 1.60668203 | 0.64833286 | 2.4781746 | 0.01320565 | 0.36723795 |
| GAR1 | 15.6846675 | -0.7306492 | 0.41750535 | -1.7500356 | 0.08011217 | NA |
| EGF | 226.576007 | 0.32343162 | 0.54024008 | 0.59868128 | 5.49E-01 | 9.12E-01 |
| ELOVL6 | 90.941087 | 0.19775747 | 0.22121351 | 0.89396652 | 0.3713398 | 0.8491102 |
| ENPEP | 13.2029037 | -0.9905773 | 0.51522441 | -1.9226133 | 0.05452863 | NA |
| FAM241A | 42.2000405 | -0.1065565 | 0.29428344 | -0.362088 | 0.71728624 | 0.94925929 |
| AP1AR | 41.9054509 | -0.4472358 | 0.37080532 | -1.2061202 | 0.22777114 | 0.77185154 |
| ALPK1 | 164.15321 | 0.14256042 | 0.35653992 | 0.39984421 | 0.68927126 | 0.94210755 |
| LARP7 | 60.3203709 | -0.0728534 | 0.21977341 | -0.331493 | 0.74027212 | 0.9544376 |
| ANK2 | 12.8460158 | -0.9008225 | 0.71819406 | -1.2542884 | 0.20973721 | NA |
| CAMK2D | 117.212703 | -0.1246607 | 0.18896773 | -0.6596932 | 0.50945071 | 0.89963034 |
| ARSJ | 85.3158462 | 0.44611567 | 0.30196985 | 1.47735171 | 0.13958138 | 0.69054855 |
| UGT8 | 96.6747899 | 0.23682985 | 0.29112705 | 0.81349314 | 0.41593538 | 0.85867915 |
| SNHG8 | 21.5871713 | 0.3807182 | 0.44645068 | 0.85276653 | 0.39378879 | 0.85605713 |
| METTL14 | 43.8766172 | -0.13265 | 0.28316906 | -0.4684481 | 0.63946415 | 0.93328692 |
| SEC24D | 63.9955599 | -0.6424019 | 0.31126761 | -2.0638252 | 0.0390343 | 0.49975976 |
| USP53 | 346.507859 | -0.207326 | 0.20767037 | -0.9983419 | 0.3181136 | 0.82354084 |
| C4orf3 | 68.3882032 | -0.6643059 | 0.50524973 | -1.3148071 | 0.18857474 | 0.74279353 |
| PDE5A | 128.40563 | -0.3271688 | 0.44834582 | -0.7297242 | 0.46555876 | 0.87981788 |
| AC108866.1 | 53.7464596 | -0.2725556 | 0.40888545 | -0.6665819 | 0.50503925 | 0.89723255 |
| PRDM5 | 54.1007011 | -0.0945486 | 0.27317218 | -0.3461135 | 0.72925739 | 0.95279138 |
| ANXA5 | 92.6668243 | 0.06772048 | 0.27897209 | 0.24275003 | 0.80819905 | 0.97257655 |
| EXOSC9 | 12.365456 | 0.05481793 | 0.43559509 | 0.12584606 | 0.89985378 | NA |
| BBS7 | 18.923913 | -0.2194808 | 0.37142878 | -0.5909094 | 0.55458113 | 0.9137961 |
| KIAA1109 | 332.120714 | -0.0469811 | 0.19597773 | -0.2397267 | 0.81054214 | 0.97257655 |
| FGF2 | 20.2822676 | -1.4484259 | 0.71496137 | -2.02588 | 0.04277708 | 0.51526789 |
| NUDT6 | 11.293461 | 0.83503868 | 0.57543236 | 1.45115003 | 0.14673809 | NA |
| SPATA5 | 135.016186 | -0.1965514 | 0.18112802 | -1.0851516 | 0.2778545 | 0.80193223 |
| SPRY1 | 22.0280612 | 1.21204184 | 0.5034921 | 2.40727084 | 1.61E-02 | 0.39748373 |
| LINC01091 | 20.7504585 | 0.54514113 | 0.62188277 | 0.87659791 | 0.38070507 | 0.85432452 |

| | | | | | | |
|---|---|---|---|---|---|---|
| ANKRD50 | 24.5544114 | 0.17516384 | 0.33613188 | 0.52111641 | 0.60228568 | 0.92407937 |
| FAT4 | 81.20589 | 0.6524468 | 0.40575246 | 1.60799223 | 1.08E-01 | 0.65986731 |
| INTU | 67.8878386 | -0.20749 | 0.30405083 | -0.6824189 | 0.4949741 | 0.89230917 |
| HSPA4L | 99.9540623 | 0.33143507 | 0.46429664 | 0.71384335 | 0.47532405 | 0.88495683 |
| MFSD8 | 46.4679763 | -0.2288222 | 0.26873056 | -0.851493 | 0.39449555 | 0.85605713 |
| ABHD18 | 161.715536 | -0.3056402 | 0.27520863 | -1.1105764 | 0.26675073 | 0.79666376 |
| LARP1B | 135.512793 | 0.11772229 | 0.26357483 | 0.44663705 | 0.65513714 | 0.93548446 |
| PGRMC2 | 44.9924979 | -0.0407269 | 0.31827287 | -0.1279622 | 0.89817892 | 0.98531749 |
| AC108062.1 | 47.0253618 | -0.3440948 | 0.3605602 | -0.9543337 | 0.33991473 | 0.83111572 |
| JADE1 | 146.894338 | 0.29632467 | 0.38476739 | 0.77013977 | 0.44121699 | 0.86925451 |
| SCLT1 | 121.727779 | -0.2634975 | 0.27543789 | -0.9566496 | 0.33874416 | 0.83066816 |
| C4orf33 | 18.7507897 | -0.250586 | 0.5136009 | -0.4879002 | 0.62562051 | 0.92995158 |
| AC096711.2 | 74.6536264 | -0.0275364 | 0.37361521 | -0.0737025 | 0.94124714 | 0.99043681 |
| NOCT | 11.6826269 | -0.1610951 | 0.41268084 | -0.3903625 | 0.69626848 | NA |
| ELF2 | 364.936871 | 0.10391264 | 0.21963779 | 0.47310912 | 0.63613533 | 0.93251095 |
| NDUFC1 | 62.7239137 | -0.3895631 | 0.4677107 | -0.8329147 | 0.40489285 | 0.85782098 |
| NAA15 | 126.959825 | -0.0587592 | 0.25086723 | -0.2342242 | 0.81481096 | 0.97284589 |
| AC097376.2 | 151.907695 | -0.5671948 | 0.40810897 | -1.3898121 | 0.16458596 | 0.71446159 |
| SETD7 | 21.0724339 | 0.2434199 | 0.33802201 | 0.72013031 | 0.47144477 | 0.88267439 |
| MGST2 | 45.2365879 | 0.31670675 | 0.35084529 | 0.90269633 | 0.36668708 | 0.84741115 |
| MAML3 | 353.916175 | 0.25311605 | 0.24889784 | 1.01694757 | 0.30917837 | 0.81568231 |
| SCOC | 105.87415 | -0.232997 | 0.3085874 | -0.7550439 | 0.45022263 | 0.87346041 |
| SCOC-AS1 | 14.6161808 | 0.52362152 | 0.60501175 | 0.86547329 | 0.38677907 | NA |
| ELMOD2 | 39.0573296 | -0.1027422 | 0.31111306 | -0.3302408 | 0.741218 | 0.95462754 |
| TBC1D9 | 470.722308 | -0.3501069 | 0.30194459 | -1.1595072 | 0.24624949 | 0.78583441 |
| RNF150 | 252.168144 | 0.66328714 | 0.32438599 | 2.04474654 | 0.04087985 | 0.50629837 |
| ZNF330 | 26.0333538 | 0.30386879 | 0.32428864 | 0.93703185 | 0.34874218 | 0.83687845 |
| IL15 | 25.0243915 | -0.2571527 | 0.50167658 | -0.5125867 | 0.60824047 | 0.92573951 |
| INPP4B | 106.299165 | -0.4679593 | 0.44589984 | -1.0494719 | 0.293961 | 0.80990527 |
| AC104596.1 | 29.8837225 | -0.2062052 | 0.63365605 | -0.3254213 | 0.74486222 | 0.95563105 |
| USP38 | 33.6789991 | 0.21122621 | 0.29436158 | 0.71757398 | 0.47302001 | 0.88344299 |
| GAB1 | 363.462945 | -0.5469084 | 0.29006434 | -1.8854726 | 0.05936607 | 0.57442774 |
| SMARCA5 | 89.9445524 | -0.1240635 | 0.19550462 | -0.634581 | 0.52570174 | 0.90487731 |
| ANAPC10 | 98.6163564 | -0.0328833 | 0.27296831 | -0.1204655 | 0.90411438 | 0.98531749 |
| ABCE1 | 36.6052707 | -0.13357 | 0.35312648 | -0.3782498 | 0.70524501 | 0.94548439 |
| OTUD4 | 91.0491501 | -0.1693346 | 0.21903755 | -0.773085 | 0.43947207 | 0.86895963 |
| SMAD1 | 133.003655 | 0.3147029 | 0.25931151 | 1.21360945 | 0.2248969 | 0.76940612 |
| MMAA | 23.5845058 | -0.3239088 | 0.33901403 | -0.9554436 | 0.33935341 | 0.83066816 |
| ZNF827 | 213.941726 | 0.29076126 | 0.23757203 | 1.22388679 | 0.22099494 | 0.76680432 |
| LSM6 | 27.2138549 | 0.34715775 | 0.34906292 | 0.99454205 | 0.31995904 | 0.82449332 |
| SLC10A7 | 107.339823 | 0.04321288 | 0.24655414 | 0.1752673 | 0.8608696 | 0.97736293 |
| TMEM184C | 48.5879202 | 0.06339343 | 0.36676948 | 0.17284272 | 0.86277506 | 0.97788853 |
| PRMT9 | 65.6251953 | 0.03841311 | 0.62853537 | 0.06111527 | 0.95126741 | 0.99105535 |
| ARHGAP10 | 32.0452475 | 0.22262941 | 0.46836003 | 0.4753382 | 0.63454593 | 0.93251095 |
| NR3C2 | 1684.6065 | 0.45830505 | 0.17294684 | 2.64997641 | 8.05E-03 | 0.30976355 |
| AC069272.1 | 22.2359178 | 0.6208858 | 0.68951441 | 0.90046819 | 0.36787114 | 0.84777627 |
| DCLK2 | 22.3311458 | -0.6473948 | 0.63834773 | -1.0141727 | 0.31050037 | 0.81690315 |
| LRBA | 1248.87031 | -0.2299445 | 0.17250996 | -1.3329345 | 0.1825533 | 0.73398872 |
| RPS3A | 68.2688847 | 0.42039219 | 0.43062991 | 0.97622616 | 0.3289524 | 0.82781307 |
| SH3D19 | 148.575242 | -0.380802 | 0.26234184 | -1.4515487 | 0.14662713 | 0.69731483 |
| FAM160A1 | 273.873225 | -0.6367111 | 0.34951098 | -1.8217198 | 0.06849752 | 0.60244724 |
| GATB | 30.0710791 | 0.36330312 | 0.31728274 | 1.14504533 | 0.25219038 | 0.7902944 |
| FBXW7 | 273.24818 | -0.1867657 | 0.26744624 | -0.69833 | 0.48497086 | 0.88838332 |
| ARFIP1 | 169.738737 | -0.04791 | 0.20825029 | -0.2300599 | 0.81804525 | 0.97288403 |

| | | | | | | |
|---|---|---|---|---|---|---|
| FHDC1 | 14.7025667 | -0.2069309 | 0.42464427 | -0.4873042 | 6.26E-01 | NA |
| TRIM2 | 352.546651 | -0.2144098 | 0.16723707 | -1.2820709 | 0.19981777 | 0.75605362 |
| TMEM131L | 25.3994727 | 0.43378879 | 0.4376974 | 0.99107007 | 0.32165137 | 0.82503937 |
| PLRG1 | 25.9269027 | -0.0907674 | 0.34932382 | -0.2598374 | 0.79498917 | 0.96974891 |
| FGB | 10.8385322 | 1.62261878 | 1.759574 | 0.92216569 | 0.35644215 | NA |
| MAP9 | 33.5200739 | 0.11299144 | 0.33633051 | 0.33595358 | 0.73690587 | 0.95387069 |
| GUCY1A1 | 17.3765326 | 0.4059968 | 0.45164331 | 0.89893239 | 0.36868867 | 0.84837538 |
| GUCY1B1 | 19.6502763 | -0.4296521 | 0.50080811 | -0.8579176 | 0.39093795 | 0.85605713 |
| CTSO | 19.6391236 | -0.3166074 | 0.34150861 | -0.9270847 | 0.35388256 | 0.83979358 |
| PDGFC | 130.001943 | -0.2445106 | 0.42256239 | -0.578638 | 0.56283349 | 0.91624774 |
| TMEM144 | 12.1426282 | -0.6691145 | 0.45985629 | -1.4550513 | 0.14565506 | NA |
| ETFDH | 79.7653453 | 0.15018898 | 0.32611132 | 0.46054514 | 0.64512498 | 0.93328692 |
| PPID | 35.7283436 | -0.1823981 | 0.31974161 | -0.5704547 | 0.56836932 | 0.91698355 |
| FNIP2 | 623.487183 | -1.3773087 | 0.50630756 | -2.7203005 | 0.00652226 | 0.29935381 |
| C4orf45 | 17.8347226 | 0.00489753 | 0.6753146 | 0.00725222 | 9.94E-01 | 0.99921857 |
| RAPGEF2 | 350.646554 | -0.1380891 | 0.24227713 | -0.5699635 | 0.56870249 | 0.91698355 |
| NAF1 | 38.934115 | 0.17183057 | 0.39145417 | 0.4389545 | 0.66069451 | 0.93729379 |
| TMA16 | 19.6068124 | -0.0012329 | 0.35841919 | -0.0034399 | 0.99725539 | 0.99953287 |
| TMEM192 | 43.2452704 | -0.3593431 | 0.37032234 | -0.9703523 | 0.33187092 | 0.82890589 |
| KLHL2 | 173.897214 | -0.5988648 | 0.31570659 | -1.8969029 | 0.05784075 | 0.57141408 |
| MSMO1 | 112.818938 | -0.6747511 | 0.50126018 | -1.3461096 | 0.17826719 | 0.72841249 |
| DDX60 | 53.2606708 | -1.7568771 | 1.17989102 | -1.4890164 | 0.13648305 | 0.68406472 |
| DDX60L | 59.1162151 | -3.0566763 | 1.22551285 | -2.4942017 | 0.01262408 | 0.3648158 |
| PALLD | 119.831058 | -0.0234506 | 0.40642777 | -0.0576994 | 0.95398807 | 0.99148072 |
| CBR4 | 326.405041 | 0.30103722 | 0.26286784 | 1.14520371 | 0.25212478 | 0.7902944 |
| SH3RF1 | 656.283134 | 0.30650256 | 0.55794446 | 0.54934242 | 0.58277048 | 0.92034873 |
| NEK1 | 214.293627 | -0.0062253 | 0.33787912 | -0.0184248 | 0.9853 | 0.99748524 |
| CLCN3 | 211.01363 | 0.04043575 | 0.47837654 | 0.08452704 | 0.9326374 | 0.98960607 |
| HPF1 | 19.544748 | -0.0197462 | 0.34983766 | -0.0564439 | 0.95498815 | 0.99148072 |
| MFAP3L | 59.7656171 | 0.18021603 | 0.33297461 | 0.54123055 | 0.58834868 | 0.92175048 |
| GALNT7 | 121.209358 | 0.38446416 | 0.26342757 | 1.45946823 | 0.14443628 | 0.69512617 |
| HMGB2 | 31.3466358 | 0.36677348 | 0.42847702 | 0.85599337 | 0.39200145 | 0.85605713 |
| FBXO8 | 23.5564469 | 0.23360245 | 0.33735493 | 0.69245306 | 0.48865286 | 0.88920066 |
| CEP44 | 42.3966695 | -0.05529 | 0.34084038 | -0.1622166 | 0.87113527 | 0.97860207 |
| HPGD | 22.7262341 | 0.24257348 | 0.74112596 | 0.32730399 | 0.74343798 | 0.95563105 |
| WDR17 | 25.3449471 | 0.22462609 | 0.50111565 | 0.44825199 | 0.65397135 | 0.93548446 |
| SPCS3 | 61.984593 | 0.16886962 | 0.26366633 | 0.64046715 | 0.52186894 | 0.90451015 |
| VEGFC | 65.2818722 | 0.5119219 | 0.41076168 | 1.24627472 | 2.13E-01 | 7.64E-01 |
| AC078881.1 | 55.9224933 | 0.06544114 | 0.40864512 | 0.16014174 | 0.87276942 | 0.97866716 |
| LINC01098 | 180.359362 | 1.83059903 | 1.01738499 | 1.79931791 | 0.07196841 | 0.60821941 |
| LINC01099 | 697.907665 | 1.28202607 | 0.54796692 | 2.33960486 | 0.01930415 | 0.41544117 |
| DCTD | 21.7950806 | -0.0932391 | 0.43066674 | -0.2164995 | 0.82859841 | 0.97567084 |
| WWC2 | 212.710312 | -0.1750185 | 0.23889587 | -0.7326142 | 0.46379375 | 0.87824803 |
| CDKN2AIP | 17.9725891 | -0.554218 | 0.68246857 | -0.8120784 | 0.41674662 | 0.85867915 |
| RWDD4 | 18.964774 | 0.04184699 | 0.37726157 | 0.11092301 | 0.9116774 | 0.9858978 |
| TRAPPC11 | 66.9428565 | 0.01527794 | 0.23118402 | 0.0660856 | 0.94730967 | 0.99083449 |
| STOX2 | 332.770364 | 0.35982052 | 0.25951317 | 1.38652123 | 0.1655878 | 0.71675975 |
| IRF2 | 66.8649453 | 0.0259103 | 0.25479105 | 0.10169236 | 0.91900086 | 0.98749988 |
| CASP3 | 27.2021748 | 0.0704206 | 0.47920248 | 0.14695376 | 0.88316852 | 0.98140267 |
| PRIMPOL | 27.8265455 | -0.091895 | 0.49774101 | -0.184624 | 0.85352393 | 0.97612951 |
| ACSL1 | 172.920158 | -0.2090475 | 0.3710571 | -0.5633836 | 0.57317373 | 0.91813957 |
| SLC25A4 | 43.7112666 | 0.83987029 | 0.37659147 | 2.23018934 | 0.02573488 | 0.4436829 |
| CFAP97 | 59.9533838 | 0.11701727 | 0.27693854 | 0.42253876 | 0.67263182 | 0.93878847 |
| SNX25 | 144.013379 | -0.5756936 | 0.26502454 | -2.1722277 | 0.02983849 | 0.46381645 |

| | | | | | | |
|---|---|---|---|---|---|---|
| LRP2BP | 15.5515632 | -0.0492354 | 0.52642516 | -0.0935278 | 0.92548423 | NA |
| AC112722.1 | 19.0133275 | -0.3522097 | 0.40878455 | -0.8616024 | 0.38890637 | 0.85605713 |
| ANKRD37 | 37.325802 | -2.3242342 | 0.73805434 | -3.149137 | 0.00163753 | 0.16939173 |
| UFSP2 | 48.9693139 | 0.21638928 | 0.25987462 | 0.83266799 | 0.405032 | 0.85782098 |
| C4orf47 | 52.683581 | -0.5771708 | 0.57090116 | -1.010982 | 0.31202504 | 0.81701824 |
| SORBS2 | 319.47729 | 0.82662407 | 0.39665384 | 2.08399363 | 0.03716075 | 0.49537143 |
| TLR3 | 12.0550114 | -0.4258058 | 0.57441926 | -0.7412804 | 0.45852343 | NA |
| FAM149A | 54.5904618 | 0.19169884 | 0.26781194 | 0.71579645 | 0.47411704 | 0.88403192 |
| CYP4V2 | 37.971758 | -0.2547195 | 0.29747987 | -0.8562579 | 0.39185513 | 0.85605713 |
| KLKB1 | 22.7698839 | -0.030159 | 0.50704708 | -0.0594797 | 0.95257004 | 0.99141043 |
| F11 | 10.7288993 | 0.00799715 | 0.78056143 | 0.01024539 | 0.99182551 | NA |
| FAT1 | 226.896946 | -0.3367837 | 0.35986868 | -0.9358516 | 0.34934959 | 0.83687845 |
| FRG1-DT | 42.1733647 | -0.2249916 | 0.58110941 | -0.3871759 | 0.69862596 | 0.94395238 |
| FRG1 | 18.0203337 | 0.08739608 | 0.35719085 | 0.24467615 | 0.80670719 | 0.97216031 |
| SDHA | 106.75558 | -0.0266977 | 0.21983461 | -0.1214443 | 0.90333911 | 0.98531749 |
| PDCD6 | 53.6261493 | -0.3551201 | 0.28511576 | -1.2455295 | 0.21293717 | 0.7643572 |
| EXOC3 | 51.1438344 | -0.2438906 | 0.27302152 | -0.8933016 | 0.37169567 | 0.8491102 |
| SLC9A3 | 12.3363679 | 0.05173731 | 0.42716543 | 0.12111773 | 0.90359779 | NA |
| ZDHHC11B | 28.2998597 | 1.31149387 | 0.58717359 | 2.23357092 | 0.02551132 | 0.44138123 |
| BRD9 | 28.7672926 | -0.4479237 | 0.36337529 | -1.2326753 | 0.21769694 | 0.76666749 |
| SLC12A7 | 48.5807786 | -0.5574218 | 0.33050114 | -1.6865956 | 9.17E-02 | 0.64396102 |
| CLPTM1L | 30.8598997 | -0.1296403 | 0.35371753 | -0.366508 | 0.71398604 | 0.94841799 |
| LPCAT1 | 20.2380755 | -0.2414608 | 0.36311673 | -0.6649675 | 0.50607128 | 0.89723255 |
| NDUFS6 | 52.0638074 | -0.2490603 | 0.44856617 | -0.5552365 | 0.57873291 | 0.91940762 |
| AC022424.1 | 42.9311334 | 0.97690001 | 0.60585466 | 1.61243293 | 0.10686776 | 0.65986731 |
| ADAMTS16 | 369.372624 | 0.68830505 | 0.48674969 | 1.41408419 | 0.15733718 | 0.7098097 |
| AC022424.2 | 17.2001676 | 1.49815235 | 0.64943207 | 2.30686538 | 0.02106233 | 0.4212027 |
| ICE1 | 88.8985022 | -0.0940718 | 0.20928225 | -0.4494973 | 0.65307299 | 0.93532021 |
| MED10 | 25.3659418 | -0.3230004 | 0.3283663 | -0.9836587 | 0.32528333 | 0.82567895 |
| NSUN2 | 45.1451363 | -0.4051543 | 0.3086074 | -1.312847 | 0.18923451 | 0.74410443 |
| SRD5A1 | 11.4992658 | -0.2824347 | 0.44190035 | -0.6391366 | 0.52273406 | NA |
| TENT4A | 67.8003518 | -0.5098238 | 0.24867548 | -2.0501571 | 0.04034911 | 0.50399977 |
| MTRR | 39.4971977 | -0.1618935 | 0.34126326 | -0.4743947 | 0.63521848 | 0.93251095 |
| SEMA5A | 36.6694004 | -0.2717605 | 0.53671617 | -0.5063393 | 0.6126185 | 0.92647422 |
| FAM173B | 35.075938 | -0.065815 | 0.3104712 | -0.2119841 | 0.83211943 | 0.97600352 |
| CCT5 | 38.4377333 | 0.60491262 | 0.32183246 | 1.8795886 | 0.06016417 | 0.57733412 |
| CMBL | 34.6391659 | 0.97227173 | 0.52599853 | 1.8484305 | 0.06454009 | 0.58883813 |
| 6-Mar | 256.755735 | -0.238841 | 0.21735225 | -1.0988658 | 0.27182661 | 0.79666376 |
| DAP | 104.342792 | -0.193112 | 0.30768359 | -0.6276318 | 0.53024515 | 0.9052447 |
| DNAH5 | 14.1631001 | 1.03082585 | 0.73445769 | 1.40351972 | 0.16046192 | NA |
| TRIO | 488.886789 | -0.3053794 | 0.2740277 | -1.1144107 | 0.26510302 | 0.79666376 |
| OTULINL | 11.1187364 | -0.3662337 | 0.69057376 | -0.5303325 | 0.5958814 | NA |
| OTULIN | 36.9219862 | 0.31744348 | 0.31853139 | 0.99658462 | 0.31896618 | 0.82354084 |
| ANKH | 21.7148645 | -0.0495011 | 0.36073355 | -0.1372233 | 0.89085427 | 0.98296447 |
| FBXL7 | 24.8478013 | 0.22999203 | 0.36243643 | 0.63457207 | 0.52570755 | 0.90487731 |
| RETREG1 | 123.094319 | 0.19622444 | 0.4033442 | 0.48649376 | 6.27E-01 | 0.93032781 |
| MYO10 | 429.828417 | -0.3643445 | 0.31103037 | -1.1714113 | 0.24143348 | 0.78323911 |
| LINC02241 | 37.4886037 | -0.1902704 | 0.29661808 | -0.6414661 | 0.52121989 | 0.90451015 |
| CDH12 | 14.5999199 | 0.14507754 | 0.57232435 | 0.25348832 | 0.79989088 | NA |
| C5orf17 | 50.7945364 | 0.20590363 | 0.82332932 | 0.2500866 | 0.80252038 | 0.97106779 |
| CDH6 | 17.6691281 | 1.18408555 | 0.80488668 | 1.47112081 | 0.14125845 | 0.69282094 |
| DROSHA | 79.713451 | -0.2749514 | 0.20563978 | -1.3370537 | 0.18120509 | 0.73398872 |
| C5orf22 | 22.0490862 | 0.19430934 | 0.33568449 | 0.57884516 | 0.56269366 | 0.91624774 |
| PDZD2 | 37.7084348 | 0.27083236 | 0.48189046 | 0.56202059 | 0.57410199 | 0.91813957 |

| | | | | | | |
|---|---|---|---|---|---|---|
| GOLPH3 | 128.457689 | -0.3307579 | 0.23964309 | -1.3802106 | 0.16752182 | 0.71965603 |
| MTMR12 | 86.051751 | -0.2971972 | 0.22046154 | -1.3480684 | 0.1776364 | 0.72841249 |
| ZFR | 217.551701 | -0.139429 | 0.14929827 | -0.9338957 | 0.35035767 | 0.83708817 |
| SUB1 | 96.4399307 | 0.53644182 | 0.26339883 | 2.03661426 | 0.04168871 | 0.51069022 |
| NPR3 | 33.7978189 | 0.87354401 | 0.55695619 | 1.56842499 | 1.17E-01 | 6.67E-01 |
| TARS | 36.8945449 | -0.1443654 | 0.31547751 | -0.457609 | 0.64723335 | 0.93328692 |
| C1QTNF3-AN | 111.574115 | -0.1317281 | 0.17730339 | -0.742953 | 0.45751015 | 0.87420547 |
| RAI14 | 167.762023 | -1.1249176 | 0.42822294 | -2.6269438 | 0.00861556 | 0.31119527 |
| RAD1 | 16.949126 | -0.0964286 | 0.38030713 | -0.2535546 | 0.79983971 | 0.97019967 |
| BRIX1 | 17.1395132 | 0.34282004 | 0.3731526 | 0.91871272 | 0.35824585 | 0.84204868 |
| DNAJC21 | 54.058739 | 0.18693178 | 0.2448206 | 0.76354595 | 0.44513787 | 0.87129815 |
| PRLR | 80.9835817 | 0.22242305 | 0.4242715 | 0.52424697 | 6.00E-01 | 0.92338945 |
| SPEF2 | 55.8371415 | 0.40219635 | 0.47725389 | 0.84273038 | 0.39937925 | 0.85778581 |
| UGT3A1 | 31.0334809 | -1.0220916 | 0.80570756 | -1.268564 | 0.2045966 | 0.76173533 |
| LMBRD2 | 45.0992694 | 0.34187249 | 0.2760448 | 1.2384674 | 0.2155428 | 0.7652442 |
| SKP2 | 11.6249496 | -0.3913445 | 0.44624313 | -0.8769759 | 0.38049975 | NA |
| NADK2 | 42.2956859 | 0.00166599 | 0.35131531 | 0.00474216 | 0.99621632 | 0.99927028 |
| RANBP3L | 24.6152127 | 0.85833258 | 0.86792422 | 0.98894876 | 0.32268821 | 0.82558302 |
| NIPBL | 503.451502 | -0.2296058 | 0.16942476 | -1.3552081 | 0.17535126 | 0.72841249 |
| CPLANE1 | 202.075495 | -0.3069419 | 0.1960781 | -1.5654064 | 0.11748766 | 0.6668162 |
| NUP155 | 32.7384644 | -0.4370724 | 0.29218885 | -1.4958557 | 0.13469126 | 0.68328989 |
| WDR70 | 271.969239 | -0.1988903 | 0.1929369 | -1.0308568 | 0.30260799 | 0.81568231 |
| LIFR | 212.140239 | 0.09841946 | 0.21463566 | 0.45854195 | 6.47E-01 | 9.33E-01 |
| LIFR-AS1 | 12.458543 | 0.37776708 | 0.59005922 | 0.64021893 | 0.52203028 | NA |
| OSMR-AS1 | 171.758088 | -0.0046425 | 0.30934868 | -0.0150074 | 9.88E-01 | 0.99814944 |
| OSMR | 212.413901 | -0.014626 | 0.39040213 | -0.037464 | 9.70E-01 | 9.94E-01 |
| RICTOR | 267.659278 | -0.4225289 | 0.25278839 | -1.6714728 | 0.09462833 | 0.64732984 |
| FYB1 | 36.2305199 | -0.1111248 | 0.58273732 | -0.1906945 | 8.49E-01 | 0.97612951 |
| DAB2 | 11.4530153 | -0.3858091 | 0.63090159 | -0.6115202 | 0.54085526 | NA |
| TTC33 | 46.4629377 | 0.11180188 | 0.26851042 | 0.4163782 | 0.67713328 | 0.94034237 |
| PRKAA1 | 67.7520015 | -0.0071501 | 0.23575059 | -0.0303289 | 0.97580472 | 0.99440179 |
| RPL37 | 79.5350095 | 0.32975529 | 0.34222196 | 0.9635714 | 0.33526085 | 0.82974237 |
| OXCT1 | 33.3266807 | -0.596539 | 0.38619201 | -1.5446696 | 0.12242621 | 0.66929384 |
| C5orf51 | 16.9262696 | -0.3139381 | 0.36453371 | -0.8612046 | 0.38912539 | 0.85605713 |
| FBXO4 | 11.6367351 | 0.34489468 | 0.52931753 | 0.65158372 | 0.51466975 | NA |
| GHR | 162.511646 | 0.20132975 | 0.38619548 | 0.52131565 | 6.02E-01 | 0.92407937 |
| SELENOP | 60.8049997 | 0.16216963 | 0.41225664 | 0.39337057 | 0.69404581 | 0.94395238 |
| ZNF131 | 111.057598 | -0.0198603 | 0.25657449 | -0.0774056 | 0.9383009 | 0.99012269 |
| HMGCS1 | 134.154922 | -0.3634958 | 0.45759249 | -0.7943657 | 0.42698257 | 0.86239557 |
| TMEM267 | 49.6379384 | 0.23047556 | 0.26674094 | 0.86404268 | 0.38756444 | 0.85605713 |
| PAIP1 | 51.8220551 | -0.3803801 | 0.24768467 | -1.5357433 | 0.12460135 | 0.66929384 |
| NNT-AS1 | 89.8179986 | 0.35298423 | 0.226621 | 1.55759715 | 0.11932877 | 0.6668162 |
| NNT | 221.496617 | 0.15481982 | 0.20413113 | 0.75843316 | 0.44819171 | 0.87307707 |
| MRPS30-DT | 31.9143831 | 0.01556428 | 0.45560468 | 0.03416181 | 0.97274812 | 0.99431961 |
| MRPS30 | 23.2783688 | 0.43209558 | 0.36719134 | 1.17675863 | 0.23929186 | 0.78098673 |
| AC093297.2 | 10.091392 | 0.00882353 | 0.48864615 | 0.0180571 | 0.9855933 | NA |
| PARP8 | 122.140371 | 0.04715432 | 0.52442757 | 0.08991579 | 0.92835413 | 0.98960607 |
| ITGA1 | 71.6343279 | -0.1911214 | 0.33477438 | -0.5708961 | 0.56807005 | 0.91698355 |
| ITGA2 | 749.109461 | -1.7073116 | 0.47464541 | -3.5970255 | 0.00032188 | 0.0763464 |
| MOCS2 | 29.3410571 | 0.48125948 | 0.31126484 | 1.54614146 | 0.12207041 | 0.66929384 |
| NDUFS4 | 143.724621 | -0.0890597 | 0.22573963 | -0.3945238 | 0.69319433 | 0.94394214 |
| ARL15 | 2358.65772 | -0.0648541 | 0.29601588 | -0.2190901 | 8.27E-01 | 9.76E-01 |
| SNX18 | 54.5786784 | 0.45113403 | 0.39356447 | 1.14627732 | 0.25168042 | 0.79001617 |
| DHX29 | 63.0508558 | 0.12658865 | 0.25601293 | 0.49446192 | 0.62098 | 0.92805465 |

| | | | | | | |
|---|---|---|---|---|---|---|
| MTREX | 118.676624 | 0.18432071 | 0.22470871 | 0.82026508 | 0.41206501 | 0.85867915 |
| PLPP1 | 194.262864 | -0.0567173 | 0.20645849 | -0.2747154 | 0.78353493 | 0.96717296 |
| SLC38A9 | 66.2084427 | 0.19271277 | 0.27874881 | 0.6913492 | 0.48934612 | 0.88930178 |
| IL6ST | 148.568593 | 0.22962391 | 0.23054075 | 0.99602307 | 0.31923894 | 0.82370522 |
| AC008892.1 | 23.4642893 | 0.81025822 | 0.59224921 | 1.3681035 | 0.17127968 | 0.72463203 |
| LINC01948 | 20.9278694 | 0.54458506 | 0.49627077 | 1.09735472 | 0.27248636 | 0.79721781 |
| MAP3K1 | 254.103896 | 0.07315897 | 0.30751535 | 0.23790347 | 0.81195597 | 0.97257655 |
| MIER3 | 26.3773701 | 0.34712312 | 0.45058577 | 0.77038189 | 0.44107339 | 0.86925451 |
| GPBP1 | 257.960168 | -0.3586124 | 0.26439206 | -1.3563661 | 0.1749827 | 0.72841249 |
| PDE4D | 8929.63419 | -0.3852036 | 0.29491899 | -1.3061334 | 1.92E-01 | 7.50E-01 |
| DEPDC1B | 43.2209905 | -1.7971553 | 0.58680721 | -3.0625993 | 0.00219424 | 0.18826595 |
| ELOVL7 | 191.83662 | -0.7730085 | 0.36439582 | -2.1213428 | 0.03389296 | 0.4908133 |
| ERCC8 | 59.489553 | 0.19767785 | 0.30387171 | 0.65053062 | 0.51534953 | 0.90174188 |
| NDUFAF2 | 100.386675 | -0.4349723 | 0.33520795 | -1.2976193 | 0.1944182 | 0.75182195 |
| SMIM15 | 19.140116 | 0.07846644 | 0.41855251 | 0.18747097 | 0.85129138 | 0.97612951 |
| ZSWIM6 | 543.432255 | -0.5244759 | 0.46198753 | -1.13526 | 0.25626641 | 0.7940072 |
| KIF2A | 77.0240828 | -0.1739635 | 0.28471481 | -0.6110095 | 0.5411933 | 0.90783815 |
| DIMT1 | 12.7779249 | -0.3678945 | 0.39616731 | -0.9286342 | 0.35307869 | NA |
| IPO11 | 50.1302645 | -0.3513775 | 0.28989183 | -1.2120985 | 0.22547467 | 0.77074574 |
| RNF180 | 154.535161 | -0.0599087 | 0.27681164 | -0.216424 | 0.82865725 | 0.97567084 |
| SREK1IP1 | 48.5932462 | -0.2827708 | 0.2782143 | -1.0163778 | 0.30944951 | 0.81568231 |
| CWC27 | 131.653457 | -0.2746634 | 0.21906398 | -1.2538044 | 0.20991311 | 0.76344019 |
| PPWD1 | 48.7952255 | 0.02465908 | 0.32102211 | 0.07681427 | 0.9387713 | 0.99012269 |
| TRIM23 | 35.5966631 | 0.01757985 | 0.29680124 | 0.05923105 | 0.95276808 | 0.99143208 |
| TRAPPC13 | 51.525258 | 0.07641949 | 0.2663272 | 0.28693837 | 0.77415951 | 0.96435639 |
| NLN | 35.3765223 | -0.6366801 | 0.31784583 | -2.0031098 | 0.0451655 | 0.52675468 |
| ERBIN | 450.347616 | -0.1523903 | 0.16615035 | -0.917183 | 0.35904675 | 0.84204868 |
| SREK1 | 129.836531 | -0.0825111 | 0.23005304 | -0.3586614 | 0.71984842 | 0.94995363 |
| MAST4 | 389.653237 | -0.9555941 | 0.49457185 | -1.9321644 | 0.05333922 | 0.55938585 |
| PIK3R1 | 95.794406 | 0.10095771 | 0.29973727 | 0.33682066 | 0.73625209 | 0.95376039 |
| SLC30A5 | 59.7380674 | -0.0954148 | 0.28788167 | -0.3314377 | 0.7403139 | 0.9544376 |
| MRPS36 | 42.7715202 | 0.16614437 | 0.27024284 | 0.61479657 | 0.53868907 | 0.90763121 |
| CDK7 | 39.2671808 | -0.1432436 | 0.29088489 | -0.4924409 | 0.62240773 | 0.92897508 |
| CCDC125 | 68.7250647 | 0.12332056 | 0.38930594 | 0.31677031 | 0.75141789 | 0.9574688 |
| AK6 | 23.2443139 | -0.0627226 | 0.35387333 | -0.1772458 | 0.85931531 | 0.97691173 |
| RAD17 | 49.3512488 | -0.2529229 | 0.25482638 | -0.9925304 | 0.32093885 | 0.82464962 |
| MARVELD2 | 32.7175195 | -0.0453235 | 0.28928404 | -0.1566748 | 0.87550117 | 0.97918624 |
| OCLN | 94.6213676 | 0.13115063 | 0.23018908 | 0.56975172 | 0.5688461 | 0.91698355 |
| GTF2H2C | 29.9291862 | 0.00221798 | 0.33960299 | 0.0065311 | 0.99478897 | 0.99927028 |
| NAIP | 107.730655 | 0.08692602 | 0.40654172 | 0.21381821 | 0.83068883 | 0.97567084 |
| GTF2H2 | 27.9671197 | -0.1115633 | 0.45905392 | -0.2430288 | 0.80798308 | 0.97257655 |
| BDP1 | 173.419713 | -0.3388467 | 0.2202519 | -1.5384505 | 0.1239385 | 0.66929384 |
| MCCC2 | 84.4625431 | -0.1357149 | 0.21089074 | -0.643532 | 0.51987896 | 0.90368693 |
| MRPS27 | 72.3356673 | 0.06366027 | 0.21972204 | 0.28973094 | 0.77202208 | 0.96402154 |
| PTCD2 | 16.9991095 | 0.47750079 | 0.42154132 | 1.13274965 | 0.2573194 | 0.79532468 |
| TNPO1 | 189.459405 | -0.0994348 | 0.20499225 | -0.4850662 | 0.62762941 | 0.93061955 |
| FCHO2 | 145.377022 | -0.1884026 | 0.1806176 | -1.0431021 | 0.29690102 | 0.81137495 |
| BTF3 | 60.7281739 | 0.44148148 | 0.34606977 | 1.27570082 | 0.2020613 | 0.76038198 |
| ANKRA2 | 25.6063801 | 0.22930522 | 0.38952727 | 0.58867565 | 0.55607888 | 0.91398748 |
| ARHGEF28 | 305.300116 | 0.19710964 | 0.32044569 | 0.61511089 | 0.53848149 | 0.90763121 |
| HEXB | 23.0106438 | -0.4243519 | 0.3530039 | -1.2021167 | 0.2293183 | 0.77185154 |
| GFM2 | 40.7155065 | -0.0629704 | 0.32161899 | -0.1957921 | 0.84477292 | 0.97612951 |
| NSA2 | 30.230981 | 0.67173694 | 0.33810403 | 1.98677592 | 0.04694723 | 0.5320597 |
| FAM169A | 73.4077958 | -0.0744606 | 0.33294022 | -0.2236455 | 0.82303316 | 0.97518529 |

| | | | | | | |
|---|---|---|---|---|---|---|
| HMGCR | 95.7920905 | -0.243419 | 0.47564929 | -0.5117615 | 6.09E-01 | 0.92573951 |
| COL4A3BP | 182.341272 | -0.2869209 | 0.26840232 | -1.0689957 | 0.28507162 | 0.80431986 |
| POLK | 117.368546 | 0.03181517 | 0.24727665 | 0.12866223 | 0.89762492 | 0.98531749 |
| ANKDD1B | 25.6896279 | 0.56178726 | 0.65998475 | 0.85121248 | 0.39465133 | 0.85605713 |
| POC5 | 22.6266567 | 0.57843794 | 0.31500708 | 1.83626966 | 0.06631778 | 0.59594352 |
| IQGAP2 | 175.006488 | 0.98738512 | 0.42371618 | 2.33029834 | 0.01979039 | 0.41544117 |
| F2RL1 | 21.8926875 | 0.6942105 | 0.36313086 | 1.91173645 | 0.05591001 | 0.56884484 |
| ZBED3 | 27.0193563 | 0.53454392 | 0.35439186 | 1.50834141 | 0.13146717 | 0.67540912 |
| AC022414.1 | 21.8691241 | 1.59625095 | 0.53010849 | 3.01117788 | 0.00260236 | 0.20599404 |
| PDE8B | 15.6086598 | 0.61704304 | 0.53537054 | 1.15255322 | 0.24909381 | NA |
| WDR41 | 85.5804009 | -0.2008048 | 0.2296003 | -0.8745842 | 0.38180018 | 0.85477791 |
| TBCA | 159.414632 | -0.1640115 | 0.2296647 | -0.7141345 | 0.475144 | 0.88486821 |
| AP3B1 | 310.032282 | -0.0424843 | 0.18321788 | -0.2318787 | 0.81663223 | 0.97288403 |
| SCAMP1 | 80.0388797 | -0.0248161 | 0.24150242 | -0.1027571 | 0.91815578 | 0.98735642 |
| LHFPL2 | 12.6453533 | 0.00553143 | 0.434372 | 0.01273432 | 0.98983976 | NA |
| ARSB | 24.1812205 | 0.30323891 | 0.51715516 | 0.58635963 | 0.55763387 | 0.91425998 |
| DMGDH | 110.292512 | 0.39399248 | 0.3307996 | 1.19103069 | 0.23364154 | 0.77650198 |
| BHMT2 | 96.1290965 | 0.82284681 | 0.49080395 | 1.67652851 | 0.09363472 | 0.64572229 |
| BHMT | 27.6676763 | 0.35678063 | 0.43110161 | 0.82760219 | 0.40789583 | 0.85867915 |
| JMY | 89.7759782 | 0.19807773 | 0.28395622 | 0.69756431 | 0.48544971 | 0.88838332 |
| HOMER1 | 145.616272 | -0.9719464 | 1.09458793 | -0.8879565 | 0.37456417 | 0.85092528 |
| TENT2 | 318.088922 | -0.1363999 | 0.18896563 | -0.7218238 | 0.47040283 | 0.88260202 |
| CMYA5 | 159.086513 | -0.2295438 | 0.34493739 | -0.6654651 | 0.50575307 | 0.89723255 |
| MTX3 | 22.3076294 | 0.38808074 | 0.34744356 | 1.11696052 | 0.26401121 | 0.79666376 |
| THBS4 | 24.7352153 | 0.37895683 | 0.56681774 | 0.66856911 | 0.50377038 | 0.89624537 |
| SERINC5 | 97.3649447 | 0.03290345 | 0.22359739 | 0.1471549 | 0.88300976 | 0.98140267 |
| ZFYVE16 | 181.82668 | -0.1177202 | 0.24907291 | -0.4726334 | 0.63647478 | 0.93251095 |
| AC008771.1 | 16.2586159 | 0.60632736 | 0.51364371 | 1.18044346 | 0.23782389 | 0.77930556 |
| FAM151B | 37.2439579 | -0.3399989 | 0.3705103 | -0.9176505 | 0.35880189 | 0.84204868 |
| ANKRD34B | 16.4868037 | 0.39270939 | 1.11072543 | 0.35356117 | 0.72366778 | 0.95093722 |
| DHFR | 379.975814 | -0.5831405 | 0.49605088 | -1.1755659 | 0.23976838 | 0.78128405 |
| MSH3 | 164.382175 | -0.0446926 | 0.17852353 | -0.2503457 | 0.80231999 | 0.97095439 |
| RASGRF2 | 140.873303 | -0.3536545 | 0.24071112 | -1.469207 | 0.14177666 | 0.69328517 |
| CKMT2-AS1 | 83.3553292 | -0.0848034 | 0.26004377 | -0.3261119 | 0.74433971 | 0.95563105 |
| CKMT2 | 98.3811097 | 0.48019775 | 0.36812856 | 1.30442952 | 0.19208718 | 0.75061775 |
| ZCCHC9 | 23.6346739 | 0.4201687 | 0.31090472 | 1.35143879 | 0.17655492 | 0.72841249 |
| SSBP2 | 294.135867 | 0.05661122 | 0.20100547 | 0.2816402 | 0.77821941 | 0.96518573 |
| ATG10 | 163.358284 | -0.2107325 | 0.20532937 | -1.0263146 | 0.30474334 | 0.81568231 |
| RPS23 | 204.621105 | 0.08596842 | 0.4469287 | 0.19235377 | 0.8474651 | 0.97612951 |
| ATP6AP1L | 62.2364396 | 0.33009452 | 0.42467686 | 0.77728399 | 0.43699124 | 0.86759679 |
| TMEM167A | 42.5547821 | -0.0354683 | 0.28305775 | -0.125304 | 0.90028288 | 0.98531749 |
| XRCC4 | 72.4659506 | -0.2416849 | 0.29721438 | -0.813167 | 0.41612229 | 0.85867915 |
| COX7C | 124.875032 | 0.5468782 | 0.35996885 | 1.51923757 | 0.12870271 | 0.66929384 |
| RASA1 | 214.969634 | -0.0177045 | 0.25148889 | -0.0703987 | 0.94387635 | 0.99069421 |
| CCNH | 260.680976 | 0.5143943 | 0.35580392 | 1.44572408 | 0.14825461 | 0.69945736 |
| TMEM161B | 87.8389361 | -0.23347 | 0.23257785 | -1.0038361 | 3.15E-01 | 0.82084199 |
| TMEM161B-/ | 286.300737 | -0.3943329 | 0.25247466 | -1.5618711 | 0.11831836 | 0.6668162 |
| MEF2C | 22.5938433 | 1.54269452 | 0.76587236 | 2.01429716 | 0.04397834 | 0.52126936 |
| CETN3 | 10.6303116 | 1.03254066 | 0.62582239 | 1.64989409 | 0.0989646 | NA |
| LYSMD3 | 17.6493413 | 0.35877375 | 0.46354804 | 0.77397317 | 0.43894666 | 0.86879768 |
| ARRDC3 | 94.3519739 | -0.3799895 | 0.38710561 | -0.9816172 | 0.32628848 | 0.82567895 |
| ARRDC3-AS1 | 12.5660915 | 0.08958065 | 0.60906558 | 0.14707883 | 0.8830698 | NA |
| FAM172A | 367.874483 | -0.0199605 | 0.17732276 | -0.1125659 | 0.91037471 | 0.98556715 |
| KIAA0825 | 114.514624 | 0.27393658 | 0.36562503 | 0.74922821 | 0.45371967 | 0.87378486 |

| | | | | | | |
|---|---|---|---|---|---|---|
| SLF1 | 17.8980465 | -0.0431181 | 0.38017657 | -0.1134159 | 0.90970083 | 0.98556715 |
| MCTP1 | 69.1933661 | -0.3646982 | 0.45735397 | -0.7974091 | 0.42521347 | 0.86207532 |
| TTC37 | 141.970097 | -0.2526351 | 0.19631216 | -1.2869053 | 0.19812731 | 0.75559912 |
| ARSK | 19.1042471 | -0.4413823 | 0.4468023 | -0.9878694 | 0.32321663 | 0.82558302 |
| RHOBTB3 | 562.885777 | 0.30119421 | 0.25854962 | 1.1649377 | 2.44E-01 | 7.86E-01 |
| GLRX | 72.5614514 | 0.6378665 | 0.55857336 | 1.14195653 | 0.25347209 | 0.7912221 |
| ELL2 | 112.743307 | -1.2309638 | 0.69048524 | -1.7827517 | 0.07462673 | 0.61020315 |
| AC104123.1 | 55.1598472 | -0.1530777 | 0.30561692 | -0.500881 | 6.16E-01 | 0.92662475 |
| CAST | 191.836319 | -0.1102913 | 0.19839876 | -0.5559071 | 0.57827437 | 0.91883952 |
| ERAP1 | 44.256156 | -0.0958853 | 0.39924189 | -0.2401684 | 0.81019967 | 0.97257655 |
| AC009126.1 | 12.8764666 | 0.48223949 | 0.62739854 | 0.76863343 | 0.44211095 | NA |
| ERAP2 | 26.2524382 | 0.75088174 | 0.66493475 | 1.12925626 | 0.25878975 | 0.79532468 |
| LNPEP | 169.099214 | 0.36834748 | 0.24994907 | 1.47369014 | 0.14056504 | 0.69227547 |
| LIX1-AS1 | 38.4878488 | 0.30059048 | 0.3278345 | 0.91689702 | 0.35919661 | 0.84204868 |
| RIOK2 | 14.0673889 | -0.0143046 | 0.41107452 | -0.0347981 | 0.97224071 | NA |
| RGMB | 12.1027721 | 0.42560906 | 0.45735454 | 0.9305889 | 0.35206626 | NA |
| CHD1 | 162.866483 | -0.1416703 | 0.32638176 | -0.4340633 | 0.66424248 | 0.93860542 |
| LINC02062 | 43.6094019 | 1.15368538 | 0.55043163 | 2.09596491 | 0.0360853 | 0.49396168 |
| AC021086.1 | 12.8628223 | 0.35480934 | 0.63932949 | 0.55497102 | 0.57891448 | NA |
| FAM174A | 27.3810286 | 0.16321669 | 0.32460804 | 0.5028116 | 0.61509674 | 0.92662475 |
| SLCO4C1 | 47.6228914 | 0.20600558 | 0.40364283 | 0.510366 | 6.10E-01 | 0.92573951 |
| PAM | 365.598204 | -0.4555718 | 0.32891407 | -1.3850786 | 0.16602844 | 0.71756194 |
| GIN1 | 17.9289541 | 0.06515678 | 0.3493866 | 0.18648905 | 0.85206127 | 0.97612951 |
| PPIP5K2 | 110.445425 | -0.0436362 | 0.21669339 | -0.2013732 | 0.84040677 | 0.97600352 |
| C5orf30 | 15.3234952 | 1.09748276 | 0.44826396 | 2.44829576 | 0.01435338 | NA |
| NUDT12 | 21.8687976 | 0.10217217 | 0.37624557 | 0.27155713 | 0.78596257 | 0.96767512 |
| EFNA5 | 1053.92066 | -0.155496 | 0.28723899 | -0.5413472 | 0.58826832 | 0.92175048 |
| FBXL17 | 694.927732 | 0.40464017 | 0.16166337 | 2.5029799 | 0.01231525 | 0.36046953 |
| FER | 419.256578 | -0.3736823 | 0.19799256 | -1.8873552 | 0.05911257 | 0.57428141 |
| PJA2 | 134.029189 | 0.01639921 | 0.19187435 | 0.08546849 | 0.93188895 | 0.98960607 |
| MAN2A1 | 287.131473 | -0.0474265 | 0.19820563 | -0.2392792 | 0.8108891 | 0.97257655 |
| TMEM232 | 147.091144 | 0.0786014 | 0.47892681 | 0.16411987 | 0.86963678 | 0.97854266 |
| SLC25A46 | 63.3906347 | -0.1126266 | 0.23019918 | -0.4892572 | 0.6246596 | 0.92981013 |
| WDR36 | 48.7356833 | -0.165441 | 0.2623271 | -0.6306669 | 0.52825834 | 0.90487731 |
| NREP | 55.8032534 | -1.1777524 | 0.48755333 | -2.4156381 | 0.01570767 | 0.39416332 |
| EPB41L4A | 234.218897 | -0.2753525 | 0.19375956 | -1.421104 | 0.15528652 | 0.70572801 |
| APC | 90.5330762 | -0.4086142 | 0.30210053 | -1.352577 | 0.17619082 | 0.72841249 |
| SRP19 | 32.6325668 | -0.0974381 | 0.31043549 | -0.3138757 | 0.75361547 | 0.95826617 |
| REEP5 | 154.586247 | 0.0392792 | 0.24195402 | 0.16234159 | 0.87103687 | 0.97860207 |
| DCP2 | 106.868011 | -0.5141418 | 0.26548768 | -1.9365936 | 0.05279504 | 0.55624218 |
| MCC | 126.967816 | 0.02354234 | 0.255058 | 0.09230192 | 0.92645817 | 0.98925753 |
| YTHDC2 | 127.252159 | 0.41624589 | 0.25630488 | 1.62402638 | 0.10437018 | 0.65946648 |
| PGGT1B | 44.1765245 | 0.18016996 | 0.2412775 | 0.74673337 | 0.45522452 | 0.8739446 |
| CCDC112 | 13.4313364 | 0.43751314 | 0.41093367 | 1.06468069 | 0.28702045 | NA |
| FEM1C | 30.5973525 | -0.9010269 | 0.42480081 | -2.1210573 | 0.03391698 | 0.4908133 |
| TMED7-TICA | 16.043585 | 0.18274185 | 0.49936852 | 0.36594588 | 0.71440546 | NA |
| ATG12 | 39.5185849 | 0.12009886 | 0.30596128 | 0.3925296 | 0.69466694 | 0.94395238 |
| AP3S1 | 68.1215475 | 0.03335339 | 0.27166898 | 0.12277218 | 0.90228751 | 0.98531749 |
| COMMD10 | 281.370848 | -0.3246079 | 0.1893998 | -1.7138764 | 0.08655142 | 0.63466198 |
| SEMA6A | 89.5401589 | -0.206202 | 0.75724548 | -0.2723054 | 0.78538723 | 0.96767512 |
| SEMA6A-AS1 | 36.1708847 | -0.2475558 | 0.36377089 | -0.6805266 | 0.4961711 | 0.89244133 |
| DTWD2 | 82.3044848 | -0.1282495 | 0.19717363 | -0.6504394 | 0.51540844 | 0.90174188 |
| AC008629.1 | 38.7959656 | 0.68082675 | 0.512475 | 1.32850726 | 0.18401059 | 0.73705812 |
| DMXL1 | 235.388329 | -0.1538414 | 0.21625971 | -0.7113734 | 0.4768529 | 0.88495683 |

| | | | | | | |
|---|---|---|---|---|---|---|
| TNFAIP8 | 72.7256088 | -0.650948 | 0.39154353 | -1.6625175 | 9.64E-02 | 6.51E-01 |
| HSD17B4 | 120.127281 | 0.18424996 | 0.31060085 | 0.59320496 | 0.553044 | 0.91322379 |
| SRFBP1 | 45.2474452 | -0.0998198 | 0.30493711 | -0.3273455 | 0.7434066 | 0.95563105 |
| SNX2 | 165.463925 | -0.2212679 | 0.33861538 | -0.653449 | 0.51346686 | 0.90156229 |
| SNX24 | 226.31803 | -0.4955456 | 0.23786214 | -2.083331 | 0.03722106 | 0.49537143 |
| PPIC | 37.9934354 | -0.0961023 | 0.30057476 | -0.3197285 | 0.74917414 | 0.95661834 |
| PRDM6 | 14.8745342 | 0.86222424 | 0.52513381 | 1.64191339 | 0.10060795 | NA |
| CEP120 | 44.0323704 | -0.3166488 | 0.26910525 | -1.1766728 | 0.23932614 | 0.78098673 |
| CSNK1G3 | 101.191892 | 0.10692159 | 0.30476911 | 0.35082817 | 0.72571726 | 0.95164932 |
| ZNF608 | 225.747944 | 0.19410795 | 0.32740655 | 0.59286521 | 0.55327136 | 0.91322379 |
| GRAMD2B | 311.470428 | 0.05112265 | 0.26333457 | 0.19413572 | 0.84606961 | 0.97612951 |
| AC093535.1 | 16.9056339 | 0.55455497 | 0.71241514 | 0.77841547 | 0.43632412 | 0.86729771 |
| ALDH7A1 | 43.8111086 | 0.43170425 | 0.38122261 | 1.13242036 | 0.25745775 | 0.79532468 |
| PHAX | 23.3920926 | 0.08249039 | 0.31348695 | 0.2631382 | 0.79244406 | 0.96944541 |
| C5orf63 | 21.4156033 | 0.04165499 | 0.30637283 | 0.13596176 | 0.89185151 | 0.98329225 |
| PRRC1 | 50.6423771 | -0.4431366 | 0.33177867 | -1.3356391 | 0.18166727 | 0.73398872 |
| LINC01184 | 115.437599 | 0.18656807 | 0.29471799 | 0.63303931 | 0.52670797 | 0.90487731 |
| SLC12A2 | 96.0999684 | 0.18042511 | 0.34501182 | 0.52295341 | 0.60100667 | 0.92352528 |
| SLC27A6 | 17.3945024 | -2.6664655 | 1.72632599 | -1.5445898 | 0.12244551 | 0.66929384 |
| ISOC1 | 14.67593 | 0.3917597 | 0.45817341 | 0.85504678 | 0.39252525 | NA |
| HINT1 | 150.223307 | 0.32716678 | 0.36155517 | 0.90488757 | 0.36552495 | 0.84720508 |
| LYRM7 | 31.0342506 | 0.13322065 | 0.33466832 | 0.39806771 | 0.69058028 | 0.94276429 |
| CDC42SE2 | 294.084159 | -0.3851296 | 0.30555432 | -1.2604292 | 0.20751457 | 0.76231289 |
| P4HA2 | 13.9249012 | 0.24271937 | 0.37032875 | 0.65541595 | 0.51219998 | NA |
| SLC22A5 | 40.4652109 | -0.1308168 | 0.35202377 | -0.3716136 | 0.71018055 | 0.94603965 |
| AC116366.3 | 113.788786 | -0.2803278 | 0.55631852 | -0.503898 | 0.61433308 | 0.92662475 |
| IRF1 | 23.179989 | 0.15312812 | 0.44327545 | 0.34544689 | 7.30E-01 | 0.95279138 |
| KIF3A | 62.2542074 | -0.0023753 | 0.20792747 | -0.0114235 | 0.99088557 | 0.99814944 |
| 8-Sep | 37.2842844 | 0.75297896 | 0.35853361 | 2.10016283 | 0.03571452 | 0.49258981 |
| SHROOM1 | 15.3427615 | -0.1268104 | 0.57651307 | -0.2199611 | 0.82590147 | NA |
| UQCRQ | 57.8842147 | 0.36904814 | 0.43399355 | 0.85035396 | 0.39512832 | 0.85605713 |
| AFF4 | 298.35784 | -0.512525 | 0.23366572 | -2.1934111 | 0.02827777 | 0.46131285 |
| ZCCHC10 | 34.9633937 | -0.2127457 | 0.29679263 | -0.7168159 | 0.47348769 | 0.88359968 |
| HSPA4 | 79.0759668 | 0.01153996 | 0.36118872 | 0.03194995 | 0.97451196 | 0.99431961 |
| FSTL4 | 233.287043 | -0.9176668 | 0.66487184 | -1.3802161 | 0.16752013 | 0.71965603 |
| C5orf15 | 75.6213238 | 0.13693336 | 0.40330096 | 0.33953145 | 0.73420941 | 0.95329896 |
| VDAC1 | 129.267604 | 0.27570599 | 0.28244111 | 0.97615387 | 0.32898822 | 0.82781307 |
| SKP1 | 136.216437 | 0.35580331 | 0.34559849 | 1.02952799 | 0.30323163 | 0.81568231 |
| CDKL3 | 42.9735158 | -0.0710552 | 0.33852447 | -0.2098968 | 0.83374821 | 0.97600352 |
| UBE2B | 121.520459 | -0.2925542 | 0.24141481 | -1.2118322 | 0.22557663 | 0.77080483 |
| JADE2 | 35.2732817 | 0.15608105 | 0.37817217 | 0.41272486 | 0.67980821 | 0.94195357 |
| SAR1B | 53.5713816 | -0.0202862 | 0.26769534 | -0.0757808 | 0.93959349 | 0.99012269 |
| SEC24A | 70.0651638 | -0.4966863 | 0.28305902 | -1.7547092 | 0.07930907 | 0.61367304 |
| CAMLG | 26.9364629 | 0.27358977 | 0.31182599 | 0.87737961 | 0.38028048 | 0.85432452 |
| DDX46 | 117.384193 | -0.1723424 | 0.27048355 | -0.6371639 | 0.52401806 | 0.90487731 |
| C5orf24 | 37.336745 | 0.09924419 | 0.37736359 | 0.26299355 | 0.79255555 | 0.96944816 |
| TXNDC15 | 27.8951442 | 0.07458184 | 0.39619907 | 0.18824336 | 0.85068589 | 0.97612951 |
| PCBD2 | 102.743271 | -0.3166528 | 0.2586004 | -1.2244868 | 0.22076864 | 0.76680432 |
| H2AFY | 80.7287561 | -0.2379336 | 0.26707495 | -0.890887 | 0.37298978 | 0.8494686 |
| CXCL14 | 19.2099919 | 0.68944669 | 0.6412441 | 1.07517042 | 0.28229844 | 0.80431986 |
| SMAD5 | 92.1060026 | 0.08978493 | 0.21961687 | 0.40882531 | 0.68266787 | 0.94205398 |
| KLHL3 | 165.079367 | 0.42064183 | 0.42369334 | 0.99279783 | 3.21E-01 | 8.25E-01 |
| HNRNPA0 | 48.3222238 | 0.37500481 | 0.36558653 | 1.02576211 | 0.30500373 | 0.81568231 |
| AC106791.1 | 29.5969191 | 0.15260776 | 0.48401902 | 0.31529291 | 0.75253927 | 0.95782508 |

| | | | | | | |
|---|---|---|---|---|---|---|
| FAM13B | 221.56271 | -0.3076188 | 0.20542438 | -1.4974794 | 0.13426856 | 0.68205729 |
| NME5 | 47.8761498 | 0.41087699 | 0.45209584 | 0.90882718 | 0.36344136 | 0.84635627 |
| BRD8 | 40.6390457 | 0.08123355 | 0.30279333 | 0.26828051 | 0.78848341 | 0.96771801 |
| CDC23 | 22.1321692 | 0.25807567 | 0.37785496 | 0.68300195 | 0.49460561 | 0.89230917 |
| FAM53C | 15.7251316 | -0.1076992 | 0.41672903 | -0.2584393 | 0.79606787 | NA |
| KDM3B | 105.435812 | 0.15161195 | 0.26810699 | 0.56549046 | 0.57174021 | 0.91791025 |
| EGR1 | 66.3805632 | -1.2888717 | 1.14359657 | -1.1270335 | 0.25972831 | 0.79535701 |
| ETF1 | 72.6147707 | -0.3488728 | 0.33885807 | -1.0295544 | 0.30321922 | 0.81568231 |
| HSPA9 | 140.28453 | -0.4022553 | 0.27206256 | -1.4785397 | 0.13926339 | 0.69054855 |
| CTNNA1 | 415.41285 | -0.4190776 | 0.23205704 | -1.8059251 | 0.07093004 | 0.60572821 |
| SIL1 | 172.94147 | -0.188733 | 0.17098073 | -1.103826 | 0.26966862 | 0.79666376 |
| PAIP2 | 75.7502658 | -0.1022655 | 0.28025891 | -0.3648965 | 0.71518866 | 0.9487373 |
| UBE2D2 | 285.986816 | -0.1842412 | 0.18933399 | -0.9731014 | 0.33050291 | 0.82867637 |
| CXXC5 | 121.488543 | -0.1659523 | 0.31289151 | -0.5303829 | 0.59584648 | 0.9229638 |
| PURA | 98.5316022 | 0.27014206 | 0.25181823 | 1.07276607 | 0.28337609 | 0.80431986 |
| IGIP | 15.3765833 | 0.86424175 | 0.54027266 | 1.59963998 | 0.10967847 | NA |
| CYSTM1 | 296.778409 | -0.093347 | 0.40683557 | -0.2294466 | 0.81852181 | 0.97288403 |
| PFDN1 | 49.1849309 | -0.4144713 | 0.34019368 | -1.2183393 | 0.22309508 | 0.76834258 |
| AC008438.1 | 12.9274138 | 0.36368986 | 0.55031655 | 0.66087392 | 0.50869317 | NA |
| ANKHD1-EIF4 | 14.2747852 | 0.0456288 | 0.47254209 | 0.09656029 | 0.92307559 | NA |
| SRA1 | 23.7264055 | 0.06379772 | 0.3968219 | 0.16077166 | 0.87227324 | 0.97866716 |
| APBB3 | 10.1213058 | 0.08659189 | 0.50576478 | 0.17120981 | 0.86405879 | NA |
| SLC35A4 | 24.7800999 | 0.22646738 | 0.47536498 | 0.47640736 | 0.63378419 | 0.93251095 |
| NDUFA2 | 40.8978592 | 0.43879666 | 0.41750336 | 1.05100152 | 0.29325789 | 0.80935782 |
| IK | 19.0465437 | 0.30668839 | 0.52739309 | 0.58151766 | 0.56089162 | 0.91584721 |
| WDR55 | 16.3305557 | -0.2209974 | 0.39454739 | -0.5601288 | 0.57539159 | 0.91854298 |
| HARS | 30.1986808 | -0.3499807 | 0.2820543 | -1.2408273 | 0.21466957 | 0.7643572 |
| ZMAT2 | 19.7571515 | -0.0248079 | 0.44936092 | -0.0552071 | 0.95597345 | 0.99148072 |
| TAF7 | 30.114146 | 0.3237132 | 0.34947682 | 0.92627948 | 0.35430076 | 0.83989578 |
| DIAPH1 | 301.609151 | -0.2182463 | 0.21978041 | -0.99302 | 0.3207002 | 0.82464962 |
| HDAC3 | 33.0046905 | 0.24648121 | 0.34333836 | 0.71789593 | 0.47282146 | 0.88343467 |
| PCDH1 | 44.0343514 | 0.73171999 | 0.36645666 | 1.99674363 | 0.04585304 | 0.52835471 |
| DELE1 | 37.3582103 | -0.1217099 | 0.26506029 | -0.4591783 | 0.64610612 | 0.93328692 |
| RNF14 | 34.1379919 | -0.3789795 | 0.36972799 | -1.0250224 | 0.30535263 | 0.81568231 |
| GNPDA1 | 27.5709634 | -0.6029401 | 0.35045775 | -1.720436 | 0.08535323 | 0.63062869 |
| NDFIP1 | 187.347031 | -0.1441496 | 0.17863261 | -0.806961 | 0.41968895 | 0.86065558 |
| SPRY4-AS1 | 49.1528983 | 1.11250015 | 0.41222547 | 2.69876615 | 0.00695971 | 0.3013833 |
| ARHGAP26 | 95.3411209 | 0.57617264 | 0.39502535 | 1.4585713 | 1.45E-01 | 0.69512617 |
| YIPF5 | 22.775642 | -0.409045 | 0.31904631 | -1.2820866 | 0.19981226 | 0.75605362 |
| PRELID2 | 107.161911 | -0.6326096 | 0.41499359 | -1.5243841 | 0.12741278 | 0.66929384 |
| LARS | 91.4835574 | -0.1290518 | 0.19774473 | -0.652618 | 0.51400257 | 0.90156229 |
| TCERG1 | 114.409666 | -0.2869384 | 0.22447563 | -1.2782609 | 0.20115745 | 0.75910235 |
| PPP2R2B | 130.473295 | 0.91338856 | 0.76979985 | 1.18652734 | 0.23541411 | 0.77848938 |
| STK32A | 39.7155949 | 0.16617828 | 0.85361283 | 0.1946764 | 0.84564628 | 0.97612951 |
| AC091948.1 | 46.2502842 | 0.0948318 | 0.46049032 | 0.20593657 | 0.83684046 | 0.97600352 |
| FBXO38 | 55.883232 | -0.3476169 | 0.27114648 | -1.2820263 | 0.19983342 | 0.75605362 |
| CSNK1A1 | 429.036832 | -0.2812401 | 0.17854048 | -1.5752174 | 0.11520628 | 0.6668162 |
| ARHGEF37 | 14.9614972 | 0.40584071 | 0.59446249 | 0.68270197 | 0.49479519 | NA |
| SLC26A2 | 26.5000705 | -0.0192191 | 0.56685957 | -0.0339045 | 0.9729533 | 0.99431961 |
| HMGXB3 | 46.0966549 | -0.1421669 | 0.24272744 | -0.585706 | 0.55807309 | 0.91425998 |
| TCOF1 | 17.0935647 | -0.6425996 | 0.41795094 | -1.5375 | 0.12417092 | 0.66929384 |
| CD74 | 316.562729 | -0.0123764 | 0.65436389 | -0.0189137 | 9.85E-01 | 0.99728981 |
| RPS14 | 180.008743 | 0.3782132 | 0.3460703 | 1.09287969 | 0.27444662 | 0.79894072 |
| NDST1 | 35.1110151 | 0.48882854 | 0.30954223 | 1.57919823 | 0.1142906 | 0.66629396 |

| | | | | | | |
|---|---|---|---|---|---|---|
| SYNPO | 15.9796269 | 0.10509365 | 0.40126742 | 0.26190427 | 0.79339525 | NA |
| RBM22 | 15.2181157 | -0.2036699 | 0.44989726 | -0.452703 | 0.6507626 | NA |
| DCTN4 | 88.1517794 | -0.1929222 | 0.18822611 | -1.0249489 | 0.30538729 | 0.81568231 |
| GPX3 | 256.000626 | -0.3515617 | 0.69403279 | -0.5065491 | 0.61247125 | 0.92644418 |
| TNIP1 | 96.9297022 | -0.1190734 | 0.3352131 | -0.3552171 | 0.72242695 | 0.95093722 |
| ANXA6 | 53.5584226 | -0.0829571 | 0.4772191 | -0.1738343 | 0.86199567 | 0.97767895 |
| ATOX1 | 40.8296703 | -0.0688108 | 0.34431149 | -0.1998505 | 0.84159752 | 0.97600352 |
| G3BP1 | 125.543679 | -0.1007069 | 0.22968786 | -0.4384509 | 0.66105942 | 0.93729379 |
| FAM114A2 | 25.5538354 | -0.1265666 | 0.32101676 | -0.3942679 | 0.69338327 | 0.94394214 |
| MFAP3 | 34.915394 | 0.1842326 | 0.31042504 | 0.59348498 | 0.55285663 | 0.91322379 |
| GALNT10 | 82.9320796 | -0.0024085 | 0.33281713 | -0.0072367 | 0.994226 | 0.99921857 |
| SAP30L-AS1 | 23.1639205 | 0.61335714 | 0.4451715 | 1.37779968 | 0.16826514 | 0.72114765 |
| SAP30L | 16.6245553 | 0.10925311 | 0.37584017 | 0.29069036 | 0.77128815 | 0.96366407 |
| LARP1 | 175.129372 | -0.450391 | 0.21861778 | -2.0601753 | 0.03938178 | 0.50068767 |
| FAXDC2 | 28.1683584 | 0.63580163 | 0.48015699 | 1.32415365 | 0.18545202 | 0.73752116 |
| CNOT8 | 26.7173348 | -0.2940415 | 0.3402713 | -0.8641385 | 0.3875118 | 0.85605713 |
| GEMIN5 | 24.4800301 | 0.30978998 | 0.33607073 | 0.92179995 | 0.35663293 | 0.84192356 |
| MRPL22 | 23.2436822 | 0.50830487 | 0.33118297 | 1.53481582 | 0.12482908 | 0.66929384 |
| SGCD | 153.176125 | -0.3440274 | 0.81785107 | -0.4206479 | 0.6740122 | 0.93901638 |
| HAVCR2 | 11.6633087 | 0.12503631 | 0.52432994 | 0.23846875 | 0.81151755 | NA |
| CYFIP2 | 200.860863 | 0.26323135 | 0.31848157 | 0.8265199 | 0.40850924 | 0.85867915 |
| CLINT1 | 277.291661 | 0.08636354 | 0.20419889 | 0.42293833 | 0.67234026 | 0.93878847 |
| RNF145 | 85.170259 | 0.31440535 | 0.35464778 | 0.88652845 | 0.37533283 | 0.85178063 |
| UBLCP1 | 29.3275208 | -0.0349243 | 0.4651484 | -0.075082 | 0.94014949 | 0.99012269 |
| AC008691.1 | 15.8458014 | 1.20657107 | 0.73545545 | 1.64057669 | 0.10088532 | NA |
| TTC1 | 64.1430353 | -0.0583401 | 0.2174227 | -0.2683258 | 0.78844855 | 0.96771801 |
| PWWP2A | 103.050171 | 0.5725032 | 0.27844247 | 2.05609155 | 0.03977369 | 0.50156625 |
| CCNJL | 26.8096708 | 0.21479667 | 0.37167888 | 0.57790927 | 0.56332538 | 0.91624774 |
| SLU7 | 20.5827002 | 0.16727499 | 0.36057021 | 0.46391793 | 0.64270655 | 0.93328692 |
| AC113414.1 | 106.686714 | 0.97564378 | 0.47256271 | 2.06458053 | 0.03896271 | 0.49975128 |
| CCNG1 | 42.1239668 | 0.37680439 | 0.36654313 | 1.02799469 | 0.30395232 | 0.81568231 |
| NUDCD2 | 15.9236062 | 0.17694365 | 0.40085728 | 0.44141311 | 0.65891396 | NA |
| MAT2B | 34.9990188 | 0.30167823 | 0.43655737 | 0.69103914 | 0.48954095 | 0.88930178 |
| WWC1 | 624.225216 | -0.3516981 | 0.44260144 | -0.7946159 | 0.42683697 | 0.86239557 |
| RARS | 53.7124399 | -0.0109585 | 0.25088388 | -0.0436796 | 0.96515982 | 0.99270395 |
| PANK3 | 58.436146 | 0.10694401 | 0.29523336 | 0.36223552 | 0.71717604 | 0.94925929 |
| KCNIP1 | 210.913843 | 1.88861996 | 0.73500299 | 2.56954051 | 0.01018335 | 0.33306084 |
| RANBP17 | 68.658964 | -0.1746266 | 0.4242208 | -0.4116408 | 0.68060274 | 0.94196488 |
| NPM1 | 75.3307619 | 0.33000261 | 0.22165812 | 1.48879099 | 0.13654242 | 0.68406472 |
| FBXW11 | 248.771074 | -0.426148 | 0.26266963 | -1.6223724 | 0.10472364 | 0.65946648 |
| STK10 | 23.2068499 | -0.0522118 | 0.31923928 | -0.1635508 | 0.87008476 | 0.97854266 |
| UBTD2 | 64.9900247 | -0.3835937 | 0.30477587 | -1.2586092 | 0.20817154 | 0.76231289 |
| DUSP1 | 145.422527 | -1.7546398 | 0.5385488 | -3.2580888 | 0.00112165 | 0.14394871 |
| ERGIC1 | 151.165685 | -0.0726331 | 0.24152878 | -0.3007225 | 0.76362608 | 0.96091323 |
| ATP6V0E1 | 234.204439 | -0.1560903 | 0.21544367 | -0.7245063 | 0.46875497 | 0.88192105 |
| CREBRF | 150.614903 | -0.0552597 | 0.29672579 | -0.1862314 | 0.8522633 | 0.97612951 |
| BOD1 | 18.7250223 | -0.0545525 | 0.36868781 | -0.1479639 | 0.88237124 | 0.98137082 |
| CPEB4 | 450.875246 | -0.3154018 | 0.3481453 | -0.9059488 | 0.36496294 | 0.8468666 |
| SFXN1 | 124.025539 | -0.1114947 | 0.22286145 | -0.5002872 | 0.6168729 | 0.92709163 |
| THOC3 | 55.7145349 | -0.1315436 | 0.37836481 | -0.3476634 | 0.72809302 | 0.95272318 |
| SIMC1 | 62.114446 | -0.0504779 | 0.28154128 | -0.1792913 | 0.857709 | 0.97691173 |
| KIAA1191 | 45.4213813 | -0.1404728 | 0.36591152 | -0.3838983 | 0.70105382 | 0.94440669 |
| ARL10 | 17.3985898 | -0.37858 | 0.40677612 | -0.9306838 | 0.35201715 | 0.83872681 |
| CLTB | 35.1170005 | -0.1707973 | 0.35821138 | -0.476806 | 0.63350028 | 0.93251095 |

| | | | | | | |
|---|---|---|---|---|---|---|
| FAF2 | 84.918048 | -0.2450197 | 0.32234528 | -0.7601158 | 0.44718537 | 0.8726106 |
| RNF44 | 24.237708 | 0.07428521 | 0.36393405 | 0.20411724 | 0.83826189 | 0.97600352 |
| TSPAN17 | 15.3579372 | -0.3591176 | 0.50830889 | -0.7064948 | 0.47988048 | NA |
| UIMC1 | 106.609716 | -0.34087 | 0.19671477 | -1.7328132 | 0.08312888 | 0.62435822 |
| ZNF346 | 57.7156359 | -0.2252937 | 0.26579363 | -0.8476264 | 0.39664604 | 0.85656568 |
| NSD1 | 297.557948 | -0.0183276 | 0.19694632 | -0.0930588 | 0.92585686 | 0.98925753 |
| PRELID1 | 28.5085325 | 0.14296191 | 0.42839073 | 0.33371848 | 0.738592 | 0.9542764 |
| LMAN2 | 60.1037878 | 0.25838238 | 0.30309588 | 0.85247736 | 0.3939492 | 0.85605713 |
| DBN1 | 25.3675591 | -0.4306175 | 0.54190741 | -0.7946329 | 0.42682705 | 0.86239557 |
| PDLIM7 | 13.3140395 | -0.9637909 | 0.52606754 | -1.8320668 | 0.06694148 | NA |
| DDX41 | 11.9258338 | -0.6034835 | 0.4615479 | -1.3075208 | 0.19103591 | NA |
| FAM193B | 61.6376307 | -0.2644552 | 0.2474786 | -1.0685982 | 0.28525075 | 0.80431986 |
| TMED9 | 60.9685367 | 0.02056758 | 0.45087373 | 0.04561716 | 0.96361539 | 0.99256424 |
| B4GALT7 | 22.1755439 | 0.31430745 | 0.48117993 | 0.6532015 | 0.51362638 | 0.90156229 |
| FAM153A | 13.9269984 | -1.3756381 | 0.95601126 | -1.4389351 | 0.15016892 | NA |
| AC106795.3 | 20.5210031 | -0.3419441 | 0.48545789 | -0.7043743 | 0.48119969 | 0.88689225 |
| RMND5B | 42.9938481 | -0.1165176 | 0.30548849 | -0.3814142 | 0.70289594 | 0.94476609 |
| NHP2 | 21.9097377 | 0.2456997 | 0.4069659 | 0.60373536 | 0.5460196 | 0.91083314 |
| HNRNPAB | 29.1640557 | 0.06623196 | 0.36234013 | 0.18278948 | 0.85496321 | 0.97612951 |
| PHYKPL | 104.811131 | 0.22961854 | 0.26232504 | 0.87532071 | 0.38139943 | 0.85432452 |
| CLK4 | 49.194204 | 0.31113058 | 0.31895891 | 0.97545661 | 0.32933381 | 0.82831342 |
| ZNF354A | 20.3757298 | 0.3601497 | 0.34467622 | 1.04489278 | 0.29607253 | 0.81105874 |
| ZNF354B | 17.7306628 | 0.26909995 | 0.39496145 | 0.68133218 | 0.49566133 | 0.89230917 |
| ZFP2 | 19.0260223 | 0.26998787 | 0.48348435 | 0.55842112 | 0.57655685 | 0.91854298 |
| ZNF354C | 13.9276469 | 0.07654415 | 0.41609992 | 0.18395618 | 0.85404783 | NA |
| RUFY1 | 86.1850825 | -0.2400159 | 0.26608699 | -0.9020204 | 0.36704601 | 0.84777627 |
| HNRNPH1 | 224.124174 | -0.0587085 | 0.26519919 | -0.2213753 | 0.82480025 | 0.97518529 |
| CANX | 225.138488 | 0.11386488 | 0.28864584 | 0.39447953 | 0.69322704 | 0.94394214 |
| MAML1 | 36.0019211 | 0.08836955 | 0.29993383 | 0.29463015 | 0.76827643 | 0.96289288 |
| MGAT4B | 51.3211215 | -0.5177429 | 0.43319305 | -1.1951783 | 0.23201737 | 0.77390779 |
| SQSTM1 | 239.273779 | -0.6335748 | 0.43658037 | -1.4512215 | 0.1467182 | 0.69731483 |
| MRNIP | 36.7439255 | -0.0545703 | 0.30331491 | -0.1799129 | 0.85722096 | 0.97675334 |
| TBC1D9B | 50.4877657 | -0.2322237 | 0.28153168 | -0.824858 | 0.40945222 | 0.85867915 |
| RNF130 | 157.385352 | -0.6130956 | 0.37339963 | -1.6419287 | 0.10060479 | 0.65863153 |
| MAPK9 | 55.128699 | -0.0604905 | 0.25309764 | -0.2390007 | 0.81110505 | 0.97257655 |
| GFPT2 | 65.8807714 | -0.2327442 | 0.54314611 | -0.4285112 | 0.66827898 | 0.93878847 |
| CNOT6 | 76.6906553 | -0.2951105 | 0.22824093 | -1.2929778 | 0.19601873 | 0.75350056 |
| MGAT1 | 36.8262123 | -0.488929 | 0.28705233 | -1.7032747 | 0.08851668 | 0.63466198 |
| ZFP62 | 16.2407099 | 0.0013743 | 0.35855447 | 0.00383288 | 0.99694181 | 0.99941676 |
| TRIM41 | 19.5322922 | -0.2196924 | 0.39419527 | -0.5573187 | 0.57730973 | 0.91859435 |
| RACK1 | 124.988838 | 0.39429875 | 0.36144668 | 1.09089052 | 0.27532106 | 0.7994854 |
| TRIM52 | 22.1369558 | 0.04095421 | 0.34755425 | 0.11783543 | 0.90619806 | 0.98531749 |
| AC138035.1 | 10.5137933 | 0.15958489 | 0.53020419 | 0.30098761 | 0.76342394 | NA |
| DUSP22 | 32.4856125 | -0.2189436 | 0.26254733 | -0.8339204 | 0.40432581 | 0.85782098 |
| EXOC2 | 120.573702 | -0.3409773 | 0.2072209 | -1.6454774 | 0.0998714 | 0.65863153 |
| GMDS | 309.359404 | 0.11040807 | 0.26048307 | 0.42385893 | 0.6716687 | 0.93878847 |
| GMDS-DT | 481.115005 | 0.1892314 | 0.26344829 | 0.71828668 | 0.47258054 | 0.88316582 |
| WRNIP1 | 34.960131 | 0.20684305 | 0.29297492 | 0.7060094 | 0.48018229 | 0.88660781 |
| SERPINB1 | 38.823405 | 0.36561379 | 0.3004019 | 1.21708215 | 2.24E-01 | 7.68E-01 |
| SERPINB6 | 70.5257216 | 0.25103238 | 0.2525088 | 0.99415301 | 0.32014838 | 0.82449332 |
| NQO2 | 44.0602848 | 0.18840989 | 0.36999115 | 0.5092281 | 0.61059235 | 0.92606167 |
| RIPK1 | 51.0097032 | -0.2666283 | 0.32700067 | -0.8153755 | 0.41485743 | 0.85867915 |
| BPHL | 19.5879418 | 0.02175256 | 0.42174718 | 0.05157725 | 0.95886555 | 0.99229276 |
| PSMG4 | 65.1159189 | 0.04861128 | 0.45651026 | 0.10648453 | 0.91519793 | 0.98691474 |

| | | | | | | |
|---|---|---|---|---|---|---|
| SLC22A23 | 122.503203 | -0.7557951 | 0.68719996 | -1.0998183 | 0.27141129 | 0.79666376 |
| PXDC1 | 13.4241914 | -0.4648341 | 0.42166945 | -1.1023661 | 0.27030252 | NA |
| PRPF4B | 128.484773 | -0.2997681 | 0.20407655 | -1.4689003 | 0.14185983 | 0.69328517 |
| ECI2 | 36.9131989 | 0.13975491 | 0.27437983 | 0.50934832 | 0.61050809 | 0.92606167 |
| CDYL | 184.876626 | -0.4328151 | 0.23310939 | -1.8567038 | 0.0633533 | 0.58688097 |
| LYRM4 | 59.598 | -0.4744955 | 0.33628879 | -1.4109762 | 0.15825163 | 0.71089007 |
| FARS2 | 251.999539 | 0.10190375 | 0.23787673 | 0.42838891 | 0.668368 | 0.93878847 |
| LY86-AS1 | 14.1991552 | 1.48948347 | 0.85713598 | 1.73774465 | 0.08225582 | NA |
| RREB1 | 198.859833 | -0.0532335 | 0.18617719 | -0.2859294 | 0.77493219 | 0.96464609 |
| SSR1 | 76.7380714 | 0.22925541 | 0.29518756 | 0.77664322 | 0.43736929 | 0.86796876 |
| RIOK1 | 18.543424 | -0.1408006 | 0.36996751 | -0.3805757 | 0.70351815 | 0.94476609 |
| DSP | 137.504656 | 0.00703193 | 0.28339727 | 0.02481299 | 0.98020413 | 0.9957881 |
| SNRNP48 | 22.4643979 | 0.36689668 | 0.38644378 | 0.94941799 | 0.34240806 | 0.83340658 |
| BLOC1S5-TXI | 24.9564038 | -0.0331275 | 0.37410946 | -0.0885502 | 0.92943941 | 0.98960607 |
| SLC35B3 | 33.778963 | 0.03217885 | 0.31188777 | 0.10317444 | 0.91782453 | 0.98735011 |
| TFAP2A | 102.709303 | 0.33938586 | 0.23910722 | 1.41938774 | 1.56E-01 | 0.70690214 |
| GCNT2 | 178.583357 | -0.6225253 | 0.26184778 | -2.3774321 | 0.01743365 | 0.4017684 |
| TMEM14C | 35.4484619 | 0.79385953 | 0.34028337 | 2.33293659 | 0.01965147 | 0.41544117 |
| MAK | 41.9080938 | 0.6721203 | 0.46342217 | 1.45034126 | 0.14696338 | 0.69749723 |
| SMIM13 | 13.3068336 | 0.25016953 | 0.40995095 | 0.61024259 | 0.54170112 | NA |
| AL139807.1 | 20.9890717 | 0.18965086 | 0.52434395 | 0.36169172 | 0.71758242 | 0.94925929 |
| NEDD9 | 58.2430164 | -0.3082954 | 0.24331656 | -1.2670546 | 0.20513577 | 0.76231289 |
| TMEM170B | 21.0295769 | 0.11633308 | 0.45913409 | 0.25337495 | 0.79997848 | 0.97019967 |
| HIVEP1 | 111.408328 | -0.0454391 | 0.37707575 | -0.1205038 | 0.90408406 | 0.98531749 |
| PHACTR1 | 91.6014063 | 0.4637275 | 0.27543777 | 1.68360174 | 0.09225866 | 0.64572229 |
| TBC1D7 | 35.4618965 | 0.12239101 | 0.47420637 | 0.25809652 | 0.79633242 | 0.96986756 |
| SIRT5 | 51.1698528 | 0.64681314 | 0.34085711 | 1.89760789 | 0.05774775 | 0.57141408 |
| NOL7 | 35.7676851 | -0.0593808 | 0.33222149 | -0.1787385 | 0.85814303 | 0.97691173 |
| RANBP9 | 152.765253 | 0.13752945 | 0.19545797 | 0.70362673 | 0.48166526 | 0.88703194 |
| MCUR1 | 29.5449382 | 0.16696432 | 0.35686549 | 0.46786344 | 0.63988225 | 0.93328692 |
| JARID2 | 218.212381 | 0.03587283 | 0.47276073 | 0.07587947 | 0.93951499 | 0.99012269 |
| DTNBP1 | 123.888747 | -0.2499694 | 0.38547834 | -0.6484656 | 0.51668385 | 0.90258551 |
| MYLIP | 32.5707324 | -0.025777 | 0.43146174 | -0.0597434 | 0.95236002 | 0.99141043 |
| GMPR | 59.910236 | 0.08234217 | 0.73194607 | 0.11249759 | 0.91042888 | 0.98556715 |
| ATXN1 | 688.374596 | 0.49581802 | 0.27906732 | 1.77669679 | 0.07561814 | 0.61020315 |
| CAP2 | 97.7785546 | 0.46039432 | 0.24739224 | 1.86098932 | 6.27E-02 | 0.58462023 |
| FAM8A1 | 17.966387 | 0.28396312 | 0.39885471 | 0.71194626 | 0.47649806 | 0.88495683 |
| NUP153 | 117.922682 | -0.2770024 | 0.2617174 | -1.0584027 | 0.28987188 | 0.80536924 |
| KIF13A | 153.773708 | 0.08633591 | 0.20989601 | 0.41132707 | 0.68083272 | 0.94196488 |
| TPMT | 28.3391766 | 0.12220165 | 0.28678466 | 0.42610945 | 0.67002809 | 0.93878847 |
| KDM1B | 22.3088395 | -0.309476 | 0.3579002 | -0.864699 | 0.387204 | 0.85605713 |
| DEK | 71.8992833 | -0.1737603 | 0.21743318 | -0.7991432 | 0.42420737 | 0.86190547 |
| RNF144B | 204.457936 | 0.06596038 | 0.41161266 | 0.16024868 | 8.73E-01 | 9.79E-01 |
| AL022068.1 | 24.6337799 | -0.7369875 | 0.66521649 | -1.1078912 | 0.26790879 | 0.79666376 |
| ID4 | 15.4547291 | -0.2355275 | 0.43246189 | -0.5446203 | 0.58601466 | NA |
| MBOAT1 | 101.863576 | 0.23687517 | 0.37523399 | 0.63127322 | 0.5278619 | 0.90487731 |
| E2F3 | 67.0161843 | 0.04280918 | 0.32273952 | 0.13264313 | 0.89447562 | 0.98421511 |
| CDKAL1 | 455.642577 | -0.3487603 | 0.16714343 | -2.0865929 | 0.03692495 | 0.49537143 |
| SOX4 | 101.044994 | -0.5764587 | 0.60111803 | -0.9589776 | 0.33757004 | 0.83055261 |
| CASC15 | 53.971436 | -0.131487 | 0.49890438 | -0.2635514 | 0.7921256 | 0.96931299 |
| DCDC2 | 2099.70376 | 0.07927153 | 0.28606136 | 0.27711372 | 0.78169279 | 0.96652357 |
| MRS2 | 57.652866 | -0.216993 | 0.25796693 | -0.8411658 | 0.40025503 | 0.85778581 |
| ALDH5A1 | 18.7786457 | 0.67839884 | 0.45749072 | 1.48286909 | 0.1381092 | 0.6888811 |
| TDP2 | 28.8557237 | 0.04577751 | 0.32819747 | 0.13948161 | 0.88906958 | 0.98270788 |

| | | | | | | |
|---|---|---|---|---|---|---|
| ACOT13 | 20.022776 | 0.55905812 | 0.44241456 | 1.26365218 | 0.2063549 | 0.76231289 |
| C6orf62 | 111.136702 | -0.3634721 | 0.27440633 | -1.3245762 | 0.18531176 | 0.73752116 |
| GMNN | 44.0979435 | -0.2842525 | 0.3772278 | -0.7535302 | 0.45113138 | 0.87346041 |
| CARMIL1 | 354.919678 | -0.4208728 | 0.36380489 | -1.156864 | 0.24732792 | 0.78583441 |
| SLC17A1 | 16.0111241 | 0.21578575 | 0.84584448 | 0.2551128 | 0.79863598 | NA |
| TRIM38 | 89.4060788 | -0.0380734 | 0.38696617 | -0.0983896 | 0.92162295 | 0.98846691 |
| HIST1H1C | 14.9883727 | -1.0666481 | 0.43983599 | -2.4251042 | 0.01530399 | NA |
| HIST1H4C | 28.5377642 | 1.43848624 | 0.90966112 | 1.58134299 | 0.11379963 | 0.6656824 |
| HIST1H2AC | 46.530773 | -0.2589492 | 0.68810669 | -0.3763213 | 0.70667802 | 0.94558664 |
| BTN3A2 | 18.8538168 | -0.246415 | 0.92975297 | -0.2650327 | 0.79098424 | 0.96909068 |
| BTN3A1 | 16.1247682 | -0.6486552 | 1.00614407 | -0.6446942 | 0.51912539 | NA |
| BTN2A1 | 18.244571 | -0.5510537 | 0.43591158 | -1.264141 | 0.20617941 | 0.76231289 |
| HMGN4 | 13.8189765 | -0.0518835 | 0.52030318 | -0.0997178 | 0.92056841 | NA |
| ZNF322 | 14.1476922 | 0.80171388 | 0.47755435 | 1.67879084 | 0.09319281 | NA |
| LINC00240 | 23.4484568 | 0.68320521 | 0.56989489 | 1.1988267 | 0.23059534 | 0.77249016 |
| ZSCAN16-AS | 19.8967794 | 0.57109384 | 0.36662774 | 1.55769403 | 0.1193058 | 0.6668162 |
| ZKSCAN8 | 20.1143503 | -0.1660025 | 0.40734711 | -0.407521 | 0.68362539 | 0.94205398 |
| ZSCAN9 | 14.8167513 | -0.2521493 | 0.41022903 | -0.6146549 | 0.53878268 | NA |
| ZSCAN23 | 11.9449694 | 0.74501297 | 0.45814075 | 1.62616614 | 0.10391432 | NA |
| TRIM27 | 52.1071922 | -0.0872165 | 0.22668895 | -0.3847408 | 0.70042947 | 0.9443142 |
| GABBR1 | 26.524296 | 0.22621921 | 0.42402375 | 0.533506 | 0.59368336 | 0.9229638 |
| HLA-F | 20.1764323 | -0.6206689 | 0.98232012 | -0.6318398 | 0.52749157 | 0.90487731 |
| HLA-A | 451.108514 | -0.3267657 | 0.61381176 | -0.5323549 | 0.59448023 | 0.9229638 |
| PPP1R11 | 18.1454275 | 0.30428669 | 0.33585921 | 0.90599478 | 0.36493862 | 0.8468666 |
| TRIM26 | 55.2109373 | 0.13523898 | 0.40153051 | 0.33680874 | 0.73626109 | 0.95376039 |
| HCG17 | 126.778943 | -0.1657214 | 0.23796967 | -0.696397 | 0.48618025 | 0.88838332 |
| HLA-E | 167.117444 | -0.1994575 | 0.52546791 | -0.3795808 | 7.04E-01 | 0.94476609 |
| GNL1 | 44.6155566 | 0.26239695 | 0.2692855 | 0.97441919 | 0.32984845 | 0.82867637 |
| PRR3 | 12.0203626 | 0.68565006 | 0.46834831 | 1.46397466 | 0.14320087 | NA |
| ABCF1 | 38.5930638 | -0.0398004 | 0.31679168 | -0.1256359 | 0.90002015 | 0.98531749 |
| PPP1R10 | 44.8132088 | 0.08548349 | 0.28961408 | 0.29516345 | 0.76786902 | 0.96283923 |
| MRPS18B | 39.0006625 | 0.10182733 | 0.37750235 | 0.2697396 | 0.78736059 | 0.96767512 |
| ATAT1 | 22.641168 | -0.5415368 | 0.36896278 | -1.4677274 | 0.1421783 | 0.69328517 |
| C6orf136 | 34.9461994 | 0.14298103 | 0.2819496 | 0.50711558 | 0.61207372 | 0.92644418 |
| DHX16 | 19.4436215 | -0.3793905 | 0.38496615 | -0.9855166 | 0.32437036 | 0.82567895 |
| MDC1 | 23.4888597 | 0.19060382 | 0.36491608 | 0.52232233 | 0.60144592 | 0.92373245 |
| TUBB | 82.1518534 | -0.5098857 | 0.50449162 | -1.0106921 | 0.31216382 | 0.8170291 |
| FLOT1 | 87.2589379 | 0.26046442 | 0.29392309 | 0.88616523 | 0.3755285 | 0.85178063 |
| IER3 | 218.535017 | -0.9311151 | 0.74930193 | -1.2426434 | 0.21399927 | 0.7643572 |
| DDR1 | 120.635465 | 0.35677554 | 0.33541059 | 1.06369791 | 0.28746557 | 0.80523851 |
| VARS2 | 12.1396653 | -0.129569 | 0.44283269 | -0.2925912 | 0.76983463 | NA |
| CCHCR1 | 17.6368499 | 0.22112138 | 0.43497619 | 0.50835284 | 0.61120592 | 0.92619266 |
| POU5F1 | 38.6992567 | -0.7938037 | 0.48650911 | -1.6316318 | 0.10275708 | 0.65946648 |
| HLA-C | 228.226222 | -0.3378669 | 0.71525631 | -0.4723718 | 0.63666146 | 0.93251095 |
| HLA-B | 373.367621 | -0.1278188 | 0.65266198 | -0.1958423 | 8.45E-01 | 0.97612951 |
| MICA | 24.2620148 | -0.0449665 | 0.49837661 | -0.0902259 | 0.92810767 | 0.98960607 |
| NFKBIL1 | 14.95118 | 0.27925551 | 0.39133984 | 0.71358824 | 0.47548183 | NA |
| PRRC2A | 54.7524527 | -0.1868417 | 0.31719124 | -0.5890506 | 0.55582736 | 0.91398748 |
| BAG6 | 28.5360785 | -0.3621999 | 0.38565886 | -0.9391717 | 0.3476426 | 0.83687845 |
| DDAH2 | 33.6871783 | 0.24596977 | 0.32890019 | 0.74785537 | 0.4545474 | 0.87378486 |
| CLIC1 | 114.245631 | 0.28064085 | 0.39950742 | 0.70246719 | 4.82E-01 | 0.88746497 |
| VARS | 13.9414722 | 0.30275214 | 0.54482009 | 0.55569195 | 0.57842146 | NA |
| LSM2 | 23.8696513 | 0.17117791 | 0.42607526 | 0.4017551 | 0.68786427 | 0.94205398 |
| HSPA1A | 95.8871017 | -0.5905136 | 0.62933625 | -0.9383117 | 0.34808422 | 0.83687845 |

| | | | | | | |
|---|---|---|---|---|---|---|
| HSPA1B | 70.1100791 | -0.2195092 | 0.66427065 | -0.3304515 | 0.74105886 | 0.95455763 |
| C6orf48 | 25.3285989 | 0.27303095 | 0.33267021 | 0.82072559 | 0.41180259 | 0.85867915 |
| NEU1 | 27.5525031 | 0.10223016 | 0.37994718 | 0.26906415 | 0.78788032 | 0.96767512 |
| SLC44A4 | 25.1413766 | 0.60046474 | 0.5377515 | 1.11662123 | 2.64E-01 | 7.97E-01 |
| EHMT2 | 22.061118 | 0.0984659 | 0.37723726 | 0.26101848 | 0.79407826 | 0.96974703 |
| NELFE | 12.3431012 | -0.3307668 | 0.4649369 | -0.7114229 | 0.47682219 | NA |
| STK19 | 19.6917571 | -0.3782919 | 0.44010114 | -0.8595568 | 0.39003342 | 0.85605713 |
| ATF6B | 36.4210544 | 0.0962523 | 0.33036412 | 0.29135216 | 0.770782 | 0.96364125 |
| AGPAT1 | 15.1651974 | -0.041641 | 0.53337028 | -0.0780715 | 0.93777117 | NA |
| RNF5 | 18.7449195 | 0.40388927 | 0.40414344 | 0.9993711 | 0.31761495 | 0.82351215 |
| PBX2 | 26.2394329 | -0.344898 | 0.35825002 | -0.9627299 | 0.33568311 | 0.82974237 |
| AL662796.1 | 21.608568 | -0.012006 | 0.56341626 | -0.0213092 | 0.98299898 | 0.99677164 |
| HLA-DRA | 95.7429591 | 1.01591652 | 0.66716472 | 1.52273717 | 0.12782447 | 0.66929384 |
| HLA-DRB5 | 23.8838559 | 1.26792355 | 0.95001952 | 1.33462895 | 0.18199782 | 0.73398872 |
| HLA-DRB1 | 58.9296258 | 0.12070951 | 0.82230642 | 0.14679384 | 0.88329475 | 0.98140267 |
| HLA-DQB1 | 17.6421015 | 1.44540651 | 0.72286311 | 1.9995577 | 0.04554805 | 0.52804951 |
| PSMB8 | 31.428674 | -0.0045081 | 0.57808892 | -0.0077983 | 0.99377795 | 0.99921857 |
| PSMB9 | 16.3904661 | -1.3701264 | 1.04087649 | -1.3163198 | 0.18806672 | 0.74207687 |
| TAP1 | 27.3134362 | -0.8921122 | 0.77083551 | -1.1573315 | 0.24713696 | 0.78583441 |
| HLA-DMA | 8.20225407 | 0.1901638 | 0.65858747 | 0.28874494 | 0.77277657 | NA |
| BRD2 | 74.7036694 | -0.2034762 | 0.28026328 | -0.7260182 | 0.46782763 | 0.88115765 |
| HLA-DPA1 | 28.4089843 | 0.72489758 | 0.90166528 | 0.80395419 | 0.42142343 | 0.86190547 |
| HLA-DPB1 | 19.5489383 | 0.47954051 | 0.83959698 | 0.5711556 | 0.56789418 | 0.91698355 |
| RXRB | 15.2763897 | 0.16995889 | 0.38882918 | 0.43710427 | 0.66203573 | NA |
| SLC39A7 | 29.8194381 | -0.3272309 | 0.59167825 | -0.5530555 | 0.58022538 | 0.92033309 |
| RING1 | 23.7255079 | -0.3520371 | 0.41658143 | -0.845062 | 0.39807623 | 0.85758974 |
| VPS52 | 27.142225 | -0.038525 | 0.36112833 | -0.1066795 | 0.91504325 | 0.98691474 |
| RPS18 | 104.600506 | -0.034643 | 0.42505674 | -0.0815022 | 0.9350426 | 0.99012269 |
| WDR46 | 12.1337824 | -0.1831955 | 0.51778095 | -0.3538089 | 0.7234821 | NA |
| PFDN6 | 15.0313585 | 0.18065936 | 0.38698264 | 0.46684098 | 0.64061365 | NA |
| RGL2 | 25.9517197 | 0.18861381 | 0.43022495 | 0.43840743 | 0.66109096 | 0.93729379 |
| TAPBP | 120.65474 | -0.2644546 | 0.40545316 | -0.6522445 | 0.51424346 | 0.90156229 |
| DAXX | 11.6965072 | -0.0178149 | 0.55991515 | -0.0318171 | 0.97461792 | NA |
| PHF1 | 28.4927773 | -0.0363119 | 0.29587318 | -0.1227278 | 0.90232262 | 0.98531749 |
| CUTA | 38.8771679 | 0.2187481 | 0.4277481 | 0.51139468 | 0.60907472 | 0.92573951 |
| SYNGAP1 | 17.6026152 | 0.06801731 | 0.40576754 | 0.1676263 | 0.86687728 | 0.97788853 |
| ITPR3 | 69.0185245 | -0.6147052 | 0.37477678 | -1.6401902 | 1.01E-01 | 6.59E-01 |
| UQCC2 | 26.3903688 | -0.1488468 | 0.34433334 | -0.4322752 | 0.66554141 | 0.93860542 |
| LEMD2 | 24.5541408 | -0.0601073 | 0.32942252 | -0.1824626 | 0.85521969 | 0.97612951 |
| HMGA1 | 30.124846 | -0.8881717 | 0.54631461 | -1.6257513 | 0.10400256 | 0.65946648 |
| SMIM29 | 17.4782234 | -0.8087634 | 0.51885144 | -1.5587572 | 0.11905385 | 0.6668162 |
| NUDT3 | 56.4824844 | -0.0432371 | 0.46942021 | -0.0921075 | 0.92661259 | 0.98925753 |
| C6orf106 | 123.165627 | -0.1984546 | 0.32048376 | -0.6192344 | 0.53576196 | 0.90763121 |
| SNRPC | 21.3726586 | 0.26256656 | 0.43771006 | 0.59986411 | 0.5485968 | 0.91208861 |
| UHRF1BP1 | 59.2668081 | -0.0303036 | 0.26774791 | -0.1131795 | 0.90988827 | 0.98556715 |
| TAF11 | 32.0850621 | -0.0902009 | 0.31156486 | -0.2895094 | 0.77219161 | 0.9640787 |
| ANKS1A | 367.959743 | -0.4026896 | 0.33796622 | -1.1915083 | 0.23345408 | 0.77650198 |
| SCUBE3 | 22.9876647 | 1.02698295 | 0.67899648 | 1.51250115 | 0.13040642 | 0.67375513 |
| ZNF76 | 53.5983709 | -0.1543691 | 0.29510517 | -0.5230987 | 0.60090554 | 0.92352528 |
| PPARD | 77.0553994 | -0.582106 | 0.33440979 | -1.7406967 | 0.08173675 | 0.61925304 |
| RPL10A | 114.413418 | 0.53544202 | 0.34351253 | 1.55872634 | 0.11906117 | 0.6668162 |
| TEAD3 | 23.97536 | 0.00475851 | 0.3408508 | 0.0139607 | 0.98886134 | 0.99814944 |
| FKBP5 | 21.4017895 | -0.7288296 | 0.86642503 | -0.8411917 | 0.40024053 | 0.85778581 |
| SRPK1 | 70.4210038 | 0.05579534 | 0.23370175 | 0.23874591 | 0.81130262 | 0.97257655 |

| | | | | | | |
|---|---|---|---|---|---|---|
| MAPK14 | 231.666944 | -0.2968699 | 0.34040392 | -0.8721107 | 0.38314799 | 0.85510589 |
| MAPK13 | 19.5119126 | 0.30368328 | 0.3802115 | 0.79872197 | 0.42445164 | 0.86190547 |
| BRPF3 | 17.3965266 | 0.0740607 | 0.36743934 | 0.20155898 | 0.84026152 | 0.97600352 |
| KCTD20 | 69.1803212 | 0.29029842 | 0.3400048 | 0.85380683 | 0.39321203 | 0.85605713 |
| STK38 | 100.775651 | 0.11680782 | 0.2148155 | 0.54375883 | 0.58660744 | 0.92076087 |
| SRSF3 | 81.6633031 | -0.1410998 | 0.27667252 | -0.5099885 | 0.61005953 | 0.92587061 |
| CDKN1A | 23.6163598 | -0.6371111 | 0.4150663 | -1.5349622 | 0.1247931 | 0.66929384 |
| CPNE5 | 35.8379623 | 0.73152253 | 0.86073317 | 0.84988305 | 0.39539011 | 0.85605713 |
| C6orf89 | 77.0931843 | 0.00988634 | 0.28550911 | 0.03462707 | 0.97237711 | 0.99431961 |
| MTCH1 | 79.9591788 | -0.4818771 | 0.2672515 | -1.8030847 | 0.07137492 | 0.60665346 |
| PIM1 | 18.7693332 | -1.7434554 | 0.60419994 | -2.8855603 | 0.00390718 | 0.24454118 |
| TBC1D22B | 58.0368513 | -0.0919978 | 0.2204469 | -0.4173241 | 0.67644134 | 0.94034237 |
| RNF8 | 35.1560006 | -0.3544851 | 0.29116121 | -1.2174875 | 0.22341877 | 0.76834258 |
| CMTR1 | 29.1904833 | -0.5579618 | 0.36459462 | -1.5303623 | 0.12592708 | 0.66929384 |
| ZFAND3 | 1053.87412 | -0.471192 | 0.23531066 | -2.0024252 | 0.04523902 | 0.52675468 |
| BTBD9 | 673.394142 | -0.5326758 | 0.25068037 | -2.1249201 | 0.03359328 | 0.4908133 |
| GLO1 | 75.7644714 | -0.0164048 | 0.22646577 | -0.0724382 | 0.94225321 | 0.99058259 |
| SAYSD1 | 11.3590798 | 0.26544756 | 0.46535755 | 0.57041635 | 0.56839534 | NA |
| KCNK5 | 19.0632552 | 0.32797751 | 0.52087526 | 0.62966613 | 0.52891305 | 0.90495622 |
| KIF6 | 17.522885 | 0.19016139 | 0.8005622 | 0.23753481 | 0.81224192 | 0.97257655 |
| MOCS1 | 21.212439 | 0.38706873 | 0.35845726 | 1.07981836 | 0.28022307 | 0.80311628 |
| UNC5CL | 29.199579 | 1.25304873 | 0.45982185 | 2.72507435 | 0.0064287 | 0.29935381 |
| OARD1 | 44.6048355 | 0.22812368 | 0.36820319 | 0.61955923 | 0.53554802 | 0.90763121 |
| NFYA | 26.381426 | -0.2764875 | 0.37394809 | -0.7393739 | 0.45967997 | 0.87607759 |
| FOXP4 | 14.3486658 | -0.3157485 | 0.37607407 | -0.8395912 | 0.40113763 | NA |
| TFEB | 83.8111538 | 0.67084636 | 0.392054 | 1.71110704 | 0.08706136 | 0.63466198 |
| USP49 | 39.7274664 | -0.0885716 | 0.34768705 | -0.2547451 | 0.79891996 | 0.97019967 |
| CCND3 | 106.102602 | -0.21071 | 0.4758746 | -0.4427847 | 0.65792148 | 0.93593674 |
| TAF8 | 32.5262262 | 0.13974532 | 0.40783833 | 0.3426488 | 0.73186268 | 0.95279138 |
| C6orf132 | 21.3389516 | 0.3302138 | 0.47335622 | 0.69760106 | 0.48542672 | 0.88838332 |
| MRPS10 | 16.9964056 | -0.3270171 | 0.36699948 | -0.8910561 | 0.37289907 | 0.8494686 |
| TRERF1 | 79.7070595 | 1.12233848 | 0.42451 | 2.6438446 | 0.00819703 | 0.30976355 |
| UBR2 | 272.636654 | -0.0196942 | 0.23131446 | -0.0851404 | 0.93214974 | 0.98960607 |
| BICRAL | 112.330926 | 0.18373035 | 0.28551434 | 0.64350655 | 0.51989547 | 0.90368693 |
| RPL7L1 | 53.9003507 | -0.0048154 | 0.29951049 | -0.0160776 | 0.98717248 | 0.99803728 |
| CNPY3 | 14.1477915 | -0.2187494 | 0.43162803 | -0.5068008 | 0.6122946 | NA |
| PEX6 | 21.7182419 | 0.09746968 | 0.41104719 | 0.23712527 | 0.81255961 | 0.97257655 |
| PPP2R5D | 14.1335723 | 0.63546447 | 0.56688109 | 1.12098372 | 0.26229479 | NA |
| KLHDC3 | 21.629431 | 0.35787289 | 0.37334064 | 0.95856933 | 0.33777575 | 0.83055261 |
| RRP36 | 28.9978202 | -0.0259224 | 0.32667647 | -0.0793518 | 0.93675277 | 0.99012269 |
| CUL7 | 16.7689642 | -0.739603 | 0.45988962 | -1.6082185 | 0.10778734 | 0.65986731 |
| KLC4 | 26.3523845 | -0.0138612 | 0.37163663 | -0.0372978 | 0.97024755 | 0.99398482 |
| MRPL2 | 18.571771 | 0.67745435 | 0.40398809 | 1.67691663 | 0.09355879 | 0.64572229 |
| PTK7 | 24.3579132 | 0.74746725 | 0.45948046 | 1.62676612 | 0.10378678 | 0.65946648 |
| CUL9 | 23.3097824 | 0.02663551 | 0.42027224 | 0.06337681 | 0.94946645 | 0.99105535 |
| DNPH1 | 27.50868 | 0.37429127 | 0.5419913 | 0.69058539 | 0.48982613 | 0.88946484 |
| ZNF318 | 22.0804201 | 0.16395343 | 0.35709055 | 0.45913686 | 0.64613589 | 0.93328692 |
| ABCC10 | 18.6159057 | -0.3151369 | 0.43773829 | -0.7199208 | 0.47157379 | 0.88267439 |
| TJAP1 | 23.5068455 | -0.406556 | 0.34882262 | -1.1655094 | 0.24381285 | 0.78544896 |
| POLR1C | 28.2339864 | 0.32135011 | 0.33194719 | 0.968076 | 0.33300642 | 0.82937275 |
| YIPF3 | 31.7899625 | 0.25016195 | 0.3401358 | 0.73547668 | 0.46204924 | 0.87689276 |
| XPO5 | 99.9204617 | -0.0634378 | 0.28656633 | -0.2213722 | 0.82480267 | 0.97518529 |
| POLH | 31.9478356 | 0.36704913 | 0.41457162 | 0.88536965 | 0.37595729 | 0.85217611 |
| GTPBP2 | 27.9722424 | -0.1476128 | 0.34787373 | -0.4243286 | 0.67132618 | 0.93878847 |

| | | | | | | |
|---|---|---|---|---|---|---|
| MRPS18A | 15.9835314 | 0.40749235 | 0.49421784 | 0.82451972 | 0.40964432 | NA |
| VEGFA | 157.234403 | -0.992992 | 0.41814794 | -2.3747384 | 0.01756139 | 0.40369024 |
| MRPL14 | 26.107473 | 0.33748855 | 0.50781912 | 0.66458418 | 0.50631649 | 0.89724484 |
| TMEM63B | 33.5709424 | -0.4213787 | 0.31106581 | -1.3546289 | 0.17553581 | 0.72841249 |
| SLC29A1 | 33.5125367 | 0.05205575 | 0.40658737 | 0.1280309 | 0.89812452 | 0.98531749 |
| HSP90AB1 | 257.671019 | 0.0557827 | 0.28370407 | 0.19662283 | 0.8441227 | 0.97612951 |
| SLC35B2 | 17.970732 | -0.0561486 | 0.47522281 | -0.1181521 | 0.90594711 | 0.98531749 |
| CDC5L | 38.704303 | 0.12473206 | 0.29736266 | 0.41946109 | 0.67487919 | 0.93936166 |
| SUPT3H | 341.967695 | -0.1789125 | 0.21613384 | -0.8277855 | 0.407792 | 0.85867915 |
| RUNX2 | 18.6775082 | -0.1174382 | 0.56401242 | -0.2082193 | 0.83505777 | 0.97600352 |
| ENPP4 | 39.1303297 | 0.86309931 | 0.40352025 | 2.13892437 | 0.03244179 | 0.48175799 |
| ENPP5 | 17.3843689 | 0.61859133 | 0.41344539 | 1.49618632 | 0.13460511 | 0.68328989 |
| RCAN2 | 76.5352793 | 1.06597594 | 0.43329591 | 2.46015694 | 0.01388763 | 0.37638271 |
| CYP39A1 | 14.8361212 | 0.70549395 | 0.61091876 | 1.15480814 | 0.248169 | NA |
| SLC25A27 | 78.5983067 | 0.4002908 | 0.39373045 | 1.01666205 | 0.30931423 | 0.81568231 |
| ADGRF1 | 128.314332 | 0.11702025 | 0.4081632 | 0.28669965 | 0.77434231 | 0.96435639 |
| TNFRSF21 | 26.2969929 | -0.4033816 | 0.44197794 | -0.9126737 | 0.3614142 | 0.84395756 |
| AL355353.1 | 11.2289793 | -0.2658516 | 0.52505624 | -0.5063298 | 0.61262512 | NA |
| CD2AP | 891.306479 | -0.2938015 | 0.26339079 | -1.1154586 | 0.26465394 | 0.79666376 |
| PTCHD4 | 30.4865123 | 3.62036366 | 0.77866526 | 4.64944803 | 3.33E-06 | 0.00378713 |
| MUT | 36.2275079 | 0.22302516 | 0.29636548 | 0.75253423 | 0.45172985 | 0.87346041 |
| PKHD1 | 2437.16735 | 0.35678325 | 0.3084611 | 1.1566556 | 0.2474131 | 0.78583441 |
| MCM3 | 19.9774646 | 0.01013655 | 0.49659351 | 0.02041217 | 0.98371457 | 0.99685229 |
| EFHC1 | 63.8206919 | 0.04348308 | 0.25791937 | 0.16859175 | 0.86611777 | 0.97788853 |
| TRAM2 | 15.5311691 | 0.48833256 | 0.40013248 | 1.2204272 | 0.22230297 | NA |
| TMEM14A | 9.90296495 | 1.03024782 | 0.49844461 | 2.06692536 | 0.03874119 | NA |
| GSTA4 | 21.6057161 | 0.21736001 | 0.38511884 | 0.56439724 | 0.57248382 | 0.91813957 |
| ICK | 22.2123717 | 0.53995559 | 0.44864984 | 1.20351227 | 0.22877815 | 0.77185154 |
| FBXO9 | 76.8649158 | -0.0963459 | 0.23064641 | -0.4177213 | 0.67615089 | 0.9402691 |
| ELOVL5 | 55.3513179 | -0.4594814 | 0.69083808 | -0.6651072 | 0.50598195 | 0.89723255 |
| GCLC | 66.75286 | -0.4115518 | 0.31072632 | -1.3244832 | 0.18534261 | 0.73752116 |
| LRRC1 | 89.8010225 | -0.2301002 | 0.33792177 | -0.6809276 | 0.49591733 | 0.8923375 |
| TINAG | 19.5983664 | 0.18276339 | 0.56197532 | 0.32521605 | 0.74501756 | 0.95563105 |
| FAM83B | 127.720833 | -0.4349665 | 0.27792124 | -1.565071 | 0.11756626 | 0.6668162 |
| DST | 160.295642 | -0.6089352 | 0.54060804 | -1.1263895 | 0.26000069 | 0.79535701 |
| ZNF451 | 107.891231 | -0.0691697 | 0.20204054 | -0.3423558 | 0.73208318 | 0.95279138 |
| ZNF451-AS1 | 14.93785 | 0.3529213 | 0.65256997 | 0.54081756 | 0.58863334 | NA |
| PRIM2 | 81.9532101 | -0.92015 | 0.31623378 | -2.9097144 | 0.00361759 | 0.23862997 |
| AL021368.2 | 13.7232923 | -0.0242325 | 0.44933949 | -0.0539291 | 0.95699164 | NA |
| PTP4A1 | 100.364811 | -1.3585085 | 1.025815 | -1.3243212 | 0.18539641 | 0.73752116 |
| PHF3 | 191.099159 | 0.32834989 | 0.209343 | 1.56847798 | 0.11676963 | 0.6668162 |
| EYS | 20.4532584 | 0.87196277 | 0.69117106 | 1.26157303 | 0.20710247 | 0.76231289 |
| LMBRD1 | 158.04552 | 0.09174513 | 0.20035926 | 0.45790314 | 0.64702203 | 0.93328692 |
| COL19A1 | 43.0391978 | 0.03630401 | 0.4098911 | 0.0885699 | 0.92942373 | 0.98960607 |
| FAM135A | 338.311579 | -0.2032907 | 0.23727448 | -0.8567746 | 0.39156947 | 0.85605713 |
| SDHAF4 | 14.9259536 | 0.41351267 | 0.38842852 | 1.06457855 | 0.28706669 | NA |
| SMAP1 | 217.717859 | -0.5990285 | 0.24120302 | -2.4835031 | 0.01300972 | 0.36664851 |
| OGFRL1 | 21.4942109 | -0.4091537 | 0.54769547 | -0.7470459 | 0.45503585 | 0.8739446 |
| AL136164.2 | 132.443814 | 0.14398682 | 0.35515115 | 0.40542407 | 0.68516582 | 0.94205398 |
| LINC00472 | 60.716645 | -0.1535616 | 0.43606777 | -0.3521509 | 0.72472512 | 0.95101237 |
| AC019205.1 | 22.0950729 | -0.3967484 | 0.393905 | -1.0072184 | 0.31382983 | 0.82005998 |
| OOEP | 106.846884 | 0.45842121 | 0.3950333 | 1.16046219 | 0.24586068 | 0.78583441 |
| MTO1 | 40.9789318 | -0.0582977 | 0.35097675 | -0.1661013 | 0.86807724 | 0.97810461 |
| EEF1A1 | 584.797577 | 0.55673559 | 0.42779334 | 1.3014125 | 0.1931173 | 0.75061775 |

| | | | | | | |
|---|---|---|---|---|---|---|
| SLC17A5 | 33.9896284 | 0.07667476 | 0.41416441 | 0.18513122 | 0.85312612 | 0.97612951 |
| COX7A2 | 68.5977002 | 0.31988851 | 0.35621783 | 0.89801376 | 0.36917821 | 0.84848455 |
| TMEM30A | 78.7064637 | 0.2922912 | 0.28760055 | 1.01630962 | 0.30948197 | 0.81568231 |
| SENP6 | 393.496172 | -0.2223837 | 0.15893368 | -1.3992231 | 0.16174609 | 0.71101785 |
| MYO6 | 777.835584 | -0.3487114 | 0.31188039 | -1.1180934 | 0.2635271 | 0.79666376 |
| MEI4 | 27.6155733 | -0.2289168 | 0.45657812 | -0.5013749 | 0.61610734 | 0.92662475 |
| IRAK1BP1 | 80.4213855 | 0.54621894 | 0.31189123 | 1.75131226 | 0.07989214 | 0.6160244 |
| PHIP | 301.439484 | -0.016237 | 0.20919216 | -0.0776178 | 0.93813208 | 0.99012269 |
| HMGN3 | 158.161217 | 0.35131225 | 0.28694763 | 1.22430788 | 0.22083611 | 0.76680432 |
| LCA5 | 26.355783 | 0.52579617 | 0.36763955 | 1.43019478 | 0.15266112 | 0.70572801 |
| SH3BGRL2 | 54.6717274 | 0.2778972 | 0.2924353 | 0.95028611 | 3.42E-01 | 0.83304756 |
| BCKDHB | 231.534726 | 0.3175731 | 0.19561983 | 1.62341973 | 0.10449971 | 0.65946648 |
| AL359715.1 | 11.04186 | 0.2770738 | 0.50652326 | 0.54701101 | 0.58437117 | NA |
| TENT5A | 27.227579 | -0.7119895 | 0.47422337 | -1.50138 | 0.13325731 | 0.67988987 |
| IBTK | 183.63343 | -0.5769774 | 0.23569462 | -2.447987 | 0.01436568 | 0.3811083 |
| TPBG | 14.2350939 | 1.09545995 | 0.44794392 | 2.44552925 | 1.45E-02 | NA |
| UBE3D | 103.93963 | 0.01940463 | 0.23843101 | 0.08138466 | 0.93513604 | 0.99012269 |
| DOPEY1 | 203.976264 | -0.0210621 | 0.21220843 | -0.0992522 | 0.92093805 | 0.98813049 |
| PGM3 | 37.3808627 | 0.2641174 | 0.33393613 | 0.7909219 | 0.42898957 | 0.8635763 |
| CYB5R4 | 51.7168458 | -0.4009374 | 0.32697223 | -1.2262123 | 0.22011879 | 0.76680432 |
| MRAP2 | 16.4118388 | -0.0568335 | 0.72556465 | -0.07833 | 0.93756558 | 0.99012269 |
| CEP162 | 78.2231608 | 0.04733064 | 0.23991122 | 0.19728397 | 0.84360532 | 0.97612951 |
| SNX14 | 133.090073 | 0.01393601 | 0.18515721 | 0.07526583 | 0.94000321 | 0.99012269 |
| SYNCRIP | 50.9398251 | 0.05912007 | 0.24970114 | 0.23676333 | 0.8128404 | 0.97269438 |
| ZNF292 | 569.321132 | -0.5826951 | 0.27931196 | -2.0861803 | 3.70E-02 | 0.49537143 |
| SMIM8 | 92.5998269 | 0.12497945 | 0.29289099 | 0.42670978 | 0.66959073 | 0.93878847 |
| RARS2 | 105.439852 | -0.4259735 | 0.25411736 | -1.6762865 | 0.09368209 | 0.64572229 |
| ORC3 | 64.7290225 | -0.4702483 | 0.28715127 | -1.6376325 | 0.10149837 | 0.65863153 |
| AKIRIN2 | 23.0187769 | 0.05364535 | 0.31107124 | 0.17245358 | 0.86308095 | 0.97788853 |
| RNGTT | 182.556774 | -0.5717489 | 0.20008989 | -2.8574602 | 0.00427046 | 0.26045571 |
| PNRC1 | 74.2073224 | 0.30202873 | 0.30562948 | 0.98821858 | 0.32304561 | 0.82558302 |
| PM20D2 | 36.8030154 | 0.03038501 | 0.36570884 | 0.08308524 | 0.93378376 | 0.98960607 |
| UBE2J1 | 39.6724517 | 0.03496419 | 0.29185976 | 0.11979793 | 0.90464322 | 0.98531749 |
| RRAGD | 73.3581314 | 0.2630509 | 0.33817639 | 0.77785116 | 0.43665676 | 0.86731104 |
| ANKRD6 | 146.695 | -0.2473031 | 0.21879859 | -1.1302776 | 0.25835928 | 0.79532468 |
| LYRM2 | 35.6673792 | 0.18693211 | 0.27995201 | 0.66772913 | 0.50430651 | 0.89662151 |
| MDN1 | 107.305298 | -0.1008855 | 0.19973347 | -0.5051004 | 0.61348833 | 0.92662475 |
| CASP8AP2 | 45.706784 | 0.00594919 | 0.29741225 | 0.02000319 | 0.98404083 | 0.99685229 |
| BACH2 | 242.706275 | 0.16228633 | 0.36626955 | 0.44307897 | 0.65770862 | 0.93593674 |
| MAP3K7 | 96.1316905 | -0.1449344 | 0.2262009 | -0.6407333 | 0.52169598 | 0.90451015 |
| MANEA | 20.2435629 | 0.16808366 | 0.3838909 | 0.43784226 | 0.66150064 | 0.93729379 |
| UFL1-AS1 | 30.8361739 | 0.24647922 | 0.45731468 | 0.53897073 | 0.58990705 | 0.92229571 |
| UFL1 | 60.604788 | 0.25365723 | 0.28366452 | 0.89421555 | 0.37120657 | 0.8491102 |
| NDUFAF4 | 21.0071624 | 0.16186494 | 0.41491718 | 0.39011386 | 0.69645235 | 0.94395238 |
| KLHL32 | 49.336428 | 0.29727242 | 0.45017195 | 0.66035305 | 0.50902729 | 0.89963034 |
| MMS22L | 62.4975879 | 0.49703045 | 0.41325427 | 1.2027231 | 0.22908349 | 0.77185154 |
| AL589740.1 | 15.201334 | -0.2989066 | 0.61301456 | -0.4876012 | 0.62583235 | NA |
| FBXL4 | 133.90743 | -0.354772 | 0.20179479 | -1.7580831 | 0.07873338 | 0.61367304 |
| FAXC | 14.7756459 | -0.7042039 | 0.42104623 | -1.6725097 | 0.09442387 | NA |
| PNISR | 408.239601 | 0.01019053 | 0.21438395 | 0.04753402 | 0.96208762 | 0.99256424 |
| AL513550.1 | 23.0428814 | 0.74202031 | 0.47904058 | 1.5489717 | 0.12138852 | 0.66771734 |
| USP45 | 72.1604313 | 0.02759327 | 0.34620221 | 0.07970275 | 0.93647367 | 0.99012269 |
| TSTD3 | 14.5975248 | -0.3865544 | 0.40780258 | -0.947896 | 0.34318241 | NA |
| CCNC | 106.242884 | 0.21429968 | 0.28323447 | 0.75661581 | 0.44928006 | 0.87314185 |

| | | | | | | |
|---|---|---|---|---|---|---|
| SIM1 | 274.251625 | -0.3126376 | 0.25224475 | -1.2394218 | 0.21518933 | 0.7643572 |
| ASCC3 | 185.622263 | -0.0675667 | 0.18478186 | -0.3656565 | 0.71462144 | 0.94841799 |
| GRIK2 | 29.6500616 | 0.54130766 | 0.51104 | 1.05922758 | 0.28949615 | 0.80536924 |
| HACE1 | 76.419824 | 0.0261414 | 0.255663 | 0.10224944 | 0.91855869 | 0.98749988 |
| PREP | 51.2794537 | 0.00632045 | 0.25336592 | 0.02494592 | 0.9800981 | 0.9957881 |
| ATG5 | 130.570148 | -0.3270549 | 0.24680337 | -1.325164 | 0.18511676 | 0.73752116 |
| CRYBG1 | 397.158481 | 0.01565805 | 0.33250733 | 0.04709085 | 0.96244082 | 0.99256424 |
| RTN4IP1 | 34.4861278 | 0.35305786 | 0.2852539 | 1.23769685 | 0.21582849 | 0.76566124 |
| QRSL1 | 53.9989929 | 0.20869804 | 0.2539864 | 0.82168983 | 0.41125345 | 0.85867915 |
| LINC02532 | 741.723447 | -0.4719889 | 0.32281796 | -1.4620899 | 0.14371657 | 0.69477001 |
| CD24 | 641.392044 | 0.34183156 | 0.31485365 | 1.08568397 | 0.27761884 | 0.80176531 |
| C6orf203 | 35.4181116 | 0.03854762 | 0.33388986 | 0.1154501 | 0.90808837 | 0.9854228 |
| PDSS2 | 238.82393 | -0.3581391 | 0.18894824 | -1.8954351 | 0.05803479 | 0.57141408 |
| SOBP | 76.9216541 | 0.43106572 | 0.48070164 | 0.89674277 | 0.36985619 | 0.84852785 |
| SEC63 | 112.687789 | -0.3929817 | 0.22461638 | -1.7495681 | 0.08019287 | 0.61685352 |
| OSTM1 | 39.7918017 | -0.2817722 | 0.3817202 | -0.7381641 | 0.46041472 | 0.87607759 |
| SNX3 | 95.2227185 | -0.2400822 | 0.19337549 | -1.2415335 | 0.21440872 | 0.7643572 |
| AFG1L | 98.8947052 | 0.01106521 | 0.25594889 | 0.04323209 | 0.96551652 | 0.99270395 |
| FOXO3 | 246.20343 | 0.20208873 | 0.20865609 | 0.96852544 | 0.33278202 | 0.82928059 |
| ARMC2 | 56.4887892 | 0.43243082 | 0.2520322 | 1.71577605 | 0.08620304 | 0.63384997 |
| SESN1 | 133.83316 | 0.37694861 | 0.20818637 | 1.8106306 | 0.07019805 | 0.60398191 |
| CEP57L1 | 22.7830756 | 0.43108561 | 0.38831268 | 1.11015075 | 0.26693407 | 0.79666376 |
| CD164 | 131.542683 | -0.1051503 | 0.24161785 | -0.4351926 | 0.66342262 | 0.93860542 |
| PPIL6 | 27.1457091 | 0.8574679 | 0.54371216 | 1.5770622 | 0.11478121 | 0.6668162 |
| MICAL1 | 11.4090086 | -0.489652 | 0.40872023 | -1.1980128 | 0.23091205 | NA |
| ZBTB24 | 12.7493361 | 0.32939436 | 0.50231443 | 0.65575333 | 0.51198285 | NA |
| AK9 | 70.9500251 | 0.07186154 | 0.3060122 | 0.23483228 | 0.81433891 | 0.97282508 |
| FIG4 | 57.3343642 | -0.270497 | 0.2857536 | -0.9466094 | 0.34383785 | 0.83465492 |
| WASF1 | 21.052254 | -0.3738861 | 0.47382723 | -0.7890769 | 0.43006709 | 0.86498027 |
| CDC40 | 36.6287336 | -0.140159 | 0.2701725 | -0.5187759 | 0.60391704 | 0.92518627 |
| CDK19 | 232.767089 | -0.6795498 | 0.29447639 | -2.3076546 | 0.02101836 | 0.4212027 |
| AMD1 | 107.586894 | -0.8289426 | 0.38192789 | -2.1704165 | 0.02997531 | 0.46405649 |
| GTF3C6 | 47.7664193 | -0.1804673 | 0.41015499 | -0.4399978 | 0.65993869 | 0.93704911 |
| RPF2 | 26.0298335 | -0.2833553 | 0.38579635 | -0.7344686 | 0.46266322 | 0.87713916 |
| SLC16A10 | 176.34085 | -0.2966394 | 0.33645529 | -0.8816606 | 3.78E-01 | 0.85310518 |
| MFSD4B | 25.3049025 | -0.31562 | 0.32743698 | -0.9639107 | 0.33509071 | 0.82974237 |
| REV3L | 282.276396 | 0.2123602 | 0.15023767 | 1.41349501 | 0.15751023 | 0.7098097 |
| TRAF3IP2-AS | 102.446467 | -0.1983262 | 0.29809257 | -0.6653174 | 0.5058475 | 0.89723255 |
| TRAF3IP2 | 33.1275774 | 0.66502587 | 0.35180426 | 1.89032919 | 0.05871395 | 0.57258816 |
| TUBE1 | 40.3919275 | 0.46956483 | 0.41941056 | 1.11958277 | 0.2628916 | 0.79666376 |
| FAM229B | 13.6197692 | 0.71963493 | 0.56157956 | 1.28144787 | 0.2000364 | NA |
| MARCKS | 25.5981806 | 0.07293922 | 0.39569022 | 0.18433415 | 0.85375133 | 0.97612951 |
| HDAC2 | 51.5779837 | 0.22499987 | 0.2561778 | 0.87829573 | 0.37978325 | 0.85431805 |
| FRK | 247.702204 | -0.046653 | 0.2611391 | -0.1786521 | 0.85821091 | 0.97691173 |
| NT5DC1 | 265.909777 | -0.0744238 | 0.20966265 | -0.3549694 | 0.72261249 | 0.95093722 |
| COL10A1 | 12.8696641 | 0.75077557 | 0.81491442 | 0.92129376 | 0.35689708 | NA |
| TSPYL4 | 16.4787549 | 0.32117994 | 0.46119578 | 0.69640693 | 0.48617402 | 0.88838332 |
| DSE | 93.2185235 | 0.71083056 | 0.37239704 | 1.90879754 | 0.05628821 | 0.56976762 |
| TSPYL1 | 50.9459359 | 0.57717708 | 0.3949052 | 1.46155859 | 0.14386221 | 0.6948824 |
| RWDD1 | 29.8319523 | 0.18111267 | 0.3410756 | 0.53100448 | 0.59541568 | 0.9229638 |
| ZUP1 | 22.1372668 | 0.09178806 | 0.4007024 | 0.22906791 | 0.81881613 | 0.97288403 |
| KPNA5 | 61.0461765 | 0.12562721 | 0.25429913 | 0.49401354 | 0.62129662 | 0.92837542 |
| DCBLD1 | 34.5646841 | -0.2702769 | 0.33827332 | -0.7989898 | 0.4242963 | 0.86190547 |
| GOPC | 49.9134921 | 0.02787187 | 0.31476167 | 0.08854914 | 0.92944023 | 0.98960607 |

| | | | | | | |
|---|---|---|---|---|---|---|
| NUS1 | 59.6811919 | 0.19193125 | 0.2834524 | 0.67711987 | 0.4983299 | 0.89378207 |
| CEP85L | 83.3300436 | 0.05636661 | 0.19618064 | 0.28731994 | 0.77386736 | 0.96435639 |
| MCM9 | 102.751926 | 0.22132999 | 0.2158534 | 1.02537182 | 0.30518778 | 0.81568231 |
| AL365275.1 | 29.9141565 | 0.21869377 | 0.51582403 | 0.42396973 | 0.67158789 | 0.93878847 |
| FAM184A | 49.4320794 | 1.12430324 | 0.36803413 | 3.05488849 | 0.00225144 | 0.18826595 |
| MAN1A1 | 443.267826 | 0.38745696 | 0.25307764 | 1.53098057 | 1.26E-01 | 0.66929384 |
| TBC1D32 | 148.330322 | 0.15857318 | 0.26206312 | 0.60509536 | 0.54511564 | 0.90999223 |
| HSF2 | 19.1206931 | -0.028641 | 0.38250157 | -0.0748782 | 0.94031164 | 0.99012269 |
| SERINC1 | 82.6595172 | 0.1547538 | 0.27524501 | 0.56224018 | 0.57395239 | 0.91813957 |
| PKIB | 17.8945353 | 0.74827338 | 0.63140995 | 1.18508328 | 0.23598452 | 0.77864547 |
| RNF217 | 101.412155 | -0.5662083 | 0.31904581 | -1.7746929 | 0.07594861 | 0.61020315 |
| TPD52L1 | 194.036681 | 0.41046632 | 0.30676671 | 1.33804062 | 0.18088319 | 0.73356278 |
| HDDC2 | 42.7231841 | 0.39536866 | 0.25613385 | 1.54360178 | 0.12268483 | 0.66929384 |
| NCOA7 | 160.242392 | 0.2626521 | 0.33235648 | 0.79027224 | 4.29E-01 | 0.86396404 |
| HINT3 | 30.7195019 | 0.31423337 | 0.34720754 | 0.90503038 | 0.36544929 | 0.84720508 |
| TRMT11 | 43.196193 | -0.2512892 | 0.28470292 | -0.8826364 | 0.37743276 | 0.85310518 |
| AC020559.1 | 16.7162829 | -0.0007186 | 0.45008992 | -0.0015965 | 0.99872618 | 0.99957534 |
| RNF146 | 32.5694678 | 0.23248014 | 0.32132933 | 0.72349492 | 0.46937588 | 0.88207878 |
| ECHDC1 | 84.1561529 | 0.04165284 | 0.20885451 | 0.19943469 | 0.84192272 | 0.97600352 |
| PTPRK | 545.50504 | -0.3655795 | 0.25316867 | -1.4440155 | 0.14873462 | 0.70006785 |
| AL356124.1 | 18.0560135 | 0.7001086 | 0.3975598 | 1.76101458 | 0.07823594 | 0.61367304 |
| ARHGAP18 | 168.241694 | 0.60345841 | 0.26553841 | 2.27258421 | 0.02305125 | 0.4299908 |
| L3MBTL3 | 35.6053945 | 0.31143959 | 0.54343929 | 0.57308994 | 0.5665838 | 0.91698355 |
| EPB41L2 | 53.4686093 | 0.00681023 | 0.38663307 | 0.01761418 | 0.98594664 | 0.99766827 |
| AKAP7 | 55.6815265 | -0.2250128 | 0.38875447 | -0.5788044 | 0.56272115 | 0.91624774 |
| MED23 | 62.6441253 | -0.3129383 | 0.22804212 | -1.372283 | 0.16997534 | 0.72416949 |
| CTGF | 13.9424581 | -0.9277384 | 0.61640967 | -1.505068 | 0.13230661 | NA |
| STX7 | 120.594136 | -0.0633372 | 0.20478776 | -0.3092821 | 0.75710697 | 0.95903525 |
| SLC18B1 | 59.2410078 | 0.05934126 | 0.35554725 | 0.1669012 | 0.86744779 | 0.97805828 |
| RPS12 | 94.0721191 | 0.35665705 | 0.44424955 | 0.80283041 | 0.42207276 | 0.86190547 |
| EYA4 | 163.904674 | -1.1045281 | 0.56931029 | -1.9401161 | 0.05236558 | 0.55403388 |
| TBPL1 | 20.164614 | 0.16685735 | 0.35629627 | 0.46831067 | 0.63956244 | 0.93328692 |
| SLC2A12 | 12.3696072 | 0.19979961 | 0.67242496 | 0.29713294 | 0.76636501 | NA |
| SGK1 | 160.527112 | -0.9806854 | 0.75887228 | -1.2922931 | 0.19625567 | 0.75350056 |
| HBS1L | 71.1359396 | -0.0110016 | 0.23808733 | -0.0462085 | 0.9631441 | 0.99256424 |
| AHI1 | 297.720493 | -0.200348 | 0.25728561 | -0.7786987 | 0.43615723 | 0.86729771 |
| LINC00271 | 123.338648 | 0.67125342 | 0.4040815 | 1.66118324 | 0.09667665 | 0.65237031 |
| PDE7B | 268.664872 | -0.6868319 | 0.44021558 | -1.5602172 | 0.11870855 | 0.6668162 |
| AL138828.1 | 44.5888006 | -0.4403311 | 0.53161672 | -0.8282868 | 4.08E-01 | 0.85867915 |
| BCLAF1 | 140.027494 | -0.0009918 | 0.18876246 | -0.0052544 | 0.99580759 | 0.99927028 |
| MAP7 | 333.217418 | -0.1841653 | 0.22854987 | -0.805799 | 0.42035874 | 0.86124817 |
| MAP3K5 | 166.367395 | 0.2637405 | 0.38722309 | 0.68110737 | 0.49580356 | 0.89230917 |
| PEX7 | 54.4223962 | -0.2677452 | 0.24747785 | -1.0818955 | 0.27929895 | 0.80219184 |
| IFNGR1 | 42.6632457 | 0.06894743 | 0.24987469 | 0.27592801 | 0.78260336 | 0.96652357 |
| PERP | 58.0295355 | 0.32154831 | 0.47647927 | 0.67484217 | 0.49977605 | 0.89433092 |
| ARFGEF3 | 57.9631293 | -0.3476556 | 0.25422569 | -1.3675076 | 1.71E-01 | 7.25E-01 |
| HEBP2 | 49.1230582 | 0.26986397 | 0.33957898 | 0.79470164 | 0.42678708 | 0.86239557 |
| NHSL1 | 169.174016 | -0.9075403 | 0.30213187 | -3.0037887 | 0.0026664 | 0.20745535 |
| CCDC28A | 24.6322997 | 0.26210074 | 0.38079436 | 0.68829997 | 0.4912639 | 0.88994534 |
| REPS1 | 136.28979 | 0.03406683 | 0.20256335 | 0.16817866 | 0.86644273 | 0.97788853 |
| HECA | 18.4346328 | -0.1799515 | 0.40614915 | -0.4430674 | 0.65771696 | 0.93593674 |
| CITED2 | 26.93539 | -0.7832297 | 0.37744464 | -2.075085 | 0.03797866 | 0.49671741 |
| VTA1 | 58.1564534 | -0.0402157 | 0.26194415 | -0.1535278 | 0.87798206 | 0.98040612 |
| ADGRG6 | 36.6630719 | -0.0247207 | 0.61911534 | -0.0399291 | 0.96814963 | 0.99358129 |

| | | | | | | |
|---|---|---|---|---|---|---|
| HIVEP2 | 307.16891 | -0.4836355 | 0.49732549 | -0.9724727 | 0.33081542 | 0.82867637 |
| AIG1 | 212.035521 | -0.0987746 | 0.15962328 | -0.6187984 | 0.53604919 | 0.90763121 |
| ADAT2 | 22.4971148 | 0.19932106 | 0.40773004 | 0.48885548 | 0.62494402 | 0.92981013 |
| PEX3 | 64.4328083 | 0.05714463 | 0.29092514 | 0.19642384 | 0.84427844 | 0.97612951 |
| FUCA2 | 27.2280034 | -0.1587381 | 0.34403756 | -0.4613977 | 0.64451329 | 0.93328692 |
| PHACTR2 | 348.981171 | -0.1138746 | 0.33546118 | -0.3394568 | 0.73426566 | 0.95329896 |
| PLAGL1 | 81.6917743 | 0.00786896 | 0.33381738 | 0.02357265 | 0.98119349 | 0.99596391 |
| SF3B5 | 46.8504934 | 1.00033249 | 0.45366791 | 2.20498841 | 0.02745491 | 0.45275729 |
| UTRN | 393.1947 | -0.1759335 | 0.19173934 | -0.9175662 | 0.35884601 | 0.84204868 |
| EPM2A | 61.8985682 | -0.032959 | 0.47282347 | -0.0697068 | 0.94442704 | 0.99069421 |
| FBXO30 | 13.7844368 | 0.62902316 | 0.48331908 | 1.3014656 | 0.19309913 | NA |
| AL356599.1 | 51.4229902 | 0.7708243 | 0.46029639 | 1.67462598 | 0.09400764 | 0.64572229 |
| SHPRH | 217.489268 | 0.36294224 | 0.20675961 | 1.75538272 | 0.07919387 | 0.61367304 |
| STXBP5-AS1 | 17.7940563 | 0.35577466 | 0.43314443 | 0.82137651 | 0.41143184 | 0.85867915 |
| STXBP5 | 199.453368 | -0.2009529 | 0.16974743 | -1.183835 | 0.2364784 | 0.77864547 |
| SAMD5 | 402.768701 | -0.1176537 | 0.42342497 | -0.2778621 | 0.78111822 | 0.96652357 |
| AL033504.1 | 259.945986 | -0.4314095 | 0.44240453 | -0.9751471 | 0.3294873 | 0.82831342 |
| AL513164.1 | 14.871533 | 0.13072022 | 0.70464427 | 0.18551236 | 0.85282719 | NA |
| SASH1 | 160.61695 | -0.3666288 | 0.23317574 | -1.5723283 | 1.16E-01 | 0.6668162 |
| UST | 78.8428242 | 0.24789804 | 0.36667421 | 0.67607166 | 4.99E-01 | 0.89378207 |
| TAB2 | 197.153296 | -0.1874781 | 0.26640715 | -0.7037277 | 0.48160235 | 0.88703194 |
| PPIL4 | 71.3571666 | -0.1313521 | 0.2586734 | -0.5077912 | 0.6115998 | 0.92630153 |
| GINM1 | 89.0811647 | 0.01065537 | 0.23289316 | 0.04575219 | 0.96350777 | 0.99256424 |
| KATNA1 | 45.2880066 | -0.2826928 | 0.31964472 | -0.8843967 | 0.37648208 | 0.85294086 |
| LATS1 | 96.2290566 | -0.0849903 | 0.27604303 | -0.3078878 | 0.7581677 | 0.95903525 |
| NUP43 | 33.6075885 | -0.0586141 | 0.33762718 | -0.1736059 | 0.86217518 | 0.97767895 |
| PCMT1 | 62.8496441 | -0.0042157 | 0.27968309 | -0.0150732 | 0.98797382 | 0.99814944 |
| LRP11 | 21.7022 | 0.4450868 | 0.40916781 | 1.08778546 | 0.27668983 | 0.80159799 |
| PPP1R14C | 31.7118963 | -1.9726215 | 0.59777344 | -3.2999484 | 9.67E-04 | 0.13754433 |
| PLEKHG1 | 67.6348757 | 0.61070415 | 0.56226807 | 1.0861441 | 0.27741524 | 0.80176531 |
| MTHFD1L | 194.682599 | -0.6651248 | 0.23823249 | -2.7919147 | 5.24E-03 | 0.28057141 |
| ZBTB2 | 11.361809 | -0.0419249 | 0.55303895 | -0.0758083 | 0.93957164 | NA |
| RMND1 | 32.2873237 | -0.2046755 | 0.36662113 | -0.5582753 | 0.57665644 | 0.91854298 |
| ARMT1 | 18.7446994 | -0.0662813 | 0.48586649 | -0.1364188 | 0.89149018 | 0.98329225 |
| SYNE1 | 424.314822 | 0.58045281 | 0.3054064 | 1.90059151 | 0.05735554 | 0.57060925 |
| MTRF1L | 22.7109593 | -0.1804506 | 0.38189799 | -0.4725099 | 0.63656287 | 0.93251095 |
| OPRM1 | 14.0856067 | 0.42651827 | 0.67349934 | 0.63328685 | 0.52654634 | NA |
| IPCEF1 | 209.751353 | 0.34023839 | 0.45508969 | 0.74762931 | 0.45468378 | 0.87378486 |
| CNKSR3 | 253.823795 | 0.08332278 | 0.31870032 | 0.26144554 | 0.79374894 | 0.96960502 |
| SCAF8 | 255.285754 | -0.1848319 | 0.2312476 | -0.7992812 | 0.42412735 | 0.86190547 |
| TIAM2 | 134.941619 | 0.23179852 | 0.29264574 | 0.79207891 | 0.42831467 | 0.86298106 |
| TFB1M | 67.1669578 | -0.1974785 | 0.2575908 | -0.7666364 | 0.4432977 | 0.87006015 |
| ARID1B | 1071.89804 | -0.1341607 | 0.112179 | -1.1959519 | 2.32E-01 | 0.77320554 |
| TMEM242 | 30.3016814 | 0.55621541 | 0.30381836 | 1.83074982 | 0.06713789 | 0.59683227 |
| ZDHHC14 | 150.660679 | -0.2612078 | 0.29903031 | -0.8735161 | 0.38238184 | 0.8548566 |
| SNX9 | 207.378476 | -0.5675793 | 0.26910998 | -2.1090979 | 0.03493613 | 0.49209205 |
| SYNJ2 | 43.6790138 | -0.7210324 | 0.39732781 | -1.8147041 | 6.96E-02 | 6.03E-01 |
| SERAC1 | 21.3882633 | -0.2847105 | 0.38776303 | -0.7342385 | 0.4628034 | 0.87713916 |
| GTF2H5 | 27.541574 | 0.32163995 | 0.4695561 | 0.68498726 | 0.49335196 | 0.89092098 |
| TULP4 | 227.395949 | -0.2042799 | 0.27376199 | -0.7461952 | 0.4555495 | 0.8739446 |
| TMEM181 | 55.1340849 | 0.30089136 | 0.24277793 | 1.23936868 | 0.215209 | 0.7643572 |
| DYNLT1 | 76.8158071 | -0.678737 | 0.39861188 | -1.7027516 | 0.08861456 | 0.63466198 |
| SYTL3 | 56.2828134 | -0.9270845 | 0.78498169 | -1.1810269 | 0.23759205 | 0.77930556 |
| EZR | 227.558111 | -0.0413948 | 0.3809053 | -0.1086747 | 0.91346048 | 0.986403 |

| | | | | | | |
|---|---|---|---|---|---|---|
| RSPH3 | 55.0491548 | -0.0592342 | 0.35292569 | -0.1678376 | 0.86671102 | 0.97788853 |
| AL035530.2 | 46.0386615 | 0.07836453 | 0.48898698 | 0.16025893 | 0.87267711 | 0.97866716 |
| SOD2.1 | 22.83831 | -0.4468333 | 0.54419033 | -0.8210975 | 0.41159073 | 0.85867915 |
| WTAP | 137.968987 | -0.6506221 | 0.24832042 | -2.6200909 | 0.00879063 | 0.31380835 |
| ACAT2 | 22.0842115 | -0.1411174 | 0.69937496 | -0.2017765 | 0.84009147 | 0.97600352 |
| TCP1 | 35.0594774 | 0.36560814 | 0.30304387 | 1.20645284 | 0.22764293 | 0.77185154 |
| MRPL18 | 22.9459868 | 0.59437899 | 0.38241287 | 1.55428606 | 0.1201162 | 0.6668162 |
| IGF2R | 246.741654 | -0.2973859 | 0.24554051 | -1.2111481 | 0.22583863 | 0.77093069 |
| SLC22A2 | 26.8803894 | 1.03889256 | 0.73093145 | 1.42132693 | 0.15522174 | 0.70572801 |
| SLC22A3 | 34.9749058 | -0.1716669 | 0.93639177 | -0.183328 | 0.85454063 | 0.97612951 |
| MAP3K4 | 103.434372 | -0.111924 | 0.20401122 | -0.5486169 | 0.58326835 | 0.92034873 |
| AGPAT4 | 41.7839352 | -0.3353508 | 0.32529475 | -1.0309136 | 0.30258134 | 0.81568231 |
| PRKN | 854.382885 | 0.16930286 | 0.24366887 | 0.6948071 | 0.4871762 | 0.88838332 |
| PACRG | 149.698716 | -0.202999 | 0.29310954 | -0.6925705 | 0.48857914 | 0.88920066 |
| QKI | 401.146623 | -0.154752 | 0.22023258 | -0.7026755 | 0.48225801 | 0.88740543 |
| PDE10A | 574.301372 | -1.7887701 | 0.70079355 | -2.5524923 | 0.01069553 | 0.33942942 |
| SFT2D1 | 43.7157281 | -0.1022752 | 0.26789421 | -0.3817745 | 0.7026286 | 0.94476609 |
| MPC1 | 173.788843 | 0.22335367 | 0.2295775 | 0.97289007 | 0.33060794 | 0.82867637 |
| RPS6KA2 | 116.186003 | -0.1340132 | 0.24764874 | -0.5411423 | 0.58840954 | 0.92175048 |
| Z94721.1 | 24.4806558 | -0.4619427 | 0.43166098 | -1.0701516 | 0.2845511 | 0.80431986 |
| FGFR1OP | 80.6819569 | -0.1448322 | 0.22403885 | -0.6464602 | 0.51798134 | 0.90346506 |
| AFDN | 352.716975 | -0.2063811 | 0.18462923 | -1.1178135 | 0.26364665 | 0.79666376 |
| WDR27 | 117.809307 | 0.13031185 | 0.26837249 | 0.48556336 | 0.62727678 | 0.93057946 |
| PHF10 | 47.9070636 | 0.64608087 | 0.34801545 | 1.85647178 | 0.06338634 | 0.58688097 |
| ERMARD | 24.5999919 | 0.06780979 | 0.46184473 | 0.14682378 | 0.88327112 | 0.98140267 |
| FAM120B | 170.937521 | -0.3134471 | 0.21453346 | -1.4610641 | 0.14399784 | 0.69502246 |
| PSMB1 | 66.5812396 | 0.55444732 | 0.28117681 | 1.97188137 | 0.04862315 | 0.5391188 |
| TBP | 23.5505575 | -0.1160675 | 0.33268716 | -0.3488788 | 0.72718029 | 0.95211709 |
| PDCD2 | 26.9783207 | 0.37955279 | 0.32143454 | 1.18080898 | 0.23767862 | 0.77930556 |
| FAM20C | 193.296063 | -0.2405276 | 0.40821554 | -0.589217 | 0.55571569 | 0.91398748 |
| PDGFA | 32.5709511 | -0.7674834 | 0.45208196 | -1.6976643 | 0.08957114 | 0.63653714 |
| PRKAR1B | 54.1610343 | -0.4500245 | 0.30558204 | -1.4726799 | 0.14083736 | 0.69227547 |
| DNAAF5 | 38.578187 | -0.2015636 | 0.34272859 | -0.5881145 | 0.55645547 | 0.9141017 |
| SUN1 | 89.7724252 | -0.0405894 | 0.20143816 | -0.2014978 | 0.84030933 | 0.97600352 |
| COX19 | 37.5160754 | 0.106489 | 0.41506706 | 0.25655855 | 0.79751957 | 0.97019967 |
| C7orf50 | 79.4691882 | -0.3786011 | 0.25420183 | -1.4893721 | 0.1363894 | 0.68406472 |
| MICALL2 | 13.1519634 | 0.40746151 | 0.49144968 | 0.82910118 | 0.40704716 | NA |
| INTS1 | 36.4372135 | -0.1023598 | 0.36989802 | -0.2767243 | 0.78199182 | 0.96652357 |
| MAFK | 32.0374284 | -1.0569517 | 0.46700329 | -2.2632639 | 0.02361943 | 0.43174223 |
| TMEM184A | 14.5989488 | -0.8132453 | 0.42626853 | -1.9078239 | 0.05641398 | NA |
| PSMG3 | 11.8963298 | 0.03246294 | 0.47895562 | 0.06777859 | 0.94596189 | NA |
| MAD1L1 | 58.4685925 | -0.1490068 | 0.3051399 | -0.488323 | 0.62532106 | 0.92981013 |
| SNX8 | 21.8329464 | -0.9996541 | 0.45107783 | -2.2161456 | 0.02668153 | 0.44895006 |
| EIF3B | 58.7322952 | -0.5987347 | 0.26205903 | -2.2847322 | 0.02232853 | 0.42433539 |
| CHST12 | 19.9231201 | -0.2034101 | 0.43865393 | -0.4637144 | 0.6428524 | 0.93328692 |
| BRAT1 | 16.3523873 | -0.0366858 | 0.40277499 | -0.0910827 | 0.9274269 | 0.9894945 |
| IQCE | 34.8364177 | -0.2653085 | 0.31868169 | -0.8325188 | 0.40511616 | 0.85782098 |
| TTYH3 | 22.0493412 | -0.4898269 | 0.37835122 | -1.2946354 | 0.19544603 | 0.75323675 |
| GNA12 | 55.97232 | -0.0855535 | 0.29154631 | -0.2934474 | 0.76918023 | 0.96289288 |
| SDK1 | 550.479436 | 0.52985355 | 0.32923613 | 1.60934206 | 0.10754157 | 0.65986731 |
| FOXK1 | 116.039163 | -0.4657343 | 0.24485445 | -1.9020862 | 0.05715989 | 0.57049536 |
| AP5Z1 | 18.7860013 | -0.2858826 | 0.37674799 | -0.7588166 | 0.44796225 | 0.87291341 |
| RBAK-RBAKI | 73.2518338 | -0.5220099 | 0.25945492 | -2.0119482 | 0.0442254 | 0.52351599 |
| WIPI2 | 57.5108558 | -0.0014701 | 0.33677761 | -0.0043651 | 0.99651719 | 0.99927028 |

| | | | | | | |
|---|---|---|---|---|---|---|
| TNRC18 | 233.331169 | 0.25559787 | 0.23761586 | 1.07567684 | 0.28207181 | 0.80431986 |
| FBXL18 | 27.6988542 | 0.17895143 | 0.32826309 | 0.54514637 | 0.58565285 | 0.92076087 |
| ACTB | 638.639471 | -0.3529048 | 0.46895649 | -0.752532 | 0.45173121 | 0.87346041 |
| RNF216 | 161.916369 | -0.2678861 | 0.24209705 | -1.1065235 | 0.26850001 | 0.79666376 |
| CCZ1 | 20.2804902 | -0.785876 | 0.37210564 | -2.1119702 | 0.03468901 | 0.49098143 |
| PMS2 | 39.4575023 | -0.1610327 | 0.3123434 | -0.5155631 | 0.60615959 | 0.92569157 |
| AIMP2 | 20.7333548 | -0.1725308 | 0.43605541 | -0.3956625 | 0.69235406 | 0.94374959 |
| EIF2AK1 | 54.1858766 | 0.11150451 | 0.24893087 | 0.44793364 | 0.65420109 | 0.93548446 |
| USP42 | 59.9277148 | -0.394441 | 0.27904724 | -1.4135276 | 0.15750065 | 0.7098097 |
| CYTH3 | 78.8431502 | -0.1270624 | 0.26541928 | -0.4787232 | 0.63213555 | 0.93251095 |
| RAC1 | 145.797656 | -0.0467426 | 0.22743491 | -0.2055207 | 0.83716529 | 0.97600352 |
| DAGLB | 19.8138668 | -0.466143 | 0.42879139 | -1.0871091 | 0.2769886 | 0.80159799 |
| ZDHHC4 | 16.4618004 | -0.0533018 | 0.39109886 | -0.1362873 | 0.89159419 | 0.98329225 |
| C7orf26 | 14.7545938 | -0.6408294 | 0.44840598 | -1.4291276 | 0.15296756 | NA |
| ZNF316 | 21.4105828 | -0.1572243 | 0.37317733 | -0.4213125 | 0.67352688 | 0.93901638 |
| AC073343.2 | 17.1709404 | 0.60101767 | 0.5903873 | 1.01800575 | 0.30867522 | 0.81568231 |
| ZNF12 | 39.748531 | -0.0513919 | 0.39245722 | -0.130949 | 0.89581568 | 0.98503595 |
| CCZ1B | 25.3240009 | 0.0792408 | 0.38890746 | 0.20375233 | 0.83854705 | 0.97600352 |
| C1GALT1 | 106.999285 | -0.3046117 | 0.36547142 | -0.8334761 | 0.40457628 | 0.85782098 |
| COL28A1 | 79.3144002 | 0.39352675 | 0.28101521 | 1.40037525 | 0.16140098 | 0.71101785 |
| AC004982.2 | 16.9227009 | 0.43983562 | 0.41436382 | 1.06147208 | 0.28847542 | 0.80523851 |
| MIOS | 61.8591268 | -0.0922718 | 0.29975076 | -0.3078285 | 0.75821285 | 0.95903525 |
| RPA3 | 31.064261 | 0.89396633 | 0.58948739 | 1.51651477 | 0.12938924 | 0.67112834 |
| UMAD1 | 315.892382 | 0.14213566 | 0.17528198 | 0.81089715 | 0.41742474 | 0.85949274 |
| GLCCI1 | 56.5965164 | 0.29743927 | 0.28645474 | 1.03834648 | 0.29910878 | 0.81301499 |
| ICA1 | 239.624503 | -0.1181811 | 0.20999249 | -0.5627873 | 0.57357971 | 0.91813957 |
| NDUFA4 | 153.262414 | 0.49994431 | 0.41179006 | 1.21407571 | 0.22471881 | 0.76931755 |
| PHF14 | 153.991776 | -0.1764441 | 0.27853829 | -0.6334644 | 0.52643044 | 0.90487731 |
| THSD7A | 1006.9009 | -0.9104203 | 0.45681421 | -1.9929771 | 0.04626395 | 0.53087707 |
| TMEM106B | 76.6140306 | -0.1836475 | 0.22800625 | -0.8054494 | 0.4205604 | 0.86146745 |
| SCIN | 444.850318 | 0.11118186 | 0.35934366 | 0.30940259 | 0.7570153 | 0.95903525 |
| AC011287.1 | 26.9280364 | -0.1813117 | 0.58772504 | -0.3084975 | 0.75770377 | 0.95903525 |
| ISPD | 81.7219407 | 0.11830705 | 0.33405071 | 0.35415894 | 0.72321977 | 0.95093722 |
| ANKMY2 | 50.9265151 | 0.1230776 | 0.2437728 | 0.5048865 | 0.61363855 | 0.92662475 |
| BZW2 | 109.21117 | 0.33572912 | 0.38196351 | 0.87895598 | 0.37942514 | 0.85429312 |
| AC073332.1 | 17.0837068 | 0.2207603 | 0.55849682 | 0.39527584 | 0.69263933 | 0.94374959 |
| SNX13 | 168.116808 | -0.0468031 | 0.21521544 | -0.2174708 | 0.82784145 | 0.97567084 |
| TWISTNB | 13.9944813 | 0.49744429 | 0.40540234 | 1.22703852 | 0.21980811 | NA |
| MACC1 | 265.368485 | -0.2287638 | 0.24429679 | -0.9364177 | 0.34905819 | 0.83687845 |
| ITGB8 | 200.644135 | 0.93464847 | 0.4064508 | 2.29953658 | 0.02147449 | 0.4212027 |
| SP4 | 49.1325363 | -0.0553457 | 0.32656312 | -0.1694795 | 0.86541953 | 0.97788853 |
| RAPGEF5 | 195.736171 | 1.5722172 | 0.51373313 | 3.0603773 | 2.21E-03 | 1.88E-01 |
| STEAP1B | 105.097429 | 0.1529299 | 0.40991944 | 0.37307306 | 0.70909407 | 0.94558664 |
| TOMM7 | 54.3505096 | 0.02165302 | 0.38421176 | 0.05635699 | 0.95505742 | 0.99148072 |
| KLHL7 | 50.9336099 | -0.1529737 | 0.24430158 | -0.6261677 | 0.53120498 | 0.90536584 |
| NUPL2 | 50.0185744 | 0.64712641 | 0.35486219 | 1.82359922 | 0.06821268 | 0.60244724 |
| AC005082.1 | 10.2192477 | 0.78115159 | 0.49751813 | 1.57009674 | 0.11639261 | NA |
| GPNMB | 29.0045673 | -0.5443019 | 0.37105944 | -1.4668859 | 1.42E-01 | 0.69328517 |
| MALSU1 | 30.4011016 | 0.36258921 | 0.3290541 | 1.10191365 | 0.27049921 | 0.79666376 |
| TRA2A | 209.784808 | -0.1273987 | 0.22285481 | -0.5716666 | 0.56754785 | 0.91698355 |
| CCDC126 | 28.2339866 | 0.15724333 | 0.4434672 | 0.35457714 | 0.72290641 | 0.95093722 |
| FAM221A | 18.1902068 | 0.24767814 | 0.41903705 | 0.59106502 | 0.55447686 | 0.9137961 |
| MPP6 | 33.0214601 | 0.22186542 | 0.60802231 | 0.36489685 | 0.71518841 | 0.9487373 |
| OSBPL3 | 423.398296 | -0.5044332 | 0.28453419 | -1.7728384 | 0.07625549 | 0.61051335 |

| | | | | | | |
|---|---|---|---|---|---|---|
| CYCS | 81.4515593 | 0.05744977 | 0.4131658 | 0.13904774 | 0.88941243 | 0.9827634 |
| NFE2L3 | 14.4610159 | 0.39730659 | 0.55398962 | 0.71717334 | 0.47326715 | NA |
| HNRNPA2B1 | 404.14164 | -0.0659458 | 0.18016136 | -0.3660374 | 0.71433717 | 0.94841799 |
| CBX3 | 76.2952545 | -0.012892 | 0.22733126 | -0.0567102 | 0.95477605 | 0.99148072 |
| SNX10 | 324.288297 | 0.02712054 | 0.37976187 | 0.07141459 | 0.9430678 | 0.99058259 |
| AC004540.1 | 71.2494012 | 0.98630828 | 0.59325373 | 1.66254037 | 0.09640448 | 0.65104726 |
| SKAP2 | 259.061609 | 0.09504667 | 0.26259734 | 0.36194834 | 0.71739064 | 0.94925929 |
| HOTAIRM1 | 40.0084743 | -0.0459306 | 0.40774875 | -0.1126444 | 0.91031249 | 0.98556715 |
| HOXA3 | 61.9452948 | 0.45340239 | 0.29551891 | 1.53425849 | 0.12496607 | 0.66929384 |
| HOXA-AS2 | 15.9947542 | 0.28084905 | 0.43430479 | 0.6466635 | 0.51784975 | NA |
| HOXA-AS3 | 9.69075047 | 1.06990691 | 0.64245217 | 1.66534872 | 0.0958432 | NA |
| HOXA7 | 10.9919257 | 1.16140356 | 0.66021297 | 1.75913472 | 0.07855463 | NA |
| HOXA9 | 52.119952 | -0.0772724 | 0.33679391 | -0.2294352 | 0.81853068 | 0.97288403 |
| HIBADH | 216.178316 | 0.12546077 | 0.19164211 | 0.6546618 | 5.13E-01 | 0.90156229 |
| TAX1BP1 | 259.638622 | -0.0353967 | 0.18171991 | -0.1947872 | 0.84555957 | 0.97612951 |
| JAZF1 | 57.1434421 | 0.66138485 | 0.53971464 | 1.22543433 | 0.22041162 | 0.76680432 |
| CREB5 | 29.8411106 | 0.52495556 | 0.85244338 | 0.61582455 | 0.53801032 | 0.90763121 |
| AC005162.3 | 27.8559453 | 0.55615617 | 0.59347645 | 0.93711582 | 0.34869898 | 0.83687845 |
| CPVL | 123.799875 | 0.46256824 | 0.28224486 | 1.63888985 | 0.1012362 | 0.65863153 |
| AC007255.1 | 29.9852572 | -0.2564877 | 0.47652501 | -0.538246 | 0.59040724 | 0.92233545 |
| PRR15 | 19.263232 | 0.18748533 | 0.36521711 | 0.51335307 | 0.60770436 | 0.92569157 |
| SCRN1 | 71.4886281 | 0.27870631 | 0.22154243 | 1.25802678 | 0.20838207 | 0.7623519 |
| FKBP14 | 20.8924218 | 0.45856771 | 0.35177805 | 1.30357114 | 0.19237985 | 0.75061775 |
| PLEKHA8 | 70.3121049 | -0.3943167 | 0.35776595 | -1.102164 | 0.27039039 | 0.79666376 |
| MTURN | 109.829793 | 0.45872394 | 0.22050533 | 2.08033038 | 0.03749524 | 0.49538341 |
| ZNRF2 | 129.409544 | 0.35166671 | 0.21848446 | 1.60957313 | 0.10749108 | 0.65986731 |
| AC006978.2 | 110.028378 | 0.3353246 | 0.28869069 | 1.1615359 | 0.24542403 | 0.78583441 |
| NOD1 | 27.5183956 | 0.60354968 | 0.60136015 | 1.00364097 | 0.3155517 | 0.82084199 |
| GGCT | 12.2271423 | 0.45526311 | 0.54652883 | 0.83300842 | 0.40483998 | NA |
| AC005154.6 | 34.1953314 | 0.47884654 | 0.34299643 | 1.39606856 | 0.16269385 | 0.71325312 |
| GARS | 55.2795517 | -0.7178488 | 0.29312413 | -2.4489584 | 0.014327 | 0.3811083 |
| AQP1 | 19.2763899 | 0.80418588 | 0.5884413 | 1.36663738 | 0.171739 | 0.72505043 |
| PDE1C | 258.60781 | -0.9652864 | 0.62628444 | -1.5412908 | 0.12324602 | 0.66929384 |
| LSM5 | 20.7312239 | 0.49255559 | 0.44850297 | 1.09822147 | 0.27210779 | 0.7968545 |
| AVL9 | 192.250203 | -0.2336146 | 0.2364505 | -0.9880064 | 0.3231495 | 0.82558302 |
| KBTBD2 | 43.8047193 | -0.2656754 | 0.24175302 | -1.0989538 | 0.27178821 | 0.79666376 |
| FKBP9 | 38.9578356 | 0.45012572 | 0.30823814 | 1.46031804 | 0.14420269 | 0.69512617 |
| NT5C3A | 74.3309346 | -0.5538629 | 0.45251738 | -1.2239593 | 0.22096758 | 0.76680432 |
| RP9 | 15.8459999 | 0.34522942 | 0.45753204 | 0.75454698 | 0.45052085 | NA |
| BBS9 | 208.137694 | 0.18294447 | 0.23409552 | 0.78149499 | 0.43451143 | 0.86729771 |
| DPY19L1 | 80.2400944 | -0.5244043 | 0.28537667 | -1.8375864 | 0.06612337 | 0.59523726 |
| HERPUD2 | 59.9828948 | 0.00378639 | 0.27463061 | 0.01378719 | 0.98899976 | 0.99814944 |
| SEPT7-AS1 | 25.1354125 | 0.66274525 | 0.52940993 | 1.25185648 | 0.21062216 | 0.76397692 |
| 7-Sep | 173.417616 | -0.4251407 | 0.28960952 | -1.467979 | 0.14210994 | 0.69328517 |
| EEPD1 | 75.3144268 | -0.9406241 | 0.39441861 | -2.384837 | 0.01708669 | 0.40063153 |
| KIAA0895 | 44.1749666 | 0.03132527 | 0.27968923 | 0.11200027 | 0.91082319 | 0.98568025 |
| AOAH | 45.3789014 | -0.8625571 | 0.62507224 | -1.3799319 | 1.68E-01 | 7.20E-01 |
| EPDR1 | 14.8516987 | 0.37186624 | 0.54020285 | 0.68838259 | 0.49121188 | NA |
| STARD3NL | 32.7629289 | 0.31932635 | 0.29801748 | 1.07150208 | 0.28394373 | 0.80431986 |
| VPS41 | 129.70348 | -0.3538148 | 0.19171468 | -1.8455277 | 0.06496081 | 0.5907475 |
| YAE1D1 | 15.5492729 | 0.50473431 | 0.44069037 | 1.14532639 | 0.25207398 | NA |
| RALA | 74.684288 | -0.3033059 | 0.25436169 | -1.1924196 | 0.23309674 | 0.77639211 |
| LINC00265 | 16.4853751 | 0.07602311 | 0.38493018 | 0.19749844 | 0.8434375 | 0.97612951 |
| CDK13 | 318.012163 | -0.16962 | 0.20351377 | -0.8334573 | 0.40458685 | 0.85782098 |

| | | | | | | |
|---|---|---|---|---|---|---|
| MPLKIP | 27.9573916 | 0.41381808 | 0.31433746 | 1.31647715 | 0.18801394 | 0.74207687 |
| SUGCT | 69.6015088 | 0.27681069 | 0.32594293 | 0.84926121 | 0.39573596 | 0.85605713 |
| MRPL32 | 23.0915051 | 0.20493063 | 0.37229383 | 0.55045399 | 0.58200802 | 0.92034873 |
| STK17A | 33.8243165 | -0.267456 | 0.46184754 | -0.5791002 | 0.56252155 | 0.91624774 |
| COA1 | 177.229837 | -0.0399697 | 0.18644685 | -0.2143761 | 0.83025381 | 0.97567084 |
| BLVRA | 24.5261549 | 0.29641519 | 0.30686872 | 0.96593486 | 0.33407677 | 0.82974237 |
| URGCP | 47.2852709 | 0.28602168 | 0.42925006 | 0.66632881 | 0.50520095 | 0.89723255 |
| UBE2D4 | 14.6452532 | 0.20781795 | 0.39493999 | 0.52620134 | 0.59874832 | NA |
| AC004951.1 | 10.6802138 | -0.4032162 | 0.52407496 | -0.7693864 | 0.44166393 | NA |
| DBNL | 26.4599255 | 0.17998028 | 0.44217353 | 0.40703539 | 0.68398199 | 0.94205398 |
| POLM | 21.6341156 | 0.45660171 | 0.42409106 | 1.0766596 | 0.28163237 | 0.80417172 |
| POLD2 | 16.1550233 | 0.52947452 | 0.41718763 | 1.269152 | 0.20438686 | 0.76173533 |
| YKT6 | 16.5232923 | -0.3202529 | 0.40372484 | -0.7932456 | 0.42763477 | 0.86239557 |
| NUDCD3 | 116.204091 | -0.5988484 | 0.18801567 | -3.1850984 | 0.00144705 | 0.16191873 |
| DDX56 | 21.848334 | 0.17269602 | 0.37070043 | 0.46586411 | 0.64131277 | 0.93328692 |
| TMED4 | 58.5774969 | 0.38206288 | 0.36479962 | 1.04732258 | 0.29495083 | 0.81018025 |
| OGDH | 281.556694 | -0.0804468 | 0.2137639 | -0.3763347 | 0.7066681 | 0.94558664 |
| ZMIZ2 | 52.9024662 | 0.1866208 | 0.26777848 | 0.69692231 | 0.48585142 | 0.88838332 |
| PPIA | 187.323883 | 0.04573148 | 0.38352009 | 0.11924142 | 0.90508409 | 0.98531749 |
| H2AFV | 80.6712587 | -0.0415295 | 0.32160216 | -0.1291331 | 0.89725234 | 0.98531749 |
| PURB | 49.2173905 | 0.17552056 | 0.29557748 | 0.5938225 | 0.55263084 | 0.91322379 |
| CCM2 | 57.4306253 | -0.0196283 | 0.31047201 | -0.0632208 | 0.94959072 | 0.99105535 |
| TBRG4 | 16.3569854 | -0.1292323 | 0.41931812 | -0.3081963 | 0.75793297 | 0.95903525 |
| ADCY1 | 21.3155562 | 0.85591163 | 0.41138297 | 2.08057139 | 3.75E-02 | 4.95E-01 |
| IGFBP3 | 127.27307 | -1.9561524 | 0.52278975 | -3.7417573 | 0.00018274 | 0.05491436 |
| AC004870.4 | 68.1735074 | 1.49070039 | 0.73321245 | 2.03310838 | 0.04204157 | 0.51232189 |
| TNS3 | 452.853889 | -0.3151301 | 0.32693387 | -0.9638957 | 0.33509823 | 0.82974237 |
| HUS1 | 29.1525239 | -0.3299287 | 0.32752569 | -1.0073368 | 0.31377293 | 0.82005998 |
| UPP1 | 99.9869357 | -1.4228583 | 0.70787211 | -2.01005 | 0.0444259 | 0.52480969 |
| GRB10 | 75.6961267 | -0.9122934 | 0.44343428 | -2.0573363 | 0.03965389 | 0.50156625 |
| COBL | 117.861134 | -0.10521 | 0.30511955 | -0.3448156 | 0.73023303 | 0.95279138 |
| SEC61G | 28.4717056 | -0.2871232 | 0.46971124 | -0.6112759 | 0.54101692 | 0.90783815 |
| EGFR | 200.738906 | 0.10393267 | 0.17329326 | 0.59975023 | 0.54867271 | 0.91208861 |
| LANCL2 | 35.9580085 | -0.306043 | 0.34547284 | -0.8858669 | 0.37568925 | 0.85178063 |
| VOPP1 | 145.397803 | 0.38771709 | 0.33618075 | 1.15329951 | 0.24878747 | 0.78729484 |
| NIPSNAP2 | 120.591606 | 0.0080268 | 0.24255176 | 0.03309315 | 0.9736003 | 0.99431961 |
| PSPH | 17.9429608 | 0.19305635 | 0.47315362 | 0.40802046 | 0.68325866 | 0.94205398 |
| CCT6A | 50.1723697 | -0.0626143 | 0.28247691 | -0.2216616 | 0.82457732 | 0.97518529 |
| SUMF2 | 55.1423421 | 0.0256392 | 0.25337229 | 0.1011918 | 0.91939821 | 0.9875219 |
| PHKG1 | 17.3720503 | -0.0270855 | 0.43335833 | -0.0625015 | 0.95016349 | 0.99105535 |
| CHCHD2 | 110.650911 | 0.35145192 | 0.36135389 | 0.9725976 | 0.33075334 | 0.82867637 |
| NUPR2 | 15.659437 | 0.26084317 | 0.47709376 | 0.54673355 | 0.5845618 | NA |
| ZNF727 | 17.4879739 | -0.1909723 | 0.42696293 | -0.4472807 | 0.65467237 | 0.93548446 |
| ZNF736 | 80.0498319 | 0.26827058 | 0.2743866 | 0.97771023 | 0.32821766 | 0.82740664 |
| ZNF680 | 100.436219 | 0.23094815 | 0.28567097 | 0.8084411 | 0.4188367 | 0.86022698 |
| ZNF107 | 25.8499569 | 0.15295083 | 0.34058837 | 0.44907826 | 0.6533752 | 0.93532021 |
| ZNF138 | 62.3373606 | 0.3444429 | 0.31270599 | 1.10149119 | 0.27068294 | 0.79666376 |
| ZNF273 | 13.4137805 | 0.28003684 | 0.62280455 | 0.44963839 | 0.6529712 | NA |
| ZNF117 | 36.0264905 | -0.2584718 | 0.3520078 | -0.7342787 | 0.4627789 | 0.87713916 |
| VKORC1L1 | 85.1842543 | -0.2940013 | 0.33386298 | -0.8806046 | 0.37853185 | 0.85354853 |
| GUSB | 21.4725376 | 0.73299642 | 0.34864368 | 2.1024228 | 0.03551626 | 0.49209205 |
| ASL | 14.5724568 | -0.4736147 | 0.41478734 | -1.1418253 | 0.25352666 | NA |
| AC068533.4 | 24.188978 | 0.10933466 | 0.49190102 | 0.22226965 | 0.82410398 | 0.97518529 |
| CRCP | 17.1103078 | -0.0363705 | 0.47517882 | -0.0765406 | 0.93898898 | 0.99012269 |

| | | | | | | |
|---|---|---|---|---|---|---|
| AC068533.3 | 16.2396778 | 0.56752496 | 0.52219315 | 1.08681041 | 0.2771206 | 0.80159799 |
| TPST1 | 461.5758 | -0.0201842 | 0.21487574 | -0.0939341 | 0.9251615 | 0.98925753 |
| AC027644.4 | 83.9195979 | -0.1371951 | 0.41041831 | -0.3342812 | 0.73816736 | 0.9542764 |
| TMEM248 | 72.7798848 | 0.04899364 | 0.25447467 | 0.19252855 | 0.84732821 | 0.97612951 |
| SBDS | 42.3949811 | 0.3618456 | 0.31152487 | 1.1615304 | 0.24542627 | 0.78583441 |
| TYW1 | 141.129811 | -0.2206931 | 0.19622778 | -1.1246781 | 0.26072548 | 0.79574359 |
| AC006480.2 | 13.2196355 | 0.14638079 | 0.45028081 | 0.32508778 | 0.74511464 | NA |
| AUTS2 | 125.800071 | -0.1237599 | 0.31741623 | -0.3898978 | 0.69661213 | 0.94395238 |
| GALNT17 | 78.7620474 | -1.125734 | 0.59590303 | -1.8891228 | 5.89E-02 | 5.73E-01 |
| CALN1 | 17.3959471 | 0.67470785 | 0.47601172 | 1.41741856 | 0.15636059 | 0.70818462 |
| TYW1B | 107.419742 | -0.083034 | 0.23468853 | -0.3538052 | 0.72348488 | 0.95093722 |
| POM121 | 105.504441 | -0.2677051 | 0.22568767 | -1.1861751 | 0.23555317 | 0.77848938 |
| NSUN5 | 18.960353 | 0.12272749 | 0.43495535 | 0.28216112 | 0.77781997 | 0.96518573 |
| BAZ1B | 105.092735 | -0.3477739 | 0.2485276 | -1.3993372 | 0.16171188 | 0.71101785 |
| BCL7B | 25.9702965 | -0.2677827 | 0.35391137 | -0.7566377 | 0.44926695 | 0.87314185 |
| MLXIPL | 23.9904955 | 0.3068513 | 0.45975434 | 0.66742448 | 0.50450103 | 0.89679221 |
| DNAJC30 | 11.3010216 | 0.40388075 | 0.55645798 | 0.72580638 | 0.46795748 | NA |
| BUD23 | 38.538894 | 0.01821304 | 0.32905106 | 0.05535019 | 0.95585948 | 0.99148072 |
| ABHD11 | 28.8757695 | -0.1021046 | 0.40596718 | -0.2515094 | 0.8014203 | 0.97045451 |
| CLDN3 | 52.9260049 | 0.07865536 | 0.48450973 | 0.16234011 | 8.71E-01 | 9.79E-01 |
| CLDN4 | 110.822499 | -0.0611435 | 0.32459381 | -0.1883693 | 8.51E-01 | 9.76E-01 |
| LIMK1 | 17.302532 | -0.5104305 | 0.46765431 | -1.0914698 | 0.27506622 | 0.7994854 |
| EIF4H | 55.9050417 | 0.23159109 | 0.23453829 | 0.98743405 | 0.3234299 | 0.82558302 |
| RFC2 | 11.0638861 | -0.2053598 | 0.47274615 | -0.4343976 | 0.66399977 | NA |
| CLIP2 | 64.184205 | -1.0648245 | 0.49540101 | -2.1494194 | 0.03160117 | 0.47705711 |
| GTF2IRD1 | 604.958066 | -1.4459446 | 0.43522223 | -3.3223132 | 0.00089274 | 0.13544418 |
| GTF2I | 356.185662 | 0.30859172 | 0.16219682 | 1.90257567 | 0.05709594 | 0.57049536 |
| GTF2IRD2 | 42.2141345 | -0.1974908 | 0.38567651 | -0.5120634 | 0.60860662 | 0.92573951 |
| CASTOR2 | 44.3858642 | -0.3010626 | 0.42409174 | -0.7098997 | 0.47776633 | 0.88545331 |
| RCC1L | 41.7827785 | 0.04019353 | 0.26514997 | 0.15158793 | 0.87951196 | 0.98091121 |
| GTF2IRD2B | 63.9511382 | -0.4398511 | 0.39142334 | -1.1237222 | 0.26113089 | 0.79589391 |
| TRIM73 | 31.0620334 | 0.71621715 | 0.5231533 | 1.36903876 | 0.17098716 | 0.72463203 |
| POM121C | 92.1896713 | -0.5111008 | 0.22280996 | -2.2938868 | 0.021797 | 0.4212027 |
| SPDYE5 | 17.335068 | 0.5591975 | 0.52103713 | 1.07323925 | 0.28316378 | 0.80431986 |
| HIP1 | 192.686904 | -0.1798801 | 0.36893641 | -0.4875639 | 0.62585881 | 0.93015391 |
| RHBDD2 | 29.0830653 | -0.5342763 | 0.42648692 | -1.2527379 | 0.21030111 | 0.76397692 |
| POR | 81.8355116 | -0.6791921 | 0.44360907 | -1.5310599 | 0.12575459 | 0.66929384 |
| TMEM120A | 12.3695776 | 0.2380533 | 0.42278101 | 0.56306526 | 0.57339045 | NA |
| STYXL1 | 42.0459455 | 0.2916704 | 0.30287119 | 0.96301799 | 0.3355385 | 0.82974237 |
| MDH2 | 45.1641638 | 0.41968287 | 0.34854537 | 1.20409824 | 0.22855161 | 0.77185154 |
| HSPB1 | 194.7878 | 0.73375062 | 0.54896179 | 1.33661512 | 0.1813483 | 0.73398872 |
| YWHAG | 61.9084376 | -0.3524705 | 0.30719183 | -1.1473953 | 0.2512183 | 0.78896325 |
| SSC4D | 45.4962927 | 0.90142305 | 0.34581573 | 2.60665712 | 0.00914309 | 0.32071089 |
| DTX2 | 33.7689766 | -0.1157303 | 0.3578863 | -0.3233718 | 0.74641368 | 0.95563105 |
| POMZP3 | 20.2595432 | 0.3387997 | 0.39989291 | 0.84722608 | 0.39686912 | 0.85656568 |
| CCDC146 | 96.4448821 | 0.36633281 | 0.2998728 | 1.22162736 | 0.22184858 | 0.76680432 |
| FGL2 | 62.4045482 | 1.49240318 | 0.75124019 | 1.98658592 | 0.0469683 | 0.5320597 |
| GSAP | 67.4280595 | 0.29637092 | 0.45402295 | 0.65276638 | 0.5139069 | 0.90156229 |
| PTPN12 | 177.430684 | -0.5338603 | 0.25002376 | -2.1352382 | 0.03274156 | 0.48384156 |
| APTR | 31.7760344 | 0.26778356 | 0.33567232 | 0.79775289 | 0.42501391 | 0.86207532 |
| RSBN1L | 104.55555 | -0.0075994 | 0.19188233 | -0.0396046 | 0.96840834 | 0.99373477 |
| TMEM60 | 10.8203395 | 0.91364308 | 0.58942433 | 1.55006 | 0.12112712 | NA |
| PHTF2 | 126.075779 | 0.25778 | 0.31151022 | 0.82751701 | 0.40794409 | 0.85867915 |
| MAGI2 | 830.089497 | -0.1294011 | 0.31155979 | -0.415333 | 0.67789812 | 0.94093842 |

| | | | | | | |
|---|---|---|---|---|---|---|
| GNAI1 | 97.3606327 | -1.571981 | 0.54330536 | -2.8933655 | 0.00381137 | 0.24454118 |
| SEMA3C | 22.7345474 | -1.4220976 | 0.64748172 | -2.1963518 | 0.02806677 | 0.45940027 |
| CACNA2D1 | 182.236255 | -0.1348994 | 0.29418193 | -0.4585576 | 0.64655188 | 0.93328692 |
| PCLO | 228.900055 | -0.0621762 | 0.42149606 | -0.1475131 | 0.88272704 | 0.98137082 |
| KIAA1324L | 28.1032362 | -0.7119182 | 0.41167376 | -1.7293261 | 0.08375075 | 0.62534311 |
| DMTF1 | 126.383897 | -0.2737464 | 0.28298579 | -0.9673503 | 0.33336897 | 0.82974237 |
| TMEM243 | 44.2667075 | 0.64118266 | 0.27809845 | 2.30559595 | 0.02113322 | 0.4212027 |
| TP53TG1 | 11.0092605 | 0.315216 | 0.48781394 | 0.64618079 | 0.51816228 | NA |
| CROT | 70.8535282 | 0.8164156 | 0.26024313 | 3.13712646 | 0.00170613 | 0.17205462 |
| ABCB1 | 99.3188021 | 0.46332331 | 0.42305119 | 1.09519445 | 0.27343146 | 0.79726027 |
| SLC25A40 | 50.4308527 | 0.67685552 | 0.31428743 | 2.15361942 | 0.03127003 | 0.4760612 |
| DBF4 | 17.504685 | -0.0284611 | 0.4449351 | -0.0639668 | 0.94899664 | 0.99105535 |
| ADAM22 | 229.561048 | 0.84949115 | 0.27626067 | 3.07496238 | 2.11E-03 | 0.18826595 |
| SRI | 46.9559128 | 0.42688792 | 0.30420831 | 1.40327503 | 0.16053484 | 0.71101785 |
| AC003991.1 | 19.8348276 | 0.76513232 | 0.57498364 | 1.33070278 | 0.18328683 | 0.73565256 |
| STEAP2 | 26.0740901 | 1.42576942 | 0.45705984 | 3.11943708 | 0.00181197 | 0.17553107 |
| CFAP69 | 48.4772598 | 0.31624307 | 0.34614052 | 0.91362626 | 0.36091326 | 0.84331775 |
| GTPBP10 | 44.4691443 | -0.2060441 | 0.32365918 | -0.6366084 | 0.52437998 | 0.90487731 |
| CLDN12 | 43.1979929 | 0.18018104 | 0.27205778 | 0.66228961 | 0.50778563 | 0.89884243 |
| CDK14 | 334.054845 | 0.0275142 | 0.21339148 | 0.12893765 | 0.89740699 | 0.98531749 |
| FZD1 | 10.530972 | 0.02196757 | 0.77169074 | 0.0284668 | 0.97728984 | NA |
| AC079760.1 | 32.3058273 | 0.23286291 | 0.5034255 | 0.46255684 | 0.64368205 | 0.93328692 |
| MTERF1 | 154.813041 | 0.58312322 | 0.41402823 | 1.40841417 | 0.15900847 | 0.71094363 |
| AKAP9 | 359.742353 | -0.102021 | 0.17189356 | -0.5935126 | 0.55283814 | 0.91322379 |
| KRIT1 | 130.347607 | 0.13388223 | 0.21326749 | 0.62776672 | 0.53015677 | 0.9052447 |
| ANKIB1 | 266.06453 | -0.2120739 | 0.16080077 | -1.3188615 | 0.18721541 | 0.74096603 |
| GATAD1 | 44.8848174 | 0.43148728 | 0.32075947 | 1.34520512 | 0.178559 | 0.72888906 |
| AC007566.1 | 33.0268167 | 0.47782778 | 0.55463643 | 0.86151532 | 0.38895429 | 0.85605713 |
| PEX1 | 51.7059828 | 0.48009541 | 0.266861 | 1.79904675 | 0.07201129 | 0.60821941 |
| RBM48 | 11.6005888 | 0.67347768 | 0.47225548 | 1.4260876 | 0.15384306 | NA |
| FAM133B | 66.1879008 | -0.1477391 | 0.25639761 | -0.5762107 | 0.56447275 | 0.91658856 |
| CDK6 | 369.025814 | 0.24208881 | 0.31192262 | 0.77611815 | 0.43767923 | 0.86820528 |
| SAMD9 | 15.004754 | -1.3063232 | 0.80668333 | -1.6193754 | 0.10536651 | NA |
| SAMD9L | 41.1112188 | -1.4782847 | 0.81077812 | -1.8232913 | 0.06825929 | 0.60244724 |
| VPS50 | 127.143835 | 0.07754437 | 0.24135588 | 0.32128645 | 0.74799333 | 0.95563105 |
| GNGT1 | 81.4190455 | -0.3648725 | 0.4956913 | -0.7360881 | 0.46167706 | 0.87676694 |
| GNG11 | 15.5652184 | 0.08579206 | 0.53270485 | 0.16104989 | 0.87205411 | NA |
| BET1 | 44.9245827 | 0.61720177 | 0.33789187 | 1.82662511 | 0.06775615 | 0.60115427 |
| CASD1 | 68.4626809 | 0.13595554 | 0.26948379 | 0.50450357 | 0.61390755 | 0.92662475 |
| SGCE | 72.1820096 | -0.1309125 | 0.51719178 | -0.2531217 | 0.80017415 | 0.97019967 |
| PPP1R9A | 440.664272 | -0.2023156 | 0.17213435 | -1.1753354 | 0.23986055 | 0.78128405 |
| AC002429.2 | 35.0023098 | 0.59029963 | 0.61443399 | 0.96072099 | 0.33669248 | 0.83037432 |
| PON2 | 82.2431756 | -0.0500776 | 0.3036502 | -0.1649186 | 0.86900803 | 0.97842672 |
| PDK4 | 334.614482 | -1.205455 | 1.19656625 | -1.0074286 | 0.31372887 | 0.82005998 |
| SLC25A13 | 223.430445 | -0.3064742 | 0.20955578 | -1.4624944 | 0.14360577 | 0.69460324 |
| SEM1 | 87.8856015 | -0.1156256 | 0.23353889 | -0.4951022 | 0.620528 | 0.92798856 |
| SDHAF3 | 24.5918628 | 0.46374639 | 0.31194525 | 1.48662748 | 0.13711323 | 0.68541554 |
| ASNS | 20.4645382 | -0.3953664 | 0.52167934 | -0.7578724 | 0.44852735 | 0.87314185 |
| LMTK2 | 96.6114407 | -0.2314702 | 0.27384723 | -0.8452529 | 0.39796968 | 0.85758974 |
| TECPR1 | 19.6032446 | -0.3595741 | 0.38081868 | -0.9442134 | 0.34506062 | 0.83562656 |
| BRI3 | 114.352213 | -0.4642252 | 0.25823535 | -1.7976827 | 0.07222731 | 0.60821941 |
| BAIAP2L1 | 227.340577 | 0.1788645 | 0.36339087 | 0.49220967 | 0.62257113 | 0.92897508 |
| TRRAP | 142.88383 | -0.1247493 | 0.1949137 | -0.640023 | 0.52215762 | 0.90451015 |
| SMURF1 | 205.842262 | -0.3289738 | 0.27438964 | -1.1989293 | 0.23055544 | 0.77249016 |

| | | | | | | |
|---|---|---|---|---|---|---|
| ARPC1B | 25.6108386 | 0.27655299 | 0.47305527 | 0.58461032 | 0.55880977 | 0.91457126 |
| PDAP1 | 30.0544244 | -0.2135944 | 0.32314078 | -0.6609947 | 0.5086157 | 0.89936455 |
| BUD31 | 21.6929067 | -0.0188582 | 0.48459676 | -0.0389152 | 0.968958 | 0.99379133 |
| CPSF4 | 15.8743283 | 0.04156024 | 0.36201137 | 0.11480369 | 0.90860072 | NA |
| ZNF789 | 32.2755996 | 0.15255911 | 0.33298345 | 0.45815824 | 0.64683876 | 0.93328692 |
| ZNF394 | 22.8322975 | 0.33925428 | 0.34803685 | 0.97476541 | 0.32967664 | 0.82856058 |
| ZKSCAN5 | 20.3541795 | 0.08913486 | 0.40866486 | 0.21811236 | 0.82734157 | 0.97567084 |
| FAM200A | 17.2276721 | 0.55236907 | 0.53553955 | 1.03142535 | 0.3023414 | 0.81568231 |
| ZSCAN25 | 12.4646291 | 0.07327156 | 0.48153959 | 0.15216103 | 0.87905993 | NA |
| TRIM4 | 41.9565334 | 0.39400454 | 0.258198 | 1.52597823 | 0.12701529 | 0.66929384 |
| ZKSCAN1 | 142.943071 | 0.16225473 | 0.21640302 | 0.74978032 | 0.45338702 | 0.87378486 |
| ZNF3 | 22.9405169 | -0.3355603 | 0.34154047 | -0.9824906 | 0.32585821 | 0.82567895 |
| COPS6 | 32.0761632 | 0.32770721 | 0.3008468 | 1.08928265 | 0.27602926 | 0.80073115 |
| MCM7 | 11.9189891 | -0.2443945 | 0.43838101 | -0.5574934 | 0.57719035 | NA |
| AP4M1 | 20.6885549 | -0.0279311 | 0.35219671 | -0.0793055 | 0.93678967 | 0.99012269 |
| TAF6 | 15.6566858 | 0.15052668 | 0.50204667 | 0.29982607 | 0.76430983 | NA |
| LAMTOR4 | 65.4374447 | 0.1821696 | 0.33916853 | 0.53710643 | 0.5911941 | 0.92254886 |
| CASTOR3 | 76.1027178 | -0.4068094 | 0.46787184 | -0.869489 | 0.38457973 | 0.85511218 |
| SPDYE3 | 33.2654943 | 0.79773025 | 0.50332998 | 1.58490509 | 0.1129879 | 0.6656824 |
| PILRB | 59.9657573 | -0.2212846 | 0.33419264 | -0.6621468 | 0.50787714 | 0.89884243 |
| PILRA | 17.1325418 | -0.6154085 | 0.72199229 | -0.8523755 | 0.39400574 | 0.85605713 |
| MEPCE | 15.3695794 | 0.23616963 | 0.52695853 | 0.44817499 | 0.65402691 | NA |
| PPP1R35 | 20.8261696 | -0.279674 | 0.35531196 | -0.7871225 | 0.43121017 | 0.86604857 |
| TSC22D4 | 36.4387801 | 0.25908157 | 0.2837981 | 0.91290807 | 0.36129089 | 0.84393918 |
| AGFG2 | 20.4421652 | 0.02420454 | 0.47954887 | 0.05047357 | 0.95974501 | 0.99245244 |
| LRCH4 | 15.0690813 | 0.24872274 | 0.4055279 | 0.61333078 | 0.53965764 | NA |
| GNB2 | 69.1218282 | 0.0692233 | 0.39066609 | 0.17719302 | 0.85935678 | 0.97691173 |
| GIGYF1 | 45.3779771 | -0.2704137 | 0.35958097 | -0.7520244 | 0.45203638 | 0.87346041 |
| EPHB4 | 16.1049012 | 0.28421978 | 0.4628377 | 0.61408089 | 0.53916188 | NA |
| SLC12A9 | 30.6352188 | 1.00045526 | 0.51241701 | 1.95242399 | 0.05088789 | 0.5488536 |
| TRIP6 | 24.5786259 | 0.09533391 | 0.38833867 | 0.24549169 | 0.80607574 | 0.97216031 |
| SRRT | 21.9459961 | -0.2139768 | 0.36679517 | -0.5833685 | 0.55964523 | 0.91511596 |
| TRIM56 | 104.677037 | -0.3752309 | 0.36420655 | -1.0302694 | 0.30288354 | 0.81568231 |
| AP1S1 | 18.6069731 | 0.44363171 | 0.49226122 | 0.90121198 | 0.36747562 | 0.84777627 |
| PLOD3 | 18.2207641 | 0.37845846 | 0.37981273 | 0.99643436 | 0.31903915 | 0.82354084 |
| ZNHIT1 | 35.3687204 | 0.25561425 | 0.34163583 | 0.74820679 | 0.45433544 | 0.87378486 |
| FIS1 | 55.9191281 | 0.38159507 | 0.38825224 | 0.98285347 | 0.32567956 | 0.82567895 |
| IFT22 | 14.1435483 | 0.37952676 | 0.47548524 | 0.79818831 | 0.42476122 | NA |
| COL26A1 | 17.0200634 | 0.30720504 | 0.62030024 | 0.49525217 | 0.62042213 | 0.92798856 |
| CUX1 | 528.091248 | -0.25115 | 0.16104938 | -1.5594595 | 0.11888766 | 0.6668162 |
| PRKRIP1 | 150.700083 | -0.2535866 | 0.18411055 | -1.3773606 | 0.16840079 | 0.72138936 |
| ORAI2 | 30.2765029 | -0.1127844 | 0.45656558 | -0.2470278 | 0.80488669 | 0.97195485 |
| ALKBH4 | 12.7716878 | 0.28640948 | 0.43648919 | 0.65616627 | 0.51171715 | NA |
| POLR2J | 18.3727557 | 0.18112481 | 0.4662792 | 0.38844712 | 0.69768518 | 0.94395238 |
| AC093668.2 | 10.1571002 | -0.2953489 | 0.70106825 | -0.4212841 | 0.67354761 | NA |
| SPDYE2 | 11.7889979 | 0.23625099 | 0.6502411 | 0.3633283 | 0.71635966 | NA |
| AC105052.1 | 10.7278601 | -1.6051472 | 0.67328875 | -2.3840399 | 0.01712375 | NA |
| AC105052.3 | 36.4809153 | 0.01982896 | 0.44323202 | 0.04473719 | 0.96431679 | 0.99264137 |
| FAM185A | 79.5739714 | 0.39342192 | 0.32648889 | 1.20500859 | 0.22819999 | 0.77185154 |
| FBXL13 | 56.8916872 | -0.1340702 | 0.38598442 | -0.347346 | 0.72833138 | 0.95272318 |
| ARMC10 | 43.0055984 | -0.0730789 | 0.27165209 | -0.2690165 | 0.78791695 | 0.96767512 |
| NAPEPLD | 70.1360571 | 0.57310163 | 0.27078421 | 2.11645146 | 0.03430643 | 0.4908133 |
| PMPCB | 40.7174765 | 0.24054751 | 0.29029535 | 0.82863023 | 0.40731368 | 0.85867915 |
| DNAJC2 | 36.0064258 | -0.0268007 | 0.3033357 | -0.0883534 | 0.92959584 | 0.98960607 |

| | | | | | | |
|---|---|---|---|---|---|---|
| PSMC2 | 24.7178782 | 0.0379381 | 0.3872954 | 0.0979565 | 0.92196683 | 0.98846691 |
| ORC5 | 55.4168813 | -0.0857745 | 0.26730423 | -0.3208871 | 0.74829596 | 0.95563105 |
| LHFPL3 | 136.402364 | 0.48934977 | 0.45653481 | 1.07187835 | 0.28377468 | 0.80431986 |
| LINC01004 | 146.877172 | -0.2198957 | 0.23018603 | -0.9552956 | 0.33942822 | 0.83066816 |
| KMT2E-AS1 | 9.23988354 | -0.9378443 | 0.56187005 | -1.669148 | 0.09508804 | NA |
| KMT2E | 341.663533 | 0.04981623 | 0.18468371 | 0.2697381 | 0.78736175 | 0.96767512 |
| SRPK2 | 478.291294 | -0.3433416 | 0.18414178 | -1.8645501 | 0.06224449 | 0.58370968 |
| PUS7 | 30.4570064 | -0.61626 | 0.34650629 | -1.7784958 | 0.07532245 | 0.61020315 |
| RINT1 | 26.9638192 | -0.3354205 | 0.46912787 | -0.7149874 | 0.4746168 | 0.88424821 |
| ATXN7L1 | 159.449347 | -0.0327029 | 0.2778677 | -0.1176923 | 0.90631144 | 0.98531749 |
| CDHR3 | 27.8000666 | 1.52564393 | 0.62902573 | 2.42540785 | 0.0152912 | 0.39253938 |
| SYPL1 | 82.5295012 | 0.08644991 | 0.34279028 | 0.25219475 | 0.80089053 | 0.97025639 |
| NAMPT | 221.711141 | 0.26800168 | 0.40620402 | 0.65977112 | 5.09E-01 | 9.00E-01 |
| HBP1 | 81.2678007 | -0.1977879 | 0.23849543 | -0.8293153 | 0.40692602 | 0.85867915 |
| COG5 | 278.969198 | -0.4189696 | 0.19583058 | -2.1394495 | 0.03239928 | 0.48175799 |
| DUS4L | 12.8377155 | -0.025116 | 0.38700697 | -0.064898 | 0.94825523 | NA |
| BCAP29 | 50.1459728 | 0.15518975 | 0.27467649 | 0.56499103 | 0.57207986 | 0.91813957 |
| CBLL1 | 16.1293287 | -0.4394592 | 0.39798697 | -1.1042051 | 0.26950419 | NA |
| SLC26A3 | 350.944498 | -1.31173 | 0.64753542 | -2.0257271 | 0.04279276 | 0.51526789 |
| DLD | 81.4817688 | 0.21935659 | 0.33766682 | 0.64962434 | 0.51593491 | 0.90213064 |
| LAMB1 | 874.803024 | -1.1840145 | 0.38513761 | -3.0742634 | 0.00211023 | 0.18826595 |
| NRCAM | 303.68516 | -0.0330784 | 0.3151007 | -0.1049771 | 9.16E-01 | 0.98710975 |
| PNPLA8 | 82.5427312 | -0.2181795 | 0.28038101 | -0.7781536 | 0.43647848 | 0.86729771 |
| DNAJB9 | 19.469242 | 0.09579073 | 0.42109552 | 0.2274798 | 0.82005067 | 0.97338914 |
| IMMP2L | 2440.1663 | -0.4159434 | 0.24492904 | -1.6982202 | 0.08946621 | 0.63653714 |
| LRRN3 | 13.5937219 | 0.84176662 | 0.72720946 | 1.15752979 | 0.24705597 | NA |
| DOCK4 | 99.7210561 | -0.4140868 | 0.38972067 | -1.0625221 | 0.28799874 | 0.80523851 |
| ZNF277 | 106.921696 | 0.06705065 | 0.20665404 | 0.32445845 | 0.74559098 | 0.95563105 |
| IFRD1 | 109.409629 | -0.0073353 | 0.56489758 | -0.0129852 | 0.98963964 | 0.99814944 |
| TMEM168 | 101.061737 | 0.21765403 | 0.27311569 | 0.7969298 | 0.4254918 | 0.86239557 |
| BMT2 | 121.437239 | -0.109982 | 0.17040104 | -0.6454302 | 0.51864844 | 0.90368693 |
| SMIM30 | 16.4526394 | -0.1115878 | 0.38461446 | -0.2901289 | 0.77171759 | 0.96377353 |
| MDFIC | 46.2085542 | 1.00161378 | 0.44612719 | 2.24513052 | 0.02475975 | 0.43948277 |
| TES | 67.2749372 | -0.5323897 | 0.40017556 | -1.3303904 | 0.18338968 | 0.73569753 |
| CAV2 | 29.3518527 | 0.67830407 | 0.44413054 | 1.52726285 | 0.12669566 | 0.66929384 |
| MET | 176.06809 | -0.8589048 | 0.67621765 | -1.2701602 | 0.20402756 | 0.76173533 |
| CAPZA2 | 148.082224 | -0.4392945 | 0.26513094 | -1.6568962 | 0.09754046 | 0.65521765 |
| ST7 | 213.336856 | -0.403364 | 0.22605205 | -1.7843855 | 0.07436105 | 0.61020315 |
| ST7-AS2 | 12.2025149 | 0.35482029 | 0.70970935 | 0.49995155 | 0.61710919 | NA |
| CFTR | 28.5964318 | 1.44512294 | 0.72752151 | 1.98636455 | 4.70E-02 | 5.32E-01 |
| CTTNBP2 | 37.4009622 | 1.45670389 | 0.76995428 | 1.89193557 | 0.05849957 | 0.57141408 |
| LSM8 | 36.3845638 | 0.13898688 | 0.29700985 | 0.46795379 | 0.63981764 | 0.93328692 |
| ANKRD7 | 57.0641482 | 0.15648446 | 0.38836556 | 0.40293085 | 0.6869991 | 0.94205398 |
| TSPAN12 | 71.2642813 | 0.66072588 | 0.42353186 | 1.56003818 | 0.11875086 | 0.6668162 |
| ING3 | 23.2397556 | 0.22293352 | 0.41118276 | 0.54217623 | 0.58769711 | 0.92148551 |
| FAM3C | 74.1510513 | -0.6130855 | 0.27282708 | -2.2471577 | 0.02462995 | 0.43948277 |
| AASS | 47.7064617 | 0.37773982 | 0.26281471 | 1.43728568 | 0.15063684 | 0.70572801 |
| CADPS2 | 2371.71549 | 0.29863736 | 0.22935512 | 1.3020741 | 1.93E-01 | 0.75061775 |
| AC004594.1 | 109.355192 | 1.34460399 | 0.50738652 | 2.65005854 | 0.00804778 | 0.30976355 |
| SLC13A1 | 15.9515852 | 2.22835711 | 0.64824588 | 3.43751833 | 0.00058707 | NA |
| IQUB | 114.376525 | 0.25185478 | 0.3811281 | 0.660814 | 0.50873161 | 0.8993948 |
| NDUFA5 | 108.484531 | 0.29349805 | 0.34585977 | 0.84860419 | 0.39610158 | 0.85605713 |
| WASL | 120.84253 | -0.1666594 | 0.20826755 | -0.8002178 | 0.42358465 | 0.86190547 |
| POT1 | 55.5243736 | -0.1284241 | 0.28105658 | -0.4569331 | 0.64771912 | 0.93328692 |

| | | | | | | |
|---|---|---|---|---|---|---|
| POT1-AS1 | 31.0563174 | 0.54666522 | 0.35159102 | 1.55483271 | 0.11998592 | 0.6668162 |
| GRM8 | 69.5145738 | -0.0127216 | 0.76911159 | -0.0165407 | 0.98680306 | 0.99803728 |
| ZNF800 | 76.5990357 | 0.24322152 | 0.31779447 | 0.76534219 | 0.44406781 | 0.8704597 |
| ARF5 | 22.8742804 | 0.22608902 | 0.60850958 | 0.37154555 | 0.71023124 | 0.94603965 |
| SND1 | 265.387858 | -0.6368286 | 0.2348301 | -2.7118696 | 0.00669049 | 0.30102597 |
| RBM28 | 45.2678233 | -0.2787366 | 0.3485604 | -0.7996795 | 0.42389648 | 0.86190547 |
| IMPDH1 | 15.1791651 | 0.28147878 | 0.40440064 | 0.6960394 | 0.48640416 | NA |
| HILPDA | 54.2162711 | -1.8238456 | 0.76043007 | -2.3984396 | 1.65E-02 | 0.39748373 |
| METTL2B | 40.0298405 | 0.05296555 | 0.31429967 | 0.16851928 | 0.86617478 | 0.97788853 |
| CALU | 67.2095527 | -0.5349087 | 0.33678426 | -1.5882828 | 0.1122224 | 0.6656824 |
| KCP | 57.9368177 | 0.40000406 | 0.3443843 | 1.16150491 | 0.24543663 | 0.78583441 |
| ATP6V1F | 81.1456317 | 0.45567146 | 0.48877998 | 0.93226294 | 0.35120065 | 0.83800638 |
| TNPO3 | 137.157495 | -0.438178 | 0.23730194 | -1.8464999 | 0.06481966 | 0.59005336 |
| TSPAN33 | 31.0178581 | 0.22513647 | 0.36562326 | 0.61576078 | 0.53805242 | 0.90763121 |
| SMO | 20.6104647 | 0.82960298 | 0.39720329 | 2.08861052 | 0.0367428 | 0.49537143 |
| AHCYL2 | 331.102938 | 0.03587061 | 0.24292065 | 0.14766389 | 0.88260803 | 0.98137082 |
| STRIP2 | 18.4973023 | -1.7190774 | 0.82471857 | -2.0844412 | 0.03712005 | 0.49537143 |
| NRF1 | 165.75809 | 0.13483124 | 0.2179603 | 0.61860458 | 0.53617688 | 0.90763121 |
| UBE2H | 654.514384 | -0.5685206 | 0.24090657 | -2.3599217 | 0.01827879 | 0.41186103 |
| ZC3HC1 | 29.4369447 | -0.6481654 | 0.31311232 | -2.0700732 | 0.03844549 | 0.49671741 |
| KLHDC10 | 174.350278 | -0.1327736 | 0.17674037 | -0.7512355 | 0.45251096 | 0.87346041 |
| TMEM209 | 25.4753831 | -0.1707012 | 0.35076901 | -0.4866484 | 0.62650753 | 0.93032781 |
| CEP41 | 12.1925016 | 0.07903099 | 0.44997432 | 0.17563445 | 0.86058113 | NA |
| MEST | 38.6770261 | -2.0459708 | 1.18900046 | -1.7207486 | 0.08529646 | 0.63062869 |
| COPG2 | 56.6492719 | -0.367794 | 0.24835828 | -1.4809008 | 0.13863301 | 0.69023184 |
| AC016831.7 | 740.142406 | -0.5939458 | 0.25190227 | -2.3578421 | 0.01838151 | 0.41315275 |
| LINC00513 | 62.0739223 | -0.0069142 | 0.59942798 | -0.0115347 | 0.99079683 | 0.99814944 |
| AC058791.1 | 14.9606006 | -0.3128602 | 0.72530473 | -0.43135 | 0.66621393 | NA |
| MKLN1 | 596.935274 | -0.5711271 | 0.21755115 | -2.6252545 | 0.00865842 | 0.31119527 |
| PODXL | 11.3891408 | 0.07399632 | 0.50273939 | 0.14718624 | 0.88298502 | NA |
| CHCHD3 | 476.361759 | -0.1297722 | 0.16882966 | -0.7686576 | 0.4420966 | 0.86925451 |
| EXOC4 | 702.349091 | -0.173281 | 0.15695839 | -1.1039929 | 0.2695962 | 0.79666376 |
| LRGUK | 31.0796459 | 0.14601848 | 0.34983147 | 0.41739664 | 0.67638831 | 0.94034237 |
| SLC35B4 | 102.61492 | 0.44346558 | 0.35426178 | 1.25180192 | 2.11E-01 | 0.76397692 |
| AKR1B1 | 54.4859526 | -0.2333204 | 0.38804584 | -0.6012702 | 0.54766005 | 0.91139843 |
| BPGM | 31.3113514 | 0.18471907 | 0.37519661 | 0.49232608 | 0.62248885 | 0.92897508 |
| CALD1 | 25.1364085 | -0.093805 | 0.48272589 | -0.1943236 | 0.84592251 | 0.97612951 |
| AGBL3 | 23.512902 | 0.19682584 | 0.3597818 | 0.54707001 | 0.58433063 | 0.92071026 |
| CYREN | 31.7450107 | 0.0461943 | 0.3180005 | 0.14526488 | 0.88450174 | 0.98154805 |
| TMEM140 | 33.1468426 | -1.497583 | 0.56737385 | -2.6394995 | 0.00830286 | 0.30976355 |
| WDR91 | 86.7050049 | 0.0282471 | 0.37053277 | 0.07623373 | 0.93923315 | 0.99012269 |
| CNOT4 | 219.364835 | -0.148644 | 0.21936448 | -0.6776119 | 0.49801779 | 0.89378207 |
| NUP205 | 52.0248312 | -0.2900035 | 0.25992458 | -1.1157218 | 0.26454123 | 0.79666376 |
| STMP1 | 76.8218725 | 0.26354593 | 0.28295951 | 0.93139096 | 0.35165136 | 0.83863829 |
| SLC13A4 | 11.4161137 | 1.96325134 | 0.61814599 | 3.17603184 | 0.00149305 | NA |
| MTPN | 69.4506717 | -0.1775099 | 0.30544133 | -0.5811588 | 0.56113346 | 0.91606849 |
| CREB3L2 | 89.4336352 | 0.15013715 | 0.20152615 | 0.74500083 | 4.56E-01 | 0.87396417 |
| TRIM24 | 124.704283 | -0.5042036 | 0.22328381 | -2.2581291 | 0.02393761 | 0.43580821 |
| SVOPL | 46.1081968 | -1.0256824 | 0.73545966 | -1.394614 | 0.16313226 | 0.71325312 |
| ATP6V0A4 | 315.103939 | 0.20964929 | 0.30523656 | 0.68684202 | 4.92E-01 | 0.89067343 |
| TMEM213 | 245.799702 | 0.90669403 | 0.27394122 | 3.30981234 | 0.00093359 | 0.13754433 |
| KIAA1549 | 46.9107893 | 0.2862682 | 0.35068363 | 0.81631469 | 0.41432017 | 0.85867915 |
| ZC3HAV1 | 91.9775174 | -0.5469868 | 0.44952313 | -1.2168156 | 0.22367437 | 0.76834258 |
| TTC26 | 28.9078627 | 0.03759094 | 0.35598435 | 0.10559716 | 0.91590198 | 0.98691474 |

| | | | | | | |
|---|---|---|---|---|---|---|
| UBN2 | 124.41921 | 0.05107032 | 0.18087161 | 0.28235676 | 0.77766996 | 0.96518573 |
| LUC7L2 | 38.44431 | 0.09600929 | 0.33834378 | 0.28376253 | 0.77659237 | 0.9649632 |
| HIPK2 | 375.165944 | -0.005974 | 0.20358227 | -0.0293445 | 0.9765898 | 0.99441127 |
| TBXAS1 | 28.0933151 | -0.3674918 | 0.43544457 | -0.8439461 | 0.39869952 | 0.85759965 |
| PARP12 | 44.7866041 | -1.4515342 | 0.6744609 | -2.1521398 | 0.03138634 | 0.47623559 |
| KDM7A | 133.244757 | -0.4911259 | 0.20640648 | -2.3794115 | 1.73E-02 | 0.40063153 |
| SLC37A3 | 64.3733172 | -0.5981631 | 0.28522806 | -2.0971396 | 0.03598122 | 0.493821 |
| MKRN1 | 62.177191 | 0.05879574 | 0.27002571 | 0.21774126 | 0.82763072 | 0.97567084 |
| DENND2A | 18.7615723 | -0.938509 | 0.57552689 | -1.6306953 | 0.10295462 | 0.65946648 |
| NDUFB2 | 90.3792809 | 0.44933733 | 0.33242211 | 1.35170712 | 0.17646903 | 0.72841249 |
| BRAF | 535.250788 | -0.3332272 | 0.30291532 | -1.1000673 | 0.27130282 | 0.79666376 |
| MRPS33 | 24.4249205 | 0.30528981 | 0.42203357 | 0.72337803 | 0.46944766 | 0.88207878 |
| TMEM178B | 80.747069 | 1.73251831 | 0.53537118 | 3.23610682 | 0.00121172 | 0.14513539 |
| AGK | 55.607315 | -0.1961274 | 0.23322692 | -0.8409295 | 0.40038743 | 0.85778581 |
| AC004918.1 | 15.3299894 | -0.0505316 | 0.4106545 | -0.1230515 | 0.90206635 | NA |
| KIAA1147 | 50.9611635 | 0.11726726 | 0.29163098 | 0.40210839 | 0.68760426 | 0.94205398 |
| SSBP1 | 68.9254792 | -0.275432 | 0.2459169 | -1.1200208 | 0.26270489 | 0.79666376 |
| TRPV6 | 50.7642594 | 0.39033793 | 0.46571245 | 0.83815224 | 0.40194521 | 0.85778581 |
| TRPV5 | 227.427665 | 2.21141178 | 0.50416882 | 4.3862526 | 1.15E-05 | 0.00954327 |
| GSTK1 | 72.8060952 | 0.65606604 | 0.34219524 | 1.91722724 | 0.05520907 | 0.5645224 |
| TMEM139 | 11.2054917 | 0.2923296 | 0.53507488 | 0.54633401 | 0.58483637 | NA |
| CASP2 | 13.9751204 | -0.1926152 | 0.38202705 | -0.5041925 | 0.61412608 | NA |
| ZYX | 17.181682 | -0.435265 | 0.4953782 | -0.8786518 | 0.37959007 | 0.85431805 |
| TCAF2 | 34.2206482 | -0.3297473 | 0.62718119 | -0.5257608 | 0.59905441 | 0.92298075 |
| TCAF1 | 67.8704038 | 0.5592369 | 0.24991787 | 2.23768276 | 0.02524175 | 0.4403385 |
| AC004889.1 | 59.1577519 | 0.03544672 | 0.34877363 | 0.10163245 | 0.91904842 | 0.98749988 |
| AC074386.1 | 78.2193038 | -0.299628 | 0.35908495 | -0.8344208 | 0.40404388 | 0.85782098 |
| ARHGEF5 | 19.9847026 | -0.2073954 | 0.36725223 | -0.5647221 | 0.57226281 | 0.91813957 |
| TPK1 | 76.8691773 | -0.3166423 | 0.23893044 | -1.3252489 | 0.18508859 | 0.73752116 |
| CNTNAP2 | 12.0326987 | -0.3714221 | 0.48322935 | -0.768625 | 0.44211598 | NA |
| CUL1 | 138.830483 | -0.0946838 | 0.20946555 | -0.4520257 | 0.65125049 | 0.93453685 |
| EZH2 | 18.7046525 | 0.46340111 | 0.48083829 | 0.96373586 | 0.33517837 | 0.82974237 |
| PDIA4 | 39.2971266 | -0.4467873 | 0.57425894 | -0.7780242 | 0.43655477 | 0.86729771 |
| ZNF786 | 18.4721592 | -0.1206958 | 0.46686454 | -0.2585242 | 0.79600239 | 0.96974891 |
| ZNF425 | 10.9991385 | 0.04748172 | 0.45049927 | 0.10539801 | 0.91606 | NA |
| ZNF398 | 42.121338 | -0.0937865 | 0.30143254 | -0.3111359 | 0.75569731 | 0.95849416 |
| ZNF282 | 22.2098093 | 0.22140741 | 0.31134384 | 0.7111347 | 0.47700077 | 0.88495683 |
| ZNF212 | 11.2457021 | 0.35288457 | 0.45578074 | 0.77424196 | 0.43878772 | NA |
| ZNF783 | 16.8123759 | -0.2398329 | 0.38209892 | -0.6276722 | 0.5302187 | 0.9052447 |
| ZNF777 | 11.6847002 | 0.2680813 | 0.40849458 | 0.65626649 | 0.51165268 | NA |
| ZNF746 | 18.003958 | -0.2461149 | 0.3819283 | -0.6444008 | 0.51931559 | 0.90368693 |
| ZNF467 | 12.6266969 | 0.52643888 | 0.45104087 | 1.16716447 | 0.24314396 | NA |
| ZNF862 | 48.889031 | 0.56733722 | 0.28228679 | 2.00979021 | 0.0444534 | 0.52480969 |
| ATP6V0E2 | 18.4785749 | 0.1753365 | 0.61204987 | 0.2864742 | 0.77451495 | 0.96435639 |
| AC092666.1 | 13.2225379 | 1.61266579 | 0.76572704 | 2.10605829 | 0.03519929 | NA |
| ACTR3C | 269.470159 | 0.20780663 | 0.23244772 | 0.89399297 | 0.37132565 | 0.8491102 |
| LRRC61 | 18.3334749 | -0.0897578 | 0.45218319 | -0.1984987 | 0.84265491 | 0.97612951 |
| TMEM176B | 69.2297038 | 1.62949004 | 0.68820326 | 2.36774533 | 0.01789685 | 0.40933424 |
| TMEM176A | 31.9551747 | 0.97260871 | 0.61366766 | 1.58491115 | 0.11298652 | 0.6656824 |
| AOC1 | 37.1836427 | 1.45672978 | 1.37053699 | 1.0628898 | 0.28783193 | 0.80523851 |
| ABCB8 | 19.861447 | 0.12954682 | 0.37895731 | 0.3418507 | 0.73246325 | 0.95279138 |
| SLC4A2 | 23.581301 | -0.6504633 | 0.37299437 | -1.7438957 | 0.08117727 | 0.61925304 |
| FASTK | 25.5826829 | -0.5990813 | 0.42685978 | -1.4034616 | 0.16047923 | 0.71101785 |
| TMUB1 | 27.0599526 | -0.5709916 | 0.45587709 | -1.2525121 | 0.2103833 | 0.76397692 |

| | | | | | | |
|---|---|---|---|---|---|---|
| AGAP3 | 21.6349859 | 0.8687192 | 0.5062214 | 1.71608547 | 0.0861464 | 0.63384997 |
| CHPF2 | 13.7218497 | -0.5764488 | 0.54177471 | -1.0640009 | 0.28732831 | NA |
| SMARCD3 | 98.2448107 | 0.72672477 | 0.27153201 | 2.67638709 | 0.00744206 | 0.30654023 |
| NUB1 | 59.4888905 | -0.5759698 | 0.32792558 | -1.7564038 | 0.07901948 | 0.61367304 |
| RHEB | 119.471264 | -0.3372673 | 0.31302195 | -1.0774556 | 0.28127677 | 0.80416687 |
| PRKAG2 | 579.651372 | -0.5622677 | 0.38171868 | -1.4729898 | 0.14075379 | 0.69227547 |
| GALNT11 | 233.985935 | -0.4274435 | 0.46337585 | -0.9224553 | 0.35629116 | 0.84192356 |
| KMT2C | 580.823139 | -0.3016723 | 0.1975725 | -1.5268942 | 0.12678732 | 0.66929384 |
| ACTR3B | 74.9105549 | -0.110313 | 0.257619 | -0.4282021 | 0.66850402 | 0.93878847 |
| DPP6 | 165.858681 | 0.18543521 | 0.24983965 | 0.74221692 | 0.45795592 | 0.87469004 |
| PAXIP1-AS2 | 23.8794233 | 0.09576196 | 0.44461775 | 0.21538041 | 0.82947074 | 0.97567084 |
| PAXIP1 | 40.3230349 | -0.3689868 | 0.33367895 | -1.105814 | 0.26880705 | 0.79666376 |
| PAXIP1-AS1 | 11.6149355 | 0.86656532 | 0.55320366 | 1.56644901 | 0.11724354 | NA |
| INSIG1 | 70.6195298 | -1.84961 | 0.74564669 | -2.4805448 | 0.01311818 | 0.36723795 |
| AC009403.1 | 24.2565595 | 0.76547653 | 0.42137683 | 1.81660803 | 0.06927715 | 0.60275677 |
| RBM33 | 193.506174 | -0.2070823 | 0.17862236 | -1.15933 | 0.24632168 | 0.78583441 |
| LINC01006 | 58.2431686 | -0.2011991 | 0.32050528 | -0.627756 | 0.53016382 | 0.9052447 |
| LMBR1 | 226.716345 | -0.0567314 | 0.26692802 | -0.2125343 | 0.83169024 | 0.97587989 |
| NOM1 | 21.6268582 | -0.2404224 | 0.34318823 | -0.7005556 | 0.48358039 | 0.88838332 |
| UBE3C | 252.791388 | -0.5116764 | 0.2685975 | -1.9049931 | 0.05678097 | 0.56976762 |
| DNAJB6 | 131.224819 | -0.2841358 | 0.35029823 | -0.8111253 | 0.4172937 | 0.85949274 |
| PTPRN2 | 11.2280392 | 1.22120797 | 0.52225758 | 2.338325 | 0.0193704 | NA |
| ESYT2 | 325.331896 | -0.3416692 | 0.68480828 | -0.4989267 | 0.61783099 | 0.92749951 |
| WDR60 | 162.121315 | -0.2545595 | 0.23686676 | -1.0746951 | 0.28251127 | 0.80431986 |
| GTPBP6 | 35.833986 | -0.3241997 | 0.3845297 | -0.8431071 | 0.39916857 | 0.85778581 |
| LINC00685 | 20.6582503 | -0.042018 | 0.72233908 | -0.0581693 | 0.95361378 | 0.99148072 |
| PPP2R3B | 15.6402727 | -0.1907145 | 0.39794575 | -0.4792474 | 0.63176263 | NA |
| IL3RA | 15.7597177 | 1.71568723 | 0.68443437 | 2.50672278 | 0.01218562 | NA |
| SLC25A6 | 147.054053 | 0.60192431 | 0.39270847 | 1.53275103 | 0.12533721 | 0.66929384 |
| LINC00106 | 50.1605333 | -0.1551469 | 0.46438495 | -0.3340911 | 0.73831084 | 0.9542764 |
| ASMTL-AS1 | 24.347216 | -0.3699758 | 0.48423426 | -0.7640429 | 0.44484166 | 0.87102466 |
| ASMTL | 27.6561702 | -0.0406592 | 0.3211861 | -0.1265909 | 0.89926422 | 0.98531749 |
| AKAP17A | 29.9068379 | 0.41873988 | 0.31655936 | 1.3227847 | 0.18590699 | 0.73804265 |
| DHRSX | 172.129665 | -0.6041885 | 0.32220766 | -1.8751526 | 0.06077174 | 0.57806178 |
| CD99 | 34.8405361 | 0.52073087 | 0.35181726 | 1.48011744 | 0.13884191 | 0.69023184 |
| ARSD | 76.8552611 | 0.55048658 | 0.30016678 | 1.83393575 | 0.06666353 | 0.59652811 |
| MXRA5 | 53.4400536 | -0.019396 | 0.41701885 | -0.0465112 | 0.96290281 | 0.99256424 |
| PRKX | 35.7478987 | -0.054871 | 0.33645188 | -0.1630872 | 0.87044982 | 0.97854266 |
| BX890604.1 | 11.8063591 | 0.11163035 | 0.48367094 | 0.23079814 | 0.81747163 | NA |
| PUDP | 38.4925274 | -0.4318231 | 0.29023707 | -1.4878288 | 0.13679606 | 0.68452453 |
| STS | 15.3897161 | -0.6571646 | 0.70727367 | -0.9291518 | 0.35281044 | NA |
| PNPLA4 | 15.4812861 | 0.14689685 | 0.44455333 | 0.33043696 | 0.74106982 | NA |
| TBL1X | 256.086788 | -0.454332 | 0.30411396 | -1.4939531 | 0.13518788 | 0.68329553 |
| SHROOM2 | 43.3825941 | 0.13094269 | 0.30953914 | 0.42302468 | 0.67227725 | 0.93878847 |
| WWC3 | 27.5728608 | -0.0955084 | 0.33854984 | -0.2821102 | 0.77785903 | 0.96518573 |
| MID1 | 592.835023 | 0.32168004 | 0.26370763 | 1.21983591 | 0.22252709 | 0.76787872 |
| AC073529.1 | 37.9221338 | -0.0215261 | 0.37255939 | -0.0577789 | 0.95392473 | 0.99148072 |
| ARHGAP6 | 829.699061 | 0.24506241 | 0.26716837 | 0.9172583 | 0.3590073 | 0.84204868 |
| PRPS2 | 48.8666613 | 0.06046875 | 0.49927969 | 0.12111198 | 0.90360234 | 0.98531749 |
| TMSB4X | 468.215026 | -0.7046623 | 0.55056281 | -1.2798945 | 0.20058225 | 0.75794945 |
| RAB9A | 45.9224221 | -0.1471702 | 0.29358663 | -0.5012836 | 0.61617153 | 0.92662475 |
| TRAPPC2 | 21.7515078 | 0.14580601 | 0.35943315 | 0.40565543 | 0.6849958 | 0.94205398 |
| OFD1 | 116.758411 | -0.4803251 | 0.26329553 | -1.8242813 | 0.06810956 | 0.60244724 |
| GPM6B | 32.0736969 | -0.7974141 | 0.53083314 | -1.5021935 | 0.13304715 | 0.67964548 |

| | | | | | | |
|---|---|---|---|---|---|---|
| GEMIN8 | 32.693359 | -0.500695 | 0.32627016 | -1.5346026 | 0.12488147 | 0.66929384 |
| MOSPD2 | 24.282358 | -0.1485734 | 0.33955929 | -0.4375479 | 0.66171407 | 0.93734459 |
| PIR | 23.2980437 | 0.77729615 | 0.79051392 | 0.98327952 | 0.32546988 | 0.82567895 |
| CLTRN | 15.9767796 | -0.2509399 | 0.85926183 | -0.2920413 | 0.77025505 | NA |
| CA5B | 51.8198056 | -0.41336 | 0.26005258 | -1.5895245 | 0.11194202 | 0.66558344 |
| ZRSR2 | 25.4795773 | -0.3381261 | 0.32474631 | -1.0412008 | 0.29778235 | 0.81214575 |
| AP1S2 | 20.0562751 | -0.4236922 | 0.55632338 | -0.7615934 | 0.44630274 | 0.87231627 |
| CTPS2 | 61.6025908 | -0.2419174 | 0.23874421 | -1.013291 | 0.31092118 | 0.81701824 |
| SYAP1 | 64.6592172 | 0.38831317 | 0.39062121 | 0.99409136 | 0.32017839 | 0.82449332 |
| TXLNG | 41.5728663 | -0.2425703 | 0.33714584 | -0.7194819 | 0.47184403 | 0.88267439 |
| RBBP7 | 53.9221018 | 0.60524237 | 0.27247591 | 2.22126925 | 0.02633273 | 0.44713242 |
| REPS2 | 105.419487 | -0.1589713 | 0.47278284 | -0.3362459 | 0.73668546 | 0.95387069 |
| NHS | 512.989023 | -0.263537 | 0.29973499 | -0.8792332 | 0.37927482 | 0.85429312 |
| SCML1 | 113.252066 | 0.25197329 | 0.31080407 | 0.81071424 | 0.4175298 | 0.85950165 |
| RAI2 | 31.3045133 | 0.65170715 | 0.36374233 | 1.79167258 | 0.07318543 | 0.60952146 |
| CDKL5 | 84.0983292 | -0.1852982 | 0.18840265 | -0.9835225 | 0.32535037 | 0.82567895 |
| PHKA2 | 34.7042052 | -0.0042253 | 0.29077388 | -0.0145313 | 0.98840607 | 0.99814944 |
| PDHA1 | 108.119931 | 0.89246173 | 0.28845804 | 3.09390483 | 0.00197541 | 0.18210466 |
| MAP3K15 | 88.1180066 | -0.347214 | 0.35686896 | -0.9729454 | 3.31E-01 | 0.82867637 |
| SH3KBP1 | 43.418386 | -0.7599251 | 0.3658562 | -2.0771141 | 0.03779103 | 0.49640946 |
| BCLAF3 | 61.3303403 | -0.4430596 | 0.21040901 | -2.1057065 | 0.03522985 | 0.49209205 |
| EIF1AX | 85.7897045 | -0.1165509 | 0.3211206 | -0.3629505 | 0.71664189 | 0.94913834 |
| RPS6KA3 | 132.03546 | -0.3450115 | 0.26432774 | -1.3052416 | 0.1918106 | 0.75034462 |
| MBTPS2 | 33.6911429 | 0.01046005 | 0.32004871 | 0.03268268 | 0.97392763 | 0.99431961 |
| SMS | 127.354075 | -0.5603647 | 0.24966823 | -2.2444374 | 0.02480428 | 0.43948277 |
| PTCHD1-AS | 54.5254407 | -0.4184746 | 0.61347436 | -0.6821388 | 0.49515121 | 0.89230917 |
| PRDX4 | 12.9037952 | 0.52814154 | 0.47094547 | 1.12144947 | 0.26209659 | NA |
| ACOT9 | 27.2905908 | -0.671928 | 0.38725419 | -1.7351084 | 0.0827216 | 0.62228971 |
| SAT1 | 638.018156 | -0.2059627 | 0.31727516 | -0.6491611 | 0.51623424 | 0.90225158 |
| APOO | 307.409531 | 0.10476354 | 0.33586163 | 0.31192469 | 0.75509775 | 0.95832511 |
| EIF2S3 | 62.2071284 | -0.1429704 | 0.27022447 | -0.5290802 | 0.59674984 | 0.9229638 |
| ZFX | 69.6777206 | -0.5180675 | 0.29675666 | -1.7457655 | 0.08085171 | 0.61925304 |
| PDK3 | 28.5107907 | -0.168622 | 0.53881924 | -0.3129472 | 0.75432078 | 0.95826617 |
| POLA1 | 88.3417594 | -0.2945416 | 0.22662648 | -1.2996787 | 0.19371111 | 0.75067401 |
| IL1RAPL1 | 125.882226 | 0.59411976 | 0.58265392 | 1.01967866 | 0.30788089 | 0.81568231 |
| GK | 32.0304937 | -0.4358091 | 0.34245163 | -1.2726152 | 0.20315462 | 0.76169832 |
| TAB3 | 80.0895662 | 0.25954935 | 0.24618599 | 1.05428156 | 0.29175403 | 0.80724527 |
| DMD | 34.4584717 | -0.0320211 | 0.35442276 | -0.0903473 | 0.92801125 | 0.98960607 |
| TMEM47 | 22.4144846 | -0.057929 | 0.61433497 | -0.0942954 | 0.92487446 | 0.98925753 |
| CFAP47 | 28.1789248 | -0.1630293 | 0.53314159 | -0.3057898 | 0.75976471 | 0.95913478 |
| AF241726.2 | 40.836714 | -0.2728441 | 0.44559371 | -0.6123159 | 0.54032878 | 0.90763121 |
| DYNLT3 | 41.5711581 | -0.7055 | 0.44891046 | -1.5715828 | 0.11604734 | 0.6668162 |
| RPGR | 33.4259638 | -0.4508464 | 0.28270433 | -1.5947631 | 0.11076522 | 0.66311481 |
| MID1IP1 | 14.0268008 | -0.0532492 | 0.48237542 | -0.1103896 | 0.91210042 | NA |
| BCOR | 67.1728876 | -0.2437242 | 0.30201398 | -0.8069964 | 0.41966858 | 0.86065558 |
| ATP6AP2 | 70.5426926 | 0.50453213 | 0.33548748 | 1.50387767 | 0.13261287 | 0.67780737 |
| CXorf38 | 29.5985818 | -0.3068732 | 0.31013377 | -0.9894866 | 0.32242513 | 0.82558302 |
| MED14 | 91.81103 | -0.3896979 | 0.284332 | -1.3705735 | 0.17050793 | 0.72463203 |
| USP9X | 240.728971 | -0.3479189 | 0.24032228 | -1.4477179 | 0.14769596 | 0.69842926 |
| DDX3X | 90.9729815 | -0.1446358 | 0.30597929 | -0.4726979 | 0.63642875 | 0.93251095 |
| CASK | 126.469009 | 0.07262559 | 0.2048778 | 0.35448246 | 0.72297735 | 0.95093722 |
| MAOA | 60.7041686 | 0.7052696 | 0.53599994 | 1.31580164 | 0.18824063 | 0.74214877 |
| MAOB | 15.4927848 | 0.24711139 | 0.74417046 | 0.33206288 | 0.73984178 | NA |
| EFHC2 | 16.4093075 | 0.24988966 | 0.49193858 | 0.50796923 | 0.61147492 | 0.92630153 |

| | | | | | | |
|---|---|---|---|---|---|---|
| FUNDC1 | 13.4112451 | -0.0530703 | 0.46892181 | -0.1131752 | 0.90989166 | NA |
| KDM6A | 269.742879 | -0.519863 | 0.36773348 | -1.4136951 | 0.15745145 | 0.7098097 |
| MIR222HG | 79.3228386 | -0.7631978 | 0.54715033 | -1.3948594 | 0.16305824 | 0.71325312 |
| KRBOX4 | 33.6728263 | -0.0791342 | 0.32145541 | -0.2461746 | 0.80554706 | 0.97214568 |
| ZNF674 | 11.4539951 | -0.0316108 | 0.55656523 | -0.0567963 | 0.95470749 | NA |
| SLC9A7 | 50.9865379 | -0.3670788 | 0.33864237 | -1.0839719 | 0.27837727 | 0.80210301 |
| RP2 | 14.7066144 | -0.1567907 | 0.35988275 | -0.4356717 | 0.66307493 | NA |
| JADE3 | 68.9642832 | -0.2812912 | 0.33310173 | -0.8444602 | 0.39841228 | 0.85758974 |
| RGN | 30.7536065 | 0.86656464 | 0.62780297 | 1.38031308 | 0.16749027 | 0.71965603 |
| NDUFB11 | 58.8219752 | 0.15869781 | 0.53526127 | 0.29648664 | 0.76685847 | 0.96283923 |
| RBM10 | 46.3886871 | 0.17606895 | 0.24562893 | 0.71680865 | 0.47349217 | 0.88359968 |
| UBA1 | 89.815165 | -0.1297193 | 0.27001776 | -0.4804101 | 0.63093579 | 0.93250784 |
| CDK16 | 44.0924202 | -0.6204773 | 0.39747261 | -1.5610569 | 0.11851034 | 0.6668162 |
| USP11 | 38.9385818 | -0.2945245 | 0.32345423 | -0.9105599 | 0.36252729 | 0.84512775 |
| ZNF41 | 21.5907914 | 0.55661789 | 0.3510759 | 1.58546315 | 0.11286114 | 0.6656824 |
| ARAF | 20.5106444 | -0.0562064 | 0.4140092 | -0.1357614 | 0.89200994 | 0.98329225 |
| TIMP1 | 55.5273876 | 0.30250274 | 0.4724753 | 0.64025091 | 5.22E-01 | 9.05E-01 |
| UXT | 14.7431306 | -0.0633543 | 0.42227638 | -0.1500304 | 0.88074067 | NA |
| ZNF81 | 59.6515311 | 0.42053248 | 0.32282696 | 1.30265599 | 0.19269223 | 0.75061775 |
| ZNF630 | 24.4900744 | 1.00594887 | 0.56189838 | 1.79026833 | 0.07341078 | 0.60981847 |
| EBP | 22.8118706 | 0.19499199 | 0.4508634 | 0.43248573 | 0.66538842 | 0.93860542 |
| TBC1D25 | 14.226997 | 0.41775153 | 0.50034179 | 0.83493231 | 0.40375582 | NA |
| RBM3 | 31.3220804 | 0.17264835 | 0.43003315 | 0.40147684 | 0.68806909 | 0.94205398 |
| WDR13 | 30.3281361 | -0.3317543 | 0.37785879 | -0.8779849 | 0.37995189 | 0.85432452 |
| HDAC6 | 23.0263223 | -0.1534885 | 0.37904753 | -0.404932 | 0.68552749 | 0.94205398 |
| PCSK1N | 104.492188 | -0.0448681 | 0.52700097 | -0.0851385 | 0.93215131 | 0.98960607 |
| TIMM17B | 16.2411326 | 0.39211891 | 0.45057634 | 0.87026076 | 0.38415792 | 0.85511218 |
| OTUD5 | 42.2529019 | 0.14276632 | 0.27173968 | 0.525379 | 0.59931975 | 0.92298075 |
| GRIPAP1 | 40.3509422 | -0.4584262 | 0.28482936 | -1.6094766 | 0.10751217 | 0.65986731 |
| TFE3 | 25.6444729 | -0.1250386 | 0.39819568 | -0.314013 | 0.75351115 | 0.95826617 |
| CCDC120 | 12.6646127 | 0.22436616 | 0.41538177 | 0.54014446 | 0.58909741 | NA |
| WDR45 | 25.3257459 | 0.1901118 | 0.37406347 | 0.50823408 | 0.61128919 | 0.92619266 |
| PLP2 | 19.666585 | 0.1134875 | 0.49119073 | 0.23104569 | 8.17E-01 | 9.73E-01 |
| PPP1R3F | 9.86386115 | 0.60699388 | 0.50357681 | 1.20536504 | 0.22806241 | NA |
| CLCN5 | 211.026039 | 0.45553851 | 0.23915937 | 1.90474871 | 0.05681274 | 0.56976762 |
| SHROOM4 | 146.437485 | -0.1918821 | 0.19789582 | -0.9696116 | 0.33224015 | 0.82890589 |
| AL158055.1 | 14.0842978 | 0.29789363 | 0.58751531 | 0.50703978 | 0.61212691 | NA |
| MAGED1 | 185.663522 | 0.0739364 | 0.1995811 | 0.37045793 | 0.71104131 | 0.94656465 |
| FAM156A | 13.166969 | 0.80879322 | 0.65653329 | 1.23191503 | 0.21798083 | NA |
| TSPYL2 | 14.3717245 | -0.8959224 | 0.55649052 | -1.609951 | 0.10740856 | NA |
| KANTR | 83.5578042 | 0.35990167 | 0.32586734 | 1.10444228 | 0.26940133 | 0.79666376 |
| KDM5C | 42.9604271 | -0.5171698 | 0.30870224 | -1.6753031 | 0.09387478 | 0.64572229 |
| IQSEC2 | 23.3874654 | 0.61200864 | 0.51504074 | 1.18827228 | 0.23472615 | 0.77743304 |
| SMC1A | 45.5612236 | -0.3087563 | 0.32837446 | -0.9402568 | 0.34708585 | 0.83687845 |
| HSD17B10 | 23.0589843 | 0.46593457 | 0.49674588 | 0.93797369 | 0.34825793 | 0.83687845 |
| HUWE1 | 229.705678 | -0.0718778 | 0.19771821 | -0.3635364 | 0.71620421 | 0.94885853 |
| PHF8 | 61.8336864 | -0.0239101 | 0.26993775 | -0.0885763 | 0.92941862 | 0.98960607 |
| FAM120C | 31.6331089 | 0.49931586 | 0.44770982 | 1.11526672 | 0.26473614 | 0.79666376 |
| WNK3 | 39.1390422 | 0.31910494 | 0.3006726 | 1.06130369 | 0.28855191 | 0.80523851 |
| TSR2 | 16.1523449 | 0.68495784 | 0.40053909 | 1.71008988 | 0.08724925 | 0.63466198 |
| FGD1 | 17.519312 | 0.45224424 | 0.39243816 | 1.15239621 | 0.2491583 | 0.78763828 |
| GNL3L | 11.9899837 | -0.3199615 | 0.46937409 | -0.681677 | 0.49544323 | NA |
| MAGED2 | 36.911041 | 0.49019372 | 0.48745916 | 1.00560983 | 0.31460329 | 0.82058274 |
| TRO | 10.5730089 | -0.0788698 | 0.5174411 | -0.1524228 | 0.87885346 | NA |

| | | | | | | |
|---|---|---|---|---|---|---|
| PFKFB1 | 47.8502759 | 2.37587716 | 0.64083766 | 3.70745558 | 0.00020935 | 0.05955414 |
| APEX2 | 10.5380877 | 0.15696736 | 0.50606966 | 0.31016948 | 0.75643208 | NA |
| FAM104B | 11.1046672 | 0.08100136 | 0.57155234 | 0.14172169 | 0.88729984 | NA |
| RRAGB | 82.1526802 | 0.12979281 | 0.27797561 | 0.46692159 | 0.64055598 | 0.93328692 |
| AL445472.1 | 97.4930262 | 0.04966361 | 0.24317055 | 0.20423367 | 0.8381709 | 0.97600352 |
| KLF8 | 14.1383225 | 0.3547066 | 0.588042 | 0.60319943 | 0.54637603 | NA |
| UBQLN2 | 20.4118951 | -0.2500242 | 0.45417568 | -0.5505011 | 0.58197574 | 0.92034873 |
| NBDY | 111.1959 | 0.04201226 | 0.200223 | 0.20982734 | 0.83380243 | 0.97600352 |
| SPIN3 | 26.5613139 | 0.09623854 | 0.38995985 | 0.24679091 | 0.80507005 | 0.97195485 |
| FAAH2 | 267.53691 | 0.3119888 | 0.31088876 | 1.00353837 | 0.31560118 | 0.82084199 |
| ARHGEF9 | 57.9076604 | 0.06080655 | 0.36960243 | 0.1645188 | 0.86932275 | 0.97854266 |
| MTMR8 | 36.2423038 | -0.0023322 | 0.30302805 | -0.0076965 | 0.99385918 | 0.99921857 |
| ZC4H2 | 66.142873 | 0.5353959 | 0.30153853 | 1.77554719 | 0.07580758 | 0.61020315 |
| ZC3H12B | 40.9814865 | 0.20131035 | 0.31952174 | 0.63003648 | 0.52867072 | 0.90487731 |
| LAS1L | 12.8538394 | -0.195669 | 0.45919017 | -0.4261175 | 0.67002223 | NA |
| MSN | 179.139647 | -0.2997927 | 0.26983031 | -1.1110414 | 0.26655052 | 0.79666376 |
| OPHN1 | 350.65656 | -0.1083862 | 0.21397085 | -0.5065467 | 6.12E-01 | 9.26E-01 |
| YIPF6 | 54.5246864 | -0.2105329 | 0.29796031 | -0.7065804 | 0.47982726 | 0.88660781 |
| STARD8 | 14.0302025 | -0.4501193 | 0.57546778 | -0.7821798 | 0.43410892 | NA |
| EFNB1 | 17.1314346 | 0.49884426 | 0.51775106 | 0.96348285 | 0.33530527 | 0.82974237 |
| EDA | 53.5027431 | -0.2113049 | 0.24736186 | -0.8542338 | 0.39297547 | 0.85605713 |
| IGBP1 | 39.4231026 | 0.23950596 | 0.28616456 | 0.83695187 | 0.40261962 | 0.85782098 |
| DLG3 | 57.0432007 | -0.050122 | 0.30363004 | -0.165076 | 0.86888419 | 0.97840831 |
| SNX12 | 14.4535009 | 0.17498386 | 0.56787265 | 0.30813927 | 0.75797636 | NA |
| MED12 | 11.4764264 | -0.0194507 | 0.51932105 | -0.037454 | 0.97012298 | NA |
| ZMYM3 | 16.0241379 | 0.43883885 | 0.51588984 | 0.85064449 | 0.39496687 | NA |
| NONO | 57.819025 | 0.25369489 | 0.27076225 | 0.93696553 | 0.34877629 | 0.83687845 |
| TAF1 | 114.406027 | -0.2308561 | 0.21783313 | -1.0597841 | 0.28924282 | 0.80536924 |
| OGT | 230.668325 | -0.3172739 | 0.3124824 | -1.0153338 | 0.30994673 | 0.81568231 |
| GCNA | 28.3018985 | 0.80342041 | 0.45927063 | 1.74933984 | 0.08023229 | 0.61685352 |
| NHSL2 | 38.02617 | -0.0639337 | 0.30481368 | -0.2097468 | 0.83386532 | 0.97600352 |
| PIN4 | 42.7363429 | -0.3882039 | 0.31062951 | -1.2497329 | 0.21139713 | 0.7643572 |
| RPS4X | 198.548918 | 0.5245393 | 0.41830019 | 1.25397815 | 0.20984995 | 0.76344019 |
| HDAC8 | 293.03415 | 0.06172733 | 0.19224407 | 0.32108836 | 0.74814343 | 0.95563105 |
| PHKA1 | 26.9080732 | 0.16514637 | 0.42151811 | 0.3917895 | 0.69521375 | 0.94395238 |
| NAP1L2 | 11.3852777 | 1.61399658 | 0.73666625 | 2.19094683 | 0.02845564 | NA |
| CHIC1 | 93.2107896 | -0.0597064 | 0.25030048 | -0.2385388 | 0.8114632 | 0.97257655 |
| XIST | 409.855067 | -8.861049 | 3.77753329 | -2.3457236 | 0.01899018 | 0.41544117 |
| JPX | 254.321206 | -0.0961449 | 0.40371891 | -0.2381482 | 0.81176617 | 0.97257655 |
| FTX | 1456.39912 | 0.1334577 | 0.24703791 | 0.54023166 | 0.58903728 | 0.92197072 |
| SLC16A2 | 15.1055909 | 0.61634885 | 0.42693504 | 1.44365956 | 0.14883476 | NA |
| RLIM | 41.9288239 | 0.18849607 | 0.30084708 | 0.62655109 | 0.53095354 | 0.90525279 |
| NEXMIF | 117.15783 | -0.0677621 | 0.33064179 | -0.2049411 | 0.83761816 | 0.97600352 |
| ABCB7 | 107.906857 | -0.1796392 | 0.19579098 | -0.9175052 | 0.35887799 | 0.84204868 |
| UPRT | 20.3876671 | 0.08541102 | 0.35282753 | 0.24207584 | 0.80872139 | 0.97257655 |
| ZDHHC15 | 23.8970858 | 0.37143902 | 0.33569136 | 1.10648966 | 0.26851464 | 0.79666376 |
| PBDC1 | 10.9138255 | 0.34219417 | 0.54611376 | 0.6265987 | 0.53092232 | NA |
| ATRX | 397.57622 | -0.0580458 | 0.18174749 | -0.3193759 | 0.74944148 | 0.95680183 |
| MAGT1 | 78.76512 | 0.07076773 | 0.27783812 | 0.25470848 | 0.79894826 | 0.97019967 |
| COX7B | 129.247349 | 0.06348752 | 0.37310623 | 0.17015936 | 0.86488481 | 0.97788853 |
| PGK1 | 185.936616 | -0.6456794 | 0.40667055 | -1.587721 | 0.11234945 | 0.6656824 |
| BRWD3 | 104.770264 | 0.07670086 | 0.29202325 | 0.26265326 | 0.79281785 | 0.96947477 |
| HMGN5 | 20.1933373 | -0.3216198 | 0.33972379 | -0.9467098 | 0.34378668 | 0.83465492 |
| SH3BGRL | 196.892612 | 0.4010672 | 0.2111127 | 1.89977767 | 0.0574623 | 0.57104732 |

| | | | | | | |
|---|---|---|---|---|---|---|
| RPS6KA6 | 153.731146 | 0.29524261 | 0.27662601 | 1.06729879 | 0.28583693 | 0.80431986 |
| HDX | 48.1026466 | -0.417615 | 0.36965633 | -1.1297386 | 0.25858637 | 0.79532468 |
| APOOL | 67.0433895 | 0.12680669 | 0.23322003 | 0.54372128 | 0.58663329 | 0.92076087 |
| ZNF711 | 17.7014451 | -0.2207 | 0.44802839 | -0.4926027 | 0.62229331 | 0.92897508 |
| CHM | 94.6987894 | -0.2754005 | 0.20689002 | -1.3311444 | 0.18314149 | 0.73539348 |
| DIAPH2 | 206.082512 | -0.4598479 | 0.20262794 | -2.26942 | 0.0232428 | 0.43085291 |
| DIAPH2-AS1 | 15.6059353 | -0.0290967 | 0.56055302 | -0.0519071 | 0.95860271 | NA |
| TSPAN6 | 20.1862581 | 0.69051612 | 0.45386515 | 1.52141253 | 0.12815635 | 0.66929384 |
| SYTL4 | 24.9739943 | 0.64276752 | 0.50462978 | 1.27374076 | 0.20275531 | 0.76141979 |
| TRMT2B | 47.4507268 | -0.2087307 | 0.33884933 | -0.6159986 | 0.53789545 | 0.90763121 |
| HNRNPH2 | 28.2042622 | -0.1212381 | 0.50153685 | -0.2417331 | 0.80898696 | 0.97257655 |
| ARMCX4 | 150.58756 | -0.0717263 | 0.35637567 | -0.2012661 | 0.84049052 | 0.97600352 |
| ARMCX6 | 15.2287999 | 0.04158173 | 0.55167504 | 0.07537359 | 0.93991747 | NA |
| ARMCX3 | 44.0541567 | 0.10133957 | 0.27003901 | 0.37527752 | 0.70745408 | 0.94558664 |
| ARMCX2 | 21.7593692 | -0.4143535 | 0.43850175 | -0.94493 | 0.34469463 | 0.83545751 |
| ZMAT1 | 122.974997 | 0.57632844 | 0.37936001 | 1.51921242 | 0.12870904 | 0.66929384 |
| GPRASP1 | 18.5445853 | 1.04003788 | 0.49208603 | 2.11352858 | 0.03455555 | 0.49098143 |
| BHLHB9 | 20.3208728 | 0.40235127 | 0.38822736 | 1.03638053 | 0.30002465 | 0.81388512 |
| LINC00630 | 61.5748211 | -0.3102164 | 0.29490749 | -1.0519108 | 0.29284046 | 0.80935782 |
| BEX4 | 40.635828 | 0.66154264 | 0.32138176 | 2.05843242 | 0.03954864 | 0.50156625 |
| TCEAL8 | 19.389114 | -0.1410175 | 0.45457811 | -0.3102162 | 0.75639652 | 0.95903253 |
| BEX2 | 34.2805068 | 0.5843791 | 0.42723479 | 1.3678172 | 0.1713693 | 0.72463203 |
| TCEAL9 | 50.3859515 | 0.26822142 | 0.38588764 | 0.69507647 | 0.48700738 | 0.88838332 |
| BEX3 | 84.9653596 | 0.62691406 | 0.36788692 | 1.70409446 | 0.08836344 | 0.63466198 |
| TCEAL4 | 59.2667354 | 0.13315697 | 0.31470472 | 0.42311716 | 0.67220978 | 0.93878847 |
| TCEAL3 | 27.6328381 | 0.33794518 | 0.34359734 | 0.98355006 | 0.32533679 | 0.82567895 |
| MORF4L2 | 92.6052981 | -0.4472047 | 0.25380546 | -1.761998 | 0.07806964 | 0.61367304 |
| SLC25A53 | 51.1194205 | -0.0224742 | 0.23968427 | -0.0937658 | 0.92529518 | 0.98925753 |
| FAM199X | 89.1619805 | -0.0601316 | 0.25849228 | -0.2326245 | 0.816053 | 0.97288403 |
| IL1RAPL2 | 52.7723002 | 0.33654947 | 0.73189135 | 0.45983529 | 0.64563445 | 0.93328692 |
| NRK | 53.8021634 | -0.0089245 | 0.34417332 | -0.0259304 | 9.79E-01 | 9.95E-01 |
| RNF128 | 74.2206418 | -0.7831649 | 0.49678924 | -1.576453 | 0.11492145 | 0.6668162 |
| TBC1D8B | 36.1282155 | 0.16061494 | 0.38706872 | 0.414952 | 0.67817704 | 0.94107403 |
| MORC4 | 28.4458446 | 0.76866514 | 0.4502531 | 1.70718455 | 0.08778776 | 0.63466198 |
| RBM41 | 94.692206 | 0.12122631 | 0.24696125 | 0.4908718 | 0.62351713 | 0.92943979 |
| NUP62CL | 28.9093063 | 0.17620153 | 0.44462314 | 0.39629411 | 0.69188807 | 0.94355912 |
| TSC22D3 | 38.4822957 | 0.27839528 | 0.44927038 | 0.61966088 | 0.53548107 | 0.90763121 |
| MID2 | 26.6503455 | -0.6662814 | 0.32274646 | -2.0644113 | 0.03897873 | 0.49975128 |
| ATG4A | 12.3163697 | -0.0342688 | 0.46526771 | -0.073654 | 0.94128567 | NA |
| COL4A6 | 94.9759048 | -0.0329803 | 0.59294654 | -0.0556211 | 0.95564369 | 0.99148072 |
| COL4A5 | 290.316715 | -0.2881516 | 0.40636978 | -0.7090872 | 0.47827036 | 0.88546214 |
| ACSL4 | 740.459127 | -0.2709823 | 0.55334539 | -0.4897164 | 0.62433461 | 0.92981013 |
| TMEM164 | 164.137215 | -0.015493 | 0.22833732 | -0.0678515 | 0.94590381 | 0.99083449 |
| AMMECR1 | 54.6063226 | 0.19856853 | 0.31517457 | 0.63002713 | 0.52867684 | 0.90487731 |
| ALG13 | 100.798249 | -0.1673852 | 0.44867901 | -0.3730623 | 0.70910211 | 0.94558664 |
| AMOT | 21.5593549 | 0.64366543 | 0.33497011 | 1.92156079 | 0.05466105 | 0.5645224 |
| PLS3-AS1 | 40.1250526 | 0.65734283 | 0.4567643 | 1.43912915 | 1.50E-01 | 0.70364941 |
| PLS3 | 193.611763 | -0.0393935 | 0.32594746 | -0.1208586 | 0.90380305 | 0.98531749 |
| DANT2 | 646.471327 | -0.3965784 | 0.24858743 | -1.5953276 | 0.110639 | 0.66311481 |
| KLHL13 | 355.806507 | -0.2080285 | 0.35420449 | -0.5873117 | 0.55699435 | 0.9141017 |
| WDR44 | 104.976573 | -0.5359212 | 0.48577154 | -1.1032372 | 0.26992417 | 0.79666376 |
| IL13RA1 | 66.4353401 | 0.15518329 | 0.27096549 | 0.57270499 | 5.67E-01 | 9.17E-01 |
| PGRMC1 | 66.4483705 | 0.20262722 | 0.37500801 | 0.54032772 | 0.58897105 | 0.92197072 |
| SLC25A43 | 29.4516113 | 0.10402841 | 0.5487565 | 0.18957116 | 0.84964519 | 0.97612951 |

| | | | | | | |
|---|---|---|---|---|---|---|
| SLC25A5 | 70.1649231 | 1.02782587 | 0.53025553 | 1.93835955 | 0.05257937 | 0.55525525 |
| CXorf56 | 11.1705034 | 0.25145483 | 0.48753585 | 0.51576684 | 0.60601727 | NA |
| UBE2A | 31.6301412 | -0.4123815 | 0.40071522 | -1.0291135 | 0.30342633 | 0.81568231 |
| NKRF | 17.0508793 | 0.75523905 | 0.40173603 | 1.87993856 | 0.06011645 | 0.57733412 |
| 6-Sep | 102.6494 | 0.57536497 | 0.29229679 | 1.96842727 | 0.0490189 | 0.54152793 |
| UPF3B | 48.8931505 | -0.6275869 | 0.49361766 | -1.2714029 | 0.20358536 | 0.76173533 |
| NDUFA1 | 66.8808737 | 0.31707034 | 0.38546276 | 0.82257063 | 0.41075221 | 0.85867915 |
| NKAP | 16.9851731 | 0.58810527 | 0.36891705 | 1.59413958 | 0.11090477 | 0.66331548 |
| RHOXF1-AS1 | 33.9887766 | 0.06332407 | 0.64682403 | 0.09789999 | 0.9220117 | 0.98846691 |
| LAMP2 | 85.0505276 | 0.1144135 | 0.30847596 | 0.37089923 | 0.71071259 | 0.94626542 |
| CUL4B | 30.743077 | -0.0902518 | 0.29728244 | -0.3035896 | 0.7614406 | 0.95989389 |
| MCTS1 | 28.8248988 | 0.18712021 | 0.31442525 | 0.59511825 | 0.55176443 | 0.91288833 |
| C1GALT1C1 | 12.3612645 | 0.08383434 | 0.45728342 | 0.18333124 | 0.85453813 | NA |
| THOC2 | 181.730676 | -0.2305538 | 0.20749013 | -1.1111554 | 0.26650145 | 0.79666376 |
| XIAP | 140.590391 | -0.0990643 | 0.18401089 | -0.5383609 | 0.59032793 | 0.92233545 |
| STAG2 | 202.569118 | -0.0729275 | 0.1770434 | -0.4119189 | 0.68039887 | 0.94196488 |
| TENM1 | 22.3973634 | 1.03760798 | 0.67688234 | 1.53292221 | 0.12529502 | 0.66929384 |
| SMARCA1 | 77.2328021 | 0.13328221 | 0.29321671 | 0.45455189 | 0.64943166 | 0.93437275 |
| OCRL | 17.9633639 | -0.2325827 | 0.37238072 | -0.624583 | 0.5322448 | 0.90612014 |
| ZDHHC9 | 25.7264501 | -0.1040255 | 0.27396887 | -0.3796983 | 7.04E-01 | 9.45E-01 |
| UTP14A | 11.1915236 | -0.2131775 | 0.58684368 | -0.3632611 | 0.71640982 | NA |
| BCORL1 | 19.1176681 | 0.69715759 | 0.41547858 | 1.67796277 | 0.09335437 | 0.64572229 |
| AIFM1 | 38.3210652 | 0.3767114 | 0.25961614 | 1.45103229 | 0.14677087 | 0.69731483 |
| ZNF280C | 18.4893058 | -0.0806743 | 0.40146605 | -0.2009492 | 0.84073826 | 0.97600352 |
| SLC25A14 | 12.8048183 | -0.2164681 | 0.46581977 | -0.4647035 | 0.6421438 | NA |
| RBMX2 | 9.79156834 | -0.0052211 | 0.48076057 | -0.0108602 | 0.991335 | NA |
| ENOX2 | 75.4271825 | -0.5586918 | 0.26946927 | -2.0733042 | 0.03814398 | 0.49671741 |
| RAP2C | 32.177412 | -0.018364 | 0.3973283 | -0.0462187 | 0.96313595 | 0.99256424 |
| RAP2C-AS1 | 89.1561424 | -0.1692187 | 0.28359912 | -0.5966829 | 0.55071913 | 0.91248567 |
| MBNL3 | 56.7388946 | 0.00047933 | 0.38598952 | 0.00124181 | 0.99900918 | 0.99957534 |
| GPC4 | 22.5779468 | -0.2866414 | 0.46806998 | -0.6123899 | 0.54027979 | 0.90763121 |
| PHF6 | 41.2153625 | -0.242264 | 0.28572569 | -0.8478901 | 0.39649916 | 0.85650971 |
| HPRT1 | 118.835297 | 0.28498866 | 0.19128425 | 1.48987 | 0.13625842 | 0.68406472 |
| FAM122B | 21.1524168 | -0.3959329 | 0.36062271 | -1.0979145 | 0.27224184 | 0.7968545 |
| FAM122C | 16.9199169 | 0.14566665 | 0.68074027 | 0.21398272 | 0.83056053 | 0.97567084 |
| RTL8C | 37.0120447 | -0.3056109 | 0.4778426 | -0.6395639 | 0.52245615 | 0.90485509 |
| RTL8A | 21.7643931 | 0.36659939 | 0.45324638 | 0.80883027 | 0.41861278 | 0.86022698 |
| ZNF75D | 53.3039588 | -0.011073 | 0.28172944 | -0.0393036 | 0.96864831 | 0.99379133 |
| INTS6L | 22.8774917 | 0.48713171 | 0.40406747 | 1.20557023 | 0.22798325 | 0.77185154 |
| MMGT1 | 12.8176245 | 0.28683128 | 0.42789099 | 0.67033727 | 0.50264281 | NA |
| SLC9A6 | 23.0822569 | 0.14578166 | 0.37400001 | 0.38979051 | 0.69669146 | 0.94395238 |
| HTATSF1 | 26.0640048 | 0.10831466 | 0.36034 | 0.30059016 | 0.76372704 | 0.96091323 |
| AL683813.1 | 24.22411 | 0.33855774 | 0.5001787 | 0.67687355 | 0.49848619 | 0.89378207 |
| RBMX | 150.484052 | 0.12317708 | 0.1993494 | 0.61789543 | 0.53664427 | 0.90763121 |
| FGF13 | 477.357124 | -0.2726052 | 0.48923086 | -0.5572118 | 0.5773827 | 0.91859435 |
| ATP11C | 82.0247416 | -0.5225227 | 0.36175292 | -1.4444188 | 0.1486212 | 0.70006785 |
| FMR1 | 66.5528823 | 0.02322759 | 0.30419379 | 0.07635787 | 0.93913439 | 0.99012269 |
| IDS | 61.1865804 | -0.3233554 | 0.3321258 | -0.9735933 | 0.33025852 | 0.82867637 |
| TMEM185A | 18.8873631 | 0.10309483 | 0.42114886 | 0.24479427 | 0.80661573 | 0.97216031 |
| LINC00894 | 130.916329 | 0.05128704 | 0.31427824 | 0.16318991 | 0.87036891 | 0.97854266 |
| MTM1 | 68.3363238 | -0.364185 | 0.23742313 | -1.5339072 | 0.12505249 | 0.66929384 |
| MTMR1 | 66.2498675 | -0.0680247 | 0.28682243 | -0.2371667 | 0.81252746 | 0.97257655 |
| CD99L2 | 43.4937603 | -0.1663661 | 0.25723257 | -0.6467537 | 0.51779136 | 0.90330678 |
| VMA21 | 28.6829288 | -0.1938149 | 0.46336303 | -0.4182787 | 0.67574336 | 0.94013324 |

| | | | | | | |
|---|---|---|---|---|---|---|
| CETN2 | 18.9020144 | 0.05238184 | 0.57248551 | 0.09149898 | 0.92709612 | 0.98925753 |
| ZNF185 | 12.6007642 | 0.84987456 | 0.65111009 | 1.30527015 | 0.19180087 | NA |
| DUSP9 | 17.0037982 | 0.21266079 | 0.80243372 | 0.26501976 | 7.91E-01 | 0.96909068 |
| SLC6A8 | 42.2506459 | -0.5924698 | 0.49030198 | -1.2083773 | 0.22690216 | 0.77185154 |
| BCAP31 | 59.231942 | 0.31353858 | 0.36941308 | 0.84874791 | 0.39602158 | 0.85605713 |
| IDH3G | 17.9248849 | 0.02261699 | 0.42550796 | 0.05315291 | 0.95761007 | 0.99170927 |
| SSR4 | 29.5212135 | 0.03426318 | 0.45838342 | 0.07474785 | 0.94041533 | 0.99012269 |
| L1CAM | 38.0611331 | 0.28857077 | 0.60594228 | 0.47623476 | 0.63390714 | 0.93251095 |
| ARHGAP4 | 13.7390875 | -0.1085236 | 0.42931547 | -0.2527829 | 0.80043602 | NA |
| HCFC1 | 22.123157 | 0.1675188 | 0.43016721 | 0.38942718 | 0.69696017 | 0.94395238 |
| IRAK1 | 20.9715188 | -0.5020865 | 0.60289279 | -0.8327957 | 0.40495998 | 0.85782098 |
| MECP2 | 117.67322 | 0.0130635 | 0.21872248 | 0.05972637 | 0.95237357 | 0.99141043 |
| FLNA | 70.7412909 | -0.1379108 | 0.62704284 | -0.2199384 | 0.82591909 | 0.97567084 |
| EMD | 39.0142832 | -0.2917856 | 0.6237563 | -0.4677879 | 0.63993629 | 0.93328692 |
| RPL10 | 230.148954 | 0.27735941 | 0.50388464 | 0.55044228 | 0.58201605 | 0.92034873 |
| DNASE1L1 | 12.3896992 | -0.3828283 | 0.54400933 | -0.7037164 | 0.48160938 | NA |
| TAZ | 25.5159751 | 0.02864319 | 0.33482272 | 0.08554732 | 0.93182628 | 0.98960607 |
| ATP6AP1 | 52.3159754 | 0.24580766 | 0.48834505 | 0.50334832 | 0.6147194 | 0.92662475 |
| GDI1 | 28.2698985 | -0.7983447 | 0.38843193 | -2.0553013 | 0.03984991 | 0.50156625 |
| FAM50A | 23.0221485 | -0.9250148 | 0.36835779 | -2.5111857 | 0.01203264 | 0.35795129 |
| PLXNA3 | 12.825278 | -0.5055251 | 0.40772589 | -1.2398651 | 0.21502528 | NA |
| FAM3A | 18.1334119 | 0.26443963 | 0.45362443 | 0.58294838 | 0.55992805 | 0.91516276 |
| IKBKG | 11.6438225 | -0.1354115 | 0.44746073 | -0.3026222 | 0.7621778 | NA |
| DKC1 | 20.9136295 | -0.2289989 | 0.39003161 | -0.5871291 | 0.55711702 | 0.9141017 |
| MPP1 | 24.3360406 | 1.34768646 | 0.42038135 | 3.20586642 | 0.00134656 | 0.15715102 |
| F8 | 16.753391 | 0.4205663 | 0.42479579 | 0.99004346 | 0.32215288 | 0.82558302 |
| FUNDC2 | 37.3526932 | -0.1884333 | 0.36335533 | -0.5185924 | 0.60404504 | 0.92522665 |
| BRCC3 | 40.7821701 | 0.16665429 | 0.29546121 | 0.56404796 | 0.57272149 | 0.91813957 |
| VBP1 | 17.6748037 | -0.0038407 | 0.38881664 | -0.009878 | 0.99211865 | 0.99814944 |
| TMLHE-AS1 | 34.9230396 | 1.29871115 | 0.65126058 | 1.99414977 | 0.04613569 | 0.53026917 |
| TMLHE | 87.7627159 | 0.34404825 | 0.26912073 | 1.27841601 | 0.20110278 | 0.75910235 |
| VAMP7 | 46.2409603 | -0.2206736 | 0.27171483 | -0.8121514 | 0.41670477 | 0.85867915 |
| ZNF596 | 20.0089576 | -0.4783471 | 0.33591882 | -1.4239961 | 0.15444761 | 0.70572801 |
| FBXO25 | 49.8532654 | -0.250592 | 0.28220069 | -0.8879922 | 0.37454492 | 0.85092528 |
| TDRP | 70.7361529 | -0.2497827 | 0.47337582 | -0.5276627 | 0.59773349 | 0.9229638 |
| ERICH1 | 139.413559 | -0.6389714 | 0.39028337 | -1.6371985 | 0.10158899 | 0.65863153 |
| AC246817.2 | 19.8121085 | 0.96188023 | 0.55658173 | 1.72819225 | 0.08395376 | 0.62590589 |
| AC246817.1 | 46.6929003 | 0.18772383 | 0.53730325 | 0.34938152 | 0.7268029 | 0.95209193 |
| CSMD1 | 52.1034862 | -1.1204675 | 1.15566144 | -0.9695465 | 0.33227261 | 0.82890589 |
| MCPH1 | 165.318347 | -0.1430666 | 0.21312299 | -0.6712865 | 0.50203803 | 0.89499455 |
| ANGPT2 | 24.0094621 | 0.41775349 | 0.75583046 | 0.55270793 | 0.58046342 | 0.92034873 |
| MCPH1-AS1 | 36.8829013 | 0.07586411 | 0.34046915 | 0.22282228 | 0.82367383 | 0.97518529 |
| AGPAT5 | 64.6748381 | -0.5657913 | 0.29900374 | -1.8922549 | 0.05845703 | 0.57141408 |
| AF233439.1 | 424.103605 | 0.36701886 | 0.75311296 | 0.48733573 | 0.62602043 | 0.93024224 |
| DEFB1 | 1095.19534 | -0.0289723 | 0.49698115 | -0.0582965 | 0.95351242 | 0.99148072 |
| FAM85B | 108.212346 | 0.19575824 | 0.39326228 | 0.49778037 | 0.61863885 | 0.92766755 |
| MFHAS1 | 661.165105 | 0.29373549 | 0.32753782 | 0.89679869 | 0.36982635 | 0.84852785 |
| ERI1 | 41.4003637 | 0.1112947 | 0.27539036 | 0.40413433 | 0.68611394 | 0.94205398 |
| PPP1R3B | 20.8114721 | -0.8335811 | 0.33348855 | -2.4995795 | 1.24E-02 | 0.36046953 |
| TNKS | 300.394129 | -0.3138734 | 0.16349524 | -1.9197708 | 0.05488686 | 0.5645224 |
| MSRA | 106.449534 | -0.3405428 | 0.34750246 | -0.9799725 | 0.32709971 | 0.82612204 |
| XKR6 | 368.628136 | -0.2260467 | 0.20261288 | -1.1156579 | 0.26456858 | 0.79666376 |
| MTMR9 | 45.4157795 | 0.06188596 | 0.2641078 | 0.23432085 | 0.8147359 | 0.97284589 |
| FDFT1 | 100.37924 | 0.01539639 | 0.38114568 | 0.04039502 | 0.9677782 | 0.99332167 |

| | | | | | | |
|---|---|---|---|---|---|---|
| CTSB | 83.4780294 | 0.2265118 | 0.3639819 | 0.62231612 | 5.34E-01 | 0.90763121 |
| FAM66D | 37.134793 | 0.05560188 | 0.77541027 | 0.07170641 | 0.94283556 | 0.99058259 |
| AC145124.1 | 10.8972218 | 1.56226119 | 0.73543117 | 2.12427926 | 0.0336468 | NA |
| LONRF1 | 57.3816853 | -0.3562038 | 0.41422358 | -0.8599313 | 0.38982693 | 0.85605713 |
| TRMT9B | 116.746782 | 0.52867433 | 0.3079393 | 1.71681344 | 0.08601326 | 0.63376931 |
| TUSC3 | 119.333984 | -0.4439883 | 0.22597046 | -1.9648067 | 0.04943662 | 0.54320302 |
| MSR1 | 274.985721 | -0.1676975 | 0.27975476 | -0.5994447 | 0.54887635 | 0.91208861 |
| AC011586.1 | 18.3557917 | 0.17735642 | 0.6215071 | 0.28536507 | 0.77536447 | 0.96469469 |
| AC068992.1 | 117.717006 | -0.3914767 | 0.32012711 | -1.222879 | 0.22137541 | 0.76680432 |
| MICU3 | 96.0736876 | -0.2862206 | 0.35835783 | -0.7987005 | 0.42446411 | 0.86190547 |
| ZDHHC2 | 133.696925 | 0.38160811 | 0.26025131 | 1.46630624 | 0.14256488 | 0.69328517 |
| CNOT7 | 37.9037477 | 0.12503517 | 0.26847989 | 0.4657152 | 0.64141937 | 0.93328692 |
| VPS37A | 59.26638 | -0.1349893 | 0.25374118 | -0.5319959 | 0.59472886 | 0.9229638 |
| MTUS1 | 271.187576 | -0.39497 | 0.14184202 | -2.7845769 | 0.00535976 | 0.28183167 |
| AC027117.2 | 29.8550103 | -0.7935709 | 0.33781844 | -2.3491049 | 0.01881861 | 0.41544117 |
| PCM1 | 317.3166 | -0.1378434 | 0.17139884 | -0.8042259 | 0.42126652 | 0.86190547 |
| ASAH1 | 110.410631 | 0.531344 | 0.34255874 | 1.55110332 | 0.12087693 | 0.66727534 |
| PSD3 | 137.900816 | -0.0958316 | 0.64892775 | -0.1476769 | 0.88259776 | 0.98137082 |
| SH2D4A | 45.5589204 | -0.0304729 | 0.40257626 | -0.0756947 | 0.93966197 | 0.99012269 |
| CSGALNACT1 | 711.108596 | -1.8888081 | 0.56124339 | -3.3653992 | 0.00076433 | 0.12591256 |
| INTS10 | 120.942868 | 0.01235348 | 0.2869827 | 0.04304609 | 0.96566479 | 0.99270995 |
| ATP6V1B2 | 41.226011 | -0.2024919 | 0.34068494 | -0.594367 | 0.55226668 | 0.91322136 |
| XPO7 | 92.7319205 | 0.02170306 | 0.23216867 | 0.09347973 | 0.92552245 | 0.98925753 |
| DMTN | 50.650354 | 0.1150071 | 0.25566868 | 0.44982865 | 0.652834 | 0.93532021 |
| FAM160B2 | 24.753015 | -0.1256715 | 0.31300844 | -0.4014956 | 0.68805529 | 0.94205398 |
| SLC39A14 | 49.1926915 | -0.789071 | 0.38583596 | -2.0450945 | 0.04084555 | 0.50629837 |
| PPP3CC | 118.568875 | -0.5495898 | 0.28847653 | -1.9051458 | 0.05676113 | 0.56976762 |
| CCAR2 | 18.8014253 | -0.2580122 | 0.37468369 | -0.6886134 | 0.49106656 | 0.88987842 |
| BIN3 | 52.0198149 | -0.488746 | 0.24589324 | -1.9876348 | 0.04685209 | 0.5320597 |
| RHOBTB2 | 12.2302718 | -1.0092388 | 0.48019883 | -2.1017103 | 0.03557866 | NA |
| TNFRSF10B | 70.4709085 | -0.3539671 | 0.32691963 | -1.0827342 | 0.27892642 | 0.80219184 |
| TNFRSF10D | 36.7190171 | -0.3028941 | 0.30162889 | -1.0041945 | 3.15E-01 | 0.82084199 |
| TNFRSF10A | 15.6691618 | -0.3343342 | 0.34740588 | -0.9623733 | 3.36E-01 | NA |
| CHMP7 | 15.6949181 | -0.0246363 | 0.43277124 | -0.0569269 | 0.95460343 | NA |
| R3HCC1 | 11.2971943 | -0.2411065 | 0.43334615 | -0.5563832 | 0.57794891 | NA |
| ENTPD4 | 46.6430512 | -0.1989733 | 0.33698073 | -0.5904591 | 0.55488289 | 0.9137961 |
| SLC25A37 | 133.48312 | 0.305947 | 0.27134878 | 1.12750461 | 0.25952919 | 0.79532468 |
| STC1 | 81.6996346 | 0.1354705 | 0.59543437 | 0.22751543 | 0.82002298 | 0.97338914 |
| DOCK5 | 100.625891 | -0.1844504 | 0.24100771 | -0.7653301 | 0.44407501 | 0.8704597 |
| KCTD9 | 49.3287781 | -0.1751106 | 0.29231416 | -0.5990494 | 0.54913994 | 0.91221128 |
| PPP2R2A | 199.080673 | -0.4423334 | 0.26055747 | -1.6976422 | 0.08957531 | 0.63653714 |
| BNIP3L | 146.908699 | -0.1910337 | 0.42015407 | -0.4546754 | 6.49E-01 | 0.93437275 |
| TRIM35 | 13.14148 | 0.42406119 | 0.44360054 | 0.95595281 | 0.33909607 | NA |
| PTK2B | 56.0490621 | 0.1754557 | 0.32075921 | 0.54700128 | 5.84E-01 | 0.92071026 |
| EPHX2 | 52.0794848 | 0.6254475 | 0.41664803 | 1.50114114 | 0.13331906 | 0.67988987 |
| CLU | 484.947623 | 0.73859322 | 0.60443315 | 1.22196014 | 0.2217227 | 0.76680432 |
| CCDC25 | 28.3846334 | 0.01906779 | 0.30939914 | 0.06162844 | 0.95085873 | 0.99105535 |
| ELP3 | 89.7523706 | -0.2758623 | 0.24829156 | -1.1110419 | 0.26655031 | 0.79666376 |
| ZNF395 | 28.697101 | -0.7211547 | 0.45454542 | -1.5865405 | 0.11261676 | 0.6656824 |
| FBXO16 | 38.6062242 | -0.4000735 | 0.36076731 | -1.1089517 | 0.26745101 | 0.79666376 |
| FZD3 | 32.6013723 | 0.65117965 | 0.30119124 | 2.16201388 | 0.03061711 | 0.47078973 |
| EXTL3 | 27.8351444 | -0.332557 | 0.42255366 | -0.7870172 | 0.43127179 | 0.86604857 |
| INTS9 | 62.1720223 | 0.11100364 | 0.21961378 | 0.50544933 | 0.61324328 | 0.92662475 |
| HMBOX1 | 177.942999 | -0.2942477 | 0.16886113 | -1.7425423 | 0.0814136 | 0.61925304 |

| | | | | | | |
|---|---|---|---|---|---|---|
| KIF13B | 407.993209 | -0.2876578 | 0.22657788 | -1.2695758 | 0.20423577 | 0.76173533 |
| AC108449.1 | 19.7954891 | 0.51579022 | 0.60631089 | 0.85070256 | 0.3949346 | 0.85605713 |
| SARAF | 98.3464993 | 0.42218571 | 0.26122744 | 1.6161614 | 0.10605942 | 0.65946648 |
| LEPROTL1 | 36.2672085 | 0.05598206 | 0.29743022 | 0.18821914 | 0.85070487 | 0.97612951 |
| DCTN6 | 15.5060355 | 0.01622785 | 0.36115914 | 0.04493268 | 0.96416097 | NA |
| RBPMS | 2091.33129 | -0.3593137 | 0.33937021 | -1.0587661 | 0.2897063 | 0.80536924 |
| GTF2E2 | 65.560234 | -0.1838661 | 0.23565022 | -0.78025 | 0.43524376 | 0.86729771 |
| GSR | 135.068847 | -0.0926374 | 0.27214875 | -0.3403924 | 0.73356103 | 0.95329896 |
| PPP2CB | 142.814494 | -0.2190994 | 0.20447848 | -1.0715037 | 0.283943 | 0.80431986 |
| WRN | 96.4442524 | -0.3269205 | 0.24553386 | -1.3314681 | 0.18303502 | 0.73529028 |
| NRG1 | 68.9267492 | 0.38462051 | 0.52796971 | 0.72848973 | 0.46631385 | 0.88049264 |
| FUT10 | 41.4429593 | 0.74317774 | 0.44017121 | 1.68838336 | 0.09133766 | 0.64297142 |
| TTI2 | 20.2174752 | -0.1621939 | 0.40456891 | -0.4009055 | 0.68848973 | 0.94205398 |
| UNC5D | 14.7281491 | -0.4623134 | 0.40965599 | -1.1285406 | 0.25909166 | NA |
| ERLIN2 | 40.6114813 | -0.0613574 | 0.36595761 | -0.1676626 | 0.86684868 | 0.97788853 |
| PLPBP | 29.6741163 | -0.0713037 | 0.34355706 | -0.2075453 | 0.83558399 | 0.97600352 |
| RAB11FIP1 | 147.635082 | 0.20293663 | 0.24934247 | 0.81388714 | 0.41570961 | 0.85867915 |
| ASH2L | 39.3829259 | -0.3790493 | 0.29253018 | -1.2957614 | 0.19505768 | 0.75237716 |
| LSM1 | 30.5633757 | -0.1045241 | 0.38886558 | -0.2687925 | 0.78808938 | 0.96768123 |
| BAG4 | 50.7946911 | 0.20085691 | 0.23073803 | 0.87049764 | 0.38402851 | 0.85511218 |
| DDHD2 | 56.5839365 | 0.16929913 | 0.28689923 | 0.59009963 | 0.55512386 | 0.9137961 |
| PLPP5 | 50.3017226 | -0.1089689 | 0.32278363 | -0.337591 | 0.7356714 | 0.9536908 |
| NSD3 | 231.939134 | -0.0961154 | 0.2156112 | -0.4457809 | 0.6557555 | 0.93567213 |
| LETM2 | 11.6490751 | 0.33579782 | 0.52452181 | 0.64019802 | 0.52204387 | NA |
| FGFR1 | 71.5238963 | -0.8904226 | 0.44775766 | -1.9886261 | 0.04674249 | 0.5320597 |
| TACC1 | 142.77477 | -0.2689308 | 0.43112974 | -0.6237816 | 0.53277104 | 0.90667691 |
| PLEKHA2 | 170.376123 | -1.1401915 | 0.47609105 | -2.3949021 | 0.0166248 | 0.39748373 |
| TM2D2 | 12.7325531 | 0.38783361 | 0.418346 | 0.92706422 | 0.35389319 | NA |
| ADAM9 | 122.978696 | -0.3243628 | 0.34150913 | -0.9497923 | 0.34221778 | 0.8331662 |
| TCIM | 54.1134617 | -0.4019351 | 0.4311407 | -0.9322598 | 0.35120228 | 0.83800638 |
| AC105999.2 | 54.59445 | 0.59285634 | 0.67235539 | 0.88176037 | 0.37790641 | 0.85310518 |
| GOLGA7 | 25.2991853 | 0.1212054 | 0.35504426 | 0.3413811 | 0.7328167 | 0.95311194 |
| GPAT4 | 43.7602559 | -0.7836518 | 0.38185825 | -2.0522061 | 0.04014963 | 0.50338084 |
| KAT6A | 239.32345 | 0.07783072 | 0.2388138 | 0.32590546 | 0.74449588 | 0.95563105 |
| AP3M2 | 21.0726423 | 0.68823797 | 0.40913025 | 1.68219771 | 0.09253051 | 0.64572229 |
| IKBKB | 84.837942 | -0.123542 | 0.2346306 | -0.5265383 | 0.59851424 | 0.92296674 |
| POLB | 21.5330499 | -0.269171 | 0.32502772 | -0.8281478 | 0.40758679 | 0.85867915 |
| VDAC3 | 65.6114856 | 0.26481669 | 0.28640707 | 0.92461645 | 0.35516545 | 0.84085075 |
| SLC20A2 | 199.338695 | -0.4023764 | 0.38048819 | -1.0575266 | 0.2902713 | 0.80559136 |
| SMIM19 | 31.1290549 | 0.05344898 | 0.34509403 | 0.15488235 | 0.87691407 | 0.97969425 |
| RNF170 | 70.7449938 | 0.03976882 | 0.26119124 | 0.1522594 | 0.87898234 | 0.98061726 |
| HOOK3 | 121.548947 | -0.1970843 | 0.29310114 | -0.6724105 | 0.5013224 | 0.89492452 |
| FNTA | 29.6096971 | -0.3852331 | 0.40196589 | -0.9583725 | 0.33787496 | 0.83055261 |
| HGSNAT | 133.29876 | -0.0989641 | 0.36971336 | -0.2676779 | 0.78894726 | 0.96815677 |
| SPIDR | 523.06855 | -0.4454753 | 0.16719571 | -2.6643943 | 0.00771271 | 0.30654023 |
| CEBPD | 152.199599 | -0.3804383 | 0.37074508 | -1.0261451 | 3.05E-01 | 0.81568231 |
| PRKDC | 146.814069 | -0.2941698 | 0.18987841 | -1.5492534 | 0.12132081 | 0.66771734 |
| MCM4 | 13.8787348 | -0.1414841 | 0.50013051 | -0.2828943 | 0.77725787 | NA |
| UBE2V2 | 55.6382696 | -0.1536793 | 0.33892075 | -0.4534372 | 0.65023398 | 0.93453685 |
| SNTG1 | 543.817179 | 0.0914897 | 0.57866793 | 0.15810397 | 8.74E-01 | 9.79E-01 |
| PXDNL | 24.9600425 | -0.1341417 | 0.39530723 | -0.3393352 | 0.73435724 | 0.95329896 |
| PCMTD1 | 278.682886 | 0.07738835 | 0.18944626 | 0.40849765 | 0.68290836 | 0.94205398 |
| AC064807.1 | 19.769055 | 0.56021213 | 0.53644285 | 1.04430906 | 0.29634242 | 0.81120086 |
| RB1CC1 | 185.604376 | -0.4479976 | 0.23093527 | -1.939927 | 0.05238856 | 0.55403388 |

| | | | | | | |
|---|---|---|---|---|---|---|
| ATP6V1H | 249.084174 | 0.02569062 | 0.19306663 | 0.13306606 | 8.94E-01 | 0.98408497 |
| TCEA1 | 67.9683568 | 0.23377159 | 0.23063639 | 1.01359371 | 0.31077666 | 0.81692172 |
| LYPLA1 | 124.852457 | -0.0448529 | 0.18378647 | -0.2440487 | 0.8071931 | 0.97216031 |
| MRPL15 | 30.1717406 | 0.06977916 | 0.29820074 | 0.23400061 | 0.8149845 | 0.97284589 |
| TMEM68 | 41.2480827 | 0.46093784 | 0.36225482 | 1.27241327 | 0.20322632 | 0.76169832 |
| TGS1 | 68.722827 | 0.39819701 | 0.39937195 | 0.99705804 | 0.31873634 | 0.82354084 |
| LYN | 103.564102 | 0.4710681 | 0.25584534 | 1.84122211 | 0.06558902 | 0.59349584 |
| RPS20 | 84.5059005 | 0.09118647 | 0.35624894 | 0.2559628 | 0.79797956 | 0.97019967 |
| PLAG1 | 43.2021352 | 0.82415002 | 0.39170616 | 2.10400066 | 0.03537839 | 0.49209205 |
| CHCHD7 | 18.4232949 | 0.07748961 | 0.36281392 | 0.21357949 | 0.83087499 | 0.97567084 |
| IMPAD1 | 50.2274092 | -0.1378573 | 0.34462477 | -0.4000214 | 0.68914072 | 0.94210755 |
| LINC01606 | 197.418823 | 0.45160427 | 0.42808106 | 1.05495036 | 0.29144803 | 0.80688911 |
| FAM110B | 326.910932 | -0.0428918 | 0.21670983 | -0.1979227 | 0.84310555 | 0.97612951 |
| AC027698.1 | 23.1709455 | 0.34541725 | 0.54065589 | 0.63888559 | 0.52289736 | 0.90487731 |
| UBXN2B | 54.7722945 | -0.1760626 | 0.24830233 | -0.7090656 | 0.47828379 | 0.88546214 |
| SDCBP | 93.5999061 | -0.1296389 | 0.24846512 | -0.5217589 | 6.02E-01 | 0.92400714 |
| NSMAF | 63.4213056 | -0.1252885 | 0.32097549 | -0.3903367 | 0.69628762 | 0.94395238 |
| TOX | 476.33688 | 0.18704979 | 0.46413616 | 0.40300629 | 0.6869436 | 0.94205398 |
| LINC01301 | 31.234044 | 0.65496389 | 0.64180895 | 1.02049665 | 0.30749298 | 0.81568231 |
| RAB2A | 246.87258 | -0.2745776 | 0.20203565 | -1.3590549 | 0.17412918 | 0.72841249 |
| CHD7 | 165.869542 | 0.06229365 | 0.2252576 | 0.27654404 | 0.78213024 | 0.96652357 |
| ASPH | 159.352303 | 0.05563671 | 0.31495355 | 0.17665051 | 0.85978291 | 0.97691173 |
| YTHDF3 | 98.3059386 | -0.0904025 | 0.2743572 | -0.3295065 | 0.74177287 | 0.95507192 |
| CYP7B1 | 109.131628 | -0.0172958 | 0.27184447 | -0.063624 | 9.49E-01 | 0.99105535 |
| ARMC1 | 33.7274155 | -0.1620651 | 0.2794555 | -0.5799318 | 0.56196062 | 0.91624774 |
| MTFR1 | 59.1154966 | -0.2250486 | 0.38235931 | -0.5885789 | 0.55614376 | 0.91398748 |
| PDE7A | 60.3892383 | 0.65228261 | 0.39217003 | 1.66326481 | 0.09625944 | 0.65104726 |
| ADHFE1 | 50.7413936 | 0.55673231 | 0.41219075 | 1.35066668 | 0.17680223 | 0.72841249 |
| VCPIP1 | 31.7611867 | -0.1132491 | 0.28599456 | -0.3959833 | 0.69211736 | 0.94373042 |
| SGK3 | 76.1996128 | 0.15321908 | 0.29748865 | 0.51504179 | 0.60652382 | 0.92569157 |
| PPP1R42 | 11.2486527 | -0.2384542 | 0.66616367 | -0.3579513 | 0.72037973 | NA |
| CSPP1 | 163.210102 | 0.01012016 | 0.21746645 | 0.04653664 | 0.96288253 | 0.99256424 |
| ARFGEF1 | 477.687128 | -0.3127575 | 0.20110032 | -1.555231 | 0.11989106 | 0.6668162 |
| NCOA2 | 664.783022 | -0.1452309 | 0.13425406 | -1.0817614 | 0.27935858 | 0.80219184 |
| TRAM1 | 131.037896 | -0.3797588 | 0.32075841 | -1.1839402 | 0.23643674 | 0.77864547 |
| LACTB2 | 43.3551196 | -0.0422838 | 0.30128933 | -0.1403427 | 8.88E-01 | 0.98262538 |
| XKR9 | 14.4263317 | -0.8522367 | 0.73286719 | -1.1628801 | 0.24487815 | NA |
| MSC-AS1 | 107.727207 | 1.5867764 | 0.68225989 | 2.32576534 | 0.02003107 | 0.41544117 |
| TERF1 | 64.7005348 | 0.11130525 | 0.29140332 | 0.38196287 | 0.70248891 | 0.94476609 |
| SBSPON | 17.6040384 | 0.26944808 | 0.60002064 | 0.44906469 | 0.65338499 | 0.93532021 |
| RPL7 | 169.587835 | 0.50583282 | 0.44743786 | 1.13050964 | 0.25826154 | 0.79532468 |
| RDH10 | 142.496273 | -1.2460503 | 0.45208298 | -2.7562424 | 0.00584696 | 0.2943676 |
| RDH10-AS1 | 44.208454 | -0.7579355 | 0.73315773 | -1.033796 | 0.30123154 | 0.81489175 |
| STAU2 | 220.942838 | 0.15838118 | 0.17530695 | 0.90345067 | 0.36628676 | 0.84741115 |
| AC027018.1 | 33.4443877 | -0.2161173 | 0.52641529 | -0.4105453 | 0.68140599 | 0.94196488 |
| ELOC | 38.5451723 | -0.2445384 | 0.27531246 | -0.8882214 | 0.37442169 | 0.85092528 |
| GDAP1 | 64.0378222 | -0.1419395 | 0.33153669 | -0.428126 | 0.66855942 | 0.93878847 |
| HNF4G | 20.7513197 | 0.18215691 | 0.7061311 | 0.25796472 | 0.79643414 | 0.96986756 |
| PEX2 | 39.5483468 | 0.27430945 | 0.28646974 | 0.95755122 | 0.33828911 | 0.83055261 |
| PKIA | 32.2406284 | 0.5279326 | 0.46827584 | 1.12739663 | 0.25957482 | 0.79532468 |
| ZC2HC1A | 67.5815886 | -0.2374927 | 0.28898683 | -0.8218115 | 0.41118418 | 0.85867915 |
| IL7 | 37.7332444 | 0.67180651 | 0.41709272 | 1.61068865 | 1.07E-01 | 0.65986731 |
| TPD52 | 139.071022 | -0.3147228 | 0.26147853 | -1.2036278 | 0.22873348 | 0.77185154 |
| ZBTB10 | 42.8567384 | 0.09975912 | 0.30702735 | 0.32491934 | 0.74524212 | 0.95563105 |

| | | | | | | |
|---|---|---|---|---|---|---|
| ZNF704 | 438.090411 | -0.0248828 | 0.23072143 | -0.1078478 | 0.91411643 | 0.98674278 |
| IMPA1 | 32.3281074 | -0.6135173 | 0.31982832 | -1.9182708 | 0.05507668 | 0.5645224 |
| ZFAND1 | 44.3051013 | 0.27307605 | 0.26384602 | 1.03498265 | 0.30067702 | 0.81411745 |
| CHMP4C | 19.3442537 | -0.4096642 | 0.36004878 | -1.137802 | 0.25520317 | 0.79341342 |
| SNX16 | 33.1340819 | -0.6746581 | 0.41145579 | -1.6396855 | 0.10107058 | 0.65863153 |
| AC015522.1 | 111.682659 | 1.62690701 | 0.56087862 | 2.9006401 | 0.00372401 | 0.24214066 |
| RALYL | 751.794036 | 0.24260051 | 0.37049717 | 0.6547972 | 0.51259833 | 0.90156229 |
| LRRCC1 | 19.7812424 | -0.412144 | 0.36114127 | -1.1412264 | 0.25377571 | 0.7912221 |
| E2F5 | 26.0777814 | 0.14828211 | 0.43206508 | 0.34319393 | 0.73145257 | 0.95279138 |
| C8orf59 | 30.6145103 | -0.3616363 | 0.39348579 | -0.9190581 | 0.3580652 | 0.84204868 |
| CA13 | 35.6126581 | 0.09932559 | 0.29139808 | 0.34085877 | 0.7332099 | 0.95329896 |
| CA3 | 49.8486906 | 0.1665971 | 0.28350443 | 0.5876349 | 0.55677738 | 0.9141017 |
| CA2 | 185.971023 | 0.25825684 | 0.39300897 | 0.6571271 | 0.51109919 | 0.90030978 |
| ATP6V0D2 | 55.392494 | -0.3627035 | 0.78058351 | -0.4646569 | 0.6421772 | 0.93328692 |
| WWP1 | 410.663246 | 0.18749336 | 0.1670922 | 1.12209525 | 0.26182194 | 0.79633795 |
| RMDN1 | 85.2947902 | 0.06237987 | 0.27458694 | 0.22717711 | 0.82028603 | 0.97354156 |
| CPNE3 | 73.1565595 | 0.25894368 | 0.21033678 | 1.23109084 | 0.21828889 | 0.76666749 |
| AF117829.1 | 230.278864 | 0.28941958 | 0.38877962 | 0.74443095 | 0.4566158 | 0.87396417 |
| RIPK2 | 24.7672404 | -0.3764868 | 0.32738779 | -1.1499721 | 0.25015538 | 0.78799766 |
| OSGIN2 | 26.1038794 | -0.0763873 | 0.39971605 | -0.1911039 | 0.84844418 | 0.97612951 |
| NBN | 54.3069074 | -0.238878 | 0.33044413 | -0.7228997 | 0.4697415 | 0.88207878 |
| DECR1 | 141.95714 | 0.05089569 | 0.28583273 | 0.17806111 | 0.85867499 | 0.97691173 |
| CALB1 | 797.947432 | 1.34280071 | 0.70435001 | 1.90643954 | 5.66E-02 | 0.56976762 |
| LINC00534 | 29.8511008 | 0.74633175 | 0.79417798 | 0.93975377 | 0.34734388 | 0.83687845 |
| TMEM64 | 21.9321164 | -0.0449557 | 0.61478737 | -0.073124 | 0.94170747 | 0.99045212 |
| PIP4P2 | 42.7552193 | -0.0703479 | 0.32394329 | -0.2171612 | 0.82808274 | 0.97567084 |
| OTUD6B-AS1 | 24.8047287 | 0.83142086 | 0.390141 | 2.13107792 | 0.03308272 | 0.48809081 |
| OTUD6B | 28.1271019 | 0.12973402 | 0.33167464 | 0.39114844 | 0.69568752 | 0.94395238 |
| LRRC69 | 44.9905274 | 0.07369707 | 0.3162132 | 0.23306134 | 0.81571378 | 0.97288403 |
| SLC26A7 | 48.5553734 | 0.60630371 | 0.66220925 | 0.91557723 | 0.35988868 | 0.84282379 |
| TRIQK | 68.8403923 | 0.03516239 | 0.28140161 | 0.12495447 | 0.9005596 | 0.98531749 |
| FAM92A | 36.9672551 | 0.53745721 | 0.42807023 | 1.25553512 | 0.20928457 | 0.7623519 |
| RBM12B | 48.3536639 | 0.17521788 | 0.31666857 | 0.55331629 | 0.58004685 | 0.92033309 |
| TMEM67 | 72.6244497 | -0.1989668 | 0.29875408 | -0.6659884 | 0.50541851 | 0.89723255 |
| PDP1 | 21.8414293 | -0.8661665 | 0.42583329 | -2.0340506 | 0.04194649 | 0.51213038 |
| VIRMA | 58.6469744 | -0.0295382 | 0.22264046 | -0.132672 | 0.8944528 | 0.98421511 |
| ESRP1 | 82.8469653 | -0.0673508 | 0.2708472 | -0.2486672 | 0.8036182 | 0.97169999 |
| DPY19L4 | 76.9405411 | -0.3095849 | 0.22997762 | -1.3461524 | 0.17825336 | 0.72841249 |
| INTS8 | 60.8976255 | 0.07485805 | 0.21820557 | 0.34306202 | 0.7315518 | 0.95279138 |
| NDUFAF6 | 50.4820707 | 0.16408077 | 0.31086755 | 0.52781568 | 0.59762727 | 0.9229638 |
| TP53INP1 | 20.7585572 | 0.22136311 | 0.3345369 | 0.66170013 | 0.50816342 | 0.89908875 |
| PLEKHF2 | 19.2370194 | 0.03046377 | 0.46322496 | 0.06576453 | 0.9475653 | 0.99088879 |
| C8orf37-AS1 | 50.0028384 | 1.19054749 | 0.51695254 | 2.3030112 | 0.02127821 | 0.4212027 |
| UQCRB | 131.059267 | 0.02139302 | 0.48036478 | 0.04453494 | 0.964478 | 0.99264137 |
| MTERF3 | 19.4935928 | 0.25569521 | 0.38074032 | 0.67157377 | 0.50185508 | 0.89499455 |
| PTDSS1 | 19.7609841 | -0.0124803 | 0.36043478 | -0.0346257 | 0.97237818 | 0.99431961 |
| SDC2 | 53.3144397 | -0.3794195 | 0.28358271 | -1.3379501 | 1.81E-01 | 7.34E-01 |
| CPQ | 148.465724 | 0.24560001 | 0.32401888 | 0.75798057 | 0.44846261 | 0.87314185 |
| MTDH | 241.567089 | 0.02288453 | 0.17015611 | 0.13449139 | 0.89301402 | 0.98377746 |
| LAPTM4B | 115.632003 | -0.1885589 | 0.26476446 | -0.7121762 | 0.47635568 | 0.88495683 |
| MATN2 | 18.623941 | -0.0802365 | 0.67706906 | -0.1185057 | 0.90566701 | 0.98531749 |
| RPL30 | 108.295811 | 0.19962065 | 0.38800014 | 0.51448603 | 0.60691223 | 0.92569157 |
| RIDA | 17.1068621 | 0.7359599 | 0.39233317 | 1.8758544 | 0.06067527 | 0.57774794 |
| POP1 | 13.563664 | -0.0256325 | 0.38774733 | -0.0661063 | 0.94729322 | NA |

| | | | | | | |
|---|---|---|---|---|---|---|
| NIPAL2 | 17.5632384 | -0.0780099 | 0.56480409 | -0.1381186 | 0.89014669 | 0.98289343 |
| STK3 | 479.64094 | -0.3438249 | 0.23960338 | -1.4349752 | 0.15129417 | 0.70572801 |
| AC104986.2 | 32.8965068 | 0.4453806 | 0.37130704 | 1.19949411 | 0.23033587 | 0.77249016 |
| VPS13B | 439.084108 | -0.1661789 | 0.15093803 | -1.1009744 | 0.27090782 | 0.79666376 |
| COX6C | 126.072169 | -0.1198455 | 0.23553045 | -0.5088322 | 0.61086986 | 0.92619266 |
| POLR2K | 22.4691215 | 0.2386291 | 0.47598121 | 0.50134143 | 0.61613085 | 0.92662475 |
| SPAG1 | 51.0447197 | 0.01355155 | 0.3591058 | 0.03773694 | 0.96989742 | 0.99398482 |
| RNF19A | 319.673381 | -0.9087542 | 0.41120661 | -2.2099698 | 0.02710726 | 0.45111036 |
| AP000424.1 | 40.9306355 | -1.5136347 | 0.43479044 | -3.4812971 | 4.99E-04 | 0.09270044 |
| ANKRD46 | 35.7724112 | 0.32727753 | 0.29043814 | 1.12684076 | 0.25980981 | 0.79535701 |
| PABPC1 | 158.154512 | 0.0562641 | 0.30312322 | 0.18561462 | 0.85274699 | 0.97612951 |
| YWHAZ | 231.901061 | -0.6391156 | 0.30583892 | -2.0897132 | 0.03664357 | 0.49537143 |
| ZNF706 | 63.5085782 | 0.02604137 | 0.23402759 | 0.11127477 | 0.91139846 | 0.98586235 |
| AP001207.3 | 10.9427642 | 0.82465574 | 0.58504711 | 1.40955443 | 0.15867129 | NA |
| GRHL2 | 244.691193 | 0.17690019 | 0.25555581 | 0.69221745 | 0.48880078 | 0.88920066 |
| RRM2B | 34.4156764 | 0.16564451 | 0.29270354 | 0.56591221 | 0.57145346 | 0.91791025 |
| UBR5 | 292.742078 | -0.1368401 | 0.20235742 | -0.6762299 | 0.4988947 | 0.89378207 |
| KLF10 | 24.393829 | -0.2419975 | 0.62547409 | -0.3869025 | 0.6988284 | 0.94395238 |
| AZIN1 | 68.4921513 | 0.40020245 | 0.28444361 | 1.40696588 | 0.15943751 | 0.71101785 |
| AZIN1-AS1 | 141.854038 | 0.02518588 | 0.27170196 | 0.09269671 | 0.92614451 | 0.98925753 |
| AP003354.2 | 33.1239686 | 0.30529646 | 0.36418127 | 0.83830906 | 0.40185715 | 0.85778581 |
| ATP6V1C1 | 73.0766044 | -0.3638395 | 0.25950529 | -1.4020505 | 0.16090017 | 0.71101785 |
| BAALC-AS1 | 26.6622533 | 0.31725316 | 0.31965723 | 0.99247921 | 0.32096381 | 0.82464962 |
| FZD6 | 73.7774557 | -0.0637369 | 0.25949051 | -0.2456232 | 0.80597395 | 0.97216031 |
| SLC25A32 | 35.136906 | 0.3464987 | 0.30179593 | 1.14812252 | 0.25091799 | 0.78892875 |
| DCAF13 | 19.3429625 | -0.145252 | 0.37477151 | -0.3875749 | 0.69833066 | 0.94395238 |
| LRP12 | 46.5644358 | -0.0642561 | 0.31583018 | -0.2034515 | 0.83878212 | 0.97600352 |
| ZFPM2 | 25.2563708 | 0.51869512 | 0.56248728 | 0.9221455 | 0.35645269 | 0.84192356 |
| AC027031.2 | 13.4845927 | -0.5586237 | 0.50559387 | -1.1048862 | 0.2692089 | NA |
| OXR1 | 489.124732 | -0.0013918 | 0.18668163 | -0.0074556 | 0.99405136 | 0.99921857 |
| AC090579.1 | 34.2335898 | 0.67051571 | 0.53088 | 1.26302688 | 0.20657952 | 0.76231289 |
| ANGPT1 | 177.168864 | 0.52913805 | 0.50552801 | 1.04670374 | 2.95E-01 | 8.10E-01 |
| EIF3E | 92.4461259 | 0.2753862 | 0.20968242 | 1.313349 | 0.18906537 | 0.74376063 |
| EMC2 | 75.838949 | 0.17617512 | 0.34234838 | 0.51460773 | 0.60682716 | 0.92569157 |
| NUDCD1 | 38.0808343 | -0.3158351 | 0.26464138 | -1.1934456 | 0.2326949 | 0.77562128 |
| ENY2 | 45.0187953 | 0.09143868 | 0.23839222 | 0.38356404 | 0.7013016 | 0.94440669 |
| EBAG9 | 28.7641238 | -0.2152831 | 0.32711889 | -0.6581189 | 0.51046175 | 0.90000646 |
| TRPS1 | 558.537338 | -0.3018746 | 0.28257939 | -1.0682823 | 0.28539318 | 0.80431986 |
| EIF3H | 122.272754 | 0.10987803 | 0.2170523 | 0.50622836 | 0.61269635 | 0.92647422 |
| UTP23 | 18.7155332 | -0.0062249 | 0.4898282 | -0.0127083 | 0.98986054 | 0.99814944 |
| RAD21 | 69.6410076 | -0.0188888 | 0.25714579 | -0.0734557 | 0.94144349 | 0.99043681 |
| MED30 | 11.2401855 | 0.92013881 | 0.47879547 | 1.92177844 | 0.05463364 | NA |
| EXT1 | 598.348043 | -0.533421 | 0.28224784 | -1.8899028 | 0.05877096 | 0.57258816 |
| SAMD12 | 745.21306 | 0.15387599 | 0.15642122 | 0.98372833 | 0.32524911 | 0.82567895 |
| AC023590.1 | 59.2243484 | 0.83494252 | 0.52204123 | 1.59938041 | 0.1097361 | 0.66110373 |
| SAMD12-AS1 | 49.658075 | 0.085026 | 0.29590606 | 0.28734119 | 7.74E-01 | 9.64E-01 |
| TNFRSF11B | 62.9512998 | 0.53799713 | 0.62807585 | 0.85657986 | 3.92E-01 | 0.85605713 |
| COLEC10 | 23.4579946 | 0.12485814 | 0.46271753 | 0.26983664 | 7.87E-01 | 0.96767512 |
| MAL2 | 159.092043 | -0.6227254 | 0.29618735 | -2.1024714 | 0.035512 | 0.49209205 |
| TAF2 | 95.4681139 | -0.3394384 | 0.23734906 | -1.430123 | 0.15268171 | 0.70572801 |
| DEPTOR | 502.676174 | -0.3263818 | 0.27261207 | -1.1972389 | 0.23121346 | 0.77260653 |
| AC091563.1 | 32.8640313 | 0.05674437 | 0.31321526 | 0.18116732 | 8.56E-01 | 9.76E-01 |
| COL14A1 | 21.1908831 | -0.1087209 | 0.3814681 | -0.2850066 | 0.77563908 | 0.96469469 |
| MRPL13 | 29.9282673 | 0.19382808 | 0.3909839 | 0.49574441 | 0.62007475 | 0.92798856 |

| | | | | | | |
|---|---|---|---|---|---|---|
| MTBP | 14.2451151 | -0.1507531 | 0.44544827 | -0.3384302 | 0.73503903 | NA |
| SNTB1 | 129.850807 | 0.44211807 | 0.35612341 | 1.24147434 | 0.21443057 | 0.7643572 |
| ZHX2 | 327.559485 | -0.2853785 | 0.27266447 | -1.0466288 | 0.29527083 | 0.81038435 |
| DERL1 | 37.3038998 | -0.2701631 | 0.32271195 | -0.8371649 | 0.40249991 | 0.85782098 |
| TBC1D31 | 19.9503163 | 0.58140271 | 0.39782643 | 1.46144817 | 0.14389249 | 0.6948824 |
| ATAD2 | 17.1198984 | -0.6737837 | 0.44904772 | -1.5004723 | 0.13349211 | 0.68039121 |
| WDYHV1 | 19.8179388 | 0.38099199 | 0.47908485 | 0.79524951 | 0.42646838 | 0.86239557 |
| FBXO32 | 23.0976841 | -0.0717165 | 0.32286757 | -0.2221237 | 0.82421757 | 0.97518529 |
| FAM91A1 | 81.2451366 | -0.1576063 | 0.24663144 | -0.6390358 | 0.52279961 | 0.90487731 |
| TMEM65 | 106.646069 | -0.0161503 | 0.2159994 | -0.0747703 | 0.94039749 | 0.99012269 |
| TATDN1 | 71.2423797 | -0.3404161 | 0.23480292 | -1.4497948 | 0.14711575 | 0.69749723 |
| NDUFB9 | 60.4154199 | 0.11298589 | 0.34764511 | 0.32500354 | 0.74517839 | 0.95563105 |
| MTSS1 | 280.039218 | 0.43200148 | 0.26471802 | 1.63193079 | 0.10269407 | 0.65946648 |
| SQLE | 60.0845387 | -0.2874764 | 0.52103102 | -0.5517453 | 0.58112288 | 0.92034873 |
| WASHC5 | 47.8951372 | -0.233273 | 0.29699523 | -0.7854435 | 0.43219356 | 0.86695857 |
| NSMCE2 | 395.52146 | -0.4879851 | 0.2658426 | -1.8356167 | 0.06641436 | 0.59622279 |
| TRIB1 | 36.3076299 | 0.1440047 | 0.45222357 | 0.31843695 | 0.75015351 | 0.95716475 |
| FAM84B | 71.0919336 | -1.2561922 | 0.45774329 | -2.7443159 | 0.00606371 | 0.29709153 |
| PCAT1 | 369.53378 | -0.6098686 | 0.30017776 | -2.0316914 | 0.04218491 | 0.51338127 |
| PVT1 | 24.8587583 | 1.56061909 | 0.65571877 | 2.38001284 | 0.01731203 | 0.40063153 |
| FAM49B | 427.715453 | -0.313258 | 0.2862002 | -1.0945416 | 0.2737175 | 0.79764696 |
| ASAP1 | 222.495343 | -0.0437887 | 0.33457346 | -0.1308792 | 0.89587088 | 0.98503595 |
| EFR3A | 187.907733 | -0.2861749 | 0.32222231 | -0.8881287 | 0.3744715 | 0.85092528 |
| LRRC6 | 61.3075333 | 0.24918761 | 0.34001983 | 0.73286198 | 0.46364262 | 0.87824803 |
| PHF20L1 | 96.0645984 | 0.14920792 | 0.30407499 | 0.49069446 | 0.62364256 | 0.92943979 |
| NDRG1 | 636.639778 | -1.0274706 | 0.41420985 | -2.4805556 | 0.01311778 | 0.36723795 |
| ST3GAL1 | 44.1563742 | -0.2432885 | 0.43180248 | -0.5634255 | 0.57314517 | 0.91813957 |
| ZFAT | 44.2875166 | -0.7050224 | 0.30719974 | -2.2949967 | 0.02173331 | 0.4212027 |
| AC083843.3 | 41.6545719 | -0.4750834 | 0.33875026 | -1.4024592 | 0.16077815 | 0.71101785 |
| KHDRBS3 | 133.100037 | -0.2885 | 0.34552571 | -0.8349595 | 0.40374054 | 0.85782098 |
| TRAPPC9 | 543.651551 | -0.1119579 | 0.18288044 | -0.6121916 | 0.54041098 | 0.90763121 |
| CHRAC1 | 12.7628661 | 0.16873697 | 0.5091184 | 0.33142971 | 0.74031992 | NA |
| AGO2 | 121.041696 | -0.3003009 | 0.33974624 | -0.8838975 | 0.37675154 | 0.85310518 |
| PTK2 | 770.148999 | -0.3157924 | 0.1595606 | -1.9791377 | 0.04780051 | 0.53653269 |
| DENND3 | 34.0110411 | 0.26627324 | 0.39577657 | 0.67278678 | 0.50108295 | 0.89492452 |
| SLC45A4 | 137.006131 | -0.8785702 | 0.53466386 | -1.6432198 | 0.10033747 | 0.65863153 |
| PTP4A3 | 12.602008 | 0.55665266 | 0.61040488 | 0.91194005 | 0.36180028 | NA |
| TSNARE1 | 39.2734863 | 0.01706719 | 0.27119163 | 0.06293404 | 0.94981903 | 0.99105535 |
| JRK | 14.3988461 | -0.072054 | 0.40655 | -0.1772327 | 0.85932562 | NA |
| LY6E | 51.2532284 | -2.0263912 | 1.26247694 | -1.6050916 | 0.10847364 | 0.65986731 |
| TOP1MT | 14.0566607 | -0.4387544 | 0.42144844 | -1.0410632 | 0.29784624 | NA |
| RHPN1 | 23.5846451 | -0.1877847 | 0.43400545 | -0.4326781 | 0.66524862 | 0.93860542 |
| ZC3H3 | 28.0643126 | 0.07586181 | 0.41245953 | 0.18392546 | 0.85407193 | 0.97612951 |
| GSDMD | 22.8021317 | 0.57125522 | 0.4760663 | 1.19994887 | 0.2301592 | 0.77249016 |
| NAPRT | 18.060239 | -0.4922517 | 0.44279691 | -1.1116873 | 0.26627262 | 0.79666376 |
| EEF1D | 108.273797 | -0.1827406 | 0.27760618 | -0.6582726 | 0.51036295 | 0.90000646 |
| TSTA3 | 13.5642282 | 0.3189492 | 0.50363208 | 0.63329802 | 0.52653905 | NA |
| ZNF623 | 18.3953301 | -0.0121094 | 0.37482389 | -0.032307 | 0.9742272 | 0.99431961 |
| FAM83H | 23.4387386 | -0.2540351 | 0.53414008 | -0.4755963 | 0.63436199 | 0.93251095 |
| IQANK1 | 22.7107778 | 0.30507042 | 0.54384008 | 0.56095612 | 0.57482745 | 0.91824492 |
| SCRIB | 16.9483779 | 0.41664609 | 0.49314561 | 0.84487439 | 0.39818098 | 0.85758974 |
| PUF60 | 22.3446037 | -0.4485517 | 0.47181042 | -0.9507032 | 0.34175505 | 0.83294501 |
| NRBP2 | 16.9063275 | 0.30133068 | 0.43094447 | 0.69923321 | 0.4844063 | 0.88838332 |
| PLEC | 50.3828441 | -0.215037 | 0.29465998 | -0.72978 | 0.46552468 | 0.87981788 |

| | | | | | | |
|---|---|---|---|---|---|---|
| PARP10 | 19.9648291 | -0.9102815 | 0.76676088 | -1.1871778 | 0.23515748 | 0.77830675 |
| GRINA | 33.0190857 | -0.1887411 | 0.48243566 | -0.3912253 | 0.69563067 | 0.94395238 |
| OPLAH | 9.43911344 | -0.1200279 | 0.71362409 | -0.1681948 | 0.86643001 | NA |
| GPAA1 | 26.6278184 | 1.27557359 | 0.5538592 | 2.30306472 | 0.0212752 | 0.4212027 |
| CYC1 | 41.8148175 | 0.72941717 | 0.44088434 | 1.65444108 | 0.09803793 | 0.65668818 |
| SHARPIN | 25.2739566 | -0.0958565 | 0.35064328 | -0.2733732 | 0.78456637 | 0.96734494 |
| MAF1 | 27.0081596 | 0.03870634 | 0.45966562 | 0.08420542 | 0.93289311 | 0.98960607 |
| MROH1 | 120.465974 | -0.5870283 | 0.47816909 | -1.2276585 | 0.2195752 | 0.76680432 |
| HSF1 | 63.7212598 | -0.1643307 | 0.28333993 | -0.5799773 | 0.5619299 | 0.91624774 |
| DGAT1 | 18.1056883 | 0.48168899 | 0.44462355 | 1.08336364 | 0.27864705 | 0.80218978 |
| CPSF1 | 15.1335917 | -0.4525989 | 0.5370313 | -0.8427794 | 0.39935182 | NA |
| VPS28 | 59.5184588 | 0.4523715 | 0.32865609 | 1.37642815 | 0.16868912 | 0.72160574 |
| CYHR1 | 39.845375 | 0.49258266 | 0.30098606 | 1.63656303 | 0.10172181 | 0.65863153 |
| KIFC2 | 17.6220086 | -0.0850082 | 0.42632096 | -0.1993995 | 0.84195025 | 0.97600352 |
| PPP1R16A | 33.5013211 | -0.0054971 | 0.33896473 | -0.0162172 | 0.9870611 | 0.99803728 |
| ARHGAP39 | 22.5070416 | 0.08735078 | 0.36492506 | 0.23936634 | 0.81082153 | 0.97257655 |
| ZNF251 | 65.2696843 | -0.1267953 | 0.24089934 | -0.5263413 | 0.59865109 | 0.92296674 |
| RPL8 | 211.050892 | 0.2574953 | 0.39165762 | 0.65745 | 0.51089161 | 0.90024126 |
| ZNF7 | 16.7415461 | 0.2047484 | 0.42487221 | 0.48190584 | 0.62987284 | 0.9318063 |
| ZNF250 | 16.2198671 | 0.20820329 | 0.51387922 | 0.40515997 | 0.68535993 | 0.94205398 |
| ZNF16 | 15.1329841 | -0.2347208 | 0.46503323 | -0.5047399 | 0.61374151 | NA |
| C8orf33 | 14.7125323 | 0.59131644 | 0.46619599 | 1.26838596 | 0.20466015 | NA |
| WASHC1 | 124.011857 | -0.0546865 | 0.27163477 | -0.2013236 | 8.40E-01 | 0.97600352 |
| CBWD1 | 28.8412946 | -0.2719933 | 0.35163362 | -0.7735134 | 0.4392186 | 0.86879768 |
| DOCK8 | 20.5159274 | -0.3095942 | 0.50309922 | -0.6153741 | 0.53830771 | 0.90763121 |
| KANK1 | 612.944627 | 0.16704525 | 0.24724486 | 0.67562681 | 4.99E-01 | 8.94E-01 |
| SMARCA2 | 304.886148 | 0.15439305 | 0.38309254 | 0.40301763 | 0.68693526 | 0.94205398 |
| VLDLR-AS1 | 27.4008938 | -0.5125016 | 0.67347122 | -0.7609852 | 0.44666593 | 0.87231627 |
| VLDLR | 33.1223518 | -0.523718 | 0.32250077 | -1.6239279 | 1.04E-01 | 6.59E-01 |
| PUM3 | 223.516888 | -0.0519423 | 0.28517322 | -0.1821431 | 0.85547045 | 0.97622508 |
| RFX3 | 211.249077 | -0.0767162 | 0.20436104 | -0.3753954 | 0.7073664 | 0.94558664 |
| RFX3-AS1 | 178.474041 | 0.29002132 | 0.41091586 | 0.70579247 | 0.4803172 | 0.88660781 |
| GLIS3 | 1583.52597 | -0.577923 | 0.4028702 | -1.434514 | 0.15142563 | 0.70572801 |
| SLC1A1 | 49.7954987 | 0.47529921 | 0.31829676 | 1.49325809 | 0.13536964 | 0.6835006 |
| SPATA6L | 108.43936 | 0.18059972 | 0.41679979 | 0.4333009 | 0.66479618 | 0.93860542 |
| CDC37L1 | 54.9404488 | -0.0500491 | 0.2768423 | -0.1807857 | 0.85653579 | 0.97646931 |
| AK3 | 292.684422 | 0.09472089 | 0.24141025 | 0.39236482 | 6.95E-01 | 9.44E-01 |
| RCL1 | 91.1052177 | -0.0474222 | 0.34476699 | -0.1375486 | 0.89059722 | 0.98296447 |
| JAK2 | 121.101327 | -0.1482297 | 0.1897237 | -0.7812927 | 0.43463036 | 0.86729771 |
| PLGRKT | 62.8013461 | -0.0806727 | 0.25128953 | -0.3210349 | 0.74818394 | 0.95563105 |
| RIC1 | 194.838704 | -0.3657845 | 0.29336915 | -1.2468405 | 0.212456 | 0.7643572 |
| ERMP1 | 207.017667 | -0.0095026 | 0.24459092 | -0.0388508 | 9.69E-01 | 0.99379133 |
| KIAA2026 | 215.850201 | -0.1771569 | 0.27143339 | -0.6526718 | 0.51396791 | 0.90156229 |
| RANBP6 | 18.3088221 | 0.20827512 | 0.38312206 | 0.54362603 | 0.58669885 | 0.92076087 |
| UHRF2 | 107.068498 | -0.4758314 | 0.24941655 | -1.9077779 | 0.05641993 | 0.56976762 |
| GLDC | 256.279775 | 0.08033371 | 0.41702374 | 0.19263582 | 8.47E-01 | 0.97612951 |
| AL162411.1 | 26.8758228 | 0.39050598 | 0.35632325 | 1.09593179 | 0.27310863 | 0.79726027 |
| KDM4C | 399.537983 | -0.3769542 | 0.17350693 | -2.1725602 | 0.02981344 | 0.46381645 |
| DMAC1 | 25.3396303 | 0.89131408 | 0.3545593 | 2.51386463 | 0.01194163 | 0.35710117 |
| PTPRD | 75.5572951 | -0.3276416 | 0.73297216 | -0.4470041 | 0.65487208 | 0.93548446 |
| LURAP1L | 75.3649063 | -1.2659903 | 0.71635163 | -1.767275 | 0.07718219 | 0.61329066 |
| MPDZ | 97.6911398 | -0.2602426 | 0.25342375 | -1.0269068 | 0.30446435 | 0.81568231 |
| NFIB | 577.058662 | -0.6035105 | 0.33250842 | -1.8150231 | 0.06952036 | 0.60275677 |
| ZDHHC21 | 261.712602 | -0.5931815 | 0.30472831 | -1.9465915 | 0.05158373 | 0.55289583 |

| | | | | | | |
|---|---|---|---|---|---|---|
| FREM1 | 361.716736 | 0.49806282 | 0.42796272 | 1.16379955 | 2.45E-01 | 7.86E-01 |
| TTC39B | 87.5209176 | -0.5632662 | 0.28679348 | -1.9640132 | 0.04952856 | 0.54320302 |
| SNAPC3 | 71.9384604 | -0.087361 | 0.21446146 | -0.4073507 | 0.68375046 | 0.94205398 |
| PSIP1 | 59.5971226 | 0.05099067 | 0.27444974 | 0.18579237 | 0.85260759 | 0.97612951 |
| CCDC171 | 165.540406 | 0.2248002 | 0.28782583 | 0.78102857 | 0.43478569 | 0.86729771 |
| BNC2 | 93.8534246 | -0.9379395 | 0.56187534 | -1.6693018 | 0.09505758 | 0.64839204 |
| CNTLN | 64.792099 | 0.042441 | 0.35100454 | 0.12091296 | 0.90375998 | 0.98531749 |
| SH3GL2 | 73.3160801 | -0.6113386 | 0.48759297 | -1.2537888 | 0.20991879 | 0.76344019 |
| RRAGA | 22.2027764 | -0.1439719 | 0.60648943 | -0.2373857 | 0.81235759 | 0.97257655 |
| HAUS6 | 15.535494 | -0.0457783 | 0.44622562 | -0.10259 | 0.91828838 | NA |
| PLIN2 | 43.2996522 | -1.3665607 | 0.56356671 | -2.4248428 | 0.01531502 | 0.39253938 |
| DENND4C | 265.710197 | -0.3021062 | 0.20865848 | -1.4478501 | 0.14765897 | 0.69842926 |
| RPS6 | 214.833606 | 0.17663657 | 0.42539737 | 0.41522722 | 0.67797558 | 0.94093842 |
| MLLT3 | 713.959011 | -0.5451789 | 0.19526457 | -2.7920012 | 0.00523832 | 0.28057141 |
| FOCAD | 228.015536 | -0.1343695 | 0.21161406 | -0.6349743 | 0.5254452 | 0.90487731 |
| KLHL9 | 23.2904331 | 0.25598837 | 0.53196328 | 0.48121436 | 0.63036415 | 0.9320416 |
| LINC01239 | 21.5162501 | 0.96862359 | 0.49090638 | 1.97313301 | 0.04848041 | 0.5391188 |
| CAAP1 | 154.695542 | -0.3070808 | 0.23312615 | -1.3172302 | 0.18776146 | 0.7419 |
| PLAA | 42.5442396 | 0.07872813 | 0.31918343 | 0.24665481 | 0.80517538 | 0.97195485 |
| IFT74 | 112.634315 | 0.1220083 | 0.26257718 | 0.46465689 | 0.64217719 | 0.93328692 |
| LRRC19 | 19.1352332 | 0.7898281 | 0.47802551 | 1.65227186 | 0.09847914 | 0.65695163 |
| MOB3B | 185.195156 | 0.17610673 | 0.32407642 | 0.54341113 | 0.58684676 | 0.92076087 |
| C9orf72 | 36.5472604 | 0.32062397 | 0.37922453 | 0.84547266 | 0.39784699 | 0.85758974 |
| LINGO2 | 102.467186 | 1.39083068 | 0.66059639 | 2.10541672 | 0.03525505 | 0.49209205 |
| ACO1 | 47.5210476 | 0.15731701 | 0.31873384 | 0.4935686 | 0.62161088 | 0.92868594 |
| DDX58 | 81.0079734 | -2.4170602 | 0.82367953 | -2.9344668 | 0.00334121 | 0.23150356 |
| TOPORS | 25.5946068 | 0.26567457 | 0.31000852 | 0.85699117 | 0.39144976 | 0.85605713 |
| NDUFB6 | 46.7645112 | -0.096581 | 0.3045475 | -0.3171294 | 0.7511454 | 0.95741436 |
| APTX | 69.0154783 | 0.5523906 | 0.31716045 | 1.74167558 | 0.08156523 | 0.61925304 |
| DNAJA1 | 61.2564197 | 0.40937268 | 0.24802605 | 1.65052291 | 0.09883603 | 0.65744285 |
| SMU1 | 38.9299839 | -0.0414464 | 0.28430415 | -0.1457818 | 0.88409364 | 0.98154805 |
| B4GALT1 | 188.992274 | -0.2344309 | 0.25582085 | -0.9163868 | 0.35946404 | 0.84204868 |
| BAG1 | 61.567315 | 0.09983275 | 0.36082759 | 0.27667716 | 0.78202801 | 0.96652357 |
| CHMP5 | 40.6364973 | -0.4273035 | 0.41840911 | -1.0212577 | 0.30713237 | 0.81568231 |
| NFX1 | 156.032084 | -0.7498651 | 0.26171255 | -2.8652241 | 0.00416714 | 0.25676573 |
| AQP3 | 186.839873 | -1.6154634 | 0.69387269 | -2.3281841 | 0.01990233 | 0.41544117 |
| UBE2R2 | 217.201037 | -0.4837695 | 0.29178553 | -1.6579627 | 0.09732499 | 0.65521765 |
| UBAP2 | 147.295178 | -0.3501213 | 0.25736157 | -1.3604258 | 0.17369521 | 0.72794035 |
| DCAF12 | 33.0078769 | -0.3487459 | 0.29876462 | -1.1672931 | 0.24309201 | 0.78495944 |
| UBAP1 | 166.29416 | -0.3835402 | 0.26524779 | -1.4459694 | 0.14818578 | 0.69945736 |
| NUDT2 | 11.7167789 | -0.0021253 | 0.44633128 | -0.0047618 | 0.99620063 | NA |
| FAM219A | 41.9917239 | -0.6537621 | 0.37280863 | -1.7536132 | 0.07949681 | 0.61431194 |
| DNAI1 | 16.5637463 | 0.51014394 | 0.58529839 | 0.87159635 | 0.38342862 | 0.85511218 |
| RPP25L | 12.6412227 | -0.1432705 | 0.51685249 | -0.277198 | 0.78162806 | NA |
| DCTN3 | 14.6893993 | 1.13238409 | 0.60040184 | 1.88604367 | 0.05928907 | NA |
| VCP | 73.3950605 | 0.20893993 | 0.24707544 | 0.84565234 | 0.39774672 | 0.85758974 |
| PIGO | 10.263438 | 0.63673163 | 0.52967753 | 1.20211184 | 0.2293202 | NA |
| STOML2 | 25.4381713 | 0.60599274 | 0.36954582 | 1.63983114 | 0.10104028 | 0.65863153 |
| FAM214B | 10.8671276 | -0.2314727 | 0.43167385 | -0.5362212 | 0.5918057 | NA |
| UNC13B | 183.991599 | -0.3120309 | 0.28734993 | -1.0858918 | 0.27752686 | 0.80176531 |
| RUSC2 | 84.3660393 | -0.2236574 | 0.32830093 | -0.6812574 | 0.49570865 | 0.89230917 |
| TESK1 | 14.4586934 | -0.4679739 | 0.41982722 | -1.1146822 | 0.26498663 | NA |
| CCDC107 | 21.3762535 | -0.4769738 | 0.41384411 | -1.1525446 | 0.24909736 | 0.78763828 |
| TLN1 | 43.1967608 | -0.208722 | 0.30296876 | -0.6889224 | 0.49087209 | 0.88987842 |

| | | | | | | |
|---|---|---|---|---|---|---|
| CREB3 | 12.7603062 | 0.46704277 | 0.49646723 | 0.94073232 | 0.34684205 | NA |
| GBA2 | 51.3947782 | -0.1951037 | 0.26351994 | -0.7403755 | 0.45907217 | 0.87574928 |
| RGP1 | 25.2994521 | -0.1458769 | 0.31851279 | -0.4579938 | 0.64695691 | 0.93328692 |
| AL133410.1 | 25.308547 | -0.1257397 | 0.40248008 | -0.3124123 | 0.75472719 | 0.95832511 |
| HINT2 | 9.80863302 | -0.1370249 | 0.69711299 | -0.1965605 | 0.84417148 | NA |
| TMEM8B | 26.283306 | -0.0736284 | 0.42685103 | -0.1724921 | 0.86305068 | 0.97788853 |
| RECK | 26.8113989 | -0.0556382 | 0.32985114 | -0.1686766 | 0.866051 | 0.97788853 |
| CLTA | 124.110859 | -0.1695325 | 0.30096324 | -0.5632997 | 0.57323079 | 0.91813957 |
| GNE | 34.5350853 | -0.5356273 | 0.34450531 | -1.5547722 | 0.12000033 | 0.6668162 |
| RNF38 | 217.823656 | -0.3374905 | 0.1958072 | -1.7235858 | 0.08478265 | 0.62880224 |
| EBLN3P | 41.3763545 | 0.33084838 | 0.26207207 | 1.26243282 | 0.20679309 | 0.76231289 |
| ZCCHC7 | 415.095479 | -0.3711233 | 0.15299084 | -2.425788 | 0.01527519 | 0.39253938 |
| GRHPR | 28.1623913 | 0.58066464 | 0.40713755 | 1.42621245 | 0.15380703 | 0.70572801 |
| ZBTB5 | 22.9618234 | -0.144028 | 0.40280654 | -0.3575612 | 0.72067175 | 0.95048937 |
| AL138752.2 | 55.2509738 | -0.1957782 | 0.31495674 | -0.6216035 | 0.53420262 | 0.90763121 |
| FRMPD1 | 14.0316131 | 1.3892313 | 0.66823668 | 2.07895096 | 0.03762186 | NA |
| TRMT10B | 45.6045227 | 0.37040469 | 0.37093066 | 0.99858202 | 0.31799721 | 0.82354084 |
| DCAF10 | 139.377701 | 0.08669005 | 0.22094654 | 0.39235757 | 0.69479403 | 0.94395238 |
| FAM95C | 21.2083209 | 0.41218431 | 0.38333939 | 1.07524643 | 0.28226441 | 0.80431986 |
| ANKRD18A | 19.7117079 | 0.661835 | 0.53845484 | 1.22913744 | 0.21902028 | 0.76666749 |
| GLIDR | 34.661622 | -0.0069949 | 0.36257208 | -0.0192924 | 0.98460788 | 0.99720578 |
| BMS1P14 | 25.9751407 | -0.0507916 | 0.54493225 | -0.0932072 | 0.92573897 | 0.98925753 |
| CBWD6 | 14.4365833 | 0.20919267 | 0.51931663 | 0.40282297 | 0.68707846 | NA |
| CNTNAP3B | 82.2215497 | 0.07832024 | 0.36098409 | 0.21696313 | 0.82823708 | 0.97567084 |
| AL391987.2 | 10.9763371 | 0.38114654 | 0.44612752 | 0.85434438 | 0.3929142 | NA |
| AL512625.3 | 13.2053559 | 0.8694183 | 0.60945764 | 1.42654427 | 0.1537113 | NA |
| CBWD5 | 38.2298712 | -0.5513674 | 0.36457631 | -1.5123511 | 0.13044456 | 0.67375513 |
| ZNF658 | 14.4722534 | 0.02066326 | 0.56184234 | 0.03677768 | 0.97066227 | NA |
| PGM5 | 82.8983848 | 0.04523904 | 0.39672488 | 0.11403125 | 0.90921302 | 0.9854228 |
| AL161457.2 | 14.0491507 | -0.2754398 | 0.42551815 | -0.6473044 | 0.51743493 | NA |
| PIP5K1B | 127.269639 | 0.30552359 | 0.50488823 | 0.60513115 | 0.54509186 | 0.90999223 |
| FAM189A2 | 20.2764718 | -0.0589364 | 0.42696672 | -0.1380351 | 0.8902127 | 0.98289343 |
| PTAR1 | 102.396054 | 0.16064791 | 0.2230667 | 0.72017881 | 0.47141491 | 0.88267439 |
| MAMDC2-AS | 55.2191877 | 0.09800728 | 0.35213916 | 0.27831973 | 0.78076693 | 0.96652357 |
| MAMDC2 | 100.96613 | 0.68208307 | 0.43086124 | 1.583069 | 0.11340574 | 0.6656824 |
| SMC5-AS1 | 82.0295036 | 0.21365107 | 0.23516698 | 0.90850796 | 0.36360991 | 0.84653222 |
| SMC5 | 192.355198 | -0.2274867 | 0.21544113 | -1.0559112 | 0.29100879 | 0.80640885 |
| KLF9 | 43.8748803 | 0.51210667 | 0.30773055 | 1.66413987 | 0.09608448 | 0.65104726 |
| TRPM3 | 922.631488 | 1.08122241 | 0.37859556 | 2.85587719 | 4.29E-03 | 2.60E-01 |
| CEMIP2 | 174.500886 | 0.15868951 | 0.30308289 | 0.52358451 | 6.01E-01 | 0.92352528 |
| ABHD17B | 25.8663696 | 0.46302935 | 0.37645936 | 1.22995838 | 0.21871269 | 0.76666749 |
| C9orf85 | 58.5136768 | 0.07027735 | 0.27463272 | 0.25589576 | 0.79803133 | 0.97019967 |
| GDA | 42.3154961 | 1.64303042 | 0.67912866 | 2.41932128 | 0.0155495 | 0.39401695 |
| ZFAND5 | 191.211504 | -0.0249376 | 0.40306174 | -0.0618705 | 0.95066598 | 0.99105535 |
| ALDH1A1 | 87.2765217 | 1.3797354 | 0.45062974 | 3.06179391 | 0.00220015 | 0.18826595 |
| TRPM6 | 278.74792 | -0.9645441 | 0.6688006 | -1.4421998 | 1.49E-01 | 0.70138697 |
| C9orf41-AS1 | 12.496636 | 0.64170603 | 0.59648549 | 1.07581164 | 0.28201151 | NA |
| CARNMT1 | 106.863456 | 0.41782348 | 0.21723529 | 1.92336831 | 5.44E-02 | 5.65E-01 |
| NMRK1 | 47.8463724 | 0.21659824 | 0.52268405 | 0.41439611 | 0.67858404 | 0.9413518 |
| OSTF1 | 37.4690003 | 0.0237559 | 0.38804504 | 0.06121944 | 0.95118445 | 0.99105535 |
| PCSK5 | 73.9105272 | 0.06792782 | 0.65657762 | 0.10345742 | 0.91759995 | 0.98722519 |
| RFK | 18.4179886 | 0.60263974 | 0.48619167 | 1.23951064 | 0.21515645 | 0.7643572 |
| VPS13A | 173.778828 | -0.1341168 | 0.21832785 | -0.6142909 | 0.5390231 | 0.90763121 |
| GNAQ | 593.215394 | -0.2291488 | 0.1787269 | -1.2821171 | 0.19980157 | 0.75605362 |

| | | | | | | |
|---|---|---|---|---|---|---|
| CEP78 | 15.1330525 | -0.3464231 | 0.37525624 | -0.923164 | 0.35592173 | NA |
| TLE4 | 20.4002551 | -0.0718514 | 0.40824467 | -0.1760009 | 0.86029327 | 0.97718459 |
| TLE1 | 21.7083523 | 0.53648362 | 0.40233354 | 1.33343002 | 0.18239073 | 0.73398872 |
| AL591368.1 | 10.5259782 | -0.109164 | 0.85456228 | -0.1277426 | 0.89835268 | NA |
| RASEF | 50.7860243 | -0.0094558 | 0.31786599 | -0.0297479 | 0.97626813 | 0.99440179 |
| FRMD3 | 148.283267 | -0.4828993 | 0.45387608 | -1.0639453 | 0.28735348 | 0.80523851 |
| IDNK | 14.6946696 | 0.18113818 | 0.42543089 | 0.42577581 | 0.67027122 | NA |
| UBQLN1 | 102.859092 | -0.0818729 | 0.20300456 | -0.4033059 | 0.68672319 | 0.94205398 |
| GKAP1 | 51.8749012 | 0.20359166 | 0.27623166 | 0.7370323 | 0.4611027 | 0.87647064 |
| KIF27 | 65.4585016 | 0.04344821 | 0.29871932 | 0.14544827 | 0.88435696 | 0.98154805 |
| HNRNPK | 174.797143 | -0.1706961 | 0.26974925 | -0.6327953 | 0.52686734 | 0.90487731 |
| RMI1 | 15.4985754 | 0.67077505 | 0.37043195 | 1.81079159 | 0.07017312 | NA |
| AGTPBP1 | 98.4976735 | 0.01922621 | 0.20626815 | 0.09320977 | 0.92573691 | 0.98925753 |
| NAA35 | 120.620402 | -0.4676349 | 0.21838404 | -2.141342 | 0.03224647 | 0.48175799 |
| GOLM1 | 55.6426822 | -0.0265783 | 0.26241376 | -0.1012841 | 0.91932491 | 0.9875219 |
| ISCA1 | 31.5970431 | 0.1868715 | 0.30447195 | 0.61375605 | 0.53937654 | 0.90763121 |
| TUT7 | 86.9027566 | -0.1518479 | 0.20311236 | -0.7476052 | 0.45469833 | 0.87378486 |
| DAPK1 | 369.976633 | -0.7813298 | 0.33406723 | -2.33884 | 0.01934372 | 0.41544117 |
| CTSL | 50.1416358 | 0.08267143 | 0.35843331 | 0.2306466 | 0.81758936 | 0.97288403 |
| SPIN1 | 292.112408 | -0.0262516 | 0.16674088 | -0.1574395 | 0.87489846 | 0.97918624 |
| SHC3 | 16.7163298 | -1.2666248 | 0.71730604 | -1.765808 | 0.07742806 | 0.61342702 |
| SECISBP2 | 141.840311 | -0.532692 | 0.21658292 | -2.4595293 | 0.01391193 | 0.37638271 |
| SEMA4D | 147.154386 | -0.5129394 | 0.3159409 | -1.6235297 | 0.10447622 | 0.65946648 |
| LINC01508 | 57.5416954 | 0.634276 | 0.53296523 | 1.19008889 | 0.23401146 | 0.7767179 |
| SYK | 122.54838 | 0.18774538 | 0.24983867 | 0.75146646 | 0.45237198 | 0.87346041 |
| AL158071.4 | 40.729314 | -0.8939758 | 0.36244301 | -2.4665278 | 0.01364301 | 0.37407336 |
| AUH | 299.493761 | -0.3232687 | 0.22821905 | -1.4164844 | 0.15663372 | 0.70821492 |
| NFIL3 | 25.5811415 | -0.538609 | 0.69285907 | -0.7773717 | 0.43693953 | 0.86759679 |
| ROR2 | 107.952294 | 0.19297046 | 0.31494475 | 0.61271211 | 0.54006671 | 0.90763121 |
| SPTLC1 | 75.5858029 | -0.140379 | 0.20980059 | -0.6691066 | 0.50342748 | 0.89612329 |
| IARS | 75.0280248 | -0.3570544 | 0.36072169 | -0.9898333 | 0.32225559 | 0.82558302 |
| NOL8 | 28.7183884 | -0.1395628 | 0.36175028 | -0.3857987 | 0.69964573 | 0.94419257 |
| CENPP | 55.6584168 | -0.7954122 | 0.36473775 | -2.1807784 | 0.02919981 | 0.46307641 |
| IPPK | 23.5834228 | -0.3309452 | 0.39562265 | -0.8365173 | 0.40286397 | 0.85782098 |
| BICD2 | 28.5236867 | 0.43485434 | 0.3599577 | 1.20807067 | 0.22702008 | 0.77185154 |
| ZNF484 | 13.7069981 | -0.5356179 | 0.40117373 | -1.3351271 | 0.18183474 | NA |
| FGD3 | 51.1432026 | 0.55315979 | 0.58641499 | 0.94329067 | 3.46E-01 | 0.83564825 |
| CARD19 | 22.3199949 | -0.3915411 | 0.47960089 | -0.8163895 | 0.4142774 | 0.85867915 |
| NINJ1 | 23.7234822 | 0.73809577 | 0.33939559 | 2.1747359 | 0.02964991 | 0.46381645 |
| FAM120AOS | 19.3846801 | 0.17001447 | 0.40125925 | 0.42370231 | 0.67178293 | 0.93878847 |
| FAM120A | 170.565217 | 0.06589617 | 0.17696218 | 0.3723743 | 0.70961419 | 0.94577129 |
| PHF2 | 50.1327341 | 0.01469295 | 0.26282256 | 0.05590445 | 0.95541792 | 0.99148072 |
| PTPDC1 | 46.2098246 | 0.04412567 | 0.2932026 | 0.15049549 | 0.88037371 | 0.98093344 |
| AL158152.2 | 63.9345695 | 0.22814955 | 0.30799689 | 0.74075277 | 0.45884336 | 0.87565012 |
| ZNF169 | 30.5870137 | 0.02819872 | 0.29516703 | 0.09553478 | 0.92389007 | 0.98925753 |
| MFSD14B | 62.024186 | -0.0994201 | 0.29006969 | -0.3427457 | 0.73178975 | 0.95279138 |
| FBP1 | 23.067904 | 0.3528246 | 0.53637604 | 0.65779337 | 0.51067092 | 0.90019019 |
| C9orf3 | 578.46971 | -0.4171707 | 0.21459578 | -1.9439838 | 0.0518974 | 0.55403388 |
| FANCC | 143.069115 | -0.1034926 | 0.2575215 | -0.4018793 | 0.68777286 | 0.94205398 |
| PTCH1 | 32.4385686 | 0.01374869 | 0.36496449 | 0.0376713 | 0.96994976 | 0.99398482 |
| LINC00476 | 74.1338184 | -0.1968162 | 0.44162285 | -0.4456658 | 0.65583866 | 0.93567213 |
| ERCC6L2 | 115.249687 | -0.1519535 | 0.23333955 | -0.651212 | 0.51490963 | 0.90156229 |
| SLC35D2 | 22.6619526 | 0.48001997 | 0.32917707 | 1.45824243 | 0.14477373 | 0.69512617 |
| HABP4 | 58.0509205 | -0.2441933 | 0.40269912 | -0.6063914 | 0.54425486 | 0.90965439 |

| | | | | | | |
|---|---|---|---|---|---|---|
| CDC14B | 142.027989 | 0.62506542 | 0.33793363 | 1.8496692 | 0.06436124 | 0.58882451 |
| AL589843.1 | 22.2445823 | 0.0637745 | 0.36747814 | 0.17354639 | 0.86222197 | 0.97767895 |
| ZNF510 | 16.4322269 | 0.30252283 | 0.40457063 | 0.74776271 | 0.4546033 | 0.87378486 |
| ZNF782 | 58.3399983 | 0.04507112 | 0.28854188 | 0.15620306 | 0.87587297 | 0.97918624 |
| MFSD14C | 144.576494 | -0.2111968 | 0.2209718 | -0.9557634 | 0.33919177 | 0.83066816 |
| CCDC180 | 14.3563025 | 0.75236273 | 0.4762253 | 1.57984621 | 0.11414209 | NA |
| TDRD7 | 30.2657669 | -0.1894896 | 0.47582931 | -0.3982303 | 0.69046045 | 0.94276429 |
| TSTD2 | 32.2909944 | -0.086528 | 0.27489296 | -0.31477 | 0.75293632 | 0.95815628 |
| NCBP1 | 38.9532281 | -0.0172886 | 0.32125854 | -0.0538152 | 0.95708239 | 0.99170927 |
| XPA | 40.2828057 | 0.4340835 | 0.28521472 | 1.52195334 | 0.12802077 | 0.66929384 |
| ANP32B | 58.0611459 | -0.0032508 | 0.27269888 | -0.0119208 | 0.99048877 | 0.99814944 |
| NANS | 14.4036844 | -0.2796094 | 0.37747472 | -0.7407367 | 0.45885309 | NA |
| TRIM14 | 20.1428878 | -0.8316827 | 0.95368336 | -0.8720743 | 0.38316785 | 0.85510589 |
| ANKS6 | 35.1734256 | 0.60968951 | 0.35517335 | 1.71659703 | 0.08605282 | 0.63376931 |
| TGFBR1 | 89.2546209 | -0.3172011 | 0.55874304 | -0.5677049 | 0.57023539 | 0.91778542 |
| ALG2 | 21.4910327 | 0.08680321 | 0.40195464 | 0.21595275 | 0.82902458 | 0.97567084 |
| SEC61B | 62.3108817 | 0.17834627 | 0.37340507 | 0.47762144 | 0.63291967 | 0.93251095 |
| AL359710.1 | 72.0958106 | 0.71137867 | 0.40318272 | 1.76440763 | 0.07766337 | 0.61367304 |
| STX17 | 62.2315906 | -0.2142139 | 0.30274009 | -0.7075837 | 0.47920383 | 0.88622003 |
| ERP44 | 104.48139 | -0.1891657 | 0.24452035 | -0.7736195 | 0.43915583 | 0.86879768 |
| INVS | 204.513273 | 0.38356978 | 0.24410467 | 1.57133323 | 0.11610527 | 0.6668162 |
| TEX10 | 31.3865749 | -0.3699093 | 0.29651557 | -1.2475207 | 0.21220665 | 0.7643572 |
| PLPPR1 | 44.5858739 | 0.28444929 | 0.50276148 | 0.56577383 | 5.72E-01 | 9.18E-01 |
| MRPL50 | 14.2854871 | 0.72574503 | 0.45316143 | 1.60151543 | 0.10926281 | NA |
| ZNF189 | 24.159816 | -0.1097522 | 0.35772622 | -0.306805 | 0.7589918 | 0.95903525 |
| ALDOB | 40.4037311 | 0.06299065 | 0.88226856 | 0.07139623 | 0.94308241 | 0.99058259 |
| RNF20 | 19.8674873 | 0.53588402 | 0.37435796 | 1.43147489 | 0.15229416 | 0.70572801 |
| SMC2 | 16.6745007 | 0.82744636 | 0.45739092 | 1.80905724 | 0.07044211 | 0.60436809 |
| NIPSNAP3A | 18.9139088 | -0.0442607 | 0.38165529 | -0.1159703 | 0.90767611 | 0.9854228 |
| ABCA1 | 36.6015098 | -0.1986916 | 0.36193172 | -0.5489753 | 0.58302242 | 0.92034873 |
| SLC44A1 | 241.639848 | -1.2852781 | 0.36908933 | -3.4822954 | 0.00049713 | 0.09270044 |
| FSD1L | 85.5572992 | 0.09304846 | 0.23149169 | 0.40195163 | 0.68771962 | 0.94205398 |
| FKTN | 57.1953865 | 0.07242714 | 0.27446228 | 0.26388741 | 0.79186668 | 0.96925674 |
| TMEM38B | 36.2415927 | -0.4755013 | 0.58456587 | -0.8134264 | 0.41597361 | 0.85867915 |
| LINC01505 | 55.8900842 | -0.9241272 | 0.71751635 | -1.2879528 | 0.1977624 | 0.75559912 |
| ZNF462 | 73.8117799 | 0.8959382 | 0.3201976 | 2.79807909 | 0.00514075 | 0.28057141 |
| RAD23B | 138.1942 | -0.2719267 | 0.20889867 | -1.3017159 | 0.19301353 | 0.75061775 |
| ELP1 | 45.2352781 | 0.1918042 | 0.25694124 | 0.74649049 | 0.45537117 | 0.8739446 |
| FAM206A | 14.9722269 | 0.30961448 | 0.40466231 | 0.76511817 | 0.44420118 | NA |
| CTNNAL1 | 81.4681379 | -0.3572053 | 0.41315473 | -0.8645798 | 0.38726944 | 0.85605713 |
| TMEM245 | 234.018633 | -0.2197451 | 0.21963606 | -1.0004965 | 0.31707028 | 0.82275575 |
| EPB41L4B | 54.9660362 | 0.39118178 | 0.28323063 | 1.38114224 | 0.16723523 | 0.71944343 |
| PTPN3 | 335.974119 | -0.5967844 | 0.22549452 | -2.6465584 | 0.00813155 | 0.30976355 |
| PALM2 | 32.3696431 | -0.2215026 | 0.46382376 | -0.4775576 | 0.63296511 | 0.93251095 |
| TXN | 65.5065453 | -0.555359 | 0.25547878 | -2.1737971 | 0.02972038 | 0.46381645 |
| ECPAS | 180.666669 | -0.4030605 | 0.25008863 | -1.6116705 | 0.10703366 | 0.65986731 |
| ZNF483 | 70.4976075 | 0.0763041 | 0.36926911 | 0.20663547 | 0.83629455 | 0.97600352 |
| PTGR1 | 299.050599 | -0.1790173 | 0.41580345 | -0.4305336 | 0.66680757 | 0.93860542 |
| C9orf84 | 44.4086758 | 0.57045187 | 0.52707974 | 1.0822876 | 0.27912475 | 0.80219184 |
| UGCG | 231.755077 | -0.6424173 | 0.27566279 | -2.3304463 | 1.98E-02 | 4.15E-01 |
| SUSD1 | 192.99957 | -0.3211313 | 0.3210238 | -1.0003349 | 0.31714846 | 0.82275575 |
| PTBP3 | 608.242271 | -0.2345664 | 0.33334972 | -0.7036645 | 0.48164173 | 0.88703194 |
| HSDL2 | 122.057681 | 0.06398604 | 0.26833362 | 0.23845705 | 0.81152662 | 0.97257655 |
| C9orf147 | 20.8404272 | 0.34954885 | 0.47393231 | 0.73755015 | 0.46078786 | 0.87626776 |

| | | | | | | |
|---|---|---|---|---|---|---|
| KIAA1958 | 328.150078 | 0.02782199 | 0.20749869 | 0.13408274 | 0.89333715 | 0.98377746 |
| INIP | 45.5226935 | -0.0772024 | 0.2856786 | -0.270242 | 0.78697408 | 0.96767512 |
| SNX30 | 43.0729415 | -0.3173691 | 0.27988479 | -1.1339278 | 0.25682484 | 0.79456034 |
| FKBP15 | 41.5438781 | 0.08327888 | 0.24783719 | 0.33602251 | 0.73685389 | 0.95387069 |
| SLC31A1 | 36.3393562 | 0.29567481 | 0.35870203 | 0.82429088 | 0.4097743 | 0.85867915 |
| CDC26 | 14.5575588 | 0.53928221 | 0.44761602 | 1.20478756 | 0.22828533 | NA |
| PRPF4 | 18.1840051 | -0.0232855 | 0.51780953 | -0.0449692 | 0.96413186 | 0.99264137 |
| WDR31 | 16.0404111 | 0.66168671 | 0.53110357 | 1.24587134 | 0.21281163 | NA |
| BSPRY | 28.9077059 | 0.15715031 | 0.35260834 | 0.44567951 | 0.65582877 | 0.93567213 |
| HDHD3 | 33.3942316 | 0.20448158 | 0.39171078 | 0.52202183 | 0.60165513 | 0.92389788 |
| ALAD | 15.6486588 | 0.62396186 | 0.46032624 | 1.35547751 | 0.17526547 | NA |
| RGS3 | 89.8644984 | 0.52290145 | 0.33600645 | 1.55622444 | 0.11965473 | 0.6668162 |
| ZNF618 | 271.517152 | 0.51329132 | 0.30966248 | 1.65758321 | 0.09740162 | 0.65521765 |
| KIF12 | 81.4267743 | 0.09248292 | 0.29714221 | 0.31124127 | 0.75561721 | 0.95849416 |
| COL27A1 | 51.1252153 | 0.13769793 | 0.36747893 | 0.37470972 | 0.70787636 | 0.94558664 |
| AKNA | 13.9203312 | -0.1028598 | 0.43909528 | -0.2342539 | 0.81478786 | NA |
| WHRN | 33.0425757 | -0.3237211 | 0.37516418 | -0.8628784 | 0.38820433 | 0.85605713 |
| ATP6V1G1 | 78.371762 | 0.20031569 | 0.35099846 | 0.57070246 | 0.56820135 | 0.91698355 |
| TMEM268 | 24.232688 | -0.4890506 | 0.41844981 | -1.16872 | 0.24251648 | 0.78464787 |
| TNFSF15 | 19.67718 | -0.750162 | 0.72093433 | -1.0405413 | 0.29808847 | 0.81214575 |
| PAPPA | 327.078961 | -1.9158139 | 1.07845005 | -1.7764512 | 0.07565857 | 0.61020315 |
| ASTN2 | 187.452541 | -0.9420309 | 0.66010446 | -1.4270936 | 0.15355291 | 0.70572801 |
| CDK5RAP2 | 74.0374751 | -0.2985045 | 0.25347403 | -1.1776534 | 0.23893483 | 0.78098673 |
| MEGF9 | 121.157805 | 0.29225078 | 0.24574934 | 1.18922302 | 0.23435192 | 0.77743304 |
| FBXW2 | 59.319732 | -0.1273671 | 0.24622151 | -0.5172865 | 0.6049562 | 0.92569157 |
| PSMD5 | 48.3833926 | 0.09839237 | 0.31349046 | 0.31386081 | 0.75362675 | 0.95826617 |
| CNTRL | 36.2483223 | 0.01872108 | 0.34885237 | 0.05366477 | 0.95720225 | 0.99170927 |
| RAB14 | 96.8304189 | 0.14868869 | 0.23574547 | 0.63071708 | 0.52822553 | 0.90487731 |
| GSN | 113.478476 | -0.377824 | 0.24876623 | -1.5187915 | 0.12881498 | 0.66929384 |
| STOM | 65.629111 | -0.5145787 | 0.44405438 | -1.1588191 | 2.47E-01 | 0.78583441 |
| AL359644.1 | 10.8819575 | 0.46812889 | 0.52429429 | 0.89287429 | 0.3719245 | NA |
| DAB2IP | 141.801269 | -0.5399596 | 0.3112608 | -1.7347499 | 0.08278511 | 0.62228971 |
| TTLL11 | 30.6484972 | -0.0383035 | 0.36675596 | -0.1044388 | 0.91682117 | 0.98710975 |
| NDUFA8 | 31.5100917 | 0.46095431 | 0.32429677 | 1.42139655 | 0.15520151 | 0.70572801 |
| RBM18 | 16.3353818 | 0.19019229 | 0.44521657 | 0.4271905 | 0.66924058 | 0.93878847 |
| MRRF | 90.430153 | -0.0871424 | 0.26316661 | -0.33113 | 0.74054628 | 0.9544376 |
| PDCL | 19.7391596 | -0.250261 | 0.3329095 | -0.7517389 | 0.4522081 | 0.87346041 |
| RC3H2 | 77.484462 | -0.1536463 | 0.21237747 | -0.7234586 | 0.46939817 | 0.88207878 |
| ZBTB26 | 13.7238227 | -0.2948631 | 0.40681424 | -0.7248102 | 0.4685685 | NA |
| RABGAP1 | 219.361071 | -0.2273837 | 0.19507254 | -1.1656364 | 0.24376149 | 0.78544896 |
| STRBP | 219.086227 | -0.1273465 | 0.18481552 | -0.6890466 | 0.49079393 | 0.88987842 |
| DENND1A | 275.657529 | 0.12940514 | 0.25217838 | 0.51314922 | 0.60784695 | 0.92573951 |
| NEK6 | 11.8610529 | 0.32982333 | 0.5913091 | 0.55778498 | 5.77E-01 | NA |
| PSMB7 | 62.6245969 | -0.3642892 | 0.2926296 | -1.2448815 | 0.21317532 | 0.7643572 |
| NR6A1 | 329.758826 | 0.06172665 | 0.21664501 | 0.2849207 | 0.7757049 | 0.96469469 |
| MIR181A2HG | 38.368002 | -0.3713134 | 0.40449748 | -0.9179621 | 0.35863868 | 0.84204868 |
| RPL35 | 93.2292701 | -0.4321084 | 0.3657768 | -1.1813446 | 0.23746585 | 0.77930556 |
| ARPC5L | 35.3215077 | -0.0026084 | 0.31642547 | -0.0082432 | 0.99342295 | 0.99921857 |
| GOLGA1 | 50.1315963 | -0.1007872 | 0.32713602 | -0.3080894 | 0.75801427 | 0.95903525 |
| SCAI | 70.6794829 | -0.0239582 | 0.26047233 | -0.0919797 | 0.92671419 | 0.98925753 |
| PPP6C | 75.8007563 | 0.09246077 | 0.26081121 | 0.35451225 | 0.72295502 | 0.95093722 |
| RABEPK | 24.8670982 | -0.0242066 | 0.32363926 | -0.0747949 | 0.94037792 | 0.99012269 |
| HSPA5 | 140.629071 | -0.4343966 | 0.62949658 | -0.6900699 | 0.49015026 | 0.88970788 |
| GAPVD1 | 107.617883 | -0.0066976 | 0.24650238 | -0.0271706 | 0.97832367 | 0.99471172 |

| | | | | | | |
|---|---|---|---|---|---|---|
| MAPKAP1 | 190.884522 | -0.4642653 | 0.21240978 | -2.1857061 | 0.02883712 | 0.46148943 |
| PBX3 | 134.064847 | 0.46068986 | 0.25620762 | 1.79811144 | 0.07215935 | 0.60821941 |
| MVB12B | 37.2319489 | -0.2021744 | 0.33080423 | -0.6111602 | 0.54109352 | 0.90783815 |
| ZBTB43 | 67.7222886 | -0.4413661 | 0.22048753 | -2.0017733 | 0.04530912 | 0.52675468 |
| ZBTB34 | 12.1078697 | -0.5165403 | 0.41206269 | -1.2535478 | 0.21000642 | NA |
| RALGPS1 | 211.090991 | 0.0150638 | 0.24673493 | 0.06105258 | 0.95131733 | 0.99105535 |
| GARNL3 | 26.1402638 | 0.28383622 | 0.31531289 | 0.90017321 | 0.36802808 | 0.84777627 |
| RPL12 | 43.3566474 | -0.4505058 | 0.40371828 | -1.1158914 | 0.26446862 | 0.79666376 |
| LRSAM1 | 65.3297772 | -0.1158049 | 0.25016537 | -0.4629132 | 0.64342659 | 0.93328692 |
| FAM129B | 74.5284447 | -0.9225277 | 0.34255376 | -2.6930888 | 0.00707934 | 0.30254194 |
| STXBP1 | 46.1781275 | -0.3015573 | 0.34223001 | -0.8811537 | 0.37823462 | 0.85330104 |
| PTRH1 | 12.9558264 | 0.43981271 | 0.46723133 | 0.94131682 | 0.34654253 | NA |
| CDK9 | 21.4800878 | -0.1540859 | 0.39664504 | -0.388473 | 0.697666 | 0.94395238 |
| FPGS | 28.4391282 | -0.1532542 | 0.33960925 | -0.4512662 | 0.65179769 | 0.93482187 |
| ENG | 21.6645597 | 0.38984526 | 0.62828481 | 0.62049129 | 0.53493438 | 0.90763121 |
| DPM2 | 36.0389367 | -1.5861585 | 0.76923626 | -2.0619913 | 0.03920856 | 0.50068767 |
| FAM102A | 177.563327 | -0.7500622 | 0.45013544 | -1.6663034 | 0.095653 | 0.65011451 |
| SLC25A25 | 57.8136537 | -0.3291025 | 0.41173346 | -0.7993096 | 0.4241109 | 0.86190547 |
| SLC25A25-AS | 23.8045108 | 0.2362327 | 0.46104869 | 0.51238124 | 0.60838422 | 0.92573951 |
| PTGES2 | 36.3656858 | 0.13809415 | 0.28887558 | 0.47804022 | 0.63262158 | 0.93251095 |
| LCN2 | 53.4263314 | 0.10840307 | 1.16311936 | 0.0932003 | 9.26E-01 | 9.89E-01 |
| C9orf16 | 55.7568667 | 0.49419823 | 0.44343969 | 1.11446547 | 0.26507955 | 0.79666376 |
| CIZ1 | 23.9392407 | -0.3088391 | 0.3321281 | -0.9298795 | 0.35243345 | 0.83872681 |
| DNM1 | 20.6943829 | 0.66498455 | 0.42834727 | 1.55244261 | 0.12055635 | 0.6668162 |
| GOLGA2 | 70.9506866 | -0.6921973 | 0.33138014 | -2.0888316 | 0.03672289 | 0.49537143 |
| SWI5 | 21.9795077 | 0.2422836 | 0.35728438 | 0.67812536 | 0.49769221 | 0.89378207 |
| TRUB2 | 17.1263979 | 0.22178038 | 0.44785139 | 0.49520976 | 0.62045206 | 0.92798856 |
| COQ4 | 67.8846121 | 0.08948632 | 0.25385769 | 0.35250586 | 0.72445893 | 0.95093722 |
| SLC27A4 | 32.226168 | 0.21526078 | 0.42796225 | 0.50299011 | 0.61497122 | 0.92662475 |
| URM1 | 20.1511766 | -0.1899637 | 0.39484473 | -0.4811098 | 0.63043843 | 0.9320416 |
| CERCAM | 18.4851113 | 0.23571449 | 0.4689887 | 0.50260164 | 0.61524437 | 0.92662475 |
| ODF2 | 36.7137324 | -1.1942788 | 0.37591912 | -3.1769567 | 0.00148829 | 0.16322802 |
| GLE1 | 32.7795619 | -0.5241602 | 0.35139945 | -1.4916365 | 0.13579448 | 0.68406472 |
| SPTAN1 | 118.36501 | -0.1628541 | 0.25595185 | -0.6362686 | 0.52460137 | 0.90487731 |
| WDR34 | 29.0068286 | 0.17183988 | 0.46027178 | 0.37334437 | 0.70889216 | 0.94558664 |
| SET | 104.620924 | -0.1786828 | 0.31128556 | -0.5740158 | 0.56595709 | 0.91698355 |
| ZDHHC12 | 16.8715285 | 0.44253258 | 0.61340281 | 0.72143879 | 0.47063959 | 0.8826189 |
| ZER1 | 96.2576126 | 0.05523263 | 0.2573306 | 0.21463684 | 0.83005047 | 0.97567084 |
| TBC1D13 | 22.07849 | 0.11647365 | 0.38256546 | 0.30445418 | 0.7607819 | 0.95956874 |
| ENDOG | 15.4244383 | 0.38893378 | 0.54869416 | 0.70883529 | 0.4784267 | NA |
| KYAT1 | 28.5537896 | 0.21944447 | 0.39989709 | 0.54875237 | 0.5831754 | 0.92034873 |
| LRRC8A | 16.0178492 | -0.0876811 | 0.4054905 | -0.2162347 | 0.82880478 | NA |
| PHYHD1 | 23.5961082 | -0.3021121 | 0.47335036 | -0.638242 | 0.52331615 | 0.90487731 |
| NUP188 | 24.1747436 | -0.3325077 | 0.38479681 | -0.8641124 | 0.38752613 | 0.85605713 |
| SH3GLB2 | 54.2665047 | -0.0878347 | 0.4127275 | -0.2128153 | 0.83147104 | 0.97587427 |
| MIGA2 | 32.4736196 | -0.2275906 | 0.34025154 | -0.6688893 | 0.50356608 | 0.89618027 |
| PTPA | 61.4635868 | -0.6619605 | 0.39503569 | -1.6756979 | 0.09379737 | 0.64572229 |
| IER5L | 38.9700987 | -0.7554517 | 0.3040029 | -2.4850149 | 1.30E-02 | 0.36664851 |
| LINC01503 | 125.64129 | -0.8971436 | 0.39780906 | -2.2552117 | 0.02412005 | 0.43641593 |
| NTMT1 | 19.9152058 | -0.1028048 | 0.34574642 | -0.2973417 | 0.76620563 | 0.96270421 |
| ASB6 | 13.9000659 | -0.2453102 | 0.4763008 | -0.515032 | 0.60653066 | NA |
| TOR1B | 17.1385469 | -0.0498171 | 0.35453755 | -0.1405128 | 0.88825485 | 0.98259617 |
| C9orf78 | 23.1656751 | 0.25115654 | 0.41383199 | 0.60690461 | 0.54391422 | 0.90965439 |
| USP20 | 30.9266095 | 0.04255346 | 0.29799477 | 0.14279935 | 0.88644865 | 0.98216235 |

| | | | | | | |
|---|---|---|---|---|---|---|
| FNBP1 | 87.4220195 | 0.0404088 | 0.24706001 | 0.16355864 | 0.87007861 | 0.97854266 |
| GPR107 | 85.1835253 | -0.3388432 | 0.22305372 | -1.5191102 | 0.12873475 | 0.66929384 |
| NCS1 | 17.4972413 | -0.7339359 | 0.46467167 | -1.5794721 | 0.11422782 | 0.66629396 |
| ASS1 | 21.1079367 | 0.15916658 | 0.34897086 | 0.4561028 | 0.64831606 | 0.93365307 |
| FUBP3 | 85.977496 | -0.2079005 | 0.19425649 | -1.0702372 | 0.28451253 | 0.80431986 |
| ABL1 | 212.234951 | -0.1938395 | 0.1815211 | -1.0678623 | 0.28558263 | 0.80431986 |
| AIF1L | 275.545618 | 0.59277387 | 0.3800037 | 1.55991604 | 0.11877972 | 0.6668162 |
| NUP214 | 207.494572 | -0.1355421 | 0.18745948 | -0.7230472 | 4.70E-01 | 0.88207878 |
| AL157938.2 | 37.9671128 | 0.84411003 | 0.63054392 | 1.3387014 | 0.1806679 | 0.73322332 |
| AL157938.3 | 17.2541582 | -0.5278965 | 0.47949772 | -1.1009364 | 0.27092434 | 0.79666376 |
| PRRC2B | 246.585514 | -0.292543 | 0.21506792 | -1.3602352 | 0.1737555 | 0.72794035 |
| POMT1 | 20.8836764 | 0.00557503 | 0.34704462 | 0.01606431 | 0.98718309 | 0.99803728 |
| RAPGEF1 | 101.317881 | -0.4112416 | 0.36666128 | -1.1215846 | 0.26203912 | 0.79633795 |
| MED27 | 108.910016 | -0.2426579 | 0.26204581 | -0.9260131 | 0.35443917 | 0.84000513 |
| SETX | 189.90258 | -0.7934967 | 0.28938074 | -2.7420509 | 0.00610569 | 0.29709153 |
| TTF1 | 28.6823002 | 0.04874984 | 0.28397217 | 0.17167119 | 0.86369603 | 0.97788853 |
| DDX31 | 36.0229225 | -0.6490585 | 0.28861566 | -2.2488678 | 0.02452091 | 0.43948277 |
| GTF3C4 | 18.4818593 | 0.17858984 | 0.35908297 | 0.49734979 | 0.61894241 | 0.92766755 |
| AK8 | 16.5550436 | 0.3102592 | 0.6036446 | 0.5139766 | 0.60726835 | 0.92569157 |
| TSC1 | 85.3782752 | -0.4582277 | 0.30594937 | -1.4977241 | 0.13420495 | 0.68205729 |
| GTF3C5 | 21.0347844 | -0.6259692 | 0.402019 | -1.5570638 | 0.11945533 | 0.6668162 |
| RALGDS | 12.7846893 | -0.4813449 | 0.44549882 | -1.0804628 | 0.27993612 | NA |
| ABO | 123.969168 | 0.51949952 | 0.29570322 | 1.7568274 | 0.07894723 | 0.61367304 |
| SURF6 | 15.1012571 | 0.22646679 | 0.45940798 | 0.49295354 | 0.62204541 | NA |
| MED22 | 12.5755724 | -0.1986792 | 0.41834436 | -0.4749179 | 0.63484552 | NA |
| RPL7A | 128.672253 | 0.11255455 | 0.44303185 | 0.25405522 | 0.7994529 | 0.97019967 |
| SURF1 | 15.2900009 | 0.32284966 | 0.47414305 | 0.68091193 | 0.49592722 | NA |
| SURF4 | 60.6637407 | -0.1884298 | 0.3342897 | -0.5636721 | 0.57297732 | 0.91813957 |
| REXO4 | 15.7181733 | -0.0331328 | 0.36376424 | -0.0910832 | 0.92742652 | NA |
| CACFD1 | 20.9913342 | -0.1758739 | 0.41060953 | -0.4283239 | 0.66841531 | 0.93878847 |
| VAV2 | 206.61365 | -1.3434824 | 0.49367045 | -2.7214154 | 0.0065003 | 0.29935381 |
| BRD3OS | 10.4445315 | 0.33297769 | 0.43646573 | 0.76289539 | 0.44552578 | NA |
| BRD3 | 40.210929 | 0.0141502 | 0.34961788 | 0.04047334 | 0.96771576 | 0.99332167 |
| WDR5 | 21.7703395 | 0.29336062 | 0.36578691 | 0.80199869 | 0.42255372 | 0.86190547 |
| RXRA | 24.8653834 | 0.44242035 | 0.50937883 | 0.86854876 | 0.38509399 | 0.85540507 |
| PPP1R26-AS1 | 31.0034045 | -0.3330615 | 0.32658631 | -1.019827 | 0.30781053 | 0.81568231 |
| PPP1R26 | 16.7406581 | 0.45191855 | 0.35877584 | 1.25961253 | 0.20780917 | 0.76231289 |
| MRPS2 | 12.7315595 | 0.41568852 | 0.62308568 | 0.66714504 | 0.50467949 | NA |
| CAMSAP1 | 49.3833155 | -0.5340122 | 0.31047108 | -1.7200061 | 0.08543132 | 0.63062869 |
| UBAC1 | 48.476221 | 0.11130282 | 0.2374288 | 0.46878399 | 0.63922404 | 0.93328692 |
| NACC2 | 37.7993458 | 0.20202914 | 0.43840664 | 0.46082591 | 0.64492351 | 0.93328692 |
| TMEM250 | 14.1622511 | -0.8689304 | 0.52760028 | -1.6469484 | 0.09956866 | NA |
| QSOX2 | 11.4273817 | 0.06145345 | 0.43357487 | 0.14173666 | 0.88728802 | NA |
| SNAPC4 | 17.5654526 | -0.2439998 | 0.40477903 | -0.6027975 | 0.54664338 | 0.91137265 |
| ENTR1 | 19.1663535 | -0.5435069 | 0.35625528 | -1.5256108 | 0.12710682 | 0.66929384 |
| PMPCA | 23.3346383 | 0.20104512 | 0.38223492 | 0.52597265 | 0.59890721 | 0.92298075 |
| INPP5E | 12.898725 | -0.1529047 | 0.41774729 | -0.366022 | 0.71434864 | NA |
| SEC16A | 54.346847 | -0.2017071 | 0.23762234 | -0.8488559 | 0.39596149 | 0.85605713 |
| NOTCH1 | 20.4989696 | 0.48932144 | 0.42328173 | 1.15601834 | 0.24767365 | 0.78593073 |
| AGPAT2 | 45.6288645 | 0.56866236 | 0.4681778 | 1.21462905 | 0.2245076 | 0.76931755 |
| SNHG7 | 35.0766674 | 0.29573832 | 0.30743431 | 0.96195613 | 0.33607164 | 0.82974237 |
| RABL6 | 68.3209728 | 0.41563886 | 0.37720348 | 1.10189561 | 0.27050706 | 0.79666376 |
| PHPT1 | 60.3191337 | 0.45237111 | 0.40948138 | 1.10474159 | 0.26927158 | 0.79666376 |
| EDF1 | 71.4105554 | -0.2022004 | 0.33626387 | -0.6013148 | 0.54763032 | 0.91139843 |

| | | | | | | |
|---|---|---|---|---|---|---|
| TRAF2 | 15.3157144 | -0.2327 | 0.38276511 | -0.6079446 | 0.54322421 | NA |
| FBXW5 | 34.8844078 | 0.13937578 | 0.37011594 | 0.37657328 | 0.70649075 | 0.94558664 |
| LCN12 | 14.7378214 | 0.1871021 | 0.55802307 | 0.33529457 | 0.73740289 | NA |
| PAXX | 13.1866146 | 0.204711 | 0.50761557 | 0.40327959 | 0.68674256 | NA |
| ABCA2 | 12.456633 | 0.98312018 | 0.75387092 | 1.30409616 | 0.1922008 | NA |
| MAN1B1 | 34.9276615 | 0.02126455 | 0.36325868 | 0.05853832 | 0.95331984 | 0.99148072 |
| DPP7 | 30.1917051 | 0.37560512 | 0.40407862 | 0.92953476 | 0.35261202 | 0.83872681 |
| SSNA1 | 22.7898619 | 0.34616028 | 0.4463533 | 0.77552977 | 0.43802668 | 0.86828562 |
| TPRN | 17.1584849 | 0.45505196 | 0.42910507 | 1.06046746 | 0.28893199 | 0.80536924 |
| TMEM203 | 15.7848628 | 0.05200414 | 0.47030583 | 0.11057516 | 0.91195324 | NA |
| NDOR1 | 10.742927 | -0.1967274 | 0.45862305 | -0.4289522 | 0.66795801 | NA |
| TUBB4B | 81.6902494 | -0.3015217 | 0.44091018 | -0.6838621 | 0.49406226 | 0.8918201 |
| NELFB | 44.3143607 | -0.1916761 | 0.2537548 | -0.7553593 | 0.45003341 | 0.87346041 |
| EXD3 | 64.1317152 | -0.3125056 | 0.25509146 | -1.2250728 | 0.22054778 | 0.76680432 |
| NOXA1 | 17.4366665 | 0.35843047 | 0.37439037 | 0.95737096 | 0.33838005 | 0.83055261 |
| NSMF | 14.1435512 | 0.17670969 | 0.5283257 | 0.33447112 | 0.73802408 | NA |
| PNPLA7 | 30.8285586 | 0.81033583 | 0.47535545 | 1.70469451 | 0.08825141 | 0.63466198 |
| MRPL41 | 77.9004334 | -0.1267231 | 0.31527843 | -0.4019402 | 0.68772802 | 0.94205398 |
| DPH7 | 22.5328387 | -0.3864699 | 0.35271973 | -1.0956854 | 0.27321648 | 0.79726027 |
| ARRDC1 | 28.0406842 | -0.3778927 | 0.42635889 | -0.8863254 | 0.37544219 | 0.85178063 |
| EHMT1 | 233.243204 | -0.1766722 | 0.22181676 | -0.7964781 | 0.42575422 | 0.86239557 |
| AC069287.3 | 9.91772888 | 0.21197148 | 0.7736561 | 0.27398669 | 0.78409485 | NA |
| BET1L | 18.8544698 | 0.02388015 | 0.354153 | 0.06742891 | 0.94624026 | 0.99083449 |
| RIC8A | 15.1137397 | -0.4117241 | 0.47225624 | -0.8718236 | 0.38330463 | NA |
| SIRT3 | 19.2501339 | 0.55149938 | 0.42897542 | 1.28562001 | 0.19857571 | 0.75570012 |
| PSMD13 | 38.6118632 | 0.01047904 | 0.38955218 | 0.02690021 | 0.97853932 | 0.99471172 |
| PGGHG | 25.1439066 | -0.0171316 | 0.51406734 | -0.0333256 | 0.97341497 | 0.99431961 |
| IFITM2 | 204.140224 | 0.14311446 | 0.46747479 | 0.3061437 | 7.59E-01 | 0.95903525 |
| IFITM3 | 267.802052 | -0.5165007 | 0.51725321 | -0.9985452 | 3.18E-01 | 8.24E-01 |
| SIGIRR | 38.3378064 | -0.1564283 | 0.264294 | -0.5918722 | 0.55393618 | 0.9137961 |
| ANO9 | 10.8872938 | 0.18267116 | 0.46930493 | 0.38923767 | 0.69710034 | NA |
| PTDSS2 | 28.8810773 | -0.2260421 | 0.31085823 | -0.727155 | 0.46713102 | 0.88090126 |
| RNH1 | 42.8124989 | 0.54803467 | 0.38562541 | 1.42115809 | 0.15527081 | 0.70572801 |
| HRAS | 21.4944654 | -0.7224157 | 0.33594198 | -2.1504181 | 0.03152216 | 0.47665483 |
| RASSF7 | 56.2064582 | -0.1566928 | 0.3498741 | -0.4478549 | 0.65425792 | 0.93548446 |
| MIR210HG | 14.0332138 | -0.6152122 | 0.49000156 | -1.2555312 | 2.09E-01 | NA |
| PHRF1 | 34.2643763 | -0.1496124 | 0.31823204 | -0.4701361 | 0.63825778 | 0.9331046 |
| IRF7 | 12.7590034 | -2.8829092 | 1.04431718 | -2.7605686 | 0.00577008 | NA |
| DEAF1 | 64.4463231 | -0.4887516 | 0.25493338 | -1.9171738 | 0.05521585 | 0.5645224 |
| AC131934.1 | 13.9645995 | 0.92144587 | 0.53573695 | 1.71995952 | 0.0854398 | NA |
| EPS8L2 | 72.7558942 | 0.0550356 | 0.25762675 | 0.21362534 | 0.83083924 | 0.97567084 |
| TALDO1 | 73.5273014 | 0.07973633 | 0.27693515 | 0.2879242 | 0.77340476 | 0.96435639 |
| GATD1 | 23.5274288 | -0.4084106 | 0.41663768 | -0.9802537 | 0.32696089 | 0.82612204 |
| AP006621.3 | 20.8660809 | 0.22294245 | 0.41646905 | 0.53531577 | 0.59243153 | 0.92278087 |
| RPLP2 | 97.6376071 | -0.1852828 | 0.42072612 | -0.4403883 | 0.65965593 | 0.93704911 |
| PNPLA2 | 31.9416374 | 0.47720518 | 0.32900013 | 1.4504711 | 0.14692719 | 0.69749723 |
| CD151 | 79.9941299 | 0.12929196 | 0.46465426 | 0.27825412 | 0.78081729 | 0.96652357 |
| POLR2L | 59.2453537 | 0.04701514 | 0.50986446 | 0.09221105 | 0.92653036 | 0.98925753 |
| TSPAN4 | 15.8855823 | 0.38767085 | 0.39312488 | 0.98612646 | 0.32407106 | NA |
| CHID1 | 25.7037657 | 0.65440512 | 0.34335653 | 1.90590559 | 0.05666246 | 0.56976762 |
| AP2A2 | 210.388171 | -0.1280106 | 0.28727612 | -0.4456013 | 0.6558853 | 0.93567213 |
| TOLLIP | 32.1953668 | 0.05413284 | 0.38010038 | 0.14241722 | 0.88675047 | 0.98216235 |
| MOB2 | 26.113365 | 0.03182089 | 0.3413062 | 0.09323268 | 0.92571871 | 0.98925753 |
| MRPL23 | 12.7618071 | 0.47325221 | 0.47959108 | 0.98678275 | 0.32374915 | NA |

| | | | | | | |
|---|---|---|---|---|---|---|
| CD81-AS1 | 15.0006505 | -0.0638596 | 0.47178524 | -0.1353573 | 0.8923294 | NA |
| CD81 | 210.775168 | 0.52232461 | 0.37518172 | 1.39219099 | 0.16386457 | 0.7133712 |
| TSSC4 | 14.712294 | -0.1312943 | 0.42039811 | -0.3123093 | 0.75480545 | NA |
| KCNQ1 | 428.7332 | 0.14498431 | 0.1934486 | 0.74947204 | 0.45357274 | 0.87378486 |
| KCNQ1OT1 | 291.16304 | -0.3332309 | 0.39247597 | -0.849048 | 0.39585461 | 0.85605713 |
| CDKN1C | 37.7330368 | -0.8298476 | 0.5101797 | -1.6265789 | 1.04E-01 | 0.65946648 |
| SLC22A18 | 26.9687102 | 0.56864628 | 0.60866988 | 0.93424415 | 0.35017797 | 0.83687845 |
| NAP1L4 | 239.74243 | 0.2099844 | 0.19679012 | 1.06704749 | 0.28595039 | 0.80431986 |
| CARS | 46.0299643 | -0.9584217 | 0.33978114 | -2.8207029 | 0.00479186 | 0.27434125 |
| OSBPL5 | 19.1628043 | 0.28329967 | 0.37416391 | 0.7571539 | 0.44895766 | 0.87314185 |
| ZNF195 | 31.4365786 | -0.1885495 | 0.27905408 | -0.6756737 | 0.49924782 | 0.89379039 |
| AC127526.1 | 28.4685518 | 0.9069211 | 0.43862579 | 2.06764197 | 0.0386737 | 0.49794443 |
| NUP98 | 143.936913 | -0.3710049 | 0.31358544 | -1.1831062 | 0.23676707 | 0.77864547 |
| PGAP2 | 26.8679164 | -0.0425714 | 0.44060599 | -0.0966201 | 0.92302807 | 0.98915831 |
| RHOG | 10.4639871 | -0.1038681 | 0.53385227 | -0.1945635 | 0.84573472 | NA |
| STIM1 | 92.6185076 | 0.0532515 | 0.26284299 | 0.20259813 | 0.83944915 | 0.97600352 |
| RRM1 | 21.0256452 | 0.01153154 | 0.38744322 | 0.02976318 | 0.97625593 | 0.99440179 |
| TRIM68 | 12.3068181 | 0.21692653 | 0.48309634 | 0.44903369 | 0.65340735 | NA |
| AC104389.4 | 51.7686561 | 0.16127954 | 0.61460273 | 0.26241266 | 0.79300331 | 0.96947477 |
| TRIM5 | 31.0597319 | 0.76952827 | 0.55248615 | 1.39284626 | 0.16366628 | 0.71332671 |
| TRIM22 | 51.8383552 | -1.1374289 | 1.11490886 | -1.020199 | 0.30763411 | 0.81568231 |
| FAM160A2 | 21.8685876 | 0.21522839 | 0.39258939 | 0.54822775 | 0.58353553 | 0.92061074 |
| SMPD1 | 17.7476482 | 0.71369134 | 0.62850673 | 1.13553492 | 0.25615126 | 0.7940072 |
| APBB1 | 20.4611794 | -0.2890549 | 0.38043242 | -0.7598061 | 0.44737048 | 0.87275002 |
| ARFIP2 | 15.9669924 | -0.3676808 | 0.36536837 | -1.0063291 | 0.31425731 | NA |
| AC084337.2 | 27.8079043 | -0.6102984 | 0.54323604 | -1.1234498 | 0.2612465 | 0.79589391 |
| DNHD1 | 41.4183597 | -0.1307305 | 0.45303921 | -0.2885634 | 0.7729155 | 0.96435639 |
| RRP8 | 9.93214976 | 0.43960307 | 0.55608671 | 0.79052971 | 0.42921848 | NA |
| ILK | 37.4935749 | 0.43738174 | 0.28458865 | 1.53689102 | 0.12432 | 0.66929384 |
| TAF10 | 19.7761622 | -0.3353545 | 0.37832342 | -0.8864228 | 0.37538976 | 0.85178063 |
| TPP1 | 29.2465589 | -0.3664427 | 0.35384062 | -1.0356152 | 0.30038168 | 0.81399689 |
| MRPL17 | 20.5598387 | -0.0868264 | 0.44347866 | -0.1957848 | 0.84477864 | 0.97612951 |
| AC087280.2 | 19.2833127 | -0.160476 | 0.82953562 | -0.1934529 | 0.84660433 | 0.97612951 |
| ZNF214 | 13.8422411 | -0.0497232 | 0.41630761 | -0.1194385 | 0.90492797 | NA |
| AC107884.1 | 22.7381071 | -0.2585082 | 0.49881037 | -0.5182495 | 0.60428421 | 0.9252816 |
| PPFIBP2 | 59.0520295 | 0.3821337 | 0.34597254 | 1.10452032 | 0.26936749 | 0.79666376 |
| OVCH2 | 23.6775772 | 0.9495423 | 0.67290983 | 1.41109887 | 0.15821546 | 0.71089007 |
| AC044810.3 | 28.1493651 | 1.83420473 | 0.69849822 | 2.62592612 | 0.00864136 | 0.31119527 |
| EIF3F | 40.9520726 | 0.21585846 | 0.3578576 | 0.60319652 | 0.54637796 | 0.91109702 |
| RIC3 | 62.8792529 | -0.3897951 | 0.3558408 | -1.09542 | 0.27333268 | 0.79726027 |
| STK33 | 363.441346 | 0.44497133 | 0.24496806 | 1.81644632 | 0.06930194 | 0.60275677 |
| TRIM66 | 73.1783472 | -0.3261051 | 0.3541361 | -0.9208467 | 0.35713047 | 0.84204868 |
| RPL27A | 115.156858 | 0.0356299 | 0.36285614 | 0.09819291 | 0.92177911 | 0.98846691 |
| ST5 | 277.12289 | -0.3798346 | 0.46568556 | -0.815646 | 4.15E-01 | 8.59E-01 |
| AC026894.1 | 20.853993 | 0.58437918 | 0.52108833 | 1.12145896 | 0.26209255 | 0.79633795 |
| AKIP1 | 11.4631067 | 0.18209392 | 0.42457001 | 0.4288902 | 0.66800313 | NA |
| TMEM9B | 34.7316498 | 0.03361167 | 0.36466015 | 0.09217259 | 0.92656092 | 0.98925753 |
| TMEM9B-AS | 22.4162456 | 0.53026075 | 0.60273867 | 0.87975234 | 0.37899349 | 0.85416631 |
| DENND5A | 101.009221 | -0.5336625 | 0.30305887 | -1.7609202 | 0.0782519 | 0.61367304 |
| TMEM41B | 96.1283962 | -1.1855694 | 0.35852891 | -3.306761 | 0.00094381 | 0.13754433 |
| IPO7 | 114.029007 | -0.2269818 | 0.31034942 | -0.731375 | 0.46455014 | 0.87896358 |
| ZNF143 | 26.7768443 | -0.4670902 | 0.38232162 | -1.2217207 | 0.22181325 | 0.76680432 |
| WEE1 | 16.38216 | -0.3711978 | 0.42517199 | -0.8730534 | 0.38263397 | 0.8548566 |
| SWAP70 | 77.8985328 | -0.2193361 | 0.26135775 | -0.8392179 | 4.01E-01 | 0.85778581 |

| | | | | | | |
|---|---|---|---|---|---|---|
| SBF2-AS1 | 41.4554667 | -0.541616 | 0.29379252 | -1.8435323 | 0.06525133 | 0.5922062 |
| SBF2 | 618.103744 | -0.1572941 | 0.18736345 | -0.8395131 | 0.40118144 | 0.85778581 |
| AC011092.2 | 24.5142559 | 0.6734248 | 0.53346519 | 1.26235942 | 0.20681949 | 0.76231289 |
| ADM | 98.65119 | -2.2481049 | 0.47478995 | -4.7349463 | 2.19E-06 | 0.00350412 |
| AMPD3 | 61.6343439 | -0.4811913 | 0.38122571 | -1.2622214 | 0.20686912 | 0.76231289 |
| MTRNR2L8 | 128.503042 | 0.46974429 | 0.72326021 | 0.64948173 | 0.51602705 | 0.90213064 |
| RNF141 | 24.6174074 | 0.47223808 | 0.34483507 | 1.36946071 | 0.17085531 | 0.72463203 |
| CTR9 | 56.1847503 | -0.0414538 | 0.28091313 | -0.1475681 | 0.88268362 | 0.98137082 |
| EIF4G2 | 134.75641 | 0.00296121 | 0.26110514 | 0.01134106 | 0.99095133 | 0.99814944 |
| ZBED5 | 49.4632736 | 0.26231115 | 0.2776398 | 0.94478944 | 0.34476639 | 0.83545751 |
| ZBED5-AS1 | 15.558445 | 0.9117933 | 0.49790633 | 1.83125468 | 0.06706254 | NA |
| GALNT18 | 246.049382 | -0.0121208 | 0.36101913 | -0.0335737 | 0.97321706 | 0.99431961 |
| USP47 | 238.27234 | 0.204905 | 0.15735415 | 1.30218997 | 0.19285145 | 0.75061775 |
| PARVA | 99.2312478 | -0.093537 | 0.36915945 | -0.2533783 | 0.79997586 | 0.97019967 |
| TEAD1 | 392.38992 | -0.8706465 | 0.33606624 | -2.5906991 | 0.00957812 | 0.32468359 |
| LINC00958 | 47.3737295 | -0.3188191 | 0.25367849 | -1.2567842 | 0.20883177 | 0.7623519 |
| ARNTL | 30.7682057 | 0.22972604 | 0.41242065 | 0.55701877 | 0.57751461 | 0.91859435 |
| BTBD10 | 69.5698936 | -0.1312562 | 0.30437857 | -0.431227 | 0.66630336 | 0.93860542 |
| FAR1 | 120.653608 | 0.05504646 | 0.23595122 | 0.23329594 | 0.81553161 | 0.97288403 |
| RRAS2 | 76.51751 | -0.2479038 | 0.30012813 | -0.8259933 | 0.40880787 | 0.85867915 |
| COPB1 | 85.3534315 | -0.3046815 | 0.2454279 | -1.2414297 | 0.21444707 | 0.7643572 |
| PSMA1 | 88.1013836 | -0.301577 | 0.22474376 | -1.3418703 | 0.17963806 | 0.73132616 |
| PDE3B | 249.324487 | -0.4901197 | 0.35838342 | -1.3675849 | 0.17144206 | 0.72463203 |
| CYP2R1 | 15.811177 | -0.1195674 | 0.4037359 | -0.2961525 | 0.76711363 | NA |
| AC073172.1 | 39.4758617 | 1.94808328 | 0.62744517 | 3.1047865 | 0.00190416 | 0.17924252 |
| SOX6 | 576.959123 | 0.42217411 | 0.32705155 | 1.29084882 | 0.1967561 | 0.75350056 |
| C11orf58 | 105.92404 | -0.0121146 | 0.1755695 | -0.0690015 | 0.94498842 | 0.99069787 |
| PLEKHA7 | 534.643606 | 0.23050019 | 0.18938526 | 1.21709675 | 0.22356741 | 0.76834258 |
| RPS13 | 91.2156311 | 0.01877455 | 0.3745602 | 0.05012424 | 0.96002338 | 0.99256424 |
| PIK3C2A | 241.068481 | -0.2997784 | 0.20561893 | -1.4579319 | 0.14485932 | 0.69512617 |
| NUCB2 | 50.231332 | 0.19862429 | 0.36429345 | 0.54523157 | 0.58559426 | 0.92076087 |
| SERGEF | 109.412482 | -0.1754892 | 0.29423749 | -0.5964202 | 0.55089456 | 0.91258455 |
| HPS5 | 28.3284487 | -0.1726913 | 0.37601841 | -0.459263 | 0.64604531 | 0.93328692 |
| GTF2H1 | 55.4941512 | -0.3377414 | 0.29274014 | -1.1537241 | 0.24861329 | 0.78729484 |
| LDHA | 216.353023 | -0.3655015 | 0.32747977 | -1.1161039 | 2.64E-01 | 7.97E-01 |
| TSG101 | 87.4558499 | -0.3946155 | 0.20311711 | -1.9427977 | 0.0520406 | 0.55403388 |
| UEVLD | 67.7683097 | -0.1277816 | 0.22905531 | -0.5578637 | 0.57693748 | 0.91859435 |
| SPTY2D1OS | 23.6921326 | 0.40694919 | 0.37181565 | 1.09449183 | 0.27373933 | 0.79764696 |
| SPTY2D1 | 23.6089307 | -0.3921877 | 0.3457378 | -1.1343501 | 0.25664773 | 0.79437752 |
| ZDHHC13 | 65.5903374 | -0.0742431 | 0.22411416 | -0.3312737 | 0.74043774 | 0.9544376 |
| NAV2 | 235.705383 | 0.14286209 | 0.2074934 | 0.68851392 | 4.91E-01 | 8.90E-01 |
| HTATIP2 | 24.1458506 | -0.121871 | 0.44914056 | -0.2713427 | 0.78612749 | 0.96767512 |
| PRMT3 | 79.7936222 | -0.1491498 | 0.2189713 | -0.6811386 | 0.49578377 | 0.89230917 |
| NELL1 | 71.8803263 | -0.744771 | 0.59705314 | -1.2474117 | 2.12E-01 | 7.64E-01 |
| ANO5 | 45.0926723 | 0.61206643 | 0.38711947 | 1.58107893 | 0.11385999 | 0.6656824 |
| GAS2 | 24.6073978 | -0.4696743 | 0.47746434 | -0.9836846 | 0.3252706 | 0.82567895 |
| SVIP | 52.9355184 | 0.40828827 | 0.27666027 | 1.47577489 | 0.14000433 | 0.6912882 |
| AC006299.1 | 15.6571145 | 0.05252025 | 0.66457559 | 0.07902826 | 0.93701015 | NA |
| SLC5A12 | 10.9401542 | -0.9597133 | 0.806039 | -1.1906537 | 0.23378956 | NA |
| FIBIN | 37.0259124 | -0.3042316 | 0.81609461 | -0.3727896 | 0.70930505 | 0.94563619 |
| BBOX1 | 48.8280561 | -0.3316269 | 0.6432476 | -0.5155509 | 6.06E-01 | 0.92569157 |
| LGR4 | 580.430291 | -0.5739416 | 0.21323943 | -2.6915362 | 0.00711238 | 0.30254194 |
| LIN7C | 31.3076263 | -0.0412435 | 0.30358018 | -0.135857 | 0.89193429 | 0.98329225 |
| BDNF-AS | 116.927603 | 0.28951641 | 0.21512351 | 1.34581485 | 0.17836224 | 0.72841249 |

| | | | | | | |
|---|---|---|---|---|---|---|
| METTL15 | 254.183954 | 0.11789259 | 0.26916047 | 0.43800113 | 0.66138546 | 0.93729379 |
| ARL14EP | 34.5790833 | 0.40031881 | 0.28070366 | 1.42612609 | 0.15383195 | 0.70572801 |
| MPPED2 | 661.009519 | 0.31858856 | 0.26861613 | 1.1860366 | 0.23560785 | 0.77848938 |
| AL136088.1 | 33.9454146 | 1.1881276 | 0.46607797 | 2.54920352 | 0.01079693 | 0.33942942 |
| DCDC1 | 95.467935 | -0.2129278 | 0.41786065 | -0.5095666 | 0.61035515 | 0.92606167 |
| DNAJC24 | 41.5127218 | -0.3897642 | 0.30280773 | -1.2871674 | 0.19803595 | 0.75559912 |
| IMMP1L | 117.022436 | -0.1153746 | 0.25516736 | -0.4521528 | 0.6511589 | 0.93453685 |
| ELP4 | 176.893543 | 0.20172273 | 0.18275114 | 1.10381101 | 0.26967513 | 0.79666376 |
| AC131571.1 | 17.3834798 | -0.0741267 | 0.69074918 | -0.1073135 | 0.91454029 | 0.98691474 |
| RCN1 | 26.1310161 | 0.36590469 | 0.45763965 | 0.79954761 | 0.42397295 | 0.86190547 |
| EIF3M | 49.3472601 | 0.18288864 | 0.29942479 | 0.61079994 | 0.54133203 | 0.90783815 |
| PRRG4 | 45.6466929 | 0.32296423 | 0.38985501 | 0.82842139 | 0.4074319 | 0.85867915 |
| QSER1 | 83.4701817 | 0.26417416 | 0.22871819 | 1.15502033 | 0.2480821 | 0.78658703 |
| TCP11L1 | 40.8130273 | -0.3869273 | 0.29024004 | -1.3331287 | 0.18248957 | 0.73398872 |
| CSTF3 | 76.3176757 | -0.0432256 | 0.23821749 | -0.1814542 | 0.85601105 | 0.97634735 |
| HIPK3 | 206.43535 | -0.1646818 | 0.20624801 | -0.798465 | 0.42460071 | 0.861935 |
| FBXO3 | 48.4700086 | 0.21715715 | 0.2512938 | 0.86415644 | 0.38750196 | 0.85605713 |
| CAPRIN1 | 128.695641 | 0.08645524 | 0.21229206 | 0.4072467 | 0.6838268 | 0.94205398 |
| NAT10 | 33.275019 | -0.1089046 | 0.28538828 | -0.3816015 | 0.70275698 | 0.94476609 |
| ABTB2 | 672.630911 | -0.8342208 | 0.53731252 | -1.5525802 | 0.12052346 | 0.6668162 |
| CAT | 80.9182455 | 0.50863585 | 0.45111473 | 1.12750884 | 0.25952741 | 0.79532468 |
| ELF5 | 21.3097699 | 1.70765758 | 0.82051274 | 2.08120789 | 0.03741488 | 0.49538341 |
| EHF | 99.2123967 | 0.48771577 | 0.50983023 | 0.95662388 | 0.33875713 | 0.83066816 |
| APIP | 45.9306132 | -1.0459608 | 0.33531503 | -3.1193378 | 0.00181258 | 0.17553107 |
| PDHX | 90.6008912 | -0.0515866 | 0.23192541 | -0.2224276 | 0.82398105 | 0.97518529 |
| CD44 | 77.840161 | -0.6956068 | 0.50706201 | -1.3718377 | 1.70E-01 | 7.24E-01 |
| TRIM44 | 203.726469 | -0.2301534 | 0.17050919 | -1.3498005 | 0.17707997 | 0.72841249 |
| COMMD9 | 18.0973661 | 0.11689272 | 0.36129788 | 0.32353558 | 0.74628967 | 0.95563105 |
| PRR5L | 37.9961378 | 0.315763 | 0.52756407 | 0.59853015 | 0.54948625 | 0.91221128 |
| TRAF6 | 34.1387014 | -0.2696285 | 0.32768085 | -0.8228387 | 0.4105997 | 0.85867915 |
| C11orf74 | 19.1456063 | 0.86859636 | 0.48119584 | 1.80507871 | 0.07106237 | 0.60573888 |
| LRRC4C | 44.5479635 | 2.63675324 | 0.55770686 | 4.72784793 | 2.27E-06 | 0.00350412 |
| API5 | 66.6176385 | -0.026981 | 0.22769316 | -0.1184973 | 0.90567362 | 0.98531749 |
| TTC17 | 337.815866 | -0.1373647 | 0.15202976 | -0.9035382 | 0.3662403 | 0.84741115 |
| HSD17B12 | 293.219173 | -0.0396214 | 0.16687892 | -0.2374257 | 0.81232652 | 0.97257655 |
| ALKBH3 | 25.5685378 | -0.2556322 | 0.34119783 | -0.7492198 | 0.45372476 | 0.87378486 |
| ACCS | 12.8957476 | 0.91213232 | 0.68666296 | 1.32835522 | 0.18406079 | NA |
| EXT2 | 84.0970445 | -0.318359 | 0.23642037 | -1.3465802 | 0.17811548 | 0.72841249 |
| CD82 | 40.1356773 | 0.93227277 | 0.47960943 | 1.94381658 | 0.05191757 | 0.55403388 |
| TSPAN18 | 17.0951247 | 1.44855774 | 0.76409096 | 1.89579227 | 0.05798752 | 0.57141408 |
| PRDM11 | 27.8165594 | -0.2926965 | 0.32935642 | -0.8886921 | 0.37416857 | 0.85087596 |
| SYT13 | 22.4149151 | 0.22675971 | 0.39177141 | 0.57880616 | 0.56271998 | 0.91624774 |
| CRY2 | 46.8899991 | -0.0750236 | 0.31431697 | -0.2386877 | 0.81134775 | 0.97257655 |
| LARGE2 | 12.9930307 | 1.43108665 | 0.5926581 | 2.41469179 | 0.01574853 | NA |
| PHF21A | 293.949445 | -0.136874 | 0.2428922 | -0.5635174 | 0.5730826 | 0.91813957 |
| CREB3L1 | 15.1229663 | -0.1782033 | 0.7041329 | -0.2530819 | 0.80020491 | NA |
| DGKZ | 17.7527996 | -0.5254252 | 0.4140443 | -1.2690073 | 0.20443847 | 0.76173533 |
| MDK | 16.8584571 | -1.6640634 | 1.00042144 | -1.6633623 | 0.09623993 | 0.65104726 |
| AMBRA1 | 367.985733 | -0.0780756 | 0.20867387 | -0.3741515 | 0.70829164 | 0.94558664 |
| ATG13 | 45.3854782 | -0.2568906 | 0.29065928 | -0.8838206 | 0.37679309 | 0.85310518 |
| ARHGAP1 | 51.0825742 | -0.2222025 | 0.27987032 | -0.793948 | 0.4272257 | 0.86239557 |
| CKAP5 | 82.3298906 | -0.2438906 | 0.2353728 | -1.0361883 | 0.30011429 | 0.81388512 |
| LRP4 | 43.0617526 | 0.50170931 | 0.55759657 | 0.89977115 | 0.36824205 | 0.84777627 |
| C11orf49 | 261.466141 | -0.1632946 | 0.20052646 | -0.8143294 | 0.41545627 | 0.85867915 |

| | | | | | | |
|---|---|---|---|---|---|---|
| ARFGAP2 | 34.4415108 | -0.25954 | 0.30634773 | -0.8472073 | 0.39687961 | 0.85656568 |
| DDB2 | 24.6170372 | 0.79259786 | 0.56931044 | 1.3922068 | 0.16385979 | 0.7133712 |
| ACP2 | 17.3756042 | 0.10358893 | 0.49147014 | 0.21077359 | 0.83306395 | 0.97600352 |
| NR1H3 | 18.8006529 | -0.0383147 | 0.45971529 | -0.0833444 | 0.93357772 | 0.98960607 |
| MADD | 29.221526 | -0.4767469 | 0.30813422 | -1.5472052 | 0.12181378 | 0.66879213 |
| SLC39A13 | 29.5690392 | -0.0481759 | 0.4134065 | -0.1165339 | 0.90722946 | 0.9854228 |
| PSMC3 | 14.1481603 | -0.119001 | 0.43237513 | -0.2752264 | 0.78314233 | NA |
| CELF1 | 302.074678 | -0.4515194 | 0.20140097 | -2.2418929 | 0.02496829 | 0.43948277 |
| NDUFS3 | 31.0871045 | 0.25199892 | 0.33478683 | 0.75271456 | 0.45162146 | 0.87346041 |
| MTCH2 | 67.2029729 | -0.1289705 | 0.29569456 | -0.4361611 | 0.66271982 | 0.93807161 |
| FNBP4 | 170.700725 | -0.2283672 | 0.22971373 | -0.9941384 | 0.3201555 | 0.82449332 |
| NUP160 | 94.2206736 | -0.5883109 | 0.21359053 | -2.7543868 | 0.00588022 | 0.2943676 |
| PTPRJ | 334.912685 | -0.3319796 | 0.32605028 | -1.0181854 | 0.30858985 | 0.81568231 |
| TNKS1BP1 | 47.6747254 | 0.08570676 | 0.29627725 | 0.2892789 | 0.77236796 | 0.9640787 |
| SSRP1 | 24.257403 | 0.43437387 | 0.47741412 | 0.90984715 | 0.36290313 | 0.84543391 |
| TIMM10 | 17.0155142 | 0.44254916 | 0.41786084 | 1.05908265 | 0.28956214 | 0.80536924 |
| UBE2L6 | 20.1976869 | -0.7471732 | 0.80401335 | -0.9293045 | 0.35273129 | 0.83872681 |
| SERPING1 | 160.419358 | -0.402817 | 0.48093908 | -0.8375635 | 4.02E-01 | 8.58E-01 |
| ZDHHC5 | 61.5347806 | -0.2813803 | 0.27371612 | -1.0280005 | 0.30394958 | 0.81568231 |
| CTNND1 | 170.606275 | -0.0935975 | 0.23137454 | -0.4045281 | 0.6858244 | 0.94205398 |
| LPXN | 30.6071506 | 0.21404351 | 0.34707249 | 0.61671124 | 0.5374252 | 0.90763121 |
| FAM111A | 42.902591 | 0.0927494 | 0.37860605 | 0.24497602 | 0.806475 | 0.97216031 |
| OSBP | 84.7144864 | -0.4743433 | 0.25563422 | -1.8555546 | 0.06351707 | 0.58688097 |
| PATL1 | 54.41333 | -0.584195 | 0.30361429 | -1.9241355 | 0.0543376 | 0.5645224 |
| STX3 | 91.5599996 | -0.477975 | 0.32285277 | -1.4804735 | 0.13874692 | 0.69023184 |
| MRPL16 | 23.4576453 | -0.0087006 | 0.36244822 | -0.024005 | 0.98084866 | 0.99592962 |
| PRPF19 | 22.3968938 | 0.36171925 | 0.39193676 | 0.92290208 | 0.35605824 | 0.84192356 |
| TMEM109 | 18.2235159 | 0.27891012 | 0.54859887 | 0.50840446 | 0.61116972 | 0.92619266 |
| VPS37C | 35.1720939 | -0.0040939 | 0.32597095 | -0.0125591 | 0.98997958 | 0.99814944 |
| DDB1 | 60.545773 | 0.11382361 | 0.35377201 | 0.32174285 | 0.74764752 | 0.95563105 |
| TKFC | 16.0905683 | 0.55078068 | 0.46895265 | 1.17449103 | 0.2401984 | NA |
| TMEM138 | 10.7037561 | 0.36034239 | 0.45405026 | 0.79361783 | 0.42741795 | NA |
| CPSF7 | 59.6076274 | -0.1650815 | 0.21889649 | -0.7541531 | 0.45075729 | 0.87346041 |
| AP003108.2 | 53.8253333 | -0.3237277 | 0.38176521 | -0.8479759 | 0.3964514 | 0.85650971 |
| SYT7 | 20.6730059 | -0.1207008 | 0.7736986 | -0.156005 | 0.87602908 | 0.97918624 |
| MYRF | 35.7262818 | -0.1734812 | 0.56293904 | -0.3081704 | 0.75795266 | 0.95903525 |
| TMEM258 | 33.7717048 | -0.162525 | 0.41424313 | -0.392342 | 0.69480554 | 0.94395238 |
| FADS1 | 16.8366013 | -1.1188717 | 0.7357988 | -1.5206218 | 0.12835477 | 0.66929384 |
| FTH1 | 575.418765 | 0.094129 | 0.45634297 | 0.20626811 | 8.37E-01 | 0.97600352 |
| ASRGL1 | 34.549164 | 0.35982595 | 0.54437006 | 0.66099512 | 0.50861544 | 0.89936455 |
| AHNAK | 183.425085 | -0.1750357 | 0.2913316 | -0.6008125 | 0.54796486 | 0.91157238 |
| B3GAT3 | 26.9561365 | 0.16436575 | 0.33977616 | 0.48374716 | 0.62856532 | 0.93128745 |
| GANAB | 75.6087321 | 0.37138731 | 0.26008968 | 1.42792021 | 0.15331483 | 0.70572801 |
| UQCC3 | 20.1901407 | 0.08377543 | 0.39298913 | 0.21317492 | 0.83119053 | 0.97567084 |
| UBXN1 | 21.5628927 | 0.43577965 | 0.4097476 | 1.06353192 | 0.2875408 | 0.80523851 |
| TTC9C | 33.0956306 | -0.1421532 | 0.36721087 | -0.387116 | 0.69867031 | 0.94395238 |
| POLR2G | 11.4835128 | 0.67314291 | 0.5067504 | 1.328352 | 0.18406185 | NA |
| TAF6L | 15.4984049 | -0.300866 | 0.51691699 | -0.5820393 | 0.56054022 | NA |
| TMEM223 | 19.8316142 | -0.202609 | 0.37696234 | -0.5374781 | 0.5909374 | 0.92254886 |
| TMEM179B | 29.8747407 | 0.61863654 | 0.32881389 | 1.88141853 | 0.05991501 | 0.57714955 |
| NXF1 | 16.3382111 | -0.1595994 | 0.41149884 | -0.387849 | 0.69812777 | 0.94395238 |
| STX5 | 61.5126343 | -0.1849634 | 0.33694133 | -0.5489483 | 0.58304095 | 0.92034873 |
| SLC3A2 | 56.6510017 | -0.9774268 | 0.35842213 | -2.7270269 | 0.00639078 | 0.29935381 |
| RARRES3 | 34.538269 | -0.0333387 | 1.17103962 | -0.0284693 | 0.97728783 | 0.99466135 |

| | | | | | | |
|---|---|---|---|---|---|---|
| PLA2G16 | 17.2486483 | 0.73149263 | 0.58916022 | 1.24158523 | 0.21438963 | 0.7643572 |
| ATL3 | 65.2261415 | -0.0715815 | 0.22093551 | -0.3239929 | 0.74594342 | 0.95563105 |
| RTN3 | 180.470052 | 0.0489532 | 0.26513844 | 0.18463259 | 0.85351722 | 0.97612951 |
| MARK2 | 98.9369366 | -0.0505168 | 0.19949059 | -0.253229 | 0.80009123 | 0.97019967 |
| NAA40 | 28.9336966 | -0.2908319 | 0.33519314 | -0.8676546 | 0.38558344 | 0.85540507 |
| OTUB1 | 29.7691457 | 0.00135601 | 0.35969907 | 0.00376986 | 0.9969921 | 0.99941676 |
| MACROD1 | 58.6635034 | 0.94993149 | 0.3876068 | 2.45076066 | 0.01425547 | 0.38054999 |
| AP000721.2 | 17.8141071 | 1.13411959 | 0.55549376 | 2.04164236 | 0.04118702 | 0.50744725 |
| STIP1 | 37.1547479 | -0.1890295 | 0.33908495 | -0.5574695 | 0.57720672 | 0.91859435 |
| TRPT1 | 11.2049651 | 0.20871093 | 0.50237418 | 0.41544916 | 0.67781312 | NA |
| NUDT22 | 23.7185853 | 0.15477762 | 0.39883991 | 0.38806955 | 0.69796457 | 0.94395238 |
| DNAJC4 | 15.8606557 | 0.14443458 | 0.53790089 | 0.26851524 | 0.78830275 | NA |
| VEGFB | 41.6607728 | -0.1402605 | 0.3318617 | -0.4226473 | 0.67255261 | 0.93878847 |
| FKBP2 | 49.7654311 | 0.27807709 | 0.3275889 | 0.84885993 | 0.39595924 | 0.85605713 |
| PPP1R14B | 34.9830647 | -0.3740177 | 0.43909273 | -0.8517965 | 0.39432704 | 0.85605713 |
| BAD | 25.4799084 | 0.5033551 | 0.34348328 | 1.46544282 | 0.14280014 | 0.69328517 |
| GPR137 | 18.8490335 | 0.08627351 | 0.43021122 | 0.20053756 | 0.84106018 | 0.97600352 |
| ESRRA | 70.5625489 | -0.3764574 | 0.35791753 | -1.0517992 | 0.2928917 | 0.80935782 |
| TRMT112 | 35.6282148 | 0.0264583 | 0.33187955 | 0.07972259 | 0.93645789 | 0.99012269 |
| PRDX5 | 62.8724083 | -0.11637 | 0.50896057 | -0.2286424 | 0.81914687 | 0.97296246 |
| RPS6KA4 | 10.604081 | -0.4075651 | 0.55822212 | -0.7301127 | 0.46532129 | NA |
| SF1 | 170.806344 | -0.6011896 | 0.29936188 | -2.0082369 | 0.04461813 | 0.52543182 |
| MAP4K2 | 10.8833888 | 0.05711665 | 0.4557818 | 0.12531578 | 0.90027356 | NA |
| MEN1 | 11.8558343 | -0.0304872 | 0.46837168 | -0.0650918 | 0.94810089 | NA |
| EHD1 | 19.4234773 | 0.06567903 | 0.42027504 | 0.15627631 | 0.87581523 | 0.97918624 |
| ATG2A | 17.9492532 | -0.5152121 | 0.49029242 | -1.0508262 | 0.29333842 | 0.80935782 |
| PPP2R5B | 11.7292007 | 0.40448427 | 0.51259792 | 0.78908684 | 0.43006125 | NA |
| VPS51 | 42.25587 | 0.48275752 | 0.29166831 | 1.65515931 | 0.09789219 | 0.65621002 |
| TM7SF2 | 14.9778717 | 0.67986381 | 0.50922953 | 1.33508323 | 0.18184911 | NA |
| FAU | 118.623652 | 0.0037834 | 0.4367225 | 0.00866317 | 0.99308788 | 0.99901414 |
| SYVN1 | 14.9715742 | 0.34855014 | 0.47761168 | 0.72977727 | 0.46552634 | NA |
| MRPL49 | 15.4399424 | 0.22561499 | 0.3617681 | 0.62364534 | 0.53286053 | NA |
| CAPN1 | 59.3292767 | 0.00332836 | 0.30279322 | 0.01099219 | 0.99122967 | 0.99814944 |
| POLA2 | 11.1173474 | -1.0665104 | 0.45777369 | -2.3297766 | 0.01981796 | NA |
| DPF2 | 30.4478261 | 0.21813528 | 0.28378236 | 0.76867102 | 0.44208863 | 0.86925451 |
| FRMD8 | 28.7671595 | -0.486668 | 0.39704443 | -1.2257268 | 0.22030152 | 0.76680432 |
| NEAT1 | 9036.78831 | -0.1365547 | 0.50260908 | -0.2716917 | 0.78585906 | 0.96767512 |
| MALAT1 | 89533.2753 | -0.3446874 | 0.37195258 | -0.9266972 | 0.35408379 | 0.83979358 |
| SCYL1 | 25.5541042 | -0.3812994 | 0.50607342 | -0.7534468 | 0.45118149 | 0.87346041 |
| LTBP3 | 54.5820518 | 0.04032291 | 0.32524857 | 0.12397567 | 0.90133454 | 0.98531749 |
| EHBP1L1 | 19.8311536 | -0.3646543 | 0.44429319 | -0.8207514 | 0.41178786 | 0.85867915 |
| MAP3K11 | 17.9377073 | -0.2087808 | 0.39634455 | -0.5267659 | 0.5983562 | 0.92296674 |
| PCNX3 | 17.628284 | -0.3981843 | 0.47199471 | -0.8436202 | 0.39888164 | 0.85778581 |
| SIPA1 | 15.1746798 | 0.41133604 | 0.39616128 | 1.03830449 | 2.99E-01 | NA |
| RELA | 23.8865871 | -0.5196833 | 0.37310964 | -1.3928434 | 0.16366714 | 0.71332671 |
| KAT5 | 25.381085 | 0.10308485 | 0.37171458 | 0.27732259 | 0.78153241 | 0.96652357 |
| RNASEH2C | 19.7739998 | 0.68539716 | 0.486665 | 1.40835516 | 0.15902593 | 0.71094363 |
| CFL1 | 123.109532 | 0.00814687 | 0.39224969 | 0.02076959 | 0.98342946 | 0.99685229 |
| MUS81 | 12.9443641 | 0.46032649 | 0.39408638 | 1.16808526 | 0.24277238 | NA |
| EFEMP2 | 14.7859619 | 0.8050487 | 0.47605161 | 1.69109542 | 0.09081858 | NA |
| CCDC85B | 11.3232017 | 0.49747911 | 0.55461638 | 0.89697875 | 0.36973025 | NA |
| DRAP1 | 20.1526385 | -0.0589734 | 0.41883679 | -0.1408029 | 0.88802566 | 0.9825428 |
| SART1 | 18.9398222 | -0.3452645 | 0.40179289 | -0.8593097 | 0.39016966 | 0.85605713 |
| BANF1 | 35.7977508 | -0.0746872 | 0.40280232 | -0.1854189 | 0.85290047 | 0.97612951 |

| | | | | | | |
|---|---|---|---|---|---|---|
| SF3B2 | 47.3965697 | 0.14617786 | 0.39139058 | 0.37348334 | 0.70878874 | 0.94558664 |
| PACS1 | 319.711703 | -0.0023569 | 0.2073777 | -0.0113653 | 0.99093201 | 0.99814944 |
| KLC2 | 13.888805 | -0.3902612 | 0.4320476 | -0.9032828 | 0.36637585 | NA |
| RAB1B | 33.4984933 | 0.12449406 | 0.4875713 | 0.25533509 | 0.7984643 | 0.97019967 |
| YIF1A | 14.6906058 | -0.66947 | 0.54909717 | -1.2192196 | 0.22276085 | NA |
| BRMS1 | 12.0006399 | -0.0326917 | 0.65984984 | -0.0495441 | 0.96048567 | NA |
| SLC29A2 | 28.2424129 | 0.18915573 | 0.32621852 | 0.57984363 | 0.56202007 | 0.91624774 |
| MRPL11 | 13.0027069 | 0.37471976 | 0.40492079 | 0.92541497 | 0.3547501 | NA |
| ZDHHC24 | 21.1851176 | 0.28870709 | 0.43788793 | 0.65931731 | 0.50969203 | 0.89963034 |
| CTSF | 22.8253115 | 1.45978982 | 0.43551644 | 3.3518593 | 0.00080271 | 0.12609798 |
| CCS | 29.5144136 | 0.75255465 | 0.36313607 | 2.07237649 | 0.03823034 | 0.49671741 |
| RBM14-RBM | 32.5986095 | 0.04611038 | 0.30556122 | 0.15090392 | 0.88005151 | 0.98093344 |
| RBM4 | 22.4349932 | -0.3133147 | 0.43560826 | -0.719258 | 0.47198196 | 0.88267439 |
| RBM4B | 25.9715946 | 0.49479003 | 0.35798528 | 1.38215186 | 0.16692508 | 0.71878854 |
| SPTBN2 | 127.550612 | 0.5606458 | 0.38582367 | 1.45311405 | 0.14619209 | 0.69713616 |
| C11orf80 | 469.954027 | 0.37008853 | 0.25867234 | 1.43072326 | 0.15250954 | 0.70572801 |
| RCE1 | 13.0704916 | 0.20347663 | 0.4742922 | 0.42901112 | 0.66791513 | NA |
| PC | 12.2508287 | -0.5577186 | 0.43476394 | -1.2828079 | 0.19955939 | NA |
| RHOD | 18.9153626 | -0.1534249 | 0.44075888 | -0.3480926 | 0.72777066 | 0.95253722 |
| KDM2A | 327.122663 | -0.4732819 | 0.16104727 | -2.9387763 | 0.00329511 | 0.23150356 |
| GRK2 | 19.2280843 | 0.09267806 | 0.38615289 | 0.24000355 | 0.81032751 | 0.97257655 |
| ANKRD13D | 24.9637733 | 0.0444717 | 0.37223972 | 0.11947058 | 0.90490255 | 0.98531749 |
| SSH3 | 17.7443793 | -0.2190773 | 0.41457931 | -0.5284329 | 0.59719892 | 0.9229638 |
| RAD9A | 130.952448 | 0.56302655 | 0.29963127 | 1.87906474 | 0.06023565 | 0.57733412 |
| PPP1CA | 38.8297303 | 0.13531545 | 0.50114782 | 0.27001105 | 0.78715175 | 0.96767512 |
| RPS6KB2 | 14.0001115 | 0.6888945 | 0.44453109 | 1.54971053 | 0.12121101 | NA |
| CORO1B | 24.9683045 | 0.13309368 | 0.33298824 | 0.39969484 | 0.6893813 | 0.94211649 |
| TMEM134 | 27.8478273 | 0.11264847 | 0.37070744 | 0.30387432 | 0.76122364 | 0.9597533 |
| AIP | 12.733279 | 0.09856702 | 0.43944011 | 0.22430138 | 0.8225228 | NA |
| PITPNM1 | 22.5284451 | -0.5868777 | 0.63580235 | -0.9230505 | 0.35598087 | 0.84192356 |
| CDK2AP2 | 41.5289912 | -0.653989 | 0.52223696 | -1.2522839 | 0.21046643 | 0.76397692 |
| GSTP1 | 270.745117 | 0.36428161 | 0.54124598 | 0.67304262 | 0.50092018 | 0.89492452 |
| NDUFV1 | 75.3228131 | 0.43020416 | 0.32471904 | 1.32485041 | 0.18522077 | 0.73752116 |
| UNC93B1 | 16.2980778 | -0.6470425 | 0.70240267 | -0.9211845 | 0.3569541 | 0.84204868 |
| NDUFS8 | 64.3960377 | 0.34983798 | 0.33658976 | 1.03936015 | 0.29863727 | 0.81246117 |
| TCIRG1 | 10.6169754 | -0.7201234 | 0.7483499 | -0.9622816 | 0.33590816 | NA |
| CHKA | 112.490446 | -0.5885283 | 0.25273224 | -2.3286634 | 0.01987691 | 0.41544117 |
| KMT5B | 113.064707 | 0.11138319 | 0.21437226 | 0.51957839 | 0.60335746 | 0.92498018 |
| LRP5 | 32.7304539 | 0.45545968 | 0.38643133 | 1.17863033 | 0.23854541 | 0.78096803 |
| PPP6R3 | 413.406665 | -0.1588659 | 0.16316109 | -0.9736754 | 0.33021772 | 0.82867637 |
| TESMIN | 11.0557294 | 0.11123368 | 0.44859042 | 0.24796267 | 0.80416329 | NA |
| CPT1A | 119.042626 | -0.0003282 | 0.30425314 | -0.0010786 | 9.99E-01 | 0.99957534 |
| MRPL21 | 13.2992326 | -0.3391995 | 0.40439673 | -0.8387791 | 0.4015933 | NA |
| IGHMBP2 | 16.8688822 | 0.232866 | 0.43558089 | 0.53461024 | 0.5929194 | 0.92278087 |
| TPCN2 | 29.0785071 | -0.378979 | 0.3744941 | -1.0119759 | 0.31154957 | 0.81701824 |
| CCND1 | 36.2443285 | 0.30715246 | 0.35882062 | 0.85600559 | 0.39199469 | 0.85605713 |
| ORAOV1 | 23.1337148 | -0.2727735 | 0.32829956 | -0.8308678 | 0.4060483 | 0.85867915 |
| PPFIA1 | 146.932339 | -0.5468376 | 0.31997798 | -1.7089851 | 0.08745372 | 0.63466198 |
| AP002336.2 | 53.6458815 | -0.29137 | 0.55065938 | -0.5291293 | 0.59671578 | 0.9229638 |
| AP000487.1 | 12.6058407 | -0.3744932 | 0.41556723 | -0.9011615 | 0.36750247 | NA |
| CTTN | 167.836458 | -0.609992 | 0.20456563 | -2.9818888 | 0.00286476 | 0.21375341 |
| SHANK2 | 469.437055 | -0.5923459 | 0.21977329 | -2.6952588 | 0.0070334 | 0.30200482 |
| DHCR7 | 14.4059373 | -0.4406893 | 0.56633971 | -0.7781359 | 0.4364889 | NA |
| NADSYN1 | 35.3150774 | -0.0172342 | 0.25795545 | -0.0668109 | 0.94673221 | 0.99083449 |

| | | | | | | |
|---|---|---|---|---|---|---|
| AP002495.2 | 64.6602455 | 0.24182359 | 0.29068878 | 0.83189862 | 0.40546617 | 0.8583057 |
| RNF121 | 77.1992962 | -0.2187118 | 0.25590495 | -0.8546604 | 0.39273916 | 0.85605713 |
| NUMA1 | 99.9569513 | -0.1777091 | 0.22536445 | -0.788541 | 0.43038031 | 0.86534658 |
| LAMTOR1 | 45.5553919 | -0.0026946 | 0.27267309 | -0.0098821 | 9.92E-01 | 0.99814944 |
| ANAPC15 | 10.6005169 | 0.32228171 | 0.5760961 | 0.55942352 | 0.57587271 | NA |
| FOLR3 | 14.2884867 | 1.59000215 | 0.54092405 | 2.93941846 | 0.00328829 | NA |
| FOLR1 | 24.5450889 | 1.43377113 | 0.51141827 | 2.80351959 | 0.00505482 | 0.28057141 |
| INPPL1 | 25.3785023 | 0.73830229 | 0.3481904 | 2.12039819 | 0.03397248 | 0.4908133 |
| CLPB | 28.2572812 | -0.0883717 | 0.34145588 | -0.2588087 | 0.79578286 | 0.96974891 |
| ARAP1 | 32.0180797 | 0.20669148 | 0.39782658 | 0.51955171 | 0.60337606 | 0.92498018 |
| ATG16L2 | 24.8670641 | 0.30694148 | 0.36649062 | 0.83751524 | 0.40230301 | 0.85782098 |
| FCHSD2 | 216.733094 | -0.1013016 | 0.21808013 | -0.4645153 | 0.6422786 | 0.93328692 |
| P2RY6 | 55.2540642 | 1.95619511 | 0.50940609 | 3.84014865 | 0.00012296 | 0.04981461 |
| ARHGEF17 | 34.4233515 | 0.27378463 | 0.27052878 | 1.01203514 | 0.31152125 | 0.81701824 |
| FAM168A | 159.12215 | 0.01853738 | 0.23353991 | 0.07937562 | 0.93673386 | 0.99012269 |
| RAB6A | 215.16023 | -0.427999 | 0.24587342 | -1.740729 | 0.08173109 | 0.61925304 |
| MRPL48 | 45.6208906 | -0.3607831 | 0.3277641 | -1.10074 | 0.27100983 | 0.79666376 |
| COA4 | 26.8480598 | 0.19984366 | 0.30549077 | 0.65417249 | 0.51300069 | 0.90156229 |
| PAAF1 | 37.9877506 | -0.1217826 | 0.32102968 | -0.37935 | 0.70442795 | 0.94480737 |
| UCP2 | 16.5706319 | -0.8243655 | 0.81431577 | -1.0123414 | 0.31137487 | 0.81701824 |
| C2CD3 | 68.9712449 | -0.213248 | 0.27254162 | -0.7824418 | 0.43395495 | 0.86729771 |
| PPME1 | 79.4460718 | 0.01274792 | 0.29064847 | 0.04386028 | 0.96501578 | 0.99270395 |
| P4HA3 | 13.0315103 | 0.217805 | 0.68325939 | 0.31877352 | 0.74989826 | NA |
| PGM2L1 | 30.4159872 | -0.3904362 | 0.37868481 | -1.0310321 | 0.30252579 | 0.81568231 |
| POLD3 | 41.6686939 | 0.23315409 | 0.37511343 | 0.62155623 | 0.53423371 | 0.90763121 |
| RNF169 | 115.69291 | -0.3821663 | 0.25836957 | -1.4791459 | 0.13910133 | 0.69054855 |
| XRRA1 | 102.317985 | -0.4936729 | 0.42299199 | -1.1670975 | 0.243171 | 0.78495944 |
| SPCS2 | 88.7028816 | -0.2251453 | 0.2450435 | -0.9187973 | 0.35820159 | 0.84204868 |
| NEU3 | 15.9739881 | 0.17384729 | 0.38279837 | 0.45414847 | 0.64972197 | NA |
| ARRB1 | 51.6239882 | -0.4026437 | 0.31590971 | -1.2745532 | 0.20246743 | 0.76065252 |
| RPS3 | 206.340973 | 0.38085107 | 0.3712389 | 1.02589212 | 0.30494244 | 0.81568231 |
| SERPINH1 | 22.7681799 | 0.16818488 | 0.5584905 | 0.30114188 | 0.7633063 | 0.96091323 |
| UVRAG | 268.246244 | -0.2157981 | 0.25415457 | -0.8490823 | 0.39583551 | 0.85605713 |
| THAP12 | 20.9264638 | -0.0785539 | 0.36363051 | -0.2160267 | 0.82896696 | 0.97567084 |
| AP002360.1 | 9.29383511 | 0.77749086 | 0.67780316 | 1.14707471 | 0.25135075 | NA |
| EMSY | 110.92496 | -0.287895 | 0.28256918 | -1.018848 | 0.30827513 | 0.81568231 |
| ACER3 | 131.356729 | 0.26818859 | 0.31529558 | 0.85059419 | 0.39499482 | 0.85605713 |
| CAPN5 | 26.3793656 | 1.08284955 | 0.57647773 | 1.87838923 | 0.06032794 | 0.57746079 |
| PAK1 | 105.72285 | -0.0860578 | 0.4580144 | -0.1878933 | 0.85096032 | 0.97612951 |
| CLNS1A | 63.4806697 | -0.1327453 | 0.24951812 | -0.5320068 | 0.5947213 | 0.9229638 |
| RSF1 | 218.102328 | -0.2538829 | 0.17745748 | -1.4306689 | 0.15252512 | 0.70572801 |
| AAMDC | 53.9627019 | 0.5134122 | 0.38441757 | 1.33555862 | 0.18169358 | 0.73398872 |
| AP002812.2 | 22.799678 | 0.04385811 | 0.3729452 | 0.11759935 | 0.90638512 | 0.98531749 |
| INTS4 | 57.5916287 | -0.5306512 | 0.27980973 | -1.8964715 | 0.05789773 | 0.57141408 |
| ALG8 | 28.6282489 | -0.2357983 | 0.30756634 | -0.7666585 | 0.44328461 | 0.87006015 |
| GAB2 | 131.613737 | -0.9643386 | 0.48117165 | -2.0041468 | 0.04505434 | 0.52648218 |
| AP003086.2 | 34.2640988 | -0.2228137 | 0.64340956 | -0.3463016 | 0.72911609 | 0.95279138 |
| NARS2 | 81.0205695 | -0.1706746 | 0.21720926 | -0.7857613 | 0.43200733 | 0.86695857 |
| PRCP | 60.4688081 | 0.39563985 | 0.28883245 | 1.36979017 | 0.17075241 | 0.72463203 |
| RAB30 | 31.0614984 | -0.1654744 | 0.34989784 | -0.4729219 | 0.63626888 | 0.93251095 |
| RAB30-AS1 | 87.8261199 | -0.0177682 | 0.35063513 | -0.0506744 | 0.95958494 | 0.99245244 |
| PCF11 | 82.6388712 | -0.0390194 | 0.29274168 | -0.1332895 | 0.8939644 | 0.98408497 |
| ANKRD42 | 38.7741184 | 0.13314029 | 0.27840079 | 0.47823245 | 0.63248477 | 0.93251095 |
| CCDC90B | 34.6151299 | -0.0884885 | 0.33745894 | -0.2622199 | 0.7931519 | 0.96947761 |

| | | | | | | |
|---|---|---|---|---|---|---|
| AP000446.1 | 26.8741619 | 0.56041117 | 0.50642999 | 1.10659159 | 0.26847054 | 0.79666376 |
| DLG2 | 636.582955 | 0.35060607 | 0.50239682 | 0.6978668 | 0.4852605 | 0.88838332 |
| TMEM126B | 21.4108442 | -0.0060863 | 0.33491858 | -0.0181723 | 0.98550138 | 0.99752207 |
| CREBZF | 71.2782477 | -0.0976507 | 0.24933788 | -0.3916399 | 0.69532427 | 0.94395238 |
| SYTL2 | 442.877718 | -0.2942364 | 0.34637196 | -0.849481 | 0.39561372 | 0.85605713 |
| PICALM | 481.706531 | -0.3226202 | 0.17390108 | -1.8551937 | 0.06356856 | 0.58688097 |
| EED | 41.902121 | -0.3763243 | 0.32228578 | -1.1676727 | 0.2429388 | 0.78495944 |
| HIKESHI | 43.1240611 | -0.2103749 | 0.2969785 | -0.7083843 | 0.47870665 | 0.88588465 |
| AP001831.1 | 37.3705955 | 0.75796383 | 0.47607235 | 1.59211898 | 0.11135797 | 0.66417829 |
| ME3 | 146.268316 | 0.01448039 | 0.25741173 | 0.05625379 | 9.55E-01 | 9.91E-01 |
| AP003059.1 | 17.7893208 | 0.31727509 | 0.53160389 | 0.59682612 | 0.55062347 | 0.91248567 |
| AP003059.2 | 13.2374498 | 1.13712008 | 0.5705437 | 1.99304643 | 0.04625636 | NA |
| PRSS23 | 92.4897773 | 1.15305545 | 0.37911869 | 3.04141021 | 0.00235473 | 0.1931089 |
| AP001528.2 | 57.5645433 | 1.53951983 | 0.56835108 | 2.70874796 | 0.00675376 | 0.30102597 |
| FZD4 | 75.1049426 | -0.5095971 | 0.51692705 | -0.9858202 | 0.32422133 | 0.82567895 |
| AP001528.1 | 17.2045398 | 0.73201859 | 0.50277887 | 1.4559454 | 0.14540771 | 0.69628954 |
| TMEM135 | 165.891928 | -0.288798 | 0.23428795 | -1.2326624 | 0.21770173 | 0.76666749 |
| RAB38 | 21.1008919 | -0.4631807 | 0.5288946 | -0.8757523 | 0.38116469 | 0.85432452 |
| CTSC | 49.1255011 | 0.31313393 | 0.38980464 | 0.80330991 | 0.42179563 | 0.86190547 |
| NOX4 | 12.6793996 | 0.18227214 | 0.62786702 | 0.29030374 | 0.77158388 | NA |
| CHORDC1 | 41.7853341 | -0.07639 | 0.44980207 | -0.1698303 | 0.86514356 | 0.97788853 |
| SLC36A4 | 106.890333 | -0.2617593 | 0.2666884 | -0.9815173 | 0.32633771 | 0.82567895 |
| SMCO4 | 48.7224768 | -0.2004185 | 0.40194644 | -0.49862 | 0.61804713 | 0.92749951 |
| CEP295 | 53.0875067 | -0.1232488 | 0.27832216 | -0.4428279 | 0.65789023 | 0.93593674 |
| TAF1D | 112.31504 | 0.26004572 | 0.22882999 | 1.1364145 | 0.25578313 | 0.7940072 |
| PANX1 | 13.2988063 | -0.0048619 | 0.39244338 | -0.0123888 | 0.99011539 | NA |
| MRE11 | 46.494938 | -0.1139877 | 0.28627311 | -0.3981782 | 0.69049881 | 0.94276429 |
| ANKRD49 | 12.0781697 | -0.1673215 | 0.46812974 | -0.3574254 | 0.72077336 | NA |
| AMOTL1 | 76.3183426 | -0.1667146 | 0.2831112 | -0.588866 | 0.55595118 | 0.91398748 |
| CWC15 | 24.3835942 | -0.0516822 | 0.29762303 | -0.1736498 | 0.86214069 | 0.97767895 |
| SRSF8 | 21.9054664 | 0.70420861 | 0.43868134 | 1.60528509 | 0.10843108 | 0.65986731 |
| ENDOD1 | 35.3558281 | -0.1463257 | 0.33249946 | -0.4400781 | 0.65988054 | 0.93704911 |
| AP000787.1 | 27.7047333 | 0.06090786 | 0.35969765 | 0.16933071 | 0.86553652 | 0.97788853 |
| SESN3 | 50.7787427 | 0.42524655 | 0.30618573 | 1.38885165 | 0.16487787 | 0.71485946 |
| FAM76B | 30.7319564 | 0.05141843 | 0.35478175 | 0.14492975 | 0.88476634 | 0.9816 |
| CEP57 | 102.707147 | 0.09485396 | 0.21503097 | 0.44111767 | 0.65912781 | 0.93677446 |
| MTMR2 | 67.1136318 | -0.3818299 | 0.28251261 | -1.3515498 | 0.17651937 | 0.72841249 |
| MAML2 | 1169.41882 | -0.1269796 | 0.27560765 | -0.4607258 | 0.64499531 | 0.93328692 |
| CCDC82 | 43.2954343 | -0.37403 | 0.28609047 | -1.3073837 | 0.19108244 | 0.74847043 |
| JRKL | 13.0679135 | -0.135531 | 0.53999507 | -0.2509857 | 0.80182519 | NA |
| CNTN5 | 11.9999257 | -0.5588448 | 0.57451085 | -0.9727316 | 0.33068674 | NA |
| ARHGAP42 | 41.2584575 | 0.00455815 | 0.3625147 | 0.01257369 | 0.98996791 | 0.99814944 |
| CEP126 | 32.8521521 | -0.02923 | 0.3993924 | -0.0731861 | 0.94165802 | 0.99045212 |
| CFAP300 | 20.4900647 | 0.72034684 | 0.5075906 | 1.41914928 | 0.1558555 | 0.70690214 |
| YAP1 | 533.798695 | -0.1463634 | 0.19813678 | -0.7386988 | 0.46008994 | 0.87607759 |
| BIRC3 | 104.841787 | 0.37466173 | 0.65855054 | 0.56891872 | 0.5694113 | 0.91756967 |
| BIRC2 | 51.8885747 | -0.2151596 | 0.24791844 | -0.8678644 | 0.38546858 | 0.85540507 |
| TMEM123 | 137.211627 | -0.0160249 | 0.27866543 | -0.0575059 | 0.95414225 | 0.99148072 |
| MMP7 | 220.484378 | 0.74425235 | 0.53065393 | 1.40251924 | 1.61E-01 | 7.11E-01 |
| DCUN1D5 | 40.1034174 | -0.0654328 | 0.32362184 | -0.202189 | 0.83976897 | 0.97600352 |
| DYNC2H1 | 397.900918 | 0.46016042 | 0.30146507 | 1.52641371 | 0.12690686 | 0.66929384 |
| PDGFD | 25.6875536 | -0.041569 | 0.55609657 | -0.0747515 | 0.94041244 | 0.99012269 |
| CASP4 | 61.5024381 | -0.1736688 | 0.33057965 | -0.5253462 | 5.99E-01 | 0.92298075 |
| MSANTD4 | 15.0863904 | 0.43803396 | 0.4103333 | 1.06750773 | 0.28574263 | NA |

| | | | | | | |
|---|---|---|---|---|---|---|
| KBTBD3 | 17.335482 | 0.65902694 | 0.42220032 | 1.56093426 | 0.11853926 | 0.6668162 |
| AASDHPPT | 31.7976212 | 0.19823156 | 0.32897286 | 0.60257724 | 0.54678996 | 0.91139843 |
| CWF19L2 | 106.42752 | -0.3733587 | 0.21266006 | -1.7556597 | 0.07914654 | 0.61367304 |
| ALKBH8 | 20.7813901 | 0.18395974 | 0.41246729 | 0.44599838 | 0.65559842 | 0.93567213 |
| SLC35F2 | 31.7016769 | 0.36425795 | 0.40850728 | 0.89168042 | 0.37256425 | 0.8491102 |
| CUL5 | 175.560967 | -0.1064245 | 0.22125102 | -0.4810123 | 0.63050776 | 0.9320416 |
| ACAT1 | 69.7997372 | 0.33151798 | 0.22796648 | 1.45424003 | 0.14587977 | 0.69679188 |
| NPAT | 53.4197498 | 0.09026139 | 0.27010254 | 0.33417454 | 0.73824786 | 0.9542764 |
| ATM | 138.055119 | -0.2660525 | 0.21087092 | -1.261684 | 0.20706251 | 0.76231289 |
| C11orf65 | 43.9783583 | 0.99231692 | 0.56848135 | 1.74555757 | 0.08088786 | 0.61925304 |
| KDELC2 | 24.447681 | 0.58261632 | 0.56510925 | 1.03097997 | 0.30255021 | 0.81568231 |
| EXPH5 | 126.521087 | -0.3496072 | 0.34807157 | -1.0044119 | 0.31518013 | 0.82084199 |
| DDX10 | 112.835154 | -0.2815484 | 0.24697463 | -1.139989 | 0.25429087 | 0.79171259 |
| RDX | 146.762431 | 0.38550489 | 0.34965697 | 1.10252313 | 0.27023431 | 0.79666376 |
| ZC3H12C | 124.838092 | 0.39444682 | 0.57892296 | 0.68134596 | 0.49565261 | 0.89230917 |
| FDX1 | 88.3934602 | 0.03896254 | 0.31742309 | 0.1227464 | 0.90230792 | 0.98531749 |
| COLCA1 | 21.3560385 | 0.50600337 | 0.47312393 | 1.06949435 | 2.85E-01 | 8.04E-01 |
| COLCA2 | 10.900664 | 0.10922826 | 0.53021892 | 0.20600597 | 0.83678625 | NA |
| SIK2 | 322.397327 | -0.4329664 | 0.52534486 | -0.8241565 | 0.40985066 | 0.85867915 |
| PPP2R1B | 49.2378967 | -0.0941236 | 0.28307802 | -0.3325005 | 0.73951133 | 0.9544376 |
| C11orf1 | 14.06023 | 0.83447589 | 0.45841055 | 1.82036798 | 0.06870299 | NA |
| CRYAB | 220.518828 | 0.11107234 | 0.63251651 | 0.17560386 | 0.86060517 | 0.97718459 |
| DIXDC1 | 54.9747754 | 0.1075067 | 0.31635216 | 0.33983235 | 0.73398278 | 0.95329896 |
| DLAT | 43.3335737 | 0.40214228 | 0.29708288 | 1.35363668 | 0.17585232 | 0.72841249 |
| NKAPD1 | 15.423902 | 0.35596008 | 0.49232147 | 0.72302367 | 0.46966534 | NA |
| TIMM8B | 24.4986796 | 0.26572819 | 0.4158869 | 0.6389434 | 0.52285975 | 0.90487731 |
| AP002884.3 | 28.5692793 | -0.622722 | 0.72866865 | -0.8546024 | 0.39277128 | 0.85605713 |
| PTS | 52.1728013 | 0.09394794 | 0.41882944 | 0.22431073 | 0.82251553 | 0.97478959 |
| AP002884.1 | 37.1858639 | -0.690623 | 0.51169771 | -1.34967 | 0.17712187 | 0.72841249 |
| TTC12 | 47.641328 | 0.10464136 | 0.32481896 | 0.32215285 | 0.7473369 | 0.95563105 |
| ZW10 | 22.3482731 | -0.4126342 | 0.35250402 | -1.1705803 | 0.24176754 | 0.78376421 |
| USP28 | 36.2738286 | 0.25668153 | 0.34177276 | 0.7510298 | 0.45263472 | 0.87346041 |
| ZBTB16 | 31.6395098 | 0.26379984 | 0.80679297 | 0.3269734 | 0.74368801 | 0.95563105 |
| NNMT | 44.0166613 | 0.52095746 | 0.80931049 | 0.6437053 | 5.20E-01 | 9.04E-01 |
| REXO2 | 34.0896364 | 0.16535979 | 0.31137696 | 0.53105983 | 0.59537732 | 0.9229638 |
| CADM1 | 189.930979 | -0.6104108 | 0.41298369 | -1.4780506 | 0.13939423 | 0.69054855 |
| BUD13 | 11.7223697 | -0.4341904 | 0.46188429 | -0.9400416 | 0.34719625 | NA |
| SIK3 | 778.494419 | -0.3014547 | 0.35278104 | -0.8545095 | 0.39282277 | 0.85605713 |
| PAFAH1B2 | 87.6143763 | -0.2479709 | 0.2199088 | -1.1276078 | 0.25948559 | 0.79532468 |
| SIDT2 | 21.0035214 | -0.029351 | 0.34751073 | -0.0844606 | 0.93269026 | 0.98960607 |
| PCSK7 | 68.1597669 | -0.1854603 | 0.25514251 | -0.7268891 | 0.4672939 | 0.88090126 |
| RNF214 | 53.1223153 | -0.6986732 | 0.26203619 | -2.6663232 | 0.00766859 | 0.30654023 |
| BACE1 | 35.6509677 | 0.83233988 | 0.27329469 | 3.04557646 | 0.00232235 | 0.19218479 |
| CEP164 | 41.9430818 | -0.1875822 | 0.29047248 | -0.6457829 | 0.51841997 | 0.90368693 |
| DSCAML1 | 243.847948 | 1.16360071 | 0.6546138 | 1.7775377 | 0.07547981 | 0.61020315 |
| FXYD2 | 68.2859308 | 0.52267475 | 0.48522358 | 1.07718333 | 0.28139837 | 0.80416687 |
| AP000757.1 | 21.4081638 | 1.0287943 | 0.54638078 | 1.88292549 | 0.05971047 | 0.57639913 |
| TMPRSS4 | 25.3173933 | 0.47490954 | 0.82851064 | 0.57320874 | 5.67E-01 | 9.17E-01 |
| MPZL3 | 21.3888946 | -0.4170075 | 0.36196801 | -1.1520561 | 0.24929801 | 0.78763828 |
| MPZL2 | 48.4805511 | 0.12004805 | 0.58267817 | 0.20602806 | 8.37E-01 | 0.97600352 |
| UBE4A | 61.2361523 | -0.1006551 | 0.24353796 | -0.4133035 | 0.67938428 | 0.94176503 |
| ATP5MG | 121.486395 | 0.11034399 | 0.37068493 | 0.29767596 | 0.7659505 | 0.96270421 |
| KMT2A | 211.714108 | -0.0780399 | 0.16561598 | -0.47121 | 0.63749081 | 0.93271601 |
| TMEM25 | 13.3435844 | 0.83942843 | 0.44910251 | 1.86912435 | 0.06160552 | NA |

| | | | | | | |
|---|---|---|---|---|---|---|
| IFT46 | 19.5937272 | -0.0933373 | 0.36823543 | -0.2534719 | 0.7999036 | 0.97019967 |
| ARCN1 | 35.7557438 | -0.0660921 | 0.32635321 | -0.2025172 | 0.83951244 | 0.97600352 |
| PHLDB1 | 47.4006978 | 0.24537498 | 0.37779112 | 0.64949905 | 0.51601586 | 0.90213064 |
| AP002954.1 | 16.2844288 | 0.13000623 | 0.42275886 | 0.30751863 | 0.75844865 | 0.95903525 |
| DDX6 | 112.860111 | 0.08904056 | 0.17865919 | 0.49838221 | 0.61821468 | 0.92757676 |
| BCL9L | 20.3888475 | -0.1400394 | 0.45833911 | -0.3055367 | 0.75995742 | 0.95913478 |
| CCDC84 | 80.1250498 | -0.4255758 | 0.310665 | -1.3698865 | 0.17072234 | 0.72463203 |
| RPS25 | 116.361488 | 0.17681998 | 0.41971708 | 0.42128374 | 0.6735479 | 0.93901638 |
| TRAPPC4 | 25.7685851 | 0.76047629 | 0.33197882 | 2.29073734 | 0.02197861 | 0.4212027 |
| HYOU1 | 23.2216063 | -0.1358141 | 0.45889732 | -0.2959574 | 0.7672626 | 0.96283923 |
| VPS11 | 14.5322574 | -0.1572017 | 0.37236183 | -0.4221745 | 0.67289768 | NA |
| HINFP | 18.7438059 | -0.0521921 | 0.34962059 | -0.1492821 | 0.88133101 | 0.98137082 |
| NLRX1 | 15.0292249 | 0.15463767 | 0.39938429 | 0.38719017 | 0.69861543 | NA |
| CBL | 73.037525 | -0.6784005 | 0.36337171 | -1.8669602 | 0.06190715 | 0.58370968 |
| USP2 | 12.4939188 | 0.31976235 | 0.62005263 | 0.51570195 | 6.06E-01 | NA |
| THY1 | 10.7774387 | -0.7081415 | 0.99659372 | -0.7105618 | 0.47735579 | NA |
| NECTIN1 | 11.4478685 | 0.06653924 | 0.58787733 | 0.11318559 | 0.90988342 | NA |
| OAF | 50.6615381 | -0.5330488 | 0.34846645 | -1.5296992 | 0.1260912 | 0.66929384 |
| TMEM136 | 13.0470438 | 0.51536052 | 0.42186662 | 1.22161958 | 0.22185152 | NA |
| ARHGEF12 | 502.322672 | -0.1055905 | 0.17805754 | -0.5930133 | 0.55317223 | 0.91322379 |
| SC5D | 25.8999575 | 0.0308138 | 0.48877032 | 0.06304352 | 0.94973185 | 0.99105535 |
| SORL1 | 190.896363 | 0.07376103 | 0.36852488 | 0.20015209 | 0.84136163 | 0.97600352 |
| UBASH3B | 23.9686417 | -0.1451657 | 0.64060074 | -0.2266086 | 0.82072808 | 0.97368535 |
| JHY | 18.0770651 | -0.0008245 | 0.69231408 | -0.0011909 | 0.9990498 | 0.99957534 |
| HSPA8 | 100.890794 | 0.58762918 | 0.46464456 | 1.26468536 | 0.20598414 | 0.76231289 |
| GRAMD1B | 41.3177994 | -0.2162264 | 0.52319195 | -0.4132831 | 0.67939921 | 0.94176503 |
| VWA5A | 26.7458067 | 0.61399381 | 0.38182377 | 1.60805549 | 0.10782303 | 0.65986731 |
| TBRG1 | 51.7758373 | 0.27262823 | 0.24617484 | 1.10745774 | 0.26809608 | 0.79666376 |
| SIAE | 43.4085945 | 0.79604316 | 0.30638319 | 2.5981946 | 0.00937154 | 0.32071089 |
| MSANTD2 | 63.8033827 | 0.46513792 | 0.32158945 | 1.44637182 | 0.14807294 | 0.69945736 |
| EI24 | 45.3354087 | 0.34052494 | 0.36025172 | 0.94524167 | 0.34453551 | 0.83545751 |
| STT3A | 30.6618519 | -0.452309 | 0.41045263 | -1.101976 | 0.27047208 | 0.79666376 |
| RPUSD4 | 11.409078 | -0.1178657 | 0.48224456 | -0.2444106 | 0.80691282 | NA |
| FAM118B | 23.1130596 | -0.0324786 | 0.4817934 | -0.0674118 | 0.94625384 | 0.99083449 |
| SRPRA | 29.5074054 | 0.02559306 | 0.33276643 | 0.07690999 | 0.93869515 | 0.99012269 |
| FOXRED1 | 21.3942 | 0.31002636 | 0.37593429 | 0.8246823 | 0.40955198 | 0.85867915 |
| TIRAP | 10.4565693 | 0.42094716 | 0.57188136 | 0.73607429 | 0.46168549 | NA |
| DCPS | 36.3962278 | -1.4147759 | 0.37116272 | -3.8117401 | 0.00013799 | 0.05233918 |
| ST3GAL4 | 37.8245396 | -1.5898052 | 0.54247293 | -2.9306627 | 0.0033824 | 0.23150356 |
| ETS1 | 87.9584439 | -0.8700377 | 0.36056315 | -2.4129969 | 0.01582196 | 0.39477603 |
| KCNJ1 | 265.479011 | 0.75328686 | 0.40518191 | 1.8591325 | 0.06300836 | 0.58586834 |
| ARHGAP32 | 225.619435 | 0.2224083 | 0.25675808 | 0.86621736 | 0.38637098 | 0.8560201 |
| BARX2 | 144.832807 | 0.46707672 | 0.4005151 | 1.16619005 | 0.24353761 | 0.78542956 |
| TMEM45B | 86.9895954 | 0.33262033 | 0.28685323 | 1.15954887 | 0.24623253 | 0.78583441 |
| NFRKB | 35.736858 | 0.16606683 | 0.34815569 | 0.47699014 | 0.63336915 | 0.93251095 |
| PRDM10 | 36.1437726 | -0.3852105 | 0.32147053 | -1.1982763 | 0.23080948 | 0.77249016 |
| APLP2 | 229.530994 | 0.60195008 | 0.3361015 | 1.79097708 | 0.07329697 | 0.60981847 |
| ST14 | 98.8921252 | 0.0210747 | 0.29951221 | 0.07036342 | 0.94390441 | 0.99069421 |
| ZBTB44 | 146.559693 | 0.40992996 | 0.20750512 | 1.97551739 | 0.04820946 | 0.5380949 |
| AP002986.1 | 24.6121646 | 0.82919027 | 0.81579695 | 1.01641747 | 0.30943063 | 0.81568231 |
| SNX19 | 56.9767297 | -0.3188265 | 0.3907499 | -0.8159349 | 0.41453735 | 0.85867915 |
| JAM3 | 18.8582805 | -0.664372 | 0.40322978 | -1.6476262 | 0.09942939 | 0.65860455 |
| NCAPD3 | 19.6086798 | -0.1456788 | 0.33892385 | -0.4298276 | 0.66732109 | 0.93878847 |
| VPS26B | 22.9555597 | 0.47643292 | 0.37284111 | 1.27784438 | 0.2013043 | 0.75910235 |

| | | | | | | |
|---|---|---|---|---|---|---|
| ACAD8 | 17.3631948 | 0.21710494 | 0.42073709 | 0.51601093 | 0.60584677 | 0.92569157 |
| ZMYND11 | 201.994088 | 0.04186849 | 0.15822939 | 0.26460631 | 0.79131276 | 0.96909997 |
| DIP2C | 391.968949 | -0.9159335 | 0.28356795 | -3.2300319 | 0.00123776 | 0.14632943 |
| LARP4B | 262.523651 | -0.3152055 | 0.194201 | -1.6230889 | 0.1045704 | 0.65946648 |
| GTPBP4 | 26.2384012 | -0.3386867 | 0.31910104 | -1.0613777 | 0.28851828 | 0.80523851 |
| IDI1 | 58.0378368 | -0.2746918 | 0.51344273 | -0.5349998 | 0.59264998 | 0.92278087 |
| WDR37 | 111.935418 | -0.0980104 | 0.25306468 | -0.387294 | 0.69853858 | 0.94395238 |
| PFKP | 144.483507 | -0.5611837 | 0.48195176 | -1.164398 | 0.24426277 | 0.78561483 |
| PITRM1 | 40.0377661 | -0.2405526 | 0.30712165 | -0.7832487 | 0.43348108 | 0.86729771 |
| KLF6 | 219.814505 | -1.2114299 | 0.6040551 | -2.0054957 | 0.04491008 | 0.52610161 |
| AKR1C3 | 20.8569453 | -0.2165592 | 0.71900234 | -0.301194 | 0.76326659 | 0.96091323 |
| NET1 | 12.7059962 | 0.19120922 | 0.46351212 | 0.4125226 | 0.67995642 | NA |
| ASB13 | 14.6249884 | 0.95006787 | 0.43054552 | 2.20666068 | 0.02733777 | NA |
| FAM208B | 169.378609 | -0.6931424 | 0.26227882 | -2.6427691 | 0.00822311 | 0.30976355 |
| GDI2 | 113.21098 | -0.1783767 | 0.19493874 | -0.9150397 | 0.36017077 | 0.84282379 |
| ANKRD16 | 17.1429343 | 0.35613283 | 0.44350223 | 0.8030012 | 0.42197404 | 0.86190547 |
| FBH1 | 80.5209238 | -0.2231141 | 0.22629212 | -0.9859559 | 0.32415474 | 0.82567895 |
| IL15RA | 15.3256437 | 0.15454453 | 0.58431134 | 0.26449004 | 0.79140234 | NA |
| RBM17 | 63.2665019 | -0.1844263 | 0.24089961 | -0.7655731 | 0.44393038 | 0.8704597 |
| PFKFB3 | 289.418912 | -0.6437682 | 0.31107262 | -2.0695111 | 0.03849816 | 0.49671741 |
| AL137145.2 | 56.6059883 | -0.3513732 | 0.36680975 | -0.9579168 | 0.3381047 | 0.83055261 |
| PRKCQ | 58.2150624 | -0.1460148 | 0.35038768 | -0.4167237 | 0.67688053 | 0.94034237 |
| SFMBT2 | 96.1796266 | -0.141362 | 0.27357309 | -0.5167247 | 0.60534834 | 0.92569157 |
| KIN | 25.0320948 | -0.2055297 | 0.31279157 | -0.6570818 | 0.5111283 | 0.90030978 |
| ATP5F1C | 146.621745 | 0.3322571 | 0.20948999 | 1.58602853 | 0.11273284 | 0.6656824 |
| TAF3 | 178.374265 | -0.349708 | 0.1968028 | -1.7769464 | 0.07557705 | 0.61020315 |
| GATA3 | 297.105486 | 0.03489127 | 0.30564275 | 0.11415704 | 0.90911331 | 0.9854228 |
| GATA3-AS1 | 34.9728391 | -0.2458509 | 0.40833569 | -0.6020803 | 0.54712066 | 0.91139843 |
| USP6NL | 104.899444 | 0.02520934 | 0.21817274 | 0.11554763 | 0.90801107 | 0.9854228 |
| ECHDC3 | 15.5146081 | 0.96241492 | 0.46849953 | 2.05424949 | 0.03995155 | NA |
| PROSER2 | 38.6952547 | 0.24578232 | 0.41896553 | 0.58664093 | 0.55744489 | 0.91425998 |
| UPF2 | 103.375295 | -0.33591 | 0.26437025 | -1.2706044 | 0.20386942 | 0.76173533 |
| DHTKD1 | 41.8144227 | -0.0387428 | 0.27662603 | -0.1400548 | 0.88861669 | 0.98270788 |
| SEC61A2 | 18.492536 | 0.2226573 | 0.36080643 | 0.61711013 | 0.53716209 | 0.90763121 |
| NUDT5 | 40.1185085 | 0.03411576 | 0.27190789 | 0.12546809 | 0.90015299 | 0.98531749 |
| CDC123 | 43.3217419 | -0.2006853 | 0.24161824 | -0.8305882 | 0.4062063 | 0.85867915 |
| OPTN | 39.2005417 | -0.5430394 | 0.38087492 | -1.4257683 | 0.15393524 | 0.70572801 |
| PHYH | 33.121866 | 0.4307521 | 0.31996712 | 1.34623862 | 0.17822557 | 0.72841249 |
| SEPHS1 | 32.7591461 | 0.15706557 | 0.34188352 | 0.45941253 | 0.64593795 | 0.93328692 |
| BEND7 | 107.52375 | -0.2108391 | 0.22605295 | -0.9326979 | 0.35097598 | 0.83797093 |
| PRPF18 | 15.7064895 | -0.1488239 | 0.39566947 | -0.3761318 | 0.7068189 | NA |
| AL157392.5 | 30.3041636 | -0.5109924 | 0.2994679 | -1.7063343 | 0.08794585 | 0.63466198 |
| FRMD4A | 144.622253 | 0.23447859 | 0.32140258 | 0.72954796 | 0.46566654 | 0.87981788 |
| FAM107B | 347.013887 | -0.305426 | 0.25421417 | -1.2014514 | 0.22957616 | 0.77201028 |
| DCLRE1C | 32.584151 | 0.09501317 | 0.2941397 | 0.32302056 | 0.74667967 | 0.95563105 |
| RPP38 | 42.7611391 | 0.13391404 | 0.35835788 | 0.37368801 | 0.70863645 | 0.94558664 |
| NMT2 | 48.1175609 | -0.1230411 | 0.35046724 | -0.3510773 | 0.72553038 | 0.95164932 |
| FAM171A1 | 389.865943 | 0.49673758 | 0.29356119 | 1.69210919 | 0.09062516 | 0.64149363 |
| MINDY3 | 75.2047174 | -0.1830998 | 0.23953327 | -0.7644024 | 0.44462748 | 0.87079259 |
| PTER | 77.5560595 | 0.01838797 | 0.30023692 | 0.06124486 | 0.9511642 | 0.99105535 |
| RSU1 | 91.0851043 | -0.5312938 | 0.31627147 | -1.6798663 | 0.09298334 | 0.64572229 |
| CUBN | 43.0226791 | 0.2758825 | 0.46269159 | 0.59625571 | 0.55100442 | 0.91258455 |
| TRDMT1 | 14.9213052 | 0.50561549 | 0.43785368 | 1.15475903 | 0.24818912 | NA |
| VIM | 32.3933776 | -0.0285652 | 0.41849302 | -0.0682574 | 9.46E-01 | 9.91E-01 |

| | | | | | | |
|---|---|---|---|---|---|---|
| HACD1 | 34.6282421 | -2.0134537 | 0.54815526 | -3.673145 | 0.00023958 | 0.06160773 |
| STAM | 105.906745 | -1.3733743 | 0.42275037 | -3.2486649 | 0.00115948 | 0.14394871 |
| CACNB2 | 13.6689376 | -0.051355 | 0.62432477 | -0.0822569 | 0.93444242 | NA |
| AL450384.2 | 22.5248964 | -0.1399499 | 0.41490489 | -0.337306 | 0.7358862 | 0.95376039 |
| NSUN6 | 68.0314673 | 0.11766469 | 0.31160914 | 0.37760346 | 0.70572519 | 0.94558664 |
| ARL5B | 34.1485105 | -1.2029003 | 0.40848417 | -2.9447905 | 0.00323173 | 0.23150356 |
| MALRD1 | 71.8034874 | -0.3439216 | 0.62995718 | -0.5459443 | 0.5851042 | 0.92076087 |
| PLXDC2 | 336.839912 | 0.23548901 | 0.28003339 | 0.8409319 | 0.40038609 | 0.85778581 |
| NEBL | 537.932021 | 0.4534351 | 0.35921537 | 1.26229318 | 0.20684331 | 0.76231289 |
| MLLT10 | 332.851488 | -0.4036483 | 0.15202204 | -2.6551958 | 0.00792624 | 0.30966764 |
| DNAJC1 | 298.571604 | -0.3157964 | 0.19102382 | -1.6531783 | 0.09829459 | 0.65695163 |
| BMI1 | 18.3497603 | 0.01995186 | 0.53440272 | 0.03733488 | 0.97021799 | 0.99398482 |
| PIP4K2A | 146.026562 | -0.0610421 | 0.25339144 | -0.2409003 | 0.80963241 | 0.97257655 |
| MSRB2 | 36.0227038 | -0.2317453 | 0.27726691 | -0.8358204 | 0.40325594 | 0.85782098 |
| OTUD1 | 19.3430196 | -0.0993327 | 0.44641919 | -0.2225098 | 0.82391703 | 0.97518529 |
| KIAA1217 | 599.521729 | -0.0636542 | 0.23212198 | -0.2742272 | 0.78391002 | 0.96727606 |
| ARHGAP21 | 174.725535 | -0.3204197 | 0.22013489 | -1.4555609 | 0.14551403 | 0.69643231 |
| PRTFDC1 | 27.6635188 | -0.3758319 | 0.36807519 | -1.0210737 | 0.30721952 | 0.81568231 |
| PDSS1 | 24.7560243 | -0.4974285 | 0.36679039 | -1.3561655 | 0.17504652 | 0.72841249 |
| ABI1 | 167.286844 | -0.2794577 | 0.21786248 | -1.2827254 | 0.19958828 | 0.75605362 |
| ANKRD26 | 157.819106 | -0.1322619 | 0.25998233 | -0.5087342 | 0.61093855 | 0.92619266 |
| YME1L1 | 109.654772 | -0.131045 | 0.22761242 | -0.5757377 | 0.5647925 | 0.9167807 |
| MASTL | 20.8029871 | -0.3819073 | 0.36529096 | -1.045488 | 0.29579749 | 0.81054924 |
| ACBD5 | 60.7842066 | -0.1567287 | 0.23444016 | -0.6685234 | 0.50379958 | 0.89624537 |
| RAB18 | 105.880304 | -0.3380878 | 0.21378882 | -1.5814099 | 0.11378434 | 0.6656824 |
| MPP7 | 121.991975 | 0.25827946 | 0.26581691 | 0.9716442 | 0.33122758 | 0.82879734 |
| WAC-AS1 | 39.7317967 | 0.1893059 | 0.50233775 | 0.37684984 | 0.7062852 | 0.94558664 |
| WAC | 375.152857 | -0.2186024 | 0.21322815 | -1.0252042 | 0.30526684 | 0.81568231 |
| SVIL | 244.801669 | -0.4189478 | 0.27208339 | -1.5397772 | 1.24E-01 | 6.69E-01 |
| MTPAP | 24.1433622 | 0.16031092 | 0.3540143 | 0.45283741 | 0.65066582 | 0.93453685 |
| MAP3K8 | 37.2106301 | -0.3691813 | 0.34384586 | -1.0736827 | 0.28296493 | 0.80431986 |
| ZNF438 | 85.8446363 | 0.03164892 | 0.19288833 | 0.16407898 | 0.86966897 | 0.97854266 |
| ZEB1 | 59.0987146 | -1.6572046 | 0.62379091 | -2.6566668 | 0.00789174 | 0.30964876 |
| ARHGAP12 | 200.199352 | -0.333208 | 0.16761555 | -1.9879304 | 0.04681939 | 0.5320597 |
| KIF5B | 115.984365 | 0.01836257 | 0.19204717 | 0.09561492 | 0.92382641 | 0.98925753 |
| EPC1 | 187.092947 | 0.05331207 | 0.19051201 | 0.27983574 | 0.77960353 | 0.96621832 |
| CCDC7 | 148.359455 | 0.28089179 | 0.30620554 | 0.91733083 | 0.35896931 | 0.84204868 |
| ITGB1 | 203.478684 | -0.1642951 | 0.35275368 | -0.4657503 | 0.64139424 | 0.93328692 |
| NRP1 | 375.972782 | 0.11005364 | 0.37501802 | 0.29346228 | 0.76916883 | 0.96289288 |
| PARD3 | 1076.21746 | -0.3192279 | 0.15295237 | -2.0871066 | 0.0368785 | 0.49537143 |
| CUL2 | 73.470813 | -0.0798171 | 0.24748284 | -0.3225158 | 0.74706199 | 0.95563105 |
| CREM | 179.102901 | -0.8097689 | 0.50929045 | -1.5899942 | 0.11183611 | 0.66538829 |
| CCNY | 347.831933 | -0.4970642 | 0.17993242 | -2.7625049 | 0.00573597 | 0.2943676 |
| AL117336.3 | 38.1690685 | 0.01121071 | 0.32923945 | 0.03405031 | 0.97283703 | 0.99431961 |
| FZD8 | 18.9561965 | -0.0121484 | 0.51131618 | -0.0237591 | 0.98104473 | 0.99592962 |
| ZNF248 | 135.709373 | 0.13783249 | 0.34469599 | 0.39986683 | 0.6892546 | 0.94210755 |
| ZNF25 | 22.5660858 | 0.36714112 | 0.32693975 | 1.12296262 | 0.26145337 | 0.79625629 |
| ZNF33A | 220.172076 | -0.0139835 | 0.2429167 | -0.0575651 | 0.95409504 | 0.99148072 |
| AL117339.5 | 19.1822127 | 0.0124248 | 0.40062493 | 0.03101354 | 0.97525874 | 0.99440004 |
| ZNF33B | 125.803802 | -0.4160279 | 0.28018915 | -1.4848108 | 0.13759396 | 0.68706409 |
| BMS1 | 60.0462003 | -0.0076331 | 0.27159146 | -0.0281052 | 0.97757822 | 0.99469237 |
| CSGALNACT2 | 39.0961259 | -0.2011833 | 0.36562626 | -0.5502431 | 0.58215264 | 0.92034873 |
| FXYD4 | 61.077251 | -0.2903079 | 0.70413903 | -0.4122878 | 0.68012848 | 0.94195357 |
| HNRNPF | 87.9065731 | -0.3390831 | 0.24111131 | -1.4063343 | 0.15962488 | 0.71101785 |

| | | | | | | |
|---|---|---|---|---|---|---|
| ZNF487 | 19.0925991 | 0.0469986 | 0.38410827 | 0.1223577 | 0.90261574 | 0.98531749 |
| ZNF32 | 23.5950417 | 0.37522817 | 0.37434513 | 1.00235888 | 0.31617029 | 0.82160953 |
| TMEM72-AS | 28.9850591 | 0.35203948 | 0.52081208 | 0.67594337 | 0.4990766 | 0.89378207 |
| TMEM72 | 116.084416 | 1.12565675 | 0.44183592 | 2.5476805 | 0.01084417 | 0.33942942 |
| RASSF4 | 25.8521835 | 1.07872888 | 0.43316624 | 2.49033462 | 0.01276229 | 0.36544853 |
| DEPP1 | 20.0394906 | 0.1432087 | 0.35764014 | 0.4004268 | 0.68884219 | 0.94205398 |
| ZNF22 | 18.2652647 | 0.43585561 | 0.45628014 | 0.95523686 | 0.33945793 | 0.83066816 |
| 8-Mar | 99.516752 | -0.1833072 | 0.23124956 | -0.7926813 | 0.42796356 | 0.86279045 |
| ZFAND4 | 28.7039548 | 0.08989166 | 0.41304961 | 0.2176292 | 0.82771803 | 0.97567084 |
| WASHC2C | 31.5244134 | -0.4110609 | 0.41975319 | -0.9792918 | 0.32743583 | 0.82612204 |
| AGAP4 | 47.874834 | 0.08164175 | 0.33399768 | 0.24443808 | 0.80689155 | 0.97216031 |
| TIMM23 | 41.7668627 | -0.2709236 | 0.35305845 | -0.767362 | 0.44286632 | 0.86991418 |
| NCOA4 | 55.0436849 | 0.42038604 | 0.32092439 | 1.30992238 | 0.1902221 | 0.74573288 |
| PTPN20 | 56.8604627 | 0.18554458 | 0.48010684 | 0.38646519 | 0.69915218 | 0.94395238 |
| AGAP9 | 45.7413114 | 1.20645913 | 0.45659817 | 2.64227765 | 0.00823505 | 0.30976355 |
| MAPK8 | 415.194759 | -0.4654854 | 0.25828243 | -1.8022342 | 0.07150857 | 0.60722248 |
| ARHGAP22 | 20.2785552 | -0.3402454 | 0.41100655 | -0.8278344 | 4.08E-01 | 8.59E-01 |
| ERCC6 | 56.2647243 | 0.06583042 | 0.29770139 | 0.22112902 | 0.82499197 | 0.97518529 |
| OGDHL | 140.299751 | 0.96979445 | 0.33853756 | 2.86465834 | 4.17E-03 | 0.25676573 |
| PARG | 129.764939 | -0.2319071 | 0.20433874 | -1.134915 | 0.25641094 | 0.79418467 |
| TIMM23B | 134.423635 | -0.7837442 | 0.43690963 | -1.793836 | 0.07283936 | 0.60942708 |
| LINC00843 | 28.0867011 | 0.30569079 | 0.3165887 | 0.96557705 | 0.33425586 | 0.82974237 |
| AGAP6 | 37.7562058 | 0.12530491 | 0.33627303 | 0.37262848 | 0.70942497 | 0.94565757 |
| WASHC2A | 33.8545184 | -0.7517976 | 0.27765888 | -2.7076303 | 0.00677655 | 0.30102597 |
| SGMS1 | 558.54914 | 0.07419956 | 0.14242371 | 0.52097756 | 0.6023824 | 0.92407937 |
| ASAH2B | 14.8829796 | -0.2209211 | 0.51471515 | -0.4292104 | 0.66777013 | NA |
| PRKG1 | 2036.17323 | -0.2393694 | 0.28790679 | -0.8314128 | 4.06E-01 | 8.59E-01 |
| IPMK | 37.1342927 | -0.1368536 | 0.35401937 | -0.3865708 | 0.69907397 | 0.94395238 |
| CISD1 | 87.1978267 | 0.07543233 | 0.269579 | 0.27981532 | 0.7796192 | 0.96621832 |
| UBE2D1 | 31.7882245 | -0.2774097 | 0.36024684 | -0.7700544 | 0.4412676 | 0.86925451 |
| TFAM | 30.8275471 | -0.2111933 | 0.33598378 | -0.6285817 | 0.52962292 | 0.9052447 |
| BICC1 | 2038.15406 | -0.2519993 | 0.36476093 | -0.6908614 | 0.48965262 | 0.88932718 |
| PHYHIPL | 18.2214419 | 0.34850863 | 0.46874177 | 0.74349814 | 0.45718015 | 0.87419345 |
| CCDC6 | 99.5737905 | 0.08662128 | 0.193011 | 0.44878933 | 0.65358364 | 0.93532021 |
| ANK3 | 1656.87893 | 0.34464272 | 0.22207401 | 1.55192727 | 1.21E-01 | 6.67E-01 |
| RHOBTB1 | 14.7734265 | -0.4567184 | 0.51169195 | -0.8925652 | 0.37209007 | NA |
| ARID5B | 523.558929 | -0.9272088 | 0.40020981 | -2.3168067 | 0.02051426 | 0.42078083 |
| NRBF2 | 12.1963728 | -0.3599653 | 0.43153296 | -0.8341548 | 0.40419378 | NA |
| JMJD1C | 823.21649 | -0.4121056 | 0.19837012 | -2.0774582 | 0.03775929 | 0.49640946 |
| REEP3 | 157.461721 | -0.4789171 | 0.37949786 | -1.2619758 | 0.20695749 | 0.76231289 |
| LINC01515 | 27.5051545 | 0.28029558 | 0.53231793 | 0.52655672 | 0.59850145 | 0.92296674 |
| CTNNA3 | 17.2071025 | -0.1386184 | 0.60104659 | -0.2306284 | 0.81760347 | 0.97288403 |
| SIRT1 | 48.4560873 | -0.7720033 | 0.36745725 | -2.1009336 | 0.03564679 | 0.49240174 |
| HERC4 | 314.18011 | -0.4043113 | 0.21618792 | -1.8701846 | 0.06145819 | 0.58094903 |
| PBLD | 55.0232115 | -0.3313856 | 0.31933795 | -1.037727 | 0.29939719 | 0.81335071 |
| HNRNPH3 | 112.27884 | -0.0580711 | 0.24818267 | -0.2339855 | 0.81499623 | 0.97284589 |
| RUFY2 | 104.920109 | -0.1345395 | 0.21198296 | -0.6346713 | 0.52564278 | 0.90487731 |
| DNA2 | 13.9530029 | -0.1470015 | 0.49064677 | -0.2996075 | 0.76447655 | NA |
| SLC25A16 | 116.70524 | -0.2429403 | 0.26476742 | -0.9175612 | 0.35884866 | 0.84204868 |
| TET1 | 38.4226877 | 0.0863453 | 0.44034393 | 0.19608605 | 8.45E-01 | 9.76E-01 |
| CCAR1 | 198.526364 | -0.4743723 | 0.29376774 | -1.6147869 | 0.10635685 | 0.65986731 |
| STOX1 | 23.8894122 | 0.1239539 | 0.49424694 | 0.25079347 | 0.80197379 | 0.97079354 |
| DDX50 | 59.9380032 | 0.06448043 | 0.2912834 | 0.22136663 | 0.82480697 | 0.97518529 |
| DDX21 | 48.665127 | -0.2540788 | 0.44216999 | -0.574618 | 0.56554969 | 0.91698355 |

| | | | | | | |
|---|---|---|---|---|---|---|
| KIF1BP | 29.3078086 | 0.48489636 | 0.30354763 | 1.59743088 | 0.11016969 | 0.66283855 |
| VPS26A | 66.1091292 | -0.1637175 | 0.33361685 | -0.4907352 | 0.62361375 | 0.92943979 |
| SUPV3L1 | 36.3401669 | -0.3043275 | 0.31159337 | -0.9766814 | 0.3287269 | 0.82781307 |
| HKDC1 | 95.6281822 | -0.3631083 | 0.75172756 | -0.4830318 | 0.62907318 | 0.9314335 |
| HK1 | 155.034897 | -0.4854224 | 0.2476339 | -1.9602423 | 0.04996748 | 0.54408368 |
| TSPAN15 | 55.851501 | 0.72057741 | 0.24741152 | 2.91246507 | 3.59E-03 | 0.23862997 |
| H2AFY2 | 44.7703895 | 0.11539842 | 0.29575986 | 0.39017607 | 0.69640635 | 0.94395238 |
| AIFM2 | 22.8919205 | 0.16983498 | 0.33055005 | 0.51379505 | 0.60739529 | 0.92569157 |
| TYSND1 | 16.087338 | 0.6100887 | 0.45267447 | 1.34774266 | 0.17774117 | NA |
| SAR1A | 46.4825931 | -0.1318625 | 0.3208674 | -0.4109562 | 0.68110463 | 0.94196488 |
| PPA1 | 76.1995636 | 0.03339628 | 0.2699809 | 0.12369868 | 9.02E-01 | 9.85E-01 |
| EIF4EBP2 | 56.5215173 | 0.12798666 | 0.27768904 | 0.46089922 | 0.64487091 | 0.93328692 |
| SGPL1 | 101.860365 | -0.1031089 | 0.21176534 | -0.4869019 | 0.62632788 | 0.93032781 |
| PCBD1 | 43.2090659 | 1.01875646 | 0.43627535 | 2.33512266 | 0.01953701 | 0.41544117 |
| PSAP | 262.222933 | 0.25079622 | 0.38648571 | 0.6489146 | 0.51639358 | 0.90225158 |
| CHST3 | 42.1820058 | -0.2504252 | 0.28486261 | -0.8791088 | 0.37934229 | 0.85429312 |
| ASCC1 | 71.2936676 | -0.2815151 | 0.28825307 | -0.976625 | 0.32875485 | 0.82781307 |
| ANAPC16 | 180.39896 | 0.0022937 | 0.2257308 | 0.01016124 | 0.99189265 | 0.99814944 |
| DDIT4 | 108.360228 | -1.7112222 | 0.46595881 | -3.6724753 | 0.00024021 | 0.06160773 |
| DNAJB12 | 21.9562095 | -0.1331669 | 0.36385193 | -0.3659921 | 0.71437099 | 0.94841799 |
| MICU1 | 208.54501 | -0.6108382 | 0.18586325 | -3.2864928 | 0.00101443 | 0.13958055 |
| MCU | 256.946938 | -0.8678222 | 0.40993157 | -2.1169929 | 0.03426045 | 0.4908133 |
| P4HA1 | 131.452453 | -0.696323 | 0.40125722 | -1.7353531 | 0.08267828 | 0.62228971 |
| ECD | 39.4204606 | -0.3999857 | 0.29686244 | -1.3473771 | 0.1778588 | 0.72841249 |
| FAM149B1 | 59.939001 | 0.00741375 | 0.2663835 | 0.02783113 | 0.97779684 | 0.99469237 |
| DNAJC9 | 45.0804199 | -0.0604136 | 0.32922206 | -0.1835041 | 0.85440248 | 0.97612951 |
| MRPS16 | 32.9169789 | 0.03106445 | 0.31713073 | 0.09795472 | 0.92196825 | 0.98846691 |
| CFAP70 | 84.174624 | 0.69047991 | 0.57681069 | 1.19706504 | 0.2312812 | 0.77260653 |
| ANXA7 | 104.549778 | -0.1274404 | 0.2122461 | -0.600437 | 0.54821503 | 0.91182193 |
| PPP3CB | 107.383223 | -0.3251803 | 0.20836056 | -1.5606615 | 0.11860364 | 0.6668162 |
| USP54 | 81.4344672 | -0.4666816 | 0.24306739 | -1.9199682 | 0.05486192 | 0.5645224 |
| AGAP5 | 18.1222979 | 0.44793885 | 0.35319346 | 1.26825353 | 0.20470743 | 0.76183635 |
| SEC24C | 31.7387006 | -0.3304679 | 0.29208843 | -1.1313969 | 0.25788806 | 0.79532468 |
| FUT11 | 34.7530908 | -0.6121503 | 0.45106595 | -1.3571193 | 1.75E-01 | 0.72841249 |
| CHCHD1 | 26.0806859 | 0.2722268 | 0.45789325 | 0.59452023 | 0.55216422 | 0.91322136 |
| ZSWIM8 | 47.4834498 | -0.3275005 | 0.30478642 | -1.0745245 | 0.28258766 | 0.80431986 |
| CAMK2G | 39.987713 | -0.4256652 | 0.32152266 | -1.3239043 | 0.18553482 | 0.73752116 |
| VCL | 243.539267 | -0.2220629 | 0.38460442 | -0.5773799 | 0.56368282 | 0.91628656 |
| AP3M1 | 25.9211261 | 0.28569212 | 0.31815372 | 0.89796884 | 0.36920216 | 0.84848455 |
| ADK | 711.463153 | -0.3511629 | 0.13066351 | -2.6875362 | 0.00719813 | 0.30476547 |
| KAT6B | 241.754421 | -0.4508779 | 0.15941785 | -2.8282773 | 0.00467993 | 0.27019594 |
| SAMD8 | 59.1645299 | -0.2429502 | 0.26178221 | -0.9280622 | 0.35337531 | 0.83945817 |
| VDAC2 | 52.1596081 | 0.38266567 | 0.29618186 | 1.29199564 | 0.19635866 | 0.75350056 |
| COMTD1 | 20.9401828 | 0.20186543 | 0.46840504 | 0.43096339 | 0.66649499 | 0.93860542 |
| LRMDA | 1721.68553 | 0.01434955 | 0.17160762 | 0.08361836 | 0.93335987 | 0.98960607 |
| DLG5 | 82.5103366 | -0.4236281 | 0.30862105 | -1.3726482 | 0.16986175 | 0.72416949 |
| POLR3A | 34.7466741 | -0.0865989 | 0.33380753 | -0.2594276 | 0.79530533 | 0.96974891 |
| RPS24 | 239.278264 | -0.0312738 | 0.44576698 | -0.0701572 | 0.94406854 | 0.99069421 |
| ZMIZ1-AS1 | 13.6514232 | -0.6653162 | 0.64136631 | -1.0373419 | 0.29957653 | NA |
| ZMIZ1 | 261.52095 | -0.2247322 | 0.38231382 | -0.5878214 | 0.55665219 | 0.9141017 |
| PPIF | 27.750695 | -0.3572414 | 0.37844274 | -0.9439773 | 0.34518123 | 0.83564825 |
| NUTM2B-AS | 208.387427 | 0.14650098 | 0.34785109 | 0.42116004 | 0.67363823 | 0.93901638 |
| TMEM254 | 18.6570629 | 0.22528646 | 0.37016037 | 0.60861853 | 0.54277732 | 0.90953689 |
| ANXA11 | 260.420138 | 0.13368429 | 0.22089229 | 0.60520123 | 0.5450453 | 0.90999223 |

| | | | | | | |
|---|---|---|---|---|---|---|
| FAM213A | 62.4955667 | 0.10528719 | 0.29755753 | 0.35383811 | 0.72346021 | 0.95093722 |
| TSPAN14 | 56.9172613 | 0.21078665 | 0.28188834 | 0.74776647 | 0.45460103 | 0.87378486 |
| NRG3 | 62.3534248 | 1.15578999 | 0.41670909 | 2.77361356 | 5.54E-03 | 2.87E-01 |
| GHITM | 72.343298 | 0.44768285 | 0.34308947 | 1.30485745 | 0.19194139 | 0.75053372 |
| CCSER2 | 180.905914 | -0.4048348 | 0.25924817 | -1.5615727 | 0.1183887 | 0.6668162 |
| WAPL | 151.424748 | -0.0918973 | 0.22364067 | -0.410915 | 0.68113488 | 0.94196488 |
| BMPR1A | 200.279928 | 0.06769078 | 0.1768133 | 0.38283761 | 0.70184017 | 0.94476609 |
| ADIRF | 17.9631243 | 0.9672914 | 0.66243202 | 1.46021231 | 0.14423173 | 0.69512617 |
| GLUD1 | 100.37472 | 0.43045656 | 0.23496128 | 1.83203191 | 0.06694667 | 0.59652811 |
| SHLD2 | 155.260183 | 0.02140346 | 0.20081108 | 0.10658505 | 0.91511818 | 0.98691474 |
| NUTM2A-AS | 219.224994 | -0.1273972 | 0.28641523 | -0.4447988 | 0.65646516 | 0.93570875 |
| LINC00864 | 22.3683205 | 1.2101647 | 0.49012029 | 2.46911773 | 0.01354466 | 0.37249875 |
| MINPP1 | 19.166287 | 0.23180054 | 0.42215977 | 0.5490825 | 0.58294884 | 0.92034873 |
| PAPSS2 | 76.4727671 | -0.3778167 | 0.41200342 | -0.9170232 | 0.3591305 | 0.84204868 |
| ATAD1 | 116.879531 | -0.4163044 | 0.28993411 | -1.4358587 | 0.15104254 | 0.70572801 |
| PTEN | 267.932607 | -0.2687625 | 0.24168722 | -1.1120261 | 0.26612693 | 0.79666376 |
| RNLS | 92.4909469 | -0.1568733 | 0.37152524 | -0.4222414 | 0.67284885 | 0.93880075 |
| STAMBPL1 | 12.1733892 | 0.38119481 | 0.46103789 | 0.82681883 | 0.40833975 | NA |
| LIPA | 42.2187618 | -0.5615606 | 0.31256373 | -1.7966276 | 0.07239475 | 0.60824637 |
| IFIT2 | 29.3865877 | -3.436068 | 1.43524566 | -2.3940627 | 0.0166629 | 0.39748373 |
| IFIT3 | 59.2979362 | -2.7907563 | 1.14219726 | -2.4433225 | 0.01455273 | 0.38288257 |
| IFIT1 | 47.6560821 | -5.2505138 | 1.42304364 | -3.6896365 | 0.00022457 | 0.06160773 |
| PANK1 | 35.5780438 | 0.52193429 | 0.44935564 | 1.16151715 | 0.24543165 | 0.78583441 |
| RPP30 | 16.0769507 | 0.20900808 | 0.44740299 | 0.46715844 | 0.64038653 | NA |
| PCGF5 | 203.110576 | -0.4166386 | 0.28307837 | -1.4718137 | 0.14107119 | 0.69227547 |
| HECTD2 | 85.6792952 | 0.0012641 | 0.26210997 | 0.0048228 | 0.99615198 | 0.99927028 |
| PPP1R3C | 15.2744841 | -0.5370623 | 0.7125685 | -0.7536992 | 0.45102984 | NA |
| TNKS2 | 159.774748 | -0.4218607 | 0.17169173 | -2.4570823 | 0.01400706 | 0.37656544 |
| BTAF1 | 150.319493 | -0.2737364 | 0.3642727 | -0.75146 | 0.45237584 | 0.87346041 |
| CPEB3 | 366.510816 | 0.34960088 | 0.26658175 | 1.31142087 | 1.90E-01 | 7.45E-01 |
| 5-Mar | 80.2999028 | -0.1437398 | 0.25346982 | -0.5670882 | 0.57065424 | 0.91778542 |
| IDE | 81.3732804 | 0.00388208 | 0.31004377 | 0.01252106 | 0.9900099 | 0.99814944 |
| EXOC6 | 107.108154 | -0.3077562 | 0.27198671 | -1.131512 | 0.25783967 | 0.79532468 |
| MYOF | 161.405033 | -0.7047525 | 0.24041014 | -2.9314589 | 0.00337374 | 0.23150356 |
| FRA10AC1 | 55.7311981 | -0.1090402 | 0.28842835 | -0.3780496 | 0.70539373 | 0.94548439 |
| SLC35G1 | 14.5849925 | 1.13173124 | 0.44182306 | 2.56150332 | 0.01042202 | NA |
| PLCE1 | 13.5574352 | 0.36180869 | 0.64102431 | 0.56442272 | 0.57246648 | NA |
| NOC3L | 20.6990876 | -0.4267123 | 0.33981324 | -1.255726 | 0.20921533 | 0.7623519 |
| TBC1D12 | 70.5878724 | 0.11738625 | 0.21971962 | 0.53425476 | 0.59316529 | 0.9229638 |
| HELLS | 58.2548234 | 0.53492392 | 0.24041545 | 2.22499808 | 0.02608137 | 0.4455707 |
| SORBS1 | 45.4288353 | -0.4515194 | 0.37289402 | -1.2108518 | 2.26E-01 | 0.77093069 |
| ALDH18A1 | 51.0037136 | -0.4562154 | 0.37005756 | -1.2328227 | 0.21764192 | 0.76666749 |
| TCTN3 | 53.417069 | 0.06029399 | 0.2823433 | 0.21354851 | 0.83089916 | 0.97567084 |
| ENTPD1-AS1 | 148.863393 | 0.48710424 | 0.33931365 | 1.43555744 | 0.15112831 | 0.70572801 |
| ZNF518A | 202.175837 | 0.11327503 | 0.2394285 | 0.47310587 | 0.63613765 | 0.93251095 |
| BLNK | 371.231628 | 0.23686016 | 0.31185705 | 0.75951517 | 0.44754444 | 0.87275002 |
| TM9SF3 | 196.019001 | 0.02481928 | 0.1624461 | 0.15278473 | 0.87856804 | 0.98061726 |
| LCOR | 330.382472 | -0.222794 | 0.23392631 | -0.9524111 | 0.34088852 | 0.8319325 |
| RRP12 | 24.9858536 | -0.1960723 | 0.43888145 | -0.4467545 | 0.65505234 | 0.93548446 |
| PGAM1 | 87.6458896 | -0.0538356 | 0.36925343 | -0.1457958 | 0.8840826 | 0.98154805 |
| EXOSC1 | 24.9751786 | 0.16956562 | 0.34503044 | 0.49145119 | 0.62310737 | 0.92925071 |
| ZDHHC16 | 18.8043509 | -0.455719 | 0.38084153 | -1.1966104 | 0.23145843 | 0.7726315 |
| MMS19 | 54.4838258 | -0.3120432 | 0.24451473 | -1.2761735 | 0.20189421 | 0.76038186 |
| UBTD1 | 38.719299 | -0.0670813 | 0.32753268 | -0.2048081 | 0.83772209 | 0.97600352 |

| | | | | | | |
|---|---|---|---|---|---|---|
| MORN4 | 22.2692574 | 0.28137671 | 0.33099801 | 0.8500858 | 0.39527738 | 0.85605713 |
| ZFYVE27 | 31.0989274 | 0.07638154 | 0.28763101 | 0.26555392 | 0.79058278 | 0.96904219 |
| R3HCC1L | 106.049173 | -0.2066992 | 0.19338649 | -1.0688401 | 0.28514173 | 0.80431986 |
| HPS1 | 19.2521565 | 0.09407268 | 0.36083069 | 0.26071142 | 0.79431506 | 0.96974891 |
| GOT1 | 163.925919 | 0.07465508 | 0.32543004 | 0.22940439 | 0.81855463 | 0.97288403 |
| AL391684.1 | 11.1769075 | -0.5564425 | 0.5595116 | -0.9945148 | 0.31997233 | NA |
| SLC25A28 | 34.9034944 | -0.2244672 | 0.54643923 | -0.4107817 | 0.68123262 | 0.94196488 |
| ENTPD7 | 11.5760294 | -0.4802454 | 0.45141216 | -1.0638734 | 0.28738605 | NA |
| CUTC | 39.1255572 | 0.24619606 | 0.27677954 | 0.88950237 | 0.37373315 | 0.85056774 |
| COX15 | 35.3169392 | 0.14131166 | 0.26024567 | 0.54299332 | 0.5871344 | 0.92086224 |
| DNMBP | 91.2251447 | -0.4085777 | 0.20510225 | -1.9920686 | 0.04636354 | 0.53087707 |
| ERLIN1 | 25.5607918 | 0.27942147 | 0.3599773 | 0.77621969 | 0.43761928 | 0.86820528 |
| CHUK | 28.8543938 | -0.2198524 | 0.4376949 | -0.502296 | 0.61545935 | 0.92662475 |
| CWF19L1 | 33.5490523 | -0.0567717 | 0.32129606 | -0.1766958 | 0.85974733 | 0.97691173 |
| BLOC1S2 | 20.9666987 | -0.2616976 | 0.34700017 | -0.7541713 | 0.45074636 | 0.87346041 |
| SEC31B | 15.898343 | 0.34849573 | 0.41322683 | 0.84335214 | 0.39903153 | NA |
| HIF1AN | 33.5496592 | -0.5669822 | 0.34926219 | -1.6233711 | 0.10451009 | 0.65946648 |
| PAX2 | 334.947742 | 0.31261586 | 0.30446192 | 1.0267815 | 0.30452336 | 0.81568231 |
| SLF2 | 76.1514629 | -0.3755679 | 0.31589971 | -1.1888835 | 0.23448552 | 0.77743304 |
| MRPL43 | 19.1933028 | 0.69260244 | 0.4220893 | 1.64089075 | 0.1008201 | 0.65863153 |
| LZTS2 | 28.3851557 | -0.4182171 | 0.32396009 | -1.2909526 | 0.1967201 | 0.75350056 |
| BTRC | 190.637611 | -0.0278242 | 0.18465739 | -0.1506803 | 0.8802279 | 0.98093344 |
| DPCD | 19.8838904 | -0.0589052 | 0.42511182 | -0.1385639 | 0.88979476 | 0.9827634 |
| POLL | 21.3274743 | 0.17738516 | 0.41847455 | 0.42388517 | 0.67164956 | 0.93878847 |
| FBXW4 | 133.29854 | -0.0780812 | 0.2112839 | -0.3695559 | 0.71171339 | 0.94676706 |
| OGA | 246.61626 | -0.0898643 | 0.18599593 | -0.483152 | 0.62898783 | 0.9314335 |
| ARMH3 | 175.084198 | -0.2011941 | 0.20283974 | -0.9918869 | 0.32125268 | 0.82490712 |
| LDB1 | 26.5098714 | 0.48153192 | 0.38175617 | 1.26135988 | 0.20717921 | 0.76231289 |
| PPRC1 | 20.4062469 | -0.7684529 | 0.4801225 | -1.6005351 | 0.10947994 | 0.66067258 |
| NOLC1 | 15.5456684 | 0.34291261 | 0.45423062 | 0.75493065 | 0.4502906 | NA |
| GBF1 | 242.495438 | -0.1812448 | 0.20774996 | -0.8724179 | 0.38298045 | 0.85510589 |
| NFKB2 | 20.0308026 | -0.4632449 | 0.41115665 | -1.126687 | 0.25987485 | 0.79535701 |
| FBXL15 | 23.7554121 | 0.62012221 | 0.41391415 | 1.49819042 | 0.13408378 | 0.68205729 |
| CUEDC2 | 20.1580938 | 0.04560114 | 0.33699138 | 0.13531843 | 0.89236012 | 0.98343271 |
| ACTR1A | 29.2003674 | -0.3066905 | 0.37386198 | -0.8203309 | 0.41202751 | 0.85867915 |
| SUFU | 65.8988794 | 0.33130776 | 0.25157401 | 1.31693953 | 0.18785889 | 0.7419 |
| TRIM8 | 66.9411877 | 0.59875112 | 0.3429731 | 1.74576697 | 0.08085145 | 0.61925304 |
| ARL3 | 45.3388672 | 0.32054995 | 0.26974183 | 1.18835835 | 0.23469226 | 0.77743304 |
| SFXN2 | 15.2732188 | 0.35406165 | 0.55120784 | 0.64233784 | 0.52065385 | NA |
| WBP1L | 64.3414875 | -0.4958947 | 0.26323892 | -1.8838196 | 0.05958938 | 0.57584087 |
| CNNM2 | 255.181949 | 0.17530605 | 0.20849188 | 0.84082915 | 4.00E-01 | 8.58E-01 |
| NT5C2 | 349.611549 | -0.4562709 | 0.25586273 | -1.7832643 | 0.07454329 | 0.61020315 |
| PCGF6 | 30.9088111 | -0.1826502 | 0.41163373 | -0.4437202 | 0.6572449 | 0.93593674 |
| ATP5MD | 41.3830422 | -0.1408356 | 0.26501807 | -0.531419 | 0.59512843 | 0.9229638 |
| PDCD11 | 26.0245442 | -0.7854575 | 0.31407585 | -2.5008528 | 0.01238947 | 0.36046953 |
| SH3PXD2A | 54.8433205 | 0.18316373 | 0.3015602 | 0.60738696 | 0.54359414 | 0.90965439 |
| STN1 | 34.9834138 | -0.7341521 | 0.38793497 | -1.8924617 | 0.05842949 | 0.57141408 |
| SLK | 112.365273 | -0.4026793 | 0.26510127 | -1.518964 | 0.12877155 | 0.66929384 |
| CFAP43 | 31.3073615 | 0.27884798 | 0.44858744 | 0.62161344 | 0.53419608 | 0.90763121 |
| GSTO1 | 43.215608 | -0.1262443 | 0.2750308 | -0.4590187 | 0.64622072 | 0.93328692 |
| GSTO2 | 83.5236135 | 0.23492495 | 0.34967002 | 0.67184756 | 0.50168074 | 0.89492452 |
| LINC01435 | 70.7325829 | -1.1720715 | 0.86718536 | -1.3515812 | 0.17650931 | 0.72841249 |
| XPNPEP1 | 44.9907787 | -0.1896444 | 0.3427078 | -0.5533707 | 0.58000962 | 0.92033309 |
| ADD3 | 215.917973 | 0.16431614 | 0.28939699 | 0.56778802 | 0.57017891 | 0.91778542 |

| | | | | | | |
|---|---|---|---|---|---|---|
| MXI1 | 61.3413874 | 0.14774858 | 0.32954354 | 0.44834311 | 0.65390559 | 0.93548446 |
| SMNDC1 | 27.5300849 | 0.025329 | 0.34457975 | 0.07350694 | 0.94140272 | 0.99043681 |
| SMC3 | 46.8489347 | -0.126155 | 0.27911053 | -0.4519894 | 0.65127661 | 0.93453685 |
| PDCD4 | 60.9130952 | 0.25404194 | 0.44428113 | 0.57180447 | 0.56745445 | 0.91698355 |
| BBIP1 | 35.8011463 | -0.3198099 | 0.3097415 | -1.0325056 | 0.30183531 | 0.81568231 |
| SHOC2 | 119.510923 | -0.1593628 | 0.24285345 | -0.6562098 | 0.51168914 | 0.90094899 |
| GPAM | 24.8592495 | 0.39994467 | 0.37783526 | 1.05851602 | 0.28982025 | 0.80536924 |
| ZDHHC6 | 29.5825572 | 0.04601693 | 0.28071025 | 0.16393035 | 0.86978597 | 0.97854266 |
| VTI1A | 412.93118 | -0.1511413 | 0.16432663 | -0.9197615 | 0.35769739 | 0.84204868 |
| TCF7L2 | 350.899908 | -0.2556287 | 0.19339104 | -1.3218231 | 0.18622707 | 0.73831062 |
| CASP7 | 30.2375465 | -0.2809511 | 0.3209764 | -0.8753014 | 0.38140996 | 0.85432452 |
| NHLRC2 | 84.5351084 | 0.18571611 | 0.21211928 | 0.87552679 | 0.38128734 | 0.85432452 |
| CCDC186 | 82.3130642 | -0.0029535 | 0.2231508 | -0.0132357 | 0.98943978 | 0.99814944 |
| AFAP1L2 | 105.981536 | 0.79304994 | 0.3292532 | 2.40863241 | 0.01601242 | 0.39716901 |
| ABLIM1 | 480.616253 | 0.17575239 | 0.29371497 | 0.59837737 | 0.54958816 | 0.91221128 |
| FAM160B1 | 48.4447289 | 0.04499667 | 0.3002851 | 0.14984648 | 0.88088574 | 0.98112111 |
| TRUB1 | 27.3683928 | 0.23621318 | 0.39676562 | 0.5953469 | 0.55161161 | 0.9128014 |
| HSPA12A | 148.543648 | 0.32202903 | 0.28224591 | 1.14095199 | 0.2538899 | 0.7912221 |
| SHTN1 | 98.5102295 | -0.2942896 | 0.31306151 | -0.9400377 | 0.34719824 | 0.83687845 |
| PDZD8 | 120.956431 | -0.4372085 | 0.27075512 | -1.6147746 | 0.10635951 | 0.65986731 |
| AC005871.2 | 14.8805775 | 0.55674128 | 0.57871131 | 0.96203629 | 0.33603138 | NA |
| EMX2OS | 315.537225 | 0.1885091 | 0.23515745 | 0.8016293 | 0.42276743 | 0.86190547 |
| EMX2 | 118.928434 | 0.14018477 | 0.25992034 | 0.53933744 | 0.58965404 | 0.92227161 |
| RAB11FIP2 | 217.177383 | 0.14165947 | 0.29131686 | 0.48627281 | 0.62677375 | 0.93032781 |
| AC022395.1 | 17.5715524 | 0.78299255 | 0.63471177 | 1.23361908 | 0.21734488 | 0.76666749 |
| CASC2 | 103.93494 | 0.76820075 | 0.44568581 | 1.72363747 | 0.08477331 | 0.62880224 |
| FAM204A | 61.4798904 | -0.0263087 | 0.21411497 | -0.1228717 | 0.90220871 | 0.98531749 |
| CACUL1 | 137.342019 | -0.5287182 | 0.22467903 | -2.3532157 | 0.01861183 | 0.41322807 |
| EIF3A | 148.214774 | -0.2514508 | 0.29586476 | -0.8498841 | 0.39538953 | 0.85605713 |
| FAM45A | 25.2054894 | 0.17231693 | 0.31857002 | 0.54090756 | 0.5885713 | 0.92184525 |
| SFXN4 | 45.795003 | 0.09150279 | 0.38346874 | 0.23861865 | 0.8114013 | 0.97257655 |
| PRDX3 | 60.3555861 | 0.5296307 | 0.34035112 | 1.55613031 | 0.11967711 | 0.6668162 |
| TIAL1 | 78.464444 | 0.12516885 | 0.22679933 | 0.55189248 | 0.58102202 | 0.92034873 |
| BAG3 | 35.9882661 | 0.26582191 | 0.32241062 | 0.82448248 | 0.40966546 | 0.85867915 |
| INPP5F | 62.6117642 | 0.34144443 | 0.23902348 | 1.42849744 | 0.15314873 | 0.70572801 |
| MCMBP | 47.6512639 | 0.21060621 | 0.25654741 | 0.82092511 | 0.41168892 | 0.85867915 |
| SEC23IP | 76.6061106 | -0.5897732 | 0.284873 | -2.0703023 | 0.03842405 | 0.49671741 |
| PLPP4 | 103.908616 | 1.16418001 | 0.40361829 | 2.88435889 | 3.92E-03 | 2.45E-01 |
| WDR11-AS1 | 22.5021854 | 1.12304067 | 0.42794246 | 2.6242796 | 0.00868325 | 0.31119527 |
| WDR11 | 108.796893 | 0.17852035 | 0.18123 | 0.98504853 | 0.32460023 | 0.82567895 |
| FGFR2 | 160.642912 | -0.0256014 | 0.34871875 | -0.0734157 | 0.94147531 | 0.99043681 |
| ATE1 | 155.782857 | -0.008904 | 0.20788879 | -0.0428306 | 0.96583661 | 0.99277446 |
| NSMCE4A | 18.4272236 | -0.0322704 | 0.36476691 | -0.0884686 | 0.92950422 | 0.98960607 |
| TACC2 | 159.374415 | -0.3108985 | 0.28888564 | -1.0761992 | 0.28183817 | 0.80425482 |
| PLEKHA1 | 387.196401 | -0.7588357 | 0.27707915 | -2.7386965 | 6.17E-03 | 0.29709153 |
| C10orf88 | 18.7055887 | -0.0261516 | 0.36664215 | -0.0713272 | 0.94313735 | 0.99058259 |
| PSTK | 12.9130253 | 0.1931364 | 0.45792067 | 0.42176825 | 0.67319418 | NA |
| IKZF5 | 22.8414368 | 0.08558144 | 0.3557036 | 0.24059763 | 0.80986698 | 0.97257655 |
| ACADSB | 51.5025994 | 0.01116146 | 0.25783134 | 0.04328978 | 0.96547054 | 0.99270395 |
| BUB3 | 31.0610517 | -0.3569163 | 0.35822729 | -0.9963404 | 0.31908476 | 0.82354084 |
| CPXM2 | 71.8027106 | -0.3175407 | 0.69845373 | -0.4546338 | 0.6493727 | 0.93437275 |
| OAT | 31.4094546 | 0.32826038 | 0.39661513 | 0.8276547 | 0.40786608 | 0.85867915 |
| LHPP | 38.2820089 | 0.20644743 | 0.3028299 | 0.68172735 | 0.49541137 | 0.89230917 |
| ABRAXAS2 | 32.1169956 | -0.362485 | 0.31746412 | -1.1418141 | 0.2535313 | 0.7912221 |

| | | | | | | |
|---|---|---|---|---|---|---|
| ZRANB1 | 70.1475269 | -0.2378198 | 0.26295092 | -0.9044264 | 0.36576932 | 0.84741115 |
| CTBP2 | 391.32175 | -0.4667103 | 0.22085823 | -2.1131669 | 0.03458649 | 0.49098143 |
| EDRF1 | 48.7333936 | -0.1005414 | 0.23750016 | -0.4233318 | 0.67205318 | 0.93878847 |
| UROS | 26.5848105 | -0.0219884 | 0.34798352 | -0.0631879 | 0.94961685 | 0.99105535 |
| BCCIP | 44.8393436 | -0.0863179 | 0.27044567 | -0.319169 | 0.74959837 | 0.95680183 |
| DHX32 | 108.564769 | -0.3753036 | 0.19194317 | -1.9552848 | 0.05054947 | 0.54649857 |
| FANK1 | 55.8131953 | -0.4593519 | 0.37539165 | -1.2236605 | 0.22108034 | 0.76680432 |
| DOCK1 | 590.154248 | -0.2799502 | 0.24751294 | -1.1310527 | 0.25803291 | 0.79532468 |
| FAM196A | 22.1937777 | 0.5993918 | 0.4872402 | 1.23017725 | 0.21863074 | 0.76666749 |
| MGMT | 151.104686 | -0.0950377 | 0.22442201 | -0.4234778 | 0.67194669 | 0.93878847 |
| AL157832.3 | 23.1906095 | -0.4296068 | 0.38593489 | -1.1131587 | 0.26564029 | 0.79666376 |
| C10orf143 | 19.4361554 | -0.4217879 | 0.35131133 | -1.2006101 | 0.22990248 | 0.77225177 |
| GLRX3 | 67.119138 | -0.5972515 | 0.24661944 | -2.4217536 | 0.01544582 | 0.39384675 |
| PPP2R2D | 139.230621 | -0.3496891 | 0.25058479 | -1.395492 | 0.16286752 | 0.71325312 |
| BNIP3 | 83.048924 | -0.7977491 | 0.55301359 | -1.4425487 | 0.14914763 | 0.70128662 |
| STK32C | 22.3941565 | -0.4126023 | 0.3147979 | -1.3106896 | 0.18996267 | 0.74535784 |
| LRRC27 | 21.0115552 | -0.3302831 | 0.37663388 | -0.8769341 | 0.38052244 | 0.85432452 |
| PWWP2B | 16.9166842 | 0.18675946 | 0.49310638 | 0.3787407 | 0.70488042 | 0.94527497 |
| INPP5A | 85.3577111 | 0.06698257 | 0.26726721 | 0.25062023 | 0.80210774 | 0.97082659 |
| TUBGCP2 | 60.0313616 | -0.1370031 | 0.35467566 | -0.3862772 | 0.69929139 | 0.94395238 |
| ECHS1 | 58.3246041 | 0.66417627 | 0.45667832 | 1.45436348 | 0.14584556 | 0.69679188 |
| SCART1 | 18.081803 | -0.2712333 | 0.53991854 | -0.5023596 | 0.61541457 | 0.92662475 |
| SLC6A12 | 44.7425446 | -0.6275508 | 0.42406108 | -1.4798594 | 0.13891077 | 0.69023184 |
| KDM5A | 150.367581 | -0.0997972 | 0.18412681 | -0.5420026 | 0.58781671 | 0.92148551 |
| CCDC77 | 13.718907 | -0.3846871 | 0.42174138 | -0.9121398 | 0.36169515 | NA |
| B4GALNT3 | 286.576064 | -0.2425357 | 0.4220974 | -0.5745965 | 5.66E-01 | 0.91698355 |
| WNK1 | 996.846815 | 0.86214489 | 0.3652137 | 2.3606587 | 1.82E-02 | 4.12E-01 |
| RAD52 | 42.430973 | -0.0337414 | 0.30276008 | -0.111446 | 0.91126265 | 0.98586235 |
| ERC1 | 663.497734 | -0.2081938 | 0.19599803 | -1.0622241 | 0.28813395 | 0.80523851 |
| FBXL14 | 20.025739 | 0.01989504 | 0.42169474 | 0.04717878 | 0.96237074 | 0.99256424 |
| ADIPOR2 | 115.395692 | -0.0997665 | 0.33853233 | -0.2947029 | 0.76822082 | 0.96289288 |
| DCP1B | 44.6438453 | 0.2739752 | 0.31826217 | 0.86084752 | 0.38932203 | 0.85605713 |
| FKBP4 | 34.8444307 | 0.52504111 | 0.51673232 | 1.01607949 | 0.30959153 | 0.81568231 |
| ITFG2 | 68.610574 | -0.3180708 | 0.3860471 | -0.823917 | 0.40998672 | 0.85867915 |
| TULP3 | 91.9749641 | -0.20454 | 0.28702373 | -0.712624 | 0.47607844 | 0.88495683 |
| TEAD4 | 42.6868012 | -0.5936263 | 0.43758507 | -1.3565962 | 1.75E-01 | 0.72841249 |
| TSPAN9 | 318.623415 | -0.1931586 | 0.18984202 | -1.01747 | 0.30892989 | 0.81568231 |
| PARP11 | 24.3970768 | -0.3584049 | 0.32624984 | -1.0985596 | 0.27196021 | 0.79679877 |
| TIGAR | 27.4953104 | -0.4484059 | 0.55989194 | -0.8008793 | 0.42320153 | 0.86190547 |
| C12orf4 | 31.8685801 | -0.1541287 | 0.4187343 | -0.3680824 | 0.71281177 | 0.94781267 |
| CD9 | 134.30131 | 0.44213831 | 0.2489867 | 1.7757507 | 0.07577402 | 0.61020315 |
| TNFRSF1A | 54.7522338 | 0.25744434 | 0.38519576 | 0.66834677 | 0.50391227 | 0.89627069 |
| SCNN1A | 255.017249 | 0.29273048 | 0.18896264 | 1.54914478 | 1.21E-01 | 6.68E-01 |
| LTBR | 29.6633139 | -0.734963 | 0.33549214 | -2.1907011 | 0.02847343 | 0.46148943 |
| TAPBPL | 17.0056896 | 0.26811887 | 0.50643444 | 0.52942463 | 0.59651091 | 0.9229638 |
| MRPL51 | 36.641415 | 0.10725667 | 0.47139073 | 0.22753241 | 0.82000977 | 0.97338914 |
| NCAPD2 | 21.0840264 | 0.15218136 | 0.46445284 | 0.3276573 | 0.7431708 | 0.95563105 |
| GAPDH | 386.464706 | -0.0294551 | 0.49101027 | -0.0599887 | 0.95216464 | 0.99141043 |
| CHD4 | 35.888682 | 0.31770389 | 0.31184579 | 1.01878524 | 0.30830493 | 0.81568231 |
| AC006064.2 | 14.2796721 | 0.82644614 | 0.60789643 | 1.35951801 | 0.1739825 | NA |
| ING4 | 24.3257976 | -0.1250353 | 0.32522747 | -0.3844551 | 0.7006412 | 0.94440669 |
| ZNF384 | 16.9210528 | 0.2050566 | 0.37529716 | 0.54638462 | 0.58480158 | 0.92076087 |
| COPS7A | 29.3406467 | 0.5291219 | 0.41766209 | 1.26686598 | 0.20520322 | 0.76231289 |
| MLF2 | 44.3366241 | 0.30671849 | 0.36389902 | 0.84286704 | 0.39930281 | 0.85778581 |

| | | | | | | |
|---|---|---|---|---|---|---|
| PTMS | 89.6111351 | 0.68602274 | 0.40297836 | 1.70238108 | 0.08868395 | 0.63466198 |
| USP5 | 19.4882654 | -0.0188174 | 0.58714592 | -0.0320489 | 0.97443308 | 0.99431961 |
| TPI1 | 116.471048 | -0.1942448 | 0.5233515 | -0.3711554 | 0.71052177 | 0.94614974 |
| LRRC23 | 24.0553937 | -0.0161063 | 0.33624513 | -0.0479006 | 0.96179549 | 0.99256424 |
| ATN1 | 52.8868261 | -0.5132065 | 0.2930696 | -1.7511421 | 0.07992143 | 0.6160244 |
| C12orf57 | 29.2969802 | 0.46248893 | 0.51030615 | 0.906297 | 0.36477867 | 0.8468666 |
| PTPN6 | 18.5926821 | 0.4464721 | 0.40051068 | 1.11475705 | 0.26495455 | 0.79666376 |
| PHB2 | 41.6436737 | 0.61371824 | 0.34406242 | 1.78374097 | 0.07446577 | 0.61020315 |
| LPCAT3 | 93.4130333 | 0.07136025 | 0.21356392 | 0.33414002 | 0.7382739 | 0.9542764 |
| C1S | 34.2964775 | -0.3525737 | 0.53053796 | -0.6645588 | 0.50633272 | 0.89724484 |
| C1R | 20.3771596 | -0.1720996 | 0.74001646 | -0.2325619 | 0.81610164 | 0.97288403 |
| C1RL | 67.2015367 | 0.03925971 | 0.23565859 | 0.16659571 | 8.68E-01 | 9.78E-01 |
| C1RL-AS1 | 11.2636443 | -0.9359278 | 0.56868885 | -1.6457643 | 0.09981229 | NA |
| CLSTN3 | 24.5898426 | 0.10392068 | 0.41106345 | 0.25280935 | 0.80041554 | 0.97019967 |
| PEX5 | 15.2483003 | -0.4896793 | 0.36136816 | -1.3550705 | 0.1753951 | NA |
| FOXJ2 | 18.0434135 | 0.07493308 | 0.37977223 | 0.19731058 | 0.8435845 | 0.97612951 |
| NECAP1 | 23.9128205 | -0.5189773 | 0.37062567 | -1.4002734 | 0.16143145 | 0.71101785 |
| FAM66C | 15.3627036 | -0.1839253 | 0.46968521 | -0.3915928 | 0.69535912 | NA |
| LINC00937 | 26.9890695 | 0.39415275 | 0.31120857 | 1.26652283 | 0.20532596 | 0.76231289 |
| RIMKLB | 125.698432 | -1.5836941 | 0.30342357 | -5.2194168 | 1.79E-07 | 0.00081694 |
| A2ML1-AS1 | 15.3996098 | -0.958391 | 0.56520481 | -1.6956527 | 0.08995168 | NA |
| PHC1 | 33.1678009 | 0.03876101 | 0.31066674 | 0.12476717 | 0.90070788 | 0.98531749 |
| M6PR | 53.1594234 | 0.13621665 | 0.29749469 | 0.45787928 | 0.64703918 | 0.93328692 |
| KLRG1 | 11.5394083 | 0.43203115 | 0.51884006 | 0.83268658 | 0.40502152 | NA |
| CLEC2D | 36.3552387 | 0.00885374 | 0.37275516 | 0.02375216 | 0.9810503 | 0.99592962 |
| TMEM52B | 237.2275 | 1.63997569 | 0.61454444 | 2.6686039 | 0.00761672 | 0.30654023 |
| GABARAPL1 | 170.06882 | -0.2670561 | 0.34581949 | -0.7722412 | 0.43997158 | 0.86895963 |
| KLRD1 | 59.5735849 | 0.40553422 | 0.46850813 | 0.86558629 | 0.38671708 | 0.85605713 |
| MAGOHB | 14.3558619 | 0.46132347 | 0.46698544 | 0.98787549 | 0.32321363 | NA |
| YBX3 | 215.123281 | -0.582518 | 0.29809981 | -1.954104 | 5.07E-02 | 0.54735613 |
| PRH1 | 13.4503617 | 1.03737949 | 0.54009051 | 1.9207512 | 0.05476308 | NA |
| SMIM10L1 | 17.9726664 | 0.30909777 | 0.36860362 | 0.83856412 | 0.40171395 | 0.85778581 |
| AC244131.2 | 25.3220326 | 0.80576526 | 0.74002029 | 1.08884211 | 0.27622352 | 0.8010394 |
| ETV6 | 390.698342 | -0.0862249 | 0.31708379 | -0.271931 | 0.78567509 | 0.96767512 |
| BCL2L14 | 42.7104384 | 0.48262075 | 0.42450417 | 1.13690462 | 0.25557816 | 0.7940072 |
| LRP6 | 147.674293 | 0.19903618 | 0.22564738 | 0.88206733 | 0.3777404 | 0.85310518 |
| MANSC1 | 16.4288025 | 0.18996481 | 0.42450031 | 0.44750216 | 0.65451253 | 0.93548446 |
| BORCS5 | 53.6243856 | -0.0611807 | 0.2509316 | -0.2438142 | 0.80737471 | 0.97216031 |
| DUSP16 | 87.6546008 | -0.0913576 | 0.20839315 | -0.4383908 | 0.66110301 | 0.93729379 |
| CREBL2 | 35.4619584 | -0.0896879 | 0.31835042 | -0.2817271 | 0.7781528 | 0.96518573 |
| CDKN1B | 23.6557716 | 0.92928546 | 0.47384587 | 1.96115556 | 0.04986088 | 0.54357315 |
| GPRC5A | 50.1416517 | -0.7178235 | 0.6231691 | -1.1518921 | 0.24936543 | 0.78763828 |
| GPRC5D-AS1 | 49.2050774 | -0.5742193 | 0.37932405 | -1.5137961 | 0.13007755 | 0.67331904 |
| HEBP1 | 37.6755969 | 0.45816285 | 0.29761576 | 1.53944421 | 0.12369589 | 0.66929384 |
| FAM234B | 14.7292006 | 0.06347508 | 0.42365592 | 0.14982697 | 0.88090113 | NA |
| EMP1 | 30.5458685 | -0.4737836 | 0.31168423 | -1.5200756 | 1.28E-01 | 6.69E-01 |
| ATF7IP | 229.136528 | -0.098362 | 0.23588001 | -0.4170002 | 0.67667826 | 0.94034237 |
| PLBD1 | 22.0474169 | 0.27898577 | 0.49505802 | 0.56354156 | 0.57306617 | 0.91813957 |
| AC010168.1 | 21.0012901 | 0.26014772 | 0.64162896 | 0.40544884 | 0.68514762 | 0.94205398 |
| H2AFJ | 53.0470356 | 0.46869533 | 0.32323825 | 1.44999959 | 0.14705863 | 0.69749723 |
| WBP11 | 35.9959311 | 0.49215323 | 0.29446192 | 1.67136463 | 0.09464968 | 0.64732984 |
| C12orf60 | 28.0477397 | 0.55541062 | 0.39571761 | 1.40355296 | 0.16045201 | 0.71101785 |
| MGP | 69.4695152 | -0.1889416 | 0.63613839 | -0.2970134 | 0.76645627 | 0.96270421 |
| ERP27 | 11.652194 | 2.85132572 | 0.78589663 | 3.62811801 | 0.00028549 | NA |

| | | | | | | |
|---|---|---|---|---|---|---|
| ARHGDIB | 14.0942528 | 1.34525062 | 0.6531894 | 2.05951079 | 0.03944533 | NA |
| RERG | 47.0649765 | -0.0523516 | 0.44111049 | -0.1186814 | 0.90552776 | 0.98531749 |
| EPS8 | 164.657641 | -0.6334364 | 0.25005084 | -2.5332303 | 0.01130167 | 0.34756443 |
| STRAP | 52.7005681 | 0.32334718 | 0.28806197 | 1.12249173 | 0.26165342 | 0.79633269 |
| DERA | 131.068257 | -0.2955428 | 0.18074174 | -1.6351662 | 0.1020142 | 0.65946373 |
| MGST1 | 52.1187808 | 0.18023103 | 0.68287159 | 0.26393107 | 0.79183304 | 0.96925674 |
| RERGL | 24.9306045 | -0.1574294 | 0.63121925 | -0.2494053 | 0.80304731 | 0.97131805 |
| PIK3C2G | 921.676957 | 1.35895003 | 0.46266459 | 2.93722506 | 0.00331164 | 0.23150356 |
| PLCZ1 | 13.851797 | 1.31684463 | 0.83112248 | 1.58441706 | 0.11309884 | NA |
| AC087242.1 | 35.7649663 | 1.49884918 | 0.54343811 | 2.75808623 | 0.00581409 | 0.2943676 |
| PLEKHA5 | 889.229147 | -0.4799232 | 0.24715758 | -1.9417701 | 0.05216493 | 0.55403388 |
| AEBP2 | 245.070897 | 0.46358503 | 0.23278602 | 1.99146421 | 0.04642988 | 0.5309688 |
| PYROXD1 | 29.3995228 | 0.15870318 | 0.33707881 | 0.47081921 | 0.63776984 | 0.93277938 |
| RECQL | 24.7314149 | -0.2335405 | 0.3480412 | -0.671014 | 0.5022116 | 0.89499455 |
| GOLT1B | 31.06351 | -0.2966117 | 0.29314082 | -1.0118402 | 0.31161447 | 0.81701824 |
| LDHB | 330.56483 | 0.77060346 | 0.37240945 | 2.06923716 | 0.03852384 | 0.49671741 |
| AC010197.1 | 147.321999 | 0.98146978 | 0.49761896 | 1.97233196 | 0.04857172 | 0.5391188 |
| AC010185.1 | 19.9355068 | 1.66332879 | 0.56650334 | 2.93613235 | 0.00332332 | 0.23150356 |
| ABCC9 | 75.9076485 | 2.53425739 | 1.09393915 | 2.31663471 | 0.02052364 | 0.42078083 |
| AC008250.1 | 134.233911 | 1.88686281 | 0.51527727 | 3.66183982 | 0.00025041 | 0.06160773 |
| CMAS | 28.1562007 | 0.05070512 | 0.27174242 | 0.18659257 | 0.85198009 | 0.97612951 |
| C2CD5 | 120.256165 | -0.3541816 | 0.2022144 | -1.7515154 | 0.07985718 | 0.6160244 |
| ETNK1 | 272.011962 | -0.2893946 | 0.33337511 | -0.868075 | 0.38535327 | 0.85540507 |
| SOX5 | 912.873468 | -0.0972173 | 0.42399104 | -0.2292909 | 0.81864281 | 0.97288403 |
| CASC1 | 31.8213813 | 0.51855108 | 0.65352546 | 0.7934673 | 0.42750561 | 0.86239557 |
| ETFRF1 | 30.279462 | 0.79700031 | 0.33841486 | 2.35509846 | 0.0185178 | 0.41315568 |
| KRAS | 89.655665 | -0.2751271 | 0.28670033 | -0.959633 | 0.33723995 | 0.83055261 |
| RASSF8-AS1 | 26.9399613 | -0.5558476 | 0.32148553 | -1.7289973 | 0.08380958 | 0.62534311 |
| RASSF8 | 160.431242 | -0.4816359 | 0.31627368 | -1.5228452 | 0.12779743 | 0.66929384 |
| AC024145.1 | 13.0783103 | 0.64729199 | 0.75559972 | 0.85665992 | 0.39163286 | NA |
| ITPR2 | 782.447772 | -0.4276218 | 0.23907017 | -1.7886875 | 0.07366515 | 0.60985192 |
| AC023051.1 | 17.4064433 | 0.51916262 | 0.57899048 | 0.89666866 | 0.36989575 | 0.84852785 |
| INTS13 | 20.4602798 | -0.0692805 | 0.33201663 | -0.2086657 | 0.83470921 | 0.97600352 |
| FGFR1OP2 | 58.0447206 | 0.25471463 | 0.29775604 | 0.85544742 | 0.3923035 | 0.85605713 |
| TM7SF3 | 66.9520708 | 0.26896937 | 0.29029438 | 0.92654006 | 0.35416539 | 0.83979358 |
| MED21 | 24.0117136 | -0.1493173 | 0.32715755 | -0.4564078 | 0.64809678 | 0.93359698 |
| STK38L | 48.5223753 | 0.05433298 | 0.28647672 | 0.18965931 | 0.84957611 | 0.97612951 |
| ARNTL2 | 17.8270039 | -0.2435699 | 0.7213206 | -0.3376721 | 0.73561028 | 0.9536908 |
| PPFIBP1 | 382.187434 | -0.014316 | 0.30048849 | -0.0476423 | 0.96200135 | 0.99256424 |
| MRPS35 | 48.0529803 | 0.13509674 | 0.28770733 | 0.46956307 | 0.63866722 | 0.9331046 |
| KLHL42 | 38.6257946 | 0.73874319 | 0.33732919 | 2.18997706 | 0.0285259 | 0.46148943 |
| CCDC91 | 300.334791 | -0.1922679 | 0.14548106 | -1.3216007 | 0.18630116 | 0.73831062 |
| FAR2 | 33.1385603 | -0.0702399 | 0.49577189 | -0.1416778 | 0.88733452 | 0.98229432 |
| AC009318.1 | 19.0216054 | -0.5261109 | 0.54244788 | -0.9698829 | 0.33210488 | 0.82890589 |
| ERGIC2 | 69.1231605 | 0.04098486 | 0.25215606 | 0.16253767 | 0.87088246 | 0.97860207 |
| TMTC1 | 23.3999574 | -1.9415155 | 0.51906404 | -3.740416 | 0.00018372 | 0.05491436 |
| IPO8 | 77.3841656 | 0.38875696 | 0.23574765 | 1.6490385 | 0.09913975 | 0.65777633 |
| CAPRIN2 | 56.4980132 | -0.1223259 | 0.24350394 | -0.5023568 | 0.61541654 | 0.92662475 |
| DDX11 | 11.7286476 | -1.0322794 | 0.53268833 | -1.9378675 | 0.05263939 | NA |
| SINHCAF | 106.356611 | -0.2580459 | 0.17680468 | -1.4594969 | 0.14442841 | 0.69512617 |
| DENND5B | 239.663403 | -0.3467639 | 0.32446033 | -1.0687404 | 0.28518667 | 0.80431986 |
| AMN1 | 58.6909171 | -0.230516 | 0.26494538 | -0.870051 | 0.38427256 | 0.85511218 |
| KIAA1551 | 90.3853408 | -0.0630401 | 0.24624369 | -0.2560071 | 0.79794535 | 0.97019967 |
| BICD1 | 449.345742 | 0.05995365 | 0.29573284 | 0.20272911 | 0.83934677 | 0.97600352 |

| | | | | | | |
|---|---|---|---|---|---|---|
| FGD4 | 955.020134 | -0.2869828 | 0.17887967 | -1.6043343 | 0.10864039 | 0.65986731 |
| DNM1L | 141.963997 | -0.1042329 | 0.16509442 | -0.6313535 | 0.52780941 | 0.90487731 |
| YARS2 | 42.4358222 | 0.19333221 | 0.30581895 | 0.63217865 | 0.52727014 | 0.90487731 |
| PKP2 | 14.8607119 | -0.1397667 | 0.4159106 | -0.3360498 | 0.73683335 | NA |
| ALG10B | 14.543675 | 0.57121227 | 0.4137237 | 1.38066122 | 0.16738315 | NA |
| CPNE8 | 203.500112 | -0.718497 | 0.42842436 | -1.6770685 | 0.09352909 | 0.64572229 |
| KIF21A | 56.3750668 | 0.18778285 | 0.35212386 | 0.53328636 | 0.59383537 | 0.9229638 |
| SLC2A13 | 78.3945307 | 0.00473011 | 0.35758143 | 0.01322806 | 9.89E-01 | 9.98E-01 |
| LRRK2 | 140.749034 | -0.4231633 | 0.51153284 | -0.8272456 | 0.40809787 | 0.85867915 |
| GXYLT1 | 55.1286721 | 0.45815351 | 0.22348995 | 2.04999604 | 0.04036482 | 0.50399977 |
| YAF2 | 119.608833 | -0.0883598 | 0.22354764 | -0.3952617 | 0.69264979 | 0.94374959 |
| PPHLN1 | 135.954497 | 0.0037382 | 0.18815991 | 0.01986715 | 0.98414935 | 0.99685229 |
| ZCRB1 | 24.6638681 | 0.06599158 | 0.35290123 | 0.18699732 | 0.85166274 | 0.97612951 |
| PRICKLE1 | 75.8939319 | 0.93193224 | 0.75702314 | 1.23104855 | 2.18E-01 | 0.76666749 |
| PUS7L | 44.0762473 | 0.35823739 | 0.30970609 | 1.15670115 | 0.24739448 | 0.78583441 |
| IRAK4 | 42.1545299 | 0.35516216 | 0.37467094 | 0.94793089 | 0.34316464 | 0.83426365 |
| TWF1 | 32.5213892 | 0.00891908 | 0.3052361 | 0.02922025 | 0.97668893 | 0.99441127 |
| TMEM117 | 382.666561 | 0.18413948 | 0.20091796 | 0.91649087 | 0.35940949 | 0.84204868 |
| ANO6 | 202.09161 | -0.4246311 | 0.26199481 | -1.6207613 | 0.10506884 | 0.65946648 |
| AC008124.1 | 16.7863179 | 0.9095388 | 0.3821993 | 2.37975004 | 0.01732438 | 0.40063153 |
| ARID2 | 283.559434 | 0.00121601 | 0.23742272 | 0.00512172 | 0.99591347 | 0.99927028 |
| SCAF11 | 275.652082 | 0.08271192 | 0.14862967 | 0.55649667 | 0.57787137 | 0.91878391 |
| SLC38A1 | 366.317632 | -0.5450606 | 0.48843802 | -1.1159259 | 0.26445385 | 0.79666376 |
| SLC38A2 | 190.275893 | -0.1306135 | 0.20672309 | -0.6318281 | 0.52749923 | 0.90487731 |
| AC008014.1 | 508.506433 | 0.34279737 | 0.27730328 | 1.23618216 | 0.21639086 | 0.76616336 |
| SLC38A4 | 26.2194865 | 0.57370451 | 0.79771434 | 0.71918541 | 0.47202669 | 0.88267439 |
| PCED1B | 122.790825 | -1.6592627 | 0.48755445 | -3.4032356 | 0.00066593 | 0.11225824 |
| RPAP3 | 27.401129 | -0.3781499 | 0.30629399 | -1.2345978 | 0.21698022 | 0.76666749 |
| RAPGEF3 | 109.660199 | 0.04094705 | 0.25314368 | 0.16175418 | 8.71E-01 | 0.97860207 |
| SLC48A1 | 41.8526486 | -0.2887329 | 0.31302632 | -0.9223918 | 0.35632422 | 0.84192356 |
| HDAC7 | 108.362797 | -0.7873377 | 0.38938938 | -2.0219803 | 0.04317839 | 0.51638379 |
| VDR | 35.2900484 | 0.26024807 | 0.54186698 | 0.48028037 | 0.63102805 | 0.93250784 |
| TMEM106C | 24.8743962 | 0.55738808 | 0.45318845 | 1.22992562 | 0.21872496 | 0.76666749 |
| SENP1 | 36.9474596 | 0.17514558 | 0.35802806 | 0.48919512 | 0.62470357 | 0.92981013 |
| PFKM | 24.3150659 | 0.94873956 | 0.3549693 | 2.67273696 | 0.00752352 | 0.30654023 |
| ASB8 | 24.3947351 | 0.35226259 | 0.34574495 | 1.01885099 | 0.30827371 | 0.81568231 |
| ZNF641 | 14.4719222 | -0.1652583 | 0.43800441 | -0.3772982 | 0.70595199 | NA |
| AC090115.1 | 16.5732922 | 1.08159465 | 0.59354285 | 1.82226886 | 0.0684142 | 0.60244724 |
| KANSL2 | 22.5029237 | 0.47873228 | 0.33413585 | 1.43274742 | 0.15193003 | 0.70572801 |
| CCNT1 | 48.057109 | -0.0607015 | 0.23664672 | -0.2565066 | 0.79755966 | 0.97019967 |
| ADCY6 | 43.6583434 | 0.11668137 | 0.2698513 | 0.43239134 | 0.66545701 | 0.93860542 |
| DDX23 | 24.3332219 | 0.07087772 | 0.42632631 | 0.16625228 | 0.86795842 | 0.97810461 |
| CCDC65 | 23.3643159 | 0.27469286 | 0.43958102 | 0.624897 | 0.53203866 | 0.90610813 |
| PRKAG1 | 31.1498185 | -0.2153361 | 0.27923208 | -0.7711726 | 0.44060463 | 0.86895963 |
| KMT2D | 31.8763511 | 0.01426421 | 0.34078717 | 0.04185665 | 0.96661297 | 0.99301184 |
| LMBR1L | 36.7055104 | 0.05098665 | 0.35536511 | 0.1434768 | 0.88591363 | 0.98216235 |
| AC011603.2 | 15.322265 | 0.351369 | 0.40755966 | 0.86212901 | 0.38861653 | NA |
| TUBA1B | 89.5113411 | -0.0626214 | 0.56111706 | -0.1116014 | 0.91113946 | 0.98586235 |
| TUBA1A | 38.803284 | -0.3242362 | 0.56218937 | -0.5767384 | 0.56411621 | 0.91650701 |
| TUBA1C | 64.1733397 | -0.1502183 | 0.29354607 | -0.5117365 | 0.60883541 | 0.92573951 |
| SPATS2 | 268.137882 | -0.1129217 | 0.34458136 | -0.3277071 | 0.74313317 | 0.95563105 |
| MCRS1 | 16.1660806 | 0.169768 | 0.44256644 | 0.3835989 | 0.70127575 | 0.94440669 |
| PRPF40B | 87.2082743 | 0.17560794 | 0.23181421 | 0.75753744 | 0.44872794 | 0.87314185 |
| FMNL3 | 12.8518197 | 1.44708353 | 0.59120627 | 2.44767959 | 0.01437795 | NA |

| | | | | | | |
|---|---|---|---|---|---|---|
| TMBIM6 | 294.251233 | -0.0501733 | 0.3966808 | -0.1264827 | 0.89934981 | 0.98531749 |
| NCKAP5L | 24.2166076 | 0.50407896 | 0.36429975 | 1.38369283 | 0.16645253 | 0.718453 |
| AQP2 | 591.177486 | -1.2099309 | 0.73963391 | -1.635851 | 0.10187077 | 0.6590829 |
| SMARCD1 | 24.2902174 | -0.1288545 | 0.32394189 | -0.3977705 | 0.69079937 | 0.94292198 |
| AC074032.1 | 79.5010428 | -0.0371573 | 0.30241635 | -0.1228679 | 0.90221171 | 0.98531749 |
| CERS5 | 60.2986283 | 0.0251809 | 0.29447558 | 0.08551101 | 0.93185514 | 0.98960607 |
| LIMA1 | 266.649594 | -0.4200997 | 0.35455247 | -1.1848731 | 0.23606761 | 0.77864547 |
| LARP4 | 166.618703 | 0.02764239 | 0.24138924 | 0.11451377 | 0.90883053 | 0.9854228 |
| DIP2B | 235.260002 | -0.5269508 | 0.20162195 | -2.6135585 | 0.00896047 | 0.31797637 |
| ATF1 | 70.1423612 | -0.1061547 | 0.22987678 | -0.4617896 | 0.64423223 | 0.93328692 |
| METTL7A | 51.2068722 | 0.39041902 | 0.51718649 | 0.75489021 | 0.45031486 | 0.87346041 |
| SLC11A2 | 232.240716 | -0.0553376 | 0.28362439 | -0.1951087 | 0.84530788 | 0.97612951 |
| LETMD1 | 92.446289 | 0.1699555 | 0.26357103 | 0.64481857 | 0.51904476 | 0.90368693 |
| CSRNP2 | 26.4895707 | -0.1117467 | 0.32341197 | -0.3455244 | 0.72970017 | 0.95279138 |
| TFCP2 | 171.829886 | 0.1011639 | 0.22917945 | 0.44141786 | 0.65891052 | 0.93664697 |
| POU6F1 | 24.5901395 | 0.05515467 | 0.3576913 | 0.15419628 | 0.87745498 | 0.980058 |
| DAZAP2 | 127.224273 | 0.00566508 | 0.32539435 | 0.01740989 | 0.98610962 | 0.99766827 |
| SMAGP | 16.0712097 | 0.09014939 | 0.53511757 | 0.16846651 | 0.86621629 | NA |
| ACVR1B | 248.208086 | -0.1354558 | 0.17032099 | -0.7952973 | 0.42644059 | 0.86239557 |
| NR4A1 | 109.622728 | -1.0191464 | 1.09500428 | -0.9307236 | 3.52E-01 | 0.83872681 |
| ATG101 | 14.9861461 | 0.23389735 | 0.61426114 | 0.38077836 | 0.70336772 | NA |
| AC078864.1 | 31.6426144 | -0.7515016 | 0.50852922 | -1.4777944 | 0.13946282 | 0.69054855 |
| KRT7 | 61.5621236 | 0.28753023 | 0.63944923 | 0.44965294 | 0.65296071 | 0.93532021 |
| KRT8 | 144.830037 | 0.28396465 | 0.42502299 | 0.66811596 | 0.50405957 | 0.89635755 |
| KRT18 | 152.879784 | 0.10407034 | 0.46720015 | 0.22275322 | 0.82372758 | 0.97518529 |
| EIF4B | 121.320031 | 0.04706669 | 0.24675991 | 0.19073881 | 0.84873023 | 0.97612951 |
| AC068888.1 | 40.6517349 | 0.44785308 | 0.33324855 | 1.34390106 | 0.17898038 | 0.72924345 |
| TNS2 | 29.4525239 | 0.30938624 | 0.45350987 | 0.68220398 | 0.49510998 | 0.89230917 |
| CSAD | 85.6911381 | 0.08429298 | 0.19931875 | 0.42290541 | 6.72E-01 | 0.93878847 |
| ZNF740 | 15.5418699 | 0.32929875 | 0.41193741 | 0.79939027 | 0.42406415 | NA |
| RARG | 16.205395 | 0.65136507 | 0.48835156 | 1.3338036 | 0.18226823 | 0.73398872 |
| PFDN5 | 79.3472078 | 0.26379977 | 0.4307183 | 0.61246474 | 0.54023032 | 0.90763121 |
| C12orf10 | 11.2219619 | 0.86437413 | 0.48019756 | 1.80003856 | 0.07185455 | NA |
| AAAS | 14.9260634 | -0.2406273 | 0.39514585 | -0.6089583 | 0.54255209 | NA |
| SP1 | 77.9279777 | 0.39683481 | 0.2579675 | 1.53831321 | 0.12397205 | 0.66929384 |
| PRR13 | 70.0921825 | 0.51398883 | 0.36047339 | 1.42587175 | 0.15390537 | 0.70572801 |
| PCBP2 | 179.47173 | -0.2020234 | 0.18519019 | -1.0908971 | 0.27531818 | 0.7994854 |
| MAP3K12 | 18.15695 | 0.07486494 | 0.41016879 | 0.18252226 | 0.85517288 | 0.97612951 |
| ATP5MC2 | 122.473806 | -0.1596709 | 0.24922305 | -0.6406746 | 0.52173409 | 0.90451015 |
| CALCOCO1 | 32.6135353 | 0.26735119 | 0.41944864 | 0.63738719 | 0.52387267 | 0.90487731 |
| HOXC4 | 13.1807205 | -0.0657788 | 0.4397868 | -0.1495698 | 0.88110401 | NA |
| SMUG1 | 23.6990418 | 0.35715309 | 0.32406011 | 1.10211988 | 0.27040956 | 0.79666376 |
| LINC02381 | 35.1903273 | -0.0842671 | 0.350633 | -0.2403285 | 0.81007561 | 0.97257655 |
| CBX5 | 108.268745 | 0.55008588 | 0.22438878 | 2.45148569 | 0.01422678 | 0.38054999 |
| HNRNPA1 | 66.1563637 | 0.24128504 | 0.24625463 | 0.9798193 | 0.32717532 | 0.82612204 |
| COPZ1 | 51.9193337 | -0.1095338 | 0.24123286 | -0.4540585 | 0.64978674 | 0.93453685 |
| CD63 | 128.044465 | -0.1027432 | 0.41885587 | -0.2452949 | 0.80622813 | 0.97216031 |
| ORMDL2 | 12.8778016 | -0.0647542 | 0.4360107 | -0.1485151 | 0.88193626 | NA |
| RAB5B | 48.1963537 | -0.0615734 | 0.27208048 | -0.2263058 | 0.82096358 | 0.97383783 |
| SUOX | 18.8540128 | 0.71321171 | 0.47874916 | 1.48973988 | 0.13629265 | 0.68406472 |
| IKZF4 | 14.3737801 | 0.58368764 | 0.45689611 | 1.27750626 | 0.20142357 | NA |
| RPS26 | 21.8389242 | -0.0177001 | 0.43830133 | -0.0403833 | 0.96778753 | 0.99332167 |
| ERBB3 | 22.2282112 | 0.11934074 | 0.46377831 | 0.25732283 | 0.79692957 | 0.97010563 |
| PA2G4 | 23.5123438 | 0.58062259 | 0.39446166 | 1.47193667 | 0.14103798 | 0.69227547 |

| | | | | | | |
|---|---|---|---|---|---|---|
| RPL41 | 338.234232 | 0.05991975 | 0.39247156 | 0.15267285 | 0.87865627 | 0.98061726 |
| ESYT1 | 22.324144 | 0.01200507 | 0.32232974 | 0.03724468 | 0.97028991 | 0.99398482 |
| MYL6B | 14.2379571 | 0.08899972 | 0.45259398 | 0.19664362 | 0.84410643 | NA |
| MYL6 | 166.4437 | -0.1187727 | 0.40054711 | -0.2965261 | 0.76682837 | 0.96283923 |
| SMARCC2 | 72.9513788 | 0.12307393 | 0.27212979 | 0.45226187 | 0.65108034 | 0.93453685 |
| RNF41 | 18.6810275 | 0.19494563 | 0.32277374 | 0.60396993 | 0.54586363 | 0.91073986 |
| ANKRD52 | 26.3065472 | 0.19486776 | 0.30571409 | 0.63741831 | 0.5238524 | 0.90487731 |
| COQ10A | 8.53385393 | 1.00711625 | 0.66774765 | 1.50822882 | 0.13149597 | NA |
| CS | 48.2982773 | -0.1283556 | 0.31154509 | -0.4119967 | 0.68034183 | 0.94196488 |
| AC073896.2 | 13.240312 | -0.1174576 | 0.39212753 | -0.2995393 | 0.76452856 | NA |
| PAN2 | 30.5830961 | 0.13838723 | 0.31384293 | 0.44094424 | 0.65925337 | 0.93680665 |
| STAT2 | 48.9689338 | -0.8790252 | 0.43130624 | -2.0380535 | 0.04154459 | 0.51036486 |
| SPRYD4 | 13.2197035 | -0.1700655 | 0.45861865 | -0.3708211 | 0.71077078 | NA |
| RBMS2 | 89.0276396 | -0.0875546 | 0.35490436 | -0.246699 | 0.80514115 | 0.97195485 |
| BAZ2A | 75.7275416 | -0.1984358 | 0.21551469 | -0.9207531 | 0.35717933 | 0.84204868 |
| ATP5F1B | 156.950944 | 0.50519963 | 0.40889572 | 1.23552192 | 0.21663632 | 0.76658236 |
| PTGES3 | 195.538347 | -0.0856701 | 0.28139978 | -0.3044428 | 0.76079054 | 0.95956874 |
| NACA | 74.4339448 | 0.50727295 | 0.25107067 | 2.02043891 | 0.04333788 | 0.51699786 |
| NEMP1 | 13.8880623 | -0.5177157 | 0.41247316 | -1.2551502 | 0.20942426 | NA |
| STAT6 | 77.6661021 | -0.1598234 | 0.2648944 | -0.6033476 | 0.54627746 | 0.91109632 |
| R3HDM2 | 257.706643 | -0.2237138 | 0.23571954 | -0.9490679 | 0.34258609 | 0.833617 |
| MARS | 31.9386086 | -1.1735295 | 0.44067516 | -2.6630261 | 0.00774414 | 0.30654023 |
| DDIT3 | 50.5460416 | -1.7853046 | 0.70770053 | -2.5226837 | 0.01164631 | 0.35575956 |
| MBD6 | 19.3173169 | -0.3525155 | 0.44432841 | -0.7933669 | 0.42756408 | 0.86239557 |
| DCTN2 | 47.2555997 | -0.1462379 | 0.27541307 | -0.5309767 | 0.5954349 | 0.9229638 |
| PIP4K2C | 34.2576619 | 0.09365379 | 0.33164824 | 0.28238891 | 0.77764531 | 0.96518573 |
| DTX3 | 15.2825511 | -0.4438477 | 0.46591635 | -0.9526339 | 0.34077558 | NA |
| OS9 | 65.7832581 | 0.53252777 | 0.35678388 | 1.49257798 | 0.13554769 | 0.6835006 |
| TSPAN31 | 19.1816598 | 0.37970921 | 0.45128271 | 0.84139986 | 0.40012397 | 0.85778581 |
| CDK4 | 16.8201527 | -0.1058752 | 0.41394754 | -0.2557695 | 0.79812881 | 0.97019967 |
| 9-Mar | 18.1960105 | 0.46274343 | 0.33917611 | 1.36431611 | 0.1724681 | 0.72649757 |
| TSFM | 23.7110601 | 0.51358576 | 0.43749532 | 1.17392285 | 0.24042593 | 0.78192112 |
| AVIL | 35.9955869 | 0.44551958 | 0.3490521 | 1.27636986 | 0.2018248 | 0.76038186 |
| CTDSP2 | 72.2426381 | 0.05288799 | 0.19082813 | 0.27714987 | 0.78166503 | 0.96652357 |
| LRIG3 | 18.4745695 | 0.34204459 | 0.55802328 | 0.61295756 | 0.5399044 | 0.90763121 |
| SLC16A7 | 370.101239 | 0.12602487 | 0.46213873 | 0.27269921 | 0.78508444 | 0.9674911 |
| FAM19A2 | 68.8708543 | 0.0436425 | 0.42983384 | 0.10153341 | 0.91912704 | 0.98749988 |
| USP15 | 208.052708 | -0.3177477 | 0.2155401 | -1.4741928 | 0.14042967 | 0.69227547 |
| MON2 | 286.419012 | -0.244886 | 0.1689668 | -1.4493143 | 0.14724984 | 0.69776953 |
| AC048341.1 | 45.5534569 | -1.2995349 | 0.40829887 | -3.1828032 | 0.00145857 | 0.16191873 |
| PPM1H | 228.099883 | -0.4155326 | 0.25293256 | -1.6428594 | 0.10041202 | 0.65863153 |
| AVPR1A | 28.6357104 | -0.3856243 | 0.51789725 | -0.7445962 | 0.45651585 | 0.87396417 |
| AC026116.1 | 21.644983 | -0.5551301 | 0.68169578 | -0.814337 | 0.4154519 | 0.85867915 |
| DPY19L2 | 521.730721 | 0.28819779 | 0.28345384 | 1.0167362 | 3.09E-01 | 8.16E-01 |
| RXYLT1 | 26.1278898 | 0.11651839 | 0.30109116 | 0.3869871 | 0.69876576 | 0.94395238 |
| SRGAP1 | 118.802352 | -0.7228041 | 0.324917 | -2.2245807 | 0.0261094 | 0.4455707 |
| XPOT | 32.8573857 | -0.152408 | 0.39060369 | -0.3901859 | 0.69639912 | 0.94395238 |
| TBK1 | 76.3196246 | -0.5387986 | 0.25470809 | -2.1153571 | 0.03439953 | 0.4908133 |
| RASSF3 | 28.4990996 | -0.546521 | 0.46189577 | -1.1832128 | 0.23672481 | 0.77864547 |
| GNS | 135.165036 | 0.12931674 | 0.27799677 | 0.46517351 | 0.64180721 | 0.93328692 |
| TBC1D30 | 50.9879872 | 0.42823369 | 0.23847015 | 1.79575383 | 0.07253368 | 0.60824637 |
| LINC02389 | 12.5656671 | 0.26586169 | 0.45022377 | 0.59051012 | 0.5548487 | NA |
| LEMD3 | 100.699013 | -0.2479751 | 0.23617833 | -1.0499487 | 0.29374172 | 0.8097379 |
| HMGA2 | 26.3618493 | -0.6468988 | 0.76894686 | -0.841279 | 0.40019167 | 0.85778581 |

| | | | | | | |
|---|---|---|---|---|---|---|
| AC090673.1 | 41.1849592 | 0.63567135 | 0.75034859 | 0.84716804 | 0.39690147 | 0.85656568 |
| LLPH | 10.7248702 | 0.071048 | 0.48543071 | 0.14636074 | 0.88363662 | NA |
| HELB | 40.6332568 | 0.00731968 | 0.35651381 | 0.02053127 | 0.98361957 | 0.99685229 |
| GRIP1 | 1232.33117 | 0.116655 | 0.18831667 | 0.61946187 | 5.36E-01 | 0.90763121 |
| CAND1 | 99.1342327 | -0.1672681 | 0.22387613 | -0.7471458 | 0.45497558 | 0.8739446 |
| DYRK2 | 23.2693453 | 0.48901934 | 0.51833259 | 0.94344702 | 3.45E-01 | 0.83564825 |
| MDM1 | 13.3555578 | -0.2091524 | 0.47344359 | -0.4417683 | 0.65865685 | NA |
| RAP1B | 318.502845 | -0.2084165 | 0.27014418 | -0.7715011 | 4.40E-01 | 0.86895963 |
| NUP107 | 54.1006497 | -0.1596013 | 0.39132558 | -0.4078478 | 0.68338542 | 0.94205398 |
| SLC35E3 | 46.0541486 | 0.04717057 | 0.39334723 | 0.11992095 | 0.90454577 | 0.98531749 |
| MDM2 | 82.3552542 | 0.19475976 | 0.32337542 | 0.60227138 | 0.54699351 | 0.91139843 |
| CPM | 75.4041933 | -1.0850275 | 0.6801732 | -1.5952223 | 0.11066253 | 0.66311481 |
| CPSF6 | 151.272744 | -0.2086811 | 0.25686206 | -0.8124248 | 0.41654792 | 0.85867915 |
| YEATS4 | 20.5050803 | -0.4311082 | 0.46036206 | -0.9364547 | 0.34903914 | 0.83687845 |
| FRS2 | 134.633974 | -0.3992872 | 0.20969713 | -1.9041136 | 0.05689539 | 0.56976762 |
| CCT2 | 39.3219184 | 0.05868197 | 0.36286863 | 0.16171685 | 0.87152884 | 0.97860207 |
| RAB3IP | 136.899333 | 0.06283335 | 0.24540892 | 0.25603532 | 0.79792357 | 0.97019967 |
| MYRFL | 61.2373515 | 1.56102987 | 0.5496111 | 2.84024442 | 0.0045079 | 0.26474449 |
| AC025159.1 | 43.9617017 | 0.87617975 | 0.61950765 | 1.41431628 | 0.15726906 | 0.7098097 |
| CNOT2 | 189.264151 | -0.1157918 | 0.21879872 | -0.5292162 | 0.59665549 | 0.9229638 |
| KCNMB4 | 17.1313895 | 0.83902467 | 0.52363223 | 1.60231672 | 0.10908559 | 0.6606663 |
| ZFC3H1 | 139.798606 | -0.2903597 | 0.27301045 | -1.0635479 | 0.28753355 | 0.80523851 |
| RAB21 | 124.143569 | -0.2400092 | 0.23586069 | -1.017589 | 0.30887332 | 0.81568231 |
| TBC1D15 | 106.952692 | -0.2901202 | 0.24944907 | -1.1630439 | 0.24481169 | 0.78583441 |
| TRHDE | 368.078712 | 0.46912671 | 0.22828473 | 2.05500695 | 3.99E-02 | 0.50156625 |
| ATXN7L3B | 42.8111297 | 0.47804076 | 0.29852267 | 1.60135496 | 0.10929832 | 0.66067258 |
| CAPS2 | 112.24987 | 1.04238476 | 0.52580047 | 1.98247208 | 0.04742643 | 0.53530393 |
| KRR1 | 39.7830391 | -0.1648849 | 0.25050363 | -0.6582135 | 0.51040093 | 0.90000646 |
| PHLDA1 | 20.1748437 | -0.3910524 | 0.61980136 | -0.6309318 | 0.52808514 | 0.90487731 |
| NAP1L1 | 132.084727 | 0.33674429 | 0.19979578 | 1.68544244 | 0.09190324 | 0.64452635 |
| OSBPL8 | 204.360819 | -0.1724609 | 0.2193192 | -0.7863468 | 0.43166435 | 0.8664698 |
| ZDHHC17 | 97.1020782 | -0.1830821 | 0.26722881 | -0.6851136 | 0.49327226 | 0.89092098 |
| SYT1 | 25.2423822 | -1.5840022 | 0.63914697 | -2.4783067 | 0.01320076 | 0.36723795 |
| AC027288.3 | 27.3974647 | 0.34017352 | 0.64581155 | 0.52673806 | 0.5983755 | 0.92296674 |
| PAWR | 302.030631 | -0.0707523 | 0.23347682 | -0.3030377 | 0.76186115 | 0.96015811 |
| AC073569.2 | 18.7147124 | 1.07787763 | 0.47510878 | 2.26869651 | 0.02328679 | 0.43085291 |
| PPP1R12A | 269.205851 | -0.253575 | 0.16680834 | -1.5201576 | 0.12847137 | 0.66929384 |
| LIN7A | 127.627842 | 0.4436615 | 0.39984177 | 1.10959269 | 0.26717458 | 0.79666376 |
| ACSS3 | 127.127937 | 1.12093224 | 0.53687979 | 2.08786446 | 0.03681006 | 0.49537143 |
| PPFIA2 | 60.826963 | 2.14518331 | 0.73697539 | 2.91079367 | 0.00360512 | 0.23862997 |
| CCDC59 | 21.3039095 | 1.12091765 | 0.42685221 | 2.62600881 | 0.00863926 | 0.31119527 |
| METTL25 | 91.6350115 | -0.0213055 | 0.22111685 | -0.0963538 | 0.92323956 | 0.98915831 |
| TMTC2 | 1135.40106 | 0.85970701 | 0.27247805 | 3.15514225 | 0.0016042 | 0.16785081 |
| LRRIQ1 | 47.8685624 | -0.0072905 | 0.58887335 | -0.0123804 | 0.99012209 | 0.99814944 |
| RASSF9 | 54.7805007 | 0.16732921 | 0.38561222 | 0.43393129 | 0.66433834 | 0.93860542 |
| CEP290 | 90.0047544 | 0.14477278 | 0.22681078 | 0.63829762 | 0.52327996 | 0.90487731 |
| TMTC3 | 69.6248761 | -0.0002242 | 0.22741024 | -0.000986 | 0.99921328 | 0.99957534 |
| KITLG | 1053.51218 | 0.05456878 | 0.50285538 | 0.10851783 | 0.91358493 | 0.986403 |
| AC024941.2 | 9.19893786 | 0.56335429 | 0.8483346 | 0.66407086 | 0.50664496 | NA |
| DUSP6 | 127.889886 | -0.859973 | 0.70407136 | -1.2214288 | 0.22192372 | 0.76680432 |
| POC1B | 64.4519354 | 0.01409295 | 0.22724645 | 0.06201616 | 0.95054996 | 0.99105535 |
| POC1B-AS1 | 25.5680316 | -0.2581462 | 0.41003611 | -0.6295693 | 0.52897639 | 0.90495622 |
| ATP2B1 | 106.003789 | 0.02421625 | 0.36709936 | 0.06596646 | 0.94740452 | 0.99083449 |
| BTG1 | 83.5431483 | 0.12983381 | 0.38236563 | 0.33955408 | 0.73419236 | 0.95329896 |

| | | | | | | |
|---|---|---|---|---|---|---|
| LINC02391 | 30.8633644 | 0.63690267 | 0.47041368 | 1.35392039 | 0.17576178 | 0.72841249 |
| PLEKHG7 | 14.3879124 | -1.4836612 | 0.87185902 | -1.7017214 | 0.0888076 | NA |
| EEA1 | 132.527475 | -0.2863462 | 0.29963641 | -0.9556455 | 0.33925136 | 0.83066816 |
| AC138123.1 | 146.3178 | 0.14320244 | 0.40427732 | 0.35421833 | 7.23E-01 | 9.51E-01 |
| AC124947.1 | 42.3292354 | -0.3269382 | 0.52672699 | -0.6206977 | 0.53479855 | 0.90763121 |
| NUDT4 | 209.722025 | 0.64986549 | 0.30986887 | 2.09722743 | 3.60E-02 | 4.94E-01 |
| UBE2N | 63.1268662 | 0.03529029 | 0.23796592 | 0.14829979 | 0.88210618 | 0.98137082 |
| MRPL42 | 63.2753524 | -0.0661047 | 0.22646295 | -0.2919005 | 0.77036268 | 0.96342652 |
| SOCS2 | 30.9381303 | 0.67663803 | 0.37298298 | 1.81412575 | 0.06965837 | 0.60275677 |
| CRADD | 361.425878 | 0.30876244 | 0.28547451 | 1.08157621 | 0.27944088 | 0.80219184 |
| AC012085.2 | 25.4610714 | 1.26116208 | 0.68882378 | 1.83089219 | 0.06711664 | 0.59683227 |
| CEP83 | 77.4326458 | -0.190903 | 0.26198329 | -0.7286838 | 0.46619508 | 0.880451 |
| TMCC3 | 42.2935355 | 0.46902301 | 0.37675167 | 1.24491288 | 0.21316378 | 0.7643572 |
| NDUFA12 | 50.8979805 | 0.37717249 | 0.25734673 | 1.46561992 | 0.14275186 | 0.69328517 |
| NR2C1 | 80.295004 | 0.00770073 | 0.21396954 | 0.03598983 | 0.97129047 | 0.99403127 |
| FGD6 | 86.7585528 | 0.35272034 | 0.33785156 | 1.04400979 | 0.29648086 | 0.81120086 |
| VEZT | 187.258584 | 0.03587709 | 0.22008774 | 0.16301267 | 0.87050846 | 0.97854266 |
| METAP2 | 204.697506 | 1.44764965 | 0.30022392 | 4.82189985 | 1.42E-06 | 0.00350412 |
| USP44 | 13.0696031 | 1.27216726 | 0.41869612 | 3.03840228 | 0.00237836 | NA |
| NTN4 | 189.747219 | 0.98704752 | 0.39899486 | 2.47383515 | 0.01336714 | 0.36985119 |
| SNRPF | 19.2774275 | 0.21913921 | 0.39042143 | 0.56128888 | 0.57460062 | 0.91824492 |
| LTA4H | 38.0886096 | -0.2531673 | 0.3579289 | -0.7073118 | 0.47937275 | 0.88622003 |
| ELK3 | 30.0699017 | -0.5766069 | 0.31686271 | -1.8197375 | 6.88E-02 | 6.03E-01 |
| CDK17 | 92.3981903 | -0.6106925 | 0.27222237 | -2.2433589 | 0.02487368 | 0.43948277 |
| CFAP54 | 14.8136026 | -0.6594329 | 0.68381934 | -0.9643379 | 0.33487655 | NA |
| NEDD1 | 39.3223605 | -0.1599349 | 0.26801502 | -0.5967385 | 0.55068198 | 0.91248567 |
| TMPO | 52.5710196 | 0.37583558 | 0.23958128 | 1.56871847 | 0.11671355 | 0.6668162 |
| SLC25A3 | 131.239313 | 0.65626611 | 0.39110699 | 1.67797079 | 0.0933528 | 0.64572229 |
| IKBIP | 14.7565418 | -0.5911637 | 0.4866594 | -1.214738 | 0.22446603 | NA |
| APAF1 | 38.6670186 | -0.2258011 | 0.29420024 | -0.7675081 | 0.44277948 | 0.86991418 |
| ANKS1B | 91.1701086 | 1.29292679 | 0.67210199 | 1.92370624 | 0.05439142 | 0.5645224 |
| UHRF1BP1L | 62.8503588 | -0.0622237 | 0.26406523 | -0.2356376 | 0.81371387 | 0.97282508 |
| ACTR6 | 26.5165922 | 0.47402728 | 0.34862923 | 1.35968886 | 0.1739284 | 0.72794035 |
| SCYL2 | 116.581234 | -0.3215692 | 0.32011828 | -1.0045324 | 0.31512208 | 0.82084199 |
| NR1H4 | 169.556969 | -0.3191768 | 0.54667493 | -0.5838511 | 5.59E-01 | 0.91477029 |
| GAS2L3 | 21.7439171 | -1.2206295 | 0.49402082 | -2.4708057 | 1.35E-02 | 0.37186867 |
| UTP20 | 44.4422048 | -0.2478072 | 0.25153368 | -0.9851848 | 0.32453331 | 0.82567895 |
| ARL1 | 47.0102286 | 0.30981501 | 0.31455161 | 0.98494175 | 0.32465269 | 0.82567895 |
| CHPT1 | 221.827463 | 0.37636016 | 0.29106126 | 1.29306168 | 0.19598973 | 0.75350056 |
| GNPTAB | 93.2446297 | -0.145353 | 0.25166554 | -0.5775643 | 0.56355829 | 0.91624774 |
| DRAM1 | 44.6905541 | -0.3329533 | 0.40659495 | -0.818882 | 4.13E-01 | 0.85867915 |
| WASHC3 | 53.2959952 | -0.2203458 | 0.30338624 | -0.7262881 | 0.46766219 | 0.88108588 |
| NUP37 | 21.2545716 | 0.16593101 | 0.37697745 | 0.44016164 | 0.65982004 | 0.93704911 |
| NT5DC3 | 63.9220583 | -0.8509421 | 0.33503343 | -2.5398721 | 0.0110893 | 0.34397433 |
| HSP90B1 | 182.179483 | -0.232574 | 0.54732668 | -0.4249273 | 0.67088968 | 0.93878847 |
| C12orf73 | 11.7764628 | 0.33110543 | 0.41252922 | 0.80262299 | 0.42219268 | NA |
| TDG | 47.3967412 | 0.04428077 | 0.37575074 | 0.11784612 | 0.90618958 | 0.98531749 |
| HCFC2 | 29.3034006 | 0.32579095 | 0.32218189 | 1.0112019 | 0.3119198 | 0.81701824 |
| NFYB | 18.2955398 | 0.69059047 | 0.36703226 | 1.88155249 | 0.0598968 | 0.57714955 |
| TXNRD1 | 226.551831 | -0.0850706 | 0.25975302 | -0.3275055 | 0.74328555 | 0.95563105 |
| CHST11 | 14.5365382 | 0.6955941 | 0.49861923 | 1.39504068 | 0.16300357 | NA |
| SLC41A2 | 32.763836 | 0.00380434 | 0.62637611 | 0.00607357 | 0.99515402 | 0.99927028 |
| C12orf45 | 11.4647916 | -0.5738949 | 0.47078601 | -1.2190143 | 0.22283875 | NA |
| WASHC4 | 147.552279 | -0.4293968 | 0.23788705 | -1.8050447 | 0.07106768 | 0.60573888 |

| | | | | | | |
|---|---|---|---|---|---|---|
| APPL2 | 129.203813 | -0.2414045 | 0.30927348 | -0.7805534 | 0.43506518 | 0.86729771 |
| C12orf75 | 193.244447 | 0.16543346 | 0.52123435 | 0.31738787 | 7.51E-01 | 9.57E-01 |
| NUAK1 | 17.0665647 | 0.22193325 | 0.62483404 | 0.35518751 | 0.72244912 | 0.95093722 |
| CKAP4 | 20.4516273 | 0.62773728 | 0.48409584 | 1.29672106 | 0.19472718 | 0.75191699 |
| TCP11L2 | 30.9536245 | 0.15814131 | 0.38878965 | 0.40675289 | 0.68418949 | 0.94205398 |
| POLR3B | 30.1961479 | -0.3683842 | 0.31807056 | -1.158184 | 0.24678894 | 0.78583441 |
| RIC8B | 71.723022 | -0.001168 | 0.26130416 | -0.0044699 | 0.99643351 | 0.99927028 |
| TMEM263 | 54.3453312 | -0.0727204 | 0.35954653 | -0.202256 | 0.8397166 | 0.97600352 |
| MTERF2 | 19.1443946 | 1.41865958 | 0.47856754 | 2.96438738 | 0.00303286 | 0.22086516 |
| CRY1 | 74.645757 | -0.8032412 | 0.61782622 | -1.3001086 | 0.19356374 | 0.75067401 |
| BTBD11 | 115.239743 | -0.0291001 | 0.67335412 | -0.0432167 | 0.96552881 | 0.99270395 |
| PWP1 | 30.3544146 | -0.095485 | 0.31670332 | -0.3014968 | 0.76303567 | 0.96091323 |
| PRDM4 | 46.3520645 | -0.1002933 | 0.30253208 | -0.331513 | 0.74025703 | 0.9544376 |
| SART3 | 37.8436696 | 0.08829118 | 0.27790256 | 0.31770551 | 0.75070833 | 0.95736872 |
| ISCU | 83.014425 | 0.71530939 | 0.33237803 | 2.15209592 | 0.0313898 | 0.47623559 |
| CORO1C | 29.4714078 | -0.7464553 | 0.59153424 | -1.2618971 | 0.20698582 | 0.76231289 |
| SSH1 | 137.687622 | -0.2553374 | 0.31882771 | -0.8008631 | 0.4232109 | 0.86190547 |
| USP30 | 22.2110446 | -0.2949506 | 0.35826717 | -0.8232699 | 0.41035455 | 0.85867915 |
| UNG | 17.4426124 | 0.59688422 | 0.41725196 | 1.4305127 | 0.15256992 | 0.70572801 |
| ACACB | 246.002303 | 0.10269367 | 0.29674526 | 0.34606677 | 0.72929253 | 0.95279138 |
| KCTD10 | 23.7571136 | -0.1185076 | 0.43783504 | -0.2706672 | 0.78664702 | 0.96767512 |
| UBE3B | 41.1749058 | 0.05645429 | 0.31920705 | 0.1768579 | 0.85962001 | 0.97691173 |
| MMAB | 34.2715796 | -0.0974758 | 0.38783213 | -0.2513349 | 0.80155517 | 0.97045451 |
| MVK | 26.9045558 | 0.07250316 | 0.292521 | 0.24785625 | 0.80424563 | 0.97175786 |
| FAM222A | 23.1056061 | 0.51097686 | 0.43137939 | 1.1845185 | 0.23620788 | 0.77864547 |
| TRPV4 | 134.400421 | 0.58401211 | 0.41737399 | 1.39925372 | 0.16173691 | 0.71101785 |
| GLTP | 50.2802977 | 0.13846425 | 0.34481593 | 0.4015599 | 0.68800795 | 0.94205398 |
| TCHP | 39.497782 | -0.2197687 | 0.38978034 | -0.5638271 | 0.57287181 | 0.91813957 |
| GIT2 | 90.023566 | -0.347796 | 0.26924705 | -1.2917355 | 0.19644874 | 0.75350056 |
| ANKRD13A | 132.064059 | -0.4052706 | 0.2905646 | -1.3947694 | 0.1630854 | 0.71325312 |
| C12orf76 | 19.235241 | 0.21381197 | 0.42414919 | 0.50409614 | 0.61419382 | 0.92662475 |
| IFT81 | 55.9494982 | 0.01274988 | 0.26397185 | 0.04830013 | 0.96147705 | 0.99256424 |
| ATP2A2 | 138.674651 | -0.2037328 | 0.37861255 | -0.5381037 | 0.59050544 | 0.92233545 |
| ANAPC7 | 45.1863134 | -0.8101388 | 0.32618902 | -2.4836482 | 0.01300442 | 0.36664851 |
| ARPC3 | 111.312191 | 0.21018829 | 0.21768871 | 0.96554519 | 0.33427181 | 0.82974237 |
| GPN3 | 12.6942448 | 0.39673587 | 0.47528042 | 0.83474062 | 0.40386376 | NA |
| VPS29 | 58.1446648 | 0.35715632 | 0.2090945 | 1.70810954 | 0.08761602 | 0.63466198 |
| PPTC7 | 69.4555395 | 0.20691238 | 0.28276476 | 0.7317474 | 0.46432276 | 0.87877814 |
| TCTN1 | 31.5034058 | 0.76868035 | 0.40478661 | 1.89897673 | 0.05756754 | 0.57118657 |
| PPP1CC | 32.9883988 | 0.37981437 | 0.31396105 | 1.20974998 | 0.22637484 | 0.77169045 |
| SH2B3 | 17.5217951 | 0.01641296 | 0.36884898 | 0.04449777 | 0.96450763 | 0.99264137 |
| ATXN2 | 322.773026 | -0.3715616 | 0.21084022 | -1.7622898 | 0.07802035 | 0.61367304 |
| BRAP | 42.6234419 | -0.058967 | 0.29367482 | -0.2007903 | 0.84086257 | 0.97600352 |
| ACAD10 | 53.7312356 | -0.5480166 | 0.39399572 | -1.3909201 | 0.16424964 | 0.71397803 |
| ALDH2 | 91.7341067 | 0.63037227 | 0.32656046 | 1.93033866 | 0.05356489 | 0.56110609 |
| MAPKAPK5 | 45.1493624 | -0.0823364 | 0.29004955 | -0.2838702 | 0.77650989 | 0.9649632 |
| TMEM116 | 155.246244 | 0.25100927 | 0.22548856 | 1.11317961 | 2.66E-01 | 7.97E-01 |
| ERP29 | 86.7554676 | 0.68045546 | 0.29175628 | 2.33227359 | 0.0196863 | 0.41544117 |
| NAA25 | 97.1424671 | -0.476086 | 0.27052957 | -1.7598298 | 0.07843667 | 0.61367304 |
| TRAFD1 | 43.42206 | 0.02884928 | 0.3274576 | 0.08810082 | 0.92979655 | 0.98960607 |
| HECTD4 | 257.834783 | -0.165407 | 0.19674159 | -0.8407325 | 0.40049781 | 0.85778581 |
| RPL6 | 137.676391 | 0.68471006 | 0.39389348 | 1.73831276 | 0.08215572 | 0.61960523 |
| PTPN11 | 88.2133194 | -0.1238256 | 0.25843936 | -0.4791283 | 0.63184738 | 0.93251095 |
| OAS1 | 19.9455526 | -6.6072401 | 2.83306844 | -2.3321851 | 0.01969095 | 0.41544117 |

| | | | | | | |
|---|---|---|---|---|---|---|
| OAS3 | 16.1805679 | -7.4625845 | 2.48413538 | -3.0040973 | 0.0026637 | 0.20745535 |
| DTX1 | 11.2119867 | -0.5228548 | 0.51675418 | -1.0118056 | 0.31163099 | NA |
| RASAL1 | 105.554023 | -1.1700321 | 0.47887137 | -2.4433119 | 0.01455316 | 0.38288257 |
| DDX54 | 20.3059866 | -0.1145821 | 0.43131205 | -0.2656594 | 0.7905015 | 0.96904219 |
| TPCN1 | 148.752526 | -0.1120807 | 0.29079951 | -0.3854226 | 0.69992436 | 0.94419257 |
| SLC8B1 | 45.1047188 | 0.01808398 | 0.2733925 | 0.06614659 | 0.94726112 | 0.99083449 |
| PLBD2 | 17.4246523 | -0.4502347 | 0.3943878 | -1.1416039 | 0.25361871 | 0.7912221 |
| SDSL | 20.315851 | -0.3208719 | 0.51290237 | -0.6256004 | 0.53157705 | 0.90583039 |
| RBM19 | 40.5997775 | -0.3153404 | 0.28612597 | -1.1021035 | 0.27041668 | 0.79666376 |
| MED13L | 1227.94164 | -0.3671131 | 0.24831715 | -1.478404 | 0.13929968 | 0.69054855 |
| MAP1LC3B2 | 15.4672302 | 0.12202724 | 0.40464717 | 0.30156454 | 0.76298405 | NA |
| C12orf49 | 48.1352302 | -0.6781752 | 0.43666196 | -1.5530897 | 0.1204017 | 0.6668162 |
| RNFT2 | 12.0328966 | -0.483464 | 0.52724005 | -0.9169713 | 0.35915766 | NA |
| FBXW8 | 77.6215198 | -0.0917074 | 0.27854103 | -0.329242 | 0.74197281 | 0.95519424 |
| TESC | 30.8022756 | 0.91715307 | 0.57957165 | 1.58246711 | 1.14E-01 | 0.6656824 |
| FBXO21 | 74.5768069 | 0.22100923 | 0.23198202 | 0.9526998 | 0.34074219 | 0.8319325 |
| NOS1 | 27.2139135 | 0.86733935 | 0.69778055 | 1.24299731 | 2.14E-01 | 7.64E-01 |
| KSR2 | 183.913985 | -1.4248472 | 0.64229409 | -2.218372 | 0.02652948 | 0.44763538 |
| WSB2 | 33.4088215 | -0.4134095 | 0.32798118 | -1.2604671 | 0.2075009 | 0.76231289 |
| VSIG10 | 56.3479524 | 0.25276519 | 0.39955317 | 0.63261964 | 0.52698206 | 0.90487731 |
| PEBP1 | 237.027918 | 0.88242299 | 0.40974845 | 2.15357251 | 0.03127371 | 0.4760612 |
| TAOK3 | 216.707064 | 0.00797019 | 0.19155712 | 0.04160738 | 0.96681169 | 0.99308116 |
| SUDS3 | 49.7472306 | -0.1620672 | 0.26723382 | -0.6064622 | 0.54420787 | 0.90965439 |
| AC002070.1 | 68.2983286 | 0.58827129 | 0.53513319 | 1.09929884 | 0.27163774 | 0.79666376 |
| PRKAB1 | 59.7157108 | 0.3914827 | 0.28650049 | 1.36642943 | 0.17180422 | 0.72505043 |
| CIT | 45.7398169 | -0.1111375 | 0.47500304 | -0.2339721 | 0.81500661 | 0.97284589 |
| BICDL1 | 86.1555303 | 0.0616806 | 0.33265405 | 0.18541965 | 0.8528999 | 0.97612951 |
| RAB35 | 24.9352207 | -0.5418596 | 0.39723794 | -1.364068 | 0.17254616 | 0.72649757 |
| GCN1 | 42.9809995 | -0.3970369 | 0.28221613 | -1.4068539 | 0.15947073 | 0.71101785 |
| RPLP0 | 201.514982 | 0.36110994 | 0.4673539 | 0.77266914 | 0.43971822 | 0.86895963 |
| PXN | 69.7117365 | -0.0358413 | 0.2348302 | -0.1526265 | 0.87869285 | 0.98061726 |
| GATC | 18.1681352 | 0.29134254 | 0.36359193 | 0.8012899 | 0.42296384 | 0.86190547 |
| SRSF9 | 59.5473563 | 0.38956924 | 0.32108995 | 1.21327138 | 0.22502608 | 0.76950128 |
| DYNLL1 | 96.9898006 | 0.32028372 | 0.58232661 | 0.55000701 | 0.58231457 | 0.92034873 |
| COQ5 | 25.4539385 | -0.2745289 | 0.35387136 | -0.7757872 | 0.43787467 | 0.86821458 |
| RNF10 | 187.808986 | -0.1321244 | 0.19605794 | -0.6739051 | 5.00E-01 | 0.89469323 |
| POP5 | 20.180581 | 0.68082103 | 0.39326203 | 1.73121476 | 0.08341347 | 0.62487197 |
| MLEC | 107.711383 | 0.71070329 | 0.34902244 | 2.03626818 | 0.04172343 | 0.51069022 |
| UNC119B | 26.1196959 | 0.611528 | 0.3386271 | 1.80590392 | 0.07093335 | 0.60572821 |
| ACADS | 32.628942 | -0.0226397 | 0.28942335 | -0.0782234 | 0.93765035 | 0.99012269 |
| SPPL3 | 312.815575 | -0.1780665 | 0.2156483 | -0.8257265 | 0.40895926 | 0.85867915 |
| P2RX4 | 36.3740279 | 0.01947762 | 0.38200265 | 0.05098818 | 0.95933494 | 0.99245244 |
| CAMKK2 | 23.8929104 | -0.2326144 | 0.39736344 | -0.5853946 | 0.55828241 | 0.91425998 |
| ANAPC5 | 82.8864439 | 0.03487147 | 0.20567458 | 0.16954679 | 0.86536657 | 0.97788853 |
| RNF34 | 42.8624926 | -0.1554975 | 0.28868559 | -0.5386396 | 0.59013558 | 0.92233545 |
| KDM2B | 74.4787839 | -0.0947425 | 0.3001757 | -0.3156234 | 0.7522884 | 0.95782508 |
| TMEM120B | 96.8241893 | 0.1284959 | 0.2845067 | 0.45164457 | 6.52E-01 | 0.93472538 |
| RHOF | 19.0681066 | -0.2540859 | 0.48226304 | -0.5268617 | 0.59828963 | 0.92296674 |
| SETD1B | 26.3225171 | 0.06819383 | 0.33879722 | 0.20128213 | 0.84047797 | 0.97600352 |
| WDR66 | 31.0578797 | -0.9671439 | 0.42161878 | -2.2938824 | 0.02179725 | 0.4212027 |
| BCL7A | 71.5905777 | 0.00658194 | 0.36411157 | 0.0180767 | 0.98557766 | 0.99752207 |
| MLXIP | 98.4134178 | -0.3349205 | 0.29491588 | -1.1356474 | 0.25610416 | 0.7940072 |
| CLIP1 | 228.738771 | -0.5885344 | 0.36302755 | -1.6211839 | 0.1049782 | 0.65946648 |
| ZCCHC8 | 57.3984055 | 0.02057693 | 0.30074267 | 0.06842038 | 0.945451 | 0.99083449 |

| | | | | | | |
|---|---|---|---|---|---|---|
| RSRC2 | 85.590246 | 0.01266041 | 0.21712981 | 0.05830803 | 0.95350327 | 0.99148072 |
| KNTC1 | 15.5708723 | -0.4070517 | 0.40601931 | -1.0025428 | 0.31608153 | NA |
| DENR | 88.3004328 | 0.01905535 | 0.2287487 | 0.08330255 | 0.93361097 | 0.98960607 |
| HIP1R | 142.71101 | 0.38349323 | 0.31293068 | 1.22548939 | 0.22039088 | 0.76680432 |
| VPS37B | 198.349735 | -0.1987232 | 0.52576296 | -0.3779711 | 0.70545205 | 0.94548439 |
| ABCB9 | 29.3420898 | 1.13143473 | 0.50287847 | 2.24991684 | 0.02445422 | 0.43948277 |
| ARL6IP4 | 63.5726983 | 0.12869296 | 0.30808072 | 0.41772481 | 0.67614833 | 0.9402691 |
| PITPNM2 | 149.31132 | 0.25850157 | 0.23520162 | 1.09906371 | 0.27174028 | 0.79666376 |
| MPHOSPH9 | 81.5534865 | -0.27648 | 0.275507 | -1.0035317 | 0.31560441 | 0.82084199 |
| C12orf65 | 37.3271178 | 0.20365395 | 0.30373928 | 0.67048934 | 0.5025459 | 0.89523979 |
| SBNO1 | 95.3489972 | -0.310341 | 0.22001473 | -1.4105463 | 0.15837844 | 0.71089007 |
| KMT5A | 22.2240066 | 0.51758035 | 0.3604889 | 1.43577334 | 0.15106685 | 0.70572801 |
| RILPL2 | 28.5556141 | 0.29311342 | 0.28776338 | 1.01859183 | 0.30839679 | 0.81568231 |
| SNRNP35 | 27.9488461 | 0.31179544 | 0.28792362 | 1.08291027 | 0.27884825 | 0.80219184 |
| RILPL1 | 73.3685973 | 0.09981001 | 0.23465413 | 0.42534949 | 0.67058192 | 0.93878847 |
| TMED2 | 44.9000419 | -0.0241562 | 0.29037061 | -0.0831909 | 0.93369975 | 0.98960607 |
| DDX55 | 16.9751759 | -0.1964624 | 0.37239572 | -0.5275635 | 0.59780232 | 0.9229638 |
| EIF2B1 | 23.4797052 | -0.1932203 | 0.32134842 | -0.6012798 | 0.54765362 | 0.91139843 |
| GTF2H3 | 25.1959475 | 0.20471408 | 0.3185454 | 0.64265276 | 0.52044944 | 0.90427107 |
| TCTN2 | 19.8167068 | -0.4047977 | 0.4430845 | -0.9135902 | 0.36093222 | 0.84331775 |
| ATP6V0A2 | 29.9228884 | -0.1510577 | 0.36792307 | -0.4105686 | 0.68138891 | 0.94196488 |
| CCDC92 | 75.1008007 | -0.1666656 | 0.20459535 | -0.8146106 | 0.41529522 | 0.85867915 |
| RFLNA | 16.7043713 | -0.6392897 | 0.49682227 | -1.2867573 | 0.19817889 | 0.75559912 |
| NCOR2 | 127.575085 | -0.2827567 | 0.26971005 | -1.048373 | 0.29446679 | 0.81012446 |
| UBC | 311.711781 | -0.4271174 | 0.2992124 | -1.4274721 | 0.15344386 | 0.70572801 |
| DHX37 | 13.4262038 | -0.4899316 | 0.46153834 | -1.0615187 | 0.28845423 | NA |
| BRI3BP | 27.0528816 | 0.94948199 | 0.45781788 | 2.07392945 | 0.03808586 | 0.49671741 |
| AACS | 29.1508814 | 0.30113267 | 0.34686003 | 0.86816769 | 0.38530254 | 0.85540507 |
| SLC15A4 | 39.7324198 | -0.4464982 | 0.3100975 | -1.439864 | 0.14990587 | 0.70336675 |
| RIMBP2 | 156.080166 | -0.0282523 | 0.38760838 | -0.0728887 | 0.94189472 | 0.99053461 |
| STX2 | 39.5628419 | -0.2099626 | 0.36840205 | -0.5699279 | 0.56872661 | 0.91698355 |
| RAN | 52.4305859 | 0.24329099 | 0.46049768 | 0.52832186 | 0.59727596 | 0.9229638 |
| SFSWAP | 101.587882 | -0.2279002 | 0.21055641 | -1.0823713 | 0.2790876 | 0.80219184 |
| ULK1 | 52.662924 | 0.16644345 | 0.2442039 | 0.6815757 | 0.49550728 | 0.89230917 |
| PUS1 | 16.0433892 | -0.4355425 | 0.41168083 | -1.0579615 | 0.29007298 | NA |
| EP400 | 99.3923229 | 0.07139764 | 0.26185027 | 0.27266591 | 0.78511005 | 0.9674911 |
| DDX51 | 12.6344327 | 0.15811542 | 0.50916924 | 0.31053609 | 0.75615332 | NA |
| FBRSL1 | 49.8088419 | -0.448187 | 0.30485082 | -1.4701846 | 0.14151175 | 0.69328517 |
| POLE | 9.60802261 | -0.1623454 | 0.53379155 | -0.3041363 | 0.76102403 | NA |
| ANKLE2 | 61.7900404 | -0.8637435 | 0.33221796 | -2.5999301 | 0.00932427 | 0.32071089 |
| GOLGA3 | 66.9602867 | -0.758503 | 0.3032512 | -2.5012367 | 0.01237604 | 0.36046953 |
| CHFR | 73.4697684 | -0.3822174 | 0.31599533 | -1.2095667 | 0.22644521 | 0.77169045 |
| ZNF26 | 46.3364821 | -0.3201045 | 0.36315329 | -0.8814582 | 0.37806985 | 0.85314077 |
| ZNF84 | 65.1990031 | -0.1580598 | 0.28364213 | -0.5572509 | 0.57735603 | 0.91859435 |
| ZNF140 | 31.2127561 | -0.3386372 | 0.31596071 | -1.0717699 | 0.28382339 | 0.80431986 |
| ZNF891 | 31.7031069 | 0.36802318 | 0.42295973 | 0.87011399 | 0.38423811 | 0.85511218 |
| ZNF268 | 14.0628162 | -0.3625127 | 0.46331149 | -0.7824383 | 0.43395702 | NA |
| MPHOSPH8 | 160.446294 | 0.42093423 | 0.25032826 | 1.68152899 | 0.09266021 | 0.64572229 |
| PSPC1 | 217.30765 | -0.3332186 | 0.28779868 | -1.1578184 | 0.24693816 | 0.78583441 |
| ZMYM5 | 65.2202344 | 0.3117041 | 0.26675333 | 1.16851064 | 0.24260085 | 0.78464787 |
| ZMYM2 | 415.996695 | -0.1869229 | 0.26932646 | -0.6940382 | 0.48765824 | 0.88838332 |
| CRYL1 | 115.660815 | 0.36585128 | 0.27107028 | 1.34965468 | 0.17712678 | 0.72841249 |
| IFT88 | 165.570836 | -0.2994083 | 0.22415922 | -1.3356949 | 0.18164901 | 0.73398872 |
| EEF1AKMT1 | 19.3352831 | 0.53376085 | 0.54465426 | 0.97999941 | 0.32708641 | 0.82612204 |

| | | | | | | |
|---|---|---|---|---|---|---|
| XPO4 | 133.330822 | -0.198568 | 0.21267772 | -0.9336568 | 0.35048098 | 0.83716304 |
| LATS2 | 108.079458 | -0.2348809 | 0.35769667 | -0.6566483 | 0.51140711 | 0.9006266 |
| SAP18 | 53.6966837 | 0.41946795 | 0.38906108 | 1.07815449 | 0.28096482 | 0.80377836 |
| MRPL57 | 43.1833209 | 0.39661355 | 0.34474638 | 1.15045021 | 0.24995849 | 0.78799766 |
| ZDHHC20 | 125.785204 | -0.0505721 | 0.26718993 | -0.1892741 | 0.84987802 | 0.97612951 |
| MICU2 | 200.686651 | 0.02686062 | 0.19008429 | 0.14130898 | 0.88762585 | 0.98237789 |
| FGF9 | 60.9034959 | 0.83662324 | 0.37404859 | 2.23666998 | 0.02530792 | 0.4403385 |
| AL136962.1 | 59.2425598 | 0.74505271 | 0.67329402 | 1.10657853 | 0.26847619 | 0.79666376 |
| TNFRSF19 | 84.2549311 | 0.08452938 | 0.33012625 | 0.25605167 | 0.79791094 | 0.97019967 |
| MIPEP | 119.536222 | -0.3355108 | 0.21153568 | -1.586072 | 0.11272299 | 0.6656824 |
| PARP4 | 103.488707 | -0.1919885 | 0.33871992 | -0.566806 | 0.57084596 | 0.91791025 |
| ATP12A | 29.8317898 | 0.35626313 | 0.88994817 | 0.40031897 | 6.89E-01 | 9.42E-01 |
| MTMR6 | 33.8069946 | 0.00279833 | 0.27732194 | 0.01009056 | 0.99194903 | 0.99814944 |
| NUP58 | 77.3072997 | -0.6071037 | 0.34526718 | -1.7583591 | 0.07868642 | 0.61367304 |
| RNF6 | 14.132045 | 0.74471487 | 0.43901907 | 1.69631555 | 0.08982614 | NA |
| CDK8 | 226.151489 | -0.2182753 | 0.2404734 | -0.90769 | 0.36404203 | 0.8466721 |
| WASF3 | 39.2264372 | -0.1080371 | 0.24697738 | -0.4374372 | 0.66179432 | 0.93734459 |
| USP12 | 157.147181 | -0.2597111 | 0.27908484 | -0.9305812 | 0.35207026 | 0.83872681 |
| LINC02340 | 26.5236748 | 0.08722134 | 0.52605698 | 0.16580207 | 0.86831271 | 0.97812778 |
| RPL21 | 72.0002721 | 0.28395328 | 0.43576711 | 0.65161706 | 0.51464824 | 0.90156229 |
| RASL11A | 16.3847689 | 0.77394218 | 0.58445482 | 1.32421216 | 0.18543259 | 0.73752116 |
| GTF3A | 24.3679479 | 0.1358846 | 0.35056354 | 0.38761761 | 0.69829904 | 0.94395238 |
| MTIF3 | 48.0365733 | 0.02678292 | 0.26919203 | 0.09949375 | 0.92074625 | 0.98804116 |
| LNX2 | 63.0792587 | 0.02936349 | 0.27315209 | 0.10749867 | 0.91439338 | 0.98691474 |
| POLR1D | 44.1357548 | 0.31257881 | 0.24638518 | 1.26865915 | 0.20456266 | 0.76173533 |
| PAN3 | 439.172815 | -0.1350443 | 0.19243558 | -0.7017638 | 0.48282649 | 0.88781758 |
| POMP | 40.1976172 | -0.1928477 | 0.30507893 | -0.6321241 | 0.52730582 | 0.90487731 |
| SLC46A3 | 17.5779191 | 1.09985676 | 0.47291599 | 2.32569162 | 0.02003501 | 0.41544117 |
| SLC7A1 | 124.704427 | -0.5458847 | 0.56450435 | -0.9670158 | 0.33353611 | 0.82974237 |
| UBL3 | 176.407822 | -0.0199404 | 0.26173409 | -0.0761857 | 0.93927139 | 0.99012269 |
| KATNAL1 | 26.3739235 | -0.0263848 | 0.34110287 | -0.0773515 | 0.93834395 | 0.99012269 |
| HMGB1 | 208.425886 | 0.33110817 | 0.23479692 | 1.41018957 | 0.15848371 | 0.71089007 |
| USPL1 | 39.4092148 | -0.2463153 | 0.33095841 | -0.7442486 | 0.45672607 | 0.87399147 |
| HSPH1 | 87.0188595 | -0.0585275 | 0.52633764 | -0.1111976 | 0.91145969 | 0.98586235 |
| B3GLCT | 45.2386369 | 0.30559009 | 0.31417797 | 0.97266556 | 0.33071955 | 0.82867637 |
| FRY | 330.891869 | -0.2095978 | 0.38422211 | -0.5455121 | 0.58540135 | 0.92076087 |
| N4BP2L1 | 29.4697091 | 0.33393967 | 0.47391889 | 0.70463465 | 0.48103763 | 0.88689225 |
| N4BP2L2 | 719.599435 | -0.1976782 | 0.22786887 | -0.8675086 | 0.38566341 | 0.85540507 |
| PDS5B | 264.399177 | -0.1795274 | 0.20659635 | -0.8689768 | 0.38485984 | 0.85531718 |
| KL | 148.227061 | -0.1077165 | 0.43623123 | -0.2469253 | 8.05E-01 | 9.72E-01 |
| STARD13 | 475.966685 | -0.4983537 | 0.38399479 | -1.2978138 | 0.19435132 | 0.75182195 |
| AL139383.1 | 81.1672999 | 0.68733429 | 0.31954921 | 2.15094972 | 0.03148017 | 0.47665483 |
| RFC3 | 30.4235486 | -0.0044217 | 0.40707958 | -0.0108619 | 0.99133364 | 0.99814944 |
| LINC02343 | 179.577502 | 0.87384322 | 0.45691872 | 1.91246971 | 0.05581597 | 0.56884484 |
| NBEA | 427.388545 | -0.7157984 | 0.29867868 | -2.39655 | 0.01655024 | 0.39748373 |
| SPART | 84.4943859 | 0.24320808 | 0.30689193 | 0.79248769 | 0.42807637 | 0.86279045 |
| RFXAP | 15.1605389 | 1.36491065 | 0.55042816 | 2.47972531 | 0.01314836 | NA |
| ALG5 | 49.8548711 | -0.1582488 | 0.24276383 | -0.6518633 | 0.51448938 | 0.90156229 |
| EXOSC8 | 22.722818 | -0.2520582 | 0.3346537 | -0.7531911 | 0.45133508 | 0.87346041 |
| SUPT20H | 107.607744 | 0.02894542 | 0.23574903 | 0.12278063 | 0.90228081 | 0.98531749 |
| LINC00571 | 32.6702586 | -0.0749711 | 0.38976491 | -0.1923495 | 0.84746849 | 0.97612951 |
| UFM1 | 61.5339743 | -0.0272582 | 0.27666971 | -0.0985225 | 0.9215174 | 0.98846691 |
| FREM2 | 195.775382 | 0.3940305 | 0.33562224 | 1.17402977 | 0.2403831 | 0.78192112 |
| PROSER1 | 35.9793289 | -0.1080344 | 0.38728371 | -0.2789542 | 0.78027999 | 0.96650724 |

| | | | | | | |
|---|---|---|---|---|---|---|
| NHLRC3 | 58.6117438 | 0.72248655 | 0.34357175 | 2.10286951 | 0.03547718 | 0.49209205 |
| COG6 | 107.150231 | -0.3045046 | 0.28584669 | -1.0652724 | 0.28675269 | 0.80515414 |
| LINC00598 | 23.0678815 | -0.3518193 | 0.40627485 | -0.8659638 | 0.38651001 | 0.8560201 |
| FOXO1 | 343.10403 | -0.9185825 | 0.45886031 | -2.0018783 | 0.04529783 | 0.52675468 |
| MRPS31 | 75.9063452 | 0.20736796 | 0.29221868 | 0.70963278 | 0.47793189 | 0.88545331 |
| ELF1 | 272.117221 | -0.1970288 | 0.14600255 | -1.3494884 | 0.17718013 | 0.72841249 |
| WBP4 | 39.050885 | -0.313428 | 0.33139211 | -0.9457919 | 0.34425474 | 0.83522146 |
| AL354696.2 | 18.5285517 | -0.1215757 | 0.48079717 | -0.2528627 | 0.80037432 | 0.97019967 |
| MTRF1 | 20.6203335 | -0.0302664 | 0.3831641 | -0.0789908 | 0.93703998 | 0.99012269 |
| NAA16 | 59.6674023 | 0.36212494 | 0.26580188 | 1.36238669 | 0.17307588 | 0.72721271 |
| VWA8 | 321.880545 | 0.2589852 | 0.20138838 | 1.28599873 | 1.98E-01 | 7.56E-01 |
| DGKH | 155.088294 | 0.18393218 | 0.48459672 | 0.37955721 | 0.70427413 | 0.94476609 |
| AKAP11 | 89.2944776 | 0.40274042 | 0.22648956 | 1.77818536 | 0.07537342 | 0.61020315 |
| LINC02341 | 32.2277789 | 0.08347829 | 0.86715467 | 0.0962669 | 0.9233086 | 0.98915831 |
| EPSTI1 | 73.4146355 | -2.194889 | 1.19445356 | -1.8375674 | 0.06612617 | 0.59523726 |
| DNAJC15 | 75.5993291 | 0.03515809 | 0.35282985 | 0.099646 | 0.92062537 | 0.98804116 |
| ENOX1 | 19.4522318 | 1.25946683 | 0.71300192 | 1.76642838 | 0.07732401 | 0.61342702 |
| CCDC122 | 59.3662128 | 0.4824375 | 0.40518788 | 1.19065134 | 0.23379049 | 0.77671344 |
| SMIM2-AS1 | 16.2736423 | 0.78334784 | 0.66121291 | 1.18471346 | 0.23613076 | 0.77864547 |
| SERP2 | 13.4038374 | -0.1934509 | 0.43612512 | -0.4435674 | 0.65735539 | NA |
| TSC22D1 | 594.752793 | 0.12744295 | 0.18414995 | 0.69206074 | 0.48889919 | 0.88920066 |
| NUFIP1 | 15.34593 | -0.0834204 | 0.40981457 | -0.2035564 | 0.8387002 | NA |
| GPALPP1 | 49.1878729 | 0.36773298 | 0.28682469 | 1.28208272 | 0.19981363 | 0.75605362 |
| GTF2F2 | 109.820195 | -0.3574978 | 0.30494375 | -1.1723403 | 0.24106046 | 0.78310048 |
| TPT1 | 395.009249 | 0.19636737 | 0.43358272 | 0.45289481 | 0.65062449 | 0.93453685 |
| SLC25A30 | 67.6404138 | 0.11221894 | 0.35821345 | 0.31327394 | 0.75407254 | 0.95826617 |
| COG3 | 73.4748387 | -0.2496085 | 0.25475314 | -0.9798056 | 0.3271821 | 0.82612204 |
| LINC01055 | 44.0678095 | 1.78525289 | 0.55102509 | 3.23987587 | 0.00119582 | 0.14513539 |
| ZC3H13 | 116.384478 | 0.0238059 | 0.22337479 | 0.10657381 | 0.9151271 | 0.98691474 |
| AL138686.1 | 25.4678757 | -0.6696882 | 0.74838604 | -0.8948433 | 0.37087087 | 0.84889051 |
| LRCH1 | 444.555262 | -0.3557389 | 0.21729648 | -1.6371132 | 0.10160681 | 0.65863153 |
| ESD | 46.1844429 | -0.1169071 | 0.3445292 | -0.3393241 | 0.73436556 | 0.95329896 |
| SUCLA2 | 112.641994 | -0.2966968 | 0.26645052 | -1.1135154 | 0.26548715 | 0.79666376 |
| MED4 | 64.5712352 | -0.1108808 | 0.2767425 | -0.4006643 | 0.68866728 | 0.94205398 |
| ITM2B | 669.951422 | 0.54304539 | 0.3546492 | 1.53121842 | 0.12571542 | 0.66929384 |
| RB1 | 244.607503 | -0.0537573 | 0.25397771 | -0.2116616 | 0.83237107 | 0.97600352 |
| LPAR6 | 13.1615506 | 0.2857006 | 0.60107975 | 0.4753123 | 0.63456439 | NA |
| RCBTB2 | 30.2114152 | 0.4157275 | 0.33082204 | 1.25664995 | 0.20888042 | 0.7623519 |
| FNDC3A | 518.079979 | -0.876432 | 0.56305787 | -1.5565575 | 0.11957558 | 0.6668162 |
| CDADC1 | 32.6184197 | -0.0326068 | 0.38541646 | -0.0846015 | 0.93257822 | 0.98960607 |
| CAB39L | 64.2151644 | -0.3751173 | 0.41552746 | -0.9027496 | 0.36665879 | 0.84741115 |
| SETDB2 | 72.1179558 | -0.0984233 | 0.35455867 | -0.2775937 | 0.78132427 | 0.96652357 |
| PHF11 | 46.9395083 | -0.5917927 | 0.34700789 | -1.7054156 | 0.08811693 | 0.63466198 |
| RCBTB1 | 19.8531702 | -0.6141311 | 0.37883982 | -1.6210838 | 0.10499966 | 0.65946648 |
| EBPL | 21.5658708 | 0.49807879 | 0.44642733 | 1.1156996 | 0.26455074 | 0.79666376 |
| KPNA3 | 87.6088598 | -0.1070256 | 0.2526222 | -0.4236589 | 0.67181461 | 0.93878847 |
| SPRYD7 | 17.1930007 | 0.46660051 | 0.40877957 | 1.14144773 | 0.25368365 | 0.7912221 |
| DLEU2 | 104.610384 | 0.16864255 | 0.31777507 | 0.53069787 | 5.96E-01 | 9.23E-01 |
| TRIM13 | 40.5915253 | 0.11451615 | 0.33593675 | 0.34088605 | 0.73318937 | 0.95329896 |
| RNASEH2B | 43.8370713 | 0.03309304 | 0.27183336 | 0.12174018 | 0.90310479 | 0.98531749 |
| INTS6 | 191.401678 | -0.2324923 | 0.31605556 | -0.7356058 | 0.4619706 | 0.87689276 |
| INTS6-AS1 | 20.2662922 | 1.12586695 | 0.46008633 | 2.44707758 | 0.01440198 | 0.3811083 |
| WDFY2 | 229.542954 | -0.1075837 | 0.23395858 | -0.4598409 | 6.46E-01 | 9.33E-01 |
| ATP7B | 20.3472869 | 0.41667659 | 0.49074827 | 0.84906379 | 0.3958458 | 0.85605713 |

| | | | | | | |
|---|---|---|---|---|---|---|
| ALG11 | 13.465081 | -0.0065495 | 0.4765287 | -0.0137441 | 0.98903411 | NA |
| UTP14C | 18.3544082 | 0.08377586 | 0.46882079 | 0.17869484 | 0.85817732 | 0.97691173 |
| NEK5 | 10.567407 | 0.1703843 | 0.58792061 | 0.28980834 | 0.77196287 | NA |
| NEK3 | 42.1439715 | -0.1464645 | 0.34288474 | -0.4271538 | 0.66926731 | 0.93878847 |
| VPS36 | 98.3303109 | 0.13401031 | 0.20490267 | 0.65401937 | 0.51309932 | 0.90156229 |
| SUGT1 | 62.3035154 | -0.0475364 | 0.29474053 | -0.1612822 | 0.87187117 | 0.97866716 |
| OLFM4 | 674.312208 | -0.3153449 | 0.77240708 | -0.4082626 | 6.83E-01 | 0.94205398 |
| DIAPH3 | 25.7467522 | 0.27166778 | 0.47896238 | 0.56720066 | 0.57057786 | 0.91778542 |
| TDRD3 | 215.958267 | 0.49556533 | 0.16541285 | 2.99593013 | 0.00273609 | 0.2110732 |
| PCDH9 | 437.890693 | 1.3401067 | 0.56945468 | 2.35331579 | 0.01860682 | 0.41322807 |
| DACH1 | 1025.13921 | 0.12756501 | 0.44619738 | 0.28589368 | 0.77495956 | 0.96464609 |
| MZT1 | 12.1907443 | 0.40759133 | 0.4294861 | 0.94902101 | 0.34260993 | NA |
| DIS3 | 36.854921 | -0.2727422 | 0.26641048 | -1.0237667 | 0.30594546 | 0.81568231 |
| PIBF1 | 240.676825 | -0.3335886 | 0.16791509 | -1.9866507 | 0.04696112 | 0.5320597 |
| KLF5 | 67.9863136 | -1.0370687 | 0.38349162 | -2.70428 | 6.85E-03 | 0.30102597 |
| KLF12 | 314.628518 | -0.282992 | 0.1948351 | -1.452469 | 1.46E-01 | 0.69713616 |
| TBC1D4 | 375.398096 | 0.18191161 | 0.28203865 | 0.64498822 | 0.51893482 | 0.90368693 |
| COMMD6 | 51.9145913 | 0.16871811 | 0.48843718 | 0.34542439 | 0.72977534 | 0.95279138 |
| UCHL3 | 56.6578131 | -0.3432388 | 0.29011664 | -1.1831063 | 0.23676702 | 0.77864547 |
| LMO7-AS1 | 12.7578108 | 1.45633565 | 0.52373792 | 2.78065724 | 0.0054249 | NA |
| LMO7 | 395.22379 | 0.03192887 | 0.36039387 | 0.08859437 | 0.92940428 | 0.98960607 |
| KCTD12 | 59.0919557 | 0.88216583 | 0.60095377 | 1.46794291 | 0.14211973 | 0.69328517 |
| AC001226.2 | 19.9748037 | 0.87733621 | 0.58257473 | 1.50596337 | 0.13207658 | 0.6765887 |
| FBXL3 | 58.9237357 | 0.39310736 | 0.31091509 | 1.264356 | 0.20610228 | 0.76231289 |
| MYCBP2 | 264.384275 | -0.3390803 | 0.22134869 | -1.5318829 | 0.12555134 | 0.66929384 |
| SLAIN1 | 27.9091098 | 0.1596636 | 0.39505281 | 0.40415761 | 0.68609683 | 0.94205398 |
| EDNRB | 41.6206378 | -0.1728828 | 0.5714922 | -0.3025113 | 0.76226234 | 0.96053075 |
| RNF219 | 22.0043391 | -0.8324386 | 0.34928499 | -2.3832647 | 0.01715985 | 0.40063153 |
| RBM26 | 220.069645 | -0.1236186 | 0.16519472 | -0.7483204 | 0.45426691 | 0.87378486 |
| NDFIP2 | 125.223638 | -0.3144324 | 0.31352213 | -1.0029032 | 0.31590757 | 0.82116122 |
| GPC5 | 824.53645 | 1.09613856 | 1.10547026 | 0.99155862 | 3.21E-01 | 8.25E-01 |
| GPC6 | 99.3662503 | -1.9245015 | 0.707739 | -2.7192249 | 0.00654351 | 0.29935381 |
| TGDS | 28.7980864 | -0.5137405 | 0.41683252 | -1.2324865 | 0.21776739 | 0.76666749 |
| GPR180 | 24.846761 | -0.4171667 | 0.44480063 | -0.9378734 | 0.3483095 | 0.83687845 |
| ABCC4 | 66.7600927 | 0.89776858 | 0.49914801 | 1.79860196 | 0.07208167 | 0.60821941 |
| CLDN10 | 107.225923 | 0.45124695 | 0.40464618 | 1.11516425 | 0.26478004 | 0.79666376 |
| CLDN10-AS1 | 22.835977 | 0.25138286 | 0.58272867 | 0.43138922 | 0.66618539 | 0.93860542 |
| DZIP1 | 24.1694925 | -0.0699854 | 0.39213297 | -0.1784736 | 0.85835104 | 0.97691173 |
| DNAJC3-DT | 17.2427775 | 0.51201689 | 0.50371577 | 1.01647976 | 0.30940098 | 0.81568231 |
| DNAJC3 | 119.269747 | -0.1890057 | 0.29805056 | -0.6341398 | 0.52598957 | 0.90487731 |
| UGGT2 | 228.809299 | -0.3250907 | 0.17918089 | -1.8143158 | 0.06962913 | 0.60275677 |
| MBNL2 | 235.426831 | -0.2034399 | 0.28628862 | -0.7106113 | 0.47732514 | 0.88512746 |
| RAP2A | 15.6632105 | 0.56235742 | 0.47031829 | 1.19569542 | 0.23181544 | NA |
| IPO5 | 68.9877359 | 0.13988298 | 0.23325401 | 0.59970237 | 0.54870461 | 0.91208861 |
| FARP1 | 637.01564 | -0.2339831 | 0.18243405 | -1.2825626 | 0.19964535 | 0.75605362 |
| STK24 | 292.644222 | -0.3406111 | 0.30795985 | -1.1060243 | 0.26871599 | 0.79666376 |
| DOCK9 | 143.631662 | 0.08083417 | 0.24623249 | 0.32828393 | 0.742697 | 0.95563105 |
| UBAC2 | 296.417434 | -0.4048943 | 0.20168579 | -2.0075497 | 0.04469116 | 0.52549539 |
| TM9SF2 | 122.392429 | -0.2035668 | 0.18501968 | -1.1002439 | 0.27122585 | 0.79666376 |
| CLYBL | 185.079286 | -0.3505112 | 0.23518747 | -1.4903481 | 0.13613273 | 0.68406472 |
| AL355338.1 | 20.3725046 | -0.1959586 | 0.40284127 | -0.4864413 | 0.62665428 | 0.93032781 |
| PCCA | 1012.06105 | -0.2135934 | 0.21624849 | -0.9877222 | 0.32328874 | 0.82558302 |
| GGACT | 30.7181009 | 0.20549533 | 0.4759899 | 0.43172204 | 0.66594344 | 0.93860542 |
| TMTC4 | 28.8697018 | 0.32696212 | 0.37414659 | 0.87388776 | 0.38217938 | 0.8548566 |

| | | | | | | |
|---|---|---|---|---|---|---|
| FGF14 | 52.2236988 | -0.0337084 | 0.66661775 | -0.0505663 | 0.95967115 | 0.99245244 |
| TPP2 | 83.3347495 | -0.2276324 | 0.24949719 | -0.9123644 | 0.3615769 | 0.84395756 |
| EFNB2 | 32.668901 | 1.87440708 | 0.48883784 | 3.83441485 | 0.00012586 | 0.04981461 |
| ARGLU1 | 211.519339 | 0.22712992 | 0.23246964 | 0.97703048 | 0.32855406 | 0.82781307 |
| FAM155A | 610.910333 | -1.1207706 | 1.02623603 | -1.0921177 | 0.27478136 | 0.7994854 |
| LIG4 | 17.7616936 | -0.1597243 | 0.35750828 | -0.4467708 | 0.65504057 | 0.93548446 |
| ABHD13 | 27.4742861 | -0.0023479 | 0.35824444 | -0.006554 | 0.9947707 | 0.99927028 |
| TNFSF13B | 35.3986111 | 0.01278631 | 0.41725395 | 0.03064396 | 0.97555349 | 0.99440179 |
| MYO16 | 28.6474023 | 2.06968883 | 0.75194112 | 2.75246131 | 0.00591491 | 0.2943676 |
| IRS2 | 64.5958608 | -0.5514698 | 0.42972233 | -1.2833165 | 0.19938121 | 0.75605362 |
| COL4A1 | 140.375329 | 0.29872781 | 0.30148088 | 0.99086819 | 0.32174995 | 0.82503937 |
| COL4A2 | 174.778099 | 0.63503206 | 0.25544488 | 2.48598468 | 0.01291935 | 0.36664851 |
| RAB20 | 34.0432397 | -0.1201702 | 0.32153396 | -0.3737404 | 0.70859749 | 0.94558664 |
| NAXD | 33.0223487 | 0.25370415 | 0.28935721 | 0.8767853 | 0.38060326 | 0.85432452 |
| CARS2 | 61.0646785 | -0.0662913 | 0.23977593 | -0.276472 | 0.78218553 | 0.96652357 |
| ANKRD10 | 302.458732 | 0.01749912 | 0.20824067 | 0.08403315 | 0.93303007 | 0.98960607 |
| ARHGEF7 | 182.621743 | -0.2038548 | 0.16370906 | -1.2452264 | 0.21304853 | 0.7643572 |
| TUBGCP3 | 68.2342697 | 0.03505745 | 0.38161503 | 0.09186601 | 0.92680449 | 0.98925753 |
| ATP11A | 62.1917278 | 0.09950638 | 0.63362335 | 0.15704343 | 8.75E-01 | 0.97918624 |
| MCF2L | 61.9801021 | -0.008884 | 0.29654982 | -0.029958 | 0.97610055 | 0.99440179 |
| PCID2 | 44.8464934 | 0.43222075 | 0.30393643 | 1.42207613 | 0.15500415 | 0.70572801 |
| CUL4A | 131.948543 | -0.1247153 | 0.21184266 | -0.5887166 | 0.55605137 | 0.91398748 |
| LAMP1 | 112.254041 | 0.17914774 | 0.3143047 | 0.56998111 | 0.56869051 | 0.91698355 |
| GRTP1 | 38.151721 | 0.26589051 | 0.31526208 | 0.84339514 | 0.39900749 | 0.85778581 |
| DCUN1D2 | 42.8228264 | 0.14496784 | 0.39536639 | 0.36666709 | 0.71386735 | 0.94841799 |
| TMCO3 | 50.6459422 | -0.0528326 | 0.26683777 | -0.1979951 | 0.84304893 | 0.97612951 |
| TFDP1 | 44.9388756 | -0.7516875 | 0.36081263 | -2.0833181 | 0.03722224 | 0.49537143 |
| GAS6 | 70.5452086 | 0.5900758 | 0.28702063 | 2.05586549 | 0.03979548 | 0.50156625 |
| RASA3 | 33.7860771 | -0.701076 | 0.61182786 | -1.1458713 | 0.25184841 | 0.79027098 |
| CDC16 | 63.9184469 | -0.3231256 | 0.25113758 | -1.2866476 | 0.19821715 | 0.75559912 |
| UPF3A | 67.1159364 | 0.25442908 | 0.28273112 | 0.89989768 | 0.3681747 | 0.84777627 |
| CHAMP1 | 15.7109005 | 0.03856155 | 0.39384068 | 0.09791154 | 0.92200253 | NA |
| TTC5 | 24.0601373 | -0.0376336 | 0.36055616 | -0.1043765 | 0.91687057 | 0.98710975 |
| CCNB1IP1 | 52.4430805 | -0.2271531 | 0.30836929 | -0.7366267 | 4.61E-01 | 0.87657347 |
| PARP2 | 12.358428 | 0.59131577 | 0.49011432 | 1.20648539 | 0.22763039 | NA |
| TEP1 | 25.2025002 | -0.1825026 | 0.31384293 | -0.5815092 | 0.56089731 | 0.91584721 |
| OSGEP | 21.815539 | -0.2138303 | 0.38676802 | -0.5528645 | 0.5803562 | 0.92034873 |
| APEX1 | 27.6559736 | 0.54045106 | 0.38861918 | 1.39069579 | 0.16431769 | 0.71397803 |
| PIP4P1 | 14.8803316 | 0.05782414 | 0.45350715 | 0.12750436 | 0.89854123 | NA |
| PNP | 49.2039752 | 0.00277304 | 0.53231123 | 0.00520944 | 0.99584349 | 0.99927028 |
| METTL17 | 22.0415898 | -0.3895526 | 0.37162742 | -1.0482344 | 0.29453061 | 0.81012446 |
| NDRG2 | 34.8360453 | 0.62075949 | 0.38727624 | 1.60288555 | 0.10895992 | 0.6603659 |
| ARHGEF40 | 16.4857513 | 0.48970949 | 0.39763093 | 1.23156788 | 0.21811054 | 0.76666749 |
| HNRNPC | 427.349919 | -0.3577162 | 0.21844213 | -1.6375788 | 0.10150958 | 0.65863153 |
| SUPT16H | 58.2371339 | 0.04790026 | 0.29890866 | 0.1602505 | 0.87268375 | 0.97866716 |
| CHD8 | 94.395609 | -0.292715 | 0.21577422 | -1.3565801 | 0.17491468 | 0.72841249 |
| RAB2B | 35.6892055 | 0.05300284 | 0.41963939 | 0.12630569 | 0.89948995 | 0.98531749 |
| TOX4 | 40.594625 | 0.2229277 | 0.2860686 | 0.77928055 | 0.43581447 | 0.86729771 |
| METTL3 | 14.5454278 | 0.05430993 | 0.45368712 | 0.11970789 | 0.90471455 | NA |
| DAD1 | 72.0066448 | -0.2831193 | 0.34913524 | -0.8109158 | 0.41741401 | 0.85949274 |
| ABHD4 | 26.0071547 | -0.2274048 | 0.31967404 | -0.7113647 | 0.47685826 | 0.88495683 |
| OXA1L | 22.4777554 | 0.25307039 | 0.3409568 | 0.74223595 | 0.45794439 | 0.87469004 |
| MRPL52 | 19.6571722 | 0.37593613 | 0.34040642 | 1.10437438 | 0.26943077 | 0.79666376 |
| MMP14 | 18.5977916 | -0.2298618 | 0.66728223 | -0.3444747 | 0.73048935 | 0.95279138 |

| | | | | | | |
|---|---|---|---|---|---|---|
| LRP10 | 72.3770544 | 0.03651232 | 0.30944834 | 0.11799165 | 9.06E-01 | 9.85E-01 |
| RBM23 | 57.2251159 | -0.1934037 | 0.28539302 | -0.6776749 | 0.49797783 | 0.89378207 |
| PRMT5 | 11.1324893 | 0.23998462 | 0.47555025 | 0.5046462 | 0.61380735 | NA |
| AL132780.1 | 26.1018947 | 1.1245897 | 0.65250301 | 1.72350118 | 0.08479793 | 0.62880224 |
| C14orf93 | 14.0504849 | 0.63013365 | 0.55472858 | 1.13593148 | 0.25598524 | NA |
| PSMB5 | 41.0077655 | 0.45417484 | 0.29091417 | 1.56119877 | 0.11847686 | 0.6668162 |
| ACIN1 | 117.069708 | -0.2321156 | 0.18860026 | -1.2307281 | 0.21842459 | 0.76666749 |
| C14orf119 | 17.250505 | 0.0078023 | 0.4241066 | 0.01839703 | 0.98532212 | 0.99748524 |
| SLC7A8 | 71.92336 | -0.9653388 | 0.56419049 | -1.7110158 | 0.08707819 | 0.63466198 |
| HOMEZ | 15.0858627 | -0.1880529 | 0.40205933 | -0.4677243 | 0.63998176 | NA |
| PPP1R3E | 14.8853092 | 0.32201575 | 0.62110963 | 0.51845236 | 0.60414269 | NA |
| BCL2L2-PABI | 19.951915 | 0.35767395 | 0.37064585 | 0.96500192 | 0.33454385 | 0.82974237 |
| PABPN1 | 70.5279624 | -0.5691228 | 0.39374099 | -1.4454244 | 0.14833872 | 0.69945736 |
| SLC22A17 | 14.2869205 | -0.44778 | 0.52117117 | -0.8591803 | 0.39024104 | NA |
| NGDN | 20.1572513 | 0.26686621 | 0.39401803 | 0.67729441 | 0.49821918 | 0.89378207 |
| ZFHX2 | 26.4186099 | 0.35014124 | 0.4596638 | 0.76173333 | 0.44621918 | 0.87231627 |
| AP1G2 | 21.0936552 | -0.6811857 | 0.37043946 | -1.8388585 | 0.06593601 | 0.59523726 |
| DHRS4-AS1 | 21.1263038 | 0.97169581 | 0.52097738 | 1.86514012 | 0.06216177 | 0.58370968 |
| DHRS4L2 | 17.5209483 | 0.82679135 | 0.44409076 | 1.8617621 | 0.06263663 | 0.5842021 |
| NRL | 10.8887271 | -0.0416828 | 0.48217283 | -0.0864479 | 0.93111036 | NA |
| PSME1 | 40.1957557 | 0.09613325 | 0.4400417 | 0.21846397 | 0.82706763 | 0.97567084 |
| PSME2 | 11.6102584 | -0.6908763 | 0.49622721 | -1.392258 | 0.16384427 | NA |
| AL136295.1 | 23.7742595 | -0.4603603 | 0.40212994 | -1.144805 | 0.25228996 | 0.7902944 |
| GMPR2 | 14.3919917 | 1.21095892 | 0.4542787 | 2.665674 | 0.00768342 | NA |
| TINF2 | 13.2586827 | -0.2904856 | 0.44110036 | -0.6585476 | 0.5101863 | NA |
| DHRS1 | 17.5324266 | 0.15690646 | 0.43735554 | 0.35876179 | 0.7197733 | 0.94995363 |
| KHNYN | 18.6507444 | -0.2622035 | 0.32368087 | -0.8100679 | 0.41790116 | 0.85950165 |
| SDR39U1 | 22.0606985 | 0.3150555 | 0.35094272 | 0.89774051 | 0.36932391 | 0.84852785 |
| NOVA1 | 101.276804 | 0.34468197 | 0.22617156 | 1.52398456 | 0.12751257 | 0.66929384 |
| LINC00645 | 63.235476 | 0.75081124 | 0.57151883 | 1.31371218 | 0.18894308 | 0.7436009 |
| PRKD1 | 526.097654 | -0.2096365 | 0.22495556 | -0.9319018 | 0.35138726 | 0.83822804 |
| G2E3 | 37.4536365 | 0.10667645 | 0.29337889 | 0.36361325 | 0.71614683 | 0.94885853 |
| SCFD1 | 152.322589 | -0.4362837 | 0.20427211 | -2.1357966 | 0.032696 | 0.48384156 |
| COCH | 11.9183615 | 0.06483405 | 0.44525754 | 0.14561023 | 0.88422909 | NA |
| STRN3 | 375.295989 | -0.4673544 | 0.33702605 | -1.3867011 | 0.16553294 | 0.71675975 |
| AP4S1 | 26.3277289 | -0.5044773 | 0.35196365 | -1.433322 | 0.15176584 | 0.70572801 |
| HECTD1 | 293.544902 | -0.8585231 | 0.31249292 | -2.7473362 | 0.00600815 | 0.29709153 |
| HEATR5A | 43.499863 | -0.0464784 | 0.33011796 | -0.1407933 | 0.8880332 | 0.9825428 |
| AL139353.1 | 11.0285075 | 0.50319166 | 0.43484309 | 1.15717985 | 0.24719889 | NA |
| NUBPL | 146.80839 | -0.2364905 | 0.2468498 | -0.958034 | 0.33804559 | 0.83055261 |
| ARHGAP5 | 360.875005 | -0.30445 | 0.20052283 | -1.5182808 | 0.12894364 | 0.66958011 |
| AKAP6 | 107.154844 | 0.40065803 | 0.34722012 | 1.15390211 | 0.2485403 | 0.78729484 |
| NPAS3 | 106.795781 | 0.67858749 | 0.63961259 | 1.06093517 | 0.28871937 | 0.80536924 |
| EGLN3 | 37.9685779 | -0.3727268 | 0.36264003 | -1.0278147 | 0.304037 | 0.81568231 |
| EAPP | 49.3578682 | 0.32448744 | 0.34700491 | 0.93510908 | 0.34973209 | 0.83687845 |
| SNX6 | 152.716799 | -0.3788617 | 0.20787961 | -1.8225052 | 0.06837836 | 0.60244724 |
| CFL2 | 12.0038317 | 1.02890237 | 0.48471948 | 2.12267593 | 0.03378102 | NA |
| BAZ1A | 63.20223 | -0.5743098 | 0.35457002 | -1.6197359 | 0.10528903 | 0.65946648 |
| SRP54 | 88.5119275 | -0.309741 | 0.29254878 | -1.058767 | 0.28970593 | 0.80536924 |
| FAM177A1 | 62.0223052 | 0.13437586 | 0.26744212 | 0.50244838 | 0.61535216 | 0.92662475 |
| PPP2R3C | 34.5215739 | -0.2907284 | 0.30349901 | -0.9579221 | 0.33810203 | 0.83055261 |
| NFKBIA | 43.3752324 | 0.3887359 | 0.4892422 | 0.79456738 | 0.4268652 | 0.86239557 |
| RALGAPA1 | 354.703194 | -0.1311112 | 0.30362792 | -0.4318154 | 0.66587559 | 0.93860542 |
| BRMS1L | 21.682951 | 0.74253417 | 0.42318294 | 1.75464109 | 0.07932072 | 0.61367304 |

| | | | | | | |
|---|---|---|---|---|---|---|
| PTCSC3 | 11.6680229 | 0.41327298 | 0.84859611 | 0.48700786 | 0.62625276 | NA |
| MBIP | 53.1801503 | -0.0435689 | 0.33470965 | -0.1301693 | 0.89643245 | 0.98531749 |
| MIPOL1 | 132.186143 | 0.20684586 | 0.23650561 | 0.87459178 | 0.38179606 | 0.85477791 |
| SEC23A | 115.323141 | -0.3178569 | 0.34218859 | -0.9288939 | 0.35294408 | 0.83886421 |
| GEMIN2 | 13.7372148 | 0.091634 | 0.46294489 | 0.19793717 | 0.84309422 | NA |
| TRAPPC6B | 65.2709396 | -0.0890165 | 0.30859406 | -0.2884584 | 0.7729959 | 0.96435639 |
| PNN | 99.5995748 | -0.2719031 | 0.24676243 | -1.101882 | 0.27051298 | 0.79666376 |
| MIA2 | 190.744083 | -0.0728599 | 0.2199679 | -0.3312296 | 0.7404711 | 0.9544376 |
| FBXO33 | 71.339867 | 0.06154193 | 0.30196724 | 0.20380333 | 0.83850719 | 0.97600352 |
| AL049875.1 | 22.6046126 | 0.02987333 | 0.44814824 | 0.06665949 | 0.94685279 | 0.99083449 |
| KLHL28 | 59.2825988 | 0.01006743 | 0.32226946 | 0.03123918 | 0.9750788 | 0.99440004 |
| TOGARAM1 | 107.053441 | -0.1720843 | 0.25286493 | -0.6805382 | 0.49616373 | 0.89244133 |
| PRPF39 | 46.2213463 | -0.3218733 | 0.2922589 | -1.1013293 | 0.27075336 | 0.79666376 |
| FKBP3 | 35.6597444 | 0.21724014 | 0.25589147 | 0.8489542 | 0.39590678 | 0.85605713 |
| FANCM | 26.0665274 | -0.1862584 | 0.36279462 | -0.513399 | 0.60767222 | 0.92569157 |
| MIS18BP1 | 32.4462781 | 0.55431769 | 0.35120493 | 1.57833116 | 0.11448955 | 0.66679359 |
| RPS29 | 92.4915968 | 0.16326825 | 0.36622456 | 0.44581459 | 0.65573118 | 0.93567213 |
| RPL36AL | 90.2163681 | 0.28090321 | 0.39370711 | 0.7134827 | 0.47554711 | 0.88495683 |
| DNAAF2 | 16.0300304 | 0.38079748 | 0.38645437 | 0.98536207 | 0.32444625 | NA |
| POLE2 | 14.9135468 | 0.59246336 | 0.57671997 | 1.02729816 | 0.30428009 | NA |
| KLHDC1 | 47.3091277 | 0.61611761 | 0.38719562 | 1.59123084 | 0.11155764 | 0.6645905 |
| KLHDC2 | 47.5264293 | 0.1104287 | 0.29280711 | 0.37713803 | 0.70607103 | 0.94558664 |
| NEMF | 213.297411 | -0.1963341 | 0.25059681 | -0.7834661 | 0.43335346 | 0.86729771 |
| ARF6 | 39.7997963 | -0.6264426 | 0.55514271 | -1.1284352 | 0.25913619 | 0.79532468 |
| VCPKMT | 25.8496199 | -0.219492 | 0.41507692 | -0.5287983 | 0.59694535 | 0.9229638 |
| SOS2 | 300.27586 | -0.1078568 | 0.21778712 | -0.4952396 | 0.62043097 | 0.92798856 |
| L2HGDH | 108.545888 | -0.0405896 | 0.21651279 | -0.1874697 | 0.85129241 | 0.97612951 |
| ATP5S | 46.3905809 | 0.14789334 | 0.26360418 | 0.56104322 | 0.57476807 | 0.91824492 |
| CDKL1 | 135.177133 | -0.0625983 | 0.36021586 | -0.17378 | 0.86203835 | 0.97767895 |
| MAP4K5 | 193.760995 | -0.2660712 | 0.25439832 | -1.0458842 | 0.29561451 | 0.81054924 |
| ATL1 | 20.4495864 | -0.0855831 | 0.52192954 | -0.1639744 | 0.86975126 | 0.97854266 |
| SAV1 | 44.7001575 | -0.205833 | 0.25561571 | -0.805244 | 0.42067887 | 0.86148657 |
| NIN | 60.1478631 | 0.30795105 | 0.24501559 | 1.25686307 | 0.20880322 | 0.7623519 |
| PYGL | 24.8019376 | 0.67826994 | 0.37899294 | 1.78966378 | 0.07350798 | 0.60985192 |
| TMX1 | 38.3635973 | -0.140736 | 0.29579109 | -0.4757952 | 0.63422028 | 0.93251095 |
| FRMD6 | 18.2053033 | 0.65926967 | 0.50104867 | 1.3157797 | 1.88E-01 | 0.74214877 |
| RTRAF | 40.3717136 | -0.2535302 | 0.43293758 | -0.5856045 | 0.55814131 | 0.91425998 |
| TXNDC16 | 29.7608289 | 0.02706346 | 0.31703965 | 0.08536302 | 0.93197279 | 0.98960607 |
| GPR137C | 16.7720252 | 0.53354396 | 0.44379292 | 1.20223629 | 0.22927199 | 0.77185154 |
| ERO1A | 198.517333 | -0.9108109 | 0.49167685 | -1.8524583 | 6.40E-02 | 5.88E-01 |
| AL133453.1 | 25.8956807 | -0.2341174 | 0.70112722 | -0.3339157 | 7.38E-01 | 9.54E-01 |
| PSMC6 | 74.3916208 | -0.5441176 | 0.23249659 | -2.3403248 | 0.01926697 | 0.41544117 |
| STYX | 33.2501991 | 0.10272292 | 0.31643357 | 0.32462713 | 0.74546329 | 0.95563105 |
| GNPNAT1 | 22.4084418 | 0.04892943 | 0.46769355 | 0.10461858 | 0.91667847 | 0.98710975 |
| FERMT2 | 242.342756 | -0.2759055 | 0.2266781 | -1.2171687 | 0.22354002 | 0.76834258 |
| DDHD1 | 66.80456 | 0.0946392 | 0.28846658 | 0.32807683 | 0.74285357 | 0.95563105 |
| CNIH1 | 29.5882343 | 0.21711151 | 0.32296163 | 0.67225172 | 0.50142346 | 0.89492452 |
| GMFB | 24.9739996 | -0.0883498 | 0.34628419 | -0.2551367 | 0.79861752 | 0.97019967 |
| CGRRF1 | 17.132666 | 0.31730314 | 0.40677603 | 0.78004385 | 0.43536506 | 0.86729771 |
| SAMD4A | 152.638045 | -0.2346218 | 0.47892522 | -0.4898923 | 0.62421009 | 0.92981013 |
| GCH1 | 15.8171482 | -0.4178793 | 0.65764182 | -0.6354208 | 0.52515401 | NA |
| WDHD1 | 14.600021 | -0.7420568 | 0.45319068 | -1.6374053 | 0.10154581 | NA |
| SOCS4 | 28.9645499 | 0.45753768 | 0.36240218 | 1.26251359 | 0.20676404 | 0.76231289 |
| MAPK1IP1L | 126.897751 | -0.2800681 | 0.29774085 | -0.9406437 | 0.34688748 | 0.83687845 |

| | | | | | | |
|---|---|---|---|---|---|---|
| LGALS3 | 156.771366 | -0.4899074 | 0.35379459 | -1.3847227 | 1.66E-01 | 0.71756194 |
| FBXO34 | 191.505584 | -0.1491789 | 0.24327143 | -0.6132198 | 0.53973101 | 0.90763121 |
| AL158801.3 | 13.752609 | 0.28121977 | 0.552251 | 0.50922456 | 0.61059483 | NA |
| ATG14 | 38.9634112 | 0.02043132 | 0.29421231 | 0.06944414 | 0.94463609 | 0.99069421 |
| KTN1-AS1 | 11.5885884 | -0.4178733 | 0.62291666 | -0.6708333 | 0.5023267 | NA |
| KTN1 | 317.872866 | -0.1418755 | 0.22284473 | -0.6366565 | 0.52434861 | 0.90487731 |
| PELI2 | 226.43173 | -0.1367145 | 0.28171789 | -0.4852885 | 0.62747168 | 0.93057946 |
| TMEM260 | 122.139731 | 0.14644 | 0.18760436 | 0.78057887 | 0.43505022 | 0.86729771 |
| EXOC5 | 87.4312433 | -0.0392337 | 0.19657775 | -0.1995836 | 0.84180624 | 0.97600352 |
| AP5M1 | 77.1454916 | 0.21146351 | 0.27898635 | 0.7579708 | 0.44846846 | 0.87314185 |
| NAA30 | 31.5685388 | -0.1633144 | 0.30391795 | -0.5373633 | 0.59101669 | 0.92254886 |
| CCDC198 | 268.62169 | 0.4861891 | 0.47092584 | 1.03241118 | 0.30187953 | 0.81568231 |
| ARMH4 | 57.678407 | 0.02790007 | 0.30479173 | 0.09153815 | 0.92706499 | 0.98925753 |
| ACTR10 | 70.1202221 | -0.2646187 | 0.22082657 | -1.1983099 | 0.23079638 | 0.77249016 |
| PSMA3 | 61.6191015 | -0.4401206 | 0.32248588 | -1.3647749 | 0.1723238 | 0.72649757 |
| ARID4A | 98.8726587 | 0.09089094 | 0.2331924 | 0.38976801 | 0.6967081 | 0.94395238 |
| TIMM9 | 27.6053158 | -0.0262552 | 0.3345824 | -0.0784716 | 0.93745292 | 0.99012269 |
| KIAA0586 | 75.6598139 | 0.00568474 | 0.25251675 | 0.02251235 | 0.98203926 | 0.99626066 |
| DAAM1 | 119.03782 | -0.3571537 | 0.23403665 | -1.5260589 | 0.12699519 | 0.66929384 |
| GPR135 | 23.5914908 | -0.0240348 | 0.40466781 | -0.0593939 | 0.95263837 | 0.99141043 |
| L3HYPDH | 22.6109388 | -0.1875122 | 0.44002441 | -0.4261405 | 0.67000545 | 0.93878847 |
| JKAMP | 35.6157284 | 0.55501001 | 0.35822215 | 1.54934589 | 0.12129859 | 0.66771734 |
| AL157911.1 | 12.3049963 | 0.2919229 | 0.58086563 | 0.50256529 | 0.61526994 | NA |
| PCNX4 | 142.840821 | -0.3493662 | 0.28389386 | -1.2306227 | 0.21846401 | 0.76666749 |
| DHRS7 | 32.0692996 | 0.71534842 | 0.33760294 | 2.11890461 | 0.03409853 | 0.4908133 |
| PPM1A | 86.7865062 | -0.2651373 | 0.250535 | -1.0582843 | 0.28992585 | 0.80536924 |
| MNAT1 | 267.354765 | -0.0791175 | 0.19622207 | -0.4032038 | 0.6867983 | 0.94205398 |
| SLC38A6 | 45.5583552 | -0.2204789 | 0.31642593 | -0.696779 | 0.48594112 | 0.88838332 |
| PRKCH | 73.110405 | -0.5330129 | 0.28801228 | -1.8506603 | 0.06421844 | 0.58870134 |
| TMEM30B | 35.7581626 | 0.15257902 | 0.45612231 | 0.33451338 | 0.73799219 | 0.9542764 |
| HIF1A | 1095.66565 | -0.627927 | 0.3793781 | -1.6551481 | 0.09789445 | 0.65621002 |
| HIF1A-AS2 | 164.891486 | -0.2350073 | 0.5416056 | -0.4339086 | 6.64E-01 | 0.93860542 |
| SNAPC1 | 98.396044 | -0.4676891 | 0.36004141 | -1.298987 | 0.19394837 | 0.75067401 |
| SYT16 | 44.0062881 | -0.5907367 | 0.82454944 | -0.7164357 | 0.47372233 | 0.88366688 |
| PPP2R5E | 269.017507 | -0.5304602 | 0.26270676 | -2.0192103 | 0.04346537 | 0.51720946 |
| WDR89 | 41.9176777 | -0.2261014 | 0.33388014 | -0.6771934 | 0.49828327 | 0.89378207 |
| SGPP1 | 311.30871 | -0.9906135 | 0.34976853 | -2.8321973 | 0.00462293 | 0.26975991 |
| SYNE2 | 947.851962 | -0.1452336 | 0.22169232 | -0.6551132 | 5.12E-01 | 0.9014941 |
| ESR2 | 55.964918 | 0.55920455 | 0.52515018 | 1.06484692 | 0.28694521 | 0.80523851 |
| MTHFD1 | 27.2810642 | 0.00971841 | 0.3520003 | 0.02760908 | 0.97797394 | 0.99469237 |
| AL122035.1 | 23.6313848 | -0.5483884 | 0.34785074 | -1.5765049 | 0.1149095 | 0.6668162 |
| ZBTB25 | 40.8251776 | -0.0173327 | 0.36428789 | -0.0475797 | 0.96205125 | 0.99256424 |
| ZBTB1 | 102.988754 | 0.27949296 | 0.23748611 | 1.17688128 | 0.2392429 | 0.78098673 |
| AL049869.3 | 11.0258018 | 1.0174596 | 0.68811893 | 1.47861012 | 0.13924455 | NA |
| HSPA2 | 23.3993718 | 1.05904007 | 0.35579353 | 2.97655793 | 0.00291504 | 0.21399685 |
| PLEKHG3 | 84.779319 | 0.19014462 | 0.3476633 | 0.54692176 | 0.58443248 | 0.92071026 |
| CHURC1-FNT | 40.4610995 | 0.05920787 | 0.42908481 | 0.1379864 | 0.89025117 | 0.98289343 |
| GPX2 | 36.6499462 | 1.68825778 | 0.74494556 | 2.26628343 | 0.02343404 | 0.43088993 |
| RAB15 | 50.2814957 | 0.53764195 | 0.30424818 | 1.76711642 | 0.07720873 | 0.61329066 |
| MAX | 76.8874174 | -0.41271 | 0.29869837 | -1.3816948 | 0.16706545 | 0.71905284 |
| FUT8 | 74.264571 | 0.54463213 | 0.2832979 | 1.92247149 | 0.05454646 | 0.5645224 |
| AL359232.1 | 67.3588078 | -0.0505809 | 0.44819095 | -0.1128556 | 0.910145 | 0.98556715 |
| GPHN | 1135.07581 | -0.1259361 | 0.23930996 | -0.5262468 | 5.99E-01 | 9.23E-01 |
| MPP5 | 115.898225 | -0.0896093 | 0.19175882 | -0.4673021 | 0.64028373 | 0.93328692 |

| | | | | | | |
|---|---|---|---|---|---|---|
| ATP6V1D | 60.6519281 | -0.3983046 | 0.31603113 | -1.2603334 | 0.20754913 | 0.76231289 |
| EIF2S1 | 33.5435756 | -0.0525618 | 0.26944993 | -0.1950708 | 0.84533753 | 0.97612951 |
| PLEKHH1 | 39.2913866 | -0.1138067 | 0.29613644 | -0.3843049 | 0.70075246 | 0.94440669 |
| PIGH | 24.6844066 | 0.28361843 | 0.50754944 | 0.5587996 | 0.57629849 | 0.91854298 |
| ARG2 | 111.293575 | 0.11129561 | 0.58661962 | 0.18972364 | 0.84952569 | 0.97612951 |
| VTI1B | 35.0122023 | -0.1866555 | 0.37926876 | -0.4921456 | 0.62261639 | 0.92897508 |
| ZFYVE26 | 30.4768826 | -0.9237912 | 0.34574363 | -2.6718966 | 0.00754239 | 0.30654023 |
| RAD51B | 1041.3788 | 0.06786269 | 0.2938946 | 0.23090825 | 8.17E-01 | 0.97288403 |
| ZFP36L1 | 423.025798 | -0.5293692 | 0.28173665 | -1.8789503 | 0.06025128 | 0.57733412 |
| ACTN1 | 49.4152586 | 0.17455798 | 0.33889754 | 0.51507598 | 0.60649992 | 0.92569157 |
| DCAF5 | 177.026756 | 0.14696461 | 0.19900822 | 0.7384851 | 0.46021972 | 0.87607759 |
| EXD2 | 42.184757 | 0.22436605 | 0.31358388 | 0.71548973 | 0.47430648 | 0.88403192 |
| GALNT16 | 39.9324636 | 2.52828948 | 0.53875646 | 4.69282441 | 2.69E-06 | 0.00350412 |
| ERH | 51.5005404 | 0.07278164 | 0.22819243 | 0.31894852 | 0.74976555 | 0.95683664 |
| SLC39A9 | 69.065041 | -0.0959135 | 0.22468567 | -0.4268787 | 0.66946768 | 0.93878847 |
| PLEKHD1 | 19.3723658 | -0.5203265 | 1.02517035 | -0.5075513 | 0.61176808 | 0.92630153 |
| SUSD6 | 113.7882 | -0.0019992 | 0.17952583 | -0.0111361 | 0.99111483 | 0.99814944 |
| SRSF5 | 230.188485 | 0.07204081 | 0.24230072 | 0.29731983 | 7.66E-01 | 0.96270421 |
| AL357153.2 | 42.0818621 | 0.71157715 | 0.3694434 | 1.92607895 | 0.05409452 | 0.5642154 |
| MED6 | 32.18043 | 0.08649123 | 0.30695705 | 0.28176982 | 0.77812001 | 0.96518573 |
| MAP3K9 | 42.5949054 | 0.02141873 | 0.40734576 | 0.05258119 | 0.9580656 | 0.99183169 |
| PCNX1 | 250.304676 | -0.3224005 | 0.22491913 | -1.4334061 | 0.15174181 | 0.70572801 |
| SIPA1L1 | 487.879476 | -0.4061542 | 0.20018733 | -2.0288707 | 0.04247146 | 0.51343649 |
| RGS6 | 699.719603 | -0.4554941 | 1.05591706 | -0.431373 | 0.66619716 | 0.93860542 |
| DPF3 | 14.8931507 | 0.52362203 | 0.50845286 | 1.02983399 | 0.30308794 | NA |
| ZFYVE1 | 42.7171423 | -0.5526788 | 0.31249872 | -1.7685794 | 0.07696409 | 0.61295198 |
| RBM25 | 158.309226 | -0.2192629 | 0.17407219 | -1.2596089 | 0.20781048 | 0.76231289 |
| PSEN1 | 114.212847 | -0.4572975 | 0.31233479 | -1.464126 | 0.14315953 | 0.69428142 |
| PAPLN | 20.7253016 | 0.02366078 | 0.38228088 | 0.06189369 | 0.95064749 | 0.99105535 |
| NUMB | 280.210335 | -0.1394745 | 0.21434144 | -0.6507116 | 0.51523266 | 0.90174188 |
| HEATR4 | 13.5491129 | 1.10387136 | 0.57961003 | 1.904507 | 0.05684419 | NA |
| DNAL1 | 42.90299 | 0.19331645 | 0.25796172 | 0.74939977 | 0.45361629 | 0.87378486 |
| PNMA1 | 15.11972 | -0.0170923 | 0.46957187 | -0.0363999 | 0.97096353 | NA |
| ELMSAN1 | 51.5703511 | -0.5743816 | 0.26433718 | -2.1729127 | 0.02978689 | 0.46381645 |
| AC005480.1 | 24.7275292 | 0.26429201 | 0.44734973 | 0.59079506 | 0.55465775 | 0.9137961 |
| COQ6 | 12.0801058 | 0.07716676 | 0.45826697 | 0.16838822 | 0.86627788 | NA |
| ENTPD5 | 45.7233138 | -0.3217346 | 0.30193227 | -1.0655852 | 0.28661118 | 0.80500512 |
| BBOF1 | 40.1745449 | 0.30768961 | 0.26386293 | 1.16609636 | 0.24357549 | 0.78542956 |
| ALDH6A1 | 42.1214946 | 0.9143964 | 0.42370796 | 2.15808172 | 0.03092148 | 0.47307264 |
| LIN52 | 63.5735114 | -0.287512 | 0.30942442 | -0.9291832 | 0.35279413 | 0.83872681 |
| ABCD4 | 64.5997498 | 0.35733971 | 0.30385469 | 1.17602169 | 0.23958621 | 0.78114372 |
| NPC2 | 60.5535611 | 0.54385659 | 0.29974414 | 1.81440275 | 0.06961574 | 0.60275677 |
| ISCA2 | 16.8951972 | 0.25819326 | 0.42535616 | 0.60700487 | 0.54384768 | 0.90965439 |
| AREL1 | 57.2306156 | -0.1798229 | 0.25799786 | -0.6969939 | 0.48580661 | 0.88838332 |
| FCF1 | 23.7167845 | -0.2010665 | 0.31685023 | -0.634579 | 0.52570306 | 0.90487731 |
| YLPM1 | 131.996766 | 0.2014187 | 0.18467271 | 1.0906793 | 0.27541402 | 0.7994854 |
| DLST | 54.381636 | -0.0358342 | 0.26166187 | -0.1369487 | 0.89107138 | 0.98296447 |
| EIF2B2 | 12.9523328 | 0.55078424 | 0.39132223 | 1.40749541 | 0.15928054 | NA |
| MLH3 | 89.9262341 | 0.21723621 | 0.23236559 | 0.93488974 | 0.34984512 | 0.83687845 |
| ACYP1 | 50.1547354 | 0.2007381 | 0.29850568 | 0.67247665 | 0.5012803 | 0.89492452 |
| ZC2HC1C | 15.9043346 | 0.50072356 | 0.59482169 | 0.84180447 | 0.39989741 | NA |
| NEK9 | 64.591549 | -0.3543902 | 0.22095369 | -1.6039118 | 0.10873349 | 0.65986731 |
| TMED10 | 118.42142 | 0.29985499 | 0.22731863 | 1.31909556 | 0.18713717 | 0.74096603 |
| FOS | 46.5033122 | -0.9080507 | 0.91091065 | -0.9968603 | 0.31883233 | 0.82354084 |

| | | | | | | |
|---|---|---|---|---|---|---|
| JDP2 | 11.9030321 | -0.1598479 | 0.4325632 | -0.3695365 | 0.7117279 | NA |
| TTLL5 | 151.639411 | -0.1593468 | 0.20032853 | -0.7954273 | 0.42636497 | 0.86239557 |
| ERG28 | 18.1298529 | -0.0262265 | 0.53982298 | -0.0485835 | 0.96125119 | 0.99256424 |
| IFT43 | 104.614369 | -0.0332933 | 0.20575824 | -0.1618077 | 0.87145725 | 0.97860207 |
| GPATCH2L | 146.377915 | -0.3393799 | 0.27513648 | -1.2334966 | 0.21739054 | 0.76666749 |
| ESRRB | 97.9480675 | -1.2821666 | 0.47927435 | -2.6752249 | 0.00746791 | 0.30654023 |
| ANGEL1 | 19.9882503 | -0.4081656 | 0.35331192 | -1.1552558 | 0.24798569 | 0.78655532 |
| IRF2BPL | 41.1221367 | 0.14004349 | 0.37436035 | 0.37408739 | 0.7083393 | 0.94558664 |
| POMT2 | 18.3346425 | -0.5213851 | 0.43167972 | -1.2078054 | 0.22712211 | 0.77185154 |
| GSTZ1 | 10.8604949 | 0.74029195 | 0.45108874 | 1.64112264 | 0.10077196 | NA |
| TMED8 | 57.0289839 | -0.1344695 | 0.24758717 | -0.5431197 | 0.5870474 | 0.92086224 |
| VIPAS39 | 23.4587827 | -0.004371 | 0.36381106 | -0.0120144 | 0.99041411 | 0.99814944 |
| AHSA1 | 25.1484591 | 0.74992382 | 0.33423317 | 2.24371457 | 0.02485077 | 0.43948277 |
| SPTLC2 | 127.995096 | -0.4204554 | 0.27652529 | -1.5204953 | 0.12838654 | 0.66929384 |
| ALKBH1 | 34.7678201 | -0.2986506 | 0.33997567 | -0.8784469 | 0.37970124 | 0.85431805 |
| SLIRP | 40.8319899 | 0.57852711 | 0.36134523 | 1.60103708 | 0.10936871 | 0.66067258 |
| SNW1 | 43.5208833 | -0.1873896 | 0.28331815 | -0.6614104 | 0.50834917 | 0.89924262 |
| ADCK1 | 22.870662 | -0.8965635 | 0.40538604 | -2.211629 | 0.02699231 | 0.45084589 |
| NRXN3 | 14.1897781 | 0.53329118 | 0.63950837 | 0.83390806 | 0.4043328 | NA |
| CEP128 | 50.628563 | -0.1392547 | 0.32723195 | -0.4255535 | 0.67043325 | 0.93878847 |
| AC007262.2 | 29.0441913 | 1.20581757 | 0.52399036 | 2.30122089 | 0.02137915 | 0.4212027 |
| GTF2A1 | 43.4487244 | 0.56706726 | 0.39692451 | 1.42865267 | 0.15310409 | 0.70572801 |
| STON2 | 139.57353 | -0.0820335 | 0.32624811 | -0.2514452 | 0.80146991 | 0.97045451 |
| SEL1L | 98.2243197 | -0.1755652 | 0.35618798 | -0.4929005 | 0.62208287 | 0.92897508 |
| FLRT2 | 14.4887406 | -1.2618815 | 0.77317015 | -1.6320878 | 0.102661 | NA |
| GALC | 26.7589936 | 0.4055343 | 0.44139077 | 0.91876478 | 0.35821862 | 0.84204868 |
| SPATA7 | 65.5139879 | 0.03264239 | 0.24318926 | 0.13422628 | 0.89322364 | 0.98377746 |
| PTPN21 | 78.0107052 | -0.4637595 | 0.29755513 | -1.5585667 | 0.11909897 | 0.6668162 |
| ZC3H14 | 113.287343 | -0.1311161 | 0.2783308 | -0.47108 | 0.63758358 | 0.93271601 |
| EML5 | 62.8463416 | -1.6261886 | 0.7009863 | -2.3198579 | 0.02034857 | 0.41907916 |
| TTC8 | 49.5885271 | 0.02080284 | 0.2461342 | 0.08451827 | 0.93264437 | 0.98960607 |
| FOXN3 | 246.902734 | -0.0342214 | 0.19709611 | -0.1736279 | 8.62E-01 | 9.78E-01 |
| EFCAB11 | 51.4181026 | 0.43081934 | 0.38812571 | 1.10999948 | 0.26699925 | 0.79666376 |
| TDP1 | 26.6156919 | -0.1675932 | 0.35523413 | -0.4717824 | 0.63708215 | 0.93259128 |
| KCNK13 | 117.843799 | -0.6118758 | 0.46249964 | -1.3229757 | 0.18584346 | 0.73804265 |
| PSMC1 | 37.5959462 | 0.16143742 | 0.28924809 | 0.55812788 | 0.57675706 | 0.91854298 |
| NRDE2 | 47.6682866 | 0.00913679 | 0.28821971 | 0.03170079 | 0.97471067 | 0.99431961 |
| CALM1 | 113.626245 | -0.0165162 | 0.3386759 | -0.0487669 | 0.96110509 | 0.99256424 |
| TTC7B | 150.697585 | -0.3733353 | 0.32689157 | -1.1420767 | 0.25342212 | 0.7912221 |
| RPS6KA5 | 62.453162 | -0.8868504 | 0.51831712 | -1.711019 | 0.0870776 | 0.63466198 |
| DGLUCY | 173.759497 | 0.05259083 | 0.40310641 | 0.13046388 | 0.89619943 | 0.98527819 |
| CCDC88C | 44.3736348 | 0.19668856 | 0.42483473 | 0.46297665 | 0.6433811 | 0.93328692 |
| PPP4R3A | 144.197899 | -0.0033977 | 0.23459509 | -0.0144834 | 0.9884443 | 0.99814944 |
| CATSPERB | 52.4738939 | 0.03557845 | 0.42830001 | 0.08306899 | 0.93379669 | 0.98960607 |
| TC2N | 60.0545772 | -0.0964715 | 0.3719999 | -0.2593321 | 0.79537903 | 0.96974891 |
| TRIP11 | 148.205743 | -0.0906521 | 0.15817649 | -0.5731072 | 0.56657213 | 0.91698355 |
| ATXN3 | 87.1418037 | -0.2084305 | 0.32208064 | -0.6471376 | 0.5175429 | 0.90320951 |
| NDUFB1 | 53.3684989 | -0.0118491 | 0.27227045 | -0.0435195 | 0.96528741 | 0.99270395 |
| CPSF2 | 48.3198694 | 0.14785925 | 0.28189802 | 0.52451327 | 0.59992159 | 0.92326057 |
| LGMN | 39.4137707 | -0.6694112 | 0.43529767 | -1.5378239 | 0.12409168 | 0.66929384 |
| GOLGA5 | 35.4720688 | -0.6308635 | 0.292125 | -2.1595668 | 0.03080622 | 0.47270799 |
| ITPK1 | 98.5382645 | 0.29766176 | 0.2087576 | 1.42587266 | 0.1539051 | 0.70572801 |
| MOAP1 | 18.561823 | 0.02931761 | 0.50006938 | 0.05862709 | 0.95324913 | 0.99148072 |
| UBR7 | 14.4448969 | 0.29188314 | 0.4530364 | 0.64428187 | 0.51939267 | NA |

| | | | | | | |
|---|---|---|---|---|---|---|
| BTBD7 | 393.051945 | 0.1184213 | 0.17189815 | 0.68890388 | 0.49088376 | 0.88987842 |
| UNC79 | 35.4743334 | 1.09187824 | 0.61186438 | 1.78451021 | 0.0743408 | 0.61020315 |
| DDX24 | 94.096417 | 0.55760854 | 0.23299797 | 2.39319056 | 0.01670256 | 0.39748373 |
| IFI27 | 26.9498754 | -4.0849102 | 1.33691045 | -3.0554852 | 0.00224697 | 0.18826595 |
| SERPINA1 | 42.7345444 | 0.21503106 | 0.38011303 | 0.56570296 | 5.72E-01 | 9.18E-01 |
| DICER1 | 233.513239 | 0.04656306 | 0.21701352 | 0.21456297 | 0.83010807 | 0.97567084 |
| DICER1-AS1 | 22.8227599 | 0.34555995 | 0.5660903 | 0.61043254 | 0.54157532 | 0.90791162 |
| CLMN | 485.607177 | 0.53560369 | 0.24334731 | 2.2009846 | 0.02773711 | 0.456584 |
| GLRX5 | 38.7488558 | 0.21119558 | 0.39127455 | 0.53976314 | 0.58936039 | 0.92197072 |
| ATG2B | 103.214814 | 0.01729798 | 0.22771155 | 0.07596442 | 0.9394474 | 0.99012269 |
| GSKIP | 22.8789328 | 0.26901125 | 0.34360571 | 0.78290682 | 0.43368183 | 0.86729771 |
| AK7 | 21.8268625 | 0.25739264 | 0.33077961 | 0.77813938 | 0.43648685 | 0.86729771 |
| PAPOLA | 185.003636 | -0.0065835 | 0.22586871 | -0.0291473 | 0.97674709 | 0.99441127 |
| VRK1 | 15.8634988 | -0.1679648 | 0.42135711 | -0.3986282 | 0.69016717 | NA |
| LINC01550 | 11.6729843 | 0.9115988 | 0.72017131 | 1.26580827 | 0.20558173 | NA |
| SETD3 | 120.044424 | 0.14808633 | 0.21841102 | 0.67801676 | 0.49776107 | 0.89378207 |
| CCNK | 36.1265798 | -0.2391305 | 0.28474359 | -0.83981 | 0.40101494 | 0.85778581 |
| CCDC85C | 21.1962138 | 0.00920372 | 0.32881953 | 0.0279902 | 0.97766997 | 0.99469237 |
| EVL | 122.858577 | 0.21171504 | 0.34800577 | 0.60836644 | 0.54294446 | 0.90953689 |
| YY1 | 122.037251 | -0.0560841 | 0.20528333 | -0.2732033 | 0.78469693 | 0.96737489 |
| SLC25A29 | 292.325315 | -0.6216383 | 0.37735236 | -1.6473682 | 0.0994824 | 0.65860455 |
| WARS | 81.9893858 | -1.473617 | 0.51732415 | -2.848537 | 0.00439207 | 0.26131408 |
| WDR25 | 40.059955 | -0.3852436 | 0.30665536 | -1.2562754 | 0.20901612 | 0.7623519 |
| PPP2R5C | 154.734955 | 0.08637772 | 0.20952744 | 0.41225015 | 0.68015608 | 0.94195357 |
| DYNC1H1 | 121.926697 | -0.2375376 | 0.23432845 | -1.013695 | 0.31072833 | 0.81692172 |
| HSP90AA1 | 430.457893 | 0.11065163 | 0.55188136 | 0.20049895 | 0.84109038 | 0.97600352 |
| WDR20 | 123.697351 | -0.1843596 | 0.2030692 | -0.907866 | 0.36394903 | 0.8466721 |
| MOK | 72.246902 | -0.0705667 | 0.38562652 | -0.1829924 | 0.85480401 | 0.97612951 |
| ZNF839 | 23.5441225 | 0.22628903 | 0.40219443 | 0.56263591 | 0.57368284 | 0.91813957 |
| CINP | 14.2854014 | 0.11034458 | 0.45643031 | 0.24175559 | 0.80896955 | NA |
| TECPR2 | 67.5793298 | -0.1250558 | 0.2886975 | -0.4331723 | 0.6648896 | 0.93860542 |
| ANKRD9 | 41.9294897 | 0.02359144 | 0.33916455 | 0.06955751 | 0.94454586 | 0.99069421 |
| RCOR1 | 180.205673 | -0.4775679 | 0.35415676 | -1.3484648 | 0.17750894 | 0.72841249 |
| TRAF3 | 114.460199 | 0.01403466 | 0.28270435 | 0.04964429 | 0.96040586 | 0.99256424 |
| CDC42BPB | 201.678195 | -0.6977558 | 0.27728913 | -2.5163475 | 0.01185782 | 0.35710117 |
| TNFAIP2 | 23.5766398 | 2.23289613 | 0.92323613 | 2.41855367 | 0.01558235 | 0.39401695 |
| EIF5 | 91.4791284 | 0.10348539 | 0.27720661 | 0.37331501 | 0.70891401 | 0.94558664 |
| MARK3 | 381.734439 | -0.2541264 | 0.17007556 | -1.4941968 | 0.1351242 | 0.68329553 |
| CKB | 164.836425 | 0.43927021 | 0.45104757 | 0.97388886 | 0.33011173 | 0.82867637 |
| BAG5 | 41.9875347 | 0.67280743 | 0.31696441 | 2.12265925 | 0.03378242 | 0.4908133 |
| KLC1 | 75.0512248 | -0.4231305 | 0.22134964 | -1.9115932 | 0.05592839 | 0.56884484 |
| ZFYVE21 | 29.5396189 | -0.0486557 | 0.35488954 | -0.1371011 | 0.89095089 | 0.98296447 |
| PPP1R13B | 64.6872259 | 0.07961345 | 0.25480435 | 0.31244932 | 0.75469907 | 0.95832511 |
| ATP5MPL | 55.3314697 | 0.16675817 | 0.30258071 | 0.55111962 | 0.58155168 | 0.92034873 |
| INF2 | 14.1380465 | -0.0801242 | 0.4891983 | -0.1637867 | 0.86989905 | NA |
| SIVA1 | 24.3555772 | 0.09465544 | 0.398168 | 0.23772738 | 0.81209254 | 0.97257655 |
| AKT1 | 34.9217204 | -0.2489224 | 0.29700572 | -0.8381063 | 0.40197101 | 0.85778581 |
| CEP170B | 23.1061404 | -0.781144 | 0.33554659 | -2.3279747 | 0.01991345 | 0.41544117 |
| CLBA1 | 13.3663726 | -0.0804714 | 0.45215874 | -0.1779717 | 0.85874524 | NA |
| GPR132 | 25.7759118 | 0.33251782 | 0.89481952 | 0.37160322 | 0.71018829 | 0.94603965 |
| NUDT14 | 12.6917049 | 0.70171617 | 0.51626553 | 1.35921561 | 0.17407828 | NA |
| BRF1 | 43.8144947 | -0.2107395 | 0.29386966 | -0.7171189 | 0.47330073 | 0.88359968 |
| BTBD6 | 15.509852 | -0.3016747 | 0.54418425 | -0.5543614 | 0.57933155 | NA |
| PACS2 | 67.1398388 | -0.1177124 | 0.2975837 | -0.3955607 | 0.69242914 | 0.94374959 |

| | | | | | | |
|---|---|---|---|---|---|---|
| TEX22 | 10.6888247 | 0.17830718 | 0.53856655 | 0.33107734 | 0.74058607 | NA |
| MTA1 | 33.6781834 | -0.0800769 | 0.34490278 | -0.2321723 | 0.81640416 | 0.97288403 |
| CRIP2 | 32.6280999 | -0.1710291 | 0.55252504 | -0.3095408 | 0.75691014 | 0.95903525 |
| AC087386.1 | 12.3022447 | -0.0439035 | 0.60794937 | -0.0722158 | 0.94243018 | NA |
| NIPA1 | 45.6915044 | -0.0522878 | 0.31065491 | -0.1683147 | 0.86633567 | 0.97788853 |
| NIPA2 | 48.7482869 | -0.0265127 | 0.25580132 | -0.1036457 | 0.91745055 | 0.98718112 |
| CYFIP1 | 103.624032 | 0.1443411 | 0.22935508 | 0.62933468 | 0.52912997 | 0.90504888 |
| TUBGCP5 | 40.5524867 | -0.2138088 | 0.28425709 | -0.7521669 | 0.45195072 | 0.87346041 |
| AC087463.4 | 11.6967863 | 1.44069984 | 0.76690022 | 1.87860141 | 0.06029894 | NA |
| PWRN3 | 39.6113823 | 0.73710783 | 0.83601849 | 0.88168844 | 0.37794532 | 0.85310518 |
| PWRN1 | 312.498638 | 0.64194985 | 0.42249361 | 1.51943092 | 1.29E-01 | 0.66929384 |
| NPAP1 | 9.15892915 | -0.2676722 | 0.84678379 | -0.3161046 | 0.75192313 | NA |
| SNRPN | 33.3217944 | 1.02582562 | 0.63215332 | 1.62274812 | 0.10464326 | 0.65946648 |
| AC124312.1 | 319.708406 | 0.09701982 | 0.24047706 | 0.40344729 | 0.6866192 | 0.94205398 |
| PWAR6 | 80.95726 | 0.26556261 | 0.31088618 | 0.8542117 | 0.3929877 | 0.85605713 |
| UBE3A | 213.0888 | -0.1357742 | 0.26350342 | -0.5152655 | 0.60636753 | 0.92569157 |
| GABRB3 | 95.2694941 | 0.46378035 | 0.37957202 | 1.22185072 | 0.22176408 | 0.76680432 |
| HERC2 | 221.406419 | 0.01010643 | 0.25776026 | 0.03920863 | 0.96872405 | 0.99379133 |
| FAM189A1 | 99.808694 | 1.16721486 | 0.32712639 | 3.56808525 | 0.0003596 | 0.0763464 |
| TJP1 | 247.84526 | -0.3658245 | 0.27759389 | -1.3178407 | 0.18755699 | 0.74167302 |
| FAN1 | 54.8155373 | 0.29942355 | 0.34427369 | 0.86972533 | 0.38445052 | 0.85511218 |
| MTMR10 | 328.876076 | 0.12763369 | 0.2787728 | 0.45784126 | 0.64706649 | 0.93328692 |
| KLF13 | 62.8398402 | 0.43367168 | 0.31302117 | 1.38543882 | 0.16591833 | 0.71756194 |
| OTUD7A | 87.6613013 | 0.38155403 | 0.32681218 | 1.16750246 | 0.24300752 | 0.78495944 |
| LINC02256 | 128.655111 | 0.29841369 | 0.27991984 | 1.06606837 | 0.28639274 | 0.80463985 |
| AC123768.3 | 18.5590676 | 0.07022092 | 0.44501032 | 0.15779615 | 0.87461743 | 0.97918624 |
| GOLGA8N | 15.6422523 | 0.32240149 | 0.4819025 | 0.6690181 | 0.50348393 | NA |
| FMN1 | 266.54555 | 0.44476991 | 0.35822638 | 1.24158895 | 0.21438826 | 0.7643572 |
| AVEN | 51.5080667 | -0.0322801 | 0.28832786 | -0.1119561 | 0.9108582 | 0.98568025 |
| EMC7 | 28.522336 | 0.05751307 | 0.36396331 | 0.15801886 | 0.87444194 | 0.97918624 |
| KATNBL1 | 118.153455 | -0.0378284 | 0.21207801 | -0.1783703 | 0.85843214 | 0.97691173 |
| EMC4 | 12.2039168 | 0.46788123 | 0.51809993 | 0.9030714 | 0.366488 | NA |
| SLC12A6 | 52.1839738 | -0.0245402 | 0.45282788 | -0.0541932 | 0.9567812 | 0.99170927 |
| NOP10 | 35.3927018 | -0.3827305 | 0.31406916 | -1.2186187 | 0.22298896 | 0.76834258 |
| LPCAT4 | 11.5689152 | 0.13201959 | 0.46487587 | 0.28398892 | 0.77641887 | NA |
| GOLGA8A | 73.0443569 | 0.13605638 | 0.34242611 | 0.39733061 | 0.69112368 | 0.94293839 |
| GOLGA8B | 70.5902352 | 0.66609263 | 0.326191 | 2.04203255 | 0.0411483 | 0.50744725 |
| AQR | 91.9807326 | -0.1619254 | 0.22826705 | -0.7093684 | 0.47809586 | 0.88546214 |
| ZNF770 | 65.0782713 | 0.41742158 | 0.33241996 | 1.25570552 | 0.20922276 | 0.7623519 |
| AC021231.1 | 87.5412534 | 0.68771538 | 0.29768121 | 2.31024112 | 0.02087481 | 0.4212027 |
| DPH6 | 280.662306 | 0.35040905 | 0.21606919 | 1.62174466 | 0.10485803 | 0.65946648 |
| DPH6-DT | 246.863855 | 0.56608963 | 0.60550654 | 0.93490258 | 0.34983851 | 0.83687845 |
| C15orf41 | 89.6002191 | 0.01427586 | 0.24900807 | 0.05733091 | 0.95428159 | 0.99148072 |
| SPRED1 | 126.963701 | -0.4092925 | 0.36662712 | -1.1163727 | 0.26426261 | 0.79666376 |
| FAM98B | 38.5875185 | -0.0359543 | 0.28170817 | -0.1276298 | 0.89844198 | 0.98531749 |
| RASGRP1 | 107.254617 | 0.20323239 | 0.29162886 | 0.69688711 | 0.48587345 | 0.88838332 |
| C15orf53 | 247.889599 | 1.25198888 | 0.53815208 | 2.32645924 | 2.00E-02 | 4.15E-01 |
| AC013652.1 | 73.8636806 | 0.46569607 | 0.56518053 | 0.82397755 | 0.40995231 | 0.85867915 |
| THBS1 | 67.0075906 | 0.71629871 | 0.60286261 | 1.18816245 | 0.23476941 | 0.77743304 |
| FSIP1 | 23.3724126 | 0.9836335 | 0.41607276 | 2.36409013 | 0.01807441 | 0.41119297 |
| EIF2AK4 | 56.8804692 | -0.0045958 | 0.22959036 | -0.0200172 | 0.98402962 | 0.99685229 |
| SRP14 | 191.475447 | 0.41738727 | 0.37108133 | 1.1247865 | 0.26067952 | 0.79574359 |
| SRP14-AS1 | 17.4122974 | 0.22498054 | 0.48301333 | 0.46578536 | 0.64136915 | 0.93328692 |
| IVD | 60.3627179 | 0.39215935 | 0.35742701 | 1.09717326 | 0.27256566 | 0.79721781 |

| | | | | | | |
|---|---|---|---|---|---|---|
| BAHD1 | 20.8578932 | 0.3208071 | 0.34875911 | 0.91985296 | 0.3576496 | 0.84204868 |
| CCDC32 | 46.5489961 | 0.00514992 | 0.35120328 | 0.01466365 | 0.98830052 | 0.99814944 |
| RMDN3 | 26.8145708 | 0.29041629 | 0.38550701 | 0.75333596 | 0.45124806 | 0.87346041 |
| GCHFR | 13.1154175 | 0.83747305 | 0.69717915 | 1.20123078 | 0.22966169 | NA |
| DNAJC17 | 41.2028763 | -0.1515024 | 0.24979644 | -0.6065036 | 0.54418041 | 0.90965439 |
| ZFYVE19 | 18.0678551 | -0.1785144 | 0.41061563 | -0.4347482 | 0.6637452 | 0.93860542 |
| SPINT1-AS1 | 31.7829291 | 0.79185118 | 0.43187947 | 1.83350041 | 0.06672819 | 0.59652811 |
| SPINT1 | 71.5721925 | -0.257236 | 0.39318401 | -0.6542382 | 0.51295833 | 0.90156229 |
| VPS18 | 13.8244706 | -0.5459336 | 0.46918403 | -1.163581 | 0.24459386 | NA |
| INO80 | 135.923633 | -0.6636539 | 0.27346323 | -2.4268486 | 0.01523061 | 0.39253938 |
| CHP1 | 83.0794254 | -0.0480051 | 0.47152326 | -0.1018085 | 0.91890868 | 0.98749988 |
| NDUFAF1 | 14.7506736 | 0.54526425 | 0.40167473 | 1.35747713 | 0.17462966 | NA |
| RTF1 | 70.6607021 | 0.19076802 | 0.34534321 | 0.55240124 | 0.58067348 | 0.92034873 |
| RPAP1 | 15.4781308 | 0.31691761 | 0.40704183 | 0.77858734 | 0.43622284 | NA |
| MGA | 113.041901 | -0.2108076 | 0.24971746 | -0.8441844 | 0.39856637 | 0.85758979 |
| MAPKBP1 | 66.0828895 | -0.5760756 | 0.35251492 | -1.6341878 | 0.10221941 | 0.65946373 |
| EHD4 | 60.0816693 | -0.1688891 | 0.39337601 | -0.4293324 | 0.66768134 | 0.93878847 |
| PLA2G4F | 23.2921461 | 0.78252648 | 0.6574186 | 1.19030171 | 0.23392783 | 0.7767179 |
| VPS39 | 73.9217362 | -0.0285698 | 0.22993323 | -0.1242524 | 0.90111543 | 0.98531749 |
| TMEM87A | 118.379245 | -0.1195441 | 0.18678022 | -0.6400256 | 0.52215599 | 0.90451015 |
| GANC | 50.5084411 | -0.0413662 | 0.35184897 | -0.1175682 | 0.90640983 | 0.98531749 |
| ZNF106 | 92.7061744 | -0.1309999 | 0.23816836 | -0.5500305 | 0.58229843 | 0.92034873 |
| SNAP23 | 68.3085266 | -0.3156562 | 0.25341398 | -1.2456146 | 0.21290591 | 0.7643572 |
| LRRC57 | 12.3805022 | -0.2538823 | 0.43711035 | -0.5808196 | 0.56136203 | NA |
| HAUS2 | 18.09665 | -0.2858382 | 0.36399891 | -0.7852722 | 0.43229397 | 0.86696894 |
| TTBK2 | 122.733568 | 0.1663181 | 0.27015869 | 0.61563114 | 0.53813799 | 0.90763121 |
| UBR1 | 180.960341 | -0.2324052 | 0.20312248 | -1.144163 | 0.25255606 | 0.79056088 |
| AC068724.3 | 26.4906121 | 0.53052436 | 0.53242299 | 0.99643398 | 0.31903933 | 0.82354084 |
| TMEM62 | 50.1368124 | -0.4386889 | 0.38134272 | -1.1503796 | 0.24998756 | 0.78799766 |
| CCNDBP1 | 19.8371555 | 1.00033237 | 0.36237122 | 2.76051823 | 0.00577097 | 0.2943676 |
| ADAL | 37.6909019 | 0.22407822 | 0.40579047 | 0.55220179 | 0.58081011 | 0.92034873 |
| ZSCAN29 | 11.0296218 | 0.44197621 | 0.45957 | 0.96171686 | 0.33619185 | NA |
| TUBGCP4 | 26.4379048 | 0.18087753 | 0.35448323 | 0.51025696 | 0.60987145 | 0.92573951 |
| TP53BP1 | 153.272156 | -0.2225429 | 0.21382175 | -1.0407871 | 0.29797438 | 0.81214575 |
| PPIP5K1 | 58.8894673 | -0.4174604 | 0.22709276 | -1.8382815 | 0.06602094 | 0.59523726 |
| CATSPER2 | 68.5618401 | -0.2170971 | 0.36265947 | -0.5986252 | 0.54942283 | 0.91221128 |
| PDIA3 | 138.935593 | -0.0755048 | 0.44045372 | -0.1714251 | 0.8638895 | 0.97788853 |
| SERF2 | 358.507382 | -0.2271888 | 0.4155947 | -0.5466596 | 0.58461264 | 0.92071433 |
| MFAP1 | 25.9033062 | 0.27282169 | 0.36461962 | 0.74823645 | 0.45431755 | 0.87378486 |
| FRMD5 | 13.1269685 | 1.06343932 | 0.50546671 | 2.10387607 | 0.03538926 | NA |
| CASC4 | 249.238914 | -0.1121304 | 0.14273305 | -0.785595 | 0.43210481 | 0.86695857 |
| CTDSPL2 | 134.306462 | -0.1316124 | 0.22756497 | -0.578351 | 0.56302719 | 0.91624774 |
| EIF3J-DT | 34.2908014 | 0.47484408 | 0.36227061 | 1.31074415 | 0.18994422 | 0.74535784 |
| EIF3J | 67.259915 | -0.0144228 | 0.21775168 | -0.066235 | 0.94719072 | 0.99083449 |
| SPG11 | 195.722794 | -0.2857693 | 0.1823237 | -1.5673732 | 0.11702748 | 0.6668162 |
| B2M | 644.519896 | 0.16755043 | 0.44258553 | 0.37857187 | 7.05E-01 | 0.94530386 |
| TRIM69 | 19.493062 | -0.4306567 | 0.40929362 | -1.052195 | 0.29271008 | 0.80935782 |
| SORD | 19.4227597 | 0.57048499 | 0.52451242 | 1.0876482 | 0.27675044 | 0.80159799 |
| DUOX1 | 15.630118 | 0.13487228 | 0.40078275 | 0.33652216 | 0.73647715 | NA |
| GATM | 18.3505943 | 0.17235811 | 0.61070221 | 0.28222938 | 0.77776763 | 0.96518573 |
| SPATA5L1 | 23.139131 | -0.0646715 | 0.36298086 | -0.1781679 | 0.85859114 | 0.97691173 |
| SLC30A4 | 12.8927215 | 0.71665101 | 0.4998204 | 1.43381706 | 0.15162447 | NA |
| SQOR | 17.4882757 | -0.1610032 | 0.38640038 | -0.4166746 | 0.67691643 | 0.94034237 |
| SEMA6D | 37.5716559 | 1.07133001 | 0.44109937 | 2.4287725 | 0.01515004 | 0.39253938 |

| | | | | | | |
|---|---|---|---|---|---|---|
| MYEF2 | 89.44782 | -0.0468061 | 0.24853223 | -0.1883301 | 0.85061786 | 0.97612951 |
| SLC12A1 | 46.0800454 | 0.53098741 | 0.7065255 | 0.75154741 | 4.52E-01 | 8.73E-01 |
| DUT | 41.3529988 | 0.50385561 | 0.33377958 | 1.50954593 | 0.13115933 | 0.67492558 |
| CEP152 | 40.7752729 | -0.0417594 | 0.32286664 | -0.1293395 | 0.89708903 | 0.98531749 |
| EID1 | 76.2455629 | 0.46220856 | 0.41063245 | 1.12560165 | 0.26033417 | 0.79574359 |
| SECISBP2L | 176.715883 | 0.18937037 | 0.16207598 | 1.16840485 | 0.2426435 | 0.78464787 |
| COPS2 | 43.2872876 | -0.192056 | 0.35308445 | -0.5439378 | 0.58648424 | 0.92076087 |
| GALK2 | 107.689232 | -0.0294161 | 0.26946337 | -0.1091655 | 0.91307122 | 0.98639439 |
| FAM227B | 70.0897932 | 0.55401399 | 0.45407902 | 1.22008277 | 0.2224335 | 0.76784685 |
| DTWD1 | 59.3439991 | -0.1785097 | 0.28387415 | -0.628834 | 0.52945777 | 0.9052447 |
| SLC27A2 | 12.0943189 | 1.53396571 | 0.65018978 | 2.35925842 | 0.0183115 | NA |
| GABPB1 | 51.0441139 | -0.6497793 | 0.27579468 | -2.3560255 | 0.01847165 | 0.41315568 |
| GABPB1-AS1 | 95.1728424 | 0.71581157 | 0.35676345 | 2.00640385 | 0.04481318 | 0.52568861 |
| USP8 | 133.642926 | -0.0266803 | 0.22642256 | -0.1178342 | 0.90619905 | 0.98531749 |
| USP50 | 13.5192907 | -0.4798698 | 0.54425509 | -0.8817002 | 0.37793895 | NA |
| TRPM7 | 228.229664 | -0.1932979 | 0.20059001 | -0.9636469 | 0.33522297 | 0.82974237 |
| SPPL2A | 158.389748 | -0.0948641 | 0.30398708 | -0.3120662 | 0.75499018 | 0.95832511 |
| AP4E1 | 50.2010719 | 0.01780864 | 0.28858827 | 0.0617095 | 0.95079417 | 0.99105535 |
| DMXL2 | 102.909918 | -0.2114808 | 0.25793422 | -0.819902 | 0.41227198 | 0.85867915 |
| TMOD2 | 15.4243405 | 0.32940025 | 0.42621806 | 0.77284442 | 0.43961446 | NA |
| TMOD3 | 122.635026 | -0.3054191 | 0.18308197 | -1.6682095 | 0.09527415 | 0.64916213 |
| LEO1 | 18.4396575 | 0.0316401 | 0.37190464 | 0.08507585 | 0.93220109 | 0.98960607 |
| MAPK6 | 75.7522774 | -0.6227537 | 0.3180728 | -1.9578969 | 0.05024211 | 0.54473367 |
| GNB5 | 74.5957177 | 0.01022725 | 0.32419967 | 0.03154613 | 0.974834 | 0.99431961 |
| MYO5C | 116.352962 | 0.29069504 | 0.31872149 | 0.91206601 | 0.36173397 | 0.84398658 |
| MYO5A | 276.042882 | -0.5492898 | 0.31233906 | -1.7586331 | 0.07863984 | 0.61367304 |
| ARPP19 | 102.025133 | -0.0889217 | 0.27198751 | -0.3269332 | 0.74371844 | 0.95563105 |
| FAM214A | 141.698787 | -0.1138029 | 0.22885207 | -0.497277 | 0.61899371 | 0.92766755 |
| WDR72 | 562.490766 | 0.14002204 | 0.41778165 | 0.33515604 | 0.73750738 | 0.9542764 |
| UNC13C | 78.9607151 | -0.8814668 | 0.70020772 | -1.2588647 | 0.20807922 | 0.76231289 |
| RSL24D1 | 27.7701021 | 0.86811433 | 0.30908926 | 2.80862019 | 0.00497543 | 0.27957621 |
| RAB27A | 69.3942706 | 0.12039344 | 0.2251309 | 0.53477085 | 5.93E-01 | 0.92278087 |
| PIGB | 31.5307008 | 0.07761636 | 0.3216615 | 0.24129826 | 0.80932396 | 0.97257655 |
| CCPG1 | 123.650651 | 0.24729844 | 0.20000833 | 1.23644073 | 0.21629478 | 0.76616336 |
| DNAAF4 | 56.1376444 | -0.0765573 | 0.32842083 | -0.2331075 | 0.81567796 | 0.97288403 |
| PYGO1 | 23.2046587 | -0.5524737 | 0.426706 | -1.2947408 | 0.19540965 | 0.75323675 |
| PRTG | 116.730409 | 1.22995681 | 0.5627004 | 2.18581114 | 0.02882943 | 0.46148943 |
| NEDD4 | 47.4404112 | 0.1835248 | 0.33624447 | 0.54580764 | 0.58519817 | 0.92076087 |
| RFX7 | 157.559513 | -0.0279028 | 0.20012229 | -0.1394285 | 0.88911151 | 0.98270788 |
| TEX9 | 102.288424 | 0.4871643 | 0.30765383 | 1.58348196 | 0.11331165 | 0.6656824 |
| MNS1 | 35.2974844 | 0.63772845 | 0.46951535 | 1.35826964 | 0.17437814 | 0.72841249 |
| ZNF280D | 65.6684022 | 0.30526376 | 0.35164798 | 0.86809475 | 0.38534247 | 0.85540507 |
| AC090517.4 | 39.1602609 | 0.17142427 | 0.40203294 | 0.4263936 | 0.66982107 | 0.93878847 |
| TCF12 | 762.207935 | -0.1767443 | 0.19575602 | -0.9028804 | 0.36658936 | 0.84741115 |
| CGNL1 | 321.598667 | 0.02509452 | 0.38254476 | 0.06559892 | 0.94769715 | 0.99091283 |
| POLR2M | 17.8199903 | -0.4554104 | 0.36991726 | -1.2311143 | 0.21828011 | 0.76666749 |
| ALDH1A2 | 146.145692 | -0.9714187 | 0.61994755 | -1.5669368 | 0.11712945 | 0.6668162 |
| LIPC | 55.7698851 | -0.1664711 | 0.50888761 | -0.3271274 | 0.74357151 | 0.95563105 |
| ADAM10 | 474.314251 | -0.6302595 | 0.33790473 | -1.8651989 | 0.06215353 | 0.58370968 |
| MINDY2 | 257.790773 | -0.6994411 | 0.19090137 | -3.6638871 | 0.00024842 | 0.06160773 |
| RNF111 | 191.509722 | -0.13273 | 0.28518694 | -0.4654141 | 0.64163497 | 0.93328692 |
| SLTM | 238.564096 | -0.2153635 | 0.18324595 | -1.17527 | 0.23988673 | 0.78128405 |
| MYO1E | 631.136795 | -0.6899833 | 0.43503609 | -1.5860369 | 0.11273094 | 0.6656824 |
| FAM81A | 75.6233569 | -0.1047197 | 0.30816469 | -0.3398173 | 0.73399409 | 0.95329896 |

| | | | | | | |
|---|---|---|---|---|---|---|
| GCNT3 | 51.3620161 | 1.03147898 | 0.46359501 | 2.22495705 | 2.61E-02 | 4.46E-01 |
| GTF2A2 | 43.745665 | 0.52183654 | 0.32534292 | 1.60395849 | 0.1087232 | 0.65986731 |
| BNIP2 | 60.9103194 | -0.1521706 | 0.2842304 | -0.5353777 | 0.5923887 | 0.92278087 |
| ANXA2 | 159.178375 | -0.4818959 | 0.47129552 | -1.022492 | 3.07E-01 | 0.81568231 |
| ICE2 | 93.0609883 | 0.16501432 | 0.24355859 | 0.67751386 | 0.49807998 | 0.89378207 |
| RORA-AS1 | 58.688051 | 0.54959353 | 0.41238688 | 1.33271343 | 0.18262586 | 0.73398872 |
| RORA | 1126.35285 | 0.48698551 | 0.38107205 | 1.27793552 | 0.20127216 | 0.75910235 |
| AC012404.1 | 31.8005279 | 0.74536144 | 0.62215802 | 1.19802593 | 0.23090692 | 0.77249016 |
| AC009554.1 | 43.8180595 | 0.15784523 | 0.57193268 | 0.27598569 | 0.78255906 | 0.96652357 |
| VPS13C | 369.323139 | -0.4155564 | 0.1963454 | -2.1164561 | 0.03430603 | 0.4908133 |
| C2CD4A | 33.719514 | -0.9130961 | 0.60382718 | -1.5121812 | 1.30E-01 | 6.74E-01 |
| TLN2 | 312.422339 | 0.21017764 | 0.34423363 | 0.61056686 | 5.41E-01 | 9.08E-01 |
| TPM1 | 115.587942 | 0.77671009 | 0.3249812 | 2.39001544 | 0.01684766 | 0.39748373 |
| LACTB | 27.721331 | -0.0164703 | 0.39959844 | -0.0412172 | 0.96712271 | 0.9930872 |
| RPS27L | 43.1768834 | 0.65057265 | 0.33122434 | 1.96414506 | 0.04951327 | 0.54320302 |
| RAB8B | 50.9647054 | -0.2344123 | 0.26489066 | -0.8849398 | 0.37618908 | 0.85248921 |
| APH1B | 25.0850698 | 0.00535212 | 0.40757535 | 0.01313161 | 0.98952279 | 0.99814944 |
| CA12 | 1006.50152 | 0.38004279 | 0.28614756 | 1.32813571 | 0.18413328 | 0.73705812 |
| LINC02568 | 51.2542801 | 0.69890968 | 0.47158562 | 1.48204197 | 0.13832913 | 0.68960025 |
| USP3 | 240.345257 | -0.1797486 | 0.20660294 | -0.8700196 | 0.38428967 | 0.85511218 |
| USP3-AS1 | 42.0183462 | 0.24719425 | 0.48416974 | 0.51055287 | 0.60966418 | 0.92573951 |
| HERC1 | 295.300141 | -0.7280594 | 0.34440995 | -2.1139325 | 0.03452103 | 0.49098143 |
| FAM96A | 40.958433 | 0.17823079 | 0.33752954 | 0.52804502 | 0.59746809 | 0.9229638 |
| SNX1 | 91.8804595 | -0.1338513 | 0.21554314 | -0.6209953 | 0.5346027 | 0.90763121 |
| PPIB | 107.686413 | 0.50752413 | 0.26234782 | 1.93454673 | 5.30E-02 | 0.55759493 |
| CSNK1G1 | 29.8400323 | 0.20014639 | 0.35781954 | 0.55935008 | 0.57592283 | 0.91854298 |
| TRIP4 | 61.7021499 | -0.7572798 | 0.28645458 | -2.6436296 | 0.00820224 | 0.30976355 |
| ZNF609 | 304.218947 | -0.4388026 | 0.19827905 | -2.2130558 | 0.0268938 | 0.45084589 |
| OAZ2 | 36.3768433 | 0.41922055 | 0.36150182 | 1.15966373 | 0.24618574 | 0.78583441 |
| RBPMS2 | 13.6647566 | 0.04119563 | 0.60645675 | 0.06792839 | 0.94584264 | NA |
| SPG21 | 50.4707054 | -0.2947588 | 0.29919651 | -0.9851679 | 0.3245416 | 0.82567895 |
| MTFMT | 36.8541389 | -0.1549257 | 0.32974678 | -0.4698323 | 0.63847482 | 0.9331046 |
| UBAP1L | 19.5261096 | 0.89401088 | 0.36881932 | 2.42398062 | 0.01535142 | 0.39253938 |
| PDCD7 | 35.0827056 | -0.0263065 | 0.28569535 | -0.092079 | 0.92663527 | 0.98925753 |
| CLPX | 62.6299823 | 0.08481954 | 0.20812998 | 0.4075316 | 0.68361759 | 0.94205398 |
| PARP16 | 24.8989598 | 0.08292163 | 0.34626994 | 0.23947106 | 0.81074033 | 0.97257655 |
| DPP8 | 114.517398 | -0.2544583 | 0.18937181 | -1.3436971 | 0.17904637 | 0.72924345 |
| HACD3 | 135.660558 | 0.33315612 | 0.26848118 | 1.24089189 | 0.21464569 | 0.7643572 |
| INTS14 | 30.5382044 | -0.2579377 | 0.31692847 | -0.8138672 | 0.41572101 | 0.85867915 |
| SLC24A1 | 18.6962277 | 0.85728458 | 0.52448909 | 1.63451366 | 0.10215104 | 0.65946373 |
| DENND4A | 227.531517 | 0.12609707 | 0.22130601 | 0.569786 | 0.56882285 | 0.91698355 |
| RAB11A | 144.167454 | 0.37671537 | 0.34364541 | 1.09623279 | 0.27297691 | 0.79726027 |
| MEGF11 | 258.574597 | -0.1824261 | 0.45519307 | -0.4007664 | 6.89E-01 | 9.42E-01 |
| DIS3L | 29.8028676 | 0.72722939 | 0.31347886 | 2.31986739 | 0.02034805 | 0.41907916 |
| TIPIN | 18.8461349 | 0.52624311 | 0.53354413 | 0.986316 | 0.32397807 | 0.82567895 |
| MAP2K1 | 102.099552 | -0.6399715 | 0.23383195 | -2.7368863 | 0.00620237 | 0.2971589 |
| SNAPC5 | 15.7493718 | -0.0521035 | 0.38697749 | -0.1346421 | 0.89289482 | NA |
| RPL4 | 126.699569 | 0.28692644 | 0.45017767 | 0.63736266 | 0.52388865 | 0.90487731 |
| SMAD6 | 24.2778644 | 0.25356096 | 0.41881466 | 0.60542523 | 0.54489649 | 0.90999223 |
| SMAD3 | 488.302997 | -1.5894421 | 0.29244704 | -5.434974 | 5.48E-08 | 0.00049888 |
| AAGAB | 102.085001 | -0.2254407 | 0.24548325 | -0.9183548 | 0.35843317 | 0.84204868 |
| IQCH | 110.201631 | 0.21810164 | 0.35972314 | 0.60630418 | 0.54431278 | 0.90965439 |
| IQCH-AS1 | 58.825074 | 0.271072 | 0.40098387 | 0.67601723 | 0.49902971 | 0.89378207 |
| MAP2K5 | 188.227705 | 0.22515954 | 0.26005232 | 0.865824 | 0.38658669 | 0.8560201 |

| | | | | | | |
|---|---|---|---|---|---|---|
| PIAS1 | 286.574826 | -0.1890314 | 0.18314904 | -1.0321179 | 0.30201687 | 0.81568231 |
| FEM1B | 36.9594022 | -0.0292517 | 0.30365736 | -0.0963311 | 0.92325758 | 0.98915831 |
| CORO2B | 23.3019026 | -0.7058864 | 0.56493666 | -1.2494966 | 0.21148351 | 0.7643572 |
| ANP32A | 64.522962 | 0.2173882 | 0.26560299 | 0.81847046 | 0.4130886 | 0.85867915 |
| SPESP1 | 14.9484137 | 0.74675443 | 0.52625596 | 1.41899473 | 0.15590055 | NA |
| GLCE | 107.019449 | -0.0547134 | 0.26448774 | -0.2068655 | 0.8361149 | 0.97600352 |
| AC026992.2 | 15.3720322 | 0.52938844 | 0.64764937 | 0.81739976 | 0.41370001 | NA |
| PAQR5 | 391.645324 | -1.1381061 | 0.38113182 | -2.9861219 | 0.0028254 | 0.21255882 |
| KIF23 | 32.1769433 | -1.5167827 | 0.53993128 | -2.8092143 | 0.00496626 | 0.27957621 |
| RPLP1 | 242.030981 | 0.00981726 | 0.44130984 | 0.02224573 | 0.98225194 | 0.99626066 |
| TLE3 | 60.4928121 | 0.02093727 | 0.30828668 | 0.06791494 | 0.94585334 | 0.99083449 |
| UACA | 176.367968 | 0.0901334 | 0.25068471 | 0.35954885 | 0.71918454 | 0.94959812 |
| LARP6 | 23.290905 | -0.0531839 | 0.39177016 | -0.1357529 | 0.89201663 | 0.98329225 |
| LRRC49 | 53.1705772 | -0.0992747 | 0.31320379 | -0.3169652 | 0.75127 | 0.95741436 |
| THSD4 | 206.743749 | -0.7199446 | 0.33893366 | -2.124146 | 0.03365794 | 0.4908133 |
| THSD4-AS1 | 72.0766383 | 0.04363846 | 0.27615098 | 0.15802393 | 0.87443795 | 0.97918624 |
| MYO9A | 468.195838 | 0.05307068 | 0.22245206 | 0.23857129 | 0.81143803 | 0.97257655 |
| AC022872.1 | 16.8782513 | -0.6948187 | 0.67675189 | -1.0266963 | 0.30456349 | 0.81568231 |
| SENP8 | 14.2219052 | 0.51636836 | 0.47048529 | 1.09752287 | 0.27241289 | NA |
| PKM | 465.381823 | -0.2049725 | 0.26923147 | -0.7613243 | 4.46E-01 | 8.72E-01 |
| PARP6 | 46.1121354 | -0.3893965 | 0.24599886 | -1.58292 | 0.11343969 | 0.6656824 |
| AC009690.2 | 21.2978892 | 0.53104579 | 0.66073103 | 0.80372461 | 0.42155604 | 0.86190547 |
| AC009690.1 | 76.8238839 | 0.08491634 | 0.36148991 | 0.23490653 | 0.81428128 | 0.97282508 |
| ARIH1 | 405.226015 | -0.5000596 | 0.3073854 | -1.6268162 | 0.10377614 | 0.65946648 |
| BBS4 | 63.174858 | 0.10924441 | 0.23893782 | 0.45720852 | 0.64752117 | 0.93328692 |
| ADPGK | 43.1889644 | -0.1090736 | 0.26560909 | -0.4106545 | 0.68132586 | 0.94196488 |
| NEO1 | 117.520801 | 0.39879867 | 0.21754673 | 1.83316324 | 0.0667783 | 0.59652811 |
| NPTN | 150.227233 | -0.1158158 | 0.27416828 | -0.4224259 | 0.67271418 | 0.93878847 |
| CD276 | 18.4789275 | -0.8251742 | 0.39572462 | -2.0852233 | 0.03704904 | 0.49537143 |
| LOXL1 | 12.2041584 | 0.92114621 | 0.458719 | 2.00808384 | 0.04463438 | NA |
| PML | 32.5948542 | -0.1750203 | 0.49555481 | -0.3531806 | 0.72395307 | 0.95093722 |
| ARID3B | 25.0473028 | -0.1971431 | 0.33227123 | -0.5933199 | 0.55296707 | 0.91322379 |
| CLK3 | 28.0888705 | -0.043675 | 0.36023199 | -0.1212413 | 0.90349988 | 0.98531749 |
| EDC3 | 33.5691824 | -0.1826653 | 0.33405588 | -0.5468105 | 0.58450891 | 0.92071026 |
| CSK | 12.9955053 | 0.14458562 | 0.51170364 | 0.28255734 | 0.77751618 | NA |
| ULK3 | 20.0792374 | 0.35932184 | 0.4301515 | 0.83533788 | 0.4035275 | 0.85782098 |
| SCAMP2 | 42.4199198 | -0.6276328 | 0.39380701 | -1.5937572 | 0.11099042 | 0.66339185 |
| FAM219B | 18.1986613 | -0.2925533 | 0.38467104 | -0.7605286 | 0.44693871 | 0.87231627 |
| COX5A | 72.160218 | 0.06107381 | 0.23478264 | 0.26012917 | 0.79476414 | 0.96974891 |
| SCAMP5 | 40.7896009 | 0.17102948 | 0.33159152 | 0.51578363 | 0.60600554 | 0.92569157 |
| PPCDC | 65.2384714 | -0.2531902 | 0.39974789 | -0.6333746 | 0.52648903 | 0.90487731 |
| COMMD4 | 22.8840135 | -0.5169845 | 0.37374638 | -1.3832494 | 0.1665884 | 0.71869865 |
| NEIL1 | 22.7660184 | 0.41678244 | 0.40711142 | 1.02375522 | 0.30595091 | 0.81568231 |
| MAN2C1 | 42.5779377 | -0.3149158 | 0.33045193 | -0.9529852 | 0.34059759 | 0.8319325 |
| SIN3A | 80.1871144 | 0.06325764 | 0.2847125 | 0.22218077 | 0.82417316 | 0.97518529 |
| PTPN9 | 86.5277678 | -0.3794827 | 0.2765527 | -1.3721896 | 0.17000442 | 0.72416949 |
| SNUPN | 18.2324515 | -0.1320738 | 0.4205112 | -0.3140792 | 0.75346091 | 0.95826617 |
| IMP3 | 14.5157038 | 0.61370885 | 0.37678851 | 1.62878866 | 0.10335777 | NA |
| UBE2Q2 | 38.2065931 | 0.3072226 | 0.27741582 | 1.1074444 | 2.68E-01 | 0.79666376 |
| FBXO22 | 64.9893362 | 0.10806576 | 0.23984197 | 0.45057069 | 0.65229899 | 0.93524614 |
| ETFA | 95.9693486 | 0.18657599 | 0.21983529 | 0.84870808 | 0.39604375 | 0.85605713 |
| SCAPER | 307.983565 | 0.40957174 | 0.23111467 | 1.77215813 | 0.07636831 | 0.61087939 |
| RCN2 | 63.6970927 | 0.28535423 | 0.23240415 | 1.22783623 | 0.21950845 | 0.76680432 |
| TSPAN3 | 66.0319246 | 0.14058547 | 0.2708382 | 0.51907548 | 0.60370811 | 0.92508698 |

| | | | | | | |
|---|---|---|---|---|---|---|
| PEAK1 | 141.02551 | -0.2814826 | 0.19385532 | -1.452024 | 0.14649492 | 0.69713616 |
| HMG20A | 95.4481625 | 0.0571695 | 0.20256382 | 0.28222957 | 0.77776749 | 0.96518573 |
| LINGO1 | 589.740485 | -1.0343043 | 0.57852105 | -1.7878421 | 0.07380149 | 0.60985192 |
| TBC1D2B | 36.3479298 | -0.6004566 | 0.27513619 | -2.1823978 | 0.02908018 | 0.46268871 |
| IDH3A | 67.5384988 | 0.01125841 | 0.29675879 | 0.03793791 | 0.96973718 | 0.99398482 |
| DNAJA4 | 16.8657534 | -0.6475004 | 0.63250917 | -1.0237011 | 0.30597646 | 0.81568231 |
| WDR61 | 38.1549183 | 0.66751746 | 0.29264368 | 2.28099056 | 0.02254901 | 0.42564995 |
| IREB2 | 93.8011598 | -0.1938999 | 0.1998716 | -0.9701225 | 0.33198542 | 0.82890589 |
| PSMA4 | 43.5914352 | -0.1114066 | 0.35403923 | -0.3146729 | 0.75301 | 0.95815628 |
| MORF4L1 | 141.965959 | -0.0334365 | 0.16165947 | -0.206833 | 0.83614032 | 0.97600352 |
| CTSH | 139.020324 | -0.2867164 | 0.38563873 | -0.7434843 | 0.4571885 | 0.87419345 |
| TMED3 | 33.2320481 | -0.1696497 | 0.45185637 | -0.3754504 | 0.70732553 | 0.94558664 |
| ZFAND6 | 211.224867 | -0.0269126 | 0.25725483 | -0.1046145 | 0.91668167 | 0.98710975 |
| FAH | 11.6138928 | 0.3822688 | 0.46962559 | 0.81398631 | 0.4156528 | NA |
| AC016705.2 | 44.8468755 | -0.9348723 | 0.46527768 | -2.0092782 | 0.04450764 | 0.52480969 |
| ARNT2 | 150.591423 | -0.2178334 | 0.34293526 | -0.6352027 | 0.5252962 | 0.90487731 |
| ABHD17C | 62.4112132 | 0.03712947 | 0.30997735 | 0.11978124 | 0.90465645 | 0.98531749 |
| CEMIP | 13.9899339 | 1.58999561 | 0.82725275 | 1.92201913 | 0.05460335 | NA |
| AC027808.1 | 21.8328799 | 0.51483285 | 0.42545635 | 1.21007207 | 0.22625124 | 0.77166168 |
| MESD | 44.7161072 | 0.52395819 | 0.26643845 | 1.96652618 | 0.04923786 | 0.54262984 |
| EFL1 | 55.8073394 | -0.5338721 | 0.2446938 | -2.1817968 | 0.02912453 | 0.46268871 |
| WHAMM | 35.9081185 | -0.3736509 | 0.27411898 | -1.3630977 | 0.17285172 | 0.72711145 |
| HOMER2 | 30.6057845 | -0.0725034 | 0.30055672 | -0.2412302 | 0.80937668 | 0.97257655 |
| C15orf40 | 27.0621416 | 0.1653813 | 0.31605127 | 0.52327365 | 0.60078384 | 0.92352528 |
| BTBD1 | 73.7622576 | -0.1640977 | 0.23038075 | -0.7122892 | 0.47628573 | 0.88495683 |
| HDGFL3 | 82.6977909 | 0.08238544 | 0.29572653 | 0.27858657 | 0.78056212 | 0.96652357 |
| ADAMTSL3 | 99.7910212 | 0.05701751 | 0.49428975 | 0.11535239 | 0.90816581 | 0.9854228 |
| WDR73 | 60.7705859 | 0.07992702 | 0.25498447 | 0.31345839 | 0.75393243 | 0.95826617 |
| SEC11A | 124.96647 | -0.1016512 | 0.24652124 | -0.4123426 | 0.68008835 | 0.94195357 |
| ZNF592 | 61.019411 | -0.4133078 | 0.29325606 | -1.4093752 | 0.15872424 | 0.71094363 |
| ALPK3 | 62.8945767 | -0.4332385 | 0.51997995 | -0.8331831 | 0.40474147 | 0.85782098 |
| PDE8A | 360.015991 | -0.1987439 | 0.22899355 | -0.8679016 | 0.38544819 | 0.85540507 |
| AKAP13 | 447.173826 | -0.5143527 | 0.22944181 | -2.2417566 | 0.02497711 | 0.43948277 |
| KLHL25 | 22.7278904 | -0.6899846 | 0.34930798 | -1.97529 | 0.04823525 | 0.5380949 |
| MRPL46 | 15.1430302 | 0.27136532 | 0.36595034 | 0.74153591 | 0.45836857 | NA |
| MRPS11 | 25.3341157 | 0.48681013 | 0.37090927 | 1.31247766 | 0.18935902 | 0.74427251 |
| DET1 | 23.8292046 | -0.0889299 | 0.47291274 | -0.1880472 | 0.85083967 | 0.97612951 |
| ABHD2 | 90.9255309 | -0.0444361 | 0.40746542 | -0.1090549 | 0.91315891 | 0.98639439 |
| POLG | 18.5919112 | -0.013085 | 0.34878182 | -0.0375162 | 0.97007344 | 0.99398482 |
| RHCG | 216.49326 | 0.92982716 | 0.512964 | 1.81265578 | 6.99E-02 | 6.03E-01 |
| PEX11A | 11.3887628 | 0.05814772 | 0.4189641 | 0.13878926 | 0.88961668 | NA |
| ANPEP | 15.1784524 | 0.0149305 | 0.47333908 | 0.03154292 | 0.97483656 | NA |
| ZNF710 | 54.4272117 | 0.10432282 | 0.33860944 | 0.3080919 | 0.75801241 | 0.95903525 |
| ZNF710-AS1 | 38.9753541 | -0.1071258 | 0.49621764 | -0.2158848 | 0.82907755 | 0.97567084 |
| IDH2 | 238.706825 | 0.10762218 | 0.29847651 | 0.36057171 | 0.71841964 | 0.94925929 |
| IDH2-DT | 31.9105953 | -0.0667969 | 0.52138196 | -0.1281152 | 0.89805784 | 0.98531749 |
| SEMA4B | 240.454793 | -0.6304916 | 0.24887665 | -2.5333498 | 1.13E-02 | 3.48E-01 |
| CIB1 | 98.5683015 | -0.2284111 | 0.47398764 | -0.4818925 | 0.62988234 | 0.9318063 |
| AC091167.6 | 9.50415294 | 0.02800298 | 0.60188484 | 0.04652548 | 0.96289142 | NA |
| ZNF774 | 10.0943542 | 0.2200276 | 0.48542423 | 0.45326868 | 0.65035528 | NA |
| IQGAP1 | 250.227178 | -0.0153618 | 0.17301496 | -0.0887889 | 0.92924967 | 0.98960607 |
| CRTC3 | 175.883449 | 0.17306983 | 0.18403592 | 0.94041331 | 0.3470056 | 0.83687845 |
| CRTC3-AS1 | 24.5885649 | -0.32213 | 0.41367199 | -0.7787087 | 0.43615134 | 0.86729771 |
| FURIN | 31.2934795 | -0.1945526 | 0.51918823 | -0.3747246 | 0.70786526 | 0.94558664 |

| | | | | | | |
|---|---|---|---|---|---|---|
| MAN2A2 | 17.8645258 | 0.21640359 | 0.37834701 | 0.57197119 | 0.56734149 | 0.91698355 |
| UNC45A | 31.5981258 | 0.3330766 | 0.32018806 | 1.04025301 | 0.29822237 | 0.81214575 |
| AC104035.1 | 60.9586968 | -0.6190286 | 0.85306415 | -0.7256531 | 0.46805146 | 0.88139687 |
| SLCO3A1 | 93.2710931 | 1.07967032 | 0.5113443 | 2.11143515 | 3.47E-02 | 0.49098143 |
| FAM174B | 34.5138716 | -0.403098 | 0.31444887 | -1.2819189 | 0.19987108 | 0.75605362 |
| AC013394.1 | 131.210186 | -0.1086113 | 0.25146999 | -0.4319058 | 0.66580991 | 0.93860542 |
| CHD2 | 258.314677 | -0.4722507 | 0.3213744 | -1.4694722 | 0.14170476 | 0.69328517 |
| MCTP2 | 51.4196333 | 0.35805597 | 0.50651719 | 0.70689796 | 0.47962989 | 0.88651186 |
| AC012409.2 | 49.2289805 | -0.2727085 | 0.65123671 | -0.4187547 | 0.67539539 | 0.93993644 |
| NR2F2-AS1 | 327.302746 | -0.4545086 | 0.34366492 | -1.3225342 | 0.18599031 | 0.73804265 |
| NR2F2 | 97.1085105 | -0.5363877 | 0.28216385 | -1.9009796 | 0.05730468 | 0.57060925 |
| ARRDC4 | 40.3184107 | 0.08405575 | 0.4939052 | 0.17018601 | 0.86486385 | 0.97788853 |
| FAM169B | 14.8679117 | -0.4583484 | 0.68629496 | -0.6678592 | 5.04E-01 | NA |
| IGF1R | 550.901399 | -0.3460494 | 0.2114531 | -1.6365303 | 0.10172865 | 0.65863153 |
| TTC23 | 63.7399921 | -0.2267774 | 0.24239601 | -0.9355659 | 0.34949674 | 0.83687845 |
| LRRC28 | 132.875535 | 0.0835289 | 0.2593125 | 0.32211675 | 0.74736425 | 0.95563105 |
| MEF2A | 380.124585 | 0.0017509 | 0.17354358 | 0.01008912 | 0.99195018 | 0.99814944 |
| AC090825.1 | 25.8377143 | 0.38427185 | 0.46648412 | 0.82376192 | 0.41007485 | 0.85867915 |
| ADAMTS17 | 53.692837 | -0.6350858 | 0.47363022 | -1.3408896 | 0.17995629 | 0.73229421 |
| LINS1 | 24.8617248 | -0.0437202 | 0.35088614 | -0.1245994 | 0.90084068 | 0.98531749 |
| ASB7 | 31.0029305 | 0.24593757 | 0.33852555 | 0.72649633 | 0.46753456 | 0.88108588 |
| CHSY1 | 257.170126 | -0.2383584 | 0.49194085 | -0.4845265 | 0.62801229 | 0.93061955 |
| SELENOS | 53.0937145 | 0.05276378 | 0.28409978 | 0.18572272 | 0.85266221 | 0.97612951 |
| SNRPA1 | 32.8897596 | -0.3020376 | 0.31392392 | -0.9621365 | 0.33598106 | 0.82974237 |
| PCSK6 | 25.7190964 | 0.06066005 | 0.43431145 | 0.13966948 | 0.88892115 | 0.98270788 |
| TM2D3 | 38.9745265 | 0.32382353 | 0.27254514 | 1.18814643 | 0.23477573 | 0.77743304 |
| TARSL2 | 71.2430969 | -0.1093361 | 0.35463709 | -0.3083042 | 0.75785089 | 0.95903525 |
| SNRNP25 | 11.1877749 | 0.32316529 | 0.50156935 | 0.6443083 | 0.51937553 | NA |
| MPG | 30.5132076 | -0.0542345 | 0.2887111 | -0.1878503 | 0.85099403 | 0.97612951 |
| NPRL3 | 64.4621978 | 0.05953297 | 0.29882618 | 0.19922276 | 0.8420885 | 0.97600352 |
| LUC7L | 135.170543 | -0.090934 | 0.26347617 | -0.3451319 | 0.7299952 | 0.95279138 |
| FAM234A | 22.390201 | -0.4164724 | 0.47894042 | -0.8695703 | 0.38453528 | 0.85511218 |
| AXIN1 | 26.1490453 | -0.5759302 | 0.38589267 | -1.4924621 | 0.13557805 | 0.6835006 |
| MRPL28 | 21.8474294 | 0.35711631 | 0.45665904 | 0.78201957 | 0.43420307 | 0.86729771 |
| TMEM8A | 60.7872729 | 0.03789826 | 0.30147076 | 0.12571122 | 0.89996052 | 0.98531749 |
| NME4 | 14.7536453 | 0.47425788 | 0.41889352 | 1.13216811 | 0.25756377 | NA |
| RAB11FIP3 | 183.200535 | 0.10800623 | 0.2470871 | 0.43711803 | 0.66202575 | 0.93752651 |
| CAPN15 | 29.6269171 | -0.5624586 | 0.40250952 | -1.3973797 | 0.16229942 | 0.71269251 |
| PRR35 | 17.5240668 | 1.28662869 | 0.76785544 | 1.67561318 | 0.09381398 | 0.64572229 |
| PIGQ | 73.3634626 | 0.40639331 | 0.39555334 | 1.02740457 | 0.30423 | 0.81568231 |
| RAB40C | 78.1379746 | -0.4120598 | 0.30901984 | -1.3334414 | 0.18238699 | 0.73398872 |
| METTL26 | 12.8202423 | -0.6797331 | 0.58227679 | -1.1673711 | 0.24306052 | NA |
| MCRIP2 | 50.4794461 | -0.2593088 | 0.3326832 | -0.7794466 | 0.43571666 | 0.86729771 |
| RHOT2 | 40.115664 | -0.2889977 | 0.29146127 | -0.9915476 | 0.32141828 | 0.82490712 |
| STUB1 | 35.2594335 | 0.21754235 | 0.40687778 | 0.53466265 | 0.59288316 | 0.92278087 |
| JMJD8 | 17.5962907 | 0.36620118 | 0.49861755 | 0.73443299 | 0.46268488 | 0.87713916 |
| METRN | 23.9835554 | 1.16642347 | 0.43234935 | 2.69787263 | 0.00697841 | 0.3013833 |
| FAM173A | 24.5757204 | -0.1188279 | 0.53158621 | -0.2235347 | 0.82311941 | 0.97518529 |
| NARFL | 15.6844959 | -0.111836 | 0.38079043 | -0.2936942 | 0.76899155 | NA |
| LMF1 | 58.2698299 | -0.1537342 | 0.26678546 | -0.5762466 | 0.56444849 | 0.91658856 |
| CACNA1H | 18.247143 | -1.4600458 | 0.77388306 | -1.8866491 | 0.05920754 | 0.57442774 |
| UBE2I | 87.9663285 | -0.4684873 | 0.3000427 | -1.5614019 | 0.11842895 | 0.6668162 |
| BAIAP3 | 10.3410387 | -0.617122 | 0.51259728 | -1.203912 | 0.22862361 | NA |
| TSR3 | 21.2920732 | -0.315634 | 0.51266295 | -0.6156755 | 0.53810872 | 0.90763121 |

| | | | | | | |
|---|---|---|---|---|---|---|
| GNPTG | 38.2143936 | -0.3286601 | 0.33859874 | -0.9706478 | 0.33172369 | 0.82890589 |
| UNKL | 25.2352587 | -0.0481126 | 0.42109042 | -0.1142573 | 0.90903386 | 0.9854228 |
| CLCN7 | 43.2277811 | -0.5844811 | 0.32838487 | -1.7798663 | 0.07509785 | 0.61020315 |
| TELO2 | 11.6344707 | 0.27252901 | 0.56823886 | 0.4796029 | 0.63150978 | NA |
| IFT140 | 45.3937555 | 0.21311578 | 0.27878863 | 0.76443498 | 0.44460807 | 0.87079259 |
| JPT2 | 31.3131632 | -0.0243762 | 0.34162694 | -0.0713532 | 0.94311663 | 0.99058259 |
| MAPK8IP3 | 71.2298014 | -0.0419117 | 0.19572611 | -0.2141343 | 0.83044233 | 0.97567084 |
| NME3 | 64.0270161 | 0.63516309 | 0.40163251 | 1.58145337 | 0.11377441 | 0.6656824 |
| MRPS34 | 29.5534316 | 0.03565198 | 0.47889115 | 0.07444694 | 0.94065476 | 0.99026024 |
| SPSB3 | 39.0608691 | 0.47082786 | 0.28119878 | 1.67435957 | 0.09405995 | 0.64572229 |
| NUBP2 | 11.3620592 | 0.00444501 | 0.4872448 | 0.00912274 | 0.9927212 | NA |
| HAGH | 53.4646205 | 0.0462031 | 0.36284859 | 0.12733436 | 0.89867576 | 0.98531749 |
| FAHD1 | 19.6729534 | 0.40111634 | 0.40559914 | 0.98894771 | 0.32268872 | 0.82558302 |
| NDUFB10 | 42.5938521 | 0.52276999 | 0.46335384 | 1.12823062 | 0.25922254 | 0.79532468 |
| RPS2 | 145.218784 | 0.03000839 | 0.42575414 | 0.07048291 | 0.9438093 | 0.99069421 |
| TSC2 | 55.908746 | -0.4219938 | 0.33243268 | -1.2694112 | 0.20429443 | 0.76173533 |
| PKD1 | 33.0924283 | -0.1728393 | 0.32470629 | -0.5322943 | 0.59452215 | 0.9229638 |
| SNHG19 | 17.4167252 | -0.4050434 | 0.39046689 | -1.037331 | 0.29958163 | 0.81357148 |
| TRAF7 | 16.3521437 | -0.097616 | 0.4248506 | -0.2297655 | 0.81827397 | 0.97288403 |
| PGP | 43.84617 | 0.49668118 | 0.38557588 | 1.28815418 | 0.1976923 | 0.75559912 |
| E4F1 | 17.2713459 | -0.1144259 | 0.37492338 | -0.3051982 | 0.7602152 | 0.95923591 |
| ECI1 | 96.1894664 | 0.19021443 | 0.20427933 | 0.93114868 | 0.35177666 | 0.83871737 |
| RNPS1 | 51.2945509 | 0.07711927 | 0.24625937 | 0.31316278 | 0.75415699 | 0.95826617 |
| ABCA3 | 22.7344352 | 0.56806127 | 0.34747646 | 1.63481945 | 0.1020869 | 0.65946373 |
| AMDHD2 | 11.2675951 | -1.0925499 | 0.44036523 | -2.4810086 | 0.01310112 | NA |
| PDPK1 | 103.871652 | 0.16109046 | 0.26128119 | 0.61654058 | 0.53753779 | 0.90763121 |
| AC093525.4 | 12.5469028 | -0.2863608 | 0.45201606 | -0.6335191 | 0.5263947 | NA |
| KCTD5 | 12.0278391 | -0.2567618 | 0.51382201 | -0.4997097 | 0.61727953 | NA |
| SRRM2 | 288.070936 | -0.223283 | 0.23845162 | -0.936387 | 0.34907398 | 0.83687845 |
| ELOB | 60.6524955 | -0.0812127 | 0.40173571 | -0.2021546 | 0.83979588 | 0.97600352 |
| PRSS22 | 22.1857896 | -0.1527573 | 0.45101775 | -0.3386946 | 0.73483981 | 0.95329896 |
| FLYWCH2 | 18.412197 | -0.1498868 | 0.43820356 | -0.3420484 | 0.73231444 | 0.95279138 |
| FLYWCH1 | 39.669732 | 0.10133456 | 0.30558804 | 0.33160511 | 0.74018746 | 0.9544376 |
| TNFRSF12A | 78.5108784 | -1.2841975 | 0.69299063 | -1.853124 | 0.06386459 | 0.58781448 |
| HCFC1R1 | 31.3191169 | -0.0589838 | 0.51501631 | -0.114528 | 0.90881923 | 0.9854228 |
| BICDL2 | 17.0478504 | 0.38995316 | 0.48296861 | 0.80740891 | 0.41943094 | 0.86065558 |
| IL32 | 16.1983826 | 0.75206854 | 0.42812602 | 1.75665227 | 0.0789771 | 0.61367304 |
| ZNF213-AS1 | 15.5727491 | -0.0430618 | 0.55329294 | -0.0778282 | 0.93796472 | NA |
| ZNF213 | 33.0510456 | 0.54076328 | 0.44201388 | 1.22340792 | 0.22117567 | 0.76680432 |
| ZNF263 | 21.1045588 | -0.4339492 | 0.40646579 | -1.0676155 | 0.28569397 | 0.80431986 |
| TIGD7 | 17.7674975 | -0.4529737 | 0.3633881 | -1.2465286 | 0.21257039 | 0.7643572 |
| ZNF75A | 55.4393923 | -0.5117915 | 0.30845295 | -1.6592208 | 0.09707131 | 0.65442315 |
| AC025283.3 | 22.7108589 | -0.6317333 | 0.34737201 | -1.8186074 | 0.06897135 | 0.60275677 |
| ZSCAN32 | 10.4019801 | 0.6073912 | 0.51750772 | 1.1736853 | 0.2405211 | NA |
| NAA60 | 27.0399987 | -0.176088 | 0.43481944 | -0.4049682 | 0.6855009 | 0.94205398 |
| CLUAP1 | 49.513225 | -0.0918408 | 0.32249549 | -0.2847817 | 0.77581136 | 0.96469469 |
| DNASE1 | 84.4661328 | 0.39571583 | 0.37278814 | 1.06150326 | 0.28846125 | 0.80523851 |
| TRAP1 | 52.5096649 | 0.05859876 | 0.27247376 | 0.21506203 | 0.82971896 | 0.97567084 |
| CREBBP | 210.858594 | -0.1221541 | 0.24657679 | -0.4953999 | 0.6203179 | 0.92798856 |
| ADCY9 | 98.9617564 | 0.0415638 | 0.26548327 | 0.15655902 | 0.8755924 | 0.97918624 |
| GLIS2 | 43.5973358 | -0.0993867 | 0.3435943 | -0.2892559 | 0.77238558 | 0.9640787 |
| VASN | 58.5625559 | 0.45924202 | 0.3394602 | 1.35285969 | 0.17610046 | 0.72841249 |
| DNAJA3 | 38.2024769 | 0.06900383 | 0.30134046 | 0.22898959 | 0.81887701 | 0.97288403 |
| NMRAL1 | 29.4476224 | 0.15424826 | 0.27401091 | 0.56292744 | 0.57348429 | 0.91813957 |

| | | | | | | |
|---|---|---|---|---|---|---|
| HMOX2 | 49.6105997 | -0.0843182 | 0.28778761 | -0.2929877 | 0.76953155 | 0.96289288 |
| CDIP1 | 26.3598678 | 0.31871722 | 0.30633963 | 1.04040478 | 0.29815188 | 0.81214575 |
| UBALD1 | 11.6262198 | -0.0048763 | 0.45558981 | -0.0107033 | 0.99146014 | NA |
| MGRN1 | 64.3143916 | -0.1332142 | 0.21551343 | -0.618125 | 0.53649294 | 0.90763121 |
| NUDT16L1 | 16.0232334 | 1.04842054 | 0.52822173 | 1.98481146 | 0.04716545 | NA |
| ANKS3 | 42.4726371 | -0.4677557 | 0.260573 | -1.7951043 | 0.07263708 | 0.60829377 |
| ZNF500 | 16.558892 | 0.37333276 | 0.41683511 | 0.89563657 | 0.3704469 | 0.84852785 |
| ROGDI | 11.7918372 | 1.03709129 | 0.68504021 | 1.51391302 | 0.1300479 | NA |
| GLYR1 | 113.976903 | -0.0403112 | 0.21140649 | -0.190681 | 0.84877556 | 0.97612951 |
| UBN1 | 42.4571403 | -0.0545783 | 0.24969032 | -0.218584 | 0.8269741 | 0.97567084 |
| PPL | 44.0691507 | -0.3853974 | 0.25439324 | -1.5149673 | 0.12978068 | 0.67277537 |
| ALG1 | 16.5191255 | 0.48666266 | 0.40034449 | 1.21560974 | 0.22413362 | 0.76875975 |
| RBFOX1 | 20.7082543 | -0.2116775 | 0.43586065 | -0.4856541 | 0.62721247 | 0.93057946 |
| METTL22 | 16.7264719 | 0.05674937 | 0.44471066 | 0.12760965 | 0.89845789 | 0.98531749 |
| ABAT | 20.0704436 | 0.62070026 | 0.37102272 | 1.67294408 | 0.09433831 | 0.64665783 |
| PMM2 | 72.7500507 | -0.979113 | 0.29391004 | -3.3313358 | 0.0008643 | 0.13335165 |
| CARHSP1 | 41.8228933 | -0.0753956 | 0.32683152 | -0.2306864 | 8.18E-01 | 0.97288403 |
| USP7 | 68.7566308 | 0.2154541 | 0.20586443 | 1.04658245 | 0.2952922 | 0.81038435 |
| C16orf72 | 71.5417808 | -0.2018757 | 0.23541232 | -0.8575408 | 0.39114606 | 0.85605713 |
| ATF7IP2 | 81.0628319 | 0.26280019 | 0.30944148 | 0.84927267 | 0.39572958 | 0.85605713 |
| NUBP1 | 14.4393104 | -0.2472991 | 0.40982824 | -0.6034213 | 0.54622845 | NA |
| CIITA | 32.2474155 | 0.83270613 | 0.57598653 | 1.44570418 | 0.14826019 | 0.69945736 |
| DEXI | 23.0377555 | 0.15115724 | 0.34801181 | 0.43434515 | 0.66403783 | 0.93860542 |
| CLEC16A | 267.759406 | 0.18161963 | 0.2891451 | 0.62812624 | 0.52992124 | 0.9052447 |
| AC007014.1 | 15.8458045 | 0.32898728 | 0.68684095 | 0.47898612 | 0.6319485 | NA |
| AC007220.1 | 17.5565953 | 0.42998868 | 0.45146549 | 0.95242868 | 0.34087962 | 0.8319325 |
| SOCS1 | 59.9883319 | -0.3738087 | 0.48887417 | -0.7646317 | 4.44E-01 | 8.71E-01 |
| AC099489.1 | 13.8798475 | -0.7071438 | 0.52861957 | -1.3377178 | 0.18098843 | NA |
| LITAF | 128.502338 | 0.07444091 | 0.19489635 | 0.3819513 | 7.02E-01 | 0.94476609 |
| TXNDC11 | 99.3279935 | -0.1692049 | 0.19508052 | -0.8673594 | 0.38574512 | 0.85540507 |
| ZC3H7A | 97.3931731 | -0.0360677 | 0.22704069 | -0.1588602 | 0.87377905 | 0.97918624 |
| RSL1D1 | 22.9204351 | 0.02217782 | 0.32138656 | 0.06900669 | 0.94498429 | 0.99069787 |
| AC007216.4 | 43.9042423 | 0.41929619 | 0.39402073 | 1.06414752 | 0.28726187 | 0.80523851 |
| GSPT1 | 172.102806 | -0.1583911 | 0.2466202 | -0.642247 | 0.52071279 | 0.90441454 |
| SNX29 | 1001.74751 | -0.1450866 | 0.20216032 | -0.7176809 | 0.47295406 | 0.88344299 |
| CPPED1 | 131.861733 | -1.2667705 | 0.3358073 | -3.7723137 | 0.00016174 | 0.05491436 |
| SHISA9 | 61.6980483 | -2.0696971 | 0.48978018 | -4.2257675 | 2.38E-05 | 0.01667444 |
| U91319.1 | 69.0830613 | -2.0977967 | 0.63900137 | -3.2829299 | 0.00102734 | 0.13958055 |
| ERCC4 | 22.2673864 | 0.70626496 | 0.32711727 | 2.15905739 | 0.03084572 | 0.47270799 |
| MKL2 | 260.779683 | -0.2220744 | 0.19356286 | -1.1472984 | 0.25125832 | 0.78896325 |
| PARN | 114.854614 | -0.1700944 | 0.19458593 | -0.8741353 | 0.38204457 | 0.8548566 |
| BFAR | 33.5977105 | -0.0461405 | 0.28132626 | -0.1640106 | 0.86972281 | 0.97854266 |
| NOMO1 | 15.8344661 | 0.23696542 | 0.49469082 | 0.47901721 | 0.63192639 | NA |
| PDXDC1 | 236.606376 | -0.3446433 | 0.23629375 | -1.4585375 | 0.14469245 | 0.69512617 |
| NTAN1 | 12.777131 | -0.4534097 | 0.39246693 | -1.1552813 | 0.24797524 | NA |
| RRN3 | 32.6381472 | -0.2326535 | 0.30363429 | -0.7662294 | 0.44353979 | 0.87034764 |
| C16orf45 | 76.2651407 | -0.1568739 | 0.27911071 | -0.562049 | 0.57408266 | 0.91813957 |
| MARF1 | 50.771609 | -0.3967488 | 0.44387419 | -0.8938316 | 0.371412 | 0.8491102 |
| NDE1 | 25.2440273 | 0.24865364 | 0.39767902 | 0.62526216 | 0.53179901 | 0.906039 |
| FOPNL | 20.1843917 | 0.46959429 | 0.36372195 | 1.29108044 | 0.19667578 | 0.75350056 |
| ABCC1 | 75.9518043 | -0.3002115 | 0.29988097 | -1.0011023 | 0.31677735 | 0.82275575 |
| XYLT1 | 19.1066973 | -0.6228965 | 0.49335372 | -1.2625759 | 0.20674164 | 0.76231289 |
| RPS15A | 24.2922911 | 0.27480918 | 0.58891952 | 0.46663284 | 0.64076259 | 0.93328692 |
| SMG1 | 216.418747 | -0.4179553 | 0.20550244 | -2.0338215 | 0.0419696 | 0.51213038 |

| | | | | | | |
|---|---|---|---|---|---|---|
| COQ7 | 20.0615707 | 0.19230145 | 0.37433881 | 0.51370964 | 0.60745501 | 0.92569157 |
| GDE1 | 25.8921578 | 0.08388743 | 0.33908104 | 0.2473964 | 0.80460146 | 0.97195485 |
| CCP110 | 28.8749979 | 0.03097461 | 0.39187995 | 0.07904107 | 0.93699996 | 0.99012269 |
| VPS35L | 114.627729 | -0.1193321 | 0.21433661 | -0.556751 | 0.57769757 | 0.91872484 |
| KNOP1 | 24.4292557 | -0.4064063 | 0.48668864 | -0.8350437 | 0.40369312 | 0.85782098 |
| IQCK | 173.060028 | -0.0462777 | 0.21214969 | -0.2181368 | 0.82732249 | 0.97567084 |
| GPRC5B | 118.98469 | -0.0998858 | 0.31783162 | -0.3142726 | 0.75331397 | 0.95826617 |
| UMOD | 63.0895026 | 0.15351599 | 0.55689614 | 0.27566359 | 0.78280646 | 0.96661519 |
| ACSM2B | 12.5247033 | 0.71291741 | 0.93411075 | 0.76320437 | 0.44534152 | NA |
| ACSM3 | 51.2499397 | 1.17275162 | 0.43475183 | 2.6975197 | 0.00698582 | 0.3013833 |
| THUMPD1 | 50.5497184 | 0.63219431 | 0.31297854 | 2.01992863 | 0.04339079 | 0.51699786 |
| ERI2 | 23.0597765 | 0.66981921 | 0.49221331 | 1.36083116 | 0.17356705 | 0.72794035 |
| LYRM1 | 24.6353509 | 0.17532144 | 0.36610697 | 0.47888036 | 0.63202374 | 0.93251095 |
| TMEM159 | 16.9420689 | 0.34833645 | 0.46075345 | 0.75601485 | 0.44964029 | 0.87328261 |
| METTL9 | 144.916486 | -0.4221218 | 0.21514036 | -1.9620761 | 0.04975362 | 0.54357315 |
| NPIPB4 | 30.8907156 | -1.1107016 | 0.40504936 | -2.7421388 | 0.00610405 | 0.29709153 |
| UQCRC2 | 134.481672 | 0.00640624 | 0.22285823 | 0.0287458 | 0.97706733 | 0.99454813 |
| MOSMO | 83.4742881 | -0.0649543 | 0.24835302 | -0.2615403 | 0.79367585 | 0.96960502 |
| EEF2K | 25.4637656 | 0.47039665 | 0.37362742 | 1.25899925 | 0.2080306 | 0.76231289 |
| POLR3E | 37.8090868 | 0.1083188 | 0.37228151 | 0.2909594 | 0.77108237 | 0.96364125 |
| CDR2 | 201.311703 | -0.4317539 | 0.26703699 | -1.616832 | 0.10591456 | 0.65946648 |
| NPIPB5 | 82.2736766 | -0.1540173 | 0.30800928 | -0.5000412 | 0.61704605 | 0.92711833 |
| USP31 | 40.7554629 | 0.13761385 | 0.35733482 | 0.3851118 | 0.70015459 | 0.94424267 |
| SCNN1G | 365.584483 | 0.0238874 | 0.38972403 | 0.06129312 | 9.51E-01 | 9.91E-01 |
| AC099482.1 | 35.1803331 | -0.2088051 | 0.71176123 | -0.2933639 | 0.76924399 | 0.96289288 |
| SCNN1B | 185.855852 | 0.23249362 | 0.46618005 | 0.49872065 | 6.18E-01 | 0.92749951 |
| COG7 | 94.3635826 | -0.2068998 | 0.18401084 | -1.1243892 | 0.26084795 | 0.79574359 |
| GGA2 | 110.756809 | -0.2484584 | 0.23703264 | -1.0482035 | 0.29454488 | 0.81012446 |
| EARS2 | 16.6256243 | -0.5089332 | 0.39179945 | -1.2989636 | 0.19395642 | 0.75067401 |
| UBFD1 | 20.1807728 | -0.5918419 | 0.51746236 | -1.143739 | 0.2527319 | 0.79059056 |
| NDUFAB1 | 38.9980814 | -0.168725 | 0.31423276 | -0.5369427 | 0.59130722 | 0.92254886 |
| PALB2 | 20.5779914 | -0.5128477 | 0.42787853 | -1.1985824 | 0.23069036 | 0.77249016 |
| DCTN5 | 49.3047097 | -0.0519623 | 0.26768022 | -0.1941206 | 0.84608142 | 0.97612951 |
| RBBP6 | 88.3042953 | 0.14903381 | 0.24297063 | 0.61338202 | 0.53962377 | 0.90763121 |
| TNRC6A | 468.122968 | -0.3478189 | 0.23168045 | -1.5012872 | 0.13328131 | 0.67988987 |
| AC008731.1 | 12.7866853 | 0.50954603 | 0.58303806 | 0.87394985 | 0.38214557 | NA |
| ARHGAP17 | 93.7372127 | -0.1296277 | 0.26896889 | -0.4819431 | 0.62984638 | 0.9318063 |
| LCMT1 | 57.5166739 | -0.0363254 | 0.30752029 | -0.1181235 | 0.90596979 | 0.98531749 |
| AC008741.2 | 11.2936422 | 0.03269278 | 0.46846036 | 0.06978772 | 0.94436262 | NA |
| NSMCE1 | 28.2691812 | 0.06127733 | 0.33268527 | 0.1841901 | 0.85386433 | 0.97612951 |
| IL4R | 28.1883401 | 0.69050999 | 0.76174228 | 0.90648768 | 3.65E-01 | 0.8468666 |
| GTF3C1 | 57.2305489 | -0.3644861 | 0.28829949 | -1.2642621 | 0.20613596 | 0.76231289 |
| KIAA0556 | 87.7443014 | -0.3909471 | 0.30713137 | -1.2728985 | 0.20305407 | 0.76169832 |
| XPO6 | 99.6463476 | -0.5726451 | 0.24210397 | -2.3652858 | 0.01801616 | 0.41103033 |
| NUPR1 | 44.5111112 | -1.5850693 | 0.89485372 | -1.7713166 | 0.07650806 | 0.61119868 |
| SGF29 | 16.1727889 | -0.4896059 | 0.39631016 | -1.2354109 | 0.21667762 | 0.76658236 |
| SULT1A1 | 29.5480891 | 0.455842 | 0.45569136 | 1.00033056 | 0.31715056 | 0.82275575 |
| ATXN2L | 48.5441723 | -0.3235607 | 0.31519746 | -1.0265333 | 0.30464028 | 0.81568231 |
| TUFM | 74.7348237 | 0.29416462 | 0.43089744 | 0.68267897 | 0.49480972 | 0.89230917 |
| SH2B1 | 52.1487776 | 0.29879423 | 0.27919987 | 1.07018041 | 0.28453811 | 0.80431986 |
| RABEP2 | 57.7529082 | -0.3942509 | 0.35561492 | -1.1086457 | 0.26758307 | 0.79666376 |
| NFATC2IP | 91.8059429 | -0.0705448 | 0.19944481 | -0.353706 | 0.72355925 | 0.95093722 |
| SPNS1 | 18.2446414 | -0.1432893 | 0.40647838 | -0.352514 | 0.72445284 | 0.95093722 |
| QPRT | 29.8172294 | 0.59981176 | 0.53390219 | 1.12344876 | 0.26124695 | 0.79589391 |

| | | | | | | |
|---|---|---|---|---|---|---|
| KIF22 | 13.6580728 | 0.03623449 | 0.37430424 | 0.09680491 | 0.92288132 | NA |
| MAZ | 20.3984503 | 0.20634891 | 0.42798817 | 0.48213695 | 0.62970866 | 0.9318063 |
| AC009133.1 | 16.61536 | -0.5910117 | 0.46098118 | -1.2820734 | 2.00E-01 | 0.75605362 |
| MVP | 84.7084806 | -0.0059557 | 0.28661266 | -0.0207796 | 9.83E-01 | 9.97E-01 |
| CDIPT | 30.2790318 | 0.03039179 | 0.31903351 | 0.09526205 | 0.92410668 | 0.98925753 |
| KCTD13 | 32.0579535 | -0.4501721 | 0.30680723 | -1.46728 | 0.14229989 | 0.69328517 |
| TMEM219 | 53.1159391 | -0.2165122 | 0.4425931 | -0.4891903 | 0.624707 | 0.92981013 |
| TAOK2 | 17.2188834 | 0.03001894 | 0.55783248 | 0.05381354 | 0.95708372 | 0.99170927 |
| INO80E | 21.4638798 | 0.20024326 | 0.3822866 | 0.52380404 | 0.60041484 | 0.92352528 |
| PPP4C | 35.5246801 | 0.10819153 | 0.39552711 | 0.27353758 | 0.78444001 | 0.96732016 |
| YPEL3 | 36.6879199 | 0.53133681 | 0.29130669 | 1.82397735 | 0.0681555 | 0.60244724 |
| MAPK3 | 12.1909119 | -0.1022796 | 0.49030304 | -0.2086048 | 0.83475678 | NA |
| NPIPB13 | 20.3946876 | -0.8434242 | 0.49253413 | -1.7124177 | 0.08681971 | 0.63466198 |
| CD2BP2 | 20.7243684 | -0.0295674 | 0.44424611 | -0.0665564 | 0.94693487 | 0.99083449 |
| TBC1D10B | 14.0840753 | -0.3100807 | 0.51501711 | -0.6020785 | 0.54712188 | NA |
| DCTPP1 | 13.5299916 | 0.76133784 | 0.53482086 | 1.42353803 | 0.15458024 | NA |
| SEPHS2 | 47.9194947 | 0.2103529 | 0.48042902 | 0.43784386 | 0.66149947 | 0.93729379 |
| ZNF768 | 19.247647 | -0.5836758 | 0.37972784 | -1.5370898 | 0.12427133 | 0.66929384 |
| ZNF785 | 23.5734282 | -0.1605165 | 0.32413246 | -0.4952187 | 0.62044573 | 0.92798856 |
| PRR14 | 13.2748817 | -0.0855147 | 0.44865512 | -0.1906023 | 0.84883721 | NA |
| FBRS | 26.7978944 | -0.0448182 | 0.31012472 | -0.1445167 | 0.88509244 | 0.98184212 |
| PHKG2 | 16.3094209 | 0.17768011 | 0.42093075 | 0.42211244 | 0.67294295 | 0.93880075 |
| RNF40 | 34.2280986 | -0.3193395 | 0.29191356 | -1.0939523 | 0.2739759 | 0.79808085 |
| ZNF629 | 13.4698961 | -0.1881104 | 0.48278001 | -0.3896399 | 0.69680281 | NA |
| BCL7C | 67.4843715 | 0.02871943 | 0.26243125 | 0.10943603 | 0.91285666 | 0.98639439 |
| MIR762HG | 14.1104586 | 0.06710573 | 0.41817817 | 0.16047163 | 0.87250957 | NA |
| FBXL19 | 20.3326587 | -0.3407224 | 0.49302445 | -0.6910863 | 0.48951132 | 0.88930178 |
| SETD1A | 14.2123392 | -0.4777239 | 0.40466903 | -1.1805299 | 0.23778951 | NA |
| STX4 | 17.9659911 | -0.2953052 | 0.34672285 | -0.8517038 | 0.39437852 | 0.85605713 |
| BCKDK | 21.746183 | -0.2981076 | 0.50435843 | -0.591063 | 0.55447824 | 0.9137961 |
| KAT8 | 31.958726 | -0.4202578 | 0.2932147 | -1.4332766 | 0.15177881 | 0.70572801 |
| PRSS8 | 42.9380632 | -0.8684544 | 0.44868687 | -1.9355468 | 0.05292324 | 0.55694824 |
| FUS | 145.247901 | 0.13863338 | 0.21271242 | 0.6517409 | 0.51456833 | 0.90156229 |
| AC026471.4 | 19.4591928 | 0.10445536 | 0.47441386 | 0.22017772 | 0.82573275 | 0.97567084 |
| C16orf58 | 26.1208341 | -0.2268701 | 0.39199217 | -0.5787618 | 0.56274988 | 0.91624774 |
| ZNF720 | 53.8295292 | 0.54710079 | 0.26739334 | 2.04605239 | 0.04075122 | 0.50616193 |
| ZNF267 | 12.5559062 | 0.12967732 | 0.4133644 | 0.31371187 | 0.75373989 | NA |
| SHCBP1 | 20.8178678 | 0.52239312 | 0.4621182 | 1.13043183 | 2.58E-01 | 7.95E-01 |
| VPS35 | 62.9476501 | -0.0481006 | 0.27847566 | -0.1727283 | 0.86286497 | 0.97788853 |
| C16orf87 | 16.9354845 | 0.7285554 | 0.40925324 | 1.78020681 | 0.07504212 | 0.61020315 |
| GPT2 | 15.8867623 | -0.919769 | 0.39187176 | -2.3471174 | 0.01891929 | NA |
| DNAJA2 | 27.9326251 | 0.19598255 | 0.31111123 | 0.62994365 | 0.52873145 | 0.90487731 |
| ITFG1 | 388.474303 | -0.2516493 | 0.16528241 | -1.522541 | 0.12787358 | 0.66929384 |
| PHKB | 262.958439 | 0.03292569 | 0.15098584 | 0.21807135 | 0.82737352 | 0.97567084 |
| LONP2 | 175.948342 | 0.17206333 | 0.17597017 | 0.97779825 | 0.32817412 | 0.82740664 |
| SIAH1 | 35.9170325 | -0.0912941 | 0.33875918 | -0.2694956 | 0.78754829 | 0.96767512 |
| N4BP1 | 88.5989545 | -0.2009023 | 0.32327471 | -0.6214601 | 0.53429691 | 0.90763121 |
| CNEP1R1 | 13.7465754 | 0.02912601 | 0.42900129 | 0.06789259 | 0.94587113 | NA |
| HEATR3 | 19.0255335 | 0.1413047 | 0.35290125 | 0.40040861 | 0.68885558 | 0.94205398 |
| TENT4B | 86.0222116 | -0.0003069 | 0.27954226 | -0.001098 | 0.99912395 | 0.99957534 |
| BRD7 | 57.1227378 | -0.1863131 | 0.28523319 | -0.6531958 | 0.51363005 | 0.90156229 |
| CYLD | 42.8789785 | -0.1877951 | 0.29490996 | -0.6367881 | 0.52426288 | 0.90487731 |
| SALL1 | 26.7367625 | -0.3503036 | 0.29052841 | -1.2057464 | 0.22791528 | 0.77185154 |
| AC087564.1 | 90.4409645 | -0.078987 | 0.36949147 | -0.2137722 | 0.8307247 | 0.97567084 |

| | | | | | | |
|---|---|---|---|---|---|---|
| TOX3 | 773.937192 | -0.3586043 | 0.41917694 | -0.8554963 | 0.39227644 | 0.85605713 |
| AC007906.2 | 11.4524923 | 2.00162922 | 0.57588439 | 3.47574837 | 0.00050943 | NA |
| CHD9 | 553.310804 | 0.01929578 | 0.22685239 | 0.08505873 | 0.9322147 | 0.98960607 |
| RBL2 | 107.792419 | -0.0515982 | 0.24495961 | -0.2106396 | 0.8331685 | 0.97600352 |
| AKTIP | 22.601537 | -0.0362993 | 0.36291242 | -0.1000223 | 0.92032661 | 0.98794023 |
| RPGRIP1L | 22.7952922 | -0.2543249 | 0.44787797 | -0.5678441 | 0.57014085 | 0.91778542 |
| FTO | 301.729189 | 0.06149236 | 0.15255001 | 0.40309641 | 0.6868773 | 0.94205398 |
| LPCAT2 | 85.6249849 | -0.1611169 | 0.34419962 | -0.4680913 | 0.63971928 | 0.93328692 |
| AMFR | 124.012513 | 0.1457933 | 0.36380175 | 0.4007493 | 0.68860471 | 0.94205398 |
| NUDT21 | 71.3813809 | 0.08969207 | 0.25340505 | 0.35394745 | 0.72337826 | 0.95093722 |
| OGFOD1 | 25.5718764 | 0.11431103 | 0.32307223 | 0.35382501 | 0.72347003 | 0.95093722 |
| BBS2 | 56.0234735 | 0.00744851 | 0.24919493 | 0.0298903 | 0.97615454 | 0.99440179 |
| MT2A | 234.385598 | -1.4816426 | 0.60254825 | -2.458961 | 0.01393398 | 0.37638271 |
| MT1E | 92.6283809 | -1.4824128 | 0.70545523 | -2.1013563 | 0.0356097 | 0.49240174 |
| MT1F | 19.9110896 | -0.5023762 | 0.66814017 | -0.7519024 | 0.45210974 | 0.87346041 |
| MT1G | 150.930429 | -1.8341254 | 0.75433965 | -2.4314318 | 0.01503928 | 0.3922709 |
| MT1H | 45.2062286 | -2.0544084 | 1.17776924 | -1.7443217 | 0.08110302 | 0.61925304 |
| MT1X | 48.0171658 | -0.7443757 | 0.68798863 | -1.0819593 | 0.27927061 | 0.80219184 |
| NUP93 | 40.0781042 | 0.05786186 | 0.30363606 | 0.19056322 | 0.84886781 | 0.97612951 |
| SLC12A3 | 256.297294 | 0.5610777 | 0.68958939 | 0.81364027 | 4.16E-01 | 8.59E-01 |
| HERPUD1 | 60.589277 | -0.695487 | 0.32713092 | -2.1260205 | 0.03350155 | 0.4908133 |
| NLRC5 | 24.3732611 | -2.2616159 | 0.54691965 | -4.1351886 | 3.55E-05 | 0.02152333 |
| FAM192A | 56.3360175 | -0.3633204 | 0.22086203 | -1.6450106 | 0.09996763 | 0.65863153 |
| RSPRY1 | 78.1424542 | -0.2469244 | 0.25816965 | -0.9564424 | 0.33884878 | 0.83066816 |
| ARL2BP | 40.2746134 | -0.1323021 | 0.31445717 | -0.4207317 | 0.67395102 | 0.93901638 |
| PLLP | 61.7682247 | 1.35906004 | 0.5686774 | 2.38986118 | 0.01685474 | 0.39748373 |
| CX3CL1 | 22.645322 | 0.11447213 | 0.48294774 | 0.23702797 | 0.81263509 | 0.97257655 |
| CIAPIN1 | 40.9141814 | -0.2383639 | 0.29282021 | -0.8140283 | 0.41562876 | 0.85867915 |
| COQ9 | 77.0610835 | -0.0030453 | 0.23282747 | -0.0130797 | 0.98956424 | 0.99814944 |
| POLR2C | 20.0774002 | -0.1385745 | 0.46038663 | -0.300996 | 0.76341758 | 0.96091323 |
| DOK4 | 23.615816 | 0.53037824 | 0.51022229 | 1.03950425 | 0.29857028 | 0.81246117 |
| ADGRG1 | 231.219729 | 0.08904424 | 0.30985699 | 0.28737205 | 0.77382746 | 0.96435639 |
| KIFC3 | 189.33426 | 0.00498485 | 0.22741155 | 0.02191996 | 9.83E-01 | 0.99641321 |
| USB1 | 14.7197695 | 0.09898984 | 0.40114533 | 0.24676802 | 0.80508776 | NA |
| MMP15 | 12.816733 | 0.81408745 | 0.61276244 | 1.32855312 | 0.18399545 | NA |
| CSNK2A2 | 53.1040264 | -0.3698266 | 0.23694055 | -1.5608412 | 0.11856122 | 0.6668162 |
| SETD6 | 11.4950421 | -0.257517 | 0.45394176 | -0.5672908 | 0.57051662 | NA |
| CNOT1 | 169.234496 | -0.0745542 | 0.2132414 | -0.3496234 | 0.72662137 | 0.95209193 |
| SLC38A7 | 22.4142917 | -0.6438705 | 0.4769425 | -1.3499961 | 0.17701725 | 0.72841249 |
| GOT2 | 76.4139343 | 0.48181965 | 0.27141799 | 1.77519423 | 0.07586582 | 0.61020315 |
| AC092378.1 | 22.0426968 | 2.11113773 | 0.5334824 | 3.9572772 | 7.58E-05 | 0.03632046 |
| AC138305.1 | 695.657996 | 0.18744224 | 0.30962465 | 0.6053854 | 5.45E-01 | 9.10E-01 |
| AC010542.3 | 21.4069254 | -0.002991 | 0.49785769 | -0.0060077 | 0.99520661 | 0.99927028 |
| TK2 | 20.7614391 | 0.07941375 | 0.34660032 | 0.22912197 | 0.81877412 | 0.97288403 |
| CMTM4 | 236.635502 | 0.19851642 | 0.23779987 | 0.83480455 | 4.04E-01 | 0.85782098 |
| DYNC1LI2 | 116.856977 | -0.2066044 | 0.27505363 | -0.7511423 | 0.45256702 | 0.87346041 |
| NAE1 | 42.0139776 | -0.1455166 | 0.32810724 | -0.4435031 | 0.65740188 | 0.93593674 |
| PDP2 | 25.9728034 | -0.1477551 | 0.49111962 | -0.3008535 | 0.76352619 | 0.96091323 |
| CDH16 | 545.065085 | 0.40200056 | 0.38265086 | 1.0505675 | 0.29345727 | 0.80935782 |
| RRAD | 208.084361 | 0.04025107 | 0.72897339 | 0.0552161 | 0.9559663 | 0.99148072 |
| FAM96B | 42.8293466 | -0.2177745 | 0.35128707 | -0.6199331 | 0.53530184 | 0.90763121 |
| CES2 | 33.5997921 | 0.21825943 | 0.35417173 | 0.61625311 | 0.53772748 | 0.90763121 |
| CBFB | 164.703201 | 0.04668791 | 0.27610796 | 0.16909294 | 0.86572354 | 0.97788853 |
| C16orf70 | 23.5978189 | -0.3118403 | 0.41956993 | -0.743238 | 0.45733757 | 0.87420547 |

| | | | | | | |
|---|---|---|---|---|---|---|
| TRADD | 15.243634 | 0.07267392 | 0.45105841 | 0.16111864 | 0.87199996 | NA |
| NOL3 | 26.4198358 | 0.19218439 | 0.4902747 | 0.39199329 | 0.69506317 | 0.94395238 |
| KIAA0895L | 20.2367541 | -0.601321 | 0.34740619 | -1.7308874 | 0.08347186 | 0.62487197 |
| E2F4 | 19.3569212 | 0.07050259 | 0.35923827 | 0.19625578 | 0.84440997 | 0.97612951 |
| ELMO3 | 14.1135004 | 0.06482709 | 0.39409631 | 0.16449555 | 0.86934105 | NA |
| LRRC29 | 14.5163137 | -0.2384014 | 0.41572107 | -0.5734649 | 0.56632998 | NA |
| TMEM208 | 26.7833339 | 0.56456446 | 0.3646961 | 1.54804083 | 0.12161247 | 0.66809793 |
| ZDHHC1 | 31.140216 | 0.30802936 | 0.30459352 | 1.0112801 | 0.31188239 | 0.81701824 |
| HSD11B2 | 724.825019 | 1.26633522 | 0.57289185 | 2.2104263 | 2.71E-02 | 0.45111036 |
| ATP6V0D1 | 68.4918433 | 0.13333059 | 0.30693926 | 0.43438753 | 0.66400705 | 0.93860542 |
| AC009061.2 | 46.1947681 | 0.15320514 | 0.40338043 | 0.37980311 | 0.70409158 | 0.94476609 |
| AC027682.6 | 9.87796682 | 0.90327252 | 0.68197205 | 1.32450079 | 0.18533678 | NA |
| RIPOR1 | 45.9177952 | 0.10335412 | 0.33726572 | 0.30644715 | 0.75926421 | 0.95903525 |
| AC027682.4 | 12.0768596 | 0.31687409 | 0.57168767 | 0.55427834 | 0.57938837 | NA |
| CTCF | 80.0228728 | -0.113536 | 0.31825707 | -0.3567432 | 0.72128409 | 0.95088329 |
| ENKD1 | 14.2452424 | 0.19060845 | 0.45909504 | 0.41518299 | 0.67800795 | NA |
| GFOD2 | 25.4999513 | -0.0559501 | 0.35038247 | -0.159683 | 0.87313081 | 0.97895181 |
| RANBP10 | 89.8906601 | -0.33388 | 0.27451066 | -1.2162733 | 0.22388083 | 0.76840496 |
| NUTF2 | 16.5194665 | -0.3937136 | 0.36554511 | -1.0770588 | 0.28145399 | 0.80416687 |
| PSKH1 | 25.4243817 | 0.39157869 | 0.3309126 | 1.18332967 | 0.23667852 | 0.77864547 |
| DUS2 | 18.5923753 | -0.1870181 | 0.42830379 | -0.4366483 | 0.6623664 | 0.93786301 |
| NFATC3 | 136.619082 | 0.06432576 | 0.17888993 | 0.35958289 | 0.71915908 | 0.94959812 |
| ESRP2 | 11.0926451 | 0.2466864 | 0.53364834 | 0.46226397 | 0.64389203 | NA |
| PLA2G15 | 23.1436655 | 0.27489608 | 0.37529099 | 0.73248783 | 0.46387087 | 0.87824803 |
| SLC7A6 | 57.2400583 | -0.0761215 | 0.34534035 | -0.2204246 | 0.82554052 | 0.97567084 |
| SLC7A6OS | 29.1102458 | -0.3040289 | 0.43458739 | -0.6995807 | 0.48418921 | 0.88838332 |
| PRMT7 | 37.2631874 | -0.2078526 | 0.38599982 | -0.5384785 | 0.59024676 | 0.92233545 |
| ZFP90 | 43.3820299 | -0.0828896 | 0.27687789 | -0.2993723 | 0.76465601 | 0.9616833 |
| CDH3 | 19.4326303 | 0.57016328 | 0.59362391 | 0.96047897 | 0.33681421 | 0.83044955 |
| CDH1 | 257.934294 | -0.1893874 | 0.27396055 | -0.6912946 | 0.48938043 | 0.88930178 |
| TANGO6 | 77.0562124 | 0.16285687 | 0.2447827 | 0.66531202 | 0.50585094 | 0.89723255 |
| CHTF8 | 33.4660385 | 0.32884854 | 0.40583769 | 0.81029571 | 0.41777024 | 0.85950165 |
| UTP4 | 23.2120453 | -0.2707117 | 0.33846154 | -0.7998299 | 0.42380935 | 0.86190547 |
| SNTB2 | 40.082923 | 0.06140936 | 0.4067911 | 0.15096044 | 0.88000692 | 0.98093344 |
| VPS4A | 24.9483287 | 0.04666928 | 0.35878583 | 0.13007559 | 0.89650662 | 0.98531749 |
| COG8 | 13.1464167 | 0.33029785 | 0.45397968 | 0.72756087 | 0.46688244 | NA |
| TERF2 | 36.1263437 | 0.03159626 | 0.26690905 | 0.11837837 | 0.90576787 | 0.98531749 |
| CYB5B | 120.151086 | -0.3980993 | 0.18050468 | -2.205479 | 0.0274205 | 0.45275729 |
| NFAT5 | 1080.79479 | -0.1349841 | 0.31035653 | -0.4349324 | 0.66361152 | 0.93860542 |
| NQO1 | 38.0859157 | 0.25289653 | 0.56303683 | 0.44916517 | 0.65331251 | 0.93532021 |
| WWP2 | 181.92459 | -0.3705196 | 0.23850322 | -1.5535204 | 0.12029885 | 0.6668162 |
| PDPR | 58.8934741 | -0.5792414 | 0.32634606 | -1.7749299 | 0.07590946 | 0.61020315 |
| AC009060.1 | 15.9028382 | 0.46309794 | 0.55453253 | 0.8351141 | 0.40365347 | NA |
| EXOSC6 | 24.0459309 | -0.1881791 | 0.3624099 | -0.5192438 | 0.60359076 | 0.92508698 |
| AARS | 45.2905681 | -0.8012776 | 0.4081337 | -1.9632723 | 0.04961454 | 0.54349113 |
| ST3GAL2 | 39.0922899 | 0.06179304 | 0.29827537 | 0.20716776 | 0.83587885 | 0.97600352 |
| FUK | 21.6810312 | 0.11484974 | 0.40334514 | 0.2847431 | 0.77584097 | 0.96469469 |
| COG4 | 41.4927081 | -0.0125788 | 0.27311533 | -0.0460567 | 0.96326508 | 0.99256424 |
| SF3B3 | 72.915129 | -0.0828452 | 0.20250659 | -0.4090989 | 0.68246706 | 0.94205398 |
| MTSS1L | 27.0204171 | -1.2522052 | 0.49904899 | -2.5091828 | 0.01210108 | 0.35881486 |
| VAC14 | 43.7528137 | -0.3688565 | 0.39197444 | -0.9410218 | 0.34669368 | 0.83687845 |
| HYDIN | 127.829883 | -0.0375506 | 0.40088146 | -0.0936702 | 0.92537118 | 0.98925753 |
| CMTR2 | 39.0334144 | -0.5738138 | 0.4150496 | -1.3825185 | 0.16681255 | 0.71878854 |
| ZNF19 | 29.5147697 | 0.35521508 | 0.34889127 | 1.01812546 | 0.30861834 | 0.81568231 |

| | | | | | | |
|---|---|---|---|---|---|---|
| MARVELD3 | 16.7779951 | 0.09808794 | 0.36197789 | 0.27097771 | 0.78640818 | 0.96767512 |
| PHLPP2 | 40.1623187 | -0.0378466 | 0.31699611 | -0.1193915 | 0.90496518 | 0.98531749 |
| AP1G1 | 184.804611 | -0.4023353 | 0.27903057 | -1.4419039 | 0.14932948 | 0.70141707 |
| IST1 | 106.498912 | -0.0587729 | 0.19905178 | -0.2952646 | 0.76779174 | 0.96283923 |
| ZNF821 | 35.0341744 | 0.19403234 | 0.3318534 | 0.58469295 | 0.5587542 | 0.91457126 |
| TXNL4B | 19.4254271 | -0.5543946 | 0.35804612 | -1.5483889 | 0.12152868 | 0.66804082 |
| DHX38 | 17.574351 | -0.5111544 | 0.35858322 | -1.4254833 | 0.15401754 | 0.70572801 |
| LINC01572 | 55.7858036 | 0.31142265 | 0.45353888 | 0.68665038 | 0.49230307 | 0.89067343 |
| AC004943.2 | 33.7441431 | 0.14677496 | 0.42627404 | 0.34432067 | 0.73060514 | 0.95279138 |
| ZFHX3 | 276.503039 | -0.4848097 | 0.19243452 | -2.5193487 | 0.01175722 | 0.35687659 |
| AC138627.1 | 24.2608901 | 0.23074548 | 0.76562893 | 0.30138031 | 0.76312451 | 0.96091323 |
| PSMD7 | 42.710095 | -0.0885171 | 0.2727674 | -0.3245149 | 0.74554828 | 0.95563105 |
| NPIPB15 | 14.8013042 | 0.98220594 | 0.92190123 | 1.06541342 | 0.28668889 | NA |
| GLG1 | 296.862635 | -0.1963991 | 0.16239549 | -1.2093875 | 0.22651399 | 0.77169045 |
| RFWD3 | 43.7216022 | -0.6380724 | 0.37417033 | -1.7052992 | 0.08813864 | 0.63466198 |
| FA2H | 39.3838132 | 0.25106823 | 0.3744386 | 0.67051909 | 5.03E-01 | 0.89523979 |
| WDR59 | 130.434138 | -0.036897 | 0.18752591 | -0.196757 | 0.84401767 | 0.97612951 |
| ZNRF1 | 118.703566 | -0.9739291 | 0.39998979 | -2.4348849 | 0.01489653 | 0.389664 |
| ZFP1 | 13.283947 | -0.4559396 | 0.3966239 | -1.1495515 | 0.25032865 | NA |
| BCAR1 | 70.4666236 | -0.3840996 | 0.31830598 | -1.2066994 | 0.22754794 | 0.77185154 |
| CFDP1 | 184.330585 | -0.702555 | 0.24314889 | -2.8894026 | 0.00385975 | 0.24454118 |
| GABARAPL2 | 55.9013057 | 0.55192645 | 0.37557079 | 1.46956703 | 0.14167906 | 0.69328517 |
| ADAT1 | 19.5944323 | 0.04239547 | 0.40782854 | 0.10395414 | 0.91720574 | 0.98710975 |
| KARS | 29.2099788 | -0.0089104 | 0.34553162 | -0.0257874 | 0.97942693 | 0.99539168 |
| TERF2IP | 135.01974 | 0.42538108 | 0.35746417 | 1.18999639 | 0.23404781 | 0.7767179 |
| NUDT7 | 12.067569 | 0.38001148 | 0.54905211 | 0.69212279 | 0.48886022 | NA |
| WWOX | 1212.5945 | 0.32786871 | 0.2169195 | 1.51147642 | 0.13066711 | 0.6742986 |
| MAF | 11.9374332 | 0.34068113 | 0.49555378 | 0.6874756 | 0.49178307 | NA |
| CDYL2 | 32.3840763 | 1.25966307 | 0.36688558 | 3.43339486 | 0.00059607 | 0.10434722 |
| CMC2 | 80.8724562 | 0.17599557 | 0.29696835 | 0.59264083 | 0.55342155 | 0.9132498 |
| ATMIN | 24.2704887 | 0.20544356 | 0.34781033 | 0.59067701 | 0.55473686 | 0.9137961 |
| GAN | 48.5317493 | -0.4102504 | 0.279648 | -1.4670244 | 0.14236941 | 0.69328517 |
| CMIP | 251.368236 | -0.3085224 | 0.23518002 | -1.3118564 | 0.18956859 | 0.74477466 |
| PLCG2 | 233.096341 | -0.1645236 | 0.24004827 | -0.6853772 | 0.49310594 | 0.89092098 |
| SDR42E1 | 11.8987617 | 0.02356417 | 0.50726224 | 0.04645363 | 0.96294869 | NA |
| AC092142.1 | 16.8107323 | -0.1083955 | 0.59206002 | -0.183082 | 0.85473367 | 0.97612951 |
| MPHOSPH6 | 37.440134 | -0.1938919 | 0.32437539 | -0.5977393 | 0.55001388 | 0.91248567 |
| HSBP1 | 48.9241729 | -0.0548508 | 0.30943505 | -0.177261 | 0.85930339 | 0.97691173 |
| MBTPS1 | 83.2743643 | 0.1806698 | 0.21237905 | 0.850695 | 0.3949388 | 0.85605713 |
| HSDL1 | 16.5181329 | -0.49685 | 0.42982042 | -1.155948 | 0.24770244 | 0.78593073 |
| TAF1C | 18.0732519 | -0.0907689 | 0.39603749 | -0.2291928 | 0.8187191 | 0.97288403 |
| TLDC1 | 29.2637963 | 0.04938369 | 0.39221881 | 0.12590852 | 0.89980434 | 0.98531749 |
| COTL1 | 23.2700385 | -0.7384264 | 0.58722994 | -1.257474 | 0.20858204 | 0.7623519 |
| KLHL36 | 44.85967 | 0.36657229 | 0.28328019 | 1.29402726 | 0.195656 | 0.75350056 |
| USP10 | 89.1224623 | -0.5244738 | 0.19893445 | -2.6364149 | 0.00837872 | 0.31004673 |
| ZDHHC7 | 62.2373341 | -0.4850431 | 0.29041414 | -1.6701772 | 0.09488431 | 0.64796091 |
| KIAA0513 | 17.2082803 | -0.6263564 | 0.54459132 | -1.1501403 | 0.25008609 | 0.78799766 |
| GSE1 | 135.309726 | -0.0701561 | 0.24838771 | -0.2824458 | 0.77760173 | 0.96518573 |
| EMC8 | 23.3464656 | -0.5012173 | 0.37525992 | -1.3356536 | 0.18166251 | 0.73398872 |
| COX4I1 | 194.705816 | 0.47443859 | 0.37016087 | 1.28170918 | 0.19994468 | 0.75605362 |
| MTHFSD | 15.6248106 | -0.3621096 | 0.39760013 | -0.910738 | 0.36243342 | NA |
| FBXO31 | 22.179377 | -0.2043032 | 0.34316042 | -0.5953578 | 0.55160431 | 0.9128014 |
| MAP1LC3B | 49.1002881 | -0.25109 | 0.31481944 | -0.7975683 | 0.42512106 | 0.86207532 |
| ZCCHC14 | 162.444987 | -1.0157021 | 0.48003972 | -2.1158709 | 0.03435579 | 0.4908133 |

| | | | | | | |
|---|---|---|---|---|---|---|
| KLHDC4 | 51.0848135 | -0.1610641 | 0.37784033 | -0.4262756 | 0.66990705 | 0.93878847 |
| BANP | 80.4145107 | -0.5786155 | 0.51142936 | -1.1313694 | 0.25789964 | 0.79532468 |
| ZFPM1 | 34.6371211 | 0.49605316 | 0.39498818 | 1.25586837 | 0.2091637 | 0.7623519 |
| ZC3H18 | 42.256957 | -0.0360407 | 0.31595155 | -0.1140704 | 0.90918202 | 0.9854228 |
| CYBA | 56.0715247 | 0.72591828 | 0.60129862 | 1.20725087 | 0.22733555 | 0.77185154 |
| MVD | 29.4763078 | -0.6517438 | 0.48303539 | -1.3492672 | 0.17725115 | 0.72841249 |
| PIEZO1 | 75.2807256 | -0.2387075 | 0.33167976 | -0.7196927 | 0.47171423 | 0.88267439 |
| APRT | 38.7245082 | 0.58535641 | 0.59157312 | 0.98949123 | 0.32242286 | 0.82558302 |
| GALNS | 22.8333927 | -0.0266197 | 0.36775281 | -0.0723847 | 0.94229572 | 0.99058259 |
| TRAPPC2L | 12.2765328 | -0.3549279 | 0.48845787 | -0.7266295 | 0.46745294 | NA |
| ACSF3 | 32.1419993 | 0.0635914 | 0.31907573 | 0.19929877 | 0.84202904 | 0.97600352 |
| ZNF778 | 12.9879912 | -0.1653031 | 0.44579341 | -0.3708066 | 0.71078159 | NA |
| ANKRD11 | 379.387702 | -0.5857801 | 0.25713987 | -2.2780601 | 0.022723 | 0.42648962 |
| SPG7 | 134.809855 | -0.7171041 | 0.28149654 | -2.5474704 | 0.0108507 | 0.33942942 |
| RPL13 | 328.681532 | 0.04691906 | 0.43069617 | 0.10893773 | 0.91325188 | 0.98639439 |
| CHMP1A | 33.8700575 | -0.2398706 | 0.33276924 | -0.7208316 | 0.4710131 | 0.88267439 |
| SPATA33 | 11.3464463 | 0.1236123 | 0.43174891 | 0.28630599 | 0.77464377 | NA |
| CDK10 | 35.7039049 | -0.0039812 | 0.28400296 | -0.0140181 | 0.98881557 | 0.99814944 |
| VPS9D1 | 36.8658881 | -0.2159553 | 0.28094553 | -0.7686731 | 0.44208739 | 0.86925451 |
| ZNF276 | 20.3326316 | -0.179406 | 0.33857613 | -0.5298836 | 0.59619261 | 0.9229638 |
| SPIRE2 | 22.195877 | 0.35322896 | 0.47833036 | 0.73846234 | 0.46023354 | 0.87607759 |
| TCF25 | 215.928298 | -0.3257548 | 0.26669611 | -1.2214455 | 0.22191739 | 0.76680432 |
| DEF8 | 19.4673274 | 0.01740048 | 0.32366695 | 0.05376046 | 0.95712601 | 0.99170927 |
| GAS8 | 43.9273502 | -0.3115638 | 0.28400685 | -1.097029 | 0.2726287 | 0.79721781 |
| FAM157C | 58.478495 | -0.098341 | 0.45120114 | -0.2179539 | 0.82746505 | 0.97567084 |
| LINC02193 | 10.8206639 | -0.8592599 | 0.48024322 | -1.7892181 | 0.0735797 | NA |
| DOC2B | 51.1079246 | -0.2066191 | 0.35054002 | -0.5894309 | 5.56E-01 | 0.91398748 |
| RPH3AL | 117.912851 | -0.2497263 | 0.32825078 | -0.760779 | 0.44678909 | 0.87231627 |
| AC129507.2 | 16.818488 | 0.54368199 | 0.59157255 | 0.91904533 | 0.35807186 | 0.84204868 |
| C17orf97 | 9.3968333 | -0.3842373 | 0.55012643 | -0.6984527 | 0.48489411 | NA |
| VPS53 | 186.868076 | -0.4258364 | 0.16627179 | -2.5610865 | 0.01043454 | 0.33556978 |
| GLOD4 | 35.0234445 | 0.35994056 | 0.3745643 | 0.96095798 | 0.3365733 | 0.83037432 |
| NXN | 22.7470195 | 0.94066608 | 0.60566942 | 1.55310148 | 0.12039889 | 0.6668162 |
| TIMM22 | 12.3460478 | 0.49143042 | 0.42632718 | 1.15270722 | 0.24903058 | NA |
| ABR | 39.2505595 | 0.20510227 | 0.23856103 | 0.85974757 | 3.90E-01 | 8.56E-01 |
| YWHAE | 230.20413 | -0.183683 | 0.20856937 | -0.8806806 | 0.37849072 | 0.85354853 |
| CRK | 81.4883792 | -0.0236956 | 0.30014992 | -0.0789458 | 0.93707577 | 0.99012269 |
| MYO1C | 79.5382793 | 0.0871186 | 0.27568819 | 0.31600411 | 0.75199939 | 0.95769549 |
| INPP5K | 13.8173366 | -0.0749013 | 0.4224614 | -0.1772975 | 0.85927474 | NA |
| PITPNA | 113.803789 | 0.01767092 | 0.23943537 | 0.07380245 | 0.94116757 | 0.99043681 |
| SLC43A2 | 111.74944 | 0.76545279 | 0.40780414 | 1.87701082 | 0.06051661 | 0.57774794 |
| SCARF1 | 16.9765767 | 0.58751237 | 0.58466852 | 1.00486403 | 0.31496233 | 0.82084199 |
| RILP | 11.0670384 | 0.31752242 | 0.530241 | 0.59882661 | 0.54928851 | NA |
| PRPF8 | 90.042239 | 0.05968432 | 0.24768723 | 0.24096648 | 0.8095811 | 0.97257655 |
| MIR22HG | 25.8501486 | -0.7668183 | 0.69410932 | -1.1047514 | 0.26926732 | 0.79666376 |
| WDR81 | 14.1875575 | -0.1021938 | 0.59155206 | -0.1727553 | 0.86284374 | NA |
| SERPINF2 | 13.6633067 | 0.86980267 | 0.50603124 | 1.71887147 | 0.08563778 | NA |
| SMYD4 | 21.6824661 | 0.55325346 | 0.65705957 | 0.84201416 | 0.39978003 | 0.85778581 |
| RPA1 | 40.1720727 | -0.0484821 | 0.27213616 | -0.1781538 | 0.8586022 | 0.97691173 |
| SMG6 | 227.218216 | -0.351224 | 0.21059086 | -1.6678026 | 0.09535492 | 0.64922651 |
| TSR1 | 21.2080263 | -0.3022907 | 0.37186258 | -0.8129097 | 0.41626983 | 0.85867915 |
| SGSM2 | 44.3962096 | -0.3055128 | 0.25302638 | -1.2074344 | 0.22726489 | 0.77185154 |
| MNT | 18.6687949 | -0.273127 | 0.43871803 | -0.6225571 | 0.5335756 | 0.90760867 |
| METTL16 | 82.2601617 | -0.3461154 | 0.19250569 | -1.7979491 | 0.07218508 | 0.60821941 |

| | | | | | | |
|---|---|---|---|---|---|---|
| PAFAH1B1 | 302.272814 | -0.2070295 | 0.2256228 | -0.9175912 | 0.35883294 | 0.84204868 |
| CLUH | 65.4764497 | -0.5348783 | 0.33038591 | -1.6189502 | 0.10545798 | 0.65946648 |
| RAP1GAP2 | 143.028474 | -0.1685992 | 0.29791255 | -0.5659353 | 0.57143778 | 0.91791025 |
| SPATA22 | 12.6916851 | 1.4552695 | 0.86712355 | 1.67827237 | 0.09329394 | NA |
| ASPA | 41.7300737 | 0.05266675 | 0.6308087 | 0.08349084 | 0.93346126 | 0.98960607 |
| CTNS | 19.9810052 | -0.4544208 | 0.52008134 | -0.8737495 | 0.38225466 | 0.8548566 |
| TAX1BP3 | 20.7132364 | -0.113505 | 0.54369575 | -0.2087657 | 0.83463115 | 0.97600352 |
| EMC6 | 18.2048086 | 0.55000089 | 0.35931556 | 1.53069043 | 0.12584592 | 0.66929384 |
| ITGAE | 17.9628127 | 0.60792019 | 0.36997185 | 1.64315256 | 0.10035137 | 0.65863153 |
| NCBP3 | 65.8157023 | -0.2726299 | 0.23133204 | -1.1785219 | 0.23858861 | 0.78096803 |
| CAMKK1 | 10.6561496 | 1.05258015 | 0.49533652 | 2.12497992 | 0.03358829 | NA |
| ZZEF1 | 108.569303 | -0.3397012 | 0.23210389 | -1.4635738 | 0.14331042 | 0.69428142 |
| CYB5D2 | 38.232471 | 0.01765099 | 0.48452543 | 0.03642944 | 0.97093994 | 0.99403127 |
| ANKFY1 | 100.564318 | -0.3987374 | 0.30752563 | -1.2965989 | 0.19476922 | 0.75191699 |
| UBE2G1 | 141.665789 | -0.308349 | 0.26471747 | -1.1648232 | 0.24409057 | 0.78561483 |
| SPNS2 | 31.3318714 | 0.12542718 | 0.63958873 | 0.19610599 | 0.84452721 | 0.97612951 |
| MYBBP1A | 11.4199802 | -0.1569239 | 0.5436313 | -0.2886587 | 0.77284259 | NA |
| GGT6 | 16.6898356 | 1.07090842 | 0.6224385 | 1.72050479 | 0.08534072 | 0.63062869 |
| SMTNL2 | 18.892344 | 1.31274924 | 0.50419359 | 2.60366111 | 0.00922339 | 0.32071089 |
| PELP1 | 25.4359861 | -0.1689353 | 0.34151688 | -0.4946617 | 0.62083894 | 0.92805465 |
| MED11 | 11.5066864 | 0.70972408 | 0.44386542 | 1.59896229 | 0.10982898 | NA |
| CXCL16 | 15.9598345 | 0.24546531 | 0.59145388 | 0.41502021 | 0.67812711 | NA |
| PSMB6 | 30.3906433 | -0.1573906 | 0.43604409 | -0.3609511 | 0.71813597 | 0.94925929 |
| MINK1 | 59.1163547 | -0.2004364 | 0.22874735 | -0.8762348 | 0.3809024 | 0.85432452 |
| SLC25A11 | 25.0090457 | 1.21819725 | 0.4686995 | 2.59910081 | 0.00934683 | 0.32071089 |
| RNF167 | 26.6331611 | -0.2394427 | 0.36831468 | -0.6501037 | 0.51562522 | 0.90194781 |
| PFN1 | 106.948532 | -0.2865877 | 0.38726352 | -0.7400327 | 0.45928014 | 0.87574928 |
| SPAG7 | 39.1246959 | -0.3621493 | 0.38007786 | -0.9528293 | 0.34067656 | 0.8319325 |
| CAMTA2 | 19.7792668 | 0.2435563 | 0.4048481 | 0.60159922 | 0.54744095 | 0.91139843 |
| KIF1C | 31.8457092 | 0.22240494 | 0.39006873 | 0.5701686 | 0.56856335 | 0.91698355 |
| ZNF594 | 23.1403245 | 0.10121333 | 0.35327008 | 0.28650412 | 0.77449204 | 0.96435639 |
| AC087500.1 | 50.987622 | 0.3692328 | 0.33327253 | 1.10790049 | 0.26790479 | 0.79666376 |
| RABEP1 | 170.457712 | -0.2818639 | 0.31083682 | -0.9067906 | 0.36451754 | 0.8468666 |
| NUP88 | 110.398805 | -0.1784279 | 0.2875992 | -0.6204047 | 0.53499136 | 0.90763121 |
| RPAIN | 53.9411799 | 0.20837558 | 0.27208873 | 0.76583687 | 0.44377337 | 0.8704597 |
| C1QBP | 34.9362393 | 0.80730715 | 0.65262823 | 1.23700923 | 0.21608365 | 0.76616336 |
| DERL2 | 25.4336476 | 0.2379282 | 0.34664091 | 0.68638235 | 0.49247203 | 0.89071585 |
| MIS12 | 13.9304421 | 0.09787074 | 0.44980067 | 0.21758691 | 0.82775098 | NA |
| KIAA0753 | 58.3110685 | -0.0937401 | 0.25098309 | -0.3734915 | 0.70878264 | 0.94558664 |
| TXNDC17 | 16.7114495 | -0.1575822 | 0.40778731 | -0.3864324 | 0.69917644 | 0.94395238 |
| MED31 | 13.7907511 | 0.2070843 | 0.49276321 | 0.42025114 | 0.674302 | NA |
| XAF1 | 57.2567996 | -4.0268664 | 1.68351906 | -2.391934 | 0.01675986 | 0.39748373 |
| ALOX12-AS1 | 42.1821259 | 1.02350372 | 0.6305688 | 1.62314363 | 0.10455871 | 0.65946648 |
| DLG4 | 11.7511573 | 0.3275115 | 0.66746584 | 0.49067904 | 0.62365347 | NA |
| ACADVL | 179.168982 | 0.12610472 | 0.32381325 | 0.38943657 | 0.69695322 | 0.94395238 |
| DVL2 | 14.0790552 | -0.709708 | 0.40455222 | -1.754305 | 0.07937825 | NA |
| ELP5 | 18.0692068 | 0.17411068 | 0.42316042 | 0.41145313 | 0.68074031 | 0.94196488 |
| EIF5A | 30.7680394 | 0.12176739 | 0.50714362 | 0.24010436 | 0.81024936 | 0.97257655 |
| GPS2 | 25.0006619 | -0.3890042 | 0.45686235 | -0.8514691 | 0.39450881 | 0.85605713 |
| ZBTB4 | 79.3655059 | 0.37065198 | 0.21794098 | 1.70069885 | 0.08899955 | 0.63542189 |
| POLR2A | 90.91011 | -0.0949529 | 0.26289872 | -0.3611768 | 0.71796726 | 0.94925929 |
| EIF4A1 | 32.7859458 | -0.1247925 | 0.43974357 | -0.2837848 | 0.7765753 | 0.9649632 |
| MPDU1 | 20.7351336 | -0.1232577 | 0.41193398 | -0.2992172 | 0.76477433 | 0.96169923 |
| FXR2 | 21.2262024 | -0.0689221 | 0.40510068 | -0.1701358 | 0.86490338 | 0.97788853 |

| | | | | | | |
|---|---|---|---|---|---|---|
| SAT2 | 44.5619193 | 0.43258938 | 0.33529219 | 1.29018624 | 0.196986 | 0.75406374 |
| TP53 | 23.8108933 | 0.44592774 | 0.34229571 | 1.30275584 | 0.19265813 | 0.75061775 |
| KDM6B | 32.5696305 | -0.7109145 | 0.43914373 | -1.6188651 | 0.10547628 | 0.65946648 |
| NAA38 | 71.4753395 | 0.60306303 | 0.26178906 | 2.30362198 | 0.02124387 | 0.4212027 |
| CYB5D1 | 10.867662 | -0.010198 | 0.51198667 | -0.0199186 | 0.98410834 | NA |
| CHD3 | 48.968764 | 0.12550517 | 0.40464404 | 0.31016192 | 0.75643783 | 0.95903253 |
| TRAPPC1 | 13.2298758 | 0.49064597 | 0.60522825 | 0.81067923 | 0.4175499 | NA |
| CNTROB | 16.2457372 | -0.3253414 | 0.40602061 | -0.8012928 | 0.42296215 | 0.86190547 |
| PER1 | 43.7073583 | -0.9885343 | 0.5198336 | -1.9016361 | 0.05721875 | 0.57049536 |
| TMEM107 | 11.2848653 | 0.38744683 | 0.54804434 | 0.70696257 | 0.47958974 | NA |
| CTC1 | 20.236635 | 0.37774463 | 0.37120436 | 1.01761904 | 0.30885903 | 0.81568231 |
| SLC25A35 | 10.7747913 | 0.35918873 | 0.66003444 | 0.54419696 | 0.58630595 | NA |
| NDEL1 | 70.8787329 | -0.1257363 | 0.36486294 | -0.3446126 | 0.73038567 | 0.95279138 |
| MYH10 | 276.114639 | 0.31363322 | 0.25337416 | 1.23782639 | 0.21578044 | 0.76566124 |
| NTN1 | 25.6508304 | 0.5047075 | 0.36324592 | 1.38943749 | 0.16469976 | 0.71461481 |
| STX8 | 206.626712 | -0.2604478 | 0.2727601 | -0.9548603 | 0.33964837 | 0.83071861 |
| USP43 | 44.4804039 | 0.01182131 | 0.38308172 | 0.03085847 | 0.97538241 | 0.99440004 |
| SCO1 | 30.6668731 | -0.2934218 | 0.28947945 | -1.0136188 | 0.3107647 | 0.81692172 |
| TMEM220 | 34.4581369 | -0.2721988 | 0.42575252 | -0.6393357 | 0.52260458 | 0.90487731 |
| TMEM220-A: | 22.0663305 | -0.1419037 | 0.33250846 | -0.4267673 | 0.66954882 | 0.93878847 |
| SHISA6 | 25.2352904 | 1.2014655 | 0.66109721 | 1.817381 | 0.0691588 | 0.60275677 |
| DNAH9 | 12.6863334 | 0.35358613 | 0.45337278 | 0.77990154 | 0.43544883 | NA |
| MAP2K4 | 238.702928 | -0.2536688 | 0.26284163 | -0.9651011 | 0.33449418 | 0.82974237 |
| ARHGAP44 | 113.611752 | 0.01810272 | 0.27394758 | 0.06608096 | 0.94731337 | 0.99083449 |
| ELAC2 | 20.8813625 | -0.0597232 | 0.33061916 | -0.1806403 | 0.85664988 | 0.97646931 |
| COX10 | 79.8947675 | -0.1666769 | 0.3154485 | -0.5283807 | 0.59723513 | 0.9229638 |
| TRIM16 | 22.404584 | 0.41179378 | 0.40170197 | 1.02512262 | 0.30530533 | 0.81568231 |
| ZSWIM7 | 48.0475708 | 0.08541918 | 0.26105011 | 0.32721374 | 0.74350623 | 0.95563105 |
| TTC19 | 128.642223 | 0.37323014 | 0.3236446 | 1.15320985 | 0.24882426 | 0.78729484 |
| NCOR1 | 406.127482 | 0.01452782 | 0.13952784 | 0.1041213 | 9.17E-01 | 0.98710975 |
| PIGL | 102.468423 | 0.06453859 | 0.30220715 | 0.21355745 | 0.83089218 | 0.97567084 |
| UBB | 174.107246 | 0.74470464 | 0.40652973 | 1.83185775 | 0.06697262 | 0.59652811 |
| LRRC75A | 15.8447312 | -0.0310731 | 0.39964293 | -0.0777522 | 0.93802518 | NA |
| AC098850.3 | 50.429587 | -0.3306781 | 0.74289682 | -0.4451198 | 0.65623319 | 0.93570875 |
| MPRIP | 177.923388 | -0.5803142 | 0.25289781 | -2.2946588 | 0.02175269 | 0.4212027 |
| COPS3 | 27.1889997 | 0.17021736 | 0.34199804 | 0.49771444 | 0.61868533 | 0.92766755 |
| NT5M | 18.7030686 | -0.3134107 | 0.40214448 | -0.7793484 | 0.43577451 | 0.86729771 |
| MED9 | 46.198882 | -0.3464374 | 0.30153469 | -1.1489139 | 0.2505915 | 0.78877402 |
| RASD1 | 201.885142 | -0.1818574 | 0.39480503 | -0.4606257 | 0.64506715 | 0.93328692 |
| PEMT | 22.9318098 | -0.330871 | 0.36445181 | -0.9078595 | 0.36395245 | 0.8466721 |
| RAI1 | 111.917872 | -0.0347504 | 0.17907383 | -0.1940565 | 0.84613166 | 0.97612951 |
| SREBF1 | 57.1086498 | -1.1606513 | 0.37985334 | -3.0555247 | 0.00224667 | 0.18826595 |
| TOM1L2 | 176.891945 | -0.1507725 | 0.27255189 | -0.5531883 | 0.58013447 | 0.92033309 |
| DRC3 | 16.6438793 | 0.33519643 | 0.52652417 | 0.63662116 | 0.52437164 | 0.90487731 |
| ATPAF2 | 17.80748 | -0.2000972 | 0.41269394 | -0.4848561 | 0.62777843 | 0.93061955 |
| GID4 | 14.1652401 | 0.52763745 | 0.40762674 | 1.29441322 | 0.19552273 | NA |
| DRG2 | 11.180171 | -0.0404074 | 0.50113687 | -0.0806314 | 0.93573507 | NA |
| ALKBH5 | 52.5217325 | -0.1565358 | 0.26417681 | -0.5925419 | 0.55348777 | 0.9132498 |
| FLII | 37.7717858 | -0.0423247 | 0.3830884 | -0.1104829 | 0.9120264 | 0.9859256 |
| MIEF2 | 13.9593662 | -0.1831357 | 0.3626735 | -0.5049601 | 0.61358685 | NA |
| TOP3A | 27.1026802 | -0.3325288 | 0.34241756 | -0.9711207 | 0.33148817 | 0.82890589 |
| SMCR8 | 13.9949257 | 0.52014564 | 0.38357682 | 1.35604034 | 0.17508633 | NA |
| SHMT1 | 51.1352193 | 0.7753354 | 0.42777889 | 1.81246764 | 0.06991397 | 0.60302096 |
| TRIM16L | 11.7566529 | 0.12818002 | 0.46312135 | 0.27677415 | 0.78195353 | NA |

| | | | | | | |
|---|---|---|---|---|---|---|
| TVP23B | 25.6893321 | -0.024495 | 0.35761557 | -0.0684953 | 0.94539139 | 0.99083449 |
| PRPSAP2 | 65.9942985 | -0.0194134 | 0.31447459 | -0.061733 | 0.95077549 | 0.99105535 |
| EPN2 | 150.032874 | -0.154577 | 0.24568023 | -0.6291797 | 0.52923139 | 0.90505228 |
| B9D1 | 26.9954424 | -0.4326791 | 0.30910125 | -1.3997974 | 0.161574 | 0.71101785 |
| AC004448.2 | 15.4430185 | 0.32635089 | 0.43262733 | 0.75434644 | 0.45064122 | NA |
| SLC47A1 | 21.8023943 | -0.3356696 | 0.40300825 | -0.83291 | 0.4048955 | 0.85782098 |
| ALDH3A2 | 47.7766664 | -0.1622669 | 0.36614745 | -0.4431737 | 0.65764009 | 0.93593674 |
| ULK2 | 57.7561239 | 0.55792361 | 0.26929951 | 2.07175875 | 0.03828795 | 0.49671741 |
| AKAP10 | 141.750417 | -0.3226516 | 0.21312213 | -1.5139283 | 0.13004401 | 0.67331904 |
| SPECC1 | 18.0928984 | 0.30467781 | 0.41846499 | 0.72808433 | 0.46656197 | 0.88059581 |
| CCDC144NL-/ | 22.2039212 | 0.5645013 | 0.39672503 | 1.42290316 | 0.15476423 | 0.70572801 |
| USP22 | 127.937763 | -0.0140542 | 0.22650082 | -0.0620494 | 0.95052346 | 0.99105535 |
| DHRS7B | 86.6939021 | 0.28820171 | 0.27085565 | 1.06404171 | 0.2873098 | 0.80523851 |
| TMEM11 | 12.4905564 | 0.04844901 | 0.44962604 | 0.10775401 | 0.91419082 | NA |
| AC087294.1 | 14.4842542 | 0.00344572 | 0.4797285 | 0.00718264 | 0.99426913 | NA |
| MAP2K3 | 41.188356 | -1.0268199 | 0.44497203 | -2.3076055 | 0.02102109 | 0.4212027 |
| WSB1 | 1084.92906 | -0.6386412 | 0.38105855 | -1.6759662 | 0.09374482 | 0.64572229 |
| KSR1 | 91.8663516 | 0.3031806 | 0.24900842 | 1.21755158 | 0.22339442 | 0.76834258 |
| NLK | 433.108672 | -0.3498608 | 0.38823081 | -0.901167 | 0.36749954 | 0.84777627 |
| IFT20 | 15.7142915 | -0.1349352 | 0.4796186 | -0.2813386 | 0.77845073 | NA |
| TNFAIP1 | 30.1730416 | 0.07127077 | 0.53236189 | 0.13387655 | 0.8935002 | 0.98377746 |
| POLDIP2 | 30.678096 | -0.1522548 | 0.40909841 | -0.3721716 | 0.70976512 | 0.94583397 |
| SARM1 | 26.5938125 | 0.19908708 | 0.33954252 | 0.58633917 | 0.55764762 | 0.91425998 |
| AC005726.1 | 19.9255883 | 0.36679428 | 0.40959352 | 0.89550802 | 0.37051558 | 0.84852785 |
| KIAA0100 | 54.8873183 | -0.1748653 | 0.30878262 | -0.5663055 | 0.57118614 | 0.91791025 |
| SDF2 | 25.5215268 | -0.0097405 | 0.36081874 | -0.0269957 | 0.97846319 | 0.99471172 |
| SUPT6H | 71.5395688 | -0.2911982 | 0.24647537 | -1.1814494 | 0.23742423 | 0.77930556 |
| RAB34 | 38.2111819 | 0.49508222 | 0.39308008 | 1.25949455 | 0.20785176 | 0.76231289 |
| RPL23A | 115.136952 | 0.67645761 | 0.27013023 | 2.50419075 | 0.01227318 | 0.36046953 |
| NEK8 | 20.0893643 | -0.3284756 | 0.40007191 | -0.8210414 | 0.41162268 | 0.85867915 |
| TRAF4 | 57.5038033 | -0.0045205 | 0.340875 | -0.0132614 | 0.98941926 | 0.99814944 |
| FAM222B | 198.437171 | -0.4277528 | 0.26887243 | -1.5909136 | 0.11162901 | 0.6645905 |
| ERAL1 | 12.0773486 | 0.22713 | 0.42093819 | 0.5395804 | 0.58948644 | NA |
| FLOT2 | 89.4679552 | -0.0319263 | 0.31591442 | -0.1010598 | 0.91950298 | 0.9875219 |
| DHRS13 | 14.3438051 | 0.07594103 | 0.54069408 | 0.14045101 | 0.88830366 | NA |
| PHF12 | 59.900731 | -0.0699662 | 0.23329262 | -0.2999076 | 0.76424761 | 0.96130247 |
| MYO18A | 43.7653421 | -0.496087 | 0.45078607 | -1.1004933 | 0.27111725 | 0.79666376 |
| NUFIP2 | 150.99397 | -0.1138215 | 0.1804176 | -0.6308778 | 0.5281204 | 0.90487731 |
| TAOK1 | 269.728019 | -0.3266236 | 0.22424511 | -1.4565471 | 0.14524144 | 0.69606446 |
| ABHD15-AS1 | 30.143293 | 0.16238216 | 0.52189357 | 0.31114037 | 0.75569391 | 0.95849416 |
| GIT1 | 20.3866062 | -0.1931159 | 0.36756586 | -0.5253912 | 0.59931126 | 0.92298075 |
| SSH2 | 575.440729 | -0.2098611 | 0.21499986 | -0.9760989 | 0.32901546 | 0.82781307 |
| AC023389.1 | 18.5700504 | -0.0784004 | 0.42725773 | -0.1834968 | 0.85440826 | 0.97612951 |
| EFCAB5 | 18.5693097 | 0.17971189 | 0.5084683 | 0.35343775 | 0.72376029 | 0.95093722 |
| NSRP1 | 169.23043 | 0.05455524 | 0.18452172 | 0.29565756 | 0.76749161 | 0.96283923 |
| AC104984.4 | 19.0886526 | 1.33146507 | 0.67781247 | 1.96435611 | 0.04948881 | 0.54320302 |
| BLMH | 43.6315752 | 0.69891085 | 0.30224309 | 2.312413 | 0.02075493 | 0.4212027 |
| CPD | 122.345276 | 0.09664048 | 0.37215648 | 0.25967701 | 0.79511293 | 0.96974891 |
| GOSR1 | 96.6036884 | -0.223692 | 0.1914054 | -1.1686817 | 0.24253189 | 0.78464787 |
| CRLF3 | 65.6334512 | 0.07617871 | 0.32367919 | 0.23535251 | 0.81393514 | 0.97282508 |
| ATAD5 | 15.2786923 | -0.5352769 | 0.47401932 | -1.1292302 | 0.25880075 | NA |
| TEFM | 31.1664214 | 0.43668727 | 0.35655433 | 1.22474257 | 0.22067223 | 0.76680432 |
| RNF135 | 21.3613267 | -0.2527637 | 0.36695859 | -0.6888072 | 0.49094462 | 0.88987842 |
| NF1 | 531.388191 | -0.2905939 | 0.18445328 | -1.5754337 | 0.11515636 | 0.6668162 |

| | | | | | | |
|---|---|---|---|---|---|---|
| RAB11FIP4 | 40.226954 | 0.20755334 | 0.34108778 | 0.60850416 | 0.54285315 | 0.90953689 |
| COPRS | 27.6589112 | -0.017775 | 0.41705286 | -0.0426206 | 0.966004 | 0.99283442 |
| UTP6 | 33.041179 | -0.8692605 | 0.31727152 | -2.7397998 | 0.00614766 | 0.29709153 |
| SUZ12 | 130.173128 | -0.129869 | 0.20259815 | -0.6410177 | 0.52151117 | 0.90451015 |
| LRRC37B | 38.8005138 | -0.3828583 | 0.39016263 | -0.9812787 | 0.32645533 | 0.82570794 |
| RHOT1 | 114.026239 | 0.09912025 | 0.19778997 | 0.50113892 | 0.61627336 | 0.92662475 |
| C17orf75 | 16.0507029 | 0.08138851 | 0.40888727 | 0.19904878 | 0.84222459 | NA |
| ZNF207 | 187.507869 | 0.11585361 | 0.19705905 | 0.58791319 | 0.55659056 | 0.9141017 |
| PSMD11 | 50.417544 | -0.5154581 | 0.30153753 | -1.7094326 | 0.08737084 | 0.63466198 |
| MYO1D | 239.387438 | -0.1195852 | 0.20014499 | -0.5974927 | 0.55017846 | 0.91248567 |
| ASIC2 | 26.9658986 | 0.37004868 | 0.53088463 | 0.69704161 | 0.48577675 | 0.88838332 |
| LIG3 | 16.1207505 | -0.161052 | 0.37843956 | -0.4255685 | 0.67042228 | NA |
| SLFN5 | 52.9541889 | 0.29707487 | 0.54249819 | 0.54760528 | 5.84E-01 | 9.21E-01 |
| LINC02001 | 16.760869 | -0.0643847 | 0.36785324 | -0.1750283 | 0.86105742 | 0.97745426 |
| AP2B1 | 133.507397 | -0.0588497 | 0.19085669 | -0.3083449 | 0.75781993 | 0.95903525 |
| TAF15 | 214.919065 | -0.0818085 | 0.17549975 | -0.4661457 | 0.64111118 | 0.93328692 |
| AC015849.3 | 12.325295 | 0.19115857 | 0.59132325 | 0.32327254 | 0.74648885 | NA |
| ZNHIT3 | 14.8657852 | 0.30826987 | 0.418694 | 0.73626531 | 0.46156925 | NA |
| MYO19 | 83.8423279 | -0.2003956 | 0.19959493 | -1.0040115 | 0.31537306 | 0.82084199 |
| GGNBP2 | 105.138037 | -0.3772548 | 0.25626716 | -1.4721152 | 0.14098976 | 0.69227547 |
| DHRS11 | 14.6747755 | 0.58121844 | 0.47103383 | 1.2339208 | 0.21723242 | NA |
| LHX1-DT | 158.673896 | 0.29498282 | 0.38095992 | 0.77431458 | 0.43874478 | 0.86879768 |
| LHX1 | 225.100109 | 0.50228525 | 0.32848545 | 1.52909435 | 0.12624106 | 0.66929384 |
| AATF | 121.190918 | -0.1740032 | 0.24747732 | -0.7031077 | 0.48198861 | 0.88708902 |
| ACACA | 194.235804 | -0.2013931 | 0.2171946 | -0.9272471 | 0.35379824 | 0.83979358 |
| TADA2A | 46.3724345 | -0.4453301 | 0.2448237 | -1.8189828 | 0.06891406 | 0.60275677 |
| SYNRG | 113.878649 | -0.2398539 | 0.21106322 | -1.1364078 | 0.25578595 | 0.7940072 |
| DDX52 | 61.0798209 | -0.677428 | 0.28321565 | -2.3919158 | 0.01676069 | 0.39748373 |
| HNF1B | 276.931115 | -0.8794559 | 0.21644431 | -4.063197 | 4.84E-05 | 0.0275395 |
| MRPL45 | 33.3000149 | -0.1068361 | 0.27753059 | -0.3849524 | 0.70027269 | 0.94424267 |
| SOCS7 | 33.608607 | 0.08196723 | 0.3122534 | 0.26250229 | 0.79293422 | 0.96947477 |
| MLLT6 | 117.132158 | 0.06981162 | 0.23822826 | 0.29304507 | 0.7694877 | 0.96289288 |
| CISD3 | 18.9994209 | 0.56663236 | 0.43341173 | 1.30737661 | 0.19108484 | 0.74847043 |
| PCGF2 | 29.1803872 | -0.149572 | 0.32490587 | -0.460355 | 0.64526143 | 0.93328692 |
| PSMB3 | 35.4650701 | 0.2127683 | 0.3663075 | 0.58084615 | 0.56134415 | 0.91608386 |
| PIP4K2B | 39.611928 | -0.1389058 | 0.27547415 | -0.5042425 | 0.61409097 | 0.92662475 |
| CWC25 | 26.9549588 | -0.497654 | 0.33389503 | -1.4904504 | 0.13610585 | 0.68406472 |
| RPL23 | 91.3205546 | 0.55120491 | 0.36042202 | 1.52933195 | 0.12618217 | 0.66929384 |
| LASP1 | 58.4689853 | -0.2525131 | 0.31035771 | -0.8136195 | 0.41586298 | 0.85867915 |
| RPL19 | 161.964914 | 0.0955789 | 0.45591673 | 0.20964112 | 0.83394778 | 0.97600352 |
| FBXL20 | 181.380788 | -0.144798 | 0.3192342 | -0.4535791 | 0.65013177 | 0.93453685 |
| MED1 | 37.1650573 | 0.06193579 | 0.30847554 | 0.20078024 | 0.84087041 | 0.97600352 |
| CDK12 | 118.456053 | 0.11808503 | 0.24638147 | 0.47927725 | 0.63174141 | 0.93251095 |
| PPP1R1B | 14.0836341 | 1.66440883 | 0.55517331 | 2.99799865 | 0.00271759 | NA |
| STARD3 | 26.0838056 | 0.36991388 | 0.33156653 | 1.11565508 | 0.2645698 | 0.79666376 |
| PGAP3 | 19.9707073 | 1.11055972 | 0.43617222 | 2.54614961 | 0.01089185 | 0.33954968 |
| ERBB2 | 133.877067 | 0.60028621 | 0.27998571 | 2.14398873 | 0.03203379 | 0.48175799 |
| MIEN1 | 18.1969676 | -0.2406806 | 0.39808409 | -0.6045974 | 0.54544657 | 0.91037772 |
| GRB7 | 73.4830026 | -0.3065268 | 0.24915014 | -1.2302895 | 0.2185887 | 0.76666749 |
| ORMDL3 | 15.270611 | 0.26177308 | 0.46114658 | 0.56765698 | 0.57026791 | NA |
| PSMD3 | 27.5179246 | -0.2629051 | 0.33742661 | -0.7791474 | 0.4358929 | 0.86729771 |
| MED24 | 38.094028 | -0.1600917 | 0.27599083 | -0.5800618 | 0.56187294 | 0.91624774 |
| THRA | 32.7518285 | 0.40692909 | 0.29925542 | 1.35980525 | 0.17389156 | 0.72794035 |
| NR1D1 | 15.4963542 | -1.0292185 | 0.69324823 | -1.4846321 | 0.13764132 | NA |

| | | | | | | |
|---|---|---|---|---|---|---|
| MSL1 | 39.9886162 | -0.2032997 | 0.28910958 | -0.7031925 | 0.48193577 | 0.88708902 |
| CASC3 | 50.1956938 | -0.048929 | 0.28091869 | -0.174175 | 0.8617279 | 0.97767895 |
| WIPF2 | 136.110603 | -0.5725798 | 0.23307231 | -2.4566615 | 0.01402347 | 0.37656544 |
| RARA | 24.8981028 | -0.3159223 | 0.43281056 | -0.729932 | 0.46543173 | 0.87981788 |
| IGFBP4 | 14.0530776 | -0.5040167 | 0.48935903 | -1.0299529 | 0.30303212 | NA |
| TNS4 | 11.0914713 | 4.23561186 | 1.31023014 | 3.23272359 | 0.00122616 | NA |
| KRT10 | 30.3584445 | -0.1998489 | 0.401281 | -0.4980274 | 0.61846475 | 0.92766755 |
| KRT19 | 590.811324 | 0.2935981 | 0.44097881 | 0.66578731 | 0.50554706 | 0.89723255 |
| EIF1 | 201.229621 | 0.012578 | 0.34999252 | 0.03593791 | 0.97133186 | 0.99403127 |
| JUP | 170.063875 | -0.3301426 | 0.32639184 | -1.0114917 | 0.31178117 | 0.81701824 |
| P3H4 | 14.8608136 | 0.01416215 | 0.42890785 | 0.03301909 | 0.97365937 | NA |
| NT5C3B | 24.0008291 | -0.1860284 | 0.30630187 | -0.6073367 | 0.54362749 | 0.90965439 |
| ACLY | 38.9736838 | -0.3288469 | 0.34147699 | -0.9630134 | 0.3355408 | 0.82974237 |
| TTC25 | 11.4193269 | 0.11040408 | 0.4809733 | 0.22954305 | 0.81844687 | NA |
| CNP | 15.801734 | -0.0874099 | 0.60239517 | -0.1451039 | 0.88462887 | NA |
| DNAJC7 | 83.2923907 | -0.179726 | 0.23934397 | -0.7509107 | 0.45270638 | 0.87346041 |
| NKIRAS2 | 11.8987259 | 0.05005818 | 0.46397737 | 0.10788926 | 0.91408354 | NA |
| KAT2A | 20.2592407 | 0.57018573 | 0.42317376 | 1.34740331 | 0.17785037 | 0.72841249 |
| STAT5B | 114.179594 | -0.04104 | 0.22501683 | -0.1823864 | 0.85527947 | 0.97612951 |
| STAT5A | 16.5228316 | -0.1633867 | 0.45283386 | -0.3608094 | 0.71824195 | 0.94925929 |
| STAT3 | 475.103643 | -0.0502288 | 0.17881975 | -0.2808909 | 7.79E-01 | 0.96559006 |
| ATP6V0A1 | 90.099885 | -0.3332656 | 0.20455019 | -1.6292608 | 0.10325782 | 0.65946648 |
| NAGLU | 24.946384 | -0.4727595 | 0.44875566 | -1.0534897 | 0.2921166 | 0.80800285 |
| COASY | 28.1278127 | 0.47870683 | 0.42516103 | 1.1259424 | 0.2601899 | 0.79560251 |
| MLX | 20.2727591 | 0.0857509 | 0.36417648 | 0.23546523 | 0.81384766 | 0.97282508 |
| RETREG3 | 61.6704383 | -0.0909341 | 0.26827181 | -0.3389624 | 0.73463804 | 0.95329896 |
| TUBG2 | 26.3085549 | -0.5843513 | 0.32731839 | -1.7852688 | 0.07421772 | 0.61020315 |
| PLEKHH3 | 12.5894687 | 0.63313771 | 0.42111542 | 1.50347785 | 0.13271586 | NA |
| EZH1 | 66.2649858 | -0.290404 | 0.33980582 | -0.8546176 | 0.39276286 | 0.85605713 |
| VPS25 | 24.9041562 | 0.03882501 | 0.40930648 | 0.09485559 | 0.92442953 | 0.98925753 |
| WNK4 | 62.9904246 | -0.4758145 | 0.51676323 | -0.9207593 | 3.57E-01 | 0.84204868 |
| COA3 | 60.7333219 | 0.68807194 | 0.42545623 | 1.6172567 | 0.10582288 | 0.65946648 |
| BECN1 | 38.3642594 | -0.0519381 | 0.28188695 | -0.1842517 | 0.853816 | 0.97612951 |
| PSME3 | 26.7151016 | -0.0582778 | 0.31275332 | -0.1863379 | 0.85217981 | 0.97612951 |
| RUNDC1 | 22.760013 | 0.1246099 | 0.35295294 | 0.35304961 | 0.72405125 | 0.95093722 |
| RPL27 | 61.169945 | 0.11470725 | 0.4178429 | 0.27452244 | 0.78368316 | 0.96717296 |
| IFI35 | 16.7739377 | -1.3311028 | 1.05489002 | -1.2618404 | 0.20700622 | 0.76231289 |
| VAT1 | 18.5320844 | -0.6083418 | 0.3563462 | -1.707165 | 8.78E-02 | 0.63466198 |
| BRCA1 | 21.407671 | 0.63136113 | 0.46798305 | 1.3491111 | 0.17730128 | 0.72841249 |
| NBR1 | 70.9462054 | 0.3545225 | 0.25242231 | 1.40448163 | 0.16017548 | 0.71101785 |
| LINC00854 | 36.1261617 | -1.4144837 | 0.54633456 | -2.5890431 | 0.0096243 | 0.32468359 |
| ARL4D | 54.3288991 | 0.53321044 | 0.46765837 | 1.14017084 | 0.25421513 | 0.79171259 |
| DHX8 | 29.6922716 | 0.46520192 | 0.32708707 | 1.42225715 | 0.15495161 | 0.70572801 |
| DUSP3 | 46.2310322 | -0.0355773 | 0.30875731 | -0.1152273 | 0.908265 | 0.9854228 |
| TMEM101 | 126.062976 | 0.97370003 | 0.57405113 | 1.69619041 | 0.08984983 | 0.63749257 |
| LSM12 | 43.9737275 | -0.6129714 | 0.30991324 | -1.9778807 | 0.04794216 | 0.53707392 |
| G6PC3 | 20.4145252 | -0.3010968 | 0.6232964 | -0.4830717 | 0.62904485 | 0.9314335 |
| HDAC5 | 44.0614034 | -0.04786 | 0.34914291 | -0.1370786 | 0.89096865 | 0.98296447 |
| ASB16-AS1 | 11.3943724 | 0.05521212 | 0.43524881 | 0.12685185 | 0.89905765 | NA |
| TMUB2 | 24.9106756 | 0.05965995 | 0.41983926 | 0.14210189 | 0.88699953 | 0.98216235 |
| ATXN7L3 | 16.3370712 | -0.2977911 | 0.40520705 | -0.734911 | 0.46239368 | 0.87713916 |
| UBTF | 49.6843054 | -0.2373447 | 0.3060918 | -0.7754035 | 0.43810127 | 0.86828562 |
| SLC25A39 | 40.1670634 | 0.12993152 | 0.35323678 | 0.36783123 | 0.71299908 | 0.94792328 |
| GRN | 85.924677 | 0.11189522 | 0.43137213 | 0.25939371 | 0.79533148 | 0.96974891 |

| | | | | | | |
|---|---|---|---|---|---|---|
| GPATCH8 | 333.805133 | -0.1970156 | 0.13970762 | -1.4101992 | 0.15848086 | 0.71089007 |
| DBF4B | 15.0886996 | -0.1911072 | 0.48919519 | -0.3906564 | 0.69605126 | NA |
| EFTUD2 | 35.1860027 | -0.1124864 | 0.35001082 | -0.3213798 | 0.74792257 | 0.95563105 |
| DCAKD | 50.0826465 | -0.0556282 | 0.30208973 | -0.1841445 | 0.85390009 | 0.97612951 |
| NMT1 | 76.8112011 | -0.3692738 | 0.20155493 | -1.8321249 | 0.06693281 | 0.59652811 |
| ACBD4 | 14.3286197 | -0.2635542 | 0.40775137 | -0.6463601 | 0.51804619 | NA |
| HEXIM1 | 19.1714565 | -0.4970826 | 0.38621488 | -1.2870622 | 1.98E-01 | 0.75559912 |
| AC138150.1 | 50.7458306 | 0.43939652 | 0.47024884 | 0.93439151 | 0.35010197 | 0.83687845 |
| MAP3K14 | 86.5715459 | 0.04366794 | 0.47367275 | 0.0921901 | 0.926547 | 0.98925753 |
| ARHGAP27 | 18.6575487 | -0.1356442 | 0.35120682 | -0.386223 | 0.69933152 | 0.94395238 |
| PLEKHM1 | 34.4971549 | -0.7799132 | 0.47278685 | -1.6496086 | 0.09902302 | 0.65748109 |
| LINC02210-C | 33.0273942 | -0.3473249 | 0.42007391 | -0.8268186 | 0.40833988 | 0.85867915 |
| KANSL1 | 608.119325 | -0.0077505 | 0.25126456 | -0.0308458 | 0.97539251 | 0.99440004 |
| ARL17B | 131.221057 | 1.04002472 | 1.07292501 | 0.9693359 | 0.33237763 | 0.82894069 |
| LRRC37A2 | 35.596559 | -0.247386 | 0.39253795 | -0.6302218 | 0.52854947 | 0.90487731 |
| ARL17A | 38.2058734 | -0.06109 | 0.54211124 | -0.1126891 | 0.91027701 | 0.98556715 |
| NSF | 173.123097 | -0.2142334 | 0.35249697 | -0.6077594 | 0.54334704 | 0.90965439 |
| AC005670.2 | 28.608801 | 0.20861415 | 0.37322113 | 0.55895589 | 0.57619183 | 0.91854298 |
| CDC27 | 162.298527 | -0.4669479 | 0.27006339 | -1.7290308 | 0.08380359 | 0.62534311 |
| AC068234.1 | 35.8752844 | -0.052941 | 0.73440089 | -0.0720874 | 9.43E-01 | 9.91E-01 |
| EFCAB13 | 55.653552 | 0.08402308 | 0.37214845 | 0.22577838 | 0.82137379 | 0.97396757 |
| NPEPPS | 389.545906 | -0.6122194 | 0.17703439 | -3.458195 | 0.00054381 | 0.09900562 |
| AC025682.1 | 34.6095053 | 0.18408844 | 0.36002763 | 0.51131754 | 0.60912872 | 0.92573951 |
| KPNB1 | 137.704851 | -0.1768443 | 0.23349792 | -0.7573699 | 0.44882828 | 0.87314185 |
| MRPL10 | 13.0154566 | 0.00454663 | 0.43565671 | 0.01043628 | 0.99167321 | NA |
| SCRN2 | 17.6127457 | -0.0364413 | 0.38298437 | -0.0951509 | 0.92419498 | 0.98925753 |
| SP2 | 14.5186271 | -0.1328039 | 0.40442512 | -0.328377 | 0.74262662 | NA |
| PNPO | 22.5775224 | 1.04691065 | 0.43793358 | 2.39056951 | 0.01682226 | 0.39748373 |
| PRR15L | 11.0483914 | 1.84882532 | 0.70639738 | 2.61725959 | 0.00886389 | NA |
| CDK5RAP3 | 36.198564 | 0.3851869 | 0.28844 | 1.33541432 | 0.18174078 | 0.73398872 |
| NFE2L1 | 86.3780453 | -0.0586661 | 0.26040714 | -0.2252862 | 0.82175667 | 0.97427076 |
| CBX1 | 40.8640756 | 0.38426789 | 0.43183613 | 0.88984655 | 0.37354829 | 0.85052778 |
| SNX11 | 15.1525652 | -0.2178856 | 0.39164084 | -0.5563404 | 0.5779782 | NA |
| SKAP1 | 390.872936 | 0.14732624 | 0.21498596 | 0.68528307 | 0.49316531 | 0.89092098 |
| HOXB2 | 21.140311 | -0.0061961 | 0.45686252 | -0.0135623 | 0.98917921 | 0.99814944 |
| HOXB-AS1 | 15.6635896 | 0.14431092 | 0.40464542 | 0.3566355 | 0.7213647 | NA |
| HOXB3 | 235.314652 | 0.30518924 | 0.27287245 | 1.11843185 | 0.2633826 | 0.79666376 |
| HOXB-AS3 | 57.8625327 | 0.14895018 | 0.46811181 | 0.3181936 | 0.75033809 | 0.95716475 |
| HOXB7 | 54.3367865 | 0.40824901 | 0.36419754 | 1.12095486 | 0.26230708 | 0.79666376 |
| AC103702.2 | 24.1234391 | -0.1249838 | 0.34876512 | -0.358361 | 0.72007319 | 0.95011251 |
| LINC02086 | 59.1354759 | -0.3941139 | 0.38615454 | -1.0206119 | 0.30743834 | 0.81568231 |
| TTLL6 | 11.2367465 | 0.9574225 | 0.67667881 | 1.41488472 | 0.1571023 | NA |
| CALCOCO2 | 58.6546229 | -0.1726137 | 0.29036337 | -0.5944747 | 0.55219463 | 0.91322136 |
| ATP5MC1 | 70.1902783 | 0.47564877 | 0.42799015 | 1.11135447 | 0.2664158 | 0.79666376 |
| UBE2Z | 90.2416067 | -0.2942506 | 0.21112401 | -1.3937335 | 0.1633981 | 0.71332671 |
| SNF8 | 30.9011726 | -0.320635 | 0.28578842 | -1.1219314 | 0.26189159 | 0.79633795 |
| ZNF652 | 138.162044 | -0.0441422 | 0.20357888 | -0.2168307 | 0.82834025 | 0.97567084 |
| PHB | 52.4243874 | 0.56052411 | 0.48285855 | 1.16084535 | 0.2457048 | 0.78583441 |
| SPOP | 83.1512415 | -0.0331877 | 0.19922357 | -0.166585 | 0.86769657 | 0.97805828 |
| SLC35B1 | 12.7491825 | -0.3932597 | 0.4866156 | -0.8081526 | 0.41900274 | NA |
| FAM117A | 10.5063604 | 0.08405401 | 0.48412032 | 0.17362216 | 0.86216242 | NA |
| KAT7 | 39.8648061 | -0.2308806 | 0.28022457 | -0.8239129 | 0.40998907 | 0.85867915 |
| ITGA3 | 257.239984 | -0.8884289 | 0.55650808 | -1.5964349 | 1.10E-01 | 0.66311481 |
| PDK2 | 21.451403 | 0.79289996 | 0.3964944 | 1.99977592 | 0.04552447 | 0.52804951 |

| | | | | | | |
|---|---|---|---|---|---|---|
| PPP1R9B | 12.9891397 | -0.3807868 | 0.48899464 | -0.7787137 | 0.43614839 | NA |
| XYLT2 | 21.1912772 | 0.23317158 | 0.41929347 | 0.55610593 | 0.57813844 | 0.91878391 |
| MRPL27 | 34.4565827 | -0.1694188 | 0.32706034 | -0.5180047 | 0.60445498 | 0.92531731 |
| LRRC59 | 15.230327 | -0.0927111 | 0.37518733 | -0.2471062 | 0.80482606 | NA |
| ACSF2 | 22.5838001 | 1.26483763 | 0.49260149 | 2.56766911 | 0.01023848 | 0.33306084 |
| RSAD1 | 19.0802362 | 0.39057164 | 0.39266386 | 0.99467172 | 0.31989595 | 0.82449332 |
| EPN3 | 17.1551373 | 1.34873693 | 0.59823384 | 2.25453132 | 0.02416276 | 0.43641593 |
| SPATA20 | 16.2057666 | 0.39785185 | 0.45108563 | 0.88198743 | 0.37778361 | 0.85310518 |
| ABCC3 | 85.3774105 | -0.3772635 | 0.78704229 | -0.4793434 | 0.63169435 | 0.93251095 |
| ANKRD40 | 21.4387343 | 0.10275557 | 0.33529691 | 0.30646142 | 0.75925335 | 0.95903525 |
| LUC7L3 | 226.69458 | -0.1169953 | 0.21836824 | -0.5357708 | 0.592117 | 0.92278087 |
| TOB1 | 66.5885164 | -0.0010509 | 0.35709591 | -0.002943 | 0.99765186 | 0.99956021 |
| SPAG9 | 273.222602 | -0.8233073 | 0.30277639 | -2.7191925 | 0.00654415 | 0.29935381 |
| AC005920.2 | 23.2637142 | -0.1418067 | 0.68370628 | -0.2074087 | 0.83569066 | 0.97600352 |
| MBTD1 | 130.559057 | 0.0627552 | 0.23759682 | 0.26412477 | 0.79168378 | 0.96925674 |
| UTP18 | 30.4289116 | -0.6720508 | 0.29392274 | -2.2864881 | 0.02222572 | 0.4232651 |
| CA10 | 84.6581135 | -0.3817417 | 0.78623499 | -0.4855314 | 0.62729948 | 0.93057946 |
| TOM1L1 | 219.193163 | 0.01401481 | 0.17937669 | 0.07813063 | 0.93772414 | 0.99012269 |
| COX11 | 41.4369468 | 0.22554354 | 0.2934424 | 0.76861264 | 0.4421233 | 0.86925451 |
| STXBP4 | 241.630792 | 0.06933297 | 0.25813138 | 0.26859568 | 0.78824084 | 0.96768123 |
| HLF | 134.830657 | -0.242558 | 0.41144607 | -0.5895256 | 0.55550877 | 0.91398748 |
| MMD | 20.7454284 | 0.3714682 | 0.43554947 | 0.85287257 | 0.39372997 | 0.85605713 |
| PCTP | 19.8754072 | 0.36129135 | 0.3795081 | 0.95199905 | 0.34109746 | 0.8322193 |
| DGKE | 35.4143685 | 0.36231306 | 0.38727149 | 0.93555314 | 0.34950331 | 0.83687845 |
| TRIM25 | 36.7495319 | -0.4052225 | 0.48524971 | -0.8350803 | 0.4036725 | 0.85782098 |
| COIL | 23.1978898 | -0.1744768 | 0.30721748 | -0.5679259 | 0.5700853 | 0.91778542 |
| SCPEP1 | 151.99763 | 0.68166586 | 0.31490065 | 2.16470135 | 0.03041056 | 0.46919892 |
| AKAP1 | 61.7852357 | 0.62648919 | 0.30102423 | 2.08119192 | 0.03741634 | 0.49538341 |
| MSI2 | 1187.41282 | 0.02093764 | 0.15620241 | 0.13404171 | 0.89336959 | 0.98377746 |
| MRPS23 | 21.6117875 | -0.0072594 | 0.36315241 | -0.0199898 | 0.98405148 | 0.99685229 |
| CUEDC1 | 110.84524 | -0.3829448 | 0.41192957 | -0.9296365 | 0.35255932 | 0.83872681 |
| VEZF1 | 65.9792639 | 0.04988243 | 0.22655377 | 0.22017922 | 0.82573158 | 0.97567084 |
| AC015813.2 | 31.386521 | 0.26585705 | 0.34841527 | 0.76304647 | 0.44543568 | 0.87162533 |
| SRSF1 | 14.9581941 | 0.41009991 | 0.55082177 | 0.74452379 | 0.45655965 | NA |
| DYNLL2 | 44.4477536 | 0.27178579 | 0.37599039 | 0.72285304 | 0.46977018 | 0.88207878 |
| MKS1 | 13.1154444 | -0.3667247 | 0.44891425 | -0.8169148 | 0.4139771 | NA |
| TSPOAP1 | 28.8666974 | 0.98506916 | 0.48513336 | 2.0305121 | 0.04230451 | 0.51343649 |
| TSPOAP1-AS | 11.5313852 | 1.09861994 | 0.6865172 | 1.60028028 | 0.10953642 | NA |
| SUPT4H1 | 16.0959986 | 0.13233522 | 0.413217 | 0.32025598 | 0.74877429 | NA |
| RNF43 | 76.724433 | 0.64475923 | 0.42668433 | 1.5110919 | 0.13076504 | 0.67442162 |
| MTMR4 | 29.1931432 | 0.14134978 | 0.37037508 | 0.38163957 | 0.70272873 | 0.94476609 |
| RAD51C | 19.1152265 | 0.17204868 | 0.42460509 | 0.40519693 | 0.68533277 | 0.94205398 |
| TRIM37 | 55.9782669 | -0.3632806 | 0.30867972 | -1.1768852 | 0.23924131 | 0.78098673 |
| SKA2 | 26.274969 | 0.28382105 | 0.36727937 | 0.77276613 | 0.4396608 | 0.86895963 |
| GDPD1 | 19.854771 | 0.35265135 | 0.45267495 | 0.77903881 | 0.43595686 | 0.86729771 |
| YPEL2 | 145.643667 | 0.18889762 | 0.40393745 | 0.46764077 | 0.64004151 | 0.93328692 |
| DHX40 | 53.8253676 | -0.069711 | 0.304908 | -0.2286296 | 0.8191568 | 0.97296246 |
| CLTC | 249.255619 | -0.1681747 | 0.217334 | -0.7738078 | 0.43904446 | 0.86879768 |
| PTRH2 | 33.9365835 | -0.1256605 | 0.32641338 | -0.3849736 | 0.70025694 | 0.94424267 |
| VMP1 | 507.576981 | -0.9294257 | 0.39018863 | -2.3819908 | 0.01721933 | 0.40063153 |
| TUBD1 | 34.3416124 | 0.63413739 | 0.41335239 | 1.53413264 | 0.12499703 | 0.66929384 |
| RPS6KB1 | 50.9931195 | 0.17802883 | 0.3079466 | 0.57811591 | 0.56318586 | 0.91624774 |
| RNFT1 | 11.7391635 | 0.20022187 | 0.45572081 | 0.43935205 | 0.66040647 | NA |
| HEATR6 | 15.7772898 | 0.17193087 | 0.42625037 | 0.40335652 | 0.68668597 | NA |

| | | | | | | |
|---|---|---|---|---|---|---|
| USP32 | 199.21357 | -0.5260084 | 0.25765076 | -2.041556 | 0.0411956 | 0.50744725 |
| APPBP2 | 133.954577 | -0.0233568 | 0.17988957 | -0.1298396 | 0.89669331 | 0.98531749 |
| AC011921.1 | 10.8910187 | 1.55378588 | 0.5869285 | 2.64731714 | 0.00811332 | NA |
| PPM1D | 56.3849953 | 0.3803063 | 0.38573613 | 0.98592347 | 0.32417067 | 0.82567895 |
| BCAS3 | 682.156833 | -0.0187369 | 0.18370017 | -0.101997 | 0.91875908 | 0.98749988 |
| AC005746.2 | 25.3830229 | 0.90096398 | 0.57056094 | 1.57908457 | 0.11431666 | 0.66629396 |
| TBX2 | 164.05819 | -0.0595098 | 0.32921477 | -0.1807629 | 8.57E-01 | 0.97646931 |
| INTS2 | 33.5498444 | -0.1113155 | 0.30746862 | -0.3620385 | 0.71732329 | 0.94925929 |
| MED13 | 312.508608 | -0.3233279 | 0.22579869 | -1.4319297 | 0.15216394 | 0.70572801 |
| METTL2A | 17.3213192 | 0.32523693 | 0.38696462 | 0.84048233 | 0.400638 | 0.85778581 |
| TLK2 | 156.196301 | -0.2079084 | 0.21588542 | -0.9630496 | 0.33552264 | 0.82974237 |
| TANC2 | 491.192728 | -0.3367784 | 0.22000706 | -1.5307616 | 0.12582832 | 0.66929384 |
| AC015923.1 | 30.3362087 | 0.20676634 | 0.67265046 | 0.30739047 | 0.75854618 | 0.95903525 |
| CYB561 | 23.6183011 | 0.12798725 | 0.43256611 | 0.29587906 | 0.76732244 | 0.96283923 |
| DCAF7 | 116.623773 | 0.06967433 | 0.23465457 | 0.29692295 | 0.76652533 | 0.96270421 |
| MAP3K3 | 98.5577994 | -0.3498591 | 0.30403449 | -1.1507217 | 0.24984675 | 0.78799766 |
| CCDC47 | 23.969349 | -0.1405344 | 0.32026952 | -0.4388005 | 0.66080613 | 0.93729379 |
| DDX42 | 87.3268703 | -0.2533946 | 0.26669835 | -0.9501169 | 0.34205287 | 0.83304756 |
| FTSJ3 | 16.4783054 | -0.1775026 | 0.38601141 | -0.4598377 | 0.64563272 | 0.93328692 |
| PSMC5 | 17.9044767 | 0.31675405 | 0.4994008 | 0.6342682 | 0.5259058 | 0.90487731 |
| SMARCD2 | 27.3063383 | 0.35238682 | 0.42483229 | 0.82947277 | 0.40683694 | 0.85867915 |
| ERN1 | 60.2599882 | -0.7916668 | 0.35594058 | -2.2241544 | 2.61E-02 | 0.4455707 |
| SNHG25 | 19.7360377 | -0.3417839 | 0.50389337 | -0.6782862 | 0.49759022 | 0.89378207 |
| TEX2 | 73.0783394 | -0.8622311 | 0.27779097 | -3.1038847 | 0.00190998 | 0.17924252 |
| POLG2 | 26.3415605 | -0.3345325 | 0.33876009 | -0.9875205 | 0.32338754 | 0.82558302 |
| DDX5 | 360.863728 | 0.15869046 | 0.2360119 | 0.67238328 | 0.50133972 | 0.89492452 |
| CEP95 | 126.182523 | -0.1345218 | 0.28184552 | -0.4772892 | 0.63315619 | 0.93251095 |
| SMURF2 | 130.970477 | -0.3184681 | 0.23131833 | -1.3767527 | 0.16858873 | 0.72151539 |
| LRRC37A3 | 134.876373 | 0.04376356 | 0.31587914 | 0.13854525 | 0.88980951 | 0.9827634 |
| GNA13 | 100.882159 | -0.9773803 | 0.32281445 | -3.0276845 | 0.00246435 | 0.19678071 |
| CEP112 | 158.406986 | 0.43671115 | 0.37731792 | 1.15740899 | 0.2471053 | 0.78583441 |
| PRKCA | 836.401913 | -0.4217804 | 0.31113827 | -1.3556044 | 0.17522506 | 0.72841249 |
| PRKCA-AS1 | 34.0161454 | -0.3069312 | 0.64802669 | -0.4736397 | 0.63575688 | 0.93251095 |
| HELZ | 248.959375 | -0.302874 | 0.17359674 | -1.7446985 | 0.08103737 | 0.61925304 |
| PSMD12 | 47.6523592 | 0.00108362 | 0.24887246 | 0.0043541 | 0.99652594 | 0.99927028 |
| PITPNC1 | 184.654253 | -0.4835996 | 0.37287944 | -1.2969329 | 0.19465427 | 0.75191699 |
| NOL11 | 28.2132073 | -0.6212306 | 0.32556676 | -1.9081513 | 0.05637167 | 0.56976762 |
| BPTF | 349.869325 | -0.1269642 | 0.19273995 | -0.6587334 | 0.51006699 | 0.8998333 |
| AC005332.7 | 14.3096104 | -0.4957744 | 0.39963296 | -1.2405743 | 0.21476305 | NA |
| AMZ2 | 13.2762029 | -0.4659029 | 0.44097731 | -1.0565235 | 0.29072913 | NA |
| ARSG | 50.8507025 | -0.2087222 | 0.42183828 | -0.494792 | 0.62074696 | 0.92805465 |
| WIPI1 | 40.0717236 | -0.233893 | 0.4178389 | -0.5597683 | 0.5756375 | 0.91854298 |
| PRKAR1A | 197.734694 | -0.0951044 | 0.32964241 | -0.2885078 | 0.77295805 | 0.96435639 |
| FAM20A | 192.530542 | 0.20834789 | 0.42537336 | 0.48980004 | 6.24E-01 | 9.30E-01 |
| LINC01482 | 147.87783 | 1.40041119 | 0.60932742 | 2.29829011 | 0.02154528 | 0.4212027 |
| ABCA10 | 48.5264819 | -0.2157399 | 0.37086547 | -0.5817202 | 0.56075516 | 0.91584721 |
| ABCA5 | 160.273387 | 0.11236994 | 0.28623664 | 0.39257705 | 0.69463189 | 0.94395238 |
| MAP2K6 | 18.2601093 | -0.3834604 | 0.79094822 | -0.484811 | 0.6278104 | 0.93061955 |
| AC005208.1 | 246.613788 | 0.87784495 | 0.64643484 | 1.35797902 | 0.17447034 | 0.72841249 |
| KCNJ16 | 863.984139 | 0.19299877 | 0.33057243 | 0.58383201 | 0.55933334 | 0.91477029 |
| SOX9 | 70.1093218 | 0.72023349 | 0.52690943 | 1.36690188 | 1.72E-01 | 7.25E-01 |
| LINC00511 | 92.1495568 | 0.67925388 | 0.78057143 | 0.8702008 | 3.84E-01 | 8.55E-01 |
| SLC39A11 | 150.944001 | -0.3239836 | 0.26252685 | -1.2340971 | 0.21716673 | 0.76666749 |
| COG1 | 18.761246 | 0.00419055 | 0.36611096 | 0.01144611 | 0.99086752 | 0.99814944 |

| | | | | | | |
|---|---|---|---|---|---|---|
| FAM104A | 22.0994222 | 0.69904345 | 0.33725463 | 2.0727468 | 0.03819585 | 0.49671741 |
| C17orf80 | 16.8505891 | 0.25963019 | 0.38044763 | 0.68243345 | 0.49496491 | 0.89230917 |
| CDC42EP4 | 26.6240793 | 0.63377134 | 0.34406852 | 1.84199162 | 0.06547637 | 0.59306611 |
| RPL38 | 46.554864 | 0.20035076 | 0.44566486 | 0.44955475 | 0.65303152 | 0.93532021 |
| GPRC5C | 155.615083 | -0.1555921 | 0.23390024 | -0.6652071 | 0.50591805 | 0.89723255 |
| SLC9A3R1 | 23.3049188 | 0.07258511 | 0.45952724 | 0.15795606 | 0.87449142 | 0.97918624 |
| NAT9 | 13.3786151 | 0.21142173 | 0.43832 | 0.48234561 | 0.62956045 | NA |
| TMEM104 | 11.5224074 | -0.7239661 | 0.43827941 | -1.6518369 | 0.09856781 | NA |
| CDR2L | 10.3621763 | 0.19946609 | 0.460006 | 0.43361628 | 0.66456711 | NA |
| KCTD2 | 30.2985187 | -0.3757544 | 0.27802712 | -1.3515026 | 0.17653448 | 0.72841249 |
| ATP5PD | 40.6664053 | 0.05120578 | 0.36854615 | 0.13893993 | 0.88949761 | 0.9827634 |
| SLC16A5 | 113.38139 | 0.01905112 | 0.40938305 | 0.04653617 | 0.9628829 | 0.99256424 |
| ARMC7 | 31.6451914 | 0.76536889 | 0.34393107 | 2.22535546 | 0.02605738 | 0.4455707 |
| NT5C | 32.2666178 | 0.31800328 | 0.38144035 | 0.83369073 | 0.40445528 | 0.85782098 |
| JPT1 | 37.2423472 | -0.3358113 | 0.43610422 | -0.7700254 | 0.44128485 | 0.86925451 |
| SUMO2 | 27.2763187 | -0.5595145 | 0.30268143 | -1.8485259 | 0.0645263 | 0.58883813 |
| NUP85 | 31.3357962 | -0.2307603 | 0.30413874 | -0.7587338 | 0.4480118 | 0.87291341 |
| GGA3 | 35.0166192 | -0.067409 | 0.3272708 | -0.2059731 | 0.8368119 | 0.97600352 |
| MRPS7 | 25.6807366 | 0.30555925 | 0.42141465 | 0.72507981 | 0.46840307 | 0.88170434 |
| GRB2 | 109.795308 | -0.2774865 | 0.21453495 | -1.2934326 | 0.19586147 | 0.75350056 |
| AC011933.4 | 13.802925 | 0.23837067 | 0.61245653 | 0.38920422 | 0.69712508 | NA |
| TMEM94 | 29.716313 | 0.09326726 | 0.42118961 | 0.22143771 | 0.82475163 | 0.97518529 |
| CASKIN2 | 17.0855709 | 0.13546314 | 0.42029019 | 0.32230858 | 0.74721893 | 0.95563105 |
| TSEN54 | 23.7249207 | 0.283777 | 0.30956966 | 0.91668219 | 0.3593092 | 0.84204868 |
| LLGL2 | 371.785717 | 0.30824802 | 0.23950348 | 1.28702941 | 0.19808404 | 0.75559912 |
| MYO15B | 44.8340018 | -0.3853622 | 0.34265834 | -1.1246251 | 0.26074793 | 0.79574359 |
| RECQL5 | 41.4620356 | -0.3269788 | 0.34162189 | -0.9571366 | 0.33849832 | 0.83055261 |
| SMIM5 | 48.0578122 | -0.4643443 | 0.53572518 | -0.8667584 | 3.86E-01 | 0.85586379 |
| SAP30BP | 48.5474908 | -0.5722843 | 0.30548843 | -1.8733421 | 0.06102116 | 0.57922376 |
| ITGB4 | 16.6147732 | 0.95944826 | 0.56914788 | 1.68576269 | 0.09184152 | 0.64452635 |
| H3F3B | 212.101713 | 0.07448818 | 0.43436402 | 0.17148791 | 0.86384013 | 0.97788853 |
| UNK | 70.4445852 | -0.156482 | 0.30548363 | -0.5122434 | 0.60848068 | 0.92573951 |
| WBP2 | 54.6333324 | -0.0672671 | 0.32530725 | -0.2067801 | 0.83618162 | 0.97600352 |
| TRIM47 | 17.3744225 | -0.0217487 | 0.52582072 | -0.0413615 | 0.96700773 | 0.99308116 |
| TRIM65 | 11.7121547 | -0.1582666 | 0.40801346 | -0.3878955 | 0.69809337 | NA |
| FBF1 | 8.96380413 | 0.35866669 | 0.56894497 | 0.63040664 | 0.52842857 | NA |
| ACOX1 | 59.4342259 | 0.20777478 | 0.2329868 | 0.89178777 | 0.3725067 | 0.8491102 |
| EVPL | 19.4333821 | 0.41843788 | 0.40623801 | 1.03003135 | 0.30299529 | 0.81568231 |
| SRP68 | 31.9469962 | -0.1253273 | 0.30092528 | -0.4164733 | 0.67706369 | 0.94034237 |
| EXOC7 | 74.420555 | -0.0854077 | 0.32169019 | -0.2654967 | 0.79062687 | 0.96904219 |
| RNF157 | 10.8777914 | 0.30847679 | 0.50554601 | 0.61018538 | 0.54173901 | NA |
| UBALD2 | 30.4551543 | -0.4112237 | 0.38942609 | -1.0559737 | 0.29098024 | 0.80640885 |
| PRPSAP1 | 41.2992884 | 0.22552618 | 0.30517251 | 0.73901212 | 0.45989963 | 0.87607759 |
| UBE2O | 51.5790864 | -0.1011759 | 0.25406963 | -0.3982211 | 0.69046719 | 0.94276429 |
| RHBDF2 | 16.791004 | 0.38228522 | 0.48277735 | 0.7918458 | 0.42845059 | 0.86306389 |
| MXRA7 | 24.2008685 | 0.56105299 | 0.41858397 | 1.34035947 | 0.1801285 | 0.73244853 |
| JMJD6 | 15.6151721 | -0.0471228 | 0.4842059 | -0.0973197 | 0.92247248 | NA |
| SRSF2 | 75.9145362 | 0.16606818 | 0.29755979 | 0.55810019 | 0.57677597 | 0.91854298 |
| MFSD11 | 46.7821637 | -0.1167465 | 0.26561963 | -0.4395252 | 0.660281 | 0.93728744 |
| SEC14L1 | 96.3509023 | -0.270795 | 0.26731446 | -1.0130203 | 0.31105049 | 0.81701824 |
| 9-Sep | 370.079058 | -0.4433088 | 0.32514303 | -1.3634269 | 0.172748 | 0.72701113 |
| TNRC6C | 89.6905658 | -0.102889 | 0.33012449 | -0.3116674 | 0.75529329 | 0.95838233 |
| TMC6 | 15.6936583 | 0.26116454 | 0.44011526 | 0.59340033 | 0.55291327 | NA |
| SYNGR2 | 41.7801641 | -0.1713184 | 0.38285077 | -0.4474808 | 0.65452796 | 0.93548446 |

| | | | | | | |
|---|---|---|---|---|---|---|
| AFMID | 57.5292093 | 0.07073249 | 0.30457508 | 0.23223336 | 8.16E-01 | 0.97288403 |
| SOCS3 | 38.3825536 | 0.20901846 | 0.45644375 | 0.45792818 | 6.47E-01 | 9.33E-01 |
| PGS1 | 40.8980034 | -0.0970093 | 0.31980485 | -0.303339 | 0.76163156 | 0.96000167 |
| CYTH1 | 107.17323 | -0.1244053 | 0.27109568 | -0.4588979 | 0.64630745 | 0.93328692 |
| USP36 | 90.5711008 | -0.7946699 | 0.56489467 | -1.4067576 | 0.15949928 | 0.71101785 |
| TIMP2 | 46.4569696 | -0.1656171 | 0.45283757 | -0.3657318 | 0.71456523 | 0.94841799 |
| LGALS3BP | 95.521811 | -0.1049957 | 0.65034894 | -0.1614452 | 0.87174274 | 0.97866716 |
| CANT1 | 25.4459567 | 0.34049894 | 0.48256413 | 0.7056035 | 0.48043474 | 0.88660781 |
| ENGASE | 14.6305717 | 0.36176156 | 0.47397408 | 0.76325177 | 0.44531326 | NA |
| TBC1D16 | 126.335966 | 0.17908591 | 0.33912931 | 0.52807559 | 0.59744688 | 0.9229638 |
| CCDC40 | 40.2293705 | -0.0172581 | 0.38738811 | -0.0445498 | 0.96446612 | 0.99264137 |
| GAA | 47.7768065 | 0.45477792 | 0.43597538 | 1.04312752 | 0.29688923 | 0.81137495 |
| EIF4A3 | 20.3402475 | 0.26551884 | 0.37606289 | 0.70604903 | 0.48015765 | 0.88660781 |
| AC087741.1 | 17.2314947 | -0.0986785 | 0.51179509 | -0.1928086 | 0.84710889 | 0.97612951 |
| SGSH | 13.3869434 | 0.14756868 | 0.42295043 | 0.34890301 | 0.72716212 | NA |
| SLC26A11 | 37.6912592 | 0.11180687 | 0.31567377 | 0.35418485 | 0.72320035 | 0.95093722 |
| RNF213 | 633.065969 | -0.3941347 | 0.51380973 | -0.7670829 | 4.43E-01 | 0.86991418 |
| AC124319.2 | 139.95584 | 0.02198427 | 0.34862285 | 0.06306033 | 0.94971846 | 0.99105535 |
| ENDOV | 19.5685268 | -0.0680806 | 0.37425786 | -0.1819083 | 0.85565469 | 0.97631294 |
| RPTOR | 146.921598 | -0.4452749 | 0.23224599 | -1.9172556 | 0.05520547 | 0.5645224 |
| CHMP6 | 11.0253994 | 0.02763517 | 0.46935159 | 0.05887947 | 0.95304811 | NA |
| BAIAP2-DT | 11.0108507 | 0.66061499 | 0.56485213 | 1.16953616 | 0.24218768 | NA |
| BAIAP2 | 25.9473328 | -0.6225524 | 0.34607822 | -1.7988777 | 0.07203803 | 0.60821941 |
| TEPSIN | 11.3288058 | 0.7858038 | 0.49249037 | 1.59557191 | 0.1105844 | NA |
| NDUFAF8 | 26.3694735 | 0.0767966 | 0.41695964 | 0.18418232 | 0.85387043 | 0.97612951 |
| SLC38A10 | 69.372487 | 0.30667117 | 0.21471716 | 1.42825648 | 0.15321805 | 0.70572801 |
| BAHCC1 | 16.8174948 | 0.39326711 | 0.44092322 | 0.89191744 | 0.37243719 | 0.8491102 |
| ACTG1 | 435.72274 | -0.3089672 | 0.54142025 | -0.5706607 | 0.56822968 | 0.91698355 |
| FAAP100 | 10.8923029 | -0.3905262 | 0.47655112 | -0.8194843 | 0.41251016 | NA |
| NPLOC4 | 103.361736 | -0.4638673 | 0.20906662 | -2.2187535 | 0.0265035 | 0.44763538 |
| OXLD1 | 16.294178 | -0.0208933 | 0.36294839 | -0.0575656 | 0.95409465 | 0.99148072 |
| CCDC137 | 21.5640273 | -0.5911293 | 0.4383027 | -1.3486783 | 0.17744032 | 0.72841249 |
| ARL16 | 19.9616382 | 0.08631088 | 0.3935214 | 0.21932958 | 0.82639332 | 0.97567084 |
| HGS | 43.8814086 | -0.7775898 | 0.28588107 | -2.7199765 | 0.00652866 | 0.29935381 |
| GCGR | 31.4129979 | 1.11425173 | 0.51451398 | 2.16563936 | 0.03033876 | 0.46888573 |
| MCRIP1 | 26.6861136 | -0.3037012 | 0.33122083 | -0.9169145 | 0.35918746 | 0.84204868 |
| P4HB | 113.69438 | -0.0812318 | 0.24250211 | -0.3349736 | 0.73764499 | 0.9542764 |
| ARHGDIA | 50.918287 | -0.5096472 | 0.47485506 | -1.0732689 | 0.28315049 | 0.80431986 |
| ANAPC11 | 35.7979312 | 0.70048134 | 0.40224631 | 1.74142392 | 0.0816093 | 0.61925304 |
| PCYT2 | 13.9100638 | 0.29535817 | 0.69328868 | 0.4260248 | 0.67008977 | NA |
| SIRT7 | 16.9055489 | 0.04370504 | 0.42624947 | 0.10253395 | 0.91833287 | 0.9874302 |
| MAFG | 15.7161012 | -0.5656865 | 0.34643964 | -1.6328572 | 0.10249904 | NA |
| ASPSCR1 | 25.9506788 | -0.2927173 | 0.37396105 | -0.782748 | 0.43377511 | 0.86729771 |
| CENPX | 12.9666564 | 0.31122875 | 0.62002917 | 0.50195824 | 0.6156969 | NA |
| DCXR | 19.7167195 | -0.2483345 | 0.45546404 | -0.5452341 | 0.58559254 | 0.92076087 |
| RFNG | 18.4702613 | 0.23776387 | 0.44823146 | 0.53044886 | 0.59580076 | 0.9229638 |
| GPS1 | 20.71524 | 0.04929051 | 0.43311889 | 0.11380365 | 0.90939345 | 0.98550102 |
| DUS1L | 23.4093686 | -0.0067789 | 0.50608515 | -0.0133947 | 0.98931289 | 0.99814944 |
| CCDC57 | 122.271278 | 0.04897469 | 0.27211662 | 0.17997686 | 0.85717074 | 0.97675334 |
| SLC16A3 | 44.597908 | -0.6367457 | 0.48599499 | -1.3101898 | 0.19013164 | 0.74569941 |
| CSNK1D | 86.1545697 | -0.541211 | 0.30356761 | -1.7828351 | 0.07461315 | 0.61020315 |
| OGFOD3 | 26.3808197 | -0.0744493 | 0.35898095 | -0.2073905 | 0.83570488 | 0.97600352 |
| HEXDC | 40.7579582 | -0.3352084 | 0.40899695 | -0.8195865 | 0.41245187 | 0.85867915 |
| CYBC1 | 22.2604025 | 0.18806659 | 0.37903104 | 0.49617727 | 0.61976935 | 0.92798856 |

| | | | | | | |
|---|---|---|---|---|---|---|
| NARF | 37.2267213 | 0.01906458 | 0.3558008 | 0.05358218 | 0.95726805 | 0.99170927 |
| FOXK2 | 129.343934 | -0.3185791 | 0.26618482 | -1.1968344 | 0.23137111 | 0.77262335 |
| WDR45B | 81.5304441 | -0.6178001 | 0.26966983 | -2.2909502 | 0.02196629 | 0.4212027 |
| RAB40B | 52.9565122 | -1.0449475 | 0.39991572 | -2.6129192 | 0.00897725 | 0.31797637 |
| FN3KRP | 16.7652121 | -0.8283741 | 0.41093998 | -2.0158032 | 0.04382056 | 0.52007638 |
| FN3K | 41.7642445 | 0.15797626 | 0.52428313 | 0.30131861 | 0.76317155 | 0.96091323 |
| TBCD | 54.2487093 | -0.2825867 | 0.34149348 | -0.8275025 | 0.40795228 | 0.85867915 |
| METRNL | 50.358582 | -0.355428 | 0.4417668 | -0.8045603 | 0.42107346 | 0.86190547 |
| USP14 | 95.7269609 | -0.5956629 | 0.29797302 | -1.9990498 | 0.04560297 | 0.52804951 |
| THOC1 | 78.0071189 | -0.1231113 | 0.24691408 | -0.4985999 | 0.6180613 | 0.92749951 |
| ENOSF1 | 274.278614 | 0.04411615 | 0.30170213 | 0.1462242 | 0.8837444 | 0.98154805 |
| YES1 | 98.9241004 | -0.712027 | 0.2186262 | -3.2568235 | 0.00112666 | 0.14394871 |
| METTL4 | 27.9023777 | -0.2828004 | 0.3562266 | -0.7938779 | 0.42726652 | 0.86239557 |
| SMCHD1 | 217.673117 | -0.292276 | 0.32369827 | -0.9029271 | 0.36656456 | 0.84741115 |
| AP001011.1 | 30.1124712 | 0.78115864 | 0.64846965 | 1.20461865 | 0.22835055 | 0.77185154 |
| LPIN2 | 141.895007 | -0.4248662 | 0.38321717 | -1.1086826 | 0.26756711 | 0.79666376 |
| MYL12A | 96.3425313 | 0.06112196 | 0.35673988 | 0.17133482 | 0.86396051 | 0.97788853 |
| MYL12B | 202.869526 | 0.31467036 | 0.39580895 | 0.79500568 | 0.4266102 | 0.86239557 |
| TGIF1 | 42.4446809 | -1.0124695 | 0.46359786 | -2.183939 | 0.02896673 | 0.46182173 |
| DLGAP1 | 237.337582 | 0.18771126 | 0.77776223 | 0.24134787 | 8.09E-01 | 9.73E-01 |
| DLGAP1-AS1 | 21.3295744 | -0.2527981 | 0.50562392 | -0.4999726 | 0.61709436 | 0.92711833 |
| LINC00667 | 14.1748134 | 0.18260093 | 0.39031025 | 0.46783535 | 0.63990234 | NA |
| EPB41L3 | 324.649008 | -0.2450553 | 0.36450555 | -0.6722951 | 5.01E-01 | 0.89492452 |
| L3MBTL4 | 182.391401 | 0.06764478 | 0.36181345 | 0.18696038 | 0.8516917 | 0.97612951 |
| ARHGAP28 | 21.9017706 | -0.8351986 | 0.61489882 | -1.35827 | 0.17437802 | 0.72841249 |
| LAMA1 | 67.9089494 | -0.2125642 | 0.58122214 | -0.3657194 | 0.71457448 | 0.94841799 |
| PTPRM | 722.007561 | 0.1124635 | 0.16070069 | 0.69983211 | 0.48403216 | 0.88838332 |
| RAB12 | 62.0926071 | -0.0833032 | 0.25520149 | -0.3264212 | 0.74410572 | 0.95563105 |
| MTCL1 | 16.9975324 | 0.30820899 | 0.44573351 | 0.6914647 | 0.48927355 | 0.88930178 |
| NDUFV2 | 93.2545176 | -0.5365192 | 0.25621693 | -2.0940038 | 0.03625964 | 0.49485984 |
| ANKRD12 | 396.088914 | -0.3887583 | 0.16404262 | -2.3698616 | 0.01779474 | 0.40802409 |
| TWSG1 | 159.399223 | -0.0587057 | 0.36430907 | -0.1611425 | 0.87198118 | 0.97866716 |
| AP005432.2 | 12.4713295 | 2.63797269 | 1.14347446 | 2.30697999 | 0.02105594 | NA |
| RALBP1 | 391.116798 | 0.84636144 | 0.33807321 | 2.50348573 | 0.01229766 | 0.36046953 |
| PPP4R1 | 137.684269 | -0.4157713 | 0.30828433 | -1.3486617 | 0.17744566 | 0.72841249 |
| RAB31 | 53.4410244 | -0.3161346 | 0.37100467 | -0.8521042 | 0.39415628 | 0.85605713 |
| VAPA | 204.312281 | -0.2989084 | 0.24022446 | -1.2442882 | 0.21339352 | 0.7643572 |
| NAPG | 62.7958643 | -0.2420773 | 0.27706568 | -0.8737181 | 0.38227178 | 0.8548566 |
| GNAL | 28.4834939 | 0.9973305 | 0.43924032 | 2.2705805 | 0.02317238 | 0.43085291 |
| MPPE1 | 42.0302175 | -0.1864055 | 0.32333977 | -0.5765004 | 0.564277 | 0.91658856 |
| IMPA2 | 95.6700576 | -0.4153116 | 0.35365481 | -1.1743418 | 0.24025813 | 0.78192112 |
| AFG3L2 | 125.824144 | -0.2348311 | 0.26418476 | -0.8888895 | 0.37406245 | 0.8508472 |
| SPIRE1 | 680.557982 | -0.3978603 | 0.28780441 | -1.3823982 | 0.16684948 | 0.71878854 |
| PSMG2 | 98.1497027 | 0.24954043 | 0.22867563 | 1.09124191 | 0.27516645 | 0.7994854 |
| CEP76 | 23.96227 | 0.288279 | 0.49369544 | 0.58392072 | 0.55927365 | 0.91477029 |
| PTPN2 | 147.394276 | -0.6721123 | 0.28530624 | -2.3557575 | 0.01848498 | 0.41315568 |
| SEH1L | 32.895236 | -0.2943219 | 0.34583642 | -0.8510437 | 0.39474506 | 0.85605713 |
| CEP192 | 138.742506 | -0.26474 | 0.31346957 | -0.8445477 | 0.39836339 | 0.85758974 |
| LDLRAD4 | 29.9837225 | -0.1340311 | 0.61717518 | -0.2171687 | 0.82807688 | 0.97567084 |
| FAM210A | 54.5509619 | -0.7653262 | 0.33340302 | -2.2954988 | 0.02170455 | 0.4212027 |
| RNMT | 86.4887991 | -0.6190379 | 0.21892661 | -2.8276047 | 0.00468977 | 0.27019594 |
| ROCK1 | 209.177599 | -0.2915543 | 0.16393518 | -1.778473 | 0.0753262 | 0.61020315 |
| ESCO1 | 79.4062948 | -0.5415832 | 0.2494286 | -2.1712954 | 0.02990885 | 0.46381645 |
| SNRPD1 | 23.0165421 | -0.1267112 | 0.33331117 | -0.380159 | 0.70382738 | 0.94476609 |

| | | | | | | |
|---|---|---|---|---|---|---|
| ABHD3 | 84.014506 | -1.0649345 | 0.51544784 | -2.0660374 | 0.03882495 | 0.49918575 |
| MIB1 | 172.896737 | -0.4398865 | 0.2813952 | -1.5632338 | 0.11799762 | 0.6668162 |
| AC090912.1 | 25.8384198 | 0.16828631 | 0.55320065 | 0.30420483 | 0.76097184 | 0.95956874 |
| RBBP8 | 491.182402 | -0.4184653 | 0.51430228 | -0.8136562 | 0.41584192 | 0.85867915 |
| CABLES1 | 29.7418655 | 0.07467774 | 0.46541382 | 0.16045448 | 8.73E-01 | 9.79E-01 |
| TMEM241 | 31.2074475 | -0.119676 | 0.28359908 | -0.4219899 | 0.67303237 | 0.93880075 |
| RIOK3 | 103.021837 | -0.2804897 | 0.18146257 | -1.5457169 | 0.12217297 | 0.66929384 |
| RMC1 | 36.5403961 | -0.0752269 | 0.26975442 | -0.2788718 | 0.78034317 | 0.96650724 |
| NPC1 | 131.606146 | -1.235603 | 0.38611482 | -3.2000922 | 0.00137384 | 0.15830421 |
| TTC39C | 104.708443 | 0.34797764 | 0.33580559 | 1.03624731 | 0.30008678 | 0.81388512 |
| OSBPL1A | 73.0540788 | 0.37001347 | 0.29744259 | 1.24398285 | 0.21350588 | 0.7643572 |
| IMPACT | 18.2948543 | -0.0216763 | 0.40721503 | -0.0532307 | 0.95754812 | 0.99170927 |
| SS18 | 145.341949 | -0.1679782 | 0.24263587 | -0.6923056 | 0.48874543 | 0.88920066 |
| TAF4B | 21.7885199 | 0.0963141 | 0.32419703 | 0.29708508 | 0.76640155 | 0.96270421 |
| KCTD1 | 243.818488 | 0.04352013 | 0.21954756 | 0.19822644 | 0.8428679 | 0.97612951 |
| CHST9 | 24.9585615 | 0.71070982 | 0.57100584 | 1.24466297 | 0.21325566 | 0.7643572 |
| DSC2 | 14.9964547 | 0.10726924 | 0.41728829 | 0.25706266 | 0.7971304 | NA |
| DSG2 | 57.721871 | 0.38379941 | 0.34922764 | 1.09899495 | 0.27177027 | 0.79666376 |
| B4GALT6 | 22.9951767 | 0.43472206 | 0.44381632 | 0.97950897 | 0.32732856 | 0.82612204 |
| TRAPPC8 | 141.339931 | -0.250136 | 0.29469398 | -0.8487991 | 0.39599309 | 0.85605713 |
| RNF138 | 46.2313478 | -0.1307246 | 0.28355794 | -0.4610154 | 0.64478755 | 0.93328692 |
| GAREM1 | 769.873416 | -0.7898544 | 0.46890347 | -1.6844713 | 0.09209062 | 0.64534327 |
| KLHL14 | 41.5825846 | 0.81947254 | 0.49095534 | 1.66913867 | 0.0950899 | 0.64839204 |
| CCDC178 | 436.055751 | 1.18545456 | 0.44402898 | 2.66976845 | 0.00759036 | 0.30654023 |
| ASXL3 | 138.396139 | 0.30355341 | 0.31026036 | 0.97838283 | 0.32788502 | 0.82702613 |
| DTNA | 31.5752069 | 0.70467175 | 0.51499152 | 1.3683172 | 0.17121281 | 0.72463203 |
| MAPRE2 | 27.4072378 | -0.4230935 | 0.38984327 | -1.0852913 | 0.27779265 | 0.80193223 |
| ZNF397 | 49.3169813 | 0.0103273 | 0.29469111 | 0.03504448 | 0.97204427 | 0.99431961 |
| ZSCAN30 | 39.3607573 | -0.1322669 | 0.31367878 | -0.4216637 | 0.67327053 | 0.93898906 |
| ZNF24 | 46.4066756 | 0.26955692 | 0.29903438 | 0.90142451 | 0.36736265 | 0.84777627 |
| GALNT1 | 64.1365983 | 0.0104035 | 0.29858038 | 0.0348432 | 0.97220477 | 0.99431961 |
| RPRD1A | 142.048068 | -0.1915226 | 0.23359189 | -0.8199025 | 0.41227168 | 0.85867915 |
| SLC39A6 | 51.1435847 | 0.49580836 | 0.39438339 | 1.25717351 | 0.20869081 | 0.7623519 |
| ELP2 | 95.93948 | -0.0874498 | 0.22879443 | -0.38222 | 0.70229821 | 0.94476609 |
| MOCOS | 97.9713577 | -0.5628125 | 0.42401692 | -1.327335 | 0.18439789 | 0.73705812 |
| FHOD3 | 823.322571 | 0.12223628 | 0.20477366 | 0.59693358 | 5.51E-01 | 9.12E-01 |
| TPGS2 | 44.3753791 | -0.2610417 | 0.26957859 | -0.9683326 | 0.33287831 | 0.82928059 |
| KIAA1328 | 167.62801 | -0.4188895 | 0.22467591 | -1.8644166 | 0.06226322 | 0.58370968 |
| AC016205.1 | 25.4466871 | 0.60852294 | 0.47861315 | 1.27142962 | 0.20357585 | 0.76173533 |
| PIK3C3 | 153.506605 | -0.3690915 | 0.17909875 | -2.0608268 | 0.03931956 | 0.50068767 |
| LINC00907 | 32.6482771 | -0.4355446 | 0.29657117 | -1.4686005 | 0.14194118 | 0.69328517 |
| SETBP1 | 985.040337 | -0.1682341 | 0.1428746 | -1.1774949 | 2.39E-01 | 7.81E-01 |
| SLC14A2 | 161.448113 | 0.41526284 | 1.08684183 | 0.38208213 | 0.70240044 | 0.94476609 |
| EPG5 | 92.1506202 | -0.222486 | 0.21273619 | -1.0458306 | 0.29563925 | 0.81054924 |
| PSTPIP2 | 14.0582333 | -0.076846 | 0.56363431 | -0.1363402 | 0.89155238 | NA |
| ATP5F1A | 134.508675 | 0.1887205 | 0.33605742 | 0.56157218 | 0.57440754 | 0.91824492 |
| HAUS1 | 13.5441487 | -0.3734507 | 0.4857796 | -0.7687658 | 0.44203236 | NA |
| C18orf25 | 70.9715036 | -0.4114728 | 0.28134937 | -1.4624976 | 0.14360489 | 0.69460324 |
| ST8SIA5 | 21.2946135 | 1.09255337 | 0.57370796 | 1.904372 | 0.05686175 | 0.56976762 |
| PIAS2 | 195.818767 | -0.5519409 | 0.23527575 | -2.3459319 | 0.01897958 | 0.41544117 |
| KATNAL2 | 149.169047 | 0.32176598 | 0.31568471 | 1.01926375 | 3.08E-01 | 0.81568231 |
| HDHD2 | 11.0773407 | 0.10370818 | 0.50782489 | 0.20422035 | 0.83818131 | NA |
| SMAD2 | 237.125145 | -0.720064 | 0.31380532 | -2.2946202 | 0.0217549 | 0.4212027 |
| ZBTB7C | 47.3731797 | -0.4982231 | 0.76027423 | -0.6553203 | 0.51226157 | 0.9014941 |

| | | | | | | |
|---|---|---|---|---|---|---|
| CTIF | 83.5506464 | 0.02509617 | 0.32145562 | 0.07807039 | 0.93777206 | 0.99012269 |
| SMAD7 | 46.0868924 | -0.1071821 | 0.2943643 | -0.3641137 | 7.16E-01 | 9.49E-01 |
| DYM | 326.196078 | -0.2519524 | 0.20448319 | -1.2321422 | 0.21789599 | 0.76666749 |
| LIPG | 109.537878 | -0.9994753 | 0.72889679 | -1.3712166 | 0.17030746 | 0.72463203 |
| ACAA2 | 43.3790899 | 0.46706892 | 0.25638164 | 1.82177211 | 0.06848957 | 0.60244724 |
| SNHG22 | 28.621335 | 0.16227474 | 0.34212963 | 0.47430775 | 0.63528046 | 0.93251095 |
| MYO5B | 167.388954 | -0.2714791 | 0.24010529 | -1.130667 | 0.25819528 | 0.79532468 |
| MBD1 | 29.1484569 | -0.2171544 | 0.34826271 | -0.6235361 | 0.53293226 | 0.90678175 |
| CXXC1 | 18.0946209 | -0.3840109 | 0.51482868 | -0.7459003 | 0.45572765 | 0.87394768 |
| MAPK4 | 69.362746 | 0.33244759 | 0.44080894 | 0.75417616 | 4.51E-01 | 8.73E-01 |
| ME2 | 37.676768 | -0.0410012 | 0.28791239 | -0.1424087 | 0.88675719 | 0.98216235 |
| AC091551.1 | 14.7314836 | 0.51806528 | 0.54241623 | 0.95510653 | 0.33952383 | NA |
| SMAD4 | 56.0230459 | -0.3664822 | 0.26787063 | -1.3681312 | 0.17127102 | 0.72463203 |
| MEX3C | 17.9811406 | -0.0399792 | 0.39900936 | -0.100196 | 0.92018869 | 0.98790868 |
| MBD2 | 148.20318 | -0.6138064 | 0.2274126 | -2.699087 | 0.006953 | 0.3013833 |
| POLI | 54.8381154 | 0.06233789 | 0.28495976 | 0.21876033 | 0.82683675 | 0.97567084 |
| TCF4 | 75.8472443 | 0.16568131 | 0.48117307 | 0.3443279 | 0.7305997 | 0.95279138 |
| TXNL1 | 54.3211788 | 0.03002691 | 0.28339781 | 0.10595321 | 0.91561947 | 0.98691474 |
| WDR7 | 172.835404 | -0.09646 | 0.15751124 | -0.6124006 | 0.54027272 | 0.90763121 |
| FECH | 24.4495465 | 0.70343386 | 0.36633986 | 1.92016743 | 0.05483675 | 0.5645224 |
| NARS | 82.5094335 | -0.1154992 | 0.32873056 | -0.3513491 | 0.72532647 | 0.95164932 |
| AC027097.2 | 41.5123337 | -0.8049708 | 0.36158173 | -2.2262486 | 0.02599753 | 0.4455707 |
| ATP8B1 | 179.831358 | 0.21292594 | 0.27304424 | 0.77982212 | 0.43549558 | 0.86729771 |
| NEDD4L | 729.096819 | -0.390508 | 0.22793153 | -1.7132691 | 8.67E-02 | 6.35E-01 |
| MALT1 | 129.124514 | -0.2394233 | 0.41935699 | -0.5709296 | 0.56804737 | 0.91698355 |
| AC104365.1 | 14.7771027 | 0.67227514 | 0.67904409 | 0.99003165 | 0.32215865 | NA |
| ZNF532 | 127.066113 | -0.6738277 | 0.18823436 | -3.5797274 | 0.00034395 | 0.0763464 |
| SEC11C | 20.4505489 | 0.62493934 | 0.36778614 | 1.69919219 | 0.08928298 | 0.63653714 |
| LMAN1 | 93.1653843 | -0.0530513 | 0.38610643 | -0.1374008 | 0.89071401 | 0.98296447 |
| CDH20 | 24.9096079 | 0.40152343 | 0.9376285 | 0.42823296 | 0.66848152 | 0.93878847 |
| RNF152 | 103.04132 | -0.1340454 | 0.49584253 | -0.2703386 | 0.78689977 | 0.96767512 |
| PIGN | 117.88091 | -0.0939149 | 0.1886219 | -0.4979001 | 0.61855444 | 0.92766755 |
| KIAA1468 | 168.281271 | 0.03752815 | 0.1811517 | 0.2071642 | 0.83588162 | 0.97600352 |
| ZCCHC2 | 73.7044336 | -0.1006133 | 0.25993299 | -0.387074 | 0.69870141 | 0.94395238 |
| PHLPP1 | 124.031587 | -0.2335301 | 0.2494582 | -0.9361493 | 0.34919631 | 0.83687845 |
| BCL2 | 224.497629 | -0.5990399 | 0.37178968 | -1.6112332 | 0.1071289 | 0.65986731 |
| KDSR | 62.0520719 | -0.2099984 | 0.28515457 | -0.7364371 | 0.46146473 | 0.87660964 |
| VPS4B | 69.7536766 | 0.08348675 | 0.2116741 | 0.39441173 | 0.69327709 | 0.94394214 |
| LINC01924 | 164.087587 | 0.18865312 | 0.42674931 | 0.44207012 | 6.58E-01 | 9.36E-01 |
| TMX3 | 77.3402017 | -0.3295556 | 0.2177184 | -1.5136783 | 0.13010746 | 0.67331904 |
| CCDC102B | 35.8078643 | 0.55277687 | 0.55989688 | 0.98728336 | 0.32350375 | 0.82558302 |
| RTTN | 64.3108951 | -0.191266 | 0.24877041 | -0.7688454 | 0.44198508 | 0.86925451 |
| SOCS6 | 34.0240361 | 0.30389035 | 0.32517584 | 0.9345416 | 0.35002459 | 0.83687845 |
| TIMM21 | 15.4410319 | 0.62218275 | 0.42179701 | 1.47507624 | 0.14019205 | NA |
| CYB5A | 175.264421 | 0.25042601 | 0.45028728 | 0.5561472 | 0.57811023 | 0.91878391 |
| CNDP2 | 125.929486 | 0.34231491 | 0.32650696 | 1.04841538 | 0.29444727 | 0.81012446 |
| LINC00909 | 13.4980654 | 0.34549822 | 0.45157799 | 0.76509093 | 0.4442174 | NA |
| ZNF407 | 257.073687 | -0.379302 | 0.19069778 | -1.9890216 | 0.04669881 | 0.5320597 |
| ZADH2 | 33.5230048 | 0.7043099 | 0.30741141 | 2.29109874 | 0.0219577 | 0.4212027 |
| TSHZ1 | 50.6697097 | 0.40146621 | 0.3324352 | 1.20765254 | 0.22718094 | 0.77185154 |
| ZNF516 | 172.951383 | -0.3981402 | 0.27414622 | -1.4522915 | 0.14642058 | 0.69713616 |
| ZNF236-DT | 11.4424809 | 0.2246188 | 0.45744361 | 0.49103059 | 0.62340481 | NA |
| ZNF236 | 85.6378904 | -0.0935039 | 0.22550008 | -0.4146514 | 0.67839712 | 0.94123594 |
| MBP | 226.55563 | -0.1590527 | 0.21008542 | -0.7570857 | 0.44899851 | 0.87314185 |

| | | | | | | |
|---|---|---|---|---|---|---|
| LINC01896 | 25.2899729 | 0.77638495 | 0.56864999 | 1.36531251 | 0.17215485 | 0.72619351 |
| SALL3 | 33.6441844 | 0.76525556 | 0.4767441 | 1.6051705 | 0.10845629 | 0.65986731 |
| ATP9B | 281.304933 | -0.173292 | 0.20599212 | -0.8412557 | 0.4002047 | 0.85778581 |
| NFATC1 | 23.3336743 | 0.19101064 | 0.39644185 | 0.48181251 | 0.62993914 | 0.9318063 |
| CTDP1 | 28.6057932 | -0.2849021 | 0.30358892 | -0.9384468 | 0.34801482 | 0.83687845 |
| PQLC1 | 36.3977048 | 0.24749345 | 0.28068611 | 0.88174458 | 0.37791495 | 0.85310518 |
| HSBP1L1 | 43.2269611 | 0.02843646 | 0.30564589 | 0.09303728 | 0.92587394 | 0.98925753 |
| TXNL4A | 61.4464762 | -0.0942371 | 0.25480803 | -0.3698358 | 0.71150482 | 0.94676706 |
| ADNP2 | 56.5022981 | -0.1625321 | 0.31620889 | -0.5140022 | 0.60725044 | 0.92569157 |
| C20orf96 | 17.185022 | -0.0400523 | 0.466624 | -0.0858342 | 0.93159822 | 0.98960607 |
| ZCCHC3 | 15.0681947 | 0.07025546 | 0.54608041 | 0.12865405 | 0.89763139 | NA |
| NRSN2-AS1 | 94.3425123 | -0.1874098 | 0.46248849 | -0.4052206 | 0.6853154 | 0.94205398 |
| SOX12 | 19.6343624 | -0.1070151 | 0.52801176 | -0.2026756 | 0.8393886 | 0.97600352 |
| NRSN2 | 17.8303093 | -0.5229502 | 0.44243404 | -1.1819846 | 0.23721183 | 0.77930556 |
| RBCK1 | 65.8349617 | -0.2734796 | 0.33554881 | -0.815022 | 0.41505974 | 0.85867915 |
| TBC1D20 | 29.0826209 | -0.0581954 | 0.31439444 | -0.185103 | 0.85314824 | 0.97612951 |
| CSNK2A1 | 199.93846 | -0.2624 | 0.17292533 | -1.5174181 | 0.12916116 | 0.67032726 |
| SLC52A3 | 67.5501065 | 0.37789441 | 0.35519814 | 1.0638975 | 0.28737514 | 0.80523851 |
| PSMF1 | 78.5311793 | -0.1284585 | 0.2399584 | -0.5353367 | 0.59241707 | 0.92278087 |
| AL136531.2 | 37.1606554 | -0.5305714 | 0.36203574 | -1.4655221 | 0.14277854 | 0.69328517 |
| SDCBP2-AS1 | 52.9751868 | 0.19988851 | 0.25620641 | 0.78018546 | 0.43528172 | 0.86729771 |
| NSFL1C | 36.5599708 | -0.1382563 | 0.29318105 | -0.4715732 | 0.63723144 | 0.93259128 |
| SIRPA | 10.1068177 | 1.25014227 | 0.58908679 | 2.12216992 | 0.03382347 | NA |
| STK35 | 29.7917759 | -0.1196482 | 0.31807372 | -0.3761649 | 0.70679433 | 0.94558664 |
| SNRPB | 38.1764292 | -0.1366839 | 0.39872344 | -0.3428037 | 0.73174613 | 0.95279138 |
| NOP56 | 19.9780975 | 0.05709601 | 0.46082947 | 0.12389834 | 0.90139577 | 0.98531749 |
| IDH3B | 18.7906482 | 0.23715244 | 0.37342127 | 0.63508017 | 0.52537613 | 0.90487731 |
| EBF4 | 18.6247192 | 0.04086649 | 0.47518257 | 0.08600166 | 0.9314651 | 0.98960607 |
| PCED1A | 21.1061579 | 0.0435011 | 0.35941719 | 0.12103232 | 0.90366544 | 0.98531749 |
| PTPRA | 203.040318 | -0.3197266 | 0.19344495 | -1.652804 | 0.09837075 | 0.65695163 |
| MRPS26 | 30.0198425 | 0.77480594 | 0.38176788 | 2.02952101 | 0.04240525 | 0.51343649 |
| UBOX5-AS1 | 15.1645355 | 0.63575854 | 0.59682343 | 1.06523725 | 0.28676858 | NA |
| UBOX5 | 17.8693071 | -0.0915442 | 0.36923666 | -0.2479284 | 0.80418984 | 0.97175786 |
| LZTS3 | 21.0604818 | 0.57013205 | 0.32703753 | 1.74332304 | 0.0812772 | 0.61925304 |
| DDRGK1 | 30.0121066 | 0.22995176 | 0.29400525 | 0.78213487 | 0.43413531 | 0.86729771 |
| ITPA | 43.3977855 | -0.2051825 | 0.2748808 | -0.7464419 | 0.45540051 | 0.8739446 |
| C20orf194 | 438.29426 | 0.29350139 | 0.21737369 | 1.35021582 | 0.17694676 | 0.72841249 |
| ATRN | 178.959907 | 0.06797572 | 0.19496315 | 0.34865934 | 0.72734508 | 0.95211709 |
| CENPB | 26.7587738 | 0.36870328 | 0.46285992 | 0.79657638 | 0.4256971 | 0.86239557 |
| CDC25B | 12.9875392 | 0.26531674 | 0.51199721 | 0.51819958 | 0.60431902 | NA |
| MAVS | 75.3160825 | 0.18516808 | 0.31621218 | 0.58558175 | 0.55815661 | 0.91425998 |
| PANK2 | 67.7559085 | -0.1877348 | 0.2343755 | -0.8010002 | 0.42313151 | 0.86190547 |
| RNF24 | 232.280138 | -0.5463265 | 0.33588904 | -1.6265089 | 0.10384144 | 0.65946648 |
| PRNP | 21.3331775 | 0.22577048 | 0.32775829 | 0.68883224 | 0.49092885 | 0.88987842 |
| SLC23A2 | 77.3174981 | -0.2952704 | 0.31373165 | -0.941156 | 0.34662492 | 0.83687845 |
| TMEM230 | 59.3350252 | 0.04328006 | 0.26420698 | 0.16381118 | 0.86987979 | 0.97854266 |
| PCNA | 16.7173068 | 0.66296495 | 0.5134533 | 1.2911884 | 0.19663836 | 0.75350056 |
| CDS2 | 75.98684 | -0.498397 | 0.27635389 | -1.8034739 | 0.07131382 | 0.60665346 |
| GPCPD1 | 137.950997 | -0.2022767 | 0.39159223 | -0.5165492 | 0.60547086 | 0.92569157 |
| SHLD1 | 44.9589869 | -0.4977924 | 0.34931389 | -1.4250576 | 0.15414054 | 0.70572801 |
| TRMT6 | 11.806459 | -0.2118739 | 0.51419919 | -0.4120464 | 0.68030543 | NA |
| MCM8 | 13.9506019 | 0.26529411 | 0.42519103 | 0.62394098 | 0.53266635 | NA |
| CRLS1 | 71.6007112 | 0.0078609 | 0.26909977 | 0.02921184 | 0.97669564 | 0.99441127 |
| TMX4 | 48.3929518 | 0.34289556 | 0.35215196 | 0.97371478 | 0.33019818 | 0.82867637 |

| | | | | | | |
|---|---|---|---|---|---|---|
| PLCB1 | 61.5126328 | -0.2640695 | 0.64524579 | -0.4092541 | 0.68235323 | 0.94205398 |
| PLCB4 | 151.285196 | 0.28821464 | 0.4643622 | 0.62066774 | 0.53481826 | 0.90763121 |
| ANKEF1 | 10.6171885 | 0.36223213 | 0.45691181 | 0.79278347 | 0.42790399 | NA |
| SNAP25 | 22.6410234 | 0.41668264 | 0.99234172 | 0.41989834 | 0.67455972 | 0.93934789 |
| MKKS | 22.0717778 | 0.43065669 | 0.3445101 | 1.25005535 | 0.21127933 | 0.7643572 |
| SLX4IP | 147.86707 | -0.0028992 | 0.27754839 | -0.0104457 | 0.99166567 | 0.99814944 |
| JAG1 | 61.2964324 | 0.22441774 | 0.39563437 | 0.5672352 | 0.5705544 | 0.91778542 |
| AL050403.2 | 43.3166356 | 0.02430015 | 0.43067152 | 0.05642387 | 0.95500414 | 0.99148072 |
| BTBD3 | 90.6322203 | -0.2646459 | 0.21807301 | -1.2135654 | 0.22491373 | 0.76940612 |
| SPTLC3 | 91.9149631 | 0.03522686 | 0.23453828 | 0.15019664 | 0.88060947 | 0.98093344 |
| TASP1 | 184.677426 | -0.2589525 | 0.17759895 | -1.4580743 | 0.14482007 | 0.69512617 |
| ESF1 | 72.0930051 | -0.4451337 | 0.26120613 | -1.7041472 | 0.08835358 | 0.63466198 |
| NDUFAF5 | 49.985211 | -0.4653018 | 0.32685625 | -1.4235671 | 0.15457182 | 0.70572801 |
| MACROD2 | 666.319209 | 0.37626528 | 0.29984221 | 1.25487764 | 0.20952318 | 0.76291581 |
| FLRT3 | 23.3620527 | -0.1159794 | 0.43094118 | -0.2691304 | 0.7878293 | 0.96767512 |
| KIF16B | 208.975575 | -0.1407712 | 0.25542785 | -0.5511192 | 0.58155199 | 0.92034873 |
| SNRPB2 | 25.5143398 | 0.24425778 | 0.39487376 | 0.6185718 | 0.53619848 | 0.90763121 |
| DSTN | 242.968122 | 0.30502075 | 0.29233159 | 1.04340672 | 0.29675996 | 0.81137495 |
| RRBP1 | 107.925125 | -0.0734928 | 0.22114179 | -0.3323335 | 7.40E-01 | 0.9544376 |
| SNX5 | 39.619328 | 0.33109884 | 0.32601182 | 1.01560378 | 0.30981811 | 0.81568231 |
| OVOL2 | 37.5464115 | 0.51018651 | 0.41733982 | 1.22247263 | 0.22152895 | 0.76680432 |
| MGME1 | 41.2203864 | 0.15734351 | 0.293067 | 0.53688581 | 0.5913465 | 0.92254886 |
| KAT14 | 13.3679499 | 0.68030171 | 0.48128016 | 1.41352535 | 0.15750131 | NA |
| ZNF133 | 48.6165752 | 0.24155735 | 0.28816508 | 0.83826032 | 0.40188451 | 0.85778581 |
| DZANK1 | 28.6134831 | -0.1979477 | 0.3108753 | -0.6367432 | 0.52429214 | 0.90487731 |
| POLR3F | 12.2279362 | -0.3586636 | 0.42034293 | -0.8532643 | 0.39351276 | NA |
| RBBP9 | 11.0498636 | 0.99167898 | 0.59653554 | 1.66239715 | 0.09643317 | NA |
| SEC23B | 68.9807805 | 0.10182356 | 0.25196374 | 0.40411989 | 0.68612456 | 0.94205398 |
| DTD1 | 14.348339 | 0.27665788 | 0.37861742 | 0.73070562 | 0.46495898 | NA |
| SLC24A3 | 65.5198989 | -0.5700138 | 0.72835503 | -0.7826043 | 0.43385954 | 0.86729771 |
| AL121761.1 | 50.2129291 | 0.14542545 | 0.44057086 | 0.33008414 | 0.74133639 | 0.95464495 |
| RIN2 | 213.136441 | 0.4233585 | 0.21465722 | 1.97225374 | 0.04858065 | 0.5391188 |
| NAA20 | 72.4535474 | 0.45546612 | 0.26618811 | 1.71106862 | 0.08706845 | 0.63466198 |
| CRNKL1 | 18.3395851 | 0.03889413 | 0.35234938 | 0.11038514 | 0.91210394 | 0.9859256 |
| RALGAPA2 | 336.398616 | 0.1122987 | 0.20438185 | 0.54945533 | 5.83E-01 | 9.20E-01 |
| KIZ | 189.222994 | 0.09772003 | 0.34043576 | 0.28704396 | 0.77407866 | 0.96435639 |
| KIZ-AS1 | 38.2468917 | 0.08437096 | 0.49794206 | 0.16943931 | 0.86545111 | 0.97788853 |
| XRN2 | 112.969268 | -0.2593743 | 0.26824011 | -0.9669481 | 0.33357 | 0.82974237 |
| NAPB | 51.2192515 | 0.21972265 | 0.30331268 | 0.72440972 | 0.46881423 | 0.88192105 |
| CST3 | 129.446957 | 0.53026893 | 0.53009711 | 1.00032412 | 0.31715368 | 0.82275575 |
| APMAP | 33.2707268 | 0.39566575 | 0.46329783 | 0.8540203 | 0.39309374 | 0.85605713 |
| ACSS1 | 110.276711 | 0.37298552 | 0.30168172 | 1.2363544 | 0.21632686 | 0.76616336 |
| ENTPD6 | 36.7086658 | -0.2815471 | 0.27954809 | -1.0071507 | 0.31386235 | 0.82005998 |
| PYGB | 50.7879338 | -0.010673 | 0.33462957 | -0.0318949 | 0.97455584 | 0.99431961 |
| ABHD12 | 79.2521458 | -0.1920646 | 0.22534373 | -0.8523183 | 0.39403747 | 0.85605713 |
| NINL | 232.549377 | -0.3575634 | 0.3278424 | -1.0906563 | 0.27542417 | 0.7994854 |
| ZNF337-AS1 | 28.0170977 | -0.5921013 | 0.37827128 | -1.5652822 | 0.11751675 | 0.6668162 |
| ZNF337 | 34.654942 | -0.1580599 | 0.32692632 | -0.4834725 | 0.62876026 | 0.93142468 |
| FAM182B | 44.1749038 | 0.56908188 | 0.38202864 | 1.48963146 | 0.13632116 | 0.68406472 |
| HM13 | 99.4569869 | -0.3231643 | 0.2102327 | -1.5371742 | 0.12425065 | 0.66929384 |
| ID1 | 74.2034002 | -0.5703717 | 0.69700258 | -0.8183207 | 0.41317406 | 0.85867915 |
| BCL2L1 | 284.184922 | -0.6334656 | 0.30492421 | -2.0774527 | 0.0377598 | 0.49640946 |
| AL117381.1 | 49.8311751 | -0.5847495 | 0.59455866 | -0.9835018 | 0.32536054 | 0.82567895 |
| DUSP15 | 36.9574107 | 0.73414556 | 0.62367624 | 1.17712606 | 0.2391452 | 0.78098673 |

| | | | | | | |
|---|---|---|---|---|---|---|
| TTLL9 | 12.6461853 | 0.15359524 | 0.4780545 | 0.32129231 | 0.74798888 | NA |
| PDRG1 | 11.840912 | 0.63838345 | 0.44968789 | 1.41961449 | 0.15571995 | NA |
| TM9SF4 | 77.0292867 | -0.3450946 | 0.21026069 | -1.64127 | 0.10074138 | 0.65863153 |
| POFUT1 | 20.4144665 | 0.50459217 | 0.48469063 | 1.0410603 | 0.29784757 | 0.81214575 |
| KIF3B | 94.1198342 | 0.17729495 | 0.23925002 | 0.74104466 | 0.45866636 | 0.87549589 |
| ASXL1 | 215.319757 | -0.2999453 | 0.21824747 | -1.3743359 | 0.16933742 | 0.72335924 |
| NOL4L | 40.9311822 | 0.06363432 | 0.43102857 | 0.14763365 | 0.8826319 | 0.98137082 |
| MAPRE1 | 33.8260206 | -0.0928329 | 0.36117711 | -0.2570286 | 0.79715668 | 0.97019967 |
| AL035071.1 | 10.0463439 | -0.5109691 | 0.53514145 | -0.95483 | 0.33966366 | NA |
| CDK5RAP1 | 38.697338 | -0.0977959 | 0.25719529 | -0.3802399 | 0.70376732 | 0.94476609 |
| CBFA2T2 | 307.641166 | -0.2136535 | 0.19912191 | -1.0729783 | 0.28328086 | 0.80431986 |
| NECAB3 | 13.6479457 | 0.37228477 | 0.46282023 | 0.80438308 | 0.42117577 | NA |
| PXMP4 | 15.6924569 | 0.34304391 | 0.4865515 | 0.7050516 | 0.48077813 | NA |
| CHMP4B | 56.9966873 | -0.3080164 | 0.22865641 | -1.347071 | 0.17795735 | 0.72841249 |
| RALY | 103.480895 | -0.3427549 | 0.19325449 | -1.7735934 | 0.07613042 | 0.61051335 |
| EIF2S2 | 72.9631201 | -0.3414899 | 0.30075599 | -1.1354385 | 0.25619165 | 0.7940072 |
| AHCY | 16.7707008 | 0.27137127 | 0.41865361 | 0.6482 | 0.5168556 | 0.9026606 |
| ITCH | 254.684319 | -0.237401 | 0.19314636 | -1.2291249 | 0.21902496 | 0.76666749 |
| DYNLRB1 | 83.4449386 | -0.0733514 | 0.38398536 | -0.1910266 | 0.84850474 | 0.97612951 |
| MAP1LC3A | 26.5014847 | 0.62654256 | 0.58041951 | 1.07946501 | 0.28038048 | 0.80311628 |
| PIGU | 57.5095068 | -0.6083583 | 0.26860196 | -2.2649063 | 0.02351843 | 0.43088993 |
| NCOA6 | 171.060463 | -0.2505207 | 0.24613256 | -1.0178282 | 0.30875962 | 0.81568231 |
| TP53INP2 | 31.8183125 | -0.3789054 | 0.60368924 | -0.6276498 | 0.53023341 | 0.9052447 |
| GGT7 | 60.7935367 | 0.43244485 | 0.27154969 | 1.59250728 | 0.11127077 | 0.66417829 |
| ACSS2 | 66.6426511 | 0.10246296 | 0.29891977 | 0.34277746 | 0.73176589 | 0.95279138 |
| GSS | 24.4956996 | 0.04328735 | 0.29827502 | 0.14512562 | 0.88461169 | 0.98154805 |
| TRPC4AP | 142.964414 | -0.3526419 | 0.21040759 | -1.675994 | 0.09373936 | 0.64572229 |
| EDEM2 | 34.2966063 | 0.25658165 | 0.31556158 | 0.81309536 | 0.41616339 | 0.85867915 |
| PROCR | 15.4121412 | 0.20853445 | 0.64964604 | 0.32099703 | 0.74821264 | NA |
| MMP24OS | 50.2896685 | -0.1332227 | 0.34332778 | -0.3880336 | 0.69799117 | 0.94395238 |
| MMP24 | 44.0115588 | 0.27553417 | 0.40059684 | 0.68780915 | 0.49157297 | 0.8899739 |
| UQCC1 | 96.5614844 | -0.1882481 | 0.19388457 | -0.9709289 | 0.33158369 | 0.82890589 |
| CEP250 | 24.2159912 | -1.0832411 | 0.41618813 | -2.6027679 | 0.00924745 | 0.32071089 |
| ERGIC3 | 45.885273 | -0.1621879 | 0.32797359 | -0.4945151 | 0.62094244 | 0.92805465 |
| CPNE1 | 40.4251521 | 0.09691348 | 0.29497632 | 0.32854663 | 0.74249839 | 0.95563105 |
| NFS1 | 10.7498449 | -0.2805518 | 0.46643783 | -0.6014774 | 0.54752207 | NA |
| ROMO1 | 31.4118357 | -0.8041146 | 0.4521699 | -1.778346 | 0.07534704 | 0.61020315 |
| RBM39 | 334.67541 | -0.2214264 | 0.18914209 | -1.1706879 | 0.24172424 | 0.78376421 |
| PHF20 | 155.657216 | -0.4981873 | 0.2788828 | -1.7863681 | 0.07403967 | 0.61020315 |
| SCAND1 | 57.1119145 | -0.0611693 | 0.38473973 | -0.1589889 | 0.87367763 | 0.97918624 |
| NORAD | 164.986863 | -0.1092468 | 0.24790877 | -0.4406734 | 0.65944946 | 0.93693904 |
| EPB41L1 | 386.925239 | 0.10642132 | 0.24895047 | 0.4274799 | 0.66902982 | 0.93878847 |
| AAR2 | 20.1291776 | 0.21829985 | 0.41587213 | 0.52492061 | 0.59963838 | 0.92298075 |
| DLGAP4 | 82.9007798 | -0.1451286 | 0.27213944 | -0.5332878 | 0.59383439 | 0.9229638 |
| DLGAP4-AS1 | 40.674843 | -0.4500118 | 0.28938578 | -1.5550582 | 0.1199322 | 0.6668162 |
| MYL9 | 13.0481762 | 0.48889495 | 0.57379087 | 0.8520438 | 0.39418978 | NA |
| RAB5IF | 15.3739868 | -0.3677658 | 0.4414386 | -0.8331074 | 0.40478414 | NA |
| NDRG3 | 38.0079404 | 0.26749521 | 0.26589914 | 1.00600257 | 0.31441434 | 0.82040655 |
| SOGA1 | 51.36429 | -0.0279499 | 0.33166801 | -0.0842709 | 0.93284108 | 0.98960607 |
| SAMHD1 | 19.5643254 | -1.9562758 | 0.76194976 | -2.5674604 | 0.01024465 | 0.33306084 |
| RBL1 | 27.1629358 | 0.0102998 | 0.37165537 | 0.02771331 | 0.97789081 | 0.99469237 |
| RPN2 | 78.275623 | -0.1403463 | 0.33023027 | -0.4249953 | 0.67084007 | 0.93878847 |
| MANBAL | 20.6831994 | -0.5008419 | 0.32210522 | -1.5549016 | 0.1199695 | 0.6668162 |
| SRC | 32.050193 | -0.1904957 | 0.29486025 | -0.6460542 | 0.51824424 | 0.90368693 |

| | | | | | | |
|---|---|---|---|---|---|---|
| BLCAP | 87.2687843 | -0.3775975 | 0.25921615 | -1.4566895 | 0.1452021 | 0.69606446 |
| CTNNBL1 | 57.3329752 | -0.0922034 | 0.2379358 | -0.3875138 | 0.6983759 | 0.94395238 |
| TTI1 | 30.6215301 | -0.177249 | 0.37523064 | -0.4723735 | 0.63666024 | 0.93251095 |
| RPRD1B | 73.1988939 | -0.2517667 | 0.22112713 | -1.1385609 | 0.25488634 | 0.79297003 |
| TGM2 | 90.9645897 | 0.35606261 | 0.55499857 | 0.6415559 | 5.21E-01 | 9.05E-01 |
| RALGAPB | 135.736451 | -0.232603 | 0.2575595 | -0.903104 | 0.36647072 | 0.84741115 |
| DHX35 | 32.8764512 | -0.1873036 | 0.32415766 | -0.5778163 | 0.56338815 | 0.91624774 |
| TOP1 | 245.160332 | -0.3351547 | 0.27885379 | -1.2019012 | 0.22940182 | 0.77185154 |
| PLCG1 | 28.7346737 | 0.07985118 | 0.37739567 | 0.21158479 | 0.83243098 | 0.97600352 |
| ZHX3 | 211.634555 | 0.46619686 | 0.31384459 | 1.48543855 | 0.13742769 | 0.68661046 |
| LPIN3 | 35.1012707 | -0.5924505 | 0.35203961 | -1.6829088 | 0.09239275 | 0.64572229 |
| CHD6 | 212.001335 | 0.04200232 | 0.15781533 | 0.26614855 | 0.79012481 | 0.96894869 |
| SRSF6 | 51.6804349 | -0.2986527 | 0.31506051 | -0.9479216 | 0.34316935 | 0.83426365 |
| L3MBTL1 | 26.7176602 | 0.17718063 | 0.34370148 | 0.51550733 | 0.60619855 | 0.92569157 |
| SGK2 | 91.8149864 | 0.14849231 | 0.27285375 | 0.5442194 | 5.86E-01 | 9.21E-01 |
| IFT52 | 36.1253483 | 0.21421352 | 0.36533643 | 0.5863459 | 0.55764309 | 0.91425998 |
| OSER1 | 27.1173255 | 0.41114119 | 0.51971624 | 0.79108783 | 0.42889274 | 0.86357236 |
| TTPAL | 17.1317265 | -0.1785162 | 0.45658957 | -0.3909773 | 0.695814 | 0.94395238 |
| SERINC3 | 53.6680224 | 0.01383453 | 0.259568 | 0.05329828 | 0.95749425 | 0.99170927 |
| PKIG | 79.0634743 | -0.18884 | 0.22994765 | -0.8212302 | 0.41151516 | 0.85867915 |
| YWHAB | 152.252805 | 0.26714274 | 0.41576737 | 0.64252935 | 0.52052954 | 0.90427107 |
| PABPC1L | 28.597011 | 0.29118765 | 0.30342673 | 0.95966382 | 0.33722444 | 0.83055261 |
| TOMM34 | 15.9007106 | 0.31066216 | 0.41715347 | 0.7447191 | 0.45644155 | NA |
| STK4 | 65.4565122 | -0.21385 | 0.27425728 | -0.7797424 | 0.43554252 | 0.86729771 |
| SLPI | 258.303652 | 0.24567915 | 0.8008808 | 0.3067612 | 7.59E-01 | 9.59E-01 |
| SDC4 | 257.009601 | -0.1076348 | 0.36227323 | -0.2971093 | 0.76638304 | 0.96270421 |
| PIGT | 28.3743142 | 0.12155955 | 0.50789153 | 0.23934155 | 0.81084075 | 0.97257655 |
| WFDC2 | 1513.67598 | 1.15162124 | 0.52844444 | 2.17926643 | 2.93E-02 | 4.64E-01 |
| DNTTIP1 | 18.0651158 | -0.028316 | 0.38488266 | -0.0735704 | 0.94135226 | 0.99043681 |
| SNX21 | 21.0512966 | 0.26162925 | 0.42440239 | 0.61646507 | 0.53758762 | 0.90763121 |
| ACOT8 | 17.4303474 | 0.35612328 | 0.38373029 | 0.92805621 | 0.35337841 | 0.83945817 |
| CTSA | 48.6535821 | 0.23957657 | 0.3435872 | 0.69728026 | 0.48562742 | 0.88838332 |
| ZNF335 | 21.095275 | 0.3335016 | 0.42201143 | 0.79026675 | 0.429372 | 0.86396404 |
| NCOA5 | 38.2839677 | 0.12432077 | 0.3199835 | 0.38852244 | 0.69762945 | 0.94395238 |
| CD40 | 17.8343854 | 0.43667356 | 0.50751563 | 0.86041401 | 0.38956086 | 0.85605713 |
| SLC35C2 | 28.8393783 | -0.2097754 | 0.3016185 | -0.6954991 | 0.48674258 | 0.88838332 |
| ELMO2 | 28.0170044 | -0.1987611 | 0.34338421 | -0.5788301 | 0.56270379 | 0.91624774 |
| ZNF334 | 18.6227148 | 0.00271598 | 0.60386989 | 0.00449762 | 0.99641143 | 0.99927028 |
| SLC13A3 | 26.2864402 | -0.7022545 | 0.60063167 | -1.1691932 | 0.24232578 | 0.78464787 |
| EYA2 | 51.5292115 | 0.5404531 | 0.62400504 | 0.86610373 | 0.38643328 | 0.8560201 |
| ZMYND8 | 320.429698 | -0.2012129 | 0.26689604 | -0.7539 | 0.45090925 | 0.87346041 |
| NCOA3 | 178.918434 | -0.5716968 | 0.17450237 | -3.2761548 | 0.00105231 | 0.14056417 |
| SULF2 | 18.6223363 | 0.86522437 | 0.54236902 | 1.59526879 | 0.11065214 | 0.66311481 |
| ARFGEF2 | 217.991296 | -0.0786292 | 0.22790516 | -0.3450084 | 0.73008808 | 0.95279138 |
| CSE1L | 54.5142341 | 0.10060661 | 0.23202566 | 0.4336012 | 0.66457806 | 0.93860542 |
| STAU1 | 130.904165 | -0.0860025 | 0.18630046 | -0.4616335 | 0.6443442 | 0.93328692 |
| DDX27 | 23.3021536 | 0.17136001 | 0.37202548 | 0.46061364 | 0.64507582 | 0.93328692 |
| ZNFX1 | 45.0762766 | -1.1446406 | 0.42718145 | -2.6795185 | 0.00737281 | 0.30654023 |
| ZFAS1 | 42.1458449 | -0.1338558 | 0.33211981 | -0.4030346 | 0.68692275 | 0.94205398 |
| B4GALT5 | 174.586771 | -0.0845656 | 0.22127491 | -0.3821742 | 0.70233213 | 0.94476609 |
| SLC9A8 | 69.8978163 | -0.5370153 | 0.23960292 | -2.2412719 | 0.02500847 | 0.43948277 |
| RNF114 | 87.194897 | 0.04628948 | 0.40548799 | 0.11415745 | 0.90911298 | 0.9854228 |
| CEBPB | 42.5758851 | 0.01411176 | 0.41508287 | 0.03399746 | 9.73E-01 | 9.94E-01 |
| PTPN1 | 144.223 | -0.7581478 | 0.4002516 | -1.894178 | 0.05820139 | 0.57141408 |

| | | | | | | |
|---|---|---|---|---|---|---|
| RIPOR3 | 16.3711988 | 0.90504801 | 0.64668509 | 1.39951889 | 0.16165744 | 0.71101785 |
| PARD6B | 211.537405 | -0.1726013 | 0.37777901 | -0.4568843 | 0.64775423 | 0.93328692 |
| BCAS4 | 30.1167906 | 0.0935519 | 0.38679119 | 0.24186668 | 0.80888347 | 0.97257655 |
| ADNP | 244.371832 | -0.0574084 | 0.15126124 | -0.3795313 | 0.70429338 | 0.94476609 |
| DPM1 | 50.3771435 | -0.1906379 | 0.22898434 | -0.8325368 | 0.40510603 | 0.85782098 |
| ATP9A | 81.1923611 | 0.10689102 | 0.25856521 | 0.41340065 | 0.6793131 | 0.94176503 |
| ZFP64 | 41.6674767 | -0.0472104 | 0.26739601 | -0.1765561 | 0.85985709 | 0.97691173 |
| ZNF217 | 64.7073751 | -0.3177641 | 0.26889102 | -1.1817579 | 0.23730179 | 0.77930556 |
| PFDN4 | 11.2826 | 0.10807062 | 0.43741871 | 0.24706446 | 0.80485834 | NA |
| DOK5 | 36.4857173 | -0.7432381 | 0.66981879 | -1.1096107 | 0.26716684 | 0.79666376 |
| FAM210B | 19.1759675 | 0.10337852 | 0.42324024 | 0.24425494 | 0.80703338 | 0.97216031 |
| RTF2 | 34.0993349 | -0.1575987 | 0.27865066 | -0.5655781 | 0.57168061 | 0.91791025 |
| RAE1 | 18.5763065 | -0.0876984 | 0.36548007 | -0.2399541 | 0.81036584 | 0.97257655 |
| PCK1 | 42.066121 | -0.2773012 | 1.21527785 | -0.2281793 | 0.81950685 | 0.97323535 |
| PMEPA1 | 24.58607 | -0.1135538 | 0.74677619 | -0.1520587 | 0.87914067 | 0.98061726 |
| RAB22A | 65.1413878 | -0.1564831 | 0.25480589 | -0.6141267 | 0.53913162 | 0.90763121 |
| VAPB | 90.9914867 | -0.0225498 | 0.21208157 | -0.1063261 | 0.91532362 | 0.98691474 |
| APCDD1L | 55.2238838 | -1.0077403 | 0.5990467 | -1.6822399 | 0.09252233 | 0.64572229 |
| STX16-NPEPl | 10.516302 | -0.2002366 | 0.5427105 | -0.3689566 | 0.71216005 | NA |
| NPEPL1 | 28.6006688 | 0.25778934 | 0.41807884 | 0.6166046 | 0.53749556 | 0.90763121 |
| GNAS | 294.474925 | 0.1599195 | 0.28187479 | 0.56734232 | 0.57048163 | 0.91778542 |
| NELFCD | 33.6665908 | 0.11842497 | 0.29145476 | 0.40632367 | 0.6845048 | 0.94205398 |
| CTSZ | 24.1404291 | 0.46814251 | 0.40788647 | 1.14772749 | 0.25108108 | 0.78895101 |
| ATP5F1E | 129.435572 | -0.11691 | 0.54432832 | -0.2147784 | 0.8299401 | 0.97567084 |
| PRELID3B | 10.897364 | 0.33262674 | 0.48954516 | 0.67946078 | 0.49684595 | NA |
| FAM217B | 10.1963743 | 0.52154774 | 0.61706628 | 0.84520539 | 0.39799617 | NA |
| MIR646HG | 238.105609 | -0.5868582 | 0.76454352 | -0.7675929 | 0.44272907 | 0.86991418 |
| TAF4 | 51.2660304 | -0.0540724 | 0.29498296 | -0.1833068 | 0.85455729 | 0.97612951 |
| LSM14B | 12.596778 | -0.3050824 | 0.4122718 | -0.7400031 | 0.45929809 | NA |
| PSMA7 | 71.2707285 | 0.07845629 | 0.37556185 | 0.20890378 | 0.83452335 | 0.97600352 |
| SS18L1 | 46.071138 | -0.6144876 | 0.34496136 | -1.7813229 | 0.07485971 | 0.61020315 |
| MTG2 | 39.4242021 | -0.2269856 | 0.28536806 | -0.7954134 | 0.42637305 | 0.86239557 |
| OSBPL2 | 74.887579 | -0.3264265 | 0.2212938 | -1.475082 | 0.1401905 | 0.69168246 |
| ADRM1 | 10.0417671 | -0.2704228 | 0.53073709 | -0.509523 | 0.61038565 | NA |
| LAMA5 | 64.464545 | 0.25075711 | 0.29786898 | 0.84183695 | 0.39987923 | 0.85778581 |
| RPS21 | 13.4764366 | -0.4602489 | 0.53075734 | -0.867155 | 0.3858571 | NA |
| SLCO4A1 | 39.7880253 | -1.219451 | 0.59715075 | -2.0421158 | 0.04114005 | 0.50744725 |
| OGFR | 29.9793435 | -0.6485877 | 0.31475461 | -2.0606139 | 0.03933989 | 0.50068767 |
| TCFL5 | 17.9947055 | -0.1834641 | 0.453798 | -0.4042857 | 0.68600262 | 0.94205398 |
| DIDO1 | 106.82649 | -0.0925294 | 0.18473629 | -0.5008731 | 0.61646045 | 0.92662475 |
| GID8 | 37.7731818 | 0.22797053 | 0.35359709 | 0.64471834 | 0.51910973 | 0.90368693 |
| YTHDF1 | 36.9175639 | -0.1761309 | 0.27445876 | -0.6417391 | 0.52104258 | 0.90451015 |
| ARFGAP1 | 30.0921528 | -0.8014158 | 0.37316174 | -2.1476366 | 0.03174263 | 0.47839934 |
| PPDPF | 169.418328 | -0.0937686 | 0.45923284 | -0.2041853 | 0.8382087 | 0.97600352 |
| HELZ2 | 16.620562 | -2.1028035 | 0.79710157 | -2.6380622 | 0.00833813 | 0.30980404 |
| GMEB2 | 23.4525098 | -0.7473339 | 0.34090632 | -2.1921972 | 0.02836527 | 0.46148943 |
| MHENCR | 9.47785003 | 0.5165942 | 0.48219933 | 1.07132916 | 0.28402145 | NA |
| ARFRP1 | 21.1337902 | -0.2672922 | 0.3791425 | -0.7049914 | 0.48081556 | 0.88675516 |
| SLC2A4RG | 31.6449554 | 0.17182084 | 0.53123333 | 0.32343761 | 0.74636385 | 0.95563105 |
| ZBTB46 | 29.2082169 | 1.17231367 | 0.43342008 | 2.70479775 | 0.0068346 | 0.30102597 |
| TPD52L2 | 41.1957978 | -0.4352552 | 0.28549113 | -1.5245841 | 0.12736286 | 0.66929384 |
| DNAJC5 | 109.145002 | -0.5063003 | 0.27392533 | -1.848315 | 0.06455679 | 0.58883813 |
| UCKL1 | 21.9510193 | 0.08151654 | 0.36807553 | 0.22146688 | 0.82472892 | 0.97518529 |
| ZNF512B | 14.745448 | 0.12446371 | 0.41416929 | 0.30051411 | 0.76378504 | NA |

| | | | | | | |
|---|---|---|---|---|---|---|
| PRPF6 | 53.8512889 | -0.1028159 | 0.23554567 | -0.4365009 | 0.66247335 | 0.93786857 |
| TCEA2 | 11.1604846 | 0.30582081 | 0.48987409 | 0.62428452 | 0.53244075 | NA |
| PCMTD2 | 77.6218411 | 0.19750033 | 0.26466582 | 0.74622529 | 0.45553133 | 0.8739446 |
| PLPP2 | 20.7867472 | 0.47067547 | 0.47493191 | 0.9910378 | 3.22E-01 | 8.25E-01 |
| MIER2 | 17.8509204 | -0.7565744 | 0.4374031 | -1.7296961 | 0.08368459 | 0.62534311 |
| TPGS1 | 21.4340282 | 0.15470539 | 0.39339624 | 0.39325589 | 0.69413049 | 0.94395238 |
| CDC34 | 16.1390517 | 0.24999721 | 0.44704997 | 0.55921537 | 0.57601475 | NA |
| BSG | 114.883589 | 0.16717781 | 0.35545895 | 0.47031538 | 0.63812971 | 0.9331046 |
| RNF126 | 18.9710855 | -0.0566498 | 0.39568515 | -0.1431688 | 0.88615685 | 0.98216235 |
| PALM | 26.0295375 | 0.44702206 | 0.40558109 | 1.10217679 | 2.70E-01 | 7.97E-01 |
| PTBP1 | 65.4441913 | 0.09933697 | 0.2467929 | 0.40251145 | 0.68730766 | 0.94205398 |
| MED16 | 23.1161836 | -0.068893 | 0.37373556 | -0.1843363 | 0.85374966 | 0.97612951 |
| R3HDM4 | 22.100358 | -0.2383015 | 0.46010836 | -0.5179247 | 0.60451082 | 0.92531731 |
| ARID3A | 80.5732882 | -0.3011653 | 0.29871905 | -1.008189 | 0.31336373 | 0.81969828 |
| WDR18 | 16.6296286 | 0.52472367 | 0.42667975 | 1.2297834 | 0.21877823 | 0.76666749 |
| TMEM259 | 59.5024717 | 0.03581932 | 0.3529149 | 0.10149562 | 0.91915703 | 0.98749988 |
| CNN2 | 19.191035 | 0.5938026 | 0.40300468 | 1.47343847 | 0.14063284 | 0.69227547 |
| ABCA7 | 13.8627526 | 0.0957845 | 0.61749975 | 0.15511666 | 0.87672934 | NA |
| POLR2E | 26.7405175 | 0.15578184 | 0.38104926 | 0.40882336 | 0.68266929 | 0.94205398 |
| GPX4 | 107.895386 | -0.2297315 | 0.35469861 | -0.6476808 | 0.51719146 | 0.9029524 |
| SBNO2 | 83.7669762 | -0.2337326 | 0.2649667 | -0.8821207 | 3.78E-01 | 8.53E-01 |
| STK11 | 49.0879923 | -0.1428112 | 0.30153 | -0.4736219 | 0.63576955 | 0.93251095 |
| ATP5F1D | 102.222029 | 0.14625645 | 0.42316391 | 0.34562602 | 0.72962379 | 0.95279138 |
| MIDN | 126.821491 | -0.6339334 | 0.52487471 | -1.2077804 | 0.22713173 | 0.77185154 |
| CIRBP | 135.777593 | 0.23191035 | 0.28489698 | 0.81401476 | 0.4156365 | 0.85867915 |
| C19orf24 | 18.364811 | 0.16516252 | 0.50682139 | 0.32587914 | 0.7445158 | 0.95563105 |
| MUM1 | 54.6678428 | 0.05037371 | 0.30060628 | 0.16757372 | 0.86691865 | 0.97788853 |
| NDUFS7 | 34.4699446 | 0.0849247 | 0.41935692 | 0.20251175 | 0.83951668 | 0.97600352 |
| DAZAP1 | 77.3156476 | -0.2492384 | 0.28220353 | -0.8831865 | 0.3771355 | 0.85310518 |
| RPS15 | 199.056845 | 0.15432204 | 0.4478615 | 0.34457536 | 0.73041363 | 0.95279138 |
| C19orf25 | 40.2469409 | -0.1398331 | 0.30494156 | -0.4585572 | 0.6465522 | 0.93328692 |
| MBD3 | 23.6581626 | -0.3541396 | 0.4403859 | -0.8041574 | 0.42130609 | 0.86190547 |
| TCF3 | 24.3642131 | -0.4308069 | 0.43805363 | -0.9834569 | 0.3253826 | 0.82567895 |
| REXO1 | 20.5288237 | -0.6059802 | 0.37120743 | -1.632457 | 0.10258325 | 0.65946648 |
| ABHD17A | 28.2809105 | 0.26519191 | 0.39322967 | 0.67439445 | 0.50006057 | 0.89448839 |
| SCAMP4 | 46.6061024 | 0.34148658 | 0.3020128 | 1.13070234 | 0.2581804 | 0.79532468 |
| CSNK1G2 | 46.7712447 | -0.2486125 | 0.29751265 | -0.8356366 | 0.40335936 | 0.85782098 |
| BTBD2 | 49.1578291 | -0.4086438 | 0.32880288 | -1.2428229 | 0.2139331 | 0.7643572 |
| MKNK2 | 39.7116658 | 0.14752039 | 0.32429325 | 0.45489811 | 0.64918254 | 0.93437275 |
| MOB3A | 29.0111768 | -0.465391 | 0.34159671 | -1.3623988 | 0.17307205 | 0.72721271 |
| AP3D1 | 75.7366127 | -0.4712449 | 0.31023189 | -1.5190085 | 0.12876035 | 0.66929384 |
| DOT1L | 79.906053 | -0.5582104 | 0.48549894 | -1.1497664 | 0.25024011 | 0.78799766 |
| PLEKHJ1 | 52.9339204 | -0.1339532 | 0.47629469 | -0.2812401 | 0.77852624 | 0.9653895 |
| SF3A2 | 10.2289334 | -0.1901226 | 0.46175626 | -0.411738 | 0.6805315 | NA |
| LSM7 | 48.6099783 | 0.32609462 | 0.39774322 | 0.81986215 | 0.41229469 | 0.85867915 |
| SPPL2B | 45.2073076 | -0.0486504 | 0.30774134 | -0.1580888 | 0.87438685 | 0.97918624 |
| TIMM13 | 29.5331438 | 0.68661555 | 0.46292571 | 1.48320891 | 0.13801891 | 0.68880821 |
| LMNB2 | 11.2452659 | -0.0108638 | 0.4990545 | -0.0217688 | 0.98263241 | NA |
| GADD45B | 99.3053535 | -0.6723772 | 0.41055897 | -1.6377117 | 1.01E-01 | 0.65863153 |
| GNG7 | 76.7722066 | 0.31729998 | 0.30374595 | 1.0446229 | 2.96E-01 | 0.81115642 |
| SGTA | 29.0769149 | 0.00862809 | 0.34171352 | 0.02524949 | 0.97985596 | 0.99571654 |
| THOP1 | 10.9119729 | -0.3672061 | 0.56203777 | -0.6533477 | 0.51353215 | NA |
| ZNF555 | 14.3015563 | -0.004387 | 0.53638869 | -0.0081788 | 0.99347434 | NA |
| TLE2 | 46.8093932 | -0.0387533 | 0.33355626 | -0.1161822 | 0.90750814 | 0.9854228 |

| | | | | | | |
|---|---|---|---|---|---|---|
| AES | 150.695766 | -0.2771981 | 0.31594567 | -0.8773599 | 0.3802912 | 0.85432452 |
| GNA11 | 38.7245005 | -0.0186363 | 0.38979413 | -0.0478107 | 0.9618671 | 0.99256424 |
| NCLN | 23.7341073 | -0.1910101 | 0.37179349 | -0.5137532 | 0.60742457 | 0.92569157 |
| NFIC | 221.802933 | -0.1228013 | 0.28222888 | -0.4351126 | 0.6634807 | 0.93860542 |
| SMIM24 | 19.9863488 | -0.1414606 | 0.56813586 | -0.2489908 | 0.80336789 | 0.97157671 |
| FZR1 | 22.2235032 | -0.4837784 | 0.39341037 | -1.2297043 | 0.21880786 | 0.76666749 |
| MFSD12 | 36.4778134 | -0.9240854 | 0.32395052 | -2.8525511 | 0.00433698 | 0.26125499 |
| HMG20B | 23.9897065 | -0.5416802 | 0.34689528 | -1.5615093 | 0.11840363 | 0.6668162 |
| PIP5K1C | 27.3054431 | 0.02774907 | 0.33425133 | 0.08301857 | 0.93383677 | 0.98960607 |
| TJP3 | 47.5509373 | 0.21717459 | 0.31162165 | 0.69691754 | 0.4858544 | 0.88838332 |
| MRPL54 | 26.3791883 | -0.394266 | 0.54626053 | -0.7217546 | 0.47044537 | 0.88260202 |
| DAPK3 | 20.007036 | -0.2375812 | 0.32146542 | -0.7390568 | 0.45987251 | 0.87607759 |
| EEF2 | 178.425523 | 0.79600819 | 0.39659611 | 2.00710035 | 0.04473898 | 0.52549539 |
| PIAS4 | 21.3761963 | -0.5672931 | 0.38321422 | -1.480355 | 0.13877852 | 0.69023184 |
| ZBTB7A | 67.4400545 | -0.1273549 | 0.3531897 | -0.360585 | 0.71840969 | 0.94925929 |
| MAP2K2 | 49.9578478 | 0.04518412 | 0.27322365 | 0.16537413 | 0.86864952 | 0.9783491 |
| SIRT6 | 11.4802642 | 0.06667714 | 0.48124213 | 0.13855217 | 0.88980405 | NA |
| YJU2 | 12.6256539 | -0.7284612 | 0.49908452 | -1.4595949 | 0.14440144 | NA |
| STAP2 | 14.0157216 | 0.1851796 | 0.38904995 | 0.47597897 | 0.63408936 | NA |
| SH3GL1 | 43.8229123 | -0.6305014 | 0.2459442 | -2.5635953 | 0.01035943 | 0.33440378 |
| UBXN6 | 34.6160364 | -0.2896287 | 0.40706111 | -0.7115115 | 0.47676731 | 0.88495683 |
| HDGFL2 | 20.0592134 | 0.3101159 | 0.3794546 | 0.81726749 | 0.41377558 | 0.85867915 |
| PLIN5 | 56.7255708 | 1.09550494 | 0.33170485 | 3.30264974 | 0.00095776 | 0.13754433 |
| MYDGF | 30.4596789 | 0.20644141 | 0.32059821 | 0.64392564 | 0.51962365 | 0.90368693 |
| DPP9 | 33.3771867 | 0.04228975 | 0.33260979 | 0.12714524 | 0.89882544 | 0.98531749 |
| TICAM1 | 11.1140467 | 0.66179923 | 0.57574915 | 1.14945759 | 0.25036734 | NA |
| PLIN3 | 37.7332696 | -0.2609973 | 0.33531698 | -0.7783598 | 4.36E-01 | 0.86729771 |
| KDM4B | 101.020521 | -0.3568303 | 0.25474415 | -1.40074 | 0.16129183 | 0.71101785 |
| PTPRS | 57.0440817 | 0.05223775 | 0.27475109 | 0.19012754 | 0.84920919 | 0.97612951 |
| TINCR | 11.358723 | 0.16266996 | 0.50897892 | 0.31960059 | 0.74927113 | NA |
| SAFB2 | 88.3322551 | -0.2269673 | 0.24622347 | -0.9217941 | 0.35663597 | 0.84192356 |
| SAFB | 88.1100074 | -0.1824243 | 0.22738081 | -0.8022853 | 0.42238796 | 0.86190547 |
| RPL36 | 136.487862 | -0.2935293 | 0.5466118 | -0.5369978 | 0.59126912 | 0.92254886 |
| C19orf70 | 41.22627 | 0.64648139 | 0.38878363 | 1.66283079 | 0.09634631 | 0.65104726 |
| LONP1 | 49.4069085 | -0.4586839 | 0.31540731 | -1.4542589 | 0.14587454 | 0.69679188 |
| CAPS | 39.8756469 | 0.11619113 | 0.56935473 | 0.20407511 | 0.83829481 | 0.97600352 |
| RANBP3 | 35.9790663 | -0.1188815 | 0.26837886 | -0.4429616 | 0.65779351 | 0.93593674 |
| RFX2 | 34.8166242 | -0.2244179 | 0.34811992 | -0.6446569 | 0.51914955 | 0.90368693 |
| MLLT1 | 34.8427529 | -0.23922 | 0.32445334 | -0.7373017 | 0.46093889 | 0.87634226 |
| CLPP | 18.3500996 | 0.57155748 | 0.40355859 | 1.41629369 | 0.15668954 | 0.70821492 |
| ALKBH7 | 22.6515571 | 0.27171603 | 0.43999661 | 0.6175412 | 0.53687782 | 0.90763121 |
| GTF2F1 | 23.0868159 | -0.0689296 | 0.37427276 | -0.1841693 | 0.85388063 | 0.97612951 |
| KHSRP | 39.8966293 | -0.3719356 | 0.32116994 | -1.1580647 | 0.24683763 | 0.78583441 |
| SLC25A23 | 43.4979794 | 0.73974646 | 0.41372984 | 1.78799398 | 0.07377698 | 0.60985192 |
| AC010503.4 | 16.2631684 | -0.5871827 | 0.52494554 | -1.1185592 | 0.26332823 | 0.79666376 |
| C3 | 51.1813688 | 0.89771924 | 1.34715558 | 0.66638127 | 0.50516743 | 0.89723255 |
| TRIP10 | 33.5787058 | -0.4052006 | 0.28457237 | -1.4238928 | 0.1544775 | 0.70572801 |
| SH2D3A | 40.2319458 | -0.8904938 | 0.36210995 | -2.4591807 | 0.01392545 | 0.37638271 |
| ZNF557 | 11.2386983 | -0.2248372 | 0.43384163 | -0.5182472 | 0.60428583 | NA |
| INSR | 268.443661 | -0.5042854 | 0.27714902 | -1.8195462 | 0.06882815 | 0.60275677 |
| ZNF358 | 21.4559222 | -0.2001307 | 0.56207855 | -0.3560546 | 0.72179966 | 0.95093722 |
| PNPLA6 | 26.1186016 | 0.63547212 | 0.41322222 | 1.53784596 | 0.12408628 | 0.66929384 |
| CAMSAP3 | 20.4044421 | 0.63354097 | 0.37886519 | 1.67220688 | 0.09448353 | 0.64716599 |
| XAB2 | 12.2982198 | -0.3002325 | 0.45483492 | -0.6600911 | 0.5091954 | NA |

| | | | | | | |
|---|---|---|---|---|---|---|
| STXBP2 | 17.1483556 | 0.02326726 | 0.4308502 | 0.05400314 | 0.95693266 | 0.99170927 |
| EVI5L | 20.0115476 | -0.079144 | 0.51875218 | -0.1525661 | 0.87874049 | 0.98061726 |
| MAP2K7 | 43.1286972 | -0.6766918 | 0.46949472 | -1.4413192 | 0.14949452 | 0.70183012 |
| TIMM44 | 25.7640835 | -0.7120149 | 0.36332931 | -1.9596958 | 0.05003135 | 0.54412834 |
| ELAVL1 | 69.4156256 | -0.2694351 | 0.24590226 | -1.0956999 | 0.27321015 | 0.79726027 |
| CERS4 | 56.6270212 | -0.4467624 | 0.37262014 | -1.1989755 | 0.23053746 | 0.77249016 |
| AC022146.2 | 11.2709603 | 0.39889861 | 0.48271427 | 0.82636589 | 0.40859657 | NA |
| CD320 | 15.0397307 | 0.07129 | 0.67996002 | 0.1048444 | 0.91649928 | NA |
| RPS28 | 47.867176 | -0.2764466 | 0.42035588 | -0.6576489 | 0.51076374 | 0.90019019 |
| ANGPTL4 | 59.0726203 | -1.9081969 | 1.25411105 | -1.5215534 | 0.12812102 | 0.66929384 |
| RAB11B-AS1 | 13.8650454 | -0.2102891 | 0.41106951 | -0.5115659 | 0.60895485 | NA |
| RAB11B | 43.7047485 | -0.1602522 | 0.32794365 | -0.4886579 | 0.62508394 | 0.92981013 |
| 2-Mar | 16.5416331 | -0.1155075 | 0.364861 | -0.3165795 | 0.75156271 | 0.9575193 |
| HNRNPM | 140.839787 | 0.12947216 | 0.21718299 | 0.5961432 | 0.55107956 | 0.91258455 |
| ZNF558 | 14.6067091 | 0.25729656 | 0.3734997 | 0.68888023 | 0.49089864 | NA |
| ZNF317 | 14.7302729 | 0.38249705 | 0.42545597 | 0.89902853 | 0.36863747 | NA |
| ZNF559-ZNF | 23.5504631 | -0.1662735 | 0.36348501 | -0.4574424 | 0.64735307 | 0.93328692 |
| ZNF266 | 29.8413801 | 0.35986195 | 0.29239302 | 1.2307474 | 0.21841735 | 0.76666749 |
| ZNF426 | 16.7447926 | 0.6153447 | 0.3665763 | 1.67862653 | 0.09322485 | 0.64572229 |
| ZNF121 | 30.3553285 | -0.090975 | 0.34148293 | -0.2664115 | 0.78992229 | 0.96883086 |
| ZNF561 | 22.3481504 | -0.4541789 | 0.3195486 | -1.4213141 | 0.15522546 | 0.70572801 |
| ZNF562 | 55.7747839 | -0.0742322 | 0.28362065 | -0.2617306 | 0.79352913 | 0.96960502 |
| ZNF846 | 47.0737488 | 0.11760883 | 0.27641984 | 0.42547173 | 0.67049283 | 0.93878847 |
| FBXL12 | 17.6986052 | 0.15411808 | 0.39260908 | 0.39254845 | 0.69465302 | 0.94395238 |
| UBL5 | 34.4684542 | 0.3732064 | 0.41809555 | 0.89263423 | 0.37205308 | 0.8491102 |
| PIN1 | 20.6453294 | 0.54022431 | 0.37856615 | 1.4270275 | 0.15357197 | 0.70572801 |
| C19orf66 | 32.1407779 | -0.7309693 | 0.62226768 | -1.1746862 | 0.24012028 | 0.78176499 |
| EIF3G | 19.6269325 | -0.203927 | 0.41458048 | -0.4918876 | 0.6227988 | 0.92909497 |
| DNMT1 | 22.709814 | -0.5801045 | 0.34968044 | -1.6589561 | 0.09712465 | 0.65442315 |
| MRPL4 | 28.3918104 | 0.42498813 | 0.33043833 | 1.2861345 | 0.19839612 | 0.75563666 |
| AC011511.2 | 37.8164028 | -0.3621171 | 0.71423848 | -0.5069975 | 0.61215655 | 0.92644418 |
| ICAM1 | 22.1737196 | 0.12887561 | 0.48172755 | 0.267528 | 0.78906266 | 0.96816786 |
| TYK2 | 39.6258168 | -0.0143045 | 0.34492744 | -0.0414709 | 0.96692045 | 0.99308116 |
| CDC37 | 62.6801736 | -0.2162532 | 0.24316271 | -0.8893356 | 0.37382276 | 0.85056774 |
| PDE4A | 39.1347226 | 0.13065975 | 0.44456855 | 0.29390238 | 0.7688325 | 0.96289288 |
| KEAP1 | 20.4203123 | 0.15735762 | 0.49627628 | 0.31707664 | 0.75118545 | 0.95741436 |
| ATG4D | 19.4674665 | -0.1229839 | 0.36583581 | -0.3361724 | 0.73674086 | 0.95387069 |
| KRI1 | 17.875673 | -0.0447256 | 0.36906098 | -0.1211875 | 0.90354254 | 0.98531749 |
| AP1M2 | 40.4589035 | 0.21265584 | 0.41669536 | 0.51033887 | 0.60981408 | 0.92573951 |
| SLC44A2 | 52.8306397 | 0.11129742 | 0.4283568 | 0.25982411 | 0.79499945 | 0.96974891 |
| ILF3-DT | 27.0530161 | -0.0382312 | 0.40299913 | -0.0948667 | 0.92442074 | 0.98925753 |
| ILF3 | 88.5752078 | -0.1617929 | 0.24538608 | -0.6593404 | 0.50967722 | 0.89963034 |
| QTRT1 | 30.9086212 | -0.2172363 | 0.37943098 | -0.5725317 | 0.56696179 | 0.91698355 |
| DNM2 | 294.683485 | -0.1546531 | 0.23088642 | -0.6698234 | 0.5029704 | 0.89564545 |
| TMED1 | 13.679387 | 0.57677571 | 0.52309509 | 1.10262115 | 0.27019172 | NA |
| CARM1 | 24.6716992 | 0.27574713 | 0.33849206 | 0.81463399 | 0.41528185 | 0.85867915 |
| YIPF2 | 21.9404709 | -0.4483602 | 0.3347801 | -1.3392678 | 0.18048351 | 0.73303885 |
| TIMM29 | 11.9407605 | 0.56247201 | 0.57699733 | 0.97482602 | 0.32964657 | NA |
| SMARCA4 | 150.148433 | -0.2913351 | 0.28769663 | -1.0126468 | 0.31122891 | 0.81701824 |
| LDLR | 86.7137573 | -1.5532313 | 0.62366638 | -2.4904842 | 0.01275692 | 0.36544853 |
| KANK2 | 17.8691224 | 0.10238455 | 0.41187262 | 0.24858306 | 0.80368332 | 0.97169999 |
| DOCK6 | 74.5391354 | 0.12922346 | 0.25783551 | 0.50118565 | 0.61624047 | 0.92662475 |
| RAB3D | 21.5769868 | 0.46176458 | 0.46783245 | 0.98702982 | 0.32362802 | 0.82566868 |
| EPOR | 38.385921 | 0.38151882 | 0.40591878 | 0.93988954 | 0.34727422 | 0.83687845 |

| | | | | | | |
|---|---|---|---|---|---|---|
| RGL3 | 270.418966 | 0.08208955 | 0.21227379 | 0.38671545 | 0.69896688 | 0.94395238 |
| PRKCSH | 39.2206205 | 0.21453853 | 0.42029064 | 0.5104528 | 0.60973427 | 0.92573951 |
| ECSIT | 34.6490618 | -0.105024 | 0.34756665 | -0.3021693 | 0.76252302 | 0.96072624 |
| ELOF1 | 12.1879063 | -0.2817282 | 0.47847862 | -0.5887999 | 0.55599552 | NA |
| ZNF627 | 34.5006552 | -0.0570777 | 0.33417882 | -0.1707999 | 0.86438113 | 0.97788853 |
| AC008543.1 | 15.9276461 | 1.06099145 | 0.63037465 | 1.6831125 | 0.09235331 | NA |
| ZNF823 | 39.969146 | 0.23074636 | 0.31692212 | 0.72808539 | 0.46656131 | 0.88059581 |
| ZNF441 | 28.45351 | 0.49301489 | 0.32920232 | 1.49760456 | 0.13423602 | 0.68205729 |
| ZNF440 | 35.6418385 | -0.2316746 | 0.42570578 | -0.544213 | 0.58629488 | 0.92076087 |
| ZNF439 | 47.2732839 | 0.10473305 | 0.30552557 | 0.34279635 | 0.73175167 | 0.95279138 |
| ZNF69 | 59.0212565 | 0.08548844 | 0.37485228 | 0.228059 | 0.81960037 | 0.97323535 |
| ZNF433 | 13.1555268 | 0.14532654 | 0.44235819 | 0.32852684 | 0.74251335 | NA |
| ZNF844 | 26.7388839 | 0.2631989 | 0.37923768 | 0.69402096 | 0.48766906 | 0.88838332 |
| ZNF788P | 14.2635179 | 0.60951492 | 0.4695307 | 1.29813646 | 0.19424045 | NA |
| ZNF136 | 18.3647637 | -0.1900939 | 0.37628526 | -0.5051856 | 0.61342849 | 0.92662475 |
| ZNF44 | 205.320747 | -0.0005432 | 0.18805652 | -0.0028887 | 0.99769516 | 0.99956021 |
| ZNF563 | 11.2475428 | 0.60689998 | 0.66072429 | 0.91853741 | 0.35833758 | NA |
| AC008758.5 | 19.6995956 | -0.6063164 | 0.56232928 | -1.078223 | 0.28093426 | 0.80377836 |
| ZNF490 | 27.9881406 | 0.72497627 | 0.40642157 | 1.78380362 | 0.07445558 | 0.61020315 |
| ZNF791 | 48.3510036 | -0.0886119 | 0.255355 | -0.3470145 | 0.72858045 | 0.95279138 |
| MAN2B1 | 11.7699685 | -0.180125 | 0.40727721 | -0.4422663 | 0.65829651 | NA |
| DHPS | 15.1542537 | 0.45784228 | 0.41162936 | 1.11226828 | 0.26602282 | NA |
| TNPO2 | 43.4510507 | 0.02753762 | 0.3445575 | 0.0799217 | 0.93629953 | 0.99012269 |
| TRIR | 54.8543953 | 0.15071954 | 0.37469339 | 0.40224765 | 0.68750178 | 0.94205398 |
| ASNA1 | 17.2046902 | 0.59445853 | 0.42589284 | 1.39579367 | 0.16277664 | 0.71325312 |
| HOOK2 | 127.330364 | -0.1019144 | 0.35980368 | -0.2832499 | 0.77698526 | 0.96518573 |
| JUNB | 65.97608 | -0.5754264 | 0.56194397 | -1.0239924 | 0.30583885 | 0.81568231 |
| DNASE2 | 9.54337348 | 0.71042252 | 0.54458647 | 1.30451738 | 0.19205724 | NA |
| GCDH | 19.9513136 | 0.09195808 | 0.4981127 | 0.184613 | 0.85353259 | 0.97612951 |
| FARSA | 16.8739086 | 0.37910046 | 0.48576146 | 0.78042515 | 0.43514067 | 0.86729771 |
| CALR | 103.707211 | 0.20073847 | 0.55611463 | 0.36096599 | 0.71812487 | 0.94925929 |
| AC092069.1 | 17.9280309 | 0.29505159 | 0.65382503 | 0.45126995 | 0.651795 | 0.93482187 |
| RAD23A | 38.1797606 | -0.4397186 | 0.25649439 | -1.7143401 | 0.08646628 | 0.63466198 |
| GADD45GIP1 | 47.6045321 | 0.0340407 | 0.42640226 | 0.07983237 | 0.93637058 | 0.99012269 |
| NFIX | 18.863061 | -0.2084024 | 0.48331444 | -0.4311942 | 0.66632714 | 0.93860542 |
| TRMT1 | 10.3945144 | -0.2377775 | 0.51604244 | -0.4607712 | 0.64496277 | NA |
| NACC1 | 21.7346717 | -0.063258 | 0.40124405 | -0.1576548 | 0.87472884 | 0.97918624 |
| STX10 | 12.9851917 | -0.0376474 | 0.48097528 | -0.0782731 | 0.93761085 | NA |
| IER2 | 68.77771 | -0.5350331 | 0.57997985 | -0.922503 | 0.35626628 | 0.84192356 |
| CCDC130 | 61.4803544 | 0.11255288 | 0.22365907 | 0.50323414 | 0.61479966 | 0.92662475 |
| MRI1 | 25.1863891 | 0.52821949 | 0.36378925 | 1.45199314 | 0.14650351 | 0.69713616 |
| C19orf53 | 48.7785928 | -0.0246145 | 0.43132856 | -0.0570668 | 0.95449201 | 0.99148072 |
| ZSWIM4 | 22.4149763 | -0.3646653 | 0.35885407 | -1.0161939 | 0.30953706 | 0.81568231 |
| CC2D1A | 42.4124865 | -0.0461118 | 0.36963779 | -0.1247487 | 0.90072249 | 0.98531749 |
| DCAF15 | 12.3281179 | -0.4004089 | 0.40631211 | -0.9854713 | 0.32439262 | NA |
| RFX1 | 31.0791694 | -0.5171059 | 0.32748452 | -1.5790239 | 0.11433057 | 0.66629396 |
| PALM3 | 32.892755 | 0.37034931 | 0.29869448 | 1.23989337 | 0.21501484 | 0.7643572 |
| SAMD1 | 12.8430422 | 0.0820623 | 0.58811337 | 0.13953483 | 0.88902754 | NA |
| PRKACA | 33.1100659 | 0.19018637 | 0.31135838 | 0.61082783 | 0.54131356 | 0.90783815 |
| ADGRL1 | 34.208699 | 0.22890642 | 0.34534498 | 0.66283408 | 0.50743682 | 0.89851807 |
| PKN1 | 34.3789903 | 0.03829118 | 0.32394881 | 0.11820133 | 0.90590814 | 0.98531749 |
| PTGER1 | 31.8115922 | 1.39132383 | 0.824879 | 1.6867005 | 0.09166096 | 0.64396102 |
| GIPC1 | 96.3361849 | -0.4221773 | 0.33389671 | -1.264395 | 0.2060883 | 0.76231289 |
| DNAJB1 | 41.721283 | -0.2195401 | 0.687948 | -0.3191231 | 0.74963316 | 0.95680183 |

| | | | | | | |
|---|---|---|---|---|---|---|
| TECR | 37.9469195 | 0.4587707 | 0.36538529 | 1.25558066 | 0.20926805 | 0.7623519 |
| NDUFB7 | 67.4027271 | 0.07186699 | 0.42094336 | 0.17072841 | 0.86443732 | 0.97788853 |
| ZNF333 | 11.3708048 | -0.4232704 | 0.51025521 | -0.8295269 | 0.4068063 | NA |
| ILVBL | 25.7466448 | 0.01587871 | 0.35924107 | 0.0442007 | 0.96474442 | 0.99270395 |
| NOTCH3 | 48.5726925 | -0.4116117 | 0.41442719 | -0.9932064 | 0.3206094 | 0.82464962 |
| BRD4 | 218.314015 | -0.3370195 | 0.24514419 | -1.3747805 | 0.1691995 | 0.72310941 |
| AKAP8 | 45.08944 | -0.1807162 | 0.27204755 | -0.6642818 | 0.50650998 | 0.89738426 |
| AKAP8L | 123.913244 | -0.2666049 | 0.2183313 | -1.2211024 | 0.22204726 | 0.76680432 |
| WIZ | 12.6575208 | 0.37525385 | 0.41749431 | 0.89882386 | 0.36874649 | NA |
| TPM4 | 150.04238 | -0.0019286 | 0.43932627 | -0.00439 | 0.99649731 | 0.99927028 |
| RAB8A | 24.5553439 | -0.0682826 | 0.30431837 | -0.2243787 | 0.82246265 | 0.97478959 |
| FAM32A | 17.1373726 | 0.04078922 | 0.3734147 | 0.10923304 | 0.91301766 | 0.98639439 |
| AP1M1 | 18.3160838 | -0.59359 | 0.34954006 | -1.6982031 | 0.08946944 | 0.63653714 |
| EPS15L1 | 74.9719556 | -0.3703356 | 0.21938482 | -1.6880638 | 0.09139899 | 0.64297142 |
| C19orf44 | 14.6120068 | 0.55989147 | 0.44290568 | 1.26413251 | 0.20618247 | NA |
| TMEM38A | 24.8374158 | -1.0198505 | 0.48813376 | -2.0892848 | 0.03668209 | 0.49537143 |
| SIN3B | 49.0992738 | -0.3645757 | 0.28424034 | -1.2826318 | 0.19962109 | 0.75605362 |
| MYO9B | 75.3384077 | -0.7390605 | 0.32595522 | -2.2673684 | 0.02336773 | 0.43088993 |
| NR2F6 | 32.5215663 | 0.38943416 | 0.40600614 | 0.95918293 | 0.3374666 | 0.83055261 |
| ABHD8 | 15.0405686 | 0.15609167 | 0.3802187 | 0.41053128 | 0.68141626 | NA |
| MRPL34 | 33.2602636 | 0.00032457 | 0.34342948 | 0.0009451 | 0.99924592 | 0.99957534 |
| DDA1 | 15.9420633 | 0.18614493 | 0.38576059 | 0.48254004 | 0.62942236 | NA |
| BST2 | 65.789054 | -0.6218876 | 0.82441352 | -0.7543394 | 0.45064542 | 0.87346041 |
| MVB12A | 22.1046105 | 0.22838759 | 0.34403848 | 0.66384316 | 0.5067907 | 0.89770689 |
| SLC27A1 | 27.8445768 | 1.11608761 | 0.42256671 | 2.64121044 | 0.00826104 | 0.30976355 |
| AC010618.3 | 21.2701231 | 0.26079431 | 0.44772401 | 0.582489 | 0.56023735 | 0.91533003 |
| PGLS | 38.6590292 | 0.02633149 | 0.37445672 | 0.07031919 | 0.94393961 | 0.99069421 |
| COLGALT1 | 16.0349734 | -0.3203705 | 0.49355808 | -0.6491039 | 0.51627125 | NA |
| RPL18A | 12.4681678 | 0.32122596 | 0.59625266 | 0.53874135 | 0.59006534 | NA |
| CCDC124 | 18.2108679 | -0.1717754 | 0.44097463 | -0.3895359 | 0.69687978 | 0.94395238 |
| ARRDC2 | 13.7094536 | -0.0310095 | 0.51624086 | -0.0600679 | 0.95210157 | NA |
| IQCN | 11.4953672 | 0.88701217 | 0.78369939 | 1.13182705 | 0.25770715 | NA |
| JUND | 155.306836 | -1.4740141 | 0.45040462 | -3.2726442 | 0.00106546 | 0.14056417 |
| LSM4 | 50.0973379 | -0.0969236 | 0.39595173 | -0.2447865 | 0.80662175 | 0.97216031 |
| PGPEP1 | 53.0590275 | -0.2354145 | 0.36574983 | -0.6436489 | 0.51980314 | 0.90368693 |
| GDF15 | 210.885181 | -1.6192751 | 0.60634368 | -2.6705566 | 0.00757256 | 0.30654023 |
| SSBP4 | 29.5266878 | 0.10018241 | 0.39679193 | 0.25248096 | 0.80066933 | 0.97021033 |
| ISYNA1 | 14.9988479 | 0.08204896 | 0.59472785 | 0.13796052 | 0.89027162 | NA |
| ELL | 58.2308499 | 0.37455597 | 0.30043525 | 1.24671112 | 0.21250344 | 0.7643572 |
| FKBP8 | 47.865281 | -0.1083391 | 0.44714046 | -0.2422932 | 0.80855296 | 0.97257655 |
| KXD1 | 17.8342837 | -0.2230948 | 0.40218657 | -0.5547049 | 0.57909655 | 0.91976673 |
| UBA52 | 74.3303311 | -0.1242642 | 0.47876331 | -0.2595525 | 0.79520898 | 0.96974891 |
| REX1BD | 23.616595 | 0.2197025 | 0.41718397 | 0.52663218 | 0.59844904 | 0.92296674 |
| CRTC1 | 94.5560289 | -0.0691417 | 0.26488261 | -0.2610277 | 0.79407117 | 0.96974703 |
| UPF1 | 42.7843662 | 0.02027263 | 0.2711463 | 0.07476639 | 0.94040058 | 0.99012269 |
| COPE | 30.3113176 | 0.18018319 | 0.37129344 | 0.48528514 | 0.6274741 | 0.93057946 |
| DDX49 | 12.7836141 | -0.1508613 | 0.48509401 | -0.3109939 | 0.75580525 | NA |
| SUGP2 | 100.734667 | -0.0655135 | 0.23745505 | -0.2758984 | 0.78262608 | 0.96652357 |
| SLC25A42 | 24.7661421 | 0.66322152 | 0.40521931 | 1.63669772 | 0.10169364 | 0.65863153 |
| TMEM161A | 18.466375 | 0.03054543 | 0.38362605 | 0.07962293 | 0.93653716 | 0.99012269 |
| RFXANK | 12.3901687 | -0.2548997 | 0.41467058 | -0.614704 | 0.5387502 | NA |
| SUGP1 | 24.6000691 | -0.4158799 | 0.33776324 | -1.2312762 | 0.21821956 | 0.76666749 |
| MAU2 | 63.4978507 | -0.364673 | 0.25865824 | -1.4098644 | 0.15857973 | 0.71089007 |
| GATAD2A | 151.151946 | -0.0853541 | 0.23696984 | -0.3601898 | 0.71870518 | 0.94927064 |

| | | | | | | |
|---|---|---|---|---|---|---|
| PBX4 | 13.596815 | 0.10960041 | 0.42433629 | 0.25828667 | 0.79618568 | NA |
| ATP13A1 | 21.2076291 | 0.17596722 | 0.40613714 | 0.43327044 | 0.66481831 | 0.93860542 |
| ZNF101 | 10.6562684 | -0.0829582 | 0.42226482 | -0.19646 | 0.84425011 | NA |
| ZNF14 | 27.525959 | -0.2039594 | 0.33911003 | -0.6014551 | 0.54753694 | 0.91139843 |
| AC011477.4 | 11.3438139 | 0.59543008 | 0.5145671 | 1.15714758 | 0.24721207 | NA |
| ZNF253 | 37.5900376 | 0.3521956 | 0.30677417 | 1.14806147 | 0.25094319 | 0.78892875 |
| ZNF93 | 14.7754629 | -0.3767102 | 0.44717299 | -0.8424261 | 0.39954947 | NA |
| ZNF682 | 30.0149496 | -0.0461698 | 0.43614168 | -0.1058596 | 0.91569373 | 0.98691474 |
| ZNF90 | 90.6452696 | 0.30700329 | 0.26975108 | 1.13809846 | 0.25507937 | 0.79329946 |
| AC011447.3 | 50.4878999 | 0.32190287 | 0.41849733 | 0.76918738 | 0.44178208 | 0.86925451 |
| ZNF486 | 57.2673961 | -0.2599246 | 0.56322441 | -0.4614938 | 0.64444437 | 0.93328692 |
| AC008554.1 | 20.2566748 | 0.82271488 | 0.91394372 | 0.90018112 | 0.36802387 | 0.84777627 |
| ZNF737 | 40.7864925 | 0.2552858 | 0.32941083 | 0.77497693 | 0.43835329 | 0.868596 |
| AC010636.2 | 12.9920206 | 0.00133534 | 0.38583145 | 0.00346093 | 0.99723858 | NA |
| ZNF66 | 11.7643969 | 0.32340243 | 0.46807392 | 0.69092171 | 0.48961475 | NA |
| ZNF85 | 18.3301609 | 0.5427306 | 0.48445935 | 1.12028098 | 0.26259405 | 0.79666376 |
| ZNF430 | 55.1284326 | -0.3106266 | 0.32926689 | -0.9433886 | 0.34548217 | 0.83564825 |
| ZNF714 | 22.6864271 | 0.45436658 | 0.43892696 | 1.03517583 | 0.30058681 | 0.81411535 |
| ZNF431 | 90.3205435 | -0.1120693 | 0.21963045 | -0.5102631 | 0.60986712 | 0.92573951 |
| ZNF708 | 65.2092563 | 0.00061977 | 0.20876619 | 0.00296874 | 0.99763129 | 0.99956021 |
| ZNF738 | 15.5415302 | -0.5695371 | 0.42870719 | -1.328499 | 0.18401331 | NA |
| ZNF493 | 39.1430829 | -0.0246942 | 0.32132181 | -0.0768518 | 0.93874142 | 0.99012269 |
| AC010615.4 | 59.3768368 | 1.21822879 | 0.56896083 | 2.14114702 | 0.03226218 | 0.48175799 |
| ZNF429 | 61.5390388 | 0.47050969 | 0.33849399 | 1.39000899 | 0.16452615 | 0.71446159 |
| AC123912.1 | 22.5284909 | 0.09472205 | 0.61374595 | 0.1543343 | 0.87734616 | 0.9800567 |
| AC123912.4 | 13.3153223 | 0.05139546 | 0.61492796 | 0.08357964 | 0.93339066 | NA |
| ZNF100 | 70.4655298 | -0.1405077 | 0.34563945 | -0.4065151 | 0.68436416 | 0.94205398 |
| ZNF43 | 102.256722 | 0.06940414 | 0.26916239 | 0.25785228 | 0.79652091 | 0.96986756 |
| ZNF208 | 181.455218 | 0.10440266 | 0.38605332 | 0.27043585 | 0.78682496 | 0.96767512 |
| ZNF257 | 12.879406 | 0.07403085 | 0.47609214 | 0.15549689 | 0.8764296 | NA |
| ZNF676 | 27.8060414 | 0.85096314 | 0.60304941 | 1.41110019 | 0.15821508 | 0.71089007 |
| ZNF98 | 140.358803 | 0.23983559 | 0.44076985 | 0.54412883 | 0.58635282 | 0.92076087 |
| ZNF99 | 17.9712314 | 1.2477227 | 0.64784528 | 1.92595784 | 0.05410964 | 0.5642154 |
| ZNF730 | 13.3680786 | -0.9137796 | 0.43888212 | -2.0820616 | 0.03733685 | NA |
| ZNF91 | 254.814177 | -0.2700503 | 0.29506594 | -0.9152203 | 0.360076 | 0.84282379 |
| ZNF675 | 36.253809 | 0.12737817 | 0.27754177 | 0.45895134 | 0.64626911 | 0.93328692 |
| AC139769.2 | 14.315313 | 1.13840016 | 0.68782682 | 1.65506799 | 0.09791071 | NA |
| ZNF726 | 10.8116712 | 1.22701117 | 0.57777608 | 2.12367943 | 0.03369696 | NA |
| ZNF254 | 150.705066 | -0.2816423 | 0.22511029 | -1.2511304 | 0.21088689 | 0.76397692 |
| LINC00662 | 51.9066756 | -0.0050704 | 0.44802854 | -0.0113171 | 0.99097047 | 0.99814944 |
| AC006504.5 | 48.7388322 | 1.01720243 | 0.2609022 | 3.89878831 | 9.67E-05 | 0.04400174 |
| UQCRFS1 | 58.8427618 | 1.11647325 | 0.35659201 | 3.13095423 | 0.00174239 | 0.17240222 |
| C19orf12 | 23.3141743 | 0.19271156 | 0.31894234 | 0.60422069 | 0.54569693 | 0.91062862 |
| URI1 | 182.915935 | -0.017932 | 0.21362501 | -0.0839414 | 0.933103 | 0.98960607 |
| ZNF507 | 48.7435722 | -0.1764464 | 0.30044939 | -0.5872749 | 0.5570191 | 0.9141017 |
| DPY19L3 | 58.000034 | -0.5800698 | 0.27992381 | -2.0722417 | 0.03824291 | 0.49671741 |
| PDCD5 | 21.3873714 | 0.0066515 | 0.37463233 | 0.01775474 | 0.98583451 | 0.99766827 |
| ANKRD27 | 38.5636116 | -0.0186934 | 0.37915552 | -0.0493027 | 0.96067806 | 0.99256424 |
| NUDT19 | 13.8372367 | 0.16446918 | 0.4169164 | 0.39448959 | 0.69321961 | NA |
| CEP89 | 40.5764745 | -0.17453 | 0.34665291 | -0.503472 | 0.61463244 | 0.92662475 |
| RHPN2 | 135.658367 | -0.395699 | 0.37223418 | -1.0630377 | 0.28776484 | 0.80523851 |
| GPATCH1 | 20.6402278 | -0.2974925 | 0.45400138 | -0.6552678 | 0.51229536 | 0.9014941 |
| LRP3 | 12.8676999 | 0.91775848 | 0.51378816 | 1.78625852 | 0.07405741 | NA |
| PEPD | 34.8468732 | -0.2194124 | 0.33855109 | -0.6480925 | 0.51692516 | 0.9026606 |

| | | | | | | |
|---|---|---|---|---|---|---|
| LSM14A | 233.452449 | -0.1398675 | 0.17604464 | -0.7945003 | 0.42690424 | 0.86239557 |
| KIAA0355 | 117.591324 | -0.2228627 | 0.23222321 | -0.9596918 | 0.33721035 | 0.83055261 |
| GPI | 179.543438 | -0.4421529 | 0.17324418 | -2.552195 | 0.01070466 | 0.33942942 |
| UBA2 | 92.5626186 | -0.2522741 | 0.20987267 | -1.2020341 | 0.22935032 | 0.77185154 |
| WTIP | 26.0103492 | 1.04228623 | 0.43557276 | 2.39290959 | 0.01671536 | 0.39748373 |
| SCGB2B2 | 57.6410976 | -0.1856523 | 0.48512864 | -0.3826868 | 0.70195203 | 0.94476609 |
| ZNF302 | 68.1119946 | 0.33654008 | 0.35371722 | 0.9514382 | 0.34138198 | 0.8325058 |
| ZNF30 | 11.8291268 | 1.01797924 | 0.70680645 | 1.44025176 | 0.14979618 | NA |
| GRAMD1A | 40.5625385 | 0.11798125 | 0.34849194 | 0.33854801 | 7.35E-01 | 0.95329896 |
| HPN | 127.224381 | 0.67608119 | 0.29655082 | 2.27981557 | 0.02261863 | 0.42564995 |
| LSR | 47.8358816 | -0.0209446 | 0.29850518 | -0.0701648 | 0.94406247 | 0.99069421 |
| USF2 | 31.0901144 | -0.1216304 | 0.42716888 | -0.284736 | 0.77584641 | 0.96469469 |
| DMKN | 69.1957052 | 0.09593889 | 0.33782776 | 0.2839876 | 0.77641989 | 0.9649632 |
| TMEM147 | 25.2338655 | 0.40920335 | 0.41778814 | 0.97945179 | 0.3273568 | 0.82612204 |
| RBM42 | 13.1465005 | -0.0589524 | 0.5696496 | -0.103489 | 0.9175749 | NA |
| COX6B1 | 106.355337 | -0.050462 | 0.35344755 | -0.1427709 | 0.8864711 | 0.98216235 |
| KMT2B | 28.7265565 | -0.1692222 | 0.31997505 | -0.5288606 | 0.59690212 | 0.9229638 |
| PROSER3 | 15.9525311 | -0.050282 | 0.44621715 | -0.112685 | 0.91028029 | NA |
| SYNE4 | 11.297934 | 0.98789509 | 0.44767871 | 2.20670555 | 0.02733464 | NA |
| TBCB | 26.2152118 | -0.0314824 | 0.3622349 | -0.0869115 | 0.93074187 | 0.98960607 |
| CAPNS1 | 88.5720777 | 0.12167877 | 0.37906613 | 0.32099615 | 0.74821331 | 0.95563105 |
| COX7A1 | 9.31761887 | 0.23133985 | 0.82168335 | 0.28154379 | 0.77829334 | NA |
| ZNF565 | 57.6003472 | 0.43998659 | 0.43277783 | 1.01665693 | 0.30931666 | 0.81568231 |
| ZNF146 | 69.8583245 | -0.0768654 | 0.30131895 | -0.2550965 | 0.79864854 | 0.97019967 |
| LINC00665 | 32.0511194 | 0.28064615 | 0.35133544 | 0.79879828 | 0.42440739 | 0.86190547 |
| ZFP14 | 59.2463369 | 0.15636649 | 0.31990944 | 0.48878359 | 0.62499491 | 0.92981013 |
| ZFP82 | 21.5593919 | 0.37000329 | 0.39436562 | 0.938224 | 0.34812931 | 0.83687845 |
| ZNF566 | 35.1631361 | -0.2058309 | 0.28095198 | -0.7326195 | 0.46379052 | 0.87824803 |
| ZNF260 | 29.5091368 | 0.1529604 | 0.29901527 | 0.51154712 | 0.608968 | 0.92573951 |
| ZNF529 | 45.4072069 | 0.00747185 | 0.3196115 | 0.02337791 | 0.98134883 | 0.99599026 |
| ZNF529-AS1 | 41.8928241 | -0.3178926 | 0.32417467 | -0.9806213 | 0.32677952 | 0.82612204 |
| ZNF382 | 11.6672714 | 0.30984083 | 0.44016346 | 0.70392221 | 0.48148122 | NA |
| ZNF461 | 24.8218949 | -0.8149437 | 0.34891174 | -2.3356728 | 0.0195083 | 0.41544117 |
| ZNF567 | 26.0374168 | 0.02614596 | 0.38785726 | 0.0674113 | 0.94625428 | 0.99083449 |
| AC020928.1 | 47.7222684 | -0.0422993 | 0.25009443 | -0.1691334 | 0.86569173 | 0.97788853 |
| ZNF790 | 32.6243403 | 0.59920904 | 0.38342057 | 1.56279836 | 0.11810003 | 0.6668162 |
| ZNF345 | 58.7158843 | 0.37674811 | 0.41871142 | 0.89977987 | 0.36823741 | 0.84777627 |
| ZNF568 | 44.4046945 | -0.4827876 | 0.34591221 | -1.3956939 | 0.16280669 | 0.71325312 |
| ZNF420 | 85.8792757 | -0.2027527 | 0.2923775 | -0.6934619 | 0.48801971 | 0.88866642 |
| ZNF383 | 23.508214 | -0.434566 | 0.37565486 | -1.1568226 | 0.24734486 | 0.78583441 |
| LINC01535 | 15.8731859 | 0.32725195 | 0.47969052 | 0.68221474 | 0.49510317 | NA |
| HKR1 | 69.8652357 | 0.14397442 | 0.25142379 | 0.57263644 | 0.56689088 | 0.91698355 |
| ZNF569 | 23.2127363 | 0.11483394 | 0.37399129 | 0.30704977 | 0.75880549 | 0.95903525 |
| ZNF793 | 39.1676179 | 0.32710851 | 0.31932915 | 1.02436155 | 0.30566454 | 0.81568231 |
| ZNF540 | 31.3605575 | 0.50360287 | 0.50031078 | 1.00658009 | 0.31413661 | 0.82040655 |
| ZNF571 | 31.257816 | 0.41474684 | 0.40832838 | 1.01571886 | 0.30976328 | 0.81568231 |
| ZFP30 | 34.5205731 | 0.17159787 | 0.32679085 | 0.52509997 | 0.59951369 | 0.92298075 |
| ZNF573 | 40.6776273 | 0.00998951 | 0.30594053 | 0.0326518 | 0.97395226 | 0.99431961 |
| SIPA1L3 | 273.178042 | -0.2511927 | 0.25145064 | -0.9989744 | 0.31780712 | 0.82351215 |
| SPINT2 | 17.9034249 | -0.1196721 | 0.35544129 | -0.3366858 | 0.73635373 | 0.95376039 |
| CATSPERG | 19.8601528 | -0.510664 | 0.50207013 | -1.0171168 | 0.30909785 | 0.81568231 |
| PSMD8 | 38.9963686 | 0.28942107 | 0.31667688 | 0.91393178 | 0.36075269 | 0.84331775 |
| EIF3K | 72.7971436 | 0.01636593 | 0.29493014 | 0.05549087 | 0.95574741 | 0.99148072 |
| ACTN4 | 316.883899 | -0.4699138 | 0.29144124 | -1.612379 | 0.10687948 | 0.65986731 |

| | | | | | | |
|---|---|---|---|---|---|---|
| ECH1 | 40.4763642 | 0.64132144 | 0.52410415 | 1.22365268 | 0.22108328 | 0.76680432 |
| HNRNPL | 54.3217608 | -0.3287849 | 0.23498558 | -1.3991706 | 0.16176183 | 0.71101785 |
| SIRT2 | 31.6712155 | 0.55935319 | 0.37838799 | 1.47825301 | 0.13934007 | 0.69054855 |
| NFKBIB | 14.664173 | -0.1245949 | 0.37484482 | -0.3323906 | 0.73959433 | NA |
| MRPS12 | 24.5575348 | 0.566359 | 0.39974356 | 1.41680582 | 0.15653971 | 0.70821492 |
| FBXO17 | 113.583688 | 0.21464352 | 0.33484853 | 0.64101675 | 0.5215118 | 0.90451015 |
| PAK4 | 74.4418975 | 0.00788846 | 0.28859318 | 0.02733418 | 0.9781932 | 0.9946931 |
| SAMD4B | 92.2295056 | -0.0186959 | 0.32095559 | -0.0582507 | 0.95354896 | 0.99148072 |
| PAF1 | 16.176074 | 0.33893374 | 0.42098072 | 0.80510514 | 0.42075901 | 0.86148657 |
| MED29 | 21.6969144 | -0.1505045 | 0.3276419 | -0.4593566 | 0.6459781 | 0.93328692 |
| ZFP36 | 32.8132908 | -0.7236427 | 0.63661865 | -1.1366972 | 0.25566487 | 0.7940072 |
| RPS16 | 77.4821969 | 0.07829468 | 0.42128757 | 0.18584616 | 0.8525654 | 0.97612951 |
| SUPT5H | 63.0838417 | -0.1269652 | 0.25766943 | -0.4927444 | 0.6221932 | 0.92897508 |
| TIMM50 | 17.6620631 | -0.0334154 | 0.37963062 | -0.0880208 | 0.92986013 | 0.98960607 |
| DYRK1B | 10.9709601 | -0.0580667 | 0.64239961 | -0.0903903 | 0.92797709 | NA |
| FBL | 14.4222965 | 0.36644944 | 0.55588849 | 0.65921393 | 0.5097584 | NA |
| PSMC4 | 20.6517196 | 0.56359183 | 0.3677651 | 1.53247773 | 0.12540459 | 0.66929384 |
| ZNF546 | 15.9770918 | 1.07074154 | 0.61163405 | 1.75062447 | 0.08001062 | NA |
| ZNF780B | 97.4012614 | 0.01266533 | 0.2315607 | 0.0546955 | 0.95638105 | 0.99170927 |
| ZNF780A | 16.1993736 | 0.2805499 | 0.3703903 | 0.75744398 | 0.44878391 | 0.87314185 |
| MAP3K10 | 16.0025369 | 0.05981861 | 0.44771295 | 0.1336093 | 0.89371153 | NA |
| AKT2 | 80.7162081 | -0.321755 | 0.22560836 | -1.4261661 | 0.15382039 | 0.70572801 |
| PLD3 | 90.0497388 | -0.0728775 | 0.30999515 | -0.2350922 | 0.81413714 | 0.97282508 |
| SERTAD1 | 14.483126 | -0.2396626 | 0.46331333 | -0.5172797 | 0.60496092 | NA |
| BLVRB | 81.7401175 | -0.0841504 | 0.2488556 | -0.3381496 | 0.73525045 | 0.95341665 |
| SPTBN4 | 20.1571189 | 0.11404191 | 0.50516351 | 0.22575248 | 0.82139394 | 0.97396757 |
| SHKBP1 | 13.8847738 | 0.27460903 | 0.45630923 | 0.60180469 | 0.54730415 | NA |
| COQ8B | 19.0227782 | 0.25962519 | 0.4847219 | 0.5356168 | 0.59222342 | 0.92278087 |
| ITPKC | 28.5990149 | -0.3450071 | 0.31734685 | -1.087161 | 0.27696565 | 0.80159799 |
| SNRPA | 20.2661685 | -0.0531138 | 0.37213864 | -0.142726 | 0.88650661 | 0.98216235 |
| AC008537.1 | 17.899614 | -1.3422569 | 0.42520244 | -3.1567478 | 0.00159539 | 0.16785081 |
| AXL | 13.00563 | -0.3933023 | 0.67182814 | -0.5854209 | 0.55826471 | NA |
| HNRNPUL1 | 49.9002237 | 0.21825871 | 0.24803978 | 0.87993431 | 0.3788949 | 0.85415558 |
| TGFB1 | 37.316486 | 0.0981702 | 0.59176417 | 0.16589413 | 0.86824027 | 0.97812778 |
| DMAC2 | 13.8099287 | -0.0206801 | 0.38103567 | -0.0542735 | 0.95671725 | NA |
| RPS19 | 267.005342 | 0.37333842 | 0.39627874 | 0.94211064 | 0.346136 | 0.83666384 |
| ARHGEF1 | 25.6189686 | -0.6653412 | 0.33547785 | -1.9832641 | 0.04733794 | 0.53530098 |
| RABAC1 | 50.2014265 | 0.0230993 | 0.46409427 | 0.04977287 | 0.96030339 | 0.99256424 |
| GRIK5 | 41.3069285 | 0.26428746 | 0.31627045 | 0.83563754 | 0.40335884 | 0.85782098 |
| DEDD2 | 14.0324933 | -0.0605625 | 0.39758219 | -0.1523271 | 0.87892897 | NA |
| GSK3A | 12.7810143 | -0.3174623 | 0.62847216 | -0.5051334 | 0.61346518 | NA |
| CIC | 22.6640281 | -0.7390788 | 0.42872128 | -1.7239145 | 0.08472328 | 0.62880224 |
| PAFAH1B3 | 11.529169 | -0.3264022 | 0.42562194 | -0.7668829 | 0.44315113 | NA |
| MEGF8 | 30.7302733 | 0.39526466 | 0.31606551 | 1.2505783 | 0.21108837 | 0.76432085 |
| LIPE-AS1 | 34.9056478 | -0.0171919 | 0.349321 | -0.0492151 | 0.96074787 | 0.99256424 |
| PHLDB3 | 16.3329502 | -0.1523139 | 0.44474582 | -0.3424741 | 0.73199414 | 0.95279138 |
| ETHE1 | 28.9341642 | -0.3001579 | 0.41509075 | -0.7231138 | 0.46960994 | 0.88207878 |
| ZNF428 | 28.6986172 | 0.36355663 | 0.37665056 | 0.96523586 | 0.33442669 | 0.82974237 |
| SMG9 | 19.0331153 | -0.3758516 | 0.35879513 | -1.0475383 | 0.29485139 | 0.8101516 |
| ZNF283 | 26.5231941 | 0.30600928 | 0.46430719 | 0.65906642 | 0.50985312 | 0.89963034 |
| ZNF45 | 12.8487471 | 0.25448985 | 0.42926311 | 0.59285283 | 0.55327965 | NA |
| ZNF155 | 23.2744505 | 0.81362954 | 0.42960658 | 1.89389449 | 0.05823902 | 0.57141408 |
| ZNF224 | 30.827152 | -0.0193857 | 0.36709476 | -0.0528085 | 0.95788452 | 0.99183169 |
| ZNF234 | 16.4639341 | -0.1552549 | 0.41804263 | -0.3713852 | 0.71035062 | 0.94606024 |

| | | | | | | |
|---|---|---|---|---|---|---|
| ZNF226 | 29.4098772 | 0.12989697 | 0.34497244 | 0.37654303 | 0.70651323 | 0.94558664 |
| ZNF227 | 17.9915646 | -0.4065469 | 0.37971887 | -1.0706525 | 0.28432573 | 0.80431986 |
| ZNF235 | 74.0288164 | 0.84434 | 0.4064136 | 2.07753876 | 0.03775186 | 0.49640946 |
| ZNF229 | 17.1710241 | -0.638283 | 0.3401262 | -1.8766065 | 0.06057204 | 0.57774794 |
| ZNF180 | 12.0961627 | -0.0032632 | 0.48088723 | -0.0067857 | 0.99458583 | NA |
| AC243964.2 | 21.54566 | -0.168797 | 0.50571939 | -0.3337761 | 0.73854852 | 0.9542764 |
| CEACAM19 | 16.0913757 | -0.9147742 | 0.54927862 | -1.6654101 | 0.09583096 | NA |
| BCL3 | 23.2653816 | -0.3918618 | 0.46886457 | -0.8357675 | 4.03E-01 | 0.85782098 |
| CBLC | 22.3017939 | -0.2175223 | 0.48852691 | -0.4452616 | 0.65613072 | 0.93570875 |
| BCAM | 196.78794 | 0.75311301 | 0.38951984 | 1.93343941 | 0.05318209 | 0.55838128 |
| NECTIN2 | 90.3801571 | 0.12724952 | 0.33418554 | 0.38077506 | 0.70337017 | 0.94476609 |
| TOMM40 | 21.228571 | -0.2345761 | 0.39208272 | -0.5982822 | 0.54965164 | 0.91221128 |
| APOE | 57.6194269 | -1.2129265 | 0.70066378 | -1.7311106 | 0.08343204 | 0.62487197 |
| CLPTM1 | 43.5843189 | -0.1850616 | 0.39478413 | -0.4687665 | 0.63923652 | 0.93328692 |
| RELB | 48.329272 | -0.2069661 | 0.41151565 | -0.5029362 | 0.61500911 | 0.92662475 |
| CLASRP | 51.0225188 | -0.1829852 | 0.26530592 | -0.6897141 | 0.49037402 | 0.8897498 |
| GEMIN7-AS1 | 18.108645 | -0.0004892 | 0.42496939 | -0.0011511 | 0.99908154 | 0.99957534 |
| MARK4 | 55.7847406 | -0.0827823 | 0.30509118 | -0.2713363 | 0.78613237 | 0.96767512 |
| TRAPPC6A | 25.1908265 | 0.2588882 | 0.4190837 | 0.61774819 | 0.53674134 | 0.90763121 |
| ERCC2 | 12.0795961 | 0.88448652 | 0.49797618 | 1.77616231 | 0.07570617 | NA |
| PPP1R13L | 39.2383703 | -0.3181912 | 0.26886904 | -1.1834431 | 0.2366336 | 0.77864547 |
| ERCC1 | 18.931013 | -0.687282 | 0.4508263 | -1.5244941 | 0.12738533 | 0.66929384 |
| VASP | 33.6520505 | -0.2732198 | 0.38501876 | -0.7096273 | 0.4779353 | 0.88545331 |
| OPA3 | 58.6692574 | -0.3321135 | 0.44132664 | -0.7525344 | 0.45172972 | 0.87346041 |
| EML2 | 42.9686197 | 0.80081255 | 0.30483952 | 2.62699714 | 0.00861421 | 0.31119527 |
| SNRPD2 | 48.2676119 | -0.0844386 | 0.42116982 | -0.2004858 | 0.84110068 | 0.97600352 |
| FBXO46 | 20.955902 | 0.28712069 | 0.4518654 | 0.63541198 | 0.52515976 | 0.90487731 |
| DMPK | 15.9282891 | -0.2802389 | 0.45991829 | -0.6093233 | 0.54231014 | NA |
| SYMPK | 89.7365248 | -0.5205256 | 0.23255524 | -2.238288 | 0.02520228 | 0.4403385 |
| MYPOP | 18.8569985 | 0.58207017 | 0.40603626 | 1.43354234 | 0.15170291 | 0.70572801 |
| PPP5C | 49.0620931 | -0.1117732 | 0.29656401 | -0.376894 | 0.70625238 | 0.94558664 |
| CCDC8 | 13.8618037 | 0.35451322 | 0.43048423 | 0.82352198 | 0.41021122 | NA |
| PNMA8A | 10.7845083 | 0.5691715 | 0.66601379 | 0.85459417 | 0.39277585 | NA |
| PPP5D1 | 30.7923349 | -0.1345549 | 0.35953596 | -0.374246 | 0.70822129 | 0.94558664 |
| CALM3 | 41.1263559 | -0.1719149 | 0.58421948 | -0.2942642 | 0.76855602 | 0.96289288 |
| PRKD2 | 50.3150647 | -0.5036238 | 0.28435499 | -1.7711094 | 0.07654251 | 0.61119868 |
| STRN4 | 19.4309357 | -0.1119626 | 0.36056654 | -0.3105185 | 0.75616668 | 0.95895587 |
| FKRP | 18.6936049 | -0.2301529 | 0.42301596 | -0.5440761 | 0.58638909 | 0.92076087 |
| SLC1A5 | 14.4730468 | 0.05037694 | 0.5979541 | 0.08424884 | 0.93285859 | NA |
| AP2S1 | 34.1614861 | 0.01519627 | 0.32750964 | 0.04639947 | 0.96299186 | 0.99256424 |
| ARHGAP35 | 130.673938 | 0.22812832 | 0.32809256 | 0.69531694 | 0.4868567 | 0.88838332 |
| TMEM160 | 31.3360554 | 0.74459599 | 0.52395195 | 1.42111502 | 0.15528333 | 0.70572801 |
| ZC3H4 | 31.4372787 | -0.3761188 | 0.3867211 | -0.9725842 | 0.33076001 | 0.82867637 |
| SAE1 | 34.7546794 | -0.8416586 | 0.32709795 | -2.5731088 | 0.01007895 | 0.33306084 |
| CCDC9 | 11.4514388 | -0.2464964 | 0.60521898 | -0.4072846 | 0.68379895 | NA |
| INAFM1 | 12.0344152 | -0.5313881 | 0.52347932 | -1.0151081 | 0.31005432 | NA |
| C5AR1 | 42.0140159 | 0.74880879 | 0.39293527 | 1.90567975 | 0.05669177 | 0.56976762 |
| DHX34 | 26.2887099 | -0.4238941 | 0.31948649 | -1.3267982 | 0.18457544 | 0.73705812 |
| NAPA-AS1 | 11.2395005 | 0.49240898 | 0.58192419 | 0.84617376 | 0.39745582 | NA |
| NAPA | 79.9618566 | -0.2157688 | 0.26122907 | -0.8259756 | 0.40881795 | 0.85867915 |
| NOP53 | 18.4887848 | 1.08870612 | 0.50029413 | 2.1761321 | 0.02954538 | 0.46381645 |
| SELENOW | 79.3925069 | 0.06839028 | 0.40401706 | 0.16927573 | 0.86557976 | 0.97788853 |
| ZSWIM9 | 17.3621802 | 0.52095742 | 0.36560888 | 1.42490364 | 0.15418506 | 0.70572801 |
| CARD8 | 36.868843 | 0.08690092 | 0.3519435 | 0.24691725 | 0.80497226 | 0.97195485 |

| | | | | | | |
|---|---|---|---|---|---|---|
| TMEM143 | 25.0691178 | -0.0401394 | 0.45209306 | -0.0887856 | 0.92925228 | 0.98960607 |
| KDELR1 | 51.1149213 | 0.05121881 | 0.27253248 | 0.18793653 | 0.85092641 | 0.97612951 |
| CYTH2 | 25.5875881 | -0.3566708 | 0.3754159 | -0.9500683 | 0.34207753 | 0.83304756 |
| LMTK3 | 17.8536607 | 0.17073063 | 0.36729274 | 0.4648353 | 0.64204941 | 0.93328692 |
| RPL18 | 121.78239 | 0.26813841 | 0.52489059 | 0.51084628 | 0.6094587 | 0.92573951 |
| SPHK2 | 12.1481463 | 0.10486439 | 0.49176024 | 0.21324292 | 0.83113749 | NA |
| BCAT2 | 24.9178946 | 0.32029781 | 0.41507916 | 0.77165477 | 0.44031893 | 0.86895963 |
| HSD17B14 | 20.4720841 | -0.9780427 | 0.54633211 | -1.7901981 | 0.07342206 | 0.60981847 |
| PLEKHA4 | 22.1309212 | -0.5324051 | 0.73309981 | -0.7262382 | 0.46769274 | 0.88108588 |
| NUCB1 | 78.6904324 | -0.013281 | 0.29245123 | -0.0454126 | 0.96377846 | 0.99256424 |
| BAX | 24.4649038 | 0.37581712 | 0.3545387 | 1.06001722 | 0.28913677 | 0.80536924 |
| FTL | 808.076448 | -0.1104114 | 0.53408777 | -0.2067289 | 0.83622158 | 0.97600352 |
| GYS1 | 58.7411159 | -0.5310092 | 0.33002972 | -1.6089739 | 0.10762205 | 0.65986731 |
| RUVBL2 | 10.5737607 | -0.3307664 | 0.50427271 | -0.6559275 | 0.51187075 | NA |
| SNRNP70 | 176.343455 | 0.00495134 | 0.233528 | 0.02120235 | 0.98308424 | 0.99677164 |
| TRPM4 | 45.2544275 | 0.64625865 | 0.45162608 | 1.43095956 | 0.1524418 | 0.70572801 |
| SLC6A16 | 39.1172615 | -0.5339173 | 0.42612667 | -1.2529544 | 0.2102223 | 0.76397692 |
| TEAD2 | 19.4983301 | -0.9039535 | 0.5591859 | -1.6165528 | 0.10597486 | 0.65946648 |
| PIH1D1 | 20.986658 | 0.25839016 | 0.351081 | 0.73598444 | 0.46174017 | 0.87676694 |
| RPL13A | 156.075336 | 0.17742637 | 0.48769476 | 0.36380618 | 0.71600275 | 0.94885853 |
| RPS11 | 125.052364 | -0.1069135 | 0.3951706 | -0.2705502 | 0.78673701 | 0.96767512 |
| FCGRT | 43.7339327 | 0.73281663 | 0.46846811 | 1.56428285 | 0.11775116 | 0.6668162 |
| NOSIP | 30.6703326 | -0.0182448 | 0.29800558 | -0.0612232 | 0.95118147 | 0.99105535 |
| PRRG2 | 11.2901527 | 0.45958411 | 0.42806261 | 1.07363759 | 0.28298514 | NA |
| PRR12 | 29.5903622 | -0.3015203 | 0.36182371 | -0.8333349 | 0.40465588 | 0.85782098 |
| SCAF1 | 18.8046078 | -0.1449445 | 0.39723101 | -0.3648872 | 0.71519562 | 0.9487373 |
| IRF3 | 44.5691468 | 0.26349792 | 0.34170887 | 0.77111817 | 0.44063689 | 0.86895963 |
| PRMT1 | 37.4295005 | 0.50476364 | 0.31547139 | 1.60002987 | 0.10959196 | 0.66067258 |
| AP2A1 | 29.9444832 | -0.2494497 | 0.39435178 | -0.6325562 | 0.52702351 | 0.90487731 |
| MED25 | 33.0978381 | -0.0586116 | 0.33737261 | -0.1737297 | 0.86207792 | 0.97767895 |
| PTOV1 | 32.5195793 | 0.33473607 | 0.41149254 | 0.81346814 | 0.41594971 | 0.85867915 |
| PNKP | 32.0928215 | -0.4647269 | 0.39623711 | -1.1728504 | 0.2408558 | 0.78275985 |
| TBC1D17 | 29.8934772 | -0.4905525 | 0.3324288 | -1.475662 | 0.14003465 | 0.6912882 |
| IL4I1 | 17.2168822 | 0.75906406 | 0.88321549 | 0.85943247 | 0.39010196 | 0.85605713 |
| ATF5 | 15.0358562 | 0.74278556 | 0.40986158 | 1.81228393 | 0.06994234 | NA |
| VRK3 | 39.3161004 | -0.1352241 | 0.28605913 | -0.4727139 | 0.63641731 | 0.93251095 |
| MYH14 | 64.9661729 | -0.1415098 | 0.29995774 | -0.4717658 | 0.63709397 | 0.93259128 |
| KCNC3 | 33.8440723 | 0.83935888 | 0.38433679 | 2.183915 | 0.02896849 | 0.46182173 |
| NR1H2 | 41.6797229 | -0.3637302 | 0.28376931 | -1.2817812 | 0.19991943 | 0.75605362 |
| EMC10 | 72.0900337 | 0.22588702 | 0.28506555 | 0.79240376 | 0.42812529 | 0.86279045 |
| JOSD2 | 12.1675032 | 0.735496 | 0.44199211 | 1.66404782 | 0.09610287 | NA |
| CLEC11A | 16.0084522 | 0.48948019 | 0.48339809 | 1.01258197 | 0.31125988 | NA |
| C19orf48 | 15.988775 | -0.7223466 | 0.46527847 | -1.5525038 | 0.12054172 | NA |
| KLK1 | 68.5823637 | -0.6023724 | 1.47992336 | -0.4070295 | 0.68398633 | 0.94205398 |
| ETFB | 51.8126594 | 0.53903202 | 0.46511262 | 1.15892797 | 0.24648555 | 0.78583441 |
| ZNF175 | 14.0238565 | 0.09756051 | 0.45516592 | 0.21434055 | 0.8302815 | NA |
| SPACA6 | 34.2549654 | -0.3733919 | 0.43856392 | -0.8513968 | 0.39454896 | 0.85605713 |
| ZNF577 | 57.8931811 | 0.0238179 | 0.32980254 | 0.07221867 | 0.94242788 | 0.99058259 |
| ZNF613 | 13.1813422 | -0.0595396 | 0.4337312 | -0.137273 | 0.89081499 | NA |
| ZNF350-AS1 | 80.1747476 | -0.5889622 | 0.47442757 | -1.2414165 | 0.21445194 | 0.7643572 |
| ZNF350 | 33.9139092 | -0.4713637 | 0.4128037 | -1.1418593 | 0.25351253 | 0.7912221 |
| ZNF615 | 20.7098612 | -0.099892 | 0.42402159 | -0.2355825 | 0.81375669 | 0.97282508 |
| ZNF614 | 12.3973899 | -0.2418178 | 0.45736468 | -0.5287199 | 0.59699977 | NA |
| ZNF432 | 41.2721137 | -0.538802 | 0.3069094 | -1.7555735 | 0.07916126 | 0.61367304 |

| | | | | | | |
|---|---|---|---|---|---|---|
| ZNF616 | 13.8930279 | 0.26647563 | 0.53723796 | 0.49601043 | 0.61988705 | NA |
| PPP2R1A | 115.156034 | -0.1400717 | 0.37271283 | -0.3758168 | 0.7070531 | 0.94558664 |
| ZNF766 | 32.7995792 | 0.38789118 | 0.27570681 | 1.40689735 | 0.15945783 | 0.71101785 |
| ZNF480 | 31.3911284 | -0.7600851 | 0.37909643 | -2.0049913 | 0.04496397 | 0.52610161 |
| ZNF610 | 11.187004 | 0.22803766 | 0.46052098 | 0.49517324 | 0.62047784 | NA |
| ZNF880 | 12.2240614 | 0.22523406 | 0.4331712 | 0.51996545 | 0.60308765 | NA |
| ZNF528 | 25.274433 | -0.7205977 | 0.31433479 | -2.2924528 | 0.02187953 | 0.4212027 |
| ZNF808 | 26.8551223 | -0.0279737 | 0.40298603 | -0.0694161 | 0.94465843 | 0.99069421 |
| ZNF701 | 33.7448638 | 0.20997231 | 0.38357001 | 0.54741587 | 0.58409306 | 0.92071026 |
| ZNF83 | 222.910803 | -0.4342347 | 0.23164828 | -1.8745432 | 0.0608556 | 0.57825523 |
| ZNF611 | 100.514287 | -0.6546666 | 0.33583118 | -1.9493919 | 0.05124864 | 0.55209033 |
| ZNF600 | 14.3393986 | -0.3240966 | 0.37764966 | -0.8581939 | 0.39078541 | NA |
| ZNF28 | 17.8815556 | -0.3762709 | 0.35789592 | -1.0513417 | 0.29310166 | 0.80935782 |
| AC010487.3 | 14.6019851 | 0.03318778 | 0.4610275 | 0.07198656 | 0.94261261 | NA |
| ZNF160 | 78.3701717 | 0.08577793 | 0.24221256 | 0.35414319 | 0.72323158 | 0.95093722 |
| ZNF347 | 27.5857892 | 0.21000403 | 0.35855493 | 0.58569556 | 0.55808012 | 0.91425998 |
| ZNF677 | 20.6108753 | -0.1167776 | 0.35212451 | -0.3316372 | 0.74016322 | 0.9544376 |
| ZNF525 | 20.5379385 | 0.13859051 | 0.41836085 | 0.33127026 | 0.74044035 | 0.9544376 |
| ZNF765 | 22.9239714 | 0.10964986 | 0.35538436 | 0.3085388 | 0.75767238 | 0.95903525 |
| ZNF761 | 33.2226834 | -0.5148001 | 0.33562525 | -1.5338538 | 0.12506562 | 0.66929384 |
| ZNF331 | 120.174605 | -0.39961 | 1.13408105 | -0.3523645 | 0.72456489 | 0.95093919 |
| NDUFA3 | 21.7164736 | -0.03165 | 0.69585508 | -0.0454836 | 0.96372183 | 0.99256424 |
| PRPF31 | 33.0202993 | 0.04955644 | 0.3413737 | 0.14516772 | 0.88457845 | 0.98154805 |
| CNOT3 | 20.848703 | -0.4672755 | 0.33937952 | -1.3768523 | 0.16855793 | 0.72151539 |
| TMC4 | 75.4413014 | -0.2413871 | 0.2368076 | -1.0193387 | 0.30804221 | 0.81568231 |
| MBOAT7 | 69.8001082 | -0.306287 | 0.33622794 | -0.9109505 | 0.36232147 | 0.84504544 |
| TSEN34 | 24.3063895 | 0.13065745 | 0.4712507 | 0.27725678 | 0.78158295 | 0.96652357 |
| RPS9 | 251.949287 | 0.12855 | 0.36702207 | 0.35025142 | 0.72615002 | 0.9519216 |
| LENG8-AS1 | 16.234631 | -0.0211108 | 0.41001212 | -0.0514881 | 0.95893655 | 0.99229276 |
| LENG8 | 113.457248 | -0.8913052 | 0.30349194 | -2.9368331 | 0.00331582 | 0.23150356 |
| AC011476.3 | 18.4892583 | 0.79681872 | 0.68695586 | 1.1599271 | 0.24607849 | 0.78583441 |
| RDH13 | 53.2791163 | -0.2835071 | 0.32603891 | -0.86955 | 0.38454638 | 0.85511218 |
| EPS8L1 | 22.9201796 | 0.10746806 | 0.45551528 | 0.23592635 | 0.81348982 | 0.97282508 |
| PPP1R12C | 33.8682402 | -0.1161155 | 0.35096035 | -0.3308506 | 0.74075732 | 0.95455763 |
| PPP6R1 | 33.1535319 | -0.2197406 | 0.34565243 | -0.635727 | 0.52495437 | 0.90487731 |
| HSPBP1 | 18.5084233 | 0.42731138 | 0.54571893 | 0.78302466 | 0.43361263 | 0.86729771 |
| KMT5C | 10.3076743 | 0.11887637 | 0.48069361 | 0.24730175 | 0.8046747 | NA |
| TMEM238 | 24.7517584 | 0.36210908 | 0.37748785 | 0.95926022 | 0.33742767 | 0.83055261 |
| RPL28 | 185.913372 | -0.0022255 | 0.50247201 | -0.004429 | 0.99646617 | 0.99927028 |
| ISOC2 | 19.8426115 | 0.24267169 | 0.46809319 | 0.51842602 | 0.60416107 | 0.92524869 |
| NAT14 | 11.9685895 | 0.49666756 | 0.79303812 | 0.6262846 | 0.53112828 | NA |
| ZNF524 | 10.5831668 | 0.27666955 | 0.58006867 | 0.47696001 | 0.6333906 | NA |
| CCDC106 | 10.1223666 | 0.12982186 | 0.665424 | 0.19509645 | 0.84531744 | NA |
| U2AF2 | 22.9143371 | -0.1667189 | 0.32668184 | -0.5103401 | 0.60981319 | 0.92573951 |
| EPN1 | 57.6860155 | -0.3034726 | 0.28879844 | -1.0508112 | 0.2933453 | 0.80935782 |
| ZNF787 | 19.2944432 | -0.2453057 | 0.37009416 | -0.6628197 | 0.50744605 | 0.89851807 |
| ZNF444 | 29.7247452 | 0.24596559 | 0.29348959 | 0.83807261 | 0.40198992 | 0.85778581 |
| ZSCAN5A | 81.6244226 | -0.3023865 | 0.27890256 | -1.0842012 | 0.27827561 | 0.80210301 |
| ZNF583 | 11.2903976 | -0.3173407 | 0.48123754 | -0.6594264 | 0.50962201 | NA |
| ZNF667 | 22.2118532 | 0.05447879 | 0.36558733 | 0.14901718 | 0.88154007 | 0.98137082 |
| ZNF667-AS1 | 31.7418634 | -0.2177398 | 0.34837571 | -0.6250143 | 0.53196166 | 0.90610813 |
| ZNF471 | 27.6781228 | 0.30214005 | 0.35336567 | 0.85503509 | 0.39253172 | 0.85605713 |
| AC005498.3 | 13.8880277 | 0.29772938 | 0.44605633 | 0.66747036 | 0.50447174 | NA |
| ZNF264 | 37.9912218 | 0.03031814 | 0.28045711 | 0.1081026 | 0.91391431 | 0.9866416 |

| | | | | | | |
|---|---|---|---|---|---|---|
| ZNF460-AS1 | 16.0860805 | 0.87081485 | 0.5630583 | 1.54658027 | 0.12196449 | NA |
| ZNF460 | 15.5075769 | -0.493482 | 0.35830987 | -1.3772493 | 0.16843519 | NA |
| AC005261.1 | 55.3324919 | 0.10964886 | 0.31009902 | 0.35359305 | 0.72364389 | 0.95093722 |
| AC003002.2 | 10.3796603 | 0.55092554 | 0.58863132 | 0.9359433 | 0.34930238 | NA |
| AC004076.1 | 28.8738655 | -0.21423 | 0.45654605 | -0.4692408 | 0.63889751 | 0.93322914 |
| ZNF549 | 19.1879324 | -0.0025682 | 0.36596081 | -0.0070176 | 0.99440082 | 0.99923067 |
| ZNF134 | 10.4660631 | 0.60289455 | 0.51344265 | 1.17421986 | 0.24030697 | NA |
| AC003006.1 | 12.6858178 | 0.12511368 | 0.53132738 | 0.2354738 | 0.81384101 | NA |
| ZNF154 | 14.9674675 | -0.1960999 | 0.43063789 | -0.4553708 | 0.64884252 | NA |
| ZNF586 | 113.392792 | -0.1865051 | 0.22870979 | -0.8154662 | 0.41480549 | 0.85867915 |
| ZNF552 | 42.7808148 | 0.83487057 | 0.35537706 | 2.34925281 | 0.01881113 | 0.41544117 |
| ZNF814 | 75.6733514 | -0.1602051 | 0.33585085 | -0.4770127 | 0.63335308 | 0.93251095 |
| ZNF587 | 37.1728463 | 0.23244968 | 0.32664055 | 0.71163756 | 0.47668924 | 0.88495683 |
| ZNF418 | 14.1317732 | -0.2877548 | 0.42386858 | -0.6788775 | 0.49721549 | NA |
| C19orf18 | 12.0666746 | -0.1945674 | 0.47394448 | -0.4105278 | 0.68141884 | NA |
| ZNF606 | 19.1984297 | 0.41101633 | 0.35899731 | 1.14490085 | 0.25225023 | 0.7902944 |
| ZNF135 | 35.9693095 | 0.11988653 | 0.3486102 | 0.3438985 | 0.73092262 | 0.95279138 |
| ZSCAN18 | 71.5246303 | 0.42369032 | 0.2226213 | 1.90318857 | 0.05701594 | 0.57034741 |
| ZNF329 | 28.5971622 | 0.37196303 | 0.32982108 | 1.12777215 | 0.25941616 | 0.79532468 |
| ZNF274 | 37.8813801 | -0.1794794 | 0.34150904 | -0.5255479 | 0.59920237 | 0.92298075 |
| AC020915.3 | 18.8533143 | 0.08773623 | 0.41923124 | 0.20927884 | 0.83423057 | 0.97600352 |
| RPS5 | 70.5330573 | 0.12133369 | 0.42341306 | 0.28656105 | 0.77444844 | 0.96435639 |
| ZNF584 | 9.48496727 | 0.56525397 | 0.5067289 | 1.11549582 | 0.26463801 | NA |
| SLC27A5 | 11.696629 | 0.72384479 | 0.49398778 | 1.46530909 | 0.14283661 | NA |
| TRIM28 | 35.5556345 | 0.37195714 | 0.41005981 | 0.90708022 | 0.36436438 | 0.8468666 |
| CHMP2A | 33.0483578 | 0.37843199 | 0.48256585 | 0.78420797 | 0.43291809 | 0.86729771 |
| UBE2M | 26.5707672 | -0.024814 | 0.32826981 | -0.0755902 | 0.93974512 | 0.99012269 |
| MZF1-AS1 | 12.9958663 | 0.49362646 | 0.51309363 | 0.96205922 | 0.33601986 | NA |
| MZF1 | 54.520444 | -0.4380682 | 0.31238273 | -1.4023446 | 0.16081237 | 0.71101785 |
| RPS4Y1 | 66.5642906 | 0.94845792 | 2.04313692 | 0.46421652 | 0.64249262 | 0.93328692 |
| ZFY | 24.705633 | 0.37778098 | 1.85363586 | 0.20380539 | 0.83850558 | 0.97600352 |
| LINC00278 | 168.915305 | 0.1226846 | 2.5767093 | 0.0476129 | 0.96202475 | 0.99256424 |
| TBL1Y | 45.5594598 | -0.0469604 | 2.03488176 | -0.0230777 | 0.98158828 | 0.99603144 |
| TTTY15 | 34.7154845 | 1.09118501 | 2.17499183 | 0.50169614 | 0.61588128 | 0.92662475 |
| USP9Y | 127.13777 | 1.0862376 | 2.47232791 | 0.43935823 | 0.66040199 | 0.93728744 |
| DDX3Y | 36.8837691 | 0.61147055 | 2.171472 | 0.28159265 | 0.77825588 | 0.96518573 |
| UTY | 303.318669 | 0.75964782 | 2.10559119 | 0.3607765 | 0.71826653 | 0.94925929 |
| NLGN4Y | 50.4974739 | 1.67967396 | 2.03431285 | 0.82567141 | 0.40899051 | 0.85867915 |
| TTTY14 | 339.616502 | 0.59972347 | 1.16135809 | 0.51639841 | 0.60557618 | 0.92569157 |
| KDM5D | 34.562307 | 0.74955584 | 2.17052731 | 0.34533352 | 0.72984365 | 0.95279138 |
| TTTY10 | 19.6290661 | 0.29436137 | 1.59159913 | 0.18494693 | 0.85327066 | 0.97612951 |
| EIF1AY | 13.2667999 | 1.00147411 | 1.92223612 | 0.52099433 | 0.60237072 | NA |
| IL17RA | 23.4694963 | -0.3022235 | 0.34475721 | -0.876627 | 0.38068928 | 0.85432452 |
| HDHD5 | 23.7789427 | -0.188258 | 0.39234265 | -0.4798306 | 0.63134785 | 0.93251095 |
| HDHD5-AS1 | 9.57096203 | 0.52647704 | 0.66940494 | 0.78648515 | 0.43158331 | NA |
| ADA2 | 16.4428513 | 0.27406444 | 0.59216086 | 0.46282092 | 0.64349273 | 0.93328692 |
| CECR2 | 69.5398583 | -0.2994295 | 0.36703936 | -0.8157966 | 0.4146165 | 0.85867915 |
| ATP6V1E1 | 67.5899742 | -0.0786143 | 0.25414816 | -0.3093246 | 0.75707459 | 0.95903525 |
| BCL2L13 | 126.29814 | -0.1129511 | 0.20786716 | -0.5433811 | 0.58686747 | 0.92076087 |
| MICAL3 | 135.814322 | -0.3553005 | 0.32328289 | -1.0990389 | 0.27175111 | 0.79666376 |
| PEX26 | 43.9885307 | -0.2057551 | 0.27931677 | -0.7366373 | 0.46134296 | 0.87657347 |
| USP18 | 13.8238994 | -1.6193016 | 1.02510567 | -1.5796436 | 0.11418852 | NA |
| DGCR2 | 80.138799 | -0.1775145 | 0.26687019 | -0.6651717 | 0.5059407 | 0.89723255 |
| SLC25A1 | 35.3228686 | 0.01481247 | 0.4412466 | 0.0335696 | 0.97322037 | 0.99431961 |

| | | | | | | |
|---|---|---|---|---|---|---|
| CLTCL1 | 24.4688469 | -0.4310007 | 0.37130235 | -1.160781 | 0.24573099 | 0.78583441 |
| HIRA | 25.9191088 | -0.8140448 | 0.38779017 | -2.0991888 | 0.03580026 | 0.49302533 |
| MRPL40 | 21.3831743 | 0.72739348 | 0.48303565 | 1.50587948 | 0.13209812 | 0.6765887 |
| UFD1 | 27.1337177 | -0.3478286 | 0.39971962 | -0.8701814 | 0.38420129 | 0.85511218 |
| TXNRD2 | 27.7154498 | 0.0538967 | 0.41870726 | 0.12872168 | 0.89757788 | 0.98531749 |
| COMT | 82.0309196 | 0.44976862 | 0.27335301 | 1.64537649 | 0.0998922 | 0.65863153 |
| TANGO2 | 45.5974123 | -0.2358296 | 0.41259557 | -0.5715758 | 0.5676094 | 0.91698355 |
| DGCR8 | 27.6521529 | -0.7616448 | 0.43132055 | -1.7658439 | 0.07742204 | 0.61342702 |
| RANBP1 | 15.4021314 | 0.34774458 | 0.51281939 | 0.67810342 | 0.49770612 | NA |
| ZDHHC8 | 17.649663 | -0.1055807 | 0.48743843 | -0.2166031 | 0.8285177 | 0.97567084 |
| DGCR6L | 11.3823319 | 0.07799099 | 0.61355676 | 0.12711292 | 0.89885103 | NA |
| KLHL22 | 58.7617455 | 0.18938858 | 0.28078796 | 0.67448969 | 0.50000004 | 0.89448839 |
| MED15 | 92.1292958 | -0.3915302 | 0.19118019 | -2.0479641 | 0.04056351 | 0.50512951 |
| PI4KA | 125.366674 | -0.1707497 | 0.19905916 | -0.8577838 | 0.39101188 | 0.85605713 |
| SNAP29 | 27.3211119 | -0.1789602 | 0.32484959 | -0.5509016 | 0.58170113 | 0.92034873 |
| CRKL | 30.7779705 | 0.10027423 | 0.36608321 | 0.27391102 | 0.784153 | 0.96732016 |
| LZTR1 | 12.9728452 | 0.44822194 | 0.4554347 | 0.98416292 | 0.32503541 | NA |
| GGT2 | 97.0432577 | 1.31610481 | 0.7484837 | 1.75836136 | 0.07868604 | 0.61367304 |
| HIC2 | 28.0403008 | 0.14557012 | 0.39997868 | 0.3639447 | 0.7158993 | 0.94885853 |
| UBE2L3 | 85.240742 | -0.3020701 | 0.22816134 | -1.323932 | 0.18552563 | 0.73752116 |
| SDF2L1 | 21.3732533 | 0.04070848 | 0.58355385 | 0.06975959 | 0.94438501 | 0.99069421 |
| PPIL2 | 62.021236 | -0.5414588 | 0.31656292 | -1.7104302 | 0.08718634 | 0.63466198 |
| YPEL1 | 11.5316879 | -0.6945021 | 0.5993661 | -1.1587277 | 0.24656718 | NA |
| MAPK1 | 162.291098 | -0.5012045 | 0.23734722 | -2.1116932 | 0.03471277 | 0.49098143 |
| PPM1F | 20.7367506 | -0.1203546 | 0.52389695 | -0.2297295 | 0.81830199 | 0.97288403 |
| TOP3B | 13.9956632 | 0.18333688 | 0.46595714 | 0.39346296 | 0.69397758 | NA |
| AC245060.5 | 40.1173395 | -0.5856012 | 0.38821505 | -1.5084454 | 0.13144057 | 0.67540912 |
| RSPH14 | 10.6864082 | 0.0825517 | 0.51127213 | 0.16146333 | 0.87172849 | NA |
| RAB36 | 13.2417343 | 0.48023737 | 0.54010489 | 0.88915575 | 0.37391938 | NA |
| BCR | 196.476616 | -0.7632617 | 0.31438696 | -2.4277778 | 0.01519165 | 0.39253938 |
| CHCHD10 | 154.174224 | -0.1819551 | 0.37259651 | -0.4883436 | 0.62530649 | 0.92981013 |
| SMARCB1 | 47.7441994 | -0.3017328 | 0.32027085 | -0.9421177 | 0.34613237 | 0.83666384 |
| DDT | 70.7053142 | 0.08764162 | 0.55208451 | 0.15874675 | 0.8738684 | 0.97918624 |
| CABIN1 | 158.67924 | -0.1614326 | 0.21658804 | -0.7453441 | 0.45606371 | 0.87394768 |
| GUCD1 | 24.8455263 | 0.49187681 | 0.34916698 | 1.40871515 | 1.59E-01 | 0.71094363 |
| SNRPD3 | 44.6849858 | -0.0148577 | 0.26705769 | -0.0556346 | 0.95563287 | 0.99148072 |
| SGSM1 | 101.651013 | -0.2818227 | 0.41992554 | -0.6711255 | 0.50214059 | 0.89499455 |
| KIAA1671 | 152.500434 | -0.2041971 | 0.27939091 | -0.7308654 | 0.4648614 | 0.87920909 |
| LRP5L | 55.2823457 | 0.10407399 | 0.39551614 | 0.26313463 | 0.79244681 | 0.96944541 |
| GRK3 | 41.8645284 | -0.5449795 | 0.35825995 | -1.5211844 | 0.12821356 | 0.66929384 |
| HPS4 | 47.7060567 | 0.02316963 | 0.2659139 | 0.08713208 | 0.93056653 | 0.98960607 |
| TFIP11 | 14.6695025 | 0.04387824 | 0.47870787 | 0.09165974 | 0.92696838 | NA |
| TPST2 | 62.2065684 | -0.3842824 | 0.37335192 | -1.0292767 | 0.30334969 | 0.81568231 |
| MIATNB | 39.9147823 | -0.6874611 | 0.35317604 | -1.9465113 | 0.05159336 | 0.55289583 |
| PITPNB | 118.350655 | -0.1046922 | 0.25819484 | -0.4054774 | 0.68512664 | 0.94205398 |
| TTC28 | 199.930381 | 0.18666641 | 0.24116407 | 0.77402247 | 0.43891751 | 0.86879768 |
| HSCB | 39.3227894 | -0.0121345 | 0.28605346 | -0.0424205 | 0.96616349 | 0.99288623 |
| CCDC117 | 17.7167083 | 0.03345166 | 0.39750295 | 0.08415448 | 0.93293361 | 0.98960607 |
| XBP1 | 58.4653409 | 0.02903885 | 0.35646148 | 0.08146421 | 0.93507279 | 0.99012269 |
| Z93930.2 | 78.244553 | 0.42495716 | 0.39256113 | 1.08252481 | 0.2790194 | 0.80219184 |
| ZNRF3 | 221.625666 | -0.0257116 | 0.24727197 | -0.1039812 | 0.91718427 | 0.98710975 |
| KREMEN1 | 56.838151 | 0.31350425 | 0.29338178 | 1.06858802 | 0.28525535 | 0.80431986 |
| RHBDD3 | 14.8282357 | -0.3582743 | 0.38317481 | -0.9350153 | 0.34978041 | NA |
| EWSR1 | 115.50507 | -0.3838483 | 0.27058438 | -1.4185901 | 0.15601856 | 0.70728931 |

| | | | | | | |
|---|---|---|---|---|---|---|
| GAS2L1 | 9.65104979 | 0.53620363 | 0.62725783 | 0.85483768 | 0.39264101 | NA |
| AP1B1 | 46.8765142 | 0.44845718 | 0.26365044 | 1.70095361 | 0.0889517 | 0.63542189 |
| THOC5 | 15.4273486 | 0.13720623 | 0.38971884 | 0.35206465 | 0.72478978 | NA |
| NIPSNAP1 | 26.6974314 | -0.2650878 | 0.37279939 | -0.7110735 | 0.47703869 | 0.88495683 |
| NF2 | 71.6885631 | 0.2996133 | 0.33418077 | 0.89656057 | 0.36995345 | 0.84852785 |
| ZMAT5 | 11.2085499 | 0.85507815 | 0.46147258 | 1.85293382 | 0.06389184 | NA |
| UQCR10 | 116.256855 | 0.26492268 | 0.53938846 | 0.49115378 | 0.62331769 | 0.92941211 |
| ASCC2 | 38.0856425 | -0.0259141 | 0.24949552 | -0.1038659 | 0.91727578 | 0.98710975 |
| MTMR3 | 211.96325 | -0.3038278 | 0.21676349 | -1.4016556 | 0.16101812 | 0.71101785 |
| TBC1D10A | 17.843772 | -0.1678741 | 0.35149922 | -0.4775945 | 0.63293885 | 0.93251095 |
| SF3A1 | 18.6782013 | 0.11297075 | 0.42675401 | 0.264721 | 0.7912244 | 0.96909997 |
| RNF215 | 23.1084043 | -0.5761901 | 0.51323832 | -1.122656 | 0.26158361 | 0.79633269 |
| SEC14L6 | 21.8210579 | 0.87927473 | 0.54700833 | 1.60742474 | 0.10796123 | 0.65986731 |
| GAL3ST1 | 11.4948496 | 0.10935769 | 0.46100774 | 0.23721443 | 0.81249044 | NA |
| PES1 | 17.4514873 | -0.4124109 | 0.39088895 | -1.055059 | 0.29139834 | 0.80688911 |
| TCN2 | 18.1971057 | -0.0813032 | 0.71922249 | -0.1130432 | 0.90999629 | 0.98556715 |
| DUSP18 | 23.8623384 | 0.02680254 | 0.3879799 | 0.06908229 | 0.94492412 | 0.99069787 |
| OSBP2 | 62.8593329 | -0.4594177 | 0.2830035 | -1.6233641 | 0.10451159 | 0.65946648 |
| MORC2 | 29.6759836 | -0.2141216 | 0.30933255 | -0.6922051 | 0.48880852 | 0.88920066 |
| SMTN | 12.4013106 | -1.0891895 | 0.50697576 | -2.1484055 | 0.03168156 | NA |
| SELENOM | 66.6676488 | 0.75957825 | 0.55682479 | 1.36412432 | 0.17252844 | 0.72649757 |
| INPP5J | 62.1609854 | 1.04856128 | 0.46451827 | 2.257309 | 0.02398877 | 0.43586787 |
| RNF185 | 57.570324 | -0.2132677 | 0.30292047 | -0.7040384 | 0.48140883 | 0.88703194 |
| LIMK2 | 63.9653969 | 0.4177202 | 0.22910525 | 1.8232677 | 0.06826286 | 0.60244724 |
| PIK3IP1 | 25.5287144 | 0.37947881 | 0.4133358 | 0.91808843 | 0.35857257 | 0.84204868 |
| PIK3IP1-AS1 | 14.5838081 | -0.5626534 | 0.49966715 | -1.1260565 | 0.26014163 | NA |
| PATZ1 | 35.0951895 | 0.38842924 | 0.31065422 | 1.25035879 | 0.21116851 | 0.76432085 |
| DRG1 | 56.527752 | -0.2522668 | 0.30220238 | -0.834761 | 0.40385227 | 0.85782098 |
| EIF4ENIF1 | 56.3139922 | 0.01959035 | 0.25513138 | 0.07678536 | 0.9387943 | 0.99012269 |
| SFI1 | 54.3054389 | -0.9343036 | 0.24322955 | -3.841242 | 0.00012241 | 0.04981461 |
| PISD | 123.967103 | -0.2558514 | 0.25182467 | -1.0159903 | 0.309634 | 0.81568231 |
| PRR14L | 104.262018 | -0.2959668 | 0.22760571 | -1.3003486 | 0.19348152 | 0.75067401 |
| YWHAH | 27.4461606 | 0.4294721 | 0.51557639 | 0.83299413 | 0.40484804 | 0.85782098 |
| RTCB | 49.5372151 | 0.12101287 | 0.23133631 | 0.52310363 | 0.60090214 | 0.92352528 |
| FBXO7 | 55.6476353 | -0.1444493 | 0.27102361 | -0.5329769 | 0.59404958 | 0.9229638 |
| SYN3 | 19.1149782 | 0.19052486 | 0.67251377 | 0.28330255 | 0.77694492 | 0.96518573 |
| TIMP3 | 222.702024 | -0.4055884 | 0.4376336 | -0.9267762 | 0.35404277 | 0.83979358 |
| LARGE1 | 305.3289 | -0.5738698 | 0.36739084 | -1.5620145 | 0.11828458 | 0.6668162 |
| HMGXB4 | 41.8472311 | -0.0667467 | 0.32202812 | -0.2072699 | 0.83579905 | 0.97600352 |
| TOM1 | 53.3284598 | -0.3021564 | 0.26867223 | -1.1246285 | 0.2607465 | 0.79574359 |
| APOL6 | 34.0779538 | -0.711111 | 0.71064404 | -1.000657 | 0.31699265 | 0.82275575 |
| RBFOX2 | 365.423961 | 0.0558425 | 0.19567673 | 0.28538142 | 0.77535194 | 0.96469469 |
| APOL2 | 42.0030934 | -1.0059761 | 0.57337264 | -1.7544892 | 0.07934672 | 0.61367304 |
| APOL1 | 25.2330608 | -0.8805633 | 0.63225129 | -1.3927426 | 0.16369763 | 0.71332671 |
| MYH9 | 171.510663 | -0.4078959 | 0.26286031 | -1.5517591 | 0.12071988 | 0.6668162 |
| TXN2 | 35.3651655 | 0.44626107 | 0.37271598 | 1.19732206 | 0.23118105 | 0.77260653 |
| EIF3D | 40.838714 | 0.39332916 | 0.36525824 | 1.07685225 | 0.28154628 | 0.80417172 |
| IFT27 | 18.7105315 | 0.60143479 | 0.40226872 | 1.49510702 | 0.13488651 | 0.68328989 |
| PVALB | 33.3136002 | 0.50347459 | 0.50041685 | 1.00611038 | 3.14E-01 | 0.82040655 |
| TST | 21.8394809 | -0.0482759 | 0.54929962 | -0.0878862 | 0.9299671 | 0.98960607 |
| MPST | 25.2860617 | 0.05357026 | 0.37661711 | 0.14224063 | 0.88688994 | 0.98216235 |
| CARD10 | 21.4087378 | 0.15816647 | 0.47859902 | 0.33047805 | 0.74103877 | 0.95455763 |
| CDC42EP1 | 20.1634132 | 0.31899696 | 0.55468285 | 0.57509794 | 0.56522506 | 0.91698355 |
| GGA1 | 52.6300315 | 0.15633094 | 0.24700465 | 0.63290686 | 0.52679447 | 0.90487731 |

| | | | | | | |
|---|---|---|---|---|---|---|
| LGALS1 | 22.2082621 | -0.3639235 | 0.51926962 | -0.7008372 | 4.83E-01 | 0.88833419 |
| TRIOBP | 73.0226952 | 0.07712628 | 0.2751043 | 0.28035287 | 0.77920679 | 0.96597023 |
| H1F0 | 53.0319764 | -1.7536604 | 0.53980429 | -3.2486966 | 0.00115935 | 0.14394871 |
| GCAT | 11.244394 | 0.35740231 | 0.52610375 | 0.67933809 | 0.49692367 | NA |
| ANKRD54 | 15.7333369 | 0.57627625 | 0.40338164 | 1.42861297 | 0.1531155 | NA |
| EIF3L | 73.3225478 | -0.1531983 | 0.34618869 | -0.4425284 | 0.65810692 | 0.93605426 |
| MICALL1 | 14.7488897 | -0.3448708 | 0.4444983 | -0.7758654 | 0.43782848 | NA |
| POLR2F | 26.8735688 | -0.2307129 | 0.33571245 | -0.6872336 | 0.49193551 | 0.89045316 |
| PICK1 | 21.315395 | -0.1113482 | 0.36269059 | -0.307006 | 0.75883883 | 0.95903525 |
| PLA2G6 | 22.7520358 | 0.21596068 | 0.38655575 | 0.55867927 | 0.57638063 | 0.91854298 |
| MAFF | 18.6815935 | -0.8238164 | 0.65604719 | -1.2557273 | 0.20921485 | 0.7623519 |
| TMEM184B | 43.8353281 | -0.2641721 | 0.34505605 | -0.765592 | 0.44391912 | 0.8704597 |
| CSNK1E | 15.3132521 | -0.0688495 | 0.45723852 | -0.1505767 | 0.88030963 | NA |
| TPTEP2-CSNI | 113.40594 | 0.2621619 | 0.23187769 | 1.13060423 | 0.25822171 | 0.79532468 |
| DDX17 | 781.66943 | -0.0113441 | 0.12257647 | -0.0925469 | 9.26E-01 | 9.89E-01 |
| FAM227A | 24.9353074 | 0.32839354 | 0.43811083 | 0.74956726 | 0.45351537 | 0.87378486 |
| CBY1 | 32.0528459 | -0.121855 | 0.27475063 | -0.4435113 | 0.65739592 | 0.93593674 |
| TOMM22 | 13.6919185 | 0.03086957 | 0.6402703 | 0.04821335 | 0.96154621 | NA |
| JOSD1 | 34.4506477 | -0.4278505 | 0.580117 | -0.7375245 | 0.46080344 | 0.87626776 |
| GTPBP1 | 24.3542702 | -0.4271734 | 0.38757791 | -1.1021613 | 0.27039153 | 0.79666376 |
| SUN2 | 28.7071078 | 0.2949489 | 0.40710874 | 0.72449662 | 0.4687609 | 0.88192105 |
| AL021707.1 | 17.5709997 | -0.4600143 | 0.36769402 | -1.2510791 | 0.21090564 | 0.76397692 |
| CBX6 | 43.8587543 | 0.21231879 | 0.45881278 | 0.46275692 | 0.64353861 | 0.93328692 |
| CBX7 | 28.4766094 | 0.4024704 | 0.3154606 | 1.27581828 | 0.20201977 | 0.76038198 |
| RPL3 | 156.142458 | 0.21538601 | 0.50084598 | 0.43004439 | 0.66716335 | 0.93878847 |
| SYNGR1 | 26.3691161 | 0.83040543 | 0.35536248 | 2.33678419 | 0.01945041 | 0.41544117 |
| TAB1 | 24.9932606 | -0.0519681 | 0.30742958 | -0.1690408 | 0.86576454 | 0.97788853 |
| MIEF1 | 17.8869263 | 0.19220196 | 0.36155826 | 0.53159333 | 0.59500769 | 0.9229638 |
| ATF4 | 55.7509272 | -1.0413768 | 0.33752384 | -3.0853429 | 0.00203318 | 0.18508 |
| RPS19BP1 | 29.4758049 | -0.0007812 | 0.38815056 | -0.0020126 | 0.99839418 | 0.99957534 |
| FAM83F | 29.2265254 | 0.25480868 | 0.4309788 | 0.59123252 | 0.55436464 | 0.9137961 |
| TNRC6B | 689.763582 | 0.01321656 | 0.21936916 | 0.06024805 | 0.95195808 | 0.99141043 |
| MKL1 | 298.695827 | -0.5050533 | 0.32365914 | -1.5604481 | 0.11865404 | 0.6668162 |
| SLC25A17 | 54.2381538 | 0.00404886 | 0.23341625 | 0.0173461 | 0.98616051 | 0.99766827 |
| ST13 | 100.858919 | 0.34138601 | 0.21209615 | 1.60958134 | 0.10748928 | 0.65986731 |
| XPNPEP3 | 70.5060883 | 0.00570927 | 0.24539337 | 0.02326577 | 0.98143827 | 0.99599026 |
| RBX1 | 53.8729493 | -0.2521642 | 0.30064022 | -0.8387575 | 0.40160541 | 0.85778581 |
| EP300 | 164.435797 | -0.2421879 | 0.2114336 | -1.1454562 | 0.25202024 | 0.7902944 |
| L3MBTL2 | 18.2616631 | -0.1547728 | 0.38130165 | -0.4059065 | 0.68481134 | 0.94205398 |
| RANGAP1 | 29.7242927 | -0.2532908 | 0.2938522 | -0.8619665 | 0.38870595 | 0.85605713 |
| ZC3H7B | 60.5995692 | -0.2636267 | 0.30711206 | -0.8584056 | 0.39066852 | 0.85605713 |
| TEF | 23.7599774 | -0.2901389 | 0.51271741 | -0.5658847 | 0.57147218 | 0.91791025 |
| TOB2 | 42.4883541 | 0.24653979 | 0.46320631 | 0.53224617 | 0.59455551 | 0.9229638 |
| ACO2 | 150.752358 | -0.3179168 | 0.23941269 | -1.3279028 | 0.18421022 | 0.73705812 |
| POLR3H | 13.2558733 | -0.2843243 | 0.42667802 | -0.6663673 | 0.50517637 | NA |
| PMM1 | 13.7011918 | 0.50953283 | 0.4312316 | 1.18157581 | 0.23737406 | NA |
| DESI1 | 41.2457598 | -0.3956752 | 0.44174067 | -0.8957183 | 0.37040322 | 0.84852785 |
| XRCC6 | 104.074699 | 0.18229311 | 0.2598846 | 0.70143868 | 0.48302929 | 0.88792723 |
| SNU13 | 80.327318 | 0.14582616 | 0.27910894 | 0.5224704 | 0.60134285 | 0.92373 |
| CCDC134 | 13.6638902 | 0.01141711 | 0.45980106 | 0.02483053 | 0.98019014 | NA |
| SREBF2 | 354.165301 | -0.4709123 | 0.33552448 | -1.4035111 | 0.16046447 | 0.71101785 |
| WBP2NL | 35.9307167 | 0.39654689 | 0.57062668 | 0.69493226 | 0.48709775 | 0.88838332 |
| SMDT1 | 28.2684195 | 1.11923988 | 0.31646329 | 3.53671321 | 0.00040514 | 0.07958059 |
| NDUFA6 | 42.4222205 | -0.2622094 | 0.50448206 | -0.5197597 | 0.60323106 | 0.92498018 |

| | | | | | | |
|---|---|---|---|---|---|---|
| TCF20 | 132.602732 | -0.5294711 | 0.22764949 | -2.3258174 | 0.0200283 | 0.41544117 |
| RRP7A | 28.039002 | -0.3316006 | 0.38948043 | -0.8513922 | 0.39455155 | 0.85605713 |
| SERHL2 | 17.5343319 | 0.32981701 | 0.39019429 | 0.84526355 | 0.39796371 | 0.85758974 |
| POLDIP3 | 31.7818319 | 0.03327835 | 0.289446 | 0.11497256 | 0.90846687 | 0.9854228 |
| CYB5R3 | 83.0948733 | -0.2442911 | 0.29681744 | -0.8230347 | 0.41048824 | 0.85867915 |
| A4GALT | 88.2389009 | 0.26930034 | 0.35452811 | 0.75960222 | 0.44749239 | 0.87275002 |
| ARFGAP3 | 146.7914 | -0.0973868 | 0.19438375 | -0.501003 | 6.16E-01 | 9.27E-01 |
| PACSIN2 | 213.750998 | -0.3861693 | 0.27558336 | -1.4012795 | 0.16113052 | 0.71101785 |
| TTLL1 | 10.9499924 | -0.3210951 | 0.55371281 | -0.5798947 | 0.56198566 | NA |
| MCAT | 12.7039804 | 0.53732209 | 0.48239287 | 1.11386823 | 0.26533572 | NA |
| TSPO | 66.4037203 | 0.41837782 | 0.36093361 | 1.1591545 | 0.24639322 | 0.78583441 |
| EFCAB6 | 106.561345 | 0.00959162 | 0.30373093 | 0.03157932 | 0.97480753 | 0.99431961 |
| SAMM50 | 16.7353024 | -0.0958142 | 0.40561611 | -0.2362189 | 0.81326283 | 0.97282508 |
| PARVB | 13.122584 | -0.7922564 | 0.91402447 | -0.8667781 | 0.38606363 | NA |
| NUP50-DT | 63.6624126 | 0.16379273 | 0.29827377 | 0.54913553 | 0.58291244 | 0.92034873 |
| NUP50 | 32.137708 | -0.1986739 | 0.27866285 | -0.7129543 | 0.47587401 | 0.88495683 |
| KIAA0930 | 23.3603094 | -0.535906 | 0.45664513 | -1.1735721 | 0.24056646 | 0.78209874 |
| FAM118A | 24.559923 | -0.7171394 | 0.44337451 | -1.6174575 | 0.10577955 | 0.65946648 |
| ATXN10 | 202.862392 | -0.1506947 | 0.17537805 | -0.8592564 | 0.39019908 | 0.85605713 |
| MIRLET7BHG | 83.2629086 | 0.05845949 | 0.2805409 | 0.20838133 | 0.83493123 | 0.97600352 |
| PPARA | 181.148459 | -0.168629 | 0.27516246 | -0.6128342 | 0.53998597 | 0.90763121 |
| TTC38 | 29.5968482 | 0.54280809 | 0.34317936 | 1.58170375 | 0.11371722 | 0.6656824 |
| TRMU | 38.7858722 | -0.0524446 | 0.30536327 | -0.171745 | 0.86363799 | 0.97788853 |
| CELSR1 | 50.0145531 | 0.07764491 | 0.26677791 | 0.29104701 | 0.77101537 | 0.96364125 |
| GRAMD4 | 26.0558311 | -0.2230987 | 0.38982275 | -0.5723082 | 0.56711323 | 0.91698355 |
| CERK | 32.7027506 | 0.35937451 | 0.30977248 | 1.16012409 | 0.24599829 | 0.78583441 |
| TBC1D22A | 270.556293 | -0.5472174 | 0.27380146 | -1.9985919 | 0.04565253 | 0.52804951 |
| BRD1 | 56.3717619 | -0.343904 | 0.31066603 | -1.1069894 | 0.2682985 | 0.79666376 |
| ZBED4 | 29.0369712 | -0.6234647 | 0.33553887 | -1.8580997 | 0.06315485 | 0.58663125 |
| ALG12 | 13.6442543 | -0.0374921 | 0.42449892 | -0.0883209 | 0.92962163 | NA |
| PIM3 | 87.3212395 | -0.6712823 | 0.40676889 | -1.6502794 | 0.0988858 | 0.65744285 |
| TRABD | 35.9289843 | 0.10786687 | 0.38805854 | 0.27796546 | 0.78103887 | 0.96652357 |
| SELENOO | 54.0691403 | -0.0836415 | 0.23659017 | -0.3535291 | 0.72369185 | 0.95093722 |
| TUBGCP6 | 16.1926647 | -0.2989975 | 0.39780907 | -0.7516105 | 0.45228531 | 0.87346041 |
| PLXNB2 | 108.379275 | 0.09214793 | 0.31013579 | 0.29712125 | 0.76637394 | 0.96270421 |
| DENND6B | 17.6466854 | 0.18019761 | 0.4513852 | 0.39921028 | 0.68973827 | 0.94246285 |
| PPP6R2 | 218.007551 | -0.2540294 | 0.20466088 | -1.2412213 | 0.21452402 | 0.7643572 |
| SBF1 | 11.3935242 | -0.2199299 | 0.41987963 | -0.5237927 | 0.60042272 | NA |
| MIOX | 14.3195222 | -0.8090279 | 0.89345625 | -0.9055037 | 0.36519861 | NA |
| LMF2 | 16.7497903 | -0.228876 | 0.48311999 | -0.4737457 | 0.63568125 | 0.93251095 |
| KLHDC7B | 23.2817378 | -0.4222563 | 0.59331364 | -0.7116915 | 4.77E-01 | 8.85E-01 |
| ARSA | 14.1700595 | 0.16124989 | 0.47870419 | 0.33684663 | 0.73623252 | NA |
| RABL2B | 24.736866 | -0.4979813 | 0.34494213 | -1.4436663 | 0.14883286 | 0.70016822 |
| CU633967.1 | 32.3438182 | 0.65207905 | 0.36069353 | 1.80784792 | 0.07063017 | 0.60513619 |
| FP671120.1 | 506.551147 | -0.9406973 | 0.43919658 | -2.1418593 | 0.03220481 | 0.48175799 |
| FP236383.1 | 290.726029 | -0.9227568 | 0.40087727 | -2.3018436 | 0.02134399 | 0.4212027 |
| CR381653.1 | 42.3289359 | -0.5655063 | 0.47099242 | -1.2006696 | 0.22987939 | 0.77225177 |
| RBM11 | 11.5962983 | -0.5789259 | 0.62663641 | -0.9238625 | 0.35555791 | NA |
| HSPA13 | 14.5837407 | -0.3002706 | 0.46647431 | -0.6437024 | 0.51976846 | NA |
| NRIP1 | 94.498655 | -0.5175505 | 0.27972773 | -1.8501938 | 0.06428562 | 0.58872433 |
| USP25 | 316.40353 | -0.7317369 | 0.19448721 | -3.7623911 | 0.0001683 | 0.05491436 |
| MIR99AHG | 125.921461 | -0.2699834 | 0.35707969 | -0.7560872 | 0.44959689 | 0.87328261 |
| CXADR | 72.0278688 | 0.24313293 | 0.28481108 | 0.85366387 | 0.39329126 | 0.85605713 |
| BTG3 | 52.735998 | 0.37372492 | 0.45942838 | 0.81345633 | 0.41595647 | 0.85867915 |

| | | | | | | |
|---|---|---|---|---|---|---|
| C21orf91 | 16.86096 | 0.43298522 | 0.37982911 | 1.13994742 | 0.25430821 | 0.79171259 |
| CHODL | 55.4894768 | 1.45370389 | 0.60520139 | 2.40201676 | 0.01630496 | 0.39748373 |
| MRPL39 | 22.9095172 | -0.1384098 | 0.33654565 | -0.4112659 | 0.68087755 | 0.94196488 |
| ATP5PF | 89.6371589 | 0.27366721 | 0.39770925 | 0.68810874 | 0.49138431 | 0.88995517 |
| GABPA | 32.5024849 | -0.8003651 | 0.3184671 | -2.5131799 | 0.01196483 | 0.35710117 |
| APP | 804.421499 | 0.18268904 | 0.17729902 | 1.03040072 | 0.30282194 | 0.81568231 |
| CYYR1 | 51.7792722 | 1.52481202 | 0.35931616 | 4.24365 | 2.20E-05 | 0.01667444 |
| ADAMTS1 | 376.204314 | 0.08504638 | 0.49563325 | 0.17159135 | 8.64E-01 | 9.78E-01 |
| ADAMTS5 | 14.997443 | -0.168696 | 0.72165793 | -0.2337618 | 0.81516994 | NA |
| AF165147.1 | 20.1850293 | -0.5677629 | 0.62717851 | -0.9052653 | 0.36532487 | 0.84720508 |
| N6AMT1 | 12.3173717 | 0.0544581 | 0.49975926 | 0.10896867 | 0.91322734 | NA |
| LTN1 | 78.0951909 | -0.0944592 | 0.24505767 | -0.3854572 | 0.69989874 | 0.94419257 |
| USP16 | 46.2804914 | -0.4460987 | 0.29231266 | -1.5261013 | 0.12698463 | 0.66929384 |
| CCT8 | 35.1209489 | 0.19300318 | 0.31500609 | 0.61269667 | 0.54007693 | 0.90763121 |
| MAP3K7CL | 36.8237754 | -0.0660625 | 0.37614618 | -0.1756298 | 0.86058482 | 0.97718459 |
| BACH1 | 378.961619 | -0.4661653 | 0.22150487 | -2.1045373 | 3.53E-02 | 0.49209205 |
| GRIK1 | 21.3309227 | 1.15886081 | 0.61919661 | 1.87155547 | 0.06126813 | 0.58024409 |
| CLDN8 | 71.290138 | 1.2169403 | 0.54401177 | 2.23697419 | 0.02528803 | 0.4403385 |
| SOD1 | 92.3267835 | 0.58498754 | 0.4629267 | 1.26367207 | 0.20634775 | 0.76231289 |
| SCAF4 | 71.8335777 | -0.4028515 | 0.26708845 | -1.5083074 | 0.13147588 | 0.67540912 |
| HUNK | 178.163962 | 0.02649248 | 0.42551372 | 0.06226 | 0.95035578 | 0.99105535 |
| URB1 | 27.4812846 | -0.467491 | 0.39190289 | -1.1928746 | 0.2329185 | 0.7760824 |
| EVA1C | 53.8618163 | 0.87418303 | 0.38825447 | 2.25157234 | 0.02434931 | 0.4389144 |
| SYNJ1 | 75.6235121 | 0.1695713 | 0.24881059 | 0.68152765 | 0.49553768 | 0.89230917 |
| PAXBP1 | 108.941097 | 0.05485022 | 0.21164984 | 0.25915549 | 0.79551527 | 0.96974891 |
| C21orf62-AS | 31.055202 | -0.2024733 | 0.60134471 | -0.3367009 | 0.7363424 | 0.95376039 |
| AP000282.1 | 17.1256973 | 0.66810095 | 0.61270942 | 1.09040424 | 0.27553511 | 0.79955249 |
| IFNAR1 | 69.8377482 | -0.0106035 | 0.23342667 | -0.0454253 | 0.96376833 | 0.99256424 |
| IFNGR2 | 96.065609 | 0.22781384 | 0.38378995 | 0.59358989 | 0.55278644 | 0.91322379 |
| TMEM50B | 94.6899898 | -0.2831031 | 0.34349283 | -0.8241892 | 0.40983209 | 0.85867915 |
| GART | 43.926542 | -0.2221815 | 0.27752968 | -0.8005684 | 0.42338152 | 0.86190547 |
| SON | 186.03528 | 0.10040156 | 0.26688584 | 0.37619666 | 0.7067707 | 0.94558664 |
| AP000311.1 | 57.9087747 | 0.11596139 | 0.47084573 | 0.24628318 | 0.80546303 | 0.97214568 |
| ITSN1 | 233.376388 | -0.3579974 | 0.24065643 | -1.4875872 | 0.13685979 | 0.68452453 |
| MRPS6 | 145.508372 | -0.6943273 | 0.36750614 | -1.8892943 | 0.0588524 | 0.57258816 |
| LINC00310 | 16.5718293 | -0.3581788 | 0.60001858 | -0.5969462 | 0.5505433 | 0.91248567 |
| AP000317.1 | 25.1179483 | -0.699209 | 0.35745269 | -1.9560882 | 0.05045477 | 0.54612337 |
| RCAN1 | 76.6354303 | 0.00141561 | 0.5707572 | 0.00248023 | 0.99802106 | 0.99957534 |
| CLIC6 | 26.4076721 | 0.10210006 | 0.44424394 | 0.22982882 | 0.81822479 | 0.97288403 |
| RUNX1 | 580.998556 | -1.0817566 | 0.67584882 | -1.6005896 | 0.10946784 | 0.66067258 |
| SETD4 | 32.6774513 | -0.132993 | 0.35543324 | -0.3741714 | 0.70827677 | 0.94558664 |
| CBR1 | 81.5883103 | 0.53723905 | 0.34399842 | 1.56174859 | 0.11834723 | 0.6668162 |
| AP000688.2 | 14.4973849 | 0.06780658 | 0.49994405 | 0.13562834 | 0.8921151 | NA |
| CBR3 | 42.6202501 | 0.14716667 | 0.45675568 | 0.32219997 | 0.74730121 | 0.95563105 |
| DOPEY2 | 66.4790335 | -0.1943759 | 0.30988036 | -0.627261 | 0.53048817 | 0.9052447 |
| MORC3 | 103.939084 | -0.4842978 | 0.17838879 | -2.7148446 | 0.00663069 | 0.30102597 |
| CLDN14 | 29.7059083 | -1.0113687 | 0.50620353 | -1.9979488 | 4.57E-02 | 5.28E-01 |
| HLCS | 211.044456 | 0.08182067 | 0.18215017 | 0.4491935 | 0.65329208 | 0.93532021 |
| PIGP | 14.0303909 | 1.0284532 | 0.63312931 | 1.62439677 | 0.10429116 | NA |
| TTC3 | 284.845006 | -0.2677256 | 0.22106785 | -1.211056 | 0.22587394 | 0.77093069 |
| VPS26C | 54.4320213 | -0.1865856 | 0.27215641 | -0.6855824 | 0.49297648 | 0.89092098 |
| DYRK1A | 412.060967 | -0.3002967 | 0.21356966 | -1.4060832 | 0.15969943 | 0.71101785 |
| KCNJ15 | 357.791543 | 0.77239257 | 0.39215898 | 1.96959044 | 0.04888533 | 0.54070856 |
| ETS2 | 59.8704757 | -1.2384581 | 0.44639795 | -2.7743365 | 5.53E-03 | 0.28673151 |

| | | | | | | |
|---|---|---|---|---|---|---|
| BRWD1 | 309.175862 | -0.0763212 | 0.16153936 | -0.4724618 | 0.63659723 | 0.93251095 |
| HMGN1 | 92.9850147 | 0.17347532 | 0.26627271 | 0.65149492 | 0.51472706 | 0.90156229 |
| WRB | 27.3450447 | -0.1331501 | 0.33830576 | -0.3935792 | 0.69389177 | 0.94395238 |
| LCA5L | 14.9056484 | 0.22096563 | 0.44686593 | 0.49447858 | 0.62096823 | NA |
| SH3BGR | 18.5754767 | 0.13633284 | 0.64121805 | 0.21261541 | 0.83162695 | 0.97587989 |
| BACE2 | 442.369906 | -1.342829 | 0.5559453 | -2.4153977 | 0.01571804 | 0.39416332 |
| FAM3B | 41.0320957 | 0.16402388 | 0.31503071 | 0.52065996 | 0.60260367 | 0.92426305 |
| MX1 | 101.504519 | -2.8820937 | 1.14457136 | -2.518055 | 0.01180049 | 0.35687659 |
| TMPRSS2 | 268.082725 | 0.3694095 | 0.39510159 | 0.93497348 | 3.50E-01 | 8.37E-01 |
| RIPK4 | 25.4373997 | -0.599765 | 0.30483418 | -1.9675123 | 0.04912418 | 0.54203319 |
| PRDM15 | 15.8472625 | 0.02836793 | 0.59228468 | 0.04789577 | 0.96179931 | NA |
| C2CD2 | 32.97353 | -0.1201326 | 0.39473319 | -0.3043388 | 0.76086976 | 0.95956874 |
| ZBTB21 | 28.1303162 | -0.2545331 | 0.40333468 | -0.6310716 | 0.52799368 | 0.90487731 |
| RSPH1 | 15.7735643 | 0.61738453 | 0.489965 | 1.26005843 | 0.20764828 | NA |
| SLC37A1 | 49.5208663 | 0.24970132 | 0.21843338 | 1.14314633 | 2.53E-01 | 7.91E-01 |
| LINC01671 | 18.335095 | -0.023024 | 0.47706028 | -0.0482623 | 0.96150716 | 0.99256424 |
| PDE9A | 166.227287 | 0.06284188 | 0.30325061 | 0.20722755 | 0.83583216 | 0.97600352 |
| WDR4 | 20.0620337 | -0.5390579 | 0.44517117 | -1.2109003 | 0.22593362 | 0.77093069 |
| NDUFV3 | 68.8057685 | 0.02561543 | 0.21033197 | 0.12178574 | 0.90306871 | 0.98531749 |
| PKNOX1 | 71.5510019 | 0.28874536 | 0.26975789 | 1.0703871 | 0.2844451 | 0.80431986 |
| RRP1B | 27.0038798 | -0.1914403 | 0.3481591 | -0.5498646 | 0.58241228 | 0.92034873 |
| PDXK | 135.562761 | -0.5040536 | 0.30669588 | -1.6434964 | 0.10028026 | 0.65863153 |
| CSTB | 112.087862 | -0.2151171 | 0.35196295 | -0.6111924 | 5.41E-01 | 0.90783815 |
| AGPAT3 | 119.275164 | 0.09514491 | 0.19846487 | 0.47940427 | 0.63165105 | 0.93251095 |
| TRAPPC10 | 105.004641 | -0.2337046 | 0.2468563 | -0.9467231 | 0.34377989 | 0.83465492 |
| PFKL | 147.180414 | -0.4594203 | 0.27586498 | -1.6653811 | 0.09583674 | 0.65056069 |
| C21orf2 | 16.659501 | 0.36061966 | 0.56920307 | 0.63355185 | 0.52637333 | 0.90487731 |
| UBE2G2 | 91.2202524 | 0.24465723 | 0.21808438 | 1.12184662 | 0.26192766 | 0.79633795 |
| SUMO3 | 33.1650297 | -0.3295189 | 0.44053244 | -0.7480015 | 0.45445924 | 0.87378486 |
| PTTG1IP | 135.650853 | -0.080137 | 0.22435935 | -0.3571815 | 7.21E-01 | 9.51E-01 |
| FAM207A | 13.2438718 | 0.10224879 | 0.50874685 | 0.20098167 | 0.84071291 | NA |
| ADARB1 | 35.0134461 | 0.5583978 | 0.3470512 | 1.60897815 | 0.10762112 | 0.65986731 |
| POFUT2 | 20.0814728 | -0.6893635 | 0.37933785 | -1.8172812 | 0.06917407 | 0.60275677 |
| COL18A1 | 274.767156 | 0.29540502 | 0.29024194 | 1.0177889 | 0.30877829 | 0.81568231 |
| COL6A1 | 56.3747359 | 0.3288795 | 0.43242729 | 0.7605429 | 0.44693014 | 0.87231627 |
| LSS | 29.7565058 | -0.2518951 | 0.51778047 | -0.4864901 | 0.6266197 | 0.93032781 |
| MCM3AP-AS | 14.4500417 | -0.1852302 | 0.41139082 | -0.4502535 | 0.65252766 | NA |
| MCM3AP | 65.0221379 | -0.2730209 | 0.24667826 | -1.1067895 | 0.26838493 | 0.79666376 |
| YBEY | 11.8429736 | 0.09281924 | 0.55342721 | 0.16771717 | 0.86680579 | NA |
| PCNT | 33.4970516 | -0.2186068 | 0.44757988 | -0.4884197 | 0.62525262 | 0.92981013 |
| DIP2A | 111.486224 | -0.2694229 | 0.20154755 | -1.336771 | 0.1812974 | 0.73398872 |
| PRMT2 | 104.158109 | -0.0073944 | 0.23479919 | -0.0314926 | 0.97487669 | 0.99431961 |
| AC011043.1 | 33.765574 | 0.42472823 | 0.46785969 | 0.90781111 | 0.36397803 | 0.8466721 |
| AL592183.1 | 68.9485541 | 0.25344348 | 0.5036436 | 0.5032199 | 0.61480967 | 0.92662475 |

**Collecting duct principal cells**

*This means that positive fold changes imply higher mean value in non-COVID AKI than in COVID AKI and vice versa.

| | baseMean | log2 fold cha | lfcSE | stat | pvalue | padj |
|---|---|---|---|---|---|---|
| AL669831.5 | 117.625093 | 0.26115562 | 0.41103427 | 0.63536216 | 0.52519224 | 0.91924275 |
| AL645608.1 | 37.838824 | 0.3492314 | 0.62262773 | 0.56089921 | 0.57486625 | 0.93275827 |
| SAMD11 | 118.20368 | 0.53204097 | 0.58738155 | 0.90578427 | 0.36505005 | 0.85866868 |
| NOC2L | 80.3916088 | 0.33799722 | 0.23344842 | 1.44784537 | 0.1476603 | 0.69834914 |
| HES4 | 131.461973 | -3.7360064 | 1.02802902 | -3.6341449 | 0.0002789 | 0.03764581 |
| ISG15 | 634.877613 | -5.5190753 | 1.6169775 | -3.4132048 | 0.00064204 | 0.05952104 |
| AGRN | 470.478214 | 0.11385887 | 0.3073928 | 0.37040188 | 0.71108307 | 0.95773687 |
| C1orf159 | 129.910862 | -0.4990123 | 0.24293676 | -2.054083 | 0.03996766 | 0.45827251 |
| LINC01342 | 43.7069962 | -0.0309782 | 0.4045115 | -0.0765817 | 0.93895631 | 0.99243654 |
| AL390719.2 | 38.6331177 | -0.0563877 | 0.44643129 | -0.1263077 | 0.89948839 | 0.98660836 |
| SDF4 | 194.743158 | 0.20899629 | 0.29111672 | 0.71791235 | 0.47281133 | 0.90213021 |
| UBE2J2 | 120.117691 | -0.2180935 | 0.2307703 | -0.9450674 | 0.34462447 | 0.85148675 |
| INTS11 | 123.392404 | -0.1689354 | 0.21967937 | -0.7690089 | 0.44188804 | 0.88997055 |
| DVL1 | 44.9676192 | -0.1573632 | 0.36826414 | -0.4273106 | 0.66915314 | 0.95304061 |
| AURKAIP1 | 240.795271 | 0.22733256 | 0.50942935 | 0.44624943 | 0.65541708 | 0.95100227 |
| CCNL2 | 274.326611 | -0.6263531 | 0.25441992 | -2.4618872 | 0.01382081 | 0.28408502 |
| MRPL20 | 244.574701 | 0.33198501 | 0.32241601 | 1.02967904 | 0.3031607 | 0.83709431 |
| VWA1 | 109.481148 | 0.5911299 | 0.51165199 | 1.15533587 | 0.24795291 | 0.80097144 |
| ATAD3C | 29.2610394 | 0.78797191 | 0.43890809 | 1.79530048 | 0.07260584 | 0.56657423 |
| ATAD3B | 71.7650765 | -0.4461546 | 0.30997347 | -1.4393317 | 0.15005656 | 0.7029344 |
| ATAD3A | 37.646709 | -0.0542919 | 0.45675449 | -0.1188646 | 0.90538262 | 0.9875066 |
| SSU72 | 251.688729 | -0.0697105 | 0.35212907 | -0.1979687 | 0.84306953 | 0.97560439 |
| MIB2 | 64.2940999 | -0.1107688 | 0.35671867 | -0.3105215 | 0.75616439 | 0.96120879 |
| CDK11B | 102.200513 | -0.0508488 | 0.54699201 | -0.0929608 | 0.92593472 | 0.99051393 |
| SLC35E2B | 85.2218091 | -0.4320924 | 0.39489495 | -1.0941957 | 0.27386916 | 0.81980828 |
| CDK11A | 45.9637821 | -0.0982355 | 0.37175969 | -0.2642448 | 0.79159133 | 0.9653598 |
| SLC35E2A | 95.2218604 | -0.4074722 | 0.38947832 | -1.0462 | 0.29546869 | 0.83396494 |
| NADK | 134.31432 | -0.5461615 | 0.25483534 | -2.1431938 | 0.03209754 | 0.41517597 |
| GNB1 | 948.793386 | -0.4510298 | 0.23613192 | -1.9100755 | 0.05612349 | 0.52189372 |
| PRKCZ | 179.441755 | 0.0515207 | 0.23783749 | 0.21662142 | 0.82850339 | 0.97117144 |
| FAAP20 | 177.778053 | -0.1805288 | 0.30818749 | -0.585776 | 0.55802608 | 0.92901206 |
| SKI | 241.762857 | 0.19242333 | 0.40446943 | 0.47574259 | 0.63425778 | 0.94775679 |
| MORN1 | 122.211515 | -0.2213254 | 0.23412729 | -0.9453207 | 0.34449517 | 0.85148675 |
| RER1 | 305.643726 | -0.1456741 | 0.34689293 | -0.4199397 | 0.67452949 | 0.95524793 |
| PEX10 | 76.0813871 | -0.1361705 | 0.40287305 | -0.3379986 | 0.73536426 | 0.96078945 |
| AL139246.5 | 78.1323222 | -1.5061057 | 0.69129875 | -2.1786611 | 0.02935685 | 0.40623517 |
| TNFRSF14-A | 103.219763 | -1.2200534 | 0.59955296 | -2.0349386 | 0.04185706 | 0.46765666 |
| TNFRSF14 | 258.973109 | -1.1120881 | 0.40471581 | -2.7478248 | 0.00599921 | 0.19123755 |
| FAM213B | 117.683282 | -0.5033534 | 0.4979301 | -1.0108917 | 0.31206826 | 0.84279957 |
| MMEL1 | 37.0811475 | -0.5860127 | 0.57063719 | -1.0269445 | 0.30444658 | 0.83709431 |
| LINC00982 | 377.940021 | -0.3016943 | 0.433129 | -0.696546 | 0.48608695 | 0.90695174 |
| PRDM16 | 1017.52816 | -0.3664253 | 0.3720472 | -0.9848892 | 0.32467852 | 0.84285425 |
| ARHGEF16 | 89.1986175 | -0.7405119 | 0.4162765 | -1.7788944 | 0.07525708 | 0.57072562 |
| TPRG1L | 81.2447232 | -0.0471202 | 0.28665233 | -0.1643809 | 0.86943129 | 0.9820194 |
| WRAP73 | 42.0303651 | -0.0859353 | 0.2873924 | -0.2990173 | 0.76492682 | 0.96285313 |
| LRRC47 | 67.7882402 | -0.0779108 | 0.3218831 | -0.2420468 | 0.80874386 | 0.96775927 |
| CEP104 | 234.865246 | -0.5786314 | 0.32079849 | -1.8037223 | 0.07127485 | 0.5633527 |
| C1orf174 | 50.188868 | -0.4181679 | 0.43928942 | -0.9519189 | 0.34113809 | 0.85042291 |
| NPHP4 | 69.6948276 | 0.2426268 | 0.27415951 | 0.88498407 | 0.37616522 | 0.86357183 |
| RPL22 | 190.042941 | 0.01599301 | 0.33469484 | 0.04778386 | 0.9618885 | 0.99635733 |
| RNF207 | 81.4088126 | 0.28829845 | 0.23073008 | 1.24950522 | 0.21148034 | 0.77683631 |
| ICMT | 99.8263651 | 0.03651852 | 0.28053156 | 0.13017616 | 0.89642705 | 0.98597539 |

| | | | | | | |
|---|---|---|---|---|---|---|
| ESPN | 86.5684673 | 0.10482449 | 0.62580477 | 0.16750351 | 0.86697389 | 0.98087945 |
| NOL9 | 93.1733938 | 0.24288252 | 0.29290543 | 0.82921822 | 0.40698094 | 0.87948803 |
| ZBTB48 | 47.724122 | -0.070919 | 0.31889047 | -0.222393 | 0.82400797 | 0.97018919 |
| KLHL21 | 58.7267779 | -0.0178513 | 0.41562438 | -0.0429506 | 0.96574088 | 0.99637506 |
| PHF13 | 47.3358921 | 0.34117548 | 0.26958069 | 1.26557834 | 0.20566409 | 0.77193931 |
| THAP3 | 40.278405 | 0.02775465 | 0.26170698 | 0.10605238 | 0.91554079 | 0.98868699 |
| DNAJC11 | 175.71743 | -0.1638792 | 0.20652058 | -0.7935249 | 0.42747208 | 0.88518226 |
| CAMTA1 | 1006.88619 | -0.5440431 | 0.22728714 | -2.393638 | 0.0166822 | 0.30789747 |
| VAMP3 | 108.061993 | 0.11315167 | 0.33732509 | 0.33543805 | 0.73729467 | 0.96078945 |
| PER3 | 198.979431 | 0.12437976 | 0.58088958 | 0.21411945 | 0.83045391 | 0.97160502 |
| PARK7 | 422.227972 | 0.67127205 | 0.44616563 | 1.50453556 | 0.13244352 | 0.67466347 |
| ERRFI1 | 328.135005 | -2.4427341 | 1.19280618 | -2.0478885 | 0.04057092 | 0.46316263 |
| RERE | 2945.66642 | -0.1262475 | 0.22292347 | -0.5663265 | 0.57117187 | 0.93125376 |
| ENO1 | 1957.39888 | 0.05967848 | 0.44130075 | 0.13523312 | 0.89242756 | 0.98597539 |
| GPR157 | 36.3476524 | -0.0130871 | 0.30650492 | -0.0426979 | 0.96594232 | 0.99637506 |
| MIR34AHG | 44.3913371 | 0.6930596 | 0.78690603 | 0.88073997 | 0.37845858 | 0.86515603 |
| H6PD | 117.65147 | -0.0052386 | 0.22223835 | -0.0235721 | 0.98119393 | 0.99696835 |
| SPSB1 | 127.347321 | -0.592669 | 0.43393507 | -1.365801 | 0.17200143 | 0.72776437 |
| SLC25A33 | 147.008776 | -0.3321721 | 0.43128517 | -0.7701914 | 0.44118639 | 0.88964371 |
| PIK3CD | 37.551523 | 0.26554052 | 0.46947686 | 0.56560939 | 0.57165934 | 0.93172471 |
| CLSTN1 | 260.614526 | 0.47828484 | 0.19345104 | 2.47238178 | 0.01342161 | 0.28170334 |
| CTNNBIP1 | 91.3337699 | -0.1509955 | 0.31232584 | -0.4834551 | 0.62877266 | 0.94595714 |
| LZIC | 106.411177 | -0.1352797 | 0.24065022 | -0.5621425 | 0.57401896 | 0.93270589 |
| NMNAT1 | 119.906887 | 0.14011311 | 0.24079466 | 0.58187796 | 0.56064889 | 0.92934517 |
| UBE4B | 407.426611 | -0.3643518 | 0.2522517 | -1.4443978 | 0.14862712 | 0.70131121 |
| KIF1B | 457.333209 | -0.0671496 | 0.20014779 | -0.3355003 | 0.73724774 | 0.96078945 |
| PGD | 89.896907 | -0.1439407 | 0.35063411 | -0.4105154 | 0.68142787 | 0.95574492 |
| DFFA | 134.99283 | 0.25331295 | 0.18594354 | 1.36231112 | 0.17309972 | 0.72964461 |
| PEX14 | 333.793624 | -0.3711322 | 0.17538989 | -2.1160407 | 0.03434134 | 0.43050784 |
| CASZ1 | 639.012269 | 0.1179757 | 0.38559453 | 0.30595791 | 0.75963669 | 0.96190674 |
| SRM | 50.1581133 | -0.3280585 | 0.37459769 | -0.8757621 | 0.38115937 | 0.86593382 |
| EXOSC10 | 97.3655633 | -0.4128294 | 0.22268428 | -1.8538775 | 0.06375669 | 0.54850785 |
| MTOR | 195.624262 | -0.2114784 | 0.28705129 | -0.736727 | 0.46128841 | 0.89799005 |
| FBXO2 | 120.212283 | 0.56807824 | 0.59497158 | 0.95479895 | 0.33967938 | 0.84928129 |
| FBXO44 | 54.3543407 | 0.20942789 | 0.28592012 | 0.73246993 | 0.46388179 | 0.89801718 |
| AGTRAP | 124.79997 | -0.0934083 | 0.3949027 | -0.2365349 | 0.81301759 | 0.9680627 |
| MTHFR | 71.6660496 | 0.02089927 | 0.31653949 | 0.06602421 | 0.94735855 | 0.99314907 |
| CLCN6 | 93.0908092 | -1.0519338 | 0.35803563 | -2.9380702 | 0.00330262 | 0.14358074 |
| KIAA2013 | 86.9208264 | 0.14867356 | 0.30931661 | 0.48065172 | 0.63076405 | 0.94596555 |
| PLOD1 | 52.4796337 | 0.43848099 | 0.32819781 | 1.33602658 | 0.18154059 | 0.74315969 |
| MFN2 | 126.603999 | -0.1159596 | 0.25145368 | -0.461157 | 0.64468594 | 0.94949493 |
| VPS13D | 780.902363 | -0.1571075 | 0.15818769 | -0.9931715 | 0.32062637 | 0.84285425 |
| DHRS3 | 171.081593 | -0.7193412 | 0.38620369 | -1.8625952 | 6.25E-02 | 0.54645673 |
| PRDM2 | 537.651129 | 0.07522389 | 0.26034464 | 0.28893964 | 0.77262757 | 0.96388926 |
| KAZN | 10345.8817 | -0.8607725 | 0.34186308 | -2.5178868 | 0.01180613 | 0.26780995 |
| TMEM51-AS | 137.149302 | -0.9417014 | 0.2952141 | -3.1898931 | 0.00142325 | 0.09561351 |
| TMEM51 | 942.153407 | -0.4992647 | 0.38966235 | -1.2812751 | 0.20009705 | 0.7687013 |
| C1orf195 | 38.9416262 | -0.5271038 | 0.70932257 | -0.7431087 | 0.45741585 | 0.89705917 |
| FHAD1 | 54.0977 | 0.92169342 | 0.27863228 | 3.30792047 | 0.00093991 | 0.07350977 |
| EFHD2 | 32.3744813 | 0.37850678 | 0.35774437 | 1.05803699 | 0.29003858 | 0.83103268 |
| CASP9 | 208.132488 | -0.6296982 | 0.86852972 | -0.7250163 | 0.46844206 | 0.90026987 |
| DNAJC16 | 193.13378 | -0.1585999 | 0.19805728 | -0.8007778 | 0.42326031 | 0.88371155 |
| DDI2 | 155.113889 | -0.2572576 | 0.17308953 | -1.4862691 | 0.13720796 | 0.68129775 |
| PLEKHM2 | 153.184686 | -0.519491 | 0.22350104 | -2.3243339 | 0.0201076 | 0.33933607 |

| | | | | | | |
|---|---|---|---|---|---|---|
| FBLIM1 | 142.039183 | 0.04006903 | 0.65006872 | 0.06163815 | 0.950851 | 0.99404295 |
| UQCRHL | 65.2065987 | 0.37103043 | 0.51864577 | 0.71538312 | 0.47437234 | 0.90242252 |
| SPEN | 371.626344 | 0.21597183 | 0.19379834 | 1.11441527 | 0.26510108 | 0.81163409 |
| ZBTB17 | 45.118242 | -0.359391 | 0.31151274 | -1.1536961 | 0.24862479 | 0.80097144 |
| CLCNKA | 59.9151531 | 0.70341821 | 0.64755198 | 1.08627296 | 0.27735825 | 0.82198569 |
| CLCNKB | 85.0498752 | 0.06483705 | 0.99311055 | 0.06528684 | 0.94794562 | 0.99314907 |
| EPHA2 | 56.3072095 | -0.1450266 | 0.66421911 | -0.2183415 | 0.82716301 | 0.97069946 |
| FBXO42 | 419.64711 | -0.3419994 | 0.24995756 | -1.3682298 | 0.17124014 | 0.72776437 |
| SZRD1 | 140.50314 | 0.1509633 | 0.21940613 | 0.68805415 | 0.49141868 | 0.90730741 |
| NECAP2 | 87.1199448 | -0.3435282 | 0.2286095 | -1.5026858 | 0.13292009 | 0.67594872 |
| NBPF1 | 107.798233 | -0.0375534 | 0.2543373 | -0.1476519 | 0.88261747 | 0.98341679 |
| CROCC | 99.2842336 | 0.16237166 | 0.3269274 | 0.49665969 | 0.61942905 | 0.94300065 |
| ATP13A2 | 55.7764779 | 0.07417442 | 0.30969748 | 0.23950604 | 0.81071321 | 0.9680627 |
| SDHB | 202.678012 | -0.0716792 | 0.19979898 | -0.3587568 | 0.71977707 | 0.95796325 |
| RCC2 | 168.609561 | -0.2171562 | 0.26573794 | -0.8171818 | 0.41382456 | 0.88211689 |
| ARHGEF10L | 1114.38733 | 0.25920831 | 0.41979458 | 0.61746465 | 0.53692829 | 0.92324352 |
| KLHDC7A | 59.0736232 | 0.61016961 | 0.44385174 | 1.37471493 | 0.16921984 | 0.72596839 |
| UBR4 | 452.743077 | -0.3321867 | 0.24225355 | -1.3712355 | 0.17030154 | 0.72675731 |
| AL035413.1 | 67.0274747 | 0.23142894 | 0.2466113 | 0.93843607 | 0.34802036 | 0.85348649 |
| EMC1 | 70.0611332 | 0.28995095 | 0.26888962 | 1.07832708 | 0.28088782 | 0.82628479 |
| MRTO4 | 48.971908 | 0.07826066 | 0.30792616 | 0.25415398 | 0.7993766 | 0.96556863 |
| AKR7A2 | 121.446563 | 0.4710224 | 0.44062805 | 1.06897962 | 0.28507885 | 0.82804896 |
| CAPZB | 520.874249 | -0.257209 | 0.21473508 | -1.1977966 | 0.2309962 | 0.79130016 |
| NBL1 | 190.560919 | -0.5661183 | 0.46766677 | -1.2105165 | 0.22608078 | 0.78982769 |
| TMCO4 | 387.482238 | 0.03151042 | 0.28535639 | 0.1104248 | 0.91207248 | 0.98796069 |
| OTUD3 | 69.6283561 | -0.5283728 | 0.31359564 | -1.6848856 | 0.09201064 | 0.61120158 |
| VWA5B1 | 44.1749799 | -0.9607671 | 0.82205546 | -1.1687376 | 0.24250937 | 0.80072691 |
| CAMK2N1 | 822.682763 | 0.10976617 | 0.44872639 | 0.24461714 | 0.80675289 | 0.96752828 |
| MUL1 | 47.6634072 | 0.03258147 | 0.33569255 | 0.09705747 | 0.92268076 | 0.99051393 |
| PINK1 | 146.357573 | 0.23836989 | 0.38583739 | 0.61779883 | 0.53670796 | 0.92324352 |
| DDOST | 117.3025 | 0.47979671 | 0.26373628 | 1.81922906 | 0.06887649 | 0.55917298 |
| HP1BP3 | 621.034485 | 0.01354213 | 0.15365868 | 0.08813126 | 0.92977235 | 0.99216065 |
| EIF4G3 | 1709.10288 | -0.2065359 | 0.25453305 | -0.8114305 | 0.41711849 | 0.88242985 |
| ECE1 | 45.433246 | 0.28822788 | 0.39781158 | 0.72453366 | 0.46873817 | 0.90026987 |
| NBPF3 | 44.1343552 | 0.15212421 | 0.31964237 | 0.47592004 | 0.63413134 | 0.94775679 |
| RAP1GAP | 178.175856 | 0.42276619 | 0.37380687 | 1.13097489 | 0.25806566 | 0.80709747 |
| USP48 | 371.882542 | -0.1575539 | 0.18878698 | -0.834559 | 0.40396606 | 0.87834687 |
| HSPG2 | 378.91538 | 1.11285294 | 0.35203193 | 3.1612273 | 0.00157106 | 0.09927452 |
| CDC42 | 632.178369 | -0.0444905 | 0.25253757 | -0.1761739 | 0.86015729 | 0.97841862 |
| ZBTB40 | 105.559977 | -0.0278031 | 0.32227736 | -0.0862706 | 0.93125127 | 0.99243654 |
| EPHB2 | 155.456401 | 0.25222633 | 0.46838707 | 0.53849978 | 0.59023206 | 0.93483536 |
| KDM1A | 334.690617 | -0.0165714 | 0.17579757 | -0.0942642 | 0.92489926 | 0.99051393 |
| AL031428.1 | 76.2179286 | 0.28265706 | 0.50727199 | 0.55721006 | 0.57738391 | 0.93340377 |
| LUZP1 | 257.893076 | 0.31046505 | 0.26253922 | 1.18254733 | 0.2369886 | 0.79797745 |
| LINC01355 | 48.8025773 | 0.79886245 | 0.39670201 | 2.01375952 | 0.04403479 | 0.48009422 |
| HNRNPR | 453.180165 | 0.03128527 | 0.16671231 | 0.18766022 | 0.85114302 | 0.97718353 |
| TCEA3 | 183.846387 | 0.5244391 | 0.38470838 | 1.36321204 | 0.17281569 | 0.72940007 |
| ASAP3 | 136.635425 | 0.32533247 | 0.35790831 | 0.90898271 | 0.36335925 | 0.85694172 |
| ID3 | 260.672505 | -1.0588863 | 0.75878633 | -1.3955 | 0.16286513 | 0.71543692 |
| RPL11 | 1139.17476 | 0.19494031 | 0.5421954 | 0.35953885 | 0.71919202 | 0.95796325 |
| ELOA | 79.1258634 | 0.18863169 | 0.22298625 | 0.84593415 | 0.39758948 | 0.87462201 |
| ELOA-AS1 | 80.1749033 | 0.25339054 | 0.2705107 | 0.93671171 | 0.34890687 | 0.8535343 |
| PITHD1 | 99.538017 | 0.13904229 | 0.28308383 | 0.49117002 | 0.6233062 | 0.94456546 |
| LYPLA2 | 89.4895438 | -0.2626175 | 0.37318655 | -0.7037164 | 0.48160937 | 0.90561114 |

| | | | | | | |
|---|---|---|---|---|---|---|
| GALE | 143.289428 | -0.7134321 | 0.24080034 | -2.9627535 | 0.00304901 | 0.14138755 |
| HMGCL | 160.306143 | 0.10712778 | 0.26633071 | 0.40223592 | 0.68751041 | 0.95588883 |
| FUCA1 | 404.593578 | 0.17984008 | 0.50097158 | 0.3589826 | 0.71960811 | 0.95796325 |
| PNRC2 | 224.389965 | 0.28742729 | 0.34355508 | 0.83662653 | 0.40280253 | 0.87834687 |
| SRSF10 | 356.534344 | 0.274253 | 0.22133268 | 1.23909853 | 0.21530901 | 0.77894575 |
| IFNLR1 | 61.8167618 | -0.2552077 | 0.34033489 | -0.7498724 | 0.45333156 | 0.89544675 |
| STPG1 | 81.8260301 | -0.0808353 | 0.23193872 | -0.3485199 | 0.72744979 | 0.95979778 |
| NIPAL3 | 255.000562 | 0.46717444 | 0.27408177 | 1.70450757 | 0.0882863 | 0.60651125 |
| RCAN3 | 131.189247 | 0.03252344 | 0.34915585 | 0.09314878 | 0.92578536 | 0.99051393 |
| NCMAP | 109.424766 | 0.13368042 | 0.79139878 | 0.16891664 | 8.66E-01 | 0.98023062 |
| SRRM1 | 367.70621 | 0.02537764 | 0.19672338 | 0.12900167 | 0.89735633 | 0.98612679 |
| CLIC4 | 117.072269 | -1.4643636 | 0.5001228 | -2.928008 | 0.00341141 | 0.14358074 |
| SYF2 | 164.675969 | 0.50173789 | 0.40293685 | 1.2452023 | 0.2130574 | 0.77799356 |
| RSRP1 | 628.684243 | 0.22961276 | 0.3291538 | 0.69758503 | 0.48543675 | 0.90695174 |
| TMEM50A | 221.659194 | 0.08357793 | 0.23648529 | 0.35341704 | 0.72377582 | 0.95963742 |
| MACO1 | 203.295896 | -0.349871 | 0.20868087 | -1.676584 | 0.09362386 | 0.61317397 |
| LDLRAP1 | 52.403616 | -0.176283 | 0.27739273 | -0.6354996 | 0.52510266 | 0.91924275 |
| MAN1C1 | 534.034854 | 0.1243272 | 0.48781009 | 0.25486805 | 0.79882502 | 0.9653598 |
| AL031280.1 | 54.2432632 | 0.21487945 | 0.65273813 | 0.32919703 | 0.74200677 | 0.96086423 |
| MTFR1L | 116.670478 | 0.3022023 | 0.38714339 | 0.78059527 | 0.43504058 | 0.88785137 |
| AL033528.2 | 66.1860173 | -0.039072 | 0.50323624 | -0.0776415 | 0.93811319 | 0.99243654 |
| PAQR7 | 142.561257 | -0.0789892 | 0.32201393 | -0.2452976 | 0.80622604 | 0.96752828 |
| STMN1 | 175.725479 | -0.4632067 | 0.69160019 | -0.6697608 | 0.50301029 | 0.90964098 |
| PAFAH2 | 92.4337536 | 0.24056773 | 0.21601396 | 1.11366752 | 0.26542185 | 0.81191505 |
| PDIK1L | 33.7890669 | 0.37786925 | 0.29115021 | 1.29784979 | 0.19433896 | 0.76162393 |
| ZNF593 | 90.1074409 | 0.23314352 | 0.7585071 | 0.30737158 | 0.75856056 | 0.96180522 |
| CNKSR1 | 78.7576155 | -0.3768321 | 0.27253479 | -1.3826935 | 0.16675887 | 0.72097071 |
| CEP85 | 34.9747699 | 0.10120712 | 0.35255858 | 0.2870647 | 0.77406278 | 0.96388926 |
| SH3BGRL3 | 224.632146 | -0.4850382 | 0.45126703 | -1.0748364 | 0.28244798 | 0.82666614 |
| UBXN11 | 47.5477272 | -0.5414954 | 0.33233448 | -1.6293687 | 0.10323499 | 0.62635087 |
| DHDDS | 116.96175 | -0.5162494 | 0.2437858 | -2.1176351 | 0.03420598 | 0.43050784 |
| HMGN2 | 285.687663 | 0.33038604 | 0.40690399 | 0.81195086 | 0.41681983 | 0.88242985 |
| RPS6KA1 | 131.294394 | 0.39808351 | 0.23764449 | 1.675122 | 0.0939103 | 0.61323531 |
| ARID1A | 519.847675 | -0.0292675 | 0.32578355 | -0.0898373 | 0.92841647 | 0.99202739 |
| ZDHHC18 | 38.4641641 | -0.0925796 | 0.39324014 | -0.2354276 | 0.81387686 | 0.96820092 |
| GPN2 | 77.0192507 | -0.0676831 | 0.21475149 | -0.3151692 | 0.75263315 | 0.96120879 |
| NUDC | 220.276074 | -0.2106498 | 0.31372656 | -0.6714439 | 0.50193782 | 0.90945856 |
| NR0B2 | 97.365884 | -1.5927938 | 0.65830226 | -2.4195477 | 0.01553982 | 0.30047476 |
| KDF1 | 37.9525618 | -0.0213273 | 0.32859257 | -0.0649049 | 0.94824969 | 0.99332441 |
| TRNP1 | 94.2973391 | 0.70239171 | 0.44924804 | 1.56348306 | 0.11793902 | 0.65447949 |
| SLC9A1 | 115.052503 | -0.1899915 | 0.34206652 | -0.5554228 | 0.57860552 | 0.93345492 |
| WDTC1 | 169.958294 | 0.24827433 | 0.22753373 | 1.09115398 | 0.27520514 | 0.8202929 |
| TMEM222 | 71.0179272 | -0.2832004 | 0.26388269 | -1.0732058 | 0.28317881 | 0.82666614 |
| WASF2 | 435.698649 | -0.30259 | 0.18196755 | -1.6628787 | 0.09633671 | 0.61440337 |
| AHDC1 | 115.047607 | 1.23605882 | 0.41335166 | 2.99033227 | 0.00278674 | 0.13678792 |
| IFI6 | 506.653485 | -3.7472795 | 1.41908842 | -2.6406244 | 0.00827534 | 0.22277119 |
| FAM76A | 88.085855 | -0.1502018 | 0.32975213 | -0.4554992 | 0.64875011 | 0.95061388 |
| STX12 | 291.854552 | -0.0881181 | 0.25886091 | -0.340407 | 0.73355007 | 0.96035038 |
| PPP1R8 | 66.648626 | 0.12905817 | 0.26334547 | 0.49007173 | 0.62408314 | 0.94457465 |
| RPA2 | 116.571271 | 0.02868169 | 0.35189357 | 0.08150671 | 0.93503899 | 0.99243654 |
| EYA3 | 1663.55735 | 0.85820848 | 0.37525864 | 2.2869786 | 0.02219707 | 0.35843654 |
| PTAFR | 49.2511279 | 0.62424133 | 0.72824289 | 0.85718837 | 0.39134079 | 0.87015937 |
| DNAJC8 | 165.55849 | -0.1171464 | 0.22916219 | -0.5111945 | 0.60921488 | 0.94003018 |
| ATP5IF1 | 714.682592 | 0.67343595 | 0.4171744 | 1.61427918 | 0.10646688 | 0.63461115 |

| | | | | | | |
|---|---|---|---|---|---|---|
| SESN2 | 39.6966167 | 0.35782867 | 0.30434778 | 1.17572296 | 0.2397056 | 0.80019197 |
| PHACTR4 | 540.120061 | -0.0387751 | 0.20081823 | -0.1930854 | 0.84689205 | 0.97656375 |
| RCC1 | 86.7078187 | -0.23675 | 0.31981905 | -0.7402624 | 0.45914079 | 0.89753337 |
| TRNAU1AP | 147.244612 | -0.3536208 | 0.25552232 | -1.3839134 | 0.16638498 | 0.72054851 |
| TAF12 | 83.487837 | 0.07368124 | 0.37148998 | 0.19833977 | 0.84277924 | 0.97560439 |
| GMEB1 | 87.2502021 | -0.3674934 | 0.27296808 | -1.3462872 | 0.17820991 | 0.73737861 |
| YTHDF2 | 218.538744 | 0.11240466 | 0.21044553 | 0.53412708 | 0.59325361 | 0.93491607 |
| EPB41 | 391.934336 | 0.08217841 | 0.18806387 | 0.43697072 | 0.66213258 | 0.95142083 |
| SRSF4 | 622.392479 | -0.1907736 | 0.1971718 | -0.9675499 | 0.33326919 | 0.84442294 |
| MECR | 76.1403365 | -0.0126221 | 0.2829459 | -0.0446095 | 0.96441855 | 0.99637506 |
| PTPRU | 42.3054309 | 0.1650155 | 0.37062141 | 0.44524007 | 0.65614627 | 0.95100227 |
| SDC3 | 74.155773 | -0.2462994 | 0.41490149 | -0.5936335 | 5.53E-01 | 0.92872652 |
| PUM1 | 1206.23581 | 0.028338 | 0.24435871 | 0.11596884 | 0.90767724 | 0.9875066 |
| SNRNP40 | 122.29513 | -0.0322978 | 0.22506453 | -0.1435044 | 0.88589182 | 0.98486779 |
| ZCCHC17 | 185.706914 | 0.15451945 | 0.18760725 | 0.82363263 | 0.41014833 | 0.8801764 |
| SERINC2 | 470.38466 | -0.2594199 | 0.29685765 | -0.8738865 | 0.38218005 | 0.86593382 |
| TINAGL1 | 141.719441 | -0.077985 | 0.36869817 | -0.2115144 | 0.83248592 | 0.97208613 |
| PEF1 | 127.740521 | -0.0970301 | 0.35954701 | -0.2698676 | 0.78726209 | 0.96513523 |
| PTP4A2 | 390.329501 | -1.1330657 | 0.61331099 | -1.847457 | 0.06468094 | 0.55065495 |
| KHDRBS1 | 483.386738 | -0.4126363 | 0.24126475 | -1.710305 | 0.08720948 | 0.60607118 |
| TMEM39B | 41.5369332 | -0.0065048 | 0.33327862 | -0.0195176 | 0.98442817 | 0.99756277 |
| KPNA6 | 387.836786 | -0.0743148 | 0.16129489 | -0.4607387 | 0.64498611 | 0.94949493 |
| TXLNA | 113.643447 | -0.074897 | 0.2018425 | -0.3710664 | 0.71058811 | 0.95756248 |
| TMEM234 | 42.5973776 | 0.01359792 | 0.3149175 | 0.0431793 | 0.96555861 | 0.99637506 |
| EIF3I | 141.603637 | 0.52491565 | 0.4786478 | 1.09666367 | 0.27278844 | 0.81922962 |
| HDAC1 | 228.359656 | 0.32490837 | 0.25342693 | 1.28205937 | 0.19982182 | 0.7687013 |
| MARCKSL1 | 103.850744 | -2.2295559 | 0.69633233 | -3.2018561 | 0.00136545 | 0.0943274 |
| BSDC1 | 163.179509 | -0.2402532 | 0.20842646 | -1.1527003 | 0.24903343 | 0.80097144 |
| ZBTB8OS | 230.57967 | 0.08261732 | 0.22957654 | 0.35986831 | 0.71894562 | 0.95796325 |
| RBBP4 | 297.062143 | -0.1686896 | 0.18247656 | -0.9244452 | 0.35525456 | 0.8557799 |
| SYNC | 79.3060396 | -0.1149763 | 0.31038172 | -0.3704351 | 0.71105835 | 0.95773687 |
| KIAA1522 | 461.930942 | -0.1669705 | 0.19745002 | -0.845634 | 0.39775696 | 0.87462201 |
| YARS | 105.501448 | -0.7788584 | 0.29934893 | -2.6018414 | 0.00927247 | 0.23300339 |
| S100PBP | 219.212996 | -0.109447 | 0.17859997 | -0.612805 | 0.54000525 | 0.92414041 |
| TMEM54 | 96.8300194 | 0.08450909 | 0.45485454 | 0.18579365 | 0.85260658 | 0.97718353 |
| RNF19B | 104.671708 | -1.1952282 | 0.73083019 | -1.635439 | 0.10195705 | 0.62351102 |
| AL020995.1 | 63.8883558 | 0.16839505 | 0.63201215 | 0.26644274 | 0.78989825 | 0.96522573 |
| AK2 | 345.087233 | 0.15834522 | 0.25493477 | 0.62112054 | 0.53452031 | 0.92303275 |
| TRIM62 | 36.0729333 | -0.0830259 | 0.31737822 | -0.2615991 | 7.94E-01 | 0.9653598 |
| ZNF362 | 100.78087 | 0.09617279 | 0.40555149 | 0.23714077 | 0.81254759 | 0.9680627 |
| AL513327.1 | 58.6898798 | -0.2711741 | 0.65383788 | -0.4147421 | 0.67833074 | 0.95574492 |
| PHC2 | 345.877088 | -0.4892304 | 0.25952699 | -1.8850848 | 0.05941838 | 0.53405203 |
| SMIM12 | 109.238792 | 0.07862882 | 0.34246622 | 0.22959584 | 0.81840584 | 0.96883031 |
| AC114490.3 | 30.6738716 | 0.52528355 | 0.48553073 | 1.08187499 | 0.27930808 | 0.82501371 |
| ZMYM1 | 143.404268 | 0.28975645 | 0.22010862 | 1.31642486 | 0.18803148 | 0.7526909 |
| SFPQ | 565.883419 | -0.189765 | 0.21332841 | -0.8895439 | 0.37371086 | 0.86209328 |
| ZMYM4 | 1080.5385 | -0.0575407 | 0.15386876 | -0.3739597 | 0.7084343 | 0.95752169 |
| KIAA0319L | 604.871011 | 0.36745271 | 0.259158 | 1.4178714 | 0.15622831 | 0.7109898 |
| PSMB2 | 152.969838 | 0.11747088 | 0.24396645 | 0.48150423 | 0.63015817 | 0.94596555 |
| AGO4 | 249.834588 | -0.0762352 | 0.2039493 | -0.3737947 | 0.70855709 | 0.95752169 |
| AGO1 | 96.0591099 | -0.1188297 | 0.26417833 | -0.4498087 | 0.6528484 | 0.95100227 |
| AGO3 | 656.799001 | -0.2495369 | 0.14585865 | -1.7108134 | 0.08711557 | 0.60599181 |
| AL138787.2 | 34.2457536 | -0.3192517 | 0.43332974 | -0.7367407 | 0.46128006 | 0.89799005 |
| ADPRHL2 | 88.4757974 | 0.01201672 | 0.40232424 | 0.02986825 | 0.97617213 | 0.99656026 |

| | | | | | | |
|---|---|---|---|---|---|---|
| TRAPPC3 | 103.32709 | 0.0714935 | 0.2596677 | 0.27532689 | 0.7830651 | 0.96513523 |
| MAP7D1 | 62.4961369 | -0.1199409 | 0.26070667 | -0.4600606 | 0.64547275 | 0.94973439 |
| THRAP3 | 530.052005 | -0.1462274 | 0.16992775 | -0.8605272 | 0.38949851 | 0.87004477 |
| EVA1B | 51.7255514 | 0.46899838 | 0.32046229 | 1.46350569 | 0.14332906 | 0.69023197 |
| STK40 | 125.467527 | -0.4643529 | 0.37882664 | -1.2257661 | 0.22028669 | 0.78633738 |
| LSM10 | 50.1135247 | 0.34927528 | 0.60608981 | 0.57627644 | 0.56442834 | 0.93094256 |
| OSCP1 | 46.5342461 | 0.22042243 | 0.28488667 | 0.77371971 | 0.43909656 | 0.88917548 |
| MRPS15 | 73.6851179 | -0.0462239 | 0.46708982 | -0.0989615 | 0.92116887 | 0.9904507 |
| LINC01137 | 42.3246439 | -0.0463532 | 0.4179948 | -0.1108942 | 0.91170025 | 0.98788932 |
| ZC3H12A | 104.234178 | -0.4216505 | 0.87715087 | -0.4807047 | 0.6307264 | 0.94596555 |
| MEAF6 | 145.188655 | 0.20589152 | 0.28998915 | 0.70999732 | 0.4777058 | 0.90245446 |
| SNIP1 | 54.8561217 | -0.2671079 | 0.2920054 | -0.9147362 | 0.36033016 | 0.85608464 |
| DNALI1 | 58.044687 | 0.30853135 | 0.38270219 | 0.80619176 | 0.42013229 | 0.8828928 |
| GNL2 | 44.0021222 | -0.6212959 | 0.35141791 | -1.7679687 | 0.07706613 | 0.57405253 |
| C1orf122 | 162.034531 | 0.42745274 | 0.36763159 | 1.16272038 | 0.24494297 | 0.80072691 |
| MTF1 | 82.1399954 | -0.1086931 | 0.20875939 | -0.5206623 | 0.60260203 | 0.93804805 |
| AL929472.2 | 29.7127187 | 0.34043427 | 0.6290487 | 0.54118906 | 0.58837728 | 0.93483536 |
| INPP5B | 220.186984 | 0.3869771 | 0.34288867 | 1.12857943 | 0.25907529 | 0.80709747 |
| SF3A3 | 100.127115 | -0.0318466 | 0.24626795 | -0.1293167 | 0.89710705 | 0.98601802 |
| UTP11 | 32.1309271 | 0.04863869 | 0.35488464 | 0.13705493 | 0.89098737 | 0.98597539 |
| RRAGC | 101.636007 | -1.1281354 | 0.2541817 | -4.4383031 | 9.07E-06 | 0.00339368 |
| AL139260.1 | 116.180628 | -1.103751 | 0.49940396 | -2.2101367 | 0.02709567 | 0.40076654 |
| MYCBP | 33.0828475 | 0.91427097 | 0.46372089 | 1.97159757 | 0.04865556 | 0.48777647 |
| AKIRIN1 | 194.451125 | 0.10410332 | 0.32752875 | 0.31784484 | 0.75060263 | 0.96109466 |
| NDUFS5 | 486.227944 | -0.1240068 | 0.35492483 | -0.3493889 | 0.72679734 | 0.95979351 |
| MACF1 | 2184.10625 | 0.07454768 | 0.23823881 | 0.31291158 | 0.75434784 | 0.96120879 |
| PABPC4 | 157.179917 | -0.3508098 | 0.45387728 | -0.7729177 | 0.43957106 | 0.88917548 |
| PPIE | 79.5246979 | 0.37022775 | 0.29067987 | 1.27366144 | 0.20278343 | 0.76996624 |
| TRIT1 | 185.40328 | -0.479074 | 0.28206702 | -1.6984403 | 0.08942468 | 0.60739841 |
| MFSD2A | 221.101566 | -5.2494611 | 1.44884099 | -3.6232141 | 0.00029096 | 0.03764581 |
| CAP1 | 317.031895 | -0.1901858 | 0.32300498 | -0.5888016 | 0.5559944 | 0.92872652 |
| PPT1 | 111.092582 | 0.31145862 | 0.26592574 | 1.17122402 | 0.24150874 | 0.80072691 |
| RLF | 675.565037 | -0.1851585 | 0.37113432 | -0.4988989 | 0.61785064 | 0.94244117 |
| ZMPSTE24 | 217.363795 | -0.108411 | 0.20542356 | -0.527744 | 0.59767706 | 0.9367387 |
| COL9A2 | 158.449471 | 1.14592772 | 0.42452477 | 2.69931889 | 0.00694816 | 0.20503022 |
| SMAP2 | 86.9380736 | -0.2107766 | 0.28273464 | -0.7454926 | 0.455974 | 0.89626521 |
| AL603839.3 | 68.2766557 | 0.49413707 | 0.42351368 | 1.16675587 | 0.24330897 | 0.80072691 |
| ZNF684 | 124.571708 | 0.81855781 | 0.37198425 | 2.20051738 | 0.02777021 | 0.40380659 |
| NFYC | 306.69665 | 0.40504648 | 0.35476286 | 1.14173869 | 0.25356265 | 0.80414079 |
| SLFNL1-AS1 | 40.9777881 | 0.0937247 | 0.50606492 | 0.18520291 | 0.85306989 | 0.97718353 |
| SCMH1 | 1323.74787 | 0.00843357 | 0.30284598 | 0.02784772 | 0.97778361 | 0.99696835 |
| AC093151.3 | 33.4345019 | 0.08950264 | 0.80454294 | 0.11124657 | 0.91142082 | 0.98788932 |
| HIVEP3 | 127.687032 | 0.74308836 | 0.40853133 | 1.81892625 | 0.06892268 | 0.55917298 |
| FOXJ3 | 524.9843 | -0.2117391 | 0.20554373 | -1.0301414 | 0.30294366 | 0.83709431 |
| RIMKLA | 36.7382704 | 0.29582878 | 0.62560363 | 0.47286934 | 0.6363064 | 0.94821838 |
| PPCS | 118.343045 | 0.51473526 | 0.32771982 | 1.57065649 | 0.11626246 | 0.65257994 |
| CCDC30 | 486.914119 | 0.30525202 | 0.26454821 | 1.15386161 | 0.24855691 | 0.80097144 |
| PPIH | 61.3547535 | -0.1518599 | 0.24873615 | -0.6105262 | 0.54151331 | 0.92443671 |
| YBX1 | 446.06213 | -0.0843694 | 0.49655861 | -0.1699082 | 0.86508232 | 0.98000894 |
| SVBP | 41.6130875 | 0.08017593 | 0.3085108 | 0.25988045 | 0.79495599 | 0.9653598 |
| ERMAP | 67.5376608 | 0.13228348 | 0.34876088 | 0.37929564 | 0.70446834 | 0.95707594 |
| SLC2A1 | 675.977665 | -1.5624983 | 0.83376548 | -1.8740261 | 0.06092683 | 0.5418629 |
| SLC2A1-AS1 | 75.3709917 | -0.7779436 | 0.68295596 | -1.1390832 | 0.25466847 | 0.80414079 |
| EBNA1BP2 | 90.7587039 | 0.01376125 | 0.2806931 | 0.04902597 | 0.9608986 | 0.99635733 |

| | | | | | | |
|---|---|---|---|---|---|---|
| TMEM125 | 50.1898641 | 0.62561648 | 0.37523473 | 1.66726699 | 0.09546133 | 0.61367241 |
| C1orf210 | 36.6032379 | 0.43196777 | 0.4301253 | 1.00428357 | 0.31524195 | 0.84285425 |
| ELOVL1 | 76.7984395 | 0.21010903 | 0.27643439 | 0.76006834 | 0.44721373 | 0.89187439 |
| MED8 | 40.6126208 | -0.1533362 | 0.34154978 | -0.4489426 | 0.65347307 | 0.95100227 |
| SZT2 | 117.89245 | -0.3009971 | 0.25237467 | -1.1926599 | 0.2330026 | 0.7926597 |
| HYI | 43.6187533 | 0.20051392 | 0.37298413 | 0.53759371 | 0.59085757 | 0.93483536 |
| PTPRF | 592.603403 | 0.28875574 | 0.30026332 | 0.96167502 | 0.33621287 | 0.84565588 |
| IPO13 | 33.427407 | 0.11379136 | 0.37086896 | 0.30682364 | 0.75897761 | 0.96180522 |
| ATP6V0B | 366.475306 | 0.4299323 | 0.35484525 | 1.21160503 | 0.22566361 | 0.78982769 |
| B4GALT2 | 44.8637001 | 0.04326025 | 0.44080531 | 0.09813912 | 0.92182182 | 0.99051393 |
| CCDC24 | 106.546895 | 0.49544405 | 0.24647866 | 2.01008903 | 0.04442177 | 0.48009422 |
| DMAP1 | 63.1790279 | -0.0497075 | 0.25437659 | -0.195409 | 0.84507279 | 0.97590775 |
| ERI3 | 258.768972 | 0.3484265 | 0.28577602 | 1.2192293 | 0.22275718 | 0.78821341 |
| RNF220 | 76.2152161 | 0.10507224 | 0.27250957 | 0.38557267 | 0.69981317 | 0.95672887 |
| TMEM53 | 45.4225601 | -0.0264704 | 0.34597782 | -0.076509 | 0.93901413 | 0.99243654 |
| ARMH1 | 70.6893567 | 0.26648898 | 0.45184171 | 0.58978393 | 0.55533552 | 0.92872652 |
| RPS8 | 771.956117 | 0.17477358 | 0.44493794 | 0.39280439 | 0.69446396 | 0.95672887 |
| EIF2B3 | 140.597869 | -0.1336434 | 0.31259957 | -0.4275226 | 0.66899869 | 0.95298441 |
| HECTD3 | 38.3156741 | -0.1673677 | 0.27802023 | -0.6019984 | 0.54717522 | 0.92639682 |
| UROD | 44.1961927 | -0.0001199 | 0.42502599 | -0.0002821 | 0.99977495 | 0.9999633 |
| ZSWIM5 | 221.230743 | -0.1753324 | 0.50461042 | -0.3474609 | 0.72824512 | 0.95985088 |
| TESK2 | 73.7245193 | -0.3719263 | 0.40111735 | -0.9272258 | 0.35380932 | 0.8557799 |
| PRDX1 | 1214.13835 | -0.0212866 | 0.60951443 | -0.0349239 | 0.97214039 | 0.99656026 |
| AKR1A1 | 78.4198128 | -0.5819251 | 0.33600628 | -1.7318875 | 0.0832936 | 0.59231889 |
| NASP | 283.002052 | -0.6256611 | 0.31391184 | -1.9931108 | 0.04624932 | 0.48600281 |
| GPBP1L1 | 693.222781 | -0.2685374 | 0.17735641 | -1.5141118 | 0.12999748 | 0.67275878 |
| TMEM69 | 50.9990297 | 0.02785268 | 0.33420017 | 0.08334132 | 0.93358015 | 0.99243654 |
| IPP | 141.832707 | -0.1662289 | 0.238999 | -0.6955212 | 0.48672872 | 0.9070379 |
| MAST2 | 164.04168 | -0.1278238 | 0.21447078 | -0.5959962 | 0.55117778 | 0.92837402 |
| PIK3R3 | 33.9505263 | -0.054113 | 0.5111599 | -0.1058631 | 0.91569097 | 0.98868699 |
| TSPAN1 | 1304.56006 | 0.34476399 | 0.34549418 | 0.99788653 | 0.31833439 | 0.84285425 |
| POMGNT1 | 133.678431 | -0.0016532 | 0.29519267 | -0.0056004 | 0.99553156 | 0.99963311 |
| LURAP1 | 140.360903 | 0.21350032 | 0.2177028 | 0.98069625 | 0.32674255 | 0.84381657 |
| LRRC41 | 146.991187 | 0.17807388 | 0.27768388 | 0.64128273 | 0.52133901 | 0.91787515 |
| UQCRH | 611.542952 | 0.15469947 | 0.47212129 | 0.32766891 | 0.74316201 | 0.96105014 |
| NSUN4 | 98.0989004 | -0.376165 | 0.20571033 | -1.8286148 | 0.06745733 | 0.55848603 |
| FAAH | 49.5651106 | -0.138738 | 0.2879097 | -0.4818803 | 0.62989095 | 0.94596555 |
| MKNK1 | 116.041428 | 0.34546244 | 0.24951605 | 1.38452992 | 0.16619625 | 0.72054851 |
| ATPAF1 | 155.470703 | 0.44871264 | 0.19349659 | 2.31896924 | 0.0203967 | 0.34310989 |
| EFCAB14-AS | 43.6837064 | 1.14412324 | 0.67701553 | 1.68995125 | 0.09103728 | 0.60962984 |
| EFCAB14 | 566.61901 | 0.29371928 | 0.30000848 | 0.9790366 | 0.3275619 | 0.84381657 |
| PDZK1IP1 | 162.05664 | 0.2091228 | 0.77237594 | 0.27075261 | 0.78658131 | 0.96513523 |
| CMPK1 | 228.748195 | 0.23853767 | 0.23725556 | 1.00540392 | 0.31470239 | 0.84285425 |
| TRABD2B | 746.205175 | 0.59162452 | 0.57206521 | 1.03419069 | 0.30104702 | 0.83709431 |
| SPATA6 | 474.78239 | 0.10478115 | 0.2500282 | 0.41907732 | 0.67515963 | 0.95524793 |
| AGBL4 | 2350.09715 | 0.80240725 | 0.351558 | 2.28243206 | 0.02246384 | 0.35887361 |
| FAF1 | 1284.76968 | -0.1481776 | 0.16940564 | -0.8746908 | 0.38174215 | 0.86593382 |
| RNF11 | 268.201759 | -0.12841 | 0.17503753 | -0.7336141 | 0.46318395 | 0.89801718 |
| TTC39A | 157.249802 | -0.8115263 | 0.35350122 | -2.2956816 | 2.17E-02 | 0.35579666 |
| EPS15 | 648.44047 | -0.3364123 | 0.1852029 | -1.8164527 | 0.06930096 | 0.55917298 |
| OSBPL9 | 592.962565 | -0.1590255 | 0.15943284 | -0.9974452 | 0.31854846 | 0.84285425 |
| NRDC | 502.024415 | -0.3652059 | 0.21003887 | -1.7387538 | 0.08207808 | 0.59202053 |
| TXNDC12 | 110.351929 | 0.14560614 | 0.23957179 | 0.60777667 | 0.54333561 | 0.92482794 |
| BTF3L4 | 211.674133 | 0.07248331 | 0.20046063 | 0.36158375 | 0.71766311 | 0.95796325 |

| | | | | | | |
|---|---|---|---|---|---|---|
| ZFYVE9 | 364.354859 | 0.17227701 | 0.21937857 | 0.78529553 | 0.43228032 | 0.88707611 |
| CC2D1B | 74.0899405 | -0.8277302 | 0.36537335 | -2.2654368 | 0.02348589 | 0.36567438 |
| PRPF38A | 82.230589 | 0.0417629 | 0.25690701 | 0.16256038 | 0.87086459 | 0.98208511 |
| TUT4 | 633.014317 | -0.0335501 | 0.22382979 | -0.1498912 | 0.88085049 | 0.98333507 |
| ZYG11B | 270.619629 | -0.1336691 | 0.21284422 | -0.6280139 | 0.52999484 | 0.92181514 |
| ECHDC2 | 597.828245 | 0.49117565 | 0.29219155 | 1.68100568 | 0.09276181 | 0.61218122 |
| SCP2 | 499.574198 | 0.21049966 | 0.18956719 | 1.11042241 | 0.26681704 | 0.81351996 |
| CPT2 | 67.1596139 | 0.69424124 | 0.27725097 | 2.50401738 | 0.0122792 | 0.27034878 |
| C1orf123 | 53.1150534 | 0.34272894 | 0.38504146 | 0.8901092 | 0.37340726 | 0.86201082 |
| MAGOH | 89.701129 | 0.21104596 | 0.35690195 | 0.59132757 | 0.55430096 | 0.92872652 |
| LRP8 | 58.1671802 | -0.0792026 | 0.48204168 | -0.1643066 | 0.86948981 | 0.9820194 |
| NDC1 | 100.952854 | 0.07472489 | 0.31523897 | 0.23704204 | 0.81262418 | 0.9680627 |
| YIPF1 | 56.8358815 | 0.01825159 | 0.279027 | 0.06541157 | 0.94784631 | 0.99314907 |
| HSPB11 | 112.95165 | -0.1523063 | 0.25434211 | -0.5988246 | 0.54928984 | 0.9274143 |
| LRRC42 | 59.8658634 | 0.09961204 | 0.31167846 | 0.31959873 | 0.74927254 | 0.96109466 |
| TMEM59 | 535.176843 | 0.70497766 | 0.34511 | 2.04276219 | 0.04107599 | 0.46430138 |
| TCEANC2 | 76.0253457 | 0.33835157 | 0.29332757 | 1.15349392 | 0.24870771 | 0.80097144 |
| MRPL37 | 53.7681798 | -0.0965694 | 0.31168841 | -0.3098266 | 0.75669278 | 0.96133693 |
| SSBP3 | 464.006012 | -0.4110348 | 0.51452939 | -0.7988558 | 0.42437401 | 0.88391318 |
| ACOT11 | 534.401997 | -0.0356042 | 0.40813168 | -0.0872371 | 0.93048308 | 0.99231329 |
| MROH7-TTC4 | 47.881913 | -0.2821025 | 0.52769752 | -0.5345913 | 0.59293252 | 0.93491607 |
| TTC22 | 31.9160757 | -0.1257019 | 0.39555262 | -0.3177881 | 0.75064571 | 0.96109466 |
| DHCR24 | 149.888359 | -0.1467531 | 0.63036405 | -0.2328069 | 0.81591139 | 0.96844493 |
| USP24 | 713.964448 | -0.340204 | 0.21790697 | -1.5612351 | 0.11846828 | 0.65447949 |
| PLPP3 | 235.205851 | -0.3405393 | 0.31043556 | -1.0969725 | 0.27265341 | 0.81922962 |
| PRKAA2 | 703.901679 | 0.39532064 | 0.24373827 | 1.62190632 | 0.10482341 | 0.6317386 |
| FYB2 | 1496.87844 | 0.48491007 | 0.47797486 | 1.01450956 | 0.31033967 | 0.84213522 |
| DAB1 | 463.746613 | 0.48080881 | 0.80121492 | 0.60009966 | 0.54843982 | 0.92704021 |
| TACSTD2 | 659.512064 | 0.44741088 | 0.67654735 | 0.66131496 | 0.50841035 | 0.91250907 |
| MYSM1 | 290.686547 | -0.2836301 | 0.19156458 | -1.480598 | 0.13871373 | 0.68156449 |
| JUN | 302.306899 | -0.5230594 | 0.67568516 | -0.7741171 | 0.43886156 | 0.88909126 |
| LINC01135 | 126.977163 | 0.98949389 | 0.35366978 | 2.79779031 | 0.00514535 | 0.1779646 |
| FGGY | 468.51233 | -0.1994269 | 0.31829363 | -0.6265501 | 0.53095418 | 0.92211604 |
| HOOK1 | 354.135637 | 0.40412752 | 0.2820673 | 1.43273438 | 0.15193376 | 0.7036084 |
| NFIA | 816.862283 | 0.42399631 | 0.40158589 | 1.05580481 | 0.2910574 | 0.83183872 |
| TM2D1 | 278.020816 | -0.3449957 | 0.16047411 | -2.1498528 | 0.03156686 | 0.41477661 |
| PATJ | 3650.1527 | 0.17531362 | 0.19345639 | 0.90621775 | 0.36482061 | 0.85866868 |
| USP1 | 77.816446 | -0.0094884 | 0.22614255 | -0.0419578 | 0.96653236 | 0.99637506 |
| DOCK7 | 1123.02204 | -0.1362806 | 0.2332514 | -0.5842651 | 0.55904198 | 0.92901206 |
| AC103923.1 | 48.3170134 | 0.67262866 | 0.43303049 | 1.55330556 | 0.12035015 | 0.65691125 |
| ATG4C | 138.527406 | 0.1407362 | 0.30235695 | 0.46546375 | 0.64159939 | 0.94935885 |
| ALG6 | 59.3838369 | -0.1541997 | 0.2467228 | -0.6249917 | 0.53197649 | 0.92211604 |
| ITGB3BP | 200.177768 | 0.51257553 | 0.44328425 | 1.15631343 | 0.24755298 | 0.80072691 |
| EFCAB7 | 117.151134 | 0.14043698 | 0.26427586 | 0.531403 | 0.59513954 | 0.9353279 |
| PGM1 | 162.36331 | 0.1517188 | 0.37618125 | 0.40331305 | 0.68671795 | 0.9558749 |
| ROR1 | 100.65168 | -0.9813763 | 0.74218055 | -1.3222877 | 0.18607235 | 0.74938192 |
| CACHD1 | 199.829066 | 0.36255846 | 0.66552256 | 0.54477261 | 0.58590992 | 0.93483536 |
| RAVER2 | 163.106843 | -0.078866 | 0.28613257 | -0.2756276 | 0.78283412 | 0.96513523 |
| JAK1 | 846.864383 | 0.2057128 | 0.30233558 | 0.68041214 | 0.49624354 | 0.90839719 |
| AK4 | 134.859561 | -1.0113429 | 0.71631764 | -1.4118638 | 0.15799007 | 0.71264724 |
| DNAJC6 | 52.3023997 | -0.2579023 | 0.40473196 | -0.6372176 | 0.52398313 | 0.91871245 |
| LEPR | 234.24322 | -0.406926 | 0.39233995 | -1.0371771 | 0.29965333 | 0.83618229 |
| PDE4B | 123.38708 | -1.8623902 | 1.32010155 | -1.4107932 | 0.15830561 | 0.71264724 |
| WDR78 | 211.696445 | 0.46061928 | 0.35679912 | 1.2909765 | 0.19671183 | 0.76478825 |

| | | | | | | |
|---|---|---|---|---|---|---|
| MIER1 | 435.708116 | -0.1214666 | 0.18760301 | -0.6474661 | 0.51733033 | 0.91751084 |
| SLC35D1 | 135.018543 | -0.004464 | 0.26843319 | -0.0166298 | 0.98673193 | 0.99791717 |
| SERBP1 | 489.1971 | 0.40107973 | 0.28371728 | 1.41365983 | 0.1574618 | 0.71160548 |
| GADD45A | 102.858025 | 0.18154837 | 0.32937693 | 0.55118727 | 0.58150531 | 0.93386308 |
| GNG12 | 682.259091 | -0.1357972 | 0.15129649 | -0.8975569 | 0.36942181 | 0.86020443 |
| GNG12-AS1 | 224.50603 | -0.3254904 | 0.3176619 | -1.0246442 | 0.30553109 | 0.83709431 |
| WLS | 304.941344 | 0.39054597 | 0.32011106 | 1.22003272 | 0.22245247 | 0.78760199 |
| LRRC40 | 133.813838 | -0.3681321 | 0.20766293 | -1.7727386 | 0.07627203 | 0.57145422 |
| SRSF11 | 689.456573 | 0.05142436 | 0.1947421 | 0.26406391 | 0.79173067 | 0.9653598 |
| ANKRD13C | 398.887495 | -0.1950412 | 0.33873594 | -0.5757912 | 0.56475632 | 0.93104392 |
| HHLA3 | 42.345526 | 0.30430305 | 0.41033431 | 0.74159787 | 0.45833101 | 0.89747927 |
| LINC01788 | 163.892487 | 0.40120901 | 0.65683856 | 0.61081829 | 0.54131988 | 0.92443671 |
| PTGER3 | 884.465057 | 0.43280977 | 0.40885123 | 1.05859966 | 0.28978214 | 0.83103268 |
| ZRANB2-AS1 | 46.6181173 | 0.62397267 | 0.64289891 | 0.97056109 | 0.33176689 | 0.84442294 |
| ZRANB2 | 563.890686 | -0.2262508 | 0.17846013 | -1.2677946 | 0.2048713 | 0.77193931 |
| NEGR1 | 46.8095705 | 0.7170773 | 0.47791628 | 1.50042449 | 0.13350448 | 0.67640892 |
| LRRIQ3 | 42.0070767 | -0.092216 | 0.56943024 | -0.1619444 | 0.87134967 | 0.98208511 |
| CRYZ | 404.303801 | 0.55788939 | 0.20400774 | 2.73464813 | 0.0062447 | 0.19425932 |
| TYW3 | 92.9211729 | 0.20496068 | 0.20593392 | 0.995274 | 0.31960302 | 0.84285425 |
| SLC44A5 | 97.5813723 | -1.3665651 | 1.27758091 | -1.0696505 | 0.28477663 | 0.82781987 |
| ACADM | 290.414273 | 0.42863126 | 0.17578399 | 2.43839755 | 0.01475254 | 0.29134933 |
| RABGGTB | 65.7560337 | -0.0881829 | 0.29256581 | -0.3014121 | 0.76310027 | 0.96285313 |
| ST6GALNAC3 | 55.233953 | 0.10239335 | 0.44064028 | 0.23237403 | 0.81624752 | 0.96844493 |
| PIGK | 173.062297 | 0.18559369 | 0.18213053 | 1.01901474 | 0.30819597 | 0.83980597 |
| AC118549.1 | 729.749476 | -0.1888043 | 0.2625992 | -0.7189828 | 0.47215153 | 0.90188689 |
| USP33 | 467.633953 | -0.188307 | 0.20922987 | -0.9000007 | 0.36811987 | 0.85902833 |
| MIGA1 | 341.68976 | -0.0286548 | 0.18550962 | -0.1544653 | 0.87724285 | 0.98277085 |
| NEXN | 36.8179886 | 0.73788519 | 0.42075387 | 1.75372168 | 0.07947821 | 0.58461232 |
| FUBP1 | 370.803236 | -0.0694184 | 0.18414674 | -0.3769732 | 0.70619352 | 0.95730303 |
| PTGFR | 315.240855 | 0.96561912 | 0.42470523 | 2.27362193 | 0.02298873 | 0.36283414 |
| IFI44L | 360.955907 | -4.285652 | 1.62290846 | -2.6407232 | 0.00827293 | 0.22277119 |
| IFI44 | 123.186765 | -2.5148728 | 0.86750005 | -2.8989887 | 0.00374368 | 0.14917797 |
| AL136234.1 | 104.297624 | 0.27413967 | 0.64067803 | 0.42788991 | 0.66873127 | 0.95298441 |
| LINC01781 | 73.0825915 | 0.45758942 | 0.66896771 | 0.68402319 | 0.49396052 | 0.90839719 |
| ADGRL2 | 961.92353 | 0.27058303 | 0.26917925 | 1.00521504 | 0.31479331 | 0.84285425 |
| LINC01362 | 53.0282692 | 0.66150359 | 0.74181914 | 0.89173164 | 0.37253679 | 0.86201082 |
| LINC01725 | 732.977474 | 0.02336382 | 0.41126473 | 0.05680969 | 0.95469679 | 0.99537617 |
| PRKACB | 350.858025 | 0.42976511 | 0.25138406 | 1.70959574 | 0.08734065 | 0.60617884 |
| SAMD13 | 40.4538776 | 0.88032325 | 0.52457281 | 1.67817171 | 0.09331358 | 0.61317397 |
| RPF1 | 55.0169987 | 0.47586379 | 0.2919843 | 1.62975815 | 0.10315262 | 0.62635087 |
| GNG5 | 292.930632 | 0.11543196 | 0.37219198 | 0.31014091 | 0.7564538 | 0.96127511 |
| SPATA1 | 61.407825 | 0.49426341 | 0.52784182 | 0.93638548 | 0.34907475 | 0.8535343 |
| CTBS | 100.944422 | 0.02614863 | 0.3308819 | 0.07902709 | 0.93701107 | 0.99243654 |
| SSX2IP | 64.8020427 | 0.01795302 | 0.27154935 | 0.06611329 | 0.94728763 | 0.99314907 |
| WDR63 | 58.5709384 | -7.72E-05 | 0.46633067 | -0.0001656 | 0.99986789 | 0.9999633 |
| MCOLN3 | 56.0659469 | -0.6697931 | 0.55605528 | -1.2045441 | 0.22837935 | 0.79042788 |
| SYDE2 | 62.3358545 | 0.1657338 | 0.26859179 | 0.61704716 | 0.53720362 | 0.92324352 |
| C1orf52 | 110.751507 | -0.2372588 | 0.27896518 | -0.850496 | 0.3950494 | 0.87344256 |
| BCL10 | 124.082712 | 0.13510929 | 0.36861514 | 0.36653212 | 0.71396805 | 0.95791799 |
| AL078459.1 | 234.143691 | 0.74320644 | 0.45221258 | 1.64348906 | 0.10028179 | 0.61891077 |
| DDAH1 | 921.460836 | 0.14826928 | 0.197438 | 0.75096629 | 0.45267295 | 0.89520141 |
| CYR61 | 51.9484265 | 1.44878227 | 0.85808282 | 1.68839445 | 0.09133554 | 0.6104569 |
| ZNHIT6 | 204.275066 | -0.0230191 | 0.28182581 | -0.0816786 | 0.93490231 | 0.99243654 |
| AL136382.1 | 41.2595926 | 0.22593096 | 0.64557118 | 0.34997064 | 0.72636073 | 0.95979351 |

| | | | | | | |
|---|---|---|---|---|---|---|
| ODF2L | 542.036972 | -0.8067612 | 0.2092873 | -3.8548026 | 0.00011582 | 0.02194494 |
| SH3GLB1 | 345.744949 | 0.14911348 | 0.17717364 | 0.84162338 | 0.3999988 | 0.87661803 |
| SELENOF | 345.822168 | -0.004468 | 0.16600956 | -0.0269143 | 0.97852811 | 0.99696835 |
| HS2ST1 | 225.409101 | -0.2071043 | 0.25988778 | -0.796899 | 0.4255097 | 0.88403732 |
| LMO4 | 174.429873 | 0.67507762 | 0.35511047 | 1.90103554 | 0.05729736 | 0.5267336 |
| PKN2-AS1 | 60.0801645 | 0.79806114 | 0.44076321 | 1.81063466 | 0.07019742 | 0.5621116 |
| PKN2 | 812.031449 | -0.2102007 | 0.16624328 | -1.2644165 | 0.20608057 | 0.77193931 |
| GTF2B | 100.904313 | 0.0936726 | 0.25514077 | 0.36714084 | 0.71351396 | 0.95791799 |
| KYAT3 | 179.332074 | 0.19209404 | 0.22768892 | 0.84366881 | 0.3988545 | 0.87575847 |
| GBP3 | 125.157947 | 0.46276141 | 0.37558162 | 1.23211942 | 0.21790448 | 0.78394745 |
| GBP2 | 88.8645168 | -0.6120824 | 0.56948919 | -1.074792 | 2.82E-01 | 0.82666614 |
| LRRC8B | 202.127395 | -0.1531641 | 0.23548837 | -0.6504107 | 0.51542699 | 0.91612355 |
| LRRC8D | 36.3676859 | -0.0485185 | 0.34640938 | -0.1400612 | 0.88861163 | 0.98551241 |
| ZNF326 | 217.368938 | -0.1865876 | 0.18206824 | -1.0248224 | 0.30544699 | 0.83709431 |
| ZNF644 | 581.254403 | -0.1248181 | 0.16267163 | -0.7673008 | 0.44290272 | 0.89005186 |
| HFM1 | 44.8428067 | 0.39573381 | 0.38315203 | 1.03283756 | 0.30167992 | 0.83709431 |
| TGFBR3 | 240.03009 | -0.7376467 | 0.64271939 | -1.1476963 | 0.25109396 | 0.80240987 |
| BTBD8 | 129.865689 | 0.09356251 | 0.22474863 | 0.41629848 | 0.6771916 | 0.95536622 |
| GLMN | 121.637575 | -0.2685122 | 0.25982468 | -1.0334359 | 0.30139993 | 0.83709431 |
| RPAP2 | 366.897953 | 0.00567057 | 0.18795124 | 0.03017044 | 0.97593113 | 0.99656026 |
| EVI5 | 800.152032 | -0.1087017 | 0.23114442 | -0.4702759 | 0.6381579 | 0.94863756 |
| RPL5 | 544.570179 | 0.34945378 | 0.33965025 | 1.0288636 | 0.30354378 | 0.83709431 |
| FAM69A | 83.2111835 | 0.27821318 | 0.31131138 | 0.89368138 | 0.37149239 | 0.86201082 |
| MTF2 | 456.165574 | -0.1839966 | 0.21342608 | -0.8621094 | 0.3886273 | 0.86970192 |
| TMED5 | 449.557184 | 0.20627708 | 0.36191631 | 0.56995795 | 0.56870622 | 0.93122667 |
| CCDC18 | 116.253877 | -0.4303646 | 0.41171562 | -1.0452958 | 0.29588626 | 0.83444322 |
| DR1 | 113.680114 | 0.33474701 | 0.24489696 | 1.36688921 | 0.17166004 | 0.72776437 |
| FNBP1L | 1483.92658 | -0.1230473 | 0.32813303 | -0.3749921 | 0.70766637 | 0.95752169 |
| BCAR3 | 732.260826 | 0.28631015 | 0.42258004 | 0.6775288 | 0.4980705 | 0.90839719 |
| DNTTIP2 | 50.1179814 | -0.2319827 | 0.44212897 | -0.5246947 | 0.59979543 | 0.93728325 |
| GCLM | 136.349628 | -0.1973368 | 0.33722941 | -0.5851708 | 5.58E-01 | 0.92901206 |
| ABCA4 | 88.3571112 | 1.02081923 | 0.71825255 | 1.42125389 | 0.15524296 | 0.70907003 |
| ARHGAP29 | 2060.67737 | 0.72342995 | 0.40730182 | 1.77615202 | 0.07570786 | 0.57072562 |
| ABCD3 | 565.260758 | 0.27422124 | 0.22822017 | 1.20156446 | 0.22953232 | 0.79047378 |
| SLC44A3 | 257.175071 | 0.4215671 | 0.37108906 | 1.13602676 | 0.25594537 | 0.80525591 |
| CNN3 | 533.100164 | -0.0131012 | 0.34830961 | -0.0376136 | 0.96999577 | 0.9965254 |
| AC105942.1 | 51.4473948 | -0.1064181 | 0.33460695 | -0.318039 | 0.75045535 | 0.96109466 |
| ALG14 | 109.447745 | -0.2369829 | 0.30334983 | -0.7812198 | 0.43467325 | 0.88785137 |
| TMEM56-RW | 33.2226697 | -0.0951401 | 0.43445501 | -0.2189872 | 0.82666002 | 0.97068874 |
| AC092802.1 | 59.4015085 | 0.58534272 | 0.50486802 | 1.15939749 | 0.2462942 | 0.80072691 |
| PTBP2 | 683.057643 | -0.2341789 | 0.21216782 | -1.1037435 | 0.2697044 | 0.81817548 |
| DPYD | 556.313727 | 0.03015129 | 0.2376574 | 0.12686873 | 0.8990443 | 0.98660836 |
| SNX7 | 147.560779 | -0.0670378 | 0.23624773 | -0.2837608 | 0.77659368 | 0.96452972 |
| FRRS1 | 114.439575 | -0.6938566 | 0.44558284 | -1.5571889 | 0.11942564 | 0.65527787 |
| AGL | 257.337226 | -0.563749 | 0.23353553 | -2.4139751 | 0.01577954 | 0.30067202 |
| SASS6 | 37.4204082 | -0.2652601 | 0.35020724 | -0.7574375 | 0.44878781 | 0.89314398 |
| TRMT13 | 147.239516 | 0.01831263 | 0.25617155 | 0.07148581 | 0.94301111 | 0.99265578 |
| DBT | 200.80657 | 0.27306451 | 0.18852975 | 1.44838949 | 0.14750815 | 0.69834914 |
| RTCA | 108.695249 | -0.0034977 | 0.27502732 | -0.0127176 | 0.98985309 | 0.99861959 |
| CDC14A | 111.525575 | 0.54223708 | 0.35454618 | 1.52938349 | 0.1261694 | 0.66795428 |
| EXTL2 | 85.206148 | -0.0949641 | 0.29325874 | -0.3238236 | 0.74607159 | 0.96109466 |
| SLC30A7 | 274.990959 | -0.330781 | 0.21553702 | -1.5346832 | 0.12486167 | 0.66626798 |
| DPH5 | 64.5952301 | 0.09692231 | 0.23110255 | 0.41939094 | 0.67493045 | 0.95524793 |
| AC093157.1 | 90.0533258 | 0.30060901 | 0.26113767 | 1.15115146 | 0.24966993 | 0.80149394 |

| | | | | | | |
|---|---|---|---|---|---|---|
| RNPC3 | 356.64996 | -0.2622864 | 0.29149995 | -0.899782 | 0.36823626 | 0.85910864 |
| AMY2B | 95.5061295 | -0.3577624 | 0.34028266 | -1.0513683 | 0.29308948 | 0.83336457 |
| PRMT6 | 31.6889994 | 0.41639396 | 0.35700083 | 1.16636692 | 0.24346613 | 0.80072691 |
| VAV3 | 1724.28945 | 0.32791346 | 0.40838929 | 0.80294332 | 0.42200749 | 0.88371155 |
| SLC25A24 | 152.393984 | 0.41180897 | 0.20709582 | 1.98849481 | 0.04675699 | 0.48600281 |
| FAM102B | 82.8298527 | -1.04304 | 0.48860336 | -2.1347376 | 0.03278245 | 0.42102945 |
| PRPF38B | 246.458005 | -0.0787268 | 0.28239667 | -0.278781 | 0.78041293 | 0.96513523 |
| STXBP3 | 293.255599 | 0.04568045 | 0.17782114 | 0.25688987 | 7.97E-01 | 0.9653598 |
| AL449266.1 | 42.8219428 | -0.6266764 | 0.36523847 | -1.7158008 | 0.08619851 | 0.60143834 |
| GPSM2 | 65.0829203 | 0.16874731 | 0.45307463 | 0.37244926 | 0.70955838 | 0.95752169 |
| WDR47 | 71.3303453 | -0.2342624 | 0.25522474 | -0.9178671 | 0.35868846 | 0.8557799 |
| TAF13 | 56.9810052 | 0.01358341 | 0.36530499 | 0.03718374 | 0.9703385 | 0.99656026 |
| TMEM167B | 87.1325128 | 0.0107384 | 0.27850696 | 0.03855702 | 0.96924357 | 0.99645289 |
| SARS | 183.061209 | 0.10492601 | 0.17965248 | 0.58404989 | 0.55918674 | 0.92901206 |
| CELSR2 | 143.874565 | 0.37271463 | 0.35675294 | 1.04474156 | 0.29614243 | 0.83451806 |
| SORT1 | 341.391809 | 0.35027354 | 0.17722805 | 1.9764002 | 0.04810947 | 0.48661187 |
| PSMA5 | 153.973659 | -0.2720708 | 0.2783019 | -0.9776101 | 0.32826719 | 0.84381657 |
| GNAI3 | 271.555535 | -0.2828992 | 0.22205565 | -1.2740014 | 0.20266293 | 0.76996624 |
| AMPD2 | 53.910004 | -0.0010012 | 0.39420371 | -0.0025398 | 0.99797357 | 0.99967893 |
| GSTM4 | 70.8293538 | 0.38176658 | 0.33183441 | 1.15047314 | 0.24994905 | 0.80149394 |
| GSTM3 | 637.994537 | 0.18133063 | 0.53235843 | 0.34061756 | 0.73339151 | 0.96035038 |
| AHCYL1 | 490.272107 | 0.37811745 | 0.21573154 | 1.75272214 | 0.07964971 | 0.58461232 |
| STRIP1 | 72.8918776 | 0.20607983 | 0.24094673 | 0.85529209 | 0.39238946 | 0.87123763 |
| KCNC4 | 182.659882 | 0.02839418 | 0.47055886 | 0.06034141 | 0.95188372 | 0.99419388 |
| RBM15 | 86.2073164 | 0.37166284 | 0.29573973 | 1.25672276 | 0.20885404 | 0.77436433 |
| SLC16A4 | 512.67367 | -0.3148245 | 0.40332302 | -0.7805765 | 0.4350516 | 0.88785137 |
| AL355488.1 | 38.5583839 | 0.18556576 | 0.33278175 | 0.55762 | 0.5771039 | 0.93319686 |
| LAMTOR5 | 487.422757 | 0.36170641 | 0.47490874 | 0.76163351 | 0.44627877 | 0.8918683 |
| LRIF1 | 48.8936316 | 0.5034394 | 0.46771079 | 1.07639039 | 0.2817527 | 0.82661559 |
| DRAM2 | 135.060528 | 0.08811941 | 0.1710967 | 0.51502696 | 0.60653418 | 0.93919596 |
| CEPT1 | 156.270568 | -0.0974875 | 0.23129371 | -0.421488 | 0.67339874 | 0.95471033 |
| DENND2D | 101.221173 | -0.018637 | 0.351771 | -0.0529804 | 0.95774755 | 0.99579587 |
| WDR77 | 45.3452344 | -0.0794764 | 0.41610814 | -0.1909993 | 0.84852615 | 0.97656375 |
| ATP5PB | 310.354266 | 0.68736566 | 0.43402139 | 1.58371378 | 0.11325887 | 0.64757575 |
| RAP1A | 437.816402 | 0.00184375 | 0.16514689 | 0.01116428 | 0.99109238 | 0.99889703 |
| CTTNBP2NL | 186.150856 | 0.01060048 | 0.29259466 | 0.03622924 | 0.97109957 | 0.99656026 |
| ST7L | 191.043689 | -0.0073597 | 0.20323053 | -0.0362137 | 0.97111193 | 0.99656026 |
| CAPZA1 | 377.554835 | -0.1319131 | 0.23043271 | -0.5724581 | 0.56701169 | 0.93122667 |
| MOV10 | 103.246369 | 0.06312302 | 0.32894249 | 0.19189683 | 0.84782302 | 0.97656375 |
| RHOC | 252.177597 | -0.010435 | 0.498467 | -0.0209342 | 0.98329811 | 0.9971769 |
| LRIG2 | 179.508628 | -0.2253702 | 0.23528831 | -0.9578469 | 0.33813995 | 0.84744028 |
| MAGI3 | 1234.11247 | 0.15394922 | 0.33302814 | 0.46227092 | 0.64388705 | 0.94949493 |
| PHTF1 | 158.333284 | -0.1235778 | 0.32268411 | -0.3829683 | 0.70174327 | 0.95672887 |
| RSBN1 | 199.964561 | 0.26033757 | 0.1670776 | 1.55818356 | 0.11918974 | 0.65527787 |
| HIPK1 | 248.552295 | 0.36097116 | 0.16521637 | 2.18483899 | 0.02890065 | 0.40623517 |
| TRIM33 | 572.224699 | -0.2501456 | 0.24753616 | -1.0105417 | 0.31223582 | 0.84279957 |
| BCAS2 | 89.9198533 | 0.0408501 | 0.5131658 | 0.0796041 | 0.93655214 | 0.99243654 |
| NRAS | 69.7578055 | -0.6760901 | 0.31337661 | -2.1574365 | 0.03097167 | 0.41259458 |
| CSDE1 | 620.706592 | 0.10131711 | 0.2101972 | 0.48200981 | 0.62979898 | 0.94596555 |
| SIKE1 | 89.5671357 | -0.1916294 | 0.24891402 | -0.7698616 | 0.44138197 | 0.88964371 |
| VANGL1 | 35.545361 | -0.0415634 | 0.37030419 | -0.1122413 | 0.91063209 | 0.98788932 |
| SLC22A15 | 244.908474 | 0.10425783 | 0.36581704 | 0.28499993 | 0.7756442 | 0.9641503 |
| ATP1A1 | 2055.60026 | 0.51349758 | 0.27986247 | 1.83482116 | 0.06653219 | 0.55602662 |
| LINC01762 | 319.798884 | 2.93397569 | 0.62836153 | 4.66924779 | 3.02E-06 | 0.00176561 |

| | | | | | | |
|---|---|---|---|---|---|---|
| CD58 | 110.199924 | 0.10195938 | 0.23287663 | 0.43782576 | 0.6615126 | 0.95137258 |
| IGSF3 | 151.79031 | 0.24093168 | 0.40068587 | 0.60129817 | 0.54764141 | 0.92673696 |
| PTGFRN | 65.4090099 | 0.00204946 | 0.37985234 | 0.0053954 | 0.99569511 | 0.99967893 |
| TTF2 | 76.8123376 | -0.2220982 | 0.29457586 | -0.7539594 | 0.45087362 | 0.893393 |
| VTCN1 | 290.794849 | 4.09839899 | 0.6470679 | 6.33380054 | 2.39E-10 | 6.27E-07 |
| MAN1A2 | 573.051448 | -0.1177426 | 0.31208259 | -0.3772803 | 0.70596534 | 0.95730303 |
| GDAP2 | 195.125213 | -0.2613669 | 0.21692104 | -1.2048941 | 0.22824419 | 0.79042788 |
| WDR3 | 53.7378582 | 0.22160872 | 0.26782047 | 0.82745251 | 0.40798063 | 0.87948803 |
| SPAG17 | 445.663514 | 0.07628227 | 0.2438125 | 0.31287268 | 0.75437739 | 0.96120879 |
| WARS2 | 159.761388 | -0.4213149 | 0.19926116 | -2.1143854 | 0.03448236 | 0.43072125 |
| AL359915.2 | 345.796083 | -0.9988366 | 0.3978745 | -2.5104314 | 0.01205837 | 0.26883805 |
| PHGDH | 53.7026597 | -0.0114555 | 0.69995387 | -0.0163661 | 0.98694235 | 0.99791717 |
| HMGCS2 | 612.409901 | 0.97050214 | 0.59087203 | 1.64249124 | 0.10048824 | 0.61911626 |
| NOTCH2 | 198.435468 | 0.07562714 | 0.22209681 | 0.34051432 | 0.73346924 | 0.96035038 |
| SEC22B | 204.376146 | -0.2470408 | 0.21092588 | -1.1712207 | 0.24151008 | 0.80072691 |
| NBPF26 | 42.7122275 | -0.255237 | 0.4460625 | -0.5722 | 0.56718647 | 0.93122667 |
| LINC00623 | 499.366113 | 0.23520717 | 0.37136256 | 0.63336263 | 0.52649686 | 0.91974278 |
| SRGAP2C | 113.161826 | 0.52109775 | 0.3335286 | 1.56237799 | 0.11819897 | 0.65447949 |
| AC239800.3 | 85.2034806 | -0.0778054 | 0.41361447 | -0.188111 | 0.85078966 | 0.97718353 |
| NBPF15 | 73.2113098 | -0.0198266 | 0.34013348 | -0.0582905 | 0.95351723 | 0.99514888 |
| FP700111.1 | 66.2046797 | -0.7266645 | 0.30993322 | -2.344584 | 0.01904832 | 0.33090117 |
| SRGAP2B | 123.000387 | 0.43008627 | 0.26632476 | 1.61489403 | 0.10633364 | 0.63461115 |
| GPR89A | 50.4331264 | 0.16476948 | 0.28347422 | 0.58125031 | 0.56107177 | 0.92934517 |
| PDZK1 | 157.620061 | 1.9118012 | 0.46187723 | 4.13919772 | 3.49E-05 | 0.00869647 |
| RNF115 | 486.500655 | 0.2172671 | 0.3300354 | 0.65831453 | 0.51033604 | 0.91334187 |
| POLR3C | 70.4655552 | -0.026398 | 0.28777533 | -0.0917312 | 0.9269116 | 0.99092457 |
| PIAS3 | 34.202358 | 0.13778089 | 0.31512009 | 0.43723297 | 0.6619424 | 0.95142083 |
| PEX11B | 59.8327465 | -0.0751347 | 0.28902155 | -0.2599622 | 0.79489292 | 0.9653598 |
| RBM8A | 139.896246 | 0.25030939 | 0.41210346 | 0.60739454 | 0.54358911 | 0.92482794 |
| POLR3GL | 38.5022252 | 0.38102165 | 0.40955105 | 0.93033982 | 0.35219517 | 0.85536253 |
| TXNIP | 2661.63171 | 1.40247688 | 0.58329301 | 2.40441229 | 0.01619849 | 0.30477595 |
| NBPF12 | 107.685347 | -0.1176845 | 0.26995081 | -0.435948 | 0.66287444 | 0.9515075 |
| PRKAB2 | 97.3375914 | 0.61392373 | 0.40030968 | 1.53362197 | 0.12512268 | 0.6663037 |
| AC242426.2 | 209.913406 | 0.24988254 | 0.29126693 | 0.85791593 | 0.39093888 | 0.87004477 |
| FMO5 | 149.164883 | 0.63463611 | 0.39777609 | 1.59546068 | 0.11060926 | 0.64136059 |
| CHD1L | 146.603318 | 0.18496518 | 0.27631851 | 0.66939121 | 0.50324596 | 0.90964098 |
| BCL9 | 191.448064 | 0.1592009 | 0.38646908 | 0.41193696 | 0.68038563 | 0.95574492 |
| ACP6 | 189.821715 | 0.3876018 | 0.24807453 | 1.5624409 | 0.11818416 | 0.65447949 |
| GPR89B | 63.0748305 | -0.2604726 | 0.34625277 | -0.7522616 | 0.45189378 | 0.89473773 |
| NBPF11 | 101.927454 | -0.0750104 | 0.32073266 | -0.2338721 | 0.81508429 | 0.96839471 |
| AC239809.3 | 72.9340394 | -0.3830015 | 0.4943405 | -0.7747726 | 0.43847404 | 0.88864101 |
| NBPF14 | 49.4595219 | 0.24135995 | 0.32981119 | 0.73181247 | 0.46428303 | 0.89801718 |
| PDE4DIP | 432.267491 | -0.0038588 | 0.42402823 | -0.0091004 | 0.992739 | 0.99919666 |
| NBPF9 | 45.7550977 | -0.314214 | 0.26155444 | -1.2013331 | 0.22962201 | 0.79047378 |
| AC245297.3 | 567.09025 | 0.09607458 | 0.3001432 | 0.3200958 | 0.74889571 | 0.96109466 |
| NBPF19 | 73.685457 | -0.1618197 | 0.63067381 | -0.2565823 | 0.79750126 | 0.9653598 |
| HIST2H2BF | 65.2363784 | 0.28897047 | 0.84243499 | 0.34301813 | 0.73158482 | 0.96008876 |
| HIST2H2BE | 70.7998257 | 0.63659848 | 0.74295659 | 0.85684478 | 0.39153067 | 0.87015937 |
| SF3B4 | 57.2255286 | 0.04581991 | 0.42258216 | 0.10842841 | 0.91365587 | 0.98830345 |
| OTUD7B | 261.362741 | -0.3010928 | 0.24555474 | -1.226174 | 0.22013319 | 0.78633738 |
| VPS45 | 288.328347 | -0.2552555 | 0.19313272 | -1.3216583 | 0.18628196 | 0.74938192 |
| ANP32E | 223.025386 | 0.27790058 | 0.26018746 | 1.06807829 | 0.2854852 | 0.82810567 |
| APH1A | 197.286495 | 0.34979907 | 0.27943988 | 1.2517865 | 0.21064767 | 0.77610028 |
| C1orf54 | 38.1873017 | 1.320402 | 0.47754575 | 2.76497489 | 0.00569272 | 0.187252 |

| | | | | | | |
|---|---|---|---|---|---|---|
| MRPS21 | 282.838576 | 0.1318428 | 0.3406426 | 0.38704142 | 0.69872554 | 0.95672887 |
| PRPF3 | 155.886357 | -0.3926669 | 0.23542483 | -1.6679077 | 0.09533404 | 0.61367241 |
| RPRD2 | 885.573085 | 0.12393224 | 0.33802313 | 0.36663835 | 0.7138888 | 0.95791799 |
| TARS2 | 57.0182196 | 0.29850982 | 0.28393448 | 1.05133344 | 0.29310547 | 0.83336457 |
| MCL1 | 237.057571 | 0.06753866 | 0.33067249 | 0.20424639 | 0.83816096 | 0.97385715 |
| ENSA | 442.411354 | -0.3197049 | 0.17573224 | -1.8192731 | 0.06886978 | 0.55917298 |
| GOLPH3L | 178.844829 | -0.1572048 | 0.22781718 | -0.6900482 | 0.49016386 | 0.90730741 |
| CTSS | 72.4613685 | -0.6316354 | 0.30093593 | -2.0989032 | 0.03582544 | 0.4362094 |
| ARNT | 249.552899 | -0.400286 | 0.20797681 | -1.9246666 | 0.05427108 | 0.51425036 |
| SETDB1 | 147.827143 | -0.3003033 | 0.17263121 | -1.7395653 | 0.08193539 | 0.59202053 |
| CERS2 | 125.822965 | 0.18424404 | 0.3316183 | 0.55559069 | 0.5784907 | 0.93345492 |
| ANXA9 | 79.8732817 | 2.12194364 | 0.38325816 | 5.53659093 | 3.08E-08 | 4.04E-05 |
| MINDY1 | 91.236805 | 0.47736661 | 0.21642104 | 2.20573109 | 0.02740283 | 0.40165272 |
| PRUNE1 | 98.7293897 | 0.13439523 | 0.21864941 | 0.61466085 | 0.53877872 | 0.92365631 |
| BNIPL | 78.0746614 | 1.35392754 | 0.55455291 | 2.44147584 | 0.01462737 | 0.28992884 |
| C1orf56 | 61.2650168 | -0.0991091 | 0.3957443 | -0.2504371 | 0.80224933 | 0.9661728 |
| CDC42SE1 | 142.605824 | -0.0240626 | 0.25517072 | -0.0943 | 0.92487088 | 0.99051393 |
| GABPB2 | 211.297327 | 0.08721917 | 0.25468438 | 0.34245981 | 0.73200488 | 0.96008876 |
| SCNM1 | 45.8408588 | 0.13677704 | 0.41345498 | 0.33081483 | 0.74078436 | 0.96084411 |
| VPS72 | 53.6024799 | -0.2537532 | 0.27039378 | -0.9384579 | 0.34800916 | 0.85348649 |
| PIP5K1A | 593.929692 | 0.1227595 | 0.26299148 | 0.46678129 | 0.64065636 | 0.94935885 |
| PSMD4 | 156.534136 | 0.12414485 | 0.30507552 | 0.40693154 | 0.68405827 | 0.95574492 |
| ZNF687 | 34.8082028 | 0.52292813 | 0.28173431 | 1.85610384 | 0.06343876 | 0.54850785 |
| PI4KB | 162.264538 | 0.0407474 | 0.25012095 | 0.16291079 | 0.87058867 | 0.98208511 |
| SELENBP1 | 51.0686497 | 0.53317587 | 0.62470173 | 0.8534887 | 0.39338835 | 0.87223214 |
| PSMB4 | 120.663915 | -0.0337841 | 0.3805699 | -0.0887723 | 0.92926288 | 0.99216065 |
| POGZ | 918.219102 | 0.00573049 | 0.19014843 | 0.03013695 | 0.97595783 | 0.99656026 |
| CGN | 136.988392 | 0.32019592 | 0.27695743 | 1.15611965 | 0.24763222 | 0.80073609 |
| TUFT1 | 151.993985 | -0.127931 | 0.45285719 | -0.2824974 | 0.77756211 | 0.96472142 |
| SNX27 | 256.100119 | -0.270159 | 0.15155128 | -1.7826241 | 0.07464752 | 0.57072562 |
| MRPL9 | 41.3367863 | -0.0911362 | 0.49425308 | -0.1843918 | 0.85370612 | 0.97718353 |
| TDRKH | 31.8455878 | 0.00544401 | 0.39012728 | 0.01395443 | 0.98886633 | 0.99824159 |
| RORC | 92.9607202 | -1.0734332 | 0.38272351 | -2.8047225 | 0.00503599 | 0.17602704 |
| THEM4 | 252.088539 | 0.79043037 | 0.52320706 | 1.51074102 | 0.13085445 | 0.67355334 |
| S100A10 | 1328.25283 | -0.1464341 | 0.56210462 | -0.2605104 | 0.7944701 | 0.9653598 |
| S100A11 | 3245.8857 | -0.640912 | 0.54752134 | -1.1705698 | 0.24177173 | 0.80072691 |
| FLG-AS1 | 355.202572 | 0.25478774 | 0.38669599 | 0.65888383 | 0.50997037 | 0.91334187 |
| S100A6 | 4301.60372 | -0.6814217 | 0.67853341 | -1.0042566 | 0.31525494 | 0.84285425 |
| S100A4 | 193.239316 | -0.1279832 | 1.30590143 | -0.0980037 | 0.92192933 | 0.99051393 |
| S100A2 | 818.295799 | -1.611678 | 0.78457039 | -2.0542172 | 0.03995468 | 0.45827251 |
| S100A16 | 196.116123 | -0.1896483 | 0.61000208 | -0.3108977 | 0.75587838 | 0.96120879 |
| S100A14 | 124.679602 | -0.5122503 | 0.6706609 | -0.7637992 | 0.4449869 | 0.89133461 |
| S100A13 | 359.530208 | 0.48337392 | 0.56732747 | 0.85201924 | 0.39420341 | 0.87253562 |
| S100A1 | 185.669373 | -1.1414339 | 0.93453235 | -1.2213958 | 0.22193621 | 0.78760199 |
| CHTOP | 147.378416 | 0.08052934 | 0.20156441 | 0.39952163 | 0.68950889 | 0.95645972 |
| SNAPIN | 36.8957756 | 0.17023986 | 0.32479948 | 0.52413832 | 0.60018233 | 0.93728325 |
| ILF2 | 133.750343 | 0.08394621 | 0.41740428 | 0.20111488 | 0.84060875 | 0.97505032 |
| INTS3 | 223.03248 | -0.036405 | 0.20993087 | -0.1734141 | 0.86232595 | 0.97910899 |
| GATAD2B | 604.773002 | -0.0347609 | 0.26192835 | -0.1327114 | 0.89442166 | 0.98597539 |
| DENND4B | 81.6995273 | -0.3696453 | 0.21194152 | -1.7440912 | 0.0811432 | 0.59202053 |
| CRTC2 | 51.9977489 | 0.15802439 | 0.27204259 | 0.58088106 | 0.56132062 | 0.92934517 |
| SLC39A1 | 129.733837 | -0.1043955 | 0.34238867 | -0.3049034 | 0.76043975 | 0.96191672 |
| JTB | 158.407666 | 0.37813247 | 0.41032154 | 0.9215516 | 0.35676251 | 0.8557799 |
| RAB13 | 254.256411 | 0.33218517 | 0.46454143 | 0.71508191 | 0.47455843 | 0.90242252 |

| | | | | | | |
|---|---|---|---|---|---|---|
| RPS27 | 225.980462 | -0.0676464 | 0.31858802 | -0.2123319 | 0.83184808 | 0.97177214 |
| NUP210L | 45.0837913 | 0.28312533 | 0.98147964 | 0.28846786 | 0.77298863 | 0.96388926 |
| TPM3 | 275.47415 | -0.1268431 | 0.35041506 | -0.3619796 | 0.7173673 | 0.95791799 |
| C1orf43 | 188.741126 | -0.097366 | 0.19598574 | -0.4968014 | 0.61932907 | 0.94300065 |
| UBAP2L | 238.388835 | -0.5127274 | 0.25752705 | -1.9909651 | 0.04648472 | 0.48600281 |
| HAX1 | 65.6744717 | 0.47726433 | 0.44840145 | 1.06436841 | 0.28716184 | 0.82973699 |
| IL6R | 38.9614489 | -0.7689297 | 0.48849182 | -1.5740891 | 0.11546684 | 0.65156815 |
| UBE2Q1 | 85.6343903 | -0.0480929 | 0.35812568 | -0.1342906 | 0.8931728 | 0.98597539 |
| ADAR | 390.753053 | -0.3333681 | 0.30718681 | -1.0852293 | 0.27782011 | 0.82270551 |
| PMVK | 124.376449 | 0.56554401 | 0.42443939 | 1.33244941 | 0.18271256 | 0.74315969 |
| PBXIP1 | 127.621888 | 0.7230872 | 0.21939096 | 3.29588425 | 0.00098112 | 0.07616432 |
| PYGO2 | 47.7386737 | 0.16208289 | 0.29519396 | 0.54907252 | 0.58295568 | 0.93415528 |
| FLAD1 | 33.20632 | 0.16952308 | 0.31033345 | 0.54626107 | 0.58488649 | 0.93483536 |
| ZBTB7B | 54.0272713 | 0.44272147 | 0.31157824 | 1.42089985 | 0.15534588 | 0.70907003 |
| EFNA1 | 139.410547 | -0.8243372 | 0.40101333 | -2.0556355 | 0.03981767 | 0.45805615 |
| SLC50A1 | 56.6655161 | -0.1840969 | 0.40309091 | -0.4567131 | 0.64787728 | 0.95061388 |
| DPM3 | 137.863669 | 0.03886471 | 0.27087205 | 0.14347997 | 0.88591113 | 0.98486779 |
| MUC1 | 69.6195062 | -0.3265502 | 0.34652791 | -0.9423488 | 0.34601408 | 0.8523814 |
| THBS3 | 44.3707727 | -0.1040497 | 0.3235644 | -0.3215733 | 0.74777594 | 0.96109466 |
| MTX1 | 48.4057963 | 0.00355435 | 0.41142786 | 0.00863905 | 0.99310712 | 0.99919666 |
| GBA | 57.803082 | 0.09396861 | 0.38501974 | 0.2440618 | 0.80718295 | 0.96752828 |
| SCAMP3 | 95.5794928 | -0.0782346 | 0.24574051 | -0.3183626 | 0.75020989 | 0.96109466 |
| CLK2 | 66.9318458 | -0.1241883 | 0.3211743 | -0.3866694 | 0.69900098 | 0.95672887 |
| FDPS | 237.105507 | -0.1396201 | 0.54442785 | -0.256453 | 0.7976011 | 0.9653598 |
| RUSC1 | 39.8810695 | -0.0278593 | 0.28068529 | -0.0992547 | 0.92093603 | 0.9904507 |
| ASH1L | 1188.97471 | 0.08366476 | 0.15464176 | 0.54102309 | 0.58849167 | 0.93483536 |
| YY1AP1 | 220.720083 | -0.0793746 | 0.19854659 | -0.3997781 | 0.68931998 | 0.95632425 |
| DAP3 | 235.063691 | -0.0506455 | 0.19556814 | -0.2589658 | 0.79566164 | 0.9653598 |
| GON4L | 300.279487 | -0.2120467 | 0.2187039 | -0.9695607 | 0.33226551 | 0.84442294 |
| SYT11 | 38.8919344 | 1.14740034 | 0.46600376 | 2.46221264 | 0.01380828 | 0.28408502 |
| RIT1 | 121.324413 | -0.6046996 | 0.40095197 | -1.5081598 | 0.13151363 | 0.67390608 |
| KHDC4 | 276.380659 | 0.04206485 | 0.21039966 | 0.19992831 | 0.84153665 | 0.975446 |
| ARHGEF2 | 97.7337995 | 0.27935579 | 0.28470599 | 0.98120799 | 0.32649019 | 0.84381657 |
| SSR2 | 237.532269 | 0.15716933 | 0.3828561 | 0.41051803 | 0.68142598 | 0.95574492 |
| UBQLN4 | 77.7628717 | 0.1602251 | 0.28777131 | 0.55677925 | 0.57767826 | 0.93345492 |
| LAMTOR2 | 91.1586569 | -0.0070522 | 0.42783217 | -0.0164836 | 0.98684859 | 0.99791717 |
| RAB25 | 157.704009 | 0.49820164 | 0.37460045 | 1.32995474 | 0.18353318 | 0.74504275 |
| LMNA | 482.848967 | -0.187414 | 0.49420391 | -0.3792241 | 0.70452148 | 0.95707594 |
| SEMA4A | 94.9116122 | 0.47961688 | 0.39863525 | 1.20314717 | 0.22891937 | 0.79047378 |
| SLC25A44 | 52.4359847 | 0.17360992 | 0.25559742 | 0.6792319 | 0.49699094 | 0.90839719 |
| SMG5 | 154.745071 | -0.5745182 | 0.22473334 | -2.5564438 | 0.01057481 | 0.25038862 |
| GLMP | 62.6355806 | -0.0780043 | 0.37322392 | -0.2090013 | 0.83444724 | 0.97232209 |
| CCT3 | 242.371796 | 0.38541335 | 0.24228283 | 1.590758 | 0.11166405 | 0.64353508 |
| RHBG | 200.065166 | 0.99887774 | 0.57250377 | 1.74475313 | 0.08102785 | 0.5917574 |
| MEF2D | 126.787235 | -0.5834031 | 0.29716131 | -1.9632539 | 0.04961668 | 0.49381084 |
| NAXE | 88.359842 | 0.06985514 | 0.38208839 | 0.18282455 | 0.85493568 | 0.97718353 |
| GPATCH4 | 45.1380628 | -0.0302037 | 0.35957202 | -0.083999 | 0.93305725 | 0.99243654 |
| CRABP2 | 47.1994763 | 0.62528089 | 0.6859645 | 0.91153535 | 0.36201337 | 0.85690129 |
| ISG20L2 | 63.6261365 | 0.10226735 | 0.36869528 | 0.27737634 | 0.78149115 | 0.96513523 |
| RRNAD1 | 64.8630836 | 0.06792328 | 0.29948738 | 0.22679847 | 0.82058046 | 0.96886922 |
| MRPL24 | 47.9327475 | 0.32769498 | 0.30368529 | 1.07906111 | 0.28056049 | 0.82628479 |
| HDGF | 262.780255 | -0.0936622 | 0.30695045 | -0.3051378 | 0.76026123 | 0.96191449 |
| PRCC | 153.119889 | 0.08437048 | 0.20883975 | 0.40399627 | 0.68621546 | 0.9558749 |
| ARHGEF11 | 301.370543 | 0.11712744 | 0.28548533 | 0.41027481 | 0.68160437 | 0.95574492 |

| | | | | | | |
|---|---|---|---|---|---|---|
| ETV3 | 48.5224023 | 0.71789901 | 0.34622251 | 2.07351918 | 0.03812399 | 0.44791264 |
| KIRREL1 | 64.8954283 | 0.56971308 | 0.64930758 | 0.87741633 | 0.38026055 | 0.86593382 |
| IFI16 | 59.6801299 | -1.0730553 | 0.62453844 | -1.7181573 | 0.08576792 | 0.60075714 |
| DUSP23 | 131.274898 | -0.576143 | 0.43552136 | -1.3228811 | 0.18587491 | 0.74914489 |
| TAGLN2 | 472.707934 | -1.2356753 | 0.51829999 | -2.3840928 | 0.01712128 | 0.3131432 |
| IGSF8 | 68.2009579 | 0.18673463 | 0.36039371 | 0.51814066 | 0.60436013 | 0.93818608 |
| PEA15 | 52.2768746 | -0.1454385 | 0.38998615 | -0.3729324 | 0.70919877 | 0.95752169 |
| DCAF8 | 101.369344 | 0.12874183 | 0.26658949 | 0.48292163 | 0.62915138 | 0.94596555 |
| COPA | 470.903069 | -0.1938093 | 0.26045533 | -0.7441173 | 0.45680552 | 0.89667014 |
| NCSTN | 121.402268 | 0.39026668 | 0.25616336 | 1.52350704 | 0.1276319 | 0.67051794 |
| VANGL2 | 31.7008338 | 0.71183269 | 0.54549131 | 1.30493866 | 0.19191374 | 0.7597012 |
| F11R | 404.88785 | -0.026849 | 0.30872926 | -0.086966 | 0.93069852 | 0.99243654 |
| AL591806.3 | 74.9974784 | 0.52987705 | 0.39724957 | 1.33386438 | 0.18224831 | 0.74315969 |
| USF1 | 32.4455948 | -0.4917269 | 0.39599975 | -1.2417355 | 0.21433416 | 0.77890374 |
| NECTIN4 | 122.403778 | -1.3837137 | 0.42476393 | -3.2576064 | 0.00112356 | 0.08298537 |
| PFDN2 | 185.063312 | 0.29209514 | 0.51417467 | 0.56808544 | 0.56997695 | 0.93122667 |
| NIT1 | 121.414157 | 0.4216922 | 0.35000076 | 1.20483225 | 0.22826807 | 0.79042788 |
| DEDD | 44.3539013 | 0.20360919 | 0.30914795 | 0.65861408 | 0.51014362 | 0.91334187 |
| UFC1 | 166.953438 | 0.60302188 | 0.41738665 | 1.44475602 | 0.14852643 | 0.70115179 |
| PPOX | 32.0903013 | 0.20933799 | 0.3844182 | 0.54455795 | 0.58605758 | 0.93483536 |
| B4GALT3 | 52.5659315 | 0.10902016 | 0.39694571 | 0.27464753 | 0.78358704 | 0.96513523 |
| NDUFS2 | 162.576373 | 0.00440624 | 0.30895863 | 0.0142616 | 0.98862128 | 0.99814557 |
| SDHC | 314.595515 | 0.27361291 | 0.20232889 | 1.35231757 | 0.17627375 | 0.7345557 |
| AL592295.4 | 40.9625429 | 0.21766822 | 0.2967638 | 0.73347297 | 0.46327001 | 0.89801718 |
| DUSP12 | 57.5919096 | -0.670171 | 0.40210592 | -1.6666529 | 0.09558345 | 0.61367241 |
| ATF6 | 855.749944 | -0.4843039 | 0.35428242 | -1.3669996 | 0.17162544 | 0.72776437 |
| NOS1AP | 1112.49666 | -0.8874676 | 0.51029487 | -1.7391269 | 0.08201244 | 0.59202053 |
| UHMK1 | 320.980805 | -0.155956 | 0.2175608 | -0.7168389 | 0.47347351 | 0.90242252 |
| UAP1 | 90.7677711 | -0.0257161 | 0.67327484 | -0.0381955 | 0.96953179 | 0.99645289 |
| HSD17B7 | 70.3110138 | -0.2971918 | 0.35295329 | -0.8420147 | 0.39977974 | 0.87650454 |
| PBX1 | 610.494478 | 0.28953779 | 0.2512073 | 1.15258511 | 0.24908072 | 0.80097144 |
| MGST3 | 1091.14978 | -0.3496368 | 0.35452251 | -0.986219 | 0.32402567 | 0.84285425 |
| ALDH9A1 | 188.303017 | 0.61130264 | 0.29707957 | 2.05770674 | 0.03961829 | 0.45799149 |
| TMCO1 | 266.263064 | 0.35233533 | 0.30755184 | 1.14561281 | 0.25195539 | 0.80268905 |
| UCK2 | 119.792179 | 0.29875983 | 0.30204587 | 0.98912071 | 0.32260408 | 0.84285425 |
| POGK | 129.073962 | 0.03783926 | 0.31582931 | 0.11980921 | 0.90463429 | 0.9875066 |
| TADA1 | 52.36667 | -0.0013962 | 0.33924486 | -0.0041157 | 0.99671615 | 0.99967893 |
| POU2F1 | 601.468531 | 0.13116634 | 0.25299282 | 0.51845875 | 0.60413823 | 0.93804805 |
| CREG1 | 122.219194 | 0.25104043 | 0.35151785 | 0.71416125 | 0.47512748 | 0.90242252 |
| MPZL1 | 199.990735 | -0.2721978 | 0.29515702 | -0.9222136 | 0.35641715 | 0.8557799 |
| MPC2 | 202.382291 | 0.66062791 | 0.2910905 | 2.26949317 | 0.02323835 | 0.36457774 |
| DCAF6 | 779.941669 | -0.2552063 | 0.26191254 | -0.9743953 | 0.32986032 | 0.84442294 |
| TIPRL | 102.490113 | 0.32158486 | 0.22806092 | 1.41008315 | 0.15851513 | 0.71297794 |
| SFT2D2 | 190.259831 | 0.0316798 | 0.28362286 | 0.11169692 | 0.91106372 | 0.98788932 |
| TBX19 | 34.9510003 | -0.0696849 | 0.32397832 | -0.2150913 | 0.82969615 | 0.97139806 |
| LINC00970 | 165.535836 | 0.42761148 | 0.61089963 | 0.6999701 | 0.48394597 | 0.90647968 |
| ATP1B1 | 4443.92114 | 0.87310771 | 0.37063602 | 2.35570118 | 0.01848778 | 0.32618176 |
| NME7 | 678.960494 | 0.27964763 | 0.17164628 | 1.62920881 | 0.10326882 | 0.62635087 |
| BLZF1 | 135.572738 | -0.0888331 | 0.21713783 | -0.4091091 | 0.68245958 | 0.95574492 |
| CCDC181 | 107.84852 | 0.69140892 | 0.24548599 | 2.81649038 | 0.00485515 | 0.17248125 |
| SLC19A2 | 338.342772 | -1.5407129 | 0.56463578 | -2.7286845 | 0.00635875 | 0.19542438 |
| C1orf112 | 37.9159551 | -0.5982547 | 0.39413161 | -1.5179058 | 0.12903815 | 0.67206108 |
| SCYL3 | 127.519931 | 0.25391807 | 0.29710003 | 0.85465515 | 0.39274208 | 0.87165122 |
| KIFAP3 | 291.373103 | -0.125446 | 0.15350946 | -0.8171874 | 0.41382133 | 0.88211689 |

| | | | | | | |
|---|---|---|---|---|---|---|
| GORAB | 48.6908346 | -0.3091538 | 0.31486144 | -0.9818725 | 0.32616265 | 0.84381657 |
| FMO4 | 87.4043552 | 0.46813315 | 0.33431905 | 1.40025868 | 0.16143587 | 0.71416435 |
| PRRC2C | 884.655718 | -0.3098681 | 0.1655682 | -1.8715434 | 0.0612698 | 0.5418629 |
| VAMP4 | 76.2215287 | -0.1402036 | 0.27032596 | -0.5186464 | 0.60400732 | 0.93804805 |
| METTL13 | 38.5948368 | -0.0743094 | 0.30557445 | -0.2431793 | 0.80786653 | 0.96752828 |
| DNM3 | 132.472285 | -0.0532363 | 0.40947318 | -0.1300118 | 0.89655711 | 0.98597539 |
| PIGC | 77.9906777 | 0.15077107 | 0.31056038 | 0.48548069 | 0.62733541 | 0.94511671 |
| SUCO | 260.313964 | -0.1533915 | 0.24073496 | -0.6371802 | 0.52400747 | 0.91871245 |
| AL645568.1 | 84.8161877 | 0.6298486 | 0.41374791 | 1.52230038 | 0.12793383 | 0.67070864 |
| PRDX6 | 434.182427 | 0.75868941 | 0.42513961 | 1.78456533 | 0.07433185 | 0.5698716 |
| ANKRD45 | 43.3352507 | 0.47383965 | 0.36055167 | 1.31420734 | 0.18877644 | 0.75510575 |
| KLHL20 | 165.651394 | -0.0562266 | 0.29426596 | -0.191074 | 0.8484676 | 0.97656375 |
| DARS2 | 47.7091827 | -0.0866292 | 0.34013666 | -0.2546895 | 0.79896295 | 0.9653598 |
| ZBTB37 | 143.531683 | -0.2545545 | 0.25062139 | -1.0156936 | 0.30977531 | 0.84194359 |
| RC3H1 | 379.720196 | 0.1339188 | 0.22017922 | 0.6082263 | 0.54303739 | 0.92482794 |
| RABGAP1L | 2249.92191 | -0.3226845 | 0.2051263 | -1.5731015 | 0.11569532 | 0.65187472 |
| CACYBP | 74.151909 | 0.11326841 | 0.35813962 | 0.31626885 | 0.75179845 | 0.96120879 |
| MRPS14 | 89.2683708 | 0.20183363 | 0.39833385 | 0.50669466 | 0.61236908 | 0.94003018 |
| KIAA0040 | 66.7386802 | -0.0792192 | 0.32583045 | -0.2431302 | 0.80790457 | 0.96752828 |
| COP1 | 893.267675 | -0.291745 | 0.20336441 | -1.4345925 | 0.15140326 | 0.7029344 |
| RASAL2 | 602.105719 | 0.06904383 | 0.31095989 | 0.22203452 | 0.82428701 | 0.97018919 |
| RALGPS2 | 743.86808 | -0.1118394 | 0.2569958 | -0.4351798 | 0.6634319 | 0.95187756 |
| ANGPTL1 | 63.905346 | -0.2430245 | 0.42141407 | -0.5766881 | 0.56415018 | 0.93077674 |
| FAM20B | 139.106307 | -0.1633934 | 0.16792179 | -0.9730328 | 0.33053698 | 0.84442294 |
| TOR3A | 48.8038175 | 0.51666119 | 0.40665937 | 1.27050113 | 0.20390618 | 0.77173592 |
| ABL2 | 374.928488 | 0.15185458 | 0.25836851 | 0.58774417 | 0.55670402 | 0.92872652 |
| SOAT1 | 200.420481 | 0.4889551 | 0.24464987 | 1.99859125 | 0.0456526 | 0.48339878 |
| TDRD5 | 57.7897681 | 1.42781476 | 0.52783733 | 2.70502803 | 0.00682986 | 0.20224502 |
| TOR1AIP2 | 395.855729 | -0.0685631 | 0.22819797 | -0.3004543 | 0.76383066 | 0.96285313 |
| TOR1AIP1 | 128.963623 | -0.0515665 | 0.21141039 | -0.2439165 | 0.80729545 | 0.96752828 |
| CEP350 | 561.555453 | -0.0206771 | 0.16392184 | -0.12614 | 0.89962114 | 0.98660836 |
| QSOX1 | 202.656991 | 0.12979855 | 0.27891698 | 0.46536627 | 0.64166919 | 0.94935885 |
| ACBD6 | 523.690956 | -0.0256994 | 0.19859239 | -0.129408 | 0.89703484 | 0.98601802 |
| XPR1 | 412.08286 | 0.00952484 | 0.24153313 | 0.03943493 | 0.96854363 | 0.9964399 |
| KIAA1614 | 85.5575486 | -0.1263504 | 0.34152899 | -0.369955 | 0.71141601 | 0.957816 |
| STX6 | 117.307589 | -0.1392591 | 0.30511652 | -0.456413 | 0.64809303 | 0.95061388 |
| MR1 | 83.5171623 | 0.49189017 | 0.2148968 | 2.28895993 | 0.02208168 | 0.35843654 |
| GLUL | 190.45199 | 0.12572494 | 0.31545671 | 0.39854895 | 0.69022558 | 0.95659154 |
| RNASEL | 35.2850391 | 0.08769561 | 0.33456235 | 0.26212037 | 0.79322864 | 0.9653598 |
| DHX9 | 403.785325 | -0.1775017 | 0.30578084 | -0.5804867 | 0.56158645 | 0.92947347 |
| LAMC1 | 848.105189 | -0.1753428 | 0.36707452 | -0.4776763 | 0.63288059 | 0.94683634 |
| LAMC2 | 157.731713 | -1.0961479 | 0.61953655 | -1.769303 | 0.07684332 | 0.57318004 |
| SMG7 | 383.478089 | -0.0479899 | 0.21865143 | -0.2194811 | 0.82627527 | 0.97058548 |
| ARPC5 | 202.809329 | 0.13498872 | 0.33004672 | 0.40899882 | 0.68254053 | 0.95574492 |
| RGL1 | 124.414127 | 0.43864273 | 0.3175779 | 1.38121302 | 0.16721347 | 0.72204253 |
| TSEN15 | 112.502836 | 0.08027446 | 0.27774519 | 0.28902195 | 0.77256458 | 0.96388926 |
| EDEM3 | 255.757775 | -0.1418111 | 0.21650187 | -0.6550111 | 0.51246063 | 0.91445384 |
| RNF2 | 111.949135 | 0.0372725 | 0.28391379 | 0.13128106 | 0.89555297 | 0.98597539 |
| TRMT1L | 130.514047 | 0.34860885 | 0.30114549 | 1.1576094 | 0.24702347 | 0.80072691 |
| SWT1 | 159.393478 | -0.2925156 | 0.28971847 | -1.0096545 | 0.31266085 | 0.84279957 |
| IVNS1ABP | 1792.3199 | -1.1448046 | 0.63287845 | -1.8088854 | 0.07046881 | 0.5621116 |
| HMCN1 | 64.7775178 | 1.84774362 | 0.50200575 | 3.68072205 | 0.00023257 | 0.0343293 |
| TPR | 401.775455 | -0.1863585 | 0.17577417 | -1.0602152 | 0.28904669 | 0.83060304 |
| ODR4 | 210.466969 | -0.1178205 | 0.18829911 | -0.6257095 | 0.53150547 | 0.92211604 |

| | | | | | | |
|---|---|---|---|---|---|---|
| BRINP3 | 334.657962 | -0.617591 | 0.67302048 | -0.9176408 | 0.35880695 | 0.8557799 |
| AL390957.1 | 57.0902677 | -0.5432389 | 0.46972247 | -1.1565105 | 0.24747242 | 0.80072691 |
| AL136454.1 | 282.591721 | -0.1315511 | 0.50755317 | -0.2591868 | 0.79549115 | 0.9653598 |
| RGS2 | 194.725036 | -2.7866942 | 1.389832 | -2.0050583 | 0.04495681 | 0.48262669 |
| UCHL5 | 137.395793 | -0.0996536 | 0.20600942 | -0.4837335 | 0.62857505 | 0.94593144 |
| TROVE2 | 273.648502 | 0.01212568 | 0.17288692 | 0.07013648 | 0.94408503 | 0.99265578 |
| GLRX2 | 33.7258035 | 0.07259329 | 0.42998628 | 0.16882698 | 0.86593273 | 0.98023062 |
| CDC73 | 487.118272 | -0.2852736 | 0.25044234 | -1.1390791 | 0.25467016 | 0.80414079 |
| ZBTB41 | 154.843088 | -0.0138832 | 0.24749069 | -0.056096 | 0.9552653 | 0.99538167 |
| CRB1 | 74.6625975 | 1.18638214 | 0.74202121 | 1.59885205 | 0.10985348 | 0.63959138 |
| DENND1B | 731.840528 | -0.3304334 | 0.18289687 | -1.806665 | 0.07081454 | 0.56265074 |
| NEK7 | 673.109842 | 0.08371184 | 0.19025383 | 0.44000079 | 0.65993653 | 0.95127957 |
| ZNF281 | 62.3197897 | 0.17034431 | 0.27466802 | 0.62018254 | 0.53513761 | 0.92303275 |
| DDX59 | 61.2042264 | -0.0237592 | 0.24725368 | -0.0960923 | 0.92344725 | 0.99051393 |
| CAMSAP2 | 420.755846 | -0.0569728 | 0.1988454 | -0.286518 | 0.77448141 | 0.96388926 |
| INAVA | 32.5165257 | -0.7213528 | 0.36360425 | -1.9838954 | 0.0472675 | 0.48600281 |
| TMEM9 | 123.303529 | 0.59583921 | 0.35157807 | 1.69475645 | 0.09012163 | 0.60739841 |
| LAD1 | 83.5476669 | 0.39070429 | 0.22039455 | 1.77274929 | 0.07627026 | 0.57145422 |
| PHLDA3 | 44.8931307 | 1.20464179 | 0.3500002 | 3.44183173 | 0.00057779 | 0.05789336 |
| CSRP1 | 148.550338 | -0.2768041 | 0.4485556 | -0.6171009 | 0.53716818 | 0.92324352 |
| AC096677.1 | 94.0129456 | 0.10801218 | 0.36788628 | 0.29360208 | 0.76906199 | 0.96345263 |
| NAV1 | 122.15627 | 0.22902211 | 0.28238759 | 0.81102044 | 4.17E-01 | 0.88242985 |
| IPO9-AS1 | 44.1458569 | 0.57004243 | 0.4845483 | 1.1764409 | 0.23941874 | 0.79984328 |
| IPO9 | 211.98666 | -0.2621499 | 0.23257069 | -1.1271836 | 0.25966485 | 0.80709747 |
| TIMM17A | 82.0449648 | -0.1266273 | 0.3698154 | -0.3424069 | 0.73204469 | 0.96008876 |
| RNPEP | 133.807287 | -0.0748867 | 0.22633312 | -0.3308695 | 0.7407431 | 0.96084411 |
| ELF3 | 1160.00513 | -0.4341273 | 0.50462344 | -0.8602994 | 0.38962402 | 0.87004477 |
| ARL8A | 107.216545 | -0.3428669 | 0.22780408 | -1.5050954 | 0.13229956 | 0.67466347 |
| PPP1R12B | 610.675119 | -0.3327741 | 0.1882435 | -1.7677852 | 0.07709682 | 0.57405253 |
| KDM5B | 633.077272 | -0.2076287 | 0.25240314 | -0.8226073 | 0.41073137 | 0.88043868 |
| KLHL12 | 86.5635563 | -0.1905384 | 0.23361137 | -0.8156214 | 0.41471668 | 0.88242985 |
| ADIPOR1 | 78.278407 | -0.0600854 | 0.36141664 | -0.1662498 | 0.8679604 | 0.9815717 |
| CYB5R1 | 66.5657765 | 0.67823086 | 0.33751623 | 2.00947625 | 0.04448665 | 0.48014429 |
| TMEM183A | 45.5672774 | -0.2554253 | 0.27737533 | -0.9208652 | 0.35712081 | 0.8557799 |
| BTG2 | 1321.48329 | -0.6879566 | 0.50845388 | -1.3530363 | 0.17604402 | 0.73444173 |
| ATP2B4 | 170.258957 | 0.32776969 | 0.33521526 | 0.97778871 | 0.32817884 | 0.84381657 |
| SNRPE | 73.9447588 | 0.0848619 | 0.34991023 | 0.24252479 | 0.80837355 | 0.96775815 |
| SOX13 | 240.437183 | 1.01668306 | 0.48566383 | 2.09338845 | 0.03631449 | 0.43729328 |
| ETNK2 | 36.2734322 | 0.3056387 | 0.40067153 | 0.76281611 | 0.44557307 | 0.8918683 |
| GOLT1A | 47.4166024 | 0.12254981 | 0.30388285 | 0.40327978 | 0.68674241 | 0.9558749 |
| PLEKHA6 | 573.066613 | -0.7760243 | 0.26503479 | -2.9280091 | 0.0034114 | 0.14358074 |
| PPP1R15B | 111.239245 | 0.26521953 | 0.36786556 | 0.72096863 | 0.47092882 | 0.90123338 |
| PIK3C2B | 62.2556443 | 0.37108166 | 0.4739665 | 0.78292803 | 0.43366938 | 0.887667 |
| MDM4 | 467.223078 | -0.0665584 | 0.2080113 | -0.3199748 | 0.74898747 | 0.96109466 |
| LRRN2 | 131.723419 | 0.94556524 | 0.63326706 | 1.493154 | 0.13539688 | 0.67980087 |
| RBBP5 | 99.8323828 | -0.0524262 | 0.21862483 | -0.2397997 | 0.81048552 | 0.9680627 |
| DSTYK | 201.467276 | 0.14236356 | 0.31913168 | 0.44609662 | 0.65552746 | 0.95100227 |
| NUAK2 | 401.741455 | 0.22031642 | 0.44698141 | 0.49289838 | 0.62208439 | 0.94418789 |
| CDK18 | 428.945093 | -1.2411655 | 0.54452632 | -2.279349 | 0.02264633 | 0.35959765 |
| MFSD4A | 54.6940133 | -0.0915195 | 0.65731185 | -0.1392329 | 0.88926609 | 0.98551241 |
| ELK4 | 222.876884 | -0.8151555 | 0.45503961 | -1.7913946 | 0.07322999 | 0.56668778 |
| NUCKS1 | 1050.15491 | 0.2238573 | 0.16788703 | 1.33338055 | 0.18240695 | 0.74315969 |
| RAB29 | 100.72905 | -0.0561308 | 0.31802562 | -0.1764979 | 0.85990283 | 0.97841862 |
| SLC41A1 | 52.4965196 | 0.13552747 | 0.33616126 | 0.40316208 | 0.68682899 | 0.9558749 |

| | | | | | | |
|---|---|---|---|---|---|---|
| RHEX | 561.005933 | 0.17600109 | 0.39719103 | 0.44311446 | 0.65768295 | 0.95100227 |
| SRGAP2 | 196.129585 | 0.31906294 | 0.28711048 | 1.11128976 | 0.26644365 | 0.81314193 |
| EIF2D | 66.7557175 | -0.1221711 | 0.22594346 | -0.5407152 | 0.58870388 | 0.93483536 |
| DYRK3 | 47.3837411 | 0.04843886 | 0.3111776 | 0.15566306 | 0.87629862 | 0.98269442 |
| MAPKAPK2 | 341.781193 | -0.1655463 | 0.33054999 | -0.5008209 | 0.61649718 | 0.94206626 |
| PIGR | 791.063809 | 1.57617047 | 0.58087151 | 2.71345803 | 0.0066585 | 0.20224502 |
| C1orf116 | 38.8347453 | 0.7106252 | 0.37200812 | 1.9102411 | 0.05610218 | 0.52189372 |
| PFKFB2 | 164.479742 | -1.2144899 | 0.46015698 | -2.6392947 | 0.00830787 | 0.22277119 |
| YOD1 | 61.4569273 | -1.7685937 | 0.5955149 | -2.9698563 | 0.00297939 | 0.14126521 |
| CD55 | 299.655594 | -0.7529834 | 0.75023487 | -1.0036636 | 0.31554079 | 0.84285425 |
| CD46 | 689.28181 | 0.13106745 | 0.19628553 | 0.66773873 | 0.50430038 | 0.9107831 |
| MIR29B2CHG | 558.911946 | 0.17622084 | 0.48791435 | 0.36117168 | 0.71797111 | 0.95796325 |
| IRF6 | 98.7085751 | 0.3379246 | 0.47535248 | 0.71089269 | 0.47715073 | 0.90242252 |
| UTP25 | 40.6450665 | -0.1114882 | 0.27251964 | -0.4091015 | 0.68246516 | 0.95574492 |
| HHAT | 170.19601 | -0.0269862 | 0.33467719 | -0.0806336 | 0.93573334 | 0.99243654 |
| RCOR3 | 504.929892 | 0.13935265 | 0.23909835 | 0.58282563 | 0.56001069 | 0.92934517 |
| TRAF5 | 159.159388 | 0.22503548 | 0.34971145 | 0.64348903 | 0.51990683 | 0.91751084 |
| LINC00467 | 92.8903478 | -0.012151 | 0.38477127 | -0.0315798 | 0.97480715 | 0.99656026 |
| SLC30A1 | 59.6670747 | 0.49011179 | 0.35404622 | 1.38431586 | 0.16626176 | 0.72054851 |
| LPGAT1 | 372.865827 | -0.1141089 | 0.21239893 | -0.5372388 | 0.59110268 | 0.93483997 |
| INTS7 | 85.1339249 | -0.3134218 | 0.4572497 | -0.68545 | 0.49306002 | 0.90804281 |
| PPP2R5A | 393.945373 | -0.108612 | 0.24907004 | -0.4360701 | 0.66278585 | 0.9515075 |
| AL360091.3 | 58.3034416 | 0.4714641 | 0.63992642 | 0.73674736 | 0.461276 | 0.89799005 |
| TMEM206 | 48.8832556 | 0.04011522 | 0.48180016 | 0.08326111 | 0.93364392 | 0.99243654 |
| NENF | 210.914574 | 0.06802576 | 0.49956154 | 0.13617092 | 0.89168616 | 0.98597539 |
| ATF3 | 394.581128 | -3.0575912 | 1.25964299 | -2.4273475 | 0.01520968 | 0.29684131 |
| NSL1 | 154.064064 | -0.1860218 | 0.20407473 | -0.9115378 | 0.36201206 | 0.85690129 |
| TATDN3 | 118.893902 | -0.1130903 | 0.36728891 | -0.3079056 | 0.75815415 | 0.96180522 |
| FLVCR1 | 117.404913 | -0.6269703 | 0.23801197 | -2.6341966 | 0.00843366 | 0.22489769 |
| ANGEL2 | 127.74713 | -0.1919511 | 0.19189862 | -1.0002733 | 0.31717825 | 0.84285425 |
| RPS6KC1 | 345.273261 | -0.337376 | 0.22612861 | -1.4919653 | 0.13570824 | 0.67994379 |
| SMYD2 | 104.021163 | -0.2613076 | 0.25469252 | -1.0259728 | 0.30490442 | 0.83709431 |
| PTPN14 | 802.214698 | -0.1321095 | 0.29166861 | -0.4529438 | 0.65058924 | 0.95061388 |
| KCTD3 | 232.168673 | 0.07951955 | 0.18484615 | 0.43019315 | 0.66705514 | 0.9527406 |
| ESRRG | 4685.15676 | 0.45739887 | 0.28903134 | 1.58252347 | 0.11353012 | 0.6483668 |
| GPATCH2 | 558.422677 | -0.2367053 | 0.24174423 | -0.9791559 | 0.32750294 | 0.84381657 |
| SPATA17 | 150.326628 | -0.0874885 | 0.30556559 | -0.2863165 | 0.77463572 | 0.96388926 |
| RRP15 | 76.7199502 | -0.3548266 | 0.22395398 | -1.5843727 | 0.11310893 | 0.64739574 |
| LYPLAL1 | 492.767449 | 0.12719398 | 0.28698658 | 0.44320533 | 0.65761722 | 0.95100227 |
| EPRS | 222.561454 | -0.1867125 | 0.16467684 | -1.1338113 | 0.25687371 | 0.80676096 |
| BPNT1 | 66.4151381 | 0.42659928 | 0.26638093 | 1.60146326 | 0.10927435 | 0.63854503 |
| IARS2 | 216.283733 | 0.06584358 | 0.18827606 | 0.34971827 | 0.72655014 | 0.95979351 |
| RAB3GAP2 | 561.035997 | -0.3668061 | 0.1870338 | -1.9611753 | 0.04985857 | 0.49497258 |
| MARK1 | 302.956524 | 0.20651709 | 0.32874528 | 0.62819787 | 0.52987432 | 0.92181514 |
| C1orf115 | 51.2018643 | 0.29417302 | 0.41061251 | 0.71642487 | 0.47372904 | 0.90242252 |
| 2-Mar | 197.155226 | 0.37715714 | 0.26329881 | 1.43243008 | 0.15202077 | 0.7037004 |
| 1-Mar | 61.1186637 | -0.6229975 | 0.48499979 | -1.2845316 | 0.19895603 | 0.76762045 |
| AL513314.2 | 45.5209922 | 0.12627524 | 0.36996035 | 0.341321 | 0.73286194 | 0.96035038 |
| MIA3 | 145.510025 | 0.04642231 | 0.24363496 | 0.19054044 | 0.84888566 | 0.97656375 |
| AIDA | 79.568725 | -0.3934494 | 0.28889175 | -1.3619268 | 0.17322097 | 0.72964461 |
| BROX | 187.601335 | 0.08914425 | 0.23454277 | 0.38007672 | 0.70388847 | 0.95672887 |
| FAM177B | 86.1211772 | -1.1460263 | 0.62530343 | -1.8327524 | 0.06683941 | 0.55637569 |
| AC093152.1 | 125.048443 | 0.08993329 | 0.29437356 | 0.30550738 | 0.75997975 | 0.96190674 |
| TLR5 | 55.5003259 | 0.46270158 | 0.47466146 | 0.97480335 | 0.32965781 | 0.84442294 |

| | | | | | | |
|---|---|---|---|---|---|---|
| SUSD4 | 197.316278 | 0.06211444 | 0.4451172 | 0.13954626 | 0.8890185 | 0.98551241 |
| CAPN2 | 292.743663 | 0.12537243 | 0.18731063 | 0.66932899 | 0.50328564 | 0.90964098 |
| TP53BP2 | 150.541406 | -0.4372923 | 0.28047753 | -1.5590994 | 0.11897286 | 0.65485063 |
| FBXO28 | 199.200685 | -0.052968 | 0.19383212 | -0.2732676 | 0.78464751 | 0.96513523 |
| NVL | 200.934739 | -0.1780758 | 0.24821841 | -0.7174157 | 0.47311765 | 0.90216029 |
| CNIH4 | 167.264923 | -0.2452685 | 0.23508107 | -1.0433359 | 0.29679276 | 0.83536682 |
| WDR26 | 454.330915 | -0.167398 | 0.19106983 | -0.876109 | 0.38097075 | 0.86593382 |
| CNIH3 | 154.381813 | 0.6083339 | 0.33056112 | 1.84030689 | 0.06572319 | 0.55444578 |
| DNAH14 | 711.985737 | 0.04180132 | 0.29521533 | 0.14159605 | 0.88739909 | 0.98551241 |
| LBR | 169.919962 | -0.1198105 | 0.1953631 | -0.6132711 | 0.53969708 | 0.92403616 |
| ENAH | 277.18518 | 0.14951358 | 0.18981127 | 0.78769604 | 0.43087452 | 0.88627379 |
| SRP9 | 284.377017 | 0.24874381 | 0.32645673 | 0.76195033 | 0.44608965 | 0.8918683 |
| EPHX1 | 77.6988403 | 0.12381366 | 0.29638743 | 0.41774262 | 0.67613531 | 0.95529325 |
| AL591895.1 | 86.2907283 | -0.0228762 | 0.24607177 | -0.0929655 | 0.92593095 | 0.99051393 |
| TMEM63A | 183.735207 | -0.2385991 | 0.23182353 | -1.0292272 | 0.30337291 | 0.83709431 |
| SDE2 | 39.2140526 | -0.076876 | 0.33232584 | -0.2313271 | 0.81706071 | 0.96878439 |
| H3F3A | 63.5071434 | 0.28978175 | 0.31555249 | 0.91833137 | 0.35844541 | 0.8557799 |
| ACBD3 | 239.454657 | -0.1393355 | 0.15711682 | -0.8868272 | 0.37517193 | 0.8635629 |
| PARP1 | 119.963401 | 0.47612463 | 0.30562784 | 1.55785751 | 0.11926703 | 0.65527787 |
| ITPKB | 185.812836 | -0.0639801 | 0.39556204 | -0.1617448 | 0.87150683 | 0.98208511 |
| COQ8A | 82.4677882 | 0.29280731 | 0.35527338 | 0.82417463 | 0.40984034 | 0.88016834 |
| CDC42BPA | 1411.4819 | 0.24027405 | 0.18751234 | 1.28137728 | 0.20006118 | 0.7687013 |
| ZNF678 | 167.453912 | -0.296977 | 0.24823768 | -1.1963412 | 0.23156342 | 0.79200039 |
| SNAP47 | 70.4914553 | 0.30032694 | 0.25132097 | 1.19499355 | 0.23208955 | 0.7922741 |
| ARF1 | 385.657952 | -0.1499844 | 0.23206216 | -0.6463115 | 0.51807765 | 0.91751084 |
| C1orf35 | 50.5621818 | -0.1399984 | 0.30561356 | -0.4580896 | 0.64688811 | 0.95061388 |
| MRPL55 | 72.5436738 | 0.01846486 | 0.38073192 | 0.04849832 | 0.9613191 | 0.99635733 |
| GUK1 | 447.230002 | -0.1363913 | 0.54852291 | -0.2486521 | 0.80362994 | 0.96678163 |
| OBSCN | 210.848264 | 0.36695725 | 0.26331719 | 1.39359399 | 0.16344025 | 0.71642393 |
| RNF187 | 117.760415 | 0.28586153 | 0.31908311 | 0.89588424 | 0.37031459 | 0.86127318 |
| RAB4A | 125.947613 | 0.16780664 | 0.23381243 | 0.71769768 | 0.47294371 | 0.90213021 |
| NUP133 | 165.651686 | -0.3590568 | 0.18079539 | -1.9859843 | 0.04703507 | 0.48600281 |
| ABCB10 | 78.3367057 | -0.1874218 | 0.27159856 | -0.6900693 | 0.49015058 | 0.90730741 |
| TAF5L | 41.2854042 | -0.1547083 | 0.25923801 | -0.5967808 | 0.55065372 | 0.92793873 |
| GALNT2 | 259.612447 | -0.1602696 | 0.38837697 | -0.4126651 | 0.67985197 | 0.95574492 |
| COG2 | 93.2829256 | 0.05105097 | 0.23506625 | 0.21717697 | 0.82807043 | 0.97117144 |
| C1orf198 | 56.8823011 | -0.0121293 | 0.26288822 | -0.0461387 | 0.9631997 | 0.99637506 |
| TTC13 | 123.925533 | -0.1118363 | 0.26847593 | -0.41656 | 0.67700028 | 0.95529325 |
| ARV1 | 70.3488804 | 0.27826963 | 0.23540605 | 1.18208361 | 0.23717253 | 0.7980661 |
| C1orf131 | 51.1111752 | 0.02926615 | 0.37209072 | 0.07865327 | 0.93730841 | 0.99243654 |
| GNPAT | 148.330109 | -0.3450404 | 0.17834098 | -1.934723 | 0.05302429 | 0.50981156 |
| EGLN1 | 416.727644 | -0.4724731 | 0.4023864 | -1.1741775 | 0.24032394 | 0.80057052 |
| DISC1 | 51.3692416 | -0.0104927 | 0.59684714 | -0.0175803 | 0.98597371 | 0.99768314 |
| SIPA1L2 | 120.213234 | 0.89438439 | 0.48522296 | 1.84324415 | 0.06529337 | 0.55362017 |
| NTPCR | 104.424939 | -0.0276381 | 0.25514399 | -0.1083235 | 0.91373911 | 0.98830345 |
| PCNX2 | 549.392048 | -0.2088492 | 0.19189147 | -1.0883714 | 0.27643119 | 0.82184366 |
| MAP3K21 | 1286.20522 | 1.41410971 | 0.50488412 | 2.80085993 | 0.00509666 | 0.17686435 |
| KCNK1 | 166.646431 | 0.03661969 | 0.51248245 | 0.0714555 | 0.94303524 | 0.99265578 |
| COA6 | 73.2049892 | 0.43368047 | 0.39726154 | 1.09167495 | 0.274976 | 0.8202929 |
| TARBP1 | 193.228017 | -0.3008874 | 0.27713368 | -1.0857121 | 0.27760639 | 0.82230498 |
| IRF2BP2 | 258.697261 | 0.31221811 | 0.48178342 | 0.64804661 | 0.51695481 | 0.91751084 |
| TOMM20 | 310.137666 | 0.42279103 | 0.33122118 | 1.27646133 | 0.20179249 | 0.76957251 |
| ARID4B | 1066.43615 | -0.1179978 | 0.24570976 | -0.4802325 | 0.63106205 | 0.94622065 |
| B3GALNT2 | 134.475882 | 0.11321955 | 0.26876142 | 0.42126417 | 0.67356219 | 0.9548129 |

| | | | | | | |
|---|---|---|---|---|---|---|
| LYST | 310.827602 | -0.1316073 | 0.27064048 | -0.4862811 | 0.62676789 | 0.94511671 |
| GPR137B | 66.960005 | -0.4814937 | 0.44635262 | -1.0787295 | 0.28070834 | 0.82628479 |
| ERO1B | 89.4106164 | 0.60270191 | 0.43961863 | 1.37096536 | 0.17038576 | 0.72675731 |
| EDARADD | 32.0990232 | -0.5060923 | 0.42371475 | -1.1944174 | 0.23231472 | 0.7922741 |
| LGALS8 | 93.3263875 | -0.0451298 | 0.30608537 | -0.147442 | 0.88278315 | 0.98347692 |
| HEATR1 | 142.579237 | -0.5871765 | 0.28326903 | -2.072858 | 0.0381855 | 0.44791264 |
| MTR | 1862.55103 | 0.55324427 | 0.30587058 | 1.80875279 | 0.07048942 | 0.5621116 |
| CHRM3 | 225.076653 | -1.1621993 | 0.69297033 | -1.6771271 | 0.09351763 | 0.61317397 |
| FH | 60.2117379 | 0.46390829 | 0.32104186 | 1.44500875 | 0.14845543 | 0.70113244 |
| OPN3 | 30.2697442 | 0.3217316 | 0.36198005 | 0.88881031 | 0.37410503 | 0.8625432 |
| CEP170 | 466.5905 | -1.4434551 | 0.48073212 | -3.0026183 | 0.00267668 | 0.13421817 |
| SDCCAG8 | 1179.93166 | -0.7490015 | 0.268435 | -2.7902528 | 0.00526669 | 0.17920428 |
| AKT3 | 2882.06145 | -0.0496702 | 0.21295553 | -0.2332423 | 0.81557328 | 0.96844493 |
| ZBTB18 | 66.0607946 | 0.29083116 | 0.30671833 | 0.94820274 | 0.34302625 | 0.85120525 |
| AL645465.1 | 40.2001952 | 0.08541494 | 0.5835995 | 0.14635883 | 0.88363812 | 0.98369742 |
| ADSS | 241.290104 | -0.0183758 | 0.27573276 | -0.0666435 | 0.94686553 | 0.99314907 |
| CATSPERE | 180.753478 | 0.00060016 | 0.36295682 | 0.00165353 | 0.99868067 | 0.99967893 |
| DESI2 | 173.107999 | -0.1074154 | 0.23006631 | -0.4668887 | 0.64057948 | 0.94935885 |
| COX20 | 316.804384 | 0.18463111 | 0.36720206 | 0.50280523 | 0.61510122 | 0.94169251 |
| HNRNPU | 1158.38699 | -0.0820633 | 0.24128004 | -0.3401165 | 0.73376882 | 0.9603968 |
| EFCAB2 | 312.536954 | -0.1084024 | 0.38198885 | -0.2837843 | 0.77657569 | 0.96452972 |
| KIF26B | 513.04736 | 0.83417252 | 0.56683558 | 1.47163049 | 0.14112069 | 0.68601316 |
| SMYD3 | 2044.03155 | -0.1796478 | 0.19353972 | -0.928222 | 0.35329241 | 0.8557799 |
| TFB2M | 43.0214804 | -0.1216588 | 0.36447435 | -0.3337926 | 0.73853606 | 0.96078945 |
| CNST | 278.505986 | 0.05788786 | 0.19847479 | 0.29166354 | 0.77054389 | 0.96352463 |
| SCCPDH | 670.846096 | -0.2099796 | 0.53021597 | -0.3960265 | 0.69208546 | 0.95672887 |
| AHCTF1 | 288.606022 | -0.043911 | 0.16441696 | -0.267071 | 0.78941446 | 0.96515649 |
| ZNF124 | 169.932374 | -0.0167239 | 0.33620135 | -0.0497437 | 0.96032661 | 0.99635733 |
| AL390728.6 | 60.6287968 | -0.3271163 | 0.40278769 | -0.8121308 | 0.4167166 | 0.88242985 |
| ZNF496 | 65.5755634 | 0.16413283 | 0.34277493 | 0.47883558 | 0.63205561 | 0.94654542 |
| ZNF672 | 55.2640072 | 0.44391939 | 0.29105736 | 1.52519555 | 0.12721033 | 0.67016545 |
| ZNF692 | 48.6661866 | 0.12124183 | 0.30214057 | 0.40127622 | 0.68821677 | 0.95597173 |
| FAM110C | 133.121822 | -0.603316 | 0.61921003 | -0.9743317 | 0.32989186 | 0.84442294 |
| SH3YL1 | 346.593695 | 0.25848555 | 0.20286908 | 1.2741496 | 0.2026104 | 0.76996624 |
| ACP1 | 129.653585 | 0.0483215 | 0.19313848 | 0.25019093 | 0.8024397 | 0.9661728 |
| TMEM18 | 38.8683924 | 0.02924143 | 0.41929961 | 0.06973874 | 0.94440161 | 0.99265578 |
| EIPR1 | 343.487914 | -0.208868 | 0.32583582 | -0.6410222 | 0.52150826 | 0.91787515 |
| TRAPPC12 | 363.036849 | -0.0988287 | 0.25836289 | -0.3825189 | 0.70207654 | 0.95672887 |
| ADI1 | 219.393011 | 0.40069779 | 0.46874299 | 0.85483474 | 0.39264264 | 0.87161511 |
| AC108488.2 | 33.7045294 | -0.4629134 | 0.33111018 | -1.3980646 | 0.16209367 | 0.71482182 |
| RPS7 | 560.381465 | 0.01765933 | 0.49570175 | 0.0356249 | 0.97158145 | 0.99656026 |
| RNF144A | 150.901845 | -0.5022741 | 0.34701361 | -1.447419 | 0.14777962 | 0.69834914 |
| ID2 | 408.763862 | -0.2804413 | 0.66396045 | -0.4223765 | 0.67275019 | 0.9544944 |
| KIDINS220 | 415.923807 | -0.266464 | 0.20162806 | -1.3215622 | 0.18631397 | 0.74938192 |
| MBOAT2 | 702.519878 | 0.25960682 | 0.36815299 | 0.70516017 | 0.48071057 | 0.90478569 |
| ASAP2 | 1844.45995 | 0.46470005 | 0.23211239 | 2.00204757 | 0.04527962 | 0.48339878 |
| ITGB1BP1 | 195.502782 | -0.0261229 | 0.38924849 | -0.0671111 | 0.94649329 | 0.99307065 |
| CPSF3 | 71.9430117 | -0.3306905 | 0.33228109 | -0.9952132 | 0.31963259 | 0.84285425 |
| IAH1 | 149.749581 | 0.11688812 | 0.29867553 | 0.39135486 | 0.69553496 | 0.95672887 |
| ADAM17 | 352.974644 | -0.3703567 | 0.23320194 | -1.5881374 | 0.11225527 | 0.64471804 |
| YWHAQ | 331.789698 | 0.09466924 | 0.25249943 | 0.37492853 | 0.70771362 | 0.95752169 |
| TAF1B | 331.749766 | 0.12585299 | 0.2904905 | 0.43324305 | 0.66483821 | 0.9520866 |
| GRHL1 | 87.3929267 | 0.11029963 | 0.37668096 | 0.29281977 | 0.76965992 | 0.96345263 |
| KLF11 | 66.4234491 | 0.0902101 | 0.31618637 | 0.28530673 | 0.77540916 | 0.96408684 |

| | | | | | | |
|---|---|---|---|---|---|---|
| CYS1 | 812.255191 | 0.88764266 | 0.30474122 | 2.91277521 | 0.00358232 | 0.14665138 |
| HPCAL1 | 296.922102 | -0.3395518 | 0.47400624 | -0.7163447 | 0.47377854 | 0.90242252 |
| ODC1 | 178.501633 | 0.22200751 | 0.50243062 | 0.441867 | 0.65858545 | 0.95113765 |
| NOL10 | 254.920506 | -0.2231685 | 0.20219532 | -1.1037273 | 0.26971145 | 0.81817548 |
| PDIA6 | 347.153349 | 0.60086162 | 0.27545696 | 2.1813267 | 0.02915926 | 0.40623517 |
| PQLC3 | 109.500371 | 0.2052323 | 0.46224948 | 0.443986 | 0.65705271 | 0.95100227 |
| ROCK2 | 907.415106 | 0.08463416 | 0.16754871 | 0.50513168 | 6.13E-01 | 9.40E-01 |
| E2F6 | 41.8185768 | 0.06088429 | 0.29849532 | 0.20397065 | 0.83837643 | 0.97397018 |
| LPIN1 | 181.419379 | -0.2900032 | 0.46933975 | -0.6178961 | 0.5366438 | 0.92324352 |
| MIR3681HG | 119.977215 | 0.10475896 | 0.52929251 | 0.19792261 | 0.84310561 | 0.97560439 |
| TRIB2 | 163.631163 | -0.2173609 | 0.44772564 | -0.4854778 | 0.62733746 | 0.94511671 |
| FAM84A | 220.260482 | -0.0559619 | 0.51134939 | -0.1094396 | 0.91285381 | 0.98830345 |
| NBAS | 703.367171 | -0.4912142 | 0.19424882 | -2.5287885 | 0.0114457 | 0.26305028 |
| DDX1 | 205.144906 | -0.0065353 | 0.17107837 | -0.0382005 | 0.96952786 | 0.99645289 |
| SMC6 | 202.154444 | -0.0330633 | 0.30472142 | -0.1085033 | 0.91359643 | 0.98830345 |
| GEN1 | 65.0476542 | -0.3485953 | 0.47160261 | -0.7391716 | 0.45980281 | 0.89753337 |
| KCNS3 | 61.9815187 | 1.0582807 | 0.42814519 | 2.47177997 | 0.01344422 | 0.28170334 |
| RDH14 | 59.5334146 | 0.12773325 | 0.36074938 | 0.35407753 | 0.72328078 | 0.95963742 |
| LINC01376 | 193.322838 | 0.3074874 | 0.35177987 | 0.87409038 | 0.38206904 | 0.86593382 |
| TTC32 | 52.0934663 | -0.2769876 | 0.29804226 | -0.9293567 | 0.35270427 | 0.85536253 |
| WDR35 | 221.214424 | -0.177596 | 0.19322551 | -0.9191124 | 0.35803677 | 0.8557799 |
| LAPTM4A | 565.043441 | 0.4683292 | 0.34612965 | 1.3530456 | 0.17604106 | 0.73444173 |
| SDC1 | 86.4506542 | -0.7861331 | 0.48877242 | -1.6083827 | 0.10775138 | 0.63520805 |
| PUM2 | 922.247787 | -0.210789 | 0.16895902 | -1.2475746 | 0.21218688 | 0.77709975 |
| RHOB | 198.024956 | -1.4902231 | 0.56701447 | -2.6281924 | 0.008584 | 0.22546434 |
| HS1BP3 | 55.6431537 | -0.1951341 | 0.24196859 | -0.806444 | 0.41998686 | 0.8828928 |
| GDF7 | 50.657202 | 1.36748177 | 0.43981684 | 3.10920738 | 0.0018759 | 0.1093833 |
| LDAH | 287.239887 | -0.0215027 | 0.31927298 | -0.0673488 | 0.946304 | 0.99307065 |
| AC018742.1 | 203.226907 | -0.8441654 | 0.71589135 | -1.1791808 | 0.23832618 | 0.79848413 |
| KLHL29 | 64.1653855 | -1.1278047 | 0.41463746 | -2.7199778 | 0.00652863 | 0.19947532 |
| ATAD2B | 520.743538 | -0.0003028 | 0.26590454 | -0.0011389 | 0.99909127 | 0.99968413 |
| UBXN2A | 202.261665 | 0.02211406 | 0.26384222 | 0.08381546 | 0.93320316 | 0.99243654 |
| SF3B6 | 196.997961 | 0.35570808 | 0.50994065 | 0.69754801 | 0.48545991 | 0.90695174 |
| FAM228B | 57.0890769 | 0.0173362 | 0.47229073 | 0.03670662 | 0.97071893 | 0.99656026 |
| TP53I3 | 111.330729 | 0.00875766 | 0.40541846 | 0.02160154 | 0.9827658 | 0.9971769 |
| ITSN2 | 587.044892 | -0.1383302 | 0.14194284 | -0.9745489 | 0.32978408 | 0.84442294 |
| NCOA1 | 1185.22248 | 0.04885783 | 0.23886369 | 0.20454272 | 0.83792941 | 0.97385715 |
| PTRHD1 | 38.903839 | 0.08897505 | 0.53737729 | 0.16557278 | 0.86849317 | 0.98185636 |
| DNAJC27 | 94.9726916 | -0.0338394 | 0.25162834 | -0.1344818 | 0.89302158 | 0.98597539 |
| DNAJC27-AS | 59.3268205 | 0.66399054 | 0.43894334 | 1.51270217 | 0.13035533 | 0.67329908 |
| DNMT3A | 300.569476 | 0.10193519 | 0.29799374 | 0.34207157 | 0.73229704 | 0.96008876 |
| DTNB | 759.464198 | 0.39245556 | 0.30341374 | 1.29346664 | 0.19584971 | 0.76403539 |
| ASXL2 | 418.872992 | -0.2760994 | 0.21806915 | -1.2661093 | 0.20547397 | 0.77193931 |
| KIF3C | 34.6863794 | 0.01227773 | 0.51251493 | 0.02395584 | 0.98088783 | 0.99696835 |
| RAB10 | 917.195203 | -0.3185628 | 0.19776251 | -1.610835 | 0.1072157 | 0.63477557 |
| HADHA | 401.55045 | 0.18730347 | 0.23145755 | 0.80923466 | 0.41838018 | 0.88242985 |
| HADHB | 451.423601 | 0.19494401 | 0.17977121 | 1.08440063 | 0.2781872 | 0.82310625 |
| SELENOI | 189.569965 | -0.4053302 | 0.19662446 | -2.0614433 | 0.03926077 | 0.45565104 |
| MAPRE3 | 76.0245339 | 0.28846588 | 0.30170164 | 0.95612964 | 0.33900674 | 0.84863112 |
| TMEM214 | 59.0746425 | -0.2022395 | 0.3203435 | -0.6313209 | 0.52783074 | 0.92102334 |
| AGBL5 | 94.9200919 | 0.1200188 | 0.31482855 | 0.38121954 | 0.70304035 | 0.95672887 |
| OST4 | 313.078856 | 0.07757098 | 0.40773417 | 0.19024892 | 0.84911408 | 0.97659302 |
| PREB | 30.4483897 | 0.63016544 | 0.37205107 | 1.69376061 | 0.09031078 | 0.60779802 |
| ATRAID | 198.979152 | 0.17157516 | 0.44020196 | 0.38976464 | 0.69671059 | 0.95672887 |

| | | | | | | |
|---|---|---|---|---|---|---|
| CAD | 35.5804407 | -0.0833769 | 0.35157319 | -0.2371538 | 0.81253745 | 0.9680627 |
| MPV17 | 63.0971635 | -0.0602586 | 0.25159333 | -0.2395079 | 0.81071179 | 0.9680627 |
| GTF3C2 | 155.718962 | -0.1232036 | 0.26119915 | -0.4716844 | 0.63715209 | 0.94821838 |
| EIF2B4 | 43.6039707 | -0.0011361 | 0.25083143 | -0.0045295 | 0.99638599 | 0.99967893 |
| SNX17 | 90.8550613 | 0.2251632 | 0.42557702 | 0.52907743 | 0.59675174 | 0.93617523 |
| PPM1G | 255.204172 | -0.0933553 | 0.33114862 | -0.2819136 | 0.77800978 | 0.96472201 |
| NRBP1 | 132.963189 | -0.0915026 | 0.23548713 | -0.3885671 | 0.69759639 | 0.95672887 |
| KRTCAP3 | 53.2773305 | -0.7368001 | 0.31733012 | -2.3218725 | 0.0202398 | 0.34101791 |
| IFT172 | 165.045077 | -0.1063916 | 0.2226606 | -0.4778198 | 0.63277847 | 0.94683634 |
| ZNF512 | 187.813493 | -0.2925434 | 0.24257782 | -1.2059777 | 0.22782611 | 0.79042788 |
| GPN1 | 45.5448134 | -0.0028596 | 0.31542497 | -0.0090657 | 0.99276671 | 0.99919666 |
| SUPT7L | 60.2810088 | -0.0421413 | 0.25761892 | -0.1635798 | 0.87006191 | 0.98202612 |
| SLC4A1AP | 80.7254782 | -0.0364722 | 0.23030053 | -0.1583678 | 0.87416699 | 0.98246825 |
| MRPL33 | 605.699512 | 0.46527594 | 0.32316825 | 1.43973283 | 0.149943 | 0.7029344 |
| RBKS | 451.381183 | -1.1005711 | 0.62274447 | -1.7672917 | 0.07717939 | 0.57405253 |
| BABAM2 | 1635.23234 | 0.12403801 | 0.31750556 | 0.39066405 | 0.69604557 | 0.95672887 |
| FOSL2 | 1205.35802 | -1.1143246 | 0.7393037 | -1.5072624 | 0.13174342 | 0.67390608 |
| PPP1CB | 548.608627 | -0.3549292 | 0.69319375 | -0.5120202 | 0.60863685 | 0.94003018 |
| TRMT61B | 99.0293199 | 0.21263706 | 0.2751815 | 0.7727157 | 0.43969065 | 0.88917548 |
| WDR43 | 209.980761 | -0.367453 | 0.30419016 | -1.2079712 | 0.22705835 | 0.79029275 |
| CLIP4 | 358.622512 | -0.180577 | 0.43902026 | -0.4113181 | 0.68083927 | 0.95574492 |
| YPEL5 | 334.217048 | 0.55333841 | 0.54879184 | 1.00828468 | 0.31331782 | 0.84285425 |
| LCLAT1 | 278.239452 | -0.1727789 | 0.25113357 | -0.687996 | 0.49145533 | 0.90730741 |
| GALNT14 | 372.468318 | 0.86399979 | 0.60481771 | 1.42852925 | 0.15313958 | 0.70700565 |
| MEMO1 | 33.9052865 | -0.2000967 | 0.34410375 | -0.581501 | 0.56090285 | 0.92934517 |
| DPY30 | 193.302449 | -0.1986488 | 0.38642517 | -0.514068 | 0.60720444 | 0.93940102 |
| SPAST | 203.279813 | -0.0164062 | 0.29791759 | -0.0550695 | 0.95608307 | 0.99565888 |
| SLC30A6 | 167.055032 | -0.2039684 | 0.22268051 | -0.9159688 | 0.35968324 | 0.85608464 |
| YIPF4 | 264.006948 | 0.06902399 | 0.18940673 | 0.36442204 | 0.71554288 | 0.95791799 |
| BIRC6 | 1493.89129 | -0.2740332 | 0.21150112 | -1.2956585 | 0.19509316 | 0.76233115 |
| TTC27 | 215.55311 | -0.3174364 | 0.21377175 | -1.4849314 | 0.13756199 | 0.68146162 |
| LINC00486 | 300.432025 | -0.397126 | 0.20600831 | -1.9277182 | 0.05389019 | 0.51332378 |
| LTBP1 | 68.1630228 | 1.33089724 | 0.67965079 | 1.95820745 | 0.05020568 | 0.49590526 |
| FAM98A | 60.6222576 | 0.19833212 | 0.26333286 | 0.75316132 | 0.45135298 | 0.89400477 |
| LINC01317 | 56.428322 | 1.35507229 | 0.52760074 | 2.56836694 | 1.02E-02 | 0.24616901 |
| AC068599.1 | 473.166867 | 2.3341506 | 0.56882362 | 4.10346987 | 4.07E-05 | 0.00969399 |
| LINC01320 | 700.908043 | -0.168701 | 0.43769396 | -0.3854313 | 0.69991788 | 0.95672887 |
| AC012593.1 | 116.816291 | 1.20269852 | 1.00283007 | 1.1993044 | 0.2304096 | 0.79075638 |
| CRIM1 | 962.397628 | -0.0389803 | 0.30899822 | -0.1261506 | 0.89961274 | 0.98660836 |
| FEZ2 | 328.864526 | 0.15971034 | 0.29419538 | 0.54287168 | 0.58721815 | 0.93483536 |
| STRN | 484.899194 | -0.1822892 | 0.15407046 | -1.1831546 | 2.37E-01 | 0.79797745 |
| HEATR5B | 217.473122 | -0.3303974 | 0.27170456 | -1.2160171 | 0.2239784 | 0.78960314 |
| EIF2AK2 | 333.125301 | -0.6288146 | 0.4418 | -1.4233016 | 0.15464874 | 0.70839173 |
| CEBPZOS | 170.707623 | 0.13202385 | 0.32680233 | 0.40398685 | 0.68622239 | 0.9558749 |
| CEBPZ | 180.683849 | 0.07947836 | 0.29069604 | 0.27340709 | 0.78454031 | 0.96513523 |
| NDUFAF7 | 103.338157 | 0.17071489 | 0.2524767 | 0.67616097 | 0.49893845 | 0.90839719 |
| PRKD3 | 287.833726 | 0.09674935 | 0.1951521 | 0.49576385 | 0.62006103 | 0.94319304 |
| CDC42EP3 | 51.5844807 | 0.26020608 | 0.31990716 | 0.81337997 | 0.41600024 | 0.88242985 |
| RMDN2 | 138.853622 | 0.35071014 | 0.28461625 | 1.23222105 | 0.21786652 | 0.78394745 |
| CYP1B1-AS1 | 43.5411835 | 1.11011804 | 0.53966378 | 2.05705493 | 0.03968094 | 0.45799149 |
| ATL2 | 233.18124 | -0.0425417 | 0.21951275 | -0.1938004 | 0.84633218 | 0.97656375 |
| HNRNPLL | 184.64837 | 0.10310763 | 0.22665372 | 0.4549126 | 0.64917212 | 0.95061388 |
| GALM | 188.597175 | 0.48762937 | 0.31745406 | 1.53606279 | 0.12452298 | 0.66615663 |
| SRSF7 | 206.787556 | 0.52539863 | 0.36207545 | 1.45107498 | 0.14675898 | 0.69720496 |

| | | | | | | |
|---|---|---|---|---|---|---|
| DHX57 | 161.697901 | -0.0784263 | 0.29641155 | -0.2645859 | 0.79132852 | 0.9653598 |
| MORN2 | 90.5599003 | 0.57478502 | 0.44584611 | 1.28920049 | 0.19732839 | 0.76534786 |
| SOS1 | 810.416268 | -0.0537194 | 0.26159004 | -0.2053572 | 0.83729309 | 0.97357501 |
| MAP4K3 | 1681.99623 | 0.4813391 | 0.24806954 | 1.94033939 | 0.05233846 | 0.50740706 |
| AC007388.1 | 812.826622 | 0.80573372 | 0.27886464 | 2.88933631 | 0.00386056 | 0.15207852 |
| TMEM178A | 133.974867 | 2.32804661 | 0.35251968 | 6.60401885 | 4.00E-11 | 2.10E-07 |
| THUMPD2 | 168.405922 | -0.1648859 | 0.34863159 | -0.4729518 | 0.63624758 | 0.94821838 |
| SLC8A1 | 845.096328 | 1.57178475 | 0.75829759 | 2.07278089 | 0.03819268 | 0.44791264 |
| EML4 | 894.184628 | 0.06340004 | 0.26627606 | 0.2380989 | 0.81180439 | 0.9680627 |
| COX7A2L | 276.704032 | 0.09582673 | 0.26335559 | 0.36386821 | 0.71595642 | 0.95791799 |
| MTA3 | 468.401578 | 0.0066713 | 0.22797931 | 0.02926274 | 0.97665504 | 0.99656026 |
| ZFP36L2 | 757.336618 | 0.90034987 | 0.38172446 | 2.35863814 | 0.01834213 | 0.32472005 |
| THADA | 659.662634 | -0.1580584 | 0.25367635 | -0.6230711 | 0.53323783 | 0.92277617 |
| PLEKHH2 | 536.997015 | 0.14642789 | 0.38979033 | 0.3756581 | 0.70717109 | 0.95752169 |
| DYNC2LI1 | 469.439462 | 0.14739256 | 0.19332763 | 0.7623978 | 0.44582262 | 0.8918683 |
| LRPPRC | 447.344024 | -0.2208095 | 0.18037502 | -1.2241692 | 0.2208884 | 0.78752408 |
| PPM1B | 415.13433 | 0.3087886 | 0.25105244 | 1.22997649 | 0.21870591 | 0.78557122 |
| SLC3A1 | 225.634202 | 2.0221617 | 0.64512114 | 3.13454569 | 0.0017212 | 0.10426718 |
| PREPL | 533.869504 | 0.2569583 | 0.30877608 | 0.83218329 | 0.40530549 | 0.87905661 |
| CAMKMT | 1366.58143 | -0.1524335 | 0.21970237 | -0.6938182 | 0.48779624 | 0.90730741 |
| SRBD1 | 316.190003 | -0.3666758 | 0.26104555 | -1.4046429 | 0.16012751 | 0.71409437 |
| PRKCE | 1317.67186 | 0.09272959 | 0.29794049 | 0.31123525 | 0.75562178 | 0.96120879 |
| EPAS1 | 70.8983863 | -1.2525366 | 0.43614341 | -2.8718457 | 0.00408082 | 0.15545856 |
| RHOQ | 285.507879 | -0.5770356 | 0.34188638 | -1.6877994 | 0.09144973 | 0.61047264 |
| AC018682.1 | 49.7560376 | -0.8222718 | 0.66889756 | -1.2292941 | 0.21896157 | 0.78597171 |
| PIGF | 119.735088 | -0.250575 | 0.19621411 | -1.2770488 | 0.20158503 | 0.76957251 |
| CRIPT | 61.2643707 | 0.50205004 | 0.31769861 | 1.58027145 | 0.11404472 | 0.6492062 |
| SOCS5 | 131.731066 | -0.0004353 | 0.27243506 | -0.0015979 | 0.99872504 | 0.99967893 |
| MCFD2 | 194.883748 | -0.1545201 | 0.23055074 | -0.6702215 | 0.50271658 | 0.90962743 |
| TTC7A | 201.190574 | 0.07721793 | 0.32637148 | 0.23659521 | 0.81297084 | 0.9680627 |
| AC106869.1 | 48.2540411 | 0.32333937 | 0.41150163 | 0.78575476 | 0.43201118 | 0.88697335 |
| EPCAM | 1419.62174 | 0.83375607 | 0.43216711 | 1.92924463 | 0.0537005 | 0.51316717 |
| MSH2 | 218.673766 | 0.34050856 | 0.28849461 | 1.18029436 | 0.23788316 | 0.79810098 |
| MSH6 | 78.3851895 | 0.27910927 | 0.31303098 | 0.89163465 | 0.37258879 | 0.86201082 |
| FBXO11 | 765.165809 | -0.0591777 | 0.19763209 | -0.2994335 | 0.7646093 | 0.96285313 |
| FOXN2 | 107.433606 | -0.1902878 | 0.24962348 | -0.7622993 | 0.44588141 | 0.8918683 |
| PPP1R21 | 614.758555 | -0.2577857 | 0.18386537 | -1.4020354 | 0.16090469 | 0.71409437 |
| GPR75-ASB3 | 107.513759 | 0.2464613 | 0.34841808 | 0.70737231 | 0.47933514 | 0.90300777 |
| ERLEC1 | 112.236619 | 0.11483308 | 0.18369982 | 0.62511266 | 0.53189712 | 0.92211604 |
| PSME4 | 381.969003 | -0.0135706 | 0.25659452 | -0.0528872 | 0.95782176 | 0.99579587 |
| ACYP2 | 670.723793 | -0.1647023 | 0.31763783 | -0.5185223 | 0.60409391 | 0.93804805 |
| SPTBN1 | 1738.60311 | -0.8963766 | 0.21980148 | -4.0781189 | 4.54E-05 | 0.01041317 |
| AC093110.1 | 86.7018687 | -0.4531919 | 0.6213693 | -0.7293438 | 0.46579136 | 0.89883965 |
| EML6 | 812.950245 | 1.05776138 | 0.36845321 | 2.87081601 | 0.00409414 | 0.15545856 |
| RTN4 | 2369.52103 | 0.24137632 | 0.24726486 | 0.97618531 | 0.32897264 | 0.84409527 |
| CLHC1 | 158.447722 | 0.27628314 | 0.20517075 | 1.34660101 | 0.17810877 | 0.73737861 |
| RPS27A | 1293.27595 | -0.0274125 | 0.49781773 | -0.0550654 | 0.95608635 | 0.99565888 |
| MTIF2 | 100.970518 | 0.12272143 | 0.22912543 | 0.53560807 | 0.59222945 | 0.93491607 |
| AC012358.3 | 101.027733 | 0.14919801 | 0.44815967 | 0.33291261 | 0.73920025 | 0.96078945 |
| CCDC88A | 154.735193 | -0.2187485 | 0.35862075 | -0.6099718 | 0.5418805 | 0.92479556 |
| CFAP36 | 236.085207 | -0.0075255 | 0.28964433 | -0.0259818 | 0.97927188 | 0.99696835 |
| PPP4R3B | 410.574084 | -0.2800919 | 0.2199436 | -1.2734713 | 0.20285084 | 0.76996624 |
| PNPT1 | 159.821105 | -0.7993854 | 0.36930557 | -2.1645635 | 0.03042113 | 0.41080114 |
| EFEMP1 | 151.432865 | 1.05474578 | 0.73543297 | 1.43418343 | 0.15151992 | 0.7029344 |

| | | | | | | |
|---|---|---|---|---|---|---|
| VRK2 | 268.449411 | -0.1690053 | 0.26305589 | -0.6424693 | 0.52056854 | 0.91751084 |
| FANCL | 179.05181 | -0.0669589 | 0.2682836 | -0.2495826 | 0.80291019 | 0.9665172 |
| PAPOLG | 107.277796 | -0.3831585 | 0.24293656 | -1.5771959 | 0.11475047 | 0.65008305 |
| REL | 345.438024 | -0.3626079 | 0.53161778 | -0.6820839 | 0.49518589 | 0.90839719 |
| PUS10 | 74.7475413 | 0.33751897 | 0.32087739 | 1.05186273 | 0.29286253 | 0.83336457 |
| PEX13 | 106.951046 | -0.1596078 | 0.30552818 | -0.5223998 | 0.60139202 | 0.93802476 |
| KIAA1841 | 78.5721397 | -0.5881197 | 0.30068167 | -1.9559547 | 0.05047049 | 0.49596288 |
| C2orf74 | 61.6724883 | 0.07680358 | 0.47464831 | 0.16181156 | 0.87145425 | 0.98208511 |
| USP34 | 1601.24102 | -0.3004834 | 0.20572418 | -1.4606131 | 0.14412164 | 0.69093999 |
| AC016727.1 | 38.6398269 | 0.25713096 | 0.36938399 | 0.69610748 | 0.48636152 | 0.90695174 |
| XPO1 | 730.685419 | 0.02878547 | 0.34125086 | 0.08435281 | 0.93277592 | 0.99243654 |
| FAM161A | 56.3912533 | 0.47342032 | 0.27965354 | 1.69288155 | 0.09047802 | 0.60782667 |
| CCT4 | 191.045494 | 0.07603935 | 0.22166738 | 0.34303354 | 0.73157323 | 0.96008876 |
| COMMD1 | 559.141738 | -0.1842985 | 0.22257505 | -0.8280287 | 0.40765427 | 0.87948803 |
| AC018462.1 | 38.6489386 | 0.17003325 | 0.47503956 | 0.35793492 | 0.72039202 | 0.95796325 |
| B3GNT2 | 98.3700242 | 0.14715045 | 0.26265305 | 0.56024648 | 0.57531133 | 0.93275827 |
| EHBP1 | 697.353487 | 0.10413141 | 0.16091538 | 0.64711904 | 0.5175549 | 0.91751084 |
| WDPCP | 800.714608 | -0.2676647 | 0.205227 | -1.3042372 | 0.19215273 | 0.7597012 |
| MDH1 | 259.216819 | 0.5856117 | 0.32438232 | 1.80531325 | 0.07102568 | 0.56333976 |
| UGP2 | 304.183298 | 0.09187801 | 0.25665424 | 0.35798361 | 0.72035558 | 0.95796325 |
| VPS54 | 507.685534 | 0.1515314 | 0.23540723 | 0.64369901 | 0.51977063 | 0.91751084 |
| PELI1 | 138.98807 | -0.1541771 | 0.43071035 | -0.35796 | 0.72037325 | 0.95796325 |
| AC008074.1 | 52.371359 | 0.36438603 | 0.3892097 | 0.9362203 | 0.34915977 | 0.8535343 |
| AFTPH | 366.684155 | -0.1078214 | 0.23861844 | -0.4518571 | 0.65137196 | 0.95061388 |
| SERTAD2 | 116.779304 | 0.16028774 | 0.53572831 | 0.29919595 | 0.76479054 | 0.96285313 |
| CEP68 | 88.8015043 | 0.06576883 | 0.30147727 | 0.2181552 | 0.82730819 | 0.97069946 |
| RAB1A | 688.44015 | -0.467046 | 0.25393788 | -1.8392137 | 0.06588376 | 0.55444578 |
| ACTR2 | 528.007567 | 0.01665933 | 0.24744236 | 0.06732611 | 0.94632209 | 0.99307065 |
| SPRED2 | 390.155343 | 0.54489393 | 0.23166884 | 2.35203805 | 0.01867087 | 0.32830649 |
| ETAA1 | 67.4551322 | 0.08356136 | 0.24071734 | 0.34713478 | 0.72849006 | 0.95985088 |
| C1D | 122.077549 | -0.0905218 | 0.28326131 | -0.3195698 | 0.74929444 | 0.96109466 |
| PNO1 | 39.6956252 | -0.1382848 | 0.31200063 | -0.4432197 | 0.65760681 | 0.95100227 |
| APLF | 172.738028 | -0.1326624 | 0.19793315 | -0.6702384 | 0.50270584 | 0.90962743 |
| ANTXR1 | 64.987032 | 0.57144937 | 0.44384364 | 1.28750155 | 0.19791953 | 0.76582694 |
| GFPT1 | 394.918782 | -0.1250286 | 0.26379855 | -0.473955 | 0.63553198 | 0.94807428 |
| NFU1 | 150.853841 | 0.13069986 | 0.32281924 | 0.40487011 | 0.68557299 | 0.9558749 |
| AAK1 | 1835.02893 | 0.01064446 | 0.39982196 | 0.02662299 | 0.97876044 | 0.99696835 |
| ANXA4 | 785.419559 | -0.0527907 | 0.37610675 | -0.1403609 | 0.88837486 | 0.98551241 |
| GMCL1 | 206.06264 | -0.5223882 | 0.19414338 | -2.6907341 | 0.0071295 | 0.20697274 |
| MXD1 | 92.1258825 | -0.9137988 | 0.5360832 | -1.7045839 | 0.08827206 | 0.60651125 |
| PCBP1 | 319.726851 | -0.2820883 | 0.37852478 | -0.7452308 | 0.45613218 | 0.89626521 |
| C2orf42 | 56.0611426 | 0.05046583 | 0.38304448 | 0.13174926 | 0.89518262 | 0.98597539 |
| PCYOX1 | 293.505707 | 0.89389364 | 0.33491514 | 2.66901531 | 0.0076074 | 0.21455924 |
| SNRPG | 122.548333 | -0.0425163 | 0.36294399 | -0.1171429 | 0.9067468 | 0.9875066 |
| FAM136A | 88.4667065 | 0.14818617 | 0.26938501 | 0.55009063 | 0.58225721 | 0.93402467 |
| TGFA | 240.444538 | 0.62809115 | 0.27297181 | 2.30093781 | 0.02139515 | 0.35310415 |
| NAGK | 111.269254 | -0.7553773 | 0.3729854 | -2.0252194 | 0.04284484 | 0.47339997 |
| MCEE | 87.3639232 | -0.2227436 | 0.32769264 | -0.6797332 | 0.49667338 | 0.90839719 |
| MPHOSPH10 | 73.565617 | 0.30526248 | 0.25545055 | 1.19499638 | 0.23208845 | 0.7922741 |
| PAIP2B | 48.6113931 | 0.10831045 | 0.36153754 | 0.29958286 | 0.76449536 | 0.96285313 |
| ZNF638 | 1304.00868 | -0.1791505 | 0.2445143 | -0.732679 | 0.46375421 | 0.89801718 |
| EXOC6B | 1152.98202 | -0.3492024 | 0.22279301 | -1.5673849 | 0.11702475 | 0.65289383 |
| SPR | 69.9415091 | 0.82515282 | 0.42135518 | 1.9583308 | 0.05019121 | 0.49590526 |
| EMX1 | 759.045274 | 1.45983431 | 0.4017523 | 3.63366757 | 0.00027942 | 0.03764581 |

| | | | | | | |
|---|---|---|---|---|---|---|
| SFXN5 | 450.887724 | 0.46376512 | 0.40901439 | 1.13386017 | 0.25685321 | 0.80676096 |
| RAB11FIP5 | 98.0244257 | 0.56942156 | 0.47906659 | 1.18860629 | 0.23459463 | 0.79543722 |
| CCT7 | 182.996257 | 0.50592882 | 0.40420073 | 1.25167715 | 0.21068753 | 0.77610028 |
| ALMS1 | 271.084976 | -0.090361 | 0.21095132 | -0.4283499 | 0.66839637 | 0.95294349 |
| TPRKB | 34.7429325 | -0.3066924 | 0.34043177 | -0.9008924 | 0.36764552 | 0.85902833 |
| DUSP11 | 61.5870284 | -0.3211306 | 0.26636287 | -1.2056131 | 0.22796669 | 0.79042788 |
| STAMBP | 131.71573 | -0.2143333 | 0.22920146 | -0.9351306 | 0.34972102 | 0.8535343 |
| DGUOK | 159.689026 | -0.190888 | 0.35370101 | -0.5396873 | 0.58941268 | 0.93483536 |
| TET3 | 158.652411 | -0.5710323 | 0.33934979 | -1.6827246 | 0.09242842 | 0.61120158 |
| BOLA3 | 53.9829385 | 0.45358615 | 0.44967569 | 1.00869617 | 0.31312037 | 0.84285425 |
| MOB1A | 304.326036 | 0.19590457 | 0.26673164 | 0.73446319 | 0.46266648 | 0.89801718 |
| AC006030.1 | 76.4236705 | -0.5926888 | 0.27994031 | -2.1171971 | 0.03424312 | 0.43050784 |
| WDR54 | 48.6914874 | 0.2548612 | 0.42131988 | 0.6049114 | 0.54523787 | 0.92579581 |
| RTKN | 78.8058759 | -0.2747827 | 0.28662683 | -0.9586774 | 0.33772128 | 0.84672703 |
| MOGS | 77.7898831 | 0.19642461 | 0.30181129 | 0.65081929 | 0.51516315 | 0.91599162 |
| TTC31 | 35.5002423 | -0.4918343 | 0.32894972 | -1.4951657 | 0.13487121 | 0.6785647 |
| AUP1 | 56.1191184 | -0.3775904 | 0.35016305 | -1.0783273 | 0.28088772 | 0.82628479 |
| M1AP | 105.778573 | 0.01306678 | 0.44347117 | 0.02946478 | 0.9764939 | 0.99656026 |
| HK2 | 90.9363955 | -5.0174955 | 1.5606468 | -3.2150103 | 0.0013044 | 0.09113398 |
| POLE4 | 83.6385061 | 0.31815808 | 0.42632569 | 0.74627939 | 0.45549866 | 0.89626521 |
| TACR1 | 51.876616 | 0.91477306 | 0.571878 | 1.59959478 | 0.10968851 | 0.63926535 |
| AC007681.1 | 57.6716004 | -0.154301 | 0.46484242 | -0.3319426 | 0.73993257 | 0.96084411 |
| EVA1A | 77.6558336 | 0.5549403 | 0.42995824 | 1.2906842 | 0.19681321 | 0.76478825 |
| MRPL19 | 295.164355 | 0.39386354 | 0.34667659 | 1.13611232 | 0.25590956 | 0.80525591 |
| GCFC2 | 330.858178 | 0.36532993 | 0.23516277 | 1.55351944 | 0.12029909 | 0.65691125 |
| CTNNA2 | 39.9922541 | -0.026241 | 0.55843578 | -0.0469901 | 0.96252111 | 0.99637506 |
| SUCLG1 | 292.393534 | 0.262925 | 0.2040907 | 1.28827524 | 0.19765017 | 0.76534786 |
| DNAH6 | 203.897415 | 0.06691177 | 0.44719734 | 0.14962471 | 0.88106071 | 0.98333507 |
| TMSB10 | 1182.22193 | -1.2541306 | 0.32611993 | -3.8456117 | 0.00012025 | 0.02194494 |
| KCMF1 | 331.684171 | -0.0982549 | 0.2418787 | -0.4062155 | 0.68458429 | 0.9558749 |
| TCF7L1 | 974.132364 | 0.26887659 | 0.35563248 | 0.75605183 | 0.44961812 | 0.89324553 |
| TGOLN2 | 691.902252 | 0.3462351 | 0.27228688 | 1.27158202 | 0.20352167 | 0.77083739 |
| RETSAT | 69.6239264 | 0.02816795 | 0.29500185 | 0.09548397 | 0.92393042 | 0.99051393 |
| ELMOD3 | 161.358428 | -0.0303425 | 0.21616148 | -0.1403697 | 0.88836786 | 0.98551241 |
| CAPG | 206.925873 | -0.1398071 | 0.47442227 | -0.2946892 | 0.76823133 | 0.96345263 |
| MAT2A | 268.350063 | -1.1250578 | 0.41906261 | -2.6847011 | 0.00725947 | 0.2101379 |
| GGCX | 93.2569261 | -0.0949224 | 0.26649114 | -0.3561933 | 0.72169578 | 0.95868461 |
| VAMP8 | 372.172125 | 0.25529891 | 0.43654018 | 0.58482338 | 0.55866648 | 0.92901206 |
| RNF181 | 53.6787812 | -0.056594 | 0.35701372 | -0.1585206 | 0.87404662 | 0.98246825 |
| USP39 | 171.325309 | -0.020527 | 0.20419357 | -0.1005274 | 0.91992565 | 0.99014946 |
| C2orf68 | 65.6156472 | -0.1670784 | 0.30358408 | -0.550353 | 0.58207726 | 0.93402467 |
| POLR1A | 88.8364203 | 0.16931076 | 0.26363964 | 0.64220524 | 0.52073993 | 0.91751084 |
| PTCD3 | 218.94066 | 0.11429641 | 0.18586979 | 0.61492732 | 0.53860272 | 0.92365631 |
| IMMT | 306.16983 | 0.08252487 | 0.20623089 | 0.40015767 | 0.68904039 | 0.95606293 |
| MRPL35 | 77.6798687 | 0.44194003 | 0.37196564 | 1.18812055 | 0.23478592 | 0.79578151 |
| KDM3A | 263.73855 | -0.4511753 | 0.44535524 | -1.0130684 | 0.31102748 | 0.84248332 |
| CHMP3 | 162.476921 | 0.07078738 | 0.31474851 | 0.22490139 | 0.82205599 | 0.96995572 |
| RMND5A | 218.387356 | 0.08942326 | 0.25631779 | 0.34887654 | 0.727182 | 0.95979351 |
| PLGLB1 | 56.6790579 | 0.64299706 | 0.66554753 | 0.96611742 | 0.33398543 | 0.8445213 |
| CYTOR | 214.472161 | 0.15477266 | 0.68008756 | 0.22757755 | 0.81997468 | 0.96886922 |
| KRCC1 | 198.840115 | 0.28460097 | 0.24364451 | 1.16809928 | 0.24276672 | 0.80072691 |
| THNSL2 | 78.7870685 | -0.1585179 | 0.83182312 | -0.1905668 | 0.848865 | 0.97656375 |
| EIF2AK3 | 526.814096 | -1.9472736 | 0.79354501 | -2.4538919 | 0.01413195 | 0.28646578 |
| RPIA | 101.310794 | -0.0025376 | 0.26138718 | -0.0097084 | 0.99225397 | 0.99919666 |

| | | | | | | |
|---|---|---|---|---|---|---|
| MAL | 1109.5094 | 1.0970577 | 0.40510123 | 2.70810754 | 0.00676681 | 0.20224502 |
| MRPS5 | 287.368158 | -0.1540812 | 0.19312861 | -0.7978166 | 0.42497695 | 0.88392184 |
| AC092835.1 | 46.4881044 | 0.2245754 | 0.36437046 | 0.6163381 | 0.5376714 | 0.92324352 |
| PROM2 | 161.266745 | 0.42176677 | 0.42116365 | 1.00143205 | 0.31661797 | 0.84285425 |
| KCNIP3 | 47.8360945 | -0.1698871 | 0.63559531 | -0.2672882 | 0.7892473 | 0.96515649 |
| FAHD2A | 53.8570624 | 0.09894013 | 0.27633648 | 0.35804224 | 0.72031171 | 0.95796325 |
| ANKRD36C | 173.04571 | -0.1517373 | 0.29813462 | -0.5089555 | 0.61078338 | 0.94003018 |
| STARD7 | 220.138031 | 0.4342601 | 0.33951532 | 1.27905895 | 0.2008763 | 0.76957251 |
| TMEM127 | 126.856997 | 0.05253205 | 0.28303592 | 0.18560206 | 0.85275684 | 0.97718353 |
| CIAO1 | 115.011838 | 0.20979371 | 0.24749652 | 0.84766326 | 0.39662553 | 0.87462201 |
| SNRNP200 | 233.846176 | -0.02908 | 0.2232906 | -0.1302339 | 0.89638139 | 0.98597539 |
| NEURL3 | 71.3208945 | -0.2789839 | 0.71651237 | -0.3893637 | 0.6970071 | 0.95672887 |
| KANSL3 | 261.743984 | -0.1471856 | 0.20016221 | -0.7353317 | 0.46213748 | 0.89801718 |
| FER1L5 | 59.285176 | 0.02941028 | 0.2951784 | 0.09963561 | 0.92063362 | 0.9904507 |
| LMAN2L | 100.002324 | 0.04294924 | 0.23840918 | 0.18014928 | 0.85703538 | 0.97718353 |
| CNNM4 | 74.5471376 | -0.0346138 | 0.42828581 | -0.0808195 | 0.9355855 | 0.99243654 |
| ANKRD23 | 56.3434021 | 0.1696319 | 0.35392145 | 0.47929251 | 0.63173055 | 0.94654542 |
| ANKRD36 | 245.049268 | -0.2763482 | 0.28293223 | -0.9767293 | 0.32870317 | 0.84409527 |
| AC092683.1 | 195.707461 | 0.5666221 | 0.41444812 | 1.36717258 | 0.17157122 | 0.72776437 |
| ANKRD36B | 69.3218223 | -0.4290223 | 0.40176486 | -1.0678442 | 0.28559079 | 0.82816588 |
| COX5B | 464.990953 | 0.32143444 | 0.46147922 | 0.69653069 | 0.48609654 | 0.90695174 |
| ACTR1B | 86.7263249 | -0.0532919 | 0.27179391 | -0.1960745 | 0.84455183 | 0.97573093 |
| TMEM131 | 1065.46296 | -0.0339005 | 0.22148695 | -0.1530587 | 0.87835196 | 0.98300919 |
| INPP4A | 156.753417 | -0.133287 | 0.23189819 | -0.574765 | 0.56545027 | 0.93114846 |
| COA5 | 102.924755 | 0.35677826 | 0.35080719 | 1.01702095 | 0.30914346 | 0.84136316 |
| UNC50 | 106.084912 | 0.20967509 | 0.28367972 | 0.73912611 | 0.45983042 | 0.89753337 |
| MGAT4A | 461.390015 | 0.19262891 | 0.45501644 | 0.42334495 | 0.6720436 | 0.9544944 |
| KIAA1211L | 124.796201 | 0.28861304 | 0.29989976 | 0.96236503 | 0.33586627 | 0.84552074 |
| TSGA10 | 481.88565 | 0.21165995 | 0.27670316 | 0.76493506 | 0.44431021 | 0.8909978 |
| AC079447.1 | 138.325538 | -0.2168964 | 0.23862757 | -0.9089327 | 0.36338566 | 0.85694172 |
| MITD1 | 112.422831 | -0.1820224 | 0.30589661 | -0.5950456 | 0.55181302 | 0.92872652 |
| TXNDC9 | 87.2342192 | -0.2596761 | 0.27938792 | -0.9294464 | 0.35265778 | 0.85536253 |
| EIF5B | 576.088184 | -0.1957787 | 0.19820835 | -0.9877418 | 0.32327913 | 0.84285425 |
| REV1 | 635.113339 | -0.0392135 | 0.1587767 | -0.2469728 | 0.80492924 | 0.96704419 |
| LONRF2 | 54.9820512 | 1.64025509 | 0.60004199 | 2.7335672 | 0.00626523 | 0.19425932 |
| NPAS2 | 1265.31355 | -0.8420914 | 0.36148916 | -2.3295066 | 0.01983225 | 0.33831338 |
| RPL31 | 285.404807 | 0.2756083 | 0.33345421 | 0.82652518 | 0.40850624 | 0.87948803 |
| TBC1D8 | 668.465358 | -0.129926 | 0.28385559 | -0.4577188 | 0.64715446 | 0.95061388 |
| CNOT11 | 143.101672 | 0.17818718 | 0.20152846 | 0.88417872 | 0.37659974 | 0.86400292 |
| RNF149 | 841.784237 | -0.5440524 | 0.31605136 | -1.7214051 | 0.08517733 | 0.59845478 |
| MAP4K4 | 403.755735 | -0.2874014 | 0.32200961 | -0.8925243 | 0.372112 | 0.86201082 |
| IL1R1 | 237.421858 | -0.5149421 | 0.35557437 | -1.448198 | 0.14756168 | 0.69834914 |
| SLC9A4 | 325.240523 | 2.06098687 | 0.46245127 | 4.45665742 | 8.32E-06 | 0.00323124 |
| SLC9A2 | 307.389552 | 0.19089054 | 0.51770323 | 0.3687258 | 0.71233212 | 0.95791799 |
| MFSD9 | 51.3371137 | 0.09728185 | 0.3511495 | 0.27703828 | 0.78175072 | 0.96513523 |
| TMEM182 | 86.9651002 | -0.1968431 | 0.31472805 | -0.6254387 | 0.53168317 | 0.92211604 |
| POU3F3 | 525.30728 | -0.4458444 | 0.26637978 | -1.6737171 | 0.09418621 | 0.61330068 |
| MRPS9 | 173.583994 | 0.15243 | 0.15428968 | 0.98794682 | 0.3231787 | 0.84285425 |
| TGFBRAP1 | 76.9468271 | -0.0728171 | 0.23976937 | -0.3036965 | 0.7613591 | 0.96218738 |
| C2orf49 | 68.512796 | 0.36853925 | 0.28793049 | 1.2799591 | 0.20055952 | 0.7694065 |
| FHL2 | 67.9243543 | -0.6555493 | 1.34542224 | -0.4872443 | 0.62608522 | 0.94511671 |
| NCK2 | 327.535943 | -0.000937 | 0.24902473 | -0.0037626 | 0.99699792 | 0.99967893 |
| UXS1 | 285.846523 | 0.62825144 | 0.32566751 | 1.92911919 | 0.05371607 | 0.51316717 |
| SULT1C2 | 372.83687 | -0.0368065 | 0.48376559 | -0.0760832 | 0.93935288 | 0.99243654 |

| | | | | | | |
|---|---|---|---|---|---|---|
| GCC2 | 820.058825 | 0.17597131 | 0.24398038 | 0.7212519 | 0.47075455 | 0.90123338 |
| GCC2-AS1 | 74.3238336 | -0.0898674 | 0.44865826 | -0.2003027 | 0.84124389 | 0.975446 |
| LIMS1 | 775.600699 | -0.8635753 | 0.25409974 | -3.3985682 | 0.0006774 | 0.05965636 |
| RANBP2 | 1106.65406 | -0.189837 | 0.76360471 | -0.2486063 | 0.8036653 | 0.96678163 |
| CCDC138 | 133.233175 | -0.6051259 | 0.34136729 | -1.7726534 | 0.07628616 | 0.57145422 |
| 10-Sep | 670.76311 | -1.0530207 | 0.40991707 | -2.5688627 | 0.01020329 | 0.24616901 |
| SOWAHC | 119.534201 | -1.0360352 | 0.77890535 | -1.3301169 | 0.18347976 | 0.74504275 |
| NPHP1 | 160.728225 | 0.25697325 | 0.29171837 | 0.880895 | 0.37837466 | 0.86515603 |
| SMIM37 | 109.858744 | 0.36274612 | 0.37102432 | 0.97768827 | 0.32822852 | 0.84381657 |
| LINC01106 | 38.0014621 | -1.0245658 | 0.42483377 | -2.4116863 | 0.01587894 | 0.30146967 |
| MIR4435-2H | 829.364086 | 0.41314712 | 0.39842414 | 1.03695303 | 3.00E-01 | 0.83618229 |
| BCL2L11 | 91.1521748 | 0.14441106 | 0.47557353 | 0.30365664 | 0.7613895 | 0.96218738 |
| AC017002.5 | 69.0101794 | -0.3980743 | 0.91151461 | -0.4367174 | 0.66231631 | 0.95142083 |
| ANAPC1 | 265.636794 | -0.1944345 | 0.15886513 | -1.2238966 | 0.22099125 | 0.78752408 |
| MERTK | 129.311738 | 0.4345368 | 0.40406697 | 1.07540786 | 0.28219217 | 0.82666614 |
| TMEM87B | 229.766214 | -0.3953936 | 0.17910468 | -2.2076117 | 0.02727135 | 0.40084681 |
| ZC3H8 | 126.62768 | 0.14513011 | 0.26755176 | 0.5424375 | 0.58751715 | 0.93483536 |
| ZC3H6 | 263.610888 | -0.1922261 | 0.319419 | -0.6017992 | 0.54730783 | 0.92647167 |
| TTL | 79.6900358 | -0.2961744 | 0.38270911 | -0.773889 | 0.43899641 | 0.88917548 |
| POLR1B | 44.9248214 | 0.03079642 | 0.32859724 | 0.09372088 | 0.92533088 | 0.99051393 |
| CHCHD5 | 80.1097348 | 0.34617665 | 0.40120164 | 0.86284954 | 0.3882202 | 0.86953359 |
| SLC20A1 | 68.9370511 | -0.4857066 | 0.53942421 | -0.9004168 | 0.36789848 | 0.85902833 |
| PSD4 | 76.0486855 | -0.4660881 | 0.27830908 | -1.674714 | 0.09399036 | 0.61323531 |
| PAX8 | 4625.23234 | 0.2739728 | 0.23927732 | 1.14500112 | 0.25220869 | 0.8029001 |
| CBWD2 | 102.296564 | -0.6297903 | 0.25306863 | -2.4886147 | 0.01282418 | 0.27834029 |
| RABL2A | 80.3499791 | 0.11969405 | 0.32093678 | 0.37295211 | 0.70918409 | 0.95752169 |
| SLC35F5 | 726.182942 | -0.0272677 | 0.23252232 | -0.117269 | 0.90664687 | 0.9875066 |
| ACTR3 | 508.980569 | -0.2630827 | 0.2477569 | -1.0618581 | 0.2883001 | 0.83060304 |
| DDX18 | 168.886852 | -0.2442229 | 0.33532377 | -0.7283197 | 0.46641788 | 0.89883965 |
| CCDC93 | 248.018848 | -0.4429996 | 0.17528454 | -2.527317 | 0.01149377 | 0.26357703 |
| INSIG2 | 274.463432 | -0.4913305 | 0.51482328 | -0.9543674 | 0.33989772 | 0.84954164 |
| C2orf76 | 74.7955925 | 0.10369398 | 0.29816416 | 0.34777481 | 0.72800931 | 0.9598317 |
| DBI | 546.424155 | -0.1021713 | 0.4843984 | -0.2109242 | 0.83294642 | 0.97208613 |
| TMEM37 | 64.0857816 | 0.81008822 | 0.56355427 | 1.43746266 | 0.15058657 | 0.7029344 |
| CFAP221 | 545.485075 | 0.94517626 | 0.4482827 | 2.10843795 | 0.03499312 | 0.43353051 |
| PTPN4 | 441.06628 | -0.0606065 | 0.24701379 | -0.2453567 | 0.80618026 | 0.96752828 |
| EPB41L5 | 1164.40565 | 0.00086135 | 0.29798606 | 0.00289057 | 0.99769366 | 0.99967893 |
| RALB | 101.399587 | -0.061926 | 0.27001841 | -0.22934 | 0.81860466 | 0.96883031 |
| TFCP2L1 | 1502.28279 | 0.70546624 | 0.39097192 | 1.80439106 | 0.07117002 | 0.56333976 |
| CLASP1 | 1021.3717 | 8.81E-05 | 0.25433846 | 0.00034625 | 0.99972373 | 0.9999633 |
| NIFK-AS1 | 71.6047874 | 0.08036627 | 0.38121122 | 0.21081822 | 0.83302912 | 0.97208613 |
| NIFK | 50.9068367 | -0.0316029 | 0.36771033 | -0.0859452 | 0.93150997 | 0.99243654 |
| TSN | 124.865419 | 0.16706575 | 0.29566178 | 0.56505699 | 0.572035 | 0.93200748 |
| ERCC3 | 110.556859 | 0.02652298 | 0.31796915 | 0.0834137 | 0.93352259 | 0.99243654 |
| MAP3K2 | 576.573891 | -0.1757379 | 0.16597571 | -1.058817 | 0.28968314 | 0.83103268 |
| IWS1 | 395.845208 | -0.8839336 | 0.29877588 | -2.9585173 | 0.00309123 | 0.14271399 |
| AC010976.1 | 39.2158188 | -0.4425331 | 0.57040177 | -0.7758271 | 0.43785108 | 0.88825695 |
| WDR33 | 425.620069 | -0.2878008 | 0.2234153 | -1.2881873 | 0.19768077 | 0.76534786 |
| POLR2D | 43.5818669 | 0.17917388 | 0.25826515 | 0.69375945 | 0.48783308 | 0.90730741 |
| AMMECR1L | 58.5814055 | -0.2187674 | 0.23697497 | -0.9231667 | 0.35592036 | 0.8557799 |
| SAP130 | 127.319872 | -0.3060115 | 0.23168132 | -1.3208292 | 0.18655832 | 0.74995444 |
| UGGT1 | 148.101033 | -0.2064231 | 0.24456621 | -0.8440377 | 0.39864835 | 0.87557619 |
| HS6ST1 | 1190.32929 | 1.06503588 | 0.38836824 | 2.74233515 | 0.00610041 | 0.19256704 |
| SMPD4 | 118.581442 | -0.4627819 | 0.20297896 | -2.2799501 | 0.02261065 | 0.35959765 |

| | | | | | | |
|---|---|---|---|---|---|---|
| MZT2B | 250.103674 | 0.36186366 | 0.46007537 | 0.78653125 | 0.43155632 | 0.88650713 |
| CCDC115 | 101.34767 | 0.74267822 | 0.37377864 | 1.98694666 | 0.04692831 | 0.48600281 |
| IMP4 | 51.9526313 | 0.1160568 | 0.34721431 | 0.3342512 | 0.73819001 | 0.96078945 |
| PTPN18 | 125.569197 | 0.13764731 | 0.26967309 | 0.51042287 | 0.60975524 | 0.94003018 |
| FAM168B | 333.451223 | -0.2421348 | 0.23495831 | -1.0305437 | 0.30275485 | 0.83709431 |
| PLEKHB2 | 412.323027 | -0.0203322 | 0.27103495 | -0.0750167 | 0.94020139 | 0.99243654 |
| MZT2A | 174.554198 | 0.21788297 | 0.33156041 | 0.65714411 | 0.51108826 | 0.91390025 |
| GPR39 | 398.642992 | 0.20123947 | 0.26934603 | 0.74714104 | 0.45497843 | 0.89626521 |
| NCKAP5 | 293.035599 | 0.78687123 | 0.38210784 | 2.05929097 | 3.95E-02 | 4.57E-01 |
| MGAT5 | 209.479202 | -0.0816319 | 0.29087958 | -0.2806382 | 0.77898788 | 0.96512745 |
| CCNT2-AS1 | 89.8036212 | -0.629459 | 0.50567968 | -1.2447782 | 0.21321329 | 0.77821377 |
| CCNT2 | 318.229745 | -0.0936458 | 0.19255357 | -0.4863362 | 0.62672881 | 0.94511671 |
| RAB3GAP1 | 619.358277 | -0.1435627 | 0.18190535 | -0.7892166 | 0.42998542 | 0.88579433 |
| ZRANB3 | 258.196627 | -0.1365192 | 0.24462003 | -0.5580868 | 0.57678512 | 0.93319686 |
| R3HDM1 | 473.194605 | -0.0560614 | 0.23529656 | -0.2382584 | 0.8116807 | 0.9680627 |
| UBXN4 | 512.482837 | -0.055969 | 0.23242951 | -0.2407998 | 0.80971029 | 0.9679755 |
| DARS | 479.309964 | -0.4239475 | 0.29576629 | -1.4333869 | 0.15174731 | 0.70307801 |
| HNMT | 222.367928 | 0.46691384 | 0.30007824 | 1.5559737 | 0.11971435 | 0.65651824 |
| AC114763.1 | 123.858247 | -0.6695064 | 0.37408941 | -1.7896964 | 0.07350274 | 0.56723762 |
| SPOPL | 217.607872 | -0.0450779 | 0.24467088 | -0.1842389 | 0.85382604 | 0.97718353 |
| LRP1B | 842.682545 | -0.4997017 | 0.52172242 | -0.9577923 | 3.38E-01 | 0.84744028 |
| GTDC1 | 265.504496 | -0.3289537 | 0.33635879 | -0.9779845 | 0.32808199 | 0.84381657 |
| TEX41 | 1492.53726 | 0.39715238 | 0.77477396 | 0.51260419 | 0.60822822 | 0.94003018 |
| ACVR2A | 306.291782 | 0.29047047 | 0.44290023 | 0.65583725 | 0.51192885 | 0.91408994 |
| ORC4 | 429.516296 | -0.1054219 | 0.2958173 | -0.3563751 | 0.72155971 | 0.95868461 |
| MBD5 | 2411.65952 | -0.1374668 | 0.22299841 | -0.6164476 | 0.53759912 | 0.92324352 |
| EPC2 | 419.631757 | -0.1098472 | 0.22832746 | -0.481095 | 0.63044896 | 0.94596555 |
| AC105402.3 | 45.5580766 | -0.7822845 | 1.05975912 | -0.738172 | 0.46040995 | 0.89785937 |
| KIF5C | 255.436649 | -0.8186907 | 0.65605078 | -1.2479075 | 0.21206494 | 0.77709975 |
| LYPD6B | 693.790863 | 0.01972859 | 0.3099438 | 0.06365214 | 0.94924721 | 0.99361356 |
| MMADHC | 91.5309511 | 0.18931237 | 0.25845805 | 0.73246848 | 0.46388268 | 0.89801718 |
| RBM43 | 42.8115377 | 0.13539487 | 0.41178722 | 0.32879814 | 0.74230827 | 0.96086911 |
| NMI | 71.9536005 | -0.4482578 | 0.50453542 | -0.8884566 | 0.37429517 | 0.86268163 |
| RIF1 | 335.434157 | -0.2212839 | 0.1749528 | -1.2648206 | 0.20593564 | 0.77193931 |
| ARL5A | 118.92029 | 0.22537195 | 0.19531556 | 1.15388631 | 0.24854678 | 0.80097144 |
| CACNB4 | 82.3134031 | 0.95152791 | 0.50829334 | 1.87200547 | 0.06120585 | 0.5418629 |
| STAM2 | 280.094544 | -0.2159492 | 0.23497756 | -0.9190207 | 0.35808475 | 0.8557799 |
| FMNL2 | 1464.0061 | -0.3761791 | 0.2648036 | -1.4205967 | 0.15543403 | 0.70916354 |
| PRPF40A | 378.951115 | -0.0768646 | 0.13532004 | -0.5680207 | 0.5700209 | 0.93122667 |
| ARL6IP6 | 130.227716 | -0.3313031 | 0.25062752 | -1.3218944 | 0.18620332 | 0.74938192 |
| KCNJ3 | 1223.97177 | 0.24645203 | 0.3527331 | 0.69869267 | 0.48474411 | 0.90689776 |
| LINC01876 | 362.034491 | -0.4631154 | 0.61276691 | -0.7557775 | 0.4497826 | 0.89324553 |
| NR4A2 | 431.712541 | -1.245793 | 1.40786572 | -0.8848806 | 0.37622105 | 0.86357183 |
| GPD2 | 425.87309 | 0.03544838 | 0.26995956 | 0.13130996 | 0.89553011 | 0.98597539 |
| ACVR1 | 219.560094 | 0.07259123 | 0.47162546 | 0.15391711 | 0.8776751 | 0.98279964 |
| UPP2 | 87.1797168 | 0.67013473 | 0.61605052 | 1.08779184 | 0.27668701 | 0.82198569 |
| CCDC148 | 512.172438 | 0.6031409 | 0.34700979 | 1.73810914 | 0.08219158 | 0.59202053 |
| PKP4 | 1522.2614 | -0.1283128 | 0.21706155 | -0.5911357 | 0.55442947 | 0.92872652 |
| PKP4-AS1 | 54.2796845 | 0.27409377 | 0.54976519 | 0.49856517 | 0.61808575 | 0.94246161 |
| TANC1 | 859.994405 | 0.00252084 | 0.2310495 | 0.0109104 | 0.99129494 | 0.99889703 |
| WDSUB1 | 309.902458 | -0.3749145 | 0.2984853 | -1.2560567 | 0.20909541 | 0.77436433 |
| BAZ2B | 1993.7951 | -0.1671369 | 0.24729122 | -0.6758708 | 0.49912267 | 0.90839719 |
| AC009961.1 | 144.676258 | -0.0020011 | 0.47058071 | -0.0042525 | 0.99660704 | 0.99967893 |
| 7-Mar | 714.508005 | -0.1523909 | 0.24904394 | -0.6119038 | 0.54060144 | 0.92422563 |

| | | | | | | |
|---|---|---|---|---|---|---|
| PLA2R1 | 85.8755346 | 1.10578331 | 0.34658218 | 3.19053709 | 0.00142009 | 0.09561351 |
| ITGB6 | 1056.35284 | -0.128423 | 0.8025298 | -0.1600227 | 0.87286321 | 0.98246494 |
| RBMS1 | 1033.24959 | -0.0400466 | 0.26666672 | -0.1501747 | 0.88062681 | 0.98333507 |
| TANK | 377.567401 | -0.2283788 | 0.20761252 | -1.100024 | 0.27132165 | 0.81922962 |
| PSMD14 | 192.547492 | -0.1714679 | 0.21368504 | -0.802433 | 0.42230255 | 0.88371155 |
| IFIH1 | 73.9125154 | -1.4226363 | 0.70179757 | -2.0271319 | 0.04264892 | 0.47247434 |
| GCA | 151.443927 | -0.2482996 | 0.24756607 | -1.0029629 | 0.31587876 | 0.84285425 |
| FIGN | 85.3974017 | 1.11308149 | 0.37201323 | 2.99204816 | 0.00277113 | 0.13678792 |
| GRB14 | 140.768852 | 0.57334371 | 0.57329589 | 1.00008343 | 0.31727014 | 0.84285425 |
| COBLL1 | 9286.99595 | 0.66604088 | 0.31773384 | 2.09622267 | 0.03606244 | 0.4362094 |
| AC019197.1 | 13234.3471 | 0.88454253 | 0.35323373 | 2.50412814 | 0.01227536 | 0.27034878 |
| SLC38A11 | 576.883719 | 0.45399894 | 0.521878 | 0.8699331 | 0.38433697 | 0.86725322 |
| SCN3A | 632.036451 | 1.29754071 | 0.72843644 | 1.78126824 | 0.07486864 | 0.57072562 |
| SCN2A | 1790.8389 | 0.76355387 | 0.41703005 | 1.8309325 | 0.06711062 | 0.55774725 |
| CSRNP3 | 410.738715 | 0.52019277 | 0.35357711 | 1.47122866 | 0.14122929 | 0.68617662 |
| GALNT3 | 220.378303 | -0.0468048 | 0.27039879 | -0.1730956 | 0.86257626 | 0.979268 |
| TTC21B | 202.163243 | 0.18035038 | 0.22185805 | 0.81290891 | 0.41627028 | 0.88242985 |
| STK39 | 387.307735 | 0.26120089 | 0.45155949 | 0.57844181 | 0.56296587 | 0.9306324 |
| CERS6 | 125.568047 | 0.19583647 | 0.37255738 | 0.52565452 | 0.59912827 | 0.93728325 |
| LRP2 | 90.0603257 | 0.31313976 | 0.76112949 | 0.41141457 | 0.68076857 | 0.95574492 |
| FASTKD1 | 173.564939 | 0.38736317 | 0.26419976 | 1.46617531 | 0.14260053 | 0.6890058 |
| PPIG | 440.913135 | -0.0869462 | 0.20047016 | -0.4337112 | 0.66449814 | 0.95200828 |
| KLHL23 | 102.880776 | -0.2235729 | 0.32570919 | -0.6864187 | 0.49244914 | 0.90780423 |
| SSB | 192.084979 | 0.05334435 | 0.27818977 | 0.19175523 | 0.84793394 | 0.97656375 |
| METTL5 | 69.6124053 | -0.3670849 | 0.268458 | -1.3673831 | 0.17150524 | 0.72776437 |
| UBR3 | 662.982482 | -0.0680837 | 0.18652737 | -0.3650066 | 0.71510646 | 0.95791799 |
| MYO3B | 950.056337 | 1.53911939 | 0.35601324 | 4.32320834 | 1.54E-05 | 0.00519863 |
| GORASP2 | 168.28883 | 0.05697211 | 0.20176125 | 0.28237389 | 0.77765683 | 0.96472142 |
| TLK1 | 1105.26139 | -0.1286261 | 0.28427718 | -0.4524672 | 0.65093242 | 0.95061388 |
| METTL8 | 140.186715 | -0.2503875 | 0.26944887 | -0.9292579 | 0.35275543 | 0.85536253 |
| DCAF17 | 102.389187 | 0.12592427 | 0.26421289 | 0.47660156 | 0.63364587 | 0.94744025 |
| CYBRD1 | 164.925015 | 0.90114969 | 0.61680864 | 1.46098747 | 0.14401888 | 0.69093999 |
| DYNC1I2 | 521.743891 | 0.04072692 | 0.13340715 | 0.3052829 | 0.7601507 | 0.96191449 |
| SLC25A12 | 364.765437 | 0.12084826 | 0.26564116 | 0.45493047 | 0.64915926 | 0.95061388 |
| HAT1 | 162.172926 | -0.2788887 | 0.24102276 | -1.1571051 | 0.24722942 | 0.80072691 |
| METAP1D | 78.1513573 | -0.0842814 | 0.26604093 | -0.3167986 | 0.7513964 | 0.96120879 |
| ITGA6 | 382.041604 | 0.39017978 | 0.2249914 | 1.73419863 | 0.08288285 | 0.59202053 |
| PDK1 | 258.524086 | -1.5242802 | 0.84616902 | -1.8013897 | 0.07164148 | 0.56343001 |
| MAP3K20 | 91.8096036 | -0.4972684 | 0.32833539 | -1.5145137 | 0.12989561 | 0.672582 |
| SP3 | 284.997915 | 0.1062965 | 0.18096902 | 0.58737403 | 0.55695253 | 0.92872652 |
| OLA1 | 482.326317 | -0.1066361 | 0.1975699 | -0.5397387 | 0.58937728 | 0.93483536 |
| CIR1 | 196.950743 | 0.00191525 | 0.23399702 | 0.00818494 | 0.99346944 | 0.99919666 |
| SCRN3 | 70.2518056 | 0.09700344 | 0.2774658 | 0.34960504 | 0.72663513 | 0.95979351 |
| GPR155 | 110.77174 | 0.09883944 | 0.38097796 | 0.25943612 | 0.79529876 | 0.9653598 |
| AC010894.3 | 518.222464 | 0.46946244 | 0.50411999 | 0.93125138 | 0.35172354 | 0.85536253 |
| WIPF1 | 114.568098 | -0.1757301 | 0.36213674 | -0.4852591 | 0.62749257 | 0.94511671 |
| ATF2 | 327.883167 | -0.1997657 | 0.1941336 | -1.0290116 | 0.30347425 | 0.83709431 |
| ATP5MC3 | 541.602713 | 0.6753853 | 0.52396424 | 1.2889912 | 0.19740114 | 0.76534786 |
| LNPK | 164.525995 | -0.2468098 | 0.27206076 | -0.9071864 | 0.36430826 | 0.85796641 |
| HOXD10 | 367.997358 | 0.08591228 | 0.55252454 | 0.15549043 | 0.87643469 | 0.98269442 |
| HOXD-AS2 | 97.8750665 | -0.3629134 | 0.61792071 | -0.5873139 | 0.55699291 | 0.92872652 |
| HOXD9 | 156.077738 | -0.02076 | 0.50764101 | -0.0408951 | 0.96737954 | 0.99637506 |
| HOXD8 | 278.751419 | -0.1274393 | 0.52623109 | -0.2421736 | 0.80864565 | 0.96775927 |
| HOXD3 | 403.126584 | 0.13412422 | 0.39094464 | 0.34307727 | 0.73154033 | 0.96008876 |

| | | | | | | |
|---|---|---|---|---|---|---|
| HAGLR | 65.9503915 | -0.0842451 | 0.31016162 | -0.2716168 | 0.78591664 | 0.96513523 |
| MTX2 | 616.080712 | -0.0588018 | 0.20027323 | -0.293608 | 0.7690575 | 0.96345263 |
| LINC01116 | 132.305668 | 0.60796624 | 0.34304946 | 1.77224074 | 0.0763546 | 0.57145422 |
| LINC01117 | 62.1182697 | -0.0215764 | 0.42839693 | -0.0503654 | 0.95983121 | 0.99609409 |
| HNRNPA3 | 694.304409 | -0.2597077 | 0.24967892 | -1.0401666 | 0.29826251 | 0.83556453 |
| NFE2L2 | 1195.5949 | 0.27242972 | 0.24848581 | 1.09635925 | 0.27292159 | 0.81922962 |
| AGPS | 260.267427 | 0.08069569 | 0.2074377 | 0.38901169 | 0.6972675 | 0.95672887 |
| RBM45 | 67.1386073 | -0.5308258 | 0.29513691 | -1.7985749 | 0.07208595 | 0.5644261 |
| PRKRA | 159.580674 | -0.0654471 | 0.18522885 | -0.3533312 | 0.72384015 | 0.95963742 |
| PJVK | 35.890042 | 0.57574732 | 0.3213198 | 1.79182023 | 0.07316177 | 0.56668778 |
| PLEKHA3 | 128.225839 | -0.0083893 | 0.24887741 | -0.0337085 | 0.97310958 | 0.99656026 |
| TTN-AS1 | 388.963974 | -0.0078251 | 0.34032482 | -0.0229932 | 0.98165572 | 0.99696835 |
| TTN | 46.1856391 | 0.68308646 | 0.70682075 | 0.96642106 | 0.33383353 | 0.8445213 |
| SESTD1 | 784.622494 | -0.8599073 | 0.47093711 | -1.8259492 | 0.06785791 | 0.55864172 |
| ZNF385B | 81.7035559 | -0.3112856 | 0.60514254 | -0.5144004 | 0.60697207 | 0.93925224 |
| CWC22 | 158.513244 | -0.2352074 | 0.23626343 | -0.9955302 | 0.31947844 | 0.84285425 |
| UBE2E3 | 443.31037 | -0.0233834 | 0.24010649 | -0.0973876 | 0.92241859 | 0.99051393 |
| CERKL | 42.1369746 | 0.91484727 | 0.65417255 | 1.39848007 | 0.16196895 | 0.71470931 |
| SSFA2 | 134.443657 | -0.3766975 | 0.35041606 | -1.0750008 | 0.28237439 | 0.82666614 |
| PDE1A | 3339.95871 | -0.3976574 | 0.24826357 | -1.6017552 | 0.10920976 | 0.63854503 |
| DNAJC10 | 257.198347 | -0.5085899 | 0.27747193 | -1.8329418 | 0.06681123 | 0.55637569 |
| NCKAP1 | 798.730188 | -0.3087371 | 0.25426411 | -1.2142377 | 0.22465697 | 0.78980378 |
| NUP35 | 74.109932 | -0.0824469 | 0.25725616 | -0.3204857 | 0.74860018 | 0.96109466 |
| FSIP2 | 50.7067098 | -0.0862235 | 0.66084602 | -0.1304743 | 0.89619115 | 0.98597539 |
| FSIP2-AS1 | 37.6338229 | 0.35160261 | 0.3542813 | 0.99243909 | 0.32098338 | 0.84285425 |
| LINC01473 | 103.085373 | 0.12051017 | 0.26103164 | 0.46166881 | 0.64431884 | 0.94949493 |
| ZC3H15 | 222.88347 | -0.010067 | 0.22499205 | -0.0447438 | 0.96431148 | 0.99637506 |
| ITGAV | 952.07375 | 0.90941544 | 0.3817624 | 2.38215037 | 0.01721187 | 0.313705 |
| AC017101.1 | 109.882729 | 1.50586762 | 0.6347071 | 2.37253943 | 0.01766628 | 0.31976267 |
| AC007319.1 | 264.818725 | -0.0955799 | 0.88393106 | -0.1081305 | 0.9138922 | 0.98830345 |
| TFPI | 204.845951 | -1.8581066 | 0.77722108 | -2.3907053 | 0.01681604 | 0.30900364 |
| GULP1 | 420.062448 | -0.3960862 | 0.43054878 | -0.9199567 | 0.35759539 | 0.8557799 |
| WDR75 | 100.104202 | -0.2460374 | 0.20280391 | -1.2131787 | 0.22506151 | 0.78982769 |
| SLC40A1 | 66.7935776 | 0.10532328 | 0.23680116 | 0.44477519 | 0.65648223 | 0.95100227 |
| ASNSD1 | 56.1404911 | 0.33121381 | 0.28772966 | 1.15112849 | 0.24967938 | 0.80149394 |
| ANKAR | 216.410849 | 0.14352422 | 0.30192888 | 0.47535771 | 0.63453203 | 0.94775679 |
| ORMDL1 | 166.474338 | 0.29054883 | 0.2682402 | 1.08316659 | 0.27873449 | 0.82401619 |
| PMS1 | 180.815447 | -0.0445298 | 0.24771291 | -0.1797637 | 0.85733809 | 0.97718353 |
| C2orf88 | 85.3237196 | 0.74147507 | 0.40800035 | 1.81733929 | 0.06916518 | 0.55917298 |
| HIBCH | 582.862397 | 0.37328505 | 0.23484236 | 1.58951331 | 0.11194455 | 0.64424981 |
| INPP1 | 101.9883 | 0.02283793 | 0.32063011 | 0.07122827 | 0.94321608 | 0.99265578 |
| MFSD6 | 197.757008 | -0.0920863 | 0.22478127 | -0.4096706 | 0.68204757 | 0.95574492 |
| NAB1 | 102.177231 | 0.06993971 | 0.38717434 | 0.18064139 | 0.85664907 | 0.97718353 |
| GLS | 3668.9224 | 1.02362459 | 0.36222896 | 2.82590491 | 0.00471473 | 0.16921345 |
| STAT1 | 1025.9541 | -0.4289174 | 0.81430261 | -0.5267297 | 0.5983813 | 0.93728325 |
| MYO1B | 1326.4769 | -0.3005399 | 0.41169416 | -0.7300077 | 0.4653855 | 0.89883965 |
| NABP1 | 46.9612065 | -0.7015984 | 0.74768869 | -0.9383564 | 3.48E-01 | 0.85348649 |
| SLC39A10 | 216.236138 | 0.84358152 | 0.2502977 | 3.37031268 | 0.00075083 | 0.06345719 |
| DNAH7 | 487.162815 | -0.129764 | 0.24928518 | -0.5205446 | 0.60268408 | 0.93804805 |
| STK17B | 42.625696 | -0.0761306 | 0.65393176 | -0.1164198 | 0.90731983 | 0.9875066 |
| GTF3C3 | 107.129846 | 0.0911253 | 0.22328016 | 0.40812089 | 0.68318492 | 0.95574492 |
| PGAP1 | 450.436456 | 0.52087159 | 0.50005682 | 1.04162482 | 0.29758566 | 0.83556453 |
| ANKRD44 | 376.788487 | 0.03464715 | 0.29350408 | 0.11804657 | 0.90603076 | 0.9875066 |
| SF3B1 | 437.729735 | 0.12403368 | 0.15192203 | 0.8164299 | 0.4142543 | 0.88239535 |

| | | | | | | |
|---|---|---|---|---|---|---|
| COQ10B | 93.9738639 | -0.1964295 | 0.28739652 | -0.6834791 | 0.49430415 | 0.90839719 |
| HSPD1 | 333.213574 | 0.33334733 | 0.27493876 | 1.21244215 | 0.22534318 | 0.78982769 |
| MOB4 | 61.3099176 | 0.31210982 | 0.35305585 | 0.88402391 | 0.3766833 | 0.86400548 |
| PLCL1 | 315.071646 | 1.49840029 | 0.49597402 | 3.02112655 | 0.00251836 | 0.13065555 |
| FTCDNL1 | 87.1208125 | 0.71845749 | 0.48156742 | 1.49191463 | 0.13572153 | 0.67994379 |
| C2orf69 | 59.048889 | -0.0439117 | 0.41045754 | -0.1069823 | 0.91480305 | 0.98868699 |
| TYW5 | 90.126488 | 0.08314483 | 0.21944636 | 0.37888452 | 0.70477362 | 0.95723529 |
| MAIP1 | 51.6931805 | 0.0762021 | 0.29136337 | 0.26153629 | 0.79367897 | 0.9653598 |
| SPATS2L | 849.123895 | -0.0962314 | 0.40786235 | -0.2359408 | 0.81347859 | 0.96819339 |
| KCTD18 | 52.8278746 | 0.04102925 | 0.29825586 | 0.13756394 | 0.89058506 | 0.98597539 |
| BZW1 | 106.297038 | -0.5013744 | 0.39770917 | -1.2606558 | 0.20743288 | 0.77307845 |
| CLK1 | 275.021284 | -0.7210322 | 0.33254227 | -2.1682423 | 0.03014026 | 0.40968855 |
| PPIL3 | 84.1044893 | -0.3089795 | 0.31972914 | -0.966379 | 0.33385456 | 0.8445213 |
| NIF3L1 | 51.3424596 | -0.012851 | 0.29094708 | -0.0441694 | 0.96476934 | 0.99637506 |
| ORC2 | 300.397274 | -0.4384697 | 0.24319087 | -1.802986 | 0.07139042 | 0.56343001 |
| AC005037.1 | 65.5154438 | -0.1269638 | 0.32788656 | -0.3872187 | 0.69859432 | 0.95672887 |
| FAM126B | 265.166215 | -0.3913274 | 0.21290391 | -1.8380471 | 0.06605547 | 0.55444578 |
| NDUFB3 | 247.651752 | 0.46671845 | 0.43723625 | 1.06742854 | 0.28577837 | 0.82848058 |
| CFLAR | 524.231459 | -0.2237417 | 0.17728168 | -1.2620689 | 0.20692399 | 0.77200549 |
| CFLAR-AS1 | 44.4551193 | -0.0823268 | 0.59077638 | -0.1393535 | 0.88917078 | 0.98551241 |
| CASP10 | 121.732125 | -0.0720101 | 0.35413023 | -0.2033436 | 0.83886651 | 0.97430756 |
| CASP8 | 150.068996 | 0.15824416 | 0.28158374 | 0.56197902 | 0.57413032 | 0.93270589 |
| TRAK2 | 358.449157 | -0.1525888 | 0.21966951 | -0.6946289 | 0.48728789 | 0.90730741 |
| STRADB | 167.889135 | -0.0604437 | 0.37183037 | -0.1625572 | 0.87086713 | 0.98208511 |
| TMEM237 | 40.4102636 | -0.1928084 | 0.35573982 | -0.5419928 | 0.58782348 | 0.93483536 |
| ALS2 | 185.324039 | -0.2565402 | 0.30279256 | -0.8472472 | 0.39685733 | 0.87462201 |
| SUMO1 | 279.083033 | 0.35192079 | 0.3871915 | 0.90890628 | 0.3633996 | 0.85694172 |
| NOP58 | 217.112247 | -0.1898849 | 0.22396999 | -0.8478139 | 0.3965416 | 0.87462201 |
| BMPR2 | 990.959521 | 0.14928537 | 0.24027086 | 0.62132117 | 0.53438833 | 0.92303275 |
| FAM117B | 513.487808 | -0.2129764 | 0.26233559 | -0.8118471 | 0.41687935 | 0.88242985 |
| ICA1L | 270.885092 | -0.1269011 | 0.28887192 | -0.4392988 | 0.66044504 | 0.95127957 |
| WDR12 | 116.46342 | 0.09276735 | 0.34616409 | 0.26798665 | 0.78870959 | 0.96515649 |
| CARF | 196.327301 | 0.31981462 | 0.33646935 | 0.95050149 | 0.34185749 | 0.85092302 |
| NBEAL1 | 1837.89291 | -0.2906147 | 0.226068 | -1.2855188 | 0.19861104 | 0.76668122 |
| CYP20A1 | 305.173097 | -0.1620837 | 0.2850856 | -0.568544 | 0.56966563 | 0.93122667 |
| ABI2 | 617.89375 | 0.19857978 | 0.27915561 | 0.71135875 | 0.47686195 | 0.90242252 |
| RAPH1 | 920.53565 | -0.2931629 | 0.43490254 | -0.6740888 | 0.50025489 | 0.90839719 |
| PARD3B | 1913.41926 | 0.03208538 | 0.13015735 | 0.24651222 | 8.05E-01 | 9.67E-01 |
| INO80D | 686.911254 | -0.8084754 | 0.64860952 | -1.2464747 | 0.21259017 | 0.77709975 |
| NDUFS1 | 431.909574 | 0.05862102 | 0.18189908 | 0.32227219 | 0.7472465 | 0.96109466 |
| EEF1B2 | 191.704489 | 0.2352452 | 0.5085796 | 0.46255335 | 0.64368455 | 0.94949493 |
| ZDBF2 | 178.090511 | 0.06415575 | 0.71173664 | 0.09013973 | 0.92817618 | 0.99197291 |
| ADAM23 | 122.27126 | 0.23545582 | 0.47834224 | 0.49223297 | 0.62255466 | 0.94419288 |
| FASTKD2 | 94.7530893 | -0.3320881 | 0.23654197 | -1.4039289 | 0.16034004 | 0.71409437 |
| CREB1 | 316.57669 | -0.0965564 | 0.15789779 | -0.6115118 | 0.54086082 | 0.92443671 |
| METTL21A | 81.8403209 | 0.08291034 | 0.34862023 | 0.23782424 | 0.81201742 | 0.9680627 |
| CCNYL1 | 197.491258 | -0.2232229 | 0.45000136 | -0.4960493 | 0.61985964 | 0.94319304 |
| PLEKHM3 | 182.451012 | -0.0054649 | 0.20764829 | -0.0263179 | 0.97900379 | 0.99696835 |
| IDH1 | 73.7795411 | 0.03378222 | 0.41130106 | 0.08213503 | 0.93453934 | 0.99243654 |
| PIKFYVE | 329.912343 | -0.1444002 | 0.19947199 | -0.7239122 | 0.46911963 | 0.90026987 |
| PTH2R | 3246.7206 | -0.3979188 | 0.30838741 | -1.2903211 | 0.19693918 | 0.76498244 |
| RPE | 213.038556 | -0.3623093 | 0.31506296 | -1.1499586 | 0.25016094 | 0.80149394 |
| KANSL1L | 1668.6619 | 0.58955738 | 0.31428087 | 1.87589334 | 0.06066993 | 0.54112412 |
| AC007038.1 | 169.073168 | -0.053558 | 0.32322467 | -0.165699 | 0.86839387 | 0.98185001 |

| | | | | | | |
|---|---|---|---|---|---|---|
| ACADL | 253.921639 | -0.0178904 | 0.38385553 | -0.0466071 | 0.96282635 | 0.99637506 |
| LANCL1 | 207.971432 | -0.1396616 | 0.19889828 | -0.7021758 | 0.48256957 | 0.90614725 |
| ERBB4 | 4148.00411 | 0.10247732 | 0.26995981 | 0.37960213 | 0.70424078 | 0.9570077 |
| IKZF2 | 421.028177 | 0.67874729 | 0.39879062 | 1.70201418 | 0.08875271 | 0.60673738 |
| SPAG16 | 1828.96363 | 0.89451157 | 0.2648439 | 3.37750489 | 0.00073147 | 0.06232333 |
| AC068051.1 | 412.426308 | 0.59863055 | 0.4438845 | 1.34861781 | 0.17745976 | 0.73683766 |
| VWC2L | 38.2387144 | 1.15388717 | 0.62219341 | 1.85454738 | 0.06366089 | 0.54850785 |
| BARD1 | 73.4349262 | 0.95140476 | 0.31489671 | 3.02132328 | 0.00251672 | 0.13065555 |
| AC016708.1 | 119.35412 | 0.10866736 | 0.24990781 | 0.43482978 | 0.66368599 | 0.95187756 |
| ATIC | 131.150803 | 0.25933473 | 0.24842836 | 1.04390148 | 0.29653097 | 0.83516382 |
| MREG | 78.3175492 | -0.1543303 | 0.23984118 | -0.6434688 | 0.51991996 | 0.91751084 |
| PECR | 173.756107 | 0.15631138 | 0.29373089 | 0.53215848 | 0.59461624 | 0.935202 |
| XRCC5 | 761.362706 | -0.2221473 | 0.19957061 | -1.1131264 | 0.26565416 | 0.8121456 |
| SMARCAL1 | 103.763313 | -0.0536306 | 0.24310888 | -0.2206032 | 0.82540141 | 0.97018919 |
| RPL37A | 530.26165 | -0.1286514 | 0.28537841 | -0.4508099 | 0.65212654 | 0.95079106 |
| IGFBP2 | 387.581537 | 0.9649914 | 0.64401248 | 1.49840483 | 0.1340281 | 0.67790276 |
| IGFBP5 | 97.5383664 | -1.2792906 | 0.4228883 | -3.0251266 | 0.00248529 | 0.13022925 |
| DIRC3 | 143.686529 | -0.739636 | 0.76692752 | -0.9644145 | 0.33483817 | 0.84521974 |
| TNS1 | 405.698307 | 0.9494799 | 0.36866795 | 2.57543378 | 0.01001144 | 0.24399985 |
| ARPC2 | 558.000775 | -0.2100276 | 0.31519236 | -0.6663473 | 0.50518913 | 0.9107831 |
| AAMP | 116.091984 | 0.53927249 | 0.39268406 | 1.37329864 | 0.16965952 | 0.72632018 |
| PNKD | 447.266161 | 0.48667287 | 0.32994634 | 1.47500611 | 0.1402109 | 0.68408298 |
| TMBIM1 | 253.478084 | -0.2249728 | 0.31554787 | -0.7129593 | 0.47587092 | 0.90242252 |
| CATIP-AS2 | 62.4531756 | 0.41788468 | 0.50643133 | 0.82515567 | 0.40928322 | 0.87985398 |
| CATIP | 46.7747168 | 0.43657436 | 0.66522373 | 0.65628201 | 0.51164269 | 0.91408377 |
| CTDSP1 | 212.033844 | -0.1772544 | 0.22322846 | -0.7940494 | 0.42716668 | 0.8850745 |
| USP37 | 313.171132 | -0.835251 | 0.52518809 | -1.5903845 | 0.11174815 | 0.64353508 |
| CNOT9 | 140.384524 | -0.5104607 | 0.26625511 | -1.9171864 | 0.05521426 | 0.51757193 |
| ZNF142 | 48.3162615 | 0.00653591 | 0.24625362 | 0.02654138 | 0.97882553 | 0.99696835 |
| STK36 | 62.4707143 | -0.0527365 | 0.27459427 | -0.1920523 | 0.84770127 | 0.97656375 |
| TTLL4 | 43.8679468 | 0.02156107 | 0.32967517 | 0.06540096 | 0.94785476 | 0.99314907 |
| CYP27A1 | 45.409806 | 1.40274168 | 0.53318629 | 2.63086598 | 0.00851676 | 0.22546434 |
| CNPPD1 | 107.833879 | 0.45109339 | 0.3245077 | 1.39008533 | 0.16450297 | 0.71715698 |
| RETREG2 | 135.619146 | -0.0267286 | 0.26042365 | -0.1026352 | 0.9182525 | 0.99014946 |
| ZFAND2B | 50.1749877 | -0.1522442 | 0.34440474 | -0.4420504 | 0.65845275 | 0.95113765 |
| ANKZF1 | 106.467963 | -0.5122888 | 0.35908474 | -1.4266517 | 0.15368033 | 0.70747602 |
| STK16 | 56.1930789 | -0.1053121 | 0.28874789 | -0.3647199 | 0.71532048 | 0.95791799 |
| TUBA4A | 189.01939 | -0.4808405 | 0.75888082 | -0.633618 | 0.52633018 | 0.91974278 |
| DNAJB2 | 129.801564 | -0.0573283 | 0.31321846 | -0.1830297 | 0.85477474 | 0.97718353 |
| DNPEP | 69.616461 | -0.0072468 | 0.28590217 | -0.0253472 | 0.97977806 | 0.99696835 |
| CHPF | 63.6986503 | -0.3916919 | 0.48111008 | -0.8141419 | 0.41556368 | 0.88242985 |
| OBSL1 | 152.222607 | 0.23284083 | 0.20357388 | 1.14376579 | 0.25272078 | 0.80350545 |
| STK11IP | 31.7968961 | -0.2801014 | 0.30952161 | -0.9049494 | 0.36549217 | 0.85866868 |
| SGPP2 | 649.945548 | 0.17610942 | 0.43486559 | 0.40497438 | 0.68549634 | 0.9558749 |
| FARSB | 104.272345 | -0.3451791 | 0.29567034 | -1.1674458 | 0.24303039 | 0.80072691 |
| ACSL3 | 450.863357 | -0.6667108 | 0.36458096 | -1.8287044 | 0.0674439 | 0.55848603 |
| AP1S3 | 67.2338071 | -0.1409927 | 0.41147392 | -0.3426527 | 0.73185975 | 0.96008876 |
| WDFY1 | 195.851449 | -0.5499505 | 0.18764698 | -2.9307719 | 0.00338121 | 0.14358074 |
| MRPL44 | 50.4032612 | 0.00424778 | 0.34955133 | 0.0121521 | 0.99030427 | 0.99882205 |
| CUL3 | 715.585178 | 0.31428094 | 0.14996392 | 2.09571036 | 0.03610789 | 0.4362094 |
| DOCK10 | 198.355787 | -0.6667013 | 0.57536596 | -1.158743 | 0.24656096 | 0.80072691 |
| IRS1 | 51.508641 | -0.3863464 | 0.6123892 | -0.6308837 | 0.52811658 | 0.92106733 |
| RHBDD1 | 336.162122 | -0.2552942 | 0.20548434 | -1.242402 | 0.21408827 | 0.77890374 |
| COL4A4 | 198.473437 | -0.0676072 | 0.94811216 | -0.0713072 | 0.94315326 | 0.99265578 |

| | | | | | | |
|---|---|---|---|---|---|---|
| COL4A3 | 195.396082 | 0.082415 | 0.81644422 | 0.10094383 | 0.91959505 | 0.99014946 |
| AC097662.1 | 55.8429113 | -0.1001477 | 0.45241427 | -0.2213628 | 0.82480996 | 0.97018919 |
| MFF | 390.955655 | 0.41451766 | 0.24087166 | 1.72090674 | 0.08526775 | 0.59845478 |
| AGFG1 | 527.286677 | -0.1636363 | 0.26558827 | -0.6161277 | 0.53781027 | 0.92324352 |
| TRIP12 | 852.239884 | -0.1403004 | 0.18888081 | -0.7427988 | 0.45760348 | 0.89706032 |
| FBXO36 | 100.851837 | 0.1416726 | 0.24997637 | 0.56674397 | 0.57088814 | 0.93125087 |
| SLC16A14 | 47.4715605 | 0.75518549 | 0.69641379 | 1.08439192 | 0.27819106 | 0.82310625 |
| SP110 | 153.711536 | -0.5510271 | 0.450844 | -1.2222123 | 0.22162734 | 0.78752765 |
| SP140L | 167.211188 | -0.427145 | 0.31030531 | -1.3765314 | 0.16865716 | 0.72499057 |
| SP100 | 433.786753 | -0.7384609 | 0.42899341 | -1.7213805 | 0.08518179 | 0.59845478 |
| CAB39 | 788.004298 | 0.10676313 | 0.16088335 | 0.6636058 | 5.07E-01 | 0.91163608 |
| ITM2C | 577.416849 | 1.46357876 | 0.46043546 | 3.17868383 | 0.00147945 | 0.09875589 |
| PSMD1 | 328.425288 | -0.394531 | 0.26752643 | -1.4747367 | 0.14028335 | 0.68411798 |
| ARMC9 | 477.779714 | 0.08125428 | 0.21881738 | 0.37133373 | 0.71038898 | 0.95756248 |
| NCL | 370.571876 | 0.22033618 | 0.3303932 | 0.66689077 | 0.50484191 | 0.9107831 |
| PTMA | 2137.96196 | -0.1847128 | 0.33342586 | -0.5539846 | 0.5795894 | 0.93345492 |
| PDE6D | 85.2402472 | -0.0342786 | 0.29888162 | -0.1146896 | 0.90869118 | 0.98758338 |
| COPS7B | 89.5048371 | -0.2752793 | 0.27491725 | -1.0013171 | 0.31667354 | 0.84285425 |
| DIS3L2 | 578.287404 | -0.1737772 | 0.23993802 | -0.7242589 | 0.46890681 | 0.90026987 |
| EIF4E2 | 157.5624 | 0.32949524 | 0.1602847 | 2.05568747 | 0.03981265 | 0.45805615 |
| EFHD1 | 221.202613 | 1.25992184 | 0.36726861 | 3.43051876 | 0.00060243 | 0.05792155 |
| GIGYF2 | 717.875931 | -0.3102695 | 0.22281721 | -1.3924843 | 0.16377578 | 0.71715698 |
| ATG16L1 | 55.9259851 | -0.3114359 | 0.26868256 | -1.1591221 | 0.24640642 | 0.80072691 |
| DGKD | 82.1579064 | -0.2684966 | 0.31652964 | -0.8482511 | 0.39629813 | 0.87462201 |
| USP40 | 461.551687 | 0.02838568 | 0.33202947 | 0.08549144 | 0.9318707 | 0.99243654 |
| AC114812.2 | 192.785358 | 0.74211331 | 0.96299638 | 0.77062939 | 0.44092664 | 0.88964371 |
| SH3BP4 | 165.300016 | -0.6416715 | 0.35193508 | -1.8232666 | 0.06826302 | 0.55917298 |
| AGAP1 | 1220.12964 | -0.5452482 | 0.31299884 | -1.7420135 | 0.08150607 | 0.59202053 |
| IQCA1 | 317.634146 | -0.2061302 | 0.21199879 | -0.972318 | 0.33089237 | 0.84442294 |
| ACKR3 | 65.9500219 | 0.91433093 | 0.65725651 | 1.39113257 | 0.16418523 | 0.71715698 |
| AC105760.2 | 41.0345866 | -0.013364 | 0.39983471 | -0.0334238 | 0.97333661 | 0.99656026 |
| COPS8 | 83.1935106 | 0.23222998 | 0.30168084 | 0.76978697 | 0.44142627 | 0.88964371 |
| RAB17 | 374.100533 | 0.15221825 | 0.48058242 | 0.31673704 | 0.75144314 | 0.96120879 |
| LRRFIP1 | 2507.12995 | -0.4742697 | 0.40461883 | -1.1721395 | 0.24114105 | 0.80072691 |
| ILKAP | 208.892852 | -0.0992336 | 0.17996793 | -0.5513961 | 0.58136216 | 0.93386308 |
| PER2 | 92.0147979 | -0.4410122 | 0.36458278 | -1.2096354 | 0.22641884 | 0.78982769 |
| TRAF3IP1 | 147.9627 | -0.1177376 | 0.22858804 | -0.5150645 | 0.60650798 | 0.93919596 |
| ASB1 | 44.4608943 | 0.23290257 | 0.26289334 | 0.88592036 | 0.37566044 | 0.86357183 |
| HDAC4 | 1313.43307 | -0.799376 | 0.40549803 | -1.9713438 | 0.04868456 | 0.48777647 |
| NDUFA10 | 469.939162 | 0.18957569 | 0.234042 | 0.81000712 | 0.41793609 | 0.88242985 |
| COPS9 | 144.137492 | -0.0032742 | 0.42152631 | -0.0077675 | 0.99380252 | 0.99919666 |
| GPC1 | 89.3886211 | -0.6207779 | 0.32999254 | -1.8811877 | 0.05994639 | 0.53787511 |
| ANKMY1 | 129.976148 | -0.0459124 | 0.35662472 | -0.1287414 | 0.89756224 | 0.98612679 |
| RNPEPL1 | 102.437945 | -0.2881413 | 0.30711049 | -0.9382333 | 0.34812451 | 0.85348649 |
| CAPN10 | 41.9838526 | -0.2956025 | 0.3590678 | -0.8232497 | 0.41036601 | 0.88019562 |
| SNED1 | 251.01957 | -1.4454576 | 0.38965117 | -3.7096195 | 0.00020757 | 0.03246782 |
| AC005237.1 | 107.399125 | -1.6014851 | 0.54495773 | -2.9387327 | 0.00329557 | 0.14358074 |
| MTERF4 | 126.210932 | -0.1874498 | 0.29143913 | -0.6431867 | 0.52010299 | 0.91751084 |
| PASK | 34.5477525 | -0.1199046 | 0.33400365 | -0.3589918 | 0.71960123 | 0.95796325 |
| PPP1R7 | 122.923872 | 0.20084847 | 0.23027472 | 0.87221241 | 0.38309251 | 0.86638702 |
| HDLBP | 611.803897 | -0.1391888 | 0.18014402 | -0.7726526 | 0.43972799 | 0.88917548 |
| 2-Sep | 818.831876 | 0.04131785 | 0.17376273 | 0.23778314 | 0.81204929 | 0.9680627 |
| FARP2 | 292.873424 | -0.1087783 | 0.20673848 | -0.5261638 | 0.59877437 | 0.93728325 |
| STK25 | 127.03663 | -0.1642723 | 0.26218445 | -0.6265526 | 0.53095254 | 0.92211604 |

| | | | | | | |
|---|---|---|---|---|---|---|
| THAP4 | 168.5505 | 0.06192846 | 0.23193669 | 0.26700589 | 0.78946463 | 0.96515649 |
| ATG4B | 219.85591 | -0.371053 | 0.23220172 | -1.5979769 | 0.11004811 | 0.64001342 |
| DTYMK | 59.3866596 | 0.21526737 | 0.41446272 | 0.51938897 | 0.60348952 | 0.93804805 |
| ING5 | 120.378054 | -0.5715734 | 0.27628325 | -2.068795 | 0.03856533 | 0.45064107 |
| D2HGDH | 173.726574 | 0.27016824 | 0.26255403 | 1.02900056 | 0.30347941 | 0.83709431 |
| CHL1 | 95.7319262 | 1.42802099 | 0.87636905 | 1.62947447 | 0.10321262 | 0.62635087 |
| LINC01266 | 565.761094 | 0.42384683 | 0.48841058 | 0.86780845 | 0.3854992 | 0.86751807 |
| CNTN4 | 198.183022 | -0.0206406 | 0.47612195 | -0.0433516 | 0.96542129 | 0.99637506 |
| TRNT1 | 127.659712 | 0.01664632 | 0.18607074 | 0.08946229 | 0.92871452 | 0.99216065 |
| CRBN | 156.619486 | 0.16877698 | 0.25055808 | 0.67360423 | 0.50056297 | 0.90839719 |
| AC034195.1 | 237.913968 | 0.3619012 | 0.51892109 | 0.69741085 | 0.48554571 | 0.90695174 |
| SUMF1 | 1198.19665 | -0.4434561 | 0.35746175 | -1.2405694 | 0.21476485 | 0.77890374 |
| SETMAR | 35.3342327 | 0.47699058 | 0.39000209 | 1.22304622 | 0.22131224 | 0.78752408 |
| ITPR1 | 491.509398 | 0.08375175 | 0.33839879 | 0.24749425 | 0.80452574 | 0.96679622 |
| EGOT | 114.942212 | -0.1531978 | 0.83653288 | -0.1831343 | 0.85469266 | 0.97718353 |
| AC018816.1 | 63.8522581 | -0.4688202 | 0.31458494 | -1.4902817 | 0.13615018 | 0.67998335 |
| BHLHE40-AS | 68.811835 | 0.13460467 | 0.59217155 | 0.22730688 | 0.82018512 | 0.96886922 |
| BHLHE40 | 1164.36445 | -1.336596 | 0.65487472 | -2.040995 | 0.04125132 | 0.4648536 |
| ARL8B | 372.018752 | -0.7417371 | 0.25252033 | -2.9373363 | 0.00331045 | 0.14358074 |
| AC026202.2 | 68.7829322 | -0.8184927 | 0.47866803 | -1.7099382 | 0.08727731 | 0.60614063 |
| EDEM1 | 95.5129926 | -0.2185928 | 0.34414988 | -0.6351675 | 0.52531916 | 0.91924275 |
| LMCD1-AS1 | 310.907736 | 0.99863623 | 0.38658315 | 2.58323785 | 0.00978778 | 0.24135519 |
| LMCD1 | 40.666427 | 1.16638137 | 0.53303898 | 2.18817275 | 0.02865702 | 0.40623517 |
| RAD18 | 76.9126873 | -0.0811299 | 0.33162918 | -0.2446404 | 0.80673485 | 0.96752828 |
| SRGAP3 | 937.076841 | 0.72133689 | 0.29114112 | 2.47761936 | 0.01322622 | 0.28170334 |
| THUMPD3-A: | 264.202691 | -0.0070824 | 0.21240013 | -0.0333447 | 0.97339967 | 0.99656026 |
| THUMPD3 | 92.8573891 | 0.0217391 | 0.214232 | 0.10147457 | 0.91917374 | 0.99014946 |
| SETD5 | 923.894402 | -0.042367 | 0.16408065 | -0.2582081 | 0.79624628 | 0.9653598 |
| MTMR14 | 156.46338 | 0.02219252 | 0.3835386 | 0.05786255 | 0.95385811 | 0.99514888 |
| OGG1 | 60.148999 | -0.0749202 | 0.32654515 | -0.2294329 | 0.81853246 | 0.96883031 |
| CAMK1 | 46.8856648 | 0.37418913 | 0.27286756 | 1.37132144 | 0.17027477 | 0.72675731 |
| TADA3 | 90.4981537 | 0.00159221 | 0.23767295 | 0.00669918 | 0.99465487 | 0.99919666 |
| RPUSD3 | 50.0907971 | 0.00190704 | 0.25184299 | 0.00757235 | 0.9939582 | 0.99919666 |
| JAGN1 | 64.2516252 | 0.36121448 | 0.42706297 | 0.84581081 | 0.39765829 | 0.87462201 |
| IL17RE | 55.3790469 | 0.68261406 | 0.32501037 | 2.10028394 | 0.03570387 | 0.4362094 |
| IL17RC | 49.6468701 | 0.4799945 | 0.34121785 | 1.40670981 | 0.15951345 | 0.71409437 |
| CRELD1 | 80.3260401 | 0.08523377 | 0.34227334 | 0.24902251 | 0.80334337 | 0.96678163 |
| EMC3 | 146.312828 | 0.17579792 | 0.21425288 | 0.82051604 | 0.41192199 | 0.88098902 |
| BRK1 | 219.093909 | -0.0048685 | 0.44995602 | -0.0108199 | 0.99136718 | 0.99889703 |
| VHL | 90.9062772 | -0.0045183 | 0.23652993 | -0.0191025 | 0.98475932 | 0.99762072 |
| SEC13 | 154.187161 | -0.1262803 | 0.30405366 | -0.4153223 | 0.67790596 | 0.9556705 |
| ATG7 | 567.924371 | -0.409519 | 0.21323848 | -1.9204742 | 0.05479803 | 0.51597783 |
| VGLL4 | 596.513848 | 0.08970452 | 0.23582521 | 0.38038562 | 0.70365919 | 0.95672887 |
| TAMM41 | 173.901078 | -0.0577339 | 0.26796482 | -0.2154534 | 0.82941382 | 0.97131041 |
| PPARG | 1505.56936 | -0.3102134 | 0.43730942 | -0.7093681 | 0.47809606 | 0.90262056 |
| TSEN2 | 218.353432 | 0.07432384 | 0.27230761 | 0.27294074 | 0.78489877 | 0.96513523 |
| MKRN2OS | 40.6826461 | 0.11113702 | 0.4885673 | 0.22747536 | 0.82005413 | 0.96886922 |
| MKRN2 | 48.0674004 | 0.06782159 | 0.28517306 | 0.23782607 | 0.812016 | 0.9680627 |
| RAF1 | 431.997897 | -0.0848864 | 0.17512517 | -0.4847185 | 6.28E-01 | 0.94555838 |
| RPL32 | 844.374661 | -0.0287472 | 0.46855835 | -0.0613524 | 0.95107858 | 0.99404743 |
| IQSEC1 | 100.236477 | 0.77323892 | 0.24387228 | 3.17067164 | 0.00152087 | 0.09927452 |
| NUP210 | 96.9177639 | 0.39516608 | 0.35833657 | 1.1027791 | 0.27012311 | 0.81817548 |
| HDAC11 | 68.9688822 | 0.24810155 | 0.24973415 | 0.99346267 | 0.32048453 | 0.84285425 |
| XPC | 186.538518 | 0.21379729 | 0.28315423 | 0.75505596 | 0.4502154 | 0.893271 |

| | | | | | | |
|---|---|---|---|---|---|---|
| LSM3 | 185.363791 | 0.12220956 | 0.5442305 | 0.22455478 | 0.82232564 | 0.97010249 |
| SLC6A6 | 248.491399 | -0.5225078 | 0.65935074 | -0.792458 | 0.42809365 | 0.88535099 |
| CCDC174 | 96.6084104 | -0.081707 | 0.20968403 | -0.3896671 | 0.69678272 | 0.95672887 |
| FGD5-AS1 | 295.66451 | 0.09421556 | 0.2753239 | 0.342199 | 0.73220114 | 0.96008876 |
| NR2C2 | 294.826773 | -0.2398682 | 0.27292615 | -0.8788758 | 0.37946864 | 0.86584614 |
| MRPS25 | 285.811136 | -0.1416313 | 0.162923 | -0.8693145 | 0.38467512 | 0.86725322 |
| RBSN | 102.12909 | -0.3099475 | 0.25705916 | -1.2057439 | 0.22791626 | 0.79042788 |
| CAPN7 | 418.75116 | -0.0947274 | 0.20824491 | -0.4548846 | 0.64919226 | 0.95061388 |
| SH3BP5-AS1 | 36.6609034 | -0.2572713 | 0.42819718 | -0.6008244 | 0.54795698 | 0.92682201 |
| SH3BP5 | 459.514087 | -2.5528178 | 0.88028344 | -2.8999952 | 0.00373168 | 0.14917797 |
| METTL6 | 91.4247232 | -0.3787912 | 0.25734255 | -1.4719339 | 0.14103873 | 0.68601316 |
| EAF1 | 48.2840016 | 0.35318866 | 0.39062299 | 0.90416761 | 0.36590653 | 0.8588355 |
| COLQ | 54.621641 | -0.0080992 | 0.30592155 | -0.0264747 | 0.9788787 | 0.99696835 |
| HACL1 | 174.030356 | -0.0287037 | 0.25871092 | -0.1109488 | 0.91165697 | 0.98788932 |
| BTD | 72.1385746 | 0.58683473 | 0.28591817 | 2.05245697 | 0.04012527 | 0.4595769 |
| ANKRD28 | 838.62589 | 0.03522425 | 0.21483298 | 0.16396107 | 0.86976179 | 0.98202612 |
| DPH3 | 38.3970177 | -0.4968596 | 0.46964717 | -1.0579423 | 0.29008173 | 0.83103268 |
| OXNAD1 | 211.62511 | -0.1162386 | 0.37727678 | -0.3080991 | 0.75800695 | 0.96180522 |
| RFTN1 | 39.2082546 | 0.87219271 | 0.41596484 | 2.09679431 | 0.03601179 | 0.4362094 |
| PLCL2 | 2591.27144 | -0.0831337 | 0.40393906 | -0.2058075 | 0.8369413 | 0.97353666 |
| TBC1D5 | 1752.93411 | -0.0405904 | 0.17019042 | -0.2384997 | 0.81149357 | 0.9680627 |
| RAB5A | 315.69223 | -0.1903167 | 0.18796217 | -1.0125266 | 0.31128632 | 0.84279403 |
| KAT2B | 236.02665 | 0.16088954 | 0.30923853 | 0.5202765 | 0.60287087 | 0.93804805 |
| ZNF385D | 53.8558268 | -0.2102965 | 0.50733372 | -0.4145132 | 0.67849832 | 0.95574492 |
| UBE2E2 | 793.837475 | 0.08435748 | 0.33697396 | 0.25033827 | 0.80232576 | 0.9661728 |
| UBE2E1 | 572.965901 | -0.117729 | 0.23629996 | -0.4982183 | 0.61833018 | 0.94256005 |
| NKIRAS1 | 61.5288036 | 0.28598307 | 0.29625203 | 0.9653371 | 0.334376 | 0.84480724 |
| RPL15 | 1478.5133 | 0.5368814 | 0.44901845 | 1.19567781 | 0.23182231 | 0.79222614 |
| NR1D2 | 163.725299 | 0.20853684 | 0.33085256 | 0.63030143 | 0.52849739 | 0.92121123 |
| THRB | 1791.69811 | 0.4842846 | 0.2184184 | 2.21723355 | 2.66E-02 | 0.39496145 |
| RARB | 251.884691 | -0.3378624 | 0.32150114 | -1.0508903 | 0.29330896 | 0.83336457 |
| TOP2B | 526.377907 | -0.1402327 | 0.20261943 | -0.6920988 | 0.48887528 | 0.90730741 |
| NGLY1 | 345.43345 | -1.0337158 | 0.61605517 | -1.6779598 | 0.09335496 | 0.61317397 |
| OXSM | 44.4344443 | -0.0078157 | 0.37299113 | -0.020954 | 0.98328234 | 0.9971769 |
| NEK10 | 381.700248 | 0.57336325 | 0.30119848 | 1.90360605 | 0.05696151 | 0.52502782 |
| SLC4A7 | 305.268932 | -0.8634526 | 0.63194212 | -1.3663476 | 0.17182988 | 0.72776437 |
| CMC1 | 313.768321 | 0.08965051 | 0.18061916 | 0.49635103 | 0.61964677 | 0.94313269 |
| AZI2 | 244.754583 | -0.2364098 | 0.25622993 | -0.9226472 | 0.35619109 | 0.8557799 |
| TGFBR2 | 326.911867 | -0.1782967 | 0.45337789 | -0.3932629 | 0.69412534 | 0.95672887 |
| STT3B | 318.912124 | -0.0056364 | 0.23172836 | -0.0243233 | 0.98059473 | 0.99696835 |
| OSBPL10 | 110.566128 | -0.5465038 | 0.53300985 | -1.0253165 | 0.30521387 | 0.83709431 |
| GPD1L | 36.1646103 | 0.67687152 | 0.30652102 | 2.20823852 | 0.02722765 | 0.40076654 |
| CMTM8 | 56.410003 | 0.939999 | 0.43193231 | 2.1762646 | 0.02953548 | 0.40623517 |
| CMTM7 | 153.836752 | 1.08200588 | 0.50723578 | 2.13314188 | 0.03291309 | 0.42193013 |
| CMTM6 | 202.379482 | -0.0612349 | 0.25888533 | -0.2365328 | 0.8130193 | 0.9680627 |
| DYNC1LI1 | 89.1859915 | -0.1641661 | 0.23003622 | -0.7136535 | 0.47544148 | 0.90242252 |
| CNOT10 | 265.492627 | -0.5404152 | 0.20316846 | -2.6599368 | 0.00781553 | 0.217192 |
| GLB1 | 235.287875 | 0.44755719 | 0.24542881 | 1.8235723 | 0.06821676 | 0.55917298 |
| CRTAP | 135.370921 | 0.17997371 | 0.2886764 | 0.62344448 | 0.53299248 | 0.92268523 |
| FBXL2 | 524.223079 | -0.480079 | 0.20604771 | -2.3299412 | 0.01980926 | 0.33831338 |
| UBP1 | 163.287607 | 0.10281721 | 0.29566084 | 0.3477539 | 0.72802502 | 0.9598317 |
| CLASP2 | 786.524792 | -0.1268928 | 0.27865624 | -0.4553741 | 0.64884014 | 0.95061388 |
| PDCD6IP | 451.098163 | 0.08542773 | 0.15005613 | 0.56930517 | 0.56914906 | 0.93122667 |
| STAC | 181.748257 | -0.5190601 | 0.52746368 | -0.9840679 | 0.32508215 | 0.84285425 |

| | | | | | | |
|---|---|---|---|---|---|---|
| EPM2AIP1 | 151.167897 | 0.2750595 | 0.21116793 | 1.30256284 | 0.19272405 | 0.7597012 |
| MLH1 | 95.1141704 | -0.2024382 | 0.23624546 | -0.8568977 | 0.39150143 | 0.87015937 |
| LRRFIP2 | 740.129556 | -0.1377555 | 0.24141564 | -0.5706156 | 0.56826021 | 0.93122667 |
| GOLGA4 | 1251.1376 | -0.3557726 | 0.20425699 | -1.7417892 | 0.08154534 | 0.59202053 |
| ITGA9 | 69.5473024 | -0.679922 | 0.44787565 | -1.5181043 | 0.12898811 | 0.67206108 |
| CTDSPL | 551.781073 | -0.0958379 | 0.28470792 | -0.3366182 | 0.73640473 | 0.96078945 |
| PLCD1 | 49.5337436 | 0.41898165 | 0.42768911 | 0.97964069 | 0.32726351 | 0.84381657 |
| ACAA1 | 145.588168 | 0.70192641 | 0.43486257 | 1.61413389 | 0.10649839 | 0.63461115 |
| OXSR1 | 646.329861 | -0.3215362 | 0.51386423 | -0.6257221 | 0.53149722 | 0.92211604 |
| ACVR2B | 69.4550421 | 0.13027051 | 0.29813871 | 0.43694597 | 0.66215053 | 0.95142083 |
| EXOG | 54.0804094 | 0.18323705 | 0.2864661 | 0.63964656 | 0.52240241 | 0.91845051 |
| WDR48 | 196.409716 | -0.0249481 | 0.22766926 | -0.1095803 | 0.91274225 | 0.98830345 |
| GORASP1 | 62.2634128 | -0.0885847 | 0.2527786 | -0.3504438 | 0.72600564 | 0.95979351 |
| TTC21A | 62.3819314 | 0.50062957 | 0.38021393 | 1.31670498 | 0.18793753 | 0.7526909 |
| CSRNP1 | 78.0875936 | -1.3749199 | 0.76977419 | -1.7861341 | 0.07407755 | 0.56915891 |
| SLC25A38 | 53.9844736 | 0.62466366 | 0.31102959 | 2.00837375 | 0.04460359 | 0.48091114 |
| RPSA | 566.747636 | -0.0891536 | 0.40506995 | -0.2200942 | 0.82579777 | 0.97043739 |
| MYRIP | 60.095204 | 0.16683436 | 0.52079844 | 0.32034343 | 0.74870801 | 0.96109466 |
| EIF1B | 145.913303 | 0.5825843 | 0.47744773 | 1.2202054 | 0.22238702 | 0.78760199 |
| ENTPD3-AS1 | 84.4607322 | -0.0319822 | 0.57179754 | -0.0559327 | 0.95539544 | 0.99538167 |
| RPL14 | 962.514007 | 0.12746424 | 0.56117546 | 0.22713794 | 0.82031649 | 0.96886922 |
| ZNF621 | 129.175161 | -0.4055236 | 0.23572333 | -1.7203372 | 0.08537117 | 0.59845478 |
| CTNNB1 | 380.944811 | -0.0817328 | 0.17287585 | -0.4727832 | 0.63636786 | 0.94821838 |
| ULK4 | 1027.54577 | 0.4359116 | 0.30675163 | 1.42105716 | 0.15530014 | 0.70907003 |
| TRAK1 | 1140.64711 | 0.26109555 | 0.37505093 | 0.69616025 | 0.48632848 | 0.90695174 |
| SEC22C | 96.9850628 | -0.3137884 | 0.32149112 | -0.9760407 | 0.32904429 | 0.84409527 |
| SS18L2 | 63.1296276 | -0.3054904 | 0.4124704 | -0.740636 | 0.4589142 | 0.89753337 |
| NKTR | 732.769937 | -0.2175992 | 0.1462302 | -1.4880596 | 0.13673518 | 0.68107639 |
| AC006059.1 | 110.984423 | 0.1793671 | 0.50030641 | 0.3585145 | 0.71995832 | 0.95796325 |
| CCDC13 | 36.265746 | 0.27805947 | 0.37229138 | 0.74688666 | 0.45513198 | 0.89626521 |
| AC099329.3 | 39.4004537 | 0.18041049 | 0.39770349 | 0.45363063 | 0.65009471 | 0.95061388 |
| POMGNT2 | 37.8644752 | 0.48110368 | 0.29838889 | 1.61233782 | 0.10688845 | 0.63477557 |
| SNRK | 284.982527 | -0.0331191 | 0.47541859 | -0.0696631 | 0.94446184 | 0.99265578 |
| ANO10 | 604.723295 | -0.0356985 | 0.34029329 | -0.1049052 | 0.91645102 | 0.98905249 |
| ABHD5 | 74.1997716 | -0.3794111 | 0.31390637 | -1.2086759 | 0.22678738 | 0.78990116 |
| TCAIM | 548.653208 | 0.09902209 | 0.25384921 | 0.39008232 | 0.69647568 | 0.95672887 |
| ZNF445 | 153.455067 | -0.2407679 | 0.28379781 | -0.8483783 | 0.3962273 | 0.87462201 |
| ZNF852 | 38.0886034 | -0.1177108 | 0.26491687 | -0.4443312 | 0.65680313 | 0.95100227 |
| AC099669.1 | 178.190271 | 0.2661654 | 0.37209608 | 0.71531364 | 0.47441526 | 0.90242252 |
| ZNF197 | 95.717252 | 0.40817871 | 0.33581912 | 1.2154719 | 0.22418616 | 0.78960314 |
| KIAA1143 | 76.2403643 | 0.11185162 | 0.36411891 | 0.30718431 | 0.75870308 | 0.96180522 |
| ZDHHC3 | 303.494781 | 0.30556728 | 0.18829768 | 1.6227884 | 0.10463465 | 0.63166537 |
| EXOSC7 | 95.2602009 | -0.0536845 | 0.28523136 | -0.1882139 | 0.85070894 | 0.97718353 |
| CDCP1 | 204.851737 | 0.22925801 | 0.4247614 | 0.53973363 | 0.58938075 | 0.93483536 |
| LARS2 | 206.461204 | -0.0506403 | 0.25460593 | -0.1988968 | 0.84234344 | 0.97560439 |
| LIMD1 | 148.735213 | 0.34076901 | 0.2860841 | 1.19114977 | 0.2335948 | 0.79354083 |
| SACM1L | 227.122905 | 0.00758138 | 0.21541915 | 0.0351936 | 0.97192536 | 0.99656026 |
| SLC6A20 | 43.5895795 | 2.72344458 | 0.63851746 | 4.26526252 | 2.00E-05 | 0.00614104 |
| LZTFL1 | 70.6868679 | 0.1865573 | 0.23874281 | 0.78141535 | 0.43455825 | 0.88785137 |
| FYCO1 | 113.225462 | -0.0755686 | 0.18530041 | -0.4078165 | 0.68340843 | 0.95574492 |
| LTF | 896.092408 | 1.2321325 | 1.33766573 | 0.92110642 | 0.35699487 | 0.8557799 |
| ALS2CL | 35.5537642 | -0.0737091 | 0.26821191 | -0.2748165 | 7.83E-01 | 0.96513523 |
| CCDC12 | 316.284078 | -0.1013209 | 0.24475999 | -0.4139602 | 0.67890324 | 0.95574492 |
| NBEAL2 | 200.239995 | -1.0709165 | 0.2764054 | -3.8744411 | 0.00010687 | 0.02190188 |

| | | | | | | |
|---|---|---|---|---|---|---|
| SETD2 | 918.377898 | -0.0579649 | 0.17953239 | -0.3228662 | 0.74679654 | 0.96109466 |
| KIF9-AS1 | 241.688533 | 0.34792654 | 0.26797781 | 1.29834086 | 0.19417023 | 0.76162393 |
| KIF9 | 53.210615 | 0.55551356 | 0.38147743 | 1.45621605 | 0.14533291 | 0.69325556 |
| KLHL18 | 101.085008 | 0.16966186 | 0.22378503 | 0.75814662 | 0.44836321 | 0.89297728 |
| PTPN23 | 75.4942687 | -0.0409007 | 0.25252255 | -0.1619684 | 0.87133074 | 0.98208511 |
| SCAP | 158.065056 | 0.04079136 | 0.2506323 | 0.1627538 | 0.87071228 | 0.98208511 |
| ELP6 | 79.5399996 | 0.18610064 | 0.28884446 | 0.64429361 | 0.51938506 | 0.91751084 |
| SMARCC1 | 800.37027 | -0.3177648 | 0.26451603 | -1.2013065 | 0.22963232 | 0.79047378 |
| DHX30 | 164.675145 | -0.0771505 | 0.22931962 | -0.3364323 | 0.73654486 | 0.96078945 |
| MAP4 | 1647.04582 | -0.083742 | 0.20997266 | -0.3988235 | 0.69002327 | 0.95659154 |
| ZNF589 | 65.0167272 | 0.10317466 | 0.28159445 | 0.36639453 | 0.7140707 | 0.95791799 |
| NME6 | 64.1493465 | 0.19290605 | 0.27517749 | 0.70102409 | 0.48328798 | 0.90631121 |
| PLXNB1 | 370.316853 | -0.1272851 | 0.27499115 | -0.4628697 | 0.64345774 | 0.94949493 |
| TMA7 | 376.272148 | 0.18959239 | 0.49680759 | 0.38162137 | 0.70274223 | 0.95672887 |
| SHISA5 | 287.087954 | -0.5649473 | 0.36792483 | -1.5354964 | 0.12466194 | 0.66626798 |
| PFKFB4 | 199.882039 | -1.5872241 | 0.78180114 | -2.0302147 | 0.04233472 | 0.46998719 |
| UQCRC1 | 165.025059 | 0.19663207 | 0.28121407 | 0.69922558 | 0.48441107 | 0.90659463 |
| NCKIPSD | 49.4030793 | -0.1132367 | 0.39822465 | -0.2843539 | 0.77613919 | 0.96437053 |
| IP6K2 | 219.914004 | -0.0505757 | 0.16191452 | -0.3123605 | 0.75476655 | 0.96120879 |
| PRKAR2A | 383.373607 | -0.1580869 | 0.21181919 | -0.7463296 | 0.45546831 | 0.89626521 |
| SLC25A20 | 34.7265744 | -0.1325485 | 0.30062924 | -0.4409035 | 0.65928286 | 0.95127957 |
| ARIH2 | 314.723224 | -0.2046742 | 0.1929509 | -1.0607578 | 0.2888 | 0.83060304 |
| P4HTM | 112.489035 | 0.50907316 | 0.23226271 | 2.19179895 | 0.02839403 | 0.40459506 |
| WDR6 | 204.046466 | -0.3897458 | 0.21638428 | -1.8011745 | 0.07167538 | 0.56343001 |
| DALRD3 | 59.2538256 | 0.17640647 | 0.3729896 | 0.47295278 | 0.63624687 | 0.94821838 |
| NDUFAF3 | 137.544509 | 0.34105963 | 0.55291139 | 0.6168432 | 0.53733815 | 0.92324352 |
| IMPDH2 | 56.1574682 | 0.10876414 | 0.31044529 | 0.35034882 | 0.72607693 | 0.95979351 |
| QRICH1 | 202.502279 | -0.1389242 | 0.16587405 | -0.8375283 | 0.40229568 | 0.87834687 |
| QARS | 77.238538 | -0.1380449 | 0.24178132 | -0.5709492 | 0.56803406 | 0.93122667 |
| USP19 | 48.755571 | 0.05492223 | 0.29407281 | 0.18676404 | 0.85184564 | 0.97718353 |
| LAMB2 | 122.830412 | 0.74460732 | 0.34314312 | 2.16996138 | 0.03000977 | 0.40897583 |
| KLHDC8B | 52.4952796 | 1.03434024 | 0.3008217 | 3.43838304 | 0.0005852 | 0.05789336 |
| C3orf62 | 35.0254852 | 0.53028516 | 0.30626052 | 1.7314839 | 0.08336549 | 0.59231889 |
| USP4 | 199.046849 | -0.3984916 | 0.27929964 | -1.4267529 | 0.15365113 | 0.70747602 |
| RHOA | 569.489337 | 0.11012859 | 0.23304565 | 0.47256232 | 0.63652547 | 0.94821838 |
| TCTA | 57.7255601 | 0.22682541 | 0.3353316 | 0.67642122 | 0.49877325 | 0.90839719 |
| DAG1 | 229.384616 | 0.42605938 | 0.24630949 | 1.72977249 | 0.08367093 | 0.59328239 |
| APEH | 98.1398248 | 0.4363355 | 0.35750112 | 1.22051507 | 0.22226968 | 0.78760199 |
| MST1 | 76.1597838 | -0.0530618 | 0.33323113 | -0.1592343 | 0.87348428 | 0.98246494 |
| RNF123 | 57.9327589 | 0.19788018 | 0.22457339 | 0.88113818 | 0.37824304 | 0.86512157 |
| IP6K1 | 249.445479 | -0.1510605 | 0.2607744 | -0.5792764 | 0.56240268 | 0.93008996 |
| RBM6 | 1104.37516 | -0.2384775 | 0.24182386 | -0.9861618 | 0.32405374 | 0.84285425 |
| RBM5 | 401.814096 | -0.3111676 | 0.14335144 | -2.1706624 | 0.0299567 | 0.40893584 |
| SEMA3F-AS1 | 54.4035796 | -1.4169468 | 0.50930868 | -2.7820983 | 0.00540087 | 0.18165004 |
| SEMA3F | 217.878339 | -0.2182711 | 0.35540927 | -0.6141402 | 0.53912267 | 0.92365631 |
| GNAI2 | 248.824433 | -0.0418687 | 0.27175436 | -0.1540682 | 0.87755595 | 0.98279964 |
| SEMA3B | 356.751866 | -1.1328448 | 0.64762027 | -1.7492424 | 0.08024914 | 0.58729814 |
| IFRD2 | 52.2585143 | -0.2706817 | 0.3523913 | -0.7681281 | 0.4424111 | 0.89005186 |
| HYAL1 | 48.1073109 | 0.44446594 | 0.44819185 | 0.9916868 | 0.32135033 | 0.84285425 |
| HYAL2 | 54.8112093 | 0.33037352 | 0.2856779 | 1.15645459 | 0.24749526 | 0.80072691 |
| TUSC2 | 76.711823 | 0.2993478 | 0.47189894 | 0.63434725 | 0.52585423 | 0.91956488 |
| RASSF1 | 41.3647808 | -0.910065 | 0.41863669 | -2.1738777 | 0.02971432 | 0.40653537 |
| TMEM115 | 54.8386121 | 0.42581605 | 0.41068597 | 1.036841 | 0.29980996 | 0.83618229 |
| CACNA2D2 | 89.8670547 | 1.49996516 | 0.72941893 | 2.05638365 | 0.03974555 | 0.45805615 |

| | | | | | | |
|---|---|---|---|---|---|---|
| HEMK1 | 51.7330867 | -0.0328456 | 0.3236895 | -0.1014727 | 0.91917525 | 0.99014946 |
| CISH | 77.9282485 | 0.18508269 | 0.35045888 | 0.52811527 | 0.59741933 | 0.93662454 |
| MAPKAPK3 | 97.2461093 | -0.2779788 | 0.29904768 | -0.9295466 | 0.35260587 | 0.85536253 |
| DOCK3 | 157.196907 | 0.66063044 | 0.63365991 | 1.04256309 | 0.29715069 | 0.83556453 |
| MANF | 84.9299355 | 0.4393511 | 0.41390977 | 1.06146589 | 0.28847823 | 0.83060304 |
| RBM15B | 50.2671663 | -0.6545295 | 0.5632262 | -1.1621077 | 0.24519172 | 0.80072691 |
| DCAF1 | 162.74177 | -0.6732633 | 0.26399744 | -2.5502644 | 0.01076412 | 0.25256429 |
| RAD54L2 | 153.808306 | -0.0148715 | 0.3077972 | -0.0483157 | 0.9614646 | 0.99635733 |
| TEX264 | 162.195114 | 0.40215046 | 0.35777092 | 1.12404458 | 0.26099411 | 0.80805101 |
| RPL29 | 1258.11328 | 0.2775086 | 0.51435757 | 0.53952468 | 0.58952487 | 0.93483536 |
| ALAS1 | 280.626107 | 0.58782713 | 0.41893885 | 1.40313348 | 0.16057704 | 0.71409437 |
| WDR82 | 195.943891 | 0.09639806 | 0.19909489 | 0.4841815 | 0.62825707 | 0.94572452 |
| DNAH1 | 36.7499772 | -0.1043602 | 0.40983418 | -0.2546402 | 0.79900104 | 0.9653598 |
| BAP1 | 56.7365308 | -0.1602359 | 0.35440864 | -0.4521218 | 0.65118125 | 0.95061388 |
| PHF7 | 34.8432096 | 0.28505795 | 0.43410246 | 0.65666052 | 0.51139922 | 0.91403647 |
| NISCH | 215.089732 | -0.0719538 | 0.2263092 | -0.3179445 | 0.75052704 | 0.96109466 |
| SMIM4 | 116.801818 | -0.3415609 | 0.24559446 | -1.3907516 | 0.16430076 | 0.71715698 |
| PBRM1 | 538.81353 | -0.4798136 | 0.18958876 | -2.5308122 | 0.01137988 | 0.26211233 |
| GNL3 | 92.4525512 | -0.3961336 | 0.45859163 | -0.8638046 | 0.38769524 | 0.86910078 |
| GLT8D1 | 80.1022967 | 0.62387302 | 0.2524599 | 2.47117672 | 0.01346692 | 0.28170334 |
| SPCS1 | 361.887495 | 0.49721795 | 0.46443339 | 1.07059045 | 0.28435362 | 0.82778499 |
| NEK4 | 152.288952 | -0.1236721 | 0.16418516 | -0.7532476 | 0.45130112 | 0.89400477 |
| SFMBT1 | 87.8031725 | 0.17116797 | 0.23242685 | 0.73643802 | 0.46146417 | 0.89799005 |
| PRKCD | 64.8924528 | 0.2229196 | 0.30833263 | 0.72298413 | 0.46968963 | 0.90053922 |
| TKT | 224.827064 | -0.1209662 | 0.37509795 | -0.3224923 | 7.47E-01 | 0.96109466 |
| DCP1A | 285.222627 | -0.2287554 | 0.194145 | -1.1782711 | 0.23868855 | 0.79867687 |
| CACNA1D | 77.2256565 | 0.01040343 | 0.3936197 | 0.02643017 | 9.79E-01 | 9.97E-01 |
| CHDH | 49.4150599 | 0.05702143 | 0.41647104 | 0.13691572 | 0.89109742 | 0.98597539 |
| ACTR8 | 35.137569 | 0.15937413 | 0.33431404 | 0.47671983 | 0.63356164 | 0.94744025 |
| SELENOK | 125.353647 | -0.0868351 | 0.67462555 | -0.128716 | 0.89758239 | 0.98612679 |
| CACNA2D3 | 60.4256208 | -0.4434451 | 0.40625822 | -1.091535 | 0.27503755 | 0.8202929 |
| ERC2 | 215.097114 | -0.1816266 | 0.30113114 | -0.6031477 | 0.5464104 | 0.92597739 |
| CCDC66 | 309.24526 | -0.1249487 | 0.19323107 | -0.6466287 | 0.51787229 | 0.91751084 |
| FAM208A | 372.13904 | -0.2060015 | 0.20833406 | -0.9888039 | 0.32275908 | 0.84285425 |
| ARHGEF3 | 1057.11433 | -0.5280162 | 0.33583272 | -1.5722595 | 0.11589039 | 0.65192493 |
| APPL1 | 241.306544 | -0.0542131 | 0.16219352 | -0.3342495 | 0.7381913 | 0.96078945 |
| PDE12 | 64.8197196 | 0.01529572 | 0.35373918 | 0.04324011 | 0.96551013 | 0.99637506 |
| ARF4 | 263.573591 | -0.0798523 | 0.28304522 | -0.2821185 | 0.77785267 | 0.96472142 |
| DENND6A | 284.857553 | -0.1948037 | 0.21133599 | -0.9217726 | 0.3566472 | 0.8557799 |
| DENND6A-AS | 65.7906743 | -0.2406795 | 0.3788256 | -0.6353306 | 0.52521284 | 0.91924275 |
| SLMAP | 765.183752 | -0.0859829 | 0.27206211 | -0.3160416 | 0.75197093 | 0.96120879 |
| FLNB | 768.735975 | 0.02962043 | 0.19459249 | 0.15221774 | 0.8790152 | 0.98311774 |
| RPP14 | 41.4971039 | 0.29757375 | 0.30314193 | 0.98163176 | 0.32628129 | 0.84381657 |
| PXK | 216.558168 | 0.02940303 | 0.37958922 | 0.07746014 | 0.9382575 | 0.99243654 |
| PDHB | 104.359722 | 0.42942622 | 0.36711456 | 1.16973356 | 0.24210821 | 0.80072691 |
| KCTD6 | 51.2385773 | 0.71464308 | 0.50939719 | 1.40291918 | 0.16064094 | 0.71409437 |
| FAM107A | 93.2145681 | 1.85738544 | 0.62927787 | 2.95161413 | 0.00316118 | 0.14358074 |
| C3orf67 | 64.9461286 | 0.20409148 | 0.34096331 | 0.59857314 | 0.54945757 | 0.9274143 |
| FHIT | 4610.70647 | 0.18166595 | 0.25012867 | 0.72628999 | 0.46766102 | 0.89976199 |
| PTPRG | 1509.98122 | 0.41826056 | 0.4634214 | 0.90254908 | 0.36676526 | 0.85902833 |
| THOC7 | 124.036318 | 0.10431807 | 0.27957742 | 0.37312765 | 0.70905344 | 0.95752169 |
| ATXN7 | 57.5541445 | 0.28891186 | 0.23668084 | 1.22068122 | 0.22220674 | 0.78760199 |
| PSMD6-AS2 | 45.0057026 | -0.7483044 | 0.38240641 | -1.9568301 | 0.05036745 | 0.49596288 |
| PSMD6 | 64.1020803 | 0.27897062 | 0.2577757 | 1.08222234 | 0.27915374 | 0.82479031 |

| | | | | | | |
|---|---|---|---|---|---|---|
| PRICKLE2 | 629.602764 | 0.45013414 | 0.46479777 | 0.9684516 | 0.33281888 | 0.84442294 |
| ADAMTS9 | 1146.30309 | -0.5035371 | 0.5016341 | -1.0037936 | 0.3154781 | 0.84285425 |
| ADAMTS9-A! | 99.7558157 | -0.2662519 | 0.53391581 | -0.4986776 | 0.61800653 | 0.94246161 |
| ADAMTS9-A! | 4499.34738 | 0.99452426 | 0.34684192 | 2.86737042 | 0.00413898 | 0.15603071 |
| MAGI1 | 4163.72096 | -0.0584874 | 0.12691666 | -0.4608332 | 6.45E-01 | 0.94949493 |
| MAGI1-AS1 | 71.3016291 | 0.33656857 | 0.64734542 | 0.51992115 | 0.60311854 | 0.93804805 |
| SLC25A26 | 264.366938 | -0.4355171 | 0.25721409 | -1.6932087 | 0.09041575 | 0.60779802 |
| LRIG1 | 38.1414179 | 0.78395764 | 0.45827257 | 1.71067982 | 0.08714023 | 0.60599181 |
| SUCLG2 | 685.568242 | -0.0201057 | 0.22758225 | -0.0883446 | 0.92960282 | 0.99216065 |
| SUCLG2-AS1 | 348.071751 | 0.59756427 | 0.35867837 | 1.66601704 | 0.09571002 | 0.61367241 |
| FAM19A1 | 62.0396522 | 0.1426143 | 0.55199775 | 0.2583603 | 0.79612885 | 0.9653598 |
| EOGT | 73.9618167 | -0.2010228 | 0.22684364 | -0.8861732 | 0.3755242 | 0.86357183 |
| AC109587.1 | 38.7316319 | 0.10334449 | 0.7291135 | 0.14173993 | 0.88728544 | 0.98551241 |
| TMF1 | 322.890986 | -0.1181023 | 0.25966426 | -0.454827 | 0.64923368 | 0.95061388 |
| UBA3 | 201.120355 | -0.1332868 | 0.17615272 | -0.756655 | 0.44925658 | 0.89324553 |
| ARL6IP5 | 290.42072 | 0.13917953 | 0.43619126 | 0.31907914 | 0.7496665 | 0.96109466 |
| FRMD4B | 89.9345197 | 0.13546556 | 0.34324049 | 0.39466662 | 0.69308894 | 0.95672887 |
| MITF | 1035.327 | 0.50720081 | 0.21414421 | 2.36850117 | 0.01786032 | 0.32105694 |
| FOXP1 | 1792.95274 | -0.093227 | 0.21548619 | -0.4326355 | 0.6652796 | 0.9520866 |
| EIF4E3 | 84.9775451 | 0.15852603 | 0.30334952 | 0.52258538 | 0.60126281 | 0.93802476 |
| RYBP | 505.490748 | -0.5264157 | 0.47360635 | -1.1115046 | 0.26635119 | 0.81309656 |
| SHQ1 | 172.087473 | 0.17399496 | 0.31569016 | 0.55115736 | 0.58152581 | 0.93386308 |
| GXYLT2 | 83.3474569 | -0.3129062 | 0.43789962 | -0.7145615 | 0.47488004 | 0.90242252 |
| PPP4R2 | 349.239488 | -0.0482246 | 0.20896169 | -0.230782 | 0.8174842 | 0.96883031 |
| PDZRN3 | 412.975415 | 1.08683898 | 0.57567527 | 1.88793757 | 0.05903433 | 0.53286516 |
| CNTN3 | 252.183686 | -0.2072237 | 0.26164356 | -0.7920078 | 0.42835616 | 0.88535099 |
| ZNF717 | 198.395744 | -0.0264806 | 0.26227293 | -0.1009657 | 0.91957771 | 0.99014946 |
| ROBO2 | 1195.50779 | -2.4390167 | 0.93757382 | -2.601413 | 0.00928406 | 0.23300339 |
| ROBO1 | 114.96236 | 1.23192177 | 0.56309451 | 2.18777088 | 0.0286863 | 0.40623517 |
| GBE1 | 1191.98923 | -1.2611167 | 0.59502466 | -2.119436 | 0.03405363 | 0.43049709 |
| VGLL3 | 156.933399 | -0.0427124 | 0.37616297 | -0.1135476 | 0.90959644 | 0.98779844 |
| CHMP2B | 217.474858 | -0.4249001 | 0.26030357 | -1.6323255 | 0.10261094 | 0.62630326 |
| CGGBP1 | 321.749308 | 0.07472232 | 0.23509775 | 0.31783511 | 0.75061002 | 0.96109466 |
| ZNF654 | 390.441403 | -0.0516482 | 0.277234 | -0.1862981 | 0.85221102 | 0.97718353 |
| C3orf38 | 83.9922527 | -0.0570624 | 0.18341525 | -0.3111102 | 0.75571686 | 0.96120879 |
| ARL13B | 165.573988 | -0.003218 | 0.20706968 | -0.0155408 | 0.98760076 | 0.99791717 |
| NSUN3 | 113.906755 | -0.6034425 | 0.24394003 | -2.473733 | 0.01337096 | 0.28170334 |
| MTRNR2L12 | 767.994788 | -0.1931519 | 0.57167506 | -0.3378701 | 0.73546106 | 0.96078945 |
| ARL6 | 124.578136 | -0.1102218 | 0.20601023 | -0.5350307 | 0.59262862 | 0.93491607 |
| CRYBG3 | 302.270978 | -0.2555434 | 0.26558247 | -0.9621998 | 0.33594927 | 0.84552074 |
| RIOX2 | 95.3499393 | -0.0198434 | 0.24477581 | -0.0810677 | 0.93538812 | 0.99243654 |
| CLDND1 | 86.163137 | -0.2912877 | 0.41574117 | -0.7006468 | 0.48352344 | 0.90631121 |
| DCBLD2 | 144.799325 | 0.00312064 | 0.33206541 | 0.00939766 | 0.99250186 | 0.99919666 |
| CMSS1 | 640.960263 | 0.06451205 | 0.41666773 | 0.15482852 | 0.87695651 | 0.98277085 |
| FILIP1L | 189.15376 | 0.77239302 | 0.70057053 | 1.10251999 | 0.27023567 | 0.81817548 |
| TBC1D23 | 200.024197 | -0.1459021 | 0.17037918 | -0.8563376 | 0.39181109 | 0.87030305 |
| NIT2 | 63.5744395 | -0.2531999 | 0.27295651 | -0.9276199 | 0.35360479 | 0.8557799 |
| TOMM70 | 158.904023 | 0.28109104 | 0.22188926 | 1.26680777 | 0.20522403 | 0.77193931 |
| TFG | 316.37336 | -0.2114449 | 0.15341696 | -1.3782366 | 0.16813026 | 0.72391336 |
| ABI3BP | 54.250466 | -0.8568608 | 0.67937803 | -1.2612431 | 0.20722128 | 0.77256457 |
| SENP7 | 432.842912 | -0.1792519 | 0.24060634 | -0.7450008 | 0.45627126 | 0.89626521 |
| TRMT10C | 65.796392 | 0.37317233 | 0.42251701 | 0.88321256 | 0.37712144 | 0.8642538 |
| PCNP | 278.786728 | -0.1902328 | 0.22130101 | -0.8596111 | 0.39000348 | 0.87004477 |
| ZBTB11 | 99.080838 | -0.4243311 | 0.19729466 | -2.1507482 | 0.03149608 | 0.41477661 |

| | | | | | | |
|---|---|---|---|---|---|---|
| RPL24 | 438.638102 | 0.03313679 | 0.42609556 | 0.07776845 | 0.93801224 | 0.99243654 |
| CEP97 | 70.9068166 | 0.04591051 | 0.21762189 | 0.2109646 | 0.8329149 | 0.97208613 |
| NFKBIZ | 685.82635 | -0.5244393 | 0.79577224 | -0.6590319 | 0.50987526 | 0.91334187 |
| ZPLD1 | 571.530324 | -2.1848106 | 1.57325803 | -1.3887173 | 0.16491873 | 0.71715698 |
| ALCAM | 237.13238 | -0.0554113 | 0.43861508 | -0.1263323 | 0.89946889 | 0.98660836 |
| CBLB | 593.374278 | 0.13400272 | 0.30216294 | 0.44347832 | 0.6574198 | 0.95100227 |
| LINC00882 | 278.960289 | 0.00692845 | 0.38060986 | 0.01820356 | 0.98547646 | 0.99762072 |
| DUBR | 51.5500039 | -0.3099548 | 0.37183852 | -0.8335736 | 0.4045213 | 0.87834687 |
| BBX | 1334.83899 | 0.29026431 | 0.29377926 | 0.9880354 | 0.32313531 | 0.84285425 |
| CD47 | 250.668021 | -0.086148 | 0.27873323 | -0.3090698 | 0.7572684 | 0.96149417 |
| IFT57 | 164.458159 | 0.33480794 | 0.21688485 | 1.54371286 | 0.12265791 | 0.66466126 |
| CIP2A | 38.9093563 | -0.5430921 | 0.48163001 | -1.1276127 | 0.25948352 | 0.80709747 |
| DZIP3 | 517.165478 | 0.67927053 | 0.40825766 | 1.66382801 | 0.0961468 | 0.61407447 |
| NECTIN3 | 72.1467905 | -0.0108485 | 0.2730052 | -0.0397373 | 0.96830256 | 0.9964399 |
| PHLDB2 | 909.442911 | -0.1399563 | 0.16065223 | -0.8711755 | 0.38365836 | 0.8665075 |
| ABHD10 | 74.6485749 | 0.53476184 | 0.25522099 | 2.09528941 | 0.03614527 | 0.4362094 |
| C3orf52 | 258.441721 | 0.26262101 | 0.43353264 | 0.60576986 | 0.54466759 | 0.92561018 |
| AC128688.2 | 38.9097092 | 0.33330207 | 0.57068472 | 0.58403888 | 0.55919415 | 0.92901206 |
| SLC9C1 | 214.28726 | 1.66086873 | 0.9285607 | 1.78864853 | 0.07367144 | 0.56770343 |
| ATG3 | 78.8836447 | -0.173697 | 0.24084776 | -0.7211898 | 0.47079274 | 0.90123338 |
| SLC35A5 | 54.7112079 | -0.4248306 | 0.26275314 | -1.6168432 | 0.10591213 | 0.63461115 |
| GTPBP8 | 41.5745418 | 0.0790068 | 0.38420481 | 0.2056372 | 0.83707432 | 0.97353666 |
| NEPRO | 97.846333 | 0.00268245 | 0.23693902 | 0.01132128 | 0.99096712 | 0.99889703 |
| CFAP44 | 226.185299 | 0.00228001 | 0.22841302 | 0.00998197 | 0.99203567 | 0.99919666 |
| SPICE1 | 167.022215 | -0.242809 | 0.2392265 | -1.0149754 | 0.31011755 | 0.84213522 |
| SIDT1 | 97.6297134 | -0.2331635 | 0.85627291 | -0.2723005 | 0.78539101 | 0.96513523 |
| USF3 | 177.823686 | 0.25754226 | 0.26186183 | 0.98350438 | 0.32535926 | 0.84285425 |
| NAA50 | 504.530113 | -0.3104924 | 0.23734327 | -1.3081996 | 0.19080561 | 0.75802398 |
| ATP6V1A | 822.711293 | -0.3357247 | 0.30447054 | -1.1026507 | 0.27017887 | 0.81817548 |
| GRAMD1C | 523.601705 | -0.5644418 | 0.26580333 | -2.1235319 | 0.0337093 | 0.42838847 |
| ZDHHC23 | 67.3092636 | -0.2497388 | 0.31954231 | -0.7815516 | 0.43447816 | 0.88785137 |
| CCDC191 | 246.689596 | -0.0488887 | 0.32529244 | -0.1502915 | 0.88053467 | 0.98333507 |
| QTRT2 | 90.2197298 | -0.2707355 | 0.35049786 | -0.7724311 | 0.43985911 | 0.88917548 |
| ZBTB20 | 8194.12004 | -0.0571276 | 0.20420543 | -0.2797557 | 0.77966495 | 0.96513523 |
| ZBTB20-AS1 | 41.2293648 | -0.1336345 | 0.51431884 | -0.2598281 | 0.79499636 | 0.9653598 |
| ZBTB20-AS5 | 173.608363 | 0.20028157 | 0.60008057 | 0.3337578 | 0.73856233 | 0.96078945 |
| ZBTB20-AS2 | 39.5617992 | 0.07901038 | 0.56363975 | 0.14017887 | 0.88851867 | 0.98551241 |
| LSAMP | 242.053193 | -0.1204838 | 0.80241271 | -0.1501519 | 0.88064481 | 0.98333507 |
| AC068633.1 | 314.501437 | -0.7193408 | 0.82159008 | -0.8755471 | 0.38127627 | 0.86593382 |
| IGSF11 | 53.0546198 | 1.16197259 | 0.4166732 | 2.78869048 | 0.00529216 | 0.1794882 |
| B4GALT4 | 62.6926045 | 0.52013436 | 0.23088544 | 2.25278108 | 0.02427296 | 0.37408908 |
| TMEM39A | 79.5394177 | -0.6024881 | 0.27938022 | -2.1565166 | 0.03104335 | 0.41286083 |
| POGLUT1 | 57.0919883 | 0.37231654 | 0.32656999 | 1.1400819 | 0.25425218 | 0.80414079 |
| TIMMDC1 | 158.298325 | 0.23735198 | 0.18956613 | 1.25208013 | 0.21054067 | 0.77610028 |
| COX17 | 123.900887 | 0.12977641 | 0.32813818 | 0.39549317 | 0.69247896 | 0.95672887 |
| MAATS1 | 174.13772 | 0.25590597 | 0.27266557 | 0.93853423 | 0.34796993 | 0.85348649 |
| GSK3B | 853.848243 | -0.1536746 | 0.25307847 | -0.6072211 | 0.54370422 | 0.92482794 |
| LRRC58 | 152.910253 | -0.1334748 | 0.24783232 | -0.5385692 | 0.59018417 | 0.93483536 |
| FSTL1 | 40.7392558 | 0.80339265 | 0.61305602 | 1.31047184 | 0.19003626 | 0.75619436 |
| NDUFB4 | 194.478777 | -0.0041794 | 0.28339815 | -0.0147474 | 0.98823367 | 0.99814557 |
| HGD | 216.631502 | -1.3119978 | 0.55838216 | -2.3496414 | 0.0187915 | 0.32850268 |
| RABL3 | 98.7207018 | -0.2614763 | 0.20899214 | -1.2511296 | 0.21088718 | 0.77656279 |
| GOLGB1 | 723.28774 | 0.05270738 | 0.19495411 | 0.2703579 | 0.78688492 | 0.96513523 |
| IQCB1 | 235.203742 | -0.0692475 | 0.163564 | -0.4233665 | 0.67202785 | 0.9544944 |

| | | | | | | |
|---|---|---|---|---|---|---|
| SLC15A2 | 47.8278653 | 0.54056161 | 0.55805263 | 0.96865704 | 0.33271634 | 0.84442294 |
| ILDR1 | 59.3482849 | -0.1925582 | 0.30681671 | -0.6276002 | 0.53026585 | 0.9218955 |
| CCDC58 | 58.8617298 | -0.4255811 | 0.28143966 | -1.5121577 | 0.13049376 | 0.67339292 |
| FAM162A | 439.742829 | -0.4278606 | 0.54529237 | -0.7846444 | 0.43266209 | 0.88722778 |
| AC083798.2 | 96.6984702 | 0.62628682 | 0.39760421 | 1.57515137 | 0.11522151 | 0.65130605 |
| KPNA1 | 337.664831 | -0.1534588 | 0.2474697 | -0.6201114 | 0.53518447 | 0.92303275 |
| PARP9 | 185.977084 | -0.8796 | 0.58728285 | -1.4977451 | 0.1341995 | 0.67832119 |
| DTX3L | 82.882122 | -0.7248371 | 0.48169734 | -1.5047562 | 0.13238676 | 0.67466347 |
| PARP14 | 555.580294 | -1.4076558 | 0.63467919 | -2.2179014 | 0.02656156 | 0.39484417 |
| HSPBAP1 | 96.5497458 | -0.1063456 | 0.38090101 | -0.2791948 | 0.78009535 | 0.96513523 |
| DIRC2 | 299.439487 | -0.277448 | 0.5971416 | -0.4646269 | 0.64219869 | 0.94949493 |
| PDIA5 | 43.4959712 | -0.3102186 | 0.37309243 | -0.8314793 | 0.40570293 | 0.87937203 |
| SEC22A | 240.539977 | -0.0743816 | 0.20080169 | -0.3704233 | 0.7110671 | 0.95773687 |
| ADCY5 | 51.0682238 | 0.28020434 | 0.78174998 | 0.35843217 | 0.72001992 | 0.95796325 |
| HACD2 | 161.219346 | 0.2241257 | 0.1721381 | 1.30201098 | 0.19291263 | 0.7597012 |
| MYLK-AS1 | 41.6737738 | 0.14717581 | 0.37009828 | 0.39766682 | 0.6908758 | 0.95659154 |
| MYLK | 39.9581598 | 0.35293119 | 0.61999707 | 0.56924655 | 0.56918883 | 0.93122667 |
| CCDC14 | 266.47929 | -0.0164696 | 0.16191655 | -0.1017164 | 0.91898177 | 0.99014946 |
| KALRN | 473.381185 | 0.47589723 | 0.48325295 | 0.98477873 | 0.32473277 | 0.84285425 |
| UMPS | 58.7645334 | 0.31333667 | 0.27032407 | 1.15911492 | 0.24640934 | 0.80072691 |
| ITGB5 | 330.174136 | 0.04058956 | 0.26870716 | 0.15105498 | 0.87993234 | 0.98333507 |
| SLC12A8 | 70.0875304 | -0.0660432 | 0.57545997 | -0.1147659 | 0.90863071 | 0.98758338 |
| ZNF148 | 933.853488 | -0.0255443 | 0.22699279 | -0.1125334 | 0.91040052 | 0.98788932 |
| SNX4 | 345.687406 | -0.3026944 | 0.1603222 | -1.8880378 | 0.05902087 | 0.53286516 |
| OSBPL11 | 131.827294 | -0.0783024 | 0.24645307 | -0.3177174 | 0.7506993 | 0.96109466 |
| SLC41A3 | 78.8058983 | -0.1830011 | 0.25390924 | -0.7207344 | 0.47107297 | 0.90123338 |
| ALDH1L1 | 140.01634 | 1.25131031 | 0.5208785 | 2.40230746 | 0.01629201 | 0.30598611 |
| ZXDC | 328.46019 | -0.1762348 | 0.2169353 | -0.8123841 | 0.41657127 | 0.88242985 |
| TXNRD3NB | 43.2365244 | 0.03182994 | 0.62648367 | 0.05080729 | 0.95947908 | 0.99609409 |
| TXNRD3 | 53.2336898 | -0.1661077 | 0.39976359 | -0.4155148 | 0.6777651 | 0.95560047 |
| CHCHD6 | 135.64595 | 0.01165131 | 0.24998893 | 0.04660731 | 0.96282621 | 0.99637506 |
| PLXNA1 | 40.4529607 | -0.1121791 | 0.33286142 | -0.3370143 | 0.73610609 | 0.96078945 |
| TPRA1 | 52.2699344 | 0.25910333 | 0.33391582 | 0.77595405 | 0.43777612 | 0.88825695 |
| ABTB1 | 46.1029953 | 0.5084234 | 0.3103165 | 1.63840272 | 0.10133771 | 0.62142728 |
| MGLL | 503.722367 | 0.21248483 | 0.30521581 | 0.69617897 | 4.86E-01 | 0.90695174 |
| SEC61A1 | 190.167253 | -0.1026708 | 0.30531021 | -0.3362836 | 0.73665698 | 0.96078945 |
| RUVBL1 | 129.075215 | 0.20795475 | 0.24232881 | 0.8581512 | 0.39080898 | 0.87004477 |
| EEFSEC | 203.781915 | -0.0616064 | 0.20897525 | -0.2948024 | 0.76814483 | 0.96345263 |
| GATA2 | 535.069708 | 1.05708429 | 0.53147089 | 1.98897872 | 0.04670355 | 0.48600281 |
| GATA2-AS1 | 118.489296 | -1.1999292 | 0.39945074 | -3.0039479 | 0.00266501 | 0.13421817 |
| RPN1 | 161.967536 | 0.20362895 | 0.301889 | 0.67451597 | 0.49998333 | 0.90839719 |
| RAB7A | 654.358429 | -0.6672849 | 0.2420942 | -2.7563025 | 0.00584589 | 0.19023403 |
| ACAD9 | 144.456747 | -0.135115 | 0.24558886 | -0.5501674 | 0.5822046 | 0.93402467 |
| CNBP | 384.397577 | 0.60221937 | 0.33204338 | 1.81367677 | 0.0697275 | 0.55992041 |
| COPG1 | 100.920077 | -0.401064 | 0.25167891 | -1.5935544 | 0.11103587 | 0.64219424 |
| HMCES | 103.435353 | 0.14488485 | 0.3144075 | 0.46081869 | 0.64492869 | 0.94949493 |
| H1FX | 236.762812 | 0.22889799 | 0.47768026 | 0.47918663 | 0.63180587 | 0.94654542 |
| MBD4 | 116.027721 | 0.01723674 | 0.20737759 | 0.08311767 | 0.93375797 | 0.99243654 |
| IFT122 | 174.664863 | 0.11147444 | 0.2457255 | 0.45365435 | 0.65007763 | 0.95061388 |
| TMCC1 | 1049.46768 | 0.03043114 | 0.27514279 | 0.11060127 | 0.91193254 | 0.98791121 |
| PIK3R4 | 127.316179 | -0.1913361 | 0.21363984 | -0.8956011 | 0.37046587 | 0.8613284 |
| ATP2C1 | 460.267042 | -0.0903523 | 0.24966102 | -0.3618999 | 0.71742684 | 0.95791799 |
| NEK11 | 461.158184 | -0.097519 | 0.30372286 | -0.3210788 | 0.7481507 | 0.96109466 |
| NUDT16 | 96.004867 | 0.38480433 | 0.30351215 | 1.26783831 | 0.2048557 | 0.77193931 |

| | | | | | | |
|---|---|---|---|---|---|---|
| MRPL3 | 147.662451 | 0.35240771 | 0.22333578 | 1.57792766 | 0.11458223 | 0.65008305 |
| DNAJC13 | 333.595932 | -0.5806542 | 0.26352631 | -2.2034011 | 0.02756649 | 0.40348721 |
| UBA5 | 293.844825 | -0.0619759 | 0.32933331 | -0.188186 | 0.85073088 | 0.97718353 |
| CDV3 | 326.621263 | -0.6463246 | 0.4830483 | -1.3380125 | 0.18089238 | 0.74315969 |
| TOPBP1 | 95.79904 | -0.3939961 | 0.19641043 | -2.0059833 | 0.04485802 | 0.48216625 |
| SRPRB | 56.7053482 | -0.0643856 | 0.33538719 | -0.1919738 | 0.8477627 | 0.97656375 |
| RAB6B | 65.4690525 | 0.4015147 | 0.37796855 | 1.06229659 | 0.28810106 | 0.8303903 |
| SLCO2A1 | 118.924191 | 0.66468021 | 0.79342687 | 0.83773343 | 0.40218044 | 0.87834687 |
| RYK | 457.548425 | 0.12057267 | 0.14935154 | 0.80730783 | 0.41948916 | 0.88278039 |
| AMOTL2 | 108.466716 | 1.64626605 | 0.61955909 | 2.65715744 | 0.00788026 | 0.21790279 |
| ANAPC13 | 95.6300981 | 0.39990216 | 0.29795847 | 1.3421406 | 0.17955041 | 0.74111395 |
| CEP63 | 163.682845 | 0.19915466 | 0.28995998 | 0.68683499 | 0.49218672 | 0.90769984 |
| EPHB1 | 147.708789 | 0.44221982 | 0.57210464 | 0.77297017 | 0.43954003 | 0.88917548 |
| PPP2R3A | 2003.57004 | 0.13893774 | 0.15002883 | 0.92607359 | 0.35440774 | 0.8557799 |
| MSL2 | 127.500159 | 0.37004969 | 0.27135478 | 1.36371171 | 0.17265832 | 0.72940007 |
| PCCB | 142.904329 | 0.17275764 | 0.28079421 | 0.61524646 | 0.53839197 | 0.92346119 |
| STAG1 | 1548.34725 | -0.3694087 | 0.22441404 | -1.6461036 | 0.09974244 | 0.61822465 |
| NCK1 | 111.187214 | 0.11190611 | 0.31188841 | 0.35880175 | 0.71974341 | 0.95796325 |
| DZIP1L | 87.2462189 | 1.0199007 | 0.38635466 | 2.63980432 | 0.00829539 | 0.22277119 |
| DBR1 | 49.9006776 | -0.5242691 | 0.24062362 | -2.1787933 | 0.02934703 | 0.40623517 |
| ARMC8 | 349.791477 | -0.2786528 | 0.17704896 | -1.5738741 | 0.11551657 | 0.65156815 |
| NME9 | 71.4514222 | 0.12730304 | 0.35565532 | 0.35793938 | 0.72038868 | 0.95796325 |
| MRAS | 148.315721 | -0.1277231 | 0.3180592 | -0.4015703 | 0.68800032 | 0.95597173 |
| CEP70 | 417.971519 | -0.1035187 | 0.26390512 | -0.3922571 | 0.69486825 | 0.95672887 |
| FAIM | 67.5581237 | 0.20000735 | 0.24120131 | 0.82921338 | 0.40698368 | 0.87948803 |
| PIK3CB | 564.927244 | 0.15448583 | 0.28079599 | 0.55017107 | 0.58220204 | 0.93402467 |
| MRPS22 | 42.8153175 | 0.08138665 | 0.28438285 | 0.28618692 | 0.77473496 | 0.96388926 |
| COPB2 | 240.38433 | -0.1167813 | 0.21660363 | -0.5391473 | 0.58978522 | 0.93483536 |
| AC097103.2 | 285.065758 | -0.3657962 | 0.3303923 | -1.107157 | 0.26822603 | 0.8165043 |
| NMNAT3 | 123.265987 | 0.12256214 | 0.32367785 | 0.37865472 | 0.70494428 | 0.95730303 |
| SLC25A36 | 622.086315 | -0.3182528 | 0.35340561 | -0.9005312 | 0.36783762 | 0.85902833 |
| PXYLP1 | 48.6888607 | 0.67917671 | 0.36545966 | 1.85841775 | 0.06310971 | 0.54798979 |
| ZBTB38 | 440.649123 | -0.7793421 | 0.29244341 | -2.6649332 | 0.00770036 | 0.21577483 |
| RASA2 | 602.460204 | -0.2992834 | 0.16147317 | -1.8534559 | 0.06381704 | 0.54850785 |
| RNF7 | 168.946187 | -0.0145686 | 0.47181123 | -0.0308781 | 0.97536677 | 0.99656026 |
| ATP1B3 | 1792.54498 | -0.889002 | 0.97245687 | -0.9141814 | 0.36062153 | 0.85608464 |
| TFDP2 | 988.146646 | 0.25298296 | 0.26350218 | 0.9600792 | 0.33701536 | 0.84636904 |
| GK5 | 406.180708 | -0.243207 | 0.24755131 | -0.9824507 | 0.32587786 | 0.84367588 |
| XRN1 | 713.130047 | -0.434685 | 0.35259684 | -1.2328102 | 0.2176466 | 0.78382693 |
| ATR | 393.620291 | -0.2780405 | 0.24088019 | -1.1542688 | 0.24839 | 0.80097144 |
| PLS1 | 215.207227 | 0.3862449 | 0.32006959 | 1.20675289 | 0.22752733 | 0.79042788 |
| TRPC1 | 261.973906 | 0.43528558 | 0.28378093 | 1.5338789 | 0.12505945 | 0.6663037 |
| PCOLCE2 | 36.8071554 | 0.27399204 | 0.48906683 | 0.56023435 | 0.5753196 | 0.93275827 |
| U2SURP | 409.541857 | -0.2569988 | 0.20431267 | -1.2578703 | 0.20843866 | 0.77434851 |
| C3orf58 | 70.9021565 | 0.4403515 | 0.2892586 | 1.5223454 | 0.12792256 | 0.67070864 |
| AC107021.1 | 60.9088826 | -0.4773054 | 0.70642167 | -0.6756665 | 0.49925242 | 0.90839719 |
| PLOD2 | 485.896991 | -0.3939359 | 0.42749167 | -0.9215054 | 0.35678662 | 0.8557799 |
| PLSCR4 | 113.131988 | 0.38726985 | 0.25863946 | 1.49733475 | 0.13430618 | 0.67832119 |
| PLSCR2 | 178.87523 | 0.28279671 | 0.30766725 | 0.91916415 | 0.35800972 | 0.8557799 |
| PLSCR1 | 999.639496 | -0.3399982 | 0.30113287 | -1.1290639 | 0.25887089 | 0.80709747 |
| GYG1 | 45.8184346 | 0.2412364 | 0.36468696 | 0.66148897 | 0.50829879 | 0.91250907 |
| HLTF | 119.146979 | 0.70262284 | 0.25501675 | 2.75520272 | 0.00586558 | 0.19023403 |
| HPS3 | 109.442863 | -0.0338759 | 0.25984477 | -0.1303696 | 0.89627402 | 0.98597539 |
| WWTR1 | 1892.07107 | -0.8458359 | 0.46947619 | -1.8016588 | 0.0715991 | 0.56343001 |

| | | | | | | |
|---|---|---|---|---|---|---|
| COMMD2 | 96.859152 | -0.0111935 | 0.31345626 | -0.0357098 | 0.97151373 | 0.99656026 |
| ANKUB1 | 152.628288 | 0.1528309 | 0.45486076 | 0.33599491 | 0.73687471 | 0.96078945 |
| RNF13 | 323.660443 | -0.084812 | 0.18903653 | -0.4486541 | 0.65368118 | 0.95100227 |
| TSC22D2 | 633.188532 | -0.0513315 | 0.31992329 | -0.1604495 | 0.87252702 | 0.98238968 |
| SERP1 | 446.050087 | 0.38789582 | 0.38403674 | 1.01004872 | 0.31247195 | 0.84279957 |
| EIF2A | 147.61664 | -0.3327586 | 0.2874473 | -1.1576333 | 0.24701373 | 0.80072691 |
| SELENOT | 223.160178 | 0.42651652 | 0.2023514 | 2.10780123 | 0.03504819 | 0.43353051 |
| SIAH2 | 79.2729491 | -0.3461371 | 0.514167 | -0.6731997 | 0.50082026 | 0.90857526 |
| MBNL1 | 1345.0721 | 0.06982323 | 0.22481301 | 0.3105836 | 0.7561172 | 0.96120879 |
| ARHGEF26 | 51.9638214 | 0.86664593 | 0.52857143 | 1.6396004 | 0.10108828 | 0.62105083 |
| DHX36 | 406.911818 | -0.2559454 | 0.18101569 | -1.4139403 | 0.15737945 | 0.71160548 |
| PLCH1 | 257.164874 | -0.3840733 | 0.57284627 | -0.6704648 | 0.50256151 | 0.90962743 |
| GMPS | 126.102907 | 0.05173251 | 0.18513668 | 0.27942875 | 0.77991581 | 0.96513523 |
| SSR3 | 142.902723 | 0.16354539 | 0.3223141 | 0.50740999 | 0.61186718 | 0.94003018 |
| TIPARP | 112.739891 | -1.021346 | 0.97024313 | -1.0526702 | 0.29249217 | 0.83336457 |
| LINC00886 | 132.460621 | 1.01097467 | 0.4161756 | 2.42920218 | 0.01513209 | 0.29641929 |
| LEKR1 | 124.247713 | 0.26308095 | 0.36527958 | 0.72021806 | 0.47139075 | 0.90123338 |
| CCNL1 | 527.480532 | -0.7174493 | 0.4764144 | -1.5059353 | 0.13208379 | 0.67466347 |
| VEPH1 | 682.341096 | 0.71705186 | 0.57820483 | 1.24013467 | 0.21492559 | 0.77890374 |
| PQLC2L | 134.239692 | -0.1264259 | 0.313317 | -0.403508 | 0.68657459 | 0.9558749 |
| RSRC1 | 511.074587 | -0.2341611 | 0.20577326 | -1.1379567 | 0.25513855 | 0.80489222 |
| MLF1 | 115.665537 | 0.66460679 | 0.28177766 | 2.35862129 | 0.01834296 | 0.32472005 |
| GFM1 | 145.987817 | -0.1907987 | 0.29757642 | -0.6411754 | 0.52140873 | 0.91787515 |
| RARRES1 | 35.5918188 | -0.0286597 | 0.59832975 | -0.0478995 | 0.96179634 | 0.99635733 |
| MFSD1 | 71.9928281 | -0.1182354 | 0.28453887 | -0.4155335 | 0.67775139 | 0.95560047 |
| IQCJ-SCHIP1 | 287.735687 | -0.5353709 | 0.93799042 | -0.5707637 | 0.56815982 | 0.93122667 |
| SMC4 | 143.395036 | -0.3416477 | 0.35939884 | -0.9506088 | 0.341803 | 0.85092302 |
| KPNA4 | 308.732787 | -0.181448 | 0.26610101 | -0.6818765 | 0.49531704 | 0.90839719 |
| PPM1L | 77.6731845 | 0.19979936 | 0.35004336 | 0.57078458 | 0.56814567 | 0.93122667 |
| NMD3 | 194.466915 | 0.24118443 | 0.31803005 | 0.75836994 | 0.44822955 | 0.89297728 |
| LINC01322 | 50.3940228 | -0.5176727 | 0.34829439 | -1.486308 | 0.13719767 | 0.68129775 |
| PDCD10 | 229.970312 | 0.05120988 | 0.22921682 | 0.22341238 | 0.82321457 | 0.97018919 |
| GOLIM4 | 239.233905 | 0.29855733 | 0.30927673 | 0.96534041 | 0.33437434 | 0.84480724 |
| MECOM | 23218.6649 | -0.5624509 | 0.33872196 | -1.6605091 | 0.09681209 | 0.61564967 |
| AC074033.1 | 106.595785 | -0.1054079 | 0.63192199 | -0.1668052 | 0.86752332 | 0.98135671 |
| MYNN | 60.7456558 | 0.04139127 | 0.29870187 | 0.13857053 | 0.88978954 | 0.98581358 |
| LRRC34 | 45.4982107 | -0.0305999 | 0.38722779 | -0.0790231 | 0.93701426 | 0.99243654 |
| SEC62 | 695.3026 | 0.17211663 | 0.29063175 | 0.5922155 | 0.55370629 | 0.92872652 |
| GPR160 | 498.268591 | 0.20148523 | 0.27047887 | 0.74492042 | 0.45631983 | 0.89626521 |
| PHC3 | 748.05664 | -0.0486219 | 0.2327797 | -0.2088754 | 0.83454553 | 0.97232209 |
| PRKCI | 588.550795 | -0.5280179 | 0.3778563 | -1.3974041 | 0.1622921 | 0.71482182 |
| SKIL | 317.960408 | -0.0892843 | 0.45589841 | -0.1958425 | 0.84473347 | 0.97573093 |
| RPL22L1 | 40.9793326 | 0.03045681 | 0.42825877 | 0.07111777 | 0.94330402 | 0.99265578 |
| TNIK | 203.876735 | 0.69348183 | 0.69129951 | 1.00315684 | 0.31578519 | 0.84285425 |
| PLD1 | 115.734011 | -0.0971672 | 0.40493776 | -0.2399559 | 0.81036447 | 0.9680627 |
| FNDC3B | 2708.86165 | -0.1668641 | 0.28331345 | -0.5889735 | 0.55587906 | 0.92872652 |
| TNFSF10 | 257.449112 | -1.0425349 | 0.62607144 | -1.6652011 | 0.09587264 | 0.61367241 |
| NCEH1 | 292.686737 | 0.0483802 | 0.51054931 | 0.09476107 | 0.92450461 | 0.99051393 |
| ECT2 | 72.0702911 | 0.13640831 | 0.37875078 | 0.36015322 | 0.71873255 | 0.95796325 |
| NLGN1 | 2841.82305 | -0.8998335 | 0.73978008 | -1.2163528 | 0.22385057 | 0.78960314 |
| NAALADL2 | 4050.16914 | -0.7905479 | 0.19836165 | -3.9853868 | 6.74E-05 | 0.01470918 |
| TBL1XR1 | 2153.28742 | -0.4173201 | 0.41649855 | -1.0019726 | 0.31635685 | 0.84285425 |
| LINC00578 | 61.2976523 | -0.7751622 | 0.47822093 | -1.6209291 | 0.10503284 | 0.6326116 |
| ZMAT3 | 139.80389 | 0.18065543 | 0.39530366 | 0.45700419 | 0.64766804 | 0.95061388 |

| | | | | | | |
|---|---|---|---|---|---|---|
| PIK3CA | 363.140472 | -0.2731622 | 0.16907761 | -1.6156023 | 0.10618033 | 0.63461115 |
| KCNMB3 | 55.6554431 | -0.1679203 | 0.34097531 | -0.4924704 | 0.62238682 | 0.94418789 |
| ZNF639 | 64.0285892 | 0.12735949 | 0.27813948 | 0.45789791 | 0.64702579 | 0.95061388 |
| MFN1 | 407.112891 | -0.0991831 | 0.21933107 | -0.4522073 | 0.65111962 | 0.95061388 |
| GNB4 | 51.6246015 | 0.01659225 | 0.37443174 | 0.04431315 | 0.96465479 | 0.99637506 |
| ACTL6A | 73.5939835 | -0.4017486 | 0.32718732 | -1.2278855 | 0.21948997 | 0.78630025 |
| MRPL47 | 77.680458 | -0.0306905 | 0.36042349 | -0.0851513 | 0.93214108 | 0.99243654 |
| NDUFB5 | 211.766083 | 0.73086476 | 0.32821855 | 2.22676251 | 0.02596315 | 0.3903785 |
| TTC14 | 335.113733 | -0.1357004 | 0.16595654 | -0.8176866 | 0.41353614 | 0.88211689 |
| FXR1 | 400.553686 | -0.0153657 | 0.20013895 | -0.0767749 | 0.9388026 | 0.99243654 |
| DNAJC19 | 78.8024999 | -0.1152496 | 0.29927255 | -0.3850992 | 0.70016394 | 0.95672887 |
| ATP11B | 638.322181 | -0.1354103 | 0.32949513 | -0.4109629 | 0.68109971 | 0.95574492 |
| DCUN1D1 | 653.776073 | -0.3163776 | 0.52281401 | -0.6051436 | 0.54508358 | 0.92579581 |
| MCCC1 | 602.499551 | -0.3687603 | 0.35315707 | -1.044182 | 0.29640121 | 0.83502276 |
| MCF2L2 | 55.547504 | -0.101224 | 0.62467469 | -0.1620428 | 0.87127215 | 0.98208511 |
| B3GNT5 | 51.0057272 | 0.04093797 | 0.54298701 | 0.07539402 | 0.93990122 | 0.99243654 |
| KLHL24 | 521.478054 | -0.2084036 | 0.28599332 | -0.7287008 | 0.4661847 | 0.89883965 |
| YEATS2 | 325.234883 | -0.7132147 | 0.36736678 | -1.941424 | 0.05220687 | 0.5070695 |
| ABCC5 | 583.851903 | -0.1769536 | 0.27050054 | -0.6541709 | 0.51300168 | 0.91495194 |
| EIF2B5 | 66.4421819 | -0.0926646 | 0.26704265 | -0.347003 | 0.7285891 | 0.95985088 |
| DVL3 | 113.522213 | -0.0366848 | 0.23414832 | -0.1566732 | 0.87550241 | 0.98267808 |
| AP2M1 | 283.693755 | 0.1420261 | 0.36594183 | 0.38811115 | 0.69793378 | 0.95672887 |
| ABCF3 | 33.9063009 | -0.1264954 | 0.39148785 | -0.3231144 | 0.74660857 | 0.96109466 |
| PSMD2 | 121.814356 | -0.0861512 | 0.32195137 | -0.2675906 | 0.78901446 | 0.96515649 |
| EIF4G1 | 266.33382 | -0.0734898 | 0.26189222 | -0.2806109 | 0.77900888 | 0.96512745 |
| FAM131A | 44.3967777 | -0.2765718 | 0.48346666 | -0.5720597 | 0.56728154 | 0.93122667 |
| POLR2H | 151.55007 | 0.14377486 | 0.27330717 | 0.52605593 | 0.59884934 | 0.93728325 |
| MAGEF1 | 82.9656107 | 0.47944914 | 0.32322731 | 1.48331877 | 0.13798974 | 0.68146162 |
| VPS8 | 399.172021 | -0.5407203 | 0.26563796 | -2.0355537 | 0.0417952 | 0.46746387 |
| EHHADH | 164.326176 | 0.4380009 | 0.29985519 | 1.46070807 | 0.14409557 | 0.69093999 |
| MAP3K13 | 465.753609 | 0.27999869 | 0.20005294 | 1.39962301 | 0.16162624 | 0.71416435 |
| TMEM41A | 105.887283 | -0.4079363 | 0.22966235 | -1.7762436 | 0.07569277 | 0.57072562 |
| LIPH | 236.479356 | -0.0802227 | 0.37679295 | -0.2129094 | 0.83139765 | 0.97160502 |
| SENP2 | 205.040112 | -0.3000533 | 0.23121869 | -1.2977034 | 0.19438929 | 0.76162393 |
| IGF2BP2 | 467.407036 | 0.08147956 | 0.2727361 | 0.29874871 | 0.76513179 | 0.96285313 |
| TRA2B | 253.888703 | 0.10501818 | 0.26235249 | 0.4002942 | 0.68893984 | 0.95606293 |
| AC068631.2 | 52.5041984 | 0.28180968 | 0.39535146 | 0.712808 | 0.47596457 | 0.90242252 |
| EIF4A2 | 379.408106 | 0.35547086 | 0.32586759 | 1.09084448 | 0.27534132 | 0.82046546 |
| ST6GAL1 | 3796.77654 | -0.524334 | 0.52632329 | -0.9962203 | 0.31914311 | 0.84285425 |
| BCL6 | 223.719663 | -0.87217 | 0.41590182 | -2.0970575 | 0.03598849 | 0.4362094 |
| LPP | 3264.74055 | -0.4740243 | 0.20928966 | -2.2649197 | 0.02351761 | 0.36567438 |
| P3H2 | 699.366388 | 0.84799476 | 0.24716203 | 3.43092652 | 0.00060152 | 0.05792155 |
| CCDC50 | 282.696068 | 0.0551139 | 0.1517643 | 0.36315459 | 0.71648941 | 0.95791799 |
| FGF12 | 532.005682 | 0.77207832 | 0.88283362 | 0.87454567 | 0.38182116 | 0.86593382 |
| MB21D2 | 91.8596843 | 0.62472186 | 0.44636582 | 1.39957369 | 0.16164102 | 0.71416435 |
| OPA1 | 380.671823 | -0.1049483 | 0.14173448 | -0.740457 | 0.45902272 | 0.89753337 |
| HES1 | 787.482102 | -0.7148508 | 0.53516888 | -1.3357481 | 0.18163161 | 0.74315969 |
| ATP13A3 | 696.942929 | -0.4452132 | 0.42201873 | -1.0549607 | 0.29144332 | 0.8320147 |
| LINC00884 | 49.8988747 | 0.01925286 | 0.32671863 | 0.05892795 | 0.9530095 | 0.99507219 |
| LSG1 | 87.0287109 | -0.2617361 | 0.29066661 | -0.9004684 | 0.36787103 | 0.85902833 |
| XXYLT1 | 89.1066205 | -0.0406146 | 0.30044785 | -0.1351803 | 0.89246935 | 0.98597539 |
| ACAP2 | 578.418628 | -0.2231334 | 0.21515226 | -1.0370954 | 0.29969142 | 0.83618229 |
| PPP1R2 | 179.076899 | -0.1502131 | 0.20093042 | -0.7475877 | 0.45470891 | 0.89626521 |
| MUC20-OT1 | 1528.37035 | -0.1663967 | 0.4597443 | -0.3619332 | 0.71740196 | 0.95791799 |

| | | | | | | |
|---|---|---|---|---|---|---|
| LINC01983 | 77.2255987 | 0.9867466 | 0.54164931 | 1.82174442 | 0.06849378 | 0.55917298 |
| TNK2 | 44.7853307 | 0.70281939 | 0.26994914 | 2.60352524 | 0.00922705 | 0.23300339 |
| TFRC | 257.798086 | -1.138946 | 0.50887165 | -2.2381793 | 0.02520936 | 0.38344578 |
| PCYT1A | 108.300455 | -0.109023 | 0.22196077 | -0.4911814 | 0.62329817 | 0.94456546 |
| UBXN7 | 221.507514 | 0.2015118 | 0.17306767 | 1.16435263 | 0.24428114 | 0.80072691 |
| RNF168 | 185.742924 | -0.2149459 | 0.22950487 | -0.9365635 | 0.34898312 | 0.8535343 |
| PIGX | 81.1018111 | 0.14990033 | 0.21738915 | 0.68954835 | 0.49047826 | 0.90730741 |
| PAK2 | 527.016892 | 0.13738856 | 0.17656127 | 0.77813534 | 0.43648923 | 0.88785137 |
| SENP5 | 385.289145 | -0.2523064 | 0.20283992 | -1.2438694 | 0.21354764 | 0.77869841 |
| NCBP2 | 104.392155 | 0.40911975 | 0.30140645 | 1.3573689 | 0.17466402 | 0.73219159 |
| NCBP2-AS2 | 66.3787952 | 0.2643113 | 0.51132131 | 0.51691822 | 0.60521325 | 0.93909311 |
| DLG1 | 1313.92372 | -0.2638208 | 0.22647619 | -1.164894 | 0.24406189 | 0.80072691 |
| BDH1 | 113.579409 | 0.03479225 | 0.39635949 | 0.08777953 | 0.93005191 | 0.99216065 |
| RUBCN | 108.948713 | -0.3811899 | 0.29310996 | -1.3005014 | 0.19342918 | 0.76043322 |
| FYTTD1 | 154.73654 | 0.20536734 | 0.37751418 | 0.54399902 | 0.58644215 | 0.93483536 |
| LRCH3 | 424.792329 | -0.2787615 | 0.19753731 | -1.4111841 | 0.15819034 | 0.71264724 |
| IQCG | 97.7196401 | -0.2757496 | 0.28362027 | -0.9722493 | 0.33092655 | 0.84442294 |
| RPL35A | 581.553756 | 0.1386942 | 0.42647721 | 0.32520894 | 0.74502294 | 0.96109466 |
| LMLN | 138.382746 | 0.09083853 | 0.27948577 | 0.32502023 | 0.74516576 | 0.96109466 |
| ZNF595 | 84.4382769 | 0.15585948 | 0.25150943 | 0.61969636 | 0.53545771 | 0.92303275 |
| ZNF718 | 261.274616 | -0.1004487 | 0.25578276 | -0.392711 | 0.69453295 | 0.95672887 |
| ZNF141 | 417.97102 | -0.1186949 | 0.72606338 | -0.1634773 | 0.87014261 | 0.98202612 |
| AC092574.2 | 39.189185 | 0.7137863 | 0.5725402 | 1.24670074 | 0.21250725 | 0.77709975 |
| ZNF721 | 562.80835 | 0.32885855 | 0.26007708 | 1.2644657 | 0.20606293 | 0.77193931 |
| PIGG | 123.554757 | -0.2739747 | 0.19268165 | -1.4219037 | 0.15505422 | 0.70897393 |
| ATP5ME | 141.071402 | -0.002049 | 0.2807161 | -0.007299 | 0.99417628 | 0.99919666 |
| PCGF3 | 191.117279 | -0.3639912 | 0.26206353 | -1.3889426 | 0.16485022 | 0.71715698 |
| AC139887.2 | 277.077055 | 0.30524175 | 0.34300035 | 0.88991672 | 0.3735106 | 0.86201082 |
| GAK | 250.996138 | -0.4644739 | 0.25558136 | -1.817323 | 0.06916766 | 0.55917298 |
| TMEM175 | 37.6877389 | 0.26168248 | 0.32671936 | 0.80093962 | 0.4231666 | 0.88371155 |
| IDUA | 39.2068422 | 0.399382 | 0.40556542 | 0.98475359 | 0.32474512 | 0.84285425 |
| FGFRL1 | 43.6319602 | -0.3403701 | 0.26135832 | -1.3023119 | 0.19280977 | 0.7597012 |
| RNF212 | 46.3197518 | -0.0274183 | 0.52403102 | -0.052322 | 0.95827215 | 0.99579587 |
| AC092535.3 | 161.598187 | 0.08025621 | 0.44061428 | 0.18214619 | 0.855468 | 0.97718353 |
| CTBP1-AS | 59.0279201 | -0.3269386 | 0.46520908 | -0.7027777 | 0.48219427 | 0.90595437 |
| CTBP1 | 154.213986 | -0.2462686 | 0.22828421 | -1.0787806 | 0.28068554 | 0.82628479 |
| CTBP1-DT | 35.2714685 | 0.6101945 | 0.26936142 | 2.26533736 | 0.02349198 | 0.36567438 |
| MAEA | 200.280789 | 0.06020108 | 0.23570494 | 0.25540864 | 0.7984075 | 0.9653598 |
| UVSSA | 142.897941 | -0.5325376 | 0.31396644 | -1.6961608 | 0.08985544 | 0.60739841 |
| SLBP | 223.685555 | -0.0105687 | 0.31334804 | -0.0337282 | 0.97309385 | 0.99656026 |
| TMEM129 | 84.0599311 | 0.22369968 | 0.33183027 | 0.67413887 | 0.50022303 | 0.90839719 |
| FGFR3 | 65.6691053 | 2.50489576 | 0.46204908 | 5.42127639 | 5.92E-08 | 5.64E-05 |
| LETM1 | 93.7125048 | 0.04981005 | 0.32200616 | 0.15468665 | 0.87706835 | 0.98277085 |
| NSD2 | 267.614149 | -0.174658 | 0.22827206 | -0.7651308 | 0.44419366 | 0.89096469 |
| NELFA | 237.904608 | -0.0546119 | 0.28633583 | -0.1907268 | 0.84873966 | 0.97656375 |
| C4orf48 | 63.8104006 | 0.05737877 | 0.57093881 | 0.10049899 | 0.91994818 | 0.99014946 |
| NAT8L | 34.5694601 | 1.01216442 | 0.5318434 | 1.9031249 | 0.05702425 | 0.52514423 |
| POLN | 51.0961623 | -0.3288699 | 0.48728748 | -0.6748991 | 0.49973988 | 0.90839719 |
| MXD4 | 284.459425 | -0.1642778 | 0.3026367 | -0.5428216 | 0.58725261 | 0.93483536 |
| ZFYVE28 | 176.217693 | 0.09441819 | 0.32935128 | 0.28667928 | 0.77435791 | 0.96388926 |
| RNF4 | 190.99752 | -0.3167459 | 0.19268171 | -1.6438813 | 0.10020073 | 0.61891077 |
| FAM193A | 372.960747 | -0.3754769 | 0.26466836 | -1.4186693 | 0.15599545 | 0.71066638 |
| TNIP2 | 85.9068314 | -0.1721907 | 0.36668198 | -0.4695914 | 0.63864699 | 0.94887688 |
| SH3BP2 | 279.118754 | -0.5853765 | 0.25176163 | -2.3251218 | 2.01E-02 | 0.33933607 |

| | | | | | | |
|---|---|---|---|---|---|---|
| ADD1 | 572.700364 | -0.0591863 | 0.16502023 | -0.3586609 | 0.7198488 | 0.95796325 |
| MFSD10 | 147.537602 | -0.2520778 | 0.35194753 | -0.7162368 | 0.47384516 | 0.90242252 |
| NOP14-AS1 | 35.3142208 | 0.02579667 | 0.35820877 | 0.07201574 | 0.94258938 | 0.99265578 |
| NOP14 | 52.764545 | -0.2377324 | 0.27266781 | -0.8718756 | 0.38327622 | 0.86638702 |
| GRK4 | 65.4881234 | 0.21282436 | 0.35230898 | 0.60408441 | 0.54578752 | 0.92597739 |
| HTT | 398.1139 | -0.2040687 | 0.1952794 | -1.0450087 | 0.29601893 | 0.83444322 |
| RGS12 | 125.880037 | -0.2014801 | 0.35976369 | -0.5600346 | 0.57545585 | 0.93283484 |
| LRPAP1 | 386.149957 | 1.13285807 | 0.45826208 | 2.47207462 | 0.01343315 | 0.28170334 |
| ADRA2C | 35.5710033 | 0.15677997 | 0.75033353 | 0.20894704 | 0.83448959 | 0.97232209 |
| TMEM128 | 62.0561209 | -0.0729888 | 0.26180397 | -0.2787917 | 0.78040471 | 0.96513523 |
| STX18 | 431.084186 | 0.23566092 | 0.2900074 | 0.81260312 | 0.41644564 | 0.88242985 |
| STX18-AS1 | 251.62187 | 0.08385337 | 0.35570098 | 0.23574118 | 0.81363351 | 0.96819339 |
| EVC2 | 138.682474 | -0.0191436 | 0.28535939 | -0.0670859 | 0.9465133 | 0.99307065 |
| EVC | 171.615528 | 0.13385693 | 0.42776612 | 0.31292083 | 0.75434081 | 0.96120879 |
| CRMP1 | 54.5656984 | 0.29488672 | 0.62920001 | 0.46866929 | 0.63930604 | 0.94935885 |
| WFS1 | 54.6943127 | 0.84626383 | 0.38489584 | 2.1986827 | 0.02790049 | 0.40459506 |
| MAN2B2 | 94.2693531 | 0.10237818 | 0.36411496 | 0.28116993 | 0.77858007 | 0.96505253 |
| MRFAP1 | 291.336835 | 0.59427578 | 0.43039203 | 1.38077782 | 0.16734728 | 0.72232271 |
| LINC02482 | 32.1810845 | -0.1511286 | 0.33928464 | -0.4454331 | 0.6560068 | 0.95100227 |
| AC093323.1 | 31.1045093 | 0.56225247 | 0.36352987 | 1.54664725 | 0.12194833 | 0.66181735 |
| MRFAP1L1 | 69.4583647 | 0.09057156 | 0.30413905 | 0.29779654 | 0.76585846 | 0.96285313 |
| BLOC1S4 | 54.9223479 | 0.41481911 | 0.35582872 | 1.16578309 | 0.24370216 | 0.80072691 |
| KIAA0232 | 474.592747 | -0.0474945 | 0.17605884 | -0.2697647 | 0.78734126 | 0.96513523 |
| TBC1D14 | 205.961508 | -0.2635507 | 0.21784765 | -1.2097937 | 0.22635805 | 0.78982769 |
| TADA2B | 54.2730883 | 0.32391428 | 0.28491494 | 1.13688065 | 0.25558818 | 0.80525591 |
| GRPEL1 | 64.4809411 | 0.28109734 | 0.39052794 | 0.71978804 | 0.47165551 | 0.9015046 |
| SORCS2 | 695.497825 | -0.2717203 | 0.4092694 | -0.6639155 | 0.50674443 | 0.91163608 |
| AFAP1 | 258.245794 | -0.8890943 | 0.55458423 | -1.6031727 | 0.10889654 | 0.63791825 |
| SH3TC1 | 302.922198 | -0.6262825 | 0.45178038 | -1.3862544 | 0.16566924 | 0.71952493 |
| ACOX3 | 95.8474457 | -0.0781876 | 0.23504529 | -0.3326489 | 0.73939934 | 0.96078945 |
| TRMT44 | 76.6782427 | -0.2225252 | 0.2710791 | -0.8208865 | 0.41171091 | 0.88086809 |
| WDR1 | 335.163854 | 0.14120152 | 0.27697587 | 0.5097972 | 0.61019355 | 0.94003018 |
| ZNF518B | 51.8921747 | -0.0769479 | 0.26926603 | -0.2857692 | 0.77505492 | 0.96388926 |
| HS3ST1 | 101.772771 | -0.083273 | 0.74754502 | -0.1113953 | 0.9113029 | 0.98788932 |
| AC005699.1 | 92.8786962 | -1.5790187 | 0.54183104 | -2.9142271 | 0.0035657 | 0.14654346 |
| RAB28 | 464.212268 | -0.2043616 | 0.19941667 | -1.0247971 | 0.30545891 | 0.83709431 |
| BOD1L1 | 344.220821 | -0.3244039 | 0.19314528 | -1.6795847 | 0.09303814 | 0.61317397 |
| LINC01182 | 55.8734586 | -1.8719838 | 0.4592066 | -4.0765611 | 4.57E-05 | 0.01041317 |
| CPEB2 | 377.306177 | -0.4841912 | 0.32109651 | -1.5079305 | 0.13157232 | 0.67390608 |
| AC098829.1 | 148.939748 | -0.5932371 | 0.63795591 | -0.929903 | 0.35242133 | 0.85536253 |
| CC2D2A | 180.233021 | -0.4073398 | 0.38598844 | -1.0553161 | 0.2912808 | 0.8320147 |
| FBXL5 | 381.511418 | -0.2555168 | 0.16292905 | -1.5682702 | 0.11681808 | 0.65289383 |
| FAM200B | 125.688543 | -0.1003244 | 0.26375492 | -0.3803699 | 0.70367084 | 0.95672887 |
| PROM1 | 492.301186 | 0.98033858 | 0.57831776 | 1.69515557 | 0.09004591 | 0.60739841 |
| TAPT1 | 258.201567 | 0.2291613 | 0.23223346 | 0.98677124 | 0.3237548 | 0.84285425 |
| TAPT1-AS1 | 449.734367 | 0.85837983 | 0.41061377 | 2.0904799 | 0.03657471 | 0.43729328 |
| AC097515.1 | 52.6267762 | 0.47209951 | 0.84592739 | 0.55808515 | 0.57678624 | 0.93319686 |
| QDPR | 74.9284889 | 0.35189327 | 0.31864803 | 1.10433215 | 0.26944908 | 0.81817548 |
| LAP3 | 105.625351 | -0.3491509 | 0.72498357 | -0.4815983 | 0.6300913 | 0.94596555 |
| MED28 | 167.492336 | -0.0923719 | 0.16272448 | -0.5676581 | 0.57026712 | 0.93125087 |
| DCAF16 | 149.736628 | -0.1444448 | 0.21191191 | -0.6816266 | 0.49547506 | 0.90839719 |
| LCORL | 732.338182 | 0.11378348 | 0.28883816 | 0.39393507 | 0.69362898 | 0.95672887 |
| AC024230.1 | 109.54322 | 0.49521252 | 0.44892217 | 1.10311441 | 0.26997748 | 0.81817548 |
| SLIT2 | 242.554833 | -0.2099198 | 0.62376846 | -0.3365348 | 0.73646762 | 0.96078945 |

| | | | | | | |
|---|---|---|---|---|---|---|
| PACRGL | 175.303546 | -0.2778096 | 0.26496852 | -1.0484628 | 0.29442545 | 0.8337535 |
| KCNIP4 | 12670.1722 | 0.3138443 | 0.66535392 | 0.47169528 | 0.63714431 | 0.94821838 |
| ADGRA3 | 257.734824 | 0.41633586 | 0.41981247 | 0.99171866 | 0.32133479 | 0.84285425 |
| PPARGC1A | 1609.08413 | -0.462871 | 0.37877423 | -1.2220235 | 0.22169872 | 0.78752765 |
| DHX15 | 414.328038 | -0.4210216 | 0.22557712 | -1.8664197 | 0.06198266 | 0.54480406 |
| CCDC149 | 70.5042669 | -0.5625701 | 0.37758936 | -1.4898992 | 0.13625074 | 0.67998335 |
| SEPSECS | 90.4122689 | -0.236579 | 0.28494536 | -0.830261 | 0.40639121 | 0.87948803 |
| ZCCHC4 | 54.9854381 | -0.1926072 | 0.32564071 | -0.5914716 | 0.55420446 | 0.92872652 |
| ANAPC4 | 145.637437 | -0.1362067 | 0.20483055 | -0.6649725 | 0.50606807 | 0.91163608 |
| SMIM20 | 70.0156908 | 0.40855787 | 0.26549113 | 1.5388758 | 0.12383462 | 0.66519057 |
| RBPJ | 499.248028 | -0.1666401 | 0.25562411 | -0.6518953 | 0.51446873 | 0.91544604 |
| TBC1D19 | 389.069143 | 0.07640933 | 0.24669599 | 0.30973074 | 0.75676572 | 0.96133693 |
| STIM2 | 1244.48145 | 0.71442846 | 0.51796806 | 1.37929057 | 0.16780518 | 0.72370301 |
| PCDH7 | 969.123822 | 0.8932598 | 0.59874566 | 1.49188522 | 0.13572924 | 0.67994379 |
| ARAP2 | 355.230538 | -0.000946 | 0.35941251 | -0.0026321 | 0.99789987 | 0.99967893 |
| C4orf19 | 1632.27974 | 0.25197943 | 0.3315142 | 0.76008637 | 0.44720296 | 0.89187439 |
| RELL1 | 147.517649 | -0.0678675 | 0.39713614 | -0.1708923 | 0.86430846 | 0.97942001 |
| PGM2 | 84.2001998 | -0.6749287 | 0.35897467 | -1.8801568 | 0.06008671 | 0.53867303 |
| TBC1D1 | 1360.20325 | 0.12106617 | 0.23092462 | 0.52426707 | 0.6000928 | 0.93728325 |
| KLF3 | 323.084495 | -0.5713219 | 0.44904758 | -1.2722971 | 0.20326759 | 0.7702733 |
| FAM114A1 | 73.1830723 | -0.6490313 | 0.37304878 | -1.7398028 | 0.08189365 | 0.59202053 |
| KLHL5 | 320.641601 | 0.71909267 | 0.35391016 | 2.03185087 | 0.04216876 | 0.4691386 |
| WDR19 | 223.031034 | -0.4506971 | 0.20553652 | -2.1927834 | 0.02832299 | 0.40459506 |
| RFC1 | 253.392649 | -0.1373162 | 0.24130024 | -0.5690678 | 0.56931012 | 0.93122667 |
| RPL9 | 56.6943847 | -0.0580504 | 0.2724166 | -0.2130942 | 0.83125349 | 0.97160502 |
| LIAS | 78.1125039 | 0.30162337 | 0.3342619 | 0.90235641 | 0.36686757 | 0.85902833 |
| UGDH | 127.872348 | 0.24434714 | 0.44998502 | 0.54301171 | 0.58712174 | 0.93483536 |
| UGDH-AS1 | 158.559653 | 0.35989323 | 0.28570022 | 1.25968835 | 0.20778181 | 0.77327889 |
| SMIM14 | 600.345693 | 0.18650019 | 0.19695167 | 0.94693378 | 0.34367252 | 0.85126545 |
| UBE2K | 1018.39714 | -0.1768067 | 0.25788328 | -0.6856074 | 0.4929607 | 0.90804281 |
| PDS5A | 740.904687 | -0.2598734 | 0.2053474 | -1.2655305 | 0.20568123 | 0.77193931 |
| N4BP2 | 147.859391 | 0.00395374 | 0.30042001 | 0.01316072 | 0.98949957 | 0.99835905 |
| RBM47 | 2968.33198 | -8.05E-01 | 0.41287197 | -1.95E+00 | 0.05124929 | 0.50101917 |
| NSUN7 | 228.401138 | 0.04457679 | 0.2722175 | 0.16375433 | 0.86992454 | 0.98202612 |
| APBB2 | 1059.97438 | 0.10106454 | 0.19622781 | 0.51503681 | 6.07E-01 | 0.93919596 |
| UCHL1 | 280.37801 | 0.30674339 | 0.44059882 | 0.69619657 | 0.48630573 | 0.90695174 |
| LIMCH1 | 2683.32324 | 1.13190338 | 0.42190095 | 2.68286521 | 0.00729944 | 0.2101379 |
| TMEM33 | 169.629901 | -0.1264705 | 0.28754844 | -0.4398232 | 0.66006514 | 0.95127957 |
| SLC30A9 | 608.531156 | -0.025291 | 0.15168924 | -0.1667294 | 0.86758301 | 0.98135671 |
| ATP8A1 | 251.975413 | -0.1520345 | 0.27032947 | -0.5624046 | 0.57384036 | 0.93270589 |
| KCTD8 | 1255.41717 | 0.26005078 | 0.4526507 | 0.57450653 | 0.56562508 | 0.93114846 |
| GUF1 | 79.0353377 | -0.0725229 | 0.24715896 | -0.2934263 | 0.76919631 | 0.96345263 |
| GNPDA2 | 77.5392657 | 0.01070649 | 0.25770238 | 0.04154594 | 0.96686067 | 0.99637506 |
| GABRA2 | 313.043536 | 0.48323696 | 0.56558292 | 0.85440515 | 0.39288054 | 0.87177389 |
| COMMD8 | 55.042733 | 0.53432509 | 0.45852948 | 1.16530149 | 0.24389698 | 0.80072691 |
| ATP10D | 261.199411 | 0.30009813 | 0.23221635 | 1.2923213 | 0.1962459 | 0.76469312 |
| AC107068.2 | 78.5005087 | 0.4111565 | 0.52146244 | 0.78846811 | 0.43042295 | 0.88596084 |
| NFXL1 | 248.183906 | 0.25784143 | 0.37737786 | 0.68324472 | 0.49445222 | 0.90839719 |
| NIPAL1 | 363.997679 | 0.43989528 | 0.3901794 | 1.12741801 | 0.25956579 | 0.80709747 |
| CNGA1 | 419.782641 | 0.30261399 | 0.39675451 | 0.76272349 | 0.44562832 | 0.8918683 |
| TXK | 67.9199489 | 0.09820064 | 0.54309082 | 0.18081808 | 0.85651038 | 0.97718353 |
| TEC | 163.162941 | -0.4882208 | 0.27030032 | -1.8062161 | 0.07088459 | 0.56278072 |
| SLAIN2 | 288.515351 | 0.00135433 | 0.1522579 | 0.00889494 | 0.99290295 | 0.99919666 |
| FRYL | 819.892136 | -0.1703488 | 0.2387832 | -0.7134037 | 0.47559597 | 0.90242252 |

| | | | | | | |
|---|---|---|---|---|---|---|
| OCIAD1 | 493.955522 | 0.01289224 | 0.15962187 | 0.08076737 | 0.93562696 | 0.99243654 |
| OCIAD2 | 109.378661 | -0.3268693 | 0.36987398 | -0.8837316 | 0.3768411 | 0.86417827 |
| CWH43 | 142.508055 | 0.52762831 | 0.52366653 | 1.00756546 | 0.31366312 | 0.84285425 |
| DCUN1D4 | 244.444358 | -0.0096298 | 0.28257542 | -0.0340787 | 0.97281436 | 0.99656026 |
| SGCB | 60.8151155 | 0.66585798 | 0.37944785 | 1.75480764 | 0.07929222 | 0.58437583 |
| USP46 | 119.762349 | 0.24455701 | 0.21733451 | 1.12525622 | 0.26048047 | 0.80805101 |
| ERVMER34-1 | 47.7409533 | 1.28971617 | 0.44866096 | 2.8745897 | 0.00404553 | 0.15545856 |
| RASL11B | 80.017932 | 1.05733737 | 0.47119972 | 2.24392614 | 2.48E-02 | 0.37943641 |
| SCFD2 | 253.665713 | -0.2436563 | 0.18808644 | -1.2954487 | 0.1951655 | 0.76233115 |
| AC058822.1 | 168.619421 | 0.14914506 | 0.48466758 | 0.3077265 | 0.75829045 | 0.96180522 |
| LNX1 | 249.770152 | 0.39753207 | 0.39076016 | 1.0173301 | 0.30899642 | 0.84136316 |
| CHIC2 | 245.791167 | -0.2517917 | 0.21090164 | -1.1938824 | 0.23252398 | 0.79255697 |
| TMEM165 | 475.793199 | -0.2343258 | 0.17587222 | -1.3323637 | 0.1827407 | 0.74315969 |
| CLOCK | 312.189695 | 0.03301128 | 0.19989343 | 0.16514442 | 0.86883031 | 0.9820194 |
| EXOC1 | 337.443645 | -0.3441388 | 0.19974388 | -1.7229004 | 0.08490655 | 0.59839042 |
| CEP135 | 125.70019 | -0.2235382 | 0.24354378 | -0.9178565 | 0.35869401 | 0.8557799 |
| AASDH | 157.049245 | 0.06879599 | 0.27783531 | 0.24761427 | 0.80443287 | 0.96679546 |
| PPAT | 41.5734804 | -0.3469637 | 0.39897458 | -0.8696387 | 0.38449787 | 0.86725322 |
| PAICS | 91.1888714 | 0.06612364 | 0.45859346 | 0.14418792 | 0.88535208 | 0.9845556 |
| SRP72 | 154.699345 | 0.04761232 | 0.20263363 | 0.23496753 | 0.81423393 | 0.96820092 |
| REST | 270.86016 | 0.34454029 | 0.1838792 | 1.87373172 | 0.06096741 | 0.5418629 |
| NOA1 | 40.893993 | 0.66970102 | 0.30967278 | 2.16260861 | 0.0305713 | 0.41141619 |
| POLR2B | 213.549656 | -0.092034 | 0.27533766 | -0.3342586 | 0.73818441 | 0.96078945 |
| IGFBP7 | 2812.04252 | 0.61844374 | 0.32441159 | 1.90635524 | 0.05660413 | 0.52346922 |
| AC104806.2 | 38.6296114 | 0.45886134 | 0.46150673 | 0.99426793 | 0.32009244 | 0.84285425 |
| CENPC | 283.753693 | -0.3240307 | 0.29634953 | -1.093407 | 0.27421514 | 0.81986798 |
| UBA6 | 451.992439 | -0.6196415 | 0.33236697 | -1.8643294 | 0.06227546 | 0.54569134 |
| UBA6-AS1 | 624.846624 | -0.1717014 | 0.30320087 | -0.5662959 | 0.57119267 | 0.93125376 |
| YTHDC1 | 603.796195 | -0.0643007 | 0.19347305 | -0.3323497 | 0.73962522 | 0.96078945 |
| UGT2B7 | 239.099374 | 0.11547381 | 0.5702768 | 0.2024873 | 0.83953578 | 0.97452864 |
| UTP3 | 43.8981976 | 0.08742531 | 0.35644412 | 0.24527074 | 0.8062468 | 0.96752828 |
| RUFY3 | 825.528616 | -0.3599727 | 0.32452759 | -1.1092205 | 0.26733506 | 0.81467618 |
| GRSF1 | 206.296901 | -0.206636 | 0.20921195 | -0.9876872 | 0.32330585 | 0.84285425 |
| MOB1B | 1331.46939 | -0.1414335 | 0.34998278 | -0.4041157 | 0.68612765 | 0.9558749 |
| DCK | 39.5464274 | -0.4213684 | 0.32468531 | -1.2977748 | 0.19436475 | 0.76162393 |
| SLC4A4 | 1661.00365 | -0.968446 | 0.34569258 | -2.8014661 | 0.0050871 | 0.17686435 |
| ADAMTS3 | 339.179326 | 0.94172469 | 0.50545106 | 1.86313723 | 0.06244296 | 0.5462456 |
| COX18 | 45.6474577 | -0.0242358 | 0.41415612 | -0.0585185 | 0.9533356 | 0.99512909 |
| ANKRD17 | 1583.84071 | -0.1204795 | 0.22398594 | -0.5378885 | 0.59065401 | 0.93483536 |
| AC053527.1 | 226.72021 | 0.55104341 | 0.63044527 | 0.87405431 | 0.38208868 | 0.86593382 |
| RASSF6 | 273.72114 | -0.8270624 | 0.35083018 | -2.3574437 | 0.01840125 | 0.32520246 |
| CXCL2 | 106.544056 | -0.7216428 | 1.34091313 | -0.5381727 | 0.59045784 | 0.93483536 |
| MTHFD2L | 238.519848 | -1.0877739 | 0.30087214 | -3.6154025 | 0.00029988 | 0.0383263 |
| BTC | 108.949713 | -0.0070056 | 0.36111743 | -0.0193999 | 0.98452212 | 0.99756277 |
| PARM1 | 79.7177485 | 0.93841561 | 0.27567802 | 3.40402768 | 0.000664 | 0.05965636 |
| RCHY1 | 138.620358 | 0.01562918 | 0.26845539 | 0.05821891 | 0.95357426 | 0.99514888 |
| THAP6 | 86.785795 | -0.0546091 | 0.27114635 | -0.2014007 | 0.84038528 | 0.97489902 |
| CDKL2 | 206.909709 | -0.1035268 | 0.20436473 | -0.5065784 | 0.61245063 | 0.94003018 |
| G3BP2 | 306.685011 | -0.1500128 | 0.29939536 | -0.5010524 | 0.61633426 | 0.94206626 |
| USO1 | 383.31823 | -1.92E-01 | 0.19392075 | -0.9917246 | 0.32133187 | 0.84285425 |
| NAAA | 48.3102568 | 1.62260093 | 0.48735745 | 3.32938575 | 0.00087038 | 0.07013468 |
| SDAD1 | 130.264116 | -0.3356295 | 0.266927 | -1.2573832 | 0.20861492 | 0.77436433 |
| NUP54 | 181.503231 | -0.1627021 | 0.19101758 | -0.8517652 | 0.39434443 | 0.87253562 |
| SCARB2 | 482.07626 | 0.18460575 | 0.24695119 | 0.74753946 | 0.454738 | 0.89626521 |

| | | | | | | |
|---|---|---|---|---|---|---|
| SHROOM3 | 1324.83118 | 0.88733691 | 0.42266282 | 2.09939663 | 0.03578195 | 0.4362094 |
| 11-Sep | 905.795268 | -0.0962906 | 0.27016763 | -0.3564107 | 0.72153304 | 0.95868461 |
| CCNI | 862.16023 | 0.26992115 | 0.20766092 | 1.2998168 | 0.19366377 | 0.76043322 |
| CCNG2 | 428.738863 | -0.1040598 | 0.36156149 | -0.2878065 | 0.77349484 | 0.96388926 |
| CNOT6L | 541.216256 | 0.07137201 | 0.18782033 | 0.3800015 | 7.04E-01 | 0.95672887 |
| MRPL1 | 234.244864 | 0.04577707 | 0.17225781 | 0.26574743 | 7.90E-01 | 0.96524651 |
| FRAS1 | 3774.90282 | 0.41372718 | 0.40361245 | 1.02506049 | 0.30533465 | 0.83709431 |
| ANXA3 | 201.39205 | 0.18749876 | 0.45756713 | 0.40977324 | 0.6819723 | 0.95574492 |
| BMP2K | 198.841313 | -0.4836304 | 0.39351138 | -1.2290125 | 0.21906711 | 0.78597171 |
| PAQR3 | 61.7727664 | -0.1932889 | 0.33157384 | -0.5829436 | 0.55993126 | 0.92934517 |
| CFAP299 | 64.2676119 | -0.5459238 | 0.72528993 | -0.7526974 | 0.4516318 | 0.894388 |
| RASGEF1B | 1445.19302 | -0.8114132 | 0.58676278 | -1.3828641 | 0.16670655 | 0.72097071 |
| HNRNPD | 455.082227 | -0.0987542 | 0.17181653 | -0.5747655 | 0.56544993 | 0.93114846 |
| HNRNPDL | 473.794923 | -0.5245089 | 0.18447128 | -2.8433092 | 0.00446477 | 0.16424479 |
| ENOPH1 | 88.6179764 | 0.06186845 | 0.24102645 | 0.25668737 | 0.79742012 | 0.9653598 |
| TMEM150C | 66.4110228 | 0.32147401 | 0.55817925 | 0.57593329 | 0.56466027 | 0.93104392 |
| SCD5 | 93.7317518 | 2.45316114 | 0.5083041 | 4.82616827 | 1.39E-06 | 0.00091166 |
| SEC31A | 564.375087 | -0.2904301 | 0.20584656 | -1.4109057 | 0.15827243 | 0.71264724 |
| LIN54 | 89.4566084 | -0.1590053 | 0.19926542 | -0.7979572 | 0.42489531 | 0.88392184 |
| COPS4 | 120.772014 | 0.10412242 | 0.19477672 | 0.53457324 | 0.59294499 | 0.93491607 |
| HELQ | 109.535835 | 0.34091725 | 0.30359158 | 1.12294697 | 0.26146002 | 0.80805101 |
| MRPS18C | 58.4823332 | 0.47861743 | 0.33322397 | 1.43632352 | 0.1509103 | 0.7029344 |
| ABRAXAS1 | 72.9695367 | 0.03436109 | 0.28819821 | 0.11922727 | 0.9050953 | 0.9875066 |
| GPAT3 | 200.875855 | 0.06155612 | 0.75870842 | 0.08113277 | 0.93533636 | 0.99243654 |
| CDS1 | 268.256728 | 0.09876578 | 0.14259748 | 0.6926194 | 0.48854843 | 0.90730741 |
| WDFY3 | 2075.62131 | 0.16338031 | 0.23836734 | 0.68541399 | 0.49308272 | 0.90804281 |
| ARHGAP24 | 807.61499 | 0.90242813 | 0.49371103 | 1.82784681 | 0.06757255 | 0.55848603 |
| MAPK10 | 3160.16107 | 0.28268692 | 0.29072941 | 0.97233685 | 0.330883 | 0.84442294 |
| AC104827.1 | 67.4222003 | 0.02924813 | 0.57690229 | 0.05069859 | 0.9595657 | 0.99609409 |
| PTPN13 | 2829.25022 | 0.42786866 | 0.30465082 | 1.40445596 | 0.16018312 | 0.71409437 |
| AC093827.5 | 89.0073817 | -0.2685502 | 0.30232098 | -0.8882948 | 0.37438218 | 0.86269244 |
| AFF1 | 4713.87615 | -0.3312175 | 0.22517334 | -1.4709446 | 0.14130611 | 0.68623171 |
| KLHL8 | 455.006131 | 0.07062361 | 0.30869483 | 0.2287813 | 0.8190389 | 0.96884581 |
| HSD17B11 | 95.3707051 | -0.2664991 | 0.42512685 | -0.6268696 | 0.5307447 | 0.92211604 |
| NUDT9 | 165.219588 | 0.07868454 | 0.21385813 | 0.36792866 | 0.71292643 | 0.95791799 |
| SPP1 | 2952.15867 | -0.9169452 | 0.71770853 | -1.2776011 | 2.01E-01 | 0.76957251 |
| PKD2 | 1034.34026 | -0.1185996 | 0.32933227 | -0.3601213 | 0.71875643 | 0.95796325 |
| PPM1K | 367.400585 | -0.3025688 | 0.43915334 | -0.688982 | 0.49083457 | 0.90730741 |
| HERC6 | 132.933495 | -1.7090989 | 0.74084319 | -2.3069645 | 0.0210568 | 0.35027818 |
| HERC5 | 97.9852641 | -1.3932247 | 0.64660354 | -2.1546815 | 0.03118677 | 0.41399324 |
| PYURF | 101.670435 | 0.44905104 | 0.47970667 | 0.93609506 | 0.34922425 | 0.8535343 |
| HERC3 | 324.436429 | 0.28856127 | 0.34897777 | 0.82687579 | 0.40830747 | 0.87948803 |
| FAM13A-AS1 | 95.6123953 | 0.00528393 | 0.25375515 | 0.02082296 | 0.98338688 | 0.9971769 |
| FAM13A | 750.652624 | -0.8322359 | 0.5619701 | -1.4809255 | 0.13862642 | 0.68156449 |
| GPRIN3 | 420.398449 | -1.3189879 | 0.4521059 | -2.9174313 | 0.00352927 | 0.14561728 |
| CCSER1 | 7766.5275 | 0.28059321 | 0.34509883 | 0.81308074 | 0.41617177 | 0.88242985 |
| SMARCAD1 | 307.118403 | -0.2508809 | 0.1606323 | -1.5618337 | 0.11832718 | 0.65447949 |
| PDLIM5 | 664.675028 | -0.1147565 | 0.23194585 | -0.4947557 | 0.6207726 | 0.94369714 |
| BMPR1B | 4439.78201 | 0.35821488 | 0.37885107 | 0.94552954 | 0.34438859 | 0.85148675 |
| STPG2 | 539.910632 | -0.4243007 | 0.39674199 | -1.0694625 | 0.28486132 | 0.82781987 |
| RAP1GDS1 | 511.082348 | -0.0839392 | 0.2226057 | -0.3770756 | 0.7061174 | 0.95730303 |
| TSPAN5 | 1213.06577 | -1.822441 | 0.31114252 | -5.8572547 | 4.71E-09 | 8.22E-06 |
| EIF4E | 252.451048 | -0.2533588 | 0.2415984 | -1.0486777 | 0.29432649 | 0.8337535 |
| METAP1 | 211.340772 | 0.11895664 | 0.25182383 | 0.47238039 | 0.6366553 | 0.94821838 |

| | | | | | | |
|---|---|---|---|---|---|---|
| ADH5 | 249.702516 | 0.34549822 | 0.31494616 | 1.09700725 | 0.27263822 | 0.81922962 |
| AP002026.1 | 52.4829088 | 0.52664044 | 0.68832151 | 0.76510822 | 0.44420711 | 0.89096469 |
| TRMT10A | 44.3632353 | 0.40915133 | 0.35570504 | 1.15025453 | 0.25003905 | 0.80149394 |
| AC083902.1 | 46.9785215 | 0.98122986 | 0.73091574 | 1.34246644 | 0.1794448 | 0.74096988 |
| C4orf54 | 93.6887533 | 1.58237213 | 0.61550097 | 2.57086863 | 0.01014438 | 0.2455268 |
| LAMTOR3 | 121.975953 | 0.00837716 | 0.26683309 | 0.03139477 | 0.97495472 | 0.99656026 |
| DNAJB14 | 382.669642 | 0.14256559 | 0.15276744 | 0.93321973 | 0.35070653 | 0.85477058 |
| H2AFZ | 245.327374 | 0.26325212 | 0.63126074 | 0.41702596 | 0.67665942 | 0.95529325 |
| PPP3CA | 513.103682 | -0.1362283 | 0.28862551 | -0.4719898 | 0.63693407 | 0.94821838 |
| BANK1 | 157.121493 | -0.3369846 | 0.43012245 | -0.783462 | 0.43335584 | 0.88736205 |
| AP002075.1 | 49.8550219 | 0.54848353 | 0.69414058 | 0.79016203 | 0.42943315 | 0.88579433 |
| SLC39A8 | 57.9277882 | 0.36109808 | 0.52040907 | 0.69387355 | 4.88E-01 | 0.90730741 |
| NFKB1 | 337.434426 | -0.3816897 | 0.3754448 | -1.0166332 | 0.30932794 | 0.84136316 |
| MANBA | 238.260418 | -0.1377958 | 0.3007301 | -0.4582044 | 0.64680562 | 0.95061388 |
| UBE2D3 | 949.916326 | -0.0466261 | 0.19054294 | -0.2447012 | 0.80668778 | 0.96752828 |
| CISD2 | 113.440855 | 0.16057727 | 0.26283532 | 0.61094252 | 0.54123764 | 0.92443671 |
| SLC9B1 | 44.9203267 | 0.86286531 | 0.56904465 | 1.51634026 | 0.12943334 | 0.67207756 |
| SLC9B2 | 35.3077294 | 0.12833558 | 0.27547067 | 0.46587747 | 0.64130321 | 0.94935885 |
| TET2 | 682.978307 | -0.1991241 | 0.37690661 | -0.5283115 | 0.59728314 | 0.93662454 |
| PPA2 | 616.4955 | 0.08803532 | 0.19932974 | 0.44165672 | 0.65873763 | 0.95116704 |
| ARHGEF38 | 911.09103 | -1.1858299 | 0.34863808 | -3.4013207 | 0.00067061 | 0.05965636 |
| INTS12 | 168.617419 | -0.4626791 | 0.24939295 | -1.8552213 | 0.06356464 | 0.54850785 |
| GSTCD | 271.086832 | -0.0842183 | 0.2644991 | -0.3184067 | 0.75017646 | 0.96109466 |
| NPNT | 617.249906 | 0.3466538 | 0.43763914 | 0.79209963 | 0.42830258 | 0.88535099 |
| TBCK | 889.393832 | 0.26408639 | 0.30100693 | 0.87734321 | 0.38030025 | 0.86593382 |
| AIMP1 | 156.189122 | 0.30146976 | 0.19878278 | 1.51657888 | 0.12937304 | 0.67207756 |
| PAPSS1 | 445.212601 | -0.0638648 | 0.19824068 | -0.3221581 | 7.47E-01 | 0.96109466 |
| SGMS2 | 158.497805 | 0.07134867 | 0.30186582 | 0.23635889 | 0.81315419 | 0.9680627 |
| HADH | 100.534293 | 0.60228197 | 0.26924377 | 2.23693928 | 0.02529031 | 0.38387372 |
| RPL34-AS1 | 73.9663874 | 0.6156108 | 0.51877282 | 1.18666741 | 0.23535884 | 0.79599484 |
| RPL34 | 558.811811 | 0.10721781 | 0.41397941 | 0.2589931 | 0.79564056 | 0.9653598 |
| OSTC | 127.573197 | 0.00532831 | 0.40112769 | 0.01328333 | 0.98940175 | 0.99835647 |
| COL25A1 | 116.639123 | -2.2626729 | 1.2217299 | -1.8520238 | 0.0640224 | 0.54860648 |
| SEC24B-AS1 | 40.0396106 | 0.43522186 | 0.51610596 | 0.84328006 | 0.39907183 | 0.87596764 |
| SEC24B | 554.266659 | -0.2665984 | 0.21233531 | -1.2555539 | 0.20927776 | 0.77436433 |
| CASP6 | 66.2814227 | 0.06345864 | 0.27014321 | 0.2349074 | 8.14E-01 | 0.96820092 |
| PLA2G12A | 100.914245 | 0.21157999 | 0.21094767 | 1.00299749 | 0.31586207 | 0.84285425 |
| AC126283.2 | 57.0221799 | 1.11990014 | 0.40358996 | 2.77484634 | 0.00552278 | 0.18370315 |
| GAR1 | 50.0709148 | 0.06046908 | 0.38881306 | 0.15552224 | 0.87640962 | 0.98269442 |
| ELOVL6 | 292.344619 | 0.33106577 | 0.28700962 | 1.15350059 | 0.24870497 | 0.80097144 |
| FAM241A | 293.477333 | -0.0568399 | 0.35286662 | -0.1610805 | 0.87203003 | 0.98223939 |
| AP1AR | 200.144467 | -0.5616467 | 0.30481856 | -1.8425605 | 0.0653932 | 0.55401837 |
| TIFA | 31.601756 | 0.21907608 | 0.33893575 | 0.64636463 | 0.51804324 | 0.91751084 |
| ALPK1 | 404.262768 | -0.597877 | 0.30424656 | -1.965107 | 0.04940186 | 0.49260844 |
| LARP7 | 214.375545 | -0.0911643 | 0.22449314 | -0.4060896 | 0.68467677 | 0.9558749 |
| ANK2 | 79.2600153 | -1.3765672 | 0.62963906 | -2.1862799 | 0.02879513 | 0.40623517 |
| CAMK2D | 424.535409 | -0.3840828 | 0.21027568 | -1.8265678 | 0.06776479 | 0.55864172 |
| ARSJ | 341.630393 | 0.53055634 | 0.34568877 | 1.53478037 | 0.12483779 | 0.66626798 |
| UGT8 | 278.8101 | -0.0774193 | 0.26053333 | -0.2971569 | 0.76634671 | 0.96285313 |
| SNHG8 | 93.9156242 | 0.311944 | 0.50144866 | 0.62208562 | 0.53388557 | 0.92289732 |
| METTL14 | 126.707766 | -0.1109147 | 0.19508695 | -0.5685396 | 0.5696686 | 0.93122667 |
| SEC24D | 148.991448 | -0.9534042 | 0.40582021 | -2.3493265 | 0.01880741 | 0.32850268 |
| USP53 | 1200.83333 | 0.21316424 | 0.24547244 | 0.86838361 | 0.38518437 | 0.86749643 |
| C4orf3 | 385.583612 | -0.064501 | 0.49004191 | -0.1316234 | 0.89528216 | 0.98597539 |

| | | | | | | |
|---|---|---|---|---|---|---|
| PDE5A | 392.137198 | -0.0598227 | 0.34843318 | -0.1716906 | 0.86368074 | 0.979268 |
| AC108866.1 | 179.329102 | -0.294644 | 0.36973759 | -0.7969002 | 0.42550899 | 0.88403732 |
| PRDM5 | 317.451188 | 0.02406837 | 0.33304353 | 0.07226795 | 0.94238867 | 0.99265578 |
| ANXA5 | 329.288146 | 0.21115083 | 0.39038808 | 0.54087419 | 0.5885943 | 0.93483536 |
| EXOSC9 | 35.7652657 | -0.5991592 | 0.27288704 | -2.1956307 | 0.02811838 | 0.40459506 |
| BBS7 | 70.3709504 | -0.088519 | 0.32400574 | -0.2732019 | 0.78469803 | 0.96513523 |
| KIAA1109 | 1431.63416 | -0.1373428 | 0.22525673 | -0.6097166 | 0.54204956 | 0.92481499 |
| FGF2 | 91.7231339 | -0.9122557 | 0.57456352 | -1.5877369 | 0.11234585 | 0.64478889 |
| NUDT6 | 49.3064145 | 0.79058149 | 0.28323398 | 2.79126643 | 0.00525022 | 0.17920428 |
| SPATA5 | 458.727617 | -0.384624 | 0.21347755 | -1.8017069 | 0.07159153 | 0.56343001 |
| SPRY1 | 119.854554 | 1.4298652 | 0.41835908 | 3.41779409 | 0.00063131 | 0.05952104 |
| LINC01091 | 140.944967 | 0.76965646 | 0.52491492 | 1.46624993 | 0.14258021 | 0.6890058 |
| ANKRD50 | 76.3730449 | 0.65606906 | 0.52344398 | 1.25337015 | 0.21007103 | 0.77551367 |
| FAT4 | 148.142416 | 0.59538666 | 0.45452701 | 1.30990381 | 0.19022838 | 0.75658194 |
| INTU | 262.624163 | -0.4539249 | 0.3516956 | -1.2906755 | 0.19681621 | 0.76478825 |
| HSPA4L | 214.674562 | 0.35011872 | 0.59574144 | 0.58770247 | 0.55673202 | 0.92872652 |
| MFSD8 | 176.130924 | -0.4234411 | 0.3474035 | -1.2188741 | 0.22289199 | 0.78835912 |
| ABHD18 | 477.654729 | -0.3580842 | 0.23698028 | -1.5110295 | 0.13078092 | 0.6735057 |
| LARP1B | 287.749095 | 0.18383828 | 0.23861399 | 0.77044217 | 0.44103765 | 0.88964371 |
| PGRMC2 | 187.876063 | 0.11683128 | 0.26610582 | 0.43904069 | 0.66063206 | 0.95127957 |
| AC108062.1 | 145.246092 | 0.05484865 | 0.34085495 | 0.16091493 | 0.8721604 | 0.98223939 |
| JADE1 | 413.341927 | 0.90145359 | 0.26785291 | 3.36547992 | 0.00076411 | 0.06406267 |
| SCLT1 | 354.225238 | -0.2811331 | 0.3063021 | -0.9178296 | 0.35870811 | 0.8557799 |
| C4orf33 | 70.93439 | -0.4649759 | 0.57036137 | -0.8152303 | 0.41494051 | 0.88242985 |
| AC096711.2 | 236.166225 | -0.2016534 | 0.36958237 | -0.5456249 | 0.58532378 | 0.93483536 |
| ELF2 | 868.669321 | -0.1058635 | 0.27854125 | -0.3800641 | 0.70389783 | 0.95672887 |
| NDUFC1 | 231.329064 | 0.21918501 | 0.36173499 | 0.60592704 | 0.54456321 | 0.92561018 |
| NAA15 | 324.569631 | -0.2721754 | 0.237456 | -1.1462141 | 0.25170659 | 0.80252057 |
| AC097376.2 | 350.110197 | -0.5248448 | 0.30251719 | -1.7349255 | 0.082754 | 0.59202053 |
| SETD7 | 67.4044287 | 0.07622983 | 0.39200344 | 0.19446215 | 0.84581403 | 0.97643646 |
| MGST2 | 110.200735 | 0.02437615 | 0.25854371 | 0.0942825 | 0.92488475 | 0.99051393 |
| MAML3 | 1357.77111 | 0.25496595 | 0.33192845 | 0.76813527 | 0.44240683 | 0.89005186 |
| SCOC | 414.85506 | 0.15726815 | 0.21838001 | 0.72015817 | 0.47142762 | 0.90123338 |
| SCOC-AS1 | 39.0302714 | 0.52744064 | 0.54738587 | 0.96356276 | 0.33526518 | 0.84544327 |
| ELMOD2 | 126.492853 | -0.2906662 | 0.20821365 | -1.3959996 | 0.1627146 | 0.71543692 |
| TBC1D9 | 824.771215 | -0.08217 | 0.24853103 | -0.3306228 | 0.74092942 | 0.96084411 |
| RNF150 | 555.12123 | 0.87352903 | 0.47136131 | 1.85320478 | 0.06385301 | 0.54850785 |
| ZNF330 | 80.1263448 | -0.3226657 | 0.36904629 | -0.874323 | 0.38194237 | 0.86593382 |
| IL15 | 78.7333079 | -1.0925322 | 0.3880967 | -2.8151029 | 0.00487616 | 0.17264247 |
| INPP4B | 251.117772 | -1.2681413 | 0.45589111 | -2.7816759 | 0.0054079 | 0.18165004 |
| AC104596.1 | 151.157128 | -0.3224014 | 0.63237963 | -0.5098226 | 0.61017578 | 0.94003018 |
| USP38 | 102.004784 | -0.1033781 | 0.24931517 | -0.4146481 | 0.67839953 | 0.95574492 |
| GAB1 | 1113.54709 | -0.6441918 | 0.20013833 | -3.218733 | 0.00128758 | 0.09056289 |
| SMARCA5 | 324.973778 | 0.10369477 | 0.17305243 | 0.59921016 | 0.54903275 | 0.9274143 |
| ANAPC10 | 231.917929 | -0.0991803 | 0.19307103 | -0.5136987 | 0.60746267 | 0.93952313 |
| ABCE1 | 99.9412445 | -0.2253464 | 0.23155007 | -0.9732079 | 0.33044998 | 0.84442294 |
| OTUD4 | 287.843122 | 0.14480803 | 0.18274614 | 0.79239991 | 0.42812753 | 0.88535099 |
| SMAD1 | 383.891983 | 0.50633272 | 0.36824804 | 1.37497735 | 0.16913847 | 0.72596839 |
| MMAA | 76.4364274 | 0.12468759 | 0.23534734 | 0.52980239 | 0.59624895 | 0.93593903 |
| ZNF827 | 697.996351 | 0.74597499 | 0.345651 | 2.15817396 | 0.03091431 | 0.41259458 |
| LSM6 | 94.5157418 | 0.30620412 | 0.26198077 | 1.16880376 | 0.2424827 | 0.80072691 |
| SLC10A7 | 256.696801 | -0.0589322 | 0.18100172 | -0.3255895 | 0.74473499 | 0.96109466 |
| TMEM184C | 173.571884 | 0.50878458 | 0.51360051 | 0.9906232 | 0.32186961 | 0.84285425 |
| PRMT9 | 285.103589 | 0.05366739 | 0.74879336 | 0.07167183 | 0.94286307 | 0.99265578 |

| | | | | | | |
|---|---|---|---|---|---|---|
| ARHGAP10 | 64.204251 | 0.45822006 | 0.60391463 | 0.75874974 | 0.44800228 | 0.89297728 |
| NR3C2 | 4193.69502 | 0.48141508 | 0.27138048 | 1.77394882 | 0.07607162 | 0.57145422 |
| AC069272.1 | 58.0144212 | 0.89745638 | 0.62334571 | 1.439741 | 0.14994069 | 0.7029344 |
| DCLK2 | 52.8860444 | 0.28970459 | 0.42462333 | 0.68226253 | 0.49507296 | 0.90839719 |
| LRBA | 3362.20315 | -0.2392331 | 0.29877944 | -0.8007013 | 0.42330458 | 0.88371155 |
| RPS3A | 393.989469 | 0.21799598 | 0.48764099 | 0.44704195 | 0.65484477 | 0.95100227 |
| SH3D19 | 343.251253 | 0.28985338 | 0.30401401 | 0.95342114 | 0.34037674 | 0.84975445 |
| FAM160A1 | 541.096645 | -0.54393 | 0.35338542 | -1.5391978 | 0.12375603 | 0.66519057 |
| GATB | 84.5110917 | -0.0808486 | 0.33492864 | -0.2413905 | 0.80925245 | 0.9679755 |
| FBXW7 | 685.458702 | -0.0518678 | 0.21525071 | -0.2409644 | 0.80958268 | 0.9679755 |
| TMEM154 | 37.6984548 | -0.6774795 | 0.62065829 | -1.09155 | 0.27503096 | 0.8202929 |
| ARFIP1 | 507.912867 | -0.1032723 | 0.2000087 | -0.5163391 | 0.60561761 | 0.93919596 |
| FHDC1 | 70.0489953 | -0.3517647 | 0.32604545 | -1.0788823 | 2.81E-01 | 0.82628479 |
| TRIM2 | 1038.97587 | 0.09400714 | 0.36055658 | 0.26072785 | 0.79430239 | 0.9653598 |
| TMEM131L | 97.9732393 | 0.15119478 | 0.4207839 | 0.35931693 | 0.71935801 | 0.95796325 |
| PLRG1 | 81.164015 | 0.23867991 | 0.26263057 | 0.90880473 | 0.36345321 | 0.85694172 |
| MAP9 | 152.527271 | 0.41505884 | 0.24450771 | 1.69752863 | 0.08959676 | 0.60739841 |
| GUCY1A1 | 128.051153 | 0.34032521 | 0.42796275 | 0.79522157 | 0.42648463 | 0.88477675 |
| GUCY1B1 | 132.060161 | 0.06523021 | 0.34981609 | 0.18647001 | 0.8520762 | 0.97718353 |
| CTSO | 73.6124213 | -0.0439797 | 0.38759968 | -0.1134667 | 0.90966057 | 0.98779844 |
| PDGFC | 347.461982 | 0.24138981 | 0.36198762 | 0.6668455 | 0.50487082 | 0.9107831 |
| ETFDH | 202.267622 | -0.2245447 | 0.21816523 | -1.0292415 | 0.30336619 | 0.83709431 |
| PPID | 108.434414 | -0.2963758 | 0.22725495 | -1.3041555 | 0.19218055 | 0.7597012 |
| FNIP2 | 1329.64569 | -1.2316003 | 0.47461523 | -2.5949447 | 0.00946061 | 0.23606486 |
| C4orf45 | 38.3509391 | -0.1540474 | 0.64236991 | -0.2398111 | 0.81047673 | 0.9680627 |
| RAPGEF2 | 898.822193 | 0.04609253 | 0.1620171 | 0.28449175 | 0.77603356 | 0.96437053 |
| NAF1 | 146.338943 | -0.1703138 | 0.49499148 | -0.3440743 | 0.73079044 | 0.96008876 |
| TMA16 | 75.2162032 | -0.1693939 | 0.24621265 | -0.6879983 | 0.49145387 | 0.90730741 |
| 1-Mar | 39.977174 | -0.5789081 | 0.35093323 | -1.6496247 | 0.09901972 | 0.61707826 |
| TMEM192 | 156.673511 | 0.09689698 | 0.38074511 | 0.25449303 | 0.79911469 | 0.96538581 |
| KLHL2 | 567.999783 | -0.4275951 | 0.33292017 | -1.2843772 | 0.19901 | 0.76762045 |
| MSMO1 | 364.916026 | -0.6866614 | 0.60262235 | -1.1394556 | 0.25451318 | 0.80414079 |
| DDX60 | 187.449365 | -1.9608348 | 0.83803 | -2.3398145 | 0.01929332 | 0.33290134 |
| DDX60L | 135.390964 | -1.9501037 | 0.82244198 | -2.371114 | 0.01773456 | 0.31989361 |
| PALLD | 551.78751 | -0.2026628 | 0.23212849 | -0.873063 | 0.38262874 | 0.86593382 |
| CBR4 | 943.505842 | -0.0010273 | 0.25673621 | -0.0040015 | 0.99680725 | 0.99967893 |
| SH3RF1 | 1663.55366 | -0.1630999 | 0.37697648 | -0.4326528 | 0.66526702 | 0.9520866 |
| AC096741.1 | 42.4519703 | 0.02289515 | 0.72030625 | 0.0317853 | 0.97464327 | 0.99656026 |
| NEK1 | 658.002958 | -0.4725875 | 0.34696001 | -1.3620807 | 0.17317242 | 0.72964461 |
| CLCN3 | 774.879528 | 0.10092549 | 0.44694411 | 0.22581233 | 0.82134739 | 0.96955628 |
| HPF1 | 69.2980719 | 0.44643637 | 0.30016522 | 1.48730211 | 0.13693504 | 0.68129775 |
| MFAP3L | 182.605616 | 0.77478664 | 0.33138783 | 2.33800571 | 0.01938695 | 0.33362111 |
| AC105285.1 | 42.1898194 | 0.45304956 | 0.44695947 | 1.01362559 | 0.31076144 | 0.84248332 |
| GALNT7 | 523.691073 | 0.29607421 | 0.24483122 | 1.20929924 | 0.2265479 | 0.78982769 |
| HMGB2 | 144.914196 | 0.4493039 | 0.37579334 | 1.19561433 | 0.2318471 | 0.79222614 |
| FBXO8 | 91.0939967 | 0.15723346 | 0.29058744 | 0.54108828 | 0.58844674 | 0.93483536 |
| CEP44 | 141.947427 | -0.1907484 | 0.29522715 | -0.6461072 | 0.51820994 | 0.91751084 |
| HPGD | 141.832577 | 0.55476228 | 0.57208678 | 0.969717 | 0.33218757 | 0.84442294 |
| WDR17 | 111.200366 | -0.1601387 | 0.32295531 | -0.4958539 | 0.61999746 | 0.94319304 |
| SPCS3 | 169.340288 | -0.1632855 | 0.1904529 | -0.8573539 | 0.39124933 | 0.87015937 |
| AGA | 36.7537212 | 0.45812459 | 0.40935577 | 1.11913553 | 0.26308232 | 0.8096287 |
| AC078881.1 | 149.586132 | 1.32687268 | 0.54020274 | 2.45624944 | 0.01403957 | 0.2862542 |
| DCTD | 88.4358853 | -0.0156111 | 0.27895527 | -0.0559627 | 0.95537152 | 0.99538167 |
| WWC2 | 820.16334 | -0.3192958 | 0.26390955 | -1.2098684 | 0.2263294 | 0.78982769 |

| | | | | | | |
|---|---|---|---|---|---|---|
| CDKN2AIP | 113.132819 | -0.030667 | 0.7442183 | -0.041207 | 0.96713084 | 0.99637506 |
| ING2 | 39.9947963 | 0.45752708 | 0.44002923 | 1.03976521 | 0.29844899 | 0.83556453 |
| RWDD4 | 72.9860634 | -0.0491024 | 0.25501911 | -0.1925441 | 0.84731606 | 0.97656375 |
| TRAPPC11 | 210.372147 | -0.0482703 | 0.2338433 | -0.2064216 | 0.8364616 | 0.97347225 |
| STOX2 | 889.996558 | 0.4558372 | 0.26821011 | 1.69955263 | 0.08921511 | 0.60739841 |
| IRF2 | 192.72968 | 0.16118823 | 0.27406178 | 0.58814558 | 0.55643458 | 0.92872652 |
| CASP3 | 70.4183869 | -0.1398704 | 0.33408028 | -0.418673 | 0.6754551 | 0.95529325 |
| PRIMPOL | 97.6064568 | 0.19053811 | 0.37630386 | 0.5063411 | 0.61261721 | 0.94003018 |
| ACSL1 | 304.004918 | -0.278992 | 0.42621916 | -0.654574 | 0.51274208 | 0.9146446 |
| SLC25A4 | 138.464237 | 0.71141572 | 0.50093705 | 1.42016991 | 0.15555822 | 0.70942131 |
| CFAP97 | 216.501536 | 0.20082313 | 0.2879986 | 0.69730592 | 0.48561136 | 0.90695174 |
| SNX25 | 347.181847 | -0.2559215 | 0.37916411 | -0.6749624 | 0.49969968 | 0.90839719 |
| LRP2BP | 39.788453 | -0.2502331 | 0.51749048 | -0.483551 | 0.62870454 | 0.94595714 |
| ANKRD37 | 547.444763 | -4.821622 | 1.46818791 | -3.2840633 | 0.00102322 | 0.07827256 |
| UFSP2 | 162.832403 | -0.2105148 | 0.31320302 | -0.6721353 | 0.50149755 | 0.90900452 |
| C4orf47 | 190.345375 | -1.1319734 | 0.65611107 | -1.7252772 | 0.08447754 | 0.59698218 |
| SORBS2 | 861.232673 | 0.47528963 | 0.34723864 | 1.36876941 | 0.17107136 | 0.72776437 |
| TLR3 | 58.5052682 | 0.3998414 | 0.42960039 | 0.93072867 | 0.35199393 | 0.85536253 |
| FAM149A | 366.456627 | 0.35882299 | 0.25609387 | 1.40113854 | 0.16117265 | 0.71409437 |
| CYP4V2 | 94.2908382 | -0.2236749 | 0.22431182 | -0.9971604 | 0.31868667 | 0.84285425 |
| KLKB1 | 78.8302159 | 0.09519512 | 0.44296625 | 0.21490378 | 0.82984234 | 0.97139806 |
| FAT1 | 609.547128 | 0.26199503 | 0.34321237 | 0.76336126 | 0.44524798 | 0.89168714 |
| FRG1-DT | 126.141132 | -0.366198 | 0.51560382 | -0.7102315 | 0.47756062 | 0.90242252 |
| FRG1 | 59.3623112 | 0.24758301 | 0.23569003 | 1.05046027 | 0.29350655 | 0.83336457 |
| PLEKHG4B | 66.0140424 | -0.0113885 | 0.81910408 | -0.0139036 | 0.98890689 | 0.99824159 |
| CCDC127 | 37.3757189 | 0.01736785 | 0.28474485 | 0.06099443 | 0.95136365 | 0.99404743 |
| SDHA | 371.453173 | 0.0369396 | 0.21092854 | 0.17512852 | 0.86097865 | 0.97841862 |
| PDCD6 | 143.197705 | -0.5393562 | 0.25201368 | -2.1401863 | 0.03233972 | 0.41687605 |
| EXOC3 | 145.839874 | -0.2783644 | 0.21799788 | -1.2769135 | 0.20163279 | 0.76957251 |
| SLC9A3 | 37.8313569 | 0.26885268 | 0.3375609 | 0.79645683 | 0.42576656 | 0.8840962 |
| BRD9 | 79.7123632 | 0.05707724 | 0.30628336 | 0.18635436 | 0.85216688 | 0.97718353 |
| SLC12A7 | 189.23044 | -1.7776441 | 0.50113314 | -3.5472492 | 0.00038928 | 0.04340016 |
| CLPTM1L | 100.607526 | -0.0962056 | 0.25818802 | -0.3726186 | 0.70943236 | 0.95752169 |
| LPCAT1 | 42.3442231 | -0.0432611 | 0.4654408 | -0.0929466 | 0.92594598 | 0.99051393 |
| MRPL36 | 50.2931851 | 0.49055358 | 0.49471467 | 0.99158891 | 0.3213981 | 0.84285425 |
| NDUFS6 | 196.971929 | -0.0176066 | 0.34498543 | -0.0510358 | 0.95929701 | 0.99609409 |
| AC022424.1 | 60.6968761 | 2.24266169 | 0.78846013 | 2.84435649 | 0.00445012 | 0.16424479 |
| ADAMTS16 | 514.741841 | 2.22246807 | 0.78381286 | 2.83545754 | 0.00457601 | 0.16593973 |
| ICE1 | 257.005998 | 0.03569791 | 0.19796709 | 0.18032247 | 0.85689941 | 0.97718353 |
| MED10 | 94.6874614 | -0.2406224 | 0.29460024 | -0.8167761 | 0.41405641 | 0.88215311 |
| NSUN2 | 109.075406 | -0.14155 | 0.24334476 | -0.5816849 | 0.56077894 | 0.92934517 |
| SRD5A1 | 36.4551565 | -0.1730515 | 0.2962481 | -0.5841438 | 0.55912358 | 0.92901206 |
| TENT4A | 161.328364 | -0.0656819 | 0.23472858 | -0.2798208 | 0.77961497 | 0.96513523 |
| MTRR | 89.9883943 | -0.1972517 | 0.22469237 | -0.8778743 | 0.38001193 | 0.86593382 |
| SEMA5A | 55.3968281 | -0.4982058 | 0.51692118 | -0.9637945 | 0.33514896 | 0.84544327 |
| FAM173B | 97.721459 | 0.04488096 | 0.30796735 | 0.14573285 | 0.88413229 | 0.98403849 |
| CCT5 | 144.13889 | 0.24925722 | 0.40134418 | 0.62105601 | 0.53456277 | 0.92303275 |
| CMBL | 102.872281 | 1.06416523 | 0.43252052 | 2.4603809 | 0.01387896 | 0.28408502 |
| 6-Mar | 671.573132 | -0.3195135 | 0.1727112 | -1.849987 | 0.06431542 | 0.54860648 |
| DAP | 488.50891 | -0.1583893 | 0.52131091 | -0.3038289 | 0.76125823 | 0.96218738 |
| DNAH5 | 141.71548 | 0.64433825 | 0.42408453 | 1.51936279 | 0.1286712 | 0.67206108 |
| TRIO | 1233.78242 | -0.2807298 | 0.27739982 | -1.0120044 | 0.31153596 | 0.84279403 |
| OTULINL | 82.0933738 | 0.11650753 | 0.39505847 | 0.29491211 | 0.76806102 | 0.96345263 |
| OTULIN | 96.5889649 | -0.1528814 | 0.22587404 | -0.6768438 | 0.4985051 | 0.90839719 |

| | | | | | | |
|---|---|---|---|---|---|---|
| ANKH | 90.5358442 | 0.33389122 | 0.55189943 | 0.60498562 | 0.54518855 | 0.92579581 |
| FBXL7 | 94.8189114 | 0.97485016 | 0.58550059 | 1.66498578 | 0.09591559 | 0.61367241 |
| ZNF622 | 35.0687396 | 0.33883736 | 0.36051775 | 0.9398632 | 0.34728774 | 0.85348649 |
| RETREG1 | 323.36588 | 0.1889269 | 0.30588852 | 0.61763317 | 0.53681717 | 0.92324352 |
| MYO10 | 2177.13278 | -1.1108602 | 0.37869758 | -2.9333702 | 0.00335304 | 0.14358074 |
| LINC02241 | 94.6928599 | 0.11337142 | 0.25435575 | 0.4457199 | 0.65579959 | 0.95100227 |
| CDH12 | 42.645191 | -0.0357526 | 0.40143163 | -0.0890628 | 0.92903202 | 0.99216065 |
| C5orf17 | 177.491562 | -1.0547367 | 1.44907879 | -0.727867 | 0.46669498 | 0.89883965 |
| DROSHA | 224.124632 | -0.33739 | 0.20279297 | -1.6637165 | 0.0961691 | 0.61407447 |
| C5orf22 | 57.1750667 | -0.1959908 | 0.24418047 | -0.8026473 | 0.42217861 | 0.88371155 |
| PDZD2 | 149.567645 | -0.0535895 | 0.40467408 | -0.1324263 | 0.89464709 | 0.98597539 |
| GOLPH3 | 329.914852 | -0.1676586 | 0.22710004 | -0.7382587 | 0.46035727 | 0.89785937 |
| MTMR12 | 265.925063 | 0.15216102 | 0.21399771 | 0.7110404 | 0.4770592 | 0.90242252 |
| ZFR | 709.539656 | -0.0257648 | 0.15006256 | -0.1716939 | 0.86367817 | 0.979268 |
| SUB1 | 434.567545 | 0.25175863 | 0.32041436 | 0.78572829 | 0.43202669 | 0.88697335 |
| TARS | 95.8893786 | -0.1082619 | 0.19455798 | -0.5564504 | 0.57790299 | 0.93345492 |
| C1QTNF3-AN | 291.368332 | -0.3597042 | 0.20666462 | -1.7405216 | 0.08176748 | 0.59202053 |
| RAI14 | 663.711772 | -0.7479648 | 0.31442134 | -2.3788616 | 0.01736619 | 0.31596825 |
| TTC23L | 45.1359533 | -0.0466742 | 0.4917805 | -0.0949085 | 0.92438749 | 0.99051393 |
| RAD1 | 42.8369041 | 0.07778732 | 0.32674899 | 0.23806447 | 0.81183109 | 0.9680627 |
| BRIX1 | 56.4985202 | 0.04750929 | 0.34712979 | 0.1368632 | 0.89113893 | 0.98597539 |
| DNAJC21 | 154.089955 | 0.18364591 | 0.19334218 | 0.94984918 | 0.34218889 | 0.85093239 |
| SPEF2 | 199.461306 | 0.15954169 | 0.42202393 | 0.37803944 | 0.70540129 | 0.95730303 |
| UGT3A1 | 423.170439 | -0.5301977 | 0.44755554 | -1.1846523 | 0.23615496 | 0.79736951 |
| LMBRD2 | 121.569745 | 0.24516692 | 0.23600148 | 1.03883634 | 0.29888086 | 0.83556453 |
| SKP2 | 33.8664936 | 0.08158474 | 0.33916505 | 0.24054584 | 0.80990713 | 0.96804593 |
| NADK2 | 157.278778 | -0.1271141 | 0.33724866 | -0.3769151 | 0.7062367 | 0.95730303 |
| NIPBL | 1385.0864 | -0.1943314 | 0.19609445 | -0.991009 | 0.32168117 | 0.84285425 |
| CPLANE1 | 646.20285 | -0.1025485 | 0.24546954 | -0.4177645 | 0.6761193 | 0.95529325 |
| NUP155 | 96.0921481 | -0.2162159 | 0.19363018 | -1.1166436 | 0.26414674 | 0.81061724 |
| WDR70 | 703.245631 | -0.2176956 | 0.20530908 | -1.0603311 | 0.28899402 | 0.83060304 |
| LIFR | 500.853882 | 0.30531921 | 0.27007928 | 1.13047993 | 0.25827405 | 0.80709747 |
| LIFR-AS1 | 37.4394704 | 0.44526827 | 0.54784935 | 0.81275677 | 0.41635753 | 0.88242985 |
| OSMR-AS1 | 416.681579 | -0.6867031 | 0.45721285 | -1.501933 | 0.13311442 | 0.67594872 |
| OSMR | 525.720535 | -1.0550609 | 0.60010072 | -1.7581397 | 0.07872375 | 0.58223351 |
| RICTOR | 638.46856 | -0.3905635 | 0.18549523 | -2.1055178 | 0.03524626 | 0.43442748 |
| DAB2 | 37.4201052 | 0.13616909 | 0.5164624 | 0.26365732 | 7.92E-01 | 0.9653598 |
| TTC33 | 153.325591 | 0.13526557 | 0.2555746 | 0.52926062 | 0.59662467 | 0.93617523 |
| PRKAA1 | 248.344539 | -0.0628045 | 0.20929277 | -0.3000797 | 0.76411638 | 0.96285313 |
| RPL37 | 364.283066 | -0.0688468 | 0.3265924 | -0.2108035 | 0.83304063 | 0.97208613 |
| CARD6 | 53.004368 | -0.8419605 | 0.33252505 | -2.5320213 | 0.01134071 | 0.26178556 |
| OXCT1 | 50.2202723 | 0.55894063 | 0.3924511 | 1.42423001 | 0.1543799 | 0.70805312 |
| C5orf51 | 55.1966394 | -0.3616636 | 0.30231277 | -1.1963225 | 0.23157074 | 0.79200039 |
| FBXO4 | 38.2243157 | 0.31989309 | 0.39394186 | 0.81203124 | 0.41677371 | 0.88242985 |
| GHR | 443.375426 | 0.2394261 | 0.34691733 | 0.69015319 | 0.49009786 | 0.90730741 |
| SELENOP | 347.284099 | 0.20079787 | 0.36897974 | 0.54419755 | 0.58630554 | 0.93483536 |
| ZNF131 | 337.506167 | -0.0408527 | 0.18927717 | -0.2158353 | 0.82911612 | 0.97131041 |
| HMGCS1 | 448.546705 | -1.0891766 | 0.65781062 | -1.6557602 | 0.09777039 | 0.61685067 |
| CCL28 | 70.0332648 | -0.9393 | 0.56963283 | -1.6489569 | 0.09915647 | 0.61707826 |
| TMEM267 | 137.495214 | 0.11740634 | 0.26792732 | 0.43820221 | 0.66123971 | 0.9513366 |
| PAIP1 | 154.158338 | -0.2312716 | 0.2442776 | -0.9467572 | 0.34376254 | 0.85126545 |
| NNT-AS1 | 179.136959 | 0.00773443 | 0.28008318 | 0.02761475 | 0.97796942 | 0.99696835 |
| NNT | 367.121164 | 0.26919909 | 0.23857302 | 1.1283719 | 0.25916289 | 0.80709747 |
| MRPS30-DT | 92.4142388 | -0.1555976 | 0.36657789 | -0.4244599 | 0.67123042 | 0.95430867 |

| | | | | | | |
|---|---|---|---|---|---|---|
| MRPS30 | 72.9485641 | -0.0918057 | 0.29667489 | -0.3094489 | 0.7569801 | 0.96133693 |
| PARP8 | 655.070571 | -0.5111896 | 0.39433135 | -1.2963453 | 0.19485655 | 0.76233115 |
| ITGA1 | 101.439695 | -0.0059511 | 0.37481822 | -0.0158772 | 0.98733235 | 0.99791717 |
| ITGA2 | 1347.63525 | -1.2417073 | 0.51384699 | -2.4164923 | 0.01567086 | 0.30067202 |
| MOCS2 | 105.493114 | 0.32353471 | 0.23636278 | 1.36880567 | 0.17106003 | 0.72776437 |
| NDUFS4 | 367.714842 | -0.191621 | 0.19344129 | -0.9905898 | 0.32188594 | 0.84285425 |
| ARL15 | 4975.36033 | 0.09083506 | 0.23191284 | 0.39167757 | 6.95E-01 | 0.95672887 |
| SNX18 | 363.807916 | 0.40818353 | 0.51350953 | 0.79488989 | 0.42667756 | 0.88477675 |
| CCNO | 47.2450106 | -0.170697 | 0.50807721 | -0.3359666 | 0.73689607 | 0.96078945 |
| AC026704.1 | 99.2081446 | 0.54870276 | 0.42028129 | 1.30556077 | 0.19170197 | 0.7597012 |
| DHX29 | 246.216007 | 0.15414614 | 0.19235348 | 0.80136914 | 0.42291798 | 0.88371155 |
| MTREX | 408.492454 | -0.1647505 | 0.23650111 | -0.6966162 | 0.486043 | 0.90695174 |
| PLPP1 | 488.019824 | 0.23633689 | 0.33215147 | 0.71153349 | 0.4767537 | 0.90242252 |
| SLC38A9 | 235.227341 | -0.3745527 | 0.19642054 | -1.9068915 | 0.05653464 | 0.52346922 |
| IL6ST | 454.341986 | 0.50867777 | 0.24973829 | 2.03684334 | 0.04166574 | 0.46746387 |
| LINC01948 | 38.3451511 | 0.78825734 | 0.32629174 | 2.41580537 | 0.01570045 | 0.30067202 |
| MAP3K1 | 575.712206 | 0.56054972 | 0.35994296 | 1.55732929 | 0.11939232 | 0.65527787 |
| MIER3 | 76.6613108 | 0.01858504 | 0.24556762 | 0.07568199 | 0.93967211 | 0.99243654 |
| GPBP1 | 741.789626 | -0.1697054 | 0.20946482 | -0.8101857 | 0.41783345 | 0.88242985 |
| PDE4D | 26196.0279 | -0.2633951 | 0.42673945 | -0.617227 | 0.53708501 | 0.92324352 |
| AC092343.1 | 38.5766198 | -1.7037128 | 0.62810685 | -2.712457 | 0.00667864 | 0.20224502 |
| ELOVL7 | 283.976057 | -0.4229468 | 0.33915548 | -1.2470588 | 0.21237596 | 0.77709975 |
| ERCC8 | 154.154727 | -0.0998425 | 0.27576853 | -0.3620518 | 0.71731335 | 0.95791799 |
| NDUFAF2 | 243.895699 | -0.0418223 | 0.225003 | -0.1858746 | 0.8525431 | 0.97718353 |
| SMIM15 | 103.586844 | 0.14307463 | 0.32300897 | 0.4429432 | 0.65780682 | 0.95100227 |
| ZSWIM6 | 1524.22094 | -0.1257378 | 0.53528264 | -0.2348998 | 0.81428654 | 0.96820092 |
| KIF2A | 196.323017 | 0.06142351 | 0.27359708 | 0.22450355 | 0.82236551 | 0.97010249 |
| DIMT1 | 37.5814758 | 0.04275056 | 0.36192768 | 0.11811906 | 0.90597332 | 0.9875066 |
| IPO11 | 142.379832 | -0.2512094 | 0.19948942 | -1.2592617 | 0.20793581 | 0.77357731 |
| RNF180 | 391.579894 | -0.0844902 | 0.21371791 | -0.3953351 | 0.69259556 | 0.95672887 |
| SREK1IP1 | 165.727399 | -0.1634156 | 0.19185441 | -0.8517689 | 0.39434235 | 0.87253562 |
| CWC27 | 331.174505 | -0.1298247 | 0.21193133 | -0.6125791 | 0.54015465 | 0.92414041 |
| PPWD1 | 127.16546 | 0.01436559 | 0.27510124 | 0.05221929 | 0.95835397 | 0.99579587 |
| TRIM23 | 97.7649997 | 0.0693197 | 0.33804968 | 0.20505772 | 0.83752703 | 0.97369018 |
| TRAPPC13 | 159.690069 | 0.05755953 | 0.23909314 | 0.24074103 | 0.80975584 | 0.9679755 |
| NLN | 119.000731 | -0.6686447 | 0.34467396 | -1.9399338 | 0.05238774 | 0.50741541 |
| ERBIN | 1350.85021 | 0.00914309 | 0.29395809 | 0.03110337 | 0.9751871 | 0.99656026 |
| SREK1 | 393.857059 | -0.3621965 | 0.18488697 | -1.9590159 | 0.05011093 | 0.49590526 |
| MAST4 | 261.695301 | -0.5844727 | 0.41358297 | -1.4131933 | 0.15759888 | 0.71160548 |
| PIK3R1 | 160.901369 | 0.53069527 | 0.32918072 | 1.61216999 | 0.10692495 | 0.63477557 |
| SLC30A5 | 149.875868 | -0.1834482 | 0.25325943 | -0.724349 | 0.46885152 | 0.90026987 |
| MRPS36 | 143.491483 | -0.003374 | 0.28966177 | -0.0116481 | 0.99070635 | 0.99889703 |
| CDK7 | 117.321971 | -0.1233238 | 0.27891968 | -0.4421482 | 0.65838199 | 0.95113765 |
| CCDC125 | 187.280403 | -0.0758659 | 0.34042322 | -0.2228577 | 0.82364624 | 0.97018919 |
| AK6 | 87.576498 | 0.12897742 | 0.37162459 | 0.34706373 | 0.72854344 | 0.95985088 |
| RAD17 | 157.919723 | -0.08755 | 0.1865141 | -0.4694016 | 0.63878258 | 0.94889318 |
| MARVELD2 | 115.617663 | 0.04872294 | 0.26866625 | 0.18135116 | 0.85609195 | 0.97718353 |
| OCLN | 311.16926 | 0.3229471 | 0.23808599 | 1.35643051 | 0.17496223 | 0.73227004 |
| GTF2H2C | 86.5242053 | -0.037891 | 0.28259423 | -0.1340827 | 0.89333721 | 0.98597539 |
| NAIP | 281.85368 | -0.1797323 | 0.43750312 | -0.4108137 | 0.68120912 | 0.95574492 |
| GTF2H2 | 71.8766682 | -0.4182713 | 0.506724 | -0.8254421 | 0.40912065 | 0.87982122 |
| BDP1 | 515.038851 | -0.2911921 | 0.18637179 | -1.5624258 | 0.11818771 | 0.65447949 |
| MCCC2 | 237.716695 | 0.20198146 | 0.20682063 | 0.9766021 | 0.32876618 | 0.84409527 |
| MRPS27 | 216.674387 | -0.0748377 | 0.18321137 | -0.4084773 | 0.68292332 | 0.95574492 |

| | | | | | | |
|---|---|---|---|---|---|---|
| PTCD2 | 39.7281481 | 0.4070898 | 0.36541033 | 1.11406209 | 0.26525255 | 0.81163409 |
| TNPO1 | 542.806262 | 0.15741451 | 0.22728412 | 0.69258914 | 0.48856743 | 0.90730741 |
| FCHO2 | 477.621143 | -0.2138677 | 0.22128131 | -0.9664967 | 0.33379569 | 0.8445213 |
| BTF3 | 335.77621 | 0.49254183 | 0.49007839 | 1.00502663 | 0.31488403 | 0.84285425 |
| ANKRA2 | 71.3935981 | -0.1337735 | 0.25474761 | -0.5251215 | 0.59949871 | 0.93728325 |
| ARHGEF28 | 799.198378 | 0.4366055 | 0.25129943 | 1.7373915 | 0.08231809 | 0.59202053 |
| HEXB | 73.4840337 | -0.2726631 | 0.35177246 | -0.7751121 | 0.43827342 | 0.88844154 |
| GFM2 | 102.566564 | -0.0894421 | 0.23503691 | -0.3805451 | 0.70354081 | 0.95672887 |
| NSA2 | 133.558524 | 0.4314805 | 0.38365135 | 1.12466827 | 0.26072963 | 0.80805101 |
| FAM169A | 241.747857 | 0.6413383 | 0.46025588 | 1.39343858 | 0.16348721 | 0.71642393 |
| AC116337.3 | 59.8872605 | 1.78913465 | 0.48924103 | 3.65695953 | 0.00025522 | 0.03693578 |
| GCNT4 | 69.8533301 | 1.9897082 | 0.43226222 | 4.60301205 | 4.16E-06 | 0.00229691 |
| LINC01336 | 35.6918988 | 1.87872926 | 0.59611991 | 3.15159626 | 0.00162381 | 0.10129458 |
| HMGCR | 206.345229 | -0.3751561 | 0.55034071 | -0.6816796 | 0.49544154 | 0.90839719 |
| COL4A3BP | 557.541195 | -0.3593723 | 0.217432 | -1.6528031 | 0.09837095 | 0.61690569 |
| POLK | 438.708192 | -0.0329391 | 0.27164045 | -0.1212598 | 0.90348524 | 0.9875066 |
| ANKDD1B | 101.254572 | 0.42638462 | 0.46529215 | 0.91638043 | 0.3594674 | 0.85608464 |
| POC5 | 72.5895839 | -0.1557215 | 0.28110265 | -0.5539667 | 0.57960161 | 0.93345492 |
| IQGAP2 | 730.866795 | 0.593115 | 0.45511055 | 1.30323282 | 0.19249529 | 0.7597012 |
| F2RL1 | 101.534744 | 0.5940757 | 0.24542844 | 2.42056585 | 0.01549637 | 0.30047476 |
| AC008581.2 | 40.866331 | 0.22928082 | 0.45341261 | 0.50567808 | 0.61308266 | 0.94003018 |
| ZBED3 | 110.659919 | -0.3135552 | 0.31125776 | -1.0073811 | 0.31375167 | 0.84285425 |
| AC022414.1 | 47.0225109 | -0.0120836 | 0.44410884 | -0.0272087 | 0.9782933 | 0.99696835 |
| PDE8B | 53.8394532 | 0.43329313 | 0.48479022 | 0.89377448 | 0.37144257 | 0.86201082 |
| WDR41 | 252.496696 | -0.20704 | 0.23753981 | -0.8716014 | 0.38342586 | 0.86638702 |
| TBCA | 586.175268 | -0.1242678 | 0.1948943 | -0.6376164 | 0.52372339 | 0.91871245 |
| AP3B1 | 812.018946 | 0.08934687 | 0.30553375 | 0.29242879 | 0.7699588 | 0.96345263 |
| SCAMP1 | 316.272298 | -0.1060237 | 0.21605769 | -0.4907195 | 0.62362488 | 0.94457465 |
| LHFPL2 | 42.6903048 | 0.49538235 | 0.40901729 | 1.21115258 | 0.22583693 | 0.78982769 |
| ARSB | 134.642334 | 1.0229244 | 0.54994649 | 1.86004349 | 0.06287937 | 0.54732212 |
| DMGDH | 330.589434 | 0.30130928 | 0.23995176 | 1.25570774 | 0.20922195 | 0.77436433 |
| BHMT2 | 310.553533 | 0.89669077 | 0.41631346 | 2.15388369 | 0.03124929 | 0.41402352 |
| BHMT | 55.8736602 | 0.93619651 | 0.57369676 | 1.63186648 | 0.10270762 | 0.62635087 |
| JMY | 383.799582 | -0.1284207 | 0.49130199 | -0.2613886 | 0.79379284 | 0.9653598 |
| HOMER1 | 761.821788 | -0.2110923 | 0.768614 | -0.2746402 | 7.84E-01 | 0.96513523 |
| TENT2 | 824.980562 | -0.2310644 | 0.19959333 | -1.1576761 | 0.24699622 | 0.80072691 |
| CMYA5 | 398.455079 | 0.51530891 | 0.47893514 | 1.07594717 | 0.28195088 | 0.82661559 |
| MTX3 | 83.8247065 | -0.1663284 | 0.2257591 | -0.7367517 | 0.46127336 | 0.89799005 |
| THBS4 | 55.4559436 | -0.501592 | 0.54988661 | -0.9121736 | 0.36167734 | 0.85690129 |
| SERINC5 | 300.988969 | -0.2171505 | 0.23292643 | -0.9322707 | 0.35119663 | 0.85504372 |
| ZFYVE16 | 583.743633 | -0.2158111 | 0.33679704 | -0.640775 | 0.52166889 | 0.91791303 |
| AC008771.1 | 59.143821 | 0.54993835 | 0.35245684 | 1.56029986 | 0.11868904 | 0.65456964 |
| FAM151B | 89.4828319 | -0.3157076 | 0.3421755 | -0.9226482 | 0.35619057 | 0.8557799 |
| DHFR | 1121.93966 | -0.1409103 | 0.46538412 | -0.3027829 | 0.76205533 | 0.96218738 |
| MSH3 | 466.325958 | -0.0099681 | 0.23571937 | -0.0422881 | 0.96626901 | 0.99637506 |
| RASGRF2 | 367.512119 | 0.0218661 | 0.40235119 | 0.05434581 | 0.95665966 | 0.99565888 |
| CKMT2-AS1 | 264.418514 | -0.2475149 | 0.27712131 | -0.8931644 | 0.37176915 | 0.86201082 |
| CKMT2 | 178.374119 | -0.5240759 | 0.39917832 | -1.3128867 | 0.18922112 | 0.75619145 |
| ZCCHC9 | 112.00006 | 0.22524526 | 0.22336925 | 1.00839871 | 0.31326309 | 0.84285425 |
| ACOT12 | 64.1637569 | 1.35733654 | 0.57994611 | 2.3404529 | 0.01926037 | 0.33290134 |
| SSBP2 | 1367.08197 | -0.1085565 | 0.2881479 | -0.3767387 | 0.70636781 | 0.95730303 |
| ATG10 | 654.325871 | -0.0723885 | 0.23233571 | -0.3115687 | 0.75536835 | 0.96120879 |
| RPS23 | 1251.57151 | 0.08879267 | 0.50702899 | 0.17512346 | 0.86098262 | 0.97841862 |
| ATP6AP1L | 150.304854 | 0.1947982 | 0.53172255 | 0.36635309 | 0.71410162 | 0.95791799 |

| | | | | | | |
|---|---|---|---|---|---|---|
| TMEM167A | 152.386879 | -0.282569 | 0.28051431 | -1.0073247 | 0.31377876 | 0.84285425 |
| XRCC4 | 192.251052 | -0.3250895 | 0.2385096 | -1.3630038 | 0.17288131 | 0.72940007 |
| COX7C | 569.048625 | 0.46036384 | 0.35515429 | 1.29623618 | 0.19489413 | 0.76233115 |
| RASA1 | 593.042915 | -0.1328216 | 0.17241825 | -0.7703453 | 0.44109508 | 0.88964371 |
| CCNH | 698.231323 | 0.01312134 | 0.43701116 | 0.03002519 | 0.97604696 | 0.99656026 |
| TMEM161B | 224.217542 | 0.09492563 | 0.21860045 | 0.4342426 | 0.66411228 | 0.95195406 |
| TMEM161B-/ | 850.223817 | 0.06890381 | 0.2857541 | 0.24112975 | 0.80945456 | 0.9679755 |
| LINC00461 | 68.558947 | 0.87972929 | 0.66344113 | 1.32600957 | 0.18483653 | 0.7484938 |
| CETN3 | 50.9742601 | 0.12598794 | 0.33096951 | 0.38066328 | 0.70345312 | 0.95672887 |
| LYSMD3 | 59.3764514 | 0.15679935 | 0.4464002 | 0.35125287 | 0.72539865 | 0.95963742 |
| ARRDC3 | 539.076436 | -1.1072038 | 0.62153792 | -1.7813937 | 0.07484815 | 0.57072562 |
| ARRDC3-AS1 | 38.7845257 | 0.08139025 | 0.35613742 | 0.22853609 | 0.8192295 | 0.96884581 |
| FAM172A | 1041.3053 | -0.1257398 | 0.1617806 | -0.777224 | 0.43702664 | 0.88785137 |
| KIAA0825 | 424.667758 | -0.2659955 | 0.18989309 | -1.4007646 | 0.16128448 | 0.71409437 |
| SLF1 | 59.6036244 | -0.4200908 | 0.3390961 | -1.2388547 | 0.21539931 | 0.77894575 |
| MCTP1 | 530.13326 | -0.6915928 | 0.32275088 | -2.142807 | 0.0321286 | 0.41517597 |
| FAM81B | 47.1009579 | -0.1053664 | 0.59052702 | -0.1784277 | 0.85838707 | 0.97786699 |
| TTC37 | 490.490595 | -0.4721576 | 0.2202471 | -2.1437629 | 0.03205189 | 0.41517597 |
| ARSK | 85.9984463 | -0.0781659 | 0.3166977 | -0.2468155 | 0.80505103 | 0.96704419 |
| RHOBTB3 | 1165.51151 | 0.05341535 | 0.37840191 | 0.14116037 | 0.88774325 | 0.98551241 |
| GLRX | 269.640375 | 0.32459152 | 0.52827348 | 0.61443842 | 0.53892566 | 0.92365631 |
| ELL2 | 627.329785 | -1.3714759 | 0.74965431 | -1.8294778 | 0.06732807 | 0.55848603 |
| AC104123.1 | 240.885661 | -0.1830884 | 0.27049691 | -0.6768593 | 0.49849522 | 0.90839719 |
| CAST | 695.151262 | -0.1382769 | 0.19367043 | -0.7139806 | 0.47523916 | 0.90242252 |
| ERAP1 | 144.719949 | -0.2874946 | 0.31859604 | -0.9023798 | 0.36685516 | 0.85902833 |
| AC009126.1 | 58.2558605 | -0.029521 | 0.34108536 | -0.0865502 | 0.93102907 | 0.99243654 |
| ERAP2 | 93.659909 | 0.60803932 | 0.47591362 | 1.27762538 | 0.20138155 | 0.76957251 |
| LNPEP | 452.03445 | -0.0392326 | 0.16677317 | -0.2352452 | 0.81401841 | 0.96820092 |
| LIX1-AS1 | 88.8310066 | 0.39696154 | 0.37243014 | 1.06586845 | 0.28648311 | 0.82937652 |
| RIOK2 | 42.2588186 | 0.463623 | 0.39365278 | 1.17774603 | 0.23889787 | 0.79912215 |
| RGMB | 57.3118025 | -0.3698089 | 0.33804795 | -1.0939538 | 0.27397523 | 0.81986798 |
| CHD1 | 491.517575 | -0.2338526 | 0.40713665 | -0.5743836 | 0.56570823 | 0.93114846 |
| LINC02062 | 127.141074 | 0.85368985 | 0.47642808 | 1.7918546 | 0.07315626 | 0.56668778 |
| FAM174A | 79.1478865 | 0.27822851 | 0.34696602 | 0.80188981 | 0.4226167 | 0.88371155 |
| SLCO4C1 | 128.59842 | -0.2162824 | 0.33853854 | -0.6388708 | 5.23E-01 | 0.91854926 |
| PAM | 932.220981 | 0.03622581 | 0.34043927 | 0.10640902 | 0.91525783 | 0.98868699 |
| GIN1 | 55.5843912 | 0.30488582 | 0.26625221 | 1.14510153 | 0.2521671 | 0.8029001 |
| PPIP5K2 | 331.575207 | -0.1710608 | 0.17074433 | -1.0018536 | 0.31641432 | 0.84285425 |
| NUDT12 | 75.838246 | 0.06292418 | 0.31660898 | 0.19874414 | 0.84246289 | 0.97560439 |
| EFNA5 | 2275.27865 | -0.3163033 | 0.41383576 | -0.7643208 | 0.44467608 | 0.89105265 |
| FBXL17 | 1532.21245 | 0.07437878 | 0.21747473 | 0.34201114 | 0.73234251 | 0.96008876 |
| FER | 1638.80967 | -0.4219218 | 0.26678243 | -1.58152 | 0.11375918 | 0.6483668 |
| PJA2 | 474.953875 | 0.07616967 | 0.28539334 | 0.26689363 | 0.78955106 | 0.96515649 |
| MAN2A1 | 736.702288 | 0.42379545 | 0.23451987 | 1.80707693 | 0.07075029 | 0.56256676 |
| TMEM232 | 481.129651 | 0.01914577 | 0.42794663 | 0.04473869 | 0.96431559 | 0.99637506 |
| SLC25A46 | 198.050623 | 0.01398378 | 0.21096596 | 0.06628453 | 0.9471513 | 0.99314907 |
| WDR36 | 124.339891 | -0.0066792 | 0.21309696 | -0.0313435 | 0.97499557 | 0.99656026 |
| NREP | 114.259401 | -1.103767 | 0.39274482 | -2.8103922 | 0.00494812 | 0.17460017 |
| EPB41L4A | 975.536384 | 0.21931783 | 0.28487383 | 0.76987708 | 0.44137281 | 0.88964371 |
| EPB41L4A-A! | 36.1676445 | -0.113657 | 0.32910327 | -0.3453537 | 0.72982852 | 0.96008876 |
| APC | 205.787239 | -0.1753258 | 0.26197635 | -0.669243 | 0.50334046 | 0.90964098 |
| SRP19 | 111.828468 | 0.13120839 | 0.32039659 | 0.40951867 | 0.68215907 | 0.95574492 |
| REEP5 | 539.944904 | 0.2952791 | 0.24971437 | 1.18246739 | 0.2370203 | 0.79797745 |
| DCP2 | 237.285354 | -0.0375031 | 0.16341951 | -0.2294895 | 0.81848847 | 0.96883031 |

| | | | | | | |
|---|---|---|---|---|---|---|
| MCC | 509.460163 | 0.77712265 | 0.42790764 | 1.81609904 | 0.06935518 | 0.55917298 |
| YTHDC2 | 366.508836 | 0.06062618 | 0.171509 | 0.35348687 | 7.24E-01 | 0.95963742 |
| PGGT1B | 149.180905 | -0.1929288 | 0.17616182 | -1.0951795 | 0.27343801 | 0.81922962 |
| CCDC112 | 65.3270599 | -0.3765625 | 0.31676235 | -1.1887854 | 0.23452412 | 0.79543722 |
| FEM1C | 94.1687324 | -0.679642 | 0.50363241 | -1.3494802 | 0.17718276 | 0.73633806 |
| TMED7-TICA | 57.7394586 | 0.04965055 | 0.27847638 | 0.17829357 | 0.85849243 | 0.97786699 |
| ATG12 | 128.618248 | -0.1946871 | 0.22376889 | -0.8700367 | 0.38428036 | 0.86725322 |
| AP3S1 | 256.065126 | -0.2407392 | 0.30486495 | -0.7896585 | 0.42972722 | 0.88579433 |
| COMMD10 | 739.98867 | -0.5259422 | 0.17717365 | -2.9685124 | 0.00299245 | 0.14126521 |
| SEMA6A | 384.846099 | -0.4678815 | 0.8305058 | -0.5633693 | 0.57318343 | 0.93258748 |
| SEMA6A-AS1 | 123.795287 | -0.8007883 | 0.44955457 | -1.7812927 | 0.07486465 | 0.57072562 |
| DTWD2 | 221.480574 | -0.2262369 | 0.21708571 | -1.0421545 | 0.29734006 | 0.83556453 |
| AC008629.1 | 97.5471402 | 0.37668153 | 0.61066904 | 0.61683417 | 0.53734411 | 0.92324352 |
| DMXL1 | 759.617852 | -0.0525367 | 0.20411351 | -0.2573894 | 0.79687818 | 0.9653598 |
| TNFAIP8 | 216.212087 | -0.7474685 | 0.36064258 | -2.0726019 | 0.03820934 | 0.44791264 |
| HSD17B4 | 375.126151 | 0.25220256 | 0.17692138 | 1.4255064 | 0.15401087 | 0.7079031 |
| SRFBP1 | 127.531404 | -0.2614082 | 0.19058673 | -1.3715971 | 0.17018891 | 0.72675731 |
| SNX2 | 599.140185 | -0.1399845 | 0.42332437 | -0.3306791 | 0.74088691 | 0.96084411 |
| SNX24 | 532.280157 | 0.05648557 | 0.31512196 | 0.17924987 | 0.85774151 | 0.97750446 |
| PPIC | 132.837067 | 0.34349432 | 0.39952198 | 0.85976324 | 0.38991957 | 0.87004477 |
| CEP120 | 154.146755 | -0.0855278 | 0.28195074 | -0.3033431 | 0.76162843 | 0.96218738 |
| CSNK1G3 | 345.340819 | 0.08192937 | 0.23521506 | 0.34831685 | 0.72760224 | 0.9598317 |
| ZNF608 | 566.511827 | 0.8277888 | 0.5444956 | 1.52028557 | 0.12843922 | 0.67206108 |
| GRAMD2B | 1006.19448 | -0.1852456 | 0.1892468 | -0.978857 | 0.32765063 | 0.84381657 |
| AC093535.1 | 56.9275236 | 0.15071178 | 0.56779932 | 0.26543143 | 0.79067713 | 0.96527296 |
| ALDH7A1 | 153.639939 | 0.14674739 | 0.31145334 | 0.47116976 | 0.63751951 | 0.94849581 |
| PHAX | 86.0119446 | 0.24457396 | 0.2834123 | 0.86296171 | 0.38815852 | 0.86953359 |
| C5orf63 | 68.5932445 | 0.15617959 | 0.29521526 | 0.52903631 | 0.59678026 | 0.93617523 |
| PRRC1 | 147.40806 | -0.2728881 | 0.250582 | -1.0890171 | 0.27614636 | 0.82167981 |
| LINC01184 | 301.815539 | 0.06946744 | 0.33986531 | 0.20439696 | 0.83804331 | 0.97385715 |
| SLC12A2 | 96.5206685 | 0.04142218 | 0.3075065 | 0.13470342 | 0.89284637 | 0.98597539 |
| SLC27A6 | 53.2478693 | -1.4732181 | 1.55523538 | -0.9472637 | 0.34350441 | 0.85126545 |
| ISOC1 | 44.6297846 | 0.67133808 | 0.37326289 | 1.79856638 | 0.0720873 | 0.5644261 |
| HINT1 | 812.973088 | 0.43452112 | 0.47741342 | 0.91015689 | 0.36273977 | 0.85694172 |
| LYRM7 | 85.5598126 | 0.3506278 | 0.2054481 | 1.70664905 | 0.0878873 | 0.60651125 |
| CDC42SE2 | 977.958626 | -0.1080024 | 0.38694707 | -0.2791142 | 0.78015722 | 0.96513523 |
| P4HA2 | 50.4581791 | -0.2745943 | 0.40316788 | -0.6810916 | 0.49581354 | 0.90839719 |
| SLC22A4 | 55.9763908 | 0.41641158 | 0.48870742 | 0.85206722 | 0.39417678 | 0.87253562 |
| SLC22A5 | 224.476069 | -0.087923 | 0.33449986 | -0.2628492 | 0.79266685 | 0.9653598 |
| AC116366.3 | 283.129129 | -0.1767065 | 0.49446714 | -0.3573675 | 0.72081673 | 0.95828483 |
| IRF1 | 71.7915026 | -0.4059071 | 0.39042958 | -1.0396423 | 0.2985061 | 0.83556453 |
| AC004039.1 | 44.8221869 | 0.60715397 | 0.65978924 | 0.92022411 | 0.35745566 | 0.8557799 |
| KIF3A | 251.864726 | 0.21594503 | 0.2151798 | 1.00355625 | 0.31559255 | 0.84285425 |
| 8-Sep | 148.509762 | 0.38692396 | 0.29689703 | 1.3032261 | 0.19249758 | 0.7597012 |
| SOWAHA | 34.6776175 | 0.82267814 | 0.56637546 | 1.45253139 | 0.14635391 | 0.6955959 |
| SHROOM1 | 94.1288329 | -0.4231344 | 0.5562717 | -0.7606613 | 0.44685939 | 0.8918683 |
| UQCRQ | 277.368064 | 0.15485291 | 0.39222627 | 0.39480503 | 0.69298678 | 0.95672887 |
| AFF4 | 959.983481 | -0.533567 | 0.28085271 | -1.8998107 | 0.05745796 | 0.52728498 |
| ZCCHC10 | 93.1220596 | -0.0042176 | 0.24320664 | -0.0173416 | 0.98616409 | 0.99768314 |
| HSPA4 | 231.318808 | -0.0512496 | 0.26802896 | -0.1912093 | 0.84836165 | 0.97656375 |
| FSTL4 | 221.492103 | 0.31370171 | 0.94978822 | 0.33028595 | 0.7411839 | 0.96084411 |
| C5orf15 | 278.918483 | 0.39564513 | 0.31869592 | 1.24145025 | 0.21443947 | 0.77890374 |
| VDAC1 | 381.886915 | 0.43395318 | 0.29398685 | 1.47609724 | 0.13991779 | 0.68408298 |
| SKP1 | 739.19565 | 0.66165196 | 0.45214098 | 1.46337534 | 0.1433647 | 0.69023197 |

| | | | | | | |
|---|---|---|---|---|---|---|
| CDKL3 | 124.274474 | -0.1318928 | 0.32328722 | -0.407974 | 0.68329278 | 0.95574492 |
| UBE2B | 362.61157 | -0.5642709 | 0.26612824 | -2.1202971 | 0.033981 | 0.43009768 |
| JADE2 | 127.788469 | 0.13360606 | 0.28160189 | 0.47445015 | 0.63517894 | 0.94792975 |
| SAR1B | 164.296992 | -0.1446463 | 0.21729809 | -0.6656585 | 0.50562939 | 0.91126329 |
| SEC24A | 170.112393 | -0.7439289 | 0.39640231 | -1.8767017 | 0.06055898 | 0.54068346 |
| CAMLG | 112.087655 | 0.52886963 | 0.30999232 | 1.70607334 | 0.08799443 | 0.60651125 |
| DDX46 | 342.612566 | -0.0757621 | 0.2672958 | -0.2834393 | 0.7768401 | 0.96454202 |
| C5orf24 | 275.271637 | -0.0951765 | 0.46202883 | -0.2059969 | 0.83679334 | 0.97353666 |
| TXNDC15 | 103.135501 | 0.21270428 | 0.20567902 | 1.0341564 | 0.30106305 | 0.83709431 |
| PCBD2 | 264.061858 | 0.02555331 | 0.35185396 | 0.07262477 | 0.94210471 | 0.99265578 |
| H2AFY | 255.467696 | -0.1783393 | 0.23415265 | -0.7616368 | 0.44627678 | 0.8918683 |
| CXCL14 | 48.6430069 | 0.80769735 | 0.71435739 | 1.13066283 | 0.25819703 | 0.80709747 |
| SMAD5 | 293.40913 | 0.16835383 | 0.14990451 | 1.12307381 | 0.26140615 | 0.80805101 |
| KLHL3 | 140.836433 | 0.40203539 | 0.41689037 | 0.96436719 | 0.33486187 | 0.84521974 |
| HNRNPA0 | 212.535823 | 0.17979859 | 0.34174 | 0.52612683 | 0.59880009 | 0.93728325 |
| AC106791.1 | 109.388332 | -0.07618 | 0.44290556 | -0.1720005 | 0.86343714 | 0.979268 |
| FAM13B | 660.363108 | -0.2282634 | 0.18058209 | -1.2640423 | 0.20621486 | 0.77193956 |
| NME5 | 150.876114 | 0.04161621 | 0.34944298 | 0.11909299 | 0.90520168 | 0.9875066 |
| BRD8 | 132.970903 | -0.0921009 | 0.22984895 | -0.4007019 | 0.68863965 | 0.95604695 |
| CDC23 | 71.6610368 | -0.07584 | 0.21751646 | -0.3486632 | 0.72734216 | 0.95979351 |
| FAM53C | 60.1221428 | -0.1202598 | 0.31567635 | -0.3809593 | 0.70323345 | 0.95672887 |
| KDM3B | 314.31743 | -0.1129346 | 0.16349534 | -0.6907512 | 0.48972194 | 0.90730741 |
| EGR1 | 373.69177 | -3.535938 | 1.59004368 | -2.2237993 | 0.02616195 | 0.39168177 |
| ETF1 | 264.927265 | -0.2427963 | 0.33499591 | -0.7247739 | 0.46859075 | 0.90026987 |
| HSPA9 | 429.349523 | -0.379401 | 0.31367874 | -1.2095208 | 0.22646284 | 0.78982769 |
| CTNNA1 | 1378.55555 | -0.2588991 | 0.24306557 | -1.0651412 | 0.28681205 | 0.82943317 |
| SIL1 | 485.44343 | 0.08282501 | 0.2410024 | 0.3436688 | 0.73109538 | 0.96008876 |
| PAIP2 | 249.971415 | 0.12371718 | 0.18987655 | 0.65156641 | 0.51468092 | 0.91557155 |
| TMEM173 | 32.3125504 | 0.96819909 | 0.40389568 | 2.39715138 | 0.0165231 | 0.30723893 |
| UBE2D2 | 763.103175 | -0.0726519 | 0.18015874 | -0.403266 | 0.68675258 | 0.9558749 |
| CXXC5 | 517.494716 | 0.18140542 | 0.29185026 | 0.62157017 | 0.53422454 | 0.92303275 |
| PURA | 383.40347 | 0.34760662 | 0.17476168 | 1.98903223 | 0.04669765 | 0.48600281 |
| IGIP | 66.7335846 | 0.47869912 | 0.33246759 | 1.43983695 | 0.14991353 | 0.7029344 |
| CYSTM1 | 1545.39421 | -0.2156481 | 0.5513246 | -0.3911455 | 0.69568972 | 0.95672887 |
| PFDN1 | 175.362367 | -0.3967265 | 0.26773313 | -1.4817981 | 0.13839402 | 0.68146162 |
| ANKHD1-EIF4 | 41.0769349 | -0.1165322 | 0.42419143 | -0.2747159 | 0.78353448 | 0.96513523 |
| SRA1 | 90.3665465 | 0.12965007 | 0.37788669 | 0.34309246 | 0.7315289 | 0.96008876 |
| SLC35A4 | 91.4828925 | 0.11166884 | 0.32767089 | 0.34079572 | 0.73325737 | 0.96035038 |
| NDUFA2 | 198.290395 | 0.2592049 | 0.3563426 | 0.72740362 | 0.46697874 | 0.8989598 |
| IK | 87.3881852 | -0.0041599 | 0.31149984 | -0.0133543 | 0.98934515 | 0.99835647 |
| WDR55 | 54.079677 | 0.11348221 | 0.43469881 | 0.2610594 | 0.7940467 | 0.9653598 |
| HARS | 96.3219174 | -0.0749608 | 0.19154981 | -0.3913385 | 0.69554701 | 0.95672887 |
| ZMAT2 | 94.3564211 | 0.11759595 | 0.44413938 | 0.26477263 | 0.79118462 | 0.9653598 |
| TAF7 | 128.011653 | 0.49204953 | 0.3245508 | 1.51609401 | 0.12949558 | 0.67207756 |
| DIAPH1 | 613.369959 | -0.229252 | 0.2554738 | -0.8973602 | 0.36952675 | 0.86020443 |
| HDAC3 | 106.849428 | -0.1481747 | 0.38065432 | -0.3892632 | 0.69708144 | 0.95672887 |
| PCDH1 | 143.00079 | 0.87569944 | 0.52947508 | 1.65390114 | 0.0981476 | 0.61690569 |
| DELE1 | 102.719067 | -0.0536136 | 0.3366075 | -0.1592763 | 0.87345117 | 0.98246494 |
| RNF14 | 109.046946 | -0.2477755 | 0.28178913 | -0.8792941 | 0.37924183 | 0.86565578 |
| GNPDA1 | 96.7290686 | -0.5227834 | 0.33307542 | -1.5695646 | 0.11651644 | 0.65257994 |
| NDFIP1 | 633.019973 | -0.1462185 | 0.16631199 | -0.879182 | 0.37930261 | 0.86565578 |
| SPRY4-AS1 | 539.769373 | 0.66280041 | 0.45290939 | 1.46342828 | 0.14335022 | 0.69023197 |
| FGF1 | 96.6421623 | 1.17819075 | 0.6076179 | 1.93903233 | 0.0524974 | 0.5080081 |
| ARHGAP26 | 123.606041 | -0.2814648 | 0.3624548 | -0.7765514 | 0.43742349 | 0.88806628 |

| | | | | | | |
|---|---|---|---|---|---|---|
| YIPF5 | 71.8699603 | -0.2219034 | 0.22001882 | -1.0085657 | 0.31318298 | 0.84285425 |
| PRELID2 | 346.662282 | -0.7466251 | 0.46298236 | -1.6126427 | 0.10682215 | 0.63477557 |
| LARS | 286.042147 | 0.11341676 | 0.21099795 | 0.5375254 | 0.59090475 | 0.93483536 |
| TCERG1 | 335.161974 | -0.3883635 | 0.17604284 | -2.2060738 | 0.02737883 | 0.40165272 |
| PPP2R2B | 130.208708 | 2.28149139 | 0.67467688 | 3.38160601 | 0.00072063 | 0.06190366 |
| STK32A | 41.4056512 | 1.39372281 | 0.83625904 | 1.66661615 | 0.09559076 | 0.61367241 |
| AC091948.1 | 84.0131596 | 0.41249288 | 0.45034974 | 0.91593899 | 0.3596989 | 0.85608464 |
| FBXO38 | 157.08987 | -0.1081318 | 0.18891915 | -0.5723706 | 0.56707093 | 0.93122667 |
| CSNK1A1 | 1293.17334 | -0.2239914 | 0.18986463 | -1.1797426 | 0.23810262 | 0.79810098 |
| ARHGEF37 | 77.7290334 | 0.26557355 | 0.25667515 | 1.03466796 | 0.30082401 | 0.83709431 |
| SLC26A2 | 56.8813341 | 0.67379455 | 0.33909919 | 1.98701316 | 0.04692094 | 0.48600281 |
| HMGXB3 | 130.09529 | -0.6084733 | 0.19839983 | -3.0669044 | 0.00216288 | 0.11990273 |
| TCOF1 | 59.0632145 | -0.3631871 | 0.29550767 | -1.2290278 | 0.21906139 | 0.78597171 |
| CD74 | 813.710538 | -0.231184 | 0.6376581 | -0.3625516 | 0.71693987 | 0.95791799 |
| RPS14 | 969.174859 | 0.11197055 | 0.46405533 | 0.24128707 | 0.80933263 | 0.9679755 |
| NDST1 | 127.392559 | 0.13931949 | 0.24286567 | 0.57364836 | 0.56620578 | 0.93122667 |
| SYNPO | 45.2807751 | 0.08472089 | 0.43198859 | 0.19611835 | 0.84451754 | 0.97573093 |
| RBM22 | 57.8728608 | 0.04125639 | 0.28456956 | 0.14497823 | 0.88472806 | 0.98438795 |
| DCTN4 | 308.626978 | -0.3192351 | 0.23232596 | -1.3740828 | 0.16941597 | 0.72596839 |
| GPX3 | 677.603539 | -0.5582919 | 0.75817644 | -0.7363615 | 0.46151073 | 0.89799005 |
| TNIP1 | 248.88547 | -0.2627859 | 0.26319643 | -0.99844 | 3.18E-01 | 0.84285425 |
| ANXA6 | 76.6015023 | 0.83231125 | 0.44117353 | 1.88658473 | 0.0592162 | 0.53314931 |
| ATOX1 | 186.403633 | 0.03319192 | 0.22595454 | 0.14689644 | 0.88321376 | 0.9835847 |
| G3BP1 | 371.573631 | 0.07430947 | 0.25520641 | 0.291174 | 0.77091824 | 0.96387774 |
| FAM114A2 | 87.6579823 | -0.6468695 | 0.22910093 | -2.8235131 | 0.00475005 | 0.16989933 |
| MFAP3 | 106.900829 | -0.167068 | 0.25475833 | -0.6557903 | 0.51195903 | 0.91408994 |
| GALNT10 | 125.924919 | -0.0105802 | 0.33503497 | -0.0315794 | 0.97480744 | 0.99656026 |
| SAP30L-AS1 | 57.1879863 | 0.12178336 | 0.31945924 | 0.38121723 | 0.70304207 | 0.95672887 |
| SAP30L | 51.1973561 | -0.0018453 | 0.51129091 | -0.0036091 | 0.99712039 | 0.99967893 |
| LARP1 | 503.506834 | -0.3784608 | 0.23451871 | -1.6137765 | 0.10657592 | 0.63461115 |
| FAXDC2 | 96.8825087 | 0.21887183 | 0.42801006 | 0.51137076 | 0.60909147 | 0.94003018 |
| CNOT8 | 108.826917 | -0.1552744 | 0.19802153 | -0.7841289 | 0.4329645 | 0.88722778 |
| GEMIN5 | 66.7927835 | -0.1100998 | 0.34505191 | -0.319082 | 0.74966435 | 0.96109466 |
| MRPL22 | 96.9475527 | 0.10435444 | 0.23471334 | 0.44460378 | 0.65660612 | 0.95100227 |
| SGCD | 401.108047 | 1.54503613 | 0.87944821 | 1.75682446 | 0.07894773 | 0.5826987 |
| HAVCR2 | 37.1204823 | -0.2772952 | 0.32584549 | -0.8510022 | 0.39476815 | 0.87300489 |
| CYFIP2 | 1371.94276 | -0.0354497 | 0.40630059 | -0.08725 | 0.93047283 | 0.99231329 |
| ADAM19 | 64.5273458 | 0.35838805 | 0.4013655 | 0.89292192 | 0.37189899 | 0.86201082 |
| CLINT1 | 751.015732 | 0.16634358 | 0.15794622 | 1.05316597 | 0.29226493 | 0.83322538 |
| RNF145 | 224.385156 | 0.1307465 | 0.24451117 | 0.53472609 | 0.59283927 | 0.93491607 |
| UBLCP1 | 80.3794031 | -0.0056901 | 0.31889644 | -0.0178431 | 0.98576405 | 0.99762072 |
| TTC1 | 214.622283 | -0.180376 | 0.17451775 | -1.033568 | 0.30133815 | 0.83709431 |
| PWWP2A | 298.711072 | 0.35075486 | 0.35319737 | 0.99308458 | 0.32066874 | 0.84285425 |
| SLU7 | 70.3174487 | -0.1930897 | 0.29335577 | -0.6582099 | 0.51040326 | 0.91334187 |
| AC113414.1 | 252.221605 | 0.42288874 | 0.4576701 | 0.92400343 | 0.35548453 | 0.8557799 |
| CCNG1 | 143.539962 | 0.49921318 | 0.32115286 | 1.55444102 | 0.12007925 | 0.65679551 |
| NUDCD2 | 73.4275509 | 0.46749223 | 0.45190535 | 1.03449147 | 0.30090646 | 0.83709431 |
| MAT2B | 168.29816 | 0.19360432 | 0.28742155 | 0.67359014 | 0.50057192 | 0.90839719 |
| WWC1 | 1874.09253 | 0.3936176 | 0.36720705 | 1.07192276 | 0.28375472 | 0.82696038 |
| RARS | 146.619851 | -0.2390509 | 0.23848574 | -1.0023699 | 0.316165 | 0.84285425 |
| PANK3 | 226.521644 | -0.2908472 | 0.27796473 | -1.0463456 | 0.29540151 | 0.83396494 |
| KCNIP1 | 93.5476576 | 0.88043191 | 0.96411056 | 0.91320637 | 0.36113401 | 0.8565748 |
| RANBP17 | 332.182904 | 0.25724544 | 0.36998888 | 0.69527885 | 0.48688056 | 0.9070379 |
| NPM1 | 316.660153 | 0.28442941 | 0.32329469 | 0.87978372 | 0.37897649 | 0.86528836 |

| | | | | | | |
|---|---|---|---|---|---|---|
| FBXW11 | 745.233665 | 0.22414078 | 0.24423836 | 0.91771325 | 0.35876901 | 0.8557799 |
| STK10 | 57.5671873 | 0.13597008 | 0.32040274 | 0.42437242 | 0.67129423 | 0.95430867 |
| UBTD2 | 165.304881 | 0.07089699 | 0.21902178 | 0.32369834 | 0.74616643 | 0.96109466 |
| SH3PXD2B | 44.3193452 | -0.9890642 | 0.47210555 | -2.0950065 | 0.03617042 | 0.4362094 |
| AC022217.3 | 56.5086369 | -0.8442343 | 0.74012266 | -1.1406681 | 0.25400805 | 0.80414079 |
| DUSP1 | 1337.87601 | -2.068897 | 0.76765562 | -2.6950848 | 0.00703707 | 0.20555027 |
| ERGIC1 | 573.973789 | -0.4492519 | 0.31921683 | -1.4073565 | 0.15932172 | 0.71409437 |
| RPL26L1 | 51.2118544 | 0.05029038 | 0.37971708 | 0.13244172 | 0.89463492 | 0.98597539 |
| ATP6V0E1 | 1072.08163 | -0.0794383 | 0.22768613 | -0.3488937 | 0.72716911 | 0.95979351 |
| CREBRF | 555.365394 | -0.296595 | 0.22187136 | -1.3367881 | 0.18129181 | 0.74315969 |
| BNIP1 | 38.3632377 | -0.1703143 | 0.36563025 | -0.4658101 | 0.64135141 | 0.94935885 |
| BOD1 | 76.5417916 | 0.15645659 | 0.3876365 | 0.40361677 | 0.68649455 | 0.9558749 |
| CPEB4 | 562.187342 | 0.38649329 | 0.31800338 | 1.21537478 | 0.22422318 | 0.78960314 |
| SFXN1 | 374.716465 | -0.1186576 | 0.21592265 | -0.5495377 | 0.5826365 | 0.93407228 |
| THOC3 | 171.378221 | -0.5522018 | 0.43308946 | -1.2750293 | 0.20229886 | 0.76996624 |
| SIMC1 | 270.54088 | -0.2002002 | 0.30174832 | -0.6634674 | 0.50703122 | 0.91163608 |
| KIAA1191 | 183.651761 | 0.12108999 | 0.26015937 | 0.46544544 | 0.6416125 | 0.94935885 |
| ARL10 | 63.1685869 | 0.29666512 | 0.42243973 | 0.70226613 | 0.48251321 | 0.90614725 |
| CLTB | 129.976575 | -0.0559995 | 0.41832831 | -0.1338649 | 0.8935094 | 0.98597539 |
| FAF2 | 242.751292 | -0.4559263 | 0.16359937 | -2.7868461 | 0.00532237 | 0.1799306 |
| RNF44 | 80.3423746 | 0.23954272 | 0.22636784 | 1.05820117 | 0.28996374 | 0.83103268 |
| TSPAN17 | 53.1442474 | -0.3721598 | 0.32622841 | -1.1407952 | 0.25395518 | 0.80414079 |
| UIMC1 | 313.931168 | -0.0084422 | 0.22122528 | -0.038161 | 0.96955928 | 0.99645289 |
| ZNF346 | 150.466877 | -0.2467443 | 0.29691234 | -0.8310341 | 0.40595438 | 0.87937203 |
| NSD1 | 807.232616 | 0.17780755 | 0.30491766 | 0.58313301 | 0.55980376 | 0.92934517 |
| PRELID1 | 149.0939 | 0.03030477 | 0.53213855 | 0.05694902 | 0.95458581 | 0.99537617 |
| LMAN2 | 221.152608 | 0.07715102 | 0.27491322 | 0.28063772 | 0.77898828 | 0.96512745 |
| DBN1 | 93.0908528 | -0.351435 | 0.35498146 | -0.9900094 | 0.32216954 | 0.84285425 |
| DDX41 | 42.8285263 | -0.1179373 | 0.32373407 | -0.3643032 | 0.71563163 | 0.95791799 |
| FAM193B | 163.961177 | -0.2357643 | 0.28127556 | -0.8381969 | 0.40192012 | 0.87834687 |
| TMED9 | 226.823506 | 0.2246953 | 0.40502559 | 0.55476817 | 0.57905324 | 0.93345492 |
| B4GALT7 | 73.4281462 | 0.12158793 | 0.31712898 | 0.38340214 | 0.70142162 | 0.95672887 |
| FAM153A | 56.8931363 | -1.1293645 | 1.0063317 | -1.1222587 | 0.26175246 | 0.80847797 |
| AC106795.3 | 63.4020082 | -0.1915728 | 0.37679792 | -0.5084233 | 0.61115651 | 0.94003018 |
| RMND5B | 118.986654 | -0.0849864 | 0.28323758 | -0.3000535 | 0.76413631 | 0.96285313 |
| NHP2 | 117.257457 | 0.14607214 | 0.47218557 | 0.30935324 | 0.75705283 | 0.96133693 |
| HNRNPAB | 149.164159 | -0.2639427 | 0.33125498 | -0.796796 | 0.42556949 | 0.88403732 |
| PHYKPL | 304.830584 | 0.21963467 | 0.30481227 | 0.72055718 | 0.471182 | 0.90123338 |
| CLK4 | 118.992443 | 0.10160407 | 0.25076269 | 0.40518016 | 0.68534509 | 0.9558749 |
| ZNF354A | 74.8982148 | 0.47752923 | 0.31004516 | 1.54019248 | 0.12351344 | 0.66519057 |
| ZNF354B | 47.2548949 | -0.1786313 | 0.35237539 | -0.5069347 | 0.61220063 | 0.94003018 |
| ZFP2 | 53.6604667 | 0.93561379 | 0.3609029 | 2.59242526 | 0.00953019 | 0.23684368 |
| ZNF354C | 48.5218594 | -0.0303126 | 0.32286088 | -0.0938873 | 0.92519865 | 0.99051393 |
| RUFY1 | 242.755145 | -0.0831288 | 0.23397186 | -0.3552939 | 0.72236942 | 0.95913233 |
| HNRNPH1 | 718.5327 | -0.3654076 | 0.19443624 | -1.8793184 | 0.06020103 | 0.53892231 |
| CANX | 730.302777 | 0.26070658 | 0.19709209 | 1.32276531 | 0.18591344 | 0.74914489 |
| MAML1 | 87.0218969 | -0.018272 | 0.21470715 | -0.0851021 | 0.93218025 | 0.99243654 |
| MGAT4B | 173.484464 | -1.2370631 | 0.42235962 | -2.9289332 | 0.00340128 | 0.14358074 |
| SQSTM1 | 1162.24988 | -1.1858428 | 0.57057559 | -2.0783272 | 0.03767923 | 0.44445707 |
| MRNIP | 133.450156 | -0.081079 | 0.2786967 | -0.290922 | 0.77111101 | 0.96388926 |
| TBC1D9B | 169.313637 | -0.176152 | 0.20161845 | -0.8736897 | 0.38228726 | 0.86593382 |
| RNF130 | 462.338562 | -0.1768832 | 0.33821217 | -0.5229947 | 0.60097796 | 0.93795727 |
| MAPK9 | 193.712571 | 0.11219204 | 0.18455313 | 0.6079119 | 0.5432459 | 0.92482794 |
| GFPT2 | 174.694211 | -2.1718008 | 0.8629097 | -2.5168344 | 0.01184145 | 0.26803102 |

| | | | | | | |
|---|---|---|---|---|---|---|
| CNOT6 | 223.593089 | -0.1412943 | 0.2768463 | -0.5103708 | 0.60979168 | 0.94003018 |
| MGAT1 | 92.8619843 | -0.1269887 | 0.28081535 | -0.4522143 | 0.65111458 | 0.95061388 |
| ZFP62 | 46.2776172 | 0.01007282 | 0.27283696 | 0.03691882 | 0.97054974 | 0.99656026 |
| TRIM41 | 63.3579246 | -0.0795883 | 0.30006033 | -0.265241 | 0.79082382 | 0.96527296 |
| RACK1 | 552.650257 | 0.21086634 | 0.36957735 | 0.57056079 | 0.56829741 | 0.93122667 |
| TRIM52 | 64.5613556 | -0.1871046 | 0.25613547 | -0.7304909 | 0.4650902 | 0.89883965 |
| AC138035.1 | 42.2975099 | 0.04476877 | 0.37965313 | 0.11792019 | 0.9061309 | 0.9875066 |
| DUSP22 | 87.7415336 | -0.2910758 | 0.24735859 | -1.1767361 | 0.23930084 | 0.79984328 |
| EXOC2 | 341.801281 | -0.5152888 | 0.24180461 | -2.1310132 | 0.03308805 | 0.42324075 |
| FOXQ1 | 74.1744335 | -0.7941778 | 1.27369871 | -0.623521 | 0.53294223 | 0.92268523 |
| GMDS | 974.868461 | -0.0833485 | 0.27044974 | -0.3081847 | 0.75794177 | 0.96180522 |
| GMDS-DT | 1267.42186 | 0.00448573 | 0.27022636 | 0.01659992 | 0.98675579 | 0.99791717 |
| WRNIP1 | 120.591467 | -0.090794 | 0.26397449 | -0.34395 | 0.73088387 | 0.96008876 |
| SERPINB1 | 115.20738 | -0.072231 | 0.35845056 | -0.201509 | 0.84030059 | 0.97489902 |
| SERPINB6 | 248.630438 | 0.18562623 | 0.23451689 | 0.79152606 | 0.42863708 | 0.88535099 |
| NQO2 | 91.7834327 | 0.22380646 | 0.33577506 | 0.66653688 | 0.505068 | 0.9107831 |
| RIPK1 | 122.749068 | -0.2598271 | 0.31413382 | -0.8271223 | 0.40816773 | 0.87948803 |
| BPHL | 91.4191875 | -0.0809331 | 0.30619389 | -0.2643199 | 0.79153345 | 0.9653598 |
| PSMG4 | 130.790803 | 0.35215662 | 0.32347298 | 1.08867399 | 0.27629768 | 0.82167981 |
| SLC22A23 | 177.494314 | -0.1118321 | 0.35659922 | -0.3136073 | 0.75381934 | 0.96120879 |
| PXDC1 | 34.6132437 | 1.02898551 | 0.48870287 | 2.10554423 | 0.03524396 | 0.43442748 |
| PRPF4B | 370.93988 | -0.0305774 | 0.19023153 | -0.1607379 | 0.87229981 | 0.98223939 |
| ECI2 | 140.674576 | 0.44550498 | 0.28763715 | 1.54884368 | 0.1214193 | 0.66068239 |
| CDYL | 507.469744 | -0.0701666 | 0.28318572 | -0.2477759 | 0.8043078 | 0.96679546 |
| LYRM4 | 173.731905 | -0.5954055 | 0.25611028 | -2.3248013 | 0.02008259 | 0.33933607 |
| FARS2 | 716.430152 | -0.2391268 | 0.18933754 | -1.2629654 | 0.20660161 | 0.77200549 |
| RREB1 | 758.18852 | -0.1008886 | 0.30346137 | -0.3324596 | 0.73954225 | 0.96078945 |
| SSR1 | 250.245308 | 0.12658366 | 0.20993726 | 0.6029595 | 0.54653563 | 0.92597739 |
| AL139095.4 | 46.8734024 | 0.50134286 | 0.56740066 | 0.8835782 | 0.37692395 | 0.86417917 |
| RIOK1 | 44.9240529 | -0.3936958 | 0.30234649 | -1.3021343 | 0.19287046 | 0.7597012 |
| DSP | 518.518748 | -0.2139976 | 0.3169799 | -0.6751141 | 0.49960327 | 0.90839719 |
| SNRNP48 | 67.9945602 | -0.0711452 | 0.31891799 | -0.223083 | 0.82347087 | 0.97018919 |
| BLOC1S5-TXI | 85.9965943 | 0.56431895 | 0.41531373 | 1.35877751 | 0.1742171 | 0.73178166 |
| SLC35B3 | 127.747398 | 0.35088618 | 0.40370384 | 0.8691673 | 0.38475563 | 0.86725322 |
| TFAP2A | 281.705203 | -0.0878942 | 0.32724423 | -0.268589 | 0.788246 | 0.96515649 |
| TFAP2A-AS1 | 32.9000437 | -0.0150532 | 0.34723414 | -0.0433518 | 0.96542112 | 0.99637506 |
| GCNT2 | 142.884102 | -0.6264099 | 0.47005066 | -1.3326434 | 0.18264887 | 0.74315969 |
| TMEM14C | 176.539107 | 0.34785091 | 0.42616157 | 0.81624185 | 0.41436182 | 0.88242985 |
| MAK | 133.59252 | 0.16770925 | 0.54629863 | 0.30699189 | 0.75884955 | 0.96180522 |
| SMIM13 | 32.7725749 | 0.33441192 | 0.31469346 | 1.06265925 | 0.28793651 | 0.8303903 |
| AL139807.1 | 61.793219 | 0.70907806 | 0.52546861 | 1.34942039 | 0.17720198 | 0.73633806 |
| NEDD9 | 157.699672 | 0.28299391 | 0.38656719 | 0.73206913 | 0.46412638 | 0.89801718 |
| TMEM170B | 41.8218565 | 0.29792908 | 0.3155098 | 0.9442784 | 0.3450274 | 0.85220061 |
| HIVEP1 | 250.571018 | -0.1818817 | 0.31366458 | -0.5798605 | 0.56200873 | 0.92973188 |
| EDN1 | 62.211549 | 2.44752144 | 0.57548492 | 4.2529723 | 2.11E-05 | 0.00614104 |
| PHACTR1 | 540.782573 | -0.05011 | 0.27881138 | -0.1797273 | 0.85736666 | 0.97718353 |
| TBC1D7 | 166.297015 | 0.26476728 | 0.43429356 | 0.60965048 | 0.54209337 | 0.92481499 |
| SIRT5 | 114.705958 | 0.03442634 | 0.28589786 | 0.12041481 | 0.90415456 | 0.9875066 |
| NOL7 | 165.108758 | 0.02942935 | 0.42752555 | 0.06883647 | 0.94511979 | 0.99276891 |
| RANBP9 | 461.331322 | 0.07249693 | 0.16292774 | 0.44496369 | 0.656346 | 0.95100227 |
| MCUR1 | 111.087193 | 0.21103153 | 0.40668149 | 0.51891108 | 0.60382275 | 0.93804805 |
| JARID2 | 829.597261 | 0.01780813 | 0.5265177 | 0.03382248 | 9.73E-01 | 0.99656026 |
| DTNBP1 | 280.570259 | -0.3073347 | 0.25376313 | -1.2111085 | 0.22585383 | 0.78982769 |
| MYLIP | 83.5844155 | 1.21895439 | 0.38884833 | 3.13478107 | 0.00171982 | 0.10426718 |

| | | | | | | |
|---|---|---|---|---|---|---|
| GMPR | 111.52382 | -0.1891011 | 0.78527013 | -0.2408102 | 0.80970221 | 0.9679755 |
| ATXN1 | 1434.37092 | 0.45278153 | 0.35692096 | 1.26857647 | 0.20459216 | 0.77193931 |
| CAP2 | 147.153279 | 0.94310613 | 0.37970533 | 2.48378428 | 0.01299945 | 0.27974182 |
| FAM8A1 | 60.4012245 | 0.26750329 | 0.35052159 | 0.76315782 | 0.44536928 | 0.89175966 |
| NUP153 | 347.77379 | -0.0669539 | 0.23202942 | -0.2885578 | 0.77291983 | 0.96388926 |
| KIF13A | 535.749272 | -0.1747594 | 0.32629965 | -0.5355795 | 0.59224923 | 0.93491607 |
| TPMT | 74.9615861 | 0.3713494 | 0.27810039 | 1.33530701 | 0.18177588 | 0.74315969 |
| KDM1B | 67.2387733 | -0.4071896 | 0.27442327 | -1.4838012 | 0.13786166 | 0.68146162 |
| DEK | 228.265225 | 0.08308878 | 0.25667651 | 0.32371011 | 0.74615752 | 0.96109466 |
| RNF144B | 614.492087 | -0.2077819 | 0.38727591 | -0.5365215 | 0.59159817 | 0.93491607 |
| AL022068.1 | 387.850832 | 0.39461493 | 0.45196954 | 0.87310073 | 0.38260818 | 0.86593382 |
| ID4 | 282.548997 | 0.22606212 | 0.44313354 | 0.51014446 | 0.60995026 | 0.94003018 |
| MBOAT1 | 242.557609 | 0.80421018 | 0.66951755 | 1.20117864 | 0.22968191 | 0.79047378 |
| E2F3 | 133.30516 | 0.30443654 | 0.47609172 | 0.63944934 | 0.52253066 | 0.91845051 |
| CDKAL1 | 1282.24026 | -0.8520208 | 0.23982059 | -3.5527425 | 0.00038124 | 0.04296095 |
| SOX4 | 496.356773 | -0.4802268 | 0.68840222 | -0.6975962 | 0.48542973 | 0.90695174 |
| CASC15 | 86.6369761 | 0.28044027 | 0.59606008 | 0.47048993 | 0.63800502 | 0.94854485 |
| DCDC2 | 4916.3203 | 0.61458094 | 0.40570669 | 1.51484055 | 0.12981279 | 0.67256577 |
| MRS2 | 122.585343 | -0.1417522 | 0.236466 | -0.5994612 | 0.54886535 | 0.9274143 |
| GPLD1 | 40.6471069 | 0.69226273 | 0.46615299 | 1.4850548 | 0.13752931 | 0.68146162 |
| ALDH5A1 | 79.8366635 | 0.64368161 | 0.29328279 | 2.19474726 | 0.02818173 | 0.40459506 |
| TDP2 | 92.7073686 | 0.21959053 | 0.22414582 | 0.9796771 | 0.32724553 | 0.84381657 |
| ACOT13 | 76.4850989 | 0.53947139 | 0.26126887 | 2.06481314 | 0.03894068 | 0.4535395 |
| C6orf62 | 409.998298 | -0.3290202 | 0.28411104 | -1.1580689 | 0.24683593 | 0.80072691 |
| GMNN | 220.398225 | -0.7100033 | 0.56359687 | -1.2597715 | 0.2077518 | 0.77327889 |
| CARMIL1 | 1453.43022 | -0.5578264 | 0.40328681 | -1.3832003 | 0.16660347 | 0.72089361 |
| TRIM38 | 267.136682 | -0.2695491 | 0.36379094 | -0.7409449 | 0.45872682 | 0.89753337 |
| HIST1H1C | 158.3701 | 0.68645541 | 0.60369698 | 1.13708603 | 0.25550232 | 0.80525591 |
| HFE | 31.7629771 | 0.63053112 | 0.37587889 | 1.67748478 | 0.09344772 | 0.61317397 |
| HIST1H4C | 191.593773 | 1.31207748 | 0.60194241 | 2.17973924 | 2.93E-02 | 0.40623517 |
| HIST1H2BC | 108.485296 | -0.376823 | 0.68133679 | -0.5530642 | 5.80E-01 | 9.33E-01 |
| HIST1H2AC | 1211.29043 | 0.44900247 | 0.59442063 | 0.75536152 | 4.50E-01 | 0.89324553 |
| HIST1H1E | 97.9109089 | 0.87729733 | 0.61551665 | 1.42530237 | 0.15406982 | 0.7079031 |
| HIST1H2BD | 184.916599 | 0.67075711 | 0.51400239 | 1.30496887 | 0.19190345 | 0.7597012 |
| AL031777.3 | 59.2740957 | 0.90088533 | 0.76455386 | 1.17831505 | 2.39E-01 | 0.79867687 |
| BTN3A2 | 55.7506793 | -0.4646255 | 0.82002711 | -0.5665977 | 0.57098754 | 0.93125087 |
| BTN3A1 | 52.2668311 | -0.7655543 | 0.6333931 | -1.2086558 | 0.2267951 | 0.78990116 |
| BTN2A1 | 51.8622384 | -0.191919 | 0.25963542 | -0.7391866 | 0.45979373 | 0.89753337 |
| HCG11 | 37.631277 | 0.66008693 | 0.39808252 | 1.65816606 | 0.09728395 | 0.61662137 |
| HMGN4 | 54.9814686 | 0.43866951 | 0.40524935 | 1.08246812 | 0.27904458 | 0.82470027 |
| ABT1 | 39.2659935 | 0.0055695 | 0.49707955 | 0.01120445 | 0.99106033 | 0.99889703 |
| ZNF322 | 45.2878185 | 0.31193484 | 0.37370328 | 0.8347126 | 0.40387954 | 0.87834687 |
| LINC00240 | 51.7236004 | 0.32856452 | 0.55434456 | 0.59270812 | 0.55337651 | 0.92872652 |
| ZSCAN16-AS | 77.8199984 | 0.24890208 | 0.36258544 | 0.68646464 | 0.49242016 | 0.90780423 |
| ZKSCAN8 | 67.9835336 | 0.04767963 | 0.28623152 | 0.16657714 | 0.86770278 | 0.98138627 |
| ZSCAN9 | 43.3515297 | -0.3914021 | 0.28548165 | -1.3710238 | 0.17036754 | 0.72675731 |
| ZSCAN23 | 38.9962909 | 0.18005043 | 0.34479426 | 0.52219672 | 0.60153337 | 0.93804805 |
| TRIM27 | 145.767025 | -0.0798384 | 0.24159483 | -0.330464 | 0.74104941 | 0.96084411 |
| GABBR1 | 130.867512 | 0.27568997 | 0.28616768 | 0.96338613 | 0.33535379 | 0.84544327 |
| HLA-F | 57.6250082 | -1.3964192 | 0.80983756 | -1.7243201 | 0.08465006 | 0.59720183 |
| HLA-A | 1378.48429 | -0.3641861 | 0.56984933 | -0.639092 | 0.5227631 | 0.91845051 |
| ZNRD1 | 51.7252194 | 0.19525821 | 0.70176137 | 0.27824019 | 0.78082799 | 0.96513523 |
| PPP1R11 | 75.178808 | 0.08568283 | 0.32841557 | 0.26089758 | 0.79417149 | 0.9653598 |
| TRIM26 | 166.168844 | -0.200238 | 0.31894933 | -0.627805 | 0.53013166 | 0.92181514 |

| | | | | | | |
|---|---|---|---|---|---|---|
| HCG17 | 313.697445 | 0.01424202 | 0.26341701 | 0.05406645 | 0.95688222 | 0.99565888 |
| HLA-E | 557.920367 | -0.4781336 | 0.4830496 | -0.9898231 | 0.32226061 | 0.84285425 |
| GNL1 | 163.0327 | 0.07689266 | 0.25845187 | 0.29751248 | 0.76607528 | 0.96285313 |
| PRR3 | 39.3218836 | 0.39531948 | 0.43787469 | 0.90281418 | 0.36662452 | 0.85902833 |
| ABCF1 | 146.47149 | -0.0274118 | 0.27928722 | -0.0981491 | 0.92181388 | 0.99051393 |
| PPP1R10 | 182.157576 | -0.3933097 | 0.33928565 | -1.1592288 | 0.24636293 | 0.80072691 |
| MRPS18B | 143.482656 | 0.31244679 | 0.26099521 | 1.19713611 | 0.23125351 | 0.79148816 |
| ATAT1 | 55.9498553 | -0.4614637 | 0.35203812 | -1.3108344 | 0.18991373 | 0.75619145 |
| C6orf136 | 95.6699062 | 0.4128483 | 0.23906201 | 1.7269507 | 0.08417652 | 0.59525634 |
| DHX16 | 62.7303081 | -0.4022859 | 0.31817014 | -1.2643735 | 0.20609601 | 0.77193931 |
| MDC1 | 87.3550991 | 0.2023006 | 0.22172304 | 0.91240221 | 0.36155703 | 0.85690129 |
| TUBB | 439.133775 | -0.6656527 | 0.62422053 | -1.0663743 | 0.2862545 | 0.8291345 |
| FLOT1 | 354.317003 | 0.62790769 | 0.31039875 | 2.02290662 | 0.04308278 | 0.47527104 |
| IER3 | 1262.10078 | -1.5988743 | 0.8427263 | -1.897264 | 0.0577931 | 0.52943333 |
| DDR1 | 416.249562 | -0.068286 | 0.21938771 | -0.3112574 | 0.75560497 | 0.96120879 |
| VARS2 | 33.9695765 | 0.00132399 | 0.33587144 | 0.00394196 | 0.99685478 | 0.99967893 |
| CCHCR1 | 72.6870586 | 0.23636726 | 0.21418651 | 1.10355808 | 0.26978489 | 0.81817548 |
| TCF19 | 37.6998712 | 0.13763229 | 0.61971677 | 0.22208901 | 0.82424459 | 0.97018919 |
| POU5F1 | 320.388956 | -0.8413603 | 0.35110813 | -2.3962996 | 0.01656155 | 0.30723893 |
| HLA-C | 899.143165 | -0.5958374 | 0.65968622 | -0.9032133 | 0.36641269 | 0.85902833 |
| HLA-B | 1351.27791 | -0.2306387 | 0.65897665 | -0.3499952 | 0.72634226 | 0.95979351 |
| MICA | 64.9182135 | 0.4904182 | 0.290113 | 1.69043858 | 0.09094408 | 0.60942442 |
| NFKBIL1 | 50.969583 | 0.29749331 | 0.30641995 | 0.97086795 | 0.33161404 | 0.84442294 |
| PRRC2A | 192.672748 | 0.13401644 | 0.19368453 | 0.69193159 | 0.48898029 | 0.90730741 |
| BAG6 | 95.917251 | -0.3270683 | 0.24538886 | -1.3328571 | 0.1825787 | 0.74315969 |
| C6orf47 | 39.287563 | 0.18289228 | 0.40851771 | 0.44769731 | 0.65437166 | 0.95100227 |
| GPANK1 | 37.1421694 | 0.30460465 | 0.29231638 | 1.0420376 | 0.29739425 | 0.83556453 |
| DDAH2 | 158.872248 | -0.0646444 | 0.27222474 | -0.2374669 | 0.81229456 | 0.9680627 |
| CLIC1 | 501.810502 | -0.0087811 | 0.3692158 | -0.0237832 | 0.98102558 | 0.99696835 |
| VARS | 42.3769479 | -0.2104385 | 0.38089631 | -0.5524823 | 0.58061794 | 0.93354125 |
| LSM2 | 103.174911 | 0.14447171 | 0.31676966 | 0.4560781 | 0.64833382 | 0.95061388 |
| HSPA1A | 469.176785 | 0.31217343 | 0.43854532 | 0.71183848 | 0.4765648 | 0.90242252 |
| HSPA1B | 274.467515 | 0.48953245 | 0.52208521 | 0.93764857 | 0.34842504 | 0.8535343 |
| C6orf48 | 87.0119629 | 0.42611894 | 0.44663079 | 0.95407427 | 0.34004605 | 0.84970973 |
| NEU1 | 95.6017845 | 0.05169362 | 0.40386324 | 0.12799784 | 0.89815069 | 0.9863394 |
| SLC44A4 | 144.813663 | -0.4293239 | 0.53960904 | -0.7956202 | 0.42625281 | 0.88458009 |
| EHMT2 | 64.2351746 | 0.03914944 | 0.29131351 | 0.13438936 | 0.8930947 | 0.98597539 |
| NELFE | 51.6654822 | -0.3271832 | 0.38952576 | -0.8399525 | 0.40093499 | 0.87775198 |
| STK19 | 73.0370909 | -0.0159693 | 0.50494002 | -0.0316261 | 0.97477021 | 0.99656026 |
| ATF6B | 126.825492 | -0.0436871 | 0.21238826 | -0.2056946 | 0.83702949 | 0.97353666 |
| AGPAT1 | 63.3659143 | -0.1564378 | 0.36222516 | -0.43188 | 0.66582865 | 0.95243327 |
| RNF5 | 114.921342 | -0.0160224 | 0.51395449 | -0.0311747 | 0.97513026 | 0.99656026 |
| PBX2 | 82.142906 | 0.18388443 | 0.23610318 | 0.77883081 | 0.43607939 | 0.88785137 |
| AL662796.1 | 86.5302745 | 0.12545109 | 0.71147273 | 0.17632593 | 0.86003789 | 0.97841862 |
| HLA-DRA | 402.005642 | 0.85113105 | 0.64272343 | 1.32425708 | 0.18541768 | 0.74895057 |
| HLA-DRB5 | 65.4841813 | 0.6146725 | 0.86300014 | 0.71225076 | 0.47630951 | 0.90242252 |
| HLA-DRB1 | 250.775874 | 0.48949813 | 0.98771268 | 0.49558758 | 0.62018542 | 0.94319304 |
| HLA-DQB1 | 70.6845642 | 0.15707733 | 1.41684088 | 0.11086448 | 0.91172381 | 0.98788932 |
| PSMB8 | 122.488861 | 0.18010332 | 0.55458602 | 0.32475272 | 0.74536823 | 0.96109466 |
| TAP1 | 67.6844114 | -1.3140807 | 0.83555666 | -1.5727009 | 0.11578809 | 0.65192493 |
| BRD2 | 260.124856 | -0.123696 | 0.16400717 | -0.7542108 | 0.45072264 | 0.89333031 |
| HLA-DPA1 | 73.541158 | -0.0259831 | 0.87722483 | -0.0296197 | 0.97637036 | 0.99656026 |
| HLA-DPB1 | 82.8702073 | 0.76085553 | 0.77126362 | 0.98650514 | 0.32388529 | 0.84285425 |
| RXRB | 54.2316518 | 0.20847839 | 0.24729068 | 0.84304993 | 0.39920052 | 0.87596764 |

| | | | | | | |
|---|---|---|---|---|---|---|
| SLC39A7 | 89.0599268 | 0.20072046 | 0.42501369 | 0.47226823 | 0.63673534 | 0.94821838 |
| RING1 | 75.6017638 | -0.1533087 | 0.21755967 | -0.7046744 | 0.48101289 | 0.90519215 |
| VPS52 | 82.6105669 | -0.0995693 | 0.22183179 | -0.4488505 | 0.65353951 | 0.95100227 |
| RPS18 | 518.752546 | -0.0990608 | 0.44193777 | -0.224151 | 0.82263983 | 0.97018919 |
| B3GALT4 | 33.1307452 | 0.22009418 | 0.29597304 | 0.74362915 | 0.45710086 | 0.8969884 |
| WDR46 | 43.1995083 | 0.45184526 | 0.30912504 | 1.46169091 | 0.14382593 | 0.69093999 |
| PFDN6 | 72.7270522 | 0.10121181 | 0.31209196 | 0.32430124 | 0.74570998 | 0.96109466 |
| RGL2 | 127.310323 | -0.0107681 | 0.2137703 | -0.0503723 | 0.95982568 | 0.99609409 |
| TAPBP | 341.435843 | -0.4111274 | 0.36582841 | -1.1238259 | 0.26108689 | 0.80805101 |
| DAXX | 45.2958065 | 0.20322223 | 0.38977871 | 0.52137847 | 0.60210315 | 0.93804805 |
| PHF1 | 101.10297 | -0.3507907 | 0.28320389 | -1.2386508 | 0.21547485 | 0.77895013 |
| CUTA | 187.678483 | -0.0373637 | 0.48436552 | -0.0771395 | 0.93851256 | 0.99243654 |
| SYNGAP1 | 58.0568225 | -0.0400643 | 0.24910108 | -0.1608355 | 0.87222298 | 0.98223939 |
| BAK1 | 46.0390744 | -0.0819362 | 0.44302316 | -0.184948 | 0.85326986 | 0.97718353 |
| ITPR3 | 300.284641 | -0.8801183 | 0.48826709 | -1.8025345 | 0.07146136 | 0.56343001 |
| UQCC2 | 79.8671521 | 0.16391043 | 0.2917505 | 0.56181712 | 0.57424063 | 0.93274051 |
| LEMD2 | 82.7119426 | -0.5428487 | 0.27379203 | -1.9827045 | 0.04740045 | 0.48600281 |
| HMGA1 | 108.619218 | -0.8215303 | 0.41125257 | -1.9976296 | 0.04575683 | 0.48339878 |
| SMIM29 | 48.8744807 | -0.3167096 | 0.3385101 | -0.9355985 | 0.34947992 | 0.8535343 |
| NUDT3 | 227.058555 | 0.15465783 | 0.38432952 | 0.40240944 | 0.68738272 | 0.95588883 |
| C6orf106 | 409.340949 | -0.1347939 | 0.34665377 | -0.3888429 | 0.69739235 | 0.95672887 |
| SNRPC | 90.9457735 | 0.07708602 | 0.42001667 | 0.18353085 | 0.85438152 | 0.97718353 |
| UHRF1BP1 | 144.855245 | -0.009038 | 0.22637212 | -0.0399254 | 0.96815261 | 0.9964399 |
| TAF11 | 113.479095 | -0.3023349 | 0.25507302 | -1.1852876 | 0.23590376 | 0.79724974 |
| ANKS1A | 724.38962 | -0.8583201 | 0.43265579 | -1.9838406 | 0.04727362 | 0.48600281 |
| SCUBE3 | 44.1714063 | 0.23043602 | 0.59131989 | 0.38969773 | 0.69676007 | 0.95672887 |
| ZNF76 | 162.351856 | -0.1259863 | 0.28273558 | -0.4455975 | 0.65588803 | 0.95100227 |
| PPARD | 198.793181 | -0.2893178 | 0.27311779 | -1.0593151 | 0.2894563 | 0.83103268 |
| RPL10A | 622.279298 | 0.28664744 | 0.50227016 | 0.57070371 | 0.5682005 | 0.93122667 |
| TEAD3 | 74.7643368 | 0.11428843 | 0.30409406 | 0.37583251 | 0.70704142 | 0.95752169 |
| SRPK1 | 193.289811 | -0.2323124 | 0.26753964 | -0.868329 | 0.38521423 | 0.86749643 |
| MAPK14 | 650.364842 | -0.8676023 | 0.2878301 | -3.0142863 | 0.00257585 | 0.13156951 |
| MAPK13 | 63.4962841 | 0.16987355 | 0.24536936 | 0.69231769 | 0.48873784 | 0.90730741 |
| BRPF3 | 42.6681903 | -0.1559711 | 0.32019513 | -0.4871126 | 0.62617855 | 0.94511671 |
| KCTD20 | 238.488303 | -0.145599 | 0.41650544 | -0.349573 | 0.72665919 | 0.95979351 |
| STK38 | 379.063902 | -0.0229478 | 0.19813972 | -0.115816 | 0.90779834 | 0.9875066 |
| SRSF3 | 352.382152 | -0.1017851 | 0.38273537 | -0.2659412 | 0.79028451 | 0.96524651 |
| CDKN1A | 124.270842 | -1.3011275 | 0.82566529 | -1.5758534 | 0.11505958 | 0.65074172 |
| C6orf89 | 222.641678 | -0.124187 | 0.24895178 | -0.4988395 | 0.61789249 | 0.94244117 |
| MTCH1 | 265.83923 | -0.264289 | 0.27133385 | -0.9740362 | 0.33003859 | 0.84442294 |
| PIM1 | 57.9290561 | -1.0233256 | 0.49800063 | -2.054868 | 0.03989175 | 0.45827251 |
| TBC1D22B | 143.619647 | -0.2498528 | 0.21291673 | -1.1734766 | 0.24060473 | 0.80057149 |
| RNF8 | 110.088444 | 0.55095717 | 0.26704641 | 2.06315135 | 0.03909825 | 0.45477211 |
| CMTR1 | 88.2070323 | -0.1501257 | 0.2546404 | -0.5895595 | 0.55548599 | 0.92872652 |
| ZFAND3 | 3112.5367 | -0.2786179 | 0.24228088 | -1.1499791 | 0.25015248 | 0.80149394 |
| BTBD9 | 1783.20776 | -0.4322167 | 0.18665979 | -2.3155321 | 0.02058383 | 0.34432775 |
| GLO1 | 340.736474 | 0.18958891 | 0.19829114 | 0.95611384 | 0.33901472 | 0.84863112 |
| SAYSD1 | 48.4526542 | -0.2456214 | 0.41581658 | -0.5906965 | 0.55472382 | 0.92872652 |
| KCNK5 | 40.8711611 | 0.80795058 | 0.4299442 | 1.87919868 | 0.06021737 | 0.53892231 |
| KIF6 | 82.7005009 | 0.54369685 | 0.54767994 | 0.99272735 | 0.32084284 | 0.84285425 |
| MOCS1 | 50.9690111 | 0.01123661 | 0.34083049 | 0.03296832 | 0.97369985 | 0.99656026 |
| UNC5CL | 104.4415 | 0.15864577 | 0.64886031 | 0.24449912 | 0.80684429 | 0.96752828 |
| OARD1 | 163.423614 | -0.0274931 | 0.27711242 | -0.0992127 | 0.92096937 | 0.9904507 |
| NFYA | 93.9386223 | -0.0247136 | 0.34140365 | -0.0723883 | 0.94229286 | 0.99265578 |

| | | | | | | |
|---|---|---|---|---|---|---|
| FOXP4 | 36.4824831 | 0.09371758 | 0.3637917 | 0.2576133 | 0.79670536 | 0.9653598 |
| TFEB | 132.300204 | 0.98285444 | 0.53026624 | 1.85351126 | 0.06380911 | 0.54850785 |
| USP49 | 89.8868701 | -0.0103482 | 0.27235515 | -0.0379952 | 0.96969154 | 0.99645289 |
| CCND3 | 253.564351 | 0.10261617 | 0.47400029 | 0.21648969 | 0.82860606 | 0.97117144 |
| TAF8 | 94.1253299 | -0.0846187 | 0.3484669 | -0.2428315 | 0.80813595 | 0.96769478 |
| C6orf132 | 85.442489 | 0.41793438 | 0.34630375 | 1.20684334 | 0.22749248 | 0.79042788 |
| MRPS10 | 61.162782 | 0.10871621 | 0.27784984 | 0.39127684 | 0.69559262 | 0.95672887 |
| TRERF1 | 407.658839 | 0.69414243 | 0.44371437 | 1.56439024 | 0.11772595 | 0.65447949 |
| UBR2 | 701.839416 | -0.2283153 | 0.23214648 | -0.9834968 | 0.32536297 | 0.84285425 |
| BICRAL | 333.65748 | 0.30318005 | 0.35129534 | 0.86303465 | 0.38811842 | 0.86953359 |
| RPL7L1 | 198.293041 | 0.42339384 | 0.27488874 | 1.5402371 | 0.12350257 | 0.66519057 |
| C6orf226 | 34.5428638 | -0.1265586 | 0.38220935 | -0.3311237 | 0.74055104 | 0.96084411 |
| CNPY3 | 50.0572851 | -0.2803231 | 0.28448635 | -0.9853658 | 0.32444444 | 0.84285425 |
| PEX6 | 74.7761668 | 0.48397122 | 0.33712766 | 1.43557257 | 0.151124 | 0.7029344 |
| PPP2R5D | 40.3597724 | -0.1372561 | 0.27372399 | -0.5014397 | 0.61606171 | 0.94190661 |
| MEA1 | 56.0749669 | -0.0253358 | 0.55128544 | -0.0459578 | 0.96334392 | 0.99637506 |
| KLHDC3 | 98.9300556 | -0.0142207 | 0.24762922 | -0.0574274 | 0.95420475 | 0.99522948 |
| RRP36 | 108.687688 | -0.0607885 | 0.32759784 | -0.1855583 | 0.85279117 | 0.97718353 |
| CUL7 | 50.0421931 | -0.2028568 | 0.31932015 | -0.6352771 | 0.52524772 | 0.91924275 |
| KLC4 | 71.4686265 | 0.30662257 | 0.26337465 | 1.16420684 | 0.2443402 | 0.80072691 |
| MRPL2 | 72.3577034 | 0.29765023 | 0.37440403 | 0.79499739 | 0.42661502 | 0.88477675 |
| CUL9 | 73.1932252 | -0.2190316 | 0.29532123 | -0.7416723 | 0.45828589 | 0.89747927 |
| DNPH1 | 146.276341 | 0.55186105 | 0.48919311 | 1.12810471 | 0.2592757 | 0.80709747 |
| ZNF318 | 64.2234155 | 0.29765636 | 0.33460025 | 0.88958799 | 0.37368716 | 0.86209328 |
| ABCC10 | 61.5084902 | -0.6187072 | 0.36997529 | -1.6722934 | 0.09446648 | 0.61338829 |
| TJAP1 | 52.3970953 | -0.0114817 | 0.28100519 | -0.0408596 | 0.96740786 | 0.99637506 |
| POLR1C | 92.3671117 | -0.2181215 | 0.23762974 | -0.9179049 | 0.35866864 | 0.8557799 |
| YIPF3 | 129.309496 | 0.15611138 | 0.31877282 | 0.48972612 | 0.62432771 | 0.94457465 |
| XPO5 | 341.984295 | -0.2900258 | 0.23788579 | -1.2191808 | 0.22277558 | 0.78821341 |
| POLH | 102.336971 | 0.33911387 | 0.27121531 | 1.25034931 | 0.21117197 | 0.7767562 |
| GTPBP2 | 76.5645391 | -0.4464885 | 0.29499565 | -1.5135427 | 0.13014186 | 0.67285975 |
| MRPS18A | 67.7441201 | 0.22012082 | 0.43453328 | 0.50656838 | 0.6124577 | 0.94003018 |
| VEGFA | 884.575293 | -1.4715277 | 0.76443731 | -1.9249815 | 0.05423167 | 0.51425036 |
| MRPL14 | 110.500499 | 0.30369802 | 0.4275926 | 0.71025087 | 0.47754858 | 0.90242252 |
| TMEM63B | 81.1050278 | -0.2599947 | 0.26956441 | -0.9644992 | 0.33479571 | 0.84521974 |
| SLC29A1 | 126.068227 | -0.3553703 | 0.37816191 | -0.9397305 | 0.3473558 | 0.85348649 |
| HSP90AB1 | 1056.49456 | 0.15297538 | 0.22642419 | 0.67561413 | 0.49928567 | 0.90839719 |
| SLC35B2 | 56.7334072 | 0.07746187 | 0.5009472 | 0.1546308 | 0.87711238 | 0.98277085 |
| CDC5L | 133.599342 | -0.2248371 | 0.24018821 | -0.9360871 | 0.34922834 | 0.8535343 |
| SUPT3H | 869.03028 | -0.3160062 | 0.26120542 | -1.2097996 | 0.22635579 | 0.78982769 |
| ENPP4 | 109.088592 | 0.29126961 | 0.37776971 | 0.77102425 | 0.44069256 | 0.88956997 |
| ENPP5 | 77.7837416 | 0.58685006 | 0.33946158 | 1.72876726 | 0.08385076 | 0.59375402 |
| RCAN2 | 167.729883 | 1.84309441 | 0.47407154 | 3.88779804 | 0.00010116 | 0.02120266 |
| CYP39A1 | 51.5788481 | 1.33130977 | 0.42082464 | 3.16357375 | 0.00155845 | 0.09927452 |
| SLC25A27 | 200.286296 | 0.82778489 | 0.41616354 | 1.98908555 | 0.04669176 | 0.48600281 |
| ADGRF1 | 495.276723 | 0.41323336 | 0.4074684 | 1.01414822 | 0.31051203 | 0.84217549 |
| TNFRSF21 | 52.6242727 | -0.7446579 | 0.72541225 | -1.0265306 | 0.30464154 | 0.83709431 |
| CD2AP | 2070.65024 | 0.29822032 | 0.30416232 | 0.98046437 | 0.32685695 | 0.84381657 |
| MUT | 124.812537 | 0.10833553 | 0.30094153 | 0.35998864 | 0.71885563 | 0.95796325 |
| TFAP2B | 210.254727 | -0.4269657 | 0.52782599 | -0.8089137 | 0.41856477 | 0.88242985 |
| PKHD1 | 7003.51155 | 0.4656629 | 0.26846333 | 1.73454937 | 0.08282065 | 0.59202053 |
| MCM3 | 86.5062388 | -0.057531 | 0.33307694 | -0.1727258 | 0.86286694 | 0.979268 |
| EFHC1 | 184.288594 | 0.06600358 | 0.21255961 | 0.31051795 | 0.75616711 | 0.96120879 |
| TRAM2 | 53.6106891 | 0.85400939 | 0.41791595 | 2.04349556 | 0.04100341 | 0.46430138 |

| | | | | | | |
|---|---|---|---|---|---|---|
| TMEM14A | 58.9452828 | 0.10443006 | 0.34658549 | 0.30131111 | 0.76317727 | 0.96285313 |
| GSTA4 | 81.4650924 | 0.37975114 | 0.34856153 | 1.08948092 | 0.27594186 | 0.82155419 |
| ICK | 71.3323437 | 0.18437527 | 0.31346567 | 0.58818328 | 0.55640928 | 0.92872652 |
| FBXO9 | 215.19846 | -0.0150306 | 0.24363757 | -0.0616924 | 0.95080777 | 0.99404295 |
| ELOVL5 | 111.911068 | 0.16374217 | 0.5034809 | 0.32522022 | 0.7450144 | 0.96109466 |
| GCLC | 115.027073 | -0.1637852 | 0.40698264 | -0.4024378 | 0.68736186 | 0.95588883 |
| LRRC1 | 390.500201 | -0.2325989 | 0.43275691 | -0.5374816 | 0.59093498 | 0.93483536 |
| FAM83B | 339.547571 | -0.5578976 | 0.41255462 | -1.3523 | 0.17627935 | 0.7345557 |
| DST | 559.208721 | -0.1553064 | 0.38531086 | -0.4030679 | 0.68689827 | 0.9558749 |
| ZNF451 | 323.162196 | 0.07192873 | 0.14690526 | 0.48962661 | 0.62439814 | 0.94457465 |
| ZNF451-AS1 | 42.2740003 | 0.37307261 | 0.61903249 | 0.60267048 | 0.54672792 | 0.92597739 |
| PRIM2 | 300.592513 | -0.633347 | 0.26957521 | -2.349426 | 0.01880238 | 0.32850268 |
| AL021368.2 | 39.5411637 | -0.0009474 | 0.26848845 | -0.0035287 | 0.99718453 | 0.99967893 |
| PTP4A1 | 680.466116 | -2.0372343 | 0.86174098 | -2.3640912 | 0.01807436 | 0.32379373 |
| PHF3 | 631.113904 | 0.32043884 | 0.18667848 | 1.71652802 | 0.08606544 | 0.60140155 |
| EYS | 55.0979432 | 1.17430899 | 0.66115408 | 1.77615025 | 0.07570816 | 0.57072562 |
| LMBRD1 | 473.484187 | 0.02773377 | 0.16203463 | 0.17115954 | 0.86409832 | 0.97939196 |
| COL19A1 | 109.398385 | -0.2467673 | 0.35522949 | -0.69467 | 0.4872621 | 0.90730741 |
| FAM135A | 453.221931 | 0.11504055 | 0.29310248 | 0.39249259 | 0.69469429 | 0.95672887 |
| SDHAF4 | 104.335052 | 0.07858197 | 0.4506997 | 0.1743555 | 0.86158609 | 0.97848095 |
| SMAP1 | 844.568502 | -0.4850763 | 0.35224185 | -1.3771114 | 0.16847781 | 0.72499026 |
| OGFRL1 | 58.0650118 | -0.3937423 | 0.43795183 | -0.899054 | 0.36862391 | 0.85931545 |
| AL136164.2 | 330.786573 | 0.90593042 | 0.42528301 | 2.13018248 | 0.03315655 | 0.42324075 |
| LINC00472 | 117.955877 | 0.47119138 | 0.45981303 | 1.02474562 | 0.30548323 | 0.83709431 |
| AC019205.1 | 77.8355313 | 0.0832024 | 0.32980601 | 0.25227678 | 0.80082714 | 0.96611815 |
| OOEP | 583.097435 | 0.08846683 | 0.4573838 | 0.19341924 | 0.84663065 | 0.97656375 |
| MTO1 | 98.809049 | 0.07115687 | 0.26608562 | 0.26742094 | 0.78914508 | 0.96515649 |
| EEF1A1 | 3240.38349 | 0.37628903 | 0.45586316 | 0.82544295 | 0.40912016 | 0.87982122 |
| SLC17A5 | 117.460118 | -0.3613689 | 0.26804125 | -1.3481838 | 0.17759926 | 0.73685825 |
| COX7A2 | 266.905879 | 0.16446096 | 0.363064 | 0.45298063 | 0.65056269 | 0.95061388 |
| TMEM30A | 237.715691 | 0.55201534 | 0.22863766 | 2.41436757 | 0.01576255 | 0.30067202 |
| SENP6 | 1049.66499 | -0.0112359 | 0.21086091 | -0.0532858 | 0.95750419 | 0.99579587 |
| MYO6 | 2330.19756 | 0.33267744 | 0.32392719 | 1.02701301 | 0.30441434 | 0.83709431 |
| MEI4 | 145.409418 | -0.0752774 | 0.2628955 | -0.2863397 | 0.77461792 | 0.96388926 |
| IRAK1BP1 | 245.397344 | 0.66925906 | 0.36492402 | 1.83396823 | 0.06665871 | 0.55619688 |
| PHIP | 998.745758 | -0.1980148 | 0.18025053 | -1.098553 | 0.27196307 | 0.81922962 |
| HMGN3 | 582.633102 | 0.3559916 | 0.2325753 | 1.53065095 | 0.12585568 | 0.6674937 |
| LCA5 | 120.683831 | 0.37980082 | 0.26392624 | 1.43904152 | 0.15013876 | 0.7029344 |
| SH3BGRL2 | 187.974205 | 0.25945608 | 0.19245129 | 1.34816494 | 0.17760534 | 0.73685825 |
| BCKDHB | 613.576134 | -0.2616905 | 0.2609903 | -1.0026827 | 0.31601396 | 0.84285425 |
| TENT5A | 132.14608 | -0.8795426 | 0.52303532 | -1.6816122 | 0.09264407 | 0.61178944 |
| IBTK | 533.364681 | -0.8014294 | 0.24485028 | -3.2731408 | 0.00106359 | 0.08019046 |
| TPBG | 42.0088486 | 0.19529352 | 0.5546752 | 0.35208627 | 0.72477357 | 0.95963742 |
| UBE3D | 290.196937 | 0.15430542 | 0.23968414 | 0.64378653 | 0.51971387 | 0.91751084 |
| DOPEY1 | 589.246112 | 0.73421938 | 0.31703453 | 2.31589722 | 0.02056388 | 0.34432775 |
| PGM3 | 105.906882 | -0.4336754 | 0.28727651 | -1.5096096 | 0.13114306 | 0.67390608 |
| ME1 | 46.689117 | 1.36810976 | 0.45337648 | 3.01760198 | 0.00254783 | 0.13153345 |
| CYB5R4 | 147.594126 | -0.4936456 | 0.29539384 | -1.6711437 | 0.0946933 | 0.61367241 |
| CEP162 | 252.502516 | 0.12917957 | 0.2223918 | 0.5808648 | 0.56133158 | 0.92934517 |
| SNX14 | 405.997074 | -0.1609735 | 0.15179832 | -1.0604435 | 0.28894289 | 0.83060304 |
| SYNCRIP | 139.779391 | -0.0481044 | 0.18317963 | -0.2626079 | 0.7928528 | 0.9653598 |
| ZNF292 | 1531.6656 | -0.994441 | 0.31063698 | -3.2012964 | 0.00136811 | 0.0943274 |
| SMIM8 | 218.369411 | 0.05757067 | 0.39140631 | 0.14708671 | 0.88306358 | 0.98358022 |
| SLC35A1 | 33.442596 | 0.54425561 | 0.55154203 | 0.98678901 | 0.32374608 | 0.84285425 |

| | | | | | | |
|---|---|---|---|---|---|---|
| RARS2 | 279.497076 | -0.3600007 | 0.22362992 | -1.6098057 | 0.10744027 | 0.63477557 |
| ORC3 | 192.489778 | -0.1415067 | 0.21380168 | -0.6618594 | 0.50806131 | 0.91235137 |
| AKIRIN2 | 62.6266183 | 0.12416842 | 0.30955887 | 0.40111409 | 0.68833613 | 0.95597173 |
| RNGTT | 492.302251 | -0.3659755 | 0.24682785 | -1.4827155 | 0.13815002 | 0.68146162 |
| PNRC1 | 311.833245 | 0.04173815 | 0.37525759 | 0.11122534 | 0.91143766 | 0.98788932 |
| PM20D2 | 111.42854 | 0.18858333 | 0.21469135 | 0.8783928 | 0.37973059 | 0.86587828 |
| UBE2J1 | 140.921241 | 0.07870611 | 0.25999712 | 0.30271918 | 0.76210388 | 0.96218738 |
| RRAGD | 92.9830235 | 0.03757759 | 0.50125267 | 0.07496737 | 0.94024067 | 0.99243654 |
| ANKRD6 | 356.818138 | 0.04985618 | 0.2082984 | 0.2393498 | 0.81083435 | 0.9680627 |
| LYRM2 | 139.055121 | 0.34309252 | 0.31974291 | 1.0730262 | 0.28325936 | 0.82666614 |
| MDN1 | 312.247734 | -0.2781125 | 0.22890356 | -1.2149767 | 0.22437497 | 0.78966267 |
| CASP8AP2 | 142.875892 | -0.2869249 | 0.22654598 | -1.2665197 | 0.2053271 | 0.77193931 |
| BACH2 | 715.295586 | 0.00906079 | 0.63102238 | 0.0143589 | 0.98854365 | 0.99814557 |
| MAP3K7 | 317.403628 | -0.1851686 | 0.24638234 | -0.75155 | 0.45232174 | 0.89473986 |
| MANEA | 70.5676054 | 0.14143011 | 0.23111398 | 0.61194962 | 0.54057109 | 0.92422563 |
| UFL1-AS1 | 99.3308094 | 0.64007023 | 0.43996041 | 1.45483597 | 0.14571468 | 0.69350129 |
| UFL1 | 196.906623 | 0.11434168 | 0.17492878 | 0.65364701 | 0.51333925 | 0.91512427 |
| NDUFAF4 | 51.4114665 | 0.36177475 | 0.50718906 | 0.71329368 | 0.47566405 | 0.90242252 |
| KLHL32 | 126.264293 | -0.0269531 | 0.49383417 | -0.0545793 | 0.95647361 | 0.99565888 |
| MMS22L | 254.1414 | 0.63669145 | 0.41219962 | 1.54461921 | 0.1224384 | 0.66381501 |
| AL589740.1 | 60.8537199 | 0.44587247 | 0.53873862 | 0.82762299 | 0.40788405 | 0.87948803 |
| FBXL4 | 369.006691 | -0.1657273 | 0.3159286 | -0.524572 | 0.59988076 | 0.93728325 |
| PNISR | 1032.76889 | -0.007609 | 0.20054943 | -0.0379406 | 0.96973502 | 0.99645289 |
| AL513550.1 | 58.7164843 | -0.257538 | 0.50054009 | -0.5145202 | 0.60688831 | 0.93925224 |
| USP45 | 205.570842 | 0.01865016 | 0.342075 | 0.05452068 | 0.95652033 | 0.99565888 |
| TSTD3 | 47.4523557 | -0.4240801 | 0.35036954 | -1.2103794 | 0.22613333 | 0.78982769 |
| CCNC | 303.309127 | -0.0650034 | 0.2829964 | -0.2296969 | 0.81832731 | 0.96883031 |
| SIM1 | 725.462292 | -0.0806562 | 0.36153899 | -0.2230912 | 0.82346454 | 0.97018919 |
| ASCC3 | 339.960792 | -0.1168023 | 0.20389079 | -0.5728669 | 5.67E-01 | 0.93122667 |
| GRIK2 | 153.368133 | 1.26459377 | 0.55708895 | 2.27000333 | 0.02320738 | 0.36457774 |
| HACE1 | 197.284276 | 0.02431802 | 0.29368209 | 0.08280388 | 0.93400748 | 0.99243654 |
| PREP | 134.255413 | -0.0167638 | 0.23978317 | -0.0699124 | 0.94426337 | 0.99265578 |
| ATG5 | 393.014223 | -0.0552445 | 0.21560401 | -0.2562315 | 0.79777207 | 0.9653598 |
| CRYBG1 | 1600.4954 | 0.49927001 | 0.39283583 | 1.27093807 | 0.20375068 | 0.77142599 |
| RTN4IP1 | 82.2373424 | -0.1738729 | 0.38388195 | -0.4529333 | 0.65059674 | 0.95061388 |
| QRSL1 | 135.740654 | -0.116668 | 0.20430622 | -0.5710447 | 0.56796933 | 0.93122667 |
| LINC02532 | 3118.4703 | -1.1530028 | 0.21238112 | -5.4289327 | 5.67E-08 | 5.64E-05 |
| CD24 | 3068.37802 | -0.1096105 | 0.29354946 | -0.3733972 | 0.70885285 | 0.95752169 |
| C6orf203 | 75.3657877 | 0.36829677 | 0.31363139 | 1.17429819 | 0.24027561 | 0.80057052 |
| PDSS2 | 709.094622 | -0.2906568 | 0.21403998 | -1.3579556 | 0.17447776 | 0.73199638 |
| SOBP | 434.35978 | 0.71900808 | 0.6214694 | 1.15694848 | 0.24729341 | 0.80072691 |
| SCML4 | 58.3238506 | -0.3421737 | 0.45112152 | -0.7584955 | 0.44815439 | 0.89297728 |
| SEC63 | 382.472171 | -0.2036604 | 0.17247969 | -1.1807792 | 0.23769043 | 0.79810098 |
| OSTM1 | 122.397043 | -0.1982717 | 0.28336258 | -0.6997103 | 0.48410823 | 0.90656031 |
| SNX3 | 363.979179 | -0.1647186 | 0.3256021 | -0.5058891 | 0.61293449 | 0.94003018 |
| AFG1L | 247.01512 | -0.3278704 | 0.3042308 | -1.0777028 | 0.28116639 | 0.82628479 |
| FOXO3 | 658.850834 | 0.19172464 | 0.26968852 | 0.71091137 | 0.47713916 | 0.90242252 |
| ARMC2 | 186.555636 | -0.0424445 | 0.26985248 | -0.1572876 | 0.87501819 | 0.98267808 |
| SESN1 | 330.815969 | -0.0538945 | 0.27734479 | -0.194323 | 0.84592299 | 0.97643646 |
| CEP57L1 | 93.0668038 | -0.3054312 | 0.3678257 | -0.8303694 | 0.40632994 | 0.87948803 |
| CD164 | 551.519243 | 0.04128169 | 0.26001468 | 0.15876677 | 0.87385263 | 0.98246825 |
| PPIL6 | 125.706731 | 0.66220179 | 0.34748955 | 1.90567399 | 0.05669252 | 0.52346922 |
| MICAL1 | 34.0059309 | -0.1173768 | 0.32275919 | -0.3636669 | 0.71610678 | 0.95791799 |
| ZBTB24 | 38.5536957 | 0.70168628 | 0.36661297 | 1.91397015 | 0.05562397 | 0.51955366 |

| | | | | | | |
|---|---|---|---|---|---|---|
| AK9 | 198.006622 | -0.0213703 | 0.24345158 | -0.0877805 | 0.93005114 | 0.99216065 |
| FIG4 | 179.03236 | -0.336878 | 0.24806885 | -1.358002 | 0.17446304 | 0.73199638 |
| WASF1 | 40.984607 | -0.0575557 | 0.31605369 | -0.1821074 | 0.85549845 | 0.97718353 |
| CDC40 | 101.127286 | 0.14674303 | 0.24720286 | 0.59361382 | 0.55277043 | 0.92872652 |
| CDK19 | 582.178194 | -0.2944942 | 0.33079823 | -0.8902533 | 0.3733299 | 0.86201082 |
| AMD1 | 262.966278 | -0.1239643 | 0.34221645 | -0.3622394 | 0.71717316 | 0.95791799 |
| GTF3C6 | 201.406498 | -0.0577765 | 0.46751356 | -0.1235826 | 0.90164579 | 0.9875066 |
| RPF2 | 76.5522086 | -0.3863235 | 0.22853755 | -1.6904157 | 0.09094845 | 0.60942442 |
| SLC16A10 | 450.132783 | -0.0585487 | 0.36657228 | -0.1597194 | 0.87310214 | 0.98246494 |
| MFSD4B | 100.003856 | 0.17593112 | 0.25588458 | 0.68754091 | 0.49174193 | 0.90730741 |
| REV3L | 929.339904 | 0.18268433 | 0.28325583 | 0.64494465 | 0.51896305 | 0.91751084 |
| TRAF3IP2-AS | 290.266769 | -0.0396314 | 0.33506675 | -0.1182791 | 0.90584649 | 0.9875066 |
| TRAF3IP2 | 105.785082 | 0.50613492 | 0.34201859 | 1.47984623 | 0.13891429 | 0.68156449 |
| TUBE1 | 107.48822 | 0.16095735 | 0.24835168 | 0.64810252 | 0.51691864 | 0.91751084 |
| FAM229B | 59.7904443 | 0.67257365 | 0.33646929 | 1.99891542 | 0.04561751 | 0.48339878 |
| MARCKS | 155.561469 | -0.4322255 | 0.29232621 | -1.4785725 | 0.1392546 | 0.68227594 |
| HDAC2 | 175.395678 | 0.11470133 | 0.16955127 | 0.6764994 | 0.49872362 | 0.90839719 |
| FRK | 635.326715 | 0.16959845 | 0.24835893 | 0.68287639 | 0.49468496 | 0.90839719 |
| NT5DC1 | 637.842634 | -0.0981115 | 0.21972462 | -0.4465202 | 0.65522155 | 0.95100227 |
| TSPYL4 | 58.6317339 | 0.16611548 | 0.35887396 | 0.46287973 | 0.64345058 | 0.94949493 |
| DSE | 313.160108 | 0.50478248 | 0.25807194 | 1.95597583 | 0.050468 | 0.49596288 |
| TSPYL1 | 308.710282 | 0.68537408 | 0.34609311 | 1.98031707 | 0.04766791 | 0.48600281 |
| RWDD1 | 104.120208 | 0.04800304 | 0.40499785 | 0.11852666 | 0.90565037 | 0.9875066 |
| ZUP1 | 49.0685669 | -0.325432 | 0.28464866 | -1.1432761 | 0.25292395 | 0.80350545 |
| KPNA5 | 155.041907 | 0.12342499 | 0.25791964 | 0.4785405 | 0.63226556 | 0.94659186 |
| DCBLD1 | 139.614433 | -0.0662372 | 0.29746815 | -0.2226699 | 0.8237924 | 0.97018919 |
| GOPC | 166.907504 | 0.3775047 | 0.25785191 | 1.4640369 | 0.14318387 | 0.69023197 |
| NUS1 | 183.669764 | 0.21354689 | 0.16917196 | 1.26230659 | 0.20683848 | 0.77200549 |
| SLC35F1 | 46.6194522 | 0.18773412 | 0.29484723 | 0.63671658 | 0.52430947 | 0.91885674 |
| CEP85L | 266.678195 | 0.38756896 | 0.33484589 | 1.15745471 | 0.24708663 | 0.80072691 |
| MCM9 | 240.562749 | -0.0116484 | 0.27613927 | -0.0421831 | 0.96635276 | 0.99637506 |
| ASF1A | 39.4121774 | 0.31090379 | 0.31035322 | 1.00177401 | 0.31645275 | 0.84285425 |
| AL365275.1 | 67.1245852 | 0.33229393 | 0.65849154 | 0.504629 | 0.61381944 | 0.94048939 |
| FAM184A | 72.1874647 | 1.04487614 | 0.49401551 | 2.11506749 | 0.0344242 | 0.43050784 |
| MAN1A1 | 2158.07567 | 0.59679271 | 0.32246672 | 1.85071099 | 0.06421114 | 0.54860648 |
| TBC1D32 | 507.283107 | 0.15433545 | 0.26131867 | 0.59060249 | 0.5547868 | 0.92872652 |
| HSF2 | 60.944035 | -0.2373847 | 0.22199274 | -1.0693355 | 0.28491854 | 0.82781987 |
| SERINC1 | 348.956307 | 0.08996303 | 0.32945401 | 0.27306702 | 0.7848017 | 0.96513523 |
| PKIB | 52.942757 | 0.64453732 | 0.49731791 | 1.29602678 | 0.19496626 | 0.76233115 |
| RNF217 | 266.001141 | 0.14919855 | 0.39093652 | 0.38164392 | 0.7027255 | 0.95672887 |
| TPD52L1 | 396.937549 | 0.45089816 | 0.36172418 | 1.2465248 | 0.21257179 | 0.77709975 |
| HDDC2 | 168.158007 | 0.28838536 | 0.31723584 | 0.9090567 | 0.3633202 | 0.85694172 |
| NCOA7 | 689.483031 | -0.4135388 | 0.24773361 | -1.6692881 | 0.0950603 | 0.61367241 |
| HINT3 | 141.975697 | -0.198885 | 0.28294271 | -0.7029161 | 0.48210805 | 0.90595437 |
| TRMT11 | 139.612397 | -0.359272 | 0.23558548 | -1.5250176 | 0.1272547 | 0.67016545 |
| RNF146 | 89.826986 | 0.27490105 | 0.25222606 | 1.08989948 | 0.27575743 | 0.82123837 |
| ECHDC1 | 222.168234 | -0.0390531 | 0.2771534 | -0.1409079 | 0.88794266 | 0.98551241 |
| PTPRK | 1473.44996 | -0.4596973 | 0.30116749 | -1.5263841 | 0.12691423 | 0.66971859 |
| ARHGAP18 | 297.212279 | 0.78088017 | 0.31559289 | 2.47432749 | 0.01334873 | 0.28170334 |
| L3MBTL3 | 96.0163804 | 0.36220643 | 0.44230689 | 0.81890297 | 0.41284177 | 0.88153663 |
| EPB41L2 | 71.6817193 | 0.39569035 | 0.42975304 | 0.92073892 | 0.35718675 | 0.8557799 |
| AKAP7 | 139.493989 | 0.16883589 | 0.43604215 | 0.38720085 | 0.69860752 | 0.95672887 |
| MED23 | 178.791097 | -0.1620009 | 0.17980601 | -0.9009761 | 0.36760103 | 0.85902833 |
| STX7 | 476.562699 | -0.0168494 | 0.27681465 | -0.060869 | 0.95146357 | 0.99404743 |

| | | | | | | |
|---|---|---|---|---|---|---|
| SLC18B1 | 181.691736 | -0.2601683 | 0.41992283 | -0.6195623 | 0.53554602 | 0.92303275 |
| RPS12 | 499.539189 | 0.19315652 | 0.39442036 | 0.4897225 | 0.62433028 | 0.94457465 |
| EYA4 | 2129.66239 | 0.10445955 | 0.33133034 | 0.31527311 | 7.53E-01 | 0.96120879 |
| TARID | 55.6600659 | 0.48483388 | 0.63735717 | 0.76069416 | 0.44683976 | 0.8918683 |
| TBPL1 | 56.1180904 | 0.33490306 | 0.29048701 | 1.15290203 | 0.2489506 | 0.80097144 |
| SGK1 | 198.832939 | 0.40388825 | 0.72906231 | 0.55398318 | 0.57959036 | 0.93345492 |
| HBS1L | 204.124683 | -0.2089706 | 0.2211321 | -0.9450034 | 0.34465714 | 0.85148675 |
| AHI1 | 1105.49054 | -0.3413019 | 0.23958524 | -1.424553 | 0.15428646 | 0.70805312 |
| LINC00271 | 363.429699 | 0.52382884 | 0.46616155 | 1.12370668 | 0.26113748 | 0.80805101 |
| PDE7B | 1959.59449 | -0.7833817 | 0.4418268 | -1.7730516 | 0.07622015 | 0.57145422 |
| AL138828.1 | 252.833254 | -0.7093995 | 0.4678224 | -1.5163863 | 0.12942169 | 0.67207756 |
| BCLAF1 | 456.898232 | 0.10099198 | 0.14580814 | 0.69263606 | 0.48853797 | 0.90730741 |
| MAP7 | 920.815313 | -0.022023 | 0.17644654 | -0.1248139 | 0.90067087 | 0.9875066 |
| MAP3K5 | 459.41119 | 0.64622253 | 0.45300158 | 1.42653484 | 0.15371402 | 0.70747602 |
| PEX7 | 166.627413 | 0.04408237 | 0.20604383 | 0.21394659 | 0.83058871 | 0.97160502 |
| IFNGR1 | 158.027757 | 0.30005802 | 0.39714913 | 0.75552984 | 0.44993114 | 0.89324553 |
| PERP | 256.596157 | -0.0038956 | 0.35060109 | -0.0111111 | 0.99113478 | 0.99889703 |
| ARFGEF3 | 204.654468 | -0.8566275 | 0.34409924 | -2.4894779 | 0.01279309 | 0.27834029 |
| HEBP2 | 235.340133 | -0.1595135 | 0.39338051 | -0.4054941 | 0.68511433 | 0.9558749 |
| NHSL1 | 693.210837 | -0.926612 | 0.35050274 | -2.6436656 | 0.00820136 | 0.22277119 |
| CCDC28A | 94.2970194 | 0.73706243 | 0.3616166 | 2.03824279 | 0.04152566 | 0.46694087 |
| REPS1 | 324.356715 | 0.02622171 | 0.20774055 | 0.12622338 | 0.89955511 | 0.98660836 |
| HECA | 74.1678998 | -0.1771296 | 0.54932905 | -0.3224471 | 0.74711399 | 0.96109466 |
| CITED2 | 79.1864937 | 0.03923471 | 0.37122657 | 0.10568939 | 0.9158288 | 0.98868699 |
| VTA1 | 156.248948 | -0.0631315 | 0.24900682 | -0.2535333 | 0.79985614 | 0.96568323 |
| ADGRG6 | 100.170578 | 0.26238287 | 0.6284377 | 0.41751612 | 0.67630094 | 0.95529325 |
| HIVEP2 | 907.045303 | -0.6361614 | 0.51809891 | -1.2278764 | 0.21949336 | 0.78630025 |
| AIG1 | 618.719664 | 0.07577501 | 0.13050947 | 0.5806093 | 0.5615038 | 0.92947347 |
| ADAT2 | 63.1434075 | -0.1675328 | 0.27089446 | -0.6184431 | 0.53628329 | 0.92324352 |
| PEX3 | 185.139956 | 0.05034786 | 0.2674818 | 0.1882291 | 0.85069707 | 0.97718353 |
| FUCA2 | 77.4179547 | 0.06381059 | 0.22589986 | 0.28247289 | 0.77758093 | 0.96472142 |
| PHACTR2 | 377.380165 | 0.7748883 | 0.4876904 | 1.58889389 | 0.11208435 | 0.64470032 |
| PLAGL1 | 166.691381 | 0.40938506 | 0.57000734 | 0.71821015 | 0.47262772 | 0.90213021 |
| SF3B5 | 213.931391 | 0.46213096 | 0.40740754 | 1.13432108 | 0.25665989 | 0.80676096 |
| UTRN | 1246.70008 | 0.01675219 | 0.22379053 | 0.07485657 | 0.94032883 | 0.99243654 |
| EPM2A | 104.727346 | -0.0248567 | 0.36439267 | -0.0682139 | 0.94561534 | 0.99299085 |
| FBXO30 | 37.2840953 | 0.29308979 | 0.35439822 | 0.82700694 | 0.40823313 | 0.87948803 |
| AL356599.1 | 139.861638 | 0.43991218 | 0.39557443 | 1.11208447 | 0.26610183 | 0.81257203 |
| SHPRH | 623.863666 | -0.0600843 | 0.23868757 | -0.2517278 | 0.80125145 | 0.9661728 |
| RAB32 | 41.6884725 | 0.03747807 | 0.36350014 | 0.10310332 | 0.91788097 | 0.99005687 |
| STXBP5 | 869.700403 | -0.0840545 | 0.18014286 | -0.466599 | 0.64078682 | 0.94935885 |
| SAMD5 | 770.128883 | -0.6459646 | 0.28244672 | -2.2870318 | 0.02219397 | 0.35843654 |
| AL033504.1 | 480.448606 | -0.8886183 | 0.28033693 | -3.1698224 | 0.00152532 | 0.09927452 |
| SASH1 | 547.665738 | -1.0297523 | 0.39725716 | -2.5921554 | 0.00953767 | 0.23684368 |
| TAB2 | 518.589448 | 0.03748753 | 0.19013913 | 0.19715842 | 0.84370357 | 0.97561662 |
| PPIL4 | 237.066849 | 0.11043783 | 0.23508988 | 0.46976854 | 0.63852039 | 0.94887688 |
| GINM1 | 349.733992 | 0.56226182 | 0.14781972 | 3.80369976 | 0.00014255 | 0.02411295 |
| KATNA1 | 144.423053 | -0.1535229 | 0.20349963 | -0.7544139 | 0.45060073 | 0.89333031 |
| LATS1 | 250.001389 | 0.01819448 | 0.2694587 | 0.06752234 | 0.94616588 | 0.99307065 |
| NUP43 | 102.303412 | -0.0716097 | 0.2473762 | -0.2894768 | 0.7722165 | 0.96388926 |
| PCMT1 | 233.592797 | -0.0655154 | 0.23451671 | -0.2793635 | 0.77996587 | 0.96513523 |
| LRP11 | 58.2500656 | 0.98502871 | 0.45002402 | 2.18883586 | 0.02860877 | 0.40623517 |
| PPP1R14C | 216.684624 | -0.6927171 | 0.39194242 | -1.767395 | 0.07716209 | 0.57405253 |
| PLEKHG1 | 269.749613 | -0.0585798 | 0.33487095 | -0.1749325 | 0.86113271 | 0.97841862 |

| | | | | | | |
|---|---|---|---|---|---|---|
| MTHFD1L | 521.037637 | -0.4132091 | 0.48806545 | -0.8466264 | 0.39720342 | 0.87462201 |
| AKAP12 | 109.753079 | -0.107103 | 0.74849347 | -0.1430914 | 0.88621799 | 0.98489549 |
| RMND1 | 98.8131921 | -0.0534382 | 0.29338231 | -0.1821454 | 0.85546864 | 0.97718353 |
| ARMT1 | 59.7820491 | 0.35277042 | 0.37599072 | 0.93824236 | 0.34811987 | 0.85348649 |
| SYNE1 | 1557.42088 | 1.48609318 | 0.32606243 | 4.55769515 | 5.17E-06 | 0.00257078 |
| MTRF1L | 58.7492777 | 0.0033701 | 0.35402335 | 0.00951942 | 0.99240472 | 0.99919666 |
| OPRM1 | 47.5646245 | -0.278558 | 0.83039587 | -0.3354521 | 0.73728407 | 0.96078945 |
| IPCEF1 | 471.333886 | 0.0150361 | 0.65258287 | 0.02304091 | 0.98161764 | 0.99696835 |
| CNKSR3 | 568.981084 | 0.40841682 | 0.33905673 | 1.2045678 | 0.22837019 | 0.79042788 |
| SCAF8 | 673.662711 | -0.0731173 | 0.23222005 | -0.314862 | 0.75286647 | 0.96120879 |
| TIAM2 | 362.275049 | -0.104773 | 0.33701071 | -0.3108894 | 0.75588473 | 0.96120879 |
| TFB1M | 192.521051 | -0.3310583 | 0.26687657 | -1.2404921 | 0.21479345 | 0.77890374 |
| ARID1B | 3251.88329 | -0.2065185 | 0.22117098 | -0.9337504 | 0.35043266 | 0.85447517 |
| TMEM242 | 87.3266976 | 0.0971086 | 0.2363696 | 0.41083372 | 0.68119447 | 0.95574492 |
| ZDHHC14 | 290.275109 | 0.60217781 | 0.3579338 | 1.682372 | 0.09249673 | 0.61120158 |
| SNX9 | 667.529645 | -0.7799285 | 0.39428679 | -1.9780741 | 0.04792035 | 0.48616194 |
| SYNJ2 | 186.869828 | -2.0225572 | 0.56372216 | -3.5878618 | 0.0003334 | 0.04110637 |
| SERAC1 | 58.5143634 | -0.1655042 | 0.30625398 | -0.540415 | 0.58891086 | 0.93483536 |
| GTF2H5 | 155.513972 | 0.03503402 | 0.43336261 | 0.08084227 | 0.93556739 | 0.99243654 |
| TULP4 | 520.772389 | -0.2187955 | 0.23064204 | -0.9486367 | 0.3428054 | 0.85093239 |
| TMEM181 | 163.22806 | -0.3312818 | 0.2357701 | -1.4051053 | 0.15998998 | 0.71409437 |
| DYNLT1 | 354.052983 | -0.4701169 | 0.56810707 | -0.8275146 | 0.40794543 | 0.87948803 |
| SYTL3 | 296.633044 | -2.7587935 | 0.93782032 | -2.941708 | 0.00326408 | 0.14358074 |
| EZR | 819.57587 | -0.1755245 | 0.47001647 | -0.3734433 | 0.7088185 | 0.95752169 |
| RSPH3 | 180.409313 | 0.18684118 | 0.26371053 | 0.7085086 | 0.47862947 | 0.90265195 |
| AL035530.2 | 125.649968 | 0.34370867 | 0.45356093 | 0.75780043 | 0.44857046 | 0.89301266 |
| SOD2 | 33.8775365 | -1.0928812 | 0.56242262 | -1.9431672 | 0.05199596 | 0.5054895 |
| SOD2.1 | 90.1358494 | -0.9587518 | 0.45483117 | -2.107929 | 0.03503713 | 0.43353051 |
| WTAP | 378.015048 | -0.6164773 | 0.31561089 | -1.9532828 | 0.05078609 | 0.49835041 |
| ACAT2 | 83.1544011 | -0.5379564 | 0.67299752 | -0.7993439 | 0.42409105 | 0.88391318 |
| TCP1 | 139.201902 | 0.15673366 | 0.32497506 | 0.48229441 | 0.62959682 | 0.94596555 |
| MRPL18 | 106.854751 | 0.3867086 | 0.47847738 | 0.80820665 | 0.41897163 | 0.88261006 |
| IGF2R | 583.142703 | -0.3394602 | 0.26781169 | -1.2675332 | 0.20496469 | 0.77193931 |
| MAP3K4 | 312.742524 | -0.1784612 | 0.20172655 | -0.8846689 | 0.37633525 | 0.86358517 |
| AGPAT4 | 149.844798 | -0.4471784 | 0.24108972 | -1.8548215 | 0.06362171 | 0.54850785 |
| PRKN | 2904.81475 | -0.1927778 | 0.18950637 | -1.0172626 | 0.3090285 | 0.84136316 |
| PACRG | 584.103346 | -0.165633 | 0.28717369 | -0.5767693 | 0.56409533 | 0.93077674 |
| QKI | 1021.18887 | -0.0605819 | 0.18865818 | -0.3211199 | 0.7481195 | 0.96109466 |
| PDE10A | 4570.11467 | -2.6128756 | 0.76603449 | -3.4109112 | 0.00064746 | 0.05952104 |
| LINC00473 | 260.90232 | -5.28829 | 1.35568654 | -3.9008206 | 9.59E-05 | 0.02050383 |
| SFT2D1 | 156.543667 | -0.0041805 | 0.22327454 | -0.0187236 | 0.98506156 | 0.99762072 |
| MPC1 | 418.514648 | 0.29161283 | 0.2215205 | 1.31641466 | 0.18803491 | 0.7526909 |
| RPS6KA2 | 364.035089 | 0.12760533 | 0.25074252 | 0.50890984 | 0.61081542 | 0.94003018 |
| Z94721.1 | 89.7361994 | -0.6323307 | 0.43183714 | -1.4642805 | 0.14311732 | 0.69023197 |
| FGFR1OP | 249.359619 | -0.4135484 | 0.19328916 | -2.1395323 | 0.03239259 | 0.41704459 |
| AFDN | 1278.32155 | -0.4241718 | 0.19922999 | -2.1290562 | 0.03324961 | 0.42373931 |
| WDR27 | 277.837022 | 0.12611255 | 0.20153531 | 0.62575909 | 0.53147297 | 0.92211604 |
| C6orf120 | 40.3061726 | 0.42680948 | 0.29889995 | 1.42793426 | 0.15331078 | 0.70717297 |
| PHF10 | 142.93927 | 0.28686756 | 0.2797024 | 1.02561707 | 0.30507212 | 0.83709431 |
| ERMARD | 80.2963022 | 0.03337599 | 0.27149483 | 0.12293416 | 0.90215923 | 0.9875066 |
| FAM120B | 446.165156 | -0.2473732 | 0.24832984 | -0.9961476 | 0.31917845 | 0.84285425 |
| PSMB1 | 278.34928 | 0.2661638 | 0.38461584 | 0.69202505 | 0.4889216 | 0.90730741 |
| TBP | 61.5037146 | -0.3079327 | 0.26060618 | -1.1816018 | 0.23736374 | 0.7980661 |
| PDCD2 | 93.1545939 | -0.0578892 | 0.24793433 | -0.2334861 | 0.81538398 | 0.96844493 |

| | | | | | | |
|---|---|---|---|---|---|---|
| FAM20C | 797.758847 | -0.6801777 | 0.38426127 | -1.7700917 | 0.07671187 | 0.57260711 |
| PDGFA | 53.9602868 | -1.2186856 | 0.39628716 | -3.075259 | 0.0021032 | 0.11724213 |
| PRKAR1B | 134.429704 | -0.0137393 | 0.26135889 | -0.0525686 | 0.95807561 | 0.99579587 |
| DNAAF5 | 108.366631 | -0.2838679 | 0.26561555 | -1.0687172 | 0.28519714 | 0.82804896 |
| SUN1 | 250.807525 | -0.1026731 | 0.17406381 | -0.5898592 | 0.55528504 | 0.92872652 |
| ADAP1 | 44.342013 | 0.24942189 | 0.43269381 | 0.57643968 | 0.56431803 | 0.93090712 |
| COX19 | 107.910716 | -0.221729 | 0.26886162 | -0.8246958 | 0.40954434 | 0.88008904 |
| C7orf50 | 326.583443 | -0.2887595 | 0.32574859 | -0.8864489 | 0.37537567 | 0.86357183 |
| MICALL2 | 49.1878605 | -0.2101668 | 0.35688977 | -0.5888842 | 0.55593894 | 0.92872652 |
| INTS1 | 129.461697 | 0.04180026 | 0.29287921 | 0.14272183 | 0.88650988 | 0.98511542 |
| MAFK | 143.211914 | -0.5824899 | 0.56749376 | -1.0264251 | 0.30469124 | 0.83709431 |
| TMEM184A | 53.8701286 | -0.3346318 | 0.41820586 | -0.8001605 | 0.4236178 | 0.88391318 |
| PSMG3 | 53.6812529 | 0.39634736 | 0.31029407 | 1.27732817 | 0.20148641 | 0.76957251 |
| MAD1L1 | 159.065332 | -0.4308263 | 0.28081109 | -1.534221 | 0.1249753 | 0.6663037 |
| MRM2 | 39.2205033 | 0.33523899 | 0.35835788 | 0.93548656 | 0.3495376 | 0.8535343 |
| SNX8 | 81.1811385 | -0.1012875 | 0.39867781 | -0.2540586 | 0.79945029 | 0.96556863 |
| EIF3B | 198.775556 | -0.1747272 | 0.22747164 | -0.7681275 | 0.44241143 | 0.89005186 |
| CHST12 | 50.6870445 | -0.4005391 | 0.38222074 | -1.0479261 | 0.29467268 | 0.8337535 |
| BRAT1 | 49.4904702 | 0.22540164 | 0.27842152 | 0.80956975 | 0.4181875 | 0.88242985 |
| IQCE | 127.680644 | -0.2335135 | 0.25752722 | -0.9067528 | 0.36453752 | 0.85831347 |
| TTYH3 | 99.860459 | -0.5321861 | 0.51630305 | -1.030763 | 0.30265196 | 0.83709431 |
| GNA12 | 160.464887 | 0.03259679 | 0.26854545 | 0.12138278 | 0.90338785 | 0.9875066 |
| SDK1 | 2224.78755 | 0.2989813 | 0.31507428 | 0.94892323 | 0.34265966 | 0.85093239 |
| FOXK1 | 310.274388 | -0.2070379 | 0.26105981 | -0.7930671 | 0.42773872 | 0.88535099 |
| AP5Z1 | 51.0760532 | -0.5812207 | 0.35022205 | -1.6595778 | 0.09699941 | 0.61609322 |
| RBAK-RBAKI | 171.529503 | -0.273041 | 0.18509809 | -1.475115 | 0.14018162 | 0.68408298 |
| WIPI2 | 212.841273 | -0.2195856 | 0.17950627 | -1.2232756 | 0.22122562 | 0.78752408 |
| TNRC18 | 781.264254 | -0.097896 | 0.26112436 | -0.3749017 | 0.70773356 | 0.95752169 |
| FBXL18 | 62.6400967 | 0.03759772 | 0.41213458 | 0.0912268 | 0.92731238 | 0.99125191 |
| ACTB | 2795.32504 | -0.0765333 | 0.38296242 | -0.1998455 | 0.84160141 | 0.975446 |
| RNF216 | 365.282063 | -0.1529039 | 0.20690342 | -0.7390112 | 0.45990021 | 0.89753337 |
| CCZ1 | 65.833248 | -0.5984056 | 0.30291307 | -1.9755027 | 0.04821113 | 0.48661187 |
| PMS2 | 108.289039 | 0.15211803 | 0.32670771 | 0.46560892 | 0.64149545 | 0.94935885 |
| AIMP2 | 65.7090925 | 0.20541428 | 0.35758914 | 0.57444218 | 0.56566862 | 0.93114846 |
| EIF2AK1 | 209.690604 | 0.16521521 | 0.18072845 | 0.9141627 | 0.36063136 | 0.85608464 |
| USP42 | 168.910193 | -0.1294744 | 0.29415878 | -0.4401512 | 0.65982758 | 0.95127957 |
| CYTH3 | 172.66848 | 0.04909857 | 0.3047753 | 0.1610976 | 0.87201653 | 0.98223939 |
| RAC1 | 633.68506 | -0.0161953 | 0.26684167 | -0.0606924 | 0.95160416 | 0.99404743 |
| DAGLB | 46.403502 | -0.3160233 | 0.35789731 | -0.883 | 0.37723626 | 0.86432794 |
| ZDHHC4 | 60.0722951 | -0.314214 | 0.2409924 | -1.3038336 | 0.19229032 | 0.7597012 |
| C7orf26 | 42.5895903 | 0.04702486 | 0.29544223 | 0.15916771 | 0.87353674 | 0.98246494 |
| ZNF316 | 60.20809 | -0.2747203 | 0.30705253 | -0.8947014 | 0.37094671 | 0.86188219 |
| AC073343.2 | 50.6335637 | 0.15899546 | 0.4647254 | 0.34212776 | 0.73225475 | 0.96008876 |
| ZNF12 | 110.827071 | 0.12473802 | 0.28083421 | 0.44416961 | 0.65691996 | 0.95100227 |
| CCZ1B | 77.3422047 | -0.6174817 | 0.31117189 | -1.9843749 | 0.04721407 | 0.48600281 |
| C1GALT1 | 327.668798 | -0.4860679 | 0.36417612 | -1.3347055 | 0.18197276 | 0.74315969 |
| COL28A1 | 264.023639 | 0.88223068 | 0.37195525 | 2.37187317 | 0.01769816 | 0.31978753 |
| AC004982.2 | 57.9092211 | 0.83599439 | 0.31064273 | 2.69117638 | 0.00712005 | 0.20697274 |
| MIOS | 163.527634 | 0.28812948 | 0.23287445 | 1.23727389 | 0.21598542 | 0.78025756 |
| RPA3 | 134.298061 | 0.46225208 | 0.46631465 | 0.99128792 | 0.32154501 | 0.84285425 |
| UMAD1 | 1020.16366 | -0.0991511 | 0.25624277 | -0.3869421 | 0.6987991 | 0.95672887 |
| GLCCI1 | 174.334644 | 0.27752114 | 0.31364261 | 0.88483238 | 0.37624704 | 0.86357183 |
| ICA1 | 622.243555 | -0.2029272 | 0.3599641 | -0.563743 | 0.57292904 | 0.93248895 |
| NDUFA4 | 755.276147 | 0.69467363 | 0.38775302 | 1.79153636 | 0.07320727 | 0.56668778 |

| | | | | | | |
|---|---|---|---|---|---|---|
| PHF14 | 529.253456 | -0.46801 | 0.20791854 | -2.2509298 | 0.02438999 | 0.37424167 |
| THSD7A | 6303.15089 | -1.1931183 | 0.55518749 | -2.1490366 | 0.0316315 | 0.41489121 |
| TMEM106B | 255.195424 | -0.1454735 | 0.17830854 | -0.8158528 | 0.41458434 | 0.88242985 |
| VWDE | 47.6157968 | 1.46518302 | 0.48426061 | 3.02560851 | 0.00248133 | 0.13022925 |
| AC013470.2 | 52.6471789 | 0.86288514 | 0.5425802 | 1.59033659 | 0.11175895 | 0.64353508 |
| SCIN | 2169.03331 | 0.27604638 | 0.2956194 | 0.93378981 | 0.35041233 | 0.85447517 |
| ARL4A | 34.4491057 | 0.46350212 | 0.71933306 | 0.64434981 | 0.51934862 | 0.91751084 |
| AC011287.1 | 65.2063937 | -0.3916221 | 0.45937878 | -0.8525037 | 0.39393461 | 0.87253562 |
| AGMO | 251.914964 | -0.1936908 | 0.58636824 | -0.3303228 | 0.74115608 | 0.96084411 |
| ISPD | 144.318997 | -0.2408035 | 0.32783442 | -0.7345277 | 0.46262718 | 0.89801718 |
| ANKMY2 | 155.901087 | -0.2760816 | 0.25032403 | -1.1028971 | 0.27007187 | 0.81817548 |
| BZW2 | 292.00027 | -0.9985774 | 0.4658457 | -2.1435798 | 0.03206657 | 0.41517597 |
| TSPAN13 | 48.823807 | 0.05177807 | 0.33783949 | 0.15326233 | 0.87819141 | 0.98300919 |
| AGR2 | 143.086866 | 1.37878127 | 0.38503877 | 3.58088942 | 3.42E-04 | 0.04172826 |
| AC073332.1 | 52.2954245 | 0.32770072 | 0.42143277 | 0.77758718 | 0.43681242 | 0.88785137 |
| SNX13 | 450.865325 | -0.180654 | 0.22628363 | -0.7983519 | 0.42466631 | 0.88391318 |
| HDAC9 | 69.3094969 | -0.8264954 | 0.54685257 | -1.5113678 | 0.13069477 | 0.67339292 |
| TWISTNB | 53.8983601 | 0.44919369 | 0.45297339 | 0.99165582 | 0.32136545 | 0.84285425 |
| MACC1 | 1713.24472 | -0.0315406 | 0.25308136 | -0.1246263 | 0.90081944 | 0.9875066 |
| MACC1-AS1 | 39.102979 | 0.33915053 | 0.67858725 | 0.49978912 | 0.61722357 | 0.94233994 |
| ITGB8 | 967.250693 | 0.50627169 | 0.30090177 | 1.68251482 | 0.09246905 | 0.61120158 |
| SP4 | 167.601182 | 0.25016972 | 0.29373533 | 0.85168413 | 0.39438943 | 0.87253562 |
| DNAH11 | 140.137155 | -3.8592956 | 1.28789457 | -2.9965928 | 0.00273015 | 0.13560183 |
| RAPGEF5 | 68.5468706 | 1.96228242 | 0.56062316 | 3.50018081 | 0.00046494 | 0.05075624 |
| STEAP1B | 552.447921 | 0.29113258 | 0.41673909 | 0.69859677 | 0.48480406 | 0.90689776 |
| TOMM7 | 300.801368 | 0.34616334 | 0.3965001 | 0.87304729 | 0.3826373 | 0.86593382 |
| KLHL7 | 148.953332 | -0.2187483 | 0.28482346 | -0.7680136 | 0.44247908 | 0.89005186 |
| NUPL2 | 133.684426 | -0.1221326 | 0.30156625 | -0.4049943 | 0.68548169 | 0.9558749 |
| AC005082.1 | 55.7301231 | 0.29355623 | 0.32440996 | 0.90489276 | 0.3655222 | 0.85866868 |
| GPNMB | 55.6451156 | -0.2794905 | 0.68231516 | -0.4096208 | 6.82E-01 | 0.95574492 |
| MALSU1 | 112.406469 | 0.00876188 | 0.24640638 | 0.03555864 | 0.97163428 | 0.99656026 |
| TRA2A | 644.520471 | 0.01475928 | 0.23192792 | 0.06363736 | 0.94925898 | 0.99361356 |
| CCDC126 | 68.9977784 | 0.34406235 | 0.32213603 | 1.0680654 | 0.28549101 | 0.82810567 |
| FAM221A | 59.19666 | -0.2209515 | 0.28793514 | -0.7673657 | 0.44286414 | 0.89005186 |
| OSBPL3 | 873.188569 | 0.13948761 | 0.27359743 | 0.50982792 | 0.61017203 | 0.94003018 |
| CYCS | 331.533549 | 0.36691471 | 0.50537622 | 0.72602291 | 0.46782472 | 0.89976199 |
| HNRNPA2B1 | 1222.20395 | -0.1040026 | 0.2552624 | -0.407434 | 0.68368929 | 0.95574492 |
| CBX3 | 221.070726 | 0.02175388 | 0.26102868 | 0.08333906 | 0.93358194 | 0.99243654 |
| SNX10 | 160.788137 | -0.2245447 | 0.40177577 | -0.5588807 | 0.57624311 | 0.9331615 |
| AC004540.2 | 57.6809575 | 0.17604857 | 0.34683915 | 0.50757985 | 0.61174802 | 0.94003018 |
| AC004540.1 | 348.270242 | 0.68908469 | 0.49313934 | 1.39734277 | 0.16231052 | 0.71482182 |
| SKAP2 | 159.510795 | 0.22537184 | 0.5900062 | 0.38198217 | 0.70247459 | 0.95672887 |
| HIBADH | 700.017248 | 0.13029177 | 0.25669769 | 0.50756893 | 0.61175569 | 0.94003018 |
| TAX1BP1 | 898.537091 | 0.18366597 | 0.22020462 | 0.83406958 | 0.40424177 | 0.87834687 |
| JAZF1 | 199.342337 | 1.18209784 | 0.48024745 | 2.46143491 | 0.01383825 | 0.28408502 |
| AC005162.3 | 77.6331086 | 1.33715174 | 0.44378386 | 3.01306975 | 0.00258619 | 0.13156951 |
| CPVL | 558.636406 | 0.90205074 | 0.28842494 | 3.12750601 | 0.00176296 | 0.10618301 |
| AC007255.1 | 76.2903464 | -0.5687943 | 0.45371273 | -1.2536441 | 0.20997139 | 0.77551367 |
| PRR15 | 88.2596085 | -0.7287676 | 0.52825966 | -1.3795632 | 0.16772117 | 0.72363848 |
| SCRN1 | 279.402219 | 0.49937902 | 0.17179335 | 2.9068588 | 0.00365078 | 0.14773661 |
| FKBP14 | 63.4703872 | 0.23253565 | 0.27846624 | 0.83505866 | 0.40368468 | 0.87834687 |
| PLEKHA8 | 152.944103 | -0.3759163 | 0.22449532 | -1.6744952 | 0.09403332 | 0.61323531 |
| MTURN | 358.796738 | 0.66826367 | 0.26861944 | 2.48777101 | 0.01285465 | 0.27834029 |
| ZNRF2 | 368.263186 | 0.58421888 | 0.40817964 | 1.43127886 | 0.15235031 | 0.70429257 |

| | | | | | | |
|---|---|---|---|---|---|---|
| AC006978.2 | 336.842808 | 0.49447345 | 0.25874409 | 1.91105219 | 0.05599787 | 0.52165131 |
| NOD1 | 128.144166 | 0.08583167 | 0.37721927 | 0.22753786 | 0.82000553 | 0.96886922 |
| GGCT | 83.4851044 | 0.53364439 | 0.52141631 | 1.02345167 | 0.30609434 | 0.83709431 |
| AC005154.6 | 90.635987 | -0.134073 | 0.27849603 | -0.4814179 | 0.63021955 | 0.94596555 |
| GARS | 182.1536 | -0.6012828 | 0.32927961 | -1.8260553 | 0.06784193 | 0.55864172 |
| AQP1 | 51.9255984 | 0.07299462 | 0.51909872 | 0.14061799 | 0.88817174 | 0.98551241 |
| LSM5 | 94.2393704 | 0.01485545 | 0.37554879 | 0.03955664 | 0.96844659 | 0.9964399 |
| AVL9 | 505.340331 | -0.2043096 | 0.2610918 | -0.7825203 | 0.43390886 | 0.88785137 |
| KBTBD2 | 119.358056 | -0.1774603 | 0.2003014 | -0.8859661 | 0.37563579 | 0.86357183 |
| FKBP9 | 128.571628 | 0.31190491 | 0.24073149 | 1.29565484 | 0.19509443 | 0.76233115 |
| NT5C3A | 191.472294 | -0.5807303 | 0.38026427 | -1.5271755 | 0.12671738 | 0.66960383 |
| RP9 | 43.9697682 | 0.09776197 | 0.30717922 | 0.31825709 | 0.75028993 | 0.96109466 |
| BBS9 | 584.125217 | 0.20700757 | 0.1631218 | 1.26903686 | 0.20442792 | 0.77193931 |
| DPY19L1 | 133.397891 | -0.520673 | 0.2996084 | -1.737845 | 0.08223813 | 0.59202053 |
| HERPUD2 | 169.693851 | -0.2716956 | 0.22370274 | -1.2145386 | 0.22454211 | 0.78980378 |
| SEPT7-AS1 | 59.0806824 | 0.63972956 | 0.39221925 | 1.63105092 | 0.10287958 | 0.62635087 |
| 7-Sep | 580.382497 | -0.2061414 | 0.26285004 | -0.7842549 | 0.43289053 | 0.88722778 |
| EEPD1 | 130.867434 | -0.1251062 | 0.60083927 | -0.208219 | 0.83505797 | 0.97246185 |
| KIAA0895 | 149.330715 | 0.12691968 | 0.24953076 | 0.50863342 | 0.6110092 | 0.94003018 |
| AOAH | 82.067537 | -1.9425288 | 0.66551669 | -2.9188281 | 3.51E-03 | 0.14553941 |
| STARD3NL | 95.1928116 | 0.06875273 | 0.25011194 | 0.27488783 | 0.78340242 | 0.96513523 |
| VPS41 | 456.11122 | -0.2829266 | 0.24425536 | -1.1583232 | 0.24673217 | 0.80072691 |
| YAE1D1 | 48.9782868 | 0.49338461 | 0.28985501 | 1.70217729 | 0.08872214 | 0.60673738 |
| RALA | 197.935747 | 0.12507239 | 0.32437462 | 0.38558007 | 0.6998077 | 0.95672887 |
| LINC00265 | 35.40831 | 0.18270821 | 0.45907985 | 0.39798786 | 0.69063913 | 0.95659154 |
| CDK13 | 785.533825 | -0.1335371 | 0.22567443 | -0.5917247 | 0.55403495 | 0.92872652 |
| MPLKIP | 109.57671 | 0.66762801 | 0.31831764 | 2.09736417 | 0.03596135 | 0.4362094 |
| SUGCT | 94.6244775 | 0.38237443 | 0.36051016 | 1.06064813 | 0.28884984 | 0.83060304 |
| MRPL32 | 90.6074159 | 0.30268118 | 0.35902131 | 0.84307302 | 0.39918761 | 0.87596764 |
| STK17A | 67.8806584 | 0.22486178 | 0.48290782 | 0.4656412 | 0.64147235 | 0.94935885 |
| COA1 | 527.393144 | -0.1061141 | 0.18104064 | -0.5861343 | 0.55778526 | 0.92901206 |
| BLVRA | 91.5173131 | 0.16292053 | 0.32727895 | 0.49780326 | 0.61862272 | 0.94286883 |
| URGCP | 113.636408 | 0.233635 | 0.3852464 | 0.60645602 | 0.54421198 | 0.92526631 |
| UBE2D4 | 50.7113395 | -0.3173363 | 0.27302105 | -1.1623145 | 0.24510774 | 0.80072691 |
| DBNL | 102.074965 | 0.03489567 | 0.26089902 | 0.13375163 | 0.89359898 | 0.98597539 |
| POLM | 61.3466473 | 0.0929787 | 0.32115934 | 0.28950957 | 0.77219146 | 0.96388926 |
| POLD2 | 72.1114509 | 0.46334085 | 0.32287171 | 1.43506179 | 0.15126949 | 0.7029344 |
| YKT6 | 50.5520182 | -0.302225 | 0.34459998 | -0.8770315 | 0.38046954 | 0.86593382 |
| NUDCD3 | 325.964336 | -0.24826 | 0.21763573 | -1.1407136 | 0.25398912 | 0.80414079 |
| DDX56 | 78.0712568 | 0.24052003 | 0.2389923 | 1.00639239 | 0.31422685 | 0.84285425 |
| TMED4 | 278.81233 | 0.56837184 | 0.33356758 | 1.70391809 | 0.08839639 | 0.60667595 |
| OGDH | 635.834937 | 0.00534914 | 0.25859431 | 0.02068546 | 0.98349657 | 0.9971769 |
| ZMIZ2 | 158.615802 | -0.0811661 | 0.27913392 | -0.2907782 | 0.77122098 | 0.96388926 |
| PPIA | 917.126129 | 0.10700957 | 0.48005094 | 0.22291294 | 0.82360327 | 0.97018919 |
| H2AFV | 283.547365 | -0.0096052 | 0.3286762 | -0.0292238 | 0.97668611 | 0.99656026 |
| PURB | 166.767254 | 0.15816912 | 0.17131412 | 0.92326963 | 0.35586671 | 0.8557799 |
| CCM2 | 176.504458 | -0.2724051 | 0.26724071 | -1.0193249 | 0.30804875 | 0.83980597 |
| TBRG4 | 50.8194852 | 0.25017332 | 0.25996838 | 0.96232213 | 0.33588782 | 0.84552074 |
| ADCY1 | 40.8003994 | 0.29562714 | 0.59568431 | 0.49628157 | 0.61969577 | 0.94313269 |
| IGFBP3 | 260.575085 | -1.8613506 | 0.86588679 | -2.1496466 | 0.03158318 | 0.41477661 |
| AC004870.4 | 320.750001 | 1.3761357 | 0.80688858 | 1.70548417 | 0.08810416 | 0.60651125 |
| TNS3 | 1017.74514 | -0.1409305 | 0.29983379 | -0.4700287 | 0.63833449 | 0.94876549 |
| HUS1 | 82.0764648 | -0.0445015 | 0.24261492 | -0.1834246 | 0.85446486 | 0.97718353 |
| UPP1 | 192.46331 | -1.6832542 | 0.94916388 | -1.7734074 | 0.07616123 | 0.57145422 |

| | | | | | | |
|---|---|---|---|---|---|---|
| GRB10 | 136.511754 | -1.1163489 | 0.39322654 | -2.8389461 | 0.00452628 | 0.16528022 |
| COBL | 500.941011 | 0.00974254 | 0.22622101 | 0.04306645 | 0.96564856 | 0.99637506 |
| SEC61G | 144.284175 | -0.1035491 | 0.3695765 | -0.2801832 | 0.77933695 | 0.96513523 |
| EGFR | 844.377475 | 0.27621087 | 0.23784544 | 1.16130405 | 0.24551827 | 0.80072691 |
| LANCL2 | 124.80556 | -0.1264247 | 0.3503837 | -0.3608178 | 0.71823564 | 0.95796325 |
| VOPP1 | 410.068798 | 0.23249302 | 0.23663229 | 0.98250759 | 0.32584985 | 0.84367588 |
| NIPSNAP2 | 427.172141 | 0.29153584 | 0.24713558 | 1.17965951 | 0.23813566 | 0.79810098 |
| PSPH | 49.2480263 | 0.47964167 | 0.41148811 | 1.16562703 | 0.24376527 | 0.80072691 |
| CCT6A | 151.974824 | 0.02617601 | 0.32717837 | 0.08000533 | 0.93623302 | 0.99243654 |
| SUMF2 | 215.180808 | -0.0201773 | 0.21445894 | -0.0940848 | 0.9250418 | 0.99051393 |
| PHKG1 | 60.6745363 | -0.1006102 | 0.33936722 | -0.2964642 | 0.7668756 | 0.96285313 |
| CHCHD2 | 604.866974 | 0.34907013 | 0.50737671 | 0.68799006 | 0.49145904 | 0.90730741 |
| NUPR2 | 92.486461 | 0.89045595 | 0.59628683 | 1.49333494 | 0.13534953 | 0.67980087 |
| ZNF727 | 78.9062033 | -0.5489741 | 0.43038528 | -1.2755411 | 0.20211778 | 0.76996624 |
| ZNF736 | 294.550087 | 0.15534145 | 0.26520691 | 0.58573681 | 0.5580524 | 0.92901206 |
| ZNF680 | 273.437233 | 0.169866 | 0.30692261 | 0.55344896 | 0.57995602 | 0.93345492 |
| ZNF107 | 114.888927 | -0.3978926 | 0.3637822 | -1.0937659 | 0.27405767 | 0.81986798 |
| ZNF138 | 211.947052 | -0.0200988 | 0.30719493 | -0.0654267 | 0.94783423 | 0.99314907 |
| ZNF273 | 49.1903406 | 0.30786765 | 0.3968628 | 0.77575336 | 0.43789463 | 0.88825695 |
| ZNF117 | 136.109797 | 0.15354758 | 0.27467029 | 0.55902507 | 0.57614461 | 0.9331615 |
| VKORC1L1 | 208.73695 | -0.0493845 | 0.25782511 | -0.1915427 | 0.8481004 | 0.97656375 |
| GUSB | 79.6749061 | 0.5397773 | 0.24995398 | 2.15950672 | 0.03081088 | 0.41259458 |
| ASL | 41.4021891 | -0.4747955 | 0.35388562 | -1.3416637 | 0.17970507 | 0.74130581 |
| AC068533.4 | 55.507312 | -0.191525 | 0.4009159 | -0.4777187 | 0.63285044 | 0.94683634 |
| CRCP | 63.072304 | -0.2557704 | 0.33322037 | -0.7675714 | 0.44274183 | 0.89005186 |
| AC068533.3 | 45.5933357 | 0.12744616 | 0.45950475 | 0.27735549 | 0.78150716 | 0.96513523 |
| TPST1 | 1420.25482 | -0.1206156 | 0.26374847 | -0.4573128 | 0.64744622 | 0.95061388 |
| AC027644.4 | 348.67097 | 0.31477663 | 0.29792871 | 1.05655016 | 0.29071694 | 0.83156124 |
| AC027644.3 | 51.370311 | 0.00328316 | 0.45579342 | 0.00720317 | 0.99425275 | 0.99919666 |
| TMEM248 | 248.855978 | 0.0174426 | 0.2449753 | 0.07120145 | 0.94323743 | 0.99265578 |
| SBDS | 198.386469 | 0.76219265 | 0.49174963 | 1.54996078 | 0.12115093 | 0.65972015 |
| TYW1 | 415.86816 | -0.2047678 | 0.2405484 | -0.851254 | 0.39462828 | 0.87287977 |
| AUTS2 | 285.729752 | -1.0094985 | 0.5675653 | -1.7786474 | 0.07529759 | 0.57072562 |
| TYW1B | 294.35724 | -0.0303167 | 0.25019655 | -0.1211717 | 0.90355507 | 0.9875066 |
| POM121 | 273.34483 | -0.327008 | 0.23992822 | -1.3629408 | 0.17290116 | 0.72940007 |
| NSUN5 | 54.3127248 | -0.1710744 | 0.37667495 | -0.4541699 | 0.64970651 | 0.95061388 |
| BAZ1B | 304.335133 | 0.01609851 | 0.23942238 | 0.06723894 | 0.94639149 | 0.99307065 |
| BCL7B | 83.7218368 | -0.1851328 | 0.27219903 | -0.6801377 | 0.49641728 | 0.90839719 |
| TBL2 | 34.2938167 | -0.2349303 | 0.30996602 | -0.7579228 | 0.44849721 | 0.89301266 |
| MLXIPL | 47.1298668 | 0.60640097 | 0.65451212 | 0.92649311 | 0.35418978 | 0.8557799 |
| VPS37D | 48.3985511 | 0.7226973 | 0.36148376 | 1.99925246 | 0.04558104 | 0.48339878 |
| DNAJC30 | 48.8878252 | 0.79654299 | 0.27243276 | 2.92381496 | 0.0034577 | 0.14494688 |
| BUD23 | 145.067922 | -0.0433764 | 0.26344585 | -0.16465 | 0.86921947 | 0.9820194 |
| ABHD11 | 67.9470496 | 0.00246209 | 0.38565558 | 0.00638416 | 0.99490622 | 0.99919666 |
| CLDN3 | 185.424393 | 0.33012408 | 0.40084961 | 0.82356094 | 4.10E-01 | 0.8801764 |
| CLDN4 | 405.305314 | 0.18816312 | 0.37578489 | 0.5007203 | 0.61656798 | 0.94206626 |
| LIMK1 | 51.1922262 | -0.3921556 | 0.34955202 | -1.1218806 | 0.26191323 | 0.8087362 |
| EIF4H | 283.926173 | 0.0116669 | 0.44733028 | 0.02608117 | 0.9791926 | 0.99696835 |
| CLIP2 | 189.348934 | -0.221592 | 0.47057076 | -0.4709004 | 0.63771186 | 0.94851267 |
| GTF2IRD1 | 1185.14719 | -0.4776512 | 0.33763173 | -1.4147106 | 0.15715336 | 0.71160548 |
| GTF2I | 1143.51532 | 0.13429778 | 0.19654049 | 0.68330846 | 0.49441195 | 0.90839719 |
| GTF2IRD2 | 123.77853 | -0.7392291 | 0.29505705 | -2.505377 | 0.01223209 | 0.27034878 |
| CASTOR2 | 86.0861946 | 0.14858064 | 0.54744371 | 0.27140807 | 0.78607719 | 0.96513523 |
| RCC1L | 141.236627 | 0.12073059 | 0.21825263 | 0.55316899 | 0.5801477 | 0.93345492 |

| | | | | | | |
|---|---|---|---|---|---|---|
| GTF2IRD2B | 179.860729 | -0.6212643 | 0.23823999 | -2.6077246 | 0.00911462 | 0.23300339 |
| TRIM73 | 78.3839986 | 0.27239904 | 0.36508763 | 0.74611959 | 0.45559517 | 0.89626521 |
| POM121C | 231.318306 | -0.3880716 | 0.25350604 | -1.5308179 | 0.1258144 | 0.6674937 |
| SPDYE5 | 55.714431 | -0.199041 | 0.40402131 | -0.4926497 | 0.62226009 | 0.94418789 |
| HIP1 | 737.281204 | -0.7130285 | 0.46441642 | -1.5353216 | 0.12470486 | 0.66626798 |
| RHBDD2 | 125.178204 | -0.6674703 | 0.53392728 | -1.2501147 | 0.21125766 | 0.7767562 |
| POR | 455.680159 | -1.3522516 | 0.50110782 | -2.6985242 | 0.00696477 | 0.20503022 |
| TMEM120A | 55.4405258 | -0.8057656 | 0.33135832 | -2.4317046 | 0.01502796 | 0.29493072 |
| STYXL1 | 126.143468 | 0.2008082 | 0.36109233 | 0.55611316 | 0.5781335 | 0.93345492 |
| MDH2 | 206.522283 | 0.53580841 | 0.40111553 | 1.33579575 | 0.18161604 | 0.74315969 |
| HSPB1 | 704.744436 | 0.89692695 | 0.44591383 | 2.01143562 | 0.04427946 | 0.48009422 |
| YWHAG | 203.8976 | 0.01875469 | 0.30878532 | 0.060737 | 0.95156867 | 0.99404743 |
| SSC4D | 80.3096166 | 1.19628081 | 0.48918415 | 2.44546109 | 0.01446671 | 0.28823399 |
| DTX2 | 82.4385335 | -0.1799291 | 0.35397465 | -0.5083107 | 0.61123545 | 0.94003018 |
| POMZP3 | 58.9688355 | 0.09305035 | 0.33969989 | 0.27391928 | 0.78414666 | 0.96513523 |
| CCDC146 | 293.174348 | 0.22376875 | 0.29887005 | 0.74871588 | 0.45402847 | 0.8959176 |
| FGL2 | 583.705792 | 1.55498269 | 0.53544231 | 2.90410875 | 0.003683 | 0.14788453 |
| GSAP | 171.846491 | -0.1042469 | 0.30595012 | -0.3407316 | 0.73330566 | 0.96035038 |
| PTPN12 | 521.211489 | -0.7226725 | 0.29226416 | -2.4726689 | 0.01341083 | 0.28170334 |
| APTR | 95.0830342 | 0.40408231 | 0.28634722 | 1.41116196 | 0.15819686 | 0.71264724 |
| RSBN1L | 348.324506 | 0.1853531 | 0.18981202 | 0.97650879 | 0.3288124 | 0.84409527 |
| TMEM60 | 52.3450029 | 0.188588 | 0.57383154 | 0.32864697 | 0.74242254 | 0.96086911 |
| PHTF2 | 607.937073 | 0.52011662 | 0.31219125 | 1.66601919 | 9.57E-02 | 0.61367241 |
| MAGI2 | 1296.44291 | -0.0792437 | 0.31962002 | -0.247931 | 0.80418778 | 0.96679546 |
| GNAI1 | 531.261588 | -1.3414884 | 0.66677131 | -2.0119168 | 0.04422871 | 0.48009422 |
| SEMA3C | 59.5581822 | -0.6216429 | 0.7491018 | -0.829851 | 0.40662302 | 0.87948803 |
| CACNA2D1 | 844.147766 | 0.19443058 | 0.3119921 | 0.62319072 | 0.5331592 | 0.92277617 |
| PCLO | 766.034908 | -0.4133719 | 0.48423875 | -0.8536531 | 0.39329722 | 0.87223214 |
| KIAA1324L | 80.3994996 | 0.18178937 | 0.63260229 | 0.28736755 | 0.7738309 | 0.96388926 |
| DMTF1 | 383.117446 | -0.0642349 | 0.25539969 | -0.2515075 | 0.80142177 | 0.9661728 |
| TMEM243 | 148.503609 | 0.81557756 | 0.30008409 | 2.71783002 | 0.00657116 | 0.20019111 |
| TP53TG1 | 59.3577986 | 0.55632108 | 0.28193329 | 1.97323659 | 0.04846861 | 0.48661187 |
| CROT | 314.730861 | 0.88952164 | 0.33895591 | 2.62429896 | 0.00868275 | 0.22635636 |
| ABCB1 | 764.996177 | 1.28212228 | 0.26101781 | 4.91201061 | 9.01E-07 | 0.00070934 |
| SLC25A40 | 129.950102 | 0.19369051 | 0.21823509 | 0.88753146 | 0.37479284 | 0.86306942 |
| DBF4 | 42.6801158 | -0.34875 | 0.27109891 | -1.2864309 | 0.19829271 | 0.76654652 |
| ADAM22 | 571.174526 | 1.06198246 | 0.42254195 | 2.51331842 | 0.01196013 | 0.26883805 |
| SRI | 277.716321 | 0.3642984 | 0.40207613 | 0.90604334 | 0.36491291 | 0.85866868 |
| STEAP2 | 101.999841 | 0.85318793 | 0.29609719 | 2.88144553 | 0.00395856 | 0.15365064 |
| CFAP69 | 188.372912 | 0.35177777 | 0.1924238 | 1.82814061 | 0.06752845 | 0.55848603 |
| GTPBP10 | 124.476829 | 0.07137671 | 0.2796316 | 0.25525266 | 0.79852796 | 0.9653598 |
| CLDN12 | 121.476445 | 0.18187031 | 0.24243833 | 0.75017144 | 0.45315146 | 0.89544675 |
| CDK14 | 687.054271 | -0.1080313 | 0.28910989 | -0.3736686 | 0.70865086 | 0.95752169 |
| FZD1 | 42.8848311 | 0.37788043 | 0.40543562 | 0.93203559 | 0.35131813 | 0.85504372 |
| AC079760.1 | 82.4747946 | -0.2019289 | 0.66173096 | -0.3051525 | 0.76024998 | 0.96191449 |
| MTERF1 | 359.537428 | 0.07685024 | 0.42980582 | 0.17880223 | 0.85809299 | 0.97779869 |
| AKAP9 | 1089.1838 | 0.06040138 | 0.14932567 | 0.40449431 | 0.68584926 | 0.9558749 |
| KRIT1 | 367.496795 | 0.04340661 | 0.21705516 | 0.19997962 | 0.84149652 | 0.975446 |
| ANKIB1 | 796.993393 | -0.3428174 | 0.14175444 | -2.4183894 | 0.01558938 | 0.30067202 |
| GATAD1 | 133.842956 | 0.12125752 | 0.24284765 | 0.4993152 | 0.61755735 | 0.94244117 |
| AC007566.1 | 86.4498321 | 0.09589384 | 0.43702903 | 0.21942213 | 0.82632124 | 0.97058548 |
| PEX1 | 155.393933 | 0.2072048 | 0.2074371 | 0.99888016 | 0.31785275 | 0.84285425 |
| RBM48 | 33.9233757 | 0.64930786 | 0.37828383 | 1.71645681 | 0.08607847 | 0.60140155 |
| FAM133B | 189.508577 | -0.0950524 | 0.22948309 | -0.4142021 | 0.67872611 | 0.95574492 |

| | | | | | | |
|---|---|---|---|---|---|---|
| CDK6 | 1062.57092 | 0.2056674 | 0.25738912 | 0.79905242 | 4.24E-01 | 0.88391318 |
| SAMD9 | 72.4499237 | -0.5273104 | 0.64084209 | -0.8228399 | 0.41059905 | 0.88036458 |
| SAMD9L | 165.804253 | -0.8540018 | 0.72297058 | -1.18124 | 0.2375074 | 0.79810098 |
| VPS50 | 372.118407 | -0.1474748 | 0.27044995 | -0.5452943 | 0.58555111 | 0.93483536 |
| GNGT1 | 59.6224909 | 0.46598027 | 0.67346759 | 0.69191195 | 0.48899263 | 0.90730741 |
| TFPI2 | 62.2223104 | -1.0645679 | 1.00715531 | -1.0570047 | 0.29050944 | 0.83138693 |
| GNG11 | 134.968711 | 0.53625227 | 0.55338362 | 0.96904253 | 0.33252397 | 0.84442294 |
| BET1 | 152.865985 | 0.03291011 | 0.31111841 | 0.10578 | 0.9157569 | 0.98868699 |
| CASD1 | 213.005552 | 0.13783306 | 0.27964482 | 0.49288617 | 0.62209302 | 0.94418789 |
| SGCE | 291.355738 | 0.40600849 | 0.30828765 | 1.31697943 | 0.18784552 | 0.7526909 |
| PEG10 | 36.1966726 | 0.75391139 | 0.65313566 | 1.15429524 | 0.24837915 | 0.80097144 |
| PPP1R9A | 1880.6393 | -0.1235117 | 0.17355521 | -0.7116564 | 0.4766776 | 0.90242252 |
| AC002429.2 | 165.634289 | 0.09743332 | 0.51908421 | 0.18770234 | 0.85111 | 0.97718353 |
| PON3 | 38.0438964 | -0.4805135 | 0.42037343 | -1.1430635 | 0.25301221 | 0.80350545 |
| PON2 | 241.72705 | -0.3843272 | 0.23590307 | -1.6291741 | 0.10327617 | 0.62635087 |
| PDK4 | 3412.80273 | -2.9802227 | 1.23740855 | -2.4084388 | 0.01602091 | 0.30351249 |
| SLC25A13 | 513.393326 | -0.658907 | 0.27026702 | -2.4379854 | 0.01476937 | 0.29134933 |
| SEM1 | 291.466121 | -0.0253086 | 0.13871875 | -0.1824454 | 0.85523319 | 0.97718353 |
| SDHAF3 | 78.6836336 | -0.1563684 | 0.22350336 | -0.6996242 | 0.48416203 | 0.90656031 |
| ASNS | 46.5664721 | -0.2085067 | 0.3164325 | -0.6589294 | 0.50994113 | 0.91334187 |
| LMTK2 | 277.774404 | -0.3254771 | 0.36310251 | -0.8963779 | 0.37005099 | 0.86085113 |
| TECPR1 | 50.3599891 | -0.4593274 | 0.32730055 | -1.4033812 | 0.1605032 | 0.71409437 |
| BRI3 | 448.610748 | -0.6996494 | 0.35020767 | -1.9978129 | 0.04573694 | 0.48339878 |
| BAIAP2L1 | 1070.92166 | 0.1942857 | 0.6155848 | 0.3156116 | 0.75229733 | 0.96120879 |
| TRRAP | 341.180579 | -0.1009261 | 0.17887625 | -0.5642231 | 0.57260231 | 0.93218128 |
| SMURF1 | 422.868293 | -0.3032443 | 0.29965107 | -1.0119914 | 0.31154218 | 0.84279403 |
| ARPC1B | 97.989595 | -0.0332397 | 0.52609345 | -0.0631821 | 0.94962151 | 0.99361356 |
| PDAP1 | 126.990308 | 0.05572047 | 0.37666083 | 0.14793273 | 0.88239585 | 0.98335905 |
| BUD31 | 95.2954769 | 0.35867095 | 0.30654795 | 1.17003215 | 0.24198803 | 0.80072691 |
| CPSF4 | 47.6991049 | 0.06062354 | 0.24927158 | 0.24320278 | 0.80784831 | 0.96752828 |
| ZNF789 | 98.6411929 | 0.15029065 | 0.33272806 | 0.45169215 | 0.65149078 | 0.95061388 |
| ZNF394 | 77.0853014 | 0.29540677 | 0.24510362 | 1.2052322 | 0.22811368 | 0.79042788 |
| ZKSCAN5 | 56.5638842 | 0.12896369 | 0.343052 | 0.37593045 | 0.7069686 | 0.95752169 |
| FAM200A | 48.521818 | 0.51811148 | 0.4155679 | 1.24675529 | 0.21248724 | 0.77709975 |
| ZSCAN25 | 40.4130549 | 0.18672015 | 0.33765244 | 0.55299511 | 0.58026676 | 0.93345492 |
| TRIM4 | 123.776546 | 0.19912768 | 0.21632787 | 0.92049018 | 0.35731666 | 0.8557799 |
| ZKSCAN1 | 424.758342 | 0.00418106 | 0.19532156 | 0.02140604 | 0.98292175 | 0.9971769 |
| ZSCAN21 | 34.0012893 | 0.07919072 | 0.36588205 | 0.21643785 | 0.82864647 | 0.97117144 |
| ZNF3 | 71.8385747 | -0.294089 | 0.35622857 | -0.8255626 | 0.40905224 | 0.87982122 |
| COPS6 | 117.339004 | 0.17855845 | 0.30355807 | 0.58821843 | 0.55638569 | 0.92872652 |
| MCM7 | 39.3917872 | 0.08750572 | 0.4255147 | 0.20564675 | 0.83706686 | 0.97353666 |
| AP4M1 | 61.8579227 | -0.1105257 | 0.3008557 | -0.3673713 | 0.71334208 | 0.95791799 |
| TAF6 | 66.9531523 | 0.1780588 | 0.21713185 | 0.82004919 | 0.41218807 | 0.88103833 |
| LAMTOR4 | 313.701419 | 0.20953698 | 0.45419163 | 0.46134047 | 0.64455435 | 0.94949493 |
| CASTOR3 | 158.600199 | -0.2693516 | 0.47317876 | -0.5692387 | 0.56919419 | 0.93122667 |
| SPDYE3 | 118.888761 | 0.29140024 | 0.44692337 | 0.65201388 | 0.51439222 | 0.91544604 |
| PILRB | 191.635901 | -0.7348095 | 0.39489607 | -1.8607669 | 0.0627771 | 0.54720847 |
| PILRA | 36.355978 | -0.5946777 | 0.43683032 | -1.3613472 | 0.17340399 | 0.73012208 |
| MEPCE | 67.5437774 | -0.1961647 | 0.48367403 | -0.4055721 | 0.68505705 | 0.9558749 |
| PPP1R35 | 97.8541896 | -0.7716825 | 0.35967503 | -2.1454991 | 0.03191296 | 0.41517597 |
| TSC22D4 | 146.760313 | -0.2982192 | 0.38191245 | -0.7808576 | 0.43488626 | 0.88785137 |
| AGFG2 | 122.300893 | -0.2865438 | 0.48624883 | -0.5892946 | 0.55566369 | 0.92872652 |
| LRCH4 | 47.4210766 | -0.1234587 | 0.41398984 | -0.2982166 | 0.76553783 | 0.96285313 |
| GNB2 | 315.421954 | 0.15975518 | 0.36335109 | 0.43967167 | 0.66017492 | 0.95127957 |

| | | | | | | |
|---|---|---|---|---|---|---|
| GIGYF1 | 128.334234 | -0.3297464 | 0.29584685 | -1.1145848 | 0.26502837 | 0.81163409 |
| POP7 | 38.2878731 | 0.01504251 | 0.50669887 | 0.02968727 | 0.97631647 | 0.99656026 |
| EPHB4 | 47.3140139 | 0.1980419 | 0.36752003 | 0.53886016 | 0.58998335 | 0.93483536 |
| SLC12A9 | 96.7299454 | 0.58604323 | 0.37047702 | 1.58186121 | 0.11368126 | 0.6483668 |
| TRIP6 | 148.898503 | 0.17624208 | 0.40619216 | 0.43388843 | 0.66436946 | 0.95195406 |
| SRRT | 77.1959564 | -0.0755944 | 0.3064011 | -0.2467172 | 0.8051271 | 0.96704419 |
| TRIM56 | 369.393067 | -0.1093206 | 0.29673557 | -0.3684109 | 0.71256686 | 0.95791799 |
| AP1S1 | 95.3467602 | 0.44518975 | 0.47238751 | 0.94242489 | 0.34597515 | 0.8523814 |
| PLOD3 | 57.3138187 | -0.0304702 | 0.31527449 | -0.0966467 | 0.92300696 | 0.99051393 |
| ZNHIT1 | 162.701533 | 0.15413744 | 0.38078601 | 0.40478757 | 0.68563366 | 0.9558749 |
| CLDN15 | 38.3961943 | 0.10099673 | 0.27105265 | 0.37260926 | 0.70943928 | 0.95752169 |
| FIS1 | 314.59783 | 0.36805649 | 0.45651739 | 0.80622666 | 0.42011217 | 0.8828928 |
| IFT22 | 85.4813849 | 0.67364835 | 0.437196 | 1.5408383 | 0.12335614 | 0.66519057 |
| COL26A1 | 89.9263576 | 0.18815804 | 0.58048999 | 0.32413658 | 0.74583463 | 0.96109466 |
| CUX1 | 1463.02968 | -0.0973443 | 0.16777149 | -0.5802193 | 0.56176675 | 0.92959451 |
| PRKRIP1 | 401.355165 | -0.4091318 | 0.25102466 | -1.6298472 | 0.10313379 | 0.62635087 |
| ORAI2 | 161.014486 | 0.18811085 | 0.27722817 | 0.6785416 | 0.49742836 | 0.90839719 |
| ALKBH4 | 39.7254097 | 0.44313713 | 0.29362795 | 1.50917899 | 0.13125305 | 0.67390608 |
| POLR2J | 98.203159 | 0.12401306 | 0.46721273 | 0.26543169 | 0.79067693 | 0.96527296 |
| SPDYE2 | 43.4721056 | -0.2599209 | 0.51342795 | -0.5062461 | 0.6126839 | 0.94003018 |
| AC105052.3 | 101.122939 | 0.03967154 | 0.47760072 | 0.08306424 | 0.93380046 | 0.99243654 |
| FAM185A | 237.736102 | 0.01265946 | 0.32111733 | 0.03942315 | 0.96855302 | 0.9964399 |
| FBXL13 | 182.938043 | -0.0108356 | 0.30645293 | -0.0353582 | 0.97179414 | 0.99656026 |
| ARMC10 | 140.494855 | -0.3628166 | 0.25852401 | -1.4034153 | 0.16049303 | 0.71409437 |
| NAPEPLD | 257.013688 | -0.2060356 | 0.26568976 | -0.7754742 | 0.4380595 | 0.88832499 |
| PMPCB | 124.995796 | -0.0509087 | 0.23899009 | -0.213016 | 0.8313145 | 0.97160502 |
| DNAJC2 | 103.474328 | -0.0552466 | 0.25579253 | -0.2159821 | 0.82900168 | 0.97131041 |
| PSMC2 | 97.6722282 | -0.1458668 | 0.20813209 | -0.7008378 | 0.48340427 | 0.90631121 |
| ORC5 | 156.563179 | -0.0166119 | 0.27636832 | -0.0601079 | 0.95206969 | 0.99428903 |
| LHFPL3 | 310.967075 | 0.2404068 | 0.50581191 | 0.47528893 | 0.63458105 | 0.94775679 |
| LINC01004 | 404.997178 | 0.07646293 | 0.34554853 | 0.22127985 | 0.82487454 | 0.97018919 |
| KMT2E | 1188.68748 | -0.0085219 | 0.28764723 | -0.0296263 | 0.97636506 | 0.99656026 |
| SRPK2 | 1216.72821 | -0.0920293 | 0.22080591 | -0.416788 | 0.67683347 | 0.95529325 |
| PUS7 | 93.2142115 | -0.4514971 | 0.34201102 | -1.3201244 | 0.18679347 | 0.75061181 |
| RINT1 | 65.6579048 | -0.6067398 | 0.29197213 | -2.0780743 | 0.03770251 | 0.44445707 |
| EFCAB10 | 43.958892 | -0.2412414 | 0.56225211 | -0.4290627 | 0.6678776 | 0.95294349 |
| ATXN7L1 | 431.63914 | -0.0886193 | 0.21905548 | -0.4045519 | 0.68580694 | 0.9558749 |
| CDHR3 | 100.774048 | 0.91696326 | 0.49360714 | 1.85767828 | 0.06321472 | 0.54841905 |
| SYPL1 | 386.1813 | 0.45203778 | 0.3235699 | 1.39703283 | 0.1624037 | 0.71482182 |
| NAMPT | 834.552059 | -1.7120115 | 0.81460898 | -2.101636 | 0.03558517 | 0.43592878 |
| PRKAR2B | 75.9312175 | 0.72839869 | 0.47367433 | 1.53776264 | 0.12410666 | 0.66528789 |
| HBP1 | 293.399157 | -0.3556785 | 0.31456545 | -1.130698 | 0.25818224 | 0.80709747 |
| COG5 | 741.035728 | -0.2802695 | 0.14580069 | -1.9222782 | 0.05457076 | 0.51597783 |
| DUS4L | 41.4542585 | -0.346462 | 0.32131976 | -1.0782467 | 0.28092368 | 0.82628479 |
| BCAP29 | 178.737415 | -0.2442168 | 0.27078185 | -0.901895 | 0.36711263 | 0.85902833 |
| CBLL1 | 47.1076114 | 0.09019076 | 0.2457395 | 0.36701778 | 0.71360576 | 0.95791799 |
| SLC26A3 | 1144.35256 | -0.9514266 | 0.57883958 | -1.6436793 | 0.10024246 | 0.61891077 |
| DLD | 274.232066 | 0.27528105 | 0.27521338 | 1.00024586 | 0.31719154 | 0.84285425 |
| LAMB1 | 2398.95964 | -1.2579658 | 0.30080104 | -4.1820525 | 2.89E-05 | 0.00776297 |
| NRCAM | 810.948114 | 0.28236469 | 0.54793464 | 0.51532549 | 0.60632559 | 0.93919596 |
| PNPLA8 | 264.88972 | -0.5058909 | 0.4000429 | -1.2645917 | 0.20601773 | 0.77193931 |
| DNAJB9 | 68.2698845 | -0.5639075 | 0.42136337 | -1.3382926 | 0.18080109 | 0.74315969 |
| IMMP2L | 7403.74566 | -0.2748904 | 0.20092869 | -1.3680995 | 0.17128092 | 0.72776437 |
| LRRN3 | 49.4436396 | 0.55647382 | 0.73202707 | 0.76018202 | 0.44714579 | 0.89187439 |

| | | | | | | |
|---|---|---|---|---|---|---|
| DOCK4 | 75.4282718 | 0.13626108 | 0.46052066 | 0.29588484 | 0.76731803 | 0.96305305 |
| ZNF277 | 228.017597 | -0.0899678 | 0.20508025 | -0.4386957 | 0.66088201 | 0.95127957 |
| IFRD1 | 297.469206 | -0.5198998 | 0.62119364 | -0.8369368 | 0.40262809 | 0.87834687 |
| TMEM168 | 271.200241 | -0.0453427 | 0.2494946 | -0.1817382 | 0.85578816 | 0.97718353 |
| BMT2 | 355.49198 | 0.13747052 | 0.22640066 | 0.60720019 | 0.54371806 | 0.92482794 |
| SMIM30 | 80.966934 | 0.10101322 | 0.31342901 | 0.3222842 | 0.7472374 | 0.96109466 |
| FOXP2 | 178.42868 | -2.3301598 | 1.52370705 | -1.5292702 | 0.12619747 | 0.66795428 |
| MDFIC | 355.135372 | 0.43572214 | 0.32286408 | 1.34955287 | 0.17715945 | 0.73633806 |
| TES | 243.351268 | -0.6009022 | 0.43422342 | -1.3838549 | 0.1664029 | 0.72054851 |
| CAV2 | 108.69592 | 1.14518074 | 0.59348998 | 1.92957048 | 0.05366008 | 0.51316717 |
| MET | 336.362484 | -1.1628282 | 0.76237385 | -1.525273 | 0.12719102 | 0.67016545 |
| CAPZA2 | 522.024494 | -0.4031925 | 0.25031781 | -1.6107222 | 0.10724028 | 0.63477557 |
| ST7 | 725.671957 | -0.4120484 | 0.26302729 | -1.5665613 | 0.11721728 | 0.65342396 |
| ST7-AS2 | 56.0138249 | -0.3608936 | 0.54495668 | -0.6622427 | 0.50781566 | 0.91214298 |
| LSM8 | 131.993564 | -0.1353054 | 0.20121645 | -0.672437 | 0.50130556 | 0.90900452 |
| ANKRD7 | 153.184915 | 0.26376374 | 0.31703364 | 0.83197399 | 0.40542363 | 0.8791309 |
| TSPAN12 | 227.493721 | 0.81165729 | 0.26049758 | 3.11579587 | 0.00183449 | 0.10848841 |
| ING3 | 59.684752 | 0.18274558 | 0.25134051 | 0.72708366 | 0.46717471 | 0.89917189 |
| FAM3C | 185.509642 | -0.2192864 | 0.28704125 | -0.7639542 | 0.44489452 | 0.89131994 |
| AASS | 132.924522 | -0.0385407 | 0.33283538 | -0.1157952 | 0.90781483 | 0.9875066 |
| CADPS2 | 6641.07028 | 1.13516853 | 0.3126151 | 3.63120188 | 0.0002821 | 0.03764581 |
| AC004594.1 | 404.846818 | 1.29472586 | 0.55852264 | 2.318126 | 0.02044247 | 0.34332873 |
| IQUB | 488.419978 | 1.11179256 | 0.45105858 | 2.46485181 | 0.01370699 | 0.28408502 |
| NDUFA5 | 500.457139 | 0.55377282 | 0.38032895 | 1.45603644 | 0.14538255 | 0.69325556 |
| WASL | 414.034279 | 0.15503937 | 0.16751583 | 0.9255207 | 0.35469512 | 0.8557799 |
| POT1 | 177.554905 | -0.3072087 | 0.14691532 | -2.0910596 | 0.03652272 | 0.43729328 |
| POT1-AS1 | 106.106509 | 0.06751199 | 0.22076056 | 0.30581543 | 0.75974518 | 0.96190674 |
| ZNF800 | 205.877634 | 0.04183296 | 0.17263347 | 0.24232239 | 0.80853036 | 0.96775927 |
| ARF5 | 131.538756 | -0.1244849 | 0.41905077 | -0.2970639 | 0.76641771 | 0.96285313 |
| SND1 | 610.619583 | -0.3518216 | 0.25744865 | -1.3665699 | 0.17176016 | 0.72776437 |
| RBM28 | 123.944943 | -0.254642 | 0.19927575 | -1.2778374 | 0.20130676 | 0.76957251 |
| IMPDH1 | 35.4041283 | 0.38306073 | 0.47809582 | 0.80122168 | 0.42300332 | 0.88371155 |
| HILPDA | 366.542934 | -3.1968343 | 1.26662152 | -2.5239065 | 0.01160588 | 0.26383866 |
| METTL2B | 104.865203 | -0.0674491 | 0.24861373 | -0.2713007 | 0.7861598 | 0.96513523 |
| CALU | 178.879166 | -0.3052343 | 0.25334823 | -1.2048014 | 0.22827999 | 0.79042788 |
| KCP | 379.033239 | -0.1852527 | 0.36479794 | -0.5078227 | 0.61157769 | 0.94003018 |
| ATP6V1F | 453.548002 | 0.2100352 | 0.48018063 | 0.43740873 | 0.66181495 | 0.95142083 |
| IRF5 | 42.6820304 | 0.05076139 | 0.32584043 | 0.15578605 | 0.87620167 | 0.98269442 |
| TNPO3 | 400.899253 | -0.281735 | 0.27115818 | -1.0390059 | 0.29880198 | 0.83556453 |
| TSPAN33 | 53.4353632 | 0.29748094 | 0.31447571 | 0.94595839 | 0.34416981 | 0.85148675 |
| SMO | 34.9757339 | 0.80795212 | 0.40414469 | 1.99916547 | 0.04559045 | 0.48339878 |
| AHCYL2 | 647.24603 | -0.0188947 | 0.30325168 | -0.0623071 | 0.95031826 | 0.99394565 |
| AC078846.1 | 33.5862687 | -0.1121976 | 0.32179248 | -0.3486644 | 0.72734124 | 0.95979351 |
| NRF1 | 454.581628 | 0.16701063 | 0.23798201 | 0.70177837 | 0.48281739 | 0.90614725 |
| UBE2H | 1626.3455 | -0.472599 | 0.23627922 | -2.0001716 | 0.04548174 | 0.48339878 |
| ZC3HC1 | 67.2500941 | -0.1673381 | 0.26108108 | -0.6409429 | 0.52155978 | 0.91787515 |
| KLHDC10 | 465.343557 | 0.24417069 | 0.22136662 | 1.10301492 | 0.27002068 | 0.81817548 |
| TMEM209 | 87.6981774 | -0.162488 | 0.27538356 | -0.5900425 | 0.55516214 | 0.92872652 |
| CEP41 | 49.4598764 | -0.3896317 | 0.32810541 | -1.1875199 | 0.23502261 | 0.79599484 |
| MEST | 115.115849 | -0.062553 | 0.75196068 | -0.0831865 | 0.93370325 | 0.99243654 |
| COPG2 | 174.074602 | -0.3770773 | 0.2651595 | -1.4220772 | 0.15500383 | 0.70897393 |
| AC016831.7 | 2631.27207 | -0.7178366 | 0.47846502 | -1.5002907 | 0.13353913 | 0.67640892 |
| LINC00513 | 182.402953 | -0.6801113 | 0.63395871 | -1.0728007 | 0.28336056 | 0.82673124 |
| AC058791.1 | 49.8658702 | -0.9174572 | 0.69098519 | -1.3277523 | 0.18425996 | 0.74701911 |

| | | | | | | |
|---|---|---|---|---|---|---|
| MKLN1 | 1744.37313 | -0.6203945 | 0.20623139 | -3.0082449 | 0.00262761 | 0.13303085 |
| CHCHD3 | 1293.06002 | 0.07960955 | 0.22815457 | 0.34892814 | 0.72714326 | 0.95979351 |
| EXOC4 | 1987.60185 | -0.2838271 | 0.15931153 | -1.7815853 | 0.07481688 | 0.57072562 |
| LRGUK | 95.7609402 | -0.1541286 | 0.30528125 | -0.5048741 | 0.61364726 | 0.94048308 |
| SLC35B4 | 190.472617 | 0.49914496 | 0.37695258 | 1.32415851 | 0.18545041 | 0.74895057 |
| AKR1B1 | 579.221354 | -0.2650752 | 0.77060306 | -0.3439841 | 0.73085821 | 0.96008876 |
| BPGM | 102.695129 | 0.66538059 | 0.50033288 | 1.3298758 | 0.1835592 | 0.74504275 |
| CALD1 | 71.0706126 | -0.1557179 | 0.45009185 | -0.3459692 | 0.72936588 | 0.96008876 |
| AGBL3 | 87.0792135 | -0.179378 | 0.43788634 | -0.409645 | 0.68206637 | 0.95574492 |
| CYREN | 117.919923 | 0.08380157 | 0.23038523 | 0.36374542 | 0.71604812 | 0.95791799 |
| TMEM140 | 105.366932 | -2.0768045 | 0.45834908 | -4.5310542 | 5.87E-06 | 0.00257078 |
| WDR91 | 142.160524 | -0.3644961 | 0.34762484 | -1.0485328 | 0.29439318 | 0.8337535 |
| CNOT4 | 545.939797 | -0.0893736 | 0.19361758 | -0.4615984 | 0.64436931 | 0.94949493 |
| NUP205 | 134.384433 | -0.4568104 | 0.17748881 | -2.5737422 | 0.01006052 | 0.24462705 |
| STMP1 | 246.496667 | 0.10269706 | 0.26561116 | 0.38664435 | 0.69901952 | 0.95672887 |
| MTPN | 247.328575 | -0.0357867 | 0.30716544 | -0.1165062 | 0.90725134 | 0.9875066 |
| CREB3L2 | 218.497313 | -0.0441071 | 0.24087339 | -0.1831132 | 0.85470923 | 0.97718353 |
| TRIM24 | 374.834586 | -0.7535547 | 0.25286874 | -2.9800233 | 0.00288226 | 0.13856025 |
| SVOPL | 143.600402 | 0.12049842 | 0.58654053 | 0.20543921 | 0.83722899 | 0.97357501 |
| ATP6V0A4 | 562.686261 | 0.75641257 | 0.33324038 | 2.26987067 | 0.02321543 | 0.36457774 |
| TMEM213 | 440.912996 | 0.94315943 | 0.29790616 | 3.16596148 | 0.00154571 | 0.09927452 |
| KIAA1549 | 299.088298 | -0.2075394 | 0.30718659 | -0.6756136 | 0.49928604 | 0.90839719 |
| ZC3HAV1 | 741.517621 | -1.1508205 | 0.51117984 | -2.2513026 | 0.02436638 | 0.37424167 |
| TTC26 | 155.129767 | -0.1473547 | 0.18873139 | -0.780764 | 0.43494134 | 0.88785137 |
| UBN2 | 380.59672 | 0.02295359 | 0.19878208 | 0.1154711 | 0.90807172 | 0.98758267 |
| LUC7L2 | 137.494569 | 0.10618595 | 0.25870138 | 0.4104576 | 0.6814703 | 0.95574492 |
| HIPK2 | 1058.65016 | 0.55823403 | 0.19443832 | 2.87100823 | 0.00409165 | 0.15545856 |
| TBXAS1 | 46.8661677 | -0.2487681 | 0.42602534 | -0.5839279 | 0.55926881 | 0.92901206 |
| PARP12 | 131.410486 | -1.1254285 | 0.61205397 | -1.8387734 | 0.06594852 | 0.55444578 |
| KDM7A | 425.45851 | -0.2924149 | 0.33031586 | -0.8852585 | 0.37601721 | 0.86357183 |
| SLC37A3 | 160.645309 | -0.5365311 | 0.16474191 | -3.2567976 | 0.00112677 | 0.08298537 |
| MKRN1 | 217.492977 | 0.00786592 | 0.17848734 | 0.0440699 | 0.96484868 | 0.99637506 |
| NDUFB2 | 436.674996 | 0.27544305 | 0.40175751 | 0.68559527 | 0.49296836 | 0.90804281 |
| BRAF | 1535.65409 | -0.1197724 | 0.23735514 | -0.5046127 | 0.61383086 | 0.94048939 |
| MRPS33 | 113.060395 | 0.50731243 | 0.48777786 | 1.04004808 | 0.29831756 | 0.83556453 |
| AGK | 171.084141 | -0.4149474 | 0.25767002 | -1.6103831 | 0.10731425 | 0.63477557 |
| AC004918.1 | 47.8387971 | -0.3493892 | 0.43590507 | -0.801526 | 0.4228272 | 0.88371155 |
| KIAA1147 | 187.973598 | -0.4693527 | 0.23419761 | -2.0040884 | 0.04506059 | 0.48262669 |
| WEE2-AS1 | 37.4704222 | -0.3648008 | 0.41880609 | -0.8710493 | 0.38372723 | 0.8665075 |
| SSBP1 | 226.566078 | -0.3035669 | 0.25179021 | -1.2056344 | 0.22795848 | 0.79042788 |
| EPHB6 | 51.3780354 | 0.84178806 | 0.32484359 | 2.59136422 | 0.00955963 | 0.23684368 |
| TRPV6 | 158.124681 | 0.65149637 | 0.4327551 | 1.50546203 | 0.13220533 | 0.67466347 |
| TRPV5 | 115.30927 | 1.66663086 | 0.6028531 | 2.76457211 | 5.70E-03 | 0.187252 |
| GSTK1 | 262.054913 | 0.35168075 | 0.29280077 | 1.20109229 | 0.22971541 | 0.79047378 |
| AC073342.1 | 34.1559277 | 0.61680792 | 0.45799751 | 1.34674951 | 0.17806092 | 0.73737861 |
| TMEM139 | 72.5654783 | 1.76514902 | 0.53906063 | 3.27449071 | 0.00105853 | 0.08019046 |
| CASP2 | 55.3212956 | -0.3710893 | 0.27208372 | -1.3638792 | 0.17260558 | 0.72940007 |
| ZYX | 71.8992527 | -0.3570387 | 0.46280829 | -0.7714614 | 0.44043349 | 0.88956997 |
| TCAF2 | 122.418399 | 0.27354292 | 0.56359924 | 0.48535006 | 0.62742805 | 0.94511671 |
| TCAF1 | 258.01517 | 0.23620108 | 0.24703036 | 0.95616215 | 0.33899032 | 0.84863112 |
| AC004889.1 | 217.204657 | 0.20963651 | 0.34228925 | 0.61245426 | 0.54023725 | 0.92414041 |
| AC074386.1 | 276.872473 | -0.2685642 | 0.32488969 | -0.826632 | 0.40844568 | 0.87948803 |
| ARHGEF5 | 76.8804983 | -0.2814707 | 0.38625307 | -0.728721 | 0.46617236 | 0.89883965 |
| TPK1 | 203.681604 | -0.6774608 | 0.23398148 | -2.8953607 | 0.00378723 | 0.14977426 |

| | | | | | | |
|---|---|---|---|---|---|---|
| CNTNAP2 | 44.4297586 | -0.3405346 | 0.38554909 | -0.8832459 | 0.37710345 | 0.8642538 |
| CUL1 | 328.458485 | 0.12143587 | 0.17975691 | 0.67555608 | 0.49932254 | 0.90839719 |
| PDIA4 | 121.738925 | 0.32794344 | 0.34570216 | 0.94863 | 0.34280883 | 0.85093239 |
| ZNF786 | 42.23227 | 0.19852399 | 0.48957963 | 0.40549888 | 0.68511084 | 0.9558749 |
| ZNF398 | 120.600697 | -0.0651897 | 0.34525308 | -0.188817 | 0.85023622 | 0.97713298 |
| ZNF282 | 68.3740768 | 0.06918626 | 0.27286899 | 0.2535512 | 0.7998423 | 0.96568323 |
| ZNF212 | 32.2941756 | -0.1139102 | 0.3424055 | -0.3326763 | 0.73937864 | 0.96078945 |
| ZNF783 | 62.2086912 | 0.27751408 | 0.28331921 | 0.97951028 | 0.32732791 | 0.84381657 |
| ZNF777 | 38.5623423 | 0.1946651 | 0.34888672 | 0.55796076 | 0.57687118 | 0.93319686 |
| ZNF746 | 43.856302 | 0.21020051 | 0.27189949 | 0.77308164 | 0.43947407 | 0.88917548 |
| ZNF467 | 33.41682 | 0.71861428 | 0.73307262 | 0.98027706 | 0.32694937 | 0.84381657 |
| ZNF862 | 143.600166 | 0.34261864 | 0.33273216 | 1.02971302 | 0.30314474 | 0.83709431 |
| ATP6V0E2 | 51.1027014 | 0.65007973 | 0.37488153 | 1.73409377 | 0.08290145 | 0.59202053 |
| ACTR3C | 457.096878 | 0.29287042 | 0.25877458 | 1.13175885 | 0.25773583 | 0.80709747 |
| LRRC61 | 51.5568876 | 0.6070053 | 0.35802474 | 1.69542835 | 0.08999419 | 0.60739841 |
| TMEM176B | 349.565449 | 0.90320598 | 0.62395578 | 1.44754806 | 0.14774349 | 0.69834914 |
| TMEM176A | 168.517329 | 0.5941381 | 0.64498847 | 0.92116081 | 0.35696648 | 0.8557799 |
| AOC1 | 284.794248 | 0.49868227 | 0.84446458 | 0.59053071 | 0.5548349 | 0.92872652 |
| ABCB8 | 72.7950548 | -0.2960063 | 0.2356963 | -1.25588 | 0.2091595 | 0.77436433 |
| SLC4A2 | 59.4047949 | -0.2321119 | 0.3246136 | -0.7150405 | 0.47458403 | 0.90242252 |
| FASTK | 78.5908871 | -0.3276338 | 0.30619715 | -1.0700095 | 0.28461505 | 0.82781987 |
| TMUB1 | 95.947015 | -0.0370339 | 0.43761159 | -0.0846273 | 0.93255771 | 0.99243654 |
| AGAP3 | 61.1097951 | 0.16297962 | 0.49960102 | 0.32621956 | 0.74425824 | 0.96109466 |
| CHPF2 | 45.7094161 | -0.1096662 | 0.5265185 | -0.2082855 | 0.83500607 | 0.97246185 |
| SMARCD3 | 177.113695 | 0.32755561 | 0.33837798 | 0.96801692 | 3.33E-01 | 0.84442294 |
| NUB1 | 182.724547 | -0.2400315 | 0.32948383 | -0.7285078 | 0.4663028 | 0.89883965 |
| RHEB | 320.956854 | 0.19993974 | 0.3352733 | 0.59634853 | 0.55094242 | 0.92816739 |
| PRKAG2 | 4993.40831 | -1.31337 | 0.52231702 | -2.5145074 | 0.01191988 | 0.26883805 |
| PRKAG2-AS1 | 63.7517734 | -0.4879928 | 0.85971296 | -0.567623 | 0.57029101 | 0.93125087 |
| GALNT11 | 886.143011 | -0.3342328 | 0.45379913 | -0.7365214 | 0.46141343 | 0.89799005 |
| KMT2C | 1760.14377 | -0.3678144 | 0.21845223 | -1.6837292 | 0.09223401 | 0.61120158 |
| ACTR3B | 163.44233 | 0.12795513 | 0.23540354 | 0.54355652 | 0.58674669 | 0.93483536 |
| DPP6 | 289.627629 | 0.20063167 | 0.30570694 | 0.65628759 | 0.5116391 | 0.91408377 |
| PAXIP1-AS2 | 65.6985786 | 0.07009785 | 0.33208583 | 0.21108354 | 0.83282208 | 0.97208613 |
| PAXIP1 | 102.805716 | 0.11213231 | 0.26247049 | 0.42721872 | 0.66922003 | 0.95304061 |
| PAXIP1-AS1 | 47.8562 | 0.18187497 | 0.46303178 | 0.39279154 | 0.69447346 | 0.95672887 |
| INSIG1 | 281.683368 | -1.8481891 | 0.93359491 | -1.9796478 | 0.04774311 | 0.48600281 |
| AC009403.1 | 64.0330162 | -0.1305271 | 0.37187223 | -0.3509999 | 0.72558844 | 0.95963742 |
| RBM33 | 517.914279 | -0.3161547 | 0.23924456 | -1.3214709 | 0.1863444 | 0.74938192 |
| LINC01006 | 201.86677 | 0.1343762 | 0.39791887 | 0.33769748 | 0.73559118 | 0.96078945 |
| LMBR1 | 486.910264 | 0.00336233 | 0.2101581 | 0.01599907 | 0.98723513 | 0.99791717 |
| NOM1 | 50.8377971 | -0.0538342 | 0.4049576 | -0.1329378 | 0.89424259 | 0.98597539 |
| UBE3C | 651.613283 | -0.3596265 | 0.25079974 | -1.433919 | 0.15159537 | 0.70297324 |
| DNAJB6 | 516.440915 | -0.1874486 | 0.31073364 | -0.6032454 | 0.54634546 | 0.92597739 |
| PTPRN2 | 50.1401678 | 0.83671884 | 0.41097202 | 2.03595087 | 0.04175529 | 0.46746387 |
| NCAPG2 | 44.4089303 | -0.1025812 | 0.36767503 | -0.2789996 | 0.78024513 | 0.96513523 |
| ESYT2 | 2331.76522 | -0.2652793 | 0.82034752 | -0.3233742 | 0.74641184 | 0.96109466 |
| WDR60 | 608.198757 | -0.1362969 | 0.25049223 | -0.5441162 | 0.58636149 | 0.93483536 |
| GTPBP6 | 127.766686 | -0.172147 | 0.32746575 | -0.5256947 | 0.59910037 | 0.93728325 |
| LINC00685 | 58.0919269 | 0.50896936 | 0.62773368 | 0.8108046 | 0.41747789 | 0.88242985 |
| PPP2R3B | 50.149927 | -0.0894814 | 0.37423349 | -0.2391059 | 0.81102344 | 0.9680627 |
| SLC25A6 | 556.365284 | 0.46797943 | 0.43497283 | 1.075882 | 0.28198003 | 0.82661559 |
| LINC00106 | 108.017606 | 0.29227923 | 0.40130379 | 0.72832412 | 0.4664152 | 0.89883965 |
| ASMTL-AS1 | 54.7814216 | 0.40791909 | 0.33671256 | 1.21147571 | 0.22571314 | 0.78982769 |

| | | | | | | |
|---|---|---|---|---|---|---|
| ASMTL | 82.4235873 | 0.12509691 | 0.33395487 | 0.37459226 | 0.70796373 | 0.95752169 |
| AKAP17A | 82.0852298 | -0.0805806 | 0.29550121 | -0.2726914 | 0.78509046 | 0.96513523 |
| DHRSX | 526.35023 | -0.1941671 | 0.29840087 | -0.6506921 | 0.51524529 | 0.91599162 |
| CD99 | 134.267094 | 0.34518558 | 0.34982944 | 0.98672536 | 0.3237773 | 0.84285425 |
| ARSD | 290.749109 | -0.0587057 | 0.24884023 | -0.2359172 | 0.81349695 | 0.96819339 |
| MXRA5 | 237.372739 | -1.0484826 | 0.39067454 | -2.6837752 | 0.0072796 | 0.2101379 |
| PRKX | 74.9772145 | -0.210636 | 0.28659133 | -0.7349698 | 0.4623579 | 0.89801718 |
| BX890604.1 | 35.2408115 | 0.01835769 | 0.34898729 | 0.05260275 | 0.95804842 | 0.99579587 |
| PUDP | 97.9057616 | 0.04200891 | 0.35003513 | 0.12001342 | 0.90447252 | 0.9875066 |
| STS | 46.0089792 | -0.4856632 | 0.56609667 | -0.8579157 | 0.39093901 | 0.87004477 |
| PNPLA4 | 53.6963564 | -0.0539431 | 0.25199187 | -0.2140668 | 0.83049497 | 0.97160502 |
| TBL1X | 912.434414 | 0.12270919 | 0.37835484 | 0.32432304 | 0.74569348 | 0.96109466 |
| SHROOM2 | 125.314886 | 0.48570856 | 0.33544233 | 1.44796443 | 0.147627 | 0.69834914 |
| WWC3 | 95.2898755 | 0.57513267 | 0.20214019 | 2.84521682 | 0.00443812 | 0.16424479 |
| MID1 | 1365.60404 | -0.5970386 | 0.3627732 | -1.6457628 | 0.0998126 | 0.61822465 |
| AC073529.1 | 82.4376486 | 0.23669174 | 0.32457379 | 0.7292386 | 0.46585573 | 0.89883965 |
| HCCS | 31.1723237 | 0.35426207 | 0.2759364 | 1.28385408 | 0.19919302 | 0.76804372 |
| ARHGAP6 | 1754.16313 | -0.1464928 | 0.33441945 | -0.438051 | 0.6613493 | 0.9513366 |
| MSL3 | 40.0183892 | -0.2065961 | 0.37808435 | -0.5464285 | 0.58477143 | 0.93483536 |
| PRPS2 | 212.567006 | -0.0015794 | 0.41934689 | -0.0037662 | 0.99699498 | 0.99967893 |
| TMSB4X | 4320.04348 | -0.6467748 | 0.55809458 | -1.1588982 | 0.24649767 | 0.80072691 |
| RAB9A | 146.622302 | -0.2361079 | 0.44203598 | -0.5341374 | 0.59324651 | 0.93491607 |
| TRAPPC2 | 72.2977212 | 0.15121391 | 0.33816877 | 0.44715515 | 0.65476304 | 0.95100227 |
| OFD1 | 360.25933 | -0.5852215 | 0.35326568 | -1.6566046 | 0.09759945 | 0.61685067 |
| GPM6B | 306.659407 | -0.559352 | 0.48504498 | -1.153196 | 0.24882993 | 0.80097144 |
| GEMIN8 | 129.833652 | -0.1229634 | 0.30982234 | -0.3968835 | 0.69145335 | 0.95672887 |
| MOSPD2 | 91.1713028 | -0.4012453 | 0.26199122 | -1.5315221 | 0.12564042 | 0.66745103 |
| PIR | 41.0188203 | 0.58317391 | 0.68995976 | 0.84522887 | 0.39798307 | 0.87462201 |
| CLTRN | 42.1896018 | -0.7269647 | 0.71969873 | -1.0100958 | 0.31244938 | 0.84279957 |
| CA5B | 154.253094 | -0.6280607 | 0.28430017 | -2.2091464 | 0.02716446 | 0.40076654 |
| ZRSR2 | 75.9714494 | -0.1712659 | 0.29165735 | -0.587216 | 0.55705867 | 0.92872652 |
| CTPS2 | 192.946082 | -0.6355792 | 0.20483719 | -3.1028505 | 0.00191666 | 0.10976307 |
| SYAP1 | 261.204001 | 0.00074539 | 0.43285779 | 0.00172203 | 0.99862602 | 0.99967893 |
| TXLNG | 127.615572 | -0.1790152 | 0.26669675 | -0.6712313 | 0.50207316 | 0.90954654 |
| RBBP7 | 202.349341 | 0.48813952 | 0.20421939 | 2.39027021 | 0.01683598 | 0.30900364 |
| REPS2 | 582.435872 | 0.21890683 | 0.35687522 | 0.61339881 | 0.53961267 | 0.92403616 |
| NHS | 299.857893 | 0.56563671 | 0.58405398 | 0.96846651 | 0.33281144 | 0.84442294 |
| SCML1 | 397.247174 | 0.67414172 | 0.34880359 | 1.93272588 | 0.05326998 | 0.51123567 |
| RAI2 | 124.566766 | 0.39562385 | 0.20828815 | 1.89940639 | 0.05751106 | 0.52731054 |
| CDKL5 | 318.940431 | -0.0998727 | 0.19242549 | -0.51902 | 0.60374681 | 0.93804805 |
| PHKA2 | 118.565337 | -0.3667516 | 0.22060885 | -1.6624519 | 0.0964222 | 0.61440337 |
| PDHA1 | 503.615329 | 0.94347198 | 0.32693 | 2.88585318 | 0.00390354 | 0.15207852 |
| MAP3K15 | 471.538265 | -1.133154 | 0.49299005 | -2.2985332 | 0.02153146 | 0.35423813 |
| SH3KBP1 | 207.023157 | -0.3497632 | 0.33886713 | -1.0321545 | 0.30199973 | 0.83709431 |
| BCLAF3 | 191.475873 | -0.4085377 | 0.19565984 | -2.088 | 0.03679784 | 0.4382288 |
| EIF1AX | 371.990138 | -0.0531959 | 0.35441503 | -0.150095 | 0.88068969 | 0.98333507 |
| RPS6KA3 | 350.803619 | -0.2310551 | 0.21974389 | -1.0514744 | 0.29304076 | 0.83336457 |
| MBTPS2 | 99.9224081 | 0.17361886 | 0.29718684 | 0.58420778 | 0.55908053 | 0.92901206 |
| SMS | 472.943409 | -0.2483227 | 0.41553406 | -0.5975988 | 0.55010765 | 0.92780338 |
| PTCHD1-AS | 208.540637 | -2.6905195 | 0.59463231 | -4.5246777 | 6.05E-06 | 0.00257078 |
| PRDX4 | 60.3878766 | -0.0567047 | 0.36171203 | -0.1567676 | 0.87542798 | 0.98267808 |
| ACOT9 | 92.6115817 | -0.3702977 | 0.42765783 | -0.8658739 | 0.38655932 | 0.86852426 |
| SAT1 | 1694.81145 | -0.2208041 | 0.37265086 | -0.5925227 | 0.55350062 | 0.92872652 |
| APOO | 1503.42118 | 0.0704792 | 0.44597906 | 0.15803254 | 0.87443116 | 0.98258673 |

| | | | | | | |
|---|---|---|---|---|---|---|
| EIF2S3 | 193.50081 | -0.3541568 | 0.30877702 | -1.146966 | 0.25139566 | 0.80240987 |
| ZFX | 210.718317 | -0.2264447 | 0.27475862 | -0.8241587 | 0.40984938 | 0.88016834 |
| PDK3 | 132.736781 | -1.2311172 | 0.71083511 | -1.7319308 | 0.08328589 | 0.59231889 |
| POLA1 | 262.13951 | 0.05544287 | 0.24436328 | 0.22688707 | 0.82051155 | 0.96886922 |
| IL1RAPL1 | 72.825927 | -1.0961824 | 0.70531131 | -1.5541823 | 0.12014093 | 0.65679551 |
| GK | 80.0676586 | -0.1242181 | 0.33406054 | -0.3718431 | 0.71000964 | 0.95752169 |
| TAB3 | 259.087835 | 0.3552556 | 0.25775177 | 1.37828578 | 0.16811507 | 0.72391336 |
| DMD | 149.287621 | -0.4401378 | 0.26913924 | -1.6353534 | 0.10197499 | 0.62351102 |
| CFAP47 | 91.535345 | 0.01561635 | 0.51669286 | 0.03022366 | 0.97588868 | 0.99656026 |
| AF241726.2 | 172.302165 | 0.62140079 | 0.41578434 | 1.49452667 | 0.13503802 | 0.67875224 |
| DYNLT3 | 234.47998 | -0.0393193 | 0.42068064 | -0.093466 | 0.92553338 | 0.99051393 |
| RPGR | 128.847111 | -0.0553634 | 0.28072324 | -0.197217 | 0.84365776 | 0.97561662 |
| MID1IP1 | 41.1713456 | -0.0962023 | 0.3698545 | -0.2601086 | 0.79478 | 0.9653598 |
| BCOR | 280.801684 | -0.0273024 | 0.28372519 | -0.0962283 | 0.92333926 | 0.99051393 |
| ATP6AP2 | 300.432573 | 0.56810303 | 0.31834428 | 1.78455552 | 0.07433344 | 0.5698716 |
| CXorf38 | 116.2235 | 0.03461413 | 0.21204935 | 0.16323618 | 0.87033248 | 0.98208511 |
| MED14 | 244.876384 | -0.3049246 | 0.27661512 | -1.1023424 | 0.27031285 | 0.81817548 |
| USP9X | 884.021033 | -0.3607788 | 0.34106603 | -1.0577975 | 0.29014777 | 0.83103268 |
| DDX3X | 337.148019 | -0.0887906 | 0.34276242 | -0.2590441 | 0.79560123 | 0.9653598 |
| CASK | 298.984255 | -0.0721479 | 0.22178767 | -0.3253014 | 0.74495294 | 0.96109466 |
| MAOA | 195.133532 | 0.81319422 | 0.50071801 | 1.62405625 | 0.1043638 | 0.63075702 |
| EFHC2 | 109.900911 | 0.2893165 | 0.3156438 | 0.91659176 | 0.3593566 | 0.85608464 |
| FUNDC1 | 53.8237392 | 0.33902658 | 0.3629741 | 0.93402418 | 0.35029142 | 0.85447517 |
| KDM6A | 756.184899 | -0.489435 | 0.43804781 | -1.1173095 | 0.26386201 | 0.81021794 |
| MIR222HG | 79.830422 | -0.6576307 | 0.82523551 | -0.7969006 | 0.42550876 | 0.88403732 |
| KRBOX4 | 94.7823441 | -0.0124294 | 0.3068616 | -0.0405049 | 0.96769062 | 0.9964399 |
| SLC9A7 | 93.7744339 | 0.10930983 | 0.2714735 | 0.40265377 | 0.68720294 | 0.95588883 |
| RP2 | 52.4202959 | -0.7128123 | 0.30532222 | -2.334623 | 0.01956312 | 0.33555077 |
| JADE3 | 157.045371 | -0.026375 | 0.38124582 | -0.069181 | 0.94484553 | 0.99276891 |
| RGN | 117.878691 | 1.28061541 | 0.6785785 | 1.88720304 | 0.05913302 | 0.53286516 |
| NDUFB11 | 265.220171 | 0.31324863 | 0.48776238 | 0.64221563 | 0.52073318 | 0.91751084 |
| RBM10 | 118.695532 | 0.14032688 | 0.22586935 | 0.62127456 | 0.53441899 | 0.92303275 |
| UBA1 | 306.253202 | -0.1666333 | 0.21602672 | -0.7713551 | 0.4404965 | 0.88956997 |
| CDK16 | 133.959896 | -0.3295336 | 0.30141033 | -1.0933056 | 0.27425963 | 0.81986798 |
| USP11 | 170.608238 | -0.1055681 | 0.28331233 | -0.3726208 | 0.7094307 | 0.95752169 |
| ZNF41 | 75.4179647 | 0.09007297 | 0.3063711 | 0.29399958 | 0.76875823 | 0.96345263 |
| ARAF | 70.7863799 | 0.21018244 | 0.28664429 | 0.7332518 | 0.46340486 | 0.89801718 |
| TIMP1 | 66.9413126 | -0.5521656 | 0.6199256 | -0.8906966 | 3.73E-01 | 0.86201082 |
| ELK1 | 34.8086777 | 0.19133676 | 0.34262896 | 0.55843721 | 0.57654587 | 0.9331615 |
| UXT | 50.8729536 | -0.0444925 | 0.2851018 | -0.1560583 | 0.8759871 | 0.98269442 |
| ZNF81 | 164.320138 | 0.18105484 | 0.31394183 | 0.5767146 | 0.56413227 | 0.93077674 |
| ZNF182 | 35.6645066 | 0.08299013 | 0.30346882 | 0.2734717 | 0.78449065 | 0.96513523 |
| ZNF630 | 72.3052341 | 0.51182636 | 0.43618472 | 1.17341653 | 0.24062881 | 0.80057149 |
| EBP | 78.503344 | 0.20116547 | 0.62274775 | 0.32302881 | 0.74667342 | 0.96109466 |
| TBC1D25 | 40.7325417 | 0.16780572 | 0.33034234 | 0.50797522 | 0.61147072 | 0.94003018 |
| RBM3 | 158.34827 | 0.05927947 | 0.43952163 | 0.13487271 | 0.89271252 | 0.98597539 |
| WDR13 | 125.603521 | -0.1458646 | 0.27109106 | -0.538065 | 0.59053217 | 0.93483536 |
| HDAC6 | 78.4180167 | 0.10973855 | 0.22586024 | 0.48586927 | 0.62705986 | 0.94511671 |
| PCSK1N | 598.989942 | -0.0212028 | 0.41316671 | -0.0513178 | 0.95907232 | 0.99609409 |
| TIMM17B | 64.2821041 | 0.21082401 | 0.47589593 | 0.44300445 | 0.65776252 | 0.95100227 |
| SLC35A2 | 32.8586202 | 0.21412413 | 0.36177456 | 0.59187173 | 0.55393649 | 0.92872652 |
| OTUD5 | 156.979507 | -0.1989268 | 0.16535434 | -1.2030333 | 0.22896344 | 0.79047378 |
| GRIPAP1 | 126.012983 | -0.1231204 | 0.18888016 | -0.6518443 | 0.51450164 | 0.91544604 |
| TFE3 | 83.5337524 | -0.0402446 | 0.20507489 | -0.1962436 | 0.84441947 | 0.97573093 |

| | | | | | | |
|---|---|---|---|---|---|---|
| CCDC120 | 47.3601182 | -0.1598301 | 0.31578831 | -0.5061307 | 0.61276493 | 0.94003018 |
| WDR45 | 84.4465309 | 0.02522988 | 0.34749189 | 0.07260566 | 0.94211993 | 0.99265578 |
| MAGIX | 39.2496674 | 0.28334995 | 0.35040302 | 0.80864012 | 0.41872218 | 0.88258416 |
| PLP2 | 98.3670084 | -0.4615448 | 0.49110482 | -0.9398091 | 0.34731548 | 0.85348649 |
| PPP1R3F | 37.3420977 | 0.49225891 | 0.31222256 | 1.57662825 | 0.11488109 | 0.65008305 |
| CLCN5 | 567.477677 | 0.07452312 | 0.30053701 | 0.24796652 | 0.80416031 | 0.96679546 |
| SHROOM4 | 371.71897 | -0.2400903 | 0.21822743 | -1.1001839 | 0.271252 | 0.81922962 |
| MAGED1 | 366.904386 | 0.30501748 | 0.30446076 | 1.00182854 | 0.31642641 | 0.84285425 |
| FAM156A | 47.5043337 | 0.38831802 | 0.35253458 | 1.10150332 | 0.27067766 | 0.81867298 |
| GPR173 | 68.5827421 | 0.43540024 | 0.47196638 | 0.92252383 | 0.3562554 | 0.8557799 |
| TSPYL2 | 45.9447928 | -0.5632401 | 0.45772647 | -1.2305169 | 0.21850361 | 0.78529418 |
| KANTR | 243.790407 | -0.0778126 | 0.24853178 | -0.313089 | 0.75421304 | 0.96120879 |
| KDM5C | 136.926469 | -0.7233008 | 0.40312065 | -1.794254 | 0.07277264 | 0.56668778 |
| IQSEC2 | 77.5022288 | 0.74141439 | 0.32465663 | 2.28368785 | 0.02238988 | 0.35859484 |
| SMC1A | 136.17566 | -0.30788 | 0.20943441 | -1.4700547 | 0.14154695 | 0.68671506 |
| HSD17B10 | 114.755102 | 0.44638841 | 0.49765484 | 0.89698397 | 0.36972747 | 0.86048055 |
| HUWE1 | 802.028103 | 0.04983332 | 0.15136886 | 0.32921779 | 0.74199108 | 0.96086423 |
| PHF8 | 192.039165 | -0.1091242 | 0.2680326 | -0.4071302 | 0.68391237 | 0.95574492 |
| FAM120C | 101.045336 | 0.48097609 | 0.31848234 | 1.51021272 | 0.13098915 | 0.67390608 |
| WNK3 | 107.976195 | 0.28893021 | 0.3890875 | 0.74258416 | 0.45773348 | 0.89714735 |
| TSR2 | 83.4298438 | 0.32793225 | 0.34837136 | 0.94132955 | 0.346536 | 0.8525698 |
| FGD1 | 44.0579742 | 0.37822339 | 0.35270266 | 1.07235764 | 0.28355943 | 0.82685108 |
| MAGED2 | 179.551214 | 0.51767609 | 0.2953646 | 1.75266803 | 0.07965901 | 0.58461232 |
| TRO | 34.1586363 | 0.90440887 | 0.4555204 | 1.98544095 | 0.04709543 | 0.48600281 |
| APEX2 | 33.4107645 | 0.81043981 | 0.44064369 | 1.83921801 | 0.06588313 | 0.55444578 |
| FAM104B | 60.5843731 | 0.2865845 | 0.30816692 | 0.92996516 | 0.35238913 | 0.85536253 |
| RRAGB | 233.672947 | 0.17071938 | 0.2814637 | 0.60654139 | 0.54415531 | 0.92526631 |
| AL445472.1 | 343.850292 | -0.0328362 | 0.2453979 | -0.133808 | 0.89355444 | 0.98597539 |
| KLF8 | 41.399577 | 0.84284879 | 0.44170603 | 1.90816682 | 0.05636966 | 0.5228532 |
| UBQLN2 | 86.2336078 | -0.0759141 | 0.28912272 | -0.2625671 | 0.79288427 | 0.9653598 |
| NBDY | 344.558872 | -0.0458233 | 0.28279632 | -0.1620366 | 0.87127706 | 0.98208511 |
| SPIN3 | 64.6430432 | -0.0095971 | 0.27893811 | -0.0344059 | 0.97255345 | 0.99656026 |
| FAAH2 | 655.602316 | -0.7668203 | 0.31459501 | -2.437484 | 0.01478987 | 0.29134933 |
| ARHGEF9 | 131.958597 | 0.11671863 | 0.34276676 | 0.34051909 | 0.73346565 | 0.96035038 |
| MTMR8 | 108.563852 | -0.1589464 | 0.27942763 | -0.5688285 | 0.56947255 | 0.93122667 |
| ZC4H2 | 223.221779 | 0.14990649 | 0.19406479 | 0.77245589 | 0.43984446 | 0.88917548 |
| ZC3H12B | 165.136403 | 0.52865779 | 0.34966627 | 1.51189245 | 0.13056123 | 0.67339292 |
| LAS1L | 54.1883317 | -0.1588607 | 0.34749523 | -0.4571594 | 0.64755646 | 0.95061388 |
| MSN | 613.37117 | -0.0022849 | 0.48693403 | -0.0046924 | 0.99625605 | 0.99967893 |
| OPHN1 | 1038.05103 | 0.05184223 | 0.34484921 | 0.15033303 | 8.81E-01 | 0.98333507 |
| YIPF6 | 190.905158 | -0.0168325 | 0.21663851 | -0.0776985 | 0.93806786 | 0.99243654 |
| EFNB1 | 89.0963999 | 0.6229335 | 0.40379785 | 1.5426865 | 0.12290686 | 0.66475376 |
| EDA | 122.734079 | -0.3213992 | 0.20772218 | -1.547255 | 0.12180176 | 0.66173276 |
| IGBP1 | 136.683707 | 0.29984189 | 0.30941924 | 0.96904734 | 0.33252157 | 0.84442294 |
| PDZD11 | 43.8811716 | 0.21011038 | 0.49251772 | 0.42660472 | 0.66966726 | 0.9533662 |
| DLG3 | 155.249788 | 0.07378228 | 0.32224852 | 0.22896082 | 0.81889937 | 0.96884581 |
| SNX12 | 70.1092386 | -0.0186264 | 0.22916511 | -0.0812793 | 0.93521986 | 0.99243654 |
| MED12 | 37.6045916 | -0.1243806 | 0.29605988 | -0.4201198 | 0.67439797 | 0.95524793 |
| ZMYM3 | 64.9318378 | 0.12095252 | 0.27905033 | 0.43344338 | 0.66469269 | 0.9520866 |
| NONO | 218.386793 | 0.06148242 | 0.23500769 | 0.26161878 | 0.79361536 | 0.9653598 |
| TAF1 | 298.495984 | -0.3228939 | 0.24662284 | -1.309262 | 0.19044563 | 0.75715865 |
| OGT | 610.320174 | -0.2289364 | 0.3562387 | -0.6426488 | 0.52045202 | 0.91751084 |
| GCNA | 38.2631764 | 0.86180821 | 0.50193727 | 1.71696399 | 0.08598575 | 0.60140155 |
| NHSL2 | 87.3503129 | 0.18994677 | 0.2303314 | 0.82466726 | 0.40956052 | 0.88008904 |

| | | | | | | |
|---|---|---|---|---|---|---|
| PIN4 | 131.939897 | -0.459324 | 0.37625396 | -1.2207817 | 0.22216868 | 0.78760199 |
| RPS4X | 1024.55867 | 0.54957876 | 0.48643334 | 1.1298131 | 0.25855499 | 0.80709747 |
| HDAC8 | 788.36516 | -0.0046952 | 0.22932689 | -0.0204736 | 0.98366553 | 0.9971769 |
| PHKA1 | 63.8304748 | 0.47118583 | 0.35193597 | 1.33883965 | 0.18062288 | 0.7429073 |
| NAP1L2 | 37.7152719 | 1.42706247 | 0.61064874 | 2.3369613 | 0.0194412 | 0.33400613 |
| CHIC1 | 228.600477 | 0.03213013 | 0.24745678 | 0.12984137 | 0.89669193 | 0.98597539 |
| XIST | 1177.92794 | -9.5225521 | 3.87824539 | -2.4553764 | 0.01407372 | 0.28639328 |
| JPX | 688.232672 | -0.1357473 | 0.44533852 | -0.3048182 | 0.76050465 | 0.96191672 |
| FTX | 4660.79993 | 0.07718433 | 0.28245924 | 0.27325828 | 0.78465468 | 0.96513523 |
| SLC16A2 | 50.0121442 | 1.12522649 | 0.29607506 | 3.80047707 | 0.00014442 | 0.02411295 |
| RLIM | 186.940565 | 0.07811832 | 0.31470735 | 0.24822529 | 0.8039601 | 0.96678163 |
| NEXMIF | 423.807766 | 0.25298474 | 0.37475162 | 0.67507311 | 0.49962932 | 0.90839719 |
| ABCB7 | 283.071469 | -0.1340057 | 0.24186949 | -0.5540413 | 0.57955056 | 0.93345492 |
| UPRT | 60.8782775 | 0.04829325 | 0.30237781 | 0.15971161 | 0.87310825 | 0.98246494 |
| ZDHHC15 | 76.3558672 | 0.430248 | 0.3373959 | 1.27520222 | 0.20223768 | 0.76996624 |
| PBDC1 | 68.3430793 | 0.00997399 | 0.46239194 | 0.02157042 | 0.98279063 | 0.9971769 |
| ATRX | 1376.34322 | -0.0983478 | 0.17214497 | -0.5713082 | 0.56779077 | 0.93122667 |
| MAGT1 | 200.830742 | 0.22160488 | 0.22854276 | 0.96964297 | 0.33222448 | 0.84442294 |
| COX7B | 534.137458 | 0.33197983 | 0.39278582 | 0.845193 | 0.39800309 | 0.87462201 |
| PGK1 | 893.829113 | -0.5061629 | 0.55098105 | -0.9186575 | 0.35827474 | 0.8557799 |
| TAF9B | 45.5608398 | -0.0608487 | 0.41054712 | -0.1482136 | 0.8821742 | 0.98335905 |
| BRWD3 | 349.169288 | 0.02867266 | 0.36042785 | 0.07955172 | 0.93659379 | 0.99243654 |
| HMGN5 | 135.301804 | 0.66116584 | 0.39987364 | 1.65343691 | 0.09824198 | 0.61690569 |
| SH3BGRL | 720.072991 | 0.50614722 | 0.16013931 | 3.16066822 | 0.00157408 | 0.09927452 |
| POU3F4 | 57.421268 | 1.05629074 | 0.42488465 | 2.4860647 | 0.01291645 | 0.27852752 |
| RPS6KA6 | 374.867101 | 0.18607299 | 0.32824438 | 0.56687333 | 0.57080025 | 0.93125087 |
| HDX | 94.0486357 | -0.0514731 | 0.3433948 | -0.1498947 | 0.88084771 | 0.98333507 |
| APOOL | 174.592239 | -0.1288926 | 0.23695593 | -0.5439519 | 0.5864746 | 0.93483536 |
| ZNF711 | 61.0612009 | 0.21887328 | 0.35170465 | 0.62232126 | 0.53373065 | 0.92289732 |
| POF1B | 104.987383 | 0.11573942 | 0.36356518 | 0.31834573 | 0.7502227 | 0.96109466 |
| CHM | 292.935971 | -0.2402449 | 0.24703997 | -0.972494 | 0.33080486 | 0.84442294 |
| PCDH11X | 42.1030528 | 0.24299457 | 0.45678477 | 0.53196733 | 0.59474862 | 0.935202 |
| DIAPH2 | 683.479111 | -0.1585292 | 0.18461678 | -0.8586935 | 0.39050961 | 0.87004477 |
| DIAPH2-AS1 | 54.2324571 | 0.42735201 | 0.61180957 | 0.69850495 | 0.48486146 | 0.90689776 |
| TSPAN6 | 105.81857 | 0.6231793 | 0.31236685 | 1.99502377 | 0.04604029 | 0.48492685 |
| SYTL4 | 63.6098208 | 0.14344046 | 0.54136347 | 0.26496147 | 0.79103915 | 0.9653598 |
| TRMT2B | 113.760522 | -0.5381432 | 0.30566531 | -1.7605636 | 0.07831229 | 0.58000906 |
| GLA | 38.9403345 | -0.5468085 | 0.5190086 | -1.0535635 | 0.29208282 | 0.83315947 |
| HNRNPH2 | 122.923811 | -0.2112689 | 0.38270831 | -0.5520362 | 0.58092353 | 0.93375439 |
| ARMCX4 | 410.032146 | 0.08111562 | 0.35731228 | 0.22701605 | 0.82041126 | 0.96886922 |
| ARMCX1 | 38.3960292 | -0.0850451 | 0.43135376 | -0.1971586 | 0.84370347 | 0.97561662 |
| ARMCX6 | 78.8869162 | -0.3003734 | 0.33091809 | -0.9076972 | 0.36403824 | 0.85752322 |
| ARMCX3 | 217.676552 | -0.4606644 | 0.34917403 | -1.3192973 | 0.18706974 | 0.75114593 |
| ARMCX2 | 170.992103 | -0.6673053 | 0.52343166 | -1.2748661 | 0.20235663 | 0.76996624 |
| ZMAT1 | 343.154138 | 0.25995228 | 0.3014971 | 0.86220491 | 0.38857477 | 0.86970192 |
| TCEAL2 | 66.6181897 | 1.56863713 | 0.61789223 | 2.53869049 | 0.01112682 | 0.25855673 |
| BEX5 | 37.6452653 | 1.52032127 | 0.58744127 | 2.5880396 | 0.00965239 | 0.2385779 |
| ARMCX5 | 34.6472906 | 0.45353382 | 0.29843824 | 1.51969071 | 0.12858873 | 0.67206108 |
| GPRASP1 | 61.1926167 | 1.46476926 | 0.49404587 | 2.96484467 | 0.00302836 | 0.14138755 |
| BHLHB9 | 63.1260008 | 0.45527094 | 0.34707044 | 1.31175374 | 0.18960324 | 0.75619145 |
| LINC00630 | 178.763639 | -0.0413181 | 0.28185748 | -0.1465923 | 0.88345385 | 0.98359676 |
| BEX4 | 265.834532 | 0.5578137 | 0.42986011 | 1.29766332 | 0.19440305 | 0.76162393 |
| TCEAL8 | 96.6647534 | 0.34449783 | 0.37692406 | 0.91397145 | 0.36073185 | 0.85608464 |
| BEX2 | 158.057016 | 0.85444655 | 0.34896939 | 2.44848567 | 0.01434581 | 0.28746487 |

| | | | | | | |
|---|---|---|---|---|---|---|
| TCEAL9 | 384.957505 | 0.2913776 | 0.42747306 | 0.681628 | 0.49547421 | 0.90839719 |
| BEX3 | 654.640692 | 0.48654911 | 0.50362837 | 0.96608758 | 0.33400036 | 0.8445213 |
| TCEAL4 | 318.318995 | 0.43344333 | 0.35744926 | 1.21260101 | 0.2252824 | 0.78982769 |
| TCEAL3 | 142.662099 | 0.29346303 | 0.3958509 | 0.7413474 | 0.45848283 | 0.89753337 |
| TCEAL1 | 32.0731125 | 0.44744448 | 0.49256366 | 0.90839928 | 0.36366731 | 0.85694172 |
| MORF4L2 | 343.112351 | -0.3352066 | 0.31727317 | -1.0565236 | 0.29072905 | 0.83156124 |
| SLC25A53 | 145.071668 | -0.1070598 | 0.25013724 | -0.4280041 | 0.66864813 | 0.95298441 |
| FAM199X | 324.126228 | 0.0545302 | 0.23210044 | 0.23494226 | 0.81425355 | 0.96820092 |
| IL1RAPL2 | 902.942044 | 0.71533495 | 0.56205966 | 1.27270288 | 0.2031235 | 0.7702733 |
| RNF128 | 208.542185 | 0.07894884 | 0.30728011 | 0.25692793 | 0.79723441 | 0.9653598 |
| TBC1D8B | 122.311678 | 0.12033953 | 0.35660575 | 0.3374582 | 0.73577152 | 0.96078945 |
| MORC4 | 100.867633 | 0.63146134 | 0.44583135 | 1.41636818 | 0.15666774 | 0.71126849 |
| RBM41 | 222.54259 | 0.30647719 | 0.29587405 | 1.03583667 | 0.30027835 | 0.83694603 |
| NUP62CL | 86.6415049 | 0.67864883 | 0.4472305 | 1.51744756 | 0.12915372 | 0.67206108 |
| PRPS1 | 40.3182296 | -0.4904787 | 0.35951807 | -1.3642673 | 0.17248347 | 0.72940007 |
| TSC22D3 | 126.687688 | 0.14150187 | 0.31037127 | 0.45591163 | 0.64845353 | 0.95061388 |
| MID2 | 65.2997236 | -0.1279421 | 0.2920687 | -0.4380548 | 0.66134655 | 0.9513366 |
| PSMD10 | 34.4975715 | 0.22661967 | 0.33696764 | 0.6725265 | 0.50124858 | 0.90900452 |
| ATG4A | 35.6536734 | -0.1985611 | 0.32242809 | -0.6158307 | 0.53800627 | 0.92324744 |
| COL4A6 | 569.887617 | 0.94400664 | 0.36335475 | 2.59803027 | 9.38E-03 | 0.23451245 |
| COL4A5 | 1402.04403 | -0.1153388 | 0.29407398 | -0.3922101 | 6.95E-01 | 0.95672887 |
| ACSL4 | 1214.28609 | -0.0602836 | 0.60858044 | -0.0990561 | 0.92109375 | 0.9904507 |
| TMEM164 | 306.969926 | 0.42148341 | 0.37098794 | 1.13611079 | 0.2559102 | 0.80525591 |
| AMMECR1 | 156.190254 | -0.0183732 | 0.25627791 | -0.0716925 | 0.9428466 | 0.99265578 |
| ALG13 | 412.158543 | -0.0190173 | 0.51550103 | -0.0368909 | 0.97057202 | 0.99656026 |
| AMOT | 41.5177878 | 0.62837717 | 0.33918732 | 1.8525963 | 0.06394024 | 0.54860648 |
| PLS3-AS1 | 97.0571927 | 1.17597337 | 0.48712788 | 2.4140958 | 0.01577431 | 0.30067202 |
| PLS3 | 873.822724 | 0.53896518 | 0.29125598 | 1.85048627 | 0.0642435 | 0.54860648 |
| DANT2 | 1442.11006 | -0.1018674 | 0.29368835 | -0.3468553 | 0.72870003 | 0.95987635 |
| KLHL13 | 498.287297 | -0.7811588 | 0.36490671 | -2.1407081 | 0.03229758 | 0.41684566 |
| WDR44 | 319.759012 | -0.3092527 | 0.46142956 | -0.6702056 | 0.50272673 | 0.90962743 |
| IL13RA1 | 153.621481 | 0.19039151 | 0.30526796 | 0.62368653 | 0.53283347 | 0.92268523 |
| PGRMC1 | 253.119424 | 0.49280511 | 0.20539957 | 2.39925088 | 0.01642865 | 0.30723893 |
| SLC25A43 | 102.93804 | -0.3179903 | 0.44560104 | -0.713621 | 0.47546157 | 0.90242252 |
| SLC25A5 | 309.90276 | 0.59790695 | 0.58794598 | 1.01694198 | 0.30918103 | 0.84136316 |
| CXorf56 | 36.1587554 | -0.0818632 | 0.31553575 | -0.2594419 | 0.7952943 | 0.9653598 |
| UBE2A | 112.507204 | -0.091976 | 0.31024286 | -0.2964646 | 0.76687531 | 0.96285313 |
| NKRF | 52.6789112 | 0.65906655 | 0.34308424 | 1.92100505 | 0.05473107 | 0.51597783 |
| 6-Sep | 264.227439 | 0.1249707 | 0.27374867 | 0.45651621 | 0.64801882 | 0.95061388 |
| RPL39 | 38.6508123 | -0.172501 | 0.41773487 | -0.4129436 | 0.67964791 | 0.95574492 |
| UPF3B | 117.566265 | -0.4999481 | 0.3038324 | -1.6454734 | 0.09987222 | 0.61822853 |
| NDUFA1 | 288.077608 | 0.25522088 | 0.3524401 | 0.72415391 | 0.46897125 | 0.90026987 |
| NKAP | 62.3447511 | -0.064045 | 0.25035569 | -0.2558161 | 0.79809282 | 0.9653598 |
| RHOXF1-AS1 | 76.7219528 | 0.04856878 | 0.53341573 | 0.0910524 | 0.92745095 | 0.99129893 |
| LAMP2 | 289.199955 | -0.0265325 | 0.22802722 | -0.1163567 | 0.90736985 | 0.9875066 |
| CUL4B | 105.654958 | -0.0872268 | 0.21627909 | -0.4033065 | 0.68672273 | 0.9558749 |
| MCTS1 | 91.1931907 | -0.1064324 | 0.23527782 | -0.4523691 | 0.65100311 | 0.95061388 |
| C1GALT1C1 | 54.3670905 | 0.31410456 | 0.42664329 | 0.7362229 | 0.46159506 | 0.89799005 |
| THOC2 | 573.113784 | -0.3569325 | 0.20626599 | -1.7304476 | 0.08355033 | 0.59322999 |
| XIAP | 509.30518 | -0.2032148 | 0.18294752 | -1.110782 | 0.2666622 | 0.81333524 |
| STAG2 | 720.625269 | -0.0352488 | 0.19100817 | -0.1845407 | 0.85358929 | 0.97718353 |
| SMARCA1 | 294.546106 | 0.02048431 | 0.25881571 | 0.07914633 | 0.93691623 | 0.99243654 |
| OCRL | 71.5803353 | -0.2421973 | 0.31179162 | -0.7767921 | 0.43728144 | 0.88794991 |
| ZDHHC9 | 106.07939 | -0.5907302 | 0.22469536 | -2.6290272 | 0.00856295 | 0.22546434 |

| | | | | | | |
|---|---|---|---|---|---|---|
| UTP14A | 40.3652438 | -0.1615517 | 0.26774913 | -0.6033695 | 0.54626289 | 0.92597739 |
| BCORL1 | 66.6624964 | 0.73324104 | 0.34539977 | 2.12287646 | 0.03376421 | 0.42838847 |
| ELF4 | 36.0618923 | -0.2399044 | 0.39605357 | -0.6057372 | 0.54468931 | 0.92561018 |
| AIFM1 | 110.811423 | 0.32061394 | 0.21707134 | 1.47699803 | 0.13967617 | 0.6834438 |
| ZNF280C | 44.1305204 | 0.35299997 | 0.26409852 | 1.33662231 | 0.18134595 | 0.74315969 |
| SLC25A14 | 33.2915341 | -0.3801522 | 0.30034201 | -1.2657311 | 0.20560938 | 0.77193931 |
| RBMX2 | 38.4572635 | 0.00792399 | 0.27415917 | 0.02890289 | 0.97694204 | 0.99672436 |
| ENOX2 | 201.745475 | -0.5186664 | 0.18877156 | -2.7475877 | 0.00600355 | 0.19123755 |
| RAP2C | 72.1606697 | -0.3280458 | 0.331012 | -0.991039 | 0.32166654 | 0.84285425 |
| RAP2C-AS1 | 258.09991 | 0.12764416 | 0.2966251 | 0.43032151 | 0.66696178 | 0.9527406 |
| MBNL3 | 219.736559 | 0.47924623 | 0.29199076 | 1.64130613 | 0.10073388 | 0.61990083 |
| GPC4 | 112.371734 | 0.33333853 | 0.28649197 | 1.1635179 | 0.24461944 | 0.80072691 |
| GPC3 | 43.6823585 | -0.4247865 | 0.72000075 | -0.5899807 | 0.55520361 | 0.92872652 |
| PHF6 | 137.170512 | -0.092698 | 0.20761106 | -0.4464982 | 0.65523743 | 0.95100227 |
| HPRT1 | 243.231424 | 0.14189616 | 0.33630515 | 0.4219268 | 0.67307845 | 0.9545145 |
| FAM122B | 57.0414313 | -0.3076523 | 0.29684005 | -1.0364244 | 0.30000419 | 0.83640433 |
| FAM122C | 53.634948 | -0.3709196 | 0.44515135 | -0.8332439 | 0.40470719 | 0.87834687 |
| MOSPD1 | 33.471731 | 0.01277218 | 0.31112456 | 0.04105167 | 0.9672547 | 0.99637506 |
| RTL8C | 150.054006 | -0.0804703 | 0.38369534 | -0.2097246 | 0.83388264 | 0.97232209 |
| RTL8A | 97.6236185 | 0.13070815 | 0.31364348 | 0.41674116 | 0.67686775 | 0.95529325 |
| ZNF75D | 132.476704 | -0.0301388 | 0.35020368 | -0.0860608 | 0.93141808 | 0.99243654 |
| ZNF449 | 37.0266132 | -0.1677321 | 0.36806053 | -0.4557189 | 0.64859215 | 0.95061388 |
| INTS6L | 68.8261875 | -0.1185152 | 0.43901965 | -0.2699542 | 0.78719553 | 0.96513523 |
| MMGT1 | 48.9649528 | 0.09364135 | 0.31500431 | 0.29727004 | 0.76626035 | 0.96285313 |
| SLC9A6 | 87.3653005 | 0.01514823 | 0.33753788 | 0.04487862 | 0.96420406 | 0.99637506 |
| FHL1 | 149.499166 | -0.0908976 | 0.91875626 | -0.0989355 | 0.92118949 | 0.9904507 |
| HTATSF1 | 144.625567 | 0.53793897 | 0.41517832 | 1.29568172 | 0.19508516 | 0.76233115 |
| AL683813.1 | 78.829577 | 0.16120976 | 0.36709048 | 0.43915538 | 0.66054895 | 0.95127957 |
| RBMX | 453.225003 | -0.2829476 | 0.2081736 | -1.3591905 | 0.17408624 | 0.73178166 |
| ATP11C | 415.839642 | -0.9646081 | 0.57640132 | -1.6735009 | 0.09422873 | 0.61330068 |
| LDOC1 | 38.7527757 | 0.65198979 | 0.39220688 | 1.66236195 | 0.09644022 | 0.61440337 |
| FMR1 | 221.757363 | -0.1806932 | 0.21546921 | -0.8386033 | 0.40169195 | 0.87834687 |
| IDS | 257.129648 | -0.2549296 | 0.3601208 | -0.7079002 | 0.47900721 | 0.90291126 |
| TMEM185A | 61.4730347 | 0.10127409 | 0.28489536 | 0.35547822 | 0.72223136 | 0.95907053 |
| LINC00894 | 355.454778 | -0.0563722 | 0.37298612 | -0.1511376 | 0.87986721 | 0.98333507 |
| MTM1 | 168.817877 | -0.2522603 | 0.23115428 | -1.0913071 | 0.27513776 | 0.8202929 |
| MTMR1 | 161.010644 | -0.246667 | 0.16246287 | -1.5182976 | 0.1289394 | 0.67206108 |
| CD99L2 | 127.038414 | -0.0453315 | 0.22726888 | -0.1994618 | 0.84190151 | 0.97557804 |
| HMGB3 | 47.0079842 | -0.8891967 | 0.56234898 | -1.5812186 | 0.11382805 | 0.6483668 |
| VMA21 | 117.57327 | -0.4512523 | 0.43791853 | -1.030448 | 0.30279974 | 0.83709431 |
| CETN2 | 120.411695 | 0.19939098 | 0.43121969 | 0.46238839 | 0.64380282 | 0.94949493 |
| ZNF185 | 63.2110924 | 1.13718779 | 0.59801369 | 1.90160827 | 0.05722239 | 0.52650623 |
| BCAP31 | 197.998036 | 0.1092543 | 0.35402434 | 0.30860676 | 0.75762068 | 0.96180522 |
| IDH3G | 66.9447731 | 0.27332738 | 0.37447028 | 0.72990406 | 0.46544883 | 0.89883965 |
| SSR4 | 143.248679 | 0.19059664 | 0.42701499 | 0.44634648 | 0.65534698 | 0.95100227 |
| L1CAM | 314.622781 | 0.20674911 | 0.44376645 | 0.46589621 | 0.64128979 | 0.94935885 |
| AVPR2 | 118.60423 | 0.78485871 | 0.609287 | 1.28815929 | 0.19769052 | 0.76534786 |
| HCFC1 | 78.4859478 | -0.0720754 | 0.2275655 | -0.3167236 | 0.75145333 | 0.96120879 |
| IRAK1 | 65.222452 | -0.5289655 | 0.33719326 | -1.5687308 | 0.11671067 | 0.65289383 |
| MECP2 | 318.233316 | -0.1793109 | 0.20702131 | -0.8661472 | 0.38640944 | 0.86844754 |
| FLNA | 296.162638 | 0.04687389 | 0.3958169 | 0.11842317 | 0.90573237 | 0.9875066 |
| EMD | 187.886274 | -0.0941248 | 0.53613855 | -0.1755606 | 0.86063919 | 0.97841862 |
| RPL10 | 1364.34063 | 0.20861932 | 0.56847828 | 0.36697853 | 0.71363503 | 0.95791799 |
| DNASE1L1 | 39.4739106 | 0.12719245 | 0.29696254 | 0.42831144 | 0.66842439 | 0.95294349 |

| | | | | | | |
|---|---|---|---|---|---|---|
| TAZ | 91.8297492 | -0.3635391 | 0.24404628 | -1.4896318 | 0.13632109 | 0.67998335 |
| ATP6AP1 | 235.306037 | 0.62077432 | 0.34766785 | 1.78553846 | 0.07417402 | 0.56948258 |
| GDI1 | 101.325827 | -0.5402469 | 0.28059543 | -1.9253587 | 0.0541845 | 0.51425036 |
| FAM50A | 81.4161533 | -0.5797508 | 0.40786346 | -1.4214336 | 0.15519073 | 0.70907003 |
| PLXNA3 | 52.0423147 | -0.557953 | 0.43268858 | -1.2895025 | 0.19722345 | 0.76534786 |
| LAGE3 | 35.7085192 | 0.66817435 | 0.4148143 | 1.61077944 | 0.1072278 | 0.63477557 |
| UBL4A | 40.8475878 | 0.45378991 | 0.41972122 | 1.08116981 | 0.27962159 | 0.82547443 |
| FAM3A | 64.4529201 | -0.1509373 | 0.23835333 | -0.6332502 | 0.52657029 | 0.91974278 |
| IKBKG | 35.188024 | -0.578581 | 0.35042818 | -1.6510686 | 0.09872457 | 0.61707826 |
| DKC1 | 69.1520524 | 0.17899688 | 0.23519837 | 0.76104642 | 0.44662934 | 0.8918683 |
| MPP1 | 71.4306085 | 0.69610444 | 0.44400276 | 1.56779305 | 0.11692944 | 0.65289383 |
| F8 | 52.6951122 | 0.51598748 | 0.37019953 | 1.3938091 | 0.16337526 | 0.71642393 |
| FUNDC2 | 161.712404 | 0.00163313 | 0.37462899 | 0.00435933 | 0.99652177 | 0.99967893 |
| BRCC3 | 131.777629 | -0.0755875 | 0.20383788 | -0.3708217 | 0.71077035 | 0.95768492 |
| VBP1 | 59.0689875 | 0.47405661 | 0.2920067 | 1.62344426 | 0.10449447 | 0.63118273 |
| TMLHE-AS1 | 124.186808 | 0.69819652 | 0.6721999 | 1.03867393 | 0.29895641 | 0.83556453 |
| TMLHE | 259.286757 | 0.29788281 | 0.26889756 | 1.10779289 | 0.26795127 | 0.81607943 |
| VAMP7 | 139.792948 | 0.14072854 | 0.30516615 | 0.46115383 | 0.64468824 | 0.94949493 |
| ZNF596 | 53.9548076 | 0.33230067 | 0.27315032 | 1.2165487 | 0.22377597 | 0.78960314 |
| FBXO25 | 212.800859 | -0.3493117 | 0.4006335 | -0.8718984 | 0.38326383 | 0.86638702 |
| TDRP | 263.366322 | 0.13457797 | 0.26579307 | 0.50632609 | 0.61262775 | 0.94003018 |
| ERICH1 | 311.145266 | -0.5432756 | 0.34653939 | -1.5677168 | 0.11694725 | 0.65289383 |
| AC100810.1 | 49.553699 | 0.21118087 | 0.39250566 | 0.53803267 | 0.59055449 | 0.93483536 |
| AC246817.2 | 50.3613341 | 0.56807528 | 0.66133808 | 0.85897863 | 0.3903523 | 0.87004477 |
| AC246817.1 | 91.5428001 | 0.46495945 | 0.57810673 | 0.8042796 | 0.42123552 | 0.88360543 |
| CSMD1 | 175.27034 | -0.5929188 | 0.9476614 | -0.6256653 | 0.53153453 | 0.92211604 |
| MCPH1 | 427.462222 | 0.04935909 | 0.22367987 | 0.22066846 | 0.82535059 | 0.97018919 |
| ANGPT2 | 67.9304238 | 0.514678 | 0.6335497 | 0.81237194 | 0.41657824 | 0.88242985 |
| MCPH1-AS1 | 154.347206 | 0.29561682 | 0.29448636 | 1.00383876 | 0.31545634 | 0.84285425 |
| AGPAT5 | 405.815959 | -0.2297336 | 0.24667022 | -0.931339 | 0.35167822 | 0.85536253 |
| AF233439.1 | 703.175892 | -1.6811252 | 0.77285874 | -2.1752037 | 0.02961485 | 0.40623517 |
| DEFB1 | 2065.50345 | -0.693142 | 0.49612021 | -1.397125 | 0.16237597 | 0.71482182 |
| FAM85B | 417.242069 | 0.44482352 | 0.43893435 | 1.01341697 | 0.31086104 | 0.84248332 |
| PRAG1 | 48.1843386 | 0.56883357 | 0.55742648 | 1.02046385 | 0.30750853 | 0.83924202 |
| MFHAS1 | 1196.75838 | 1.04117505 | 0.33511419 | 3.10692615 | 0.00189044 | 0.1093833 |
| ERI1 | 117.952488 | -0.1017237 | 0.31703643 | -0.3208581 | 0.74831796 | 0.96109466 |
| PPP1R3B | 112.098081 | -0.5728281 | 0.40889727 | -1.4009096 | 0.1612411 | 0.71409437 |
| TNKS | 856.625126 | -0.2426892 | 0.24361253 | -0.99621 | 0.31914815 | 0.84285425 |
| MSRA | 226.885975 | -0.1930008 | 0.28712449 | -0.672185 | 0.50146591 | 0.90900452 |
| XKR6 | 689.382506 | 0.16576523 | 0.33752444 | 0.49112067 | 0.6233411 | 0.94456546 |
| MTMR9 | 118.816892 | -0.3639278 | 0.23685023 | -1.5365314 | 0.12440811 | 0.665882 |
| FAM167A | 45.6926844 | 0.41026261 | 0.43110174 | 0.95166076 | 0.34126906 | 0.85042291 |
| NEIL2 | 30.0701917 | 0.05583201 | 0.46986368 | 0.11882597 | 0.90541323 | 0.9875066 |
| FDFT1 | 352.8773 | -0.1639985 | 0.5497054 | -0.2983389 | 0.76544455 | 0.96285313 |
| CTSB | 245.653839 | 0.19778977 | 0.31983228 | 0.61841715 | 0.53630039 | 0.92324352 |
| FAM66D | 111.709968 | 0.46438624 | 0.61943907 | 0.74968833 | 0.45344244 | 0.89544675 |
| AC145124.1 | 48.2787777 | 0.80727741 | 0.6171233 | 1.30812985 | 0.19082927 | 0.75802398 |
| LONRF1 | 129.369612 | -0.6561314 | 0.27804687 | -2.3597872 | 0.01828542 | 0.32472005 |
| TRMT9B | 112.422359 | 0.05834889 | 0.46827482 | 0.12460394 | 0.90083711 | 0.9875066 |
| TUSC3 | 364.98135 | 0.1736476 | 0.26079929 | 0.66582851 | 0.50552072 | 0.91122414 |
| MSR1 | 537.818406 | -0.1814133 | 0.50078643 | -0.3622569 | 0.71716007 | 0.95791799 |
| AC068992.1 | 225.746464 | -0.1359344 | 0.41562394 | -0.327061 | 0.74362172 | 0.96105014 |
| MICU3 | 268.476839 | 0.11880527 | 0.26465164 | 0.44891192 | 0.6534952 | 0.95100227 |
| ZDHHC2 | 386.721963 | 0.14443778 | 0.19216635 | 0.75162885 | 0.45227429 | 0.89473986 |

| | | | | | | |
|---|---|---|---|---|---|---|
| CNOT7 | 124.297046 | 0.23460989 | 0.28893217 | 0.81198952 | 0.41679765 | 0.88242985 |
| VPS37A | 152.843088 | -0.3365768 | 0.26593711 | -1.2656256 | 0.20564717 | 0.77193931 |
| SLC7A2 | 180.728749 | -1.9222028 | 1.56574544 | -1.2276598 | 0.21957469 | 0.78630025 |
| MTUS1 | 1046.57871 | -0.3906132 | 0.37827937 | -1.032605 | 0.30178878 | 0.83709431 |
| AC027117.2 | 130.862799 | 0.26809222 | 0.40417966 | 0.66329963 | 0.50713867 | 0.91163608 |
| PCM1 | 1327.67818 | -0.1054146 | 0.18538221 | -0.5686336 | 0.5696048 | 0.93122667 |
| ASAH1 | 899.730123 | 0.07713253 | 0.30717635 | 0.2511018 | 0.80173541 | 0.9661728 |
| PSD3 | 675.972565 | -1.3954408 | 0.68662267 | -2.0323256 | 0.04212071 | 0.46910202 |
| SH2D4A | 175.833282 | 0.30233259 | 0.35098412 | 0.86138537 | 0.38902583 | 0.87004477 |
| CSGALNACT1 | 1266.42798 | -2.9192418 | 0.9523208 | -3.0653975 | 0.00217381 | 0.11990273 |
| INTS10 | 257.203723 | -0.435115 | 0.19700359 | -2.2086652 | 0.02719794 | 0.40076654 |
| ATP6V1B2 | 159.338581 | 0.12698675 | 0.26338194 | 0.48213919 | 0.62970708 | 0.94596555 |
| XPO7 | 262.812465 | -0.1990328 | 0.18227114 | -1.0919602 | 0.27485062 | 0.8202929 |
| DMTN | 128.010536 | -0.1705466 | 0.26826206 | -0.6357461 | 0.5249419 | 0.91924275 |
| FAM160B2 | 69.0305377 | 0.09703798 | 0.23479418 | 0.41328956 | 0.67939448 | 0.95574492 |
| SLC39A14 | 203.314447 | -1.5592785 | 0.59405697 | -2.6247962 | 0.00867009 | 0.22635636 |
| PPP3CC | 292.808386 | -0.6565981 | 0.27648208 | -2.3748306 | 0.01755701 | 0.31833463 |
| CCAR2 | 68.1777036 | -0.0878918 | 0.30824531 | -0.2851359 | 0.77554003 | 0.96413517 |
| BIN3 | 131.702493 | -0.4692081 | 0.21069357 | -2.2269692 | 0.02594933 | 0.3903785 |
| TNFRSF10B | 110.326762 | -0.2173801 | 0.55257093 | -0.3933977 | 0.69402576 | 0.95672887 |
| TNFRSF10D | 84.9572786 | -0.5962012 | 0.55494566 | -1.0743416 | 0.28266962 | 0.82666614 |
| TNFRSF10A | 61.9944701 | -0.5909455 | 0.35998509 | -1.6415832 | 0.1006764 | 0.61990083 |
| CHMP7 | 62.6235825 | -0.08824 | 0.28166612 | -0.3132787 | 0.75406895 | 0.96120879 |
| R3HCC1 | 32.9679211 | 0.06919369 | 0.45736215 | 0.15128863 | 0.87974804 | 0.98333507 |
| ENTPD4 | 94.438475 | -0.1352836 | 0.26622394 | -0.5081571 | 0.61134316 | 0.94003018 |
| SLC25A37 | 273.677426 | 0.2461295 | 0.23506528 | 1.04706869 | 0.2950679 | 0.83396494 |
| STC1 | 296.987457 | 0.48351938 | 0.53075344 | 0.91100564 | 0.3622924 | 0.85694172 |
| DOCK5 | 371.370578 | -0.3277599 | 0.3044954 | -1.0764035 | 0.28174683 | 0.82661559 |
| KCTD9 | 174.856957 | 0.02825461 | 0.24296673 | 0.11629004 | 0.90742268 | 0.9875066 |
| PPP2R2A | 547.869689 | -0.22922 | 0.20271068 | -1.130774 | 0.25815022 | 0.80709747 |
| BNIP3L | 544.485587 | -0.5205899 | 0.50441526 | -1.0320661 | 0.30204117 | 0.83709431 |
| PTK2B | 180.623138 | 0.23758446 | 0.35456581 | 0.67007154 | 0.50281219 | 0.90962743 |
| EPHX2 | 121.011716 | 0.58156809 | 0.38156039 | 1.52418358 | 0.12746286 | 0.67051794 |
| CLU | 1870.04194 | 0.61419342 | 0.54037225 | 1.13661168 | 0.25570066 | 0.80525591 |
| SCARA3 | 48.0394906 | 0.9905572 | 0.96184088 | 1.02985558 | 0.30307781 | 0.83709431 |
| CCDC25 | 128.964622 | 0.36645105 | 0.19651435 | 1.86475465 | 0.0622158 | 0.54562478 |
| ELP3 | 229.272627 | -0.2255548 | 0.26192017 | -0.8611588 | 0.38915061 | 0.87004477 |
| ZNF395 | 103.8567 | -0.4701222 | 0.48727735 | -0.9647938 | 0.33464808 | 0.84521974 |
| FBXO16 | 123.356911 | -0.4359045 | 0.42595825 | -1.0233502 | 0.30614229 | 0.83709431 |
| FZD3 | 104.060297 | 0.23326908 | 0.26945081 | 0.86572048 | 0.38664347 | 0.86852426 |
| EXTL3 | 54.9989418 | -0.0128468 | 0.37718281 | -0.0340599 | 0.97282935 | 0.99656026 |
| INTS9 | 184.879479 | -0.4376917 | 0.18547225 | -2.3598769 | 0.018281 | 0.32472005 |
| HMBOX1 | 556.162662 | -0.3959906 | 0.18875806 | -2.0978738 | 0.03591629 | 0.4362094 |
| KIF13B | 1612.98219 | -0.1395954 | 0.44864388 | -0.3111498 | 0.75568677 | 0.96120879 |
| AC108449.1 | 67.7533713 | -0.0197446 | 0.56441947 | -0.0349822 | 0.97209391 | 0.99656026 |
| SARAF | 425.189089 | 0.37325437 | 0.37917805 | 0.98437759 | 0.32492989 | 0.84285425 |
| LEPROTL1 | 143.406119 | 0.23366497 | 0.24563484 | 0.95126963 | 0.34146752 | 0.85042291 |
| DCTN6 | 78.7919165 | 0.15196727 | 0.42114871 | 0.36083993 | 0.71821911 | 0.95796325 |
| RBPMS | 7726.74508 | -0.4239662 | 0.24529537 | -1.7283906 | 0.08391823 | 0.59383053 |
| GTF2E2 | 229.204597 | -0.2458546 | 0.22844104 | -1.0762279 | 0.28182533 | 0.82661559 |
| GSR | 425.395473 | -0.0364113 | 0.3750278 | -0.0970895 | 0.9226553 | 0.99051393 |
| UBXN8 | 40.3899545 | 0.34661432 | 0.33784093 | 1.02596902 | 0.30490619 | 0.83709431 |
| PPP2CB | 557.501205 | 0.01588 | 0.16210744 | 0.09795971 | 0.92196428 | 0.99051393 |
| WRN | 275.470065 | -0.2609429 | 0.22310438 | -1.1696001 | 0.24216194 | 0.80072691 |

| | | | | | | |
|---|---|---|---|---|---|---|
| NRG1 | 68.438469 | 0.78158731 | 0.71768222 | 1.08904372 | 0.27613461 | 0.82167981 |
| FUT10 | 114.026276 | 0.35725778 | 0.4075306 | 0.87664037 | 0.380682 | 0.86593382 |
| TTI2 | 46.0297743 | 0.09051153 | 0.33247478 | 0.27223579 | 0.78544073 | 0.96513523 |
| UNC5D | 53.7611685 | -0.1743683 | 0.39733041 | -0.4388497 | 0.66077045 | 0.95127957 |
| ERLIN2 | 152.510048 | 0.37508314 | 0.20667865 | 1.81481316 | 0.06955263 | 0.55983989 |
| PLPBP | 104.067921 | 0.18694012 | 0.31557928 | 0.59237132 | 0.55360197 | 0.92872652 |
| RAB11FIP1 | 478.17657 | 0.14115728 | 0.24521136 | 0.57565556 | 0.56484802 | 0.93104864 |
| EIF4EBP1 | 41.1424191 | -0.0111076 | 0.53841302 | -0.0206302 | 0.98354065 | 0.9971769 |
| ASH2L | 128.567157 | -0.1911253 | 0.19651277 | -0.9725848 | 0.33075969 | 0.84442294 |
| LSM1 | 134.752989 | 0.30967026 | 0.43541381 | 0.7112091 | 0.47695467 | 0.90242252 |
| BAG4 | 166.022559 | 0.25191716 | 0.25058174 | 1.0053293 | 0.31473831 | 0.84285425 |
| DDHD2 | 227.536722 | -0.1377848 | 0.24423374 | -0.5641515 | 0.57265106 | 0.93218128 |
| PLPP5 | 187.906177 | -0.1704321 | 0.42376393 | -0.4021865 | 0.68754675 | 0.95588883 |
| NSD3 | 752.192712 | 0.02629616 | 0.14351365 | 0.1832311 | 0.85461669 | 0.97718353 |
| LETM2 | 48.9927798 | 0.24922176 | 0.47457526 | 0.52514697 | 0.59948103 | 0.93728325 |
| FGFR1 | 275.435487 | -0.5884054 | 0.56653071 | -1.0386116 | 0.2989854 | 0.83556453 |
| TACC1 | 374.812639 | -0.9381856 | 0.63202469 | -1.4844129 | 0.1376994 | 0.68146162 |
| PLEKHA2 | 1399.2522 | -1.64752 | 0.52415221 | -3.1432092 | 0.00167106 | 0.10301616 |
| AC108863.1 | 58.1705022 | -1.8597458 | 0.82466694 | -2.2551477 | 0.02412406 | 0.37234186 |
| TM2D2 | 73.5170363 | 0.513864 | 0.58852733 | 0.87313532 | 0.38258932 | 0.86593382 |
| ADAM9 | 510.126564 | -0.2415153 | 0.54328095 | -0.4445496 | 0.65664529 | 0.95100227 |
| TCIM | 128.238277 | -1.2723661 | 0.80175146 | -1.5869832 | 0.11251644 | 0.64541452 |
| GOLGA7 | 84.0765187 | 0.21349867 | 0.34758985 | 0.61422583 | 0.53906611 | 0.92365631 |
| GPAT4 | 97.700508 | -0.4719964 | 0.41263687 | -1.1438541 | 0.25268417 | 0.80350545 |
| KAT6A | 515.696939 | -0.020173 | 0.22963416 | -0.0878486 | 0.92999703 | 0.99216065 |
| AP3M2 | 67.0413108 | 0.10746132 | 0.28524973 | 0.37672717 | 0.70637637 | 0.95730303 |
| IKBKB | 258.699822 | -0.2159866 | 0.22308427 | -0.9681838 | 0.33295257 | 0.84442294 |
| POLB | 105.868771 | -0.3377551 | 0.66643286 | -0.5068104 | 0.61228783 | 0.94003018 |
| VDAC3 | 239.862125 | 0.0958045 | 0.27277118 | 0.35122662 | 0.72541834 | 0.95963742 |
| SLC20A2 | 987.339934 | -0.426874 | 0.44859011 | -0.9515904 | 0.34130476 | 0.85042291 |
| SMIM19 | 146.096777 | -0.2973307 | 0.38181066 | -0.7787386 | 0.43613373 | 0.88785137 |
| RNF170 | 237.149324 | -0.0717612 | 0.28605374 | -0.2508663 | 0.80191751 | 0.9661728 |
| HOOK3 | 482.323201 | -0.2952961 | 0.54518767 | -0.5416412 | 0.58806568 | 0.93483536 |
| FNTA | 104.716101 | -0.0864951 | 0.27051377 | -0.3197438 | 0.74916254 | 0.96109466 |
| HGSNAT | 579.909076 | -0.6993871 | 0.58190934 | -1.2018833 | 0.22940875 | 0.79047378 |
| SPIDR | 1435.48935 | -0.4279676 | 0.18406454 | -2.3250952 | 0.02006687 | 0.33933607 |
| CEBPD | 769.05717 | -1.053156 | 0.71970562 | -1.463315 | 0.1433812 | 0.69023197 |
| PRKDC | 434.744755 | -0.1839888 | 0.18383671 | -1.0008275 | 0.31691021 | 0.84285425 |
| MCM4 | 37.5658969 | -0.0268433 | 0.45884506 | -0.0585019 | 0.95334886 | 0.99512909 |
| UBE2V2 | 195.030723 | -0.5242671 | 0.34645288 | -1.5132421 | 0.13021817 | 0.67292228 |
| SNTG1 | 310.563397 | 1.83785148 | 0.59009753 | 3.11448766 | 0.00184265 | 0.10848841 |
| PXDNL | 62.2151952 | 1.19E-01 | 0.38697561 | 0.30642922 | 0.75927787 | 0.96182318 |
| PCMTD1 | 889.078914 | 0.14287364 | 0.19777091 | 0.72241991 | 0.47003635 | 0.90087435 |
| AC064807.1 | 50.2384557 | -0.2541187 | 0.47468731 | -0.5353392 | 0.59241533 | 0.93491607 |
| RB1CC1 | 481.162109 | -0.3450968 | 0.22646484 | -1.5238427 | 0.12754802 | 0.67051794 |
| ATP6V1H | 551.019528 | 0.19013885 | 0.3003422 | 0.63307403 | 0.5266853 | 0.91979036 |
| TCEA1 | 219.76576 | -0.0398742 | 0.18270043 | -0.218249 | 0.82723509 | 0.97069946 |
| LYPLA1 | 328.214033 | -0.0724367 | 0.22666414 | -0.3195772 | 0.7492889 | 0.96109466 |
| MRPL15 | 104.47952 | 0.07742463 | 0.43386391 | 0.17845373 | 0.85836666 | 0.97786699 |
| TMEM68 | 169.472101 | 0.0635351 | 0.26842287 | 0.23669779 | 0.81289125 | 0.9680627 |
| TGS1 | 343.977606 | 0.62529544 | 0.39223737 | 1.5941761 | 0.1108966 | 0.64209743 |
| LYN | 578.080477 | 0.14564373 | 0.33940221 | 0.4291184 | 0.66783706 | 0.95294349 |
| RPS20 | 421.67466 | 0.01994486 | 0.37842626 | 0.05270475 | 0.95796715 | 0.99579587 |
| PLAG1 | 118.763511 | 0.57352143 | 0.35729375 | 1.6051818 | 0.1084538 | 0.63603573 |

| | | | | | | |
|---|---|---|---|---|---|---|
| CHCHD7 | 72.6079072 | 0.09915232 | 0.26524626 | 0.37381231 | 0.70854396 | 0.95752169 |
| IMPAD1 | 169.464288 | 0.13616985 | 0.29073053 | 0.46837137 | 0.63951904 | 0.94935885 |
| LINC01606 | 255.505391 | 0.03238743 | 0.73079769 | 0.04431791 | 0.96465099 | 0.99637506 |
| FAM110B | 920.896009 | 0.17013504 | 0.3914711 | 0.43460434 | 0.66384965 | 0.95187756 |
| AC027698.1 | 67.7388069 | 0.70028292 | 0.59184858 | 1.18321298 | 0.23672475 | 0.79797745 |
| UBXN2B | 177.798513 | -0.1701594 | 0.16584078 | -1.0260407 | 0.3048724 | 0.83709431 |
| SDCBP | 505.978531 | -0.2742886 | 0.35582585 | -0.7708506 | 0.4407955 | 0.88956997 |
| NSMAF | 174.352508 | -0.1472779 | 0.25255799 | -0.5831447 | 0.55979589 | 0.92934517 |
| TOX | 3955.20487 | -0.3342268 | 0.49439453 | -0.6760326 | 0.49901997 | 0.90839719 |
| LINC01301 | 103.624603 | 0.86325293 | 0.53974268 | 1.59937869 | 0.10973648 | 0.63926535 |
| RAB2A | 731.974987 | -0.0587326 | 0.18039912 | -0.3255705 | 0.74474934 | 0.96109466 |
| CHD7 | 398.158986 | 0.20401464 | 0.27989052 | 0.72890872 | 0.4660575 | 0.89883965 |
| ASPH | 497.178411 | -0.0414971 | 0.21375376 | -0.1941352 | 0.84607004 | 0.97643646 |
| YTHDF3 | 279.079855 | -0.0577147 | 0.34086727 | -0.1693173 | 0.86554703 | 0.98018684 |
| CYP7B1 | 320.391673 | 0.33390965 | 0.30501131 | 1.09474515 | 0.2736283 | 0.81932131 |
| ARMC1 | 103.803769 | -0.2599568 | 0.23914997 | -1.0870031 | 0.27703542 | 0.82198569 |
| MTFR1 | 126.976773 | -0.3750174 | 0.36250716 | -1.0345103 | 0.30089765 | 0.83709431 |
| PDE7A | 121.256429 | 0.03376884 | 0.42995712 | 0.07854001 | 0.93739851 | 0.99243654 |
| ADHFE1 | 102.009024 | 0.43476003 | 0.50250905 | 0.86517851 | 0.38694082 | 0.86852426 |
| VCPIP1 | 98.3903433 | -0.0812062 | 0.28821275 | -0.2817577 | 0.7781293 | 0.96472201 |
| SGK3 | 166.718163 | -0.1092745 | 0.2485978 | -0.4395636 | 0.66025321 | 0.95127957 |
| PPP1R42 | 59.6481637 | 0.62536075 | 0.3755517 | 1.66517886 | 0.09587708 | 0.61367241 |
| CSPP1 | 443.292032 | -0.2330423 | 0.28576732 | -0.8154969 | 0.41478797 | 0.88242985 |
| ARFGEF1 | 1485.22398 | -0.0149276 | 0.17388325 | -0.0858486 | 0.93158675 | 0.99243654 |
| NCOA2 | 1785.67129 | 0.02327737 | 0.13627881 | 0.17080698 | 0.86437554 | 0.97942001 |
| TRAM1 | 379.323258 | 0.05016959 | 0.21440117 | 0.23399868 | 0.814986 | 0.96839471 |
| LACTB2 | 69.6586089 | 0.03523909 | 0.32402279 | 0.10875499 | 0.91339683 | 0.98830345 |
| MSC-AS1 | 434.866312 | 0.3707242 | 0.89399381 | 0.41468318 | 0.67837385 | 0.95574492 |
| TERF1 | 200.718993 | -0.0631087 | 0.1733742 | -0.3640027 | 0.71585601 | 0.95791799 |
| SBSPON | 115.967499 | 0.64571697 | 0.69281386 | 0.93202086 | 0.35132574 | 0.85504372 |
| RPL7 | 972.146465 | 0.26764177 | 0.52036591 | 0.51433378 | 0.60701865 | 0.93925224 |
| RDH10 | 906.156206 | -1.9086815 | 0.55315717 | -3.4505229 | 0.0005595 | 0.05789336 |
| RDH10-AS1 | 302.569083 | -1.5404479 | 0.75424105 | -2.0423814 | 0.04111371 | 0.46430138 |
| STAU2 | 653.000366 | -0.3516144 | 0.23921776 | -1.4698506 | 0.14160222 | 0.68671506 |
| AC027018.1 | 97.8465963 | -0.2307234 | 0.47526446 | -0.4854633 | 0.62734775 | 0.94511671 |
| ELOC | 142.490292 | -0.1419029 | 0.34510369 | -0.4111891 | 0.68093388 | 0.95574492 |
| TMEM70 | 42.6842144 | 0.24013712 | 0.47829433 | 0.50206976 | 0.61561845 | 0.94190661 |
| GDAP1 | 155.324306 | 0.04970466 | 0.38242883 | 0.12997101 | 0.89658936 | 0.98597539 |
| HNF4G | 220.894328 | -0.647478 | 0.34229616 | -1.8915726 | 0.05854796 | 0.53216182 |
| ZFHX4 | 48.3933537 | -1.7293292 | 1.07614678 | -1.6069641 | 0.10806225 | 0.63533473 |
| PEX2 | 154.866996 | 0.21270936 | 0.24662336 | 0.86248667 | 0.38841977 | 0.86970192 |
| PKIA-AS1 | 50.0825663 | 0.27630777 | 0.59805287 | 0.46201228 | 0.64407251 | 0.94949493 |
| PKIA | 611.026227 | -0.2786864 | 0.42491913 | -0.6558576 | 0.51191575 | 0.91408994 |
| ZC2HC1A | 372.387764 | -0.210916 | 0.19690002 | -1.0711832 | 0.28408704 | 0.82746865 |
| IL7 | 144.664781 | 0.23831168 | 0.43361541 | 0.54959226 | 0.58259907 | 0.93407228 |
| TPD52 | 457.361445 | -0.3135512 | 0.41240376 | -0.7603015 | 0.4470744 | 0.89187439 |
| ZBTB10 | 131.709364 | 0.17053162 | 0.37584881 | 0.45372397 | 0.65002751 | 0.95061388 |
| ZNF704 | 1120.75772 | 0.48774524 | 0.2919236 | 1.67079753 | 0.09476168 | 0.61367241 |
| PAG1 | 92.4141329 | -0.1826066 | 0.48305958 | -0.3780209 | 0.70541506 | 0.95730303 |
| IMPA1 | 74.9066441 | -0.2802716 | 0.23990385 | -1.1682664 | 0.24269933 | 0.80072691 |
| ZFAND1 | 140.56002 | -0.0415643 | 0.20895236 | -0.1989175 | 0.84232729 | 0.97560439 |
| CHMP4C | 52.7010152 | 0.053558 | 0.33949666 | 0.15775708 | 0.87464822 | 0.98266529 |
| SNX16 | 110.234234 | -0.6494116 | 0.28359983 | -2.289887 | 0.02202787 | 0.35843654 |
| AC015522.1 | 229.29151 | 1.37914445 | 1.4881192 | 0.92677014 | 0.35404589 | 0.8557799 |

| | | | | | | |
|---|---|---|---|---|---|---|
| RALYL | 2412.03299 | 0.84802775 | 0.28735385 | 2.95116194 | 0.00316581 | 0.14358074 |
| LRRCC1 | 64.1130758 | -0.2104815 | 0.28359887 | -0.7421805 | 0.45797799 | 0.89736194 |
| E2F5 | 81.7045756 | -0.2555105 | 0.31118685 | -0.821084 | 0.41159842 | 0.88086809 |
| C8orf59 | 111.595455 | -0.1188663 | 0.26318019 | -0.4516536 | 0.65151857 | 0.95061388 |
| CA13 | 106.675294 | -0.1182038 | 0.33435812 | -0.3535245 | 0.72369526 | 0.95963742 |
| CA1 | 38.3134012 | -0.0329402 | 0.62854936 | -0.0524067 | 0.95820463 | 0.99579587 |
| CA3 | 119.902805 | -0.2737414 | 0.28553362 | -0.9587011 | 0.33770936 | 0.84672703 |
| CA2 | 823.521408 | 0.67238778 | 0.52448879 | 1.28198695 | 0.19984722 | 0.7687013 |
| WWP1 | 1152.58764 | 0.14985513 | 0.24356638 | 0.61525377 | 0.53838714 | 0.92346119 |
| RMDN1 | 235.911663 | 0.00849741 | 0.27804219 | 0.03056159 | 0.97561918 | 0.99656026 |
| CPNE3 | 232.649245 | -0.0780585 | 0.16807588 | -0.4644241 | 0.64234395 | 0.94949493 |
| CNGB3 | 44.8898322 | 0.31185497 | 0.59213615 | 0.52666092 | 0.59842908 | 0.93728325 |
| AF117829.1 | 671.525865 | 0.57538418 | 0.31344235 | 1.8356938 | 0.06640296 | 0.55583305 |
| RIPK2 | 97.4509702 | -0.3426621 | 0.29589984 | -1.158034 | 0.24685014 | 0.80072691 |
| OSGIN2 | 101.873164 | 0.17327733 | 0.33096125 | 0.52355775 | 0.60058617 | 0.93760511 |
| NBN | 182.504179 | -0.0779178 | 0.22981693 | -0.339043 | 0.7345773 | 0.96061519 |
| DECR1 | 438.884643 | 0.3281295 | 0.24172508 | 1.35744911 | 0.17463855 | 0.73219159 |
| CALB1 | 432.990686 | 1.6347803 | 0.82425334 | 1.98334689 | 0.04732869 | 0.48600281 |
| TMEM64 | 119.518224 | 0.71133101 | 0.32383543 | 2.1965818 | 0.02805033 | 0.40459506 |
| PIP4P2 | 133.222573 | -0.2330368 | 0.32217957 | -0.7233135 | 0.46948732 | 0.90053922 |
| OTUD6B-AS1 | 99.9881358 | 0.45547114 | 0.39317344 | 1.15844839 | 0.2466811 | 0.80072691 |
| OTUD6B | 99.4262093 | 0.0574196 | 0.30294786 | 0.18953624 | 0.84967256 | 0.97671331 |
| LRRC69 | 141.505593 | -0.0331752 | 0.28931662 | -0.1146675 | 0.90870864 | 0.98758338 |
| SLC26A7 | 235.24274 | 1.42702775 | 0.54451889 | 2.62071301 | 0.00877461 | 0.22761858 |
| AC117834.1 | 62.3127884 | 0.24366557 | 0.56518329 | 0.43112663 | 0.6663763 | 0.9527406 |
| TRIQK | 294.342632 | -0.0737521 | 0.16572618 | -0.4450241 | 0.65630234 | 0.95100227 |
| FAM92A | 96.108735 | 0.55504573 | 0.27595922 | 2.01133247 | 0.04429035 | 0.48009422 |
| RBM12B | 124.138812 | -0.1705245 | 0.24233343 | -0.7036771 | 0.48163386 | 0.90561114 |
| TMEM67 | 250.737304 | -0.0927967 | 0.19096903 | -0.4859252 | 0.6270202 | 0.94511671 |
| PDP1 | 76.0415572 | 0.10570598 | 0.42225435 | 0.25033721 | 0.80232658 | 0.9661728 |
| VIRMA | 189.790889 | -0.0334204 | 0.21106986 | -0.158338 | 0.8741905 | 0.98246825 |
| ESRP1 | 268.783659 | -0.0735271 | 0.26926252 | -0.2730686 | 0.7848005 | 0.96513523 |
| DPY19L4 | 225.613776 | -0.0918323 | 0.17309151 | -0.5305419 | 0.59573628 | 0.93574883 |
| INTS8 | 152.801859 | -0.1418084 | 0.29051141 | -0.4881338 | 0.62545508 | 0.94508035 |
| NDUFAF6 | 150.17164 | 0.00117351 | 0.32004347 | 0.00366672 | 0.99707438 | 0.99967893 |
| TP53INP1 | 69.9655644 | -0.1949738 | 0.24408818 | -0.7987843 | 0.4244155 | 0.88391318 |
| PLEKHF2 | 68.1958017 | 0.49259496 | 0.41709742 | 1.18100697 | 0.23759995 | 0.79810098 |
| C8orf37-AS1 | 173.865744 | 0.73602351 | 0.31601082 | 2.32910859 | 0.01985331 | 0.33831338 |
| UQCRB | 560.583501 | 0.3396536 | 0.42034598 | 0.80803342 | 0.41907134 | 0.88261006 |
| MTERF3 | 39.6592936 | 0.05062136 | 0.2945086 | 0.17188416 | 0.8635286 | 0.979268 |
| PTDSS1 | 60.3118488 | -0.0296055 | 0.38697115 | -0.0765058 | 0.93901668 | 0.99243654 |
| SDC2 | 99.9161433 | -0.0742511 | 0.52755731 | -0.140745 | 0.88807136 | 0.98551241 |
| CPQ | 559.351425 | 0.0350235 | 0.26777588 | 0.13079407 | 0.89593822 | 0.98597539 |
| TSPYL5 | 52.7290184 | 0.28419376 | 0.38759013 | 0.73323272 | 0.4634165 | 0.89801718 |
| MTDH | 695.996461 | 0.00529736 | 0.13490171 | 0.03926828 | 0.9686765 | 0.9964399 |
| LAPTM4B | 432.948051 | -0.0735175 | 0.25071752 | -0.2932285 | 0.7693475 | 0.96345263 |
| MATN2 | 100.546017 | 0.084553 | 0.48363196 | 0.17482921 | 0.86121383 | 0.97841862 |
| RPL30 | 701.987599 | -0.1972803 | 0.48681181 | -0.4052497 | 0.68529399 | 0.9558749 |
| RIDA | 58.8724242 | 0.37196366 | 0.34401039 | 1.08125705 | 0.27958279 | 0.82547443 |
| POP1 | 35.5489972 | 0.27507766 | 0.34224954 | 0.80373422 | 0.42155049 | 0.88360543 |
| NIPAL2 | 44.8136327 | -1.0341729 | 0.3638105 | -2.8426141 | 0.00447452 | 0.16424479 |
| STK3 | 1325.59488 | -0.2102833 | 0.26158741 | -0.803874 | 0.42146975 | 0.88360543 |
| AC104986.2 | 89.8355278 | 0.45917232 | 0.31151369 | 1.47400365 | 0.14048061 | 0.6847613 |
| VPS13B | 1276.1304 | -0.2020719 | 0.21597526 | -0.9356253 | 0.34946616 | 0.8535343 |

| | | | | | | |
|---|---|---|---|---|---|---|
| COX6C | 466.224506 | -0.3229795 | 0.29702244 | -1.0873908 | 0.27686412 | 0.82198569 |
| POLR2K | 115.968331 | 0.16420352 | 0.52046868 | 0.31549164 | 0.7523884 | 0.96120879 |
| SPAG1 | 176.41171 | 0.03460364 | 0.33920675 | 0.10201341 | 0.91874603 | 0.99014946 |
| RNF19A | 1406.77501 | -1.032502 | 0.39621759 | -2.6058965 | 0.00916342 | 0.23300339 |
| AP000424.1 | 235.690655 | -1.3320037 | 0.52580797 | -2.5332513 | 0.01130099 | 0.26144451 |
| ANKRD46 | 146.398824 | 0.39698281 | 0.25796917 | 1.53887694 | 0.12383434 | 0.66519057 |
| PABPC1 | 640.436902 | -0.0800651 | 0.36439736 | -0.2197193 | 0.82608978 | 0.97058548 |
| YWHAZ | 906.533269 | -0.2578932 | 0.34843631 | -0.7401445 | 0.45921232 | 0.89753337 |
| ZNF706 | 243.660884 | 0.01962377 | 0.24389686 | 0.0804593 | 0.93587196 | 0.99243654 |
| AP001207.3 | 42.7051627 | 0.50973445 | 0.57213699 | 0.89093078 | 0.37296631 | 0.86201082 |
| GRHL2 | 830.925684 | 0.15038951 | 0.28703743 | 0.52393693 | 0.60032241 | 0.93733296 |
| NCALD | 67.9365815 | -2.5610282 | 0.62210283 | -4.1167281 | 3.84E-05 | 0.00936593 |
| RRM2B | 152.44683 | 0.22052683 | 0.30003791 | 0.73499656 | 0.46234159 | 0.89801718 |
| UBR5 | 875.981148 | -0.0627206 | 0.20333621 | -0.3084577 | 0.75773409 | 0.96180522 |
| KLF10 | 173.040168 | -1.0272306 | 0.81916571 | -1.253996 | 0.20984345 | 0.77551367 |
| AZIN1 | 256.189782 | -0.0087159 | 0.29786945 | -0.0292608 | 0.97665661 | 0.99656026 |
| AZIN1-AS1 | 382.416283 | -0.0379268 | 0.35946148 | -0.1055101 | 0.91597107 | 0.98870912 |
| AP003354.2 | 75.8851322 | -0.2931123 | 0.41307814 | -0.7095808 | 0.47796414 | 0.90262056 |
| ATP6V1C1 | 258.137178 | -0.4291148 | 0.25499758 | -1.6828191 | 0.09241012 | 0.61120158 |
| BAALC-AS1 | 68.9810223 | -0.1877049 | 0.37939902 | -0.4947428 | 0.62078167 | 0.94369714 |
| FZD6 | 244.924995 | 0.05146676 | 0.17593514 | 0.29253254 | 0.76987948 | 0.96345263 |
| SLC25A32 | 76.6981862 | -0.0762603 | 0.23116483 | -0.3298956 | 0.74147886 | 0.96084411 |
| DCAF13 | 55.5947282 | 0.1026515 | 0.32052681 | 0.32025872 | 0.74877221 | 0.96109466 |
| LRP12 | 130.545631 | 0.3578648 | 0.23265885 | 1.53815255 | 0.12401132 | 0.66528789 |
| OXR1 | 985.616974 | 0.43661233 | 0.37865827 | 1.15305111 | 0.24888941 | 0.80097144 |
| AC090579.1 | 81.3423451 | 0.7795956 | 0.4644762 | 1.67844039 | 0.09326116 | 0.61317397 |
| ANGPT1 | 85.1815908 | 0.30815628 | 0.64703433 | 0.47625956 | 0.63388947 | 0.94753411 |
| EIF3E | 307.800447 | 0.18991444 | 0.19138326 | 0.99232526 | 0.32103889 | 0.84285425 |
| EMC2 | 264.925375 | -0.3524932 | 0.35660186 | -0.9884783 | 0.32291843 | 0.84285425 |
| NUDCD1 | 109.37163 | -0.0121701 | 0.2385036 | -0.0510269 | 0.95930405 | 0.99609409 |
| ENY2 | 193.67305 | 0.12570659 | 0.27030849 | 0.46504863 | 0.64189663 | 0.94935885 |
| EBAG9 | 100.67872 | 0.00673562 | 0.22224388 | 0.03030735 | 0.97582194 | 0.99656026 |
| SYBU | 84.1366676 | 0.75149122 | 0.38416607 | 1.9561624 | 0.05044603 | 0.49596288 |
| TRPS1 | 2811.38665 | -0.4531402 | 0.53573881 | -0.845823 | 0.3976515 | 0.87462201 |
| EIF3H | 419.102567 | -0.0718505 | 0.22073702 | -0.3255026 | 0.74480075 | 0.96109466 |
| UTP23 | 54.1156007 | -0.0675316 | 0.244551 | -0.2761452 | 0.78243652 | 0.96513523 |
| RAD21 | 271.241555 | 0.07845639 | 0.24600003 | 0.31892837 | 0.74978083 | 0.96109466 |
| MED30 | 43.4053947 | 0.16177565 | 0.29933662 | 0.54044724 | 0.58888863 | 0.93483536 |
| EXT1 | 1452.00162 | 0.06860665 | 0.32140133 | 0.213461 | 0.83096741 | 0.97160502 |
| SAMD12 | 2693.27401 | 0.01848616 | 0.18354806 | 0.10071564 | 0.91977619 | 0.99014946 |
| AC023590.1 | 278.966799 | 0.25474889 | 0.63314735 | 0.40235324 | 0.68742407 | 0.95588883 |
| SAMD12-AS1 | 184.450021 | -0.9299218 | 0.26109528 | -3.5616185 | 0.00036858 | 0.04296095 |
| TNFRSF11B | 327.357974 | 0.59786348 | 0.59206033 | 1.00980161 | 0.31259035 | 0.84279957 |
| COLEC10 | 96.334981 | 0.2142134 | 0.42065309 | 0.50924005 | 0.61058397 | 0.94003018 |
| MAL2 | 1136.24785 | -0.4700829 | 0.38039719 | -1.2357687 | 0.21654456 | 0.78093153 |
| MAL2-AS1 | 74.7549785 | 0.21009248 | 0.72814663 | 0.28853045 | 0.77294073 | 0.96388926 |
| NOV | 106.62949 | -1.9883026 | 0.68895718 | -2.8859596 | 0.00390222 | 0.15207852 |
| TAF2 | 280.326646 | -0.2578507 | 0.21213029 | -1.2155301 | 0.22416399 | 0.78960314 |
| DEPTOR | 605.598762 | -0.0813197 | 0.47534208 | -0.1710762 | 0.86416388 | 0.97939196 |
| AC091563.1 | 60.6777062 | -0.1721476 | 0.43514281 | -0.3956117 | 0.6923915 | 0.95672887 |
| MRPL13 | 116.403928 | 0.16957843 | 0.39402619 | 0.43037351 | 0.66692397 | 0.9527406 |
| MTBP | 57.427399 | -0.1968868 | 0.32517124 | -0.6054864 | 0.54485584 | 0.92561018 |
| SNTB1 | 172.282796 | 0.49305071 | 0.42055037 | 1.17239396 | 0.24103892 | 0.80072691 |
| ZHX2 | 1015.40152 | -0.6124611 | 0.32792547 | -1.8676839 | 0.06180615 | 0.54419361 |

| | | | | | | |
|---|---|---|---|---|---|---|
| AC104316.1 | 42.3209481 | -0.3057523 | 0.45298427 | -0.6749733 | 0.49969275 | 0.90839719 |
| DERL1 | 125.890017 | -0.2516201 | 0.2556631 | -0.9841862 | 0.32502396 | 0.84285425 |
| TBC1D31 | 54.2137132 | 0.09817089 | 0.32519039 | 0.30188742 | 0.76273788 | 0.96272347 |
| ATAD2 | 78.2479493 | 0.09360188 | 0.3818598 | 0.24512106 | 0.8063627 | 0.96752828 |
| WDYHV1 | 54.0799063 | -0.2864369 | 0.27744844 | -1.0323967 | 0.30188632 | 0.83709431 |
| FBXO32 | 89.1872372 | -0.7270749 | 0.64454317 | -1.1280469 | 0.2593001 | 0.80709747 |
| FAM91A1 | 260.942962 | -0.3781209 | 0.18993647 | -1.9907756 | 0.04650556 | 0.48600281 |
| TMEM65 | 273.64325 | -0.0701642 | 0.33550091 | -0.2091327 | 0.83434466 | 0.97232209 |
| RNF139 | 40.7741897 | 0.13575048 | 0.44842327 | 0.30272846 | 0.76209681 | 0.96218738 |
| TATDN1 | 186.228604 | -0.3534075 | 0.18740449 | -1.8858007 | 0.05932182 | 0.53364175 |
| NDUFB9 | 204.990054 | 0.29007696 | 0.36499994 | 0.79473151 | 0.4267697 | 0.88477675 |
| MTSS1 | 600.144937 | 0.7261352 | 0.38402334 | 1.89086216 | 0.05864275 | 0.53256151 |
| SQLE | 200.843979 | -0.0161544 | 0.66801082 | -0.0241829 | 0.98070674 | 0.99696835 |
| WASHC5 | 136.331849 | -0.4802484 | 0.22701449 | -2.1154968 | 0.03438763 | 0.43050784 |
| NSMCE2 | 895.967565 | -0.3880248 | 0.21285622 | -1.8229432 | 0.06831199 | 0.55917298 |
| TRIB1 | 87.4964509 | 0.80788595 | 0.43673126 | 1.84984685 | 0.06433563 | 0.54860648 |
| FAM84B | 209.610466 | -0.859924 | 0.35572625 | -2.4173757 | 0.01563287 | 0.30067202 |
| PCAT1 | 1161.15295 | -0.6248378 | 0.25235571 | -2.4760202 | 0.01328561 | 0.28170334 |
| PVT1 | 163.335202 | 0.94890122 | 0.71094943 | 1.3346958 | 0.18197593 | 0.74315969 |
| FAM49B | 1290.36985 | -0.213403 | 0.25219456 | -0.8461841 | 0.39745005 | 0.87462201 |
| ASAP1 | 407.644081 | 0.28247609 | 0.38801791 | 0.72799756 | 0.46661508 | 0.89883965 |
| EFR3A | 754.241251 | 0.31628546 | 0.27337623 | 1.15696036 | 0.24728856 | 0.80072691 |
| LRRC6 | 203.584288 | 0.06468056 | 0.23922812 | 0.2703719 | 0.78687416 | 0.96513523 |
| PHF20L1 | 236.946806 | -0.2673305 | 0.23491247 | -1.1380005 | 0.25512027 | 0.80489222 |
| NDRG1 | 4661.83604 | -0.7957309 | 0.52890317 | -1.5044926 | 0.13245457 | 0.67466347 |
| ZFAT | 98.5452952 | -0.4247803 | 0.37216818 | -1.1413664 | 0.25371746 | 0.80414079 |
| AC083843.3 | 78.442458 | -0.2288071 | 0.43843311 | -0.5218745 | 0.60175772 | 0.93804805 |
| KHDRBS3 | 97.3409384 | 0.23344024 | 0.43331165 | 0.53873521 | 0.59006958 | 0.93483536 |
| COL22A1 | 95.4975039 | 0.39492053 | 0.94134746 | 0.41952685 | 0.67483114 | 0.95524793 |
| TRAPPC9 | 1268.20196 | -0.0849043 | 0.20752386 | -0.4091302 | 0.68244412 | 0.95574492 |
| CHRAC1 | 72.6458887 | -0.3146533 | 0.42833282 | -0.7346 | 0.46258311 | 0.89801718 |
| AGO2 | 345.081534 | -0.3376794 | 0.28077416 | -1.2026725 | 0.22910309 | 0.79047378 |
| PTK2 | 2630.17004 | -0.2668976 | 0.2146934 | -1.243157 | 0.21380997 | 0.77890374 |
| DENND3 | 142.21838 | -0.8165013 | 0.52323395 | -1.5604899 | 0.11864415 | 0.65456964 |
| SLC45A4 | 1174.04603 | -2.7322262 | 0.83865775 | -3.257856 | 0.00112257 | 0.08298537 |
| TSNARE1 | 103.307696 | -0.6014772 | 0.33492322 | -1.795866 | 0.07251583 | 0.56629348 |
| JRK | 44.8747785 | 0.07436259 | 0.3211171 | 0.23157469 | 0.81686837 | 0.96874284 |
| THEM6 | 40.8298605 | 0.50735709 | 0.44793634 | 1.13265446 | 0.25735939 | 0.80709747 |
| LY6E | 146.173729 | -1.711969 | 1.3006729 | -1.3162179 | 0.1881009 | 0.7526909 |
| GLI4 | 37.6882713 | -0.0074132 | 0.39849111 | -0.0186032 | 0.98515765 | 0.99762072 |
| TOP1MT | 54.5885941 | -0.4801048 | 0.32975671 | -1.4559363 | 0.14541023 | 0.69325556 |
| RHPN1 | 94.9730563 | -0.4842197 | 0.39196806 | -1.2353549 | 0.21669845 | 0.78121766 |
| ZC3H3 | 79.5605791 | -0.1456606 | 0.33430487 | -0.4357118 | 0.66304583 | 0.95159487 |
| GSDMD | 100.708695 | 0.42740338 | 0.42340811 | 1.00943599 | 0.31276559 | 0.84283449 |
| NAPRT | 83.991955 | -0.0804933 | 0.54238255 | -0.1484069 | 0.88202166 | 0.98335905 |
| EEF1D | 425.55393 | -0.1700833 | 0.40202099 | -0.4230707 | 0.6722437 | 0.9544944 |
| TSTA3 | 54.4740809 | 0.22784523 | 0.66065832 | 0.34487604 | 0.73018756 | 0.96008876 |
| ZNF623 | 51.8901919 | -0.1688702 | 0.28593857 | -0.590582 | 0.55480053 | 0.92872652 |
| FAM83H | 106.005099 | -0.0713952 | 0.33978246 | -0.2101204 | 0.8335737 | 0.97232209 |
| IQANK1 | 99.7432499 | 0.24788898 | 0.37057495 | 0.66893075 | 0.50353965 | 0.90984406 |
| SCRIB | 67.37657 | -0.1843492 | 0.36741402 | -0.5017479 | 0.61584483 | 0.94190661 |
| PUF60 | 105.766061 | 0.26580368 | 0.3984629 | 0.66707259 | 0.50472577 | 0.9107831 |
| NRBP2 | 71.5557411 | -0.2740684 | 0.31820141 | -0.8613048 | 0.38907018 | 0.87004477 |
| EPPK1 | 35.5422656 | -0.532747 | 0.48588825 | -1.0964394 | 0.27288654 | 0.81922962 |

| | | | | | | |
|---|---|---|---|---|---|---|
| PLEC | 174.074302 | -0.5071363 | 0.25354432 | -2.0001882 | 0.04547995 | 0.48339878 |
| PARP10 | 68.7125641 | -1.2401035 | 0.55710907 | -2.2259617 | 0.02601675 | 0.39062391 |
| GRINA | 142.948949 | -0.3697065 | 0.35287141 | -1.047709 | 0.29477272 | 0.83379706 |
| EXOSC4 | 35.3687248 | 0.30865554 | 0.54478882 | 0.56655997 | 0.57101318 | 0.93125087 |
| GPAA1 | 94.2100871 | 0.58312307 | 0.41519383 | 1.40445985 | 0.16018196 | 0.71409437 |
| CYC1 | 168.941387 | 0.36988597 | 0.55179023 | 0.67033802 | 0.50264234 | 0.90962743 |
| SHARPIN | 80.0530419 | -0.1070306 | 0.23070269 | -0.4639328 | 0.64269586 | 0.94949493 |
| MAF1 | 124.981391 | 0.04464839 | 0.32691315 | 0.13657571 | 0.89136618 | 0.98597539 |
| MROH1 | 273.026199 | -0.8448873 | 0.39538581 | -2.1368681 | 0.03260872 | 0.41931218 |
| HSF1 | 196.768827 | -0.0713842 | 0.24991459 | -0.2856342 | 0.77515827 | 0.96388926 |
| DGAT1 | 61.8957743 | 0.25096944 | 0.36218948 | 0.69292306 | 0.48835784 | 0.90730741 |
| SLC52A2 | 43.4649457 | 0.09615572 | 0.41088628 | 0.23402028 | 0.81496923 | 0.96839471 |
| CPSF1 | 44.2456584 | -0.1585186 | 0.41166103 | -0.3850706 | 0.70018512 | 0.95672887 |
| SLC39A4 | 43.9774383 | 0.12253787 | 0.41170699 | 0.29763369 | 0.76598276 | 0.96285313 |
| VPS28 | 321.631394 | 0.02424024 | 0.38069979 | 0.06367286 | 0.94923072 | 0.99361356 |
| CYHR1 | 168.767825 | 0.70237442 | 0.3318967 | 2.11624409 | 0.03432405 | 0.43050784 |
| KIFC2 | 62.058143 | 0.20779708 | 0.36081016 | 0.57591804 | 0.56467058 | 0.93104392 |
| PPP1R16A | 125.01867 | -0.2323686 | 0.23546388 | -0.9868545 | 0.32371396 | 0.84285425 |
| MFSD3 | 31.1731029 | 0.16187933 | 0.5570556 | 0.29059816 | 0.77135867 | 0.96388926 |
| LRRC14 | 31.5203112 | 0.2806006 | 0.39759991 | 0.70573608 | 0.48035228 | 0.9044362 |
| ARHGAP39 | 53.8017416 | 0.05179472 | 0.34034563 | 0.15218271 | 0.87904283 | 0.98311774 |
| ZNF251 | 165.982064 | 0.1082181 | 0.30057439 | 0.36003767 | 0.71881896 | 0.95796325 |
| RPL8 | 1106.11385 | 0.05488705 | 0.50845925 | 0.10794778 | 0.91403712 | 0.98830345 |
| ZNF7 | 64.7501584 | 0.05329585 | 0.27449873 | 0.194157 | 0.84605295 | 0.97643646 |
| COMMD5 | 34.5835414 | -0.2681882 | 0.37850362 | -0.7085487 | 0.47860458 | 0.90265195 |
| ZNF250 | 56.5507119 | -0.1052921 | 0.33307699 | -0.3161193 | 0.75191194 | 0.96120879 |
| ZNF16 | 43.1251335 | -0.0235617 | 0.35624426 | -0.0661393 | 0.94726695 | 0.99314907 |
| C8orf33 | 45.5894598 | 0.33933889 | 0.38182818 | 0.88872144 | 0.37415281 | 0.8625432 |
| WASHC1 | 355.473354 | 0.21859156 | 0.29305792 | 0.74589883 | 0.45572853 | 0.89626521 |
| CBWD1 | 85.0879344 | -0.3422318 | 0.26280573 | -1.3022236 | 0.19283996 | 0.7597012 |
| DOCK8 | 128.674376 | -2.1976607 | 0.83827088 | -2.6216593 | 0.00875029 | 0.22755084 |
| KANK1 | 1668.41327 | 0.81543813 | 0.34773804 | 2.34497822 | 0.01902819 | 0.33090117 |
| SMARCA2 | 1270.07307 | 0.11906368 | 0.52812797 | 0.22544476 | 0.8216333 | 0.96967533 |
| VLDLR-AS1 | 46.9324815 | -0.797532 | 0.61789479 | -1.2907246 | 0.1967992 | 0.76478825 |
| VLDLR | 75.9151142 | -0.3234568 | 0.43868793 | -0.7373278 | 4.61E-01 | 8.98E-01 |
| PUM3 | 394.079128 | -0.2504326 | 0.31155199 | -0.8038229 | 0.42149924 | 0.88360543 |
| RFX3 | 625.514624 | 0.0871589 | 0.20257221 | 0.43026092 | 0.66700585 | 0.9527406 |
| RFX3-AS1 | 534.361585 | 0.47335409 | 0.43528023 | 1.08746976 | 0.27682926 | 0.82198569 |
| GLIS3 | 5184.86761 | -0.6636512 | 0.41118041 | -1.6140147 | 0.10652424 | 0.63461115 |
| SLC1A1 | 133.770968 | -0.0349979 | 0.37352427 | -0.0936965 | 0.92535022 | 0.99051393 |
| SPATA6L | 395.92495 | -0.0852391 | 0.37252065 | -0.2288171 | 0.81901105 | 0.96884581 |
| CDC37L1 | 158.214193 | 0.13362282 | 0.27479475 | 0.48626411 | 0.62677992 | 0.94511671 |
| AK3 | 603.364537 | 0.26941332 | 0.22156071 | 1.21597967 | 0.22399267 | 0.78960314 |
| RCL1 | 178.968005 | -0.0163398 | 0.31140554 | -0.052471 | 0.95815339 | 0.99579587 |
| JAK2 | 350.114116 | -0.2029284 | 0.19671587 | -1.0315812 | 0.30226835 | 0.83709431 |
| PLGRKT | 179.744847 | 0.07635979 | 0.21849414 | 0.34948208 | 0.72672742 | 0.95979351 |
| RIC1 | 585.711035 | -0.3399357 | 0.16964044 | -2.0038599 | 4.51E-02 | 0.48262669 |
| ERMP1 | 443.261682 | 0.44054413 | 0.24546058 | 1.79476528 | 0.07269111 | 0.56668778 |
| KIAA2026 | 527.547992 | 0.010874 | 0.22788156 | 0.04771778 | 0.96194117 | 0.99635733 |
| RANBP6 | 56.1896995 | 0.10152986 | 0.29553065 | 0.34355105 | 0.73118394 | 0.96008876 |
| UHRF2 | 292.599053 | -0.2699169 | 0.223185 | -1.2093864 | 0.22651443 | 0.78982769 |
| GLDC | 685.820922 | 1.15591553 | 0.43318267 | 2.66842517 | 0.00762078 | 0.21455924 |
| AL162411.1 | 83.0507824 | 0.49267122 | 0.53014485 | 0.92931436 | 0.3527262 | 0.85536253 |
| KDM4C | 1053.53094 | -0.4219387 | 0.26263086 | -1.6065844 | 0.10814556 | 0.63533473 |

| | | | | | | |
|---|---|---|---|---|---|---|
| DMAC1 | 110.720742 | 0.54864623 | 0.34452779 | 1.59245857 | 0.11128171 | 0.64290644 |
| PTPRD | 435.337356 | -2.6193345 | 0.72575703 | -3.6091066 | 0.00030725 | 0.03833351 |
| MPDZ | 247.605687 | -0.1496167 | 0.21680781 | -0.6900892 | 0.4901381 | 0.90730741 |
| NFIB | 2640.38716 | -0.2761757 | 0.3425294 | -0.8062832 | 0.42007956 | 0.8828928 |
| ZDHHC21 | 957.758899 | -0.3265598 | 0.23294576 | -1.4018707 | 0.16095386 | 0.71409437 |
| FREM1 | 815.607774 | -0.1744917 | 0.31948818 | -0.5461601 | 5.85E-01 | 0.93483536 |
| TTC39B | 246.908506 | -0.288647 | 0.38045023 | -0.7586984 | 0.44803297 | 0.89297728 |
| SNAPC3 | 237.569108 | -0.2397359 | 0.24894952 | -0.9629899 | 0.33555259 | 0.84552074 |
| PSIP1 | 225.407208 | 0.1998499 | 0.20407689 | 0.97928723 | 0.32743808 | 0.84381657 |
| CCDC171 | 458.379344 | 0.03226291 | 0.28276696 | 0.11409716 | 0.90916078 | 0.98776746 |
| BNC2 | 501.968903 | -0.8075032 | 0.40063645 | -2.0155509 | 0.04384696 | 0.48009422 |
| CNTLN | 170.212866 | 0.05901964 | 0.28573802 | 0.20655157 | 0.83636009 | 0.97346221 |
| SH3GL2 | 229.673301 | -0.54363 | 0.62641348 | -0.8678453 | 0.385479 | 0.86751807 |
| RRAGA | 85.1647975 | 0.56349544 | 0.47214765 | 1.19347294 | 0.2326842 | 0.7926597 |
| HAUS6 | 63.183519 | -0.5323356 | 0.25437758 | -2.0926987 | 0.03637605 | 0.43729328 |
| PLIN2 | 224.15008 | -0.7134651 | 0.54311013 | -1.3136656 | 0.18895876 | 0.75554668 |
| DENND4C | 820.621107 | -0.4267071 | 0.29704729 | -1.4364956 | 0.15086136 | 0.7029344 |
| RPS6 | 1111.89188 | 0.07812577 | 0.51113127 | 0.15284874 | 0.87851756 | 0.98300919 |
| ACER2 | 40.3314149 | -0.5357015 | 0.47122386 | -1.1368301 | 0.25560932 | 0.80525591 |
| MLLT3 | 1485.48113 | -0.1215843 | 0.26136997 | -0.465181 | 0.64180187 | 0.94935885 |
| FOCAD | 610.326301 | 0.09035998 | 0.30139434 | 0.2998065 | 0.76432476 | 0.96285313 |
| KLHL9 | 97.1127898 | 0.09174362 | 0.2682115 | 0.342057 | 0.73230799 | 0.96008876 |
| LINC01239 | 76.7648083 | 0.81443224 | 0.52863381 | 1.54063593 | 0.12340542 | 0.66519057 |
| AL391117.1 | 70.8703503 | 0.71983643 | 0.53405711 | 1.34786415 | 0.17770208 | 0.73696787 |
| TUSC1 | 37.6888037 | 0.0981563 | 0.29481194 | 0.33294548 | 0.73917544 | 0.96078945 |
| CAAP1 | 418.264272 | -0.0353349 | 0.22583753 | -0.1564616 | 0.87566921 | 0.98268059 |
| PLAA | 139.227175 | -0.0488489 | 0.24459128 | -0.1997163 | 0.8417025 | 0.97545529 |
| IFT74 | 379.602028 | 0.34577212 | 0.23185854 | 1.49130641 | 0.13588107 | 0.67998335 |
| LRRC19 | 87.5442245 | 1.26072794 | 0.49306364 | 2.55692742 | 0.01056012 | 0.25038862 |
| MOB3B | 340.029187 | 0.84990639 | 0.30172984 | 2.81677935 | 0.00485078 | 0.17248125 |
| C9orf72 | 183.063783 | -0.2174289 | 0.63977221 | -0.3398535 | 0.73396682 | 0.96041607 |
| LINGO2 | 112.305023 | 2.52710759 | 0.80360156 | 3.14472709 | 0.00166242 | 0.10301616 |
| ACO1 | 157.261589 | 0.16689626 | 0.20461318 | 0.81566719 | 0.41469051 | 0.88242985 |
| DDX58 | 184.369669 | -1.8036352 | 0.77903178 | -2.3152268 | 0.02060052 | 0.34432775 |
| TOPORS | 78.0953178 | 0.31604278 | 0.29313595 | 1.07814408 | 0.28096946 | 0.82628479 |
| NDUFB6 | 135.886721 | 0.01424519 | 0.28668498 | 0.04968935 | 0.96036994 | 0.99635733 |
| APTX | 229.991527 | 0.09294226 | 0.25626479 | 0.36268057 | 0.71684351 | 0.95791799 |
| DNAJA1 | 337.838872 | 0.77823068 | 0.39792076 | 1.95574282 | 0.05049546 | 0.49596288 |
| SMU1 | 136.771498 | 0.01344968 | 0.19474897 | 0.06906162 | 0.94494057 | 0.99276891 |
| B4GALT1 | 943.179809 | -0.5799659 | 0.31831913 | -1.8219637 | 0.0684605 | 0.55917298 |
| BAG1 | 483.247498 | -0.4217385 | 0.62085574 | -0.6792857 | 0.49695683 | 0.90839719 |
| CHMP5 | 184.217517 | -0.1510695 | 0.43427414 | -0.3478667 | 0.72794029 | 0.9598317 |
| NFX1 | 490.671689 | -0.5494243 | 0.25244701 | -2.1763947 | 0.02952576 | 0.40623517 |
| AQP7 | 123.283988 | -1.0404583 | 0.54188226 | -1.9200819 | 0.05484755 | 0.51598058 |
| AQP3 | 3438.93743 | -0.8999158 | 0.68023668 | -1.3229451 | 0.18585365 | 0.74914489 |
| PRSS3 | 52.7886919 | -0.0791188 | 0.74557789 | -0.1061175 | 0.91548914 | 0.98868699 |
| UBE2R2 | 543.532496 | -0.0370989 | 0.29167801 | -0.1271913 | 0.89878902 | 0.98660836 |
| UBAP2 | 418.065773 | -0.4790075 | 0.25704923 | -1.8634854 | 0.06239401 | 0.5462456 |
| DCAF12 | 92.3620103 | 0.08131108 | 0.28856478 | 0.28177757 | 0.77811407 | 0.96472201 |
| UBAP1 | 462.290494 | -0.4443945 | 0.29510583 | -1.5058818 | 0.13209751 | 0.67466347 |
| KIF24 | 37.1083414 | 0.39754186 | 0.38933176 | 1.02108769 | 0.3072129 | 0.83865359 |
| NUDT2 | 38.7312362 | 0.19463255 | 0.36134418 | 0.53863481 | 0.59013887 | 0.93483536 |
| FAM219A | 94.1518795 | -0.7832391 | 0.38316961 | -2.0441055 | 0.04094313 | 0.46430138 |
| DNAI1 | 74.6499215 | 1.256024 | 0.37469949 | 3.35208355 | 0.00080206 | 0.06566851 |

| | | | | | | |
|---|---|---|---|---|---|---|
| RPP25L | 43.197209 | 0.28848407 | 0.36995674 | 0.77977785 | 0.43552165 | 0.88785137 |
| DCTN3 | 96.1120108 | 0.72885491 | 0.55743555 | 1.30751424 | 0.19103812 | 0.75807631 |
| VCP | 287.459247 | 0.04904991 | 0.25798859 | 0.19012434 | 0.8492117 | 0.97659811 |
| PIGO | 43.6290758 | 0.57199284 | 0.28961478 | 1.97501261 | 0.04826672 | 0.48661187 |
| STOML2 | 121.688598 | 0.43170421 | 0.4152827 | 1.03954297 | 0.29855228 | 0.83556453 |
| FAM214B | 37.4123954 | -0.2027616 | 0.28484122 | -0.7118408 | 0.47656336 | 0.90242252 |
| UNC13B | 530.783875 | -0.2486182 | 0.3028487 | -0.8209321 | 0.41168493 | 0.88086809 |
| RUSC2 | 220.365866 | 0.71502664 | 0.38425541 | 1.86081084 | 0.06277089 | 0.54720847 |
| TESK1 | 41.1045554 | 0.00670469 | 0.31019473 | 0.02161446 | 0.98275549 | 0.9971769 |
| CD72 | 38.9062632 | -0.0423101 | 0.39852223 | -0.1061674 | 0.91544952 | 0.98868699 |
| CCDC107 | 128.961253 | -0.7946811 | 0.61761695 | -1.2866893 | 0.19820259 | 0.76648087 |
| TLN1 | 183.130332 | -0.0745789 | 0.21081379 | -0.3537668 | 0.72351367 | 0.95963742 |
| CREB3 | 64.7540139 | 0.41169887 | 0.36085166 | 1.14090888 | 0.25390784 | 0.80414079 |
| GBA2 | 144.357707 | -0.3558987 | 0.27378068 | -1.2999408 | 0.19362125 | 0.76043322 |
| RGP1 | 83.1537211 | 0.2624804 | 0.22586768 | 1.16209808 | 0.24519563 | 0.80072691 |
| AL133410.1 | 91.8726444 | 0.23825035 | 0.30388507 | 0.78401468 | 0.4330315 | 0.88722778 |
| HINT2 | 54.7480906 | -0.0530743 | 0.43048861 | -0.1232886 | 0.90187858 | 0.9875066 |
| TMEM8B | 87.2207323 | 0.25837483 | 0.40427346 | 0.63910905 | 0.52275199 | 0.91845051 |
| RECK | 70.1815836 | 0.17443839 | 0.33675998 | 0.51799028 | 0.60446505 | 0.93820997 |
| CLTA | 429.313156 | -0.157691 | 0.32760204 | -0.4813494 | 0.63026819 | 0.94596555 |
| GNE | 70.0669373 | 0.11381956 | 0.37770883 | 0.30134207 | 0.76315366 | 0.96285313 |
| RNF38 | 625.664552 | -0.2397341 | 0.20714349 | -1.1573333 | 0.24713621 | 0.80072691 |
| EBLN3P | 166.545739 | 0.45481105 | 0.26229862 | 1.73394373 | 0.08292807 | 0.59202053 |
| ZCCHC7 | 1225.35562 | -0.3284982 | 0.2282412 | -1.4392591 | 0.15007713 | 0.7029344 |
| GRHPR | 108.658534 | 0.57522011 | 0.2904915 | 1.98016161 | 0.04768537 | 0.48600281 |
| ZBTB5 | 58.4612916 | -0.1234662 | 0.32088982 | -0.3847621 | 0.70041371 | 0.95672887 |
| AL138752.2 | 186.123814 | -0.1266228 | 0.26693113 | -0.4743651 | 0.63523957 | 0.94792975 |
| TRMT10B | 128.754352 | 0.05907917 | 0.31615013 | 0.18687062 | 0.85176207 | 0.97718353 |
| DCAF10 | 395.530155 | -0.0577478 | 0.21257536 | -0.2716581 | 0.78588491 | 0.96513523 |
| FAM95C | 55.5033409 | -0.5134463 | 0.44625113 | -1.150577 | 0.24990628 | 0.80149394 |
| ANKRD18A | 61.9602621 | -0.2263014 | 0.44625243 | -0.5071152 | 0.61207401 | 0.94003018 |
| CNTNAP3 | 41.7413073 | 0.0429088 | 0.33869734 | 0.12668775 | 0.89918754 | 0.98660836 |
| GLIDR | 138.481892 | 0.20756402 | 0.43080719 | 0.48180258 | 0.6299462 | 0.94596555 |
| BMS1P14 | 83.1867503 | -0.5865975 | 0.4700607 | -1.2479187 | 0.21206083 | 0.77709975 |
| CBWD6 | 53.9893237 | -0.1993153 | 0.42345425 | -0.4706892 | 0.63786272 | 0.94854485 |
| CNTNAP3B | 838.282311 | 0.55596319 | 0.38117255 | 1.4585604 | 0.14468614 | 0.69206332 |
| FAM27C | 33.9630263 | -0.0191739 | 0.51558142 | -0.0371889 | 0.97033442 | 0.99656026 |
| AL512625.3 | 46.1562595 | 0.4385705 | 0.46892934 | 0.93525924 | 0.34965471 | 0.8535343 |
| CBWD5 | 164.162982 | -0.6511129 | 0.2747303 | -2.3700077 | 0.01778772 | 0.32030113 |
| ZNF658 | 48.7968853 | 0.7000003 | 0.39449241 | 1.77443287 | 0.07599158 | 0.57145422 |
| CBWD3 | 47.6018508 | 0.00225452 | 0.34988025 | 0.00644368 | 0.99485872 | 0.99919666 |
| PGM5 | 393.837569 | 0.58380194 | 0.25515078 | 2.2880664 | 0.02213365 | 0.35843654 |
| AL161457.2 | 49.4733796 | 0.61969119 | 0.44538172 | 1.39137096 | 0.16411297 | 0.71715698 |
| PIP5K1B | 89.7339503 | -0.7284888 | 0.5804029 | -1.2551433 | 0.20942676 | 0.77445039 |
| FAM189A2 | 39.6776642 | 0.03086615 | 0.54780436 | 0.0563452 | 0.95506681 | 0.99537617 |
| APBA1 | 119.000617 | -0.0310806 | 0.37141936 | -0.0836806 | 0.93331035 | 0.99243654 |
| PTAR1 | 293.007172 | -0.0040143 | 0.17546303 | -0.0228782 | 0.98174746 | 0.99696835 |
| MAMDC2-AS | 119.913945 | -0.0831934 | 0.33717912 | -0.2467335 | 0.80511451 | 0.96704419 |
| MAMDC2 | 529.437754 | 0.3184765 | 0.48973299 | 0.65030639 | 0.51549433 | 0.91612355 |
| SMC5-AS1 | 190.818402 | -0.091388 | 0.24557482 | -0.3721393 | 0.70978914 | 0.95752169 |
| SMC5 | 547.851997 | -0.2052196 | 0.23018185 | -0.8915541 | 0.37263198 | 0.86201082 |
| KLF9 | 136.91644 | 0.38605309 | 0.34467114 | 1.12006215 | 0.26268728 | 0.8096287 |
| TRPM3 | 3768.17106 | 1.59085657 | 0.3501841 | 4.5429149 | 5.55E-06 | 0.00257078 |
| CEMIP2 | 495.180835 | 0.43125364 | 0.29137534 | 1.48006226 | 0.13885663 | 0.68156449 |

| | | | | | | |
|---|---|---|---|---|---|---|
| ABHD17B | 64.5933552 | 0.47332946 | 0.3012025 | 1.57146593 | 0.11607446 | 0.65192493 |
| C9orf85 | 149.522788 | 0.14497673 | 0.26788532 | 0.54118951 | 0.58837697 | 0.93483536 |
| GDA | 60.5925684 | 0.74653564 | 0.54215469 | 1.37697904 | 0.16851873 | 0.72499026 |
| ZFAND5 | 725.795732 | 0.40416144 | 0.53493562 | 0.75553286 | 0.44992932 | 0.89324553 |
| ALDH1A1 | 349.618784 | 1.16104363 | 0.4188221 | 2.77216419 | 0.00556849 | 0.18409406 |
| CARNMT1 | 230.753325 | 0.11999861 | 0.20639118 | 0.58141347 | 0.56096182 | 0.92934517 |
| NMRK1 | 159.891472 | 0.14903909 | 0.34213911 | 0.43560964 | 0.66311998 | 0.95159487 |
| OSTF1 | 117.189609 | -0.0536142 | 0.44220959 | -0.1212416 | 0.90349969 | 0.9875066 |
| PCSK5 | 186.755606 | -0.4793818 | 0.70946468 | -0.6756951 | 0.49923427 | 0.90839719 |
| RFK | 76.6409356 | 0.66754068 | 0.33675574 | 1.98226962 | 0.04744907 | 0.48600281 |
| VPS13A | 523.022719 | -0.1864061 | 0.19752693 | -0.9436997 | 0.34532312 | 0.8523814 |
| GNAQ | 1856.84582 | 0.00726794 | 0.198914 | 0.0365381 | 0.9708533 | 0.99656026 |
| CEP78 | 60.4639739 | -0.2549258 | 0.35735793 | -0.7133627 | 0.47562134 | 0.90242252 |
| TLE4 | 67.5851438 | 0.1468463 | 0.40532713 | 0.36229083 | 0.71713471 | 0.95791799 |
| TLE1 | 85.8890885 | 0.35910128 | 0.63092515 | 0.56916621 | 0.56924335 | 0.93122667 |
| AL591368.1 | 32.9057827 | 0.3084687 | 0.81655103 | 0.37777027 | 0.70560126 | 0.95730303 |
| RASEF | 206.122069 | -0.2136119 | 0.2393445 | -0.8924873 | 0.37213179 | 0.86201082 |
| FRMD3 | 206.144612 | 0.14154374 | 0.27949479 | 0.50642709 | 6.13E-01 | 0.94003018 |
| IDNK | 51.7112614 | 0.16873952 | 0.3852628 | 0.43798549 | 0.6613968 | 0.9513366 |
| UBQLN1 | 294.260365 | -0.0302179 | 0.15838685 | -0.1907853 | 0.84869384 | 0.97656375 |
| GKAP1 | 139.509995 | -0.0433957 | 0.22680671 | -0.1913334 | 0.84826441 | 0.97656375 |
| KIF27 | 198.504379 | -0.0062091 | 0.26518435 | -0.0234144 | 0.98131969 | 0.99696835 |
| HNRNPK | 764.472642 | -0.1695956 | 0.33945068 | -0.4996179 | 0.61734416 | 0.94233994 |
| RMI1 | 66.4516598 | 0.49237524 | 0.26591467 | 1.85162872 | 0.06407916 | 0.54860648 |
| AGTPBP1 | 249.338356 | -0.0967304 | 0.32974248 | -0.2933513 | 0.76925363 | 0.96345263 |
| NAA35 | 314.811589 | -0.2196948 | 0.23835463 | -0.9217142 | 0.35667769 | 0.8557799 |
| GOLM1 | 77.6429028 | 0.61216734 | 0.30644126 | 1.9976662 | 0.04575286 | 0.48339878 |
| ISCA1 | 113.196711 | 0.02819741 | 0.36055661 | 0.07820523 | 0.9376648 | 0.99243654 |
| TUT7 | 275.491221 | -0.2067826 | 0.18731282 | -1.1039425 | 0.26961809 | 0.81817548 |
| DAPK1 | 1988.92862 | -0.7043261 | 0.56960066 | -1.2365262 | 0.21626305 | 0.78072228 |
| CTSL | 207.00918 | 0.02887535 | 0.39889584 | 0.07238819 | 0.94229298 | 0.99265578 |
| SPIN1 | 813.193115 | 0.23231729 | 0.2133854 | 1.08872161 | 0.27627667 | 0.82167981 |
| SECISBP2 | 379.935284 | -0.3333966 | 0.21210264 | -1.5718646 | 0.11598196 | 0.65192493 |
| SEMA4D | 506.062329 | -0.1894078 | 0.30382097 | -0.623419 | 0.5330092 | 0.92268523 |
| LINC01508 | 213.256008 | 1.28827669 | 0.51270895 | 2.51268619 | 0.01198159 | 0.26883805 |
| DIRAS2 | 37.5058596 | -0.1853724 | 0.37489472 | -0.4944653 | 0.62097761 | 0.94371308 |
| SYK | 667.98879 | 0.10236698 | 0.27930806 | 0.36650207 | 0.71399047 | 0.95791799 |
| AL158071.4 | 64.706502 | -0.4308299 | 0.37079045 | -1.1619228 | 0.24526681 | 0.80072691 |
| AUH | 716.387899 | -0.4525808 | 0.20485311 | -2.2092943 | 0.02715417 | 0.40076654 |
| NFIL3 | 126.765468 | -1.7198174 | 0.98975132 | -1.7376258 | 0.08227678 | 0.59202053 |
| ROR2 | 756.841973 | 0.00682134 | 0.2445546 | 0.02789291 | 0.97774756 | 0.99696835 |
| SPTLC1 | 245.321414 | -0.0480831 | 0.17346415 | -0.2771931 | 0.78163185 | 0.96513523 |
| IARS | 198.857188 | -0.3132347 | 0.19683281 | -1.5913744 | 0.11152533 | 0.64353508 |
| NOL8 | 74.9318206 | -0.1305301 | 0.24455062 | -0.5337549 | 0.59351113 | 0.93491607 |
| CENPP | 155.726452 | -1.0231872 | 0.33429935 | -3.0606914 | 0.00220827 | 0.12116555 |
| IPPK | 47.5556783 | -0.4147156 | 0.33249206 | -1.2472947 | 0.21228946 | 0.77709975 |
| BICD2 | 71.6877234 | 0.24595431 | 0.25318554 | 0.97143901 | 0.33132971 | 0.84442294 |
| ZNF484 | 38.9844942 | -0.2080355 | 0.26673482 | -0.7799339 | 0.43542976 | 0.88785137 |
| FGD3 | 228.589675 | 1.1194455 | 0.51575099 | 2.17051548 | 0.02996782 | 0.40893584 |
| CARD19 | 65.3908699 | -0.5965755 | 0.34326886 | -1.7379248 | 0.08222406 | 0.59202053 |
| NINJ1 | 77.4116302 | 0.42483676 | 0.23258692 | 1.82657201 | 0.06776414 | 0.55864172 |
| FAM120AOS | 68.4404452 | 0.16490173 | 0.29783336 | 0.55367113 | 0.57980394 | 0.93345492 |
| FAM120A | 525.164763 | 0.1165987 | 0.18921418 | 0.616226 | 0.53774537 | 0.92324352 |
| PHF2 | 117.982825 | 0.10989367 | 0.28874393 | 0.38059215 | 0.70350591 | 0.95672887 |

| | | | | | | |
|---|---|---|---|---|---|---|
| PTPDC1 | 104.342451 | -0.3739635 | 0.23548399 | -1.5880635 | 0.11227199 | 0.64471804 |
| AL158152.2 | 136.771076 | 0.23774085 | 0.30349866 | 0.78333412 | 0.43343093 | 0.88736205 |
| ZNF169 | 66.2115564 | -0.1304077 | 0.36286982 | -0.3593788 | 0.71931172 | 0.95796325 |
| MFSD14B | 183.09002 | -0.1521967 | 0.22107426 | -0.6884414 | 0.49117487 | 0.90730741 |
| FBP1 | 164.82255 | 0.69714757 | 0.45280026 | 1.53963598 | 0.12364911 | 0.66519057 |
| C9orf3 | 1149.76024 | -0.0766088 | 0.29013189 | -0.2640481 | 0.79174286 | 0.9653598 |
| FANCC | 447.407728 | -0.3746006 | 0.24013832 | -1.5599367 | 0.11877485 | 0.65456964 |
| PTCH1 | 69.064994 | 0.308427 | 0.32971576 | 0.93543299 | 3.50E-01 | 0.8535343 |
| LINC00476 | 169.525054 | -0.1816074 | 0.40864523 | -0.4444133 | 0.6567438 | 0.95100227 |
| ERCC6L2 | 291.948299 | -0.1112422 | 0.24139964 | -0.4608217 | 0.6449265 | 0.94949493 |
| SLC35D2 | 47.4258618 | 0.27895857 | 0.35330897 | 0.78955982 | 0.42978488 | 0.88579433 |
| HABP4 | 137.8256 | 0.23660191 | 0.31540096 | 0.75016231 | 0.45315696 | 0.89544675 |
| CDC14B | 387.22337 | 0.46519537 | 0.19696762 | 2.36178598 | 0.01818714 | 0.32470394 |
| AL589843.1 | 60.3386466 | 0.6664836 | 0.30891954 | 2.15746663 | 0.03096932 | 0.41259458 |
| ZNF510 | 37.8637447 | 0.40292503 | 0.31485691 | 1.27970841 | 0.20064771 | 0.7694065 |
| ZNF782 | 137.989509 | -0.2885879 | 0.24244653 | -1.1903157 | 0.23392235 | 0.79439604 |
| MFSD14C | 386.775648 | -0.0873593 | 0.19042445 | -0.4587607 | 0.64640603 | 0.95048567 |
| CCDC180 | 38.1380002 | 0.42568046 | 0.46480865 | 0.91581872 | 0.35976199 | 0.85608464 |
| TDRD7 | 78.6169372 | -0.3197662 | 0.45995985 | -0.6952047 | 0.48692703 | 0.9070379 |
| TSTD2 | 110.625745 | -0.044861 | 0.25481239 | -0.1760552 | 0.86025059 | 0.97841862 |
| NCBP1 | 99.8293825 | -0.2099077 | 0.24255886 | -0.8653888 | 0.38682542 | 0.86852426 |
| XPA | 96.61703 | 0.52221904 | 0.23048102 | 2.26577895 | 0.02346492 | 0.36567438 |
| ANP32B | 196.430629 | 0.32931893 | 0.21063025 | 1.56349301 | 0.11793668 | 0.65447949 |
| NANS | 38.0490828 | 0.15187801 | 0.38943495 | 0.38999584 | 0.69653963 | 0.95672887 |
| TRIM14 | 56.5989667 | -0.921522 | 0.8015691 | -1.1496476 | 0.25028904 | 0.80165927 |
| ANKS6 | 114.846497 | 0.51326989 | 0.36813561 | 1.39424136 | 0.16324473 | 0.71642393 |
| GALNT12 | 44.8234315 | 0.04231558 | 0.53161903 | 0.07959756 | 0.93655734 | 0.99243654 |
| TGFBR1 | 696.54011 | -1.4580212 | 0.55906268 | -2.6079745 | 0.00910798 | 0.23300339 |
| ALG2 | 74.4600291 | 0.20795843 | 0.2996384 | 0.69403129 | 0.48766259 | 0.90730741 |
| SEC61B | 260.331387 | 0.12007633 | 0.39252827 | 0.30590491 | 0.75967705 | 0.96190674 |
| AL359710.1 | 220.666289 | 0.72284702 | 0.33728859 | 2.14311141 | 0.03210415 | 0.41517597 |
| NR4A3 | 174.524968 | -2.4749155 | 1.75002529 | -1.414217 | 0.1572982 | 0.71160548 |
| STX17 | 192.383759 | 0.24709362 | 0.27027801 | 0.91422022 | 0.36060114 | 0.85608464 |
| ERP44 | 267.180581 | 0.03951864 | 0.22651669 | 0.17446239 | 0.86150209 | 0.97848095 |
| INVS | 566.834633 | 0.19924282 | 0.16078492 | 1.23918844 | 0.21527572 | 0.77894575 |
| TEX10 | 97.3811374 | -0.1089691 | 0.28222576 | -0.3861061 | 0.69941811 | 0.95672887 |
| MRPL50 | 68.1879055 | 0.36426855 | 0.43340095 | 0.84048859 | 0.4006345 | 0.87775198 |
| ZNF189 | 58.7956595 | 0.37137236 | 0.3085316 | 1.20367689 | 0.22871449 | 0.79047378 |
| ALDOB | 110.236773 | 0.14478965 | 0.9644341 | 0.15012913 | 0.88066274 | 0.98333507 |
| TMEM246 | 43.8540377 | 0.38346058 | 0.28972863 | 1.32351635 | 0.18566372 | 0.74914489 |
| RNF20 | 81.6217629 | 0.38879624 | 0.33888376 | 1.14728494 | 0.25126388 | 0.80240987 |
| SMC2 | 59.0945513 | 0.36158668 | 0.31742188 | 1.13913597 | 0.25464645 | 0.80414079 |
| NIPSNAP3A | 83.5854802 | -0.1730355 | 0.29093302 | -0.5947606 | 0.55200348 | 0.92872652 |
| ABCA1 | 157.636979 | 0.62326149 | 0.42436551 | 1.46869024 | 0.14191683 | 0.68792245 |
| SLC44A1 | 1490.3124 | -1.6852984 | 0.49011706 | -3.438563 | 0.00058481 | 0.05789336 |
| FSD1L | 256.548746 | -0.3928399 | 0.17365073 | -2.2622417 | 0.02368248 | 0.36714846 |
| FKTN | 178.300946 | 0.1130967 | 0.22126393 | 0.51113937 | 0.60925347 | 0.94003018 |
| TMEM38B | 71.9226399 | -0.3596925 | 0.39352987 | -0.9140158 | 0.36070855 | 0.85608464 |
| LINC01505 | 70.9875608 | -0.4596086 | 0.54706074 | -0.8401418 | 0.40082891 | 0.87775198 |
| ZNF462 | 94.5206284 | 1.08089028 | 0.40719687 | 2.65446608 | 0.0079434 | 0.21849569 |
| RAD23B | 429.106295 | -0.0814897 | 0.18771232 | -0.4341201 | 0.66420126 | 0.95195406 |
| ELP1 | 142.165226 | -0.199067 | 0.19320447 | -1.0303439 | 0.30284862 | 0.83709431 |
| FAM206A | 54.4523583 | 0.19242089 | 0.30773497 | 0.6252812 | 0.53178652 | 0.92211604 |
| CTNNAL1 | 219.465484 | 0.87974711 | 0.18147425 | 4.84777939 | 1.25E-06 | 0.00087229 |

| | | | | | | |
|---|---|---|---|---|---|---|
| TMEM245 | 1029.55571 | -0.3881044 | 0.29478225 | -1.31658 | 0.18797945 | 0.7526909 |
| EPB41L4B | 209.42693 | 0.40315356 | 0.29355528 | 1.37334802 | 0.16964418 | 0.72632018 |
| PTPN3 | 966.674468 | 0.05267558 | 0.29021601 | 0.18150472 | 0.85597143 | 0.97718353 |
| PALM2 | 148.867284 | 0.26697769 | 0.44507074 | 0.5998545 | 0.54860321 | 0.92716685 |
| PALM2-AKAF | 40.4183532 | 0.24478709 | 0.50311615 | 0.48654189 | 0.62658301 | 0.94511671 |
| TXN | 243.741466 | -0.223228 | 0.33671098 | -0.6629662 | 0.50735221 | 0.91185923 |
| ECPAS | 449.964841 | -0.3862019 | 0.20002887 | -1.9307309 | 0.05351634 | 0.51313014 |
| ZNF483 | 255.4643 | -0.0136113 | 0.28178102 | -0.0483045 | 0.96147359 | 0.99635733 |
| PTGR1 | 969.51936 | 0.58740672 | 0.49232022 | 1.19313953 | 0.23281473 | 0.7926597 |
| C9orf84 | 98.4766797 | 0.488996 | 0.56593439 | 0.8640507 | 0.38756004 | 0.86910078 |
| UGCG | 288.441768 | -0.405167 | 0.33123408 | -1.2232044 | 0.22125249 | 0.78752408 |
| SUSD1 | 632.119147 | 0.4290633 | 0.35730327 | 1.20083789 | 0.22981409 | 0.79047378 |
| PTBP3 | 2262.31299 | 0.10718263 | 0.3667877 | 0.2922198 | 0.77011857 | 0.96345263 |
| HSDL2 | 366.705772 | 0.07986161 | 0.28616204 | 0.27907827 | 0.78018476 | 0.96513523 |
| C9orf147 | 50.5941267 | 0.32880577 | 0.28025657 | 1.17323127 | 0.24070308 | 0.80057149 |
| KIAA1958 | 598.206929 | 0.00158149 | 0.2082368 | 0.00759467 | 0.99394039 | 0.99919666 |
| INIP | 105.546822 | -0.1330902 | 0.22148267 | -0.6009059 | 0.54790268 | 0.92682201 |
| SNX30 | 76.6074805 | 0.01398737 | 0.26034485 | 0.05372633 | 0.9571532 | 0.99579587 |
| FKBP15 | 105.991323 | 0.11503653 | 0.18670476 | 0.61614139 | 0.53780121 | 0.92324352 |
| SLC31A1 | 86.9876456 | 0.11503251 | 0.24305611 | 0.47327552 | 0.63601663 | 0.94821838 |
| CDC26 | 79.9776011 | 0.10158592 | 0.44459553 | 0.22849065 | 0.81926482 | 0.96884581 |
| PRPF4 | 69.308401 | -0.5111035 | 0.43503562 | -1.1748543 | 0.240053 | 0.80056465 |
| WDR31 | 57.2054063 | 0.39891254 | 0.41980431 | 0.95023451 | 0.34199311 | 0.85092302 |
| BSPRY | 101.274874 | -0.1606064 | 0.28077691 | -0.5720073 | 0.56731705 | 0.93122667 |
| HDHD3 | 141.511903 | -0.3218612 | 0.30294046 | -1.062457 | 0.28802827 | 0.8303903 |
| ALAD | 69.7757853 | -0.2569215 | 0.21985868 | -1.1685757 | 0.24257461 | 0.80072691 |
| RGS3 | 138.563203 | 0.04489078 | 0.51212823 | 0.08765535 | 0.93015061 | 0.99216065 |
| ZNF618 | 802.801796 | 0.90349527 | 0.2409605 | 3.7495576 | 0.00017715 | 0.02856152 |
| KIF12 | 320.137869 | 0.2894079 | 0.29238006 | 0.98983457 | 0.32225498 | 0.84285425 |
| COL27A1 | 130.064239 | -0.3969333 | 0.60250429 | -0.6588058 | 0.51002046 | 0.91334187 |
| AKNA | 43.2427387 | -0.3182196 | 0.33364475 | -0.9537676 | 0.3402013 | 0.84975445 |
| WHRN | 67.3991064 | -0.2391752 | 0.27434787 | -0.8717952 | 0.38332011 | 0.86638702 |
| ATP6V1G1 | 407.002157 | 0.28693362 | 0.47875471 | 0.59933325 | 0.54895068 | 0.9274143 |
| TMEM268 | 51.0320544 | -0.0945855 | 0.41460906 | -0.2281319 | 0.81954371 | 0.96884581 |
| PAPPA | 3461.5327 | -2.419028 | 0.75743162 | -3.1937246 | 0.0014045 | 0.09557897 |
| AL137024.1 | 126.354687 | -2.5093759 | 0.85214227 | -2.9447852 | 0.00323179 | 0.14358074 |
| ASTN2 | 377.892156 | -0.8779366 | 0.66673375 | -1.3167725 | 0.18791491 | 0.7526909 |
| CDK5RAP2 | 244.105938 | -0.3437347 | 0.20378877 | -1.6867206 | 0.09165709 | 0.61104726 |
| MEGF9 | 406.583681 | 0.33507524 | 0.36467505 | 0.91883238 | 0.35818326 | 0.8557799 |
| FBXW2 | 171.462249 | -0.2696607 | 0.18070252 | -1.4922906 | 0.13562299 | 0.67994379 |
| PSMD5 | 148.069046 | -0.0310296 | 0.23416969 | -0.1325091 | 0.89458162 | 0.98597539 |
| TRAF1 | 35.1376588 | 0.71090815 | 0.44050042 | 1.61386488 | 0.10655674 | 0.63461115 |
| CNTRL | 133.601592 | 0.13534361 | 0.30569146 | 0.44274579 | 0.65794962 | 0.95107752 |
| RAB14 | 393.72945 | 0.23059488 | 0.19839455 | 1.16230448 | 0.24511181 | 0.80072691 |
| GSN | 294.654232 | -0.1422054 | 0.19815007 | -0.7176653 | 4.73E-01 | 0.90213021 |
| STOM | 96.4993973 | 0.09131805 | 0.36325014 | 0.25139165 | 0.80151132 | 0.9661728 |
| DAB2IP | 298.016442 | 0.13649121 | 0.28004822 | 0.48738467 | 0.62598576 | 0.94511671 |
| TTLL11 | 67.7031163 | 0.16420511 | 0.3142578 | 0.5225172 | 0.60131027 | 0.93802476 |
| NDUFA8 | 135.69595 | 0.2710876 | 0.46981033 | 0.577015 | 0.56392932 | 0.93077674 |
| RBM18 | 57.4566297 | 0.09409585 | 0.29498939 | 0.31898045 | 0.74974134 | 0.96109466 |
| MRRF | 225.31215 | -0.1101121 | 0.23508328 | -0.4683963 | 0.6395012 | 0.94935885 |
| PTGS1 | 119.319234 | 0.0336257 | 0.83269372 | 0.04038183 | 0.96778871 | 0.9964399 |
| PDCL | 76.4030943 | 0.32642061 | 0.23816125 | 1.37058654 | 0.17050388 | 0.72686344 |
| RC3H2 | 259.691335 | 0.00764973 | 0.20236368 | 0.03780189 | 0.96984564 | 0.99646885 |

| | | | | | | |
|---|---|---|---|---|---|---|
| ZBTB26 | 32.825053 | 0.33664781 | 0.35734872 | 0.94207083 | 0.34615638 | 0.8523814 |
| RABGAP1 | 657.153707 | -0.1144312 | 0.18036545 | -0.6344408 | 0.52579322 | 0.91956488 |
| STRBP | 588.407444 | -0.0810251 | 0.29708105 | -0.2727374 | 0.78505509 | 0.96513523 |
| DENND1A | 928.0985 | 0.37351855 | 0.28001354 | 1.33393032 | 0.18222669 | 0.74315969 |
| PSMB7 | 209.88261 | -0.2500111 | 0.26508978 | -0.9431188 | 0.34562016 | 0.8523814 |
| NR6A1 | 804.269667 | -0.0018101 | 0.20751034 | -0.0087228 | 0.9930403 | 0.99919666 |
| MIR181A2HC | 118.512159 | 0.28898327 | 0.31069255 | 0.93012617 | 0.35230576 | 0.85536253 |
| RPL35 | 421.82524 | -0.2072839 | 0.37336175 | -0.5551824 | 0.57876987 | 0.93345492 |
| ARPC5L | 146.660511 | 0.08225035 | 0.44664371 | 0.18415203 | 0.85389419 | 0.97718353 |
| GOLGA1 | 136.540138 | -0.5589763 | 0.18904739 | -2.9568051 | 0.00310845 | 0.14287942 |
| SCAI | 256.980204 | -0.2574065 | 0.30674035 | -0.8391673 | 0.40137546 | 0.87816594 |
| PPP6C | 195.691573 | 0.08621395 | 0.15775167 | 0.54651686 | 0.5847107 | 0.93483536 |
| RABEPK | 62.3872166 | -0.2763977 | 0.29917956 | -0.9238521 | 0.35556331 | 0.8557799 |
| HSPA5 | 436.139176 | 0.20790682 | 0.45928831 | 0.45267171 | 0.65078515 | 0.95061388 |
| GAPVD1 | 320.398041 | -0.2088897 | 0.21521628 | -0.9706034 | 0.33174581 | 0.84442294 |
| MAPKAP1 | 474.502485 | -0.1139635 | 0.21503315 | -0.529981 | 0.59612507 | 0.93593903 |
| PBX3 | 370.303098 | 0.72810379 | 0.37651167 | 1.93381467 | 0.05313592 | 0.51041652 |
| MVB12B | 106.283246 | 0.37702061 | 0.32537581 | 1.15872354 | 0.24656889 | 0.80072691 |
| ZBTB43 | 130.589586 | -0.1046742 | 0.24183476 | -0.4328335 | 0.66513577 | 0.9520866 |
| ZBTB34 | 33.2715789 | -0.2054059 | 0.30681028 | -0.6694883 | 0.50318407 | 0.90964098 |
| RALGPS1 | 538.467529 | 0.01394529 | 0.257861 | 0.05408064 | 0.95687092 | 0.99565888 |
| GARNL3 | 89.2155692 | 0.18212994 | 0.31688376 | 0.57475314 | 0.56545826 | 0.93114846 |
| SLC2A8 | 31.7758476 | -0.2100573 | 0.28770872 | -0.7301039 | 0.46532669 | 0.89883965 |
| RPL12 | 157.252194 | -0.1909882 | 0.34412221 | -0.5550011 | 0.57889392 | 0.93345492 |
| LRSAM1 | 187.573384 | -0.1989056 | 0.23462535 | -0.8477583 | 0.39657259 | 0.87462201 |
| FAM129B | 304.451781 | -0.8262086 | 0.37631084 | -2.1955482 | 0.02812429 | 0.40459506 |
| STXBP1 | 117.231754 | 0.00174361 | 0.19993639 | 0.00872084 | 0.99304187 | 0.99919666 |
| PTRH1 | 68.1929266 | 0.30174645 | 0.40185801 | 0.7508783 | 0.4527259 | 0.89520141 |
| CDK9 | 81.7139297 | -0.025662 | 0.31332885 | -0.081901 | 0.93472541 | 0.99243654 |
| FPGS | 93.811504 | -0.2732985 | 0.34168091 | -0.7998646 | 0.42378924 | 0.88391318 |
| ENG | 63.4086527 | -0.1993138 | 0.47537872 | -0.4192737 | 0.67501611 | 0.95524793 |
| DPM2 | 114.867263 | -1.049005 | 0.47804429 | -2.1943679 | 0.02820897 | 0.40459506 |
| FAM102A | 695.822352 | 0.0261217 | 0.55914244 | 0.04671743 | 0.96273844 | 0.99637506 |
| SLC25A25 | 176.239672 | -0.2149814 | 0.49980432 | -0.4301312 | 0.66710024 | 0.9527406 |
| SLC25A25-AS | 70.4266547 | 0.20893989 | 0.45419485 | 0.46002258 | 0.64550001 | 0.94973439 |
| PTGES2 | 118.78739 | 0.28198875 | 0.28462111 | 0.99075135 | 0.32180701 | 0.84285425 |
| LCN2 | 97.4290056 | -0.3434283 | 0.68277714 | -0.5029875 | 6.15E-01 | 9.42E-01 |
| C9orf16 | 275.401313 | 0.21536239 | 0.4792829 | 0.44934295 | 0.65318428 | 0.95100227 |
| CIZ1 | 99.9103976 | -0.0380872 | 0.2523466 | -0.1509322 | 0.88002921 | 0.98333507 |
| DNM1 | 61.273769 | 0.87913783 | 0.44247561 | 1.98686167 | 0.04693773 | 0.48600281 |
| GOLGA2 | 188.75176 | -0.2947811 | 0.27178552 | -1.0846095 | 0.27809466 | 0.82310625 |
| SWI5 | 101.593306 | -0.0077863 | 0.3344351 | -0.0232819 | 0.9814254 | 0.99696835 |
| TRUB2 | 68.0085889 | 0.48151921 | 0.31539087 | 1.52673795 | 0.12682619 | 0.66960383 |
| COQ4 | 223.502591 | 0.14068153 | 0.18933108 | 0.74304513 | 0.45745436 | 0.89705917 |
| SLC27A4 | 124.285034 | -0.2309535 | 0.31563613 | -0.7317079 | 0.46434686 | 0.89801718 |
| URM1 | 80.4457973 | -0.482041 | 0.26198481 | -1.8399578 | 0.06577444 | 0.55444578 |
| CERCAM | 84.1095326 | 0.38588388 | 0.30018366 | 1.28549265 | 0.19862018 | 0.76668122 |
| ODF2 | 80.9131822 | -0.2460659 | 0.31623559 | -0.7781095 | 0.43650449 | 0.88785137 |
| GLE1 | 95.3305369 | -0.0787062 | 0.25134955 | -0.3131344 | 0.75417858 | 0.96120879 |
| SPTAN1 | 473.387185 | -0.2959578 | 0.21298663 | -1.3895605 | 0.16466237 | 0.71715698 |
| WDR34 | 122.308659 | 0.41336278 | 0.33358992 | 1.23913451 | 0.21529569 | 0.77894575 |
| SET | 496.532731 | 0.12896072 | 0.30418333 | 0.42395721 | 0.67159702 | 0.95444808 |
| ZDHHC12 | 70.0331796 | -0.2214571 | 0.35674437 | -0.6207726 | 0.53474922 | 0.92303275 |
| ZER1 | 256.05289 | 0.18801365 | 0.2729977 | 0.68870049 | 0.49101177 | 0.90730741 |

| | | | | | | |
|---|---|---|---|---|---|---|
| TBC1D13 | 75.0815403 | -0.2481814 | 0.26765801 | -0.9272334 | 0.35380538 | 0.8557799 |
| ENDOG | 73.1052891 | 0.22334773 | 0.39732682 | 0.56212599 | 0.57403018 | 0.93270589 |
| SPOUT1 | 34.2638525 | 0.04267452 | 0.32848328 | 0.12991383 | 0.8966346 | 0.98597539 |
| KYAT1 | 80.1327815 | 0.33395124 | 0.31790389 | 1.05047861 | 0.29349812 | 0.83336457 |
| LRRC8A | 41.9732514 | -0.0992494 | 0.2585153 | -0.3839207 | 0.70103721 | 0.95672887 |
| PHYHD1 | 60.8799109 | -0.0555017 | 0.46173464 | -0.1202025 | 0.90432271 | 0.9875066 |
| NUP188 | 58.5052835 | 0.0177929 | 0.25076593 | 0.07095423 | 0.94343418 | 0.99265578 |
| SH3GLB2 | 245.487094 | -0.1460214 | 0.30001035 | -0.4867213 | 0.62645582 | 0.94511671 |
| MIGA2 | 65.2982174 | -0.2289738 | 0.30681131 | -0.7463018 | 0.45548514 | 0.89626521 |
| PTPA | 155.255067 | 0.16869848 | 0.29027339 | 0.58117101 | 0.5611252 | 0.92934517 |
| IER5L | 271.651809 | -0.8204905 | 0.45371919 | -1.8083663 | 0.07054951 | 0.5621116 |
| LINC01503 | 767.402659 | 0.1926 | 0.50095258 | 0.38446752 | 0.70063196 | 0.95672887 |
| NTMT1 | 70.7113629 | 0.47293932 | 0.37842178 | 1.24976771 | 0.21138442 | 0.7767562 |
| ASB6 | 51.07482 | 0.05064538 | 0.23501729 | 0.21549641 | 0.82938031 | 0.97131041 |
| TOR1B | 39.2035093 | -0.0637981 | 0.31240796 | -0.2042141 | 0.83818621 | 0.97385715 |
| TOR1A | 37.0348226 | 0.03991586 | 0.28658103 | 0.13928298 | 0.88922654 | 0.98551241 |
| C9orf78 | 107.089014 | 0.5094779 | 0.38610153 | 1.31954386 | 0.18698736 | 0.75110291 |
| USP20 | 78.9140537 | -0.1022404 | 0.27726416 | -0.3687472 | 0.71231617 | 0.95791799 |
| FNBP1 | 242.972754 | 0.29712592 | 0.31161485 | 0.95350372 | 0.34033491 | 0.84975445 |
| GPR107 | 257.206334 | -0.1330374 | 0.15702947 | -0.8472127 | 0.3968766 | 0.87462201 |
| NCS1 | 101.422835 | -0.1057902 | 0.33626675 | -0.3146022 | 0.75306374 | 0.96120879 |
| ASS1 | 77.8813294 | 0.95151058 | 0.4159991 | 2.28729001 | 0.0221789 | 0.35843654 |
| FUBP3 | 236.908841 | -0.3401827 | 0.18003408 | -1.8895463 | 0.05881867 | 0.53286516 |
| ABL1 | 481.777267 | -0.1417868 | 0.24696663 | -0.5741131 | 0.56589129 | 0.93122667 |
| AIF1L | 1619.04972 | 1.25176754 | 0.36407483 | 3.43821497 | 0.00058556 | 0.05789336 |
| NUP214 | 759.673468 | 0.31440072 | 0.21260066 | 1.47883232 | 0.13918513 | 0.68225454 |
| AL157938.2 | 204.130346 | 1.08120237 | 0.65066867 | 1.66167885 | 0.09657718 | 0.61485924 |
| AL157938.3 | 55.5559174 | 0.41336062 | 0.5401369 | 0.76528861 | 0.44409971 | 0.89096469 |
| PRRC2B | 800.491517 | -0.0512507 | 0.19008467 | -0.2696205 | 0.78745219 | 0.96513523 |
| POMT1 | 65.5732988 | 0.23019457 | 0.23929795 | 0.96195798 | 0.33607071 | 0.8456233 |
| UCK1 | 39.5405263 | 0.40892498 | 0.43087586 | 0.94905521 | 0.34259253 | 0.85093239 |
| RAPGEF1 | 234.945607 | -0.1858005 | 0.29776188 | -0.6239902 | 0.53263403 | 0.92268523 |
| MED27 | 287.491845 | -0.2130956 | 0.22637682 | -0.9413313 | 0.3465351 | 0.8525698 |
| SETX | 470.854772 | -0.2761767 | 0.18128156 | -1.5234685 | 0.12764154 | 0.67051794 |
| TTF1 | 97.6422799 | -0.177075 | 0.24285417 | -0.7291413 | 0.46591525 | 0.89883965 |
| DDX31 | 90.4689006 | -0.2099023 | 0.25316301 | -0.829119 | 0.40703706 | 0.87948803 |
| GTF3C4 | 63.5153812 | 0.24135661 | 0.26476761 | 0.91157906 | 0.36199035 | 0.85690129 |
| AK8 | 74.7244864 | 0.14828112 | 0.36552468 | 0.4056665 | 0.68498766 | 0.9558749 |
| SPACA9 | 36.7371444 | 0.19378887 | 0.36303544 | 0.53380153 | 0.59347886 | 0.93491607 |
| TSC1 | 249.844004 | -0.7161954 | 0.26829947 | -2.669388 | 0.00759896 | 0.21455924 |
| GTF3C5 | 59.3924053 | -0.5153097 | 0.36659537 | -1.4056633 | 0.15982413 | 0.71409437 |
| RALGDS | 44.3799728 | -0.3987516 | 0.32261874 | -1.2359841 | 0.21646446 | 0.78091138 |
| ABO | 383.074303 | 0.316401 | 0.26542321 | 1.1920623 | 0.23323682 | 0.7926597 |
| SURF6 | 67.0690701 | -0.0541986 | 0.25458857 | -0.2128871 | 0.83141501 | 0.97160502 |
| MED22 | 45.0336832 | -0.4265418 | 0.27576496 | -1.5467584 | 0.12192152 | 0.66181735 |
| RPL7A | 664.239902 | 0.02848976 | 0.49131685 | 0.05798652 | 0.95375936 | 0.99514888 |
| SURF1 | 61.7654338 | 0.09262964 | 0.28612833 | 0.32373459 | 0.74613898 | 0.96109466 |
| SURF2 | 43.4324177 | 0.16574498 | 0.54680814 | 0.3031136 | 0.76180329 | 0.96218738 |
| SURF4 | 190.09179 | -0.1699071 | 0.46612426 | -0.3645102 | 0.71547705 | 0.95791799 |
| REXO4 | 60.0930901 | 0.43370129 | 0.2520772 | 1.72050983 | 0.08533981 | 0.59845478 |
| ADAMTS13 | 35.4752124 | 0.30885661 | 0.37022758 | 0.83423448 | 0.40414887 | 0.87834687 |
| CACFD1 | 55.1637755 | -0.3116062 | 0.39146825 | -0.7959934 | 0.42603585 | 0.88430496 |
| VAV2 | 930.343974 | -1.9377224 | 0.64258337 | -3.0155191 | 0.0025654 | 0.13156951 |
| BRD3OS | 54.9837827 | 0.71945597 | 0.42430336 | 1.69561694 | 0.08995845 | 0.60739841 |

| | | | | | | |
|---|---|---|---|---|---|---|
| BRD3 | 141.145179 | 0.16063688 | 0.28901994 | 0.55579858 | 0.57834855 | 0.93345492 |
| WDR5 | 63.3260155 | -0.0496213 | 0.32116909 | -0.154502 | 0.8772139 | 0.98277085 |
| RXRA | 95.1647106 | 0.19105665 | 0.40403958 | 0.47286618 | 0.63630866 | 0.94821838 |
| PPP1R26-AS1 | 77.2824342 | 0.03932201 | 0.43765138 | 0.08984779 | 0.92840817 | 0.99202739 |
| PPP1R26 | 58.0877126 | 0.23199658 | 0.25168802 | 0.92176252 | 0.35665246 | 0.8557799 |
| MRPS2 | 66.7754819 | 0.38530779 | 0.34405054 | 1.11991624 | 0.26274946 | 0.8096287 |
| CAMSAP1 | 97.7170283 | -0.2718604 | 0.33142922 | -0.8202669 | 0.41206399 | 0.88098902 |
| UBAC1 | 116.72837 | 0.07450557 | 0.33740264 | 0.22082096 | 0.82523184 | 0.97018919 |
| NACC2 | 87.2127338 | -0.0267395 | 0.24785026 | -0.1078857 | 0.91408639 | 0.98830345 |
| TMEM250 | 41.9156423 | -0.1023937 | 0.34598251 | -0.2959506 | 0.76726782 | 0.96305305 |
| QSOX2 | 41.3045684 | -0.0802076 | 0.3122068 | -0.2569055 | 0.79725174 | 0.9653598 |
| SNAPC4 | 51.8081886 | -0.1410671 | 0.37091618 | -0.3803208 | 0.7037073 | 0.95672887 |
| ENTR1 | 54.7309534 | -0.6707909 | 0.33731424 | -1.9886232 | 0.04674281 | 0.48600281 |
| PMPCA | 80.2772955 | 0.17765744 | 0.25453301 | 0.69797406 | 0.48519342 | 0.90695174 |
| INPP5E | 39.1744846 | -0.4902825 | 0.31389073 | -1.5619529 | 0.11829909 | 0.65447949 |
| SEC16A | 154.539858 | -0.4294865 | 0.22083501 | -1.9448298 | 0.05179546 | 0.50451388 |
| NOTCH1 | 43.2712468 | 0.63716784 | 0.3630015 | 1.75527606 | 0.0792121 | 0.5841962 |
| AGPAT2 | 141.81678 | 0.44937608 | 0.33519227 | 1.34065168 | 0.18003357 | 0.74223125 |
| SNHG7 | 118.193628 | 0.68135185 | 0.31558846 | 2.15898846 | 0.03085106 | 0.41259458 |
| RABL6 | 212.575259 | 0.12820266 | 0.26557002 | 0.48274522 | 0.62927665 | 0.94596555 |
| PHPT1 | 228.38733 | 0.65911583 | 0.50268058 | 1.31120209 | 0.1897895 | 0.75619145 |
| EDF1 | 357.806725 | -0.025526 | 0.45341685 | -0.056297 | 0.95510523 | 0.99537617 |
| TRAF2 | 38.9484847 | 0.1558395 | 0.39571446 | 0.39381807 | 0.69371537 | 0.95672887 |
| FBXW5 | 110.200152 | 0.07724793 | 0.31683498 | 0.24381126 | 0.807377 | 0.96752828 |
| LCN12 | 91.8616333 | 0.71196244 | 0.43198724 | 1.64810987 | 0.09933013 | 0.61779213 |
| PAXX | 69.4509047 | -0.0200608 | 0.36085312 | -0.0555928 | 0.9556662 | 0.99556479 |
| ABCA2 | 33.0452626 | 1.49449306 | 0.59515756 | 2.51108808 | 0.01203597 | 0.26883805 |
| MAN1B1 | 110.507855 | 0.01356901 | 0.21655333 | 0.06265899 | 0.95003806 | 0.99385095 |
| DPP7 | 119.042546 | 0.32152414 | 0.37161239 | 0.86521373 | 0.38692149 | 0.86852426 |
| SSNA1 | 120.584513 | -0.1626195 | 0.44469511 | -0.3656876 | 0.71459823 | 0.95791799 |
| TPRN | 44.3165985 | 0.16603659 | 0.33321752 | 0.4982829 | 0.61828466 | 0.94256005 |
| TMEM203 | 62.872876 | 0.43356431 | 0.48295864 | 0.89772555 | 0.36933189 | 0.86020443 |
| RNF208 | 33.549932 | 0.68216902 | 0.44354432 | 1.53799518 | 0.12404979 | 0.66528789 |
| TUBB4B | 378.910348 | -0.1079565 | 0.59933732 | -0.1801265 | 0.8570533 | 0.97718353 |
| NELFB | 135.194179 | -0.0333907 | 0.26784471 | -0.1246645 | 0.90078917 | 0.9875066 |
| TOR4A | 43.6502878 | -0.0997188 | 0.29930589 | -0.3331668 | 0.73900839 | 0.96078945 |
| EXD3 | 180.439044 | -0.3455955 | 0.25222563 | -1.370184 | 0.17062947 | 0.72690927 |
| NOXA1 | 76.7437867 | -0.0265993 | 0.2795322 | -0.0951564 | 0.9241906 | 0.99051393 |
| NSMF | 38.8227217 | -0.1863568 | 0.31402292 | -0.5934496 | 0.55288029 | 0.92872652 |
| PNPLA7 | 105.434096 | 0.81269605 | 0.4466362 | 1.81959287 | 0.06882103 | 0.55917298 |
| MRPL41 | 316.340791 | 0.02132677 | 0.43017913 | 0.04957648 | 0.96045989 | 0.99635733 |
| DPH7 | 64.4647334 | -0.0748108 | 0.24373186 | -0.3069388 | 0.75888995 | 0.96180522 |
| ARRDC1 | 105.394303 | -0.0624464 | 0.23049131 | -0.2709274 | 0.78644685 | 0.96513523 |
| ARRDC1-AS1 | 31.3819182 | 0.2333588 | 0.3611435 | 0.64616642 | 0.51817159 | 0.91751084 |
| EHMT1 | 627.39191 | -0.1404206 | 0.2080418 | -0.6749635 | 0.49969896 | 0.90839719 |
| AC069287.3 | 44.2212623 | 0.57337311 | 0.68044334 | 0.84264637 | 0.39942625 | 0.87617225 |
| BET1L | 66.6179047 | -0.2663411 | 0.29891676 | -0.891021 | 0.37291793 | 0.86201082 |
| RIC8A | 65.0825855 | -0.1198256 | 0.39118145 | -0.3063173 | 0.75936307 | 0.96182318 |
| SIRT3 | 69.2556685 | 0.12043898 | 0.27987491 | 0.43033146 | 0.66695455 | 0.9527406 |
| PSMD13 | 126.970639 | -0.1927046 | 0.26398278 | -0.7299895 | 0.46539663 | 0.89883965 |
| AC136475.3 | 38.1994481 | -0.5875153 | 1.01594837 | -0.5782925 | 0.56306665 | 0.9306324 |
| PGGHG | 246.063759 | -0.6778877 | 0.65384809 | -1.0367664 | 0.29984476 | 0.83618229 |
| IFITM2 | 651.158259 | -0.0786983 | 0.39869742 | -0.1973886 | 8.44E-01 | 0.97561662 |
| IFITM3 | 1112.39215 | -0.9418832 | 0.56526786 | -1.66626 | 0.09566164 | 0.61367241 |

| | | | | | | |
|---|---|---|---|---|---|---|
| PKP3 | 53.5440523 | -0.2028992 | 0.30252869 | -0.6706776 | 0.50242595 | 0.90962743 |
| SIGIRR | 99.2448245 | -0.3468227 | 0.29215042 | -1.1871373 | 0.23517346 | 0.79599484 |
| ANO9 | 38.7573661 | -0.4876357 | 0.35532751 | -1.3723557 | 0.16995274 | 0.72675731 |
| PTDSS2 | 79.133099 | -0.336758 | 0.25855801 | -1.3024468 | 0.19276371 | 0.7597012 |
| RNH1 | 177.109333 | 0.05806297 | 0.36532159 | 0.15893661 | 0.87371882 | 0.98246494 |
| HRAS | 97.1299798 | -0.5828716 | 0.4003995 | -1.4557252 | 0.14546861 | 0.69325556 |
| AP006284.1 | 45.5352068 | -0.614157 | 0.62930615 | -0.9759273 | 0.3291005 | 0.84409527 |
| RASSF7 | 263.284834 | 0.18673465 | 0.3530165 | 0.52896861 | 0.59682723 | 0.93617523 |
| MIR210HG | 83.8403415 | -0.1770248 | 0.66458846 | -0.2663675 | 0.78995621 | 0.96522573 |
| PHRF1 | 108.138277 | -0.4643283 | 0.23525013 | -1.9737644 | 0.04840854 | 0.48661187 |
| DEAF1 | 198.214173 | 0.02642544 | 0.20623888 | 0.12813023 | 0.89804591 | 0.9863394 |
| AC131934.1 | 35.5665652 | 0.07924769 | 0.40086874 | 0.19768988 | 0.84328771 | 0.97560439 |
| EPS8L2 | 467.741274 | 0.93148994 | 0.28812394 | 3.2329488 | 0.0012252 | 0.08794552 |
| AP006621.4 | 65.1828093 | 0.8301692 | 0.59222798 | 1.40177301 | 0.16098304 | 0.71409437 |
| TALDO1 | 273.146939 | 0.05342608 | 0.42043642 | 0.12707291 | 0.89888269 | 0.98660836 |
| GATD1 | 68.2908658 | -0.3442596 | 0.27317281 | -1.2602264 | 0.2075877 | 0.77317759 |
| AP006621.3 | 85.6722259 | -0.258071 | 0.32397293 | -0.796582 | 0.42569385 | 0.8840962 |
| RPLP2 | 458.584995 | -0.1448696 | 0.35150032 | -0.4121463 | 0.6802322 | 0.95574492 |
| PNPLA2 | 155.948992 | -0.248564 | 0.33227622 | -0.7480645 | 0.45442125 | 0.89626521 |
| CD151 | 355.850665 | 0.21828157 | 0.33859568 | 0.64466733 | 0.51914279 | 0.91751084 |
| POLR2L | 316.233674 | 0.37428924 | 0.49544585 | 0.75545943 | 0.44997337 | 0.89324553 |
| TSPAN4 | 43.7496413 | 0.96301547 | 0.35548686 | 2.70900439 | 0.00674854 | 0.20224502 |
| CHID1 | 96.5820291 | 0.68019619 | 0.26442157 | 2.57239303 | 0.01009982 | 0.24501406 |
| AP2A2 | 642.654688 | 0.01261465 | 0.32320326 | 0.03903009 | 0.9688664 | 0.99645289 |
| MUC6 | 82.6057908 | -0.5741256 | 0.89247699 | -0.6432946 | 0.52003299 | 0.91751084 |
| TOLLIP | 96.2288976 | -0.2621295 | 0.25084857 | -1.0449713 | 0.29603625 | 0.83444322 |
| MOB2 | 74.1856073 | 0.27639927 | 0.24387447 | 1.13336696 | 0.25706018 | 0.80706732 |
| MRPL23 | 41.8786936 | 1.31670756 | 0.51003037 | 2.58162582 | 0.00983361 | 0.24191614 |
| CD81-AS1 | 41.1741237 | 0.60536602 | 0.33781229 | 1.79201895 | 0.07312993 | 0.56668778 |
| CD81 | 1179.15295 | 0.26335363 | 0.30676482 | 0.85848707 | 0.39062357 | 0.87004477 |
| TSSC4 | 57.7318225 | -0.1838291 | 0.37614667 | -0.4887164 | 0.62504246 | 0.9448702 |
| KCNQ1 | 1183.1197 | -0.0536681 | 0.23795633 | -0.2255378 | 0.82156091 | 0.96967533 |
| KCNQ1OT1 | 1165.96078 | -0.062239 | 0.41823985 | -0.1488117 | 0.88170224 | 0.98335905 |
| CDKN1C | 507.853436 | -2.0113145 | 0.72393268 | -2.778317 | 0.00546413 | 0.18295228 |
| SLC22A18 | 111.857186 | -0.8733662 | 0.38139017 | -2.2899545 | 0.02202395 | 0.35843654 |
| PHLDA2 | 134.179809 | -2.196612 | 1.21727212 | -1.8045365 | 0.07114724 | 0.56333976 |
| NAP1L4 | 694.968777 | 0.21965271 | 0.21823692 | 1.00648741 | 0.31418116 | 0.84285425 |
| CARS | 106.117455 | -0.6525003 | 0.26813135 | -2.43351 | 0.01495322 | 0.2940146 |
| OSBPL5 | 53.3326278 | -0.1069354 | 0.39324104 | -0.2719335 | 0.78567313 | 0.96513523 |
| ZNF195 | 93.72594 | 0.09457838 | 0.34394252 | 0.27498309 | 0.78332923 | 0.96513523 |
| AC127526.1 | 59.3690936 | 0.27662128 | 0.43224732 | 0.63996065 | 0.52219818 | 0.91845051 |
| NUP98 | 452.840345 | -0.2504849 | 0.28394 | -0.8821754 | 0.37768198 | 0.86489308 |
| PGAP2 | 93.4953221 | 0.3877303 | 0.33613378 | 1.15349996 | 0.24870523 | 0.80097144 |
| RHOG | 44.7082379 | -0.1252864 | 0.38292795 | -0.3271801 | 0.74353165 | 0.96105014 |
| STIM1 | 215.280882 | 0.07728426 | 0.30883052 | 0.25024812 | 0.80239547 | 0.9661728 |
| RRM1 | 78.0175643 | 0.13562711 | 0.28331728 | 0.47871103 | 0.63214422 | 0.94654542 |
| TRIM68 | 28.4810399 | -0.0484867 | 0.40655909 | -0.1192611 | 0.90506849 | 0.9875066 |
| AC104389.4 | 181.843855 | -0.1717602 | 0.43189592 | -0.3976887 | 0.69085963 | 0.95659154 |
| TRIM5 | 74.5564131 | -0.8819893 | 0.49241516 | -1.7911497 | 0.07326927 | 0.56668778 |
| TRIM22 | 148.173748 | -0.9781878 | 0.75165524 | -1.3013782 | 0.19312902 | 0.7597568 |
| FAM160A2 | 70.3910495 | -0.4543123 | 0.30940812 | -1.468327 | 0.14201543 | 0.68808217 |
| SMPD1 | 49.3267085 | 0.90124851 | 0.46761877 | 1.92731464 | 0.05394043 | 0.51332378 |
| APBB1 | 67.5746921 | 0.08494863 | 0.2650529 | 0.3204969 | 0.74859168 | 0.96109466 |
| ARFIP2 | 59.9844327 | -0.1751224 | 0.32937055 | -0.5316882 | 0.59494196 | 0.935202 |

| | | | | | | |
|---|---|---|---|---|---|---|
| AC084337.2 | 58.7276715 | -0.7010588 | 0.47439937 | -1.4777819 | 0.13946616 | 0.68299314 |
| DNHD1 | 95.2219765 | -0.2700636 | 0.44794119 | -0.6028997 | 0.5465754 | 0.92597739 |
| RRP8 | 31.7719418 | 0.1442458 | 0.31449113 | 0.45866415 | 0.64647537 | 0.95048567 |
| ILK | 153.144525 | 0.21939064 | 0.29783538 | 0.73661711 | 0.46135523 | 0.89799005 |
| TAF10 | 91.7688046 | -0.38413 | 0.37206258 | -1.0324338 | 0.30186893 | 0.83709431 |
| TPP1 | 79.2322838 | -0.2577192 | 0.25524604 | -1.0096894 | 0.31264412 | 0.84279957 |
| MRPL17 | 103.889828 | 0.37622975 | 0.43320612 | 0.86847746 | 0.38513301 | 0.86749643 |
| AC087280.2 | 90.2477361 | 0.17055699 | 0.5886999 | 0.28971806 | 0.77203194 | 0.96388926 |
| ZNF215 | 46.9519981 | 0.58907714 | 0.50770654 | 1.16027093 | 0.24593851 | 0.80072691 |
| ZNF214 | 41.1152172 | 0.5778585 | 0.3991588 | 1.44769074 | 0.14770356 | 0.69834914 |
| SYT9 | 166.518954 | 1.08077315 | 0.57937372 | 1.86541626 | 0.06212308 | 0.5452679 |
| AC107884.1 | 73.0929232 | 0.24248637 | 0.49542011 | 0.48945604 | 0.62451887 | 0.94457465 |
| PPFIBP2 | 230.33918 | 0.11412131 | 0.28880645 | 0.39514806 | 0.69273362 | 0.95672887 |
| OVCH2 | 97.1602346 | 1.16570592 | 0.45243936 | 2.57649097 | 0.00998088 | 0.24399985 |
| AC044810.3 | 206.022039 | 2.81660744 | 0.63127151 | 4.46180035 | 8.13E-06 | 0.00323124 |
| EIF3F | 191.977584 | 0.31402633 | 0.30005343 | 1.04656806 | 0.29529884 | 0.83396494 |
| TUB | 34.2315877 | 1.06906802 | 0.59316964 | 1.80229726 | 0.07149865 | 0.56343001 |
| RIC3 | 134.905538 | -0.0884414 | 0.41862226 | -0.2112677 | 0.8326784 | 0.97208613 |
| STK33 | 1145.68823 | 0.55027761 | 0.20548328 | 2.67796776 | 0.00740703 | 0.21151424 |
| TRIM66 | 199.907867 | 0.04376265 | 0.30273322 | 0.14455847 | 0.88505949 | 0.98450356 |
| RPL27A | 518.230893 | -0.0737607 | 0.34286303 | -0.2151317 | 0.82966465 | 0.97139806 |
| ST5 | 409.758737 | -0.1363503 | 0.34755358 | -0.3923145 | 6.95E-01 | 0.95672887 |
| AC026894.1 | 39.1158964 | -0.2375982 | 0.56780513 | -0.4184503 | 0.67561792 | 0.95529325 |
| AKIP1 | 42.7818488 | 0.0886244 | 0.35992943 | 0.24622715 | 0.8055064 | 0.96719834 |
| TMEM9B | 161.121771 | -0.0834329 | 0.34267036 | -0.2434787 | 0.80763462 | 0.96752828 |
| TMEM9B-AS | 53.7107897 | 0.28848116 | 0.43701032 | 0.66012437 | 0.50917402 | 0.91334187 |
| DENND5A | 247.593023 | -0.5741293 | 0.34698552 | -1.6546202 | 0.09800156 | 0.61685067 |
| TMEM41B | 269.927209 | -0.689012 | 0.32977571 | -2.0893352 | 0.03667756 | 0.43729328 |
| IPO7 | 332.73799 | -0.1201889 | 0.31185436 | -0.3854008 | 0.69994047 | 0.95672887 |
| ZNF143 | 92.5560567 | -0.4022767 | 0.40252997 | -0.9993709 | 0.31761506 | 0.84285425 |
| WEE1 | 81.8484863 | 0.40881477 | 0.31178902 | 1.3111904 | 0.18979344 | 0.75619145 |
| SWAP70 | 349.568178 | -0.116076 | 0.26444351 | -0.4389442 | 0.66070193 | 0.95127957 |
| SBF2-AS1 | 107.806654 | -0.306836 | 0.3513638 | -0.8732716 | 0.38251504 | 0.86593382 |
| SBF2 | 1774.48941 | -0.2053593 | 0.2370405 | -0.8663469 | 0.38629996 | 0.86844754 |
| AC011092.2 | 84.9058383 | 0.04861197 | 0.47238504 | 0.10290752 | 0.91803637 | 0.99012259 |
| ADM | 233.181281 | -3.1662223 | 1.27212856 | -2.4889169 | 0.01281329 | 0.27834029 |
| AMPD3 | 95.8626138 | -0.6070601 | 0.66808187 | -0.9086613 | 0.36352895 | 0.85694172 |
| MTRNR2L8 | 274.671879 | 0.39425054 | 0.5831543 | 0.67606557 | 0.49899902 | 0.90839719 |
| RNF141 | 94.9310413 | 0.21596788 | 0.23981615 | 0.90055603 | 0.36782442 | 0.85902833 |
| CTR9 | 150.676887 | 0.19260372 | 0.18467611 | 1.04292707 | 0.29698207 | 0.83553613 |
| EIF4G2 | 567.063012 | 0.09998313 | 0.31894011 | 0.31348558 | 0.75391177 | 0.96120879 |
| ZBED5 | 132.799068 | 0.31157101 | 0.18263366 | 1.70598902 | 0.08801012 | 0.60651125 |
| ZBED5-AS1 | 49.8957404 | 0.55761125 | 0.44690194 | 1.24772617 | 0.21213135 | 0.77709975 |
| GALNT18 | 1035.79625 | 0.22503628 | 0.45588882 | 0.49362096 | 0.62157389 | 0.94413291 |
| USP47 | 691.923734 | -0.0809768 | 0.1977943 | -0.4093992 | 0.68224669 | 0.95574492 |
| PARVA | 255.996129 | 0.10457252 | 0.24952867 | 0.41908019 | 0.67515753 | 0.95524793 |
| TEAD1 | 1439.22824 | -0.3640468 | 0.27968497 | -1.3016315 | 0.1930424 | 0.7597012 |
| LINC00958 | 109.606203 | 0.00416908 | 0.22840488 | 0.01825305 | 0.98543698 | 0.99762072 |
| ARNTL | 84.2170048 | 0.19312686 | 0.36348848 | 0.53131494 | 0.59520055 | 0.9353279 |
| BTBD10 | 179.652119 | 0.12883237 | 0.26131987 | 0.49300641 | 0.62200806 | 0.94418789 |
| FAR1 | 420.506334 | -0.1371915 | 0.15686285 | -0.8745951 | 0.38179427 | 0.86593382 |
| RRAS2 | 204.26874 | -0.057671 | 0.31748662 | -0.1816487 | 0.85585846 | 0.97718353 |
| COPB1 | 268.642233 | -0.2473271 | 0.26061953 | -0.9489966 | 0.34262233 | 0.85093239 |
| PSMA1 | 119.385473 | -0.5966938 | 0.72842892 | -0.8191517 | 0.41269986 | 0.88153663 |

| | | | | | | |
|---|---|---|---|---|---|---|
| PDE3B | 473.163542 | -1.6773576 | 0.7635732 | -2.1967214 | 0.02804035 | 0.40459506 |
| AC073172.2 | 46.9564675 | 2.93258213 | 0.85966822 | 3.41129529 | 0.00064655 | 0.05952104 |
| AC073172.1 | 324.51854 | 2.93840258 | 0.62939648 | 4.66860343 | 3.03E-06 | 0.00176561 |
| AC087379.2 | 546.335085 | 2.60820978 | 0.83986872 | 3.10549699 | 0.0018996 | 0.1093833 |
| SOX6 | 2427.26555 | 0.40291409 | 0.37090288 | 1.08630617 | 0.27734356 | 0.82198569 |
| C11orf58 | 404.317035 | 0.04911498 | 0.19655884 | 0.2498742 | 0.80268464 | 0.96635669 |
| PLEKHA7 | 1710.05719 | -0.0146588 | 0.21772395 | -0.0673273 | 0.94632118 | 0.99307065 |
| RPS13 | 464.423299 | 0.01137384 | 0.54617198 | 0.02082465 | 0.98338554 | 0.9971769 |
| PIK3C2A | 644.999137 | -0.2259013 | 0.19114005 | -1.1818625 | 0.23726026 | 0.7980661 |
| NUCB2 | 165.665552 | 0.20587953 | 0.23786949 | 0.86551468 | 0.38675636 | 0.86852426 |
| SERGEF | 219.678937 | -0.2777981 | 0.24895467 | -1.1158581 | 0.26448287 | 0.81108996 |
| HPS5 | 102.232056 | -0.8636588 | 0.4285651 | -2.0152337 | 0.04388017 | 0.48009422 |
| GTF2H1 | 159.523578 | -0.5003206 | 0.24231822 | -2.0647253 | 0.038949 | 0.4535395 |
| LDHA | 990.259918 | -0.3761971 | 0.55834961 | -0.6737662 | 0.50045998 | 0.90839719 |
| TSG101 | 242.297093 | 0.0903981 | 0.21670524 | 0.41714774 | 0.67657035 | 0.95529325 |
| UEVLD | 194.462827 | -0.1676599 | 0.25231976 | -0.6644739 | 0.50638705 | 0.91163608 |
| SPTY2D1OS | 61.679994 | 0.41819743 | 0.26906837 | 1.55424224 | 0.12012665 | 0.65679551 |
| SPTY2D1 | 94.876844 | -0.3252469 | 0.30940128 | -1.0512138 | 0.29316039 | 0.83336457 |
| ZDHHC13 | 182.409903 | -0.2095248 | 0.17284251 | -1.2122294 | 0.22542458 | 0.78982769 |
| NAV2 | 769.044816 | 0.22957142 | 0.49152408 | 0.46706037 | 0.64045669 | 0.94935885 |
| HTATIP2 | 114.847721 | -0.4191264 | 0.54309279 | -0.77174 | 0.44026843 | 0.88956997 |
| PRMT3 | 242.461581 | -0.4883588 | 0.39415327 | -1.2390073 | 0.21534281 | 0.77894575 |
| ANO5 | 82.7415628 | 0.81541231 | 0.31866672 | 2.55882481 | 0.01050267 | 0.25038862 |
| FANCF | 32.5796836 | 0.24451402 | 0.2883317 | 0.8480303 | 0.3964211 | 0.87462201 |
| GAS2 | 75.186458 | -0.4263664 | 0.43135012 | -0.9884461 | 0.3229342 | 0.84285425 |
| SVIP | 229.542146 | 0.16369699 | 0.28785409 | 0.56868043 | 0.56957303 | 0.93122667 |
| MUC15 | 72.6937926 | 1.27481058 | 0.61622687 | 2.06873578 | 0.03857089 | 0.45064107 |
| FIBIN | 51.5727056 | -0.3644711 | 0.83478547 | -0.4366045 | 0.66239822 | 0.95142083 |
| BBOX1 | 89.9161365 | -0.6010567 | 0.53712522 | -1.1190253 | 0.26312933 | 0.8096287 |
| CCDC34 | 53.3128234 | 0.86524133 | 0.40360708 | 2.14377143 | 0.03205121 | 0.41517597 |
| LGR4 | 761.076012 | 0.06895921 | 0.35525814 | 0.19411014 | 0.84608964 | 0.97643646 |
| LIN7C | 104.875295 | 0.08129614 | 0.34862975 | 0.2331876 | 0.81561573 | 0.96844493 |
| BDNF-AS | 415.379849 | -0.0849858 | 0.18641688 | -0.4558913 | 0.64846812 | 0.95061388 |
| METTL15 | 861.397104 | -0.166067 | 0.31898846 | -0.5206049 | 0.60264203 | 0.93804805 |
| ARL14EP | 106.246657 | 0.35826734 | 0.24284747 | 1.47527717 | 0.14013804 | 0.68408298 |
| MPPED2 | 1723.36919 | 0.48916831 | 0.35182667 | 1.39036733 | 0.16441736 | 0.71715698 |
| AL136088.1 | 50.2149934 | 0.72752396 | 0.43552248 | 1.67046246 | 0.0948279 | 0.61367241 |
| DCDC1 | 410.172021 | -0.1964212 | 0.25251822 | -0.7778495 | 0.43665773 | 0.88785137 |
| DNAJC24 | 107.779403 | -0.3872729 | 0.27349922 | -1.4159925 | 0.15677769 | 0.71126849 |
| IMMP1L | 263.434763 | -0.0947156 | 0.25085556 | -0.3775701 | 0.70575 | 0.95730303 |
| ELP4 | 444.78749 | 0.04024264 | 0.21995354 | 0.18295975 | 0.8548296 | 0.97718353 |
| AC131571.1 | 57.4231018 | 0.61200681 | 0.6641166 | 0.92153517 | 0.35677109 | 0.8557799 |
| RCN1 | 99.2217096 | 0.59077097 | 0.25727804 | 2.2962355 | 0.02166242 | 0.35579666 |
| EIF3M | 146.911079 | 0.07233973 | 0.19609952 | 0.36889297 | 0.7122075 | 0.95791799 |
| PRRG4 | 142.646502 | 0.31151605 | 0.37559368 | 0.8293964 | 0.40688014 | 0.87948803 |
| QSER1 | 230.200228 | 0.24745867 | 0.24182238 | 1.02330756 | 0.30616245 | 0.83709431 |
| TCP11L1 | 105.804851 | -0.1593435 | 0.35612185 | -0.4474408 | 0.65455679 | 0.95100227 |
| CSTF3 | 213.466602 | 0.07770824 | 0.24913912 | 0.311907 | 0.7551112 | 0.96120879 |
| HIPK3 | 595.290402 | 0.17496871 | 0.14092227 | 1.24159733 | 0.21438517 | 0.77890374 |
| FBXO3 | 142.663666 | 0.21093087 | 0.26419841 | 0.79838056 | 0.42464968 | 0.88391318 |
| CAPRIN1 | 430.04368 | -0.1649462 | 0.17371924 | -0.9494988 | 0.34236699 | 0.85093239 |
| NAT10 | 94.9966037 | -0.1047953 | 0.22715634 | -0.4613356 | 0.64455781 | 0.94949493 |
| ABTB2 | 3134.42333 | -0.9256523 | 0.52881356 | -1.7504323 | 0.08004375 | 0.58661433 |
| CAT | 328.284641 | 0.66596894 | 0.33530266 | 1.98617257 | 0.04701417 | 0.48600281 |

| | | | | | | |
|---|---|---|---|---|---|---|
| ELF5 | 323.172577 | 0.46258455 | 0.650002 | 0.71166634 | 0.47667141 | 0.90242252 |
| EHF | 664.125793 | 0.60031872 | 0.36311738 | 1.653236 | 0.09828284 | 0.61690569 |
| APIP | 163.11281 | -0.9275891 | 0.32637872 | -2.8420637 | 0.00448225 | 0.16424479 |
| PDHX | 251.216044 | -0.4255488 | 0.22205996 | -1.9163687 | 0.05531818 | 0.51808266 |
| CD44 | 121.482935 | -3.1640531 | 0.87199484 | -3.6285227 | 2.85E-04 | 0.03764581 |
| TRIM44 | 506.888303 | -0.1129917 | 0.16415079 | -0.6883409 | 0.49123812 | 0.90730741 |
| COMMD9 | 58.4429387 | 0.06190205 | 0.23121441 | 0.26772576 | 0.78891042 | 0.96515649 |
| PRR5L | 162.238007 | 0.62240025 | 0.40903591 | 1.52162739 | 0.12810247 | 0.67092149 |
| TRAF6 | 76.8563068 | -0.0766781 | 0.27242007 | -0.28147 | 0.77834992 | 0.96488138 |
| C11orf74 | 74.9488782 | 0.41356557 | 0.24031822 | 1.72090808 | 0.08526751 | 0.59845478 |
| LRRC4C | 85.4616137 | 1.36762785 | 0.67913098 | 2.01379098 | 0.04403148 | 0.48009422 |
| API5 | 214.838718 | -0.0403955 | 0.21204703 | -0.1905024 | 0.84891549 | 0.97656375 |
| TTC17 | 939.633607 | -0.2024602 | 0.18486618 | -1.0951719 | 0.27344134 | 0.81922962 |
| HSD17B12 | 830.811828 | 0.09201566 | 0.22489806 | 0.40914389 | 0.68243407 | 0.95574492 |
| ALKBH3 | 68.3208524 | 0.02533272 | 0.28584037 | 0.08862542 | 0.9293796 | 0.99216065 |
| ACCS | 43.288646 | 0.59765774 | 0.35173447 | 1.69917308 | 0.08928658 | 0.60739841 |
| EXT2 | 216.220113 | -0.0373911 | 0.24190945 | -0.1545666 | 0.87716298 | 0.98277085 |
| CD82 | 281.941312 | 0.67132754 | 0.47998564 | 1.39864089 | 0.1619207 | 0.71470931 |
| TSPAN18 | 55.6227555 | 0.96181652 | 0.58374984 | 1.64765189 | 0.09942413 | 0.61800999 |
| TP53I11 | 57.8248884 | 0.29784729 | 0.52430104 | 0.56808448 | 0.5699776 | 0.93122667 |
| PRDM11 | 83.9227207 | 0.19269343 | 0.25079832 | 0.76832026 | 0.44229694 | 0.89005186 |
| CRY2 | 125.733261 | 0.31405429 | 0.33068352 | 0.94971257 | 0.34225832 | 0.85093239 |
| LARGE2 | 33.0587196 | 0.95957166 | 0.37996413 | 2.52542697 | 0.01155578 | 0.26383866 |
| PHF21A | 872.318133 | 0.05779049 | 0.26814933 | 0.21551608 | 0.82936498 | 0.97131041 |
| CREB3L1 | 117.716732 | -1.5109724 | 1.01799674 | -1.4842605 | 0.13773982 | 0.68146162 |
| DGKZ | 49.2873976 | -0.1738452 | 0.28757643 | -0.6045182 | 0.54549916 | 0.92595257 |
| AMBRA1 | 1464.06065 | 0.13528497 | 0.2674522 | 0.50582858 | 0.612977 | 0.94003018 |
| ATG13 | 145.160423 | -0.1850802 | 0.21689057 | -0.8533346 | 0.3934738 | 0.87223214 |
| ARHGAP1 | 161.932861 | -0.4354045 | 0.21381206 | -2.0363889 | 0.04171132 | 0.46746387 |
| CKAP5 | 210.137472 | -0.1376128 | 0.18371544 | -0.749054 | 0.45382466 | 0.89568407 |
| LRP4 | 164.409668 | -0.0851763 | 0.45530677 | -0.1870745 | 0.85160222 | 0.97718353 |
| C11orf49 | 856.603633 | -0.0242907 | 0.21389315 | -0.1135644 | 0.9095831 | 0.98779844 |
| ARFGAP2 | 112.272017 | -0.1001819 | 0.27159991 | -0.3688585 | 0.7122332 | 0.95791799 |
| DDB2 | 57.5803033 | 0.04176359 | 0.42042493 | 0.09933662 | 0.920871 | 0.9904507 |
| ACP2 | 58.3503359 | 0.33389527 | 0.29332704 | 1.13830372 | 0.25499368 | 0.80489222 |
| NR1H3 | 42.1718625 | -0.2512126 | 0.31333884 | -0.8017283 | 0.42271016 | 0.88371155 |
| MADD | 77.3692957 | -0.3389132 | 0.22474846 | -1.5079668 | 0.13156301 | 0.67390608 |
| SLC39A13 | 91.3805655 | -0.2064278 | 0.24772598 | -0.833291 | 0.40468064 | 0.87834687 |
| PSMC3 | 51.9044585 | -0.2097161 | 0.54002743 | -0.3883434 | 0.69776196 | 0.95672887 |
| CELF1 | 832.391924 | -0.243239 | 0.24393792 | -0.9971348 | 0.3186991 | 0.84285425 |
| NDUFS3 | 124.789298 | 0.35708946 | 0.44331586 | 0.80549669 | 0.42053312 | 0.88320382 |
| MTCH2 | 193.877188 | -0.0564679 | 0.25483944 | -0.2215821 | 0.8246392 | 0.97018919 |
| FNBP4 | 459.500975 | -0.1434119 | 0.1729661 | -0.8291329 | 0.40702919 | 0.87948803 |
| NUP160 | 274.691983 | -0.4096856 | 0.16537503 | -2.4773122 | 0.0132376 | 0.28170334 |
| PTPRJ | 1227.68697 | 0.28736272 | 0.33557376 | 0.85633251 | 0.39181388 | 0.87030305 |
| TNKS1BP1 | 176.924607 | -0.0466323 | 0.21807933 | -0.2138318 | 0.83067825 | 0.97160502 |
| SSRP1 | 98.6348984 | 0.44589053 | 0.30980179 | 1.43927683 | 0.1500721 | 0.7029344 |
| TIMM10 | 80.2285765 | 0.28936698 | 0.42351228 | 0.68325522 | 0.49444558 | 0.90839719 |
| UBE2L6 | 94.9547934 | -0.5362456 | 0.6364296 | -0.8425843 | 0.39946097 | 0.87617225 |
| SERPING1 | 522.025256 | -0.3953645 | 0.36196117 | -1.0922843 | 2.75E-01 | 0.8202929 |
| ZDHHC5 | 161.816573 | -0.3207661 | 0.19410362 | -1.6525506 | 0.09842236 | 0.61690569 |
| MED19 | 39.6211768 | -0.1731743 | 0.43853928 | -0.3948888 | 0.69292495 | 0.95672887 |
| CTNND1 | 624.600448 | -0.1793524 | 0.15164514 | -1.1827108 | 0.23692378 | 0.79797745 |
| LPXN | 97.6049595 | 0.23000301 | 0.36494718 | 0.63023643 | 0.52853991 | 0.92121123 |

| | | | | | | |
|---|---|---|---|---|---|---|
| FAM111A | 153.174088 | -0.0248737 | 0.25731728 | -0.0966654 | 0.92299213 | 0.99051393 |
| OSBP | 295.966239 | 0.03424557 | 0.17901775 | 0.19129704 | 0.84829288 | 0.97656375 |
| PATL1 | 168.563182 | -0.5036924 | 0.31889434 | -1.5794962 | 0.11422228 | 0.64951142 |
| STX3 | 194.308674 | -0.0809938 | 0.4308711 | -0.1879768 | 0.85089481 | 0.97718353 |
| MRPL16 | 98.2217093 | 0.46136814 | 0.35879664 | 1.28587642 | 0.19848619 | 0.76668122 |
| PRPF19 | 87.377977 | -0.1625683 | 0.35684039 | -0.4555772 | 0.64869407 | 0.95061388 |
| TMEM109 | 60.0985466 | 0.400494 | 0.38162313 | 1.04944896 | 0.29397153 | 0.8337535 |
| VPS37C | 92.9551359 | -0.0129778 | 0.27031362 | -0.04801 | 0.96170827 | 0.99635733 |
| DDB1 | 231.755365 | 0.0895082 | 0.2267579 | 0.39473025 | 0.69304197 | 0.95672887 |
| TKFC | 50.6567595 | 0.32529267 | 0.25783785 | 1.2616172 | 0.20708656 | 0.77233708 |
| CYB561A3 | 31.3151376 | -0.4431631 | 0.33805027 | -1.3109386 | 0.18987851 | 0.75619145 |
| TMEM138 | 35.8998962 | 0.00909287 | 0.36237665 | 0.02509231 | 0.97998133 | 0.99696835 |
| CPSF7 | 169.249267 | -0.0521437 | 0.17260082 | -0.3021059 | 0.76257137 | 0.96262924 |
| AP003108.2 | 133.119064 | 0.21055801 | 0.27904744 | 0.7545599 | 0.45051309 | 0.89333031 |
| SYT7 | 40.4954521 | -1.5513844 | 0.81304553 | -1.9081151 | 0.05637635 | 0.5228532 |
| MYRF | 170.171025 | -0.185969 | 0.39376736 | -0.4722813 | 0.63672599 | 0.94821838 |
| TMEM258 | 154.239831 | 0.06980805 | 0.2335102 | 0.29895077 | 0.76497761 | 0.96285313 |
| FADS2 | 54.7298511 | -1.4888247 | 0.90395508 | -1.6470119 | 0.09955561 | 0.61822465 |
| FADS1 | 73.5171536 | -0.9787775 | 0.83788104 | -1.1681581 | 0.24274301 | 0.80072691 |
| FTH1 | 2515.18135 | -0.4736246 | 0.51769494 | -0.914872 | 0.36025883 | 0.85608464 |
| ASRGL1 | 87.7585163 | 0.30358695 | 0.28744823 | 1.05614478 | 0.29090207 | 0.83182912 |
| AHNAK | 822.694267 | 0.43830715 | 0.31143977 | 1.40735769 | 0.15932135 | 0.71409437 |
| B3GAT3 | 87.7555348 | 0.06569929 | 0.3523009 | 0.18648631 | 0.85206341 | 0.97718353 |
| GANAB | 262.533585 | 0.46738498 | 0.26661696 | 1.75302046 | 0.0795985 | 0.58461232 |
| UQCC3 | 86.4532355 | -0.1290939 | 0.34119333 | -0.3783599 | 0.70516327 | 0.95730303 |
| UBXN1 | 144.705994 | 0.34623695 | 0.42707696 | 0.81071325 | 0.41753036 | 0.88242985 |
| TTC9C | 88.587008 | -0.0688657 | 0.25727061 | -0.2676782 | 0.78894705 | 0.96515649 |
| POLR2G | 56.7975599 | 0.21467469 | 0.36626902 | 0.58611205 | 0.55780022 | 0.92901206 |
| TAF6L | 40.827168 | -0.0652737 | 0.38930387 | -0.1676677 | 0.86684468 | 0.98083915 |
| TMEM223 | 88.7392795 | 0.0938592 | 0.31902603 | 0.29420547 | 0.7686009 | 0.96345263 |
| TMEM179B | 124.92911 | 0.49353856 | 0.37742733 | 1.30763862 | 0.19099591 | 0.75807631 |
| NXF1 | 63.7178831 | 0.07651615 | 0.28151001 | 0.27180614 | 0.78577108 | 0.96513523 |
| STX5 | 169.767517 | -0.2857335 | 0.22760156 | -1.255411 | 0.20932959 | 0.77436433 |
| SLC3A2 | 163.609417 | -0.9484955 | 0.30023145 | -3.1592144 | 0.00158195 | 0.09927452 |
| RARRES3 | 134.382666 | -0.6018233 | 0.79428699 | -0.75769 | 0.44863661 | 0.89301266 |
| PLA2G16 | 60.25525 | 0.74518406 | 0.59022452 | 1.26254337 | 0.20675333 | 0.77200549 |
| ATL3 | 248.623087 | 0.09452934 | 0.1592371 | 0.59363895 | 0.55275362 | 0.92872652 |
| RTN3 | 782.01783 | 0.2668635 | 0.23185585 | 1.1509889 | 0.2497368 | 0.80149394 |
| MARK2 | 280.366073 | 0.04447397 | 0.20073666 | 0.22155378 | 0.82466127 | 0.97018919 |
| NAA40 | 93.935446 | 0.35420515 | 0.23581946 | 1.50201832 | 0.13309238 | 0.67594872 |
| OTUB1 | 119.351988 | -0.0592526 | 0.26826994 | -0.2208694 | 0.82519409 | 0.97018919 |
| MACROD1 | 169.764855 | 0.55939514 | 0.32804095 | 1.70526008 | 0.08814593 | 0.60651125 |
| AP000721.2 | 47.6889049 | 1.0071393 | 0.56633069 | 1.77835904 | 0.0753449 | 0.57072562 |
| STIP1 | 105.414879 | 0.28626604 | 0.2452643 | 1.16717368 | 0.24314024 | 0.80072691 |
| TRPT1 | 37.9672638 | -0.337826 | 0.44054909 | -0.7668293 | 0.44318301 | 0.89022532 |
| NUDT22 | 78.9254131 | 0.00248516 | 0.28907887 | 0.00859684 | 0.9931408 | 0.99919666 |
| DNAJC4 | 68.1679777 | 0.00831463 | 0.34191292 | 0.02431797 | 0.98059898 | 0.99696835 |
| VEGFB | 186.929384 | -0.5198306 | 0.32166989 | -1.6160376 | 0.10608619 | 0.63461115 |
| FKBP2 | 209.29935 | 0.41463984 | 0.38469847 | 1.07783075 | 0.28110929 | 0.82628479 |
| PPP1R14B | 146.805245 | -0.4882537 | 0.51303345 | -0.9516995 | 0.34124939 | 0.85042291 |
| PLCB3 | 29.8169693 | -0.0973125 | 0.42380997 | -0.2296136 | 0.81839201 | 0.96883031 |
| BAD | 127.490127 | 0.34219911 | 0.43420737 | 0.78810065 | 0.43063784 | 0.88596084 |
| GPR137 | 54.7525843 | 0.47765558 | 0.28490723 | 1.67653019 | 0.09363439 | 0.61317397 |
| ESRRA | 161.634853 | -0.1863622 | 0.28767593 | -0.64782 | 0.5171014 | 0.91751084 |

| | | | | | | |
|---|---|---|---|---|---|---|
| TRMT112 | 228.435233 | -0.1722902 | 0.60074822 | -0.2867927 | 0.77427105 | 0.96388926 |
| PRDX5 | 340.44528 | -0.084679 | 0.55578845 | -0.1523583 | 0.87890438 | 0.98311774 |
| SF1 | 506.217239 | -0.3743292 | 0.19985425 | -1.8730111 | 0.06106685 | 0.5418629 |
| MAP4K2 | 38.9373877 | 0.55397144 | 0.32667687 | 1.69577799 | 0.08992793 | 0.60739841 |
| MEN1 | 38.2092933 | -0.1096027 | 0.36731351 | -0.29839 | 0.76540552 | 0.96285313 |
| EHD1 | 59.5541969 | 0.30951212 | 0.32321003 | 0.95761916 | 0.33825483 | 0.84745652 |
| ATG2A | 62.0808585 | -0.027184 | 0.30704337 | -0.0885348 | 0.92945161 | 0.99216065 |
| PPP2R5B | 33.8907685 | -0.2163121 | 0.35417838 | -0.6107433 | 0.54136955 | 0.92443671 |
| ZFPL1 | 40.7434383 | -0.3649654 | 0.29428018 | -1.240197 | 0.21490254 | 0.77890374 |
| VPS51 | 155.164645 | -0.06278 | 0.23252191 | -0.2699962 | 0.78716318 | 0.96513523 |
| TM7SF2 | 73.578837 | 0.31232797 | 0.5665508 | 0.55127974 | 0.58144193 | 0.93386308 |
| FAU | 624.935209 | -0.0656886 | 0.42536975 | -0.1544271 | 0.87727303 | 0.98277085 |
| SYVN1 | 56.2037345 | -0.2693276 | 0.33287868 | -0.8090864 | 0.41846546 | 0.88242985 |
| MRPL49 | 72.1091832 | 0.29120023 | 0.3744365 | 0.77770256 | 0.43674438 | 0.88785137 |
| CAPN1 | 251.786977 | -0.076328 | 0.28953059 | -0.2636266 | 0.79206767 | 0.9653598 |
| CDC42EP2 | 47.7923626 | 0.44584543 | 0.38303559 | 1.16397913 | 0.24443247 | 0.80072691 |
| DPF2 | 99.606968 | 0.02526584 | 0.19967567 | 0.12653442 | 0.89930891 | 0.98660836 |
| FRMD8 | 61.546989 | -0.5511277 | 0.32893221 | -1.6755053 | 0.09383514 | 0.61323531 |
| NEAT1 | 18827.8969 | -1.3337724 | 0.43728858 | -3.0500966 | 0.00228768 | 0.12358178 |
| MALAT1 | 224543.617 | -0.5571025 | 0.37250054 | -1.495575 | 0.13476444 | 0.6785647 |
| SCYL1 | 72.8052854 | 0.36470348 | 0.28198573 | 1.29334019 | 0.19589342 | 0.76403539 |
| LTBP3 | 236.141351 | 0.32396854 | 0.28647261 | 1.13088836 | 0.25810208 | 0.80709747 |
| SSSCA1 | 55.0013222 | 0.37777358 | 0.44631785 | 0.84642274 | 0.39731696 | 0.87462201 |
| FAM89B | 44.1169399 | -0.5141462 | 0.48770168 | -1.0542228 | 0.29178093 | 0.83275168 |
| EHBP1L1 | 40.8531187 | -0.0731055 | 0.54819695 | -0.1333563 | 0.89391162 | 0.98597539 |
| MAP3K11 | 73.7873818 | -0.2273052 | 0.28495715 | -0.7976822 | 0.42505494 | 0.88392184 |
| PCNX3 | 58.3388758 | -0.078524 | 0.2950969 | -0.2660955 | 0.79016566 | 0.96524651 |
| SIPA1 | 59.3129637 | 0.10705088 | 0.51017316 | 0.20983245 | 0.83379845 | 0.97232209 |
| RELA | 95.1858308 | -0.271387 | 0.30564519 | -0.8879152 | 0.37458638 | 0.86295837 |
| KAT5 | 97.4307568 | -0.3295803 | 0.23127735 | -1.4250438 | 0.15414455 | 0.7079031 |
| RNASEH2C | 82.3983814 | 1.05469148 | 0.39898504 | 2.64343617 | 0.00820692 | 0.22277119 |
| OVOL1 | 34.9653243 | 0.59424495 | 0.47877499 | 1.24117792 | 0.21454003 | 0.77890374 |
| CFL1 | 623.560423 | -0.0842099 | 0.48831222 | -0.1724509 | 0.86308307 | 0.979268 |
| MUS81 | 51.5745846 | -0.3463315 | 0.24642097 | -1.4054467 | 0.15988849 | 0.71409437 |
| EFEMP2 | 83.8897973 | 0.75324091 | 0.28310002 | 2.66068826 | 0.00779811 | 0.217192 |
| FIBP | 37.9924586 | 0.33157446 | 0.42230723 | 0.78514986 | 0.43236571 | 0.88707765 |
| CCDC85B | 46.2585636 | 0.8155267 | 0.44817492 | 1.81966162 | 0.06881055 | 0.55917298 |
| C11orf68 | 38.7737309 | 0.24058454 | 0.46359754 | 0.51895128 | 0.60379471 | 0.93804805 |
| DRAP1 | 116.013683 | 0.09055655 | 0.50332702 | 0.17991593 | 0.85721857 | 0.97718353 |
| SART1 | 52.8870014 | -0.1605177 | 0.27818971 | -0.5770081 | 0.56393398 | 0.93077674 |
| BANF1 | 161.182147 | 0.11050473 | 0.4711003 | 0.23456731 | 0.81454459 | 0.96839471 |
| SF3B2 | 185.248602 | -0.0687107 | 0.18842558 | -0.364657 | 0.71536747 | 0.95791799 |
| PACS1 | 640.559957 | -0.1576241 | 0.1599241 | -0.9856184 | 0.32432042 | 0.84285425 |
| KLC2 | 46.8852697 | -0.0390303 | 0.27464771 | -0.1421105 | 0.88699276 | 0.98544303 |
| RAB1B | 124.802807 | 0.066515 | 0.30718236 | 0.21653263 | 0.82857259 | 0.97117144 |
| YIF1A | 51.6629999 | -0.5694192 | 0.49650559 | -1.1468535 | 0.25144218 | 0.80240987 |
| BRMS1 | 50.3605467 | 0.33199098 | 0.42071759 | 0.78910649 | 0.43004977 | 0.88579433 |
| B4GAT1 | 38.9705039 | 0.9914259 | 0.36768473 | 2.69640213 | 0.0070093 | 0.20555027 |
| SLC29A2 | 101.634922 | 0.14001379 | 0.32347544 | 0.43284212 | 0.66512947 | 0.9520866 |
| MRPL11 | 66.4760952 | 0.53205004 | 0.4543505 | 1.17101233 | 0.24159382 | 0.80072691 |
| ZDHHC24 | 64.2083033 | 0.40075249 | 0.2450554 | 1.63535462 | 0.10197473 | 0.62351102 |
| CTSF | 96.2682079 | 1.05671429 | 0.35347259 | 2.98952255 | 0.00279414 | 0.13678792 |
| CCS | 117.2377 | 0.52726216 | 0.31927947 | 1.65141267 | 0.09865434 | 0.61707826 |
| RBM14-RBM | 67.6079391 | -0.1127438 | 0.36147575 | -0.3118987 | 0.7551175 | 0.96120879 |

| | | | | | | |
|---|---|---|---|---|---|---|
| RBM4 | 57.3957572 | 0.24549159 | 0.39255555 | 0.62536777 | 0.53172971 | 0.92211604 |
| RBM4B | 92.1269623 | -0.0325036 | 0.27605539 | -0.1177429 | 0.9062714 | 0.9875066 |
| SPTBN2 | 988.286561 | 0.84850275 | 0.45401883 | 1.86887128 | 0.06164073 | 0.54376667 |
| C11orf80 | 2380.26282 | 0.58849292 | 0.37768362 | 1.55816372 | 0.11919444 | 0.65527787 |
| RCE1 | 85.6413523 | 0.16062182 | 0.39459084 | 0.40705917 | 0.68396453 | 0.95574492 |
| LRFN4 | 29.967122 | -0.4861334 | 0.59949221 | -0.8109086 | 0.41741817 | 0.88242985 |
| RHOD | 84.6532079 | 0.46800929 | 0.35955251 | 1.30164377 | 0.19303819 | 0.7597012 |
| KDM2A | 919.638631 | -0.403734 | 0.23971179 | -1.6842477 | 0.0921338 | 0.61120158 |
| GRK2 | 73.8476833 | -0.1617223 | 0.2476524 | -0.6530214 | 0.51374246 | 0.91544604 |
| ANKRD13D | 94.8092464 | -0.2770223 | 0.21770087 | -1.2724906 | 0.20319888 | 0.7702733 |
| SSH3 | 89.4411122 | -0.2825399 | 0.2938684 | -0.9614504 | 0.33632573 | 0.84565588 |
| RAD9A | 399.135377 | 0.25450511 | 0.21174364 | 1.20194924 | 0.2293832 | 0.79047378 |
| PPP1CA | 171.17088 | 0.33305258 | 0.44310025 | 0.75164159 | 0.45226662 | 0.89473986 |
| RPS6KB2 | 49.4296768 | 0.13741024 | 0.38312908 | 0.35865258 | 0.71985501 | 0.95796325 |
| CORO1B | 112.808922 | -0.3204023 | 0.44916359 | -0.713331 | 0.47564094 | 0.90242252 |
| TMEM134 | 144.619786 | -0.110653 | 0.3351811 | -0.3301289 | 0.74130254 | 0.96084411 |
| AIP | 81.1832803 | 0.28523387 | 0.4804409 | 0.59369189 | 0.55271821 | 0.92872652 |
| PITPNM1 | 94.9937908 | -0.5980367 | 0.51629271 | -1.1583288 | 0.24672987 | 0.80072691 |
| CDK2AP2 | 216.573843 | -0.6171821 | 0.6222977 | -0.9917796 | 0.32130507 | 0.84285425 |
| GSTP1 | 1591.21378 | -0.0894871 | 0.53088383 | -0.1685625 | 0.86614076 | 0.98036021 |
| NDUFV1 | 228.64654 | 0.32995893 | 0.23752222 | 1.38917079 | 0.16478082 | 0.71715698 |
| NUDT8 | 41.8155364 | 0.35840207 | 0.43338189 | 0.82698904 | 0.40824328 | 0.87948803 |
| UNC93B1 | 64.1387187 | -0.6156008 | 0.46490562 | -1.3241415 | 0.18545605 | 0.74895057 |
| NDUFS8 | 253.77333 | 0.41895869 | 0.41614419 | 1.0067633 | 0.31404854 | 0.84285425 |
| TCIRG1 | 53.5873239 | -0.4347293 | 0.33923982 | -1.2814806 | 0.20002491 | 0.7687013 |
| CHKA | 352.364449 | -0.2874791 | 0.25892464 | -1.1102809 | 0.26687802 | 0.81351996 |
| KMT5B | 350.976802 | 0.21306041 | 0.24210225 | 0.88004307 | 0.37883598 | 0.86515603 |
| LRP5 | 94.1972517 | 0.50979676 | 0.33654853 | 1.51477933 | 0.1298283 | 0.67256577 |
| PPP6R3 | 1240.39828 | -0.1419159 | 0.24376843 | -0.582175 | 0.56044882 | 0.92934517 |
| TESMIN | 39.4784559 | 0.57452777 | 0.42258471 | 1.35955646 | 0.17397033 | 0.7316248 |
| CPT1A | 329.892351 | 0.32039322 | 0.27623525 | 1.15985638 | 0.24610728 | 0.80072691 |
| MRPL21 | 57.4974695 | 0.25566533 | 0.30661494 | 0.83383194 | 0.4043757 | 0.87834687 |
| IGHMBP2 | 47.1086879 | -0.222119 | 0.34385973 | -0.6459583 | 0.51830636 | 0.91751084 |
| TPCN2 | 75.3105553 | -0.1881216 | 0.27530127 | -0.6833299 | 0.49439839 | 0.90839719 |
| CCND1 | 148.202442 | 1.05647567 | 0.43906487 | 2.40619493 | 0.01611966 | 0.30396236 |
| ORAOV1 | 68.3023656 | -0.0724492 | 0.29852145 | -0.2426936 | 0.8082428 | 0.96771216 |
| FADD | 37.8164569 | -0.1516453 | 0.35955578 | -0.4217574 | 0.67320213 | 0.95456072 |
| PPFIA1 | 513.924925 | -0.4651341 | 0.23550724 | -1.9750311 | 0.04826462 | 0.48661187 |
| AP002336.2 | 167.244788 | -0.2538701 | 0.42522924 | -0.5970195 | 0.55049437 | 0.92781939 |
| AP000487.1 | 44.51998 | 0.11447312 | 0.39448778 | 0.29018168 | 0.77167725 | 0.96388926 |
| CTTN | 658.795311 | -0.7528422 | 0.20850647 | -3.6106421 | 0.00030544 | 0.03833351 |
| SHANK2 | 1517.45476 | -0.2928774 | 0.28133889 | -1.0410128 | 0.29786959 | 0.83556453 |
| DHCR7 | 54.7748678 | -0.1346399 | 0.68685116 | -0.1960248 | 0.84459076 | 0.97573093 |
| NADSYN1 | 94.3022612 | -0.3120541 | 0.20504857 | -1.5218545 | 0.12804555 | 0.67092149 |
| AP002495.2 | 259.071023 | 0.079057 | 0.23846268 | 0.33152774 | 0.74024589 | 0.96084411 |
| RNF121 | 225.623866 | -0.0072003 | 0.23087812 | -0.0311866 | 0.97512069 | 0.99656026 |
| NUMA1 | 394.888538 | -0.0137212 | 0.25028204 | -0.0548229 | 0.95627957 | 0.99565888 |
| LAMTOR1 | 242.609422 | -0.1393565 | 0.41079097 | -0.3392394 | 0.73442936 | 0.96054158 |
| ANAPC15 | 49.8191832 | 0.20434005 | 0.48927003 | 0.41764269 | 0.67620838 | 0.95529325 |
| FOLR1 | 308.611435 | 1.09160932 | 0.6279491 | 1.7383723 | 0.08214523 | 0.59202053 |
| INPPL1 | 96.9726136 | 0.37604451 | 0.30508336 | 1.23259593 | 0.21772656 | 0.78384552 |
| CLPB | 85.5452889 | -0.0206011 | 0.27529287 | -0.0748334 | 0.94034728 | 0.99243654 |
| ARAP1 | 82.9669603 | 0.02484084 | 0.26992013 | 0.09203034 | 0.92667393 | 0.99087265 |
| ATG16L2 | 63.6293881 | 0.33352356 | 0.3389271 | 0.98405692 | 0.32508753 | 0.84285425 |

| | | | | | | |
|---|---|---|---|---|---|---|
| FCHSD2 | 402.214446 | -0.1326208 | 0.21661377 | -0.6122457 | 0.54037523 | 0.92414041 |
| ARHGEF17 | 98.574831 | 0.6516413 | 0.28768274 | 2.26513867 | 0.02350417 | 0.36567438 |
| FAM168A | 455.990974 | -0.1008852 | 0.27761236 | -0.3634032 | 0.7163037 | 0.95791799 |
| PLEKHB1 | 45.1846953 | 1.18285564 | 0.56771637 | 2.08353272 | 0.03720269 | 0.44129343 |
| RAB6A | 836.254837 | -0.3226937 | 0.19336598 | -1.6688238 | 0.09515231 | 0.61367241 |
| MRPL48 | 131.568792 | -0.4389452 | 0.34662388 | -1.2663445 | 0.20538979 | 0.77193931 |
| COA4 | 122.956393 | 0.57809656 | 0.4899915 | 1.17980937 | 0.23807604 | 0.79810098 |
| PAAF1 | 125.923986 | 0.0714654 | 0.34932924 | 0.20457892 | 0.83790113 | 0.97385715 |
| UCP2 | 242.621086 | -0.0966039 | 0.43217838 | -0.2235278 | 0.82312473 | 0.97018919 |
| C2CD3 | 182.425209 | -0.1384284 | 0.17810833 | -0.7772147 | 0.43703211 | 0.88785137 |
| PPME1 | 279.530021 | -0.3485005 | 0.45781509 | -0.7612255 | 0.44652241 | 0.8918683 |
| P4HA3 | 55.0866041 | -0.2291945 | 0.63770781 | -0.3594036 | 0.71929321 | 0.95796325 |
| PGM2L1 | 86.8026897 | -0.6422522 | 0.46591628 | -1.3784712 | 0.16805785 | 0.72391336 |
| POLD3 | 148.156689 | 0.11757496 | 0.29715953 | 0.39566277 | 0.69235382 | 0.95672887 |
| RNF169 | 271.704808 | -0.2412091 | 0.22809574 | -1.0574907 | 0.29028768 | 0.83103998 |
| XRRA1 | 272.94439 | -0.5084618 | 0.47972534 | -1.0599019 | 0.28918923 | 0.83078485 |
| SPCS2 | 307.576468 | 0.2765363 | 0.31927021 | 0.86615126 | 0.38640722 | 0.86844754 |
| NEU3 | 39.7413074 | -0.018689 | 0.28993243 | -0.0644597 | 0.94860418 | 0.99336519 |
| ARRB1 | 103.357512 | 0.01696268 | 0.36946959 | 0.04591091 | 0.96338126 | 0.99637506 |
| RPS3 | 1048.10816 | 0.37576317 | 0.45331837 | 0.82891671 | 0.40715154 | 0.87948803 |
| SERPINH1 | 65.0310611 | 0.76964993 | 0.42258711 | 1.82128111 | 0.06856414 | 0.55917298 |
| MAP6 | 42.9410958 | 1.81802547 | 0.31969939 | 5.68667177 | 1.30E-08 | 1.94E-05 |
| UVRAG | 594.39936 | -0.2039199 | 0.35949781 | -0.5672356 | 0.57055414 | 0.93125087 |
| THAP12 | 66.9416968 | 0.11582773 | 0.2480519 | 0.46694959 | 0.64053595 | 0.94935885 |
| EMSY | 244.45519 | -0.2507676 | 0.22396241 | -1.1196859 | 0.26284761 | 0.8096287 |
| ACER3 | 275.916858 | -0.0424259 | 0.30433659 | -0.1394046 | 0.88913045 | 0.98551241 |
| CAPN5 | 101.222374 | 0.43971523 | 0.34694739 | 1.26738304 | 0.20501836 | 0.77193931 |
| PAK1 | 328.221599 | -0.3175559 | 0.44967415 | -0.7061911 | 0.48006934 | 0.90406588 |
| CLNS1A | 220.030507 | 0.20687205 | 0.19860689 | 1.0416157 | 0.29758988 | 0.83556453 |
| RSF1 | 687.403339 | 0.00484077 | 0.18000967 | 0.0268917 | 0.97854612 | 0.99696835 |
| AAMDC | 183.06409 | 0.06904177 | 0.27080329 | 0.25495173 | 0.79876038 | 0.9653598 |
| AP002812.2 | 65.8200269 | 0.12753502 | 0.47726273 | 0.26722183 | 0.78929837 | 0.96515649 |
| INTS4 | 152.850736 | 0.01086699 | 0.24009494 | 0.04526122 | 0.9638991 | 0.99637506 |
| ALG8 | 82.893568 | -0.390788 | 0.26230976 | -1.4897958 | 0.13627793 | 0.67998335 |
| GAB2 | 327.164242 | -1.068994 | 0.5303746 | -2.0155453 | 0.04384755 | 0.48009422 |
| AP003086.2 | 103.420599 | -0.6629838 | 0.79909512 | -0.8296682 | 0.40672641 | 0.87948803 |
| NARS2 | 210.751347 | -0.1428617 | 0.20929378 | -0.6825893 | 0.49486637 | 0.90839719 |
| PRCP | 141.415427 | 0.45547204 | 0.21672883 | 2.10157571 | 0.03559046 | 0.43592878 |
| RAB30 | 125.802258 | -0.5020732 | 0.37667627 | -1.3329036 | 0.18256345 | 0.74315969 |
| RAB30-AS1 | 251.542512 | 0.03614955 | 0.32168389 | 0.11237601 | 0.91052527 | 0.98788932 |
| PCF11 | 218.13641 | -0.2397507 | 0.25167292 | -0.9526279 | 0.34077863 | 0.85032381 |
| ANKRD42 | 121.849794 | 0.1874155 | 0.21390247 | 0.87617269 | 0.38093616 | 0.86593382 |
| CCDC90B | 108.398668 | -0.0890179 | 0.25321514 | -0.3515505 | 0.72517536 | 0.95963742 |
| AP000446.1 | 64.2039768 | -0.1651798 | 0.30938448 | -0.5338982 | 0.59341193 | 0.93491607 |
| DLG2 | 2470.36525 | 0.28274972 | 0.53544698 | 0.52806296 | 0.59745563 | 0.93662454 |
| TMEM126B | 75.8287177 | -0.5157785 | 0.26063003 | -1.9789679 | 0.04781962 | 0.48600281 |
| TMEM126A | 42.5607351 | -0.0666415 | 0.44537694 | -0.1496293 | 0.88105705 | 0.98333507 |
| CREBZF | 190.41495 | -0.0126988 | 0.23843377 | -0.0532591 | 0.95752543 | 0.99579587 |
| SYTL2 | 1274.72603 | -0.1250368 | 0.54910032 | -0.2277121 | 0.81987009 | 0.96886922 |
| PICALM | 1571.23473 | -0.0755656 | 0.23924809 | -0.3158464 | 0.75211912 | 0.96120879 |
| EED | 94.1927131 | -0.3204522 | 0.30498448 | -1.0507164 | 0.29338884 | 0.83336457 |
| HIKESHI | 110.707779 | 0.30558979 | 0.20506532 | 1.49020709 | 0.13616979 | 0.67998335 |
| AP001831.1 | 102.297165 | 1.71045966 | 0.34810244 | 4.91366754 | 8.94E-07 | 0.00070934 |
| ME3 | 234.673569 | 0.74568854 | 0.31822859 | 2.34324811 | 0.01911667 | 0.33114489 |

| | | | | | | |
|---|---|---|---|---|---|---|
| AP003059.1 | 35.7223638 | 1.24599037 | 0.63705426 | 1.95586225 | 0.05048138 | 0.49596288 |
| PRSS23 | 264.027381 | 0.48618568 | 0.45932644 | 1.05847529 | 0.28983881 | 0.83103268 |
| AP001528.2 | 250.32665 | 0.21531512 | 0.56555604 | 0.38071404 | 0.70341545 | 0.95672887 |
| FZD4 | 268.708683 | -0.996395 | 0.45790995 | -2.1759628 | 0.02955804 | 0.40623517 |
| AP001528.1 | 45.6401468 | -0.4173141 | 0.42899554 | -0.9727702 | 0.33066754 | 0.84442294 |
| TMEM135 | 457.344751 | -0.3048893 | 0.21300101 | -1.4313983 | 0.1523161 | 0.70429257 |
| RAB38 | 55.6154427 | 0.27011301 | 0.45201171 | 0.59757967 | 0.55012043 | 0.92780338 |
| CTSC | 168.113204 | 0.67077969 | 0.25648036 | 2.61532571 | 0.00891424 | 0.23010158 |
| NOX4 | 98.9245513 | 0.07044724 | 0.58728081 | 0.11995494 | 0.90451884 | 0.9875066 |
| CHORDC1 | 87.5376401 | 0.15274832 | 0.31832999 | 0.4798427 | 0.63133925 | 0.94628651 |
| SLC36A4 | 232.554241 | -0.4331085 | 0.28490985 | -1.5201597 | 0.12847084 | 0.67206108 |
| SMCO4 | 181.294364 | -0.3362672 | 0.37428589 | -0.8984233 | 0.36895993 | 0.8598399 |
| CEP295 | 151.055061 | -0.2097852 | 0.28829169 | -0.727684 | 0.46680706 | 0.89883965 |
| TAF1D | 299.128454 | -0.2091982 | 0.31039821 | -0.6739671 | 0.50033226 | 0.90839719 |
| PANX1 | 39.4229649 | 0.09503416 | 0.34237804 | 0.27757085 | 0.78134181 | 0.96513523 |
| MRE11 | 179.160796 | -0.18421 | 0.21782095 | -0.8456945 | 0.39772318 | 0.87462201 |
| ANKRD49 | 35.8311677 | -0.0083369 | 0.42273976 | -0.0197212 | 0.98426578 | 0.99749593 |
| AMOTL1 | 137.012411 | 0.33319947 | 0.52820759 | 0.63081159 | 0.52816372 | 0.92106733 |
| CWC15 | 91.1778147 | 0.08657925 | 0.29302387 | 0.29546826 | 0.7676362 | 0.96322167 |
| SRSF8 | 76.0391114 | 0.54318938 | 0.38871963 | 1.39738088 | 0.16229907 | 0.71482182 |
| ENDOD1 | 51.9154382 | 0.02051558 | 0.29490492 | 0.06956676 | 0.94453849 | 0.99265578 |
| AP000787.1 | 134.45786 | -0.0791551 | 0.57390804 | -0.137923 | 0.89030123 | 0.98597539 |
| SESN3 | 348.450253 | -0.3837381 | 0.69150923 | -0.5549283 | 0.57894368 | 0.93345492 |
| FAM76B | 85.9257215 | -0.3792186 | 0.27207096 | -1.3938226 | 0.16337118 | 0.71642393 |
| CEP57 | 313.55144 | 0.00296594 | 0.1580192 | 0.01876949 | 0.98502499 | 0.99762072 |
| MTMR2 | 206.670052 | -0.0339878 | 0.24823541 | -0.1369175 | 0.89109601 | 0.98597539 |
| MAML2 | 2870.0755 | 0.50768392 | 0.31704279 | 1.60131041 | 0.10930818 | 0.63854503 |
| CCDC82 | 166.746381 | -0.5397029 | 0.26652075 | -2.0249939 | 0.04286799 | 0.47339997 |
| JRKL | 52.1733259 | 0.11259914 | 0.33784078 | 0.33329055 | 0.73891498 | 0.96078945 |
| CNTN5 | 69.1506959 | -1.1364547 | 0.617601 | -1.8401114 | 0.06575188 | 0.55444578 |
| ARHGAP42 | 146.396084 | -0.9794434 | 0.91251002 | -1.0733508 | 0.28311373 | 0.82666614 |
| CEP126 | 104.734017 | 0.37399628 | 0.37384895 | 1.00039409 | 0.31711983 | 0.84285425 |
| CFAP300 | 93.4926174 | 0.74748987 | 0.4399591 | 1.69899855 | 0.08931946 | 0.60739841 |
| YAP1 | 1428.90631 | 0.09243955 | 0.31194374 | 0.29633404 | 0.76697499 | 0.96285313 |
| BIRC3 | 183.216604 | -0.1027204 | 0.92121454 | -0.1115054 | 0.91121556 | 0.98788932 |
| BIRC2 | 164.165388 | -0.4460547 | 0.31243002 | -1.4276947 | 0.15337974 | 0.70717982 |
| TMEM123 | 519.25002 | 0.01499756 | 0.16950665 | 0.08847769 | 0.92949702 | 0.99216065 |
| MMP7 | 447.803752 | 0.44646475 | 0.74700059 | 0.59767657 | 0.55005576 | 0.92780338 |
| DCUN1D5 | 97.9689193 | -0.0688028 | 0.19938353 | -0.3450777 | 0.73003595 | 0.96008876 |
| DYNC2H1 | 731.83034 | -0.0334568 | 0.27551751 | -0.1214325 | 0.90334844 | 0.9875066 |
| PDGFD | 199.139512 | -0.4454978 | 0.5884316 | -0.7570936 | 0.44899381 | 0.89324553 |
| CASP4 | 158.480091 | -0.2198516 | 0.34507527 | -0.6371121 | 0.52405182 | 0.91871245 |
| MSANTD4 | 49.7417872 | 0.23341689 | 0.26120645 | 0.89361073 | 0.3715302 | 0.86201082 |
| KBTBD3 | 64.0318587 | 0.31477587 | 0.24574491 | 1.28090495 | 0.20022705 | 0.76890831 |
| AASDHPPT | 116.23223 | 0.19637098 | 0.3046524 | 0.64457389 | 0.51920336 | 0.91751084 |
| CWF19L2 | 339.631128 | 0.02758222 | 0.23622262 | 0.11676366 | 0.90704734 | 0.9875066 |
| ALKBH8 | 55.4000799 | -0.0413897 | 0.26201634 | -0.1579662 | 0.87448344 | 0.98258673 |
| SLC35F2 | 35.0067634 | 0.37614144 | 0.3469792 | 1.08404606 | 0.27834438 | 0.82332743 |
| CUL5 | 559.461897 | -0.032593 | 0.19259262 | -0.1692329 | 0.86561347 | 0.98018684 |
| ACAT1 | 229.747277 | -0.0396032 | 0.29588625 | -0.1338461 | 0.89352428 | 0.98597539 |
| NPAT | 171.158569 | -0.1275471 | 0.17966248 | -0.7099265 | 0.47774975 | 0.90245446 |
| ATM | 374.559224 | -0.2744624 | 0.23393872 | -1.1732235 | 0.24070618 | 0.80057149 |
| C11orf65 | 116.120523 | 0.16685738 | 0.44553235 | 0.37451238 | 0.70802315 | 0.95752169 |
| KDELC2 | 70.6315679 | 0.85358901 | 0.32739697 | 2.60719887 | 0.00912863 | 0.23300339 |

| | | | | | | |
|---|---|---|---|---|---|---|
| EXPH5 | 273.147655 | -0.0096648 | 0.37125087 | -0.0260332 | 0.97923087 | 0.99696835 |
| DDX10 | 272.961586 | -0.1282102 | 0.31484443 | -0.4072176 | 0.68384817 | 0.95574492 |
| RDX | 500.799086 | 0.06411619 | 0.41940615 | 0.15287375 | 0.87849783 | 0.98300919 |
| ZC3H12C | 313.639648 | 0.52059333 | 0.61109792 | 0.85189838 | 0.39427049 | 0.87253562 |
| FDX1 | 322.250276 | 0.2109761 | 0.37171088 | 0.56758117 | 0.5703194 | 0.93125087 |
| COLCA1 | 73.9167686 | 0.39633183 | 0.49266833 | 0.80445972 | 0.42113153 | 0.88360543 |
| COLCA2 | 59.4676875 | -0.317991 | 0.31243904 | -1.0177696 | 0.30878745 | 0.84119898 |
| SIK2 | 1051.21931 | -0.5524809 | 0.54946069 | -1.0054967 | 0.31465772 | 0.84285425 |
| PPP2R1B | 150.687711 | -0.5027192 | 0.2305405 | -2.1806111 | 0.02921219 | 0.40623517 |
| C11orf1 | 47.5946676 | 1.08786737 | 0.46213739 | 2.35399122 | 0.01857305 | 0.32713539 |
| CRYAB | 863.436759 | -0.8326875 | 0.88766816 | -0.9380617 | 0.34821268 | 0.85348649 |
| DIXDC1 | 189.749746 | -0.042884 | 0.27997876 | -0.1531686 | 0.87826532 | 0.98300919 |
| DLAT | 109.12129 | -0.0703715 | 0.26727051 | -0.2632971 | 0.7923216 | 0.9653598 |
| NKAPD1 | 58.7058833 | 0.35290877 | 0.26398725 | 1.3368402 | 0.1812748 | 0.74315969 |
| TIMM8B | 95.6979541 | 0.03827175 | 0.3847963 | 0.09945976 | 0.92077323 | 0.9904507 |
| AP002884.3 | 55.805209 | -0.8239152 | 0.63999169 | -1.2873842 | 0.19796042 | 0.76582694 |
| PTS | 198.794858 | -0.1591861 | 0.38620639 | -0.4121788 | 0.68020834 | 0.95574492 |
| AP002884.1 | 115.960833 | -0.8085472 | 0.50233681 | -1.6095718 | 0.10749136 | 0.63477557 |
| TTC12 | 163.496672 | 0.18937042 | 0.28749556 | 0.65868991 | 0.51009491 | 0.91334187 |
| ZW10 | 57.9641941 | -0.1545031 | 0.23908584 | -0.6462245 | 0.51813399 | 0.91751084 |
| USP28 | 106.387366 | -0.1450971 | 0.21060976 | -0.688938 | 0.49086227 | 0.90730741 |
| ZBTB16 | 129.819137 | 1.19191557 | 0.7995614 | 1.49071174 | 0.13603719 | 0.67998335 |
| NNMT | 64.405721 | -0.0545023 | 0.87331839 | -0.0624083 | 9.50E-01 | 0.99394565 |
| REXO2 | 117.543914 | 0.27205359 | 0.42068057 | 0.64669874 | 0.51782694 | 0.91751084 |
| CADM1 | 549.829778 | -0.459194 | 0.52811501 | -0.8694963 | 0.38457574 | 0.86725322 |
| BUD13 | 37.5108492 | 0.14328718 | 0.29846949 | 0.48007311 | 0.63117541 | 0.94622065 |
| ZPR1 | 36.939489 | -0.3986508 | 0.30922097 | -1.2892101 | 0.19732506 | 0.76534786 |
| SIK3 | 2603.0962 | -0.4949234 | 0.28693831 | -1.7248426 | 0.08455583 | 0.59713285 |
| PAFAH1B2 | 315.961112 | -0.0102907 | 0.21913133 | -0.0469613 | 0.96254409 | 0.99637506 |
| SIDT2 | 74.7886947 | 0.12551496 | 0.24482425 | 0.51267372 | 0.60817958 | 0.94003018 |
| PCSK7 | 153.169325 | -0.3555434 | 0.3055824 | -1.1634943 | 0.24462901 | 0.80072691 |
| RNF214 | 138.997973 | -0.0890886 | 0.28908781 | -0.3081714 | 0.75795191 | 0.96180522 |
| BACE1 | 84.556561 | 0.51052462 | 0.30637585 | 1.66633441 | 0.09564683 | 0.61367241 |
| CEP164 | 128.122268 | -0.1307813 | 0.28255898 | -0.4628459 | 0.64347484 | 0.94949493 |
| FXYD2 | 47.4998657 | -0.6610688 | 0.65548834 | -1.0085135 | 0.31320801 | 0.84285425 |
| MPZL3 | 71.3333676 | -0.5766812 | 0.27252552 | -2.1160632 | 0.03433943 | 0.43050784 |
| MPZL2 | 194.723634 | -0.3997603 | 0.34091162 | -1.1726217 | 2.41E-01 | 0.80072691 |
| UBE4A | 177.259462 | -0.1093205 | 0.19677134 | -0.5555714 | 0.57850389 | 0.93345492 |
| ATP5MG | 500.247101 | 0.27819681 | 0.38620779 | 0.72032937 | 0.47132223 | 0.90123338 |
| KMT2A | 629.226354 | 0.0701004 | 0.19072996 | 0.3675374 | 0.7132182 | 0.95791799 |
| TMEM25 | 51.8907413 | 0.4883588 | 0.29343844 | 1.66426324 | 0.09605983 | 0.61407447 |
| IFT46 | 73.5434508 | -0.0219723 | 0.35062306 | -0.0626664 | 0.95003215 | 0.99385095 |
| ARCN1 | 123.847445 | -0.1571481 | 0.32348249 | -0.485801 | 0.62710829 | 0.94511671 |
| PHLDB1 | 157.673901 | 0.82962426 | 0.2909963 | 2.85097872 | 0.00435849 | 0.163132 |
| AP002954.1 | 71.7136977 | -0.0306432 | 0.47260779 | -0.0648385 | 0.94830255 | 0.99332441 |
| DDX6 | 377.607007 | -0.058207 | 0.15291718 | -0.3806438 | 0.70346761 | 0.95672887 |
| BCL9L | 67.4811834 | -0.1326512 | 0.3662436 | -0.3621938 | 0.7172072 | 0.95791799 |
| CCDC84 | 184.848588 | 0.24975089 | 0.28874087 | 0.86496549 | 0.38705773 | 0.86860065 |
| RPS25 | 624.950265 | -0.1603111 | 0.49300014 | -0.3251746 | 0.7450489 | 0.96109466 |
| TRAPPC4 | 146.611483 | 0.76007559 | 0.46008872 | 1.65201961 | 0.09853055 | 0.61707826 |
| HYOU1 | 64.0574497 | 0.13171047 | 0.3193582 | 0.41242238 | 0.68002986 | 0.95574492 |
| VPS11 | 58.689245 | -0.1530422 | 0.2735748 | -0.5594162 | 0.5758777 | 0.93294146 |
| H2AFX | 34.2059393 | 0.44915629 | 0.43244945 | 1.03863304 | 0.29897544 | 0.83556453 |
| DPAGT1 | 36.3628258 | 0.46499453 | 0.39314094 | 1.18276802 | 0.2369011 | 0.79797745 |

| | | | | | | |
|---|---|---|---|---|---|---|
| HINFP | 51.0666698 | 0.25169187 | 0.26226478 | 0.95968612 | 0.33721321 | 0.84636904 |
| NLRX1 | 50.8538302 | 0.48395359 | 0.33410661 | 1.44850048 | 0.14747713 | 0.69834914 |
| CBL | 191.852584 | -0.3172435 | 0.33065301 | -0.9594454 | 0.33733444 | 0.84636904 |
| RNF26 | 39.4762661 | 0.09629876 | 0.27256691 | 0.35330321 | 0.72386114 | 0.95963742 |
| NECTIN1 | 34.3881947 | 0.29302302 | 0.43482662 | 0.67388474 | 0.50038459 | 0.90839719 |
| OAF | 164.378073 | -1.2244498 | 0.54745386 | -2.2366266 | 0.02531076 | 0.38387372 |
| TMEM136 | 64.4765532 | -0.0584841 | 0.394782 | -0.1481427 | 0.88223012 | 0.98335905 |
| ARHGEF12 | 1313.66969 | 0.01050687 | 0.23424581 | 0.04485404 | 0.96422365 | 0.99637506 |
| SC5D | 77.5136632 | -0.0398559 | 0.35263775 | -0.1130223 | 0.9100129 | 0.98788661 |
| SORL1 | 602.998718 | 0.41231339 | 0.29591915 | 1.3933312 | 0.16351966 | 0.71642393 |
| JHY | 70.5165624 | 0.28171768 | 0.34636924 | 0.81334495 | 0.41602031 | 0.88242985 |
| HSPA8 | 460.815413 | 0.14847694 | 0.4839249 | 0.30681815 | 0.75898179 | 0.96180522 |
| VWA5A | 86.7006699 | 0.2264554 | 0.23430008 | 0.96651868 | 0.33378471 | 0.8445213 |
| TBRG1 | 138.188761 | 0.0271743 | 0.27494989 | 0.09883363 | 0.92127037 | 0.9904507 |
| SIAE | 142.085814 | 0.90179863 | 0.24211418 | 3.72468327 | 0.00019556 | 0.03105265 |
| SPA17 | 44.0823678 | -0.0444813 | 0.33053514 | -0.1345736 | 0.89294904 | 0.98597539 |
| MSANTD2 | 190.619906 | 0.38190933 | 0.23077667 | 1.65488713 | 0.0979474 | 0.61685067 |
| TMEM218 | 40.723017 | 0.54668549 | 0.32128318 | 1.70156897 | 0.0888362 | 0.60691223 |
| EI24 | 188.433268 | 0.10109078 | 0.30520876 | 0.33121848 | 0.74047946 | 0.96084411 |
| STT3A | 86.8030252 | -0.3088494 | 0.2690146 | -1.1480767 | 0.25093689 | 0.80240987 |
| FAM118B | 66.0993527 | -0.32101 | 0.30192146 | -1.0632235 | 0.28768062 | 0.8303903 |
| SRPRA | 118.389849 | 0.00687995 | 0.30671907 | 0.0224308 | 0.98210431 | 0.99713749 |
| FOXRED1 | 65.1982595 | -0.1840975 | 0.34486275 | -0.5338283 | 0.5934603 | 0.93491607 |
| TIRAP | 39.7175287 | 0.68169841 | 0.29225199 | 2.33257066 | 0.01967069 | 0.3368445 |
| DCPS | 329.275457 | -1.7515932 | 0.58891759 | -2.9742587 | 0.00293697 | 0.13990675 |
| ST3GAL4 | 485.308978 | -2.482947 | 0.72014695 | -3.4478338 | 0.0005651 | 0.05789336 |
| ETS1 | 260.169387 | -0.0722857 | 0.54821183 | -0.1318573 | 0.89509719 | 0.98597539 |
| KCNJ1 | 597.333512 | 1.14563029 | 0.53415074 | 2.14476962 | 0.03197127 | 0.41517597 |
| ARHGAP32 | 582.591592 | -0.047719 | 0.23512857 | -0.2029484 | 0.8391754 | 0.97446628 |
| BARX2 | 100.894809 | 0.96739442 | 0.84714095 | 1.14195213 | 0.25347392 | 0.80414079 |
| TMEM45B | 381.786012 | 0.96394136 | 0.42216861 | 2.28330893 | 0.02241218 | 0.35859484 |
| NFRKB | 101.800922 | 0.0667756 | 0.21419291 | 0.3117545 | 0.7552271 | 0.96120879 |
| PRDM10 | 111.819916 | -0.2995233 | 0.32103236 | -0.9330004 | 0.35081978 | 0.85477058 |
| APLP2 | 843.818097 | 0.57009779 | 0.22209147 | 2.56695046 | 0.01025973 | 0.24660995 |
| ST14 | 270.33243 | -0.0583229 | 0.2522062 | -0.2312509 | 0.81711987 | 0.96878439 |
| ZBTB44 | 510.992683 | 0.38301627 | 0.26855088 | 1.42623355 | 0.15380094 | 0.70756533 |
| AP002986.1 | 44.321867 | 1.24053528 | 0.64855653 | 1.91276353 | 0.05577833 | 0.52053155 |
| SNX19 | 178.302856 | -0.2121813 | 0.35536263 | -0.5970839 | 0.55045135 | 0.92781939 |
| JAM3 | 177.049755 | 0.02298357 | 0.31833696 | 0.07219887 | 0.94244364 | 0.99265578 |
| NCAPD3 | 69.974597 | 0.23140017 | 0.28395013 | 0.81493242 | 0.41511099 | 0.88242985 |
| VPS26B | 112.357964 | -0.1678938 | 0.33576419 | -0.5000349 | 0.61705048 | 0.94233994 |
| THYN1 | 36.9191514 | 0.55401999 | 0.31481273 | 1.75983982 | 0.07843497 | 0.5805074 |
| ACAD8 | 67.9377749 | 0.80054976 | 0.27006134 | 2.96432568 | 0.00303347 | 0.14138755 |
| ZMYND11 | 600.617512 | 0.41822934 | 0.21500567 | 1.94520141 | 0.05175074 | 0.50451388 |
| DIP2C | 1290.97161 | -1.0288808 | 0.36276885 | -2.8361883 | 0.00456555 | 0.16593973 |
| LARP4B | 953.146223 | -0.1136642 | 0.20260274 | -0.5610201 | 0.57478384 | 0.93275827 |
| GTPBP4 | 81.3476043 | -0.2353816 | 0.2345393 | -1.0035914 | 0.31557561 | 0.84285425 |
| IDI1 | 251.98914 | -0.6787007 | 0.66873559 | -1.0149015 | 0.3101528 | 0.84213522 |
| WDR37 | 263.220433 | -0.2252216 | 0.24069794 | -0.9357024 | 0.34942645 | 0.8535343 |
| PFKP | 640.266538 | -1.6036185 | 0.73955009 | -2.1683703 | 0.03013052 | 0.40968855 |
| PITRM1 | 94.9404253 | -0.5944609 | 0.20450494 | -2.9068293 | 0.00365112 | 0.14773661 |
| KLF6 | 1108.94764 | -1.4887711 | 0.5840121 | -2.5492127 | 0.01079664 | 0.25256429 |
| AKR1C1 | 169.834971 | -1.0586798 | 1.49695206 | -0.7072236 | 0.47942756 | 0.90301956 |
| AKR1C3 | 120.514467 | -0.0883782 | 0.65795385 | -0.1343228 | 0.89314729 | 0.98597539 |

| | | | | | | |
|---|---|---|---|---|---|---|
| NET1 | 48.3799193 | 0.72790143 | 0.28642154 | 2.54136411 | 0.01104209 | 0.25715792 |
| ASB13 | 43.3780838 | 0.23730455 | 0.40713221 | 0.58286853 | 0.55998181 | 0.92934517 |
| FAM208B | 416.107007 | -0.1420322 | 0.22614145 | -0.6280679 | 0.52995944 | 0.92181514 |
| GDI2 | 322.452808 | -0.1421448 | 0.2094309 | -0.6787192 | 0.49731581 | 0.90839719 |
| ANKRD16 | 42.3599019 | -0.1990135 | 0.34230398 | -0.5813941 | 0.56097488 | 0.92934517 |
| FBH1 | 210.127434 | -0.0351763 | 0.19747333 | -0.1781318 | 0.85861947 | 0.97786699 |
| IL15RA | 52.4774455 | -0.2566969 | 0.42024778 | -0.6108227 | 0.54131697 | 0.92443671 |
| RBM17 | 194.525334 | -0.088708 | 0.23735369 | -0.3737374 | 0.7085997 | 0.95752169 |
| PFKFB3 | 2322.21366 | -0.1475382 | 0.29857103 | -0.4941479 | 0.62120174 | 0.94377998 |
| AL137145.2 | 380.657653 | -0.1879342 | 0.49167917 | -0.3822293 | 0.70229126 | 0.95672887 |
| AL137145.1 | 43.0030958 | 0.51500392 | 0.57789428 | 0.89117325 | 0.37283623 | 0.86201082 |
| PRKCQ | 160.720933 | 0.20278561 | 0.46962644 | 0.43180195 | 0.66588536 | 0.95243327 |
| SFMBT2 | 272.087861 | 0.13804661 | 0.35126308 | 0.39300063 | 0.69431902 | 0.95672887 |
| KIN | 81.0227603 | -0.1757258 | 0.21607924 | -0.8132472 | 0.41607637 | 0.88242985 |
| ATP5F1C | 389.053221 | 0.26286451 | 0.17818373 | 1.47524422 | 0.1401469 | 0.68408298 |
| TAF3 | 470.925771 | -0.0775408 | 0.17835203 | -0.4347626 | 0.66373477 | 0.95187756 |
| GATA3 | 2027.51207 | -0.0043743 | 0.27314301 | -0.0160148 | 0.98722259 | 0.99791717 |
| GATA3-AS1 | 238.936081 | -0.6488514 | 0.4513621 | -1.4375407 | 0.15056441 | 0.7029344 |
| USP6NL | 386.331819 | 0.15673688 | 0.14361441 | 1.09137294 | 0.27510881 | 0.8202929 |
| ECHDC3 | 87.9688896 | 1.40214246 | 0.39405352 | 3.55825385 | 0.00037333 | 0.04296095 |
| PROSER2 | 157.986669 | 0.66075095 | 0.35648709 | 1.853506 | 0.06380987 | 0.54850785 |
| UPF2 | 334.986443 | -0.1882102 | 0.18252483 | -1.0311484 | 0.30247125 | 0.83709431 |
| DHTKD1 | 130.084938 | -0.0490223 | 0.22472533 | -0.2181432 | 0.82731752 | 0.97069946 |
| SEC61A2 | 57.7890785 | -0.0259695 | 0.26867823 | -0.0966565 | 0.92299922 | 0.99051393 |
| NUDT5 | 133.240616 | -0.1465665 | 0.23624047 | -0.6204122 | 0.53498643 | 0.92303275 |
| CDC123 | 136.734322 | -0.0174302 | 0.21517345 | -0.0810053 | 0.93543773 | 0.99243654 |
| CAMK1D | 100.244939 | 0.24067157 | 0.81050506 | 0.29694024 | 0.76651213 | 0.96285313 |
| OPTN | 136.597028 | -0.4194244 | 0.36692889 | -1.1430672 | 0.25301069 | 0.80350545 |
| PHYH | 109.2179 | 0.51125027 | 0.37073134 | 1.3790317 | 0.16788498 | 0.72374931 |
| SEPHS1 | 116.123137 | 0.06207072 | 0.24583448 | 0.25248989 | 0.80066243 | 0.96603066 |
| BEND7 | 307.2374 | 0.21086286 | 0.21660143 | 0.97350628 | 0.33030174 | 0.84442294 |
| PRPF18 | 39.770121 | -0.0006984 | 0.36976469 | -0.0018888 | 0.99849299 | 0.99967893 |
| AL157392.5 | 93.4287454 | -0.1157573 | 0.3174503 | -0.364647 | 0.71537491 | 0.95791799 |
| FRMD4A | 347.695138 | 0.59379226 | 0.43636638 | 1.36076538 | 0.17358785 | 0.73030936 |
| FAM107B | 1914.25123 | 0.04612357 | 0.42406854 | 0.10876441 | 0.91338935 | 0.98830345 |
| DCLRE1C | 103.552469 | -0.3179118 | 0.28085847 | -1.1319287 | 0.25766442 | 0.80709747 |
| RPP38 | 152.737965 | 0.00976665 | 0.23238948 | 0.04202706 | 0.96647713 | 0.99637506 |
| NMT2 | 200.526034 | -0.7120129 | 0.33170952 | -2.1464951 | 0.0318335 | 0.41517597 |
| FAM171A1 | 805.499663 | 0.23781065 | 0.40555468 | 0.58638369 | 0.5576177 | 0.92901206 |
| MINDY3 | 220.313317 | -0.1170164 | 0.21354329 | -0.5479748 | 0.58370919 | 0.93473293 |
| PTER | 274.563172 | -0.0642242 | 0.21520441 | -0.2984333 | 0.76537244 | 0.96285313 |
| RSU1 | 253.44206 | -0.3800982 | 0.33355269 | -1.1395447 | 0.25447602 | 0.80414079 |
| CUBN | 114.15157 | 0.84063828 | 0.51638421 | 1.6279318 | 0.10353935 | 0.62722102 |
| TRDMT1 | 57.4648111 | 0.1079373 | 0.31541687 | 0.34220522 | 0.73219646 | 0.96008876 |
| VIM-AS1 | 36.1911455 | 0.56771398 | 0.56987031 | 0.9962161 | 0.31914516 | 0.84285425 |
| VIM | 169.390309 | -0.8524537 | 0.61983214 | -1.3752976 | 0.16903919 | 0.72596839 |
| HACD1 | 312.586652 | -0.90799 | 0.51546114 | -1.7615101 | 0.0781521 | 0.5795765 |
| STAM | 807.762315 | -1.3420414 | 0.50534315 | -2.6557032 | 0.00791432 | 0.21826871 |
| AL450384.2 | 63.6309485 | -0.1899943 | 0.44427547 | -0.4276498 | 0.66890608 | 0.95298441 |
| NSUN6 | 212.901292 | -0.3096662 | 0.28239476 | -1.0965721 | 0.2728285 | 0.81922962 |
| ARL5B | 104.803556 | -1.5933293 | 0.62333513 | -2.556136 | 0.01058418 | 0.25038862 |
| MALRD1 | 164.9398 | -0.0644044 | 0.55100344 | -0.1168857 | 0.9069506 | 0.9875066 |
| PLXDC2 | 857.703511 | 0.10314885 | 0.27726931 | 0.37201683 | 0.70988032 | 0.95752169 |
| NEBL | 1355.26891 | 0.37528245 | 0.45565223 | 0.82361597 | 0.4101578 | 0.8801764 |

| | | | | | | |
|---|---|---|---|---|---|---|
| MLLT10 | 855.646491 | -0.3746208 | 0.25246967 | -1.4838248 | 0.13785541 | 0.68146162 |
| DNAJC1 | 650.620743 | -0.1612517 | 0.18299523 | -0.8811796 | 0.37822062 | 0.86512157 |
| BMI1 | 48.4528218 | 0.09382898 | 0.35748535 | 0.26246943 | 0.79295955 | 0.9653598 |
| PIP4K2A | 488.982773 | 0.02211973 | 0.29386366 | 0.07527208 | 0.93999824 | 0.99243654 |
| MSRB2 | 147.374194 | 0.03832238 | 0.29940523 | 0.12799503 | 0.89815291 | 0.9863394 |
| OTUD1 | 63.1224998 | -0.1236202 | 0.32735988 | -0.377628 | 0.70570699 | 0.95730303 |
| KIAA1217 | 2355.90023 | 0.18124039 | 0.23231983 | 0.78013312 | 0.43531253 | 0.88785137 |
| ARHGAP21 | 452.597883 | -0.3790965 | 0.32717312 | -1.158703 | 0.24657727 | 0.80072691 |
| PRTFDC1 | 87.0208177 | 0.62695979 | 0.36819858 | 1.70277623 | 0.08860995 | 0.60673738 |
| PDSS1 | 46.9566742 | -0.4162445 | 0.47210068 | -0.8816859 | 0.37794668 | 0.86501008 |
| ABI1 | 425.355121 | -0.1253435 | 0.20867737 | -0.600657 | 0.54806849 | 0.92684569 |
| ANKRD26 | 383.527279 | -0.0975039 | 0.26642302 | -0.3659741 | 0.71438437 | 0.95791799 |
| YME1L1 | 361.270146 | -0.1430191 | 0.16891758 | -0.8466797 | 0.39717365 | 0.87462201 |
| MASTL | 52.5029706 | -0.4564673 | 0.25117957 | -1.8172946 | 0.06917202 | 0.55917298 |
| ACBD5 | 257.227264 | 0.07198485 | 0.17602404 | 0.40894897 | 0.68257711 | 0.95574492 |
| RAB18 | 341.4094 | -0.1360772 | 0.19761104 | -0.6886115 | 0.49106778 | 0.90730741 |
| MPP7 | 451.813573 | -0.1000474 | 0.26251708 | -0.3811082 | 0.70312299 | 0.95672887 |
| AL358234.1 | 76.6627022 | 0.04653537 | 0.68179854 | 0.06825384 | 0.94558357 | 0.99299085 |
| WAC-AS1 | 209.394094 | -0.1588207 | 0.36524083 | -0.4348382 | 0.66367986 | 0.95187756 |
| WAC | 1255.18474 | -0.3113523 | 0.171445 | -1.8160477 | 0.06936306 | 0.55917298 |
| SVIL | 468.781859 | -0.6888757 | 0.25685319 | -2.6819823 | 0.00731873 | 0.2101379 |
| MTPAP | 71.0401687 | -0.0101492 | 0.23558284 | -0.0430813 | 0.96563674 | 0.99637506 |
| MAP3K8 | 134.510732 | -0.5781837 | 0.32248693 | -1.7928902 | 0.07299048 | 0.56668778 |
| ZNF438 | 324.411759 | -0.190178 | 0.25545642 | -0.7444635 | 0.45659611 | 0.89642698 |
| ZEB1 | 60.555923 | -1.9378987 | 0.51303197 | -3.7773449 | 0.00015851 | 0.02595588 |
| ARHGAP12 | 682.96247 | -0.2761871 | 0.1373633 | -2.0106322 | 0.04436432 | 0.48009422 |
| KIF5B | 442.700171 | 0.02800992 | 0.17115996 | 0.16364763 | 0.87000855 | 0.98202612 |
| EPC1 | 585.654168 | 0.12357044 | 0.24208411 | 0.51044424 | 0.60974027 | 0.94003018 |
| CCDC7 | 477.911157 | -0.1546649 | 0.33126695 | -0.4668889 | 0.64057939 | 0.94935885 |
| ITGB1 | 663.554374 | 0.02145608 | 0.30543456 | 0.0702477 | 0.94399651 | 0.99265578 |
| NRP1 | 399.589457 | 0.28313972 | 0.64045742 | 0.44208984 | 6.58E-01 | 0.95113765 |
| PARD3 | 2953.60663 | 0.10752212 | 0.22050029 | 0.48762801 | 0.62581335 | 0.94511671 |
| CUL2 | 182.424622 | -0.0682348 | 0.17754395 | -0.3843264 | 0.70073655 | 0.95672887 |
| CREM | 978.880953 | -1.1390125 | 0.69062272 | -1.6492543 | 0.09909554 | 0.61707826 |
| CCNY | 1183.92754 | -0.1541274 | 0.21274452 | -0.7244717 | 0.46877618 | 0.90026987 |
| AL117336.3 | 113.22244 | -0.0160204 | 0.3316365 | -0.048307 | 0.96147156 | 0.99635733 |
| ZNF248 | 338.035266 | 0.06059131 | 0.2634442 | 0.22999676 | 0.81809429 | 0.96883031 |
| ZNF25 | 62.6457103 | 0.30938134 | 0.36031386 | 0.858644 | 0.39053695 | 0.87004477 |
| ZNF33A | 603.829349 | -0.2671167 | 0.26728707 | -0.9993626 | 0.31761905 | 0.84285425 |
| AL117339.5 | 38.0528339 | -0.086345 | 0.30480934 | -0.2832753 | 0.7769658 | 0.96454202 |
| ZNF33B | 420.832909 | -0.0670149 | 0.26159028 | -0.2561828 | 0.79780968 | 0.9653598 |
| BMS1 | 154.720673 | -0.2895997 | 0.185488 | -1.5612854 | 0.11845642 | 0.65447949 |
| CSGALNACT2 | 125.194137 | 0.02945677 | 0.27590957 | 0.10676239 | 0.91497748 | 0.98868699 |
| RASGEF1A | 49.3389884 | -0.5060502 | 0.44176035 | -1.1455311 | 0.25198922 | 0.80268905 |
| FXYD4 | 1628.65458 | 0.29821831 | 0.65277919 | 0.45684408 | 0.64778312 | 0.95061388 |
| HNRNPF | 368.81537 | -0.1934215 | 0.2924409 | -0.6614037 | 0.50835346 | 0.91250907 |
| ZNF487 | 83.2117134 | -0.441494 | 0.41717982 | -1.0582823 | 0.28992678 | 0.83103268 |
| ZNF32 | 94.5394837 | 0.1884994 | 0.27582348 | 0.68340593 | 0.49435037 | 0.90839719 |
| TMEM72-AS | 113.88352 | 0.7507068 | 0.59476636 | 1.26218772 | 0.20688125 | 0.77200549 |
| TMEM72 | 345.66373 | 2.16421228 | 0.65042634 | 3.32737488 | 0.00087668 | 0.07013468 |
| RASSF4 | 40.7960944 | 1.0989075 | 0.52581134 | 2.08992733 | 0.03662433 | 0.43729328 |
| DEPP1 | 52.1170845 | 1.39296342 | 0.58071141 | 2.39871888 | 0.01645254 | 0.30723893 |
| ZNF22 | 85.8568746 | 0.58408201 | 0.29519598 | 1.97862456 | 0.04785829 | 0.48600281 |
| 8-Mar | 288.460638 | 0.08492786 | 0.24140063 | 0.3518129 | 0.72497859 | 0.95963742 |

| | | | | | | |
|---|---|---|---|---|---|---|
| ZFAND4 | 79.9859849 | 0.37599521 | 0.23404512 | 1.60650735 | 0.10816248 | 0.63533473 |
| WASHC2C | 106.815641 | -0.4110595 | 0.34783354 | -1.1817706 | 0.23729672 | 0.7980661 |
| AGAP4 | 137.965163 | 0.22765006 | 0.33726957 | 0.67497953 | 0.49968877 | 0.90839719 |
| TIMM23 | 124.27734 | -0.1299348 | 0.26608513 | -0.4883203 | 0.62532296 | 0.94508035 |
| NCOA4 | 196.634786 | 0.4047867 | 0.27467131 | 1.4737131 | 0.14055885 | 0.68482417 |
| PTPN20 | 230.566259 | 0.54091249 | 0.40492498 | 1.33583381 | 0.1816036 | 0.74315969 |
| AGAP9 | 152.123752 | 0.80610687 | 0.27493986 | 2.93193891 | 0.00336853 | 0.14358074 |
| MAPK8 | 945.038763 | -0.4747749 | 0.27063177 | -1.7543206 | 0.07937559 | 0.58457919 |
| ARHGAP22 | 44.4851941 | -1.7906389 | 0.41650064 | -4.2992465 | 1.71E-05 | 0.00544261 |
| ERCC6 | 142.463876 | 0.20795063 | 0.24156638 | 0.86084258 | 0.38932475 | 0.87004477 |
| OGDHL | 231.012834 | 1.23164961 | 0.39756511 | 3.09798212 | 0.00194843 | 0.11037603 |
| PARG | 357.673184 | -0.1507364 | 0.20447976 | -0.7371705 | 0.46101869 | 0.89799005 |
| TIMM23B | 266.117324 | -0.3565777 | 0.38385546 | -0.9289374 | 0.35292153 | 0.85545893 |
| LINC00843 | 73.5656572 | 0.09179194 | 0.35625633 | 0.25765702 | 0.79667162 | 0.9653598 |
| AGAP6 | 111.857384 | -0.1607395 | 0.23915661 | -0.6721099 | 0.50151375 | 0.90900452 |
| WASHC2A | 98.2664831 | -0.6221378 | 0.29375563 | -2.1178754 | 0.03418562 | 0.43050784 |
| SGMS1 | 1455.35676 | 0.46791759 | 0.2204785 | 2.12228217 | 0.03381405 | 0.4285021 |
| SGMS1-AS1 | 37.3245096 | 0.2493175 | 0.3679935 | 0.67750518 | 0.49808548 | 0.90839719 |
| ASAH2B | 51.9670686 | -0.2320095 | 0.36415333 | -0.6371204 | 0.52404641 | 0.91871245 |
| PRKG1 | 1636.88218 | 0.46422168 | 0.38323889 | 1.21131151 | 0.22577604 | 0.78982769 |
| CSTF2T | 33.6005989 | 0.16710192 | 0.35264827 | 0.47384868 | 0.63560782 | 0.94807428 |
| IPMK | 111.96959 | -0.3114179 | 0.36496498 | -0.8532815 | 0.3935032 | 0.87223214 |
| CISD1 | 244.660974 | 0.41416601 | 0.24318725 | 1.7030745 | 0.08855413 | 0.60673738 |
| UBE2D1 | 83.6314169 | -0.2737834 | 0.30074155 | -0.910361 | 0.36263215 | 0.85694172 |
| TFAM | 93.0994337 | 0.13284079 | 0.24766966 | 0.5363628 | 0.59170784 | 0.93491607 |
| BICC1 | 4333.08272 | 1.03234057 | 0.36995172 | 2.79047379 | 0.0052631 | 0.17920428 |
| PHYHIPL | 41.2720206 | 0.80554747 | 0.65918438 | 1.22203664 | 0.22169377 | 0.78752765 |
| CCDC6 | 240.703793 | 0.14871027 | 0.23318243 | 0.63774218 | 0.52364152 | 0.91871245 |
| ANK3 | 8417.15534 | 0.50475184 | 0.14770543 | 3.41728693 | 6.32E-04 | 5.95E-02 |
| RHOBTB1 | 64.0100402 | -0.8483426 | 0.4269741 | -1.9868714 | 0.04693665 | 0.48600281 |
| CABCOCO1 | 68.6059043 | 0.93301575 | 0.36770954 | 2.53737164 | 0.01116883 | 0.25895876 |
| ARID5B | 1146.41567 | -0.0812153 | 0.57879303 | -0.1403184 | 0.88840839 | 0.98551241 |
| ADO | 33.522221 | 0.67412457 | 0.40296118 | 1.67292684 | 0.0943417 | 0.61330068 |
| NRBF2 | 36.5323641 | -0.0109412 | 0.31932923 | -0.0342632 | 0.97266729 | 0.99656026 |
| JMJD1C | 2830.41985 | -0.3086131 | 0.25614661 | -1.2048299 | 0.22826896 | 0.79042788 |
| REEP3 | 629.241619 | -0.4958473 | 0.50397713 | -0.9838686 | 0.32518011 | 0.84285425 |
| LINC01515 | 195.275907 | -0.1130753 | 0.39935122 | -0.2831475 | 0.77706377 | 0.96454202 |
| CTNNA3 | 49.7712739 | -0.1179505 | 0.49344264 | -0.2390359 | 0.81107774 | 0.9680627 |
| SIRT1 | 118.563195 | -0.2062716 | 0.25679754 | -0.803246 | 0.42183258 | 0.88371155 |
| HERC4 | 848.65544 | -0.4457885 | 0.20327546 | -2.1930267 | 0.02830546 | 0.40459506 |
| PBLD | 212.005811 | -0.3945829 | 0.25770845 | -1.5311211 | 0.12573946 | 0.6674937 |
| HNRNPH3 | 365.085919 | 0.00886094 | 0.19231815 | 0.04607438 | 0.96325097 | 0.99637506 |
| RUFY2 | 322.878104 | -0.1644191 | 0.20323828 | -0.8089969 | 0.41851693 | 0.88242985 |
| DNA2 | 37.5167011 | 0.03444614 | 0.42366822 | 0.08130452 | 0.93519978 | 0.99243654 |
| SLC25A16 | 252.013617 | -0.2765968 | 0.30576335 | -0.9046108 | 0.36567159 | 0.85866868 |
| TET1 | 158.735759 | 0.2599867 | 0.44229528 | 0.58781252 | 0.55665814 | 0.92872652 |
| CCAR1 | 542.217791 | -0.1672176 | 0.24409191 | -0.6850598 | 0.49330616 | 0.90826574 |
| STOX1 | 76.8773213 | 0.34892987 | 0.49228376 | 0.70879825 | 0.47844969 | 0.90263775 |
| DDX50 | 207.12499 | -0.0409742 | 0.18596004 | -0.2203389 | 0.8256072 | 0.97032224 |
| DDX21 | 200.58915 | -0.3914114 | 0.45086814 | -0.8681284 | 0.38532403 | 0.86749643 |
| KIF1BP | 117.246979 | 0.12557869 | 0.25820618 | 0.48635044 | 0.62671872 | 0.94511671 |
| VPS26A | 224.013342 | -0.6111322 | 0.2706747 | -2.2578104 | 0.02395748 | 0.37077504 |
| SUPV3L1 | 109.910235 | -0.1867807 | 0.29605341 | -0.6309021 | 0.52810452 | 0.92106733 |
| HKDC1 | 409.8574 | -0.31913 | 0.93594759 | -0.34097 | 0.7331262 | 0.96035038 |

| | | | | | | |
|---|---|---|---|---|---|---|
| HK1 | 499.55359 | -0.1905533 | 0.3851621 | -0.4947352 | 0.62078703 | 0.94369714 |
| TSPAN15 | 219.115146 | 0.27679235 | 0.27710667 | 0.99886572 | 0.31785975 | 0.84285425 |
| FAM241B | 31.6918811 | 0.04356904 | 0.34071114 | 0.12787677 | 0.89824649 | 0.9863394 |
| H2AFY2 | 105.506025 | 0.42141523 | 0.39222647 | 1.07441812 | 0.28263532 | 0.82666614 |
| AIFM2 | 72.2724425 | 0.04234153 | 0.32721406 | 0.12940009 | 0.89704108 | 0.98601802 |
| TYSND1 | 50.0754081 | 0.31209623 | 0.2935985 | 1.06300351 | 0.28778036 | 0.8303903 |
| SAR1A | 160.507953 | -0.1919897 | 0.22144623 | -0.8669811 | 0.38595234 | 0.86831175 |
| PPA1 | 247.620761 | -0.3650558 | 0.40555291 | -0.9001434 | 0.36804394 | 0.85902833 |
| EIF4EBP2 | 228.033812 | 0.0913734 | 0.22269215 | 0.41031261 | 0.68157664 | 0.95574492 |
| SGPL1 | 356.470321 | -0.4219683 | 0.30394929 | -1.3882853 | 0.16505019 | 0.71743095 |
| PCBD1 | 234.305626 | 0.77593515 | 0.54786395 | 1.41629167 | 0.15669013 | 0.71126849 |
| UNC5B | 34.8627213 | 0.43376995 | 0.75027724 | 0.57814622 | 0.56316541 | 0.9306324 |
| PSAP | 1130.34439 | 0.18378336 | 0.30131678 | 0.60993406 | 0.54190549 | 0.92479556 |
| CHST3 | 112.696437 | 0.65058946 | 0.42221439 | 1.54089837 | 0.12334152 | 0.66519057 |
| ASCC1 | 212.044497 | -0.2634955 | 0.26719775 | -0.9861441 | 0.32406241 | 0.84285425 |
| ANAPC16 | 604.967561 | 0.1374413 | 0.22506092 | 0.61068488 | 0.54140822 | 0.92443671 |
| DDIT4 | 1153.15052 | -1.475806 | 0.69505604 | -2.1232906 | 0.0337295 | 0.42838847 |
| DNAJB12 | 76.2102083 | -0.2240082 | 0.30423395 | -0.7363023 | 0.46154674 | 0.89799005 |
| MICU1 | 766.691103 | -0.3029854 | 0.17464746 | -1.7348398 | 0.08276918 | 0.59202053 |
| MCU | 1123.12458 | -0.2485756 | 0.33629404 | -0.7391614 | 4.60E-01 | 0.89753337 |
| P4HA1 | 554.942878 | -0.8518807 | 0.6710223 | -1.2695267 | 0.20425327 | 0.77193931 |
| ECD | 113.095572 | -0.070511 | 0.30280885 | -0.2328565 | 0.81587281 | 0.96844493 |
| FAM149B1 | 173.920221 | 0.15379367 | 0.23568825 | 0.65253008 | 0.51405927 | 0.91544604 |
| DNAJC9 | 140.748033 | 0.00449756 | 0.25822622 | 0.01741713 | 0.98610385 | 0.99768314 |
| MRPS16 | 127.867153 | 0.21703658 | 0.27004514 | 0.80370481 | 0.42156747 | 0.88360543 |
| CFAP70 | 245.361711 | 0.56026777 | 0.41956685 | 1.33534803 | 0.18176246 | 0.74315969 |
| ANXA7 | 326.385681 | -0.0408403 | 0.17463806 | -0.2338565 | 0.81509635 | 0.96839471 |
| PPP3CB | 342.224155 | -0.2442711 | 0.21763834 | -1.1223717 | 0.26170445 | 0.80847797 |
| USP54 | 212.9767 | -0.0608393 | 0.23721832 | -0.2564698 | 0.79758811 | 0.9653598 |
| AGAP5 | 40.3777779 | -0.3613324 | 0.27083097 | -1.3341621 | 0.18215075 | 0.74315969 |
| SEC24C | 99.6560093 | -0.4449133 | 0.23092884 | -1.9266253 | 0.05402635 | 0.51332378 |
| FUT11 | 197.398817 | -1.1265733 | 0.67982975 | -1.6571404 | 0.09749109 | 0.61685067 |
| CHCHD1 | 130.816131 | 0.26013737 | 0.4546426 | 0.57217992 | 0.56720009 | 0.93122667 |
| ZSWIM8 | 146.35581 | -0.0972319 | 0.20223208 | -0.4807934 | 0.63066331 | 0.94596555 |
| CAMK2G | 104.058165 | -0.1689349 | 0.3461127 | -0.4880923 | 0.62548447 | 0.94508035 |
| VCL | 600.41398 | 0.05260882 | 0.26548979 | 0.19815761 | 0.84292175 | 0.97560439 |
| AP3M1 | 80.1457372 | -0.146559 | 0.24845167 | -0.5898893 | 0.55526489 | 0.92872652 |
| ADK | 2198.71613 | 0.00384242 | 0.16466629 | 0.02333461 | 0.98138336 | 0.99696835 |
| KAT6B | 604.69284 | -0.2928942 | 0.27132487 | -1.0794962 | 0.2803666 | 0.82628479 |
| SAMD8 | 182.017954 | 0.08019574 | 0.22595392 | 0.35492076 | 0.72264895 | 0.95926626 |
| VDAC2 | 215.574237 | 0.51052109 | 0.39307154 | 1.29879943 | 0.19401277 | 0.76151829 |
| COMTD1 | 76.9165615 | 0.2083149 | 0.39539538 | 0.52685213 | 0.59829628 | 0.93728325 |
| ZNF503-AS1 | 50.487867 | 0.25488057 | 0.41090408 | 0.62029215 | 0.53506547 | 0.92303275 |
| AC010997.3 | 73.2058347 | 0.47148357 | 0.6545317 | 0.72033726 | 0.47131737 | 0.90123338 |
| ZNF503-AS2 | 45.3771467 | -0.3723942 | 0.4263617 | -0.8734231 | 0.38243248 | 0.86593382 |
| LRMDA | 3673.47355 | -0.1306228 | 0.28996659 | -0.4504752 | 0.65236783 | 0.95092279 |
| DLG5 | 180.703316 | -0.4360187 | 0.28741197 | -1.5170512 | 0.12925376 | 0.67207756 |
| POLR3A | 93.908421 | -0.3390076 | 0.25438224 | -1.3326699 | 0.18264015 | 0.74315969 |
| RPS24 | 1136.32392 | 0.07134817 | 0.47831671 | 0.14916512 | 0.88142334 | 0.98335905 |
| ZMIZ1 | 542.085489 | 0.12306447 | 0.42727176 | 0.28802387 | 0.77332847 | 0.96388926 |
| PPIF | 71.1115501 | 0.16332146 | 0.49388848 | 0.33068489 | 0.74088252 | 0.96084411 |
| NUTM2B-AS | 544.379128 | -0.0882446 | 0.36808726 | -0.2397383 | 0.81053315 | 0.9680627 |
| TMEM254 | 66.0044474 | 0.15053014 | 0.30131825 | 0.49957193 | 0.61737653 | 0.94233994 |
| ANXA11 | 660.16523 | 0.31010356 | 0.24946435 | 1.24307763 | 0.21383923 | 0.77890374 |

| | | | | | | |
|---|---|---|---|---|---|---|
| FAM213A | 277.454743 | -0.3751301 | 0.4129937 | -0.9083192 | 0.36370962 | 0.85694172 |
| TSPAN14 | 173.05157 | -0.1543624 | 0.22375565 | -0.6898703 | 4.90E-01 | 0.90730741 |
| NRG3 | 125.628403 | -0.9883975 | 0.90862141 | -1.0877991 | 0.27668383 | 0.82198569 |
| GHITM | 256.668218 | 0.37697284 | 0.36591536 | 1.03021869 | 0.30290736 | 0.83709431 |
| CCSER2 | 447.898692 | -0.3303657 | 0.25594083 | -1.2907894 | 0.19677671 | 0.76478825 |
| WAPL | 505.638135 | -0.0012582 | 0.19070151 | -0.0065977 | 0.99473584 | 0.99919666 |
| BMPR1A | 567.376027 | 0.04470356 | 0.15420609 | 0.2898949 | 0.77189665 | 0.96388926 |
| ADIRF | 51.7087352 | -0.5079828 | 0.56906167 | -0.8926674 | 0.37203531 | 0.86201082 |
| GLUD1 | 272.520355 | 0.44903277 | 0.22822808 | 1.96747384 | 0.04912861 | 0.49081776 |
| SHLD2 | 407.502794 | 0.22734766 | 0.28532296 | 0.79680813 | 0.42556247 | 0.88403732 |
| NUTM2A-AS | 562.757403 | -0.468559 | 0.19576148 | -2.39352 | 0.01668757 | 0.30789747 |
| MINPP1 | 59.6069686 | 0.38985183 | 0.29715202 | 1.31196089 | 0.18953333 | 0.75619145 |
| PAPSS2 | 429.388384 | 0.63746586 | 0.4887675 | 1.3042313 | 0.19215473 | 0.7597012 |
| ATAD1 | 282.331378 | 0.17250637 | 0.20875876 | 0.82634314 | 0.40860947 | 0.87948803 |
| PTEN | 882.193648 | -0.0188882 | 0.22596577 | -0.0835888 | 0.93338339 | 0.99243654 |
| RNLS | 200.15984 | -0.4657624 | 0.17613978 | -2.6442773 | 0.00818656 | 0.22277119 |
| LIPA | 116.65149 | -0.2426121 | 0.3973873 | -0.6105179 | 0.54151879 | 0.92443671 |
| IFIT2 | 95.6682629 | -2.6595385 | 1.00568942 | -2.6444929 | 0.00818134 | 0.22277119 |
| IFIT3 | 178.340804 | -2.6403489 | 0.92744508 | -2.8469059 | 0.00441464 | 0.16424479 |
| PANK1 | 73.6053751 | 0.56989012 | 0.36108706 | 1.57826239 | 0.11450534 | 0.65008305 |
| KIF20B | 55.2690233 | -0.0789381 | 0.24046316 | -0.3282753 | 0.74270355 | 0.96086911 |
| RPP30 | 53.6250862 | 0.07540375 | 0.29411396 | 0.25637596 | 0.79766055 | 0.9653598 |
| PCGF5 | 869.173068 | -0.8313182 | 0.47910067 | -1.735164 | 0.08271177 | 0.59202053 |
| HECTD2 | 237.794378 | 0.26194718 | 0.17259403 | 1.51770707 | 0.12908826 | 0.67206108 |
| PPP1R3C | 136.998192 | -1.5491384 | 0.86579125 | -1.7892747 | 0.07357059 | 0.56734347 |
| TNKS2 | 492.593373 | -0.2627759 | 0.14300331 | -1.8375511 | 0.06612857 | 0.55444578 |
| BTAF1 | 446.999132 | -0.5481658 | 0.40806206 | -1.3433392 | 0.17916217 | 0.74042867 |
| CPEB3 | 711.507879 | 0.44462373 | 0.29653493 | 1.49939746 | 0.13377055 | 0.67725381 |
| 5-Mar | 228.490296 | -0.0392824 | 0.24760634 | -0.1586485 | 0.87394582 | 0.98246825 |
| IDE | 180.913869 | -0.0461646 | 0.24715193 | -0.1867863 | 0.85182821 | 0.97718353 |
| EXOC6 | 242.631105 | -0.4219033 | 0.26968936 | -1.5644048 | 0.11772253 | 0.65447949 |
| MYOF | 490.593387 | -0.1973979 | 0.43754294 | -0.4511508 | 0.65188083 | 0.95079106 |
| FRA10AC1 | 173.860427 | -0.327287 | 0.23325762 | -1.4031137 | 0.16058293 | 0.71409437 |
| SLC35G1 | 36.4288178 | 0.67120031 | 0.38500542 | 1.7433529 | 0.08127199 | 0.59202053 |
| NOC3L | 76.613269 | -0.0797108 | 0.24214863 | -0.3291815 | 0.74201853 | 0.96086423 |
| TBC1D12 | 312.485286 | 0.02887072 | 0.24951418 | 0.11570774 | 0.90788417 | 0.9875066 |
| HELLS | 351.833168 | 0.327423 | 0.34742627 | 0.94242442 | 0.34597539 | 0.8523814 |
| PDLIM1 | 59.3740615 | 0.56717049 | 0.38807346 | 1.46150293 | 0.14387747 | 0.69093999 |
| SORBS1 | 231.32613 | 0.36901671 | 0.3766449 | 0.97974701 | 0.32721102 | 0.84381657 |
| ALDH18A1 | 149.44701 | -0.1583509 | 0.37517607 | -0.422071 | 0.6729732 | 0.9544944 |
| TCTN3 | 145.653103 | -0.0179301 | 0.23004399 | -0.0779419 | 0.93787429 | 0.99243654 |
| ENTPD1-AS1 | 484.327032 | 0.29093946 | 0.34247112 | 0.84952992 | 0.39558649 | 0.87426116 |
| AL365273.2 | 38.395824 | 0.09388031 | 0.52694871 | 0.17815835 | 0.85859862 | 0.97786699 |
| CC2D2B | 44.4797779 | 0.2310743 | 0.60559772 | 0.38156401 | 0.70278478 | 0.95672887 |
| ZNF518A | 465.995159 | -0.0751316 | 0.27932155 | -0.268979 | 0.78794583 | 0.96515649 |
| BLNK | 1049.73766 | 0.60731372 | 0.29835064 | 2.03557037 | 0.04179352 | 0.46746387 |
| TM9SF3 | 624.576814 | -0.0365689 | 0.1677561 | -0.2179882 | 0.82743829 | 0.97073248 |
| LCOR | 930.893146 | 0.06316495 | 0.2271617 | 0.27806161 | 0.78096506 | 0.96513523 |
| RRP12 | 87.9328665 | -0.5034393 | 0.29853799 | -1.6863493 | 0.09172855 | 0.61113488 |
| PGAM1 | 390.08918 | 0.11339342 | 0.36940743 | 0.30696032 | 0.75887358 | 0.96180522 |
| EXOSC1 | 84.2595589 | 0.02115259 | 0.22520838 | 0.09392455 | 0.9251691 | 0.99051393 |
| ZDHHC16 | 56.4893816 | 0.11025233 | 0.25137196 | 0.43860234 | 0.6609497 | 0.95127957 |
| MMS19 | 187.511149 | -0.4220433 | 0.22041379 | -1.9147772 | 0.05552092 | 0.51905372 |
| UBTD1 | 131.962959 | 0.54902803 | 0.32252156 | 1.70229872 | 0.08869938 | 0.60673738 |

| | | | | | | |
|---|---|---|---|---|---|---|
| AL355315.1 | 32.9021923 | 0.86565387 | 0.43011297 | 2.01261979 | 0.04415464 | 0.48009422 |
| MORN4 | 71.2573363 | 0.11691394 | 0.32068655 | 0.36457388 | 0.71542952 | 0.95791799 |
| AVPI1 | 62.3303626 | -0.7531 | 0.93948045 | -0.8016133 | 0.42277667 | 0.88371155 |
| ZFYVE27 | 82.768116 | 0.08562287 | 0.23730229 | 0.36081769 | 0.71823573 | 0.95796325 |
| CRTAC1 | 62.1727138 | 1.09822859 | 0.9336741 | 1.17624403 | 0.23949737 | 0.79985101 |
| R3HCC1L | 284.408302 | -0.2884576 | 0.19353912 | -1.4904358 | 0.13610969 | 0.67998335 |
| HPS1 | 71.0352919 | -0.2988624 | 0.30388523 | -0.9834714 | 0.32537551 | 0.84285425 |
| GOT1 | 359.394171 | 0.32272527 | 0.54284792 | 0.59450402 | 0.55217505 | 0.92872652 |
| SLC25A28 | 102.536886 | -0.2449266 | 0.4062707 | -0.6028655 | 0.54659819 | 0.92597739 |
| ENTPD7 | 39.9264682 | -0.1733255 | 0.47738081 | -0.3630759 | 0.71654816 | 0.95791799 |
| CUTC | 135.192861 | -0.0962421 | 0.23551448 | -0.4086462 | 0.68279933 | 0.95574492 |
| COX15 | 124.121655 | -0.0538458 | 0.26603166 | -0.2024038 | 0.83960106 | 0.97452864 |
| DNMBP | 260.267965 | 0.1878195 | 0.33130676 | 0.56690511 | 0.57077866 | 0.93125087 |
| ERLIN1 | 82.2865111 | -0.0040967 | 0.22826798 | -0.0179467 | 0.98568135 | 0.99762072 |
| CHUK | 81.7215704 | -0.1041366 | 0.31214201 | -0.3336193 | 0.73866683 | 0.96078945 |
| CWF19L1 | 96.6328013 | -0.1344455 | 0.27304957 | -0.4923849 | 0.62244728 | 0.94418789 |
| BLOC1S2 | 62.1646302 | 0.21118724 | 0.33403425 | 0.63223231 | 0.52723509 | 0.92044374 |
| SEC31B | 50.9857858 | -0.0374523 | 0.28335649 | -0.1321737 | 0.89484693 | 0.98597539 |
| HIF1AN | 104.923921 | -0.1212278 | 0.20540264 | -0.5901958 | 0.55505939 | 0.92872652 |
| PAX2 | 1473.71141 | 0.80171608 | 0.36403503 | 2.20230473 | 0.02764379 | 0.40349154 |
| SLF2 | 196.140317 | -0.4042089 | 0.25042649 | -1.6140818 | 0.10650967 | 0.63461115 |
| MRPL43 | 102.10886 | 0.42771307 | 0.36105487 | 1.18462068 | 0.23616746 | 0.79736951 |
| LZTS2 | 109.181255 | -0.5384191 | 0.29232758 | -1.8418346 | 0.06549934 | 0.55444578 |
| BTRC | 511.031932 | 0.1121532 | 0.22553546 | 0.49727524 | 0.61899497 | 0.94292825 |
| DPCD | 92.8858543 | 0.02651789 | 0.37768094 | 0.07021241 | 0.9440246 | 0.99265578 |
| POLL | 70.1361919 | 0.12889221 | 0.25290729 | 0.50964214 | 0.61030219 | 0.94003018 |
| FBXW4 | 422.073514 | -0.0798445 | 0.20921855 | -0.3816321 | 0.70273426 | 0.95672887 |
| OGA | 823.766282 | -0.3737023 | 0.23892413 | -1.5641047 | 0.11779298 | 0.65447949 |
| ARMH3 | 469.271373 | -0.0767695 | 0.22101645 | -0.3473476 | 0.72833019 | 0.95985088 |
| LDB1 | 83.6508577 | 0.92721935 | 0.29827749 | 3.10857976 | 0.00187989 | 0.1093833 |
| PPRC1 | 43.2740341 | -0.6288722 | 0.3671057 | -1.7130549 | 0.08670243 | 0.60414994 |
| NOLC1 | 58.829126 | 0.17188788 | 0.34604218 | 0.49672522 | 0.61938284 | 0.94300065 |
| GBF1 | 651.56545 | -0.058649 | 0.16620141 | -0.3528792 | 0.72417902 | 0.95963742 |
| NFKB2 | 46.6447223 | -0.4309074 | 0.39662401 | -1.086438 | 0.27728526 | 0.82198569 |
| FBXL15 | 90.1638365 | 0.19108222 | 0.4331776 | 0.4411175 | 0.65912794 | 0.95127957 |
| CUEDC2 | 84.5528087 | 0.31777773 | 0.33571985 | 0.94655626 | 0.34386494 | 0.85126545 |
| ACTR1A | 92.8538475 | -0.4989194 | 0.23586858 | -2.115243 | 0.03440924 | 0.43050784 |
| SUFU | 161.980189 | 0.44950421 | 0.37614728 | 1.19502184 | 0.2320785 | 0.7922741 |
| TRIM8 | 320.812776 | 0.74869291 | 0.31863819 | 2.34966474 | 0.01879033 | 0.32850268 |
| ARL3 | 230.513625 | 0.62735424 | 0.30732921 | 2.04131015 | 0.04122001 | 0.4648536 |
| SFXN2 | 42.9234361 | 0.037765 | 0.36520308 | 0.10340822 | 0.91763899 | 0.98999965 |
| WBP1L | 197.86303 | -0.3077045 | 0.21592603 | -1.425046 | 0.15414391 | 0.7079031 |
| CNNM2 | 401.334242 | -0.2082787 | 0.39157034 | -0.5319061 | 0.59479102 | 0.935202 |
| NT5C2 | 1048.13202 | -0.634081 | 0.2880605 | -2.2012077 | 0.02772133 | 0.40380659 |
| PCGF6 | 81.8964939 | -0.2160563 | 0.3783572 | -0.5710379 | 0.56797392 | 0.93122667 |
| ATP5MD | 168.822507 | -0.28148 | 0.31483942 | -0.894043 | 0.3712989 | 0.86201082 |
| PDCD11 | 80.4762258 | -0.3696004 | 0.25383003 | -1.4560941 | 0.14536661 | 0.69325556 |
| SH3PXD2A | 76.7699205 | 0.27583385 | 0.34746456 | 0.79384744 | 0.42728425 | 0.88510458 |
| STN1 | 79.2464869 | -0.2515077 | 0.34868745 | -0.7212983 | 0.47072599 | 0.90123338 |
| SLK | 278.581875 | -0.1379194 | 0.15318759 | -0.90033 | 0.36794464 | 0.85902833 |
| CFAP43 | 138.197361 | 0.78481558 | 0.41485575 | 1.89177944 | 0.05852037 | 0.53216182 |
| GSTO1 | 145.529131 | -0.061099 | 0.22668931 | -0.2695275 | 0.78752379 | 0.96513523 |
| GSTO2 | 296.792479 | 0.02159841 | 0.22470674 | 0.0961182 | 0.9234267 | 0.99051393 |
| XPNPEP1 | 131.174247 | -0.3046267 | 0.27168789 | -1.1212378 | 0.26218665 | 0.80886551 |

| | | | | | | |
|---|---|---|---|---|---|---|
| ADD3 | 710.528739 | 0.42888576 | 0.26360425 | 1.62700626 | 0.10373577 | 0.62768525 |
| MXI1 | 243.156386 | -0.404464 | 0.51423418 | -0.7865367 | 0.43155315 | 0.88650713 |
| SMNDC1 | 89.3470383 | -0.007495 | 0.27656458 | -0.0271003 | 0.97837973 | 0.99696835 |
| SMC3 | 153.182409 | -0.1413382 | 0.26080227 | -0.5419362 | 0.58786245 | 0.93483536 |
| PDCD4 | 217.088522 | -0.1838874 | 0.24055304 | -0.7644361 | 0.44460739 | 0.89105265 |
| BBIP1 | 121.071919 | -0.1513319 | 0.24917952 | -0.6073209 | 0.54363799 | 0.92482794 |
| SHOC2 | 332.821751 | -0.0658648 | 0.19149379 | -0.3439528 | 0.73088176 | 0.96008876 |
| GPAM | 62.4002252 | 0.16972607 | 0.39421063 | 0.43054667 | 0.66679802 | 0.9527406 |
| ACSL5 | 33.265527 | 0.47795905 | 0.3401412 | 1.40517837 | 0.15996825 | 0.71409437 |
| ZDHHC6 | 104.543946 | -0.2021382 | 0.22115487 | -0.9140119 | 0.36071058 | 0.85608464 |
| VTI1A | 1042.24401 | -0.2474408 | 0.18274748 | -1.3540041 | 0.17573505 | 0.73403881 |
| TCF7L2 | 1336.39533 | -0.4069209 | 0.33931672 | -1.1992363 | 0.23043607 | 0.79075638 |
| CASP7 | 100.838909 | -0.1669043 | 0.26800655 | -0.6227619 | 0.53344101 | 0.92282301 |
| NHLRC2 | 260.159371 | 0.05864289 | 0.23613777 | 0.24834185 | 0.80386992 | 0.96678163 |
| CCDC186 | 323.414232 | 0.18670846 | 0.22416987 | 0.8328883 | 0.40490773 | 0.8783757 |
| AFAP1L2 | 453.745299 | 0.99319372 | 0.35984558 | 2.76005537 | 0.00577916 | 0.18867776 |
| ABLIM1 | 1353.19725 | 0.70452699 | 0.32728885 | 2.15261532 | 0.03134892 | 0.41423957 |
| FAM160B1 | 135.727374 | 0.0351219 | 0.18526686 | 0.18957464 | 0.84964246 | 0.97671331 |
| TRUB1 | 83.4251139 | -0.0897872 | 0.20082365 | -0.4470948 | 0.65480658 | 0.95100227 |
| HSPA12A | 363.742281 | 0.413612 | 0.26838525 | 1.54111301 | 0.12328928 | 0.66519057 |
| SHTN1 | 227.808899 | -0.6376553 | 0.34464088 | -1.8502022 | 0.06428441 | 0.54860648 |
| PDZD8 | 259.868612 | -0.1910597 | 0.22297125 | -0.8568803 | 0.39151103 | 0.87015937 |
| EMX2OS | 645.604666 | 0.37853007 | 0.27211895 | 1.39104635 | 0.16421137 | 0.71715698 |
| EMX2 | 285.494285 | 0.4533159 | 0.32718477 | 1.38550428 | 0.16589832 | 0.72022139 |
| RAB11FIP2 | 518.278259 | 0.256586 | 0.20302487 | 1.26381561 | 0.20629622 | 0.77193956 |
| AC022395.1 | 45.0986665 | 0.77417129 | 0.63795234 | 1.21352527 | 0.22492906 | 0.78982769 |
| CASC2 | 348.204849 | 0.2712948 | 0.42190684 | 0.64302062 | 0.52021072 | 0.91751084 |
| FAM204A | 187.863741 | -0.030627 | 0.19578418 | -0.1564324 | 0.87569218 | 0.98268059 |
| CACUL1 | 389.863686 | -0.3625199 | 0.25187748 | -1.4392706 | 0.15007387 | 0.7029344 |
| EIF3A | 485.966979 | -0.2163935 | 0.31463503 | -0.6877604 | 0.4916037 | 0.90730741 |
| FAM45A | 104.930764 | -0.1739208 | 0.27095782 | -0.6418741 | 0.52095496 | 0.91769212 |
| SFXN4 | 87.9497228 | 0.17792118 | 0.38426238 | 0.46302003 | 0.64335001 | 0.94949493 |
| PRDX3 | 245.739621 | 0.56331052 | 0.41673427 | 1.35172592 | 0.17646301 | 0.73502875 |
| TIAL1 | 216.828206 | -0.1320541 | 0.17730199 | -0.7447976 | 0.45639411 | 0.89626521 |
| BAG3 | 65.0828619 | 0.31628329 | 0.31326628 | 1.00963081 | 0.31267221 | 0.84279957 |
| INPP5F | 148.297063 | 0.0873598 | 0.25402689 | 0.34389981 | 0.73092164 | 0.96008876 |
| MCMBP | 141.817355 | -0.0007706 | 0.17049696 | -0.0045199 | 0.99639363 | 0.99967893 |
| SEC23IP | 219.285641 | -0.2541819 | 0.24829673 | -1.0237023 | 0.3059759 | 0.83709431 |
| PLPP4 | 156.35539 | 3.01354582 | 0.47505701 | 6.34354565 | 2.25E-10 | 6.27E-07 |
| WDR11-AS1 | 53.3027372 | 0.61313284 | 0.45188783 | 1.35682529 | 0.17483673 | 0.73227004 |
| WDR11 | 304.893845 | -0.0456022 | 0.17278106 | -0.2639308 | 0.79183326 | 0.9653598 |
| FGFR2 | 529.703732 | 0.49455771 | 0.43984941 | 1.12437962 | 0.26085201 | 0.80805101 |
| ATE1 | 465.577815 | -0.0125014 | 0.1769112 | -0.0706648 | 0.94366454 | 0.99265578 |
| NSMCE4A | 61.639604 | -0.5697068 | 0.36914989 | -1.5432939 | 0.12275949 | 0.66474886 |
| TACC2 | 375.759513 | -0.7952296 | 0.46164705 | -1.7225922 | 0.0849623 | 0.59839042 |
| PLEKHA1 | 917.046695 | -0.7633625 | 0.27782117 | -2.7476759 | 0.00600193 | 0.19123755 |
| C10orf88 | 51.443043 | 0.30455435 | 0.27114127 | 1.12323126 | 0.26133929 | 0.80805101 |
| PSTK | 40.9782422 | 0.31508333 | 0.40392348 | 0.78005699 | 0.43535733 | 0.88785137 |
| IKZF5 | 75.7563425 | 0.10129106 | 0.22592238 | 0.44834453 | 0.65390457 | 0.95100227 |
| ACADSB | 141.598166 | 0.3209427 | 0.26719524 | 1.20115425 | 0.22969137 | 0.79047378 |
| BUB3 | 104.061987 | 0.05852372 | 0.26348732 | 0.2221121 | 0.82422662 | 0.97018919 |
| OAT | 147.836263 | 0.35378322 | 0.32196194 | 1.09883551 | 0.27183982 | 0.81922962 |
| LHPP | 126.460218 | 0.20138837 | 0.24836879 | 0.81084412 | 0.41745519 | 0.88242985 |
| ABRAXAS2 | 81.7460769 | -0.4121984 | 0.23786572 | -1.7329037 | 0.08311279 | 0.59231889 |

| | | | | | | |
|---|---|---|---|---|---|---|
| ZRANB1 | 209.699219 | -0.1627037 | 0.21098453 | -0.7711642 | 0.44060961 | 0.88956997 |
| CTBP2 | 1041.25562 | -0.1069145 | 0.20624082 | -0.5183964 | 0.60418171 | 0.93804805 |
| EDRF1 | 147.450645 | -0.4438984 | 0.18974665 | -2.3394267 | 0.01931336 | 0.33290134 |
| UROS | 114.073779 | -0.1118239 | 0.21708592 | -0.5151137 | 0.60647356 | 0.93919596 |
| BCCIP | 147.988473 | 0.08697202 | 0.23588216 | 0.3687096 | 0.71234419 | 0.95791799 |
| DHX32 | 342.358692 | -0.1675756 | 0.20735546 | -0.8081563 | 0.41900059 | 0.88261006 |
| FANK1 | 270.690415 | -0.2542657 | 0.25735346 | -0.9880017 | 0.32315179 | 0.84285425 |
| DOCK1 | 1572.32422 | -0.2811272 | 0.17574332 | -1.5996463 | 0.10967707 | 0.63926535 |
| FAM196A | 68.79994 | -0.0036858 | 0.48657446 | -0.007575 | 0.99395611 | 0.99919666 |
| MGMT | 766.546107 | 0.01306077 | 0.20270458 | 0.06443251 | 0.94862584 | 0.99336519 |
| AL157832.3 | 120.022711 | -0.4511997 | 0.45880569 | -0.9834223 | 0.32539965 | 0.84285425 |
| C10orf143 | 48.8400287 | 0.30323065 | 0.40886066 | 0.7416479 | 0.4583007 | 0.89747927 |
| GLRX3 | 171.320406 | -0.3757512 | 0.25692356 | -1.4625019 | 0.14360373 | 0.69093999 |
| PPP2R2D | 308.038051 | -0.0955764 | 0.22287761 | -0.4288289 | 0.66804774 | 0.95294349 |
| BNIP3 | 412.466808 | -0.4265077 | 0.64237835 | -0.6639509 | 0.50672174 | 0.91163608 |
| STK32C | 85.0999311 | 0.04389556 | 0.28191667 | 0.15570403 | 0.87626632 | 0.98269442 |
| LRRC27 | 81.4231945 | 0.05452459 | 0.30213824 | 0.1804624 | 0.85678957 | 0.97718353 |
| PWWP2B | 68.4796858 | -0.1263836 | 0.43574548 | -0.29004 | 0.7717856 | 0.96388926 |
| INPP5A | 186.039915 | 0.02692934 | 0.28230323 | 0.09539153 | 0.92400384 | 0.99051393 |
| CFAP46 | 81.2616226 | -1.2139807 | 0.57584402 | -2.1081763 | 0.03501575 | 0.43353051 |
| TUBGCP2 | 191.255325 | 0.04325933 | 0.23066578 | 0.18754115 | 0.85123636 | 0.97718353 |
| ECHS1 | 311.353484 | 0.82509687 | 0.44902452 | 1.83753188 | 0.06613141 | 0.55444578 |
| SCART1 | 43.6048628 | 0.10850799 | 0.40198035 | 0.26993358 | 0.78721135 | 0.96513523 |
| SLC6A12 | 165.065896 | -0.3226745 | 0.65732534 | -0.4908901 | 0.62350419 | 0.94457465 |
| SLC6A13 | 42.675542 | 0.56688403 | 0.48940499 | 1.15831273 | 2.47E-01 | 8.01E-01 |
| KDM5A | 469.809089 | 0.07951322 | 0.1823572 | 0.43603004 | 0.66281494 | 0.9515075 |
| CCDC77 | 36.9232053 | -0.553788 | 0.32829067 | -1.6868832 | 0.09162581 | 0.61104726 |
| B4GALNT3 | 199.106339 | -0.2493563 | 0.31674473 | -0.7872469 | 0.43113734 | 0.88646642 |
| WNK1 | 907.769012 | 1.27111963 | 0.38593227 | 3.29363394 | 0.00098901 | 0.07621211 |
| RAD52 | 137.494117 | -0.0171256 | 0.29600617 | -0.0578555 | 0.9538637 | 0.99514888 |
| ERC1 | 3535.34324 | 0.31090677 | 0.25117021 | 1.23783299 | 0.21577799 | 0.77977703 |
| FBXL14 | 50.3399038 | -0.0668056 | 0.33674395 | -0.198387 | 0.84274226 | 0.97560439 |
| ADIPOR2 | 326.474309 | -0.3351858 | 0.242609 | -1.3815887 | 0.16709804 | 0.72184149 |
| DCP1B | 107.715471 | 0.25995637 | 0.32799959 | 0.79255088 | 0.42803953 | 0.88535099 |
| FKBP4 | 165.992504 | 0.53914528 | 0.43926906 | 1.22736912 | 0.21968389 | 0.78630025 |
| ITFG2 | 157.46848 | -0.3238273 | 0.3591259 | -0.9017096 | 0.36721117 | 0.85902833 |
| TULP3 | 248.539946 | -0.1659777 | 0.20771509 | -0.7990644 | 0.42425308 | 0.88391318 |
| TEAD4 | 70.4738687 | -0.2800378 | 0.52864939 | -0.5297231 | 0.5963039 | 0.93593903 |
| TSPAN9 | 780.006489 | 0.21882112 | 0.24085766 | 0.90850805 | 0.36360986 | 0.85694172 |
| PARP11 | 88.0476789 | -0.1775761 | 0.28220651 | -0.6292415 | 0.52919094 | 0.92155551 |
| CCND2 | 135.297648 | -0.2919214 | 0.40398942 | -0.7225966 | 0.46992774 | 0.90083093 |
| TIGAR | 76.5145322 | 0.24941778 | 0.48107201 | 0.51846247 | 0.60413564 | 0.93804805 |
| C12orf4 | 92.7471932 | -0.2216509 | 0.29065164 | -0.7625998 | 0.44570208 | 0.8918683 |
| ANO2 | 66.0914876 | 0.85280677 | 0.40621787 | 2.0993827 | 0.03578318 | 0.4362094 |
| CD9 | 1283.8246 | 0.09256799 | 0.62467721 | 0.14818532 | 0.88219652 | 0.98335905 |
| TNFRSF1A | 173.799067 | -0.0314752 | 0.3703075 | -0.0849975 | 0.93226339 | 0.99243654 |
| SCNN1A | 810.930009 | 0.34648437 | 0.32390591 | 1.06970684 | 0.28475129 | 0.82781987 |
| LTBR | 102.261028 | -0.3240026 | 0.30484875 | -1.0628307 | 0.28785872 | 0.8303903 |
| TAPBPL | 59.1318741 | -0.0367273 | 0.395117 | -0.092953 | 0.92594086 | 0.99051393 |
| MRPL51 | 190.766624 | 0.13930433 | 0.51197812 | 0.2720904 | 0.78555251 | 0.96513523 |
| NCAPD2 | 64.8131398 | -0.1082584 | 0.40007588 | -0.2705948 | 0.78670271 | 0.96513523 |
| GAPDH | 2169.01156 | -0.2562242 | 0.60467539 | -0.4237384 | 0.67175658 | 0.95444808 |
| CHD4 | 97.1831164 | 0.42982662 | 0.30730852 | 1.39868111 | 0.16190863 | 0.71470931 |
| AC006064.2 | 42.8714332 | 0.23793925 | 0.34568732 | 0.68830771 | 0.49125903 | 0.90730741 |

| | | | | | | |
|---|---|---|---|---|---|---|
| ING4 | 87.0826366 | -0.0061579 | 0.25724538 | -0.0239379 | 0.98090216 | 0.99696835 |
| ZNF384 | 64.4720752 | -0.1030895 | 0.26098276 | -0.3950049 | 0.69283929 | 0.95672887 |
| COPS7A | 99.3304652 | 0.10809595 | 0.32910083 | 0.32845847 | 0.74256504 | 0.96086911 |
| MLF2 | 187.558099 | 0.25259129 | 0.33805004 | 0.74720089 | 0.45494231 | 0.89626521 |
| PTMS | 446.576367 | 0.34724904 | 0.43769358 | 0.79336105 | 0.4275675 | 0.88520493 |
| USP5 | 69.2932838 | -0.0041605 | 0.30939534 | -0.0134473 | 0.9892709 | 0.99835647 |
| TPI1 | 571.585601 | 0.05773997 | 0.48956974 | 0.11794023 | 0.90611501 | 0.9875066 |
| LRRC23 | 84.9323462 | -0.2037257 | 0.28322642 | -0.7193035 | 0.47195395 | 0.90186595 |
| ENO2 | 75.4899392 | -2.5382217 | 0.78616595 | -3.2286081 | 0.00124394 | 0.08868379 |
| ATN1 | 206.473705 | -0.3455242 | 0.23018766 | -1.5010545 | 0.13334147 | 0.67606123 |
| C12orf57 | 129.684104 | 0.24443663 | 0.52620629 | 0.46452624 | 0.64227076 | 0.94949493 |
| PTPN6 | 66.2216212 | -0.1395322 | 0.33859517 | -0.4120916 | 0.68027232 | 0.95574492 |
| PHB2 | 176.830098 | 0.23243112 | 0.39207534 | 0.59282258 | 0.55329989 | 0.92872652 |
| LPCAT3 | 254.106399 | 0.05886311 | 0.27393243 | 0.21488185 | 0.82985943 | 0.97139806 |
| C1S | 90.2526436 | -0.3919318 | 0.37441827 | -1.0467752 | 0.29520328 | 0.83396494 |
| C1R | 43.6277424 | -0.7042662 | 0.67742283 | -1.0396258 | 0.29851379 | 0.83556453 |
| C1RL | 210.95572 | -0.2087279 | 0.25925392 | -0.8051099 | 0.42075627 | 0.88331845 |
| CLSTN3 | 64.3713901 | -0.2365813 | 0.32002048 | -0.7392694 | 0.45974344 | 0.89753337 |
| PEX5 | 51.854655 | -0.1562133 | 0.30235995 | -0.5166467 | 0.60540282 | 0.93919596 |
| CD163L1 | 38.5327373 | 0.49024196 | 0.47504626 | 1.03198783 | 0.30207782 | 0.83709431 |
| FOXJ2 | 48.2022153 | 0.24828247 | 0.28354215 | 0.87564572 | 0.38122266 | 0.86593382 |
| NECAP1 | 56.8747516 | -0.2879047 | 0.27515546 | -1.0463348 | 0.2954065 | 0.83396494 |
| FAM66C | 47.6417875 | 0.85351568 | 0.43305696 | 1.97090858 | 0.04873433 | 0.4878088 |
| LINC00937 | 89.660974 | 0.77581698 | 0.31542818 | 2.45956772 | 0.01391044 | 0.28417439 |
| RIMKLB | 268.103472 | -1.7195991 | 0.39145092 | -4.3928856 | 1.12E-05 | 0.00404224 |
| PHC1 | 123.802772 | -0.2164888 | 0.22401671 | -0.9663958 | 0.33384619 | 0.8445213 |
| M6PR | 172.878567 | 0.02947512 | 0.24349585 | 0.12104977 | 0.90365161 | 0.9875066 |
| CLEC2D | 147.333305 | -0.0782794 | 0.29146305 | -0.2685741 | 0.78825746 | 0.96515649 |
| GABARAPL1 | 566.738693 | -0.1842581 | 0.48123289 | -0.3828876 | 0.70180309 | 0.95672887 |
| KLRD1 | 164.033303 | 0.49835923 | 0.50384892 | 0.9891045 | 0.32261202 | 0.84285425 |
| MAGOHB | 52.4233798 | -0.035268 | 0.29289334 | -0.1204124 | 0.90415647 | 0.9875066 |
| STYK1 | 49.1630661 | -0.9038094 | 0.58760869 | -1.5381144 | 0.12402065 | 0.66528789 |
| YBX3 | 914.921923 | -1.0256737 | 0.51980841 | -1.9731765 | 0.04847546 | 0.48661187 |
| SMIM10L1 | 59.7032597 | 0.26184102 | 0.3436403 | 0.76196247 | 0.4460824 | 0.8918683 |
| AC244131.2 | 66.4952235 | 1.15319824 | 0.68436442 | 1.68506458 | 0.09197611 | 0.61120158 |
| ETV6 | 940.267926 | -0.0744683 | 0.22739974 | -0.3274774 | 0.7433068 | 0.96105014 |
| BCL2L14 | 146.087536 | 0.42634003 | 0.32992836 | 1.29222001 | 0.19628096 | 0.76469312 |
| LRP6 | 359.94435 | 0.47234414 | 0.27140654 | 1.74035654 | 0.08179643 | 0.59202053 |
| MANSC1 | 105.681832 | 0.49448407 | 0.26447784 | 1.86966165 | 0.06153082 | 0.54371246 |
| BORCS5 | 165.83022 | -0.0773987 | 0.2430317 | -0.3184716 | 0.75012726 | 0.96109466 |
| DUSP16 | 275.00522 | 0.00945369 | 0.16758587 | 0.05641103 | 0.95501437 | 0.99537617 |
| CREBL2 | 155.578727 | 0.29470877 | 0.2429401 | 1.21309232 | 0.22509453 | 0.78982769 |
| CDKN1B | 129.508073 | 0.64624996 | 0.39906389 | 1.61941477 | 0.10535805 | 0.63347812 |
| GPRC5A | 524.510886 | -1.1663771 | 0.94834307 | -1.2299105 | 0.21873061 | 0.78557122 |
| GPRC5D-AS1 | 165.065272 | -0.2233299 | 0.43988109 | -0.5077052 | 0.6116601 | 0.94003018 |
| HEBP1 | 175.99334 | 0.51443908 | 0.4705557 | 1.09325863 | 0.27428026 | 0.81986798 |
| FAM234B | 73.736048 | 0.40143377 | 0.31121454 | 1.28989401 | 0.19708746 | 0.76527476 |
| GRIN2B | 128.06352 | -0.9120716 | 0.68635008 | -1.3288723 | 0.18389011 | 0.74609694 |
| ATF7IP | 788.211328 | 0.23071945 | 0.323711 | 0.71273281 | 0.47601111 | 0.90242252 |
| PLBD1 | 100.452384 | 0.9544147 | 0.4285004 | 2.22733676 | 0.02592478 | 0.3903785 |
| AC010168.1 | 61.701819 | 0.60488931 | 0.54136395 | 1.11734317 | 0.26384763 | 0.81021794 |
| H2AFJ | 267.606952 | 0.51063328 | 0.42051287 | 1.21431073 | 0.22462909 | 0.78980378 |
| WBP11 | 154.042277 | -0.0051377 | 0.28236611 | -0.0181952 | 0.98548313 | 0.99762072 |
| C12orf60 | 79.7209376 | 0.17701935 | 0.38961542 | 0.45434379 | 0.6495814 | 0.95061388 |

| | | | | | | |
|---|---|---|---|---|---|---|
| MGP | 279.281754 | -0.1945363 | 0.77569562 | -0.2507895 | 0.80197684 | 0.9661728 |
| ERP27 | 229.998626 | 2.76449205 | 0.50850655 | 5.43649248 | 5.43E-08 | 5.64E-05 |
| ARHGDIB | 99.9471095 | 0.41086248 | 0.6758519 | 0.60791792 | 0.54324191 | 0.92482794 |
| RERG | 94.0092113 | 0.54394933 | 0.61808896 | 0.88005023 | 0.3788321 | 0.86515603 |
| EPS8 | 604.110956 | -0.2847952 | 0.1922489 | -1.4813881 | 0.13850317 | 0.68146162 |
| STRAP | 178.862953 | 0.15433015 | 0.34422526 | 0.44834057 | 0.65390743 | 0.95100227 |
| DERA | 344.974051 | -0.490465 | 0.21629156 | -2.2676104 | 0.02335296 | 0.36567438 |
| MGST1 | 365.371746 | 0.23305131 | 0.3981564 | 0.58532605 | 0.5583285 | 0.92901206 |
| LMO3 | 260.188563 | 0.41516757 | 0.39587642 | 1.04873024 | 0.29430229 | 0.8337535 |
| RERGL | 93.9297361 | 1.64313265 | 0.69640374 | 2.35945409 | 0.01830185 | 0.32472005 |
| PIK3C2G | 2909.47011 | 0.73757677 | 0.4116097 | 1.79193241 | 0.07314379 | 0.56668778 |
| PLCZ1 | 60.2488241 | 1.92403796 | 0.88345268 | 2.17786192 | 0.02941632 | 0.40623517 |
| AC087242.1 | 86.1737009 | 1.17298891 | 0.61132216 | 1.91877375 | 0.05501297 | 0.51660929 |
| PLEKHA5 | 2322.07794 | -0.336649 | 0.30465136 | -1.1050303 | 0.26914646 | 0.81817548 |
| AEBP2 | 623.465089 | 0.25971303 | 0.25867772 | 1.00400232 | 0.3153775 | 0.84285425 |
| AC024901.1 | 76.2546094 | 1.3816002 | 0.63767223 | 2.16663065 | 0.03026303 | 0.41080114 |
| LINC02398 | 60.2139907 | 1.15825369 | 0.79590763 | 1.45526144 | 0.1455969 | 0.69325556 |
| PDE3A | 59.6894747 | 0.55152743 | 0.70980014 | 0.77701793 | 0.43714819 | 0.88785137 |
| PYROXD1 | 102.515395 | -0.137039 | 0.28818044 | -0.4755319 | 0.63440788 | 0.94775679 |
| RECQL | 68.4183901 | -0.4327839 | 0.30533961 | -1.4173852 | 0.15637033 | 0.71120384 |
| GOLT1B | 76.9536309 | -0.4360288 | 0.24245498 | -1.7983906 | 0.07211513 | 0.5644261 |
| LDHB | 1077.14508 | 0.4929657 | 0.39021192 | 1.26332813 | 0.20647128 | 0.77200549 |
| CMAS | 114.398281 | -0.1779305 | 0.18787157 | -0.9470859 | 0.34359502 | 0.85126545 |
| C2CD5 | 349.547875 | -0.2459304 | 0.27025566 | -0.9099918 | 0.36282681 | 0.85694172 |
| ETNK1 | 713.957318 | -0.2380074 | 0.24549106 | -0.9695155 | 0.33228803 | 0.84442294 |
| AC084816.1 | 30.9728802 | -0.8614943 | 0.81849392 | -1.052536 | 0.29255372 | 0.83336457 |
| SOX5 | 2688.08229 | -0.2077278 | 0.47220168 | -0.4399134 | 0.65999984 | 0.95127957 |
| BCAT1 | 66.1239749 | 1.99304302 | 1.33231199 | 1.49592816 | 0.13467238 | 0.6785647 |
| CASC1 | 153.771758 | 0.55728321 | 0.48120427 | 1.15810112 | 0.24682277 | 0.80072691 |
| ETFRF1 | 96.4643067 | 0.51231022 | 0.25858345 | 1.98121815 | 0.04756681 | 0.48600281 |
| KRAS | 276.11438 | -0.084035 | 0.26318613 | -0.3192987 | 0.7495 | 0.96109466 |
| RASSF8-AS1 | 98.1895643 | -1.1033183 | 0.32147547 | -3.4320451 | 0.00059905 | 0.05792155 |
| RASSF8 | 651.642557 | -0.9315112 | 0.34917715 | -2.6677323 | 0.00763651 | 0.21455924 |
| BHLHE41 | 64.6359536 | -0.0777374 | 0.67096555 | -0.1158589 | 0.90776434 | 0.9875066 |
| SSPN | 85.6608814 | 0.67327375 | 0.38850918 | 1.73296741 | 0.08310146 | 0.59231889 |
| AC024145.1 | 37.3878944 | -0.1970051 | 0.63453382 | -0.3104723 | 0.75620186 | 0.96120879 |
| ITPR2 | 1377.00116 | -0.2588334 | 0.23151744 | -1.1179864 | 0.26357278 | 0.81004186 |
| AC023051.1 | 40.1227672 | 0.58892416 | 0.74467032 | 0.7908522 | 0.42903025 | 0.88570555 |
| INTS13 | 63.5983994 | -0.5436161 | 0.2956857 | -1.8384931 | 0.06598977 | 0.55444578 |
| FGFR1OP2 | 181.270756 | -0.0254987 | 0.29865476 | -0.0853785 | 0.93196046 | 0.99243654 |
| TM7SF3 | 326.904646 | -0.0581972 | 0.32919798 | -0.1767848 | 0.85967747 | 0.97837316 |
| MED21 | 128.650046 | 0.10516353 | 0.38014242 | 0.27664244 | 0.78205467 | 0.96513523 |
| STK38L | 154.327614 | 0.11516641 | 0.34384631 | 0.33493572 | 0.73767358 | 0.96078945 |
| PPFIBP1 | 945.206451 | 0.49576612 | 0.47305328 | 1.04801328 | 0.29463249 | 0.8337535 |
| MRPS35 | 162.23055 | 0.31847116 | 0.16075125 | 1.9811427 | 0.04757527 | 0.48600281 |
| KLHL42 | 105.694155 | 0.29580159 | 0.20161783 | 1.46714003 | 0.14233797 | 0.6890058 |
| CCDC91 | 964.597135 | -0.1183363 | 0.19219737 | -0.6157021 | 0.53809116 | 0.92324744 |
| FAR2 | 58.1164572 | -0.1501399 | 0.57687483 | -0.2602643 | 0.79465993 | 0.9653598 |
| AC009318.1 | 44.0273028 | -0.1976802 | 0.57167278 | -0.3457925 | 0.72949864 | 0.96008876 |
| ERGIC2 | 221.164723 | -0.2163905 | 0.18647106 | -1.160451 | 0.24586522 | 0.80072691 |
| OVCH1-AS1 | 45.0420922 | -1.0469544 | 0.98243439 | -1.0656737 | 0.28657119 | 0.8294024 |
| TMTC1 | 172.743733 | -1.4706254 | 0.86507455 | -1.6999985 | 0.08913122 | 0.60739841 |
| IPO8 | 226.688455 | 0.24860741 | 0.19121862 | 1.30012136 | 0.19355938 | 0.76043322 |
| CAPRIN2 | 147.895556 | -0.0394358 | 0.21350473 | -0.1847067 | 0.85345911 | 0.97718353 |

| | | | | | | |
|---|---|---|---|---|---|---|
| DDX11 | 35.0666636 | -0.4344063 | 0.34516788 | -1.2585362 | 0.20819789 | 0.77419992 |
| SINHCAF | 326.466017 | 0.18917257 | 0.20933959 | 0.90366362 | 0.36617379 | 0.85902833 |
| DENND5B | 390.012938 | -0.3749075 | 0.35082858 | -1.0686344 | 0.28523442 | 0.82804896 |
| AMN1 | 204.11519 | -0.394214 | 0.32430809 | -1.2155539 | 0.22415491 | 0.78960314 |
| KIAA1551 | 215.811785 | 0.33154876 | 0.30759536 | 1.0778731 | 0.28109039 | 0.82628479 |
| BICD1 | 1543.0822 | 0.69905517 | 0.46822667 | 1.49298451 | 0.13544124 | 0.67980087 |
| FGD4 | 2216.96119 | 0.26612513 | 0.31471727 | 0.84560066 | 0.39777556 | 0.87462201 |
| DNM1L | 434.086498 | 0.12894037 | 0.19033858 | 0.67742633 | 0.4981355 | 0.90839719 |
| YARS2 | 110.093248 | 0.09983178 | 0.25070821 | 0.39819908 | 0.69048345 | 0.95659154 |
| PKP2 | 42.8640794 | 0.51737436 | 0.47037792 | 1.0999121 | 0.27137042 | 0.81922962 |
| ALG10B | 47.3875159 | -0.1426194 | 0.32159872 | -0.4434702 | 0.65742569 | 0.95100227 |
| CPNE8 | 731.130206 | -1.2564141 | 0.48303814 | -2.601066 | 0.00929346 | 0.23300339 |
| KIF21A | 147.784052 | -0.4504371 | 0.38051419 | -1.1837589 | 0.23650852 | 0.79797745 |
| SLC2A13 | 242.906397 | -1.0259345 | 0.24337334 | -4.2154759 | 2.49E-05 | 0.00687411 |
| LRRK2 | 612.609038 | -0.5280343 | 0.26074944 | -2.025064 | 0.04286079 | 0.47339997 |
| CNTN1 | 125.222335 | 1.47921945 | 0.47125415 | 3.13889954 | 0.00169584 | 0.10393192 |
| GXYLT1 | 192.209619 | 0.18473239 | 0.1864795 | 0.9906311 | 0.32186575 | 0.84285425 |
| YAF2 | 303.53204 | -0.0806227 | 0.22639808 | -0.3561105 | 0.72175778 | 0.95868461 |
| PPHLN1 | 438.541736 | -0.0307258 | 0.19955988 | -0.1539679 | 0.87763502 | 0.98279964 |
| ZCRB1 | 127.535613 | -0.058245 | 0.33317516 | -0.1748179 | 0.86122274 | 0.97841862 |
| PRICKLE1 | 176.668374 | 2.19090872 | 0.5915907 | 3.7034198 | 0.00021271 | 0.03247931 |
| PUS7L | 117.545269 | -0.0922771 | 0.25406496 | -0.3632027 | 0.71645346 | 0.95791799 |
| IRAK4 | 105.352288 | 0.52209797 | 0.27665257 | 1.88719728 | 0.0591338 | 0.53286516 |
| TWF1 | 123.213648 | -0.077506 | 0.28296635 | -0.2739054 | 0.78415729 | 0.96513523 |
| TMEM117 | 661.184365 | 0.17869879 | 0.2691551 | 0.66392497 | 0.50673833 | 0.91163608 |
| ANO6 | 448.464044 | 0.1547256 | 0.37796444 | 0.40936548 | 0.68227147 | 0.95574492 |
| AC008124.1 | 50.5810192 | 0.3446789 | 0.38404881 | 0.89748723 | 0.36945898 | 0.86020443 |
| ARID2 | 804.120725 | -0.0042983 | 0.17769431 | -0.0241891 | 0.98070177 | 0.99696835 |
| SCAF11 | 809.588481 | 0.02393089 | 0.13053668 | 0.18332695 | 0.85454149 | 0.97718353 |
| SLC38A1 | 1945.66568 | -0.4419358 | 0.37623548 | -1.1746254 | 0.24014461 | 0.80056465 |
| SLC38A2 | 529.338882 | -0.0912017 | 0.29842413 | -0.3056109 | 0.75990092 | 0.96190674 |
| AC008014.1 | 1259.29803 | 0.42838502 | 0.29686652 | 1.44302233 | 0.14901418 | 0.70215315 |
| SLC38A4 | 136.158824 | 0.69473859 | 0.42560139 | 1.63236918 | 0.10260174 | 0.62630326 |
| PCED1B | 166.742873 | -2.1077643 | 0.63583291 | -3.3149658 | 9.17E-04 | 0.07236181 |
| RPAP3 | 85.2358332 | -0.0445021 | 0.3348121 | -0.1329168 | 0.89425922 | 0.98597539 |
| RAPGEF3 | 502.329676 | 1.37968461 | 0.30056676 | 4.59027675 | 4.43E-06 | 0.00231953 |
| SLC48A1 | 187.957078 | 0.97310132 | 0.32564778 | 2.98820194 | 0.00280624 | 0.13678792 |
| HDAC7 | 306.436577 | 0.41235674 | 0.33730227 | 1.22251399 | 0.22151332 | 0.78752765 |
| VDR | 39.6798855 | -0.5969337 | 0.38978166 | -1.5314566 | 0.12565657 | 0.66745103 |
| TMEM106C | 98.8483337 | 0.58385424 | 0.37848798 | 1.54259653 | 0.1229287 | 0.66475376 |
| SENP1 | 100.427891 | -0.2340564 | 0.20241586 | -1.1563145 | 0.24755255 | 0.80072691 |
| PFKM | 70.6309347 | 0.16550825 | 0.28076157 | 0.58949753 | 0.55552757 | 0.92872652 |
| ASB8 | 88.783444 | 0.31645271 | 0.31961363 | 0.99011019 | 0.32212026 | 0.84285425 |
| ZNF641 | 50.9247928 | 0.12154947 | 0.30607149 | 0.39712772 | 0.69127328 | 0.95672887 |
| AC090115.1 | 38.9487884 | 0.51685845 | 0.49143249 | 1.05173844 | 0.29291957 | 0.83336457 |
| KANSL2 | 51.7978809 | 0.07584918 | 0.262828 | 0.28858865 | 0.77289618 | 0.96388926 |
| CCNT1 | 139.422899 | -0.1940387 | 0.22305124 | -0.869929 | 0.38433922 | 0.86725322 |
| ADCY6 | 170.833657 | 0.2790325 | 0.22572243 | 1.23617534 | 0.21639339 | 0.78091138 |
| DDX23 | 89.8742391 | -0.1705789 | 0.28075575 | -0.6075704 | 0.54347242 | 0.92482794 |
| CCDC65 | 76.8284043 | -0.0026469 | 0.3434199 | -0.0077073 | 0.9938505 | 0.99919666 |
| PRKAG1 | 100.417926 | -0.2559703 | 0.25306447 | -1.0114825 | 0.31178554 | 0.84279403 |
| KMT2D | 97.1374617 | -0.2014387 | 0.27516487 | -0.7320654 | 0.46412867 | 0.89801718 |
| LMBR1L | 89.5702932 | -0.2348993 | 0.20307973 | -1.1566853 | 0.24740096 | 0.80072691 |
| AC011603.2 | 34.6474032 | 0.31282559 | 0.44601347 | 0.70138147 | 0.48306498 | 0.90628733 |

| | | | | | | |
|---|---|---|---|---|---|---|
| TUBA1B | 457.176001 | -0.8258945 | 0.61129811 | -1.3510502 | 0.17667934 | 0.73554204 |
| TUBA1A | 227.182226 | -1.5264002 | 0.73431587 | -2.0786697 | 0.03764772 | 0.44445707 |
| TUBA1C | 245.360141 | -0.5044582 | 0.36250384 | -1.3915941 | 0.16404535 | 0.71715698 |
| DNAJC22 | 32.1404299 | 1.1287889 | 0.54655727 | 2.06527104 | 0.03889736 | 0.4535395 |
| SPATS2 | 776.658629 | 0.18764342 | 0.30462955 | 0.61597248 | 0.53791268 | 0.92324352 |
| MCRS1 | 55.8586398 | -0.0687213 | 0.2899675 | -0.2369965 | 0.81265953 | 0.9680627 |
| PRPF40B | 228.396552 | 0.31818534 | 0.23414965 | 1.3588974 | 0.1741791 | 0.73178166 |
| FMNL3 | 69.3358981 | 0.48066412 | 0.53609498 | 0.89660254 | 0.36993104 | 0.86076317 |
| TMBIM6 | 1131.3004 | -0.0977519 | 0.31487087 | -0.3104508 | 0.75621818 | 0.96120879 |
| NCKAP5L | 63.1826084 | -0.0531899 | 0.32388391 | -0.1642252 | 0.86955385 | 0.9820194 |
| AQP2 | 9762.38265 | -0.3916488 | 0.60476371 | -0.6476064 | 0.51723958 | 0.91751084 |
| AC025154.2 | 178.193546 | -0.4053217 | 0.70932442 | -0.5714194 | 0.56771541 | 0.93122667 |
| SMARCD1 | 79.53272 | -0.1290357 | 0.28418766 | -0.4540509 | 0.64979221 | 0.95061388 |
| AC074032.1 | 186.696158 | -0.2131621 | 0.25490274 | -0.8362489 | 0.40301491 | 0.87834687 |
| CERS5 | 154.110796 | -0.0004799 | 0.22173688 | -0.0021641 | 0.9982733 | 0.99967893 |
| LIMA1 | 1008.83766 | -0.5822093 | 0.32753855 | -1.7775291 | 0.07548122 | 0.57072562 |
| LARP4 | 388.282794 | -0.0180474 | 0.16724892 | -0.1079077 | 0.91406892 | 0.98830345 |
| DIP2B | 557.380542 | -0.0807248 | 0.29289885 | -0.2756063 | 0.78285046 | 0.96513523 |
| ATF1 | 167.355925 | -0.070711 | 0.22416248 | -0.3154452 | 0.75242368 | 0.96120879 |
| METTL7A | 217.480357 | 0.72627818 | 0.38111323 | 1.90567557 | 0.05669231 | 0.52346922 |
| SLC11A2 | 1009.6862 | -0.5548169 | 0.26384938 | -2.102779 | 0.03548509 | 0.43592878 |
| LETMD1 | 225.688989 | -0.0369576 | 0.19562651 | -0.1889189 | 0.85015635 | 0.97713298 |
| CSRNP2 | 68.2796078 | -0.0989587 | 0.35813747 | -0.2763147 | 0.78230634 | 0.96513523 |
| TFCP2 | 451.227678 | 0.08005001 | 0.2074936 | 0.38579508 | 0.69964844 | 0.95672887 |
| POU6F1 | 102.389391 | -0.6965069 | 0.28530242 | -2.4412932 | 0.01463477 | 0.28992884 |
| DAZAP2 | 565.055585 | 0.18286667 | 0.30792294 | 0.59387152 | 0.55259805 | 0.92872652 |
| SMAGP | 37.9979067 | -0.0994789 | 0.39945125 | -0.2490388 | 0.80333075 | 0.96678163 |
| FIGNL2 | 51.4678417 | -0.2304395 | 0.42991592 | -0.5360105 | 0.59195127 | 0.93491607 |
| ACVR1B | 602.916339 | 0.61677646 | 0.37197626 | 1.65810703 | 0.09729587 | 0.61662137 |
| NR4A1 | 651.622634 | -1.5526794 | 0.98168948 | -1.58164 | 1.14E-01 | 0.6483668 |
| ATG101 | 62.7952863 | 0.21622367 | 0.4188919 | 0.51618011 | 0.60572862 | 0.93919596 |
| AC078864.2 | 68.0691921 | -0.4223158 | 0.75922559 | -0.5562455 | 0.57804307 | 0.93345492 |
| AC078864.1 | 166.783883 | -0.620411 | 0.49552937 | -1.2520166 | 0.21056381 | 0.77610028 |
| KRT7 | 227.278188 | 0.10117084 | 0.58791476 | 0.17208419 | 0.86337134 | 0.979268 |
| KRT8 | 722.124091 | 0.0178881 | 0.5572208 | 0.03210235 | 0.97439043 | 0.99656026 |
| KRT18 | 834.989409 | 0.3091279 | 0.60002287 | 0.51519352 | 0.6064178 | 0.93919596 |
| EIF4B | 427.342437 | -0.0003647 | 0.27909203 | -0.0013067 | 0.99895744 | 0.99968413 |
| AC068888.1 | 146.599711 | -0.2013031 | 0.27807446 | -0.7239181 | 0.46911599 | 0.90026987 |
| TNS2 | 59.6449924 | 0.46521786 | 0.45508541 | 1.02226493 | 0.30665553 | 0.8375684 |
| SPRYD3 | 40.9667994 | 0.21632041 | 0.33933102 | 0.63749083 | 0.52380518 | 0.91871245 |
| CSAD | 251.06643 | -0.2924433 | 0.28122502 | -1.0398908 | 0.29839066 | 0.83556453 |
| ZNF740 | 52.1403052 | -0.1847671 | 0.31616588 | -0.5843994 | 0.55895163 | 0.92901206 |
| RARG | 79.7518273 | 0.37264694 | 0.36880694 | 1.01041197 | 0.31229796 | 0.84279957 |
| MFSD5 | 36.7989491 | -0.2092719 | 0.39804062 | -0.5257551 | 0.59905836 | 0.93728325 |
| PFDN5 | 440.933131 | 0.17519756 | 0.46468897 | 0.37702112 | 0.70615791 | 0.95730303 |
| C12orf10 | 47.3596472 | 0.46742588 | 0.376497 | 1.24151291 | 0.21441633 | 0.77890374 |
| AAAS | 42.8081747 | -0.0465118 | 0.36346242 | -0.1279687 | 0.89817371 | 0.9863394 |
| SP1 | 201.803557 | -0.3346683 | 0.21871753 | -1.5301392 | 0.12598229 | 0.66763141 |
| PRR13 | 235.894336 | 0.67829294 | 0.31578185 | 2.1479795 | 0.03171538 | 0.41517597 |
| PCBP2 | 566.013632 | -0.2241799 | 0.19651048 | -1.1408035 | 0.25395169 | 0.80414079 |
| MAP3K12 | 58.0594322 | 0.56312424 | 0.33188559 | 1.69674207 | 0.08974543 | 0.60739841 |
| ATP5MC2 | 532.60026 | 0.04173419 | 0.38273715 | 0.10904138 | 0.91316967 | 0.98830345 |
| CALCOCO1 | 108.875426 | 0.27840267 | 0.27865126 | 0.99910788 | 0.31774243 | 0.84285425 |
| SMUG1 | 58.0997965 | 0.51352771 | 0.28139327 | 1.82494666 | 0.06800908 | 0.55917298 |

| | | | | | | |
|---|---|---|---|---|---|---|
| LINC02381 | 89.1206424 | 0.75414701 | 0.261227 | 2.88694125 | 0.00389007 | 0.15207852 |
| CBX5 | 469.201012 | 0.35420467 | 0.17590076 | 2.01366196 | 0.04404504 | 0.48009422 |
| HNRNPA1 | 319.58324 | -0.2263838 | 0.35547227 | -0.6368535 | 0.52422025 | 0.91885402 |
| COPZ1 | 189.792507 | -0.0568378 | 0.23328398 | -0.2436419 | 0.80750817 | 0.96752828 |
| CD63 | 596.493001 | -0.3338737 | 0.47928164 | -0.6966128 | 0.48604512 | 0.90695174 |
| ORMDL2 | 63.6871063 | -0.3107445 | 0.39817035 | -0.780431 | 0.43513721 | 0.88785137 |
| PYM1 | 35.5394179 | 0.06726147 | 0.26019234 | 0.25850673 | 0.79601585 | 0.9653598 |
| RAB5B | 180.483371 | -0.2210547 | 0.23659823 | -0.9343041 | 0.35014705 | 0.85437511 |
| SUOX | 87.1475216 | 0.70350278 | 0.34494817 | 2.0394449 | 0.04140565 | 0.46609147 |
| IKZF4 | 57.9945298 | 0.04011603 | 0.31481166 | 0.12742868 | 0.89860112 | 0.98660836 |
| RPS26 | 104.936399 | -0.4631117 | 0.40023544 | -1.1570982 | 0.24723224 | 0.80072691 |
| ERBB3 | 69.3689298 | -0.4169854 | 0.36356279 | -1.146942 | 0.25140561 | 0.80240987 |
| PA2G4 | 83.2556182 | 0.05607414 | 0.35698341 | 0.15707773 | 0.87518359 | 0.98267808 |
| RPL41 | 2120.69781 | 0.02307013 | 0.41807651 | 0.0551816 | 0.95599379 | 0.99565888 |
| ESYT1 | 102.656721 | -0.4806297 | 0.23910246 | -2.0101411 | 0.04441626 | 0.48009422 |
| MYL6B | 64.4171982 | 0.19912073 | 0.43012267 | 0.46293938 | 0.64340782 | 0.94949493 |
| MYL6 | 927.382018 | -0.1429068 | 0.45966277 | -0.3108949 | 0.75588052 | 0.96120879 |
| SMARCC2 | 237.41906 | 0.0028302 | 0.23426955 | 0.01208094 | 0.99036104 | 0.99882205 |
| RNF41 | 66.0992722 | 0.10739715 | 0.27551856 | 0.38980004 | 0.69668442 | 0.95672887 |
| NABP2 | 37.0404256 | -0.0796993 | 0.30205784 | -0.2638545 | 0.79189204 | 0.9653598 |
| ANKRD52 | 79.1459125 | 0.35730972 | 0.27955829 | 1.27812243 | 0.20120626 | 0.76957251 |
| CS | 124.088126 | -0.0220429 | 0.30450442 | -0.0723893 | 0.94229209 | 0.99265578 |
| PAN2 | 78.786446 | -0.0790125 | 0.2375187 | -0.3326579 | 0.7393925 | 0.96078945 |
| STAT2 | 138.059741 | -0.6329109 | 0.34452339 | -1.8370622 | 0.06620071 | 0.55458309 |
| SPRYD4 | 46.9506708 | -0.000725 | 0.33158423 | -0.0021863 | 0.99825555 | 0.99967893 |
| RBMS2 | 326.940196 | 0.01249048 | 0.17703202 | 0.0705549 | 9.44E-01 | 0.99265578 |
| BAZ2A | 214.713401 | -0.14602 | 0.16758494 | -0.8713194 | 0.38357976 | 0.8665075 |
| ATP5F1B | 638.878877 | 0.47473104 | 0.39744532 | 1.19445623 | 0.23229955 | 0.7922741 |
| PTGES3 | 795.022672 | 0.129697 | 0.23024044 | 0.56331112 | 0.57322305 | 0.93258748 |
| NACA | 463.688074 | 0.32082994 | 0.49085507 | 0.65361439 | 0.51336027 | 0.91512427 |
| NEMP1 | 45.9171928 | -0.3393566 | 0.25262761 | -1.3433075 | 0.17917243 | 0.74042867 |
| STAT6 | 299.202064 | -0.4273405 | 0.21582574 | -1.9800257 | 0.04770064 | 0.48600281 |
| LRP1 | 45.2105827 | 0.38527413 | 0.32304627 | 1.19262831 | 0.23301497 | 0.7926597 |
| R3HDM2 | 619.562483 | -0.1589449 | 0.21803495 | -0.7289882 | 0.46600885 | 0.89883965 |
| MARS | 80.3737665 | -0.5978928 | 0.36095524 | -1.6564182 | 0.09763716 | 0.61685067 |
| DDIT3 | 223.257429 | -1.7815179 | 0.82310639 | -2.1643835 | 0.03043493 | 0.41080114 |
| MBD6 | 86.213692 | 0.13573216 | 0.24886652 | 0.54540143 | 0.58547746 | 0.93483536 |
| DCTN2 | 181.127194 | -0.2329199 | 0.32721075 | -0.7118345 | 0.4765673 | 0.90242252 |
| PIP4K2C | 114.682443 | -0.0883434 | 0.23114052 | -0.3822063 | 0.70230837 | 0.95672887 |
| DTX3 | 61.7346269 | -0.277315 | 0.29249717 | -0.9480948 | 0.3430812 | 0.85120525 |
| OS9 | 258.983612 | 0.2851837 | 0.21398979 | 1.33269769 | 0.18263103 | 0.74315969 |
| TSPAN31 | 70.1464777 | -0.0988759 | 0.31208176 | -0.3168269 | 0.75137496 | 0.96120879 |
| CDK4 | 70.2116364 | -0.2210403 | 0.33885262 | -0.6523199 | 0.5141948 | 0.91544604 |
| 9-Mar | 61.5709187 | 0.11032938 | 0.32871496 | 0.33563846 | 0.73714352 | 0.96078945 |
| TSFM | 88.2503758 | 0.89233893 | 0.39621605 | 2.25215239 | 0.02431264 | 0.37415052 |
| AVIL | 47.3403499 | 0.94336766 | 0.5356284 | 1.76123531 | 0.07819858 | 0.5795765 |
| CTDSP2 | 255.071225 | 0.20217578 | 0.24362059 | 0.8298797 | 0.4066068 | 0.87948803 |
| LRIG3 | 75.7107555 | 0.48105729 | 0.39228122 | 1.22630721 | 0.22008309 | 0.78633738 |
| SLC16A7 | 497.954629 | 0.11777172 | 0.31421339 | 0.37481446 | 0.70779846 | 0.95752169 |
| FAM19A2 | 214.135782 | -0.1863739 | 0.4128048 | -0.451482 | 0.6516422 | 0.9506139 |
| USP15 | 575.810717 | -0.3382912 | 0.23587323 | -1.4342076 | 0.15151303 | 0.7029344 |
| MON2 | 1047.31256 | -0.1733936 | 0.18989451 | -0.9131051 | 0.36118729 | 0.8565748 |
| AC048341.1 | 58.9012419 | -0.4318104 | 0.47328664 | -0.9123654 | 0.36157638 | 0.85690129 |
| PPM1H | 485.56644 | -0.3917554 | 0.26559081 | -1.4750336 | 0.14020351 | 0.68408298 |

| | | | | | | |
|---|---|---|---|---|---|---|
| DPY19L2 | 665.799316 | -0.1277792 | 0.3004156 | -0.4253415 | 0.67058777 | 0.95395138 |
| RXYLT1 | 69.8638304 | 0.26778404 | 0.26097357 | 1.02609642 | 0.30484614 | 0.83709431 |
| SRGAP1 | 150.152576 | 0.00162951 | 0.5126384 | 0.00317867 | 0.99746379 | 0.99967893 |
| XPOT | 91.056251 | -0.031088 | 0.26166938 | -0.1188066 | 0.90542862 | 0.9875066 |
| TBK1 | 228.741775 | -0.4652905 | 0.23084094 | -2.0156323 | 0.04383844 | 0.48009422 |
| RASSF3 | 77.350566 | -0.5407177 | 0.40603216 | -1.3317114 | 0.18295504 | 0.74374275 |
| GNS | 455.075481 | -0.0487478 | 0.27385058 | -0.1780089 | 0.85871602 | 0.97787068 |
| TBC1D30 | 167.605737 | -0.0836356 | 0.26729226 | -0.3128994 | 0.75435713 | 0.96120879 |
| LEMD3 | 269.312142 | -0.3184395 | 0.2129594 | -1.4953063 | 0.13483453 | 0.6785647 |
| AC090673.1 | 65.9976936 | 1.92623821 | 0.87890275 | 2.19163975 | 0.02840553 | 0.40459506 |
| LLPH | 50.2775489 | -0.0087655 | 0.35914046 | -0.024407 | 0.98052799 | 0.99696835 |
| HELB | 92.5791583 | 0.0130485 | 0.31411458 | 0.04154058 | 0.96686494 | 0.99637506 |
| GRIP1 | 4720.01277 | 0.37427071 | 0.23225549 | 1.61146123 | 1.07E-01 | 0.63477557 |
| CAND1 | 320.319933 | 0.11444555 | 0.16750364 | 0.68324214 | 0.49445385 | 0.90839719 |
| DYRK2 | 46.2915855 | 1.0933203 | 0.3214871 | 3.40082172 | 0.00067184 | 0.05965636 |
| MDM1 | 48.5998799 | -0.2381031 | 0.2666754 | -0.8928573 | 0.3719336 | 0.86201082 |
| RAP1B | 1611.14686 | -1.1861707 | 0.44945171 | -2.6391506 | 0.00831141 | 0.22277119 |
| NUP107 | 174.310087 | -0.5819761 | 0.23789335 | -2.4463739 | 0.01443013 | 0.28805285 |
| SLC35E3 | 136.03039 | -0.8155365 | 0.30262374 | -2.6948859 | 0.00704127 | 0.20555027 |
| MDM2 | 274.658345 | -0.2934603 | 0.32946871 | -0.8907078 | 0.37308597 | 0.86201082 |
| CPM | 50.0561963 | -0.9080151 | 0.48074134 | -1.888781 | 0.05892118 | 0.53286516 |
| CPSF6 | 415.114311 | -0.1295435 | 0.20020139 | -0.6470659 | 0.51758927 | 0.91751084 |
| YEATS4 | 82.9664752 | -0.3706242 | 0.40296358 | -0.9197461 | 0.35770545 | 0.8557799 |
| FRS2 | 439.162562 | -0.1032464 | 0.18421638 | -0.5604624 | 0.57516409 | 0.93275827 |
| CCT2 | 152.033865 | 0.24702443 | 0.29714675 | 0.83132133 | 0.40579213 | 0.87937203 |
| RAB3IP | 271.102213 | -0.043538 | 0.1969939 | -0.221012 | 0.8250831 | 0.97018919 |
| MYRFL | 74.2000275 | 1.19576992 | 0.65604756 | 1.82268786 | 0.06835068 | 0.55917298 |
| AC025159.1 | 186.317266 | 0.41326211 | 0.5759102 | 0.71758081 | 0.47301579 | 0.90213021 |
| CNOT2 | 586.916366 | -0.2024015 | 0.23148781 | -0.8743505 | 0.38192743 | 0.86593382 |
| KCNMB4 | 65.1669968 | -0.7145336 | 0.91953811 | -0.7770571 | 0.43712507 | 0.88785137 |
| ZFC3H1 | 366.679759 | -0.3127124 | 0.20738225 | -1.5079035 | 0.13157923 | 0.67390608 |
| TMEM19 | 35.9833711 | 0.17710697 | 0.32988496 | 0.53687495 | 0.591354 | 0.93489061 |
| RAB21 | 383.59546 | -0.4048677 | 0.25789261 | -1.5699082 | 0.11643647 | 0.65257994 |
| TBC1D15 | 307.127457 | -0.5808915 | 0.3212872 | -1.8080133 | 0.07060443 | 0.5621116 |
| TRHDE | 721.982401 | 0.43429233 | 0.29009092 | 1.49709042 | 0.13436973 | 0.67832119 |
| ATXN7L3B | 187.358762 | 0.37959259 | 0.35357221 | 1.07359284 | 0.28300521 | 0.82666614 |
| CAPS2 | 293.602314 | 0.48076628 | 0.27699707 | 1.73563666 | 0.0826281 | 0.59202053 |
| KRR1 | 141.35752 | 0.15294051 | 0.26865918 | 0.56927336 | 0.56917064 | 0.93122667 |
| PHLDA1 | 188.369895 | 1.21027557 | 0.49418922 | 2.44901248 | 0.01432485 | 0.28746487 |
| NAP1L1 | 456.745083 | 0.63702502 | 0.18819141 | 3.38498455 | 0.00071182 | 0.06165207 |
| OSBPL8 | 541.636279 | -0.481968 | 0.19522592 | -2.4687708 | 0.0135578 | 0.28303942 |
| ZDHHC17 | 329.797688 | -0.2248333 | 0.22421408 | -1.0027616 | 0.31597591 | 0.84285425 |
| AC027288.3 | 78.0846591 | -0.5948541 | 0.61156875 | -0.9726692 | 0.33071776 | 0.84442294 |
| PAWR | 928.838127 | 6.40E-05 | 0.27753807 | 0.00023072 | 0.99981591 | 0.9999633 |
| AC073569.2 | 54.7441024 | 0.85461726 | 0.4514828 | 1.8929121 | 0.05836956 | 0.53192434 |
| PPP1R12A | 725.768026 | -0.0623653 | 0.17472909 | -0.3569256 | 0.72114752 | 0.95848142 |
| LIN7A | 327.971724 | 0.66028787 | 0.44160455 | 1.49520167 | 0.13486182 | 0.6785647 |
| ACSS3 | 254.829568 | 0.84816865 | 0.47752698 | 1.77616908 | 0.07570505 | 0.57072562 |
| PPFIA2 | 183.717702 | 1.29377851 | 0.76791888 | 1.68478538 | 0.09202999 | 0.61120158 |
| CCDC59 | 75.0064686 | 0.55208954 | 0.43594588 | 1.26641759 | 0.20536362 | 0.77193931 |
| METTL25 | 195.10881 | -0.0401833 | 0.30485241 | -0.1318122 | 0.89513284 | 0.98597539 |
| TMTC2 | 3622.43225 | 0.55310949 | 0.32551646 | 1.69917518 | 0.08928618 | 0.60739841 |
| LRRIQ1 | 245.973315 | -0.4088702 | 0.23490096 | -1.7406066 | 0.08175255 | 0.59202053 |
| RASSF9 | 104.756076 | 0.58014903 | 0.31931861 | 1.8168344 | 0.06924247 | 0.55917298 |

| | | | | | | |
|---|---|---|---|---|---|---|
| C12orf29 | 39.6772812 | 0.25277256 | 0.35657072 | 0.70889881 | 0.47838727 | 0.90263775 |
| CEP290 | 288.828607 | -0.0788064 | 0.21171067 | -0.3722363 | 0.70971693 | 0.95752169 |
| TMTC3 | 187.360352 | 0.14768807 | 0.26549877 | 0.5562665 | 0.57802868 | 0.93345492 |
| KITLG | 2086.92693 | 0.43482627 | 0.44412615 | 0.97906027 | 0.3275502 | 0.84381657 |
| DUSP6 | 607.397 | -1.3223047 | 0.81148946 | -1.6294786 | 0.10321174 | 0.62635087 |
| POC1B | 202.476506 | 0.14781272 | 0.25710733 | 0.57490668 | 0.56535441 | 0.93114846 |
| POC1B-AS1 | 72.5463574 | -0.9276371 | 0.31157712 | -2.9772312 | 0.00290865 | 0.13919 |
| ATP2B1 | 329.022434 | 0.05077129 | 0.34905648 | 0.14545292 | 0.88435328 | 0.98407543 |
| BTG1 | 398.840983 | -0.1487883 | 0.54253706 | -0.2742454 | 0.78389604 | 0.96513523 |
| PLEKHG7 | 53.3061108 | -2.3204429 | 0.53585117 | -4.3303868 | 1.49E-05 | 0.00519863 |
| EEA1 | 296.513733 | -0.2437216 | 0.19275502 | -1.2644113 | 0.20608245 | 0.77193931 |
| AC138123.1 | 218.161786 | -0.3012582 | 0.3422176 | -0.8803118 | 0.37869043 | 0.86515603 |
| AC124947.1 | 64.6069698 | -0.2170733 | 0.49727278 | -0.4365276 | 0.66245399 | 0.95142083 |
| NUDT4 | 294.719801 | -0.0154687 | 0.38378422 | -0.0403057 | 0.96784937 | 0.9964399 |
| UBE2N | 195.604131 | 0.15858941 | 0.17743012 | 0.89381335 | 0.37142177 | 0.86201082 |
| MRPL42 | 194.728089 | -0.2219971 | 0.20191624 | -1.0994515 | 0.27157116 | 0.81922962 |
| SOCS2 | 116.382347 | 0.93300068 | 0.46497441 | 2.0065635 | 0.04479616 | 0.48199565 |
| CRADD | 947.805531 | -0.0126013 | 0.26039457 | -0.048393 | 0.96140305 | 0.99635733 |
| AC012085.2 | 83.8629143 | 0.36858775 | 0.59497761 | 0.61949853 | 0.53558799 | 0.92303275 |
| CEP83 | 228.665231 | -0.0121862 | 0.18337785 | -0.066454 | 0.94701637 | 0.99314907 |
| TMCC3 | 149.02779 | 0.25760043 | 0.3285029 | 0.78416486 | 0.43294338 | 0.88722778 |
| NDUFA12 | 189.637171 | 0.07164775 | 0.31241102 | 0.2293381 | 0.81860614 | 0.96883031 |
| NR2C1 | 195.81098 | -0.136486 | 0.23868944 | -0.5718141 | 0.56744792 | 0.93122667 |
| FGD6 | 186.324525 | 0.59722034 | 0.35645917 | 1.67542425 | 0.09385102 | 0.61323531 |
| VEZT | 640.979465 | 0.25964668 | 0.27289905 | 0.95143857 | 0.34138179 | 0.85042291 |
| METAP2 | 1011.19976 | 1.37034724 | 0.37813286 | 3.62398345 | 0.0002901 | 0.03764581 |
| USP44 | 78.3164204 | 0.50599867 | 0.37487274 | 1.34978785 | 0.17708404 | 0.73633806 |
| NTN4 | 1492.13477 | 1.46184817 | 0.39487258 | 3.7020757 | 0.00021384 | 0.03247931 |
| SNRPF | 78.4557018 | -0.2798418 | 0.38184162 | -0.7328739 | 0.46363533 | 0.89801718 |
| LTA4H | 223.984521 | -0.8309051 | 0.37861867 | -2.1945698 | 0.02819447 | 0.40459506 |
| ELK3 | 87.1530459 | -0.3062373 | 0.39831157 | -0.7688385 | 0.44198916 | 0.88997055 |
| CDK17 | 183.890731 | -0.2322501 | 0.29036554 | -0.7998544 | 0.42379517 | 0.88391318 |
| CFAP54 | 104.429981 | -0.0563275 | 0.50641823 | -0.1112272 | 0.91143615 | 0.98788932 |
| NEDD1 | 160.536622 | 0.12688472 | 0.22550931 | 0.56265844 | 0.57366749 | 0.93270589 |
| AC007656.1 | 55.1944004 | 0.48502542 | 0.64171431 | 0.75582766 | 0.44975253 | 0.89324553 |
| TMPO | 225.671426 | 0.51385899 | 0.21608251 | 2.37806836 | 0.0174036 | 0.31610003 |
| SLC25A3 | 538.77688 | 0.56038754 | 0.47631308 | 1.17651093 | 0.23939077 | 0.79984328 |
| IKBIP | 46.3548067 | -0.2819356 | 0.32096663 | -0.8783954 | 0.37972917 | 0.86587828 |
| APAF1 | 148.555268 | -0.015832 | 0.25057387 | -0.0631829 | 0.94962086 | 0.99361356 |
| ANKS1B | 252.610823 | 0.71274055 | 0.56398518 | 1.26375758 | 0.20631705 | 0.77193956 |
| UHRF1BP1L | 151.88945 | -0.1228388 | 0.22718432 | -0.5407012 | 0.58871354 | 0.93483536 |
| ACTR6 | 106.088285 | -0.0423203 | 0.33267232 | -0.1272131 | 0.89877174 | 0.98660836 |
| DEPDC4 | 41.7530483 | -0.0957599 | 0.40501562 | -0.236435 | 0.81309517 | 0.9680627 |
| SCYL2 | 357.32192 | -0.235187 | 0.21632645 | -1.0871857 | 0.27695477 | 0.82198569 |
| NR1H4 | 542.053839 | -1.664048 | 0.50206298 | -3.3144207 | 0.00091833 | 0.07236181 |
| UTP20 | 115.305475 | -0.4938652 | 0.26384444 | -1.8718043 | 0.06123368 | 0.5418629 |
| ARL1 | 197.86086 | 0.16935837 | 0.28346878 | 0.59744981 | 0.55020711 | 0.92780338 |
| CHPT1 | 614.317615 | 0.51120151 | 0.31615642 | 1.61692591 | 0.10589428 | 0.63461115 |
| GNPTAB | 244.054282 | -0.0707346 | 0.21948743 | -0.3222718 | 0.74724682 | 0.96109466 |
| DRAM1 | 160.035144 | -0.4200574 | 0.34904894 | -1.2034341 | 0.22880838 | 0.79047378 |
| WASHC3 | 195.339981 | -0.1880542 | 0.23145001 | -0.8125046 | 0.41650213 | 0.88242985 |
| NUP37 | 80.0487833 | 0.0946684 | 0.2148138 | 0.44069981 | 0.65943034 | 0.95127957 |
| NT5DC3 | 51.8269232 | -0.6033262 | 0.37024165 | -1.6295471 | 0.10319726 | 0.62635087 |
| HSP90B1 | 567.973933 | 0.05778523 | 0.40332717 | 0.14327137 | 0.88607587 | 0.98489549 |

| | | | | | | |
|---|---|---|---|---|---|---|
| C12orf73 | 41.1423556 | 0.39862467 | 0.27056068 | 1.4733282 | 0.14066256 | 0.68501098 |
| TDG | 110.678291 | -0.0418291 | 0.19986387 | -0.2092881 | 0.83422334 | 0.97232209 |
| HCFC2 | 82.1936221 | -0.0865738 | 0.21038079 | -0.4115102 | 0.68069849 | 0.95574492 |
| NFYB | 81.6611482 | 0.0644122 | 0.2428157 | 0.26527196 | 0.79079996 | 0.96527296 |
| TXNRD1 | 958.776245 | -0.124885 | 0.46808495 | -0.2667999 | 0.78962326 | 0.96515649 |
| SLC41A2 | 64.367712 | 0.12915955 | 0.50442076 | 0.25605519 | 0.79790823 | 0.9653598 |
| C12orf45 | 42.2004361 | -0.2718538 | 0.30535473 | -0.8902885 | 0.37331102 | 0.86201082 |
| WASHC4 | 437.423449 | -0.3524544 | 0.21784169 | -1.6179383 | 0.10567589 | 0.63429746 |
| APPL2 | 294.127435 | -0.3947779 | 0.28048683 | -1.4074739 | 0.15928692 | 0.71409437 |
| C12orf75 | 500.003373 | -0.1879577 | 0.50781495 | -0.3701303 | 0.71128539 | 0.957816 |
| NUAK1 | 41.2479664 | 0.23765224 | 0.50921274 | 0.46670522 | 0.64071079 | 0.94935885 |
| CKAP4 | 84.9340048 | 0.29682734 | 0.32023735 | 0.92689793 | 0.35397953 | 0.8557799 |
| TCP11L2 | 83.0825045 | 0.01448175 | 0.33236853 | 0.04357135 | 0.96524609 | 0.99637506 |
| POLR3B | 88.4399354 | -0.0740087 | 0.31140419 | -0.2376612 | 0.81214388 | 0.9680627 |
| RIC8B | 188.409687 | 0.22822591 | 0.21425055 | 1.06522908 | 0.28677228 | 0.82943317 |
| TMEM263 | 195.328387 | -0.022354 | 0.27701902 | -0.0806949 | 0.93568456 | 0.99243654 |
| MTERF2 | 64.6507594 | 0.16335348 | 0.32727723 | 0.49912877 | 0.61768867 | 0.94244117 |
| CRY1 | 173.028095 | -0.5982455 | 0.50974058 | -1.1736274 | 0.24054429 | 0.80057149 |
| PWP1 | 92.6832726 | 0.24277819 | 0.23511738 | 1.03258293 | 0.30179912 | 0.83709431 |
| PRDM4 | 141.586511 | -0.11097 | 0.28449394 | -0.3900612 | 0.69649132 | 0.95672887 |
| SART3 | 107.47794 | 0.07432493 | 0.17716669 | 0.41951976 | 0.67483631 | 0.95524793 |
| ISCU | 377.050539 | 0.76534158 | 0.45579676 | 1.67912907 | 0.09312689 | 0.61317397 |
| CORO1C | 83.140349 | -0.9192656 | 0.40500187 | -2.2697811 | 0.02322087 | 0.36457774 |
| SSH1 | 314.373383 | -0.1569381 | 0.24050677 | -0.6525309 | 0.51405872 | 0.91544604 |
| USP30 | 69.0138912 | -0.3774081 | 0.21565883 | -1.750024 | 0.08011417 | 0.58672016 |
| UNG | 109.301288 | -0.3792085 | 0.52114452 | -0.7276456 | 0.46683055 | 0.89883965 |
| ACACB | 899.799482 | -1.0619234 | 0.40292379 | -2.6355441 | 0.00840025 | 0.22457816 |
| MYO1H | 36.9735335 | 0.80431244 | 0.62763092 | 1.28150544 | 0.20001619 | 0.7687013 |
| KCTD10 | 85.0930729 | 0.1969486 | 0.26271059 | 0.74967896 | 0.45344808 | 0.89544675 |
| UBE3B | 142.644962 | 0.04390807 | 0.22185668 | 0.19791184 | 0.84311404 | 0.97560439 |
| MMAB | 132.978215 | 0.20054472 | 0.36848849 | 0.54423605 | 0.58627905 | 0.93483536 |
| MVK | 108.267569 | -0.0152107 | 0.42103533 | -0.0361268 | 0.97118124 | 0.99656026 |
| TRPV4 | 445.170552 | 0.62094001 | 0.34543128 | 1.79757898 | 0.07224375 | 0.56501083 |
| GLTP | 197.609169 | 0.51702762 | 0.32300329 | 1.60068844 | 0.10944594 | 0.63899359 |
| TCHP | 141.012141 | -0.187132 | 0.34731369 | -0.5387984 | 0.59002599 | 0.93483536 |
| GIT2 | 340.666388 | -0.1541034 | 0.45572344 | -0.3381512 | 0.73524924 | 0.96078945 |
| ANKRD13A | 661.947182 | -0.3970105 | 0.49363645 | -0.8042569 | 0.42124862 | 0.88360543 |
| C12orf76 | 89.961479 | 0.08736107 | 0.53204363 | 0.16419907 | 0.86957443 | 0.9820194 |
| IFT81 | 224.134628 | -0.1529849 | 0.18091093 | -0.8456363 | 0.39775565 | 0.87462201 |
| ATP2A2 | 291.381182 | -0.3643476 | 0.31244457 | -1.1661191 | 0.2435663 | 0.80072691 |
| ANAPC7 | 123.853983 | -0.5004239 | 0.30231211 | -1.6553222 | 0.09785916 | 0.61685067 |
| ARPC3 | 563.448177 | 0.12581044 | 0.45553507 | 0.27618168 | 0.78240852 | 0.96513523 |
| GPN3 | 55.7737972 | 0.48883093 | 0.34579066 | 1.41366145 | 0.15746133 | 0.71160548 |
| VPS29 | 232.781353 | 0.22560699 | 0.31642362 | 0.71299036 | 0.47585172 | 0.90242252 |
| PPTC7 | 116.865141 | 0.16824437 | 0.26096734 | 0.64469511 | 0.51912479 | 0.91751084 |
| TCTN1 | 102.205138 | 0.65548675 | 0.32582698 | 2.01176327 | 0.0442449 | 0.48009422 |
| PPP1CC | 117.864951 | 0.10346734 | 0.27653474 | 0.37415675 | 0.7082877 | 0.95752169 |
| SH2B3 | 57.0922596 | 0.71608541 | 0.3531776 | 2.0275505 | 0.04260615 | 0.47247434 |
| ATXN2 | 902.224375 | -0.227375 | 0.27201391 | -0.8358948 | 0.40321408 | 0.87834687 |
| BRAP | 119.238577 | 0.00884555 | 0.26574881 | 0.03328537 | 0.97344702 | 0.99656026 |
| ACAD10 | 135.150577 | -0.1090893 | 0.35941408 | -0.3035198 | 0.76149375 | 0.96218738 |
| ALDH2 | 371.312175 | -0.0267467 | 0.34983473 | -0.0764552 | 0.93905699 | 0.99243654 |
| MAPKAPK5 | 131.733365 | -0.1386375 | 0.17627352 | -0.7864907 | 0.43158009 | 0.88650713 |
| TMEM116 | 370.293579 | 0.12207409 | 0.22100464 | 0.55235982 | 0.58070185 | 0.93354125 |

| | | | | | | |
|---|---|---|---|---|---|---|
| ERP29 | 456.206875 | 0.65997043 | 0.39874731 | 1.65510943 | 0.0979023 | 0.61685067 |
| NAA25 | 220.949283 | -0.4184412 | 0.24110823 | -1.7354911 | 0.08265385 | 0.59202053 |
| TRAFD1 | 110.489079 | 0.16294204 | 0.2788219 | 0.58439468 | 0.5589548 | 0.92901206 |
| HECTD4 | 780.476154 | -0.1266486 | 0.21687814 | -0.583962 | 0.55924587 | 0.92901206 |
| RPL6 | 633.958003 | 0.44142469 | 0.46801935 | 0.94317616 | 0.34559081 | 0.8523814 |
| PTPN11 | 269.411659 | 0.03895832 | 0.26313882 | 0.14805236 | 0.88230145 | 0.98335905 |
| RASAL1 | 535.947066 | -1.1282693 | 0.40984115 | -2.7529429 | 0.00590622 | 0.19045289 |
| DDX54 | 90.0294536 | 0.07766713 | 0.32466038 | 0.23922577 | 0.81093052 | 0.9680627 |
| IQCD | 42.4037316 | -0.0109406 | 0.26829441 | -0.0407784 | 0.96747258 | 0.99637506 |
| TPCN1 | 686.699745 | -0.0585203 | 0.25160855 | -0.2325848 | 0.81608381 | 0.96844493 |
| SLC8B1 | 137.886408 | -0.0618835 | 0.31242858 | -0.1980726 | 0.84298829 | 0.97560439 |
| PLBD2 | 47.4737983 | -0.2652627 | 0.27555865 | -0.9626362 | 0.33573012 | 0.84552074 |
| SDSL | 77.3714235 | 1.03857441 | 0.4657966 | 2.22967365 | 0.02576912 | 0.38969746 |
| RBM19 | 105.663141 | -0.6020048 | 0.26308664 | -2.2882377 | 0.02212368 | 0.35843654 |
| TBX3 | 162.08484 | 0.4543542 | 0.61903098 | 0.73397652 | 0.46296305 | 0.89801718 |
| MED13L | 3118.31309 | -0.4199121 | 0.2629048 | -1.597202 | 0.11022068 | 0.64066154 |
| C12orf49 | 85.0369402 | 0.286686 | 0.39650035 | 0.72304098 | 0.46965471 | 0.90053922 |
| FBXW8 | 206.833452 | -0.0865822 | 0.25964687 | -0.3334613 | 0.73878613 | 0.96078945 |
| FBXO21 | 254.993312 | 0.32299807 | 0.19358311 | 1.66852404 | 0.09521174 | 0.61367241 |
| KSR2 | 1867.7967 | -1.9588378 | 0.66689868 | -2.9372346 | 0.00331154 | 0.14358074 |
| RFC5 | 42.4200769 | -0.6022944 | 0.39030876 | -1.543123 | 0.12280093 | 0.66474886 |
| WSB2 | 139.168784 | -0.6098475 | 0.38744013 | -1.5740431 | 0.11547747 | 0.65156815 |
| VSIG10 | 198.015635 | -0.0135066 | 0.32840386 | -0.0411281 | 0.96719378 | 0.99637506 |
| PEBP1 | 1626.9796 | 0.33656005 | 0.55327089 | 0.60830969 | 0.5429821 | 0.92482794 |
| TAOK3 | 679.772473 | -0.054951 | 0.23302236 | -0.2358188 | 0.81357331 | 0.96819339 |
| SUDS3 | 174.966462 | 0.00416286 | 0.17538742 | 0.02373523 | 0.98106381 | 0.99696835 |
| AC002070.1 | 156.147628 | 0.76667265 | 0.53339514 | 1.43734463 | 0.15062009 | 0.7029344 |
| PRKAB1 | 220.65313 | 0.44394897 | 0.35954866 | 1.2347396 | 0.21692743 | 0.7815055 |
| CIT | 154.227566 | 1.38418721 | 0.5025499 | 2.75432791 | 0.00588128 | 0.19023403 |
| BICDL1 | 89.1091587 | -0.4477381 | 0.33818571 | -1.3239415 | 0.18552249 | 0.74895057 |
| RAB35 | 65.3102304 | 0.06979597 | 0.35616843 | 0.19596339 | 0.84463882 | 0.97573093 |
| GCN1 | 122.989356 | -0.1922063 | 0.22045564 | -0.8718591 | 0.38328524 | 0.86638702 |
| RPLP0 | 846.278295 | -0.0462978 | 0.47063272 | -0.0983735 | 0.92163575 | 0.99051393 |
| PXN | 184.018413 | -0.0557839 | 0.15936254 | -0.3500438 | 0.72630586 | 0.95979351 |
| GATC | 56.9182016 | -0.0482052 | 0.26730479 | -0.180338 | 0.85688719 | 0.97718353 |
| SRSF9 | 319.018242 | 0.26468456 | 0.42272938 | 0.62613239 | 0.53122811 | 0.92211604 |
| DYNLL1 | 506.656271 | 1.08085973 | 0.55790728 | 1.93734654 | 0.052703 | 0.50904874 |
| COQ5 | 67.3774623 | 0.0145074 | 0.23898103 | 0.06070526 | 0.95159395 | 0.99404743 |
| RNF10 | 485.550096 | 0.41295648 | 0.1744804 | 2.36677861 | 0.01794366 | 0.32200272 |
| POP5 | 80.1259349 | 0.20567822 | 0.34430994 | 0.59736359 | 0.55026466 | 0.92780338 |
| MLEC | 320.983334 | 0.18041449 | 0.21647064 | 0.83343632 | 0.4045987 | 0.87834687 |
| UNC119B | 80.8252343 | 0.49150961 | 0.28643334 | 1.71596507 | 0.08616844 | 0.60143834 |
| ACADS | 89.662606 | 0.61180386 | 0.29709054 | 2.05931784 | 0.0394638 | 0.45702496 |
| SPPL3 | 871.241996 | -0.2378533 | 0.22103473 | -1.0760902 | 0.2818869 | 0.82661559 |
| C12orf43 | 37.7798939 | -0.2459018 | 0.4151357 | -0.5923407 | 0.55362248 | 0.92872652 |
| P2RX4 | 93.9654064 | 0.76161484 | 0.35219555 | 2.16247719 | 0.03058141 | 0.41141619 |
| CAMKK2 | 62.6729394 | -0.3062407 | 0.30704103 | -0.9973933 | 0.31857364 | 0.84285425 |
| ANAPC5 | 231.498637 | -0.0853008 | 0.20359523 | -0.4189724 | 0.67523632 | 0.95524793 |
| RNF34 | 105.193422 | -0.1363303 | 0.2514463 | -0.5421847 | 0.58769129 | 0.93483536 |
| KDM2B | 209.462658 | -0.2992172 | 0.3024972 | -0.989157 | 0.32258632 | 0.84285425 |
| ORAI1 | 36.3008872 | 0.33467329 | 0.30523385 | 1.09644881 | 0.27288241 | 0.81922962 |
| MORN3 | 36.8209958 | 0.72159837 | 0.42951332 | 1.68003722 | 0.09295007 | 0.61303762 |
| TMEM120B | 158.522462 | -0.0911141 | 0.31128485 | -0.2927033 | 0.76974893 | 0.96345263 |
| RHOF | 73.3763823 | -0.183011 | 0.69664173 | -0.2627046 | 0.79277824 | 0.9653598 |

| | | | | | | |
|---|---|---|---|---|---|---|
| SETD1B | 83.5051758 | 0.38850867 | 0.33906627 | 1.14581928 | 0.25186994 | 0.80268905 |
| WDR66 | 123.018202 | -0.3419976 | 0.39860607 | -0.8579839 | 0.39090137 | 0.87004477 |
| BCL7A | 241.541677 | 0.12772459 | 0.30082891 | 0.42457551 | 0.67114615 | 0.95430867 |
| MLXIP | 199.490668 | -0.088581 | 0.27058487 | -0.3273685 | 0.74338922 | 0.96105014 |
| CLIP1 | 414.007607 | -0.6941592 | 0.26528193 | -2.6166847 | 0.00887883 | 0.22975346 |
| ZCCHC8 | 137.084771 | -0.2430683 | 0.2023799 | -1.2010498 | 0.2297319 | 0.79047378 |
| RSRC2 | 279.892011 | -0.0368757 | 0.27984136 | -0.1317737 | 0.89516329 | 0.98597539 |
| KNTC1 | 46.9474156 | -0.3602471 | 0.30560791 | -1.1787884 | 0.23848244 | 0.7985324 |
| DENR | 239.315761 | 0.16791152 | 0.2015881 | 0.83294364 | 0.40487652 | 0.8783757 |
| HIP1R | 492.622837 | 0.14240383 | 0.29043415 | 0.49031365 | 0.62391197 | 0.94457465 |
| VPS37B | 612.33814 | -0.1103763 | 0.57970642 | -0.1904003 | 0.84899545 | 0.97656375 |
| ABCB9 | 42.2722554 | 0.20516701 | 0.35281006 | 0.58152258 | 0.56088831 | 0.92934517 |
| ARL6IP4 | 260.00454 | -0.0860147 | 0.39733763 | -0.2164777 | 0.8286154 | 0.97117144 |
| PITPNM2 | 262.832733 | -0.618583 | 0.24989296 | -2.4753919 | 0.013309 | 0.28170334 |
| MPHOSPH9 | 168.394284 | -0.2266275 | 0.20660702 | -1.0969015 | 0.27268447 | 0.81922962 |
| C12orf65 | 108.25951 | 0.52389913 | 0.33338518 | 1.57145295 | 0.11607747 | 0.65192493 |
| SBNO1 | 282.330514 | -0.3632102 | 0.17284248 | -2.101394 | 0.0356064 | 0.43592878 |
| KMT5A | 82.0936567 | -0.0672964 | 0.24802245 | -0.271332 | 0.78613571 | 0.96513523 |
| RILPL2 | 145.509692 | 0.88980511 | 0.29298927 | 3.03698868 | 0.00238954 | 0.1271189 |
| SNRNP35 | 105.48264 | 0.09625043 | 0.24596523 | 0.3913172 | 0.69556278 | 0.95672887 |
| RILPL1 | 193.540678 | 0.03856492 | 0.33179058 | 0.11623271 | 0.90746811 | 0.9875066 |
| TMED2 | 177.413931 | -0.0680771 | 0.25092885 | -0.2713006 | 0.78615988 | 0.96513523 |
| DDX55 | 52.8517523 | 0.00094739 | 0.44799891 | 0.00211471 | 0.99831271 | 0.99967893 |
| EIF2B1 | 79.1014697 | -0.2520281 | 0.25190191 | -1.000501 | 0.31706812 | 0.84285425 |
| GTF2H3 | 82.3696966 | -0.1231272 | 0.21906073 | -0.5620689 | 0.57406906 | 0.93270589 |
| TCTN2 | 81.1720851 | -0.0857009 | 0.25390223 | -0.3375352 | 0.73571351 | 0.96078945 |
| ATP6V0A2 | 68.7536955 | -0.5206256 | 0.45678071 | -1.1397714 | 0.25438157 | 0.80414079 |
| CCDC92 | 251.490078 | -0.0672643 | 0.17312888 | -0.3885218 | 0.69762992 | 0.95672887 |
| RFLNA | 46.4169526 | 0.59044159 | 0.43188872 | 1.36711509 | 0.17158923 | 0.72776437 |
| NCOR2 | 303.080367 | -0.0876035 | 0.24013692 | -0.3648065 | 0.71525583 | 0.95791799 |
| UBC | 1643.7255 | -0.491287 | 0.38813377 | -1.2657671 | 0.20559647 | 0.77193931 |
| DHX37 | 35.4751188 | -0.1472552 | 0.38195792 | -0.3855274 | 0.69984671 | 0.95672887 |
| BRI3BP | 77.0941436 | 0.24968744 | 0.26920363 | 0.92750397 | 0.35366493 | 0.8557799 |
| AACS | 83.285733 | 0.12382276 | 0.39550047 | 0.31307865 | 0.75422091 | 0.96120879 |
| SLC15A4 | 128.00908 | -0.3397719 | 0.28796776 | -1.1798958 | 0.23804167 | 0.79810098 |
| RIMBP2 | 492.677205 | -0.4827325 | 0.35344202 | -1.3658039 | 0.17200051 | 0.72776437 |
| STX2 | 170.562747 | -0.0638862 | 0.36342463 | -0.1757894 | 0.86045942 | 0.97841862 |
| RAN | 278.098796 | 0.53206878 | 0.51477706 | 1.0335907 | 0.30132755 | 0.83709431 |
| SFSWAP | 255.543422 | -0.5098479 | 0.21357541 | -2.3872031 | 0.01697711 | 0.3110491 |
| ULK1 | 149.806587 | -0.3585969 | 0.29236942 | -1.2265198 | 0.22000314 | 0.78633738 |
| PUS1 | 42.4096671 | -0.3933317 | 0.41198064 | -0.9547334 | 0.33971252 | 0.84928129 |
| EP400 | 255.748914 | -0.1795278 | 0.2225235 | -0.8067815 | 0.41979238 | 0.8828928 |
| DDX51 | 37.3175291 | -0.2713029 | 0.41599208 | -0.6521828 | 0.51428328 | 0.91544604 |
| FBRSL1 | 135.972497 | -0.4854163 | 0.28449018 | -1.7062674 | 0.0879583 | 0.60651125 |
| ANKLE2 | 167.054547 | -0.4383751 | 0.31219204 | -1.4041841 | 0.16026402 | 0.71409437 |
| GOLGA3 | 163.617517 | -0.4053212 | 0.31122515 | -1.3023406 | 0.19279996 | 0.7597012 |
| CHFR | 181.521579 | -0.2725144 | 0.26183761 | -1.0407764 | 0.29797933 | 0.83556453 |
| ZNF26 | 120.020117 | -0.3295721 | 0.25533094 | -1.2907646 | 0.19678532 | 0.76478825 |
| ZNF84 | 195.682842 | -0.2619322 | 0.25494345 | -1.0274131 | 0.30422599 | 0.83709431 |
| ZNF140 | 99.8154048 | 0.03322611 | 0.2708351 | 0.12268022 | 0.90236033 | 0.9875066 |
| ZNF891 | 110.530342 | 0.17745071 | 0.54555572 | 0.32526597 | 0.74497979 | 0.96109466 |
| ZNF268 | 52.4280818 | 0.6670199 | 0.33227203 | 2.00745126 | 0.04470164 | 0.48147291 |
| TPTE2 | 46.0147155 | -0.4933426 | 0.42961758 | -1.1483297 | 0.25083247 | 0.80240987 |
| MPHOSPH8 | 555.339125 | 0.25628853 | 0.24925719 | 1.02820918 | 0.30385144 | 0.83709431 |

| | | | | | | |
|---|---|---|---|---|---|---|
| PSPC1 | 564.471038 | 0.01225161 | 0.2863157 | 0.04279056 | 0.96586849 | 0.99637506 |
| ZMYM5 | 166.049491 | 0.27622194 | 0.22261137 | 1.24082588 | 0.21467008 | 0.77890374 |
| ZMYM2 | 1430.67888 | -0.1083856 | 0.292924 | -0.3700126 | 0.71137308 | 0.957816 |
| CRYL1 | 436.655152 | 0.08410065 | 0.26472282 | 0.31769323 | 0.75071764 | 0.96109466 |
| IFT88 | 603.270023 | -0.2344161 | 0.22666331 | -1.0342041 | 0.30104076 | 0.83709431 |
| EEF1AKMT1 | 62.9319318 | 0.24711909 | 0.32189172 | 0.76770874 | 0.44266024 | 0.89005186 |
| XPO4 | 325.721006 | -0.1491679 | 0.24855607 | -0.6001376 | 0.54841451 | 0.92704021 |
| LATS2 | 239.859694 | 0.49171623 | 0.25283669 | 1.94479774 | 0.05179933 | 0.50451388 |
| SAP18 | 339.29321 | 0.27018734 | 0.50443111 | 0.53562783 | 0.59221579 | 0.93491607 |
| MRPL57 | 224.602393 | 0.23094984 | 0.41993113 | 0.54997076 | 0.58233943 | 0.93402467 |
| ZDHHC20 | 383.182021 | -0.1481244 | 0.26239326 | -0.5645131 | 0.57240497 | 0.93218128 |
| MICU2 | 623.363501 | 0.00216123 | 0.19894662 | 0.01086337 | 0.99133246 | 0.99889703 |
| FGF9 | 76.0261139 | 0.97566826 | 0.55070356 | 1.77167598 | 0.07644836 | 0.57145422 |
| AL136962.1 | 73.6110028 | 2.25336511 | 1.03078724 | 2.18606228 | 0.02881105 | 0.40623517 |
| TNFRSF19 | 160.58187 | 0.76458962 | 0.51599677 | 1.48177211 | 0.13840094 | 0.68146162 |
| MIPEP | 340.115971 | -0.5323421 | 0.28612773 | -1.8605051 | 0.0628141 | 0.54720847 |
| PARP4 | 420.262053 | -0.5583183 | 0.24004871 | -2.3258541 | 0.02002634 | 0.33933607 |
| ATP12A | 78.4487913 | 1.96075764 | 0.83594722 | 2.34555197 | 1.90E-02 | 0.33090117 |
| MTMR6 | 103.636832 | 0.34914363 | 0.19942368 | 1.75076314 | 0.07998672 | 0.58660658 |
| NUP58 | 277.213598 | -0.7723862 | 0.45861274 | -1.6841795 | 0.09214699 | 0.61120158 |
| RNF6 | 46.1819655 | 0.15444962 | 0.33307068 | 0.46371424 | 0.6428525 | 0.94949493 |
| CDK8 | 420.726867 | -0.1535024 | 0.25235167 | -0.6082875 | 0.5429968 | 0.92482794 |
| WASF3 | 120.26111 | 0.03087829 | 0.23694962 | 0.13031584 | 0.89631655 | 0.98597539 |
| USP12 | 520.937698 | -0.2315044 | 0.22329262 | -1.0367756 | 0.29984043 | 0.83618229 |
| LINC02340 | 76.0363849 | -0.4727455 | 0.57114172 | -0.8277201 | 0.40782902 | 0.87948803 |
| RPL21 | 394.019541 | 0.33939793 | 0.46378675 | 0.73179738 | 0.46429225 | 0.89801718 |
| RASL11A | 143.583417 | 0.38602484 | 0.61284513 | 0.6298897 | 0.52876675 | 0.92136681 |
| GTF3A | 77.3734787 | -0.3205867 | 0.28432379 | -1.1275408 | 0.2595139 | 0.80709747 |
| MTIF3 | 175.040713 | 0.20845925 | 0.27480212 | 0.75857946 | 0.44810416 | 0.89297728 |
| LNX2 | 182.105169 | -0.1090722 | 0.21168569 | -0.5152552 | 0.6063747 | 0.93919596 |
| POLR1D | 162.51203 | -0.0686842 | 0.20672328 | -0.3322519 | 0.73969904 | 0.96078945 |
| PAN3 | 1322.70683 | -0.1157633 | 0.21122207 | -0.5480646 | 0.58364758 | 0.93473293 |
| POMP | 167.897915 | -0.11165 | 0.32439402 | -0.3441803 | 0.73071067 | 0.96008876 |
| SLC7A1 | 629.687498 | -1.1002987 | 0.66959785 | -1.6432231 | 0.10033677 | 0.61891077 |
| UBL3 | 737.776494 | -0.2278368 | 0.42049028 | -0.5418361 | 0.58793145 | 0.93483536 |
| KATNAL1 | 129.101114 | -0.5612391 | 0.26835364 | -2.0914159 | 0.0364908 | 0.43729328 |
| HMGB1 | 901.696171 | 0.02622947 | 0.24825626 | 0.10565482 | 0.91585623 | 0.98868699 |
| USPL1 | 95.5449998 | -0.0181991 | 0.28338214 | -0.0642212 | 0.94879412 | 0.99344214 |
| HSPH1 | 227.702836 | 0.40561257 | 0.43667492 | 0.92886619 | 0.35295844 | 0.85545893 |
| B3GLCT | 136.400377 | 0.27672448 | 0.29348976 | 0.94287612 | 0.34574427 | 0.8523814 |
| FRY | 1912.48795 | -0.6283978 | 0.42415576 | -1.4815261 | 0.13846645 | 0.68146162 |
| N4BP2L1 | 41.9015266 | 1.05961968 | 0.53131137 | 1.99434784 | 0.04611405 | 0.48521615 |
| N4BP2L2 | 1872.18608 | -0.2920904 | 0.18945051 | -1.5417768 | 0.12312785 | 0.66519057 |
| PDS5B | 705.454846 | -0.0412892 | 0.26557698 | -0.1554697 | 0.87645102 | 0.98269442 |
| KL | 112.136416 | 0.31783893 | 0.32770009 | 0.96990798 | 3.32E-01 | 8.44E-01 |
| STARD13 | 1577.76858 | 0.32464628 | 0.2262217 | 1.43508018 | 1.51E-01 | 0.7029344 |
| AL139383.1 | 132.73045 | -0.1217549 | 0.43111484 | -0.2824187 | 0.77762251 | 0.96472142 |
| RFC3 | 98.7186951 | -0.1611585 | 0.38602867 | -0.4174782 | 0.67632868 | 0.95529325 |
| LINC02343 | 357.525064 | 1.27280849 | 0.58493657 | 2.17597694 | 0.02955698 | 0.40623517 |
| NBEA | 1345.16719 | -0.6661492 | 0.21069109 | -3.1617343 | 0.00156833 | 0.09927452 |
| SPART | 301.699875 | -0.1895714 | 0.34661547 | -0.5469214 | 0.58443272 | 0.93483536 |
| RFXAP | 42.6748799 | -0.1006426 | 0.4017055 | -0.2505382 | 0.80217114 | 0.9661728 |
| SMAD9 | 45.6607397 | -0.2358336 | 0.60877209 | -0.3873923 | 0.69846581 | 0.95672887 |
| ALG5 | 113.119631 | 0.06636224 | 0.21243049 | 0.31239508 | 0.75474029 | 0.96120879 |

| | | | | | | |
|---|---|---|---|---|---|---|
| EXOSC8 | 86.2396063 | -0.0049355 | 0.31841247 | -0.0155003 | 0.98763303 | 0.99791717 |
| SUPT20H | 362.157628 | -0.0955748 | 0.207761 | -0.4600227 | 0.64549994 | 0.94973439 |
| LINC00571 | 116.653971 | -0.2163816 | 0.34010398 | -0.6362219 | 0.5246318 | 0.91923576 |
| UFM1 | 226.478617 | -0.1694419 | 0.2584536 | -0.6555989 | 0.51208226 | 0.91408994 |
| FREM2 | 529.25617 | 0.23199239 | 0.25241728 | 0.91908284 | 0.35805225 | 0.8557799 |
| PROSER1 | 84.9675629 | -0.5183259 | 0.34388588 | -1.5072614 | 0.13174367 | 0.67390608 |
| NHLRC3 | 171.059324 | -0.1181801 | 0.34364905 | -0.3438978 | 0.73092317 | 0.96008876 |
| COG6 | 227.386494 | -0.1787005 | 0.22850194 | -0.7820523 | 0.43418385 | 0.88785137 |
| LINC00598 | 60.574015 | -0.7087141 | 0.36922242 | -1.9194773 | 0.05492395 | 0.51623588 |
| FOXO1 | 1303.99617 | -1.4675222 | 0.52293796 | -2.8063027 | 0.00501136 | 0.17602704 |
| MRPS31 | 211.235656 | 0.12292371 | 0.256729 | 0.47880725 | 0.63207576 | 0.94654542 |
| ELF1 | 730.964487 | -0.1514693 | 0.16348481 | -0.9265035 | 0.35418438 | 0.8557799 |
| WBP4 | 122.590746 | -0.221557 | 0.22583223 | -0.9810691 | 0.32655867 | 0.84381657 |
| AL354696.2 | 56.5742345 | -0.4187203 | 0.42389421 | -0.9877944 | 0.32325336 | 0.84285425 |
| MTRF1 | 55.6489324 | -0.4129119 | 0.27909929 | -1.4794446 | 0.13902153 | 0.68177147 |
| NAA16 | 150.828978 | 0.09138742 | 0.22717475 | 0.40227808 | 0.68747938 | 0.95588883 |
| VWA8 | 740.358479 | 0.07725159 | 0.24641227 | 0.31350547 | 0.75389667 | 0.96120879 |
| DGKH | 348.730325 | 0.35019589 | 0.50728989 | 0.69032697 | 0.48998859 | 0.90730741 |
| AKAP11 | 383.071374 | 0.7592033 | 0.19762069 | 3.84171963 | 0.00012218 | 0.02194494 |
| EPSTI1 | 248.753441 | -1.653501 | 0.73853852 | -2.2388825 | 0.02516356 | 0.3833054 |
| DNAJC15 | 305.465707 | -0.2429098 | 0.23709801 | -1.0245121 | 0.30559344 | 0.83709431 |
| CCDC122 | 131.723219 | 0.44712945 | 0.37681171 | 1.18661242 | 0.23538054 | 0.79599484 |
| SERP2 | 56.7504231 | -0.0028385 | 0.27416424 | -0.0103533 | 0.99173943 | 0.99917604 |
| TSC22D1 | 1676.4044 | 0.14419082 | 0.32180797 | 0.4480648 | 0.65410643 | 0.95100227 |
| NUFIP1 | 44.0004604 | -0.4076091 | 0.32449936 | -1.2561167 | 0.20907365 | 0.77436433 |
| AL359706.1 | 40.2093632 | 0.32830419 | 0.53159741 | 0.61758049 | 0.53685191 | 0.92324352 |
| GPALPP1 | 151.319775 | -0.1481009 | 0.19375603 | -0.7643677 | 0.44464813 | 0.89105265 |
| GTF2F2 | 244.509095 | -0.1086545 | 0.26710022 | -0.4067928 | 0.68416015 | 0.95574492 |
| TPT1 | 1827.85903 | 0.06977487 | 0.52986107 | 0.13168523 | 0.89523327 | 0.98597539 |
| SLC25A30 | 285.683407 | 0.94306099 | 0.30634891 | 3.0783886 | 0.00208123 | 0.1166381 |
| COG3 | 218.516612 | -0.1958134 | 0.18385141 | -1.0650631 | 0.28684737 | 0.82943317 |
| ZC3H13 | 363.136597 | -0.0490794 | 0.16310136 | -0.3009137 | 0.76348027 | 0.96285313 |
| AL138686.1 | 111.548856 | 0.43574404 | 0.65787205 | 0.66235378 | 0.50774452 | 0.91214298 |
| LRCH1 | 992.759824 | 0.11276695 | 0.25776068 | 0.43748701 | 0.66175819 | 0.95142083 |
| ESD | 178.770144 | 0.16060578 | 0.27708843 | 0.57961924 | 0.56217142 | 0.92985424 |
| SUCLA2 | 271.805612 | -0.1865383 | 0.25608376 | -0.7284268 | 0.46635235 | 0.89883965 |
| MED4 | 195.057289 | -0.0658096 | 0.23621245 | -0.2786033 | 0.78054927 | 0.96513523 |
| ITM2B | 2677.27816 | 0.50388794 | 0.33667572 | 1.4966566 | 0.13448263 | 0.67856427 |
| RB1 | 778.896408 | -0.1496205 | 0.21037692 | -0.7112019 | 0.47695915 | 0.90242252 |
| LPAR6 | 41.3331321 | -0.1999345 | 0.47066331 | -0.4247931 | 0.67098748 | 0.95426093 |
| RCBTB2 | 150.738364 | -0.5187721 | 0.31649089 | -1.6391377 | 0.10118459 | 0.62105083 |
| CYSLTR2 | 76.3281243 | -1.3230254 | 0.31077518 | -4.2571785 | 2.07E-05 | 0.00614104 |
| FNDC3A | 2236.98222 | -1.116579 | 0.68827541 | -1.6222852 | 0.1047423 | 0.6317386 |
| CDADC1 | 150.59431 | -0.3412376 | 0.76696022 | -0.4449222 | 0.65637599 | 0.95100227 |
| CAB39L | 293.609798 | -0.6529906 | 0.62313319 | -1.047915 | 0.29467776 | 0.8337535 |
| SETDB2 | 193.869372 | -0.473729 | 0.37693995 | -1.2567759 | 0.20883479 | 0.77436433 |
| PHF11 | 136.199861 | -0.9090873 | 0.30867062 | -2.9451697 | 0.00322778 | 0.14358074 |
| RCBTB1 | 74.5454059 | -0.8834304 | 0.24232504 | -3.645642 | 0.00026673 | 0.03764581 |
| EBPL | 76.9514974 | 0.16654257 | 0.35799457 | 0.46520976 | 0.64178125 | 0.94935885 |
| KPNA3 | 264.808017 | -0.0964597 | 0.29038313 | -0.3321806 | 0.73975287 | 0.96078945 |
| SPRYD7 | 54.8040574 | 0.15649586 | 0.34179738 | 0.4578615 | 0.64705194 | 0.95061388 |
| DLEU2 | 480.557704 | -0.3661584 | 0.34803397 | -1.0520768 | 0.29276434 | 0.83336457 |
| TRIM13 | 137.309181 | -0.0549296 | 0.24764423 | -0.2218086 | 0.82446287 | 0.97018919 |
| RNASEH2B | 113.425869 | 0.0740593 | 0.2191002 | 0.33801568 | 0.73535137 | 0.96078945 |

| | | | | | | |
|---|---|---|---|---|---|---|
| FAM124A | 115.969252 | -1.2562786 | 0.35874346 | -3.5018856 | 0.00046198 | 0.05075624 |
| INTS6 | 555.592124 | -0.0890107 | 0.33801293 | -0.2633353 | 0.79229215 | 0.9653598 |
| INTS6-AS1 | 56.8337084 | 0.51277783 | 0.37677582 | 1.36096269 | 0.17352548 | 0.73030936 |
| WDFY2 | 921.14649 | 0.500393 | 0.31042195 | 1.61197685 | 0.10696698 | 0.63477557 |
| ATP7B | 70.2373108 | 0.14278346 | 0.36582459 | 0.39030581 | 0.69631043 | 0.95672887 |
| UTP14C | 80.4532195 | -0.1741549 | 0.25665991 | -0.6785434 | 0.4974272 | 0.90839719 |
| NEK5 | 89.206893 | -0.0696928 | 0.56447784 | -0.1234642 | 0.90173956 | 0.9875066 |
| NEK3 | 142.848706 | -0.0763223 | 0.44336573 | -0.172143 | 0.86332515 | 0.979268 |
| VPS36 | 274.028616 | -0.2592306 | 0.16698165 | -1.5524498 | 0.12055463 | 0.6576848 |
| CKAP2 | 33.3618398 | 0.83286176 | 0.4780919 | 1.74205371 | 0.08149905 | 0.59202053 |
| SUGT1 | 273.328956 | -0.2072914 | 0.40805859 | -0.5079942 | 0.61145739 | 0.94003018 |
| OLFM4 | 141.863187 | -0.6572013 | 0.80673053 | -0.8146479 | 4.15E-01 | 8.82E-01 |
| PCDH17 | 71.4574262 | 1.13963102 | 0.9440375 | 1.2071883 | 0.22735963 | 0.79042788 |
| TDRD3 | 565.674159 | 0.21933407 | 0.27789082 | 0.78928143 | 0.42994753 | 0.88579433 |
| PCDH9 | 222.97549 | 0.3674818 | 0.63125576 | 0.58214407 | 0.56046964 | 0.92934517 |
| DACH1 | 789.339609 | 1.40966776 | 0.68362038 | 2.06206222 | 0.03920181 | 0.45547117 |
| MZT1 | 58.4350691 | 0.08789324 | 0.37817784 | 0.23241245 | 0.81621768 | 0.96844493 |
| DIS3 | 108.946388 | -0.2683271 | 0.17866234 | -1.5018673 | 0.13313139 | 0.67594872 |
| PIBF1 | 617.87971 | -0.2655694 | 0.17938484 | -1.4804453 | 0.13875444 | 0.68156449 |
| KLF5 | 229.999878 | -0.2741951 | 0.51846234 | -0.5288622 | 0.59690104 | 0.93617523 |
| KLF12 | 601.041563 | -0.211636 | 0.2866076 | -0.7384173 | 0.46026093 | 0.89785937 |
| TBC1D4 | 723.148086 | 0.04218994 | 0.23725734 | 0.17782352 | 0.85886157 | 0.97788846 |
| COMMD6 | 266.952291 | 0.30686582 | 0.37729571 | 0.81332975 | 0.41602902 | 0.88242985 |
| UCHL3 | 140.718068 | -0.2015784 | 0.22759518 | -0.8856885 | 0.37578538 | 0.86357183 |
| LMO7-AS1 | 62.2903644 | 1.02569335 | 0.42344715 | 2.42224643 | 0.01542489 | 0.30046991 |
| LMO7 | 1018.19608 | 0.30731908 | 0.30728007 | 1.00012695 | 0.31724907 | 0.84285425 |
| KCTD12 | 426.839992 | 2.48499253 | 0.69779515 | 3.56120635 | 0.00036915 | 0.04296095 |
| AC001226.2 | 85.4699145 | 0.89519463 | 0.60831576 | 1.47159532 | 0.14113019 | 0.68601316 |
| FBXL3 | 209.695548 | 0.57403851 | 0.34551077 | 1.66142002 | 0.09662912 | 0.61485924 |
| MYCBP2 | 696.23987 | -0.1522104 | 0.16625324 | -0.9155336 | 0.35991155 | 0.85608464 |
| SLAIN1 | 72.1591663 | 0.46376577 | 0.39905913 | 1.16214802 | 0.24517535 | 0.80072691 |
| EDNRB | 56.2834412 | -0.5523216 | 0.6984587 | -0.7907721 | 0.42907701 | 0.88570555 |
| RNF219 | 61.0085908 | 0.11888899 | 0.25615206 | 0.46413441 | 0.64255145 | 0.94949493 |
| RBM26 | 599.514608 | -0.1345368 | 0.21080181 | -0.6382148 | 0.52333388 | 0.91871245 |
| NDFIP2 | 285.967962 | -0.0252272 | 0.16082976 | -0.1568565 | 0.87535792 | 0.98267808 |
| GPC5 | 116.610583 | 1.12242826 | 0.5994876 | 1.87231272 | 0.06116335 | 0.5418629 |
| GPC6 | 325.19696 | -0.877985 | 0.93149943 | -0.9425502 | 0.34591103 | 0.8523814 |
| TGDS | 70.8700378 | -0.2471115 | 0.25520897 | -0.9682713 | 0.33290889 | 0.84442294 |
| GPR180 | 117.754598 | -0.5664491 | 0.3132075 | -1.8085425 | 0.0705221 | 0.5621116 |
| ABCC4 | 73.6138148 | 0.27213439 | 0.46786562 | 0.58165076 | 0.56080195 | 0.92934517 |
| CLDN10 | 156.519743 | 0.52002649 | 0.48238008 | 1.07804304 | 0.28101455 | 0.82628479 |
| DNAJC3-DT | 44.0332121 | -0.1930596 | 0.50271491 | -0.384034 | 0.70095322 | 0.95672887 |
| DNAJC3 | 306.960789 | -0.0363343 | 0.17236485 | -0.2107986 | 0.83304442 | 0.97208613 |
| UGGT2 | 702.525812 | -0.4372962 | 0.14883321 | -2.9381628 | 0.00330164 | 0.14358074 |
| MBNL2 | 694.929756 | 0.49396807 | 0.22974306 | 2.15008919 | 0.03154816 | 0.41477661 |
| RAP2A | 55.5524672 | -0.0796869 | 0.25566585 | -0.311684 | 0.7552807 | 0.96120879 |
| IPO5 | 209.625222 | 0.09227539 | 0.19980844 | 0.46181929 | 0.64421092 | 0.94949493 |
| FARP1 | 1569.7818 | 0.17624967 | 0.17798842 | 0.99023113 | 0.32206116 | 0.84285425 |
| STK24 | 705.960921 | -0.3600439 | 0.40817342 | -0.8820856 | 0.3777305 | 0.86489308 |
| SLC15A1 | 297.13111 | -6.084511 | 1.7041817 | -3.5703417 | 0.00035652 | 0.04294582 |
| DOCK9 | 544.531723 | 0.14587494 | 0.22194937 | 0.65724421 | 0.5110239 | 0.91390025 |
| UBAC2 | 803.84153 | -0.265146 | 0.12360744 | -2.1450651 | 0.03194764 | 0.41517597 |
| TM9SF2 | 407.258824 | 0.14427901 | 0.18336856 | 0.78682522 | 0.43138419 | 0.88650713 |
| CLYBL | 462.063435 | -0.2124931 | 0.29602305 | -0.7178263 | 0.4728644 | 0.90213021 |

| | | | | | | |
|---|---|---|---|---|---|---|
| AL355338.1 | 46.1168216 | 0.53889461 | 0.34170609 | 1.57707056 | 0.11477929 | 0.65008305 |
| PCCA | 2387.30086 | 0.08975906 | 0.30063267 | 0.29856722 | 0.76527028 | 0.96285313 |
| GGACT | 145.668035 | -0.1266573 | 0.34568509 | -0.366395 | 0.71407034 | 0.95791799 |
| TMTC4 | 89.085065 | -0.2615722 | 0.37600232 | -0.6956664 | 0.48663779 | 0.9070379 |
| FGF14 | 201.198396 | 0.09070571 | 0.63940749 | 0.141859 | 0.88719138 | 0.98551241 |
| TPP2 | 227.074853 | -0.2246569 | 0.22926358 | -0.9799067 | 0.32713217 | 0.84381657 |
| EFNB2 | 84.9803445 | 1.1985392 | 0.30967101 | 3.87036291 | 0.00010867 | 0.02190188 |
| ARGLU1 | 594.255505 | -0.1478327 | 0.17648457 | -0.8376525 | 0.40222591 | 0.87834687 |
| FAM155A | 1061.09161 | -1.2669486 | 0.91314862 | -1.3874506 | 0.16530439 | 0.71823799 |
| LIG4 | 55.6620841 | -0.0731002 | 0.33663678 | -0.2171485 | 0.82809262 | 0.97117144 |
| ABHD13 | 81.2055018 | 0.05962798 | 0.27165426 | 0.21949952 | 0.82626096 | 0.97058548 |
| TNFSF13B | 63.1217859 | 0.3112242 | 0.45316214 | 0.68678331 | 0.49221929 | 0.90769984 |
| MYO16 | 49.3748785 | 2.19328873 | 0.60529936 | 3.6234777 | 0.00029067 | 0.03764581 |
| IRS2 | 266.123167 | 0.08739049 | 0.38215827 | 0.22867617 | 0.81912062 | 0.96884581 |
| COL4A1 | 456.133857 | 0.05400159 | 0.44526083 | 0.1212808 | 0.90346863 | 0.9875066 |
| COL4A2 | 540.78759 | 0.32113258 | 0.40177591 | 0.79928282 | 0.42412644 | 0.88391318 |
| RAB20 | 117.419004 | 0.13095686 | 0.48112078 | 0.27219124 | 0.78547498 | 0.96513523 |
| NAXD | 116.498391 | -0.2009493 | 0.30255751 | -0.6641689 | 0.50658224 | 0.91163608 |
| CARS2 | 183.370712 | -0.1269328 | 0.22119756 | -0.5738437 | 0.56607356 | 0.93122667 |
| ING1 | 36.6579108 | 0.28789235 | 0.57586346 | 0.49993161 | 0.61712323 | 0.94233994 |
| ANKRD10 | 883.167844 | -0.339167 | 0.27835025 | -1.2184899 | 0.22303785 | 0.78860888 |
| ARHGEF7 | 491.838432 | -0.2271005 | 0.23569644 | -0.9635298 | 0.33528172 | 0.84544327 |
| TUBGCP3 | 263.506801 | -0.347875 | 0.44661057 | -0.7789225 | 0.43602536 | 0.88785137 |
| ATP11A | 349.383039 | -0.8521904 | 0.56733498 | -1.5020939 | 0.13307287 | 0.67594872 |
| MCF2L | 232.22734 | -0.1336847 | 0.31130412 | -0.4294345 | 0.66760706 | 0.95294349 |
| PCID2 | 149.811803 | -0.0470925 | 0.2260922 | -0.2082889 | 0.83500339 | 0.97246185 |
| CUL4A | 387.076538 | -0.4386157 | 0.1681783 | -2.60804 | 0.00910623 | 0.23300339 |
| LAMP1 | 504.055126 | 0.42572361 | 0.25223953 | 1.68777518 | 0.09145439 | 0.61047264 |
| GRTP1 | 83.3306621 | -0.0701734 | 0.27134291 | -0.2586153 | 0.79593211 | 0.9653598 |
| DCUN1D2 | 106.406344 | -0.0873918 | 0.24845525 | -0.3517405 | 0.72503285 | 0.95963742 |
| TMCO3 | 179.872799 | 0.06648693 | 0.28273632 | 0.23515525 | 0.81408824 | 0.96820092 |
| TFDP1 | 248.158709 | -1.2426097 | 0.61497408 | -2.0205887 | 0.04332236 | 0.47741153 |
| GAS6 | 39.7951853 | 0.63591164 | 0.56207251 | 1.13136941 | 0.25789964 | 0.80709747 |
| RASA3 | 39.9168406 | -0.7327992 | 0.73296142 | -0.9997787 | 0.31741762 | 0.84285425 |
| CDC16 | 208.21558 | -0.4348522 | 0.17718834 | -2.4541806 | 0.0141206 | 0.28646578 |
| UPF3A | 222.891937 | -0.0610768 | 0.19390429 | -0.314984 | 0.75277379 | 0.96120879 |
| CHAMP1 | 51.0068111 | 0.33472327 | 0.35011106 | 0.95604882 | 0.33904757 | 0.84863112 |
| TTC5 | 86.9248305 | -0.1674231 | 0.24736308 | -0.6768315 | 0.49851288 | 0.90839719 |
| CCNB1IP1 | 116.088224 | 0.35089704 | 0.29224777 | 1.2006834 | 0.22987404 | 0.79047378 |
| PARP2 | 47.6168912 | -0.161167 | 0.28902447 | -0.5576241 | 0.57710113 | 0.93319686 |
| TEP1 | 64.7863938 | 0.16995675 | 0.35787823 | 0.47490107 | 0.63485749 | 0.94776446 |
| OSGEP | 72.609086 | -0.363152 | 0.25919447 | -1.4010791 | 0.16119041 | 0.71409437 |
| APEX1 | 123.747318 | 0.26943859 | 0.48060351 | 0.5606255 | 0.57505286 | 0.93275827 |
| PIP4P1 | 56.8720752 | -0.3113727 | 0.42526466 | -0.7321857 | 0.46405525 | 0.89801718 |
| PNP | 331.622985 | -0.3299135 | 0.64635993 | -0.5104176 | 0.60975892 | 0.94003018 |
| METTL17 | 59.8809527 | -0.2095676 | 0.26914951 | -0.778629 | 0.43619829 | 0.88785137 |
| NDRG2 | 102.001372 | 0.73991152 | 0.27320206 | 2.7082941 | 0.00676301 | 0.20224502 |
| ARHGEF40 | 34.9438115 | 0.53407616 | 0.51317842 | 1.04072218 | 0.29800451 | 0.83556453 |
| HNRNPC | 1168.01829 | -0.2527732 | 0.19836025 | -1.2743137 | 0.20255227 | 0.76996624 |
| SUPT16H | 210.130377 | 0.15335397 | 0.19123042 | 0.80193294 | 0.42259175 | 0.88371155 |
| CHD8 | 288.794165 | -0.2457691 | 0.22883916 | -1.0739818 | 0.28283085 | 0.82666614 |
| RAB2B | 101.377701 | -0.4237469 | 0.19888414 | -2.130622 | 0.0331203 | 0.42324075 |
| TOX4 | 156.599859 | 0.01126304 | 0.23145648 | 0.04866158 | 0.96118899 | 0.99635733 |
| METTL3 | 52.3358413 | -0.3479356 | 0.30786463 | -1.1301579 | 0.25840969 | 0.80709747 |

| | | | | | | |
|---|---|---|---|---|---|---|
| DAD1 | 305.183069 | -0.0952299 | 0.42690471 | -0.2230707 | 0.82348049 | 0.97018919 |
| ABHD4 | 80.205 | 0.35368296 | 0.30928075 | 1.14356604 | 0.25280365 | 0.80350545 |
| OXA1L | 77.0897192 | 0.08827373 | 0.36873588 | 0.23939556 | 0.81079887 | 0.9680627 |
| MRPL52 | 75.7738044 | -0.039158 | 0.26688063 | -0.1467249 | 0.88334916 | 0.9835847 |
| MMP14 | 57.9058307 | -0.3550322 | 0.35710282 | -0.9942015 | 0.32012477 | 0.84285425 |
| LRP10 | 249.714515 | 0.17306332 | 0.26801766 | 0.6457161 | 0.51846323 | 0.91751084 |
| RBM23 | 161.130434 | -0.1852606 | 0.17723487 | -1.0452829 | 0.29589222 | 0.83444322 |
| PRMT5 | 45.9063962 | 0.15509956 | 0.25748092 | 0.60237305 | 0.54692585 | 0.92612423 |
| AL132780.1 | 104.23397 | 1.28088128 | 0.52399781 | 2.44444014 | 0.01450772 | 0.28850257 |
| C14orf93 | 44.1867853 | 0.51078956 | 0.33740589 | 1.51387267 | 0.13005814 | 0.67275878 |
| PSMB5 | 189.409381 | 0.26717616 | 0.42300557 | 0.63161381 | 0.52763926 | 0.92084254 |
| ACIN1 | 327.549624 | -0.2866293 | 0.12896431 | -2.2225474 | 0.02624634 | 0.39193403 |
| C14orf119 | 81.2651854 | 0.14408331 | 0.43580047 | 0.33061761 | 0.74093334 | 0.96084411 |
| SLC7A8 | 147.702189 | -1.9840075 | 0.46846013 | -4.2351682 | 2.28E-05 | 0.00646873 |
| HOMEZ | 50.2611621 | -0.1166453 | 0.29294007 | -0.3981883 | 0.69049142 | 0.95659154 |
| PPP1R3E | 45.5707688 | -0.4027959 | 0.42988699 | -0.9369808 | 0.34876843 | 0.8535343 |
| BCL2L2-PABI | 56.4222829 | -0.3955472 | 0.3221985 | -1.2276505 | 0.21957818 | 0.78630025 |
| PABPN1 | 232.533162 | -0.5956261 | 0.3210722 | -1.8551159 | 0.06357968 | 0.54850785 |
| SLC22A17 | 58.3790921 | 0.48654605 | 0.46424581 | 1.04803541 | 0.29462229 | 0.8337535 |
| NGDN | 60.0742697 | 0.00946694 | 0.28251407 | 0.03350961 | 0.9732682 | 0.99656026 |
| ZFHX2 | 97.7501131 | 0.54357822 | 0.42408233 | 1.28177522 | 0.19992151 | 0.7687013 |
| AP1G2 | 63.736377 | -0.7169159 | 0.26218178 | -2.7344231 | 0.00624897 | 0.19425932 |
| DHRS4-AS1 | 84.4471821 | 0.30396671 | 0.36400401 | 0.83506418 | 0.40368158 | 0.87834687 |
| DHRS4L2 | 77.9252234 | 0.29652064 | 0.39719542 | 0.7465359 | 0.45534376 | 0.89626521 |
| PCK2 | 37.1369237 | 0.16083909 | 0.2783312 | 0.57786942 | 0.56335228 | 0.9306324 |
| PSME1 | 235.37395 | 0.24709355 | 0.50822002 | 0.48619405 | 0.62682959 | 0.94511671 |
| EMC9 | 46.07675 | 0.49381746 | 0.37893766 | 1.3031628 | 0.19251919 | 0.7597012 |
| REC8 | 64.6221736 | -0.8478442 | 0.37244112 | -2.2764515 | 0.022819 | 0.36178988 |
| AL136295.1 | 61.4252418 | -0.5900923 | 0.34930591 | -1.6893282 | 0.09115655 | 0.60989481 |
| GMPR2 | 77.3890446 | 0.86681521 | 0.41646779 | 2.08134993 | 0.03740189 | 0.44240608 |
| TINF2 | 41.8570001 | -0.1851706 | 0.41305605 | -0.448294 | 0.65394101 | 0.95100227 |
| DHRS1 | 63.910255 | -0.5109638 | 0.34489012 | -1.4815263 | 0.13846638 | 0.68146162 |
| NFATC4 | 48.3539755 | -0.0750177 | 0.46319947 | -0.1619555 | 0.87134091 | 0.98208511 |
| KHNYN | 66.6992157 | -0.5721958 | 0.23234125 | -2.4627385 | 0.01378804 | 0.28408502 |
| SDR39U1 | 68.8865127 | 0.1046955 | 0.31035195 | 0.33734442 | 0.73585728 | 0.96078945 |
| STXBP6 | 315.541426 | -0.0814579 | 0.62176443 | -0.1310109 | 8.96E-01 | 0.98597539 |
| NOVA1 | 203.471465 | -0.6357906 | 0.50330709 | -1.263226 | 0.20650797 | 0.77200549 |
| LINC00645 | 74.6764619 | -0.4027289 | 0.91274428 | -0.4412286 | 0.65904752 | 0.95127957 |
| PRKD1 | 1578.75929 | -0.20174 | 0.1984728 | -1.0164615 | 0.3094097 | 0.84136316 |
| G2E3 | 132.102801 | -0.1746148 | 0.24209105 | -0.7212774 | 0.47073886 | 0.90123338 |
| SCFD1 | 409.782646 | -0.5061958 | 0.17905756 | -2.8270003 | 0.00469863 | 0.16921345 |
| COCH | 45.0082176 | -0.2163769 | 0.44677215 | -0.4843115 | 0.62816481 | 0.94572238 |
| STRN3 | 863.219518 | -0.4591799 | 0.3067099 | -1.4971146 | 0.13436345 | 0.67832119 |
| AP4S1 | 74.0242395 | 0.05335027 | 0.41374053 | 0.12894619 | 0.89740022 | 0.98612679 |
| HECTD1 | 911.77546 | -0.5650775 | 0.35480449 | -1.5926446 | 0.11123993 | 0.64290644 |
| HEATR5A | 106.659452 | -0.216234 | 0.30481095 | -0.7094037 | 0.478074 | 0.90262056 |
| AL139353.1 | 32.511718 | 0.6319962 | 0.45447942 | 1.39059365 | 0.16434868 | 0.71715698 |
| NUBPL | 487.482155 | -0.2402411 | 0.2685384 | -0.8946249 | 0.37098765 | 0.86188219 |
| ARHGAP5 | 926.804643 | 0.15921845 | 0.16096894 | 0.98912531 | 0.32260183 | 0.84285425 |
| AKAP6 | 223.163741 | -0.2307078 | 0.24818319 | -0.9295866 | 0.35258517 | 0.85536253 |
| NPAS3 | 1082.74482 | 0.19883949 | 0.52572235 | 0.37822149 | 0.70526607 | 0.95730303 |
| EGLN3 | 146.27035 | -0.4485361 | 0.6282308 | -0.7139671 | 0.47524751 | 0.90242252 |
| EAPP | 182.539306 | 0.1946177 | 0.23023539 | 0.84529881 | 0.39794402 | 0.87462201 |
| SNX6 | 439.3222 | -0.2622133 | 0.18361387 | -1.4280693 | 0.1532719 | 0.70717297 |

| | | | | | | |
|---|---|---|---|---|---|---|
| CFL2 | 63.2349758 | 0.19868443 | 0.3080893 | 0.64489233 | 0.51899696 | 0.91751084 |
| BAZ1A | 147.691668 | -0.2678763 | 0.28049224 | -0.9550222 | 0.33956646 | 0.84928129 |
| SRP54 | 296.046131 | -0.301617 | 0.20316037 | -1.4846251 | 0.13764317 | 0.68146162 |
| FAM177A1 | 316.805647 | -0.0530019 | 0.38083027 | -0.1391747 | 0.88931211 | 0.98551241 |
| PPP2R3C | 126.880778 | -0.417691 | 0.33230055 | -1.2569675 | 0.2087654 | 0.77436433 |
| NFKBIA | 262.942099 | 0.03713352 | 0.70212587 | 0.05288726 | 0.95782173 | 0.99579587 |
| RALGAPA1 | 1133.07387 | -0.3741649 | 0.33142793 | -1.128948 | 0.25891977 | 0.80709747 |
| BRMS1L | 61.7203975 | 0.04527877 | 0.25084289 | 0.18050647 | 0.85675497 | 0.97718353 |
| LINC00609 | 60.7968524 | 1.60191663 | 0.6547597 | 2.44657183 | 0.01442221 | 0.28805285 |
| PTCSC3 | 229.703533 | 2.5935612 | 0.64576045 | 4.01628997 | 5.91E-05 | 0.01318284 |
| MBIP | 195.721455 | 0.22087094 | 0.29899624 | 0.73870811 | 0.46008426 | 0.89772539 |
| MIPOL1 | 446.228417 | 0.22042079 | 0.22287207 | 0.98900144 | 0.32266244 | 0.84285425 |
| SEC23A | 338.426463 | -0.4578563 | 0.25445 | -1.7993961 | 0.07195604 | 0.5644261 |
| GEMIN2 | 36.8348167 | -0.1246403 | 0.28837588 | -0.4322147 | 0.66558538 | 0.95238247 |
| TRAPPC6B | 151.371846 | -0.027316 | 0.20794848 | -0.1313593 | 0.89549105 | 0.98597539 |
| PNN | 313.724751 | -0.2866724 | 0.19969228 | -1.4355707 | 0.15112454 | 0.7029344 |
| MIA2 | 426.061662 | -0.26559 | 0.23694598 | -1.1208885 | 0.26233533 | 0.809086 |
| FBXO33 | 223.786553 | -0.201779 | 0.27249398 | -0.7404897 | 0.45900294 | 0.89753337 |
| AL049875.1 | 64.0320957 | -0.1708416 | 0.3312283 | -0.515782 | 0.60600665 | 0.93919596 |
| KLHL28 | 162.05845 | 0.15789216 | 0.29341929 | 0.53811105 | 0.59050039 | 0.93483536 |
| TOGARAM1 | 299.144172 | -0.0041182 | 0.27907211 | -0.0147567 | 0.98822626 | 0.99814557 |
| PRPF39 | 132.893646 | -0.3838535 | 0.30907902 | -1.2419267 | 0.21426362 | 0.77890374 |
| FKBP3 | 148.366553 | 0.20116129 | 0.28554095 | 0.70449191 | 0.48112649 | 0.90524337 |
| FANCM | 69.5264671 | -0.4582596 | 0.34533876 | -1.3269857 | 0.18451341 | 0.74775735 |
| MIS18BP1 | 92.9573364 | 0.11930909 | 0.46597552 | 0.25604155 | 0.79791876 | 0.9653598 |
| RPS29 | 328.453786 | 0.1296948 | 0.26442788 | 0.49047327 | 0.62379904 | 0.94457465 |
| RPL36AL | 484.506431 | 0.10592516 | 0.45820811 | 0.23117259 | 0.81718072 | 0.96878439 |
| DNAAF2 | 44.6935119 | 0.53569626 | 0.27317036 | 1.96103364 | 0.0498751 | 0.49497258 |
| POLE2 | 44.0890959 | 0.11808385 | 0.35879889 | 0.32910874 | 0.7420735 | 0.96086423 |
| KLHDC1 | 140.956873 | 0.27630507 | 0.33825902 | 0.81684466 | 0.4140172 | 0.88215311 |
| KLHDC2 | 192.71911 | 0.30052917 | 0.25189619 | 1.19306754 | 0.23284292 | 0.7926597 |
| NEMF | 645.985896 | -0.2264402 | 0.2034594 | -1.1129505 | 0.2657297 | 0.8121456 |
| ARF6 | 221.926527 | 0.16927365 | 0.40631975 | 0.41660207 | 0.6769695 | 0.95529325 |
| LINC01588 | 51.6358684 | 0.51198672 | 0.27666655 | 1.85055521 | 0.06423357 | 0.54860648 |
| VCPKMT | 72.9927219 | -0.03071 | 0.22607606 | -0.1358391 | 0.8919485 | 0.98597539 |
| SOS2 | 859.333518 | 0.22381315 | 0.28799975 | 0.77712967 | 0.43708227 | 0.88785137 |
| L2HGDH | 384.218946 | 0.0962293 | 0.26288378 | 0.36605263 | 0.7143258 | 0.95791799 |
| ATP5S | 164.117811 | 0.27551567 | 0.17424615 | 1.58118654 | 0.11383539 | 0.6483668 |
| AL359397.2 | 37.2671268 | -0.1948085 | 0.35632224 | -0.5467201 | 0.58457104 | 0.93483536 |
| CDKL1 | 644.591154 | -0.0453827 | 0.37217668 | -0.1219386 | 0.90294762 | 0.9875066 |
| MAP4K5 | 936.660405 | -0.2624025 | 0.31418505 | -0.8351845 | 0.40361384 | 0.87834687 |
| ATL1 | 101.116963 | 0.03857983 | 0.36894858 | 0.10456696 | 0.91671943 | 0.98921125 |
| SAV1 | 148.121361 | 0.04662314 | 0.23858889 | 0.19541204 | 0.84507039 | 0.97590775 |
| NIN | 127.413043 | -0.2636564 | 0.29458296 | -0.8950159 | 0.37077858 | 0.86177855 |
| PYGL | 56.6486229 | 1.05693642 | 0.53377262 | 1.98012482 | 0.04768951 | 0.48600281 |
| TMX1 | 124.947369 | 0.02423753 | 0.24885829 | 0.09739492 | 0.92241278 | 0.99051393 |
| RTRAF | 187.921839 | -0.0464396 | 0.42385093 | -0.1095659 | 0.91275363 | 0.98830345 |
| NID2 | 38.4503393 | -1.0416232 | 0.99347528 | -1.0484641 | 0.29442483 | 0.8337535 |
| TXNDC16 | 71.8613851 | -0.2759304 | 0.23738963 | -1.1623524 | 0.24509237 | 0.80072691 |
| GPR137C | 42.5084293 | -0.2968301 | 0.6501167 | -0.4565797 | 0.6479732 | 0.95061388 |
| ERO1A | 1133.75371 | -2.4147066 | 0.880136 | -2.7435608 | 0.00607768 | 0.19244777 |
| AL133453.1 | 104.279616 | -0.6242891 | 0.8094042 | -0.7712947 | 0.44053229 | 0.88956997 |
| PSMC6 | 201.309845 | -0.1175562 | 0.26475849 | -0.4440128 | 0.65703331 | 0.95100227 |
| STYX | 78.9705564 | -0.0392524 | 0.38940225 | -0.1008017 | 0.91970786 | 0.99014946 |

| | | | | | | |
|---|---|---|---|---|---|---|
| GNPNAT1 | 89.6990379 | -0.1689962 | 0.30796696 | -0.548748 | 0.58317842 | 0.93436933 |
| FERMT2 | 438.090622 | 0.2557956 | 0.2628462 | 0.97317595 | 0.33046586 | 0.84442294 |
| DDHD1 | 134.183829 | 0.25080926 | 0.27376304 | 0.91615459 | 0.35958582 | 0.85608464 |
| CNIH1 | 112.708954 | 0.11038286 | 0.28191626 | 0.39154485 | 0.69539454 | 0.95672887 |
| GMFB | 120.458247 | -0.1783804 | 0.23722954 | -0.7519318 | 0.45209207 | 0.89473986 |
| CGRRF1 | 60.661146 | 0.07253648 | 0.34638027 | 0.20941285 | 0.83412596 | 0.97232209 |
| SAMD4A | 504.33887 | -0.2779798 | 0.54660135 | -0.5085604 | 0.61106038 | 0.94003018 |
| GCH1 | 44.8397964 | -0.5687774 | 0.54574451 | -1.0422044 | 0.2973169 | 0.83556453 |
| WDHD1 | 45.4267174 | -0.7433972 | 0.42704584 | -1.74079 | 0.08172039 | 0.59202053 |
| SOCS4 | 99.3523533 | -0.0652264 | 0.36224958 | -0.1800594 | 0.85710596 | 0.97718353 |
| MAPK1IP1L | 428.588226 | -0.3219668 | 0.33070803 | -0.9735683 | 0.33027095 | 0.84442294 |
| LGALS3 | 581.999307 | -0.9417514 | 0.55326597 | -1.7021676 | 0.08872395 | 0.60673738 |
| FBXO34 | 630.736551 | -0.041158 | 0.27379422 | -0.1503245 | 0.88050857 | 0.98333507 |
| AL158801.3 | 44.2601526 | 0.06499171 | 0.49437496 | 0.13146238 | 0.89540954 | 0.98597539 |
| ATG14 | 132.091295 | -0.0320916 | 0.30850957 | -0.1040215 | 0.91715228 | 0.98957643 |
| KTN1-AS1 | 42.7616455 | -0.2763532 | 0.44198731 | -0.6252514 | 0.53180608 | 0.92211604 |
| KTN1 | 1158.9562 | 0.1565786 | 0.2462146 | 0.63594362 | 0.52481317 | 0.91924275 |
| PELI2 | 958.194675 | -0.2878594 | 0.30291406 | -0.9503006 | 0.34195951 | 0.85092302 |
| TMEM260 | 386.46363 | -0.0143655 | 0.26578943 | -0.0540484 | 0.95689658 | 0.99565888 |
| EXOC5 | 260.578556 | -0.019448 | 0.16550975 | -0.1175036 | 0.90646099 | 0.9875066 |
| AP5M1 | 179.608707 | 0.18465004 | 0.26871347 | 0.68716332 | 0.49197981 | 0.90757762 |
| NAA30 | 86.9679686 | 0.12564683 | 0.20832474 | 0.60312968 | 0.54642242 | 0.92597739 |
| CCDC198 | 803.787015 | 1.07589044 | 0.40925761 | 2.62888317 | 0.00856658 | 0.22546434 |
| ARMH4 | 137.128281 | -0.0848893 | 0.28892714 | -0.2938088 | 0.76890404 | 0.96345263 |
| ACTR10 | 226.306213 | -0.1821221 | 0.17665892 | -1.0309249 | 0.30257606 | 0.83709431 |
| PSMA3 | 174.649969 | -0.0050429 | 0.27756534 | -0.0181682 | 0.9855047 | 0.99762072 |
| ARID4A | 349.199592 | 0.00782899 | 0.22969799 | 0.03408385 | 0.97281029 | 0.99656026 |
| AL139021.1 | 38.3658745 | 0.22158523 | 0.48217714 | 0.45955151 | 0.64583817 | 0.94994864 |
| TIMM9 | 79.8705354 | 0.06567912 | 0.22963808 | 0.28601147 | 0.77486934 | 0.96388926 |
| KIAA0586 | 179.957869 | -0.2117319 | 0.17678379 | -1.1976886 | 0.23103827 | 0.79130016 |
| DAAM1 | 272.929859 | 0.40982351 | 0.21879737 | 1.87307335 | 0.06105826 | 0.5418629 |
| GPR135 | 49.3435781 | -0.1640405 | 0.52392994 | -0.3130962 | 0.75420754 | 0.96120879 |
| L3HYPDH | 50.1261098 | -0.3944353 | 0.33775719 | -1.1678074 | 0.24288447 | 0.80072691 |
| JKAMP | 115.529631 | 0.12182352 | 0.314559 | 0.38728354 | 0.69854631 | 0.95672887 |
| CCDC175 | 82.5739556 | -0.4158114 | 0.50241692 | -0.8276222 | 0.40788447 | 0.87948803 |
| PCNX4 | 472.31091 | -0.4658078 | 0.47638988 | -0.977787 | 0.32817969 | 0.84381657 |
| DHRS7 | 193.742635 | 0.23393369 | 0.59786036 | 0.39128483 | 0.69558671 | 0.95672887 |
| PPM1A | 324.572145 | -0.2910272 | 0.33068734 | -0.8800677 | 0.37882264 | 0.86515603 |
| MNAT1 | 751.739804 | 0.09493668 | 0.23844005 | 0.39815743 | 0.69051415 | 0.95659154 |
| SLC38A6 | 160.205963 | -0.2760276 | 0.21518669 | -1.2827356 | 0.19958472 | 0.7687013 |
| PRKCH | 325.244137 | 0.22243716 | 0.48034815 | 0.46307488 | 0.6433107 | 0.94949493 |
| TMEM30B | 176.602353 | 0.2954019 | 0.33763451 | 0.87491619 | 0.38161951 | 0.86593382 |
| AL359220.1 | 35.9729989 | 1.02616603 | 0.67146441 | 1.52825081 | 0.12645028 | 0.66895453 |
| HIF1A | 3115.05291 | -0.5859118 | 0.45162455 | -1.2973428 | 0.19451327 | 0.76177095 |
| HIF1A-AS2 | 872.421697 | -1.1850585 | 0.69935904 | -1.6944923 | 0.09017178 | 0.60739841 |
| SNAPC1 | 433.523945 | -0.7661802 | 0.48756002 | -1.5714582 | 0.11607626 | 0.65192493 |
| SYT16 | 45.0306683 | 0.10510994 | 0.59864227 | 0.17558055 | 0.86062348 | 0.97841862 |
| PPP2R5E | 677.30136 | -0.2518965 | 0.25913709 | -0.9720588 | 0.33102129 | 0.84442294 |
| WDR89 | 148.17765 | -0.4016951 | 0.33971831 | -1.1824359 | 0.2370328 | 0.79797745 |
| SGPP1 | 1265.58723 | -0.8571846 | 0.38518586 | -2.2253793 | 0.02605579 | 0.39065042 |
| SYNE2 | 2764.83313 | -0.1653926 | 0.20630346 | -0.8016955 | 0.42272913 | 0.88371155 |
| ESR2 | 221.63662 | -0.2150287 | 0.63339833 | -0.3394841 | 0.73424507 | 0.96042436 |
| MTHFD1 | 66.3919687 | -0.2095725 | 0.22783234 | -0.919854 | 0.35764905 | 0.8557799 |
| AL122035.1 | 70.0202636 | -0.6927533 | 0.37062761 | -1.8691357 | 0.06160394 | 0.54376667 |

| | | | | | | |
|---|---|---|---|---|---|---|
| ZBTB25 | 160.493046 | -0.4703704 | 0.34226798 | -1.3742751 | 0.16935629 | 0.72596839 |
| ZBTB1 | 314.488891 | -0.2015343 | 0.28769489 | -0.7005139 | 0.48360641 | 0.90631121 |
| AL049869.3 | 38.5974472 | 0.05739477 | 0.71434034 | 0.08034653 | 0.93596165 | 0.99243654 |
| HSPA2 | 155.613944 | 0.90412025 | 0.51302358 | 1.76233664 | 0.07801244 | 0.57901582 |
| PPP1R36 | 39.154808 | 0.20115338 | 0.45753549 | 0.43964542 | 0.66019394 | 0.95127957 |
| PLEKHG3 | 385.869959 | -0.0083193 | 0.40577268 | -0.0205023 | 0.98364267 | 0.9971769 |
| SPTB | 35.1116152 | 0.80695231 | 0.55321646 | 1.45865563 | 0.14465991 | 0.69206332 |
| CHURC1-FNT | 93.6845949 | -0.2701366 | 0.39377987 | -0.6860091 | 0.49270734 | 0.90804281 |
| RAB15 | 135.412247 | 0.21727277 | 0.41702968 | 0.52100073 | 0.60236626 | 0.93804805 |
| MAX | 241.549489 | -0.2933822 | 0.25488849 | -1.151022 | 0.2497232 | 0.80149394 |
| FUT8 | 229.396509 | 0.67327273 | 0.4107872 | 1.63898174 | 0.10121706 | 0.62105083 |
| AL359232.1 | 109.946292 | 0.00952733 | 0.41105989 | 0.02317749 | 0.9815087 | 0.99696835 |
| GPHN | 1957.08997 | 0.22674589 | 0.33469608 | 0.67746803 | 0.49810905 | 0.90839719 |
| MPP5 | 389.387593 | -0.1759488 | 0.13626049 | -1.2912678 | 0.19661085 | 0.76478825 |
| ATP6V1D | 220.393398 | -0.1356012 | 0.22501345 | -0.6026361 | 0.54675077 | 0.92597739 |
| EIF2S1 | 119.558549 | -0.0687921 | 0.25559873 | -0.2691411 | 0.7878211 | 0.96515649 |
| PLEKHH1 | 131.594637 | -0.2894078 | 0.24142504 | -1.1987482 | 0.23062589 | 0.7911487 |
| PIGH | 88.1181447 | -0.1453158 | 0.62399172 | -0.232881 | 0.81585382 | 0.96844493 |
| ARG2 | 855.487349 | -0.4482833 | 0.81386453 | -0.5508083 | 0.58176509 | 0.93402467 |
| VTI1B | 141.036407 | 0.09352492 | 0.40935923 | 0.22846662 | 0.8192835 | 0.96884581 |
| ZFYVE26 | 95.5564769 | -0.8439175 | 0.27755269 | -3.040567 | 0.00236133 | 0.12625894 |
| RAD51B | 2428.82614 | 0.29014689 | 0.33651962 | 0.86219903 | 0.38857801 | 0.86970192 |
| ZFP36L1 | 2127.96204 | -0.7094919 | 0.35204232 | -2.0153596 | 0.04386698 | 0.48009422 |
| ACTN1 | 139.247971 | -0.3233891 | 0.42284417 | -0.764795 | 0.44439365 | 0.8909978 |
| DCAF5 | 475.194937 | 0.08031978 | 0.23549135 | 0.34107315 | 0.73304852 | 0.96035038 |
| EXD2 | 115.617866 | 0.0892185 | 0.21115134 | 0.42253344 | 0.6726357 | 0.9544944 |
| ERH | 212.333511 | 0.113186 | 0.3541257 | 0.31962097 | 0.74925568 | 0.96109466 |
| SLC39A9 | 193.089862 | -0.1007103 | 0.15994424 | -0.6296585 | 0.52891801 | 0.92136681 |
| SUSD6 | 185.875125 | 0.45237944 | 0.25530645 | 1.77190762 | 0.07640989 | 0.57145422 |
| SRSF5 | 640.127204 | 0.03096731 | 0.21027877 | 0.14726788 | 0.88292058 | 0.98352548 |
| AL357153.2 | 109.779295 | 0.36649484 | 0.38484046 | 0.95232928 | 0.34093001 | 0.85042291 |
| MED6 | 99.0849082 | 0.07508166 | 0.2573641 | 0.29173322 | 0.77049061 | 0.96352463 |
| MAP3K9 | 106.752477 | 0.39890042 | 0.40474072 | 0.98557029 | 0.32434402 | 0.84285425 |
| PCNX1 | 657.290321 | 0.01311572 | 0.25633897 | 0.05116552 | 0.95919362 | 0.99609409 |
| SIPA1L1 | 1092.60059 | -0.3266784 | 0.21258818 | -1.5366724 | 0.12437356 | 0.665882 |
| ZFYVE1 | 98.1212552 | -0.259449 | 0.327635 | -0.7918843 | 0.42842817 | 0.88535099 |
| RBM25 | 493.015498 | -0.4315592 | 0.14854544 | -2.9052337 | 0.00366979 | 0.14788453 |
| PSEN1 | 321.662456 | -0.362405 | 0.21240772 | -1.706176 | 0.08797532 | 0.60651125 |
| PAPLN | 88.773131 | 0.02831894 | 0.73695776 | 0.03842681 | 0.96934738 | 0.99645289 |
| NUMB | 924.841601 | -0.1438485 | 0.23490307 | -0.6123739 | 0.5402904 | 0.92414041 |
| HEATR4 | 43.3521893 | 0.51246733 | 0.36481208 | 1.40474331 | 1.60E-01 | 0.71409437 |
| DNAL1 | 150.122087 | 0.222096 | 0.26561357 | 0.83616211 | 0.40306371 | 0.87834687 |
| PNMA1 | 68.3290215 | 0.49346059 | 0.32337548 | 1.52596785 | 0.12701787 | 0.66992818 |
| ELMSAN1 | 219.980858 | -0.8254234 | 0.27965448 | -2.9515829 | 0.0031615 | 0.14358074 |
| AC005480.1 | 89.0342427 | -0.14072 | 0.47171748 | -0.2983142 | 0.7654634 | 0.96285313 |
| COQ6 | 39.19258 | 0.04163542 | 0.30438804 | 0.13678402 | 0.89120151 | 0.98597539 |
| ENTPD5 | 124.034177 | -0.368663 | 0.30647606 | -1.2029095 | 0.22901134 | 0.79047378 |
| BBOF1 | 156.21987 | 0.20999489 | 0.27155985 | 0.7732914 | 0.43934995 | 0.88917548 |
| ALDH6A1 | 167.812999 | 0.80976432 | 0.45511898 | 1.77923656 | 0.07520099 | 0.57072562 |
| LIN52 | 159.422797 | -0.0845477 | 0.33035566 | -0.2559294 | 0.79800533 | 0.9653598 |
| ABCD4 | 193.222375 | 0.02494167 | 0.2218639 | 0.11241877 | 0.91049138 | 0.98788932 |
| NPC2 | 350.545772 | 0.25133383 | 0.4615409 | 0.54455375 | 0.58606047 | 0.93483536 |
| ISCA2 | 88.8830892 | 0.40788478 | 0.52481637 | 0.77719522 | 0.4370436 | 0.88785137 |
| AREL1 | 139.083838 | -0.0071856 | 0.23827474 | -0.030157 | 0.97594188 | 0.99656026 |

| | | | | | | |
|---|---|---|---|---|---|---|
| FCF1 | 73.7295574 | -0.0960584 | 0.24155996 | -0.3976587 | 0.69088181 | 0.95659154 |
| YLPM1 | 352.713113 | 0.23558866 | 0.23176658 | 1.01649106 | 0.3093956 | 0.84136316 |
| DLST | 156.498382 | -0.1467895 | 0.17960638 | -0.8172846 | 0.41376578 | 0.88211689 |
| EIF2B2 | 46.4927692 | -0.0577831 | 0.38797317 | -0.1489358 | 0.88160432 | 0.98335905 |
| MLH3 | 278.247206 | 0.01585509 | 0.17219158 | 0.09207819 | 0.92663591 | 0.99087265 |
| ACYP1 | 194.643918 | -0.1753533 | 0.40361902 | -0.4344526 | 0.66395979 | 0.95188763 |
| ZC2HC1C | 40.4677777 | -0.0485065 | 0.47764089 | -0.1015543 | 0.91911049 | 0.99014946 |
| NEK9 | 211.603854 | -0.0367339 | 0.18754474 | -0.1958675 | 0.84471384 | 0.97573093 |
| AL049780.2 | 44.8529445 | 1.1368374 | 0.61505621 | 1.84834715 | 0.06455214 | 0.55000521 |
| TMED10 | 517.961065 | 0.14111487 | 0.25438684 | 0.55472552 | 0.57908242 | 0.93345492 |
| FOS | 526.506955 | -2.3348362 | 1.37531551 | -1.6976731 | 0.08956947 | 0.60739841 |
| JDP2 | 56.5808086 | -0.0142127 | 0.59905849 | -0.023725 | 0.98107196 | 0.99696835 |
| TTLL5 | 448.162606 | -0.2091847 | 0.25879592 | -0.8082997 | 0.41891809 | 0.88261006 |
| ERG28 | 83.5183836 | -0.2002581 | 0.35321835 | -0.5669528 | 0.57074627 | 0.93125087 |
| IFT43 | 310.480364 | 0.10492299 | 0.21309997 | 0.49236509 | 0.62246127 | 0.94418789 |
| GPATCH2L | 318.057293 | -0.1365447 | 0.25001069 | -0.5461556 | 0.58495898 | 0.93483536 |
| ANGEL1 | 46.112947 | 0.33827147 | 0.41934966 | 0.80665732 | 0.41986394 | 0.8828928 |
| IRF2BPL | 203.817786 | 0.20048764 | 0.37992694 | 0.52770051 | 0.59770722 | 0.9367387 |
| POMT2 | 57.6173598 | -0.5114006 | 0.32691754 | -1.5643108 | 0.11774459 | 0.65447949 |
| GSTZ1 | 41.5830766 | 0.38180802 | 0.34621224 | 1.10281491 | 0.27010755 | 0.81817548 |
| TMED8 | 206.119004 | -0.1477726 | 0.16230974 | -0.9104356 | 0.36259281 | 0.85694172 |
| VIPAS39 | 73.490448 | 0.02949089 | 0.23808161 | 0.12386881 | 0.90141915 | 0.9875066 |
| AHSA1 | 94.5718995 | 0.08009186 | 0.4084542 | 0.19608529 | 0.84454341 | 0.97573093 |
| SPTLC2 | 612.474361 | 0.22920303 | 0.2991543 | 0.76616994 | 0.44357519 | 0.89037885 |
| ALKBH1 | 88.9301097 | -0.0415536 | 0.23645935 | -0.1757326 | 0.86050403 | 0.97841862 |
| SLIRP | 129.432979 | 0.33907026 | 0.23761746 | 1.42695856 | 0.15359184 | 0.70747602 |
| SNW1 | 147.524358 | 0.01157258 | 0.23641651 | 0.04894995 | 0.96095918 | 0.99635733 |
| ADCK1 | 56.3234144 | -0.4558629 | 0.44533782 | -1.023634 | 0.30600819 | 0.83709431 |
| CEP128 | 206.180143 | -0.6767864 | 0.28051123 | -2.4126891 | 0.01583532 | 0.30118722 |
| AC007262.2 | 65.3392466 | 0.83788829 | 0.53936515 | 1.55347132 | 0.12031058 | 0.65691125 |
| GTF2A1 | 141.893513 | 0.19676617 | 0.27549758 | 0.71422105 | 0.4750905 | 0.90242252 |
| STON2 | 448.209745 | 0.1427473 | 0.31415922 | 0.45437883 | 0.64955618 | 0.95061388 |
| SEL1L | 314.409855 | -0.0013045 | 0.19400248 | -0.0067241 | 0.99463495 | 0.99919666 |
| GALC | 80.6667013 | -0.3132306 | 0.19477013 | -1.6082064 | 0.10778999 | 0.63520805 |
| SPATA7 | 229.237525 | -0.3342262 | 0.32534784 | -1.0272887 | 0.30428452 | 0.83709431 |
| PTPN21 | 312.99644 | 0.15272783 | 0.36173674 | 0.42220712 | 0.67287384 | 0.9544944 |
| ZC3H14 | 340.632249 | 0.12357104 | 0.22743379 | 0.54332754 | 0.58690431 | 0.93483536 |
| EML5 | 90.9553767 | -0.708658 | 0.46239996 | -1.532565 | 0.12538308 | 0.66667411 |
| TTC8 | 180.280493 | 0.13062461 | 0.28826981 | 0.45313317 | 0.65045285 | 0.95061388 |
| FOXN3 | 675.702735 | -0.138789 | 0.25645801 | -0.5411763 | 0.58838608 | 0.93483536 |
| EFCAB11 | 180.600182 | 0.46977332 | 0.36304458 | 1.29398247 | 0.19567148 | 0.7637382 |
| TDP1 | 94.1590421 | 0.30887338 | 0.25210918 | 1.22515724 | 0.22051598 | 0.78685989 |
| KCNK13 | 455.971689 | 0.88965343 | 0.55082518 | 1.61512847 | 0.10628288 | 0.63461115 |
| PSMC1 | 132.889725 | 0.05392299 | 0.21525908 | 0.25050276 | 0.80219857 | 0.9661728 |
| NRDE2 | 127.177774 | -0.1104169 | 0.31374926 | -0.3519273 | 0.72489279 | 0.95963742 |
| CALM1 | 694.441038 | -0.158624 | 0.5005138 | -0.3169222 | 0.7513026 | 0.96120879 |
| TTC7B | 415.883162 | -0.4260299 | 0.28184842 | -1.5115566 | 0.13064669 | 0.67339292 |
| RPS6KA5 | 198.684627 | -0.4237555 | 0.3419948 | -1.2390699 | 0.21531961 | 0.77894575 |
| DGLUCY | 421.423218 | -0.3630325 | 0.35441764 | -1.0243071 | 0.30569025 | 0.83709431 |
| CCDC88C | 300.945543 | -0.1268503 | 0.46091023 | -0.275217 | 0.7831495 | 0.96513523 |
| PPP4R3A | 465.747588 | -0.1628473 | 0.38722794 | -0.4205464 | 0.67408638 | 0.95511203 |
| CATSPERB | 255.282211 | -1.1192884 | 0.46710065 | -2.3962468 | 0.01656393 | 0.30723893 |
| TC2N | 526.850572 | -0.985567 | 0.70328268 | -1.4013811 | 0.16110014 | 0.71409437 |
| TRIP11 | 483.500847 | 0.25291047 | 0.17395882 | 1.45385255 | 0.14598719 | 0.69448286 |

| | | | | | | |
|---|---|---|---|---|---|---|
| ATXN3 | 265.376523 | -0.1996003 | 0.22896838 | -0.8717375 | 0.3833516 | 0.86638702 |
| NDUFB1 | 156.662623 | 0.07431756 | 0.26558448 | 0.27982642 | 0.77961068 | 0.96513523 |
| CPSF2 | 147.754774 | -0.0111246 | 0.19753815 | -0.0563161 | 0.95509003 | 0.99537617 |
| LGMN | 118.992457 | -0.1355701 | 0.4694111 | -0.2888089 | 0.7727276 | 0.96388926 |
| GOLGA5 | 107.263681 | -0.3889817 | 0.2034109 | -1.9122955 | 0.05583831 | 0.52062762 |
| ITPK1 | 415.811069 | -0.1308498 | 0.31705437 | -0.4127045 | 0.67982312 | 0.95574492 |
| MOAP1 | 117.309851 | 0.25082579 | 0.40009332 | 0.62691821 | 0.53071285 | 0.92211604 |
| GON7 | 48.6700632 | 0.27770625 | 0.46493679 | 0.59729895 | 0.55030781 | 0.92780338 |
| UBR7 | 47.5582962 | -0.093984 | 0.30181234 | -0.3113987 | 0.75549756 | 0.96120879 |
| BTBD7 | 932.307852 | 0.31537607 | 0.23305087 | 1.35324993 | 0.1759758 | 0.73444173 |
| UNC79 | 47.3320138 | 1.08269854 | 0.57150005 | 1.89448547 | 0.0581606 | 0.53192434 |
| DDX24 | 425.920846 | 0.36684126 | 0.30172595 | 1.21580948 | 0.22405751 | 0.78960314 |
| IFI27L1 | 39.5552547 | 0.06285561 | 0.2927768 | 0.2146878 | 0.83001074 | 0.97146666 |
| IFI27L2 | 59.3402784 | 0.01079715 | 0.32666167 | 0.03305301 | 0.97363231 | 0.99656026 |
| SERPINA1 | 84.4563915 | -0.6332677 | 0.75937111 | -0.8339371 | 0.40431643 | 0.87834687 |
| DICER1 | 652.857139 | 0.01228092 | 0.15710517 | 0.07817004 | 0.9376928 | 0.99243654 |
| DICER1-AS1 | 60.677484 | 0.21817735 | 0.46762582 | 0.46656394 | 0.64081189 | 0.94935885 |
| CLMN | 1300.99712 | 0.5304465 | 0.28935034 | 1.83323267 | 0.06676798 | 0.55637569 |
| GLRX5 | 213.78954 | -0.0200795 | 0.55249431 | -0.0363433 | 0.97100861 | 0.99656026 |
| ATG2B | 268.80644 | -0.0187659 | 0.25113267 | -0.0747252 | 0.94043335 | 0.99243654 |
| GSKIP | 60.7249234 | -0.0550228 | 0.30209235 | -0.1821389 | 0.85547376 | 0.97718353 |
| AK7 | 80.9062657 | -0.1968841 | 0.29946016 | -0.6574635 | 0.51088293 | 0.91390025 |
| PAPOLA | 592.466936 | -0.1180046 | 0.16101929 | -0.73286 | 0.46364385 | 0.89801718 |
| VRK1 | 69.2037427 | 0.22721445 | 0.25114778 | 0.90470422 | 0.36562211 | 0.85866868 |
| LINC01550 | 60.1585732 | 0.83433373 | 0.5004191 | 1.66726996 | 0.09546074 | 0.61367241 |
| SETD3 | 385.615106 | 0.01665943 | 0.16512591 | 0.10088927 | 0.91963836 | 0.99014946 |
| CCNK | 109.902377 | -0.0825276 | 0.19018138 | -0.4339415 | 0.6643309 | 0.95195406 |
| CCDC85C | 59.5244894 | -0.0193457 | 0.25860659 | -0.0748075 | 0.94036791 | 0.99243654 |
| EML1 | 46.1083431 | 0.05751881 | 0.37319846 | 0.15412391 | 0.87751203 | 0.98279964 |
| EVL | 228.71569 | -0.1102662 | 0.38795551 | -0.2842238 | 0.77623892 | 0.96437053 |
| DEGS2 | 69.0749299 | 1.29758191 | 0.71228525 | 1.82171667 | 0.06849799 | 0.55917298 |
| YY1 | 469.768244 | -0.1792971 | 0.26635007 | -0.6731632 | 0.50084344 | 0.90857526 |
| SLC25A29 | 1635.69472 | -1.4413417 | 0.51393276 | -2.8045337 | 0.00503894 | 0.17602704 |
| WARS | 220.032154 | -1.9733031 | 0.51747799 | -3.8133083 | 0.00013712 | 0.0239501 |
| WDR25 | 128.206554 | -0.4707062 | 0.25648417 | -1.8352251 | 0.06647234 | 0.55596979 |
| PPP2R5C | 470.307195 | -0.1164165 | 0.16545299 | -0.703623 | 0.4816676 | 0.90561114 |
| DYNC1H1 | 409.796666 | -0.3358687 | 0.17305138 | -1.9408613 | 0.05227511 | 0.50726213 |
| HSP90AA1 | 1573.49229 | 0.15723109 | 0.4266919 | 0.36848857 | 0.71250896 | 0.95791799 |
| WDR20 | 316.443997 | -0.0766914 | 0.21626541 | -0.3546172 | 0.72287639 | 0.95944081 |
| MOK | 268.693899 | 0.12833503 | 0.2655922 | 0.48320331 | 0.62895139 | 0.94596555 |
| ZNF839 | 77.582291 | 0.04341669 | 0.30110672 | 0.14419038 | 0.88535014 | 0.9845556 |
| CINP | 62.0537057 | -0.0505376 | 0.24653392 | -0.2049925 | 0.83757795 | 0.97369018 |
| TECPR2 | 170.672383 | -0.0345883 | 0.24812032 | -0.1394013 | 0.88913303 | 0.98551241 |
| ANKRD9 | 99.0713812 | -0.1068628 | 0.38490534 | -0.2776341 | 0.78129325 | 0.96513523 |
| RCOR1 | 436.109288 | -0.0493605 | 0.30410355 | -0.1623147 | 0.87105807 | 0.98208511 |
| TRAF3 | 302.677254 | 0.63892983 | 0.37486591 | 1.70442232 | 0.08830222 | 0.60651125 |
| CDC42BPB | 875.310045 | -0.7369229 | 0.28798195 | -2.5589205 | 0.01049977 | 0.25038862 |
| TNFAIP2 | 54.5442756 | -0.5378644 | 0.6572855 | -0.8183117 | 0.41317921 | 0.88189778 |
| EIF5 | 320.418666 | -0.0915336 | 0.34931811 | -0.262035 | 0.79329445 | 0.9653598 |
| MARK3 | 1153.95913 | -0.1875471 | 0.18401489 | -1.019195 | 0.30811039 | 0.83980597 |
| CKB | 494.462254 | -0.1073764 | 0.56107006 | -0.1913778 | 0.84822961 | 0.97656375 |
| BAG5 | 136.685313 | 0.09752579 | 0.21807551 | 0.4472111 | 0.65472265 | 0.95100227 |
| KLC1 | 228.568296 | -0.5164369 | 0.20808175 | -2.4818942 | 0.01306861 | 0.28065372 |
| ZFYVE21 | 149.638128 | 0.38406827 | 0.37814063 | 1.01567574 | 0.30978382 | 0.84194359 |

| | | | | | | |
|---|---|---|---|---|---|---|
| PPP1R13B | 277.580326 | -0.0246901 | 0.21134249 | -0.1168253 | 0.90699852 | 0.9875066 |
| ATP5MPL | 207.469871 | 0.23376556 | 0.26351153 | 0.88711698 | 0.37501593 | 0.86339343 |
| INF2 | 60.1087475 | -0.2855971 | 0.29702208 | -0.9615348 | 0.33628331 | 0.84565588 |
| SIVA1 | 126.206245 | 0.30352801 | 0.47420226 | 0.64008132 | 0.52211973 | 0.91845051 |
| AKT1 | 105.095376 | -0.1080554 | 0.33352977 | -0.3239752 | 0.7459568 | 0.96109466 |
| CEP170B | 74.2132204 | -0.5609009 | 0.31497684 | -1.7807688 | 0.07495023 | 0.57072562 |
| CLBA1 | 53.2993471 | 0.17033645 | 0.24567611 | 0.69333745 | 0.48809781 | 0.90730741 |
| GPR132 | 72.1477299 | 0.98797967 | 1.26485329 | 0.78110218 | 0.4347424 | 0.88785137 |
| NUDT14 | 45.8618483 | -0.3742159 | 0.35224945 | -1.0623604 | 0.28807211 | 0.8303903 |
| BRF1 | 124.070168 | -0.2198463 | 0.25558662 | -0.8601635 | 0.38969894 | 0.87004477 |
| BTBD6 | 72.1933696 | 0.10636796 | 0.382426 | 0.27813998 | 0.78090491 | 0.96513523 |
| PACS2 | 217.158929 | -0.5124907 | 0.3062867 | -1.6732385 | 0.09428035 | 0.61330068 |
| TEX22 | 34.0127707 | 0.58205443 | 0.39837267 | 1.46108023 | 0.14399342 | 0.69093999 |
| MTA1 | 143.844116 | -0.3752345 | 0.21460348 | -1.7485016 | 0.08037721 | 0.58782498 |
| CRIP2 | 291.563798 | -0.2993048 | 0.41343769 | -0.7239418 | 0.46910148 | 0.90026987 |
| NIPA1 | 105.721018 | -0.0155242 | 0.20208085 | -0.0768216 | 0.93876545 | 0.99243654 |
| NIPA2 | 144.627023 | -0.0361936 | 0.31758015 | -0.1139668 | 0.90926409 | 0.9877773 |
| CYFIP1 | 289.854453 | -0.1761453 | 0.20495866 | -0.8594188 | 0.39010953 | 0.87004477 |
| TUBGCP5 | 117.684371 | -0.4050814 | 0.25124396 | -1.612303 | 0.10689603 | 0.63477557 |
| NDN | 34.2999914 | 0.5161454 | 0.45285045 | 1.13977008 | 0.2543821 | 0.80414079 |
| AC087463.4 | 122.997134 | 0.42217951 | 0.62270591 | 0.67797576 | 4.98E-01 | 0.90839719 |
| PWRN3 | 387.976753 | 0.52146359 | 0.94531988 | 0.5516266 | 0.58120421 | 0.93386308 |
| PWRN1 | 3243.20903 | 0.44755299 | 0.49457376 | 0.90492667 | 3.66E-01 | 0.85866868 |
| NPAP1 | 103.61937 | 0.30154274 | 0.56659565 | 0.53220095 | 5.95E-01 | 0.935202 |
| SNRPN | 184.165342 | 1.14027401 | 0.56545857 | 2.01654742 | 0.04374276 | 0.48009422 |
| AC124312.1 | 945.491539 | 0.24995332 | 0.22780083 | 1.09724495 | 0.27253433 | 0.81922962 |
| PWAR6 | 232.035092 | 0.25740833 | 0.22657895 | 1.13606462 | 0.25592952 | 0.80525591 |
| UBE3A | 539.098116 | -0.056994 | 0.16058489 | -0.3549149 | 0.72265335 | 0.95926626 |
| GABRB3 | 94.2331414 | 0.67129401 | 0.5565957 | 1.20607114 | 0.22779006 | 0.79042788 |
| HERC2 | 577.222752 | -0.2767976 | 0.22782913 | -1.2149353 | 0.22439078 | 0.78966267 |
| FAM189A1 | 56.8139376 | 0.65769566 | 0.35243163 | 1.86616526 | 0.06201825 | 0.54480406 |
| TJP1 | 536.403586 | -0.1805885 | 0.26697473 | -0.6764256 | 0.49877045 | 0.90839719 |
| FAN1 | 155.649452 | -0.0025005 | 0.27150144 | -0.0092097 | 0.9926518 | 0.99919666 |
| MTMR10 | 1123.36812 | 0.90565075 | 0.41560445 | 2.17911706 | 0.02932297 | 0.40623517 |
| KLF13 | 222.630328 | -0.2154844 | 0.36661719 | -0.587764 | 0.55669074 | 0.92872652 |
| OTUD7A | 198.476007 | -0.1133228 | 0.26619711 | -0.42571 | 0.67031919 | 0.95382825 |
| LINC02256 | 376.607683 | 0.02270511 | 0.29474137 | 0.077034 | 0.9385965 | 0.99243654 |
| AC123768.3 | 61.6223982 | 0.40527828 | 0.3938905 | 1.02891103 | 0.30352148 | 0.83709431 |
| GOLGA8N | 44.1623628 | 0.0512493 | 0.30991362 | 0.16536641 | 0.86865559 | 0.98193405 |
| FMN1 | 103.591261 | 0.60292836 | 0.51671236 | 1.16685494 | 0.24326896 | 0.80072691 |
| AVEN | 122.379618 | -0.2318997 | 0.28972295 | -0.800419 | 0.42346809 | 0.88387684 |
| EMC7 | 111.213242 | 0.3995152 | 0.34737468 | 1.15009877 | 0.25010319 | 0.80149394 |
| KATNBL1 | 397.464346 | -0.2789525 | 0.16474437 | -1.6932444 | 0.09040896 | 0.60779802 |
| EMC4 | 67.0198684 | -0.0302073 | 0.38501992 | -0.0784564 | 0.93746505 | 0.99243654 |
| SLC12A6 | 116.924946 | -0.1770497 | 0.35110795 | -0.5042599 | 0.61407877 | 0.94073169 |
| NOP10 | 191.836099 | -0.1498246 | 0.54418479 | -0.2753193 | 0.78307096 | 0.96513523 |
| LPCAT4 | 35.626357 | -0.2858571 | 0.29019822 | -0.9850407 | 0.32460407 | 0.84285425 |
| GOLGA8A | 180.228499 | -0.105527 | 0.28821714 | -0.366137 | 0.71426281 | 0.95791799 |
| GOLGA8B | 194.642239 | 0.4521783 | 0.35664165 | 1.26787856 | 0.20484132 | 0.77193931 |
| AQR | 288.018795 | -0.143402 | 0.17726068 | -0.8089895 | 0.41852117 | 0.88242985 |
| ZNF770 | 217.844081 | 0.33784768 | 0.25224302 | 1.33937376 | 0.18044903 | 0.74280139 |
| AC021231.1 | 296.23874 | 0.49114876 | 0.43890349 | 1.11903589 | 0.26312483 | 0.8096287 |
| DPH6 | 840.938414 | 0.08293579 | 0.32116867 | 0.25823127 | 0.79622843 | 0.9653598 |
| DPH6-DT | 569.628415 | 1.12147396 | 0.60826266 | 1.84373303 | 0.06522206 | 0.55362017 |

| | | | | | | |
|---|---|---|---|---|---|---|
| C15orf41 | 252.299265 | -0.3452129 | 0.22165036 | -1.557466 | 0.11935988 | 0.65527787 |
| SPRED1 | 287.565162 | 0.15673589 | 0.40964665 | 0.3826124 | 0.70200717 | 0.95672887 |
| FAM98B | 124.960328 | -0.1291615 | 0.18946628 | -0.6817121 | 0.49542104 | 0.90839719 |
| RASGRP1 | 408.157657 | -0.3546622 | 0.31757186 | -1.1167936 | 0.26408258 | 0.81061724 |
| C15orf53 | 74.1956982 | 1.11342058 | 0.50177671 | 2.21895627 | 0.0264897 | 0.3943353 |
| AC013652.1 | 69.0492967 | 0.69538147 | 0.31719685 | 2.19227106 | 0.02835994 | 0.40459506 |
| THBS1 | 167.722063 | 1.96630409 | 0.55136026 | 3.56627825 | 0.00036209 | 0.04296095 |
| FSIP1 | 128.710677 | 0.76372654 | 0.29017436 | 2.63195735 | 0.00848945 | 0.22546434 |
| EIF2AK4 | 193.529711 | 0.02436247 | 0.18362962 | 0.13267176 | 0.89445297 | 0.98597539 |
| SRP14 | 1192.79645 | 0.29995308 | 0.47944912 | 0.62562025 | 0.53156405 | 0.92211604 |
| SRP14-AS1 | 45.8655785 | 0.75253461 | 0.38315202 | 1.96406274 | 0.04952282 | 0.49334517 |
| BMF | 54.1089125 | 1.08975688 | 0.68272881 | 1.59617826 | 0.110449 | 0.6412773 |
| PAK6 | 33.8989286 | 0.55866204 | 0.49449493 | 1.12976293 | 0.25857613 | 0.80709747 |
| IVD | 217.397224 | 0.72117786 | 0.2029215 | 3.55397467 | 0.00037946 | 0.04296095 |
| BAHD1 | 70.405815 | 0.3147098 | 0.27153702 | 1.15899409 | 0.24645859 | 0.80072691 |
| CCDC32 | 142.091446 | 0.11825263 | 0.31587586 | 0.37436426 | 0.70813333 | 0.95752169 |
| RMDN3 | 88.0670917 | -0.2310909 | 0.21598002 | -1.0699641 | 0.28463548 | 0.82781987 |
| GCHFR | 50.9446309 | 0.04080664 | 0.70645172 | 0.05776282 | 0.95393756 | 0.99514888 |
| DNAJC17 | 125.874132 | -0.2545065 | 0.21253673 | -1.1974709 | 0.23112307 | 0.79130016 |
| ZFYVE19 | 53.6066808 | -0.1019955 | 0.32718952 | -0.3117322 | 0.75524408 | 0.96120879 |
| SPINT1-AS1 | 115.948989 | 0.65317738 | 0.39723992 | 1.64428934 | 0.10011645 | 0.61891077 |
| SPINT1 | 259.044325 | 0.23346152 | 0.314587 | 0.74212069 | 0.45801422 | 0.89736194 |
| RHOV | 53.3205692 | -2.4037485 | 0.87717652 | -2.7403247 | 0.00613785 | 0.19316723 |
| VPS18 | 56.9046037 | -0.4745081 | 0.39536699 | -1.2001713 | 0.23007283 | 0.79054532 |
| INO80 | 408.962766 | -0.7048163 | 0.24244236 | -2.9071501 | 0.00364738 | 0.14773661 |
| CHP1 | 323.462868 | -0.4367779 | 0.48270532 | -0.9048542 | 0.36554262 | 0.85866868 |
| NDUFAF1 | 41.150476 | -0.1338077 | 0.25721811 | -0.5202109 | 0.60291656 | 0.93804805 |
| RTF1 | 274.278524 | 0.11899197 | 0.29795186 | 0.39936644 | 0.68962322 | 0.9564917 |
| RPAP1 | 44.2893619 | -0.1558584 | 0.30062726 | -0.518444 | 0.60414855 | 0.93804805 |
| MGA | 329.22879 | 0.18831105 | 0.28568984 | 0.65914508 | 0.50980261 | 0.91334187 |
| MAPKBP1 | 173.915333 | -0.3746439 | 0.37036668 | -1.0115486 | 0.31175393 | 0.84279403 |
| EHD4 | 289.508648 | -0.1958317 | 0.34995476 | -0.5595916 | 0.57575805 | 0.93294146 |
| PLA2G4F | 67.7834412 | 0.30826654 | 0.50600112 | 0.60922107 | 0.54237792 | 0.92482794 |
| VPS39 | 205.705503 | -0.1041139 | 0.27175166 | -0.3831216 | 0.70162963 | 0.95672887 |
| TMEM87A | 353.053936 | -0.4180524 | 0.16857972 | -2.4798498 | 0.01314377 | 0.28169069 |
| GANC | 141.852607 | -0.0769813 | 0.32377231 | -0.2377638 | 0.81206427 | 0.9680627 |
| ZNF106 | 292.612752 | 0.30169573 | 0.19718276 | 1.53003099 | 0.12600906 | 0.66763141 |
| SNAP23 | 209.992004 | -0.1030366 | 0.20228622 | -0.5093604 | 0.61049966 | 0.94003018 |
| LRRC57 | 47.7889584 | 0.06010627 | 0.27816647 | 0.2160802 | 0.82892523 | 0.97131041 |
| HAUS2 | 48.1351725 | 0.22333764 | 0.33324159 | 0.67019738 | 0.50273198 | 0.90962743 |
| TTBK2 | 303.332084 | -0.0958509 | 0.21150025 | -0.453195 | 0.65040832 | 0.95061388 |
| UBR1 | 563.071936 | -0.4363299 | 0.22049236 | -1.9788888 | 0.04782852 | 0.48600281 |
| AC068724.3 | 123.965962 | 0.15291423 | 0.389614 | 0.39247622 | 0.69470638 | 0.95672887 |
| TMEM62 | 157.714016 | -0.4099812 | 0.29963482 | -1.3682696 | 0.1712277 | 0.72776437 |
| CCNDBP1 | 83.795662 | 0.75264051 | 0.39244149 | 1.9178413 | 0.05513114 | 0.51725543 |
| ADAL | 91.9444377 | 0.37301627 | 0.31148165 | 1.19755457 | 0.23109047 | 0.79130016 |
| ZSCAN29 | 33.746607 | 0.16418309 | 0.31319346 | 0.5242226 | 0.60012372 | 0.93728325 |
| TUBGCP4 | 85.23533 | 0.04198107 | 0.26708172 | 0.15718437 | 0.87509954 | 0.98267808 |
| TP53BP1 | 455.10523 | 0.08195653 | 0.16886906 | 0.48532595 | 0.62744515 | 0.94511671 |
| PPIP5K1 | 123.328566 | 0.25810572 | 0.27257889 | 0.94690283 | 0.34368829 | 0.85126545 |
| CATSPER2 | 148.663998 | 0.04153945 | 0.40226678 | 0.10326344 | 0.91775389 | 0.9900217 |
| PDIA3 | 439.608098 | 0.10123882 | 0.19736192 | 0.51296026 | 0.60797912 | 0.94003018 |
| SERF2 | 1586.81829 | 0.06138545 | 0.40729921 | 0.15071341 | 0.88020179 | 0.98333507 |
| MFAP1 | 98.0381595 | 0.23116986 | 0.27597435 | 0.83764981 | 0.40222741 | 0.87834687 |

| | | | | | | |
|---|---|---|---|---|---|---|
| FRMD5 | 49.0659834 | -0.0061686 | 0.39536596 | -0.0156021 | 0.9875518 | 0.99791717 |
| CASC4 | 779.125995 | -0.1460589 | 0.23668862 | -0.6170932 | 0.53717323 | 0.92324352 |
| AC025043.1 | 40.1583118 | 0.12202649 | 0.41722272 | 0.29247327 | 0.76992479 | 0.96345263 |
| CTDSPL2 | 403.45643 | -0.3169061 | 0.22903052 | -1.3836849 | 0.16645496 | 0.72054851 |
| EIF3J-DT | 123.635777 | 0.35441971 | 0.37360866 | 0.94863893 | 0.34280428 | 0.85093239 |
| EIF3J | 223.9642 | 0.31553378 | 0.26470872 | 1.19200373 | 0.23325978 | 0.7926597 |
| SPG11 | 585.278687 | -0.3095042 | 0.21946721 | -1.4102527 | 0.15846507 | 0.71297794 |
| B2M | 2851.24279 | -0.1776176 | 0.52710112 | -0.3369706 | 0.73613905 | 0.96078945 |
| TRIM69 | 68.4733634 | -0.5304758 | 0.48099546 | -1.1028706 | 0.27008336 | 0.81817548 |
| SORD | 135.197373 | 0.5434107 | 0.31322661 | 1.73488036 | 0.082762 | 0.59202053 |
| DUOX1 | 47.9971772 | -0.0084648 | 0.40193934 | -0.0210598 | 0.98319792 | 0.9971769 |
| GATM | 108.58395 | 0.15829303 | 0.32226217 | 0.49119333 | 0.62328971 | 0.94456546 |
| SPATA5L1 | 74.8078931 | 0.0585509 | 0.27966946 | 0.20935752 | 0.83416915 | 0.97232209 |
| SLC30A4 | 39.295068 | 0.3488049 | 0.28564514 | 1.22111269 | 0.22204336 | 0.78760199 |
| SQOR | 48.9360039 | -0.2257297 | 0.31650973 | -0.713184 | 0.47573194 | 0.90242252 |
| SEMA6D | 153.5313 | -0.160536 | 0.29024175 | -0.5531115 | 0.58018709 | 0.93345492 |
| MYEF2 | 296.806612 | -0.2115538 | 0.23570842 | -0.8975233 | 0.36943977 | 0.86020443 |
| SLC12A1 | 110.504699 | 0.51242886 | 0.68522389 | 0.74782691 | 4.55E-01 | 0.89626521 |
| DUT | 164.681575 | 0.41788269 | 0.37047546 | 1.12796322 | 0.25933545 | 0.80709747 |
| FBN1 | 46.9035943 | 1.22214437 | 0.68278066 | 1.78995165 | 0.07346168 | 0.56723762 |
| CEP152 | 100.286436 | -0.3273802 | 0.30409918 | -1.0765573 | 0.2816781 | 0.82661559 |
| EID1 | 480.680463 | 0.39383294 | 0.41046773 | 0.95947357 | 0.33732023 | 0.84636904 |
| SECISBP2L | 598.318065 | 0.08884893 | 0.14459772 | 0.61445595 | 0.53891407 | 0.92365631 |
| COPS2 | 141.215523 | -0.0046739 | 0.26084266 | -0.0179184 | 0.98570397 | 0.99762072 |
| GALK2 | 282.050046 | -0.0012019 | 0.24499152 | -0.0049061 | 0.99608555 | 0.99967893 |
| FAM227B | 167.345472 | 0.24460359 | 0.41846436 | 0.5845267 | 0.55886601 | 0.92901206 |
| DTWD1 | 202.066932 | -0.077687 | 0.23500103 | -0.3305816 | 0.74096053 | 0.96084411 |
| GABPB1 | 150.605447 | -0.5212069 | 0.2894151 | -1.8008974 | 0.07171905 | 0.56343001 |
| GABPB1-AS1 | 252.007388 | 0.4617732 | 0.39419377 | 1.17143709 | 0.24142313 | 0.80072691 |
| USP8 | 389.398717 | 0.08827452 | 0.19580679 | 0.45082462 | 0.65211595 | 0.95079106 |
| TRPM7 | 573.921312 | -0.3128785 | 0.1900566 | -1.6462385 | 0.09971467 | 0.61822465 |
| SPPL2A | 382.110784 | -0.0773663 | 0.18961825 | -0.4080106 | 0.68326588 | 0.95574492 |
| AP4E1 | 142.057698 | -0.0785537 | 0.26601356 | -0.2952994 | 0.76776514 | 0.96326812 |
| DMXL2 | 343.999605 | -0.04787 | 0.26452471 | -0.1809659 | 0.85639432 | 0.97718353 |
| LYSMD2 | 43.3743173 | 0.46626974 | 0.47770965 | 0.97605259 | 0.32903841 | 0.84409527 |
| TMOD2 | 53.7676833 | 0.20773574 | 0.3042749 | 0.68272388 | 0.49478133 | 0.90839719 |
| TMOD3 | 402.747806 | -0.18824 | 0.1765475 | -1.0662284 | 0.2863204 | 0.8291345 |
| LEO1 | 57.8600522 | -0.0757013 | 0.2737897 | -0.2764943 | 0.78216845 | 0.96513523 |
| MAPK6 | 246.084567 | -0.7439431 | 0.38732479 | -1.9207216 | 0.05476682 | 0.51597783 |
| GNB5 | 286.613767 | 0.13229078 | 0.45483976 | 0.29085139 | 0.77116498 | 0.96388926 |
| CERNA1 | 36.2590488 | 0.23359283 | 0.52731578 | 0.44298471 | 0.6577768 | 0.95100227 |
| MYO5C | 309.508107 | -0.3180289 | 0.15543748 | -2.0460248 | 0.04075393 | 0.46430138 |
| MYO5A | 935.167916 | -0.4794716 | 0.33377217 | -1.4365236 | 0.15085342 | 0.7029344 |
| ARPP19 | 413.108212 | -0.2115242 | 0.26570903 | -0.7960745 | 0.42598871 | 0.88430496 |
| FAM214A | 371.518501 | -0.1110504 | 0.17418094 | -0.6375576 | 0.5237617 | 0.91871245 |
| WDR72 | 1820.17789 | -0.315322 | 0.38143382 | -0.8266756 | 0.40842095 | 0.87948803 |
| UNC13C | 343.434638 | -0.5363096 | 0.79177146 | -0.677354 | 0.49818137 | 0.90839719 |
| RSL24D1 | 121.919092 | 0.34300005 | 0.47438988 | 0.72303407 | 0.46965895 | 0.90053922 |
| RAB27A | 312.277697 | -0.4920157 | 0.32745639 | -1.5025381 | 0.13295821 | 0.67594872 |
| PIGB | 153.293881 | -0.1729519 | 0.47778705 | -0.3619854 | 0.71736297 | 0.95791799 |
| CCPG1 | 573.818837 | 0.4728282 | 0.27709883 | 1.7063522 | 0.08794252 | 0.60651125 |
| C15orf65 | 41.839754 | 0.41479423 | 0.36721375 | 1.12957161 | 0.25865678 | 0.80709747 |
| DNAAF4 | 227.036603 | 0.08930377 | 0.25425282 | 0.35124001 | 0.72540829 | 0.95963742 |
| PYGO1 | 106.271021 | 0.23898915 | 0.31392659 | 0.76128992 | 4.46E-01 | 0.8918683 |

| | | | | | | |
|---|---|---|---|---|---|---|
| PRTG | 409.711011 | 0.99743851 | 0.60159219 | 1.65799778 | 0.09731791 | 0.61662137 |
| NEDD4 | 55.3008408 | 0.16906877 | 0.45541378 | 0.3712421 | 7.10E-01 | 0.95756248 |
| RFX7 | 510.863769 | 0.11476339 | 0.28040415 | 0.40927851 | 0.68233529 | 0.95574492 |
| TEX9 | 310.341798 | -0.0469853 | 0.25695357 | -0.1828553 | 0.85491152 | 0.97718353 |
| MNS1 | 341.818092 | 0.75425235 | 0.40243879 | 1.87420391 | 0.06090233 | 0.5418629 |
| ZNF280D | 170.179359 | -0.0487216 | 0.2325205 | -0.2095368 | 0.83402921 | 0.97232209 |
| AC090517.4 | 107.804103 | -0.336061 | 0.3372663 | -0.9964263 | 0.31904306 | 0.84285425 |
| TCF12 | 1969.32938 | -0.0355091 | 0.23995697 | -0.147981 | 0.88235778 | 0.98335905 |
| CGNL1 | 648.033903 | -0.5224383 | 0.50824866 | -1.0279187 | 0.30398807 | 0.83709431 |
| POLR2M | 52.4647553 | -0.3634194 | 0.32362216 | -1.1229744 | 0.26144837 | 0.80805101 |
| ALDH1A2 | 280.916103 | -1.3114198 | 0.41954713 | -3.1257986 | 0.00177323 | 0.10619113 |
| LIPC | 205.71047 | -0.9386873 | 0.37424231 | -2.5082341 | 0.01213363 | 0.26883805 |
| ADAM10 | 1321.57511 | -0.2607252 | 0.37758368 | -0.6905097 | 0.48987373 | 0.90730741 |
| MINDY2 | 830.261493 | -0.4329428 | 0.324833 | -1.3328166 | 0.18259199 | 0.74315969 |
| RNF111 | 562.572565 | -0.1801131 | 0.26167022 | -0.6883211 | 0.49125061 | 0.90730741 |
| SLTM | 751.899499 | -0.1871965 | 0.17997019 | -1.040153 | 0.29826885 | 0.83556453 |
| MYO1E | 2528.70519 | -0.4672324 | 0.51268863 | -0.9113376 | 0.36211753 | 0.85690129 |
| FAM81A | 475.645451 | 0.34100629 | 0.27574578 | 1.23666915 | 0.21620994 | 0.78072228 |
| GCNT3 | 179.08929 | 1.67606076 | 0.73045024 | 2.29455843 | 0.02175844 | 0.3562945 |
| GTF2A2 | 179.651676 | 0.37501984 | 0.39742655 | 0.9436205 | 0.34536361 | 0.8523814 |
| BNIP2 | 203.603871 | -0.2557984 | 0.16690085 | -1.5326372 | 0.12536528 | 0.66667411 |
| ANXA2 | 711.871371 | -0.1305312 | 0.42902758 | -0.304249 | 0.76093819 | 0.96218738 |
| ICE2 | 281.470803 | 0.11512778 | 0.22962198 | 0.50137961 | 0.61610399 | 0.94190661 |
| RORA-AS1 | 185.276582 | 0.166307 | 0.48534606 | 0.34265653 | 0.73185687 | 0.96008876 |
| RORA | 2134.01178 | 0.43753342 | 0.43443074 | 1.00714193 | 0.31386657 | 0.84285425 |
| AC012404.1 | 70.6292638 | 0.95763982 | 0.68792949 | 1.39206102 | 0.16390392 | 0.71715698 |
| AC009554.1 | 180.721257 | 0.17956778 | 0.57983352 | 0.30968852 | 0.75679783 | 0.96133693 |
| VPS13C | 1233.49382 | -0.4382881 | 0.2594746 | -1.6891367 | 0.09119324 | 0.60989481 |
| TLN2 | 985.947792 | 0.15076822 | 0.244021 | 0.61784936 | 0.53667464 | 0.92324352 |
| TPM1 | 426.890142 | 0.35471238 | 0.43298175 | 0.81923173 | 0.41265422 | 0.88153663 |
| LACTB | 98.5949192 | -0.3688895 | 0.37090059 | -0.9945778 | 0.31994166 | 0.84285425 |
| RPS27L | 195.878089 | -0.0541387 | 0.24373082 | -0.2221251 | 0.82421651 | 0.97018919 |
| RAB8B | 143.036551 | -0.2227232 | 0.20300548 | -1.0971289 | 0.27258506 | 0.81922962 |
| APH1B | 86.1572108 | 0.21282688 | 0.29942716 | 0.71078014 | 0.47722049 | 0.90242252 |
| CA12 | 1203.18607 | 0.18139472 | 0.45811411 | 0.39595969 | 0.69213477 | 0.95672887 |
| LINC02568 | 98.1825754 | 0.38114404 | 0.50783676 | 0.75052471 | 4.53E-01 | 0.89544675 |
| USP3 | 704.931078 | -0.0822164 | 0.27063259 | -0.3037934 | 0.76128526 | 0.96218738 |
| USP3-AS1 | 158.185152 | 0.33585484 | 0.43103 | 0.77919134 | 0.43586701 | 0.88785137 |
| HERC1 | 708.790961 | -0.4750765 | 0.21166029 | -2.2445235 | 0.02479874 | 0.37940263 |
| AC073167.1 | 36.5664443 | -0.055651 | 0.63222105 | -0.0880246 | 0.92985711 | 0.99216065 |
| DAPK2 | 76.6121281 | -1.9267781 | 0.70388757 | -2.7373379 | 0.00619386 | 0.19425932 |
| FAM96A | 104.926229 | 0.0633719 | 0.22667102 | 0.27957653 | 0.77980242 | 0.96513523 |
| SNX1 | 271.487295 | -0.0208975 | 0.15405964 | -0.1356453 | 0.89210167 | 0.98597539 |
| PPIB | 438.577096 | 0.39500151 | 0.35095924 | 1.12549115 | 0.26038097 | 0.80805101 |
| CSNK1G1 | 107.640389 | 0.03873777 | 0.31708744 | 0.12216747 | 0.90276639 | 0.9875066 |
| TRIP4 | 162.532542 | -0.3910496 | 0.25994261 | -1.5043689 | 0.1324864 | 0.67466347 |
| ZNF609 | 814.616347 | -0.1410705 | 0.23896615 | -0.5903368 | 0.55496485 | 0.92872652 |
| OAZ2 | 133.824764 | 0.31308133 | 0.27820276 | 1.12537104 | 0.26043184 | 0.80805101 |
| SPG21 | 175.624383 | -0.0927534 | 0.25467127 | -0.3642085 | 0.71570235 | 0.95791799 |
| MTFMT | 95.8656211 | 0.01886923 | 0.25142164 | 0.07505013 | 0.94017483 | 0.99243654 |
| UBAP1L | 44.8672636 | -0.3045597 | 0.33359711 | -0.9129567 | 0.36126533 | 0.8565748 |
| PDCD7 | 94.4509088 | 0.36259686 | 0.25480413 | 1.42304156 | 0.15472411 | 0.70839173 |
| CLPX | 200.989863 | 0.07080299 | 0.16690372 | 0.42421459 | 0.67140932 | 0.95434283 |
| PARP16 | 52.5942269 | -0.0753433 | 0.36051395 | -0.2089887 | 0.8344571 | 0.97232209 |

| | | | | | | |
|---|---|---|---|---|---|---|
| DPP8 | 330.291923 | -0.1621262 | 0.20635413 | -0.7856698 | 0.43206097 | 0.88697335 |
| HACD3 | 514.559071 | 0.43787693 | 0.25744806 | 1.70083598 | 0.08897379 | 0.60739841 |
| INTS14 | 93.6491244 | 0.0856694 | 0.19507721 | 0.43915638 | 0.66054823 | 0.95127957 |
| SLC24A1 | 61.1710278 | 0.18243991 | 0.34107058 | 0.53490368 | 0.59271646 | 0.93491607 |
| DENND4A | 706.142081 | -0.2587127 | 0.28721671 | -0.9007579 | 0.36771705 | 0.85902833 |
| RAB11A | 542.306947 | -0.0322406 | 0.36703545 | -0.0878407 | 0.93000329 | 0.99216065 |
| MEGF11 | 126.70834 | -1.140176 | 0.63827708 | -1.7863339 | 7.40E-02 | 5.69E-01 |
| DIS3L | 84.421569 | 0.31062402 | 0.26566767 | 1.16922027 | 0.2423149 | 0.80072691 |
| TIPIN | 76.2925701 | 0.10342896 | 0.43458284 | 0.23799596 | 0.81188423 | 0.9680627 |
| MAP2K1 | 291.981374 | -0.1640809 | 0.2357752 | -0.6959209 | 0.48647838 | 0.90700827 |
| SNAPC5 | 59.240706 | 0.32414449 | 0.34776216 | 0.93208671 | 0.35129171 | 0.85504372 |
| RPL4 | 533.259186 | 0.16547107 | 0.41211656 | 0.40151522 | 0.68804084 | 0.95597173 |
| ZWILCH | 35.4266692 | -0.690867 | 0.30361835 | -2.2754456 | 0.02287921 | 0.3621966 |
| SMAD6 | 105.915789 | 0.3033028 | 0.43202294 | 0.70205252 | 0.48264641 | 0.90614725 |
| SMAD3 | 1613.12777 | -1.3855044 | 0.53265161 | -2.6011457 | 0.0092913 | 0.23300339 |
| AAGAB | 359.840626 | -0.1233414 | 0.23584511 | -0.5229763 | 0.60099075 | 0.93795727 |
| IQCH | 360.143139 | 0.12352374 | 0.2945211 | 0.41940541 | 0.67491987 | 0.95524793 |
| IQCH-AS1 | 159.517655 | -0.2094105 | 0.32412323 | -0.646083 | 0.51822561 | 0.91751084 |
| MAP2K5 | 576.668468 | -0.191779 | 0.23260598 | -0.8244799 | 0.40966693 | 0.88013723 |
| PIAS1 | 739.439097 | -0.1719456 | 0.241012 | -0.7134319 | 0.47557857 | 0.90242252 |
| FEM1B | 117.474692 | 0.28686434 | 0.18370944 | 1.56151118 | 0.11840319 | 0.65447949 |
| CORO2B | 215.178363 | -0.6175837 | 0.38461933 | -1.605701 | 0.10833962 | 0.63572183 |
| ANP32A | 287.79363 | 0.10138085 | 0.19957683 | 0.50797904 | 0.61146804 | 0.94003018 |
| SPESP1 | 56.47211 | 1.23488262 | 0.45608891 | 2.70754799 | 0.00677823 | 0.20224502 |
| NOX5 | 33.7044407 | 1.62618878 | 0.74203752 | 2.19151826 | 0.02841431 | 0.40459506 |
| GLCE | 231.28595 | -0.322806 | 0.30826369 | -1.047175 | 0.29501886 | 0.83396494 |
| PAQR5 | 420.720664 | -0.8133648 | 0.28778006 | -2.8263417 | 0.0047083 | 0.16921345 |
| RPLP1 | 1089.08306 | -0.0073163 | 0.45230551 | -0.0161755 | 0.98709437 | 0.99791717 |
| TLE3 | 240.905548 | 0.10160329 | 0.4009416 | 0.25341169 | 0.7999501 | 0.96568323 |
| UACA | 816.765497 | -0.116238 | 0.26980101 | -0.4308286 | 0.66659299 | 0.9527406 |
| LARP6 | 59.88171 | 0.28163059 | 0.27351434 | 1.02967395 | 0.30316309 | 0.83709431 |
| LRRC49 | 147.659301 | 0.35018335 | 0.30073804 | 1.16441321 | 0.2442566 | 0.80072691 |
| THSD4 | 1733.70207 | -0.9599788 | 0.46964949 | -2.0440324 | 0.04095034 | 0.46430138 |
| THSD4-AS1 | 390.238685 | -0.154369 | 0.36521572 | -0.4226791 | 0.67252943 | 0.9544944 |
| MYO9A | 1592.4132 | -0.0711378 | 0.21295207 | -0.3340553 | 0.73833787 | 0.96078945 |
| AC022872.1 | 58.6312907 | -0.0412305 | 0.49139095 | -0.0839057 | 0.93313142 | 0.99243654 |
| SENP8 | 42.8823977 | 0.10726828 | 0.44999071 | 0.23837888 | 0.81158725 | 0.9680627 |
| PKM | 1655.17477 | -0.1754939 | 0.26298792 | -0.6673077 | 5.05E-01 | 0.9107831 |
| PARP6 | 143.914163 | -0.4100218 | 0.18846609 | -2.1755735 | 0.02958717 | 0.40623517 |
| AC009690.2 | 63.311166 | 0.22567264 | 0.58439546 | 0.38616427 | 0.69937501 | 0.95672887 |
| AC009690.1 | 201.075 | -0.0435815 | 0.36538426 | -0.1192759 | 0.9050568 | 0.9875066 |
| ARIH1 | 1192.41948 | -0.3422791 | 0.30428186 | -1.1248752 | 0.26064191 | 0.80805101 |
| BBS4 | 215.244422 | -0.2580423 | 0.21643819 | -1.1922215 | 0.23317442 | 0.7926597 |
| ADPGK | 124.133059 | -0.2800686 | 0.25561029 | -1.0956859 | 0.27321625 | 0.81922962 |
| NEO1 | 418.5561 | 0.16142187 | 0.20468703 | 0.78862778 | 0.43032959 | 0.88596084 |
| NPTN | 626.108948 | 0.11489135 | 0.17867042 | 0.64303506 | 0.52020135 | 0.91751084 |
| CD276 | 52.9904039 | -0.4991952 | 0.47122521 | -1.0593559 | 0.28943774 | 0.83103268 |
| LOXL1-AS1 | 31.0634135 | 0.64807289 | 0.44196468 | 1.46634543 | 0.1425542 | 0.6890058 |
| LOXL1 | 37.3500601 | 0.81401544 | 0.40040604 | 2.03297491 | 0.04205506 | 0.46886915 |
| PML | 107.358342 | -0.0834537 | 0.36565534 | -0.2282306 | 0.81946698 | 0.96884581 |
| ARID3B | 78.3883146 | 0.02974532 | 0.42212649 | 0.07046542 | 0.94382322 | 0.99265578 |
| CLK3 | 91.7424588 | -0.1747582 | 0.25093955 | -0.6964155 | 0.48616864 | 0.90695174 |
| EDC3 | 97.1690355 | 0.11573951 | 0.22962462 | 0.50403791 | 0.61423473 | 0.94083309 |
| CSK | 40.1479114 | -0.2908074 | 0.36785755 | -0.7905435 | 0.42921044 | 0.88579433 |

| | | | | | | |
|---|---|---|---|---|---|---|
| ULK3 | 56.0771648 | 0.05578662 | 0.3237337 | 0.17232255 | 0.86318396 | 0.979268 |
| SCAMP2 | 170.523554 | -0.2412661 | 0.25890336 | -0.9318769 | 0.35140012 | 0.85504372 |
| MPI | 45.0948261 | 0.29149941 | 0.37197845 | 0.78364595 | 0.43324789 | 0.88736205 |
| FAM219B | 83.2326466 | -0.0182519 | 0.24345052 | -0.0749716 | 0.94023727 | 0.99243654 |
| COX5A | 253.279228 | 0.18477332 | 0.4312801 | 0.42842997 | 0.66833811 | 0.95294349 |
| SCAMP5 | 243.84427 | 0.06612929 | 0.43302642 | 0.15271421 | 0.87862366 | 0.98302295 |
| PPCDC | 205.655037 | -0.5243967 | 0.37334263 | -1.404599 | 0.16014056 | 0.71409437 |
| C15orf39 | 48.796486 | 0.25150964 | 0.38473999 | 0.65371329 | 0.51329654 | 0.91512427 |
| COMMD4 | 102.475787 | -0.0270235 | 0.43513166 | -0.0621041 | 0.95047989 | 0.9940155 |
| NEIL1 | 80.817034 | -0.0881754 | 0.25614095 | -0.3442454 | 0.73066172 | 0.96008876 |
| MAN2C1 | 121.689371 | -0.5360098 | 0.283065 | -1.8935925 | 0.05827912 | 0.53192434 |
| SIN3A | 245.003642 | -0.0608299 | 0.16767769 | -0.3627785 | 0.71677037 | 0.95791799 |
| PTPN9 | 237.286459 | -0.1868734 | 0.23938084 | -0.7806533 | 0.43500642 | 0.88785137 |
| SNUPN | 76.6465523 | 0.13252312 | 0.22362735 | 0.59260693 | 0.55344424 | 0.92872652 |
| IMP3 | 92.7849438 | 0.5859353 | 0.45247467 | 1.29495711 | 0.19533502 | 0.76270904 |
| UBE2Q2 | 138.065333 | 0.51272731 | 0.25860099 | 1.98269659 | 0.04740133 | 0.48600281 |
| FBXO22 | 192.403289 | 0.06174568 | 0.2074967 | 0.29757429 | 0.7660281 | 0.96285313 |
| NRG4 | 124.325718 | 0.05670434 | 0.60831827 | 0.09321493 | 0.92573282 | 0.99051393 |
| ETFA | 281.515256 | 0.138906 | 0.14459055 | 0.96068514 | 0.3367105 | 0.84621728 |
| SCAPER | 973.369311 | 0.11132265 | 0.22406414 | 0.49683384 | 0.61930623 | 0.94300065 |
| RCN2 | 215.060609 | 0.26368932 | 0.20568012 | 1.28203602 | 0.19983001 | 0.7687013 |
| TSPAN3 | 268.54976 | 0.29781155 | 0.20337026 | 1.46438103 | 0.14308987 | 0.69023197 |
| PEAK1 | 478.156176 | 0.20779677 | 0.17006395 | 1.22187431 | 0.22175516 | 0.78752765 |
| HMG20A | 266.137749 | -0.0283958 | 0.23295544 | -0.1218935 | 0.90298338 | 0.9875066 |
| LINGO1 | 2068.12934 | -0.8393973 | 0.55908021 | -1.5013898 | 0.13325478 | 0.67594872 |
| TBC1D2B | 120.279886 | -1.1818493 | 0.37267658 | -3.1712465 | 0.00151786 | 0.09927452 |
| IDH3A | 122.443672 | 0.21373902 | 0.2258317 | 0.94645269 | 0.34391774 | 0.85126545 |
| DNAJA4 | 41.9833767 | 0.19828629 | 0.39534472 | 0.5015529 | 0.61598206 | 0.94190661 |
| WDR61 | 128.661673 | 0.24554382 | 0.2071078 | 1.18558462 | 0.23578638 | 0.79711008 |
| IREB2 | 278.03199 | 0.06534579 | 0.16885088 | 0.38700298 | 0.698754 | 0.95672887 |
| PSMA4 | 164.914913 | 0.06547217 | 0.29454102 | 0.2222854 | 0.82409171 | 0.97018919 |
| MORF4L1 | 601.617615 | 0.16286142 | 0.26144622 | 0.62292511 | 0.53333373 | 0.92278975 |
| CTSH | 1162.13192 | -0.2768342 | 0.3016671 | -0.9176812 | 0.35878578 | 0.8557799 |
| TMED3 | 99.9080902 | -0.2425816 | 0.32166061 | -0.754154 | 0.45075675 | 0.89333031 |
| ZFAND6 | 610.017468 | -0.2231454 | 0.22965018 | -0.9716752 | 0.33121217 | 0.84442294 |
| AC016705.2 | 66.5520752 | 0.08670551 | 0.44896807 | 0.19312177 | 0.8468636 | 0.97656375 |
| ARNT2 | 221.452077 | 0.28933631 | 0.49604022 | 0.58329205 | 0.55969672 | 0.92934517 |
| ABHD17C | 91.1734859 | 0.12603702 | 0.30969058 | 0.40697725 | 0.6840247 | 0.95574492 |
| AC027808.1 | 53.7433683 | -0.0654119 | 0.43886153 | -0.1490491 | 0.88151485 | 0.98335905 |
| MESD | 184.361789 | 0.35211993 | 0.29672693 | 1.18668005 | 0.23535385 | 0.79599484 |
| EFL1 | 122.57824 | -0.2814732 | 0.34208537 | -0.8228157 | 0.41061279 | 0.88036458 |
| RPS17 | 47.0764224 | -0.0454802 | 0.40961067 | -0.1110328 | 0.91159037 | 0.98788932 |
| WHAMM | 102.657192 | -0.0877987 | 0.22594565 | -0.3885832 | 0.69758449 | 0.95672887 |
| HOMER2 | 52.2792119 | 0.12564 | 0.4234774 | 0.29668642 | 0.76670592 | 0.96285313 |
| C15orf40 | 103.189696 | -0.0331615 | 0.28130861 | -0.117883 | 0.90616035 | 0.9875066 |
| BTBD1 | 198.856655 | -0.0885917 | 0.22080448 | -0.4012225 | 0.6882563 | 0.95597173 |
| HDGFL3 | 247.655552 | 0.24441754 | 0.2043333 | 1.19617087 | 0.23162988 | 0.79200039 |
| AC103876.1 | 37.0439173 | 0.09419276 | 0.48308486 | 0.19498181 | 0.84540719 | 0.97618636 |
| ADAMTSL3 | 673.697953 | 0.63753221 | 0.49813454 | 1.2798394 | 0.20060162 | 0.7694065 |
| WDR73 | 193.069021 | -0.0108439 | 0.22496585 | -0.0482022 | 0.96155507 | 0.99635733 |
| NMB | 38.6251348 | 0.02342277 | 0.61081208 | 0.03834693 | 0.96941107 | 0.99645289 |
| SEC11A | 409.191773 | 0.20069791 | 0.28355906 | 0.70778171 | 0.47908083 | 0.90291126 |
| ZNF592 | 138.270037 | 0.1940337 | 0.30854785 | 0.62886097 | 0.52944008 | 0.92164157 |
| ALPK3 | 100.486559 | -0.3302603 | 0.60783793 | -0.5433361 | 0.58689841 | 0.93483536 |

| | | | | | | |
|---|---|---|---|---|---|---|
| PDE8A | 1002.18721 | -0.3768787 | 0.28130353 | -1.3397583 | 0.18032394 | 0.74280139 |
| AKAP13 | 1035.33135 | -0.2883211 | 0.30821553 | -0.9354529 | 0.34955495 | 0.8535343 |
| AC087286.2 | 48.4492079 | -0.3508811 | 0.57113072 | -0.6143622 | 0.53897604 | 0.92365631 |
| KLHL25 | 43.1283723 | -0.1452656 | 0.38205971 | -0.3802171 | 0.70378426 | 0.95672887 |
| MRPL46 | 64.8647998 | 0.05556135 | 0.32415632 | 0.17140295 | 0.86390693 | 0.97939196 |
| MRPS11 | 90.8157611 | 0.02521646 | 0.2930042 | 0.08606178 | 0.9314173 | 0.99243654 |
| DET1 | 70.9207678 | 0.07640598 | 0.46401809 | 0.16466164 | 0.86921031 | 0.9820194 |
| ABHD2 | 241.235718 | 0.18210218 | 0.35672741 | 0.51047991 | 0.60971529 | 0.94003018 |
| POLG | 61.1045875 | -0.1912059 | 0.22263687 | -0.8588242 | 0.39043749 | 0.87004477 |
| PEX11A | 51.6362014 | 0.36124342 | 0.30149288 | 1.19818224 | 0.23084608 | 0.79130016 |
| AP3S2 | 38.8051793 | -0.0887585 | 0.31059747 | -0.2857671 | 0.77505653 | 0.96388926 |
| ZNF710 | 192.540649 | 0.03140051 | 0.30834217 | 0.10183657 | 0.9188864 | 0.99014946 |
| ZNF710-AS1 | 75.3994895 | -0.3247318 | 0.35713692 | -0.9092642 | 0.36321071 | 0.85694172 |
| IDH2 | 484.317039 | -0.1011368 | 0.22963676 | -0.4404209 | 6.60E-01 | 0.95127957 |
| SEMA4B | 825.484961 | -1.3599046 | 0.49567711 | -2.7435292 | 0.00607826 | 0.19244777 |
| CIB1 | 412.033843 | -0.5719258 | 0.45951313 | -1.2446343 | 0.21326622 | 0.77821377 |
| IQGAP1 | 1051.09897 | -0.1711068 | 0.16495302 | -1.0373062 | 0.29959318 | 0.83618229 |
| CRTC3 | 747.098999 | -0.1889732 | 0.1837695 | -1.0283164 | 0.303801 | 0.83709431 |
| CRTC3-AS1 | 67.4710106 | -0.1076111 | 0.46452907 | -0.2316564 | 0.81680491 | 0.96874284 |
| FURIN | 96.1485041 | -0.2378703 | 0.35285558 | -0.6741293 | 0.50022912 | 0.90839719 |
| MAN2A2 | 54.8735269 | -0.1123616 | 0.31977022 | -0.3513824 | 0.72530145 | 0.95963742 |
| HDDC3 | 46.4996782 | 0.25690976 | 0.31298681 | 0.82083255 | 0.41174166 | 0.88086809 |
| UNC45A | 110.689379 | -0.0192301 | 0.25740409 | -0.0747077 | 0.94044726 | 0.99243654 |
| RCCD1 | 33.3727201 | 0.14818949 | 0.34585786 | 0.42846934 | 0.66830945 | 0.95294349 |
| AC104035.1 | 351.273843 | -0.5547818 | 0.77029895 | -0.7202162 | 0.47139191 | 0.90123338 |
| SLCO3A1 | 833.728373 | -0.0100752 | 0.42705211 | -0.0235924 | 0.98117772 | 0.99696835 |
| FAM174B | 126.783199 | -0.5342302 | 0.36022289 | -1.4830544 | 0.13805995 | 0.68146162 |
| AC013394.1 | 437.490464 | -0.2253484 | 0.30475199 | -0.7394486 | 0.45963467 | 0.89753337 |
| CHD2 | 801.814887 | -0.595591 | 0.36386174 | -1.6368607 | 0.10165958 | 0.62303648 |
| MCTP2 | 211.311745 | 0.16099625 | 0.30182173 | 0.53341506 | 0.5937463 | 0.93514596 |
| AC012409.2 | 245.346537 | -1.2314236 | 0.66954449 | -1.839196 | 0.06588636 | 0.55444578 |
| NR2F2-AS1 | 1390.19254 | -1.320684 | 0.44557637 | -2.9639903 | 0.00303678 | 0.14138755 |
| NR2F2 | 650.361112 | -1.2007199 | 0.54964533 | -2.1845358 | 0.0289229 | 0.40623517 |
| AC087477.2 | 42.9000311 | -0.9612617 | 0.45492039 | -2.1130328 | 0.03459796 | 0.43165078 |
| LINC00923 | 180.071782 | 0.21117701 | 0.58373395 | 0.36176929 | 0.71752445 | 0.95791799 |
| ARRDC4 | 127.094005 | 1.09777684 | 0.37326461 | 2.94101507 | 0.00327139 | 0.14358074 |
| IGF1R | 1168.59772 | -0.4377967 | 0.20320891 | -2.1544167 | 0.03120751 | 0.41399324 |
| TTC23 | 194.750078 | -0.2559808 | 0.22843205 | -1.1205994 | 0.2624584 | 0.80922744 |
| LRRC28 | 452.480693 | 0.21515977 | 0.26782664 | 0.80335464 | 0.42176978 | 0.88371155 |
| MEF2A | 1342.93967 | 0.14394542 | 0.24248904 | 0.59361618 | 0.55276885 | 0.92872652 |
| AC090825.1 | 101.038471 | 0.63157774 | 0.3753115 | 1.68280945 | 0.09241198 | 0.61120158 |
| LINS1 | 56.8289088 | 0.13476205 | 0.29620816 | 0.45495726 | 0.64913999 | 0.95061388 |
| ASB7 | 92.4443493 | -0.0343204 | 0.26248935 | -0.1307498 | 0.89597323 | 0.98597539 |
| AC015712.1 | 34.6430037 | 0.19638396 | 0.51917552 | 0.37826121 | 0.70523656 | 0.95730303 |
| ALDH1A3 | 48.7929581 | -2.4683625 | 1.27543679 | -1.9353076 | 0.05295257 | 0.50958943 |
| CHSY1 | 530.852771 | -0.3917047 | 0.4689721 | -0.8352409 | 0.4035821 | 0.87834687 |
| SELENOS | 181.606836 | 0.0014667 | 0.42757166 | 0.00343029 | 0.99726303 | 0.99967893 |
| SNRPA1 | 111.114016 | -0.4297061 | 0.19850801 | -2.1646791 | 0.03041227 | 0.41080114 |
| TM2D3 | 129.275311 | 0.14232297 | 0.28901588 | 0.49243996 | 0.62240836 | 0.94418789 |
| TARSL2 | 165.405435 | 0.03854579 | 0.28673127 | 0.13443175 | 0.89306118 | 0.98597539 |
| SNRNP25 | 61.8048654 | 0.07369498 | 0.48441793 | 0.152131 | 0.87908362 | 0.98311774 |
| MPG | 111.381288 | 0.0691573 | 0.23967391 | 0.28854747 | 0.7729277 | 0.96388926 |
| NPRL3 | 185.239486 | -0.0828337 | 0.2787356 | -0.2971767 | 0.76633162 | 0.96285313 |
| LUC7L | 329.48958 | -0.3640011 | 0.19328177 | -1.8832668 | 0.05966422 | 0.53580211 |

| | | | | | | |
|---|---|---|---|---|---|---|
| FAM234A | 43.5459749 | 0.51365213 | 0.5040421 | 1.01906593 | 0.30817167 | 0.83980597 |
| AXIN1 | 76.1049796 | -0.2626305 | 0.31422946 | -0.835792 | 0.40327195 | 0.87834687 |
| MRPL28 | 113.242664 | 0.62558655 | 0.39652564 | 1.57766985 | 0.11464148 | 0.65008305 |
| TMEM8A | 166.452225 | -0.0519643 | 0.2777539 | -0.1870875 | 0.851592 | 0.97718353 |
| NME4 | 89.114036 | -0.2747786 | 0.49689477 | -0.5529916 | 0.58026916 | 0.93345492 |
| RAB11FIP3 | 385.113049 | 0.40871155 | 0.35394499 | 1.15473182 | 0.24820027 | 0.80097144 |
| CAPN15 | 80.7897483 | -0.3292729 | 0.30751692 | -1.0707473 | 0.28428309 | 0.82778499 |
| PRR35 | 135.58046 | 2.31356516 | 0.80507343 | 2.87373186 | 0.00405653 | 0.15545856 |
| PIGQ | 433.389218 | 0.96325568 | 0.63434421 | 1.51850631 | 0.12888681 | 0.67206108 |
| RAB40C | 332.941933 | -0.3449814 | 0.31954184 | -1.0796127 | 0.28031467 | 0.82628479 |
| METTL26 | 62.1425838 | -0.0389118 | 0.52701807 | -0.0738339 | 0.94114252 | 0.99265578 |
| MCRIP2 | 151.742605 | -0.045821 | 0.33830648 | -0.1354421 | 0.89226231 | 0.98597539 |
| WDR90 | 47.1894921 | 0.18856861 | 0.34206008 | 0.55127336 | 0.5814463 | 0.93386308 |
| RHOT2 | 113.751832 | -0.0854516 | 0.22063725 | -0.3872945 | 0.69853823 | 0.95672887 |
| STUB1 | 206.757262 | 0.26024053 | 0.44107983 | 0.59000777 | 0.55518544 | 0.92872652 |
| JMJD8 | 78.8285602 | 0.2883098 | 0.45849605 | 0.62881631 | 0.52946932 | 0.92164157 |
| METRN | 85.9478704 | 0.02082877 | 0.5300886 | 0.03929299 | 0.96865679 | 0.9964399 |
| FAM173A | 89.3546707 | 0.1715034 | 0.40317797 | 0.4253789 | 0.67056048 | 0.95395138 |
| NARFL | 53.485907 | -0.5682086 | 0.3748949 | -1.5156478 | 0.12960844 | 0.67209127 |
| LMF1 | 167.147391 | -0.3427102 | 0.30338516 | -1.1296208 | 0.25863603 | 0.80709747 |
| UBE2I | 302.197459 | -0.5443122 | 0.24279629 | -2.2418472 | 0.02497125 | 0.38092969 |
| TSR3 | 98.5347523 | 0.06134387 | 0.46538971 | 0.13181183 | 0.89513313 | 0.98597539 |
| GNPTG | 109.954656 | -0.2614732 | 0.29071023 | -0.8994288 | 0.36842429 | 0.8591648 |
| UNKL | 70.7062988 | 0.09650678 | 0.29382167 | 0.32845358 | 0.74256873 | 0.96086911 |
| CLCN7 | 122.424167 | -0.6392869 | 0.34857779 | -1.8339864 | 0.06665602 | 0.55619688 |
| TELO2 | 40.7035936 | -0.2963923 | 0.37159679 | -0.797618 | 0.42509218 | 0.88392184 |
| IFT140 | 145.737497 | 0.16591814 | 0.28365592 | 0.58492746 | 0.5585965 | 0.92901206 |
| JPT2 | 143.079488 | -0.6771138 | 0.37330608 | -1.8138302 | 0.06970387 | 0.55992041 |
| MAPK8IP3 | 260.473181 | -0.4705824 | 0.39201283 | -1.200426 | 0.22997394 | 0.79047378 |
| NME3 | 257.851761 | 0.56060924 | 0.38872386 | 1.44217865 | 0.14925198 | 0.70236227 |
| MRPS34 | 139.025228 | 0.0882764 | 0.56074588 | 0.15742674 | 0.87490854 | 0.98267808 |
| SPSB3 | 151.211275 | -0.0216586 | 0.27734161 | -0.0780936 | 0.93775356 | 0.99243654 |
| NUBP2 | 59.6340716 | -0.206986 | 0.41364715 | -0.5003926 | 0.61679867 | 0.94228135 |
| HAGH | 163.805382 | -0.1338234 | 0.25658978 | -0.5215461 | 0.60198637 | 0.93804805 |
| FAHD1 | 67.9940143 | 0.11680416 | 0.38856326 | 0.30060527 | 0.76371551 | 0.96285313 |
| MSRB1 | 41.4069324 | 0.07245011 | 0.36642475 | 0.19772166 | 0.84326284 | 0.97560439 |
| NDUFB10 | 220.214287 | 0.34718755 | 0.5300121 | 0.6550559 | 0.51243177 | 0.91445384 |
| RPS2 | 614.382214 | -0.1602612 | 0.39898204 | -0.4016752 | 0.6879231 | 0.95597173 |
| SNHG9 | 34.7232007 | -0.4249369 | 0.29511243 | -1.4399153 | 0.14989137 | 0.7029344 |
| TSC2 | 163.376356 | -0.3022543 | 0.27573226 | -1.0961877 | 0.27299664 | 0.81922962 |
| PKD1 | 94.3550475 | -0.2699424 | 0.22074479 | -1.222871 | 0.22137843 | 0.78752408 |
| SNHG19 | 50.7991489 | 0.11132696 | 0.35132934 | 0.31687351 | 0.75133959 | 0.96120879 |
| TRAF7 | 51.8340785 | -0.0340524 | 0.30453953 | -0.111816 | 0.91096928 | 0.98788932 |
| MLST8 | 34.4951873 | -0.3079376 | 0.36794818 | -0.8369048 | 0.40264608 | 0.87834687 |
| PGP | 144.755545 | 0.11115769 | 0.32848899 | 0.33839093 | 0.73506861 | 0.96078945 |
| E4F1 | 49.0233043 | -0.3852613 | 0.30135273 | -1.2784398 | 0.20109441 | 0.76957251 |
| ECI1 | 311.74507 | 0.14286703 | 0.29782712 | 0.47969786 | 0.63144225 | 0.94630556 |
| RNPS1 | 172.190798 | 0.08145834 | 0.20326047 | 0.40075837 | 0.68859803 | 0.95604695 |
| ABCA3 | 81.3370274 | 0.01566611 | 0.3503604 | 0.04471426 | 0.96433507 | 0.99637506 |
| PDPK1 | 266.678036 | 0.03557253 | 0.20127736 | 0.17673391 | 0.85971741 | 0.97837316 |
| KCTD5 | 42.431574 | -0.191509 | 0.36503308 | -0.5246346 | 0.59983721 | 0.93728325 |
| SRRM2 | 870.451883 | -0.3544635 | 0.20757603 | -1.7076319 | 0.08770466 | 0.60651125 |
| ELOB | 284.4664 | 0.18647082 | 0.48187722 | 0.38696749 | 0.69878028 | 0.95672887 |
| PRSS22 | 73.5310029 | -0.6423084 | 0.37975505 | -1.6913755 | 0.0907651 | 0.60936468 |

| | | | | | | |
|---|---|---|---|---|---|---|
| FLYWCH2 | 88.0044919 | 0.36545219 | 0.56606588 | 0.64560011 | 0.51853837 | 0.91751084 |
| FLYWCH1 | 147.364691 | -0.126856 | 0.44150126 | -0.2873287 | 0.77386067 | 0.96388926 |
| TNFRSF12A | 198.698107 | 0.71758895 | 0.74112149 | 0.96824739 | 0.33292083 | 0.84442294 |
| HCFC1R1 | 191.324635 | 1.08949175 | 0.54532473 | 1.99787704 | 0.04572999 | 0.48339878 |
| BICDL2 | 88.9804967 | 0.29231735 | 0.27148154 | 1.07674855 | 0.28159262 | 0.82661559 |
| ZNF213-AS1 | 61.1077243 | -0.3483357 | 0.25504687 | -1.3657714 | 0.17201072 | 0.72776437 |
| ZNF205 | 32.4258622 | -0.1533304 | 0.32311774 | -0.4745343 | 0.63511896 | 0.94792975 |
| ZNF213 | 101.68056 | 0.1327163 | 0.40239166 | 0.32981872 | 0.74153694 | 0.96084411 |
| ZNF263 | 54.3162535 | -0.3363909 | 0.248393 | -1.3542688 | 0.17565065 | 0.73403881 |
| TIGD7 | 44.7828099 | -0.1323446 | 0.30315075 | -0.4365637 | 0.66242777 | 0.95142083 |
| ZNF75A | 156.730675 | -0.1594802 | 0.38997409 | -0.4089507 | 0.68257582 | 0.95574492 |
| AC025283.3 | 59.9970254 | -0.2591651 | 0.53849087 | -0.4812804 | 0.6303172 | 0.94596555 |
| ZSCAN32 | 34.9696023 | 0.47067089 | 0.3278211 | 1.43575534 | 0.15107197 | 0.7029344 |
| NAA60 | 93.5934947 | 0.09775623 | 0.27654968 | 0.35348524 | 0.72372469 | 0.95963742 |
| CLUAP1 | 184.356943 | 0.08615148 | 0.22579167 | 0.38155296 | 0.70279298 | 0.95672887 |
| DNASE1 | 169.714616 | 0.3381116 | 0.31950402 | 1.05823896 | 0.28994651 | 0.83103268 |
| TRAP1 | 138.597374 | 0.11055784 | 0.2163495 | 0.51101502 | 0.60934054 | 0.94003018 |
| CREBBP | 548.053827 | -0.1328765 | 0.21066612 | -0.6307446 | 0.52820751 | 0.92106733 |
| ADCY9 | 610.372705 | -0.1418035 | 0.44970996 | -0.3153222 | 0.75251702 | 0.96120879 |
| GLIS2 | 133.741658 | 0.52699375 | 0.49914638 | 1.05578998 | 0.29106418 | 0.83183872 |
| VASN | 251.412941 | 1.2261641 | 0.42739177 | 2.86894643 | 0.00411842 | 0.15581586 |
| DNAJA3 | 111.987047 | -0.0558463 | 0.20507662 | -0.2723194 | 0.78537641 | 0.96513523 |
| NMRAL1 | 93.0304736 | -0.0424075 | 0.2505755 | -0.1692405 | 0.86560746 | 0.98018684 |
| HMOX2 | 104.982964 | -0.0993849 | 0.34070621 | -0.2917025 | 0.77051409 | 0.96352463 |
| CDIP1 | 85.4748042 | 0.64253951 | 0.30469341 | 2.10880672 | 0.03496127 | 0.43353051 |
| UBALD1 | 50.3322849 | -0.0136934 | 0.52167643 | -0.0262488 | 0.97905893 | 0.99696835 |
| MGRN1 | 193.066713 | -0.1655174 | 0.15087977 | -1.0970149 | 0.27263488 | 0.81922962 |
| NUDT16L1 | 89.8387829 | 0.72473067 | 0.46294052 | 1.56549414 | 0.11746709 | 0.65446842 |
| ANKS3 | 131.897071 | -0.1849085 | 0.21519838 | -0.8592468 | 0.39020434 | 0.87004477 |
| ZNF500 | 52.2952732 | -0.0009642 | 0.38668721 | -0.0024934 | 0.99801055 | 0.99967893 |
| SMIM22 | 114.022488 | 0.10958829 | 0.41134949 | 0.26641164 | 0.7899222 | 0.96522573 |
| ROGDI | 69.6057935 | 0.48828668 | 0.5031675 | 0.97042572 | 0.33183434 | 0.84442294 |
| GLYR1 | 367.887474 | -0.0762953 | 0.1992779 | -0.3828588 | 0.70182445 | 0.95672887 |
| UBN1 | 130.306144 | -0.1756211 | 0.29351934 | -0.5983289 | 0.54962047 | 0.92753985 |
| PPL | 224.852135 | 0.21529241 | 0.20543523 | 1.04798194 | 0.29464693 | 0.8337535 |
| ALG1 | 34.9613904 | -0.1771633 | 0.33798235 | -0.524179 | 0.60015403 | 0.93728325 |
| RBFOX1 | 51.5066585 | -0.0408313 | 0.44453293 | -0.0918521 | 0.92681558 | 0.99092301 |
| METTL22 | 48.5010275 | -0.4716913 | 0.3361421 | -1.4032498 | 0.16054236 | 0.71409437 |
| ABAT | 53.0542429 | -0.0334239 | 0.30187216 | -0.110722 | 0.91183678 | 0.98790958 |
| PMM2 | 156.200077 | -0.5392623 | 0.34362265 | -1.5693444 | 0.11656771 | 0.65257994 |
| CARHSP1 | 226.861192 | 0.19882669 | 0.46479022 | 0.42777728 | 0.66881328 | 0.95298441 |
| USP7 | 213.065392 | 0.01346216 | 0.16082824 | 0.08370518 | 0.93329083 | 0.99243654 |
| C16orf72 | 227.591556 | -0.0447062 | 0.17364401 | -0.2574587 | 0.79682473 | 0.9653598 |
| AC087190.1 | 42.5929762 | 0.109233 | 0.55728952 | 0.19600764 | 0.84460419 | 0.97573093 |
| ATF7IP2 | 185.612758 | 0.82945647 | 0.37120832 | 2.23447706 | 0.0254517 | 0.3854535 |
| EMP2 | 168.560416 | 0.39472388 | 0.49878991 | 0.791363 | 0.42873219 | 0.88535099 |
| NUBP1 | 55.7249595 | -0.1406901 | 0.34122618 | -0.4123074 | 0.68011411 | 0.95574492 |
| CIITA | 72.6264195 | 0.12896289 | 0.50540536 | 0.25516725 | 0.79859393 | 0.9653598 |
| DEXI | 82.6064026 | 0.40637325 | 0.23612146 | 1.72103481 | 0.08524451 | 0.59845478 |
| CLEC16A | 699.178712 | 0.17650139 | 0.23204232 | 0.7606431 | 0.44687027 | 0.8918683 |
| AC007014.1 | 36.1181734 | 0.2418337 | 0.48176234 | 0.50197717 | 0.61568358 | 0.94190661 |
| SOCS1 | 397.479005 | -0.6808894 | 0.38582849 | -1.7647464 | 7.76E-02 | 5.76E-01 |
| LITAF | 452.471308 | 0.16201022 | 0.3110343 | 0.52087576 | 0.60245332 | 0.93804805 |
| SNN | 44.6394594 | -0.7527033 | 0.59835511 | -1.2579542 | 0.20840831 | 0.77434851 |

| | | | | | | |
|---|---|---|---|---|---|---|
| TXNDC11 | 290.420705 | -0.0637536 | 0.21315956 | -0.2990885 | 0.76487253 | 0.96285313 |
| ZC3H7A | 316.264644 | 0.02721502 | 0.20397866 | 0.1334209 | 0.89386052 | 0.98597539 |
| RSL1D1 | 93.6813866 | -0.1625084 | 0.29654178 | -0.5480119 | 0.58368373 | 0.93473293 |
| AC007216.4 | 133.566615 | 0.12033583 | 0.32264674 | 0.37296466 | 0.70917475 | 0.95752169 |
| GSPT1 | 437.844023 | -0.1786109 | 0.21371201 | -0.8357549 | 0.4032928 | 0.87834687 |
| SNX29 | 1952.71874 | 0.11302871 | 0.31240138 | 0.36180606 | 0.71749696 | 0.95791799 |
| CPPED1 | 197.300714 | -0.6508846 | 0.43704662 | -1.4892795 | 0.13641379 | 0.68012205 |
| U91319.1 | 53.9532224 | 0.92991348 | 0.51251218 | 1.8144222 | 0.06961275 | 0.5598938 |
| ERCC4 | 62.448225 | -0.1954503 | 0.24766372 | -0.7891763 | 0.43000897 | 0.88579433 |
| MKL2 | 782.4171 | -0.205365 | 0.20991682 | -0.978316 | 0.32791807 | 0.84381657 |
| PARN | 331.942455 | -0.1780393 | 0.21095535 | -0.8439668 | 0.39868794 | 0.87557619 |
| BFAR | 111.014581 | -0.255176 | 0.20412594 | -1.2500908 | 0.21126639 | 0.7767562 |
| NOMO1 | 52.1460928 | -0.1707588 | 0.31315015 | -0.5452937 | 0.58555154 | 0.93483536 |
| PDXDC1 | 637.606089 | -0.1997859 | 0.15080445 | -1.3248013 | 0.18523706 | 0.74895057 |
| NTAN1 | 51.503669 | -0.4434984 | 0.24736758 | -1.7928719 | 0.07299341 | 0.56668778 |
| RRN3 | 107.336871 | -0.0894207 | 0.23583556 | -0.3791655 | 0.70456497 | 0.95707594 |
| C16orf45 | 155.126753 | 0.67642169 | 0.33302804 | 2.03112534 | 0.04224228 | 0.46945823 |
| MARF1 | 182.987429 | -0.2242591 | 0.2380476 | -0.9420767 | 0.34615338 | 0.8523814 |
| NDE1 | 58.2504474 | 0.14818648 | 0.30231059 | 0.49017959 | 0.62400683 | 0.94457465 |
| FOPNL | 80.1701129 | 0.27734845 | 0.24673794 | 1.12406081 | 0.26098723 | 0.80805101 |
| ABCC1 | 350.09668 | -0.0162492 | 0.38448042 | -0.0422627 | 0.96628926 | 0.99637506 |
| XYLT1 | 182.799012 | -0.3497774 | 0.42663675 | -0.8198484 | 0.41230256 | 0.88110334 |
| RPS15A | 134.489006 | 0.00429682 | 0.39482129 | 0.01088295 | 0.99131684 | 0.99889703 |
| SMG1 | 684.292297 | -0.2791564 | 0.14640975 | -1.906679 | 0.05656216 | 0.52346922 |
| COQ7 | 73.207073 | -0.1936216 | 0.35036427 | -0.5526295 | 0.58051713 | 0.93353069 |
| SYT17 | 158.550094 | 0.4204006 | 0.38397518 | 1.094864 | 0.27357622 | 0.81932131 |
| GDE1 | 74.8704895 | 0.07522563 | 0.2827451 | 0.26605459 | 0.79019716 | 0.96524651 |
| CCP110 | 101.588449 | -0.0089307 | 0.35704992 | -0.0250126 | 0.98004493 | 0.99696835 |
| VPS35L | 326.876307 | -0.3227039 | 0.228727 | -1.4108693 | 0.15828317 | 0.71264724 |
| KNOP1 | 81.886891 | -0.4942977 | 0.34118233 | -1.4487787 | 0.14739938 | 0.69834914 |
| IQCK | 654.717174 | 0.42888353 | 0.37974825 | 1.12938909 | 0.25873373 | 0.80709747 |
| GPRC5B | 421.951623 | 0.58742552 | 0.3419433 | 1.71790328 | 0.08581426 | 0.60075714 |
| UMOD | 171.570971 | -0.4329458 | 0.50517766 | -0.8570168 | 0.39143557 | 0.87015937 |
| ACSM3 | 239.246449 | 0.30625812 | 0.33395087 | 0.91707536 | 0.35910315 | 0.85608464 |
| THUMPD1 | 218.574096 | 0.3425797 | 0.20503265 | 1.67085436 | 0.09475045 | 0.61367241 |
| ERI2 | 75.1883004 | 0.28024319 | 0.38853152 | 0.72128817 | 0.47073223 | 0.90123338 |
| LYRM1 | 76.9301191 | 0.60513961 | 0.2796275 | 2.16409188 | 0.0304573 | 0.41080114 |
| TMEM159 | 48.1946242 | 0.22364994 | 0.35939675 | 0.62229261 | 0.53374948 | 0.92289732 |
| CRYM | 33.1752178 | 0.98825185 | 0.4777086 | 2.06873361 | 0.03857109 | 0.45064107 |
| METTL9 | 470.504439 | 0.09367398 | 0.20704144 | 0.45244074 | 0.6509515 | 0.95061388 |
| NPIPB4 | 88.0940159 | -0.70964 | 0.35832022 | -1.9804632 | 0.04765151 | 0.48600281 |
| UQCRC2 | 380.282389 | 0.02200957 | 0.17776642 | 0.12381175 | 0.90146433 | 0.9875066 |
| MOSMO | 217.696593 | 0.17377062 | 0.23714182 | 0.73277089 | 0.46369818 | 0.89801718 |
| EEF2K | 201.805476 | -0.3889551 | 0.55730135 | -0.697926 | 0.48522346 | 0.90695174 |
| POLR3E | 115.137878 | -0.2558611 | 0.46709773 | -0.5477679 | 0.58385131 | 0.93473293 |
| CDR2 | 647.844031 | -0.0765363 | 0.24286547 | -0.3151388 | 0.75265623 | 0.96120879 |
| NPIPB5 | 242.094699 | -0.0439129 | 0.22877217 | -0.1919504 | 0.84778102 | 0.97656375 |
| USP31 | 98.259963 | -0.5145287 | 0.27670974 | -1.8594526 | 0.06296301 | 0.54759529 |
| SCNN1G | 2133.53593 | -0.5210979 | 0.44365243 | -1.1745635 | 0.2401694 | 0.80056465 |
| AC099482.1 | 249.001529 | -0.6959851 | 0.63514749 | -1.0957851 | 0.27317284 | 0.81922962 |
| SCNN1B | 898.792127 | 0.16978824 | 0.43234481 | 0.39271487 | 0.69453009 | 0.95672887 |
| COG7 | 330.306634 | -0.6649478 | 0.15440327 | -4.3065651 | 1.66E-05 | 0.00543025 |
| GGA2 | 293.57614 | -0.3154211 | 0.26293857 | -1.1996001 | 0.23029469 | 0.79075638 |
| EARS2 | 49.1513166 | -0.1947045 | 0.26344276 | -0.7390772 | 0.45986013 | 0.89753337 |

| | | | | | | |
|---|---|---|---|---|---|---|
| UBFD1 | 83.835962 | -0.0208114 | 0.23377407 | -0.0890234 | 0.92906329 | 0.99216065 |
| NDUFAB1 | 144.943542 | 0.02003584 | 0.4193599 | 0.0477772 | 0.96189381 | 0.99635733 |
| PALB2 | 49.2678433 | -0.5422222 | 0.36580879 | -1.4822558 | 0.13827223 | 0.68146162 |
| DCTN5 | 147.880502 | 0.0275547 | 0.23999767 | 0.11481237 | 0.90859384 | 0.98758338 |
| RBBP6 | 235.496454 | 0.02694255 | 0.2502804 | 0.10764948 | 0.91427375 | 0.98840405 |
| TNRC6A | 1048.42298 | 0.01641152 | 0.22889475 | 0.07169898 | 0.94284147 | 0.99265578 |
| AC008731.1 | 37.5286602 | 0.44030822 | 0.41135883 | 1.07037503 | 0.28445053 | 0.82781987 |
| ARHGAP17 | 241.160171 | -0.2161697 | 0.20232858 | -1.0684093 | 0.28533591 | 0.82810567 |
| LCMT1 | 170.804732 | 0.02838652 | 0.20499583 | 0.13847367 | 0.88986608 | 0.98581358 |
| NSMCE1 | 105.878493 | 0.39557049 | 0.28864472 | 1.37044077 | 0.17054935 | 0.72686344 |
| GTF3C1 | 156.028903 | -0.2254907 | 0.20107059 | -1.1214506 | 0.2620961 | 0.80882423 |
| KIAA0556 | 268.823565 | -0.2446856 | 0.19626373 | -1.2467184 | 0.21250077 | 0.77709975 |
| XPO6 | 274.876984 | -0.3430645 | 0.29659232 | -1.156687 | 0.24740026 | 0.80072691 |
| NUPR1 | 510.324641 | -1.3285316 | 0.93646907 | -1.4186605 | 0.15599803 | 0.71066638 |
| SGF29 | 49.1753776 | -0.0035073 | 0.45798271 | -0.0076581 | 0.99388981 | 0.99919666 |
| SULT1A1 | 48.6152454 | 0.79403538 | 0.55569936 | 1.42889382 | 0.15303475 | 0.70683307 |
| ATXN2L | 142.995765 | -0.3655356 | 0.21805656 | -1.6763337 | 0.09367286 | 0.61317397 |
| TUFM | 306.593058 | 0.64732453 | 0.39486054 | 1.63937508 | 0.10113517 | 0.62105083 |
| SH2B1 | 153.53659 | 0.00635313 | 0.26528886 | 0.02394796 | 0.98089411 | 0.99696835 |
| RABEP2 | 116.122104 | -0.3099169 | 0.28290158 | -1.0954937 | 0.2733004 | 0.81922962 |
| NFATC2IP | 253.22771 | -0.0108981 | 0.26721833 | -0.0407836 | 0.96746843 | 0.99637506 |
| SPNS1 | 60.5089427 | -0.6360973 | 0.30535731 | -2.0831245 | 0.03723988 | 0.44129343 |
| QPRT | 215.30371 | 0.77847765 | 0.40787741 | 1.90860695 | 0.05631281 | 0.5228532 |
| KIF22 | 40.2635946 | -0.2078764 | 0.42033218 | -0.4945526 | 0.62091595 | 0.94371308 |
| MAZ | 60.9498958 | -0.3243941 | 0.38613203 | -0.8401118 | 0.40084572 | 0.87775198 |
| AC009133.1 | 43.1423209 | -0.5007133 | 0.40210812 | -1.2452206 | 0.21305069 | 0.77799356 |
| MVP | 272.453028 | -0.6382246 | 0.24779368 | -2.5756288 | 1.00E-02 | 0.24399985 |
| CDIPT | 114.063136 | 0.57889034 | 0.265845 | 2.17754831 | 0.02943968 | 0.40623517 |
| KCTD13 | 77.8879029 | -0.2737952 | 0.23580689 | -1.1610991 | 0.24560162 | 0.80072691 |
| TMEM219 | 246.626963 | -0.2340911 | 0.35652313 | -0.6565944 | 0.51144178 | 0.91403647 |
| TAOK2 | 53.4878532 | -0.3215848 | 0.27909206 | -1.1522535 | 0.24921692 | 0.8011636 |
| INO80E | 75.9494136 | -0.00232 | 0.27122225 | -0.0085538 | 0.99317517 | 0.99919666 |
| PPP4C | 145.854529 | -0.0499929 | 0.31362542 | -0.1594032 | 0.87335125 | 0.98246494 |
| YPEL3 | 152.172177 | 0.23196839 | 0.43623371 | 0.53175256 | 0.59489739 | 0.935202 |
| MAPK3 | 63.0539974 | 0.27601304 | 0.35436226 | 0.77890077 | 0.43603817 | 0.88785137 |
| NPIPB13 | 51.3713982 | -0.965988 | 0.3825667 | -2.5250184 | 0.01156922 | 0.26383866 |
| CD2BP2 | 85.6123154 | -0.043964 | 0.38903573 | -0.1130076 | 0.91002456 | 0.98788661 |
| TBC1D10B | 51.5502094 | -0.0441131 | 0.2560046 | -0.1723135 | 0.86319105 | 0.979268 |
| DCTPP1 | 66.8900973 | 0.41994449 | 0.40963439 | 1.02516902 | 0.30528344 | 0.83709431 |
| SEPHS2 | 230.591907 | -0.2921569 | 0.4287657 | -0.6813906 | 0.49562437 | 0.90839719 |
| ZNF768 | 63.5150094 | 0.0224895 | 0.36512027 | 0.06159476 | 0.95088555 | 0.99404295 |
| ZNF688 | 32.6627434 | -0.0227581 | 0.32260557 | -0.0705447 | 0.94376016 | 0.99265578 |
| ZNF785 | 64.8695625 | 0.23689456 | 0.31738318 | 0.74639923 | 0.45542628 | 0.89626521 |
| PRR14 | 40.7538739 | -0.2265702 | 0.29525815 | -0.7673632 | 0.44286558 | 0.89005186 |
| FBRS | 91.7520585 | -0.0059493 | 0.21453987 | -0.0277306 | 0.97787702 | 0.99696835 |
| PHKG2 | 60.0095455 | 0.06681356 | 0.30506788 | 0.21901211 | 0.82664061 | 0.97068874 |
| RNF40 | 109.890035 | -0.2411154 | 0.2874859 | -0.8387034 | 0.40163575 | 0.87834687 |
| ZNF629 | 44.7048363 | -0.0927047 | 0.40395608 | -0.229492 | 0.81848653 | 0.96883031 |
| BCL7C | 252.197988 | 0.52543932 | 0.27138922 | 1.9361098 | 0.05285425 | 0.50911084 |
| MIR762HG | 44.6321821 | -0.0430994 | 0.35178663 | -0.1225158 | 0.9024905 | 0.9875066 |
| FBXL19 | 67.9477814 | -0.0216896 | 0.37685312 | -0.0575545 | 0.95410347 | 0.99522289 |
| ORAI3 | 33.1685755 | 0.62356372 | 0.31980325 | 1.94983546 | 0.05119573 | 0.50096292 |
| SETD1A | 48.8666469 | -0.28034 | 0.28976453 | -0.9674754 | 0.33330646 | 0.84442294 |
| STX4 | 67.4366077 | -0.2088048 | 0.37320536 | -0.5594905 | 0.57582703 | 0.93294146 |

| | | | | | | |
|---|---|---|---|---|---|---|
| BCKDK | 100.061718 | -0.0556783 | 0.39176683 | -0.142121 | 0.88698441 | 0.98544303 |
| KAT8 | 106.016046 | -0.4367199 | 0.25557071 | -1.7088028 | 0.08748749 | 0.60651125 |
| PRSS8 | 220.454732 | -0.1719969 | 0.37867773 | -0.4542038 | 0.64968213 | 0.95061388 |
| FUS | 414.85916 | 0.08249271 | 0.18810712 | 0.43854112 | 0.66099407 | 0.95127957 |
| PYCARD | 37.9322197 | 0.78530201 | 0.58610466 | 1.33986652 | 0.18028874 | 0.74280139 |
| AC026471.4 | 55.8002338 | 0.21871209 | 0.36855422 | 0.5934326 | 0.55289168 | 0.92872652 |
| C16orf58 | 77.5622184 | -0.3942712 | 0.26517556 | -1.4868308 | 0.13705951 | 0.68129775 |
| ZNF720 | 154.600336 | 0.23824682 | 0.25327329 | 0.9406709 | 0.34687353 | 0.85295795 |
| ZNF267 | 40.5727722 | 0.18736916 | 0.29762802 | 0.62954141 | 0.52899467 | 0.92136681 |
| SHCBP1 | 172.044097 | 0.39157289 | 0.35598234 | 1.09997843 | 2.71E-01 | 8.19E-01 |
| VPS35 | 299.555802 | 0.15758253 | 0.25127855 | 0.62712288 | 0.53057869 | 0.92211604 |
| C16orf87 | 113.802643 | 0.47663476 | 0.21060542 | 2.26316474 | 0.02362554 | 0.36680832 |
| GPT2 | 65.9626501 | -0.1514335 | 0.46990998 | -0.3222607 | 0.7472552 | 0.96109466 |
| DNAJA2 | 118.469151 | 0.19471682 | 0.27801155 | 0.70039113 | 0.48368307 | 0.90631121 |
| ITFG1 | 1179.78254 | -0.2620628 | 0.15735754 | -1.6653973 | 0.09583351 | 0.61367241 |
| PHKB | 890.08298 | 0.14413793 | 0.16142817 | 0.89289204 | 0.37191499 | 0.86201082 |
| LONP2 | 558.144194 | -0.0597465 | 0.1506338 | -0.3966339 | 0.69163745 | 0.95672887 |
| SIAH1 | 122.751499 | -0.242034 | 0.28875894 | -0.8381871 | 0.40192561 | 0.87834687 |
| N4BP1 | 255.556412 | -0.1088856 | 0.27366958 | -0.3978724 | 0.69072428 | 0.95659154 |
| CNEP1R1 | 34.7624321 | -0.075675 | 0.25893732 | -0.2922523 | 0.77009371 | 0.96345263 |
| HEATR3 | 55.3046749 | -0.3294723 | 0.27381487 | -1.2032668 | 0.2288731 | 0.79047378 |
| TENT4B | 255.728302 | 0.40209119 | 0.24598926 | 1.63458841 | 0.10213536 | 0.62412744 |
| BRD7 | 191.280602 | -0.0883485 | 0.29394905 | -0.3005573 | 0.76375208 | 0.96285313 |
| CYLD | 125.521432 | -0.165417 | 0.19741973 | -0.8378949 | 0.40208971 | 0.87834687 |
| HNRNPA1P4 | 58.2627096 | -0.2432892 | 0.48100794 | -0.5057905 | 0.61300373 | 0.94003018 |
| TOX3 | 1508.04781 | 0.33720121 | 0.30650639 | 1.10014414 | 0.27126933 | 0.81922962 |
| AC007906.2 | 72.7057415 | 2.97221675 | 0.43326679 | 6.86001514 | 6.89E-12 | 7.22E-08 |
| CHD9 | 1450.0366 | 0.05987675 | 0.19172636 | 0.31230317 | 0.75481013 | 0.96120879 |
| RBL2 | 415.639566 | -0.0759284 | 0.17185756 | -0.4418102 | 0.65862652 | 0.95113765 |
| AKTIP | 73.1849879 | 0.17821747 | 0.34157392 | 0.52175373 | 0.60184181 | 0.93804805 |
| RPGRIP1L | 85.9773996 | 0.49683712 | 0.27545521 | 1.80369479 | 0.07127917 | 0.5633527 |
| FTO | 952.135781 | 0.03060345 | 0.19817064 | 0.15442981 | 0.87727085 | 0.98277085 |
| MMP2 | 42.2483845 | 0.25584715 | 0.68475573 | 0.37363272 | 0.70867759 | 0.95752169 |
| LPCAT2 | 591.382502 | 0.16924888 | 0.37358586 | 0.45303877 | 0.65052083 | 0.95061388 |
| AMFR | 514.998664 | 0.30873698 | 0.38870731 | 0.79426596 | 0.42704062 | 0.88498827 |
| NUDT21 | 273.089594 | 0.29945452 | 0.22425923 | 1.33530523 | 0.18177647 | 0.74315969 |
| OGFOD1 | 97.4564113 | 0.06244962 | 0.24391254 | 0.25603286 | 0.79792547 | 0.9653598 |
| BBS2 | 171.235188 | -0.1997171 | 0.20450427 | -0.9765913 | 0.3287715 | 0.84409527 |
| MT2A | 663.235469 | -1.6716363 | 0.91818786 | -1.820582 | 0.06867041 | 0.55917298 |
| MT1E | 310.004878 | -1.5920323 | 0.95897152 | -1.6601456 | 0.09688517 | 0.61574078 |
| MT1F | 68.4012928 | -0.4434242 | 0.78001551 | -0.5684812 | 0.56970826 | 0.93122667 |
| MT1G | 451.539241 | -2.5790873 | 0.94273264 | -2.7357568 | 0.0062237 | 0.19425932 |
| MT1H | 132.639091 | -3.3996508 | 1.01091773 | -3.3629352 | 0.00077118 | 0.06414299 |
| MT1X | 160.166167 | -1.0579151 | 0.9246738 | -1.1440955 | 0.25258403 | 0.80350545 |
| NUP93 | 139.937932 | -0.4623867 | 0.31241956 | -1.4800185 | 0.13886831 | 0.68156449 |
| SLC12A3 | 69.0748464 | 0.346051 | 0.43164031 | 0.80171151 | 4.23E-01 | 8.84E-01 |
| HERPUD1 | 193.170276 | -0.6791124 | 0.55508109 | -1.2234472 | 0.22116084 | 0.78752408 |
| FAM192A | 173.887668 | -0.2296974 | 0.21573645 | -1.0647128 | 0.2870059 | 0.82951513 |
| RSPRY1 | 217.422925 | -0.1750605 | 0.21247591 | -0.8239076 | 0.40999209 | 0.8801764 |
| ARL2BP | 150.58516 | 0.23819759 | 0.19463388 | 1.22382387 | 0.22101868 | 0.78752408 |
| PLLP | 269.077543 | 1.33667124 | 0.50906509 | 2.62573738 | 0.00864615 | 0.22635636 |
| CIAPIN1 | 109.835535 | -0.2358938 | 0.24885756 | -0.9479069 | 0.34317685 | 0.85124103 |
| COQ9 | 227.259534 | 0.09547103 | 0.28628005 | 0.33348823 | 0.73876577 | 0.96078945 |
| POLR2C | 62.7989838 | 0.41689857 | 0.36973572 | 1.12755829 | 0.25950651 | 0.80709747 |

| | | | | | | |
|---|---|---|---|---|---|---|
| DOK4 | 54.1833619 | 0.57941225 | 0.37951236 | 1.5267283 | 0.12682859 | 0.66960383 |
| ADGRG1 | 640.193277 | -0.0631258 | 0.28285835 | -0.2231712 | 0.82340228 | 0.97018919 |
| KIFC3 | 886.793631 | 0.76403007 | 0.29845079 | 2.55998675 | 0.01046762 | 0.25038862 |
| USB1 | 38.4567851 | 0.37430652 | 0.31153384 | 1.20149555 | 0.22955903 | 0.79047378 |
| MMP15 | 39.4829973 | 0.01283007 | 0.36833752 | 0.03483237 | 0.97221341 | 0.99656026 |
| CSNK2A2 | 149.056851 | -0.1929723 | 0.23160135 | -0.8332091 | 0.40472681 | 0.87834687 |
| CCDC113 | 39.5166936 | -0.4921248 | 0.27352698 | -1.7991821 | 0.07198989 | 0.5644261 |
| SETD6 | 41.1499522 | -0.2139327 | 0.27456885 | -0.7791587 | 0.43588624 | 0.88785137 |
| CNOT1 | 465.194656 | -0.010849 | 0.16795323 | -0.0645954 | 0.94849611 | 0.99336519 |
| SLC38A7 | 82.8124262 | -0.3368106 | 0.43955595 | -0.7662521 | 0.44352634 | 0.89037885 |
| GOT2 | 286.94093 | 0.64591048 | 0.24265987 | 2.66179354 | 0.00777255 | 0.217192 |
| AC092378.1 | 66.7647461 | 0.77806646 | 0.39874517 | 1.95128751 | 0.05102285 | 0.49973786 |
| AC138305.1 | 615.796582 | 0.39375519 | 0.39844426 | 0.98823154 | 0.32303927 | 0.84285425 |
| AC010542.3 | 39.2919462 | 0.22699803 | 0.58198675 | 0.39003986 | 0.69650707 | 0.95672887 |
| TK2 | 55.9675285 | 0.21432131 | 0.32467009 | 0.66012027 | 0.50917665 | 0.91334187 |
| CMTM4 | 475.142426 | 0.17031231 | 0.20604448 | 0.82658034 | 0.40847496 | 0.87948803 |
| DYNC1LI2 | 329.445572 | -0.207381 | 0.25266359 | -0.8207789 | 0.41177221 | 0.88086809 |
| NAE1 | 124.493545 | -0.1470322 | 0.25413919 | -0.5785499 | 0.56289295 | 0.9306324 |
| PDP2 | 82.3686408 | 0.2047643 | 0.46112797 | 0.44405092 | 0.65700577 | 0.95100227 |
| CDH16 | 1495.63039 | 0.67551609 | 0.37976307 | 1.77878299 | 0.07527535 | 0.57072562 |
| RRAD | 287.689451 | -1.0348777 | 0.72071761 | -1.435899 | 0.15103109 | 0.7029344 |
| FAM96B | 210.992063 | -0.4863361 | 0.44256522 | -1.0989027 | 0.27181051 | 0.81922962 |
| CES2 | 152.43788 | -0.7512163 | 0.26241004 | -2.8627575 | 0.00419972 | 0.15775288 |
| CBFB | 492.201771 | 0.06943569 | 0.21309866 | 0.32583823 | 0.74454675 | 0.96109466 |
| C16orf70 | 56.2500168 | 0.03239995 | 0.33008513 | 0.09815633 | 0.92180816 | 0.99051393 |
| TRADD | 55.7466901 | 0.21818318 | 0.3412334 | 0.63939573 | 0.52256553 | 0.91845051 |
| NOL3 | 49.7533674 | 0.20497175 | 0.6120867 | 0.33487373 | 0.73772034 | 0.96078945 |
| KIAA0895L | 43.8599779 | -0.8557481 | 0.39282986 | -2.1784191 | 0.02937485 | 0.40623517 |
| E2F4 | 60.7625496 | -0.2333791 | 0.38114923 | -0.6123037 | 0.54033684 | 0.92414041 |
| ELMO3 | 55.1187451 | 0.26794714 | 0.43859094 | 0.61092722 | 0.54124776 | 0.92443671 |
| LRRC29 | 46.3028284 | -0.4397172 | 0.362533 | -1.2129024 | 0.22516713 | 0.78982769 |
| TMEM208 | 93.6240153 | 0.14410893 | 0.3684381 | 0.3911347 | 0.69569767 | 0.95672887 |
| ZDHHC1 | 103.133827 | 0.22482252 | 0.21985766 | 1.02258217 | 0.30650544 | 0.8373767 |
| HSD11B2 | 2291.87214 | 0.82017045 | 0.74423286 | 1.10203472 | 0.27044658 | 0.8182102 |
| ATP6V0D1 | 208.770896 | 0.20784822 | 0.2937396 | 0.70759347 | 0.47919775 | 0.90291126 |
| AC009061.2 | 98.4288273 | 0.12631359 | 0.45881095 | 0.2753064 | 0.78308084 | 0.96513523 |
| RIPOR1 | 120.225219 | 0.29220401 | 0.26775757 | 1.09130062 | 0.27514063 | 0.8202929 |
| AC027682.4 | 40.1453921 | 0.1249299 | 0.48920887 | 0.2553713 | 0.79843634 | 0.9653598 |
| CTCF | 256.356405 | -0.2112042 | 0.24866437 | -0.8493546 | 0.39568402 | 0.87429232 |
| ENKD1 | 61.4482953 | 0.24680903 | 0.37420988 | 0.65954708 | 0.50954453 | 0.91334187 |
| GFOD2 | 84.083065 | -0.144311 | 0.27641862 | -0.5220742 | 0.60161866 | 0.93804805 |
| RANBP10 | 222.086796 | -0.301096 | 0.23019935 | -1.3079795 | 0.19088027 | 0.75802398 |
| CENPT | 33.6363999 | -0.0268006 | 0.28068564 | -0.0954825 | 0.92393156 | 0.99051393 |
| THAP11 | 39.9922649 | 0.40933687 | 0.47359474 | 0.86431887 | 0.38741275 | 0.86910078 |
| NUTF2 | 51.2134515 | -0.2884203 | 0.29965271 | -0.9625151 | 0.33579092 | 0.84552074 |
| PSKH1 | 63.171998 | -0.1539867 | 0.27976275 | -0.5504188 | 0.58203212 | 0.93402467 |
| DUS2 | 51.3391191 | -0.1679412 | 0.30255026 | -0.5550854 | 0.57883628 | 0.93345492 |
| NFATC3 | 393.713574 | 0.09394434 | 0.17164384 | 0.54732137 | 0.58415797 | 0.93483536 |
| AC020978.5 | 33.7286827 | 0.2453169 | 0.35466237 | 0.69169137 | 0.48913117 | 0.90730741 |
| ESRP2 | 56.7238459 | 0.04695578 | 0.34656513 | 0.13548904 | 0.89222523 | 0.98597539 |
| PLA2G15 | 87.962995 | -0.0239041 | 0.21490516 | -0.1112309 | 0.91143324 | 0.98788932 |
| SLC7A6 | 196.86345 | 0.1977784 | 0.27511081 | 0.71890452 | 0.47219975 | 0.90188689 |
| SLC7A6OS | 104.520396 | -0.1356254 | 0.39895922 | -0.3399481 | 0.73389563 | 0.96041607 |
| PRMT7 | 137.106857 | -0.2447508 | 0.29313485 | -0.8349428 | 0.40374991 | 0.87834687 |

| | | | | | | |
|---|---|---|---|---|---|---|
| ZFP90 | 161.869323 | 0.22467013 | 0.19807609 | 1.13426174 | 0.25668477 | 0.80676096 |
| CDH3 | 55.1906103 | 0.98402445 | 0.3546705 | 2.77447501 | 0.00552909 | 0.18370315 |
| CDH1 | 1035.58982 | 0.02976116 | 0.29614398 | 0.10049559 | 0.91995088 | 0.99014946 |
| TANGO6 | 187.577336 | -0.1602805 | 0.29953777 | -0.5350927 | 0.59258573 | 0.93491607 |
| CHTF8 | 121.255039 | 0.26799992 | 0.35770068 | 0.74922956 | 0.45371885 | 0.89564392 |
| UTP4 | 69.4626912 | -0.2073905 | 0.28869858 | -0.7183634 | 0.47253325 | 0.90213021 |
| SNTB2 | 127.492847 | 0.34899553 | 0.34405904 | 1.01434779 | 0.31041683 | 0.84213522 |
| VPS4A | 99.8891216 | -0.0887922 | 0.24107629 | -0.3683157 | 0.71263784 | 0.95791799 |
| COG8 | 49.1708851 | 0.1573313 | 0.28209408 | 0.55772633 | 0.57703127 | 0.93319686 |
| NIP7 | 35.4734695 | 0.14921081 | 0.55560376 | 0.26855616 | 0.78827125 | 0.96515649 |
| TERF2 | 133.107045 | -0.1998281 | 0.28324325 | -0.7054998 | 0.48049923 | 0.90455038 |
| CYB5B | 369.66994 | -0.2598433 | 0.19931111 | -1.3037071 | 0.19233345 | 0.7597012 |
| NFAT5 | 4177.16644 | 0.28099731 | 0.28675835 | 0.9799098 | 0.32713065 | 0.84381657 |
| NQO1 | 158.83977 | 0.19965338 | 0.60656739 | 0.32915285 | 0.74204016 | 0.96086423 |
| NOB1 | 37.3597057 | -0.1520361 | 0.35994716 | -0.4223845 | 0.67274439 | 0.9544944 |
| WWP2 | 551.517528 | -0.5497893 | 0.22453925 | -2.4485221 | 0.01434436 | 0.28746487 |
| PDPR | 150.90518 | -0.1846945 | 0.30835931 | -0.5989587 | 0.54920045 | 0.9274143 |
| AC009060.1 | 46.6408355 | -0.2709609 | 0.39647659 | -0.6834223 | 0.49434004 | 0.90839719 |
| EXOSC6 | 86.3222061 | -0.6732055 | 0.27326142 | -2.4635949 | 0.01375515 | 0.28408502 |
| AARS | 124.654929 | -0.3544625 | 0.25517786 | -1.3890802 | 0.16480838 | 0.71715698 |
| DDX19A | 37.4145478 | -0.2087887 | 0.30674411 | -0.6806609 | 0.49608611 | 0.90839719 |
| ST3GAL2 | 118.005349 | 0.54768512 | 0.27861895 | 1.96571381 | 0.04933168 | 0.49237714 |
| FUK | 65.298963 | 0.04448093 | 0.2358133 | 0.18862775 | 0.85038458 | 0.97718353 |
| COG4 | 123.064263 | -0.1454999 | 0.2716611 | -0.5355934 | 0.59223962 | 0.93491607 |
| SF3B3 | 237.003709 | 0.04313328 | 0.16712197 | 0.25809459 | 0.79633391 | 0.9653598 |
| MTSS1L | 45.4980103 | -0.4631991 | 0.41441424 | -1.1177199 | 0.26368662 | 0.81015415 |
| VAC14 | 120.12421 | -0.737111 | 0.37127355 | -1.9853583 | 0.04710462 | 0.48600281 |
| HYDIN | 429.515705 | 0.09316578 | 0.31086179 | 0.29970161 | 0.76440477 | 0.96285313 |
| CMTR2 | 87.8747216 | -0.3930653 | 0.34943661 | -1.1248543 | 0.2606508 | 0.80805101 |
| ZNF19 | 89.4626526 | 0.2136677 | 0.36463316 | 0.58597988 | 0.55788903 | 0.92901206 |
| MARVELD3 | 52.7317953 | -0.2914829 | 0.28481135 | -1.0234244 | 0.30610722 | 0.83709431 |
| PHLPP2 | 102.292698 | 0.52206891 | 0.30189688 | 1.72929547 | 0.08375623 | 0.59348564 |
| AP1G1 | 630.2974 | -0.2058222 | 0.20186511 | -1.0196025 | 0.307917 | 0.83980597 |
| IST1 | 351.370779 | -0.1537854 | 0.13665975 | -1.1253156 | 0.26045531 | 0.80805101 |
| ZNF821 | 120.92653 | 0.30599203 | 0.40885021 | 0.74842087 | 0.45420634 | 0.89609985 |
| TXNL4B | 53.769151 | 0.01633217 | 0.30065083 | 0.05432271 | 0.95667805 | 0.99565888 |
| DHX38 | 60.298105 | -0.1765494 | 0.25956016 | -0.6801869 | 0.49638612 | 0.90839719 |
| LINC01572 | 204.520776 | 0.20501965 | 0.32694255 | 0.62708159 | 0.53060576 | 0.92211604 |
| AC004943.2 | 156.503906 | 0.12566858 | 0.26241207 | 0.47889786 | 0.63201129 | 0.94654542 |
| ZFHX3 | 1509.32034 | -0.571439 | 0.26065324 | -2.1923341 | 0.02835539 | 0.40459506 |
| AC138627.1 | 46.0457108 | 0.77006342 | 0.64504221 | 1.19381863 | 0.23254892 | 0.79255697 |
| PSMD7 | 145.069571 | 0.28130854 | 0.32958989 | 0.85351084 | 0.39337608 | 0.87223214 |
| NPIPB15 | 39.7491995 | 0.54143807 | 0.87080814 | 0.62176506 | 0.53409636 | 0.92303275 |
| GLG1 | 762.372825 | -0.1494305 | 0.18958957 | -0.7881789 | 0.43059205 | 0.88596084 |
| RFWD3 | 90.5392777 | -0.1036484 | 0.23227095 | -0.4462391 | 0.65542455 | 0.95100227 |
| WDR59 | 404.723163 | -0.2008506 | 0.22188711 | -0.9051929 | 0.36536322 | 0.85866868 |
| ZNRF1 | 410.207774 | -0.6081525 | 0.39662388 | -1.533323 | 0.1251963 | 0.66635715 |
| ZFP1 | 34.0397565 | 0.15648066 | 0.32438864 | 0.48238639 | 0.62953149 | 0.94596555 |
| BCAR1 | 201.35192 | -0.3183086 | 0.2047082 | -1.5549384 | 0.11996074 | 0.65679551 |
| CFDP1 | 574.482214 | -0.5597587 | 0.18350732 | -3.0503347 | 0.00228586 | 0.12358178 |
| GABARAPL2 | 306.587516 | 0.86952144 | 0.51316271 | 1.69443615 | 0.09018244 | 0.60739841 |
| ADAT1 | 75.806757 | 0.04621267 | 0.26544017 | 0.17409826 | 0.86178824 | 0.97860448 |
| KARS | 99.961551 | -0.1753944 | 0.31032679 | -0.5651927 | 0.57194272 | 0.93200748 |
| TERF2IP | 479.05469 | 0.65585994 | 0.39210621 | 1.67265889 | 0.09439447 | 0.61330068 |

| | | | | | | |
|---|---|---|---|---|---|---|
| AC010528.1 | 33.8724668 | -0.0322295 | 0.54937519 | -0.0586658 | 0.95321832 | 0.99512909 |
| NUDT7 | 40.9431633 | 0.48697624 | 0.4040838 | 1.20513674 | 0.22815052 | 0.79042788 |
| WWOX | 3224.61534 | -0.1739149 | 0.26863606 | -0.6473996 | 0.51737333 | 0.91751084 |
| CDYL2 | 99.2196669 | 0.04545598 | 0.36590731 | 0.12422812 | 0.90113466 | 0.9875066 |
| CMC2 | 197.297967 | 0.24079507 | 0.26178973 | 0.91980333 | 0.35767554 | 0.8557799 |
| ATMIN | 97.3824613 | 0.20290154 | 0.3110971 | 0.65221289 | 0.51426384 | 0.91544604 |
| GAN | 117.884891 | -0.1445298 | 0.21777035 | -0.6636797 | 0.50689531 | 0.91163608 |
| CMIP | 578.57046 | -0.0625355 | 0.24764138 | -0.2525246 | 0.80063561 | 0.96603066 |
| PLCG2 | 770.296458 | -0.1281653 | 0.27873881 | -0.4598042 | 0.64565675 | 0.9498151 |
| MPHOSPH6 | 86.7872566 | -0.2321964 | 0.2835052 | -0.81902 | 0.41277503 | 0.88153663 |
| HSBP1 | 219.15001 | 0.23776446 | 0.41701411 | 0.57015927 | 0.56856968 | 0.93122667 |
| MBTPS1 | 258.861022 | -0.1371122 | 0.15405384 | -0.890028 | 0.37345084 | 0.86201082 |
| HSDL1 | 45.4551469 | -0.3988265 | 0.50397866 | -0.791356 | 0.42873629 | 0.88535099 |
| TAF1C | 56.0831674 | -0.3652725 | 0.24499534 | -1.4909366 | 0.13597814 | 0.67998335 |
| TLDC1 | 105.522852 | -0.0320024 | 0.34131075 | -0.0937631 | 0.92529735 | 0.99051393 |
| COTL1 | 103.801481 | -0.4532144 | 0.54128922 | -0.837287 | 0.40243125 | 0.87834687 |
| KLHL36 | 131.627731 | 0.07114104 | 0.19349628 | 0.36766099 | 0.71312603 | 0.95791799 |
| USP10 | 255.906699 | -0.1624312 | 0.1997774 | -0.8130611 | 0.41618304 | 0.88242985 |
| ZDHHC7 | 183.050137 | -0.2815645 | 0.32111678 | -0.8768291 | 0.38057949 | 0.86593382 |
| KIAA0513 | 47.082174 | -0.4958081 | 0.56026671 | -0.8849501 | 0.37618353 | 0.86357183 |
| GSE1 | 388.958235 | 0.06087845 | 0.2717952 | 0.22398646 | 0.82276784 | 0.97018919 |
| EMC8 | 70.4206776 | -0.1707598 | 0.29156424 | -0.5856677 | 0.55809884 | 0.92901206 |
| COX4I1 | 880.499527 | 0.29688581 | 0.49226268 | 0.60310444 | 0.54643921 | 0.92597739 |
| MTHFSD | 51.2761226 | -0.3507441 | 0.32259528 | -1.0872572 | 0.27692314 | 0.82198569 |
| FBXO31 | 46.3532624 | -0.1335617 | 0.32476956 | -0.4112506 | 0.68088876 | 0.95574492 |
| MAP1LC3B | 228.551781 | -0.2889696 | 0.42419658 | -0.6812161 | 0.49573473 | 0.90839719 |
| ZCCHC14 | 239.872099 | -0.2654099 | 0.30539106 | -0.8690821 | 0.38480224 | 0.86725322 |
| KLHDC4 | 123.139763 | 0.12047725 | 0.35736421 | 0.33712734 | 0.73602091 | 0.96078945 |
| BANP | 440.026984 | -1.7730467 | 0.68818554 | -2.576408 | 0.00998328 | 0.24399985 |
| ZFPM1 | 119.049023 | 0.20337695 | 0.27944024 | 0.72780122 | 0.46673528 | 0.89883965 |
| ZC3H18 | 115.027988 | -0.066721 | 0.26348979 | -0.2532203 | 0.80009802 | 0.96568323 |
| CYBA | 209.557502 | 0.27709981 | 0.45817517 | 0.60479011 | 0.54531847 | 0.92579581 |
| MVD | 89.5781015 | -0.5219809 | 0.53348084 | -0.9784436 | 0.32785497 | 0.84381657 |
| PIEZO1 | 349.642089 | -0.0217503 | 0.46796078 | -0.046479 | 0.9629285 | 0.99637506 |
| APRT | 189.377901 | 0.3575098 | 0.48767211 | 0.73309463 | 0.46350071 | 0.89801718 |
| GALNS | 62.0688061 | -0.1305442 | 0.30211508 | -0.4321009 | 0.66566809 | 0.95238247 |
| TRAPPC2L | 60.8254146 | 0.02289852 | 0.40577583 | 0.05643145 | 0.95499811 | 0.99537617 |
| ACSF3 | 84.8968902 | -0.0912712 | 0.31944672 | -0.2857165 | 0.77509523 | 0.96388926 |
| ZNF778 | 41.8634868 | -0.2419933 | 0.37915764 | -0.6382394 | 0.52331787 | 0.91871245 |
| ANKRD11 | 1253.13571 | -0.3528534 | 0.29463388 | -1.1975995 | 0.23107297 | 0.79130016 |
| SPG7 | 386.512496 | -0.5073186 | 0.33387454 | -1.5194888 | 0.12863951 | 0.67206108 |
| RPL13 | 1740.41321 | 0.06088531 | 0.44357169 | 0.13726148 | 0.89082411 | 0.98597539 |
| CHMP1A | 127.442302 | -0.1206882 | 0.253222 | -0.4766101 | 0.63363978 | 0.94744025 |
| SPATA33 | 43.6857876 | 0.11822348 | 0.33771721 | 0.35006649 | 0.7262888 | 0.95979351 |
| CDK10 | 115.180093 | -0.1135319 | 0.24332021 | -0.4665945 | 0.64079003 | 0.94935885 |
| VPS9D1 | 92.7242038 | -0.2946566 | 0.36925585 | -0.7979741 | 0.42488552 | 0.88392184 |
| ZNF276 | 58.7772684 | -0.0866139 | 0.26509513 | -0.3267278 | 0.74387378 | 0.96109466 |
| SPIRE2 | 98.5848933 | 0.84022759 | 0.52697097 | 1.59444757 | 0.11083583 | 0.64209743 |
| TCF25 | 667.446292 | -0.247432 | 0.18371918 | -1.3467946 | 0.17804639 | 0.73737861 |
| DEF8 | 63.1341629 | -0.3623471 | 0.21362647 | -1.6961712 | 0.08985347 | 0.60739841 |
| GAS8 | 126.510268 | -0.1289071 | 0.2812584 | -0.4583227 | 0.64672059 | 0.95061388 |
| FAM157C | 192.603144 | 0.1258466 | 0.39513713 | 0.31848841 | 0.75011448 | 0.96109466 |
| AC133919.2 | 36.6124485 | 0.49630768 | 0.69564357 | 0.71345111 | 0.47556665 | 0.90242252 |
| DOC2B | 295.498107 | -0.0411605 | 0.50911315 | -0.0808475 | 0.93556322 | 0.99243654 |

| | | | | | | |
|---|---|---|---|---|---|---|
| RPH3AL | 483.353042 | -0.7590855 | 0.41174532 | -1.8435801 | 0.06524435 | 0.55362017 |
| AC129507.2 | 68.2978587 | 0.26035191 | 0.48303141 | 0.53899581 | 0.58988974 | 0.93483536 |
| C17orf97 | 42.3103836 | -0.315179 | 0.40995552 | -0.7688127 | 0.44200449 | 0.88997055 |
| VPS53 | 558.088295 | -0.5075736 | 0.182983 | -2.7738841 | 0.00553914 | 0.18370315 |
| FAM57A | 41.7817972 | -0.6742235 | 0.4673597 | -1.4426223 | 0.1491269 | 0.70215315 |
| GLOD4 | 148.290385 | 0.09394839 | 0.32065176 | 0.29299198 | 0.76952829 | 0.96345263 |
| NXN | 87.0364446 | 1.54206942 | 0.49773485 | 3.0981745 | 0.00194717 | 0.11037603 |
| TIMM22 | 44.7376932 | 0.21199901 | 0.35008692 | 0.60556106 | 0.54480626 | 0.92561018 |
| ABR | 96.5591962 | -0.0110681 | 0.27832238 | -0.0397673 | 0.96827862 | 0.9964399 |
| YWHAE | 892.494076 | -0.029662 | 0.23974777 | -0.1237217 | 0.90153562 | 0.9875066 |
| CRK | 203.72587 | -0.1284612 | 0.28001215 | -0.45877 | 0.64639936 | 0.95048567 |
| MYO1C | 294.773947 | -0.006214 | 0.21614467 | -0.0287492 | 0.97706458 | 0.99675234 |
| INPP5K | 51.0945053 | -0.2355274 | 0.28054671 | -0.83953 | 0.40117195 | 0.87808732 |
| PITPNA | 284.123572 | -0.0275702 | 0.25225014 | -0.1092969 | 0.91296703 | 0.98830345 |
| SLC43A2 | 228.186393 | 0.40675772 | 0.42663984 | 0.95339834 | 0.34038828 | 0.84975445 |
| PRPF8 | 280.986155 | -0.0502491 | 0.18573711 | -0.2705388 | 0.78674575 | 0.96513523 |
| MIR22HG | 58.3080918 | -0.2843323 | 0.65719099 | -0.4326479 | 0.66527062 | 0.9520866 |
| WDR81 | 42.0274378 | -0.5059292 | 0.4405868 | -1.1483076 | 0.25084162 | 0.80240987 |
| SMYD4 | 66.9617305 | 0.01380889 | 0.44513047 | 0.03102212 | 0.9752519 | 0.99656026 |
| RPA1 | 146.893465 | 0.04623047 | 0.2166029 | 0.21343421 | 0.8309883 | 0.97160502 |
| SMG6 | 620.935109 | -0.2669031 | 0.20245994 | -1.3183006 | 0.18740303 | 0.75219601 |
| TSR1 | 69.7264964 | -0.1391255 | 0.26128952 | -0.5324572 | 0.59440939 | 0.935202 |
| SGSM2 | 139.221148 | -0.5001146 | 0.2224464 | -2.2482476 | 0.02456041 | 0.37630563 |
| MNT | 76.0023868 | -0.8169025 | 0.43155058 | -1.8929473 | 0.05836487 | 0.53192434 |
| METTL16 | 234.732744 | -0.2651641 | 0.16763279 | -1.5818153 | 0.11369175 | 0.6483668 |
| PAFAH1B1 | 868.284527 | -0.2849456 | 0.15572926 | -1.8297501 | 0.06728731 | 0.55848603 |
| CLUH | 158.544676 | -0.5196142 | 0.27435466 | -1.8939506 | 0.05823156 | 0.53192434 |
| RAP1GAP2 | 343.417471 | 0.30106369 | 0.39785697 | 0.75671336 | 0.44922161 | 0.89324553 |
| SPATA22 | 89.7952515 | 1.19335003 | 0.7400035 | 1.61262755 | 0.10682545 | 0.63477557 |
| ASPA | 226.995608 | 0.4357912 | 0.41377498 | 1.05320821 | 0.29224558 | 0.83322538 |
| TRPV1 | 36.4301977 | -0.3111862 | 0.31328912 | -0.9932876 | 0.32056983 | 0.84285425 |
| CTNS | 70.5114158 | -0.3431219 | 0.41288451 | -0.8310361 | 0.40595325 | 0.87937203 |
| TAX1BP3 | 110.19279 | 0.39586584 | 0.45549449 | 0.86909029 | 0.38479775 | 0.86725322 |
| EMC6 | 83.1144787 | 0.33453672 | 0.56946988 | 0.58745287 | 0.55689959 | 0.92872652 |
| ITGAE | 67.153934 | -0.0755873 | 0.28072982 | -0.2692529 | 0.78773509 | 0.96515649 |
| NCBP3 | 187.560656 | -0.0598756 | 0.22693936 | -0.2638397 | 0.79190341 | 0.9653598 |
| ZZEF1 | 270.684187 | -0.3113201 | 0.21967251 | -1.4172011 | 0.15642413 | 0.71120384 |
| CYB5D2 | 121.181475 | 0.7113194 | 0.32653994 | 2.17835342 | 0.02937973 | 0.40623517 |
| ANKFY1 | 267.645575 | -0.3336032 | 0.2662159 | -1.2531303 | 0.21015828 | 0.77551367 |
| UBE2G1 | 377.709119 | 0.05856805 | 0.28462205 | 0.20577483 | 0.83696681 | 0.97353666 |
| SPNS2 | 258.634069 | 0.98578018 | 0.73941505 | 1.33318922 | 0.18246972 | 0.74315969 |
| MYBBP1A | 39.4715635 | -0.3292837 | 0.34091391 | -0.9658852 | 0.33410165 | 0.8445213 |
| GGT6 | 65.1863565 | 1.70106785 | 0.44310378 | 3.83898297 | 0.00012354 | 0.02194494 |
| SMTNL2 | 75.7319186 | 0.97961334 | 0.44837741 | 2.18479637 | 0.02890378 | 0.40623517 |
| PELP1 | 82.3218875 | -0.2121508 | 0.22792501 | -0.9307921 | 0.35196114 | 0.85536253 |
| MED11 | 49.5545005 | -0.1218981 | 0.36521928 | -0.333767 | 0.7385554 | 0.96078945 |
| CXCL16 | 47.6428279 | -0.0253971 | 0.54262119 | -0.0468044 | 0.96266912 | 0.99637506 |
| PSMB6 | 136.580815 | 0.08126399 | 0.56145501 | 0.1447382 | 0.88491758 | 0.98449429 |
| MINK1 | 159.59627 | -0.1360546 | 0.20311922 | -0.6698266 | 0.50296836 | 0.90964098 |
| SLC25A11 | 126.433042 | 0.85236704 | 0.43511432 | 1.95894962 | 0.05011869 | 0.49590526 |
| RNF167 | 82.7368625 | 0.20718683 | 0.33997989 | 0.6094091 | 0.54225331 | 0.92482794 |
| PFN1 | 572.042999 | -0.1610337 | 0.49840216 | -0.3230999 | 0.74661958 | 0.96109466 |
| SPAG7 | 161.634915 | 0.1313542 | 0.38000149 | 0.34566759 | 0.72959254 | 0.96008876 |
| CAMTA2 | 67.9750728 | 0.07125705 | 0.25253299 | 0.28216925 | 0.77781373 | 0.96472142 |

| | | | | | | |
|---|---|---|---|---|---|---|
| KIF1C | 128.311209 | 0.17361079 | 0.3304943 | 0.52530644 | 0.59937018 | 0.93728325 |
| ZNF594 | 67.0004516 | 0.2604982 | 0.30056547 | 0.86669369 | 0.38610986 | 0.86833291 |
| AC087500.1 | 145.376923 | 0.15711421 | 0.41742115 | 0.37639254 | 0.70662509 | 0.95751629 |
| RABEP1 | 403.751782 | -0.0657279 | 0.2290558 | -0.2869516 | 0.77414939 | 0.96388926 |
| NUP88 | 273.037489 | 0.40416283 | 0.23851832 | 1.6944729 | 0.09017546 | 0.60739841 |
| RPAIN | 154.87332 | 0.17214267 | 0.26155619 | 0.65814795 | 0.51044306 | 0.91334187 |
| C1QBP | 161.804962 | 0.67950166 | 0.41171186 | 1.65043013 | 0.09885499 | 0.61707826 |
| DERL2 | 103.191392 | 0.20075669 | 0.31281375 | 0.64177706 | 0.52101795 | 0.91769212 |
| MIS12 | 46.0060697 | 0.40011052 | 0.4396021 | 0.91016516 | 0.36273542 | 0.85694172 |
| PITPNM3 | 36.2875478 | 0.66823153 | 0.45093007 | 1.48189615 | 0.13836793 | 0.68146162 |
| KIAA0753 | 179.422495 | 0.01499263 | 0.19657416 | 0.07626961 | 0.9392046 | 0.99243654 |
| TXNDC17 | 73.7341931 | -0.186138 | 0.33798955 | -0.5507211 | 0.58182485 | 0.93402467 |
| MED31 | 54.7087923 | -0.1230643 | 0.32848093 | -0.3746466 | 0.70792333 | 0.95752169 |
| XAF1 | 171.329676 | -4.0734055 | 1.50591037 | -2.7049455 | 0.00683156 | 0.20224502 |
| ALOX12-AS1 | 127.555435 | 0.41703155 | 0.55195732 | 0.75555036 | 0.44991883 | 0.89324553 |
| DLG4 | 44.9243858 | -0.3383891 | 0.41177309 | -0.8217854 | 0.41119904 | 0.88078476 |
| ACADVL | 657.849211 | -0.2486781 | 0.2837991 | -0.8762469 | 0.38089583 | 0.86593382 |
| DVL2 | 59.2252735 | -0.3318468 | 0.26548916 | -1.2499449 | 0.21131966 | 0.7767562 |
| PHF23 | 64.4075705 | 0.22688385 | 0.25640573 | 0.88486261 | 0.37623074 | 0.86357183 |
| ELP5 | 69.7992062 | 0.39666025 | 0.29362608 | 1.35090268 | 0.17672661 | 0.73554204 |
| CLDN7 | 42.3940771 | 0.3511327 | 0.4988253 | 0.70391918 | 0.48148311 | 0.90561114 |
| EIF5A | 150.698317 | -0.5091743 | 0.52593356 | -0.9681343 | 0.3329773 | 0.84442294 |
| GPS2 | 104.350495 | -0.3348187 | 0.36771374 | -0.9105416 | 0.36253695 | 0.85694172 |
| ZBTB4 | 324.495451 | 0.12664589 | 0.21471099 | 0.58984351 | 0.55529557 | 0.92872652 |
| POLR2A | 342.588586 | 0.40373731 | 0.36091152 | 1.11866009 | 0.26328518 | 0.80987046 |
| EIF4A1 | 134.015605 | -0.4243844 | 0.49565862 | -0.856203 | 0.39188551 | 0.87030305 |
| MPDU1 | 71.6112497 | -0.4209394 | 0.29577818 | -1.4231591 | 0.15469003 | 0.70839173 |
| FXR2 | 65.8451627 | -0.396087 | 0.25613409 | -1.546405 | 0.12200679 | 0.66181735 |
| SAT2 | 150.3662 | 0.60404973 | 0.37060776 | 1.62988958 | 0.10312484 | 0.62635087 |
| TP53 | 67.3749535 | 0.55789655 | 0.48348476 | 1.15390722 | 0.24853821 | 0.80097144 |
| KDM6B | 116.073208 | -1.0332453 | 0.29778147 | -3.4698107 | 0.00052083 | 0.05569641 |
| NAA38 | 296.652631 | 0.08024802 | 0.23565978 | 0.3405249 | 0.73346127 | 0.96035038 |
| CYB5D1 | 41.0688645 | 0.12007401 | 0.33109817 | 0.3626538 | 0.71686351 | 0.95791799 |
| CHD3 | 388.034331 | -0.1282235 | 0.33464136 | -0.3831668 | 0.70159606 | 0.95672887 |
| TRAPPC1 | 92.8491537 | -0.0519911 | 0.51016343 | -0.1019107 | 0.91882752 | 0.99014946 |
| CNTROB | 39.5003042 | 0.04179668 | 0.36262068 | 0.11526281 | 0.90823681 | 0.98758267 |
| PER1 | 226.413538 | -1.6851292 | 0.68012236 | -2.4776854 | 0.01322377 | 0.28170334 |
| TMEM107 | 62.3084864 | 0.15463174 | 0.32554628 | 0.47499157 | 0.63479298 | 0.94776446 |
| CTC1 | 77.526932 | -0.1063147 | 0.24351246 | -0.4365883 | 0.66240995 | 0.95142083 |
| PFAS | 33.5233805 | -0.2976205 | 0.3221348 | -0.9239005 | 0.35553811 | 0.8557799 |
| SLC25A35 | 43.9616595 | 0.43459529 | 0.49283446 | 0.88182814 | 0.37786975 | 0.86501008 |
| RANGRF | 39.5249556 | 0.06642148 | 0.317275 | 0.20934986 | 0.83417513 | 0.97232209 |
| NDEL1 | 194.73047 | -0.2503567 | 0.38429845 | -0.6514642 | 0.5147469 | 0.91557155 |
| MYH10 | 1734.58036 | 0.33628328 | 0.225983 | 1.48809101 | 0.13672689 | 0.68107639 |
| STX8 | 554.045719 | -0.4794854 | 0.19753588 | -2.4273331 | 0.01521028 | 0.29684131 |
| USP43 | 80.0308531 | -0.4389572 | 0.28363112 | -1.5476341 | 0.12171041 | 0.66157942 |
| SCO1 | 105.008656 | 0.16130523 | 0.25595257 | 0.63021529 | 0.52855374 | 0.92121123 |
| TMEM220 | 147.872665 | -0.3810101 | 0.41632196 | -0.9151815 | 0.36009637 | 0.85608464 |
| TMEM220-A! | 98.6111663 | -1.164202 | 0.30640451 | -3.799559 | 0.00014495 | 0.02411295 |
| TMEM238L | 108.453998 | -3.16152 | 1.25935865 | -2.5104207 | 0.01205874 | 0.26883805 |
| SHISA6 | 80.7002628 | 0.29899109 | 0.60112363 | 0.49738702 | 0.61891616 | 0.94292825 |
| DNAH9 | 73.3722016 | 0.56146408 | 0.30922872 | 1.81569187 | 0.06941765 | 0.55918293 |
| MAP2K4 | 669.669608 | -0.0533374 | 0.21222035 | -0.2513301 | 0.80155887 | 0.9661728 |
| ARHGAP44 | 475.014569 | -0.1980082 | 0.32549911 | -0.6083217 | 0.54297411 | 0.92482794 |

| | | | | | | |
|---|---|---|---|---|---|---|
| ELAC2 | 79.5786651 | 0.02547072 | 0.25263943 | 0.10081848 | 0.91969456 | 0.99014946 |
| COX10 | 159.870663 | 0.04849416 | 0.26346606 | 0.18406225 | 0.85396462 | 0.97718353 |
| HS3ST3B1 | 44.6772238 | 0.848247 | 0.78559637 | 1.07974913 | 0.28025391 | 0.82628479 |
| TRIM16 | 58.1946101 | -0.0900433 | 0.37341591 | -0.241134 | 0.8094513 | 0.9679755 |
| AC015922.4 | 36.3187845 | 0.33285324 | 0.42768492 | 0.77826743 | 0.43641137 | 0.88785137 |
| ZSWIM7 | 173.50217 | 0.1962617 | 0.26655253 | 0.7362965 | 0.46155027 | 0.89799005 |
| TTC19 | 472.040892 | 0.19010701 | 0.48103405 | 0.3952049 | 0.69269168 | 0.95672887 |
| NCOR1 | 1457.86596 | 0.1316998 | 0.24733146 | 0.53248301 | 0.59439151 | 0.935202 |
| PIGL | 307.53343 | -0.3746742 | 0.29396959 | -1.2745338 | 0.20247429 | 0.76996624 |
| CENPV | 45.9869017 | 0.4228213 | 0.33884548 | 1.24782926 | 0.21209359 | 0.77709975 |
| UBB | 1006.28674 | 0.56063374 | 0.4635848 | 1.20934455 | 0.2265305 | 0.78982769 |
| LRRC75A | 49.2142423 | 0.03680095 | 0.44968287 | 0.08183756 | 0.93477588 | 0.99243654 |
| AC098850.3 | 90.4183456 | -0.2169955 | 0.67870337 | -0.3197207 | 0.7491801 | 0.96109466 |
| MPRIP | 513.969871 | -0.3192046 | 0.17689254 | -1.8045117 | 0.07115113 | 0.56333976 |
| COPS3 | 61.8447074 | -0.0053142 | 0.25430665 | -0.020897 | 0.98332782 | 0.9971769 |
| NT5M | 41.6581467 | 0.04961041 | 0.3225649 | 0.15379977 | 0.87776761 | 0.98279964 |
| MED9 | 101.896544 | -0.0288482 | 0.29512142 | -0.0977502 | 0.92213065 | 0.99051393 |
| RASD1 | 1098.29402 | -0.019892 | 0.74002858 | -0.02688 | 0.97855543 | 0.99696835 |
| PEMT | 48.8204594 | -0.142473 | 0.33111799 | -0.4302788 | 0.66699286 | 0.9527406 |
| RAI1 | 237.896884 | 0.16193976 | 0.18848063 | 0.85918512 | 0.39023839 | 0.87004477 |
| SREBF1 | 213.806562 | -1.3217039 | 0.4323896 | -3.056743 | 0.00223756 | 0.12213349 |
| TOM1L2 | 321.718597 | 0.14654689 | 0.22442234 | 0.65299598 | 0.51375887 | 0.91544604 |
| DRC3 | 61.1619194 | 0.35383289 | 0.38490037 | 0.91928437 | 0.35794685 | 0.8557799 |
| ATPAF2 | 59.7460449 | 0.25636495 | 0.23040324 | 1.11267947 | 0.26584611 | 0.81226451 |
| GID4 | 36.8051159 | 0.29784057 | 0.29392662 | 1.01331607 | 0.31090921 | 0.84248332 |
| DRG2 | 46.7512671 | -0.0994477 | 0.32052235 | -0.3102677 | 0.75635738 | 0.96126914 |
| ALKBH5 | 193.836028 | -0.029523 | 0.23979058 | -0.1231198 | 0.90201222 | 0.9875066 |
| FLII | 147.389221 | -0.1653019 | 0.2673445 | -0.6183103 | 0.53637079 | 0.92324352 |
| MIEF2 | 41.9473778 | 0.34046841 | 0.33223515 | 1.02478141 | 0.30546634 | 0.83709431 |
| TOP3A | 65.7170034 | -0.4456615 | 0.27016129 | -1.6496126 | 0.09902219 | 0.61707826 |
| SMCR8 | 51.7994171 | -0.2790488 | 0.26618883 | -1.0483116 | 0.29449509 | 0.8337535 |
| SHMT1 | 257.545301 | 0.50415125 | 0.33257296 | 1.51591176 | 0.12954167 | 0.67207756 |
| TVP23B | 70.4561078 | -0.1845576 | 0.33353088 | -0.5533448 | 0.58002736 | 0.93345492 |
| PRPSAP2 | 176.452311 | -0.1281817 | 0.21052212 | -0.6088751 | 0.54260721 | 0.92482794 |
| EPN2 | 480.048968 | -0.2005776 | 0.21046298 | -0.9530302 | 0.34057477 | 0.85001754 |
| B9D1 | 116.199056 | 0.34260054 | 0.21075889 | 1.62555679 | 0.10404397 | 0.62918687 |
| AC004448.2 | 59.102014 | -0.3806452 | 0.48234082 | -0.7891623 | 0.43001718 | 0.88579433 |
| SLC47A1 | 92.3946071 | 0.03593689 | 0.27420124 | 0.13106026 | 8.96E-01 | 0.98597539 |
| ALDH3A2 | 199.046753 | 0.21645118 | 0.34114153 | 0.63449086 | 0.52576053 | 0.91956488 |
| ULK2 | 183.491215 | -0.0598485 | 0.24921864 | -0.2401446 | 0.81021814 | 0.9680627 |
| AKAP10 | 414.425072 | -0.4366016 | 0.28445032 | -1.5348957 | 0.12480946 | 0.66626798 |
| CCDC144NL-/ | 108.879766 | 0.05859531 | 0.5155627 | 0.11365313 | 0.90951277 | 0.98779844 |
| USP22 | 412.266276 | -0.0807155 | 0.20959455 | -0.3851029 | 0.70016118 | 0.95672887 |
| DHRS7B | 242.244262 | 0.04328332 | 0.23378895 | 0.18513843 | 0.85312046 | 0.97718353 |
| TMEM11 | 45.4292512 | 0.33235987 | 0.36463363 | 0.91148989 | 0.36203731 | 0.85690129 |
| AC087294.1 | 38.6692405 | -0.1858483 | 0.29518732 | -0.6295945 | 0.52895994 | 0.92136681 |
| MAP2K3 | 128.485811 | -1.0427848 | 0.54113217 | -1.9270427 | 0.05397431 | 0.51332378 |
| WSB1 | 3662.47465 | -0.9590573 | 0.54004464 | -1.7758852 | 0.07575185 | 0.57072562 |
| KSR1 | 234.221206 | -0.1265934 | 0.31634152 | -0.4001795 | 0.68902431 | 0.95606293 |
| NLK | 1590.88877 | 0.09945926 | 0.30342362 | 0.32779009 | 0.74307038 | 0.96105014 |
| ERVE-1 | 34.5873982 | 0.69975543 | 0.74494511 | 0.93933824 | 0.34755711 | 0.85348649 |
| IFT20 | 75.8484462 | -0.1321889 | 0.32889812 | -0.4019145 | 0.68774697 | 0.95597173 |
| TNFAIP1 | 126.583342 | -0.5169989 | 0.37146925 | -1.3917676 | 0.16399277 | 0.71715698 |
| POLDIP2 | 119.458365 | 0.22675992 | 0.21293004 | 1.06495035 | 0.2868984 | 0.82943317 |

| | | | | | | |
|---|---|---|---|---|---|---|
| TMEM199 | 33.4657841 | -0.1028744 | 0.35436609 | -0.2903055 | 0.77158253 | 0.96388926 |
| SARM1 | 64.4420186 | -0.3748767 | 0.38299906 | -0.9787928 | 0.32768236 | 0.84381657 |
| AC005726.1 | 49.6077325 | -0.5323629 | 0.32002622 | -1.6634979 | 0.09621281 | 0.61407447 |
| KIAA0100 | 161.986287 | -0.1636862 | 0.25455587 | -0.6430266 | 0.52020683 | 0.91751084 |
| SDF2 | 105.024281 | 0.36388858 | 0.29817132 | 1.22040101 | 0.22231289 | 0.78760199 |
| SUPT6H | 219.602362 | -0.1420337 | 0.20932697 | -0.6785257 | 0.49743844 | 0.90839719 |
| RAB34 | 139.423276 | 0.5054692 | 0.33528046 | 1.50760112 | 0.13165665 | 0.67390608 |
| RPL23A | 559.72769 | 0.21957492 | 0.45741775 | 0.48003147 | 0.63120501 | 0.94622065 |
| NEK8 | 65.1595737 | -0.3399036 | 0.45457536 | -0.7477386 | 0.45461783 | 0.89626521 |
| TRAF4 | 163.555892 | -0.2159698 | 0.33620338 | -0.6423783 | 0.52062757 | 0.91751084 |
| FAM222B | 506.937081 | -0.2587683 | 0.2581012 | -1.0025847 | 0.31606127 | 0.84285425 |
| ERAL1 | 42.762552 | 0.1371892 | 0.31659684 | 0.43332461 | 0.66477896 | 0.9520866 |
| FLOT2 | 334.97431 | -0.1109096 | 0.2652574 | -0.4181208 | 0.67585883 | 0.95529325 |
| DHRS13 | 56.4924857 | -0.1150907 | 0.47261093 | -0.243521 | 0.80760185 | 0.96752828 |
| PHF12 | 157.998076 | 0.14309278 | 0.28512822 | 0.50185414 | 0.61577013 | 0.94190661 |
| MYO18A | 109.916731 | -0.3276342 | 0.3203066 | -1.0228769 | 0.30636605 | 0.8372264 |
| NUFIP2 | 414.450572 | -0.1134599 | 0.20879416 | -0.5434057 | 0.58685052 | 0.93483536 |
| TAOK1 | 864.253383 | -0.1953913 | 0.22506553 | -0.8681531 | 0.3853105 | 0.86749643 |
| ABHD15-AS1 | 119.665148 | -0.0675943 | 0.47231664 | -0.1431123 | 0.88620152 | 0.98489549 |
| GIT1 | 61.399037 | 0.21999406 | 0.37371282 | 0.58867143 | 0.55608171 | 0.92872652 |
| SSH2 | 2020.49578 | -0.1145837 | 0.20865934 | -0.5491427 | 0.58290755 | 0.93415528 |
| AC023389.1 | 74.7785846 | -0.1401658 | 0.35392546 | -0.3960319 | 0.69208146 | 0.95672887 |
| EFCAB5 | 91.4728291 | -0.3849076 | 0.43905512 | -0.8766726 | 0.38066449 | 0.86593382 |
| NSRP1 | 465.646863 | -0.1811425 | 0.17724632 | -1.0219815 | 0.30678963 | 0.83771635 |
| AC104984.4 | 92.0421799 | 2.06382654 | 0.66259058 | 3.11478403 | 0.0018408 | 0.10848841 |
| BLMH | 127.579768 | 0.42215625 | 0.21909145 | 1.9268495 | 0.05399839 | 0.51332378 |
| CPD | 259.128015 | 0.0537462 | 0.23642014 | 0.22733344 | 0.82016448 | 0.96886922 |
| GOSR1 | 267.334007 | -0.0013872 | 0.17032601 | -0.0081443 | 0.99350185 | 0.99919666 |
| CRLF3 | 201.964402 | 0.07207161 | 0.28494917 | 0.25292796 | 0.80032389 | 0.96584458 |
| ATAD5 | 43.6412519 | -0.0219312 | 0.49558848 | -0.0442528 | 0.96470292 | 0.99637506 |
| TEFM | 89.417981 | 0.15430045 | 0.2559927 | 0.60275334 | 0.54667279 | 0.92597739 |
| RNF135 | 70.5292971 | 0.34092765 | 0.23349753 | 1.46009104 | 0.14426505 | 0.69131128 |
| NF1 | 1476.51781 | -0.3036588 | 0.18902596 | -1.6064396 | 0.10817735 | 0.63533473 |
| OMG | 35.5972169 | -0.1835092 | 0.65625075 | -0.2796328 | 0.77975925 | 0.96513523 |
| RAB11FIP4 | 119.831752 | 0.32247187 | 0.33765642 | 0.9550296 | 0.33956273 | 0.84928129 |
| COPRS | 129.393766 | -0.1414468 | 0.57963518 | -0.2440273 | 0.80720965 | 0.96752828 |
| UTP6 | 99.3740662 | -0.3757815 | 0.19620711 | -1.9152286 | 0.05546335 | 0.51897853 |
| SUZ12 | 401.39175 | 0.0181821 | 0.19720552 | 0.09219876 | 0.92654013 | 0.99087265 |
| LRRC37B | 109.476972 | -0.4423155 | 0.31997076 | -1.3823624 | 0.16686046 | 0.72111243 |
| RHOT1 | 304.81643 | -0.1747508 | 0.19013352 | -0.9190953 | 0.35804571 | 0.8557799 |
| C17orf75 | 47.6206991 | -0.0046918 | 0.30632651 | -0.0153165 | 0.98777971 | 0.9979689 |
| ZNF207 | 544.618592 | -0.1196314 | 0.1647354 | -0.7262032 | 0.46771418 | 0.89976199 |
| PSMD11 | 144.066075 | -0.1901644 | 0.23041005 | -0.8253304 | 0.40918403 | 0.87982122 |
| MYO1D | 967.956324 | -0.0210866 | 0.26726832 | -0.0788966 | 0.93711485 | 0.99243654 |
| TMEM98 | 36.3781729 | 1.4176597 | 0.62013305 | 2.28605734 | 0.0222509 | 0.35844607 |
| ASIC2 | 103.881204 | -0.3833758 | 0.68863836 | -0.5567157 | 0.57772166 | 0.93345492 |
| ZNF830 | 38.4414615 | 0.41228811 | 0.43898957 | 0.93917519 | 0.34764081 | 0.85348649 |
| LIG3 | 53.4517875 | -0.2014054 | 0.23423113 | -0.8598574 | 0.38986768 | 0.87004477 |
| SLFN5 | 198.202666 | -0.3047322 | 0.41753075 | -0.7298436 | 4.65E-01 | 0.89883965 |
| LINC02001 | 60.3607588 | 0.15020237 | 0.26897047 | 0.55843442 | 0.57654778 | 0.9331615 |
| AP2B1 | 388.636345 | -0.1260136 | 0.2708412 | -0.4652673 | 0.64174006 | 0.94935885 |
| TAF15 | 695.758744 | -0.2199418 | 0.12637971 | -1.7403251 | 0.08180195 | 0.59202053 |
| AC015849.3 | 44.5642619 | -0.2709776 | 0.48108622 | -0.5632619 | 0.57325654 | 0.93258748 |
| ZNHIT3 | 66.1344448 | 0.41030477 | 0.40043928 | 1.02463666 | 0.30553466 | 0.83709431 |

| | | | | | | |
|---|---|---|---|---|---|---|
| MYO19 | 184.212775 | -0.1159095 | 0.28579234 | -0.4055723 | 0.68505687 | 0.9558749 |
| GGNBP2 | 315.986886 | -0.2928244 | 0.20059589 | -1.4597727 | 0.14435255 | 0.69141441 |
| DHRS11 | 34.1711499 | 1.97704698 | 0.54093044 | 3.65490055 | 0.00025728 | 0.03693578 |
| LHX1-DT | 516.167417 | 1.15977811 | 0.42020084 | 2.76005663 | 0.00577913 | 0.18867776 |
| LHX1 | 600.099274 | 1.76588025 | 0.39052892 | 4.5217656 | 6.13E-06 | 0.00257078 |
| AATF | 286.510864 | -0.1945074 | 0.22514079 | -0.8639369 | 0.38762256 | 0.86910078 |
| ACACA | 617.132511 | -0.400445 | 0.2692179 | -1.4874383 | 0.1368991 | 0.68129775 |
| TADA2A | 124.799365 | -0.3231276 | 0.22685132 | -1.4244025 | 0.15433 | 0.70805312 |
| SYNRG | 350.05855 | -0.1172852 | 0.18672598 | -0.628114 | 0.52992925 | 0.92181514 |
| DDX52 | 170.922652 | -0.1968601 | 0.18323228 | -1.0743745 | 0.28265485 | 0.82666614 |
| HNF1B | 642.215938 | 0.05891791 | 0.29477846 | 0.19987184 | 0.84158082 | 0.975446 |
| MRPL45 | 100.686727 | -0.0385797 | 0.1995026 | -0.1933792 | 0.84666201 | 0.97656375 |
| SOCS7 | 91.0158429 | -0.130043 | 0.31813927 | -0.4087612 | 0.68271493 | 0.95574492 |
| MLLT6 | 356.046923 | 0.17383782 | 0.20805981 | 0.83551851 | 0.40342583 | 0.87834687 |
| CISD3 | 81.1671904 | 0.23770957 | 0.44638467 | 0.53252179 | 0.59436465 | 0.935202 |
| PCGF2 | 121.897853 | 0.0575026 | 0.2456711 | 0.23406334 | 0.81493581 | 0.96839471 |
| PSMB3 | 162.458648 | -0.1562503 | 0.32892162 | -0.4750382 | 0.63475978 | 0.94776446 |
| PIP4K2B | 120.460573 | -0.0901506 | 0.20679411 | -0.4359438 | 0.66287751 | 0.9515075 |
| CWC25 | 86.9278765 | -0.3613419 | 0.2032662 | -1.7776784 | 0.07545669 | 0.57072562 |
| RPL23 | 432.195856 | 0.30765169 | 0.38730459 | 0.79434042 | 0.42699728 | 0.88498827 |
| LASP1 | 257.798481 | 0.04821607 | 0.26676382 | 0.18074442 | 0.85656819 | 0.97718353 |
| RPL19 | 959.405894 | 0.02567542 | 0.55880317 | 0.04594716 | 0.96335236 | 0.99637506 |
| STAC2 | 100.74122 | 0.08018811 | 0.86377335 | 0.09283467 | 9.26E-01 | 0.99051393 |
| FBXL20 | 543.094164 | -0.1732341 | 0.25322646 | -0.6841074 | 0.49390738 | 0.90839719 |
| MED1 | 124.133022 | 0.07092609 | 0.19315688 | 0.36719422 | 0.71347415 | 0.95791799 |
| CDK12 | 327.037105 | 0.06311537 | 0.20835124 | 0.30292774 | 0.76194493 | 0.96218738 |
| STARD3 | 68.4285485 | -0.2199147 | 0.24621571 | -0.8931788 | 0.37176141 | 0.86201082 |
| PGAP3 | 56.6455528 | 0.21080999 | 0.33764103 | 0.62436129 | 0.53239034 | 0.9226808 |
| ERBB2 | 646.240507 | 0.56417124 | 0.25611131 | 2.20283614 | 0.0276063 | 0.40349154 |
| MIEN1 | 88.7136812 | -0.2744927 | 0.54235318 | -0.5061143 | 0.6127764 | 0.94003018 |
| GRB7 | 233.764442 | -0.0840009 | 0.27490212 | -0.3055667 | 0.75993461 | 0.96190674 |
| ORMDL3 | 63.0405451 | 0.56203417 | 0.41144771 | 1.36599176 | 0.17194155 | 0.72776437 |
| PSMD3 | 95.3453168 | -0.1178997 | 0.28363682 | -0.4156714 | 0.67765047 | 0.95560047 |
| MED24 | 111.662191 | -0.139644 | 0.24874769 | -0.5613882 | 0.57453291 | 0.93275827 |
| THRA | 175.109107 | 0.4555457 | 0.29381268 | 1.55046306 | 0.12103041 | 0.65959373 |
| NR1D1 | 48.0797188 | -1.3835671 | 0.86997574 | -1.5903513 | 0.11175564 | 0.64353508 |
| MSL1 | 177.341143 | -0.3686578 | 0.18341618 | -2.0099526 | 0.0444362 | 0.48009422 |
| CASC3 | 169.866035 | -0.1965024 | 0.26039727 | -0.7546255 | 0.45047373 | 0.89333031 |
| WIPF2 | 415.799756 | -0.4641771 | 0.25839712 | -1.7963711 | 0.07243552 | 0.56608818 |
| RARA | 90.788251 | -0.2896995 | 0.64605718 | -0.4484115 | 0.65385625 | 0.95100227 |
| KRT10 | 157.128585 | -0.254632 | 0.59112516 | -0.4307581 | 0.66664426 | 0.9527406 |
| TMEM99 | 36.3719505 | -0.2538263 | 0.47444842 | -0.5349924 | 0.5926551 | 0.93491607 |
| KRT19 | 3220.16233 | 0.16719743 | 0.40013089 | 0.41785684 | 0.67605179 | 0.95529325 |
| EIF1 | 958.329245 | 0.2455008 | 0.43653378 | 0.56238671 | 0.57385257 | 0.93270589 |
| JUP | 856.814574 | -0.2923311 | 0.3042596 | -0.9607949 | 0.33665528 | 0.84621728 |
| P3H4 | 37.2078906 | 0.23099619 | 0.34058091 | 0.67824174 | 0.49761843 | 0.90839719 |
| NT5C3B | 64.7991617 | 0.14882331 | 0.27691038 | 0.53744215 | 0.59096224 | 0.93483536 |
| ACLY | 144.02162 | -0.2543885 | 0.27007758 | -0.9419089 | 0.34623928 | 0.8523814 |
| TTC25 | 43.1778316 | 0.13277195 | 0.27139539 | 0.48921963 | 0.62468622 | 0.94469142 |
| CNP | 83.1343201 | 0.07619148 | 0.29756022 | 0.25605398 | 0.79790916 | 0.9653598 |
| DNAJC7 | 271.37319 | -0.1267915 | 0.17270931 | -0.7341325 | 0.46286797 | 0.89801718 |
| NKIRAS2 | 44.8726041 | -0.1190826 | 0.31786274 | -0.3746354 | 0.70793165 | 0.95752169 |
| DHX58 | 33.76011 | -0.2464907 | 0.4957203 | -0.4972375 | 0.6190216 | 0.94292825 |
| KAT2A | 64.6461638 | -0.200227 | 0.42435643 | -0.4718368 | 0.63704328 | 0.94821838 |

| | | | | | | |
|---|---|---|---|---|---|---|
| GHDC | 33.2926697 | 0.2930076 | 0.30243363 | 0.96883271 | 0.33262866 | 0.84442294 |
| STAT5B | 262.730614 | 0.23664269 | 0.27524411 | 0.85975571 | 0.38992372 | 0.87004477 |
| STAT5A | 50.6143794 | -0.129113 | 0.35268188 | -0.3660892 | 0.71429853 | 0.95791799 |
| STAT3 | 1514.89265 | -0.6870599 | 0.31597231 | -2.1744306 | 0.02967281 | 0.40649813 |
| CAVIN1 | 33.4306493 | 0.65703403 | 0.58706787 | 1.119179 | 0.26306378 | 0.8096287 |
| ATP6V0A1 | 253.370345 | -0.4837216 | 0.2516862 | -1.9219236 | 0.05461537 | 0.51597783 |
| NAGLU | 78.437669 | -0.2991581 | 0.52951907 | -0.564962 | 0.57209963 | 0.93200748 |
| COASY | 95.3415224 | 0.27717653 | 0.29964469 | 0.92501733 | 0.35495689 | 0.8557799 |
| MLX | 67.1921033 | -0.0004492 | 0.40353219 | -0.0011132 | 0.99911179 | 0.99968413 |
| RETREG3 | 181.720121 | 0.10013882 | 0.22429581 | 0.44645873 | 0.65526592 | 0.95100227 |
| TUBG2 | 68.4002707 | -0.2294838 | 0.26566576 | -0.8638063 | 0.38769431 | 0.86910078 |
| PLEKHH3 | 48.6313948 | 0.42920891 | 0.30257199 | 1.41853485 | 0.15603467 | 0.71066638 |
| EZH1 | 207.543414 | -0.3429751 | 0.24375427 | -1.4070527 | 0.15941177 | 0.71409437 |
| VPS25 | 97.3054269 | -0.257832 | 0.28906447 | -0.8919534 | 0.37241791 | 0.86201082 |
| WNK4 | 67.8476704 | -0.223311 | 0.46698827 | -0.4781939 | 0.63251218 | 0.94669061 |
| COA3 | 329.577444 | 0.68471839 | 0.45428023 | 1.50725994 | 0.13174404 | 0.67390608 |
| BECN1 | 103.884973 | 0.05861166 | 0.19495052 | 0.30064892 | 0.76368223 | 0.96285313 |
| PSME3 | 72.9734233 | -0.0005146 | 0.25655947 | -0.0020057 | 0.9983997 | 0.99967893 |
| RUNDC1 | 91.0221949 | -0.3644872 | 0.38971472 | -0.9352666 | 0.34965093 | 0.8535343 |
| RPL27 | 279.803917 | -0.0301356 | 0.37585544 | -0.0801788 | 0.93609504 | 0.99243654 |
| IFI35 | 62.8881921 | -1.2517325 | 0.7125108 | -1.7567909 | 0.07895345 | 0.5826987 |
| VAT1 | 95.1578662 | -0.6857861 | 0.31863313 | -2.152275 | 0.0313757 | 0.41423957 |
| BRCA1 | 90.8268815 | 0.30797118 | 0.40783503 | 0.75513666 | 0.45016699 | 0.893271 |
| NBR1 | 262.996148 | 0.40289304 | 0.20003465 | 2.01411627 | 0.04399733 | 0.48009422 |
| LINC00854 | 98.4278971 | -0.9381611 | 0.59205345 | -1.5845885 | 0.11305985 | 0.64739574 |
| ARL4D | 353.29955 | -0.4024422 | 0.68449178 | -0.587943 | 0.55657053 | 0.92872652 |
| DHX8 | 92.2701601 | -0.0016996 | 0.2614495 | -0.0065008 | 0.99481313 | 0.99919666 |
| DUSP3 | 149.640189 | -0.4684281 | 0.28462672 | -1.6457629 | 0.09981259 | 0.61822465 |
| LINC01976 | 54.5570547 | 1.82574051 | 1.01888099 | 1.79190752 | 0.07314778 | 0.56668778 |
| TMEM101 | 427.05998 | 0.85534957 | 0.66535652 | 1.28555076 | 0.19859989 | 0.76668122 |
| LSM12 | 145.28627 | -0.2537539 | 0.27580632 | -0.9200438 | 0.35754988 | 0.8557799 |
| G6PC3 | 75.5647644 | 0.321268 | 0.4127078 | 0.77843936 | 0.43631004 | 0.88785137 |
| HDAC5 | 104.952016 | 0.88429728 | 0.25960432 | 3.40632734 | 0.00065843 | 0.05965636 |
| TMUB2 | 85.6174112 | -0.307443 | 0.31763384 | -0.9679165 | 0.33308609 | 0.84442294 |
| ATXN7L3 | 55.9915113 | -0.1739684 | 0.2904113 | -0.5990416 | 0.54914517 | 0.9274143 |
| UBTF | 193.851499 | 0.05798754 | 0.27255798 | 0.21275303 | 0.8315196 | 0.97160502 |
| SLC25A39 | 123.281643 | -0.0959182 | 0.30524525 | -0.3142331 | 0.75334399 | 0.96120879 |
| GRN | 416.373522 | 0.19811615 | 0.40796862 | 0.48561615 | 0.62723934 | 0.94511671 |
| GPATCH8 | 855.596906 | 0.03386425 | 0.20038877 | 0.16899275 | 0.86580234 | 0.98023062 |
| CCDC43 | 35.2735581 | -0.2271732 | 0.35946033 | -0.631984 | 0.52739734 | 0.92057364 |
| DBF4B | 46.5418455 | 0.21058741 | 0.30725237 | 0.68538907 | 0.49309844 | 0.90804281 |
| EFTUD2 | 109.703285 | -0.1756103 | 0.23756554 | -0.739208 | 0.4597807 | 0.89753337 |
| DCAKD | 138.457159 | 0.26798797 | 0.37066409 | 0.72299415 | 0.46968348 | 0.90053922 |
| NMT1 | 203.255735 | 0.01321675 | 0.20900264 | 0.06323724 | 0.94957759 | 0.99361356 |
| ACBD4 | 56.3581066 | 0.08829822 | 0.2440088 | 0.3618649 | 0.71745299 | 0.95791799 |
| HEXIM1 | 95.846955 | 0.24020381 | 0.58105712 | 0.41339104 | 0.67932014 | 0.95574492 |
| AC138150.1 | 136.212261 | 0.48541408 | 0.4965252 | 0.97762226 | 0.32826118 | 0.84381657 |
| MAP3K14 | 217.378447 | 0.22231221 | 0.43189711 | 0.51473419 | 0.60673878 | 0.93925224 |
| ARHGAP27 | 58.0865479 | -0.1423257 | 0.28371576 | -0.501649 | 0.61591448 | 0.94190661 |
| PLEKHM1 | 106.547787 | -0.8295512 | 0.21568574 | -3.8461107 | 0.00012001 | 0.02194494 |
| LINC02210-C | 94.3759704 | -0.191701 | 0.4282984 | -0.4475874 | 0.65445099 | 0.95100227 |
| KANSL1 | 1770.28817 | 0.05911879 | 0.37186996 | 0.15897706 | 0.87368694 | 0.98246494 |
| ARL17B | 386.087638 | 0.97745854 | 0.97480586 | 1.00272123 | 0.31599538 | 0.84285425 |
| LRRC37A2 | 105.240406 | -0.0870473 | 0.31352809 | -0.2776379 | 0.78129034 | 0.96513523 |

| | | | | | | |
|---|---|---|---|---|---|---|
| ARL17A | 119.338759 | -0.2533324 | 0.48335608 | -0.5241113 | 0.60020113 | 0.93728325 |
| NSF | 470.986215 | -0.0741428 | 0.32231047 | -0.2300352 | 0.81806438 | 0.96883031 |
| WNT9B | 123.162769 | 1.73185389 | 0.99077172 | 1.74798478 | 0.08046666 | 0.58806877 |
| AC005670.2 | 73.9160633 | 0.29497409 | 0.35650661 | 0.82740147 | 0.40800955 | 0.87948803 |
| CDC27 | 486.486242 | -0.3886958 | 0.24323923 | -1.5979979 | 0.11004345 | 0.64001342 |
| AC068234.1 | 83.4950252 | 0.11101618 | 0.937389 | 0.11843128 | 0.90572594 | 0.9875066 |
| EFCAB13 | 183.907035 | 0.46759919 | 0.34748096 | 1.34568289 | 0.17840481 | 0.73783835 |
| AC040934.1 | 34.1886232 | -0.7013445 | 0.46021388 | -1.5239535 | 0.12752033 | 0.67051794 |
| NPEPPS | 989.692958 | -0.2350516 | 0.1492682 | -1.5746931 | 0.1153273 | 0.6515526 |
| AC025682.1 | 109.719044 | 0.15824087 | 0.29417304 | 0.53791764 | 0.59063391 | 0.93483536 |
| KPNB1 | 439.2991 | -0.1481793 | 0.18702209 | -0.792309 | 0.42818052 | 0.88535099 |
| MRPL10 | 53.0693598 | 0.15568448 | 0.40763516 | 0.38192114 | 0.70251986 | 0.95672887 |
| SCRN2 | 67.209666 | 0.36766459 | 0.32482176 | 1.13189644 | 0.25767798 | 0.80709747 |
| SP2 | 48.3077584 | 0.00602121 | 0.29679346 | 0.02028756 | 0.98381398 | 0.99723092 |
| PNPO | 67.6317358 | 0.93239055 | 0.32189975 | 2.89652456 | 0.00377321 | 0.14977426 |
| CDK5RAP3 | 114.486789 | -0.0673162 | 0.2617817 | -0.2571464 | 0.79706578 | 0.9653598 |
| NFE2L1 | 322.934723 | 0.15801667 | 0.22879659 | 0.69064262 | 0.48979016 | 0.90730741 |
| CBX1 | 174.764173 | 0.29140513 | 0.30835236 | 0.94503939 | 0.34463877 | 0.85148675 |
| SNX11 | 43.4265739 | -0.1124984 | 0.29397325 | -0.3826825 | 0.70195517 | 0.95672887 |
| SKAP1 | 1282.40932 | 0.0580395 | 0.19776752 | 0.29347338 | 0.76916035 | 0.96345263 |
| HOXB2 | 138.867956 | 0.58636149 | 0.4351817 | 1.34739465 | 0.17785316 | 0.73730267 |
| HOXB-AS1 | 35.7445148 | 0.3988713 | 0.40282394 | 0.99018767 | 0.3220824 | 0.84285425 |
| HOXB3 | 907.151453 | 0.15917741 | 0.24038524 | 0.66217631 | 0.50785824 | 0.91214298 |
| HOXB-AS3 | 213.418486 | 0.5687671 | 0.37725326 | 1.50765327 | 0.13164329 | 0.67390608 |
| HOXB7 | 314.250955 | 0.44111798 | 0.33277062 | 1.32559171 | 0.18497497 | 0.74875926 |
| AC103702.2 | 81.0779424 | -0.2177852 | 0.44064866 | -0.4942377 | 0.62113834 | 0.94377998 |
| LINC02086 | 153.170667 | -0.3670388 | 0.41874429 | -0.8765225 | 0.38074604 | 0.86593382 |
| CALCOCO2 | 204.682244 | 0.04724402 | 0.2192205 | 0.21550915 | 0.82937038 | 0.97131041 |
| ATP5MC1 | 253.702771 | 0.49114895 | 0.40939096 | 1.19970641 | 0.23025338 | 0.79075638 |
| UBE2Z | 285.808768 | -0.2111937 | 0.19599511 | -1.0775459 | 0.28123644 | 0.82628479 |
| SNF8 | 120.966298 | -0.1705727 | 0.33220581 | -0.5134549 | 0.60763318 | 0.93964819 |
| B4GALNT2 | 160.064734 | -0.6718943 | 0.8045436 | -0.8351247 | 0.40364748 | 0.87834687 |
| ZNF652 | 421.078833 | 0.12869564 | 0.19598503 | 0.65666057 | 0.51139919 | 0.91403647 |
| AC091180.5 | 46.7709908 | 0.31068667 | 0.50119487 | 0.61989197 | 0.53532891 | 0.92303275 |
| PHB | 230.828747 | 0.46308963 | 0.43216966 | 1.07154589 | 0.28392405 | 0.8272238 |
| SPOP | 196.507136 | 0.13459956 | 0.17039316 | 0.78993526 | 0.42956558 | 0.88579433 |
| SLC35B1 | 42.1320537 | -0.3308492 | 0.33469097 | -0.9885215 | 0.3228973 | 0.84285425 |
| FAM117A | 41.8799451 | -0.4529874 | 0.32430242 | -1.3968054 | 0.16247209 | 0.71482262 |
| KAT7 | 120.889351 | 0.25378658 | 0.26553952 | 0.9557394 | 0.33920391 | 0.84881972 |
| ITGA3 | 553.19487 | -1.3019232 | 0.65954346 | -1.9739763 | 0.04838443 | 0.48661187 |
| PDK2 | 72.9473033 | 0.63754415 | 0.29231607 | 2.18100957 | 0.02918271 | 0.40623517 |
| PPP1R9B | 42.3193158 | -0.2345143 | 0.30607209 | -0.7662062 | 0.44355363 | 0.89037885 |
| XYLT2 | 71.1722102 | 0.31376533 | 0.26708626 | 1.17477152 | 0.24008614 | 0.80056465 |
| MRPL27 | 132.633859 | 0.11813421 | 0.31278861 | 0.37768066 | 0.70566784 | 0.95730303 |
| LRRC59 | 52.4117321 | -0.2838087 | 0.32458003 | -0.8743875 | 0.38190726 | 0.86593382 |
| ACSF2 | 65.618969 | 0.89182219 | 0.28157669 | 3.16724434 | 0.00153891 | 0.09927452 |
| RSAD1 | 64.0397287 | 0.10848666 | 0.24008153 | 0.45187422 | 0.6513596 | 0.95061388 |
| EPN3 | 56.9897658 | 0.44949618 | 0.49324883 | 0.91129699 | 0.36213891 | 0.85690129 |
| SPATA20 | 66.7053141 | 0.04568634 | 0.39717873 | 0.11502716 | 0.9084236 | 0.98758338 |
| ABCC3 | 236.983123 | 0.55794202 | 0.81818086 | 0.68192993 | 0.49528326 | 0.90839719 |
| ANKRD40 | 75.2380698 | -0.2222139 | 0.3649756 | -0.608846 | 0.54262653 | 0.92482794 |
| LUC7L3 | 683.569853 | -0.1661632 | 0.13235331 | -1.2554519 | 0.20931477 | 0.77436433 |
| TOB1 | 357.589654 | -0.4185164 | 0.47633003 | -0.878627 | 0.37960354 | 0.86587828 |
| SPAG9 | 885.59692 | -0.4651072 | 0.26034737 | -1.7864872 | 0.07402041 | 0.56915891 |

| | | | | | | |
|---|---|---|---|---|---|---|
| AC005920.2 | 89.2696896 | -0.6477193 | 0.6125328 | -1.0574442 | 0.29030891 | 0.83103998 |
| MBTD1 | 370.814849 | -0.2178333 | 0.22012706 | -0.9895798 | 0.32237957 | 0.84285425 |
| UTP18 | 96.6782134 | -0.5020006 | 0.22594775 | -2.2217554 | 0.02629984 | 0.39206594 |
| TOM1L1 | 633.594893 | 0.17808346 | 0.19261026 | 0.9245793 | 0.35518479 | 0.8557799 |
| COX11 | 128.854895 | 0.00775536 | 0.22020957 | 0.03521809 | 0.97190583 | 0.99656026 |
| STXBP4 | 469.934093 | 0.14293056 | 0.23969074 | 0.59631241 | 0.55096654 | 0.92816739 |
| HLF | 111.513983 | 0.28953106 | 0.67880558 | 0.42653018 | 0.66972156 | 0.9533662 |
| MMD | 50.6531984 | 0.55683342 | 0.56337046 | 0.98839654 | 0.32295848 | 0.84285425 |
| PCTP | 53.3082178 | 0.54729172 | 0.25429795 | 2.15216728 | 0.03138418 | 0.41423957 |
| DGKE | 71.9978711 | 0.676214 | 0.27590316 | 2.45091067 | 0.01424953 | 0.28746487 |
| TRIM25 | 80.2874996 | -0.3428476 | 0.31063582 | -1.1036964 | 0.26972484 | 0.81817548 |
| COIL | 80.2098731 | -0.0712327 | 0.22295113 | -0.3194994 | 0.74934788 | 0.96109466 |
| SCPEP1 | 353.946473 | 0.61943512 | 0.23559667 | 2.62921855 | 0.00855813 | 0.22546434 |
| AKAP1 | 166.526344 | 0.18072021 | 0.2514926 | 0.71859058 | 0.47239322 | 0.90209201 |
| MSI2 | 2132.88633 | 0.0821965 | 0.20706541 | 0.39695911 | 0.69139761 | 0.95672887 |
| MRPS23 | 100.979298 | 0.301239 | 0.3292075 | 0.91504292 | 0.3601691 | 0.85608464 |
| CUEDC1 | 280.64232 | 0.19287455 | 0.39333687 | 0.49035462 | 0.62388299 | 0.94457465 |
| VEZF1 | 189.358847 | 0.3101452 | 0.23640224 | 1.31193853 | 0.18954088 | 0.75619145 |
| AC015813.2 | 76.7588064 | -0.1766341 | 0.24096189 | -0.7330373 | 0.46353567 | 0.89801718 |
| SRSF1 | 42.4416839 | -0.3813797 | 0.36634829 | -1.0410303 | 0.2978615 | 0.83556453 |
| DYNLL2 | 167.360996 | 0.39666348 | 0.25213701 | 1.57320613 | 0.1156711 | 0.65187472 |
| MKS1 | 39.1628551 | -0.3767082 | 0.33291371 | -1.131549 | 0.25782409 | 0.80709747 |
| TSPOAP1 | 109.114917 | 0.28795404 | 0.41865704 | 0.68780412 | 0.49157614 | 0.90730741 |
| TSPOAP1-AS | 35.4856295 | 0.63683934 | 0.56947456 | 1.11829287 | 0.26344193 | 0.80987722 |
| SUPT4H1 | 76.8486166 | 0.09102771 | 0.43179145 | 0.21081407 | 0.83303236 | 0.97208613 |
| RNF43 | 206.670356 | 0.16220001 | 0.48782734 | 0.33249471 | 0.73951573 | 0.96078945 |
| MTMR4 | 68.7773607 | 0.00046667 | 0.27050187 | 0.00172522 | 0.99862348 | 0.99967893 |
| RAD51C | 78.0254796 | -0.0668903 | 0.3330432 | -0.2008456 | 0.8408193 | 0.97514319 |
| TRIM37 | 179.413792 | -0.3610502 | 0.25493539 | -1.416242 | 0.15670465 | 0.71126849 |
| SKA2 | 81.887359 | 0.72269763 | 0.27730076 | 2.60618696 | 0.00915565 | 0.23300339 |
| GDPD1 | 45.2356943 | 0.23045142 | 0.37586366 | 0.61312504 | 0.53979367 | 0.92405057 |
| YPEL2 | 518.026254 | 0.65415454 | 0.45574365 | 1.43535633 | 0.15118558 | 0.7029344 |
| DHX40 | 207.084585 | 0.26227144 | 0.25904744 | 1.01244563 | 0.31132503 | 0.84279403 |
| CLTC | 718.284503 | -0.1201611 | 0.21093329 | -0.5696641 | 0.56890557 | 0.93122667 |
| PTRH2 | 95.7867532 | -0.0863258 | 0.34091191 | -0.2532202 | 0.80009804 | 0.96568323 |
| VMP1 | 972.281599 | -0.9334288 | 0.43268811 | -2.1572786 | 0.03098396 | 0.41259458 |
| TUBD1 | 58.9773394 | 0.36224353 | 0.28517051 | 1.27026997 | 0.20398848 | 0.77176869 |
| RPS6KB1 | 136.567917 | -0.2702974 | 0.26637974 | -1.014707 | 0.31024551 | 0.84213522 |
| HEATR6 | 50.1890464 | 0.26298019 | 0.29491948 | 0.89170165 | 0.37255287 | 0.86201082 |
| USP32 | 483.105332 | -0.4272726 | 0.31035335 | -1.3767294 | 0.16859592 | 0.72499057 |
| APPBP2 | 388.727671 | -0.1121539 | 0.19095889 | -0.5873197 | 0.556989 | 0.92872652 |
| PPM1D | 131.950944 | 0.10312779 | 0.19919726 | 0.5177169 | 0.6046558 | 0.93836707 |
| BCAS3 | 1727.76718 | 0.09249772 | 0.17324226 | 0.53392126 | 0.59339602 | 0.93491607 |
| AC005746.2 | 78.9563209 | 0.89325596 | 0.46883195 | 1.90527963 | 0.05674373 | 0.52348091 |
| TBX2 | 549.397843 | 0.0007318 | 0.44999004 | 0.00162627 | 0.99870243 | 0.99967893 |
| NACA2 | 69.3134989 | 0.64113274 | 0.59725351 | 1.07346835 | 0.28306103 | 0.82666614 |
| INTS2 | 109.471757 | -0.3639291 | 0.24321665 | -1.4963166 | 0.13457116 | 0.6785647 |
| MED13 | 867.291771 | -0.2146767 | 0.24589856 | -0.8730297 | 0.3826469 | 0.86593382 |
| METTL2A | 56.0286317 | -0.144581 | 0.32138135 | -0.4498737 | 0.65280153 | 0.95100227 |
| TLK2 | 452.931454 | -0.1716203 | 0.26762659 | -0.6412679 | 0.52134862 | 0.91787515 |
| 10-Mar | 34.9447144 | 0.76889229 | 0.49301266 | 1.5595792 | 0.11885935 | 0.65456964 |
| TANC2 | 1737.86277 | 0.0848499 | 0.22151951 | 0.38303576 | 0.70169325 | 0.95672887 |
| AC015923.1 | 142.063389 | 0.24106981 | 0.62348965 | 0.38664605 | 0.69901826 | 0.95672887 |
| CYB561 | 107.802903 | 0.03496145 | 0.32197623 | 0.10858396 | 0.91353249 | 0.98830345 |

| | | | | | | |
|---|---|---|---|---|---|---|
| DCAF7 | 388.126425 | 0.03393745 | 0.21629227 | 0.15690552 | 0.87531931 | 0.98267808 |
| TACO1 | 35.8442247 | 0.46894086 | 0.42685986 | 1.09858271 | 0.27195012 | 0.81922962 |
| MAP3K3 | 253.847368 | -0.5550996 | 0.23998029 | -2.3131052 | 0.02071686 | 0.34572082 |
| CCDC47 | 104.417813 | 0.06263692 | 0.40290015 | 0.15546512 | 0.87645465 | 0.98269442 |
| DDX42 | 267.38093 | -0.09787 | 0.19373542 | -0.5051733 | 0.61343714 | 0.94034333 |
| FTSJ3 | 54.0238594 | -0.2392581 | 0.2792557 | -0.8567705 | 0.39157172 | 0.87015937 |
| PSMC5 | 80.1192558 | 0.18937436 | 0.36024539 | 0.52568158 | 0.59910947 | 0.93728325 |
| SMARCD2 | 79.1955621 | 0.12692905 | 0.24389382 | 0.5204275 | 0.60276565 | 0.93804805 |
| ERN1 | 188.407885 | -1.1366345 | 0.38734279 | -2.934441 | 0.00334149 | 0.14358074 |
| SNHG25 | 51.7075862 | -0.5565864 | 0.4536106 | -1.2270137 | 0.21981744 | 0.78633738 |
| TEX2 | 126.474122 | -0.0807812 | 0.28755386 | -0.2809255 | 0.77876752 | 0.96512745 |
| POLG2 | 66.5304483 | -0.3156166 | 0.27040606 | -1.1671949 | 0.24313167 | 0.80072691 |
| DDX5 | 1063.92328 | -0.0420332 | 0.28639534 | -0.1467665 | 0.88331632 | 0.9835847 |
| CEP95 | 340.82121 | -0.1661561 | 0.18262432 | -0.9098245 | 0.36291508 | 0.85694172 |
| SMURF2 | 341.746175 | -0.1427967 | 0.24850483 | -0.5746234 | 0.56554604 | 0.93114846 |
| LRRC37A3 | 467.981258 | 0.12553727 | 0.32347242 | 0.38809264 | 0.69794748 | 0.95672887 |
| GNA13 | 421.900784 | -0.8478792 | 0.59022503 | -1.4365355 | 0.15085002 | 0.7029344 |
| CEP112 | 567.291227 | 0.30621379 | 0.25173499 | 1.21641333 | 0.22382751 | 0.78960314 |
| PRKCA | 3531.77386 | -0.1865908 | 0.26365206 | -0.707716 | 0.47912161 | 0.90291126 |
| PRKCA-AS1 | 164.377907 | -0.1649324 | 0.47233417 | -0.3491858 | 0.72694979 | 0.95979351 |
| HELZ | 661.842702 | -0.2713123 | 0.230785 | -1.1756065 | 0.23975217 | 0.80019197 |
| PSMD12 | 142.814792 | -0.226934 | 0.20816131 | -1.0901836 | 0.27563228 | 0.82109899 |
| PITPNC1 | 273.25357 | -1.0552517 | 0.50831512 | -2.0759793 | 0.03789587 | 0.44623447 |
| NOL11 | 80.7419835 | 0.07847525 | 0.22356155 | 0.35102303 | 0.72557107 | 0.95963742 |
| BPTF | 1099.06 | -0.1965435 | 0.23755624 | -0.8273556 | 0.40803555 | 0.87948803 |
| AC005332.7 | 44.1441267 | -0.1402728 | 0.31795437 | -0.4411728 | 0.65908793 | 0.95127957 |
| AMZ2 | 57.0083814 | -0.0401371 | 0.29954069 | -0.1339955 | 0.89340616 | 0.98597539 |
| ARSG | 127.48531 | -0.3100976 | 0.45717265 | -0.6782943 | 0.49758514 | 0.90839719 |
| WIPI1 | 86.816679 | -0.1386455 | 0.40181765 | -0.3450458 | 0.73005997 | 0.96008876 |
| PRKAR1A | 967.785635 | -0.2921233 | 0.41104552 | -0.7106835 | 0.47728039 | 0.90242252 |
| FAM20A | 1212.28652 | -0.2117191 | 0.38936996 | -0.5437479 | 0.58661498 | 0.93483536 |
| AC079210.1 | 49.9741979 | 0.1594062 | 0.52663965 | 0.30268553 | 0.76212953 | 0.96218738 |
| LINC01482 | 1058.0531 | 0.83347183 | 0.53440118 | 1.55963695 | 0.1188457 | 0.65456964 |
| ABCA10 | 69.9216559 | 0.28374644 | 0.48142481 | 0.58938891 | 0.55560041 | 0.92872652 |
| ABCA5 | 324.631129 | -0.1395683 | 0.27556191 | -0.5064863 | 0.61251534 | 0.94003018 |
| AC005208.1 | 878.213906 | -0.1176226 | 0.66186288 | -0.1777145 | 0.85894722 | 0.97788846 |
| KCNJ16 | 2926.83949 | -0.3016684 | 0.40454833 | -0.7456918 | 0.4558536 | 0.89626521 |
| SOX9 | 109.24278 | 0.90986071 | 0.57034115 | 1.59529206 | 1.11E-01 | 6.41E-01 |
| SLC39A11 | 455.063011 | -0.3580113 | 0.19803687 | -1.8078014 | 0.07063742 | 0.5621116 |
| COG1 | 53.522171 | -0.0841866 | 0.31524559 | -0.267051 | 0.78942992 | 0.96515649 |
| FAM104A | 72.9928106 | 0.31986072 | 0.2526697 | 1.26592434 | 0.20554017 | 0.77193931 |
| C17orf80 | 51.7238018 | 0.24110588 | 0.29920143 | 0.80583131 | 0.42034012 | 0.88315246 |
| CDC42EP4 | 138.375777 | 0.08147525 | 0.32112726 | 0.25371641 | 0.79971466 | 0.96568323 |
| RPL38 | 221.875476 | 0.12634409 | 0.31564731 | 0.4002698 | 0.68895781 | 0.95606293 |
| GPRC5C | 762.150097 | 0.26666556 | 0.15905901 | 1.67651967 | 0.09363645 | 0.61317397 |
| SLC9A3R1 | 45.2679193 | 0.22507779 | 0.55314066 | 0.40690877 | 0.684075 | 0.95574492 |
| NAT9 | 44.1229701 | -0.3706551 | 0.32581594 | -1.137621 | 0.25527877 | 0.80509223 |
| TMEM104 | 34.7253726 | -0.3859422 | 0.39336609 | -0.9811274 | 0.32652993 | 0.84381657 |
| MRPL58 | 40.224783 | 0.29380582 | 0.42393699 | 0.69304123 | 0.48828368 | 0.90730741 |
| KCTD2 | 79.4488568 | -0.106205 | 0.2172377 | -0.4888885 | 0.62492064 | 0.9448702 |
| ATP5PD | 177.277764 | 0.08125281 | 0.44628593 | 0.18206447 | 0.85553213 | 0.97718353 |
| SLC16A5 | 607.534667 | 0.36262458 | 0.39391113 | 0.92057462 | 0.35727256 | 0.8557799 |
| ARMC7 | 147.337198 | 0.41393774 | 0.35493424 | 1.16623784 | 0.2435183 | 0.80072691 |
| NT5C | 162.754621 | -0.1504181 | 0.34018112 | -0.4421705 | 0.65836582 | 0.95113765 |

| | | | | | | |
|---|---|---|---|---|---|---|
| JPT1 | 337.630363 | -0.6687931 | 0.67532142 | -0.9903331 | 0.32201134 | 0.84285425 |
| SUMO2 | 115.339237 | 0.11770683 | 0.34913343 | 0.33713997 | 0.73601139 | 0.96078945 |
| NUP85 | 83.5807709 | -0.4257733 | 0.23479804 | -1.8133598 | 0.06977634 | 0.55992041 |
| GGA3 | 101.347166 | -0.0371015 | 0.19992573 | -0.1855762 | 0.85277708 | 0.97718353 |
| MRPS7 | 121.817025 | 0.28935499 | 0.36746427 | 0.78743707 | 0.43102605 | 0.88641151 |
| MIF4GD | 35.8676186 | 0.34989757 | 0.36830154 | 0.95003015 | 0.34209693 | 0.85093239 |
| GRB2 | 362.305808 | -0.1647779 | 0.18719895 | -0.8802289 | 0.3787353 | 0.86515603 |
| AC011933.4 | 50.9531827 | -0.1967528 | 0.42046929 | -0.4679361 | 0.63983025 | 0.94935885 |
| TMEM94 | 100.102505 | 0.07835951 | 0.18921074 | 0.41413882 | 0.67877245 | 0.95574492 |
| CASKIN2 | 53.4231718 | -0.075834 | 0.29555631 | -0.2565804 | 0.7975027 | 0.9653598 |
| TSEN54 | 60.607911 | 0.16103457 | 0.2997779 | 0.53717961 | 0.59114355 | 0.93483997 |
| LLGL2 | 1146.0293 | -0.1261163 | 0.26130081 | -0.4826481 | 0.62934565 | 0.94596555 |
| MYO15B | 169.345517 | -0.0163907 | 0.34211556 | -0.0479098 | 0.96178814 | 0.99635733 |
| RECQL5 | 169.051469 | -0.4025368 | 0.24819047 | -1.6218867 | 0.10482762 | 0.6317386 |
| SMIM5 | 377.52308 | -1.0449032 | 0.65051242 | -1.6062772 | 0.10821302 | 0.63533473 |
| SMIM6 | 68.1568043 | -1.0390073 | 0.73503203 | -1.4135537 | 0.15749298 | 0.71160548 |
| SAP30BP | 145.426292 | -0.4778673 | 0.20754626 | -2.3024617 | 0.02130915 | 0.35223957 |
| ITGB4 | 83.6794475 | 0.42893799 | 0.37916478 | 1.13127064 | 0.2579412 | 0.80709747 |
| H3F3B | 1690.00429 | 0.25719865 | 0.6162878 | 0.4173353 | 0.67643317 | 0.95529325 |
| UNK | 218.359976 | -0.3565169 | 0.24462705 | -1.4573895 | 0.14500889 | 0.69321483 |
| WBP2 | 188.026044 | -0.4538131 | 0.2177045 | -2.084537 | 0.03711135 | 0.44129343 |
| TRIM47 | 40.7916774 | -0.0994751 | 0.40448315 | -0.2459314 | 8.06E-01 | 0.96736244 |
| TRIM65 | 35.3683634 | 0.24408819 | 0.32993783 | 0.73980057 | 0.45942101 | 0.89753337 |
| FBF1 | 38.7082125 | 0.36934827 | 0.44861313 | 0.82331133 | 0.41033097 | 0.88019562 |
| ACOX1 | 190.298046 | 0.20183428 | 0.16180676 | 1.24737859 | 0.21225871 | 0.77709975 |
| TEN1 | 40.7292374 | 0.44455751 | 0.46346969 | 0.95919434 | 0.33746085 | 0.84647911 |
| EVPL | 51.7217131 | 0.36417539 | 0.26583805 | 1.36991443 | 0.17071361 | 0.72697224 |
| SRP68 | 95.3156274 | 0.10804652 | 0.19343512 | 0.55856721 | 0.57645713 | 0.9331615 |
| EXOC7 | 228.669991 | 0.09627801 | 0.27893802 | 0.34515915 | 0.72997473 | 0.96008876 |
| UBALD2 | 114.346063 | -0.0492611 | 0.32212623 | -0.1529248 | 0.87845757 | 0.98300919 |
| PRPSAP1 | 94.1945617 | -0.0584043 | 0.2811155 | -0.207759 | 0.83541712 | 0.97268875 |
| UBE2O | 140.400585 | 0.10053578 | 0.21083607 | 0.47684336 | 0.63347366 | 0.94744025 |
| MXRA7 | 111.803402 | 0.00776728 | 0.36134788 | 0.02149531 | 0.98285054 | 0.9971769 |
| JMJD6 | 54.4848665 | -0.5513602 | 0.41500796 | -1.3285532 | 0.18399541 | 0.74623524 |
| SRSF2 | 242.794983 | -0.11501 | 0.34853094 | -0.3299851 | 0.7414112 | 0.96084411 |
| MFSD11 | 126.841355 | -0.022405 | 0.26203711 | -0.0855032 | 0.93186132 | 0.99243654 |
| SEC14L1 | 295.151717 | -1.0835815 | 0.46884627 | -2.3111658 | 0.0208237 | 0.3469513 |
| 9-Sep | 1352.22855 | -0.9220276 | 0.54734164 | -1.6845559 | 0.09207428 | 0.61120158 |
| TNRC6C | 186.432473 | 0.15135793 | 0.40247441 | 0.37606846 | 0.706866 | 0.95752169 |
| TMC6 | 70.9618949 | -0.0173208 | 0.37886476 | -0.0457175 | 0.96353538 | 0.99637506 |
| SYNGR2 | 171.229835 | -0.2247553 | 0.39432253 | -0.5699784 | 0.56869235 | 0.93122667 |
| AFMID | 266.203752 | -0.6892053 | 0.44139982 | -1.5614082 | 0.11842746 | 0.65447949 |
| SOCS3 | 226.655715 | -0.6469905 | 0.93455142 | -0.6923006 | 0.48874857 | 0.90730741 |
| PGS1 | 94.4076501 | -0.3766525 | 0.27638684 | -1.3627728 | 0.17295412 | 0.72940007 |
| CYTH1 | 273.482706 | -0.2255743 | 0.24047993 | -0.9380173 | 0.34823552 | 0.85348649 |
| USP36 | 387.246517 | -2.047767 | 0.85933721 | -2.3829609 | 0.01717402 | 0.31356046 |
| TIMP2 | 207.66613 | -0.4059422 | 0.36938747 | -1.0989604 | 0.27178532 | 0.81922962 |
| LGALS3BP | 417.830292 | -0.0411311 | 0.58081572 | -0.0708161 | 0.94354413 | 0.99265578 |
| CANT1 | 68.1421954 | 0.24489184 | 0.27105316 | 0.9034827 | 0.36626977 | 0.85902833 |
| ENGASE | 34.4832288 | 0.26125348 | 0.45244471 | 0.57742631 | 0.5636515 | 0.93077674 |
| CBX4 | 32.9630876 | -0.124677 | 0.60115259 | -0.2073966 | 0.83570012 | 0.97280209 |
| TBC1D16 | 483.790452 | 0.73231369 | 0.38107334 | 1.92171324 | 0.05464185 | 0.51597783 |
| CCDC40 | 137.216598 | 1.40424143 | 0.54890808 | 2.55824514 | 0.01052019 | 0.25038862 |
| GAA | 181.778987 | 1.04040529 | 0.28153881 | 3.69542413 | 0.00021952 | 0.03286528 |

| | | | | | | |
|---|---|---|---|---|---|---|
| EIF4A3 | 109.650986 | 0.17412985 | 0.42760529 | 0.40722099 | 0.68384568 | 0.95574492 |
| AC087741.1 | 47.3240928 | -0.0756528 | 0.40798695 | -0.1854295 | 0.85289214 | 0.97718353 |
| SGSH | 53.1525252 | -0.0054895 | 0.3802693 | -0.0144358 | 0.98848229 | 0.99814557 |
| SLC26A11 | 128.28708 | 0.07788479 | 0.25032197 | 0.31113844 | 0.75569538 | 0.96120879 |
| RNF213 | 1822.34254 | -0.4326925 | 0.5201827 | -0.8318088 | 0.40551688 | 0.87915121 |
| AC124319.2 | 382.264203 | -0.0372581 | 0.28809155 | -0.1293274 | 0.89709856 | 0.98601802 |
| ENDOV | 80.0696601 | -0.1695843 | 0.26744991 | -0.6340787 | 0.52602944 | 0.9197178 |
| RPTOR | 358.379854 | -0.3646216 | 0.25927728 | -1.4062999 | 0.15963508 | 0.71409437 |
| CHMP6 | 43.6005051 | -0.1519365 | 0.30653998 | -0.4956498 | 0.62014149 | 0.94319304 |
| BAIAP2-DT | 43.0505187 | 0.63188335 | 0.33820431 | 1.86834805 | 0.06171357 | 0.54395143 |
| BAIAP2 | 136.42731 | -0.5656381 | 0.55261103 | -1.0235737 | 0.30603666 | 0.83709431 |
| TEPSIN | 30.4549176 | 0.00470075 | 0.32632705 | 0.01440503 | 0.98850685 | 0.99814557 |
| NDUFAF8 | 102.552333 | 0.05545713 | 0.38052991 | 0.14573658 | 0.88412934 | 0.98403849 |
| SLC38A10 | 201.671764 | 0.37497202 | 0.19434477 | 1.92941662 | 0.05367916 | 0.51316717 |
| AC110285.6 | 38.0791264 | -0.3171968 | 0.56781008 | -0.5586318 | 0.57641301 | 0.9331615 |
| BAHCC1 | 58.8634029 | 0.66532218 | 0.42451985 | 1.56723455 | 0.11705988 | 0.65289383 |
| ACTG1 | 2411.33955 | 0.01932098 | 0.440718 | 0.04383979 | 0.96503211 | 0.99637506 |
| FAAP100 | 35.3533942 | -0.099031 | 0.33633237 | -0.2944438 | 0.76841881 | 0.96345263 |
| NPLOC4 | 309.174681 | -0.4007571 | 0.24341897 | -1.6463678 | 0.09968806 | 0.61822465 |
| OXLD1 | 79.8233701 | 0.28372507 | 0.27110294 | 1.04655843 | 0.29530328 | 0.83396494 |
| CCDC137 | 70.8320184 | -0.2661357 | 0.26299164 | -1.0119551 | 0.31155954 | 0.84279403 |
| ARL16 | 85.9425479 | 0.58642758 | 0.43166903 | 1.35851205 | 0.17430126 | 0.73184183 |
| HGS | 155.466533 | -0.5914693 | 0.27730785 | -2.132898 | 0.0329331 | 0.42193013 |
| MCRIP1 | 122.123658 | -0.4760851 | 0.34388075 | -1.3844484 | 0.16622121 | 0.72054851 |
| P4HB | 392.058954 | -0.4562032 | 0.32269205 | -1.4137418 | 0.15743773 | 0.71160548 |
| ARHGDIA | 222.224582 | -0.5863052 | 0.31504556 | -1.8610171 | 0.06274177 | 0.54720847 |
| ANAPC11 | 186.242694 | 0.31601253 | 0.52574203 | 0.60107908 | 0.54778732 | 0.92682201 |
| PCYT2 | 43.8779314 | -0.0391244 | 0.5677942 | -0.0689059 | 0.94506448 | 0.99276891 |
| SIRT7 | 47.821198 | 0.10741338 | 0.30530092 | 0.35182789 | 0.72496734 | 0.95963742 |
| MAFG | 43.3401454 | -0.6005814 | 0.37853423 | -1.5865974 | 0.11260386 | 0.64556262 |
| ASPSCR1 | 63.9462927 | -0.3861698 | 0.32760624 | -1.178762 | 0.23849298 | 0.7985324 |
| CENPX | 57.6540857 | 0.29344073 | 0.57556591 | 0.50982992 | 0.61017062 | 0.94003018 |
| DCXR | 58.7689274 | -0.1070664 | 0.34940757 | -0.3064226 | 0.75928293 | 0.96182318 |
| RFNG | 66.1280813 | -0.267572 | 0.23395396 | -1.1436952 | 0.25275007 | 0.80350545 |
| GPS1 | 83.7857403 | 0.02750835 | 0.37836357 | 0.07270348 | 0.94204208 | 0.99265578 |
| DUS1L | 83.1789512 | -0.2498176 | 0.30822937 | -0.8104926 | 0.41765712 | 0.88242985 |
| FASN | 41.0966027 | -0.3728911 | 0.82279646 | -0.4531997 | 0.65040492 | 0.95061388 |
| CCDC57 | 356.02673 | -0.15539 | 0.24292462 | -0.6396633 | 0.52239154 | 0.91845051 |
| SLC16A3 | 201.190775 | -0.5942498 | 0.51836648 | -1.1463893 | 0.2516341 | 0.80252057 |
| CSNK1D | 264.056827 | -0.5863332 | 0.25672362 | -2.2839084 | 0.02237691 | 0.35859484 |
| OGFOD3 | 86.9766161 | -0.2372908 | 0.2919815 | -0.8126911 | 0.41639517 | 0.88242985 |
| HEXDC | 96.0156284 | -0.5287852 | 0.27131662 | -1.94896 | 0.05130021 | 0.50104955 |
| CYBC1 | 67.9175004 | -0.2007229 | 0.31016755 | -0.6471435 | 0.51753909 | 0.91751084 |
| NARF | 100.614251 | -0.4442493 | 0.28700499 | -1.54788 | 0.1216512 | 0.66157942 |
| FOXK2 | 390.275043 | -0.3228617 | 0.39290354 | -0.8217328 | 0.41122899 | 0.88078476 |
| WDR45B | 305.390255 | -0.6559915 | 0.53850506 | -1.2181715 | 0.2231588 | 0.78877038 |
| RAB40B | 149.166346 | -0.3408178 | 0.3457631 | -0.9856976 | 0.32428155 | 0.84285425 |
| FN3KRP | 61.8696206 | -0.1855809 | 0.39160691 | -0.4738958 | 0.63557423 | 0.94807428 |
| FN3K | 228.887253 | -1.0924512 | 0.7892692 | -1.3841301 | 0.16631863 | 0.72054851 |
| TBCD | 178.580711 | -0.3172318 | 0.23564181 | -1.3462457 | 0.17822328 | 0.73737861 |
| METRNL | 202.182091 | 0.03571848 | 0.37655254 | 0.09485656 | 0.92442876 | 0.99051393 |
| AC144831.1 | 43.3066365 | 0.03500003 | 0.51987932 | 0.06732337 | 0.94632427 | 0.99307065 |
| USP14 | 257.76768 | -0.4058651 | 0.17651421 | -2.2993338 | 0.02148599 | 0.35404595 |
| THOC1 | 193.693789 | -0.3650193 | 0.18359656 | -1.98816 | 0.046794 | 0.48600281 |

| | | | | | | |
|---|---|---|---|---|---|---|
| ENOSF1 | 742.029463 | -0.1757228 | 0.31761293 | -0.5532608 | 0.58008484 | 0.93345492 |
| YES1 | 291.460946 | -0.2720975 | 0.19214578 | -1.4160994 | 0.1567464 | 0.71126849 |
| METTL4 | 80.3050374 | -0.5215755 | 0.22646877 | -2.303079 | 0.0212744 | 0.35222066 |
| SMCHD1 | 589.088031 | -0.065328 | 0.29892801 | -0.2185411 | 0.82700757 | 0.97069946 |
| AP001011.1 | 100.890877 | 0.30940153 | 0.63211347 | 0.4894715 | 0.62450793 | 0.94457465 |
| LPIN2 | 840.07737 | 0.30072119 | 0.41412875 | 0.72615386 | 0.46774445 | 0.89976199 |
| AP000919.2 | 81.5642283 | 0.09421478 | 0.36494663 | 0.25816044 | 0.79628309 | 0.9653598 |
| MYOM1 | 49.16541 | 0.26593729 | 0.59743097 | 0.44513476 | 0.65622237 | 0.95100227 |
| MYL12A | 587.986297 | 0.34884368 | 0.49096529 | 0.71052616 | 0.47737792 | 0.90242252 |
| MYL12B | 1068.07224 | 0.43380735 | 0.51277201 | 0.84600435 | 0.39755032 | 0.87462201 |
| TGIF1 | 188.548009 | -0.5682076 | 0.54530459 | -1.0420004 | 0.29741148 | 0.83556453 |
| DLGAP1 | 383.003119 | -0.2818372 | 0.90047811 | -0.3129862 | 0.75429118 | 0.96120879 |
| DLGAP1-AS1 | 105.615094 | -1.0658807 | 0.67471535 | -1.5797487 | 0.11416442 | 0.64951142 |
| DLGAP1-AS2 | 43.5511284 | -1.173708 | 0.66008489 | -1.7781167 | 0.07538469 | 0.57072562 |
| LINC00667 | 60.2844868 | -0.2724269 | 0.33273767 | -0.8187438 | 0.41293258 | 0.88155092 |
| EPB41L3 | 296.915659 | -1.1580687 | 0.54905027 | -2.1092215 | 0.03492546 | 0.43353051 |
| L3MBTL4 | 443.199401 | -0.3935185 | 0.34290378 | -1.1476062 | 0.25113116 | 0.80240987 |
| AP001021.2 | 37.399076 | -0.47058 | 0.65860839 | -0.7145065 | 0.47491403 | 0.90242252 |
| ARHGAP28 | 109.411566 | -0.2897503 | 0.44023473 | -0.6581722 | 0.51042747 | 0.91334187 |
| PTPRM | 1646.10398 | 0.41174981 | 0.39078415 | 1.05365024 | 0.29204308 | 0.83315947 |
| RAB12 | 162.947747 | -0.2344715 | 0.21849983 | -1.0730968 | 0.28322769 | 0.82666614 |
| MTCL1 | 56.9477479 | 0.25712028 | 0.74791553 | 0.34378252 | 0.73100985 | 0.96008876 |
| NDUFV2 | 299.148565 | -0.1373286 | 0.22183714 | -0.6190516 | 0.5358824 | 0.92324352 |
| ANKRD12 | 1369.10183 | -0.3357139 | 0.1961276 | -1.7117115 | 0.08694984 | 0.60547132 |
| TWSG1 | 604.958536 | 0.35713175 | 0.34368823 | 1.03911544 | 0.29875105 | 0.83556453 |
| AP005432.2 | 57.5286614 | 2.18280114 | 1.01036468 | 2.1604092 | 0.03074101 | 0.41259458 |
| RALBP1 | 1593.3708 | 0.65559234 | 0.34629457 | 1.89316381 | 0.05833609 | 0.53192434 |
| PPP4R1 | 342.700512 | -0.3867181 | 0.3252228 | -1.1890867 | 0.23440555 | 0.79543722 |
| RAB31 | 61.8180248 | 0.3028359 | 0.43680723 | 0.69329414 | 0.48812498 | 0.90730741 |
| VAPA | 530.060498 | -0.164201 | 0.21900162 | -0.7497707 | 0.45339279 | 0.89544675 |
| NAPG | 165.726405 | -0.3160452 | 0.27711539 | -1.1404823 | 0.25408542 | 0.80414079 |
| MPPE1 | 119.763256 | -0.289006 | 0.26608708 | -1.086133 | 0.27742017 | 0.82198569 |
| IMPA2 | 160.217242 | 0.22786215 | 0.36751198 | 0.62001284 | 0.53524934 | 0.92303275 |
| AFG3L2 | 377.148681 | -0.0646045 | 0.24190153 | -0.2670694 | 0.78941572 | 0.96515649 |
| SPIRE1 | 1580.98745 | 0.04395246 | 0.31396656 | 0.13999089 | 0.88866719 | 0.98551241 |
| PSMG2 | 260.281325 | 0.27800331 | 0.18822777 | 1.47695161 | 0.13968861 | 0.6834438 |
| CEP76 | 55.8436717 | 0.70876202 | 0.3549344 | 1.99688172 | 0.04583803 | 0.48376897 |
| PTPN2 | 339.299769 | -0.3780801 | 0.26066871 | -1.450424 | 0.14694032 | 0.69743411 |
| SEH1L | 85.6805644 | -0.3859087 | 0.25066896 | -1.5395152 | 0.12367857 | 0.66519057 |
| CEP192 | 342.429811 | -0.2424973 | 0.29888018 | -0.8113528 | 0.41716308 | 0.88242985 |
| LDLRAD4 | 440.18779 | -0.3747878 | 0.4952294 | -0.7567964 | 0.44917186 | 0.89324553 |
| FAM210A | 158.911575 | -0.752093 | 0.45120028 | -1.6668716 | 0.09553993 | 0.61367241 |
| RNMT | 345.604604 | -0.9599469 | 0.48540611 | -1.9776161 | 0.04797203 | 0.48621556 |
| ZNF519 | 43.7842932 | -0.1880216 | 0.39309621 | -0.4783093 | 0.63243011 | 0.94669061 |
| ROCK1 | 562.511948 | -0.152675 | 0.1936716 | -0.788319 | 0.43051013 | 0.88596084 |
| GREB1L | 391.175758 | -1.5621195 | 0.96732515 | -1.6148856 | 0.10633546 | 0.63461115 |
| ESCO1 | 225.266607 | -0.2200831 | 0.26220837 | -0.8393444 | 0.40127607 | 0.8781318 |
| SNRPD1 | 79.4186729 | -0.0125311 | 0.39015581 | -0.0321181 | 0.97437788 | 0.99656026 |
| ABHD3 | 192.85036 | -1.4642178 | 0.45231005 | -3.2371994 | 0.00120709 | 0.08724348 |
| MIB1 | 502.037668 | -0.4955114 | 0.26404331 | -1.8766292 | 0.06056893 | 0.54068346 |
| AC090912.1 | 76.0026976 | -0.282662 | 0.54360825 | -0.5199736 | 0.60308196 | 0.93804805 |
| RBBP8 | 1303.84118 | -0.5730698 | 0.49911991 | -1.1481606 | 0.25090229 | 0.80240987 |
| CABLES1 | 147.401249 | -0.8267717 | 0.55312223 | -1.494736 | 0.13498335 | 0.67875224 |
| TMEM241 | 106.327438 | -0.5137275 | 0.23263096 | -2.2083369 | 0.0272208 | 0.40076654 |

| | | | | | | |
|---|---|---|---|---|---|---|
| RIOK3 | 333.689046 | -0.4465246 | 0.35788691 | -1.2476696 | 0.21215208 | 0.77709975 |
| RMC1 | 76.8043362 | -0.3527773 | 0.27097068 | -1.3019022 | 0.19294983 | 0.7597012 |
| NPC1 | 243.291184 | -1.015742 | 0.34505582 | -2.9437033 | 0.00324311 | 0.14358074 |
| TTC39C | 176.224871 | -0.4625538 | 0.26963471 | -1.715483 | 0.0862567 | 0.60144393 |
| OSBPL1A | 128.036866 | -0.0133522 | 0.40574756 | -0.0329077 | 0.97374817 | 0.99656026 |
| IMPACT | 74.2791773 | 0.28201879 | 0.25706977 | 1.09705153 | 0.27261887 | 0.81922962 |
| SS18 | 397.389065 | 0.04659422 | 0.18551879 | 0.25115636 | 0.80169323 | 0.9661728 |
| TAF4B | 45.20201 | 0.30959813 | 0.28347866 | 1.0921391 | 0.27477197 | 0.8202929 |
| KCTD1 | 721.109027 | -0.6621313 | 0.30091877 | -2.2003656 | 0.02778097 | 0.40380659 |
| AQP4-AS1 | 65.0851716 | 0.28842624 | 0.35942269 | 0.80247088 | 4.22E-01 | 0.88371155 |
| CHST9 | 109.738366 | 1.98329956 | 0.56926409 | 3.48397096 | 0.00049403 | 0.05337595 |
| DSG2 | 99.8638295 | 0.60909843 | 0.47768898 | 1.27509415 | 0.20227592 | 0.76996624 |
| TRAPPC8 | 373.568583 | -0.1498784 | 0.26749331 | -0.5603071 | 0.57527002 | 0.93275827 |
| RNF125 | 65.4102782 | -1.4678183 | 0.7047193 | -2.082841 | 0.03726571 | 0.44129343 |
| RNF138 | 137.809085 | -0.3512031 | 0.35334661 | -0.9939338 | 0.32025511 | 0.84285425 |
| GAREM1 | 2695.956 | -0.0882519 | 0.42304589 | -0.2086108 | 0.83475211 | 0.97245466 |
| AC025887.2 | 58.6015972 | 0.49775157 | 0.45945783 | 1.0833455 | 0.2786551 | 0.82401394 |
| KLHL14 | 213.892342 | 0.99161735 | 0.57314618 | 1.7301299 | 0.08360707 | 0.59323093 |
| CCDC178 | 362.144145 | 0.71878107 | 0.47364193 | 1.51756216 | 0.12912481 | 0.67206108 |
| ASXL3 | 313.886898 | 0.2166455 | 0.38661437 | 0.56036587 | 0.5752299 | 0.93275827 |
| MAPRE2 | 50.827116 | -0.2649052 | 0.40830321 | -0.6487954 | 0.51647064 | 0.91751084 |
| ZNF397 | 141.178275 | -0.0443049 | 0.25397624 | -0.174445 | 0.86151577 | 0.97848095 |
| ZSCAN30 | 101.355853 | 0.23512861 | 0.24943081 | 0.94266068 | 0.34585449 | 0.8523814 |
| ZNF24 | 140.118645 | 0.16919188 | 0.20903572 | 0.8093922 | 0.41828959 | 0.88242985 |
| GALNT1 | 166.347862 | 0.05683021 | 0.24376054 | 0.2331395 | 0.81565308 | 0.96844493 |
| RPRD1A | 348.930048 | -0.2748226 | 0.25907955 | -1.0607653 | 0.28879659 | 0.83060304 |
| SLC39A6 | 366.920071 | 0.6759021 | 0.43323953 | 1.5601118 | 0.11873346 | 0.65456964 |
| ELP2 | 350.496244 | -0.3235706 | 0.34462668 | -0.9389017 | 0.34778119 | 0.85348649 |
| MOCOS | 220.438913 | -0.2387263 | 0.39686091 | -0.6015364 | 0.54748279 | 0.92661816 |
| FHOD3 | 1782.91728 | 0.66024482 | 0.38892286 | 1.69762409 | 0.08957873 | 0.60739841 |
| TPGS2 | 155.873645 | -0.4247715 | 0.29870148 | -1.4220601 | 0.15500881 | 0.70897393 |
| KIAA1328 | 465.020167 | -0.3925561 | 0.27386936 | -1.4333699 | 0.15175214 | 0.70307801 |
| AC016205.1 | 154.428684 | 0.24554314 | 0.52048215 | 0.47176093 | 0.63709744 | 0.94821838 |
| PIK3C3 | 377.985728 | -0.4625217 | 0.20489726 | -2.2573348 | 0.02398716 | 0.37077504 |
| LINC00907 | 70.0538867 | -0.2912678 | 0.49663627 | -0.5864812 | 0.55755222 | 0.92901206 |
| SETBP1 | 2144.48856 | 0.49847085 | 0.3190774 | 1.56222551 | 0.11823487 | 0.65447949 |
| SLC14A2 | 742.310479 | 0.05801057 | 1.26788068 | 0.04575397 | 0.96350635 | 0.99637506 |
| EPG5 | 234.146233 | -0.183687 | 0.27153931 | -0.6764656 | 0.49874507 | 0.90839719 |
| ATP5F1A | 424.610975 | 0.4135451 | 0.32506092 | 1.27220799 | 0.20329923 | 0.7702733 |
| HAUS1 | 33.4979641 | -0.4688644 | 0.46852093 | -1.0007331 | 0.31695588 | 0.84285425 |
| C18orf25 | 147.467985 | -0.0917573 | 0.21392243 | -0.4289279 | 0.66797573 | 0.95294349 |
| PIAS2 | 440.335013 | -0.3570985 | 0.32383717 | -1.1027102 | 0.27015305 | 0.81817548 |
| KATNAL2 | 236.809542 | 0.22877074 | 0.43560043 | 0.52518482 | 0.59945472 | 0.93728325 |
| HDHD2 | 39.6447421 | 0.18719566 | 0.35326428 | 0.5299026 | 0.59617946 | 0.93593903 |
| SMAD2 | 1078.22121 | -1.4023962 | 0.46794974 | -2.996895 | 0.00272745 | 0.13560183 |
| ZBTB7C | 94.6287785 | 0.53035782 | 0.79553501 | 0.66666811 | 0.50498415 | 0.9107831 |
| CTIF | 273.095846 | 0.42027356 | 0.33096055 | 1.26985998 | 0.20413451 | 0.77193931 |
| SMAD7 | 116.152285 | 0.55283156 | 0.30997557 | 1.78346813 | 0.07451013 | 0.57072562 |
| DYM | 787.222834 | -0.1618493 | 0.18819253 | -0.8600199 | 0.38977805 | 0.87004477 |
| LIPG | 169.218326 | -2.0464454 | 0.89743103 | -2.2803372 | 0.0225877 | 0.35959765 |
| ACAA2 | 171.68114 | 0.26348034 | 0.31204291 | 0.84437212 | 0.3984615 | 0.8754458 |
| SNHG22 | 76.700141 | 0.25173109 | 0.35435085 | 0.71040069 | 0.4774557 | 0.90242252 |
| MYO5B | 455.969277 | 0.22543366 | 0.30631571 | 0.735952 | 0.46175991 | 0.89799005 |
| CFAP53 | 35.3914547 | 0.61308299 | 0.38420655 | 1.59571198 | 0.11055311 | 0.64136059 |

| | | | | | | |
|---|---|---|---|---|---|---|
| MBD1 | 96.6437382 | -0.2748843 | 0.24094449 | -1.1408616 | 0.25392752 | 0.80414079 |
| CXXC1 | 70.7589721 | -0.2049917 | 0.31884476 | -0.6429202 | 0.52027587 | 0.91751084 |
| MAPK4 | 184.168175 | -0.1825723 | 0.51794561 | -0.3524933 | 0.72446836 | 0.95963742 |
| ME2 | 106.86988 | 0.06972825 | 0.22981445 | 0.30341107 | 0.76157661 | 0.96218738 |
| AC091551.1 | 49.6901828 | 0.26850161 | 0.35490739 | 0.75653993 | 0.44932554 | 0.89324553 |
| SMAD4 | 184.535979 | -0.1428128 | 0.23259376 | -0.6140009 | 0.53921476 | 0.92366307 |
| MEX3C | 51.7871238 | 0.28610589 | 0.25717446 | 1.11249726 | 0.26592441 | 0.81226691 |
| MBD2 | 446.707977 | -0.3222812 | 0.23455533 | -1.3740093 | 0.1694388 | 0.72596839 |
| POLI | 164.127506 | 0.21244866 | 0.27052571 | 0.78531781 | 0.43226726 | 0.88707611 |
| TCF4 | 260.795714 | -0.9607984 | 0.49049517 | -1.9588336 | 0.05013228 | 0.49590526 |
| TXNL1 | 173.700846 | -0.0321865 | 0.19919986 | -0.1615791 | 0.87163735 | 0.98209302 |
| WDR7 | 399.018196 | -0.1706876 | 0.21862732 | -0.7807242 | 0.43496473 | 0.88785137 |
| FECH | 64.7286007 | 0.20152945 | 0.32398839 | 0.62202675 | 0.53392428 | 0.92289732 |
| NARS | 247.55551 | -0.0228609 | 0.29814669 | -0.0766768 | 0.93888066 | 0.99243654 |
| AC027097.2 | 152.527745 | 0.03060656 | 0.44460458 | 0.06883995 | 0.94511702 | 0.99276891 |
| ATP8B1 | 1204.37303 | 0.60116712 | 0.49729743 | 1.20886834 | 0.22671343 | 0.78990116 |
| NEDD4L | 2661.7568 | 0.07231758 | 0.30037599 | 0.24075684 | 8.10E-01 | 0.9679755 |
| MALT1 | 319.263847 | -0.0644508 | 0.4460583 | -0.1444897 | 0.88511379 | 0.98450356 |
| AC104365.1 | 50.2882625 | 0.49542572 | 0.60285628 | 0.8217974 | 0.41119221 | 0.88078476 |
| ZNF532 | 282.62227 | -0.2269025 | 0.43429104 | -0.5224664 | 0.60134565 | 0.93802476 |
| SEC11C | 61.3492345 | 0.21956683 | 0.36678022 | 0.59863323 | 0.54941749 | 0.9274143 |
| LMAN1 | 314.504632 | 0.09289858 | 0.32686913 | 0.28420727 | 0.77625154 | 0.96437053 |
| RNF152 | 253.678287 | -1.1015522 | 0.55744054 | -1.9760891 | 0.04814469 | 0.48661187 |
| PIGN | 344.263644 | -0.0731673 | 0.18356008 | -0.3986013 | 0.69018701 | 0.95659154 |
| KIAA1468 | 530.799048 | -0.3005072 | 0.22379623 | -1.3427715 | 0.17934597 | 0.74085367 |
| ZCCHC2 | 228.531933 | -0.2082284 | 0.35338695 | -0.5892362 | 0.55570282 | 0.92872652 |
| PHLPP1 | 383.249464 | 0.27801035 | 0.21198777 | 1.31144524 | 0.18970738 | 0.75619145 |
| BCL2 | 811.460611 | -0.1534049 | 0.30063235 | -0.5102742 | 0.60985936 | 0.94003018 |
| KDSR | 148.030857 | -0.1231619 | 0.29820224 | -0.4130147 | 0.67959582 | 0.95574492 |
| VPS4B | 211.511881 | -0.0178983 | 0.18012756 | -0.0993647 | 0.92084868 | 0.9904507 |
| LINC01924 | 519.698105 | -1.5106606 | 0.73437235 | -2.0570771 | 0.03967881 | 0.45799149 |
| TMX3 | 190.013374 | -0.4168395 | 0.23999816 | -1.7368446 | 0.0824146 | 0.59202053 |
| CCDC102B | 87.0095581 | 0.51284 | 0.40124028 | 1.27813689 | 0.20120116 | 0.76957251 |
| RTTN | 181.881882 | -0.5858801 | 0.35359802 | -1.6569099 | 0.0975377 | 0.61685067 |
| SOCS6 | 104.130037 | 0.02241696 | 0.30410872 | 0.07371363 | 0.94123825 | 0.99265578 |
| FBXO15 | 36.3154841 | 0.54679986 | 0.49079103 | 1.1141195 | 0.26522792 | 0.81163409 |
| TIMM21 | 44.0645838 | -0.1737171 | 0.26230642 | -0.6622678 | 0.50779963 | 0.91214298 |
| CYB5A | 1159.06398 | 1.09297451 | 0.43514466 | 2.51174979 | 0.01201342 | 0.26883805 |
| CNDP2 | 496.373131 | -0.0218002 | 0.25806286 | -0.0844762 | 0.93267783 | 0.99243654 |
| LINC00909 | 42.423181 | 0.21308252 | 0.37494185 | 0.56830818 | 0.56982572 | 0.93122667 |
| ZNF407 | 649.589187 | -0.4577137 | 0.2303386 | -1.9871344 | 0.04690751 | 0.48600281 |
| ZADH2 | 79.942865 | 0.54623795 | 0.23882065 | 2.28723083 | 0.02218235 | 0.35843654 |
| TSHZ1 | 68.820972 | 0.96395773 | 0.35598954 | 2.707826 | 0.00677255 | 0.20224502 |
| ZNF516 | 449.569489 | -0.5728201 | 0.34680697 | -1.6516972 | 0.0985963 | 0.61707826 |
| ZNF236-DT | 45.9185892 | 0.7049158 | 0.4971805 | 1.41782671 | 0.15624136 | 0.7109898 |
| ZNF236 | 316.319908 | 0.29183897 | 0.47373974 | 0.61603228 | 0.53787321 | 0.92324352 |
| MBP | 962.377646 | 0.12452318 | 0.34777235 | 0.35805946 | 0.72029882 | 0.95796325 |
| ATP9B | 602.439696 | -0.1335614 | 0.20609087 | -0.6480706 | 0.51693928 | 0.91751084 |
| NFATC1 | 78.479445 | 0.5514402 | 0.38405651 | 1.43583092 | 0.15105046 | 0.7029344 |
| CTDP1 | 84.1105281 | -0.722075 | 0.2904191 | -2.4863206 | 0.01290716 | 0.27852752 |
| PQLC1 | 110.224906 | -0.0665346 | 0.19653493 | -0.3385384 | 0.73495747 | 0.96078945 |
| HSBP1L1 | 140.786489 | 0.25067697 | 0.38147839 | 0.65711971 | 0.51110395 | 0.91390025 |
| TXNL4A | 186.486905 | 0.02230253 | 0.35227569 | 0.06330986 | 0.94951976 | 0.99361356 |
| ADNP2 | 146.550033 | 0.04046531 | 0.30796656 | 0.13139514 | 0.89546273 | 0.98597539 |

| | | | | | | |
|---|---|---|---|---|---|---|
| C20orf96 | 62.0332208 | 0.30387883 | 0.3006967 | 1.01058255 | 0.31221627 | 0.84279957 |
| ZCCHC3 | 54.2673113 | 0.45807955 | 0.27683832 | 1.6546826 | 0.0979889 | 0.61685067 |
| NRSN2-AS1 | 193.070032 | -0.1379132 | 0.40202609 | -0.3430454 | 0.73156428 | 0.96008876 |
| SOX12 | 67.2993008 | 0.57386105 | 0.40132646 | 1.42991082 | 0.15274262 | 0.70579481 |
| NRSN2 | 62.9033982 | 0.06897685 | 0.32337016 | 0.21330618 | 0.83108815 | 0.97160502 |
| TRIB3 | 35.7749199 | 0.03633252 | 0.37647082 | 0.09650819 | 0.92311697 | 0.99051393 |
| RBCK1 | 215.953821 | 0.03195423 | 0.26313036 | 0.12143879 | 0.90334349 | 0.9875066 |
| TBC1D20 | 94.198619 | 0.16280708 | 0.23482158 | 0.69332249 | 0.48810719 | 0.90730741 |
| CSNK2A1 | 522.16283 | 0.18411456 | 0.20450863 | 0.90027774 | 0.36797247 | 0.85902833 |
| SLC52A3 | 210.811225 | 0.09796773 | 0.39985694 | 0.24500696 | 0.80645104 | 0.96752828 |
| PSMF1 | 221.783616 | -0.1757563 | 0.19416872 | -0.9051731 | 0.36537367 | 0.85866868 |
| AL136531.2 | 94.6131337 | -0.2678534 | 0.38749478 | -0.6912439 | 0.48941226 | 0.90730741 |
| SDCBP2-AS1 | 152.41371 | 0.13912675 | 0.29202087 | 0.47642742 | 0.63376991 | 0.94749053 |
| NSFL1C | 130.838003 | -0.0263948 | 0.21859773 | -0.120746 | 0.90389223 | 0.9875066 |
| STK35 | 70.2785644 | -0.0927107 | 0.24790693 | -0.3739739 | 0.70842375 | 0.95752169 |
| SNRPB | 164.728898 | 0.13503772 | 0.46796164 | 0.28856579 | 0.77291368 | 0.96388926 |
| NOP56 | 55.4280587 | 0.0606089 | 0.33577998 | 0.18050181 | 0.85675863 | 0.97718353 |
| IDH3B | 66.0862937 | 0.104713 | 0.3476688 | 0.30118608 | 0.7632726 | 0.96285313 |
| EBF4 | 43.3242377 | 1.06044445 | 0.40705144 | 2.60518534 | 0.00918246 | 0.23300339 |
| PCED1A | 54.0911085 | 0.17037111 | 0.28766592 | 0.59225338 | 0.55368093 | 0.92872652 |
| PTPRA | 522.65348 | -0.1190048 | 0.18926811 | -0.6287631 | 0.52950419 | 0.92164157 |
| MRPS26 | 161.264859 | 0.67154413 | 0.52130897 | 1.28818833 | 0.19768041 | 0.76534786 |
| UBOX5-AS1 | 49.4781526 | 0.53803665 | 0.51564855 | 1.04341736 | 0.29675504 | 0.83536682 |
| UBOX5 | 67.9893243 | 0.55723136 | 0.37607072 | 1.48171961 | 0.13841492 | 0.68146162 |
| LZTS3 | 88.4086576 | -0.0007483 | 0.2960419 | -0.0025276 | 0.9979833 | 0.99967893 |
| DDRGK1 | 118.161136 | 0.10408236 | 0.39547913 | 0.2631804 | 0.79241154 | 0.9653598 |
| ITPA | 124.58247 | 0.00514528 | 0.35517411 | 0.01448664 | 0.98844174 | 0.99814557 |
| C20orf194 | 1234.11826 | -0.0944496 | 0.19857794 | -0.4756301 | 0.63433791 | 0.94775679 |
| ATRN | 492.131826 | -0.2349135 | 0.23763319 | -0.9885553 | 0.32288078 | 0.84285425 |
| CENPB | 109.706087 | -0.114567 | 0.35341618 | -0.3241703 | 0.74580914 | 0.96109466 |
| CDC25B | 38.6893008 | 0.45262731 | 0.34134655 | 1.32600521 | 0.18483797 | 0.7484938 |
| MAVS | 203.936537 | -0.1004192 | 0.23776771 | -0.4223418 | 0.67277559 | 0.9544944 |
| PANK2 | 284.555633 | -0.0173182 | 0.25829939 | -0.0670472 | 0.94654415 | 0.99307065 |
| RNF24 | 928.073311 | -0.5840765 | 0.23472729 | -2.4883196 | 0.01283483 | 0.27834029 |
| SMOX | 40.4531253 | -1.2266753 | 0.6127398 | -2.0019514 | 0.04528996 | 0.48339878 |
| PRNP | 84.2169301 | -0.1782403 | 0.38460705 | -0.4634348 | 0.64305275 | 0.94949493 |
| SLC23A2 | 450.575469 | -0.1201949 | 0.27130633 | -0.4430228 | 0.65774925 | 0.95100227 |
| TMEM230 | 252.892907 | 0.02177824 | 0.28319128 | 0.07690293 | 0.93870077 | 0.99243654 |
| PCNA | 86.1692971 | 0.47649538 | 0.39694082 | 1.20041919 | 0.22997658 | 0.79047378 |
| CDS2 | 332.979887 | -0.5491388 | 0.28591368 | -1.9206454 | 0.05477643 | 0.51597783 |
| GPCPD1 | 756.273777 | -0.7053853 | 0.52666071 | -1.3393542 | 0.18045538 | 0.74280139 |
| SHLD1 | 142.300135 | -0.2227096 | 0.30603054 | -0.7277365 | 0.46677492 | 0.89883965 |
| CHGB | 315.287755 | -0.7813435 | 1.28478547 | -0.608151 | 0.54308734 | 0.92482794 |
| TRMT6 | 82.2769137 | 0.17807407 | 0.404422 | 0.44031747 | 0.65970719 | 0.95127957 |
| MCM8 | 78.4111755 | -0.0365076 | 0.25339532 | -0.1440735 | 0.88544242 | 0.9845556 |
| CRLS1 | 242.571089 | 0.09903648 | 0.3334287 | 0.29702445 | 0.76644784 | 0.96285313 |
| TMX4 | 155.123914 | 0.05327426 | 0.33234098 | 0.16030001 | 0.87264475 | 0.98241669 |
| PLCB1 | 130.303616 | -0.2333555 | 0.5845092 | -0.3992333 | 0.68972134 | 0.95650121 |
| PLCB4 | 169.385072 | 0.86029894 | 0.53715917 | 1.60157172 | 0.10925035 | 0.63854503 |
| ANKEF1 | 40.7076403 | -0.071983 | 0.25078751 | -0.2870279 | 0.77409094 | 0.96388926 |
| MKKS | 58.0556183 | 0.45610569 | 0.45755806 | 0.99682581 | 0.31884907 | 0.84285425 |
| SLX4IP | 399.849817 | 0.07784412 | 0.30948508 | 0.25152785 | 0.80140603 | 0.9661728 |
| JAG1 | 177.017493 | 1.00619431 | 0.31195955 | 3.22539997 | 0.00125797 | 0.08907772 |
| AL050403.2 | 99.5230541 | 0.09390525 | 0.43039283 | 0.21818498 | 0.82728499 | 0.97069946 |

| | | | | | | |
|---|---|---|---|---|---|---|
| BTBD3 | 186.371483 | 0.07104914 | 0.21181953 | 0.33542298 | 0.73730603 | 0.96078945 |
| SPTLC3 | 244.120525 | 0.06274881 | 0.34186957 | 0.18354605 | 0.85436959 | 0.97718353 |
| TASP1 | 633.827127 | -0.0770266 | 0.17951859 | -0.429073 | 0.6678701 | 0.95294349 |
| ESF1 | 196.271173 | -0.2949935 | 0.23964818 | -1.2309442 | 0.21834374 | 0.78525821 |
| NDUFAF5 | 107.863676 | -0.1568478 | 0.36807198 | -0.4261334 | 0.67001063 | 0.95351867 |
| MACROD2 | 1716.94239 | 0.26947849 | 0.16631363 | 1.62030311 | 0.10516718 | 0.6330569 |
| FLRT3 | 66.3045266 | -0.7058167 | 0.50268169 | -1.4041026 | 0.1602883 | 0.71409437 |
| KIF16B | 572.142837 | -0.0482611 | 0.40998255 | -0.1177151 | 0.90629343 | 0.9875066 |
| SNRPB2 | 100.052253 | 0.3514012 | 0.25182202 | 1.39543475 | 0.16288478 | 0.71543692 |
| DSTN | 1070.72382 | 0.80807862 | 0.31692809 | 2.54972228 | 0.01078088 | 0.25256429 |
| RRBP1 | 265.122876 | 0.22602339 | 0.24581771 | 0.91947558 | 0.35784687 | 0.8557799 |
| SNX5 | 172.944731 | 0.72350164 | 0.23369618 | 3.09590698 | 0.00196212 | 0.1105538 |
| OVOL2 | 127.582564 | -0.3183579 | 0.41174099 | -0.7731995 | 0.43940434 | 0.88917548 |
| MGME1 | 130.199845 | 0.51543255 | 0.26509628 | 1.94432206 | 0.05185663 | 0.50460301 |
| KAT14 | 45.6235634 | 0.16267627 | 0.39911009 | 0.40759748 | 0.68356921 | 0.95574492 |
| ZNF133 | 154.939174 | 0.37540345 | 0.25603859 | 1.46619872 | 0.14259416 | 0.6890058 |
| DZANK1 | 110.886695 | -0.1223344 | 0.26348225 | -0.4642984 | 0.64243396 | 0.94949493 |
| POLR3F | 38.4233936 | 0.0200719 | 0.33877587 | 0.05924832 | 0.95275433 | 0.99490488 |
| RBBP9 | 51.3606493 | 0.48877757 | 0.32609028 | 1.4989026 | 0.1338989 | 0.67757629 |
| SEC23B | 173.435858 | -0.0045414 | 0.2291652 | -0.0198172 | 0.98418917 | 0.99749593 |
| DTD1 | 66.9381166 | 0.11538078 | 0.3605605 | 0.32000394 | 0.74896534 | 0.96109466 |
| SLC24A3 | 98.951366 | -0.4266875 | 1.01393693 | -0.4208226 | 0.67388465 | 0.95501165 |
| AL121761.1 | 142.015122 | 0.15948108 | 0.45901998 | 0.34743821 | 0.72826213 | 0.95985088 |
| RIN2 | 425.190147 | 0.20985944 | 0.23269918 | 0.90184863 | 0.36713728 | 0.85902833 |
| NAA20 | 271.209317 | 0.74472711 | 0.31510996 | 2.36338805 | 0.0181087 | 0.32385519 |
| CRNKL1 | 67.9303505 | 0.26240129 | 0.26134297 | 1.00404955 | 0.31535473 | 0.84285425 |
| CFAP61 | 39.0542355 | 0.52434283 | 0.32897596 | 1.59386367 | 0.11096657 | 0.64214781 |
| RALGAPA2 | 878.224843 | -0.1773422 | 0.27390689 | -0.6474544 | 0.51733788 | 0.91751084 |
| KIZ | 800.228981 | 1.03510177 | 0.47787045 | 2.16607194 | 0.03030569 | 0.41080114 |
| KIZ-AS1 | 191.094177 | 1.13585616 | 0.55378623 | 2.05107331 | 0.04025981 | 0.4606144 |
| XRN2 | 412.577631 | -0.3494743 | 0.20828449 | -1.67787 | 0.09337248 | 0.61317397 |
| NXT1 | 35.4005482 | -0.1675853 | 0.52534768 | -0.3189988 | 0.74972742 | 0.96109466 |
| GZF1 | 59.0660798 | -0.4174547 | 0.80407167 | -0.519176 | 0.603638 | 0.93804805 |
| NAPB | 173.00583 | -0.2098636 | 0.24934926 | -0.8416451 | 0.39998662 | 0.87661803 |
| CST3 | 618.001326 | 0.42142438 | 0.37814137 | 1.11446252 | 0.26508082 | 0.81163409 |
| AL035661.1 | 52.4474397 | 0.8750109 | 0.52758688 | 1.65851527 | 0.09721351 | 0.61662137 |
| APMAP | 136.300331 | 0.44263643 | 0.33143025 | 1.33553418 | 0.18170158 | 0.74315969 |
| ACSS1 | 168.732674 | 0.15510371 | 0.29019648 | 0.53447827 | 0.59301068 | 0.93491607 |
| ENTPD6 | 97.9138011 | -0.1278145 | 0.24036932 | -0.5317423 | 0.59490447 | 0.935202 |
| PYGB | 133.364896 | 0.21025113 | 0.32898413 | 0.63909203 | 0.52276307 | 0.91845051 |
| ABHD12 | 227.192776 | -0.1258221 | 0.28434231 | -0.4425022 | 0.65812585 | 0.95113765 |
| NINL | 390.534571 | 0.10535536 | 0.38072363 | 0.27672399 | 0.78199205 | 0.96513523 |
| ZNF337-AS1 | 79.7540579 | 0.26098489 | 0.39325717 | 0.66364941 | 0.50691473 | 0.91163608 |
| ZNF337 | 101.101702 | 0.01925753 | 0.29429406 | 0.06543636 | 0.94782657 | 0.99314907 |
| FAM182B | 126.2486 | 0.50435445 | 0.3393657 | 1.486168 | 0.13723469 | 0.68129775 |
| HM13 | 248.872611 | 0.02321538 | 0.20006988 | 0.11603634 | 0.90762374 | 0.9875066 |
| ID1 | 1215.32897 | 0.4242227 | 0.77221815 | 0.54935603 | 0.58276115 | 0.93412921 |
| BCL2L1 | 345.215872 | -0.0243199 | 0.35617723 | -0.0682803 | 0.94556249 | 0.99299085 |
| AL117381.1 | 58.7054991 | -0.1068105 | 0.57016611 | -0.1873323 | 0.85140009 | 0.97718353 |
| DUSP15 | 74.6735588 | 0.90886009 | 0.59488784 | 1.52778394 | 0.12656619 | 0.6692299 |
| TTLL9 | 35.3784016 | 0.13457385 | 0.46118923 | 0.29179747 | 0.77044148 | 0.96352463 |
| PDRG1 | 42.3351619 | -0.0164197 | 0.33288684 | -0.0493252 | 0.96066017 | 0.99635733 |
| TM9SF4 | 202.774198 | -0.2991773 | 0.20739133 | -1.4425739 | 0.14914055 | 0.70215315 |
| POFUT1 | 96.7533813 | 0.41233243 | 0.26036161 | 1.58369139 | 0.11326396 | 0.64757575 |

| | | | | | | |
|---|---|---|---|---|---|---|
| KIF3B | 335.494697 | 0.45794067 | 0.1911073 | 2.39624899 | 0.01656383 | 0.30723893 |
| ASXL1 | 561.269783 | 0.03326135 | 0.2544724 | 0.1307071 | 0.89600702 | 0.98597539 |
| NOL4L | 183.393799 | 0.46186158 | 0.3627523 | 1.27321475 | 0.20294185 | 0.77003279 |
| MAPRE1 | 156.055903 | 0.04169877 | 0.3551862 | 0.11739976 | 0.90654327 | 0.9875066 |
| AL035071.2 | 35.1451816 | -0.4531201 | 0.45920529 | -0.9867485 | 0.32376596 | 0.84285425 |
| AL035071.1 | 32.9942211 | -0.6520288 | 0.39517227 | -1.6499863 | 0.09894573 | 0.61707826 |
| CDK5RAP1 | 112.256274 | -0.1211007 | 0.23836961 | -0.5080374 | 0.61142709 | 0.94003018 |
| CBFA2T2 | 711.467334 | -0.2210838 | 0.27382416 | -0.8073935 | 0.41943982 | 0.88278039 |
| PXMP4 | 49.8367675 | 0.78011369 | 0.38989163 | 2.00084748 | 0.04540883 | 0.48339878 |
| CHMP4B | 197.138182 | -0.1662743 | 0.29249087 | -0.5684769 | 0.56971121 | 0.93122667 |
| RALY | 247.540695 | -0.0574779 | 0.29677871 | -0.1936726 | 0.84643227 | 0.97656375 |
| EIF2S2 | 255.470081 | 0.16804835 | 0.36271325 | 0.46330909 | 0.64314283 | 0.94949493 |
| AHCY | 65.2656887 | 0.28949775 | 0.35031674 | 0.82638856 | 0.40858371 | 0.87948803 |
| ITCH | 701.920078 | -0.077585 | 0.2160623 | -0.3590862 | 0.71953059 | 0.95796325 |
| DYNLRB1 | 432.391182 | 0.27313777 | 0.41099953 | 0.66456956 | 0.50632584 | 0.91163608 |
| MAP1LC3A | 123.073972 | 0.88151518 | 0.49240399 | 1.79022755 | 0.07341734 | 0.56723762 |
| PIGU | 188.770743 | -0.1356282 | 0.24533514 | -0.5528284 | 0.58038094 | 0.93345492 |
| NCOA6 | 435.897277 | -0.0229254 | 0.23989582 | -0.095564 | 0.92386682 | 0.99051393 |
| TP53INP2 | 96.384839 | -0.3475546 | 0.53086519 | -0.6546947 | 0.51266433 | 0.9146446 |
| GGT7 | 146.504022 | 0.4768301 | 0.18887463 | 2.52458526 | 0.01158349 | 0.26383866 |
| ACSS2 | 150.103192 | -0.0855844 | 0.39323563 | -0.2176415 | 0.82770841 | 0.9709407 |
| GSS | 77.7780833 | 0.51852898 | 0.35996017 | 1.44051766 | 0.149721 | 0.7029344 |
| TRPC4AP | 461.85497 | -0.4003231 | 0.1880571 | -2.1287316 | 0.03327647 | 0.42373931 |
| EDEM2 | 85.2385596 | 0.07891158 | 0.26388595 | 0.29903668 | 0.76491206 | 0.96285313 |
| PROCR | 55.6688629 | 0.16419025 | 0.4657623 | 0.3525194 | 0.72444877 | 0.95963742 |
| MMP24OS | 140.739708 | -0.218451 | 0.25457498 | -0.8581009 | 0.39083675 | 0.87004477 |
| MMP24 | 106.840936 | 0.34112879 | 0.38940088 | 0.876035 | 0.38101101 | 0.86593382 |
| UQCC1 | 253.410866 | -0.3406156 | 0.21720856 | -1.5681498 | 0.11684618 | 0.65289383 |
| CEP250 | 65.5851633 | -0.4211062 | 0.2889817 | -1.4572073 | 0.14505917 | 0.69321483 |
| ERGIC3 | 143.90442 | -0.0886102 | 0.27477314 | -0.3224849 | 0.7470854 | 0.96109466 |
| CPNE1 | 123.409779 | 0.06850003 | 0.31647105 | 0.2164496 | 0.82863731 | 0.97117144 |
| ROMO1 | 152.208573 | -0.3246288 | 0.50502997 | -0.6427912 | 0.52035962 | 0.91751084 |
| RBM39 | 915.473265 | -0.0503679 | 0.15664136 | -0.3215491 | 0.74779433 | 0.96109466 |
| PHF20 | 457.522113 | -0.2064394 | 0.2514797 | -0.8208986 | 0.411704 | 0.88086809 |
| SCAND1 | 229.147071 | 0.15748486 | 0.43327414 | 0.36347626 | 0.71624915 | 0.95791799 |
| NORAD | 608.672782 | -0.138838 | 0.15160284 | -0.915801 | 0.35977126 | 0.85608464 |
| EPB41L1 | 1045.11239 | 0.40704269 | 0.34246386 | 1.18857122 | 0.23460844 | 0.79543722 |
| AAR2 | 51.4794892 | 0.22403293 | 0.25235353 | 0.88777412 | 0.37466227 | 0.86295837 |
| DLGAP4 | 224.811239 | -0.3223962 | 0.22786883 | -1.4148324 | 0.15711765 | 0.71160548 |
| DLGAP4-AS1 | 142.88053 | 0.30204079 | 0.28320498 | 1.06650944 | 0.28619341 | 0.8291345 |
| MYL9 | 98.6822381 | 0.73354489 | 0.35213303 | 2.08314706 | 0.03723782 | 0.44129343 |
| RAB5IF | 79.7960732 | -0.152077 | 0.51980312 | -0.2925666 | 0.76985348 | 0.96345263 |
| NDRG3 | 90.1587583 | -0.0409158 | 0.34720428 | -0.1178436 | 0.90619159 | 0.9875066 |
| SOGA1 | 230.729709 | 0.02181525 | 0.38239762 | 0.05704861 | 0.95450647 | 0.99537617 |
| SAMHD1 | 58.5972613 | -0.9119021 | 0.63702191 | -1.4315083 | 0.1522846 | 0.70429257 |
| RBL1 | 64.6564638 | 0.42606683 | 0.32951789 | 1.29300058 | 0.19601086 | 0.764209 |
| RPN2 | 244.480756 | 0.02402939 | 0.25389667 | 0.0946424 | 0.92459887 | 0.99051393 |
| MANBAL | 84.7255886 | -0.0915167 | 0.26854869 | -0.3407827 | 0.7332672 | 0.96035038 |
| SRC | 121.097719 | 0.08224614 | 0.24598886 | 0.33434906 | 0.73811618 | 0.96078945 |
| BLCAP | 235.802334 | 0.09315612 | 0.23748935 | 0.39225388 | 0.69487064 | 0.95672887 |
| CTNNBL1 | 191.087308 | 0.14241579 | 0.24267594 | 0.58685581 | 0.55730055 | 0.92898199 |
| TTI1 | 90.4289504 | -0.2735346 | 0.30300119 | -0.9027509 | 0.36665812 | 0.85902833 |
| RPRD1B | 179.760358 | -0.1080655 | 0.22114187 | -0.4886706 | 0.62507491 | 0.9448702 |
| TGM2 | 68.4941402 | -0.782328 | 0.59701225 | -1.3104053 | 0.19005877 | 0.75619436 |

| | | | | | | |
|---|---|---|---|---|---|---|
| RALGAPB | 397.769371 | -0.1346113 | 0.17377356 | -0.7746364 | 0.43855451 | 0.88864101 |
| DHX35 | 88.3010857 | -0.4032462 | 0.32174748 | -1.2533003 | 0.21009646 | 0.77551367 |
| TOP1 | 834.695407 | -0.2937852 | 0.35142657 | -0.835979 | 0.40316669 | 0.87834687 |
| PLCG1 | 89.4558494 | -0.2251831 | 0.29943316 | -0.7520313 | 0.45203224 | 0.89473986 |
| ZHX3 | 567.181717 | 0.49907474 | 0.3631939 | 1.37412751 | 0.1694021 | 0.72596839 |
| LPIN3 | 101.29379 | -0.001836 | 0.26041857 | -0.0070502 | 0.99437478 | 0.99919666 |
| CHD6 | 621.747212 | -0.0152566 | 0.23183532 | -0.0658079 | 0.94753079 | 0.99314907 |
| SRSF6 | 148.445761 | -0.0675947 | 0.35263934 | -0.1916823 | 0.84799111 | 0.97656375 |
| L3MBTL1 | 76.8371631 | 0.55180388 | 0.29223683 | 1.88820788 | 0.05899805 | 0.53286516 |
| SGK2 | 215.27606 | 0.02931289 | 0.35786032 | 0.08191155 | 0.93471705 | 0.99243654 |
| IFT52 | 138.981876 | 0.03068541 | 0.26843831 | 0.11431083 | 0.90899139 | 0.98768583 |
| OSER1 | 74.7358527 | 0.31038918 | 0.44222771 | 0.70187637 | 0.48275627 | 0.90614725 |
| FITM2 | 32.5806751 | 0.07041199 | 0.33920317 | 0.20758059 | 0.83555646 | 0.97274291 |
| TTPAL | 48.6633398 | 0.0532915 | 0.35599083 | 0.14969907 | 0.88100204 | 0.98333507 |
| SERINC3 | 159.277531 | -0.0926615 | 0.32178871 | -0.2879576 | 0.77337922 | 0.96388926 |
| PKIG | 285.969329 | 0.00139934 | 0.18525272 | 0.00755368 | 0.9939731 | 0.99919666 |
| YWHAB | 697.523445 | 0.2211001 | 0.3603038 | 0.61364911 | 0.53944723 | 0.92391027 |
| PABPC1L | 84.4699116 | 0.03655779 | 0.26152667 | 0.13978608 | 0.88882901 | 0.98551241 |
| TOMM34 | 46.2947012 | 0.26908825 | 0.27815237 | 0.96741309 | 0.33333757 | 0.84442294 |
| STK4 | 199.886467 | -0.4360944 | 0.17383429 | -2.508679 | 0.01211835 | 0.26883805 |
| SLPI | 2334.70845 | 1.51405389 | 1.2980049 | 1.16644697 | 0.24343378 | 0.80072691 |
| SDC4 | 644.189925 | 0.52305334 | 0.45600955 | 1.14702278 | 0.25137221 | 0.80240987 |
| PIGT | 112.763575 | 0.40665245 | 0.35917623 | 1.13218084 | 0.25755841 | 0.80709747 |
| WFDC2 | 6791.59647 | 1.12573197 | 0.42609511 | 2.64197344 | 0.00824245 | 0.22277119 |
| DNTTIP1 | 58.1457439 | 0.01858759 | 0.30356528 | 0.06123094 | 0.95117529 | 0.99404743 |
| SNX21 | 60.0430657 | 0.73899753 | 0.40820369 | 1.81036464 | 0.07023926 | 0.5621116 |
| ACOT8 | 45.4908403 | -0.0232891 | 0.27012916 | -0.0862147 | 0.93129571 | 0.99243654 |
| CTSA | 161.473596 | 0.0171767 | 0.33992214 | 0.05053128 | 0.95969902 | 0.99609409 |
| ZNF335 | 62.0306526 | -0.413592 | 0.2387771 | -1.732126 | 0.08325114 | 0.59231889 |
| NCOA5 | 101.489667 | -0.148641 | 0.28327786 | -0.524718 | 0.59977922 | 0.93728325 |
| CD40 | 71.6893032 | 0.24280077 | 0.53763624 | 0.45160789 | 0.65155148 | 0.95061388 |
| SLC35C2 | 79.1831012 | 0.07817446 | 0.27069954 | 0.28878684 | 0.77274451 | 0.96388926 |
| ELMO2 | 83.3027083 | -0.020185 | 0.30451648 | -0.0662854 | 0.94715064 | 0.99314907 |
| ZNF334 | 43.1901822 | -0.1957389 | 0.57826957 | -0.3384908 | 0.73499339 | 0.96078945 |
| SLC13A3 | 65.1119499 | -0.719421 | 0.64479208 | -1.1157411 | 0.26453297 | 0.81108996 |
| ZMYND8 | 976.926235 | 0.03788147 | 0.37295093 | 0.10157225 | 0.9190962 | 0.99014946 |
| NCOA3 | 477.140443 | -0.3192441 | 0.24095789 | -1.3248958 | 0.18520572 | 0.74895057 |
| ARFGEF2 | 556.534808 | -0.2458051 | 0.17544915 | -1.4010048 | 0.16121264 | 0.71409437 |
| CSE1L | 193.342346 | -0.1724422 | 0.21170591 | -0.8145367 | 0.41533756 | 0.88242985 |
| STAU1 | 397.460777 | 0.06614995 | 0.20024938 | 0.33033785 | 0.7411447 | 0.96084411 |
| DDX27 | 93.6194354 | -0.045763 | 0.42945662 | -0.1065603 | 0.91513785 | 0.98868699 |
| ZNFX1 | 140.46099 | -0.4389733 | 0.47933339 | -0.9157996 | 0.35977201 | 0.85608464 |
| ZFAS1 | 145.996879 | -0.3445582 | 0.22231833 | -1.5498417 | 0.12117951 | 0.65972015 |
| B4GALT5 | 460.099313 | 0.03937618 | 0.28394978 | 0.13867305 | 0.88970852 | 0.98581358 |
| SLC9A8 | 168.974179 | -0.1022535 | 0.2385738 | -0.4286032 | 0.66821205 | 0.95294349 |
| RNF114 | 317.089285 | 0.08825284 | 0.31153142 | 0.28328713 | 0.77695674 | 0.96454202 |
| CEBPB | 123.662788 | 0.31555127 | 0.61861157 | 0.51009597 | 6.10E-01 | 0.94003018 |
| PTPN1 | 280.274899 | -0.1686487 | 0.23718579 | -0.7110404 | 0.47705921 | 0.90242252 |
| RIPOR3 | 61.6393003 | 1.79729418 | 0.56251077 | 3.19512845 | 0.00139769 | 0.09557897 |
| PARD6B | 764.625983 | 0.8719773 | 0.4154995 | 2.09862417 | 0.03585004 | 0.4362094 |
| BCAS4 | 111.141348 | -0.5728839 | 0.33603596 | -1.704829 | 0.08822631 | 0.60651125 |
| ADNP | 757.864146 | -0.148118 | 0.28497803 | -0.5197524 | 0.60323617 | 0.93804805 |
| DPM1 | 170.807908 | -0.1414308 | 0.21974857 | -0.6436028 | 0.51983304 | 0.91751084 |
| ATP9A | 298.081573 | -0.0220765 | 0.25106479 | -0.0879317 | 0.92993098 | 0.99216065 |

| | | | | | | |
|---|---|---|---|---|---|---|
| ZFP64 | 108.318364 | 0.03078763 | 0.39184854 | 0.07857023 | 0.93737447 | 0.99243654 |
| ZNF217 | 261.170622 | 0.35617831 | 0.34821552 | 1.02286742 | 0.30637054 | 0.8372264 |
| PFDN4 | 53.0014081 | -0.0089427 | 0.36060917 | -0.0247989 | 0.98021533 | 0.99696835 |
| DOK5 | 115.424493 | 1.11421444 | 0.56773546 | 1.96255918 | 0.04969742 | 0.49414511 |
| FAM210B | 79.2892651 | 0.39333046 | 0.29074768 | 1.35282408 | 0.17611184 | 0.73444173 |
| CSTF1 | 42.011984 | 0.16490627 | 0.3546082 | 0.46503795 | 0.64190428 | 0.94935885 |
| RTF2 | 94.6092178 | -0.2022008 | 0.21856649 | -0.9251226 | 0.35490213 | 0.8557799 |
| RAE1 | 54.5971836 | 0.2839438 | 0.25893954 | 1.09656408 | 0.272832 | 0.81922962 |
| PCK1 | 178.34614 | -1.2772744 | 1.26839009 | -1.0070044 | 0.31393264 | 0.84285425 |
| PMEPA1 | 101.700994 | 0.48738152 | 0.73926848 | 0.6592754 | 0.50971894 | 0.91334187 |
| RAB22A | 188.839785 | -0.243232 | 0.24219239 | -1.0042926 | 0.31523761 | 0.84285425 |
| VAPB | 268.613873 | 0.00331928 | 0.17888887 | 0.01855496 | 0.98519614 | 0.99762072 |
| APCDD1L | 112.847528 | 0.59736186 | 0.58194023 | 1.02650036 | 0.30465579 | 0.83709431 |
| NPEPL1 | 94.4183509 | 0.23398707 | 0.24222198 | 0.96600266 | 0.33404285 | 0.8445213 |
| GNAS | 1019.84681 | 0.18205748 | 0.26180897 | 0.69538287 | 0.48681539 | 0.9070379 |
| NELFCD | 123.728506 | 0.10965705 | 0.19528134 | 0.5615337 | 0.57443376 | 0.93275827 |
| CTSZ | 132.534707 | 0.23579966 | 0.32083616 | 0.73495349 | 0.46236782 | 0.89801718 |
| ATP5F1E | 543.092586 | 0.03654239 | 0.45769492 | 0.07984005 | 0.93636447 | 0.99243654 |
| PRELID3B | 64.4485073 | 0.44557517 | 0.36973904 | 1.20510717 | 0.22816193 | 0.79042788 |
| FAM217B | 31.9054984 | 0.12656044 | 0.4088204 | 0.30957468 | 0.75688441 | 0.96133693 |
| MIR646HG | 198.044832 | 1.79359483 | 0.95698419 | 1.87421573 | 0.0609007 | 0.5418629 |
| TAF4 | 125.894127 | 0.08498104 | 0.26657496 | 0.31878852 | 0.74988688 | 0.96109466 |
| LSM14B | 40.6400833 | 0.0807465 | 0.31372335 | 0.25738124 | 0.79688449 | 0.9653598 |
| PSMA7 | 333.950501 | 0.23676382 | 0.46075437 | 0.51386126 | 0.60734899 | 0.93948597 |
| SS18L1 | 101.57399 | -0.411451 | 0.2124214 | -1.9369565 | 0.05275066 | 0.50904874 |
| MTG2 | 107.285465 | 0.37631836 | 0.29684062 | 1.2677455 | 0.20488885 | 0.77193931 |
| OSBPL2 | 203.759642 | -0.1474088 | 0.17300648 | -0.852042 | 0.39419078 | 0.87253562 |
| ADRM1 | 46.1934825 | -0.1919597 | 0.42980632 | -0.4466191 | 0.65515009 | 0.95100227 |
| LAMA5 | 227.778986 | 0.02610431 | 0.32534859 | 0.08023488 | 0.93605045 | 0.99243654 |
| RPS21 | 49.7247394 | -0.5821727 | 0.278212 | -2.0925507 | 0.03638929 | 0.43729328 |
| SLCO4A1 | 177.88349 | -2.0487795 | 1.18322904 | -1.7315156 | 0.08335985 | 0.59231889 |
| NTSR1 | 42.4217955 | -0.993973 | 0.53575391 | -1.8552791 | 0.06355638 | 0.54850785 |
| MRGBP | 46.3358005 | -0.2937104 | 0.31830445 | -0.9227341 | 0.35614578 | 0.8557799 |
| OGFR | 119.789309 | -0.3830618 | 0.32973246 | -1.1617352 | 0.24534303 | 0.80072691 |
| TCFL5 | 76.8386274 | 0.00047777 | 0.32471864 | 0.00147135 | 0.99882604 | 0.99968413 |
| DIDO1 | 298.352878 | -0.1692689 | 0.14447065 | -1.171649 | 0.241338 | 0.80072691 |
| GID8 | 132.18796 | 0.15048118 | 0.22655292 | 0.66422086 | 0.50654896 | 0.91163608 |
| YTHDF1 | 96.7378227 | 0.11747773 | 0.23370244 | 0.5026808 | 0.61518871 | 0.94169251 |
| ARFGAP1 | 89.3558488 | -0.5821064 | 0.32005365 | -1.8187777 | 0.06894535 | 0.55917298 |
| PPDPF | 941.663333 | -0.2637157 | 0.44526667 | -0.5922646 | 0.55367343 | 0.92872652 |
| GMEB2 | 66.8778559 | -0.614042 | 0.22977269 | -2.6723889 | 0.00753133 | 0.2144792 |
| ARFRP1 | 76.0310023 | -0.0613722 | 0.35866884 | -0.171111 | 0.86413647 | 0.97939196 |
| SLC2A4RG | 144.329158 | -0.0523779 | 0.3598835 | -0.1455414 | 0.88428347 | 0.98407543 |
| ZBTB46 | 88.9046496 | 0.82247603 | 0.27024804 | 3.0434117 | 0.00233912 | 0.12571275 |
| TPD52L2 | 131.20813 | -0.2996445 | 0.25763401 | -1.1630625 | 0.24480414 | 0.80072691 |
| DNAJC5 | 308.45155 | -0.1151796 | 0.24556388 | -0.4690413 | 0.63904008 | 0.94914116 |
| UCKL1 | 72.6346848 | -0.038349 | 0.28138786 | -0.1362852 | 0.89159585 | 0.98597539 |
| ZNF512B | 53.4986063 | 0.41495766 | 0.4105484 | 1.01073992 | 0.31214092 | 0.84279957 |
| PRPF6 | 171.093024 | 0.15136709 | 0.20296552 | 0.74577737 | 0.45580191 | 0.89626521 |
| TCEA2 | 61.135082 | 0.35294383 | 0.33452166 | 1.05507019 | 0.29139323 | 0.8320147 |
| PCMTD2 | 234.906202 | 0.52549924 | 0.26333464 | 1.99555686 | 0.04598218 | 0.48480206 |
| PLPP2 | 42.2336446 | 0.74551613 | 0.55570794 | 1.34156104 | 1.80E-01 | 0.74130581 |
| MIER2 | 44.4184983 | -0.0670268 | 0.30705476 | -0.2182893 | 0.82720369 | 0.97069946 |
| TPGS1 | 74.1577767 | 0.06346939 | 0.46841774 | 0.13549741 | 0.89221861 | 0.98597539 |

| | | | | | | |
|---|---|---|---|---|---|---|
| CDC34 | 72.3584044 | -0.3447543 | 0.2963862 | -1.1631927 | 0.24475132 | 0.80072691 |
| BSG | 408.367711 | 0.12167321 | 0.36622657 | 0.3322348 | 0.73971197 | 0.96078945 |
| RNF126 | 66.2485765 | -0.6243284 | 0.28092025 | -2.2224398 | 0.0262536 | 0.39193403 |
| PTBP1 | 224.852949 | -0.1287433 | 0.27987261 | -0.4600069 | 0.64551127 | 0.94973439 |
| MED16 | 93.0463996 | -0.3513832 | 0.31419684 | -1.1183536 | 0.26341598 | 0.80987722 |
| R3HDM4 | 80.5975558 | -0.4066936 | 0.32795464 | -1.2400909 | 0.21494176 | 0.77890374 |
| ARID3A | 305.940393 | -0.3392282 | 0.47900177 | -0.7081982 | 0.47882216 | 0.90285287 |
| WDR18 | 53.98206 | 0.05684495 | 0.40267245 | 0.14116921 | 0.88773627 | 0.98551241 |
| TMEM259 | 193.6131 | -0.2034185 | 0.28283935 | -0.7192017 | 0.47201667 | 0.90186595 |
| CNN2 | 61.6716184 | 0.82679632 | 0.44489806 | 1.85839497 | 0.06311295 | 0.54798979 |
| ABCA7 | 34.5967597 | 0.63124281 | 0.43613753 | 1.44734806 | 0.14779946 | 0.69834914 |
| POLR2E | 131.409707 | 0.14371193 | 0.39145935 | 0.36711839 | 0.71353071 | 0.95791799 |
| GPX4 | 587.685435 | -0.3662387 | 0.45275101 | -0.8089185 | 0.41856204 | 0.88242985 |
| SBNO2 | 191.262112 | -0.7514217 | 0.3563545 | -2.1086354 | 0.03497607 | 0.43353051 |
| STK11 | 160.244359 | -0.0889975 | 0.21058198 | -0.4226263 | 0.67256793 | 0.9544944 |
| ATP5F1D | 481.686315 | 0.23317438 | 0.48509105 | 0.48068168 | 0.63074275 | 0.94596555 |
| MIDN | 574.993342 | -1.0109544 | 0.71034398 | -1.4231899 | 0.15468111 | 0.70839173 |
| CIRBP | 564.974638 | 0.69052221 | 0.27524035 | 2.50879718 | 0.0121143 | 0.26883805 |
| C19orf24 | 85.6833985 | -0.1492807 | 0.37698684 | -0.3959839 | 0.69211694 | 0.95672887 |
| MUM1 | 117.330299 | -0.1477176 | 0.22629812 | -0.6527566 | 0.51391322 | 0.91544604 |
| NDUFS7 | 129.984867 | 0.10057686 | 0.43786842 | 0.22969654 | 0.81832758 | 0.96883031 |
| GAMT | 51.2149778 | 1.03169273 | 0.56922278 | 1.8124586 | 0.06991537 | 0.56060676 |
| DAZAP1 | 213.389155 | -0.202062 | 0.22809212 | -0.885879 | 0.37568271 | 0.86357183 |
| RPS15 | 1061.01694 | -0.019257 | 0.46218428 | -0.0416652 | 0.96676559 | 0.99637506 |
| C19orf25 | 108.526249 | 0.00771595 | 0.22816776 | 0.03381703 | 0.97302306 | 0.99656026 |
| MBD3 | 83.9225815 | -0.4934869 | 0.39483871 | -1.2498443 | 0.21135643 | 0.7767562 |
| TCF3 | 69.1137384 | -0.4111202 | 0.40506065 | -1.0149596 | 0.31012509 | 0.84213522 |
| REXO1 | 54.6582796 | -0.4266688 | 0.30477666 | -1.3999391 | 0.16153156 | 0.71416435 |
| ABHD17A | 107.952386 | -0.2784422 | 0.36595088 | -0.7608732 | 0.44673283 | 0.8918683 |
| SCAMP4 | 169.57804 | 0.39192155 | 0.25732466 | 1.52306257 | 0.12774305 | 0.67059156 |
| CSNK1G2 | 149.159554 | -0.2718602 | 0.29326554 | -0.9270103 | 0.35392118 | 0.8557799 |
| BTBD2 | 131.545577 | -0.1678312 | 0.34022144 | -0.4933 | 0.62180061 | 0.94418789 |
| MKNK2 | 146.403758 | -0.750696 | 0.36622739 | -2.0498083 | 0.04038314 | 0.46152157 |
| MOB3A | 82.6925469 | -0.4394913 | 0.33355481 | -1.3175985 | 0.1876381 | 0.7526909 |
| AP3D1 | 205.010965 | -0.2331378 | 0.21032996 | -1.1084384 | 0.26767252 | 0.81546744 |
| DOT1L | 139.685005 | -0.4030408 | 0.51393496 | -0.7842252 | 0.43290796 | 0.88722778 |
| PLEKHJ1 | 185.506039 | -0.0573612 | 0.36387577 | -0.1576395 | 0.87474088 | 0.98266529 |
| SF3A2 | 33.0275086 | -0.282926 | 0.2993542 | -0.9451212 | 0.34459702 | 0.85148675 |
| LSM7 | 196.91554 | 0.30407319 | 0.36989105 | 0.82206151 | 0.41104189 | 0.88078476 |
| SPPL2B | 145.804628 | -0.15939 | 0.23167693 | -0.687984 | 0.49146284 | 0.90730741 |
| TIMM13 | 155.524023 | 0.36044207 | 0.4444986 | 0.81089585 | 0.41742549 | 0.88242985 |
| LMNB2 | 36.7854901 | -0.0329559 | 0.44694627 | -0.0737357 | 0.94122072 | 0.99265578 |
| GADD45B | 309.396575 | -0.8887546 | 0.56329716 | -1.5777721 | 0.11461797 | 0.65008305 |
| GNG7 | 245.044747 | 0.21168602 | 0.39808873 | 0.53175587 | 0.59489509 | 0.935202 |
| SGTA | 94.0185142 | -0.1996958 | 0.19709797 | -1.0131803 | 0.31097403 | 0.84248332 |
| THOP1 | 44.0671287 | -0.4745025 | 0.39802777 | -1.1921343 | 0.2332086 | 0.7926597 |
| ZNF555 | 44.8963569 | 0.2008897 | 0.44549684 | 0.45093407 | 0.65203707 | 0.95079106 |
| TLE2 | 288.340321 | -0.0725055 | 0.40845034 | -0.1775137 | 0.85910487 | 0.97788846 |
| AES | 656.039932 | -0.2013448 | 0.29149728 | -0.690726 | 0.48973772 | 0.90730741 |
| GNA11 | 164.024053 | 0.09257536 | 0.30138903 | 0.30716235 | 0.7587198 | 0.96180522 |
| NCLN | 76.0238381 | 0.09634594 | 0.2704256 | 0.35627523 | 0.72163446 | 0.95868461 |
| NFIC | 617.941706 | -0.0052484 | 0.2339807 | -0.0224309 | 0.98210423 | 0.99713749 |
| SMIM24 | 57.9904972 | 0.32451799 | 0.46414118 | 0.69917947 | 0.48443988 | 0.90659463 |
| FZR1 | 72.245423 | -0.1722146 | 0.22400489 | -0.7687983 | 0.44201305 | 0.88997055 |

| | | | | | | |
|---|---|---|---|---|---|---|
| MFSD12 | 135.349756 | -1.5242544 | 0.31093131 | -4.9022223 | 9.48E-07 | 0.00070934 |
| HMG20B | 90.7488399 | -0.2847075 | 0.31972363 | -0.89048 | 0.37320819 | 0.86201082 |
| PIP5K1C | 88.3374865 | -0.2538475 | 0.27507891 | -0.9228171 | 0.35610252 | 0.8557799 |
| TJP3 | 144.351751 | -0.3059036 | 0.44428114 | -0.6885361 | 0.49111528 | 0.90730741 |
| MRPL54 | 102.627238 | -0.0128277 | 0.50285317 | -0.0255099 | 0.97964829 | 0.99696835 |
| DAPK3 | 54.2523804 | -0.5023855 | 0.30341744 | -1.6557568 | 0.09777108 | 0.61685067 |
| EEF2 | 765.589234 | 0.20126531 | 0.36982309 | 0.54422052 | 0.58628973 | 0.93483536 |
| PIAS4 | 57.6902661 | -0.1848876 | 0.29187672 | -0.6334442 | 0.52644358 | 0.91974278 |
| ZBTB7A | 197.660188 | 0.24839214 | 0.35674742 | 0.69626891 | 0.48626044 | 0.90695174 |
| MAP2K2 | 192.205356 | 0.07256347 | 0.29428168 | 0.24657828 | 0.80523461 | 0.96704419 |
| SIRT6 | 33.8948093 | -0.2837868 | 0.30127218 | -0.9419614 | 0.34621242 | 0.8523814 |
| YJU2 | 37.9746872 | -0.1190394 | 0.30935721 | -0.3847961 | 0.70038851 | 0.95672887 |
| STAP2 | 48.0207799 | -0.1439093 | 0.37034765 | -0.388579 | 0.69758759 | 0.95672887 |
| SH3GL1 | 174.263605 | -0.6515526 | 0.37157831 | -1.7534732 | 0.07952082 | 0.58461232 |
| UBXN6 | 145.440305 | 0.325127 | 0.38998443 | 0.83369225 | 0.40445443 | 0.87834687 |
| HDGFL2 | 86.4614461 | 0.25803076 | 0.27284545 | 0.945703 | 0.34430009 | 0.85148675 |
| PLIN5 | 59.5062394 | 0.77418823 | 0.3995799 | 1.93750544 | 0.05268359 | 0.50904874 |
| MYDGF | 150.876381 | -0.0957932 | 0.39571256 | -0.2420778 | 0.80871991 | 0.96775927 |
| DPP9 | 88.5447748 | -0.1148947 | 0.28133795 | -0.4083869 | 0.68298963 | 0.95574492 |
| PLIN3 | 93.7733943 | 0.2956008 | 0.28158943 | 1.04975814 | 0.29382932 | 0.8337535 |
| KDM4B | 258.133659 | -0.1644316 | 0.23682817 | -0.6943075 | 0.4874894 | 0.90730741 |
| PTPRS | 220.26987 | 0.10803598 | 0.16475613 | 0.65573266 | 0.51199615 | 0.91408994 |
| TINCR | 58.5170942 | 0.01758719 | 0.3411323 | 0.05155533 | 0.95888301 | 0.99609409 |
| SAFB2 | 222.261669 | -0.1691832 | 0.24059884 | -0.7031755 | 0.48194638 | 0.90595437 |
| SAFB | 268.122266 | -0.1402559 | 0.15431771 | -0.9088773 | 0.36341492 | 0.85694172 |
| RPL36 | 663.983445 | -0.3556036 | 0.40580032 | -0.8763019 | 0.38086594 | 0.86593382 |
| C19orf70 | 190.667961 | 0.19047674 | 0.3890223 | 0.48962937 | 0.62439619 | 0.94457465 |
| LONP1 | 155.198964 | -0.6630879 | 0.2429554 | -2.7292578 | 0.00634771 | 0.19542438 |
| CAPS | 331.818379 | 0.41254787 | 0.58780327 | 0.70184684 | 0.48277468 | 0.90614725 |
| RANBP3 | 97.991845 | -0.243373 | 0.21810762 | -1.1158392 | 0.26449099 | 0.81108996 |
| RFX2 | 179.433268 | -0.8992254 | 0.6150762 | -1.461974 | 0.14374833 | 0.69093999 |
| MLLT1 | 162.334159 | 0.37351495 | 0.3637933 | 1.02672299 | 0.30455092 | 0.83709431 |
| CLPP | 95.4748495 | 0.58240893 | 0.41424014 | 1.40596933 | 0.15973323 | 0.71409437 |
| ALKBH7 | 133.030598 | 0.41618782 | 0.55978347 | 0.74348 | 0.45719113 | 0.8969884 |
| GTF2F1 | 99.852908 | 0.13090646 | 0.25119767 | 0.52112926 | 0.60227673 | 0.93804805 |
| KHSRP | 121.375057 | -0.0200927 | 0.325832 | -0.0616659 | 0.95082889 | 0.99404295 |
| SLC25A23 | 127.942519 | 0.31777258 | 0.28517706 | 1.11429925 | 0.26515083 | 0.81163409 |
| CRB3 | 52.5242488 | -0.1005408 | 0.40467989 | -0.2484454 | 0.80378983 | 0.96678163 |
| AC010503.4 | 61.1446588 | -0.4323783 | 0.31132023 | -1.3888539 | 0.16487717 | 0.71715698 |
| TRIP10 | 118.261459 | -0.0585604 | 0.32135866 | -0.1822277 | 0.85540404 | 0.97718353 |
| SH2D3A | 154.404399 | -0.5171956 | 0.45616808 | -1.1337829 | 0.25688562 | 0.80676096 |
| ZNF557 | 34.1527769 | -0.2013099 | 0.33374602 | -0.6031831 | 0.54638691 | 0.92597739 |
| INSR | 859.006138 | -0.7234853 | 0.21556588 | -3.3562143 | 0.00079017 | 0.06520484 |
| PEX11G | 30.8526406 | 0.55344173 | 0.31501736 | 1.75686105 | 0.0789415 | 0.5826987 |
| ZNF358 | 69.4748036 | 0.15139752 | 0.42281578 | 0.35806971 | 0.72029115 | 0.95796325 |
| PNPLA6 | 105.619689 | 0.16646078 | 0.28226531 | 0.58973162 | 0.55537059 | 0.92872652 |
| CAMSAP3 | 81.7793093 | -0.1924313 | 0.23835743 | -0.8073223 | 0.41948084 | 0.88278039 |
| XAB2 | 41.5615416 | 0.12294891 | 0.34879433 | 0.35249688 | 0.72446565 | 0.95963742 |
| STXBP2 | 61.764148 | -0.4812493 | 0.38966908 | -1.2350205 | 0.21682289 | 0.78139747 |
| EVI5L | 51.7905372 | 0.0680806 | 0.29329202 | 0.23212564 | 0.81644043 | 0.96856416 |
| MAP2K7 | 111.118356 | -0.3683384 | 0.29611589 | -1.2438995 | 0.21353656 | 0.77869841 |
| CTXN1 | 31.7768154 | -0.3667209 | 0.66092527 | -0.5548598 | 0.57899055 | 0.93345492 |
| TIMM44 | 67.9205556 | -0.412156 | 0.23763486 | -1.734409 | 0.08284554 | 0.59202053 |
| ELAVL1 | 230.637981 | -0.0153751 | 0.21778062 | -0.0705991 | 0.94371683 | 0.99265578 |

| | | | | | | |
|---|---|---|---|---|---|---|
| CERS4 | 127.202819 | 0.22395828 | 0.3589286 | 0.62396332 | 0.53265168 | 0.92268523 |
| CD320 | 53.0095202 | 0.78977269 | 0.38842261 | 2.03328198 | 0.04202404 | 0.46886915 |
| RPS28 | 206.11585 | -0.0653207 | 0.33907838 | -0.192642 | 0.84723937 | 0.97656375 |
| ANGPTL4 | 498.450023 | -2.4806804 | 1.45311918 | -1.7071417 | 0.08779571 | 0.60651125 |
| RAB11B | 171.802889 | 0.08798263 | 0.34533211 | 0.25477685 | 0.79889546 | 0.9653598 |
| 2-Mar | 62.4949263 | -0.5765254 | 0.25293441 | -2.2793475 | 0.02264642 | 0.35959765 |
| HNRNPM | 508.849956 | -0.3085713 | 0.23507663 | -1.3126414 | 0.18930382 | 0.75619145 |
| ZNF558 | 36.331586 | -0.3552276 | 0.30024744 | -1.1831161 | 0.23676316 | 0.79797745 |
| ZNF317 | 46.5474129 | -0.233274 | 0.25537504 | -0.9134565 | 0.36100248 | 0.85653294 |
| ZNF559-ZNF | 57.8969215 | 0.78136068 | 0.33906867 | 2.30443197 | 0.02119841 | 0.35151789 |
| ZNF266 | 94.0573034 | 0.12980827 | 0.21615336 | 0.60053782 | 0.54814786 | 0.92684569 |
| ZNF426 | 63.9775832 | 0.37216186 | 0.23385319 | 1.59143373 | 0.111512 | 0.64353508 |
| ZNF121 | 102.618979 | -0.2373819 | 0.25642416 | -0.9257391 | 0.35458157 | 0.8557799 |
| ZNF561 | 64.9622467 | 0.37779657 | 0.27066857 | 1.39579032 | 0.16277764 | 0.71543692 |
| ZNF562 | 170.886391 | -0.2081036 | 0.22620147 | -0.9199921 | 0.35757687 | 0.8557799 |
| ZNF846 | 171.297409 | 0.23342119 | 0.29215236 | 0.79897074 | 0.42430738 | 0.88391318 |
| FBXL12 | 57.5117495 | 0.15057608 | 0.2371892 | 0.6348353 | 0.52553584 | 0.91946838 |
| UBL5 | 180.658814 | 0.13378535 | 0.49132724 | 0.27229378 | 0.78539614 | 0.96513523 |
| PIN1 | 108.025128 | 0.2927745 | 0.43933302 | 0.66640677 | 0.50515114 | 0.9107831 |
| C19orf66 | 137.575512 | -0.6992216 | 0.4369888 | -1.6000904 | 0.10957854 | 0.63926535 |
| EIF3G | 98.9213141 | 0.22492116 | 0.38910242 | 0.5780513 | 0.56322948 | 0.9306324 |
| DNMT1 | 64.8301968 | -0.2155921 | 0.39355514 | -0.5478066 | 0.58382468 | 0.93473293 |
| MRPL4 | 96.0690422 | 0.52364266 | 0.31922857 | 1.64033771 | 0.10093497 | 0.62077375 |
| TYK2 | 69.3573204 | -0.5777544 | 0.22639192 | -2.5520099 | 0.01071035 | 0.25223467 |
| CDC37 | 227.165814 | -0.3705419 | 0.30223474 | -1.226007 | 0.22019603 | 0.78633738 |
| PDE4A | 68.8699036 | 0.20936003 | 0.56408059 | 0.3711527 | 0.7105238 | 0.95756248 |
| KEAP1 | 57.9511695 | 0.36803657 | 0.39545529 | 0.93066544 | 0.35202665 | 0.85536253 |
| ATG4D | 59.5728649 | 0.10331891 | 0.25491322 | 0.40531011 | 0.68524957 | 0.9558749 |
| KRI1 | 53.5714029 | -0.1453782 | 0.25845027 | -0.5624998 | 0.57377556 | 0.93270589 |
| AP1M2 | 179.677942 | 0.28915861 | 0.29448055 | 0.9819277 | 0.32613547 | 0.84381657 |
| SLC44A2 | 270.283013 | -0.044712 | 0.41997855 | -0.1064625 | 0.91521539 | 0.98868699 |
| ILF3-DT | 93.9168249 | -0.0693149 | 0.42281839 | -0.1639353 | 0.86978207 | 0.98202612 |
| ILF3 | 287.275242 | -0.1859019 | 0.27532773 | -0.6752024 | 0.49954719 | 0.90839719 |
| QTRT1 | 81.4303853 | -0.0535955 | 0.26547554 | -0.2018848 | 0.84000679 | 0.9746807 |
| DNM2 | 1069.98461 | 0.16383074 | 0.39856455 | 0.41105196 | 0.68103443 | 0.95574492 |
| TMED1 | 74.7801215 | 0.16996857 | 0.3619867 | 0.46954369 | 0.63868106 | 0.94887688 |
| CARM1 | 66.6625978 | -0.0859548 | 0.32900508 | -0.2612569 | 0.79389441 | 0.9653598 |
| YIPF2 | 61.3473741 | 0.03397215 | 0.29157475 | 0.11651265 | 0.90724626 | 0.9875066 |
| TIMM29 | 53.0476405 | 0.34185657 | 0.31072017 | 1.10020721 | 0.27124185 | 0.81922962 |
| SMARCA4 | 433.645834 | 0.06219959 | 0.34335747 | 0.18115113 | 0.85624895 | 0.97718353 |
| LDLR | 193.87897 | -0.5579651 | 0.84466153 | -0.6605784 | 0.50888274 | 0.91320053 |
| KANK2 | 107.929433 | 0.070879 | 0.38877127 | 0.18231544 | 0.85533518 | 0.97718353 |
| DOCK6 | 250.752562 | -0.278562 | 0.43292639 | -0.6434397 | 0.51993885 | 0.91751084 |
| RAB3D | 97.3903784 | -0.0240533 | 0.31905987 | -0.075388 | 0.939906 | 0.99243654 |
| EPOR | 182.321885 | 0.77520649 | 0.42015944 | 1.84502933 | 0.06503327 | 0.55275644 |
| RGL3 | 780.235815 | 0.26293134 | 0.34292694 | 0.76672697 | 0.44324387 | 0.89022532 |
| CCDC151 | 37.6388413 | 0.54865925 | 0.47568528 | 1.15340809 | 0.24874292 | 0.80097144 |
| PRKCSH | 176.891035 | 0.24809534 | 0.28619933 | 0.86686205 | 0.3860176 | 0.86831175 |
| ECSIT | 94.5044761 | -0.2382534 | 0.3036833 | -0.7845456 | 0.43272002 | 0.88722778 |
| ELOF1 | 53.8087159 | -0.3047758 | 0.41127594 | -0.7410494 | 0.4586635 | 0.89753337 |
| ZNF627 | 86.0644333 | 0.07495892 | 0.39007355 | 0.19216613 | 0.84761208 | 0.97656375 |
| AC008543.1 | 42.9292633 | 0.71759067 | 0.57090136 | 1.25694337 | 0.20877414 | 0.77436433 |
| ZNF823 | 108.298804 | 0.32664217 | 0.28444744 | 1.14833929 | 0.25082853 | 0.80240987 |
| ZNF441 | 83.8086777 | 0.36415586 | 0.4066363 | 0.8955321 | 0.37050271 | 0.8613284 |

| | | | | | | |
|---|---|---|---|---|---|---|
| ZNF440 | 118.73986 | -0.6509259 | 0.36619017 | -1.7775624 | 0.07547575 | 0.57072562 |
| ZNF439 | 118.931611 | 0.22451576 | 0.4156783 | 0.54011902 | 0.58911496 | 0.93483536 |
| ZNF69 | 207.487639 | 0.11654396 | 0.4751402 | 0.24528331 | 0.80623707 | 0.96752828 |
| ZNF433 | 40.0905839 | 0.3707703 | 0.3061487 | 1.21107911 | 0.22586509 | 0.78982769 |
| ZNF844 | 77.3562755 | 0.59466792 | 0.37502699 | 1.58566701 | 0.11281487 | 0.64641869 |
| ZNF788P | 44.5671313 | 0.60285299 | 0.37258213 | 1.61804054 | 0.10565386 | 0.63429746 |
| ZNF136 | 60.7442576 | -0.1234596 | 0.25390547 | -0.4862424 | 0.62679531 | 0.94511671 |
| ZNF44 | 580.826324 | 0.17723037 | 0.22328676 | 0.79373432 | 0.42735011 | 0.88510458 |
| ZNF563 | 37.9188054 | 0.01012285 | 0.48986246 | 0.02066468 | 0.98351315 | 0.9971769 |
| AC008758.5 | 43.3065511 | -0.6807462 | 0.50455129 | -1.3492111 | 0.17726917 | 0.73633806 |
| ZNF490 | 81.0432803 | 0.00249168 | 0.37302247 | 0.00667971 | 0.9946704 | 0.99919666 |
| ZNF791 | 143.335801 | -0.2146215 | 0.19415468 | -1.1054152 | 0.26897972 | 0.81817548 |
| DHPS | 47.4051704 | 0.19726144 | 0.40162907 | 0.49115328 | 0.62331804 | 0.94456546 |
| TNPO2 | 126.206527 | -0.0002089 | 0.2651957 | -0.0007878 | 0.99937139 | 0.99984842 |
| TRIR | 292.75303 | 0.17367538 | 0.47764308 | 0.36360912 | 0.71614991 | 0.95791799 |
| ASNA1 | 82.1842504 | 0.4180274 | 0.45467903 | 0.9193901 | 0.35789156 | 0.8557799 |
| HOOK2 | 373.776022 | -0.2916349 | 0.24028043 | -1.2137273 | 0.22485189 | 0.78982769 |
| JUNB | 553.91758 | -1.1896714 | 0.89790384 | -1.3249429 | 0.18519008 | 0.74895057 |
| DNASE2 | 51.4830939 | 0.77526469 | 0.35912719 | 2.15874682 | 0.03086981 | 0.41259458 |
| GCDH | 61.8681584 | 0.23242795 | 0.34610618 | 0.67155101 | 0.50186958 | 0.90945856 |
| FARSA | 64.2548348 | 0.22412625 | 0.31951065 | 0.70146724 | 0.48301147 | 0.90628733 |
| CALR | 424.309794 | 0.27581116 | 0.3752615 | 0.73498391 | 0.46234929 | 0.89801718 |
| AC092069.1 | 77.6175592 | -0.0315005 | 0.4367364 | -0.072127 | 0.94250085 | 0.99265578 |
| RAD23A | 127.315943 | -0.2917385 | 0.33789876 | -0.8633902 | 0.38792294 | 0.86942525 |
| GADD45GIP1 | 256.197798 | 0.26416136 | 0.49759531 | 0.53087589 | 0.59550479 | 0.93566569 |
| NFIX | 83.178793 | 0.00792128 | 0.4455934 | 0.01777693 | 0.98581681 | 0.99762072 |
| NACC1 | 67.14007 | 0.05203735 | 0.30548748 | 0.17034201 | 0.86474118 | 0.97972838 |
| STX10 | 53.9018217 | -0.2113665 | 0.31684347 | -0.6671006 | 0.50470787 | 0.9107831 |
| IER2 | 429.434472 | -0.5341224 | 0.74093825 | -0.720873 | 0.47098765 | 0.90123338 |
| CCDC130 | 189.921044 | -0.0756752 | 0.27114153 | -0.2790984 | 0.78016928 | 0.96513523 |
| MRI1 | 108.175728 | 0.61782999 | 0.31910071 | 1.93615989 | 0.05284812 | 0.50911084 |
| C19orf53 | 265.409163 | 0.36278138 | 0.51047638 | 0.71067222 | 0.47728738 | 0.90242252 |
| ZSWIM4 | 68.9535811 | -0.0079086 | 0.30949284 | -0.0255535 | 0.9796135 | 0.99696835 |
| AC020916.1 | 86.0581369 | -0.7835456 | 0.7828659 | -1.0008683 | 0.31689049 | 0.84285425 |
| CC2D1A | 120.667269 | -0.1078302 | 0.23372871 | -0.4613478 | 0.64454907 | 0.94949493 |
| DCAF15 | 44.7735112 | -0.5761351 | 0.30852157 | -1.8674061 | 0.0618449 | 0.54419361 |
| RFX1 | 87.3813873 | 0.15407085 | 0.28518384 | 0.540251 | 0.58902395 | 0.93483536 |
| PALM3 | 115.083604 | 0.18081179 | 0.20112556 | 0.89899957 | 0.36865289 | 0.85931545 |
| MISP3 | 31.8516344 | 0.16633134 | 0.46752122 | 0.35577282 | 0.7220107 | 0.95889902 |
| SAMD1 | 56.1819039 | 0.25697101 | 0.40288734 | 0.6378235 | 0.52358858 | 0.91871245 |
| PRKACA | 97.2939448 | 0.20471783 | 0.2478901 | 0.82584107 | 0.40889425 | 0.87982122 |
| ADGRL1 | 104.598006 | 0.40279577 | 0.2672474 | 1.50720184 | 0.13175893 | 0.67390608 |
| DDX39A | 37.0361627 | -1.0180462 | 0.34865643 | -2.9199122 | 0.0035013 | 0.14553941 |
| PKN1 | 187.200852 | -0.077819 | 0.33888276 | -0.2296339 | 0.81837627 | 0.96883031 |
| PTGER1 | 656.749057 | 1.86001066 | 0.68231075 | 2.72604624 | 0.0064098 | 0.19641732 |
| GIPC1 | 491.973442 | 0.14547249 | 0.32122981 | 0.45286113 | 0.65064874 | 0.95061388 |
| DNAJB1 | 137.146491 | 0.36413048 | 0.37764825 | 0.9642054 | 0.33494296 | 0.84522086 |
| TECR | 133.263068 | 0.16641885 | 0.24205208 | 0.68753323 | 0.49174677 | 0.90730741 |
| NDUFB7 | 296.28826 | -0.0166655 | 0.39493054 | -0.0421986 | 0.96634037 | 0.99637506 |
| ADGRE3 | 31.5133238 | 0.15887527 | 0.35432993 | 0.44838229 | 0.65387732 | 0.95100227 |
| ZNF333 | 43.2397857 | -0.3192181 | 0.32986625 | -0.9677199 | 0.33318426 | 0.84442294 |
| ILVBL | 106.884786 | 0.08314444 | 0.28895068 | 0.28774613 | 0.77354108 | 0.96388926 |
| NOTCH3 | 170.73236 | -0.7338172 | 0.41072517 | -1.7866379 | 0.07399603 | 0.56915891 |
| BRD4 | 550.414365 | -0.1753243 | 0.19477077 | -0.9001573 | 0.36803655 | 0.85902833 |

| | | | | | | |
|---|---|---|---|---|---|---|
| AKAP8 | 139.84579 | -0.1583346 | 0.18729336 | -0.8453829 | 0.39789707 | 0.87462201 |
| AKAP8L | 373.827475 | -0.0556152 | 0.28014227 | -0.1985247 | 0.84263455 | 0.97560439 |
| WIZ | 47.447396 | 0.07461151 | 0.28805659 | 0.25901685 | 0.79562224 | 0.9653598 |
| TPM4 | 1061.9397 | 0.29796128 | 0.55113774 | 0.54062943 | 0.58876303 | 0.93483536 |
| RAB8A | 93.8120328 | -0.0275783 | 0.29622156 | -0.0931004 | 0.9258238 | 0.99051393 |
| FAM32A | 70.6286326 | 0.15699421 | 0.45556961 | 0.34461081 | 0.73038698 | 0.96008876 |
| AP1M1 | 55.50099 | -0.1248389 | 0.232209 | -0.5376142 | 0.5908434 | 0.93483536 |
| EPS15L1 | 173.284308 | -0.3364946 | 0.28809102 | -1.1680148 | 0.24280078 | 0.80072691 |
| C19orf44 | 62.442834 | 0.24404943 | 0.30558578 | 0.79862823 | 0.42450601 | 0.88391318 |
| TMEM38A | 39.7433699 | -0.1621912 | 0.42436366 | -0.3821985 | 0.70231413 | 0.95672887 |
| SIN3B | 132.254395 | -0.2102741 | 0.21689013 | -0.9694958 | 0.33229785 | 0.84442294 |
| CPAMD8 | 391.58682 | 1.02427511 | 0.62354696 | 1.64265914 | 0.10045348 | 0.61911626 |
| MYO9B | 209.03692 | -0.3596269 | 0.28166213 | -1.2768025 | 0.201672 | 0.76957251 |
| USE1 | 37.2554688 | -0.195398 | 0.44397853 | -0.4401069 | 0.65985967 | 0.95127957 |
| OCEL1 | 41.822116 | -0.1138893 | 0.43917323 | -0.2593266 | 0.79538326 | 0.9653598 |
| NR2F6 | 135.777163 | 0.29817417 | 0.37667388 | 0.79159767 | 0.4285953 | 0.88535099 |
| ABHD8 | 50.50281 | 0.52894718 | 0.34732871 | 1.5229008 | 0.12778352 | 0.67059156 |
| MRPL34 | 141.428931 | 0.63124534 | 0.43206073 | 1.46101067 | 0.14401251 | 0.69093999 |
| DDA1 | 50.8010178 | 0.04387756 | 0.36078444 | 0.12161712 | 0.90320226 | 0.9875066 |
| BST2 | 112.242384 | -2.238639 | 1.02671118 | -2.180398 | 0.02922797 | 0.40623517 |
| MVB12A | 98.5947995 | 0.17344388 | 0.32916445 | 0.52692167 | 0.59824799 | 0.93728325 |
| SLC27A1 | 63.7853262 | 0.53331818 | 0.33299063 | 1.60160118 | 0.10924383 | 0.63854503 |
| AC010618.3 | 61.2844691 | -0.1544602 | 0.36593003 | -0.422103 | 0.67294986 | 0.9544944 |
| PGLS | 163.829346 | 0.11789767 | 0.41488689 | 0.28416822 | 0.77628147 | 0.96437053 |
| COLGALT1 | 49.7204291 | -0.9557798 | 0.32631236 | -2.9290333 | 0.00340018 | 0.14358074 |
| RPL18A | 60.5826442 | -0.3184276 | 0.53479805 | -0.5954164 | 0.55156513 | 0.92872652 |
| CCDC124 | 78.4675552 | 0.17099953 | 0.47880391 | 0.35713897 | 0.72098777 | 0.95839064 |
| ARRDC2 | 46.7100074 | -0.9062465 | 0.33173695 | -2.7318227 | 0.0062985 | 0.19471476 |
| MAST3 | 47.2132053 | 0.08151103 | 0.34911138 | 0.23348145 | 0.81538757 | 0.96844493 |
| IQCN | 50.2602984 | 0.34906025 | 0.60451032 | 0.57742644 | 0.56365142 | 0.93077674 |
| JUND | 1059.24472 | -1.8776977 | 0.54314226 | -3.4571011 | 0.00054602 | 0.0578009 |
| LSM4 | 271.190071 | -0.0969203 | 0.43412298 | -0.2232555 | 0.82333666 | 0.97018919 |
| PGPEP1 | 204.129667 | -0.4217635 | 0.26035341 | -1.6199653 | 0.10523973 | 0.63312997 |
| GDF15 | 1591.01861 | -1.8363895 | 0.68531293 | -2.6796365 | 0.00737022 | 0.21103786 |
| SSBP4 | 186.484475 | 0.20751278 | 0.26700611 | 0.77718366 | 0.43705042 | 0.88785137 |
| ISYNA1 | 75.2781587 | 0.29546465 | 0.65143398 | 0.45356039 | 0.65014526 | 0.95061388 |
| ELL | 148.019923 | 0.23141011 | 0.25603678 | 0.90381588 | 0.36609304 | 0.85902833 |
| FKBP8 | 204.306119 | 0.05994352 | 0.32546489 | 0.18417815 | 0.8538737 | 0.97718353 |
| KXD1 | 76.6464239 | -0.5075998 | 0.24828999 | -2.0443828 | 0.04091575 | 0.46430138 |
| UBA52 | 357.955943 | -0.1198598 | 0.44211825 | -0.2711036 | 0.78631137 | 0.96513523 |
| REX1BD | 94.5575026 | 0.14681857 | 0.44684937 | 0.32856391 | 0.74248533 | 0.96086911 |
| CRTC1 | 350.728885 | 0.00917003 | 0.34016945 | 0.02695725 | 0.97849383 | 0.99696835 |
| UPF1 | 164.308099 | 0.26480406 | 0.24691613 | 1.07244537 | 0.28352004 | 0.82685108 |
| COPE | 127.589227 | -0.2045377 | 0.4226058 | -0.4839917 | 0.62839175 | 0.9457914 |
| DDX49 | 48.2324309 | -0.2452921 | 0.30313465 | -0.8091853 | 0.41840855 | 0.88242985 |
| SUGP2 | 316.51293 | -0.403313 | 0.16125795 | -2.5010427 | 0.01238282 | 0.27205868 |
| SLC25A42 | 59.4907342 | 0.47615474 | 0.27432574 | 1.73572753 | 0.08261202 | 0.59202053 |
| TMEM161A | 50.6562461 | 0.42124342 | 0.30041427 | 1.40220846 | 0.16085301 | 0.71409437 |
| RFXANK | 50.1283359 | 0.04333681 | 0.33010337 | 0.13128256 | 0.89555179 | 0.98597539 |
| SUGP1 | 66.7277579 | 0.02995787 | 0.29522471 | 0.1014748 | 0.91917356 | 0.99014946 |
| MAU2 | 181.542581 | -0.1720799 | 0.21161071 | -0.8131907 | 0.41610871 | 0.88242985 |
| GATAD2A | 398.574386 | -0.1274036 | 0.1586248 | -0.803176 | 0.42187301 | 0.88371155 |
| PBX4 | 64.6214595 | 0.14642737 | 0.29349185 | 0.49891462 | 0.61783953 | 0.94244117 |
| ATP13A1 | 60.6412705 | 0.20251916 | 0.29897132 | 0.67738658 | 0.49816071 | 0.90839719 |

| | | | | | | |
|---|---|---|---|---|---|---|
| ZNF14 | 87.1045286 | 0.29893181 | 0.32572709 | 0.91773704 | 0.35875655 | 0.8557799 |
| AC011477.4 | 35.8987937 | -0.2066186 | 0.37894468 | -0.5452474 | 0.5855834 | 0.93483536 |
| ZNF253 | 93.8157612 | 0.18111047 | 0.34165214 | 0.53010196 | 0.59604124 | 0.93593903 |
| ZNF93 | 55.6738144 | -0.2918895 | 0.35722602 | -0.8171004 | 0.41387106 | 0.88211689 |
| ZNF682 | 85.8410635 | 0.01971984 | 0.47966873 | 0.04111136 | 0.96720711 | 0.99637506 |
| ZNF90 | 368.985419 | 0.24312831 | 0.31538207 | 0.77090085 | 0.4407657 | 0.88956997 |
| AC011447.3 | 281.762512 | 0.46587544 | 0.50207342 | 0.92790302 | 0.35345788 | 0.8557799 |
| ZNF486 | 230.744654 | 0.49906036 | 0.48117698 | 1.03716591 | 0.29965854 | 0.83618229 |
| AC008554.1 | 253.143424 | 1.48489119 | 0.51701842 | 2.87202762 | 0.00407847 | 0.15545856 |
| ZNF737 | 127.967858 | 0.25809898 | 0.32038401 | 0.80559257 | 0.42047781 | 0.88320382 |
| AC010636.2 | 58.5818706 | -0.5155886 | 0.42391478 | -1.2162553 | 0.2238877 | 0.78960314 |
| AC010329.1 | 43.0868877 | 0.10475613 | 0.42300798 | 0.24764575 | 0.80440851 | 0.96679546 |
| ZNF66 | 46.3957365 | -0.3594144 | 0.32802729 | -1.0956846 | 0.27321682 | 0.81922962 |
| ZNF85 | 50.0869781 | 0.13699201 | 0.55062416 | 0.24879405 | 0.8035201 | 0.96678163 |
| ZNF430 | 148.729107 | -0.2427112 | 0.21572097 | -1.1251164 | 0.26053972 | 0.80805101 |
| ZNF714 | 77.9180324 | -0.5353806 | 0.30242748 | -1.7702776 | 0.07668091 | 0.57260711 |
| ZNF431 | 270.90396 | 0.19453006 | 0.2509774 | 0.77508993 | 0.43828652 | 0.88844154 |
| ZNF708 | 177.311737 | 0.2369278 | 0.29111479 | 0.81386384 | 0.41572296 | 0.88242985 |
| ZNF738 | 50.3513358 | -0.3234074 | 0.40048146 | -0.8075465 | 0.4193517 | 0.88278039 |
| ZNF493 | 82.6309207 | 0.54991384 | 0.29269617 | 1.87878725 | 0.06027355 | 0.53896484 |
| AC010615.4 | 153.06871 | 0.95881161 | 0.67732834 | 1.41557876 | 0.15689887 | 0.71151025 |
| ZNF429 | 221.210309 | 0.31691243 | 0.31982961 | 0.99087897 | 0.32174468 | 0.84285425 |
| AC123912.1 | 67.0575637 | 0.36139047 | 0.57560293 | 0.62784683 | 0.53010428 | 0.92181514 |
| ZNF100 | 201.416543 | 0.43596723 | 0.51901518 | 0.83998936 | 0.40091435 | 0.87775198 |
| ZNF43 | 306.188032 | 0.06221309 | 0.30610326 | 0.20324218 | 0.83894575 | 0.97430756 |
| ZNF208 | 612.562079 | 0.32587406 | 0.34763619 | 0.9373997 | 0.34855299 | 0.8535343 |
| ZNF257 | 38.0725541 | 0.08213852 | 0.31712031 | 0.25901374 | 0.79562464 | 0.9653598 |
| ZNF676 | 110.618827 | 0.37638772 | 0.41118416 | 0.91537503 | 0.35999478 | 0.85608464 |
| ZNF98 | 415.289861 | 0.10976905 | 0.35747115 | 0.30707108 | 0.75878927 | 0.96180522 |
| ZNF99 | 46.1054249 | 0.67687587 | 0.50885167 | 1.3302027 | 0.1834515 | 0.74504275 |
| ZNF728 | 42.4913045 | 0.23302426 | 0.54656126 | 0.4263461 | 0.66985567 | 0.9534276 |
| ZNF730 | 69.1943551 | -0.1757755 | 0.60568762 | -0.2902082 | 0.77165695 | 0.96388926 |
| ZNF91 | 782.145089 | -0.4619711 | 0.20030888 | -2.3062937 | 0.02109423 | 0.35034471 |
| ZNF675 | 182.356566 | 0.28866313 | 0.46591086 | 0.61956729 | 0.53554271 | 0.92303275 |
| AC139769.2 | 68.6632351 | 0.41266456 | 0.58046446 | 0.71092133 | 0.47713299 | 0.90242252 |
| ZNF726 | 41.236394 | 0.59502284 | 0.50407802 | 1.18041813 | 0.23783395 | 0.79810098 |
| ZNF254 | 457.235576 | -0.2266298 | 0.25177667 | -0.9001225 | 0.36805506 | 0.85902833 |
| LINC00662 | 139.412509 | -0.1136734 | 0.33408035 | -0.3402576 | 0.73366252 | 0.96037761 |
| AC006504.5 | 141.706986 | 1.0065229 | 0.29693203 | 3.38974174 | 0.00069958 | 0.06109709 |
| UQCRFS1 | 237.927345 | 0.72336029 | 0.44991889 | 1.60775709 | 0.10788839 | 0.63520805 |
| POP4 | 39.039043 | -0.0311186 | 0.32333381 | -0.0962429 | 0.92332764 | 0.99051393 |
| C19orf12 | 97.2889856 | 0.09907376 | 0.2398236 | 0.41311095 | 0.67952533 | 0.95574492 |
| URI1 | 627.17358 | 0.1864951 | 0.17477769 | 1.06704175 | 0.28595298 | 0.82875754 |
| ZNF507 | 120.079184 | -0.0107771 | 0.2699353 | -0.0399246 | 0.96815323 | 0.9964399 |
| DPY19L3 | 158.722639 | -0.1492243 | 0.19491151 | -0.7656001 | 0.44391427 | 0.89088885 |
| PDCD5 | 66.1803186 | -0.2786328 | 0.25000618 | -1.1145036 | 0.2650632 | 0.81163409 |
| ANKRD27 | 128.991573 | -0.3134173 | 0.24998484 | -1.2537452 | 0.20993462 | 0.77551367 |
| NUDT19 | 38.3655647 | 0.00191477 | 0.32183774 | 0.00594949 | 0.99525302 | 0.99944919 |
| CEP89 | 150.509552 | -0.0241199 | 0.22242324 | -0.1084414 | 0.9136456 | 0.98830345 |
| RHPN2 | 373.96021 | -0.552785 | 0.28324325 | -1.9516263 | 0.05098259 | 0.49973786 |
| GPATCH1 | 51.6929855 | -1.21E-05 | 0.32491604 | -3.72E-05 | 0.99997035 | 0.99997035 |
| LRP3 | 91.3126842 | -0.0632376 | 0.31995098 | -0.1976478 | 0.84332063 | 0.97560439 |
| PEPD | 122.340202 | -0.1739977 | 0.33481986 | -0.5196756 | 0.60328966 | 0.93804805 |
| LSM14A | 677.347922 | 0.06045399 | 0.21151386 | 0.28581574 | 0.77501925 | 0.96388926 |

| | | | | | | |
|---|---|---|---|---|---|---|
| KIAA0355 | 337.384632 | -0.1531235 | 0.20559295 | -0.7447898 | 0.45639881 | 0.89626521 |
| GPI | 437.482786 | -0.556451 | 0.39373517 | -1.413262 | 0.1575787 | 0.71160548 |
| UBA2 | 271.609291 | -0.3408676 | 0.21177281 | -1.609591 | 0.10748717 | 0.63477557 |
| WTIP | 66.2629696 | 1.63914798 | 0.42609684 | 3.84689075 | 0.00011963 | 0.02194494 |
| SCGB2B2 | 126.362265 | 0.06245117 | 0.41709426 | 0.14972915 | 0.88097831 | 0.98333507 |
| ZNF302 | 195.97989 | 0.01418201 | 0.27694872 | 0.05120807 | 0.95915972 | 0.99609409 |
| GRAMD1A | 105.395696 | -0.297893 | 0.27742247 | -1.0737885 | 0.28291749 | 0.82666614 |
| HPN | 243.184338 | 0.12085617 | 0.40740069 | 0.29665185 | 0.76673232 | 0.96285313 |
| FXYD3 | 59.9409219 | -0.3038468 | 0.56016609 | -0.5424228 | 0.58752731 | 0.93483536 |
| FXYD5 | 58.5556003 | 0.57954575 | 0.60391192 | 0.95965277 | 0.33723 | 0.84636904 |
| LSR | 157.956993 | -0.1037575 | 0.31702609 | -0.3272838 | 0.74345328 | 0.96105014 |
| USF2 | 127.874458 | -0.0268955 | 0.33275358 | -0.0808272 | 0.93557937 | 0.99243654 |
| DMKN | 348.596673 | 0.06064306 | 0.26713677 | 0.22701127 | 0.82041498 | 0.96886922 |
| TMEM147 | 93.0705089 | 0.05518237 | 0.44402426 | 0.12427782 | 0.90109531 | 0.9875066 |
| HAUS5 | 32.0485245 | -0.2497123 | 0.35671369 | -0.7000357 | 0.48390499 | 0.90647968 |
| RBM42 | 67.1055473 | 0.05851006 | 0.35467653 | 0.1649674 | 0.86896964 | 0.9820194 |
| COX6B1 | 457.607154 | 0.25033669 | 0.36746971 | 0.68124442 | 0.49571684 | 0.90839719 |
| KMT2B | 87.6118241 | -0.3622972 | 0.29578858 | -1.2248517 | 0.22063109 | 0.78700265 |
| PROSER3 | 51.5807342 | -0.5033212 | 0.27140256 | -1.854519 | 0.06366495 | 0.54850785 |
| SDHAF1 | 39.4083618 | -0.0663055 | 0.3022984 | -0.2193379 | 0.82638686 | 0.97058548 |
| TBCB | 106.46824 | 0.27657364 | 0.38996478 | 0.70922724 | 0.47818347 | 0.90262298 |
| CAPNS1 | 421.651705 | 0.04080873 | 0.35625859 | 0.11454806 | 0.90880335 | 0.9875839 |
| ZNF565 | 183.233826 | -0.3667816 | 0.43362687 | -0.8458461 | 0.39763861 | 0.87462201 |
| ZNF146 | 218.559734 | -0.2248198 | 0.19402097 | -1.1587397 | 0.24656232 | 0.80072691 |
| LINC00665 | 105.821064 | 0.193571 | 0.27234407 | 0.71075901 | 0.47723359 | 0.90242252 |
| ZFP14 | 162.642727 | -0.1655372 | 0.30598289 | -0.5410015 | 0.58850656 | 0.93483536 |
| ZFP82 | 67.8217699 | 0.19078426 | 0.25274219 | 0.75485721 | 0.45033466 | 0.89333031 |
| ZNF566 | 77.4043352 | -0.3014231 | 0.25866115 | -1.1653203 | 0.24388935 | 0.80072691 |
| ZNF260 | 88.9703151 | 0.03876216 | 0.19137936 | 0.20254094 | 0.83949386 | 0.97452864 |
| ZNF529 | 139.186939 | -0.2672627 | 0.2263553 | -1.1807219 | 0.23771322 | 0.79810098 |
| ZNF529-AS1 | 124.348582 | -0.2297639 | 0.28994515 | -0.7924393 | 0.42810458 | 0.88535099 |
| ZNF461 | 64.7735641 | -0.1732666 | 0.2568309 | -0.6746332 | 0.49990885 | 0.90839719 |
| ZNF567 | 57.9922017 | 0.57090961 | 0.23621383 | 2.41691859 | 0.01565252 | 0.30067202 |
| AC020928.1 | 124.668046 | 0.06705295 | 0.32959675 | 0.20343934 | 0.83879165 | 0.97430756 |
| ZNF790 | 94.2923688 | 0.42017598 | 0.33290311 | 1.26215698 | 0.20689231 | 0.77200549 |
| ZNF345 | 148.513967 | 0.042737 | 0.36043703 | 0.11856995 | 0.90561608 | 0.9875066 |
| ZNF568 | 103.571572 | 0.18506474 | 0.32055074 | 0.57733367 | 0.56371407 | 0.93077674 |
| ZNF420 | 211.33662 | 0.40448929 | 0.52747318 | 0.76684333 | 0.44317467 | 0.89022532 |
| ZNF383 | 56.456271 | 0.00140379 | 0.30304462 | 0.00463229 | 0.99630398 | 0.99967893 |
| LINC01535 | 38.2317816 | -0.2747538 | 0.33096601 | -0.830157 | 0.40645001 | 0.87948803 |
| HKR1 | 225.977894 | -0.0440079 | 0.16320365 | -0.2696499 | 0.78742959 | 0.96513523 |
| ZNF569 | 67.5802043 | 0.46390305 | 0.37791971 | 1.22751747 | 0.21962816 | 0.78630025 |
| ZNF570 | 35.1873978 | 0.25564982 | 0.28958853 | 0.88280368 | 0.37734235 | 0.86438205 |
| ZNF793 | 102.755692 | 0.27809299 | 0.29563682 | 0.94065747 | 0.34688042 | 0.85295795 |
| ZNF540 | 84.0586151 | 0.80018711 | 0.51451761 | 1.5552181 | 0.11989413 | 0.65679551 |
| ZNF571 | 93.8346608 | 0.47047483 | 0.60653641 | 0.77567452 | 0.43794119 | 0.88825695 |
| ZFP30 | 98.8303029 | 0.43116483 | 0.29911722 | 1.44145773 | 0.14945541 | 0.7029344 |
| ZNF573 | 119.44649 | 0.14496385 | 0.33984547 | 0.42655812 | 0.66970121 | 0.9533662 |
| SIPA1L3 | 1054.53761 | -0.8319061 | 0.35488757 | -2.3441398 | 0.01907102 | 0.33090117 |
| SPINT2 | 58.6866271 | 0.38067797 | 0.30618867 | 1.24327908 | 0.21376501 | 0.77890374 |
| CATSPERG | 48.8153122 | -0.2850723 | 0.42721685 | -0.6672778 | 0.50459471 | 0.9107831 |
| PSMD8 | 165.451513 | 0.19782863 | 0.35777328 | 0.55294413 | 0.58030167 | 0.93345492 |
| FAM98C | 34.7701411 | 0.07055147 | 0.37226244 | 0.18952079 | 0.84968466 | 0.97671331 |
| EIF3K | 231.163256 | 0.18606938 | 0.32197 | 0.57790907 | 0.56332551 | 0.9306324 |

| | | | | | | |
|---|---|---|---|---|---|---|
| ACTN4 | 1029.48805 | 0.09085653 | 0.24482634 | 0.37110604 | 0.71055855 | 0.95756248 |
| CAPN12 | 40.7962165 | 0.04150009 | 0.35988302 | 0.1153155 | 0.90819506 | 0.98758267 |
| ECH1 | 227.961169 | 0.64226699 | 0.50311337 | 1.27658501 | 0.2017488 | 0.76957251 |
| HNRNPL | 156.379203 | -0.305513 | 0.27039158 | -1.129891 | 0.25852217 | 0.80709747 |
| SIRT2 | 101.2619 | 0.04309135 | 0.28411027 | 0.1516712 | 0.87944627 | 0.98333507 |
| NFKBIB | 42.415857 | -0.111073 | 0.2877206 | -0.3860446 | 0.69946365 | 0.95672887 |
| MRPS12 | 110.894234 | 0.80508186 | 0.43786675 | 1.83864582 | 0.0659673 | 0.55444578 |
| FBXO17 | 209.70617 | 0.6760648 | 0.53724605 | 1.25838954 | 0.20825091 | 0.77419992 |
| PAK4 | 194.549507 | 0.1024785 | 0.33775685 | 0.3034091 | 0.76157811 | 0.96218738 |
| SAMD4B | 275.764736 | -0.1297778 | 0.23074878 | -0.5624204 | 0.57382965 | 0.93270589 |
| PAF1 | 63.079207 | 0.3517335 | 0.38828041 | 0.90587495 | 0.36500205 | 0.85866868 |
| MED29 | 89.7577594 | 0.21003042 | 0.37207835 | 0.56447902 | 0.57242817 | 0.93218128 |
| ZFP36 | 240.719757 | -1.4576049 | 0.80258776 | -1.8161315 | 0.06935021 | 0.55917298 |
| RPS16 | 366.781519 | 0.11473597 | 0.39083233 | 0.29356826 | 0.76908783 | 0.96345263 |
| SUPT5H | 206.319997 | -0.2853786 | 0.15768452 | -1.8098072 | 0.07032569 | 0.5621116 |
| TIMM50 | 46.3827037 | -0.267353 | 0.27557295 | -0.9701716 | 0.33196099 | 0.84442294 |
| DYRK1B | 41.9980207 | -0.0572372 | 0.33295847 | -0.1719048 | 0.86351236 | 0.979268 |
| FBL | 73.2063644 | -0.1977117 | 0.38099961 | -0.518929 | 0.60381025 | 0.93804805 |
| FCGBP | 42.0797023 | -0.8294605 | 0.33326225 | -2.4889122 | 0.01281346 | 0.27834029 |
| PSMC4 | 74.4143438 | 0.30938354 | 0.27455137 | 1.1268694 | 0.2597977 | 0.80719831 |
| ZNF546 | 46.3919375 | 0.56468316 | 0.41055861 | 1.37540205 | 0.16900684 | 0.72596839 |
| ZNF780B | 256.997639 | 0.42983915 | 0.25988021 | 1.65398954 | 0.09812964 | 0.61690569 |
| ZNF780A | 55.5115602 | -0.0747093 | 0.29295313 | -0.2550213 | 0.79870666 | 0.9653598 |
| MAP3K10 | 44.9796755 | -0.159876 | 0.3316449 | -0.4820697 | 0.62975645 | 0.94596555 |
| AKT2 | 298.440935 | -0.9026916 | 0.29801209 | -3.0290434 | 0.00245329 | 0.12985113 |
| PLD3 | 293.828316 | 0.10114051 | 0.24147689 | 0.41884138 | 0.67533206 | 0.95525442 |
| SERTAD1 | 58.5206011 | 0.15342401 | 0.62852621 | 0.24410122 | 0.80715243 | 0.96752828 |
| SERTAD3 | 44.9016694 | 0.07608342 | 0.35599323 | 0.21372154 | 0.83076421 | 0.97160502 |
| BLVRB | 375.600937 | -0.1092575 | 0.40547644 | -0.2694547 | 0.78757982 | 0.96513523 |
| SPTBN4 | 43.861736 | 0.26589999 | 0.59437936 | 0.44735737 | 0.65461705 | 0.95100227 |
| SHKBP1 | 44.9873645 | -0.2084765 | 0.37203414 | -0.5603694 | 0.57522752 | 0.93275827 |
| LTBP4 | 83.1646452 | -0.2679274 | 0.27013704 | -0.9918204 | 0.32128512 | 0.84285425 |
| COQ8B | 54.0026685 | 0.92126142 | 0.28329673 | 3.25193096 | 0.00114624 | 0.0834207 |
| ITPKC | 70.3533723 | -0.3383498 | 0.33444894 | -1.0116634 | 0.311699 | 0.84279403 |
| SNRPA | 72.0763436 | -0.3853958 | 0.26007575 | -1.4818599 | 0.13837758 | 0.68146162 |
| AC008537.1 | 72.4154963 | -0.7884937 | 0.44440068 | -1.7742856 | 0.07601592 | 0.57145422 |
| HNRNPUL1 | 163.257767 | 0.04244709 | 0.24280529 | 0.17481945 | 0.8612215 | 0.97841862 |
| TGFB1 | 148.866583 | 0.57676262 | 0.36581595 | 1.57664701 | 0.11487677 | 0.65008305 |
| CCDC97 | 37.8869118 | -0.4432885 | 0.50003329 | -0.886518 | 0.37533845 | 0.86357183 |
| B3GNT8 | 32.7710652 | 1.58877471 | 0.66032607 | 2.4060457 | 0.01612625 | 0.30396236 |
| DMAC2 | 45.7134174 | -0.0831782 | 0.30006322 | -0.2772024 | 0.78162472 | 0.96513523 |
| RPS19 | 1220.62199 | -0.2198403 | 0.38587635 | -0.5697169 | 0.56886974 | 0.93122667 |
| ARHGEF1 | 77.9518102 | -0.5207775 | 0.26447939 | -1.9690666 | 0.04894544 | 0.48945442 |
| RABAC1 | 262.841767 | 0.09527933 | 0.48361879 | 0.1970133 | 0.84381713 | 0.97564028 |
| GRIK5 | 132.794683 | 0.31012588 | 0.30730244 | 1.0091878 | 0.31288458 | 0.84285425 |
| DEDD2 | 42.7646228 | -0.2283983 | 0.25252954 | -0.904442 | 0.36576107 | 0.85868639 |
| GSK3A | 59.5628339 | -0.1793572 | 0.32190646 | -0.5571719 | 0.57740998 | 0.93340377 |
| CIC | 79.1402561 | -0.3395952 | 0.20866277 | -1.6274835 | 0.10363445 | 0.62743442 |
| PAFAH1B3 | 53.5067795 | -0.0573526 | 0.50837519 | -0.1128156 | 0.91017677 | 0.98788932 |
| MEGF8 | 101.214861 | 0.38548576 | 0.25674491 | 1.50143485 | 0.13324313 | 0.67594872 |
| LIPE-AS1 | 75.1331446 | 0.6599401 | 0.28389627 | 2.32458177 | 0.02009433 | 0.33933607 |
| AC004784.1 | 53.1817189 | -1.2512453 | 0.97513446 | -1.2831515 | 1.99E-01 | 0.7687013 |
| PHLDB3 | 52.0169148 | 0.623101 | 0.58066399 | 1.07308358 | 0.28323362 | 0.82666614 |
| ETHE1 | 77.2822023 | -0.5268363 | 0.33385576 | -1.5780356 | 0.11455742 | 0.65008305 |

| | | | | | | |
|---|---|---|---|---|---|---|
| XRCC1 | 33.5556646 | -0.1328922 | 0.3185369 | -0.4171957 | 0.6765353 | 0.95529325 |
| PINLYP | 34.7938626 | 0.11876494 | 0.29768423 | 0.39896282 | 0.6899206 | 0.95659154 |
| ZNF428 | 137.755164 | 0.26609389 | 0.47395943 | 0.56142756 | 0.5745061 | 0.93275827 |
| PLAUR | 72.2624294 | -3.1594494 | 0.91487677 | -3.4534153 | 0.00055354 | 0.05789336 |
| SMG9 | 48.4162625 | -0.5219747 | 0.34666335 | -1.5057107 | 0.13214145 | 0.67466347 |
| ZNF283 | 54.8006129 | -0.1389901 | 0.34621033 | -0.4014615 | 0.68808039 | 0.95597173 |
| ZNF45 | 35.5140907 | 0.08565405 | 0.38819317 | 0.22064801 | 0.82536652 | 0.97018919 |
| ZNF155 | 75.7019546 | 0.98644642 | 0.40761551 | 2.42004141 | 0.01551874 | 0.30047476 |
| ZNF224 | 89.0971454 | -0.0588952 | 0.2905566 | -0.2026977 | 0.83937131 | 0.97452864 |
| ZNF234 | 39.4007091 | 0.1508665 | 0.40656448 | 0.37107645 | 0.71058059 | 0.95756248 |
| ZNF226 | 84.0319278 | 0.25245786 | 0.25309257 | 0.9974922 | 0.31852566 | 0.84285425 |
| ZNF227 | 45.33576 | 0.50691376 | 0.29742239 | 1.70435642 | 0.08831452 | 0.60651125 |
| ZNF235 | 179.650475 | 0.43891936 | 0.39134571 | 1.12156423 | 0.26204777 | 0.80882423 |
| ZNF229 | 42.8584657 | 0.51670414 | 0.40408404 | 1.27870465 | 0.20100108 | 0.76957251 |
| AC243964.2 | 42.3610636 | -0.1443913 | 0.37257402 | -0.3875506 | 0.69834866 | 0.95672887 |
| BCL3 | 66.9891495 | -0.2605889 | 0.60937885 | -0.4276304 | 0.66892021 | 0.95298441 |
| CBLC | 75.1884272 | -0.0459601 | 0.43825199 | -0.1048713 | 0.91647794 | 0.98905249 |
| BCAM | 677.203121 | 1.04331761 | 0.43632945 | 2.3911235 | 0.0167969 | 0.30900364 |
| NECTIN2 | 249.100137 | 0.21284957 | 0.22184729 | 0.95944183 | 0.33733621 | 0.84636904 |
| TOMM40 | 62.8339641 | -0.0107516 | 0.29568167 | -0.036362 | 0.97099368 | 0.99656026 |
| APOE | 156.231858 | -1.661545 | 0.81323634 | -2.0431269 | 0.04103988 | 0.46430138 |
| CLPTM1 | 138.8901 | 0.05262947 | 0.2472801 | 0.21283342 | 0.8314569 | 0.97160502 |
| RELB | 73.1950776 | -0.5376808 | 0.63251998 | -0.8500613 | 0.39529101 | 0.8737924 |
| CLASRP | 129.288845 | -0.2784515 | 0.24547023 | -1.1343594 | 0.25664383 | 0.80676096 |
| GEMIN7-AS1 | 55.4125734 | -0.272273 | 0.32556492 | -0.8363094 | 0.40298089 | 0.87834687 |
| MARK4 | 152.697134 | 0.11312802 | 0.26760571 | 0.42274142 | 0.67248393 | 0.9544944 |
| TRAPPC6A | 109.04673 | 0.42914 | 0.36222272 | 1.18474071 | 0.23611998 | 0.79736951 |
| ERCC2 | 38.4446085 | 0.2966067 | 0.43118049 | 0.68789452 | 0.49151921 | 0.90730741 |
| PPP1R13L | 122.976751 | 0.05534136 | 0.32879453 | 0.16831593 | 0.86633474 | 0.98040284 |
| ERCC1 | 57.9423684 | 0.09897502 | 0.3495336 | 0.28316309 | 0.77705182 | 0.96454202 |
| VASP | 122.808455 | -0.2229258 | 0.39165479 | -0.5691895 | 0.56922752 | 0.93122667 |
| OPA3 | 156.975177 | -0.3111855 | 0.26955202 | -1.1544544 | 0.24831392 | 0.80097144 |
| EML2 | 122.625274 | 0.78203881 | 0.24022838 | 3.25539726 | 0.00113234 | 0.08298537 |
| SNRPD2 | 237.718801 | 0.074835 | 0.44081853 | 0.16976374 | 0.86519595 | 0.98003173 |
| FBXO46 | 54.3579945 | -0.2121434 | 0.31983215 | -0.6632959 | 0.50714106 | 0.91163608 |
| DMPK | 51.3605427 | -0.4076331 | 0.41958819 | -0.9715077 | 0.33129554 | 0.84442294 |
| SYMPK | 265.817365 | -0.3779053 | 0.22354402 | -1.6905187 | 0.09092877 | 0.60942442 |
| MYPOP | 39.0828137 | 0.24569118 | 0.38172271 | 0.64363785 | 0.5198103 | 0.91751084 |
| HIF3A | 36.4901228 | 0.97959131 | 0.508166 | 1.92769946 | 0.05389252 | 0.51332378 |
| PPP5C | 146.686228 | 0.15251191 | 0.18106735 | 0.84229379 | 0.39962353 | 0.87634538 |
| CCDC8 | 62.2015426 | 0.52119717 | 0.3823495 | 1.36314333 | 0.17283734 | 0.72940007 |
| PNMA8A | 63.8005393 | 0.69241356 | 0.48149587 | 1.43804673 | 0.1504208 | 0.7029344 |
| PPP5D1 | 80.6918457 | -0.1255193 | 0.36047298 | -0.3482071 | 0.72768465 | 0.9598317 |
| CALM3 | 216.319175 | 0.30588484 | 0.30656359 | 0.99778594 | 0.31838317 | 0.84285425 |
| PRKD2 | 130.849718 | -0.3272305 | 0.27570688 | -1.1868783 | 0.23527562 | 0.79599484 |
| STRN4 | 54.1172341 | 0.06540769 | 0.22339077 | 0.29279494 | 0.7696789 | 0.96345263 |
| FKRP | 39.2052693 | 0.21648046 | 0.38070156 | 0.5686356 | 0.56960346 | 0.93122667 |
| AP2S1 | 159.517602 | 0.198438 | 0.38527849 | 0.51505082 | 0.60651751 | 0.93919596 |
| ARHGAP35 | 466.557206 | 0.5411791 | 0.36739995 | 1.47299721 | 0.14075179 | 0.68512714 |
| TMEM160 | 183.213681 | 0.52742304 | 0.4211288 | 1.25240318 | 0.21042299 | 0.77610028 |
| ZC3H4 | 103.086907 | -0.0435854 | 0.27294521 | -0.1596857 | 0.87312869 | 0.98246494 |
| SAE1 | 107.893607 | -0.2025015 | 0.31953905 | -0.6337301 | 0.52625698 | 0.91974278 |
| INAFM1 | 35.5345541 | -0.4463905 | 0.41978539 | -1.063378 | 0.28761057 | 0.8303903 |
| C5AR1 | 144.244665 | 1.35998178 | 0.48733563 | 2.79064715 | 0.00526028 | 0.17920428 |

| | | | | | | |
|---|---|---|---|---|---|---|
| DHX34 | 62.5729392 | -0.1734472 | 0.36822368 | -0.4710375 | 0.63761395 | 0.94850165 |
| NAPA-AS1 | 50.1822568 | -0.1927695 | 0.55910999 | -0.3447793 | 0.73026031 | 0.96008876 |
| NAPA | 338.402661 | -0.1485768 | 0.25576654 | -0.5809077 | 0.56130266 | 0.92934517 |
| NOP53 | 85.0190601 | 0.10915673 | 0.32512828 | 0.33573434 | 0.73707121 | 0.96078945 |
| SELENOW | 352.719879 | -0.2339658 | 0.45692693 | -0.5120421 | 0.60862152 | 0.94003018 |
| LIG1 | 36.3971218 | 0.20207313 | 0.51130201 | 0.39521287 | 0.6926858 | 0.95672887 |
| ZSWIM9 | 60.8603112 | -0.0188472 | 0.3746758 | -0.0503028 | 0.95988113 | 0.99609409 |
| CARD8 | 96.3646595 | -0.0091457 | 0.30023673 | -0.0304618 | 0.97569876 | 0.99656026 |
| TMEM143 | 44.3611737 | -0.0724781 | 0.3642303 | -0.1989899 | 0.84227066 | 0.97560439 |
| KDELR1 | 187.755449 | 0.46386011 | 0.23141409 | 2.00445928 | 0.04502088 | 0.48262669 |
| CYTH2 | 82.4125852 | -0.4547621 | 0.43482243 | -1.045857 | 0.29562708 | 0.83418733 |
| LMTK3 | 38.8629486 | -0.9339871 | 0.45654586 | -2.0457683 | 0.04077917 | 0.46430138 |
| RPL18 | 628.362633 | 0.20207901 | 0.48059581 | 0.42047601 | 0.67413775 | 0.95511203 |
| SPHK2 | 32.3464297 | -0.1083786 | 0.37683952 | -0.2875988 | 0.7736539 | 0.96388926 |
| BCAT2 | 76.2167471 | 0.10513974 | 0.30970812 | 0.3394801 | 0.73424809 | 0.96042436 |
| HSD17B14 | 40.2580605 | 0.11774935 | 0.58308935 | 0.20194049 | 0.83996325 | 0.9746807 |
| PLEKHA4 | 49.345312 | -0.2017882 | 0.75925126 | -0.2657726 | 0.79041436 | 0.96524651 |
| PPP1R15A | 45.4773276 | -0.0523437 | 0.47663128 | -0.1098201 | 0.91255205 | 0.98830345 |
| NUCB1 | 242.457583 | 0.42012628 | 0.27012293 | 1.5553151 | 0.11987104 | 0.65679551 |
| BAX | 113.357333 | 0.3170381 | 0.40473329 | 0.78332596 | 0.43343572 | 0.88736205 |
| FTL | 3207.66535 | -0.2295247 | 0.63270314 | -0.3627684 | 0.71677787 | 0.95791799 |
| GYS1 | 140.049261 | -0.4342268 | 0.41330202 | -1.0506282 | 0.29342938 | 0.83336457 |
| RUVBL2 | 36.899945 | -0.2287157 | 0.28936993 | -0.790392 | 0.42929887 | 0.88579433 |
| SNRNP70 | 490.470956 | 0.00369737 | 0.27032598 | 0.01367745 | 0.98908732 | 0.99832756 |
| TRPM4 | 125.682337 | 0.28386795 | 0.34608034 | 0.82023714 | 0.41208093 | 0.88098902 |
| SLC6A16 | 94.642618 | -0.4202896 | 0.4123279 | -1.0193092 | 0.3080562 | 0.83980597 |
| TEAD2 | 50.8469408 | -0.8066004 | 0.57475445 | -1.4033826 | 0.16050279 | 0.71409437 |
| PIH1D1 | 82.1821273 | 0.16948043 | 0.37773552 | 0.44867487 | 0.65366622 | 0.95100227 |
| RPL13A | 784.356689 | -0.0335499 | 0.43265026 | -0.0775451 | 0.93818994 | 0.99243654 |
| RPS11 | 573.543719 | -0.0448876 | 0.41122037 | -0.1091569 | 0.91307802 | 0.98830345 |
| FCGRT | 172.066061 | 0.7412928 | 0.30219338 | 2.4530412 | 0.01416541 | 0.28658979 |
| NOSIP | 119.068862 | 0.01043452 | 0.27297274 | 0.03822549 | 0.9695079 | 0.99645289 |
| PRRG2 | 48.1151451 | 0.19339286 | 0.33226602 | 0.58204225 | 0.56053822 | 0.92934517 |
| PRR12 | 82.3006452 | -0.0730809 | 0.45251904 | -0.1614979 | 0.87170127 | 0.98209302 |
| RRAS | 74.3599433 | 0.3235691 | 0.64378971 | 0.50260061 | 0.6152451 | 0.94169251 |
| SCAF1 | 62.2286798 | 0.02074926 | 0.26885161 | 0.07717735 | 0.93848246 | 0.99243654 |
| IRF3 | 142.481839 | -0.055476 | 0.22050574 | -0.2515853 | 0.80136165 | 0.9661728 |
| PRMT1 | 149.632584 | 0.19588216 | 0.29361797 | 0.66713275 | 0.50468734 | 0.9107831 |
| AP2A1 | 87.1700123 | -0.1800289 | 0.23193959 | -0.7761889 | 0.43763743 | 0.88825695 |
| MED25 | 121.748117 | -0.127122 | 0.25871626 | -0.4913567 | 0.62317421 | 0.94456546 |
| PTOV1 | 115.705778 | -0.2140882 | 0.2318814 | -0.923266 | 0.3558686 | 0.8557799 |
| PNKP | 91.0964833 | -0.1326467 | 0.31301458 | -0.4237716 | 0.67173237 | 0.95444808 |
| TBC1D17 | 87.8023377 | -0.4203084 | 0.24935652 | -1.6855722 | 0.09187823 | 0.61120158 |
| ATF5 | 57.9816968 | 0.48035156 | 0.36866094 | 1.30296299 | 0.1925874 | 0.7597012 |
| VRK3 | 103.799198 | -0.1556355 | 0.21955992 | -0.7088521 | 0.47841627 | 0.90263775 |
| MYH14 | 245.194142 | -0.2238888 | 0.31956175 | -0.7006121 | 0.4835451 | 0.90631121 |
| KCNC3 | 197.635976 | 1.71125717 | 0.28115709 | 6.08648065 | 1.15E-09 | 2.42E-06 |
| NR1H2 | 157.164062 | 0.12363308 | 0.27663852 | 0.44691201 | 0.65493859 | 0.95100227 |
| EMC10 | 209.42477 | 0.22086947 | 0.33139655 | 0.66648089 | 0.50510377 | 0.9107831 |
| JOSD2 | 54.8018985 | 0.57665965 | 0.39961051 | 1.44305425 | 0.14900519 | 0.70215315 |
| CLEC11A | 57.7085493 | -0.0485439 | 0.32617831 | -0.1488263 | 0.88169072 | 0.98335905 |
| C19orf48 | 47.8094181 | -0.4836884 | 0.31470135 | -1.536976 | 0.12429918 | 0.665882 |
| CTU1 | 35.713269 | 0.1159312 | 0.38038427 | 0.30477391 | 0.76053835 | 0.96191672 |
| ETFB | 215.225426 | 0.61387771 | 0.34486305 | 1.78006227 | 0.07506577 | 0.57072562 |

| | | | | | | |
|---|---|---|---|---|---|---|
| ZNF175 | 48.4094956 | -0.0901739 | 0.42639484 | -0.2114799 | 0.83251283 | 0.97208613 |
| SPACA6 | 68.4837595 | 0.03254117 | 0.34764564 | 0.09360442 | 0.9254234 | 0.99051393 |
| ZNF577 | 145.142716 | 0.2501053 | 0.43572441 | 0.57399883 | 0.56596859 | 0.93122667 |
| ZNF613 | 39.1076524 | -0.2892836 | 0.45752697 | -0.6322767 | 0.52720608 | 0.92044374 |
| ZNF350-AS1 | 227.336879 | -1.404348 | 0.7732724 | -1.8161104 | 0.06935344 | 0.55917298 |
| ZNF350 | 97.2989393 | -0.9089841 | 0.54443185 | -1.6696013 | 0.09499828 | 0.61367241 |
| ZNF615 | 66.8477873 | -0.2011007 | 0.42739952 | -0.4705216 | 0.63798238 | 0.94854485 |
| ZNF614 | 38.1547649 | -0.1645955 | 0.47311681 | -0.347896 | 0.72791828 | 0.9598317 |
| ZNF432 | 118.88017 | -0.3926463 | 0.28636962 | -1.3711172 | 0.17033843 | 0.72675731 |
| ZNF616 | 41.2564108 | -0.8198775 | 0.50360075 | -1.6280308 | 0.10351835 | 0.62722102 |
| AC011468.3 | 53.740638 | 0.2562996 | 0.41553883 | 0.61678856 | 0.5373742 | 0.92324352 |
| ZNF836 | 67.3101992 | -1.2084342 | 0.77930368 | -1.5506589 | 0.12098345 | 0.65959373 |
| PPP2R1A | 548.949266 | -0.3827447 | 0.40229889 | -0.951394 | 0.3414044 | 0.85042291 |
| ZNF766 | 100.099966 | -0.1434768 | 0.2651967 | -0.5410202 | 0.58849367 | 0.93483536 |
| ZNF480 | 76.58877 | -0.3154215 | 0.33310115 | -0.9469242 | 0.3436774 | 0.85126545 |
| ZNF610 | 34.1773688 | 0.5010872 | 0.5147548 | 0.97344833 | 0.33033053 | 0.84442294 |
| ZNF880 | 37.599271 | 0.71473708 | 0.3621029 | 1.97385078 | 0.04839871 | 0.48661187 |
| ZNF528 | 74.5388846 | 0.19543035 | 0.34639241 | 0.56418776 | 0.57262636 | 0.93218128 |
| ZNF808 | 55.1811536 | 0.14034921 | 0.31403778 | 0.44691822 | 0.65493411 | 0.95100227 |
| ZNF701 | 86.9178981 | 0.29158756 | 0.23879638 | 1.2210719 | 0.2220588 | 0.78760199 |
| ZNF83 | 621.414182 | -0.5143447 | 0.37026742 | -1.3891169 | 0.16479721 | 0.71715698 |
| ZNF611 | 273.290335 | -0.1233007 | 0.29576401 | -0.4168888 | 0.67675977 | 0.95529325 |
| ZNF600 | 35.6476965 | -0.7696303 | 0.32153414 | -2.3936191 | 0.01668306 | 0.30789747 |
| ZNF28 | 58.3258937 | -0.1283436 | 0.2355068 | -0.5449676 | 0.58577581 | 0.93483536 |
| AC010487.3 | 66.2529163 | 0.2325206 | 0.38270468 | 0.60757186 | 0.54347147 | 0.92482794 |
| ZNF160 | 237.972633 | -0.2950448 | 0.27151635 | -1.0866558 | 0.27718894 | 0.82198569 |
| ZNF347 | 73.7276256 | -0.0737469 | 0.31514899 | -0.2340064 | 0.81498003 | 0.96839471 |
| ZNF677 | 74.612736 | 0.0665434 | 0.30107591 | 0.22101867 | 0.8250779 | 0.97018919 |
| ZNF525 | 48.350818 | 0.10658207 | 0.39542215 | 0.26953995 | 0.7875142 | 0.96513523 |
| ZNF765 | 60.0645007 | 0.2961586 | 0.32924337 | 0.89951273 | 0.36837962 | 0.8591648 |
| ZNF761 | 88.2910421 | -0.3186183 | 0.36395488 | -0.8754335 | 0.3813381 | 0.86593382 |
| ZNF331 | 1301.54198 | -3.2977046 | 1.47955612 | -2.2288472 | 0.02582407 | 0.3899658 |
| MYADM | 58.9958069 | 0.05532453 | 0.3359828 | 0.16466477 | 0.86920785 | 0.9820194 |
| NDUFA3 | 89.2705209 | 0.33991609 | 0.43207185 | 0.78671195 | 0.43145051 | 0.88650713 |
| TFPT | 38.1497137 | -0.1549666 | 0.38676978 | -0.4006688 | 0.68866397 | 0.95604695 |
| PRPF31 | 120.712119 | 0.39299524 | 0.31185822 | 1.26017279 | 0.20760704 | 0.77317759 |
| CNOT3 | 59.8575745 | -0.2088643 | 0.25877984 | -0.807112 | 0.41960193 | 0.88284044 |
| TMC4 | 274.210744 | -0.0716573 | 0.20308968 | -0.3528358 | 0.72421153 | 0.95963742 |
| MBOAT7 | 220.087998 | -0.3138437 | 0.28427594 | -1.1040106 | 0.26958852 | 0.81817548 |
| TSEN34 | 89.1842259 | 0.67081105 | 0.40676184 | 1.64914942 | 0.09911703 | 0.61707826 |
| RPS9 | 1317.89095 | 0.10865576 | 0.46694945 | 0.23269276 | 0.81599999 | 0.96844493 |
| LENG8-AS1 | 50.9665406 | 0.14272785 | 0.3323767 | 0.42941592 | 0.66762057 | 0.95294349 |
| LENG8 | 297.625911 | -0.6283433 | 0.36443514 | -1.7241566 | 0.08467957 | 0.59720183 |
| AC011476.3 | 68.6853006 | -0.3395108 | 0.63833651 | -0.5318681 | 0.59481736 | 0.935202 |
| RDH13 | 182.365397 | -0.652418 | 0.38955326 | -1.6747851 | 0.0939764 | 0.61323531 |
| EPS8L1 | 141.354323 | 0.59773115 | 0.42626428 | 1.40225483 | 0.16083916 | 0.71409437 |
| PPP1R12C | 104.638071 | -0.2879573 | 0.22150986 | -1.2999751 | 0.19360951 | 0.76043322 |
| PPP6R1 | 103.400244 | -0.0095666 | 0.2720894 | -0.0351598 | 0.97195235 | 0.99656026 |
| HSPBP1 | 63.2178337 | 0.0932005 | 0.33748506 | 0.27616187 | 0.78242375 | 0.96513523 |
| TMEM238 | 96.6689778 | -0.6919431 | 0.43557378 | -1.5885784 | 0.1121556 | 0.64471804 |
| RPL28 | 946.436941 | -0.3145085 | 0.4090645 | -0.7688483 | 0.44198339 | 0.88997055 |
| ISOC2 | 107.659561 | 0.10244484 | 0.53071162 | 0.19303297 | 0.84693315 | 0.97656375 |
| NAT14 | 65.1965578 | 0.62605959 | 0.48884022 | 1.28070393 | 0.20029768 | 0.76890831 |
| ZNF579 | 36.2335258 | 0.15389329 | 0.40291546 | 0.38194932 | 0.70249896 | 0.95672887 |

| | | | | | | |
|---|---|---|---|---|---|---|
| ZNF524 | 37.551933 | 0.15723593 | 0.40230804 | 0.39083468 | 0.69591944 | 0.95672887 |
| CCDC106 | 51.7633823 | -0.0730372 | 0.39027092 | -0.1871448 | 0.85154714 | 0.97718353 |
| U2AF2 | 92.1529705 | -0.3099839 | 0.22890158 | -1.3542237 | 0.17566502 | 0.73403881 |
| EPN1 | 284.214673 | 0.12997379 | 0.27481289 | 0.47295374 | 0.63624618 | 0.94821838 |
| ZNF787 | 59.569634 | -0.0899014 | 0.30418386 | -0.2955497 | 0.767574 | 0.96322167 |
| ZNF444 | 82.6442409 | 0.23224647 | 0.20351062 | 1.14120078 | 0.25378638 | 0.80414079 |
| ZSCAN5A | 182.723344 | -0.3406921 | 0.32928265 | -1.0346495 | 0.30083264 | 0.83709431 |
| ZNF667 | 73.7701004 | 0.07606497 | 0.4663914 | 0.16309258 | 0.87044555 | 0.98208511 |
| ZNF667-AS1 | 102.473062 | 0.33506393 | 0.28469131 | 1.1769377 | 0.23922037 | 0.79984328 |
| ZNF471 | 73.379451 | 0.25266373 | 0.29655988 | 0.85198217 | 0.39422399 | 0.87253562 |
| AC005498.3 | 36.0259436 | 0.38205231 | 0.46718802 | 0.81776992 | 0.41348858 | 0.88211689 |
| ZNF470 | 36.0262921 | -0.241418 | 0.48510119 | -0.4976652 | 0.61872001 | 0.94287999 |
| ZNF264 | 114.268217 | -0.2097727 | 0.19881368 | -1.0551218 | 0.29136962 | 0.8320147 |
| ZNF460-AS1 | 34.6121704 | 0.27225442 | 0.33309151 | 0.81735623 | 0.41372488 | 0.88211689 |
| ZNF460 | 45.5859966 | -0.926049 | 0.42545907 | -2.1765878 | 0.02951134 | 0.40623517 |
| AC005261.1 | 203.826032 | -0.9075169 | 0.27159983 | -3.3413752 | 0.00083364 | 0.06772557 |
| AC004076.1 | 75.8165902 | -0.0314543 | 0.43388766 | -0.0724942 | 0.94220861 | 0.99265578 |
| ZNF549 | 56.1615594 | 0.59920437 | 0.26240997 | 2.28346648 | 0.02240291 | 0.35859484 |
| ZNF134 | 36.4285853 | -0.0258625 | 0.35739561 | -0.0723638 | 0.94231235 | 0.99265578 |
| ZNF154 | 43.3960731 | -0.1520785 | 0.34993758 | -0.4345874 | 0.66386193 | 0.95187756 |
| ZNF776 | 34.3093356 | -0.1136994 | 0.32233624 | -0.3527353 | 0.72428692 | 0.95963742 |
| ZNF586 | 311.375981 | -0.1323469 | 0.25659889 | -0.5157736 | 0.60601257 | 0.93919596 |
| ZNF552 | 132.553507 | 0.26491956 | 0.28912656 | 0.91627542 | 0.35952246 | 0.85608464 |
| ZNF814 | 195.601407 | 0.01490665 | 0.29485187 | 0.05055641 | 0.959679 | 0.99609409 |
| ZNF587 | 103.126636 | -0.337256 | 0.20197216 | -1.6698141 | 0.09495615 | 0.61367241 |
| ZNF418 | 34.2395057 | -0.0879094 | 0.40992884 | -0.2144504 | 0.83019582 | 0.97157478 |
| ZNF606 | 63.3855267 | 0.04482315 | 0.43818409 | 0.10229298 | 0.91852413 | 0.99014946 |
| ZNF135 | 92.7809073 | -0.0389175 | 0.3503021 | -0.111097 | 0.91153947 | 0.98788932 |
| ZSCAN18 | 314.426259 | 0.10334751 | 0.28591814 | 0.36145838 | 0.71775681 | 0.95796325 |
| ZNF329 | 81.5533957 | 0.52124717 | 0.32648294 | 1.59655254 | 0.11036549 | 0.64114761 |
| ZNF274 | 120.401665 | 0.15781334 | 0.25368303 | 0.6220887 | 0.53388355 | 0.92289732 |
| AC020915.3 | 56.3327607 | 0.07014262 | 0.34936021 | 0.2007745 | 0.84087491 | 0.97514319 |
| RPS5 | 338.880118 | -0.1065987 | 0.52803158 | -0.2018793 | 0.84001107 | 0.9746807 |
| ZNF584 | 38.132509 | 0.16714828 | 0.39698766 | 0.42104151 | 0.67372477 | 0.9549142 |
| ZNF446 | 34.5229498 | 0.11521159 | 0.30978714 | 0.37190564 | 0.70996311 | 0.95752169 |
| SLC27A5 | 55.5292242 | 0.0154286 | 0.34261805 | 0.0450315 | 0.9640822 | 0.99637506 |
| TRIM28 | 130.947994 | 0.03787373 | 0.37923425 | 0.09986896 | 0.92044836 | 0.9904507 |
| CHMP2A | 166.876349 | 0.07685054 | 0.51439268 | 0.14940053 | 0.88123759 | 0.98335905 |
| UBE2M | 115.659874 | 0.0925002 | 0.44002653 | 0.21021504 | 0.83349984 | 0.97232209 |
| MZF1-AS1 | 42.4692228 | 0.22074066 | 0.45578318 | 0.48431067 | 0.62816541 | 0.94572238 |
| MZF1 | 144.045014 | -0.3272649 | 0.21264178 | -1.539043 | 0.12379379 | 0.66519057 |
| RPS4Y1 | 274.00365 | 0.68719318 | 2.187041 | 0.31421138 | 0.7533605 | 0.96120879 |
| ZFY | 77.3103475 | 0.43739585 | 2.10157688 | 0.20812746 | 0.83512945 | 0.97246185 |
| LINC00278 | 649.214111 | 0.08223656 | 2.60699268 | 0.03154461 | 0.97483522 | 0.99656026 |
| PCDH11Y | 123.55098 | 0.2092092 | 0.92643618 | 0.22582149 | 0.82134026 | 0.96955628 |
| TBL1Y | 167.479896 | 0.02307842 | 1.83651391 | 0.01256643 | 0.98997371 | 0.99864515 |
| TTTY15 | 90.8886887 | 0.42727099 | 1.72093611 | 0.24827824 | 0.80391913 | 0.96678163 |
| USP9Y | 333.606469 | 0.46414759 | 2.43682974 | 0.1904719 | 0.84893936 | 0.97656375 |
| DDX3Y | 130.905573 | 0.87591984 | 2.5347333 | 0.34556686 | 0.72966825 | 0.96008876 |
| UTY | 764.361165 | 0.63527271 | 2.12293724 | 0.29924234 | 0.76475515 | 0.96285313 |
| NLGN4Y | 123.718279 | 1.34264037 | 1.50822831 | 0.89021029 | 0.37335298 | 0.86201082 |
| TTTY14 | 1078.81855 | 0.27129246 | 0.90101401 | 0.30109683 | 0.76334066 | 0.96285313 |
| KDM5D | 98.8400507 | 0.52822776 | 2.48469331 | 0.21259274 | 0.83164464 | 0.97164278 |
| TTTY10 | 69.7109273 | 0.41792784 | 1.58972899 | 0.26289251 | 0.79263343 | 0.9653598 |

| | | | | | | |
|---|---|---|---|---|---|---|
| EIF1AY | 61.488158 | 0.72932923 | 2.37939147 | 0.30651923 | 0.75920934 | 0.96182318 |
| IL17RA | 64.7642947 | 0.19721674 | 0.28195822 | 0.69945378 | 0.48426849 | 0.90659463 |
| HDHD5 | 78.5590332 | 0.12624924 | 0.21952251 | 0.57510842 | 0.56521798 | 0.93114846 |
| HDHD5-AS1 | 33.0246642 | 0.39958325 | 0.5069047 | 0.78828083 | 0.43053246 | 0.88596084 |
| ATP6V1E1 | 276.42159 | -0.0397407 | 0.21873468 | -0.1816847 | 0.85583019 | 0.97718353 |
| BCL2L13 | 387.108436 | -0.006869 | 0.22095956 | -0.0310873 | 0.97519988 | 0.99656026 |
| MICAL3 | 250.31445 | 0.29575734 | 0.40796642 | 0.72495511 | 0.46847957 | 0.90026987 |
| PEX26 | 124.066044 | -0.0038681 | 0.24955546 | -0.0155 | 0.98763329 | 0.99791717 |
| DGCR2 | 210.744035 | -0.0745448 | 0.22267432 | -0.3347707 | 0.7377981 | 0.96078945 |
| SLC25A1 | 144.557601 | 0.23452541 | 0.37807682 | 0.62031151 | 0.53505272 | 0.92303275 |
| CLTCL1 | 89.8462246 | -0.3693959 | 0.27239573 | -1.3561003 | 0.17506724 | 0.73241705 |
| HIRA | 80.0978902 | -0.5063707 | 0.30943942 | -1.6364131 | 0.10175315 | 0.62324547 |
| MRPL40 | 115.879162 | 0.58797734 | 0.55433849 | 1.06068288 | 0.28883404 | 0.83060304 |
| UFD1 | 68.3812432 | -0.2778141 | 0.31524839 | -0.8812547 | 0.37817998 | 0.86512157 |
| TXNRD2 | 84.0190625 | -0.1522733 | 0.26248801 | -0.5801151 | 0.56183699 | 0.92959451 |
| COMT | 349.816245 | 0.43292511 | 0.40076904 | 1.08023592 | 0.28003714 | 0.82628479 |
| TANGO2 | 105.817102 | -0.7049092 | 0.33733633 | -2.0896333 | 0.03665075 | 0.43729328 |
| DGCR8 | 77.7511122 | -0.2652361 | 0.30889838 | -0.8586516 | 0.39053276 | 0.87004477 |
| RANBP1 | 79.8204569 | 0.39902171 | 0.47163565 | 0.84603807 | 0.39753151 | 0.87462201 |
| ZDHHC8 | 52.5935503 | -0.5269646 | 0.36124351 | -1.4587517 | 0.14463347 | 0.69206332 |
| DGCR6L | 65.3009584 | 0.31531883 | 0.40827158 | 0.77232618 | 0.43992126 | 0.88917548 |
| KLHL22 | 135.954163 | 0.32138712 | 0.3053417 | 1.05254907 | 0.29254771 | 0.83336457 |
| MED15 | 281.85942 | -0.353537 | 0.1677567 | -2.1074387 | 0.03507957 | 0.43353051 |
| PI4KA | 315.283965 | 0.10966629 | 0.18063648 | 0.6071104 | 0.54377765 | 0.92482794 |
| SNAP29 | 83.6515925 | 0.07842602 | 0.30098587 | 0.26056381 | 0.7944289 | 0.9653598 |
| CRKL | 89.3460292 | 0.10950018 | 0.21212403 | 0.51620829 | 0.60570894 | 0.93919596 |
| LZTR1 | 48.1067408 | 0.10990934 | 0.26405274 | 0.41624011 | 0.67723432 | 0.95536622 |
| THAP7 | 40.6473919 | 0.12204585 | 0.38909255 | 0.31366793 | 0.75377326 | 0.96120879 |
| HIC2 | 64.0570323 | 0.31578079 | 0.39001742 | 0.80965816 | 0.41813667 | 0.88242985 |
| UBE2L3 | 294.537559 | -0.1509233 | 0.22320151 | -0.6761749 | 0.49892961 | 0.90839719 |
| YDJC | 29.9426695 | 0.60231825 | 0.37553913 | 1.60387614 | 0.10874136 | 0.63736545 |
| SDF2L1 | 70.7258067 | 0.14885838 | 0.43823246 | 0.33967903 | 0.73409826 | 0.96042436 |
| PPIL2 | 160.504604 | -0.6577098 | 0.24653646 | -2.6677995 | 0.00763498 | 0.21455924 |
| YPEL1 | 38.1884953 | -0.5301293 | 0.52246732 | -1.014665 | 0.31026554 | 0.84213522 |
| MAPK1 | 524.039032 | -0.2982778 | 0.23419314 | -1.2736403 | 0.20279092 | 0.76996624 |
| PPM1F | 53.3119765 | -0.027399 | 0.41762268 | -0.065607 | 0.94769072 | 0.99314907 |
| TOP3B | 41.2181029 | -0.0333261 | 0.39538935 | -0.0842868 | 0.93282842 | 0.99243654 |
| AC245060.5 | 94.1421505 | -0.2269238 | 0.45976578 | -0.4935639 | 0.62161423 | 0.94413291 |
| RSPH14 | 35.3731343 | 0.44428187 | 0.33588469 | 1.32272141 | 0.18592804 | 0.74914489 |
| GNAZ | 44.7698391 | 0.15026354 | 0.41018941 | 0.36632721 | 0.71412093 | 0.95791799 |
| RAB36 | 57.0922959 | 0.43451566 | 0.29556982 | 1.47009478 | 0.14153609 | 0.68671506 |
| BCR | 324.807303 | 0.24943358 | 0.2597821 | 0.96016459 | 0.33697239 | 0.84636904 |
| CHCHD10 | 655.020538 | 0.1991084 | 0.42663317 | 0.46669696 | 0.6407167 | 0.94935885 |
| SMARCB1 | 137.008579 | 0.05173339 | 0.29452296 | 0.17565145 | 0.86056778 | 0.97841862 |
| DDT | 315.800858 | 0.28979767 | 0.45732657 | 0.63367776 | 0.52629114 | 0.91974278 |
| CABIN1 | 557.474682 | 0.1481302 | 0.27422328 | 0.54018097 | 0.58907223 | 0.93483536 |
| GUCD1 | 100.937048 | 0.07199963 | 0.22625976 | 0.31821666 | 0.7503206 | 0.96109466 |
| SNRPD3 | 219.471433 | 0.14586105 | 0.31519174 | 0.46276926 | 0.64352977 | 0.94949493 |
| SGSM1 | 173.292806 | 0.58651423 | 0.40061252 | 1.46404369 | 0.14318201 | 0.69023197 |
| KIAA1671 | 356.835574 | -0.278459 | 0.27924653 | -0.9971798 | 0.31867722 | 0.84285425 |
| LRP5L | 107.695431 | -0.0151948 | 0.34002012 | -0.0446881 | 0.96435593 | 0.99637506 |
| GRK3 | 115.041657 | 0.17697782 | 0.37796851 | 0.46823428 | 0.63961706 | 0.94935885 |
| ASPHD2 | 35.9021839 | 0.14326294 | 0.64907012 | 0.22072028 | 0.82531024 | 0.97018919 |
| HPS4 | 134.217286 | -0.3400173 | 0.20681701 | -1.644049 | 0.10016608 | 0.61891077 |

| | | | | | | |
|---|---|---|---|---|---|---|
| TFIP11 | 48.0520953 | -0.0234522 | 0.26742297 | -0.087697 | 0.93011751 | 0.99216065 |
| TPST2 | 84.159154 | 0.26390793 | 0.28424985 | 0.92843646 | 0.35318121 | 0.8557799 |
| MIATNB | 87.4095428 | -0.3231311 | 0.43329005 | -0.7457617 | 0.45581136 | 0.89626521 |
| PITPNB | 286.291168 | -0.1499709 | 0.32158102 | -0.4663548 | 0.64096155 | 0.94935885 |
| TTC28 | 502.583028 | 0.12045482 | 0.21282874 | 0.56597067 | 0.57141372 | 0.93146924 |
| HSCB | 110.444772 | -0.0851589 | 0.25945962 | -0.3282163 | 0.74274816 | 0.96086911 |
| CCDC117 | 60.6255596 | 0.33048789 | 0.29751457 | 1.11082927 | 0.26664185 | 0.81333524 |
| XBP1 | 174.631586 | 0.03243617 | 0.26835098 | 0.1208722 | 0.90379226 | 0.9875066 |
| Z93930.2 | 205.543457 | 0.04631382 | 0.47335409 | 0.0978418 | 0.92205792 | 0.99051393 |
| ZNRF3 | 362.434209 | -0.3994642 | 0.37637267 | -1.0613528 | 0.2885296 | 0.83060304 |
| KREMEN1 | 90.5996368 | 0.15891386 | 0.29390374 | 0.54070037 | 0.58871412 | 0.93483536 |
| RHBDD3 | 46.0205079 | -0.3451565 | 0.34838928 | -0.9907207 | 0.32182198 | 0.84285425 |
| EWSR1 | 338.318039 | -0.2543986 | 0.24766477 | -1.0271892 | 0.3043314 | 0.83709431 |
| AP1B1 | 146.092607 | 0.17658892 | 0.26584384 | 0.66425808 | 0.50652515 | 0.91163608 |
| THOC5 | 48.5936347 | -0.1798123 | 0.31351065 | -0.5735445 | 0.56627607 | 0.93122667 |
| NIPSNAP1 | 90.2741676 | 0.11676168 | 0.28490883 | 0.40982121 | 0.68193711 | 0.95574492 |
| NF2 | 175.589332 | 0.51273743 | 0.35218801 | 1.45586281 | 0.14543055 | 0.69325556 |
| ZMAT5 | 48.0799965 | -0.1326731 | 0.24937633 | -0.5320195 | 0.59471249 | 0.935202 |
| UQCR10 | 532.436366 | 0.32555295 | 0.44526298 | 0.73114757 | 0.46468902 | 0.89851309 |
| ASCC2 | 105.075444 | -0.1537828 | 0.24850491 | -0.618832 | 0.53602704 | 0.92324352 |
| MTMR3 | 505.986066 | -0.0726528 | 0.1869716 | -0.3885767 | 0.69758931 | 0.95672887 |
| TBC1D10A | 47.3700245 | 0.11838524 | 0.45996286 | 0.25738 | 0.79688544 | 0.9653598 |
| SF3A1 | 87.2169645 | 0.02895023 | 0.30526307 | 0.09483698 | 0.92444431 | 0.99051393 |
| RNF215 | 70.617564 | -0.3340383 | 0.31428416 | -1.0628544 | 0.28784797 | 0.8303903 |
| SEC14L6 | 88.4385878 | 0.44048124 | 0.34566737 | 1.27429222 | 0.20255987 | 0.76996624 |
| GAL3ST1 | 65.83879 | 0.13922854 | 0.32747915 | 0.42515239 | 0.67072559 | 0.95401794 |
| PES1 | 64.1200205 | -0.2704893 | 0.27920122 | -0.9687969 | 0.33264653 | 0.84442294 |
| TCN2 | 49.617492 | 0.15315115 | 0.46788752 | 0.32732471 | 0.7434223 | 0.96105014 |
| DUSP18 | 81.0430312 | -0.0797749 | 0.26914394 | -0.2964024 | 0.76692278 | 0.96285313 |
| OSBP2 | 171.65056 | 0.03308667 | 0.27455968 | 0.12050811 | 0.90408065 | 0.9875066 |
| MORC2 | 77.3355493 | 0.00960239 | 0.24983992 | 0.03843418 | 0.96934151 | 0.99645289 |
| SELENOM | 410.303019 | 0.54869546 | 0.37826536 | 1.45055699 | 0.14690326 | 0.69743411 |
| INPP5J | 200.196783 | 0.69082623 | 0.51016709 | 1.35411758 | 0.17569887 | 0.73403881 |
| RNF185 | 184.594378 | 0.09712195 | 0.24079854 | 0.40333281 | 0.68670341 | 0.9558749 |
| LIMK2 | 237.511951 | 0.02435716 | 0.22807207 | 0.10679588 | 0.91495092 | 0.98868699 |
| PIK3IP1 | 169.162874 | 0.49220452 | 0.31343019 | 1.57038005 | 0.11632672 | 0.65257994 |
| PIK3IP1-AS1 | 54.883795 | -0.3212985 | 0.42479047 | -0.7563694 | 0.44942774 | 0.89324553 |
| PATZ1 | 114.907621 | -0.1080726 | 0.37509683 | -0.2881192 | 0.77325548 | 0.96388926 |
| DRG1 | 148.337495 | 0.07575663 | 0.27887466 | 0.27165118 | 0.78589024 | 0.96513523 |
| EIF4ENIF1 | 181.184574 | -0.1014033 | 0.2150886 | -0.471449 | 0.63732012 | 0.94833378 |
| SFI1 | 164.361374 | -0.6234897 | 0.2526569 | -2.4677329 | 0.01359717 | 0.28329699 |
| PISD | 389.167033 | -0.3136984 | 0.42196382 | -0.743425 | 0.45722443 | 0.8969884 |
| PRR14L | 308.892018 | -0.1145103 | 0.180015 | -0.6361154 | 0.52470118 | 0.91923576 |
| YWHAH | 99.5605996 | 0.06303866 | 0.522349 | 0.12068303 | 0.9039421 | 0.9875066 |
| RTCB | 162.596414 | 0.20083169 | 0.25073239 | 0.80098021 | 0.4231431 | 0.88371155 |
| FBXO7 | 189.571736 | -0.0942103 | 0.24261187 | -0.3883168 | 0.69778158 | 0.95672887 |
| TIMP3 | 332.235116 | 0.02169276 | 0.7676775 | 0.02825765 | 0.97745666 | 0.99696835 |
| LARGE1 | 553.484148 | -0.2308518 | 0.31072692 | -0.7429412 | 0.4575173 | 0.89705917 |
| HMGXB4 | 117.186016 | -0.342719 | 0.30787123 | -1.1131895 | 0.26562703 | 0.8121456 |
| TOM1 | 118.152518 | -0.3290841 | 0.18506102 | -1.7782465 | 0.07536338 | 0.57072562 |
| APOL6 | 97.6987026 | -0.7104942 | 0.66275397 | -1.0720331 | 0.28370516 | 0.82696038 |
| RBFOX2 | 953.688973 | 0.1061341 | 0.20851742 | 0.50899393 | 0.61075648 | 0.94003018 |
| APOL2 | 119.18544 | -0.9953507 | 0.43759793 | -2.2745782 | 0.02293125 | 0.36247279 |
| APOL1 | 110.597002 | -0.690011 | 0.41308037 | -1.6704038 | 0.0948395 | 0.61367241 |

| | | | | | | |
|---|---|---|---|---|---|---|
| MYH9 | 557.632777 | -0.3175158 | 0.19748402 | -1.6078051 | 0.10787788 | 0.63520805 |
| TXN2 | 146.112936 | 0.47268991 | 0.3484179 | 1.35667516 | 0.17488445 | 0.73227004 |
| EIF3D | 162.46874 | 0.10312629 | 0.26245731 | 0.39292596 | 0.69437417 | 0.95672887 |
| IFT27 | 80.2105165 | 0.05644002 | 0.23860168 | 0.23654493 | 0.81300985 | 0.9680627 |
| TST | 93.9726885 | 0.11014461 | 0.40064161 | 0.27492055 | 0.78337728 | 0.96513523 |
| MPST | 96.1363376 | -0.223076 | 0.35761723 | -0.6237843 | 0.53276927 | 0.92268523 |
| CARD10 | 46.3127511 | 0.9760559 | 0.40535521 | 2.40790267 | 0.01604446 | 0.30351249 |
| CDC42EP1 | 66.4587737 | 0.2417689 | 0.40745467 | 0.59336391 | 0.55293763 | 0.92872652 |
| GGA1 | 144.069047 | -0.0537254 | 0.2384522 | -0.2253088 | 0.82173905 | 0.96969094 |
| LGALS1 | 145.994595 | -1.112739 | 0.37279063 | -2.9848898 | 0.0028368 | 0.1372855 |
| TRIOBP | 247.74448 | -0.2825849 | 0.18692905 | -1.5117229 | 0.13060436 | 0.67339292 |
| H1F0 | 304.713019 | -0.3274546 | 0.67967072 | -0.4817841 | 0.62995931 | 0.94596555 |
| GCAT | 35.1390126 | 0.14210702 | 0.42470118 | 0.33460471 | 0.73792329 | 0.96078945 |
| ANKRD54 | 53.0596918 | 0.31799858 | 0.28723318 | 1.10710949 | 0.26824659 | 0.8165043 |
| EIF3L | 216.511776 | -0.043832 | 0.29472863 | -0.14872 | 0.8817746 | 0.98335905 |
| MICALL1 | 39.7920736 | 0.13502772 | 0.33420929 | 0.40402143 | 0.68619696 | 0.9558749 |
| POLR2F | 87.0104288 | -0.3495492 | 0.32930497 | -1.0614755 | 0.28847385 | 0.83060304 |
| PICK1 | 71.0616145 | -0.3139944 | 0.25298388 | -1.2411638 | 0.21454524 | 0.77890374 |
| PLA2G6 | 61.4556893 | 0.21532761 | 0.28913079 | 0.74474119 | 0.45642819 | 0.89626521 |
| MAFF | 119.013122 | -1.4989868 | 0.95501096 | -1.5696017 | 0.11650781 | 0.65257994 |
| TMEM184B | 144.333796 | -0.2086332 | 0.20967698 | -0.9950221 | 0.31972554 | 0.84285425 |
| CSNK1E | 58.78246 | -0.5293143 | 0.29279659 | -1.8077886 | 0.07063941 | 0.5621116 |
| TPTEP2-CSNI | 489.995925 | 0.34765037 | 0.28033903 | 1.24010688 | 0.21493586 | 0.77890374 |
| DDX17 | 2381.56801 | 0.0858996 | 0.19070457 | 0.45043285 | 0.65239836 | 0.95092279 |
| FAM227A | 98.1498455 | 0.21428162 | 0.28170987 | 0.76064649 | 0.44686825 | 0.8918683 |
| CBY1 | 117.46588 | 0.50754217 | 0.26714442 | 1.89987936 | 0.05744895 | 0.52728498 |
| TOMM22 | 57.5693304 | 0.42056673 | 0.45082443 | 0.93288362 | 0.35088006 | 0.85477058 |
| JOSD1 | 134.772865 | -0.8864024 | 0.69880538 | -1.2684539 | 0.20463592 | 0.77193931 |
| GTPBP1 | 91.4665406 | -0.4163204 | 0.36313508 | -1.1464616 | 0.25160421 | 0.80252057 |
| SUN2 | 84.5605582 | 0.18675486 | 0.2735719 | 0.68265365 | 0.49482573 | 0.90839719 |
| AL021707.1 | 40.4662084 | -0.4720899 | 0.32905976 | -1.434663 | 0.15138315 | 0.7029344 |
| CBX6 | 182.799423 | 0.28656995 | 0.36874878 | 0.77714143 | 0.43707533 | 0.88785137 |
| CBX7 | 103.988952 | 0.66135355 | 0.30335102 | 2.18015928 | 0.02924566 | 0.40623517 |
| RPL3 | 804.368483 | 0.31405727 | 0.42730316 | 0.7349753 | 0.46235454 | 0.89801718 |
| SYNGR1 | 86.7333857 | 0.73161793 | 0.27513237 | 2.65914888 | 0.00783383 | 0.217192 |
| TAB1 | 69.7815472 | -0.3442445 | 0.28954012 | -1.1889354 | 0.23446507 | 0.79543722 |
| MIEF1 | 58.5967715 | 0.17023199 | 0.3076355 | 0.55335614 | 0.58001957 | 0.93345492 |
| ATF4 | 159.718496 | -0.9089435 | 0.35672004 | -2.5480583 | 0.01083244 | 0.25283731 |
| RPS19BP1 | 170.475381 | 0.0330816 | 0.47483608 | 0.06966952 | 0.94445671 | 0.99265578 |
| FAM83F | 73.2511305 | -0.0706726 | 0.50675881 | -0.13946 | 0.88908664 | 0.98551241 |
| TNRC6B | 1827.66149 | -0.0576094 | 0.23605925 | -0.2440463 | 0.80719495 | 0.96752828 |
| MKL1 | 978.448517 | -0.3061878 | 0.25250337 | -1.2126086 | 0.2252795 | 0.78982769 |
| SLC25A17 | 154.083731 | -0.1644066 | 0.27268019 | -0.6029282 | 0.54655648 | 0.92597739 |
| ST13 | 396.061882 | 0.25290699 | 0.24695009 | 1.02412189 | 0.30577771 | 0.83709431 |
| XPNPEP3 | 222.627798 | -0.1505934 | 0.22859644 | -0.6587742 | 0.51004076 | 0.91334187 |
| RBX1 | 231.824164 | -0.3863459 | 0.23878807 | -1.6179446 | 0.10567454 | 0.63429746 |
| EP300 | 503.201864 | 0.02628951 | 0.19989499 | 0.13151658 | 0.89536667 | 0.98597539 |
| L3MBTL2 | 64.8261152 | -0.0401583 | 0.28743878 | -0.139711 | 0.88888837 | 0.98551241 |
| RANGAP1 | 73.0982043 | -0.3621458 | 0.38473642 | -0.941283 | 0.34655986 | 0.8525698 |
| ZC3H7B | 172.547841 | 0.04033752 | 0.24324759 | 0.16582907 | 0.86829147 | 0.98184016 |
| TEF | 63.1546153 | 0.43222307 | 0.39213128 | 1.10224073 | 0.27035703 | 0.81817548 |
| TOB2 | 277.293555 | -0.6016508 | 0.63552995 | -0.9466915 | 0.34379603 | 0.85126545 |
| PHF5A | 49.7835376 | -0.0446849 | 0.48146897 | -0.0928095 | 0.92605491 | 0.99051393 |
| ACO2 | 379.19431 | 0.07935158 | 0.20074089 | 0.39529356 | 0.69262625 | 0.95672887 |

| | | | | | | |
|---|---|---|---|---|---|---|
| POLR3H | 54.0120274 | 0.12045242 | 0.35140912 | 0.34276976 | 0.73177168 | 0.96008876 |
| PMM1 | 62.8346765 | 0.07137258 | 0.31261557 | 0.22830782 | 0.81940695 | 0.96884581 |
| DESI1 | 108.15741 | -0.1952847 | 0.50641376 | -0.3856228 | 0.69977605 | 0.95672887 |
| XRCC6 | 394.74532 | -0.004252 | 0.35467793 | -0.0119882 | 0.99043499 | 0.99882205 |
| SNU13 | 346.144405 | 0.37390416 | 0.4147647 | 0.90148501 | 0.3673305 | 0.85902833 |
| CCDC134 | 37.2323892 | -0.0470924 | 0.38865761 | -0.1211668 | 0.90355891 | 0.9875066 |
| SREBF2 | 955.544973 | -0.2023436 | 0.35702669 | -0.5667463 | 0.57088655 | 0.93125087 |
| WBP2NL | 86.3109713 | 0.1782843 | 0.51136171 | 0.34864616 | 0.72735497 | 0.95979351 |
| NAGA | 37.9905549 | 0.20116498 | 0.39086568 | 0.51466523 | 0.60678697 | 0.93925224 |
| SMDT1 | 178.154758 | 0.61237143 | 0.49912053 | 1.22690089 | 0.21985984 | 0.78633738 |
| NDUFA6 | 185.912218 | 0.15211358 | 0.42953899 | 0.35413217 | 0.72323983 | 0.95963742 |
| TCF20 | 370.220564 | -0.0715278 | 0.20000837 | -0.3576242 | 0.7206246 | 0.95815095 |
| RRP7A | 96.8937825 | -0.0298499 | 0.36407907 | -0.0819874 | 0.9346567 | 0.99243654 |
| SERHL2 | 73.6849889 | -0.4099937 | 0.33927065 | -1.2084561 | 0.22687186 | 0.78990601 |
| POLDIP3 | 124.412854 | -0.2262263 | 0.23265167 | -0.9723818 | 0.33086062 | 0.84442294 |
| CYB5R3 | 315.013898 | -0.4624877 | 0.31523543 | -1.4671185 | 0.14234383 | 0.6890058 |
| A4GALT | 130.400298 | 0.35575606 | 0.44521952 | 0.79905765 | 4.24E-01 | 8.84E-01 |
| ARFGAP3 | 383.951982 | -0.4921991 | 0.29838497 | -1.6495438 | 0.09903628 | 0.61707826 |
| PACSIN2 | 616.257945 | -0.457728 | 0.25050848 | -1.8271955 | 0.06767038 | 0.55864172 |
| TTLL1 | 41.8095365 | -0.3028722 | 0.39947559 | -0.7581745 | 0.44834651 | 0.89297728 |
| MCAT | 39.287372 | 0.15130611 | 0.36408297 | 0.4155814 | 0.67771634 | 0.95560047 |
| TSPO | 405.319195 | -0.0764407 | 0.47909731 | -0.1595515 | 0.87323437 | 0.98246494 |
| EFCAB6 | 335.186777 | -0.1112187 | 0.33797866 | -0.3290703 | 0.74210258 | 0.96086423 |
| SAMM50 | 52.2944859 | -0.2633956 | 0.27365539 | -0.9625082 | 0.33579439 | 0.84552074 |
| RTL6 | 38.1292537 | 0.04565553 | 0.39162392 | 0.11658004 | 0.90719285 | 0.9875066 |
| NUP50-DT | 259.608915 | -0.1836887 | 0.34303567 | -0.5354799 | 0.59231807 | 0.93491607 |
| NUP50 | 145.902474 | -0.2750976 | 0.36707544 | -0.7494307 | 0.45359762 | 0.8955733 |
| KIAA0930 | 84.1001549 | -0.8187217 | 0.44714393 | -1.8310026 | 0.06710015 | 0.55774725 |
| FAM118A | 71.1465193 | -0.8178707 | 0.56841259 | -1.438868 | 0.15018794 | 0.7029344 |
| ATXN10 | 709.676283 | 0.08720247 | 0.17455057 | 0.49958285 | 0.61736884 | 0.94233994 |
| MIRLET7BHG | 360.063234 | 0.1803632 | 0.35799391 | 0.5038164 | 0.6143904 | 0.94093401 |
| PPARA | 340.50714 | 0.00529432 | 0.31074371 | 0.01703759 | 0.98640663 | 0.99783219 |
| TTC38 | 82.3158725 | 0.61263556 | 0.32382145 | 1.89189307 | 0.05850523 | 0.53216182 |
| TRMU | 104.445242 | -0.1674687 | 0.20828912 | -0.8040204 | 0.4213852 | 0.88360543 |
| CELSR1 | 166.958266 | 0.40635761 | 0.35299561 | 1.15116904 | 0.2496627 | 0.80149394 |
| GRAMD4 | 50.3196308 | -0.0578061 | 0.34351797 | -0.1682768 | 0.86636552 | 0.98040284 |
| CERK | 135.222893 | -0.024235 | 0.39967552 | -0.0606366 | 0.95164865 | 0.99404743 |
| TBC1D22A | 767.599847 | -0.4876022 | 0.1979717 | -2.4629896 | 0.01377839 | 0.28408502 |
| BRD1 | 153.495991 | -0.2310426 | 0.28692563 | -0.8052351 | 0.42068403 | 0.88331845 |
| ZBED4 | 79.1077116 | -0.1734847 | 0.44532928 | -0.3895649 | 0.69685829 | 0.95672887 |
| ALG12 | 43.5966288 | 0.11617535 | 0.35805646 | 0.32446098 | 0.74558906 | 0.96109466 |
| CRELD2 | 37.8447552 | -0.095213 | 0.29245531 | -0.3255644 | 0.74475395 | 0.96109466 |
| PIM3 | 564.481997 | -1.9373839 | 0.715983 | -2.7059077 | 0.0068118 | 0.20224502 |
| TRABD | 107.353383 | 0.10251966 | 0.35660476 | 0.28748821 | 0.77373853 | 0.96388926 |
| SELENOO | 146.490371 | -0.3906248 | 0.21363752 | -1.8284466 | 0.06748256 | 0.55848603 |
| TUBGCP6 | 45.1781366 | -0.3590451 | 0.27366715 | -1.3119775 | 0.18952774 | 0.75619145 |
| PLXNB2 | 593.569235 | 0.50543951 | 0.30194493 | 1.67394601 | 0.09414121 | 0.61330068 |
| DENND6B | 74.4799549 | 0.35711178 | 0.30062246 | 1.18790785 | 0.23486972 | 0.79580816 |
| PPP6R2 | 576.158748 | -0.1054707 | 0.2833805 | -0.3721877 | 0.70975314 | 0.95752169 |
| SBF1 | 42.8889231 | -0.1067584 | 0.27335235 | -0.3905523 | 0.69612822 | 0.95672887 |
| LMF2 | 53.108038 | 0.24080672 | 0.30294958 | 0.79487394 | 0.42668684 | 0.88477675 |
| ARSA | 42.2967794 | -0.0811855 | 0.29252214 | -0.2775361 | 0.78136847 | 0.96513523 |
| RABL2B | 80.132067 | -0.3451556 | 0.23909217 | -1.443609 | 0.148849 | 0.70204209 |
| CU633967.1 | 66.4813652 | 0.89647608 | 0.58450194 | 1.53374354 | 0.12509276 | 0.6663037 |

| | | | | | | |
|---|---|---|---|---|---|---|
| FP671120.1 | 1488.62304 | -0.4598777 | 0.37518205 | -1.2257454 | 0.22029452 | 0.78633738 |
| FP236383.1 | 915.378242 | -0.5727472 | 0.38515643 | -1.4870509 | 0.13700137 | 0.68129775 |
| CR381653.1 | 97.2669494 | -0.8176872 | 0.58411815 | -1.3998662 | 0.1615534 | 0.71416435 |
| AP001347.1 | 86.962113 | -0.341271 | 0.38379358 | -0.8892045 | 0.37389316 | 0.86232402 |
| HSPA13 | 56.4496489 | -0.0085022 | 0.365365 | -0.0232706 | 0.98143445 | 0.99696835 |
| SAMSN1 | 222.037666 | -0.0225853 | 0.69896008 | -0.0323128 | 0.97422262 | 0.99656026 |
| NRIP1 | 301.249482 | -0.3905382 | 0.37435519 | -1.0432291 | 0.29684218 | 0.83536682 |
| USP25 | 814.424747 | -0.2766755 | 0.28127166 | -0.9836595 | 0.32528296 | 0.84285425 |
| MIR99AHG | 2261.71451 | -0.2336362 | 0.3546814 | -0.6587214 | 0.51007472 | 0.91334187 |
| CXADR | 379.989847 | 0.1801396 | 0.26012393 | 0.69251451 | 0.48861428 | 0.90730741 |
| BTG3 | 277.406056 | 0.45936355 | 0.24876133 | 1.84660354 | 0.06480462 | 0.55126009 |
| C21orf91 | 38.859694 | 0.45769811 | 0.33740265 | 1.35653383 | 0.17492938 | 0.73227004 |
| CHODL | 154.796164 | 0.70244328 | 0.48331234 | 1.45339406 | 0.14611437 | 0.69477252 |
| NCAM2 | 80.3758188 | -0.022736 | 0.52392553 | -0.0433956 | 0.96538622 | 0.99637506 |
| MRPL39 | 65.5985906 | -0.1538921 | 0.28997118 | -0.5307151 | 0.59561625 | 0.93570054 |
| ATP5PF | 365.288455 | 0.36999907 | 0.4735707 | 0.78129637 | 0.4346282 | 0.88785137 |
| GABPA | 115.330821 | -0.0396423 | 0.22681021 | -0.1747818 | 0.86125113 | 0.97841862 |
| APP | 2458.10458 | 0.46331525 | 0.20269298 | 2.28579819 | 0.02226607 | 0.35844607 |
| CYYR1 | 95.4167177 | 0.84352353 | 0.55580117 | 1.51767138 | 0.12909726 | 0.67206108 |
| ADAMTS1 | 250.577812 | 0.36586497 | 0.64090619 | 0.57085573 | 5.68E-01 | 0.93122667 |
| ADAMTS5 | 92.7343816 | 0.82101273 | 0.51011151 | 1.60947698 | 0.10751208 | 0.63477557 |
| N6AMT1 | 38.8535238 | 0.14193529 | 0.28345201 | 0.50073835 | 0.61655528 | 0.94206626 |
| LTN1 | 209.674612 | -0.3242209 | 0.19070824 | -1.7000884 | 0.08911429 | 0.60739841 |
| RWDD2B | 31.4727471 | 1.15804175 | 0.42115149 | 2.74970354 | 0.00596492 | 0.19123755 |
| USP16 | 148.043288 | -0.4330047 | 0.23739847 | -1.8239573 | 0.06815852 | 0.55917298 |
| CCT8 | 127.338592 | 0.11913841 | 0.2128152 | 0.55982097 | 0.57560156 | 0.93292674 |
| MAP3K7CL | 102.98955 | 0.38128942 | 0.51809277 | 0.73594815 | 0.46176225 | 0.89799005 |
| BACH1 | 1056.23484 | -0.4596698 | 0.36892572 | -1.2459685 | 0.21277597 | 0.77750772 |
| GRIK1 | 70.9021455 | 0.27921692 | 0.59834325 | 0.46665007 | 0.64075026 | 0.94935885 |
| CLDN8 | 352.473494 | 2.42991749 | 0.58283649 | 4.16912378 | 3.06E-05 | 0.00781585 |
| TIAM1 | 80.8250119 | 0.43452484 | 0.3869341 | 1.12299444 | 0.26143985 | 0.80805101 |
| SOD1 | 472.982456 | 0.72136907 | 0.43646378 | 1.65275817 | 0.09838009 | 0.61690569 |
| SCAF4 | 180.026118 | 0.02048072 | 0.19347378 | 0.10585788 | 0.91569512 | 0.98868699 |
| HUNK | 650.161629 | 0.70942577 | 0.3979545 | 1.7826806 | 0.07463831 | 0.57072562 |
| URB1 | 77.6016286 | 0.18592348 | 0.27550009 | 0.67485816 | 0.49976588 | 0.90839719 |
| EVA1C | 294.986768 | 0.22292349 | 0.51971581 | 0.42893344 | 0.66797166 | 0.95294349 |
| SYNJ1 | 229.115864 | 0.48600495 | 0.23775599 | 2.04413332 | 0.04094038 | 0.46430138 |
| PAXBP1 | 339.297896 | 0.01784912 | 0.25012232 | 0.07136155 | 0.94311001 | 0.99265578 |
| C21orf62-AS | 78.8122397 | 0.20499119 | 0.51918205 | 0.39483489 | 0.69296475 | 0.95672887 |
| IFNAR1 | 280.261854 | 0.51880444 | 0.18577294 | 2.79268043 | 0.00522733 | 0.17920428 |
| IFNGR2 | 382.757323 | 0.27768407 | 0.39512511 | 0.70277506 | 0.48219594 | 0.90595437 |
| TMEM50B | 476.437846 | 0.04740427 | 0.32110342 | 0.14762929 | 0.88263534 | 0.98341679 |
| GART | 120.088139 | -0.1841283 | 0.19904122 | -0.9250763 | 0.35492623 | 0.8557799 |
| SON | 661.187192 | 0.0420432 | 0.22131274 | 0.18997187 | 0.84933118 | 0.97662834 |
| AP000311.1 | 122.394 | 0.08177412 | 0.46051233 | 0.17757205 | 0.85905908 | 0.97788846 |
| ITSN1 | 456.687307 | -0.0913221 | 0.34083947 | -0.2679328 | 0.78875102 | 0.96515649 |
| MRPS6 | 810.965615 | -0.0494924 | 0.66684118 | -0.0742191 | 0.94083603 | 0.99265578 |
| LINC00310 | 69.4634575 | 0.09125204 | 0.68295153 | 0.13361422 | 0.89370764 | 0.98597539 |
| AP000317.1 | 85.5821482 | -0.0569404 | 0.50307106 | -0.1131855 | 0.90988347 | 0.98788661 |
| KCNE1 | 66.079562 | -0.1848146 | 0.79734964 | -0.2317861 | 0.81670413 | 0.96874284 |
| RCAN1 | 654.959155 | 0.57834155 | 0.88969496 | 0.65004477 | 0.5156633 | 0.91626847 |
| CLIC6 | 95.8364513 | 1.19061189 | 0.48383881 | 2.46076146 | 0.01386425 | 0.28408502 |
| RUNX1 | 261.458892 | -0.5877795 | 0.6019211 | -0.9765059 | 0.32881381 | 0.84409527 |
| SETD4 | 81.478231 | -0.1335361 | 0.24288012 | -0.5498025 | 0.58245482 | 0.93406679 |

| | | | | | | |
|---|---|---|---|---|---|---|
| CBR1 | 178.319339 | 0.66140468 | 0.41121535 | 1.60841435 | 0.10774446 | 0.63520805 |
| CBR3 | 88.6534658 | -0.2221298 | 0.49633646 | -0.4475388 | 0.65448608 | 0.95100227 |
| DOPEY2 | 174.87757 | -0.1551724 | 0.23843588 | -0.6507928 | 0.51518024 | 0.91599162 |
| MORC3 | 312.011377 | -0.2320522 | 0.18143125 | -1.279009 | 0.20089389 | 0.76957251 |
| CLDN14 | 100.767056 | -1.8817422 | 0.63055565 | -2.9842603 | 0.00284265 | 0.1372855 |
| HLCS | 567.042731 | 0.16206897 | 0.25435009 | 0.63718858 | 0.52400202 | 0.91871245 |
| PIGP | 71.3028666 | 0.16387804 | 0.46522097 | 0.3522585 | 0.72464441 | 0.95963742 |
| TTC3 | 1111.31462 | -0.2589337 | 0.37919262 | -0.6828554 | 0.4946982 | 0.90839719 |
| VPS26C | 148.16936 | -0.4051001 | 0.18892527 | -2.1442348 | 0.03201408 | 0.41517597 |
| DYRK1A | 1108.47364 | -0.2916552 | 0.24411114 | -1.194764 | 0.23217924 | 0.7922741 |
| KCNJ15 | 1435.57648 | 1.48910847 | 0.58276321 | 2.55525475 | 0.01061101 | 0.25045809 |
| ETS2 | 206.921898 | -2.2778847 | 0.59328982 | -3.839413 | 0.00012333 | 0.02194494 |
| PSMG1 | 37.9501106 | -0.4041357 | 0.30229706 | -1.3368827 | 0.18126092 | 0.74315969 |
| BRWD1 | 978.741863 | -0.0784248 | 0.14605368 | -0.5369589 | 0.59129599 | 0.93489061 |
| HMGN1 | 465.4897 | 0.16495884 | 0.27811837 | 0.59312455 | 0.5530978 | 0.92872652 |
| WRB | 101.466608 | -0.3344391 | 0.22980775 | -1.4552995 | 0.14558637 | 0.69325556 |
| LCA5L | 43.0106943 | 0.08275362 | 0.35833407 | 0.23093985 | 0.81736153 | 0.96883031 |
| SH3BGR | 40.3669241 | -0.0219355 | 0.40485283 | -0.0541814 | 0.95679061 | 0.99565888 |
| PCP4 | 35.2721234 | 0.53488865 | 0.5732238 | 0.93312359 | 0.35075616 | 0.85477058 |
| BACE2 | 1545.98152 | -2.2611527 | 0.54212085 | -4.1709384 | 3.03E-05 | 0.00781585 |
| FAM3B | 210.082354 | 0.34516156 | 0.28103365 | 1.22818587 | 0.2193772 | 0.78630025 |
| MX1 | 352.600715 | -2.6474889 | 0.906352 | -2.9210383 | 0.00348867 | 0.14553941 |
| TMPRSS2 | 311.075576 | 1.03540093 | 0.42251752 | 2.4505515 | 0.01426376 | 0.28746487 |
| RIPK4 | 91.4505149 | -0.1670572 | 0.36160838 | -0.4619837 | 0.64409302 | 0.94949493 |
| PRDM15 | 46.3841185 | -0.3434007 | 0.3134229 | -1.0956466 | 0.27323347 | 0.81922962 |
| C2CD2 | 116.574391 | 0.10315434 | 0.24610837 | 0.41914194 | 0.6751124 | 0.95524793 |
| ZBTB21 | 91.1867848 | -0.099345 | 0.43252933 | -0.2296839 | 0.81833739 | 0.96883031 |
| RSPH1 | 81.3855767 | 0.7740265 | 0.43832367 | 1.76587885 | 0.07741618 | 0.57540534 |
| SLC37A1 | 134.492279 | -0.0880279 | 0.32830474 | -0.2681287 | 0.78860027 | 0.96515649 |
| LINC01671 | 77.6745951 | -1.5785511 | 0.65189646 | -2.4214752 | 0.01545765 | 0.30047476 |
| PDE9A | 605.013503 | -0.9034675 | 0.4294513 | -2.1037717 | 0.03539837 | 0.43541651 |
| WDR4 | 45.304616 | -0.4542953 | 0.40305552 | -1.1271283 | 0.25968823 | 0.80709747 |
| NDUFV3 | 203.134592 | 0.25440934 | 0.18546483 | 1.37173897 | 0.17014471 | 0.72675731 |
| PKNOX1 | 247.617794 | 0.3820905 | 0.20215761 | 1.89006238 | 0.05874962 | 0.53286516 |
| RRP1B | 77.8437804 | -0.0831354 | 0.29450785 | -0.2822859 | 0.77772429 | 0.96472142 |
| PDXK | 421.625362 | -0.0846285 | 0.31576128 | -0.2680143 | 0.78868829 | 0.96515649 |
| CSTB | 616.643106 | -0.3827744 | 0.52832461 | -0.724506 | 0.46875514 | 0.90026987 |
| RRP1 | 33.3458837 | -0.2187564 | 0.40104141 | -0.545471 | 0.58542965 | 0.93483536 |
| AGPAT3 | 300.319028 | 0.00201921 | 0.20614529 | 0.00979508 | 0.99218478 | 0.99919666 |
| TRAPPC10 | 246.925268 | -0.3231246 | 0.22873591 | -1.4126537 | 0.15775756 | 0.71201518 |
| PFKL | 592.122244 | -0.6164009 | 0.33935912 | -1.8163677 | 0.06931399 | 0.55917298 |
| C21orf2 | 66.762933 | 0.59274781 | 0.44269737 | 1.33894585 | 0.1805883 | 0.7429073 |
| UBE2G2 | 279.306907 | -0.0548213 | 0.24813963 | -0.2209293 | 0.8251475 | 0.97018919 |
| SUMO3 | 165.531249 | -0.2981375 | 0.40654574 | -0.7333431 | 0.46334918 | 0.89801718 |
| PTTG1IP | 447.799622 | 0.00942926 | 0.21953042 | 0.04295195 | 0.96573984 | 0.99637506 |
| FAM207A | 49.2314787 | 0.07061811 | 0.40993433 | 0.17226688 | 0.86322772 | 0.979268 |
| ADARB1 | 123.461102 | 0.25968329 | 0.31315036 | 0.82926072 | 0.40695689 | 0.87948803 |
| POFUT2 | 76.3334372 | -0.3874274 | 0.28237307 | -1.372041 | 0.17005068 | 0.72675731 |
| COL18A1 | 827.424487 | 0.45051467 | 0.23853765 | 1.88865223 | 0.05893844 | 0.53286516 |
| PCBP3 | 198.872135 | -1.6941327 | 0.67841179 | -2.4972041 | 0.01251769 | 0.27444642 |
| COL6A1 | 46.5944288 | 0.01865249 | 0.59129525 | 0.03154513 | 0.9748348 | 0.99656026 |
| LSS | 105.902491 | -0.7086908 | 0.57584021 | -1.2307073 | 0.21843235 | 0.78529418 |
| MCM3AP-AS | 51.44215 | 0.28343478 | 0.40445652 | 0.70077934 | 0.48344073 | 0.90631121 |
| MCM3AP | 183.56963 | 0.02202192 | 0.26012543 | 0.08465886 | 0.9325326 | 0.99243654 |

| | | | | | | |
|---|---|---|---|---|---|---|
| YBEY | 43.5688778 | -0.7705251 | 0.32144331 | -2.3970791 | 0.01652635 | 0.30723893 |
| PCNT | 102.70895 | -0.3726194 | 0.23057331 | -1.6160559 | 0.10608223 | 0.63461115 |
| DIP2A | 295.757193 | -0.3616993 | 0.17181477 | -2.1051701 | 0.03527651 | 0.43442748 |
| PRMT2 | 329.61481 | -0.1751761 | 0.2706268 | -0.6472978 | 0.5174392 | 0.91751084 |
| AC011043.1 | 103.39831 | 0.83509677 | 0.39928842 | 2.09146252 | 0.03648662 | 0.43729328 |
| AL592183.1 | 188.551408 | -0.0238771 | 0.45729623 | -0.0522136 | 0.95835851 | 0.99579587 |

**Collecting duct intercalated cells type A**

*This means that positive fold changes imply higher mean value in non-COVID AKI than in COVID AKI and vice versa.

| | baseMean | log2 fold cha | lfcSE | stat | pvalue | padj |
|---|---|---|---|---|---|---|
| AL669831.5 | 23.9983634 | -0.1490466 | 0.41785619 | -0.3566936 | 0.72132116 | 0.99937574 |
| NOC2L | 21.9691869 | 0.61664624 | 0.42418536 | 1.45371881 | 0.14602428 | 0.99937574 |
| ISG15 | 175.761502 | -5.3933889 | 1.66970332 | -3.2301481 | 0.00123726 | 0.33941289 |
| AGRN | 45.9738332 | -0.3253298 | 0.47172928 | -0.6896537 | 0.49041201 | 0.99937574 |
| C1orf159 | 23.2007535 | 0.01006581 | 0.45770359 | 0.02199199 | 0.98245434 | 0.99937574 |
| AL390719.2 | 15.8393265 | -0.1218592 | 0.47866829 | -0.2545797 | 0.79904778 | 0.99937574 |
| SDF4 | 62.3658481 | 0.21385755 | 0.33362074 | 0.64101994 | 0.52150973 | 0.99937574 |
| UBE2J2 | 49.111668 | 0.08665142 | 0.3241212 | 0.26734263 | 0.78920536 | 0.99937574 |
| ACAP3 | 14.4411895 | -0.6624194 | 0.57118433 | -1.1597297 | 0.24615888 | 0.99937574 |
| INTS11 | 38.8687937 | 0.11226707 | 0.35613509 | 0.3152373 | 0.75258149 | 0.99937574 |
| CPTP | 15.173372 | -0.4263135 | 0.71315662 | -0.5977838 | 0.5499842 | 0.99937574 |
| DVL1 | 25.6079257 | 0.32236239 | 0.42151969 | 0.76476234 | 0.44441307 | 0.99937574 |
| AURKAIP1 | 87.6826146 | -0.2791276 | 0.47081776 | -0.5928571 | 0.55327679 | 0.99937574 |
| CCNL2 | 78.1415717 | 0.01397417 | 0.32343963 | 0.04320488 | 0.96553822 | 0.99937574 |
| MRPL20 | 60.6860931 | -0.2627883 | 0.34824788 | -0.7546013 | 0.45048823 | 0.99937574 |
| VWA1 | 14.0596711 | 0.81947946 | 0.68435802 | 1.19744263 | 0.23113408 | 0.99937574 |
| ATAD3C | 10.3198 | -0.0663414 | 0.70398204 | -0.0942373 | 0.92492066 | 0.99937574 |
| ATAD3B | 26.8393954 | -0.3497627 | 0.43486796 | -0.8042963 | 0.42122588 | 0.99937574 |
| ATAD3A | 17.6943416 | 0.56963117 | 0.46373507 | 1.22835473 | 0.21931383 | 0.99937574 |
| SSU72 | 79.5260158 | -0.4499686 | 0.38992798 | -1.1539787 | 0.24850889 | 0.99937574 |
| MIB2 | 16.8975317 | 0.18826859 | 0.49071578 | 0.38366117 | 0.70122959 | 0.99937574 |
| CDK11B | 29.3520728 | -0.2881508 | 0.5154858 | -0.5589888 | 0.57616933 | 0.99937574 |
| SLC35E2B | 25.2846838 | -0.5639447 | 0.58330945 | -0.9668019 | 0.33364308 | 0.99937574 |
| CDK11A | 12.9060735 | 0.40672796 | 0.57461873 | 0.70782232 | 0.4790556 | 0.99937574 |
| SLC35E2A | 21.7280457 | 0.11764669 | 0.56874087 | 0.20685464 | 0.83612339 | 0.99937574 |
| NADK | 50.1602058 | -0.4072188 | 0.34962971 | -1.1647144 | 0.24413461 | 0.99937574 |
| GNB1 | 312.9607 | -0.1336086 | 0.29894355 | -0.4469357 | 0.65492146 | 0.99937574 |
| PRKCZ | 92.9317922 | -0.640584 | 0.49835831 | -1.2853885 | 0.19865657 | 0.99937574 |
| FAAP20 | 48.9496007 | -0.1751887 | 0.43378235 | -0.4038632 | 0.68631334 | 0.99937574 |
| SKI | 34.711009 | -0.1288507 | 0.33317258 | -0.3867386 | 0.69894977 | 0.99937574 |
| MORN1 | 15.1996098 | -0.0106116 | 0.61153554 | -0.0173524 | 0.9861555 | 0.99937574 |
| RER1 | 85.0233206 | 0.24856365 | 0.23879018 | 1.04092911 | 0.29790845 | 0.99937574 |
| PEX10 | 17.5634942 | -0.0016287 | 0.52882963 | -0.0030798 | 0.99754271 | 0.99937574 |
| PANK4 | 13.3661344 | 0.2779649 | 0.52964573 | 0.52481289 | 0.59971327 | 0.99937574 |
| TNFRSF14-A | 17.1347379 | -0.7020553 | 0.601287 | -1.1675877 | 0.24297311 | 0.99937574 |
| TNFRSF14 | 45.9019014 | -0.3604278 | 0.5290057 | -0.6813306 | 0.49566231 | 0.99937574 |
| LINC00982 | 49.9927911 | 1.03638594 | 0.65793172 | 1.57521807 | 0.11520612 | 0.99937574 |
| PRDM16 | 245.376301 | 0.16315815 | 0.3710366 | 0.43973599 | 0.66012834 | 0.99937574 |
| ARHGEF16 | 40.0161729 | -0.5366472 | 0.40707856 | -1.318289 | 0.18740693 | 0.99937574 |
| TPRG1L | 25.4803409 | 0.24225416 | 0.4814961 | 0.50312797 | 0.6148743 | 0.99937574 |
| WRAP73 | 11.7268162 | -0.138797 | 0.51180501 | -0.2711912 | 0.78624395 | 0.99937574 |
| LRRC47 | 21.8441422 | 0.67299773 | 0.54413592 | 1.23681915 | 0.21615423 | 0.99937574 |
| CEP104 | 48.8020517 | -0.2085804 | 0.39804785 | -0.5240084 | 0.6002727 | 0.99937574 |
| C1orf174 | 11.2212296 | -0.5744316 | 0.53207028 | -1.079616 | 0.28031323 | 0.99937574 |
| NPHP4 | 23.0466631 | -0.1563118 | 0.50747115 | -0.3080211 | 0.75806627 | 0.99937574 |
| RPL22 | 44.7341461 | -0.1571352 | 0.47118678 | -0.3334881 | 0.73876585 | 0.99937574 |
| RNF207 | 21.3646298 | -0.4730755 | 0.47590394 | -0.9940567 | 0.32019527 | 0.99937574 |
| ICMT | 25.514415 | 0.03482289 | 0.43292776 | 0.0804358 | 0.93589066 | 0.99937574 |
| NOL9 | 26.6235704 | -0.3584237 | 0.54197455 | -0.6613294 | 0.50840111 | 0.99937574 |
| ZBTB48 | 15.5919373 | -0.0234474 | 0.52435954 | -0.0447163 | 0.96433344 | 0.99937574 |
| KLHL21 | 25.9406754 | -1.4808818 | 0.69243399 | -2.1386613 | 0.03246311 | 0.99937574 |
| DNAJC11 | 72.1303552 | -0.1637567 | 0.33869375 | -0.4834948 | 0.62874448 | 0.99937574 |

| | | | | | | |
|---|---|---|---|---|---|---|
| CAMTA1 | 380.132492 | 0.12280505 | 0.30877299 | 0.39771954 | 0.69083694 | 0.99937574 |
| VAMP3 | 30.2765187 | -0.1915146 | 0.44940516 | -0.4261513 | 0.66999763 | 0.99937574 |
| PER3 | 118.051848 | -0.9020625 | 0.45415481 | -1.9862445 | 0.04700618 | 0.99937574 |
| PARK7 | 118.135707 | 0.5525305 | 0.45014979 | 1.22743699 | 0.21965839 | 0.99937574 |
| ERRFI1 | 21.4685093 | -0.5537533 | 0.79296186 | -0.6983353 | 0.4849675 | 0.99937574 |
| RERE | 1223.53237 | 0.23277085 | 0.29703168 | 0.78365663 | 0.43324162 | 0.99937574 |
| ENO1 | 260.217417 | -0.1479519 | 0.461923 | -0.3202956 | 0.74874426 | 0.99937574 |
| GPR157 | 24.1032696 | 0.24942945 | 0.63634358 | 0.39197291 | 0.69507823 | 0.99937574 |
| H6PD | 47.0041557 | -0.0087283 | 0.3656733 | -0.0238692 | 0.98095698 | 0.99937574 |
| SPSB1 | 16.6124766 | -2.7101145 | 0.69288375 | -3.9113552 | 9.18E-05 | 0.08697803 |
| SLC25A33 | 39.4475162 | -0.2596956 | 0.49523061 | -0.5243932 | 0.60000506 | 0.99937574 |
| PIK3CD | 11.3799688 | -0.1221765 | 0.58068439 | -0.2104008 | 0.83335488 | 0.99937574 |
| CLSTN1 | 65.3058163 | 0.25029388 | 0.38583119 | 0.64871344 | 0.51652362 | 0.99937574 |
| CTNNBIP1 | 23.2544558 | -0.0827082 | 0.40854561 | -0.2024455 | 0.83956847 | 0.99937574 |
| LZIC | 28.7232656 | -0.0595945 | 0.36994993 | -0.1610881 | 0.87202404 | 0.99937574 |
| NMNAT1 | 40.9502545 | 0.31236496 | 0.46092954 | 0.67768483 | 0.49797155 | 0.99937574 |
| RBP7 | 11.8498178 | -0.1762256 | 0.7814999 | -0.2254966 | 0.82159296 | 0.99937574 |
| UBE4B | 186.937593 | -0.0121335 | 0.26326969 | -0.0460878 | 0.96324027 | 0.99937574 |
| KIF1B | 398.451509 | 0.15097388 | 0.3428606 | 0.44033604 | 0.65969374 | 0.99937574 |
| PGD | 47.7708866 | -0.3752986 | 0.43421903 | -0.8643071 | 0.38741923 | 0.99937574 |
| DFFA | 54.9936222 | -0.2852249 | 0.47262847 | -0.6034866 | 0.54618506 | 0.99937574 |
| PEX14 | 119.637217 | -0.3710189 | 0.24019602 | -1.5446506 | 0.12243081 | 0.99937574 |
| CASZ1 | 273.702332 | 0.79107512 | 0.5123289 | 1.54407671 | 0.12256975 | 0.99937574 |
| AL139423.1 | 12.6022701 | 1.00347862 | 0.87345724 | 1.14885832 | 0.25061441 | 0.99937574 |
| SRM | 14.0198955 | -0.1086064 | 0.57156566 | -0.1900157 | 0.84929686 | 0.99937574 |
| EXOSC10 | 32.3016013 | -0.5367761 | 0.37422443 | -1.4343694 | 0.15146688 | 0.99937574 |
| MTOR | 127.608258 | -0.2538139 | 0.28690056 | -0.8846755 | 0.37633169 | 0.99937574 |
| UBIAD1 | 12.5945827 | -0.6132511 | 0.72662742 | -0.8439691 | 0.39868668 | 0.99937574 |
| FBXO2 | 18.383391 | -0.236653 | 0.56779221 | -0.4167951 | 0.67682827 | 0.99937574 |
| FBXO44 | 17.8601803 | 0.06813245 | 0.56546926 | 0.12048834 | 0.90409631 | 0.99937574 |
| AGTRAP | 20.4507002 | -0.5188372 | 0.5402025 | -0.9604495 | 0.33682904 | 0.99937574 |
| MTHFR | 30.5191057 | 0.33498276 | 0.59058254 | 0.56720735 | 0.57057331 | 0.99937574 |
| CLCN6 | 32.9856089 | -0.5417162 | 0.39030437 | -1.3879328 | 0.1651575 | 0.99937574 |
| KIAA2013 | 26.0253803 | 0.32830429 | 0.47813096 | 0.68664092 | 0.49230903 | 0.99937574 |
| MFN2 | 59.7695047 | -0.2927084 | 0.37558311 | -0.7793439 | 0.43577717 | 0.99937574 |
| VPS13D | 343.01607 | -0.1963304 | 0.20182472 | -0.9727769 | 0.33066421 | 0.99937574 |
| DHRS3 | 115.573632 | -0.9416042 | 0.49192407 | -1.9141252 | 0.05560416 | 0.99937574 |
| PRDM2 | 232.797862 | 0.00365023 | 0.32634564 | 0.01118518 | 0.9910757 | 0.99937574 |
| KAZN | 992.31998 | 0.60642689 | 0.56439527 | 1.07447196 | 0.28261121 | 0.99937574 |
| TMEM51-AS | 52.8140427 | -0.5629853 | 0.52436643 | -1.0736486 | 0.28298019 | 0.99937574 |
| TMEM51 | 433.819024 | -0.6634088 | 0.4884585 | -1.3581682 | 0.17441031 | 0.99937574 |
| C1orf195 | 17.938098 | 0.22171509 | 0.53252013 | 0.41635063 | 0.67715345 | 0.99937574 |
| CASP9 | 16.6300124 | -0.0226656 | 0.63200103 | -0.0358632 | 0.97139145 | 0.99937574 |
| DNAJC16 | 60.2943817 | 0.06254028 | 0.31530757 | 0.19834689 | 0.84277367 | 0.99937574 |
| DDI2 | 62.169987 | -0.11266 | 0.30248755 | -0.3724449 | 0.7095616 | 0.99937574 |
| PLEKHM2 | 62.3262527 | -0.270306 | 0.41092861 | -0.6577931 | 0.51067106 | 0.99937574 |
| UQCRHL | 16.4537425 | 0.08651461 | 0.76275205 | 0.11342429 | 0.90969418 | 0.99937574 |
| SPEN | 116.657918 | -0.5299989 | 0.23777757 | -2.2289694 | 0.02581594 | 0.96546668 |
| ZBTB17 | 21.1938301 | -0.0689738 | 0.42525142 | -0.1621953 | 0.87115204 | 0.99937574 |
| SRARP | 32.6270528 | 1.38272982 | 0.59961564 | 2.30602692 | 0.02110913 | 0.96427895 |
| CLCNKA | 201.227552 | 1.34478688 | 0.62271837 | 2.15954264 | 0.03080809 | 0.99049468 |
| CLCNKB | 336.39926 | 1.37085048 | 0.72114117 | 1.90094607 | 0.05730908 | 0.99937574 |
| FAM131C | 18.2227334 | 0.63792171 | 1.02654586 | 0.62142544 | 0.53431974 | 0.99937574 |
| EPHA2 | 12.7350112 | -0.6499078 | 0.98496753 | -0.6598266 | 0.50936513 | 0.99937574 |

| | | | | | | |
|---|---|---|---|---|---|---|
| FBXO42 | 114.544861 | -0.1492454 | 0.2732012 | -0.5462838 | 0.58487089 | 0.99937574 |
| SZRD1 | 61.9413711 | 0.01969817 | 0.37533232 | 0.05248195 | 0.95814468 | 0.99937574 |
| NECAP2 | 20.7324652 | -0.4020827 | 0.42187961 | -0.9530746 | 0.34055226 | 0.99937574 |
| NBPF1 | 29.3109943 | 0.07975841 | 0.45256485 | 0.17623642 | 0.86010821 | 0.99937574 |
| CROCC | 26.3791058 | 0.37175066 | 0.42858621 | 0.86738831 | 0.38572929 | 0.99937574 |
| ATP13A2 | 18.8221856 | 0.37411652 | 0.49796661 | 0.75128837 | 0.45247913 | 0.99937574 |
| SDHB | 84.2649204 | 0.16035883 | 0.31811043 | 0.50409799 | 0.61419252 | 0.99937574 |
| RCC2 | 18.6532187 | -0.4209647 | 0.45950586 | -0.9161249 | 0.35960141 | 0.99937574 |
| ARHGEF10L | 127.309384 | -0.1356041 | 0.4955454 | -0.2736463 | 0.78435649 | 0.99937574 |
| KLHDC7A | 13.4925687 | -0.8973317 | 1.36606203 | -0.6568748 | 0.51126143 | 0.99937574 |
| UBR4 | 131.634152 | -0.0625783 | 0.28094279 | -0.222744 | 0.82373472 | 0.99937574 |
| AL035413.1 | 18.3873377 | 1.06786435 | 0.60271138 | 1.77176736 | 0.07643318 | 0.99937574 |
| EMC1 | 20.02347 | 0.28700627 | 0.58291611 | 0.49236291 | 0.62246281 | 0.99937574 |
| MRTO4 | 12.776665 | -0.5092563 | 0.55404019 | -0.9191684 | 0.35800748 | 0.99937574 |
| AKR7A2 | 20.1338855 | 0.37140277 | 0.62609001 | 0.59320987 | 0.55304071 | 0.99937574 |
| CAPZB | 204.922011 | -0.0402354 | 0.32599516 | -0.1234233 | 0.90177194 | 0.99937574 |
| NBL1 | 16.0745965 | 0.14977999 | 0.60218095 | 0.24872921 | 0.80357025 | 0.99937574 |
| TMCO4 | 223.10674 | 0.26776585 | 0.47690927 | 0.56146078 | 0.57448346 | 0.99937574 |
| OTUD3 | 25.2840262 | -0.0145844 | 0.49601443 | -0.0294032 | 0.97654301 | 0.99937574 |
| VWA5B1 | 220.719665 | -0.118322 | 0.76037971 | -0.1556091 | 0.87634114 | 0.99937574 |
| CAMK2N1 | 250.152909 | 0.06214122 | 0.28822959 | 0.21559625 | 0.82930248 | 0.99937574 |
| MUL1 | 19.0175074 | -0.3440386 | 0.50423284 | -0.682301 | 0.49504864 | 0.99937574 |
| CDA | 40.6734775 | -0.8175235 | 0.60009439 | -1.3623249 | 0.17309538 | 0.99937574 |
| PINK1 | 61.817769 | 0.47350786 | 0.49273738 | 0.96097409 | 0.33656519 | 0.99937574 |
| DDOST | 25.0116331 | 0.20037595 | 0.47781879 | 0.41935553 | 0.67495632 | 0.99937574 |
| HP1BP3 | 220.693694 | 0.09963739 | 0.25484832 | 0.39096741 | 0.69582133 | 0.99937574 |
| EIF4G3 | 509.462222 | -0.2794368 | 0.3676535 | -0.7600548 | 0.44722185 | 0.99937574 |
| ECE1 | 130.31039 | 0.67394403 | 0.45562846 | 1.47915261 | 0.13909953 | 0.99937574 |
| NBPF3 | 25.1203345 | 0.69661888 | 0.50995664 | 1.36603552 | 0.17192782 | 0.99937574 |
| RAP1GAP | 197.575599 | 0.59764383 | 0.49631029 | 1.20417377 | 0.22852242 | 0.99937574 |
| USP48 | 141.074019 | 0.23164426 | 0.27908559 | 0.83001155 | 0.40653225 | 0.99937574 |
| HSPG2 | 36.3732965 | 0.57464739 | 0.65561069 | 0.87650705 | 0.38075444 | 0.99937574 |
| CDC42 | 188.461102 | -0.1966541 | 0.25605952 | -0.7680017 | 0.44248617 | 0.99937574 |
| ZBTB40 | 56.1108881 | 0.05373039 | 0.41693029 | 0.12887139 | 0.89745941 | 0.99937574 |
| EPHB2 | 19.8508388 | -1.5293215 | 0.80480175 | -1.9002463 | 0.05740081 | 0.99937574 |
| KDM1A | 128.144554 | -0.0544331 | 0.34975451 | -0.1556322 | 0.87632294 | 0.99937574 |
| AL031428.1 | 28.070482 | 0.16698674 | 0.56268838 | 0.29676594 | 0.76664521 | 0.99937574 |
| LUZP1 | 140.698394 | 0.00506635 | 0.22447437 | 0.02256984 | 0.9819934 | 0.99937574 |
| LINC01355 | 12.0548772 | 0.49366696 | 0.67341477 | 0.73308009 | 0.46350958 | 0.99937574 |
| HNRNPR | 120.998681 | -0.1969874 | 0.23580087 | -0.8353972 | 0.4034941 | 0.99937574 |
| TCEA3 | 33.0076746 | 1.06535726 | 0.65229418 | 1.63324662 | 0.10241714 | 0.99937574 |
| ASAP3 | 51.9194 | 0.36154161 | 0.46952838 | 0.77001013 | 0.44129388 | 0.99937574 |
| ID3 | 29.8817794 | -1.0170127 | 0.59612512 | -1.706039 | 0.08800081 | 0.99937574 |
| RPL11 | 348.952635 | -0.1356737 | 0.55583935 | -0.2440881 | 0.80716262 | 0.99937574 |
| ELOA | 27.7869937 | -0.1270062 | 0.47442338 | -0.2677064 | 0.78892535 | 0.99937574 |
| ELOA-AS1 | 21.4079812 | 0.77902791 | 0.65470821 | 1.18988565 | 0.23409134 | 0.99937574 |
| PITHD1 | 28.0890999 | -0.6828836 | 0.43084031 | -1.585004 | 0.11296543 | 0.99937574 |
| LYPLA2 | 30.757692 | -0.5041564 | 0.48741131 | -1.0343552 | 0.30097014 | 0.99937574 |
| GALE | 39.9441778 | -0.6111141 | 0.35405281 | -1.7260536 | 8.43E-02 | 0.99937574 |
| HMGCL | 47.5642846 | 0.24180588 | 0.34357398 | 0.70379568 | 0.48156003 | 0.99937574 |
| FUCA1 | 59.4963331 | 0.48332392 | 0.33382546 | 1.44783423 | 0.14766342 | 0.99937574 |
| PNRC2 | 63.3867833 | -0.471172 | 0.41779284 | -1.1277646 | 0.25941936 | 0.99937574 |
| SRSF10 | 103.645114 | 0.05629605 | 0.33059038 | 0.17028942 | 0.86478253 | 0.99937574 |
| IFNLR1 | 19.5459155 | -0.0072069 | 0.51150243 | -0.0140896 | 0.98875847 | 0.99937574 |

| | | | | | | |
|---|---|---|---|---|---|---|
| STPG1 | 16.3573775 | -0.0289365 | 0.51148824 | -0.0565732 | 0.95488516 | 0.99937574 |
| NIPAL3 | 78.7056864 | 0.00780815 | 0.40205917 | 0.0194204 | 0.98450574 | 0.99937574 |
| RCAN3 | 55.7277606 | -0.57082 | 0.48230534 | -1.1835242 | 0.23660148 | 0.99937574 |
| SRRM1 | 117.763204 | -0.1094181 | 0.28929791 | -0.3782196 | 0.70526749 | 0.99937574 |
| CLIC4 | 21.3220876 | -1.4012913 | 0.64977826 | -2.1565685 | 0.0310393 | 0.99049468 |
| SYF2 | 47.4779667 | -0.03577 | 0.42434592 | -0.0842945 | 0.93282231 | 0.99937574 |
| RSRP1 | 258.866914 | 0.38033255 | 0.3764259 | 1.01037827 | 0.31231409 | 0.99937574 |
| TMEM50A | 77.1227292 | 0.12789343 | 0.37578279 | 0.3403387 | 0.73360148 | 0.99937574 |
| MACO1 | 73.7200718 | -0.3637145 | 0.39809526 | -0.9136368 | 0.36090775 | 0.99937574 |
| LDLRAP1 | 23.0087878 | 0.18720275 | 0.52335503 | 0.35769742 | 0.72056976 | 0.99937574 |
| MAN1C1 | 326.860318 | 0.47589104 | 0.45248659 | 1.05172409 | 0.29292616 | 0.99937574 |
| AL031280.1 | 29.5523108 | 0.00394017 | 0.54068891 | 0.00728732 | 0.99418561 | 0.99937574 |
| MTFR1L | 38.0550042 | 0.32280049 | 0.49463186 | 0.65260757 | 0.5140093 | 0.99937574 |
| PAQR7 | 10.8645823 | 0.23562624 | 0.59264875 | 0.3975816 | 0.69093862 | 0.99937574 |
| STMN1 | 13.4964445 | -0.5458321 | 0.87745233 | -0.6220646 | 0.53389937 | 0.99937574 |
| PAFAH2 | 28.5905567 | 0.40571787 | 0.42200956 | 0.96139498 | 0.33635361 | 0.99937574 |
| PDIK1L | 14.2478128 | -0.1762613 | 0.55407139 | -0.3181202 | 0.75039374 | 0.99937574 |
| ZNF593 | 24.9190626 | -0.103463 | 0.49947147 | -0.207145 | 0.83589659 | 0.99937574 |
| CNKSR1 | 27.4204385 | -0.6550735 | 0.42792482 | -1.5308145 | 0.12581524 | 0.99937574 |
| CEP85 | 99.1872962 | 0.15119254 | 0.38278448 | 0.39498086 | 6.93E-01 | 0.99937574 |
| SH3BGRL3 | 57.1116077 | -0.321622 | 0.3990833 | -0.805902 | 0.42029936 | 0.99937574 |
| DHDDS | 38.0679391 | -0.1945391 | 0.41655784 | -0.4670158 | 0.64048856 | 0.99937574 |
| HMGN2 | 43.9609475 | -0.6010809 | 0.35378679 | -1.6989921 | 0.08932068 | 0.99937574 |
| RPS6KA1 | 40.7943899 | -0.6214672 | 0.49085482 | -1.2660917 | 0.20548026 | 0.99937574 |
| ARID1A | 144.316335 | 0.12115562 | 0.30952324 | 0.39142657 | 0.69548196 | 0.99937574 |
| ZDHHC18 | 15.5912539 | 0.33812015 | 0.47592965 | 0.71044146 | 0.47743042 | 0.99937574 |
| GPN2 | 20.6579375 | -0.1390766 | 0.53470791 | -0.2600982 | 0.79478802 | 0.99937574 |
| NUDC | 52.1447005 | -0.4078858 | 0.39414759 | -1.0348555 | 0.30073639 | 0.99937574 |
| NR0B2 | 14.9197282 | -0.8900161 | 0.96700975 | -0.9203797 | 0.35737438 | 0.99937574 |
| KDF1 | 9.96923888 | 0.08921733 | 0.69633865 | 0.12812348 | 0.89805126 | 0.99937574 |
| TRNP1 | 27.6948409 | 0.30507822 | 0.50334971 | 0.60609594 | 0.54445105 | 0.99937574 |
| SLC9A1 | 34.7700708 | -0.8113051 | 0.54218621 | -1.4963588 | 0.13456018 | 0.99937574 |
| WDTC1 | 53.892314 | -0.361426 | 0.30024955 | -1.2037521 | 0.22868543 | 0.99937574 |
| TMEM222 | 21.9451615 | -0.0685823 | 0.49452353 | -0.1386837 | 0.88970013 | 0.99937574 |
| WASF2 | 119.116841 | 0.11079808 | 0.32621694 | 0.33964539 | 0.73412359 | 0.99937574 |
| AHDC1 | 17.7318173 | 0.09390423 | 0.51130775 | 0.18365501 | 0.85428411 | 0.99937574 |
| IFI6 | 181.637526 | -4.7224454 | 1.66184739 | -2.8416842 | 0.00448759 | 0.60810972 |
| FAM76A | 24.4826784 | -0.2019124 | 0.42498336 | -0.4751065 | 0.63471106 | 0.99937574 |
| STX12 | 131.443262 | -0.3140761 | 0.37779975 | -0.8313296 | 0.40578747 | 0.99937574 |
| PPP1R8 | 21.3988322 | -0.3508512 | 0.47257906 | -0.742418 | 0.45783412 | 0.99937574 |
| RPA2 | 23.8225872 | 0.02734912 | 0.44660924 | 0.06123724 | 0.95117027 | 0.99937574 |
| EYA3 | 282.001285 | -0.4199875 | 0.31232493 | -1.3447132 | 0.17871786 | 0.99937574 |
| DNAJC8 | 45.9072028 | -0.1534078 | 0.41971559 | -0.3655041 | 0.71473514 | 0.99937574 |
| ATP5IF1 | 135.433229 | 0.42483841 | 0.47238035 | 0.89935664 | 0.36846273 | 0.99937574 |
| SESN2 | 12.1313557 | -0.4244131 | 0.82979426 | -0.5114679 | 0.60902349 | 0.99937574 |
| PHACTR4 | 215.740261 | -0.2889282 | 0.25861669 | -1.1172065 | 0.26390606 | 0.99937574 |
| RCC1 | 30.0834967 | -0.0384615 | 0.40763334 | -0.0943532 | 0.92482858 | 0.99937574 |
| TRNAU1AP | 59.332435 | -0.3797609 | 0.30149609 | -1.259588 | 0.20781803 | 0.99937574 |
| SNHG12 | 12.8242921 | -0.0172576 | 0.61013599 | -0.0282849 | 0.97743495 | 0.99937574 |
| TAF12 | 25.1069917 | -0.0313671 | 0.5012629 | -0.0625761 | 0.95010406 | 0.99937574 |
| GMEB1 | 26.7185532 | -1.2365797 | 0.62388724 | -1.9820564 | 0.04747293 | 0.99937574 |
| YTHDF2 | 91.3780734 | -0.1235183 | 0.33316263 | -0.3707448 | 0.71082764 | 0.99937574 |
| EPB41 | 257.660586 | 0.09227428 | 0.39983291 | 0.2307821 | 0.81748409 | 0.99937574 |
| SRSF4 | 245.375654 | -0.3163306 | 0.25185164 | -1.2560194 | 0.20910892 | 0.99937574 |

| | | | | | | |
|---|---|---|---|---|---|---|
| MECR | 26.5901407 | 0.56589311 | 0.47901036 | 1.1813797 | 0.23745192 | 0.99937574 |
| PTPRU | 11.2849979 | 0.52403535 | 0.59925898 | 0.87447225 | 0.38186113 | 0.99937574 |
| LINC01648 | 18.8656871 | 1.08435566 | 1.96347531 | 0.55226345 | 5.81E-01 | 0.99937574 |
| PUM1 | 327.861926 | -0.0120272 | 0.21335071 | -0.0563729 | 0.95504474 | 0.99937574 |
| SNRNP40 | 44.0428847 | -0.1047521 | 0.33795961 | -0.3099544 | 0.7565956 | 0.99937574 |
| ZCCHC17 | 54.2659223 | -0.3028347 | 0.31912184 | -0.9489627 | 0.34263958 | 0.99937574 |
| SERINC2 | 118.030869 | -0.2949354 | 0.37583234 | -0.7847525 | 0.43259869 | 0.99937574 |
| TINAGL1 | 32.3380522 | -1.1232302 | 0.509544 | -2.2043831 | 0.02749741 | 0.98755826 |
| PEF1 | 31.3981212 | -0.413679 | 0.34329707 | -1.2050176 | 0.22819651 | 0.99937574 |
| PTP4A2 | 64.3151693 | -0.2394729 | 0.42460989 | -0.5639834 | 0.57276542 | 0.99937574 |
| KHDRBS1 | 136.5728 | -0.4412284 | 0.26634996 | -1.6565741 | 0.09760562 | 0.99937574 |
| TMEM39B | 13.2117413 | 0.67942949 | 0.6591331 | 1.03079256 | 0.30263811 | 0.99937574 |
| KPNA6 | 143.676094 | 0.09842406 | 0.23534349 | 0.41821453 | 0.67579028 | 0.99937574 |
| TXLNA | 38.2790628 | -0.4820622 | 0.39449714 | -1.2219664 | 0.22172035 | 0.99937574 |
| EIF3I | 38.3882493 | 0.46573483 | 0.32213449 | 1.44577761 | 0.14823959 | 0.99937574 |
| HDAC1 | 72.3963594 | 0.36567371 | 0.36673131 | 0.99711614 | 0.31870814 | 0.99937574 |
| MARCKSL1 | 49.4972918 | 0.08679841 | 0.73576135 | 0.11797087 | 0.90609074 | 0.99937574 |
| FAM229A | 13.0909113 | 0.70863844 | 0.50538087 | 1.4021869 | 0.16085944 | 0.99937574 |
| BSDC1 | 55.478729 | -0.2442396 | 0.35589178 | -0.6862748 | 0.49253983 | 0.99937574 |
| ZBTB8OS | 77.5998273 | 0.12722865 | 0.24203074 | 0.52567146 | 0.5991165 | 0.99937574 |
| RBBP4 | 73.8071335 | 0.08152033 | 0.29361332 | 0.27764522 | 0.78128472 | 0.99937574 |
| SYNC | 17.8591545 | 0.46696266 | 0.53995185 | 0.86482279 | 0.38713606 | 0.99937574 |
| KIAA1522 | 156.299673 | -0.2782569 | 0.33473193 | -0.8312827 | 0.40581397 | 0.99937574 |
| YARS | 64.3130863 | -0.6586681 | 0.37239255 | -1.7687466 | 0.07693617 | 0.99937574 |
| S100PBP | 68.7500215 | -0.0218114 | 0.365649 | -0.0596513 | 0.95243334 | 0.99937574 |
| TMEM54 | 33.7981424 | 0.0905589 | 0.52755529 | 0.17165765 | 0.86370669 | 0.99937574 |
| RNF19B | 36.0557501 | -1.4083765 | 0.47332742 | -2.9754805 | 0.0029253 | 0.51793204 |
| AL020995.1 | 29.9225966 | 0.33567427 | 0.7829852 | 0.42871088 | 0.66813364 | 0.99937574 |
| AK2 | 132.389044 | -0.097618 | 0.29940836 | -0.3260364 | 0.7443968 | 0.99937574 |
| ZNF362 | 14.2826409 | 0.8514955 | 0.68928099 | 1.23533872 | 0.21670447 | 0.99937574 |
| AL513327.1 | 22.2297266 | 0.37097468 | 0.82455782 | 0.44990742 | 0.6527772 | 0.99937574 |
| PHC2 | 122.300226 | -0.3567789 | 0.34388402 | -1.0374978 | 0.29950393 | 0.99937574 |
| SMIM12 | 32.7490789 | -0.3076261 | 0.47910619 | -0.6420834 | 0.52081902 | 0.99937574 |
| ZMYM1 | 45.8208819 | -0.0395433 | 0.36948929 | -0.1070214 | 0.91477201 | 0.99937574 |
| SFPQ | 206.290411 | -0.270376 | 0.26224973 | -1.0309867 | 0.30254708 | 0.99937574 |
| ZMYM4 | 315.856387 | 0.09195646 | 0.211458 | 0.43486866 | 0.66365776 | 0.99937574 |
| KIAA0319L | 206.429286 | -0.1776403 | 0.3052604 | -0.5819303 | 0.56061362 | 0.99937574 |
| PSMB2 | 48.0799642 | -0.0988586 | 0.37114263 | -0.2663629 | 0.78995975 | 0.99937574 |
| AGO4 | 97.5481493 | -0.0620373 | 0.33537744 | -0.1849775 | 0.85324665 | 0.99937574 |
| AGO1 | 24.4235746 | -0.0969947 | 0.41838548 | -0.2318309 | 0.81666933 | 0.99937574 |
| AGO3 | 225.917826 | -0.1753711 | 0.27506021 | -0.6375737 | 0.52375124 | 0.99937574 |
| ADPRHL2 | 17.8451982 | -1.1701342 | 0.5433102 | -2.1537129 | 0.03126269 | 0.99049468 |
| TRAPPC3 | 32.0519794 | -0.087552 | 0.38667188 | -0.2264244 | 0.82087134 | 0.99937574 |
| MAP7D1 | 16.4020039 | 0.0353383 | 0.5762461 | 0.06132501 | 0.95110038 | 0.99937574 |
| THRAP3 | 150.956946 | -0.2026568 | 0.37553227 | -0.5396521 | 0.58943696 | 0.99937574 |
| EVA1B | 22.1367435 | 0.00519723 | 0.49502857 | 0.01049884 | 0.99162329 | 0.99937574 |
| STK40 | 69.2763342 | -0.4085631 | 0.47974264 | -0.8516298 | 0.39441959 | 0.99937574 |
| LSM10 | 16.9016712 | 0.26622156 | 0.62450081 | 0.42629498 | 0.66989292 | 0.99937574 |
| MRPS15 | 28.6139964 | -0.0952073 | 0.53905946 | -0.1766174 | 0.85980892 | 0.99937574 |
| MEAF6 | 33.124331 | -0.1439713 | 0.3341022 | -0.43092 | 0.66652657 | 0.99937574 |
| SNIP1 | 15.5709844 | -0.0359957 | 0.47536802 | -0.0757217 | 0.93964055 | 0.99937574 |
| GNL2 | 14.5935614 | -1.2353769 | 0.53217593 | -2.3213695 | 0.02026691 | 0.96427895 |
| C1orf122 | 49.3303793 | 0.36347996 | 0.4753831 | 0.76460429 | 0.44450721 | 0.99937574 |
| MTF1 | 34.9584524 | -0.1963624 | 0.52437763 | -0.3744676 | 0.70805647 | 0.99937574 |

| | | | | | | |
|---|---|---|---|---|---|---|
| AL929472.2 | 13.8336167 | 0.38997612 | 0.63740772 | 0.6118158 | 0.54065964 | 0.99937574 |
| INPP5B | 99.384039 | 0.15347634 | 0.36984597 | 0.41497368 | 0.67816117 | 0.99937574 |
| SF3A3 | 29.2613836 | -0.3261007 | 0.3633824 | -0.8974036 | 0.36950359 | 0.99937574 |
| UTP11 | 10.5232274 | -0.4386233 | 0.60526895 | -0.7246751 | 0.46865137 | 0.99937574 |
| RRAGC | 40.2981918 | -0.8144092 | 0.61489487 | -1.3244691 | 0.18534731 | 0.99937574 |
| AL139260.1 | 78.8955372 | -0.7828365 | 0.73690845 | -1.0623253 | 0.28808804 | 0.99937574 |
| RHBDL2 | 23.461865 | 0.11308665 | 0.77115752 | 0.14664534 | 0.88341197 | 0.99937574 |
| AKIRIN1 | 89.841048 | -0.0395847 | 0.40549176 | -0.0976215 | 0.92223281 | 0.99937574 |
| NDUFS5 | 223.601401 | 0.27774898 | 0.36178115 | 0.76772652 | 0.44264968 | 0.99937574 |
| MACF1 | 1210.32324 | -0.3352739 | 0.29434277 | -1.1390594 | 0.2546784 | 0.99937574 |
| PABPC4 | 37.2536393 | -0.4119138 | 0.35215437 | -1.1696966 | 0.24212311 | 0.99937574 |
| PPIE | 21.9451397 | -0.1990427 | 0.44753934 | -0.4447491 | 0.65650106 | 0.99937574 |
| TRIT1 | 33.2897561 | 0.49160548 | 0.39918312 | 1.23152874 | 0.21812518 | 0.99937574 |
| CAP1 | 85.8625161 | -0.190979 | 0.30021094 | -0.6361493 | 0.52467911 | 0.99937574 |
| PPT1 | 22.801651 | -0.1081325 | 0.53105698 | -0.2036175 | 0.83865239 | 0.99937574 |
| RLF | 162.317225 | -0.0685644 | 0.28074906 | -0.2442196 | 0.80706071 | 0.99937574 |
| ZMPSTE24 | 168.753818 | 0.06903079 | 0.36785948 | 0.18765532 | 0.85114685 | 0.99937574 |
| COL9A2 | 50.7580362 | 0.88738752 | 0.93697361 | 0.94707845 | 0.3435988 | 0.99937574 |
| SMAP2 | 36.095012 | -0.0021686 | 0.42629911 | -0.0050871 | 0.99594112 | 0.99937574 |
| AL603839.3 | 14.7141143 | 0.63272572 | 0.55962764 | 1.13061914 | 0.25821543 | 0.99937574 |
| ZNF684 | 40.686732 | 0.5625046 | 0.54717823 | 1.02800982 | 0.30394521 | 0.99937574 |
| NFYC | 167.905015 | 0.63222832 | 0.28673631 | 2.20491197 | 0.02746027 | 0.98755826 |
| CITED4 | 95.0889676 | -0.342152 | 0.6682432 | -0.5120172 | 0.60863899 | 0.99937574 |
| CTPS1 | 26.3164105 | 0.19951335 | 0.53957389 | 0.36976094 | 0.71156062 | 0.99937574 |
| SLFNL1-AS1 | 21.8780985 | 0.67422448 | 0.59286098 | 1.13723875 | 0.25543849 | 0.99937574 |
| SCMH1 | 491.661835 | -0.3878351 | 0.27570162 | -1.4067204 | 0.1595103 | 0.99937574 |
| HIVEP3 | 34.1796528 | 0.60700223 | 0.44393201 | 1.36733152 | 0.17152142 | 0.99937574 |
| FOXJ3 | 173.198557 | -0.4199805 | 0.21523279 | -1.9512851 | 0.05102314 | 0.99937574 |
| PPCS | 37.9807388 | 0.2165129 | 0.42933058 | 0.50430346 | 0.61404815 | 0.99937574 |
| CCDC30 | 49.0099995 | 0.63301597 | 0.52011749 | 1.21706342 | 0.22358009 | 0.99937574 |
| PPIH | 15.6029761 | -0.3257711 | 0.50597585 | -0.6438471 | 0.51967461 | 0.99937574 |
| YBX1 | 116.861617 | -0.5277977 | 0.49752925 | -1.0608376 | 0.28876373 | 0.99937574 |
| SVBP | 19.9431761 | -0.3546651 | 0.54752109 | -0.6477652 | 0.51713684 | 0.99937574 |
| ERMAP | 29.1541755 | 0.32000874 | 0.36751697 | 0.87073187 | 0.38390057 | 0.99937574 |
| SLC2A1 | 27.4708722 | -0.8018747 | 0.70697789 | -1.1342288 | 0.25669858 | 0.99937574 |
| EBNA1BP2 | 22.8198049 | -0.4235545 | 0.4975946 | -0.851204 | 0.39465601 | 0.99937574 |
| TMEM125 | 11.5005534 | 0.0511868 | 0.76653757 | 0.06677664 | 0.94675952 | 0.99937574 |
| ELOVL1 | 26.0463493 | 0.01797194 | 0.44561063 | 0.04033103 | 0.96782922 | 0.99937574 |
| MED8 | 14.9248813 | -0.5124872 | 0.56732151 | -0.9033453 | 0.36634264 | 0.99937574 |
| SZT2 | 38.6740412 | -0.2552334 | 0.33576965 | -0.7601444 | 0.44716828 | 0.99937574 |
| PTPRF | 127.604024 | 0.09863441 | 0.36328682 | 0.27150561 | 0.78600219 | 0.99937574 |
| IPO13 | 15.3074422 | -0.1981972 | 0.5551497 | -0.3570157 | 0.72108002 | 0.99937574 |
| ATP6V0B | 531.68551 | 0.65029184 | 0.40797945 | 1.59393283 | 0.11095108 | 0.99937574 |
| B4GALT2 | 22.5393472 | -0.5532171 | 0.47733092 | -1.1589802 | 0.24646425 | 0.99937574 |
| CCDC24 | 24.7889306 | 0.12070774 | 0.49572815 | 0.24349584 | 0.80762131 | 0.99937574 |
| DMAP1 | 16.3952519 | 0.98884843 | 0.53847924 | 1.83637244 | 0.06630259 | 0.99937574 |
| ERI3 | 64.4545965 | 0.43104018 | 0.33737616 | 1.27762488 | 0.20138172 | 0.99937574 |
| RNF220 | 29.0040649 | -0.0503047 | 0.35414425 | -0.1420459 | 0.88704377 | 0.99937574 |
| TMEM53 | 46.5340829 | 0.68769101 | 0.44240593 | 1.55443443 | 0.12008083 | 0.99937574 |
| ARMH1 | 12.8470632 | 0.33273194 | 0.56805033 | 0.58574376 | 0.55804772 | 0.99937574 |
| RPS8 | 246.794105 | 0.07896027 | 0.52172569 | 0.15134442 | 0.87970403 | 0.99937574 |
| EIF2B3 | 47.2876845 | 0.00910129 | 0.3946759 | 0.02306016 | 0.98160229 | 0.99937574 |
| HECTD3 | 13.2433242 | -0.4971193 | 0.69234403 | -0.7180235 | 0.47274283 | 0.99937574 |
| UROD | 13.543191 | 0.4322407 | 0.59349079 | 0.72830229 | 0.46642856 | 0.99937574 |

| | | | | | | |
|---|---|---|---|---|---|---|
| ZSWIM5 | 116.338998 | 0.2055773 | 0.33419846 | 0.6151354 | 0.53846531 | 0.99937574 |
| TESK2 | 14.1774966 | -0.6663089 | 0.59351677 | -1.1226455 | 0.26158809 | 0.99937574 |
| PRDX1 | 297.807616 | -0.4161348 | 0.66504427 | -0.625725 | 0.53149532 | 0.99937574 |
| AKR1A1 | 18.8744774 | 0.02515442 | 0.46700328 | 0.05386348 | 0.95704393 | 0.99937574 |
| NASP | 74.2717054 | 0.15590746 | 0.39720922 | 0.39250715 | 0.69468353 | 0.99937574 |
| GPBP1L1 | 213.943532 | 0.09763613 | 0.24615269 | 0.39664864 | 0.69162658 | 0.99937574 |
| TMEM69 | 13.1441783 | 0.56280371 | 0.50238596 | 1.12026162 | 0.26260229 | 0.99937574 |
| IPP | 44.3657729 | 0.21087891 | 0.41813429 | 0.50433298 | 0.6140274 | 0.99937574 |
| MAST2 | 55.9034701 | -0.2194076 | 0.36596349 | -0.599534 | 0.54881683 | 0.99937574 |
| TSPAN1 | 309.937517 | 0.5928528 | 0.61120336 | 0.96997635 | 0.33205828 | 0.99937574 |
| POMGNT1 | 63.3393096 | 0.29884201 | 0.32941376 | 0.90719348 | 0.36430449 | 0.99937574 |
| LURAP1 | 48.6882202 | 0.42446337 | 0.47404443 | 0.8954084 | 0.37056881 | 0.99937574 |
| LRRC41 | 43.661885 | -0.3489539 | 0.30564028 | -1.1417145 | 0.25357272 | 0.99937574 |
| UQCRH | 157.174575 | -0.1625131 | 0.45880548 | -0.3542092 | 0.72318213 | 0.99937574 |
| NSUN4 | 42.5739297 | 0.00880645 | 0.33469808 | 0.02631161 | 0.97900879 | 0.99937574 |
| FAAH | 18.8190814 | 0.53759254 | 0.50444598 | 1.06570885 | 0.28655528 | 0.99937574 |
| MKNK1 | 80.6818358 | 0.39508374 | 0.2859923 | 1.38144889 | 0.16714098 | 0.99937574 |
| ATPAF1 | 61.0008257 | 0.18154064 | 0.33163979 | 0.54740306 | 0.58410185 | 0.99937574 |
| EFCAB14 | 125.384313 | -0.1358166 | 0.24910633 | -0.5452155 | 0.58560531 | 0.99937574 |
| PDZK1IP1 | 28.80599 | -0.0419408 | 0.90386865 | -0.0464015 | 0.96299025 | 0.99937574 |
| CMPK1 | 73.6281342 | -0.2985461 | 0.4076324 | -0.7323905 | 0.46393024 | 0.99937574 |
| TRABD2B | 139.339693 | 0.37016451 | 0.87346472 | 0.42378874 | 0.67171989 | 0.99937574 |
| SPATA6 | 113.47569 | 0.48592075 | 0.39341266 | 1.23514264 | 0.21677742 | 0.99937574 |
| AGBL4 | 165.87497 | 0.9539108 | 0.65011642 | 1.46729227 | 0.14229657 | 0.99937574 |
| FAF1 | 569.431393 | 0.1199772 | 0.24817402 | 0.48343979 | 0.6287835 | 0.99937574 |
| RNF11 | 113.716088 | -0.2119441 | 0.28742103 | -0.7373996 | 0.46087939 | 0.99937574 |
| TTC39A | 38.323504 | -0.5540335 | 0.53864094 | -1.0285766 | 3.04E-01 | 0.99937574 |
| EPS15 | 350.184419 | -0.2069763 | 0.30313966 | -0.6827754 | 0.49474879 | 0.99937574 |
| OSBPL9 | 246.144869 | -0.2017326 | 0.22260729 | -0.9062266 | 0.36481595 | 0.99937574 |
| NRDC | 566.567795 | 0.46809498 | 0.43322193 | 1.08049695 | 0.27992095 | 0.99937574 |
| RAB3B | 49.3007467 | 0.17963263 | 0.66538384 | 0.26996843 | 0.78718454 | 0.99937574 |
| TXNDC12 | 39.9870979 | 0.0944446 | 0.32418123 | 0.29133272 | 0.77079686 | 0.99937574 |
| BTF3L4 | 69.4679026 | -0.0103615 | 0.26589263 | -0.0389686 | 0.96891544 | 0.99937574 |
| ZFYVE9 | 92.6397773 | -0.2785561 | 0.31296885 | -0.8900441 | 0.37344223 | 0.99937574 |
| CC2D1B | 36.9761157 | -0.6157146 | 0.50193535 | -1.2266811 | 0.21994247 | 0.99937574 |
| PRPF38A | 22.0621891 | -0.3398068 | 0.53644757 | -0.6334389 | 0.52644707 | 0.99937574 |
| TUT4 | 231.293641 | -0.0817873 | 0.24288301 | -0.3367353 | 0.73631642 | 0.99937574 |
| ZYG11B | 129.2276 | -0.3157818 | 0.3666067 | -0.8613639 | 0.38903764 | 0.99937574 |
| ECHDC2 | 185.917337 | 0.14081631 | 0.27689044 | 0.50856328 | 0.61105837 | 0.99937574 |
| SCP2 | 162.542733 | 0.40072975 | 0.34682669 | 1.15541785 | 0.24791936 | 0.99937574 |
| CPT2 | 28.5505532 | 0.10530248 | 0.44140563 | 0.2385617 | 0.81144547 | 0.99937574 |
| C1orf123 | 12.2189638 | -0.436199 | 0.6435079 | -0.6778456 | 0.49786961 | 0.99937574 |
| MAGOH | 19.7983921 | -0.7886214 | 0.42366391 | -1.8614316 | 0.06268326 | 0.99937574 |
| NDC1 | 25.4649657 | 0.31305635 | 0.38247605 | 0.81849921 | 0.41307219 | 0.99937574 |
| YIPF1 | 24.2341432 | 0.18623605 | 0.40978948 | 0.45446762 | 0.64949229 | 0.99937574 |
| HSPB11 | 32.6654844 | -0.1121628 | 0.44709665 | -0.2508693 | 0.8019152 | 0.99937574 |
| LRRC42 | 16.2154744 | 0.03524674 | 0.48813674 | 0.0722067 | 0.94243741 | 0.99937574 |
| TMEM59 | 208.399496 | 0.44144686 | 0.36469587 | 1.21045204 | 0.22610548 | 0.99937574 |
| TCEANC2 | 22.366409 | 0.26110899 | 0.42620554 | 0.61263631 | 0.54011684 | 0.99937574 |
| MRPL37 | 23.60467 | 0.60702936 | 0.38969993 | 1.55768402 | 0.11930817 | 0.99937574 |
| SSBP3 | 115.652335 | 0.28745382 | 0.49886972 | 0.57621019 | 0.56447312 | 0.99937574 |
| ACOT11 | 91.9218544 | -1.5187373 | 0.4290632 | -3.5396587 | 0.00040064 | 0.22462818 |
| DHCR24 | 73.5879981 | -0.0936696 | 0.6536201 | -0.1433089 | 0.88604621 | 0.99937574 |
| TMEM61 | 129.573638 | 1.13516739 | 0.39131836 | 2.90087943 | 0.00372117 | 0.54426159 |

| | | | | | | |
|---|---|---|---|---|---|---|
| BSND | 87.5344687 | 0.97657092 | 0.42242947 | 2.31179641 | 0.02078891 | 0.96427895 |
| USP24 | 244.709084 | -0.1067755 | 0.24145325 | -0.4422201 | 0.65832997 | 0.99937574 |
| PLPP3 | 108.928072 | -0.4568181 | 0.45748888 | -0.9985338 | 0.31802057 | 0.99937574 |
| PRKAA2 | 429.250301 | 0.09924861 | 0.19182507 | 0.51739123 | 0.60488308 | 0.99937574 |
| FYB2 | 1025.82632 | 0.20715332 | 0.57224656 | 0.36200011 | 0.71735195 | 0.99937574 |
| DAB1 | 118.96525 | 0.20183207 | 0.74158606 | 0.2721627 | 0.78549692 | 0.99937574 |
| TACSTD2 | 66.8800593 | -0.6052424 | 0.88737249 | -0.6820612 | 0.49520022 | 0.99937574 |
| MYSM1 | 127.41908 | -0.2463157 | 0.28850114 | -0.8537771 | 0.39322853 | 0.99937574 |
| JUN | 68.8402241 | -1.0144274 | 0.51735147 | -1.9608089 | 0.04990132 | 0.99937574 |
| LINC01135 | 66.7847068 | 0.47732216 | 0.57648199 | 0.82799145 | 0.40767534 | 0.99937574 |
| FGGY | 116.185459 | -0.192439 | 0.42855295 | -0.4490438 | 0.65340006 | 0.99937574 |
| HOOK1 | 81.2955108 | -0.0722301 | 0.40970138 | -0.1762995 | 0.86005869 | 0.99937574 |
| NFIA | 230.604398 | 0.68940907 | 0.53901033 | 1.27902758 | 0.20088734 | 0.99937574 |
| TM2D1 | 96.0275696 | -0.2713559 | 0.2371441 | -1.1442658 | 0.25251343 | 0.99937574 |
| PATJ | 947.684313 | -0.1406462 | 0.20454965 | -0.6875895 | 0.4917113 | 0.99937574 |
| USP1 | 22.607273 | 0.05021315 | 0.50951065 | 0.09855171 | 0.92149421 | 0.99937574 |
| DOCK7 | 247.391786 | -0.0930512 | 0.41285594 | -0.2253841 | 0.82168047 | 0.99937574 |
| ATG4C | 41.2545826 | 0.80111845 | 0.59211741 | 1.35297229 | 0.17606448 | 0.99937574 |
| LINC01739 | 28.1845605 | 1.86892111 | 0.82871454 | 2.25520492 | 0.02412047 | 0.96546668 |
| ALG6 | 18.0053026 | 0.23306699 | 0.47123751 | 0.49458497 | 0.62089311 | 0.99937574 |
| ITGB3BP | 54.2933861 | 0.23435163 | 0.54566728 | 0.42947716 | 0.66757601 | 0.99937574 |
| EFCAB7 | 22.8116028 | 0.41660768 | 0.45647179 | 0.91266906 | 0.36141662 | 0.99937574 |
| PGM1 | 126.984264 | 0.44340435 | 0.29548553 | 1.50059581 | 0.13346014 | 0.99937574 |
| CACHD1 | 40.2094262 | 0.28161069 | 0.79727446 | 0.35321675 | 0.72392595 | 0.99937574 |
| RAVER2 | 32.6971025 | 0.74849016 | 0.50454413 | 1.48349791 | 0.13794217 | 0.99937574 |
| JAK1 | 286.678911 | 0.31766717 | 0.29202939 | 1.08779178 | 2.77E-01 | 0.99937574 |
| AK4 | 18.4468464 | -0.4307269 | 0.82686503 | -0.5209156 | 0.60242556 | 0.99937574 |
| DNAJC6 | 12.4259551 | -0.5794629 | 0.66866044 | -0.8666026 | 0.38615979 | 0.99937574 |
| LEPR | 72.9014727 | 0.46915212 | 0.52373269 | 0.89578544 | 0.37036737 | 0.99937574 |
| WDR78 | 23.3129927 | 0.30959503 | 0.5699538 | 0.54319321 | 0.58699678 | 0.99937574 |
| MIER1 | 120.850914 | -0.3399149 | 0.24054752 | -1.4130883 | 0.15762977 | 0.99937574 |
| SLC35D1 | 32.3168149 | 0.59952757 | 0.39220371 | 1.52861271 | 0.12636048 | 0.99937574 |
| SERBP1 | 116.912488 | 0.08089142 | 0.30941132 | 0.26143652 | 0.7937559 | 0.99937574 |
| GADD45A | 74.6254915 | -1.00455 | 0.47353319 | -2.1213931 | 0.03388874 | 0.99937574 |
| GNG12 | 408.765594 | 0.07818904 | 0.51468901 | 0.15191511 | 0.87925389 | 0.99937574 |
| GNG12-AS1 | 151.765287 | 0.93454019 | 0.45437501 | 2.05675965 | 0.03970935 | 0.99937574 |
| WLS | 224.753255 | 0.3256568 | 0.41193695 | 0.79055011 | 0.42920657 | 0.99937574 |
| LRRC40 | 52.6531858 | 0.27876045 | 0.34696128 | 0.80343388 | 0.421724 | 0.99937574 |
| SRSF11 | 214.234112 | -0.0835899 | 0.24453724 | -0.341829 | 0.7324796 | 0.99937574 |
| ANKRD13C | 102.254289 | -0.3241332 | 0.2694157 | -1.203097 | 0.22893879 | 0.99937574 |
| CTH | 15.4504603 | -1.1753781 | 0.86438791 | -1.3597808 | 0.17389929 | 0.99937574 |
| PTGER3 | 3332.00678 | 0.74688776 | 0.3997879 | 1.86821004 | 0.0617328 | 0.99937574 |
| ZRANB2-AS1 | 22.7509993 | 0.97404695 | 0.62781241 | 1.55149362 | 0.12078343 | 0.99937574 |
| ZRANB2 | 202.482207 | -0.1288585 | 0.28148547 | -0.4577804 | 0.6471102 | 0.99937574 |
| CRYZ | 122.301025 | -0.3515302 | 0.42926151 | -0.8189185 | 0.41283293 | 0.99937574 |
| TYW3 | 35.6067655 | -0.9188947 | 0.50608596 | -1.815689 | 0.06941809 | 0.99937574 |
| ACADM | 99.7046179 | 0.13932308 | 0.34426407 | 0.40469827 | 0.68569931 | 0.99937574 |
| RABGGTB | 26.8482441 | -0.5391504 | 0.53438777 | -1.0089123 | 0.31301668 | 0.99937574 |
| PIGK | 45.4252858 | 0.23168321 | 0.33308985 | 0.69555772 | 0.48670585 | 0.99937574 |
| AC118549.1 | 222.016114 | -0.1768108 | 0.19727945 | -0.8962453 | 0.37012179 | 0.99937574 |
| USP33 | 224.364838 | -0.0048612 | 0.27479401 | -0.0176904 | 0.98588584 | 0.99937574 |
| MIGA1 | 113.971006 | 0.25765241 | 0.25596159 | 1.00660578 | 0.31412426 | 0.99937574 |
| NEXN | 37.2867615 | 0.75509246 | 0.41830961 | 1.80510428 | 0.07105837 | 0.99937574 |
| FUBP1 | 135.216674 | 0.19079148 | 0.26599528 | 0.71727392 | 0.47320509 | 0.99937574 |

| | | | | | | |
|---|---|---|---|---|---|---|
| GIPC2 | 27.8512075 | 0.13372254 | 0.72055697 | 0.18558219 | 0.85277242 | 0.99937574 |
| PTGFR | 15.4134734 | -0.7604974 | 1.27749724 | -0.5953026 | 0.55164125 | 0.99937574 |
| IFI44L | 101.769402 | -7.6223721 | 2.87430213 | -2.6519036 | 0.00800394 | 0.74742128 |
| IFI44 | 44.1598777 | -3.9511928 | 1.50246977 | -2.6297985 | 0.00854355 | 0.74742128 |
| ADGRL2 | 236.764378 | 0.69061903 | 0.34160385 | 2.02169566 | 0.0432078 | 0.99937574 |
| LINC01725 | 106.11703 | 0.00788402 | 0.40290986 | 0.01956769 | 0.98438824 | 0.99937574 |
| PRKACB | 102.355385 | 0.63880157 | 0.51288279 | 1.2455118 | 0.21294368 | 0.99937574 |
| RPF1 | 20.1519276 | -0.1880647 | 0.43126941 | -0.4360724 | 0.66278421 | 0.99937574 |
| GNG5 | 76.9225415 | -0.0647827 | 0.3667256 | -0.1766518 | 0.85978194 | 0.99937574 |
| SPATA1 | 21.3041834 | -0.7610223 | 0.76096231 | -1.0000789 | 0.31727233 | 0.99937574 |
| CTBS | 46.8725198 | 0.05231826 | 0.35862333 | 0.14588637 | 0.88401109 | 0.99937574 |
| SSX2IP | 25.8437464 | 0.00287271 | 0.57927503 | 0.00495915 | 0.99604319 | 0.99937574 |
| WDR63 | 23.7910404 | 1.69569789 | 0.66619484 | 2.54534826 | 0.01091688 | 0.76274616 |
| MCOLN3 | 29.4985358 | 0.37576191 | 0.79650182 | 0.47176529 | 0.63709433 | 0.99937574 |
| SYDE2 | 19.6444307 | -0.1621665 | 0.55960925 | -0.2897852 | 0.77198059 | 0.99937574 |
| C1orf52 | 31.7468447 | 0.08237693 | 0.3738478 | 0.22034883 | 0.82559949 | 0.99937574 |
| BCL10 | 36.0707076 | -0.2651574 | 0.41795962 | -0.6344091 | 0.52581388 | 0.99937574 |
| AL078459.1 | 35.1802386 | -0.1567102 | 0.51335546 | -0.3052664 | 0.76016329 | 0.99937574 |
| DDAH1 | 98.5332609 | 0.03746691 | 0.53970083 | 0.06942163 | 0.94465401 | 0.99937574 |
| ZNHIT6 | 48.6973964 | 0.13068977 | 0.37740195 | 0.34628801 | 0.72912628 | 0.99937574 |
| COL24A1 | 180.167199 | 0.45330404 | 0.45732447 | 0.99120878 | 0.32158364 | 0.99937574 |
| ODF2L | 96.3081347 | -0.4055411 | 0.3534483 | -1.1473844 | 0.25122279 | 0.99937574 |
| SH3GLB1 | 104.044169 | 0.0214006 | 0.24925869 | 0.08585697 | 0.93158012 | 0.99937574 |
| SELENOF | 113.89338 | 0.10435891 | 0.26258602 | 0.39742751 | 0.69105223 | 0.99937574 |
| HS2ST1 | 51.9423835 | -0.1503307 | 0.35852571 | -0.4193024 | 0.67499513 | 0.99937574 |
| LMO4 | 32.2939283 | 0.86103974 | 0.49528321 | 1.73847956 | 0.08212635 | 0.99937574 |
| PKN2-AS1 | 14.6976123 | -0.3275 | 0.75148756 | -0.4358023 | 0.66298019 | 0.99937574 |
| PKN2 | 258.206381 | -0.1911774 | 0.22227419 | -0.8600972 | 0.38973544 | 0.99937574 |
| GTF2B | 33.6451364 | -0.8085418 | 0.43123108 | -1.8749618 | 0.06079797 | 0.99937574 |
| KYAT3 | 64.8753985 | -0.0557044 | 0.34497516 | -0.1614737 | 0.8717203 | 0.99937574 |
| GBP3 | 23.1982353 | -0.4212451 | 0.6168485 | -0.6828989 | 0.49467076 | 0.99937574 |
| GBP2 | 38.6761885 | -0.9569404 | 0.63856241 | -1.4985855 | 0.13398119 | 0.99937574 |
| LRRC8B | 86.6488411 | -0.0541332 | 0.35113532 | -0.1541662 | 0.87747873 | 0.99937574 |
| ZNF326 | 69.2505991 | -0.2452416 | 0.29460603 | -0.8324392 | 0.40516106 | 0.99937574 |
| ZNF644 | 253.804081 | 0.01829991 | 0.23109228 | 0.07918875 | 0.9368825 | 0.99937574 |
| HFM1 | 19.9248046 | -0.2331308 | 0.82959906 | -0.2810162 | 0.77869797 | 0.99937574 |
| BTBD8 | 41.687509 | 0.11059102 | 0.62968221 | 0.17562989 | 0.86058472 | 0.99937574 |
| GLMN | 44.8821826 | 0.23800108 | 0.4220179 | 0.56395968 | 0.57278157 | 0.99937574 |
| RPAP2 | 107.513089 | -0.0026834 | 0.21873501 | -0.0122678 | 0.99021199 | 0.99937574 |
| EVI5 | 275.783268 | 0.01225736 | 0.27937424 | 0.04387432 | 0.96500458 | 0.99937574 |
| RPL5 | 165.915756 | -0.0061769 | 0.4397405 | -0.0140467 | 0.98879276 | 0.99937574 |
| FAM69A | 131.015681 | -0.3734731 | 0.38777022 | -0.96313 | 0.3354823 | 0.99937574 |
| MTF2 | 144.815653 | -0.0992047 | 0.33347115 | -0.2974911 | 0.76609157 | 0.99937574 |
| TMED5 | 120.953103 | -0.1055384 | 0.38495469 | -0.274158 | 0.78396318 | 0.99937574 |
| CCDC18 | 24.481434 | -0.5321796 | 0.56013862 | -0.9500856 | 0.34206878 | 0.99937574 |
| DR1 | 40.7669639 | -0.1932818 | 0.33371955 | -0.5791745 | 0.56247147 | 0.99937574 |
| FNBP1L | 525.442488 | 0.08838973 | 0.27899466 | 0.31681512 | 0.75138389 | 0.99937574 |
| BCAR3 | 94.048058 | 0.12044458 | 0.48783224 | 0.24689753 | 0.80498753 | 0.99937574 |
| DNTTIP2 | 20.7874551 | -0.6092415 | 0.44783683 | -1.3604095 | 0.17370037 | 0.99937574 |
| GCLM | 57.8828359 | -0.42491 | 0.40373525 | -1.0524472 | 2.93E-01 | 0.99937574 |
| ARHGAP29 | 99.5513966 | 0.12916944 | 0.37932437 | 0.34052501 | 0.73346119 | 0.99937574 |
| ABCD3 | 101.019766 | 0.1810668 | 0.26430128 | 0.68507726 | 0.49329517 | 0.99937574 |
| SLC44A3 | 82.1746876 | 0.24623597 | 0.31746745 | 0.7756259 | 0.4379699 | 0.99937574 |
| CNN3 | 232.817388 | -0.0018548 | 0.37292535 | -0.0049738 | 0.99603154 | 0.99937574 |

| | | | | | | |
|---|---|---|---|---|---|---|
| AC105942.1 | 32.8914254 | 0.75819593 | 0.38907671 | 1.94870553 | 0.0513306 | 0.99937574 |
| ALG14 | 56.1098538 | 0.16475525 | 0.38120409 | 0.43219696 | 0.66559827 | 0.99937574 |
| PTBP2 | 199.477627 | 0.05398967 | 0.29523106 | 0.1828726 | 0.85489798 | 0.99937574 |
| DPYD | 270.937678 | 0.07216851 | 0.30098316 | 0.2397759 | 0.81050399 | 0.99937574 |
| DPYD-AS1 | 12.7485962 | 1.02601593 | 0.64846458 | 1.58222356 | 0.11359854 | 0.99937574 |
| SNX7 | 49.9283449 | -0.4512641 | 0.39441418 | -1.1441376 | 0.25256657 | 0.99937574 |
| FRRS1 | 23.2791093 | -0.6454981 | 0.6930672 | -0.9313643 | 0.35166513 | 0.99937574 |
| AGL | 98.8892087 | 0.24138292 | 0.24617545 | 0.98053205 | 0.32682355 | 0.99937574 |
| SASS6 | 11.5258485 | 0.05689335 | 0.58396377 | 0.09742617 | 0.92238797 | 0.99937574 |
| TRMT13 | 51.0684927 | -0.0663571 | 0.38299995 | -0.1732563 | 0.86245 | 0.99937574 |
| DBT | 68.1387014 | 0.78814131 | 0.34557646 | 2.2806568 | 0.02256876 | 0.96427895 |
| RTCA | 35.3537264 | 0.04084979 | 0.36144303 | 0.11301861 | 0.9100158 | 0.99937574 |
| CDC14A | 71.5132926 | 0.11738209 | 0.41310706 | 0.28414448 | 0.77629966 | 0.99937574 |
| EXTL2 | 57.954085 | 0.39542629 | 0.47694327 | 0.82908453 | 4.07E-01 | 0.99937574 |
| SLC30A7 | 88.6753435 | -0.1133015 | 0.28387904 | -0.3991191 | 0.68980547 | 0.99937574 |
| DPH5 | 20.8342211 | 0.74390674 | 0.46518703 | 1.59915626 | 0.10978589 | 0.99937574 |
| AC093157.1 | 20.0133503 | 0.5831181 | 0.49206657 | 1.18503905 | 0.23600201 | 0.99937574 |
| RNPC3 | 107.821716 | -0.1074297 | 0.29054121 | -0.3697572 | 0.7115634 | 0.99937574 |
| VAV3 | 276.985566 | 0.09400707 | 0.42431129 | 0.22155213 | 0.82466255 | 0.99937574 |
| SLC25A24 | 61.2439993 | -0.1446421 | 0.38865017 | -0.3721654 | 0.70976971 | 0.99937574 |
| PRPF38B | 72.8697362 | -0.3837627 | 0.41631449 | -0.9218096 | 0.35662791 | 0.99937574 |
| STXBP3 | 108.874606 | -0.0757303 | 0.28831059 | -0.2626693 | 0.79280551 | 0.99937574 |
| AL449266.1 | 21.059978 | -0.3215995 | 0.48502422 | -0.6630587 | 0.50729294 | 0.99937574 |
| GPSM2 | 56.3581426 | -0.4422401 | 0.55647671 | -0.7947145 | 0.42677961 | 0.99937574 |
| WDR47 | 17.1018108 | 0.3825183 | 0.54614586 | 0.70039585 | 0.48368013 | 0.99937574 |
| TAF13 | 20.5103556 | -0.0884762 | 0.50036473 | -0.1768233 | 0.85964716 | 0.99937574 |
| TMEM167B | 31.5026591 | 0.17992249 | 0.41494832 | 0.43360216 | 0.66457737 | 0.99937574 |
| SARS | 61.0803528 | -0.2595973 | 0.27801705 | -0.933746 | 0.35043491 | 0.99937574 |
| SORT1 | 88.7367542 | 0.22198935 | 0.26359479 | 0.84216139 | 0.39969762 | 0.99937574 |
| PSMA5 | 63.0819878 | -0.3887164 | 0.34120798 | -1.139236 | 0.25460473 | 0.99937574 |
| SYPL2 | 21.0518958 | 0.03787554 | 0.52199154 | 0.07255968 | 0.94215652 | 0.99937574 |
| CYB561D1 | 11.798385 | -0.5923735 | 0.62879288 | -0.9420805 | 0.34615142 | 0.99937574 |
| GNAI3 | 74.2856711 | -0.2916133 | 0.39991007 | -0.7291971 | 0.46588108 | 0.99937574 |
| GSTM3 | 20.1391418 | -0.7807226 | 0.82664785 | -0.944444 | 0.34494279 | 0.99937574 |
| LINC01768 | 70.6327151 | -0.164241 | 0.80932734 | -0.2029352 | 0.83918566 | 0.99937574 |
| AHCYL1 | 174.801454 | 0.16411901 | 0.31980954 | 0.51317733 | 0.60782728 | 0.99937574 |
| STRIP1 | 35.0389791 | 0.11436367 | 0.34502344 | 0.33146639 | 0.74029223 | 0.99937574 |
| KCNC4 | 96.1269627 | -0.0739187 | 0.44390683 | -0.1665185 | 0.86774894 | 0.99937574 |
| RBM15 | 34.6854443 | -0.1840681 | 0.42808403 | -0.4299812 | 0.66720931 | 0.99937574 |
| SLC16A4 | 58.8630638 | -0.3352146 | 0.76909047 | -0.4358585 | 0.66293938 | 0.99937574 |
| AL355488.1 | 12.3328107 | -0.274566 | 0.82307347 | -0.3335863 | 0.73869175 | 0.99937574 |
| LAMTOR5 | 100.276293 | -0.0607542 | 0.44245919 | -0.1373103 | 0.89078555 | 0.99937574 |
| LRIF1 | 24.7351969 | -0.1784286 | 0.43553691 | -0.409675 | 0.68204437 | 0.99937574 |
| DRAM2 | 68.7242713 | 0.56438018 | 0.40838912 | 1.38196671 | 0.16698192 | 0.99937574 |
| CEPT1 | 101.041118 | 0.193015 | 0.33295932 | 0.57969543 | 0.56212002 | 0.99937574 |
| DENND2D | 48.1945503 | 0.00255052 | 0.32142196 | 0.00793511 | 0.99366877 | 0.99937574 |
| WDR77 | 17.9469493 | -0.2953968 | 0.44317251 | -0.6665504 | 0.50505936 | 0.99937574 |
| ATP5PB | 138.467191 | 0.73059551 | 0.39465162 | 1.85124164 | 0.0641348 | 0.99937574 |
| RAP1A | 192.007247 | -0.0598857 | 0.27660437 | -0.2165032 | 0.82859549 | 0.99937574 |
| KCND3 | 28.7584006 | -0.0243942 | 0.93887654 | -0.0259823 | 0.97927146 | 0.99937574 |
| CTTNBP2NL | 57.4225416 | 0.11964156 | 0.30438586 | 0.39305887 | 0.694276 | 0.99937574 |
| WNT2B | 23.4634566 | 0.9313518 | 0.78234311 | 1.19046463 | 0.23386382 | 0.99937574 |
| ST7L | 59.0533918 | -0.0578842 | 0.33095181 | -0.1749021 | 0.86115657 | 0.99937574 |
| CAPZA1 | 125.95098 | -0.1810771 | 0.22581707 | -0.8018753 | 0.42262507 | 0.99937574 |

| | | | | | | |
|---|---|---|---|---|---|---|
| MOV10 | 30.9914886 | -0.2739001 | 0.52771012 | -0.5190351 | 0.60373627 | 0.99937574 |
| RHOC | 43.7105819 | -0.144164 | 0.46809368 | -0.3079811 | 0.75809674 | 0.99937574 |
| LRIG2 | 63.5334768 | 0.00963854 | 0.30232507 | 0.03188139 | 0.97456664 | 0.99937574 |
| MAGI3 | 1021.22249 | 0.41619704 | 0.3244452 | 1.28279611 | 0.19956352 | 0.99937574 |
| PHTF1 | 47.8685645 | -0.2115512 | 0.30850253 | -0.6857358 | 0.49287974 | 0.99937574 |
| RSBN1 | 68.8793045 | -0.2195694 | 0.29331319 | -0.7485834 | 0.45410835 | 0.99937574 |
| HIPK1 | 104.097403 | 0.03296355 | 0.31996153 | 0.10302346 | 0.91794435 | 0.99937574 |
| TRIM33 | 181.213853 | -0.2164879 | 0.22912535 | -0.9448448 | 0.34473813 | 0.99937574 |
| BCAS2 | 22.8615595 | -0.0890766 | 0.46428985 | -0.1918556 | 0.84785531 | 0.99937574 |
| DENND2C | 14.866962 | -0.7525341 | 0.64069536 | -1.1745584 | 0.24017145 | 0.99937574 |
| NRAS | 19.0036621 | -1.1467616 | 0.601565 | -1.906297 | 0.05661168 | 0.99937574 |
| CSDE1 | 184.620897 | -0.0320466 | 0.25127545 | -0.1275357 | 0.89851639 | 0.99937574 |
| SIKE1 | 32.3986199 | 0.5054106 | 0.4418314 | 1.14389922 | 0.25266543 | 0.99937574 |
| SLC22A15 | 130.225673 | -0.0171141 | 0.35959882 | -0.0475922 | 0.96204124 | 0.99937574 |
| MAB21L3 | 48.7819396 | 0.80713679 | 0.8222132 | 0.98166362 | 0.32626559 | 0.99937574 |
| ATP1A1 | 1291.35166 | 0.44730213 | 0.33003534 | 1.35531588 | 0.17531694 | 0.99937574 |
| LINC01762 | 99.6121782 | -0.1317256 | 1.38970987 | -0.0947864 | 0.92448452 | 0.99937574 |
| CD58 | 48.9228206 | -0.3146657 | 0.3402812 | -0.9247225 | 0.35511025 | 0.99937574 |
| IGSF3 | 99.2750893 | 0.53524525 | 0.38087007 | 1.40532244 | 0.15992542 | 0.99937574 |
| PTGFRN | 20.3033413 | -0.2127499 | 0.69097158 | -0.3078997 | 0.75815865 | 0.99937574 |
| TTF2 | 45.4931253 | 0.30977243 | 0.36744029 | 0.84305515 | 0.3991976 | 0.99937574 |
| MAN1A2 | 230.918626 | -0.0991389 | 0.27689501 | -0.3580379 | 0.72031492 | 0.99937574 |
| GDAP2 | 61.3837597 | 0.05968653 | 0.32509902 | 0.18359492 | 0.85433125 | 0.99937574 |
| WDR3 | 13.1112088 | 0.34261745 | 0.55819049 | 0.61380022 | 0.53934735 | 0.99937574 |
| SPAG17 | 56.4988032 | 0.71331681 | 0.72507823 | 0.98377911 | 0.32522413 | 0.99937574 |
| WARS2 | 50.0063284 | -0.3786197 | 0.36571267 | -1.0352927 | 0.30053223 | 0.99937574 |
| AL359915.2 | 178.500118 | 0.55971573 | 0.46539532 | 1.20266728 | 0.2291051 | 0.99937574 |
| HMGCS2 | 62.0341246 | -2.4150236 | 1.96581219 | -1.2285119 | 0.21925487 | 0.99937574 |
| SEC22B | 67.2545195 | 0.2285467 | 0.27514464 | 0.83064201 | 0.40617589 | 0.99937574 |
| LINC00623 | 167.825164 | -0.0405908 | 0.35753454 | -0.1135297 | 0.90961062 | 0.99937574 |
| AC239800.3 | 19.0597247 | -0.0132413 | 0.45114306 | -0.0293504 | 0.97658509 | 0.99937574 |
| NBPF15 | 15.3089346 | 0.7286967 | 0.6649602 | 1.09585009 | 0.27314438 | 0.99937574 |
| FP700111.1 | 12.0353628 | 0.48734684 | 0.55599814 | 0.87652603 | 0.38074413 | 0.99937574 |
| SRGAP2B | 13.919997 | 0.34776438 | 0.55068596 | 0.63151126 | 0.52770629 | 0.99937574 |
| GPR89A | 23.858661 | -0.2170108 | 0.39596825 | -0.548051 | 0.58365686 | 0.99937574 |
| PDZK1 | 18.7657885 | 0.221656 | 0.63799138 | 0.34742789 | 0.72826988 | 0.99937574 |
| RNF115 | 173.897227 | -0.320321 | 0.25479771 | -1.257158 | 0.20869644 | 0.99937574 |
| POLR3C | 20.4629791 | -0.4138104 | 0.52684118 | -0.7854556 | 0.43218649 | 0.99937574 |
| PEX11B | 15.2454145 | -0.3690714 | 0.49409988 | -0.7469571 | 0.45508948 | 0.99937574 |
| RBM8A | 41.2784834 | -0.4017762 | 0.43063073 | -0.9329946 | 0.35082275 | 0.99937574 |
| TXNIP | 1254.00711 | 0.10356446 | 0.49077365 | 0.21102286 | 0.83286943 | 0.99937574 |
| NBPF12 | 32.5888928 | 0.22216983 | 0.37216306 | 0.59696905 | 0.55052804 | 0.99937574 |
| PRKAB2 | 44.6453632 | 0.14804312 | 0.53786451 | 0.27524239 | 0.78313002 | 0.99937574 |
| AC242426.2 | 63.1143543 | -0.0623317 | 0.35310844 | -0.1765227 | 0.85988329 | 0.99937574 |
| FMO5 | 45.2939498 | -0.0594019 | 0.58044004 | -0.1023394 | 0.91848724 | 0.99937574 |
| CHD1L | 90.2867744 | 0.56935886 | 0.47274504 | 1.2043677 | 0.22844749 | 0.99937574 |
| LINC00624 | 23.9805634 | -1.4467224 | 0.571828 | -2.5299957 | 0.01140639 | 0.76274616 |
| BCL9 | 101.501392 | 0.74075114 | 0.47179084 | 1.57008377 | 0.11639562 | 0.99937574 |
| ACP6 | 37.7077325 | 0.36952089 | 0.41180036 | 0.89733016 | 0.36954277 | 0.99937574 |
| GPR89B | 29.0767424 | -0.0116344 | 0.39794573 | -0.029236 | 0.97667635 | 0.99937574 |
| NBPF11 | 31.8235001 | 0.24350282 | 0.42943075 | 0.5670363 | 0.57068952 | 0.99937574 |
| AC239809.3 | 26.4235632 | -0.68371 | 0.52334113 | -1.3064327 | 0.19140545 | 0.99937574 |
| PDE4DIP | 126.907256 | 0.4502239 | 0.58886796 | 0.76455832 | 0.4445346 | 0.99937574 |
| AC245297.3 | 196.134703 | -0.0606366 | 0.46620062 | -0.1300655 | 0.89651462 | 0.99937574 |

| | | | | | | |
|---|---|---|---|---|---|---|
| SF3B4 | 14.6413064 | -0.4228364 | 0.54142418 | -0.7809706 | 0.43481976 | 0.99937574 |
| OTUD7B | 76.5123973 | -0.1086525 | 0.34364533 | -0.3161764 | 0.75186863 | 0.99937574 |
| VPS45 | 106.193395 | 0.12239467 | 0.26650979 | 0.45925018 | 0.64605452 | 0.99937574 |
| ANP32E | 28.6030404 | 0.26577558 | 0.41913118 | 0.63411075 | 0.52600855 | 0.99937574 |
| APH1A | 40.7614336 | -0.4982627 | 0.46369032 | -1.0745592 | 0.28257212 | 0.99937574 |
| MRPS21 | 67.6880252 | -0.244224 | 0.31888341 | -0.7658725 | 0.44375216 | 0.99937574 |
| PRPF3 | 40.2364188 | -0.539738 | 0.40125837 | -1.3451134 | 0.17858863 | 0.99937574 |
| RPRD2 | 158.567734 | -0.0027431 | 0.23158751 | -0.0118449 | 0.99054936 | 0.99937574 |
| TARS2 | 16.0078622 | -0.1928669 | 0.59689934 | -0.3231146 | 0.74660846 | 0.99937574 |
| MCL1 | 69.6012673 | -0.4791437 | 0.48987776 | -0.9780883 | 0.32803068 | 0.99937574 |
| ENSA | 140.921739 | 0.01462186 | 0.21961696 | 0.0665789 | 0.94691695 | 0.99937574 |
| GOLPH3L | 49.6740117 | -0.204744 | 0.31423683 | -0.6515594 | 0.51468544 | 0.99937574 |
| CTSS | 24.0151775 | -0.8379053 | 0.64515971 | -1.2987563 | 0.19402756 | 0.99937574 |
| ARNT | 56.171635 | -0.164794 | 0.33019707 | -0.4990776 | 0.61772469 | 0.99937574 |
| SETDB1 | 42.7556148 | -0.3759907 | 0.41769551 | -0.900155 | 0.36803776 | 0.99937574 |
| CERS2 | 43.1568467 | 0.25067623 | 0.37697845 | 0.66496171 | 0.50607498 | 0.99937574 |
| MINDY1 | 84.1360813 | 0.6115745 | 0.33808487 | 1.80893782 | 0.07046066 | 0.99937574 |
| PRUNE1 | 43.1853572 | -0.3269861 | 0.47563367 | -0.6874746 | 0.4917837 | 0.99937574 |
| C1orf56 | 23.1917273 | 0.16407844 | 0.60043919 | 0.27326404 | 0.78465026 | 0.99937574 |
| CDC42SE1 | 62.4437155 | 0.06731549 | 0.28857807 | 0.23326613 | 0.81555475 | 0.99937574 |
| GABPB2 | 66.4734409 | 0.25217121 | 0.4325432 | 0.5829966 | 0.55989559 | 0.99937574 |
| SCNM1 | 15.2918559 | 0.14739181 | 0.53765016 | 0.27414074 | 0.78397646 | 0.99937574 |
| VPS72 | 21.9670561 | 0.48398304 | 0.39432119 | 1.22738278 | 0.21967876 | 0.99937574 |
| PIP5K1A | 195.496061 | -0.0820606 | 0.25172844 | -0.3259886 | 0.74443299 | 0.99937574 |
| PSMD4 | 46.9371035 | -0.2848007 | 0.29188304 | -0.9757356 | 0.32919548 | 0.99937574 |
| ZNF687 | 10.5386166 | -0.2068622 | 0.54556857 | -0.3791681 | 0.70456307 | 0.99937574 |
| PI4KB | 60.3131855 | -0.2628043 | 0.31109254 | -0.8447786 | 0.39823449 | 0.99937574 |
| SELENBP1 | 37.4158324 | 1.36902502 | 0.81071133 | 1.68867138 | 0.09128242 | 0.99937574 |
| PSMB4 | 41.9734105 | -0.3341293 | 0.44345154 | -0.7534742 | 0.45116503 | 0.99937574 |
| POGZ | 257.321609 | -0.0832414 | 0.22784208 | -0.3653468 | 0.71485255 | 0.99937574 |
| CGN | 19.943396 | 0.02610464 | 0.55275461 | 0.04722645 | 0.96233274 | 0.99937574 |
| TUFT1 | 39.4228909 | -0.8961184 | 0.48821555 | -1.8354975 | 0.06643201 | 0.99937574 |
| SNX27 | 81.5472617 | -0.2249941 | 0.30581396 | -0.735722 | 0.4618999 | 0.99937574 |
| RORC | 42.8931162 | -1.2313952 | 0.48720048 | -2.5274918 | 0.01148805 | 0.76274616 |
| AL450992.2 | 11.3569996 | -0.1260402 | 0.6342647 | -0.1987187 | 0.84248282 | 0.99937574 |
| THEM4 | 121.541482 | 0.08220824 | 0.33305413 | 0.24683146 | 0.80503866 | 0.99937574 |
| S100A10 | 411.034825 | -0.4026118 | 0.7041043 | -0.571807 | 0.56745271 | 0.99937574 |
| S100A11 | 629.921078 | -0.3570213 | 0.65703885 | -0.5433794 | 0.58686864 | 0.99937574 |
| FLG-AS1 | 52.0745225 | 0.58898127 | 0.52367109 | 1.12471604 | 0.26070938 | 0.99937574 |
| S100A6 | 585.229974 | -0.9397331 | 0.73559422 | -1.2775156 | 0.20142026 | 0.99937574 |
| S100A2 | 29.2176525 | -0.8462521 | 0.81853891 | -1.0338568 | 0.30120309 | 0.99937574 |
| S100A16 | 19.0364371 | -0.1136263 | 0.46177796 | -0.2460626 | 0.80563377 | 0.99937574 |
| S100A13 | 65.5535237 | -0.4772571 | 0.49362668 | -0.9668382 | 0.33362491 | 0.99937574 |
| S100A1 | 27.9635069 | -0.2364805 | 1.15011361 | -0.2056149 | 0.83709175 | 0.99937574 |
| CHTOP | 45.5501151 | 0.09854962 | 0.33468436 | 0.2944554 | 0.76840994 | 0.99937574 |
| SNAPIN | 12.3583175 | 0.52557986 | 0.65284183 | 0.80506463 | 0.42078239 | 0.99937574 |
| ILF2 | 33.6743339 | 0.78305418 | 0.37894397 | 2.06641153 | 0.03878964 | 0.99937574 |
| INTS3 | 65.1949787 | 0.08781137 | 0.46555996 | 0.18861452 | 0.85039495 | 0.99937574 |
| GATAD2B | 205.512727 | -0.1938611 | 0.284373 | -0.6817143 | 0.49541962 | 0.99937574 |
| DENND4B | 25.101714 | -0.2781504 | 0.54926302 | -0.5064065 | 0.61257128 | 0.99937574 |
| CRTC2 | 18.4091905 | -0.6720991 | 0.46217385 | -1.4542128 | 0.14588731 | 0.99937574 |
| SLC39A1 | 44.0977272 | -0.2575436 | 0.5550309 | -0.4640166 | 0.64263583 | 0.99937574 |
| JTB | 53.6862184 | 0.28311681 | 0.51697883 | 0.54763715 | 0.58394107 | 0.99937574 |
| RAB13 | 56.5064491 | 0.25666798 | 0.47217923 | 0.54358169 | 0.58672936 | 0.99937574 |

| | | | | | | |
|---|---|---|---|---|---|---|
| RPS27 | 89.5702669 | -0.0242403 | 0.39871248 | -0.0607963 | 0.9515214 | 0.99937574 |
| TPM3 | 71.1379628 | -0.1643468 | 0.42424707 | -0.3873846 | 0.69847152 | 0.99937574 |
| C1orf43 | 70.524264 | -0.0558573 | 0.3811755 | -0.1465396 | 0.8834954 | 0.99937574 |
| UBAP2L | 84.8463036 | -0.2398644 | 0.29228767 | -0.8206449 | 0.41184858 | 0.99937574 |
| HAX1 | 24.455763 | -0.1027088 | 0.45389678 | -0.2262823 | 0.82098183 | 0.99937574 |
| IL6R | 28.4738942 | -0.1161553 | 0.70803878 | -0.1640522 | 0.86969008 | 0.99937574 |
| UBE2Q1 | 26.9497252 | -0.4249905 | 0.53540067 | -0.7937804 | 0.42732329 | 0.99937574 |
| ADAR | 126.143327 | -0.5913906 | 0.4518023 | -1.3089587 | 0.19054834 | 0.99937574 |
| KCNN3 | 26.0281648 | 1.25651971 | 1.00184995 | 1.25419951 | 0.2097695 | 0.99937574 |
| PMVK | 44.9582238 | 0.67393277 | 0.39376031 | 1.71153044 | 0.08698324 | 0.99937574 |
| PBXIP1 | 42.7450808 | 0.89329433 | 0.33871735 | 2.63728544 | 0.00835725 | 0.74742128 |
| PYGO2 | 13.0013571 | -0.0397339 | 0.56132242 | -0.0707862 | 0.94356795 | 0.99937574 |
| FLAD1 | 13.2988596 | 0.19143412 | 0.58596798 | 0.32669723 | 0.74389689 | 0.99937574 |
| ZBTB7B | 15.2336594 | -0.2627841 | 0.60027745 | -0.437771 | 0.6615523 | 0.99937574 |
| EFNA1 | 72.5508792 | -0.9302881 | 0.40154171 | -2.3167907 | 0.02051514 | 0.96427895 |
| SLC50A1 | 24.5928476 | -0.0395728 | 0.43230061 | -0.0915401 | 0.92706348 | 0.99937574 |
| DPM3 | 40.221026 | 0.29631254 | 0.4826265 | 0.61395828 | 0.5392429 | 0.99937574 |
| MUC1 | 17.4445967 | 0.04521208 | 0.68664649 | 0.06584477 | 0.94750142 | 0.99937574 |
| MTX1 | 14.2510495 | -0.0861775 | 0.53253823 | -0.1618241 | 0.87144436 | 0.99937574 |
| GBA | 12.2304211 | -0.7697717 | 0.63191352 | -1.2181599 | 0.2231632 | 0.99937574 |
| SCAMP3 | 27.0441025 | -0.0498626 | 0.48749353 | -0.1022835 | 0.91853163 | 0.99937574 |
| CLK2 | 20.5959351 | -0.1676388 | 0.51815409 | -0.3235308 | 0.74629332 | 0.99937574 |
| FDPS | 106.672682 | -0.0255672 | 0.54860737 | -0.0466037 | 0.96282905 | 0.99937574 |
| RUSC1 | 12.5423353 | 0.05160585 | 0.49104681 | 0.10509354 | 0.91630159 | 0.99937574 |
| ASH1L | 466.941529 | 0.04800215 | 0.1927094 | 0.24909086 | 0.8032905 | 0.99937574 |
| YY1AP1 | 90.1023565 | -0.1360426 | 0.29760773 | -0.4571205 | 0.64758444 | 0.99937574 |
| DAP3 | 105.548897 | -0.0637319 | 0.24565904 | -0.2594322 | 0.79530182 | 0.99937574 |
| GON4L | 76.8175821 | -0.3419935 | 0.39071963 | -0.8752914 | 0.38141538 | 0.99937574 |
| RIT1 | 27.8811866 | -1.153642 | 0.46193914 | -2.4973897 | 0.01251114 | 0.77939762 |
| KHDC4 | 83.4511184 | -0.1037369 | 0.26833164 | -0.3865998 | 0.69905254 | 0.99937574 |
| ARHGEF2 | 36.0781979 | 0.06412303 | 0.36540208 | 0.17548624 | 0.86069759 | 0.99937574 |
| SSR2 | 53.0421067 | -0.035554 | 0.34998215 | -0.1015881 | 0.91908364 | 0.99937574 |
| UBQLN4 | 17.4033593 | -0.278122 | 0.54158462 | -0.5135338 | 0.60757795 | 0.99937574 |
| LAMTOR2 | 25.5442936 | 0.11379423 | 0.53795016 | 0.21153303 | 0.83247136 | 0.99937574 |
| RAB25 | 36.7536297 | 0.78001062 | 0.42556022 | 1.83290304 | 0.06681699 | 0.99937574 |
| LMNA | 103.236995 | 0.20426753 | 0.34597246 | 0.59041559 | 0.55491206 | 0.99937574 |
| SEMA4A | 15.6803505 | 0.62663049 | 0.53730392 | 1.1662496 | 0.24351354 | 0.99937574 |
| SLC25A44 | 16.3270441 | 0.14914676 | 0.53434998 | 0.27911812 | 0.78015418 | 0.99937574 |
| SMG5 | 60.8332846 | -0.5033689 | 0.38167233 | -1.3188509 | 0.18721895 | 0.99937574 |
| GLMP | 15.8586185 | -0.2683765 | 0.72891375 | -0.3681869 | 0.7127339 | 0.99937574 |
| CCT3 | 66.5779644 | 0.14148706 | 0.37725852 | 0.37504006 | 0.70763068 | 0.99937574 |
| RHBG | 286.814666 | 0.94306982 | 0.64138363 | 1.47036777 | 0.14146218 | 0.99937574 |
| MEF2D | 43.1994622 | -0.3003539 | 0.51189559 | -0.5867484 | 0.5573727 | 0.99937574 |
| NAXE | 27.8353025 | 0.07698776 | 0.66727947 | 0.11537558 | 0.90814743 | 0.99937574 |
| GPATCH4 | 13.7796366 | -0.251972 | 0.61304021 | -0.4110204 | 0.68105757 | 0.99937574 |
| ISG20L2 | 15.8790188 | -0.3272644 | 0.54626341 | -0.5990963 | 0.54910867 | 0.99937574 |
| RRNAD1 | 22.5593155 | 0.2322231 | 0.41248338 | 0.56298778 | 0.5734432 | 0.99937574 |
| MRPL24 | 11.3585564 | 0.64017014 | 0.60828195 | 1.05242337 | 0.29260535 | 0.99937574 |
| HDGF | 49.2985111 | -0.3151446 | 0.37943095 | -0.8305718 | 0.40621558 | 0.99937574 |
| PRCC | 43.1024005 | -0.2597174 | 0.31670536 | -0.8200601 | 0.41218182 | 0.99937574 |
| ARHGEF11 | 109.776999 | 0.42215332 | 0.33201634 | 1.27148358 | 0.20355666 | 0.99937574 |
| ETV3 | 11.3142876 | -0.8123618 | 0.5749361 | -1.4129601 | 0.15766747 | 0.99937574 |
| DUSP23 | 38.1086315 | -0.1629953 | 0.53671189 | -0.3036923 | 0.76136234 | 0.99937574 |
| TAGLN2 | 70.2776761 | -0.6994452 | 0.45192358 | -1.5477068 | 0.12169291 | 0.99937574 |

| | | | | | | |
|---|---|---|---|---|---|---|
| IGSF8 | 18.8013129 | 0.02709278 | 0.52169477 | 0.05193224 | 0.95858269 | 0.99937574 |
| PEA15 | 17.5388836 | -0.7414933 | 0.64265762 | -1.1537921 | 0.24858543 | 0.99937574 |
| DCAF8 | 34.3560513 | -0.1574357 | 0.32962002 | -0.477628 | 0.63291503 | 0.99937574 |
| COPA | 143.374073 | 0.16295838 | 0.25427547 | 0.64087338 | 0.52160496 | 0.99937574 |
| NCSTN | 29.5440897 | -0.0921452 | 0.47244718 | -0.1950382 | 0.84536307 | 0.99937574 |
| F11R | 55.8826765 | 0.33432779 | 0.39193998 | 0.85300763 | 0.39365507 | 0.99937574 |
| AL591806.3 | 12.0971562 | 0.05811852 | 0.73500397 | 0.07907238 | 0.93697505 | 0.99937574 |
| PFDN2 | 61.2480635 | -0.4137705 | 0.506638 | -0.8166984 | 0.41410078 | 0.99937574 |
| NIT1 | 35.7219803 | 0.48617306 | 0.46493252 | 1.04568523 | 0.29570639 | 0.99937574 |
| DEDD | 14.7597304 | -0.4440854 | 0.50236865 | -0.8839831 | 0.37670531 | 0.99937574 |
| UFC1 | 39.6103104 | 0.14777905 | 0.52524753 | 0.28135126 | 0.778441 | 0.99937574 |
| B4GALT3 | 18.1770109 | -0.1131508 | 0.46537393 | -0.2431396 | 0.80789729 | 0.99937574 |
| NDUFS2 | 90.8389044 | 0.42278726 | 0.36567149 | 1.15619421 | 0.24760173 | 0.99937574 |
| SDHC | 111.508912 | 0.23886406 | 0.24924259 | 0.95835974 | 0.33788139 | 0.99937574 |
| AL592295.4 | 10.6160377 | 0.78733249 | 0.56839843 | 1.38517712 | 0.16599832 | 0.99937574 |
| FCGR2A | 15.6057437 | 1.2258392 | 0.69484678 | 1.76418634 | 0.07770061 | 0.99937574 |
| DUSP12 | 15.5146722 | 0.53759164 | 0.48390507 | 1.11094441 | 0.26659228 | 0.99937574 |
| ATF6 | 195.462812 | -0.0140602 | 0.24080411 | -0.0583887 | 0.95343899 | 0.99937574 |
| NOS1AP | 316.317956 | -0.4013907 | 0.44095393 | -0.910278 | 0.3626759 | 0.99937574 |
| C1orf226 | 12.2284818 | -0.9093936 | 0.59724834 | -1.5226389 | 0.12784906 | 0.99937574 |
| UHMK1 | 286.341464 | -0.0924133 | 0.31584538 | -0.2925903 | 0.7698353 | 0.99937574 |
| UAP1 | 248.470107 | -0.3641389 | 0.39618463 | -0.9191142 | 0.35803587 | 0.99937574 |
| DDR2 | 46.3800492 | -0.3630747 | 0.58794902 | -0.6175275 | 0.53688684 | 0.99937574 |
| HSD17B7 | 37.2921696 | -0.1415521 | 0.55336086 | -0.2558044 | 0.7981019 | 0.99937574 |
| PBX1 | 160.226152 | 0.61818544 | 0.47109576 | 1.31222883 | 0.18944294 | 0.99937574 |
| MGST3 | 176.258596 | -0.3218784 | 0.36466433 | -0.8826703 | 0.37741443 | 0.99937574 |
| ALDH9A1 | 37.3260049 | 0.65302249 | 0.42955544 | 1.52022867 | 0.12845351 | 0.99937574 |
| TMCO1 | 77.1306034 | -0.057094 | 0.25588468 | -0.2231238 | 0.82343915 | 0.99937574 |
| UCK2 | 12.7471963 | 0.43538246 | 0.61131335 | 0.71220833 | 0.47633578 | 0.99937574 |
| POGK | 36.4837269 | -0.1207822 | 0.56682189 | -0.2130866 | 0.83125939 | 0.99937574 |
| TADA1 | 20.2100222 | -0.1994858 | 0.46919577 | -0.4251653 | 0.67071619 | 0.99937574 |
| POU2F1 | 199.187204 | 0.07601067 | 0.28165756 | 0.2698691 | 0.78726096 | 0.99937574 |
| CREG1 | 26.5379682 | 0.59130997 | 0.42114741 | 1.40404513 | 0.16030541 | 0.99937574 |
| MPZL1 | 102.980531 | -0.1204707 | 0.4072093 | -0.2958446 | 0.76734874 | 0.99937574 |
| MPC2 | 56.6817598 | 0.82739104 | 0.52254546 | 1.58338576 | 0.11333357 | 0.99937574 |
| DCAF6 | 257.087523 | 0.10610146 | 0.28301135 | 0.37490177 | 0.70773352 | 0.99937574 |
| TIPRL | 30.2822411 | 0.0146216 | 0.35318858 | 0.04139885 | 0.96697793 | 0.99937574 |
| SFT2D2 | 57.4372951 | -0.4268922 | 0.37416807 | -1.1409103 | 0.25390726 | 0.99937574 |
| TBX19 | 11.4099139 | -0.0363674 | 0.651379 | -0.0558314 | 0.95547615 | 0.99937574 |
| LINC00970 | 54.0480212 | 0.58587355 | 0.90356317 | 0.64840353 | 0.51672399 | 0.99937574 |
| ATP1B1 | 1151.22078 | 0.45077116 | 0.33670233 | 1.33878241 | 0.18064152 | 0.99937574 |
| NME7 | 117.131199 | 0.46491553 | 0.28884315 | 1.60957784 | 0.10749005 | 0.99937574 |
| BLZF1 | 42.8543534 | -0.5633407 | 0.33709736 | -1.6711513 | 0.0946918 | 0.99937574 |
| CCDC181 | 11.8901313 | -0.5277699 | 0.64919877 | -0.8129558 | 0.41624339 | 0.99937574 |
| SLC19A2 | 163.4661 | -0.3927479 | 0.49668515 | -0.7907381 | 0.42909682 | 0.99937574 |
| SCYL3 | 36.0683142 | 0.03350725 | 0.38268883 | 0.08755744 | 0.93022843 | 0.99937574 |
| KIFAP3 | 74.6651374 | 0.36522209 | 0.39984413 | 0.91341117 | 0.36102633 | 0.99937574 |
| GORAB | 14.5983729 | -0.4854474 | 0.54476749 | -0.8911093 | 0.37287054 | 0.99937574 |
| FMO4 | 19.063729 | -0.1306899 | 0.65637879 | -0.1991074 | 0.84217875 | 0.99937574 |
| PRRC2C | 276.929532 | -0.2555458 | 0.23890357 | -1.069661 | 0.28477191 | 0.99937574 |
| VAMP4 | 21.1566347 | 0.00148565 | 0.41761495 | 0.00355747 | 0.99716155 | 0.99937574 |
| METTL13 | 12.7784954 | -0.0686116 | 0.53660252 | -0.1278631 | 0.89825734 | 0.99937574 |
| DNM3 | 23.0341409 | -1.278833 | 0.50466105 | -2.5340433 | 0.01127548 | 0.76274616 |
| PIGC | 21.5408469 | 0.23743832 | 0.4973389 | 0.47741755 | 0.63306482 | 0.99937574 |

| | | | | | | |
|---|---|---|---|---|---|---|
| SUCO | 99.248634 | -0.3238281 | 0.33407408 | -0.9693302 | 0.33238047 | 0.99937574 |
| AL645568.1 | 25.471815 | 0.2192475 | 0.6173113 | 0.35516522 | 0.72246582 | 0.99937574 |
| PRDX6 | 147.515041 | 0.39657746 | 0.39251416 | 1.010352 | 0.31232667 | 0.99937574 |
| KLHL20 | 57.3036378 | 0.29494643 | 0.3428201 | 0.86035336 | 0.38959428 | 0.99937574 |
| DARS2 | 19.0010986 | 0.43083833 | 0.42444735 | 1.01505718 | 0.31007857 | 0.99937574 |
| ZBTB37 | 28.92072 | 0.13598052 | 0.55423192 | 0.24534949 | 0.80618584 | 0.99937574 |
| RC3H1 | 166.216206 | -0.2097443 | 0.26433637 | -0.7934749 | 0.4275012 | 0.99937574 |
| RABGAP1L | 659.045356 | -0.0400225 | 0.27523994 | -0.1454096 | 0.88438745 | 0.99937574 |
| CACYBP | 23.8912166 | -0.0037337 | 0.44481326 | -0.0083939 | 0.99330268 | 0.99937574 |
| MRPS14 | 25.0592891 | 0.06749778 | 0.47061827 | 0.14342364 | 0.88595562 | 0.99937574 |
| KIAA0040 | 74.7168657 | -0.6023152 | 0.42860281 | -1.4052993 | 0.15993228 | 0.99937574 |
| COP1 | 281.010437 | 0.13644698 | 0.19745584 | 0.69102532 | 0.48954963 | 0.99937574 |
| PAPPA2 | 47.7592759 | 1.46090509 | 0.82913831 | 1.76195584 | 0.07807676 | 0.99937574 |
| SEC16B | 19.2615235 | -0.0007612 | 0.87115229 | -0.0008738 | 0.99930284 | 0.99979721 |
| RASAL2 | 137.908374 | 0.19466849 | 0.36588912 | 0.53204229 | 0.5946967 | 0.99937574 |
| RALGPS2 | 418.897909 | 0.21831069 | 0.35445944 | 0.61589752 | 0.53796216 | 0.99937574 |
| ANGPTL1 | 53.4543119 | -0.1408874 | 0.66759084 | -0.2110386 | 0.83285718 | 0.99937574 |
| FAM20B | 50.6973091 | 0.12822604 | 0.30637122 | 0.4185316 | 0.67555849 | 0.99937574 |
| TOR3A | 15.3877687 | 0.09174453 | 0.63472925 | 0.14454121 | 0.88507312 | 0.99937574 |
| ABL2 | 175.875498 | 0.21280156 | 0.2930988 | 0.72604035 | 0.46781403 | 0.99937574 |
| SOAT1 | 113.685241 | -0.2919287 | 0.34819578 | -0.8384038 | 0.40180398 | 0.99937574 |
| TOR1AIP2 | 109.802825 | 0.16815112 | 0.30206062 | 0.55668004 | 0.57774606 | 0.99937574 |
| TOR1AIP1 | 57.6342449 | -0.5989844 | 0.3430312 | -1.7461514 | 0.08078465 | 0.99937574 |
| CEP350 | 196.210315 | 0.08891591 | 0.20135876 | 0.44157957 | 0.65879347 | 0.99937574 |
| QSOX1 | 84.0035059 | 0.2696602 | 0.35468575 | 0.76027921 | 0.44708771 | 0.99937574 |
| ACBD6 | 173.439361 | -0.1351621 | 0.29365006 | -0.4602828 | 0.64531327 | 0.99937574 |
| XPR1 | 277.583514 | 0.24542999 | 0.32604161 | 0.75275665 | 0.45159616 | 0.99937574 |
| STX6 | 56.7998212 | -0.0649247 | 0.51999676 | -0.124856 | 0.90063754 | 0.99937574 |
| MR1 | 116.475299 | 0.66142934 | 0.38121175 | 1.73507068 | 0.08272829 | 0.99937574 |
| CACNA1E | 23.7131143 | -0.8001763 | 2.04610698 | -0.3910725 | 0.69574361 | 0.99937574 |
| GLUL | 115.601994 | -0.089874 | 0.37348941 | -0.2406332 | 0.80983939 | 0.99937574 |
| RNASEL | 29.9852395 | 0.84748262 | 0.48956902 | 1.73107892 | 0.08343769 | 0.99937574 |
| RGS16 | 29.594964 | -0.4386249 | 0.92057195 | -0.47647 | 0.6337396 | 0.99937574 |
| DHX9 | 112.194165 | -0.1671037 | 0.27689616 | -0.6034887 | 0.54618365 | 0.99937574 |
| LAMC1 | 103.74101 | 0.18366846 | 0.34285056 | 0.53570995 | 0.59215902 | 0.99937574 |
| SMG7 | 187.449714 | -0.1365764 | 0.32559089 | -0.4194725 | 0.67487083 | 0.99937574 |
| ARPC5 | 65.1006292 | -0.1925032 | 0.37348095 | -0.5154299 | 0.60625265 | 0.99937574 |
| RGL1 | 16.9275161 | -0.8687517 | 0.65957401 | -1.3171406 | 0.18779149 | 0.99937574 |
| TSEN15 | 33.1132284 | 0.63279138 | 0.41757157 | 1.5154082 | 0.12966906 | 0.99937574 |
| C1orf21 | 14.3259238 | 0.31201431 | 0.66845274 | 0.46677093 | 0.64066378 | 0.99937574 |
| EDEM3 | 92.1340577 | 0.54564142 | 0.44808561 | 1.21771691 | 0.22333157 | 0.99937574 |
| RNF2 | 33.1560157 | -0.2598636 | 0.38718793 | -0.6711563 | 0.50212095 | 0.99937574 |
| TRMT1L | 35.0801078 | -0.0542885 | 0.40598555 | -0.1337203 | 0.89362372 | 0.99937574 |
| SWT1 | 56.2657792 | -0.6279718 | 0.31009428 | -2.0250997 | 0.04285713 | 0.99937574 |
| IVNS1ABP | 112.845698 | -0.6799219 | 0.47382382 | -1.4349677 | 0.15129631 | 0.99937574 |
| TPR | 132.760049 | -0.0867043 | 0.24383215 | -0.3555902 | 0.72214748 | 0.99937574 |
| ODR4 | 74.0256775 | -0.0380063 | 0.25847776 | -0.1470391 | 0.88310112 | 0.99937574 |
| PLA2G4A | 57.3849597 | 0.19207323 | 0.55494201 | 0.34611406 | 0.729257 | 0.99937574 |
| BRINP3 | 37.158075 | -0.9228275 | 0.93977149 | -0.9819701 | 0.32611459 | 0.99937574 |
| AL390957.1 | 73.9218806 | 0.30612731 | 0.88609242 | 0.34548011 | 0.72973346 | 0.99937574 |
| AL136454.1 | 71.1719748 | -0.0407537 | 0.4311099 | -0.0945321 | 0.92468651 | 0.99937574 |
| UCHL5 | 46.21622 | 0.36553603 | 0.34110818 | 1.07161322 | 0.28389379 | 0.99937574 |
| TROVE2 | 87.8474288 | 0.17190186 | 0.26624867 | 0.64564402 | 0.51850992 | 0.99937574 |
| GLRX2 | 11.4596479 | -0.0414381 | 0.55821606 | -0.0742332 | 0.94082487 | 0.99937574 |

| | | | | | | |
|---|---|---|---|---|---|---|
| CDC73 | 137.721736 | -0.0931741 | 0.26711288 | -0.348819 | 0.72722518 | 0.99937574 |
| ZBTB41 | 41.2470155 | 0.10326632 | 0.3415541 | 0.3023425 | 0.76239099 | 0.99937574 |
| DENND1B | 252.722117 | 0.07675313 | 0.34424033 | 0.2229638 | 0.82356368 | 0.99937574 |
| NEK7 | 209.715766 | -0.2740989 | 0.27677454 | -0.9903326 | 0.32201157 | 0.99937574 |
| ATP6V1G3 | 200.728887 | 1.58463124 | 0.7123641 | 2.22446813 | 0.02611696 | 0.96546668 |
| PTPRC | 20.1804007 | 1.76728981 | 0.69804151 | 2.53178326 | 0.01134841 | 0.76274616 |
| ZNF281 | 17.6875325 | 0.16621284 | 0.51514172 | 0.3226546 | 0.74695684 | 0.99937574 |
| DDX59 | 35.2018543 | 0.32950669 | 0.35268104 | 0.93429089 | 0.35015386 | 0.99937574 |
| CAMSAP2 | 382.657111 | 0.24151078 | 0.30770663 | 0.78487351 | 0.43252773 | 0.99937574 |
| TMEM9 | 29.7946218 | 0.91271215 | 0.46879257 | 1.94694242 | 0.05154164 | 0.99937574 |
| LAD1 | 15.1475792 | -0.4173577 | 0.5086227 | -0.8205644 | 0.41189443 | 0.99937574 |
| CSRP1 | 19.7810749 | -0.6221346 | 0.64477184 | -0.9648911 | 0.33459935 | 0.99937574 |
| AC096677.1 | 20.5090107 | -0.1069166 | 0.56479183 | -0.1893027 | 0.84985562 | 0.99937574 |
| NAV1 | 18.9733751 | -0.950549 | 0.89505755 | -1.0619976 | 0.28823678 | 0.99937574 |
| IPO9 | 71.7532566 | -0.2821101 | 0.35242395 | -0.800485 | 0.42342985 | 0.99937574 |
| TIMM17A | 40.2709475 | -0.3562571 | 0.44080244 | -0.8082013 | 0.41897473 | 0.99937574 |
| RNPEP | 46.9698796 | -0.2614828 | 0.30079355 | -0.8693098 | 0.38467768 | 0.99937574 |
| ELF3 | 211.348752 | -0.4775273 | 0.37758117 | -1.264701 | 0.20597855 | 0.99937574 |
| ARL8A | 30.2615882 | -0.1282016 | 0.45648595 | -0.2808447 | 0.77882954 | 0.99937574 |
| PPP1R12B | 521.106898 | 0.81556722 | 0.29475646 | 2.76691886 | 0.00565888 | 0.65033075 |
| KDM5B | 159.358407 | 0.09820004 | 0.26453251 | 0.37122105 | 0.71047289 | 0.99937574 |
| RABIF | 9.94999756 | -0.968329 | 0.77593245 | -1.2479553 | 0.21204742 | 0.99937574 |
| KLHL12 | 38.6711105 | -0.1439451 | 0.33813196 | -0.425707 | 0.67032139 | 0.99937574 |
| ADIPOR1 | 24.8024526 | 0.03684441 | 0.41336189 | 0.08913355 | 0.92897578 | 0.99937574 |
| CYB5R1 | 19.047876 | 0.26526634 | 0.58800279 | 0.45113108 | 0.65189508 | 0.99937574 |
| TMEM183A | 14.0748976 | -1.0023703 | 0.57342178 | -1.7480507 | 0.08045525 | 0.99937574 |
| BTG2 | 101.368425 | -0.831439 | 0.820503 | -1.0133284 | 0.3109033 | 0.99937574 |
| ATP2B4 | 66.2691825 | 1.2427584 | 0.47459642 | 2.6185583 | 0.00883022 | 0.74742128 |
| SNRPE | 28.4614595 | 0.42954689 | 0.37546894 | 1.14402775 | 0.25261213 | 0.99937574 |
| SOX13 | 26.8166779 | -0.0650605 | 0.60919813 | -0.106797 | 0.91495004 | 0.99937574 |
| GOLT1A | 28.5918614 | -0.4663629 | 0.48185672 | -0.9678457 | 0.33312145 | 0.99937574 |
| PLEKHA6 | 124.174524 | -0.6494339 | 0.35250631 | -1.8423328 | 0.06542649 | 0.99937574 |
| PPP1R15B | 37.7333395 | -0.3637524 | 0.4302933 | -0.8453591 | 0.39791036 | 0.99937574 |
| MDM4 | 136.547928 | -0.3213796 | 0.25211723 | -1.2747229 | 0.20240736 | 0.99937574 |
| LRRN2 | 95.5991802 | 0.80059073 | 0.53671907 | 1.49163832 | 0.13579399 | 0.99937574 |
| RBBP5 | 43.6168006 | 0.17533165 | 0.35363003 | 0.49580532 | 0.62003177 | 0.99937574 |
| DSTYK | 56.9497127 | 0.03300086 | 0.41641934 | 0.0792491 | 0.93683449 | 0.99937574 |
| NUAK2 | 49.8777091 | -0.279565 | 0.65499798 | -0.4268181 | 0.66951184 | 0.99937574 |
| CDK18 | 17.8185356 | 0.11892203 | 0.4781865 | 0.24869382 | 0.80359763 | 0.99937574 |
| MFSD4A | 17.8885704 | 0.97541069 | 0.58818625 | 1.65833644 | 0.09724958 | 0.99937574 |
| ELK4 | 148.390215 | 0.07111497 | 0.481682 | 0.14763883 | 0.88262781 | 0.99937574 |
| NUCKS1 | 205.132484 | -0.0045378 | 0.1932217 | -0.0234847 | 0.98126362 | 0.99937574 |
| RAB29 | 12.6012714 | -0.5319344 | 0.65999285 | -0.80597 | 0.42026017 | 0.99937574 |
| SLC41A1 | 16.6436472 | -0.3988255 | 0.46383086 | -0.8598512 | 0.38987108 | 0.99937574 |
| RHEX | 18.8473102 | -2.0007391 | 0.87369595 | -2.2899718 | 0.02202295 | 0.96427895 |
| SRGAP2 | 12.5119452 | -0.961223 | 0.60083889 | -1.5998016 | 0.10964261 | 0.99937574 |
| EIF2D | 20.0412192 | 0.12566255 | 0.46035988 | 0.2729659 | 0.78487943 | 0.99937574 |
| MAPKAPK2 | 129.25088 | -0.287198 | 0.27028568 | -1.0625721 | 0.28797605 | 0.99937574 |
| PIGR | 46.8061186 | 1.18719543 | 0.65097617 | 1.82371567 | 0.06819507 | 0.99937574 |
| PFKFB2 | 107.362073 | -0.2284033 | 0.37288862 | -0.6125241 | 0.54019104 | 0.99937574 |
| YOD1 | 15.4481175 | -0.3710536 | 0.6353589 | -0.5840064 | 0.55921603 | 0.99937574 |
| CD55 | 86.6038922 | -0.4628777 | 0.56568029 | -0.8182674 | 0.41320453 | 0.99937574 |
| CR1L | 97.9691751 | -0.8155016 | 0.76372071 | -1.0678008 | 0.28561039 | 0.99937574 |
| CD46 | 324.730856 | 0.40797946 | 0.2644355 | 1.54283161 | 0.12287164 | 0.99937574 |

| | | | | | | |
|---|---|---|---|---|---|---|
| MIR29B2CH( | 187.754128 | 0.08728099 | 0.37480514 | 0.23287032 | 0.81586211 | 0.99937574 |
| PLXNA2 | 56.3010084 | 0.93317768 | 0.56724048 | 1.64511827 | 0.09994542 | 0.99937574 |
| G0S2 | 134.399858 | 0.26268892 | 1.22781444 | 0.21394839 | 0.8305873 | 0.99937574 |
| IRF6 | 38.5124171 | 0.07954466 | 0.46221032 | 0.17209625 | 0.86336186 | 0.99937574 |
| HHAT | 48.1639963 | 0.39604978 | 0.6374373 | 0.62131566 | 0.53439195 | 0.99937574 |
| RCOR3 | 163.455435 | -0.0281361 | 0.26425605 | -0.1064728 | 0.91520724 | 0.99937574 |
| LINC00467 | 33.5065624 | -0.551477 | 0.46280342 | -1.191601 | 0.23341775 | 0.99937574 |
| SLC30A1 | 14.5572249 | -0.7424388 | 0.69537487 | -1.0676813 | 0.28566428 | 0.99937574 |
| LPGAT1 | 261.582417 | -0.1356229 | 0.36112697 | -0.3755545 | 0.7072481 | 0.99937574 |
| INTS7 | 62.3142407 | 0.20084247 | 0.37450734 | 0.53628447 | 0.59176196 | 0.99937574 |
| PPP2R5A | 551.035396 | 0.0039496 | 0.38405499 | 0.01028396 | 0.99179473 | 0.99937574 |
| AL360091.3 | 76.1966827 | 0.26145257 | 0.71318427 | 0.3665989 | 0.71391823 | 0.99937574 |
| NENF | 33.5351096 | -0.1942867 | 0.38861972 | -0.4999405 | 0.61711696 | 0.99937574 |
| NSL1 | 45.3980347 | 0.29346145 | 0.34062207 | 0.86154559 | 0.38893762 | 0.99937574 |
| TATDN3 | 46.6662431 | -0.1946905 | 0.42679253 | -0.4561714 | 0.64826675 | 0.99937574 |
| FLVCR1 | 32.8029817 | -0.3309694 | 0.62039221 | -0.5334842 | 0.59369848 | 0.99937574 |
| ANGEL2 | 36.3930902 | -0.0311627 | 0.35846332 | -0.0869341 | 0.93072388 | 0.99937574 |
| RPS6KC1 | 140.56877 | -0.4509365 | 0.22678031 | -1.9884288 | 0.04676428 | 0.99937574 |
| SMYD2 | 36.2747197 | -0.1861614 | 0.44290857 | -0.4203156 | 0.67425494 | 0.99937574 |
| PTPN14 | 117.434823 | -0.2328753 | 0.29317016 | -0.794335 | 0.42700044 | 0.99937574 |
| KCTD3 | 72.8508916 | 0.12027489 | 0.28299628 | 0.4250052 | 0.67083289 | 0.99937574 |
| ESRRG | 3023.32315 | 0.58564662 | 0.39572906 | 1.47991816 | 0.13889509 | 0.99937574 |
| GPATCH2 | 153.908973 | 0.02382612 | 0.26858637 | 0.08870935 | 0.9293129 | 0.99937574 |
| RRP15 | 27.4319967 | -0.1886389 | 0.3916126 | -0.4816978 | 0.63002065 | 0.99937574 |
| LYPLAL1 | 209.836564 | 0.10736931 | 0.31282841 | 0.34322109 | 0.73143214 | 0.99937574 |
| ZC3H11B | 10.4481885 | -0.2655512 | 0.71780907 | -0.3699468 | 0.71142211 | 0.99937574 |
| EPRS | 56.6937783 | 0.07403964 | 0.2824129 | 0.26216804 | 0.79319189 | 0.99937574 |
| BPNT1 | 24.0412326 | 0.35949246 | 0.45738063 | 0.78598095 | 0.43187865 | 0.99937574 |
| IARS2 | 71.4183988 | 0.10490877 | 0.28963541 | 0.36220974 | 0.7171953 | 0.99937574 |
| RAB3GAP2 | 194.417958 | 0.11988337 | 0.29117116 | 0.41172817 | 0.68053868 | 0.99937574 |
| MARK1 | 80.7175584 | 0.14280952 | 0.43292604 | 0.32987049 | 0.74149783 | 0.99937574 |
| 2-Mar | 55.1330895 | 0.50560136 | 0.39000616 | 1.29639328 | 0.19484003 | 0.99937574 |
| AL513314.2 | 16.9660664 | -0.1842796 | 0.65810232 | -0.2800167 | 0.77946469 | 0.99937574 |
| MIA3 | 42.67005 | -0.1923074 | 0.32594908 | -0.589992 | 0.55519598 | 0.99937574 |
| AIDA | 18.4505439 | 0.30626883 | 0.43816415 | 0.69898195 | 0.48456331 | 0.99937574 |
| BROX | 56.2137846 | 0.04017656 | 0.31521033 | 0.12745954 | 0.8985767 | 0.99937574 |
| FAM177B | 27.6966519 | -1.190954 | 0.79345419 | -1.5009739 | 0.13336232 | 0.99937574 |
| AC093152.1 | 23.3649729 | -0.4065552 | 0.41219952 | -0.9863067 | 0.32398262 | 0.99937574 |
| SUSD4 | 11.3998305 | 0.61494214 | 0.83231413 | 0.7388342 | 0.46000768 | 0.99937574 |
| CAPN2 | 248.173629 | 0.33649079 | 0.25284574 | 1.33081456 | 0.18325004 | 0.99937574 |
| TP53BP2 | 52.2668678 | -0.479375 | 0.39508301 | -1.2133527 | 0.22499501 | 0.99937574 |
| FBXO28 | 86.5151069 | -0.4320236 | 0.29355461 | -1.4716976 | 0.14110256 | 0.99937574 |
| DEGS1 | 13.4334769 | -0.5971476 | 0.90274172 | -0.6614823 | 0.50830309 | 0.99937574 |
| NVL | 68.0811393 | -0.0533897 | 0.29158443 | -0.1831019 | 0.85471803 | 0.99937574 |
| CNIH4 | 61.2461446 | -0.4349608 | 0.372846 | -1.1665963 | 0.24337343 | 0.99937574 |
| WDR26 | 168.879664 | -0.1176063 | 0.34562549 | -0.3402709 | 0.73365256 | 0.99937574 |
| CNIH3 | 43.0295348 | 0.55524008 | 0.41223857 | 1.34689018 | 0.17801561 | 0.99937574 |
| DNAH14 | 254.228899 | -0.0943019 | 0.32985263 | -0.2858909 | 0.7749617 | 0.99937574 |
| LBR | 52.7016555 | -0.3887483 | 0.42502177 | -0.9146551 | 0.3603727 | 0.99937574 |
| ENAH | 33.1840993 | 0.05925637 | 0.43562005 | 0.13602764 | 0.89179943 | 0.99937574 |
| SRP9 | 100.345026 | 0.12474157 | 0.41374147 | 0.30149641 | 0.76303599 | 0.99937574 |
| EPHX1 | 21.139246 | -0.796458 | 0.62360527 | -1.2771829 | 0.2015377 | 0.99937574 |
| AL591895.1 | 23.1163699 | 0.19321334 | 0.50718179 | 0.38095481 | 0.70323678 | 0.99937574 |
| TMEM63A | 64.7521073 | -0.1622202 | 0.43002971 | -0.3772302 | 0.70600253 | 0.99937574 |

| | | | | | | |
|---|---|---|---|---|---|---|
| SDE2 | 17.3667492 | -0.4020738 | 0.5526534 | -0.7275334 | 0.46689927 | 0.99937574 |
| H3F3A | 24.931393 | 0.32461623 | 0.42583586 | 0.76230365 | 0.4458788 | 0.99937574 |
| ACBD3 | 103.040916 | -0.1404567 | 0.28027552 | -0.5011378 | 0.61627417 | 0.99937574 |
| LIN9 | 14.605825 | 0.70957172 | 0.53836256 | 1.31801832 | 0.18749751 | 0.99937574 |
| PARP1 | 52.3358032 | 0.13887429 | 0.44720832 | 0.31053603 | 0.75615337 | 0.99937574 |
| STUM | 106.557595 | -0.1069473 | 0.53585357 | -0.199583 | 0.84180669 | 0.99937574 |
| ITPKB | 50.4833383 | 0.17412198 | 0.4251747 | 0.40953044 | 0.68215043 | 0.99937574 |
| PSEN2 | 46.461462 | -0.5370981 | 0.49826352 | -1.0779399 | 0.28106058 | 0.99937574 |
| COQ8A | 42.4268475 | -0.213411 | 0.54358129 | -0.3926018 | 0.69461361 | 0.99937574 |
| CDC42BPA | 450.782517 | 0.13854476 | 0.29269031 | 0.47334931 | 0.63596398 | 0.99937574 |
| ZNF678 | 73.8892878 | 0.37133811 | 0.32879606 | 1.12938733 | 0.25873447 | 0.99937574 |
| SNAP47 | 19.1443132 | 0.15790125 | 0.42727276 | 0.36955608 | 0.71171328 | 0.99937574 |
| ARF1 | 136.103725 | 0.04714547 | 0.29675346 | 0.15887082 | 0.87377065 | 0.99937574 |
| MRPL55 | 18.5361676 | 0.46564237 | 0.50328004 | 0.92521524 | 0.35485396 | 0.99937574 |
| GUK1 | 128.243799 | -0.1031632 | 0.34399304 | -0.2998991 | 0.76425414 | 0.99937574 |
| IBA57 | 13.5023886 | 0.02981876 | 0.83664092 | 0.03564104 | 0.97156858 | 0.99937574 |
| OBSCN | 80.5307159 | 0.54656243 | 0.3071815 | 1.77928175 | 0.07519358 | 0.99937574 |
| TRIM11 | 11.9330403 | -0.0976611 | 0.64840905 | -0.1506166 | 0.8802782 | 0.99937574 |
| RNF187 | 27.9934814 | -0.1033616 | 0.4819936 | -0.2144461 | 0.83019922 | 0.99937574 |
| RHOU | 24.0880491 | -0.4699712 | 0.51361619 | -0.9150241 | 3.60E-01 | 0.99937574 |
| RAB4A | 43.6529163 | 0.07174088 | 0.35135414 | 0.20418396 | 0.83820975 | 0.99937574 |
| NUP133 | 50.760794 | 0.06128559 | 0.30574235 | 0.20044847 | 0.84112985 | 0.99937574 |
| ABCB10 | 26.2009398 | 0.41878033 | 0.42493468 | 0.98551694 | 0.32437021 | 0.99937574 |
| TAF5L | 17.7813993 | -0.6535942 | 0.58306335 | -1.120966 | 0.26230235 | 0.99937574 |
| GALNT2 | 48.5781396 | -0.306944 | 0.36923844 | -0.8312893 | 0.40581022 | 0.99937574 |
| COG2 | 33.3132375 | 0.04375056 | 0.3999202 | 0.10939823 | 0.91288664 | 0.99937574 |
| C1orf198 | 17.1156901 | -0.2316775 | 0.51598708 | -0.4489986 | 0.65343265 | 0.99937574 |
| TTC13 | 25.2204729 | 0.23072393 | 0.5896274 | 0.39130462 | 0.69557209 | 0.99937574 |
| ARV1 | 23.8635367 | 0.19899334 | 0.46400988 | 0.42885582 | 0.66802815 | 0.99937574 |
| C1orf131 | 18.6098672 | -0.1959574 | 0.4567557 | -0.4290202 | 0.66790854 | 0.99937574 |
| GNPAT | 59.7262557 | 0.47075952 | 0.30447106 | 1.54615521 | 0.12206709 | 0.99937574 |
| EXOC8 | 11.791216 | -0.2041778 | 0.5634758 | -0.3623541 | 0.71708741 | 0.99937574 |
| SPRTN | 15.3224501 | 0.4560736 | 0.64921992 | 0.70249478 | 0.48237066 | 0.99937574 |
| EGLN1 | 238.939835 | -0.0901399 | 0.3266855 | -0.2759225 | 0.78260761 | 0.99937574 |
| DISC1 | 52.185804 | -0.3253302 | 0.74310797 | -0.4377966 | 0.6615337 | 0.99937574 |
| SIPA1L2 | 106.961805 | 0.6065053 | 0.50790626 | 1.19412842 | 0.23242774 | 0.99937574 |
| NTPCR | 26.7405479 | -0.1830411 | 0.4076862 | -0.4489754 | 0.65344941 | 0.99937574 |
| PCNX2 | 146.967564 | -0.4426002 | 0.34917217 | -1.26757 | 0.20495157 | 0.99937574 |
| MAP3K21 | 130.032987 | 0.0799868 | 0.28390688 | 0.28173605 | 0.77814591 | 0.99937574 |
| KCNK1 | 193.430889 | 0.25597044 | 0.39426386 | 0.64923638 | 0.5161856 | 0.99937574 |
| SLC35F3 | 106.575839 | -0.2468178 | 0.52252737 | -0.4723538 | 0.63667431 | 0.99937574 |
| COA6 | 21.767948 | -0.5582225 | 0.6185235 | -0.9025082 | 0.36678699 | 0.99937574 |
| TARBP1 | 59.3376465 | 0.31727863 | 0.54930425 | 0.57760091 | 0.5635336 | 0.99937574 |
| IRF2BP2 | 59.9609771 | 0.4268746 | 0.38805602 | 1.10003345 | 0.27131755 | 0.99937574 |
| TOMM20 | 92.2399121 | 0.03167156 | 0.39148938 | 0.08090017 | 0.93552134 | 0.99937574 |
| ARID4B | 368.158288 | -0.1430249 | 0.3217081 | -0.4445797 | 0.65662354 | 0.99937574 |
| B3GALNT2 | 54.1888424 | -0.3427272 | 0.38081417 | -0.8999854 | 0.36812802 | 0.99937574 |
| LYST | 137.358917 | 0.13136424 | 0.32423166 | 0.4051555 | 0.68536321 | 0.99937574 |
| GPR137B | 81.0392155 | -0.4896706 | 0.32234336 | -1.5190961 | 0.1287383 | 0.99937574 |
| ERO1B | 146.725005 | -0.3127166 | 0.65420284 | -0.4780117 | 0.63264188 | 0.99937574 |
| EDARADD | 140.186563 | 0.27771209 | 0.56349329 | 0.49284009 | 0.62212558 | 0.99937574 |
| LGALS8 | 45.3874262 | -0.3872706 | 0.31324258 | -1.2363282 | 0.21633658 | 0.99937574 |
| HEATR1 | 35.7598486 | -0.0612206 | 0.40759191 | -0.1502008 | 0.88060622 | 0.99937574 |
| MTR | 695.705029 | -0.0500497 | 0.51347082 | -0.0974733 | 0.92235053 | 0.99937574 |

| | | | | | | |
|---|---|---|---|---|---|---|
| CHRM3 | 18.4078542 | 0.46950267 | 1.08053436 | 0.43450971 | 0.66391834 | 0.99937574 |
| FH | 34.1399224 | 0.60734515 | 0.36298223 | 1.67320903 | 0.09428615 | 0.99937574 |
| OPN3 | 21.0671509 | 0.00453265 | 0.47815808 | 0.00947941 | 0.99243664 | 0.99937574 |
| CEP170 | 54.1942024 | 0.47826849 | 0.42147368 | 1.13475294 | 0.25647885 | 0.99937574 |
| SDCCAG8 | 261.873517 | 0.09617256 | 0.26926256 | 0.3571702 | 0.72096439 | 0.99937574 |
| AKT3 | 378.952892 | 0.08190066 | 0.27593204 | 0.29681462 | 0.76660804 | 0.99937574 |
| ZBTB18 | 44.1516442 | -0.1072015 | 0.41728128 | -0.2569046 | 0.79725239 | 0.99937574 |
| AL645465.1 | 15.3597603 | 0.14885449 | 0.62272238 | 0.23903828 | 0.8110759 | 0.99937574 |
| ADSS | 55.7937226 | -0.373086 | 0.42766841 | -0.8723722 | 0.38300538 | 0.99937574 |
| CATSPERE | 35.6016241 | -0.3303741 | 0.58313363 | -0.5665495 | 0.57102027 | 0.99937574 |
| DESI2 | 68.6727324 | -0.0658583 | 0.36115322 | -0.1823556 | 0.85530363 | 0.99937574 |
| COX20 | 81.9067869 | -0.0148459 | 0.39597662 | -0.0374919 | 0.97009282 | 0.99937574 |
| HNRNPU | 263.328632 | -0.0706572 | 0.27323133 | -0.2585985 | 0.79594501 | 0.99937574 |
| EFCAB2 | 101.528491 | -0.9179555 | 0.52622707 | -1.7444096 | 0.0810877 | 0.99937574 |
| KIF26B | 18.3022415 | -0.183974 | 0.57717128 | -0.3187511 | 0.74991525 | 0.99937574 |
| SMYD3 | 351.996689 | -0.3005196 | 0.43031711 | -0.6983677 | 0.48494728 | 0.99937574 |
| TFB2M | 11.3509915 | 0.23049966 | 0.57529737 | 0.40066176 | 0.68866917 | 0.99937574 |
| CNST | 85.9905028 | -0.2187716 | 0.26526124 | -0.8247403 | 0.40951903 | 0.99937574 |
| SCCPDH | 128.800395 | -0.3481993 | 0.62928267 | -0.5533274 | 0.58003924 | 0.99937574 |
| AHCTF1 | 99.7347564 | 0.22275269 | 0.29038969 | 0.76708196 | 0.44303279 | 0.99937574 |
| ZNF124 | 32.0854441 | -0.0205684 | 0.37735758 | -0.0545065 | 0.95653161 | 0.99937574 |
| AL390728.6 | 21.9113058 | -0.2319494 | 0.53193505 | -0.4360484 | 0.6628016 | 0.99937574 |
| ZNF496 | 20.7613237 | 0.32744563 | 0.61120587 | 0.53573705 | 0.59214029 | 0.99937574 |
| SH3BP5L | 10.741168 | -0.4100845 | 0.60590703 | -0.6768109 | 0.49852594 | 0.99937574 |
| ZNF672 | 32.7342144 | -0.2290595 | 0.36227538 | -0.6322802 | 0.52720383 | 0.99937574 |
| ZNF692 | 15.9771473 | 0.41901607 | 0.51063908 | 0.82057188 | 0.41189017 | 0.99937574 |
| FAM110C | 12.8325646 | 0.29382107 | 0.63969229 | 0.45931625 | 6.46E-01 | 0.99937574 |
| SH3YL1 | 116.414303 | 0.26051605 | 0.35240429 | 0.73925333 | 0.45975318 | 0.99937574 |
| ACP1 | 45.9317881 | 0.12065381 | 0.36425759 | 0.33123212 | 0.74046916 | 0.99937574 |
| TMEM18 | 12.8045445 | 0.14190347 | 0.62094374 | 0.22852871 | 0.81923524 | 0.99937574 |
| EIPR1 | 101.739432 | 0.00375228 | 0.493137 | 0.007609 | 0.99392895 | 0.99937574 |
| TRAPPC12 | 125.853887 | 0.12503202 | 0.39699203 | 0.31494844 | 0.7528008 | 0.99937574 |
| ADI1 | 91.7243452 | 0.69194495 | 0.54576308 | 1.26784859 | 0.20485202 | 0.99937574 |
| AC108488.2 | 11.1912791 | 0.00400414 | 0.59145197 | 0.00677002 | 0.99459835 | 0.99937574 |
| RPS7 | 179.534943 | 0.00671468 | 0.59046099 | 0.01137193 | 0.99092671 | 0.99937574 |
| RNF144A | 28.9637705 | -0.1554866 | 0.59266108 | -0.2623533 | 0.79304904 | 0.99937574 |
| ID2 | 20.9131645 | -0.4922851 | 0.63233383 | -0.7785209 | 0.43626198 | 0.99937574 |
| KIDINS220 | 278.688723 | 0.14163131 | 0.23366765 | 0.60612289 | 0.54443315 | 0.99937574 |
| MBOAT2 | 241.470627 | -0.7554943 | 0.73699854 | -1.025096 | 0.30531787 | 0.99937574 |
| ASAP2 | 224.497767 | -0.0930079 | 0.449014 | -0.2071381 | 0.83590201 | 0.99937574 |
| ITGB1BP1 | 68.2014416 | -0.0798054 | 0.40042883 | -0.1992997 | 0.8420283 | 0.99937574 |
| CPSF3 | 25.9638641 | -0.2601995 | 0.43392931 | -0.5996356 | 0.54874909 | 0.99937574 |
| IAH1 | 58.3429357 | 0.04521036 | 0.40793432 | 0.11082754 | 0.91175311 | 0.99937574 |
| ADAM17 | 123.05057 | -0.3690563 | 0.29091 | -1.268627 | 0.20457412 | 0.99937574 |
| YWHAQ | 147.534 | 0.01780608 | 0.2236127 | 0.07962912 | 0.93653224 | 0.99937574 |
| TAF1B | 82.8814696 | -0.103765 | 0.35086465 | -0.2957408 | 0.76742805 | 0.99937574 |
| GRHL1 | 39.5306247 | 0.20529095 | 0.36507344 | 0.56232781 | 0.5738927 | 0.99937574 |
| KLF11 | 22.9526235 | -1.2242572 | 0.50425472 | -2.4278547 | 0.01518843 | 0.86113259 |
| CYS1 | 33.9354087 | 0.43855741 | 0.53481009 | 0.82002456 | 0.41220211 | 0.99937574 |
| HPCAL1 | 49.1993385 | 0.38176397 | 0.5616702 | 0.67969419 | 0.49669811 | 0.99937574 |
| ODC1 | 21.1837657 | 0.01977962 | 0.48547755 | 0.0407426 | 0.9675011 | 0.99937574 |
| NOL10 | 98.7599373 | -0.3267561 | 0.3093243 | -1.0563545 | 0.29080629 | 0.99937574 |
| ATP6V1C2 | 550.290121 | 0.55232602 | 0.49965732 | 1.10540964 | 0.26898213 | 0.99937574 |
| PDIA6 | 113.455001 | 0.21883713 | 0.3729731 | 0.58673703 | 0.55738033 | 0.99937574 |

| | | | | | | |
|---|---|---|---|---|---|---|
| PQLC3 | 43.1391364 | 0.74832753 | 0.48133858 | 1.55468015 | 0.12002226 | 0.99937574 |
| ROCK2 | 189.30584 | -0.1705422 | 0.26115859 | -0.6530215 | 0.51374243 | 0.99937574 |
| E2F6 | 18.757128 | -0.4269654 | 0.46135357 | -0.9254625 | 0.35472537 | 0.99937574 |
| LPIN1 | 93.9896303 | -0.3642484 | 0.5140566 | -0.7085765 | 0.4785873 | 0.99937574 |
| MIR3681HG | 163.296787 | 0.40839558 | 0.5273827 | 0.77438182 | 0.43870503 | 0.99937574 |
| TRIB2 | 43.7188106 | -0.2896229 | 0.4809495 | -0.6021898 | 0.54704783 | 0.99937574 |
| NBAS | 240.644136 | -0.004073 | 0.34169507 | -0.01192 | 0.99048941 | 0.99937574 |
| DDX1 | 85.8505536 | 0.40203944 | 0.29827886 | 1.34786435 | 0.17770201 | 0.99937574 |
| SMC6 | 41.1091186 | -0.300016 | 0.43634655 | -0.6875635 | 0.49172771 | 0.99937574 |
| GEN1 | 17.0939029 | -0.7826738 | 0.69447093 | -1.1270072 | 0.25973942 | 0.99937574 |
| RDH14 | 25.1905699 | 0.52121751 | 0.54561411 | 0.95528598 | 0.3394331 | 0.99937574 |
| LINC01376 | 58.7860609 | -0.0759271 | 0.38190311 | -0.1988124 | 0.84240946 | 0.99937574 |
| TTC32 | 20.9639887 | -0.493757 | 0.48040121 | -1.0278014 | 0.30404325 | 0.99937574 |
| WDR35 | 45.1212363 | -0.0169343 | 0.36803671 | -0.0460126 | 0.96330022 | 0.99937574 |
| LAPTM4A | 173.497271 | 0.21169447 | 0.36823756 | 0.57488558 | 0.56536868 | 0.99937574 |
| SDC1 | 29.801949 | -0.4541017 | 0.56986403 | -0.7968597 | 0.42553254 | 0.99937574 |
| PUM2 | 292.590532 | -0.152325 | 0.17767709 | -0.8573138 | 0.39127148 | 0.99937574 |
| RHOB | 23.3858221 | 0.04470027 | 0.6359222 | 0.07029204 | 0.94396122 | 0.99937574 |
| HS1BP3 | 11.9116018 | -0.5762145 | 0.61955088 | -0.9300519 | 0.3523442 | 0.99937574 |
| LDAH | 74.2996913 | 0.1462298 | 0.29761577 | 0.49133753 | 0.62318774 | 0.99937574 |
| KLHL29 | 12.7455147 | -0.8720952 | 0.86264847 | -1.0109509 | 0.31203994 | 0.99937574 |
| ATAD2B | 221.300874 | -0.0133391 | 0.34887539 | -0.0382347 | 0.96950058 | 0.99937574 |
| UBXN2A | 77.7329625 | -0.1670442 | 0.32165311 | -0.5193303 | 0.60353046 | 0.99937574 |
| SF3B6 | 37.8488751 | -0.09696 | 0.55469533 | -0.1747987 | 0.86123778 | 0.99937574 |
| FAM228B | 17.3206558 | 0.26696163 | 0.49858665 | 0.53543678 | 0.59234786 | 0.99937574 |
| TP53I3 | 26.6013495 | -0.1550322 | 0.58161646 | -0.266554 | 0.78981254 | 0.99937574 |
| ITSN2 | 349.29837 | -0.1155921 | 0.23010576 | -0.5023432 | 0.61542614 | 0.99937574 |
| NCOA1 | 469.333775 | 0.11149949 | 0.2388141 | 0.46688824 | 0.64057984 | 0.99937574 |
| PTRHD1 | 12.6150849 | 0.05824278 | 0.82002332 | 0.07102576 | 0.94337726 | 0.99937574 |
| DNAJC27 | 34.0647406 | -0.1006791 | 0.39497425 | -0.2549003 | 0.79880007 | 0.99937574 |
| DNMT3A | 92.4770651 | 0.21242358 | 0.44474495 | 0.4776301 | 0.6329135 | 0.99937574 |
| DTNB | 153.424277 | 0.48801944 | 0.33967073 | 1.43674268 | 0.15079112 | 0.99937574 |
| ASXL2 | 154.431532 | -0.5020927 | 0.38170076 | -1.3154091 | 0.18837244 | 0.99937574 |
| RAB10 | 288.696717 | -0.301454 | 0.28839515 | -1.045281 | 0.2958931 | 0.99937574 |
| HADHA | 208.90849 | 0.28963823 | 0.223583 | 1.29543939 | 0.1951687 | 0.99937574 |
| HADHB | 319.807886 | 0.81873103 | 0.33698504 | 2.42957678 | 0.01511646 | 0.86113259 |
| SELENOI | 178.915814 | -0.0913071 | 0.24789748 | -0.3683261 | 0.71263011 | 0.99937574 |
| AC013472.2 | 73.7897044 | -0.676267 | 0.74196788 | -0.9114504 | 3.62E-01 | 0.99937574 |
| MAPRE3 | 103.611356 | 0.03690775 | 0.40201963 | 0.09180584 | 0.92685231 | 0.99937574 |
| TMEM214 | 22.810919 | -0.2476801 | 0.48791094 | -0.5076339 | 0.61171009 | 0.99937574 |
| AGBL5 | 24.3165802 | 0.2104486 | 0.44347562 | 0.47454378 | 0.63511218 | 0.99937574 |
| OST4 | 73.6289535 | 0.25531636 | 0.53513068 | 0.47711031 | 0.63328357 | 0.99937574 |
| ATRAID | 56.7474196 | 0.13895284 | 0.44385034 | 0.31306237 | 0.75423327 | 0.99937574 |
| MPV17 | 19.8618907 | 0.00153374 | 0.47929678 | 0.00319998 | 0.99744679 | 0.99937574 |
| GTF3C2 | 46.7694232 | 0.04902695 | 0.35805089 | 0.13692732 | 0.89108825 | 0.99937574 |
| EIF2B4 | 10.5843879 | -0.1356914 | 0.57935522 | -0.2342111 | 0.81482114 | 0.99937574 |
| SNX17 | 24.6885083 | 0.45271142 | 0.39833084 | 1.13652113 | 0.25573853 | 0.99937574 |
| ZNF513 | 16.4519162 | 0.4096468 | 0.47595631 | 0.86068152 | 0.38941347 | 0.99937574 |
| PPM1G | 92.6128714 | -0.0159133 | 0.41691724 | -0.038169 | 0.9695529 | 0.99937574 |
| NRBP1 | 33.6167576 | -0.1889657 | 0.40020645 | -0.4721706 | 0.63680503 | 0.99937574 |
| KRTCAP3 | 22.3278249 | 0.32259654 | 0.59356956 | 0.54348565 | 0.58679547 | 0.99937574 |
| IFT172 | 28.9089837 | 0.85707175 | 0.56517746 | 1.51646483 | 0.12940186 | 0.99937574 |
| ZNF512 | 55.0626993 | 0.06747229 | 0.49308688 | 0.13683652 | 0.89116001 | 0.99937574 |
| GPN1 | 12.2459956 | 0.3644471 | 0.60385866 | 0.60353047 | 0.54615585 | 0.99937574 |

| | | | | | | |
|---|---|---|---|---|---|---|
| SUPT7L | 23.5969219 | 0.05638844 | 0.40805354 | 0.13818881 | 0.89009119 | 0.99937574 |
| SLC4A1AP | 23.1756091 | 0.03777426 | 0.40330852 | 0.09366095 | 0.92537849 | 0.99937574 |
| MRPL33 | 84.5220593 | 0.0160942 | 0.43792203 | 0.0367513 | 0.9706833 | 0.99937574 |
| RBKS | 47.9655851 | 0.181649 | 0.45263423 | 0.40131521 | 0.68818807 | 0.99937574 |
| BABAM2 | 447.465842 | -0.0761922 | 0.41308156 | -0.1844483 | 0.8536618 | 0.99937574 |
| FOSL2 | 121.902912 | -0.3975654 | 0.44415359 | -0.895108 | 0.37072938 | 0.99937574 |
| PPP1CB | 84.8484955 | -0.0490633 | 0.31769437 | -0.1544355 | 0.87726634 | 0.99937574 |
| TRMT61B | 30.3903999 | -0.2756586 | 0.41162597 | -0.6696821 | 0.50306044 | 0.99937574 |
| WDR43 | 56.8914501 | -0.2182719 | 0.29160968 | -0.7485071 | 0.45415433 | 0.99937574 |
| CLIP4 | 150.077929 | 0.02158727 | 0.31644805 | 0.06821743 | 0.94561255 | 0.99937574 |
| YPEL5 | 72.8911126 | -0.1588707 | 0.33755521 | -0.4706509 | 0.63789004 | 0.99937574 |
| LBH | 17.7638597 | -1.8463737 | 0.84980242 | -2.1727094 | 0.0298022 | 0.99049468 |
| LCLAT1 | 106.482364 | -0.224629 | 0.48377579 | -0.4643246 | 0.64241524 | 0.99937574 |
| CAPN13 | 45.4153964 | 1.11223264 | 0.66292627 | 1.6777622 | 0.09339353 | 0.99937574 |
| MEMO1 | 19.5567699 | 0.47369609 | 0.49449967 | 0.95793004 | 0.33809804 | 0.99937574 |
| DPY30 | 44.0171964 | -0.2716526 | 0.44720837 | -0.6074409 | 0.54355838 | 0.99937574 |
| SPAST | 48.1664194 | -0.033443 | 0.34699893 | -0.0963778 | 0.92322054 | 0.99937574 |
| SLC30A6 | 61.1052892 | -0.0154555 | 0.25652034 | -0.0602504 | 0.95195621 | 0.99937574 |
| YIPF4 | 115.848894 | -0.3995083 | 0.26045314 | -1.533897 | 0.12505499 | 0.99937574 |
| BIRC6 | 546.157275 | 0.00630384 | 0.24019671 | 0.02624448 | 0.97906233 | 0.99937574 |
| TTC27 | 65.0987826 | -0.5491761 | 0.40803502 | -1.3459044 | 0.17833336 | 0.99937574 |
| LINC00486 | 97.4282287 | -0.1080297 | 0.29344027 | -0.3681488 | 0.71276231 | 0.99937574 |
| LTBP1 | 15.3701799 | 0.44616756 | 1.00364182 | 0.44454859 | 0.656646 | 0.99937574 |
| RASGRP3 | 25.4078689 | -1.867147 | 0.50236801 | -3.7166917 | 0.00020185 | 0.13580369 |
| FAM98A | 19.7265299 | -0.1545394 | 0.47734545 | -0.3237475 | 0.74612918 | 0.99937574 |
| LINC01317 | 13.6519823 | 0.60075131 | 1.02204437 | 0.58779376 | 0.55667073 | 0.99937574 |
| AC068599.1 | 72.1513171 | -0.375717 | 1.06279764 | -0.3535169 | 0.72370093 | 0.99937574 |
| LINC01320 | 129.09247 | -0.1000968 | 0.66114549 | -0.1513991 | 0.87966088 | 0.99937574 |
| AC012593.1 | 19.6761973 | 1.63141578 | 0.88031844 | 1.85321096 | 0.06385213 | 0.99937574 |
| CRIM1 | 317.649176 | -0.1379998 | 0.52598023 | -0.262367 | 0.79303854 | 0.99937574 |
| FEZ2 | 141.58688 | 0.39954582 | 0.30426364 | 1.31315663 | 0.18913018 | 0.99937574 |
| STRN | 222.950911 | -0.3744101 | 0.31281747 | -1.1968963 | 0.23134699 | 0.99937574 |
| HEATR5B | 70.6102611 | -0.0151243 | 0.28076903 | -0.0538676 | 0.95704068 | 0.99937574 |
| EIF2AK2 | 108.629646 | -0.9724686 | 0.63251114 | -1.5374726 | 0.12417761 | 0.99937574 |
| CEBPZOS | 51.1304937 | 0.3179403 | 0.45819049 | 0.69390419 | 0.4877423 | 0.99937574 |
| CEBPZ | 60.7312432 | 0.05711798 | 0.32320586 | 0.17672322 | 0.8597258 | 0.99937574 |
| NDUFAF7 | 39.8197228 | 0.01163825 | 0.356642 | 0.03263287 | 0.97396736 | 0.99937574 |
| PRKD3 | 73.9102123 | -0.1095582 | 0.51627406 | -0.2122094 | 0.8319437 | 0.99937574 |
| CDC42EP3 | 22.2515092 | -0.102927 | 0.50990674 | -0.2018546 | 0.84003036 | 0.99937574 |
| RMDN2 | 28.4540093 | 0.24249124 | 0.49321483 | 0.4916544 | 0.62296368 | 0.99937574 |
| ATL2 | 212.198437 | 0.34524659 | 0.2854519 | 1.2094738 | 0.22648087 | 0.99937574 |
| HNRNPLL | 54.7874518 | -0.2246943 | 0.35100765 | -0.6401408 | 0.52208107 | 0.99937574 |
| GALM | 35.8050135 | -0.0301832 | 0.52267948 | -0.0577471 | 0.95395005 | 0.99937574 |
| SRSF7 | 71.7739596 | -0.1920375 | 0.49119284 | -0.3909615 | 0.69582568 | 0.99937574 |
| DHX57 | 36.3926163 | -0.6303225 | 0.45697277 | -1.3793436 | 0.16778884 | 0.99937574 |
| SOS1 | 320.821645 | 0.00786815 | 0.26826726 | 0.02932953 | 0.97660178 | 0.99937574 |
| MAP4K3 | 731.793332 | 0.11458391 | 0.22441279 | 0.51059439 | 0.60963511 | 0.99937574 |
| AC007388.1 | 386.667995 | 0.39250091 | 0.30707384 | 1.27819715 | 0.20117992 | 0.99937574 |
| TMEM178A | 25.0992963 | 0.58572103 | 0.47886522 | 1.22314381 | 0.22127538 | 0.99937574 |
| THUMPD2 | 51.1742582 | -0.4034893 | 0.29641526 | -1.36123 | 0.17344101 | 0.99937574 |
| SLC8A1 | 83.1480985 | 1.63757471 | 0.67200666 | 2.43684297 | 0.01481611 | 0.85933448 |
| PKDCC | 12.5279806 | 0.34034346 | 0.59357682 | 0.57337729 | 0.56638927 | 0.99937574 |
| EML4 | 244.569056 | -0.3395702 | 0.35665999 | -0.9520839 | 0.34105446 | 0.99937574 |
| COX7A2L | 136.184341 | -0.2830496 | 0.45748326 | -0.6187104 | 0.53610716 | 0.99937574 |

| | | | | | | |
|---|---|---|---|---|---|---|
| KCNG3 | 31.6516161 | -0.7214124 | 0.93706946 | -0.7698601 | 0.44138288 | 0.99937574 |
| MTA3 | 118.388748 | -0.3991277 | 0.29636237 | -1.3467558 | 0.1780589 | 0.99937574 |
| ZFP36L2 | 96.5968665 | -0.6205841 | 0.52671693 | -1.1782117 | 0.23871221 | 0.99937574 |
| THADA | 158.411485 | -0.140287 | 0.34505107 | -0.4065689 | 0.68432465 | 0.99937574 |
| PLEKHH2 | 88.7315253 | -0.7196299 | 0.5115569 | -1.4067446 | 0.15950313 | 0.99937574 |
| DYNC2LI1 | 80.8237775 | -0.0909133 | 0.42702373 | -0.2129 | 0.83140498 | 0.99937574 |
| LRPPRC | 151.832215 | -0.1291987 | 0.24739098 | -0.5222448 | 0.6014999 | 0.99937574 |
| PPM1B | 195.221845 | 0.04960667 | 0.27495864 | 0.18041503 | 0.85682675 | 0.99937574 |
| SLC3A1 | 130.568029 | 0.82530825 | 0.3790781 | 2.17714569 | 0.0294697 | 0.99049468 |
| PREPL | 144.148648 | 0.13223647 | 0.37311608 | 0.35441109 | 0.72303083 | 0.99937574 |
| CAMKMT | 332.488747 | -0.1961253 | 0.23766002 | -0.8252345 | 0.40923844 | 0.99937574 |
| SRBD1 | 93.2802937 | -0.1088727 | 0.26682975 | -0.4080231 | 0.6832567 | 0.99937574 |
| PRKCE | 581.871993 | 0.11963346 | 0.21300401 | 0.56164885 | 0.57435529 | 0.99937574 |
| EPAS1 | 130.55453 | 0.33506172 | 0.29970907 | 1.11795654 | 0.26358556 | 0.99937574 |
| RHOQ | 95.6387782 | -0.7200258 | 0.36583978 | -1.9681452 | 0.04905133 | 0.99937574 |
| AC018682.1 | 16.9849341 | 0.4534496 | 0.88817617 | 0.51054015 | 0.60967309 | 0.99937574 |
| PIGF | 43.771876 | 0.17614287 | 0.38294012 | 0.45997496 | 0.64553419 | 0.99937574 |
| CRIPT | 14.0984543 | 0.29160545 | 0.54174625 | 0.53826943 | 0.59039106 | 0.99937574 |
| SOCS5 | 50.5013358 | 0.13631812 | 0.33313423 | 0.40919879 | 0.68239378 | 0.99937574 |
| MCFD2 | 67.9493256 | 0.4668961 | 0.39833991 | 1.17210476 | 0.241155 | 0.99937574 |
| TTC7A | 56.9558899 | 0.40064362 | 0.43804644 | 0.91461449 | 0.36039405 | 0.99937574 |
| AC106869.1 | 31.8586988 | 0.92961639 | 0.63969706 | 1.45321348 | 0.14616449 | 0.99937574 |
| EPCAM | 357.082781 | -0.0917062 | 0.4451374 | -0.2060178 | 0.83677702 | 0.99937574 |
| MSH2 | 42.8260025 | -0.0123162 | 0.3963595 | -0.0310733 | 0.97521108 | 0.99937574 |
| MSH6 | 23.5911814 | 0.40308853 | 0.5629061 | 0.71608485 | 0.47393896 | 0.99937574 |
| FBXO11 | 250.138029 | -0.1885348 | 0.18341792 | -1.0278972 | 0.30399818 | 0.99937574 |
| FOXN2 | 51.5625225 | -0.0504134 | 0.4368942 | -0.1153904 | 0.90813566 | 0.99937574 |
| PPP1R21 | 101.363221 | -0.3141026 | 0.41586146 | -0.7553059 | 0.45006546 | 0.99937574 |
| GPR75-ASB3 | 44.7084243 | 0.92311177 | 0.52161384 | 1.76972254 | 0.07677337 | 0.99937574 |
| ERLEC1 | 44.5896067 | 0.13475235 | 0.31492755 | 0.42788364 | 0.66873584 | 0.99937574 |
| PSME4 | 212.790032 | -0.249222 | 0.27611283 | -0.9026091 | 0.36673339 | 0.99937574 |
| ACYP2 | 206.677661 | -0.0283541 | 0.24626699 | -0.1151356 | 0.90833765 | 0.99937574 |
| SPTBN1 | 583.022791 | -0.4723199 | 0.36914945 | -1.2794817 | 0.20072749 | 0.99937574 |
| AC093110.1 | 24.2591934 | -0.3964541 | 0.72880158 | -0.5439808 | 0.5864547 | 0.99937574 |
| EML6 | 39.6791609 | -1.3392208 | 0.84581261 | -1.5833541 | 0.11334079 | 0.99937574 |
| RTN4 | 1013.86352 | 0.63584111 | 0.31787099 | 2.00031184 | 0.0454666 | 0.99937574 |
| CLHC1 | 36.7161566 | 0.06542628 | 0.46011535 | 0.14219538 | 0.88692568 | 0.99937574 |
| RPS27A | 331.313438 | 0.16963712 | 0.53638 | 0.31626294 | 0.75180294 | 0.99937574 |
| MTIF2 | 38.7332112 | -0.169681 | 0.40300183 | -0.4210427 | 0.67372391 | 0.99937574 |
| AC012358.3 | 30.0442409 | 0.07630967 | 0.51973799 | 0.14682335 | 0.88327146 | 0.99937574 |
| CCDC88A | 38.6839442 | 0.14315881 | 0.47169925 | 0.30349595 | 0.76151193 | 0.99937574 |
| CFAP36 | 35.0814148 | 0.50666943 | 0.36062731 | 1.4049669 | 0.16003112 | 0.99937574 |
| PPP4R3B | 146.162908 | -0.0944249 | 0.28348984 | -0.3330805 | 0.7390735 | 0.99937574 |
| PNPT1 | 66.2868284 | -1.4504275 | 0.71352869 | -2.0327529 | 0.04207749 | 0.99937574 |
| EFEMP1 | 103.928625 | 0.84063096 | 0.76229226 | 1.10276727 | 0.27012824 | 0.99937574 |
| VRK2 | 119.40192 | -0.1314196 | 0.34309954 | -0.3830365 | 0.70169272 | 0.99937574 |
| FANCL | 49.9125522 | -0.2605328 | 0.42478839 | -0.6133238 | 0.53966225 | 0.99937574 |
| PAPOLG | 31.8936341 | 0.18002973 | 0.37280412 | 0.48290703 | 0.62916175 | 0.99937574 |
| REL | 63.308748 | -0.1432237 | 0.51362723 | -0.2788476 | 0.78036181 | 0.99937574 |
| PEX13 | 37.29939 | -0.0791116 | 0.43010736 | -0.1839346 | 0.85406475 | 0.99937574 |
| KIAA1841 | 19.7088333 | -0.7688574 | 0.53047172 | -1.4493842 | 0.14723031 | 0.99937574 |
| C2orf74 | 14.0508103 | -0.6067259 | 0.77891702 | -0.7789352 | 0.4360179 | 0.99937574 |
| USP34 | 687.341247 | -0.2993454 | 0.25476462 | -1.1749883 | 0.2399994 | 0.99937574 |
| AC016727.1 | 15.9384518 | 0.02961642 | 0.47033267 | 0.06296908 | 0.94979112 | 0.99937574 |

| | | | | | | |
|---|---|---|---|---|---|---|
| XPO1 | 211.364048 | -0.4403211 | 0.31903358 | -1.3801718 | 0.16753376 | 0.99937574 |
| FAM161A | 13.414897 | -0.3420699 | 0.74468043 | -0.4593512 | 0.64598196 | 0.99937574 |
| CCT4 | 73.9164908 | 0.33317223 | 0.30211698 | 1.10279216 | 0.27011743 | 0.99937574 |
| COMMD1 | 218.265504 | -0.2657699 | 0.27243184 | -0.9755464 | 0.32928928 | 0.99937574 |
| AC018462.1 | 14.6181133 | -0.587651 | 0.73989499 | -0.7942356 | 0.42705828 | 0.99937574 |
| B3GNT2 | 128.823164 | -0.0901926 | 0.3854162 | -0.2340136 | 0.81497439 | 0.99937574 |
| EHBP1 | 124.374907 | -0.1608826 | 0.30931142 | -0.5201315 | 0.60297192 | 0.99937574 |
| WDPCP | 177.229487 | -0.0571942 | 0.29280783 | -0.19533 | 0.84513458 | 0.99937574 |
| MDH1 | 103.832701 | 0.30785536 | 0.41934141 | 0.73414012 | 0.46286334 | 0.99937574 |
| UGP2 | 131.221274 | 0.19842758 | 0.27334211 | 0.72593125 | 0.46788092 | 0.99937574 |
| VPS54 | 171.181873 | -0.1759919 | 0.33751045 | -0.5214413 | 0.60205936 | 0.99937574 |
| PELI1 | 41.1208232 | -1.3453133 | 0.73187165 | -1.838182 | 0.0660356 | 0.99937574 |
| AFTPH | 82.472937 | -0.23693 | 0.28140786 | -0.8419453 | 0.39981857 | 0.99937574 |
| SERTAD2 | 22.7733347 | 0.52914701 | 0.52952134 | 0.99929308 | 0.31765273 | 0.99937574 |
| CEP68 | 43.2099334 | 0.31727366 | 0.38069368 | 0.83340932 | 0.40461392 | 0.99937574 |
| RAB1A | 252.10034 | -0.0135481 | 0.23525644 | -0.0575886 | 0.95407634 | 0.99937574 |
| ACTR2 | 193.977229 | 0.04698154 | 0.26268032 | 0.17885443 | 0.85805201 | 0.99937574 |
| SPRED2 | 252.130353 | 0.11411856 | 0.34017067 | 0.33547442 | 0.73726724 | 0.99937574 |
| ETAA1 | 18.4207869 | -0.1972335 | 0.48256997 | -0.4087149 | 0.68274892 | 0.99937574 |
| C1D | 40.3684157 | -0.2630498 | 0.39640192 | -0.6635937 | 0.50695036 | 0.99937574 |
| PNO1 | 14.2141236 | -0.2235028 | 0.58192771 | -0.3840732 | 0.70092421 | 0.99937574 |
| APLF | 42.8773614 | 0.077814 | 0.42528663 | 0.18296836 | 0.85482285 | 0.99937574 |
| GFPT1 | 116.458961 | -0.0755101 | 0.29684532 | -0.2543752 | 0.79920573 | 0.99937574 |
| NFU1 | 49.6539599 | -0.2303789 | 0.33137614 | -0.6952188 | 0.48691817 | 0.99937574 |
| AAK1 | 354.026112 | -0.3054017 | 0.22917345 | -1.3326227 | 0.18265566 | 0.99937574 |
| ANXA4 | 520.6698 | 0.41394369 | 0.32160224 | 1.28712939 | 0.19804919 | 0.99937574 |
| GMCL1 | 64.3362781 | -0.2248029 | 0.31830996 | -0.7062388 | 0.48003963 | 0.99937574 |
| MXD1 | 31.8431758 | -0.3935667 | 0.39467902 | -0.9971817 | 0.3186763 | 0.99937574 |
| PCBP1 | 83.5234228 | -0.3095135 | 0.41813242 | -0.7402285 | 0.45916137 | 0.99937574 |
| C2orf42 | 20.1633236 | -0.3937435 | 0.51298345 | -0.767556 | 0.44275103 | 0.99937574 |
| PCYOX1 | 66.9811046 | 1.11141027 | 0.46616039 | 2.38417998 | 0.01711723 | 0.89506255 |
| SNRPG | 38.3724133 | -0.0474779 | 0.50108576 | -0.0947501 | 0.92451334 | 0.99937574 |
| FAM136A | 19.840325 | -0.0855195 | 0.45640921 | -0.1873747 | 0.85136687 | 0.99937574 |
| TGFA | 27.8910428 | -0.0713487 | 0.66291366 | -0.1076289 | 0.91429006 | 0.99937574 |
| ATP6V1B1 | 39.6545437 | 1.63763757 | 0.89961578 | 1.82037443 | 0.068702 | 0.99937574 |
| AC007040.2 | 119.997313 | 0.20931416 | 1.32810939 | 0.1576031 | 0.87476956 | 0.99937574 |
| NAGK | 24.3086559 | 0.03710529 | 0.64402241 | 0.0576149 | 0.95405538 | 0.99937574 |
| MCEE | 38.8211769 | 0.44061204 | 0.39627468 | 1.11188542 | 0.26618742 | 0.99937574 |
| MPHOSPH10 | 27.7886923 | -0.4306657 | 0.43693379 | -0.9856543 | 0.32430279 | 0.99937574 |
| ZNF638 | 446.046585 | 0.13819911 | 0.23760412 | 0.58163598 | 0.5608119 | 0.99937574 |
| EXOC6B | 399.625312 | -0.5406524 | 0.31562048 | -1.7129825 | 0.08671576 | 0.99937574 |
| SPR | 16.1413345 | 1.07625737 | 0.69691886 | 1.54430799 | 0.12251374 | 0.99937574 |
| EMX1 | 18.1276969 | -0.8631649 | 0.82884253 | -1.04141 | 0.29768529 | 0.99937574 |
| SFXN5 | 46.0761288 | -0.4021662 | 0.62095624 | -0.6476563 | 0.51720726 | 0.99937574 |
| RAB11FIP5 | 17.1793199 | -0.6899003 | 0.56146505 | -1.2287503 | 0.21916546 | 0.99937574 |
| SMYD5 | 12.4182901 | 0.09353188 | 0.59542162 | 0.15708513 | 0.87517776 | 0.99937574 |
| PRADC1 | 13.2877947 | 0.22663107 | 0.79503812 | 0.28505686 | 0.77560058 | 0.99937574 |
| CCT7 | 69.578205 | 0.36168128 | 0.38185207 | 0.94717644 | 0.34354888 | 0.99937574 |
| ALMS1 | 105.287499 | -0.2331022 | 0.33174247 | -0.7026602 | 0.48226755 | 0.99937574 |
| DUSP11 | 17.5913743 | 0.16084288 | 0.49491924 | 0.32498813 | 0.74519006 | 0.99937574 |
| STAMBP | 42.6869102 | -0.2593674 | 0.35295751 | -0.7348402 | 0.46243684 | 0.99937574 |
| DGUOK | 45.8395992 | -0.4383764 | 0.37347361 | -1.1737814 | 0.2404826 | 0.99937574 |
| TET3 | 51.5610588 | -0.4724361 | 0.48549705 | -0.9730978 | 0.33050468 | 0.99937574 |
| BOLA3 | 20.3333795 | 0.75402854 | 0.7144651 | 1.05537491 | 0.29125389 | 0.99937574 |

| | | | | | | |
|---|---|---|---|---|---|---|
| MOB1A | 115.46267 | -0.2479094 | 0.3307059 | -0.749637 | 0.45347338 | 0.99937574 |
| AC006030.1 | 31.297854 | 0.50138901 | 0.38831697 | 1.29118493 | 0.19663956 | 0.99937574 |
| DCTN1 | 18.8944269 | -0.2273894 | 0.58898698 | -0.3860686 | 0.69944585 | 0.99937574 |
| RTKN | 32.5664094 | -0.0015279 | 0.39179501 | -0.0038997 | 0.99688846 | 0.99937574 |
| MOGS | 26.8854269 | -0.058264 | 0.53004841 | -0.1099219 | 0.91247128 | 0.99937574 |
| AUP1 | 26.6398056 | -0.1541372 | 0.58419871 | -0.2638437 | 0.79190033 | 0.99937574 |
| M1AP | 26.0046744 | -1.1795283 | 0.85705025 | -1.376265 | 0.16873959 | 0.99937574 |
| POLE4 | 29.5756893 | -0.0799971 | 0.42872766 | -0.1865918 | 0.85198066 | 0.99937574 |
| TACR1 | 16.8273232 | 0.31379417 | 0.72459558 | 0.43306111 | 0.66497037 | 0.99937574 |
| AC007681.1 | 13.9107694 | 0.25041683 | 0.76686343 | 0.32654683 | 0.74401066 | 0.99937574 |
| MRPL19 | 68.7974648 | 0.57537776 | 0.39543092 | 1.45506516 | 0.14565122 | 0.99937574 |
| GCFC2 | 87.6233385 | 0.57295709 | 0.39038408 | 1.46767535 | 0.14219243 | 0.99937574 |
| SUCLG1 | 119.692204 | 0.27278258 | 0.27666434 | 0.98596942 | 0.32414812 | 0.99937574 |
| DNAH6 | 45.5528212 | 0.74090722 | 0.8778021 | 0.84404813 | 0.39864251 | 0.99937574 |
| TMSB10 | 223.133561 | -0.8041206 | 0.50448828 | -1.5939332 | 0.110951 | 0.99937574 |
| KCMF1 | 119.91063 | -0.2803711 | 0.30672018 | -0.9140941 | 0.3606674 | 0.99937574 |
| TCF7L1 | 159.530529 | 0.03072504 | 0.42076718 | 0.07302149 | 0.94178902 | 0.99937574 |
| TGOLN2 | 181.109194 | 0.41610221 | 0.37353874 | 1.11394661 | 0.26530209 | 0.99937574 |
| RETSAT | 30.1895995 | 0.05355056 | 0.4696578 | 0.11402039 | 0.90922163 | 0.99937574 |
| ELMOD3 | 47.9896043 | 0.23163855 | 0.37145276 | 0.62360165 | 0.53288923 | 0.99937574 |
| CAPG | 55.7774191 | -0.2069834 | 0.54739647 | -0.3781233 | 0.70533901 | 0.99937574 |
| MAT2A | 51.0953156 | -0.4205771 | 0.37980199 | -1.1073587 | 0.26813885 | 0.99937574 |
| GGCX | 21.8702965 | 0.53454261 | 0.42984838 | 1.24356083 | 0.21366124 | 0.99937574 |
| VAMP8 | 126.007269 | -0.1666489 | 0.49659807 | -0.3355811 | 0.7371868 | 0.99937574 |
| VAMP5 | 10.9938864 | -0.6781433 | 0.58322234 | -1.1627526 | 0.24492991 | 0.99937574 |
| RNF181 | 17.1885435 | -0.2084729 | 0.51769755 | -0.4026924 | 0.68717455 | 0.99937574 |
| USP39 | 50.8532771 | 0.14258139 | 0.32506281 | 0.43862718 | 0.6609317 | 0.99937574 |
| C2orf68 | 20.1328335 | -0.1686755 | 0.42459256 | -0.3972643 | 0.69117257 | 0.99937574 |
| ST3GAL5 | 11.3571549 | 0.86479633 | 0.73460545 | 1.17722559 | 0.23910548 | 0.99937574 |
| POLR1A | 41.4301001 | -0.3154354 | 0.37968749 | -0.8307765 | 0.40609991 | 0.99937574 |
| PTCD3 | 85.9167946 | 0.01772917 | 0.30613736 | 0.05791248 | 0.95381834 | 0.99937574 |
| IMMT | 167.99172 | 0.36456448 | 0.240443 | 1.51621996 | 0.12946374 | 0.99937574 |
| MRPL35 | 35.6826465 | 0.46515865 | 0.44933189 | 1.03522288 | 0.30056484 | 0.99937574 |
| REEP1 | 10.9544044 | 0.93209269 | 0.85925727 | 1.08476556 | 0.2780255 | 0.99937574 |
| KDM3A | 114.75045 | -0.2624527 | 0.42176115 | -0.6222781 | 0.53375901 | 0.99937574 |
| CHMP3 | 45.7702292 | -0.0038871 | 0.39413439 | -0.0098624 | 0.99213108 | 0.99937574 |
| RNF103-CHN | 19.2477838 | -3.1471753 | 1.22107617 | -2.5773783 | 0.00995529 | 0.76274616 |
| RMND5A | 65.269176 | -0.391683 | 0.37827405 | -1.0354478 | 0.30045982 | 0.99937574 |
| CYTOR | 13.6909659 | -0.4866871 | 0.78724186 | -0.618218 | 0.53643164 | 0.99937574 |
| KRCC1 | 63.229798 | 0.38305392 | 0.29636327 | 1.29251484 | 0.19617891 | 0.99937574 |
| THNSL2 | 19.4720374 | -0.1198426 | 0.86454613 | -0.138619 | 0.8897512 | 0.99937574 |
| EIF2AK3 | 101.461049 | 0.12065785 | 0.43380901 | 0.27813588 | 0.78090806 | 0.99937574 |
| RPIA | 26.2623233 | 0.21311662 | 0.51920555 | 0.41046675 | 0.68146359 | 0.99937574 |
| MAL | 370.049254 | 0.64519962 | 0.42542293 | 1.51660754 | 0.1293658 | 0.99937574 |
| MRPS5 | 153.56079 | 0.33774599 | 0.25524439 | 1.32322588 | 0.18576027 | 0.99937574 |
| ZNF514 | 10.1907726 | 0.14068717 | 0.55293513 | 0.25443703 | 0.79915795 | 0.99937574 |
| AC092835.1 | 19.4364772 | 0.86133576 | 0.56587034 | 1.52214333 | 0.12797317 | 0.99937574 |
| PROM2 | 20.7748113 | 0.92601637 | 0.6356866 | 1.45671841 | 0.14519413 | 0.99937574 |
| KCNIP3 | 24.2131724 | 0.42464833 | 0.69335578 | 0.61245373 | 0.5402376 | 0.99937574 |
| FAHD2A | 19.3877345 | 0.46746858 | 0.4871976 | 0.95950509 | 0.33730436 | 0.99937574 |
| ANKRD36C | 30.9258677 | -0.3854001 | 0.45624914 | -0.8447142 | 0.39827045 | 0.99937574 |
| STARD7 | 107.544926 | 0.07039104 | 0.24692303 | 0.28507282 | 0.77558836 | 0.99937574 |
| TMEM127 | 51.9279137 | -0.3615875 | 0.45841683 | -0.7887745 | 0.43024382 | 0.99937574 |
| CIAO1 | 40.9616241 | -0.0754135 | 0.34732873 | -0.2171242 | 0.82811156 | 0.99937574 |

| | | | | | | |
|---|---|---|---|---|---|---|
| SNRNP200 | 65.214764 | -0.0517153 | 0.36512481 | -0.1416373 | 0.88736654 | 0.99937574 |
| KANSL3 | 102.014666 | -0.3269482 | 0.23114382 | -1.4144794 | 0.15722119 | 0.99937574 |
| LMAN2L | 24.8495056 | 0.25528111 | 0.42322266 | 0.60318393 | 0.54638634 | 0.99937574 |
| CNNM4 | 31.3657037 | -1.4532548 | 0.65031 | -2.2347108 | 0.02543634 | 0.96546668 |
| ANKRD23 | 14.1131075 | -0.186824 | 0.56005779 | -0.3335798 | 0.73869669 | 0.99937574 |
| ANKRD36 | 35.4635883 | -0.1724448 | 0.48258811 | -0.3573333 | 0.72084234 | 0.99937574 |
| AC092683.1 | 14.2980811 | -0.2449473 | 0.57092986 | -0.4290322 | 0.66789981 | 0.99937574 |
| ANKRD36B | 15.6875725 | -0.1650364 | 0.67822496 | -0.2433359 | 0.80774523 | 0.99937574 |
| COX5B | 149.307523 | 0.36892525 | 0.58047199 | 0.63556081 | 0.52506272 | 0.99937574 |
| ACTR1B | 22.3900845 | 0.05795839 | 0.44067632 | 0.13152144 | 0.89536282 | 0.99937574 |
| TMEM131 | 186.494379 | 4.89E-05 | 0.23635078 | 0.00020685 | 0.99983495 | 0.99993404 |
| INPP4A | 386.803831 | 0.09391191 | 0.3754647 | 0.25012181 | 0.80249315 | 0.99937574 |
| COA5 | 23.9477843 | 0.06310184 | 0.5612498 | 0.11243094 | 0.91048173 | 0.99937574 |
| UNC50 | 44.5019645 | 0.39066558 | 0.38631214 | 1.01126921 | 0.3118876 | 0.99937574 |
| MGAT4A | 63.3717854 | 0.55681422 | 0.34517725 | 1.61312547 | 0.10671725 | 0.99937574 |
| KIAA1211L | 19.2728912 | -0.5606158 | 0.83826247 | -0.6687832 | 0.5036338 | 0.99937574 |
| TSGA10 | 102.797837 | -0.010192 | 0.37215338 | -0.0273866 | 0.97815141 | 0.99937574 |
| AC079447.1 | 33.5127712 | 0.56493475 | 0.50664836 | 1.11504308 | 0.26483196 | 0.99937574 |
| MITD1 | 35.0371669 | -0.2118051 | 0.32319258 | -0.6553526 | 0.51224078 | 0.99937574 |
| TXNDC9 | 29.0816411 | 0.18881041 | 0.45283287 | 0.41695386 | 0.67671216 | 0.99937574 |
| EIF5B | 145.694034 | -0.2032687 | 0.33676479 | -0.6035925 | 0.5461146 | 0.99937574 |
| REV1 | 245.731699 | 0.12972326 | 0.21970457 | 0.59044404 | 0.55489299 | 0.99937574 |
| LONRF2 | 160.164085 | -0.3089182 | 0.53904803 | -0.5730809 | 0.56658989 | 0.99937574 |
| PDCL3 | 18.4789129 | -0.9403752 | 0.51919139 | -1.8112304 | 0.07010519 | 0.99937574 |
| RPL31 | 89.9858127 | 0.38674022 | 0.34166547 | 1.13192658 | 0.2576653 | 0.99937574 |
| TBC1D8 | 66.8048465 | -0.675046 | 0.41499166 | -1.6266494 | 0.10381157 | 0.99937574 |
| CNOT11 | 38.7511928 | -0.4391952 | 0.32948701 | -1.3329666 | 0.18254277 | 0.99937574 |
| RNF149 | 179.367072 | -0.8076815 | 0.24435803 | -3.3053202 | 0.00094868 | 0.29012345 |
| MAP4K4 | 155.744728 | -0.0193907 | 0.44048707 | -0.0440211 | 0.96488757 | 0.99937574 |
| IL1R1 | 113.613473 | -0.4214594 | 0.46118618 | -0.9138596 | 0.36079064 | 0.99937574 |
| IL1RL2 | 37.7311097 | -0.5581728 | 0.52698985 | -1.0591718 | 0.28952153 | 0.99937574 |
| SLC9A4 | 19.0539636 | 1.01922729 | 0.73760438 | 1.38180753 | 0.16703081 | 0.99937574 |
| SLC9A2 | 62.017599 | 0.81006597 | 0.58952274 | 1.3741047 | 0.16940918 | 0.99937574 |
| MFSD9 | 13.8375037 | 0.61003153 | 0.48787989 | 1.25037237 | 0.21116355 | 0.99937574 |
| TMEM182 | 21.2910021 | -0.3903736 | 0.44836011 | -0.8706697 | 0.38393451 | 0.99937574 |
| POU3F3 | 54.7897237 | -0.098068 | 0.42602276 | -0.2301942 | 0.81794086 | 0.99937574 |
| MRPS9 | 53.0238032 | 0.15882815 | 0.37947971 | 0.41854188 | 0.67555097 | 0.99937574 |
| TGFBRAP1 | 26.8820081 | -0.459973 | 0.46595593 | -0.9871599 | 0.32356423 | 0.99937574 |
| C2orf49 | 23.9303294 | 0.06386285 | 0.39060864 | 0.16349574 | 0.87012813 | 0.99937574 |
| NCK2 | 58.057098 | 0.29915777 | 0.38349243 | 0.78008782 | 0.43533918 | 0.99937574 |
| UXS1 | 47.3715915 | -0.0877475 | 0.38466959 | -0.2281113 | 0.81955973 | 0.99937574 |
| LINC01594 | 14.1986754 | 0.23745845 | 0.83156028 | 0.28555771 | 0.77521689 | 0.99937574 |
| SULT1C2 | 262.146355 | -0.0524086 | 0.43285341 | -0.1210769 | 0.90363009 | 0.99937574 |
| GCC2 | 240.724995 | -0.2292363 | 0.28449045 | -0.8057784 | 0.42037065 | 0.99937574 |
| GCC2-AS1 | 44.9993136 | 0.66422303 | 0.40194415 | 1.65252567 | 0.09842744 | 0.99937574 |
| LIMS1 | 1241.07222 | -0.7012566 | 0.27667853 | -2.5345538 | 1.13E-02 | 7.63E-01 |
| RANBP2 | 208.842814 | 0.12446449 | 0.21678288 | 0.57414355 | 0.56587066 | 0.99937574 |
| CCDC138 | 35.8481036 | 0.09929884 | 0.38882603 | 0.25538115 | 0.79842873 | 0.99937574 |
| 10-Sep | 193.781005 | 0.3059574 | 0.38209844 | 0.80072925 | 4.23E-01 | 0.99937574 |
| SOWAHC | 37.4375054 | 0.51138619 | 0.41949385 | 1.21905528 | 2.23E-01 | 0.99937574 |
| AC013271.1 | 99.7327844 | -0.9762548 | 0.73422731 | -1.3296357 | 0.18363834 | 0.99937574 |
| NPHP1 | 22.2469795 | 0.44399545 | 0.56504899 | 0.78576452 | 0.43200546 | 0.99937574 |
| SMIM37 | 87.1822752 | 0.28412688 | 0.66451093 | 0.42757293 | 0.66896208 | 0.99937574 |
| LINC01106 | 11.8683803 | 0.32658077 | 0.64055615 | 0.50983941 | 0.61016397 | 0.99937574 |

| | | | | | | |
|---|---|---|---|---|---|---|
| AC112229.3 | 203.998796 | 0.18767366 | 0.67284615 | 0.27892506 | 0.78030233 | 0.99937574 |
| MIR4435-2H | 83.4850519 | 0.02571653 | 0.38717047 | 0.06642172 | 0.94704208 | 0.99937574 |
| BCL2L11 | 35.6645343 | -0.2088406 | 0.43885363 | -0.4758775 | 0.63416166 | 0.99937574 |
| ANAPC1 | 69.2295536 | -0.2193519 | 0.27017796 | -0.8118791 | 0.41686099 | 0.99937574 |
| MERTK | 62.9244854 | 0.03288557 | 0.44919323 | 0.0732103 | 0.94163877 | 0.99937574 |
| TMEM87B | 85.8172898 | -0.4938351 | 0.29748452 | -1.6600363 | 0.09690714 | 0.99937574 |
| ZC3H8 | 38.8302159 | -0.2515588 | 0.34306551 | -0.7332675 | 0.46339529 | 0.99937574 |
| ZC3H6 | 79.8526608 | 0.18799446 | 0.41878212 | 0.44890755 | 0.65349835 | 0.99937574 |
| RGPD8 | 11.0009153 | -1.0151687 | 0.62086124 | -1.6350976 | 0.10202858 | 0.99937574 |
| TTL | 45.7874172 | -0.0467197 | 0.42155549 | -0.1108269 | 0.91175361 | 0.99937574 |
| POLR1B | 11.1787049 | -0.7502532 | 0.61077895 | -1.2283548 | 0.21931382 | 0.99937574 |
| CHCHD5 | 21.7908913 | 0.00497627 | 0.52778972 | 0.0094285 | 0.99247726 | 0.99937574 |
| SLC20A1 | 90.4865593 | -0.6755462 | 0.51368707 | -1.3150929 | 0.18847869 | 0.99937574 |
| PSD4 | 41.4823585 | -0.278232 | 0.4211388 | -0.6606658 | 0.50882669 | 0.99937574 |
| PAX8 | 1677.22527 | 0.2027296 | 0.42129943 | 0.48120074 | 0.63037384 | 0.99937574 |
| CBWD2 | 31.7909251 | -0.6732613 | 0.45255288 | -1.4876965 | 0.13683095 | 0.99937574 |
| SLC35F5 | 252.216112 | -0.2895111 | 0.27006028 | -1.0720241 | 0.2837092 | 0.99937574 |
| ACTR3 | 233.079915 | -0.3065122 | 0.29178702 | -1.0504654 | 0.2935042 | 0.99937574 |
| DPP10 | 51.5067365 | 1.60203976 | 0.6081994 | 2.63406995 | 0.00843681 | 0.74742128 |
| DDX18 | 53.7102272 | -0.0260662 | 0.32762781 | -0.0795604 | 0.93658692 | 0.99937574 |
| CCDC93 | 114.703937 | -0.2697624 | 0.32971534 | -0.8181677 | 0.41326146 | 0.99937574 |
| INSIG2 | 90.5059254 | -0.1395753 | 0.38767231 | -0.3600343 | 0.71882149 | 0.99937574 |
| C2orf76 | 19.3295433 | -0.1718266 | 0.5405068 | -0.3178991 | 0.75056146 | 0.99937574 |
| DBI | 191.83145 | 0.13987119 | 0.50270956 | 0.27823459 | 0.78083229 | 0.99937574 |
| TMEM37 | 19.3108456 | 0.09463103 | 0.67919615 | 0.13932799 | 0.88919097 | 0.99937574 |
| SCTR | 66.4079207 | -0.7165896 | 1.305614 | -0.5488525 | 0.58310665 | 0.99937574 |
| CFAP221 | 91.6702039 | 0.18176257 | 0.79903836 | 0.22747664 | 0.82005313 | 0.99937574 |
| PTPN4 | 168.142293 | 0.37709112 | 0.30792329 | 1.22462682 | 0.22071586 | 0.99937574 |
| EPB41L5 | 336.11128 | 0.09171061 | 0.23910321 | 0.38356076 | 0.70130403 | 0.99937574 |
| RALB | 31.7190994 | -0.1849723 | 0.38875082 | -0.4758121 | 0.63420826 | 0.99937574 |
| TFCP2L1 | 1106.23603 | 0.44943473 | 0.47805249 | 0.94013679 | 0.3471474 | 0.99937574 |
| CLASP1 | 298.879291 | 0.2108702 | 0.33596925 | 0.62764731 | 0.53023501 | 0.99937574 |
| NIFK-AS1 | 22.2245636 | 0.37618576 | 0.55097843 | 0.6827595 | 0.49475882 | 0.99937574 |
| NIFK | 16.3627258 | -0.4638542 | 0.43935916 | -1.0557516 | 0.2910817 | 0.99937574 |
| TSN | 24.5101754 | -0.7470333 | 0.44264507 | -1.6876575 | 0.09147699 | 0.99937574 |
| CNTNAP5 | 283.033813 | -2.2108378 | 0.87425941 | -2.5288122 | 0.01144493 | 0.76274616 |
| BIN1 | 18.0032348 | 0.9912041 | 0.60212439 | 1.6461783 | 0.09972706 | 0.99937574 |
| ERCC3 | 35.0656862 | -0.002402 | 0.47862627 | -0.0050186 | 0.99599574 | 0.99937574 |
| MAP3K2 | 188.459629 | -0.3031168 | 0.24589491 | -1.2327087 | 0.21768447 | 0.99937574 |
| IWS1 | 118.907588 | 0.25387654 | 0.23505746 | 1.08006163 | 0.28011474 | 0.99937574 |
| AC010976.1 | 12.083965 | -0.8623014 | 0.8401658 | -1.0263467 | 0.3047282 | 0.99937574 |
| WDR33 | 170.668419 | -0.0411529 | 0.25493331 | -0.1614263 | 0.87175769 | 0.99937574 |
| POLR2D | 12.2766819 | -0.8148018 | 0.57265458 | -1.4228504 | 0.15477951 | 0.99937574 |
| AMMECR1L | 22.0646257 | -0.4162671 | 0.42928571 | -0.9696739 | 0.33220907 | 0.99937574 |
| SAP130 | 42.2956771 | -0.1479305 | 0.35171032 | -0.4206031 | 0.67404493 | 0.99937574 |
| UGGT1 | 65.6589591 | -0.0397336 | 0.30369562 | -0.1308335 | 0.895907 | 0.99937574 |
| HS6ST1 | 104.917502 | 0.24750324 | 0.44878022 | 0.55150212 | 0.58128952 | 0.99937574 |
| SMPD4 | 35.4713797 | 0.09018155 | 0.44708556 | 0.20170982 | 0.84014359 | 0.99937574 |
| MZT2B | 49.9011223 | 0.26244484 | 0.35753146 | 0.73404683 | 0.4629202 | 0.99937574 |
| CCDC115 | 17.1216425 | 1.06820813 | 0.45030059 | 2.37221124 | 0.01768198 | 0.91044145 |
| IMP4 | 18.1358263 | 0.33467568 | 0.54735136 | 0.61144579 | 0.54090449 | 0.99937574 |
| PTPN18 | 44.1196307 | 0.45151598 | 0.37898732 | 1.1913749 | 0.23350644 | 0.99937574 |
| FAM168B | 131.911395 | -0.3123858 | 0.30771484 | -1.0151796 | 0.31002024 | 0.99937574 |
| PLEKHB2 | 272.915857 | -0.117521 | 0.42936793 | -0.2737069 | 0.78430986 | 0.99937574 |

| | | | | | | |
|---|---|---|---|---|---|---|
| MZT2A | 24.3644612 | -0.1462866 | 0.48047618 | -0.3044616 | 0.76077621 | 0.99937574 |
| GPR39 | 23.7256362 | -0.5954425 | 0.70722279 | -0.8419448 | 0.39981887 | 0.99937574 |
| NCKAP5 | 164.335649 | -0.7100486 | 0.91396943 | -0.7768844 | 0.43722698 | 0.99937574 |
| MGAT5 | 152.568939 | 0.66424133 | 0.48061535 | 1.38206431 | 0.16695195 | 0.99937574 |
| TMEM163 | 86.7810103 | -1.2769991 | 0.65287774 | -1.9559544 | 0.05047053 | 0.99937574 |
| CCNT2-AS1 | 18.5696794 | -0.0804429 | 0.70749517 | -0.113701 | 0.90947482 | 0.99937574 |
| CCNT2 | 201.278907 | 0.27110538 | 0.42658214 | 0.63552914 | 0.52508336 | 0.99937574 |
| RAB3GAP1 | 226.171484 | -0.0819978 | 0.22903729 | -0.3580106 | 0.72033538 | 0.99937574 |
| ZRANB3 | 89.9084192 | 0.04126647 | 0.31159612 | 0.13243578 | 0.89463962 | 0.99937574 |
| R3HDM1 | 138.893497 | 0.52719167 | 0.23988548 | 2.19768067 | 0.02797187 | 0.99049468 |
| UBXN4 | 186.32567 | -0.1988475 | 0.26823691 | -0.741313 | 0.45850371 | 0.99937574 |
| MCM6 | 11.1237697 | -0.052273 | 0.62517468 | -0.0836134 | 0.93336378 | 0.99937574 |
| DARS | 173.668406 | -0.3219441 | 0.40647682 | -0.7920356 | 0.42833991 | 0.99937574 |
| THSD7B | 164.748557 | 1.24701704 | 0.83041652 | 1.50167659 | 0.13318065 | 0.99937574 |
| HNMT | 53.7417534 | 0.68185881 | 0.36271897 | 1.87985428 | 0.06012794 | 0.99937574 |
| AC114763.1 | 45.3004749 | -0.04813 | 0.61708702 | -0.0779955 | 0.93783166 | 0.99937574 |
| SPOPL | 170.630092 | 0.31158531 | 0.44333369 | 0.70282344 | 0.48216579 | 0.99937574 |
| NXPH2 | 417.983523 | 0.33087485 | 0.70859339 | 0.46694599 | 0.64053852 | 0.99937574 |
| LRP1B | 26.4502921 | -0.4521998 | 0.83379171 | -0.5423415 | 0.58758329 | 0.99937574 |
| KYNU | 46.9593979 | 0.25255113 | 0.61766857 | 0.40887806 | 0.68262915 | 0.99937574 |
| GTDC1 | 69.953453 | -0.0548739 | 0.35047134 | -0.1565718 | 0.8755823 | 0.99937574 |
| ACVR2A | 62.4075362 | 0.26789442 | 0.34317869 | 0.78062662 | 0.43502213 | 0.99937574 |
| ORC4 | 136.88463 | 0.19367112 | 0.32527285 | 0.59541129 | 0.55156858 | 0.99937574 |
| MBD5 | 668.083498 | 0.36206742 | 0.2564987 | 1.41157605 | 0.15807483 | 0.99937574 |
| EPC2 | 186.576828 | -0.0961795 | 0.29031717 | -0.3312911 | 0.74042463 | 0.99937574 |
| AC105402.3 | 18.5305982 | -0.7564903 | 1.64634619 | -0.4594965 | 0.64587768 | 0.99937574 |
| LYPD6B | 30.0530844 | 1.09650059 | 0.74380941 | 1.47416875 | 1.40E-01 | 0.99937574 |
| MMADHC | 34.0551997 | 0.14431278 | 0.38477347 | 0.37505907 | 0.70761653 | 0.99937574 |
| NMI | 20.16201 | -1.4907092 | 0.70300222 | -2.1204901 | 0.03396474 | 0.99937574 |
| RIF1 | 113.580851 | -0.1530607 | 0.30374125 | -0.5039179 | 0.61431906 | 0.99937574 |
| ARL5A | 35.0443114 | 0.305625 | 0.34661141 | 0.88175112 | 0.37791141 | 0.99937574 |
| STAM2 | 91.2004481 | -0.0118673 | 0.24039911 | -0.0493651 | 0.96062831 | 0.99937574 |
| FMNL2 | 499.763142 | -0.5814673 | 0.25069111 | -2.3194573 | 0.02037025 | 0.96427895 |
| PRPF40A | 113.982011 | 0.00415704 | 0.23968137 | 0.01734403 | 0.98616216 | 0.99937574 |
| ARL6IP6 | 43.9404076 | -0.1524119 | 0.4350817 | -0.3503065 | 0.7261087 | 0.99937574 |
| KCNJ3 | 309.104635 | 0.18285246 | 0.65547586 | 0.27896139 | 0.78027445 | 0.99937574 |
| LINC01876 | 110.194188 | -1.076288 | 0.81527735 | -1.3201495 | 0.18678509 | 0.99937574 |
| NR4A2 | 25.7750381 | -0.5618303 | 0.63970808 | -0.8782604 | 0.37980244 | 0.99937574 |
| GPD2 | 234.324604 | 0.34161663 | 0.39920251 | 0.85574769 | 0.39213736 | 0.99937574 |
| CYTIP | 44.7217408 | -0.109858 | 0.82766103 | -0.132733 | 0.89440452 | 0.99937574 |
| ACVR1C | 17.1799065 | 0.48810729 | 0.90964269 | 0.53659233 | 0.59154924 | 0.99937574 |
| ACVR1 | 96.2119087 | -0.0830664 | 0.41975938 | -0.1978906 | 0.84313064 | 0.99937574 |
| UPP2 | 14.2458129 | 0.58732593 | 0.74200455 | 0.79153952 | 0.42862922 | 0.99937574 |
| CCDC148 | 42.5885931 | -0.6279373 | 0.58892439 | -1.0662444 | 0.28631319 | 0.99937574 |
| PKP4 | 274.485131 | -0.4647792 | 0.31773214 | -1.4628019 | 0.14352159 | 0.99937574 |
| DAPL1 | 31.6857664 | 1.35106431 | 0.94639232 | 1.42759434 | 1.53E-01 | 0.99937574 |
| TANC1 | 200.46546 | 0.02844423 | 0.46982746 | 0.06054187 | 0.95172407 | 0.99937574 |
| WDSUB1 | 90.8373955 | -0.0964629 | 0.30258279 | -0.3187983 | 0.7498795 | 0.99937574 |
| BAZ2B | 443.073797 | 0.17873051 | 0.4010098 | 0.4457011 | 0.65581317 | 0.99937574 |
| AC009961.1 | 16.2840558 | 0.42110492 | 0.55403356 | 0.76007115 | 0.44721206 | 0.99937574 |
| 7-Mar | 189.539779 | -0.106107 | 0.2703889 | -0.3924238 | 0.69474514 | 0.99937574 |
| PLA2R1 | 12.3720803 | -0.4988423 | 0.79605495 | -0.626643 | 0.53089325 | 0.99937574 |
| ITGB6 | 82.4212593 | -0.6261601 | 0.54600952 | -1.1467933 | 0.25146705 | 0.99937574 |
| RBMS1 | 164.97372 | 0.10070537 | 0.43498096 | 0.23151672 | 0.8169134 | 0.99937574 |

| | | | | | | |
|---|---|---|---|---|---|---|
| TANK | 129.769729 | -0.2490515 | 0.30500421 | -0.816551 | 0.41418509 | 0.99937574 |
| PSMD14 | 77.737245 | -0.0865818 | 0.31402883 | -0.2757129 | 0.78276855 | 0.99937574 |
| IFIH1 | 30.6343987 | -0.9988339 | 0.68390775 | -1.4604805 | 0.14415806 | 0.99937574 |
| GCA | 37.9245247 | -0.0023591 | 0.39648981 | -0.00595 | 0.99525264 | 0.99937574 |
| FIGN | 34.5641714 | 1.14261751 | 0.74194307 | 1.54003394 | 0.12355208 | 0.99937574 |
| GRB14 | 670.937814 | 0.58859216 | 0.34291798 | 1.71642255 | 0.08608473 | 0.99937574 |
| COBLL1 | 1965.30172 | -0.4236223 | 0.28171335 | -1.5037353 | 0.13264954 | 0.99937574 |
| AC019197.1 | 2807.52072 | 1.00278546 | 0.39214773 | 2.55716249 | 0.01055299 | 0.76274616 |
| SLC38A11 | 96.163377 | -1.1917381 | 0.76556442 | -1.5566791 | 0.1195467 | 0.99937574 |
| SCN3A | 127.362681 | 0.60645032 | 0.85648946 | 0.70806513 | 0.47890481 | 0.99937574 |
| SCN2A | 411.850455 | 1.52518314 | 0.51603297 | 2.95559241 | 0.00312069 | 0.52695149 |
| CSRNP3 | 54.1352494 | 0.60394122 | 0.56928502 | 1.06087672 | 0.28874593 | 0.99937574 |
| GALNT3 | 71.0545276 | -0.2041353 | 0.4538303 | -0.4498053 | 0.65285085 | 0.99937574 |
| TTC21B | 77.1499768 | 0.28395382 | 0.3602187 | 0.78828172 | 0.43053194 | 0.99937574 |
| SCN9A | 26.0751825 | -0.4682533 | 0.73290131 | -0.6389036 | 0.52288563 | 0.99937574 |
| STK39 | 496.727536 | 0.65439699 | 0.53696127 | 1.21870427 | 0.22295646 | 0.99937574 |
| CERS6 | 87.2107676 | 0.32867741 | 0.45755776 | 0.71832986 | 0.47255392 | 0.99937574 |
| LRP2 | 25.6994891 | 0.81802794 | 0.96063738 | 0.85154707 | 0.39446553 | 0.99937574 |
| FASTKD1 | 99.7147129 | 0.12217662 | 0.37704135 | 0.32404037 | 0.74590747 | 0.99937574 |
| PPIG | 142.590592 | -0.2935555 | 0.22759406 | -1.2898205 | 0.19711298 | 0.99937574 |
| KLHL23 | 39.4901197 | 0.26045906 | 0.35376643 | 0.73624583 | 0.4615811 | 0.99937574 |
| SSB | 50.9712561 | -0.0251117 | 0.31065285 | -0.0808352 | 0.935573 | 0.99937574 |
| METTL5 | 30.0834992 | -0.3218606 | 0.48373178 | -0.66537 | 0.50581385 | 0.99937574 |
| UBR3 | 558.592638 | 0.08658383 | 0.27608434 | 0.31361368 | 0.75381447 | 0.99937574 |
| MYO3B | 191.924398 | 0.52867619 | 0.62051817 | 0.85199148 | 0.39421882 | 0.99937574 |
| AC007405.1 | 13.6668701 | 0.48846672 | 0.74697209 | 0.65392902 | 0.51315754 | 0.99937574 |
| GAD1 | 58.6036333 | 0.36991384 | 0.9060409 | 0.40827499 | 0.6830718 | 0.99937574 |
| GORASP2 | 67.0587797 | -0.2826264 | 0.39103204 | -0.7227705 | 0.46982089 | 0.99937574 |
| TLK1 | 261.340787 | -0.2813318 | 0.19237307 | -1.4624282 | 0.1436239 | 0.99937574 |
| METTL8 | 21.1701619 | -0.7676968 | 0.5056288 | -1.5183013 | 0.12893847 | 0.99937574 |
| DCAF17 | 25.9721787 | 0.02034801 | 0.4156106 | 0.04895931 | 0.96095172 | 0.99937574 |
| DYNC1I2 | 133.413109 | -0.1687492 | 0.25269002 | -0.6678112 | 0.50425412 | 0.99937574 |
| SLC25A12 | 153.244509 | 0.04011925 | 0.42299779 | 0.09484505 | 0.9244379 | 0.99937574 |
| HAT1 | 38.2152713 | -0.2258837 | 0.34593827 | -0.6529595 | 0.51378236 | 0.99937574 |
| METAP1D | 18.751087 | 0.24238765 | 0.45136687 | 0.53700805 | 0.59126205 | 0.99937574 |
| ITGA6 | 1079.88322 | 0.61710606 | 0.36612667 | 1.68549879 | 0.09189238 | 0.99937574 |
| AC078883.1 | 43.6160276 | 0.23400486 | 0.6958867 | 0.33626861 | 0.73666832 | 0.99937574 |
| PDK1 | 74.4575379 | -0.515532 | 0.52623169 | -0.9796674 | 0.32725033 | 0.99937574 |
| MAP3K20 | 61.4945602 | 0.46457507 | 0.63853782 | 0.72756077 | 0.4668825 | 0.99937574 |
| SP3 | 95.6902541 | -0.0468583 | 0.26342906 | -0.1778782 | 0.85881864 | 0.99937574 |
| OLA1 | 151.973563 | -0.0479252 | 0.26205152 | -0.1828847 | 0.85488852 | 0.99937574 |
| CIR1 | 54.290893 | 0.1381002 | 0.2781535 | 0.49648915 | 0.61954934 | 0.99937574 |
| SCRN3 | 22.8770764 | 0.34219524 | 0.49614572 | 0.68970713 | 0.49037838 | 0.99937574 |
| GPR155 | 41.6789737 | 0.81857235 | 0.70837819 | 1.15555838 | 0.24786184 | 0.99937574 |
| AC010894.3 | 158.954819 | 0.32409497 | 0.44961362 | 0.72082997 | 0.47101414 | 0.99937574 |
| WIPF1 | 142.794301 | -0.1357431 | 0.52895143 | -0.2566267 | 0.79746697 | 0.99937574 |
| CHN1 | 12.9896525 | -0.9046641 | 0.73170633 | -1.2363759 | 0.21631888 | 0.99937574 |
| ATF2 | 122.673944 | -0.0661433 | 0.30573786 | -0.2163398 | 0.82872292 | 0.99937574 |
| ATP5MC3 | 203.418308 | 0.75566169 | 0.53659701 | 1.40824804 | 0.15905763 | 0.99937574 |
| LNPK | 75.4412208 | 0.28140298 | 0.34583034 | 0.81370241 | 0.41581545 | 0.99937574 |
| HOXD10 | 14.3541556 | 0.58668433 | 0.92969944 | 0.63104732 | 0.52800959 | 0.99937574 |
| HOXD-AS2 | 21.2314149 | 0.64031506 | 0.53516071 | 1.19649118 | 0.23150494 | 0.99937574 |
| HOXD8 | 38.9239969 | 0.9038001 | 0.41914008 | 2.15631991 | 0.03105869 | 0.99049468 |
| HOXD3 | 49.9535729 | 0.61139025 | 0.5057959 | 1.20876869 | 0.22675172 | 0.99937574 |

| | | | | | | |
|---|---|---|---|---|---|---|
| HAGLR | 43.402256 | 0.27322935 | 0.34010561 | 0.8033662 | 0.4217631 | 0.99937574 |
| HOXD1 | 15.543455 | 0.76174208 | 0.62361536 | 1.22149345 | 0.22189924 | 0.99937574 |
| MTX2 | 249.891147 | 0.16309132 | 0.31709326 | 0.51433235 | 0.60701965 | 0.99937574 |
| LINC01116 | 15.2777675 | -0.0911658 | 0.61596918 | -0.1480038 | 0.88233977 | 0.99937574 |
| AC074286.1 | 27.0405632 | -1.3261688 | 1.14205879 | -1.1612089 | 0.24555697 | 0.99937574 |
| HNRNPA3 | 194.128769 | -0.1641697 | 0.26645822 | -0.6161179 | 0.53781673 | 0.99937574 |
| NFE2L2 | 428.889409 | -0.0309938 | 0.25765788 | -0.1202906 | 0.90425297 | 0.99937574 |
| AGPS | 75.7041055 | -0.0463353 | 0.25030252 | -0.1851173 | 0.853137 | 0.99937574 |
| AC012499.1 | 16.1792967 | 1.00300116 | 0.76890554 | 1.30445302 | 1.92E-01 | 0.99937574 |
| RBM45 | 23.3536201 | -0.0934147 | 0.48601805 | -0.1922042 | 0.84758229 | 0.99937574 |
| AC009948.1 | 37.8825015 | -0.6413881 | 0.70777631 | -0.9062016 | 0.36482913 | 0.99937574 |
| PRKRA | 85.9817509 | -0.0111294 | 0.28786653 | -0.0386616 | 0.96916017 | 0.99937574 |
| PLEKHA3 | 52.9870615 | -0.1387303 | 0.2994493 | -0.4632847 | 0.6431603 | 0.99937574 |
| TTN-AS1 | 125.100511 | 0.37209069 | 0.32943078 | 1.12949583 | 0.25868873 | 0.99937574 |
| SESTD1 | 234.077598 | -0.6984035 | 0.41488532 | -1.6833652 | 0.09230442 | 0.99937574 |
| ZNF385B | 67.3737023 | 0.50742747 | 0.57318989 | 0.88526941 | 0.37601135 | 0.99937574 |
| CWC22 | 46.6357351 | -0.3887925 | 0.334525 | -1.1622226 | 0.24514505 | 0.99937574 |
| UBE2E3 | 123.951892 | 0.12630051 | 0.28026769 | 0.45064244 | 0.65224727 | 0.99937574 |
| SSFA2 | 65.302693 | -0.5512421 | 0.45928255 | -1.2002244 | 0.23005222 | 0.99937574 |
| PPP1R1C | 21.3982337 | 0.04602362 | 0.92940997 | 0.04951918 | 0.96050555 | 0.99937574 |
| PDE1A | 1099.90465 | -0.504325 | 0.27133748 | -1.8586634 | 6.31E-02 | 9.99E-01 |
| DNAJC10 | 70.3509188 | -0.2715684 | 0.28599543 | -0.949555 | 0.34233844 | 0.99937574 |
| NCKAP1 | 269.642378 | -0.0639159 | 0.33222201 | -0.192389 | 0.8474375 | 0.99937574 |
| NUP35 | 22.7563128 | -0.232491 | 0.39094369 | -0.5946917 | 0.55204959 | 0.99937574 |
| LINC01473 | 29.3895197 | 0.76479652 | 0.4199347 | 1.82122726 | 0.06857232 | 0.99937574 |
| ZC3H15 | 72.4393566 | -0.084888 | 0.25900167 | -0.327751 | 0.74309997 | 0.99937574 |
| ITGAV | 313.041804 | -0.4577483 | 0.55158276 | -0.8298815 | 0.4066058 | 0.99937574 |
| AC017101.1 | 26.5661352 | 0.00367219 | 0.7941379 | 0.00462412 | 0.9963105 | 0.99937574 |
| AC007319.1 | 33.2415757 | 0.92396805 | 1.0084638 | 0.91621341 | 0.35955497 | 0.99937574 |
| TFPI | 24.9986264 | -0.4828708 | 0.6202946 | -0.7784541 | 0.43630138 | 0.99937574 |
| LINC01090 | 29.5479397 | -0.020248 | 0.57451579 | -0.0352436 | 0.97188547 | 0.99937574 |
| GULP1 | 703.110847 | -0.7028835 | 0.66172919 | -1.0621921 | 0.28814848 | 0.99937574 |
| WDR75 | 38.3387026 | -0.4015049 | 0.40404098 | -0.9937231 | 0.32035767 | 0.99937574 |
| SLC40A1 | 49.3999251 | 0.29950585 | 0.29748268 | 1.00680095 | 0.31403044 | 0.99937574 |
| ASNSD1 | 16.5281848 | 0.3655553 | 0.78510232 | 0.46561485 | 0.64149121 | 0.99937574 |
| ANKAR | 59.9450292 | -0.5193758 | 0.32468215 | -1.5996439 | 0.1096776 | 0.99937574 |
| ORMDL1 | 47.0978526 | -0.0451539 | 0.42551685 | -0.1061155 | 0.91549075 | 0.99937574 |
| PMS1 | 41.3489923 | -0.0550127 | 0.33000607 | -0.1667022 | 0.86760439 | 0.99937574 |
| C2orf88 | 38.2139701 | 0.22827598 | 0.39548401 | 0.57720659 | 0.56379991 | 0.99937574 |
| HIBCH | 175.889163 | 0.62823376 | 0.37647692 | 1.66871786 | 0.09517331 | 0.99937574 |
| INPP1 | 62.0558366 | -0.2721269 | 0.45100777 | -0.6033752 | 0.54625914 | 0.99937574 |
| MFSD6 | 64.3753428 | -0.3928293 | 0.38186017 | -1.0287255 | 0.30360868 | 0.99937574 |
| NAB1 | 28.3319249 | 0.52728341 | 0.42647456 | 1.23637717 | 0.2163184 | 0.99937574 |
| GLS | 484.794586 | -0.1995675 | 0.51396601 | -0.3882893 | 0.69780197 | 0.99937574 |
| STAT1 | 163.65537 | -0.7966713 | 0.73318689 | -1.0865869 | 0.27721943 | 0.99937574 |
| SLC39A10 | 18.0123109 | 0.88461561 | 0.45806751 | 1.93119047 | 0.0534595 | 0.99937574 |
| DNAH7 | 24.0844656 | 0.26529574 | 0.54801157 | 0.4841061 | 0.62831057 | 0.99937574 |
| GTF3C3 | 49.1354906 | 0.0574561 | 0.33434317 | 0.1718477 | 0.86355727 | 0.99937574 |
| PGAP1 | 50.2135052 | -0.5651839 | 0.40290907 | -1.402758 | 0.16068901 | 0.99937574 |
| ANKRD44 | 314.152441 | -0.1779195 | 0.50600943 | -0.3516129 | 0.72512858 | 0.99937574 |
| SF3B1 | 170.002391 | 0.29075385 | 0.24503162 | 1.18659727 | 0.23538651 | 0.99937574 |
| COQ10B | 51.4501528 | 0.17088778 | 0.29584729 | 0.57762159 | 0.56351963 | 0.99937574 |
| HSPD1 | 163.74233 | -0.1499931 | 0.3530435 | -0.4248574 | 0.67094064 | 0.99937574 |
| MOB4 | 21.3719938 | -0.0215906 | 0.50493216 | -0.0427594 | 0.96589334 | 0.99937574 |

| | | | | | | |
|---|---|---|---|---|---|---|
| PLCL1 | 1571.2662 | 1.12897444 | 0.39885267 | 2.83055507 | 0.00464673 | 0.60897109 |
| FTCDNL1 | 20.4727804 | -0.2212049 | 0.80160911 | -0.2759511 | 0.78258565 | 0.99937574 |
| C2orf69 | 24.5579897 | 0.56108293 | 0.49224396 | 1.13984724 | 0.25434995 | 0.99937574 |
| TYW5 | 43.1428183 | -0.1854365 | 0.36312982 | -0.5106618 | 0.60958789 | 0.99937574 |
| MAIP1 | 16.569839 | 0.26403852 | 0.49012427 | 0.5387175 | 0.5900818 | 0.99937574 |
| SPATS2L | 116.363329 | -1.6433171 | 0.61606104 | -2.6674582 | 0.00764274 | 0.74742128 |
| KCTD18 | 14.4587033 | 0.01898806 | 0.56445532 | 0.03363961 | 0.97316453 | 0.99937574 |
| BZW1 | 28.9713272 | -0.2320553 | 0.51264449 | -0.4526633 | 0.65079121 | 0.99937574 |
| CLK1 | 102.630674 | -0.3086034 | 0.29662955 | -1.0403665 | 0.29816967 | 0.99937574 |
| PPIL3 | 21.6074782 | -0.0087443 | 0.59929441 | -0.0145909 | 0.98835853 | 0.99937574 |
| NIF3L1 | 17.7612453 | 0.24599549 | 0.45968879 | 0.53513485 | 0.59255661 | 0.99937574 |
| ORC2 | 66.8990462 | -0.1287414 | 0.40325022 | -0.3192594 | 0.74952978 | 0.99937574 |
| AC005037.1 | 16.8198694 | -0.4257738 | 0.63725676 | -0.6681355 | 0.50404713 | 0.99937574 |
| FAM126B | 106.297989 | -0.1130123 | 0.2979609 | -0.3792857 | 0.70447568 | 0.99937574 |
| NDUFB3 | 78.8930029 | 0.42388026 | 0.68948402 | 0.61477894 | 0.53870072 | 0.99937574 |
| CFLAR | 244.031949 | -0.5539617 | 0.32584717 | -1.7000661 | 0.08911848 | 0.99937574 |
| CFLAR-AS1 | 25.9444357 | -0.0121832 | 0.7850271 | -0.0155195 | 0.98761775 | 0.99937574 |
| CASP10 | 40.1695566 | -0.0720491 | 0.52693818 | -0.1367317 | 0.89124288 | 0.99937574 |
| CASP8 | 32.9052236 | -0.2580978 | 0.3705392 | -0.6965466 | 0.48608659 | 0.99937574 |
| TRAK2 | 118.817114 | -0.7118099 | 0.33583534 | -2.1195205 | 0.0340465 | 0.99937574 |
| STRADB | 45.1285117 | -0.4381307 | 0.51306188 | -0.853953 | 0.39313103 | 0.99937574 |
| TMEM237 | 11.0533801 | 0.07529885 | 0.63624694 | 0.11834846 | 0.90579156 | 0.99937574 |
| ALS2 | 80.7597702 | 0.43233287 | 0.27055342 | 1.59795753 | 0.11005243 | 0.99937574 |
| FZD7 | 96.2843625 | 0.93682164 | 0.44072824 | 2.12562202 | 0.03353474 | 0.99937574 |
| SUMO1 | 57.001044 | -0.3765794 | 0.33433268 | -1.1263613 | 0.26001262 | 0.99937574 |
| NOP58 | 74.2878678 | -0.8638446 | 0.27058988 | -3.1924498 | 0.00141071 | 0.34724223 |
| BMPR2 | 315.660001 | -0.128402 | 0.34248946 | -0.3749078 | 0.70772901 | 0.99937574 |
| FAM117B | 108.682806 | -0.0475528 | 0.44312582 | -0.1073122 | 0.91454134 | 0.99937574 |
| ICA1L | 32.296111 | -1.3074783 | 0.51212865 | -2.553027 | 0.01067912 | 0.76274616 |
| WDR12 | 28.420698 | 0.11277887 | 0.53645973 | 0.21022802 | 0.83348971 | 0.99937574 |
| CARF | 39.6489118 | -0.1195708 | 0.53864329 | -0.2219852 | 0.82432539 | 0.99937574 |
| NBEAL1 | 474.564633 | 0.05876373 | 0.25487847 | 0.23055589 | 0.81765984 | 0.99937574 |
| CYP20A1 | 93.4192207 | -0.5601235 | 0.45392065 | -1.2339678 | 0.21721489 | 0.99937574 |
| ABI2 | 148.978366 | -0.0651305 | 0.36822449 | -0.1768773 | 0.85960479 | 0.99937574 |
| RAPH1 | 216.445906 | 0.07436384 | 0.48215983 | 0.15423068 | 0.87742785 | 0.99937574 |
| PARD3B | 537.646668 | 0.0838613 | 0.25653178 | 0.32690414 | 7.44E-01 | 0.99937574 |
| INO80D | 286.259602 | -0.2496257 | 0.54903551 | -0.4546621 | 0.64935234 | 0.99937574 |
| NDUFS1 | 219.823891 | 0.23142269 | 0.21017677 | 1.10108596 | 0.27085925 | 0.99937574 |
| EEF1B2 | 71.4435492 | -0.3567525 | 0.58093321 | -0.6141024 | 0.53914767 | 0.99937574 |
| ZDBF2 | 36.4587713 | -0.1268607 | 0.40082984 | -0.3164951 | 0.75162674 | 0.99937574 |
| FASTKD2 | 43.8507999 | -0.2741582 | 0.43163918 | -0.6351561 | 0.52532662 | 0.99937574 |
| KLF7 | 80.0396337 | -0.5492125 | 0.48473418 | -1.1330179 | 0.25720672 | 0.99937574 |
| CREB1 | 118.665193 | 0.13925898 | 0.28243306 | 0.49306899 | 0.62196384 | 0.99937574 |
| METTL21A | 34.4738718 | 0.25601558 | 0.45445196 | 0.56335015 | 0.57319647 | 0.99937574 |
| CCNYL1 | 39.9494376 | -0.3674614 | 0.36532088 | -1.0058592 | 0.31448332 | 0.99937574 |
| PLEKHM3 | 46.8806145 | 0.12511203 | 0.43143014 | 0.28999371 | 0.77182105 | 0.99937574 |
| IDH1 | 13.9261964 | 0.07909948 | 0.59482763 | 0.13297882 | 0.89421013 | 0.99937574 |
| PIKFYVE | 128.084668 | -0.0263973 | 0.28324759 | -0.0931951 | 0.92574856 | 0.99937574 |
| PTH2R | 76.3101922 | -1.1327165 | 0.63503948 | -1.7836947 | 0.07447329 | 0.99937574 |
| MAP2 | 111.324979 | -0.7179078 | 0.46278727 | -1.5512696 | 0.12083708 | 0.99937574 |
| UNC80 | 21.1649831 | 2.28356539 | 0.64903864 | 3.51838129 | 0.00043419 | 0.2306224 |
| RPE | 48.0992101 | -0.2322809 | 0.36539574 | -0.6356967 | 0.5249741 | 0.99937574 |
| KANSL1L | 494.382009 | 0.466712 | 0.30891539 | 1.51080849 | 0.13083725 | 0.99937574 |
| AC007038.1 | 49.8475392 | 0.03509225 | 0.54700189 | 0.0641538 | 0.94884777 | 0.99937574 |

| | | | | | | |
|---|---|---|---|---|---|---|
| ACADL | 106.276917 | 0.23200137 | 0.3115553 | 0.74465553 | 0.45647999 | 0.99937574 |
| LANCL1 | 51.0586827 | 0.14432793 | 0.32352224 | 0.44611442 | 0.6555146 | 0.99937574 |
| ERBB4 | 1516.42934 | 0.62794472 | 0.32355768 | 1.94075051 | 0.05228855 | 0.99937574 |
| IKZF2 | 48.7174235 | 0.29923128 | 0.70505005 | 0.4244114 | 0.67126581 | 0.99937574 |
| SPAG16 | 177.943562 | 0.60107986 | 0.4447567 | 1.35148016 | 0.17654167 | 0.99937574 |
| BARD1 | 16.0979133 | 0.19598757 | 0.63772506 | 0.30732299 | 0.75859754 | 0.99937574 |
| AC016708.1 | 16.8238527 | -0.091871 | 0.67980371 | -0.1351434 | 0.89249849 | 0.99937574 |
| ATIC | 23.2513184 | 0.11750609 | 0.43979422 | 0.26718426 | 0.78932729 | 0.99937574 |
| MREG | 28.2323578 | -0.6469171 | 0.60267062 | -1.0734174 | 0.28308389 | 0.99937574 |
| PECR | 14.8295755 | 1.2217489 | 0.63672294 | 1.91880774 | 0.05500867 | 0.99937574 |
| XRCC5 | 210.80627 | -0.1433476 | 0.21569927 | -0.6645717 | 0.50632448 | 0.99937574 |
| SMARCAL1 | 23.9617756 | -0.4796615 | 0.50229125 | -0.9549469 | 0.33960456 | 0.99937574 |
| RPL37A | 218.430535 | 0.0464693 | 0.45669433 | 0.10175143 | 0.91895398 | 0.99937574 |
| IGFBP5 | 1368.9944 | -0.374635 | 0.54531406 | -0.6870078 | 0.49207781 | 0.99937574 |
| AC007563.2 | 84.993138 | 0.05562135 | 0.69247292 | 0.08032278 | 0.93598054 | 0.99937574 |
| DIRC3 | 29.7217442 | -0.0904234 | 0.68154552 | -0.132674 | 0.89445117 | 0.99937574 |
| TNS1 | 139.468069 | 0.66925029 | 0.50116986 | 1.33537617 | 0.18175326 | 0.99937574 |
| ARPC2 | 127.739626 | -0.4011532 | 0.28237174 | -1.4206563 | 0.15541669 | 0.99937574 |
| AAMP | 25.5625911 | 0.01166983 | 0.47255347 | 0.02469525 | 0.98029804 | 0.99937574 |
| PNKD | 94.9173947 | -0.4199914 | 0.47286409 | -0.8881863 | 0.37444055 | 0.99937574 |
| TMBIM1 | 68.2922416 | -0.5364529 | 0.50222687 | -1.0681485 | 0.28545353 | 0.99937574 |
| CATIP-AS2 | 12.4977046 | -0.3260417 | 0.78729661 | -0.4141281 | 0.67878027 | 0.99937574 |
| CTDSP1 | 40.4601862 | -0.4394823 | 0.436889 | -1.0059358 | 0.31444645 | 0.99937574 |
| USP37 | 74.6048212 | -0.147478 | 0.38944049 | -0.378692 | 0.70491661 | 0.99937574 |
| CNOT9 | 49.5619359 | -0.2871567 | 0.36648496 | -0.783543 | 0.43330829 | 0.99937574 |
| PLCD4 | 11.6475732 | 1.44568559 | 0.87740963 | 1.64767464 | 0.09941945 | 0.99937574 |
| ZNF142 | 15.0757774 | -0.5445931 | 0.51449828 | -1.0584935 | 0.28983049 | 0.99937574 |
| TTLL4 | 15.0685033 | 0.2650723 | 0.52581864 | 0.50411355 | 0.61418159 | 0.99937574 |
| CYP27A1 | 24.109583 | 0.22400083 | 0.39101056 | 0.57287666 | 0.56672821 | 0.99937574 |
| CNPPD1 | 58.4301335 | -0.0458412 | 0.32369776 | -0.1416172 | 0.88738234 | 0.99937574 |
| RETREG2 | 51.2148927 | -0.5568544 | 0.47394645 | -1.174931 | 0.24002231 | 0.99937574 |
| ZFAND2B | 15.1307188 | -0.8278769 | 0.48342494 | -1.7125241 | 0.08680013 | 0.99937574 |
| ANKZF1 | 25.3413618 | 0.19879137 | 0.47316322 | 0.42013276 | 0.67438847 | 0.99937574 |
| STK16 | 14.7196327 | -0.3560993 | 0.55583746 | -0.6406536 | 0.52174775 | 0.99937574 |
| TUBA4A | 72.2542579 | 0.88917012 | 0.40455491 | 2.19789722 | 0.02795643 | 0.99049468 |
| DNAJB2 | 35.2050558 | -0.3958341 | 0.42212571 | -0.9377162 | 0.34839026 | 0.99937574 |
| DNPEP | 22.9545752 | -0.4446283 | 0.43459794 | -1.0230796 | 0.30627023 | 0.99937574 |
| CHPF | 20.2193098 | 0.46633769 | 0.51771752 | 0.90075702 | 0.36771752 | 0.99937574 |
| OBSL1 | 14.7280784 | 0.91415283 | 0.56094537 | 1.62966463 | 0.1031724 | 0.99937574 |
| SGPP2 | 243.513866 | -0.1454671 | 0.27816295 | -0.5229563 | 0.60100467 | 0.99937574 |
| FARSB | 40.9702292 | -0.2031622 | 0.42538619 | -0.4775947 | 0.63293872 | 0.99937574 |
| ACSL3 | 199.523743 | 0.27195627 | 0.38667051 | 0.70332818 | 0.48185125 | 0.99937574 |
| AP1S3 | 25.8515795 | 0.44975737 | 0.57777811 | 0.77842578 | 0.43631805 | 0.99937574 |
| WDFY1 | 64.6601316 | -0.6560109 | 0.41137472 | -1.5946796 | 0.11078391 | 0.99937574 |
| MRPL44 | 18.6296353 | -0.2593285 | 0.45495521 | -0.5700089 | 0.56867166 | 0.99937574 |
| CUL3 | 203.857319 | -0.2708503 | 0.24612375 | -1.100464 | 0.27113001 | 0.99937574 |
| DOCK10 | 281.151282 | 0.14991558 | 0.8683561 | 0.17264297 | 0.86293207 | 0.99937574 |
| IRS1 | 107.096248 | -0.0118092 | 0.6385209 | -0.0184946 | 0.98524432 | 0.99937574 |
| RHBDD1 | 191.047531 | -0.2473871 | 0.40038452 | -0.6178738 | 0.5366585 | 0.99937574 |
| COL4A4 | 294.068355 | 0.02655558 | 0.37690772 | 0.07045645 | 0.94383036 | 0.99937574 |
| COL4A3 | 282.496502 | 0.32738604 | 0.35180371 | 0.9305929 | 0.35206419 | 0.99937574 |
| AC097662.1 | 43.8835026 | 0.6336238 | 0.65358426 | 0.96946001 | 0.33231572 | 0.99937574 |
| MFF | 99.2194005 | 0.12513692 | 0.44843232 | 0.27905419 | 0.78020324 | 0.99937574 |
| AGFG1 | 185.795853 | -0.3774278 | 0.28973727 | -1.3026554 | 0.19269242 | 0.99937574 |

| | | | | | | |
|---|---|---|---|---|---|---|
| TRIP12 | 283.96057 | -0.3438942 | 0.19892011 | -1.7288056 | 0.0838439 | 0.99937574 |
| FBXO36 | 20.4236792 | -0.7430775 | 0.43230876 | -1.7188583 | 0.08564018 | 0.99937574 |
| SP110 | 47.9264008 | -0.8743976 | 0.62797149 | -1.392416 | 0.16379646 | 0.99937574 |
| SP140L | 40.6594906 | -1.2970545 | 0.57455002 | -2.2575136 | 0.023976 | 0.96546668 |
| SP100 | 135.202254 | -1.0673671 | 0.43307971 | -2.4645973 | 0.01371673 | 0.83391111 |
| CAB39 | 271.436903 | 0.17385967 | 0.34152856 | 0.50906335 | 0.61070782 | 0.99937574 |
| ITM2C | 49.1185991 | 0.12752952 | 0.38865453 | 0.3281308 | 0.74281277 | 0.99937574 |
| C2orf72 | 12.0705712 | -0.4441064 | 0.83839587 | -0.5297096 | 0.59631327 | 0.99937574 |
| PSMD1 | 136.785766 | -0.2872198 | 0.25050107 | -1.1465811 | 0.25155478 | 0.99937574 |
| ARMC9 | 101.088793 | 0.25864895 | 0.29863678 | 0.86609878 | 0.386436 | 0.99937574 |
| NCL | 93.6147492 | -0.1918397 | 0.50203371 | -0.3821252 | 0.70236851 | 0.99937574 |
| PTMA | 256.895035 | -0.4075269 | 0.38847156 | -1.0490522 | 0.29415409 | 0.99937574 |
| PDE6D | 34.4813789 | -0.0674599 | 0.46295386 | -0.1457164 | 0.88414531 | 0.99937574 |
| COPS7B | 33.7363734 | 0.12617811 | 0.35896107 | 0.35150918 | 0.72520638 | 0.99937574 |
| DIS3L2 | 162.176465 | -0.0499905 | 0.34252068 | -0.1459488 | 0.88396182 | 0.99937574 |
| EIF4E2 | 45.8077926 | -0.4612175 | 0.36567352 | -1.2612823 | 0.20720715 | 0.99937574 |
| EFHD1 | 293.337177 | 0.28964408 | 0.39232963 | 0.73826714 | 0.46035213 | 0.99937574 |
| GIGYF2 | 408.921577 | -0.0120694 | 0.23647959 | -0.0510378 | 0.9592954 | 0.99937574 |
| KCNJ13 | 13.1552583 | 0.46539853 | 0.90033281 | 0.51691833 | 0.60521317 | 0.99937574 |
| SNORC | 11.9406874 | 1.37323171 | 0.58187253 | 2.36002155 | 0.01827387 | 0.93502451 |
| ATG16L1 | 18.1009589 | -0.4330738 | 0.46615182 | -0.9290402 | 0.35286824 | 0.99937574 |
| DGKD | 22.5175863 | -1.0399317 | 0.53027934 | -1.9611017 | 0.04986716 | 0.99937574 |
| USP40 | 110.947764 | -0.308062 | 0.38983885 | -0.7902291 | 0.42939399 | 0.99937574 |
| SH3BP4 | 213.465623 | -0.3513565 | 0.47126523 | -0.7455599 | 0.4559333 | 0.99937574 |
| AGAP1 | 368.934924 | -0.1422021 | 0.58814279 | -0.2417816 | 0.8089494 | 0.99937574 |
| IQCA1 | 32.2575006 | -1.5983036 | 0.70602679 | -2.2638002 | 0.0235864 | 0.96546668 |
| COPS8 | 26.7023361 | -0.031483 | 0.41059338 | -0.0766767 | 0.93888071 | 0.99937574 |
| RAB17 | 97.3429201 | -0.7073023 | 0.58356189 | -1.2120434 | 0.22549578 | 0.99937574 |
| LRRFIP1 | 457.018197 | -0.4838906 | 0.28748067 | -1.6832109 | 0.09233427 | 0.99937574 |
| ILKAP | 61.737328 | -0.1831645 | 0.31679673 | -0.5781767 | 0.5631448 | 0.99937574 |
| PER2 | 26.8594522 | -0.5720918 | 0.49946058 | -1.1454193 | 0.25203553 | 0.99937574 |
| TRAF3IP1 | 41.5601143 | 0.54726607 | 0.42547508 | 1.28624704 | 0.19835686 | 0.99937574 |
| ASB1 | 28.701229 | -0.126228 | 0.6180862 | -0.204224 | 0.83817847 | 0.99937574 |
| HDAC4 | 134.536876 | 0.07823001 | 0.35146961 | 0.22257972 | 0.82386262 | 0.99937574 |
| NDUFA10 | 153.626056 | 0.38594714 | 0.27655631 | 1.39554632 | 0.16285116 | 0.99937574 |
| COPS9 | 39.6995764 | 0.34750094 | 0.52998862 | 0.65567622 | 0.51203247 | 0.99937574 |
| GPC1 | 9.98539265 | -0.4707671 | 0.77163011 | -0.6100942 | 0.54179941 | 0.99937574 |
| ANKMY1 | 33.2719838 | 0.03368937 | 0.38100532 | 0.08842231 | 0.92954103 | 0.99937574 |
| RNPEPL1 | 42.9286615 | -0.008883 | 0.44953044 | -0.0197607 | 0.98423429 | 0.99937574 |
| CAPN10 | 22.7549278 | 0.38240803 | 0.43536338 | 0.87836516 | 0.37974558 | 0.99937574 |
| SNED1 | 44.3590441 | 0.2265786 | 0.64903404 | 0.34910126 | 0.72701329 | 0.99937574 |
| AC005237.1 | 12.2145558 | 0.66348716 | 0.71632641 | 0.9262358 | 0.35432345 | 0.99937574 |
| MTERF4 | 37.0744872 | 0.0820293 | 0.32474957 | 0.25259248 | 0.80058315 | 0.99937574 |
| PPP1R7 | 44.9288056 | 0.1876159 | 0.31323093 | 0.59896991 | 0.54919295 | 0.99937574 |
| HDLBP | 180.542159 | -0.20343 | 0.23905056 | -0.8509916 | 0.39477404 | 0.99937574 |
| 2-Sep | 200.896761 | 0.2512424 | 0.24946071 | 1.00714216 | 0.31386646 | 0.99937574 |
| FARP2 | 114.972981 | 0.04979603 | 0.26527252 | 0.18771649 | 0.8510989 | 0.99937574 |
| STK25 | 38.1840571 | -0.3443257 | 0.40292599 | -0.8545632 | 0.39279303 | 0.99937574 |
| BOK | 20.0053338 | -0.6316242 | 0.52847635 | -1.1951797 | 0.23201684 | 0.99937574 |
| THAP4 | 64.7865975 | 0.05143487 | 0.27087226 | 0.18988608 | 0.8493984 | 0.99937574 |
| ATG4B | 76.9912498 | -0.0688761 | 0.33600936 | -0.2049828 | 0.83758556 | 0.99937574 |
| DTYMK | 16.8456489 | -0.3214527 | 0.60451173 | -0.5317559 | 0.59489506 | 0.99937574 |
| ING5 | 48.9930033 | -0.3314635 | 0.4286078 | -0.7733493 | 0.43931568 | 0.99937574 |
| D2HGDH | 32.9816641 | 0.49412212 | 0.42673299 | 1.15791874 | 0.2468972 | 0.99937574 |

| | | | | | | |
|---|---|---|---|---|---|---|
| CHL1 | 366.110422 | -0.1266045 | 0.85850307 | -0.1474713 | 0.88276007 | 0.99937574 |
| LINC01266 | 36.1962732 | -0.280465 | 0.57440287 | -0.4882722 | 0.62535704 | 0.99937574 |
| CNTN4 | 17.6339776 | 0.59028403 | 0.79703138 | 0.74060326 | 0.45893403 | 0.99937574 |
| TRNT1 | 41.3284673 | -0.0341861 | 0.33595931 | -0.1017566 | 0.91894988 | 0.99937574 |
| CRBN | 36.8363946 | -0.1979718 | 0.41457489 | -0.4775297 | 0.63298499 | 0.99937574 |
| AC034195.1 | 75.2510761 | 0.56761097 | 0.53224151 | 1.06645378 | 0.28621856 | 0.99937574 |
| SUMF1 | 390.95825 | 0.00044876 | 0.39946736 | 0.0011234 | 0.99910365 | 0.99979721 |
| SETMAR | 24.5172781 | 0.54416637 | 0.4549399 | 1.19612803 | 0.2316466 | 0.99937574 |
| ITPR1 | 509.346086 | 0.39731183 | 0.40938103 | 0.97051843 | 0.33178814 | 0.99937574 |
| EGOT | 17.5717947 | -0.5667157 | 0.83475995 | -0.6788966 | 0.49720336 | 0.99937574 |
| AC018816.1 | 20.6613629 | 0.88991548 | 0.49000147 | 1.81614857 | 0.06934758 | 0.99937574 |
| BHLHE40-AS | 15.1758351 | -0.0235797 | 0.90287152 | -0.0261163 | 0.97916458 | 0.99937574 |
| BHLHE40 | 171.271498 | -0.6676964 | 0.61993628 | -1.0770404 | 0.28146221 | 0.99937574 |
| ARL8B | 116.924152 | -0.4883866 | 0.29270416 | -1.6685331 | 0.09520994 | 0.99937574 |
| AC026202.2 | 20.9622226 | 0.26111079 | 0.57367164 | 0.45515723 | 0.64899613 | 0.99937574 |
| EDEM1 | 33.0901279 | -0.0361099 | 0.39313609 | -0.091851 | 0.92681641 | 0.99937574 |
| LMCD1-AS1 | 71.4192478 | -0.0334248 | 0.50040034 | -0.0667961 | 0.94674404 | 0.99937574 |
| RAD18 | 14.5092567 | -0.5379194 | 0.50280752 | -1.0698316 | 0.28469509 | 0.99937574 |
| SRGAP3 | 139.232208 | -0.1407791 | 0.82956558 | -0.1697022 | 0.86524432 | 0.99937574 |
| THUMPD3-A: | 97.6030354 | -0.2446422 | 0.39952068 | -0.6123392 | 0.54031338 | 0.99937574 |
| THUMPD3 | 32.3744019 | -0.1111031 | 0.35331284 | -0.3144611 | 0.75317086 | 0.99937574 |
| SETD5 | 301.14251 | 0.06359716 | 0.26308531 | 0.2417359 | 0.80898481 | 0.99937574 |
| MTMR14 | 28.3599124 | -0.4907183 | 0.40163066 | -1.2218149 | 0.22177764 | 0.99937574 |
| OGG1 | 15.6886022 | 0.41024935 | 0.48362503 | 0.84827981 | 0.39628216 | 0.99937574 |
| CAMK1 | 11.7573308 | -0.152033 | 0.69540587 | -0.2186249 | 0.82694226 | 0.99937574 |
| TADA3 | 29.7840129 | -0.3249325 | 0.33697462 | -0.964264 | 0.33491359 | 0.99937574 |
| JAGN1 | 15.6748376 | -0.0165815 | 0.64119259 | -0.0258604 | 0.97936867 | 0.99937574 |
| IL17RC | 12.5239514 | 0.67072895 | 0.50591658 | 1.32576987 | 0.18491594 | 0.99937574 |
| CRELD1 | 26.4303943 | 0.40107419 | 0.48427656 | 0.82819245 | 0.40756152 | 0.99937574 |
| EMC3 | 49.2438152 | 0.03917203 | 0.38803353 | 0.10095012 | 0.91959005 | 0.99937574 |
| BRK1 | 43.8215617 | -0.3876176 | 0.49214777 | -0.787604 | 0.43092835 | 0.99937574 |
| VHL | 26.2500397 | -0.5560174 | 0.49113749 | -1.1321014 | 0.25759181 | 0.99937574 |
| SEC13 | 38.0356703 | -0.1544763 | 0.45235419 | -0.341494 | 0.73273169 | 0.99937574 |
| ATG7 | 215.894626 | -0.4103959 | 0.25777225 | -1.5920872 | 0.11136511 | 0.99937574 |
| VGLL4 | 263.516358 | -0.1947576 | 0.28584454 | -0.6813409 | 0.4956558 | 0.99937574 |
| TAMM41 | 58.9934191 | 0.1065774 | 0.30856623 | 0.34539554 | 0.72979703 | 0.99937574 |
| PPARG | 283.521728 | 0.26179703 | 0.37006709 | 0.70743126 | 0.47929852 | 0.99937574 |
| TSEN2 | 44.4005185 | 0.24401557 | 0.4484792 | 0.54409562 | 0.58637567 | 0.99937574 |
| MKRN2 | 15.4864254 | 0.27556407 | 0.49944238 | 0.55174346 | 0.58112413 | 0.99937574 |
| RAF1 | 121.460211 | -0.3641725 | 0.28812759 | -1.263928 | 0.20625588 | 0.99937574 |
| RPL32 | 231.066049 | -0.0456467 | 0.57498685 | -0.0793874 | 0.93672447 | 0.99937574 |
| IQSEC1 | 100.550171 | 0.80105351 | 0.54922337 | 1.45852045 | 0.14469714 | 0.99937574 |
| NUP210 | 31.8440788 | 0.11815197 | 0.59243639 | 0.19943402 | 0.84192325 | 0.99937574 |
| HDAC11 | 17.7520416 | 0.84766791 | 0.59780196 | 1.41797447 | 0.15619822 | 0.99937574 |
| XPC | 48.7804718 | 0.00466507 | 0.32729899 | 0.01425324 | 0.98862795 | 0.99937574 |
| LSM3 | 36.8231951 | 0.05810137 | 0.50594739 | 0.11483678 | 0.90857449 | 0.99937574 |
| SLC6A6 | 78.2420864 | -0.3479816 | 0.61703896 | -0.5639541 | 5.73E-01 | 0.99937574 |
| GRIP2 | 54.8071733 | 0.86930013 | 0.93685776 | 0.92788913 | 0.35346508 | 0.99937574 |
| CCDC174 | 40.2712718 | 0.10913502 | 0.30862874 | 0.35361264 | 0.7236292 | 0.99937574 |
| FGD5-AS1 | 80.1562547 | -0.142974 | 0.30604747 | -0.4671629 | 0.64038337 | 0.99937574 |
| NR2C2 | 92.130876 | -0.4297957 | 0.33027125 | -1.3013415 | 0.19314159 | 0.99937574 |
| MRPS25 | 99.4061224 | 0.15863267 | 0.29971628 | 0.52927613 | 0.59661391 | 0.99937574 |
| RBSN | 31.7782742 | -0.1797792 | 0.41309297 | -0.4352027 | 0.66341533 | 0.99937574 |
| CAPN7 | 170.870771 | -0.2825356 | 0.26844978 | -1.0524708 | 0.2925836 | 0.99937574 |

| | | | | | | |
|---|---|---|---|---|---|---|
| SH3BP5-AS1 | 11.6283122 | 0.40705761 | 0.68261735 | 0.59631887 | 0.55096223 | 0.99937574 |
| METTL6 | 18.4092619 | -0.1244314 | 0.48831739 | -0.2548166 | 0.79886473 | 0.99937574 |
| EAF1 | 15.9643856 | -0.7331104 | 0.54133692 | -1.3542591 | 0.17565374 | 0.99937574 |
| COLQ | 14.4343223 | -0.4620522 | 0.84540178 | -0.5465475 | 0.58468966 | 0.99937574 |
| HACL1 | 45.9216113 | 0.18973184 | 0.30706048 | 0.61789729 | 0.53664304 | 0.99937574 |
| BTD | 19.3476376 | 0.06063361 | 0.44910152 | 0.13501092 | 0.89260324 | 0.99937574 |
| ANKRD28 | 300.848349 | -0.1415484 | 0.30602645 | -0.4625366 | 0.6436966 | 0.99937574 |
| DPH3 | 11.8073761 | 0.41554257 | 0.78412526 | 0.52994411 | 0.59615068 | 0.99937574 |
| OXNAD1 | 99.217619 | 0.03390275 | 0.37668416 | 0.09000311 | 0.92828474 | 0.99937574 |
| PLCL2 | 310.642758 | -0.6605225 | 0.54054202 | -1.2219633 | 0.22172151 | 0.99937574 |
| TBC1D5 | 549.155362 | 0.11640521 | 0.38478692 | 0.30251863 | 0.76225673 | 0.99937574 |
| RAB5A | 96.6638695 | -0.2536691 | 0.25188447 | -1.0070851 | 0.31389389 | 0.99937574 |
| KAT2B | 71.6146062 | 0.224302 | 0.49837712 | 0.4500648 | 0.65266372 | 0.99937574 |
| UBE2E2 | 272.439775 | 0.28603652 | 0.54672751 | 0.5231793 | 0.60084948 | 0.99937574 |
| UBE2E1 | 214.331442 | -0.3173344 | 0.27817358 | -1.1407783 | 0.25396219 | 0.99937574 |
| NKIRAS1 | 21.0081998 | 0.23237408 | 0.44548567 | 0.52161965 | 0.60193518 | 0.99937574 |
| RPL15 | 422.323987 | 0.19098925 | 0.46973174 | 0.40659217 | 0.68430755 | 0.99937574 |
| NR1D2 | 104.509563 | 0.09682855 | 0.38127374 | 0.25396071 | 0.79952591 | 0.99937574 |
| THRB | 577.656259 | 0.07950016 | 0.21372923 | 0.37196672 | 0.70991763 | 0.99937574 |
| AC092422.1 | 499.248652 | 1.7842391 | 0.68130003 | 2.61887425 | 0.00882205 | 0.74742128 |
| RARB | 344.532632 | 1.76620213 | 0.48221852 | 3.6626593 | 0.00024961 | 0.15744178 |
| TOP2B | 131.415899 | 0.15716159 | 0.30726535 | 0.51148491 | 0.60901155 | 0.99937574 |
| NGLY1 | 80.5840398 | -0.2564529 | 0.33029075 | -0.7764459 | 0.43748577 | 0.99937574 |
| OXSM | 15.7948561 | -0.2221235 | 0.62030644 | -0.3580867 | 0.72027843 | 0.99937574 |
| NEK10 | 63.7340518 | -0.432877 | 0.62021901 | -0.6979421 | 0.48521338 | 0.99937574 |
| SLC4A7 | 72.3430767 | -0.6130184 | 0.77606986 | -0.789901 | 0.42958557 | 0.99937574 |
| CMC1 | 149.288779 | 0.09022764 | 0.35458789 | 0.25445777 | 0.79914193 | 0.99937574 |
| AZI2 | 53.0592918 | -0.566171 | 0.40969904 | -1.3819193 | 0.16699648 | 0.99937574 |
| TGFBR2 | 50.5253464 | -0.2206629 | 0.4050872 | -0.5447294 | 0.58593963 | 0.99937574 |
| STT3B | 165.841472 | 0.4282881 | 0.26552516 | 1.61298499 | 0.10674777 | 0.99937574 |
| OSBPL10 | 191.184075 | 0.79529142 | 0.62637516 | 1.26967266 | 0.20420126 | 0.99937574 |
| GPD1L | 62.6619001 | 0.67145565 | 0.3460277 | 1.94046792 | 0.05232285 | 0.99937574 |
| CMTM8 | 23.8612243 | 0.30249231 | 0.42690406 | 0.7085721 | 0.47859005 | 0.99937574 |
| CMTM6 | 73.1100669 | 0.05344296 | 0.4020457 | 0.13292758 | 0.89425066 | 0.99937574 |
| DYNC1LI1 | 41.5047242 | 0.08758904 | 0.45285656 | 0.19341454 | 0.84663433 | 0.99937574 |
| CNOT10 | 65.5856418 | 0.04954288 | 0.31446678 | 0.15754567 | 0.87481482 | 0.99937574 |
| GLB1 | 97.3402288 | 0.3986481 | 0.21506345 | 1.8536302 | 0.06379208 | 0.99937574 |
| CRTAP | 57.5400706 | 0.09067786 | 0.40741624 | 0.22256811 | 0.82387166 | 0.99937574 |
| FBXL2 | 117.749125 | -0.1056191 | 0.32565801 | -0.3243251 | 0.74569192 | 0.99937574 |
| UBP1 | 55.1040505 | -0.0753219 | 0.33075364 | -0.2277281 | 0.81985759 | 0.99937574 |
| CLASP2 | 226.892239 | -0.1805625 | 0.26108435 | -0.6915868 | 0.48919683 | 0.99937574 |
| PDCD6IP | 202.341181 | -0.145664 | 0.22495766 | -0.6475174 | 0.51729717 | 0.99937574 |
| TRANK1 | 15.2775871 | -1.9939359 | 0.96841087 | -2.0589772 | 0.03949642 | 0.99937574 |
| EPM2AIP1 | 44.0039997 | 0.48283423 | 0.38112774 | 1.26685671 | 0.20520653 | 0.99937574 |
| MLH1 | 32.7223404 | 0.20701636 | 0.33713796 | 0.61404051 | 0.53918856 | 0.99937574 |
| LRRFIP2 | 448.505001 | 0.00504844 | 0.40350333 | 0.01251152 | 0.99001752 | 0.99937574 |
| GOLGA4 | 442.4556 | -0.3350304 | 0.25862883 | -1.29541 | 0.19517885 | 0.99937574 |
| ITGA9 | 34.8433501 | -1.0444162 | 0.57784964 | -1.8074185 | 0.07069705 | 0.99937574 |
| CTDSPL | 357.751328 | -0.416419 | 0.39759357 | -1.0473485 | 0.29493886 | 0.99937574 |
| ACAA1 | 38.718684 | 0.45783672 | 0.40716949 | 1.12443769 | 0.26082739 | 0.99937574 |
| OXSR1 | 117.549462 | -0.2392546 | 0.36145869 | -0.6619141 | 0.50802629 | 0.99937574 |
| ACVR2B | 19.400995 | 0.33134855 | 0.50659746 | 0.65406674 | 0.51306881 | 0.99937574 |
| EXOG | 31.4002052 | 0.01672623 | 0.39357624 | 0.04249806 | 0.96610166 | 0.99937574 |
| WDR48 | 46.5671664 | -0.1756984 | 0.36450222 | -0.4820228 | 0.62978975 | 0.99937574 |

| | | | | | | |
|---|---|---|---|---|---|---|
| GORASP1 | 23.1995924 | 0.17596685 | 0.40385325 | 0.43571977 | 0.66304006 | 0.99937574 |
| CSRNP1 | 15.2552195 | -0.834426 | 0.67449677 | -1.237109 | 0.21604662 | 0.99937574 |
| SLC25A38 | 15.8614764 | -0.4083737 | 0.49796843 | -0.8200796 | 0.41217074 | 0.99937574 |
| RPSA | 226.907394 | 0.16455972 | 0.40304146 | 0.40829477 | 0.68305728 | 0.99937574 |
| MYRIP | 143.718783 | -0.9653119 | 0.38499616 | -2.5073287 | 0.01216475 | 0.77519564 |
| EIF1B | 52.8586597 | -0.2366394 | 0.53463621 | -0.4426176 | 0.65804237 | 0.99937574 |
| ENTPD3-AS1 | 18.9370938 | 0.15834586 | 0.57501263 | 0.27537805 | 0.78302581 | 0.99937574 |
| RPL14 | 291.299755 | -0.0325727 | 0.66754709 | -0.0487946 | 0.96108295 | 0.99937574 |
| ZNF621 | 35.0603825 | -0.2397332 | 0.3331588 | -0.7195764 | 0.47178586 | 0.99937574 |
| CTNNB1 | 118.433628 | 0.05305291 | 0.30545775 | 0.17368332 | 0.86211435 | 0.99937574 |
| ULK4 | 85.2035755 | -0.6325796 | 0.33639088 | -1.8804898 | 0.06004135 | 0.99937574 |
| TRAK1 | 312.005255 | -0.0552748 | 0.28109481 | -0.1966411 | 0.84410842 | 0.99937574 |
| SEC22C | 42.073614 | 0.12156965 | 0.31475272 | 0.38623862 | 0.69931996 | 0.99937574 |
| SS18L2 | 16.9722882 | -0.8229427 | 0.62981103 | -1.3066502 | 0.19133153 | 0.99937574 |
| NKTR | 287.95176 | -0.1020769 | 0.1996191 | -0.5113584 | 0.60910013 | 0.99937574 |
| AC006059.1 | 41.6714501 | 0.53996022 | 0.62012983 | 0.87072126 | 0.38390637 | 0.99937574 |
| AC099329.3 | 16.4854825 | 0.86867858 | 0.60561891 | 1.43436502 | 0.15146812 | 0.99937574 |
| SNRK | 91.700839 | 0.09002274 | 0.33539003 | 0.26841208 | 0.78838214 | 0.99937574 |
| ANO10 | 132.419777 | -0.00822 | 0.40127361 | -0.0204848 | 0.98365662 | 0.99937574 |
| ABHD5 | 32.8726571 | -0.5501194 | 0.49654363 | -1.1078974 | 0.26790613 | 0.99937574 |
| TCAIM | 102.786225 | 0.43387688 | 0.31138856 | 1.39336164 | 0.16351046 | 0.99937574 |
| ZNF445 | 39.889596 | 0.47785955 | 0.40056712 | 1.19295752 | 0.23288601 | 0.99937574 |
| AC099669.1 | 53.0165536 | 0.77187284 | 0.5107372 | 1.51129161 | 0.13071417 | 0.99937574 |
| ZNF197 | 36.2571127 | 0.49696884 | 0.36825262 | 1.34953239 | 0.17716602 | 0.99937574 |
| KIAA1143 | 21.9291421 | -0.1893257 | 0.54872086 | -0.3450309 | 0.73007112 | 0.99937574 |
| ZDHHC3 | 278.24964 | 0.58309197 | 0.28349707 | 2.05678304 | 0.0397071 | 0.99937574 |
| EXOSC7 | 203.74562 | 0.45399602 | 0.45055649 | 1.00763396 | 0.31363022 | 0.99937574 |
| CLEC3B | 12.1577343 | 1.18748924 | 1.04807633 | 1.13301789 | 0.25720674 | 0.99937574 |
| CDCP1 | 68.6321449 | 0.26337042 | 0.61416148 | 0.42882926 | 0.66804748 | 0.99937574 |
| LARS2 | 69.8385381 | 0.36228288 | 0.37703197 | 0.96088104 | 0.33661198 | 0.99937574 |
| LIMD1 | 50.1324862 | -0.2154247 | 0.47810006 | -0.4505849 | 0.65228873 | 0.99937574 |
| SACM1L | 119.024477 | 0.25333757 | 0.3338717 | 0.75878719 | 0.44797987 | 0.99937574 |
| LZTFL1 | 20.7292311 | 0.61531404 | 0.56114461 | 1.09653381 | 0.27284523 | 0.99937574 |
| FYCO1 | 71.3760041 | 0.20628918 | 0.43048557 | 0.47920116 | 0.63179553 | 0.99937574 |
| LTF | 22.4861723 | 0.8468686 | 1.09240687 | 0.77523185 | 4.38E-01 | 0.99937574 |
| LRRC2 | 13.8813896 | 0.67213204 | 0.67841901 | 0.99073291 | 0.32181602 | 0.99937574 |
| ALS2CL | 70.6007835 | 0.3344481 | 0.53817996 | 0.62144287 | 0.53430828 | 0.99937574 |
| CCDC12 | 97.0363021 | -0.2155855 | 0.27019618 | -0.797885 | 0.42493724 | 0.99937574 |
| NBEAL2 | 45.2263332 | -0.9245591 | 0.50615295 | -1.8266398 | 0.06775395 | 0.99937574 |
| SETD2 | 372.430907 | -0.0234539 | 0.23101843 | -0.1015239 | 0.91913456 | 0.99937574 |
| KIF9-AS1 | 155.136237 | 0.36106274 | 0.38913555 | 0.92785852 | 0.35348096 | 0.99937574 |
| KIF9 | 11.3656515 | 0.13241784 | 0.51634816 | 0.25645069 | 0.79760285 | 0.99937574 |
| KLHL18 | 52.4391008 | -0.3817702 | 0.47285437 | -0.8073736 | 0.41945125 | 0.99937574 |
| PTPN23 | 30.0802258 | -0.197858 | 0.52292804 | -0.3783656 | 0.705159 | 0.99937574 |
| SCAP | 70.3598122 | 0.28120496 | 0.29146811 | 0.96478808 | 0.33465097 | 0.99937574 |
| ELP6 | 24.6489778 | 0.03083161 | 0.45155892 | 0.06827816 | 0.94556421 | 0.99937574 |
| SMARCC1 | 244.736547 | -0.2100396 | 0.28195312 | -0.7449452 | 0.45630482 | 0.99937574 |
| DHX30 | 50.1809766 | 0.07350771 | 0.31475497 | 0.23353948 | 0.81534251 | 0.99937574 |
| MAP4 | 362.142175 | -0.078061 | 0.26387898 | -0.2958214 | 0.7673665 | 0.99937574 |
| ZNF589 | 12.5425134 | -0.5888302 | 0.62222916 | -0.9463237 | 0.34398353 | 0.99937574 |
| NME6 | 16.6997031 | -0.6923821 | 0.48669204 | -1.4226288 | 0.1548438 | 0.99937574 |
| PLXNB1 | 55.9105402 | -0.6707798 | 0.30741935 | -2.1819701 | 0.02911174 | 0.99049468 |
| TMA7 | 82.7886614 | 0.57192832 | 0.53159499 | 1.07587228 | 0.28198438 | 0.99937574 |
| SHISA5 | 57.1336208 | -0.3579713 | 0.47258134 | -0.7574809 | 0.44876181 | 0.99937574 |

| | | | | | | |
|---|---|---|---|---|---|---|
| PFKFB4 | 17.2544483 | -1.5340379 | 0.95069544 | -1.6135955 | 0.10661518 | 0.99937574 |
| UQCRC1 | 83.9271467 | 0.45670129 | 0.40305641 | 1.13309524 | 0.25717426 | 0.99937574 |
| IP6K2 | 67.7429342 | 0.01389721 | 0.28773138 | 0.04829924 | 0.96147776 | 0.99937574 |
| PRKAR2A | 115.942027 | -0.1967537 | 0.37786267 | -0.5207015 | 0.60257472 | 0.99937574 |
| SLC25A20 | 15.958315 | 0.36647185 | 0.54374745 | 0.67397439 | 0.50032759 | 0.99937574 |
| ARIH2 | 133.64606 | -0.1688473 | 0.24352837 | -0.6933371 | 0.48809802 | 0.99937574 |
| P4HTM | 31.4906239 | 0.98544553 | 0.39340016 | 2.50494441 | 0.01224706 | 0.77519564 |
| WDR6 | 58.4639012 | -0.2293762 | 0.48110946 | -0.4767652 | 0.63352934 | 0.99937574 |
| DALRD3 | 13.9139235 | 0.60130939 | 0.65200439 | 0.92224745 | 0.35639952 | 0.99937574 |
| NDUFAF3 | 29.1436735 | -0.1037707 | 0.59711535 | -0.1737867 | 0.86203311 | 0.99937574 |
| IMPDH2 | 19.3470459 | 0.38981406 | 0.51696652 | 0.75404122 | 0.45082447 | 0.99937574 |
| QRICH1 | 64.2489658 | -0.2489336 | 0.2953476 | -0.8428497 | 0.39931252 | 0.99937574 |
| QARS | 27.6963237 | 0.45331376 | 0.3935498 | 1.1518587 | 0.24937914 | 0.99937574 |
| USP19 | 13.0669238 | -0.9014243 | 0.57801124 | -1.5595273 | 0.11887163 | 0.99937574 |
| LAMB2 | 27.4176655 | 0.14007262 | 0.38706861 | 0.36188059 | 0.71744126 | 0.99937574 |
| C3orf62 | 11.7030281 | -0.5123731 | 0.6627807 | -0.7730658 | 0.43948341 | 0.99937574 |
| USP4 | 82.8246935 | -0.1081946 | 0.3936497 | -0.2748501 | 0.78343144 | 0.99937574 |
| RHOA | 178.13228 | 0.04678515 | 0.31494779 | 0.14854891 | 0.88190959 | 0.99937574 |
| TCTA | 16.2301608 | 1.27253255 | 0.59373113 | 2.14328084 | 0.03209055 | 0.99937574 |
| DAG1 | 71.3647185 | 0.26020637 | 0.32663406 | 0.79662962 | 0.42566617 | 0.99937574 |
| APEH | 37.4975758 | 0.66452067 | 0.5103384 | 1.30211771 | 0.19287615 | 0.99937574 |
| MST1 | 14.5414148 | -0.1686702 | 0.57018157 | -0.2958184 | 0.76736879 | 0.99937574 |
| RNF123 | 19.4521507 | -0.0598833 | 0.42143692 | -0.1420931 | 0.88700647 | 0.99937574 |
| IP6K1 | 103.850058 | -0.2536412 | 0.32236901 | -0.7868039 | 0.43139664 | 0.99937574 |
| RBM6 | 420.369082 | -0.1150765 | 0.24392712 | -0.4717659 | 0.63709391 | 0.99937574 |
| RBM5 | 148.951582 | 0.0667229 | 0.21925329 | 0.30431881 | 0.76088502 | 0.99937574 |
| SEMA3F | 63.3379298 | 0.095914 | 0.64372385 | 0.14899868 | 0.88155468 | 0.99937574 |
| GNAI2 | 47.5291611 | 0.00956406 | 0.43322371 | 0.0220765 | 0.98238693 | 0.99937574 |
| SEMA3B | 20.825429 | 0.50873165 | 0.53224377 | 0.95582452 | 0.3391609 | 0.99937574 |
| IFRD2 | 15.6071519 | -0.128825 | 0.5508157 | -0.2338804 | 0.81507782 | 0.99937574 |
| HYAL1 | 22.2272764 | 1.29992517 | 0.50729819 | 2.5624479 | 0.01039372 | 0.76274616 |
| HYAL2 | 16.6343433 | 0.61017298 | 0.5721886 | 1.06638437 | 0.28624992 | 0.99937574 |
| TUSC2 | 21.3151044 | 0.42309679 | 0.47042618 | 0.89939039 | 0.36844476 | 0.99937574 |
| TMEM115 | 12.8161095 | -0.2699448 | 0.5153022 | -0.5238572 | 0.60037786 | 0.99937574 |
| CACNA2D2 | 20.5172042 | 1.04149529 | 0.98725009 | 1.05494575 | 0.29145014 | 0.99937574 |
| HEMK1 | 16.5249551 | -0.4205157 | 0.52866324 | -0.7954321 | 0.4263622 | 0.99937574 |
| CISH | 37.4023336 | 0.06365642 | 0.40600217 | 0.15678836 | 0.87541164 | 0.99937574 |
| MAPKAPK3 | 35.6963518 | -0.0947576 | 0.35136323 | -0.2696856 | 0.78740211 | 0.99937574 |
| DOCK3 | 19.6298827 | 0.08918349 | 0.93269789 | 0.09561884 | 0.9238233 | 0.99937574 |
| MANF | 41.4653747 | -0.054203 | 0.41898307 | -0.1293681 | 0.89706641 | 0.99937574 |
| RBM15B | 16.1728697 | -0.1842263 | 0.69657358 | -0.264475 | 0.79141394 | 0.99937574 |
| DCAF1 | 74.8546973 | -0.2559393 | 0.35096951 | -0.7292351 | 0.46585789 | 0.99937574 |
| RAD54L2 | 137.879135 | 0.26004265 | 0.37603194 | 0.69154405 | 0.48922371 | 0.99937574 |
| TEX264 | 66.6900554 | 0.57024268 | 0.41683252 | 1.3680379 | 0.17130021 | 0.99937574 |
| RPL29 | 287.770548 | 0.01768097 | 0.54977667 | 0.03216028 | 0.97434423 | 0.99937574 |
| ALAS1 | 25.2582139 | 0.70169879 | 0.42582383 | 1.64786172 | 0.09938105 | 0.99937574 |
| WDR82 | 66.4560655 | -0.0626522 | 0.30006008 | -0.2087987 | 0.83460538 | 0.99937574 |
| DNAH1 | 12.6613384 | -0.1678699 | 0.52442333 | -0.3201037 | 0.7488897 | 0.99937574 |
| BAP1 | 22.7078916 | -0.22857 | 0.56424587 | -0.4050893 | 0.68541184 | 0.99937574 |
| PHF7 | 13.4807043 | 0.10350145 | 0.57909364 | 0.17873008 | 0.85814965 | 0.99937574 |
| NISCH | 53.7333227 | 0.05966859 | 0.3278161 | 0.1820185 | 0.85556821 | 0.99937574 |
| SMIM4 | 33.9252082 | 0.46954516 | 0.43436665 | 1.08098804 | 0.27970243 | 0.99937574 |
| PBRM1 | 181.583095 | -0.331402 | 0.27594381 | -1.2009762 | 0.22976043 | 0.99937574 |
| GNL3 | 27.3800329 | -0.2609454 | 0.51843581 | -0.5033322 | 0.61473074 | 0.99937574 |

| | | | | | | |
|---|---|---|---|---|---|---|
| GLT8D1 | 19.0621216 | 0.60326198 | 0.44675078 | 1.35033223 | 0.17690944 | 0.99937574 |
| SPCS1 | 94.6022757 | 0.51128692 | 0.51874989 | 0.98561353 | 0.32432279 | 0.99937574 |
| NEK4 | 40.6761663 | 0.03638409 | 0.41954699 | 0.08672233 | 0.93089223 | 0.99937574 |
| SFMBT1 | 44.3469821 | 0.00053325 | 0.31604788 | 0.00168723 | 0.99865378 | 0.99979721 |
| PRKCD | 68.3980243 | 0.48693661 | 0.50963528 | 0.95546095 | 0.33934464 | 0.99937574 |
| TKT | 87.25246 | 0.1493458 | 0.33383386 | 0.44736563 | 0.65461109 | 0.99937574 |
| DCP1A | 93.6552059 | -0.1322677 | 0.26539945 | -0.498372 | 0.61822183 | 0.99937574 |
| CACNA1D | 390.422863 | 0.56413722 | 0.67956265 | 0.83014748 | 0.40645541 | 0.99937574 |
| CHDH | 38.0292816 | -0.3144172 | 0.5458839 | -0.5759782 | 0.56462993 | 0.99937574 |
| ACTR8 | 13.5831502 | 0.80480965 | 0.53071508 | 1.51646274 | 0.12940238 | 0.99937574 |
| SELENOK | 35.1265095 | 0.01892652 | 0.40797792 | 0.04639104 | 0.96299858 | 0.99937574 |
| CACNA2D3 | 18.3002045 | -1.2021081 | 0.67219305 | -1.7883376 | 0.07372156 | 0.99937574 |
| ERC2 | 194.8531 | 0.66460809 | 0.46191118 | 1.43882226 | 0.15020089 | 0.99937574 |
| CCDC66 | 102.217412 | 0.3117295 | 0.3112712 | 1.00147237 | 0.31659849 | 0.99937574 |
| FAM208A | 111.216806 | 0.20848192 | 0.26777969 | 0.77855762 | 0.43624035 | 0.99937574 |
| ARHGEF3 | 396.634292 | -0.1301513 | 0.42909199 | -0.3033179 | 0.7616476 | 0.99937574 |
| APPL1 | 64.1469713 | -0.176391 | 0.2990935 | -0.589752 | 0.55535695 | 0.99937574 |
| PDE12 | 28.506945 | 0.03996406 | 0.40604008 | 0.09842394 | 0.92159567 | 0.99937574 |
| ARF4 | 68.9523292 | 0.11910262 | 0.44908431 | 0.26521216 | 0.79084603 | 0.99937574 |
| DENND6A | 72.6091835 | -0.2180128 | 0.32526929 | -0.6702534 | 0.50269627 | 0.99937574 |
| DENND6A-A! | 17.6951257 | 0.12880239 | 0.54171113 | 0.23776952 | 0.81205986 | 0.99937574 |
| SLMAP | 226.634895 | -0.1298137 | 0.22784341 | -0.5697496 | 0.56884756 | 0.99937574 |
| FLNB | 361.305353 | 0.22466498 | 0.30360088 | 0.74000107 | 0.45929935 | 0.99937574 |
| ABHD6 | 38.418873 | 0.79937205 | 0.44722286 | 1.78741319 | 0.07387073 | 0.99937574 |
| RPP14 | 13.4682302 | -0.3030109 | 0.5166805 | -0.586457 | 0.55756842 | 0.99937574 |
| PXK | 315.309949 | -0.0113378 | 0.37588847 | -0.0301626 | 0.9759374 | 0.99937574 |
| PDHB | 51.6117461 | 0.57470162 | 0.38692157 | 1.48531811 | 0.13745958 | 0.99937574 |
| KCTD6 | 17.7981614 | 0.9321332 | 0.52480117 | 1.77616447 | 0.07570581 | 0.99937574 |
| FHIT | 1353.80637 | 0.49952703 | 0.29481087 | 1.69439826 | 0.09018963 | 0.99937574 |
| PTPRG | 1055.67056 | 0.65761582 | 0.78614791 | 0.83650393 | 0.40287147 | 0.99937574 |
| THOC7 | 33.2137246 | 0.37584017 | 0.34117595 | 1.10160215 | 0.27063467 | 0.99937574 |
| ATXN7 | 23.9847341 | -0.0134822 | 0.43560998 | -0.0309503 | 0.9753092 | 0.99937574 |
| PSMD6-AS2 | 14.2064105 | 0.86466909 | 0.68798619 | 1.2568117 | 0.20882183 | 0.99937574 |
| PSMD6 | 22.7503113 | 0.14437228 | 0.51869348 | 0.27833832 | 0.78075266 | 0.99937574 |
| ADAMTS9-A! | 51.2408371 | -0.5547671 | 0.60993404 | -0.9095526 | 0.36305848 | 0.99937574 |
| MAGI1 | 9871.07419 | -0.0655633 | 0.34013198 | -0.1927583 | 0.84714824 | 0.99937574 |
| MAGI1-AS1 | 175.233841 | -0.0116076 | 0.72899148 | -0.0159228 | 0.98729601 | 0.99937574 |
| SLC25A26 | 70.0400877 | -0.6571487 | 0.30458317 | -2.1575344 | 0.03096405 | 0.99049468 |
| LRIG1 | 22.6168957 | 0.76288454 | 0.84731422 | 0.90035611 | 0.36793077 | 0.99937574 |
| SUCLG2 | 238.425999 | -0.0257854 | 0.30344995 | -0.0849743 | 0.93228182 | 0.99937574 |
| SUCLG2-AS1 | 74.292053 | 0.46548298 | 0.39727804 | 1.17168062 | 0.2413253 | 0.99937574 |
| FAM19A1 | 18.2872449 | 1.14829134 | 0.63739119 | 1.80154882 | 0.07161642 | 0.99937574 |
| EOGT | 22.784742 | -0.1540805 | 0.51266002 | -0.300551 | 0.76375689 | 0.99937574 |
| TMF1 | 85.2621799 | -0.1829438 | 0.32432918 | -0.5640682 | 0.5727077 | 0.99937574 |
| UBA3 | 84.4395228 | 0.40980468 | 0.30510906 | 1.34314164 | 0.17922611 | 0.99937574 |
| ARL6IP5 | 102.180524 | 0.04942468 | 0.4146377 | 0.11919967 | 0.90511717 | 0.99937574 |
| FRMD4B | 16.7201331 | -0.5738055 | 0.56093397 | -1.0229465 | 0.30633313 | 0.99937574 |
| MITF | 338.770354 | 0.43870536 | 0.24944968 | 1.75869284 | 0.07862969 | 0.99937574 |
| FOXP1 | 1878.42348 | 0.04047777 | 0.31191901 | 0.12977012 | 0.8967483 | 0.99937574 |
| EIF4E3 | 19.7052615 | -0.3709035 | 0.48900525 | -0.7584858 | 0.4481602 | 0.99937574 |
| RYBP | 109.87254 | -0.0086247 | 0.2795501 | -0.0308519 | 0.97538762 | 0.99937574 |
| SHQ1 | 45.3136595 | -0.5744749 | 0.37188315 | -1.5447727 | 0.12240125 | 0.99937574 |
| GXYLT2 | 23.8269586 | -0.8183168 | 0.64169638 | -1.2752399 | 0.20222434 | 0.99937574 |
| PPP4R2 | 125.281965 | 0.26608957 | 0.27547578 | 0.96592726 | 0.33408058 | 0.99937574 |

| | | | | | | |
|---|---|---|---|---|---|---|
| PDZRN3 | 35.3316284 | -1.1323826 | 0.81036239 | -1.397378 | 0.16229994 | 0.99937574 |
| CNTN3 | 79.6928348 | 0.36202939 | 0.34547667 | 1.04791269 | 0.29467884 | 0.99937574 |
| ZNF717 | 50.0409278 | 0.19322475 | 0.42522423 | 0.45440674 | 0.6495361 | 0.99937574 |
| GBE1 | 107.991641 | -1.4094126 | 0.44034626 | -3.2006916 | 0.00137098 | 0.34724223 |
| CHMP2B | 47.7674533 | -0.0114871 | 0.44056646 | -0.0260735 | 0.97919871 | 0.99937574 |
| CGGBP1 | 126.757361 | -0.0780453 | 0.32996828 | -0.2365236 | 0.81302641 | 0.99937574 |
| ZNF654 | 126.222272 | -0.1584593 | 0.31368564 | -0.5051531 | 0.61345134 | 0.99937574 |
| C3orf38 | 23.91381 | -0.0769242 | 0.39189948 | -0.1962855 | 0.84438671 | 0.99937574 |
| ARL13B | 36.3696388 | -0.331895 | 0.35626988 | -0.9315832 | 0.35155199 | 0.99937574 |
| NSUN3 | 48.1891315 | -0.3702301 | 0.30867839 | -1.1994041 | 0.23037086 | 0.99937574 |
| MTRNR2L12 | 317.465453 | -0.3442099 | 0.58462311 | -0.5887723 | 0.55601406 | 0.99937574 |
| ARL6 | 87.6494661 | -0.0130514 | 0.37133445 | -0.0351473 | 0.97196227 | 0.99937574 |
| CRYBG3 | 399.200316 | -0.2741386 | 0.4029704 | -0.6802947 | 0.4963179 | 0.99937574 |
| AC110491.2 | 13.1335178 | -0.3226798 | 0.94153691 | -0.3427161 | 0.73181208 | 0.99937574 |
| RIOX2 | 86.7869359 | -0.1335072 | 0.51001757 | -0.2617699 | 0.79349886 | 0.99937574 |
| CLDND1 | 29.2528774 | -0.6956977 | 0.44796748 | -1.5530093 | 0.1204209 | 0.99937574 |
| ST3GAL6 | 333.248007 | 0.49977726 | 0.47789896 | 1.04578017 | 0.29566254 | 0.99937574 |
| DCBLD2 | 56.2400085 | 0.23722198 | 0.3126116 | 0.75883932 | 0.44794868 | 0.99937574 |
| CMSS1 | 139.144215 | -0.0149297 | 0.35810261 | -0.0416912 | 0.96674484 | 0.99937574 |
| FILIP1L | 35.2426838 | 0.27323664 | 0.56709719 | 0.48181625 | 0.62993648 | 0.99937574 |
| TBC1D23 | 69.0272979 | 0.0630265 | 0.34552891 | 0.18240586 | 0.85526422 | 0.99937574 |
| NIT2 | 18.094055 | -0.390165 | 0.46077989 | -0.8467491 | 0.39713496 | 0.99937574 |
| TOMM70 | 47.9747591 | 0.16858761 | 0.29412386 | 0.57318578 | 0.56651891 | 0.99937574 |
| TFG | 136.285821 | 0.13947961 | 0.3044068 | 0.45820135 | 0.64680779 | 0.99937574 |
| SENP7 | 121.39304 | 0.04300831 | 0.38897822 | 0.11056741 | 0.9119594 | 0.99937574 |
| TRMT10C | 25.8469485 | 0.30885609 | 0.59967908 | 0.51503564 | 0.60652812 | 0.99937574 |
| PCNP | 104.238898 | -0.5018679 | 0.25395789 | -1.9761855 | 0.04813377 | 0.99937574 |
| ZBTB11 | 46.1497197 | -0.2370485 | 0.38684063 | -0.6127808 | 0.54002131 | 0.99937574 |
| RPL24 | 138.361794 | -0.0458121 | 0.51200717 | -0.0894755 | 0.92870405 | 0.99937574 |
| CEP97 | 20.2862002 | -0.2310309 | 0.60712204 | -0.3805345 | 0.70354866 | 0.99937574 |
| NFKBIZ | 31.3455995 | -0.1275261 | 0.69701723 | -0.1829597 | 0.85482962 | 0.99937574 |
| ZPLD1 | 80.3561597 | -0.6378458 | 0.7598076 | -0.8394833 | 4.01E-01 | 9.99E-01 |
| ALCAM | 38.1509306 | -0.0897958 | 0.84603473 | -0.1061373 | 0.91547346 | 0.99937574 |
| CBLB | 143.685951 | 0.41232527 | 0.34986793 | 1.1785169 | 0.2385906 | 0.99937574 |
| BBX | 414.980437 | 0.28443935 | 0.32994875 | 0.86207132 | 0.38864827 | 0.99937574 |
| CD47 | 95.9949234 | -0.217045 | 0.41615392 | -0.5215497 | 0.60198386 | 0.99937574 |
| IFT57 | 48.5647115 | 0.49582829 | 0.51506381 | 0.96265412 | 0.33572113 | 0.99937574 |
| DZIP3 | 73.0911513 | 0.34561485 | 0.65439029 | 0.52814788 | 0.5973967 | 0.99937574 |
| NECTIN3 | 36.1436108 | 0.46371482 | 0.39611053 | 1.17067027 | 0.24173134 | 0.99937574 |
| PHLDB2 | 393.040178 | -0.3134027 | 0.2895597 | -1.0823421 | 0.27910053 | 0.99937574 |
| ABHD10 | 37.284201 | 0.67871517 | 0.39721318 | 1.70869247 | 0.08750793 | 0.99937574 |
| C3orf52 | 31.7013465 | -1.3486378 | 0.71699545 | -1.8809573 | 0.05997773 | 0.99937574 |
| SLC9C1 | 44.7377831 | -0.5179885 | 0.9344603 | -0.5543184 | 0.57936097 | 0.99937574 |
| CD200 | 27.0000434 | 1.29931414 | 0.96458734 | 1.34701555 | 0.17797522 | 0.99937574 |
| AC092894.1 | 69.5716593 | 0.16278713 | 0.59045022 | 0.27570001 | 0.78277849 | 0.99937574 |
| ATG3 | 35.7541707 | -0.3759068 | 0.40854038 | -0.9201214 | 0.35750931 | 0.99937574 |
| SLC35A5 | 24.6613409 | -0.4067364 | 0.46881449 | -0.8675849 | 0.38562163 | 0.99937574 |
| GTPBP8 | 21.9174477 | -0.4498297 | 0.50194892 | -0.8961662 | 0.370164 | 0.99937574 |
| NEPRO | 38.9517375 | 0.14889729 | 0.35124562 | 0.42391217 | 0.67162987 | 0.99937574 |
| CFAP44 | 35.7397665 | 0.27319549 | 0.3849626 | 0.70966763 | 0.47791027 | 0.99937574 |
| SPICE1 | 42.4138201 | -0.2680948 | 0.36633667 | -0.7318262 | 0.46427468 | 0.99937574 |
| SIDT1 | 14.7293892 | -0.9723529 | 0.98699995 | -0.98516 | 0.32454546 | 0.99937574 |
| USF3 | 80.1744054 | 0.03292081 | 0.37808971 | 0.08707142 | 0.93061474 | 0.99937574 |
| NAA50 | 142.214257 | -0.3114893 | 0.27392505 | -1.1371332 | 0.25548259 | 0.99937574 |

| | | | | | | |
|---|---|---|---|---|---|---|
| ATP6V1A | 889.306067 | 0.12829824 | 0.34045929 | 0.37683871 | 0.70629347 | 0.99937574 |
| GRAMD1C | 220.721783 | 0.0919983 | 0.30215432 | 0.30447455 | 0.76076638 | 0.99937574 |
| ZDHHC23 | 35.6600293 | -0.3948527 | 0.38262771 | -1.0319502 | 0.30209546 | 0.99937574 |
| CCDC191 | 42.7557422 | -0.1793224 | 0.50534821 | -0.3548492 | 0.72270253 | 0.99937574 |
| QTRT2 | 29.4202039 | -0.6471153 | 0.37742317 | -1.7145617 | 0.08642561 | 0.99937574 |
| ZBTB20 | 4900.96819 | 0.15124364 | 0.36411849 | 0.4153693 | 0.67787158 | 0.99937574 |
| ZBTB20-AS1 | 24.0149501 | 0.23783003 | 0.83502964 | 0.28481627 | 0.77578491 | 0.99937574 |
| ZBTB20-AS5 | 102.96786 | -0.0747858 | 0.8121861 | -0.0920797 | 0.92663472 | 0.99937574 |
| ZBTB20-AS2 | 30.6284389 | -0.407932 | 0.82583405 | -0.4939637 | 0.62133185 | 0.99937574 |
| LSAMP | 20.38397 | 0.17418519 | 0.54472131 | 0.31976937 | 0.74914317 | 0.99937574 |
| B4GALT4 | 20.4459059 | 0.18138727 | 0.46621968 | 0.38905967 | 0.69723201 | 0.99937574 |
| ARHGAP31 | 21.9494162 | -0.0597075 | 0.68880089 | -0.0866832 | 0.93092331 | 0.99937574 |
| TMEM39A | 27.2961357 | -0.2597811 | 0.40594768 | -0.6399373 | 0.52221334 | 0.99937574 |
| POGLUT1 | 15.1264471 | 0.88955224 | 0.54250051 | 1.6397261 | 0.10106213 | 0.99937574 |
| TIMMDC1 | 48.6670858 | 0.40946626 | 0.36492111 | 1.12206788 | 0.26183358 | 0.99937574 |
| COX17 | 40.9280254 | 0.36244716 | 0.41875398 | 0.86553723 | 0.38674399 | 0.99937574 |
| GSK3B | 295.848522 | 0.01644453 | 0.26176367 | 0.06282204 | 0.94990821 | 0.99937574 |
| LRRC58 | 36.1362213 | -0.4414968 | 0.53605322 | -0.8236064 | 0.41016325 | 0.99937574 |
| NDUFB4 | 62.769505 | 0.36332076 | 0.48575544 | 0.74794997 | 0.45449034 | 0.99937574 |
| HGD | 198.977268 | 0.38226322 | 0.70261684 | 0.54405645 | 0.58640263 | 0.99937574 |
| RABL3 | 34.8393124 | -0.391873 | 0.38430708 | -1.0196873 | 0.3078768 | 0.99937574 |
| GTF2E1 | 11.8931286 | 0.28140458 | 0.54381004 | 0.51746852 | 0.60482914 | 0.99937574 |
| GOLGB1 | 218.757235 | -0.1236954 | 0.31880254 | -0.3880001 | 0.69801598 | 0.99937574 |
| IQCB1 | 80.2967413 | -0.1788651 | 0.28166339 | -0.6350315 | 0.52540786 | 0.99937574 |
| ILDR1 | 22.8518193 | 0.79835448 | 0.51512536 | 1.54982563 | 0.12118337 | 0.99937574 |
| CCDC58 | 24.1329714 | -0.3006349 | 0.48488278 | -0.6200157 | 0.53524748 | 0.99937574 |
| FAM162A | 151.605962 | -0.2706901 | 0.49150035 | -0.5507424 | 0.58181031 | 0.99937574 |
| AC083798.2 | 20.102142 | -0.060169 | 0.56118098 | -0.1072185 | 0.91461563 | 0.99937574 |
| KPNA1 | 121.971309 | -0.3354327 | 0.26321907 | -1.2743479 | 0.20254015 | 0.99937574 |
| PARP9 | 87.1063025 | -0.7177123 | 0.75163757 | -0.9548648 | 0.33964605 | 0.99937574 |
| DTX3L | 42.5495899 | -0.3302073 | 0.51601389 | -0.6399193 | 0.52222505 | 0.99937574 |
| PARP14 | 275.347037 | -1.8237171 | 0.72101619 | -2.5293706 | 0.01142673 | 0.76274616 |
| HSPBAP1 | 27.6590849 | -0.7652849 | 0.46846103 | -1.6336148 | 0.10233976 | 0.99937574 |
| DIRC2 | 59.9068338 | -1.0048485 | 0.42017832 | -2.3914812 | 0.01678054 | 0.89506255 |
| PDIA5 | 12.8123611 | -0.2060758 | 0.50908537 | -0.4047962 | 0.6856273 | 0.99937574 |
| SEC22A | 93.9446731 | 0.21120381 | 0.30320266 | 0.69657636 | 0.48606795 | 0.99937574 |
| ADCY5 | 114.460831 | 0.70014446 | 0.55472799 | 1.26214014 | 0.20689836 | 0.99937574 |
| HACD2 | 80.3215542 | 0.16307169 | 0.38362622 | 0.42507964 | 0.67077862 | 0.99937574 |
| MYLK-AS1 | 12.2087027 | 0.05681178 | 0.5831044 | 0.09742986 | 0.92238504 | 0.99937574 |
| MYLK | 34.0409346 | -0.8433046 | 0.63691853 | -1.3240383 | 0.18549031 | 0.99937574 |
| CCDC14 | 56.8700089 | -0.0066485 | 0.44901086 | -0.014807 | 0.98818619 | 0.99937574 |
| KALRN | 24.7651797 | -0.057019 | 0.59201746 | -0.096313 | 0.92327196 | 0.99937574 |
| UMPS | 22.5844396 | -0.6361239 | 0.49085078 | -1.2959619 | 0.19498862 | 0.99937574 |
| ITGB5 | 39.5082215 | -0.2569706 | 0.4563723 | -0.5630722 | 0.57338573 | 0.99937574 |
| ZNF148 | 260.335197 | 0.02194906 | 0.21850192 | 0.10045248 | 0.91998511 | 0.99937574 |
| SNX4 | 111.314069 | -0.0733088 | 0.26525956 | -0.2763663 | 0.78226678 | 0.99937574 |
| OSBPL11 | 36.860205 | -0.3815528 | 0.412911 | -0.9240558 | 0.35545726 | 0.99937574 |
| SLC41A3 | 20.1933319 | -0.4374902 | 0.43350484 | -1.0091934 | 0.31288189 | 0.99937574 |
| ALDH1L1 | 44.0770691 | 0.84795821 | 0.70843873 | 1.19693938 | 0.23133019 | 0.99937574 |
| ZXDC | 104.503327 | -0.3362623 | 0.28766519 | -1.1689364 | 0.24242926 | 0.99937574 |
| TXNRD3NB | 12.5136203 | 0.72167862 | 0.70594077 | 1.02229345 | 0.30664203 | 0.99937574 |
| TXNRD3 | 15.6405142 | 0.06879766 | 0.46731867 | 0.14721788 | 0.88296005 | 0.99937574 |
| CHCHD6 | 29.0521893 | -0.1397933 | 0.45152169 | -0.3096047 | 0.75686155 | 0.99937574 |
| TPRA1 | 18.9539851 | -0.5663898 | 0.44765604 | -1.2652343 | 0.20578737 | 0.99937574 |

| | | | | | | |
|---|---|---|---|---|---|---|
| ABTB1 | 13.1272279 | 0.10388663 | 0.67645413 | 0.15357528 | 0.87794463 | 0.99937574 |
| MGLL | 266.491849 | -0.827435 | 0.4086081 | -2.0250087 | 0.04286647 | 0.99937574 |
| KBTBD12 | 149.177452 | 0.19710277 | 0.37632051 | 0.52376304 | 0.60044336 | 0.99937574 |
| SEC61A1 | 72.0205859 | -0.115996 | 0.45058786 | -0.2574326 | 0.79684486 | 0.99937574 |
| RUVBL1 | 37.6400491 | 0.11445958 | 0.3969633 | 0.28833794 | 0.77308807 | 0.99937574 |
| EEFSEC | 71.4278563 | -0.1506968 | 0.36853138 | -0.4089118 | 0.68260437 | 0.99937574 |
| GATA2 | 55.7307526 | 0.01926085 | 0.65631964 | 0.02934676 | 0.97658804 | 0.99937574 |
| GATA2-AS1 | 48.1172292 | 0.3648202 | 0.4173783 | 0.87407562 | 0.38207707 | 0.99937574 |
| RPN1 | 69.8891532 | -0.0684041 | 0.37612848 | -0.1818637 | 0.85568969 | 0.99937574 |
| RAB7A | 204.278724 | -0.6751084 | 0.33166384 | -2.0355201 | 0.04179857 | 0.99937574 |
| ACAD9 | 34.2811653 | 0.04431538 | 0.36333273 | 0.12196915 | 0.90292345 | 0.99937574 |
| CNBP | 113.035016 | 0.10464737 | 0.49912132 | 0.2096632 | 0.83393055 | 0.99937574 |
| COPG1 | 40.1358486 | -0.032537 | 0.40944406 | -0.0794663 | 0.93666172 | 0.99937574 |
| HMCES | 30.7123003 | -0.1423228 | 0.37086495 | -0.3837591 | 0.70115699 | 0.99937574 |
| H1FX | 59.267748 | -0.6470514 | 0.77912792 | -0.8304816 | 0.40626655 | 0.99937574 |
| MBD4 | 64.1385982 | -0.3334288 | 0.33202524 | -1.0042273 | 0.31526907 | 0.99937574 |
| IFT122 | 98.564954 | 0.20939153 | 0.52804912 | 0.39653798 | 0.6917082 | 0.99937574 |
| TMCC1 | 445.388386 | 0.06025142 | 0.31405518 | 0.19184979 | 0.84785987 | 0.99937574 |
| PIK3R4 | 38.1506014 | -0.45304 | 0.32801086 | -1.3811738 | 0.16722554 | 0.99937574 |
| ATP2C1 | 102.712523 | 0.20712273 | 0.38781126 | 0.53408128 | 0.5932853 | 0.99937574 |
| NEK11 | 51.4196997 | -0.1440412 | 0.61657746 | -0.2336141 | 0.8152846 | 0.99937574 |
| NUDT16 | 38.8374293 | 0.54881055 | 0.38932074 | 1.40966174 | 0.15863959 | 0.99937574 |
| MRPL3 | 63.0405792 | 0.4005914 | 0.34143102 | 1.17327186 | 0.2406868 | 0.99937574 |
| DNAJC13 | 93.4963164 | -0.1162842 | 0.28361967 | -0.4100005 | 0.68180556 | 0.99937574 |
| UBA5 | 58.2717763 | 0.18922873 | 0.4022853 | 0.4703844 | 0.63808041 | 0.99937574 |
| CDV3 | 65.1814234 | -0.8748849 | 0.57774576 | -1.5143078 | 0.12994779 | 0.99937574 |
| TOPBP1 | 28.992432 | -0.1165495 | 0.36728574 | -0.3173264 | 0.75099597 | 0.99937574 |
| SRPRB | 19.7628935 | 0.19941649 | 0.45199628 | 0.44119056 | 0.65907505 | 0.99937574 |
| RYK | 163.026225 | 0.08110062 | 0.2861614 | 0.28340867 | 0.77686358 | 0.99937574 |
| AMOTL2 | 26.6760718 | -0.2357034 | 0.58990077 | -0.3995645 | 0.68947734 | 0.99937574 |
| ANAPC13 | 28.8866582 | 0.4803401 | 0.40557638 | 1.18433944 | 0.23627872 | 0.99937574 |
| CEP63 | 46.4477645 | 0.03573768 | 0.31901975 | 0.11202342 | 0.91080484 | 0.99937574 |
| PPP2R3A | 500.844864 | 0.14173535 | 0.38333483 | 0.36974295 | 0.71157402 | 0.99937574 |
| MSL2 | 61.7879185 | 0.20216205 | 0.36504313 | 0.55380318 | 0.57971355 | 0.99937574 |
| PCCB | 58.8884037 | 0.47733636 | 0.33093428 | 1.4423902 | 0.14919233 | 0.99937574 |
| STAG1 | 419.306805 | -0.5514201 | 0.33222308 | -1.6597886 | 0.09695698 | 0.99937574 |
| NCK1 | 43.6458123 | -0.3837394 | 0.46136281 | -0.831752 | 0.40554896 | 0.99937574 |
| DBR1 | 12.2136416 | -0.6168322 | 0.56856272 | -1.0848975 | 0.27796708 | 0.99937574 |
| ARMC8 | 114.952407 | 0.00011911 | 0.23703897 | 0.00050248 | 0.99959908 | 0.99979721 |
| NME9 | 14.8776798 | 0.4789086 | 0.7795063 | 0.61437425 | 0.53896805 | 0.99937574 |
| MRAS | 12.3898513 | -0.3445921 | 0.69874717 | -0.4931571 | 0.62190157 | 0.99937574 |
| CEP70 | 160.74565 | 0.42607428 | 0.42918966 | 0.99274125 | 0.32083607 | 0.99937574 |
| FAIM | 17.9024741 | -0.8437708 | 0.6280606 | -1.3434545 | 0.17912485 | 0.99937574 |
| PIK3CB | 105.145887 | 0.00882947 | 0.27383847 | 0.03224335 | 0.97427799 | 0.99937574 |
| MRPS22 | 15.8840206 | 0.24115847 | 0.52039668 | 0.46341277 | 0.64306853 | 0.99937574 |
| COPB2 | 78.0602367 | 0.01172483 | 0.36875199 | 0.03179599 | 0.97463475 | 0.99937574 |
| AC097103.2 | 63.2858618 | 0.36338766 | 0.5340338 | 0.68045816 | 0.49621441 | 0.99937574 |
| NMNAT3 | 34.1053356 | 0.15030081 | 0.38980967 | 0.38557486 | 0.69981156 | 0.99937574 |
| SLC25A36 | 194.714475 | -0.0458456 | 0.32750133 | -0.1399859 | 0.88867115 | 0.99937574 |
| SPSB4 | 34.5653916 | -0.6795304 | 1.10680959 | -0.6139542 | 0.53924558 | 0.99937574 |
| PXYLP1 | 13.5998209 | -0.1762438 | 0.55782799 | -0.3159464 | 0.75204316 | 0.99937574 |
| ZBTB38 | 107.257749 | -0.3394604 | 0.2647951 | -1.2819738 | 0.19985185 | 0.99937574 |
| RASA2 | 154.277608 | -0.3843878 | 0.26176211 | -1.4684625 | 0.14197864 | 0.99937574 |
| RNF7 | 47.8662867 | -0.0110532 | 0.41457644 | -0.0266615 | 0.97872972 | 0.99937574 |

| | | | | | | |
|---|---|---|---|---|---|---|
| ATP1B3 | 77.7523687 | -1.0284372 | 0.6092732 | -1.6879737 | 0.09141627 | 0.99937574 |
| TFDP2 | 135.830626 | 0.28431593 | 0.30657757 | 0.92738659 | 0.35372585 | 0.99937574 |
| GK5 | 111.248978 | 0.03578569 | 0.27952025 | 0.12802538 | 0.89812888 | 0.99937574 |
| XRN1 | 244.195694 | -0.4137028 | 0.32405227 | -1.2766545 | 0.20172425 | 0.99937574 |
| ATR | 169.514895 | 0.05258584 | 0.28985581 | 0.18142068 | 0.85603739 | 0.99937574 |
| PLS1 | 45.2805438 | -0.1339709 | 0.48656061 | -0.2753428 | 0.78305292 | 0.99937574 |
| TRPC1 | 60.5968838 | -0.6150342 | 0.71388471 | -0.8615315 | 0.38894538 | 0.99937574 |
| PCOLCE2 | 17.5947042 | -1.1277586 | 0.69253529 | -1.6284493 | 0.10342964 | 0.99937574 |
| U2SURP | 174.031066 | 0.03342366 | 0.25311029 | 0.13205178 | 0.89494334 | 0.99937574 |
| CHST2 | 29.0264702 | -0.8698188 | 0.68443135 | -1.2708634 | 0.20377723 | 0.99937574 |
| SLC9A9 | 19.2842653 | -0.584303 | 0.65240316 | -0.8956165 | 0.37045763 | 0.99937574 |
| C3orf58 | 19.5674383 | 0.11722669 | 0.49385118 | 0.2373725 | 0.81236782 | 0.99937574 |
| AC107021.1 | 52.1360481 | -0.6563811 | 0.94172794 | -0.6969965 | 0.48580497 | 0.99937574 |
| LNCSRLR | 14.2153237 | -0.7996995 | 0.77390643 | -1.0333284 | 0.30145026 | 0.99937574 |
| PLOD2 | 384.895949 | -0.5109484 | 0.45179079 | -1.1309403 | 0.25808023 | 0.99937574 |
| PLSCR4 | 15.7474541 | 0.49615492 | 0.62904749 | 0.78874 | 0.43026398 | 0.99937574 |
| PLSCR2 | 15.770199 | 0.23023697 | 0.66032156 | 0.34867401 | 0.72733406 | 0.99937574 |
| PLSCR1 | 216.105065 | -0.80513 | 0.52510155 | -1.5332844 | 0.1252058 | 0.99937574 |
| AGTR1 | 26.2850121 | -7.0469528 | 2.7356958 | -2.575927 | 0.00999717 | 0.76274616 |
| GYG1 | 51.1516688 | -0.004401 | 0.50103182 | -0.0087839 | 0.99299154 | 0.99937574 |
| HLTF | 42.8989039 | 0.58162294 | 0.47411278 | 1.22676071 | 0.21991254 | 0.99937574 |
| HPS3 | 38.7811576 | -0.0737567 | 0.32703016 | -0.2255349 | 0.82156315 | 0.99937574 |
| WWTR1 | 224.32924 | -0.6131418 | 0.35332349 | -1.7353553 | 0.08267789 | 0.99937574 |
| COMMD2 | 28.5662401 | 0.00291959 | 0.4673768 | 0.00624677 | 0.99501583 | 0.99937574 |
| ANKUB1 | 85.7596727 | 0.54831082 | 0.81938136 | 0.66917658 | 0.50338284 | 0.99937574 |
| RNF13 | 142.422329 | 0.07504074 | 0.26878774 | 0.27918213 | 0.78010505 | 0.99937574 |
| PFN2 | 15.7421977 | 0.47105128 | 0.64405109 | 0.73138806 | 0.46454216 | 0.99937574 |
| AC117386.2 | 53.9601446 | 2.79234426 | 0.78524328 | 3.55602439 | 0.00037651 | 0.22351354 |
| TSC22D2 | 260.759841 | 0.22801144 | 0.36623268 | 0.62258626 | 0.53355644 | 0.99937574 |
| SERP1 | 112.019805 | 0.16604218 | 0.38944788 | 0.42635276 | 0.66985082 | 0.99937574 |
| EIF2A | 48.5903264 | -0.4808938 | 0.36811051 | -1.3063844 | 0.19142188 | 0.99937574 |
| SELENOT | 97.9619339 | 0.0971876 | 0.23450949 | 0.41442927 | 0.67855976 | 0.99937574 |
| SIAH2 | 25.7223222 | -0.3310544 | 0.57198527 | -0.5787813 | 0.56273673 | 0.99937574 |
| MBNL1 | 627.769484 | 0.02977375 | 0.31017659 | 0.09598967 | 0.92352878 | 0.99937574 |
| RAP2B | 28.5289967 | -0.8711035 | 0.81272605 | -1.0718292 | 0.28379677 | 0.99937574 |
| DHX36 | 152.334179 | -0.1130122 | 0.29618252 | -0.3815628 | 0.70278566 | 0.99937574 |
| PLCH1 | 164.40783 | 0.36223257 | 0.6308258 | 0.57421965 | 0.56581918 | 0.99937574 |
| GMPS | 55.9858754 | 0.00283212 | 0.32531439 | 0.0087058 | 0.99305386 | 0.99937574 |
| SSR3 | 46.1444815 | -0.0518086 | 0.41614466 | -0.1244966 | 0.90092207 | 0.99937574 |
| TIPARP | 15.9631524 | 0.24575258 | 0.44193388 | 0.55608449 | 0.57815309 | 0.99937574 |
| LINC00886 | 79.2778134 | 0.46063755 | 0.47681029 | 0.96608139 | 0.33400346 | 0.99937574 |
| LEKR1 | 20.9282174 | 0.09146535 | 0.83825793 | 0.10911361 | 0.91311238 | 0.99937574 |
| CCNL1 | 132.959794 | -0.3943335 | 0.35334693 | -1.1159951 | 0.26442421 | 0.99937574 |
| VEPH1 | 13.7661851 | 0.31025792 | 0.59373299 | 0.52255462 | 0.60128423 | 0.99937574 |
| RSRC1 | 176.423433 | -0.2120257 | 0.2150028 | -0.9861534 | 0.32405785 | 0.99937574 |
| MLF1 | 13.1911706 | 0.22394216 | 0.6979724 | 0.32084673 | 0.74832654 | 0.99937574 |
| GFM1 | 62.0944451 | -0.1682849 | 0.32251068 | -0.5217963 | 0.60181215 | 0.99937574 |
| MFSD1 | 31.4705088 | -0.2716572 | 0.46439557 | -0.5849694 | 0.5585683 | 0.99937574 |
| IQCJ-SCHIP1 | 77.8911854 | 1.45713692 | 0.86866726 | 1.67743967 | 0.09345654 | 0.99937574 |
| SMC4 | 23.9967202 | -0.4646243 | 0.55836769 | -0.8321118 | 0.40534586 | 0.99937574 |
| KPNA4 | 115.314128 | -0.4775063 | 0.36370047 | -1.3129109 | 0.18921296 | 0.99937574 |
| PPM1L | 314.740507 | -0.0052437 | 0.46497827 | -0.0112773 | 0.99100222 | 0.99937574 |
| B3GALNT1 | 12.744397 | 0.17078153 | 0.65099579 | 0.26233891 | 0.79306016 | 0.99937574 |
| NMD3 | 70.3850488 | -0.0094945 | 0.32966093 | -0.0288007 | 0.97702353 | 0.99937574 |

| | | | | | | |
|---|---|---|---|---|---|---|
| LINC01324 | 17.0393516 | 1.73670303 | 1.17998598 | 1.47179972 | 0.14107497 | 0.99937574 |
| LINC01322 | 12.8025674 | -0.2918494 | 0.65803405 | -0.4435172 | 0.65739168 | 0.99937574 |
| PDCD10 | 87.8307649 | -0.4465613 | 0.27528612 | -1.6221715 | 0.10476663 | 0.99937574 |
| GOLIM4 | 112.096905 | 0.16757049 | 0.24195988 | 0.69255484 | 4.89E-01 | 9.99E-01 |
| MECOM | 2528.44048 | 0.17306151 | 0.31411234 | 0.55095418 | 0.58166509 | 0.99937574 |
| MYNN | 25.6470238 | 0.20773402 | 0.47927955 | 0.43342976 | 0.66470259 | 0.99937574 |
| SEC62 | 162.662665 | 0.23876919 | 0.26445895 | 0.90285919 | 0.36660063 | 0.99937574 |
| GPR160 | 97.8697507 | -0.3852869 | 0.49839738 | -0.7730516 | 0.43949184 | 0.99937574 |
| PHC3 | 298.941612 | -0.0965399 | 0.31239296 | -0.3090336 | 0.75729596 | 0.99937574 |
| PRKCI | 215.151125 | -1.4964387 | 0.35285077 | -4.2409959 | 2.23E-05 | 0.0449155 |
| SKIL | 46.8283901 | -0.2871028 | 0.38270928 | -0.7501852 | 0.45314317 | 0.99937574 |
| RPL22L1 | 13.0915136 | -0.3764986 | 0.67738504 | -0.5558118 | 0.57833948 | 0.99937574 |
| TNIK | 93.1400871 | -1.0595287 | 1.03035879 | -1.0283104 | 0.30380383 | 0.99937574 |
| PLD1 | 97.9361396 | -0.2351645 | 0.62204964 | -0.3780479 | 0.70539503 | 0.99937574 |
| FNDC3B | 660.27973 | -0.3154981 | 0.31357726 | -1.0061255 | 0.31435521 | 0.99937574 |
| TNFSF10 | 57.625063 | -1.570619 | 0.73382002 | -2.1403328 | 0.03232788 | 0.99937574 |
| NCEH1 | 132.779693 | -0.0615619 | 0.41691783 | -0.1476596 | 0.88261143 | 0.99937574 |
| ECT2 | 33.336254 | 0.61168552 | 0.56871709 | 1.07555325 | 0.28212711 | 0.99937574 |
| NLGN1 | 170.770053 | 0.13617502 | 0.61861259 | 0.22012973 | 0.82577012 | 0.99937574 |
| NAALADL2 | 422.972452 | -0.4277033 | 0.4160239 | -1.0280739 | 0.30391507 | 0.99937574 |
| TBL1XR1 | 479.72771 | -0.2991378 | 0.32821005 | -0.9114218 | 0.36207315 | 0.99937574 |
| KCNMB2-AS: | 37.859971 | -1.6341247 | 1.42368577 | -1.1478128 | 0.25104586 | 0.99937574 |
| ZMAT3 | 26.9863418 | 0.23182544 | 0.44228544 | 0.52415345 | 0.60017181 | 0.99937574 |
| PIK3CA | 106.057625 | -0.2683208 | 0.2942863 | -0.9117681 | 0.36189082 | 0.99937574 |
| KCNMB3 | 16.319836 | -0.2273865 | 0.61853664 | -0.3676201 | 0.71315651 | 0.99937574 |
| ZNF639 | 20.1089758 | -0.2643726 | 0.48645497 | -0.5434678 | 0.58680773 | 0.99937574 |
| MFN1 | 88.8984668 | -0.482175 | 0.25685625 | -1.8772172 | 0.06048834 | 0.99937574 |
| ACTL6A | 20.0302535 | -0.2881455 | 0.49678426 | -0.5800213 | 0.56190022 | 0.99937574 |
| MRPL47 | 26.2025917 | -0.301959 | 0.41561711 | -0.7265318 | 0.46751282 | 0.99937574 |
| NDUFB5 | 79.2409918 | 0.85062216 | 0.45319536 | 1.87694368 | 0.06052581 | 0.99937574 |
| TTC14 | 93.5616668 | -0.4654632 | 0.28340035 | -1.6424228 | 0.10050241 | 0.99937574 |
| FXR1 | 118.986993 | 0.06021857 | 0.23563737 | 0.25555612 | 0.79829361 | 0.99937574 |
| DNAJC19 | 20.3395868 | 0.03224449 | 0.62185841 | 0.05185182 | 0.95864676 | 0.99937574 |
| ATP11B | 160.357285 | 0.08938521 | 0.22859489 | 0.39102017 | 0.69578233 | 0.99937574 |
| DCUN1D1 | 78.6139983 | 0.2171547 | 0.28554015 | 0.76050495 | 0.44695281 | 0.99937574 |
| MCCC1 | 227.341516 | -0.2846091 | 0.46165539 | -0.6164968 | 0.53756667 | 0.99937574 |
| MCF2L2 | 38.1522991 | -0.3631853 | 0.40600328 | -0.8945379 | 0.37103418 | 0.99937574 |
| B3GNT5 | 29.9246876 | -0.0058592 | 0.43711024 | -0.0134045 | 0.9893051 | 0.99937574 |
| KLHL24 | 181.617463 | -0.5578636 | 0.33364155 | -1.6720447 | 0.09451551 | 0.99937574 |
| YEATS2 | 86.0919141 | -0.2826526 | 0.43820814 | -0.6450192 | 0.51891477 | 0.99937574 |
| ABCC5 | 365.433647 | -0.1159052 | 0.33532239 | -0.3456531 | 0.72960344 | 0.99937574 |
| EIF2B5 | 25.4260061 | -0.0759097 | 0.44705338 | -0.1698002 | 0.86516729 | 0.99937574 |
| DVL3 | 35.6170933 | -0.4957683 | 0.4118468 | -1.2037688 | 0.22867895 | 0.99937574 |
| AP2M1 | 61.7586382 | 0.19696807 | 0.35515705 | 0.5545943 | 0.57917219 | 0.99937574 |
| ABCF3 | 12.0244725 | -0.0117676 | 0.6491927 | -0.0181266 | 0.9855379 | 0.99937574 |
| PSMD2 | 40.2616277 | 0.18300288 | 0.34068958 | 0.53715433 | 0.59116102 | 0.99937574 |
| EIF4G1 | 97.7329612 | -0.317721 | 0.46306534 | -0.6861257 | 0.49263384 | 0.99937574 |
| FAM131A | 14.582046 | 0.4335251 | 0.54236495 | 0.79932359 | 0.42410281 | 0.99937574 |
| POLR2H | 48.9874275 | -0.4136707 | 0.43812068 | -0.9441935 | 0.34507077 | 0.99937574 |
| MAGEF1 | 20.3613695 | -0.0525128 | 0.57252216 | -0.0917218 | 0.92691908 | 0.99937574 |
| VPS8 | 137.7107 | 0.02863907 | 0.24717889 | 0.11586375 | 0.90776053 | 0.99937574 |
| EHHADH | 31.6760802 | 0.85190337 | 0.49164522 | 1.73276042 | 0.08313826 | 0.99937574 |
| MAP3K13 | 110.583225 | 0.26374111 | 0.32603244 | 0.80894129 | 0.41854891 | 0.99937574 |
| TMEM41A | 28.3433589 | -0.5334882 | 0.3877961 | -1.3756926 | 0.16891681 | 0.99937574 |

| | | | | | | |
|---|---|---|---|---|---|---|
| LIPH | 116.381897 | -0.0457571 | 0.45557202 | -0.1004388 | 0.91999599 | 0.99937574 |
| SENP2 | 58.8767108 | -0.1406916 | 0.4130528 | -0.3406141 | 0.73339411 | 0.99937574 |
| IGF2BP2 | 196.360665 | -0.0580625 | 0.22534567 | -0.2576598 | 0.79666949 | 0.99937574 |
| TRA2B | 94.1366552 | 0.1098605 | 0.28343507 | 0.38760376 | 0.69830929 | 0.99937574 |
| TBCCD1 | 11.7055414 | -0.4454687 | 0.63303158 | -0.7037069 | 0.48161533 | 0.99937574 |
| AC068631.2 | 15.6846243 | 0.64571651 | 0.69791277 | 0.92521091 | 0.35485621 | 0.99937574 |
| EIF4A2 | 143.278384 | 0.22653701 | 0.36906625 | 0.61381125 | 0.53934006 | 0.99937574 |
| ST6GAL1 | 573.469819 | -0.3271008 | 0.24766318 | -1.3207487 | 1.87E-01 | 0.99937574 |
| AC072022.1 | 17.6308195 | 0.69766493 | 0.87663065 | 0.7958482 | 0.42612027 | 0.99937574 |
| BCL6 | 215.735107 | 0.10851757 | 0.57767333 | 0.18785282 | 0.85099203 | 0.99937574 |
| LPP | 1521.90532 | 0.14490534 | 0.31325687 | 0.46257674 | 0.64366779 | 0.99937574 |
| TPRG1 | 24.4093397 | 0.29720104 | 0.42689657 | 0.6961898 | 0.48630997 | 0.99937574 |
| P3H2 | 199.578728 | -0.6626638 | 0.79430221 | -0.8342717 | 0.40412791 | 0.99937574 |
| CCDC50 | 198.823584 | -0.2426863 | 0.30321091 | -0.8003879 | 0.42348611 | 0.99937574 |
| FGF12 | 62.7651027 | 0.14242013 | 0.45971474 | 0.30980109 | 0.75671222 | 0.99937574 |
| MB21D2 | 20.5730456 | 0.43704875 | 0.7825285 | 0.55850841 | 0.57649726 | 0.99937574 |
| OPA1 | 144.566325 | -0.1200131 | 0.23186028 | -0.5176095 | 0.60473075 | 0.99937574 |
| HES1 | 21.7157693 | 0.08732158 | 0.69409259 | 0.12580681 | 0.89988485 | 0.99937574 |
| ATP13A3 | 58.6753217 | -0.6901603 | 0.40851601 | -1.6894328 | 0.09113652 | 0.99937574 |
| LSG1 | 30.3676302 | -0.0644286 | 0.51797244 | -0.1243861 | 0.9010096 | 0.99937574 |
| XXYLT1 | 27.3935476 | 0.43292396 | 0.5409171 | 0.80035177 | 0.42350702 | 0.99937574 |
| ACAP2 | 200.098864 | -0.2793374 | 0.19565151 | -1.4277291 | 0.15336986 | 0.99937574 |
| PPP1R2 | 72.5923728 | -0.26938 | 0.31570632 | -0.8532613 | 0.39351443 | 0.99937574 |
| MUC20-OT1 | 1266.31049 | 0.14586788 | 0.48322668 | 0.30186223 | 0.76275709 | 0.99937574 |
| MUC20 | 15.5613486 | -0.3170702 | 0.51665409 | -0.6136993 | 0.53941407 | 0.99937574 |
| MUC4 | 12.7723852 | 0.078857 | 0.86421585 | 0.09124688 | 0.92729643 | 0.99937574 |
| TNK2 | 21.044615 | 0.40252925 | 0.55530734 | 0.72487652 | 0.46852779 | 0.99937574 |
| TFRC | 190.036687 | -0.2937902 | 0.41851275 | -0.7019862 | 0.48268778 | 0.99937574 |
| LINC00885 | 27.6576349 | -0.8534153 | 1.07628874 | -0.7929241 | 0.42782202 | 0.99937574 |
| PCYT1A | 35.0630616 | -0.3644259 | 0.37652369 | -0.9678698 | 0.33310939 | 0.99937574 |
| UBXN7 | 78.0205399 | 0.31936166 | 0.28737129 | 1.11132066 | 0.26643035 | 0.99937574 |
| RNF168 | 65.5302872 | -0.2017224 | 0.30795023 | -0.6550487 | 0.51243641 | 0.99937574 |
| WDR53 | 11.2948068 | -0.1880536 | 0.56639207 | -0.3320202 | 0.73987404 | 0.99937574 |
| PIGX | 19.4261299 | 0.34156835 | 0.42036666 | 0.81254862 | 0.4164769 | 0.99937574 |
| PAK2 | 133.608067 | -0.5740199 | 0.21233349 | -2.7033883 | 0.00686365 | 0.72913641 |
| SENP5 | 137.263485 | -0.5894207 | 0.30149398 | -1.955 | 0.05058307 | 0.99937574 |
| NCBP2 | 33.4406577 | -0.1648426 | 0.40628157 | -0.4057348 | 0.68493747 | 0.99937574 |
| NCBP2-AS2 | 17.315928 | -0.0125526 | 0.60275168 | -0.0208255 | 0.98338487 | 0.99937574 |
| DLG1 | 559.211007 | 0.00941422 | 0.3119701 | 0.03017669 | 0.97592614 | 0.99937574 |
| BDH1 | 32.8421481 | 0.09721145 | 0.55066568 | 0.17653442 | 0.85987411 | 0.99937574 |
| AC024560.1 | 59.4134046 | -0.5634042 | 1.8835142 | -0.2991239 | 0.7648455 | 0.99937574 |
| RUBCN | 48.0515359 | 0.06550872 | 0.42503819 | 0.15412432 | 0.87751171 | 0.99937574 |
| FYTTD1 | 67.6688035 | 0.35827782 | 0.31910028 | 1.12277501 | 0.26153306 | 0.99937574 |
| LRCH3 | 199.190361 | -0.0283815 | 0.23717623 | -0.1196644 | 0.90474902 | 0.99937574 |
| IQCG | 25.9952776 | 0.08537994 | 0.42071016 | 0.20294244 | 0.83918002 | 0.99937574 |
| RPL35A | 153.354411 | 0.28355179 | 0.500209 | 0.56686663 | 0.5708048 | 0.99937574 |
| LMLN | 26.4299815 | 0.07843446 | 0.41058484 | 0.19103107 | 0.84850125 | 0.99937574 |
| ZNF595 | 35.8363355 | 0.54119971 | 0.35202331 | 1.53739736 | 0.12419603 | 0.99937574 |
| ZNF718 | 80.9497571 | 0.37182442 | 0.38902749 | 0.95577928 | 0.33918376 | 0.99937574 |
| ZNF141 | 58.0148085 | -0.2177576 | 0.28638629 | -0.7603632 | 0.4470375 | 0.99937574 |
| AC092574.2 | 15.9310765 | 0.46291624 | 0.68688966 | 0.673931 | 0.50035518 | 0.99937574 |
| ZNF721 | 203.85264 | 0.02407646 | 0.23635724 | 0.1018647 | 0.91886407 | 0.99937574 |
| PIGG | 44.6705076 | -0.1915127 | 0.30386797 | -0.6302498 | 0.52853119 | 0.99937574 |
| ATP5ME | 45.4789136 | 0.70377925 | 0.36217807 | 1.94318568 | 0.05199373 | 0.99937574 |

| | | | | | | |
|---|---|---|---|---|---|---|
| SLC49A3 | 11.5017474 | 0.57464345 | 0.74014802 | 0.77638991 | 0.4375188 | 0.99937574 |
| PCGF3 | 59.0097412 | 0.02335935 | 0.26826331 | 0.08707621 | 0.93061094 | 0.99937574 |
| AC139887.2 | 113.516866 | 0.16831196 | 0.45135737 | 0.37290177 | 0.70922155 | 0.99937574 |
| GAK | 115.0365 | -0.0303351 | 0.36617013 | -0.0828442 | 0.93397541 | 0.99937574 |
| TMEM175 | 17.783861 | 1.09838621 | 0.45316082 | 2.42383313 | 0.01535766 | 0.86586317 |
| FGFRL1 | 26.7619236 | -0.0019632 | 0.48922782 | -0.0040128 | 0.99679828 | 0.99937574 |
| RNF212 | 28.9507327 | -0.051715 | 0.39756566 | -0.1300791 | 0.89650383 | 0.99937574 |
| AC092535.3 | 73.6317435 | 0.03991557 | 0.32182089 | 0.12403038 | 0.90129122 | 0.99937574 |
| CTBP1-AS | 19.6418997 | -0.9817891 | 0.76803869 | -1.2783068 | 0.20114125 | 0.99937574 |
| CTBP1 | 47.4675391 | -0.4759397 | 0.37344208 | -1.2744673 | 0.20249786 | 0.99937574 |
| MAEA | 77.5950226 | -0.2246025 | 0.24271076 | -0.9253915 | 0.35476228 | 0.99937574 |
| UVSSA | 53.897423 | -0.0625817 | 0.41982202 | -0.1490672 | 0.88150063 | 0.99937574 |
| SLBP | 59.677502 | -0.3027504 | 0.3526926 | -0.8583975 | 0.390673 | 0.99937574 |
| TMEM129 | 32.0286536 | -0.0913555 | 0.39865001 | -0.2291622 | 0.81874284 | 0.99937574 |
| LETM1 | 38.4154312 | -0.0489167 | 0.46784731 | -0.1045569 | 0.91672739 | 0.99937574 |
| NSD2 | 120.621644 | -0.0492424 | 0.27756589 | -0.1774079 | 0.85918799 | 0.99937574 |
| NELFA | 54.3240088 | -0.0093285 | 0.3993558 | -0.023359 | 0.98136395 | 0.99937574 |
| MXD4 | 56.9044247 | -0.2895045 | 0.42867464 | -0.6753479 | 0.49945473 | 0.99937574 |
| ZFYVE28 | 57.782282 | 0.39066183 | 0.6708736 | 0.58231808 | 0.56035245 | 0.99937574 |
| RNF4 | 58.4553112 | -0.3883697 | 0.37965717 | -1.0229484 | 0.30633223 | 0.99937574 |
| FAM193A | 120.693909 | -0.4726255 | 0.29712005 | -1.5906887 | 0.11167965 | 0.99937574 |
| TNIP2 | 25.9001773 | -0.5950267 | 0.46237259 | -1.2868987 | 0.1981296 | 0.99937574 |
| SH3BP2 | 38.0913978 | -0.8695015 | 0.42635581 | -2.03938 | 0.04141212 | 0.99937574 |
| ADD1 | 150.139171 | -0.053258 | 0.26804928 | -0.1986872 | 0.84250744 | 0.99937574 |
| MFSD10 | 50.4288448 | 0.03468148 | 0.45675445 | 0.07593026 | 0.93947458 | 0.99937574 |
| NOP14-AS1 | 10.4963181 | -0.098828 | 0.6689393 | -0.1477384 | 0.8825492 | 0.99937574 |
| NOP14 | 16.6488947 | -0.6843832 | 0.63930384 | -1.0705132 | 0.28438839 | 0.99937574 |
| GRK4 | 10.7497121 | -0.5626775 | 0.7421789 | -0.7581427 | 0.44836555 | 0.99937574 |
| HTT | 121.299875 | -0.045168 | 0.28380596 | -0.1591509 | 0.87355001 | 0.99937574 |
| RGS12 | 39.1701359 | -0.2372468 | 0.52637118 | -0.4507215 | 0.65219027 | 0.99937574 |
| LRPAP1 | 65.7884253 | 0.26921094 | 0.38525296 | 0.69879006 | 0.48468324 | 0.99937574 |
| TMEM128 | 17.792554 | -0.2268126 | 0.44911035 | -0.5050265 | 0.61354019 | 0.99937574 |
| ZBTB49 | 11.3758172 | -1.0631914 | 0.61412804 | -1.7312211 | 0.08341233 | 0.99937574 |
| STX18 | 196.89582 | 0.01082774 | 0.31614878 | 0.03424888 | 0.97267869 | 0.99937574 |
| STX18-AS1 | 66.2392636 | -0.2423773 | 0.42201847 | -0.5743287 | 0.56574538 | 0.99937574 |
| EVC2 | 55.484754 | -0.256569 | 0.46524651 | -0.5514689 | 0.5813123 | 0.99937574 |
| EVC | 45.7758305 | -0.0290912 | 0.43091931 | -0.0675096 | 0.94617603 | 0.99937574 |
| CRMP1 | 13.2160747 | -0.2048471 | 0.81188478 | -0.2523106 | 0.80080101 | 0.99937574 |
| MAN2B2 | 22.9962094 | 0.34594672 | 0.59726168 | 0.57922136 | 0.56243982 | 0.99937574 |
| MRFAP1 | 64.5603397 | 0.16077799 | 0.50095872 | 0.32094059 | 0.74825541 | 0.99937574 |
| MRFAP1L1 | 19.4558905 | -0.1903855 | 0.49121674 | -0.3875795 | 0.69832726 | 0.99937574 |
| BLOC1S4 | 16.827258 | 0.31294229 | 0.54909589 | 0.56992284 | 0.56873003 | 0.99937574 |
| KIAA0232 | 233.53541 | 0.17673643 | 0.28603881 | 0.6178757 | 0.53665728 | 0.99937574 |
| TBC1D14 | 742.538251 | 0.11530649 | 0.3400352 | 0.33910163 | 0.73453317 | 0.99937574 |
| TADA2B | 23.8222523 | -0.9491574 | 0.53429416 | -1.7764697 | 0.07565553 | 0.99937574 |
| GRPEL1 | 29.9012137 | -0.498435 | 0.46625097 | -1.0690273 | 0.28505738 | 0.99937574 |
| SORCS2 | 69.8089074 | -2.25168 | 1.38984416 | -1.6200953 | 0.10521181 | 0.99937574 |
| SH3TC1 | 124.268203 | -0.0544739 | 0.31335688 | -0.1738398 | 0.8619914 | 0.99937574 |
| ACOX3 | 27.2047997 | 0.31152224 | 0.39633502 | 0.78600735 | 0.43186318 | 0.99937574 |
| TRMT44 | 15.6015731 | -0.1117518 | 0.58530191 | -0.1909301 | 0.84858033 | 0.99937574 |
| SLC2A9 | 22.761709 | 0.75865026 | 0.52657443 | 1.44072749 | 0.14966169 | 0.99937574 |
| WDR1 | 198.792557 | 0.34069925 | 0.26697126 | 1.27616455 | 0.20189736 | 0.99937574 |
| ZNF518B | 15.5182598 | -0.2379336 | 0.5111006 | -0.4655319 | 0.6415506 | 0.99937574 |
| CLNK | 2690.97471 | 0.60055596 | 0.33202615 | 1.80876101 | 0.07048814 | 0.99937574 |

| | | | | | | |
|---|---|---|---|---|---|---|
| AC084048.1 | 50.7371784 | 0.55841621 | 0.78794452 | 0.70869991 | 0.47851072 | 0.99937574 |
| HS3ST1 | 18.1793569 | -1.3080622 | 0.75830645 | -1.7249783 | 0.08453138 | 0.99937574 |
| AC005699.1 | 27.2439196 | 0.152299 | 0.83779112 | 0.18178636 | 0.85575039 | 0.99937574 |
| RAB28 | 168.019734 | -0.0926064 | 0.25806068 | -0.358855 | 0.71970357 | 0.99937574 |
| BOD1L1 | 118.609669 | -0.0882165 | 0.26595099 | -0.3317022 | 0.74011412 | 0.99937574 |
| CPEB2 | 180.262743 | -0.4640219 | 0.46785749 | -0.9918018 | 0.3212942 | 0.99937574 |
| AC098829.1 | 51.421615 | 0.39507154 | 0.57056553 | 0.69242097 | 0.488673 | 0.99937574 |
| CC2D2A | 27.4309717 | 0.6027396 | 0.46506078 | 1.29604479 | 0.19496005 | 0.99937574 |
| FBXL5 | 114.564785 | 0.03566632 | 0.23640294 | 0.15087089 | 0.88007756 | 0.99937574 |
| FAM200B | 37.034419 | -0.0144443 | 0.49034832 | -0.0294572 | 0.97649999 | 0.99937574 |
| TAPT1 | 89.6061617 | 0.08223114 | 0.27953895 | 0.29416702 | 0.76863028 | 0.99937574 |
| TAPT1-AS1 | 168.669978 | 0.47552812 | 0.65294295 | 0.72828433 | 0.46643955 | 0.99937574 |
| AC097515.1 | 24.3472294 | -0.0766155 | 0.87026524 | -0.088037 | 0.92984728 | 0.99937574 |
| QDPR | 18.4848871 | 0.53506804 | 0.50972443 | 1.04972023 | 0.29384676 | 0.99937574 |
| LAP3 | 51.5497618 | -1.3854732 | 0.73059633 | -1.8963593 | 0.05791255 | 0.99937574 |
| MED28 | 102.768301 | -0.469433 | 0.37120286 | -1.2646266 | 0.2060052 | 0.99937574 |
| FAM184B | 156.926562 | 0.29541317 | 0.745248 | 0.3963958 | 0.69181307 | 0.99937574 |
| DCAF16 | 77.2459523 | -0.1288933 | 0.46721521 | -0.2758757 | 0.78264353 | 0.99937574 |
| LCORL | 150.566636 | -0.2702548 | 0.20894969 | -1.2933966 | 0.19587391 | 0.99937574 |
| AC024230.1 | 34.5676993 | 0.35073478 | 0.71972688 | 0.4873165 | 0.62603406 | 0.99937574 |
| SLIT2 | 1740.64149 | 0.16112906 | 0.64674979 | 0.24913663 | 0.8032551 | 0.99937574 |
| PACRGL | 55.416413 | -0.0792363 | 0.34766253 | -0.2279115 | 0.81971505 | 0.99937574 |
| KCNIP4 | 2618.39823 | 1.4137788 | 0.6702969 | 2.10918294 | 0.03492879 | 0.99937574 |
| ADGRA3 | 26.3963687 | 0.15385861 | 0.54146685 | 0.28415148 | 0.7762943 | 0.99937574 |
| PPARGC1A | 698.387475 | -0.2769199 | 0.49272625 | -0.5620157 | 0.57410532 | 0.99937574 |
| DHX15 | 123.471432 | -0.3458945 | 0.26027292 | -1.3289684 | 0.18385841 | 0.99937574 |
| CCDC149 | 31.4671256 | -0.3150177 | 0.40315996 | -0.7813715 | 0.434584 | 0.99937574 |
| SEPSECS | 31.6370157 | -1.1567617 | 0.53809639 | -2.1497295 | 0.03157661 | 0.99049468 |
| ZCCHC4 | 21.899696 | -0.4112798 | 0.47844496 | -0.8596178 | 0.38999977 | 0.99937574 |
| ANAPC4 | 87.0168927 | -0.9638816 | 0.65762259 | -1.4657064 | 0.14272828 | 0.99937574 |
| SEL1L3 | 14.9300659 | -0.8488718 | 0.83144451 | -1.0209603 | 0.30727325 | 0.99937574 |
| SMIM20 | 23.8501719 | -0.1586525 | 0.59208489 | -0.2679557 | 0.78873343 | 0.99937574 |
| RBPJ | 197.767958 | 0.26593477 | 0.29656918 | 0.896704 | 0.36987689 | 0.99937574 |
| TBC1D19 | 106.219974 | 0.51616458 | 0.48375778 | 1.06698973 | 0.28597648 | 0.99937574 |
| STIM2 | 632.966343 | 0.98609396 | 0.61545195 | 1.60222738 | 1.09E-01 | 9.99E-01 |
| PCDH7 | 114.243253 | -0.9761577 | 1.35318928 | -0.7213756 | 0.47067848 | 0.99937574 |
| LINC02506 | 42.805576 | -1.3039043 | 1.56393013 | -0.8337356 | 0.40442998 | 0.99937574 |
| ARAP2 | 114.926848 | -0.0439582 | 0.48835409 | -0.0900129 | 0.92827694 | 0.99937574 |
| C4orf19 | 195.785574 | 0.02433695 | 0.26862775 | 0.09059731 | 0.92781257 | 0.99937574 |
| RELL1 | 27.1236723 | -0.8700292 | 0.42860737 | -2.0298979 | 0.04236691 | 0.99937574 |
| PGM2 | 26.4756552 | 0.19225356 | 0.40837913 | 0.47077224 | 0.63780339 | 0.99937574 |
| TBC1D1 | 943.347652 | 0.46117203 | 0.37608383 | 1.22624798 | 0.22010537 | 0.99937574 |
| KLF3 | 62.4113747 | -0.2721972 | 0.33183052 | -0.8202897 | 0.41205099 | 0.99937574 |
| TLR1 | 13.6286716 | 0.8378194 | 0.60591177 | 1.38274159 | 0.16674411 | 0.99937574 |
| FAM114A1 | 53.0611927 | 0.61426744 | 0.46905006 | 1.3095989 | 0.19033156 | 0.99937574 |
| KLHL5 | 101.352593 | 0.173593 | 0.41088073 | 0.42249001 | 0.67266739 | 0.99937574 |
| WDR19 | 36.0448281 | -0.1368754 | 0.35554137 | -0.3849775 | 0.70025412 | 0.99937574 |
| RFC1 | 74.1394863 | 0.14104392 | 0.27470825 | 0.5134317 | 0.60764938 | 0.99937574 |
| RPL9 | 18.9997232 | -0.0391226 | 0.42543571 | -0.0919589 | 0.92673072 | 0.99937574 |
| LIAS | 26.6471065 | -0.0531674 | 0.48830362 | -0.1088818 | 0.91329621 | 0.99937574 |
| UGDH | 23.2748723 | -0.4345754 | 0.48586913 | -0.8944289 | 0.37109244 | 0.99937574 |
| UGDH-AS1 | 29.7287358 | 0.14547638 | 0.54537357 | 0.26674631 | 0.7896645 | 0.99937574 |
| SMIM14 | 94.8125953 | -0.1271125 | 0.34199098 | -0.3716837 | 0.71012836 | 0.99937574 |
| UBE2K | 282.346749 | -0.1341587 | 0.2300132 | -0.5832653 | 0.5597147 | 0.99937574 |

| | | | | | | |
|---|---|---|---|---|---|---|
| PDS5A | 215.939985 | -0.2620469 | 0.24355283 | -1.0759344 | 0.2819566 | 0.99937574 |
| N4BP2 | 48.0455592 | 0.11219062 | 0.44567685 | 0.25173086 | 0.8012491 | 0.99937574 |
| RBM47 | 876.96316 | -0.2316648 | 0.26078776 | -0.888327 | 0.37436487 | 0.99937574 |
| NSUN7 | 151.042512 | 0.41426506 | 0.4471446 | 0.92646776 | 0.35420294 | 0.99937574 |
| APBB2 | 291.868601 | -0.2823953 | 0.40841559 | -0.691441 | 0.48928845 | 0.99937574 |
| LIMCH1 | 781.205451 | 0.34686863 | 0.36419172 | 0.95243416 | 0.34087684 | 0.99937574 |
| TMEM33 | 117.208368 | 0.02938737 | 0.38043826 | 0.07724609 | 0.93842778 | 0.99937574 |
| SLC30A9 | 210.46458 | -0.124695 | 0.2373067 | -0.5254591 | 0.59926408 | 0.99937574 |
| ATP8A1 | 97.6172415 | 1.21154118 | 0.5931333 | 2.04261197 | 0.04109087 | 0.99937574 |
| GUF1 | 34.7134293 | 0.14480241 | 0.36045928 | 0.40171641 | 0.68789274 | 0.99937574 |
| GNPDA2 | 30.0645218 | 0.29247123 | 0.36329867 | 0.8050435 | 0.42079458 | 0.99937574 |
| COMMD8 | 21.2044416 | 0.01234903 | 0.44949703 | 0.027473 | 0.97808247 | 0.99937574 |
| ATP10D | 214.30463 | 0.20975464 | 0.44164778 | 0.47493647 | 0.63483226 | 0.99937574 |
| AC107068.2 | 20.0836063 | 0.2394495 | 0.8901772 | 0.26899082 | 0.78793674 | 0.99937574 |
| NFXL1 | 52.1481027 | -0.0514508 | 0.602239 | -0.0854325 | 0.93191752 | 0.99937574 |
| NIPAL1 | 55.2346162 | -0.1494413 | 0.54042043 | -0.2765278 | 0.78214268 | 0.99937574 |
| CNGA1 | 31.2466862 | 0.2443763 | 0.80765496 | 0.30257512 | 0.76221368 | 0.99937574 |
| TXK | 19.9572889 | 0.56705623 | 0.68931953 | 0.82263189 | 0.41071736 | 0.99937574 |
| TEC | 74.6147244 | -0.4161267 | 0.31303693 | -1.3293214 | 0.18374195 | 0.99937574 |
| SLAIN2 | 110.518013 | -0.1720897 | 0.27322592 | -0.629844 | 0.52879664 | 0.99937574 |
| FRYL | 275.662964 | -0.1239488 | 0.26216867 | -0.4727828 | 0.63636814 | 0.99937574 |
| OCIAD1 | 166.406534 | 0.15949121 | 0.26508809 | 0.60165362 | 0.54740473 | 0.99937574 |
| OCIAD2 | 16.2429602 | -0.3814925 | 0.58043964 | -0.6572474 | 0.51102184 | 0.99937574 |
| CWH43 | 29.9796111 | 1.90220523 | 0.90625969 | 2.09896264 | 0.03582019 | 0.99937574 |
| DCUN1D4 | 97.2495364 | -0.3338414 | 0.35472372 | -0.9411309 | 0.3466378 | 0.99937574 |
| SGCB | 11.3681691 | 0.74446919 | 0.65398234 | 1.13836283 | 0.25496901 | 0.99937574 |
| SCFD2 | 154.927325 | 0.09001304 | 0.38242171 | 0.2353764 | 0.8139166 | 0.99937574 |
| AC058822.1 | 75.2144334 | 0.54872315 | 0.58036551 | 0.94547856 | 0.34441461 | 0.99937574 |
| LNX1 | 149.776057 | 0.67348087 | 0.37402414 | 1.80063476 | 0.07176047 | 0.99937574 |
| CHIC2 | 82.1959131 | -0.0043822 | 0.26436934 | -0.0165761 | 0.98677481 | 0.99937574 |
| KIT | 833.481199 | -0.2545829 | 0.44603267 | -0.570772 | 0.56815422 | 0.99937574 |
| TMEM165 | 196.540312 | 0.25351372 | 0.28476798 | 0.89024659 | 0.37333349 | 0.99937574 |
| CLOCK | 117.024069 | 0.42002417 | 0.34395152 | 1.22117261 | 0.22202067 | 0.99937574 |
| EXOC1 | 110.06974 | -0.2295588 | 0.36989319 | -0.6206083 | 0.53485738 | 0.99937574 |
| CEP135 | 22.5833419 | -0.5225866 | 0.43081848 | -1.2130089 | 0.22512642 | 0.99937574 |
| AASDH | 54.5252608 | -0.0263901 | 0.31199745 | -0.0845843 | 0.93259188 | 0.99937574 |
| PAICS | 20.2359108 | -0.0818594 | 0.47735442 | -0.1714856 | 0.86384192 | 0.99937574 |
| SRP72 | 50.272963 | 0.06828967 | 0.30551939 | 0.22351991 | 0.82313089 | 0.99937574 |
| REST | 107.00646 | 0.0990408 | 0.34091797 | 0.29051211 | 0.77142449 | 0.99937574 |
| NOA1 | 15.2909996 | 0.09130457 | 0.50274426 | 0.18161236 | 0.85588695 | 0.99937574 |
| POLR2B | 80.0237044 | -0.0556068 | 0.29443246 | -0.1888609 | 0.8502018 | 0.99937574 |
| IGFBP7 | 1029.13987 | -0.3654216 | 0.56827494 | -0.6430366 | 0.52020035 | 0.99937574 |
| AC104806.2 | 22.6237332 | 1.39634052 | 0.7732556 | 1.80579425 | 0.07095048 | 0.99937574 |
| CENPC | 102.687881 | -0.1950804 | 0.3357848 | -0.5809685 | 0.56126169 | 0.99937574 |
| STAP1 | 245.577412 | -0.4592352 | 0.52963733 | -0.8670749 | 0.38590097 | 0.99937574 |
| UBA6 | 179.757565 | -0.2692752 | 0.33406887 | -0.8060468 | 0.42021586 | 0.99937574 |
| UBA6-AS1 | 137.972824 | -0.2151537 | 0.5033681 | -0.4274281 | 0.66906755 | 0.99937574 |
| YTHDC1 | 135.597871 | -0.4316306 | 0.26858974 | -1.6070255 | 0.10804877 | 0.99937574 |
| UGT2B7 | 50.8155375 | -0.1861641 | 0.59856033 | -0.3110198 | 0.75578558 | 0.99937574 |
| ENAM | 35.839263 | 1.37448029 | 0.60990557 | 2.25359525 | 0.02422164 | 0.96546668 |
| UTP3 | 37.9549439 | 0.12203712 | 0.36892506 | 0.33079108 | 7.41E-01 | 0.99937574 |
| RUFY3 | 714.732523 | 0.26560763 | 0.41831826 | 0.63494151 | 0.52546656 | 0.99937574 |
| GRSF1 | 90.3941541 | 0.19021344 | 0.26576455 | 0.71572165 | 0.47416324 | 0.99937574 |
| MOB1B | 889.377554 | 0.67911253 | 0.42552669 | 1.59593404 | 0.11050352 | 0.99937574 |

| | | | | | | |
|---|---|---|---|---|---|---|
| DCK | 28.2914137 | 2.00772116 | 0.67219646 | 2.98680708 | 0.00281908 | 0.5159001 |
| SLC4A4 | 13.5224984 | 0.0966497 | 0.64523769 | 0.1497893 | 0.88093085 | 0.99937574 |
| ADAMTS3 | 109.877108 | -2.5226262 | 1.45101369 | -1.7385268 | 0.08211803 | 0.99937574 |
| COX18 | 19.3896542 | 0.24174189 | 0.58235769 | 0.41510895 | 0.67806215 | 0.99937574 |
| ANKRD17 | 682.412347 | -0.091686 | 0.24032031 | -0.3815157 | 0.70282061 | 0.99937574 |
| AC053527.1 | 77.7920972 | 0.47386838 | 0.62798546 | 0.75458495 | 0.45049806 | 0.99937574 |
| RASSF6 | 48.7959219 | -0.9226011 | 0.51525423 | -1.7905745 | 0.0733616 | 0.99937574 |
| MTHFD2L | 33.8209253 | -0.3771198 | 0.35633818 | -1.0583199 | 0.28990964 | 0.99937574 |
| BTC | 13.4023326 | -1.3855194 | 0.88550646 | -1.5646632 | 0.1176619 | 0.99937574 |
| RCHY1 | 50.5710351 | 0.14834821 | 0.37786596 | 0.3925948 | 0.69461878 | 0.99937574 |
| THAP6 | 28.7001142 | -0.691932 | 0.37178354 | -1.8611152 | 0.06272792 | 0.99937574 |
| CDKL2 | 30.0745331 | -0.0601278 | 0.53417654 | -0.1125616 | 0.9103781 | 0.99937574 |
| G3BP2 | 91.6150972 | -0.1279258 | 0.35253217 | -0.3628769 | 0.71669685 | 0.99937574 |
| USO1 | 184.788279 | 0.20378591 | 0.248311 | 0.8206882 | 0.41182389 | 0.99937574 |
| SDAD1 | 41.7036802 | -0.0176444 | 0.36627999 | -0.0481718 | 0.96157936 | 0.99937574 |
| NUP54 | 58.5582369 | -0.0892354 | 0.34391049 | -0.2594728 | 0.79527044 | 0.99937574 |
| SCARB2 | 131.605 | -0.1308725 | 0.36924998 | -0.3544278 | 0.72301832 | 0.99937574 |
| SHROOM3 | 1270.09832 | -0.040445 | 0.3617547 | -0.1118024 | 0.9109801 | 0.99937574 |
| AC107072.1 | 16.0505524 | 0.99292559 | 0.85940364 | 1.15536582 | 0.24794065 | 0.99937574 |
| 11-Sep | 280.298213 | -0.1302541 | 0.40977465 | -0.3178676 | 0.75058534 | 0.99937574 |
| CCNI | 317.18141 | 0.17665566 | 0.31397014 | 0.56265115 | 0.57367245 | 0.99937574 |
| CCNG2 | 76.0622407 | -0.2945609 | 0.37883795 | -0.7775379 | 0.43684147 | 0.99937574 |
| CNOT6L | 307.057328 | 0.28370221 | 0.4884624 | 0.58080665 | 0.56137077 | 0.99937574 |
| MRPL1 | 121.157286 | -0.2100835 | 0.36238335 | -0.5797271 | 0.56209865 | 0.99937574 |
| FRAS1 | 48.0423156 | -0.409679 | 0.79193893 | -0.5173114 | 6.05E-01 | 0.99937574 |
| ANXA3 | 55.5989673 | 0.05212745 | 0.76803437 | 0.06787124 | 0.94588813 | 0.99937574 |
| BMP2K | 47.2964981 | -0.5542605 | 0.41664473 | -1.3302953 | 0.183421 | 0.99937574 |
| PAQR3 | 21.9529031 | -0.7742723 | 0.56965606 | -1.3591926 | 0.17408558 | 0.99937574 |
| RASGEF1B | 455.189422 | -0.5643982 | 0.64312612 | -0.8775856 | 0.38016863 | 0.99937574 |
| HNRNPD | 171.654729 | -0.2157904 | 0.22407134 | -0.9630431 | 0.33552589 | 0.99937574 |
| HNRNPDL | 149.928463 | -0.131419 | 0.29831805 | -0.440533 | 0.65955108 | 0.99937574 |
| ENOPH1 | 29.9603641 | 0.20539972 | 0.4559875 | 0.45045034 | 0.65238576 | 0.99937574 |
| TMEM150C | 86.8066566 | 0.24767271 | 0.36995749 | 0.66946261 | 0.50320042 | 0.99937574 |
| SCD5 | 37.4854135 | 0.95075491 | 0.60508523 | 1.57127437 | 0.11611893 | 0.99937574 |
| SEC31A | 170.261564 | -0.1478762 | 0.23602827 | -0.6265189 | 0.53097462 | 0.99937574 |
| THAP9 | 12.5800665 | -0.2149839 | 0.62009933 | -0.3466927 | 0.72882219 | 0.99937574 |
| LIN54 | 36.2522418 | -0.2269093 | 0.41260098 | -0.5499485 | 0.58235469 | 0.99937574 |
| COPS4 | 37.2125187 | 0.38227196 | 0.33258313 | 1.14940273 | 0.25038996 | 0.99937574 |
| COQ2 | 12.5028426 | 0.15793237 | 0.5592082 | 0.28242141 | 0.7776204 | 0.99937574 |
| HELQ | 27.3835832 | -0.2027199 | 0.52216839 | -0.388227 | 0.69784805 | 0.99937574 |
| MRPS18C | 15.5717603 | -0.1634698 | 0.56460808 | -0.2895279 | 0.77217743 | 0.99937574 |
| ABRAXAS1 | 16.8893405 | 0.18555983 | 0.48158084 | 0.38531397 | 0.70000481 | 0.99937574 |
| GPAT3 | 365.767108 | 0.54802105 | 0.49492561 | 1.10727964 | 0.26817304 | 0.99937574 |
| CDS1 | 93.8217891 | 0.34508949 | 0.36646198 | 0.94167881 | 0.34635711 | 0.99937574 |
| WDFY3 | 377.16992 | -0.1635248 | 0.23724468 | -0.6892663 | 0.49065572 | 0.99937574 |
| ARHGAP24 | 261.847249 | 0.19450414 | 0.46923088 | 0.41451692 | 0.67849558 | 0.99937574 |
| MAPK10 | 481.949278 | 0.36955745 | 0.542302 | 0.6814606 | 0.49558009 | 0.99937574 |
| AC104827.1 | 12.2541894 | -0.3712273 | 0.87701482 | -0.4232851 | 0.67208728 | 0.99937574 |
| PTPN13 | 323.551274 | 0.02864629 | 0.44653358 | 0.06415259 | 0.94884873 | 0.99937574 |
| AC093827.5 | 13.7271047 | 0.04307256 | 0.66426125 | 0.0648428 | 0.94829917 | 0.99937574 |
| AFF1 | 457.996273 | -0.5216961 | 0.24025744 | -2.1714044 | 0.02990062 | 0.99049468 |
| KLHL8 | 64.5986485 | 0.66038026 | 0.45790536 | 1.4421763 | 0.14925265 | 0.99937574 |
| HSD17B11 | 26.0070886 | -0.2566011 | 0.50410981 | -0.5090182 | 0.61073944 | 0.99937574 |
| NUDT9 | 65.1477701 | -0.4656875 | 0.38420784 | -1.2120719 | 0.22548487 | 0.99937574 |

| | | | | | | |
|---|---|---|---|---|---|---|
| SPARCL1 | 737.149745 | 0.09766422 | 0.63718544 | 0.1532744 | 0.87818189 | 0.99937574 |
| SPP1 | 3666.48238 | 0.61998603 | 0.76218683 | 0.81343052 | 4.16E-01 | 9.99E-01 |
| PKD2 | 142.460784 | 0.13138599 | 0.3968979 | 0.33103221 | 0.74062016 | 0.99937574 |
| PPM1K | 41.2406995 | -0.410807 | 0.66879187 | -0.6142524 | 0.53904855 | 0.99937574 |
| HERC6 | 26.5782608 | -2.2998843 | 1.03718855 | -2.2174216 | 0.02659429 | 0.97242612 |
| HERC5 | 15.2784462 | -3.2736461 | 0.96635667 | -3.3876168 | 0.00070503 | 0.29012345 |
| PYURF | 24.0090553 | 0.25346826 | 0.54694842 | 0.46342259 | 0.64306149 | 0.99937574 |
| HERC3 | 92.1227087 | -0.1384085 | 0.39319351 | -0.3520112 | 0.72482983 | 0.99937574 |
| FAM13A-AS1 | 35.3778487 | -0.0827316 | 0.42206862 | -0.1960145 | 0.84459882 | 0.99937574 |
| FAM13A | 233.422959 | 0.38681765 | 0.4699965 | 0.8230224 | 0.41049526 | 0.99937574 |
| GPRIN3 | 174.730967 | 0.3609455 | 0.35383765 | 1.02008788 | 0.30768679 | 0.99937574 |
| SNCA | 31.0653533 | -2.7731203 | 1.10858335 | -2.5014991 | 0.01236687 | 0.77519564 |
| CCSER1 | 1178.97329 | -1.3396596 | 1.21432192 | -1.1032162 | 0.26993329 | 0.99937574 |
| SMARCAD1 | 87.0156467 | -0.267046 | 0.27614539 | -0.9670486 | 0.33351973 | 0.99937574 |
| PDLIM5 | 458.677693 | 0.06090348 | 0.26325209 | 0.23135043 | 0.81704257 | 0.99937574 |
| BMPR1B | 489.270388 | -0.4450415 | 0.75513403 | -0.5893543 | 0.55562359 | 0.99937574 |
| UNC5C | 35.0065206 | 1.27937362 | 0.86807202 | 1.47381045 | 0.14053263 | 0.99937574 |
| STPG2 | 484.389812 | 0.44818671 | 0.40202405 | 1.1148256 | 2.65E-01 | 0.99937574 |
| STPG2-AS1 | 12.9445675 | 0.35358426 | 0.83959976 | 0.42113431 | 0.67365701 | 0.99937574 |
| RAP1GDS1 | 147.290675 | 0.07935728 | 0.2510391 | 0.31611523 | 0.75191505 | 0.99937574 |
| TSPAN5 | 514.494125 | -0.3739675 | 0.44833643 | -0.8341224 | 0.40421201 | 0.99937574 |
| EIF4E | 101.293304 | 0.06127877 | 0.30058566 | 0.20386459 | 0.83845932 | 0.99937574 |
| METAP1 | 61.9085297 | 0.17555706 | 0.30477838 | 0.57601548 | 0.56460472 | 0.99937574 |
| ADH5 | 52.0314734 | 0.07041017 | 0.41870729 | 0.16816084 | 0.86645675 | 0.99937574 |
| TRMT10A | 10.4805417 | 0.53687221 | 0.54917155 | 0.97760384 | 0.3282703 | 0.99937574 |
| LAMTOR3 | 32.1242263 | -0.2421172 | 0.48356931 | -0.5006877 | 0.61659095 | 0.99937574 |
| DNAJB14 | 128.846331 | -0.0155967 | 0.27190943 | -0.05736 | 0.95425846 | 0.99937574 |
| H2AFZ | 76.8030828 | 0.09313643 | 0.74877063 | 0.12438579 | 0.90100982 | 0.99937574 |
| PPP3CA | 503.162976 | 0.13732499 | 0.37622909 | 0.36500364 | 0.71510869 | 0.99937574 |
| BANK1 | 35.7955933 | -0.1599572 | 0.5609855 | -0.285136 | 0.77553993 | 0.99937574 |
| AP002075.1 | 15.4874172 | -0.0263675 | 0.81205986 | -0.0324699 | 0.97409732 | 0.99937574 |
| NFKB1 | 148.206455 | -0.4677743 | 0.33821825 | -1.3830547 | 0.1666481 | 0.99937574 |
| MANBA | 83.5545451 | -0.0283238 | 0.42767746 | -0.0662269 | 0.94719717 | 0.99937574 |
| UBE2D3 | 314.321388 | -0.0010565 | 0.22624913 | -0.0046697 | 0.9962741 | 0.99937574 |
| CISD2 | 39.5269835 | 0.31134035 | 0.41078092 | 0.7579231 | 0.44849702 | 0.99937574 |
| SLC9B1 | 13.9074952 | -0.3859181 | 0.68508729 | -0.5633123 | 0.57322224 | 0.99937574 |
| TET2 | 227.683902 | -0.2935051 | 0.27077901 | -1.0839285 | 0.27839652 | 0.99937574 |
| PPA2 | 182.985033 | 0.28012994 | 0.3438155 | 0.81476821 | 0.415205 | 0.99937574 |
| ARHGEF38 | 299.238303 | 0.08446729 | 0.39630718 | 0.2131359 | 0.83122096 | 0.99937574 |
| INTS12 | 76.7853777 | 0.2406873 | 0.38655101 | 0.62265341 | 0.53351231 | 0.99937574 |
| GSTCD | 71.2291556 | 0.23813482 | 0.30613833 | 0.77786672 | 0.43664759 | 0.99937574 |
| NPNT | 181.846279 | 0.43444189 | 0.49113866 | 0.88456057 | 0.37639368 | 0.99937574 |
| TBCK | 272.591854 | 0.06828612 | 0.38749709 | 0.17622356 | 0.86011831 | 0.99937574 |
| AIMP1 | 49.1200713 | -0.4113797 | 0.3471219 | -1.185116 | 0.2359716 | 0.99937574 |
| PAPSS1 | 46.2977778 | -0.0796133 | 0.35832901 | -0.2221793 | 0.82417432 | 0.99937574 |
| SGMS2 | 122.379366 | 0.2925598 | 0.41950742 | 0.69738887 | 0.48555946 | 0.99937574 |
| AC096564.1 | 16.0415763 | 0.24994625 | 0.73197099 | 0.34147017 | 0.73274966 | 0.99937574 |
| HADH | 25.5579581 | 0.5119565 | 0.60733354 | 0.84295774 | 0.39925208 | 0.99937574 |
| LEF1 | 454.990184 | 0.60044624 | 0.78038919 | 0.76941896 | 0.44164463 | 0.99937574 |
| RPL34-AS1 | 25.1077759 | 0.49036429 | 0.60469688 | 0.81092577 | 0.4174083 | 0.99937574 |
| RPL34 | 139.378979 | 0.01717279 | 0.48219265 | 0.03561396 | 0.97159018 | 0.99937574 |
| OSTC | 33.0132295 | 0.39164771 | 0.3599391 | 1.08809438 | 0.27655344 | 0.99937574 |
| SEC24B | 168.805109 | -0.0675166 | 0.21093351 | -0.3200847 | 0.74890411 | 0.99937574 |
| CASP6 | 26.0152245 | -0.2988007 | 0.48675374 | -0.6138641 | 0.53930512 | 0.99937574 |

| | | | | | | |
|---|---|---|---|---|---|---|
| PLA2G12A | 38.5539686 | 0.70044821 | 0.43577484 | 1.60736265 | 0.10797484 | 0.99937574 |
| GAR1 | 16.2905783 | -0.6288334 | 0.56878664 | -1.1055699 | 0.26891273 | 0.99937574 |
| EGF | 159.345597 | 0.15032592 | 0.45489365 | 0.33046389 | 0.74104948 | 0.99937574 |
| ELOVL6 | 108.28174 | -0.2282872 | 0.69643917 | -0.327792 | 0.74306895 | 0.99937574 |
| FAM241A | 39.4715362 | -0.2621814 | 0.50402549 | -0.5201749 | 0.60294168 | 0.99937574 |
| AP1AR | 37.5285404 | -0.1898593 | 0.44615852 | -0.4255422 | 0.67044149 | 0.99937574 |
| ALPK1 | 162.494497 | -0.0919142 | 0.38777236 | -0.2370313 | 8.13E-01 | 0.99937574 |
| ZGRF1 | 12.0945713 | -0.9828155 | 0.74080199 | -1.3266913 | 0.18461083 | 0.99937574 |
| LARP7 | 67.643109 | -0.336332 | 0.36628853 | -0.918216 | 0.35850579 | 0.99937574 |
| ANK2 | 34.3332949 | -0.7349734 | 0.83206686 | -0.8833105 | 0.37706853 | 0.99937574 |
| CAMK2D | 53.4430449 | -1.250666 | 0.3312679 | -3.7753914 | 0.00015976 | 0.13113484 |
| ARSJ | 188.0291 | 0.42322036 | 0.4602576 | 0.91952933 | 0.35781877 | 0.99937574 |
| UGT8 | 334.945216 | 0.28614072 | 0.33116391 | 0.86404558 | 0.38756285 | 0.99937574 |
| SNHG8 | 31.3400552 | -0.042314 | 0.60023018 | -0.0704963 | 0.94379867 | 0.99937574 |
| METTL14 | 44.680822 | -0.1904551 | 0.34265147 | -0.5558274 | 0.57832884 | 0.99937574 |
| SEC24D | 71.8644265 | 0.1239154 | 0.46422173 | 0.26693149 | 0.78952191 | 0.99937574 |
| USP53 | 255.415986 | -0.1630258 | 0.31064645 | -0.5247953 | 0.59972547 | 0.99937574 |
| C4orf3 | 75.6704058 | -0.1876737 | 0.48678254 | -0.385539 | 0.69983809 | 0.99937574 |
| PDE5A | 63.9955295 | 0.10723028 | 0.66771875 | 0.16059198 | 0.87241477 | 0.99937574 |
| AC108866.1 | 28.6734714 | -0.6888137 | 0.47736624 | -1.442946 | 0.1490357 | 0.99937574 |
| PRDM5 | 19.1863331 | 0.07652098 | 0.79301191 | 0.09649411 | 0.92312815 | 0.99937574 |
| ANXA5 | 93.2929677 | 0.3691545 | 0.41693077 | 0.88540959 | 0.37593576 | 0.99937574 |
| EXOSC9 | 13.305979 | -0.7307819 | 0.53756768 | -1.3594231 | 0.17401255 | 0.99937574 |
| BBS7 | 15.99187 | -0.638816 | 0.49097743 | -1.3011106 | 0.19322059 | 0.99937574 |
| KIAA1109 | 282.516354 | -0.2414396 | 0.24401233 | -0.9894567 | 0.32243977 | 0.99937574 |
| SPATA5 | 96.8005301 | -0.3591159 | 0.27142077 | -1.3230965 | 0.18580328 | 0.99937574 |
| SPRY1 | 28.9480404 | -0.0945609 | 0.75731616 | -0.1248632 | 0.90063185 | 0.99937574 |
| LINC01091 | 183.869358 | 0.70814612 | 0.59479239 | 1.19057697 | 0.2338197 | 0.99937574 |
| ANKRD50 | 30.3814403 | 0.56986837 | 0.47773906 | 1.19284441 | 0.23293032 | 0.99937574 |
| INTU | 64.6276316 | -0.1643749 | 0.48161878 | -0.3412967 | 0.7328802 | 0.99937574 |
| HSPA4L | 105.693749 | 0.34588146 | 0.43297574 | 0.7988472 | 0.42437902 | 0.99937574 |
| MFSD8 | 60.0074498 | -0.3069901 | 0.2970636 | -1.0334153 | 0.30140957 | 0.99937574 |
| ABHD18 | 150.953087 | -0.1571002 | 0.3260518 | -0.481826 | 0.62992958 | 0.99937574 |
| LARP1B | 135.912431 | 0.24378381 | 0.34038026 | 0.71621018 | 0.47386158 | 0.99937574 |
| PGRMC2 | 68.4880636 | -0.9039043 | 0.46530721 | -1.9425968 | 0.0520649 | 0.99937574 |
| AC108062.1 | 35.0554364 | -0.0176083 | 0.48996057 | -0.0359382 | 0.97133162 | 0.99937574 |
| AC078850.2 | 48.8199687 | 1.1841495 | 0.65101754 | 1.81892104 | 0.06892348 | 0.99937574 |
| JADE1 | 430.089017 | 0.66341738 | 0.26735726 | 2.48138909 | 0.01308714 | 0.81027883 |
| SCLT1 | 160.744442 | 0.18237964 | 0.34981644 | 0.52135811 | 0.60211733 | 0.99937574 |
| C4orf33 | 20.3700548 | -0.0763245 | 0.48826364 | -0.1563183 | 0.87578215 | 0.99937574 |
| AC096711.2 | 63.1306361 | -0.0564692 | 0.38157339 | -0.1479905 | 0.8823503 | 0.99937574 |
| AC109927.1 | 32.0188749 | 0.7764009 | 0.48604817 | 1.5973744 | 0.11018227 | 0.99937574 |
| NOCT | 21.3490975 | 0.53179585 | 0.49386024 | 1.07681447 | 0.28156316 | 0.99937574 |
| ELF2 | 475.755863 | 0.2320803 | 0.2394323 | 0.96929403 | 0.33239851 | 0.99937574 |
| NDUFC1 | 81.7285377 | 0.33091888 | 0.41135465 | 0.80446127 | 0.42113063 | 0.99937574 |
| NAA15 | 111.599412 | -0.0533052 | 0.30890355 | -0.1725625 | 0.86299532 | 0.99937574 |
| AC097376.2 | 153.577117 | 0.03749926 | 0.3511922 | 0.10677704 | 0.91496586 | 0.99937574 |
| MGST2 | 64.8638172 | -0.3922271 | 0.48393279 | -0.8104991 | 0.4176534 | 0.99937574 |
| MAML3 | 308.97824 | 0.01645399 | 0.59403602 | 0.02769865 | 0.9779025 | 0.99937574 |
| SCOC | 96.1055634 | 0.12581109 | 0.36926424 | 0.34070749 | 0.7333238 | 0.99937574 |
| ELMOD2 | 33.9120036 | -0.0997225 | 0.41030028 | -0.2430475 | 0.80796858 | 0.99937574 |
| TBC1D9 | 163.80588 | -0.4004886 | 0.33567782 | -1.1930742 | 0.2328403 | 0.99937574 |
| RNF150 | 41.411101 | 0.73498472 | 0.97319619 | 0.7552277 | 0.45011237 | 0.99937574 |
| ZNF330 | 30.2177551 | 0.00452449 | 0.47362152 | 0.00955297 | 0.99237795 | 0.99937574 |

| | | | | | | |
|---|---|---|---|---|---|---|
| IL15 | 55.1060712 | 0.03680993 | 0.52090479 | 0.07066537 | 0.94366408 | 0.99937574 |
| INPP4B | 63.6085935 | -2.2917958 | 0.57887204 | -3.9590715 | 7.52E-05 | 0.08437103 |
| USP38 | 28.6513078 | -0.0081449 | 0.39396508 | -0.0206741 | 0.98350566 | 0.99937574 |
| GAB1 | 60.522818 | -0.6626456 | 0.46024288 | -1.4397737 | 0.14993143 | 0.99937574 |
| SMARCA5 | 100.166269 | -0.1348513 | 0.25474799 | -0.5293518 | 0.59656142 | 0.99937574 |
| ANAPC10 | 99.2754725 | 0.30498514 | 0.33606377 | 0.90752163 | 0.36413102 | 0.99937574 |
| ABCE1 | 37.5984343 | 0.46023323 | 0.4122739 | 1.11632881 | 0.26428141 | 0.99937574 |
| OTUD4 | 116.687404 | 0.02801745 | 0.24934249 | 0.11236534 | 0.91053373 | 0.99937574 |
| SMAD1 | 111.592101 | 0.30792835 | 0.29793743 | 1.03353359 | 0.30135426 | 0.99937574 |
| MMAA | 18.4038133 | 0.14704285 | 0.45966232 | 0.31989319 | 0.7490493 | 0.99937574 |
| ZNF827 | 167.447856 | 0.31805354 | 0.45535688 | 0.69847093 | 0.48488273 | 0.99937574 |
| LSM6 | 18.6277379 | -0.0789991 | 0.52167305 | -0.151434 | 0.87963335 | 0.99937574 |
| SLC10A7 | 108.248023 | 0.72635515 | 0.45308919 | 1.60311738 | 0.10890874 | 0.99937574 |
| TMEM184C | 46.7612821 | 0.12650001 | 0.41647886 | 0.30373692 | 0.76132833 | 0.99937574 |
| PRMT9 | 38.4694078 | 0.44259338 | 0.35260781 | 1.25520016 | 0.20940611 | 0.99937574 |
| NR3C2 | 1206.28796 | 0.24122735 | 0.41082507 | 0.58717779 | 0.55708431 | 0.99937574 |
| AC069272.1 | 13.6395043 | -0.1775535 | 0.91542954 | -0.1939565 | 0.84620992 | 0.99937574 |
| DCLK2 | 87.8434261 | -0.143639 | 0.80876283 | -0.1776033 | 0.85903453 | 0.99937574 |
| LRBA | 1075.42828 | 0.00703174 | 0.30247984 | 0.02324699 | 0.98145326 | 0.99937574 |
| RPS3A | 109.724061 | -0.1309048 | 0.52408318 | -0.2497787 | 0.8027585 | 0.99937574 |
| SH3D19 | 59.7711313 | -0.985323 | 0.57239617 | -1.7214005 | 0.08517818 | 0.99937574 |
| FAM160A1 | 336.775051 | -0.4007916 | 0.24382994 | -1.6437342 | 0.10023112 | 0.99937574 |
| GATB | 22.5007864 | -0.2633669 | 0.49762679 | -0.5292457 | 0.59663499 | 0.99937574 |
| FBXW7 | 228.363223 | 0.11898617 | 0.24853828 | 0.47874383 | 0.63212088 | 0.99937574 |
| ARFIP1 | 178.526856 | -0.1152135 | 0.24380528 | -0.4725636 | 0.63652456 | 0.99937574 |
| FHDC1 | 40.8204929 | -0.148632 | 0.42840468 | -0.3469429 | 0.72863421 | 0.99937574 |
| TRIM2 | 649.24508 | -0.1977013 | 0.40556525 | -0.4874709 | 0.62592467 | 0.99937574 |
| MND1 | 20.8544186 | 0.10229908 | 2.02304496 | 0.05056688 | 0.95967065 | 0.99937574 |
| TMEM131L | 17.352297 | -0.114662 | 0.62209202 | -0.1843168 | 0.85376495 | 0.99937574 |
| TLR2 | 26.0184876 | -0.5282433 | 0.75984933 | -0.6951948 | 0.48693321 | 0.99937574 |
| AC079298.3 | 34.7157257 | 0.57764752 | 0.78247663 | 0.73822975 | 0.46037485 | 0.99937574 |
| PLRG1 | 21.5851522 | -0.193214 | 0.53140372 | -0.3635917 | 0.71616291 | 0.99937574 |
| MAP9 | 20.028336 | -0.0648987 | 0.42995159 | -0.1509442 | 0.88001977 | 0.99937574 |
| CTSO | 24.8642798 | -0.6941071 | 0.53885533 | -1.2881139 | 0.1977063 | 0.99937574 |
| PDGFC | 130.111023 | 0.166884 | 0.34495042 | 0.48379128 | 0.62853401 | 0.99937574 |
| TMEM144 | 12.9946841 | -0.2314066 | 0.53913007 | -0.4292222 | 0.6677615 | 0.99937574 |
| ETFDH | 132.02684 | 0.17995145 | 0.47971757 | 0.37511959 | 0.70757153 | 0.99937574 |
| PPID | 45.850265 | -0.0348816 | 0.38467341 | -0.0906785 | 0.92774807 | 0.99937574 |
| FNIP2 | 971.271805 | -0.8214942 | 0.46289182 | -1.7747002 | 7.59E-02 | 0.99937574 |
| C4orf45 | 43.795054 | 0.07825074 | 0.95257408 | 0.08214662 | 0.93453012 | 0.99937574 |
| RAPGEF2 | 563.403196 | 0.41551744 | 0.27414679 | 1.51567504 | 0.12960154 | 0.99937574 |
| NAF1 | 30.3219339 | -0.2359487 | 0.39422729 | -0.5985093 | 0.54950014 | 0.99937574 |
| TMA16 | 16.4652974 | -0.341524 | 0.4658029 | -0.7331941 | 0.46344002 | 0.99937574 |
| 1-Mar | 971.922809 | -1.474316 | 1.31637449 | -1.1199822 | 0.26272136 | 0.99937574 |
| TMEM192 | 42.6946689 | -0.6762318 | 0.52919468 | -1.2778507 | 0.20130209 | 0.99937574 |
| KLHL2 | 139.210658 | -0.0796107 | 0.49884721 | -0.1595894 | 0.87320453 | 0.99937574 |
| MSMO1 | 107.320534 | 0.1905103 | 0.72927734 | 0.26123162 | 0.7939139 | 0.99937574 |
| DDX60 | 53.2456168 | -2.9854756 | 1.32738799 | -2.2491356 | 0.02450387 | 0.96546668 |
| DDX60L | 62.2106495 | -3.6982989 | 1.09710342 | -3.3709665 | 0.00074905 | 0.29012345 |
| PALLD | 158.086308 | 0.75604967 | 0.61095443 | 1.23748945 | 0.21590543 | 0.99937574 |
| CBR4 | 153.43222 | 0.44885204 | 0.3292057 | 1.36343944 | 0.17274406 | 0.99937574 |
| SH3RF1 | 375.578247 | -0.2540467 | 0.46452953 | -0.5468903 | 0.58445408 | 0.99937574 |
| NEK1 | 129.842233 | 0.049209 | 0.24168648 | 0.20360676 | 0.83866081 | 0.99937574 |
| CLCN3 | 159.980156 | 0.13160187 | 0.24622548 | 0.53447706 | 0.59301152 | 0.99937574 |

| | | | | | | |
|---|---|---|---|---|---|---|
| HPF1 | 21.2488301 | 0.37398175 | 0.46775276 | 0.79952869 | 0.42398392 | 0.99937574 |
| MFAP3L | 24.0439678 | -0.5775875 | 0.82995898 | -0.695923 | 0.48647707 | 0.99937574 |
| AADAT | 9.59066862 | -0.060158 | 0.7323559 | -0.0821431 | 0.93453291 | 0.99937574 |
| GALNT7 | 200.48493 | -0.0229022 | 0.42589394 | -0.0537745 | 0.95711485 | 0.99937574 |
| HMGB2 | 19.923321 | -0.104795 | 0.45903473 | -0.2282942 | 0.81941751 | 0.99937574 |
| FBXO8 | 30.035864 | -0.0847493 | 0.37226407 | -0.2276592 | 0.81991118 | 0.99937574 |
| CEP44 | 42.9087223 | 0.19449689 | 0.41277075 | 0.47119833 | 0.63749911 | 0.99937574 |
| WDR17 | 14.6333877 | -0.1538513 | 0.62306973 | -0.2469246 | 0.80496654 | 0.99937574 |
| SPCS3 | 129.841722 | 0.55141153 | 0.24397363 | 2.26012754 | 0.02381334 | 0.96546668 |
| AC078881.1 | 26.4622307 | 0.12207236 | 0.50741121 | 0.24057877 | 0.80988161 | 0.99937574 |
| TENM3 | 171.15324 | -0.4455733 | 0.67402984 | -0.6610587 | 0.50857467 | 0.99937574 |
| DCTD | 33.0097919 | -0.3949679 | 0.3646991 | -1.0829966 | 0.27880994 | 0.99937574 |
| WWC2 | 171.086142 | 0.23488956 | 0.25882732 | 0.90751455 | 0.36413476 | 0.99937574 |
| CDKN2AIP | 15.7275462 | -0.0559741 | 0.51901214 | -0.1078473 | 0.91411681 | 0.99937574 |
| RWDD4 | 39.7955355 | 0.22777712 | 0.34828769 | 0.65399129 | 0.51311742 | 0.99937574 |
| TRAPPC11 | 72.6543507 | 0.36707703 | 0.40916453 | 0.89713795 | 0.36964531 | 0.99937574 |
| STOX2 | 417.01908 | -0.223263 | 0.45844092 | -0.4870049 | 0.62625485 | 0.99937574 |
| IRF2 | 80.2619092 | 0.18168258 | 0.34976249 | 0.51944557 | 0.60345006 | 0.99937574 |
| CASP3 | 25.153243 | -0.2736038 | 0.37308831 | -0.7333486 | 0.46334587 | 0.99937574 |
| PRIMPOL | 29.4773789 | 0.23883906 | 0.64228621 | 0.37185768 | 0.70999882 | 0.99937574 |
| ACSL1 | 416.514392 | 0.12744182 | 0.31447756 | 0.40524933 | 6.85E-01 | 9.99E-01 |
| SLC25A4 | 117.519764 | 0.96130118 | 0.34964834 | 2.74933713 | 0.00597159 | 0.6771384 |
| CFAP97 | 63.5660853 | -0.2425735 | 0.2810818 | -0.8629998 | 0.38813758 | 0.99937574 |
| SNX25 | 102.94155 | -0.2693974 | 0.29792269 | -0.9042529 | 0.36586133 | 0.99937574 |
| AC112722.1 | 12.5581135 | 0.58110494 | 0.57982555 | 1.00220651 | 0.31624386 | 0.99937574 |
| UFSP2 | 40.0876038 | 0.19430297 | 0.36473433 | 0.53272466 | 0.5942242 | 0.99937574 |
| SORBS2 | 188.678161 | 1.06281388 | 0.5623384 | 1.88998986 | 0.05875932 | 0.99937574 |
| FAM149A | 31.4101501 | -0.7112231 | 0.55641561 | -1.2782227 | 0.20117091 | 0.99937574 |
| CYP4V2 | 14.1125805 | -0.2150142 | 0.56313925 | -0.3818136 | 0.70259966 | 0.99937574 |
| FAT1 | 19.7900933 | 0.52028855 | 0.68747183 | 0.75681436 | 0.44916109 | 0.99937574 |
| FRG1-DT | 20.156837 | -0.7722725 | 0.62186189 | -1.2418714 | 0.21428401 | 0.99937574 |
| FRG1 | 21.0712227 | 0.11549985 | 0.41905977 | 0.27561664 | 0.78284253 | 0.99937574 |
| CCDC127 | 10.490375 | -0.3392817 | 0.59413296 | -0.5710534 | 0.56796343 | 0.99937574 |
| SDHA | 162.006676 | 0.35979886 | 0.30140643 | 1.1937332 | 0.23258235 | 0.99937574 |
| PDCD6 | 59.905699 | -0.1928215 | 0.40140667 | -0.4803643 | 0.63096835 | 0.99937574 |
| EXOC3 | 44.5102272 | 0.04396495 | 0.36033199 | 0.12201234 | 0.90288925 | 0.99937574 |
| ZDHHC11B | 21.5605928 | -0.0753478 | 0.75941209 | -0.0992186 | 0.92096472 | 0.99937574 |
| BRD9 | 29.2329852 | 0.1546184 | 0.50634357 | 0.30536263 | 0.76008998 | 0.99937574 |
| SLC12A7 | 83.3660242 | -0.2284012 | 0.39930928 | -0.5719906 | 0.56732833 | 0.99937574 |
| CLPTM1L | 39.4666626 | -0.1820753 | 0.43758505 | -0.4160912 | 0.67734328 | 0.99937574 |
| LPCAT1 | 13.0945039 | -0.2589835 | 0.6115136 | -0.4235122 | 0.6719216 | 0.99937574 |
| MRPL36 | 13.8866962 | 0.35780366 | 0.58236053 | 0.61440231 | 0.53894951 | 0.99937574 |
| NDUFS6 | 70.0657801 | -0.0454086 | 0.42012448 | -0.1080837 | 0.91392933 | 0.99937574 |
| ADAMTS16 | 15.8711151 | 1.12194069 | 0.81264803 | 1.38059855 | 0.16740243 | 0.99937574 |
| ICE1 | 90.5669859 | 0.03478883 | 0.27758336 | 0.12532752 | 0.90026427 | 0.99937574 |
| MED10 | 36.211662 | -0.1740023 | 0.35186191 | -0.4945188 | 0.62093987 | 0.99937574 |
| UBE2QL1 | 107.424685 | 0.44453311 | 0.60997989 | 0.72876684 | 0.4661443 | 0.99937574 |
| NSUN2 | 39.7374771 | -0.1463435 | 0.45146741 | -0.3241507 | 0.74582394 | 0.99937574 |
| TENT4A | 60.8012619 | -0.398573 | 0.34231378 | -1.16435 | 0.24428221 | 0.99937574 |
| MTRR | 38.4745973 | 0.24144077 | 0.37369733 | 0.64608643 | 0.51822338 | 0.99937574 |
| MIR4458HG | 12.3930996 | 0.91660033 | 0.51742225 | 1.77147451 | 0.07648183 | 0.99937574 |
| FAM173B | 24.8389611 | -0.2758035 | 0.53081241 | -0.5195875 | 0.60335108 | 0.99937574 |
| CCT5 | 57.4998496 | 0.42414702 | 0.32016975 | 1.32475669 | 0.18525187 | 0.99937574 |
| CMBL | 40.6304297 | 0.68721813 | 0.49876767 | 1.37783214 | 0.16825512 | 0.99937574 |

| | | | | | | |
|---|---|---|---|---|---|---|
| 6-Mar | 279.755284 | -0.0014287 | 0.24691565 | -0.0057863 | 0.99538319 | 0.99937574 |
| ANKRD33B | 70.7069788 | 1.07182653 | 0.84366591 | 1.27043954 | 0.20392811 | 0.99937574 |
| DAP | 187.864988 | -0.0632556 | 0.42570686 | -0.1485895 | 0.88187758 | 0.99937574 |
| DNAH5 | 140.788094 | -0.6965328 | 1.22953168 | -0.5665025 | 5.71E-01 | 0.99937574 |
| TRIO | 403.57464 | -0.2893763 | 0.53387682 | -0.5420283 | 5.88E-01 | 0.99937574 |
| OTULIN | 45.2459261 | -0.5390954 | 0.39070881 | -1.3797882 | 0.16765187 | 0.99937574 |
| ANKH | 129.869039 | -0.9099756 | 0.45663046 | -1.9928053 | 0.04628278 | 0.99937574 |
| ZNF622 | 13.4889666 | 0.31163693 | 0.63261793 | 0.49261475 | 0.62228482 | 0.99937574 |
| RETREG1 | 303.49355 | 0.24411723 | 0.34267551 | 0.71238596 | 0.4762258 | 0.99937574 |
| MYO10 | 556.253988 | -0.4006088 | 0.34805847 | -1.1509814 | 0.24973989 | 0.99937574 |
| LINC02241 | 28.9897424 | 0.450175 | 0.47414826 | 0.94943932 | 0.34239722 | 0.99937574 |
| CDH12 | 18.565626 | -0.2345293 | 0.710823 | -0.3299405 | 0.74144493 | 0.99937574 |
| C5orf17 | 60.1031334 | -0.1637509 | 1.40500382 | -0.1165484 | 0.90721797 | 0.99937574 |
| DROSHA | 73.3585366 | -0.174887 | 0.27093238 | -0.6455007 | 0.51860275 | 0.99937574 |
| C5orf22 | 23.9466677 | 0.11103581 | 0.40078071 | 0.27704878 | 0.78174265 | 0.99937574 |
| PDZD2 | 173.506115 | 0.59389572 | 0.69514036 | 0.85435368 | 0.39290905 | 0.99937574 |
| GOLPH3 | 227.939289 | -0.0363928 | 0.32066538 | -0.1134916 | 0.90964083 | 0.99937574 |
| MTMR12 | 95.4090981 | 0.33502331 | 0.31733377 | 1.05574425 | 0.29108507 | 0.99937574 |
| ZFR | 265.575716 | 0.12257522 | 0.1906531 | 0.64292279 | 0.5202742 | 0.99937574 |
| SUB1 | 122.440741 | 0.00550082 | 0.35496683 | 0.01549672 | 0.9876359 | 0.99937574 |
| TARS | 38.5688 | 0.13127904 | 0.46660626 | 0.28134864 | 0.77844301 | 0.99937574 |
| C1QTNF3-AN | 87.7179301 | -0.2610897 | 0.44907955 | -0.5813885 | 0.56097862 | 0.99937574 |
| RAI14 | 45.4753517 | -0.3545764 | 0.44424224 | -0.79816 | 0.42477767 | 0.99937574 |
| RAD1 | 14.0154589 | 0.21646915 | 0.52365479 | 0.4133814 | 0.6793272 | 0.99937574 |
| BRIX1 | 19.5876686 | -0.1873701 | 0.54601019 | -0.3431623 | 0.73147639 | 0.99937574 |
| DNAJC21 | 58.4251922 | 0.00796178 | 0.27363405 | 0.02909644 | 0.97678767 | 0.99937574 |
| PRLR | 288.396221 | 0.50178439 | 0.36249086 | 1.38426771 | 0.1662765 | 0.99937574 |
| SPEF2 | 18.5719727 | -0.2166092 | 0.66820342 | -0.3241665 | 0.74581198 | 0.99937574 |
| LMBRD2 | 41.8628226 | 0.27965539 | 0.36317181 | 0.77003607 | 0.4412785 | 0.99937574 |
| NADK2 | 50.8372617 | -0.3026969 | 0.53582841 | -0.5649139 | 0.57213231 | 0.99937574 |
| RANBP3L | 89.1376082 | 0.33893094 | 0.48875074 | 0.69346379 | 0.48801854 | 0.99937574 |
| NIPBL | 482.607684 | -0.0749777 | 0.20958176 | -0.3577493 | 0.72053096 | 0.99937574 |
| CPLANE1 | 147.299487 | -0.2240417 | 0.25211449 | -0.8886507 | 0.37419081 | 0.99937574 |
| NUP155 | 24.120242 | -0.9056035 | 0.44024611 | -2.0570392 | 0.03968246 | 0.99937574 |
| WDR70 | 205.639169 | 0.05477724 | 0.25051214 | 0.21866102 | 0.82691412 | 0.99937574 |
| LIFR | 233.261244 | 0.12661643 | 0.43397927 | 0.29175686 | 0.77047254 | 0.99937574 |
| OSMR-AS1 | 121.852879 | 0.44220994 | 0.45473553 | 0.97245521 | 0.33082414 | 0.99937574 |
| OSMR | 138.894814 | -0.0336715 | 0.50965997 | -0.0660667 | 9.47E-01 | 0.99937574 |
| RICTOR | 320.758927 | 0.24254413 | 0.31026056 | 0.78174335 | 0.43436542 | 0.99937574 |
| FYB1 | 80.5793684 | 1.25689136 | 0.99304604 | 1.26569293 | 0.20562304 | 0.99937574 |
| C9 | 24.92864 | 1.86860203 | 1.03270272 | 1.80942878 | 0.07038442 | 0.99937574 |
| TTC33 | 64.3348328 | 0.35099781 | 0.38829389 | 0.90394883 | 0.36602254 | 0.99937574 |
| PRKAA1 | 108.465332 | -0.1531576 | 0.30652195 | -0.4996628 | 0.61731251 | 0.99937574 |
| RPL37 | 123.207133 | -0.0619288 | 0.36657157 | -0.1689406 | 0.86584336 | 0.99937574 |
| CARD6 | 17.0203137 | -1.4126455 | 0.86715337 | -1.6290607 | 0.10330016 | 0.99937574 |
| C6 | 50.073041 | 2.04728485 | 1.47948226 | 1.38378466 | 0.1664244 | 0.99937574 |
| PLCXD3 | 25.2750044 | -1.2825889 | 0.92088584 | -1.3927773 | 0.16368715 | 0.99937574 |
| OXCT1 | 126.102219 | -0.5799122 | 0.57450891 | -1.0094051 | 0.31278038 | 0.99937574 |
| C5orf51 | 19.5576414 | 0.17340733 | 0.47122947 | 0.36798914 | 0.71288133 | 0.99937574 |
| FBXO4 | 15.1397059 | -0.4317466 | 0.54647463 | -0.7900579 | 0.42949394 | 0.99937574 |
| SELENOP | 125.033984 | -0.0082828 | 0.47233025 | -0.0175361 | 0.98600891 | 0.99937574 |
| ZNF131 | 128.165839 | 0.0938516 | 0.24201691 | 0.38778945 | 0.69817186 | 0.99937574 |
| HMGCS1 | 172.233105 | -0.2409917 | 0.80614762 | -0.2989424 | 0.76498397 | 0.99937574 |
| CCL28 | 14.0156576 | -1.1646174 | 0.90688173 | -1.2841999 | 0.19907202 | 0.99937574 |

| | | | | | | |
|---|---|---|---|---|---|---|
| TMEM267 | 36.3097994 | -0.0987133 | 0.38823933 | -0.2542589 | 0.79929556 | 0.99937574 |
| C5orf34 | 11.8932949 | 0.64688681 | 0.73852517 | 0.87591708 | 0.38107511 | 0.99937574 |
| PAIP1 | 54.9649343 | 0.05756829 | 0.30294314 | 0.19003002 | 0.8492856 | 0.99937574 |
| NNT-AS1 | 119.226427 | 0.65184803 | 0.31928775 | 2.04156917 | 0.04119429 | 0.99937574 |
| NNT | 396.921318 | 0.23820378 | 0.33634028 | 0.70822258 | 0.47880705 | 0.99937574 |
| MRPS30-DT | 13.06176 | -0.7598627 | 0.68574508 | -1.1080833 | 0.26782584 | 0.99937574 |
| MRPS30 | 28.0187645 | 0.03165779 | 0.40126849 | 0.07889429 | 0.9371167 | 0.99937574 |
| PARP8 | 191.152943 | -0.1487256 | 0.49562667 | -0.3000758 | 0.76411931 | 0.99937574 |
| ITGA1 | 95.7466479 | -0.344732 | 0.48111218 | -0.7165315 | 0.47366321 | 0.99937574 |
| ITGA2 | 171.409333 | -0.5072963 | 0.51872655 | -0.9779648 | 0.32809174 | 0.99937574 |
| MOCS2 | 31.9486492 | 0.40209409 | 0.34708958 | 1.15847351 | 0.24667085 | 0.99937574 |
| NDUFS4 | 153.450909 | -0.0167173 | 0.24703487 | -0.0676716 | 0.94604704 | 0.99937574 |
| ARL15 | 2430.49289 | 0.12461949 | 0.22514699 | 0.5535028 | 0.57991916 | 0.99937574 |
| SNX18 | 49.8209512 | 0.67611944 | 0.42858454 | 1.57756376 | 0.11466586 | 0.99937574 |
| AC026704.1 | 13.663009 | -0.1408561 | 0.84316332 | -0.1670568 | 0.86732538 | 0.99937574 |
| DHX29 | 76.9640649 | -0.0921922 | 0.28122931 | -0.3278188 | 0.7430487 | 0.99937574 |
| MTREX | 143.575402 | -0.0318739 | 0.27150025 | -0.1173991 | 0.90654381 | 0.99937574 |
| PLPP1 | 138.993824 | -0.5714451 | 0.47216058 | -1.2102771 | 0.22617259 | 0.99937574 |
| SLC38A9 | 77.718935 | 0.29084563 | 0.28635763 | 1.0156727 | 0.30978527 | 0.99937574 |
| IL6ST | 168.477816 | 0.04803725 | 0.27236135 | 0.17637322 | 0.86000074 | 0.99937574 |
| LINC01948 | 10.628009 | 0.7642818 | 0.9039847 | 0.84545878 | 0.39785474 | 0.99937574 |
| MAP3K1 | 156.987714 | 0.01430015 | 0.27797304 | 0.05144438 | 0.95897142 | 0.99937574 |
| MIER3 | 30.5215434 | 0.21134703 | 0.36372617 | 0.58106085 | 0.56119944 | 0.99937574 |
| GPBP1 | 278.964933 | -0.2770251 | 0.30856131 | -0.8977959 | 0.36929439 | 0.99937574 |
| PDE4D | 4382.79138 | 0.24062257 | 0.56028553 | 0.42946419 | 0.66758545 | 0.99937574 |
| PART1 | 45.556407 | 2.18454076 | 0.95102444 | 2.29703956 | 0.02161651 | 0.96427895 |
| ELOVL7 | 258.58933 | 0.39797117 | 0.57966879 | 0.68654924 | 0.49236682 | 0.99937574 |
| ERCC8 | 60.9881694 | -0.0502904 | 0.30679615 | -0.1639213 | 0.86979309 | 0.99937574 |
| NDUFAF2 | 139.150917 | -0.408955 | 0.40616762 | -1.0068626 | 0.3140008 | 0.99937574 |
| SMIM15 | 26.0674047 | -0.2114965 | 0.3757178 | -0.5629132 | 0.57349402 | 0.99937574 |
| ZSWIM6 | 980.702103 | 0.17051143 | 0.35589464 | 0.47910649 | 0.63186288 | 0.99937574 |
| KIF2A | 63.9956081 | -0.0124221 | 0.36334611 | -0.034188 | 0.97272723 | 0.99937574 |
| IPO11 | 44.8737448 | -0.2699074 | 0.39057459 | -0.6910522 | 0.48953274 | 0.99937574 |
| RNF180 | 118.429564 | 0.0207386 | 0.26692226 | 0.07769528 | 0.93807045 | 0.99937574 |
| SREK1IP1 | 43.8912865 | 0.46815898 | 0.30607961 | 1.52953335 | 0.12613228 | 0.99937574 |
| CWC27 | 97.0651157 | -0.1889581 | 0.26600595 | -0.7103527 | 0.47748542 | 0.99937574 |
| PPWD1 | 65.5214661 | -0.1529538 | 0.40693718 | -0.3758659 | 0.7070166 | 0.99937574 |
| TRIM23 | 30.9444876 | 0.09395648 | 0.38133556 | 0.24638792 | 0.80538196 | 0.99937574 |
| TRAPPC13 | 56.281182 | 0.40545704 | 0.29322131 | 1.38276799 | 0.16673602 | 0.99937574 |
| NLN | 37.5324169 | -0.5173096 | 0.57352729 | -0.9019791 | 0.36706795 | 0.99937574 |
| ERBIN | 480.186971 | -0.3136707 | 0.2566165 | -1.2223327 | 0.22158185 | 0.99937574 |
| SREK1 | 182.865638 | -0.0667514 | 0.29983012 | -0.2226307 | 0.82382291 | 0.99937574 |
| AC113378.1 | 16.7561304 | 0.54966743 | 1.03757501 | 0.52976163 | 0.59627721 | 0.99937574 |
| MAST4 | 725.533984 | 0.13062213 | 0.68145394 | 0.19168152 | 0.84799168 | 0.99937574 |
| PIK3R1 | 110.042457 | 0.63314388 | 0.52306823 | 1.21044223 | 0.22610924 | 0.99937574 |
| SLC30A5 | 54.1423928 | 0.1906734 | 0.31400207 | 0.60723611 | 0.54369423 | 0.99937574 |
| MRPS36 | 64.016107 | 0.111903 | 0.46001095 | 0.24326159 | 0.80780276 | 0.99937574 |
| CDK7 | 29.6895149 | 0.20086969 | 0.44142181 | 0.45505158 | 0.64907213 | 0.99937574 |
| CCDC125 | 42.7897737 | -0.1557188 | 0.54678067 | -0.284792 | 0.77580347 | 0.99937574 |
| AK6 | 24.4160288 | -0.0194877 | 0.48872648 | -0.0398744 | 0.96819326 | 0.99937574 |
| RAD17 | 50.0449988 | 0.17148934 | 0.30444032 | 0.56329378 | 0.57323485 | 0.99937574 |
| MARVELD2 | 39.5729943 | 0.21187807 | 0.35277395 | 0.60060578 | 0.54810259 | 0.99937574 |
| OCLN | 100.332207 | -0.3347704 | 0.38991642 | -0.8585696 | 0.39057801 | 0.99937574 |
| GTF2H2C | 62.9535363 | 0.08214627 | 0.456182 | 0.18007346 | 0.8570949 | 0.99937574 |

| | | | | | | |
|---|---|---|---|---|---|---|
| NAIP | 103.912175 | 0.17719938 | 0.41161842 | 0.43049429 | 0.66683612 | 0.99937574 |
| GTF2H2 | 53.1108517 | 0.47861231 | 0.62925166 | 0.76060556 | 0.4468927 | 0.99937574 |
| BDP1 | 211.34537 | -0.0612844 | 0.24197613 | -0.2532663 | 0.80006241 | 0.99937574 |
| MCCC2 | 86.2922928 | 0.27749421 | 0.26920629 | 1.03078651 | 0.30264095 | 0.99937574 |
| MRPS27 | 72.1235238 | 0.41835722 | 0.28148542 | 1.48624828 | 0.13721346 | 0.99937574 |
| PTCD2 | 11.3769836 | -0.265785 | 0.78931525 | -0.3367286 | 0.73632151 | 0.99937574 |
| TNPO1 | 168.692589 | 0.04267855 | 0.24678408 | 0.17293883 | 0.86269951 | 0.99937574 |
| FCHO2 | 202.407457 | 0.00522008 | 0.33864768 | 0.01541448 | 0.98770151 | 0.99937574 |
| TMEM171 | 29.3732907 | -0.6199356 | 0.65055149 | -0.9529386 | 0.34062116 | 0.99937574 |
| AC116345.3 | 21.3632928 | 1.73157087 | 0.83193259 | 2.08138361 | 0.03739881 | 0.99937574 |
| AC116345.1 | 16.6707129 | 1.45053959 | 0.80362718 | 1.8049907 | 0.07107614 | 0.99937574 |
| BTF3 | 77.3252368 | 0.2951477 | 0.55848983 | 0.52847462 | 0.59716996 | 0.99937574 |
| ANKRA2 | 25.5385147 | 0.28271264 | 0.58581218 | 0.48259945 | 0.62938017 | 0.99937574 |
| ARHGEF28 | 916.042728 | -0.4034799 | 0.27400091 | -1.4725495 | 0.14087256 | 0.99937574 |
| ENC1 | 15.9183004 | 0.50352846 | 0.95075989 | 0.52960633 | 0.5963849 | 0.99937574 |
| HEXB | 36.9600801 | -0.7132674 | 0.40078576 | -1.7796725 | 0.07512957 | 0.99937574 |
| GFM2 | 42.4823855 | 0.40182761 | 0.34806819 | 1.15445082 | 0.24831539 | 0.99937574 |
| NSA2 | 37.6107793 | -0.1357553 | 0.46237977 | -0.2936014 | 0.76906251 | 0.99937574 |
| FAM169A | 100.639557 | 0.15843059 | 0.37623603 | 0.42109363 | 0.67368672 | 0.99937574 |
| GCNT4 | 11.9438446 | 0.08896444 | 0.605581 | 0.14690758 | 0.88320497 | 0.99937574 |
| HMGCR | 128.817577 | 0.35672035 | 0.59040161 | 0.60419949 | 0.54571102 | 0.99937574 |
| COL4A3BP | 224.40394 | -0.024756 | 0.2811461 | -0.0880537 | 0.929834 | 0.99937574 |
| POLK | 142.288269 | -0.0162391 | 0.26745314 | -0.0607174 | 0.95158426 | 0.99937574 |
| ANKDD1B | 76.7633375 | 1.05089374 | 0.45817123 | 2.29367029 | 0.02180944 | 0.96427895 |
| POC5 | 26.812309 | 0.46192863 | 0.36839226 | 1.25390425 | 0.20987681 | 0.99937574 |
| IQGAP2 | 191.450213 | 0.63394083 | 0.56719138 | 1.11768418 | 0.2637019 | 0.99937574 |
| F2RL1 | 21.5668774 | -0.2807266 | 0.45682445 | -0.6145174 | 0.53887348 | 0.99937574 |
| ZBED3 | 21.492917 | 0.08449001 | 0.4452475 | 0.18975965 | 0.84949748 | 0.99937574 |
| WDR41 | 97.6152805 | -0.3721687 | 0.40951266 | -0.9088089 | 0.36345101 | 0.99937574 |
| TBCA | 179.440374 | 0.05747031 | 0.31332805 | 0.18341899 | 0.85446928 | 0.99937574 |
| AP3B1 | 295.558313 | 0.09434184 | 0.29741594 | 0.31720507 | 0.751088 | 0.99937574 |
| SCAMP1-AS1 | 10.8638167 | -0.4768698 | 0.69546173 | -0.6856881 | 0.49290981 | 0.99937574 |
| SCAMP1 | 152.151145 | -0.2763093 | 0.27454891 | -1.006412 | 0.31421743 | 0.99937574 |
| ARSB | 24.0231902 | -0.2878031 | 0.60780466 | -0.4735125 | 0.63584759 | 0.99937574 |
| DMGDH | 80.6909481 | 0.66010378 | 0.62194355 | 1.06135642 | 0.28852795 | 0.99937574 |
| BHMT2 | 42.1665386 | 0.71161164 | 0.48001352 | 1.48248249 | 0.13821196 | 0.99937574 |
| JMY | 112.362154 | 0.16894869 | 0.45639822 | 0.37017824 | 0.71124969 | 0.99937574 |
| HOMER1 | 48.3468413 | 0.53912665 | 0.52934982 | 1.0184695 | 0.30845489 | 0.99937574 |
| TENT2 | 281.991854 | -0.2631805 | 0.28448289 | -0.9251188 | 0.3549041 | 0.99937574 |
| CMYA5 | 139.820912 | -0.5689016 | 0.48783601 | -1.1661738 | 0.24354417 | 0.99937574 |
| MTX3 | 25.2397863 | 0.14858384 | 0.45141708 | 0.32914981 | 0.74204246 | 0.99937574 |
| SERINC5 | 116.298324 | -1.2172538 | 0.43110697 | -2.8235539 | 0.00474944 | 0.60897109 |
| ZFYVE16 | 178.917693 | -0.0696927 | 0.36076394 | -0.193181 | 0.84681726 | 0.99937574 |
| FAM151B | 26.9048612 | 0.11776542 | 0.81483705 | 0.14452635 | 0.88508485 | 0.99937574 |
| DHFR | 483.818527 | -0.6179949 | 0.52427299 | -1.1787655 | 0.23849156 | 0.99937574 |
| MSH3 | 107.966946 | -0.0298036 | 0.29288249 | -0.1017595 | 0.91894759 | 0.99937574 |
| RASGRF2 | 193.370972 | -0.2006957 | 0.43528576 | -0.4610665 | 0.64475088 | 0.99937574 |
| CKMT2-AS1 | 67.7548086 | -0.0338366 | 0.37793219 | -0.089531 | 0.92865993 | 0.99937574 |
| CKMT2 | 119.53857 | 0.62179742 | 0.39799186 | 1.56233703 | 0.11820861 | 0.99937574 |
| ZCCHC9 | 33.3579163 | -0.4038477 | 0.45149099 | -0.8944757 | 0.37106744 | 0.99937574 |
| SSBP2 | 47.7539067 | -1.2913776 | 0.77764385 | -1.6606286 | 0.09678807 | 0.99937574 |
| ATG10 | 117.958188 | -0.3042428 | 0.43883575 | -0.6932953 | 0.48812424 | 0.99937574 |
| RPS23 | 268.613554 | -0.1526001 | 0.55238764 | -0.2762555 | 0.7823518 | 0.99937574 |
| ATP6AP1L | 30.9351952 | 0.57069911 | 0.5174568 | 1.10289228 | 0.27007395 | 0.99937574 |

| | | | | | | |
|---|---|---|---|---|---|---|
| TMEM167A | 52.3723428 | 0.08281481 | 0.43696824 | 0.18952134 | 0.84968423 | 0.99937574 |
| XRCC4 | 53.670484 | -0.6947635 | 0.35045999 | -1.9824332 | 0.04743078 | 0.99937574 |
| COX7C | 203.519432 | 0.53354515 | 0.43954966 | 1.21384498 | 0.22480693 | 0.99937574 |
| RASA1 | 189.887054 | -0.0450906 | 0.31175013 | -0.1446371 | 0.88499742 | 0.99937574 |
| CCNH | 202.357219 | 0.38062666 | 0.36001936 | 1.05723941 | 0.29040234 | 0.99937574 |
| TMEM161B | 91.2268678 | -0.1002589 | 0.30031389 | -0.3338471 | 0.73849492 | 0.99937574 |
| TMEM161B-/ | 186.133639 | -0.0062874 | 0.55478856 | -0.0113329 | 0.99095784 | 0.99937574 |
| MEF2C | 27.9774587 | 0.88902387 | 0.79818243 | 1.11381038 | 0.26536054 | 0.99937574 |
| CETN3 | 22.2184667 | -0.2176388 | 0.59975658 | -0.3628786 | 0.7166956 | 0.99937574 |
| LYSMD3 | 38.7436211 | 0.04822203 | 0.4700908 | 0.10258025 | 0.91829612 | 0.99937574 |
| ADGRV1 | 59.0557951 | -1.0184034 | 0.62306421 | -1.6345079 | 0.10215225 | 0.99937574 |
| ARRDC3 | 48.7780022 | 0.07225791 | 0.55025958 | 0.13131604 | 0.89552531 | 0.99937574 |
| ARRDC3-AS1 | 40.5080455 | 0.72442863 | 0.39426796 | 1.83740171 | 0.06615061 | 0.99937574 |
| FAM172A | 253.19501 | -0.2097852 | 0.2928945 | -0.7162485 | 0.47383795 | 0.99937574 |
| KIAA0825 | 49.0394106 | 0.24284152 | 0.35934018 | 0.67579841 | 0.49916865 | 0.99937574 |
| SLF1 | 12.5680113 | -0.3610594 | 0.54122038 | -0.6671209 | 0.5046949 | 0.99937574 |
| TTC37 | 161.576449 | 0.0432498 | 0.24584154 | 0.17592552 | 0.86035246 | 0.99937574 |
| ARSK | 38.0847365 | 0.37730122 | 0.49372914 | 0.76418666 | 0.44475601 | 0.99937574 |
| RHOBTB3 | 252.743988 | 0.11995533 | 0.37716793 | 0.31804222 | 0.75045291 | 0.99937574 |
| GLRX | 79.3902113 | 0.1212468 | 0.58168237 | 0.20844159 | 0.83488419 | 0.99937574 |
| ELL2 | 162.072636 | -0.1120798 | 0.50346127 | -0.2226186 | 0.82383238 | 0.99937574 |
| AC104123.1 | 56.1730941 | -0.6420902 | 0.57038541 | -1.1257128 | 0.26028709 | 0.99937574 |
| CAST | 256.163604 | -0.2457984 | 0.23443071 | -1.0484905 | 0.29441268 | 0.99937574 |
| ERAP1 | 70.8283018 | -0.0514291 | 0.37409038 | -0.1374778 | 0.89065313 | 0.99937574 |
| AC009126.1 | 27.7195328 | 1.11128142 | 0.59046426 | 1.8820469 | 0.05982965 | 0.99937574 |
| ERAP2 | 32.5529022 | 0.51054622 | 0.73665418 | 0.69306092 | 0.48827132 | 0.99937574 |
| LNPEP | 275.267031 | 0.48487707 | 0.23924028 | 2.02673675 | 0.04268934 | 0.99937574 |
| LIX1-AS1 | 42.835524 | 1.06982968 | 0.67673805 | 1.58086233 | 0.11390952 | 0.99937574 |
| RIOK2 | 13.2603142 | 0.39436507 | 0.64387286 | 0.61248904 | 0.54021425 | 0.99937574 |
| RGMB | 12.9504337 | 1.10923235 | 0.56723113 | 1.95552094 | 0.05052161 | 0.99937574 |
| CHD1 | 136.386058 | -0.1778979 | 0.33723848 | -0.5275136 | 0.597837 | 0.99937574 |
| LINC02062 | 28.8230572 | 0.71818573 | 0.53200944 | 1.34994923 | 0.17703227 | 0.99937574 |
| FAM174A | 47.9351365 | 0.39418323 | 0.34431377 | 1.14483724 | 0.25227658 | 0.99937574 |
| SLCO4C1 | 33.8187867 | -1.0480316 | 1.1793041 | -0.8886865 | 0.37417159 | 0.99937574 |
| PAM | 212.488001 | -0.3899185 | 0.43425095 | -0.8979105 | 0.36923329 | 0.99937574 |
| GIN1 | 21.8431134 | -0.1365047 | 0.47032279 | -0.2902363 | 0.77163548 | 0.99937574 |
| PPIP5K2 | 78.272822 | -0.2535053 | 0.29125863 | -0.8703786 | 0.38409355 | 0.99937574 |
| C5orf30 | 12.5234446 | 0.9740138 | 0.71618372 | 1.36000551 | 0.17382818 | 0.99937574 |
| NUDT12 | 28.2278207 | -0.1898131 | 0.42746699 | -0.4440416 | 0.65701248 | 0.99937574 |
| EFNA5 | 1350.41241 | 0.427325 | 0.50173533 | 0.85169406 | 0.39438392 | 0.99937574 |
| FBXL17 | 638.854086 | 0.36336211 | 0.23699307 | 1.53321824 | 0.12522209 | 0.99937574 |
| FER | 407.956483 | -0.0038591 | 0.27893029 | -0.0138353 | 0.98896138 | 0.99937574 |
| AC008467.1 | 20.3730589 | -0.0197047 | 0.8620683 | -0.0228575 | 0.98176392 | 0.99937574 |
| PJA2 | 173.402963 | 0.15384504 | 0.21038292 | 0.73126203 | 0.46461912 | 0.99937574 |
| MAN2A1 | 336.150671 | -0.6686667 | 0.4550016 | -1.469592 | 0.14167229 | 0.99937574 |
| TMEM232 | 49.3103466 | -0.2734129 | 0.6330375 | -0.4319064 | 0.66580946 | 0.99937574 |
| SLC25A46 | 66.9049566 | -0.2154583 | 0.42698037 | -0.5046093 | 0.61383326 | 0.99937574 |
| WDR36 | 41.5090266 | 0.20067372 | 0.32542482 | 0.61665156 | 0.53746458 | 0.99937574 |
| STARD4 | 25.2586508 | 0.09562904 | 0.58784941 | 0.16267609 | 0.87077347 | 0.99937574 |
| NREP | 30.6173253 | 0.38838412 | 0.54745241 | 0.70943905 | 0.47805206 | 0.99937574 |
| EPB41L4A | 329.67599 | -0.2830757 | 0.29436787 | -0.9616394 | 0.33623079 | 0.99937574 |
| EPB41L4A-A! | 12.0166411 | -0.3023179 | 0.51383618 | -0.5883547 | 0.55629422 | 0.99937574 |
| APC | 61.8328605 | -0.4632192 | 0.28860413 | -1.6050331 | 0.10848651 | 0.99937574 |
| SRP19 | 36.0737254 | 0.46518154 | 0.49955331 | 0.931195 | 0.3517527 | 0.99937574 |

| | | | | | | |
|---|---|---|---|---|---|---|
| REEP5 | 188.814698 | 0.05934214 | 0.31945486 | 0.18576063 | 0.85263248 | 0.99937574 |
| DCP2 | 107.509752 | -0.2880861 | 0.51842617 | -0.5556936 | 0.57842034 | 0.99937574 |
| MCC | 188.391619 | -0.37195 | 0.46252751 | -0.8041683 | 0.42129979 | 0.99937574 |
| YTHDC2 | 128.912279 | 0.08563292 | 0.26906178 | 0.31826488 | 0.75028402 | 0.99937574 |
| PGGT1B | 52.2517215 | 0.39485411 | 0.39947141 | 0.98844149 | 0.32293647 | 0.99937574 |
| CCDC112 | 13.8206701 | 0.37740979 | 0.69045047 | 0.54661385 | 0.58464405 | 0.99937574 |
| FEM1C | 20.8628336 | -0.0068355 | 0.45583871 | -0.0149955 | 0.98803577 | 0.99937574 |
| TMED7-TICA | 17.1129493 | 0.82013237 | 0.49659475 | 1.65151237 | 0.098634 | 0.99937574 |
| ATG12 | 37.5105173 | 0.03694484 | 0.46451407 | 0.07953439 | 0.93660758 | 0.99937574 |
| AP3S1 | 54.1564513 | -0.5153139 | 0.35751661 | -1.4413705 | 0.14948005 | 0.99937574 |
| COMMD10 | 245.729132 | -0.4336634 | 0.21094645 | -2.0557987 | 0.03980192 | 0.99937574 |
| SEMA6A | 13.4515325 | 0.3896974 | 0.62994653 | 0.6186198 | 0.53616685 | 0.99937574 |
| SEMA6A-AS1 | 34.2109682 | 0.05015002 | 0.48726809 | 0.1029208 | 0.91802583 | 0.99937574 |
| DTWD2 | 59.751306 | -0.0611141 | 0.43076212 | -0.1418743 | 0.88717931 | 0.99937574 |
| DMXL1 | 278.769481 | -0.0167829 | 0.26522298 | -0.0632786 | 0.94954464 | 0.99937574 |
| TNFAIP8 | 52.3324953 | -0.3881941 | 0.5006611 | -0.7753629 | 0.43812524 | 0.99937574 |
| HSD17B4 | 193.610308 | 0.20149836 | 0.47179975 | 0.42708449 | 0.66931779 | 0.99937574 |
| SRFBP1 | 47.8135634 | -0.3438386 | 0.33519272 | -1.0257938 | 0.30498881 | 0.99937574 |
| AC010255.1 | 17.9801771 | -5.0973071 | 2.23916967 | -2.2764274 | 0.02282044 | 0.96427895 |
| SNX2 | 102.995115 | -0.2308472 | 0.28754224 | -0.8028289 | 0.42207363 | 0.99937574 |
| SNX24 | 250.816953 | 0.28452002 | 0.32181302 | 0.88411592 | 0.37663364 | 0.99937574 |
| PPIC | 55.5213616 | 0.33783589 | 0.32788291 | 1.0303553 | 0.30284325 | 0.99937574 |
| CEP120 | 32.230279 | 0.03578799 | 0.45593261 | 0.07849403 | 0.93743508 | 0.99937574 |
| CSNK1G3 | 111.410322 | -0.1745749 | 0.41937344 | -0.4162756 | 0.67720834 | 0.99937574 |
| ZNF608 | 123.883236 | 0.89909539 | 0.56652618 | 1.58703238 | 0.11250531 | 0.99937574 |
| GRAMD2B | 415.529507 | 0.17594258 | 0.20364965 | 0.86394736 | 0.38761681 | 0.99937574 |
| AC093535.1 | 18.4278706 | 0.46209898 | 0.84872107 | 0.54446507 | 0.58612148 | 0.99937574 |
| ALDH7A1 | 28.1058611 | 0.60081582 | 0.46449538 | 1.29348072 | 0.19584485 | 0.99937574 |
| PHAX | 28.4297497 | -0.0923193 | 0.47958654 | -0.1924977 | 0.84735239 | 0.99937574 |
| C5orf63 | 27.0822654 | -0.1509619 | 0.48745017 | -0.3096971 | 0.75679132 | 0.99937574 |
| PRRC1 | 30.6359498 | 0.00905583 | 0.37819697 | 0.02394473 | 0.98089669 | 0.99937574 |
| LINC01184 | 57.9133818 | -0.2200788 | 0.52929567 | -0.4157956 | 0.67755961 | 0.99937574 |
| SLC12A2 | 44.0785036 | 0.17357724 | 0.60391529 | 0.28741984 | 0.77379087 | 0.99937574 |
| SLC27A6 | 17.730567 | -0.5293186 | 1.70695342 | -0.3100955 | 0.75648832 | 0.99937574 |
| ISOC1 | 20.0044263 | 0.79243822 | 0.50675416 | 1.56375276 | 0.11787564 | 0.99937574 |
| HINT1 | 197.531238 | -0.0121662 | 0.4938693 | -0.0246344 | 0.98034661 | 0.99937574 |
| LYRM7 | 31.964509 | 0.4714052 | 0.386962 | 1.21822092 | 0.22314003 | 0.99937574 |
| CDC42SE2 | 448.716684 | -0.344354 | 0.41988687 | -0.8201113 | 0.41215267 | 0.99937574 |
| P4HA2 | 14.296748 | 0.33320465 | 0.57438723 | 0.58010455 | 0.56184412 | 0.99937574 |
| SLC22A4 | 13.8521868 | 0.04901857 | 0.80957845 | 0.06054826 | 0.95171898 | 0.99937574 |
| SLC22A5 | 90.633083 | 0.13399852 | 0.53554365 | 0.25021026 | 0.80242475 | 0.99937574 |
| AC116366.3 | 86.1363805 | -1.0339015 | 0.70594085 | -1.4645725 | 0.14303759 | 0.99937574 |
| IRF1 | 21.5209047 | 0.02623371 | 0.63099596 | 0.04157508 | 0.96683744 | 0.99937574 |
| KIF3A | 65.448321 | 0.38185165 | 0.30647794 | 1.24593519 | 0.21278819 | 0.99937574 |
| CCNI2 | 28.358317 | 0.04493132 | 0.87997708 | 0.05105965 | 0.95927799 | 0.99937574 |
| 8-Sep | 44.0615826 | 0.31124954 | 0.4033408 | 0.77167879 | 0.4403047 | 0.99937574 |
| SOWAHA | 15.1414701 | 0.84366614 | 0.51471695 | 1.63908754 | 0.10119503 | 0.99937574 |
| SHROOM1 | 12.8365271 | -0.2527222 | 0.56230347 | -0.449441 | 0.65311356 | 0.99937574 |
| UQCRQ | 98.2260355 | 0.35083194 | 0.46939869 | 0.74740715 | 0.45481783 | 0.99937574 |
| AFF4 | 322.917331 | -0.3704431 | 0.26375064 | -1.4045201 | 0.16016404 | 0.99937574 |
| ZCCHC10 | 47.8726134 | -0.4082446 | 0.36429459 | -1.1206441 | 0.26243938 | 0.99937574 |
| HSPA4 | 103.567337 | 0.00391547 | 0.31275742 | 0.01251919 | 0.99001139 | 0.99937574 |
| FSTL4 | 94.3025209 | 1.38168134 | 0.95281805 | 1.45009989 | 0.14703067 | 0.99937574 |
| C5orf15 | 57.7616886 | 0.09983858 | 0.29473899 | 0.33873558 | 0.73480893 | 0.99937574 |

| | | | | | | |
|---|---|---|---|---|---|---|
| VDAC1 | 178.270066 | 0.69211366 | 0.40310891 | 1.71693962 | 0.0859902 | 0.99937574 |
| TCF7 | 15.0684582 | 0.62992826 | 0.53421544 | 1.1791652 | 0.2383324 | 0.99937574 |
| SKP1 | 155.79046 | 0.20429471 | 0.49006051 | 0.4168765 | 0.67676874 | 0.99937574 |
| CDKL3 | 20.4224261 | -0.1137494 | 0.42584978 | -0.2671116 | 0.78938324 | 0.99937574 |
| UBE2B | 121.768367 | -0.3948983 | 0.31020199 | -1.2730359 | 0.2030053 | 0.99937574 |
| JADE2 | 31.3094827 | -0.5222602 | 0.3962171 | -1.3181163 | 0.18746471 | 0.99937574 |
| SAR1B | 55.7198488 | 0.07377056 | 0.32216878 | 0.22898109 | 0.81888361 | 0.99937574 |
| SEC24A | 59.5144956 | -0.1221864 | 0.34755364 | -0.3515612 | 0.72516736 | 0.99937574 |
| CAMLG | 27.7769471 | 0.2909828 | 0.48647503 | 0.5981454 | 0.54974292 | 0.99937574 |
| DDX46 | 105.173529 | 0.1929002 | 0.30490656 | 0.63265348 | 0.52695996 | 0.99937574 |
| C5orf24 | 39.87465 | -0.8396839 | 0.55400859 | -1.5156514 | 0.12960752 | 0.99937574 |
| TXNDC15 | 19.9064271 | -0.2695432 | 0.45722748 | -0.5895166 | 0.55551476 | 0.99937574 |
| PCBD2 | 95.5549506 | -0.7762059 | 0.37311851 | -2.08032 | 0.03749619 | 0.99937574 |
| H2AFY | 96.8276854 | 0.11501546 | 0.37510106 | 0.30662525 | 0.75912863 | 0.99937574 |
| CXCL14 | 15.5047917 | 1.27709307 | 0.93460566 | 1.36645125 | 0.17179737 | 0.99937574 |
| SMAD5 | 102.991751 | 0.23407434 | 0.2263965 | 1.03391326 | 0.30117671 | 0.99937574 |
| KLHL3 | 76.5487234 | 1.08622747 | 0.60537814 | 1.79429582 | 0.07276597 | 0.99937574 |
| HNRNPA0 | 70.5230849 | -0.5905139 | 0.42739157 | -1.3816695 | 0.16707322 | 0.99937574 |
| AC106791.1 | 29.7810553 | -0.1593341 | 0.79529772 | -0.2003453 | 0.84121057 | 0.99937574 |
| FAM13B | 265.733528 | -0.0256793 | 0.23927867 | -0.1073195 | 0.91453551 | 0.99937574 |
| NME5 | 14.2409984 | 0.45907136 | 0.73818398 | 0.62189287 | 0.53401231 | 0.99937574 |
| BRD8 | 41.4407132 | 0.11421419 | 0.32689114 | 0.34939519 | 0.72679265 | 0.99937574 |
| CDC23 | 24.4460781 | 0.14962756 | 0.45667161 | 0.32764803 | 0.7431778 | 0.99937574 |
| FAM53C | 23.2436053 | 0.13070001 | 0.60911319 | 0.21457426 | 0.83009926 | 0.99937574 |
| KDM3B | 113.42111 | -0.0317624 | 0.22575806 | -0.1406924 | 0.88811297 | 0.99937574 |
| EGR1 | 43.0237813 | -1.7579697 | 1.00501626 | -1.7491953 | 0.08025727 | 0.99937574 |
| ETF1 | 70.2799483 | -0.6781325 | 0.36559452 | -1.8548759 | 0.06361395 | 0.99937574 |
| HSPA9 | 180.872904 | -0.1318702 | 0.30391235 | -0.4339088 | 0.66435468 | 0.99937574 |
| CTNNA1 | 345.159927 | -0.1282252 | 0.23986873 | -0.5345641 | 0.59295129 | 0.99937574 |
| SIL1 | 142.728451 | -0.0661619 | 0.34532557 | -0.1915928 | 0.84806121 | 0.99937574 |
| PAIP2 | 76.905895 | 0.00235436 | 0.33352686 | 0.00705899 | 0.99436778 | 0.99937574 |
| TMEM173 | 25.6679144 | 0.0939207 | 0.62409184 | 0.15049179 | 0.88037663 | 0.99937574 |
| UBE2D2 | 171.431138 | -0.4104735 | 0.28779368 | -1.4262769 | 0.15378843 | 0.99937574 |
| CXXC5 | 80.129448 | 0.61497643 | 0.45587075 | 1.34901488 | 0.17733219 | 0.99937574 |
| NRG2 | 21.6582185 | 0.59259314 | 1.05914757 | 0.55950007 | 0.57582048 | 0.99937574 |
| PURA | 122.008744 | 0.22255313 | 0.2519352 | 0.88337448 | 0.37703397 | 0.99937574 |
| IGIP | 18.0196681 | 0.7835803 | 0.47398621 | 1.6531711 | 0.09829605 | 0.99937574 |
| CYSTM1 | 412.842866 | 0.0116516 | 0.43430602 | 0.02682808 | 0.97859686 | 0.99937574 |
| PFDN1 | 46.2727404 | -0.6398525 | 0.33054059 | -1.9357759 | 0.05289515 | 0.99937574 |
| SLC4A9 | 242.023824 | 0.44937555 | 0.98202041 | 0.45760306 | 0.64723764 | 0.99937574 |
| AC008438.1 | 75.989833 | -1.0939652 | 0.99379156 | -1.1007994 | 0.27098396 | 0.99937574 |
| ANKHD1-EIF4 | 12.3844752 | -0.011618 | 0.61002083 | -0.0190453 | 0.98480498 | 0.99937574 |
| SRA1 | 41.4387972 | 0.60325938 | 0.46361457 | 1.30120884 | 0.19318698 | 0.99937574 |
| SLC35A4 | 27.2364689 | 0.03100541 | 0.39150174 | 0.0791961 | 0.93687665 | 0.99937574 |
| NDUFA2 | 49.2827376 | 0.24802613 | 0.50747401 | 0.48874646 | 0.62502121 | 0.99937574 |
| IK | 21.9980648 | -0.3197959 | 0.47498616 | -0.6732741 | 0.50077295 | 0.99937574 |
| WDR55 | 16.8083469 | -0.87399 | 0.48146727 | -1.8152636 | 0.06948342 | 0.99937574 |
| HARS | 28.4606844 | 0.39864852 | 0.39402012 | 1.0117466 | 0.31165923 | 0.99937574 |
| ZMAT2 | 24.0275439 | 0.42184391 | 0.37020598 | 1.13948431 | 0.25450121 | 0.99937574 |
| TAF7 | 51.8474773 | -0.5965805 | 0.33270453 | -1.7931241 | 0.07295309 | 0.99937574 |
| DIAPH1 | 352.014012 | -0.2947601 | 0.27363563 | -1.0771993 | 0.28139124 | 0.99937574 |
| HDAC3 | 27.2239067 | 0.44711935 | 0.43602432 | 1.02544589 | 0.30515284 | 0.99937574 |
| PCDH1 | 40.9982318 | 0.60443476 | 0.45732515 | 1.32167399 | 0.18627673 | 0.99937574 |
| DELE1 | 49.3072018 | -0.2945438 | 0.30145379 | -0.9770778 | 0.32853065 | 0.99937574 |

| | | | | | | |
|---|---|---|---|---|---|---|
| RNF14 | 57.0928168 | -0.215065 | 0.35776358 | -0.6011373 | 0.54774854 | 0.99937574 |
| GNPDA1 | 23.3703933 | -0.5641884 | 0.63326349 | -0.890922 | 0.37297101 | 0.99937574 |
| NDFIP1 | 204.563489 | 0.20072338 | 0.26688725 | 0.75209054 | 0.45199661 | 0.99937574 |
| SPRY4-AS1 | 14.7668685 | -0.1289783 | 0.85831086 | -0.1502699 | 0.88055165 | 0.99937574 |
| ARHGAP26 | 30.3265876 | -0.5115821 | 0.46072421 | -1.1103869 | 0.26683236 | 0.99937574 |
| NR3C1 | 87.7357556 | -0.2433202 | 0.42616821 | -0.5709488 | 0.56803434 | 0.99937574 |
| YIPF5 | 23.973633 | -0.2515512 | 0.4436557 | -0.5669964 | 0.57071665 | 0.99937574 |
| PRELID2 | 51.6218859 | -0.2950638 | 0.61823823 | -0.4772656 | 0.63317299 | 0.99937574 |
| SH3RF2 | 16.5395312 | 1.32766473 | 0.86039455 | 1.54308825 | 0.12280937 | 0.99937574 |
| LARS | 74.4482205 | -0.0352708 | 0.33572962 | -0.1050571 | 0.91633047 | 0.99937574 |
| TCERG1 | 112.608206 | -0.2380132 | 0.24474307 | -0.9725024 | 0.33080068 | 0.99937574 |
| STK32A | 51.0774124 | 0.86305622 | 0.9637721 | 0.89549823 | 0.37052081 | 0.99937574 |
| SPINK1 | 484.331816 | 0.29187717 | 0.93664123 | 0.3116211 | 0.7553285 | 0.99937574 |
| AC091948.1 | 24.2652981 | 0.29553113 | 0.62172845 | 0.47533797 | 0.6345461 | 0.99937574 |
| FBXO38 | 58.2511143 | -0.3385139 | 0.26593661 | -1.2729119 | 0.2030493 | 0.99937574 |
| HTR4 | 18.0916314 | -0.3623619 | 1.6860334 | -0.2149197 | 0.82982989 | 0.99937574 |
| AFAP1L1 | 15.795608 | -0.4717823 | 1.10307276 | -0.4276983 | 0.6688708 | 0.99937574 |
| GRPEL2 | 21.4181623 | -0.7914012 | 0.61608749 | -1.2845597 | 0.19894617 | 0.99937574 |
| CSNK1A1 | 423.985116 | 0.00465117 | 0.25851737 | 0.01799172 | 0.98564546 | 0.99937574 |
| ARHGEF37 | 32.8719526 | 0.5696635 | 0.5664839 | 1.00561287 | 0.31460183 | 0.99937574 |
| SLC26A2 | 16.8383552 | -1.1019537 | 0.50992567 | -2.1610085 | 0.03069468 | 0.99049468 |
| HMGXB3 | 49.6147593 | -0.5720326 | 0.36988787 | -1.5465028 | 0.12198318 | 0.99937574 |
| TCOF1 | 16.1086565 | -0.9503889 | 0.67793264 | -1.4018929 | 0.16094723 | 0.99937574 |
| CD74 | 388.895084 | -0.811101 | 0.74732021 | -1.085346 | 2.78E-01 | 0.99937574 |
| RPS14 | 260.994346 | -0.0059738 | 0.4738442 | -0.012607 | 0.98994131 | 0.99937574 |
| NDST1 | 56.4166165 | -0.0206726 | 0.40752337 | -0.0507273 | 0.95954282 | 0.99937574 |
| RBM22 | 18.1655678 | -0.2289345 | 0.42732931 | -0.5357332 | 0.59214295 | 0.99937574 |
| DCTN4 | 91.5694494 | -0.288659 | 0.23757311 | -1.2150321 | 0.22435386 | 0.99937574 |
| GPX3 | 314.331175 | -0.8150663 | 0.6772003 | -1.2035824 | 0.22875104 | 0.99937574 |
| TNIP1 | 76.119994 | 0.1068232 | 0.43802486 | 0.24387474 | 0.80732783 | 0.99937574 |
| ANXA6 | 47.0881457 | 0.82635218 | 0.53681471 | 1.53936203 | 0.12371594 | 0.99937574 |
| ATOX1 | 67.8468978 | 0.30248419 | 0.40767488 | 0.74197407 | 0.45810304 | 0.99937574 |
| G3BP1 | 115.489644 | -0.0992737 | 0.29261933 | -0.3392589 | 0.73441473 | 0.99937574 |
| FAM114A2 | 27.3124537 | -0.1339438 | 0.46798149 | -0.286216 | 0.7747127 | 0.99937574 |
| MFAP3 | 41.3488 | -0.4361356 | 0.34139447 | -1.2775121 | 0.2014215 | 0.99937574 |
| GALNT10 | 188.604554 | 0.44810956 | 0.47224712 | 0.94888786 | 0.34267765 | 0.99937574 |
| SAP30L-AS1 | 25.6751194 | 0.57122728 | 0.57093587 | 1.00051041 | 0.31706356 | 0.99937574 |
| SAP30L | 14.829472 | -0.5159075 | 0.55208689 | -0.934468 | 0.35006254 | 0.99937574 |
| LARP1 | 174.070141 | -0.227113 | 0.41217691 | -0.5510086 | 0.58162778 | 0.99937574 |
| FAXDC2 | 21.5816356 | 0.98400217 | 0.57902868 | 1.69940145 | 0.08924357 | 0.99937574 |
| CNOT8 | 32.9534011 | 0.11718902 | 0.34731391 | 0.33741529 | 0.73580386 | 0.99937574 |
| GEMIN5 | 16.1729459 | 0.07872324 | 0.48000181 | 0.16400612 | 0.86972632 | 0.99937574 |
| MRPL22 | 28.4931511 | -0.1384252 | 0.393344 | -0.351919 | 0.72489898 | 0.99937574 |
| TIMD4 | 41.2891833 | 1.62379698 | 0.60881174 | 2.66715778 | 0.00764958 | 0.74742128 |
| CYFIP2 | 235.207999 | 0.90515015 | 0.45715515 | 1.97996272 | 0.04770772 | 0.99937574 |
| CLINT1 | 215.740965 | -0.2267547 | 0.29467383 | -0.7695107 | 0.44159017 | 0.99937574 |
| RNF145 | 73.1609049 | -0.0398499 | 0.28255079 | -0.1410364 | 0.88784121 | 0.99937574 |
| UBLCP1 | 27.1751918 | 0.19304118 | 0.42294774 | 0.45641853 | 0.64808904 | 0.99937574 |
| AC008691.1 | 61.9566656 | -0.4331239 | 0.5732434 | -0.7555672 | 0.44990873 | 0.99937574 |
| TTC1 | 117.270664 | -0.3153792 | 0.39391618 | -0.8006252 | 0.42334865 | 0.99937574 |
| PWWP2A | 120.121686 | 0.37718156 | 0.32496145 | 1.16069633 | 0.24576542 | 0.99937574 |
| CCNJL | 84.9157976 | -0.1332516 | 0.45910343 | -0.290243 | 0.77163035 | 0.99937574 |
| SLU7 | 19.2810389 | -0.2382926 | 0.59901111 | -0.39781 | 0.69077022 | 0.99937574 |
| AC113414.1 | 449.694888 | 1.41992802 | 0.71467735 | 1.98680988 | 0.04694347 | 0.99937574 |

| | | | | | | |
|---|---|---|---|---|---|---|
| CCNG1 | 67.4745017 | 0.31759207 | 0.4594243 | 0.6912827 | 0.4893879 | 0.99937574 |
| NUDCD2 | 18.1108993 | 0.72659137 | 0.5583064 | 1.30142045 | 0.19311458 | 0.99937574 |
| MAT2B | 45.3541064 | 0.17194005 | 0.3482344 | 0.49374804 | 0.62148413 | 0.99937574 |
| TENM2 | 11.7456499 | 0.60052993 | 0.89182535 | 0.67337167 | 0.50071087 | 0.99937574 |
| WWC1 | 411.239877 | 0.09417697 | 0.46386436 | 0.20302697 | 0.83911395 | 0.99937574 |
| RARS | 51.9477594 | 0.21374769 | 0.38311008 | 0.55792761 | 0.57689382 | 0.99937574 |
| PANK3 | 129.028858 | -0.3079919 | 0.29266038 | -1.0523868 | 0.29262212 | 0.99937574 |
| SLIT3 | 101.293398 | -2.5115149 | 1.96252721 | -1.2797351 | 0.20063832 | 0.99937574 |
| FOXI1 | 318.311598 | 0.11072135 | 0.29604819 | 0.37399774 | 0.708406 | 0.99937574 |
| LINC01187 | 611.152988 | 0.51928593 | 0.43252098 | 1.20060286 | 0.22990529 | 0.99937574 |
| RANBP17 | 69.2556967 | -0.0264327 | 0.49750506 | -0.0531305 | 0.95762791 | 0.99937574 |
| NPM1 | 80.9868682 | -0.4175145 | 0.38952072 | -1.0718672 | 0.28377968 | 0.99937574 |
| FBXW11 | 440.305063 | 0.08202216 | 0.32900455 | 0.24930405 | 0.8031256 | 0.99937574 |
| STK10 | 23.1710509 | -0.7727048 | 0.49059494 | -1.5750363 | 0.11524808 | 0.99937574 |
| UBTD2 | 71.8499243 | -0.0775649 | 0.27772874 | -0.2792829 | 0.78002775 | 0.99937574 |
| NEURL1B | 18.059383 | -2.0430139 | 0.90625233 | -2.2543544 | 0.02417388 | 0.96546668 |
| DUSP1 | 141.787467 | -0.9408977 | 0.56504457 | -1.6651743 | 0.09587799 | 0.99937574 |
| ERGIC1 | 138.268084 | 0.32587858 | 0.34494213 | 0.94473406 | 0.34479467 | 0.99937574 |
| RPL26L1 | 13.9969994 | 0.03291428 | 0.5281825 | 0.06231611 | 0.9503111 | 0.99937574 |
| ATP6V0E1 | 343.293946 | -0.1085707 | 0.27218209 | -0.3988901 | 0.68997419 | 0.99937574 |
| CREBRF | 159.861182 | -0.5623201 | 0.35249031 | -1.5952783 | 0.11065002 | 0.99937574 |
| BNIP1 | 15.0451307 | -0.7211188 | 0.56674034 | -1.2723971 | 0.20323206 | 0.99937574 |
| BOD1 | 18.8003907 | -0.3847033 | 0.53598179 | -0.7177544 | 0.47290871 | 0.99937574 |
| CPEB4 | 779.562628 | 0.09675564 | 0.230853 | 0.41912229 | 0.67512676 | 0.99937574 |
| SFXN1 | 97.0906965 | -0.0525716 | 0.29410157 | -0.1787531 | 0.8581316 | 0.99937574 |
| THOC3 | 60.3062071 | 0.39504986 | 0.63604444 | 0.62110418 | 0.53453108 | 0.99937574 |
| FAM153B | 9.89785218 | 2.24407346 | 1.07331814 | 2.09078127 | 0.03654768 | 0.99937574 |
| SIMC1 | 71.774587 | -0.220639 | 0.36494998 | -0.6045732 | 0.5454626 | 0.99937574 |
| KIAA1191 | 52.4932961 | 0.009634 | 0.39019382 | 0.02469029 | 0.980302 | 0.99937574 |
| ARL10 | 12.032242 | -0.8417654 | 0.68691768 | -1.2254239 | 0.22041553 | 0.99937574 |
| CLTB | 42.5152603 | -0.3849769 | 0.3712016 | -1.03711 | 0.2996846 | 0.99937574 |
| FAF2 | 82.5130179 | -0.2008226 | 0.3292581 | -0.6099244 | 0.54191188 | 0.99937574 |
| RNF44 | 23.0695228 | -0.256902 | 0.4620184 | -0.5560428 | 0.57818161 | 0.99937574 |
| TSPAN17 | 20.4889594 | -0.7976961 | 0.49570551 | -1.6092136 | 0.10756964 | 0.99937574 |
| UIMC1 | 103.432314 | 0.0078464 | 0.27463243 | 0.02857054 | 0.97720711 | 0.99937574 |
| ZNF346 | 51.1736679 | 1.02E-05 | 0.3051988 | 3.35E-05 | 0.99997331 | 0.99997331 |
| NSD1 | 202.780197 | 0.17161668 | 0.38302859 | 0.44805187 | 0.65411577 | 0.99937574 |
| PRELID1 | 47.11257 | -0.1262198 | 0.51342793 | -0.2458373 | 0.80580815 | 0.99937574 |
| LMAN2 | 61.1563137 | 0.19062171 | 0.34280385 | 0.55606644 | 0.57816544 | 0.99937574 |
| RGS14 | 19.9892596 | -0.0669669 | 0.59621857 | -0.1123193 | 0.9105702 | 0.99937574 |
| DBN1 | 33.1934492 | 0.12500776 | 0.48626553 | 0.25707716 | 0.79711921 | 0.99937574 |
| DDX41 | 13.3596613 | -0.2783851 | 0.52820417 | -0.5270408 | 0.59816529 | 0.99937574 |
| FAM193B | 53.571021 | 0.24147096 | 0.3017125 | 0.80033463 | 0.42351694 | 0.99937574 |
| TMED9 | 65.5918319 | -0.1878029 | 0.38271345 | -0.4907142 | 0.62362859 | 0.99937574 |
| B4GALT7 | 22.2228598 | 1.04231789 | 0.55005824 | 1.89492277 | 0.05810263 | 0.99937574 |
| FAM153A | 25.8312892 | -0.0402381 | 0.9384432 | -0.0428775 | 0.96579921 | 0.99937574 |
| AC106795.3 | 17.3632112 | 0.84771748 | 0.74741202 | 1.13420371 | 0.25670911 | 0.99937574 |
| N4BP3 | 27.1861153 | -0.0576484 | 0.5540129 | -0.1040561 | 0.91712483 | 0.99937574 |
| RMND5B | 67.076435 | 0.15100364 | 0.33589969 | 0.4495498 | 0.65303509 | 0.99937574 |
| NHP2 | 30.6646262 | 0.46376535 | 0.41895731 | 1.10695133 | 0.26831497 | 0.99937574 |
| HNRNPAB | 39.7921745 | -0.2767004 | 0.45178554 | -0.6124596 | 0.5402337 | 0.99937574 |
| PHYKPL | 104.58676 | -0.0610804 | 0.3524504 | -0.1733021 | 0.86241398 | 0.99937574 |
| CLK4 | 46.1189284 | -0.0403386 | 0.4404598 | -0.091583 | 0.92702933 | 0.99937574 |
| ZNF354A | 17.6034828 | -0.2849796 | 0.56164755 | -0.5073993 | 0.61187464 | 0.99937574 |

| | | | | | | |
|---|---|---|---|---|---|---|
| ZNF354B | 13.1355227 | -0.2239801 | 0.56184893 | -0.3986483 | 0.69015235 | 0.99937574 |
| RUFY1 | 84.8068451 | 0.06359912 | 0.28798842 | 0.22083916 | 0.82521767 | 0.99937574 |
| HNRNPH1 | 213.502388 | -0.2278688 | 0.3668421 | -0.6211632 | 0.53449228 | 0.99937574 |
| CANX | 218.177078 | 0.06831257 | 0.28750285 | 0.23760659 | 0.81218624 | 0.99937574 |
| MAML1 | 39.3269882 | -0.2111299 | 0.34014864 | -0.6206989 | 0.53479777 | 0.99937574 |
| MGAT4B | 64.0508617 | -0.3715011 | 0.58495075 | -0.635098 | 0.52536451 | 0.99937574 |
| SQSTM1 | 226.325404 | -0.8836124 | 0.46270918 | -1.9096495 | 0.05617835 | 0.99937574 |
| MRNIP | 28.3122395 | -0.3571877 | 0.48548398 | -0.7357352 | 0.46189184 | 0.99937574 |
| TBC1D9B | 39.8350025 | 0.23488927 | 0.43604144 | 0.53868566 | 0.59010377 | 0.99937574 |
| RNF130 | 199.126675 | -0.1491154 | 0.2833974 | -0.5261706 | 0.59876967 | 0.99937574 |
| MAPK9 | 57.6283099 | 0.02623946 | 0.29649916 | 0.08849759 | 0.9294812 | 0.99937574 |
| CNOT6 | 65.5751266 | 0.00155902 | 0.27885825 | 0.00559074 | 0.99553926 | 0.99937574 |
| MGAT1 | 42.5351413 | -0.516033 | 0.51583985 | -1.0003744 | 0.31712934 | 0.99937574 |
| ZFP62 | 18.1085497 | 0.74736206 | 0.53760648 | 1.39016565 | 0.16447858 | 0.99937574 |
| BTNL9 | 57.6522804 | 0.23317327 | 0.53130197 | 0.43887146 | 0.66075468 | 0.99937574 |
| TRIM41 | 17.5035663 | -0.3201168 | 0.46005461 | -0.6958235 | 0.48653934 | 0.99937574 |
| RACK1 | 152.819751 | -0.0824186 | 0.49049567 | -0.1680313 | 0.86655868 | 0.99937574 |
| TRIM52 | 23.2735025 | -0.2426038 | 0.45373962 | -0.5346763 | 0.59287373 | 0.99937574 |
| AC138035.1 | 11.0424084 | -0.9750857 | 0.71837549 | -1.3573482 | 0.17467059 | 0.99937574 |
| DUSP22 | 35.5091391 | 0.02385294 | 0.37139132 | 0.06422591 | 0.94879035 | 0.99937574 |
| EXOC2 | 135.172982 | 0.04162976 | 0.25327964 | 0.16436285 | 0.86944551 | 0.99937574 |
| GMDS | 309.300123 | -0.1666417 | 0.32769764 | -0.5085228 | 0.61108677 | 0.99937574 |
| GMDS-DT | 487.446522 | 0.10652516 | 0.437332 | 0.24357962 | 0.80755642 | 0.99937574 |
| WRNIP1 | 54.498953 | -0.408728 | 0.37752015 | -1.0826653 | 0.278957 | 0.99937574 |
| SERPINB1 | 144.743315 | 0.96603628 | 0.65364983 | 1.47791101 | 0.1394316 | 0.99937574 |
| SERPINB6 | 108.676665 | -0.0110267 | 0.32836579 | -0.0335805 | 0.9732117 | 0.99937574 |
| NQO2 | 42.0292002 | -0.4965124 | 0.36201914 | -1.3715086 | 0.17021647 | 0.99937574 |
| RIPK1 | 70.5649775 | -0.6621976 | 0.30483483 | -2.172316 | 0.02983184 | 0.99049468 |
| BPHL | 19.5008839 | -0.2875701 | 0.48325403 | -0.5950702 | 0.55179652 | 0.99937574 |
| TUBB2A | 26.1427624 | 1.01270004 | 0.50542201 | 2.00367224 | 0.04510518 | 0.99937574 |
| LINC02525 | 42.1917429 | 1.14191754 | 0.43686955 | 2.6138639 | 8.95E-03 | 0.74742128 |
| PSMG4 | 168.22518 | 0.70181052 | 0.48682431 | 1.44160944 | 0.14941258 | 0.99937574 |
| SLC22A23 | 384.383813 | 0.02652653 | 0.38047182 | 0.06972009 | 0.94441645 | 0.99937574 |
| PRPF4B | 152.159496 | -0.2984039 | 0.24054026 | -1.240557 | 0.21476944 | 0.99937574 |
| ECI2 | 51.4562343 | 0.67505872 | 0.40279622 | 1.67593111 | 0.09375169 | 0.99937574 |
| CDYL | 203.057265 | -0.4468129 | 0.39975068 | -1.1177289 | 0.26368279 | 0.99937574 |
| LYRM4 | 66.0173115 | -0.1079014 | 0.29180054 | -0.369778 | 0.71154788 | 0.99937574 |
| FARS2 | 224.018471 | 0.06698529 | 0.2831893 | 0.23653893 | 0.8130145 | 0.99937574 |
| LY86-AS1 | 280.943486 | 1.74831756 | 0.90546062 | 1.93085985 | 0.05350039 | 0.99937574 |
| LY86 | 34.5992203 | 1.26304849 | 0.97667211 | 1.2932165 | 0.19593619 | 0.99937574 |
| AL031123.1 | 72.2325399 | 1.46411828 | 0.70048283 | 2.09015585 | 0.0366038 | 0.99937574 |
| RREB1 | 234.37139 | 0.08969699 | 0.43487668 | 0.20625846 | 0.83658902 | 0.99937574 |
| SSR1 | 74.7780572 | 0.36959485 | 0.29154637 | 1.26770519 | 0.20490325 | 0.99937574 |
| AL139095.4 | 14.7123619 | 0.93369296 | 0.99323945 | 0.9400482 | 0.34719284 | 0.99937574 |
| RIOK1 | 13.7370631 | -0.1592084 | 0.54508789 | -0.2920784 | 0.77022671 | 0.99937574 |
| DSP | 299.785489 | -0.1111862 | 0.37506176 | -0.2964477 | 0.76688821 | 0.99937574 |
| SNRNP48 | 22.163312 | 0.04539304 | 0.53934561 | 0.08416318 | 0.93292669 | 0.99937574 |
| BLOC1S5-TXI | 12.4926369 | -0.1237074 | 0.62658934 | -0.1974298 | 0.84349125 | 0.99937574 |
| SLC35B3 | 63.7532725 | 0.03793198 | 0.46492831 | 0.08158673 | 0.93497536 | 0.99937574 |
| TFAP2A | 38.6265886 | 0.56349108 | 0.43295251 | 1.30150783 | 0.19308469 | 0.99937574 |
| GCNT2 | 264.198022 | 0.24244066 | 0.36890439 | 0.65719104 | 0.51105809 | 0.99937574 |
| PAK1IP1 | 11.6274625 | -0.5792155 | 0.54105927 | -1.0705213 | 0.28438472 | 0.99937574 |
| TMEM14C | 41.4850944 | 0.05691785 | 0.62229758 | 0.09146404 | 0.92712388 | 0.99937574 |
| MAK | 39.2979779 | 0.31773169 | 0.80536938 | 0.39451672 | 0.69319958 | 0.99937574 |

| | | | | | | |
|---|---|---|---|---|---|---|
| SMIM13 | 14.2159393 | -0.0162981 | 0.51024898 | -0.0319415 | 0.97451869 | 0.99937574 |
| AL139807.1 | 16.4854766 | 0.30260383 | 0.71195627 | 0.42503148 | 0.67081373 | 0.99937574 |
| NEDD9 | 41.2692935 | -0.7789023 | 0.76614548 | -1.0166506 | 0.30931967 | 0.99937574 |
| TMEM170B | 21.4345696 | 0.41242663 | 0.49666929 | 0.8303848 | 0.40632126 | 0.99937574 |
| ADTRP | 199.017884 | 0.27327088 | 0.52544148 | 0.52007863 | 0.60300878 | 0.99937574 |
| HIVEP1 | 167.734368 | -0.1363225 | 0.42038978 | -0.3242765 | 0.7457287 | 0.99937574 |
| PHACTR1 | 20.786843 | -0.0349556 | 0.84227146 | -0.0415015 | 0.96689606 | 0.99937574 |
| TBC1D7 | 11.4682578 | 0.0745642 | 0.60631331 | 0.12297965 | 0.90212321 | 0.99937574 |
| SIRT5 | 50.2038827 | 0.2190235 | 0.46846973 | 0.46752968 | 0.64012096 | 0.99937574 |
| NOL7 | 43.0684873 | -0.1635518 | 0.41734756 | -0.3918839 | 0.69514401 | 0.99937574 |
| RANBP9 | 169.602528 | 0.06817445 | 0.31751495 | 0.21471257 | 0.82999142 | 0.99937574 |
| MCUR1 | 29.9058217 | 0.6539048 | 0.5051923 | 1.29436812 | 0.1955383 | 0.99937574 |
| CD83 | 17.638919 | -0.4447349 | 0.70102871 | -0.6344032 | 0.52581773 | 0.99937574 |
| JARID2 | 183.326695 | 0.19798419 | 0.33468602 | 0.59155203 | 0.55415061 | 0.99937574 |
| DTNBP1 | 124.496886 | -0.701131 | 0.4619791 | -1.5176682 | 0.12909807 | 0.99937574 |
| MYLIP | 20.4467141 | -0.4580891 | 0.62744906 | -0.7300818 | 0.46534019 | 0.99937574 |
| GMPR | 267.508145 | 0.09344823 | 0.5352355 | 0.17459273 | 0.86139966 | 0.99937574 |
| ATXN1 | 482.463722 | 0.50251605 | 0.44548245 | 1.12802661 | 0.25930868 | 0.99937574 |
| CAP2 | 17.093558 | -0.9350306 | 0.8517827 | -1.0977337 | 0.27232077 | 0.99937574 |
| FAM8A1 | 15.9606253 | 0.26522901 | 0.47785848 | 0.55503673 | 0.57886954 | 0.99937574 |
| NUP153 | 114.02506 | 0.04684227 | 0.29961154 | 0.15634335 | 0.87576238 | 0.99937574 |
| KIF13A | 378.053004 | 0.03502591 | 0.41638476 | 0.0841191 | 0.93296174 | 0.99937574 |
| TPMT | 25.5410826 | 0.00956639 | 0.42592519 | 0.02246026 | 0.98208081 | 0.99937574 |
| KDM1B | 18.7453654 | -0.4069834 | 0.57000389 | -0.7140011 | 0.47522647 | 0.99937574 |
| DEK | 86.4026313 | -0.3142155 | 0.37655669 | -0.8344442 | 0.40403072 | 0.99937574 |
| RNF144B | 111.170138 | 0.66920663 | 0.41543114 | 1.61087258 | 0.1072075 | 0.99937574 |
| AL022068.1 | 35.9982004 | 1.89112517 | 0.70430233 | 2.68510424 | 0.00725072 | 0.74742128 |
| ID4 | 16.217886 | -0.9917111 | 0.63657827 | -1.5578777 | 0.11926225 | 0.99937574 |
| MBOAT1 | 140.124075 | -0.0403991 | 0.55449919 | -0.0728569 | 0.94191998 | 0.99937574 |
| AL158198.1 | 18.1114381 | -0.3093433 | 0.83506492 | -0.3704422 | 0.71105301 | 0.99937574 |
| E2F3 | 55.2933525 | 0.07409496 | 0.40667998 | 0.18219476 | 0.85542989 | 0.99937574 |
| CDKAL1 | 330.96406 | -0.4774946 | 0.24672732 | -1.9353129 | 0.05295191 | 0.99937574 |
| SOX4 | 71.9318852 | -0.6085735 | 0.62025454 | -0.9811673 | 0.32651026 | 0.99937574 |
| CASC15 | 164.394639 | -0.00842 | 0.42166053 | -0.0199686 | 0.98406846 | 0.99937574 |
| NBAT1 | 12.0976709 | 0.71907589 | 0.81193142 | 0.88563625 | 0.37581357 | 0.99937574 |
| DCDC2 | 170.409558 | -0.1782988 | 0.47778215 | -0.3731801 | 0.70901441 | 0.99937574 |
| MRS2 | 41.5233122 | 0.1486289 | 0.38111483 | 0.38998457 | 0.69654796 | 0.99937574 |
| ALDH5A1 | 18.4569064 | 0.21484963 | 0.57611733 | 0.37292688 | 0.70920287 | 0.99937574 |
| TDP2 | 37.3788309 | -0.0021606 | 0.41137056 | -0.0052521 | 0.99580942 | 0.99937574 |
| ACOT13 | 33.7192213 | 0.3979331 | 0.43674528 | 0.91113314 | 0.36222523 | 0.99937574 |
| C6orf62 | 105.57554 | -0.8497664 | 0.38187733 | -2.2252339 | 0.02606554 | 0.96546668 |
| GMNN | 21.1654007 | -0.3226733 | 0.6010485 | -0.5368507 | 0.59137078 | 0.99937574 |
| CARMIL1 | 380.275744 | 0.00260755 | 0.49229584 | 0.00529672 | 0.99577385 | 0.99937574 |
| TRIM38 | 84.3973557 | -0.5346824 | 0.35108741 | -1.5229323 | 0.12777564 | 0.99937574 |
| HIST1H1C | 24.1598759 | -0.2102923 | 0.50353121 | -0.417635 | 0.67621397 | 0.99937574 |
| HFE | 11.4249185 | -0.0571343 | 0.72100611 | -0.0792425 | 0.93683976 | 0.99937574 |
| HIST1H4C | 32.0534364 | 1.31687789 | 0.76327926 | 1.72528975 | 0.08447527 | 0.99937574 |
| HIST1H2AC | 36.9769788 | -0.6239557 | 0.82680906 | -0.7546551 | 0.45045597 | 0.99937574 |
| HIST1H1E | 18.4142396 | -1.004206 | 0.86728085 | -1.1578786 | 0.24691356 | 0.99937574 |
| BTN3A2 | 28.632423 | -0.1540128 | 0.99912893 | -0.1541471 | 0.87749375 | 0.99937574 |
| BTN3A1 | 17.4151215 | 0.00083897 | 0.84095613 | 0.00099764 | 0.999204 | 0.99979721 |
| BTN3A3 | 12.3273742 | 0.03170398 | 0.87463619 | 0.03624819 | 0.97108446 | 0.99937574 |
| BTN2A1 | 26.4792662 | -0.0832226 | 0.39266486 | -0.211943 | 0.83215154 | 0.99937574 |
| HCG11 | 11.8530276 | -0.173541 | 0.6212932 | -0.2793223 | 0.77999753 | 0.99937574 |

| | | | | | | |
|---|---|---|---|---|---|---|
| HMGN4 | 21.9358684 | -0.4996896 | 0.53155181 | -0.940058 | 0.34718779 | 0.99937574 |
| ZNF322 | 22.589503 | -0.1311001 | 0.45823013 | -0.2861009 | 0.77480081 | 0.99937574 |
| AL121944.1 | 11.4543003 | 0.21978218 | 0.74181364 | 0.29627681 | 0.7670187 | 0.99937574 |
| ZSCAN16-AS | 20.4653613 | 0.60310742 | 0.52073992 | 1.15817398 | 0.24679305 | 0.99937574 |
| ZKSCAN8 | 22.1342733 | -0.4198405 | 0.55048535 | -0.7626733 | 0.44565826 | 0.99937574 |
| ZSCAN9 | 11.190364 | -0.2028707 | 0.54129436 | -0.3747881 | 0.70781808 | 0.99937574 |
| ZSCAN31 | 31.6245115 | 0.92154089 | 0.66693533 | 1.3817545 | 0.16704709 | 0.99937574 |
| ZSCAN12 | 10.5439583 | 0.89660511 | 0.70914101 | 1.26435376 | 0.20610308 | 0.99937574 |
| TRIM27 | 62.5866426 | 0.03784801 | 0.28537571 | 0.13262521 | 8.94E-01 | 0.99937574 |
| GABBR1 | 22.3578918 | -0.0337992 | 0.70603097 | -0.0478721 | 0.9618182 | 0.99937574 |
| HLA-F | 58.387795 | -0.5167598 | 0.75749778 | -0.6821932 | 0.49511679 | 0.99937574 |
| HLA-A | 790.269179 | -0.3156859 | 0.53248443 | -0.5928548 | 5.53E-01 | 0.99937574 |
| ZNRD1 | 11.3587952 | -0.1772136 | 0.79866353 | -0.2218876 | 0.82440137 | 0.99937574 |
| PPP1R11 | 17.2554759 | 0.25104153 | 0.46473417 | 0.54018307 | 0.58907079 | 0.99937574 |
| TRIM26 | 61.134166 | -0.2010788 | 0.35470191 | -0.5668951 | 0.57078548 | 0.99937574 |
| HCG17 | 75.3370087 | -0.2598426 | 0.52084814 | -0.4988835 | 0.61786143 | 0.99937574 |
| HLA-E | 264.632424 | -1.0406935 | 0.49165288 | -2.116724 | 3.43E-02 | 9.99E-01 |
| GNL1 | 36.8990029 | -0.7567045 | 0.48561923 | -1.5582261 | 0.11917967 | 0.99937574 |
| ABCF1 | 43.5683993 | -0.5576408 | 0.50796763 | -1.097788 | 0.27229706 | 0.99937574 |
| PPP1R10 | 39.4544997 | 0.28983239 | 0.41102733 | 0.7051414 | 0.48072224 | 0.99937574 |
| MRPS18B | 41.3291257 | 0.48582535 | 0.43973965 | 1.10480226 | 0.26924528 | 0.99937574 |
| ATAT1 | 13.5335833 | -0.0093987 | 0.55782588 | -0.0168488 | 0.98655725 | 0.99937574 |
| C6orf136 | 32.7548817 | 0.14283288 | 0.3651049 | 0.39121054 | 0.69564162 | 0.99937574 |
| DHX16 | 16.3012496 | -0.160776 | 0.45934024 | -0.350015 | 0.72632746 | 0.99937574 |
| MDC1 | 24.8509845 | 0.1652817 | 0.42825239 | 0.38594461 | 0.69953769 | 0.99937574 |
| TUBB | 71.1660421 | 0.53145198 | 0.55816102 | 0.95214816 | 0.34102185 | 0.99937574 |
| FLOT1 | 124.935517 | 0.33135312 | 0.24685516 | 1.3422977 | 0.17949949 | 0.99937574 |
| IER3 | 126.139153 | -0.910415 | 1.30640453 | -0.696886 | 0.48587415 | 0.99937574 |
| DDR1 | 165.531172 | 0.06921056 | 0.30121919 | 0.22976809 | 0.81827198 | 0.99937574 |
| CCHCR1 | 15.4291535 | 0.47208985 | 0.54418214 | 0.86752175 | 0.3856562 | 0.99937574 |
| POU5F1 | 97.8455147 | -0.1806385 | 0.6899018 | -0.2618322 | 0.7934508 | 0.99937574 |
| HLA-C | 509.791514 | -0.1358284 | 0.68449402 | -0.1984363 | 0.84270374 | 0.99937574 |
| HLA-B | 809.05533 | -0.1594978 | 0.57278991 | -0.2784578 | 7.81E-01 | 9.99E-01 |
| MICA | 28.6692312 | 0.39974873 | 0.41666856 | 0.9593926 | 0.337361 | 0.99937574 |
| MCCD1 | 15.2545326 | 0.92928415 | 0.61697079 | 1.50620446 | 0.13201469 | 0.99937574 |
| NFKBIL1 | 12.2371651 | -0.2944198 | 0.53741292 | -0.5478465 | 0.58379733 | 0.99937574 |
| PRRC2A | 61.7679096 | 0.01914699 | 0.34079355 | 0.05618354 | 0.9551956 | 0.99937574 |
| BAG6 | 36.3874316 | -0.2885205 | 0.47348116 | -0.60936 | 0.54228582 | 0.99937574 |
| C6orf47 | 12.0151424 | -0.1184747 | 0.65942992 | -0.1796623 | 0.85741769 | 0.99937574 |
| DDAH2 | 30.4178292 | 0.53469638 | 0.4776575 | 1.11941375 | 0.26296367 | 0.99937574 |
| CLIC1 | 119.764747 | 0.11800895 | 0.43077582 | 0.27394515 | 0.78412678 | 0.99937574 |
| VARS | 14.656009 | -0.0201936 | 0.52570469 | -0.0384124 | 0.96935884 | 0.99937574 |
| LSM2 | 19.4088872 | 0.25380495 | 0.48252506 | 0.52599329 | 0.59889287 | 0.99937574 |
| HSPA1A | 137.134996 | -0.522293 | 0.49674243 | -1.0514363 | 0.29305824 | 0.99937574 |
| HSPA1B | 56.0925947 | -0.5099257 | 0.47077273 | -1.0831674 | 0.27873411 | 0.99937574 |
| C6orf48 | 35.0553143 | -0.1816282 | 0.44375343 | -0.4092997 | 0.68231971 | 0.99937574 |
| NEU1 | 34.0580102 | -0.6376941 | 0.58074134 | -1.0980691 | 0.27217433 | 0.99937574 |
| SLC44A4 | 25.6198674 | -0.5005422 | 0.88178317 | -0.5676477 | 0.57027422 | 0.99937574 |
| EHMT2 | 12.0392611 | 0.05578505 | 0.70523493 | 0.07910137 | 0.936952 | 0.99937574 |
| NELFE | 13.3928137 | -0.6593992 | 0.57547621 | -1.1458323 | 0.25186457 | 0.99937574 |
| STK19 | 16.6265562 | -0.6686603 | 0.50668175 | -1.319685 | 0.18694022 | 0.99937574 |
| ATF6B | 31.4137843 | -0.2703647 | 0.38406115 | -0.7039625 | 0.48145612 | 0.99937574 |
| AGPAT1 | 18.1580849 | 0.54896673 | 0.52863838 | 1.03845417 | 0.29905866 | 0.99937574 |
| RNF5 | 21.6709846 | 0.18861957 | 0.70375623 | 0.26801834 | 0.78868521 | 0.99937574 |

| | | | | | | |
|---|---|---|---|---|---|---|
| PBX2 | 15.504393 | 0.14369556 | 0.49322891 | 0.29133645 | 0.77079401 | 0.99937574 |
| AL662796.1 | 42.0431834 | -1.4789479 | 0.93647307 | -1.5792744 | 0.11427314 | 0.99937574 |
| HLA-DRA | 137.96498 | -1.0219938 | 0.91875984 | -1.1123623 | 0.26598241 | 0.99937574 |
| HLA-DRB5 | 16.9691447 | -0.7044208 | 0.9552577 | -0.7374144 | 0.46087035 | 0.99937574 |
| HLA-DRB1 | 83.8864186 | -2.8411152 | 1.55554236 | -1.8264467 | 0.067783 | 0.99937574 |
| HLA-DQB1 | 27.8799654 | -1.4653867 | 1.49595198 | -0.979568 | 0.3272994 | 0.99937574 |
| PSMB8 | 69.7306667 | -0.1058728 | 0.58947367 | -0.1796057 | 0.85746215 | 0.99937574 |
| PSMB9 | 29.0871113 | -0.7057256 | 0.90623464 | -0.7787449 | 0.43613 | 0.99937574 |
| TAP1 | 48.7373161 | -0.4285563 | 0.77438185 | -0.5534172 | 0.57997774 | 0.99937574 |
| HLA-DMB | 19.8191239 | -0.0047905 | 0.76517534 | -0.0062606 | 0.99500476 | 0.99937574 |
| BRD2 | 95.1157483 | -0.4141087 | 0.29993546 | -1.3806594 | 0.16738371 | 0.99937574 |
| HLA-DPA1 | 42.6416638 | -2.3900146 | 1.11837879 | -2.137035 | 0.03259514 | 0.99937574 |
| HLA-DPB1 | 34.2463693 | -3.1089939 | 1.1405814 | -2.7257975 | 0.00641463 | 0.69675365 |
| RXRB | 16.1269424 | -0.183088 | 0.45781096 | -0.3999206 | 0.689215 | 0.99937574 |
| SLC39A7 | 37.6664908 | 0.0979457 | 0.64804296 | 0.15114075 | 0.87986469 | 0.99937574 |
| RING1 | 24.2757043 | -0.5478358 | 0.52269917 | -1.04809 | 0.29459713 | 0.99937574 |
| VPS52 | 27.1861332 | -0.0960824 | 0.44483816 | -0.215994 | 0.82899245 | 0.99937574 |
| RPS18 | 151.40008 | -0.1663855 | 0.48297253 | -0.3445031 | 0.73046797 | 0.99937574 |
| WDR46 | 11.4914182 | -0.3607279 | 0.69431051 | -0.5195484 | 0.60337839 | 0.99937574 |
| PFDN6 | 19.3627623 | -0.071799 | 0.42448386 | -0.1691443 | 0.86568311 | 0.99937574 |
| RGL2 | 21.7764061 | -0.4238978 | 0.45170265 | -0.9384444 | 0.34801606 | 0.99937574 |
| TAPBP | 156.393667 | -0.0429884 | 0.40241763 | -0.1068254 | 0.91492753 | 0.99937574 |
| DAXX | 11.1486848 | -0.2937578 | 0.72937747 | -0.4027513 | 0.68713116 | 0.99937574 |
| PHF1 | 42.0952198 | -0.5489166 | 0.5365607 | -1.0230279 | 0.30629464 | 0.99937574 |
| CUTA | 45.8818729 | -0.1460876 | 0.57202679 | -0.255386 | 0.79842501 | 0.99937574 |
| SYNGAP1 | 11.6122545 | -0.7968482 | 0.64418906 | -1.2369789 | 0.21609493 | 0.99937574 |
| ITPR3 | 28.9863981 | -1.0149125 | 0.58834975 | -1.7250156 | 0.08452466 | 0.99937574 |
| UQCC2 | 31.3613918 | -0.0914781 | 0.42144526 | -0.217058 | 0.82816317 | 0.99937574 |
| LEMD2 | 31.2900533 | 0.01924359 | 0.39274577 | 0.04899758 | 0.96092123 | 0.99937574 |
| HMGA1 | 20.7199536 | -0.4309635 | 0.52009688 | -0.8286216 | 0.40731856 | 0.99937574 |
| SMIM29 | 11.4508422 | -0.567712 | 0.55691924 | -1.0193793 | 0.30802292 | 0.99937574 |
| NUDT3 | 62.6499498 | -0.1838961 | 0.6540796 | -0.2811525 | 0.77859342 | 0.99937574 |
| C6orf106 | 166.568348 | -0.1874729 | 0.21930022 | -0.8548689 | 0.39262374 | 0.99937574 |
| SNRPC | 22.1029345 | -0.055137 | 0.43425729 | -0.1269684 | 0.89896539 | 0.99937574 |
| UHRF1BP1 | 76.4227042 | -0.0767693 | 0.30761196 | -0.2495654 | 0.80292348 | 0.99937574 |
| TAF11 | 42.4905154 | 0.01939016 | 0.3865197 | 0.05016603 | 0.95999009 | 0.99937574 |
| ANKS1A | 391.87204 | 0.37810679 | 0.31692536 | 1.1930468 | 0.23285105 | 0.99937574 |
| ZNF76 | 51.3328295 | 0.09705675 | 0.35161806 | 0.2760289 | 0.78252587 | 0.99937574 |
| PPARD | 39.3148542 | 0.15539543 | 0.46741545 | 0.33245677 | 0.73954438 | 0.99937574 |
| RPL10A | 176.818023 | 0.07919321 | 0.52890438 | 0.14973067 | 0.88097711 | 0.99937574 |
| TEAD3 | 31.3402164 | 0.05906961 | 0.45406081 | 0.13009185 | 0.89649376 | 0.99937574 |
| FKBP5 | 142.42043 | -1.155864 | 0.99412701 | -1.1626925 | 0.24495429 | 0.99937574 |
| SRPK1 | 66.3170806 | 0.36916575 | 0.38936964 | 0.94811127 | 0.34307281 | 0.99937574 |
| MAPK14 | 121.822682 | -0.2882312 | 0.38396364 | -0.7506732 | 0.45284937 | 0.99937574 |
| MAPK13 | 33.5035198 | 0.59790001 | 0.38366508 | 1.55839051 | 0.11914071 | 0.99937574 |
| BRPF3 | 30.3058352 | -0.60032 | 0.3812982 | -1.5744109 | 0.11539248 | 0.99937574 |
| KCTD20 | 105.448609 | 0.10817774 | 0.28566185 | 0.37869158 | 0.70491691 | 0.99937574 |
| STK38 | 101.172473 | 0.00784333 | 0.29628236 | 0.02647248 | 0.97888049 | 0.99937574 |
| SRSF3 | 93.0676727 | -0.0413472 | 0.34713115 | -0.1191113 | 0.90518719 | 0.99937574 |
| CDKN1A | 37.2602866 | 0.32365918 | 0.6247107 | 0.5180945 | 6.04E-01 | 9.99E-01 |
| C6orf89 | 61.3153365 | -0.2568899 | 0.36015652 | -0.713273 | 0.47567687 | 0.99937574 |
| MTCH1 | 71.5645273 | -0.1664187 | 0.30089884 | -0.5530719 | 0.58021416 | 0.99937574 |
| PIM1 | 28.7250396 | -0.1222374 | 0.51493219 | -0.2373855 | 0.81235773 | 0.99937574 |
| TBC1D22B | 56.3529122 | -0.0075924 | 0.31287802 | -0.0242662 | 0.98064027 | 0.99937574 |

| | | | | | | |
|---|---|---|---|---|---|---|
| RNF8 | 37.8076856 | 0.14135858 | 0.33639651 | 0.42021417 | 0.674329 | 0.99937574 |
| CMTR1 | 28.5525762 | -0.687496 | 0.39862203 | -1.7246813 | 0.08458493 | 0.99937574 |
| ZFAND3 | 727.957361 | -0.2402673 | 0.23911306 | -1.0048274 | 0.31497997 | 0.99937574 |
| BTBD9 | 546.766135 | -0.4758663 | 0.27356161 | -1.7395214 | 0.08194309 | 0.99937574 |
| GLO1 | 80.5730573 | -0.1119411 | 0.34981417 | -0.3200016 | 0.74896713 | 0.99937574 |
| SAYSD1 | 28.2549168 | -0.051151 | 0.4233233 | -0.1208321 | 0.90382405 | 0.99937574 |
| DAAM2 | 14.9714079 | 0.7792154 | 0.81541497 | 0.95560595 | 0.33927136 | 0.99937574 |
| MOCS1 | 12.3530494 | -0.2580365 | 0.5866574 | -0.4398418 | 0.66005167 | 0.99937574 |
| OARD1 | 43.9061039 | 0.04162918 | 0.38283936 | 0.10873799 | 0.91341031 | 0.99937574 |
| NFYA | 23.4661296 | 0.07154053 | 0.39187789 | 0.18255821 | 0.85514468 | 0.99937574 |
| FOXP4 | 21.3107611 | -0.0583578 | 0.49804995 | -0.1171726 | 0.9067233 | 0.99937574 |
| TFEB | 100.139367 | 0.49641255 | 0.41821289 | 1.18698532 | 0.23523341 | 0.99937574 |
| USP49 | 44.7931861 | 0.41044106 | 0.35748988 | 1.14811938 | 0.25091928 | 0.99937574 |
| CCND3 | 64.7544721 | -1.3687585 | 0.61438174 | -2.2278633 | 0.02588963 | 0.96546668 |
| TAF8 | 33.4929522 | -0.0662699 | 0.34052556 | -0.1946107 | 0.84569775 | 0.99937574 |
| C6orf132 | 18.8440563 | -0.4974274 | 0.63224812 | -0.7867597 | 0.43142253 | 0.99937574 |
| MRPS10 | 20.0573988 | 0.35957394 | 0.49725487 | 0.72311799 | 0.4696074 | 0.99937574 |
| TRERF1 | 25.8936833 | 0.54850392 | 0.96300587 | 0.56957484 | 0.5689661 | 0.99937574 |
| UBR2 | 244.641199 | -0.1680368 | 0.311613 | -0.5392485 | 0.58971543 | 0.99937574 |
| BICRAL | 81.4088522 | 0.04898485 | 0.47005875 | 0.10421007 | 0.91700265 | 0.99937574 |
| RPL7L1 | 62.0039253 | 0.02270114 | 0.37611973 | 0.06035615 | 0.95187198 | 0.99937574 |
| CNPY3 | 13.6402668 | -0.4946595 | 0.52874411 | -0.9355366 | 0.34951183 | 0.99937574 |
| PEX6 | 25.4696027 | 0.83715677 | 0.53220535 | 1.57299578 | 0.1157198 | 0.99937574 |
| PPP2R5D | 12.4285538 | 0.03972833 | 0.49552197 | 0.08017472 | 0.9360983 | 0.99937574 |
| MEA1 | 12.4990452 | -0.1216925 | 1.04093714 | -0.1169066 | 0.90693404 | 0.99937574 |
| KLHDC3 | 20.5516049 | -0.3556501 | 0.40856389 | -0.8704883 | 0.38403363 | 0.99937574 |
| RRP36 | 33.9247927 | 0.02180507 | 0.38280434 | 0.05696139 | 0.95457595 | 0.99937574 |
| CUL7 | 18.2188598 | 0.01021884 | 0.49845531 | 0.02050102 | 0.9836437 | 0.99937574 |
| KLC4 | 21.2574336 | 0.13877239 | 0.50421116 | 0.27522673 | 0.78314205 | 0.99937574 |
| MRPL2 | 19.3517138 | 0.15970768 | 0.53561951 | 0.29817376 | 0.76557054 | 0.99937574 |
| PTK7 | 30.7971077 | 0.81893598 | 0.64036631 | 1.2788555 | 0.20094795 | 0.99937574 |
| SRF | 10.7147711 | -0.2644592 | 0.60430326 | -0.4376267 | 0.66165692 | 0.99937574 |
| CUL9 | 23.7420202 | 0.10060936 | 0.46147962 | 0.21801474 | 0.82741763 | 0.99937574 |
| DNPH1 | 26.7770363 | 0.4117915 | 0.5776117 | 0.71292098 | 0.47589466 | 0.99937574 |
| ZNF318 | 10.2385129 | 0.02283562 | 0.59124284 | 0.03862308 | 0.9691909 | 0.99937574 |
| ABCC10 | 17.0335782 | 0.01901055 | 0.44189373 | 0.04302063 | 0.96568509 | 0.99937574 |
| TJAP1 | 19.8803241 | 0.08891844 | 0.42194616 | 0.21073409 | 0.83309477 | 0.99937574 |
| POLR1C | 24.9081748 | 0.11624532 | 0.43336535 | 0.2682386 | 0.78851566 | 0.99937574 |
| YIPF3 | 40.890274 | 0.24920314 | 0.44701682 | 0.55748046 | 0.57719921 | 0.99937574 |
| XPO5 | 124.432487 | 0.01116503 | 0.31094292 | 0.03590701 | 0.97135651 | 0.99937574 |
| POLH | 20.0964919 | -0.006841 | 0.42937639 | -0.0159324 | 0.98728833 | 0.99937574 |
| GTPBP2 | 24.8495239 | -0.2197155 | 0.45292982 | -0.4850983 | 0.6276066 | 0.99937574 |
| MRPS18A | 24.4401356 | -0.1097926 | 0.45213983 | -0.2428288 | 0.80813805 | 0.99937574 |
| VEGFA | 208.280867 | -0.1603643 | 0.4130998 | -0.3881974 | 0.69786998 | 0.99937574 |
| C6orf223 | 12.9740269 | -0.5264257 | 0.81084769 | -0.6492289 | 5.16E-01 | 0.99937574 |
| MRPL14 | 50.6318978 | 0.27327027 | 0.42963419 | 0.63605335 | 0.52474165 | 0.99937574 |
| TMEM63B | 35.2494635 | 0.33539495 | 0.37681858 | 0.8900701 | 0.37342825 | 0.99937574 |
| SLC29A1 | 38.2856072 | -0.5499806 | 0.52106945 | -1.0554843 | 0.29120387 | 0.99937574 |
| HSP90AB1 | 326.408821 | 0.33527842 | 0.27635861 | 1.21320055 | 0.22505315 | 0.99937574 |
| SLC35B2 | 16.9426343 | 0.07854465 | 0.60320725 | 0.13021171 | 0.89639893 | 0.99937574 |
| CDC5L | 41.7470305 | -0.1476912 | 0.32818467 | -0.4500246 | 0.65269272 | 0.99937574 |
| SUPT3H | 267.478463 | -0.2953338 | 0.42900656 | -0.6884133 | 0.49119254 | 0.99937574 |
| RUNX2 | 15.6694082 | 0.44892519 | 0.70833192 | 0.633778 | 0.52622572 | 0.99937574 |
| ENPP4 | 49.4310969 | 0.52905548 | 0.30872679 | 1.71366885 | 0.08658956 | 0.99937574 |

| | | | | | | |
|---|---|---|---|---|---|---|
| ENPP5 | 15.6410648 | 0.20288851 | 0.57726102 | 0.35146754 | 0.72523762 | 0.99937574 |
| RCAN2 | 1550.1846 | 0.5874096 | 0.40930735 | 1.43513083 | 0.15124982 | 0.99937574 |
| CYP39A1 | 37.5551801 | 0.68825306 | 0.64426926 | 1.06826929 | 0.28539906 | 0.99937574 |
| SLC25A27 | 77.5040262 | -0.1835298 | 0.52227641 | -0.3514036 | 0.72528555 | 0.99937574 |
| ADGRF5 | 2399.6126 | 0.05498663 | 0.49918669 | 0.11015243 | 0.91228849 | 0.99937574 |
| ADGRF1 | 1225.5367 | -0.1515596 | 0.3986142 | -0.3802163 | 0.7037849 | 0.99937574 |
| TNFRSF21 | 198.631649 | -0.4538924 | 0.59610716 | -0.7614276 | 0.44640173 | 0.99937574 |
| CD2AP | 330.149102 | -0.1473819 | 0.2548287 | -0.578357 | 0.56302315 | 0.99937574 |
| MUT | 46.7013198 | 0.24024944 | 0.43022805 | 0.55842346 | 0.57655525 | 0.99937574 |
| PKHD1 | 470.176534 | -1.5604027 | 0.88984517 | -1.7535666 | 0.0795048 | 0.99937574 |
| MCM3 | 15.0938585 | 0.17487212 | 0.53197562 | 0.32872206 | 0.74236578 | 0.99937574 |
| EFHC1 | 24.2610476 | -0.0781599 | 0.46475506 | -0.1681744 | 0.86644611 | 0.99937574 |
| GSTA4 | 12.2365669 | 0.26535279 | 0.81424946 | 0.32588636 | 0.74451033 | 0.99937574 |
| ICK | 16.5811399 | 0.30552158 | 0.4875268 | 0.62667648 | 0.53087132 | 0.99937574 |
| FBXO9 | 57.4067159 | -0.2445067 | 0.29141355 | -0.839037 | 0.40144857 | 0.99937574 |
| ELOVL5 | 62.3018773 | 0.1901336 | 0.53911379 | 0.35267805 | 0.72432982 | 0.99937574 |
| GCLC | 84.1656844 | -0.3275293 | 0.42633446 | -0.768245 | 4.42E-01 | 0.99937574 |
| LRRC1 | 37.5477714 | -0.0584011 | 0.35018484 | -0.1667722 | 0.8675493 | 0.99937574 |
| FAM83B | 113.224113 | -0.1758641 | 0.27040159 | -0.6503812 | 0.51544605 | 0.99937574 |
| DST | 402.484933 | 0.00259265 | 0.44093859 | 0.00587984 | 0.99530859 | 0.99937574 |
| KIAA1586 | 10.9586124 | 0.26846855 | 0.54660732 | 0.49115433 | 0.6233173 | 0.99937574 |
| ZNF451 | 118.767894 | 0.03013169 | 0.31518625 | 0.09559965 | 0.92383855 | 0.99937574 |
| ZNF451-AS1 | 16.7232597 | 0.61691912 | 0.84985991 | 0.72590684 | 0.46789588 | 0.99937574 |
| PRIM2 | 98.6725339 | -0.4817332 | 0.33205568 | -1.4507604 | 0.14684658 | 0.99937574 |
| AL021368.2 | 11.883462 | 0.99288375 | 0.60334708 | 1.64562618 | 0.09984075 | 0.99937574 |
| PTP4A1 | 53.744079 | -0.8096308 | 0.63125651 | -1.2825703 | 0.19964267 | 0.99937574 |
| PHF3 | 168.275426 | 0.03512098 | 0.26398549 | 0.13304133 | 0.89416069 | 0.99937574 |
| EYS | 15.2022015 | 0.68832979 | 0.67397344 | 1.02130105 | 0.30711183 | 0.99937574 |
| LMBRD1 | 154.506914 | 0.20053776 | 0.34390048 | 0.58312732 | 0.55980759 | 0.99937574 |
| COL19A1 | 34.6201707 | -0.3394688 | 0.38705923 | -0.877046 | 0.38046165 | 0.99937574 |
| FAM135A | 393.60886 | 0.13039473 | 0.37495754 | 0.34775864 | 0.72802145 | 0.99937574 |
| SDHAF4 | 22.1057652 | 0.74221085 | 0.67099788 | 1.10612995 | 0.26867028 | 0.99937574 |
| SMAP1 | 221.877787 | -0.1670268 | 0.26488505 | -0.6305635 | 0.52832598 | 0.99937574 |
| OGFRL1 | 40.97823 | -1.7866532 | 0.63347516 | -2.8203998 | 0.00479638 | 0.60897109 |
| AL136164.2 | 85.655106 | 0.81876739 | 0.57565488 | 1.42232337 | 0.1549324 | 0.99937574 |
| LINC00472 | 51.4556014 | -0.062711 | 0.64244056 | -0.0976136 | 0.92223909 | 0.99937574 |
| OOEP | 173.16663 | -0.3508065 | 0.49132265 | -0.7140043 | 0.47522454 | 0.99937574 |
| MTO1 | 33.9170732 | -0.1564974 | 0.34289691 | -0.4563977 | 0.64810401 | 0.99937574 |
| EEF1A1 | 844.608995 | 0.15483844 | 0.52651025 | 0.29408438 | 0.76869343 | 0.99937574 |
| SLC17A5 | 50.5683554 | -0.9918702 | 0.50027439 | -1.9826524 | 0.04740627 | 0.99937574 |
| COX7A2 | 136.087764 | 0.30257303 | 0.43141495 | 0.70135037 | 0.48308439 | 0.99937574 |
| TMEM30A | 101.419924 | 0.24502043 | 0.27248068 | 0.89922129 | 0.3685348 | 0.99937574 |
| FILIP1 | 76.5808904 | 0.73018081 | 0.92525866 | 0.78916398 | 4.30E-01 | 9.99E-01 |
| SENP6 | 387.966711 | 0.04254311 | 0.2107223 | 0.20189183 | 0.84000129 | 0.99937574 |
| MYO6 | 740.960731 | -0.2325896 | 0.35938528 | -0.6471874 | 0.51751065 | 0.99937574 |
| IRAK1BP1 | 25.8663571 | -0.1265513 | 0.57331958 | -0.2207344 | 0.82529927 | 0.99937574 |
| PHIP | 221.521274 | -0.292853 | 0.27358864 | -1.0704136 | 0.2844332 | 0.99937574 |
| HMGN3 | 108.716634 | 0.40964555 | 0.46076257 | 0.88905997 | 0.37397085 | 0.99937574 |
| LCA5 | 15.1236132 | 0.0484155 | 0.53847354 | 0.08991249 | 0.92835675 | 0.99937574 |
| SH3BGRL2 | 16.3826416 | -0.4621576 | 0.63314236 | -0.7299426 | 0.46542528 | 0.99937574 |
| BCKDHB | 110.868729 | 1.00432364 | 0.44149054 | 2.27484746 | 0.02291508 | 0.96427895 |
| TENT5A | 65.6172414 | -0.3054681 | 0.75435368 | -0.4049401 | 0.68552153 | 0.99937574 |
| IBTK | 161.595547 | -0.3454748 | 0.21584713 | -1.6005531 | 0.10947595 | 0.99937574 |
| TPBG | 13.3161566 | 0.36941176 | 0.58824903 | 0.62798533 | 0.53001355 | 0.99937574 |

| | | | | | | |
|---|---|---|---|---|---|---|
| UBE3D | 65.9870435 | 0.09021698 | 0.36687195 | 0.24590864 | 0.80575296 | 0.99937574 |
| DOPEY1 | 140.415527 | 0.14219118 | 0.29668472 | 0.47926696 | 0.63174872 | 0.99937574 |
| PGM3 | 31.6146998 | 0.53640757 | 0.43593946 | 1.23046344 | 0.21852361 | 0.99937574 |
| CYB5R4 | 45.4194548 | -0.7602784 | 0.39130034 | -1.9429537 | 0.05202176 | 0.99937574 |
| CEP162 | 54.1064475 | -0.0890886 | 0.3262311 | -0.2730844 | 0.78478833 | 0.99937574 |
| SNX14 | 141.091609 | 0.03997867 | 0.26363453 | 0.1516443 | 0.87946749 | 0.99937574 |
| SYNCRIP | 52.3579783 | -0.2196263 | 0.43856615 | -0.5007825 | 0.61652418 | 0.99937574 |
| ZNF292 | 455.767146 | -0.8561498 | 0.32830707 | -2.6077714 | 0.00911338 | 0.74774163 |
| SMIM8 | 71.1981007 | 0.2618363 | 0.45907843 | 0.57035199 | 0.56843899 | 0.99937574 |
| RARS2 | 89.8270507 | -0.1072889 | 0.27301865 | -0.3929726 | 0.6943397 | 0.99937574 |
| ORC3 | 48.6778137 | -0.2006994 | 0.45623092 | -0.4399074 | 0.66000417 | 0.99937574 |
| AKIRIN2 | 26.2151492 | -0.1904926 | 0.3821391 | -0.4984901 | 0.61813862 | 0.99937574 |
| RNGTT | 181.488509 | -0.0032263 | 0.25152485 | -0.012827 | 0.98976581 | 0.99937574 |
| PNRC1 | 89.8957266 | 0.07339949 | 0.50071797 | 0.14658848 | 0.88345685 | 0.99937574 |
| PM20D2 | 38.0744946 | 0.04348038 | 0.36761157 | 0.11827804 | 0.90584735 | 0.99937574 |
| UBE2J1 | 43.9220479 | 0.03743878 | 0.42934086 | 0.0872006 | 0.93051206 | 0.99937574 |
| RRAGD | 80.2889287 | -0.489842 | 0.48937029 | -1.000964 | 0.31684422 | 0.99937574 |
| ANKRD6 | 51.6251868 | -0.3783794 | 0.3383126 | -1.1184313 | 0.26338282 | 0.99937574 |
| LYRM2 | 44.1433039 | -0.1821807 | 0.45939081 | -0.3965703 | 0.69168436 | 0.99937574 |
| MDN1 | 72.543856 | -0.1591887 | 0.31477993 | -0.5057142 | 0.6130573 | 0.99937574 |
| CASP8AP2 | 46.2650316 | -0.6395737 | 0.37913936 | -1.6869094 | 0.09162078 | 0.99937574 |
| BACH2 | 98.474061 | 0.55705008 | 0.7805812 | 0.71363502 | 0.4754529 | 0.99937574 |
| MAP3K7 | 94.7603619 | -0.0644773 | 0.34101455 | -0.189075 | 0.85003401 | 0.99937574 |
| MANEA | 46.0579092 | 0.12214936 | 0.37561958 | 0.32519433 | 0.745034 | 0.99937574 |
| UFL1 | 63.3424141 | 0.0002403 | 0.27781384 | 0.00086497 | 0.99930985 | 0.99979721 |
| NDUFAF4 | 34.5700647 | 0.18731081 | 0.55291588 | 0.3387691 | 0.73478368 | 0.99937574 |
| MMS22L | 25.8047461 | 0.25447366 | 0.54398622 | 0.46779432 | 0.63993168 | 0.99937574 |
| FBXL4 | 121.12491 | -0.2708102 | 0.3783336 | -0.7157974 | 0.47411647 | 0.99937574 |
| FAXC | 20.2403242 | -0.5735918 | 0.76206437 | -0.7526815 | 0.45164132 | 0.99937574 |
| COQ3 | 13.9955999 | 0.47665612 | 0.49583464 | 0.96132073 | 0.33639093 | 0.99937574 |
| PNISR | 376.192778 | 0.21878096 | 0.19036392 | 1.14927739 | 0.25044162 | 0.99937574 |
| AL513550.1 | 19.00728 | 0.68301228 | 0.5295123 | 1.28988936 | 0.19708907 | 0.99937574 |
| USP45 | 60.6524265 | 0.01067984 | 0.45752012 | 0.02334289 | 0.98137676 | 0.99937574 |
| TSTD3 | 15.8361042 | 0.0402318 | 0.55012059 | 0.0731327 | 0.94170052 | 0.99937574 |
| CCNC | 126.155552 | -0.1949983 | 0.33724556 | -0.5782085 | 0.56312333 | 0.99937574 |
| SIM1 | 323.260102 | 0.38458393 | 0.36006386 | 1.06809924 | 0.28547575 | 0.99937574 |
| ASCC3 | 216.075409 | 0.14121486 | 0.34515249 | 0.40913761 | 0.68243868 | 0.99937574 |
| HACE1 | 37.8665098 | 0.19367662 | 0.35693622 | 0.54260848 | 0.5873994 | 0.99937574 |
| PREP | 52.4538281 | 0.23121392 | 0.35670177 | 0.64819954 | 0.5168559 | 0.99937574 |
| ATG5 | 128.612029 | -0.2296902 | 0.29421257 | -0.7806948 | 0.43498201 | 0.99937574 |
| CRYBG1 | 109.968746 | -1.350703 | 0.95478505 | -1.4146671 | 0.15716611 | 0.99937574 |
| RTN4IP1 | 54.6139521 | -0.0173192 | 0.52414505 | -0.0330427 | 0.97364055 | 0.99937574 |
| QRSL1 | 69.4664218 | -0.1722718 | 0.40529596 | -0.4250518 | 0.67079888 | 0.99937574 |
| LINC02532 | 108.91333 | -0.6606478 | 0.73429656 | -0.8997016 | 0.36827905 | 0.99937574 |
| CD24 | 1021.30196 | 0.09533869 | 0.38994468 | 0.24449286 | 0.80684913 | 0.99937574 |
| C6orf203 | 26.0771368 | -0.0144785 | 0.37302042 | -0.0388142 | 0.96903852 | 0.99937574 |
| PDSS2 | 183.136009 | -0.0130618 | 0.26715225 | -0.0488926 | 0.96100487 | 0.99937574 |
| SEC63 | 93.654056 | -0.3245371 | 0.24642643 | -1.3169736 | 0.18784749 | 0.99937574 |
| OSTM1 | 75.7419203 | -0.1976044 | 0.50284658 | -0.3929716 | 0.69434043 | 0.99937574 |
| SNX3 | 119.546927 | -0.1141892 | 0.3150545 | -0.3624428 | 0.71702116 | 0.99937574 |
| AFG1L | 86.4229602 | -0.3144452 | 0.38179563 | -0.8235955 | 0.41016945 | 0.99937574 |
| FOXO3 | 313.34487 | 0.22440184 | 0.32095406 | 0.69917122 | 0.48444503 | 0.99937574 |
| ARMC2 | 23.2602134 | 0.59977046 | 0.58064011 | 1.03294699 | 0.30162871 | 0.99937574 |
| SESN1 | 112.978051 | 0.15446149 | 0.34290689 | 0.45044733 | 0.65238793 | 0.99937574 |

| | | | | | | |
|---|---|---|---|---|---|---|
| CEP57L1 | 33.8613647 | 0.3854998 | 0.39342145 | 0.97986473 | 0.32715289 | 0.99937574 |
| CD164 | 163.9972 | -0.5626153 | 0.23366325 | -2.4078041 | 0.01604879 | 0.87097377 |
| PPIL6 | 12.5918163 | 0.04131121 | 0.78463511 | 0.05265022 | 0.9580106 | 0.99937574 |
| ZBTB24 | 10.7491909 | 0.0918613 | 0.54257406 | 0.16930647 | 0.86555559 | 0.99937574 |
| AK9 | 42.0264774 | 0.39438103 | 0.61640588 | 0.63980737 | 0.52229784 | 0.99937574 |
| FIG4 | 51.7909443 | -0.7949273 | 0.32888078 | -2.4170682 | 0.01564609 | 0.86758411 |
| CDC40 | 26.9523598 | -0.205195 | 0.38498933 | -0.5329889 | 0.59404126 | 0.99937574 |
| CDK19 | 76.5592864 | -0.0161835 | 0.44208435 | -0.0366073 | 0.97079812 | 0.99937574 |
| AMD1 | 69.4247476 | -0.4638363 | 0.36976065 | -1.254423 | 0.2096883 | 0.99937574 |
| GTF3C6 | 54.2640522 | 0.23779255 | 0.50038964 | 0.47521477 | 0.6346339 | 0.99937574 |
| RPF2 | 18.3714356 | -0.5516436 | 0.47425705 | -1.1631743 | 0.24475877 | 0.99937574 |
| SLC16A10 | 212.972674 | -0.3328298 | 0.30100303 | -1.1057356 | 0.26884099 | 0.99937574 |
| MFSD4B | 16.9175247 | 0.13100574 | 0.48421566 | 0.27055246 | 0.78673526 | 0.99937574 |
| REV3L | 196.458165 | -0.1354175 | 0.26752607 | -0.5061842 | 0.61272733 | 0.99937574 |
| TRAF3IP2-AS | 66.0866359 | 0.0183165 | 0.30234 | 0.06058245 | 0.95169175 | 0.99937574 |
| TRAF3IP2 | 19.0967412 | -0.0201903 | 0.43989385 | -0.0458981 | 0.96339147 | 0.99937574 |
| TUBE1 | 29.6943756 | -0.2011747 | 0.51680468 | -0.3892664 | 0.69707906 | 0.99937574 |
| LAMA4 | 21.564648 | 1.52992773 | 1.04762871 | 1.4603721 | 0.14418784 | 0.99937574 |
| MARCKS | 129.828601 | 0.21101918 | 0.35280367 | 0.59812072 | 0.54975939 | 0.99937574 |
| HDAC2 | 45.0208543 | -0.0450847 | 0.30905708 | -0.1458783 | 0.88401746 | 0.99937574 |
| FRK | 315.886195 | 0.3109187 | 0.32742111 | 0.94959883 | 0.34231613 | 0.99937574 |
| NT5DC1 | 256.393489 | 0.0344286 | 0.30104481 | 0.11436371 | 0.90894948 | 0.99937574 |
| COL10A1 | 14.0441999 | 0.28246097 | 0.88288084 | 0.31993102 | 0.74902062 | 0.99937574 |
| TSPYL4 | 16.5922688 | 0.35994095 | 0.50919617 | 0.70688071 | 0.47964061 | 0.99937574 |
| DSE | 57.0123155 | 0.4446019 | 0.41783034 | 1.06407278 | 0.28729573 | 0.99937574 |
| TSPYL1 | 76.436559 | 0.28316477 | 0.27162252 | 1.04249372 | 0.29718284 | 0.99937574 |
| RWDD1 | 34.4078557 | 0.08891018 | 0.41663847 | 0.21339888 | 0.83101586 | 0.99937574 |
| ZUP1 | 26.2912738 | -0.0497289 | 0.41648089 | -0.1194026 | 0.90495637 | 0.99937574 |
| KPNA5 | 78.2819353 | 0.27202564 | 0.28618254 | 0.95053191 | 0.34184205 | 0.99937574 |
| DCBLD1 | 29.6946733 | -0.115525 | 0.47467829 | -0.2433754 | 0.80771461 | 0.99937574 |
| GOPC | 47.6885957 | -0.0519191 | 0.34036044 | -0.1525414 | 0.87875993 | 0.99937574 |
| NUS1 | 63.282559 | 0.14076559 | 0.31009034 | 0.45395028 | 0.64986462 | 0.99937574 |
| SLC35F1 | 12.6551197 | -0.2304682 | 0.60013479 | -0.3840274 | 0.70095812 | 0.99937574 |
| CEP85L | 51.8459905 | 0.46363559 | 0.41440251 | 1.11880499 | 0.26322334 | 0.99937574 |
| MCM9 | 109.85032 | 0.15316303 | 0.32185842 | 0.47587084 | 0.6341664 | 0.99937574 |
| AL365275.1 | 26.7218965 | -0.9633361 | 0.59906143 | -1.6080756 | 0.10781862 | 0.99937574 |
| FAM184A | 64.0302839 | 0.41986535 | 0.76100752 | 0.551723 | 0.58113815 | 0.99937574 |
| MAN1A1 | 354.693834 | -0.5976254 | 0.30789518 | -1.9410027 | 0.05225794 | 0.99937574 |
| TBC1D32 | 52.8413536 | -0.8201999 | 0.39765967 | -2.0625675 | 0.03915374 | 0.99937574 |
| HSF2 | 20.3587629 | 0.28740517 | 0.53199921 | 0.54023608 | 0.58903423 | 0.99937574 |
| SERINC1 | 153.247533 | -0.1746685 | 0.27575814 | -0.6334119 | 0.52646469 | 0.99937574 |
| PKIB | 23.4479416 | -0.6965631 | 0.64392045 | -1.0817533 | 0.27936217 | 0.99937574 |
| SMPDL3A | 19.6075634 | -0.4533219 | 0.47720389 | -0.9499543 | 0.34213547 | 0.99937574 |
| NKAIN2 | 12.359824 | -0.9620142 | 0.62770817 | -1.5325819 | 0.12537891 | 0.99937574 |
| RNF217-AS1 | 16.927139 | -0.9768674 | 0.77418738 | -1.2617971 | 0.20702182 | 0.99937574 |
| RNF217 | 247.751208 | -0.0783375 | 0.58532135 | -0.1338367 | 0.89353172 | 0.99937574 |
| TPD52L1 | 102.356131 | 0.04085745 | 0.66273855 | 0.06164943 | 0.95084201 | 0.99937574 |
| HDDC2 | 44.1471636 | -0.1250197 | 0.4518698 | -0.2766719 | 0.78203208 | 0.99937574 |
| NCOA7 | 875.971333 | 0.00785712 | 0.29271197 | 0.02684249 | 0.97858537 | 0.99937574 |
| NCOA7-AS1 | 37.7047867 | -0.3083772 | 0.71893748 | -0.4289346 | 0.6679708 | 0.99937574 |
| HINT3 | 36.9340461 | 0.16188341 | 0.34255704 | 0.47257358 | 0.63651744 | 0.99937574 |
| TRMT11 | 60.3291446 | -0.3507722 | 0.32414794 | -1.0821362 | 0.279192 | 0.99937574 |
| RNF146 | 46.8840786 | 0.30065141 | 0.44594408 | 0.67419082 | 0.50019 | 0.99937574 |
| ECHDC1 | 69.9431809 | 0.19734039 | 0.32255002 | 0.6118133 | 0.54066129 | 0.99937574 |

| | | | | | | |
|---|---|---|---|---|---|---|
| PTPRK | 641.213301 | -0.4353197 | 0.4642824 | -0.9376183 | 0.34844061 | 0.99937574 |
| LAMA2 | 249.264159 | 0.457051 | 0.68088571 | 0.67125949 | 0.50205524 | 0.99937574 |
| AL356124.1 | 218.063138 | 1.19349173 | 0.39795452 | 2.99906566 | 0.00270809 | 0.50611182 |
| ARHGAP18 | 1327.98531 | 0.22026219 | 0.21996988 | 1.00132887 | 0.31666784 | 0.99937574 |
| L3MBTL3 | 22.9158868 | -0.1140899 | 0.45203693 | -0.2523907 | 0.80073909 | 0.99937574 |
| EPB41L2 | 16.7234765 | 0.31902838 | 0.51485715 | 0.61964446 | 0.53549189 | 0.99937574 |
| AKAP7 | 44.9048811 | -1.3153134 | 0.65784389 | -1.9994309 | 0.04556175 | 0.99937574 |
| MED23 | 58.8541769 | -0.3752572 | 0.31304793 | -1.1987212 | 0.23063637 | 0.99937574 |
| STX7 | 102.05109 | -0.0667303 | 0.35060405 | -0.1903295 | 0.84905095 | 0.99937574 |
| SLC18B1 | 56.9734475 | -0.3759918 | 0.53522829 | -0.7024886 | 0.48237449 | 0.99937574 |
| RPS12 | 179.368217 | 0.07368829 | 0.47892871 | 0.15386068 | 0.87771959 | 0.99937574 |
| EYA4 | 149.198908 | -1.5689713 | 0.96556096 | -1.6249324 | 0.10417696 | 0.99937574 |
| TBPL1 | 32.8150178 | 0.23122834 | 0.49058432 | 0.4713325 | 0.63740331 | 0.99937574 |
| SLC2A12 | 27.6201955 | 1.10950981 | 0.74363239 | 1.49201382 | 0.13569552 | 0.99937574 |
| HBS1L | 61.4609874 | -0.2449825 | 0.32039945 | -0.764616 | 0.44450025 | 0.99937574 |
| AHI1 | 265.952341 | -0.0371308 | 0.26609073 | -0.139542 | 0.88902186 | 0.99937574 |
| LINC00271 | 66.6715975 | -0.2904004 | 0.65008102 | -0.4467141 | 0.6550815 | 0.99937574 |
| PDE7B | 188.040919 | 0.72378337 | 0.41988933 | 1.72374795 | 0.08475336 | 0.99937574 |
| AL138828.1 | 38.3990718 | -1.4620633 | 0.67788765 | -2.1567929 | 0.0310218 | 0.99049468 |
| BCLAF1 | 152.533603 | -0.1641975 | 0.24446959 | -0.6716478 | 0.50180797 | 0.99937574 |
| MAP7 | 277.750109 | -0.3984503 | 0.28102123 | -1.4178655 | 0.15623004 | 0.99937574 |
| MAP3K5 | 289.928692 | 0.58806549 | 0.44698616 | 1.31562348 | 0.18830045 | 0.99937574 |
| PEX7 | 58.5473766 | 0.33693166 | 0.34765232 | 0.96916271 | 0.33246401 | 0.99937574 |
| IFNGR1 | 61.7379047 | 0.40884998 | 0.36413302 | 1.12280392 | 0.26152078 | 0.99937574 |
| TNFAIP3 | 15.1840276 | -0.7137042 | 0.76828395 | -0.9289589 | 0.3529104 | 0.99937574 |
| PERP | 69.0745785 | -0.2515324 | 0.35443501 | -0.7096714 | 4.78E-01 | 0.99937574 |
| ARFGEF3 | 63.3833865 | -1.2046867 | 0.47332002 | -2.5451842 | 0.01092201 | 0.76274616 |
| HEBP2 | 52.2270597 | -0.0520903 | 0.3662712 | -0.1422178 | 0.88690801 | 0.99937574 |
| NHSL1 | 50.1353687 | -1.0645272 | 0.76213186 | -1.3967756 | 0.16248105 | 0.99937574 |
| CCDC28A | 26.8854612 | 0.25572857 | 0.51342242 | 0.49808611 | 0.61842335 | 0.99937574 |
| REPS1 | 136.558683 | 0.02978995 | 0.33737363 | 0.08829958 | 0.92963857 | 0.99937574 |
| ABRACL | 13.825136 | -1.3772536 | 0.64688016 | -2.1290707 | 0.03324841 | 0.99937574 |
| HECA | 29.6552654 | 0.00206167 | 0.54722186 | 0.00376752 | 0.99699396 | 0.99937574 |
| CITED2 | 55.1407906 | -0.0458592 | 0.64535458 | -0.0710605 | 0.94334957 | 0.99937574 |
| VTA1 | 49.8644419 | -0.1647794 | 0.31571472 | -0.5219249 | 0.60172259 | 0.99937574 |
| ADGRG6 | 13.1002402 | -0.266423 | 0.89665641 | -0.2971294 | 0.76636774 | 0.99937574 |
| HIVEP2 | 213.757847 | 0.0536664 | 0.43950288 | 0.12210706 | 0.90281423 | 0.99937574 |
| AIG1 | 189.177865 | -0.1415051 | 0.42664883 | -0.3316665 | 0.74014107 | 0.99937574 |
| ADAT2 | 16.9746826 | 0.5910595 | 0.56410836 | 1.04777653 | 0.29474158 | 0.99937574 |
| PEX3 | 52.0180333 | -0.2454822 | 0.42054643 | -0.5837221 | 0.55940732 | 0.99937574 |
| FUCA2 | 41.2058122 | -0.2561928 | 0.34932497 | -0.733394 | 0.46331818 | 0.99937574 |
| PHACTR2 | 268.336239 | -0.047002 | 0.40310913 | -0.1165986 | 0.90717813 | 0.99937574 |
| PLAGL1 | 44.4541802 | 0.41001251 | 0.76197922 | 0.53808884 | 0.59051572 | 0.99937574 |
| SF3B5 | 53.4895467 | 0.39789306 | 0.38146535 | 1.04306475 | 0.2969183 | 0.99937574 |
| UTRN | 58.2423007 | -0.4097679 | 0.69970318 | -0.585631 | 0.55812353 | 0.99937574 |
| EPM2A | 32.6111556 | -0.3497431 | 0.58270444 | -0.6002068 | 0.54836845 | 0.99937574 |
| FBXO30 | 25.6472247 | 0.53931472 | 0.53515036 | 1.00778167 | 0.31355929 | 0.99937574 |
| AL356599.1 | 44.8189738 | 0.53939952 | 0.59243876 | 0.91047304 | 0.3625731 | 0.99937574 |
| SHPRH | 157.768198 | 0.15783319 | 0.32153552 | 0.49087325 | 0.6235161 | 0.99937574 |
| STXBP5 | 167.295943 | -0.0730804 | 0.44480633 | -0.164297 | 0.86949731 | 0.99937574 |
| SAMD5 | 774.422459 | 0.41225014 | 0.39536139 | 1.04271723 | 0.29707927 | 0.99937574 |
| AL033504.1 | 425.306454 | 0.25853748 | 0.48385437 | 0.53432912 | 0.59311385 | 0.99937574 |
| AL513164.1 | 28.1456595 | 0.75465024 | 0.57347437 | 1.31592672 | 0.18819864 | 0.99937574 |
| SASH1 | 315.346489 | 0.09825193 | 0.22455114 | 0.43754812 | 0.66171389 | 0.99937574 |

| | | | | | | |
|---|---|---|---|---|---|---|
| UST | 731.579933 | -0.8438653 | 0.34159215 | -2.4703884 | 0.01349664 | 0.82550375 |
| TAB2 | 156.625583 | 0.19234082 | 0.21584813 | 0.89109327 | 0.37287914 | 0.99937574 |
| PPIL4 | 64.3596324 | -0.0676645 | 0.31442495 | -0.2152008 | 0.82961075 | 0.99937574 |
| GINM1 | 99.2957734 | 0.11380555 | 0.2886355 | 0.39428813 | 0.69336833 | 0.99937574 |
| KATNA1 | 44.6058469 | -0.2598896 | 0.33305245 | -0.7803262 | 0.43519889 | 0.99937574 |
| LATS1 | 86.3680948 | -0.2476541 | 0.28396767 | -0.8721207 | 0.38314255 | 0.99937574 |
| NUP43 | 24.4680634 | -0.0339811 | 0.39641492 | -0.0857211 | 0.93168815 | 0.99937574 |
| PCMT1 | 77.4221366 | -0.2882299 | 0.40454326 | -0.7124824 | 0.47616613 | 0.99937574 |
| LRP11 | 75.4431846 | -0.2077624 | 0.58142044 | -0.357336 | 0.72084029 | 0.99937574 |
| PLEKHG1 | 39.6160872 | -0.9317495 | 0.87016501 | -1.0707734 | 0.28427134 | 0.99937574 |
| MTHFD1L | 248.212791 | -0.223557 | 0.4037742 | -0.5536684 | 0.57980583 | 0.99937574 |
| RMND1 | 25.2024801 | -0.7744513 | 0.43936895 | -1.7626445 | 0.07796047 | 0.99937574 |
| ARMT1 | 25.5625815 | 0.13923854 | 0.51877997 | 0.26839614 | 0.78839441 | 0.99937574 |
| ESR1 | 71.9682454 | -4.4178187 | 1.46852784 | -3.0083316 | 0.00262686 | 0.50019442 |
| SYNE1 | 122.040745 | -0.3644186 | 0.77339886 | -0.471191 | 0.63750437 | 0.99937574 |
| MTRF1L | 23.395046 | 2.81E-01 | 0.5226288 | 0.53787753 | 0.5906616 | 0.99937574 |
| OPRM1 | 27.8869686 | 0.33657884 | 0.99563528 | 0.33805435 | 0.73532223 | 0.99937574 |
| IPCEF1 | 238.801486 | 0.78929714 | 0.76447462 | 1.03247003 | 0.30185198 | 0.99937574 |
| CNKSR3 | 107.673542 | -0.2272059 | 0.40184479 | -0.5654071 | 0.57179689 | 0.99937574 |
| SCAF8 | 264.795409 | -0.0079144 | 0.25374517 | -0.0311904 | 0.97511771 | 0.99937574 |
| TIAM2 | 159.079838 | 0.99150551 | 0.29763159 | 3.33131818 | 0.00086436 | 0.29012345 |
| TFB1M | 49.6375503 | -0.5425187 | 0.41950844 | -1.2932247 | 0.19593335 | 0.99937574 |
| ARID1B | 735.38085 | -0.0226909 | 0.28971747 | -0.0783207 | 0.93757292 | 0.99937574 |
| TMEM242 | 30.6741569 | -0.2810462 | 0.38039992 | -0.7388178 | 0.46001765 | 0.99937574 |
| ZDHHC14 | 112.638516 | 0.07821649 | 0.64896921 | 0.12052419 | 0.90406792 | 0.99937574 |
| SNX9 | 249.745129 | -0.4875673 | 0.23732819 | -2.0544011 | 0.03993689 | 0.99937574 |
| SYNJ2 | 32.2790785 | -0.5746198 | 0.56635608 | -1.014591 | 0.31030082 | 0.99937574 |
| SERAC1 | 25.2860828 | -0.5523744 | 0.57304016 | -0.9639366 | 0.33507769 | 0.99937574 |
| GTF2H5 | 33.1449559 | -0.1490558 | 0.5546731 | -0.2687273 | 0.78813955 | 0.99937574 |
| TULP4 | 325.485206 | 0.18057262 | 0.35266448 | 0.51202383 | 0.60863433 | 0.99937574 |
| TMEM181 | 124.827252 | 0.14222637 | 0.38260361 | 0.37173296 | 0.71009168 | 0.99937574 |
| DYNLT1 | 128.281091 | 0.08702145 | 0.46757232 | 0.18611334 | 0.85235588 | 0.99937574 |
| SYTL3 | 47.1675909 | 0.22401102 | 0.58199338 | 0.38490304 | 0.70030925 | 0.99937574 |
| EZR | 479.497246 | 0.39843122 | 0.35019491 | 1.13774132 | 0.25522851 | 0.99937574 |
| RSPH3 | 37.592712 | 0.29757164 | 0.48223892 | 0.61706267 | 0.53719339 | 0.99937574 |
| AL035530.2 | 21.2640618 | 0.41938987 | 0.53917039 | 0.77784292 | 0.43666162 | 0.99937574 |
| SOD2.1 | 27.3140782 | -0.3221375 | 0.46951966 | -0.6861002 | 0.49264992 | 0.99937574 |
| WTAP | 143.357009 | -0.7447563 | 0.37669549 | -1.9770779 | 0.04803282 | 0.99937574 |
| ACAT2 | 26.3087048 | -0.0453858 | 0.79804212 | -0.0568714 | 0.9546476 | 0.99937574 |
| TCP1 | 48.0480916 | 0.11941375 | 0.41986079 | 0.28441273 | 0.7760941 | 0.99937574 |
| MRPL18 | 29.6347207 | -0.2054573 | 0.52997128 | -0.3876764 | 0.69825556 | 0.99937574 |
| IGF2R | 245.316921 | -0.4605606 | 0.26223831 | -1.7562674 | 0.07904276 | 0.99937574 |
| MAP3K4 | 89.5761075 | 0.07921899 | 0.25264864 | 0.31355401 | 0.75385979 | 0.99937574 |
| PRKN | 1052.19368 | 0.67242933 | 0.30711541 | 2.18950044 | 0.02856049 | 0.99049468 |
| PACRG | 1892.35692 | -0.4613756 | 0.54215491 | -0.8510033 | 0.39476751 | 0.99937574 |
| PACRG-AS3 | 32.0958934 | -0.0318748 | 0.41437033 | -0.0769234 | 0.93868447 | 0.99937574 |
| QKI | 427.851018 | -0.324724 | 0.30041955 | -1.0809016 | 0.27974086 | 0.99937574 |
| PDE10A | 249.74119 | 0.48732432 | 0.57746216 | 0.84390692 | 0.39872142 | 0.99937574 |
| SFT2D1 | 58.3702648 | -0.5002942 | 0.2842293 | -1.760178 | 0.07837762 | 0.99937574 |
| MPC1 | 275.069466 | 0.5484182 | 0.35968181 | 1.52473155 | 0.12732607 | 0.99937574 |
| RPS6KA2 | 57.9231699 | -0.1990717 | 0.4251486 | -0.4682402 | 0.63961281 | 0.99937574 |
| Z94721.1 | 23.8890918 | -0.1458945 | 0.53517403 | -0.2726114 | 0.78515197 | 0.99937574 |
| FGFR1OP | 95.0961885 | 0.35350275 | 0.41148448 | 0.85909134 | 0.39029013 | 0.99937574 |
| AFDN | 351.199367 | -0.0092786 | 0.28074352 | -0.0330502 | 0.97363455 | 0.99937574 |

| | | | | | | |
|---|---|---|---|---|---|---|
| WDR27 | 65.3942815 | -0.1195053 | 0.45056595 | -0.2652337 | 0.79082944 | 0.99937574 |
| C6orf120 | 12.589484 | 0.36927126 | 0.49041088 | 0.75298342 | 0.45145988 | 0.99937574 |
| PHF10 | 50.6018849 | 0.46461054 | 0.36132283 | 1.28585991 | 0.19849196 | 0.99937574 |
| ERMARD | 21.7276281 | -0.9501655 | 0.5777022 | -1.6447324 | 0.100025 | 0.99937574 |
| DLL1 | 70.4456591 | -0.0875402 | 0.35887733 | -0.2439278 | 0.80728671 | 0.99937574 |
| FAM120B | 265.474733 | -0.1248279 | 0.32063016 | -0.3893205 | 0.69703908 | 0.99937574 |
| PSMB1 | 91.9766838 | 0.28023246 | 0.32802622 | 0.85429897 | 0.39293936 | 0.99937574 |
| TBP | 17.7441038 | 0.08925039 | 0.53926375 | 0.16550415 | 0.86854718 | 0.99937574 |
| PDCD2 | 30.8669966 | 0.46354155 | 0.42238655 | 1.09743445 | 0.27245152 | 0.99937574 |
| FAM20C | 83.354337 | 0.02103168 | 0.50255617 | 0.04184942 | 0.96661874 | 0.99937574 |
| PRKAR1B | 34.2960382 | -0.1277942 | 0.56385887 | -0.2266421 | 0.82070203 | 0.99937574 |
| DNAAF5 | 23.702499 | -0.5084575 | 0.46370705 | -1.096506 | 0.27285742 | 0.99937574 |
| SUN1 | 109.928648 | 0.06987393 | 0.22684763 | 0.30802144 | 0.75806602 | 0.99937574 |
| COX19 | 47.5892429 | 0.14981302 | 0.43022898 | 0.34821694 | 0.72767726 | 0.99937574 |
| C7orf50 | 70.739406 | -0.4505413 | 0.33776521 | -1.3338891 | 0.18224021 | 0.99937574 |
| MICALL2 | 15.2527278 | 0.32171789 | 0.51944389 | 0.61935061 | 0.53568541 | 0.99937574 |
| INTS1 | 34.2734455 | -0.2936865 | 0.50608215 | -0.5803139 | 0.56170293 | 0.99937574 |
| MAFK | 28.8520159 | -1.3076888 | 0.47070186 | -2.7781678 | 0.00546664 | 0.65033075 |
| PSMG3 | 20.6362252 | -0.2241938 | 0.55506405 | -0.4039061 | 0.68628176 | 0.99937574 |
| MAD1L1 | 48.3769589 | 0.05313946 | 0.34393832 | 0.15450286 | 0.87721325 | 0.99937574 |
| SNX8 | 15.4025773 | -1.3533971 | 0.66132962 | -2.0464789 | 0.04070928 | 0.99937574 |
| EIF3B | 50.0084648 | -0.3814211 | 0.42085147 | -0.9063081 | 0.3647728 | 0.99937574 |
| CHST12 | 19.6239698 | 0.16096449 | 0.61780834 | 0.26054114 | 0.79444638 | 0.99937574 |
| BRAT1 | 11.7970403 | 0.36470316 | 0.56916179 | 0.64077239 | 0.52167058 | 0.99937574 |
| IQCE | 23.9101493 | -0.1740257 | 0.40940967 | -0.425065 | 0.6707893 | 0.99937574 |
| GNA12 | 54.0890065 | -0.064904 | 0.35070135 | -0.185069 | 0.85317488 | 0.99937574 |
| SDK1 | 81.1407811 | 0.12938908 | 0.43550355 | 0.29710224 | 0.76638845 | 0.99937574 |
| FOXK1 | 121.437459 | -0.1467348 | 0.43781678 | -0.3351511 | 0.73751108 | 0.99937574 |
| AP5Z1 | 23.8887295 | -1.7722265 | 0.54410622 | -3.2571334 | 0.00112544 | 0.3245112 |
| RBAK-RBAKD | 50.0982879 | -0.1509434 | 0.31031482 | -0.4864203 | 0.62666919 | 0.99937574 |
| WIPI2 | 77.0451196 | -0.1738916 | 0.38850004 | -0.4475974 | 0.65444381 | 0.99937574 |
| SLC29A4 | 30.0231874 | 1.0248013 | 0.55840408 | 1.83523248 | 0.06647125 | 0.99937574 |
| TNRC18 | 167.975229 | 0.24444745 | 0.35242859 | 0.69360847 | 0.48792778 | 0.99937574 |
| FBXL18 | 23.0451654 | 0.26296443 | 0.61486201 | 0.42768039 | 0.66888383 | 0.99937574 |
| ACTB | 526.593247 | 0.53472117 | 0.50902833 | 1.05047427 | 0.29350011 | 0.99937574 |
| RNF216 | 118.055685 | 0.00535948 | 0.29658095 | 0.01807088 | 0.98558231 | 0.99937574 |
| CCZ1 | 22.2217179 | -0.6609986 | 0.50557916 | -1.3074087 | 0.19107394 | 0.99937574 |
| PMS2 | 30.1595253 | 0.32961534 | 0.37262659 | 0.88457279 | 0.37638709 | 0.99937574 |
| AIMP2 | 27.2932979 | -0.1188831 | 0.40518775 | -0.2934024 | 0.76921458 | 0.99937574 |
| EIF2AK1 | 67.2633826 | 0.11458072 | 0.30327796 | 0.37780759 | 0.70557353 | 0.99937574 |
| USP42 | 73.9725658 | -0.1884412 | 0.34310142 | -0.5492289 | 0.58284834 | 0.99937574 |
| CYTH3 | 69.0679426 | 0.24663746 | 0.4061611 | 0.60724049 | 0.54369133 | 0.99937574 |
| RAC1 | 162.307549 | -0.3097188 | 0.27635231 | -1.1207389 | 0.26239902 | 0.99937574 |
| DAGLB | 15.1471928 | -0.5926831 | 0.55847996 | -1.0612433 | 0.28857935 | 0.99937574 |
| ZDHHC4 | 18.7852048 | -0.1546217 | 0.47852838 | -0.3231193 | 0.74660491 | 0.99937574 |
| C7orf26 | 16.5666175 | -0.5572586 | 0.47606266 | -1.1705573 | 0.24177677 | 0.99937574 |
| ZNF316 | 26.6363002 | -0.5547115 | 0.62557083 | -0.8867285 | 0.37522511 | 0.99937574 |
| AC073343.2 | 15.8390772 | 0.40327296 | 0.67038883 | 0.60155083 | 0.54747316 | 0.99937574 |
| ZNF12 | 32.0987297 | 0.05727834 | 0.40744302 | 0.14058001 | 0.88820174 | 0.99937574 |
| CCZ1B | 27.323585 | -0.22901 | 0.46644675 | -0.490967 | 0.62344976 | 0.99937574 |
| C1GALT1 | 176.197669 | 0.12099435 | 0.5027515 | 0.24066433 | 0.80981529 | 0.99937574 |
| COL28A1 | 94.1428486 | 0.84755103 | 0.52467904 | 1.61537046 | 0.10623049 | 0.99937574 |
| AC004982.2 | 19.9358745 | 0.74379755 | 0.6249528 | 1.19016596 | 0.23398117 | 0.99937574 |
| MIOS | 67.8597827 | 0.38346681 | 0.39745112 | 0.96481503 | 0.33463747 | 0.99937574 |

| | | | | | | |
|---|---|---|---|---|---|---|
| RPA3 | 49.0498281 | 0.05819524 | 0.69738421 | 0.08344788 | 0.93349542 | 0.99937574 |
| UMAD1 | 475.810971 | 0.0197856 | 0.30639353 | 0.06457579 | 0.94851176 | 0.99937574 |
| GLCCI1 | 542.995483 | 0.73332683 | 0.31264759 | 2.3455381 | 0.01899964 | 0.96315543 |
| ICA1 | 251.158854 | 0.30001838 | 0.47835668 | 0.62718552 | 0.53053763 | 0.99937574 |
| NDUFA4 | 235.757176 | 0.68144317 | 0.56520148 | 1.20566416 | 0.22794701 | 0.99937574 |
| PHF14 | 117.139393 | -0.368843 | 0.2566472 | -1.4371598 | 0.1506726 | 0.99937574 |
| THSD7A | 1869.5426 | -1.5434776 | 0.53829135 | -2.8673648 | 0.00413906 | 0.57221038 |
| TMEM106B | 97.1688922 | -0.0387364 | 0.3196084 | -0.1211996 | 0.90353295 | 0.99937574 |
| VWDE | 24.0368116 | 0.33684626 | 0.59396585 | 0.56711385 | 0.57063683 | 0.99937574 |
| SCIN | 1092.41234 | -0.1416429 | 0.31603412 | -0.4481886 | 0.65401708 | 0.99937574 |
| ARL4A | 19.4626668 | -0.7042864 | 0.61599749 | -1.1433267 | 0.25290298 | 0.99937574 |
| AC011287.1 | 37.6368008 | 0.69878916 | 0.62213016 | 1.1232202 | 0.26134399 | 0.99937574 |
| ISPD | 43.4207313 | -0.2847553 | 0.39608965 | -0.7189163 | 0.47219247 | 0.99937574 |
| ANKMY2 | 42.4945599 | 0.23173617 | 0.39375774 | 0.58852475 | 0.55618013 | 0.99937574 |
| BZW2 | 34.8441454 | 0.09555357 | 0.62704529 | 0.15238703 | 0.87888169 | 0.99937574 |
| AC073332.1 | 17.196486 | -0.3816588 | 0.57112492 | -0.6682581 | 0.50396888 | 0.99937574 |
| SNX13 | 219.087034 | -0.1060421 | 0.28979134 | -0.3659257 | 0.71442055 | 0.99937574 |
| HDAC9 | 29.3558884 | -0.0014531 | 0.79399572 | -0.0018301 | 0.99853978 | 0.99979721 |
| TWISTNB | 16.2170458 | -0.2622596 | 0.51745209 | -0.5068288 | 0.61227493 | 0.99937574 |
| MACC1 | 209.723735 | -0.1802991 | 0.54468837 | -0.3310133 | 0.74063445 | 0.99937574 |
| ITGB8 | 62.9088978 | 0.51639228 | 0.60306111 | 0.85628516 | 3.92E-01 | 0.99937574 |
| SP4 | 48.0989676 | -0.2253784 | 0.33759026 | -0.6676094 | 0.50438296 | 0.99937574 |
| RAPGEF5 | 50.5731041 | -0.0374712 | 0.6704141 | -0.0558927 | 9.55E-01 | 0.99937574 |
| STEAP1B | 149.309155 | -0.0579954 | 0.47504997 | -0.1220828 | 0.90283344 | 0.99937574 |
| TOMM7 | 76.1677336 | 0.31470401 | 0.48018724 | 0.65537771 | 0.5122246 | 0.99937574 |
| FAM126A | 13.7781802 | 0.26623781 | 0.56214665 | 0.47360916 | 0.63577864 | 0.99937574 |
| KLHL7 | 42.1759568 | 0.21035377 | 0.40112345 | 0.52441156 | 0.59999232 | 0.99937574 |
| NUPL2 | 53.5199921 | -0.0398452 | 0.37316079 | -0.1067775 | 0.91496552 | 0.99937574 |
| MALSU1 | 36.4575748 | 0.07485945 | 0.38453095 | 0.19467731 | 0.84564558 | 0.99937574 |
| TRA2A | 235.382962 | 0.00621475 | 0.31558104 | 0.01969303 | 0.98428825 | 0.99937574 |
| CCDC126 | 54.7301118 | 0.2168564 | 0.28731852 | 0.75475956 | 0.45039326 | 0.99937574 |
| FAM221A | 31.1277394 | 0.04019618 | 0.41577458 | 0.09667783 | 0.92298225 | 0.99937574 |
| MPP6 | 13.9913774 | 0.77879593 | 0.66514778 | 1.17086151 | 0.24165445 | 0.99937574 |
| OSBPL3 | 540.786819 | -0.0925854 | 0.53179482 | -0.1740999 | 0.86178693 | 0.99937574 |
| CYCS | 163.50565 | 0.57097255 | 0.45388785 | 1.25795955 | 0.20840638 | 0.99937574 |
| HNRNPA2B1 | 370.297006 | -0.370011 | 0.28574831 | -1.2948843 | 0.19536014 | 0.99937574 |
| CBX3 | 71.9146312 | -0.4102511 | 0.41227645 | -0.9950874 | 0.31969378 | 0.99937574 |
| SNX10 | 84.5077468 | 0.09613709 | 0.52428481 | 0.18336807 | 0.85450923 | 0.99937574 |
| AC004540.1 | 27.5850315 | 0.55516071 | 0.84999321 | 0.65313547 | 0.51366895 | 0.99937574 |
| SKAP2 | 151.849579 | 0.62427657 | 0.49455919 | 1.26228889 | 0.20684485 | 0.99937574 |
| HOTAIRM1 | 10.9791943 | -0.234437 | 0.80682798 | -0.2905663 | 0.77138302 | 0.99937574 |
| HOXA3 | 19.8371625 | 1.26112054 | 0.83581713 | 1.50884744 | 0.13133777 | 0.99937574 |
| HOXA7 | 12.5447618 | -0.1659339 | 0.93210716 | -0.1780202 | 0.85870711 | 0.99937574 |
| HOXA9 | 16.2701228 | 0.88936038 | 0.78176215 | 1.1376355 | 0.25527271 | 0.99937574 |
| HIBADH | 253.282006 | -0.0445245 | 0.37262622 | -0.1194883 | 0.90488847 | 0.99937574 |
| TAX1BP1 | 329.51071 | 0.17857124 | 0.18593844 | 0.96037828 | 0.33686487 | 0.99937574 |
| JAZF1 | 75.9456631 | 0.52466445 | 0.43952789 | 1.19370001 | 0.23259534 | 0.99937574 |
| CPVL | 28.1813636 | -1.1262261 | 0.78450131 | -1.4355949 | 0.15111765 | 0.99937574 |
| AC007255.1 | 14.3727776 | -0.1370074 | 0.87901098 | -0.1558654 | 0.87613913 | 0.99937574 |
| PRR15 | 11.3954313 | 0.48555572 | 0.92972524 | 0.52225722 | 0.60149125 | 0.99937574 |
| WIPF3 | 15.1793338 | 0.1488952 | 0.72714413 | 0.2047671 | 0.83775409 | 0.99937574 |
| SCRN1 | 116.018014 | 0.47536198 | 0.23637848 | 2.0110206 | 0.04432328 | 0.99937574 |
| FKBP14 | 32.8243719 | 0.42948201 | 0.45724933 | 0.93927312 | 0.34759053 | 0.99937574 |
| PLEKHA8 | 164.516939 | 0.2753917 | 0.3017467 | 0.91265853 | 0.36142216 | 0.99937574 |

| | | | | | | |
|---|---|---|---|---|---|---|
| MTURN | 84.6345016 | 0.16870991 | 0.55030282 | 0.30657649 | 0.75916575 | 0.99937574 |
| ZNRF2 | 78.856805 | -0.0495803 | 0.28870977 | -0.1717307 | 0.86364922 | 0.99937574 |
| AC006978.2 | 64.4213833 | -0.0091852 | 0.33837828 | -0.0271448 | 0.97834425 | 0.99937574 |
| NOD1 | 27.5613658 | -0.3411025 | 0.54383876 | -0.6272126 | 0.53051988 | 0.99937574 |
| GGCT | 16.0442207 | -0.235911 | 0.65573365 | -0.3597665 | 0.71902179 | 0.99937574 |
| AC005154.6 | 30.0766236 | 0.87108118 | 0.42539021 | 2.04772269 | 0.04058718 | 0.99937574 |
| GARS | 50.3617727 | -0.3856501 | 0.48322937 | -0.7980685 | 0.42483076 | 0.99937574 |
| AQP1 | 14.6960426 | 0.81661681 | 0.83551487 | 0.97738154 | 0.3283803 | 0.99937574 |
| CCDC129 | 20.0698016 | 1.98484913 | 0.5337382 | 3.71876913 | 0.0002002 | 0.13580369 |
| PDE1C | 1727.28115 | 0.79014914 | 0.41842697 | 1.88838002 | 0.05897495 | 0.99937574 |
| LSM5 | 18.8357752 | -0.2174977 | 0.52906545 | -0.411098 | 0.68100071 | 0.99937574 |
| AVL9 | 219.700614 | -0.0568607 | 0.25167578 | -0.2259283 | 0.82125715 | 0.99937574 |
| KBTBD2 | 63.1835758 | -0.3021666 | 0.40077424 | -0.7539571 | 0.45087496 | 0.99937574 |
| FKBP9 | 31.492304 | 0.11745504 | 0.41483064 | 0.28313974 | 0.77706972 | 0.99937574 |
| NT5C3A | 61.0618627 | -0.6581797 | 0.43846512 | -1.5010993 | 0.13332989 | 0.99937574 |
| RP9 | 13.8902815 | 0.04484697 | 0.56106215 | 0.07993227 | 0.93629113 | 0.99937574 |
| BBS9 | 116.201357 | 0.37041446 | 0.29066988 | 1.27434759 | 0.20254026 | 0.99937574 |
| DPY19L1 | 74.7445113 | -0.3823611 | 0.45083396 | -0.8481196 | 0.39637137 | 0.99937574 |
| HERPUD2 | 50.5771387 | -0.8087881 | 0.31606976 | -2.5588911 | 0.01050066 | 0.76274616 |
| 7-Sep | 145.133826 | -0.0607379 | 0.24398301 | -0.2489431 | 0.80340477 | 0.99937574 |
| EEPD1 | 24.8439154 | 0.76040193 | 0.5916874 | 1.28514133 | 0.1987429 | 0.99937574 |
| KIAA0895 | 30.8329722 | -0.0180843 | 0.45751866 | -0.039527 | 0.96847023 | 0.99937574 |
| EPDR1 | 32.7947789 | 0.82702994 | 0.42161929 | 1.96155621 | 0.04981417 | 0.99937574 |
| STARD3NL | 38.9057949 | -0.2691489 | 0.33783675 | -0.7966834 | 0.42563493 | 0.99937574 |
| VPS41 | 135.678993 | -0.5327821 | 0.23508194 | -2.2663676 | 0.02342889 | 0.96546668 |
| YAE1D1 | 15.6361917 | 0.46829837 | 0.49458016 | 0.9468604 | 0.34370992 | 0.99937574 |
| RALA | 140.473589 | -0.7272838 | 0.54730596 | -1.3288432 | 0.18389972 | 0.99937574 |
| LINC00265 | 24.9543458 | 0.27399904 | 0.59614769 | 0.45961603 | 0.64579185 | 0.99937574 |
| CDK13 | 295.348759 | -0.0294876 | 0.23147777 | -0.1273885 | 0.89863292 | 0.99937574 |
| MPLKIP | 28.2291745 | 0.81877938 | 0.48666302 | 1.68243598 | 0.09248433 | 0.99937574 |
| SUGCT | 12.4882145 | -0.7071361 | 0.56051385 | -1.2615855 | 0.20709798 | 0.99937574 |
| MRPL32 | 27.8143934 | -0.2641085 | 0.49265538 | -0.5360917 | 0.59189515 | 0.99937574 |
| STK17A | 146.045651 | 0.50618837 | 0.36674607 | 1.38021483 | 0.16752051 | 0.99937574 |
| COA1 | 194.430446 | -0.2206296 | 0.25277627 | -0.8728256 | 0.38275816 | 0.99937574 |
| BLVRA | 36.7456583 | 0.16428843 | 0.34559438 | 0.47537935 | 0.63451661 | 0.99937574 |
| URGCP | 25.4672845 | 0.652004 | 0.5841546 | 1.11614974 | 0.26435804 | 0.99937574 |
| UBE2D4 | 15.2871775 | -0.5662078 | 0.5530438 | -1.0238028 | 0.30592842 | 0.99937574 |
| DBNL | 44.3305423 | 0.09287795 | 0.38152035 | 0.24344165 | 0.80766328 | 0.99937574 |
| POLM | 21.8071283 | 0.44469851 | 0.41980354 | 1.05930148 | 0.2894625 | 0.99937574 |
| POLD2 | 20.6784525 | 0.70987843 | 0.46671066 | 1.52102467 | 0.12825365 | 0.99937574 |
| YKT6 | 20.3220309 | -0.6044732 | 0.52005247 | -1.1623311 | 0.24510098 | 0.99937574 |
| NUDCD3 | 112.209178 | -0.4770381 | 0.27564115 | -1.730649 | 0.08351439 | 0.99937574 |
| DDX56 | 26.7907306 | 0.23876113 | 0.41382573 | 0.57696058 | 0.56396609 | 0.99937574 |
| TMED4 | 73.7650218 | 0.66449195 | 0.33993659 | 1.95475265 | 0.05061227 | 0.99937574 |
| OGDH | 301.637473 | 0.08983689 | 0.22973347 | 0.39104834 | 0.69576151 | 0.99937574 |
| ZMIZ2 | 57.862328 | -0.0009216 | 0.38798049 | -0.0023754 | 0.99810467 | 0.99968959 |
| PPIA | 251.440936 | -0.0576699 | 0.49883413 | -0.1156094 | 0.90796211 | 0.99937574 |
| H2AFV | 89.139817 | -0.054042 | 0.41981416 | -0.1287284 | 0.89757259 | 0.99937574 |
| PURB | 50.8650044 | -0.4025808 | 0.35194655 | -1.1438693 | 0.25267786 | 0.99937574 |
| SNHG15 | 11.7921116 | -0.0312028 | 0.60511547 | -0.051565 | 0.95887534 | 0.99937574 |
| CCM2 | 63.8863399 | 0.19969793 | 0.34997871 | 0.57060023 | 0.56827066 | 0.99937574 |
| TBRG4 | 23.3539222 | 0.18281027 | 0.48472548 | 0.37714185 | 0.70606819 | 0.99937574 |
| ADCY1 | 62.2136773 | 1.39752779 | 0.69382056 | 2.01424962 | 0.04398333 | 0.99937574 |
| AC004870.4 | 52.1029688 | 1.59337604 | 0.8649498 | 1.84216013 | 0.06545173 | 0.99937574 |

| | | | | | | |
|---|---|---|---|---|---|---|
| TNS3 | 695.280854 | -0.1643177 | 0.34272233 | -0.4794483 | 0.63161971 | 0.99937574 |
| HUS1 | 28.4578286 | -0.5136337 | 0.38329719 | -1.3400404 | 0.18023222 | 0.99937574 |
| UPP1 | 51.3751434 | -0.9802586 | 0.4951088 | -1.9798852 | 0.04771643 | 0.99937574 |
| GRB10 | 244.068199 | -0.0582868 | 0.40201273 | -0.1449874 | 0.88472082 | 0.99937574 |
| COBL | 120.963051 | -0.4301436 | 0.54785294 | -0.7851443 | 0.43236898 | 0.99937574 |
| SEC61G | 32.2962202 | -0.1046268 | 0.57765335 | -0.1811239 | 0.85627033 | 0.99937574 |
| EGFR | 464.200393 | 0.42352537 | 0.37732724 | 1.1224352 | 0.26167744 | 0.99937574 |
| LANCL2 | 43.2268006 | -0.669361 | 0.49458095 | -1.3533901 | 0.17593103 | 0.99937574 |
| VOPP1 | 88.8933646 | 0.03758657 | 0.34291309 | 0.10960961 | 0.91271899 | 0.99937574 |
| NIPSNAP2 | 166.703917 | 0.01680465 | 0.26289566 | 0.06392138 | 0.94903283 | 0.99937574 |
| CCT6A | 59.7188385 | 0.2120339 | 0.35007432 | 0.60568253 | 0.54472559 | 0.99937574 |
| SUMF2 | 44.0010887 | -0.2954095 | 0.29939283 | -0.9866953 | 0.32379202 | 0.99937574 |
| PHKG1 | 13.3666012 | 0.62976618 | 0.56072804 | 1.12312233 | 0.26138555 | 0.99937574 |
| CHCHD2 | 209.262959 | 0.5594656 | 0.51380567 | 1.08886615 | 0.27621292 | 0.99937574 |
| NUPR2 | 30.5083243 | 0.22694231 | 0.62849808 | 0.36108673 | 0.71803461 | 0.99937574 |
| ZNF736 | 49.9676572 | 0.32356084 | 0.35154949 | 0.9203849 | 0.35737166 | 0.99937574 |
| ZNF680 | 72.8766313 | 0.19157214 | 0.42857829 | 0.44699449 | 0.65487904 | 0.99937574 |
| ZNF107 | 41.0363929 | -0.4749288 | 0.56584307 | -0.8393296 | 0.4012844 | 0.99937574 |
| ZNF138 | 48.1539933 | 0.40917574 | 0.39576558 | 1.03388412 | 0.30119034 | 0.99937574 |
| ZNF273 | 13.2233832 | 0.05368098 | 0.51512425 | 0.10420977 | 0.91700288 | 0.99937574 |
| ZNF117 | 20.5383719 | 0.746391 | 0.42834235 | 1.74251042 | 0.08141917 | 0.99937574 |
| ZNF92 | 14.7622563 | 0.22627249 | 0.63309229 | 0.35740838 | 0.7207861 | 0.99937574 |
| VKORC1L1 | 43.0669385 | -0.4311988 | 0.37544212 | -1.1485095 | 0.25075831 | 0.99937574 |
| GUSB | 20.7980433 | 0.77900873 | 0.57480858 | 1.35524896 | 0.17533825 | 0.99937574 |
| ASL | 14.6241353 | -1.2609448 | 0.52463995 | -2.4034479 | 0.01624128 | 0.87650811 |
| AC068533.4 | 14.7199798 | 0.28108379 | 0.68950886 | 0.40765798 | 0.68352479 | 0.99937574 |
| CRCP | 20.1917478 | -0.3966089 | 0.43783992 | -0.9058308 | 0.36502544 | 0.99937574 |
| AC068533.3 | 15.7325818 | 0.25955749 | 0.64645864 | 0.40150673 | 0.68804709 | 0.99937574 |
| TPST1 | 238.271929 | 0.29808289 | 0.37560037 | 0.79361714 | 0.42741835 | 0.99937574 |
| AC027644.4 | 62.1938361 | 0.11134191 | 0.42725549 | 0.26059795 | 0.79440257 | 0.99937574 |
| AC027644.3 | 13.7377154 | 0.76664016 | 0.65123162 | 1.17721581 | 0.23910938 | 0.99937574 |
| TMEM248 | 76.4809411 | 0.06303431 | 0.26835963 | 0.23488746 | 0.81429608 | 0.99937574 |
| SBDS | 61.2815366 | -0.6128062 | 0.51136907 | -1.1983638 | 0.2307754 | 0.99937574 |
| TYW1 | 135.456159 | 0.2151592 | 0.30628082 | 0.70248998 | 0.48237365 | 0.99937574 |
| AC006480.2 | 12.8523291 | 0.55424032 | 0.50334038 | 1.10112428 | 0.27084257 | 0.99937574 |
| AUTS2 | 126.996603 | 0.2955232 | 0.47632794 | 0.62041962 | 0.53498156 | 0.99937574 |
| GALNT17 | 1230.00205 | 1.02567951 | 0.63079513 | 1.6260105 | 0.10394742 | 0.99937574 |
| CALN1 | 27.985131 | -0.9194825 | 0.73982818 | -1.2428325 | 0.21392955 | 0.99937574 |
| TYW1B | 85.2984696 | 0.12250964 | 0.33311164 | 0.36777352 | 0.71304212 | 0.99937574 |
| POM121 | 91.8282691 | -0.034468 | 0.37472583 | -0.091982 | 0.92671235 | 0.99937574 |
| NSUN5 | 33.0749717 | -0.1076512 | 0.43082862 | -0.2498701 | 0.80268783 | 0.99937574 |
| BAZ1B | 101.427936 | -0.367465 | 0.25679029 | -1.4309926 | 0.15243232 | 0.99937574 |
| BCL7B | 27.3396926 | -0.5292763 | 0.4785843 | -1.1059208 | 0.26876081 | 0.99937574 |
| MLXIPL | 20.9064138 | 1.03447834 | 0.70338816 | 1.47070763 | 0.1413702 | 0.99937574 |
| VPS37D | 15.4557422 | 0.26605625 | 0.55157863 | 0.48235417 | 0.62955438 | 0.99937574 |
| DNAJC30 | 9.94661871 | 0.03492739 | 0.61352986 | 0.05692859 | 0.95460208 | 0.99937574 |
| BUD23 | 43.5071503 | 0.11977511 | 0.41449108 | 0.28896908 | 0.77260504 | 0.99937574 |
| ABHD11 | 22.6113791 | -0.0552913 | 0.46077374 | -0.1199965 | 0.90448589 | 0.99937574 |
| CLDN3 | 25.4788487 | -0.2949039 | 0.61053318 | -0.4830269 | 0.62907664 | 0.99937574 |
| CLDN4 | 184.155138 | -0.3248406 | 0.46842205 | -0.6934784 | 0.48800936 | 0.99937574 |
| LIMK1 | 15.4053445 | -0.0757013 | 0.54486913 | -0.1389348 | 0.88950166 | 0.99937574 |
| EIF4H | 70.8854251 | -0.0335766 | 0.34356051 | -0.0977313 | 0.92214566 | 0.99937574 |
| RFC2 | 10.5250492 | -0.5872571 | 0.53921905 | -1.0890882 | 0.276115 | 0.99937574 |
| CLIP2 | 18.8428055 | -1.4486537 | 0.63355892 | -2.2865335 | 0.02222306 | 0.96427895 |

| | | | | | | |
|---|---|---|---|---|---|---|
| GTF2IRD1 | 200.545418 | -1.1725305 | 0.62356567 | -1.8803641 | 0.06005848 | 0.99937574 |
| GTF2I | 297.60378 | 0.14622331 | 0.21313986 | 0.68604391 | 0.49268542 | 0.99937574 |
| GTF2IRD2 | 47.9025886 | -0.2608782 | 0.46341533 | -0.5629469 | 0.57347106 | 0.99937574 |
| CASTOR2 | 23.6255821 | 0.38490619 | 0.6140279 | 0.62685456 | 0.53075458 | 0.99937574 |
| RCC1L | 42.3865233 | -0.1711553 | 0.35334709 | -0.4843831 | 0.62811403 | 0.99937574 |
| GTF2IRD2B | 70.742572 | -0.4799581 | 0.33610633 | -1.4279948 | 0.15329335 | 0.99937574 |
| TRIM73 | 56.9785051 | 0.14042287 | 0.45540075 | 0.30835012 | 0.75781594 | 0.99937574 |
| POM121C | 81.614518 | -0.3319016 | 0.28470965 | -1.1657548 | 0.2437136 | 0.99937574 |
| SPDYE5 | 19.0481206 | 0.68041805 | 0.73256506 | 0.92881586 | 0.35298452 | 0.99937574 |
| HIP1 | 155.401406 | -1.2529625 | 0.80462162 | -1.5572072 | 0.11942131 | 0.99937574 |
| RHBDD2 | 23.9105067 | -0.1914276 | 0.57308572 | -0.3340296 | 0.73835719 | 0.99937574 |
| POR | 49.8536916 | -0.2749147 | 0.41904002 | -0.6560584 | 0.51178657 | 0.99937574 |
| TMEM120A | 11.0052887 | 0.40487831 | 0.56825708 | 0.71249144 | 0.47616051 | 0.99937574 |
| STYXL1 | 44.2664163 | -0.1320776 | 0.36925367 | -0.3576879 | 0.72057688 | 0.99937574 |
| MDH2 | 81.6701251 | 0.38353294 | 0.45324077 | 0.84620133 | 0.39744044 | 0.99937574 |
| HSPB1 | 149.947937 | 0.21147208 | 0.48990442 | 0.43165988 | 0.66598863 | 0.99937574 |
| YWHAG | 79.6269556 | -0.3209395 | 0.31294564 | -1.0255438 | 0.30510665 | 0.99937574 |
| DTX2 | 34.3791496 | -0.4238916 | 0.41493914 | -1.0215753 | 0.30698194 | 0.99937574 |
| POMZP3 | 16.2205525 | 0.30422935 | 0.55734528 | 0.54585436 | 0.58516605 | 0.99937574 |
| CCDC146 | 55.2967849 | 0.10099362 | 0.44205999 | 0.22846134 | 0.81928761 | 0.99937574 |
| GSAP | 65.5724037 | 0.12191101 | 0.40807166 | 0.29874903 | 0.76513155 | 0.99937574 |
| PTPN12 | 131.778915 | -0.5143115 | 0.26382516 | -1.9494406 | 5.12E-02 | 0.99937574 |
| APTR | 34.3945333 | 0.46548804 | 0.4503262 | 1.03366857 | 0.30129113 | 0.99937574 |
| RSBN1L | 116.365868 | -0.2869475 | 0.30052038 | -0.9548355 | 0.33966087 | 0.99937574 |
| TMEM60 | 12.754228 | 0.07328659 | 0.55498364 | 0.13205181 | 0.89494331 | 0.99937574 |
| PHTF2 | 67.044745 | 0.05392218 | 0.40904475 | 0.13182465 | 0.89512299 | 0.99937574 |
| MAGI2 | 129.867429 | -0.5943419 | 0.88508538 | -0.671508 | 0.50189698 | 0.99937574 |
| GNAI1 | 12.8784096 | -0.0569171 | 0.79187436 | -0.0718764 | 0.94270028 | 0.99937574 |
| SEMA3C | 891.183309 | 0.22087202 | 0.4474003 | 0.49367873 | 0.62153309 | 0.99937574 |
| CACNA2D1 | 83.4866085 | -0.548325 | 0.91503647 | -0.5992385 | 0.54901388 | 0.99937574 |
| PCLO | 127.211144 | -0.7858325 | 0.71616213 | -1.0972829 | 0.27251775 | 0.99937574 |
| KIAA1324L | 38.216706 | -0.1355554 | 0.633716 | -0.2139055 | 0.83062073 | 0.99937574 |
| DMTF1 | 89.7582661 | 0.40910937 | 0.42698321 | 0.95813924 | 0.33799255 | 0.99937574 |
| TMEM243 | 47.8449898 | 0.05576239 | 0.37154104 | 0.15008406 | 0.8806983 | 0.99937574 |
| CROT | 16.5869449 | 0.83143409 | 0.52165857 | 1.59382811 | 0.11097454 | 0.99937574 |
| SLC25A40 | 64.3848915 | 0.81362558 | 0.3675653 | 2.21355381 | 0.02685949 | 0.97857744 |
| DBF4 | 35.989231 | 0.38507185 | 0.50028105 | 0.76971105 | 0.44147132 | 0.99937574 |
| ADAM22 | 221.322743 | 1.02082295 | 0.52154612 | 1.9573014 | 0.05031204 | 0.99937574 |
| SRI | 74.2648357 | 0.30118692 | 0.5272054 | 0.57128952 | 0.5678034 | 0.99937574 |
| AC003991.1 | 27.0345941 | 1.30782457 | 0.75917224 | 1.72269809 | 0.08494314 | 0.99937574 |
| CFAP69 | 21.83375 | 0.26865448 | 0.59934526 | 0.44824661 | 0.65397523 | 0.99937574 |
| GTPBP10 | 53.2267859 | -0.078509 | 0.42146382 | -0.1862769 | 0.8522276 | 0.99937574 |
| CLDN12 | 46.150891 | 0.01787121 | 0.37436586 | 0.04773729 | 0.96192561 | 0.99937574 |
| CDK14 | 229.196025 | 0.98419162 | 0.4471427 | 2.2010683 | 0.02773119 | 0.98891577 |
| AC079760.1 | 19.1763252 | 0.27979037 | 0.62425577 | 0.4481983 | 0.65401009 | 0.99937574 |
| MTERF1 | 114.579004 | 0.21615092 | 0.45155331 | 0.47868306 | 0.63216412 | 0.99937574 |
| AKAP9 | 445.506272 | -0.0466025 | 0.23370557 | -0.1994069 | 0.84194444 | 0.99937574 |
| KRIT1 | 145.107939 | 0.02721232 | 0.22525128 | 0.12080874 | 0.90384253 | 0.99937574 |
| ANKIB1 | 230.408856 | -0.2185985 | 0.21441106 | -1.0195299 | 0.30795148 | 0.99937574 |
| GATAD1 | 41.2910082 | 0.32363835 | 0.39925577 | 0.81060406 | 0.41759308 | 0.99937574 |
| AC007566.1 | 14.231572 | 0.40833802 | 0.5704053 | 0.71587347 | 0.47406948 | 0.99937574 |
| PEX1 | 46.7719007 | -0.093756 | 0.34832353 | -0.2691635 | 0.78780388 | 0.99937574 |
| RBM48 | 10.8508241 | 0.18364097 | 0.68990764 | 0.26618196 | 0.79009908 | 0.99937574 |
| FAM133B | 57.4609751 | -0.4057327 | 0.32649658 | -1.2426858 | 0.21398363 | 0.99937574 |

| | | | | | | |
|---|---|---|---|---|---|---|
| CDK6 | 60.1506761 | 1.19546567 | 0.51295161 | 2.33056229 | 0.01977645 | 0.96427895 |
| SAMD9 | 15.1244375 | -1.0746187 | 0.80786485 | -1.3301961 | 0.18345367 | 0.99937574 |
| SAMD9L | 38.0990927 | -2.472011 | 0.94585319 | -2.613525 | 0.00896135 | 0.74742128 |
| HEPACAM2 | 134.528822 | 1.48242908 | 0.71036961 | 2.08684191 | 0.03690243 | 0.99937574 |
| VPS50 | 107.081312 | -0.3014597 | 0.3156391 | -0.9550772 | 0.33953867 | 0.99937574 |
| CALCR | 302.294023 | 0.1918082 | 0.79897498 | 0.24006784 | 0.81027766 | 0.99937574 |
| GNGT1 | 113.270772 | 0.31079937 | 0.61859562 | 0.50242736 | 0.61536694 | 0.99937574 |
| BET1 | 25.7624533 | -0.2290762 | 0.61853802 | -0.370351 | 0.71112095 | 0.99937574 |
| CASD1 | 65.9193463 | 0.16467173 | 0.30352462 | 0.54253173 | 0.58745226 | 0.99937574 |
| SGCE | 30.9123024 | 0.29542781 | 0.52457158 | 0.5631792 | 0.57331286 | 0.99937574 |
| PPP1R9A | 328.839859 | -0.2329537 | 0.42438867 | -0.5489159 | 0.58306314 | 0.99937574 |
| AC002429.2 | 27.5455784 | -0.3587757 | 0.87597242 | -0.4095742 | 0.68211836 | 0.99937574 |
| PON2 | 35.5974489 | -0.0513147 | 0.3908682 | -0.1312838 | 0.89555081 | 0.99937574 |
| PDK4 | 85.7700949 | -0.9840857 | 1.28318701 | -0.7669075 | 0.44313653 | 0.99937574 |
| SLC25A13 | 168.801353 | -0.5987085 | 0.29726906 | -2.0140291 | 0.04400648 | 0.99937574 |
| SEM1 | 83.8095405 | -0.2409202 | 0.25066992 | -0.9611052 | 0.33649925 | 0.99937574 |
| SDHAF3 | 20.9780918 | 0.26763734 | 0.41963502 | 0.637786 | 0.523613 | 0.99937574 |
| ASNS | 18.8124544 | -0.2122153 | 0.56693177 | -0.3743225 | 0.70816437 | 0.99937574 |
| LMTK2 | 92.647331 | -0.1396197 | 0.430222 | -0.3245293 | 0.74553731 | 0.99937574 |
| TECPR1 | 18.1503394 | 0.45341306 | 0.5386714 | 0.84172476 | 0.39994203 | 0.99937574 |
| BRI3 | 132.888116 | -0.505839 | 0.40725617 | -1.2420659 | 0.21421223 | 0.99937574 |
| BAIAP2L1 | 491.59021 | 0.24362979 | 0.40041844 | 0.608438 | 0.54289702 | 0.99937574 |
| TRRAP | 114.645832 | -0.0482287 | 0.26213663 | -0.1839833 | 0.85402659 | 0.99937574 |
| SMURF1 | 419.570481 | 0.04572821 | 0.30198812 | 0.15142386 | 0.87964137 | 0.99937574 |
| ARPC1B | 31.6019294 | 0.07169179 | 0.50197341 | 0.1428199 | 0.88643242 | 0.99937574 |
| PDAP1 | 32.2551992 | -0.1158946 | 0.40174438 | -0.2884785 | 0.77298047 | 0.99937574 |
| BUD31 | 26.9273696 | -0.2271855 | 0.5121065 | -0.4436294 | 0.65731052 | 0.99937574 |
| CPSF4 | 14.6947877 | 0.84250856 | 0.50641263 | 1.66367998 | 0.0961764 | 0.99937574 |
| ZNF789 | 33.4405355 | 0.53574614 | 0.4382487 | 1.22247058 | 0.22152972 | 0.99937574 |
| ZNF394 | 24.0391231 | -0.1180949 | 0.4057267 | -0.29107 | 0.77099778 | 0.99937574 |
| ZKSCAN5 | 20.198447 | 0.17008 | 0.42643133 | 0.398845 | 0.69000742 | 0.99937574 |
| FAM200A | 20.184424 | 0.32718008 | 0.46422179 | 0.7047926 | 0.48093931 | 0.99937574 |
| ZSCAN25 | 14.6568413 | 0.66479174 | 0.57591275 | 1.15432718 | 0.24836606 | 0.99937574 |
| TRIM4 | 45.1889598 | 0.12655351 | 0.30896939 | 0.40959887 | 0.68210023 | 0.99937574 |
| ZKSCAN1 | 182.35651 | 0.09690249 | 0.29837533 | 0.32476711 | 0.74535734 | 0.99937574 |
| ZSCAN21 | 11.9253234 | 0.04183103 | 0.61224308 | 0.06832421 | 0.94552755 | 0.99937574 |
| ZNF3 | 14.8140412 | -0.3880519 | 0.55425756 | -0.7001292 | 0.48384665 | 0.99937574 |
| COPS6 | 30.4715771 | 0.38408523 | 0.43271687 | 0.88761326 | 0.37474882 | 0.99937574 |
| AP4M1 | 16.4471071 | 0.00980115 | 0.49638532 | 0.01974504 | 0.98424676 | 0.99937574 |
| TAF6 | 13.7512068 | 0.3849668 | 0.461498 | 0.83416786 | 0.4041864 | 0.99937574 |
| LAMTOR4 | 88.9737565 | 0.28941967 | 0.35493528 | 0.81541533 | 0.41483461 | 0.99937574 |
| CASTOR3 | 37.8891735 | 0.18153455 | 0.39415285 | 0.46056892 | 0.64510792 | 0.99937574 |
| SPDYE3 | 26.4901366 | -0.119786 | 0.60960233 | -0.1964985 | 0.84421999 | 0.99937574 |
| PILRB | 65.3599379 | -0.4600625 | 0.38323678 | -1.2004655 | 0.22995859 | 0.99937574 |
| PILRA | 12.4383063 | -0.685574 | 0.62680471 | -1.0937601 | 0.27406022 | 0.99937574 |
| MEPCE | 18.8570509 | -1.6520883 | 0.73126984 | -2.2592048 | 0.02387065 | 0.96546668 |
| PPP1R35 | 28.0997931 | 0.15190478 | 0.40328329 | 0.37667017 | 0.70641873 | 0.99937574 |
| TSC22D4 | 38.1134916 | -0.1104801 | 0.39930178 | -0.2766833 | 0.78202332 | 0.99937574 |
| AGFG2 | 17.84671 | -0.411949 | 0.7034527 | -0.5856101 | 0.55813755 | 0.99937574 |
| SAP25 | 14.7830789 | 0.22736898 | 0.596165 | 0.38138599 | 0.70291685 | 0.99937574 |
| LRCH4 | 25.0845614 | -0.4379817 | 0.45893186 | -0.9543502 | 0.33990639 | 0.99937574 |
| GNB2 | 81.1214605 | 0.10330141 | 0.43905672 | 0.23528032 | 0.81399117 | 0.99937574 |
| GIGYF1 | 41.4586706 | -0.4413286 | 0.45347719 | -0.9732102 | 0.33044885 | 0.99937574 |
| SLC12A9 | 12.0661757 | -1.2336976 | 0.60115839 | -2.0522006 | 0.04015017 | 0.99937574 |

| | | | | | | |
|---|---|---|---|---|---|---|
| TRIP6 | 12.8892057 | -0.5997839 | 0.58918549 | -1.0179883 | 0.30868353 | 0.99937574 |
| SRRT | 19.5433107 | -0.3647417 | 0.51366083 | -0.7100828 | 0.4776528 | 0.99937574 |
| TRIM56 | 80.1114146 | -0.6149935 | 0.41891972 | -1.4680462 | 0.14209166 | 0.99937574 |
| AP1S1 | 20.2847389 | 0.26423245 | 0.53861445 | 0.4905781 | 0.62372488 | 0.99937574 |
| PLOD3 | 18.5660186 | 0.2596413 | 0.55269256 | 0.46977528 | 0.63851558 | 0.99937574 |
| ZNHIT1 | 38.7763649 | -0.1320543 | 0.47438049 | -0.2783722 | 0.78072666 | 0.99937574 |
| FIS1 | 91.4177709 | 0.40219538 | 0.41278724 | 0.97434065 | 0.32988743 | 0.99937574 |
| IFT22 | 11.5849835 | 0.92707672 | 0.62071535 | 1.49356177 | 0.1352902 | 0.99937574 |
| CUX1 | 444.56853 | -0.2067484 | 0.35621977 | -0.5803957 | 0.56164778 | 0.99937574 |
| PRKRIP1 | 102.598753 | -0.0898878 | 0.29462913 | -0.3050881 | 0.76029909 | 0.99937574 |
| ORAI2 | 12.212752 | -0.9953708 | 0.96400361 | -1.0325384 | 0.30181996 | 0.99937574 |
| ALKBH4 | 11.7101726 | -0.0049775 | 0.61521962 | -0.0080907 | 0.99354465 | 0.99937574 |
| POLR2J | 21.6717529 | 0.52671407 | 0.59404563 | 0.88665591 | 0.37526418 | 0.99937574 |
| SPDYE2 | 14.7403021 | 1.06123981 | 0.67847315 | 1.56415888 | 0.11778026 | 0.99937574 |
| AC105052.3 | 29.1855179 | 0.17484235 | 0.54581404 | 0.32033319 | 0.74871577 | 0.99937574 |
| FAM185A | 54.6668057 | 0.65417793 | 0.42534841 | 1.53798136 | 0.12405317 | 0.99937574 |
| FBXL13 | 30.6258696 | -0.0815221 | 0.62377066 | -0.1306925 | 0.89601858 | 0.99937574 |
| ARMC10 | 38.6667721 | 0.16506274 | 0.40728845 | 0.40527233 | 0.68527734 | 0.99937574 |
| NAPEPLD | 66.4352459 | -0.5421549 | 0.63899537 | -0.8484489 | 0.39618803 | 0.99937574 |
| PMPCB | 71.0834498 | 0.26648002 | 0.3658708 | 0.72834459 | 0.46640267 | 0.99937574 |
| DNAJC2 | 41.4275574 | -0.4860237 | 0.42545475 | -1.1423628 | 0.25330326 | 0.99937574 |
| PSMC2 | 41.5508609 | 0.28971586 | 0.3469389 | 0.83506306 | 0.40368221 | 0.99937574 |
| SLC26A5 | 50.3464384 | 0.07517511 | 0.47427402 | 0.15850566 | 0.87405836 | 0.99937574 |
| ORC5 | 42.847881 | -0.220077 | 0.34966615 | -0.6293918 | 0.52909257 | 0.99937574 |
| LHFPL3 | 50.0053947 | -0.205527 | 0.61105579 | -0.3363474 | 0.7366089 | 0.99937574 |
| LINC01004 | 180.490732 | 0.2670641 | 0.34361941 | 0.77720901 | 0.43703547 | 0.99937574 |
| KMT2E-AS1 | 12.1937603 | 0.35232401 | 0.64383387 | 0.54722814 | 0.584222 | 0.99937574 |
| KMT2E | 422.185333 | -0.0075973 | 0.3182851 | -0.0238694 | 0.98095675 | 0.99937574 |
| SRPK2 | 507.244528 | -0.1107326 | 0.23471916 | -0.4717664 | 0.63709352 | 0.99937574 |
| PUS7 | 26.7430412 | -0.7132325 | 0.51165214 | -1.3939793 | 0.16332387 | 0.99937574 |
| RINT1 | 32.3482634 | 0.03458721 | 0.48196575 | 0.0717628 | 0.94279068 | 0.99937574 |
| ATXN7L1 | 180.884607 | -0.1539415 | 0.3143468 | -0.4897188 | 0.62433289 | 0.99937574 |
| CDHR3 | 15.941753 | 0.90988915 | 0.79834873 | 1.13971391 | 0.25440551 | 0.99937574 |
| SYPL1 | 140.130314 | 0.50233141 | 0.24414673 | 2.05749798 | 0.03963835 | 0.99937574 |
| NAMPT | 220.782267 | 0.18528448 | 0.55392234 | 0.33449541 | 7.38E-01 | 9.99E-01 |
| HBP1 | 75.3534409 | -0.3108292 | 0.33239035 | -0.9351331 | 0.34971969 | 0.99937574 |
| COG5 | 355.442357 | 0.15354589 | 0.31882995 | 0.48159181 | 0.63009594 | 0.99937574 |
| DUS4L | 12.1597868 | 0.38481106 | 0.49322092 | 0.7802002 | 0.43527305 | 0.99937574 |
| BCAP29 | 68.1218767 | 0.34931325 | 0.36619889 | 0.95388944 | 0.34013961 | 0.99937574 |
| CBLL1 | 18.8097791 | -0.2365184 | 0.48967882 | -0.4830073 | 0.62909055 | 0.99937574 |
| SLC26A3 | 385.281009 | -0.6315649 | 0.71567958 | -0.8824688 | 0.37752336 | 0.99937574 |
| DLD | 121.760528 | 0.38446454 | 0.2866517 | 1.34122538 | 0.17984729 | 0.99937574 |
| LAMB1 | 352.548738 | -0.5850005 | 0.41008335 | -1.4265406 | 0.15371236 | 0.99937574 |
| NRCAM | 114.33096 | -1.2856867 | 1.22792589 | -1.0470393 | 0.29508145 | 0.99937574 |
| PNPLA8 | 304.823869 | -0.2078537 | 0.39705202 | -0.5234923 | 0.60063172 | 0.99937574 |
| DNAJB9 | 20.7744741 | -0.3059816 | 0.49111649 | -0.6230326 | 0.53326311 | 0.99937574 |
| IMMP2L | 814.103771 | 0.26527335 | 0.38354816 | 0.69162984 | 0.48916982 | 0.99937574 |
| DOCK4 | 264.52098 | 0.08410834 | 0.4699977 | 0.1789548 | 0.85797319 | 0.99937574 |
| ZNF277 | 115.954136 | 0.20663022 | 0.28999535 | 0.71252941 | 0.47613701 | 0.99937574 |
| IFRD1 | 96.9815269 | -0.1645629 | 0.64765605 | -0.25409 | 0.79942603 | 0.99937574 |
| TMEM168 | 88.4506953 | 0.52124235 | 0.30185449 | 1.72680009 | 0.08420357 | 0.99937574 |
| BMT2 | 64.0550539 | -0.4558164 | 0.3292616 | -1.3843593 | 0.16624846 | 0.99937574 |
| SMIM30 | 13.5266645 | -0.2203828 | 0.52287164 | -0.4214856 | 0.67340056 | 0.99937574 |
| FOXP2 | 29.7503771 | 0.27849005 | 0.934005 | 0.29816762 | 0.76557523 | 0.99937574 |

| | | | | | | |
|---|---|---|---|---|---|---|
| MDFIC | 80.075138 | 0.61208136 | 0.64884584 | 0.94333865 | 0.34550771 | 0.99937574 |
| TES | 69.3539194 | 0.08324819 | 0.36024711 | 0.23108634 | 0.81724773 | 0.99937574 |
| CAV2 | 24.6675687 | 1.0899832 | 0.67038535 | 1.62590545 | 0.10396977 | 0.99937574 |
| MET | 193.300305 | -0.4790894 | 0.23644409 | -2.026227 | 0.04274153 | 0.99937574 |
| CAPZA2 | 172.005573 | -0.2298826 | 0.22318984 | -1.0299868 | 0.30301618 | 0.99937574 |
| ST7 | 290.964382 | -0.1192339 | 0.24499771 | -0.4866736 | 0.62648963 | 0.99937574 |
| ST7-AS2 | 19.8633716 | -0.7369438 | 0.78766391 | -0.935607 | 0.34947559 | 0.99937574 |
| CFTR | 496.497161 | -0.1679881 | 0.79642741 | -0.2109271 | 8.33E-01 | 9.99E-01 |
| AC000061.1 | 30.2557858 | 0.81983924 | 0.97923744 | 0.83722212 | 4.02E-01 | 0.99937574 |
| CTTNBP2 | 262.972956 | -0.4032961 | 0.35929867 | -1.1224537 | 0.2616696 | 0.99937574 |
| LSM8 | 52.3642269 | -0.0319762 | 0.31724569 | -0.1007931 | 0.91971469 | 0.99937574 |
| ANKRD7 | 17.1605695 | 0.47158078 | 0.52587463 | 0.89675515 | 0.36984958 | 0.99937574 |
| TSPAN12 | 281.166993 | 0.97636144 | 0.7508308 | 1.30037478 | 0.19347255 | 0.99937574 |
| ING3 | 42.4698137 | 0.30602848 | 0.36535009 | 0.83763077 | 0.4022381 | 0.99937574 |
| FAM3C | 171.355346 | -0.7428214 | 0.31808544 | -2.335289 | 0.01952833 | 0.96427895 |
| AASS | 34.1523708 | 0.29915256 | 0.40581556 | 0.73716384 | 0.46102272 | 0.99937574 |
| CADPS2 | 221.39756 | -0.1332954 | 0.59676147 | -0.2233646 | 0.82325173 | 0.99937574 |
| AC004594.1 | 14.0343345 | -0.8617347 | 1.14935061 | -0.7497579 | 0.45340055 | 0.99937574 |
| NDUFA5 | 137.147134 | 0.1606176 | 0.41071156 | 0.39107154 | 0.69574435 | 0.99937574 |
| WASL | 172.95697 | 0.0646517 | 0.29317351 | 0.22052368 | 0.82546333 | 0.99937574 |
| HYAL4 | 52.9321404 | 0.00088914 | 1.69935608 | 0.00052322 | 0.99958253 | 0.99979721 |
| POT1 | 53.6783085 | -0.2537317 | 0.34955812 | -0.7258642 | 0.46792201 | 0.99937574 |
| POT1-AS1 | 29.0077051 | 0.04059312 | 0.42005952 | 0.09663659 | 0.923015 | 0.99937574 |
| ZNF800 | 94.9650482 | 0.58078278 | 0.36105568 | 1.60856845 | 0.10771074 | 0.99937574 |
| ARF5 | 30.6918747 | -0.4392872 | 0.43164444 | -1.0177062 | 0.30881762 | 0.99937574 |
| SND1 | 219.749578 | -0.0480703 | 0.27129057 | -0.1771913 | 0.85935816 | 0.99937574 |
| LRRC4 | 14.8155135 | -0.3413467 | 0.55085904 | -0.6196626 | 0.53547997 | 0.99937574 |
| RBM28 | 46.8773852 | 0.14468124 | 0.42672416 | 0.33905097 | 0.73457134 | 0.99937574 |
| HILPDA | 36.3149376 | -0.2456742 | 1.61030166 | -0.1525641 | 0.87874205 | 0.99937574 |
| METTL2B | 43.7481342 | 0.21985625 | 0.33496379 | 0.65635826 | 0.51159364 | 0.99937574 |
| AC090114.2 | 12.0481679 | 0.11343919 | 0.7648324 | 0.14831902 | 0.88209101 | 0.99937574 |
| CALU | 65.1052076 | 0.29248047 | 0.3132224 | 0.93377891 | 0.35041795 | 0.99937574 |
| KCP | 31.348898 | -2.1314696 | 1.45025096 | -1.4697247 | 0.14163633 | 0.99937574 |
| ATP6V1F | 154.165501 | 0.03618269 | 0.52617064 | 0.06876607 | 0.94517583 | 0.99937574 |
| TNPO3 | 123.785468 | -0.0639505 | 0.24402039 | -0.2620702 | 0.79326734 | 0.99937574 |
| TSPAN33 | 34.8408013 | 0.43327006 | 0.53040878 | 0.81686065 | 0.41400807 | 0.99937574 |
| AHCYL2 | 705.501431 | 0.74177178 | 0.34751894 | 2.1344787 | 0.03280362 | 0.99937574 |
| NRF1 | 112.487086 | 0.5207695 | 0.26255263 | 1.98348611 | 0.04731316 | 0.99937574 |
| UBE2H | 390.922017 | -0.1826081 | 0.36600817 | -0.498918 | 0.61783714 | 0.99937574 |
| ZC3HC1 | 22.997985 | 0.11396861 | 0.38501539 | 0.29601052 | 0.76722205 | 0.99937574 |
| KLHDC10 | 111.755997 | 0.46934724 | 0.29269525 | 1.60353553 | 0.10881647 | 0.99937574 |
| TMEM209 | 23.836976 | -0.5066781 | 0.49700311 | -1.0194667 | 0.30798145 | 0.99937574 |
| COPG2 | 32.2723521 | -0.4323067 | 0.42697808 | -1.0124798 | 0.31130871 | 0.99937574 |
| AC016831.7 | 1665.25808 | -0.6472998 | 0.2992309 | -2.1632117 | 0.0305249 | 0.99049468 |
| LINC00513 | 159.314274 | -0.2086429 | 0.66013757 | -0.3160597 | 0.75195717 | 0.99937574 |
| AC058791.1 | 44.8929452 | -0.4072994 | 0.74737349 | -0.5449744 | 0.5857711 | 0.99937574 |
| MKLN1 | 755.688807 | -0.1140802 | 0.24888383 | -0.4583674 | 0.64668854 | 0.99937574 |
| PODXL | 11.9722107 | -0.4065838 | 0.5699427 | -0.7133766 | 0.47561276 | 0.99937574 |
| CHCHD3 | 446.475072 | 0.02985248 | 0.21854811 | 0.13659454 | 0.8913513 | 0.99937574 |
| EXOC4 | 553.863769 | -0.1128191 | 0.23470165 | -0.4806917 | 0.63073566 | 0.99937574 |
| SLC35B4 | 35.168753 | 0.10802129 | 0.39190921 | 0.27562836 | 0.78283353 | 0.99937574 |
| AKR1B1 | 44.4108263 | -0.223887 | 0.51080968 | -0.4382983 | 0.66117006 | 0.99937574 |
| BPGM | 43.0278895 | -1.7230065 | 0.67395078 | -2.5565761 | 0.01057079 | 0.76274616 |
| AGBL3 | 13.5487381 | -0.6804 | 0.5292996 | -1.2854723 | 0.1986273 | 0.99937574 |

| | | | | | | |
|---|---|---|---|---|---|---|
| CYREN | 32.5163689 | 0.35943627 | 0.38566233 | 0.93199734 | 0.3513379 | 0.99937574 |
| TMEM140 | 47.0311255 | -1.0349343 | 0.72560037 | -1.4263145 | 1.54E-01 | 0.99937574 |
| WDR91 | 29.4806749 | 0.55750625 | 0.63241707 | 0.88154838 | 0.37802108 | 0.99937574 |
| CNOT4 | 200.54342 | -0.1028161 | 0.23268759 | -0.4418634 | 0.65858802 | 0.99937574 |
| NUP205 | 53.3620936 | 0.224651 | 0.32109084 | 0.69964936 | 0.48414631 | 0.99937574 |
| STMP1 | 51.5213757 | 0.77935219 | 0.29394752 | 2.65133109 | 0.00801752 | 0.74742128 |
| MTPN | 89.6765708 | 0.07278145 | 0.28959307 | 0.25132319 | 0.80156425 | 0.99937574 |
| PTN | 20.7135689 | -0.0022189 | 0.92001617 | -0.0024118 | 0.99807564 | 0.99968959 |
| DGKI | 256.178157 | -0.0520251 | 0.91699785 | -0.0567341 | 0.95475698 | 0.99937574 |
| CREB3L2 | 64.9584622 | 0.80947251 | 0.31786907 | 2.54655955 | 1.09E-02 | 0.76274616 |
| TRIM24 | 70.2482673 | -0.1104974 | 0.37419269 | -0.2952953 | 0.76776829 | 0.99937574 |
| SVOPL | 128.615798 | -1.1398951 | 0.7988839 | -1.4268595 | 0.15362038 | 0.99937574 |
| ATP6V0A4 | 1316.9121 | 0.30069189 | 0.28488888 | 1.05547076 | 0.29121008 | 0.99937574 |
| TMEM213 | 1459.58914 | 0.50883439 | 0.39465958 | 1.28929949 | 0.19729399 | 0.99937574 |
| KIAA1549 | 43.0533266 | 0.50762501 | 0.42839904 | 1.18493498 | 0.23604316 | 0.99937574 |
| ZC3HAV1 | 88.8856834 | -0.9331663 | 0.4052149 | -2.3028923 | 0.0212849 | 0.96427895 |
| TTC26 | 16.044346 | -0.4396509 | 0.517966 | -0.8488026 | 0.39599114 | 0.99937574 |
| UBN2 | 150.284996 | 0.05960725 | 0.29316145 | 0.20332568 | 0.83888049 | 0.99937574 |
| LUC7L2 | 36.1511801 | -0.0980893 | 0.35375946 | -0.2772769 | 0.78156747 | 0.99937574 |
| HIPK2 | 1119.51419 | 0.14287397 | 0.43420387 | 0.32904814 | 0.7421193 | 0.99937574 |
| TBXAS1 | 269.526425 | -0.3647929 | 0.61822497 | -0.590065 | 5.55E-01 | 9.99E-01 |
| PARP12 | 54.3082348 | -1.7202404 | 0.7522745 | -2.286719 | 0.02221223 | 0.96427895 |
| KDM7A | 69.1335857 | -0.6597643 | 0.46059322 | -1.432423 | 0.1520228 | 0.99937574 |
| SLC37A3 | 67.6026069 | -0.5734516 | 0.33288791 | -1.7226567 | 0.08495063 | 0.99937574 |
| MKRN1 | 63.4756879 | -0.2253538 | 0.34563128 | -0.6520065 | 0.51439699 | 0.99937574 |
| NDUFB2 | 151.831677 | 0.56127449 | 0.41098895 | 1.36566809 | 0.17204316 | 0.99937574 |
| BRAF | 492.263668 | -0.1668828 | 0.21421011 | -0.7790612 | 0.43594369 | 0.99937574 |
| MRPS33 | 29.2412922 | 0.54077124 | 0.59288676 | 0.91209869 | 0.36171677 | 0.99937574 |
| AGK | 36.9489362 | -0.2176457 | 0.34938754 | -0.6229348 | 0.53332736 | 0.99937574 |
| KIAA1147 | 43.4186636 | 0.10894701 | 0.54280067 | 0.20071273 | 0.8409232 | 0.99937574 |
| SSBP1 | 58.211889 | 0.17026898 | 0.30366654 | 0.56071039 | 0.57499498 | 0.99937574 |
| MGAM | 37.1965901 | -0.4146182 | 0.98172507 | -0.4223364 | 0.67277951 | 0.99937574 |
| GSTK1 | 71.9700677 | 0.487757 | 0.41582759 | 1.17297892 | 0.24080426 | 0.99937574 |
| CASP2 | 13.4990446 | -0.0615078 | 0.50452064 | -0.1219133 | 0.90296765 | 0.99937574 |
| TCAF2 | 26.2447155 | -1.0942597 | 1.00886751 | -1.0846417 | 0.27808039 | 0.99937574 |
| TCAF1 | 65.282981 | 0.39561804 | 0.30578103 | 1.29379524 | 0.19573616 | 0.99937574 |
| AC004889.1 | 22.7875296 | 0.01650219 | 0.56601151 | 0.02915523 | 0.97674079 | 0.99937574 |
| AC074386.1 | 65.5521813 | -0.365769 | 0.57323214 | -0.6380818 | 0.52342045 | 0.99937574 |
| ARHGEF5 | 19.743511 | -0.1479489 | 0.63668968 | -0.2323721 | 0.816249 | 0.99937574 |
| TPK1 | 36.4979302 | -0.5134834 | 0.44257022 | -1.1602302 | 0.24595508 | 0.99937574 |
| CNTNAP2 | 14.3728905 | 0.1412479 | 0.59377929 | 0.23787947 | 0.81197458 | 0.99937574 |
| CUL1 | 135.381106 | -0.2893161 | 0.30382851 | -0.952235 | 0.34097783 | 0.99937574 |
| EZH2 | 13.3490413 | -0.4916116 | 0.55831842 | -0.8805219 | 0.37857667 | 0.99937574 |
| PDIA4 | 26.4837104 | 0.45109244 | 0.47380848 | 0.95205649 | 0.34106833 | 0.99937574 |
| ZNF786 | 12.7989086 | 0.36317657 | 0.59546549 | 0.60990365 | 0.54192564 | 0.99937574 |
| ZNF398 | 32.9376921 | 0.37845171 | 0.36683745 | 1.03166052 | 0.30223118 | 0.99937574 |
| ZNF282 | 21.9493377 | -0.3461983 | 0.58845825 | -0.5883142 | 0.55632142 | 0.99937574 |
| ZNF212 | 13.1760489 | -0.4693858 | 0.64302086 | -0.7299698 | 0.46540863 | 0.99937574 |
| ZNF783 | 23.0377407 | -0.0971303 | 0.45711643 | -0.2124848 | 0.83172882 | 0.99937574 |
| ZNF777 | 10.2971329 | 0.17849591 | 0.64841284 | 0.27528127 | 0.78310015 | 0.99937574 |
| ZNF746 | 16.7948007 | 0.06260191 | 0.49490162 | 0.12649365 | 0.89934117 | 0.99937574 |
| ZNF862 | 42.1039825 | 0.27200725 | 0.50760867 | 0.53586013 | 0.59205522 | 0.99937574 |
| ATP6V0E2 | 10.648488 | 0.93494155 | 0.69374862 | 1.34766618 | 0.17776577 | 0.99937574 |
| ACTR3C | 187.536105 | -0.2799163 | 0.33795607 | -0.8282623 | 0.40752199 | 0.99937574 |

| | | | | | | |
|---|---|---|---|---|---|---|
| LRRC61 | 19.631093 | -0.3253537 | 0.54041846 | -0.6020403 | 0.54714731 | 0.99937574 |
| RARRES2 | 30.8567302 | -0.7421833 | 0.80280959 | -0.9244824 | 0.35523523 | 0.99937574 |
| ABCB8 | 19.8210529 | 0.60923148 | 0.44982422 | 1.35437678 | 0.1756162 | 0.99937574 |
| SLC4A2 | 21.9739856 | -1.1783454 | 0.75608799 | -1.5584765 | 0.11912033 | 0.99937574 |
| FASTK | 24.0855525 | 0.4040054 | 0.41792818 | 0.96668619 | 0.33370093 | 0.99937574 |
| TMUB1 | 32.1284049 | -0.2555791 | 0.47584197 | -0.5371092 | 0.59119216 | 0.99937574 |
| AGAP3 | 30.7830399 | 0.37109628 | 0.60539593 | 0.61298114 | 0.53988881 | 0.99937574 |
| CHPF2 | 18.8272212 | -0.3886008 | 0.63748089 | -0.6095881 | 0.54213467 | 0.99937574 |
| SMARCD3 | 18.6528161 | 0.10739458 | 0.55779825 | 0.19253301 | 0.84732471 | 0.99937574 |
| NUB1 | 63.1532944 | -0.3719396 | 0.46081888 | -0.8071275 | 0.419593 | 0.99937574 |
| RHEB | 127.012761 | -0.1519349 | 0.30306276 | -0.5013315 | 0.61613782 | 0.99937574 |
| PRKAG2 | 271.896117 | 0.04860418 | 0.48127375 | 0.10099071 | 0.91955783 | 0.99937574 |
| GALNT11 | 111.661696 | 0.20868712 | 0.26594758 | 0.78469268 | 0.43263377 | 0.99937574 |
| KMT2C | 498.02938 | -0.1743135 | 0.23963442 | -0.7274143 | 0.46697217 | 0.99937574 |
| ACTR3B | 53.2222002 | -0.2361217 | 0.31146038 | -0.7581115 | 0.44838422 | 0.99937574 |
| DPP6 | 105.611213 | -0.2690028 | 0.37683748 | -0.713843 | 0.47532429 | 0.99937574 |
| PAXIP1-AS2 | 22.0757248 | 0.20298423 | 0.5754108 | 0.35276403 | 0.72426535 | 0.99937574 |
| PAXIP1 | 28.6010007 | -0.2497131 | 0.42362282 | -0.5894703 | 0.5555458 | 0.99937574 |
| INSIG1 | 105.847189 | -0.669966 | 0.75028556 | -0.8929481 | 0.37188498 | 0.99937574 |
| AC009403.1 | 12.7415891 | 0.47359158 | 0.66141671 | 0.71602603 | 0.47397528 | 0.99937574 |
| RBM33 | 175.220678 | -0.1015929 | 0.18516558 | -0.5486597 | 0.58323901 | 0.99937574 |
| LINC01006 | 35.3737844 | -0.9990669 | 0.549248 | -1.8189723 | 0.06891566 | 0.99937574 |
| LMBR1 | 180.781376 | -0.1733502 | 0.27984481 | -0.619451 | 0.53561928 | 0.99937574 |
| NOM1 | 20.1837649 | 0.00165978 | 0.55025127 | 0.0030164 | 0.99759327 | 0.99937574 |
| UBE3C | 201.257913 | -0.233778 | 0.2572207 | -0.9088616 | 0.3634232 | 0.99937574 |
| DNAJB6 | 205.536316 | -0.0759142 | 0.29439586 | -0.2578643 | 0.79651162 | 0.99937574 |
| PTPRN2 | 12.056168 | -0.1933821 | 0.75637811 | -0.2556685 | 0.79820682 | 0.99937574 |
| NCAPG2 | 15.067197 | 1.05290195 | 0.48606766 | 2.16616336 | 0.03029871 | 0.99049468 |
| ESYT2 | 225.397248 | -0.3435528 | 0.45936336 | -0.7478889 | 0.45452715 | 0.99937574 |
| WDR60 | 116.737935 | 0.43946202 | 0.30752108 | 1.42904682 | 0.15299078 | 0.99937574 |
| GTPBP6 | 39.2534743 | -0.5502042 | 0.35852012 | -1.5346537 | 0.12486891 | 0.99937574 |
| CSF2RA | 46.0183918 | -0.637346 | 0.94156603 | -0.6768999 | 0.49846946 | 0.99937574 |
| IL3RA | 29.4267304 | -0.1816665 | 0.74842628 | -0.2427313 | 0.80821358 | 0.99937574 |
| SLC25A6 | 336.307811 | 0.36613018 | 0.51418154 | 0.71206402 | 0.47642513 | 0.99937574 |
| LINC00106 | 36.1475838 | -0.1061555 | 0.59377603 | -0.1787804 | 0.85811013 | 0.99937574 |
| ASMTL-AS1 | 18.2715237 | -0.0277002 | 0.54334119 | -0.0509812 | 0.95934047 | 0.99937574 |
| ASMTL | 27.7570937 | 0.19662098 | 0.38309435 | 0.51324427 | 0.60778046 | 0.99937574 |
| AKAP17A | 23.9065876 | -0.6403793 | 0.45015503 | -1.422575 | 0.15485939 | 0.99937574 |
| DHRSX | 156.409233 | -0.163669 | 0.24220001 | -0.6757597 | 0.49919325 | 0.99937574 |
| CD99 | 23.020008 | 0.35828296 | 0.49300781 | 0.72672877 | 0.46739212 | 0.99937574 |
| ARSD | 80.3710924 | -0.1466825 | 0.45277341 | -0.3239645 | 0.74596492 | 0.99937574 |
| ARSE | 26.6883061 | 0.73307385 | 0.63199866 | 1.15992943 | 0.24607754 | 0.99937574 |
| PRKX | 92.5854418 | -1.2632564 | 0.80499286 | -1.5692765 | 0.11658352 | 0.99937574 |
| NLGN4X | 16.8726143 | -1.0764223 | 1.16606321 | -0.9231252 | 0.35594198 | 0.99937574 |
| PUDP | 38.5617498 | -0.3682965 | 0.48802923 | -0.7546608 | 0.45045251 | 0.99937574 |
| ANOS1 | 19.7135127 | -2.7858711 | 1.16590146 | -2.3894568 | 0.01687331 | 0.89506255 |
| TBL1X | 331.665108 | -0.2104235 | 0.43563806 | -0.4830237 | 0.62907892 | 0.99937574 |
| GPR143 | 15.5198578 | 1.16851958 | 0.52106465 | 2.24256159 | 0.0249251 | 0.96546668 |
| SHROOM2 | 126.954838 | 0.61161125 | 0.45492323 | 1.34442739 | 0.17881022 | 0.99937574 |
| WWC3 | 34.6631728 | 0.62403301 | 0.41308066 | 1.51068077 | 0.1308698 | 0.99937574 |
| CLCN4 | 14.0262171 | 0.97440758 | 0.53923815 | 1.80700787 | 0.07076105 | 0.99937574 |
| MID1 | 15.2930473 | 0.50195985 | 0.58588844 | 0.85674987 | 0.39158313 | 0.99937574 |
| HCCS | 12.3970283 | -0.0319131 | 0.48429216 | -0.0658964 | 0.94746029 | 0.99937574 |
| ARHGAP6 | 536.164844 | 0.47301797 | 0.60147729 | 0.78642698 | 0.43161738 | 0.99937574 |

| | | | | | | |
|---|---|---|---|---|---|---|
| PRPS2 | 22.4572784 | -0.1998952 | 0.47665611 | -0.4193699 | 0.67494579 | 0.99937574 |
| TMSB4X | 508.652312 | -0.5025019 | 0.64411443 | -0.7801438 | 0.43530623 | 0.99937574 |
| RAB9A | 79.8317139 | -0.0609085 | 0.42913772 | -0.1419322 | 0.88713353 | 0.99937574 |
| TRAPPC2 | 25.2015239 | -0.0518819 | 0.55561996 | -0.0933767 | 0.92560433 | 0.99937574 |
| OFD1 | 119.621521 | -0.1926052 | 0.36925864 | -0.5215998 | 0.60194902 | 0.99937574 |
| GEMIN8 | 22.8492854 | -0.7347453 | 0.53900501 | -1.3631511 | 0.17283488 | 0.99937574 |
| MOSPD2 | 33.535731 | 0.53717454 | 0.39749476 | 1.35140031 | 0.17656723 | 0.99937574 |
| ASB9 | 38.3857767 | 1.40371448 | 0.65801924 | 2.13324229 | 0.03290486 | 0.99937574 |
| PIGA | 29.7393743 | -0.4800086 | 0.50093246 | -0.9582303 | 0.33794665 | 0.99937574 |
| PIR | 35.7098346 | 0.11436693 | 0.52568025 | 0.21755988 | 0.82777205 | 0.99937574 |
| CLTRN | 12.9269973 | -0.1496909 | 0.77966702 | -0.1919933 | 0.84774745 | 0.99937574 |
| CA5B | 88.7562894 | -0.6447684 | 0.34541124 | -1.8666688 | 0.06194786 | 0.99937574 |
| ZRSR2 | 43.7184732 | 0.15052547 | 0.38080524 | 0.39528205 | 0.69263475 | 0.99937574 |
| AP1S2 | 72.7320197 | 0.28736384 | 0.46909518 | 0.61259174 | 0.54014632 | 0.99937574 |
| CTPS2 | 46.7166649 | 0.18791243 | 0.34551324 | 0.54386463 | 0.58653463 | 0.99937574 |
| SYAP1 | 86.8624169 | 0.08196416 | 0.36041522 | 0.22741593 | 0.82010033 | 0.99937574 |
| TXLNG | 40.6355049 | -0.4223374 | 0.40599764 | -1.0402461 | 0.2982256 | 0.99937574 |
| RBBP7 | 54.7354553 | 0.15623393 | 0.33267402 | 0.46963068 | 0.6386189 | 0.99937574 |
| REPS2 | 41.4191914 | -1.1047317 | 0.92663011 | -1.1922036 | 0.23318143 | 0.99937574 |
| NHS | 80.4139293 | 0.52094489 | 0.7503409 | 0.69427761 | 0.48750813 | 0.99937574 |
| SCML1 | 127.962966 | -0.1806341 | 0.31359775 | -0.5760058 | 0.56461125 | 0.99937574 |
| RAI2 | 30.9386266 | 0.27856373 | 0.37575967 | 0.74133483 | 0.45849045 | 0.99937574 |
| CDKL5 | 54.8003257 | 0.13271888 | 0.37865283 | 0.3505028 | 0.72596139 | 0.99937574 |
| PHKA2 | 31.549734 | 0.0369656 | 0.35229452 | 0.10492812 | 0.91643285 | 0.99937574 |
| PDHA1 | 105.213794 | 0.48575606 | 0.38025901 | 1.27743471 | 0.20144882 | 0.99937574 |
| MAP3K15 | 21.6304028 | -0.4550046 | 0.75258139 | -0.6045918 | 0.54545024 | 0.99937574 |
| BCLAF3 | 51.5871593 | -0.6563743 | 0.37792114 | -1.7368022 | 0.08242209 | 0.99937574 |
| EIF1AX | 92.6157861 | -0.1480747 | 0.33521177 | -0.4417347 | 0.65868121 | 0.99937574 |
| RPS6KA3 | 112.193756 | -0.3657027 | 0.41638365 | -0.8782832 | 0.37979008 | 0.99937574 |
| MBTPS2 | 27.584926 | 0.4647798 | 0.44909524 | 1.0349248 | 0.30070403 | 0.99937574 |
| SMS | 72.743779 | -0.3700947 | 0.34320748 | -1.078341 | 0.28088159 | 0.99937574 |
| PRDX4 | 12.8448658 | -0.0952283 | 0.64528802 | -0.1475749 | 0.88267829 | 0.99937574 |
| ACOT9 | 32.0072861 | -0.9193874 | 0.48938168 | -1.8786715 | 0.06028936 | 0.99937574 |
| SAT1 | 417.30502 | -0.0469265 | 0.42645761 | -0.1100379 | 0.91237931 | 0.99937574 |
| APOO | 487.680229 | 0.00109353 | 0.49518749 | 0.00220832 | 0.99823801 | 0.99972393 |
| KLHL15 | 11.8473233 | -0.5903595 | 0.54002955 | -1.0931985 | 0.27430666 | 0.99937574 |
| EIF2S3 | 58.8083236 | -0.378494 | 0.3883518 | -0.9746162 | 0.32975069 | 0.99937574 |
| ZFX | 90.42616 | -0.4112149 | 0.41853802 | -0.982503 | 0.32585209 | 0.99937574 |
| PDK3 | 23.975453 | -0.751176 | 0.64973089 | -1.156134 | 0.24762637 | 0.99937574 |
| POLA1 | 83.6145112 | -0.3279372 | 0.34441459 | -0.9521583 | 0.34101668 | 0.99937574 |
| GK | 59.7480441 | 0.12134111 | 0.35802152 | 0.33892128 | 0.73466903 | 0.99937574 |
| TAB3 | 69.0666897 | 0.360672 | 0.34997999 | 1.03055036 | 0.30275173 | 0.99937574 |
| DMD | 118.754388 | 0.54984871 | 0.45812964 | 1.20020332 | 0.23006039 | 0.99937574 |
| AF241726.2 | 68.9590482 | 0.104963 | 0.35702716 | 0.29399165 | 0.76876428 | 0.99937574 |
| DYNLT3 | 133.738512 | -0.524528 | 0.38932921 | -1.3472609 | 0.17789622 | 0.99937574 |
| RPGR | 35.1939243 | 0.11944923 | 0.52767176 | 0.22637033 | 0.8209134 | 0.99937574 |
| TSPAN7 | 52.4227979 | 1.61298641 | 0.70592753 | 2.28491784 | 0.02231765 | 0.96427895 |
| MID1IP1 | 28.1828301 | -0.4342323 | 0.48508146 | -0.895174 | 0.37069406 | 0.99937574 |
| BCOR | 94.6830033 | 0.11568286 | 0.53391261 | 0.21667003 | 0.8284655 | 0.99937574 |
| ATP6AP2 | 452.144033 | 0.67839926 | 0.26365273 | 2.57307878 | 0.01007983 | 0.76274616 |
| CXorf38 | 55.4781716 | 0.18515803 | 0.34881419 | 0.53082138 | 0.59554256 | 0.99937574 |
| MED14 | 115.881919 | -0.0575862 | 0.31397278 | -0.1834116 | 0.8544751 | 0.99937574 |
| USP9X | 277.330787 | -0.178124 | 0.20451139 | -0.8709733 | 0.38376874 | 0.99937574 |
| DDX3X | 129.135256 | -0.4005206 | 0.40705807 | -0.9839398 | 0.32514512 | 0.99937574 |

| | | | | | | |
|---|---|---|---|---|---|---|
| CASK | 61.4689407 | 0.23202235 | 0.3005372 | 0.77202537 | 0.4400994 | 0.99937574 |
| MAOA | 134.700663 | 0.03392274 | 0.68829599 | 0.0492851 | 0.96069209 | 0.99937574 |
| FUNDC1 | 17.2370799 | 0.26075572 | 0.59490485 | 0.43831499 | 0.66115796 | 0.99937574 |
| KDM6A | 253.983021 | -0.4375595 | 0.42338518 | -1.0334785 | 0.30138003 | 0.99937574 |
| MIR222HG | 45.3337404 | -0.1866009 | 0.61200256 | -0.3049022 | 0.76044068 | 0.99937574 |
| KRBOX4 | 31.1072912 | 0.19366786 | 0.35393709 | 0.5471816 | 0.58425398 | 0.99937574 |
| ZNF674 | 15.5971269 | 0.28232715 | 0.46676267 | 0.60486232 | 0.54527048 | 0.99937574 |
| SLC9A7 | 31.5329117 | -0.4516292 | 0.47236615 | -0.9560998 | 0.3390218 | 0.99937574 |
| RP2 | 17.6974925 | -0.5109044 | 0.55991036 | -0.9124754 | 0.36151853 | 0.99937574 |
| JADE3 | 72.9712677 | 0.09017067 | 0.39000538 | 0.23120366 | 0.81715659 | 0.99937574 |
| RGN | 16.8921804 | 1.481441 | 0.71364471 | 2.07588031 | 0.03790502 | 0.99937574 |
| NDUFB11 | 89.6140886 | -0.0438299 | 0.55299955 | -0.0792585 | 0.93682703 | 0.99937574 |
| RBM10 | 47.0897214 | -0.2249814 | 0.33701505 | -0.6675707 | 0.50440767 | 0.99937574 |
| UBA1 | 75.8446357 | -0.2676504 | 0.29743662 | -0.8998569 | 0.36819639 | 0.99937574 |
| CDK16 | 30.9263701 | -0.2400149 | 0.476607 | -0.5035909 | 0.61454889 | 0.99937574 |
| USP11 | 30.8566001 | -0.0540311 | 0.39744337 | -0.1359466 | 0.89186351 | 0.99937574 |
| ZNF41 | 25.2359286 | -0.1809065 | 0.38713876 | -0.4672911 | 0.64029165 | 0.99937574 |
| ARAF | 27.298382 | 0.11220192 | 0.44240979 | 0.25361536 | 0.79979273 | 0.99937574 |
| TIMP1 | 26.5248353 | -0.1309559 | 0.53338941 | -0.2455166 | 8.06E-01 | 9.99E-01 |
| UXT | 22.408521 | 0.12858146 | 0.39597778 | 0.32471888 | 0.74539384 | 0.99937574 |
| ZNF81 | 46.8182623 | 0.19876166 | 0.3923201 | 0.50663135 | 0.61241351 | 0.99937574 |
| ZNF630 | 19.4258651 | 0.4660742 | 0.63527532 | 0.73365702 | 0.4631578 | 0.99937574 |
| EBP | 35.3582039 | 0.56875761 | 0.44685627 | 1.27279763 | 0.20308986 | 0.99937574 |
| TBC1D25 | 13.7832087 | -0.2423694 | 0.59487808 | -0.407427 | 0.68369437 | 0.99937574 |
| RBM3 | 54.4859679 | -0.1176581 | 0.61324663 | -0.1918611 | 0.84785105 | 0.99937574 |
| WDR13 | 50.5266576 | -0.2274825 | 0.31771982 | -0.7159845 | 0.47400092 | 0.99937574 |
| HDAC6 | 24.2873292 | 0.4059689 | 0.44554835 | 0.91116688 | 0.36220745 | 0.99937574 |
| PCSK1N | 31.5239437 | -0.8547829 | 0.82147193 | -1.0405504 | 0.29808428 | 0.99937574 |
| TIMM17B | 20.8784256 | 0.15212049 | 0.47933333 | 0.31735846 | 0.75097162 | 0.99937574 |
| SLC35A2 | 9.48862791 | -0.0658668 | 0.67142088 | -0.0981007 | 0.92185234 | 0.99937574 |
| OTUD5 | 48.0146113 | -0.1740785 | 0.44087907 | -0.3948441 | 0.69295793 | 0.99937574 |
| GRIPAP1 | 38.4823631 | -0.2418357 | 0.33845597 | -0.7145264 | 0.47490172 | 0.99937574 |
| TFE3 | 18.5189233 | -0.8777447 | 0.4215923 | -2.0819753 | 0.03734473 | 0.99937574 |
| CCDC120 | 13.4922565 | -0.0497788 | 0.53369775 | -0.0932715 | 0.92568786 | 0.99937574 |
| WDR45 | 30.7331131 | -0.1120481 | 0.40468231 | -0.2768791 | 0.78187295 | 0.99937574 |
| PLP2 | 34.2521766 | -0.2272938 | 0.56960776 | -0.3990356 | 0.68986699 | 0.99937574 |
| PPP1R3F | 9.99170977 | 0.18442855 | 0.57258582 | 0.32209766 | 0.74737871 | 0.99937574 |
| CLCN5 | 116.298233 | 0.55085545 | 0.4454965 | 1.23649783 | 0.21627357 | 0.99937574 |
| MAGED1 | 149.759846 | 0.0791756 | 0.3684723 | 0.21487531 | 0.82986453 | 0.99937574 |
| FAM156A | 15.8983433 | 0.68457069 | 0.51033934 | 1.34140294 | 0.17978966 | 0.99937574 |
| TSPYL2 | 12.8448645 | -1.6935691 | 0.79764743 | -2.1232051 | 0.03373667 | 0.99937574 |
| KANTR | 55.2432209 | 0.22418721 | 0.29017507 | 0.77259293 | 0.43976333 | 0.99937574 |
| KDM5C | 46.4024659 | -0.538086 | 0.34577172 | -1.5561886 | 0.11966325 | 0.99937574 |
| IQSEC2 | 19.0383917 | 0.72882892 | 0.51054211 | 1.42755886 | 0.15341887 | 0.99937574 |
| SMC1A | 62.6898171 | -0.5409226 | 0.32284551 | -1.6754844 | 0.09383922 | 0.99937574 |
| HSD17B10 | 23.4384171 | 0.30679531 | 0.49963825 | 0.61403487 | 0.53919229 | 0.99937574 |
| HUWE1 | 204.906645 | -0.0037676 | 0.24005179 | -0.015695 | 0.98747774 | 0.99937574 |
| PHF8 | 89.7957485 | -0.1040117 | 0.47230981 | -0.2202193 | 0.82570037 | 0.99937574 |
| FAM120C | 25.5221322 | -0.1359966 | 0.41756905 | -0.3256865 | 0.74466159 | 0.99937574 |
| WNK3 | 101.925828 | 0.85367408 | 0.31325697 | 2.72515591 | 0.00642711 | 0.69675365 |
| TSR2 | 20.9576127 | 0.42527282 | 0.5755679 | 0.73887514 | 0.45998282 | 0.99937574 |
| GNL3L | 11.573193 | -0.0713012 | 0.68186924 | -0.1045672 | 0.91671923 | 0.99937574 |
| MAGED2 | 54.668713 | -0.1573014 | 0.4150294 | -0.3790126 | 0.70467849 | 0.99937574 |
| USP51 | 11.2870909 | -0.8374283 | 0.79744385 | -1.0501408 | 0.29365339 | 0.99937574 |

| | | | | | | |
|---|---|---|---|---|---|---|
| RRAGB | 53.2538556 | -0.0743885 | 0.32313812 | -0.2302065 | 0.81793128 | 0.99937574 |
| AL445472.1 | 80.1357234 | -0.367559 | 0.43195797 | -0.8509139 | 0.39481718 | 0.99937574 |
| UBQLN2 | 29.3506539 | -0.032099 | 0.43721967 | -0.0734162 | 0.94147493 | 0.99937574 |
| NBDY | 96.8719093 | 0.08523842 | 0.32160831 | 0.265038 | 0.79098019 | 0.99937574 |
| SPIN3 | 17.68986 | 0.59021241 | 0.54962433 | 1.07384695 | 0.28289128 | 0.99937574 |
| FAAH2 | 256.881596 | 0.2038228 | 0.37272179 | 0.5468497 | 0.58448199 | 0.99937574 |
| ARHGEF9 | 69.654993 | -0.0218784 | 0.43275137 | -0.0505565 | 0.95967889 | 0.99937574 |
| MTMR8 | 17.011964 | -0.1329867 | 0.53400451 | -0.2490366 | 0.80333245 | 0.99937574 |
| ZC4H2 | 35.1482058 | 0.93233622 | 0.349179 | 2.67008105 | 0.00758329 | 0.74742128 |
| LAS1L | 18.2440384 | -0.2320772 | 0.43866751 | -0.5290504 | 0.59677052 | 0.99937574 |
| MSN | 165.091527 | -0.1649506 | 0.35178534 | -0.4688956 | 0.63914424 | 0.99937574 |
| OPHN1 | 197.586612 | -0.0285554 | 0.362892 | -0.0786884 | 0.93728049 | 0.99937574 |
| YIPF6 | 69.2550106 | 0.07668945 | 0.29096339 | 0.26357079 | 0.79211067 | 0.99937574 |
| EDA | 49.5614307 | -0.2729107 | 0.56006375 | -0.4872851 | 0.6260563 | 0.99937574 |
| IGBP1 | 39.1862824 | -0.291709 | 0.38462566 | -0.7584233 | 0.44819764 | 0.99937574 |
| DLG3 | 56.4514202 | -0.0662365 | 0.39768162 | -0.1665566 | 0.86771897 | 0.99937574 |
| SNX12 | 19.6278481 | 0.00529366 | 0.51593954 | 0.01026023 | 0.99181366 | 0.99937574 |
| MED12 | 11.8263872 | 0.02999347 | 0.5371532 | 0.05583784 | 0.95547099 | 0.99937574 |
| ZMYM3 | 15.8082753 | 0.47948476 | 0.49422125 | 0.97018241 | 0.33195558 | 0.99937574 |
| NONO | 53.9833462 | -0.1777643 | 0.36324972 | -0.4893721 | 0.62457831 | 0.99937574 |
| TAF1 | 91.8550334 | 0.07039989 | 0.32216155 | 0.21852357 | 0.8270212 | 0.99937574 |
| OGT | 179.995589 | -0.1411338 | 0.37423874 | -0.3771224 | 0.70608265 | 0.99937574 |
| NHSL2 | 24.2600457 | 0.08674346 | 0.36789346 | 0.2357842 | 0.81360013 | 0.99937574 |
| PIN4 | 36.6410951 | -0.1552965 | 0.4461069 | -0.3481149 | 0.72775391 | 0.99937574 |
| RPS4X | 258.092405 | 0.07115908 | 0.5334011 | 0.13340633 | 0.89387204 | 0.99937574 |
| HDAC8 | 219.714898 | 0.13401306 | 0.35147712 | 0.38128528 | 0.70299157 | 0.99937574 |
| PHKA1 | 30.1431576 | 0.53471396 | 0.40077164 | 1.33421106 | 0.18213469 | 0.99937574 |
| CHIC1 | 100.396507 | 0.24463743 | 0.29957691 | 0.81660976 | 0.41415148 | 0.99937574 |
| XIST | 340.542559 | -14.791794 | 4.45251135 | -3.3221238 | 0.00089335 | 0.29012345 |
| JPX | 234.802382 | -0.1786765 | 0.45989488 | -0.388516 | 0.69763425 | 0.99937574 |
| FTX | 1189.56871 | 0.25351443 | 0.31024637 | 0.81713905 | 0.41384897 | 0.99937574 |
| RLIM | 43.147967 | -0.1184414 | 0.32210245 | -0.3677136 | 0.71308683 | 0.99937574 |
| ABCB7 | 94.5318127 | -0.0493509 | 0.29388257 | -0.1679273 | 0.86664048 | 0.99937574 |
| UPRT | 20.0574996 | -0.1042875 | 0.43108602 | -0.241918 | 0.80884371 | 0.99937574 |
| ZDHHC15 | 15.693532 | -0.2223868 | 0.62210451 | -0.357475 | 0.72073621 | 0.99937574 |
| PBDC1 | 29.4652059 | 0.00264059 | 0.51707844 | 0.00510675 | 0.99592542 | 0.99937574 |
| ATRX | 373.64627 | 0.04379648 | 0.20879082 | 0.20976247 | 0.83385307 | 0.99937574 |
| MAGT1 | 75.0821362 | 0.4208118 | 0.2900868 | 1.45064099 | 0.14687986 | 0.99937574 |
| COX7B | 214.00464 | 0.50812337 | 0.4655911 | 1.09135113 | 0.27511841 | 0.99937574 |
| PGK1 | 191.371499 | 0.31712526 | 0.40459997 | 0.78379951 | 0.43315776 | 0.99937574 |
| BRWD3 | 149.288742 | -0.1363862 | 0.33885614 | -0.4024899 | 0.68732351 | 0.99937574 |
| HMGN5 | 15.264439 | 0.59900593 | 0.57304799 | 1.04529802 | 0.29588525 | 0.99937574 |
| SH3BGRL | 98.3200997 | 0.05597405 | 0.27068651 | 0.20678552 | 0.83617737 | 0.99937574 |
| RPS6KA6 | 101.721223 | 0.25749217 | 0.34728838 | 0.7414362 | 0.458429 | 0.99937574 |
| HDX | 34.9263227 | -0.169455 | 0.50918785 | -0.3327946 | 0.73928933 | 0.99937574 |
| APOOL | 64.756768 | 0.18124354 | 0.28311659 | 0.64017282 | 0.52206025 | 0.99937574 |
| POF1B | 19.2549099 | 0.04249798 | 0.92279968 | 0.0460533 | 0.96326776 | 0.99937574 |
| CHM | 105.294597 | 0.19107175 | 0.27019828 | 0.70715384 | 0.47947088 | 0.99937574 |
| DIAPH2 | 107.672033 | -2.2303106 | 0.97763191 | -2.2813398 | 0.02252835 | 0.96427895 |
| TSPAN6 | 10.8559952 | 0.07155906 | 0.67491286 | 0.10602711 | 0.91556084 | 0.99937574 |
| TRMT2B | 44.8216605 | -0.1739275 | 0.38549384 | -0.4511811 | 0.65185904 | 0.99937574 |
| HNRNPH2 | 34.4089536 | -0.6485111 | 0.41777554 | -1.5522955 | 0.12059154 | 0.99937574 |
| ARMCX4 | 150.7084 | 0.15926107 | 0.37914558 | 0.42005254 | 0.67444707 | 0.99937574 |
| ARMCX6 | 10.9500992 | 0.58457082 | 0.54168607 | 1.079169 | 0.2805124 | 0.99937574 |

| | | | | | | |
|---|---|---|---|---|---|---|
| ARMCX3 | 58.673823 | 0.33191417 | 0.38795554 | 0.85554693 | 0.39224844 | 0.99937574 |
| ARMCX2 | 18.566169 | 0.05382766 | 0.48186757 | 0.11170634 | 0.91105625 | 0.99937574 |
| ZMAT1 | 74.9894002 | -0.0710664 | 0.53748096 | -0.1322212 | 0.89480933 | 0.99937574 |
| ARMCX5 | 11.0089146 | 0.28827644 | 0.54402615 | 0.52989445 | 0.59618512 | 0.99937574 |
| GPRASP1 | 15.448013 | 0.35490293 | 0.51517432 | 0.68889871 | 0.49088701 | 0.99937574 |
| BHLHB9 | 13.5035356 | 0.69934171 | 0.61551572 | 1.13618822 | 0.2558778 | 0.99937574 |
| LINC00630 | 51.7980608 | 0.18360921 | 0.33769773 | 0.54370875 | 0.58664191 | 0.99937574 |
| BEX4 | 30.2387085 | -0.0623666 | 0.40486322 | -0.1540437 | 0.87757525 | 0.99937574 |
| TCEAL8 | 18.8460782 | 0.45322249 | 0.46263869 | 0.97964673 | 0.32726053 | 0.99937574 |
| TCEAL9 | 30.6057461 | -0.0324178 | 0.54588158 | -0.0593862 | 0.95264449 | 0.99937574 |
| BEX3 | 48.3852332 | 0.23439415 | 0.43354912 | 0.54064036 | 0.5887555 | 0.99937574 |
| TCEAL4 | 56.2424371 | -0.5534234 | 0.34884843 | -1.5864294 | 0.11264193 | 0.99937574 |
| TCEAL3 | 30.0283181 | 0.29204081 | 0.45243528 | 0.64548637 | 0.51861205 | 0.99937574 |
| MORF4L2 | 92.9025067 | -0.0335346 | 0.40576814 | -0.0826447 | 0.93413406 | 0.99937574 |
| SLC25A53 | 41.9224696 | 0.15041938 | 0.41100469 | 0.36597971 | 0.71438022 | 0.99937574 |
| FAM199X | 59.0314297 | -0.4114277 | 0.25950996 | -1.5854024 | 0.11287494 | 0.99937574 |
| NRK | 23.6206122 | 1.79667157 | 0.62553992 | 2.87219332 | 0.00407634 | 0.57136631 |
| RNF128 | 17.9018837 | 0.40477372 | 0.8004961 | 0.50565359 | 0.61309986 | 0.99937574 |
| TBC1D8B | 25.8095255 | 0.27145098 | 0.55130833 | 0.49237598 | 0.62245358 | 0.99937574 |
| MORC4 | 27.4548114 | 0.4418801 | 0.47941781 | 0.92170146 | 0.35668432 | 0.99937574 |
| RBM41 | 52.5421577 | 0.14303204 | 0.28934205 | 0.49433546 | 0.62106928 | 0.99937574 |
| MID2 | 22.3237301 | 0.0083546 | 0.5418761 | 0.01541792 | 0.98769877 | 0.99937574 |
| PSMD10 | 11.1929849 | 0.36347351 | 0.52076728 | 0.69795765 | 0.48520368 | 0.99937574 |
| ATG4A | 14.8675266 | 0.4397587 | 0.54546037 | 0.80621568 | 0.4201185 | 0.99937574 |
| COL4A5 | 25.9302847 | 0.82932988 | 0.57116151 | 1.45200589 | 0.14649997 | 0.99937574 |
| ACSL4 | 119.953676 | 0.1316868 | 0.42146422 | 0.31245072 | 7.55E-01 | 0.99937574 |
| TMEM164 | 119.596498 | -0.208102 | 0.43720619 | -0.4759813 | 0.63408767 | 0.99937574 |
| AMMECR1 | 100.394187 | 0.21133129 | 0.42003496 | 0.50312787 | 0.61487437 | 0.99937574 |
| ALG13 | 93.7340589 | -0.162071 | 0.5125731 | -0.316191 | 0.75185755 | 0.99937574 |
| AMOT | 13.9255336 | 1.00650368 | 0.5364597 | 1.87619625 | 0.06062834 | 0.99937574 |
| PLS3 | 187.685971 | 0.2492323 | 0.28785766 | 0.86581786 | 0.38659005 | 0.99937574 |
| DANT2 | 223.723698 | -0.7276507 | 0.63128271 | -1.1526542 | 0.24905235 | 0.99937574 |
| KLHL13 | 50.649305 | 0.31550386 | 0.56621487 | 0.55721578 | 0.57738001 | 0.99937574 |
| WDR44 | 46.3414031 | -0.6293649 | 0.37068281 | -1.6978529 | 0.08953553 | 0.99937574 |
| IL13RA1 | 60.1473949 | 0.33396558 | 0.36319483 | 0.91952184 | 0.35782269 | 0.99937574 |
| PGRMC1 | 87.9959022 | 0.64980291 | 0.31348775 | 2.07281755 | 0.03818926 | 0.99937574 |
| SLC25A43 | 38.3337679 | 0.40909375 | 0.50851793 | 0.80448246 | 0.42111839 | 0.99937574 |
| SLC25A5 | 200.28457 | 0.7701327 | 0.50524741 | 1.52426848 | 0.12744166 | 0.99937574 |
| UBE2A | 37.4941107 | 0.1020098 | 0.45447998 | 0.22445388 | 0.82240415 | 0.99937574 |
| NKRF | 12.3504128 | -0.0205704 | 0.5212507 | -0.0394635 | 0.96852085 | 0.99937574 |
| 6-Sep | 132.846853 | 0.31996961 | 0.52488135 | 0.6096037 | 0.54212436 | 0.99937574 |
| UPF3B | 32.0372089 | -0.9404778 | 0.44737227 | -2.1022264 | 0.03553344 | 0.99937574 |
| NDUFA1 | 107.316655 | 0.37158979 | 0.45774463 | 0.81178406 | 0.41691555 | 0.99937574 |
| NKAP | 19.5822894 | 0.09777924 | 0.48130255 | 0.20315546 | 0.83901352 | 0.99937574 |
| RHOXF1-AS1 | 23.3476297 | 0.05471682 | 0.45292046 | 0.1208089 | 0.9038424 | 0.99937574 |
| ZBTB33 | 12.2518288 | -0.1966173 | 0.57113194 | -0.344259 | 0.73065154 | 0.99937574 |
| TMEM255A | 53.5030762 | -0.0426039 | 1.5299631 | -0.0278463 | 0.97778471 | 0.99937574 |
| LAMP2 | 104.266263 | -0.1140803 | 0.30135265 | -0.3785609 | 0.70501393 | 0.99937574 |
| CUL4B | 43.0399138 | 0.2063586 | 0.32136434 | 0.64213286 | 0.52078692 | 0.99937574 |
| MCTS1 | 30.9984616 | 0.10478093 | 0.38509556 | 0.27209072 | 0.78555226 | 0.99937574 |
| C1GALT1C1 | 15.2634286 | 0.86943224 | 0.46894102 | 1.85403323 | 0.06373441 | 0.99937574 |
| THOC2 | 144.136144 | -0.137441 | 0.24874751 | -0.5525322 | 0.58058378 | 0.99937574 |
| XIAP | 150.280056 | -0.2952473 | 0.21984442 | -1.3429828 | 0.17927755 | 0.99937574 |
| STAG2 | 174.548928 | -0.1228304 | 0.27649949 | -0.4442339 | 0.65687349 | 0.99937574 |

| | | | | | | |
|---|---|---|---|---|---|---|
| AL442647.1 | 18.8811068 | 1.45211775 | 0.79965075 | 1.81593995 | 0.06937958 | 0.99937574 |
| SMARCA1 | 141.433563 | 0.48494082 | 0.39724675 | 1.22075463 | 0.22217894 | 0.99937574 |
| OCRL | 31.4954247 | -0.0269656 | 0.35587487 | -0.0757727 | 0.9395999 | 0.99937574 |
| ZDHHC9 | 83.5615067 | -0.5626105 | 0.31702452 | -1.7746593 | 0.07595416 | 0.99937574 |
| UTP14A | 15.0945764 | -0.5142132 | 0.53095556 | -0.9684676 | 0.3328109 | 0.99937574 |
| BCORL1 | 15.2653902 | 0.00431284 | 0.54310425 | 0.00794109 | 0.99366399 | 0.99937574 |
| ELF4 | 17.205281 | -0.2156773 | 0.52785514 | -0.4085919 | 0.68283921 | 0.99937574 |
| AIFM1 | 62.0839554 | 0.46936753 | 0.3703222 | 1.26745716 | 0.20499187 | 0.99937574 |
| ZNF280C | 17.0703442 | 0.25147686 | 0.48312344 | 0.52052299 | 0.60269911 | 0.99937574 |
| RBMX2 | 12.5516849 | -0.2352567 | 0.5314548 | -0.4426655 | 0.65800767 | 0.99937574 |
| ENOX2 | 122.086632 | -0.3322512 | 0.44198914 | -0.751718 | 0.45222067 | 0.99937574 |
| FRMD7 | 15.3277983 | 0.38307178 | 1.05513934 | 0.36305326 | 0.7165651 | 0.99937574 |
| RAP2C | 77.7066965 | 0.31901095 | 0.41932955 | 0.7607643 | 0.44679786 | 0.99937574 |
| RAP2C-AS1 | 73.4715108 | 0.48519125 | 0.4661531 | 1.04084099 | 0.29794935 | 0.99937574 |
| MBNL3 | 54.3267585 | -0.2661003 | 0.40374801 | -0.6590752 | 0.50984746 | 0.99937574 |
| GPC4 | 16.5460887 | 0.59349307 | 0.54867635 | 1.08168151 | 0.27939407 | 0.99937574 |
| GPC3 | 20.4284728 | -0.0375583 | 0.73280202 | -0.0512529 | 9.59E-01 | 0.99937574 |
| PHF6 | 38.9455509 | 0.22507463 | 0.38393205 | 0.58623557 | 0.55771722 | 0.99937574 |
| HPRT1 | 127.427273 | 0.41447723 | 0.23435844 | 1.76856114 | 0.07696714 | 0.99937574 |
| FAM122B | 19.1216081 | -0.194755 | 0.45803919 | -0.425193 | 0.67069601 | 0.99937574 |
| FAM122C | 20.244199 | -0.668269 | 0.70470566 | -0.9482952 | 0.3429792 | 0.99937574 |
| MOSPD1 | 11.5244297 | -0.348173 | 0.58541526 | -0.5947453 | 0.55201374 | 0.99937574 |
| RTL8C | 37.4303936 | 0.22324356 | 0.36795704 | 0.60671094 | 0.54404276 | 0.99937574 |
| RTL8A | 23.5654269 | -0.2080992 | 0.40294164 | -0.51645 | 0.60554017 | 0.99937574 |
| ZNF75D | 66.4272805 | -0.0342633 | 0.30009276 | -0.1141757 | 0.90909853 | 0.99937574 |
| INTS6L | 13.851907 | 0.0437367 | 0.61726961 | 0.0708551 | 0.94351308 | 0.99937574 |
| MMGT1 | 26.8789889 | 0.39800012 | 0.35716822 | 1.11432119 | 0.26514142 | 0.99937574 |
| SLC9A6 | 19.6737172 | 0.35255304 | 0.42376426 | 0.83195559 | 0.40543401 | 0.99937574 |
| HTATSF1 | 40.7416337 | 0.07454637 | 0.54484978 | 0.13682003 | 0.89117305 | 0.99937574 |
| ARHGEF6 | 34.4137755 | 0.43061912 | 0.43316019 | 0.99413365 | 0.3201578 | 0.99937574 |
| AL683813.1 | 15.2191631 | 0.27488686 | 0.73688099 | 0.37304105 | 0.70911789 | 0.99937574 |
| RBMX | 124.36706 | -0.2580609 | 0.2892185 | -0.8922697 | 0.37224841 | 0.99937574 |
| FGF13 | 338.873609 | 0.13814498 | 0.76769847 | 0.17994692 | 0.85719424 | 0.99937574 |
| ATP11C | 53.1200872 | -0.9700163 | 0.60753337 | -1.5966469 | 0.11034443 | 0.99937574 |
| FMR1 | 71.2062604 | 0.05731981 | 0.37218779 | 0.15400776 | 0.87760362 | 0.99937574 |
| IDS | 48.5226177 | -0.1493393 | 0.34597293 | -0.4316502 | 0.66599568 | 0.99937574 |
| TMEM185A | 19.5394879 | 0.47217167 | 0.46467107 | 1.01614172 | 0.3095619 | 0.99937574 |
| LINC00894 | 117.625694 | 0.39664854 | 0.57604594 | 0.68857102 | 0.49109327 | 0.99937574 |
| MTM1 | 68.5366552 | -0.4143448 | 0.4609355 | -0.8989215 | 0.36869449 | 0.99937574 |
| MTMR1 | 73.6820337 | -0.2065453 | 0.28035723 | -0.7367217 | 0.4612916 | 0.99937574 |
| CD99L2 | 53.6321063 | -0.2249468 | 0.36261662 | -0.6203434 | 0.53503175 | 0.99937574 |
| HMGB3 | 16.3854624 | -0.2602681 | 0.46604984 | -0.5584556 | 0.57653332 | 0.99937574 |
| VMA21 | 30.7766123 | -0.2757946 | 0.43471028 | -0.634433 | 0.52579826 | 0.99937574 |
| CETN2 | 12.0523198 | 0.62823562 | 0.70694286 | 0.88866534 | 0.37418296 | 0.99937574 |
| CCNQ | 12.416022 | -0.093149 | 0.58315595 | -0.1597326 | 0.87309176 | 0.99937574 |
| SLC6A8 | 31.8121411 | -0.029962 | 0.81270409 | -0.0368671 | 0.970591 | 0.99937574 |
| BCAP31 | 66.7418526 | -0.1999106 | 0.47466184 | -0.4211643 | 0.67363509 | 0.99937574 |
| IDH3G | 21.5420619 | 0.13229888 | 0.5259039 | 0.25156475 | 0.80137751 | 0.99937574 |
| SSR4 | 46.7411497 | 0.16583127 | 0.54289418 | 0.30545781 | 0.7600175 | 0.99937574 |
| HCFC1 | 21.6147615 | -0.3381294 | 0.48804234 | -0.6928281 | 0.48841746 | 0.99937574 |
| IRAK1 | 18.8402453 | -0.7121009 | 0.8023594 | -0.8875086 | 0.37480514 | 0.99937574 |
| MECP2 | 142.765539 | -0.0242536 | 0.30066629 | -0.0806662 | 0.93570739 | 0.99937574 |
| EMD | 35.9791299 | 0.10619152 | 0.37061171 | 0.28653038 | 0.77447192 | 0.99937574 |
| RPL10 | 379.240722 | 0.12025423 | 0.61066101 | 0.19692469 | 0.84388647 | 0.99937574 |

| | | | | | | |
|---|---|---|---|---|---|---|
| DNASE1L1 | 17.4006089 | -0.420156 | 0.47601797 | -0.8826473 | 0.37742684 | 0.99937574 |
| TAZ | 25.6691545 | 0.73537648 | 0.46144971 | 1.59362215 | 0.11102069 | 0.99937574 |
| ATP6AP1 | 128.533398 | 0.79053476 | 0.39016649 | 2.02614724 | 0.0427497 | 0.99937574 |
| GDI1 | 37.2628309 | -0.1088189 | 0.43670914 | -0.2491794 | 0.80322201 | 0.99937574 |
| FAM50A | 36.8091032 | -0.6361781 | 0.55323188 | -1.1499303 | 0.25017259 | 0.99937574 |
| PLXNA3 | 17.5026517 | -0.5997343 | 0.58867719 | -1.0187829 | 0.30830603 | 0.99937574 |
| UBL4A | 13.0760619 | -0.6830363 | 0.62457264 | -1.0936059 | 0.27412784 | 0.99937574 |
| FAM3A | 26.9643231 | 0.12847819 | 0.41749253 | 0.3077377 | 0.75828193 | 0.99937574 |
| IKBKG | 13.5693305 | -0.3867547 | 0.59031398 | -0.6551678 | 0.5123597 | 0.99937574 |
| DKC1 | 18.278295 | 0.00201497 | 0.48243473 | 0.00417667 | 0.99666751 | 0.99937574 |
| MPP1 | 24.6077002 | 0.23293389 | 0.50644389 | 0.45994017 | 0.64555916 | 0.99937574 |
| FUNDC2 | 51.0414185 | -0.0269005 | 0.47355571 | -0.0568054 | 0.95470018 | 0.99937574 |
| BRCC3 | 35.9886685 | -0.3375808 | 0.36846955 | -0.91617 | 0.35957772 | 0.99937574 |
| VBP1 | 28.3432064 | -0.1605842 | 0.45733059 | -0.3511338 | 0.72548797 | 0.99937574 |
| TMLHE-AS1 | 62.6139731 | 0.42364564 | 0.79026365 | 0.53608139 | 0.5919023 | 0.99937574 |
| TMLHE | 149.861964 | 0.3660462 | 0.3506671 | 1.04385671 | 0.29655169 | 0.99937574 |
| VAMP7 | 43.7692549 | -0.3426927 | 0.40049197 | -0.8556793 | 0.39217518 | 0.99937574 |
| ZNF596 | 17.1717515 | -0.5394228 | 0.49078756 | -1.0990962 | 0.2717261 | 0.99937574 |
| FBXO25 | 60.4572094 | -0.2016671 | 0.32688241 | -0.6169407 | 0.53727385 | 0.99937574 |
| TDRP | 61.9402108 | -0.0185205 | 0.46285055 | -0.0400139 | 0.96808202 | 0.99937574 |
| ERICH1 | 102.851393 | -0.9015047 | 0.40844861 | -2.2071434 | 0.02730404 | 0.98755826 |
| AC100810.1 | 16.3160404 | 0.52864668 | 0.61778422 | 0.85571412 | 0.39215593 | 0.99937574 |
| AC246817.1 | 17.5199628 | 0.64422531 | 0.72055378 | 0.89406971 | 0.3712846 | 0.99937574 |
| CSMD1 | 52.6483055 | 0.11323427 | 1.0431387 | 0.1085515 | 0.91355823 | 0.99937574 |
| MCPH1 | 180.245427 | -0.0565971 | 0.36713429 | -0.1541591 | 0.8774843 | 0.99937574 |
| ANGPT2 | 35.2076326 | 0.7194247 | 0.83227212 | 0.86441043 | 0.38736246 | 0.99937574 |
| MCPH1-AS1 | 90.1671238 | 0.12749172 | 0.64867091 | 0.19654299 | 0.84418519 | 0.99937574 |
| AGPAT5 | 121.229813 | -0.6462855 | 0.31546617 | -2.0486683 | 0.04049456 | 0.99937574 |
| AF233439.1 | 101.486476 | -0.1285166 | 0.92978396 | -0.138222 | 8.90E-01 | 0.99937574 |
| DEFB1 | 416.205538 | 0.72546618 | 0.77454216 | 0.93663873 | 0.34894442 | 0.99937574 |
| FAM85B | 58.6693394 | 0.3925888 | 0.56865663 | 0.69037936 | 0.48995565 | 0.99937574 |
| MFHAS1 | 46.724534 | -0.2586619 | 0.7235892 | -0.3574706 | 7.21E-01 | 0.99937574 |
| ERI1 | 31.8106898 | 0.06042072 | 0.54617792 | 0.11062461 | 0.91191403 | 0.99937574 |
| TNKS | 258.020332 | -0.0809758 | 0.30213353 | -0.2680132 | 0.78868917 | 0.99937574 |
| MSRA | 108.827452 | -0.3117823 | 0.4095637 | -0.7612548 | 0.44650488 | 0.99937574 |
| XKR6 | 209.397289 | 0.0226609 | 0.52085665 | 0.04350698 | 0.9652974 | 0.99937574 |
| MTMR9 | 49.5350221 | 0.0668887 | 0.34748418 | 0.19249423 | 0.84735509 | 0.99937574 |
| FDFT1 | 168.335217 | 0.27361397 | 0.71242441 | 0.38406036 | 0.70093371 | 0.99937574 |
| CTSB | 123.907279 | -0.4864352 | 0.60023477 | -0.8104083 | 0.41770556 | 0.99937574 |
| FAM66D | 15.8966353 | 1.10715716 | 0.85753909 | 1.29108653 | 0.19667368 | 0.99937574 |
| LONRF1 | 47.2391096 | -1.2796119 | 0.3185963 | -4.0164054 | 5.91E-05 | 0.08437103 |
| TRMT9B | 156.316901 | 1.01064103 | 0.50100256 | 2.01723724 | 0.04367076 | 0.99937574 |
| DLC1 | 136.552229 | -0.316131 | 0.6406564 | -0.4934486 | 6.22E-01 | 0.99937574 |
| TUSC3 | 64.1549003 | 0.29388592 | 0.59339868 | 0.49525881 | 0.62041744 | 0.99937574 |
| MICU3 | 62.7438644 | -0.4281537 | 0.33956257 | -1.2608979 | 0.20734563 | 0.99937574 |
| ZDHHC2 | 87.4268652 | -0.0272318 | 0.35126673 | -0.0775245 | 0.93820631 | 0.99937574 |
| CNOT7 | 30.792243 | -0.0490837 | 0.46021995 | -0.1066528 | 0.91506443 | 0.99937574 |
| VPS37A | 57.0634228 | -0.4242861 | 0.35440826 | -1.1971676 | 0.23124122 | 0.99937574 |
| MTUS1 | 320.61795 | -0.4318958 | 0.28298859 | -1.526195 | 0.1269613 | 0.99937574 |
| AC027117.2 | 26.3226687 | -1.7111947 | 0.94966677 | -1.8018897 | 0.07156277 | 0.99937574 |
| PCM1 | 264.794277 | -0.2978779 | 0.26814547 | -1.1108817 | 0.26661927 | 0.99937574 |
| ASAH1 | 72.0656973 | -0.001511 | 0.42799025 | -0.0035305 | 0.9971831 | 0.99937574 |
| PSD3 | 1463.62212 | 0.04041775 | 0.58831715 | 0.06870061 | 0.94522793 | 0.99937574 |
| AC100800.1 | 18.8824355 | 0.78940072 | 0.6996392 | 1.12829687 | 0.25919456 | 0.99937574 |

| | | | | | | |
|---|---|---|---|---|---|---|
| SH2D4A | 193.488195 | -0.1066538 | 0.49694626 | -0.2146183 | 0.83006494 | 0.99937574 |
| CSGALNACT1 | 128.644598 | -0.7428271 | 0.69220391 | -1.0731333 | 0.2832113 | 0.99937574 |
| INTS10 | 96.171854 | 0.40056825 | 0.29650865 | 1.35094965 | 0.17671156 | 0.99937574 |
| ATP6V1B2 | 27.2715302 | -0.4113057 | 0.57632763 | -0.7136665 | 0.47543342 | 0.99937574 |
| XPO7 | 66.2445583 | 0.00544989 | 0.27375317 | 0.01990803 | 0.98411674 | 0.99937574 |
| DMTN | 75.0172378 | -0.2077456 | 0.28675444 | -0.724472 | 0.46877602 | 0.99937574 |
| FAM160B2 | 32.5907157 | 0.05784886 | 0.42688298 | 0.13551456 | 0.89220506 | 0.99937574 |
| SLC39A14 | 28.8408887 | 0.08849357 | 0.4467335 | 0.19809029 | 0.84297442 | 0.99937574 |
| PPP3CC | 141.723442 | -0.1065751 | 0.30643207 | -0.3477936 | 0.7279952 | 0.99937574 |
| CCAR2 | 18.6317657 | -0.0109678 | 0.5098026 | -0.0215138 | 0.98283582 | 0.99937574 |
| BIN3 | 40.8139704 | -0.2889647 | 0.43932143 | -0.6577523 | 0.51069731 | 0.99937574 |
| TNFRSF10B | 77.205804 | -0.0494934 | 0.28421569 | -0.1741402 | 0.86175531 | 0.99937574 |
| TNFRSF10D | 22.950076 | -0.257026 | 0.49230331 | -0.5220887 | 0.60160855 | 0.99937574 |
| TNFRSF10A | 18.7651318 | -0.5849169 | 0.68984299 | -0.8478986 | 0.39649446 | 0.99937574 |
| CHMP7 | 19.4260557 | 0.03933297 | 0.47365378 | 0.0830416 | 0.93381846 | 0.99937574 |
| R3HCC1 | 10.5958227 | -0.4213259 | 0.55040722 | -0.7654803 | 0.44398558 | 0.99937574 |
| ENTPD4 | 52.3392062 | 0.1951947 | 0.34464522 | 0.56636416 | 0.57114626 | 0.99937574 |
| SLC25A37 | 79.9473569 | 0.81254239 | 0.55637769 | 1.46041513 | 0.14417602 | 0.99937574 |
| DOCK5 | 268.986903 | -0.4600131 | 0.49317932 | -0.9327501 | 3.51E-01 | 0.99937574 |
| KCTD9 | 61.6748635 | -0.3504738 | 0.37790561 | -0.927411 | 0.35371318 | 0.99937574 |
| EBF2 | 115.558457 | -0.6368651 | 0.61394551 | -1.0373316 | 0.29958135 | 0.99937574 |
| PPP2R2A | 197.168296 | -0.1912634 | 0.31469224 | -0.6077791 | 0.54333398 | 0.99937574 |
| BNIP3L | 78.9544341 | -0.0950062 | 0.60920114 | -0.1559522 | 0.87607073 | 0.99937574 |
| TRIM35 | 16.6705852 | -0.9972717 | 0.49068684 | -2.0323996 | 0.04211322 | 0.99937574 |
| PTK2B | 123.06902 | 0.05676241 | 0.33785465 | 0.16800836 | 0.8665767 | 0.99937574 |
| EPHX2 | 37.3858368 | 0.49009672 | 0.60985662 | 0.80362615 | 0.42161292 | 0.99937574 |
| CLU | 115.117442 | -0.2568545 | 0.50036655 | -0.5133326 | 0.60771867 | 0.99937574 |
| CCDC25 | 33.9089478 | -0.2655808 | 0.38087362 | -0.6972938 | 0.48561897 | 0.99937574 |
| ELP3 | 276.891191 | -1.0481018 | 0.76093027 | -1.3773953 | 0.16839006 | 0.99937574 |
| ZNF395 | 26.1420173 | -0.3335947 | 0.51185839 | -0.6517324 | 0.51457385 | 0.99937574 |
| FBXO16 | 22.183318 | 0.506408 | 0.49671896 | 1.01950609 | 0.30796276 | 0.99937574 |
| FZD3 | 34.6360319 | 1.22664146 | 0.44779611 | 2.73928563 | 0.00615729 | 0.69043694 |
| EXTL3 | 26.511417 | -0.0445845 | 0.6261349 | -0.0712058 | 0.94323394 | 0.99937574 |
| INTS9 | 115.113419 | 0.2564978 | 0.28254977 | 0.90779688 | 0.36398555 | 0.99937574 |
| HMBOX1 | 323.99639 | 0.03237393 | 0.26166178 | 0.12372432 | 0.90153355 | 0.99937574 |
| KIF13B | 412.522896 | -1.2689909 | 0.4785519 | -2.651731 | 0.00800803 | 0.74742128 |
| AC108449.1 | 24.2596582 | -1.248927 | 0.92129476 | -1.3556215 | 0.17521962 | 0.99937574 |
| SARAF | 123.782852 | 0.01799073 | 0.34488907 | 0.05216382 | 0.95839816 | 0.99937574 |
| LEPROTL1 | 38.539807 | 1.01958866 | 0.3363477 | 3.03135319 | 0.0024346 | 0.48965449 |
| DCTN6 | 22.3000937 | -0.2339701 | 0.43758336 | -0.5346868 | 0.59286643 | 0.99937574 |
| RBPMS | 1589.54732 | 0.11846563 | 0.36056992 | 0.32855105 | 0.74249505 | 0.99937574 |
| GTF2E2 | 73.5676695 | -0.0536036 | 0.32148274 | -0.1667387 | 0.86757564 | 0.99937574 |
| GSR | 102.640321 | -0.0232539 | 0.35985411 | -0.0646203 | 0.94847628 | 0.99937574 |
| UBXN8 | 12.7815777 | 0.69262501 | 0.52795909 | 1.31189144 | 0.18955677 | 0.99937574 |
| PPP2CB | 139.713098 | 0.029462 | 0.26703895 | 0.11032846 | 0.91214889 | 0.99937574 |
| WRN | 106.414396 | -0.1999318 | 0.30326675 | -0.6592606 | 0.50972846 | 0.99937574 |
| NRG1 | 141.184682 | -1.1428948 | 1.21909548 | -0.9374941 | 0.34850445 | 0.99937574 |
| FUT10 | 51.1163393 | 0.73830285 | 0.34854427 | 2.11824699 | 0.03415416 | 0.99937574 |
| TTI2 | 22.0465787 | -0.2751154 | 0.48509598 | -0.5671361 | 0.57062174 | 0.99937574 |
| MAK16 | 11.4164446 | -0.7575964 | 0.58186092 | -1.3020231 | 0.19290849 | 0.99937574 |
| RNF122 | 15.6537424 | 0.2680469 | 0.68144769 | 0.39334919 | 0.69406159 | 0.99937574 |
| UNC5D | 35.0356178 | -0.1634349 | 0.39204827 | -0.4168744 | 0.67677026 | 0.99937574 |
| ERLIN2 | 37.6503684 | 0.10157041 | 0.387261 | 0.26227897 | 0.79310638 | 0.99937574 |
| PLPBP | 32.6982612 | 0.10562215 | 0.42356832 | 0.24936271 | 0.80308022 | 0.99937574 |

| | | | | | | |
|---|---|---|---|---|---|---|
| RAB11FIP1 | 81.3328717 | -0.2579147 | 0.31788981 | -0.8113337 | 0.41717409 | 0.99937574 |
| ASH2L | 42.1255112 | -0.535151 | 0.34529377 | -1.5498425 | 0.12117932 | 0.99937574 |
| LSM1 | 27.7935489 | 0.17032829 | 0.49477696 | 0.34425267 | 0.73065627 | 0.99937574 |
| BAG4 | 52.0861274 | 0.1538295 | 0.29733795 | 0.51735576 | 0.60490784 | 0.99937574 |
| DDHD2 | 64.9071687 | -0.0636691 | 0.31379041 | -0.2029034 | 0.83921054 | 0.99937574 |
| PLPP5 | 46.6047955 | -0.502692 | 0.45311926 | -1.1094032 | 0.26725627 | 0.99937574 |
| NSD3 | 192.978571 | 0.03381236 | 0.26784377 | 0.12623913 | 0.89954264 | 0.99937574 |
| FGFR1 | 77.5738881 | -0.2153738 | 0.73572433 | -0.2927371 | 0.76972315 | 0.99937574 |
| TACC1 | 124.530463 | -1.7606345 | 0.62800835 | -2.8035208 | 0.0050548 | 0.61461453 |
| PLEKHA2 | 59.7883774 | 0.01781878 | 0.42729684 | 0.04170118 | 0.96673691 | 0.99937574 |
| TM2D2 | 18.8707405 | 0.64588872 | 0.48481325 | 1.3322423 | 0.18278058 | 0.99937574 |
| ADAM9 | 93.2817002 | 0.10688716 | 0.29829411 | 0.35832809 | 0.7200978 | 0.99937574 |
| TCIM | 162.691719 | 0.69446307 | 0.79051526 | 0.87849419 | 0.37967559 | 0.99937574 |
| GOLGA7 | 23.5076691 | -0.7441907 | 0.42029454 | -1.7706408 | 0.07662045 | 0.99937574 |
| GPAT4 | 54.1106307 | -0.6244421 | 0.50876889 | -1.2273591 | 0.21968764 | 0.99937574 |
| ANK1 | 18.153455 | -1.147575 | 0.78101986 | -1.4693289 | 1.42E-01 | 0.99937574 |
| KAT6A | 223.107455 | 0.15502754 | 0.27212627 | 0.56968974 | 0.56888815 | 0.99937574 |
| AP3M2 | 18.6116754 | 0.57058416 | 0.59429229 | 0.96010696 | 0.33700138 | 0.99937574 |
| IKBKB | 79.4704052 | -0.0509734 | 0.32756796 | -0.1556116 | 0.87633916 | 0.99937574 |
| POLB | 21.6855565 | -0.4549391 | 0.6112625 | -0.7442614 | 0.45671835 | 0.99937574 |
| VDAC3 | 79.7086365 | 0.39757624 | 0.47572832 | 0.8357212 | 0.40331177 | 0.99937574 |
| SLC20A2 | 174.414094 | -0.5804226 | 0.28565456 | -2.0319039 | 0.04216338 | 0.99937574 |
| SMIM19 | 36.2020341 | -0.2562128 | 0.50650168 | -0.5058478 | 0.61296347 | 0.99937574 |
| RNF170 | 47.935323 | 0.14463035 | 0.43239692 | 0.33448516 | 0.73801349 | 0.99937574 |
| HOOK3 | 93.3886122 | 0.05237737 | 0.2782267 | 0.18825429 | 0.85067731 | 0.99937574 |
| FNTA | 33.7630173 | 0.26913262 | 0.49289531 | 0.54602389 | 0.58504951 | 0.99937574 |
| HGSNAT | 114.44584 | -0.3538104 | 0.27319031 | -1.295106 | 0.19528365 | 0.99937574 |
| SPIDR | 375.067507 | -0.4032446 | 0.32078609 | -1.2570514 | 0.20873502 | 0.99937574 |
| CEBPD | 285.414235 | -0.2121725 | 0.62778032 | -0.3379725 | 0.73538391 | 0.99937574 |
| PRKDC | 109.260993 | -0.1451106 | 0.37412271 | -0.387869 | 0.69811297 | 0.99937574 |
| UBE2V2 | 51.5702277 | 0.1280268 | 0.39652384 | 0.32287291 | 0.7467915 | 0.99937574 |
| PCMTD1 | 216.957022 | 0.01624426 | 0.25215725 | 0.06442115 | 0.94863489 | 0.99937574 |
| AC064807.1 | 12.3411628 | 0.17311466 | 0.66839622 | 0.25900005 | 0.7956352 | 0.99937574 |
| RB1CC1 | 175.138298 | -0.5472687 | 0.34241488 | -1.598262 | 0.10998469 | 0.99937574 |
| ATP6V1H | 333.750914 | 0.24565489 | 0.33597191 | 0.73117688 | 0.46467112 | 0.99937574 |
| TCEA1 | 66.717932 | -0.1400904 | 0.33147316 | -0.4226298 | 0.67256535 | 0.99937574 |
| LYPLA1 | 173.281532 | 0.11626395 | 0.23165698 | 0.50187977 | 0.6157521 | 0.99937574 |
| MRPL15 | 44.6624382 | -0.2107888 | 0.37585405 | -0.5608263 | 0.57491599 | 0.99937574 |
| TMEM68 | 36.4627052 | -0.2063218 | 0.43522155 | -0.4740615 | 0.63545605 | 0.99937574 |
| TGS1 | 55.4234299 | -0.79141 | 0.47114413 | -1.6797619 | 0.09300365 | 0.99937574 |
| LYN | 46.5111154 | 0.37810502 | 0.48064321 | 0.78666465 | 0.4314782 | 0.99937574 |
| RPS20 | 130.252492 | -0.3026906 | 0.5407529 | -0.5597577 | 0.57564475 | 0.99937574 |
| PLAG1 | 84.6608958 | 0.53604277 | 0.47620918 | 1.12564562 | 0.26031555 | 0.99937574 |
| CHCHD7 | 28.6409143 | -0.0657918 | 0.49154065 | -0.1338481 | 0.89352272 | 0.99937574 |
| IMPAD1 | 64.1030212 | 0.07551576 | 0.35622104 | 0.21199128 | 0.83211383 | 0.99937574 |
| LINC01606 | 77.8502364 | -0.1084615 | 0.61742269 | -0.1756682 | 0.86055465 | 0.99937574 |
| FAM110B | 191.484712 | 0.19518953 | 0.35669276 | 0.54722032 | 0.58422738 | 0.99937574 |
| AC027698.1 | 15.5854209 | -0.0991602 | 0.77458356 | -0.1280174 | 0.89813522 | 0.99937574 |
| UBXN2B | 55.5014372 | -0.3297823 | 0.33476077 | -0.9851283 | 0.32456106 | 0.99937574 |
| SDCBP | 71.690378 | -0.0741527 | 0.35719804 | -0.2075956 | 0.83554474 | 0.99937574 |
| NSMAF | 43.8726448 | 0.1601041 | 0.39798159 | 0.40229021 | 0.68747046 | 0.99937574 |
| TOX | 102.109945 | 2.40971244 | 0.72529179 | 3.32240415 | 0.00089245 | 0.29012345 |
| CA8 | 521.223365 | 0.13275133 | 0.75879682 | 0.17494977 | 0.8611191 | 0.99937574 |
| RAB2A | 192.191984 | -0.0925328 | 0.26536324 | -0.3487022 | 0.72731287 | 0.99937574 |

| | | | | | | |
|---|---|---|---|---|---|---|
| CHD7 | 77.6352476 | -0.5122591 | 0.4604709 | -1.1124679 | 0.26593702 | 0.99937574 |
| ASPH | 150.871161 | -0.2848073 | 0.45487338 | -0.6261243 | 0.53123343 | 0.99937574 |
| YTHDF3 | 126.125044 | -0.2887125 | 0.35226568 | -0.8195874 | 0.41245134 | 0.99937574 |
| CYP7B1 | 37.5336454 | 0.32446644 | 0.55743248 | 0.58207308 | 0.56051746 | 0.99937574 |
| ARMC1 | 34.4235176 | -0.3688322 | 0.37532543 | -0.9826996 | 0.3257553 | 0.99937574 |
| MTFR1 | 63.1171419 | 0.02697255 | 0.44191999 | 0.06103492 | 0.9513314 | 0.99937574 |
| PDE7A | 87.5886414 | 0.3772864 | 0.37603562 | 1.00332624 | 0.31570348 | 0.99937574 |
| ADHFE1 | 40.4390442 | 0.08753763 | 0.61416547 | 0.14253102 | 0.88666058 | 0.99937574 |
| VCPIP1 | 41.3834934 | -0.5683905 | 0.31791096 | -1.787892 | 0.07379343 | 0.99937574 |
| SGK3 | 151.491702 | 0.05353692 | 0.34590343 | 0.15477418 | 0.87699935 | 0.99937574 |
| CSPP1 | 112.559631 | 0.09395086 | 0.36143941 | 0.25993529 | 0.79491369 | 0.99937574 |
| ARFGEF1 | 410.861207 | -0.4054439 | 0.22364252 | -1.8129105 | 0.06984563 | 0.99937574 |
| NCOA2 | 668.03685 | 0.26909394 | 0.23777747 | 1.13170497 | 0.25775849 | 0.99937574 |
| TRAM1 | 128.208884 | -0.0810121 | 0.20909962 | -0.3874331 | 0.6984356 | 0.99937574 |
| LACTB2 | 38.8414801 | 0.26348881 | 0.40328714 | 0.65335286 | 0.51352882 | 0.99937574 |
| TERF1 | 35.4644502 | 0.43498827 | 0.34446314 | 1.26280061 | 0.20666085 | 0.99937574 |
| RPL7 | 219.280522 | 0.00652537 | 0.60369378 | 0.01080907 | 0.99137578 | 0.99937574 |
| RDH10 | 62.0398879 | -0.8471674 | 0.72340199 | -1.171088 | 0.24156341 | 0.99937574 |
| RDH10-AS1 | 25.6438193 | -0.4931883 | 0.83168319 | -0.5930002 | 0.553181 | 0.99937574 |
| STAU2 | 172.247499 | 0.51240876 | 0.26155492 | 1.95908672 | 0.05010263 | 0.99937574 |
| AC027018.1 | 30.2107824 | 0.2098305 | 0.86980656 | 0.24123813 | 0.80937056 | 0.99937574 |
| ELOC | 52.7635197 | -0.0967561 | 0.38382032 | -0.252087 | 0.80097383 | 0.99937574 |
| TMEM70 | 22.8120569 | 0.73517035 | 0.71278169 | 1.03141027 | 0.30234847 | 0.99937574 |
| JPH1 | 20.3767174 | 1.0681563 | 0.59518084 | 1.7946752 | 0.07270546 | 0.99937574 |
| GDAP1 | 39.3097846 | -0.3158348 | 0.63724696 | -0.4956238 | 0.62015985 | 0.99937574 |
| HNF4G | 51.4832233 | -2.3866999 | 0.80784886 | -2.9543892 | 0.00313289 | 0.52695149 |
| PEX2 | 50.2376886 | 0.55969825 | 0.3930865 | 1.42385517 | 0.1544884 | 0.99937574 |
| PKIA | 41.8764715 | 0.41019692 | 0.40259923 | 1.0188716 | 0.30826393 | 0.99937574 |
| ZC2HC1A | 79.1566676 | -0.4229762 | 0.37603591 | -1.1248294 | 0.26066134 | 0.99937574 |
| IL7 | 48.0280682 | 0.31940763 | 0.57699994 | 0.55356614 | 0.5798758 | 0.99937574 |
| TPD52 | 164.773605 | -0.1605161 | 0.34721611 | -0.4622946 | 0.64387005 | 0.99937574 |
| ZBTB10 | 52.4492497 | -0.1151573 | 0.37023137 | -0.3110415 | 0.75576906 | 0.99937574 |
| ZNF704 | 153.583537 | -0.0132557 | 0.4907294 | -0.0270122 | 0.97844997 | 0.99937574 |
| PAG1 | 25.4527523 | -0.9066905 | 0.83231172 | -1.089364 | 0.27599338 | 0.99937574 |
| IMPA1 | 43.5272487 | -0.4692624 | 0.35591805 | -1.3184564 | 0.18735091 | 0.99937574 |
| ZFAND1 | 49.812994 | -0.0239078 | 0.36209689 | -0.066026 | 0.94735715 | 0.99937574 |
| CHMP4C | 18.4681124 | -0.4158564 | 0.51797934 | -0.8028437 | 0.42206511 | 0.99937574 |
| SNX16 | 28.8376361 | -0.5476961 | 0.36324595 | -1.5077831 | 0.13161006 | 0.99937574 |
| LRRCC1 | 11.0027921 | -0.1935592 | 0.63251012 | -0.3060176 | 0.75959125 | 0.99937574 |
| E2F5 | 29.0019321 | 0.84001832 | 0.38275178 | 2.19468168 | 0.02818644 | 0.99049468 |
| C8orf59 | 34.1452419 | 0.09677399 | 0.46480006 | 0.20820564 | 0.83506841 | 0.99937574 |
| CA13 | 44.306518 | 0.38165454 | 0.37440028 | 1.01937569 | 0.30802465 | 0.99937574 |
| CA1 | 15.8808036 | 1.27610465 | 0.78431112 | 1.62703884 | 0.10372885 | 0.99937574 |
| CA3 | 46.7992828 | 0.82222411 | 0.46565005 | 1.76575545 | 0.07743688 | 0.99937574 |
| CA2 | 214.618002 | -0.4713049 | 0.59613545 | -0.7906003 | 0.42917727 | 0.99937574 |
| ATP6V0D2 | 1534.99309 | 0.37772265 | 0.41955984 | 0.90028314 | 0.36796959 | 0.99937574 |
| AC023194.3 | 255.238385 | 0.04235584 | 0.66541175 | 0.06365358 | 0.94924607 | 0.99937574 |
| WWP1 | 534.305048 | 0.13922045 | 0.24559873 | 0.56686143 | 0.57080833 | 0.99937574 |
| RMDN1 | 78.6165776 | 0.24800244 | 0.35517605 | 0.69825214 | 0.48501952 | 0.99937574 |
| CPNE3 | 71.3846297 | 0.44696131 | 0.28196134 | 1.58518649 | 0.11292397 | 0.99937574 |
| CNGB3 | 13.8967165 | 1.36203814 | 0.79190426 | 1.71995303 | 0.08544098 | 0.99937574 |
| AF117829.1 | 133.028192 | 0.17405735 | 0.36246341 | 0.48020668 | 0.63108044 | 0.99937574 |
| RIPK2 | 21.1835381 | -0.3555575 | 0.42884864 | -0.829098 | 0.40704898 | 0.99937574 |
| OSGIN2 | 30.88626 | -0.0336271 | 0.53985288 | -0.0622893 | 0.95033246 | 0.99937574 |

| | | | | | | |
|---|---|---|---|---|---|---|
| NBN | 40.8424286 | -0.8866362 | 0.37591855 | -2.3585859 | 0.01834471 | 0.93502451 |
| DECR1 | 140.440173 | 0.41925482 | 0.42016144 | 0.99784222 | 0.31835587 | 0.99937574 |
| CALB1 | 35.8704926 | 3.93602834 | 1.52878533 | 2.57461153 | 0.01003527 | 0.76274616 |
| PIP4P2 | 39.4481393 | -1.2645077 | 0.41000654 | -3.084116 | 0.00204158 | 0.45583974 |
| OTUD6B-AS1 | 51.8666044 | 0.39083192 | 0.45551141 | 0.85800688 | 0.39088866 | 0.99937574 |
| OTUD6B | 38.3467799 | 0.36943324 | 0.45979048 | 0.80348172 | 0.42169636 | 0.99937574 |
| LRRC69 | 49.2387987 | 1.0262696 | 0.68249952 | 1.50369278 | 0.13266049 | 0.99937574 |
| SLC26A7 | 3843.14886 | 1.92319013 | 0.83177655 | 2.31214757 | 0.02076955 | 0.96427895 |
| TRIQK | 56.1634694 | 0.52431114 | 0.40068393 | 1.30854048 | 0.19069005 | 0.99937574 |
| FAM92A | 15.379261 | 0.1484619 | 0.59198228 | 0.25078775 | 0.80197821 | 0.99937574 |
| RBM12B | 42.7317961 | -0.1879014 | 0.32819497 | -0.5725298 | 0.56696308 | 0.99937574 |
| TMEM67 | 18.8552844 | -0.1313987 | 0.47336649 | -0.2775834 | 0.78133219 | 0.99937574 |
| PDP1 | 73.2083823 | 0.17265082 | 0.45194412 | 0.38201806 | 0.70244797 | 0.99937574 |
| VIRMA | 57.129305 | -0.057469 | 0.29633849 | -0.1939301 | 0.84623058 | 0.99937574 |
| ESRP1 | 70.4507571 | -0.0752414 | 0.34450664 | -0.2184034 | 0.82711482 | 0.99937574 |
| DPY19L4 | 55.3830656 | -0.5756537 | 0.34749193 | -1.6565958 | 0.09760122 | 0.99937574 |
| INTS8 | 56.376031 | -0.2317227 | 0.44865219 | -0.5164863 | 0.60551481 | 0.99937574 |
| NDUFAF6 | 53.6898747 | 0.06431783 | 0.31278477 | 0.20562968 | 0.83708019 | 0.99937574 |
| TP53INP1 | 41.903961 | -0.0714431 | 0.39533801 | -0.1807139 | 0.85659212 | 0.99937574 |
| PLEKHF2 | 26.3912814 | -0.2461944 | 0.38516084 | -0.6391988 | 0.5226936 | 0.99937574 |
| C8orf37-AS1 | 35.7845263 | 1.15003024 | 0.55298918 | 2.07966139 | 0.0375566 | 0.99937574 |
| UQCRB | 172.169511 | 0.45782936 | 0.45425289 | 1.0078733 | 0.31351529 | 0.99937574 |
| MTERF3 | 18.1555192 | -0.2442922 | 0.45250046 | -0.5398717 | 0.58928548 | 0.99937574 |
| PTDSS1 | 18.9109825 | -0.0279777 | 0.45675283 | -0.0612535 | 0.95115729 | 0.99937574 |
| SDC2 | 220.834727 | 0.10463777 | 0.36185477 | 0.28917062 | 0.77245081 | 0.99937574 |
| CPQ | 339.329067 | 0.19556005 | 0.40028773 | 0.4885487 | 0.62516124 | 0.99937574 |
| TSPYL5 | 24.2101505 | -0.0821665 | 0.38814125 | -0.2116922 | 0.83234718 | 0.99937574 |
| MTDH | 209.604298 | -0.2951257 | 0.30750229 | -0.9597511 | 0.33718048 | 0.99937574 |
| LAPTM4B | 165.344159 | -0.4208274 | 0.42042295 | -1.000962 | 0.31684517 | 0.99937574 |
| RPL30 | 191.618218 | -0.323957 | 0.58716829 | -0.5517277 | 0.58113495 | 0.99937574 |
| RIDA | 12.8504679 | -0.0114236 | 0.73943909 | -0.015449 | 0.987674 | 0.99937574 |
| POP1 | 12.1572705 | -0.3695432 | 0.61146107 | -0.6043609 | 0.54560371 | 0.99937574 |
| NIPAL2 | 47.1145605 | -0.8144963 | 0.46383422 | -1.7560074 | 0.07908714 | 0.99937574 |
| STK3 | 820.504968 | 0.0822017 | 0.25904652 | 0.31732408 | 0.75099771 | 0.99937574 |
| AC104986.2 | 22.5449885 | 0.33951936 | 0.39752273 | 0.85408791 | 0.39305628 | 0.99937574 |
| VPS13B | 406.66571 | -0.068124 | 0.24860569 | -0.2740242 | 0.784066 | 0.99937574 |
| COX6C | 196.018179 | 0.03810834 | 0.3528145 | 0.1080124 | 0.91398585 | 0.99937574 |
| POLR2K | 32.3513768 | 0.02497519 | 0.56803217 | 0.04396792 | 0.96492998 | 0.99937574 |
| SPAG1 | 42.2791041 | 0.09848872 | 0.38950459 | 0.25285638 | 0.8003792 | 0.99937574 |
| RNF19A | 148.965634 | -0.2835316 | 0.34731221 | -0.8163595 | 0.41429453 | 0.99937574 |
| ANKRD46 | 30.458312 | 0.08111292 | 0.46455032 | 0.17460524 | 0.86138983 | 0.99937574 |
| PABPC1 | 148.103861 | -0.0856928 | 0.41502242 | -0.2064776 | 0.8364179 | 0.99937574 |
| YWHAZ | 254.9845 | 0.08801703 | 0.34206557 | 0.25731041 | 0.79693916 | 0.99937574 |
| ZNF706 | 63.6765538 | 0.26015269 | 0.4073731 | 0.63861038 | 0.52307642 | 0.99937574 |
| GRHL2 | 201.76053 | 0.02819154 | 0.43827696 | 0.06432358 | 0.94871258 | 0.99937574 |
| NCALD | 34.4905141 | 0.95679053 | 0.65142587 | 1.46876349 | 0.14189695 | 0.99937574 |
| RRM2B | 65.4856765 | 0.4169638 | 0.39096091 | 1.0665102 | 0.28619307 | 0.99937574 |
| UBR5-AS1 | 22.217675 | 0.75902073 | 0.69747287 | 1.08824411 | 0.27648736 | 0.99937574 |
| UBR5 | 306.883874 | -0.0318234 | 0.25521324 | -0.1246935 | 0.90076622 | 0.99937574 |
| KLF10 | 48.698619 | -0.2047991 | 0.52460114 | -0.39039 | 0.69624815 | 0.99937574 |
| AZIN1 | 85.4308792 | 0.42294655 | 0.35119187 | 1.20431759 | 0.22846685 | 0.99937574 |
| AZIN1-AS1 | 89.669206 | 0.07701957 | 0.46409209 | 0.16595752 | 0.86819038 | 0.99937574 |
| AP003354.2 | 14.4785235 | 0.96674043 | 0.59764702 | 1.61757758 | 0.10575366 | 0.99937574 |
| ATP6V1C1 | 79.5055918 | -0.5057989 | 0.30408877 | -1.6633263 | 0.09624714 | 0.99937574 |

| | | | | | | |
|---|---|---|---|---|---|---|
| BAALC | 89.0119144 | -0.7716304 | 0.75714828 | -1.0191272 | 0.30814259 | 0.99937574 |
| BAALC-AS1 | 32.9336919 | 0.35054733 | 0.43610592 | 0.80381237 | 0.42150535 | 0.99937574 |
| FZD6 | 107.206952 | -0.2525094 | 0.29572351 | -0.8538697 | 0.39317718 | 0.99937574 |
| SLC25A32 | 41.1328668 | 0.20082609 | 0.38368368 | 0.52341577 | 0.60068495 | 0.99937574 |
| DCAF13 | 21.2239604 | -0.5862077 | 0.50663183 | -1.1570684 | 0.24724443 | 0.99937574 |
| RIMS2 | 62.5353393 | 1.29529948 | 0.64034425 | 2.02281737 | 0.04309198 | 0.99937574 |
| LRP12 | 35.2613786 | 0.08124393 | 0.4382389 | 0.1853873 | 0.85292527 | 0.99937574 |
| ZFPM2 | 21.7553141 | 1.72135972 | 0.65088306 | 2.64465284 | 0.00817748 | 0.74742128 |
| AC027031.2 | 20.5261229 | -0.5934796 | 0.55335198 | -1.0725174 | 0.28348771 | 0.99937574 |
| OXR1 | 1151.32614 | -0.173171 | 0.38749063 | -0.4469037 | 0.65494456 | 0.99937574 |
| AC090579.1 | 110.758783 | 0.47623732 | 0.67547028 | 0.70504555 | 0.48078189 | 0.99937574 |
| ANGPT1 | 51.9406086 | 1.86651078 | 1.07309673 | 1.73936862 | 0.08196994 | 0.99937574 |
| EIF3E | 94.710159 | -0.2105503 | 0.25734105 | -0.818176 | 0.41325668 | 0.99937574 |
| EMC2 | 79.9538052 | 0.06346032 | 0.40197071 | 0.157873 | 0.87455687 | 0.99937574 |
| NUDCD1 | 33.4525578 | -0.1488071 | 0.41446223 | -0.3590365 | 0.71956782 | 0.99937574 |
| ENY2 | 50.6176141 | 0.07434281 | 0.45741391 | 0.16252853 | 0.87088967 | 0.99937574 |
| EBAG9 | 31.4649181 | -0.1277377 | 0.40952082 | -0.3119199 | 0.75510143 | 0.99937574 |
| TRPS1 | 330.106986 | -0.1073631 | 0.3165417 | -0.3391753 | 0.73447764 | 0.99937574 |
| EIF3H | 149.150389 | -0.1184378 | 0.25150671 | -0.470913 | 0.63770285 | 0.99937574 |
| UTP23 | 22.9860488 | 0.42905121 | 0.46698408 | 0.91877051 | 0.35821562 | 0.99937574 |
| RAD21 | 67.8684995 | 0.06506572 | 0.31451806 | 0.20687435 | 0.83610799 | 0.99937574 |
| MED30 | 15.5186991 | -0.1199939 | 0.56186916 | -0.2135621 | 0.83088859 | 0.99937574 |
| EXT1 | 456.825949 | 0.17730915 | 0.30937348 | 0.57312329 | 0.56656122 | 0.99937574 |
| SAMD12 | 695.659465 | -0.2965242 | 0.33821491 | -0.8767331 | 0.3806316 | 0.99937574 |
| AC023590.1 | 75.0127356 | -0.3658765 | 0.73752688 | -0.4960856 | 0.61983399 | 0.99937574 |
| SAMD12-AS1 | 49.6139028 | -0.8578063 | 0.54962525 | -1.5607113 | 0.11859189 | 0.99937574 |
| MAL2 | 71.8112962 | -0.7239264 | 0.47869082 | -1.5123047 | 0.13045637 | 0.99937574 |
| TAF2 | 89.4397792 | -0.442304 | 0.2853021 | -1.5503004 | 0.12106942 | 0.99937574 |
| DEPTOR | 135.927088 | -0.1818784 | 0.61619549 | -0.2951636 | 0.76786894 | 0.99937574 |
| MRPL13 | 46.7596481 | 0.20049243 | 0.39096251 | 0.51281754 | 0.60807896 | 0.99937574 |
| MTBP | 32.3584228 | 0.63615678 | 0.55124804 | 1.15403001 | 0.24848787 | 0.99937574 |
| SNTB1 | 1704.03353 | -0.2940539 | 0.34369785 | -0.8555594 | 0.39224152 | 0.99937574 |
| AC104958.1 | 21.7704771 | 0.60497682 | 0.83396796 | 0.72541973 | 0.46819457 | 0.99937574 |
| AC104958.2 | 18.0739148 | -0.2252839 | 0.49217785 | -0.4577286 | 6.47E-01 | 0.99937574 |
| ZHX2 | 215.843717 | -0.1051656 | 0.29911559 | -0.3515884 | 0.72514696 | 0.99937574 |
| AC104316.1 | 14.6571504 | 0.34855757 | 0.71930613 | 0.48457473 | 0.62797804 | 0.99937574 |
| DERL1 | 47.3922001 | 0.14736806 | 0.2909561 | 0.50649586 | 0.61250859 | 0.99937574 |
| TBC1D31 | 19.9857625 | 0.0131672 | 0.44918666 | 0.02931342 | 0.97661463 | 0.99937574 |
| C8orf76 | 15.928308 | 0.52961404 | 0.60092662 | 0.88132897 | 0.37813979 | 0.99937574 |
| ATAD2 | 20.7628068 | -0.6722761 | 0.56526049 | -1.1893208 | 0.23431346 | 0.99937574 |
| WDYHV1 | 24.9794673 | 0.55198399 | 0.49403659 | 1.11729374 | 0.26386875 | 0.99937574 |
| FBXO32 | 45.112587 | 0.12188721 | 0.81664167 | 0.14925421 | 0.88135305 | 0.99937574 |
| FAM91A1 | 99.8430361 | -0.183807 | 0.32962829 | -0.5576192 | 0.57710447 | 0.99937574 |
| FER1L6 | 93.8631865 | -1.256018 | 0.70319206 | -1.7861664 | 0.07407232 | 0.99937574 |
| TMEM65 | 317.22679 | 0.79006276 | 0.30143307 | 2.62102221 | 0.00876666 | 0.74742128 |
| RNF139 | 18.4774239 | 0.06001197 | 0.55503433 | 0.10812299 | 0.91389813 | 0.99937574 |
| TATDN1 | 60.6993893 | 0.20846602 | 0.34404347 | 0.60592928 | 0.54456171 | 0.99937574 |
| NDUFB9 | 92.4484396 | 0.24286346 | 0.43209467 | 0.56206077 | 0.57407461 | 0.99937574 |
| MTSS1 | 667.449224 | -0.728221 | 0.31843695 | -2.286861 | 2.22E-02 | 0.96427895 |
| SQLE | 66.8102785 | -0.2711596 | 0.79850991 | -0.3395821 | 0.73417129 | 0.99937574 |
| WASHC5 | 49.4153706 | -0.3836261 | 0.31944851 | -1.2009012 | 0.22978954 | 0.99937574 |
| NSMCE2 | 348.498493 | -0.3480949 | 0.30876144 | -1.127391 | 0.25957722 | 0.99937574 |
| TRIB1 | 54.4713853 | -0.5035682 | 0.51351892 | -0.9806225 | 0.32677892 | 0.99937574 |
| FAM84B | 23.0240238 | -0.3827516 | 0.54820035 | -0.6981967 | 0.48505422 | 0.99937574 |

| | | | | | | |
|---|---|---|---|---|---|---|
| PCAT1 | 205.238116 | -0.4492137 | 0.52818423 | -0.8504867 | 0.39505457 | 0.99937574 |
| FAM49B | 375.103583 | 0.33410658 | 0.25808288 | 1.29457087 | 0.19546831 | 0.99937574 |
| ASAP1 | 68.6255863 | -0.3590386 | 0.32768813 | -1.0956717 | 0.27322246 | 0.99937574 |
| EFR3A | 128.294721 | 0.0228181 | 0.34961166 | 0.06526697 | 0.94796144 | 0.99937574 |
| LRRC6 | 20.0974171 | 0.64978987 | 0.45829803 | 1.41783256 | 0.15623965 | 0.99937574 |
| PHF20L1 | 93.6463961 | -0.1152912 | 0.34694395 | -0.3323048 | 0.73965909 | 0.99937574 |
| NDRG1 | 286.704753 | 0.67473385 | 0.70971917 | 0.95070541 | 0.34175394 | 0.99937574 |
| ST3GAL1 | 332.96877 | 0.15608125 | 0.6485264 | 0.24067062 | 0.80981041 | 0.99937574 |
| ZFAT | 44.5608164 | -0.0842237 | 0.32913134 | -0.255897 | 0.7980304 | 0.99937574 |
| AC083843.3 | 34.3365826 | 0.72457635 | 0.46680351 | 1.55220845 | 0.12061235 | 0.99937574 |
| KHDRBS3 | 36.4531633 | -0.7284311 | 0.45486998 | -1.6014051 | 1.09E-01 | 0.99937574 |
| TRAPPC9 | 416.762054 | 0.18175166 | 0.26604845 | 0.68315251 | 0.49451048 | 0.99937574 |
| CHRAC1 | 23.5783433 | -0.5805599 | 0.47379392 | -1.2253426 | 0.22044617 | 0.99937574 |
| AGO2 | 105.831875 | -0.2106356 | 0.3914849 | -0.5380427 | 0.59054754 | 0.99937574 |
| PTK2 | 672.32205 | 0.03052981 | 0.26359681 | 0.1158201 | 0.90779512 | 0.99937574 |
| DENND3 | 38.5033463 | 0.13992855 | 0.56003948 | 0.2498548 | 0.80269964 | 0.99937574 |
| SLC45A4 | 62.6433011 | -0.0036147 | 0.4271411 | -0.0084624 | 0.99324803 | 0.99937574 |
| PTP4A3 | 78.7393298 | -0.4050193 | 0.86127411 | -0.470256 | 0.6381721 | 0.99937574 |
| TSNARE1 | 30.6661984 | -0.5613247 | 0.43610768 | -1.2871241 | 0.19805102 | 0.99937574 |
| JRK | 14.3447094 | -0.2799375 | 0.67055606 | -0.4174707 | 0.67633416 | 0.99937574 |
| THEM6 | 12.9084087 | 0.40954628 | 0.53591709 | 0.76419708 | 0.44474981 | 0.99937574 |
| LY6E | 51.5500053 | -1.2535999 | 1.68792558 | -0.7426867 | 0.45767137 | 0.99937574 |
| GLI4 | 13.3152103 | -0.369058 | 0.55691394 | -0.6626841 | 0.5075329 | 0.99937574 |
| TOP1MT | 11.3958415 | -0.0974976 | 0.55160081 | -0.176754 | 0.85970165 | 0.99937574 |
| RHPN1 | 22.2045139 | 0.46603768 | 0.54588595 | 0.85372719 | 0.39325617 | 0.99937574 |
| ZC3H3 | 31.9372714 | -0.0780174 | 0.5419622 | -0.1439537 | 0.88553706 | 0.99937574 |
| GSDMD | 30.8362658 | 0.2457701 | 0.47861792 | 0.51349959 | 0.6076019 | 0.99937574 |
| NAPRT | 11.9178965 | -0.1648862 | 0.78259671 | -0.2106911 | 0.83312829 | 0.99937574 |
| EEF1D | 160.76854 | 0.00419351 | 0.46031737 | 0.00911003 | 0.99273135 | 0.99937574 |
| TSTA3 | 12.4785426 | -0.4524056 | 0.74916349 | -0.603881 | 0.54592276 | 0.99937574 |
| ZNF623 | 16.5676022 | 0.40744955 | 0.46147014 | 0.88293806 | 0.37726974 | 0.99937574 |
| FAM83H | 21.6730713 | -0.0855494 | 0.59020238 | -0.1449493 | 0.8847509 | 0.99937574 |
| SCRIB | 27.1836548 | 0.00566155 | 0.4049072 | 0.01398233 | 0.98884408 | 0.99937574 |
| PUF60 | 34.0330992 | -0.287413 | 0.52913832 | -0.5431718 | 0.5870115 | 0.99937574 |
| NRBP2 | 19.7377696 | -0.1428241 | 0.4781546 | -0.2986986 | 0.76517005 | 0.99937574 |
| PLEC | 37.9846558 | 0.0962436 | 0.3389075 | 0.28398192 | 0.77642424 | 0.99937574 |
| PARP10 | 21.9424017 | -0.0247465 | 0.78979061 | -0.0313329 | 0.97500402 | 0.99937574 |
| GRINA | 39.7098343 | -0.0305446 | 0.53261318 | -0.0573485 | 0.95426756 | 0.99937574 |
| OPLAH | 14.2480101 | 0.40518064 | 0.62408925 | 0.64923508 | 0.51618644 | 0.99937574 |
| EXOSC4 | 13.2384886 | -0.2657565 | 0.59805317 | -0.4443694 | 0.65677554 | 0.99937574 |
| GPAA1 | 44.4430537 | 0.57140137 | 0.56949735 | 1.00334334 | 0.31569523 | 0.99937574 |
| CYC1 | 87.7506035 | 0.64435717 | 0.42383127 | 1.52031529 | 0.12843175 | 0.99937574 |
| SHARPIN | 27.2951732 | -0.1410072 | 0.49580258 | -0.284402 | 0.77610233 | 0.99937574 |
| MAF1 | 42.6132107 | -0.0589365 | 0.51204013 | -0.1151013 | 0.90836486 | 0.99937574 |
| MROH1 | 117.319934 | -0.7721945 | 0.37246396 | -2.0732059 | 0.03815312 | 0.99937574 |
| HSF1 | 70.816608 | -0.0623548 | 0.35635481 | -0.1749795 | 0.86109572 | 0.99937574 |
| DGAT1 | 18.1020633 | 0.963338 | 0.51401792 | 1.87413312 | 0.06091208 | 0.99937574 |
| SLC52A2 | 16.5351962 | -0.0135267 | 0.63543641 | -0.0212872 | 0.98301654 | 0.99937574 |
| ADCK5 | 9.75106023 | 0.75437344 | 0.78661247 | 0.95901535 | 0.33755101 | 0.99937574 |
| CPSF1 | 14.3847509 | 0.39863204 | 0.55233052 | 0.72172734 | 0.47046213 | 0.99937574 |
| SLC39A4 | 17.4298301 | 1.29893408 | 0.57720472 | 2.25038716 | 0.02442438 | 0.96546668 |
| VPS28 | 77.1336148 | 0.03588484 | 0.34366772 | 0.10441725 | 0.91683823 | 0.99937574 |
| CYHR1 | 44.1506798 | 0.42071551 | 0.32993026 | 1.27516496 | 0.20225086 | 0.99937574 |
| KIFC2 | 17.091023 | -0.2395751 | 0.57716638 | -0.4150884 | 0.67807718 | 0.99937574 |

| | | | | | | |
|---|---|---|---|---|---|---|
| PPP1R16A | 32.4530398 | -0.3062196 | 0.37819435 | -0.8096884 | 0.41811931 | 0.99937574 |
| MFSD3 | 9.14575894 | 1.68996267 | 0.7278172 | 2.32196032 | 0.02023507 | 0.96427895 |
| ZNF251 | 50.254187 | -0.3869953 | 0.40594106 | -0.9533288 | 0.3404235 | 0.99937574 |
| RPL8 | 372.596068 | -0.0239553 | 0.52012308 | -0.046057 | 0.96326483 | 0.99937574 |
| ZNF7 | 19.9922721 | -0.196473 | 0.46312145 | -0.4242364 | 0.67139345 | 0.99937574 |
| COMMD5 | 12.5454506 | -0.5789893 | 0.52226406 | -1.1086142 | 0.26759665 | 0.99937574 |
| ZNF250 | 18.8871522 | 0.34934455 | 0.50051331 | 0.69797255 | 0.48519436 | 0.99937574 |
| ZNF16 | 14.4764011 | -0.6453542 | 0.61601478 | -1.0476277 | 0.29481016 | 0.99937574 |
| C8orf33 | 16.1912616 | -0.3454994 | 0.60128766 | -0.5745992 | 0.56556239 | 0.99937574 |
| WASHC1 | 93.7380184 | 0.14716963 | 0.29364859 | 0.50117599 | 0.61624727 | 0.99937574 |
| CBWD1 | 29.1482295 | 0.35242202 | 0.45751628 | 0.77029395 | 0.44112554 | 0.99937574 |
| DOCK8 | 419.05929 | -0.0924424 | 0.37637546 | -0.2456121 | 0.80598254 | 0.99937574 |
| KANK1 | 224.795562 | 0.17502723 | 0.39791178 | 0.4398644 | 0.66003532 | 0.99937574 |
| LINC01230 | 135.932601 | 0.01872642 | 0.64384112 | 0.02908547 | 0.97679642 | 0.99937574 |
| DMRT2 | 393.336778 | 0.09019325 | 0.34349958 | 0.26257165 | 0.79288075 | 0.99937574 |
| SMARCA2 | 289.692451 | 0.54778576 | 0.29619438 | 1.84941308 | 0.06439819 | 0.99937574 |
| VLDLR-AS1 | 15.9010618 | -0.2197444 | 0.71888805 | -0.3056727 | 0.75985389 | 0.99937574 |
| VLDLR | 24.8775717 | -0.0034382 | 0.66415194 | -0.0051768 | 0.99586951 | 0.99937574 |
| PUM3 | 193.939263 | 0.56278282 | 0.37098116 | 1.51701187 | 0.12926369 | 0.99937574 |
| RFX3 | 147.847739 | 0.34518625 | 0.33343753 | 1.03523515 | 0.30055911 | 0.99937574 |
| RFX3-AS1 | 96.2418221 | 0.5180476 | 0.50611816 | 1.02357046 | 0.3060382 | 0.99937574 |
| GLIS3 | 1055.89435 | 0.19037555 | 0.41131324 | 0.4628481 | 0.64347325 | 0.99937574 |
| SLC1A1 | 17.8580085 | 0.82353797 | 0.72983317 | 1.12839209 | 0.25915437 | 0.99937574 |
| SPATA6L | 135.376232 | 0.38417143 | 0.68100293 | 0.56412597 | 0.57266841 | 0.99937574 |
| CDC37L1 | 52.4247405 | 0.07189752 | 0.34864133 | 0.20622202 | 0.83661749 | 0.99937574 |
| AK3 | 232.578282 | 0.26377207 | 0.35012431 | 0.75336689 | 0.45122947 | 0.99937574 |
| RCL1 | 49.1826411 | 0.29417016 | 0.33796186 | 0.87042413 | 0.38406867 | 0.99937574 |
| JAK2 | 120.86007 | 0.15565036 | 0.33781659 | 0.46075403 | 0.64497508 | 0.99937574 |
| PLGRKT | 62.4059174 | 0.38372126 | 0.34596151 | 1.10914437 | 0.26736791 | 0.99937574 |
| RIC1 | 202.032381 | -0.2286393 | 0.29996406 | -0.7622222 | 0.4459274 | 0.99937574 |
| ERMP1 | 148.373521 | 0.15782416 | 0.4747649 | 0.33242591 | 0.73956767 | 0.99937574 |
| KIAA2026 | 243.102332 | 0.10211224 | 0.37091944 | 0.27529491 | 0.78308967 | 0.99937574 |
| RANBP6 | 21.9188703 | 0.56676917 | 0.41615055 | 1.36193302 | 0.17321902 | 0.99937574 |
| UHRF2 | 79.218704 | -0.2099222 | 0.27472202 | -0.7641258 | 0.44479227 | 0.99937574 |
| KDM4C | 350.162487 | -0.2861054 | 0.25024012 | -1.1433234 | 0.25290436 | 0.99937574 |
| DMAC1 | 31.2365056 | 0.33672971 | 0.52108798 | 0.64620511 | 0.51814653 | 0.99937574 |
| PTPRD | 14.8475305 | -0.025211 | 0.54061959 | -0.0466336 | 0.96280524 | 0.99937574 |
| LURAP1L | 14.0594892 | -0.5799082 | 0.81058045 | -0.7154234 | 4.74E-01 | 0.99937574 |
| MPDZ | 73.695267 | 0.13125335 | 0.4364578 | 0.30072404 | 0.76362494 | 0.99937574 |
| NFIB | 291.719558 | 0.15876803 | 0.44110444 | 0.35993296 | 0.71889727 | 0.99937574 |
| ZDHHC21 | 218.006719 | 0.18553824 | 0.38157392 | 0.48624456 | 0.62679378 | 0.99937574 |
| FREM1 | 422.343172 | 0.80641933 | 0.58746544 | 1.37270939 | 0.16984271 | 0.99937574 |
| TTC39B | 352.526737 | -0.198316 | 0.34294326 | -0.5782764 | 0.56307753 | 0.99937574 |
| SNAPC3 | 93.8258284 | 0.15838756 | 0.33110006 | 0.47836765 | 0.63238855 | 0.99937574 |
| PSIP1 | 46.3394777 | -0.0237375 | 0.390531 | -0.0607827 | 0.95153225 | 0.99937574 |
| CCDC171 | 65.839903 | 0.00629448 | 0.33403347 | 0.01884386 | 0.98496566 | 0.99937574 |
| BNC2 | 102.054828 | 1.81306236 | 0.60797998 | 2.98210867 | 0.0028627 | 0.5159001 |
| CNTLN | 49.0813988 | -0.2045684 | 0.37598339 | -0.5440889 | 0.58638028 | 0.99937574 |
| SH3GL2 | 54.1625589 | -0.4990362 | 0.97617382 | -0.5112165 | 0.60919945 | 0.99937574 |
| ADAMTSL1 | 243.096461 | 1.15914453 | 0.51964375 | 2.23065231 | 2.57E-02 | 9.65E-01 |
| RRAGA | 28.4282606 | 0.45358198 | 0.60366273 | 0.7513831 | 0.45242213 | 0.99937574 |
| HAUS6 | 17.6206495 | -0.6126983 | 0.48389216 | -1.2661876 | 0.20544591 | 0.99937574 |
| PLIN2 | 44.8915011 | -0.7295745 | 0.57989767 | -1.2581091 | 0.20835231 | 0.99937574 |
| DENND4C | 219.190144 | 0.03165498 | 0.26439045 | 0.11972817 | 0.90469849 | 0.99937574 |

| | | | | | | |
|---|---|---|---|---|---|---|
| RPS6 | 346.819882 | 0.02580594 | 0.58190268 | 0.04434752 | 0.96462739 | 0.99937574 |
| SLC24A2 | 44.0330348 | 1.38379578 | 0.70432013 | 1.96472558 | 0.04944601 | 0.99937574 |
| MLLT3 | 156.67063 | -0.479776 | 0.59513344 | -0.8061654 | 0.42014749 | 0.99937574 |
| FOCAD | 245.754205 | 0.10906529 | 0.49240319 | 0.2214959 | 0.82470633 | 0.99937574 |
| KLHL9 | 36.5520889 | -0.2854702 | 0.36188375 | -0.7888451 | 0.43020255 | 0.99937574 |
| LINC01239 | 15.9893957 | 0.19832006 | 0.65646612 | 0.3021025 | 0.76257392 | 0.99937574 |
| AL391117.1 | 61.243846 | -0.4061363 | 1.06427493 | -0.3816085 | 0.70275179 | 0.99937574 |
| ELAVL2 | 40.4719593 | 1.36615017 | 0.96804785 | 1.4112424 | 0.15817315 | 0.99937574 |
| TUSC1 | 28.7789738 | 0.43992465 | 0.46917697 | 0.93765185 | 0.34842336 | 0.99937574 |
| CAAP1 | 145.441385 | 0.04167743 | 0.32279445 | 0.12911445 | 0.89726708 | 0.99937574 |
| PLAA | 51.2420951 | -0.1638398 | 0.37390979 | -0.43818 | 0.66125581 | 0.99937574 |
| IFT74 | 65.555477 | 0.18300617 | 0.37519625 | 0.48776119 | 0.625719 | 0.99937574 |
| MOB3B | 168.647029 | 0.71283601 | 0.53741195 | 1.3264238 | 0.18469936 | 0.99937574 |
| C9orf72 | 29.6765885 | -0.2436836 | 0.48763512 | -0.4997253 | 0.61726853 | 0.99937574 |
| ACO1 | 51.7865789 | 0.21242276 | 0.38109778 | 0.55739702 | 0.5772562 | 0.99937574 |
| DDX58 | 80.0233415 | -2.2844751 | 0.99650442 | -2.2924887 | 0.02187746 | 0.96427895 |
| TOPORS | 36.4569917 | -0.5109012 | 0.37875193 | -1.3489073 | 0.17736674 | 0.99937574 |
| NDUFB6 | 45.492505 | 0.29361993 | 0.43498343 | 0.67501405 | 0.49966684 | 0.99937574 |
| APTX | 69.9365457 | 0.40803317 | 0.33571408 | 1.2154187 | 0.22420644 | 0.99937574 |
| DNAJA1 | 97.7230186 | 0.37302623 | 0.45044573 | 0.82812691 | 0.40759863 | 0.99937574 |
| SMU1 | 41.9134975 | -0.0089728 | 0.29499839 | -0.0304165 | 0.9757349 | 0.99937574 |
| B4GALT1 | 118.889335 | 0.21199188 | 0.2875938 | 0.73712257 | 0.46104782 | 0.99937574 |
| BAG1 | 60.7739558 | -0.0059876 | 0.62065334 | -0.0096472 | 0.99230275 | 0.99937574 |
| CHMP5 | 59.5385754 | -0.4766687 | 0.57868268 | -0.8237135 | 0.41010238 | 0.99937574 |
| NFX1 | 142.512608 | -0.4839629 | 0.32764611 | -1.4770902 | 0.13965145 | 0.99937574 |
| AQP7 | 17.3021 | 0.40133143 | 0.81257342 | 0.49390174 | 0.62137557 | 0.99937574 |
| AQP3 | 95.2028163 | -0.3605588 | 0.94334969 | -0.3822112 | 0.70230473 | 0.99937574 |
| UBE2R2 | 179.060961 | -0.0619044 | 0.2850099 | -0.2172009 | 0.82805176 | 0.99937574 |
| UBAP2 | 102.495016 | -0.6726483 | 0.30103399 | -2.2344598 | 0.02545284 | 0.96546668 |
| DCAF12 | 27.2956211 | 0.03719422 | 0.3735768 | 0.09956245 | 0.9206917 | 0.99937574 |
| UBAP1 | 120.325539 | -0.493361 | 0.33180457 | -1.4869022 | 0.13704064 | 0.99937574 |
| NUDT2 | 13.6710149 | 0.43697461 | 0.53594155 | 0.81534004 | 0.4148777 | 0.99937574 |
| FAM219A | 45.4747306 | -0.1406349 | 0.43511012 | -0.3232168 | 0.74653105 | 0.99937574 |
| RPP25L | 12.3530217 | -0.128633 | 0.64518344 | -0.1993744 | 0.84196991 | 0.99937574 |
| DCTN3 | 21.8345804 | -0.4040518 | 0.67241206 | -0.6008992 | 0.54790715 | 0.99937574 |
| VCP | 107.99708 | -0.0721672 | 0.35118753 | -0.2054947 | 0.83718562 | 0.99937574 |
| PIGO | 16.1739664 | 0.44662036 | 0.45528133 | 0.98097665 | 0.32660425 | 0.99937574 |
| STOML2 | 29.9755872 | 0.05981595 | 0.38338025 | 0.15602251 | 0.87601528 | 0.99937574 |
| FAM214B | 10.5821385 | -0.2901452 | 0.58061627 | -0.4997194 | 0.61727265 | 0.99937574 |
| UNC13B | 132.838774 | -0.1349895 | 0.30165003 | -0.4475039 | 0.6545113 | 0.99937574 |
| RUSC2 | 57.7427817 | -0.1390728 | 0.4187998 | -0.3320746 | 0.73983295 | 0.99937574 |
| TESK1 | 15.5560022 | -0.2359522 | 0.58362739 | -0.4042857 | 0.68600264 | 0.99937574 |
| CCDC107 | 33.1858249 | -0.1128193 | 0.51927735 | -0.2172621 | 0.82800406 | 0.99937574 |
| TLN1 | 82.7021726 | 0.14444889 | 0.37777484 | 0.3823677 | 0.70218865 | 0.99937574 |
| CREB3 | 15.0132723 | 0.59576079 | 0.49027724 | 1.21515081 | 0.22430858 | 0.99937574 |
| GBA2 | 77.0415168 | -0.2639182 | 0.35509995 | -0.7432222 | 0.45734716 | 0.99937574 |
| RGP1 | 34.7442808 | 0.12188623 | 0.32256135 | 0.37786992 | 0.70552723 | 0.99937574 |
| AL133410.1 | 40.8156301 | 0.20771823 | 0.37958201 | 0.54722886 | 0.58422151 | 0.99937574 |
| HINT2 | 14.3858856 | -0.3361044 | 0.62642779 | -0.5365413 | 0.59158452 | 0.99937574 |
| TMEM8B | 25.0286764 | 0.54945017 | 0.59831905 | 0.91832303 | 0.35844977 | 0.99937574 |
| RECK | 29.6540401 | 0.22504309 | 0.47067811 | 0.47812524 | 0.63256107 | 0.99937574 |
| CLTA | 172.4023 | -0.3453153 | 0.38065902 | -0.9071513 | 0.36432679 | 0.99937574 |
| GNE | 64.3727053 | 0.17482032 | 0.48091332 | 0.36351732 | 0.71621848 | 0.99937574 |
| RNF38 | 291.156257 | -0.1754965 | 0.26207482 | -0.6696429 | 0.50308549 | 0.99937574 |

| | | | | | | |
|---|---|---|---|---|---|---|
| EBLN3P | 64.6069156 | 0.01116165 | 0.35042537 | 0.03185172 | 0.9745903 | 0.99937574 |
| ZCCHC7 | 415.434818 | -0.1356586 | 0.25430148 | -0.5334557 | 0.59371815 | 0.99937574 |
| GRHPR | 34.4633667 | 0.68104937 | 0.45969113 | 1.48153692 | 0.13846356 | 0.99937574 |
| ZBTB5 | 26.17336 | 0.11228901 | 0.45350904 | 0.24760038 | 0.80444361 | 0.99937574 |
| AL138752.2 | 49.1183532 | -0.1057407 | 0.36633176 | -0.2886475 | 0.77285117 | 0.99937574 |
| TRMT10B | 55.8529316 | 0.02440767 | 0.42342052 | 0.05764404 | 0.95403217 | 0.99937574 |
| DCAF10 | 135.251325 | 0.44636505 | 0.28908457 | 1.5440639 | 0.12257285 | 0.99937574 |
| GLIDR | 31.8101388 | -0.2218239 | 0.49870709 | -0.4447979 | 0.65646582 | 0.99937574 |
| CBWD6 | 12.2910381 | 1.08790966 | 0.71312954 | 1.52554284 | 0.12712376 | 0.99937574 |
| FAM27C | 14.2199746 | -0.3826412 | 0.62018633 | -0.6169778 | 0.5372494 | 0.99937574 |
| AL391987.2 | 11.4130228 | 1.00569418 | 0.61514183 | 1.63489807 | 0.10207042 | 0.99937574 |
| AL512625.3 | 11.695102 | 0.80226393 | 0.64550673 | 1.24284363 | 0.21392546 | 0.99937574 |
| CBWD5 | 36.4522452 | -0.5897045 | 0.38216229 | -1.5430736 | 0.12281293 | 0.99937574 |
| ZNF658 | 11.7738594 | 0.09854376 | 0.49730949 | 0.19815379 | 0.84292474 | 0.99937574 |
| PIP5K1B | 34.5919698 | -0.3411582 | 0.81962654 | -0.4162362 | 0.6772372 | 0.99937574 |
| FAM189A2 | 151.349345 | 0.09874584 | 0.4916415 | 0.20084929 | 0.84081642 | 0.99937574 |
| APBA1 | 16.1793032 | 1.15932241 | 0.60709846 | 1.90961183 | 0.05618321 | 0.99937574 |
| PTAR1 | 112.274786 | 0.14040175 | 0.27488679 | 0.51076209 | 0.60951766 | 0.99937574 |
| MAMDC2-AS | 41.1958593 | 0.61148941 | 0.37755303 | 1.61961198 | 0.10531566 | 0.99937574 |
| MAMDC2 | 151.640742 | 0.40097556 | 0.59298876 | 0.67619421 | 0.49891735 | 0.99937574 |
| SMC5-AS1 | 62.9042563 | 0.18825549 | 0.28039183 | 0.67140147 | 0.50196481 | 0.99937574 |
| SMC5 | 186.499948 | -0.4562095 | 0.29936825 | -1.5239075 | 0.12753181 | 0.99937574 |
| KLF9 | 60.0610529 | 0.62369514 | 0.37423089 | 1.66660521 | 0.09559293 | 0.99937574 |
| TRPM3 | 36.560066 | 0.36676092 | 0.869941 | 0.42159287 | 0.67332221 | 0.99937574 |
| CEMIP2 | 180.821725 | -0.8372933 | 0.72496588 | -1.1549417 | 0.24811431 | 0.99937574 |
| ABHD17B | 15.7080716 | -0.4625324 | 0.63294266 | -0.730765 | 0.4649227 | 0.99937574 |
| C9orf85 | 41.5771026 | -0.0865327 | 0.37049238 | -0.2335613 | 0.81532557 | 0.99937574 |
| LINC01504 | 20.7909861 | -0.5957891 | 0.72415228 | -0.82274 | 0.41065584 | 0.99937574 |
| ZFAND5 | 159.339943 | -0.498132 | 0.35585985 | -1.3997983 | 0.16157374 | 0.99937574 |
| ALDH1A1 | 193.78627 | 0.9768572 | 0.38771976 | 2.51949296 | 0.0117524 | 0.77408712 |
| CARNMT1 | 71.7486276 | 0.09677129 | 0.33870148 | 0.28571264 | 0.77509823 | 0.99937574 |
| NMRK1 | 55.5643226 | 0.28886903 | 0.49274815 | 0.58624071 | 0.55771377 | 0.99937574 |
| OSTF1 | 40.8878755 | 0.07155654 | 0.34697592 | 0.20622913 | 0.83661193 | 0.99937574 |
| RFK | 42.6784591 | 0.25598161 | 0.43275055 | 0.59152232 | 0.5541705 | 0.99937574 |
| GCNT1 | 21.3445154 | -0.794003 | 0.56200546 | -1.4128031 | 1.58E-01 | 0.99937574 |
| VPS13A | 188.913792 | 0.08936853 | 0.22886956 | 0.39047798 | 0.69618313 | 0.99937574 |
| GNAQ | 745.795764 | -0.0911779 | 0.22576343 | -0.4038648 | 0.68631215 | 0.99937574 |
| CEP78 | 86.2267438 | 0.13633703 | 0.6621319 | 0.20590616 | 0.83686422 | 0.99937574 |
| TLE4 | 55.7124496 | -0.6520499 | 0.60032101 | -1.0861687 | 0.27740436 | 0.99937574 |
| TLE1 | 123.190146 | -0.2924131 | 0.44257725 | -0.6607052 | 0.50880139 | 0.99937574 |
| AL591368.1 | 9.63874622 | 1.5916173 | 1.08428286 | 1.46789861 | 0.14213177 | 0.99937574 |
| RASEF | 44.1604689 | 0.06866973 | 0.59358687 | 0.11568607 | 0.90790134 | 0.99937574 |
| FRMD3 | 51.078926 | 0.33722894 | 0.35563902 | 0.94823381 | 0.34301044 | 0.99937574 |
| IDNK | 13.130595 | 0.07351803 | 0.53638342 | 0.13706246 | 0.89098143 | 0.99937574 |
| UBQLN1 | 133.786899 | 0.0298928 | 0.30655304 | 0.09751264 | 0.9223193 | 0.99937574 |
| GKAP1 | 49.3647705 | 0.47613206 | 0.39587095 | 1.20274563 | 0.22907477 | 0.99937574 |
| KIF27 | 38.5518354 | 0.1623771 | 0.44786812 | 0.36255562 | 0.71693687 | 0.99937574 |
| C9orf64 | 14.9171463 | 0.19184748 | 0.49787463 | 0.38533291 | 0.69999078 | 0.99937574 |
| HNRNPK | 195.779013 | -0.2444851 | 0.24233222 | -1.0088842 | 0.3130302 | 0.99937574 |
| RMI1 | 13.8403349 | -0.2266238 | 0.53832118 | -0.4209825 | 0.67376787 | 0.99937574 |
| AGTPBP1 | 68.718696 | -0.3213799 | 0.3163508 | -1.0158973 | 0.30967828 | 0.99937574 |
| NAA35 | 118.864122 | -0.0728122 | 0.25479829 | -0.2857639 | 0.77505893 | 0.99937574 |
| GOLM1 | 73.6689601 | -0.5578803 | 0.86934571 | -0.6417244 | 0.52105215 | 0.99937574 |
| ISCA1 | 47.9259527 | 0.0159477 | 0.49617812 | 0.03214108 | 0.97435954 | 0.99937574 |

| | | | | | | |
|---|---|---|---|---|---|---|
| TUT7 | 102.127408 | 0.02830795 | 0.25431615 | 0.11131006 | 0.91137048 | 0.99937574 |
| DAPK1 | 190.361301 | -0.4385429 | 0.55465716 | -0.7906558 | 0.42914486 | 0.99937574 |
| CTSL | 54.1367266 | -0.6509267 | 0.41738229 | -1.5595456 | 0.11886731 | 0.99937574 |
| SPIN1 | 219.281085 | -0.0251335 | 0.28740787 | -0.0874488 | 0.93031477 | 0.99937574 |
| SECISBP2 | 158.166055 | -0.3708686 | 0.23060055 | -1.608273 | 0.1077754 | 0.99937574 |
| SEMA4D | 139.219576 | -0.0355539 | 0.32546598 | -0.1092401 | 0.91301209 | 0.99937574 |
| GADD45G | 84.0340994 | -0.9602397 | 0.41623025 | -2.3069917 | 2.11E-02 | 0.96427895 |
| SYK | 157.759655 | 0.38954458 | 0.34600483 | 1.12583565 | 0.26023509 | 0.99937574 |
| AUH | 266.659527 | 0.15256986 | 0.26206937 | 0.58217357 | 0.56044977 | 0.99937574 |
| SPTLC1 | 83.8343679 | -0.2119752 | 0.27927284 | -0.7590255 | 0.44783733 | 0.99937574 |
| IARS | 73.4002012 | -0.4990176 | 0.35052736 | -1.4236194 | 0.15455666 | 0.99937574 |
| NOL8 | 25.6257267 | -0.2360668 | 0.4191317 | -0.5632283 | 0.57327947 | 0.99937574 |
| CENPP | 22.8267436 | -1.4895249 | 0.52510616 | -2.8366167 | 0.00455943 | 0.60810972 |
| IPPK | 19.7688253 | -0.1755835 | 0.53604563 | -0.3275532 | 0.7432495 | 0.99937574 |
| BICD2 | 23.6899704 | 0.27465607 | 0.46258881 | 0.59373695 | 0.55268807 | 0.99937574 |
| ZNF484 | 11.7358544 | -0.1086486 | 0.54618233 | -0.1989237 | 0.8423224 | 0.99937574 |
| SUSD3 | 12.5888764 | 0.60674876 | 0.62327131 | 0.9734906 | 0.33030953 | 0.99937574 |
| CARD19 | 24.6920719 | -0.2896799 | 0.36603623 | -0.7913969 | 0.4287124 | 0.99937574 |
| NINJ1 | 16.0500654 | 0.56730089 | 0.51331208 | 1.10517737 | 0.26908274 | 0.99937574 |
| FAM120AOS | 21.7280922 | -4.24E-01 | 0.43740091 | -9.69E-01 | 0.33236038 | 0.99937574 |
| FAM120A | 202.237621 | 0.36540592 | 0.27334876 | 1.33677548 | 0.18129594 | 0.99937574 |
| PHF2 | 44.7542555 | -0.0965372 | 0.4074595 | -0.2369245 | 0.81271534 | 0.99937574 |
| PTPDC1 | 76.8904197 | 0.00858385 | 0.27777894 | 0.03090171 | 0.97534792 | 0.99937574 |
| AL158152.2 | 57.7660757 | 0.36895074 | 0.32200756 | 1.14578288 | 0.251885 | 0.99937574 |
| ZNF169 | 19.4277482 | 0.49712983 | 0.42749269 | 1.16289667 | 0.24487143 | 0.99937574 |
| MFSD14B | 57.2116727 | -0.3303726 | 0.32006636 | -1.0322004 | 0.30197824 | 0.99937574 |
| FBP1 | 38.9339175 | -1.1383792 | 0.99712947 | -1.1416564 | 0.25359689 | 0.99937574 |
| C9orf3 | 438.607768 | -0.1483198 | 0.28422498 | -0.5218393 | 0.60178223 | 0.99937574 |
| FANCC | 163.464863 | -0.4569329 | 0.28166297 | -1.6222683 | 0.10474592 | 0.99937574 |
| PTCH1 | 30.5010855 | -0.9606757 | 0.537667 | -1.7867486 | 0.07397814 | 0.99937574 |
| LINC00476 | 35.7295466 | -0.1071676 | 0.50661149 | -0.211538 | 0.83246745 | 0.99937574 |
| ERCC6L2 | 129.538282 | -0.421916 | 0.32285922 | -1.306811 | 0.1912769 | 0.99937574 |
| SLC35D2 | 18.5098676 | -0.2232836 | 0.5092988 | -0.4384137 | 0.66108643 | 0.99937574 |
| HABP4 | 28.5962912 | -0.6010707 | 0.62173898 | -0.9667573 | 0.33366539 | 0.99937574 |
| CDC14B | 159.835636 | -0.0742559 | 0.48003774 | -0.1546876 | 0.8770676 | 0.99937574 |
| AAED1 | 16.0955327 | -0.7370234 | 0.66664723 | -1.1055673 | 0.26891384 | 0.99937574 |
| AL589843.1 | 24.5991824 | -0.0521591 | 0.7562096 | -0.0689744 | 0.94500996 | 0.99937574 |
| ZNF510 | 13.1302657 | -0.1809653 | 0.52322944 | -0.3458623 | 0.72944621 | 0.99937574 |
| ZNF782 | 46.8539345 | 0.4185297 | 0.35007617 | 1.19553896 | 0.23187652 | 0.99937574 |
| MFSD14C | 110.106142 | -0.3157974 | 0.31716616 | -0.9956844 | 0.31940349 | 0.99937574 |
| CCDC180 | 20.5376509 | 0.10568576 | 0.71781812 | 0.14723195 | 0.88294895 | 0.99937574 |
| TDRD7 | 41.4290516 | -1.0352927 | 0.48824053 | -2.1204563 | 0.03396759 | 0.99937574 |
| TSTD2 | 46.3178842 | -0.3896064 | 0.30220984 | -1.2891917 | 0.19733144 | 0.99937574 |
| NCBP1 | 33.5076478 | 0.10237201 | 0.3511102 | 0.29156662 | 0.770618 | 0.99937574 |
| XPA | 38.1265382 | 0.35848358 | 0.39943923 | 0.89746714 | 0.3694697 | 0.99937574 |
| TRMO | 12.6421438 | -0.6082258 | 0.6359381 | -0.9564229 | 0.33885863 | 0.99937574 |
| ANP32B | 58.1209528 | -0.1193789 | 0.338638 | -0.3525267 | 0.7244433 | 0.99937574 |
| NANS | 22.1692921 | -0.009372 | 0.48200108 | -0.0194439 | 0.98448702 | 0.99937574 |
| TRIM14 | 25.966337 | -1.3059762 | 0.8021623 | -1.6280698 | 0.10351009 | 0.99937574 |
| TBC1D2 | 50.9229145 | 0.38099075 | 0.50361798 | 0.75650745 | 0.44934501 | 0.99937574 |
| ANKS6 | 28.2370393 | 0.99865879 | 0.43564951 | 2.29234456 | 0.02188577 | 0.96427895 |
| TGFBR1 | 81.9225427 | -0.1199312 | 0.39155822 | -0.3062921 | 0.75938223 | 0.99937574 |
| ALG2 | 26.9412237 | 0.01488903 | 0.43953571 | 0.03387445 | 0.97297727 | 0.99937574 |
| SEC61B | 74.3090406 | 0.04027156 | 0.38739783 | 0.10395402 | 0.91720584 | 0.99937574 |

| | | | | | | |
|---|---|---|---|---|---|---|
| AL359710.1 | 65.2902188 | 0.15922537 | 0.48095581 | 0.33106029 | 0.74059895 | 0.99937574 |
| STX17 | 91.1847258 | -0.0826033 | 0.38620184 | -0.2138864 | 0.83063567 | 0.99937574 |
| ERP44 | 107.132246 | -0.3961459 | 0.25545851 | -1.5507249 | 0.12096762 | 0.99937574 |
| INVS | 173.590995 | 0.2024179 | 0.34398907 | 0.58844281 | 0.55623511 | 0.99937574 |
| TEX10 | 37.0292671 | -0.059601 | 0.39775331 | -0.149844 | 0.88088766 | 0.99937574 |
| PLPPR1 | 127.421967 | 1.55036342 | 0.53063 | 2.92174097 | 0.00348081 | 0.53256471 |
| MRPL50 | 26.0457947 | 0.69333439 | 0.51346827 | 1.35029647 | 0.1769209 | 0.99937574 |
| ZNF189 | 23.7832662 | 0.12235146 | 0.41797955 | 0.29272117 | 0.76973529 | 0.99937574 |
| ALDOB | 34.3281389 | 0.10497085 | 1.56494764 | 0.06707627 | 0.94652099 | 0.99937574 |
| RNF20 | 20.0373187 | 0.42014675 | 0.47372937 | 0.88689192 | 0.37513709 | 0.99937574 |
| SMC2 | 11.1446741 | -0.3163096 | 0.64674097 | -0.4890824 | 0.62478335 | 0.99937574 |
| NIPSNAP3A | 21.2020874 | -0.2330989 | 0.44437033 | -0.5245599 | 0.59988917 | 0.99937574 |
| ABCA1 | 20.2109315 | 0.03341299 | 0.71650902 | 0.04663303 | 0.96280571 | 0.99937574 |
| SLC44A1 | 219.576217 | -0.0155711 | 0.39778217 | -0.0391449 | 0.96877489 | 0.99937574 |
| FSD1L | 97.9712672 | -0.0635646 | 0.26051349 | -0.2439973 | 0.80723294 | 0.99937574 |
| FKTN | 53.15796 | 0.13488308 | 0.32523426 | 0.41472593 | 0.67834255 | 0.99937574 |
| TMEM38B | 29.194689 | -0.6033735 | 0.55529897 | -1.0865741 | 0.27722508 | 0.99937574 |
| ZNF462 | 20.6771231 | -1.4441888 | 0.72752429 | -1.985073 | 0.04713635 | 0.99937574 |
| RAD23B | 149.814404 | -0.3421306 | 0.23974546 | -1.4270576 | 0.1535633 | 0.99937574 |
| ELP1 | 49.6843115 | 0.53628666 | 0.36912577 | 1.4528562 | 0.14626368 | 0.99937574 |
| FAM206A | 9.96661435 | -0.0627039 | 0.54878635 | -0.1142592 | 0.90903229 | 0.99937574 |
| CTNNAL1 | 58.683025 | -0.2168555 | 0.33462681 | -0.6480519 | 0.51695138 | 0.99937574 |
| TMEM245 | 216.823187 | 0.2955164 | 0.30390427 | 0.97239962 | 0.33085178 | 0.99937574 |
| EPB41L4B | 59.0100556 | 0.74822196 | 0.52912606 | 1.41407125 | 0.15734098 | 0.99937574 |
| PTPN3 | 259.329171 | -0.4118594 | 0.25713764 | -1.601708 | 0.1092202 | 0.99937574 |
| PALM2 | 15.2663533 | -1.7124878 | 0.71812829 | -2.3846544 | 0.01709517 | 0.89506255 |
| TXN | 91.3400346 | -0.5221968 | 0.4560229 | -1.1451109 | 0.25216323 | 0.99937574 |
| SVEP1 | 61.8722841 | -1.1689235 | 0.72910804 | -1.6032239 | 0.10888522 | 0.99937574 |
| LPAR1 | 57.343815 | -0.1604713 | 0.43550446 | -0.3684722 | 0.71252117 | 0.99937574 |
| ECPAS | 241.489327 | -0.2467351 | 0.22838444 | -1.0803497 | 0.27998648 | 0.99937574 |
| ZNF483 | 83.4752665 | -0.5622973 | 0.45570852 | -1.2338968 | 0.21724138 | 0.99937574 |
| PTGR1 | 376.107047 | -1.3439022 | 0.77633454 | -1.7310864 | 0.08343635 | 0.99937574 |
| C9orf84 | 299.317969 | -0.114794 | 0.71652573 | -0.1602092 | 0.8727163 | 0.99937574 |
| UGCG | 226.011681 | -0.5807786 | 0.386204 | -1.5038131 | 0.13262951 | 0.99937574 |
| SUSD1 | 259.07698 | -0.2296057 | 0.44649243 | -0.5142433 | 0.6070819 | 0.99937574 |
| PTBP3 | 777.119894 | 0.05992102 | 0.36686513 | 0.16333256 | 0.8702566 | 0.99937574 |
| HSDL2 | 116.293862 | 0.49734121 | 0.43107698 | 1.15371784 | 0.24861587 | 0.99937574 |
| C9orf147 | 16.9421647 | 0.08791101 | 0.64310119 | 0.13669857 | 0.89126906 | 0.99937574 |
| KIAA1958 | 203.379387 | 0.30809233 | 0.26409542 | 1.16659475 | 0.24337407 | 0.99937574 |
| INIP | 37.7899862 | -0.1566478 | 0.37120323 | -0.4220001 | 0.67302494 | 0.99937574 |
| SNX30 | 33.3182681 | -1.4586987 | 0.67008511 | -2.1768857 | 0.0294891 | 0.99049468 |
| FKBP15 | 43.4961552 | 0.43146959 | 0.34787746 | 1.24029188 | 0.21486745 | 0.99937574 |
| SLC31A1 | 51.2202356 | 0.38149369 | 0.32464971 | 1.17509328 | 0.2399574 | 0.99937574 |
| CDC26 | 23.1361574 | 0.27301814 | 0.57764876 | 0.47263694 | 0.63647222 | 0.99937574 |
| PRPF4 | 15.6310924 | -1.0938217 | 0.59780062 | -1.8297434 | 0.06728832 | 0.99937574 |
| BSPRY | 26.4517725 | -0.8068075 | 0.41070223 | -1.9644586 | 0.04947694 | 0.99937574 |
| HDHD3 | 42.2168757 | -0.158883 | 0.45699903 | -0.3476659 | 0.72809112 | 0.99937574 |
| ALAD | 11.7266907 | 0.07949141 | 0.77305727 | 0.10282733 | 0.91810002 | 0.99937574 |
| RGS3 | 59.6064264 | 0.31477774 | 0.5253881 | 0.59913374 | 0.54908371 | 0.99937574 |
| ZNF618 | 172.548724 | 0.50581684 | 0.4120108 | 1.22767861 | 0.21956764 | 0.99937574 |
| KIF12 | 54.7750353 | 0.20749103 | 0.45428631 | 0.45674066 | 0.64785746 | 0.99937574 |
| AKNA | 24.6928541 | -0.3881668 | 0.68081908 | -0.5701468 | 0.56857814 | 0.99937574 |
| WHRN | 29.7512365 | -0.1914286 | 0.54458933 | -0.3515101 | 0.72520571 | 0.99937574 |
| ATP6V1G1 | 135.423485 | -0.4719843 | 0.33941708 | -1.3905733 | 0.16435485 | 0.99937574 |

| | | | | | | |
|---|---|---|---|---|---|---|
| TMEM268 | 20.422512 | -0.319766 | 0.61185457 | -0.5226176 | 0.6012404 | 0.99937574 |
| PAPPA | 12.5592231 | 0.63026941 | 0.65320666 | 0.96488515 | 0.33460234 | 0.99937574 |
| ASTN2 | 312.089181 | -0.0767846 | 0.5478796 | -0.1401486 | 0.88854257 | 0.99937574 |
| CDK5RAP2 | 102.787589 | 0.14030272 | 0.29268589 | 0.47936277 | 0.63168057 | 0.99937574 |
| MEGF9 | 232.23197 | 0.39636489 | 0.24113937 | 1.64371703 | 0.10023467 | 0.99937574 |
| FBXW2 | 57.3515478 | -0.1680915 | 0.34637117 | -0.485293 | 0.6274685 | 0.99937574 |
| PSMD5 | 52.4070265 | 0.09494138 | 0.34019282 | 0.27908107 | 0.78018261 | 0.99937574 |
| CNTRL | 42.6627579 | 0.1250697 | 0.52519505 | 0.23813953 | 0.81177287 | 0.99937574 |
| RAB14 | 120.334464 | 0.2312212 | 0.25478088 | 0.90752967 | 0.36412677 | 0.99937574 |
| GSN | 110.536162 | -0.1550247 | 0.27934259 | -0.5549626 | 0.57892022 | 0.99937574 |
| STOM | 44.277604 | -0.5891455 | 0.49368239 | -1.1933695 | 0.23272469 | 0.99937574 |
| DAB2IP | 194.904486 | -0.3063423 | 0.36006325 | -0.8508015 | 0.39487964 | 0.99937574 |
| TTLL11 | 24.4954297 | 0.91087265 | 0.3920598 | 2.32330033 | 0.02016302 | 0.96427895 |
| NDUFA8 | 50.2584383 | 0.41403075 | 0.51756311 | 0.79996186 | 0.42373289 | 0.99937574 |
| RBM18 | 23.9092719 | -0.2798677 | 0.42478946 | -0.6588387 | 0.50999937 | 0.99937574 |
| MRRF | 97.4553808 | -0.1292285 | 0.37620166 | -0.3435085 | 0.73121591 | 0.99937574 |
| PDCL | 31.4605 | 0.09933502 | 0.36458431 | 0.27246102 | 0.78526756 | 0.99937574 |
| RC3H2 | 94.2549235 | -0.2315368 | 0.33403819 | -0.6931447 | 0.48821875 | 0.99937574 |
| ZBTB26 | 11.1848373 | -0.5275588 | 0.5470928 | -0.964295 | 0.33489806 | 0.99937574 |
| RABGAP1 | 198.703065 | -0.4550911 | 0.27252529 | -1.6699041 | 0.09493834 | 0.99937574 |
| STRBP | 224.585819 | 0.16349495 | 0.29691316 | 0.55064903 | 0.58187429 | 0.99937574 |
| DENND1A | 296.648028 | 0.2688723 | 0.31004535 | 0.86720313 | 0.38583072 | 0.99937574 |
| PSMB7 | 65.9396599 | -0.3489752 | 0.29362272 | -1.1885155 | 0.23463038 | 0.99937574 |
| NR6A1 | 149.433315 | -0.0214131 | 0.48072456 | -0.0445433 | 0.9644713 | 0.99937574 |
| MIR181A2HG | 13.2480272 | 1.19001318 | 0.60805677 | 1.95707578 | 0.05033856 | 0.99937574 |
| RPL35 | 125.94597 | 0.03633369 | 0.43971035 | 0.08263097 | 0.93414497 | 0.99937574 |
| ARPC5L | 48.0357336 | 0.12398524 | 0.37748002 | 0.32845511 | 0.74256758 | 0.99937574 |
| GOLGA1 | 45.0776565 | 0.18535722 | 0.43054107 | 0.43052158 | 0.66681627 | 0.99937574 |
| SCAI | 101.127183 | 0.55694842 | 0.48534514 | 1.14753065 | 0.25116237 | 0.99937574 |
| PPP6C | 70.2838382 | -0.3790868 | 0.28982469 | -1.3079866 | 0.19087786 | 0.99937574 |
| RABEPK | 14.8607256 | 0.20344075 | 0.6231778 | 0.32645699 | 0.74407863 | 0.99937574 |
| HSPA5 | 159.748147 | -0.0667683 | 0.43452093 | -0.1536596 | 0.87787814 | 0.99937574 |
| GAPVD1 | 120.898694 | 0.0897842 | 0.24971856 | 0.35954157 | 0.71918998 | 0.99937574 |
| MAPKAP1 | 243.26161 | 0.1712853 | 0.38375642 | 0.44633859 | 0.65535268 | 0.99937574 |
| AL162584.1 | 19.8213705 | 1.55833468 | 0.76069901 | 2.0485562 | 0.04050553 | 0.99937574 |
| PBX3 | 235.598468 | 0.26334668 | 0.35641107 | 0.73888468 | 0.45997702 | 0.99937574 |
| MVB12B | 170.770225 | -1.5406959 | 0.51006788 | -3.0205702 | 0.00252299 | 0.48965449 |
| ZBTB43 | 85.2672278 | -0.2609728 | 0.41112184 | -0.6347822 | 0.52557048 | 0.99937574 |
| ZBTB34 | 14.019431 | 0.46903111 | 0.50112446 | 0.93595731 | 0.34929517 | 0.99937574 |
| RALGPS1 | 264.973641 | -0.0655066 | 0.32838826 | -0.1994791 | 0.84188803 | 0.99937574 |
| GARNL3 | 55.5768227 | 0.86249863 | 0.52565367 | 1.64081156 | 0.10083654 | 0.99937574 |
| SLC2A8 | 13.005945 | 0.45934648 | 0.52031161 | 0.88282959 | 0.37732835 | 0.99937574 |
| RPL12 | 53.9465597 | -0.3443008 | 0.40816556 | -0.8435321 | 0.39893089 | 0.99937574 |
| LRSAM1 | 79.7263832 | 0.1881985 | 0.41354386 | 0.45508716 | 0.64904653 | 0.99937574 |
| FAM129B | 58.5527852 | -0.4750742 | 0.51134763 | -0.9290631 | 0.35285636 | 0.99937574 |
| STXBP1 | 31.6764632 | -0.4285979 | 0.39670315 | -1.0803995 | 0.27996433 | 0.99937574 |
| PTRH1 | 13.1608932 | 0.00990124 | 0.49253694 | 0.02010254 | 0.98396158 | 0.99937574 |
| CDK9 | 24.5029758 | -0.0338353 | 0.42176454 | -0.0802233 | 0.93605966 | 0.99937574 |
| FPGS | 28.1424699 | -0.4976519 | 0.55604279 | -0.8949885 | 0.37079326 | 0.99937574 |
| DPM2 | 29.7407199 | -0.6558981 | 0.52877822 | -1.2404031 | 0.21482634 | 0.99937574 |
| FAM102A | 97.4660342 | -1.5834568 | 0.47319091 | -3.3463382 | 0.00081886 | 0.29012345 |
| SLC25A25 | 80.1909831 | -0.9953616 | 0.5698408 | -1.7467363 | 0.08068308 | 0.99937574 |
| SLC25A25-AS | 28.4625662 | -0.1027096 | 0.49726738 | -0.206548 | 0.83636289 | 0.99937574 |
| PTGES2 | 47.4345745 | 0.25109898 | 0.35645936 | 0.70442526 | 0.48116798 | 0.99937574 |

| | | | | | | |
|---|---|---|---|---|---|---|
| C9orf16 | 87.6718761 | -0.2445874 | 0.60444498 | -0.4046479 | 0.68573631 | 0.99937574 |
| CIZ1 | 32.4370522 | -0.2309291 | 0.38736347 | -0.5961562 | 0.55107089 | 0.99937574 |
| DNM1 | 15.694408 | 0.51227018 | 0.87853578 | 0.58309541 | 0.55982907 | 0.99937574 |
| GOLGA2 | 84.4163799 | -0.1351523 | 0.4166148 | -0.3244059 | 0.74563075 | 0.99937574 |
| SWI5 | 25.7662656 | 0.06384144 | 0.61277476 | 0.10418419 | 0.91702318 | 0.99937574 |
| TRUB2 | 21.768954 | -0.5439832 | 0.51992864 | -1.0462652 | 0.29543861 | 0.99937574 |
| COQ4 | 46.6982894 | -0.1480985 | 0.30781745 | -0.4811243 | 0.63042814 | 0.99937574 |
| SLC27A4 | 55.3983393 | 0.14593869 | 0.36433529 | 0.40056148 | 0.68874301 | 0.99937574 |
| URM1 | 23.9531231 | 0.00773729 | 0.45229035 | 0.0171069 | 0.98635133 | 0.99937574 |
| ODF2 | 20.1486019 | -1.0453113 | 0.5894038 | -1.7735063 | 0.07614485 | 0.99937574 |
| GLE1 | 29.5035794 | -0.1915804 | 0.48284694 | -0.3967725 | 0.6915352 | 0.99937574 |
| SPTAN1 | 230.076485 | 0.189451 | 0.48621434 | 0.38964503 | 0.69679905 | 0.99937574 |
| WDR34 | 24.0170716 | -0.2209293 | 0.49314562 | -0.4480001 | 0.65415316 | 0.99937574 |
| SET | 113.830761 | 0.0364615 | 0.3464512 | 0.10524282 | 0.91618313 | 0.99937574 |
| ZDHHC12 | 19.1499937 | 0.69336481 | 0.55651396 | 1.24590732 | 0.21279842 | 0.99937574 |
| ZER1 | 87.2455568 | -0.0116004 | 0.37221091 | -0.0311663 | 0.97513694 | 0.99937574 |
| TBC1D13 | 19.8484573 | 0.37958752 | 0.50176391 | 0.75650622 | 0.44934575 | 0.99937574 |
| ENDOG | 29.3740861 | 0.39738748 | 0.51258744 | 0.77525793 | 0.43818727 | 0.99937574 |
| KYAT1 | 18.1095311 | 0.34589154 | 0.66557133 | 0.51969117 | 0.60327885 | 0.99937574 |
| LRRC8A | 14.3658122 | -0.69205 | 0.52543825 | -1.3170911 | 0.1878081 | 0.99937574 |
| PHYHD1 | 14.3548603 | 0.74269002 | 0.62000074 | 1.1978857 | 0.23096152 | 0.99937574 |
| NUP188 | 22.5121251 | -0.1272572 | 0.41582348 | -0.3060366 | 0.75957674 | 0.99937574 |
| SH3GLB2 | 54.6906509 | -0.5875405 | 0.34971946 | -1.6800338 | 0.09295073 | 0.99937574 |
| MIGA2 | 26.8424809 | -0.4601982 | 0.46944402 | -0.9803047 | 0.32693576 | 0.99937574 |
| CRAT | 13.9506495 | 0.20893649 | 0.51599931 | 0.40491622 | 0.68553909 | 0.99937574 |
| PTPA | 53.6321506 | 0.09292044 | 0.43153625 | 0.21532476 | 0.82951412 | 0.99937574 |
| IER5L | 22.2602665 | -0.7965035 | 0.61327165 | -1.2987776 | 1.94E-01 | 0.99937574 |
| LINC01503 | 18.0519845 | 0.29896182 | 0.60209826 | 0.49653327 | 0.61951822 | 0.99937574 |
| NTMT1 | 22.7716548 | 0.27173986 | 0.57393933 | 0.47346443 | 0.63588187 | 0.99937574 |
| PTGES | 29.5046336 | -0.0085897 | 0.70680868 | -0.0121528 | 0.99030374 | 0.99937574 |
| TOR1B | 24.6633315 | -0.0393981 | 0.47989538 | -0.0820973 | 0.93456935 | 0.99937574 |
| C9orf78 | 30.4337715 | -0.31217 | 0.44942331 | -0.6946013 | 0.48730517 | 0.99937574 |
| USP20 | 24.4913142 | 0.33963077 | 0.44445247 | 0.76415543 | 0.44477462 | 0.99937574 |
| FNBP1 | 185.716745 | -0.1161175 | 0.32440863 | -0.3579358 | 0.72039137 | 0.99937574 |
| GPR107 | 93.2490622 | -0.046321 | 0.27378605 | -0.1691869 | 0.86564965 | 0.99937574 |
| FUBP3 | 114.307872 | -0.3910852 | 0.320849 | -1.2189073 | 0.22287936 | 0.99937574 |
| ABL1 | 161.078652 | 0.12458127 | 0.28562012 | 0.43617819 | 0.66270745 | 0.99937574 |
| AIF1L | 51.9900862 | -0.2454746 | 0.73665806 | -0.3332273 | 0.7389627 | 0.99937574 |
| NUP214 | 120.71821 | -0.4442213 | 0.26983792 | -1.6462524 | 0.0997118 | 0.99937574 |
| AL157938.2 | 36.1147542 | -0.4237378 | 0.83764169 | -0.50587 | 0.61294794 | 0.99937574 |
| PRRC2B | 258.873647 | -0.2172155 | 0.30233716 | -0.7184545 | 0.47247707 | 0.99937574 |
| POMT1 | 19.6667221 | 0.61002467 | 0.43534922 | 1.40123066 | 0.16114511 | 0.99937574 |
| UCK1 | 10.507366 | -0.8256035 | 0.66894021 | -1.2341963 | 0.21712978 | 0.99937574 |
| RAPGEF1 | 154.010026 | -0.5980658 | 0.48443378 | -1.2345667 | 0.21699181 | 0.99937574 |
| MED27 | 124.298816 | -0.0619477 | 0.27873887 | -0.2222428 | 0.82412489 | 0.99937574 |
| SETX | 193.122011 | -0.6349373 | 0.30668238 | -2.0703415 | 0.03842037 | 0.99937574 |
| TTF1 | 23.9389834 | -0.0756539 | 0.3873286 | -0.1953223 | 0.84514066 | 0.99937574 |
| DDX31 | 27.5293911 | -0.0370162 | 0.41854071 | -0.0884411 | 0.92952608 | 0.99937574 |
| GTF3C4 | 13.7969118 | -0.3184767 | 0.61703471 | -0.5161406 | 0.6057562 | 0.99937574 |
| TSC1 | 78.7173832 | -0.2041638 | 0.33178047 | -0.6153582 | 0.5383182 | 0.99937574 |
| GTF3C5 | 22.3818727 | -0.194192 | 0.5839146 | -0.3325693 | 0.73945946 | 0.99937574 |
| RALGDS | 10.1058102 | -0.7282164 | 0.64186142 | -1.1345384 | 0.25656877 | 0.99937574 |
| ABO | 85.6714368 | -0.2679197 | 0.56028318 | -0.4781861 | 0.63251772 | 0.99937574 |
| SURF6 | 16.3098628 | -0.4219344 | 0.50900718 | -0.8289361 | 0.40714055 | 0.99937574 |

| | | | | | | |
|---|---|---|---|---|---|---|
| MED22 | 12.4574338 | -0.3570647 | 0.65333914 | -0.5465227 | 0.58470671 | 0.99937574 |
| RPL7A | 201.040036 | 0.00165246 | 0.47642446 | 0.00346846 | 0.99723258 | 0.99937574 |
| SURF1 | 17.9483713 | 0.28310223 | 0.52088548 | 0.54350187 | 0.5867843 | 0.99937574 |
| SURF4 | 64.0823634 | 0.01231168 | 0.34856927 | 0.03532061 | 0.97182409 | 0.99937574 |
| REXO4 | 14.1040955 | 0.10645797 | 0.52921942 | 0.20116036 | 0.84057318 | 0.99937574 |
| CACFD1 | 16.9987233 | -0.7429448 | 0.64074331 | -1.1595045 | 0.2462506 | 0.99937574 |
| VAV2 | 103.429082 | -0.0802964 | 0.36130334 | -0.2222409 | 0.82412639 | 0.99937574 |
| BRD3 | 35.3043123 | 0.19910012 | 0.35037736 | 0.56824484 | 0.56986873 | 0.99937574 |
| WDR5 | 23.0718605 | 0.0357702 | 0.4359054 | 0.08205955 | 0.93459936 | 0.99937574 |
| RXRA | 35.1897161 | -0.2873899 | 0.54849795 | -0.523958 | 0.60030775 | 0.99937574 |
| PPP1R26-AS1 | 14.0948614 | -0.0764784 | 0.61335492 | -0.1246886 | 0.90077008 | 0.99937574 |
| PPP1R26 | 20.6301426 | 0.04765557 | 0.48258404 | 0.09875083 | 0.92133611 | 0.99937574 |
| MRPS2 | 15.5402585 | 0.18350259 | 0.67279044 | 0.2727485 | 0.78504655 | 0.99937574 |
| CAMSAP1 | 24.5734047 | -0.2872896 | 0.47887736 | -0.5999231 | 0.54855746 | 0.99937574 |
| UBAC1 | 61.4584645 | 0.41164433 | 0.28234795 | 1.45793278 | 0.14485907 | 0.99937574 |
| NACC2 | 31.1965565 | 0.17082402 | 0.57467978 | 0.29725079 | 0.76627505 | 0.99937574 |
| TMEM250 | 12.2866542 | -0.4543902 | 0.76177678 | -0.5964874 | 0.5508497 | 0.99937574 |
| SNAPC4 | 14.1855098 | 0.01100611 | 0.49104328 | 0.02241372 | 0.98211794 | 0.99937574 |
| ENTR1 | 17.1337891 | -0.7471483 | 0.50810484 | -1.470461 | 0.14143695 | 0.99937574 |
| PMPCA | 35.3265925 | 0.06392654 | 0.48366413 | 0.13217134 | 0.89484877 | 0.99937574 |
| INPP5E | 12.3194354 | 0.18063154 | 0.56311031 | 0.32077471 | 0.74838113 | 0.99937574 |
| SEC16A | 51.406114 | -0.0646064 | 0.3968438 | -0.1628006 | 0.8706754 | 0.99937574 |
| NOTCH1 | 26.0679679 | 0.33054284 | 0.53920309 | 0.61302104 | 0.53986243 | 0.99937574 |
| EGFL7 | 15.9318074 | 1.57362399 | 0.72800373 | 2.16156035 | 0.03065208 | 0.99049468 |
| AGPAT2 | 53.5969455 | 0.18942884 | 0.43707135 | 0.43340484 | 0.66472069 | 0.99937574 |
| SNHG7 | 38.8271593 | 0.03379682 | 0.55330117 | 0.06108215 | 0.95129379 | 0.99937574 |
| RABL6 | 97.1900406 | 0.01813756 | 0.33055747 | 0.05486963 | 0.95624233 | 0.99937574 |
| PHPT1 | 84.8182158 | 0.78131918 | 0.44154633 | 1.76950667 | 0.07680936 | 0.99937574 |
| EDF1 | 83.7381876 | -0.4734599 | 0.53508586 | -0.8848297 | 0.37624849 | 0.99937574 |
| TRAF2 | 19.9176037 | 0.08673005 | 0.46274185 | 0.18742642 | 0.85132631 | 0.99937574 |
| FBXW5 | 57.7900575 | -0.4963142 | 0.49719823 | -0.998222 | 0.31817173 | 0.99937574 |
| LCN12 | 10.1821381 | 0.09294245 | 0.68747551 | 0.13519383 | 0.89245863 | 0.99937574 |
| PAXX | 22.844774 | 0.31556795 | 0.48891174 | 0.64544973 | 0.51863579 | 0.99937574 |
| ABCA2 | 10.9425363 | 1.14633502 | 0.70728807 | 1.620747 | 0.10507191 | 0.99937574 |
| UAP1L1 | 14.778807 | -1.0452183 | 0.83926579 | -1.245396 | 0.21298621 | 0.99937574 |
| MAN1B1 | 29.2896859 | 0.121125 | 0.401264 | 0.30185863 | 0.76275984 | 0.99937574 |
| DPP7 | 42.3635733 | -0.1391311 | 0.50904981 | -0.2733153 | 0.78461082 | 0.99937574 |
| SSNA1 | 35.9953248 | -0.1327538 | 0.43937525 | -0.3021421 | 0.76254373 | 0.99937574 |
| TPRN | 17.8951748 | -0.3293613 | 0.46828118 | -0.7033409 | 0.48184332 | 0.99937574 |
| TMEM203 | 19.4049718 | -0.0312379 | 0.50062506 | -0.0623979 | 0.95024598 | 0.99937574 |
| TUBB4B | 132.697965 | 0.67430428 | 0.52830137 | 1.2763629 | 0.20182726 | 0.99937574 |
| NELFB | 38.0550734 | -0.2977765 | 0.41083457 | -0.7248088 | 0.46856934 | 0.99937574 |
| EXD3 | 43.3369931 | -0.0073204 | 0.34364669 | -0.021302 | 0.98300473 | 0.99937574 |
| NOXA1 | 16.7950317 | -0.2219502 | 0.55086432 | -0.4029126 | 0.68701253 | 0.99937574 |
| NSMF | 25.2412435 | 0.22207427 | 0.40005616 | 0.55510774 | 0.57882097 | 0.99937574 |
| PNPLA7 | 14.641029 | -0.3147931 | 0.80797915 | -0.3896055 | 0.69682828 | 0.99937574 |
| MRPL41 | 111.451836 | 0.05080495 | 0.49996913 | 0.10161617 | 0.91906134 | 0.99937574 |
| DPH7 | 20.7069441 | 0.17724346 | 0.41903562 | 0.42297946 | 0.67231025 | 0.99937574 |
| ARRDC1 | 30.5691774 | -0.5185562 | 0.40136702 | -1.2919752 | 0.19636575 | 0.99937574 |
| ARRDC1-AS1 | 11.9560546 | 0.38236461 | 0.52921994 | 0.72250605 | 0.46998341 | 0.99937574 |
| EHMT1 | 219.059082 | -0.0433858 | 0.23641191 | -0.1835179 | 0.85439171 | 0.99937574 |
| BET1L | 20.3688646 | -0.3846738 | 0.47432617 | -0.8109901 | 0.41737136 | 0.99937574 |
| RIC8A | 16.6435066 | -0.2314823 | 0.56274309 | -0.4113463 | 0.68081865 | 0.99937574 |
| SIRT3 | 16.9806048 | 0.1718615 | 0.4883174 | 0.35194629 | 0.72487854 | 0.99937574 |

| | | | | | | |
|---|---|---|---|---|---|---|
| PSMD13 | 40.3965241 | -0.4040698 | 0.41523984 | -0.9730998 | 0.33050368 | 0.99937574 |
| PGGHG | 19.9757088 | -1.5498099 | 0.73618012 | -2.1052047 | 0.03527349 | 0.99937574 |
| IFITM2 | 207.662065 | 0.60336177 | 0.56434084 | 1.06914427 | 2.85E-01 | 9.99E-01 |
| IFITM3 | 291.95425 | -1.0382411 | 0.62360551 | -1.6649005 | 9.59E-02 | 0.99937574 |
| SIGIRR | 38.070853 | -0.5215723 | 0.45992371 | -1.1340409 | 0.2567774 | 0.99937574 |
| ANO9 | 22.3724863 | 0.13111934 | 0.42994079 | 0.30497068 | 0.76038848 | 0.99937574 |
| PTDSS2 | 45.3491818 | -0.0651425 | 0.47490022 | -0.1371708 | 0.89089577 | 0.99937574 |
| RNH1 | 88.5313085 | -0.0046399 | 0.40657968 | -0.0114119 | 0.9908948 | 0.99937574 |
| HRAS | 20.9014158 | -1.0384453 | 0.65346248 | -1.589143 | 0.11202812 | 0.99937574 |
| RASSF7 | 39.409009 | 0.40424753 | 0.42658806 | 0.94762973 | 0.34331799 | 0.99937574 |
| MIR210HG | 13.0002066 | -1.2199093 | 0.89199395 | -1.3676206 | 0.17143087 | 0.99937574 |
| PHRF1 | 27.6968083 | -0.0149941 | 0.39826267 | -0.0376487 | 0.96996776 | 0.99937574 |
| IRF7 | 22.5083435 | -2.1311353 | 0.9135061 | -2.3329185 | 0.01965242 | 0.96427895 |
| DEAF1 | 92.9711087 | 0.29922778 | 0.28461595 | 1.05133875 | 0.29310303 | 0.99937574 |
| AC131934.1 | 16.5791249 | -0.0824225 | 0.6183292 | -0.1332988 | 0.8939571 | 0.99937574 |
| EPS8L2 | 86.3581426 | -0.2804957 | 0.44641263 | -0.6283328 | 0.52978592 | 0.99937574 |
| AP006621.4 | 13.1856512 | -0.2625661 | 0.92334221 | -0.2843649 | 0.77613078 | 0.99937574 |
| TALDO1 | 93.147711 | -0.4272148 | 0.31671689 | -1.3488853 | 1.77E-01 | 0.99937574 |
| GATD1 | 20.2651468 | -1.0081565 | 0.61982796 | -1.6265103 | 0.10384115 | 0.99937574 |
| AP006621.3 | 24.7363883 | -0.1571571 | 0.52821333 | -0.2975259 | 0.76606506 | 0.99937574 |
| RPLP2 | 137.457309 | -0.3894306 | 0.40559862 | -0.9601379 | 0.33698583 | 0.99937574 |
| PNPLA2 | 25.9416037 | -0.4025254 | 0.40291366 | -0.9990363 | 0.31777712 | 0.99937574 |
| CRACR2B | 13.3293659 | -0.692037 | 0.85303746 | -0.8112621 | 0.41721514 | 0.99937574 |
| CD151 | 84.6580494 | -0.0840283 | 0.38409558 | -0.2187692 | 0.82682985 | 0.99937574 |
| POLR2L | 83.0854186 | 0.57901417 | 0.53498684 | 1.0822961 | 0.27912098 | 0.99937574 |
| CHID1 | 22.0300107 | 0.70090521 | 0.40955345 | 1.71138886 | 0.08700935 | 0.99937574 |
| AP2A2 | 102.63137 | -0.2091593 | 0.34182009 | -0.6118988 | 0.54060473 | 0.99937574 |
| TOLLIP | 41.430001 | -0.944802 | 0.45539644 | -2.0746802 | 0.03801619 | 0.99937574 |
| MOB2 | 23.0040611 | -0.2585622 | 0.4634369 | -0.5579233 | 0.57689677 | 0.99937574 |
| DUSP8 | 12.7408758 | -1.0136062 | 0.90560938 | -1.1192532 | 0.26303215 | 0.99937574 |
| MRPL23 | 16.3121801 | 2.14412649 | 0.64763459 | 3.31070408 | 0.00093062 | 0.29012345 |
| CD81 | 283.838814 | 0.69631872 | 0.31278122 | 2.22621649 | 0.02599968 | 0.96546668 |
| TSSC4 | 22.1776807 | -0.3634836 | 0.52959121 | -0.6863475 | 0.492494 | 0.99937574 |
| KCNQ1 | 239.422168 | 0.05753734 | 0.36318866 | 0.15842273 | 0.87412371 | 0.99937574 |
| KCNQ1OT1 | 253.416363 | 0.28794194 | 0.53299634 | 0.5402325 | 0.5890367 | 0.99937574 |
| CDKN1C | 69.9432063 | 0.84543402 | 0.88630581 | 0.95388523 | 0.34014174 | 0.99937574 |
| SLC22A18 | 33.8880388 | -0.5580649 | 0.86160047 | -0.6477073 | 0.51717429 | 0.99937574 |
| NAP1L4 | 250.369438 | 0.2668655 | 0.32285342 | 0.82658407 | 0.40847285 | 0.99937574 |
| CARS | 78.4004486 | 0.1025922 | 0.55565546 | 0.18463275 | 0.8535171 | 0.99937574 |
| OSBPL5 | 18.9766479 | 0.84257406 | 0.55335653 | 1.52266039 | 0.12784369 | 0.99937574 |
| ZNF195 | 25.5544115 | -0.2604246 | 0.46848461 | -0.5558873 | 0.57828791 | 0.99937574 |
| AC127526.1 | 30.2613277 | -1.2180112 | 0.68505459 | -1.7779769 | 0.07540765 | 0.99937574 |
| NUP98 | 149.251815 | -0.1045123 | 0.32092508 | -0.3256595 | 0.744682 | 0.99937574 |
| PGAP2 | 20.6107964 | 0.47419157 | 0.61566987 | 0.77020429 | 0.44117872 | 0.99937574 |
| RHOG | 15.2777489 | -0.6233057 | 0.55795045 | -1.1171346 | 0.2639368 | 0.99937574 |
| STIM1 | 367.824772 | 0.37185771 | 0.35582823 | 1.04504834 | 0.29600063 | 0.99937574 |
| RRM1 | 17.789032 | -0.0402559 | 0.44801369 | -0.0898541 | 0.92840318 | 0.99937574 |
| AC104389.4 | 32.918653 | -1.2743795 | 0.94605246 | -1.3470496 | 0.17796427 | 0.99937574 |
| TRIM5 | 14.4417602 | -1.3265894 | 0.59804407 | -2.2182134 | 0.02654029 | 0.97242612 |
| TRIM22 | 40.052544 | -1.8268707 | 0.94649931 | -1.9301342 | 0.05359021 | 0.99937574 |
| FAM160A2 | 19.7593955 | 0.32781094 | 0.42429775 | 0.77259646 | 0.43976124 | 0.99937574 |
| SMPD1 | 18.3818365 | 0.71520326 | 0.47607868 | 1.50227954 | 0.13302493 | 0.99937574 |
| APBB1 | 20.2095014 | 0.04715729 | 0.50088925 | 0.09414714 | 0.92499227 | 0.99937574 |
| ARFIP2 | 16.9858564 | -0.2354089 | 0.46890039 | -0.5020445 | 0.61563623 | 0.99937574 |

| | | | | | | |
|---|---|---|---|---|---|---|
| AC084337.2 | 32.4359739 | -0.5114127 | 0.56393219 | -0.9068691 | 0.36447601 | 0.99937574 |
| DNHD1 | 47.8366252 | -0.2882562 | 0.56015967 | -0.5145964 | 0.60683506 | 0.99937574 |
| RRP8 | 12.0706747 | -0.1375326 | 0.55573911 | -0.2474769 | 0.80453917 | 0.99937574 |
| ILK | 33.9162288 | 0.15452166 | 0.35612609 | 0.43389592 | 0.66436402 | 0.99937574 |
| TAF10 | 30.5827382 | 0.0977056 | 0.50342348 | 0.19408232 | 0.84611143 | 0.99937574 |
| TPP1 | 36.0979205 | -0.2979284 | 0.45629761 | -0.6529255 | 0.51380429 | 0.99937574 |
| MRPL17 | 22.5580798 | -0.3885292 | 0.56345027 | -0.6895536 | 0.49047495 | 0.99937574 |
| AC087280.2 | 22.052608 | -1.2263847 | 0.64262413 | -1.9084012 | 0.05633938 | 0.99937574 |
| ZNF215 | 11.5207666 | 0.17523745 | 0.54822126 | 0.31964731 | 0.7492357 | 0.99937574 |
| AC107884.1 | 24.8494327 | 0.12368228 | 0.55969854 | 0.22098017 | 0.82510788 | 0.99937574 |
| PPFIBP2 | 198.218755 | -0.2644616 | 0.33403687 | -0.7917137 | 0.42852762 | 0.99937574 |
| EIF3F | 50.4689104 | 0.14696056 | 0.48629674 | 0.30220346 | 0.76249697 | 0.99937574 |
| RIC3 | 47.7147465 | -0.8633751 | 0.51815968 | -1.6662336 | 0.09566689 | 0.99937574 |
| STK33 | 355.270925 | 0.42941779 | 0.34862065 | 1.23176236 | 0.21803787 | 0.99937574 |
| TRIM66 | 40.7541284 | -0.2228957 | 0.58084163 | -0.3837461 | 0.70116664 | 0.99937574 |
| RPL27A | 190.287962 | -0.144182 | 0.43769479 | -0.3294122 | 0.74184418 | 0.99937574 |
| ST5 | 233.549034 | -0.0928803 | 0.51397453 | -0.1807098 | 0.85659533 | 0.99937574 |
| AC026894.1 | 16.6596267 | -0.676844 | 0.8126912 | -0.8328428 | 0.4049334 | 0.99937574 |
| AKIP1 | 25.6829554 | -0.1834834 | 0.45085203 | -0.4069704 | 0.68402974 | 0.99937574 |
| TMEM9B | 86.688912 | -0.0956352 | 0.28086661 | -0.3405005 | 0.73347961 | 0.99937574 |
| TMEM9B-AS | 26.1893728 | -0.6362897 | 0.71050126 | -0.8955505 | 0.3704929 | 0.99937574 |
| DENND5A | 86.4889162 | -0.3688499 | 0.35636384 | -1.0350373 | 0.3006515 | 0.99937574 |
| TMEM41B | 102.789541 | -0.5553991 | 0.29226139 | -1.9003506 | 0.05738712 | 0.99937574 |
| IPO7 | 131.76304 | 0.09672363 | 0.28371072 | 0.34092342 | 0.73316123 | 0.99937574 |
| ZNF143 | 33.0527032 | 0.40823307 | 0.33699337 | 1.21139791 | 0.22574294 | 0.99937574 |
| SWAP70 | 67.3342144 | -0.7167137 | 0.42140862 | -1.700757 | 0.08898862 | 0.99937574 |
| SBF2-AS1 | 88.1183859 | -0.664753 | 0.75762527 | -0.8774166 | 0.38026042 | 0.99937574 |
| SBF2 | 879.026515 | 0.44184799 | 0.30190129 | 1.46355117 | 0.14331662 | 0.99937574 |
| AC011092.2 | 36.1134884 | 0.29965164 | 0.52440072 | 0.57141729 | 0.56771681 | 0.99937574 |
| AC100763.1 | 12.3703107 | 0.9926911 | 0.87946495 | 1.12874435 | 0.25900569 | 0.99937574 |
| ADM | 40.151964 | -0.6850575 | 0.93288068 | -0.7343463 | 0.46273769 | 0.99937574 |
| AMPD3 | 112.198224 | 0.2848992 | 0.43719231 | 0.65165647 | 0.51462281 | 0.99937574 |
| MTRNR2L8 | 157.925378 | -0.3352002 | 1.21684013 | -0.2754678 | 0.78295689 | 0.99937574 |
| RNF141 | 40.9260682 | 0.06543686 | 0.32389229 | 0.20203277 | 0.83989111 | 0.99937574 |
| MRVI1 | 10.2464147 | 1.32880059 | 0.90981995 | 1.46050941 | 0.14415012 | 0.99937574 |
| CTR9 | 52.9821369 | 0.03735496 | 0.34834747 | 0.10723476 | 0.91460274 | 0.99937574 |
| EIF4G2 | 162.243561 | -0.3782353 | 0.33833505 | -1.1179313 | 0.26359635 | 0.99937574 |
| ZBED5 | 59.781115 | 0.09344552 | 0.36329164 | 0.25721903 | 0.7970097 | 0.99937574 |
| ZBED5-AS1 | 15.1790563 | 1.19595462 | 0.6663132 | 1.79488357 | 0.07267225 | 0.99937574 |
| GALNT18 | 31.2089462 | 0.13386814 | 0.76591856 | 0.17478116 | 0.86125159 | 0.99937574 |
| USP47 | 307.135888 | 0.21575008 | 0.32233171 | 0.66934178 | 0.50327748 | 0.99937574 |
| PARVA | 98.6185812 | -0.4153956 | 0.34725629 | -1.196222 | 0.23160996 | 0.99937574 |
| TEAD1 | 295.817576 | -0.9040695 | 0.4199107 | -2.1530041 | 0.03131836 | 0.99049468 |
| LINC00958 | 58.5939778 | 0.32767841 | 0.30110656 | 1.08824735 | 0.27648593 | 0.99937574 |
| ARNTL | 25.4534213 | -0.2645316 | 0.51022774 | -0.5184578 | 0.60413886 | 0.99937574 |
| BTBD10 | 68.1130872 | -0.3713646 | 0.28035607 | -1.3246178 | 0.18529795 | 0.99937574 |
| FAR1 | 168.629803 | 0.23627659 | 0.29440941 | 0.80254428 | 0.42223818 | 0.99937574 |
| RRAS2 | 52.1814795 | -0.1925627 | 0.59369902 | -0.3243439 | 0.74567765 | 0.99937574 |
| COPB1 | 87.8850552 | 0.20166481 | 0.25945296 | 0.77726927 | 0.43699992 | 0.99937574 |
| PSMA1 | 21.4089332 | 1.28782307 | 0.67694627 | 1.90240071 | 0.05711879 | 0.99937574 |
| PDE3B | 35.9183455 | 0.81277648 | 0.64692108 | 1.25637654 | 0.20897948 | 0.99937574 |
| CYP2R1 | 17.6328685 | -0.1518416 | 0.50947329 | -0.2980365 | 0.76567529 | 0.99937574 |
| CALCB | 34.4261616 | -1.8635057 | 1.58425216 | -1.1762683 | 0.23948766 | 0.99937574 |
| CALCA | 98.0121794 | -3.1430431 | 1.55362193 | -2.0230424 | 0.04306877 | 0.99937574 |

| | | | | | | |
|---|---|---|---|---|---|---|
| SOX6 | 379.961396 | -0.0533467 | 0.27722102 | -0.1924339 | 0.84740233 | 0.99937574 |
| C11orf58 | 112.334144 | -0.1715251 | 0.2837636 | -0.6044648 | 0.54553464 | 0.99937574 |
| PLEKHA7 | 660.325878 | 0.03135599 | 0.28403923 | 0.11039317 | 0.91209757 | 0.99937574 |
| RPS13 | 147.899908 | -0.1044828 | 0.60090835 | -0.1738748 | 0.86196386 | 0.99937574 |
| PIK3C2A | 279.456924 | -0.0269432 | 0.19279012 | -0.1397541 | 0.8888543 | 0.99937574 |
| NUCB2 | 69.650004 | 0.80759202 | 0.39815102 | 2.02835604 | 0.04252392 | 0.99937574 |
| NCR3LG1 | 28.3243734 | -0.7262737 | 0.60472741 | -1.2009936 | 0.22975369 | 0.99937574 |
| AC124798.1 | 11.0191251 | -1.2449946 | 0.75477766 | -1.6494853 | 0.09904826 | 0.99937574 |
| KCNC1 | 13.3143171 | 0.50861778 | 0.86647324 | 0.58699768 | 0.55720527 | 0.99937574 |
| SERGEF | 157.341559 | 0.27849052 | 0.30998461 | 0.8984011 | 0.36897175 | 0.99937574 |
| HPS5 | 32.1481666 | 0.0793421 | 0.36789623 | 0.21566435 | 0.82924939 | 0.99937574 |
| GTF2H1 | 45.3555762 | -0.4883598 | 0.34710281 | -1.40696 | 0.15943924 | 0.99937574 |
| LDHA | 345.967792 | 0.26877589 | 0.48318305 | 0.55626101 | 0.57803243 | 0.99937574 |
| TSG101 | 92.9145732 | -0.1600388 | 0.23047462 | -0.694388 | 0.48743894 | 0.99937574 |
| UEVLD | 72.1521735 | 0.05554692 | 0.27094199 | 0.20501407 | 0.83756113 | 0.99937574 |
| SPTY2D1OS | 25.0508171 | 0.40943164 | 0.47833288 | 0.85595545 | 0.39202242 | 0.99937574 |
| SPTY2D1 | 42.0886789 | -0.3955702 | 0.41643874 | -0.9498882 | 0.34216908 | 0.99937574 |
| IGSF22 | 12.1937068 | 0.28999195 | 0.84499329 | 0.34318846 | 0.73145668 | 0.99937574 |
| ZDHHC13 | 85.6724131 | -0.3575479 | 0.32825756 | -1.0892297 | 0.2760526 | 0.99937574 |
| NAV2 | 192.037261 | 0.18696811 | 0.55730874 | 0.3354839 | 0.73726009 | 0.99937574 |
| HTATIP2 | 36.5926228 | 0.27851388 | 0.46587727 | 0.59782671 | 5.50E-01 | 0.99937574 |
| PRMT3 | 51.992937 | -0.2683721 | 0.3491098 | -0.7687326 | 0.44205209 | 0.99937574 |
| NELL1 | 51.740734 | -0.697327 | 0.79789672 | -0.8739565 | 3.82E-01 | 9.99E-01 |
| ANO5 | 54.5341399 | 1.06076914 | 0.37832832 | 2.80383226 | 0.00504992 | 0.61461453 |
| FANCF | 10.3259684 | 0.36328281 | 0.66351658 | 0.54751127 | 0.58402753 | 0.99937574 |
| GAS2 | 79.5708854 | -0.5816495 | 0.65110945 | -0.8933206 | 0.3716855 | 0.99937574 |
| SVIP | 83.0414974 | 0.19239328 | 0.38671591 | 0.49750547 | 0.61883264 | 0.99937574 |
| MUC15 | 16.7914362 | 0.31333988 | 0.92275754 | 0.33956903 | 0.73418111 | 0.99937574 |
| CCDC34 | 24.1289404 | 2.51488017 | 0.47383655 | 5.3074845 | 1.11E-07 | 0.00112171 |
| LGR4 | 799.475863 | 0.33955318 | 0.38082408 | 0.89162739 | 0.37259268 | 0.99937574 |
| LIN7C | 42.3880041 | 0.03114287 | 0.44696095 | 0.06967693 | 0.94445081 | 0.99937574 |
| BDNF-AS | 122.515631 | -0.0523503 | 0.330641 | -0.1583298 | 0.87419693 | 0.99937574 |
| METTL15 | 226.426886 | 0.05219167 | 0.34167922 | 0.1527505 | 0.87859504 | 0.99937574 |
| ARL14EP | 40.0934748 | 0.55426755 | 0.36023472 | 1.5386289 | 0.12389492 | 0.99937574 |
| MPPED2 | 1229.64138 | 0.31476593 | 0.45751698 | 0.68798743 | 0.4914607 | 0.99937574 |
| AL136088.1 | 36.6328303 | 0.24588474 | 0.74750795 | 0.3289393 | 0.74220157 | 0.99937574 |
| DCDC1 | 14.2992501 | -0.138556 | 0.73942753 | -0.1873828 | 0.8513605 | 0.99937574 |
| DNAJC24 | 35.3199595 | 0.01720249 | 0.41090307 | 0.04186508 | 0.96660625 | 0.99937574 |
| IMMP1L | 96.7110599 | -0.0931887 | 0.29917689 | -0.3114835 | 0.75543311 | 0.99937574 |
| ELP4 | 116.154762 | 0.45884004 | 0.27709652 | 1.65588523 | 0.09774506 | 0.99937574 |
| AC131571.1 | 16.4203078 | 0.13408009 | 0.96264035 | 0.13928368 | 0.88922598 | 0.99937574 |
| RCN1 | 27.877219 | -0.2217206 | 0.50799959 | -0.4364583 | 0.66250428 | 0.99937574 |
| EIF3M | 50.2025706 | -0.1109179 | 0.33458311 | -0.3315108 | 0.74025868 | 0.99937574 |
| PRRG4 | 67.1074237 | -0.202524 | 0.42307016 | -0.4787007 | 0.63215154 | 0.99937574 |
| QSER1 | 64.1926041 | -0.23867 | 0.29382477 | -0.8122867 | 0.41662712 | 0.99937574 |
| TCP11L1 | 15.566413 | -0.6622876 | 0.56140419 | -1.1796984 | 0.23812021 | 0.99937574 |
| CSTF3 | 68.9630763 | 0.437687 | 0.44484097 | 0.9839179 | 0.32515588 | 0.99937574 |
| HIPK3 | 244.507364 | -0.1350337 | 0.34601851 | -0.39025 | 0.69635167 | 0.99937574 |
| AL049629.1 | 49.4296785 | 0.00757537 | 0.77254046 | 0.00980579 | 0.99217624 | 0.99937574 |
| CD59 | 11.2741662 | -0.4924728 | 0.52736031 | -0.933845 | 0.35038387 | 0.99937574 |
| FBXO3 | 61.1764584 | -0.1404147 | 0.30762151 | -0.4564529 | 0.64806434 | 0.99937574 |
| CAPRIN1 | 112.954181 | 0.25019227 | 0.22809205 | 1.09689166 | 0.27268876 | 0.99937574 |
| NAT10 | 30.1535436 | -0.1816532 | 0.40162347 | -0.4522973 | 0.65105482 | 0.99937574 |
| ABTB2 | 255.730636 | -0.2193504 | 0.58833613 | -0.3728318 | 0.70927366 | 0.99937574 |

| | | | | | | |
|---|---|---|---|---|---|---|
| CAT | 189.899674 | 0.95571715 | 0.5794244 | 1.6494251 | 0.09906058 | 0.99937574 |
| ELF5 | 22.2511979 | -5.9472331 | 2.62353809 | -2.2668751 | 0.02339786 | 0.96546668 |
| EHF | 61.3962483 | -2.1170347 | 1.05155209 | -2.0132476 | 0.04408859 | 0.99937574 |
| APIP | 66.1527488 | -0.6824707 | 0.3437192 | -1.9855472 | 0.04708363 | 0.99937574 |
| PDHX | 152.131555 | -0.0377134 | 0.32382471 | -0.1164625 | 0.90728604 | 0.99937574 |
| CD44 | 97.2192669 | 0.33062641 | 0.65519017 | 0.50462664 | 0.6138211 | 0.99937574 |
| TRIM44 | 185.85242 | -0.0907464 | 0.2868477 | -0.3163575 | 0.75173115 | 0.99937574 |
| COMMD9 | 14.9459292 | 0.07933305 | 0.51206079 | 0.15492898 | 0.87687731 | 0.99937574 |
| PRR5L | 16.9831053 | -0.6601995 | 1.11663323 | -0.5912411 | 0.55435889 | 0.99937574 |
| TRAF6 | 31.6398485 | 0.16515043 | 0.37438925 | 0.44111957 | 0.65912643 | 0.99937574 |
| C11orf74 | 16.2504106 | 0.61521744 | 0.53693735 | 1.14579 | 0.25188205 | 0.99937574 |
| API5 | 65.7195883 | -0.2602737 | 0.34935365 | -0.7450148 | 0.45626275 | 0.99937574 |
| TTC17 | 339.757628 | 0.11465635 | 0.1752063 | 0.6544077 | 0.51284918 | 0.99937574 |
| HSD17B12 | 277.271548 | 0.32753866 | 0.46286698 | 0.70763022 | 0.47917492 | 0.99937574 |
| ALKBH3 | 22.3114672 | -0.0450546 | 0.48092052 | -0.0936841 | 0.92536012 | 0.99937574 |
| ACCS | 14.2580926 | -0.1482408 | 0.70089172 | -0.2115031 | 0.83249471 | 0.99937574 |
| EXT2 | 64.3754991 | -0.5430338 | 0.27694395 | -1.9608076 | 0.04990147 | 0.99937574 |
| CD82 | 40.5932479 | -1.1231827 | 0.83612315 | -1.3433221 | 0.17916771 | 0.99937574 |
| TP53I11 | 41.7875384 | -0.4100618 | 0.40064415 | -1.0235064 | 0.30606849 | 0.99937574 |
| PRDM11 | 22.6585401 | -0.2259189 | 0.4684809 | -0.4822371 | 0.62963753 | 0.99937574 |
| SYT13 | 21.036526 | 0.70442722 | 0.72341821 | 0.97374825 | 0.33018155 | 0.99937574 |
| CRY2 | 84.5794053 | 0.08635432 | 0.4352339 | 0.19840898 | 0.84272509 | 0.99937574 |
| PEX16 | 10.5119817 | 0.20479991 | 0.55947255 | 0.3660589 | 0.71432113 | 0.99937574 |
| LARGE2 | 14.2927887 | 1.10905375 | 0.53794037 | 2.06166671 | 0.03923948 | 0.99937574 |
| PHF21A | 209.355928 | 0.18323413 | 0.30437931 | 0.60199273 | 0.54717898 | 0.99937574 |
| DGKZ | 28.6139279 | -0.3326118 | 0.60682376 | -0.5481194 | 0.58360995 | 0.99937574 |
| AMBRA1 | 452.654991 | 0.04433424 | 0.31983922 | 0.13861414 | 0.88975507 | 0.99937574 |
| ATG13 | 41.3296348 | -0.3436678 | 0.36712414 | -0.9361079 | 0.34921763 | 0.99937574 |
| ARHGAP1 | 37.0593887 | -0.6229112 | 0.3823019 | -1.6293699 | 0.10323474 | 0.99937574 |
| CKAP5 | 76.6201673 | 0.07897927 | 0.27019974 | 0.29229956 | 0.77005759 | 0.99937574 |
| LRP4 | 49.9213362 | -0.0346388 | 0.57026692 | -0.0607413 | 0.95156523 | 0.99937574 |
| C11orf49 | 164.192513 | 0.0700537 | 0.28047401 | 0.24976894 | 0.80276604 | 0.99937574 |
| ARFGAP2 | 28.7702609 | -0.3253794 | 0.46084468 | -0.70605 | 0.48015706 | 0.99937574 |
| DDB2 | 12.848043 | -0.3178356 | 0.53821833 | -0.5905329 | 0.55483342 | 0.99937574 |
| ACP2 | 13.136441 | -0.2308089 | 0.6515843 | -0.3542272 | 0.72316861 | 0.99937574 |
| NR1H3 | 19.9386946 | 0.6478843 | 0.43130997 | 1.5021315 | 0.13306315 | 0.99937574 |
| MADD | 21.3682505 | -0.695273 | 0.45140583 | -1.5402393 | 0.12350204 | 0.99937574 |
| SLC39A13 | 33.072595 | -0.4623719 | 0.46508865 | -0.9941586 | 0.32014566 | 0.99937574 |
| PSMC3 | 17.1575629 | -0.4822945 | 0.63696429 | -0.7571767 | 0.448944 | 0.99937574 |
| CELF1 | 306.809718 | -0.0158135 | 0.30541274 | -0.0517776 | 0.95870593 | 0.99937574 |
| NDUFS3 | 46.2782308 | 0.10403882 | 0.54996373 | 0.18917396 | 0.84995647 | 0.99937574 |
| MTCH2 | 71.5584532 | -0.0256151 | 0.3599638 | -0.0711602 | 0.94327024 | 0.99937574 |
| FNBP4 | 181.125433 | -0.2107951 | 0.26098305 | -0.8076965 | 0.41926534 | 0.99937574 |
| NUP160 | 130.9093 | 0.06022481 | 0.29667027 | 0.20300253 | 0.83913306 | 0.99937574 |
| PTPRJ | 1440.21174 | -0.0914974 | 0.34159607 | -0.2678526 | 0.78881276 | 0.99937574 |
| TNKS1BP1 | 50.7213586 | 0.04920262 | 0.54442281 | 0.09037575 | 0.92798863 | 0.99937574 |
| SSRP1 | 25.0769768 | -0.054254 | 0.42722802 | -0.1269907 | 0.89894775 | 0.99937574 |
| TIMM10 | 18.9981441 | 0.40963411 | 0.57330319 | 0.71451566 | 0.47490838 | 0.99937574 |
| UBE2L6 | 31.4427284 | -1.4918345 | 0.82669426 | -1.8045783 | 0.0711407 | 0.99937574 |
| SERPING1 | 284.926482 | -0.0737939 | 0.58739152 | -0.1256298 | 9.00E-01 | 9.99E-01 |
| ZDHHC5 | 85.5222303 | -0.3952854 | 0.32873406 | -1.2024474 | 0.22919022 | 0.99937574 |
| MED19 | 14.5544915 | 0.12843992 | 0.53358015 | 0.24071345 | 0.80977721 | 0.99937574 |
| CTNND1 | 183.297195 | 0.02104605 | 0.24143619 | 0.08717022 | 0.93053621 | 0.99937574 |
| LPXN | 24.0765198 | 0.1135708 | 0.52181635 | 0.21764516 | 0.82770559 | 0.99937574 |

| | | | | | | |
|---|---|---|---|---|---|---|
| FAM111A | 38.221231 | -0.2404199 | 0.3690272 | -0.6514964 | 0.51472612 | 0.99937574 |
| OSBP | 85.818634 | -0.1594379 | 0.27736482 | -0.574831 | 0.56540563 | 0.99937574 |
| PATL1 | 49.4445622 | -0.2481498 | 0.40374584 | -0.6146188 | 0.53880647 | 0.99937574 |
| STX3 | 53.8648521 | -0.6443534 | 0.57725272 | -1.1162414 | 0.26431883 | 0.99937574 |
| MRPL16 | 32.420809 | 0.4036097 | 0.4511826 | 0.89455955 | 0.37102258 | 0.99937574 |
| PRPF19 | 21.7582536 | -0.11131 | 0.54772222 | -0.2032235 | 0.83896037 | 0.99937574 |
| TMEM109 | 22.4204799 | -0.1334294 | 0.48151177 | -0.2771052 | 0.78169934 | 0.99937574 |
| VPS37C | 31.6781162 | -0.5306738 | 0.46869419 | -1.1322389 | 0.257534 | 0.99937574 |
| DDB1 | 110.707135 | 0.10391453 | 0.31531983 | 0.32955278 | 0.74173791 | 0.99937574 |
| TMEM138 | 12.2459362 | 0.09449353 | 0.53878019 | 0.17538419 | 0.86077776 | 0.99937574 |
| CPSF7 | 56.5947738 | -0.2263457 | 0.33345924 | -0.6787807 | 0.49727686 | 0.99937574 |
| AP003108.2 | 30.6645995 | -0.3270277 | 0.58539496 | -0.5586445 | 0.57640438 | 0.99937574 |
| SYT7 | 18.702871 | -0.7972557 | 1.14871533 | -0.6940412 | 0.48765639 | 0.99937574 |
| TMEM258 | 29.9935514 | -0.2205004 | 0.47445572 | -0.4647439 | 0.64211486 | 0.99937574 |
| FADS1 | 17.2156131 | -2.2667357 | 0.97856687 | -2.316383 | 0.02053737 | 0.96427895 |
| FTH1 | 985.280448 | -0.2646722 | 0.56752217 | -0.4663645 | 0.64095465 | 0.99937574 |
| ASRGL1 | 14.1547184 | -0.6298822 | 0.59469608 | -1.0591666 | 0.2895239 | 0.99937574 |
| AHNAK | 179.3537 | 0.03758095 | 0.35963387 | 0.10449782 | 0.9167743 | 0.99937574 |
| EML3 | 13.3995778 | -0.2336885 | 0.66320619 | -0.3523617 | 0.724567 | 0.99937574 |
| B3GAT3 | 33.2117855 | -0.0040709 | 0.40625904 | -0.0100206 | 0.99200487 | 0.99937574 |
| GANAB | 82.8694282 | 0.46262366 | 0.27407947 | 1.68791797 | 0.09142698 | 0.99937574 |
| UQCC3 | 27.2801999 | 0.42854337 | 0.52216359 | 0.8207071 | 0.41181312 | 0.99937574 |
| UBXN1 | 31.9818497 | 0.45761487 | 0.64736189 | 0.70689189 | 0.47963366 | 0.99937574 |
| TTC9C | 24.6181747 | -0.0248063 | 0.43105449 | -0.0575479 | 0.95410876 | 0.99937574 |
| POLR2G | 12.6748237 | 0.44303261 | 0.58193853 | 0.76130483 | 0.44647502 | 0.99937574 |
| TAF6L | 11.7350458 | -0.6839097 | 0.78960735 | -0.8661389 | 0.38641397 | 0.99937574 |
| TMEM223 | 25.8056075 | 0.09207324 | 0.40061285 | 0.22983098 | 0.81822311 | 0.99937574 |
| TMEM179B | 39.1398813 | 0.11474997 | 0.51672449 | 0.22207186 | 0.82425795 | 0.99937574 |
| NXF1 | 15.594178 | -0.2819646 | 0.49134971 | -0.5738573 | 0.56606437 | 0.99937574 |
| STX5 | 50.2290329 | -0.2243322 | 0.30861727 | -0.7268946 | 0.46729053 | 0.99937574 |
| SLC3A2 | 67.0466658 | -0.9124937 | 0.58798165 | -1.5519084 | 0.12068414 | 0.99937574 |
| RARRES3 | 333.518447 | -0.0745542 | 0.60319952 | -0.1235979 | 0.90163366 | 0.99937574 |
| PLA2G16 | 55.3803066 | -0.0859802 | 0.50010336 | -0.1719249 | 0.86349654 | 0.99937574 |
| ATL3 | 79.4771334 | -0.2213205 | 0.28923267 | -0.7651987 | 0.44415325 | 0.99937574 |
| RTN3 | 161.147546 | -0.2693576 | 0.47163565 | -0.5711138 | 0.56792251 | 0.99937574 |
| SPINDOC | 11.2260755 | 0.38216805 | 0.67527388 | 0.56594525 | 0.571431 | 0.99937574 |
| MARK2 | 92.2784791 | -0.0178614 | 0.28475697 | -0.0627252 | 0.94998531 | 0.99937574 |
| NAA40 | 27.9560086 | -0.0945464 | 0.43249331 | -0.2186077 | 0.82695563 | 0.99937574 |
| OTUB1 | 36.0103129 | 0.10865572 | 0.37994427 | 0.28597806 | 0.77489493 | 0.99937574 |
| MACROD1 | 47.1643723 | 0.32215593 | 0.40362501 | 0.79815651 | 0.42477967 | 0.99937574 |
| STIP1 | 46.2233917 | -0.4284917 | 0.33532612 | -1.2778357 | 0.20130737 | 0.99937574 |
| NUDT22 | 32.8865756 | 0.32242658 | 0.42516892 | 0.75834937 | 0.44824186 | 0.99937574 |
| DNAJC4 | 26.7731751 | 0.77309114 | 0.39899082 | 1.93761638 | 0.05267004 | 0.99937574 |
| VEGFB | 73.6222251 | -0.2821976 | 0.43445824 | -0.649539 | 0.51599004 | 0.99937574 |
| FKBP2 | 68.0565283 | 0.05553166 | 0.43684842 | 0.12711883 | 0.89884635 | 0.99937574 |
| PPP1R14B | 47.9503096 | 0.09915463 | 0.48270674 | 0.20541381 | 0.83724883 | 0.99937574 |
| BAD | 27.2019453 | 0.40800631 | 0.44820398 | 0.91031389 | 0.36265699 | 0.99937574 |
| GPR137 | 19.2764133 | 0.13283309 | 0.47430709 | 0.28005714 | 0.77943366 | 0.99937574 |
| ESRRA | 119.418281 | -0.4485493 | 0.40279777 | -1.1135843 | 0.26545758 | 0.99937574 |
| TRMT112 | 46.1073939 | -0.3430804 | 0.62751753 | -0.5467265 | 0.58456668 | 0.99937574 |
| PRDX5 | 98.8887732 | 0.08434611 | 0.56484341 | 0.14932654 | 0.88129597 | 0.99937574 |
| SF1 | 158.907672 | -0.2019196 | 0.30466346 | -0.6627627 | 0.50748254 | 0.99937574 |
| MAP4K2 | 12.5081496 | 0.3605447 | 0.55928062 | 0.64465795 | 0.51914887 | 0.99937574 |
| MEN1 | 11.6763014 | 0.45368496 | 0.57324567 | 0.79143199 | 0.42869195 | 0.99937574 |

| | | | | | | |
|---|---|---|---|---|---|---|
| EHD1 | 13.2347075 | 0.40562407 | 0.60414201 | 0.67140517 | 0.50196246 | 0.99937574 |
| ATG2A | 23.0886107 | -0.6611211 | 0.7313877 | -0.903927 | 0.36603412 | 0.99937574 |
| PPP2R5B | 12.4981186 | -0.0347839 | 0.54160415 | -0.0642239 | 0.94879195 | 0.99937574 |
| ZFPL1 | 13.0821953 | -0.2106581 | 0.58477613 | -0.3602372 | 0.71866976 | 0.99937574 |
| VPS51 | 64.4435989 | -0.1709135 | 0.36113342 | -0.4732698 | 0.63602072 | 0.99937574 |
| TM7SF2 | 18.3373053 | 0.94999287 | 0.4152259 | 2.28789407 | 0.02214369 | 0.96427895 |
| FAU | 166.675477 | -0.3212764 | 0.44425237 | -0.7231845 | 0.46956654 | 0.99937574 |
| SYVN1 | 21.0146011 | -0.1499568 | 0.4461311 | -0.3361271 | 0.73677499 | 0.99937574 |
| MRPL49 | 25.3915673 | 0.40379229 | 0.49398434 | 0.81741921 | 0.4136889 | 0.99937574 |
| CAPN1 | 80.4503871 | -0.0655672 | 0.32441987 | -0.202106 | 0.83983388 | 0.99937574 |
| DPF2 | 32.1144792 | -0.1231016 | 0.34003626 | -0.362025 | 0.71733331 | 0.99937574 |
| SLC25A45 | 15.0146194 | -0.3967971 | 0.68492514 | -0.5793291 | 0.56236714 | 0.99937574 |
| FRMD8 | 25.032029 | -0.3210034 | 0.47734134 | -0.672482 | 0.50127692 | 0.99937574 |
| NEAT1 | 5140.88093 | 0.87130596 | 0.50057558 | 1.74060818 | 0.08175228 | 0.99937574 |
| MALAT1 | 90522.2756 | 0.01655177 | 0.39969545 | 0.04141095 | 0.96696828 | 0.99937574 |
| SCYL1 | 31.5370091 | -0.0527815 | 0.47086804 | -0.1120941 | 0.91074881 | 0.99937574 |
| LTBP3 | 44.9969002 | 0.4983523 | 0.40278208 | 1.23727528 | 0.2159849 | 0.99937574 |
| SSSCA1 | 14.0317257 | 0.20270573 | 0.47092361 | 0.43044292 | 0.66687348 | 0.99937574 |
| FAM89B | 10.9292043 | -0.1286243 | 0.70079199 | -0.1835413 | 0.85437333 | 0.99937574 |
| EHBP1L1 | 18.2943254 | 0.511286 | 0.6992544 | 0.73118738 | 0.46466471 | 0.99937574 |
| MAP3K11 | 25.3722333 | -0.4646542 | 0.37171412 | -1.2500311 | 0.2112882 | 0.99937574 |
| PCNX3 | 21.2074235 | -1.0239249 | 0.57272346 | -1.7878172 | 0.0738055 | 0.99937574 |
| SIPA1 | 19.1279229 | 0.52283597 | 0.6763854 | 0.77298529 | 0.43953109 | 0.99937574 |
| RELA | 32.2751341 | -0.1170456 | 0.37858279 | -0.3091677 | 0.75719395 | 0.99937574 |
| KAT5 | 28.0825008 | -1.0555897 | 0.5106465 | -2.0671632 | 0.03871877 | 0.99937574 |
| RNASEH2C | 18.5983883 | 0.29117564 | 0.69851718 | 0.41684822 | 0.67678943 | 0.99937574 |
| AP5B1 | 10.3595638 | -0.4235203 | 0.75717208 | -0.5593449 | 0.57592637 | 0.99937574 |
| CFL1 | 169.049482 | -0.1567227 | 0.45580538 | -0.3438368 | 0.73096903 | 0.99937574 |
| MUS81 | 16.7456435 | -0.0225713 | 0.44331571 | -0.0509147 | 0.95939351 | 0.99937574 |
| EFEMP2 | 14.8071665 | 0.46044486 | 0.47257275 | 0.97433646 | 0.32988951 | 0.99937574 |
| CCDC85B | 12.2416601 | 0.11203845 | 0.49344902 | 0.22705171 | 0.82038353 | 0.99937574 |
| C11orf68 | 10.9976783 | 0.03107392 | 0.55490666 | 0.05599847 | 0.95534303 | 0.99937574 |
| DRAP1 | 26.0278204 | 0.03776815 | 0.63576198 | 0.05940612 | 0.95262864 | 0.99937574 |
| SART1 | 22.0092637 | -0.3702003 | 0.52785245 | -0.701333 | 0.48309525 | 0.99937574 |
| BANF1 | 35.0218736 | -0.1931932 | 0.46787544 | -0.4129159 | 0.67966826 | 0.99937574 |
| SF3B2 | 53.7083941 | -0.3393577 | 0.36591364 | -0.9274256 | 0.3537056 | 0.99937574 |
| PACS1 | 414.489219 | 0.01763653 | 0.27846127 | 0.06333565 | 0.94949922 | 0.99937574 |
| KLC2 | 22.5608751 | -0.1702617 | 0.45563336 | -0.3736814 | 0.7086414 | 0.99937574 |
| RAB1B | 51.335261 | 0.3068345 | 0.38022465 | 0.80698214 | 0.41967677 | 0.99937574 |
| YIF1A | 18.2290645 | 0.36875454 | 0.47225056 | 0.78084511 | 0.4348936 | 0.99937574 |
| BRMS1 | 15.8667696 | -0.1525635 | 0.46287344 | -0.3296009 | 0.74170155 | 0.99937574 |
| SLC29A2 | 41.3286933 | -0.0044452 | 0.43756423 | -0.0101589 | 0.99189448 | 0.99937574 |
| MRPL11 | 15.5072995 | 0.34776112 | 0.53521637 | 0.64975799 | 0.51584856 | 0.99937574 |
| ZDHHC24 | 15.4791279 | 0.50004909 | 0.56060686 | 0.89197819 | 0.37240462 | 0.99937574 |
| CCS | 36.3275002 | 0.48969047 | 0.43104911 | 1.13604335 | 0.25593842 | 0.99937574 |
| RBM14-RBM | 26.7717027 | 0.21154478 | 0.42995868 | 0.49201189 | 0.62271094 | 0.99937574 |
| RBM4 | 24.8422593 | -0.2163061 | 0.41217591 | -0.5247908 | 0.59972861 | 0.99937574 |
| RBM4B | 30.8866097 | 0.24283085 | 0.4776968 | 0.50833676 | 0.61121719 | 0.99937574 |
| AP001157.1 | 11.9676492 | -0.4177617 | 0.72106905 | -0.5793644 | 0.56234332 | 0.99937574 |
| SPTBN2 | 302.365386 | 0.65742965 | 0.49205507 | 1.33608957 | 0.18152 | 0.99937574 |
| C11orf80 | 541.19099 | 0.34055321 | 0.54620717 | 0.62348727 | 0.53296437 | 0.99937574 |
| RCE1 | 20.6304298 | 0.2869388 | 0.53040491 | 0.54098066 | 0.58852091 | 0.99937574 |
| PC | 12.5818994 | 0.6817061 | 0.64883715 | 1.05065824 | 0.29341558 | 0.99937574 |
| LRFN4 | 12.3812659 | -1.8336095 | 0.62400204 | -2.9384672 | 0.0032984 | 0.53256471 |

| | | | | | | |
|---|---|---|---|---|---|---|
| RHOD | 25.8819266 | -0.1452357 | 0.45664629 | -0.3180487 | 0.75044803 | 0.99937574 |
| KDM2A | 302.17532 | -0.0592364 | 0.25321485 | -0.2339372 | 0.81503371 | 0.99937574 |
| GRK2 | 20.3443233 | -0.375031 | 0.6052019 | -0.6196792 | 0.53546904 | 0.99937574 |
| ANKRD13D | 21.2633662 | 0.09346479 | 0.42963162 | 0.21754635 | 0.82778259 | 0.99937574 |
| SSH3 | 25.6018003 | 0.17354549 | 0.55143257 | 0.31471752 | 0.75297614 | 0.99937574 |
| RAD9A | 94.900514 | 0.00381752 | 0.42032618 | 0.00908227 | 0.99275349 | 0.99937574 |
| PPP1CA | 61.4944628 | 0.00530955 | 0.46996389 | 0.01129779 | 0.99098586 | 0.99937574 |
| RPS6KB2 | 26.5872931 | -0.4990399 | 0.4534094 | -1.1006387 | 0.27105394 | 0.99937574 |
| CORO1B | 30.0701652 | -0.0817475 | 0.43458067 | -0.1881067 | 0.85079303 | 0.99937574 |
| TMEM134 | 31.9024164 | -0.0687231 | 0.42844491 | -0.1604012 | 0.87256501 | 0.99937574 |
| AIP | 14.7401118 | -0.5640865 | 0.56813836 | -0.9928682 | 0.32077421 | 0.99937574 |
| PITPNM1 | 19.3649784 | -0.4199823 | 0.57079347 | -0.7357867 | 0.46186051 | 0.99937574 |
| CDK2AP2 | 29.1959196 | -0.3691377 | 0.40969241 | -0.9010117 | 0.36758208 | 0.99937574 |
| GSTP1 | 275.698176 | 0.07238037 | 0.49563146 | 0.14603667 | 0.88389244 | 0.99937574 |
| NDUFV1 | 93.2959059 | 0.64131187 | 0.34976235 | 1.83356462 | 0.06671865 | 0.99937574 |
| NUDT8 | 14.9309916 | 0.31738924 | 0.68025054 | 0.46657697 | 0.64080257 | 0.99937574 |
| UNC93B1 | 11.8619975 | -1.4810998 | 0.89499122 | -1.6548763 | 0.09794959 | 0.99937574 |
| NDUFS8 | 92.8765637 | 0.5763915 | 0.44962521 | 1.28193768 | 0.19986451 | 0.99937574 |
| TCIRG1 | 23.7213998 | -0.4222148 | 0.47085148 | -0.8967049 | 0.3698764 | 0.99937574 |
| CHKA | 140.515745 | -0.3371943 | 0.38876352 | -0.8673508 | 0.38574986 | 0.99937574 |
| KMT5B | 164.007311 | -0.2206542 | 0.35551009 | -0.6206694 | 0.53481714 | 0.99937574 |
| LRP5 | 43.1793425 | 0.5744435 | 0.46596399 | 1.23280663 | 0.21764792 | 0.99937574 |
| PPP6R3 | 393.981943 | -0.0522481 | 0.21668505 | -0.2411248 | 0.80945838 | 0.99937574 |
| TESMIN | 37.576221 | 0.1811482 | 0.50614363 | 0.3578988 | 0.72041905 | 0.99937574 |
| CPT1A | 277.792945 | 0.22314697 | 0.50555001 | 0.44139444 | 0.65892746 | 0.99937574 |
| MRPL21 | 15.7167852 | 0.47822587 | 0.46038284 | 1.03875694 | 0.29891779 | 0.99937574 |
| IGHMBP2 | 16.2163945 | -0.2024075 | 0.46866629 | -0.4318797 | 0.66582882 | 0.99937574 |
| TPCN2 | 20.3665804 | -0.0547277 | 0.60477637 | -0.0904924 | 0.92789594 | 0.99937574 |
| CCND1 | 31.9292114 | 0.42042963 | 0.43076775 | 0.97600072 | 0.32906411 | 0.99937574 |
| ORAOV1 | 18.4681504 | -0.8242114 | 0.5058715 | -1.6292901 | 0.10325162 | 0.99937574 |
| PPFIA1 | 141.518875 | -0.4957751 | 0.43069502 | -1.1511048 | 0.24968913 | 0.99937574 |
| AP002336.2 | 48.7948427 | -0.0106635 | 0.59077978 | -0.0180498 | 0.98559909 | 0.99937574 |
| CTTN | 211.326704 | -0.5120418 | 0.26427342 | -1.9375456 | 0.05267868 | 0.99937574 |
| SHANK2 | 404.908599 | 0.29437634 | 0.37002847 | 0.79555053 | 0.42629333 | 0.99937574 |
| DHCR7 | 26.424654 | 0.60477481 | 0.60119218 | 1.00595921 | 0.31443519 | 0.99937574 |
| NADSYN1 | 48.5344245 | 0.25832195 | 0.30583549 | 0.84464348 | 0.39830992 | 0.99937574 |
| KRTAP5-8 | 31.5144785 | 0.27105513 | 1.16600034 | 0.23246574 | 0.8161763 | 0.99937574 |
| AP002495.2 | 45.356211 | -0.366367 | 0.42213454 | -0.8678915 | 0.38545372 | 0.99937574 |
| RNF121 | 83.3284999 | -0.0817909 | 0.29784482 | -0.2746093 | 0.78361644 | 0.99937574 |
| NUMA1 | 122.640605 | 0.18944838 | 0.37707567 | 0.50241476 | 0.6153758 | 0.99937574 |
| LAMTOR1 | 73.4056845 | -0.1840127 | 0.3726493 | -0.4937959 | 0.62145033 | 0.99937574 |
| ANAPC15 | 17.1784731 | 0.27129567 | 0.58769562 | 0.46162616 | 0.64434943 | 0.99937574 |
| FOLR1 | 20.5532677 | 0.81268447 | 1.07931178 | 0.75296545 | 0.45147068 | 0.99937574 |
| INPPL1 | 43.1471261 | 0.14399382 | 0.36918325 | 0.39003346 | 0.69651181 | 0.99937574 |
| CLPB | 36.6155699 | 0.04820375 | 0.39974657 | 0.12058577 | 0.90401914 | 0.99937574 |
| ARAP1 | 142.91559 | -0.0436747 | 0.29551871 | -0.14779 | 0.88250852 | 0.99937574 |
| ATG16L2 | 13.3439681 | -0.1395312 | 0.53704027 | -0.2598151 | 0.79500637 | 0.99937574 |
| FCHSD2 | 88.8977327 | -0.0026819 | 0.28233613 | -0.0094989 | 0.99242105 | 0.99937574 |
| ARHGEF17 | 52.7142829 | 0.40749314 | 0.33533172 | 1.21519412 | 0.22429206 | 0.99937574 |
| FAM168A | 130.494137 | -0.0982877 | 0.32282241 | -0.3044635 | 0.76077476 | 0.99937574 |
| PLEKHB1 | 15.3237813 | 0.26605803 | 0.74318807 | 0.35799556 | 0.72034664 | 0.99937574 |
| RAB6A | 164.619548 | -0.3388162 | 0.24577928 | -1.3785387 | 0.16803702 | 0.99937574 |
| MRPL48 | 40.2720127 | 0.1342908 | 0.42139097 | 0.31868456 | 0.74996572 | 0.99937574 |
| COA4 | 31.5813579 | -0.0477818 | 0.39497117 | -0.1209754 | 0.90371055 | 0.99937574 |

| | | | | | | |
|---|---|---|---|---|---|---|
| PAAF1 | 33.0188431 | 0.77496993 | 0.49278308 | 1.57263908 | 0.11580241 | 0.99937574 |
| UCP2 | 32.2343531 | -0.7268839 | 0.85647352 | -0.8486939 | 0.39605162 | 0.99937574 |
| C2CD3 | 75.7432509 | 0.10994199 | 0.31719377 | 0.34660828 | 0.72888562 | 0.99937574 |
| PPME1 | 99.0855084 | -0.2098777 | 0.36460302 | -0.5756336 | 0.56486287 | 0.99937574 |
| P4HA3 | 19.1093123 | -0.4200345 | 0.76253309 | -0.550841 | 0.58174266 | 0.99937574 |
| POLD3 | 30.7525778 | 0.15408509 | 0.37757219 | 0.40809438 | 0.68320438 | 0.99937574 |
| RNF169 | 182.002308 | -0.041939 | 0.3471468 | -0.1208105 | 0.90384113 | 0.99937574 |
| XRRA1 | 103.225371 | -0.1589432 | 0.55566216 | -0.2860429 | 0.77484528 | 0.99937574 |
| SPCS2 | 118.382707 | 0.23866042 | 0.35458469 | 0.67307029 | 0.50090257 | 0.99937574 |
| NEU3 | 14.7445951 | 0.12084817 | 0.57420376 | 0.21046218 | 0.83330697 | 0.99937574 |
| ARRB1 | 42.8907797 | -0.4646505 | 0.56317264 | -0.8250587 | 0.40933825 | 0.99937574 |
| RPS3 | 297.895663 | 0.03568024 | 0.45286501 | 0.0787878 | 0.93720141 | 0.99937574 |
| SERPINH1 | 15.1393812 | 0.41900328 | 0.54916144 | 0.76298744 | 0.44547089 | 0.99937574 |
| UVRAG | 198.093415 | -0.1430037 | 0.26626051 | -0.537082 | 0.591211 | 0.99937574 |
| THAP12 | 28.9319559 | -0.0230086 | 0.41447064 | -0.0555133 | 0.95572953 | 0.99937574 |
| EMSY | 108.596223 | 0.15392317 | 0.27442152 | 0.56090051 | 0.57486536 | 0.99937574 |
| ACER3 | 81.2196797 | -0.1718454 | 0.31128213 | -0.5520568 | 0.58090941 | 0.99937574 |
| CAPN5 | 13.091867 | 1.2213282 | 0.71837772 | 1.70011982 | 0.08910839 | 0.99937574 |
| PAK1 | 100.165034 | 0.06831774 | 0.44069955 | 0.15502113 | 0.87680466 | 0.99937574 |
| CLNS1A | 55.3412099 | -0.1857695 | 0.31885144 | -0.5826207 | 0.56014865 | 0.99937574 |
| RSF1 | 204.527526 | 0.09245481 | 0.25257813 | 0.36604441 | 0.71433194 | 0.99937574 |
| AAMDC | 30.2977525 | 0.41869329 | 0.41253911 | 1.01491782 | 0.310145 | 0.99937574 |
| AP002812.2 | 15.4515981 | 0.07556416 | 0.7427471 | 0.10173605 | 0.91896618 | 0.99937574 |
| INTS4 | 51.1537023 | -0.1934657 | 0.29225856 | -0.6619677 | 0.50799194 | 0.99937574 |
| ALG8 | 29.4206098 | -0.1564039 | 0.42321134 | -0.3695646 | 0.71170695 | 0.99937574 |
| GAB2 | 155.140492 | -0.516687 | 0.42410148 | -1.2183098 | 0.22310625 | 0.99937574 |
| AP003086.2 | 39.9057479 | -0.2022294 | 0.57940606 | -0.3490287 | 0.72706774 | 0.99937574 |
| NARS2 | 61.7782917 | 0.04070327 | 0.31850917 | 0.12779309 | 0.89831271 | 0.99937574 |
| TENM4 | 26.177515 | 0.62749248 | 0.86961312 | 0.7215766 | 0.47055483 | 0.99937574 |
| PRCP | 39.1910916 | 0.26830039 | 0.45036444 | 0.59574061 | 0.55134852 | 0.99937574 |
| RAB30 | 21.9525368 | 0.19505853 | 0.72210489 | 0.27012493 | 0.78706415 | 0.99937574 |
| RAB30-AS1 | 47.5010168 | 0.28225358 | 0.44205765 | 0.63849947 | 0.5231486 | 0.99937574 |
| PCF11 | 62.1633882 | 0.18940808 | 0.452392 | 0.41868131 | 0.67544905 | 0.99937574 |
| ANKRD42 | 29.1864524 | 0.44275326 | 0.39196191 | 1.12958237 | 0.25865224 | 0.99937574 |
| CCDC90B | 31.1943809 | 0.10118289 | 0.41014484 | 0.24670038 | 0.80514011 | 0.99937574 |
| AP000446.1 | 26.3208328 | 0.46977837 | 0.40031992 | 1.17350735 | 0.24059241 | 0.99937574 |
| DLG2 | 63.4145482 | 0.05808877 | 0.59497311 | 0.0976326 | 0.92222404 | 0.99937574 |
| TMEM126B | 25.7965884 | -0.1966279 | 0.56864795 | -0.3457813 | 0.72950705 | 0.99937574 |
| TMEM126A | 14.1812345 | 0.28194878 | 0.60362035 | 0.46709622 | 0.64043104 | 0.99937574 |
| CREBZF | 55.0448051 | 0.14453943 | 0.29334006 | 0.49273675 | 0.62219861 | 0.99937574 |
| SYTL2 | 15.8525321 | -0.574722 | 0.83551923 | -0.687862 | 0.49153966 | 0.99937574 |
| PICALM | 1001.18224 | 0.14329071 | 0.41608609 | 0.34437756 | 0.73056236 | 0.99937574 |
| EED | 35.2182996 | 0.04637361 | 0.38927109 | 0.11912934 | 0.90517289 | 0.99937574 |
| HIKESHI | 37.2563338 | -0.0967067 | 0.39344234 | -0.2457964 | 0.80583985 | 0.99937574 |
| AP001831.1 | 48.186534 | 1.05190067 | 0.53470576 | 1.96725144 | 0.04915423 | 0.99937574 |
| ME3 | 125.995295 | 0.36936051 | 0.34836333 | 1.0602738 | 0.28902005 | 0.99937574 |
| PRSS23 | 32.6052704 | 0.3836248 | 0.65975937 | 0.58146169 | 0.56092933 | 0.99937574 |
| TMEM135 | 143.260373 | 0.0776348 | 0.32734154 | 0.23716759 | 0.81252678 | 0.99937574 |
| CTSC | 23.5389127 | 0.28208622 | 0.55753112 | 0.505956 | 0.61288754 | 0.99937574 |
| NOX4 | 53.773197 | 1.00222324 | 0.83597027 | 1.19887427 | 0.23057684 | 0.99937574 |
| CHORDC1 | 42.9590189 | -0.1874004 | 0.4698393 | -0.3988607 | 0.68999584 | 0.99937574 |
| SLC36A4 | 77.1910595 | -0.8674831 | 0.42667057 | -2.0331448 | 0.0420379 | 0.99937574 |
| CEP295 | 40.0755432 | -0.006654 | 0.39531416 | -0.0168322 | 0.98657045 | 0.99937574 |
| TAF1D | 150.033282 | 0.14907712 | 0.38671333 | 0.38549776 | 0.69986866 | 0.99937574 |

| | | | | | | |
|---|---|---|---|---|---|---|
| PANX1 | 15.2496833 | -0.0751705 | 0.46181711 | -0.1627712 | 0.87069858 | 0.99937574 |
| MRE11 | 35.3426007 | -0.3734254 | 0.39651055 | -0.9417792 | 0.34630568 | 0.99937574 |
| ANKRD49 | 14.1132366 | -0.3964956 | 0.52334001 | -0.7576253 | 0.44867532 | 0.99937574 |
| CWC15 | 24.5088325 | -0.5284913 | 0.4665566 | -1.1327486 | 0.25731985 | 0.99937574 |
| SRSF8 | 27.0309061 | -0.4034147 | 0.44156602 | -0.9136001 | 0.36092703 | 0.99937574 |
| ENDOD1 | 68.7337763 | 0.58762206 | 0.56290511 | 1.04390963 | 0.2965272 | 0.99937574 |
| AP000787.1 | 36.6059148 | -0.2183299 | 0.49007017 | -0.4455074 | 0.65595314 | 0.99937574 |
| SESN3 | 22.4814297 | 0.05778726 | 0.54577558 | 0.10588099 | 0.91567678 | 0.99937574 |
| FAM76B | 29.3873249 | 0.00896832 | 0.40506097 | 0.02214067 | 0.98233575 | 0.99937574 |
| CEP57 | 105.373235 | 0.17610412 | 0.28024568 | 0.62839192 | 0.52974723 | 0.99937574 |
| MTMR2 | 65.4774743 | -0.307563 | 0.36664477 | -0.8388582 | 0.40154891 | 0.99937574 |
| MAML2 | 36.745391 | -0.1411674 | 0.45986686 | -0.3069746 | 0.75886273 | 0.99937574 |
| CCDC82 | 37.1441079 | -0.2714224 | 0.4249646 | -0.6386941 | 0.52302193 | 0.99937574 |
| CNTN5 | 15.1422248 | -0.9957974 | 0.90023471 | -1.106153 | 0.26866029 | 0.99937574 |
| ARHGAP42 | 26.244288 | -0.0561616 | 0.56596776 | -0.0992311 | 0.92095481 | 0.99937574 |
| YAP1 | 323.205424 | -0.1725397 | 0.23672646 | -0.728857 | 0.46608913 | 0.99937574 |
| BIRC3 | 92.5408533 | 0.1627851 | 0.78482518 | 0.20741575 | 0.83568518 | 0.99937574 |
| BIRC2 | 74.2928353 | 0.16779158 | 0.31933719 | 0.525437 | 0.59927944 | 0.99937574 |
| TMEM123 | 214.09475 | 0.17714086 | 0.27992623 | 0.63281266 | 0.52685599 | 0.99937574 |
| MMP7 | 45.4807473 | 0.41959688 | 0.72540979 | 0.57842738 | 5.63E-01 | 9.99E-01 |
| DCUN1D5 | 33.6157994 | -0.070121 | 0.4152306 | -0.1688724 | 0.86589702 | 0.99937574 |
| DYNC2H1 | 63.3825483 | -0.2495405 | 0.5989244 | -0.4166478 | 0.67693607 | 0.99937574 |
| CASP4 | 45.3661186 | -0.1978027 | 0.46936598 | -0.4214253 | 0.67344454 | 0.99937574 |
| MSANTD4 | 16.1065803 | -0.1147022 | 0.48070096 | -0.2386145 | 0.81140454 | 0.99937574 |
| KBTBD3 | 18.2721863 | -0.0345758 | 0.48389026 | -0.0714538 | 0.94303656 | 0.99937574 |
| AASDHPPT | 34.8838732 | 0.47790509 | 0.42203717 | 1.13237678 | 0.25747606 | 0.99937574 |
| CWF19L2 | 93.3916555 | -0.1676801 | 0.27035447 | -0.6202231 | 0.53511093 | 0.99937574 |
| ALKBH8 | 17.0883543 | -0.0221418 | 0.44797228 | -0.0494268 | 0.96057919 | 0.99937574 |
| SLC35F2 | 14.9653744 | -0.3057535 | 0.53655423 | -0.5698464 | 0.56878187 | 0.99937574 |
| CUL5 | 361.727845 | -0.4273088 | 0.44457938 | -0.961153 | 0.33647525 | 0.99937574 |
| AP002433.1 | 15.3871825 | 0.04065299 | 1.06050134 | 0.03833374 | 0.96942159 | 0.99937574 |
| ACAT1 | 295.961025 | 0.0706604 | 0.42656564 | 0.16564953 | 0.86843277 | 0.99937574 |
| NPAT | 48.5650525 | -0.2460197 | 0.33305063 | -0.7386856 | 0.46009794 | 0.99937574 |
| ATM | 130.151105 | -0.4663986 | 0.32196976 | -1.4485787 | 0.14745526 | 0.99937574 |
| C11orf65 | 32.6782893 | -0.9433522 | 0.7323488 | -1.2881187 | 0.19770464 | 0.99937574 |
| KDELC2 | 32.1949587 | -0.0691703 | 0.73107499 | -0.0946145 | 0.92462102 | 0.99937574 |
| EXPH5 | 86.8032032 | -0.6132335 | 0.34925056 | -1.7558555 | 0.07911309 | 0.99937574 |
| DDX10 | 111.45444 | -0.2148459 | 0.28069947 | -0.7653946 | 0.4440366 | 0.99937574 |
| RDX | 148.251069 | 0.15258403 | 0.2377931 | 0.64166717 | 0.52108931 | 0.99937574 |
| ZC3H12C | 110.694021 | 0.39152186 | 0.45276432 | 0.86473655 | 0.3871834 | 0.99937574 |
| FDX1 | 54.2182391 | 0.03041007 | 0.35362795 | 0.08599454 | 0.93147076 | 0.99937574 |
| COLCA1 | 19.47675 | -0.2523944 | 0.6877703 | -0.3669749 | 0.71363774 | 0.99937574 |
| COLCA2 | 13.8442382 | -0.67062 | 0.99555726 | -0.6736126 | 0.50055761 | 0.99937574 |
| SIK2 | 353.712142 | 0.27418048 | 0.3266037 | 0.8394898 | 0.40119451 | 0.99937574 |
| PPP2R1B | 106.470637 | 0.37249763 | 0.2644287 | 1.40868836 | 0.15892734 | 0.99937574 |
| CRYAB | 200.442112 | -1.0120161 | 0.78612658 | -1.287345 | 0.19797408 | 0.99937574 |
| DIXDC1 | 22.6386593 | -0.6051212 | 0.70144839 | -0.8626739 | 0.3883168 | 0.99937574 |
| DLAT | 56.5097798 | 0.19909574 | 0.38212808 | 0.52101835 | 0.60235399 | 0.99937574 |
| NKAPD1 | 13.9843313 | 0.04284174 | 0.47649581 | 0.08991001 | 0.92835872 | 0.99937574 |
| TIMM8B | 23.8207705 | 0.52745248 | 0.50661343 | 1.04113403 | 0.29781335 | 0.99937574 |
| AP002884.3 | 35.3929025 | -0.1648178 | 0.50036608 | -0.3293943 | 0.74185764 | 0.99937574 |
| IL18 | 740.201625 | 0.59086193 | 0.28863708 | 2.04707561 | 0.04065066 | 0.99937574 |
| BCO2 | 78.1668585 | -0.3978982 | 0.62729697 | -0.6343059 | 0.52588119 | 0.99937574 |
| PTS | 72.5425575 | -0.2934786 | 0.42614353 | -0.6886848 | 0.49102167 | 0.99937574 |

| | | | | | | |
|---|---|---|---|---|---|---|
| AP002884.1 | 32.4883117 | 0.27038002 | 0.69644496 | 0.38822884 | 0.69784669 | 0.99937574 |
| TTC12 | 35.0284913 | 0.03214557 | 0.45520619 | 0.07061761 | 0.9437021 | 0.99937574 |
| DRD2 | 15.3631253 | 2.43141821 | 1.16709408 | 2.08330952 | 0.03722302 | 0.99937574 |
| ZW10 | 25.8282109 | 0.56253682 | 0.45676514 | 1.23156686 | 0.21811093 | 0.99937574 |
| USP28 | 81.5484895 | 0.41190748 | 0.35635075 | 1.15590464 | 0.24772016 | 0.99937574 |
| ZBTB16 | 247.643194 | -0.0862454 | 0.53491878 | -0.1612308 | 8.72E-01 | 9.99E-01 |
| AP002518.2 | 20.4430067 | 0.52062659 | 0.97316855 | 0.53498091 | 0.59266306 | 0.99937574 |
| NNMT | 43.6853351 | 1.04895168 | 0.69226434 | 1.51524731 | 1.30E-01 | 9.99E-01 |
| REXO2 | 21.6842664 | -0.3391281 | 0.50913297 | -0.6660895 | 0.50535388 | 0.99937574 |
| CADM1 | 31.8089638 | -1.1068437 | 0.48586114 | -2.2781071 | 0.0227202 | 0.96427895 |
| BUD13 | 12.6481933 | 0.41490821 | 0.61323157 | 0.67659304 | 0.4986642 | 0.99937574 |
| ZPR1 | 11.1307945 | -0.3192472 | 0.75564403 | -0.4224836 | 0.67267205 | 0.99937574 |
| SIK3 | 681.630758 | -0.0738977 | 0.37349382 | -0.1978552 | 0.84315839 | 0.99937574 |
| PAFAH1B2 | 131.618905 | -0.1017213 | 0.31729079 | -0.3205933 | 0.74851863 | 0.99937574 |
| SIDT2 | 39.5650923 | -0.2917999 | 0.38258911 | -0.7626979 | 0.4456436 | 0.99937574 |
| TAGLN | 14.3452241 | -0.1137736 | 0.6641359 | -0.1713106 | 0.86397951 | 0.99937574 |
| PCSK7 | 96.2270312 | 0.12920226 | 0.37318041 | 0.34621931 | 0.7291779 | 0.99937574 |
| RNF214 | 30.8206804 | -0.0241499 | 0.4154708 | -0.0581267 | 0.95364772 | 0.99937574 |
| BACE1 | 20.9071946 | 0.67323126 | 0.52745891 | 1.27636722 | 0.20182574 | 0.99937574 |
| CEP164 | 19.7333964 | -0.5479962 | 0.56137084 | -0.976175 | 0.32897776 | 0.99937574 |
| DSCAML1 | 404.239525 | 1.39757893 | 1.15681693 | 1.20812454 | 0.22699936 | 0.99937574 |
| FXYD2 | 75.7836495 | 0.68055236 | 0.66893434 | 1.01736795 | 0.30897842 | 0.99937574 |
| AP000757.1 | 29.7018452 | 1.78638527 | 0.67769479 | 2.63597317 | 0.00838964 | 0.74742128 |
| MPZL3 | 18.4458835 | -0.3654802 | 0.45246494 | -0.8077536 | 0.41923241 | 0.99937574 |
| UBE4A | 56.1221759 | 0.02344425 | 0.33514514 | 0.06995254 | 0.94423143 | 0.99937574 |
| ATP5MG | 180.707424 | 0.39837364 | 0.50555102 | 0.78799889 | 0.43069736 | 0.99937574 |
| KMT2A | 178.84555 | -0.2474811 | 0.27044589 | -0.9150855 | 0.36014674 | 0.99937574 |
| IFT46 | 12.3547004 | 0.12966928 | 0.70718588 | 0.18335954 | 0.85451592 | 0.99937574 |
| ARCN1 | 59.7826614 | 0.20669642 | 0.3253351 | 0.6353339 | 0.52521067 | 0.99937574 |
| PHLDB1 | 12.6750527 | -1.1585389 | 0.5792199 | -2.0001711 | 0.04548179 | 0.99937574 |
| AP002954.1 | 28.9162819 | 0.33252186 | 0.57053731 | 0.58282229 | 0.56001294 | 0.99937574 |
| DDX6 | 170.273532 | -0.037156 | 0.22517669 | -0.1650082 | 0.86893754 | 0.99937574 |
| BCL9L | 10.4190219 | -1.2283884 | 0.61373004 | -2.0015126 | 0.04533718 | 0.99937574 |
| CCDC84 | 45.6787423 | -0.857118 | 0.58245209 | -1.4715683 | 0.14113749 | 0.99937574 |
| RPS25 | 179.721377 | -0.1570599 | 0.48738843 | -0.322248 | 0.74726486 | 0.99937574 |
| TRAPPC4 | 48.6082815 | 0.27650605 | 0.54757124 | 0.50496817 | 0.61358119 | 0.99937574 |
| SLC37A4 | 17.032117 | 0.11488765 | 0.67398212 | 0.17046098 | 0.86464762 | 0.99937574 |
| HYOU1 | 24.2989001 | -0.4245046 | 0.4709264 | -0.9014245 | 0.36736267 | 0.99937574 |
| VPS11 | 24.1445415 | 0.28720299 | 0.41286385 | 0.69563608 | 0.48665677 | 0.99937574 |
| HINFP | 13.899138 | -0.1669594 | 0.49945205 | -0.3342852 | 0.73816434 | 0.99937574 |
| NLRX1 | 16.102041 | -0.052839 | 0.60756295 | -0.0869687 | 0.93069638 | 0.99937574 |
| CBL | 66.2120093 | -0.1873116 | 0.32274014 | -0.5803789 | 0.56165915 | 0.99937574 |
| RNF26 | 11.0877558 | -0.6689436 | 0.58658712 | -1.1403995 | 0.25411991 | 0.99937574 |
| OAF | 47.274018 | -0.1858046 | 0.65850981 | -0.2821592 | 0.77782144 | 0.99937574 |
| ARHGEF12 | 582.468682 | 0.39640391 | 0.32807587 | 1.20826901 | 0.2269438 | 0.99937574 |
| SC5D | 57.2223423 | -0.425933 | 0.48281281 | -0.8821907 | 0.3776737 | 0.99937574 |
| SORL1 | 217.31142 | 0.71069366 | 0.55679697 | 1.27639641 | 0.20181542 | 0.99937574 |
| UBASH3B | 189.028154 | -0.9589059 | 0.7592825 | -1.2629106 | 0.2066213 | 0.99937574 |
| JHY | 18.809223 | -0.1010836 | 0.53658457 | -0.1883833 | 0.8505762 | 0.99937574 |
| HSPA8 | 185.428585 | 0.48853872 | 0.4767007 | 1.02483323 | 0.30544188 | 0.99937574 |
| VWA5A | 13.8145057 | -0.1913032 | 0.70395519 | -0.2717548 | 0.78581058 | 0.99937574 |
| TBRG1 | 53.1439006 | 0.01720608 | 0.42916952 | 0.04009156 | 0.96802013 | 0.99937574 |
| SIAE | 33.983967 | 0.75310921 | 0.43865557 | 1.71685774 | 0.08600517 | 0.99937574 |
| MSANTD2 | 59.6491099 | -0.1489593 | 0.3269717 | -0.4555725 | 0.64869743 | 0.99937574 |

| | | | | | | |
|---|---|---|---|---|---|---|
| EI24 | 47.6048212 | -0.024867 | 0.37063264 | -0.0670933 | 0.94650746 | 0.99937574 |
| STT3A | 27.4472285 | -0.388323 | 0.4020839 | -0.9657761 | 0.33415623 | 0.99937574 |
| CDON | 18.1873754 | 0.80946354 | 1.09789241 | 0.73728859 | 0.46094687 | 0.99937574 |
| RPUSD4 | 12.9158769 | -0.9672022 | 0.70029693 | -1.3811315 | 0.16723852 | 0.99937574 |
| FAM118B | 30.3039014 | 0.1499089 | 0.38797518 | 0.38638786 | 0.69920944 | 0.99937574 |
| SRPRA | 52.1852857 | 0.08978809 | 0.32563345 | 0.27573363 | 0.78275266 | 0.99937574 |
| FOXRED1 | 27.9659579 | 0.79385775 | 0.40967496 | 1.93777466 | 0.05265072 | 0.99937574 |
| TIRAP | 16.4129249 | -0.2462461 | 0.52589363 | -0.4682431 | 0.63961076 | 0.99937574 |
| DCPS | 67.6485764 | -0.5474693 | 0.55990448 | -0.9777906 | 0.32817789 | 0.99937574 |
| ST3GAL4 | 56.4187198 | -0.0416982 | 0.65404256 | -0.0637546 | 0.94916564 | 0.99937574 |
| ARHGAP32 | 124.640838 | -0.0803823 | 0.36419696 | -0.2207111 | 0.82531735 | 0.99937574 |
| BARX2 | 51.5381704 | -0.1446996 | 0.64363646 | -0.2248158 | 0.82212258 | 0.99937574 |
| TMEM45B | 18.548612 | -1.7745551 | 1.05505032 | -1.6819625 | 0.09257611 | 0.99937574 |
| NFRKB | 26.012414 | 0.32806768 | 0.36213953 | 0.90591512 | 0.36498078 | 0.99937574 |
| PRDM10 | 30.152226 | 0.39141953 | 0.39404779 | 0.99333011 | 0.32054911 | 0.99937574 |
| APLP2 | 277.373277 | 0.32355529 | 0.30137661 | 1.07359124 | 0.28300592 | 0.99937574 |
| ST14 | 216.185486 | 0.09326929 | 0.32878335 | 0.28368007 | 0.77665557 | 0.99937574 |
| ZBTB44 | 112.359031 | -0.0657126 | 0.35264508 | -0.186342 | 0.85217654 | 0.99937574 |
| SNX19 | 73.2730372 | -0.5020241 | 0.48779288 | -1.0291748 | 0.30339755 | 0.99937574 |
| NCAPD3 | 16.3472916 | 0.11194109 | 0.50098778 | 0.22344076 | 0.82319248 | 0.99937574 |
| VPS26B | 11.7730909 | -0.004819 | 0.61113432 | -0.0078853 | 0.99370849 | 0.99937574 |
| ACAD8 | 25.673301 | 0.83838934 | 0.4303414 | 1.94819586 | 0.05139153 | 0.99937574 |
| AP004550.1 | 14.030086 | 1.46423361 | 0.67707153 | 2.16259811 | 0.0305721 | 0.99049468 |
| ZMYND11 | 191.669167 | 0.05374258 | 0.18683182 | 0.28765217 | 0.773613 | 0.99937574 |
| DIP2C | 765.983642 | 0.1280898 | 0.80388872 | 0.15933773 | 0.8734028 | 0.99937574 |
| LARP4B | 228.615267 | -0.4362483 | 0.23391993 | -1.8649471 | 0.06218882 | 0.99937574 |
| GTPBP4 | 28.9487194 | -0.8410143 | 0.4288825 | -1.9609433 | 0.04988563 | 0.99937574 |
| IDI1 | 91.4321415 | -0.0984237 | 0.59552656 | -0.1652717 | 0.86873011 | 0.99937574 |
| WDR37 | 90.2500865 | -0.1340073 | 0.37117711 | -0.3610333 | 0.71807453 | 0.99937574 |
| PFKP | 30.0598701 | -0.229398 | 0.49414924 | -0.4642281 | 6.42E-01 | 9.99E-01 |
| PITRM1 | 43.9471132 | -0.3043186 | 0.48992488 | -0.6211536 | 0.53449854 | 0.99937574 |
| KLF6 | 199.25586 | -0.8592605 | 0.56834405 | -1.5118668 | 0.13056775 | 0.99937574 |
| AKR1C1 | 119.782746 | -0.827976 | 0.94059389 | -0.8802693 | 0.37871341 | 0.99937574 |
| AKR1C2 | 67.9198001 | -1.8949064 | 0.93089685 | -2.0355707 | 0.04179348 | 0.99937574 |
| AKR1C3 | 59.0541199 | 0.31154379 | 0.94830103 | 0.32852836 | 0.7425122 | 0.99937574 |
| NET1 | 20.360671 | 0.14945907 | 0.44443343 | 0.33629124 | 0.73665125 | 0.99937574 |
| ASB13 | 32.2757636 | -0.1975564 | 0.40946154 | -0.4824786 | 0.62946597 | 0.99937574 |
| FAM208B | 156.268081 | -0.294675 | 0.27743427 | -1.0621434 | 0.28817058 | 0.99937574 |
| GDI2 | 97.1521341 | -0.0859802 | 0.25740643 | -0.3340252 | 0.73836051 | 0.99937574 |
| FBH1 | 100.284148 | 0.05724282 | 0.25815598 | 0.22173733 | 0.82451837 | 0.99937574 |
| IL15RA | 16.8582153 | -0.7879885 | 0.70755786 | -1.1136736 | 0.26541925 | 0.99937574 |
| RBM17 | 64.590219 | -0.1438743 | 0.3017505 | -0.4767988 | 0.63350537 | 0.99937574 |
| PFKFB3 | 161.24535 | -1.4053154 | 0.670251 | -2.0967002 | 0.03602012 | 0.99937574 |
| AL137145.2 | 49.1994498 | -1.8554431 | 0.9484037 | -1.9563854 | 0.05041978 | 0.99937574 |
| PRKCQ | 47.0867289 | 0.34889331 | 0.54214669 | 0.64354043 | 0.51987349 | 0.99937574 |
| SFMBT2 | 61.2697178 | 0.10914629 | 0.53513185 | 0.2039615 | 0.83838358 | 0.99937574 |
| ITIH5 | 396.420244 | 0.68742096 | 0.49982727 | 1.37531704 | 0.16903318 | 0.99937574 |
| KIN | 23.229342 | -0.5385649 | 0.42498316 | -1.2672618 | 0.20506168 | 0.99937574 |
| ATP5F1C | 157.545903 | 0.23839713 | 0.32023287 | 0.74444928 | 0.45660471 | 0.99937574 |
| TAF3 | 158.353525 | -0.1648714 | 0.33982752 | -0.4851619 | 0.6275615 | 0.99937574 |
| GATA3 | 64.3853992 | -1.1290486 | 0.79939826 | -1.4123732 | 0.15784012 | 0.99937574 |
| CELF2 | 675.484237 | -0.9919417 | 0.48769038 | -2.0339578 | 0.04195585 | 0.99937574 |
| USP6NL | 118.274299 | -0.0933611 | 0.28912985 | -0.3229037 | 0.74676818 | 0.99937574 |
| ECHDC3 | 35.8249943 | 0.42294609 | 0.42472866 | 0.99580303 | 0.31934586 | 0.99937574 |

| | | | | | | |
|---|---|---|---|---|---|---|
| PROSER2 | 43.4330648 | 0.24979703 | 0.45748042 | 0.54602781 | 0.58504682 | 0.99937574 |
| UPF2 | 92.6119566 | -0.2207892 | 0.26671641 | -0.8278051 | 0.40778086 | 0.99937574 |
| DHTKD1 | 43.6422382 | -0.0015087 | 0.35295664 | -0.0042744 | 0.99658956 | 0.99937574 |
| SEC61A2 | 14.3734103 | -0.0681744 | 0.61247814 | -0.1113092 | 0.91137118 | 0.99937574 |
| NUDT5 | 36.8118747 | -0.0890491 | 0.35889619 | -0.2481193 | 0.8040421 | 0.99937574 |
| CDC123 | 41.992804 | -0.2962633 | 0.34037391 | -0.8704054 | 0.38407892 | 0.99937574 |
| CAMK1D | 71.3029202 | -1.132616 | 0.70014965 | -1.6176771 | 0.1057322 | 0.99937574 |
| OPTN | 72.9521846 | -0.5688337 | 0.46050518 | -1.2352384 | 0.21674178 | 0.99937574 |
| PHYH | 38.9830188 | 0.35183902 | 0.50525454 | 0.69635994 | 0.48620344 | 0.99937574 |
| SEPHS1 | 28.5533674 | -0.0903067 | 0.41440581 | -0.2179184 | 0.82749267 | 0.99937574 |
| BEND7 | 125.111637 | -0.2854891 | 0.3089568 | -0.9240421 | 0.35546438 | 0.99937574 |
| PRPF18 | 16.0558458 | 0.20422652 | 0.60039539 | 0.34015337 | 0.73374103 | 0.99937574 |
| AL157392.5 | 38.2335024 | 0.20580627 | 0.35547643 | 0.57895896 | 0.56261687 | 0.99937574 |
| FRMD4A | 29.14053 | -0.0050235 | 0.51583286 | -0.0097387 | 0.99222979 | 0.99937574 |
| FAM107B | 567.518275 | -0.5375543 | 0.40143022 | -1.3390978 | 0.18053884 | 0.99937574 |
| DCLRE1C | 32.7960553 | -0.2911152 | 0.44985741 | -0.6471278 | 0.51754923 | 0.99937574 |
| RPP38 | 32.3210644 | 0.10652251 | 0.36190107 | 0.29434152 | 0.76849695 | 0.99937574 |
| NMT2 | 24.154773 | 0.00721386 | 0.70005064 | 0.01030477 | 0.99177813 | 0.99937574 |
| FAM171A1 | 42.5620936 | -0.674359 | 0.63094007 | -1.0688162 | 0.28515251 | 0.99937574 |
| MINDY3 | 52.4462199 | -0.1886998 | 0.32085919 | -0.5881079 | 0.55645984 | 0.99937574 |
| PTER | 125.253375 | 0.05736699 | 0.24183051 | 0.2372198 | 0.81248628 | 0.99937574 |
| RSU1 | 63.0413919 | -0.1746667 | 0.384119 | -0.4547202 | 0.64931052 | 0.99937574 |
| CUBN | 32.7271645 | 0.83292103 | 0.66987682 | 1.24339432 | 0.21372256 | 0.99937574 |
| TRDMT1 | 18.6200741 | -0.5146061 | 0.74135189 | -0.6941456 | 0.48759091 | 0.99937574 |
| VIM | 20.0075362 | -0.1145574 | 0.6767401 | -0.1692783 | 0.86557771 | 0.99937574 |
| ST8SIA6 | 123.893802 | 1.86052451 | 0.86296801 | 2.15596 | 0.03108679 | 0.99049468 |
| ST8SIA6-AS1 | 17.7721471 | 1.98023944 | 0.74734791 | 2.64968888 | 0.00805659 | 0.74742128 |
| HACD1 | 13.8356856 | 0.3971894 | 0.64430323 | 0.61646346 | 0.53758868 | 0.99937574 |
| STAM | 49.6402633 | -0.4855398 | 0.29931395 | -1.6221756 | 0.10476577 | 0.99937574 |
| CACNB2 | 665.090921 | 0.43530592 | 0.85510177 | 0.50906913 | 0.61070377 | 0.99937574 |
| AL450384.2 | 42.7774299 | 0.6554402 | 0.42937562 | 1.52649607 | 0.12688637 | 0.99937574 |
| NSUN6 | 96.1800893 | 0.37215184 | 0.33367359 | 1.11531701 | 0.2647146 | 0.99937574 |
| ARL5B | 43.8202263 | -0.5794269 | 0.53902517 | -1.0749534 | 0.28239559 | 0.99937574 |
| MALRD1 | 67.6042588 | -0.0109464 | 0.60768559 | -0.0180133 | 0.98562822 | 0.99937574 |
| PLXDC2 | 114.504698 | 0.07615022 | 0.48126453 | 0.15822945 | 0.874276 | 0.99937574 |
| NEBL | 463.529382 | -1.1845185 | 0.79489429 | -1.4901585 | 0.13618256 | 0.99937574 |
| MLLT10 | 235.297565 | -0.1425566 | 0.27997953 | -0.509168 | 0.61063446 | 0.99937574 |
| DNAJC1 | 174.015119 | -0.130973 | 0.2640886 | -0.4959433 | 0.61993443 | 0.99937574 |
| BMI1 | 21.2014926 | 0.02181027 | 0.41127306 | 0.05303111 | 0.95770712 | 0.99937574 |
| PIP4K2A | 83.0523894 | -0.540457 | 0.37154254 | -1.4546303 | 0.14577163 | 0.99937574 |
| MSRB2 | 31.2992634 | 0.04394217 | 0.48053314 | 0.09144463 | 0.9271393 | 0.99937574 |
| OTUD1 | 20.8915302 | -0.4689705 | 0.52265599 | -0.8972834 | 0.36956771 | 0.99937574 |
| KIAA1217 | 664.131694 | -0.2056811 | 0.43141875 | -0.476755 | 0.63353658 | 0.99937574 |
| ARHGAP21 | 219.224058 | 0.02371741 | 0.21380292 | 0.11093117 | 0.91167093 | 0.99937574 |
| PDSS1 | 20.660109 | 0.07344555 | 0.5134874 | 0.14303282 | 0.88626426 | 0.99937574 |
| ABI1 | 146.804845 | -0.1899122 | 0.23738053 | -0.8000326 | 0.42369192 | 0.99937574 |
| ANKRD26 | 178.42428 | 0.37727274 | 0.36635003 | 1.02981496 | 0.30309688 | 0.99937574 |
| YME1L1 | 127.358095 | -0.2105907 | 0.24720339 | -0.8518923 | 0.3942739 | 0.99937574 |
| MASTL | 21.1139754 | 0.06254564 | 0.4806275 | 0.13013329 | 0.89646097 | 0.99937574 |
| ACBD5 | 115.138219 | -0.5481314 | 0.40278031 | -1.3608693 | 0.17355498 | 0.99937574 |
| RAB18 | 124.764918 | -0.1291345 | 0.22254394 | -0.5802653 | 0.56173569 | 0.99937574 |
| MPP7 | 642.887747 | 0.38640071 | 0.36747416 | 1.05150445 | 0.29302696 | 0.99937574 |
| WAC-AS1 | 21.0914045 | -0.7717657 | 0.54366152 | -1.4195702 | 0.15573285 | 0.99937574 |
| WAC | 333.044352 | -0.2219694 | 0.2294481 | -0.9674057 | 0.33334126 | 0.99937574 |

| | | | | | | |
|---|---|---|---|---|---|---|
| SVIL | 236.37189 | -0.4918895 | 0.47089457 | -1.0445853 | 0.29621469 | 0.99937574 |
| MTPAP | 26.0051749 | -0.2563222 | 0.37261236 | -0.6879057 | 0.49151218 | 0.99937574 |
| MAP3K8 | 46.1078441 | -0.3148259 | 0.35262345 | -0.8928104 | 0.37195871 | 0.99937574 |
| ZNF438 | 91.0141804 | -0.0422324 | 0.36496104 | -0.1157175 | 0.90787642 | 0.99937574 |
| ZEB1 | 62.0097711 | -0.9346063 | 0.76455805 | -1.2224138 | 0.22155117 | 0.99937574 |
| ARHGAP12 | 218.959537 | -0.1713758 | 0.17984491 | -0.9529088 | 0.34063627 | 0.99937574 |
| KIF5B | 156.008493 | 0.00726838 | 0.25210314 | 0.02883098 | 0.97699939 | 0.99937574 |
| EPC1 | 121.43963 | -0.153341 | 0.31732031 | -0.4832372 | 0.62892732 | 0.99937574 |
| CCDC7 | 83.1895843 | -0.0988487 | 0.57174524 | -0.1728895 | 0.86273832 | 0.99937574 |
| ITGB1 | 146.03383 | 0.20994907 | 0.24793031 | 0.8468068 | 0.39710281 | 0.99937574 |
| NRP1 | 26.5987255 | 0.03568819 | 0.92613211 | 0.03853466 | 0.9692614 | 0.99937574 |
| PARD3 | 453.193479 | -0.0378102 | 0.3565855 | -0.1060341 | 0.91555533 | 0.99937574 |
| CUL2 | 67.0764605 | 0.10713646 | 0.29829068 | 0.35916797 | 0.71946944 | 0.99937574 |
| CREM | 141.793465 | -0.37784 | 0.42110742 | -0.8972533 | 0.36958379 | 0.99937574 |
| CCNY | 274.869394 | -0.3621038 | 0.27735442 | -1.3055635 | 0.19170102 | 0.99937574 |
| AL117336.3 | 23.3770123 | -0.0300071 | 0.50815036 | -0.0590516 | 0.95291098 | 0.99937574 |
| ANKRD30A | 19.4504359 | -0.7474146 | 0.83969318 | -0.8901044 | 0.37340983 | 0.99937574 |
| ZNF248 | 96.5306465 | 0.36609672 | 0.31720252 | 1.15414189 | 0.248442 | 0.99937574 |
| ZNF33A | 181.542976 | -0.1984374 | 0.25308155 | -0.784085 | 0.43299024 | 0.99937574 |
| AL117339.5 | 11.2292894 | -0.0849443 | 0.68750607 | -0.1235543 | 0.9016682 | 0.99937574 |
| ZNF33B | 103.854794 | -0.3180689 | 0.28678285 | -1.1090932 | 0.26738998 | 0.99937574 |
| BMS1 | 48.2528291 | -0.3857491 | 0.3235344 | -1.1922971 | 0.23314476 | 0.99937574 |
| CSGALNACT2 | 25.6060307 | 0.10987566 | 0.47738326 | 0.23016237 | 0.8179656 | 0.99937574 |
| FXYD4 | 30.016711 | -0.5312739 | 0.86424205 | -0.6147281 | 0.53873427 | 0.99937574 |
| HNRNPF | 147.771568 | -0.1806967 | 0.3832904 | -0.4714354 | 0.63732985 | 0.99937574 |
| ZNF487 | 16.0977731 | -0.5762808 | 0.51917561 | -1.109992 | 0.26700249 | 0.99937574 |
| ZNF32 | 19.235028 | -0.4509345 | 0.51653498 | -0.8729989 | 0.38266367 | 0.99937574 |
| TMEM72-AS | 37.0035954 | -0.0559048 | 0.62268276 | -0.0897805 | 0.92846165 | 0.99937574 |
| TMEM72 | 82.0497825 | -0.0172827 | 0.79237013 | -0.0218113 | 0.98259844 | 0.99937574 |
| AL353801.1 | 17.8935929 | -0.1816853 | 0.97554017 | -0.1862407 | 0.85225598 | 0.99937574 |
| RASSF4 | 31.6515091 | 0.73471532 | 0.53543472 | 1.37218467 | 0.17000596 | 0.99937574 |
| DEPP1 | 32.3606578 | -0.6065932 | 0.46501931 | -1.3044473 | 1.92E-01 | 9.99E-01 |
| ZNF22 | 26.4215569 | 0.24672646 | 0.45905222 | 0.53746927 | 0.59094351 | 0.99937574 |
| 8-Mar | 43.8779717 | -0.3786819 | 0.36230058 | -1.0452148 | 0.29592369 | 0.99937574 |
| ZFAND4 | 19.1938797 | 0.51279179 | 0.4985467 | 1.02857321 | 0.30368027 | 0.99937574 |
| WASHC2C | 25.5606111 | -0.9821997 | 0.51042197 | -1.9242895 | 0.0543183 | 0.99937574 |
| AGAP4 | 35.6350631 | 0.46716487 | 0.35223727 | 1.32627891 | 0.18474733 | 0.99937574 |
| TIMM23 | 39.5999532 | -0.2412761 | 0.39862468 | -0.6052712 | 0.54499879 | 0.99937574 |
| NCOA4 | 75.9807143 | 0.32628953 | 0.2694619 | 1.21089299 | 0.22593642 | 0.99937574 |
| PTPN20 | 70.2969463 | 0.19060448 | 0.54053684 | 0.35262071 | 0.72437281 | 0.99937574 |
| AGAP9 | 29.5416952 | 0.76935711 | 0.49722432 | 1.54730385 | 0.12178999 | 0.99937574 |
| MAPK8 | 312.039362 | -0.1934535 | 0.20657901 | -0.9364623 | 0.34903523 | 0.99937574 |
| ERCC6 | 56.2750077 | -0.4369646 | 0.30363387 | -1.4391169 | 0.1501174 | 0.99937574 |
| OGDHL | 196.535891 | 1.19118767 | 0.41158378 | 2.89415602 | 0.00380179 | 0.54810979 |
| PARG | 118.402705 | 0.13658067 | 0.29503686 | 0.46292749 | 0.64341635 | 0.99937574 |
| TIMM23B | 100.183539 | -0.6559233 | 0.50979142 | -1.2866504 | 0.19821616 | 0.99937574 |
| LINC00843 | 24.1320878 | 0.385071 | 0.43160088 | 0.89219234 | 0.37228985 | 0.99937574 |
| AGAP6 | 36.7399785 | 0.46281985 | 0.35044006 | 1.32068191 | 0.18660745 | 0.99937574 |
| WASHC2A | 32.8546047 | -0.9214327 | 0.50653256 | -1.8190987 | 0.06889638 | 0.99937574 |
| SGMS1 | 496.209527 | -0.0364322 | 0.32159202 | -0.1132869 | 0.90980313 | 0.99937574 |
| ASAH2B | 12.0835096 | 0.61322935 | 0.58417847 | 1.04972945 | 0.29384252 | 0.99937574 |
| PRKG1 | 1092.93782 | 0.79317831 | 0.35493337 | 2.23472455 | 0.02543544 | 0.96546668 |
| CSTF2T | 10.5782903 | 0.26738215 | 0.52416228 | 0.51011331 | 0.60997208 | 0.99937574 |
| IPMK | 43.1039497 | -0.2910673 | 0.39564714 | -0.735674 | 0.46192911 | 0.99937574 |

| | | | | | | |
|---|---|---|---|---|---|---|
| CISD1 | 123.965778 | 0.24883329 | 0.33655743 | 0.73934865 | 0.45969531 | 0.99937574 |
| UBE2D1 | 36.0994757 | -0.374581 | 0.38121018 | -0.9826102 | 0.32579933 | 0.99937574 |
| TFAM | 33.6379896 | 0.00921182 | 0.4344191 | 0.02120491 | 0.9830822 | 0.99937574 |
| BICC1 | 370.692868 | 0.12898667 | 0.53722099 | 0.24009983 | 0.81025286 | 0.99937574 |
| PHYHIPL | 12.1331821 | 0.24201563 | 0.88343847 | 0.27394735 | 0.78412508 | 0.99937574 |
| FAM13C | 180.029989 | 0.71661282 | 0.61851544 | 1.15860134 | 0.24661872 | 0.99937574 |
| CCDC6 | 60.5565659 | 0.08876097 | 0.39647702 | 0.22387418 | 0.82285521 | 0.99937574 |
| ANK3 | 671.54532 | 0.2052873 | 0.40812254 | 0.50300406 | 0.61496142 | 0.99937574 |
| RHOBTB1 | 21.9151725 | -0.0169008 | 0.57403872 | -0.029442 | 0.97651209 | 0.99937574 |
| ARID5B | 84.0011884 | -0.2763216 | 0.74493974 | -0.3709315 | 0.71068856 | 0.99937574 |
| NRBF2 | 19.6988467 | -0.1230055 | 0.59308702 | -0.2073987 | 0.83569853 | 0.99937574 |
| JMJD1C | 837.348977 | -0.1259112 | 0.22011202 | -0.5720322 | 0.56730019 | 0.99937574 |
| REEP3 | 178.546697 | 0.14420421 | 0.2193138 | 0.65752454 | 0.51084369 | 0.99937574 |
| LINC01515 | 24.7678742 | 0.14703518 | 0.44036493 | 0.33389393 | 0.7384596 | 0.99937574 |
| CTNNA3 | 17.863425 | -0.5181358 | 0.68334926 | -0.7582299 | 0.44831339 | 0.99937574 |
| SIRT1 | 39.4146238 | -0.5729251 | 0.39107407 | -1.4650041 | 0.1429198 | 0.99937574 |
| HERC4 | 244.511817 | -0.1509086 | 0.20898773 | -0.722093 | 0.47023732 | 0.99937574 |
| PBLD | 52.3568474 | -0.3790293 | 0.54503218 | -0.6954256 | 0.48678864 | 0.99937574 |
| HNRNPH3 | 109.056669 | -0.5982091 | 0.32840994 | -1.8215315 | 0.0685261 | 0.99937574 |
| RUFY2 | 63.0749086 | 0.01058489 | 0.37392817 | 0.02830728 | 0.97741708 | 0.99937574 |
| SLC25A16 | 71.2633211 | -0.0517776 | 0.38266292 | -0.1353085 | 0.89236793 | 0.99937574 |
| TET1 | 14.2230357 | 0.24791486 | 0.54031173 | 0.45883673 | 0.64635142 | 0.99937574 |
| CCAR1 | 150.561731 | -0.1147681 | 0.31288118 | -0.3668105 | 0.71376036 | 0.99937574 |
| STOX1 | 24.89805 | 0.1639615 | 0.6651556 | 0.24650097 | 0.80529445 | 0.99937574 |
| DDX50 | 62.5850352 | -0.2001664 | 0.26466741 | -0.756294 | 0.44947297 | 0.99937574 |
| DDX21 | 69.2875055 | -0.2462486 | 0.34092464 | -0.7222963 | 0.47011233 | 0.99937574 |
| KIF1BP | 41.6978608 | 0.32992528 | 0.4149043 | 0.79518404 | 0.42650646 | 0.99937574 |
| VPS26A | 75.4971082 | -0.584844 | 0.28242744 | -2.0707762 | 0.03837972 | 0.99937574 |
| SUPV3L1 | 31.8252109 | -0.0544593 | 0.35754034 | -0.1523166 | 0.87893726 | 0.99937574 |
| HKDC1 | 39.9965535 | -0.4935394 | 0.66790719 | -0.7389342 | 4.60E-01 | 0.99937574 |
| HK1 | 128.426921 | -0.1210807 | 0.32279476 | -0.3751011 | 0.70758531 | 0.99937574 |
| TSPAN15 | 37.825186 | 0.27910376 | 0.62255952 | 0.44831659 | 0.65392473 | 0.99937574 |
| H2AFY2 | 24.0719314 | 0.2862628 | 0.62204754 | 0.46019442 | 0.64537668 | 0.99937574 |
| AIFM2 | 31.4576696 | -0.6138135 | 0.42111907 | -1.4575771 | 0.14495714 | 0.99937574 |
| TYSND1 | 10.5341857 | 1.06000881 | 0.58562031 | 1.81006155 | 0.07028624 | 0.99937574 |
| SAR1A | 62.3638637 | 0.17951707 | 0.37742609 | 0.47563504 | 0.63433441 | 0.99937574 |
| PPA1 | 85.4608557 | 0.55211345 | 0.46604356 | 1.18468207 | 0.23614318 | 0.99937574 |
| EIF4EBP2 | 78.1342513 | -0.0054788 | 0.34122445 | -0.0160563 | 0.98718944 | 0.99937574 |
| SGPL1 | 97.7729295 | -0.0755053 | 0.30156975 | -0.2503742 | 0.80229797 | 0.99937574 |
| PCBD1 | 61.9774468 | 0.37493849 | 0.73725742 | 0.50855845 | 0.61106176 | 0.99937574 |
| PSAP | 251.9037 | 0.35490318 | 0.40926225 | 0.86717789 | 0.38584455 | 0.99937574 |
| CHST3 | 24.7680092 | 0.11820308 | 0.66739774 | 0.17711039 | 0.85942168 | 0.99937574 |
| ASCC1 | 62.252424 | -0.5524467 | 0.38178692 | -1.4470026 | 0.14789621 | 0.99937574 |
| ANAPC16 | 183.389236 | 0.07509611 | 0.31225685 | 0.24049469 | 0.80994678 | 0.99937574 |
| DDIT4 | 22.035714 | -1.285768 | 0.8411649 | -1.5285564 | 0.12637446 | 0.99937574 |
| DNAJB12 | 29.3075037 | -0.2230484 | 0.50121059 | -0.4450193 | 0.65630578 | 0.99937574 |
| MICU1 | 193.664687 | -0.3765139 | 0.20983148 | -1.7943633 | 0.0727552 | 0.99937574 |
| MCU | 348.718006 | -0.2283891 | 0.3450176 | -0.6619637 | 0.50799449 | 0.99937574 |
| P4HA1 | 158.165871 | -0.3943379 | 0.58322336 | -0.6761352 | 0.49895478 | 0.99937574 |
| ECD | 36.08677 | -0.1251225 | 0.34085029 | -0.3670893 | 0.7135524 | 0.99937574 |
| FAM149B1 | 53.6658264 | -0.1450711 | 0.43591427 | -0.3327973 | 0.73928732 | 0.99937574 |
| DNAJC9 | 52.8660984 | 0.5741627 | 0.38735514 | 1.4822643 | 0.13826998 | 0.99937574 |
| MRPS16 | 45.7103418 | 0.34463783 | 0.47035114 | 0.73272455 | 0.46372645 | 0.99937574 |
| CFAP70 | 34.7621251 | 0.10073023 | 0.536415 | 0.18778413 | 0.85104587 | 0.99937574 |

| | | | | | | |
|---|---|---|---|---|---|---|
| ANXA7 | 134.015583 | -0.3324404 | 0.28749013 | -1.1563543 | 0.24753628 | 0.99937574 |
| PPP3CB | 136.623664 | 0.10446951 | 0.21456039 | 0.48690029 | 0.62632899 | 0.99937574 |
| USP54 | 79.4039567 | -0.0612219 | 0.38166452 | -0.1604076 | 0.87255997 | 0.99937574 |
| AGAP5 | 14.7869353 | -0.635628 | 0.618251 | -1.0281067 | 0.30389962 | 0.99937574 |
| SEC24C | 33.3446136 | -0.4279798 | 0.40546995 | -1.0555154 | 0.29118969 | 0.99937574 |
| FUT11 | 31.5629901 | -0.2416706 | 0.57094043 | -0.4232851 | 0.6720873 | 0.99937574 |
| CHCHD1 | 31.9508261 | 0.22392279 | 0.69008076 | 0.3244878 | 0.74556876 | 0.99937574 |
| ZSWIM8 | 53.2975733 | -0.0978941 | 0.40059265 | -0.2443731 | 0.80694191 | 0.99937574 |
| CAMK2G | 26.999859 | 0.2682406 | 0.56134804 | 0.47785079 | 0.63275641 | 0.99937574 |
| VCL | 198.907726 | -0.3062674 | 0.42772916 | -0.7160311 | 0.47397217 | 0.99937574 |
| AP3M1 | 31.5977586 | -0.0407235 | 0.38369825 | -0.1061343 | 0.91547581 | 0.99937574 |
| ADK | 508.023121 | 0.14080024 | 0.24254769 | 0.58050538 | 0.56157386 | 0.99937574 |
| KAT6B | 176.310993 | -0.3823762 | 0.38427549 | -0.9950575 | 0.31970829 | 0.99937574 |
| SAMD8 | 75.27482 | -0.4636203 | 0.35542168 | -1.3044231 | 0.19208935 | 0.99937574 |
| VDAC2 | 86.385153 | 0.57220815 | 0.42217062 | 1.35539548 | 0.17529159 | 0.99937574 |
| COMTD1 | 49.7268958 | 0.09934913 | 0.540607 | 0.1837733 | 0.8541913 | 0.99937574 |
| AC010997.3 | 15.388158 | -0.3300253 | 0.8871261 | -0.3720163 | 0.70988074 | 0.99937574 |
| ZNF503-AS2 | 13.3585081 | 0.12456037 | 0.55060884 | 0.22622298 | 0.821028 | 0.99937574 |
| LRMDA | 1768.49609 | 0.0704134 | 0.3555753 | 0.1980267 | 0.84302418 | 0.99937574 |
| KCNMA1 | 79.4469988 | 0.0458755 | 1.2998263 | 0.03529357 | 0.97184565 | 0.99937574 |
| DLG5 | 160.316822 | -0.4603464 | 0.53312392 | -0.8634886 | 0.38786887 | 0.99937574 |
| POLR3A | 27.9999715 | -0.3425706 | 0.46433511 | -0.7377659 | 0.46065673 | 0.99937574 |
| RPS24 | 404.23133 | -0.1954984 | 0.50594655 | -0.3864012 | 0.69919956 | 0.99937574 |
| ZMIZ1 | 223.541767 | 0.87776679 | 0.49797139 | 1.7626852 | 0.0779536 | 0.99937574 |
| PPIF | 32.0609475 | -0.417695 | 0.63552783 | -0.6572411 | 0.5110259 | 0.99937574 |
| NUTM2B-AS | 168.440381 | 0.36850328 | 0.29668746 | 1.24205884 | 0.21421485 | 0.99937574 |
| ANXA11 | 270.488565 | 0.03319696 | 0.27886806 | 0.11904181 | 0.90524223 | 0.99937574 |
| FAM213A | 39.8249648 | -0.3622325 | 0.74361986 | -0.4871206 | 0.62617289 | 0.99937574 |
| TSPAN14 | 169.59784 | 0.38028486 | 0.3233867 | 1.17594462 | 0.23961701 | 0.99937574 |
| GHITM | 115.406799 | 0.26835687 | 0.43923882 | 0.61095891 | 0.54122678 | 0.99937574 |
| CCSER2 | 216.964151 | -0.0543879 | 0.22318984 | -0.2436844 | 0.80747523 | 0.99937574 |
| WAPL | 165.892399 | -0.5063398 | 0.23104342 | -2.1915353 | 0.02841308 | 0.99049468 |
| BMPR1A | 169.493555 | 0.10549816 | 0.24983763 | 0.42226691 | 0.67283021 | 0.99937574 |
| GLUD1 | 240.446155 | 0.09780982 | 0.37121703 | 0.26348418 | 0.79217741 | 0.99937574 |
| SHLD2 | 207.826891 | -0.2937965 | 0.36187876 | -0.8118645 | 0.41686936 | 0.99937574 |
| NUTM2A-AS | 192.351564 | 0.16122243 | 0.25245219 | 0.6386256 | 0.52306652 | 0.99937574 |
| MINPP1 | 15.2644905 | 0.10754202 | 0.56793106 | 0.18935752 | 0.84981261 | 0.99937574 |
| PAPSS2 | 56.4397047 | -2.0567296 | 1.44865755 | -1.4197487 | 0.15568087 | 0.99937574 |
| ATAD1 | 133.633586 | 0.0989736 | 0.25503527 | 0.38807807 | 0.69795826 | 0.99937574 |
| PTEN | 211.228604 | 0.50712365 | 0.26581757 | 1.90778829 | 0.05641859 | 0.99937574 |
| RNLS | 65.6034548 | -0.4459276 | 0.39794182 | -1.120585 | 0.26246456 | 0.99937574 |
| ANKRD22 | 118.933348 | 0.09223274 | 0.45692089 | 0.20185713 | 0.84002842 | 0.99937574 |
| STAMBPL1 | 18.8768378 | -0.0753315 | 0.61801923 | -0.1218919 | 0.90298467 | 0.99937574 |
| LIPA | 35.2357254 | -1.0061149 | 0.48010314 | -2.0956225 | 0.03611569 | 0.99937574 |
| IFIT2 | 22.2202995 | -3.389703 | 1.16599536 | -2.9071325 | 0.00364759 | 0.54134481 |
| IFIT3 | 47.2259311 | -3.0535981 | 0.98946933 | -3.0860967 | 0.00202803 | 0.45583974 |
| IFIT1 | 29.6360344 | -4.6650776 | 1.91989366 | -2.4298625 | 0.01510455 | 0.86113259 |
| IFIT5 | 13.6935034 | -0.8357665 | 0.86407889 | -0.967234 | 0.33342707 | 0.99937574 |
| SLC16A12 | 17.7710346 | -0.7928651 | 0.6030039 | -1.3148589 | 1.89E-01 | 0.99937574 |
| PANK1 | 25.8538172 | 1.20816756 | 0.60312999 | 2.00316281 | 0.04515982 | 0.99937574 |
| RPP30 | 16.4462149 | -0.2459793 | 0.43562583 | -0.5646574 | 0.57230684 | 0.99937574 |
| PCGF5 | 229.914719 | -0.1497974 | 0.29950342 | -0.5001527 | 0.61696756 | 0.99937574 |
| HECTD2 | 73.2761661 | -0.2415834 | 0.35677513 | -0.6771306 | 0.49832308 | 0.99937574 |
| PPP1R3C | 36.5038392 | -0.614063 | 1.02582655 | -0.5986031 | 0.54943759 | 0.99937574 |

| | | | | | | |
|---|---|---|---|---|---|---|
| TNKS2 | 165.435949 | -0.396878 | 0.28725132 | -1.3816405 | 0.16708212 | 0.99937574 |
| BTAF1 | 158.722029 | -0.1463416 | 0.35889467 | -0.4077564 | 0.68345255 | 0.99937574 |
| CPEB3 | 415.265125 | 0.33509117 | 0.38268442 | 0.87563316 | 0.38122949 | 0.99937574 |
| 5-Mar | 91.8247415 | 0.01576773 | 0.30500985 | 0.05169579 | 0.95877109 | 0.99937574 |
| IDE | 82.6403882 | -0.2890736 | 0.30157727 | -0.9585391 | 0.33779101 | 0.99937574 |
| EXOC6 | 181.640221 | -0.0925468 | 0.28903938 | -0.3201876 | 0.74882616 | 0.99937574 |
| MYOF | 169.633763 | -0.1312863 | 0.3979239 | -0.3299282 | 7.41E-01 | 0.99937574 |
| FRA10AC1 | 51.0311954 | 0.14039326 | 0.34099671 | 0.41171442 | 0.68054876 | 0.99937574 |
| PLCE1 | 15.5615295 | 0.13140275 | 0.7782217 | 0.16885002 | 0.86591461 | 0.99937574 |
| NOC3L | 18.3645774 | 0.05732978 | 0.45604059 | 0.12571202 | 0.89995989 | 0.99937574 |
| TBC1D12 | 76.4268694 | 0.44313254 | 0.26583935 | 1.66691855 | 0.0955306 | 0.99937574 |
| SORBS1 | 212.448423 | 0.16192243 | 0.47014234 | 0.34441151 | 0.73053684 | 0.99937574 |
| ALDH18A1 | 25.598745 | 0.12734986 | 0.48062984 | 0.26496454 | 0.79103678 | 0.99937574 |
| TCTN3 | 42.0459979 | -0.1979192 | 0.35144264 | -0.5631623 | 0.57332437 | 0.99937574 |
| ENTPD1-AS1 | 147.219307 | 0.64632063 | 0.43401711 | 1.48915933 | 0.13644542 | 0.99937574 |
| ZNF518A | 170.390929 | -0.2106584 | 0.30725273 | -0.6856192 | 0.49295324 | 0.99937574 |
| BLNK | 345.717857 | -0.2571174 | 0.42479686 | -0.6052715 | 5.45E-01 | 0.99937574 |
| TM9SF3 | 310.455878 | 0.14712934 | 0.25035156 | 0.58769093 | 0.55673977 | 0.99937574 |
| LCOR | 328.693364 | -0.1310207 | 0.27705955 | -0.4728972 | 0.63628654 | 0.99937574 |
| RRP12 | 21.4675982 | -0.1425442 | 0.44021429 | -0.3238064 | 0.74608458 | 0.99937574 |
| PGAM1 | 88.3012813 | 0.19960037 | 0.41791491 | 0.47761008 | 0.63292775 | 0.99937574 |
| EXOSC1 | 27.5537237 | 0.14667358 | 0.38400479 | 0.38195769 | 0.70249275 | 0.99937574 |
| ZDHHC16 | 21.9989909 | -0.4931013 | 0.47453984 | -1.0391146 | 0.29875145 | 0.99937574 |
| MMS19 | 54.2793928 | -0.0850171 | 0.29432321 | -0.2888563 | 0.77269135 | 0.99937574 |
| UBTD1 | 14.2536545 | 0.33248095 | 0.49602227 | 0.6702944 | 0.50267014 | 0.99937574 |
| MORN4 | 14.8174931 | -0.245182 | 0.55443745 | -0.4422177 | 0.65833171 | 0.99937574 |
| ZFYVE27 | 34.8494375 | -0.3559061 | 0.36990529 | -0.9621547 | 0.3359719 | 0.99937574 |
| R3HCC1L | 115.067456 | -0.3083781 | 0.29247028 | -1.0543913 | 0.29170379 | 0.99937574 |
| HPS1 | 27.0407537 | -0.3461909 | 0.48378334 | -0.7155908 | 0.47424408 | 0.99937574 |
| GOT1 | 250.060532 | 0.05805764 | 0.34736836 | 0.16713566 | 0.86726331 | 0.99937574 |
| SLC25A28 | 30.0780813 | -0.1487644 | 0.59772817 | -0.248883 | 0.80345128 | 0.99937574 |
| CUTC | 33.0578512 | -0.0260221 | 0.37549565 | -0.0693006 | 0.94475035 | 0.99937574 |
| COX15 | 36.2292822 | 0.40786067 | 0.37429404 | 1.08967985 | 0.27585419 | 0.99937574 |
| DNMBP | 104.213951 | 0.21069654 | 0.31384774 | 0.67133362 | 0.50200802 | 0.99937574 |
| ERLIN1 | 26.5407812 | -0.0285332 | 0.41431522 | -0.0688684 | 0.94509439 | 0.99937574 |
| CHUK | 37.2575256 | -0.066818 | 0.45403915 | -0.1471635 | 0.883003 | 0.99937574 |
| CWF19L1 | 26.5520182 | -0.8151825 | 0.43189687 | -1.8874471 | 0.05910022 | 0.99937574 |
| BLOC1S2 | 26.0564489 | -0.4237712 | 0.41294033 | -1.0262287 | 0.30478382 | 0.99937574 |
| SCD | 73.8196964 | -0.3838652 | 0.9333606 | -0.4112721 | 0.68087304 | 0.99937574 |
| OLMALINC | 61.7697669 | 1.60642592 | 0.62392349 | 2.57471622 | 0.01003224 | 0.76274616 |
| SEC31B | 18.2311799 | 0.67741939 | 0.44865498 | 1.50988938 | 0.13107165 | 0.99937574 |
| HIF1AN | 35.0358329 | 0.16008895 | 0.41215838 | 0.3884161 | 0.69770813 | 0.99937574 |
| PAX2 | 329.787518 | 0.2044276 | 0.57982083 | 0.35257029 | 0.72441062 | 0.99937574 |
| SLF2 | 63.0305567 | -0.3363772 | 0.32169509 | -1.0456398 | 0.29572739 | 0.99937574 |
| MRPL43 | 26.0138307 | 0.61273642 | 0.54397251 | 1.12641065 | 0.25999174 | 0.99937574 |
| LZTS2 | 34.3348206 | -0.5841337 | 0.41306402 | -1.4141481 | 0.15731843 | 0.99937574 |
| BTRC | 168.456963 | 0.04182308 | 0.30094132 | 0.1389742 | 0.88947054 | 0.99937574 |
| DPCD | 14.1437831 | 0.19760145 | 0.68349562 | 0.28910419 | 0.77250165 | 0.99937574 |
| POLL | 22.9232749 | -0.2595287 | 0.52699289 | -0.4924709 | 0.62238645 | 0.99937574 |
| FBXW4 | 156.108146 | -0.0270478 | 0.20862626 | -0.129647 | 0.89684571 | 0.99937574 |
| OGA | 239.536054 | -0.1044042 | 0.25865149 | -0.4036481 | 0.68647152 | 0.99937574 |
| ARMH3 | 169.530717 | -0.2685208 | 0.24017635 | -1.1180151 | 0.26356053 | 0.99937574 |
| LDB1 | 23.6645979 | -0.3148356 | 0.41972885 | -0.7500929 | 0.45319874 | 0.99937574 |
| PPRC1 | 15.6534738 | -0.9048133 | 0.64840434 | -1.3954461 | 0.16288135 | 0.99937574 |

| | | | | | | |
|---|---|---|---|---|---|---|
| NOLC1 | 19.8825952 | -0.4532202 | 0.49800369 | -0.910074 | 0.36278347 | 0.99937574 |
| GBF1 | 200.620577 | -0.1609137 | 0.27145089 | -0.5927914 | 0.55332078 | 0.99937574 |
| NFKB2 | 14.291756 | -0.2995473 | 0.66870467 | -0.4479516 | 0.65418813 | 0.99937574 |
| FBXL15 | 36.7966076 | 0.34446034 | 0.41712711 | 0.82579226 | 0.40892194 | 0.99937574 |
| CUEDC2 | 19.2277949 | 0.40208024 | 0.39637864 | 1.01438421 | 0.31039945 | 0.99937574 |
| ACTR1A | 33.2607974 | 0.01956658 | 0.39096949 | 0.05004631 | 0.96008549 | 0.99937574 |
| SUFU | 46.2512526 | 1.0799827 | 0.46299178 | 2.33261745 | 0.01966823 | 0.96427895 |
| TRIM8 | 135.657053 | 0.3321976 | 0.28386055 | 1.17028448 | 0.2418865 | 0.99937574 |
| ARL3 | 35.5123494 | 0.21920891 | 0.58376204 | 0.37551073 | 0.70728067 | 0.99937574 |
| SFXN2 | 10.4802063 | 1.35833478 | 0.64352724 | 2.11076502 | 0.03479251 | 0.99937574 |
| WBP1L | 153.055967 | -0.1165181 | 0.27565469 | -0.4226957 | 0.67251727 | 0.99937574 |
| CNNM2 | 80.7933157 | -0.0053989 | 0.32767419 | -0.0164764 | 0.98685432 | 0.99937574 |
| NT5C2 | 255.856836 | -0.5049895 | 0.26421931 | -1.9112513 | 0.0559723 | 0.99937574 |
| PCGF6 | 20.1130557 | -0.4555938 | 0.47484209 | -0.9594637 | 0.33732518 | 0.99937574 |
| ATP5MD | 64.4398751 | -0.0128666 | 0.46548221 | -0.0276414 | 0.97794817 | 0.99937574 |
| PDCD11 | 23.6862892 | -0.3187598 | 0.4202535 | -0.7584942 | 0.44815517 | 0.99937574 |
| NEURL1 | 99.8287157 | 1.28808124 | 0.69251508 | 1.86000462 | 0.06288487 | 0.99937574 |
| SH3PXD2A | 55.6827336 | 0.60490675 | 0.67649906 | 0.89417235 | 0.37122968 | 0.99937574 |
| STN1 | 21.3206664 | 0.02848657 | 0.53267557 | 0.05347827 | 0.95735084 | 0.99937574 |
| SLK | 98.9595181 | -0.4254558 | 0.25810572 | -1.6483782 | 0.09927509 | 0.99937574 |
| CFAP43 | 13.5491977 | -0.1477036 | 0.58369211 | -0.2530504 | 0.80022923 | 0.99937574 |
| GSTO1 | 76.6465611 | 0.16059088 | 0.3168634 | 0.50681422 | 0.61228518 | 0.99937574 |
| GSTO2 | 72.5771217 | 0.14270271 | 0.49319741 | 0.28934197 | 0.7723197 | 0.99937574 |
| LINC01435 | 42.7426514 | 0.90267005 | 0.58307616 | 1.54811688 | 0.12159416 | 0.99937574 |
| XPNPEP1 | 62.464004 | -0.1161378 | 0.34297426 | -0.3386197 | 0.73489623 | 0.99937574 |
| ADD3 | 199.426224 | 0.32891221 | 0.34167523 | 0.96264578 | 0.33572532 | 0.99937574 |
| MXI1 | 55.4416742 | -0.2197379 | 0.42582742 | -0.5160257 | 6.06E-01 | 0.99937574 |
| SMNDC1 | 28.7119064 | -0.331922 | 0.48392308 | -0.6858982 | 0.49277733 | 0.99937574 |
| SMC3 | 50.0687316 | -0.3143332 | 0.34945807 | -0.8994875 | 0.36839305 | 0.99937574 |
| PDCD4 | 84.5202016 | -0.6134018 | 0.43803274 | -1.400356 | 0.16140675 | 0.99937574 |
| BBIP1 | 55.3600761 | -0.0160136 | 0.44790055 | -0.0357525 | 0.9714797 | 0.99937574 |
| SHOC2 | 137.690282 | 0.15436325 | 0.25427308 | 0.60707667 | 0.54380004 | 0.99937574 |
| GPAM | 20.2204158 | 0.53749715 | 0.57451669 | 0.93556403 | 0.3494977 | 0.99937574 |
| ACSL5 | 36.7199356 | -0.6326611 | 0.7370726 | -0.858343 | 0.39070311 | 0.99937574 |
| ZDHHC6 | 34.3308205 | -0.5073091 | 0.35288075 | -1.4376219 | 0.15054135 | 0.99937574 |
| VTI1A | 387.032296 | -0.036992 | 0.23888137 | -0.1548551 | 0.87693553 | 0.99937574 |
| TCF7L2 | 330.517142 | 0.00568622 | 0.30808668 | 0.01845657 | 0.98527463 | 0.99937574 |
| HABP2 | 36.2939854 | -1.6588598 | 0.71236428 | -2.3286678 | 1.99E-02 | 9.64E-01 |
| CASP7 | 50.6432879 | -0.1565825 | 0.33419915 | -0.4685306 | 0.63940518 | 0.99937574 |
| PLEKHS1 | 94.2983566 | -0.3817337 | 1.60163671 | -0.2383397 | 0.8116176 | 0.99937574 |
| NHLRC2 | 185.123894 | 0.01936635 | 0.21778052 | 0.08892599 | 0.92914073 | 0.99937574 |
| ADRB1 | 18.9601241 | 0.28922798 | 1.03679027 | 0.2789648 | 0.78027184 | 0.99937574 |
| CCDC186 | 103.675967 | -0.3058906 | 0.38481725 | -0.7948985 | 0.42667256 | 0.99937574 |
| AFAP1L2 | 17.5262998 | -0.8545552 | 1.10391327 | -0.7741144 | 0.43886313 | 0.99937574 |
| ABLIM1 | 122.93992 | -0.552318 | 0.61725452 | -0.8947978 | 0.37089519 | 0.99937574 |
| FAM160B1 | 42.0976783 | -0.1472177 | 0.41191845 | -0.3573952 | 0.72079599 | 0.99937574 |
| TRUB1 | 28.0803237 | -0.0295198 | 0.42371569 | -0.0696689 | 0.94445721 | 0.99937574 |
| HSPA12A | 76.6372667 | 0.37659631 | 0.41599932 | 0.90528108 | 0.3653165 | 0.99937574 |
| ENO4 | 13.9425424 | -0.3568498 | 0.72542432 | -0.4919187 | 0.62277679 | 0.99937574 |
| SHTN1 | 199.665808 | -0.1834899 | 0.29385002 | -0.6244339 | 0.53234268 | 0.99937574 |
| PDZD8 | 201.724433 | -0.1954244 | 0.35504413 | -0.550423 | 0.58202929 | 0.99937574 |
| EMX2OS | 156.699358 | 0.47604422 | 0.35853097 | 1.32776319 | 0.18425635 | 0.99937574 |
| EMX2 | 59.1544678 | 0.35783293 | 0.32854994 | 1.08912798 | 0.27609745 | 0.99937574 |
| RAB11FIP2 | 99.5114482 | 0.31701504 | 0.34193506 | 0.92712061 | 0.35386392 | 0.99937574 |

| | | | | | | |
|---|---|---|---|---|---|---|
| CASC2 | 43.4627843 | -0.2912662 | 0.53361482 | -0.5458361 | 0.58517859 | 0.99937574 |
| FAM204A | 56.3330546 | -0.1117384 | 0.35385011 | -0.3157789 | 0.75217033 | 0.99937574 |
| CACUL1 | 154.13724 | -0.4124681 | 0.23273427 | -1.7722707 | 0.07634963 | 0.99937574 |
| EIF3A | 177.850513 | -0.4378515 | 0.40052711 | -1.0931883 | 0.27431113 | 0.99937574 |
| FAM45A | 39.5421956 | 0.09798866 | 0.33189403 | 0.2952408 | 0.76780993 | 0.99937574 |
| SFXN4 | 38.4542556 | -0.3953848 | 0.49135357 | -0.8046849 | 0.42100154 | 0.99937574 |
| PRDX3 | 71.9807652 | 0.05371636 | 0.36341966 | 0.14780808 | 0.88249423 | 0.99937574 |
| TIAL1 | 91.1116375 | -0.0202426 | 0.2842973 | -0.0712021 | 0.94323689 | 0.99937574 |
| BAG3 | 36.33018 | -0.379553 | 0.41967337 | -0.904401 | 0.36578283 | 0.99937574 |
| INPP5F | 57.7871729 | -0.3181204 | 0.4873672 | -0.6527325 | 0.51392878 | 0.99937574 |
| MCMBP | 60.2320253 | -0.1835488 | 0.31848491 | -0.5763186 | 0.56439988 | 0.99937574 |
| SEC23IP | 87.2204725 | 0.05543029 | 0.27343051 | 0.20272167 | 0.83935259 | 0.99937574 |
| PLPP4 | 96.3847358 | 0.45022369 | 0.57968225 | 0.77667323 | 0.43735158 | 0.99937574 |
| WDR11-AS1 | 13.7719618 | 0.26347297 | 0.61650416 | 0.42736608 | 0.66911271 | 0.99937574 |
| WDR11 | 124.921446 | -0.0830792 | 0.23279595 | -0.3568757 | 0.72118486 | 0.99937574 |
| FGFR2 | 72.7812235 | -0.0846909 | 0.6185348 | -0.1369219 | 0.89109254 | 0.99937574 |
| ATE1 | 231.787956 | -0.2844349 | 0.21766839 | -1.3067349 | 0.19130278 | 0.99937574 |
| NSMCE4A | 19.3538863 | 0.52784852 | 0.49703844 | 1.06198733 | 0.28824144 | 0.99937574 |
| TACC2 | 188.946638 | 0.10300341 | 0.54549619 | 0.18882517 | 0.85022984 | 0.99937574 |
| PLEKHA1 | 390.14239 | -0.2296181 | 0.426087 | -0.5388996 | 5.90E-01 | 0.99937574 |
| FAM24B | 16.5571314 | 0.64041841 | 0.67920337 | 0.94289639 | 0.3457339 | 0.99937574 |
| C10orf88 | 19.018626 | -0.4902368 | 0.49203966 | -0.9963359 | 0.31908695 | 0.99937574 |
| IKZF5 | 25.495469 | -0.2563343 | 0.40277855 | -0.636415 | 0.52450595 | 0.99937574 |
| ACADSB | 27.9939773 | 0.92602394 | 0.47316868 | 1.95706936 | 0.05033931 | 0.99937574 |
| BUB3 | 31.5977437 | -0.3468621 | 0.36252184 | -0.9568034 | 0.33866652 | 0.99937574 |
| CPXM2 | 33.9032507 | -1.1438264 | 0.82557627 | -1.3854885 | 0.16590315 | 0.99937574 |
| CHST15 | 46.2258068 | 0.03320708 | 0.71998786 | 0.04612173 | 0.96321323 | 0.99937574 |
| OAT | 41.3302365 | 0.23173714 | 0.4272274 | 0.54242105 | 0.58752848 | 0.99937574 |
| LHPP | 33.5057193 | 0.38665737 | 0.42028991 | 0.91997776 | 0.35758438 | 0.99937574 |
| ABRAXAS2 | 32.6136931 | 0.05012125 | 0.4039827 | 0.12406782 | 0.90126158 | 0.99937574 |
| ZRANB1 | 76.7823197 | -0.4092071 | 0.36962349 | -1.1070918 | 0.26825425 | 0.99937574 |
| CTBP2 | 403.361109 | -0.1154571 | 0.34211602 | -0.3374793 | 0.73575558 | 0.99937574 |
| EDRF1 | 38.3739226 | -0.3547164 | 0.47391185 | -0.748486 | 0.45416706 | 0.99937574 |
| UROS | 23.1883407 | -0.1393973 | 0.45564462 | -0.3059342 | 0.75965475 | 0.99937574 |
| BCCIP | 42.5397281 | -0.2268408 | 0.39814999 | -0.569737 | 0.5688561 | 0.99937574 |
| DHX32 | 140.136766 | -0.045008 | 0.34331233 | -0.1310994 | 0.89569671 | 0.99937574 |
| FANK1 | 37.3518605 | 0.42238243 | 0.49760751 | 0.84882647 | 0.39597786 | 0.99937574 |
| DOCK1 | 597.271276 | 0.24705119 | 0.40682405 | 0.60726792 | 0.54367312 | 0.99937574 |
| FAM196A | 25.1243818 | 0.45303762 | 0.69436537 | 0.65244847 | 0.5141119 | 0.99937574 |
| FOXI2 | 11.7624106 | 1.01881935 | 0.84771966 | 1.20183522 | 0.22942738 | 0.99937574 |
| MGMT | 125.736513 | 0.21200865 | 0.3407085 | 0.62225818 | 0.53377212 | 0.99937574 |
| C10orf143 | 12.7291421 | 0.37044007 | 0.68266571 | 0.54263759 | 0.58737935 | 0.99937574 |
| GLRX3 | 65.4359843 | -0.570322 | 0.3371252 | -1.6917215 | 0.09069908 | 0.99937574 |
| PPP2R2D | 118.126365 | -0.1471772 | 0.3083866 | -0.477249 | 0.6331848 | 0.99937574 |
| BNIP3 | 103.423752 | -0.1824513 | 0.6874142 | -0.2654168 | 0.79068837 | 0.99937574 |
| LRRC27 | 13.4822912 | -0.032833 | 0.60202608 | -0.0545375 | 0.95650693 | 0.99937574 |
| PWWP2B | 16.2559421 | -0.4432181 | 0.53648035 | -0.8261591 | 0.40871382 | 0.99937574 |
| INPP5A | 82.109123 | 0.19608945 | 0.37303074 | 0.52566565 | 0.59912053 | 0.99937574 |
| KNDC1 | 14.1233389 | 1.33734872 | 0.68288259 | 1.95838747 | 0.05018456 | 0.99937574 |
| TUBGCP2 | 58.5644169 | 0.04282397 | 0.30914298 | 0.13852481 | 0.88982566 | 0.99937574 |
| ECHS1 | 78.1795596 | 0.32738755 | 0.4889405 | 0.66958567 | 0.50312195 | 0.99937574 |
| SCART1 | 22.8720737 | -0.3406279 | 0.57879691 | -0.5885103 | 0.55618984 | 0.99937574 |
| SLC6A12 | 43.4667645 | -0.4776199 | 1.08025787 | -0.442135 | 0.65839149 | 0.99937574 |
| KDM5A | 158.133638 | -0.1041054 | 0.21560425 | -0.4828543 | 0.62919919 | 0.99937574 |

| | | | | | | |
|---|---|---|---|---|---|---|
| CCDC77 | 12.7577332 | -0.802269 | 0.60022746 | -1.3366083 | 0.18135053 | 0.99937574 |
| B4GALNT3 | 104.34988 | -1.3973884 | 0.57715205 | -2.421179 | 0.01547026 | 0.86662299 |
| WNK1 | 287.92667 | 0.33380265 | 0.26751875 | 1.24777293 | 0.21211422 | 0.99937574 |
| RAD52 | 28.1337821 | 0.91970589 | 0.44955914 | 2.04579512 | 0.04077653 | 0.99937574 |
| ERC1 | 438.100978 | 0.06689179 | 0.18955157 | 0.35289492 | 0.72416722 | 0.99937574 |
| FBXL14 | 14.8332271 | -0.4947553 | 0.6177473 | -0.8009025 | 0.42318811 | 0.99937574 |
| ADIPOR2 | 347.692237 | -0.1419721 | 0.36967817 | -0.3840425 | 0.70094696 | 0.99937574 |
| DCP1B | 50.9799702 | 0.39214646 | 0.49506013 | 0.79211885 | 0.42829138 | 0.99937574 |
| CACNA1C | 17.6009073 | 1.34905153 | 0.64237216 | 2.10010894 | 0.03571926 | 0.99937574 |
| FKBP4 | 32.1027859 | 0.2162062 | 0.48045659 | 0.45000154 | 0.65270933 | 0.99937574 |
| ITFG2 | 71.2061424 | -0.2436383 | 0.36076803 | -0.6753322 | 0.49946475 | 0.99937574 |
| TULP3 | 58.4262032 | 0.49063751 | 0.34261172 | 1.43205114 | 0.15212919 | 0.99937574 |
| TEAD4 | 36.1931391 | -0.2266063 | 0.57652966 | -0.3930524 | 0.69428081 | 0.99937574 |
| TSPAN9 | 360.115845 | 0.48911355 | 0.44784059 | 1.09215993 | 0.27476281 | 0.99937574 |
| PRMT8 | 23.6120633 | 2.73243035 | 0.93777119 | 2.91374951 | 0.00357116 | 0.53791313 |
| PARP11 | 31.1602685 | -0.0672905 | 0.40458648 | -0.1663191 | 0.8679058 | 0.99937574 |
| TIGAR | 20.5123926 | 0.17616941 | 0.49128486 | 0.35858914 | 0.71990247 | 0.99937574 |
| C12orf4 | 34.7114678 | 0.53505799 | 0.39755442 | 1.34587357 | 0.17834329 | 0.99937574 |
| ANO2 | 14.2033889 | 0.26734648 | 0.79677958 | 0.3355338 | 0.73722245 | 0.99937574 |
| VWF | 28.1727508 | 0.13791969 | 0.92167884 | 0.14963964 | 0.88104893 | 0.99937574 |
| CD9 | 446.423052 | -0.5917163 | 0.58133188 | -1.0178632 | 0.30874296 | 0.99937574 |
| TNFRSF1A | 81.7673495 | 0.05106276 | 0.37291184 | 0.13692984 | 0.89108625 | 0.99937574 |
| SCNN1A | 359.096858 | 0.23448559 | 0.39099618 | 0.59971324 | 0.54869736 | 0.99937574 |
| LTBR | 43.6993241 | -0.407992 | 0.37331066 | -1.092902 | 0.27443683 | 0.99937574 |
| TAPBPL | 16.3657032 | 0.15205422 | 0.51622815 | 0.29454849 | 0.76833882 | 0.99937574 |
| MRPL51 | 40.5243537 | -0.5730589 | 0.59775651 | -0.9586828 | 0.33771857 | 0.99937574 |
| NCAPD2 | 17.4785149 | 0.42389436 | 0.54153193 | 0.78276891 | 0.43376283 | 0.99937574 |
| GAPDH | 477.160033 | 0.18763445 | 0.57528585 | 0.32615864 | 0.74430433 | 0.99937574 |
| CHD4 | 29.7330598 | -0.2779523 | 0.49952137 | -0.5564372 | 0.57791203 | 0.99937574 |
| ING4 | 17.6224275 | 0.19122423 | 0.51850249 | 0.36880099 | 0.71227607 | 0.99937574 |
| ZNF384 | 18.2884636 | -0.4443118 | 0.45496056 | -0.9765941 | 0.32877013 | 0.99937574 |
| COPS7A | 28.5882942 | 0.30235793 | 0.45058249 | 0.6710379 | 0.50219638 | 0.99937574 |
| MLF2 | 54.3454966 | 0.08981527 | 0.31051139 | 0.28924952 | 0.77239044 | 0.99937574 |
| PTMS | 84.3218905 | 0.22406105 | 0.48645123 | 0.46060331 | 0.64508324 | 0.99937574 |
| USP5 | 21.0287632 | -0.0491303 | 0.47166849 | -0.1041628 | 0.91704013 | 0.99937574 |
| TPI1 | 148.526863 | 0.19185726 | 0.57253989 | 0.33509851 | 0.73755078 | 0.99937574 |
| LRRC23 | 26.3422823 | -0.0656156 | 0.41114681 | -0.1595916 | 0.87320277 | 0.99937574 |
| ATN1 | 57.1243504 | -0.6768777 | 0.35682112 | -1.8969663 | 0.05783238 | 0.99937574 |
| C12orf57 | 25.8533003 | -0.0799676 | 0.51178869 | -0.1562513 | 0.87583495 | 0.99937574 |
| PTPN6 | 21.3085845 | 0.16813635 | 0.45940808 | 0.36598475 | 0.71437646 | 0.99937574 |
| PHB2 | 56.7568896 | 0.13939603 | 0.45473879 | 0.30654089 | 0.75919285 | 0.99937574 |
| EMG1.1 | 12.4101921 | 0.38151026 | 0.52654871 | 0.72454884 | 0.46872885 | 0.99937574 |
| LPCAT3 | 128.538237 | -0.1246333 | 0.29650356 | -0.4203433 | 0.67423468 | 0.99937574 |
| C1S | 29.1602566 | 0.10445206 | 0.50790668 | 0.20565208 | 0.8370627 | 0.99937574 |
| C1RL | 39.3533428 | -0.2092321 | 0.39554298 | -0.5289745 | 5.97E-01 | 0.99937574 |
| CLSTN3 | 11.1465717 | 0.11569743 | 0.55973337 | 0.20670098 | 0.83624339 | 0.99937574 |
| PEX5 | 10.6485065 | -0.5040158 | 0.63181193 | -0.7977307 | 0.42502677 | 0.99937574 |
| FOXJ2 | 12.980673 | -0.1422378 | 0.55351667 | -0.2569711 | 0.79720107 | 0.99937574 |
| NECAP1 | 23.1209549 | -0.3838934 | 0.45018391 | -0.8527479 | 0.39379913 | 0.99937574 |
| LINC00937 | 11.699787 | -0.5946378 | 0.54407804 | -1.0929274 | 0.27442567 | 0.99937574 |
| RIMKLB | 113.239684 | -0.7214854 | 0.46771238 | -1.5425835 | 0.12293187 | 0.99937574 |
| A2ML1-AS1 | 17.059863 | -0.1401686 | 0.7810586 | -0.1794597 | 0.85757673 | 0.99937574 |
| PHC1 | 25.4194887 | 0.4602015 | 0.38847667 | 1.18463099 | 0.23616338 | 0.99937574 |
| M6PR | 57.4490766 | -0.1845775 | 0.40353491 | -0.4574015 | 0.64738251 | 0.99937574 |

| | | | | | | |
|---|---|---|---|---|---|---|
| CLEC2D | 20.983133 | -0.2907097 | 0.51783209 | -0.5613977 | 0.57452648 | 0.99937574 |
| TMEM52B | 48.8354216 | 1.49934916 | 1.09638318 | 1.36754119 | 0.17145573 | 0.99937574 |
| GABARAPL1 | 251.892475 | 0.21669474 | 0.3290275 | 0.65859157 | 0.51015808 | 0.99937574 |
| KLRD1 | 146.624958 | 1.85269567 | 0.66814294 | 2.77290317 | 0.00555586 | 0.65033075 |
| MAGOHB | 12.5096939 | -0.0883393 | 0.52855179 | -0.1671346 | 0.86726418 | 0.99937574 |
| YBX3 | 193.491782 | -0.1129525 | 0.36369538 | -0.3105689 | 7.56E-01 | 0.99937574 |
| SMIM10L1 | 18.3680785 | -0.5262476 | 0.63667799 | -0.8265523 | 0.40849088 | 0.99937574 |
| AC244131.2 | 24.0271813 | 0.41703444 | 0.8935598 | 0.46671128 | 0.64070646 | 0.99937574 |
| ETV6 | 260.029766 | -0.0787424 | 0.47850337 | -0.1645597 | 0.86929052 | 0.99937574 |
| BCL2L14 | 43.0457309 | 0.49921195 | 0.47723392 | 1.04605293 | 0.29553659 | 0.99937574 |
| LRP6 | 159.035961 | 0.56101849 | 0.25190272 | 2.22712358 | 2.59E-02 | 0.96546668 |
| MANSC1 | 12.9850183 | -0.0284824 | 0.51061666 | -0.0557804 | 0.95551673 | 0.99937574 |
| BORCS5 | 36.2493985 | 0.01844759 | 0.42165264 | 0.04375069 | 0.96510314 | 0.99937574 |
| DUSP16 | 69.0509127 | -0.5382266 | 0.46911639 | -1.1473199 | 0.25124942 | 0.99937574 |
| CREBL2 | 45.7252749 | -0.0105271 | 0.36605178 | -0.0287586 | 0.9770571 | 0.99937574 |
| CDKN1B | 27.3030494 | -0.0806094 | 0.47518785 | -0.169637 | 0.86529562 | 0.99937574 |
| GPRC5A | 53.4173218 | 0.9969115 | 0.63274486 | 1.57553474 | 0.11513307 | 0.99937574 |
| GPRC5D-AS1 | 44.2427954 | -0.0599558 | 0.39851566 | -0.1504477 | 0.88041139 | 0.99937574 |
| HEBP1 | 39.8965512 | 0.31373753 | 0.41802971 | 0.75051491 | 0.45294465 | 0.99937574 |
| FAM234B | 34.3913452 | 0.93632328 | 0.48707963 | 1.92232076 | 0.05456541 | 0.99937574 |
| ATF7IP | 203.5798 | 0.45185308 | 0.30019826 | 1.5051822 | 0.13227724 | 0.99937574 |
| PLBD1 | 37.3400103 | 0.14459412 | 0.55299878 | 0.26147277 | 0.79372795 | 0.99937574 |
| PLBD1-AS1 | 16.065019 | 0.68339009 | 0.69056526 | 0.98960972 | 0.32236492 | 0.99937574 |
| AC010168.1 | 24.8590472 | 0.22433435 | 0.76573759 | 0.29296505 | 0.76954887 | 0.99937574 |
| H2AFJ | 90.7457294 | -0.2770588 | 0.49011657 | -0.5652916 | 0.57187545 | 0.99937574 |
| WBP11 | 45.1992278 | -0.1775131 | 0.33938226 | -0.5230476 | 0.60094111 | 0.99937574 |
| C12orf60 | 18.7657624 | 0.49421709 | 0.69599486 | 0.71008727 | 0.47765002 | 0.99937574 |
| MGP | 23.4332803 | -0.8129723 | 0.59325074 | -1.3703689 | 0.17057178 | 0.99937574 |
| ERP27 | 214.523296 | 1.10889347 | 0.51397191 | 2.15749822 | 0.03096687 | 0.99049468 |
| RERG | 139.814269 | 0.10861642 | 0.61331897 | 0.17709614 | 8.59E-01 | 0.99937574 |
| EPS8 | 1823.87269 | -0.5412464 | 0.19843848 | -2.7275275 | 0.00638109 | 0.69675365 |
| STRAP | 73.8518425 | 0.55052398 | 0.39627888 | 1.38923373 | 0.16476169 | 0.99937574 |
| DERA | 135.982007 | -0.6216595 | 0.39357456 | -1.5795215 | 0.11421648 | 0.99937574 |
| MGST1 | 19.43716 | 0.41890672 | 0.71837543 | 0.58313063 | 0.55980536 | 0.99937574 |
| RERGL | 22.8708619 | -0.6441364 | 0.73185696 | -0.8801397 | 0.37878361 | 0.99937574 |
| PIK3C2G | 282.136029 | -0.8697002 | 0.60146328 | -1.4459738 | 0.14818454 | 0.99937574 |
| PLEKHA5 | 239.987487 | 0.02202574 | 0.34393109 | 0.06404114 | 0.94893747 | 0.99937574 |
| AEBP2 | 199.283407 | 0.48896474 | 0.3989179 | 1.22572775 | 0.22030114 | 0.99937574 |
| PYROXD1 | 39.9817379 | -0.2784799 | 0.36806542 | -0.7566043 | 0.44928697 | 0.99937574 |
| RECQL | 28.1126415 | -0.2248508 | 0.36695832 | -0.6127422 | 0.54004684 | 0.99937574 |
| GOLT1B | 44.55157 | 0.28228308 | 0.3477311 | 0.81178555 | 0.41691469 | 0.99937574 |
| SPX | 16.0358434 | 0.72060456 | 0.6123496 | 1.17678619 | 0.23928086 | 0.99937574 |
| LDHB | 286.240154 | 0.84330787 | 0.44108441 | 1.91189682 | 0.05588943 | 0.99937574 |
| AC010197.1 | 42.6912494 | 1.87292691 | 0.47980459 | 3.9035202 | 9.48E-05 | 0.08697803 |
| ABCC9 | 31.4223697 | 2.21638685 | 0.9847612 | 2.25068458 | 2.44E-02 | 0.96546668 |
| AC008250.1 | 45.5507183 | 2.16831322 | 0.74217771 | 2.92155529 | 0.00348288 | 0.53256471 |
| CMAS | 31.5303903 | -0.3003889 | 0.47801704 | -0.6284062 | 0.52973786 | 0.99937574 |
| C2CD5 | 118.243653 | -0.1025999 | 0.32170565 | -0.3189248 | 0.74978353 | 0.99937574 |
| ETNK1 | 232.125807 | -0.1448378 | 0.34175394 | -0.4238072 | 0.67170642 | 0.99937574 |
| SOX5 | 175.40036 | -2.7551087 | 0.89670661 | -3.072475 | 0.00212292 | 0.45583974 |
| LRMP | 23.5654012 | 0.21772086 | 0.85831066 | 0.25366207 | 0.79975664 | 0.99937574 |
| ETFRF1 | 43.7426919 | 0.27811339 | 0.38527412 | 0.72185847 | 0.4703815 | 0.99937574 |
| KRAS | 86.2909412 | -0.147524 | 0.30502889 | -0.4836396 | 0.62864167 | 0.99937574 |
| RASSF8-AS1 | 11.9891348 | -0.5141818 | 0.72098594 | -0.7131648 | 0.47574379 | 0.99937574 |

| | | | | | | |
|---|---|---|---|---|---|---|
| RASSF8 | 77.8830039 | -0.4735023 | 0.49270429 | -0.9610273 | 0.33653844 | 0.99937574 |
| AC022509.2 | 16.6469833 | 1.76534796 | 0.62808018 | 2.81070479 | 0.00494331 | 0.61461453 |
| AC024145.1 | 43.1768662 | -0.0687053 | 0.75097516 | -0.0914881 | 0.92710479 | 0.99937574 |
| ITPR2 | 1600.97011 | 0.2876734 | 0.40675117 | 0.70724664 | 0.47941322 | 0.99937574 |
| AC023051.1 | 50.2005681 | 0.23879443 | 0.87697096 | 0.27229457 | 0.78539553 | 0.99937574 |
| INTS13 | 18.4512867 | 0.11599311 | 0.43655549 | 0.26570074 | 0.7904697 | 0.99937574 |
| FGFR1OP2 | 68.1082756 | 0.11264399 | 0.36586521 | 0.30788386 | 0.75817071 | 0.99937574 |
| TM7SF3 | 71.5001921 | 0.12079063 | 0.50899547 | 0.23731179 | 0.81241491 | 0.99937574 |
| MED21 | 32.4659302 | -0.21625 | 0.39882721 | -0.5422147 | 0.58767061 | 0.99937574 |
| STK38L | 37.9505977 | 0.10223246 | 0.34878332 | 0.29311166 | 0.76943681 | 0.99937574 |
| PPFIBP1 | 147.209581 | -0.1725673 | 0.30624194 | -0.5634998 | 0.57309459 | 0.99937574 |
| MRPS35 | 88.6615585 | 0.32643192 | 0.32097638 | 1.01699669 | 0.30915501 | 0.99937574 |
| KLHL42 | 51.3531997 | -0.4278637 | 0.50893802 | -0.840699 | 0.40051657 | 0.99937574 |
| CCDC91 | 324.308534 | -0.2567308 | 0.27050146 | -0.9490922 | 0.34257373 | 0.99937574 |
| ERGIC2 | 64.4060099 | -0.0536898 | 0.30635784 | -0.175252 | 0.86088165 | 0.99937574 |
| TMTC1 | 127.208517 | -3.9964175 | 1.3584469 | -2.9419019 | 0.00326203 | 0.53256471 |
| IPO8 | 67.6104425 | 0.3694385 | 0.27258331 | 1.35532326 | 0.17531459 | 0.99937574 |
| CAPRIN2 | 39.0868704 | -0.0553733 | 0.39712349 | -0.1394359 | 0.88910574 | 0.99937574 |
| DDX11 | 13.553753 | -0.5857447 | 0.64950402 | -0.9018339 | 0.36714512 | 0.99937574 |
| SINHCAF | 155.50856 | -0.2272144 | 0.41282763 | -0.5503856 | 0.58205489 | 0.99937574 |
| DENND5B | 180.9846 | -0.1220891 | 0.35216866 | -0.346678 | 0.72883322 | 0.99937574 |
| AMN1 | 29.4195876 | 0.23771515 | 0.37162131 | 0.63967041 | 0.5223869 | 0.99937574 |
| KIAA1551 | 32.3818642 | -0.1961578 | 0.43964422 | -0.446174 | 0.65547155 | 0.99937574 |
| BICD1 | 34.4319205 | -0.0802833 | 0.63409354 | -0.1266112 | 0.89924813 | 0.99937574 |
| FGD4 | 848.182164 | 0.09011361 | 0.25743872 | 0.35003905 | 0.72630939 | 0.99937574 |
| DNM1L | 163.648005 | -0.0443912 | 0.19046315 | -0.2330698 | 0.81570717 | 0.99937574 |
| YARS2 | 33.41035 | 0.33170513 | 0.32814339 | 1.01085423 | 0.3120862 | 0.99937574 |
| PKP2 | 53.9213868 | -0.2878464 | 0.69223183 | -0.4158238 | 0.67753897 | 0.99937574 |
| SYT10 | 103.379121 | 1.37880406 | 0.43650855 | 3.1587103 | 0.00158469 | 0.3807782 |
| ALG10B | 11.045397 | -0.2684533 | 0.60046061 | -0.4470789 | 0.65481807 | 0.99937574 |
| CPNE8 | 27.7618984 | -1.0168502 | 0.62972814 | -1.6147447 | 0.106366 | 0.99937574 |
| KIF21A | 398.10529 | -0.2319129 | 0.44648259 | -0.519422 | 0.6034665 | 0.99937574 |
| SLC2A13 | 12.75886 | -0.6774815 | 0.51001935 | -1.3283448 | 0.18406423 | 0.99937574 |
| LRRK2 | 107.358278 | -2.5004245 | 1.01783356 | -2.4566143 | 0.01402532 | 0.84756595 |
| GXYLT1 | 58.2480121 | -0.0442859 | 0.34983273 | -0.1265917 | 0.89926355 | 0.99937574 |
| YAF2 | 229.769947 | 0.27391712 | 0.2144133 | 1.27751928 | 0.20141898 | 0.99937574 |
| PPHLN1 | 126.621944 | -0.1851021 | 0.30386724 | -0.6091544 | 0.54242212 | 0.99937574 |
| ZCRB1 | 29.6313337 | 0.07901673 | 0.41412339 | 0.1908048 | 0.84867853 | 0.99937574 |
| PUS7L | 48.2529533 | 0.64422441 | 0.36337239 | 1.77290411 | 0.0762446 | 0.99937574 |
| IRAK4 | 83.2419384 | 0.10160745 | 0.49411821 | 0.20563388 | 0.83707691 | 0.99937574 |
| TWF1 | 77.0712964 | -0.4312707 | 0.50800734 | -0.8489458 | 0.39591146 | 0.99937574 |
| TMEM117 | 1662.88394 | -0.3884205 | 0.41885481 | -0.9273392 | 0.35375046 | 0.99937574 |
| ANO6 | 99.4865337 | -0.4348534 | 0.35731708 | -1.2169959 | 0.22360579 | 0.99937574 |
| AC008124.1 | 12.3587351 | 1.25146627 | 0.61449123 | 2.03658932 | 0.04169121 | 0.99937574 |
| ARID2 | 244.864554 | 0.23383861 | 0.27755953 | 0.84248091 | 0.39951882 | 0.99937574 |
| SCAF11 | 313.932925 | -0.1364983 | 0.19701437 | -0.692834 | 0.4884137 | 0.99937574 |
| SLC38A1 | 279.710096 | -0.5683672 | 0.34995425 | -1.6241185 | 0.10435052 | 0.99937574 |
| SLC38A2 | 137.735366 | -0.5684671 | 0.72207974 | -0.7872635 | 0.4311276 | 0.99937574 |
| AC008014.1 | 340.365547 | 0.13706257 | 0.53170617 | 0.25777879 | 0.79657764 | 0.99937574 |
| SLC38A4 | 35.3004194 | 0.24585566 | 0.61317568 | 0.40095468 | 0.68845349 | 0.99937574 |
| AMIGO2 | 18.8770782 | -1.4440428 | 0.83121541 | -1.7372666 | 0.08234012 | 0.99937574 |
| PCED1B | 218.978631 | -0.3971235 | 0.31433327 | -1.2633835 | 0.20645139 | 0.99937574 |
| RPAP3 | 36.5330856 | -0.2426987 | 0.40378473 | -0.6010597 | 0.54780021 | 0.99937574 |
| RAPGEF3 | 78.4991056 | -0.8085099 | 0.56800069 | -1.4234312 | 0.15461119 | 0.99937574 |

| | | | | | | |
|---|---|---|---|---|---|---|
| SLC48A1 | 44.9268425 | -0.2809867 | 0.43651275 | -0.6437078 | 0.51976491 | 0.99937574 |
| HDAC7 | 104.526687 | -0.4764352 | 0.49358808 | -0.9652485 | 0.33442037 | 0.99937574 |
| VDR | 38.5812084 | 0.35666199 | 0.50566101 | 0.70533813 | 0.48059984 | 0.99937574 |
| TMEM106C | 109.739741 | -0.0750933 | 0.54041748 | -0.1389543 | 0.88948624 | 0.99937574 |
| SENP1 | 33.4626511 | -0.798569 | 0.36111362 | -2.2114065 | 0.0270077 | 0.98043778 |
| PFKM | 30.50071 | 0.22280422 | 0.44027959 | 0.50605166 | 0.61282038 | 0.99937574 |
| ASB8 | 29.8335684 | 0.06084387 | 0.44418329 | 0.1369792 | 0.89104724 | 0.99937574 |
| ZNF641 | 13.384439 | 0.30464106 | 0.49087154 | 0.62061259 | 0.53485456 | 0.99937574 |
| KANSL2 | 31.7306891 | -0.1648202 | 0.40660273 | -0.4053594 | 0.68521339 | 0.99937574 |
| CCNT1 | 49.4366078 | -0.165454 | 0.29652004 | -0.557986 | 0.57685396 | 0.99937574 |
| ADCY6 | 46.6746213 | 0.21377246 | 0.38124425 | 0.5607231 | 0.57498632 | 0.99937574 |
| CACNB3 | 14.6173965 | 0.2987626 | 0.48124284 | 0.62081465 | 0.53472158 | 0.99937574 |
| DDX23 | 28.8908558 | 0.02858063 | 0.40742281 | 0.0701498 | 0.94407443 | 0.99937574 |
| CCDC65 | 22.8010505 | 0.27446196 | 0.65733411 | 0.41753799 | 0.67628494 | 0.99937574 |
| PRKAG1 | 39.661683 | 0.00453639 | 0.41645681 | 0.01089282 | 0.99130896 | 0.99937574 |
| KMT2D | 37.0637162 | -0.3612724 | 0.47273397 | -0.7642193 | 0.44473658 | 0.99937574 |
| LMBR1L | 31.3745424 | -0.2335012 | 0.4269633 | -0.5468882 | 0.58445556 | 0.99937574 |
| TUBA1B | 109.233014 | 0.59288293 | 0.53113996 | 1.11624616 | 0.26431678 | 0.99937574 |
| TUBA1A | 12.4279063 | -0.34741 | 0.65167245 | -0.5331052 | 0.59396074 | 0.99937574 |
| TUBA1C | 63.4306516 | 0.27383929 | 0.43754733 | 0.62585067 | 0.53141289 | 0.99937574 |
| SPATS2 | 113.687939 | -0.0277741 | 0.32670676 | -0.0850124 | 0.9322515 | 0.99937574 |
| MCRS1 | 20.1213892 | -0.4950536 | 0.41547835 | -1.1915268 | 0.23344683 | 0.99937574 |
| PRPF40B | 68.471461 | 0.24570381 | 0.30370333 | 0.80902575 | 0.41850034 | 0.99937574 |
| FMNL3 | 35.7643212 | 1.56292975 | 0.62133879 | 2.51542278 | 0.01188897 | 0.77408712 |
| TMBIM6 | 383.854264 | 0.03689288 | 0.4289081 | 0.08601582 | 0.93145384 | 0.99937574 |
| NCKAP5L | 22.5815154 | -0.4422009 | 0.49238312 | -0.8980829 | 0.36914134 | 0.99937574 |
| BCDIN3D | 11.7793395 | 0.16331592 | 0.52186314 | 0.3129478 | 0.75432032 | 0.99937574 |
| AQP2 | 25.4946826 | -0.4680344 | 0.90177666 | -0.5190136 | 0.60375124 | 0.99937574 |
| AC025154.2 | 14.9438414 | -0.1882478 | 0.67032368 | -0.2808311 | 0.77883992 | 0.99937574 |
| AQP6 | 526.113274 | 0.62428111 | 0.69874639 | 0.89343018 | 0.37162685 | 0.99937574 |
| RACGAP1 | 70.0692513 | 0.92697335 | 0.38396177 | 2.41423349 | 0.01576835 | 0.86958595 |
| SMARCD1 | 27.0744161 | -0.2321519 | 0.45498474 | -0.510241 | 0.60988265 | 0.99937574 |
| AC074032.1 | 68.5413621 | -0.0572191 | 0.36793296 | -0.155515 | 0.87641532 | 0.99937574 |
| CERS5 | 48.1436479 | -0.2948232 | 0.29167448 | -1.0107953 | 0.31211442 | 0.99937574 |
| LIMA1 | 892.477061 | -0.2621812 | 0.37399408 | -0.7010302 | 0.48328416 | 0.99937574 |
| LARP4 | 225.771429 | 0.05184544 | 0.24123722 | 0.21491475 | 0.82983379 | 0.99937574 |
| DIP2B | 142.340744 | -0.2115319 | 0.27829128 | -0.7601096 | 0.44718909 | 0.99937574 |
| ATF1 | 49.3634849 | -0.2713617 | 0.38201709 | -0.7103391 | 0.47749388 | 0.99937574 |
| METTL7A | 74.4807964 | 0.788073 | 0.52130551 | 1.51172966 | 0.13060265 | 0.99937574 |
| SLC11A2 | 73.5309184 | -0.0864731 | 0.31785489 | -0.272052 | 0.78558201 | 0.99937574 |
| LETMD1 | 67.0230535 | 0.17988151 | 0.29658379 | 0.60651158 | 0.54417509 | 0.99937574 |
| CSRNP2 | 18.825499 | -0.4789079 | 0.49956431 | -0.9586512 | 0.33773451 | 0.99937574 |
| TFCP2 | 136.652042 | -0.1975502 | 0.39277794 | -0.5029564 | 0.61499492 | 0.99937574 |
| POU6F1 | 23.1549006 | -0.545122 | 0.39671247 | -1.3740985 | 0.1694111 | 0.99937574 |
| DAZAP2 | 151.11033 | -0.3609327 | 0.31016493 | -1.1636799 | 0.24455374 | 0.99937574 |
| SMAGP | 15.2317935 | -0.1450968 | 0.59385029 | -0.2443322 | 0.80697354 | 0.99937574 |
| ACVR1B | 146.719988 | -0.108479 | 0.38450749 | -0.2821246 | 0.77784794 | 0.99937574 |
| NR4A1 | 83.0885763 | -0.4071698 | 0.65129447 | -0.6251701 | 0.53185944 | 0.99937574 |
| ATG101 | 33.0045855 | 0.01683202 | 0.53682464 | 0.03135479 | 0.9749866 | 0.99937574 |
| AC078864.2 | 31.9062334 | 0.13189784 | 0.80596393 | 0.16365228 | 0.87000489 | 0.99937574 |
| AC078864.1 | 67.7925272 | -0.423187 | 0.33910524 | -1.247952 | 2.12E-01 | 0.99937574 |
| KRT7 | 31.2874952 | -0.5678268 | 0.8750202 | -0.6489299 | 0.51638369 | 0.99937574 |
| KRT8 | 182.540439 | -0.0612171 | 0.42433301 | -0.1442666 | 0.88528994 | 0.99937574 |
| KRT18 | 144.915628 | 0.10527621 | 0.52423983 | 0.20081689 | 0.84084176 | 0.99937574 |

| | | | | | | |
|---|---|---|---|---|---|---|
| EIF4B | 139.095703 | -0.1885162 | 0.35013245 | -0.5384141 | 0.59029119 | 0.99937574 |
| AC068888.1 | 58.5982041 | 0.32840062 | 0.50179208 | 0.65445557 | 0.51281834 | 0.99937574 |
| TNS2 | 44.352451 | 0.11693726 | 0.67915071 | 0.17218161 | 0.86329475 | 0.99937574 |
| SPRYD3 | 14.4106107 | 0.11896107 | 0.59699838 | 0.19926531 | 0.84205521 | 0.99937574 |
| CSAD | 55.2312589 | 0.15455442 | 0.46088008 | 0.33534627 | 0.73736389 | 0.99937574 |
| ZNF740 | 11.1646737 | -0.1841935 | 0.59273336 | -0.3107527 | 0.75598864 | 0.99937574 |
| MFSD5 | 11.315014 | 0.11809272 | 0.68457039 | 0.17250632 | 0.8630395 | 0.99937574 |
| PFDN5 | 122.227796 | -0.1517335 | 0.53032493 | -0.2861142 | 0.77479067 | 0.99937574 |
| C12orf10 | 13.0344612 | -0.0121116 | 0.57479712 | -0.0210711 | 0.98318895 | 0.99937574 |
| SP1 | 61.1785844 | 0.07272646 | 0.35016011 | 0.20769487 | 0.83546722 | 0.99937574 |
| PRR13 | 89.5324492 | 0.21466741 | 0.34133301 | 0.62890901 | 0.52940863 | 0.99937574 |
| PCBP2 | 183.162353 | -0.2621389 | 0.29554671 | -0.8869627 | 0.37509899 | 0.99937574 |
| MAP3K12 | 11.9601286 | 1.17170134 | 0.62160686 | 1.88495562 | 0.05943583 | 0.99937574 |
| ATP5MC2 | 144.509772 | 0.49859937 | 0.36872798 | 1.35221463 | 0.17630666 | 0.99937574 |
| CALCOCO1 | 33.1126328 | 0.20713428 | 0.45061136 | 0.45967389 | 0.64575031 | 0.99937574 |
| SMUG1 | 26.528951 | 0.50718731 | 0.41266708 | 1.22904719 | 0.21905411 | 0.99937574 |
| LINC02381 | 30.9811566 | -0.1016335 | 0.48354859 | -0.2101825 | 0.83352521 | 0.99937574 |
| CBX5 | 99.252065 | 0.32277588 | 0.37518952 | 0.86030089 | 0.3896232 | 0.99937574 |
| HNRNPA1 | 73.5155134 | -0.4835894 | 0.43709308 | -1.1063762 | 0.26856372 | 0.99937574 |
| COPZ1 | 56.6254791 | 0.07456465 | 0.2922326 | 0.25515515 | 0.79860327 | 0.99937574 |
| CD63 | 186.823862 | -0.4044964 | 0.6054733 | -0.6680665 | 0.50409112 | 0.99937574 |
| ORMDL2 | 25.6924845 | -0.4098418 | 0.47963579 | -0.8544854 | 0.39283611 | 0.99937574 |
| PYM1 | 14.7247407 | -0.1878814 | 0.49655212 | -0.3783719 | 0.70515434 | 0.99937574 |
| RAB5B | 57.1288426 | -0.1629574 | 0.36641548 | -0.4447338 | 0.65651212 | 0.99937574 |
| SUOX | 20.1646137 | 0.33059804 | 0.62652989 | 0.52766523 | 0.59773171 | 0.99937574 |
| IKZF4 | 13.9403949 | 0.32612836 | 0.56952307 | 0.57263416 | 0.56689243 | 0.99937574 |
| RPS26 | 32.7176325 | -0.6709134 | 0.38696554 | -1.7337808 | 0.08295699 | 0.99937574 |
| ERBB3 | 28.6830384 | 0.0239781 | 0.48489218 | 0.04945038 | 0.96056038 | 0.99937574 |
| PA2G4 | 23.7859856 | -0.0803149 | 0.47494264 | -0.1691045 | 0.86571445 | 0.99937574 |
| RPL41 | 521.605867 | -0.2135628 | 0.48395585 | -0.4412857 | 0.65900615 | 0.99937574 |
| ESYT1 | 23.4522761 | -0.0992224 | 0.44909306 | -0.2209395 | 0.82513951 | 0.99937574 |
| MYL6 | 181.625951 | -0.6497891 | 0.45436721 | -1.4300968 | 0.15268925 | 0.99937574 |
| SMARCC2 | 77.948499 | 0.36218804 | 0.26934923 | 1.34467819 | 0.17872918 | 0.99937574 |
| RNF41 | 20.6127837 | 0.48429599 | 0.42643468 | 1.13568622 | 0.25608791 | 0.99937574 |
| ANKRD52 | 21.3697273 | -0.0348506 | 0.42488422 | -0.0820238 | 0.93462776 | 0.99937574 |
| CS | 65.2958312 | -0.1381299 | 0.3351848 | -0.4121007 | 0.68026564 | 0.99937574 |
| PAN2 | 20.9298701 | 0.0334627 | 0.50884565 | 0.06576198 | 0.94756733 | 0.99937574 |
| STAT2 | 43.3963465 | -0.7676138 | 0.55945226 | -1.372081 | 0.17003822 | 0.99937574 |
| SPRYD4 | 10.8736429 | 0.20479591 | 0.57498261 | 0.35617757 | 0.72170759 | 0.99937574 |
| RBMS2 | 36.0070331 | -0.8112832 | 0.40048443 | -2.0257546 | 0.04278994 | 0.99937574 |
| BAZ2A | 77.3900839 | 0.05755634 | 0.3235384 | 0.17789647 | 0.85880428 | 0.99937574 |
| ATP5F1B | 298.597675 | 0.72278414 | 0.45752494 | 1.57976993 | 0.11415957 | 0.99937574 |
| PTGES3 | 185.121141 | -0.0777651 | 0.26569468 | -0.2926861 | 0.7697621 | 0.99937574 |
| NACA | 122.339865 | -0.0041161 | 0.56678808 | -0.0072621 | 0.9942057 | 0.99937574 |
| NEMP1 | 15.2848038 | -0.9420308 | 0.56418322 | -1.669725 | 0.09497379 | 0.99937574 |
| STAT6 | 113.638322 | -0.5124664 | 0.28813773 | -1.7785468 | 0.07531409 | 0.99937574 |
| R3HDM2 | 200.429085 | -0.0150049 | 0.30002995 | -0.0500112 | 0.96011344 | 0.99937574 |
| MARS | 28.5169175 | -0.9127619 | 0.44486 | -2.0517959 | 0.0401895 | 0.99937574 |
| DDIT3 | 44.6852765 | -1.4285241 | 0.73428233 | -1.9454698 | 0.05171846 | 0.99937574 |
| MBD6 | 20.7593408 | -0.0999695 | 0.53763663 | -0.1859426 | 0.85248979 | 0.99937574 |
| DCTN2 | 47.1605092 | -0.0758614 | 0.34989663 | -0.2168108 | 0.82835583 | 0.99937574 |
| PIP4K2C | 39.7248378 | 0.13480893 | 0.30712412 | 0.43893956 | 0.66070533 | 0.99937574 |
| DTX3 | 15.6889051 | -0.5811947 | 0.43878392 | -1.3245578 | 0.18531786 | 0.99937574 |
| OS9 | 77.2415536 | -0.0566409 | 0.43918985 | -0.1289667 | 0.89738396 | 0.99937574 |

| | | | | | | |
|---|---|---|---|---|---|---|
| TSPAN31 | 26.5587043 | 0.33526603 | 0.43220049 | 0.77571876 | 0.43791506 | 0.99937574 |
| CDK4 | 14.9479945 | -0.6772117 | 0.54668234 | -1.2387663 | 0.21543205 | 0.99937574 |
| 9-Mar | 25.3053053 | -0.0455278 | 0.47887359 | -0.0950727 | 0.92425705 | 0.99937574 |
| TSFM | 26.2112492 | 0.68554857 | 0.52133741 | 1.31498058 | 0.18851643 | 0.99937574 |
| AVIL | 14.2053396 | -0.8099955 | 0.850014 | -0.9529202 | 0.34063051 | 0.99937574 |
| CTDSP2 | 57.5605551 | -0.6811672 | 0.48969252 | -1.3910101 | 0.16422236 | 0.99937574 |
| LRIG3 | 16.6170576 | -0.5885382 | 0.56768138 | -1.0367404 | 0.29985687 | 0.99937574 |
| SLC16A7 | 456.688104 | -0.1910379 | 0.52449975 | -0.3642287 | 0.71568725 | 0.99937574 |
| FAM19A2 | 30.2849618 | -0.9706836 | 0.67546888 | -1.4370515 | 0.15070336 | 0.99937574 |
| USP15 | 187.58201 | -0.5021053 | 0.29225546 | -1.7180356 | 0.08579013 | 0.99937574 |
| MON2 | 359.679524 | -0.3839128 | 0.37332011 | -1.0283743 | 0.30377382 | 0.99937574 |
| AC048341.1 | 25.0241871 | 0.47590658 | 0.6653734 | 0.71524738 | 0.47445619 | 0.99937574 |
| PPM1H | 64.3667032 | -0.2657357 | 0.47666975 | -0.5574839 | 0.57719688 | 0.99937574 |
| AVPR1A | 49.5110954 | -0.0919254 | 0.7567439 | -0.1214749 | 0.9033149 | 0.99937574 |
| DPY19L2 | 457.675913 | 0.04355005 | 0.46971097 | 0.0927167 | 0.92612863 | 0.99937574 |
| RXYLT1 | 37.1254102 | 0.08907848 | 0.45098894 | 0.19751811 | 0.84342211 | 0.99937574 |
| SRGAP1 | 319.779224 | -0.4051872 | 0.45742269 | -0.8858047 | 0.37572278 | 0.99937574 |
| AC020611.2 | 17.3335114 | -0.5877009 | 0.86304687 | -0.6809606 | 0.49589643 | 0.99937574 |
| XPOT | 32.8600154 | -0.4355153 | 0.426343 | -1.0215138 | 0.30701106 | 0.99937574 |
| TBK1 | 131.271899 | -0.7199766 | 0.27045798 | -2.6620646 | 0.0077663 | 0.74742128 |
| RASSF3 | 14.08382 | -0.5018804 | 0.6555596 | -0.7655756 | 0.44392886 | 0.99937574 |
| GNS | 124.540067 | 0.17298924 | 0.37003435 | 0.46749507 | 0.64014572 | 0.99937574 |
| TBC1D30 | 45.4814896 | -0.3129707 | 0.61006711 | -0.5130103 | 0.60794413 | 0.99937574 |
| LEMD3 | 108.957992 | 0.10172345 | 0.27202075 | 0.37395476 | 0.70843798 | 0.99937574 |
| LLPH | 12.8577789 | 0.03921338 | 0.53604687 | 0.0731529 | 0.94168445 | 0.99937574 |
| HELB | 25.0577071 | -0.6244011 | 0.52406982 | -1.1914465 | 0.23347837 | 0.99937574 |
| GRIP1 | 1236.73487 | 0.35362377 | 0.27470184 | 1.28730036 | 1.98E-01 | 0.99937574 |
| CAND1 | 114.004573 | -0.0549674 | 0.30522505 | -0.1800882 | 0.8570833 | 0.99937574 |
| DYRK2 | 16.7708283 | 0.44537163 | 0.52839218 | 0.84288081 | 0.39929511 | 0.99937574 |
| MDM1 | 10.960816 | 0.2844798 | 0.5419013 | 0.52496608 | 0.59960677 | 0.99937574 |
| RAP1B | 264.948658 | -0.5148665 | 0.34485505 | -1.4929939 | 0.13543877 | 0.99937574 |
| NUP107 | 46.3875407 | 0.05199793 | 0.32602989 | 0.15948822 | 0.87328424 | 0.99937574 |
| SLC35E3 | 24.5669528 | -0.5715277 | 0.65950659 | -0.866599 | 0.38616178 | 0.99937574 |
| MDM2 | 76.0734438 | -0.6799337 | 0.35435227 | -1.9188072 | 0.05500873 | 0.99937574 |
| CPM | 119.343081 | 0.74483048 | 0.28716416 | 2.59374452 | 9.49E-03 | 0.76274616 |
| CPSF6 | 85.3570201 | -0.202453 | 0.33187377 | -0.6100301 | 0.54184188 | 0.99937574 |
| YEATS4 | 14.2945141 | 0.0856285 | 0.48394962 | 0.17693681 | 0.85955803 | 0.99937574 |
| FRS2 | 107.725832 | -0.1994626 | 0.2789687 | -0.7150001 | 0.47460901 | 0.99937574 |
| CCT2 | 47.2679428 | 0.45436858 | 0.39718744 | 1.14396514 | 0.2526381 | 0.99937574 |
| RAB3IP | 86.2331835 | -0.1039666 | 0.33248728 | -0.3126934 | 0.75451365 | 0.99937574 |
| MYRFL | 19.5016714 | 1.41416245 | 0.9971148 | 1.4182544 | 0.1561165 | 0.99937574 |
| AC025159.1 | 18.3188157 | -0.883147 | 0.48489765 | -1.8213059 | 0.06856036 | 0.99937574 |
| CNOT2 | 167.267449 | -0.2583319 | 0.28082427 | -0.9199058 | 0.35762199 | 0.99937574 |
| ZFC3H1 | 136.764273 | -0.060206 | 0.28901089 | -0.2083175 | 0.83498105 | 0.99937574 |
| TMEM19 | 18.3718213 | 0.54544631 | 0.53120458 | 1.02681026 | 0.30450982 | 0.99937574 |
| RAB21 | 126.63245 | 0.01865519 | 0.25416506 | 0.07339793 | 0.94148946 | 0.99937574 |
| TBC1D15 | 111.760388 | -0.3590633 | 0.29400008 | -1.2213035 | 0.22197112 | 0.99937574 |
| TRHDE | 41.8611504 | 0.14541808 | 0.64825447 | 0.22432253 | 0.82250635 | 0.99937574 |
| ATXN7L3B | 43.2212211 | -0.009058 | 0.37140454 | -0.0243884 | 0.98054277 | 0.99937574 |
| CAPS2 | 107.330349 | 0.34268913 | 0.58497219 | 0.58582123 | 0.55799566 | 0.99937574 |
| KRR1 | 33.9714507 | -1.0317601 | 0.39007046 | -2.6450609 | 0.00816762 | 0.74742128 |
| AC078923.1 | 128.416922 | -1.4416747 | 0.75849927 | -1.9006935 | 0.05734217 | 0.99937574 |
| PHLDA1 | 112.650975 | 0.25634426 | 0.4484952 | 0.57156523 | 0.56761656 | 0.99937574 |
| NAP1L1 | 111.863463 | 0.06155724 | 0.25595646 | 0.24049887 | 0.80994354 | 0.99937574 |

| | | | | | | |
|---|---|---|---|---|---|---|
| OSBPL8 | 186.553453 | -0.3936418 | 0.3117616 | -1.2626372 | 0.2067196 | 0.99937574 |
| ZDHHC17 | 112.907221 | 0.06509147 | 0.26325221 | 0.24725895 | 0.80470782 | 0.99937574 |
| PAWR | 181.424347 | -0.1677526 | 0.25676078 | -0.6533421 | 0.51353577 | 0.99937574 |
| PPP1R12A | 180.220194 | -0.1106833 | 0.23255191 | -0.4759509 | 0.63410935 | 0.99937574 |
| LIN7A | 316.813543 | 0.32511137 | 0.5071992 | 0.64099345 | 0.52152694 | 0.99937574 |
| ACSS3 | 113.347609 | -0.2786852 | 0.58413256 | -0.4770924 | 0.63329632 | 0.99937574 |
| CCDC59 | 20.9138033 | 0.21562815 | 0.59577522 | 0.3619287 | 0.71740531 | 0.99937574 |
| METTL25 | 61.4552522 | -0.4076373 | 0.32216262 | -1.2653154 | 0.20575829 | 0.99937574 |
| TMTC2 | 118.284864 | -0.3844614 | 0.67207051 | -0.5720552 | 0.56728459 | 0.99937574 |
| LRRIQ1 | 16.0964086 | 0.25578438 | 0.67996724 | 0.37617163 | 0.7067893 | 0.99937574 |
| C12orf29 | 14.3663077 | -0.1962612 | 0.51059904 | -0.3843745 | 0.70070093 | 0.99937574 |
| CEP290 | 78.0796158 | 0.11509712 | 0.3628884 | 0.31716947 | 0.75111501 | 0.99937574 |
| TMTC3 | 56.3680784 | 0.31615655 | 0.27256299 | 1.1599394 | 0.24607348 | 0.99937574 |
| KITLG | 28.5391935 | 0.53912328 | 0.52995677 | 1.01729671 | 0.3090123 | 0.99937574 |
| DUSP6 | 59.9793625 | 0.75452047 | 0.54533584 | 1.38358863 | 0.16648445 | 0.99937574 |
| POC1B | 53.4359796 | 0.20481726 | 0.39389551 | 0.51997866 | 0.60307845 | 0.99937574 |
| POC1B-AS1 | 22.405176 | 0.64349028 | 0.57948386 | 1.11045421 | 0.26680335 | 0.99937574 |
| ATP2B1 | 117.011496 | -0.1039393 | 0.33899197 | -0.3066128 | 0.75913807 | 0.99937574 |
| BTG1 | 206.613147 | 0.11402589 | 0.40947387 | 0.27846928 | 0.78065215 | 0.99937574 |
| EEA1 | 126.409621 | -0.6416103 | 0.37257963 | -1.7220757 | 0.08505581 | 0.99937574 |
| AC138123.1 | 16.9513614 | -0.6555548 | 0.67010742 | -0.9782832 | 0.32793426 | 0.99937574 |
| NUDT4 | 11.8360958 | 0.15747589 | 0.72620733 | 0.21684701 | 0.82832758 | 0.99937574 |
| UBE2N | 71.3151406 | 0.3821211 | 0.27509786 | 1.38903697 | 0.16482151 | 0.99937574 |
| MRPL42 | 55.5170466 | 0.10974277 | 0.29416065 | 0.37307088 | 0.70909569 | 0.99937574 |
| CRADD | 347.509674 | 0.25800062 | 0.25576164 | 1.00875418 | 0.31309254 | 0.99937574 |
| AC012085.2 | 28.7324567 | 0.94440944 | 0.8032609 | 1.17571942 | 0.23970701 | 0.99937574 |
| CEP83 | 42.4070466 | -0.2725646 | 0.35420223 | -0.7695168 | 0.44158657 | 0.99937574 |
| TMCC3 | 51.1276565 | 0.37645049 | 0.55062107 | 0.68368341 | 0.4941751 | 0.99937574 |
| NDUFA12 | 82.9647158 | 0.25773763 | 0.4418991 | 0.58324995 | 0.55972505 | 0.99937574 |
| NR2C1 | 50.7029333 | 0.20373471 | 0.32607562 | 0.62480818 | 0.53209696 | 0.99937574 |
| FGD6 | 27.9741122 | 0.27009255 | 0.56551717 | 0.47760275 | 0.63293297 | 0.99937574 |
| VEZT | 152.406093 | -0.0369657 | 0.27778994 | -0.1330706 | 0.89413758 | 0.99937574 |
| METAP2 | 79.3181405 | 0.18588957 | 0.2670361 | 0.6961215 | 0.48635274 | 0.99937574 |
| USP44 | 11.2014661 | 0.90693083 | 0.74195678 | 1.22234996 | 0.22157532 | 0.99937574 |
| NTN4 | 44.6185226 | -0.2800187 | 0.74773956 | -0.374487 | 0.70804206 | 0.99937574 |
| SNRPF | 17.7964402 | 0.13881273 | 0.57096392 | 0.24311996 | 0.80791247 | 0.99937574 |
| LTA4H | 31.9630753 | -0.0351253 | 0.41212804 | -0.085229 | 0.93207935 | 0.99937574 |
| ELK3 | 14.1684749 | -1.0389305 | 0.84070122 | -1.2357904 | 0.21653647 | 0.99937574 |
| CDK17 | 74.5937494 | -0.4599599 | 0.35761606 | -1.2861836 | 0.198379 | 0.99937574 |
| NEDD1 | 31.5523025 | 0.23714541 | 0.37369625 | 0.63459404 | 0.52569322 | 0.99937574 |
| TMPO | 76.2919812 | -0.1277519 | 0.32375939 | -0.3945891 | 0.69314616 | 0.99937574 |
| SLC25A3 | 254.25624 | 0.57916012 | 0.50876243 | 1.13837045 | 0.25496583 | 0.99937574 |
| APAF1 | 26.9851952 | 0.30015927 | 0.44954685 | 0.66769297 | 0.5043296 | 0.99937574 |
| UHRF1BP1L | 101.493807 | -0.0692551 | 0.23965243 | -0.2889815 | 0.77259551 | 0.99937574 |
| ACTR6 | 30.4214916 | 0.45604771 | 0.37520363 | 1.21546721 | 0.22418795 | 0.99937574 |
| SCYL2 | 102.131813 | -0.1216068 | 0.34216922 | -0.3553997 | 0.72229016 | 0.99937574 |
| ANO4 | 47.8124584 | 0.98692182 | 0.93349257 | 1.05723587 | 2.90E-01 | 9.99E-01 |
| UTP20 | 59.5279354 | -0.8018582 | 0.57345825 | -1.3982853 | 0.16202742 | 0.99937574 |
| ARL1 | 66.218199 | 0.58346736 | 0.41204896 | 1.41601466 | 0.15677121 | 0.99937574 |
| CHPT1 | 93.3645192 | -0.1641897 | 0.37449308 | -0.4384318 | 0.6610733 | 0.99937574 |
| GNPTAB | 106.364006 | 0.10053576 | 0.45989858 | 0.21860419 | 0.82695839 | 0.99937574 |
| DRAM1 | 143.879026 | -0.7590325 | 0.44480348 | -1.7064447 | 0.08792531 | 0.99937574 |
| WASHC3 | 119.113729 | 0.20320944 | 0.33479969 | 0.60695826 | 0.54387861 | 0.99937574 |
| NUP37 | 31.2233416 | 0.26943717 | 0.39978341 | 0.67395786 | 0.50033811 | 0.99937574 |

| | | | | | | |
|---|---|---|---|---|---|---|
| HSP90B1 | 181.072644 | 0.2260648 | 0.37049179 | 0.61017492 | 0.54174594 | 0.99937574 |
| C12orf73 | 12.6354278 | 0.02528275 | 0.53609419 | 0.04716103 | 0.96238489 | 0.99937574 |
| TDG | 53.164876 | -0.2768249 | 0.37095953 | -0.7462403 | 0.45552227 | 0.99937574 |
| HCFC2 | 46.8572203 | 0.00891287 | 0.35540293 | 0.0250782 | 0.97999259 | 0.99937574 |
| NFYB | 19.0065823 | 0.24119654 | 0.49678069 | 0.48551916 | 0.62730813 | 0.99937574 |
| TXNRD1 | 341.156309 | -0.3498316 | 0.39215089 | -0.892084 | 0.37234789 | 0.99937574 |
| SLC41A2 | 14.1104669 | -1.1148594 | 0.54995485 | -2.0271835 | 0.04264365 | 0.99937574 |
| C12orf45 | 14.2831288 | 0.15287876 | 0.61745929 | 0.24759325 | 0.80444913 | 0.99937574 |
| WASHC4 | 180.023631 | -0.040282 | 0.25601104 | -0.1573447 | 0.87497316 | 0.99937574 |
| APPL2 | 67.4370002 | 0.06353347 | 0.33053921 | 0.19221161 | 0.84757645 | 0.99937574 |
| C12orf75 | 685.515187 | 0.4217506 | 0.31509013 | 1.33850779 | 0.18073096 | 0.99937574 |
| CKAP4 | 19.918265 | 0.18831509 | 0.46186298 | 0.40772934 | 0.68347239 | 0.99937574 |
| TCP11L2 | 55.1988816 | -0.3478895 | 0.39177084 | -0.8879922 | 0.37454493 | 0.99937574 |
| POLR3B | 38.2718595 | -0.0825942 | 0.35354686 | -0.233616 | 0.81528314 | 0.99937574 |
| RIC8B | 76.8541605 | 0.3386499 | 0.28293151 | 1.19693244 | 0.23133289 | 0.99937574 |
| TMEM263 | 72.1182497 | -0.0194516 | 0.35780582 | -0.0543636 | 0.9566455 | 0.99937574 |
| MTERF2 | 17.4695137 | 0.74261682 | 0.4702067 | 1.5793412 | 0.11425782 | 0.99937574 |
| CRY1 | 66.2927514 | -0.0039505 | 0.5655914 | -0.0069848 | 0.99442701 | 0.99937574 |
| BTBD11 | 23.8937801 | 0.43109849 | 1.22898392 | 0.35077635 | 0.72575614 | 0.99937574 |
| PWP1 | 34.5496103 | 0.09179737 | 0.34911377 | 0.26294401 | 0.79259373 | 0.99937574 |
| PRDM4 | 105.08113 | 0.15539682 | 0.30637174 | 0.50721656 | 0.61200288 | 0.99937574 |
| SART3 | 35.4217064 | -0.0700591 | 0.33851344 | -0.2069611 | 0.83604028 | 0.99937574 |
| ISCU | 143.375587 | 0.36849365 | 0.44815336 | 0.82224899 | 0.4109352 | 0.99937574 |
| CORO1C | 15.1766307 | -0.4655554 | 0.54152287 | -0.8597151 | 0.38994612 | 0.99937574 |
| SSH1 | 59.6359983 | 0.00951226 | 0.32066017 | 0.0296646 | 0.97633454 | 0.99937574 |
| USP30 | 21.4114614 | 0.55422306 | 0.41448969 | 1.33712145 | 0.181183 | 0.99937574 |
| UNG | 17.6874027 | 0.53992162 | 0.44971531 | 1.20058537 | 0.22991208 | 0.99937574 |
| ACACB | 275.662553 | 0.13057937 | 0.38900279 | 0.33567719 | 0.73711431 | 0.99937574 |
| KCTD10 | 35.9047085 | 0.0162366 | 0.35952318 | 0.04516147 | 0.9639786 | 0.99937574 |
| UBE3B | 39.9765512 | 0.39586314 | 0.39543784 | 1.00107553 | 0.31679029 | 0.99937574 |
| MMAB | 46.8818231 | -0.0728233 | 0.43998756 | -0.1655122 | 0.86854085 | 0.99937574 |
| MVK | 18.2968242 | -0.239451 | 0.60053387 | -0.3987303 | 0.69009194 | 0.99937574 |
| FAM222A | 23.2642066 | 0.94746045 | 0.52759174 | 1.79582125 | 0.07252295 | 0.99937574 |
| TRPV4 | 50.2847138 | 0.77667812 | 0.68970807 | 1.1260969 | 0.26012451 | 0.99937574 |
| GLTP | 120.781012 | 0.6892959 | 0.43797745 | 1.57381594 | 0.11553001 | 0.99937574 |
| TCHP | 40.0546309 | 0.37905816 | 0.47546659 | 0.79723405 | 0.42531512 | 0.99937574 |
| GIT2 | 55.0632116 | 0.14701812 | 0.35055257 | 0.41938965 | 0.67493139 | 0.99937574 |
| ANKRD13A | 142.179008 | -0.1976238 | 0.34975426 | -0.565036 | 0.57204926 | 0.99937574 |
| C12orf76 | 18.7261526 | -0.172183 | 0.57849943 | -0.2976372 | 0.76598005 | 0.99937574 |
| IFT81 | 34.1785873 | 0.29612569 | 0.41522375 | 0.71317137 | 0.47573972 | 0.99937574 |
| ATP2A2 | 171.752872 | -0.0033774 | 0.3175711 | -0.0106351 | 0.99151454 | 0.99937574 |
| ANAPC7 | 43.831215 | -0.2976308 | 0.39822903 | -0.7473861 | 0.45483054 | 0.99937574 |
| ARPC3 | 158.400128 | 0.08273608 | 0.34971051 | 0.23658447 | 0.81297917 | 0.99937574 |
| GPN3 | 18.2865457 | -0.07634 | 0.50196359 | -0.1520828 | 0.8791216 | 0.99937574 |
| VPS29 | 64.8503687 | 0.01017579 | 0.38027052 | 0.02675935 | 0.97865168 | 0.99937574 |
| PPTC7 | 95.2675132 | -0.0147557 | 0.27862748 | -0.0529586 | 0.95776493 | 0.99937574 |
| TCTN1 | 19.900433 | -0.0088749 | 0.5351827 | -0.016583 | 0.98676927 | 0.99937574 |
| PPP1CC | 39.674439 | -0.1229464 | 0.41902961 | -0.2934075 | 0.76921071 | 0.99937574 |
| SH2B3 | 15.3981237 | -0.0534913 | 0.64285189 | -0.0832093 | 0.93368512 | 0.99937574 |
| ATXN2 | 282.510089 | -0.0238927 | 0.22416444 | -0.1065854 | 0.91511792 | 0.99937574 |
| BRAP | 36.8644894 | -0.2217784 | 0.3392963 | -0.6536424 | 0.51334224 | 0.99937574 |
| ACAD10 | 47.1465195 | 0.11439126 | 0.36631359 | 0.31227687 | 0.75483011 | 0.99937574 |
| ALDH2 | 60.7360199 | 0.49083329 | 0.49759192 | 0.98641732 | 0.32392837 | 0.99937574 |
| MAPKAPK5-A | 14.6551041 | 0.15949537 | 0.6228441 | 0.25607592 | 0.79789222 | 0.99937574 |

| | | | | | | |
|---|---|---|---|---|---|---|
| MAPKAPK5 | 47.51313 | 0.09040318 | 0.3210972 | 0.28154459 | 0.77829273 | 0.99937574 |
| TMEM116 | 297.137027 | 0.14777363 | 0.42517449 | 0.34755996 | 0.72817068 | 0.99937574 |
| ERP29 | 122.049889 | 0.23154545 | 0.32319802 | 0.71641978 | 0.47373218 | 0.99937574 |
| NAA25 | 82.5523866 | -0.0562975 | 0.2616697 | -0.2151472 | 0.82965254 | 0.99937574 |
| TRAFD1 | 36.7010968 | 0.16826013 | 0.4472692 | 0.37619431 | 0.70677244 | 0.99937574 |
| HECTD4 | 210.48952 | -0.208174 | 0.20516172 | -1.0146824 | 0.31025726 | 0.99937574 |
| RPL6 | 269.778538 | 0.09678381 | 0.53054597 | 0.18242305 | 0.85525074 | 0.99937574 |
| PTPN11 | 91.8298264 | -0.399619 | 0.35266325 | -1.1331461 | 0.25715291 | 0.99937574 |
| OAS1 | 18.725557 | -3.4535327 | 1.30279697 | -2.6508603 | 0.00802871 | 0.74742128 |
| OAS3 | 16.8415363 | -4.3660999 | 1.82232559 | -2.3958945 | 0.01657986 | 0.89002103 |
| DDX54 | 28.904015 | -0.2833302 | 0.48212205 | -0.5876732 | 0.55675165 | 0.99937574 |
| TPCN1 | 101.056312 | -0.1695679 | 0.44834292 | -0.3782102 | 0.70527442 | 0.99937574 |
| SLC8B1 | 36.2169001 | -0.7902349 | 0.51261053 | -1.5415893 | 0.12317342 | 0.99937574 |
| PLBD2 | 13.7263572 | -0.4495065 | 0.57012204 | -0.7884391 | 0.43043992 | 0.99937574 |
| SDSL | 10.1147556 | 0.99780261 | 0.82707108 | 1.20642909 | 0.22765208 | 0.99937574 |
| RBM19 | 37.4705121 | -0.7492138 | 0.40341579 | -1.8571752 | 0.06328624 | 0.99937574 |
| TBX3 | 14.3511012 | -1.7230601 | 1.34207847 | -1.2838743 | 0.19918594 | 0.99937574 |
| MED13L | 581.109222 | 0.0672044 | 0.27195188 | 0.24711873 | 0.80481634 | 0.99937574 |
| C12orf49 | 27.784177 | -1.8010802 | 0.79150595 | -2.2755107 | 0.02287531 | 0.96427895 |
| FBXW8 | 69.882033 | 0.14722834 | 0.30667536 | 0.48007879 | 0.63117137 | 0.99937574 |
| FBXO21 | 177.547614 | 0.10625226 | 0.33081154 | 0.32118667 | 0.74806893 | 0.99937574 |
| WSB2 | 48.3940294 | -0.4564277 | 0.43711502 | -1.0441821 | 0.29640113 | 0.99937574 |
| VSIG10 | 32.4280411 | -0.1737461 | 0.61896028 | -0.2807064 | 0.77893559 | 0.99937574 |
| PEBP1 | 229.304875 | -0.0020787 | 0.51155277 | -0.0040635 | 0.99675778 | 0.99937574 |
| TAOK3 | 283.502962 | -0.1078568 | 0.34766393 | -0.3102331 | 0.75638373 | 0.99937574 |
| SUDS3 | 45.4065012 | -0.0777237 | 0.33761449 | -0.2302143 | 0.81792529 | 0.99937574 |
| CCDC60 | 14.4283294 | 1.12698812 | 0.98151791 | 1.14820943 | 0.25088212 | 0.99937574 |
| AC002070.1 | 74.9927651 | 0.99762905 | 0.65947433 | 1.51276403 | 0.13033961 | 0.99937574 |
| TMEM233 | 16.5266901 | -0.601877 | 0.74816172 | -0.8044744 | 0.42112302 | 0.99937574 |
| PRKAB1 | 68.6485675 | 0.36202238 | 0.29829476 | 1.21363974 | 0.22488533 | 0.99937574 |
| CIT | 50.5747116 | -0.8165403 | 0.57775772 | -1.4132918 | 0.15756993 | 0.99937574 |
| BICDL1 | 141.971882 | 0.26846342 | 0.4227173 | 0.63508974 | 0.52536989 | 0.99937574 |
| RAB35 | 31.2513525 | -0.2393242 | 0.71614423 | -0.3341843 | 0.73824047 | 0.99937574 |
| GCN1 | 45.0259534 | -0.1567557 | 0.32944912 | -0.4758116 | 0.63420862 | 0.99937574 |
| RPLP0 | 262.429687 | 0.12587614 | 0.5021687 | 0.25066505 | 0.80207309 | 0.99937574 |
| PXN | 88.3494999 | -0.0669673 | 0.34500979 | -0.1941027 | 0.8460955 | 0.99937574 |
| GATC | 22.4165513 | -0.2441527 | 0.46886657 | -0.5207295 | 0.60255522 | 0.99937574 |
| SRSF9 | 69.923637 | 0.15228843 | 0.46017726 | 0.33093427 | 0.74069414 | 0.99937574 |
| DYNLL1 | 118.307492 | 0.75827193 | 0.60865604 | 1.24581353 | 0.21283286 | 0.99937574 |
| COQ5 | 21.2577822 | 0.3083369 | 0.44860098 | 0.68732996 | 0.49187482 | 0.99937574 |
| RNF10 | 125.967213 | 0.16995765 | 0.27590889 | 0.61599192 | 0.53789985 | 0.99937574 |
| POP5 | 24.9524103 | 0.38632612 | 0.44902824 | 0.8603604 | 0.38959041 | 0.99937574 |
| MLEC | 130.45381 | 0.20680834 | 0.29754869 | 0.69504032 | 0.48703003 | 0.99937574 |
| UNC119B | 20.3586705 | -0.1623855 | 0.43015736 | -0.3775025 | 0.70580019 | 0.99937574 |
| ACADS | 64.6134644 | 0.58777524 | 0.41227379 | 1.42569151 | 0.15395741 | 0.99937574 |
| AC069234.1 | 32.0449356 | 0.44169699 | 1.12524921 | 0.3925326 | 0.69466473 | 0.99937574 |
| SPPL3 | 359.933593 | -0.3030012 | 0.2985753 | -1.0148235 | 0.31018999 | 0.99937574 |
| C12orf43 | 10.7361195 | 0.05943246 | 0.68241049 | 0.08709194 | 0.93059843 | 0.99937574 |
| P2RX4 | 53.5271326 | -0.4258863 | 0.46926969 | -0.9075514 | 0.3641153 | 0.99937574 |
| CAMKK2 | 40.7162584 | 0.86149233 | 0.36182893 | 2.38093822 | 0.01726861 | 0.89832367 |
| ANAPC5 | 87.3922845 | 0.1373732 | 0.25797488 | 0.53250611 | 0.59437551 | 0.99937574 |
| RNF34 | 59.5913049 | 0.09281916 | 0.38347139 | 0.24204977 | 0.8087416 | 0.99937574 |
| KDM2B | 65.5969727 | -0.5304943 | 0.41228298 | -1.2867238 | 0.19819057 | 0.99937574 |
| ORAI1 | 27.4414288 | 0.11318105 | 0.41848998 | 0.27045104 | 0.78681327 | 0.99937574 |

| | | | | | | |
|---|---|---|---|---|---|---|
| TMEM120B | 110.489564 | 0.683787 | 0.42472888 | 1.60993761 | 0.10741148 | 0.99937574 |
| RHOF | 25.3408897 | 0.37154125 | 0.57889785 | 0.64180796 | 0.52099788 | 0.99937574 |
| SETD1B | 23.690589 | -0.1656367 | 0.53743449 | -0.3081988 | 0.75793107 | 0.99937574 |
| WDR66 | 12.71635 | -0.3494853 | 0.66567186 | -0.5250113 | 0.59957532 | 0.99937574 |
| BCL7A | 70.6490761 | 0.2542388 | 0.44263483 | 0.57437595 | 0.56571342 | 0.99937574 |
| MLXIP | 141.068482 | 0.05787264 | 0.40372268 | 0.14334752 | 0.88601573 | 0.99937574 |
| CLIP1 | 424.080628 | 0.16600882 | 0.33299004 | 0.49853988 | 0.61810356 | 0.99937574 |
| ZCCHC8 | 66.7621661 | -0.6050562 | 0.26340659 | -2.2970429 | 0.02161633 | 0.96427895 |
| RSRC2 | 115.491601 | -0.1651012 | 0.24994887 | -0.66054 | 0.50890734 | 0.99937574 |
| KNTC1 | 19.7295565 | 0.74799409 | 0.53742349 | 1.39181502 | 0.16397842 | 0.99937574 |
| AC026333.3 | 177.082189 | 2.49487411 | 0.57074672 | 4.37124566 | 1.24E-05 | 0.03116907 |
| HCAR1 | 18.0096238 | -0.6526896 | 0.7637429 | -0.8545934 | 0.39277629 | 0.99937574 |
| DENR | 112.460537 | -0.139257 | 0.27049747 | -0.514818 | 0.6066802 | 0.99937574 |
| HIP1R | 100.57204 | -0.071562 | 0.37526488 | -0.1906973 | 0.84876279 | 0.99937574 |
| VPS37B | 197.910161 | -0.1664474 | 0.30526635 | -0.5452531 | 0.58557947 | 0.99937574 |
| ABCB9 | 13.9860484 | 0.9749343 | 0.63756108 | 1.5291622 | 0.12622424 | 0.99937574 |
| ARL6IP4 | 75.1572227 | -0.1087731 | 0.38285006 | -0.2841142 | 0.77632287 | 0.99937574 |
| PITPNM2 | 86.4046784 | 0.51416391 | 0.28336009 | 1.81452482 | 0.06959696 | 0.99937574 |
| MPHOSPH9 | 51.0089056 | 0.28320923 | 0.36656342 | 0.77260638 | 0.43975537 | 0.99937574 |
| C12orf65 | 41.2352988 | -0.4084628 | 0.38157421 | -1.0704676 | 0.28440887 | 0.99937574 |
| SBNO1 | 94.4426519 | -0.3501727 | 0.32854405 | -1.0658319 | 0.28649965 | 0.99937574 |
| KMT5A | 17.7354978 | -0.2063153 | 0.61472037 | -0.3356246 | 0.73715398 | 0.99937574 |
| RILPL2 | 15.0859581 | -0.3549641 | 0.7020832 | -0.5055869 | 0.61314669 | 0.99937574 |
| SNRNP35 | 40.2437988 | 0.05481305 | 0.31208987 | 0.17563228 | 0.86058284 | 0.99937574 |
| RILPL1 | 46.7825947 | -0.1970983 | 0.39670064 | -0.4968439 | 0.61929911 | 0.99937574 |
| TMED2 | 52.1505869 | 0.02776803 | 0.38265218 | 0.07256728 | 0.94215047 | 0.99937574 |
| DDX55 | 19.0067586 | -0.3376543 | 0.48074051 | -0.7023629 | 0.48245285 | 0.99937574 |
| EIF2B1 | 21.3853747 | -0.0049118 | 0.41634793 | -0.0117973 | 0.99058732 | 0.99937574 |
| GTF2H3 | 25.7939144 | 0.09494283 | 0.43641514 | 0.21755163 | 0.82777847 | 0.99937574 |
| TCTN2 | 12.7247499 | -0.1942894 | 0.51932776 | -0.374117 | 0.70831726 | 0.99937574 |
| ATP6V0A2 | 27.2177191 | -0.4220399 | 0.47422601 | -0.8899552 | 0.37348996 | 0.99937574 |
| DNAH10 | 13.5927845 | -0.9848042 | 0.6836137 | -1.4405859 | 0.14970172 | 0.99937574 |
| CCDC92 | 74.7554219 | 0.15371446 | 0.31958677 | 0.4809788 | 0.63053157 | 0.99937574 |
| RFLNA | 12.0800957 | -0.7655216 | 0.66488486 | -1.1513596 | 0.24958434 | 0.99937574 |
| NCOR2 | 114.420145 | 0.04491762 | 0.49567167 | 0.09061971 | 0.92779477 | 0.99937574 |
| SCARB1 | 15.3052297 | 0.29424851 | 0.81840539 | 0.35953883 | 0.71919204 | 0.99937574 |
| UBC | 362.954877 | -0.2209013 | 0.43226629 | -0.5110305 | 0.60932967 | 0.99937574 |
| DHX37 | 12.7243636 | 0.26884312 | 0.65617905 | 0.40971001 | 0.68201869 | 0.99937574 |
| BRI3BP | 44.6502478 | 0.05001805 | 0.47873136 | 0.10448041 | 0.91678811 | 0.99937574 |
| AACS | 40.4465386 | 0.44234537 | 0.57491574 | 0.76940905 | 0.44165052 | 0.99937574 |
| SLC15A4 | 34.229272 | -0.7252279 | 0.41842955 | -1.7332139 | 0.08305766 | 0.99937574 |
| RIMBP2 | 40.9254959 | -3.2455875 | 1.09394892 | -2.9668547 | 0.00300863 | 0.52350178 |
| STX2 | 33.8812092 | 0.19691407 | 0.4835018 | 0.40726647 | 0.68381229 | 0.99937574 |
| RAN | 65.2178859 | 0.24250731 | 0.49186274 | 0.49303858 | 0.62198532 | 0.99937574 |
| SFSWAP | 79.6112365 | -0.0995475 | 0.31272307 | -0.3183249 | 0.75023852 | 0.99937574 |
| ULK1 | 94.5033967 | 0.44644498 | 0.38890991 | 1.14793933 | 0.25099361 | 0.99937574 |
| PUS1 | 15.0969983 | -0.147454 | 0.48034178 | -0.3069773 | 0.75886064 | 0.99937574 |
| EP400 | 82.8432081 | -0.069387 | 0.28679351 | -0.2419405 | 0.80882626 | 0.99937574 |
| DDX51 | 10.9844243 | -0.2198106 | 0.63287917 | -0.3473185 | 0.72835207 | 0.99937574 |
| FBRSL1 | 57.2198911 | -0.2903491 | 0.49240529 | -0.5896546 | 0.55542222 | 0.99937574 |
| ANKLE2 | 43.6748232 | -1.0194578 | 0.49372482 | -2.0648299 | 0.0389391 | 0.99937574 |
| GOLGA3 | 68.2514009 | -0.3834097 | 0.38263248 | -1.0020313 | 0.31632848 | 0.99937574 |
| CHFR | 76.797545 | -0.301959 | 0.29843216 | -1.0118179 | 0.31162513 | 0.99937574 |
| ZNF26 | 31.9623079 | -0.484904 | 0.40435261 | -1.1992109 | 0.23044596 | 0.99937574 |

| | | | | | | |
|---|---|---|---|---|---|---|
| ZNF84 | 54.0232899 | -0.2437512 | 0.33269095 | -0.7326656 | 0.46376244 | 0.99937574 |
| ZNF140 | 35.1236612 | 0.12948661 | 0.39499837 | 0.32781555 | 0.74305113 | 0.99937574 |
| ZNF891 | 18.4075574 | 0.70422396 | 0.47236853 | 1.4908359 | 0.13600458 | 0.99937574 |
| ZNF268 | 13.8398629 | -0.2396485 | 0.57900652 | -0.4138961 | 0.67895019 | 0.99937574 |
| MPHOSPH8 | 153.907623 | -0.0806112 | 0.30789102 | -0.2618174 | 0.79346225 | 0.99937574 |
| PSPC1 | 179.044509 | 0.03097084 | 0.28068905 | 0.11033861 | 0.91214084 | 0.99937574 |
| ZMYM5 | 62.5135963 | 0.21468975 | 0.30544027 | 0.70288621 | 0.48212667 | 0.99937574 |
| ZMYM2 | 406.887154 | 0.00326024 | 0.29406877 | 0.01108664 | 0.99115432 | 0.99937574 |
| CRYL1 | 213.346151 | 0.59822736 | 0.41837251 | 1.42989165 | 0.15274812 | 0.99937574 |
| IFT88 | 106.698492 | -0.7498729 | 0.3156939 | -2.3753165 | 0.01753391 | 0.90744728 |
| EEF1AKMT1 | 14.7023519 | 0.62976877 | 0.4853909 | 1.29744659 | 0.19447757 | 0.99937574 |
| XPO4 | 113.349626 | -0.2265678 | 0.28955463 | -0.78247 | 0.43393839 | 0.99937574 |
| LATS2 | 48.6156559 | -0.1757516 | 0.58027683 | -0.3028755 | 0.76198473 | 0.99937574 |
| SAP18 | 78.6770553 | -0.2024852 | 0.49867595 | -0.4060457 | 0.684709 | 0.99937574 |
| MRPL57 | 56.058074 | -0.1086064 | 0.52828557 | -0.2055828 | 0.83711679 | 0.99937574 |
| ZDHHC20 | 116.902774 | -0.6163113 | 0.29487773 | -2.0900573 | 0.03661266 | 0.99937574 |
| MICU2 | 223.207324 | -0.1670135 | 0.2844649 | -0.5871147 | 0.55712665 | 0.99937574 |
| FGF9 | 209.078865 | 0.9274727 | 0.70977296 | 1.30671743 | 0.1913087 | 0.99937574 |
| AL136962.1 | 41.069153 | 0.23647111 | 1.45476896 | 0.16254891 | 0.87087362 | 0.99937574 |
| SACS | 20.180174 | 0.24696017 | 0.57141239 | 0.43219254 | 0.66560148 | 0.99937574 |
| MIPEP | 83.2381048 | -0.1730034 | 0.3845362 | -0.4499015 | 0.65278146 | 0.99937574 |
| PARP4 | 230.269519 | -0.2805926 | 0.36116736 | -0.7769048 | 0.43721497 | 0.99937574 |
| MTMR6 | 35.5796995 | -0.4726762 | 0.36612578 | -1.2910215 | 0.19669623 | 0.99937574 |
| NUP58 | 78.3505363 | -0.4291836 | 0.41338303 | -1.0382227 | 0.29916638 | 0.99937574 |
| RNF6 | 30.9672444 | 0.05714618 | 0.61257276 | 0.09328881 | 0.92567412 | 0.99937574 |
| CDK8 | 203.98826 | 0.47764668 | 0.28797995 | 1.65861089 | 0.09719423 | 0.99937574 |
| WASF3 | 24.0389271 | 0.09162071 | 0.56433304 | 0.1623522 | 0.87102851 | 0.99937574 |
| USP12 | 136.157441 | 0.36432625 | 0.3361189 | 1.08392073 | 0.27839995 | 0.99937574 |
| LINC02340 | 13.6449857 | -0.5975231 | 0.56552441 | -1.0565824 | 0.29070222 | 0.99937574 |
| RPL21 | 124.568613 | 0.17009641 | 0.54758973 | 0.31062747 | 0.75608384 | 0.99937574 |
| GTF3A | 21.2886085 | 0.35125248 | 0.49520349 | 0.70930939 | 0.4781325 | 0.99937574 |
| MTIF3 | 66.7133791 | 0.23476578 | 0.33809143 | 0.69438548 | 0.4874405 | 0.99937574 |
| LNX2 | 84.3687477 | 0.08821858 | 0.32062986 | 0.2751415 | 0.78320753 | 0.99937574 |
| POLR1D | 58.726514 | 0.18697178 | 0.39954542 | 0.46796126 | 0.6398123 | 0.99937574 |
| PAN3 | 347.134412 | -0.3615004 | 0.23300198 | -1.5514908 | 0.1207841 | 0.99937574 |
| POMP | 46.5011467 | -0.2041211 | 0.42180846 | -0.483919 | 0.62844334 | 0.99937574 |
| SLC46A3 | 18.4085547 | 0.23432587 | 0.481977 | 0.48617647 | 0.62684206 | 0.99937574 |
| SLC7A1 | 46.5306367 | -0.7612348 | 0.35047763 | -2.1719926 | 0.02985622 | 0.99049468 |
| UBL3 | 308.532761 | 0.54704309 | 0.2960452 | 1.84783641 | 0.06462601 | 0.99937574 |
| KATNAL1 | 21.3984472 | 0.1485957 | 0.41776225 | 0.35569442 | 0.72206943 | 0.99937574 |
| HMGB1 | 194.221074 | 0.21629911 | 0.23546925 | 0.91858751 | 0.35831137 | 0.99937574 |
| USPL1 | 52.8530874 | -0.3928554 | 0.42365165 | -0.9273077 | 0.35376681 | 0.99937574 |
| HSPH1 | 105.318215 | -0.2073858 | 0.45950545 | -0.4513239 | 0.65175611 | 0.99937574 |
| B3GLCT | 55.7570626 | -0.336618 | 0.31528138 | -1.0676748 | 0.28566721 | 0.99937574 |
| FRY | 94.1681755 | -0.6535399 | 0.48627083 | -1.3439834 | 0.17895375 | 0.99937574 |
| N4BP2L1 | 35.4711 | 0.39243608 | 0.74452805 | 0.52709375 | 0.59812849 | 0.99937574 |
| N4BP2L2 | 573.449437 | -0.1572672 | 0.31519058 | -0.4989589 | 0.61780832 | 0.99937574 |
| PDS5B | 154.012612 | -0.0516941 | 0.32388234 | -0.1596075 | 0.87319025 | 0.99937574 |
| STARD13 | 87.939839 | -2.0478867 | 0.95416472 | -2.1462612 | 0.03185215 | 0.99520706 |
| AL139383.1 | 25.1940051 | 0.58305226 | 0.82077743 | 0.71036585 | 0.47747729 | 0.99937574 |
| RFC3 | 18.6569936 | 0.47611212 | 0.50118488 | 0.94997303 | 0.34212596 | 0.99937574 |
| LINC02343 | 380.851539 | 0.50756463 | 0.54558353 | 0.93031517 | 0.35220793 | 0.99937574 |
| LINC00457 | 24.8833567 | 0.72124108 | 0.89595613 | 0.80499598 | 0.420822 | 0.99937574 |
| NBEA | 147.968918 | 0.15171943 | 0.31886444 | 0.47581168 | 0.63420854 | 0.99937574 |

| | | | | | | |
|---|---|---|---|---|---|---|
| SPART | 94.8795505 | -0.1588913 | 0.38627654 | -0.4113409 | 0.6808226 | 0.99937574 |
| RFXAP | 10.7720623 | -0.0352549 | 0.61418585 | -0.057401 | 0.95422579 | 0.99937574 |
| SMAD9 | 50.5598586 | -1.6468488 | 1.46125128 | -1.1270128 | 0.25973709 | 0.99937574 |
| ALG5 | 41.5096783 | 0.33513095 | 0.33215578 | 1.00895714 | 0.31299519 | 0.99937574 |
| EXOSC8 | 20.2834346 | 0.05933106 | 0.4607123 | 0.12878114 | 0.89753083 | 0.99937574 |
| SUPT20H | 95.0971915 | 0.11841077 | 0.2729538 | 0.4338125 | 0.6644246 | 0.99937574 |
| LINC00571 | 49.3204387 | -0.6727258 | 0.39408802 | -1.7070446 | 0.08781377 | 0.99937574 |
| UFM1 | 71.8069215 | -0.2425434 | 0.36752037 | -0.6599454 | 0.5092889 | 0.99937574 |
| FREM2 | 72.2993143 | -0.0395259 | 0.5438952 | -0.0726719 | 0.94206724 | 0.99937574 |
| PROSER1 | 32.9504489 | 0.26371585 | 0.42592094 | 0.6191662 | 0.53580687 | 0.99937574 |
| NHLRC3 | 47.4367503 | 0.38785999 | 0.37846215 | 1.02483163 | 0.30544264 | 0.99937574 |
| COG6 | 79.3144168 | 0.25268903 | 0.30753282 | 0.82166526 | 0.41126744 | 0.99937574 |
| LINC00598 | 40.8078978 | 0.77146507 | 0.426239 | 1.80993541 | 0.0703058 | 0.99937574 |
| FOXO1 | 817.920906 | -0.0537917 | 0.47239223 | -0.1138709 | 0.90934015 | 0.99937574 |
| MRPS31 | 74.5269413 | 0.25143146 | 0.35340324 | 0.71145772 | 0.47680064 | 0.99937574 |
| SLC25A15 | 17.1227484 | 0.67424837 | 0.6088566 | 1.10740094 | 0.26812062 | 0.99937574 |
| ELF1 | 359.234052 | -0.1814548 | 0.36536387 | -0.4966414 | 0.61944194 | 0.99937574 |
| WBP4 | 50.8194917 | -0.1640513 | 0.3349909 | -0.4897187 | 0.62433294 | 0.99937574 |
| AL354696.2 | 13.8690793 | -0.3811019 | 0.72874953 | -0.5229533 | 0.60100678 | 0.99937574 |
| MTRF1 | 22.6006939 | -0.7616876 | 0.42066385 | -1.8106799 | 0.07019042 | 0.99937574 |
| NAA16 | 75.9002796 | -0.0052287 | 0.33368985 | -0.0156693 | 0.98749824 | 0.99937574 |
| RGCC | 82.2674265 | -0.2307381 | 1.01261482 | -0.2278637 | 0.81975222 | 0.99937574 |
| VWA8 | 412.803701 | 0.1112267 | 0.31245808 | 0.35597318 | 0.72186065 | 0.99937574 |
| DGKH | 172.052554 | -0.1493328 | 0.62990725 | -0.2370711 | 0.81260161 | 0.99937574 |
| AKAP11 | 66.5497445 | 0.05972231 | 0.33296623 | 0.17936446 | 0.85765154 | 0.99937574 |
| EPSTI1 | 65.6162357 | -3.4931679 | 1.08237743 | -3.2273103 | 0.0012496 | 0.33941289 |
| DNAJC15 | 66.3703436 | -0.1883014 | 0.3261527 | -0.5773412 | 0.56370896 | 0.99937574 |
| ENOX1 | 17.0748632 | 0.40752448 | 0.85764455 | 0.47516711 | 0.63466787 | 0.99937574 |
| CCDC122 | 64.1109957 | 0.81629937 | 0.51328424 | 1.59034567 | 0.11175691 | 0.99937574 |
| TSC22D1 | 121.81452 | -0.0053541 | 0.57812453 | -0.0092611 | 0.99261084 | 0.99937574 |
| NUFIP1 | 12.1014123 | 0.28678225 | 0.58422159 | 0.49087924 | 0.62351186 | 0.99937574 |
| AL359706.1 | 16.526806 | 0.46064348 | 0.80155816 | 0.57468504 | 0.56550433 | 0.99937574 |
| GPALPP1 | 43.838628 | -0.0118082 | 0.36820933 | -0.0320692 | 0.97441684 | 0.99937574 |
| GTF2F2 | 105.637713 | -0.0401933 | 0.31455943 | -0.1277765 | 0.89832586 | 0.99937574 |
| TPT1 | 517.608317 | 0.10208205 | 0.60271188 | 0.16937122 | 0.86550466 | 0.99937574 |
| SLC25A30 | 44.1590126 | -0.0184907 | 0.47331917 | -0.039066 | 0.96883778 | 0.99937574 |
| COG3 | 62.6772868 | -0.2954604 | 0.26047267 | -1.1343241 | 0.25665861 | 0.99937574 |
| ZC3H13 | 113.45999 | -0.1197654 | 0.28811613 | -0.4156844 | 0.67764093 | 0.99937574 |
| AL138686.1 | 43.9686975 | 0.4702568 | 0.55318389 | 0.85009129 | 0.39527434 | 0.99937574 |
| LRCH1 | 231.829225 | -0.2071268 | 0.29042697 | -0.7131804 | 0.47573412 | 0.99937574 |
| ESD | 68.8582879 | 0.08511526 | 0.29628756 | 0.28727248 | 0.77390369 | 0.99937574 |
| SUCLA2 | 112.660107 | -0.3282766 | 0.34741004 | -0.9449254 | 0.34469694 | 0.99937574 |
| NUDT15 | 11.8960297 | -0.1396143 | 0.51590521 | -0.2706201 | 0.78668323 | 0.99937574 |
| MED4 | 72.6534599 | -0.0867139 | 0.32077152 | -0.2703291 | 0.78690705 | 0.99937574 |
| ITM2B | 587.531928 | 0.00047615 | 0.36458951 | 0.00130599 | 0.99895797 | 0.99979721 |
| RB1 | 238.123 | -0.2508535 | 0.30672916 | -0.8178339 | 0.41345203 | 0.99937574 |
| LPAR6 | 12.7993341 | 0.62966519 | 0.60966952 | 1.03279755 | 0.30169865 | 0.99937574 |
| RCBTB2 | 55.1649641 | -0.159621 | 0.44134581 | -0.3616688 | 0.71759956 | 0.99937574 |
| FNDC3A | 786.430493 | -0.0012249 | 0.3999683 | -0.0030624 | 0.99755657 | 0.99937574 |
| CDADC1 | 34.048428 | -0.6323065 | 0.45053603 | -1.4034538 | 0.16048156 | 0.99937574 |
| CAB39L | 122.931903 | 0.19352257 | 0.48352857 | 0.40022986 | 0.68898722 | 0.99937574 |
| SETDB2 | 130.894533 | 0.25433451 | 0.46528196 | 0.54662448 | 0.58463675 | 0.99937574 |
| PHF11 | 70.0787402 | -0.4441257 | 0.49741538 | -0.8928669 | 0.37192847 | 0.99937574 |
| RCBTB1 | 37.9573576 | -0.1859662 | 0.41886029 | -0.4439815 | 0.65705594 | 0.99937574 |

| | | | | | | |
|---|---|---|---|---|---|---|
| EBPL | 31.8497797 | 0.1995112 | 0.55202193 | 0.36141897 | 0.71778627 | 0.99937574 |
| KPNA3 | 101.203943 | 0.03300613 | 0.26811428 | 0.12310469 | 0.90202419 | 0.99937574 |
| SPRYD7 | 18.9000725 | 0.01114581 | 0.41259322 | 0.02701405 | 0.97844853 | 0.99937574 |
| DLEU2 | 195.636483 | -0.2354426 | 0.35510111 | -0.6630297 | 0.50731152 | 0.99937574 |
| TRIM13 | 49.7618346 | -0.1480485 | 0.34789262 | -0.4255579 | 0.67043001 | 0.99937574 |
| RNASEH2B | 40.8836068 | -0.2491195 | 0.37245023 | -0.6688664 | 0.50358068 | 0.99937574 |
| INTS6 | 219.985749 | 0.62098823 | 0.36759769 | 1.68931482 | 0.09115912 | 0.99937574 |
| INTS6-AS1 | 12.1412334 | 0.00431928 | 0.5558896 | 0.00777002 | 0.99380048 | 0.99937574 |
| WDFY2 | 48.1435429 | -0.2304586 | 0.48125291 | -0.4788721 | 0.6320296 | 0.99937574 |
| ATP7B | 22.3213762 | 0.79333783 | 0.51403823 | 1.54334404 | 0.12274732 | 0.99937574 |
| ALG11 | 17.4906289 | 0.49259697 | 0.54493623 | 0.90395341 | 0.36602011 | 0.99937574 |
| UTP14C | 23.6147749 | -0.4238037 | 0.62269338 | -0.6805978 | 0.49612605 | 0.99937574 |
| NEK3 | 38.3802389 | -0.0004487 | 0.49393942 | -0.0009083 | 0.99927524 | 0.99979721 |
| VPS36 | 78.5336002 | 0.31934782 | 0.26393099 | 1.20996712 | 0.22629151 | 0.99937574 |
| SUGT1 | 55.6889592 | 0.03296768 | 0.3549726 | 0.09287386 | 0.92600377 | 0.99937574 |
| DIAPH3 | 13.7799834 | -0.870538 | 0.83298519 | -1.0450822 | 0.29598496 | 0.99937574 |
| TDRD3 | 153.126768 | 0.31105101 | 0.26178549 | 1.18819039 | 0.23475841 | 0.99937574 |
| PCDH9 | 19.5394409 | 0.42417155 | 0.87529675 | 0.48460314 | 0.62795788 | 0.99937574 |
| DACH1 | 73.177432 | 0.78572604 | 0.88877839 | 0.8840517 | 0.37666831 | 0.99937574 |
| MZT1 | 14.9804414 | 0.08592229 | 0.60588548 | 0.14181275 | 0.88722791 | 0.99937574 |
| DIS3 | 41.0817871 | 0.04247416 | 0.34048428 | 0.12474631 | 0.90072439 | 0.99937574 |
| PIBF1 | 175.691071 | -0.2074655 | 0.24682752 | -0.8405283 | 0.40061222 | 0.99937574 |
| KLF5 | 37.1205631 | -1.3368225 | 0.72004869 | -1.8565723 | 0.06337202 | 0.99937574 |
| KLF12 | 153.697691 | 0.13733079 | 0.32668117 | 0.42038173 | 0.67420661 | 0.99937574 |
| TBC1D4 | 836.041623 | -0.0735871 | 0.30659713 | -0.2400124 | 0.81032062 | 0.99937574 |
| COMMD6 | 78.9462099 | 0.92412529 | 0.51187299 | 1.80538005 | 0.07101523 | 0.99937574 |
| UCHL3 | 67.0729752 | -0.2252834 | 0.36377475 | -0.6192936 | 0.53572295 | 0.99937574 |
| LMO7-AS1 | 26.5380728 | 0.93721433 | 0.50127595 | 1.86965747 | 0.0615314 | 0.99937574 |
| LMO7 | 645.910842 | 0.09371527 | 0.49837575 | 0.18804139 | 0.85084421 | 0.99937574 |
| AL365394.1 | 34.2106665 | 2.59015678 | 0.55890906 | 4.63430809 | 3.58E-06 | 0.0120476 |
| KCTD12 | 178.491762 | 1.13082642 | 0.33848277 | 3.34086848 | 8.35E-04 | 0.29012345 |
| AC001226.2 | 30.1154121 | 0.60306161 | 0.71606113 | 0.84219292 | 0.39967998 | 0.99937574 |
| FBXL3 | 89.754666 | 0.05204078 | 0.35179817 | 0.14792794 | 0.88239964 | 0.99937574 |
| MYCBP2 | 273.293924 | -0.3980112 | 0.30392353 | -1.3095769 | 0.19033902 | 0.99937574 |
| SLAIN1 | 13.4644966 | 0.89453589 | 0.50442351 | 1.77338265 | 0.07616532 | 0.99937574 |
| RNF219 | 21.3951821 | -0.5453402 | 0.49134189 | -1.1098996 | 0.26704228 | 0.99937574 |
| RBM26 | 218.346716 | 0.18860032 | 0.29384222 | 0.64184216 | 0.52097568 | 0.99937574 |
| NDFIP2 | 97.7084628 | -0.4333131 | 0.29159454 | -1.4860124 | 0.13727583 | 0.99937574 |
| GPC5 | 20.5662929 | 0.36571783 | 0.84241682 | 0.43412931 | 0.66419454 | 0.99937574 |
| GPC6 | 64.918006 | -1.768584 | 1.42307868 | -1.2427872 | 0.21394626 | 0.99937574 |
| TGDS | 21.3161612 | -0.2793103 | 0.42987873 | -0.6497422 | 0.51585876 | 0.99937574 |
| GPR180 | 16.7695303 | -1.0452554 | 0.63282827 | -1.6517205 | 0.09859154 | 0.99937574 |
| ABCC4 | 999.375464 | -0.0852796 | 0.40157128 | -0.2123647 | 0.83182254 | 0.99937574 |
| DNAJC3-DT | 19.9584328 | 0.44068974 | 0.5678605 | 0.77605282 | 0.4377178 | 0.99937574 |
| DNAJC3 | 146.557892 | 0.06765583 | 0.34133561 | 0.19820913 | 0.84288145 | 0.99937574 |
| UGGT2 | 195.659197 | -0.2863239 | 0.24187409 | -1.1837727 | 0.23650306 | 0.99937574 |
| HS6ST3 | 804.207606 | 0.34501594 | 0.75642961 | 0.4561111 | 0.64831009 | 0.99937574 |
| OXGR1 | 30.3514311 | 0.4631762 | 0.84274471 | 0.5496044 | 0.58259074 | 0.99937574 |
| MBNL2 | 216.964338 | 0.45214652 | 0.31787664 | 1.42239618 | 0.15491127 | 0.99937574 |
| RAP2A | 13.5492103 | 0.57081161 | 0.51935503 | 1.09907784 | 0.27173411 | 0.99937574 |
| IPO5 | 60.6947058 | 0.14573566 | 0.27041819 | 0.53892697 | 0.58993725 | 0.99937574 |
| FARP1 | 417.770058 | 0.03326422 | 0.24767783 | 0.13430438 | 0.89316189 | 0.99937574 |
| STK24 | 263.244135 | -0.1135421 | 0.2483294 | -0.4572236 | 0.64751033 | 0.99937574 |
| DOCK9 | 92.6544424 | 0.15644507 | 0.30863719 | 0.50688988 | 0.61223209 | 0.99937574 |

| | | | | | | |
|---|---|---|---|---|---|---|
| UBAC2 | 279.0205 | -0.1038378 | 0.24068603 | -0.4314244 | 0.66615981 | 0.99937574 |
| TM9SF2 | 233.757176 | 0.3317232 | 0.23379586 | 1.41885833 | 0.15594032 | 0.99937574 |
| CLYBL | 347.044151 | -0.2564041 | 0.51202222 | -0.5007675 | 0.61653477 | 0.99937574 |
| PCCA | 670.866256 | 0.09007461 | 0.34204192 | 0.26334376 | 0.79228563 | 0.99937574 |
| GGACT | 41.236274 | -0.5379806 | 0.3146029 | -1.7100308 | 0.08726018 | 0.99937574 |
| TMTC4 | 40.6388952 | 0.64726242 | 0.41633451 | 1.55466915 | 0.12002489 | 0.99937574 |
| FGF14 | 38.2094105 | 0.68563778 | 0.81006456 | 0.84639893 | 0.39733024 | 0.99937574 |
| TPP2 | 115.925627 | -0.1699283 | 0.34274117 | -0.495792 | 0.62004118 | 0.99937574 |
| EFNB2 | 35.6999519 | 0.2774463 | 0.58855712 | 0.47140082 | 0.63735453 | 0.99937574 |
| ARGLU1 | 197.442636 | 0.09297327 | 0.28684722 | 0.32412121 | 0.74584627 | 0.99937574 |
| LIG4 | 24.2548555 | 0.58941488 | 0.42782264 | 1.37770849 | 0.16829331 | 0.99937574 |
| ABHD13 | 35.1756731 | 0.15934596 | 0.44201392 | 0.36049986 | 0.71847336 | 0.99937574 |
| TNFSF13B | 35.6325435 | 1.65352562 | 0.56470004 | 2.92814858 | 0.00340987 | 0.53256471 |
| IRS2 | 66.1001519 | -1.1339466 | 0.56810182 | -1.9960271 | 0.04593097 | 0.99937574 |
| AL355974.2 | 14.2686609 | -0.4795073 | 0.93285017 | -0.5140239 | 0.60723529 | 0.99937574 |
| COL4A2 | 11.8530271 | -0.3235669 | 0.7765341 | -0.4166808 | 0.67691187 | 0.99937574 |
| RAB20 | 49.506173 | 0.63129214 | 0.40223717 | 1.56945251 | 0.11654254 | 0.99937574 |
| NAXD | 42.3940051 | 0.27628174 | 0.35492041 | 0.77843295 | 0.43631382 | 0.99937574 |
| CARS2 | 74.4190468 | 0.15019384 | 0.31454792 | 0.47749114 | 0.63301243 | 0.99937574 |
| ANKRD10 | 259.487127 | -0.0154284 | 0.2819705 | -0.0547164 | 0.95636438 | 0.99937574 |
| ARHGEF7 | 228.557185 | -0.1295369 | 0.25503794 | -0.5079123 | 0.61151488 | 0.99937574 |
| TUBGCP3 | 55.8230266 | -0.4693487 | 0.31547318 | -1.4877611 | 0.13681392 | 0.99937574 |
| ATP11A | 384.588535 | -0.0212471 | 0.5492923 | -0.0386809 | 0.9691448 | 0.99937574 |
| MCF2L | 67.6737653 | 0.39113842 | 0.41879063 | 0.93397128 | 0.35031871 | 0.99937574 |
| PCID2 | 55.1869863 | 0.26296101 | 0.31102236 | 0.84547299 | 0.39784681 | 0.99937574 |
| CUL4A | 122.234004 | -0.2073412 | 0.31517532 | -0.6578598 | 0.51062821 | 0.99937574 |
| LAMP1 | 117.281628 | 0.02680578 | 0.37871807 | 0.0707803 | 0.94357261 | 0.99937574 |
| GRTP1 | 31.7697821 | 0.52510811 | 0.48577258 | 1.0809752 | 0.27970814 | 0.99937574 |
| DCUN1D2 | 46.3485367 | 0.06945788 | 0.32578931 | 0.21319878 | 0.83117192 | 0.99937574 |
| TMCO3 | 45.7098406 | 0.09503085 | 0.423377 | 0.22445918 | 0.82240003 | 0.99937574 |
| TFDP1 | 41.2573588 | 0.02700464 | 0.43247893 | 0.06244151 | 0.95021124 | 0.99937574 |
| GAS6 | 195.309998 | 0.11179191 | 0.53884052 | 0.20746753 | 0.83564475 | 0.99937574 |
| CDC16 | 65.0726763 | -0.0016892 | 0.3174337 | -0.0053215 | 0.99575409 | 0.99937574 |
| UPF3A | 77.4021017 | 0.06875454 | 0.31002525 | 0.22177077 | 0.82449234 | 0.99937574 |
| CHAMP1 | 20.2436634 | -0.4396411 | 0.47982147 | -0.9162599 | 0.35953062 | 0.99937574 |
| TTC5 | 15.032078 | -0.0934145 | 0.52156214 | -0.1791053 | 0.85785502 | 0.99937574 |
| CCNB1IP1 | 40.7633896 | -0.0170415 | 0.37987427 | -0.044861 | 0.96421809 | 0.99937574 |
| TEP1 | 36.3424625 | -0.4374934 | 0.41250434 | -1.0605789 | 0.2888813 | 0.99937574 |
| OSGEP | 16.6874475 | -0.6950964 | 0.5501037 | -1.2635734 | 0.20638318 | 0.99937574 |
| APEX1 | 19.8221998 | 0.35418308 | 0.57200873 | 0.61919173 | 0.53579006 | 0.99937574 |
| PIP4P1 | 18.7115512 | -0.6733632 | 0.54733652 | -1.2302544 | 0.21860185 | 0.99937574 |
| PNP | 19.355112 | -0.042178 | 0.43106533 | -0.097846 | 0.92205457 | 0.99937574 |
| METTL17 | 20.3126477 | 0.65235955 | 0.46820173 | 1.39333006 | 0.16352001 | 0.99937574 |
| NDRG2 | 40.9683146 | 0.78708954 | 0.48741083 | 1.61483803 | 0.10634577 | 0.99937574 |
| ZNF219 | 12.4026555 | -0.550981 | 0.78694288 | -0.7001538 | 0.48383128 | 0.99937574 |
| HNRNPC | 330.314356 | -0.225083 | 0.2716918 | -0.8284496 | 0.40741595 | 0.99937574 |
| SUPT16H | 62.4788048 | -0.1016132 | 0.33547382 | -0.3028947 | 0.76197013 | 0.99937574 |
| CHD8 | 87.5555471 | 0.00216523 | 0.36358033 | 0.00595531 | 0.99524838 | 0.99937574 |
| RAB2B | 39.9732453 | -0.3090722 | 0.36629741 | -0.843774 | 0.39879572 | 0.99937574 |
| TOX4 | 56.7694361 | -0.0010679 | 0.3088578 | -0.0034575 | 0.99724133 | 0.99937574 |
| METTL3 | 19.2444822 | 0.22351788 | 0.45804268 | 0.48798485 | 0.62556057 | 0.99937574 |
| DAD1 | 73.9897805 | 0.15116508 | 0.47612097 | 0.317493 | 0.75086955 | 0.99937574 |
| ABHD4 | 16.6562812 | -0.3707312 | 0.45971278 | -0.8064409 | 0.41998868 | 0.99937574 |
| OXA1L | 25.6305627 | 0.09942277 | 0.46217382 | 0.21511987 | 0.82967386 | 0.99937574 |

| | | | | | | |
|---|---|---|---|---|---|---|
| MRPL52 | 18.8313774 | 0.16739673 | 0.5487954 | 0.30502576 | 0.76034654 | 0.99937574 |
| MMP14 | 23.4162372 | -0.5720981 | 0.67954374 | -0.8418856 | 0.39985201 | 0.99937574 |
| LRP10 | 140.777046 | -0.456228 | 0.51800106 | -0.8807472 | 0.37845465 | 0.99937574 |
| RBM23 | 63.175821 | -0.1112934 | 0.27363205 | -0.4067266 | 0.68420879 | 0.99937574 |
| PRMT5 | 11.0208053 | 0.7850644 | 0.53699771 | 1.46195112 | 0.14375461 | 0.99937574 |
| AL132780.1 | 25.6831989 | 1.19779314 | 0.6649751 | 1.80126013 | 0.07166189 | 0.99937574 |
| PSMB5 | 62.8518561 | 0.22156795 | 0.39002338 | 0.56808888 | 0.56997462 | 0.99937574 |
| ACIN1 | 107.126899 | -0.2958446 | 0.24715326 | -1.1970087 | 0.23130317 | 0.99937574 |
| C14orf119 | 25.7700308 | 0.4705494 | 0.45649058 | 1.0307976 | 0.30263575 | 0.99937574 |
| HOMEZ | 13.2007714 | 0.47404379 | 0.53087267 | 0.89295197 | 0.3718829 | 0.99937574 |
| BCL2L2-PABI | 21.6194598 | 0.09443035 | 0.43144648 | 0.21886921 | 0.82675193 | 0.99937574 |
| PABPN1 | 63.3530095 | 0.21738638 | 0.32961026 | 0.65952552 | 0.50955836 | 0.99937574 |
| NGDN | 19.6955609 | -0.5820103 | 0.49737782 | -1.1701574 | 0.24193763 | 0.99937574 |
| ZFHX2 | 10.8552793 | 0.90830047 | 0.74326999 | 1.22203303 | 0.22169514 | 0.99937574 |
| AP1G2 | 27.0663999 | -0.0366566 | 0.46754709 | -0.0784019 | 0.93750836 | 0.99937574 |
| DHRS4-AS1 | 21.5247921 | 0.44525749 | 0.46635838 | 0.95475391 | 0.33970216 | 0.99937574 |
| DHRS4L2 | 26.1820254 | 0.89030892 | 0.55759279 | 1.59670092 | 0.11033239 | 0.99937574 |
| PSME1 | 60.5031662 | -0.0940851 | 0.51862208 | -0.1814135 | 0.856043 | 0.99937574 |
| EMC9 | 11.0950717 | -0.0416463 | 0.82255077 | -0.0506306 | 0.95961986 | 0.99937574 |
| PSME2 | 15.0548625 | 0.29474006 | 0.59361226 | 0.49651949 | 0.61952794 | 0.99937574 |
| AL136295.1 | 16.453609 | 0.15469384 | 0.46535911 | 0.3324182 | 0.7395735 | 0.99937574 |
| GMPR2 | 16.6950026 | 0.09101713 | 0.46364173 | 0.19630918 | 0.84436818 | 0.99937574 |
| TINF2 | 14.6364018 | -0.1162701 | 0.65224889 | -0.1782603 | 0.85851854 | 0.99937574 |
| DHRS1 | 12.7351915 | 0.45924712 | 0.71516312 | 0.64215716 | 0.52077115 | 0.99937574 |
| KHNYN | 19.4659048 | -0.3330517 | 0.45374732 | -0.7340027 | 0.46294712 | 0.99937574 |
| SDR39U1 | 24.6725131 | 0.04147847 | 0.38328309 | 0.10821888 | 0.91382207 | 0.99937574 |
| STXBP6 | 48.2630243 | -3.4800853 | 2.00845691 | -1.7327159 | 0.08314617 | 0.99937574 |
| NOVA1 | 92.1304304 | 0.82668774 | 0.35806645 | 2.30875508 | 0.02095718 | 0.96427895 |
| AL110292.1 | 40.7387716 | 2.10448143 | 0.69359803 | 3.03415137 | 0.00241213 | 0.48965449 |
| PRKD1 | 439.415038 | 0.14490453 | 0.54210528 | 0.26729961 | 7.89E-01 | 0.99937574 |
| G2E3 | 44.8768303 | -0.1475113 | 0.3141641 | -0.4695358 | 0.63868671 | 0.99937574 |
| SCFD1 | 145.254173 | -0.0477345 | 0.25448952 | -0.1875698 | 0.85121393 | 0.99937574 |
| COCH | 13.6373639 | -0.2758609 | 0.64132345 | -0.4301431 | 0.66709153 | 0.99937574 |
| STRN3 | 279.301909 | -0.1754473 | 0.31729672 | -0.552944 | 0.58030176 | 0.99937574 |
| AP4S1 | 13.7480363 | -0.5507794 | 0.60451302 | -0.9111126 | 0.36223607 | 0.99937574 |
| HECTD1 | 390.199864 | -0.4375014 | 0.2686377 | -1.6285929 | 0.10339924 | 0.99937574 |
| AL136418.1 | 58.9241053 | -1.8381427 | 0.97790653 | -1.8796712 | 0.06015291 | 0.99937574 |
| HEATR5A | 60.9401983 | 0.11983679 | 0.44627317 | 0.26852789 | 0.78829301 | 0.99937574 |
| NUBPL | 137.955487 | -0.2598669 | 0.48937628 | -0.5310166 | 0.59540725 | 0.99937574 |
| ARHGAP5 | 280.733311 | -0.1405228 | 0.20193947 | -0.6958661 | 0.48651268 | 0.99937574 |
| AKAP6 | 147.666417 | 0.18632465 | 0.45343674 | 0.41091651 | 0.68113375 | 0.99937574 |
| NPAS3 | 60.133948 | 0.74109418 | 0.86275403 | 0.85898664 | 0.39034789 | 0.99937574 |
| EGLN3 | 19.087463 | -1.2426615 | 0.85184092 | -1.4587953 | 0.14462145 | 0.99937574 |
| EAPP | 48.1095909 | 0.04366578 | 0.32070383 | 0.13615611 | 0.89169787 | 0.99937574 |
| SNX6 | 135.52009 | -0.2904289 | 0.22772368 | -1.2753566 | 0.20218305 | 0.99937574 |
| CFL2 | 21.9748395 | -0.0388408 | 0.4554796 | -0.0852745 | 0.9320432 | 0.99937574 |
| BAZ1A | 57.5861348 | -1.2257983 | 0.48046888 | -2.5512543 | 0.0107336 | 0.76274616 |
| SRP54 | 93.4155431 | -0.2599062 | 0.26045227 | -0.9979035 | 0.31832614 | 0.99937574 |
| FAM177A1 | 71.6477728 | -0.0240856 | 0.33459141 | -0.071985 | 0.94261384 | 0.99937574 |
| PPP2R3C | 57.2370342 | -0.0198973 | 0.42389711 | -0.0469391 | 0.96256177 | 0.99937574 |
| NFKBIA | 51.7391066 | -0.5255836 | 0.68475474 | -0.7675501 | 0.44275453 | 0.99937574 |
| RALGAPA1 | 453.614006 | 0.00261108 | 0.34774726 | 0.00750854 | 0.99400911 | 0.99937574 |
| BRMS1L | 23.3481708 | -0.2294309 | 0.49913836 | -0.4596538 | 0.6457647 | 0.99937574 |
| MBIP | 49.0115129 | -0.1280747 | 0.39210268 | -0.3266356 | 0.74394348 | 0.99937574 |

| | | | | | | |
|---|---|---|---|---|---|---|
| SLC25A21 | 21.5900354 | -0.1040062 | 0.80343231 | -0.1294523 | 0.89699974 | 0.99937574 |
| MIPOL1 | 112.520187 | 0.07446918 | 0.37944356 | 0.19625891 | 0.84440752 | 0.99937574 |
| SEC23A | 146.148197 | 0.02647905 | 0.35085759 | 0.07546952 | 0.93984115 | 0.99937574 |
| GEMIN2 | 11.2023801 | -0.3504604 | 0.58537005 | -0.598699 | 0.54937365 | 0.99937574 |
| TRAPPC6B | 100.511009 | 0.34956291 | 0.26784177 | 1.30510977 | 0.19185547 | 0.99937574 |
| PNN | 123.668304 | -0.3195708 | 0.23849206 | -1.339964 | 0.18025706 | 0.99937574 |
| MIA2 | 171.817875 | 0.0147997 | 0.26582173 | 0.05567528 | 0.9556005 | 0.99937574 |
| FBXO33 | 70.5214003 | -0.3620937 | 0.45911191 | -0.7886829 | 0.43029736 | 0.99937574 |
| AL049875.1 | 33.8151589 | -0.3863733 | 0.60266563 | -0.6411072 | 0.52145302 | 0.99937574 |
| KLHL28 | 57.5190909 | -0.0238812 | 0.35083111 | -0.0680703 | 0.94572968 | 0.99937574 |
| TOGARAM1 | 78.6947201 | -0.0194073 | 0.27305923 | -0.0710737 | 0.94333908 | 0.99937574 |
| PRPF39 | 46.1271802 | 0.00920283 | 0.36214116 | 0.02541227 | 0.97972613 | 0.99937574 |
| FKBP3 | 41.9722384 | 0.48251361 | 0.48361976 | 0.99771277 | 0.31841866 | 0.99937574 |
| FANCM | 21.3484108 | -0.9931953 | 0.61406602 | -1.617408 | 0.10579025 | 0.99937574 |
| MIS18BP1 | 33.889155 | 0.23532305 | 0.39028029 | 0.60295909 | 0.5465359 | 0.99937574 |
| RPS29 | 101.7233 | 0.28336205 | 0.35399139 | 0.80047723 | 0.42343435 | 0.99937574 |
| RPL36AL | 119.156423 | -0.3960487 | 0.50908965 | -0.7779547 | 0.43659571 | 0.99937574 |
| DNAAF2 | 14.4295716 | -0.4388261 | 0.55813526 | -0.786236 | 0.43172922 | 0.99937574 |
| KLHDC1 | 38.5911463 | 1.0269047 | 0.40949466 | 2.50773648 | 0.01215072 | 0.77519564 |
| KLHDC2 | 79.3274968 | 0.05014772 | 0.3716738 | 0.13492401 | 0.89267196 | 0.99937574 |
| NEMF | 247.484174 | -0.1251336 | 0.28391084 | -0.4407498 | 0.65939416 | 0.99937574 |
| ARF6 | 46.8784759 | 0.24738116 | 0.39595969 | 0.62476348 | 0.53212631 | 0.99937574 |
| LINC01588 | 19.0754294 | 0.48155891 | 0.59762773 | 0.80578407 | 0.42036737 | 0.99937574 |
| VCPKMT | 24.1378185 | 0.44785051 | 0.45382377 | 0.98683795 | 0.32372209 | 0.99937574 |
| SOS2 | 252.100102 | -0.0546558 | 0.2091878 | -0.2612762 | 0.79387952 | 0.99937574 |
| L2HGDH | 70.4446485 | 0.25012601 | 0.45554943 | 0.54906447 | 0.58296121 | 0.99937574 |
| ATP5S | 54.5671724 | -0.0304351 | 0.30635199 | -0.0993468 | 0.92086296 | 0.99937574 |
| CDKL1 | 121.329844 | 0.06951859 | 0.40744186 | 0.1706221 | 0.86452092 | 0.99937574 |
| MAP4K5 | 174.249797 | -0.2371807 | 0.34429075 | -0.6888964 | 0.49088847 | 0.99937574 |
| ATL1 | 20.7643496 | 0.54966037 | 0.62057973 | 0.88572079 | 0.375768 | 0.99937574 |
| SAV1 | 113.077092 | 0.03592885 | 0.41752745 | 0.08605148 | 0.93142549 | 0.99937574 |
| NIN | 19.7736071 | -1.1468294 | 0.60366176 | -1.8997881 | 0.05746093 | 0.99937574 |
| TMX1 | 25.5189161 | -0.3323724 | 0.47256974 | -0.7033299 | 0.48185017 | 0.99937574 |
| RTRAF | 52.3021284 | 0.00888483 | 0.42487951 | 0.02091142 | 0.98331632 | 0.99937574 |
| TXNDC16 | 23.1897391 | -1.2111875 | 0.53249071 | -2.2745702 | 0.02293172 | 0.96427895 |
| ERO1A | 367.490504 | -0.5153853 | 0.47054391 | -1.0952969 | 0.27338661 | 0.99937574 |
| AL133453.1 | 54.3565928 | 0.07096003 | 0.73693375 | 0.09629092 | 9.23E-01 | 0.99937574 |
| PSMC6 | 91.645845 | 0.02672768 | 0.26516096 | 0.10079795 | 0.91971086 | 0.99937574 |
| STYX | 48.4168 | 0.04052048 | 0.39692045 | 0.10208717 | 0.91868749 | 0.99937574 |
| GNPNAT1 | 92.7948523 | -0.7445138 | 0.58691174 | -1.2685276 | 0.20460959 | 0.99937574 |
| FERMT2 | 118.286643 | 0.0379809 | 0.35812396 | 0.10605519 | 0.91553857 | 0.99937574 |
| DDHD1 | 25.227115 | 0.38491111 | 0.46277245 | 0.83175028 | 0.40554992 | 0.99937574 |
| CNIH1 | 29.7118177 | 0.34653889 | 0.37690077 | 0.91944331 | 0.35786375 | 0.99937574 |
| GMFB | 34.0741689 | -0.1350936 | 0.38020603 | -0.3553168 | 0.7223523 | 0.99937574 |
| CGRRF1 | 15.782636 | 0.63748269 | 0.45443086 | 1.40281558 | 0.16067184 | 0.99937574 |
| SAMD4A | 165.261271 | 0.32366962 | 0.61898744 | 0.52290176 | 0.60104262 | 0.99937574 |
| GCH1 | 18.887184 | -1.2073132 | 0.482445 | -2.5024888 | 0.01233235 | 0.77519564 |
| SOCS4 | 38.9134395 | 0.36418676 | 0.3380394 | 1.07735003 | 0.28132392 | 0.99937574 |
| MAPK1IP1L | 132.684188 | -0.0661254 | 0.27151753 | -0.24354 | 0.80758711 | 0.99937574 |
| LGALS3 | 358.220484 | 0.23247536 | 0.41034708 | 0.56653349 | 0.57103117 | 0.99937574 |
| FBXO34 | 288.796313 | -0.3382285 | 0.33541968 | -1.0083742 | 0.31327487 | 0.99937574 |
| AL158801.3 | 14.5184863 | -0.0362002 | 0.70700992 | -0.0512019 | 0.95916464 | 0.99937574 |
| ATG14 | 38.2902471 | -0.6078329 | 0.36415881 | -1.6691425 | 0.09508915 | 0.99937574 |
| KTN1-AS1 | 12.6722584 | 0.10258209 | 0.65868154 | 0.15573853 | 0.87623913 | 0.99937574 |

| | | | | | | |
|---|---|---|---|---|---|---|
| KTN1 | 341.373004 | -0.2291161 | 0.3489049 | -0.656672 | 0.51139182 | 0.99937574 |
| PELI2 | 141.113953 | -0.1893741 | 0.38904462 | -0.4867671 | 0.62642341 | 0.99937574 |
| TMEM260 | 100.62292 | -0.0580666 | 0.34648376 | -0.1675883 | 0.86690714 | 0.99937574 |
| EXOC5 | 90.9755607 | 0.2699873 | 0.25821113 | 1.04560674 | 0.29574264 | 0.99937574 |
| AP5M1 | 69.6232744 | 0.23772005 | 0.29774 | 0.79841488 | 0.42462977 | 0.99937574 |
| NAA30 | 48.1191502 | -0.0576812 | 0.32769913 | -0.1760187 | 0.86027927 | 0.99937574 |
| CCDC198 | 190.757079 | 1.59717735 | 0.6523694 | 2.44827142 | 0.01435435 | 0.85624285 |
| ARMH4 | 62.0310204 | 0.44911104 | 0.34855355 | 1.2884994 | 0.19757218 | 0.99937574 |
| ACTR10 | 75.3625762 | -0.2403078 | 0.27296132 | -0.880373 | 0.37865729 | 0.99937574 |
| PSMA3 | 67.1156402 | -0.3595563 | 0.36386297 | -0.9881641 | 0.32307227 | 0.99937574 |
| ARID4A | 99.1975326 | -0.0668819 | 0.30635055 | -0.2183183 | 0.82718115 | 0.99937574 |
| TIMM9 | 24.2203606 | 0.46645665 | 0.38010472 | 1.22717931 | 0.2197552 | 0.99937574 |
| KIAA0586 | 72.5169077 | 0.09281306 | 0.2837417 | 0.32710406 | 0.74358918 | 0.99937574 |
| DAAM1 | 107.790471 | 0.25255434 | 0.35179624 | 0.71789949 | 0.47281926 | 0.99937574 |
| L3HYPDH | 13.9302281 | 0.18543333 | 0.55160499 | 0.3361705 | 0.73674229 | 0.99937574 |
| JKAMP | 48.3523861 | 0.19018927 | 0.31091388 | 0.61171044 | 0.54072935 | 0.99937574 |
| RTN1 | 54.2734847 | 0.86738215 | 0.59356079 | 1.46131983 | 0.14392769 | 0.99937574 |
| LRRC9 | 50.7126895 | 0.87103507 | 0.70016311 | 1.24404594 | 0.21348266 | 0.99937574 |
| AL157911.1 | 134.421196 | 0.4803767 | 0.31217948 | 1.53878372 | 1.24E-01 | 9.99E-01 |
| PCNX4 | 600.325536 | 0.17186416 | 0.44405412 | 0.38703427 | 0.69873084 | 0.99937574 |
| DHRS7 | 164.912868 | 0.35473889 | 0.42135543 | 0.84189942 | 0.39984425 | 0.99937574 |
| PPM1A | 131.564123 | -0.2209727 | 0.32052232 | -0.6894144 | 0.49056255 | 0.99937574 |
| MNAT1 | 252.528366 | -0.0738035 | 0.30146159 | -0.2448191 | 0.80659654 | 0.99937574 |
| SLC38A6 | 31.7513394 | -0.7657401 | 0.48649358 | -1.5739983 | 0.11548785 | 0.99937574 |
| PRKCH | 28.9783992 | 0.08764035 | 0.44114526 | 0.19866551 | 0.84252441 | 0.99937574 |
| TMEM30B | 38.3915602 | -0.165143 | 0.39062511 | -0.4227661 | 0.67246595 | 0.99937574 |
| HIF1A | 922.718296 | 0.09370792 | 0.44030137 | 0.21282678 | 8.31E-01 | 0.99937574 |
| HIF1A-AS2 | 148.23172 | 0.28992476 | 0.72510551 | 0.39983803 | 0.68927582 | 0.99937574 |
| SNAPC1 | 74.353814 | 0.18250962 | 0.49211606 | 0.37086702 | 0.71073658 | 0.99937574 |
| PPP2R5E | 332.320077 | -0.0816413 | 0.37888409 | -0.2154784 | 0.82939434 | 0.99937574 |
| WDR89 | 24.4288978 | -0.1095914 | 0.48861285 | -0.2242909 | 0.82253092 | 0.99937574 |
| SGPP1 | 33.6810247 | -0.1069619 | 0.40255151 | -0.2657098 | 0.79046275 | 0.99937574 |
| SYNE2 | 1355.75065 | -0.0620878 | 0.41796901 | -0.1485464 | 0.88191157 | 0.99937574 |
| ESR2 | 112.690501 | 0.11866321 | 0.99497418 | 0.1192626 | 0.90506731 | 0.99937574 |
| MTHFD1 | 15.5400926 | 0.35868386 | 0.47325615 | 0.7579064 | 0.44850702 | 0.99937574 |
| AL122035.1 | 11.9896663 | -0.5182185 | 0.5980492 | -0.8665147 | 0.38620794 | 0.99937574 |
| ZBTB25 | 32.9075064 | -0.5148983 | 0.57563251 | -0.8944913 | 0.37105906 | 0.99937574 |
| ZBTB1 | 85.0764273 | 0.14359521 | 0.37500456 | 0.3829159 | 0.70178212 | 0.99937574 |
| PLEKHG3 | 61.660703 | 0.28963623 | 0.34912457 | 0.82960712 | 0.40676095 | 0.99937574 |
| CHURC1-FNT | 34.4526845 | -0.501394 | 0.48945386 | -1.0243948 | 0.30564882 | 0.99937574 |
| GPX2 | 15.3847634 | 0.45174774 | 0.97715118 | 0.462311 | 0.64385831 | 0.99937574 |
| RAB15 | 45.2774229 | -0.8044029 | 0.70957003 | -1.1336483 | 0.25694209 | 0.99937574 |
| MAX | 84.7948783 | -0.118068 | 0.28268594 | -0.4176648 | 0.67619221 | 0.99937574 |
| FUT8 | 63.1253982 | 0.02023173 | 0.53038125 | 0.03814564 | 0.96957157 | 0.99937574 |
| AL359232.1 | 49.9198999 | 0.31419689 | 0.49405539 | 0.63595477 | 0.5248059 | 0.99937574 |
| GPHN | 1404.37506 | -0.5216634 | 0.32078159 | -1.6262262 | 0.10390154 | 0.99937574 |
| MPP5 | 146.559005 | -0.1468551 | 0.24824596 | -0.5915708 | 0.55413803 | 0.99937574 |
| ATP6V1D | 129.178684 | -0.0695799 | 0.366839 | -0.1896743 | 0.84956438 | 0.99937574 |
| EIF2S1 | 49.9467498 | -0.2188883 | 0.31606171 | -0.6925493 | 0.48859243 | 0.99937574 |
| PLEK2 | 46.5398733 | -1.3503521 | 0.5298589 | -2.5485126 | 0.01081834 | 0.76274616 |
| PLEKHH1 | 83.2151732 | 0.02847607 | 0.52419796 | 0.05432313 | 0.95667772 | 0.99937574 |
| PIGH | 38.6093216 | 0.15605485 | 0.53896685 | 0.28954444 | 0.77216478 | 0.99937574 |
| ARG2 | 79.4202035 | -0.083901 | 0.91208378 | -0.0919883 | 0.92670734 | 0.99937574 |
| VTI1B | 44.5598591 | 0.08233142 | 0.49654507 | 0.16580855 | 0.86830761 | 0.99937574 |

| | | | | | | |
|---|---|---|---|---|---|---|
| RDH11 | 11.9363085 | -0.1458977 | 0.64654917 | -0.225656 | 0.82146898 | 0.99937574 |
| ZFYVE26 | 31.0742703 | -0.8230021 | 0.42119485 | -1.9539699 | 0.05070477 | 0.99937574 |
| RAD51B | 254.031696 | -0.8543203 | 0.52531478 | -1.6263016 | 0.10388552 | 0.99937574 |
| ZFP36L1 | 127.609187 | -0.1987312 | 0.32676418 | -0.6081794 | 0.54306852 | 0.99937574 |
| ACTN1 | 17.7877627 | 0.39389245 | 0.56108887 | 0.70201436 | 0.48267021 | 0.99937574 |
| DCAF5 | 179.629551 | 0.33110557 | 0.28399471 | 1.16588639 | 0.24366038 | 0.99937574 |
| AL359317.2 | 12.8153788 | -0.1820915 | 0.73227923 | -0.248664 | 0.80362068 | 0.99937574 |
| EXD2 | 41.6654589 | 0.2554077 | 0.4073765 | 0.62695738 | 0.53068717 | 0.99937574 |
| GALNT16 | 36.0948353 | 1.30250264 | 0.72535732 | 1.79567035 | 0.07254696 | 0.99937574 |
| ERH | 51.9406018 | -0.2531828 | 0.42531995 | -0.595276 | 0.551659 | 0.99937574 |
| SLC39A9 | 71.0630667 | 0.25326655 | 0.33689644 | 0.7517638 | 0.45219311 | 0.99937574 |
| PLEKHD1 | 10.9911445 | 0.58063224 | 0.91445823 | 0.6349467 | 0.52546318 | 0.99937574 |
| SUSD6 | 148.739734 | -0.0376637 | 0.27280343 | -0.1380616 | 0.89019174 | 0.99937574 |
| SRSF5 | 273.037579 | -0.1232689 | 0.29040315 | -0.4244751 | 0.67121937 | 0.99937574 |
| SMOC1 | 75.574909 | -0.8397498 | 0.82125914 | -1.022515 | 0.30653722 | 0.99937574 |
| AL357153.2 | 33.6792591 | 0.71858296 | 0.57786314 | 1.24351756 | 0.21367717 | 0.99937574 |
| MED6 | 32.9363851 | -0.162055 | 0.43597869 | -0.371704 | 0.71011325 | 0.99937574 |
| TTC9 | 73.8684052 | -1.187623 | 0.62708962 | -1.8938649 | 5.82E-02 | 0.99937574 |
| MAP3K9 | 44.0379398 | -0.3401287 | 0.38692029 | -0.8790667 | 0.37936514 | 0.99937574 |
| PCNX1 | 321.792602 | 0.30197735 | 0.30550648 | 0.98844825 | 0.32293317 | 0.99937574 |
| SIPA1L1 | 189.361495 | -0.3340853 | 0.34074324 | -0.9804605 | 0.32685887 | 0.99937574 |
| DPF3 | 16.6910739 | 2.81593959 | 0.69005984 | 4.08071797 | 4.49E-05 | 0.07551643 |
| DCAF4 | 25.1326469 | -0.1157837 | 0.49702038 | -0.2329556 | 0.81579592 | 0.99937574 |
| ZFYVE1 | 77.1842228 | -0.2351426 | 0.32914891 | -0.7143959 | 0.47498238 | 0.99937574 |
| RBM25 | 190.189506 | -0.3679427 | 0.28318633 | -1.2992954 | 0.19384257 | 0.99937574 |
| PSEN1 | 200.891135 | 0.18906145 | 0.23006763 | 0.82176468 | 0.41121084 | 0.99937574 |
| PAPLN | 18.2084582 | 0.88257016 | 0.59616653 | 1.48040876 | 0.13876419 | 0.99937574 |
| NUMB | 488.072252 | -0.0581834 | 0.26309116 | -0.2211529 | 0.82497339 | 0.99937574 |
| HEATR4 | 23.6285126 | 0.93143971 | 0.53299622 | 1.74755405 | 8.05E-02 | 0.99937574 |
| ACOT2 | 13.0633033 | 1.23622056 | 0.78355509 | 1.57770725 | 0.11463288 | 0.99937574 |
| DNAL1 | 31.0682911 | 0.15059977 | 0.36278722 | 0.41511873 | 0.67805499 | 0.99937574 |
| PNMA1 | 13.8559956 | -0.3936547 | 0.52167441 | -0.7545985 | 0.45048993 | 0.99937574 |
| ELMSAN1 | 61.9786657 | -0.2212232 | 0.52360273 | -0.422502 | 0.67265865 | 0.99937574 |
| AC005480.1 | 18.8016095 | 0.67679939 | 0.70887333 | 0.95475364 | 0.3397023 | 0.99937574 |
| COQ6 | 15.5490852 | 0.45501729 | 0.51156982 | 0.88945296 | 0.37375969 | 0.99937574 |
| ENTPD5 | 36.4579459 | 0.24827719 | 0.42756434 | 0.58067798 | 0.56145751 | 0.99937574 |
| BBOF1 | 26.569382 | -0.0591559 | 0.55371641 | -0.1068343 | 0.91492047 | 0.99937574 |
| ALDH6A1 | 45.0980254 | 0.59654444 | 0.4494521 | 1.32727035 | 0.18441927 | 0.99937574 |
| LIN52 | 62.3998291 | -0.4070295 | 0.32615415 | -1.2479665 | 0.21204331 | 0.99937574 |
| ABCD4 | 58.2795209 | 0.15877884 | 0.31790443 | 0.49945463 | 0.61745914 | 0.99937574 |
| NPC2 | 103.661256 | -0.1391053 | 0.55986658 | -0.2484615 | 0.80377736 | 0.99937574 |
| ISCA2 | 25.7215953 | 0.14311187 | 0.56750485 | 0.25217736 | 0.80090398 | 0.99937574 |
| AREL1 | 73.8528105 | -0.047861 | 0.33381527 | -0.1433758 | 0.88599336 | 0.99937574 |
| FCF1 | 23.5641754 | 0.05262356 | 0.41319532 | 0.12735759 | 0.89865738 | 0.99937574 |
| YLPM1 | 124.613716 | 0.28023645 | 0.22978021 | 1.21958481 | 0.22262231 | 0.99937574 |
| DLST | 71.4093695 | 0.16891786 | 0.3297705 | 0.51222853 | 0.60849108 | 0.99937574 |
| EIF2B2 | 15.7148721 | 0.69636442 | 0.45551117 | 1.52875378 | 0.1263255 | 0.99937574 |
| MLH3 | 53.2757134 | -0.3141338 | 0.43924677 | -0.7151648 | 0.4745072 | 0.99937574 |
| ACYP1 | 39.1846353 | -0.0518607 | 0.54373866 | -0.0953781 | 0.92401453 | 0.99937574 |
| ZC2HC1C | 14.582453 | 0.34857089 | 0.66960833 | 0.52055937 | 0.60267376 | 0.99937574 |
| NEK9 | 66.1979551 | 0.26420129 | 0.25527581 | 1.03496409 | 0.30068568 | 0.99937574 |
| AL049780.2 | 13.4633289 | 0.88944036 | 0.75294626 | 1.18128001 | 0.23749151 | 0.99937574 |
| TMED10 | 141.77981 | 0.28855077 | 0.22534918 | 1.2804607 | 0.20038316 | 0.99937574 |
| AL691403.1 | 88.799825 | 1.98319195 | 0.39652502 | 5.00142952 | 5.69E-07 | 0.00287152 |

| | | | | | | |
|---|---|---|---|---|---|---|
| AF111167.1 | 12.9224262 | 1.7755074 | 0.85499285 | 2.0766342 | 0.03783533 | 0.99937574 |
| FOS | 43.8965323 | -0.4777534 | 0.82616115 | -0.5782811 | 0.56307433 | 0.99937574 |
| JDP2 | 14.8242118 | -0.189245 | 0.666141 | -0.2840916 | 0.77634022 | 0.99937574 |
| FLVCR2 | 51.307656 | -0.6683175 | 0.64215573 | -1.0407405 | 0.29799598 | 0.99937574 |
| TTLL5 | 129.892255 | 0.11488454 | 0.33611104 | 0.34180533 | 0.7324974 | 0.99937574 |
| ERG28 | 33.1635742 | 0.32226664 | 0.42839563 | 0.75226408 | 0.45189227 | 0.99937574 |
| IFT43 | 82.5447478 | 0.38831976 | 0.34242375 | 1.13403277 | 0.2567808 | 0.99937574 |
| GPATCH2L | 140.844125 | 0.10352435 | 0.31218223 | 0.33161513 | 0.74017989 | 0.99937574 |
| ANGEL1 | 13.6799934 | 0.18027533 | 0.51286717 | 0.35150492 | 0.72520958 | 0.99937574 |
| IRF2BPL | 46.1977443 | 0.22584373 | 0.37053268 | 0.60951096 | 0.54218581 | 0.99937574 |
| POMT2 | 14.8401188 | -0.420339 | 0.50424225 | -0.8336053 | 0.40450342 | 0.99937574 |
| GSTZ1 | 19.4325116 | 0.21171629 | 0.57182947 | 0.37024375 | 0.71120088 | 0.99937574 |
| TMED8 | 48.0136785 | 0.1613177 | 0.3595141 | 0.44871035 | 0.65364061 | 0.99937574 |
| VIPAS39 | 19.9701437 | 0.27053956 | 0.50426099 | 0.53650702 | 0.59160819 | 0.99937574 |
| AHSA1 | 40.5209846 | 0.11207952 | 0.38449152 | 0.29150062 | 0.77066847 | 0.99937574 |
| SPTLC2 | 383.820081 | -0.2387811 | 0.42714296 | -0.5590191 | 0.57614872 | 0.99937574 |
| ALKBH1 | 36.1504064 | 0.13598796 | 0.33113818 | 0.41066831 | 0.68131577 | 0.99937574 |
| SLIRP | 60.4017347 | 0.1112454 | 0.39417037 | 0.28222669 | 0.7777697 | 0.99937574 |
| SNW1 | 55.8565338 | 0.07566598 | 0.28762522 | 0.26307146 | 0.7924955 | 0.99937574 |
| ADCK1 | 27.3080865 | -0.0659624 | 0.43203848 | -0.1526772 | 0.87865284 | 0.99937574 |
| NRXN3 | 891.214485 | 0.81039469 | 0.69965972 | 1.15826976 | 0.24675397 | 0.99937574 |
| AC008056.1 | 38.6121598 | 1.50333238 | 0.78987824 | 1.90324572 | 0.05700849 | 0.99937574 |
| CEP128 | 36.1076955 | -0.6968202 | 0.40999238 | -1.699593 | 0.08920751 | 0.99937574 |
| GTF2A1 | 54.0725927 | -0.0170197 | 0.28967845 | -0.0587538 | 0.95314824 | 0.99937574 |
| STON2 | 139.834067 | 0.24693194 | 0.47969326 | 0.51477049 | 0.60671341 | 0.99937574 |
| SEL1L | 134.264132 | -0.3631959 | 0.29125677 | -1.2469954 | 0.21239917 | 0.99937574 |
| GALC | 12.025854 | -0.6442399 | 0.54169245 | -1.1893094 | 0.23431794 | 0.99937574 |
| SPATA7 | 53.4668134 | 0.37463568 | 0.39342889 | 0.95223227 | 0.3409792 | 0.99937574 |
| PTPN21 | 113.753564 | -0.0990489 | 0.31759526 | -0.3118716 | 0.75513811 | 0.99937574 |
| ZC3H14 | 111.050206 | 0.19292364 | 0.29140797 | 0.66203969 | 0.50794579 | 0.99937574 |
| EML5 | 31.801596 | -1.042172 | 0.56002956 | -1.8609232 | 0.06275502 | 0.99937574 |
| TTC8 | 43.6908195 | 0.12974586 | 0.40552862 | 0.31994255 | 0.74901189 | 0.99937574 |
| FOXN3 | 441.131026 | -0.7161574 | 0.48383594 | -1.4801658 | 0.13882901 | 0.99937574 |
| EFCAB11 | 35.4385827 | -0.1889556 | 0.46877141 | -0.4030869 | 0.6868843 | 0.99937574 |
| TDP1 | 19.2418717 | -0.04922 | 0.50394576 | -0.0976692 | 0.92219499 | 0.99937574 |
| PSMC1 | 34.9541864 | -0.1897452 | 0.43892887 | -0.4322915 | 0.66552956 | 0.99937574 |
| NRDE2 | 34.2545002 | -0.3069825 | 0.44873733 | -0.6841029 | 0.49391022 | 0.99937574 |
| CALM1 | 96.9448848 | -0.0322092 | 0.47417844 | -0.0679264 | 0.94584421 | 0.99937574 |
| TTC7B | 126.087213 | -0.1150322 | 0.37544015 | -0.306393 | 0.75930543 | 0.99937574 |
| RPS6KA5 | 39.1741597 | -0.8092086 | 0.67478563 | -1.1992085 | 0.2304469 | 0.99937574 |
| DGLUCY | 202.362745 | -0.0685553 | 0.31623001 | -0.2167893 | 0.82837252 | 0.99937574 |
| PPP4R3A | 136.483325 | -0.3638187 | 0.23869009 | -1.5242303 | 0.1274512 | 0.99937574 |
| CATSPERB | 32.0203306 | -0.1743292 | 0.5173594 | -0.3369597 | 0.73614731 | 0.99937574 |
| TC2N | 53.6159363 | -0.2980011 | 0.57233151 | -0.5206793 | 0.60259022 | 0.99937574 |
| TRIP11 | 150.231114 | 0.34876285 | 0.23528368 | 1.48230789 | 0.13825839 | 0.99937574 |
| ATXN3 | 54.2714878 | -0.2614594 | 0.30019766 | -0.8709575 | 0.38377736 | 0.99937574 |
| NDUFB1 | 63.0881963 | 0.25835201 | 0.39102045 | 0.66071227 | 0.50879685 | 0.99937574 |
| CPSF2 | 60.9936248 | 0.18351358 | 0.28893473 | 0.63513852 | 0.52533807 | 0.99937574 |
| LGMN | 72.9059523 | -0.069931 | 0.55214727 | -0.1266528 | 0.89921518 | 0.99937574 |
| GOLGA5 | 30.4428704 | -0.2657453 | 0.3531638 | -0.7524703 | 0.4517683 | 0.99937574 |
| ITPK1 | 94.0153058 | 0.06391265 | 0.48359654 | 0.1321611 | 0.89485687 | 0.99937574 |
| MOAP1 | 27.1026284 | 0.19383789 | 0.44472472 | 0.43586039 | 0.66293803 | 0.99937574 |
| UBR7 | 17.4061504 | -0.0723133 | 0.44574355 | -0.1622307 | 0.87112417 | 0.99937574 |
| BTBD7 | 205.564447 | -0.0665727 | 0.19435014 | -0.3425399 | 0.73194464 | 0.99937574 |

| | | | | | | |
|---|---|---|---|---|---|---|
| DDX24 | 155.295987 | -0.0875468 | 0.31319697 | -0.2795263 | 0.77984098 | 0.99937574 |
| IFI27L1 | 13.9739615 | 0.13664358 | 0.56799519 | 0.24057173 | 0.80988706 | 0.99937574 |
| IFI27 | 67.1230633 | -5.476684 | 1.70037296 | -3.2208722 | 0.00127801 | 0.33941289 |
| IFI27L2 | 16.0991067 | -0.1470507 | 0.6030225 | -0.2438561 | 0.80734225 | 0.99937574 |
| PPP4R4 | 18.0579945 | 0.46023964 | 0.65048706 | 0.70753082 | 0.47923667 | 0.99937574 |
| SERPINA1 | 51.436666 | -0.0732157 | 0.84698962 | -0.0864422 | 0.93111488 | 0.99937574 |
| DICER1 | 243.82189 | -0.0956492 | 0.25769707 | -0.371169 | 0.71051163 | 0.99937574 |
| DICER1-AS1 | 18.7754583 | -0.2193359 | 0.6085559 | -0.3604203 | 0.71853281 | 0.99937574 |
| CLMN | 451.893373 | 0.29805021 | 0.59681496 | 0.49940137 | 0.61749666 | 0.99937574 |
| GLRX5 | 48.5769261 | 0.08421129 | 0.49358867 | 0.17061027 | 0.86453023 | 0.99937574 |
| ATG2B | 92.0169636 | -0.4733115 | 0.29379515 | -1.6110257 | 0.10717413 | 0.99937574 |
| GSKIP | 36.6382491 | 0.06127214 | 0.4394826 | 0.1394188 | 0.88911922 | 0.99937574 |
| AK7 | 19.1605031 | 0.04787088 | 0.59561978 | 0.08037154 | 9.36E-01 | 0.99937574 |
| PAPOLA | 239.308983 | 0.32333422 | 0.26362946 | 1.22647225 | 0.22002101 | 0.99937574 |
| VRK1 | 16.7377783 | -0.7404313 | 0.56165194 | -1.3183098 | 0.18739995 | 0.99937574 |
| SETD3 | 153.58981 | -0.0651266 | 0.31006046 | -0.2100448 | 0.83363274 | 0.99937574 |
| CCNK | 43.596994 | -0.0360813 | 0.35548805 | -0.1014979 | 0.9191552 | 0.99937574 |
| CCDC85C | 41.7580095 | 0.46395259 | 0.41616369 | 1.11483197 | 0.26492244 | 0.99937574 |
| EVL | 78.915863 | 0.3274047 | 0.50047527 | 0.65418758 | 0.51299096 | 0.99937574 |
| YY1 | 133.724515 | 0.02190266 | 0.39207407 | 0.05586359 | 0.95545048 | 0.99937574 |
| SLC25A29 | 157.280906 | 0.39897178 | 0.44246325 | 0.90170604 | 0.36721304 | 0.99937574 |
| WARS | 50.4009891 | -0.4092339 | 0.55839045 | -0.7328812 | 0.4636309 | 0.99937574 |
| WDR25 | 26.7695972 | -0.829796 | 0.55688943 | -1.4900551 | 0.13620974 | 0.99937574 |
| PPP2R5C | 169.1322 | 0.23349076 | 0.27861434 | 0.83804288 | 0.40200662 | 0.99937574 |
| DYNC1H1 | 132.168087 | -0.3813593 | 0.3670285 | -1.0390454 | 0.29878361 | 0.99937574 |
| HSP90AA1 | 478.738885 | 0.04730944 | 0.38532485 | 0.12277805 | 0.90228285 | 0.99937574 |
| WDR20 | 136.121182 | 0.07504954 | 0.24740693 | 0.30334452 | 0.76162732 | 0.99937574 |
| MOK | 29.7950919 | -0.1217975 | 0.46973961 | -0.2592873 | 0.79541361 | 0.99937574 |
| ZNF839 | 23.6262917 | 0.15304858 | 0.4282544 | 0.35737771 | 0.72080906 | 0.99937574 |
| CINP | 20.9820065 | -0.0189936 | 0.48677872 | -0.0390189 | 0.96887529 | 0.99937574 |
| TECPR2 | 47.1800931 | -0.2589162 | 0.46241698 | -0.5599194 | 0.57553444 | 0.99937574 |
| ANKRD9 | 84.4504265 | -0.1249781 | 0.34999316 | -0.3570873 | 0.72102648 | 0.99937574 |
| RCOR1 | 107.360455 | 0.06846493 | 0.26715845 | 0.25627089 | 0.79774168 | 0.99937574 |
| TRAF3 | 93.4571791 | 0.04992516 | 0.43363312 | 0.11513226 | 0.90834029 | 0.99937574 |
| AMN | 48.4012983 | 0.51341199 | 0.69361434 | 0.74019807 | 0.45917982 | 0.99937574 |
| CDC42BPB | 281.097769 | -0.1772336 | 0.31625519 | -0.5604133 | 0.57519757 | 0.99937574 |
| EIF5 | 102.317889 | -0.1100172 | 0.48894266 | -0.2250105 | 0.82197111 | 0.99937574 |
| MARK3 | 372.520879 | 0.20902705 | 0.29770933 | 0.70211792 | 0.48260563 | 0.99937574 |
| CKB | 602.7937 | 1.47615716 | 0.44329962 | 3.32993105 | 0.00086867 | 0.29012345 |
| BAG5 | 63.6803304 | 0.38181222 | 0.31998054 | 1.19323574 | 0.23277706 | 0.99937574 |
| KLC1 | 107.865414 | 0.14144535 | 0.24521534 | 0.57682098 | 0.5640604 | 0.99937574 |
| AL049840.1 | 12.3282968 | 0.09090499 | 0.73318317 | 0.12398674 | 0.90132577 | 0.99937574 |
| ZFYVE21 | 55.6300164 | 0.37342795 | 0.39889127 | 0.93616476 | 0.34918836 | 0.99937574 |
| PPP1R13B | 160.760915 | -0.1639541 | 0.26259003 | -0.6243729 | 0.53238273 | 0.99937574 |
| ATP5MPL | 83.9983037 | 0.25050635 | 0.42905923 | 0.58385027 | 0.55932105 | 0.99937574 |
| TDRD9 | 15.767528 | -0.6907195 | 0.75437339 | -0.9156202 | 0.35986615 | 0.99937574 |
| INF2 | 14.2926931 | -0.5131115 | 0.60017597 | -0.8549351 | 0.39258705 | 0.99937574 |
| SIVA1 | 33.3545936 | -0.289718 | 0.4363451 | -0.6639652 | 0.50671259 | 0.99937574 |
| AKT1 | 45.6929703 | -0.1103381 | 0.40743495 | -0.2708116 | 0.7865359 | 0.99937574 |
| CEP170B | 21.466557 | -0.4942947 | 0.48314446 | -1.0230785 | 0.30627074 | 0.99937574 |
| NUDT14 | 23.6900131 | 0.15866395 | 0.47517571 | 0.33390585 | 0.73845061 | 0.99937574 |
| BRF1 | 30.4766587 | 0.11834615 | 0.45787996 | 0.25846546 | 0.79604771 | 0.99937574 |
| BTBD6 | 16.6700846 | -0.1273311 | 0.49829881 | -0.2555317 | 0.79831249 | 0.99937574 |
| PACS2 | 48.5159962 | -0.3601514 | 0.43968706 | -0.8191086 | 0.41272448 | 0.99937574 |

| | | | | | | |
|---|---|---|---|---|---|---|
| MTA1 | 29.336592 | -0.2553712 | 0.45087749 | -0.566387 | 0.57113071 | 0.99937574 |
| NIPA1 | 70.0821195 | 0.09304937 | 0.32835131 | 0.28338359 | 0.77688281 | 0.99937574 |
| NIPA2 | 52.3449674 | 0.32288861 | 0.39935001 | 0.80853537 | 0.41878245 | 0.99937574 |
| CYFIP1 | 84.3774026 | -0.0127139 | 0.31385057 | -0.0405095 | 0.96768693 | 0.99937574 |
| TUBGCP5 | 23.5202852 | 0.12570218 | 0.49215857 | 0.25540993 | 0.79840651 | 0.99937574 |
| PWRN1 | 94.0604768 | -2.5643595 | 1.77903947 | -1.4414293 | 0.14946345 | 0.99937574 |
| AC124312.1 | 208.088048 | 0.21120821 | 0.20048654 | 1.05347826 | 2.92E-01 | 0.99937574 |
| PWAR6 | 45.6026532 | 0.60714615 | 0.34957172 | 1.7368286 | 0.08241743 | 0.99937574 |
| UBE3A | 185.485897 | -0.0895092 | 0.22096552 | -0.4050821 | 0.68541719 | 0.99937574 |
| HERC2 | 194.854574 | 0.08159546 | 0.29283519 | 0.27863954 | 0.78052147 | 0.99937574 |
| TJP1 | 188.157209 | -0.0934257 | 0.25305208 | -0.3691957 | 0.71198187 | 0.99937574 |
| FAN1 | 42.4465674 | 0.39504693 | 0.42251637 | 0.93498609 | 0.34979547 | 0.99937574 |
| MTMR10 | 168.303097 | 0.04337702 | 0.30177157 | 0.14374125 | 0.88570479 | 0.99937574 |
| KLF13 | 64.6977382 | -0.1580311 | 0.56817389 | -0.2781386 | 0.78090597 | 0.99937574 |
| OTUD7A | 49.4716498 | 0.42512219 | 0.35458876 | 1.19891612 | 0.23056056 | 0.99937574 |
| LINC02256 | 151.638756 | 0.39007235 | 0.38930655 | 1.00196707 | 0.3163595 | 0.99937574 |
| AC123768.3 | 20.1031552 | 1.25021659 | 0.59596706 | 2.09779479 | 0.03592328 | 0.99937574 |
| GOLGA8N | 16.034122 | 0.76003164 | 0.47907479 | 1.58645718 | 0.11263564 | 0.99937574 |
| FMN1 | 141.15539 | 0.07002293 | 0.54464729 | 0.12856565 | 0.89770135 | 0.99937574 |
| AVEN | 58.3637924 | -0.6669516 | 0.40378905 | -1.6517328 | 0.09858903 | 0.99937574 |
| EMC7 | 47.2080414 | -0.1158214 | 0.36976678 | -0.3132283 | 0.75410725 | 0.99937574 |
| KATNBL1 | 143.102747 | -0.1821982 | 0.22745739 | -0.8010214 | 0.42311925 | 0.99937574 |
| EMC4 | 18.2860319 | -0.4933612 | 0.53427681 | -0.9234186 | 0.35578909 | 0.99937574 |
| SLC12A6 | 81.3545973 | 0.2440953 | 0.32349543 | 0.75455563 | 0.45051565 | 0.99937574 |
| NOP10 | 56.3626004 | -0.4534855 | 0.61995505 | -0.7314812 | 0.46448527 | 0.99937574 |
| LPCAT4 | 16.3284842 | 0.50873032 | 0.58498268 | 0.86965023 | 0.38449158 | 0.99937574 |
| GOLGA8A | 41.4084376 | -0.532973 | 0.40204623 | -1.3256511 | 0.18495529 | 0.99937574 |
| GOLGA8B | 56.9146658 | 0.79896409 | 0.42716557 | 1.87038505 | 0.06143037 | 0.99937574 |
| AQR | 83.1032084 | -0.0801626 | 0.26384398 | -0.3038257 | 0.76126067 | 0.99937574 |
| ZNF770 | 43.9790676 | 0.06673057 | 0.34927852 | 0.19105259 | 0.84848439 | 0.99937574 |
| AC021231.1 | 72.5860239 | 0.76732167 | 0.43064659 | 1.78178971 | 0.07478353 | 0.99937574 |
| DPH6 | 263.957518 | 0.1698672 | 0.33296648 | 0.51016307 | 0.60993723 | 0.99937574 |
| DPH6-DT | 34.633775 | 0.27649583 | 0.63464481 | 0.43567019 | 0.66307603 | 0.99937574 |
| C15orf41 | 44.5577888 | 0.28309663 | 0.43202789 | 0.65527397 | 0.51229138 | 0.99937574 |
| SPRED1 | 73.8263829 | -0.0597548 | 0.49661157 | -0.1203251 | 0.90422565 | 0.99937574 |
| FAM98B | 36.7405538 | 0.29293461 | 0.32051125 | 0.91396048 | 0.36073761 | 0.99937574 |
| RASGRP1 | 24.2157084 | 0.67043503 | 0.58606278 | 1.14396451 | 0.25263836 | 0.99937574 |
| C15orf53 | 38.2977741 | 0.97504582 | 0.95383823 | 1.02223395 | 0.30667018 | 0.99937574 |
| AC013652.1 | 56.5400224 | 0.30601944 | 0.83464877 | 0.36664457 | 0.71388416 | 0.99937574 |
| THBS1 | 17.3586275 | -1.4614749 | 0.65825692 | -2.2202195 | 0.02640387 | 0.97242612 |
| EIF2AK4 | 54.9059777 | 0.27778012 | 0.34169236 | 0.81295386 | 0.41624451 | 0.99937574 |
| SRP14 | 269.374736 | 0.09490695 | 0.51414237 | 0.18459275 | 0.85354847 | 0.99937574 |
| SRP14-AS1 | 19.8388632 | 0.55700464 | 0.69698851 | 0.79915899 | 0.42419823 | 0.99937574 |
| IVD | 73.6394053 | 0.66860039 | 0.31956975 | 2.09218922 | 0.03642159 | 0.99937574 |
| BAHD1 | 28.8752305 | -0.3573182 | 0.46100805 | -0.7750801 | 0.43829233 | 0.99937574 |
| CCDC32 | 44.8840137 | -0.2899381 | 0.39244149 | -0.738806 | 0.4600248 | 0.99937574 |
| RMDN3 | 29.5916947 | -0.3533525 | 0.51605 | -0.6847254 | 0.49351723 | 0.99937574 |
| GCHFR | 18.8766537 | 0.42293945 | 0.55313376 | 0.76462419 | 0.44449536 | 0.99937574 |
| DNAJC17 | 37.9125929 | 0.25164443 | 0.36534188 | 0.68879164 | 0.49095439 | 0.99937574 |
| ZFYVE19 | 15.8441588 | -0.0980648 | 0.48250241 | -0.2032421 | 0.8389458 | 0.99937574 |
| SPINT1-AS1 | 44.343316 | 0.42631405 | 0.43934998 | 0.97032905 | 0.3318825 | 0.99937574 |
| SPINT1 | 65.020755 | -0.0607961 | 0.32610317 | -0.186432 | 0.85210601 | 0.99937574 |
| VPS18 | 13.0154743 | -0.6615896 | 0.67110792 | -0.985817 | 0.32422294 | 0.99937574 |
| INO80 | 110.19159 | -0.6169847 | 0.3388432 | -1.8208562 | 0.06862872 | 0.99937574 |

| | | | | | | |
|---|---|---|---|---|---|---|
| CHP1 | 132.325914 | 0.32684006 | 0.37644073 | 0.86823777 | 0.38526418 | 0.99937574 |
| NDUFAF1 | 16.2825715 | -0.2179182 | 0.46142604 | -0.4722711 | 0.63673331 | 0.99937574 |
| RTF1 | 73.5298935 | -0.141831 | 0.46119794 | -0.3075274 | 0.75844194 | 0.99937574 |
| RPAP1 | 18.3077629 | 0.33400803 | 0.4635725 | 0.72050872 | 0.47121184 | 0.99937574 |
| MGA | 152.667498 | -0.0172562 | 0.29550072 | -0.0583963 | 0.95343293 | 0.99937574 |
| MAPKBP1 | 62.7035742 | 0.33663165 | 0.34053467 | 0.98853855 | 0.32288896 | 0.99937574 |
| EHD4 | 51.6000819 | -0.5332882 | 0.4159373 | -1.2821361 | 0.19979492 | 0.99937574 |
| PLA2G4F | 29.7464053 | 1.16551131 | 0.76807494 | 1.51744478 | 0.12915442 | 0.99937574 |
| VPS39 | 58.4775476 | -0.4369861 | 0.31321846 | -1.3951478 | 0.16297126 | 0.99937574 |
| TMEM87A | 196.201679 | -0.1249945 | 0.27678348 | -0.4515966 | 0.65155963 | 0.99937574 |
| GANC | 36.2257746 | 0.32799182 | 0.47961556 | 0.68386402 | 0.49406103 | 0.99937574 |
| ZNF106 | 92.2337992 | 0.09123991 | 0.25404076 | 0.35915459 | 0.71947944 | 0.99937574 |
| SNAP23 | 60.4816641 | -0.1015113 | 0.30907658 | -0.3284341 | 0.74258347 | 0.99937574 |
| LRRC57 | 19.9683259 | -0.0756257 | 0.45219565 | -0.167241 | 0.8671804 | 0.99937574 |
| HAUS2 | 17.6258132 | -0.2713865 | 0.52948642 | -0.5125467 | 0.60826842 | 0.99937574 |
| STARD9 | 27.0737757 | -1.1782129 | 0.68236185 | -1.7266687 | 0.08422717 | 0.99937574 |
| TTBK2 | 123.974806 | -0.1136264 | 0.2810814 | -0.4042473 | 0.68603088 | 0.99937574 |
| UBR1 | 159.807763 | -0.120863 | 0.27369353 | -0.4415999 | 0.65877877 | 0.99937574 |
| AC068724.3 | 35.3957023 | -0.1727209 | 0.55782636 | -0.309632 | 0.75684078 | 0.99937574 |
| TMEM62 | 49.4030476 | 0.08414774 | 0.37785715 | 0.22269723 | 0.82377116 | 0.99937574 |
| CCNDBP1 | 49.118813 | 0.23520383 | 0.38822386 | 0.60584589 | 0.54461709 | 0.99937574 |
| ADAL | 30.7326042 | 0.30400867 | 0.45779318 | 0.66407426 | 0.50664278 | 0.99937574 |
| ZSCAN29 | 13.7397354 | 0.32057462 | 0.51652319 | 0.62063936 | 0.53483694 | 0.99937574 |
| TUBGCP4 | 29.1069865 | -0.4364184 | 0.4241865 | -1.0288361 | 0.30355672 | 0.99937574 |
| TP53BP1 | 130.525372 | -0.0109179 | 0.25688875 | -0.0425004 | 0.96609978 | 0.99937574 |
| PPIP5K1 | 58.0991518 | -0.1602991 | 0.32670947 | -0.4906472 | 0.623676 | 0.99937574 |
| CATSPER2 | 60.0873366 | -0.6162728 | 0.56973404 | -1.081685 | 0.27939251 | 0.99937574 |
| PDIA3 | 170.743851 | 0.28694442 | 0.33733518 | 0.8506211 | 0.39497986 | 0.99937574 |
| SERF2 | 356.173959 | 0.09701546 | 0.41407188 | 0.23429618 | 0.81475505 | 0.99937574 |
| MFAP1 | 23.6710973 | 0.06776856 | 0.43059929 | 0.15738196 | 0.87494383 | 0.99937574 |
| CASC4 | 259.45835 | 0.06649202 | 0.29675426 | 0.22406425 | 0.82270731 | 0.99937574 |
| AC025043.1 | 13.6386342 | 1.36255987 | 0.82309406 | 1.65541209 | 0.09784093 | 0.99937574 |
| CTDSPL2 | 120.406924 | -0.1264044 | 0.28821007 | -0.4385842 | 0.66096283 | 0.99937574 |
| EIF3J-DT | 38.3289374 | -0.2206663 | 0.40037161 | -0.5511537 | 0.58152832 | 0.99937574 |
| EIF3J | 61.7483433 | -0.1883183 | 0.32188729 | -0.5850442 | 0.55851797 | 0.99937574 |
| SPG11 | 222.661712 | 0.08354287 | 0.24957496 | 0.3347406 | 0.73782077 | 0.99937574 |
| B2M | 1841.47693 | 0.11923729 | 0.49128727 | 0.2427038 | 0.80823486 | 0.99937574 |
| TRIM69 | 17.0766691 | -0.4522369 | 0.60623778 | -0.7459728 | 0.45568383 | 0.99937574 |
| SORD | 27.2018945 | -0.2415949 | 0.50503866 | -0.4783691 | 0.63238751 | 0.99937574 |
| DUOX1 | 21.1619356 | -0.5919344 | 0.5678332 | -1.0424442 | 0.29720578 | 0.99937574 |
| SPATA5L1 | 18.6218491 | -0.3951095 | 0.47763386 | -0.8272225 | 0.40811096 | 0.99937574 |
| SLC30A4 | 11.6334581 | 1.04716003 | 0.554149 | 1.88967232 | 0.0588018 | 0.99937574 |
| SQOR | 29.5543162 | 0.13121239 | 0.40855225 | 0.32116428 | 0.7480859 | 0.99937574 |
| SEMA6D | 43.1274127 | -0.969398 | 0.75856261 | -1.2779407 | 0.20127034 | 0.99937574 |
| MYEF2 | 121.771111 | 0.0713506 | 0.38845832 | 0.18367633 | 0.85426738 | 0.99937574 |
| SLC12A1 | 28.6379066 | 0.29400882 | 0.97332111 | 0.30206765 | 0.76260049 | 0.99937574 |
| DUT | 44.2123248 | 0.02769824 | 0.44095565 | 0.06281412 | 0.94991452 | 0.99937574 |
| FBN1 | 15.5856859 | 1.15847543 | 0.81056096 | 1.42922678 | 0.15293906 | 0.99937574 |
| CEP152 | 111.332439 | 0.2117854 | 0.57731989 | 0.36684237 | 0.7137366 | 0.99937574 |
| EID1 | 74.2748145 | 0.27259299 | 0.47990607 | 0.5680132 | 0.570026 | 0.99937574 |
| SECISBP2L | 219.622583 | -0.266206 | 0.29918799 | -0.8897616 | 0.37359391 | 0.99937574 |
| COPS2 | 43.4208776 | 0.02439876 | 0.33471066 | 0.07289507 | 0.94188962 | 0.99937574 |
| GALK2 | 79.3443472 | 0.32766702 | 0.30440377 | 1.07642235 | 0.28173841 | 0.99937574 |
| FAM227B | 41.3653581 | 0.5841416 | 0.4628729 | 1.26199138 | 0.20695189 | 0.99937574 |

| | | | | | | |
|---|---|---|---|---|---|---|
| FGF7 | 46.5206862 | 1.90317854 | 0.6300598 | 3.02063163 | 0.00252248 | 0.48965449 |
| DTWD1 | 47.66709 | -0.0102344 | 0.37101188 | -0.0275851 | 0.97799306 | 0.99937574 |
| GABPB1 | 62.2006573 | -0.0722738 | 0.37592432 | -0.1922564 | 0.84754141 | 0.99937574 |
| GABPB1-AS1 | 58.6678782 | 0.40088248 | 0.57232018 | 0.70045141 | 0.48364544 | 0.99937574 |
| USP8 | 119.496227 | 0.07722837 | 0.28216242 | 0.27370183 | 0.78431377 | 0.99937574 |
| TRPM7 | 196.564632 | -0.2961347 | 0.21768543 | -1.3603791 | 0.17370998 | 0.99937574 |
| SPPL2A | 182.326662 | -0.229464 | 0.24253766 | -0.9460964 | 0.3440994 | 0.99937574 |
| AC021752.1 | 21.1245502 | 0.05701513 | 0.81318774 | 0.07011312 | 0.94410362 | 0.99937574 |
| AP4E1 | 59.6657831 | -0.1872696 | 0.3348215 | -0.5593118 | 0.57594892 | 0.99937574 |
| MIR4713HG | 23.7085059 | 1.50135807 | 0.95384304 | 1.57400958 | 0.11548524 | 0.99937574 |
| GLDN | 23.813368 | 0.4913626 | 0.94830517 | 0.51814818 | 0.60435488 | 0.99937574 |
| DMXL2 | 76.0323104 | -0.0531399 | 0.4077218 | -0.1303338 | 0.89630232 | 0.99937574 |
| LYSMD2 | 13.5593434 | -0.8257552 | 0.61484341 | -1.3430334 | 0.17926117 | 0.99937574 |
| TMOD3 | 155.67113 | -0.0570831 | 0.26878451 | -0.2123748 | 0.83181466 | 0.99937574 |
| LEO1 | 18.272373 | -0.182682 | 0.49881813 | -0.3662297 | 0.71419369 | 0.99937574 |
| MAPK6 | 88.8496817 | -0.6347289 | 0.24748513 | -2.5647155 | 0.01032604 | 0.76274616 |
| GNB5 | 63.0126398 | 0.05546527 | 0.3281835 | 0.16900689 | 0.86579122 | 0.99937574 |
| MYO5C | 86.0630894 | 0.11738654 | 0.44216309 | 0.26548245 | 0.79063783 | 0.99937574 |
| MYO5A | 105.489213 | -0.3872608 | 0.26821997 | -1.4438178 | 0.14879023 | 0.99937574 |
| ARPP19 | 133.025563 | 0.06013287 | 0.3010627 | 0.19973539 | 0.84168754 | 0.99937574 |
| FAM214A | 154.457113 | -0.0904779 | 0.30351299 | -0.2981023 | 0.76562508 | 0.99937574 |
| WDR72 | 718.451508 | 0.11855859 | 0.43440085 | 0.27292439 | 0.78491133 | 0.99937574 |
| UNC13C | 25.9091533 | -0.9244629 | 1.83436653 | -0.5039685 | 0.61428355 | 0.99937574 |
| RSL24D1 | 32.9590691 | 0.11043626 | 0.43860927 | 0.25178733 | 0.80120545 | 0.99937574 |
| RAB27A | 111.467942 | -0.1693966 | 0.34589708 | -0.4897311 | 0.6243242 | 0.99937574 |
| PIGB | 51.7020516 | 0.32393817 | 0.4012557 | 0.80731106 | 0.41948729 | 0.99937574 |
| CCPG1 | 296.11298 | -0.2597994 | 0.43514056 | -0.597047 | 0.550476 | 0.99937574 |
| C15orf65 | 17.9163263 | 0.11313932 | 0.87960052 | 0.1286258 | 0.89765375 | 0.99937574 |
| DNAAF4 | 19.7740082 | 0.40355978 | 0.60196787 | 0.67040086 | 0.50260229 | 0.99937574 |
| RFX7 | 109.786801 | 0.19430049 | 0.30333601 | 0.64054542 | 0.52181807 | 0.99937574 |
| TEX9 | 39.4853315 | 0.10920586 | 0.44058247 | 0.24786701 | 0.8042373 | 0.99937574 |
| ZNF280D | 71.9508807 | 0.69217759 | 0.39246815 | 1.76365289 | 0.07779044 | 0.99937574 |
| AC090517.4 | 13.203916 | 0.0290388 | 0.58285493 | 0.04982166 | 0.96026451 | 0.99937574 |
| TCF12 | 667.446376 | -0.0695743 | 0.31580252 | -0.2203097 | 0.82562999 | 0.99937574 |
| CGNL1 | 885.857757 | 0.66865349 | 0.45148666 | 1.48100386 | 0.13860554 | 0.99937574 |
| POLR2M | 23.309976 | -0.3676772 | 0.40965247 | -0.8975346 | 0.36943374 | 0.99937574 |
| ALDH1A2 | 91.4341154 | -0.6239176 | 0.64612946 | -0.9656232 | 3.34E-01 | 0.99937574 |
| LIPC | 25.4059797 | -0.3027629 | 0.66478875 | -0.4554272 | 0.64880195 | 0.99937574 |
| ADAM10 | 207.355836 | -0.00585 | 0.23544817 | -0.0248462 | 0.98017764 | 0.99937574 |
| MINDY2 | 223.990757 | 0.19762701 | 0.21741768 | 0.90897396 | 0.36336387 | 0.99937574 |
| RNF111 | 189.622358 | 0.05967476 | 0.28945983 | 0.20615905 | 0.83666668 | 0.99937574 |
| SLTM | 232.216946 | -0.0959168 | 0.27879095 | -0.3440456 | 0.73081198 | 0.99937574 |
| MYO1E | 185.506826 | -0.467431 | 0.32107309 | -1.4558399 | 0.14543689 | 0.99937574 |
| GTF2A2 | 47.3314526 | 0.54956992 | 0.46515906 | 1.18146667 | 0.23741739 | 0.99937574 |
| BNIP2 | 66.1171195 | -0.1113131 | 0.3249647 | -0.3425392 | 0.73194518 | 0.99937574 |
| ANXA2 | 64.8446153 | -2.0148614 | 0.6557582 | -3.0725676 | 0.00212226 | 0.45583974 |
| ICE2 | 81.4642101 | 0.11251431 | 0.39105454 | 0.28772024 | 0.77356089 | 0.99937574 |
| RORA-AS1 | 59.2194672 | 0.5566557 | 0.63805627 | 0.87242416 | 0.38297702 | 0.99937574 |
| RORA | 1190.23624 | 0.29890658 | 0.64195794 | 0.46561708 | 0.64148961 | 0.99937574 |
| AC012404.1 | 43.3849886 | -0.2290428 | 0.85363121 | -0.2683159 | 0.78845616 | 0.99937574 |
| AC009554.1 | 45.1447924 | 0.08556994 | 0.63743096 | 0.1342419 | 0.89321129 | 0.99937574 |
| VPS13C | 315.731452 | -0.1987871 | 0.23416017 | -0.8489364 | 0.39591667 | 0.99937574 |
| TLN2 | 64.6642396 | -0.1209371 | 0.63011105 | -0.1919298 | 0.84779721 | 0.99937574 |
| TPM1 | 200.864428 | -0.047009 | 0.4221837 | -0.1113473 | 0.91134093 | 0.99937574 |

| | | | | | | |
|---|---|---|---|---|---|---|
| LACTB | 52.4929707 | 0.04725408 | 0.49914499 | 0.09467004 | 0.92457691 | 0.99937574 |
| RPS27L | 66.9710273 | 0.18295161 | 0.44205932 | 0.41386212 | 0.67897509 | 0.99937574 |
| RAB8B | 63.6412517 | -0.3590162 | 0.35289381 | -1.017349 | 0.30898742 | 0.99937574 |
| APH1B | 60.1339028 | -0.2182362 | 0.32188423 | -0.6779958 | 4.98E-01 | 0.99937574 |
| CA12 | 1926.50663 | -0.196535 | 0.41563838 | -0.4728509 | 0.63631958 | 0.99937574 |
| LINC02568 | 45.1268972 | 0.25475083 | 0.54809416 | 0.46479392 | 6.42E-01 | 0.99937574 |
| USP3 | 224.848526 | -0.4281606 | 0.24163314 | -1.7719448 | 0.07640373 | 0.99937574 |
| USP3-AS1 | 44.5188783 | 0.1677865 | 0.59338809 | 0.28276014 | 0.77736071 | 0.99937574 |
| HERC1 | 231.201901 | -0.1210965 | 0.23001937 | -0.526462 | 0.59856722 | 0.99937574 |
| DAPK2 | 22.0647318 | -2.1270826 | 1.11932853 | -1.9003202 | 0.05739111 | 0.99937574 |
| FAM96A | 51.1380198 | -0.0122807 | 0.44092762 | -0.0278519 | 0.97778024 | 0.99937574 |
| SNX1 | 101.761357 | 0.06007872 | 0.24374489 | 0.24648198 | 0.80530915 | 0.99937574 |
| PPIB | 126.904071 | 0.2784475 | 0.40330029 | 0.69042226 | 0.48992868 | 0.99937574 |
| CSNK1G1 | 27.489487 | 0.87591818 | 0.38082441 | 2.30005787 | 0.02144494 | 0.96427895 |
| TRIP4 | 54.2170342 | 0.08285064 | 0.35682575 | 0.23218795 | 0.81639204 | 0.99937574 |
| ZNF609 | 271.575229 | -0.1021889 | 0.31860927 | -0.3207341 | 0.74841189 | 0.99937574 |
| OAZ2 | 61.1367442 | 0.11803534 | 0.38263018 | 0.30848415 | 0.75771396 | 0.99937574 |
| RBPMS2 | 20.8868443 | 0.42623326 | 0.52228555 | 0.81609238 | 0.4144473 | 0.99937574 |
| SPG21 | 52.4753023 | -0.6514775 | 0.39209464 | -1.6615312 | 0.09660681 | 0.99937574 |
| MTFMT | 33.7882297 | -0.1885045 | 0.37194346 | -0.5068096 | 0.61228841 | 0.99937574 |
| UBAP1L | 13.2543215 | 0.15247526 | 0.73034456 | 0.20877169 | 0.83462648 | 0.99937574 |
| PDCD7 | 30.3778328 | 0.27360375 | 0.37621932 | 0.72724535 | 0.46707567 | 0.99937574 |
| CLPX | 66.0179759 | -0.1160095 | 0.32011115 | -0.3624038 | 0.71705031 | 0.99937574 |
| PARP16 | 16.6180444 | 0.42475088 | 0.63229872 | 0.67175666 | 0.50173862 | 0.99937574 |
| DPP8 | 101.902063 | -0.1693343 | 0.26123666 | -0.6482027 | 0.51685388 | 0.99937574 |
| HACD3 | 124.99066 | 0.606118 | 0.30785613 | 1.96883526 | 0.04897201 | 0.99937574 |
| INTS14 | 33.4259952 | -0.2891357 | 0.34836334 | -0.8299831 | 0.40654835 | 0.99937574 |
| SLC24A1 | 14.0704425 | -0.0859128 | 0.57985172 | -0.1481633 | 0.88221388 | 0.99937574 |
| DENND4A | 362.399381 | 0.34346346 | 0.3442645 | 0.9976732 | 0.31843785 | 0.99937574 |
| RAB11A | 258.345643 | 0.46461186 | 0.49335894 | 0.94173193 | 0.3463299 | 0.99937574 |
| DIS3L | 27.1879265 | 0.87398413 | 0.47428138 | 1.84275445 | 0.06536487 | 0.99937574 |
| TIPIN | 23.4927232 | 0.39734727 | 0.5604088 | 0.70903111 | 0.47830517 | 0.99937574 |
| MAP2K1 | 142.400345 | -0.2190117 | 0.30344358 | -0.7217542 | 0.47044562 | 0.99937574 |
| SNAPC5 | 18.3766666 | 0.54090344 | 0.55854389 | 0.96841707 | 0.33283612 | 0.99937574 |
| RPL4 | 193.125417 | -0.0466429 | 0.52737513 | -0.0884436 | 0.92952412 | 0.99937574 |
| ZWILCH | 14.1710837 | -0.6392048 | 0.56865782 | -1.1240587 | 0.26098811 | 0.99937574 |
| SMAD6 | 19.3984932 | 0.83317532 | 0.57064933 | 1.46004783 | 0.14427693 | 0.99937574 |
| SMAD3 | 386.660979 | -0.6490575 | 0.42014454 | -1.5448434 | 1.22E-01 | 0.99937574 |
| AAGAB | 113.695175 | -0.0690455 | 0.28747727 | -0.2401774 | 0.81019272 | 0.99937574 |
| IQCH | 72.4329935 | 0.31524474 | 0.55434007 | 0.56868474 | 0.56957011 | 0.99937574 |
| IQCH-AS1 | 56.614528 | 0.04016453 | 0.4650689 | 0.08636253 | 0.93117823 | 0.99937574 |
| C15orf61 | 18.0139305 | 0.35285782 | 0.51496117 | 0.68521248 | 0.49320985 | 0.99937574 |
| MAP2K5 | 194.662755 | 0.13075435 | 0.51620334 | 0.25330009 | 0.80003632 | 0.99937574 |
| PIAS1 | 295.846678 | -0.1791832 | 0.21594843 | -0.82975 | 0.40668017 | 0.99937574 |
| FEM1B | 45.6728279 | 0.06393357 | 0.3496077 | 0.18287231 | 0.85489821 | 0.99937574 |
| CORO2B | 26.8189018 | -0.7330939 | 0.50273463 | -1.4582125 | 0.14478197 | 0.99937574 |
| ANP32A | 133.026482 | 0.24536179 | 0.35460806 | 0.69192391 | 0.48898512 | 0.99937574 |
| SPESP1 | 29.9431709 | 0.95478503 | 0.54048933 | 1.76651968 | 0.0773087 | 0.99937574 |
| GLCE | 113.919986 | 0.79268077 | 0.38386733 | 2.06498627 | 0.0389243 | 0.99937574 |
| AC026992.1 | 15.1288441 | 1.85221364 | 0.68929654 | 2.68710711 | 0.00720738 | 0.74742128 |
| PAQR5 | 205.667865 | -0.2526534 | 0.36036036 | -0.7011132 | 0.48323238 | 0.99937574 |
| KIF23 | 22.5644413 | -0.0573675 | 0.95015603 | -0.0603769 | 0.95185545 | 0.99937574 |
| RPLP1 | 362.420418 | -0.2159272 | 0.50763486 | -0.4253594 | 0.67057472 | 0.99937574 |
| TLE3 | 61.5758628 | -0.3882123 | 0.37224097 | -1.0429058 | 0.29699193 | 0.99937574 |

| | | | | | | |
|---|---|---|---|---|---|---|
| UACA | 128.147296 | -0.5785308 | 0.38410344 | -1.506185 | 0.1320197 | 0.99937574 |
| LRRC49 | 34.3349958 | -0.9191346 | 0.54138676 | -1.6977412 | 0.08955663 | 0.99937574 |
| THSD4 | 211.176288 | 0.51540328 | 0.70380502 | 0.73230975 | 0.46397953 | 0.99937574 |
| THSD4-AS1 | 66.961276 | 0.05593406 | 0.36116578 | 0.15487087 | 0.87692312 | 0.99937574 |
| MYO9A | 536.851474 | 0.31069459 | 0.20158266 | 1.54127637 | 0.12324954 | 0.99937574 |
| AC022872.1 | 16.8202253 | 0.46238525 | 0.74764941 | 0.61845197 | 0.53627745 | 0.99937574 |
| SENP8 | 14.9624094 | 0.54595297 | 0.51179573 | 1.06674 | 0.28608926 | 0.99937574 |
| PKM | 463.72558 | 0.09740603 | 0.26303171 | 0.37032047 | 0.71114372 | 0.99937574 |
| PARP6 | 37.7694336 | -0.2552165 | 0.41305277 | -0.6178787 | 0.53665528 | 0.99937574 |
| AC009690.2 | 28.1763779 | 0.75770611 | 0.75621949 | 1.00196586 | 0.31636008 | 0.99937574 |
| AC009690.1 | 99.2058552 | -0.2302189 | 0.51716683 | -0.4451541 | 0.65620843 | 0.99937574 |
| ARIH1 | 370.297567 | -0.1663606 | 0.23002049 | -0.7232426 | 0.46953086 | 0.99937574 |
| BBS4 | 49.9582292 | 0.34737135 | 0.4965479 | 0.69957268 | 0.48419421 | 0.99937574 |
| ADPGK | 46.7618283 | -0.4300083 | 0.35093456 | -1.2253233 | 0.22045345 | 0.99937574 |
| NEO1 | 130.481245 | 0.23428998 | 0.43077952 | 0.54387448 | 0.58652785 | 0.99937574 |
| NPTN | 225.129825 | -0.1761693 | 0.2475717 | -0.711589 | 0.47671932 | 0.99937574 |
| CD276 | 16.592804 | 0.28187269 | 0.57065153 | 0.49394888 | 0.62134228 | 0.99937574 |
| C15orf59 | 23.2893895 | 1.05493433 | 0.6201886 | 1.70098956 | 0.08894495 | 0.99937574 |
| C15orf59-AS | 15.896818 | 2.5771673 | 0.6489776 | 3.97111902 | 7.15E-05 | 0.08437103 |
| LOXL1 | 16.8134453 | 0.27295287 | 0.53001598 | 0.51498989 | 0.60656008 | 0.99937574 |
| PML | 35.5336102 | -0.4101022 | 0.43522109 | -0.9422848 | 0.34604684 | 0.99937574 |
| CLK3 | 27.579024 | 0.30825783 | 0.3880437 | 0.79438946 | 0.42696874 | 0.99937574 |
| EDC3 | 40.1130028 | -0.0956484 | 0.36480347 | -0.2621916 | 0.79317375 | 0.99937574 |
| CSK | 14.1940044 | -1.0271031 | 0.53064365 | -1.9355798 | 0.05291919 | 0.99937574 |
| ULK3 | 16.5068386 | 0.07908394 | 0.58929821 | 0.13420021 | 0.89324426 | 0.99937574 |
| SCAMP2 | 56.3817055 | 0.020514 | 0.29470496 | 0.06960862 | 0.94450518 | 0.99937574 |
| FAM219B | 26.5904038 | -0.2631129 | 0.379953 | -0.6924881 | 0.48863083 | 0.99937574 |
| COX5A | 130.827301 | 0.31434859 | 0.46766689 | 0.67216344 | 0.50147965 | 0.99937574 |
| SCAMP5 | 28.154374 | -1.6224023 | 0.88642021 | -1.8302858 | 0.06720721 | 0.99937574 |
| PPCDC | 31.6782256 | -0.7815913 | 0.52857738 | -1.4786696 | 0.13922863 | 0.99937574 |
| COMMD4 | 28.401538 | -0.5699675 | 0.3962386 | -1.4384452 | 0.15030778 | 0.99937574 |
| NEIL1 | 12.7194115 | -0.6743294 | 0.54717353 | -1.2323867 | 0.21780467 | 0.99937574 |
| MAN2C1 | 43.768949 | -0.2463988 | 0.38689556 | -0.6368614 | 0.52421516 | 0.99937574 |
| SIN3A | 81.4439718 | 0.15083075 | 0.30233318 | 0.49888916 | 0.61785747 | 0.99937574 |
| PTPN9 | 131.339028 | -0.3545144 | 0.36599734 | -0.9686257 | 0.33273198 | 0.99937574 |
| SNUPN | 17.1750679 | -0.1956011 | 0.47280521 | -0.4137035 | 0.6790913 | 0.99937574 |
| IMP3 | 23.3354187 | 0.23896921 | 0.6106372 | 0.39134401 | 0.69554297 | 0.99937574 |
| UBE2Q2 | 45.678353 | -0.6217241 | 0.46687833 | -1.3316619 | 0.18297131 | 0.99937574 |
| FBXO22 | 73.9151545 | 0.27613544 | 0.37718641 | 0.73209276 | 0.46411195 | 0.99937574 |
| NRG4 | 19.1345042 | -0.612457 | 0.88829809 | -0.6894724 | 0.49052602 | 0.99937574 |
| ETFA | 102.897627 | 0.12398157 | 0.26666553 | 0.46493287 | 0.64197953 | 0.99937574 |
| SCAPER | 357.566701 | 0.37058948 | 0.18511661 | 2.00192448 | 0.04529285 | 0.99937574 |
| RCN2 | 76.250579 | 0.04286647 | 0.34085461 | 0.12576175 | 0.89992052 | 0.99937574 |
| TSPAN3 | 108.963289 | -0.1157808 | 0.28311312 | -0.4089558 | 0.68257206 | 0.99937574 |
| PEAK1 | 86.5200889 | -0.3230507 | 0.41727041 | -0.7742 | 0.43881255 | 0.99937574 |
| HMG20A | 91.9020704 | 0.30001291 | 0.29270904 | 1.02495268 | 0.30538552 | 0.99937574 |
| LINGO1 | 663.789751 | -0.5899247 | 0.6561856 | -0.8990211 | 0.36864143 | 0.99937574 |
| TBC1D2B | 26.3262925 | 0.12311998 | 0.5375683 | 0.22903133 | 0.81884457 | 0.99937574 |
| CIB2 | 14.5979612 | 0.12863583 | 0.59534095 | 0.21607086 | 0.82893251 | 0.99937574 |
| IDH3A | 124.541918 | 0.08077412 | 0.24754642 | 0.3262989 | 0.74419822 | 0.99937574 |
| ACSBG1 | 17.0158503 | 2.14652532 | 0.87856048 | 2.44323 | 0.01455646 | 0.85908633 |
| DNAJA4 | 51.3150122 | -1.1717225 | 0.39957275 | -2.9324385 | 0.00336311 | 0.53256471 |
| WDR61 | 48.0843056 | -0.1583146 | 0.35477784 | -0.4462357 | 0.65542697 | 0.99937574 |
| IREB2 | 114.290834 | -0.0706219 | 0.33144378 | -0.2130734 | 0.8312697 | 0.99937574 |

| | | | | | | |
|---|---|---|---|---|---|---|
| PSMA4 | 59.2916308 | -0.1987861 | 0.36961646 | -0.5378172 | 0.59070324 | 0.99937574 |
| MORF4L1 | 148.671022 | -0.0074219 | 0.26138263 | -0.0283947 | 0.97734735 | 0.99937574 |
| CTSH | 68.2946793 | 0.429029 | 0.64088971 | 0.6694272 | 0.50322301 | 0.99937574 |
| TMED3 | 29.0029288 | -0.3461239 | 0.42774227 | -0.809188 | 0.41840701 | 0.99937574 |
| ZFAND6 | 287.602896 | -0.1916578 | 0.25350559 | -0.75603 | 0.44963121 | 0.99937574 |
| FAH | 18.6655182 | 0.46720628 | 0.58662553 | 0.79643018 | 0.42578204 | 0.99937574 |
| CTXND1 | 15.092294 | 0.20624172 | 0.97156411 | 0.21227804 | 0.83189013 | 0.99937574 |
| AC016705.2 | 258.570565 | 0.24863835 | 0.34824028 | 0.71398504 | 4.75E-01 | 0.99937574 |
| ARNT2 | 309.925997 | -0.1754067 | 0.53930586 | -0.3252452 | 0.74499547 | 0.99937574 |
| ABHD17C | 52.2851911 | 0.39275525 | 0.32657665 | 1.20264337 | 0.22911436 | 0.99937574 |
| AC027808.1 | 20.9000664 | 0.4960251 | 0.4835665 | 1.02576397 | 0.30500285 | 0.99937574 |
| MESD | 80.5522617 | 0.33959609 | 0.30034801 | 1.13067536 | 0.25819176 | 0.99937574 |
| AC104041.1 | 29.6515804 | -0.0169915 | 1.07159631 | -0.0158563 | 0.98734907 | 0.99937574 |
| EFL1 | 49.4625281 | -0.3818893 | 0.35745147 | -1.0683667 | 0.28535512 | 0.99937574 |
| RPS17 | 20.6542967 | 0.04591793 | 0.75807255 | 0.06057195 | 0.95170011 | 0.99937574 |
| WHAMM | 46.7472641 | -0.7279067 | 0.32649488 | -2.2294583 | 0.02578342 | 0.96546668 |
| HOMER2 | 16.4105499 | -0.9079197 | 0.61420637 | -1.4781997 | 0.13935434 | 0.99937574 |
| C15orf40 | 22.4615953 | -0.0579342 | 0.49715425 | -0.1165316 | 0.90723128 | 0.99937574 |
| BTBD1 | 97.4829568 | -0.0209368 | 0.29765409 | -0.0703394 | 0.94392349 | 0.99937574 |
| HDGFL3 | 79.5930504 | 0.05702886 | 0.39152952 | 0.1456566 | 0.88419248 | 0.99937574 |
| UBE2Q2L | 21.1760114 | -0.9043474 | 0.56913199 | -1.5889941 | 0.11206172 | 0.99937574 |
| GOLGA6L4 | 12.3746431 | -1.3595651 | 0.63218759 | -2.1505722 | 0.03150998 | 0.99049468 |
| WDR73 | 46.6247982 | 0.06761168 | 0.32659755 | 0.20701834 | 0.83599554 | 0.99937574 |
| SEC11A | 132.993822 | 0.25259361 | 0.27718536 | 0.91128049 | 0.3621476 | 0.99937574 |
| ZNF592 | 59.5951129 | -0.6633078 | 0.31992349 | -2.0733326 | 0.03814134 | 0.99937574 |
| ALPK3 | 114.833259 | -0.5029404 | 0.71192257 | -0.7064538 | 0.47990599 | 0.99937574 |
| PDE8A | 451.173041 | 0.25365994 | 0.35882656 | 0.70691517 | 0.4796192 | 0.99937574 |
| AKAP13 | 412.946967 | 0.03758984 | 0.30364539 | 0.12379519 | 0.90147744 | 0.99937574 |
| AC087286.2 | 12.7155354 | -0.0656719 | 0.65099768 | -0.1008789 | 0.9196466 | 0.99937574 |
| KLHL25 | 20.8436005 | -0.6957753 | 0.52967368 | -1.3135924 | 0.1889834 | 0.99937574 |
| MRPL46 | 16.8316027 | -0.2003225 | 0.52679319 | -0.3802679 | 0.70374658 | 0.99937574 |
| MRPS11 | 22.5732843 | 0.24857356 | 0.45713425 | 0.5437649 | 0.58660327 | 0.99937574 |
| DET1 | 28.5478354 | -0.1811297 | 0.55585569 | -0.3258573 | 0.74453229 | 0.99937574 |
| AEN | 12.2536286 | -0.1392151 | 0.61834679 | -0.2251408 | 0.82186976 | 0.99937574 |
| ISG20 | 23.6978756 | -3.4863477 | 1.04785185 | -3.327138 | 0.00087743 | 0.29012345 |
| ABHD2 | 240.922141 | 0.9615579 | 0.35326056 | 2.7219509 | 0.00648978 | 0.69675365 |
| POLG | 28.2445267 | 0.05643124 | 0.49520179 | 0.11395604 | 0.90927264 | 0.99937574 |
| MIR9-3HG | 10.6078501 | -0.3118958 | 0.72314909 | -0.4313022 | 0.66624866 | 0.99937574 |
| RHCG | 633.886486 | 1.09850945 | 0.7480237 | 1.46854899 | 0.14195516 | 0.99937574 |
| PEX11A | 12.8557823 | 0.46554136 | 0.72924391 | 0.6383891 | 0.52322042 | 0.99937574 |
| ANPEP | 13.0175001 | -0.0422976 | 0.81190998 | -0.0520965 | 0.95845183 | 0.99937574 |
| ZNF710 | 62.9699327 | -0.0211383 | 0.4110428 | -0.0514259 | 0.95898613 | 0.99937574 |
| ZNF710-AS1 | 65.5854826 | 0.72482752 | 0.68382335 | 1.05996311 | 0.28916138 | 0.99937574 |
| IDH2 | 361.17026 | 0.6900338 | 0.30522456 | 2.26074137 | 0.02377528 | 0.96546668 |
| IDH2-DT | 20.8699496 | 0.86352786 | 0.50401788 | 1.71328814 | 0.08665954 | 0.99937574 |
| SEMA4B | 163.430482 | -0.1211125 | 0.47796264 | -0.2533933 | 8.00E-01 | 0.99937574 |
| CIB1 | 168.844472 | 0.51213942 | 0.41755643 | 1.22651544 | 0.22000477 | 0.99937574 |
| IQGAP1 | 215.948598 | -0.2045796 | 0.30918146 | -0.6616813 | 0.50817549 | 0.99937574 |
| CRTC3 | 176.774503 | -0.0214328 | 0.34318347 | -0.0624528 | 0.95020221 | 0.99937574 |
| CRTC3-AS1 | 14.4331872 | -0.4198244 | 0.6794845 | -0.6178572 | 0.53666946 | 0.99937574 |
| FURIN | 40.8838065 | -0.7977964 | 0.46474347 | -1.7166381 | 0.08604531 | 0.99937574 |
| MAN2A2 | 16.5783609 | 0.03725854 | 0.50219816 | 0.07419091 | 0.94085848 | 0.99937574 |
| HDDC3 | 13.0365197 | 0.07693312 | 0.5504157 | 0.13977276 | 0.88883954 | 0.99937574 |
| UNC45A | 33.0368692 | -0.2163888 | 0.38231653 | -0.5659938 | 0.57139799 | 0.99937574 |

| | | | | | | |
|---|---|---|---|---|---|---|
| RCCD1 | 13.7447053 | 0.45416706 | 0.68886834 | 0.65929442 | 0.50970672 | 0.99937574 |
| SLCO3A1 | 99.0373731 | -0.8260951 | 0.77442 | -1.0667275 | 0.28609491 | 0.99937574 |
| FAM174B | 70.9408173 | -0.3945204 | 0.69017054 | -0.5716274 | 0.56757446 | 0.99937574 |
| AC013394.1 | 121.630945 | -0.2810677 | 0.23692274 | -1.1863265 | 0.23549337 | 0.99937574 |
| CHD2 | 219.300211 | 0.0258651 | 0.34366024 | 0.07526357 | 0.94000501 | 0.99937574 |
| MCTP2 | 30.5698076 | 0.77969439 | 0.57492865 | 1.35615852 | 0.17504874 | 0.99937574 |
| AC012409.2 | 27.7617364 | 0.92254418 | 0.69617674 | 1.32515801 | 0.18511875 | 0.99937574 |
| NR2F2-AS1 | 146.775457 | 0.94762927 | 0.47526195 | 1.9939094 | 0.04616196 | 0.99937574 |
| NR2F2 | 64.7731268 | 0.01063986 | 0.41526267 | 0.02562201 | 0.97955883 | 0.99937574 |
| LINC00923 | 31.3623349 | -1.0263953 | 1.13073341 | -0.9077253 | 0.36402337 | 0.99937574 |
| ARRDC4 | 82.6032396 | -0.0286707 | 0.48463817 | -0.0591589 | 0.95282554 | 0.99937574 |
| FAM169B | 17.9954433 | -0.1547217 | 1.06977529 | -0.1446301 | 8.85E-01 | 0.99937574 |
| IGF1R | 562.746645 | 0.18523543 | 0.28479744 | 0.65041116 | 0.51542667 | 0.99937574 |
| AC036108.1 | 16.1133809 | 0.19061677 | 0.61715056 | 0.3088659 | 0.75742353 | 0.99937574 |
| SYNM | 13.0845504 | 0.55416167 | 1.09309207 | 0.50696706 | 0.61217793 | 0.99937574 |
| TTC23 | 63.6358285 | 0.13528263 | 0.35760184 | 0.3783052 | 0.70520388 | 0.99937574 |
| LRRC28 | 85.1951665 | 0.09444757 | 0.35876446 | 0.26325788 | 0.79235182 | 0.99937574 |
| MEF2A | 323.006563 | 0.06414564 | 0.28640941 | 0.22396485 | 0.82278465 | 0.99937574 |
| AC090825.1 | 33.4401094 | 0.69309576 | 0.43123093 | 1.60724963 | 0.10799962 | 0.99937574 |
| LINS1 | 23.3209772 | -0.0731807 | 0.45829855 | -0.159679 | 0.87313393 | 0.99937574 |
| ASB7 | 32.7795729 | -0.1835726 | 0.36729247 | -0.4997995 | 0.61721629 | 0.99937574 |
| AC015712.1 | 10.9033513 | 0.16246285 | 0.65120222 | 0.24948141 | 0.80298842 | 0.99937574 |
| ALDH1A3 | 26.0199113 | 0.95555506 | 0.6437984 | 1.48424579 | 0.13774373 | 0.99937574 |
| CHSY1 | 86.3803853 | 0.08749205 | 0.40851908 | 0.21416883 | 0.8304154 | 0.99937574 |
| SELENOS | 64.1735533 | 0.10219918 | 0.50152294 | 0.20377769 | 0.83852723 | 0.99937574 |
| SNRPA1 | 47.1183103 | 0.39722735 | 0.30552862 | 1.30013138 | 0.19355594 | 0.99937574 |
| PCSK6 | 45.552674 | 0.51133565 | 0.77173134 | 0.66258247 | 0.507598 | 0.99937574 |
| TM2D3 | 62.391094 | 0.17848811 | 0.36384365 | 0.49056265 | 0.62373581 | 0.99937574 |
| TARSL2 | 76.7296877 | -0.2552007 | 0.26507409 | -0.9627522 | 0.33567187 | 0.99937574 |
| SNRNP25 | 16.404729 | -0.2050692 | 0.50448275 | -0.4064939 | 0.68437975 | 0.99937574 |
| MPG | 36.3289777 | -0.0037302 | 0.37642753 | -0.0099096 | 0.99209341 | 0.99937574 |
| NPRL3 | 58.0112938 | 0.21582713 | 0.3229401 | 0.66831939 | 0.50392974 | 0.99937574 |
| LUC7L | 121.10685 | 0.13499573 | 0.2375955 | 0.56817461 | 0.56991641 | 0.99937574 |
| FAM234A | 17.5191385 | -0.4968932 | 0.67125288 | -0.7402475 | 0.45914985 | 0.99937574 |
| AXIN1 | 28.3933993 | -0.3143963 | 0.44545399 | -0.7057885 | 0.48031968 | 0.99937574 |
| MRPL28 | 28.100281 | 0.37127093 | 0.42943448 | 0.86455779 | 0.38728155 | 0.99937574 |
| TMEM8A | 90.2467907 | -0.1098621 | 0.40423792 | -0.2717758 | 0.78579445 | 0.99937574 |
| NME4 | 14.4138015 | 1.09488803 | 0.64594947 | 1.69500569 | 0.09007434 | 0.99937574 |
| RAB11FIP3 | 121.764921 | 0.0669171 | 0.42219802 | 0.15849694 | 0.87406523 | 0.99937574 |
| CAPN15 | 32.2465721 | -0.1057135 | 0.36505234 | -0.2895844 | 0.77213422 | 0.99937574 |
| PIGQ | 44.8501545 | 0.61115925 | 0.48544788 | 1.25895956 | 0.20804494 | 0.99937574 |
| RAB40C | 79.8758905 | 0.55399866 | 0.34271075 | 1.61651963 | 0.10598201 | 0.99937574 |
| METTL26 | 11.8264788 | -0.0160935 | 0.66742786 | -0.0241127 | 0.9807627 | 0.99937574 |
| MCRIP2 | 97.7831157 | -0.2553138 | 0.43781015 | -0.5831609 | 0.559785 | 0.99937574 |
| RHOT2 | 40.1697338 | 0.10348393 | 0.33702078 | 0.30705505 | 0.75880147 | 0.99937574 |
| STUB1 | 61.6774709 | 0.24802706 | 0.49979681 | 0.49625578 | 0.61971396 | 0.99937574 |
| JMJD8 | 18.2007931 | 0.51222006 | 0.48249279 | 1.06161185 | 0.28841193 | 0.99937574 |
| METRN | 35.0787198 | 0.89092424 | 0.3945318 | 2.25818108 | 0.02393437 | 0.96546668 |
| FAM173A | 46.0648275 | 0.3523361 | 0.30377603 | 1.15985483 | 0.24610792 | 0.99937574 |
| HAGHL | 20.2315621 | -0.1462809 | 0.71162873 | -0.2055579 | 0.83713629 | 0.99937574 |
| NARFL | 16.3567081 | 0.10035032 | 0.46698454 | 0.21489003 | 0.82985306 | 0.99937574 |
| LMF1 | 62.2868475 | 0.05723385 | 0.43737323 | 0.13085814 | 0.89588753 | 0.99937574 |
| CACNA1H | 13.292812 | 1.00323994 | 0.94212842 | 1.06486537 | 0.28693686 | 0.99937574 |
| UBE2I | 90.9585701 | -0.5962927 | 0.32502309 | -1.8346164 | 0.06656255 | 0.99937574 |

| | | | | | | |
|---|---|---|---|---|---|---|
| TSR3 | 25.1575027 | -0.3693292 | 0.4815843 | -0.7669045 | 0.44313828 | 0.99937574 |
| GNPTG | 25.4470196 | -0.6237058 | 0.4462904 | -1.3975336 | 0.16225318 | 0.99937574 |
| UNKL | 22.4784151 | -0.8886542 | 0.54326978 | -1.6357512 | 0.10189168 | 0.99937574 |
| CLCN7 | 44.5923929 | -1.0627663 | 0.49194223 | -2.1603477 | 0.03074576 | 0.99049468 |
| IFT140 | 21.0896183 | 0.56312996 | 0.43961998 | 1.28094715 | 0.20021223 | 0.99937574 |
| JPT2 | 50.0988748 | -0.284438 | 0.37881758 | -0.7508573 | 0.45273853 | 0.99937574 |
| MAPK8IP3 | 77.6870909 | 0.09411909 | 0.30648258 | 0.30709441 | 0.75877151 | 0.99937574 |
| NME3 | 56.6884006 | 0.38689413 | 0.45346542 | 0.85319433 | 0.39355154 | 0.99937574 |
| MRPS34 | 35.9724762 | -0.2281552 | 0.60478503 | -0.37725 | 0.70598781 | 0.99937574 |
| SPSB3 | 36.0436586 | 0.48469287 | 0.46285084 | 1.0471902 | 0.29501187 | 0.99937574 |
| NUBP2 | 15.6901608 | 0.21530271 | 0.47576378 | 0.45254119 | 0.65087915 | 0.99937574 |
| HAGH | 74.2565036 | -0.1663078 | 0.31692894 | -0.524748 | 0.59975836 | 0.99937574 |
| FAHD1 | 20.9886732 | -0.5747288 | 0.4751027 | -1.2096939 | 0.22639636 | 0.99937574 |
| NDUFB10 | 61.9585326 | 0.40174175 | 0.53748892 | 0.74744192 | 0.45479685 | 0.99937574 |
| RPS2 | 243.371699 | -0.0757265 | 0.48963956 | -0.1546577 | 0.87709118 | 0.99937574 |
| GFER | 12.7389232 | 0.241534 | 0.52831539 | 0.45717767 | 0.64754334 | 0.99937574 |
| ZNF598 | 12.5346561 | -0.3869327 | 0.59284543 | -0.6526705 | 0.51396875 | 0.99937574 |
| TSC2 | 41.4742228 | -0.3309389 | 0.37295098 | -0.8873524 | 0.37488922 | 0.99937574 |
| PKD1 | 39.4383305 | 0.10886499 | 0.38847407 | 0.28023746 | 0.77929533 | 0.99937574 |
| SNHG19 | 14.5140688 | -0.0698743 | 0.51109715 | -0.1367143 | 0.89125664 | 0.99937574 |
| TRAF7 | 17.6099101 | -0.0223784 | 0.49309194 | -0.0453838 | 0.9638014 | 0.99937574 |
| MLST8 | 11.3578041 | -0.3193154 | 0.80257876 | -0.3978618 | 0.69073209 | 0.99937574 |
| PGP | 61.8906752 | 0.1706539 | 0.39340254 | 0.43378952 | 0.66444129 | 0.99937574 |
| E4F1 | 19.2006943 | -0.3957349 | 0.50808409 | -0.7788767 | 0.43605234 | 0.99937574 |
| ECI1 | 161.888873 | 0.25892381 | 0.34576563 | 0.74884194 | 0.45395248 | 0.99937574 |
| RNPS1 | 60.8436358 | -0.2240759 | 0.30255701 | -0.7406072 | 0.45893166 | 0.99937574 |
| ABCA3 | 18.816568 | -0.4313741 | 0.60817385 | -0.7092941 | 0.47814201 | 0.99937574 |
| TBC1D24 | 12.0202268 | 1.26426468 | 0.78282822 | 1.61499629 | 0.1063115 | 0.99937574 |
| PDPK1 | 106.308105 | -0.0475136 | 0.29580769 | -0.1606232 | 0.87239016 | 0.99937574 |
| KCTD5 | 10.5351546 | 0.411105 | 0.75313596 | 0.54585761 | 0.58516382 | 0.99937574 |
| SRRM2 | 273.395639 | -0.3166657 | 0.27969451 | -1.1321841 | 0.25755705 | 0.99937574 |
| ELOB | 73.5095626 | 0.17205602 | 0.45176799 | 0.38085041 | 0.70331425 | 0.99937574 |
| PRSS22 | 13.3092289 | -0.6012252 | 0.65504871 | -0.9178328 | 0.35870643 | 0.99937574 |
| FLYWCH2 | 11.1020487 | 0.21361698 | 0.58310095 | 0.36634648 | 0.71410655 | 0.99937574 |
| FLYWCH1 | 30.9233004 | 0.06988316 | 0.43052699 | 0.16232004 | 0.87105383 | 0.99937574 |
| TNFRSF12A | 60.7773319 | -0.2879893 | 0.89494466 | -0.3217956 | 0.74760754 | 0.99937574 |
| HCFC1R1 | 12.8669103 | -0.4861256 | 0.59158549 | -0.8217336 | 0.41122856 | 0.99937574 |
| BICDL2 | 30.9608608 | 0.32544545 | 0.45533858 | 0.71473288 | 0.47477411 | 0.99937574 |
| IL32 | 15.1659457 | 0.83647894 | 0.54029429 | 1.54819133 | 1.22E-01 | 0.99937574 |
| ZNF213-AS1 | 14.884823 | -0.7292542 | 0.59593964 | -1.2237047 | 0.22106364 | 0.99937574 |
| ZNF213 | 22.4244455 | 0.08525266 | 0.50667788 | 0.1682581 | 0.86638023 | 0.99937574 |
| ZNF263 | 16.2861876 | -0.6518958 | 0.54778834 | -1.1900506 | 0.23402652 | 0.99937574 |
| TIGD7 | 14.4442643 | 0.3026964 | 0.47592388 | 0.63601852 | 0.52476435 | 0.99937574 |
| ZNF75A | 53.0817567 | -0.2398617 | 0.68103166 | -0.3522034 | 0.72468574 | 0.99937574 |
| AC025283.3 | 17.0924097 | -0.0124917 | 0.61060951 | -0.0204578 | 0.98367821 | 0.99937574 |
| NAA60 | 27.7432437 | -0.4295025 | 0.37239943 | -1.1533382 | 0.24877161 | 0.99937574 |
| CLUAP1 | 36.1742301 | 0.31840153 | 0.39382696 | 0.80848079 | 0.41881386 | 0.99937574 |
| SLX4 | 11.448347 | 0.48813773 | 0.60057339 | 0.81278615 | 0.41634068 | 0.99937574 |
| DNASE1 | 108.198228 | 0.31216502 | 0.28844121 | 1.08224833 | 0.2791422 | 0.99937574 |
| TRAP1 | 50.5176795 | 0.59012076 | 0.38674628 | 1.52586023 | 0.12704468 | 0.99937574 |
| CREBBP | 201.505327 | -0.3678271 | 0.29975335 | -1.2270992 | 0.21978529 | 0.99937574 |
| ADCY9 | 121.687359 | 0.00486367 | 0.49807787 | 0.00976488 | 0.99220887 | 0.99937574 |
| SRL | 22.7831384 | -3.3394086 | 2.5064769 | -1.3323117 | 0.18275777 | 0.99937574 |
| GLIS2 | 26.5332474 | 0.66961843 | 0.52159503 | 1.2837899 | 0.19921548 | 0.99937574 |

| | | | | | | |
|---|---|---|---|---|---|---|
| VASN | 28.9245421 | 0.41169962 | 0.4187526 | 0.98315716 | 0.32553009 | 0.99937574 |
| DNAJA3 | 46.6756189 | 0.64502423 | 0.47880982 | 1.34714078 | 0.1779349 | 0.99937574 |
| NMRAL1 | 29.6028786 | 0.41465575 | 0.41420265 | 1.00109389 | 0.31678142 | 0.99937574 |
| HMOX2 | 47.420567 | 0.18769879 | 0.34816676 | 0.53910602 | 0.5898137 | 0.99937574 |
| CDIP1 | 38.0185284 | -0.2319141 | 0.34872905 | -0.6650266 | 0.50603349 | 0.99937574 |
| UBALD1 | 17.7878211 | -0.4182916 | 0.46066809 | -0.9080107 | 0.36387256 | 0.99937574 |
| MGRN1 | 75.0021551 | 0.10285226 | 0.35205695 | 0.29214665 | 0.7701745 | 0.99937574 |
| NUDT16L1 | 25.6359774 | 0.43703501 | 0.50003636 | 0.87400647 | 0.38211473 | 0.99937574 |
| ANKS3 | 44.238164 | -0.2686861 | 0.41677831 | -0.6446739 | 0.51913853 | 0.99937574 |
| ZNF500 | 13.4945837 | 0.4571143 | 0.6519138 | 0.70118826 | 0.48318553 | 0.99937574 |
| SMIM22 | 18.2319134 | 0.01117205 | 0.76775017 | 0.01455168 | 0.98838985 | 0.99937574 |
| ROGDI | 15.7697931 | -0.2235175 | 0.78702221 | -0.284004 | 0.77640728 | 0.99937574 |
| GLYR1 | 121.703462 | -0.0034427 | 0.2736957 | -0.0125785 | 0.98996405 | 0.99937574 |
| UBN1 | 56.4982686 | -0.2067727 | 0.33244287 | -0.6219797 | 0.53395522 | 0.99937574 |
| PPL | 32.5776981 | -0.2278652 | 0.51558273 | -0.4419565 | 0.65852067 | 0.99937574 |
| NAGPA | 12.2363786 | -0.5768108 | 0.66413064 | -0.8685201 | 0.3851097 | 0.99937574 |
| ALG1 | 13.3146314 | -0.0437357 | 0.63571785 | -0.0687973 | 0.94515096 | 0.99937574 |
| C16orf89 | 10.2275278 | 0.98024197 | 0.8323834 | 1.17763278 | 0.23894304 | 0.99937574 |
| RBFOX1 | 46.0047992 | -0.5705983 | 0.75190492 | -0.7588703 | 0.44793016 | 0.99937574 |
| METTL22 | 15.0259194 | -0.2863992 | 0.5309765 | -0.5393821 | 0.58962321 | 0.99937574 |
| ABAT | 20.043642 | 0.40400478 | 0.46646782 | 0.86609356 | 0.38643886 | 0.99937574 |
| PMM2 | 69.9903447 | -0.0331487 | 0.36344668 | -0.0912065 | 0.9273285 | 0.99937574 |
| CARHSP1 | 40.444312 | -0.761355 | 0.60303429 | -1.2625401 | 0.2067545 | 0.99937574 |
| USP7 | 97.9837721 | 0.22274259 | 0.26601764 | 0.83732265 | 0.40241123 | 0.99937574 |
| C16orf72 | 76.0274468 | -0.1963798 | 0.29321441 | -0.6697481 | 0.5030184 | 0.99937574 |
| AC087190.1 | 12.5999277 | -0.4247372 | 0.66692227 | -0.6368617 | 0.52421495 | 0.99937574 |
| ATF7IP2 | 108.342412 | -0.0971042 | 0.4750156 | -0.2044232 | 0.83802281 | 0.99937574 |
| EMP2 | 21.5041816 | -1.381962 | 1.12100464 | -1.2327888 | 0.21765455 | 0.99937574 |
| NUBP1 | 24.9047178 | 0.10104348 | 0.4276802 | 0.23625943 | 0.81323137 | 0.99937574 |
| CIITA | 27.2121312 | -0.571784 | 0.62440874 | -0.9157208 | 0.35981337 | 0.99937574 |
| DEXI | 29.9943019 | 0.58928047 | 0.42169165 | 1.39742028 | 0.16228722 | 0.99937574 |
| CLEC16A | 398.864176 | 0.48332183 | 0.32818525 | 1.47271041 | 0.14082914 | 0.99937574 |
| AC007014.1 | 24.8512608 | 0.56343992 | 0.7120732 | 0.79126686 | 0.42878829 | 0.99937574 |
| AC007220.1 | 15.0309403 | 1.01127728 | 0.68350694 | 1.47954206 | 0.1389955 | 0.99937574 |
| SOCS1 | 81.8873988 | -0.3995902 | 0.54961659 | -0.7270344 | 4.67E-01 | 9.99E-01 |
| AC099489.1 | 80.9627784 | -0.5495165 | 0.83243728 | -0.6601296 | 0.50917066 | 0.99937574 |
| LITAF | 772.273161 | -0.1949647 | 0.37109103 | -0.5253824 | 0.59931737 | 0.99937574 |
| SNN | 14.8872602 | -1.0707224 | 0.80738364 | -1.3261631 | 0.18478567 | 0.99937574 |
| TXNDC11 | 121.386061 | -0.2666566 | 0.35886023 | -0.7430652 | 0.45744219 | 0.99937574 |
| ZC3H7A | 90.9169054 | -0.0681372 | 0.32150415 | -0.2119325 | 0.83215968 | 0.99937574 |
| RSL1D1 | 24.511903 | -0.0962784 | 0.4145548 | -0.2322453 | 0.81634746 | 0.99937574 |
| AC007216.4 | 42.511846 | 0.08198211 | 0.4241712 | 0.193276 | 0.84674282 | 0.99937574 |
| GSPT1 | 173.438304 | -0.17954 | 0.31464608 | -0.5706092 | 0.56826456 | 0.99937574 |
| SNX29 | 526.99669 | -0.3813452 | 0.38119501 | -1.0003941 | 0.31711982 | 0.99937574 |
| AC010333.2 | 69.5089485 | -0.2492548 | 0.7745091 | -0.321823 | 0.7475868 | 0.99937574 |
| CPPED1 | 289.161741 | -0.4294308 | 0.3918729 | -1.0958421 | 0.2731479 | 0.99937574 |
| AC109597.1 | 16.322704 | -0.1721327 | 0.67993112 | -0.253162 | 0.80014302 | 0.99937574 |
| ERCC4 | 21.5626753 | 0.41361069 | 0.43130535 | 0.95897417 | 0.33757176 | 0.99937574 |
| MKL2 | 317.693604 | 0.13035677 | 0.28246682 | 0.46149409 | 0.64444416 | 0.99937574 |
| MIR193BHG | 19.1814329 | -0.1208452 | 1.24312138 | -0.0972111 | 0.92255876 | 0.99937574 |
| PARN | 124.998734 | 0.20720266 | 0.37285858 | 0.55571382 | 0.57840651 | 0.99937574 |
| BFAR | 60.3516295 | -0.2230705 | 0.42483684 | -0.5250734 | 0.59953218 | 0.99937574 |
| NOMO1 | 16.2579591 | -0.3328501 | 0.54109364 | -0.6151432 | 0.53846015 | 0.99937574 |
| PDXDC1 | 201.07191 | -0.1477241 | 0.24919039 | -0.592816 | 0.55330429 | 0.99937574 |

| | | | | | | |
|---|---|---|---|---|---|---|
| NTAN1 | 16.3359635 | -0.5561733 | 0.63490001 | -0.8760013 | 0.3810293 | 0.99937574 |
| RRN3 | 35.4749642 | -0.1818143 | 0.32615665 | -0.5574448 | 0.57722355 | 0.99937574 |
| C16orf45 | 66.5346374 | 0.5672771 | 0.45288218 | 1.25259311 | 0.21035382 | 0.99937574 |
| MARF1 | 45.3635261 | -0.1692156 | 0.43425079 | -0.3896725 | 0.69677873 | 0.99937574 |
| NDE1 | 17.9645232 | -0.0401286 | 0.45111361 | -0.0889544 | 0.92911813 | 0.99937574 |
| FOPNL | 28.3602031 | 0.19445655 | 0.46078412 | 0.42201227 | 0.67301606 | 0.99937574 |
| ABCC1 | 125.502931 | -0.2047414 | 0.53292721 | -0.3841828 | 0.70084299 | 0.99937574 |
| RPS15A | 45.8515345 | -0.1150958 | 0.49815044 | -0.2310463 | 0.81727882 | 0.99937574 |
| SMG1 | 240.569967 | -0.4071998 | 0.23542758 | -1.7296183 | 0.0836985 | 0.99937574 |
| COQ7 | 24.1437385 | -0.606281 | 0.43150218 | -1.4050475 | 0.16000717 | 0.99937574 |
| AC099518.5 | 11.8084704 | 0.42048079 | 0.69309669 | 0.60666975 | 0.5440701 | 0.99937574 |
| SYT17 | 287.453977 | 0.84928727 | 0.56488518 | 1.50346885 | 0.13271818 | 0.99937574 |
| GDE1 | 40.2650446 | -0.1971706 | 0.43982326 | -0.448295 | 0.65394032 | 0.99937574 |
| CCP110 | 18.1652265 | -0.2398538 | 0.56819775 | -0.4221308 | 0.67292954 | 0.99937574 |
| VPS35L | 97.687789 | -0.4882677 | 0.32915782 | -1.4833846 | 0.13797225 | 0.99937574 |
| KNOP1 | 30.5306861 | -0.1420875 | 0.47605689 | -0.2984674 | 0.76534643 | 0.99937574 |
| IQCK | 123.05953 | 0.87451127 | 0.34516493 | 2.53360407 | 0.01128962 | 0.76274616 |
| GPRC5B | 125.761185 | 0.2015998 | 0.51381688 | 0.39235729 | 0.69479424 | 0.99937574 |
| UMOD | 60.9489349 | 0.76992888 | 0.77914398 | 0.98817279 | 0.32306803 | 0.99937574 |
| ACSM3 | 25.0236467 | 0.36459389 | 0.50675928 | 0.7194617 | 0.4718565 | 0.99937574 |
| THUMPD1 | 59.425832 | -0.1031357 | 0.37282962 | -0.2766296 | 0.78206455 | 0.99937574 |
| ERI2 | 20.4619467 | -0.2080682 | 0.57787895 | -0.360055 | 0.71880601 | 0.99937574 |
| LYRM1 | 15.8950641 | 0.30102876 | 0.6049426 | 0.4976154 | 0.61875515 | 0.99937574 |
| TMEM159 | 16.9099205 | 0.37956342 | 0.54076323 | 0.70190316 | 0.48273956 | 0.99937574 |
| METTL9 | 67.1992278 | -0.2485401 | 0.29416304 | -0.844906 | 0.3981633 | 0.99937574 |
| NPIPB4 | 18.3478988 | -1.2899626 | 0.53576077 | -2.4077213 | 0.01605243 | 0.87097377 |
| UQCRC2 | 149.60235 | 0.34624155 | 0.27121227 | 1.27664412 | 0.20172792 | 0.99937574 |
| MOSMO | 70.5960581 | -0.2820667 | 0.32549417 | -0.8665799 | 0.38617221 | 0.99937574 |
| EEF2K | 15.327878 | -0.6746045 | 0.67974243 | -0.9924414 | 0.32098227 | 0.99937574 |
| POLR3E | 41.9189191 | 0.26822667 | 0.32462815 | 0.8262582 | 0.40865764 | 0.99937574 |
| CDR2 | 142.827621 | -0.1622359 | 0.30664888 | -0.5290606 | 0.59676338 | 0.99937574 |
| NPIPB5 | 60.4990629 | -0.1093901 | 0.46864762 | -0.2334164 | 0.81543806 | 0.99937574 |
| USP31 | 24.1072185 | -0.2141381 | 0.51606921 | -0.4149407 | 0.67818531 | 0.99937574 |
| COG7 | 70.2595912 | 0.02055814 | 0.31755722 | 0.06473837 | 0.94838231 | 0.99937574 |
| GGA2 | 124.273622 | -0.3232263 | 0.32677898 | -0.9891282 | 0.32260041 | 0.99937574 |
| EARS2 | 15.3551727 | -0.2434913 | 0.48673885 | -0.5002503 | 0.61689884 | 0.99937574 |
| UBFD1 | 21.1428698 | 0.15529663 | 0.5116486 | 0.30352204 | 0.76149206 | 0.99937574 |
| NDUFAB1 | 54.3069097 | 0.18435946 | 0.4701101 | 0.39216231 | 0.69493829 | 0.99937574 |
| PALB2 | 12.4233703 | -0.6858557 | 0.63216847 | -1.0849255 | 0.27795465 | 0.99937574 |
| DCTN5 | 47.8134629 | -0.0963766 | 0.3613095 | -0.2667424 | 0.78966753 | 0.99937574 |
| RBBP6 | 106.520396 | -0.0122696 | 0.3500554 | -0.0350504 | 0.97203958 | 0.99937574 |
| TNRC6A | 579.428837 | -0.5286488 | 0.34741523 | -1.5216626 | 0.12809365 | 0.99937574 |
| AC008731.1 | 18.7808206 | 0.06470447 | 0.65684981 | 0.09850726 | 0.92152951 | 0.99937574 |
| ARHGAP17 | 74.06717 | -0.1645682 | 0.3713114 | -0.4432081 | 0.65761525 | 0.99937574 |
| LCMT1 | 49.4093256 | 0.01587665 | 0.32849761 | 0.04833111 | 0.96145236 | 0.99937574 |
| NSMCE1 | 21.1914841 | 0.84236388 | 0.4503865 | 1.87031334 | 0.06144032 | 0.99937574 |
| GTF3C1 | 40.9074685 | -0.1067422 | 0.38097974 | -0.280178 | 0.77934091 | 0.99937574 |
| KIAA0556 | 84.0409126 | 0.03891558 | 0.30373228 | 0.12812462 | 0.89805035 | 0.99937574 |
| XPO6 | 75.2267825 | -0.8389644 | 0.45965183 | -1.8252172 | 0.06796825 | 0.99937574 |
| NUPR1 | 34.4144381 | -1.9114662 | 1.0656917 | -1.793639 | 0.0728708 | 0.99937574 |
| SGF29 | 13.7375088 | -0.5512956 | 0.52889992 | -1.0423439 | 0.29725225 | 0.99937574 |
| SULT1A1 | 10.6472744 | 1.19601293 | 0.82204762 | 1.45491928 | 0.14569161 | 0.99937574 |
| ATXN2L | 52.2405942 | -0.4406939 | 0.30026886 | -1.4676644 | 0.14219539 | 0.99937574 |
| TUFM | 112.795254 | 0.68007997 | 0.44526929 | 1.52734531 | 0.12667517 | 0.99937574 |

| | | | | | | |
|---|---|---|---|---|---|---|
| SH2B1 | 53.3996768 | 0.03918221 | 0.40611031 | 0.0964817 | 0.92313801 | 0.99937574 |
| RABEP2 | 39.86709 | -0.7509306 | 0.3543888 | -2.1189455 | 0.03409507 | 0.99937574 |
| NFATC2IP | 81.8566893 | -0.3488264 | 0.38173554 | -0.9137907 | 0.36082683 | 0.99937574 |
| SPNS1 | 16.5927967 | -0.2203374 | 0.6424506 | -0.3429639 | 0.73162561 | 0.99937574 |
| QPRT | 43.8728575 | -0.9629177 | 0.64318119 | -1.4971173 | 0.13436275 | 0.99937574 |
| MAZ | 19.9634817 | -0.4513633 | 0.62898676 | -0.7176039 | 0.47300157 | 0.99937574 |
| AC009133.1 | 19.4146033 | -0.4939741 | 0.75916134 | -0.650684 | 0.51525049 | 0.99937574 |
| MVP | 97.0153846 | -0.0949059 | 0.42734198 | -0.2220843 | 8.24E-01 | 0.99937574 |
| CDIPT | 25.0842437 | 0.01934423 | 0.38913562 | 0.04971078 | 0.96035287 | 0.99937574 |
| KCTD13 | 23.3765015 | -0.2908122 | 0.44689865 | -0.6507341 | 0.51521812 | 0.99937574 |
| TMEM219 | 70.0793451 | 0.14007922 | 0.43764416 | 0.3200756 | 0.74891102 | 0.99937574 |
| TAOK2 | 17.1533181 | 0.08905578 | 0.49687304 | 0.17923246 | 0.85775518 | 0.99937574 |
| INO80E | 26.3164343 | 0.22922101 | 0.40245103 | 0.56956248 | 0.56897448 | 0.99937574 |
| PPP4C | 41.7514261 | -0.4476913 | 0.36997975 | -1.2100428 | 0.22626249 | 0.99937574 |
| YPEL3 | 34.8610196 | -0.4440362 | 0.4465489 | -0.994373 | 0.32004129 | 0.99937574 |
| MAPK3 | 28.8660423 | -0.0508578 | 0.38093324 | -0.1335083 | 0.89379137 | 0.99937574 |
| NPIPB13 | 16.0386577 | -0.3875855 | 0.65981363 | -0.5874166 | 0.55692393 | 0.99937574 |
| CD2BP2 | 23.2307689 | -0.4686634 | 0.44268807 | -1.0586763 | 0.28974724 | 0.99937574 |
| TBC1D10B | 16.6164773 | -0.1723219 | 0.5692917 | -0.3026953 | 0.76212206 | 0.99937574 |
| DCTPP1 | 14.4063585 | 0.66478833 | 0.50463015 | 1.31737733 | 0.18771218 | 0.99937574 |
| SEPHS2 | 66.4427089 | -0.335741 | 0.58363811 | -0.5752554 | 0.56511856 | 0.99937574 |
| ZNF768 | 19.4130433 | -0.2191819 | 0.60723358 | -0.3609516 | 0.71813561 | 0.99937574 |
| ZNF785 | 19.3310037 | 1.32136253 | 0.45733264 | 2.88928104 | 0.00386124 | 0.54883964 |
| PRR14 | 10.9973649 | 0.11131695 | 0.57898278 | 0.19226297 | 0.84753622 | 0.99937574 |
| FBRS | 26.0765759 | -0.2369918 | 0.43165871 | -0.549026 | 0.58298763 | 0.99937574 |
| PHKG2 | 19.2465389 | 0.43300916 | 0.4402922 | 0.98345861 | 0.32538178 | 0.99937574 |
| RNF40 | 31.7325512 | -0.1421137 | 0.39603471 | -0.3588415 | 0.71971367 | 0.99937574 |
| ZNF629 | 18.2335998 | -0.0708093 | 0.65397702 | -0.1082748 | 0.91377768 | 0.99937574 |
| BCL7C | 43.1462051 | 0.03636011 | 0.33177119 | 0.10959392 | 0.91273143 | 0.99937574 |
| FBXL19 | 22.0657104 | -0.1926565 | 0.46656652 | -0.4129239 | 0.67966237 | 0.99937574 |
| SETD1A | 18.6060393 | -0.3668026 | 0.58755617 | -0.6242852 | 0.5324403 | 0.99937574 |
| STX4 | 15.6285213 | -0.8266798 | 0.48790741 | -1.6943374 | 0.09020119 | 0.99937574 |
| BCKDK | 22.2663075 | 0.21155842 | 0.40565025 | 0.52152913 | 0.60199822 | 0.99937574 |
| KAT8 | 36.3831798 | -0.1563702 | 0.44379255 | -0.3523498 | 0.72457596 | 0.99937574 |
| PRSS8 | 61.6010947 | -0.2313681 | 0.41292982 | -0.5603085 | 0.57526901 | 0.99937574 |
| FUS | 136.900541 | -0.4639144 | 0.23017709 | -2.0154671 | 0.04385573 | 0.99937574 |
| AC026471.4 | 14.7310866 | -0.0555482 | 0.59871598 | -0.0927788 | 0.92607927 | 0.99937574 |
| C16orf58 | 25.7167297 | 0.1785916 | 0.41590797 | 0.42940173 | 0.6676309 | 0.99937574 |
| ZNF720 | 63.7317602 | 0.68182886 | 0.39984935 | 1.70521438 | 0.08815445 | 0.99937574 |
| ZNF267 | 19.2847347 | 0.11830674 | 0.52970312 | 0.22334537 | 0.82326672 | 0.99937574 |
| VPS35 | 60.8386337 | -0.0301676 | 0.37007736 | -0.0815171 | 0.9350307 | 0.99937574 |
| C16orf87 | 16.6658367 | 0.12955976 | 0.4855948 | 0.26680632 | 0.78961829 | 0.99937574 |
| GPT2 | 32.2588716 | 0.78970657 | 1.00155555 | 0.78848005 | 0.43041596 | 0.99937574 |
| DNAJA2 | 42.6895779 | -0.0252672 | 0.30572747 | -0.0826463 | 0.93413277 | 0.99937574 |
| ITFG1 | 388.192591 | -0.1409401 | 0.27543613 | -0.5116979 | 0.60886246 | 0.99937574 |
| PHKB | 348.023002 | 0.21785602 | 0.23215912 | 0.93839095 | 0.34804353 | 0.99937574 |
| LONP2 | 205.453935 | 0.13620302 | 0.22010751 | 0.61880222 | 0.53604666 | 0.99937574 |
| SIAH1 | 50.4082292 | -0.1614198 | 0.37062883 | -0.4355295 | 0.66317816 | 0.99937574 |
| N4BP1 | 135.384422 | 0.09624956 | 0.39278263 | 0.24504536 | 0.80642131 | 0.99937574 |
| CNEP1R1 | 11.8480378 | -0.172856 | 0.64959022 | -0.2661 | 0.79016217 | 0.99937574 |
| TENT4B | 44.5422647 | -0.1410509 | 0.41609263 | -0.3389893 | 0.7346178 | 0.99937574 |
| BRD7 | 56.3147448 | -0.3493927 | 0.31299023 | -1.1163054 | 0.26429145 | 0.99937574 |
| CYLD | 48.2385915 | 0.10871943 | 0.32316582 | 0.33641995 | 0.73655421 | 0.99937574 |
| SALL1 | 13.5001652 | -0.0730489 | 0.86824707 | -0.0841338 | 0.93295008 | 0.99937574 |

| | | | | | | |
|---|---|---|---|---|---|---|
| AC087564.1 | 16.5610232 | 0.55234391 | 0.73143328 | 0.75515283 | 0.45015729 | 0.99937574 |
| HNRNPA1P4 | 12.9735201 | -1.261311 | 0.6389749 | -1.9739602 | 0.04838627 | 0.99937574 |
| TOX3 | 455.838807 | 0.05715171 | 0.42623092 | 0.13408627 | 0.89333436 | 0.99937574 |
| CASC16 | 149.387499 | 1.52715795 | 0.80144903 | 1.90549604 | 0.05671562 | 0.99937574 |
| CHD9 | 532.026395 | 0.34419757 | 0.24436058 | 1.40856422 | 0.15896406 | 0.99937574 |
| RBL2 | 142.481712 | 0.10250744 | 0.4101721 | 0.24991323 | 0.80265445 | 0.99937574 |
| AKTIP | 17.4631897 | 0.36167738 | 0.52181578 | 0.69311314 | 0.48823855 | 0.99937574 |
| FTO | 225.364244 | -0.0942176 | 0.2876826 | -0.3275055 | 0.74328555 | 0.99937574 |
| LPCAT2 | 26.9101818 | -0.1048902 | 0.79305896 | -0.1322602 | 0.89477845 | 0.99937574 |
| AMFR | 284.219399 | 0.35265652 | 0.31186516 | 1.13079807 | 0.25814009 | 0.99937574 |
| AC092140.1 | 13.6098782 | -0.6873781 | 0.99993174 | -0.687425 | 0.49181493 | 0.99937574 |
| NUDT21 | 64.9728304 | 0.36632914 | 0.27223536 | 1.34563391 | 0.17842061 | 0.99937574 |
| OGFOD1 | 37.6221291 | 0.21506724 | 0.41551154 | 0.51759631 | 0.60473996 | 0.99937574 |
| BBS2 | 47.2611077 | -0.3822863 | 0.32600679 | -1.1726329 | 0.24094306 | 0.99937574 |
| MT2A | 287.069174 | -2.2571197 | 0.88202679 | -2.5590149 | 0.01049692 | 0.76274616 |
| MT1E | 69.2119825 | -2.9005972 | 0.87309354 | -3.3222067 | 0.00089309 | 0.29012345 |
| MT1G | 104.413583 | -3.0233683 | 0.90251951 | -3.3499202 | 0.00080835 | 0.29012345 |
| MT1H | 34.4190991 | -3.2052766 | 1.15871389 | -2.7662365 | 0.00567074 | 0.65033075 |
| MT1X | 135.576261 | -0.8356287 | 0.84783916 | -0.9855982 | 0.32433032 | 0.99937574 |
| NUP93 | 39.6037209 | -0.1159208 | 0.43531706 | -0.2662904 | 0.79001557 | 0.99937574 |
| SLC12A3 | 99.876341 | 1.98281598 | 0.85708218 | 2.31344908 | 0.02069796 | 0.96427895 |
| HERPUD1 | 126.418599 | -0.0812071 | 0.5251918 | -0.1546238 | 0.87711793 | 0.99937574 |
| NLRC5 | 89.7445105 | -1.2060907 | 0.85309297 | -1.4137858 | 0.1574248 | 0.99937574 |
| CPNE2 | 36.5679582 | -1.130204 | 0.75057702 | -1.5057802 | 0.13212362 | 0.99937574 |
| FAM192A | 75.142331 | -0.180931 | 0.28348347 | -0.6382419 | 0.52331625 | 0.99937574 |
| RSPRY1 | 74.2935336 | 0.08670781 | 0.26571056 | 0.32632427 | 0.74417903 | 0.99937574 |
| ARL2BP | 99.6970559 | -0.0033344 | 0.27292631 | -0.0122172 | 0.9902523 | 0.99937574 |
| PLLP | 121.68262 | 0.45946772 | 0.54180328 | 0.84803422 | 0.39641891 | 0.99937574 |
| CX3CL1 | 17.2557811 | -0.3875658 | 0.679077 | -0.5707244 | 0.56818648 | 0.99937574 |
| CIAPIN1 | 46.4318826 | -0.1469502 | 0.32643883 | -0.4501615 | 0.65259399 | 0.99937574 |
| COQ9 | 81.9136924 | 0.0694099 | 0.33837957 | 0.20512437 | 0.83747496 | 0.99937574 |
| POLR2C | 17.2818575 | 0.13622481 | 0.57173233 | 0.23826676 | 0.8116742 | 0.99937574 |
| ADGRG1 | 157.281525 | -0.3238869 | 0.53880613 | -0.6011195 | 0.54776037 | 0.99937574 |
| KIFC3 | 74.2884993 | -0.1898345 | 0.37285587 | -0.5091364 | 0.61065663 | 0.99937574 |
| USB1 | 13.3220175 | 0.2867999 | 0.53198594 | 0.5391118 | 0.58980971 | 0.99937574 |
| MMP15 | 14.7559817 | 0.23667896 | 0.62205393 | 0.38047981 | 0.70358928 | 0.99937574 |
| CFAP20 | 10.7110139 | 0.06966134 | 0.59965096 | 0.11616982 | 0.90751796 | 0.99937574 |
| CSNK2A2 | 45.229223 | -0.1895811 | 0.4048513 | -0.4682733 | 0.63958916 | 0.99937574 |
| CNOT1 | 149.212583 | -0.003275 | 0.21018758 | -0.0155814 | 0.98756837 | 0.99937574 |
| SLC38A7 | 19.6678364 | -0.5601195 | 0.70386552 | -0.7957764 | 0.42616204 | 0.99937574 |
| GOT2 | 76.5114412 | 0.33365025 | 0.30666392 | 1.08799972 | 0.27659523 | 0.99937574 |
| AC138305.1 | 686.393014 | 0.54647886 | 0.30956552 | 1.76530917 | 0.07751182 | 0.99937574 |
| AC010542.3 | 13.6337868 | 0.38116091 | 0.73141322 | 0.52112938 | 0.60227665 | 0.99937574 |
| TK2 | 16.757857 | 0.19263443 | 0.60238939 | 0.31978391 | 0.74913215 | 0.99937574 |
| CMTM4 | 294.024447 | 0.26978961 | 0.25749641 | 1.04774126 | 0.29475783 | 0.99937574 |
| DYNC1LI2 | 101.301985 | -0.0902588 | 0.26502126 | -0.3405718 | 0.73342595 | 0.99937574 |
| AC018557.1 | 15.7617748 | 1.17748421 | 0.65673383 | 1.79293979 | 0.07298255 | 0.99937574 |
| NAE1 | 37.9968807 | -0.0179299 | 0.32149428 | -0.0557707 | 0.95552451 | 0.99937574 |
| PDP2 | 26.1988403 | -0.0869474 | 0.63674801 | -0.1365491 | 0.8913872 | 0.99937574 |
| CDH16 | 267.86842 | 0.43705457 | 0.4941126 | 0.88452422 | 0.37641329 | 0.99937574 |
| RRAD | 45.0950512 | -1.178911 | 0.87702912 | -1.3442096 | 0.17888061 | 0.99937574 |
| FAM96B | 81.6857792 | 0.12848748 | 0.620806 | 0.20696881 | 0.83603422 | 0.99937574 |
| CES2 | 37.9003611 | 0.04337747 | 0.34739246 | 0.1248659 | 0.90062971 | 0.99937574 |
| CBFB | 161.265017 | -0.2425212 | 0.35043103 | -0.6920655 | 0.48889619 | 0.99937574 |

| | | | | | | |
|---|---|---|---|---|---|---|
| C16orf70 | 16.5738649 | -0.3915555 | 0.571169 | -0.6855336 | 0.49300726 | 0.99937574 |
| TRADD | 15.9884311 | 0.39272566 | 0.54111353 | 0.72577313 | 0.46797786 | 0.99937574 |
| NOL3 | 22.3628073 | 0.53180877 | 0.6895358 | 0.77125621 | 0.44055508 | 0.99937574 |
| KIAA0895L | 24.0305754 | 0.34645551 | 0.55537143 | 0.62382667 | 0.53274142 | 0.99937574 |
| E2F4 | 23.269917 | -0.3059478 | 0.44415787 | -0.6888266 | 0.49093239 | 0.99937574 |
| ELMO3 | 13.9904481 | 0.0319408 | 0.61768349 | 0.05171063 | 0.95875927 | 0.99937574 |
| TMEM208 | 34.2756288 | 0.10868782 | 0.42034798 | 0.2585663 | 0.79596989 | 0.99937574 |
| ZDHHC1 | 30.6503544 | 0.62839306 | 0.42304609 | 1.48540093 | 0.13743765 | 0.99937574 |
| HSD11B2 | 173.463102 | 1.47886629 | 0.93075102 | 1.58889569 | 0.11208394 | 0.99937574 |
| ATP6V0D1 | 51.1920065 | -0.3725178 | 0.58164296 | -0.6404578 | 0.52187499 | 0.99937574 |
| AC009061.2 | 13.0747385 | 0.68188475 | 0.79297421 | 0.85990785 | 0.38983984 | 0.99937574 |
| RIPOR1 | 29.4581463 | 0.15533788 | 0.48196636 | 0.32230025 | 0.74722524 | 0.99937574 |
| CTCF | 71.4078302 | -0.5610516 | 0.35419501 | -1.5840189 | 0.11318941 | 0.99937574 |
| ENKD1 | 10.9876799 | 0.8683286 | 0.55230838 | 1.57218076 | 0.11590864 | 0.99937574 |
| GFOD2 | 27.0228213 | 0.24024861 | 0.44001151 | 0.54600528 | 0.5850623 | 0.99937574 |
| RANBP10 | 77.7584608 | -0.5285258 | 0.420488 | -1.2569344 | 0.20877741 | 0.99937574 |
| NUTF2 | 15.9814715 | -0.735013 | 0.50253346 | -1.462615 | 0.14357276 | 0.99937574 |
| PSKH1 | 28.581252 | 0.05858128 | 0.52122151 | 0.11239229 | 0.91051237 | 0.99937574 |
| DUS2 | 18.4871619 | 0.04734492 | 0.45766797 | 0.10344819 | 0.91760727 | 0.99937574 |
| NFATC3 | 116.897242 | 0.05389837 | 0.24654528 | 0.2186145 | 0.82695036 | 0.99937574 |
| ESRP2 | 15.3511866 | 0.16627104 | 0.46066653 | 0.36093579 | 0.71814745 | 0.99937574 |
| PLA2G15 | 23.2924402 | 0.19096317 | 0.51482088 | 0.37093128 | 0.71068871 | 0.99937574 |
| SLC7A6 | 61.531682 | -0.0143738 | 0.51730906 | -0.0277857 | 0.97783306 | 0.99937574 |
| SLC7A6OS | 30.983434 | 0.1669362 | 0.43140784 | 0.3869568 | 0.69878819 | 0.99937574 |
| PRMT7 | 25.4174913 | -0.0947801 | 0.54262207 | -0.1746705 | 0.86133852 | 0.99937574 |
| ZFP90 | 21.8151684 | -0.175417 | 0.4185596 | -0.4190967 | 0.67514543 | 0.99937574 |
| CDH1 | 379.296131 | -0.0155933 | 0.23670948 | -0.0658753 | 0.94747714 | 0.99937574 |
| TANGO6 | 59.5962553 | 0.09008098 | 0.35852964 | 0.25125115 | 0.80161994 | 0.99937574 |
| CHTF8 | 41.3109081 | 0.12654799 | 0.38743403 | 0.32663105 | 0.74394696 | 0.99937574 |
| UTP4 | 17.376583 | -0.2363887 | 0.5236264 | -0.4514454 | 0.65166855 | 0.99937574 |
| SNTB2 | 48.5687117 | 0.13412979 | 0.36196692 | 0.37055813 | 0.71096667 | 0.99937574 |
| VPS4A | 31.9868838 | 0.08852551 | 0.40527669 | 0.21843227 | 0.82709233 | 0.99937574 |
| COG8 | 12.3633303 | -0.0446728 | 0.57576319 | -0.0775888 | 0.93815517 | 0.99937574 |
| TERF2 | 36.0106346 | 0.01968518 | 0.34151556 | 0.05764064 | 0.95403488 | 0.99937574 |
| CYB5B | 103.379197 | -0.32281 | 0.2831172 | -1.1401992 | 0.25420334 | 0.99937574 |
| NFAT5 | 1228.56154 | -0.1861152 | 0.30677493 | -0.6066833 | 0.54406108 | 0.99937574 |
| NQO1 | 39.1084282 | -0.6385641 | 0.54014008 | -1.1822194 | 2.37E-01 | 0.99937574 |
| WWP2 | 182.769807 | -0.2068685 | 0.46190017 | -0.447864 | 0.65425133 | 0.99937574 |
| PDPR | 58.1824931 | -0.3362524 | 0.46324731 | -0.7258594 | 0.46792497 | 0.99937574 |
| AC009060.1 | 12.5657954 | 0.25000443 | 0.64962251 | 0.3848457 | 0.70035173 | 0.99937574 |
| EXOSC6 | 26.3384221 | -0.3323184 | 0.57314648 | -0.5798141 | 0.56203997 | 0.99937574 |
| AARS | 53.2923221 | -0.0517139 | 0.36074702 | -0.1433522 | 0.88601205 | 0.99937574 |
| DDX19A | 10.1743283 | -0.146298 | 0.64670685 | -0.22622 | 0.82103034 | 0.99937574 |
| ST3GAL2 | 29.5086517 | 0.54581303 | 0.39282971 | 1.38943927 | 0.16469922 | 0.99937574 |
| FUK | 16.7167375 | 0.33919099 | 0.51767439 | 0.65522072 | 0.51232565 | 0.99937574 |
| COG4 | 33.872379 | -0.0413642 | 0.37825855 | -0.1093543 | 0.91292151 | 0.99937574 |
| SF3B3 | 71.4312152 | 0.15467438 | 0.30893696 | 0.50066646 | 0.61660588 | 0.99937574 |
| VAC14 | 40.1573561 | -0.9496806 | 0.57798432 | -1.6430906 | 0.1003642 | 0.99937574 |
| HYDIN | 48.9579916 | -0.0471218 | 0.58889011 | -0.080018 | 0.93622292 | 0.99937574 |
| CMTR2 | 31.2807117 | -0.0137429 | 0.39934815 | -0.0344134 | 0.97254749 | 0.99937574 |
| ZNF19 | 21.1589403 | 0.3905098 | 0.66923852 | 0.58351364 | 0.55954758 | 0.99937574 |
| MARVELD3 | 13.837895 | -0.7328931 | 0.5215725 | -1.4051605 | 0.15997356 | 0.99937574 |
| PHLPP2 | 37.3887304 | -0.0654543 | 0.37961104 | -0.1724247 | 0.86310369 | 0.99937574 |
| AP1G1 | 181.10527 | 0.11657301 | 0.26490795 | 0.440051 | 0.65990017 | 0.99937574 |

| | | | | | | |
|---|---|---|---|---|---|---|
| IST1 | 119.493764 | -0.2381938 | 0.25065642 | -0.95028 | 0.34197002 | 0.99937574 |
| ZNF821 | 23.968741 | 0.04537195 | 0.54791284 | 0.08280871 | 0.93400365 | 0.99937574 |
| TXNL4B | 20.1354728 | 0.39356774 | 0.41024781 | 0.95934148 | 0.33738674 | 0.99937574 |
| DHX38 | 21.9039866 | -0.8295764 | 0.49074377 | -1.6904471 | 0.09094245 | 0.99937574 |
| LINC01572 | 87.8317791 | -0.3186118 | 0.53386773 | -0.5967991 | 0.55064153 | 0.99937574 |
| AC004943.2 | 40.5128476 | 0.33945929 | 0.44185048 | 0.76826734 | 0.44232838 | 0.99937574 |
| ZFHX3 | 427.804645 | -0.3265048 | 0.40653676 | -0.8031372 | 0.42189543 | 0.99937574 |
| AC138627.1 | 299.487231 | -0.3208128 | 0.86858256 | -0.3693522 | 0.71186525 | 0.99937574 |
| PSMD7 | 60.1742571 | -0.0155232 | 0.46210733 | -0.0335923 | 0.97320227 | 0.99937574 |
| NPIPB15 | 12.7962982 | 0.32908972 | 1.11473263 | 0.29521852 | 0.76782695 | 0.99937574 |
| GLG1 | 346.618599 | 0.25431981 | 0.28831222 | 0.88209862 | 0.37772348 | 0.99937574 |
| RFWD3 | 34.1291627 | 0.27452983 | 0.40315515 | 0.68095331 | 0.49590103 | 0.99937574 |
| FA2H | 73.3652205 | 0.4989584 | 0.55671473 | 0.89625507 | 0.37011655 | 0.99937574 |
| WDR59 | 97.5137675 | 0.04819088 | 0.27942567 | 0.17246402 | 0.86307274 | 0.99937574 |
| ZNRF1 | 51.25271 | -0.2240082 | 0.55304301 | -0.4050467 | 0.68544318 | 0.99937574 |
| LDHD | 10.4310725 | 1.01051566 | 0.65108895 | 1.55203934 | 0.12065281 | 0.99937574 |
| BCAR1 | 64.9335822 | -0.0516331 | 0.38101777 | -0.1355137 | 0.89220574 | 0.99937574 |
| CFDP1 | 140.865822 | -0.3113137 | 0.25931089 | -1.2005425 | 0.22992872 | 0.99937574 |
| GABARAPL2 | 68.5655493 | 0.42381177 | 0.51904773 | 0.81651792 | 0.41420398 | 0.99937574 |
| ADAT1 | 21.5156598 | -0.0331065 | 0.46357282 | -0.0714159 | 0.94306676 | 0.99937574 |
| KARS | 28.4181036 | 0.05167409 | 0.49360927 | 0.10468623 | 0.91662479 | 0.99937574 |
| TERF2IP | 108.852585 | 0.01622614 | 0.43448594 | 0.0373456 | 0.97020945 | 0.99937574 |
| MON1B | 10.8736083 | -1.3097446 | 0.73978804 | -1.7704322 | 0.07665517 | 0.99937574 |
| ADAMTS18 | 49.7449339 | 0.2530404 | 0.41444442 | 0.61055329 | 0.54149536 | 0.99937574 |
| NUDT7 | 16.4319211 | 0.32303812 | 0.61613928 | 0.52429399 | 0.60007408 | 0.99937574 |
| VAT1L | 118.772702 | 0.2265773 | 0.55204153 | 0.41043523 | 0.6814867 | 0.99937574 |
| WWOX | 1404.75181 | 0.25057414 | 0.44826958 | 0.55898091 | 0.57617475 | 0.99937574 |
| AC046158.1 | 12.2065533 | 0.49015674 | 0.81301885 | 0.60288484 | 0.5465853 | 0.99937574 |
| MAF | 32.1255021 | 0.57031191 | 0.75004777 | 0.76036745 | 4.47E-01 | 0.99937574 |
| CDYL2 | 206.071501 | 0.67729472 | 0.446863 | 1.51566523 | 0.12960402 | 0.99937574 |
| CMC2 | 96.9910174 | -0.1195599 | 0.35484554 | -0.3369349 | 0.73616596 | 0.99937574 |
| ATMIN | 26.3194746 | -0.3328432 | 0.67564431 | -0.4926308 | 0.62227351 | 0.99937574 |
| GAN | 44.6122221 | -0.0944155 | 0.37773079 | -0.2499543 | 0.80262266 | 0.99937574 |
| CMIP | 293.312617 | 0.43516448 | 0.49071459 | 0.88679753 | 0.37518792 | 0.99937574 |
| PLCG2 | 1675.5602 | -1.4192554 | 0.58010688 | -2.4465412 | 0.01442343 | 0.85624285 |
| AC092142.1 | 16.8162735 | 0.27164449 | 0.63287088 | 0.42922577 | 0.66775893 | 0.99937574 |
| MPHOSPH6 | 62.2197888 | 0.23017758 | 0.3437295 | 0.66964745 | 0.50308256 | 0.99937574 |
| HSBP1 | 66.6131542 | 0.00968537 | 0.37311651 | 0.02595804 | 0.97929081 | 0.99937574 |
| MBTPS1 | 85.9290035 | -0.0449789 | 0.32164085 | -0.1398421 | 0.88878474 | 0.99937574 |
| HSDL1 | 19.3877541 | -0.2686393 | 0.60851275 | -0.4414687 | 0.65887372 | 0.99937574 |
| TAF1C | 12.2844008 | 0.07760278 | 0.51911756 | 0.1494898 | 0.88116716 | 0.99937574 |
| ATP2C2 | 47.6980794 | 0.49040408 | 0.95379612 | 0.51416028 | 0.60713994 | 0.99937574 |
| TLDC1 | 42.9260136 | 0.15786282 | 0.479199 | 0.32943061 | 0.74183024 | 0.99937574 |
| KLHL36 | 62.3334158 | -0.0180839 | 0.35956172 | -0.0502942 | 0.95988797 | 0.99937574 |
| USP10 | 115.964479 | -0.3179553 | 0.28153425 | -1.1293664 | 0.25874332 | 0.99937574 |
| ZDHHC7 | 59.5344553 | -0.2625768 | 0.28431388 | -0.9235455 | 0.355723 | 0.99937574 |
| KIAA0513 | 27.8433082 | -0.5939364 | 0.40351712 | -1.471899 | 0.14104816 | 0.99937574 |
| GSE1 | 341.267092 | 0.27075635 | 0.45742183 | 0.5919183 | 0.5539053 | 0.99937574 |
| C16orf74 | 23.3158995 | 1.01050449 | 0.84844576 | 1.19100658 | 0.233651 | 0.99937574 |
| EMC8 | 23.5132904 | 0.13194261 | 0.44987619 | 0.2932865 | 0.76930317 | 0.99937574 |
| COX4I1 | 422.665222 | 0.25959627 | 0.5101422 | 0.5088704 | 0.61084307 | 0.99937574 |
| MTHFSD | 15.1003289 | 0.17337802 | 0.50605008 | 0.34261039 | 0.73189158 | 0.99937574 |
| FBXO31 | 21.1850465 | -0.9453424 | 0.55570745 | -1.7011513 | 0.08891459 | 0.99937574 |
| MAP1LC3B | 98.3406303 | 0.14048819 | 0.45523143 | 0.30860828 | 0.75761952 | 0.99937574 |

| | | | | | | |
|---|---|---|---|---|---|---|
| ZCCHC14 | 184.907506 | -0.6963526 | 0.59935348 | -1.1618395 | 0.24530065 | 0.99937574 |
| KLHDC4 | 57.8492401 | 0.2402434 | 0.35529041 | 0.67618881 | 0.49892078 | 0.99937574 |
| BANP | 59.1170412 | 0.03823342 | 0.47753804 | 0.08006361 | 0.93618667 | 0.99937574 |
| ZFPM1 | 19.7381956 | 0.10799662 | 0.49066571 | 0.22010223 | 0.82579154 | 0.99937574 |
| ZC3H18 | 46.7179826 | -0.6066789 | 0.42076759 | -1.4418384 | 0.14934797 | 0.99937574 |
| CYBA | 111.331754 | 1.02582763 | 0.52846341 | 1.94115167 | 0.05223988 | 0.99937574 |
| MVD | 39.304187 | 0.0907904 | 0.68436663 | 0.13266339 | 0.8944596 | 0.99937574 |
| SNAI3-AS1 | 11.671512 | 1.41028346 | 0.60812228 | 2.31907874 | 0.02039077 | 0.96427895 |
| PIEZO1 | 25.0439314 | -0.2401012 | 0.46962255 | -0.5112642 | 0.60916609 | 0.99937574 |
| APRT | 63.8043827 | 0.27594124 | 0.48928347 | 0.56397008 | 0.57277449 | 0.99937574 |
| GALNS | 25.3346511 | 0.4300126 | 0.56390186 | 0.76256639 | 0.44572204 | 0.99937574 |
| TRAPPC2L | 16.5537709 | 0.277697 | 0.51114138 | 0.54328804 | 0.5869315 | 0.99937574 |
| ACSF3 | 27.2013344 | 0.28625083 | 0.41078577 | 0.69683724 | 0.48590466 | 0.99937574 |
| ANKRD11 | 280.199291 | -0.2127475 | 0.37109152 | -0.573302 | 0.56644026 | 0.99937574 |
| SPG7 | 112.753253 | -0.4351526 | 0.42544818 | -1.02281 | 0.30639771 | 0.99937574 |
| RPL13 | 508.140839 | -0.0620415 | 0.4536769 | -0.1367526 | 0.89122637 | 0.99937574 |
| CHMP1A | 32.0888631 | -0.422028 | 0.45443012 | -0.9286972 | 0.35304606 | 0.99937574 |
| CDK10 | 36.7840181 | 0.19452559 | 0.35877328 | 0.54219643 | 0.5876832 | 0.99937574 |
| VPS9D1 | 23.9587637 | 0.15434782 | 0.42131462 | 0.36634814 | 0.71410532 | 0.99937574 |
| ZNF276 | 19.3614287 | -0.1619582 | 0.59880064 | -0.270471 | 0.78679792 | 0.99937574 |
| SPIRE2 | 29.8369536 | 0.46048964 | 0.49114962 | 0.93757506 | 0.34846283 | 0.99937574 |
| TCF25 | 257.050176 | -0.2198809 | 0.27131986 | -0.8104121 | 0.41770337 | 0.99937574 |
| DEF8 | 20.4725942 | -0.2489667 | 0.41781516 | -0.5958777 | 0.55125692 | 0.99937574 |
| GAS8 | 29.6716136 | -0.2797098 | 0.41210926 | -0.6787272 | 0.49731072 | 0.99937574 |
| FAM157C | 47.4649535 | -0.0279636 | 0.57454746 | -0.0486707 | 0.9611817 | 0.99937574 |
| LINC02193 | 10.7205119 | 0.14459908 | 0.75205245 | 0.19227259 | 0.84752868 | 0.99937574 |
| RPH3AL | 85.4949192 | 0.03070221 | 0.55259613 | 0.05555994 | 0.95569238 | 0.99937574 |
| VPS53 | 182.730517 | -0.222141 | 0.2491722 | -0.891516 | 0.3726524 | 0.99937574 |
| GLOD4 | 43.1459993 | -0.4179882 | 0.36482616 | -1.1457189 | 0.2519115 | 0.99937574 |
| NXN | 13.1096895 | -0.0738487 | 0.54692285 | -0.1350259 | 0.8925914 | 0.99937574 |
| TIMM22 | 14.401458 | 0.73510961 | 0.49961833 | 1.47134235 | 0.14119856 | 0.99937574 |
| ABR | 178.38351 | -0.2200195 | 0.53075822 | -0.414538 | 0.67848014 | 0.99937574 |
| YWHAE | 260.132726 | 0.09889318 | 0.21885051 | 0.45187548 | 0.65135869 | 0.99937574 |
| CRK | 90.4061932 | -0.2724333 | 0.44083465 | -0.6179944 | 0.53657904 | 0.99937574 |
| MYO1C | 91.8160467 | 0.27139246 | 0.35834477 | 0.75735015 | 0.44884012 | 0.99937574 |
| INPP5K | 15.1181032 | -0.6832711 | 0.62004021 | -1.1019787 | 0.27047091 | 0.99937574 |
| PITPNA | 113.849958 | 0.13413373 | 0.26749914 | 0.50143613 | 0.61606421 | 0.99937574 |
| SLC43A2 | 407.370376 | 0.55783283 | 0.68868957 | 0.80999169 | 0.41794495 | 0.99937574 |
| SCARF1 | 16.5642111 | -0.6149992 | 0.60103325 | -1.0232366 | 0.306196 | 0.99937574 |
| RILP | 19.5473416 | -0.8450226 | 0.52382411 | -1.6131801 | 0.10670538 | 0.99937574 |
| PRPF8 | 113.574924 | 0.13907282 | 0.31122797 | 0.44685195 | 0.65498196 | 0.99937574 |
| MIR22HG | 49.6172177 | -0.930906 | 0.69745975 | -1.3347092 | 0.18197153 | 0.99937574 |
| WDR81 | 26.2170685 | -1.108802 | 0.74515136 | -1.4880225 | 0.13674496 | 0.99937574 |
| SERPINF2 | 33.4668434 | 0.17216372 | 0.60281932 | 0.28559754 | 0.77518639 | 0.99937574 |
| SMYD4 | 28.788982 | -0.4983966 | 0.4531232 | -1.0999141 | 0.27136955 | 0.99937574 |
| RPA1 | 40.9619902 | -0.5200356 | 0.36460204 | -1.4263103 | 0.15377879 | 0.99937574 |
| SMG6 | 189.260592 | -0.4584915 | 0.33006179 | -1.3891081 | 0.16479989 | 0.99937574 |
| TSR1 | 20.9740534 | -0.5130374 | 0.53077739 | -0.9665773 | 0.33375539 | 0.99937574 |
| SGSM2 | 60.4051102 | 0.07703143 | 0.29615198 | 0.26010776 | 0.79478065 | 0.99937574 |
| MNT | 24.7672742 | -0.0717647 | 0.39372121 | -0.1822728 | 0.85536863 | 0.99937574 |
| METTL16 | 95.5476179 | -0.2965145 | 0.30957601 | -0.9578083 | 0.33815945 | 0.99937574 |
| PAFAH1B1 | 354.794055 | -0.318972 | 0.24541103 | -1.2997458 | 0.19368812 | 0.99937574 |
| CLUH | 55.286945 | -0.4663103 | 0.49414234 | -0.9436761 | 0.34533518 | 0.99937574 |
| RAP1GAP2 | 62.6038143 | 0.7801374 | 0.48218645 | 1.61791646 | 0.1056806 | 0.99937574 |

| | | | | | | |
|---|---|---|---|---|---|---|
| CTNS | 26.621264 | -0.4091142 | 0.74066673 | -0.5523593 | 0.5807022 | 0.99937574 |
| TAX1BP3 | 16.1247047 | -0.6852084 | 0.61590134 | -1.1125295 | 0.26591057 | 0.99937574 |
| EMC6 | 34.52138 | 0.13264 | 0.60913485 | 0.21775145 | 0.82762277 | 0.99937574 |
| ITGAE | 17.9641446 | -0.1684595 | 0.52002975 | -0.323942 | 0.74598193 | 0.99937574 |
| NCBP3 | 80.1786546 | -0.2389124 | 0.29805206 | -0.8015793 | 0.42279638 | 0.99937574 |
| CAMKK1 | 11.5015698 | -0.5059884 | 0.73689821 | -0.6866463 | 0.49230567 | 0.99937574 |
| ZZEF1 | 122.689981 | -0.358783 | 0.30515904 | -1.1757247 | 0.23970491 | 0.99937574 |
| CYB5D2 | 37.6560673 | 0.16713958 | 0.33580303 | 0.49773102 | 0.61867364 | 0.99937574 |
| ANKFY1 | 87.299837 | -0.2904431 | 0.32978385 | -0.8807075 | 0.37847613 | 0.99937574 |
| UBE2G1 | 113.19521 | -0.2098018 | 0.32002799 | -0.6555732 | 0.51209881 | 0.99937574 |
| GGT6 | 18.9151275 | 1.38983478 | 0.71512967 | 1.9434724 | 0.05195911 | 0.99937574 |
| SMTNL2 | 16.6707708 | -0.4204201 | 0.74547244 | -0.5639647 | 0.57277815 | 0.99937574 |
| PELP1 | 27.1866936 | -0.2876101 | 0.43041551 | -0.6682149 | 0.5039964 | 0.99937574 |
| MED11 | 18.5063735 | -0.4234535 | 0.53724549 | -0.7881936 | 0.43058347 | 0.99937574 |
| PSMB6 | 46.448581 | 0.29242442 | 0.54303456 | 0.53850057 | 0.59023151 | 0.99937574 |
| MINK1 | 72.7322401 | 0.23807492 | 0.4546594 | 0.52363357 | 0.60053343 | 0.99937574 |
| SLC25A11 | 41.65353 | 0.62566684 | 0.48272197 | 1.29612256 | 0.19493326 | 0.99937574 |
| RNF167 | 23.4515378 | 0.24411117 | 0.39961866 | 0.61086028 | 0.54129208 | 0.99937574 |
| PFN1 | 121.05493 | -0.117529 | 0.46706883 | -0.251631 | 0.80132633 | 0.99937574 |
| SPAG7 | 45.3249382 | -0.2674238 | 0.35930926 | -0.7442719 | 0.45671202 | 0.99937574 |
| CAMTA2 | 26.9504418 | -0.5505506 | 0.47080582 | -1.1693793 | 0.24225085 | 0.99937574 |
| KIF1C | 32.9390739 | -0.378958 | 0.5968462 | -0.6349342 | 0.52547135 | 0.99937574 |
| ZNF594 | 16.4830772 | 0.76198382 | 0.5856114 | 1.30117656 | 0.19319803 | 0.99937574 |
| AC087500.1 | 24.4636268 | 0.95250809 | 0.44973595 | 2.11792739 | 0.03418122 | 0.99937574 |
| RABEP1 | 192.211549 | -0.2441847 | 0.32717826 | -0.7463353 | 0.45546491 | 0.99937574 |
| NUP88 | 102.467158 | -0.4619975 | 0.31540366 | -1.4647818 | 0.14298046 | 0.99937574 |
| RPAIN | 45.2114988 | -0.3688612 | 0.35307359 | -1.0447149 | 0.29615475 | 0.99937574 |
| C1QBP | 44.242673 | 0.7459373 | 0.42113219 | 1.77126642 | 0.07651641 | 0.99937574 |
| DERL2 | 35.1331786 | -0.3316148 | 0.43434026 | -0.7634909 | 0.44517071 | 0.99937574 |
| MIS12 | 13.896469 | 0.00933924 | 0.55772655 | 0.01674519 | 0.98663989 | 0.99937574 |
| PITPNM3 | 18.0362408 | 1.09487484 | 0.59005513 | 1.85554667 | 0.0635182 | 0.99937574 |
| KIAA0753 | 43.8878197 | 0.67834704 | 0.43243728 | 1.56865994 | 0.1167272 | 0.99937574 |
| TXNDC17 | 20.2627668 | 0.67233476 | 0.48943236 | 1.37370313 | 0.16953386 | 0.99937574 |
| MED31 | 15.1798309 | 0.47371034 | 0.57999913 | 0.81674319 | 0.41407521 | 0.99937574 |
| XAF1 | 46.4677716 | -4.703577 | 1.6978111 | -2.7703771 | 0.00559914 | 0.65033075 |
| ALOX12-AS1 | 39.9148152 | 0.79913909 | 0.55583618 | 1.43772413 | 0.15051234 | 0.99937574 |
| SLC16A11 | 14.3472006 | -0.73925 | 0.67296536 | -1.0984963 | 0.27198783 | 0.99937574 |
| ACADVL | 204.365364 | 0.48134605 | 0.33548849 | 1.43476175 | 0.151355 | 0.99937574 |
| DVL2 | 17.1206605 | -0.5590034 | 0.51391651 | -1.087732 | 0.27671345 | 0.99937574 |
| PHF23 | 15.5604038 | -0.5286159 | 0.59520544 | -0.8881234 | 0.37447438 | 0.99937574 |
| ELP5 | 30.488491 | 0.94390409 | 0.42343844 | 2.22914127 | 0.02580451 | 0.96546668 |
| EIF5A | 35.4255199 | -0.3601188 | 0.46251199 | -0.778615 | 0.43620651 | 0.99937574 |
| GPS2 | 27.8525511 | -0.0583168 | 0.40668577 | -0.1433952 | 0.8859781 | 0.99937574 |
| ZBTB4 | 105.321385 | 0.35125578 | 0.29910728 | 1.17434715 | 0.24025601 | 0.99937574 |
| POLR2A | 129.723006 | -0.147603 | 0.43093745 | -0.342516 | 0.73196261 | 0.99937574 |
| EIF4A1 | 39.2401029 | -0.1598374 | 0.48106427 | -0.3322579 | 0.73969455 | 0.99937574 |
| MPDU1 | 33.4141984 | -0.2802049 | 0.4358782 | -0.6428513 | 0.52032061 | 0.99937574 |
| FXR2 | 24.1813969 | 0.33482262 | 0.46574295 | 0.71890003 | 0.47220252 | 0.99937574 |
| SAT2 | 33.721901 | 0.0990596 | 0.61336887 | 0.16150085 | 0.87169894 | 0.99937574 |
| TP53 | 13.8932345 | -0.0729449 | 0.52534641 | -0.1388511 | 0.88956784 | 0.99937574 |
| KDM6B | 31.866168 | -1.2487369 | 0.54061267 | -2.3098551 | 0.02089618 | 0.96427895 |
| NAA38 | 86.8007022 | 0.3589811 | 0.42597379 | 0.84273049 | 0.39937919 | 0.99937574 |
| CHD3 | 31.3223441 | -0.0292136 | 0.4627453 | -0.063131 | 0.94966221 | 0.99937574 |
| TRAPPC1 | 17.9099522 | 0.22846658 | 0.71162065 | 0.32105108 | 0.74817168 | 0.99937574 |

| | | | | | | |
|---|---|---|---|---|---|---|
| PER1 | 50.1057909 | -0.6275658 | 0.61152333 | -1.0262335 | 0.30478151 | 0.99937574 |
| TMEM107 | 13.7092448 | 0.82148929 | 0.51745831 | 1.5875468 | 0.11238886 | 0.99937574 |
| CTC1 | 28.2406814 | -0.181611 | 0.4133744 | -0.4393378 | 0.66041679 | 0.99937574 |
| RANGRF | 14.5312312 | -0.37903 | 0.65845953 | -0.5756315 | 0.56486432 | 0.99937574 |
| NDEL1 | 61.6531319 | -0.5319598 | 0.37778306 | -1.4081092 | 0.15909874 | 0.99937574 |
| MYH10 | 87.7936174 | 0.10556443 | 0.50961888 | 0.20714387 | 0.8358975 | 0.99937574 |
| NTN1 | 250.496839 | 0.47702018 | 0.43594363 | 1.09422446 | 0.27385655 | 0.99937574 |
| STX8 | 209.464848 | -0.2362 | 0.31319564 | -0.7541612 | 0.45075241 | 0.99937574 |
| USP43 | 64.0718297 | -0.210556 | 0.7288482 | -0.2888888 | 0.77266651 | 0.99937574 |
| SCO1 | 33.9582317 | 0.21211098 | 0.4678367 | 0.45338678 | 0.65027025 | 0.99937574 |
| TMEM220 | 34.5846283 | -0.2687591 | 0.38440848 | -0.6991497 | 0.48445851 | 0.99937574 |
| TMEM220-A: | 35.9306373 | -0.2959132 | 0.37909134 | -0.7805855 | 0.43504633 | 0.99937574 |
| MAP2K4 | 219.560696 | 0.11297663 | 0.25339108 | 0.44585874 | 0.65569929 | 0.99937574 |
| ARHGAP44 | 21.291691 | 0.22565233 | 0.50773505 | 0.44442929 | 0.65673224 | 0.99937574 |
| ELAC2 | 29.7566189 | 0.10783239 | 0.40660729 | 0.26520034 | 0.79085513 | 0.99937574 |
| COX10 | 68.8410269 | 0.10510724 | 0.39478559 | 0.26623881 | 0.7900553 | 0.99937574 |
| TRIM16 | 11.8815129 | 1.07250056 | 0.54551831 | 1.9660212 | 0.04929616 | 0.99937574 |
| ZSWIM7 | 49.9678915 | -0.0904886 | 0.35993762 | -0.2514008 | 0.80150423 | 0.99937574 |
| TTC19 | 156.135873 | 0.21805429 | 0.39339961 | 0.55428191 | 0.57938593 | 0.99937574 |
| NCOR1 | 575.91848 | 0.03692189 | 0.39405293 | 0.0936978 | 0.92534922 | 0.99937574 |
| PIGL | 99.5832643 | 0.24387687 | 0.33365882 | 0.73091691 | 0.46482991 | 0.99937574 |
| CENPV | 11.6724479 | 0.16361779 | 0.63208583 | 0.25885375 | 0.79574808 | 0.99937574 |
| UBB | 230.388954 | 0.14388523 | 0.56122553 | 0.25637684 | 0.79765987 | 0.99937574 |
| LRRC75A | 25.3167651 | 0.86676795 | 0.55046148 | 1.57462056 | 0.11534406 | 0.99937574 |
| AC098850.3 | 67.3821272 | 0.0440363 | 0.72115714 | 0.0610634 | 0.95130872 | 0.99937574 |
| MPRIP | 188.289024 | 0.03055684 | 0.27580306 | 0.11079225 | 0.91178109 | 0.99937574 |
| COPS3 | 27.3010335 | -0.2081736 | 0.3822091 | -0.5446589 | 0.58598816 | 0.99937574 |
| MED9 | 53.502139 | -0.1504417 | 0.38601011 | -0.3897352 | 0.69673234 | 0.99937574 |
| RASD1 | 182.51818 | 0.11906431 | 0.5850433 | 0.20351366 | 8.39E-01 | 0.99937574 |
| PEMT | 12.5673066 | 0.55411397 | 0.69077628 | 0.80216126 | 0.42245968 | 0.99937574 |
| RAI1 | 89.8527637 | 0.24405338 | 0.46693036 | 0.52267618 | 0.60119961 | 0.99937574 |
| SREBF1 | 167.363253 | 0.23106997 | 0.4887917 | 0.47273711 | 0.63640075 | 0.99937574 |
| TOM1L2 | 125.756233 | 0.02051768 | 0.28223106 | 0.07269815 | 0.94204632 | 0.99937574 |
| ATPAF2 | 17.5371918 | -0.2571523 | 0.43059775 | -0.5971985 | 0.55037483 | 0.99937574 |
| GID4 | 11.3469183 | -0.4445276 | 0.59426661 | -0.7480272 | 0.45444374 | 0.99937574 |
| DRG2 | 14.4268427 | -0.3454768 | 0.60261402 | -0.573297 | 0.56644359 | 0.99937574 |
| ALKBH5 | 61.9858089 | -0.1349609 | 0.34195227 | -0.3946777 | 0.69308074 | 0.99937574 |
| FLII | 52.98968 | -0.0314985 | 0.33639533 | -0.0936354 | 0.92539876 | 0.99937574 |
| MIEF2 | 11.706851 | -0.2923255 | 0.52664085 | -0.5550756 | 0.57884297 | 0.99937574 |
| TOP3A | 31.6740415 | -0.0810655 | 0.3793832 | -0.2136772 | 0.83079879 | 0.99937574 |
| SMCR8 | 19.6150519 | 0.04059096 | 0.53877494 | 0.07533937 | 0.9399447 | 0.99937574 |
| SHMT1 | 27.4804583 | -0.2286158 | 0.49929884 | -0.4578736 | 0.64704325 | 0.99937574 |
| TVP23B | 23.3551445 | 0.32797391 | 0.47878167 | 0.68501767 | 0.49333277 | 0.99937574 |
| PRPSAP2 | 63.5798515 | -0.3324248 | 0.32001331 | -1.0387842 | 0.29890511 | 0.99937574 |
| FAM83G | 10.4907816 | -1.4264911 | 0.93766309 | -1.5213258 | 0.12817809 | 0.99937574 |
| EPN2 | 141.073717 | 0.36279719 | 0.33935841 | 1.06906794 | 0.28503905 | 0.99937574 |
| B9D1 | 16.430026 | -0.1794468 | 0.58739118 | -0.305498 | 0.75998689 | 0.99937574 |
| AC004448.2 | 23.654942 | -0.0464501 | 0.52956769 | -0.0877132 | 0.93010465 | 0.99937574 |
| SLC47A1 | 29.5526908 | -0.0987593 | 0.50516602 | -0.1954988 | 0.84500251 | 0.99937574 |
| ALDH3A2 | 40.1107712 | -0.315235 | 0.46134034 | -0.6833025 | 0.49441574 | 0.99937574 |
| ULK2 | 36.7472969 | 0.07266426 | 0.44993843 | 0.16149824 | 0.871701 | 0.99937574 |
| AKAP10 | 123.83025 | -0.0882424 | 0.24029581 | -0.3672242 | 0.71345176 | 0.99937574 |
| SPECC1 | 35.2478808 | -0.6678357 | 0.43779839 | -1.5254412 | 0.12714909 | 0.99937574 |
| CCDC144NL-/ | 36.465478 | 0.31089411 | 0.64212737 | 0.48416269 | 0.62827041 | 0.99937574 |

| | | | | | | |
|---|---|---|---|---|---|---|
| USP22 | 147.60874 | 0.15386308 | 0.28696483 | 0.536174 | 0.5918383 | 0.99937574 |
| DHRS7B | 95.3374772 | 0.01746225 | 0.29696967 | 0.05880147 | 0.95311024 | 0.99937574 |
| TMEM11 | 18.1161841 | 0.10913947 | 0.52408 | 0.20824964 | 0.83503405 | 0.99937574 |
| AC087294.1 | 17.1384687 | 0.09862211 | 0.54561864 | 0.18075282 | 0.8565616 | 0.99937574 |
| NATD1 | 10.6116636 | 0.16135235 | 0.65094335 | 0.24787465 | 0.8042314 | 0.99937574 |
| MAP2K3 | 21.0244496 | -0.5047782 | 0.58504299 | -0.8628053 | 0.38824453 | 0.99937574 |
| WSB1 | 598.218415 | -0.0871168 | 0.43217023 | -0.2015799 | 0.84024518 | 0.99937574 |
| KSR1 | 47.4038791 | 0.58512564 | 0.31077316 | 1.88280621 | 0.05972663 | 0.99937574 |
| NLK | 211.321572 | -0.0008185 | 0.24585585 | -0.0033294 | 0.99734356 | 0.99937574 |
| IFT20 | 14.369875 | 0.09570641 | 0.50235002 | 0.19051738 | 0.84890372 | 0.99937574 |
| TNFAIP1 | 32.5547252 | 0.08858768 | 0.4801623 | 0.18449528 | 0.85362493 | 0.99937574 |
| POLDIP2 | 34.1389765 | 0.32513399 | 0.49203097 | 0.66079986 | 0.50874067 | 0.99937574 |
| TMEM199 | 11.0059143 | -0.6401946 | 0.61424293 | -1.0422499 | 0.29729585 | 0.99937574 |
| SARM1 | 12.7336147 | -0.4959959 | 0.72745781 | -0.6818208 | 0.49535225 | 0.99937574 |
| AC005726.1 | 15.8812188 | -0.0450698 | 0.50351591 | -0.0895102 | 0.92867648 | 0.99937574 |
| KIAA0100 | 64.2897382 | -0.1465858 | 0.40778184 | -0.3594712 | 0.71924259 | 0.99937574 |
| SDF2 | 27.0854415 | -0.1960566 | 0.41277304 | -0.4749744 | 0.63480525 | 0.99937574 |
| SUPT6H | 75.6433174 | -0.4582689 | 0.37846663 | -1.2108567 | 0.22595031 | 0.99937574 |
| RAB34 | 34.5266132 | 0.20994313 | 0.43942235 | 0.47777072 | 0.63281341 | 0.99937574 |
| RPL23A | 169.068971 | 0.12581886 | 0.50094487 | 0.25116309 | 0.80168802 | 0.99937574 |
| NEK8 | 17.003977 | 0.11952696 | 0.56393422 | 0.21195196 | 0.83214451 | 0.99937574 |
| TRAF4 | 43.9161864 | 0.01751493 | 0.40411191 | 0.04334177 | 0.96542909 | 0.99937574 |
| FAM222B | 133.784342 | -0.4315564 | 0.36018478 | -1.1981529 | 0.23085749 | 0.99937574 |
| ERAL1 | 12.630577 | 0.00607756 | 0.55331632 | 0.01098388 | 0.99123631 | 0.99937574 |
| FLOT2 | 103.647477 | 0.21528392 | 0.28946511 | 0.74373012 | 0.45703976 | 0.99937574 |
| PHF12 | 55.723238 | 0.48716858 | 0.38867705 | 1.25340199 | 0.21005945 | 0.99937574 |
| MYO18A | 49.889543 | -0.6285075 | 0.49209442 | -1.2772091 | 0.20152843 | 0.99937574 |
| NUFIP2 | 170.528988 | -0.1238296 | 0.31688088 | -0.3907764 | 0.69596254 | 0.99937574 |
| TAOK1 | 245.61275 | -0.20278 | 0.22831086 | -0.8881751 | 0.37444659 | 0.99937574 |
| ABHD15-AS1 | 23.31532 | -0.6406984 | 0.69351919 | -0.9238365 | 0.35557144 | 0.99937574 |
| GIT1 | 15.2789437 | -0.2379392 | 0.59097339 | -0.4026225 | 0.68722598 | 0.99937574 |
| SSH2 | 353.616359 | -0.1220696 | 0.33907181 | -0.3600111 | 0.71883881 | 0.99937574 |
| AC023389.1 | 15.61205 | 0.04828645 | 0.55672845 | 0.0867325 | 0.93088414 | 0.99937574 |
| NSRP1 | 155.961531 | -0.102267 | 0.34819912 | -0.2937026 | 0.76898518 | 0.99937574 |
| AC104984.4 | 18.9897141 | 1.30038033 | 0.49810708 | 2.61064415 | 0.00903719 | 0.74756801 |
| BLMH | 31.2080751 | 0.09723417 | 0.40272955 | 0.24143788 | 0.80921576 | 0.99937574 |
| CPD | 151.75137 | 0.05905027 | 0.2979766 | 0.19817082 | 0.84291142 | 0.99937574 |
| GOSR1 | 99.2165608 | -0.0882832 | 0.25166045 | -0.3508029 | 0.72573618 | 0.99937574 |
| CRLF3 | 68.5484012 | 0.0734714 | 0.39740117 | 0.18487969 | 0.8533234 | 0.99937574 |
| TEFM | 24.9900177 | 0.33837937 | 0.49615016 | 0.68200999 | 0.49523263 | 0.99937574 |
| RNF135 | 17.0606924 | 0.30764999 | 0.45892687 | 0.67036821 | 0.50262309 | 0.99937574 |
| NF1 | 606.804151 | -0.0273625 | 0.29632048 | -0.0923409 | 0.92642715 | 0.99937574 |
| OMG | 16.6422253 | 0.80527754 | 0.9597348 | 0.83906256 | 0.40143421 | 0.99937574 |
| RAB11FIP4 | 42.2131428 | 0.15475537 | 0.41740037 | 0.37076001 | 0.71081629 | 0.99937574 |
| COPRS | 49.8120149 | -0.4805233 | 0.49727775 | -0.9663076 | 0.33389029 | 0.99937574 |
| UTP6 | 30.0407188 | -0.1696876 | 0.42256862 | -0.4015622 | 0.68800627 | 0.99937574 |
| SUZ12 | 164.158936 | -0.1733342 | 0.25742045 | -0.6733505 | 0.50072431 | 0.99937574 |
| LRRC37B | 45.4495422 | -0.2960345 | 0.32763747 | -0.903543 | 0.36623777 | 0.99937574 |
| RHOT1 | 82.0148441 | 0.14628145 | 0.27747128 | 0.52719492 | 0.59805824 | 0.99937574 |
| C17orf75 | 16.1121727 | 0.30898649 | 0.58479204 | 0.52836987 | 0.59724264 | 0.99937574 |
| ZNF207 | 183.895938 | 0.26098811 | 0.25108644 | 1.03943533 | 0.29860232 | 0.99937574 |
| PSMD11 | 70.5267943 | -0.2967131 | 0.26374823 | -1.124986 | 0.26059497 | 0.99937574 |
| MYO1D | 374.217954 | -0.1428761 | 0.30951878 | -0.4616072 | 0.64436306 | 0.99937574 |
| TMEM98 | 13.4952098 | 0.70832118 | 0.62741942 | 1.12894366 | 0.2589216 | 0.99937574 |

| | | | | | | |
|---|---|---|---|---|---|---|
| ASIC2 | 37.857297 | 0.28319068 | 0.81996202 | 0.34537049 | 0.72981586 | 0.99937574 |
| ZNF830 | 10.8831151 | -0.7160276 | 0.69073096 | -1.036623 | 0.29991157 | 0.99937574 |
| LIG3 | 16.4149154 | 0.38261454 | 0.44109178 | 0.86742615 | 0.38570856 | 0.99937574 |
| RFFL | 13.7836436 | -0.1025505 | 0.55704147 | -0.1840985 | 0.85393617 | 0.99937574 |
| SLFN5 | 33.7258996 | 0.07114299 | 0.55914798 | 0.12723464 | 8.99E-01 | 0.99937574 |
| SLFN13 | 16.5063444 | -0.2486648 | 0.66210091 | -0.3755693 | 0.70723709 | 0.99937574 |
| LINC02001 | 13.9988047 | -0.0740387 | 0.5122614 | -0.1445331 | 0.88507956 | 0.99937574 |
| AP2B1 | 160.382889 | 0.06383844 | 0.2862862 | 0.22298817 | 0.82354471 | 0.99937574 |
| TAF15 | 255.18003 | 0.15765763 | 0.2377995 | 0.66298555 | 0.50733981 | 0.99937574 |
| AC015849.3 | 17.8730222 | 0.66074991 | 0.624881 | 1.05740119 | 0.29032853 | 0.99937574 |
| ZNHIT3 | 18.3137194 | -0.1963893 | 0.48949794 | -0.4012055 | 0.68826884 | 0.99937574 |
| MYO19 | 171.214477 | 0.40640961 | 0.35690495 | 1.13870544 | 0.25482603 | 0.99937574 |
| GGNBP2 | 111.659615 | -0.3475112 | 0.31458814 | -1.1046545 | 0.26930932 | 0.99937574 |
| AATF | 60.9816972 | -0.3695145 | 0.35406975 | -1.0436207 | 0.2966609 | 0.99937574 |
| ACACA | 295.671659 | -0.3064651 | 0.37061575 | -0.8269079 | 0.40828924 | 0.99937574 |
| TADA2A | 40.9890635 | -0.2006369 | 0.4429584 | -0.4529475 | 0.65058653 | 0.99937574 |
| SYNRG | 152.311057 | -0.1020176 | 0.22393841 | -0.4555608 | 0.64870586 | 0.99937574 |
| DDX52 | 151.442345 | -0.0871339 | 0.39727281 | -0.21933 | 0.82639297 | 0.99937574 |
| AC243571.2 | 81.7764815 | -0.0755876 | 0.90488284 | -0.083533 | 9.33E-01 | 9.99E-01 |
| HNF1B | 154.221172 | 0.24940538 | 0.27735219 | 0.8992371 | 3.69E-01 | 0.99937574 |
| MRPL45 | 28.1214663 | -0.438 | 0.41958948 | -1.0438773 | 0.29654215 | 0.99937574 |
| SOCS7 | 26.0842504 | 0.30513149 | 0.38937299 | 0.78364832 | 0.4332465 | 0.99937574 |
| MLLT6 | 201.787266 | -0.0927059 | 0.35994286 | -0.2575572 | 0.79674866 | 0.99937574 |
| CISD3 | 26.5486906 | -0.0409506 | 0.4711852 | -0.0869097 | 0.93074331 | 0.99937574 |
| PCGF2 | 22.8420345 | -0.2888453 | 0.52317437 | -0.5521014 | 0.58087889 | 0.99937574 |
| PSMB3 | 53.8918668 | -0.0175858 | 0.35285698 | -0.0498383 | 0.96025125 | 0.99937574 |
| PIP4K2B | 26.9133784 | 0.16033188 | 0.39629656 | 0.40457551 | 0.68578956 | 0.99937574 |
| CWC25 | 28.4142664 | -0.1817322 | 0.4014994 | -0.4526339 | 0.65081238 | 0.99937574 |
| RPL23 | 146.882736 | -0.1015059 | 0.48234461 | -0.2104428 | 0.8333221 | 0.99937574 |
| LASP1 | 43.2650678 | -0.4558482 | 0.41469455 | -1.0992385 | 0.27166405 | 0.99937574 |
| RPL19 | 250.255385 | -0.1643219 | 0.58421861 | -0.2812679 | 0.77850492 | 0.99937574 |
| STAC2 | 297.181868 | -0.9029723 | 0.42952505 | -2.1022575 | 0.03553072 | 0.99937574 |
| FBXL20 | 265.874995 | 0.07922907 | 0.29762185 | 0.26620718 | 0.79007966 | 0.99937574 |
| MED1 | 49.9775158 | -0.3621356 | 0.35261921 | -1.0269878 | 0.30442619 | 0.99937574 |
| CDK12 | 104.75598 | 0.1263562 | 0.25994376 | 0.48609051 | 0.62690299 | 0.99937574 |
| STARD3 | 39.9430433 | 0.15846606 | 0.35271232 | 0.44927851 | 0.65323076 | 0.99937574 |
| PGAP3 | 28.0693788 | 1.34384588 | 0.51064563 | 2.63166041 | 0.00849687 | 0.74742128 |
| ERBB2 | 150.798708 | 0.73912762 | 0.34810806 | 2.12327064 | 0.03373118 | 0.99937574 |
| MIEN1 | 44.3741564 | 0.04820545 | 0.35253578 | 0.13673918 | 0.89123696 | 0.99937574 |
| GRB7 | 67.4091266 | -0.2209126 | 0.52662531 | -0.4194873 | 0.67486006 | 0.99937574 |
| ORMDL3 | 14.1543582 | 0.29792718 | 0.47370675 | 0.62892746 | 0.52939655 | 0.99937574 |
| PSMD3 | 29.4520961 | 0.04062954 | 0.40375631 | 0.10062887 | 0.91984508 | 0.99937574 |
| MED24 | 37.9493872 | -0.0474589 | 0.35660296 | -0.133086 | 0.89412536 | 0.99937574 |
| THRA | 26.000858 | 0.23849023 | 0.42645937 | 0.55923319 | 0.57600258 | 0.99937574 |
| MSL1 | 53.8454122 | 0.37019617 | 0.35252765 | 1.05011952 | 0.29366317 | 0.99937574 |
| CASC3 | 50.0357989 | -0.0461746 | 0.28125347 | -0.1641744 | 0.86959386 | 0.99937574 |
| WIPF2 | 108.346456 | -0.2743776 | 0.2879962 | -0.9527125 | 0.34073576 | 0.99937574 |
| RARA | 11.2126398 | -0.5046147 | 0.59123205 | -0.8534968 | 0.39338386 | 0.99937574 |
| IGFBP4 | 11.9813163 | -0.1120395 | 0.56376748 | -0.1987335 | 0.84247119 | 0.99937574 |
| TNS4 | 15.9644128 | -0.2675682 | 0.86024799 | -0.3110361 | 0.75577319 | 0.99937574 |
| KRT10 | 43.5780928 | -0.0534316 | 0.64409847 | -0.0829557 | 0.93388677 | 0.99937574 |
| KRT19 | 19.5412711 | 1.2211823 | 0.85431542 | 1.42942791 | 0.15288128 | 0.99937574 |
| EIF1 | 361.486412 | 0.14659825 | 0.47449414 | 0.30895693 | 0.75735429 | 0.99937574 |
| JUP | 143.5596 | -0.395037 | 0.39626667 | -0.9968968 | 0.31881462 | 0.99937574 |

| | | | | | | |
|---|---|---|---|---|---|---|
| NT5C3B | 24.9130751 | 0.09610874 | 0.45696784 | 0.2103184 | 0.83341918 | 0.99937574 |
| ACLY | 42.3254175 | 0.41694679 | 0.41025549 | 1.01631009 | 0.30948175 | 0.99937574 |
| CNP | 30.8571451 | -0.4822838 | 0.45746918 | -1.0542432 | 0.29177159 | 0.99937574 |
| DNAJC7 | 83.9050376 | -0.2691127 | 0.28210863 | -0.9539328 | 0.34011765 | 0.99937574 |
| NKIRAS2 | 12.1817743 | -0.91451 | 0.63993803 | -1.4290602 | 0.15298694 | 0.99937574 |
| GHDC | 10.3887804 | 0.22681997 | 0.56562174 | 0.40101 | 0.68841276 | 0.99937574 |
| STAT5B | 196.069394 | -0.3206202 | 0.51841472 | -0.6184628 | 0.53627033 | 0.99937574 |
| STAT5A | 21.4367294 | -1.0646457 | 0.72203888 | -1.4744991 | 0.14034727 | 0.99937574 |
| STAT3 | 272.4694 | -0.2405978 | 0.34967566 | -0.6880598 | 0.49141512 | 0.99937574 |
| ATP6V0A1 | 66.641234 | -0.4824149 | 0.34703137 | -1.3901191 | 0.16449273 | 0.99937574 |
| NAGLU | 27.1198363 | -1.0921812 | 0.76531442 | -1.4271013 | 0.15355071 | 0.99937574 |
| COASY | 25.765068 | 0.43275671 | 0.50582173 | 0.85555184 | 0.39224572 | 0.99937574 |
| MLX | 24.790947 | -0.0675574 | 0.58644088 | -0.115199 | 0.90828743 | 0.99937574 |
| RETREG3 | 76.7421307 | -0.2131637 | 0.32296261 | -0.6600259 | 0.5092372 | 0.99937574 |
| TUBG1 | 10.5330652 | 0.81669453 | 0.68678876 | 1.18914953 | 0.23438083 | 0.99937574 |
| TUBG2 | 15.8549163 | -0.9156336 | 0.48071614 | -1.9047282 | 0.05681542 | 0.99937574 |
| PLEKHH3 | 17.2071063 | -0.0569194 | 0.56879343 | -0.1000704 | 0.92028844 | 0.99937574 |
| EZH1 | 51.403562 | -0.5645127 | 0.31035946 | -1.8188995 | 0.06892677 | 0.99937574 |
| VPS25 | 25.6001417 | -0.3021071 | 0.38686532 | -0.7809102 | 0.43485529 | 0.99937574 |
| COA3 | 103.490922 | 0.51761852 | 0.50419127 | 1.02663127 | 0.30459413 | 0.99937574 |
| BECN1 | 40.9268387 | 0.42830868 | 0.42184674 | 1.01531821 | 0.30995417 | 0.99937574 |
| PSME3 | 27.5601843 | -0.2385968 | 0.43383139 | -0.549976 | 0.58233585 | 0.99937574 |
| LINC00671 | 32.2641761 | 0.14310728 | 0.89294342 | 0.16026467 | 0.87267259 | 0.99937574 |
| RUNDC1 | 32.365746 | 0.17841005 | 0.43368225 | 0.41138427 | 0.68079079 | 0.99937574 |
| RPL27 | 105.353607 | 0.2060735 | 0.40300816 | 0.51133828 | 0.60911421 | 0.99937574 |
| IFI35 | 26.2937053 | -1.8048863 | 0.82694058 | -2.182607 | 0.02906476 | 0.99049468 |
| VAT1 | 16.6334603 | -0.8850088 | 0.68513283 | -1.2917331 | 0.19644958 | 0.99937574 |
| BRCA1 | 21.2905778 | -0.0134865 | 0.78921469 | -0.0170885 | 0.98636601 | 0.99937574 |
| NBR1 | 96.4782554 | 0.1773257 | 0.33682502 | 0.5264624 | 0.59856697 | 0.99937574 |
| LINC00854 | 106.444489 | -1.3819127 | 0.68083526 | -2.0297314 | 0.04238385 | 0.99937574 |
| ARL4D | 105.77811 | 0.91303282 | 0.39099639 | 2.33514386 | 0.01953591 | 0.96427895 |
| DHX8 | 33.198638 | 0.06185462 | 0.4094554 | 0.15106558 | 0.87992399 | 0.99937574 |
| DUSP3 | 66.541481 | -0.2978116 | 0.44276167 | -0.6726229 | 0.50118725 | 0.99937574 |
| PYY | 125.756166 | 1.27616665 | 0.78769453 | 1.62012887 | 0.1052046 | 0.99937574 |
| TMEM101 | 980.802584 | -0.1315464 | 0.66648761 | -0.1973726 | 0.843536 | 0.99937574 |
| LSM12 | 59.8344361 | -0.4592734 | 0.35249807 | -1.3029104 | 0.19260535 | 0.99937574 |
| G6PC3 | 18.8667549 | -0.021582 | 0.50265423 | -0.0429362 | 0.96575243 | 0.99937574 |
| HDAC5 | 66.5529104 | -0.0195879 | 0.45736622 | -0.0428276 | 0.96583897 | 0.99937574 |
| ASB16-AS1 | 14.3269447 | 0.44553029 | 0.48803521 | 0.91290604 | 0.36129196 | 0.99937574 |
| TMUB2 | 26.338265 | 0.07604962 | 0.39209027 | 0.19395946 | 0.84620763 | 0.99937574 |
| ATXN7L3 | 20.386037 | -1.2963625 | 0.60126241 | -2.1560677 | 0.03107838 | 0.99049468 |
| UBTF | 60.6881955 | -0.0290655 | 0.46596792 | -0.0623767 | 0.95026287 | 0.99937574 |
| AC003102.1 | 12.3971552 | 0.15579812 | 0.65180243 | 0.23902661 | 0.81108495 | 0.99937574 |
| SLC4A1 | 876.582799 | 2.63972386 | 0.76510463 | 3.45014755 | 5.60E-04 | 0.28271741 |
| RUNDC3A-AS | 14.2501449 | 0.31098393 | 0.66173342 | 0.46995349 | 0.63838824 | 0.99937574 |
| SLC25A39 | 202.604264 | 0.37367818 | 0.34065972 | 1.09692504 | 0.27267416 | 0.99937574 |
| GRN | 85.3615813 | -0.8916804 | 0.50478607 | -1.766452 | 0.07732005 | 0.99937574 |
| GPATCH8 | 413.626582 | 0.01638744 | 0.21791446 | 0.07520124 | 0.9400546 | 0.99937574 |
| MEIOC | 21.2537582 | -0.0825319 | 0.57806868 | -0.1427718 | 0.8864704 | 0.99937574 |
| CCDC43 | 15.9969613 | -0.0405028 | 0.54118538 | -0.0748408 | 0.94034137 | 0.99937574 |
| DBF4B | 13.1994927 | 0.12140467 | 0.5316757 | 0.22834346 | 0.81937924 | 0.99937574 |
| EFTUD2 | 40.1619547 | -0.0164638 | 0.37790065 | -0.0435664 | 0.96525005 | 0.99937574 |
| DCAKD | 41.1702782 | 0.56557375 | 0.40646739 | 1.39143698 | 0.16409296 | 0.99937574 |
| NMT1 | 78.1567043 | 0.17906027 | 0.24087915 | 0.74336142 | 0.45726289 | 0.99937574 |

| | | | | | | |
|---|---|---|---|---|---|---|
| HEXIM1 | 18.4423306 | -1.401192 | 0.77326117 | -1.8120553 | 0.06997766 | 0.99937574 |
| AC138150.1 | 36.6292241 | 0.3902906 | 0.51341061 | 0.76019194 | 0.44713986 | 0.99937574 |
| MAP3K14 | 36.3182993 | -0.1082578 | 0.36896016 | -0.2934134 | 0.7692062 | 0.99937574 |
| ARHGAP27 | 26.5527479 | -0.4539877 | 0.57146755 | -0.7944243 | 0.42694846 | 0.99937574 |
| PLEKHM1 | 40.5619763 | -0.671389 | 0.42883144 | -1.5656245 | 0.11743655 | 0.99937574 |
| LINC02210-C | 24.6662324 | 0.32808963 | 0.5299532 | 0.6190917 | 0.53585595 | 0.99937574 |
| KANSL1 | 535.251765 | 0.10407546 | 0.41433711 | 0.25118546 | 0.80167072 | 0.99937574 |
| ARL17B | 115.422529 | 2.06900078 | 1.01396218 | 2.0405108 | 0.04129948 | 0.99937574 |
| LRRC37A2 | 33.104959 | 0.28626977 | 0.51018738 | 0.56110712 | 0.57472452 | 0.99937574 |
| ARL17A | 33.3539046 | -0.642648 | 0.45542126 | -1.4111068 | 0.15821312 | 0.99937574 |
| NSF | 141.835614 | -0.2167701 | 0.45891845 | -0.47235 | 0.636677 | 0.99937574 |
| AC005670.2 | 30.2512535 | 0.4753467 | 0.42955799 | 1.10659495 | 0.26846909 | 0.99937574 |
| CDC27 | 181.433167 | 0.09220835 | 0.28696526 | 0.32132234 | 0.74796613 | 0.99937574 |
| AC068234.1 | 176.278585 | -0.7669065 | 0.89404911 | -0.8577901 | 0.39100837 | 0.99937574 |
| AC068234.2 | 23.1921073 | 0.09595011 | 0.65285774 | 0.1469694 | 0.88315617 | 0.99937574 |
| EFCAB13 | 145.171847 | -0.002216 | 0.64576573 | -0.0034316 | 0.99726201 | 0.99937574 |
| NPEPPS | 316.649183 | -0.1290749 | 0.23457056 | -0.5502604 | 0.58214075 | 0.99937574 |
| AC025682.1 | 29.2703306 | 0.43466209 | 0.43530531 | 0.99852236 | 0.31802613 | 0.99937574 |
| KPNB1 | 135.954777 | -0.4256953 | 0.28337341 | -1.5022417 | 0.1330347 | 0.99937574 |
| MRPL10 | 11.6115088 | 0.00369013 | 0.58016535 | 0.00636048 | 0.9949251 | 0.99937574 |
| SCRN2 | 19.4271514 | 0.376969 | 0.55814459 | 0.67539668 | 0.49942377 | 0.99937574 |
| SP2 | 18.0964496 | -0.7071041 | 0.50872714 | -1.3899476 | 0.1645448 | 0.99937574 |
| PNPO | 23.5457136 | 0.59519912 | 0.40173115 | 1.48158568 | 0.13845057 | 0.99937574 |
| PRR15L | 12.0871341 | 0.6925057 | 0.92750301 | 0.74663445 | 0.45528424 | 0.99937574 |
| CDK5RAP3 | 33.9196012 | -0.0492454 | 0.36224849 | -0.1359436 | 0.89186584 | 0.99937574 |
| NFE2L1 | 148.15875 | -0.3257961 | 0.42440267 | -0.767658 | 0.44269037 | 0.99937574 |
| CBX1 | 33.0171752 | 0.09654188 | 0.41363351 | 0.23339957 | 0.81545115 | 0.99937574 |
| SNX11 | 16.2403889 | -0.0506933 | 0.53729397 | -0.0943494 | 0.92483164 | 0.99937574 |
| SKAP1 | 568.465237 | 0.2940906 | 0.32941441 | 0.89276788 | 0.37198149 | 0.99937574 |
| SKAP1-AS1 | 12.7123396 | 0.35110987 | 0.87611095 | 0.4007596 | 0.68859713 | 0.99937574 |
| HOXB2 | 33.175338 | 0.64267589 | 0.41949553 | 1.53202085 | 0.12551729 | 0.99937574 |
| HOXB-AS1 | 15.0917525 | -0.2635704 | 0.63908397 | -0.412419 | 0.68003232 | 0.99937574 |
| HOXB3 | 199.583885 | 0.15900098 | 0.27008309 | 0.58871134 | 0.55605493 | 0.99937574 |
| HOXB-AS3 | 61.764788 | 0.2048906 | 0.56447859 | 0.3629732 | 0.7166249 | 0.99937574 |
| HOXB7 | 54.1240031 | -0.7348909 | 0.42726806 | -1.7199763 | 0.08543676 | 0.99937574 |
| AC103702.2 | 29.8520773 | -0.4335725 | 0.43663593 | -0.9929841 | 0.32071772 | 0.99937574 |
| LINC02086 | 64.2999572 | -0.4292817 | 0.35641097 | -1.2044572 | 0.22841293 | 0.99937574 |
| CALCOCO2 | 77.7889067 | 0.025777 | 0.33873744 | 0.07609728 | 0.9393417 | 0.99937574 |
| ATP5MC1 | 98.0098365 | 0.76117934 | 0.46719008 | 1.62927121 | 0.10325562 | 0.99937574 |
| UBE2Z | 104.71963 | -0.3925551 | 0.32214425 | -1.2185692 | 0.22300775 | 0.99937574 |
| SNF8 | 32.8180145 | 0.06869054 | 0.35065663 | 0.19589118 | 0.84469534 | 0.99937574 |
| B4GALNT2 | 27.1610337 | 0.19011089 | 0.84974454 | 0.22372711 | 0.82296965 | 0.99937574 |
| ZNF652 | 189.440768 | 0.21032275 | 0.27317932 | 0.76990728 | 0.4413549 | 0.99937574 |
| AC091180.5 | 15.0668682 | 0.71948072 | 0.69931268 | 1.0288398 | 0.30355496 | 0.99937574 |
| PHB | 105.462578 | 0.55847063 | 0.36404633 | 1.53406469 | 0.12501374 | 0.99937574 |
| SPOP | 103.954283 | 0.11863766 | 0.33383445 | 0.35537872 | 0.72230589 | 0.99937574 |
| SLC35B1 | 16.7776722 | 0.01989388 | 0.45481203 | 0.04374089 | 0.96511095 | 0.99937574 |
| FAM117A | 16.4072457 | -0.7835277 | 0.56126492 | -1.3960033 | 0.16271351 | 0.99937574 |
| KAT7 | 37.5792777 | -0.1606781 | 0.34381081 | -0.4673445 | 0.64025345 | 0.99937574 |
| ITGA3 | 123.539379 | -0.5779687 | 0.5495598 | -1.051694 | 2.93E-01 | 0.99937574 |
| PDK2 | 13.5181482 | 0.91816032 | 0.50892742 | 1.80410857 | 0.07121429 | 0.99937574 |
| XYLT2 | 18.4527509 | 0.08798004 | 0.54690096 | 0.16087014 | 0.87219568 | 0.99937574 |
| MRPL27 | 54.5824511 | -0.0078148 | 0.38065795 | -0.0205297 | 0.9836208 | 0.99937574 |
| LRRC59 | 20.5515529 | -0.7835337 | 0.59790591 | -1.3104632 | 0.19003919 | 0.99937574 |

| | | | | | | |
|---|---|---|---|---|---|---|
| ACSF2 | 16.5625269 | 1.10541341 | 0.58357846 | 1.89419845 | 0.05819867 | 0.99937574 |
| RSAD1 | 22.4751411 | 0.09564545 | 0.48706447 | 0.19637123 | 0.84431961 | 0.99937574 |
| EPN3 | 13.95875 | 0.01370607 | 0.77043182 | 0.01779012 | 0.98580629 | 0.99937574 |
| SPATA20 | 22.4020382 | 0.50305238 | 0.4638693 | 1.08447009 | 0.27815642 | 0.99937574 |
| ABCC3 | 108.21528 | -0.4717041 | 0.76269019 | -0.6184741 | 0.53626286 | 0.99937574 |
| ANKRD40 | 39.3813055 | -0.4731393 | 0.43639543 | -1.0841987 | 0.27827673 | 0.99937574 |
| AC005921.2 | 11.516044 | 0.30571634 | 0.61585626 | 0.49640859 | 0.61960616 | 0.99937574 |
| LUC7L3 | 247.553516 | 0.02064421 | 0.21845402 | 0.0945014 | 0.92471087 | 0.99937574 |
| TOB1 | 54.206948 | 0.10987284 | 0.38242666 | 0.28730435 | 0.77387929 | 0.99937574 |
| SPAG9 | 466.010608 | -0.5368708 | 0.27781805 | -1.9324547 | 0.05330341 | 0.99937574 |
| AC005920.2 | 36.2851454 | -0.6250144 | 0.71488784 | -0.8742832 | 0.38196406 | 0.99937574 |
| MBTD1 | 99.4758545 | -0.3078092 | 0.31213745 | -0.9861335 | 0.32406762 | 0.99937574 |
| UTP18 | 32.3359075 | -0.5907013 | 0.3461259 | -1.7066081 | 0.08789491 | 0.99937574 |
| TOM1L1 | 195.193246 | 0.36887507 | 0.28184518 | 1.30878616 | 0.1906068 | 0.99937574 |
| COX11 | 33.5377959 | 0.22354152 | 0.5494491 | 0.40684664 | 0.68412063 | 0.99937574 |
| STXBP4 | 132.451903 | 1.03727551 | 0.42529635 | 2.43894762 | 0.0147301 | 0.85933448 |
| MMD | 15.9533836 | -0.4356333 | 0.60385593 | -0.7214193 | 0.47065156 | 0.99937574 |
| PCTP | 21.753346 | 0.71009703 | 0.52416886 | 1.35471045 | 0.17550983 | 0.99937574 |
| DGKE | 11.4056924 | 0.38678299 | 0.62161834 | 0.6222194 | 0.53379761 | 0.99937574 |
| TRIM25 | 25.9571518 | -0.9000854 | 0.52377714 | -1.7184511 | 0.08571436 | 0.99937574 |
| COIL | 20.1218447 | -0.3925968 | 0.58525083 | -0.670818 | 0.50233649 | 0.99937574 |
| SCPEP1 | 63.8057463 | 0.49651719 | 0.41181721 | 1.20567374 | 0.22794331 | 0.99937574 |
| AKAP1 | 60.2075021 | 0.44369746 | 0.3776738 | 1.17481663 | 0.24006808 | 0.99937574 |
| MSI2 | 1245.3173 | 0.25758995 | 0.33859024 | 0.76077191 | 0.44679332 | 0.99937574 |
| MRPS23 | 31.6713026 | -0.1823137 | 0.36618054 | -0.4978792 | 0.61856921 | 0.99937574 |
| CUEDC1 | 150.603079 | -0.3139808 | 0.4589054 | -0.684195 | 0.49385203 | 0.99937574 |
| VEZF1 | 67.6104055 | 0.08486877 | 0.29836945 | 0.28444188 | 0.77607177 | 0.99937574 |
| AC015813.2 | 37.6911231 | -0.2790361 | 0.45371013 | -0.6150097 | 0.53854829 | 0.99937574 |
| SRSF1 | 18.9083489 | -0.0646696 | 0.50068449 | -0.1291624 | 0.89722911 | 0.99937574 |
| DYNLL2 | 49.2320629 | 0.06793483 | 0.43243172 | 0.15709956 | 0.87516638 | 0.99937574 |
| MKS1 | 10.3543674 | 0.22174244 | 0.77623716 | 0.28566327 | 0.77513604 | 0.99937574 |
| TSPOAP1 | 12.1575953 | 0.39640174 | 0.82367362 | 0.48126069 | 0.63033123 | 0.99937574 |
| SUPT4H1 | 16.2056941 | -0.4317481 | 0.58741782 | -0.7349932 | 0.46234363 | 0.99937574 |
| RNF43 | 34.2206763 | 0.34055759 | 0.47146456 | 0.72233975 | 0.47008562 | 0.99937574 |
| MTMR4 | 24.5356041 | 0.26488893 | 0.55602738 | 0.47639548 | 0.63379265 | 0.99937574 |
| RAD51C | 16.7839947 | 0.11451347 | 0.54235653 | 0.21114057 | 0.83277758 | 0.99937574 |
| PPM1E | 113.307433 | -2.1370858 | 1.48884305 | -1.4354003 | 0.15117305 | 0.99937574 |
| TRIM37 | 58.0248003 | -0.2731301 | 0.39578803 | -0.6900918 | 0.49013646 | 0.99937574 |
| SKA2 | 29.3533583 | -0.1710317 | 0.44001929 | -0.3886914 | 0.69750448 | 0.99937574 |
| YPEL2 | 102.780025 | 0.42573973 | 0.46246821 | 0.92058161 | 0.35726891 | 0.99937574 |
| DHX40 | 36.4927761 | -0.1221509 | 0.38084376 | -0.3207376 | 0.74840927 | 0.99937574 |
| CLTC | 316.491289 | 0.3997904 | 0.15762183 | 2.53638979 | 0.0112002 | 0.76274616 |
| PTRH2 | 42.3310462 | -0.0936088 | 0.52488451 | -0.1783416 | 0.85845467 | 0.99937574 |
| VMP1 | 226.26569 | -0.4090967 | 0.28309728 | -1.4450746 | 0.14843695 | 0.99937574 |
| TUBD1 | 19.736067 | -0.3208112 | 0.46129089 | -0.6954639 | 0.48676461 | 0.99937574 |
| RPS6KB1 | 51.5435041 | -0.0613018 | 0.33253223 | -0.1843483 | 0.85374019 | 0.99937574 |
| RNFT1 | 12.5332657 | -0.1223509 | 0.5011117 | -0.244159 | 0.8071077 | 0.99937574 |
| HEATR6 | 15.5167227 | 0.5849959 | 0.57942751 | 1.00961015 | 0.31268211 | 0.99937574 |
| USP32 | 140.623127 | -0.3305115 | 0.39459902 | -0.8375883 | 0.40226197 | 0.99937574 |
| APPBP2 | 125.1666 | -0.1369061 | 0.29014117 | -0.4718604 | 0.63702647 | 0.99937574 |
| PPM1D | 41.0395308 | 0.00023837 | 0.41896031 | 0.00056895 | 0.99954605 | 0.99979721 |
| BCAS3 | 403.812354 | -0.3311861 | 0.27709656 | -1.195201 | 0.23200852 | 0.99937574 |
| TBX2 | 56.4605495 | 0.26849661 | 0.45655669 | 0.5880904 | 0.55647161 | 0.99937574 |
| NACA2 | 18.8983351 | 0.19891402 | 0.64988499 | 0.30607572 | 0.75954699 | 0.99937574 |

| | | | | | | |
|---|---|---|---|---|---|---|
| INTS2 | 34.4056096 | 0.3726724 | 0.36677708 | 1.0160733 | 0.30959448 | 0.99937574 |
| MED13 | 292.464259 | -0.2592122 | 0.23755942 | -1.0911469 | 0.27520826 | 0.99937574 |
| METTL2A | 20.0018602 | -0.1017024 | 0.49091368 | -0.2071697 | 0.83587731 | 0.99937574 |
| TLK2 | 180.655365 | -0.1145984 | 0.35027615 | -0.3271659 | 0.74354239 | 0.99937574 |
| AC005821.1 | 12.1659833 | -0.345567 | 1.15782371 | -0.2984626 | 0.76535013 | 0.99937574 |
| 10-Mar | 29.7856182 | 1.60682928 | 0.61168004 | 2.62691142 | 0.00861638 | 0.74742128 |
| TANC2 | 394.519836 | 0.36918732 | 0.25318563 | 1.45816853 | 0.1447941 | 0.99937574 |
| AC015923.1 | 31.5969317 | 0.9167165 | 0.78102406 | 1.17373657 | 0.24050056 | 0.99937574 |
| CYB561 | 20.302756 | -0.1176636 | 0.44305326 | -0.2655744 | 0.79056704 | 0.99937574 |
| DCAF7 | 117.318204 | 0.06447898 | 0.30647782 | 0.21038711 | 0.83336555 | 0.99937574 |
| TACO1 | 12.2890347 | 0.82940322 | 0.58111454 | 1.42726291 | 0.15350413 | 0.99937574 |
| MAP3K3 | 123.719487 | -0.3110237 | 0.33179742 | -0.9373903 | 0.34855783 | 0.99937574 |
| CCDC47 | 34.3223679 | -0.2282642 | 0.44086469 | -0.5177647 | 0.60462248 | 0.99937574 |
| DDX42 | 67.8518889 | -0.0707336 | 0.30100799 | -0.2349892 | 0.81421708 | 0.99937574 |
| FTSJ3 | 13.3387972 | 0.0058571 | 0.52450531 | 0.0111669 | 0.99109028 | 0.99937574 |
| PSMC5 | 23.5282653 | -0.3694543 | 0.45718028 | -0.8081152 | 0.41902426 | 0.99937574 |
| SMARCD2 | 18.5085625 | -0.174412 | 0.47869263 | -0.3643506 | 0.71559621 | 0.99937574 |
| ERN1 | 24.4742411 | -0.6157101 | 0.54240181 | -1.135155 | 0.25631039 | 0.99937574 |
| SNHG25 | 14.4857628 | -0.7228904 | 0.54300956 | -1.3312664 | 0.18310136 | 0.99937574 |
| TEX2 | 41.2655167 | 0.47474799 | 0.35838024 | 1.3247047 | 0.18526911 | 0.99937574 |
| POLG2 | 20.0815469 | 0.00924906 | 0.48951994 | 0.01889414 | 0.98492555 | 0.99937574 |
| DDX5 | 484.849612 | 0.40312432 | 0.25854787 | 1.5591864 | 0.11895227 | 0.99937574 |
| CEP95 | 101.451562 | -0.2171501 | 0.2688729 | -0.8076309 | 0.4193031 | 0.99937574 |
| SMURF2 | 121.267314 | -0.3569495 | 0.29318958 | -1.2174698 | 0.22342552 | 0.99937574 |
| LRRC37A3 | 120.941103 | 0.49075878 | 0.35612826 | 1.37803942 | 0.16819112 | 0.99937574 |
| GNA13 | 87.2510204 | -0.5356249 | 0.28285402 | -1.8936442 | 0.05827225 | 0.99937574 |
| CEP112 | 90.5503278 | 0.88668926 | 0.54363209 | 1.63104657 | 0.10288049 | 0.99937574 |
| PRKCA | 879.560335 | 0.25732679 | 0.44257937 | 0.58142518 | 0.56095393 | 0.99937574 |
| PRKCA-AS1 | 41.182393 | 0.31712139 | 0.66457957 | 0.47717595 | 0.63323683 | 0.99937574 |
| HELZ | 259.339192 | -0.3431105 | 0.23575655 | -1.4553595 | 0.14556977 | 0.99937574 |
| PSMD12 | 45.6535973 | -0.2938237 | 0.37246698 | -0.7888583 | 0.43019484 | 0.99937574 |
| PITPNC1 | 22.9654335 | -0.2513468 | 0.685986 | -0.3664022 | 0.71406496 | 0.99937574 |
| NOL11 | 25.2591574 | -0.1607543 | 0.40141997 | -0.4004642 | 0.68881465 | 0.99937574 |
| BPTF | 330.520191 | 0.0649387 | 0.30688339 | 0.2116071 | 0.83241357 | 0.99937574 |
| AC005332.7 | 15.7043892 | -0.2468475 | 0.49511117 | -0.4985699 | 0.61808239 | 0.99937574 |
| AMZ2 | 16.7115703 | -0.0627264 | 0.44423333 | -0.1412014 | 0.88771087 | 0.99937574 |
| ARSG | 36.5520209 | -0.3027773 | 0.50443337 | -0.6002325 | 0.54835127 | 0.99937574 |
| WIPI1 | 38.7821724 | -0.2932106 | 0.48833477 | -0.6004296 | 0.54821999 | 0.99937574 |
| PRKAR1A | 232.57454 | -0.0606878 | 0.23395369 | -0.2594008 | 0.79532602 | 0.99937574 |
| ABCA6 | 14.7538235 | 0.38194785 | 0.71319124 | 0.5355476 | 0.59227125 | 0.99937574 |
| ABCA10 | 87.3387078 | -0.5845389 | 0.70094448 | -0.8339304 | 0.40432018 | 0.99937574 |
| ABCA5 | 488.016245 | -0.0420954 | 0.38987457 | -0.1079717 | 0.91401814 | 0.99937574 |
| MAP2K6 | 60.7627862 | 1.8509629 | 0.9043201 | 2.04680057 | 0.04067767 | 0.99937574 |
| KCNJ16 | 19.7812153 | -0.5266214 | 0.58449923 | -0.9009787 | 0.36759962 | 0.99937574 |
| SOX9 | 20.5110951 | 1.78096547 | 0.83305445 | 2.13787404 | 0.03252697 | 0.99937574 |
| SLC39A11 | 108.910185 | -0.3654964 | 0.35280534 | -1.035972 | 0.30021522 | 0.99937574 |
| COG1 | 21.2008965 | 0.789227 | 0.40486884 | 1.94933992 | 0.05125484 | 0.99937574 |
| FAM104A | 23.5469472 | 0.18761475 | 0.42025426 | 0.44643153 | 0.65528556 | 0.99937574 |
| C17orf80 | 19.4593736 | 0.12975062 | 0.47754843 | 0.27170149 | 0.78585156 | 0.99937574 |
| CDC42EP4 | 39.3256156 | -0.1002113 | 0.45531078 | -0.2200943 | 0.82579771 | 0.99937574 |
| RPL38 | 83.7867455 | 0.1794005 | 0.4516629 | 0.39719999 | 0.69121999 | 0.99937574 |
| GPRC5C | 145.330748 | -0.0798106 | 0.35102092 | -0.2273671 | 0.82013826 | 0.99937574 |
| SLC9A3R1 | 88.0835667 | -0.0754342 | 0.36647303 | -0.2058382 | 0.8369173 | 0.99937574 |
| NAT9 | 16.8679622 | 0.17970053 | 0.44956961 | 0.39971682 | 0.6893651 | 0.99937574 |

| | | | | | | |
|---|---|---|---|---|---|---|
| TMEM104 | 17.4089249 | -0.5928062 | 0.70246848 | -0.8438902 | 0.39873079 | 0.99937574 |
| KCTD2 | 28.256191 | -0.9651103 | 0.49519479 | -1.9489509 | 0.05130129 | 0.99937574 |
| ATP5PD | 65.5399008 | 0.47744152 | 0.51472527 | 0.92756573 | 0.35363288 | 0.99937574 |
| SLC16A5 | 223.934716 | 0.25602416 | 0.67137993 | 0.3813402 | 0.70295083 | 0.99937574 |
| ARMC7 | 126.41979 | 1.26543104 | 0.58749123 | 2.15395732 | 0.03124352 | 0.99049468 |
| NT5C | 18.8515832 | -1.1862949 | 0.52933133 | -2.2411198 | 0.02501831 | 0.96546668 |
| JPT1 | 62.3043398 | -0.1010272 | 0.46935189 | -0.2152483 | 0.82957372 | 0.99937574 |
| SUMO2 | 28.5757626 | -0.3124718 | 0.53590167 | -0.5830767 | 0.55984166 | 0.99937574 |
| NUP85 | 18.9347328 | 0.20040058 | 0.46464226 | 0.43130081 | 0.66624966 | 0.99937574 |
| GGA3 | 37.4141986 | -0.0529217 | 0.39889978 | -0.1326692 | 0.89445497 | 0.99937574 |
| MRPS7 | 39.0651841 | 0.3494954 | 0.43494328 | 0.80354248 | 0.42166126 | 0.99937574 |
| GRB2 | 101.998386 | 0.05544832 | 0.26267802 | 0.21108854 | 0.83281818 | 0.99937574 |
| AC011933.4 | 12.9769796 | 0.11109941 | 0.60777874 | 0.18279582 | 0.85495823 | 0.99937574 |
| TMEM94 | 30.7303929 | 0.46795807 | 0.35875034 | 1.30441151 | 0.19209331 | 0.99937574 |
| CASKIN2 | 16.4001165 | -0.0217812 | 0.5299004 | -0.0411043 | 0.96721277 | 0.99937574 |
| TSEN54 | 22.4109551 | 0.02901301 | 0.42287611 | 0.06860876 | 0.94530105 | 0.99937574 |
| LLGL2 | 306.798947 | 0.20886584 | 0.2549745 | 0.81916363 | 0.41269306 | 0.99937574 |
| MYO15B | 38.8028342 | -0.3267862 | 0.80877013 | -0.4040533 | 0.68617352 | 0.99937574 |
| RECQL5 | 51.1365934 | -0.4484639 | 0.52293285 | -0.8575937 | 0.39111687 | 0.99937574 |
| SMIM5 | 66.3012112 | 1.25646609 | 0.57279299 | 2.19357798 | 0.02826576 | 0.99049468 |
| SMIM6 | 66.7324899 | 0.93912884 | 0.52595636 | 1.78556419 | 0.07416985 | 0.99937574 |
| SAP30BP | 45.8229007 | -0.5811777 | 0.3781912 | -1.5367299 | 0.12435946 | 0.99937574 |
| ITGB4 | 12.4894193 | 0.66468183 | 0.87270585 | 0.76163329 | 0.4462789 | 0.99937574 |
| GALK1 | 11.4232182 | 0.8424294 | 0.59854539 | 1.40746118 | 0.15929068 | 0.99937574 |
| H3F3B | 218.712086 | -0.4934506 | 0.32630111 | -1.5122553 | 0.13046893 | 0.99937574 |
| UNK | 73.1604273 | -0.0871595 | 0.31029222 | -0.280895 | 0.77879094 | 0.99937574 |
| WBP2 | 68.0369573 | -0.1125163 | 0.33688141 | -0.3339939 | 0.73838415 | 0.99937574 |
| TRIM47 | 16.7814871 | -0.247243 | 0.5882911 | -0.4202732 | 0.67428589 | 0.99937574 |
| TRIM65 | 10.7037771 | -0.237851 | 0.68882623 | -0.345299 | 0.72986959 | 0.99937574 |
| ACOX1 | 69.1540455 | 0.21793153 | 0.29255331 | 0.7449293 | 0.45631446 | 0.99937574 |
| TEN1 | 17.4904724 | -0.1185514 | 0.56726885 | -0.2089863 | 0.83445893 | 0.99937574 |
| SRP68 | 29.6844381 | 0.08655232 | 0.40425164 | 0.21410506 | 0.83046513 | 0.99937574 |
| EXOC7 | 56.6492928 | -0.2000284 | 0.41300109 | -0.4843291 | 0.62815235 | 0.99937574 |
| RNF157 | 26.9914805 | 0.01718152 | 0.45729952 | 0.0375717 | 0.97002918 | 0.99937574 |
| UBALD2 | 37.3500696 | -0.3617375 | 0.51269377 | -0.7055624 | 0.48046029 | 0.99937574 |
| PRPSAP1 | 41.1842708 | 0.08172165 | 0.35699418 | 0.22891593 | 0.81893426 | 0.99937574 |
| UBE2O | 56.4560776 | 0.08815765 | 0.33953617 | 0.2596414 | 0.7951404 | 0.99937574 |
| RHBDF2 | 26.0023791 | -0.1123791 | 0.69143887 | -0.1625293 | 0.87088906 | 0.99937574 |
| MXRA7 | 41.060379 | 0.11974862 | 0.47969822 | 0.24963325 | 0.80287098 | 0.99937574 |
| JMJD6 | 27.8712124 | -0.7046262 | 0.46114093 | -1.5280062 | 0.12651099 | 0.99937574 |
| AC005837.2 | 12.6038365 | 0.26337136 | 1.04099481 | 0.25299968 | 0.80026846 | 0.99937574 |
| SRSF2 | 97.1269821 | -0.5381738 | 0.42396346 | -1.2693872 | 0.20430301 | 0.99937574 |
| MFSD11 | 54.0940587 | -0.3017062 | 0.29375077 | -1.0270821 | 0.30438179 | 0.99937574 |
| SEC14L1 | 226.485053 | -0.1166963 | 0.38728645 | -0.3013177 | 0.76317225 | 0.99937574 |
| 9-Sep | 370.7602 | 0.05465138 | 0.45544868 | 0.1199946 | 0.90448743 | 0.99937574 |
| TNRC6C | 83.3608183 | 0.00655117 | 0.30976148 | 0.02114907 | 0.98312674 | 0.99937574 |
| TMC6 | 11.7812254 | -0.1689478 | 0.56478006 | -0.2991392 | 0.76483385 | 0.99937574 |
| SYNGR2 | 37.9126758 | -0.5497393 | 0.43020491 | -1.2778545 | 0.20130073 | 0.99937574 |
| AFMID | 29.5881571 | -0.9038488 | 0.70919665 | -1.2744685 | 0.20249745 | 0.99937574 |
| SOCS3 | 12.6068504 | 0.03443957 | 0.83083354 | 0.04145183 | 0.9669357 | 0.99937574 |
| PGS1 | 33.563348 | -0.3800864 | 0.34987439 | -1.0863509 | 0.27732378 | 0.99937574 |
| CYTH1 | 357.982491 | -0.7225212 | 0.50585306 | -1.4283223 | 1.53E-01 | 0.99937574 |
| USP36 | 83.3839226 | -1.5600554 | 0.4992044 | -3.1250834 | 0.00177755 | 0.41718606 |
| TIMP2 | 24.5605773 | -0.7701393 | 0.50282571 | -1.5316228 | 0.12561555 | 0.99937574 |

| | | | | | | |
|---|---|---|---|---|---|---|
| LGALS3BP | 93.9841366 | -0.4085585 | 0.80601855 | -0.5068848 | 0.61223566 | 0.99937574 |
| CANT1 | 19.0896814 | 0.09947212 | 0.49995841 | 0.1989608 | 0.84229341 | 0.99937574 |
| ENGASE | 16.5317954 | 0.28551902 | 0.49609103 | 0.57553757 | 0.56492779 | 0.99937574 |
| TBC1D16 | 127.425893 | 0.21953556 | 0.39069794 | 0.56190612 | 0.57417999 | 0.99937574 |
| CCDC40 | 28.9496017 | 0.43919604 | 0.50483589 | 0.86997785 | 0.38431251 | 0.99937574 |
| GAA | 29.1367655 | 0.29250502 | 0.42861091 | 0.68244885 | 0.49495518 | 0.99937574 |
| EIF4A3 | 25.797564 | 0.12174022 | 0.47243263 | 0.25768801 | 0.7966477 | 0.99937574 |
| AC087741.1 | 14.9542016 | 0.30750238 | 0.60734133 | 0.506309 | 0.61263975 | 0.99937574 |
| SGSH | 13.9302214 | -1.1438337 | 0.69251929 | -1.6516994 | 0.09859585 | 0.99937574 |
| SLC26A11 | 37.7356232 | -0.329782 | 0.52093771 | -0.6330547 | 0.52669795 | 0.99937574 |
| RNF213 | 556.128911 | -0.7844939 | 0.53066983 | -1.4783089 | 0.13932512 | 0.99937574 |
| AC124319.2 | 150.758984 | 0.14235602 | 0.36165779 | 0.39362078 | 0.69386103 | 0.99937574 |
| ENDOV | 32.4849962 | 0.28858473 | 0.49979986 | 0.57740058 | 0.56366888 | 0.99937574 |
| RPTOR | 146.911061 | -0.0242249 | 0.33542926 | -0.0722205 | 0.94242644 | 0.99937574 |
| TEPSIN | 11.2508634 | -0.4056639 | 0.72938409 | -0.5561733 | 0.57809241 | 0.99937574 |
| NDUFAF8 | 30.654312 | 0.91260889 | 0.49854071 | 1.83056043 | 0.06716618 | 0.99937574 |
| SLC38A10 | 66.0586257 | 0.15127206 | 0.27646751 | 0.54716035 | 0.58426857 | 0.99937574 |
| LINC00482 | 31.718025 | 0.11918049 | 0.45593095 | 0.2614003 | 0.79378382 | 0.99937574 |
| BAHCC1 | 13.8886343 | 0.96275708 | 0.64524477 | 1.49208041 | 0.13567807 | 0.99937574 |
| ACTG1 | 448.817749 | 0.10342328 | 0.5260975 | 0.19658577 | 0.84415171 | 0.99937574 |
| FAAP100 | 11.4866168 | -0.9234362 | 0.73081713 | -1.2635667 | 0.2063856 | 0.99937574 |
| NPLOC4 | 139.637704 | -0.2634006 | 0.35760216 | -0.7365744 | 0.46138123 | 0.99937574 |
| OXLD1 | 29.4523657 | -0.1572173 | 0.43141815 | -0.3644197 | 0.71554463 | 0.99937574 |
| CCDC137 | 32.0313733 | 0.19116677 | 0.45549966 | 0.41968587 | 0.67471495 | 0.99937574 |
| ARL16 | 17.832483 | 0.13142393 | 0.51513977 | 0.25512285 | 0.79862821 | 0.99937574 |
| HGS | 41.2716525 | -0.7225852 | 0.45293134 | -1.5953525 | 0.11063343 | 0.99937574 |
| GCGR | 36.9890836 | 1.62896329 | 0.67327108 | 2.41947611 | 0.01554288 | 0.86662299 |
| MCRIP1 | 30.8674491 | -0.1909637 | 0.55511231 | -0.3440091 | 0.73083944 | 0.99937574 |
| P4HB | 121.434826 | -0.1489922 | 0.39741536 | -0.3749031 | 0.70773257 | 0.99937574 |
| ARHGDIA | 96.9364076 | -0.5321975 | 0.6329251 | -0.8408538 | 0.40042983 | 0.99937574 |
| ANAPC11 | 63.3661062 | 0.47495422 | 0.60715997 | 0.78225549 | 0.43406444 | 0.99937574 |
| PCYT2 | 15.3281564 | 0.59839083 | 0.62968291 | 0.95030501 | 0.34195729 | 0.99937574 |
| SIRT7 | 18.1521101 | -0.444551 | 0.48859381 | -0.909858 | 0.3628974 | 0.99937574 |
| MAFG | 17.3845438 | -1.1750497 | 0.72105897 | -1.6296166 | 0.10318256 | 0.99937574 |
| ASPSCR1 | 19.0972077 | -0.2331061 | 0.63744193 | -0.3656899 | 0.71459648 | 0.99937574 |
| CENPX | 15.7386348 | -0.2159524 | 0.66434684 | -0.3250598 | 0.74513585 | 0.99937574 |
| DCXR | 29.374995 | 0.47527608 | 0.53244779 | 0.89262476 | 0.37205815 | 0.99937574 |
| RFNG | 14.1481339 | 0.4041741 | 0.52164111 | 0.77481259 | 0.43845041 | 0.99937574 |
| GPS1 | 23.163891 | 0.15457984 | 0.49817983 | 0.31028925 | 0.75634101 | 0.99937574 |
| DUS1L | 35.700971 | -0.0728509 | 0.39560222 | -0.184152 | 0.8538942 | 0.99937574 |
| FASN | 23.4683679 | -0.4183446 | 0.6389333 | -0.6547547 | 0.51262567 | 0.99937574 |
| CCDC57 | 77.2222308 | 0.29671496 | 0.31186878 | 0.95140961 | 0.34139649 | 0.99937574 |
| SLC16A3 | 99.8152106 | -0.1391 | 0.57779614 | -0.2407424 | 0.80975481 | 0.99937574 |
| CSNK1D | 94.4957835 | -0.223977 | 0.2969176 | -0.7543406 | 0.45064474 | 0.99937574 |
| OGFOD3 | 26.4957758 | 0.04370147 | 0.44520763 | 0.09815975 | 0.92180544 | 0.99937574 |
| HEXDC | 63.1181074 | -0.0473251 | 0.2924344 | -0.1618314 | 0.8714386 | 0.99937574 |
| CYBC1 | 24.5027934 | 0.08546956 | 0.41595846 | 0.20547618 | 0.83720011 | 0.99937574 |
| NARF | 39.8697229 | -0.468853 | 0.52259417 | -0.8971647 | 0.36963105 | 0.99937574 |
| FOXK2 | 169.669264 | -0.2361327 | 0.319921 | -0.7380968 | 0.46045566 | 0.99937574 |
| WDR45B | 166.446564 | -0.242368 | 0.38880705 | -0.6233632 | 0.53304589 | 0.99937574 |
| RAB40B | 159.415845 | -0.8247597 | 0.46797209 | -1.7624122 | 0.07799968 | 0.99937574 |
| FN3KRP | 11.9978456 | -0.4728464 | 0.54469074 | -0.8681007 | 0.38533922 | 0.99937574 |
| FN3K | 43.8682142 | 0.1556501 | 0.5365832 | 0.29007635 | 0.77175783 | 0.99937574 |
| TBCD | 68.3800096 | -0.2130044 | 0.43373831 | -0.4910896 | 0.6233631 | 0.99937574 |

| | | | | | | |
|---|---|---|---|---|---|---|
| METRNL | 83.5772849 | -0.7502825 | 0.40734319 | -1.8418928 | 0.06549083 | 0.99937574 |
| USP14 | 101.5274 | -0.4178402 | 0.2576335 | -1.6218398 | 0.10483766 | 0.99937574 |
| THOC1 | 73.5866451 | -0.2426223 | 0.33442055 | -0.7255006 | 0.46814496 | 0.99937574 |
| ENOSF1 | 233.68116 | -0.5000274 | 0.44016086 | -1.1360106 | 0.25595211 | 0.99937574 |
| YES1 | 148.450229 | -0.4713805 | 0.28482629 | -1.6549752 | 0.09792953 | 0.99937574 |
| METTL4 | 23.9224634 | -0.3683156 | 0.51126735 | -0.7203972 | 0.47128049 | 0.99937574 |
| SMCHD1 | 217.280293 | -0.5188068 | 0.34455096 | -1.5057477 | 0.13213195 | 0.99937574 |
| AP001011.1 | 34.3681679 | 0.0544592 | 0.66906094 | 0.08139647 | 0.93512666 | 0.99937574 |
| LPIN2 | 99.9064495 | -0.5866163 | 0.47314063 | -1.239835 | 0.21503643 | 0.99937574 |
| MYL12A | 112.202966 | -0.033733 | 0.50850995 | -0.066337 | 0.94710951 | 0.99937574 |
| MYL12B | 245.547865 | 0.08822483 | 0.51553716 | 0.17113185 | 0.86412009 | 0.99937574 |
| TGIF1 | 26.2704408 | -0.7081995 | 0.60072107 | -1.1789157 | 0.23843175 | 0.99937574 |
| DLGAP1 | 18.1359476 | 0.01817851 | 0.61792626 | 0.02941858 | 0.97653076 | 0.99937574 |
| DLGAP1-AS1 | 12.0185461 | 0.4288665 | 0.62710407 | 0.6838841 | 0.49404835 | 0.99937574 |
| LINC00667 | 20.4920517 | 0.35822417 | 0.42974943 | 0.83356521 | 0.40452604 | 0.99937574 |
| EPB41L3 | 20.0720336 | -0.210852 | 0.70819236 | -0.2977326 | 0.76590724 | 0.99937574 |
| L3MBTL4 | 427.793923 | -0.4392092 | 0.35089886 | -1.2516688 | 0.21069056 | 0.99937574 |
| AP001021.2 | 32.5483818 | -0.8519133 | 0.93117661 | -0.9148783 | 0.36025553 | 0.99937574 |
| ARHGAP28 | 76.8488904 | -0.3532808 | 0.78568775 | -0.4496453 | 0.65296622 | 0.99937574 |
| LAMA1 | 36.5222485 | -1.3674894 | 0.64770246 | -2.1112926 | 0.03474717 | 0.99937574 |
| PTPRM | 137.492253 | 0.23481043 | 0.59990917 | 0.39140998 | 0.69549422 | 0.99937574 |
| RAB12 | 60.4661053 | -0.3340338 | 0.31175439 | -1.0714647 | 0.28396054 | 0.99937574 |
| NDUFV2 | 107.546785 | 0.22409307 | 0.3671624 | 0.61033775 | 0.5416381 | 0.99937574 |
| ANKRD12 | 480.927853 | -0.3052713 | 0.24903338 | -1.2258248 | 0.22026463 | 0.99937574 |
| TWSG1 | 60.3187415 | -0.0792339 | 0.33671859 | -0.235312 | 0.81396658 | 0.99937574 |
| RALBP1 | 109.536014 | -0.368725 | 0.37066757 | -0.9947593 | 0.31985333 | 0.99937574 |
| PPP4R1 | 118.939521 | -0.4714262 | 0.3249011 | -1.4509836 | 0.14678443 | 0.99937574 |
| RAB31 | 98.7293321 | -0.4285515 | 0.66222668 | -0.6471371 | 0.51754318 | 0.99937574 |
| VAPA | 321.613511 | -0.0350955 | 0.33774229 | -0.103912 | 0.91723921 | 0.99937574 |
| NAPG | 84.7463374 | -0.1911407 | 0.28665422 | -0.6667988 | 0.50490066 | 0.99937574 |
| GNAL | 22.4671458 | 1.18349388 | 0.54600041 | 2.16756956 | 0.03019145 | 0.99049468 |
| MPPE1 | 52.8281548 | 0.40238458 | 0.39851359 | 1.00971357 | 0.31263254 | 0.99937574 |
| IMPA2 | 189.346148 | -0.1922373 | 0.44892687 | -0.4282152 | 0.66849447 | 0.99937574 |
| AFG3L2 | 142.800361 | -0.1994587 | 0.33522369 | -0.5950018 | 0.55184229 | 0.99937574 |
| SPIRE1 | 446.820595 | -0.2843574 | 0.28588435 | -0.9946588 | 0.31990222 | 0.99937574 |
| PSMG2 | 65.4637497 | -0.0465735 | 0.33464688 | -0.139172 | 0.88931425 | 0.99937574 |
| CEP76 | 15.1636589 | 0.74232637 | 0.56788692 | 1.30717286 | 0.19115401 | 0.99937574 |
| PTPN2 | 153.991537 | 0.05326042 | 0.30933028 | 0.17217979 | 0.86329618 | 0.99937574 |
| SEH1L | 32.2717791 | -0.2930151 | 0.35216964 | -0.8320283 | 0.40539295 | 0.99937574 |
| CEP192 | 152.534523 | -0.0479809 | 0.39646596 | -0.1210214 | 0.90367407 | 0.99937574 |
| LDLRAD4 | 106.50048 | -2.3625186 | 1.26438405 | -1.8685134 | 0.06169054 | 0.99937574 |
| FAM210A | 51.6373043 | -0.3040997 | 0.40879536 | -0.7438922 | 0.45694171 | 0.99937574 |
| RNMT | 89.9471865 | -0.2660204 | 0.30758582 | -0.8648658 | 0.38711246 | 0.99937574 |
| ROCK1 | 246.98667 | -0.0424938 | 0.22627725 | -0.1877952 | 0.85103716 | 0.99937574 |
| ESCO1 | 79.7223269 | -0.2976166 | 0.29479143 | -1.0095836 | 0.31269485 | 0.99937574 |
| SNRPD1 | 26.9094678 | -0.3076077 | 0.41358733 | -0.743755 | 0.45702469 | 0.99937574 |
| ABHD3 | 224.291028 | -1.3218234 | 0.53875977 | -2.4534559 | 0.01414909 | 0.84995591 |
| MIB1 | 249.903277 | -0.1094148 | 0.24599302 | -0.4447882 | 0.65647281 | 0.99937574 |
| AC090912.1 | 16.4387792 | -0.2227192 | 0.63576592 | -0.3503164 | 0.72610128 | 0.99937574 |
| RBBP8 | 280.093045 | -0.2110285 | 0.36022669 | -0.5858214 | 0.55799554 | 0.99937574 |
| CABLES1 | 30.0700874 | -0.4064373 | 0.58462434 | -0.6952111 | 0.486923 | 0.99937574 |
| TMEM241 | 24.4071156 | -0.428455 | 0.41823508 | -1.0244359 | 0.30562943 | 0.99937574 |
| RIOK3 | 106.370903 | -0.3514566 | 0.28328592 | -1.2406427 | 0.21473776 | 0.99937574 |
| RMC1 | 43.3201161 | -0.1963173 | 0.3739395 | -0.5249975 | 0.59958491 | 0.99937574 |

| | | | | | | |
|---|---|---|---|---|---|---|
| NPC1 | 267.012977 | -0.0220141 | 0.43463626 | -0.0506495 | 0.95960483 | 0.99937574 |
| ANKRD29 | 53.2197237 | 0.90282956 | 0.64915321 | 1.3907804 | 0.16429203 | 0.99937574 |
| LAMA3 | 128.163995 | -0.3550811 | 0.65789717 | -0.5397213 | 0.58938922 | 0.99937574 |
| TTC39C | 15.0230347 | 0.45136772 | 0.69133859 | 0.65288952 | 0.5138275 | 0.99937574 |
| OSBPL1A | 81.6107049 | 1.07434557 | 0.60581127 | 1.77339977 | 0.07616249 | 0.99937574 |
| IMPACT | 16.6442815 | 0.36272082 | 0.50514955 | 0.7180464 | 0.47272868 | 0.99937574 |
| SS18 | 177.194276 | 0.21717704 | 0.24529056 | 0.88538685 | 0.37594802 | 0.99937574 |
| TAF4B | 19.7127951 | -0.561646 | 0.72623474 | -0.773367 | 0.43930523 | 0.99937574 |
| KCTD1 | 262.014847 | 0.77205552 | 0.46828831 | 1.64867561 | 0.09921411 | 0.99937574 |
| DSC2 | 51.4367447 | 0.30697737 | 0.53701434 | 0.57163719 | 0.5675678 | 0.99937574 |
| DSG2 | 382.744989 | 0.60675404 | 0.48139651 | 1.26040391 | 0.20752369 | 0.99937574 |
| B4GALT6 | 63.3272715 | -0.2132499 | 0.48999415 | -0.4352091 | 0.66341066 | 0.99937574 |
| TRAPPC8 | 154.215512 | 0.04800239 | 0.27751225 | 0.17297395 | 0.8626719 | 0.99937574 |
| RNF138 | 51.9779611 | 0.0538079 | 0.31904095 | 0.16865516 | 0.86606789 | 0.99937574 |
| GAREM1 | 390.843529 | 0.04754912 | 0.43680044 | 0.10885777 | 0.9133153 | 0.99937574 |
| CCDC178 | 32.903584 | -0.0321969 | 0.56236384 | -0.0572527 | 0.95434386 | 0.99937574 |
| ASXL3 | 102.018836 | 0.03222617 | 0.49777627 | 0.06474027 | 0.9483808 | 0.99937574 |
| MAPRE2 | 190.084879 | 0.02671392 | 0.48724077 | 0.05482694 | 0.95627633 | 0.99937574 |
| ZNF397 | 43.2354531 | 0.44353739 | 0.34384821 | 1.28992208 | 0.19707771 | 0.99937574 |
| ZSCAN30 | 35.2694854 | 0.05263531 | 0.48255468 | 0.10907637 | 0.91314192 | 0.99937574 |
| ZNF24 | 64.9216833 | 0.17441588 | 0.34107499 | 0.51137107 | 0.60909125 | 0.99937574 |
| GALNT1 | 54.2729137 | -0.3584995 | 0.34889028 | -1.0275422 | 0.30416521 | 0.99937574 |
| RPRD1A | 139.310178 | 0.09957188 | 0.22509377 | 0.44235734 | 0.65823064 | 0.99937574 |
| SLC39A6 | 22.900679 | 0.01406081 | 0.54485805 | 0.02580637 | 0.97941178 | 0.99937574 |
| ELP2 | 84.3516733 | 0.06654771 | 0.2978925 | 0.22339504 | 0.82322806 | 0.99937574 |
| MOCOS | 28.7344917 | -0.9927885 | 0.61894668 | -1.6039968 | 0.10871476 | 0.99937574 |
| FHOD3 | 106.270424 | 1.13413092 | 0.73460854 | 1.54385752 | 0.12262285 | 0.99937574 |
| TPGS2 | 47.6164009 | -0.3793552 | 0.35752868 | -1.0610482 | 0.288668 | 0.99937574 |
| KIAA1328 | 274.626926 | -0.0340115 | 0.33140087 | -0.1026294 | 0.9182571 | 0.99937574 |
| AC016205.1 | 40.3604139 | 0.13990083 | 0.46168651 | 0.30302127 | 0.76187365 | 0.99937574 |
| PIK3C3 | 120.746473 | -0.0557958 | 0.30255762 | -0.1844139 | 0.8536888 | 0.99937574 |
| LINC00907 | 22.4436148 | 0.19005192 | 0.55724624 | 0.34105554 | 0.73306177 | 0.99937574 |
| SETBP1 | 687.659324 | 0.22064265 | 0.30282323 | 0.72861864 | 0.46623497 | 0.99937574 |
| SLC14A2 | 86.5031088 | -3.0918586 | 1.36111876 | -2.2715568 | 0.02311329 | 0.96546668 |
| EPG5 | 124.538191 | -0.4302039 | 0.60707004 | -0.7086562 | 0.47853787 | 0.99937574 |
| ATP5F1A | 220.920078 | 0.79066161 | 0.38577076 | 2.04956333 | 0.04040706 | 0.99937574 |
| HAUS1 | 19.7244726 | -0.8703759 | 0.5218694 | -1.6678042 | 0.09535461 | 0.99937574 |
| C18orf25 | 64.2489902 | -0.067522 | 0.31474021 | -0.2145326 | 0.83013178 | 0.99937574 |
| LOXHD1 | 45.5941992 | 1.51791939 | 1.01516795 | 1.49523967 | 0.1348519 | 0.99937574 |
| ST8SIA5 | 69.3068694 | 0.38013256 | 0.66934293 | 0.567919 | 0.57008997 | 0.99937574 |
| PIAS2 | 204.640091 | -0.2201317 | 0.34981798 | -0.6292749 | 0.52916911 | 0.99937574 |
| KATNAL2 | 88.5572543 | 0.58224949 | 0.48668138 | 1.19636689 | 0.23155342 | 0.99937574 |
| HDHD2 | 11.5034534 | -0.1599473 | 0.56838301 | -0.2814076 | 0.77839777 | 0.99937574 |
| SMAD2 | 234.095631 | 0.03443379 | 0.23218843 | 0.14830105 | 0.88210519 | 0.99937574 |
| CTIF | 42.6074696 | 0.1807545 | 0.49640806 | 0.36412484 | 0.71576479 | 0.99937574 |
| SMAD7 | 21.6146406 | 0.19444056 | 0.74202737 | 0.2620396 | 0.79329091 | 0.99937574 |
| DYM | 264.077883 | -0.0034045 | 0.24112342 | -0.0141195 | 0.98873464 | 0.99937574 |
| LIPG | 45.149696 | -1.5294476 | 0.75893741 | -2.0152486 | 0.0438786 | 0.99937574 |
| ACAA2 | 52.6381079 | 0.16047948 | 0.47578695 | 0.33729273 | 0.73589624 | 0.99937574 |
| SNHG22 | 21.8776842 | 0.50243451 | 0.60714285 | 0.8275392 | 0.40793151 | 0.99937574 |
| MYO5B | 160.273693 | -0.0098733 | 0.36557228 | -0.0270079 | 0.97845342 | 0.99937574 |
| MBD1 | 25.7177134 | -0.1606258 | 0.43250967 | -0.3713808 | 0.7103539 | 0.99937574 |
| CXXC1 | 12.2080079 | -0.5240564 | 0.62208974 | -0.8424128 | 0.39955691 | 0.99937574 |
| ME2 | 39.4936939 | 0.22320933 | 0.33811238 | 0.66016314 | 0.50914915 | 0.99937574 |

| | | | | | | |
|---|---|---|---|---|---|---|
| AC091551.1 | 12.5096021 | 0.14588362 | 0.5973303 | 0.24422605 | 0.80705576 | 0.99937574 |
| SMAD4 | 64.4028848 | -0.132188 | 0.27789202 | -0.475681 | 0.63430166 | 0.99937574 |
| MEX3C | 20.8224105 | 0.18387873 | 0.45989237 | 0.39982992 | 0.6892818 | 0.99937574 |
| MBD2 | 117.498966 | -0.3482023 | 0.36634789 | -0.9504689 | 0.34187405 | 0.99937574 |
| POLI | 42.6488215 | 0.10282519 | 0.39020548 | 0.2635155 | 0.79215328 | 0.99937574 |
| CCDC68 | 46.2533538 | 0.25359332 | 0.6265743 | 0.40472985 | 0.6856761 | 0.99937574 |
| TXNL1 | 55.9617185 | -0.492189 | 0.35050896 | -1.4042122 | 0.16025567 | 0.99937574 |
| WDR7 | 132.763786 | -0.0689465 | 0.30533442 | -0.2258066 | 0.82135187 | 0.99937574 |
| FECH | 24.7199021 | 0.47096419 | 0.45902786 | 1.0260035 | 0.30488994 | 0.99937574 |
| NARS | 88.6679974 | 0.31171713 | 0.31061555 | 1.00354642 | 0.31559729 | 0.99937574 |
| AC027097.2 | 36.1574782 | -0.5196901 | 0.44599514 | -1.1652371 | 0.24392302 | 0.99937574 |
| ATP8B1 | 125.395326 | 0.19762506 | 0.39033422 | 0.50629704 | 0.61264814 | 0.99937574 |
| NEDD4L | 2302.02208 | -0.3713417 | 0.28908943 | -1.2845217 | 0.19895946 | 0.99937574 |
| MALT1 | 42.5875671 | -0.3702272 | 0.38353326 | -0.9653067 | 0.33439121 | 0.99937574 |
| ZNF532 | 247.299882 | -0.2956172 | 0.46926216 | -0.6299616 | 0.5287197 | 0.99937574 |
| SEC11C | 29.5333683 | 0.02595104 | 0.45952747 | 0.05647332 | 0.95496475 | 0.99937574 |
| LMAN1 | 127.048269 | 0.5683529 | 0.26835471 | 2.11791666 | 0.03418213 | 0.99937574 |
| CCBE1 | 115.844069 | 0.73733747 | 0.71538686 | 1.03068356 | 0.30268924 | 0.99937574 |
| AC105094.2 | 28.2930503 | -0.6513149 | 0.82235086 | -0.7920159 | 0.42835142 | 0.99937574 |
| RNF152 | 918.346152 | -0.6703702 | 0.45443634 | -1.4751686 | 0.14016721 | 0.99937574 |
| PIGN | 102.489164 | -0.1152328 | 0.23297525 | -0.4946139 | 0.62087267 | 0.99937574 |
| KIAA1468 | 125.825806 | -0.1823565 | 0.24428505 | -0.7464908 | 0.45537101 | 0.99937574 |
| ZCCHC2 | 93.6515847 | 0.14218694 | 0.32638309 | 0.43564433 | 0.6630948 | 0.99937574 |
| PHLPP1 | 218.958846 | 0.15154232 | 0.30991353 | 0.48898259 | 0.62485402 | 0.99937574 |
| BCL2 | 345.871677 | 0.27190736 | 0.45191619 | 0.60167653 | 0.54738948 | 0.99937574 |
| KDSR | 68.7696403 | -0.4016922 | 0.43937175 | -0.9142422 | 0.3605896 | 0.99937574 |
| VPS4B | 66.7825295 | 0.01165759 | 0.29131835 | 0.04001668 | 0.96807983 | 0.99937574 |
| TMX3 | 63.5736013 | -0.2048931 | 0.2666339 | -0.7684435 | 0.44222377 | 0.99937574 |
| RTTN | 40.1675487 | -0.539432 | 0.41544344 | -1.2984487 | 0.1941332 | 0.99937574 |
| SOCS6 | 27.5059008 | 0.40789356 | 0.46699421 | 0.87344459 | 0.3824208 | 0.99937574 |
| LINC01541 | 20.1743034 | -2.9651831 | 1.05210496 | -2.818334 | 0.00482736 | 0.60897109 |
| TIMM21 | 14.7989404 | 0.47056241 | 0.64370863 | 0.73101771 | 0.46476834 | 0.99937574 |
| CYB5A | 164.548471 | 0.00290148 | 0.80680089 | 0.00359628 | 0.99713059 | 0.99937574 |
| CNDP2 | 157.406594 | 0.29504336 | 0.31426215 | 0.93884472 | 0.34781048 | 0.99937574 |
| LINC00909 | 13.6178133 | 0.83097224 | 0.50964341 | 1.63049738 | 0.10299642 | 0.99937574 |
| ZNF407 | 245.425213 | -0.2572495 | 0.22898933 | -1.1234124 | 0.26126238 | 0.99937574 |
| ZADH2 | 22.7104986 | 0.15925781 | 0.43883982 | 0.36290647 | 0.71667475 | 0.99937574 |
| TSHZ1 | 23.3684818 | 0.20833186 | 0.54887295 | 0.379563 | 0.70426983 | 0.99937574 |
| ZNF516 | 94.0692493 | 0.00173908 | 0.38836374 | 0.00447797 | 0.99642711 | 0.99937574 |
| ZNF236-DT | 10.9122567 | 0.90100947 | 0.59644121 | 1.51064256 | 0.13087955 | 0.99937574 |
| ZNF236 | 84.1937232 | -0.1245316 | 0.44587558 | -0.2792967 | 0.78001713 | 0.99937574 |
| MBP | 283.270526 | 0.12888218 | 0.32290707 | 0.39913087 | 0.68979678 | 0.99937574 |
| ATP9B | 215.903771 | 0.02744938 | 0.27071368 | 0.10139635 | 0.91923583 | 0.99937574 |
| NFATC1 | 14.8549467 | 0.31483581 | 0.67158475 | 0.46879536 | 0.63921591 | 0.99937574 |
| CTDP1 | 43.2209457 | -0.257832 | 0.3617103 | -0.7128134 | 0.47596121 | 0.99937574 |
| PQLC1 | 27.1656616 | -0.2323772 | 0.39955266 | -0.5815935 | 0.56084055 | 0.99937574 |
| HSBP1L1 | 40.6445909 | -0.2490238 | 0.52990389 | -0.4699415 | 0.63839681 | 0.99937574 |
| TXNL4A | 60.2171263 | 0.14251363 | 0.26198766 | 0.54397077 | 0.58646159 | 0.99937574 |
| ADNP2 | 89.9083317 | -0.2841311 | 0.47659299 | -0.5961714 | 0.55106076 | 0.99937574 |
| PARD6G-AS1 | 47.1011056 | 0.72296728 | 0.86089131 | 0.83978926 | 0.40102656 | 0.99937574 |
| C20orf96 | 11.0742764 | 0.03054267 | 0.67894754 | 0.04498532 | 0.96411901 | 0.99937574 |
| ZCCHC3 | 17.5005902 | -0.0111136 | 0.50210529 | -0.022134 | 0.98234104 | 0.99937574 |
| NRSN2-AS1 | 63.6659378 | -0.3752533 | 0.34105631 | -1.1002679 | 0.27121543 | 0.99937574 |
| SOX12 | 14.9949703 | -0.0985734 | 0.49681552 | -0.1984104 | 0.84272397 | 0.99937574 |

| | | | | | | |
|---|---|---|---|---|---|---|
| NRSN2 | 18.1973952 | 0.06789322 | 0.49731413 | 0.13651978 | 0.89141039 | 0.99937574 |
| RBCK1 | 68.5970259 | -0.3301505 | 0.36178066 | -0.9125708 | 0.36146829 | 0.99937574 |
| TBC1D20 | 28.2982542 | -0.0518796 | 0.40731272 | -0.1273704 | 0.89864724 | 0.99937574 |
| CSNK2A1 | 209.756514 | 0.45563562 | 0.28219235 | 1.61462783 | 0.10639132 | 0.99937574 |
| SLC52A3 | 23.2619366 | -0.1442292 | 0.63166991 | -0.2283301 | 0.81938966 | 0.99937574 |
| PSMF1 | 63.5961463 | -0.1435053 | 0.27203612 | -0.5275231 | 0.59783038 | 0.99937574 |
| SDCBP2 | 69.3129818 | 0.72809222 | 0.747456 | 0.97409374 | 3.30E-01 | 0.99937574 |
| AL136531.2 | 25.9884183 | -0.2317673 | 0.55471323 | -0.4178146 | 0.67608268 | 0.99937574 |
| SDCBP2-AS1 | 57.0822394 | -0.1731604 | 0.28648555 | -0.6044297 | 0.54555798 | 0.99937574 |
| NSFL1C | 44.7887595 | -0.2522185 | 0.34181345 | -0.7378834 | 0.46058532 | 0.99937574 |
| STK35 | 34.6123857 | 0.30813384 | 0.44681044 | 0.6896299 | 0.49042696 | 0.99937574 |
| SNRPB | 43.5102632 | -0.2104627 | 0.47989654 | -0.4385585 | 0.66098145 | 0.99937574 |
| NOP56 | 14.8197993 | -0.4603458 | 0.59556287 | -0.7729593 | 0.43954649 | 0.99937574 |
| IDH3B | 22.4466988 | 0.38057066 | 0.42922251 | 0.88665122 | 0.37526671 | 0.99937574 |
| EBF4 | 14.9345028 | 0.40460907 | 0.61414482 | 0.65881704 | 0.51001326 | 0.99937574 |
| PCED1A | 15.9868642 | 0.00981904 | 0.59157606 | 0.0165981 | 0.98675724 | 0.99937574 |
| PTPRA | 148.677416 | -0.1963855 | 0.27313142 | -0.7190147 | 0.47213183 | 0.99937574 |
| MRPS26 | 44.0420597 | 0.11786899 | 0.58863106 | 0.20024256 | 0.84129088 | 0.99937574 |
| UBOX5-AS1 | 17.0276468 | -0.4998232 | 0.68683639 | -0.7277179 | 0.46678626 | 0.99937574 |
| UBOX5 | 18.0582309 | -0.2865084 | 0.50496473 | -0.567383 | 0.57045398 | 0.99937574 |
| LZTS3 | 18.7573748 | -0.2797417 | 0.81482293 | -0.3433159 | 0.73136081 | 0.99937574 |
| DDRGK1 | 46.6931586 | 0.13066415 | 0.4814293 | 0.27140879 | 0.78607664 | 0.99937574 |
| ITPA | 35.7145445 | -0.2988083 | 0.39706698 | -0.7525387 | 0.45172717 | 0.99937574 |
| C20orf194 | 365.40099 | -0.0510308 | 0.21436288 | -0.238058 | 0.81183611 | 0.99937574 |
| ATRN | 102.151883 | 0.00276979 | 0.29676764 | 0.00933321 | 0.99255328 | 0.99937574 |
| CENPB | 31.8754845 | -0.3043146 | 0.5870687 | -0.5183628 | 0.60420519 | 0.99937574 |
| CDC25B | 15.8577809 | -0.4210132 | 0.5976291 | -0.7044723 | 0.48113867 | 0.99937574 |
| MAVS | 81.7229734 | 0.07525055 | 0.45832272 | 0.16418683 | 0.86958407 | 0.99937574 |
| PANK2 | 74.5738328 | -0.2516272 | 0.39344645 | -0.6395463 | 0.52246762 | 0.99937574 |
| RNF24 | 193.689607 | -0.6545661 | 0.77291974 | -0.8468746 | 0.39706502 | 0.99937574 |
| PRNP | 48.4667359 | 0.3838199 | 0.42409462 | 0.90503363 | 0.36544757 | 0.99937574 |
| SLC23A2 | 34.7058366 | 0.1549247 | 0.34028585 | 0.4552781 | 0.64890918 | 0.99937574 |
| TMEM230 | 53.1272706 | 0.4015416 | 0.37480461 | 1.0713358 | 0.28401847 | 0.99937574 |
| PCNA | 16.8924658 | 0.51093253 | 0.61436685 | 0.83164079 | 0.40561173 | 0.99937574 |
| CDS2 | 69.6907713 | 0.04343558 | 0.30246353 | 0.14360602 | 0.88581158 | 0.99937574 |
| GPCPD1 | 105.49157 | 0.15522007 | 0.59636698 | 0.26027609 | 0.79465081 | 0.99937574 |
| SHLD1 | 32.3145986 | -0.4194873 | 0.46759572 | -0.8971154 | 0.36965733 | 0.99937574 |
| TRMT6 | 11.1223841 | -1.025218 | 0.58878968 | -1.7412296 | 0.08164335 | 0.99937574 |
| MCM8 | 18.1775657 | -0.1600621 | 0.484803 | -0.3301591 | 0.74127978 | 0.99937574 |
| CRLS1 | 62.3708696 | -0.2308641 | 0.32545871 | -0.7093499 | 0.47810736 | 0.99937574 |
| TMX4 | 97.8160632 | 0.12948664 | 0.38616511 | 0.33531419 | 0.73738809 | 0.99937574 |
| PLCB1 | 24.2234407 | -0.9345773 | 0.73554396 | -1.2705934 | 0.20387334 | 0.99937574 |
| PLCB4 | 223.758143 | 0.73846519 | 0.54293354 | 1.3601392 | 0.17378588 | 0.99937574 |
| PAK5 | 44.7404785 | 0.67950862 | 0.59086131 | 1.15003065 | 0.25013125 | 0.99937574 |
| MKKS | 34.9052851 | 0.54480784 | 0.38698497 | 1.40782686 | 0.15918234 | 0.99937574 |
| SLX4IP | 163.752676 | 0.04416775 | 0.49073125 | 0.09000395 | 0.92828408 | 0.99937574 |
| JAG1 | 79.2796269 | 0.16025872 | 0.54712507 | 0.29291058 | 0.76959051 | 0.99937574 |
| BTBD3 | 38.2261431 | 0.1252054 | 0.36790442 | 0.34032046 | 0.73361521 | 0.99937574 |
| SPTLC3 | 150.99101 | 0.34836461 | 0.38225386 | 0.9113436 | 0.36211436 | 0.99937574 |
| TASP1 | 145.123873 | -0.0659383 | 0.29963154 | -0.2200647 | 0.82582079 | 0.99937574 |
| ESF1 | 59.7863383 | -0.282985 | 0.37125677 | -0.7622352 | 0.44591962 | 0.99937574 |
| NDUFAF5 | 49.8090253 | -0.2416186 | 0.3428099 | -0.704818 | 0.48092352 | 0.99937574 |
| MACROD2 | 676.233789 | 0.74973519 | 0.51426721 | 1.45787088 | 0.14487614 | 0.99937574 |
| KIF16B | 287.520813 | -0.0825052 | 0.3521874 | -0.234265 | 0.81477929 | 0.99937574 |

| | | | | | | |
|---|---|---|---|---|---|---|
| SNRPB2 | 29.1126445 | 0.11123237 | 0.48056929 | 0.23145959 | 0.81695778 | 0.99937574 |
| DSTN | 166.831806 | 0.01183306 | 0.31593738 | 0.03745381 | 0.97012317 | 0.99937574 |
| RRBP1 | 74.5809289 | -0.3582559 | 0.43728183 | -0.8192792 | 0.41262714 | 0.99937574 |
| SNX5 | 42.3787802 | 0.59164208 | 0.42518017 | 1.39150912 | 0.1640711 | 0.99937574 |
| OVOL2 | 24.4228178 | -0.0682384 | 0.46795717 | -0.1458218 | 0.88406203 | 0.99937574 |
| MGME1 | 30.8193532 | -0.0918756 | 0.45621541 | -0.2013864 | 0.84039649 | 0.99937574 |
| ZNF133 | 31.5284 | 0.42266882 | 0.4256409 | 0.99301742 | 0.32070147 | 0.99937574 |
| DZANK1 | 22.0682078 | 0.43852392 | 0.55757301 | 0.786487 | 0.43158223 | 0.99937574 |
| POLR3F | 13.0745215 | -0.3792316 | 0.54219297 | -0.6994402 | 0.48427697 | 0.99937574 |
| SEC23B | 113.882644 | 0.52442159 | 0.34330405 | 1.52757182 | 0.12661888 | 0.99937574 |
| DTD1 | 13.1291326 | -0.1877819 | 0.55415649 | -0.3388609 | 0.73471455 | 0.99937574 |
| AL121761.1 | 27.2408797 | 0.54786138 | 0.51588109 | 1.06199159 | 0.2882395 | 0.99937574 |
| RIN2 | 228.454472 | 0.63489394 | 0.40459053 | 1.56922593 | 0.1165953 | 0.99937574 |
| NAA20 | 40.6505512 | 0.13802465 | 0.50113098 | 0.2754263 | 0.78298874 | 0.99937574 |
| CRNKL1 | 17.8288412 | -0.0805734 | 0.48505812 | -0.1661109 | 0.8680697 | 0.99937574 |
| RALGAPA2 | 759.098426 | -0.0427865 | 0.36164712 | -0.11831 | 9.06E-01 | 9.99E-01 |
| KIZ | 50.5994091 | -0.4118894 | 0.52734124 | -0.7810681 | 0.43476244 | 0.99937574 |
| KIZ-AS1 | 20.6041169 | -0.4895451 | 0.62169374 | -0.7874377 | 0.43102566 | 0.99937574 |
| XRN2 | 139.991414 | -0.0387097 | 0.24332653 | -0.1590854 | 0.87360159 | 0.99937574 |
| GZF1 | 15.1123956 | -0.3597929 | 0.57799397 | -0.6224855 | 0.53362267 | 0.99937574 |
| NAPB | 64.0758839 | 0.05500011 | 0.41778104 | 0.13164818 | 0.89526258 | 0.99937574 |
| CST3 | 154.865849 | 0.27506346 | 0.43300802 | 0.63523873 | 0.52527273 | 0.99937574 |
| AL035661.1 | 20.7740128 | 0.29419119 | 0.52703177 | 0.55820389 | 0.57670516 | 0.99937574 |
| APMAP | 54.0416317 | 0.32361806 | 0.46355114 | 0.69812807 | 0.48509711 | 0.99937574 |
| ACSS1 | 95.6820137 | 0.51969139 | 0.45138469 | 1.15132704 | 0.24959772 | 0.99937574 |
| ENTPD6 | 50.4795961 | 0.01819033 | 0.33597661 | 0.05414167 | 0.9568223 | 0.99937574 |
| PYGB | 153.02641 | 0.4612104 | 0.57549791 | 0.80141107 | 0.42289371 | 0.99937574 |
| ABHD12 | 118.00221 | -0.1137638 | 0.39067561 | -0.2911975 | 0.77090024 | 0.99937574 |
| NINL | 196.789524 | -0.1142484 | 0.52261914 | -0.2186074 | 0.82695589 | 0.99937574 |
| ZNF337-AS1 | 21.4648652 | -0.4108898 | 0.48753905 | -0.8427833 | 0.39934963 | 0.99937574 |
| ZNF337 | 30.5245185 | 0.51640022 | 0.43439013 | 1.18879363 | 0.23452088 | 0.99937574 |
| FAM182B | 23.2839556 | 1.44523012 | 0.5097429 | 2.83521382 | 0.0045795 | 0.60810972 |
| HM13 | 60.5610977 | -0.0236001 | 0.3597186 | -0.0656072 | 0.94769058 | 0.99937574 |
| ID1 | 28.7141528 | 0.14464704 | 0.75830458 | 0.19075058 | 0.84872101 | 0.99937574 |
| BCL2L1 | 109.458385 | -0.0592523 | 0.53244012 | -0.1112845 | 0.91139073 | 0.99937574 |
| AL117381.1 | 22.5649285 | -0.4304771 | 0.61829821 | -0.6962289 | 0.4862855 | 0.99937574 |
| DUSP15 | 69.5813178 | 0.78540648 | 0.49619446 | 1.58286021 | 0.11345333 | 0.99937574 |
| PDRG1 | 11.2938359 | -1.0280199 | 0.58417311 | -1.7597865 | 0.07844401 | 0.99937574 |
| TM9SF4 | 76.7411015 | -0.1433842 | 0.34249069 | -0.4186515 | 0.67547085 | 0.99937574 |
| PLAGL2 | 30.9319883 | 0.21397686 | 0.41052807 | 0.52122345 | 0.60221112 | 0.99937574 |
| POFUT1 | 41.1203899 | 0.56155773 | 0.49339095 | 1.13815976 | 0.25505378 | 0.99937574 |
| KIF3B | 160.095045 | 0.135097 | 0.29617358 | 0.45614129 | 0.64828838 | 0.99937574 |
| ASXL1 | 457.010899 | 0.20707901 | 0.25216174 | 0.82121503 | 0.4115238 | 0.99937574 |
| NOL4L | 27.6290759 | 0.42471975 | 0.4564676 | 0.93044883 | 0.35213875 | 0.99937574 |
| MAPRE1 | 45.2148305 | 0.10169426 | 0.37147266 | 0.27375974 | 0.78426927 | 0.99937574 |
| AL035071.2 | 11.4515603 | 0.4518712 | 0.79583717 | 0.56779353 | 0.57017517 | 0.99937574 |
| CDK5RAP1 | 46.1746264 | -0.1356964 | 0.35712953 | -0.3799642 | 0.70397199 | 0.99937574 |
| SNTA1 | 18.0842771 | -0.2993308 | 0.50794302 | -0.5893 | 0.55566008 | 0.99937574 |
| CBFA2T2 | 256.368767 | -0.0943849 | 0.24765068 | -0.3811213 | 0.70311325 | 0.99937574 |
| NECAB3 | 14.0020891 | 1.14673724 | 0.52093544 | 2.20130396 | 0.02771451 | 0.98891577 |
| PXMP4 | 13.6804903 | 0.13765673 | 0.64352832 | 0.21390936 | 0.83061774 | 0.99937574 |
| CHMP4B | 58.4571875 | -0.5051516 | 0.40434904 | -1.2492959 | 0.21155686 | 0.99937574 |
| RALY | 103.471255 | -0.4101434 | 0.35595408 | -1.1522367 | 0.24922382 | 0.99937574 |
| EIF2S2 | 78.6424185 | -0.0538056 | 0.43141574 | -0.1247187 | 0.90074628 | 0.99937574 |

| | | | | | | |
|---|---|---|---|---|---|---|
| AHCY | 18.5199739 | 0.1931751 | 0.4459551 | 0.43317163 | 0.66489008 | 0.99937574 |
| ITCH | 211.423719 | -0.1020483 | 0.26057235 | -0.3916313 | 0.69533066 | 0.99937574 |
| DYNLRB1 | 127.650718 | -0.215402 | 0.47001311 | -0.4582894 | 0.64674454 | 0.99937574 |
| MAP1LC3A | 60.6272047 | 0.49084256 | 0.38407227 | 1.27799532 | 0.20125108 | 0.99937574 |
| PIGU | 72.2679208 | 0.20349682 | 0.39930806 | 0.50962362 | 0.61031517 | 0.99937574 |
| NCOA6 | 197.657178 | -0.0578094 | 0.2048311 | -0.2822295 | 0.77776752 | 0.99937574 |
| TP53INP2 | 55.7499719 | -0.7326241 | 0.48413721 | -1.5132571 | 0.13021436 | 0.99937574 |
| GGT7 | 66.7768037 | 0.06645605 | 0.33982729 | 0.19555832 | 0.84495589 | 0.99937574 |
| ACSS2 | 78.0863133 | 0.24957515 | 0.54015679 | 0.46204205 | 0.64405116 | 0.99937574 |
| GSS | 33.1010612 | 0.3892445 | 0.50133567 | 0.77641493 | 0.43750403 | 0.99937574 |
| TRPC4AP | 143.131804 | -0.2166377 | 0.30531413 | -0.7095568 | 0.47797904 | 0.99937574 |
| EDEM2 | 25.1725178 | -0.3306218 | 0.39198441 | -0.8434565 | 0.39897318 | 0.99937574 |
| PROCR | 13.8044422 | 0.2015441 | 0.69706077 | 0.28913418 | 0.7724787 | 0.99937574 |
| MMP24OS | 37.2806931 | 0.22866341 | 0.44729096 | 0.5112185 | 0.60919807 | 0.99937574 |
| MMP24 | 32.6840359 | 0.08807747 | 0.53259849 | 0.16537311 | 0.86865032 | 0.99937574 |
| UQCC1 | 93.9587914 | 0.11364816 | 0.29926522 | 0.37975731 | 0.70412557 | 0.99937574 |
| CEP250 | 13.7625792 | -0.9353399 | 0.63301835 | -1.4775873 | 0.13951827 | 0.99937574 |
| ERGIC3 | 36.9888452 | 0.14755883 | 0.47528313 | 0.31046511 | 0.75620729 | 0.99937574 |
| CPNE1 | 43.8885584 | 0.422662 | 0.42284056 | 0.9995777 | 0.31751492 | 0.99937574 |
| ROMO1 | 47.9891386 | -0.2094921 | 0.41612725 | -0.5034327 | 0.61466011 | 0.99937574 |
| RBM39 | 333.981625 | -0.1521396 | 0.21799832 | -0.6978936 | 0.48524375 | 0.99937574 |
| PHF20 | 130.198999 | -0.4834106 | 0.36513472 | -1.323924 | 0.1855283 | 0.99937574 |
| SCAND1 | 55.400476 | -0.6674864 | 0.44357788 | -1.5047783 | 0.13238108 | 0.99937574 |
| NORAD | 239.73802 | -0.6835072 | 0.57374836 | -1.1913014 | 0.2335353 | 0.99937574 |
| EPB41L1 | 243.890859 | -0.2753414 | 0.49368729 | -0.5577243 | 0.57703266 | 0.99937574 |
| AAR2 | 18.4801862 | 0.06249902 | 0.62461899 | 0.10005942 | 0.92029715 | 0.99937574 |
| DLGAP4 | 85.4034067 | 0.15659447 | 0.28471797 | 0.54999856 | 0.58232036 | 0.99937574 |
| DLGAP4-AS1 | 22.4538086 | -0.916667 | 0.57410694 | -1.5966833 | 0.11033631 | 0.99937574 |
| RAB5IF | 16.5513958 | 0.44787367 | 0.55240851 | 0.81076532 | 0.41750045 | 0.99937574 |
| NDRG3 | 23.9801778 | 0.75194786 | 0.5333963 | 1.40973582 | 0.1586177 | 0.99937574 |
| SAMHD1 | 15.6560713 | -0.9968278 | 1.07969813 | -0.9232467 | 0.35587865 | 0.99937574 |
| RBL1 | 19.0328124 | -0.979578 | 0.52608904 | -1.8620005 | 0.06260302 | 0.99937574 |
| MROH8 | 13.5380185 | 0.91317847 | 0.57322722 | 1.59304798 | 0.11114943 | 0.99937574 |
| RPN2 | 106.681436 | 0.47592186 | 0.33066128 | 1.43930327 | 0.15006462 | 0.99937574 |
| MANBAL | 26.8035216 | -0.0160302 | 0.40949311 | -0.0391465 | 0.96877361 | 0.99937574 |
| SRC | 28.3571527 | -0.4599055 | 0.39426486 | -1.1664886 | 0.24341694 | 0.99937574 |
| BLCAP | 90.3857915 | -0.3109956 | 0.28680964 | -1.0843276 | 0.27821958 | 0.99937574 |
| CTNNBL1 | 43.0760631 | -0.2096184 | 0.35967713 | -0.582796 | 0.56003062 | 0.99937574 |
| TTI1 | 32.5960977 | 0.43742178 | 0.37595541 | 1.16349378 | 0.24462922 | 0.99937574 |
| RPRD1B | 76.9930658 | -0.3442404 | 0.38147098 | -0.9024025 | 0.3668431 | 0.99937574 |
| TGM2 | 15.9483357 | -0.6601352 | 0.66004955 | -1.0001297 | 3.17E-01 | 0.99937574 |
| RALGAPB | 125.206453 | -0.16158 | 0.2491227 | -0.6485962 | 0.51659944 | 0.99937574 |
| ACTR5 | 10.9150549 | -0.8258003 | 0.62484629 | -1.3216055 | 0.18629954 | 0.99937574 |
| DHX35 | 30.8278903 | -0.0353018 | 0.36862377 | -0.0957664 | 0.92370612 | 0.99937574 |
| TOP1 | 156.158847 | -0.0360753 | 0.24312679 | -0.1483805 | 0.88204248 | 0.99937574 |
| PLCG1 | 19.7960541 | -0.208056 | 0.43282417 | -0.4806941 | 0.63073391 | 0.99937574 |
| ZHX3 | 82.2122419 | 0.39330757 | 0.48090262 | 0.81785284 | 0.41344122 | 0.99937574 |
| LPIN3 | 28.8790329 | 0.16203052 | 0.38213701 | 0.42401159 | 0.67155736 | 0.99937574 |
| CHD6 | 184.559407 | 0.39096758 | 0.24915985 | 1.56914357 | 0.11661449 | 0.99937574 |
| PTPRT | 21.6082126 | -0.3648141 | 1.25401808 | -0.2909161 | 0.77111546 | 0.99937574 |
| SRSF6 | 61.4483722 | -0.1974403 | 0.41195775 | -0.4792732 | 0.63174428 | 0.99937574 |
| L3MBTL1 | 26.8567513 | 0.2085366 | 0.39575947 | 0.52692763 | 0.59824385 | 0.99937574 |
| SGK2 | 123.61859 | -0.1882347 | 0.42100896 | -0.4471038 | 0.65480012 | 0.99937574 |
| IFT52 | 32.5361752 | 0.44394023 | 0.3529164 | 1.25791894 | 0.20842107 | 0.99937574 |

| | | | | | | |
|---|---|---|---|---|---|---|
| OSER1 | 38.6495144 | -0.1572641 | 0.53332311 | -0.2948758 | 0.76808874 | 0.99937574 |
| FITM2 | 17.1118997 | 0.15716772 | 0.5406584 | 0.2906969 | 0.77128314 | 0.99937574 |
| TTPAL | 14.0456049 | -0.5075185 | 0.56622782 | -0.8963151 | 0.37008452 | 0.99937574 |
| SERINC3 | 76.4100883 | 0.11573551 | 0.28748107 | 0.4025848 | 0.68725369 | 0.99937574 |
| PKIG | 51.0568963 | -0.1181445 | 0.33376353 | -0.3539767 | 0.72335632 | 0.99937574 |
| YWHAB | 152.608282 | 0.18084047 | 0.4135251 | 0.43731436 | 0.66188338 | 0.99937574 |
| PABPC1L | 12.544188 | -0.3242428 | 0.56245974 | -0.5764729 | 0.5642956 | 0.99937574 |
| TOMM34 | 14.6977271 | -0.0676461 | 0.55412729 | -0.1220769 | 0.90283813 | 0.99937574 |
| STK4 | 55.3814912 | -0.2621792 | 0.3355189 | -0.7814142 | 0.43455892 | 0.99937574 |
| SLPI | 49.7189 | -0.7112867 | 0.89329862 | -0.7962474 | 4.26E-01 | 9.99E-01 |
| SDC4 | 219.840978 | 0.05388004 | 0.41352455 | 0.13029467 | 0.8963333 | 0.99937574 |
| PIGT | 21.7063219 | 0.49927858 | 0.47057404 | 1.06099898 | 0.28869037 | 0.99937574 |
| WFDC2 | 712.705177 | 0.72930401 | 0.4609386 | 1.5822151 | 1.14E-01 | 9.99E-01 |
| DNTTIP1 | 15.0847738 | 0.45498282 | 0.52710738 | 0.86316913 | 0.38804448 | 0.99937574 |
| SNX21 | 15.3830619 | -0.3887687 | 0.56710262 | -0.685535 | 0.49300638 | 0.99937574 |
| ACOT8 | 18.2107725 | 0.28102437 | 0.48394753 | 0.58069183 | 0.56144817 | 0.99937574 |
| CTSA | 60.4982792 | -1.2444646 | 0.72085268 | -1.7263785 | 0.08427933 | 0.99937574 |
| ZNF335 | 23.7698293 | -0.4418552 | 0.42670115 | -1.0355145 | 0.3004287 | 0.99937574 |
| NCOA5 | 42.8474939 | -0.3501107 | 0.45956528 | -0.7618303 | 0.44616131 | 0.99937574 |
| CD40 | 12.913166 | -0.1288278 | 0.7446658 | -0.1730009 | 0.86265075 | 0.99937574 |
| SLC35C2 | 40.0762619 | 0.15077794 | 0.36937739 | 0.40819481 | 0.68313066 | 0.99937574 |
| ELMO2 | 37.328537 | 0.66596515 | 0.40198971 | 1.65667215 | 0.09758578 | 0.99937574 |
| SLC13A3 | 19.2191041 | 0.27878921 | 0.77698061 | 0.35881103 | 0.71973646 | 0.99937574 |
| ZMYND8 | 202.468892 | -0.1327061 | 0.47854594 | -0.2773111 | 0.78154124 | 0.99937574 |
| NCOA3 | 159.63209 | -0.4449614 | 0.29331212 | -1.5170235 | 0.12926076 | 0.99937574 |
| ARFGEF2 | 279.253773 | -0.4291265 | 0.23755605 | -1.806422 | 0.07085245 | 0.99937574 |
| CSE1L | 44.2222772 | 0.09183535 | 0.36998112 | 0.24821632 | 0.80396704 | 0.99937574 |
| STAU1 | 141.386441 | -0.1575804 | 0.22894149 | -0.6882998 | 0.491264 | 0.99937574 |
| DDX27 | 24.0951343 | -0.5789506 | 0.5035269 | -1.1497908 | 0.25023003 | 0.99937574 |
| ZNFX1 | 49.8234867 | -0.7992177 | 0.5524693 | -1.4466282 | 0.14800108 | 0.99937574 |
| ZFAS1 | 46.8871476 | -0.3280654 | 0.33665681 | -0.9744801 | 0.32981824 | 0.99937574 |
| B4GALT5 | 173.354026 | 0.00155374 | 0.35644808 | 0.00435895 | 0.99652207 | 0.99937574 |
| SLC9A8 | 64.135892 | -0.3780216 | 0.39075308 | -0.9674181 | 0.33333506 | 0.99937574 |
| RNF114 | 69.3690121 | -0.1561681 | 0.39330495 | -0.3970662 | 0.69131868 | 0.99937574 |
| CEBPB | 26.4620784 | -1.1330398 | 0.60364311 | -1.8770028 | 6.05E-02 | 0.99937574 |
| PTPN1 | 117.305388 | -0.860969 | 0.58164144 | -1.4802401 | 0.13880918 | 0.99937574 |
| RIPOR3 | 45.446329 | 0.36562098 | 0.82362321 | 0.44391777 | 0.65710203 | 0.99937574 |
| PARD6B | 173.04551 | -0.0370342 | 0.42653922 | -0.0868248 | 0.9308108 | 0.99937574 |
| BCAS4 | 34.5178846 | 0.55724816 | 0.54858245 | 1.01579655 | 0.30972628 | 0.99937574 |
| ADNP | 221.02045 | 0.25219445 | 0.23975649 | 1.05187748 | 0.29285577 | 0.99937574 |
| DPM1 | 58.1795577 | -0.1468112 | 0.27421853 | -0.5353804 | 0.59238686 | 0.99937574 |
| NFATC2 | 30.8609984 | 0.66435034 | 0.48124902 | 1.38047105 | 0.16744166 | 0.99937574 |
| ATP9A | 220.797653 | -0.1898824 | 0.30940576 | -0.6137004 | 0.53941332 | 0.99937574 |
| ZFP64 | 61.4909887 | -0.0698053 | 0.55847866 | -0.1249919 | 0.90052992 | 0.99937574 |
| ZNF217 | 110.005134 | -0.0140079 | 0.40361247 | -0.0347064 | 0.97231383 | 0.99937574 |
| PFDN4 | 15.293013 | 0.31460072 | 0.49545431 | 0.63497424 | 0.52544521 | 0.99937574 |
| FAM210B | 43.3511662 | 0.60966084 | 0.38706733 | 1.57507698 | 0.11523868 | 0.99937574 |
| CSTF1 | 13.3951458 | 0.76611243 | 0.52398477 | 1.46208912 | 0.14371679 | 0.99937574 |
| RTF2 | 36.287837 | -0.0892671 | 0.39891393 | -0.2237755 | 0.82293203 | 0.99937574 |
| RAE1 | 21.6612978 | -0.1961133 | 0.41238728 | -0.4755562 | 0.63439058 | 0.99937574 |
| RBM38 | 11.4486028 | -0.3268918 | 0.69598753 | -0.4696805 | 0.63858331 | 0.99937574 |
| PCK1 | 88.4638656 | -0.5284315 | 0.9436933 | -0.559961 | 0.57550602 | 0.99937574 |
| PMEPA1 | 24.0507608 | -0.2111485 | 0.59596811 | -0.354295 | 0.72311781 | 0.99937574 |
| RAB22A | 68.0226881 | -0.0620267 | 0.37495013 | -0.1654266 | 0.86860822 | 0.99937574 |

| | | | | | | |
|---|---|---|---|---|---|---|
| VAPB | 114.266834 | -0.0757682 | 0.26731492 | -0.2834416 | 0.77683833 | 0.99937574 |
| NPEPL1 | 20.6663342 | 0.54555456 | 0.46936576 | 1.16232288 | 0.24510433 | 0.99937574 |
| GNAS | 385.149301 | 0.24154857 | 0.28011586 | 0.86231668 | 0.38851328 | 0.99937574 |
| NELFCD | 37.7896887 | 0.32060662 | 0.35873407 | 0.89371666 | 0.37147351 | 0.99937574 |
| CTSZ | 59.700892 | 0.28877084 | 0.44584182 | 0.64769797 | 0.51718032 | 0.99937574 |
| ATP5F1E | 270.187488 | 0.39666259 | 0.34147908 | 1.16160145 | 0.24539739 | 0.99937574 |
| PRELID3B | 22.0673543 | 0.12285051 | 0.45177752 | 0.27192702 | 0.78567814 | 0.99937574 |
| SYCP2 | 18.9120812 | 0.67859422 | 0.7995886 | 0.84867921 | 0.39605982 | 0.99937574 |
| FAM217B | 14.1117497 | 0.46609442 | 0.59390757 | 0.78479286 | 0.43257503 | 0.99937574 |
| MIR646HG | 40.6810792 | 0.87808067 | 0.84177364 | 1.04313158 | 0.29688735 | 0.99937574 |
| TAF4 | 62.6763851 | 0.18257714 | 0.32988866 | 0.55345078 | 0.57995477 | 0.99937574 |
| LSM14B | 14.9759428 | -0.747475 | 0.48722637 | -1.5341431 | 0.12499446 | 0.99937574 |
| PSMA7 | 113.882045 | 0.00350847 | 0.42657472 | 0.00822475 | 0.99343767 | 0.99937574 |
| SS18L1 | 48.442751 | -0.3037625 | 0.37511548 | -0.8097838 | 0.41806442 | 0.99937574 |
| MTG2 | 35.2951163 | 0.0483848 | 0.44846092 | 0.10789079 | 0.91408232 | 0.99937574 |
| OSBPL2 | 83.0938805 | -0.2297157 | 0.31656171 | -0.7256584 | 0.46804818 | 0.99937574 |
| ADRM1 | 16.863718 | -0.8140016 | 0.6423577 | -1.2672092 | 0.2050805 | 0.99937574 |
| LAMA5 | 46.2896923 | 0.41090658 | 0.39145529 | 1.04968968 | 0.29386081 | 0.99937574 |
| RPS21 | 14.7500271 | -0.6464731 | 0.56564658 | -1.1428923 | 0.2530833 | 0.99937574 |
| SLCO4A1 | 45.9872273 | -0.884405 | 0.86641182 | -1.0207674 | 0.30736464 | 0.99937574 |
| MRGBP | 11.3671155 | -0.1099245 | 0.6134297 | -0.1791966 | 0.85778334 | 0.99937574 |
| OGFR | 41.6176957 | -0.2975975 | 0.46033277 | -0.6464833 | 0.51796639 | 0.99937574 |
| TCFL5 | 20.5550678 | -0.1878233 | 0.56830536 | -0.3304971 | 0.74102435 | 0.99937574 |
| DIDO1 | 102.200831 | -0.3435752 | 0.25376882 | -1.3538906 | 0.1757713 | 0.99937574 |
| GID8 | 34.6998039 | -0.2315295 | 0.39192716 | -0.5907463 | 0.55469041 | 0.99937574 |
| YTHDF1 | 40.0124909 | -0.2809527 | 0.3807599 | -0.7378736 | 0.46059125 | 0.99937574 |
| ARFGAP1 | 30.84544 | -0.0928955 | 0.46845 | -0.1983039 | 0.84280728 | 0.99937574 |
| KCNQ2 | 14.7875414 | -0.3658661 | 1.73373864 | -0.2110272 | 0.83286601 | 0.99937574 |
| EEF1A2 | 52.5894787 | 0.08117729 | 1.00364318 | 0.08088262 | 9.36E-01 | 9.99E-01 |
| PPDPF | 193.795142 | 0.02004628 | 0.51764139 | 0.0387262 | 0.96910868 | 0.99937574 |
| HELZ2 | 29.5043296 | -2.1728541 | 0.96888902 | -2.2426244 | 0.02492105 | 0.96546668 |
| GMEB2 | 23.83517 | -0.6327825 | 0.39974123 | -1.5829802 | 0.11342598 | 0.99937574 |
| MHENCR | 11.2413218 | -0.8555511 | 0.55344842 | -1.5458552 | 0.12213955 | 0.99937574 |
| ARFRP1 | 20.979633 | -0.2924112 | 0.50675892 | -0.5770223 | 0.56392436 | 0.99937574 |
| SLC2A4RG | 20.9811498 | -0.0647424 | 0.47529583 | -0.1362149 | 0.89165143 | 0.99937574 |
| ZBTB46 | 56.4112788 | -0.0720087 | 0.33895361 | -0.2124441 | 0.83176059 | 0.99937574 |
| TPD52L2 | 35.2483532 | -0.6237159 | 0.42346371 | -1.4728911 | 0.14078039 | 0.99937574 |
| DNAJC5 | 106.241838 | -0.4685252 | 0.2796515 | -1.6753895 | 0.09385783 | 0.99937574 |
| UCKL1 | 24.5640735 | -0.1909695 | 0.49895635 | -0.3827379 | 0.70191408 | 0.99937574 |
| ZNF512B | 12.5176045 | -0.6320636 | 0.72345888 | -0.873669 | 0.38229854 | 0.99937574 |
| PRPF6 | 49.1104525 | 0.02923974 | 0.37946276 | 0.07705561 | 0.93857931 | 0.99937574 |
| PCMTD2 | 110.652527 | 0.5324748 | 0.40486347 | 1.31519595 | 0.18844406 | 0.99937574 |
| MIER2 | 16.4347057 | -0.1393247 | 0.47303542 | -0.2945333 | 0.76835045 | 0.99937574 |
| SHC2 | 16.4426957 | -1.830232 | 0.81522837 | -2.2450544 | 0.02476464 | 0.96546668 |
| TPGS1 | 22.9182091 | -0.4832904 | 0.6542592 | -0.7386833 | 0.46009932 | 0.99937574 |
| CDC34 | 22.6701911 | -0.753274 | 0.42985177 | -1.7524042 | 0.07970433 | 0.99937574 |
| BSG | 350.708824 | 0.9157388 | 0.4825173 | 1.89783622 | 0.05771766 | 0.99937574 |
| HCN2 | 11.2993309 | 1.19310376 | 0.88833329 | 1.34308122 | 0.17924568 | 0.99937574 |
| RNF126 | 20.1919747 | -0.0842318 | 0.44759936 | -0.1881858 | 0.850731 | 0.99937574 |
| PALM | 37.6037844 | 0.12951531 | 0.65928462 | 0.19644825 | 0.84425933 | 0.99937574 |
| PTBP1 | 73.2492926 | -0.2639944 | 0.37534644 | -0.7033354 | 0.48184675 | 0.99937574 |
| MED16 | 29.7698532 | -0.4184648 | 0.47996277 | -0.8718693 | 0.38327969 | 0.99937574 |
| R3HDM4 | 31.9596979 | -0.7309976 | 0.52612571 | -1.3893973 | 0.16471198 | 0.99937574 |
| ARID3A | 70.4306024 | 0.1525956 | 0.46514744 | 0.32805857 | 0.74286738 | 0.99937574 |

| | | | | | | |
|---|---|---|---|---|---|---|
| WDR18 | 13.4198725 | -0.1469377 | 0.59300222 | -0.247786 | 0.80429998 | 0.99937574 |
| TMEM259 | 82.4132318 | 0.17867836 | 0.28517993 | 0.62654604 | 0.53095685 | 0.99937574 |
| CNN2 | 20.2121345 | 0.50666675 | 0.50323658 | 1.00681622 | 0.3140231 | 0.99937574 |
| ABCA7 | 12.99605 | 0.38242321 | 0.6629819 | 0.576823 | 0.56405903 | 0.99937574 |
| POLR2E | 32.6030531 | 0.13457033 | 0.35456538 | 0.37953603 | 0.70428986 | 0.99937574 |
| GPX4 | 144.41569 | -0.1896969 | 0.41568419 | -0.4563486 | 0.64813935 | 0.99937574 |
| SBNO2 | 77.1820328 | -0.4590731 | 0.54732896 | -0.8387517 | 4.02E-01 | 0.99937574 |
| STK11 | 55.9593655 | -0.0484607 | 0.34660654 | -0.1398147 | 0.88880641 | 0.99937574 |
| ATP5F1D | 176.398073 | 0.34629652 | 0.46781984 | 0.74023477 | 0.45915755 | 0.99937574 |
| MIDN | 84.1415664 | -0.8104445 | 0.58195199 | -1.3926312 | 0.16373133 | 0.99937574 |
| CIRBP | 130.864169 | -0.1437861 | 0.27918367 | -0.5150232 | 0.60653679 | 0.99937574 |
| C19orf24 | 32.2158193 | -0.3784604 | 0.41731644 | -0.9068907 | 0.36446461 | 0.99937574 |
| MUM1 | 41.6522078 | -0.3053675 | 0.35769992 | -0.8536975 | 0.3932726 | 0.99937574 |
| NDUFS7 | 56.336012 | 0.46855992 | 0.51442358 | 0.91084457 | 0.36237728 | 0.99937574 |
| DAZAP1 | 69.1113169 | -0.1140489 | 0.35360186 | -0.3225347 | 0.74704769 | 0.99937574 |
| RPS15 | 353.382758 | -0.1193324 | 0.44229126 | -0.2698051 | 0.78731022 | 0.99937574 |
| C19orf25 | 40.7032941 | -0.2450819 | 0.43404836 | -0.564642 | 0.57231728 | 0.99937574 |
| MBD3 | 31.5616025 | -0.7200032 | 0.62113687 | -1.1591699 | 0.24638694 | 0.99937574 |
| TCF3 | 52.5141045 | -0.3453197 | 0.45887002 | -0.7525437 | 0.45172419 | 0.99937574 |
| REXO1 | 23.1278453 | -1.1434175 | 0.55041335 | -2.0773797 | 0.03776653 | 0.99937574 |
| ABHD17A | 38.0888871 | -0.0893947 | 0.33810121 | -0.2644021 | 0.79147006 | 0.99937574 |
| SCAMP4 | 36.1337603 | 0.0645938 | 0.41794893 | 0.15454951 | 0.87717647 | 0.99937574 |
| CSNK1G2 | 47.3962423 | -0.3549885 | 0.48518608 | -0.7316544 | 0.46437954 | 0.99937574 |
| BTBD2 | 37.7447442 | -0.3523346 | 0.40291357 | -0.874467 | 0.38186396 | 0.99937574 |
| MKNK2 | 48.7950354 | -0.0768979 | 0.30921748 | -0.2486854 | 0.80360412 | 0.99937574 |
| MOB3A | 20.0842698 | -1.1223732 | 0.60572592 | -1.8529391 | 0.06389109 | 0.99937574 |
| AP3D1 | 68.0379949 | -0.3042072 | 0.43309149 | -0.7024086 | 0.48242439 | 0.99937574 |
| DOT1L | 60.611023 | 0.36631302 | 0.44483315 | 0.82348408 | 0.41023276 | 0.99937574 |
| PLEKHJ1 | 83.6364309 | -0.0073944 | 0.32913763 | -0.0224661 | 0.98207616 | 0.99937574 |
| SF3A2 | 12.2242223 | -0.3800329 | 0.56424402 | -0.6735258 | 0.50061281 | 0.99937574 |
| LSM7 | 49.6900553 | -0.0602903 | 0.42871846 | -0.140629 | 0.88816302 | 0.99937574 |
| SPPL2B | 33.6023217 | -0.0754689 | 0.41913209 | -0.1800599 | 0.85710552 | 0.99937574 |
| TIMM13 | 47.88331 | 0.45006968 | 0.33235239 | 1.35419422 | 0.17567442 | 0.99937574 |
| LMNB2 | 14.2975626 | -0.541663 | 0.50888383 | -1.0644138 | 0.28714129 | 0.99937574 |
| GADD45B | 61.7771144 | -0.5253736 | 0.66496136 | -0.7900814 | 4.29E-01 | 0.99937574 |
| GNG7 | 58.7096731 | 0.25357577 | 0.47126346 | 0.53807645 | 0.59052427 | 0.99937574 |
| DIRAS1 | 11.4302742 | 0.87651755 | 0.77049007 | 1.13761045 | 0.25528318 | 0.99937574 |
| SGTA | 40.0596403 | -0.2714061 | 0.48004861 | -0.5653721 | 0.5718207 | 0.99937574 |
| THOP1 | 15.7117382 | -0.0563554 | 0.58467367 | -0.0963877 | 0.92321264 | 0.99937574 |
| ZNF555 | 19.2055058 | 0.15214488 | 0.4749307 | 0.32035175 | 0.7487017 | 0.99937574 |
| TLE2 | 70.4048967 | 0.4073459 | 0.50531379 | 0.80612464 | 0.42017098 | 0.99937574 |
| AES | 230.832264 | 0.06774932 | 0.37391155 | 0.18119075 | 0.85621785 | 0.99937574 |
| GNA11 | 228.320037 | -0.0690002 | 0.48747031 | -0.1415475 | 0.88743745 | 0.99937574 |
| NCLN | 23.2785444 | -0.026127 | 0.58656168 | -0.0445427 | 0.96447184 | 0.99937574 |
| NFIC | 211.490793 | 0.04308188 | 0.36270988 | 0.11877779 | 0.90545141 | 0.99937574 |
| SMIM24 | 91.4756691 | 0.91572724 | 0.4659176 | 1.96542746 | 0.04936478 | 0.99937574 |
| DOHH | 13.7671988 | -0.4614872 | 0.56890178 | -0.8111897 | 0.41725676 | 0.99937574 |
| FZR1 | 23.9588495 | -0.4920719 | 0.41664527 | -1.1810333 | 0.2375895 | 0.99937574 |
| MFSD12 | 35.4632911 | -0.2085339 | 0.35532828 | -0.5868766 | 0.55728658 | 0.99937574 |
| HMG20B | 24.5337852 | 0.58517373 | 0.43754413 | 1.33740505 | 0.18109046 | 0.99937574 |
| PIP5K1C | 26.0516345 | 0.20727136 | 0.45601614 | 0.45452638 | 0.64945001 | 0.99937574 |
| TJP3 | 41.1574838 | -0.4158471 | 0.40958138 | -1.0152979 | 0.30996384 | 0.99937574 |
| MRPL54 | 39.615762 | -0.2361034 | 0.5282105 | -0.4469873 | 0.65488424 | 0.99937574 |
| DAPK3 | 22.6562626 | 0.03762985 | 0.46679774 | 0.08061275 | 0.93574993 | 0.99937574 |

| | | | | | | |
|---|---|---|---|---|---|---|
| EEF2 | 307.786547 | -0.0306232 | 0.50263614 | -0.0609253 | 0.95141872 | 0.99937574 |
| PIAS4 | 20.2889508 | -0.0602056 | 0.52913613 | -0.1137809 | 0.90941152 | 0.99937574 |
| ZBTB7A | 78.8958569 | -0.5154141 | 0.5310406 | -0.9705738 | 0.33176056 | 0.99937574 |
| MAP2K2 | 81.0714601 | -0.0902315 | 0.28545377 | -0.3160986 | 0.75192767 | 0.99937574 |
| SIRT6 | 12.6595326 | -0.5119788 | 0.52955374 | -0.9668118 | 0.33363811 | 0.99937574 |
| YJU2 | 10.7106296 | -0.6872785 | 0.6580338 | -1.0444426 | 0.29628067 | 0.99937574 |
| STAP2 | 22.4333809 | -0.1431597 | 0.45963449 | -0.3114642 | 0.75544774 | 0.99937574 |
| SH3GL1 | 24.2803346 | -0.8386877 | 0.51348144 | -1.6333359 | 0.10239837 | 0.99937574 |
| UBXN6 | 35.4799416 | -0.3313521 | 0.45274493 | -0.7318737 | 0.46424565 | 0.99937574 |
| HDGFL2 | 20.1342184 | 0.30336068 | 0.46353918 | 0.65444453 | 0.51282546 | 0.99937574 |
| PLIN5 | 46.8418025 | 0.3824719 | 0.38016446 | 1.00606956 | 0.31438211 | 0.99937574 |
| MYDGF | 55.1735222 | 0.18609036 | 0.3973169 | 0.4683676 | 0.63952173 | 0.99937574 |
| DPP9 | 27.851247 | -0.4408253 | 0.47627683 | -0.9255654 | 0.3546719 | 0.99937574 |
| TICAM1 | 17.9892157 | -0.0055807 | 0.44077836 | -0.0126611 | 0.98989821 | 0.99937574 |
| PLIN3 | 63.6819976 | 0.38919092 | 0.48451734 | 0.80325489 | 0.42182742 | 0.99937574 |
| KDM4B | 142.678361 | -0.0370534 | 0.38380204 | -0.0965429 | 0.92308942 | 0.99937574 |
| PTPRS | 45.6224856 | 0.07477344 | 0.51441453 | 0.1453564 | 0.88442949 | 0.99937574 |
| TINCR | 16.8980236 | -0.5706425 | 0.48165067 | -1.1847643 | 0.23611064 | 0.99937574 |
| SAFB2 | 77.0489008 | 0.12441482 | 0.27997063 | 0.44438526 | 0.65676407 | 0.99937574 |
| SAFB | 92.3522144 | -0.4623994 | 0.25257144 | -1.8307666 | 0.06713538 | 0.99937574 |
| RPL36 | 218.770255 | -0.091986 | 0.46010762 | -0.1999228 | 0.84154093 | 0.99937574 |
| C19orf70 | 70.2934819 | -0.0807073 | 0.40951035 | -0.1970825 | 0.84376301 | 0.99937574 |
| LONP1 | 61.2190389 | -0.5501651 | 0.45085418 | -1.2202728 | 0.22236149 | 0.99937574 |
| CAPS | 43.079149 | -0.2727215 | 0.79854031 | -0.3415251 | 0.73270834 | 0.99937574 |
| RANBP3 | 33.4332406 | -0.0824601 | 0.37580277 | -0.2194239 | 0.82631989 | 0.99937574 |
| RFX2 | 19.5694962 | 0.70088226 | 0.49144224 | 1.42617425 | 0.15381805 | 0.99937574 |
| MLLT1 | 29.4945178 | -0.2907483 | 0.55135457 | -0.5273346 | 0.59796125 | 0.99937574 |
| CLPP | 21.1569512 | -0.036143 | 0.50733966 | -0.0712403 | 0.94320648 | 0.99937574 |
| ALKBH7 | 30.5395796 | 0.20239819 | 0.56400676 | 0.35885775 | 0.71970151 | 0.99937574 |
| GTF2F1 | 22.9364421 | -0.6120278 | 0.53268885 | -1.1489404 | 0.25058054 | 0.99937574 |
| KHSRP | 33.3305382 | -0.596282 | 0.53891931 | -1.1064402 | 0.26853603 | 0.99937574 |
| SLC25A23 | 24.1091955 | 0.212473 | 0.47365636 | 0.44858048 | 0.65373432 | 0.99937574 |
| CRB3 | 15.9597483 | -0.5743641 | 0.69557839 | -0.825736 | 0.40895386 | 0.99937574 |
| AC010503.4 | 16.7906538 | -1.4440461 | 0.75628115 | -1.9094038 | 0.05621002 | 0.99937574 |
| TRIP10 | 20.8652537 | -0.2491107 | 0.69429153 | -0.3587984 | 0.71974591 | 0.99937574 |
| SH2D3A | 31.5557477 | -1.1778332 | 0.60531297 | -1.9458252 | 0.05167574 | 0.99937574 |
| ZNF557 | 11.3776341 | -0.3911629 | 0.60973509 | -0.6415293 | 0.52117887 | 0.99937574 |
| INSR | 453.577579 | -0.3001143 | 0.30409096 | -0.9869229 | 0.32368046 | 0.99937574 |
| ZNF358 | 13.9495751 | -0.5393365 | 0.62880717 | -0.8577136 | 0.39105063 | 0.99937574 |
| PNPLA6 | 23.4719932 | -0.0493393 | 0.59943585 | -0.0823095 | 0.93440059 | 0.99937574 |
| CAMSAP3 | 17.3574117 | -0.289296 | 0.61605361 | -0.4695955 | 0.63864404 | 0.99937574 |
| XAB2 | 13.1227421 | 0.42874938 | 0.569417 | 0.75296204 | 0.45147273 | 0.99937574 |
| STXBP2 | 23.7918227 | -0.5704223 | 0.51078606 | -1.1167539 | 0.26409956 | 0.99937574 |
| EVI5L | 54.5953195 | 0.37332323 | 0.55428201 | 0.6735258 | 0.50061284 | 0.99937574 |
| MAP2K7 | 44.0462571 | -0.3945352 | 0.51634099 | -0.7640982 | 0.44480872 | 0.99937574 |
| TIMM44 | 24.3668759 | -0.2221246 | 0.44135545 | -0.5032783 | 0.61476865 | 0.99937574 |
| ELAVL1 | 69.4090354 | -0.0609352 | 0.32068627 | -0.1900151 | 0.84929729 | 0.99937574 |
| FBN3 | 74.8897743 | 0.96369779 | 0.49449145 | 1.94886644 | 0.05131138 | 0.99937574 |
| CERS4 | 162.193609 | -0.2816298 | 0.60510978 | -0.4654194 | 0.64163114 | 0.99937574 |
| AC022146.2 | 40.5355511 | -0.1583807 | 0.57099386 | -0.2773773 | 0.7814904 | 0.99937574 |
| CD320 | 20.7862536 | 0.61109828 | 0.50421556 | 1.21197822 | 0.22552072 | 0.99937574 |
| RPS28 | 61.5211701 | -0.0402622 | 0.48596974 | -0.0828493 | 0.93397138 | 0.99937574 |
| ANGPTL4 | 16.6087511 | -1.5130581 | 1.12375006 | -1.3464365 | 0.17816178 | 0.99937574 |
| RAB11B | 48.9969333 | -0.0203499 | 0.42154375 | -0.0482747 | 0.96149728 | 0.99937574 |

| | | | | | | |
|---|---|---|---|---|---|---|
| 2-Mar | 22.4927829 | -0.0520563 | 0.54064107 | -0.0962862 | 0.92329326 | 0.99937574 |
| HNRNPM | 156.147909 | -0.2566538 | 0.27250083 | -0.9418461 | 0.34627144 | 0.99937574 |
| ZNF558 | 11.6076977 | 0.04988434 | 0.60009419 | 0.08312752 | 0.93375015 | 0.99937574 |
| ZNF317 | 18.7555724 | -0.1228474 | 0.56400978 | -0.2178108 | 0.82757655 | 0.99937574 |
| ZNF559-ZNF | 15.9466157 | 0.4979341 | 0.50831084 | 0.97958584 | 0.3272906 | 0.99937574 |
| ZNF266 | 30.4162137 | -0.6510276 | 0.43181267 | -1.507662 | 0.13164104 | 0.99937574 |
| ZNF426 | 18.0337278 | 0.14496111 | 0.47951197 | 0.30230969 | 0.76241599 | 0.99937574 |
| ZNF121 | 37.735363 | -0.5829138 | 0.39578409 | -1.4728077 | 0.14080289 | 0.99937574 |
| ZNF561 | 20.9996754 | 0.35550635 | 0.47460961 | 0.74905003 | 0.45382705 | 0.99937574 |
| ZNF562 | 51.0744965 | -0.6556205 | 0.32173835 | -2.0377444 | 0.0415755 | 0.99937574 |
| ZNF846 | 44.2499506 | -0.0215529 | 0.36857154 | -0.0584769 | 0.95336878 | 0.99937574 |
| FBXL12 | 19.5090784 | -0.6010854 | 0.45627754 | -1.317368 | 0.1877153 | 0.99937574 |
| UBL5 | 48.1520379 | 0.1652494 | 0.45648709 | 0.36200234 | 0.71735028 | 0.99937574 |
| PIN1 | 32.4221415 | -0.3863262 | 0.42765877 | -0.9033516 | 0.36633931 | 0.99937574 |
| C19orf66 | 25.6988585 | -1.2604829 | 0.75043197 | -1.6796765 | 0.09302028 | 0.99937574 |
| EIF3G | 23.0366271 | -0.1356237 | 0.50243029 | -0.2699353 | 0.78720999 | 0.99937574 |
| DNMT1 | 23.2750494 | -0.1320998 | 0.48459758 | -0.2725969 | 0.78516313 | 0.99937574 |
| MRPL4 | 31.4684062 | 0.58919265 | 0.36818806 | 1.6002492 | 0.10954331 | 0.99937574 |
| AC011511.2 | 29.3955416 | 1.64676748 | 0.7661543 | 2.14939401 | 0.03160318 | 0.99049468 |
| ICAM1 | 14.8535771 | -0.2333349 | 0.8245562 | -0.2829824 | 0.77719034 | 0.99937574 |
| TYK2 | 26.4803966 | -0.5691702 | 0.48046956 | -1.1846125 | 0.23617071 | 0.99937574 |
| CDC37 | 54.8124846 | -0.5255855 | 0.31717933 | -1.6570611 | 0.09750712 | 0.99937574 |
| PDE4A | 43.2502211 | 0.10766358 | 0.53112456 | 0.20270873 | 0.8393627 | 0.99937574 |
| KEAP1 | 23.5231422 | -0.200038 | 0.45781452 | -0.4369412 | 0.66215403 | 0.99937574 |
| ATG4D | 28.8670443 | -0.469158 | 0.44691155 | -1.0497782 | 0.29382011 | 0.99937574 |
| KRI1 | 14.4216843 | -0.2664093 | 0.58498439 | -0.4554126 | 0.6488124 | 0.99937574 |
| AP1M2 | 35.5372496 | 0.29569715 | 0.46881942 | 0.63072717 | 0.52821893 | 0.99937574 |
| SLC44A2 | 32.9457673 | -0.8391185 | 0.49176971 | -1.706324 | 0.08794777 | 0.99937574 |
| ILF3-DT | 25.820468 | -0.1574793 | 0.53664852 | -0.2934496 | 0.76917854 | 0.99937574 |
| ILF3 | 102.646434 | -0.2817192 | 0.41349638 | -0.6813099 | 0.4956754 | 0.99937574 |
| QTRT1 | 25.0933187 | 0.35641599 | 0.39343263 | 0.90591365 | 0.36498156 | 0.99937574 |
| DNM2 | 225.680589 | -0.117383 | 0.39959584 | -0.2937542 | 0.76894571 | 0.99937574 |
| TMED1 | 27.0157734 | 0.1104067 | 0.58893063 | 0.18746979 | 0.85129231 | 0.99937574 |
| CARM1 | 21.0108323 | 0.01154508 | 0.50186535 | 0.02300434 | 0.98164681 | 0.99937574 |
| YIPF2 | 21.6477796 | 0.28486485 | 0.45445622 | 0.62682571 | 0.53077349 | 0.99937574 |
| TIMM29 | 15.9129647 | -0.0398835 | 0.48098563 | -0.0829204 | 0.93391482 | 0.99937574 |
| SMARCA4 | 121.245431 | -0.042644 | 0.39762529 | -0.1072466 | 0.91459335 | 0.99937574 |
| LDLR | 179.192167 | -0.7826129 | 0.69081489 | -1.1328837 | 0.25726309 | 0.99937574 |
| KANK2 | 46.4027529 | 0.57294036 | 0.42030408 | 1.36315677 | 0.17283311 | 0.99937574 |
| DOCK6 | 62.187497 | -0.2391445 | 0.49110324 | -0.4869535 | 0.62629126 | 0.99937574 |
| CCDC159 | 12.3105648 | 0.38123637 | 0.69878668 | 0.54556903 | 0.58536222 | 0.99937574 |
| EPOR | 28.4890398 | 0.95833366 | 0.48015103 | 1.99590046 | 0.04594476 | 0.99937574 |
| RGL3 | 262.62207 | 0.39006995 | 0.32323752 | 1.20675952 | 0.22752477 | 0.99937574 |
| PRKCSH | 50.997982 | -0.0145432 | 0.40809303 | -0.0356371 | 0.97157176 | 0.99937574 |
| ECSIT | 37.7749439 | -0.1074142 | 0.34689627 | -0.3096436 | 0.75683198 | 0.99937574 |
| ELOF1 | 14.4382769 | -0.5179015 | 0.65600179 | -0.7894818 | 0.42983044 | 0.99937574 |
| ZNF627 | 26.69977 | 0.20252399 | 0.45806108 | 0.44213316 | 0.65839285 | 0.99937574 |
| AC008543.1 | 23.8745057 | 0.10146998 | 0.55159932 | 0.18395595 | 0.85404801 | 0.99937574 |
| ZNF823 | 22.7469485 | -0.4711212 | 0.43810937 | -1.0753506 | 0.28221781 | 0.99937574 |
| ZNF441 | 15.4047666 | 0.52165681 | 0.57114491 | 0.9133528 | 0.36105702 | 0.99937574 |
| ZNF440 | 41.0508245 | -0.2249974 | 0.42668881 | -0.5273104 | 0.59797806 | 0.99937574 |
| ZNF439 | 21.7844272 | 0.38506945 | 0.43862914 | 0.877893 | 0.38000179 | 0.99937574 |
| ZNF69 | 38.138675 | -0.7052297 | 0.60461564 | -1.16641 | 0.24344872 | 0.99937574 |
| ZNF433 | 10.6782032 | -0.9222947 | 0.59446245 | -1.5514767 | 0.12078748 | 0.99937574 |

| | | | | | | |
|---|---|---|---|---|---|---|
| ZNF844 | 27.8072624 | -0.0297094 | 0.43822273 | -0.0677952 | 0.94594867 | 0.99937574 |
| ZNF136 | 19.0458028 | -0.2757 | 0.52412778 | -0.5260167 | 0.5988766 | 0.99937574 |
| ZNF44 | 124.844214 | -0.1771517 | 0.28694971 | -0.6173617 | 0.53699621 | 0.99937574 |
| AC008758.5 | 17.0544262 | 0.04267937 | 0.60956807 | 0.07001576 | 0.94418111 | 0.99937574 |
| ZNF490 | 30.8499346 | 0.35543722 | 0.54086021 | 0.65717021 | 0.51107148 | 0.99937574 |
| ZNF791 | 53.2466556 | -0.1316435 | 0.34433418 | -0.3823133 | 0.70222903 | 0.99937574 |
| MAN2B1 | 11.2317711 | 0.04761006 | 0.80272932 | 0.05931023 | 0.95270501 | 0.99937574 |
| WDR83 | 12.18743 | 1.02157219 | 0.70859374 | 1.44168956 | 0.14938997 | 0.99937574 |
| DHPS | 18.9705451 | 0.85130137 | 0.50862952 | 1.673716 | 0.09418643 | 0.99937574 |
| TNPO2 | 34.7472086 | 0.37368467 | 0.46825272 | 0.79804057 | 0.42484695 | 0.99937574 |
| TRIR | 77.038096 | -0.0538657 | 0.50041469 | -0.1076421 | 0.91427962 | 0.99937574 |
| ASNA1 | 22.0622306 | -0.1855495 | 0.42855784 | -0.4329625 | 0.66504203 | 0.99937574 |
| HOOK2 | 130.368556 | -0.0344517 | 0.32267966 | -0.1067674 | 0.91497354 | 0.99937574 |
| JUNB | 66.606618 | -0.5259827 | 0.65346598 | -0.8049122 | 0.42087036 | 0.99937574 |
| DNASE2 | 20.6628234 | 0.38468762 | 0.45187696 | 0.85131053 | 0.39459688 | 0.99937574 |
| GCDH | 19.9215182 | 0.22434014 | 0.43239132 | 0.5188359 | 0.60387518 | 0.99937574 |
| FARSA | 23.4547366 | -0.1036111 | 0.48451685 | -0.2138441 | 0.83066863 | 0.99937574 |
| CALR | 179.273808 | 0.47686094 | 0.42761195 | 1.11517214 | 0.26477666 | 0.99937574 |
| AC092069.1 | 29.7285631 | -0.020767 | 0.50101475 | -0.0414499 | 0.96693723 | 0.99937574 |
| RAD23A | 38.7550072 | -0.4482129 | 0.47264208 | -0.9483136 | 0.34296982 | 0.99937574 |
| GADD45GIP1 | 67.572305 | -0.1662667 | 0.44038569 | -0.3775479 | 0.7057665 | 0.99937574 |
| NFIX | 25.5386902 | 0.35805318 | 0.62417544 | 0.57364189 | 0.56621016 | 0.99937574 |
| NACC1 | 18.9383093 | -0.6998251 | 0.53748484 | -1.3020369 | 0.19290377 | 0.99937574 |
| STX10 | 12.9879923 | -0.2014675 | 0.52223263 | -0.3857812 | 0.69965875 | 0.99937574 |
| IER2 | 56.7721618 | -0.4020452 | 0.65782748 | -0.6111712 | 0.54108624 | 0.99937574 |
| CCDC130 | 47.2733706 | -0.3250538 | 0.3681007 | -0.8830566 | 0.37720568 | 0.99937574 |
| MRI1 | 19.0285144 | 0.96350173 | 0.51790806 | 1.86037214 | 0.06283289 | 0.99937574 |
| C19orf53 | 57.2450099 | -0.2478687 | 0.46839734 | -0.5291848 | 0.59667728 | 0.99937574 |
| ZSWIM4 | 16.9598778 | -0.6765923 | 0.54195739 | -1.2484234 | 0.21187603 | 0.99937574 |
| CC2D1A | 36.5091303 | -0.0493761 | 0.33975329 | -0.1453294 | 0.88445082 | 0.99937574 |
| DCAF15 | 10.880062 | -0.2115011 | 0.643231 | -0.3288105 | 0.74229894 | 0.99937574 |
| RFX1 | 25.4422979 | 0.20592497 | 0.44307398 | 0.46476431 | 0.64210025 | 0.99937574 |
| PALM3 | 25.3156736 | -0.15842 | 0.70395992 | -0.2250412 | 0.82194725 | 0.99937574 |
| SAMD1 | 12.7106111 | -0.3914704 | 0.5730505 | -0.6831341 | 0.4945221 | 0.99937574 |
| PRKACA | 30.3080271 | 0.54551184 | 0.37305346 | 1.46228866 | 0.14366212 | 0.99937574 |
| ADGRL1 | 25.5580006 | 0.02978458 | 0.55726638 | 0.05344765 | 0.95737524 | 0.99937574 |
| DDX39A | 13.3077906 | -1.11541 | 0.79529073 | -1.4025185 | 0.16076047 | 0.99937574 |
| PKN1 | 46.4802555 | -0.9480359 | 0.55746834 | -1.7006095 | 0.08901635 | 0.99937574 |
| GIPC1 | 66.7968499 | -1.0061595 | 0.49161484 | -2.0466418 | 0.04069327 | 0.99937574 |
| DNAJB1 | 41.0789394 | -0.3601362 | 0.47045044 | -0.7655137 | 0.44396572 | 0.99937574 |
| TECR | 46.1973036 | -0.0798162 | 0.39505378 | -0.2020389 | 0.83988635 | 0.99937574 |
| NDUFB7 | 88.0326244 | 0.14414695 | 0.42062191 | 0.34269957 | 0.73182448 | 0.99937574 |
| ZNF333 | 11.6004136 | -0.7813176 | 0.59465126 | -1.313909 | 0.18887684 | 0.99937574 |
| ILVBL | 24.4993316 | 0.35890384 | 0.42963505 | 0.83536909 | 0.40350993 | 0.99937574 |
| BRD4 | 196.355625 | -0.255709 | 0.30082423 | -0.850028 | 0.3953095 | 0.99937574 |
| AKAP8 | 42.7110649 | -0.3332547 | 0.30694754 | -1.0857058 | 0.27760919 | 0.99937574 |
| AKAP8L | 116.313564 | 0.05888367 | 0.29776187 | 0.19775422 | 0.84323736 | 0.99937574 |
| WIZ | 11.711013 | 0.00519816 | 0.58505607 | 0.0088849 | 0.99291097 | 0.99937574 |
| TPM4 | 20.134245 | -2.2426367 | 1.06988222 | -2.0961529 | 0.03606863 | 0.99937574 |
| RAB8A | 23.3386115 | -0.0477505 | 0.4352609 | -0.1097054 | 0.912643 | 0.99937574 |
| FAM32A | 13.6786888 | -0.6817733 | 0.68626558 | -0.9934541 | 0.32048872 | 0.99937574 |
| AP1M1 | 17.3945995 | -0.000757 | 0.42543152 | -0.0017793 | 0.99858029 | 0.99979721 |
| EPS15L1 | 84.5824057 | 0.20607334 | 0.32488956 | 0.63428736 | 0.52589331 | 0.99937574 |
| C19orf44 | 12.7981879 | 0.34549836 | 0.52767401 | 0.6547572 | 0.51262409 | 0.99937574 |

| | | | | | | |
|---|---|---|---|---|---|---|
| TMEM38A | 96.4083062 | 0.96561201 | 0.37139461 | 2.59996237 | 0.0093234 | 0.75880433 |
| SIN3B | 47.9788171 | -0.5957729 | 0.42165916 | -1.4129253 | 0.1576777 | 0.99937574 |
| CPAMD8 | 225.126757 | 1.44973847 | 0.5432761 | 2.66851139 | 0.00761882 | 0.74742128 |
| MYO9B | 49.150495 | -0.98685 | 0.43915759 | -2.2471433 | 0.02463087 | 0.96546668 |
| OCEL1 | 13.9742052 | 0.07222104 | 0.60869542 | 0.1186489 | 0.90555352 | 0.99937574 |
| NR2F6 | 52.066762 | 0.44685666 | 0.33385999 | 1.33845525 | 0.18074808 | 0.99937574 |
| ABHD8 | 11.6025384 | 0.53931219 | 0.6231069 | 0.86552113 | 0.38675282 | 0.99937574 |
| MRPL34 | 52.4992563 | 0.14408554 | 0.54540358 | 0.26418152 | 0.79164005 | 0.99937574 |
| DDA1 | 18.4705279 | -0.1979216 | 0.56748126 | -0.348772 | 0.72726045 | 0.99937574 |
| BST2 | 46.2302923 | -1.029246 | 1.59493136 | -0.6453231 | 0.51871785 | 0.99937574 |
| MVB12A | 15.015027 | 0.00468054 | 0.60603307 | 0.00772324 | 0.99383781 | 0.99937574 |
| SLC27A1 | 18.0487294 | 0.13257398 | 0.55500253 | 0.23887095 | 0.81120565 | 0.99937574 |
| AC010618.3 | 18.7917889 | 0.73874424 | 0.55955704 | 1.32023044 | 0.18675809 | 0.99937574 |
| PGLS | 57.9473571 | 0.10861315 | 0.33703732 | 0.32225854 | 0.74725684 | 0.99937574 |
| COLGALT1 | 16.8622763 | -0.4005789 | 0.53112044 | -0.7542148 | 0.45072024 | 0.99937574 |
| RPL18A | 18.568793 | -0.5311838 | 0.50199709 | -1.0581412 | 0.28999108 | 0.99937574 |
| CCDC124 | 22.5089413 | -0.6426276 | 0.56108792 | -1.1453242 | 0.2520749 | 0.99937574 |
| ARRDC2 | 12.334949 | -0.589972 | 0.69132709 | -0.8533905 | 0.39344277 | 0.99937574 |
| MAST3 | 11.9191636 | 0.78322443 | 0.56285269 | 1.39152648 | 0.16406584 | 0.99937574 |
| JUND | 163.339654 | -1.2539375 | 0.50753104 | -2.4706618 | 0.01348633 | 0.82550375 |
| LSM4 | 65.077601 | 0.36763064 | 0.33149241 | 1.10901677 | 0.26742295 | 0.99937574 |
| PGPEP1 | 35.7611533 | 0.20953155 | 0.63155665 | 0.33176999 | 0.74006295 | 0.99937574 |
| GDF15 | 133.152067 | -1.1155558 | 0.75274968 | -1.4819744 | 1.38E-01 | 0.99937574 |
| SSBP4 | 27.2282011 | -0.0606306 | 0.38243196 | -0.1585395 | 0.87403169 | 0.99937574 |
| ISYNA1 | 12.2144223 | 0.37284911 | 0.890633 | 0.41863383 | 0.67548376 | 0.99937574 |
| ELL | 50.3005105 | -0.1134524 | 0.34609283 | -0.3278092 | 0.74305593 | 0.99937574 |
| FKBP8 | 48.563003 | -0.0529765 | 0.3676834 | -0.1440818 | 0.88543591 | 0.99937574 |
| KXD1 | 23.4181329 | -0.0811256 | 0.38281699 | -0.2119174 | 0.83217145 | 0.99937574 |
| UBA52 | 107.183817 | 0.33022898 | 0.4849783 | 0.68091496 | 0.4959253 | 0.99937574 |
| REX1BD | 29.005349 | -0.0137647 | 0.60407763 | -0.0227863 | 0.98182077 | 0.99937574 |
| KLHL26 | 14.002872 | 0.66622936 | 0.76711073 | 0.86849177 | 0.38512517 | 0.99937574 |
| CRTC1 | 114.860037 | 0.08171196 | 0.42246726 | 0.19341608 | 0.84663313 | 0.99937574 |
| UPF1 | 46.9866907 | -0.1377038 | 0.34595317 | -0.3980418 | 0.69059934 | 0.99937574 |
| COPE | 64.4935212 | -0.054372 | 0.38768127 | -0.1402492 | 0.88846313 | 0.99937574 |
| DDX49 | 11.7293169 | -0.7722286 | 0.6293864 | -1.2269547 | 0.2198396 | 0.99937574 |
| SUGP2 | 97.513652 | 0.04900183 | 0.26910152 | 0.18209421 | 0.85550879 | 0.99937574 |
| ARMC6 | 10.6905514 | -0.0833122 | 0.6885644 | -0.1209941 | 0.90369574 | 0.99937574 |
| SLC25A42 | 17.4690383 | 0.08461515 | 0.7323233 | 0.11554344 | 0.90801439 | 0.99937574 |
| TMEM161A | 15.1560776 | 0.38393811 | 0.46002889 | 0.83459565 | 0.40394541 | 0.99937574 |
| RFXANK | 10.6049126 | 0.09510708 | 0.55679055 | 0.17081303 | 0.86437078 | 0.99937574 |
| SUGP1 | 20.442616 | -0.3345329 | 0.52198454 | -0.6408866 | 0.52159636 | 0.99937574 |
| MAU2 | 66.3349777 | -0.2711022 | 0.2730283 | -0.9929453 | 0.32073659 | 0.99937574 |
| GATAD2A | 124.756784 | -0.2410237 | 0.33408451 | -0.7214452 | 0.47063564 | 0.99937574 |
| ATP13A1 | 21.6396301 | 0.062723 | 0.43615836 | 0.14380786 | 0.8856522 | 0.99937574 |
| ZNF101 | 11.0487856 | 0.60930581 | 0.53818616 | 1.13214693 | 0.25757267 | 0.99937574 |
| ZNF14 | 18.4314991 | 0.38848185 | 0.50933373 | 0.76272557 | 0.44562708 | 0.99937574 |
| ZNF253 | 30.9542279 | 0.44987285 | 0.46947105 | 0.95825472 | 0.33793433 | 0.99937574 |
| ZNF93 | 12.2069202 | 0.08360313 | 0.54761417 | 0.15266795 | 0.87866014 | 0.99937574 |
| ZNF90 | 135.580214 | -0.0429851 | 0.48495476 | -0.0886373 | 0.92937019 | 0.99937574 |
| AC011447.3 | 23.9521432 | -0.480479 | 0.59306682 | -0.8101599 | 0.41784826 | 0.99937574 |
| ZNF486 | 17.7145414 | -0.9576335 | 0.57197233 | -1.6742655 | 0.09407844 | 0.99937574 |
| ZNF737 | 33.6146293 | 0.19596311 | 0.44715823 | 0.43824108 | 0.66121153 | 0.99937574 |
| AC010636.2 | 17.7313458 | 1.12200587 | 0.6453462 | 1.7386108 | 0.08210325 | 0.99937574 |
| AC010329.1 | 15.2774886 | -0.7151145 | 0.86006117 | -0.8314693 | 0.40570855 | 0.99937574 |

| | | | | | | |
|---|---|---|---|---|---|---|
| ZNF66 | 35.2901563 | -0.2375962 | 0.43700056 | -0.5436976 | 0.58664959 | 0.99937574 |
| ZNF85 | 21.6107381 | 0.26622931 | 0.52663857 | 0.50552566 | 0.61318968 | 0.99937574 |
| ZNF430 | 42.9577217 | -0.1557909 | 0.34980767 | -0.4453617 | 0.65605839 | 0.99937574 |
| ZNF714 | 26.1869223 | -0.1143671 | 0.43151437 | -0.2650367 | 0.79098122 | 0.99937574 |
| ZNF431 | 99.3060532 | 0.11783542 | 0.27536458 | 0.42792511 | 0.66870565 | 0.99937574 |
| ZNF708 | 59.5773345 | -0.0467031 | 0.36458188 | -0.1281005 | 0.89806948 | 0.99937574 |
| ZNF738 | 12.2672349 | -0.0129165 | 0.49447342 | -0.0261218 | 0.97916021 | 0.99937574 |
| ZNF493 | 34.0609725 | 0.09439066 | 0.46469613 | 0.2031234 | 0.83903858 | 0.99937574 |
| AC010615.4 | 46.6813261 | 1.49613409 | 0.75010008 | 1.9945793 | 0.04608878 | 0.99937574 |
| ZNF429 | 62.862767 | 0.27669446 | 0.31264848 | 0.88500178 | 0.37615567 | 0.99937574 |
| AC123912.1 | 24.7886308 | 0.9624497 | 0.52123162 | 1.84649138 | 0.06482089 | 0.99937574 |
| AC123912.4 | 34.5665331 | 1.375314 | 0.52119412 | 2.63877497 | 0.00832062 | 0.74742128 |
| ZNF100 | 38.6186632 | 0.19143123 | 0.56345344 | 0.33974631 | 0.73404758 | 0.99937574 |
| ZNF43 | 44.8012871 | -0.2323314 | 0.39896904 | -0.5823294 | 0.56034485 | 0.99937574 |
| ZNF208 | 115.363864 | -0.1688519 | 0.40315392 | -0.4188273 | 0.67534234 | 0.99937574 |
| ZNF257 | 16.549018 | -0.6958768 | 0.5096765 | -1.3653303 | 0.17214925 | 0.99937574 |
| ZNF676 | 17.288943 | 0.48418377 | 0.74524983 | 0.64969323 | 0.5158904 | 0.99937574 |
| ZNF98 | 65.9096365 | -0.1039733 | 0.4377212 | -0.2375333 | 0.81224312 | 0.99937574 |
| ZNF99 | 13.7620279 | -0.4770133 | 0.7676484 | -0.6213956 | 0.53433936 | 0.99937574 |
| ZNF91 | 264.654813 | -0.2508745 | 0.29745551 | -0.8434017 | 0.3990038 | 0.99937574 |
| ZNF675 | 24.0675635 | -0.1521114 | 0.42861082 | -0.354894 | 0.72266899 | 0.99937574 |
| ZNF726 | 13.0315242 | 0.50143876 | 0.72633368 | 0.69036969 | 0.48996173 | 0.99937574 |
| ZNF254 | 165.487217 | -0.227573 | 0.26045171 | -0.8737629 | 0.38224741 | 0.99937574 |
| LINC00662 | 49.1755557 | -0.3502847 | 0.35555006 | -0.985191 | 0.32453027 | 0.99937574 |
| AC006504.5 | 77.7172112 | 0.63904698 | 0.39305729 | 1.62583677 | 0.10398438 | 0.99937574 |
| UQCRFS1 | 125.878317 | 0.85884033 | 0.50966451 | 1.68510915 | 0.09196752 | 0.99937574 |
| POP4 | 12.3951766 | 0.88464542 | 0.47556107 | 1.86021413 | 0.06285524 | 0.99937574 |
| C19orf12 | 50.6040792 | -0.249158 | 0.30953107 | -0.8049531 | 0.42084676 | 0.99937574 |
| URI1 | 140.48933 | -0.0865497 | 0.24299197 | -0.3561832 | 0.72170334 | 0.99937574 |
| ZNF507 | 42.9075006 | -0.1112645 | 0.33295275 | -0.334175 | 0.73824752 | 0.99937574 |
| DPY19L3 | 37.0103318 | -0.5182297 | 0.44505869 | -1.1644076 | 0.24425886 | 0.99937574 |
| PDCD5 | 22.9747132 | -0.3184188 | 0.47588945 | -0.6691025 | 0.50343013 | 0.99937574 |
| ANKRD27 | 44.3230883 | -0.2504163 | 0.42834647 | -0.5846115 | 0.55880895 | 0.99937574 |
| NUDT19 | 15.5126729 | 0.56202005 | 0.5575876 | 1.00794933 | 0.31347879 | 0.99937574 |
| CEP89 | 30.9626818 | -0.0098733 | 0.42445363 | -0.0232611 | 0.98144197 | 0.99937574 |
| RHPN2 | 126.187647 | -0.4040849 | 0.38043886 | -1.0621545 | 0.28816555 | 0.99937574 |
| GPATCH1 | 14.5106961 | -0.0942583 | 0.55997024 | -0.1683273 | 0.86632581 | 0.99937574 |
| CEBPG | 15.1079414 | 0.0638217 | 0.59243961 | 0.10772694 | 0.9142123 | 0.99937574 |
| PEPD | 43.9111491 | -0.3243122 | 0.48505111 | -0.6686145 | 0.50374144 | 0.99937574 |
| LSM14A | 187.342685 | -0.2598583 | 0.23053178 | -1.1272126 | 0.25965261 | 0.99937574 |
| KIAA0355 | 103.713405 | 0.02713008 | 0.23553842 | 0.11518324 | 0.90829989 | 0.99937574 |
| GPI | 135.860737 | -0.5177018 | 0.49449104 | -1.0469388 | 0.29512782 | 0.99937574 |
| UBA2 | 90.2218174 | -0.2494721 | 0.28566763 | -0.873295 | 0.38250229 | 0.99937574 |
| SCGB2B2 | 40.7027464 | -0.8628154 | 0.50986761 | -1.6922341 | 0.09060134 | 0.99937574 |
| ZNF302 | 60.996124 | -0.5825109 | 0.43510448 | -1.3387839 | 0.18064103 | 0.99937574 |
| GRAMD1A | 39.5852516 | 0.05233955 | 0.49162989 | 0.10646129 | 0.91521637 | 0.99937574 |
| HPN | 118.745567 | 0.92421775 | 0.50110749 | 1.8443503 | 0.06513211 | 0.99937574 |
| LSR | 53.3163126 | -0.7046743 | 0.46990241 | -1.4996183 | 0.1337133 | 0.99937574 |
| USF2 | 44.2423053 | -0.5970835 | 0.52148196 | -1.1449745 | 0.25221973 | 0.99937574 |
| DMKN | 88.6495643 | 0.2938329 | 0.40712929 | 0.72171889 | 0.47046733 | 0.99937574 |
| TMEM147 | 29.0907819 | -0.10899 | 0.50260703 | -0.2168492 | 0.82832583 | 0.99937574 |
| RBM42 | 16.6545305 | -0.2777317 | 0.58772064 | -0.4725573 | 0.63652905 | 0.99937574 |
| COX6B1 | 146.624485 | 0.11673355 | 0.38169814 | 0.30582688 | 0.75973646 | 0.99937574 |
| KMT2B | 28.4621199 | 0.36745913 | 0.4380914 | 0.83877274 | 0.40159685 | 0.99937574 |

| | | | | | | |
|---|---|---|---|---|---|---|
| PROSER3 | 11.252323 | -0.7440142 | 0.58451787 | -1.2728682 | 0.20306482 | 0.99937574 |
| SDHAF1 | 13.774967 | 0.22658797 | 0.5671899 | 0.39949225 | 0.68953053 | 0.99937574 |
| TBCB | 24.739272 | -0.0622075 | 0.48144986 | -0.1292087 | 0.89719255 | 0.99937574 |
| CAPNS1 | 114.415808 | -0.0390743 | 0.41468643 | -0.0942262 | 0.92492949 | 0.99937574 |
| COX7A1 | 24.5748168 | 0.78300427 | 1.08729663 | 0.72013859 | 0.47143967 | 0.99937574 |
| ZNF565 | 52.6999164 | -0.0897025 | 0.571109 | -0.1570672 | 0.87519191 | 0.99937574 |
| ZNF146 | 64.8946754 | -0.1019711 | 0.36947717 | -0.2759875 | 0.7825577 | 0.99937574 |
| LINC00665 | 20.5779856 | 1.16861158 | 0.48027653 | 2.43320566 | 0.0149658 | 0.86113259 |
| ZFP14 | 44.0770109 | 0.98629986 | 0.32378154 | 3.04618925 | 0.00231762 | 0.48727941 |
| ZFP82 | 19.3519599 | 0.31712794 | 0.47835846 | 0.66295043 | 0.5073623 | 0.99937574 |
| ZNF566 | 27.138215 | -0.4974419 | 0.44229177 | -1.1246917 | 0.26071968 | 0.99937574 |
| ZNF260 | 22.926552 | 0.42476412 | 0.42027294 | 1.01068632 | 0.31216659 | 0.99937574 |
| ZNF529 | 55.2763014 | 0.07167634 | 0.41492041 | 0.1727472 | 0.86285014 | 0.99937574 |
| ZNF529-AS1 | 32.3330118 | 0.27559093 | 0.43506928 | 0.63344148 | 0.52644539 | 0.99937574 |
| ZNF382 | 16.219139 | 0.07444426 | 0.63115782 | 0.11794872 | 0.90610829 | 0.99937574 |
| ZNF461 | 16.3745785 | 0.19501194 | 0.48740096 | 0.40010578 | 0.68907861 | 0.99937574 |
| ZNF567 | 25.8324568 | -0.0092347 | 0.40277712 | -0.0229275 | 0.98170811 | 0.99937574 |
| AC020928.1 | 37.2848096 | 0.04715029 | 0.39207234 | 0.12025915 | 0.90427787 | 0.99937574 |
| ZNF790 | 37.8051685 | 0.59762293 | 0.41132288 | 1.45292899 | 0.14624347 | 0.99937574 |
| ZNF345 | 36.9267515 | 0.35533369 | 0.43597605 | 0.8150303 | 0.41505497 | 0.99937574 |
| ZNF568 | 26.2246535 | -0.3683416 | 0.46049825 | -0.7998762 | 0.42378252 | 0.99937574 |
| ZNF420 | 74.7952236 | 0.3555843 | 0.40236363 | 0.88373868 | 0.3768373 | 0.99937574 |
| ZNF383 | 16.2709241 | -0.6041631 | 0.49445254 | -1.221883 | 0.22175188 | 0.99937574 |
| HKR1 | 80.7297287 | 0.20670393 | 0.31754479 | 0.65094416 | 0.51508253 | 0.99937574 |
| ZNF569 | 24.5856241 | 0.86855107 | 0.4315613 | 2.01257866 | 0.04415897 | 0.99937574 |
| ZNF793 | 18.0681102 | 0.66451183 | 0.56301787 | 1.18026773 | 0.23789375 | 0.99937574 |
| ZNF540 | 31.8508421 | 0.25779961 | 0.49907077 | 0.51655922 | 0.60546389 | 0.99937574 |
| ZNF571 | 18.7804124 | -0.0265876 | 0.48301482 | -0.0550452 | 0.95610245 | 0.99937574 |
| ZFP30 | 24.2436589 | 0.43106826 | 0.45451617 | 0.94841127 | 0.34292012 | 0.99937574 |
| ZNF573 | 36.004167 | 0.30478606 | 0.35693413 | 0.8539 | 0.3931604 | 0.99937574 |
| SIPA1L3 | 277.55523 | 0.37585264 | 0.33214404 | 1.1315953 | 0.25780462 | 0.99937574 |
| SPINT2 | 25.0983744 | 0.06278119 | 0.45056735 | 0.13933808 | 0.889183 | 0.99937574 |
| CATSPERG | 11.4343381 | -0.1810651 | 0.62021485 | -0.2919393 | 0.77033304 | 0.99937574 |
| PSMD8 | 67.4608131 | 0.23570934 | 0.30129162 | 0.78232956 | 0.43402092 | 0.99937574 |
| EIF3K | 89.2116615 | 0.39895181 | 0.33347422 | 1.19634979 | 0.23156009 | 0.99937574 |
| ACTN4 | 278.81953 | 0.1636577 | 0.32941778 | 0.49680895 | 0.61932378 | 0.99937574 |
| ECH1 | 53.9647234 | 0.72193413 | 0.52022093 | 1.38774527 | 0.16521463 | 0.99937574 |
| HNRNPL | 52.4333872 | -0.6943252 | 0.41165497 | -1.6866678 | 0.09166725 | 0.99937574 |
| SIRT2 | 39.0431621 | 0.00676076 | 0.38486404 | 0.01756663 | 0.98598458 | 0.99937574 |
| NFKBIB | 16.4077914 | -0.4764284 | 0.52707062 | -0.9039176 | 0.36603909 | 0.99937574 |
| MRPS12 | 52.4935954 | 0.29150408 | 0.47966227 | 0.60772777 | 0.54336804 | 0.99937574 |
| FBXO17 | 45.9277991 | 0.73892846 | 0.48164074 | 1.53419012 | 0.12498289 | 0.99937574 |
| PAK4 | 73.6288655 | -0.1923461 | 0.36291349 | -0.5300054 | 0.5961082 | 0.99937574 |
| SAMD4B | 108.962318 | -0.3283245 | 0.31289423 | -1.0493148 | 0.29403327 | 0.99937574 |
| PAF1 | 19.1749118 | -0.0638925 | 0.45138225 | -0.1415486 | 0.88743655 | 0.99937574 |
| MED29 | 22.8007144 | -0.2888567 | 0.43981483 | -0.656769 | 0.51132949 | 0.99937574 |
| ZFP36 | 39.1832375 | -0.1509868 | 0.60848731 | -0.2481347 | 0.80403019 | 0.99937574 |
| RPS16 | 158.55535 | -0.073664 | 0.4980311 | -0.1479105 | 0.88241342 | 0.99937574 |
| SUPT5H | 63.1614418 | -0.2801268 | 0.35651544 | -0.7857355 | 0.43202247 | 0.99937574 |
| TIMM50 | 16.9640341 | 0.14443868 | 0.44086932 | 0.32762243 | 0.74319716 | 0.99937574 |
| FBL | 15.3067122 | -0.1665502 | 0.53420886 | -0.3117698 | 0.75521551 | 0.99937574 |
| PSMC4 | 23.6042152 | 0.21302917 | 0.40264341 | 0.5290765 | 0.59675238 | 0.99937574 |
| ZNF546 | 14.041969 | 0.76142033 | 0.60947419 | 1.24930693 | 0.21155283 | 0.99937574 |
| ZNF780B | 64.3425712 | 0.46098566 | 0.43804975 | 1.05235915 | 0.29263481 | 0.99937574 |

| | | | | | | |
|---|---|---|---|---|---|---|
| ZNF780A | 18.4869055 | 0.03856956 | 0.4186087 | 0.09213749 | 0.9265888 | 0.99937574 |
| MAP3K10 | 15.9086194 | 0.36354029 | 0.52236957 | 0.69594461 | 0.48646352 | 0.99937574 |
| AKT2 | 79.0348462 | -0.0144204 | 0.2599155 | -0.055481 | 0.95575527 | 0.99937574 |
| PLD3 | 102.745944 | 0.23678334 | 0.29172899 | 0.81165515 | 0.41698954 | 0.99937574 |
| SERTAD1 | 14.8733927 | -0.3854851 | 0.73901126 | -0.5216227 | 0.60193304 | 0.99937574 |
| SERTAD3 | 12.4452367 | -0.4639892 | 0.60804676 | -0.7630814 | 0.44541485 | 0.99937574 |
| BLVRB | 72.4611568 | -0.3488694 | 0.32833871 | -1.062529 | 0.28799558 | 0.99937574 |
| SPTBN4 | 17.6647667 | 0.39539804 | 0.60867028 | 0.64960958 | 0.51594445 | 0.99937574 |
| SHKBP1 | 16.8748655 | -0.3253903 | 0.51547168 | -0.6312478 | 0.52787854 | 0.99937574 |
| LTBP4 | 13.0704614 | -1.0018274 | 0.68978613 | -1.4523739 | 0.14639768 | 0.99937574 |
| COQ8B | 19.4858867 | 0.91747857 | 0.4965771 | 1.84760548 | 0.06465944 | 0.99937574 |
| ITPKC | 31.4997692 | -0.5847683 | 0.42482971 | -1.3764771 | 0.16867399 | 0.99937574 |
| SNRPA | 20.2961522 | -0.1789429 | 0.47355631 | -0.3778703 | 0.70552692 | 0.99937574 |
| AC008537.1 | 21.5819505 | -1.843545 | 0.78655651 | -2.3438176 | 0.0190875 | 0.96315543 |
| HNRNPUL1 | 50.8505635 | -0.0165128 | 0.35370382 | -0.0466855 | 0.96276392 | 0.99937574 |
| TGFB1 | 60.6299088 | 0.00593911 | 0.37102327 | 0.01600737 | 0.98722851 | 0.99937574 |
| CCDC97 | 13.2829791 | -0.0618596 | 0.57953809 | -0.1067396 | 0.91499559 | 0.99937574 |
| B3GNT8 | 10.0739495 | 0.12532916 | 0.67507072 | 0.18565338 | 0.85271659 | 0.99937574 |
| DMAC2 | 18.3407777 | -0.3669281 | 0.44725772 | -0.8203953 | 0.41199077 | 0.99937574 |
| RPS19 | 394.131246 | 0.220782 | 0.34055225 | 0.64830578 | 0.5167872 | 0.99937574 |
| ARHGEF1 | 23.5485781 | -0.4945508 | 0.43953428 | -1.12517 | 0.26051699 | 0.99937574 |
| RABAC1 | 65.5270494 | -0.1581619 | 0.48790367 | -0.3241663 | 0.74581211 | 0.99937574 |
| GSK3A | 19.2452193 | 0.35210322 | 0.43391334 | 0.81145978 | 0.41710168 | 0.99937574 |
| CIC | 22.9230689 | -0.3235513 | 0.52039826 | -0.6217379 | 0.53411424 | 0.99937574 |
| MEGF8 | 30.3482321 | 0.4006193 | 0.53192114 | 0.75315544 | 0.45135651 | 0.99937574 |
| LIPE-AS1 | 29.573978 | 0.67251779 | 0.49247569 | 1.36558576 | 0.17206902 | 0.99937574 |
| ETHE1 | 11.8951434 | 0.28935667 | 0.49197316 | 0.5881554 | 0.55642799 | 0.99937574 |
| ZNF428 | 38.8926988 | -0.484797 | 0.54742348 | -0.8855978 | 0.37583429 | 0.99937574 |
| SMG9 | 15.2720945 | 0.04411792 | 0.499282 | 0.08836273 | 0.92958839 | 0.99937574 |
| ZNF283 | 14.806869 | -0.2889707 | 0.52646811 | -0.5488854 | 0.58308409 | 0.99937574 |
| ZNF45 | 11.7512036 | -0.0451909 | 0.70959288 | -0.0636856 | 0.94922053 | 0.99937574 |
| ZNF155 | 18.5342166 | 0.93888408 | 0.64079135 | 1.46519468 | 0.14286781 | 0.99937574 |
| ZNF224 | 25.7947625 | -0.0983809 | 0.45370128 | -0.2168408 | 0.82833242 | 0.99937574 |
| ZNF234 | 14.2176171 | 0.01855145 | 0.70165758 | 0.02643947 | 0.97890681 | 0.99937574 |
| ZNF226 | 27.4393752 | 0.22879849 | 0.37892162 | 0.60381483 | 0.54596676 | 0.99937574 |
| ZNF227 | 16.9683304 | 0.34893248 | 0.59194296 | 0.58946976 | 0.55554619 | 0.99937574 |
| ZNF235 | 67.4860141 | 0.80757369 | 0.40008208 | 2.01852003 | 0.04353713 | 0.99937574 |
| ZNF229 | 14.3394641 | -0.7401216 | 0.68436635 | -1.08147 | 0.27948812 | 0.99937574 |
| ZNF180 | 15.2441075 | -0.3293421 | 0.48182321 | -0.683533 | 0.49427011 | 0.99937574 |
| AC243964.2 | 34.4180651 | 0.01604594 | 0.54289432 | 0.02955628 | 0.97642093 | 0.99937574 |
| PVR | 15.9313665 | -0.2875476 | 0.58474213 | -0.4917512 | 0.62289527 | 0.99937574 |
| CEACAM19 | 19.8239962 | -0.4951475 | 0.78281422 | -0.6325223 | 0.52704561 | 0.99937574 |
| BCL3 | 17.8027664 | -0.1504653 | 0.48525241 | -0.3100764 | 7.57E-01 | 9.99E-01 |
| CBLC | 27.865875 | -0.2283557 | 0.44096791 | -0.5178512 | 0.60456211 | 0.99937574 |
| BCAM | 109.274875 | 0.99921758 | 0.46874851 | 2.13167097 | 0.0330339 | 0.99937574 |
| NECTIN2 | 111.934122 | 0.0076438 | 0.29195312 | 0.02618159 | 0.9791125 | 0.99937574 |
| TOMM40 | 26.6898959 | 0.11805913 | 0.52581519 | 0.22452589 | 0.82234812 | 0.99937574 |
| APOE | 594.469789 | -0.4217678 | 0.73338002 | -0.5751013 | 0.5652228 | 0.99937574 |
| APOC1 | 13.8706514 | 0.46152363 | 0.74516112 | 0.61936086 | 0.53567866 | 0.99937574 |
| CLPTM1 | 52.5288593 | 0.14045511 | 0.35714261 | 0.39327457 | 0.6941167 | 0.99937574 |
| RELB | 43.8555419 | -0.6161692 | 0.58774733 | -1.0483572 | 0.29447404 | 0.99937574 |
| CLASRP | 63.7070491 | 0.02901363 | 0.42159395 | 0.0688189 | 0.94513378 | 0.99937574 |
| MARK4 | 51.1667746 | 0.30629062 | 0.33472398 | 0.9150543 | 0.36016312 | 0.99937574 |
| TRAPPC6A | 25.8833153 | -0.816062 | 0.56138267 | -1.4536644 | 0.14603939 | 0.99937574 |

| | | | | | | |
|---|---|---|---|---|---|---|
| PPP1R13L | 23.2408664 | -0.3705311 | 0.41120596 | -0.901084 | 0.36754368 | 0.99937574 |
| ERCC1 | 13.3734941 | 1.07943568 | 0.59166174 | 1.82441352 | 0.06808958 | 0.99937574 |
| VASP | 31.6794301 | -0.9045837 | 0.51965512 | -1.7407386 | 0.08172941 | 0.99937574 |
| OPA3 | 80.753456 | -0.3640793 | 0.45339653 | -0.8030042 | 0.42197231 | 0.99937574 |
| EML2 | 121.75208 | 0.78507476 | 0.32596104 | 2.4084926 | 0.01601855 | 0.87097377 |
| GIPR | 32.8672229 | -0.4298489 | 0.60286916 | -0.7130053 | 0.47584249 | 0.99937574 |
| SNRPD2 | 61.2290771 | -0.0556474 | 0.50064594 | -0.1111511 | 0.91149651 | 0.99937574 |
| FBXO46 | 18.2056725 | 0.10941829 | 0.49357527 | 0.22168511 | 0.82455902 | 0.99937574 |
| DMPK | 19.0519455 | -0.1844438 | 0.67408259 | -0.2736219 | 0.78437517 | 0.99937574 |
| SYMPK | 83.1292026 | -0.1398079 | 0.32567632 | -0.4292848 | 0.66771597 | 0.99937574 |
| MYPOP | 21.3034193 | 0.80586086 | 0.53228197 | 1.51397362 | 0.13003253 | 0.99937574 |
| PPP5C | 42.0757058 | 0.33747517 | 0.30723445 | 1.09842881 | 0.27201729 | 0.99937574 |
| CCDC8 | 11.2238108 | 0.53868843 | 0.6323845 | 0.85183687 | 0.39430464 | 0.99937574 |
| CALM3 | 69.4925932 | 0.25837669 | 0.48557253 | 0.5321073 | 0.59465168 | 0.99937574 |
| PRKD2 | 47.0867907 | -0.5219587 | 0.32953581 | -1.5839212 | 0.11321166 | 0.99937574 |
| STRN4 | 22.4166071 | 0.15960277 | 0.45197034 | 0.35312666 | 0.72399349 | 0.99937574 |
| FKRP | 12.03426 | -0.051183 | 0.60172882 | -0.0850598 | 0.93221381 | 0.99937574 |
| AP2S1 | 34.7824717 | -0.0459862 | 0.41872798 | -0.1098236 | 0.91254931 | 0.99937574 |
| ARHGAP35 | 171.658788 | 0.05067517 | 0.24992022 | 0.20276537 | 0.83931842 | 0.99937574 |
| TMEM160 | 38.9586599 | -0.0094465 | 0.49922206 | -0.0189224 | 0.98490299 | 0.99937574 |
| ZC3H4 | 33.7093033 | -0.2155363 | 0.470322 | -0.4582739 | 0.6467557 | 0.99937574 |
| SAE1 | 39.9687596 | -0.470641 | 0.39965922 | -1.1776058 | 0.23895382 | 0.99937574 |
| CCDC9 | 13.8141174 | 0.08149406 | 0.56985959 | 0.14300726 | 0.88628445 | 0.99937574 |
| INAFM1 | 12.8278323 | -0.4998697 | 0.56669537 | -0.8820784 | 0.37773441 | 0.99937574 |
| C5AR2 | 10.9656865 | 2.41832408 | 0.99196793 | 2.4379055 | 0.01477264 | 0.85933448 |
| DHX34 | 24.5152174 | -0.1381375 | 0.502908 | -0.2746775 | 0.78356403 | 0.99937574 |
| NAPA-AS1 | 14.0915592 | 0.22953327 | 0.72619752 | 0.31607553 | 0.75194518 | 0.99937574 |
| NAPA | 77.6779176 | -0.607898 | 0.32181838 | -1.8889474 | 0.05889887 | 0.99937574 |
| NOP53 | 33.8767193 | 0.21061242 | 0.6321375 | 0.33317501 | 0.73900218 | 0.99937574 |
| SELENOW | 132.937827 | 0.20384251 | 0.33116381 | 0.61553377 | 0.53820228 | 0.99937574 |
| CARD8 | 30.3032204 | 0.45366116 | 0.46998406 | 0.96526924 | 0.33440997 | 0.99937574 |
| EMP3 | 15.3126195 | -0.9247898 | 0.66103107 | -1.3990111 | 0.16180965 | 0.99937574 |
| TMEM143 | 13.4355468 | 1.08508316 | 0.67749273 | 1.60161594 | 0.10924056 | 0.99937574 |
| KDELR1 | 60.2128557 | -0.2042305 | 0.31248535 | -0.6535683 | 0.51338996 | 0.99937574 |
| CYTH2 | 30.0383449 | -0.4725422 | 0.54599058 | -0.8654768 | 0.38677713 | 0.99937574 |
| LMTK3 | 15.2251921 | -0.380024 | 0.52033431 | -0.7303459 | 0.46517877 | 0.99937574 |
| RPL18 | 196.84567 | 0.04454026 | 0.34102302 | 0.13060778 | 0.89608559 | 0.99937574 |
| BCAT2 | 25.8470089 | 0.12716415 | 0.41767092 | 0.30446015 | 0.76077735 | 0.99937574 |
| PLEKHA4 | 15.4573929 | -0.027025 | 0.99452129 | -0.0271739 | 0.97832104 | 0.99937574 |
| PPP1R15A | 17.8932841 | -1.3908596 | 0.55273848 | -2.5163069 | 0.01185919 | 0.77408712 |
| NUCB1 | 70.1436419 | 0.25739021 | 0.37097437 | 0.69382208 | 0.48779379 | 0.99937574 |
| BAX | 26.6385116 | -0.1687278 | 0.4852359 | -0.3477233 | 0.72804802 | 0.99937574 |
| FTL | 1175.76333 | -0.6899318 | 0.6382055 | -1.0810495 | 0.27967508 | 0.99937574 |
| GYS1 | 38.4421192 | -0.0109343 | 0.45709517 | -0.0239214 | 0.98091532 | 0.99937574 |
| RUVBL2 | 11.4747479 | -0.0584014 | 0.60657365 | -0.0962808 | 0.92329755 | 0.99937574 |
| SNRNP70 | 173.553209 | -0.1505758 | 0.39138785 | -0.3847227 | 0.70044291 | 0.99937574 |
| TRPM4 | 47.4559953 | 0.8507506 | 0.46984061 | 1.81072172 | 0.07018394 | 0.99937574 |
| SLC6A16 | 43.726152 | -0.3322229 | 0.46731452 | -0.7109194 | 0.47713421 | 0.99937574 |
| TEAD2 | 20.2602267 | -0.085741 | 0.59724295 | -0.1435614 | 0.88584681 | 0.99937574 |
| PIH1D1 | 17.2530758 | 1.05705357 | 0.64579037 | 1.63683701 | 0.10166453 | 0.99937574 |
| RPL13A | 240.428114 | 0.0670298 | 0.41834 | 0.16022803 | 0.87270145 | 0.99937574 |
| RPS11 | 215.30327 | -0.0994839 | 0.52789127 | -0.1884553 | 0.85051975 | 0.99937574 |
| FCGRT | 50.1494654 | 0.7368688 | 0.30874108 | 2.38668853 | 0.01700089 | 0.89506255 |
| NOSIP | 33.4417843 | -0.0706123 | 0.38719591 | -0.1823683 | 0.8552937 | 0.99937574 |

| | | | | | | |
|---|---|---|---|---|---|---|
| PRRG2 | 14.9305561 | 0.15019383 | 0.50979231 | 0.29461767 | 0.76828596 | 0.99937574 |
| PRR12 | 21.1155453 | 0.27550427 | 0.5686813 | 0.48446163 | 0.62805829 | 0.99937574 |
| RRAS | 13.4231153 | 0.22115116 | 0.64960915 | 0.34043726 | 0.73352726 | 0.99937574 |
| SCAF1 | 22.3372218 | -0.5113889 | 0.47293474 | -1.0813096 | 0.27955944 | 0.99937574 |
| IRF3 | 60.9183906 | -0.1038916 | 0.34412606 | -0.3018998 | 0.76272842 | 0.99937574 |
| PRMT1 | 44.093828 | 0.26258148 | 0.35232311 | 0.74528599 | 0.45609885 | 0.99937574 |
| AP2A1 | 26.9150877 | -0.0700868 | 0.54551382 | -0.1284785 | 0.89777035 | 0.99937574 |
| MED25 | 31.3264416 | -0.1439183 | 0.37929094 | -0.3794403 | 0.70436097 | 0.99937574 |
| PTOV1 | 39.8754351 | -0.1147687 | 0.39143834 | -0.2931974 | 0.76937127 | 0.99937574 |
| PNKP | 50.3792313 | -0.2889973 | 0.49961057 | -0.5784451 | 0.56296367 | 0.99937574 |
| TBC1D17 | 20.0882237 | -0.2812089 | 0.53433465 | -0.5262787 | 0.59869459 | 0.99937574 |
| ATF5 | 18.2019844 | -0.8100464 | 0.57866762 | -1.3998474 | 0.16155902 | 0.99937574 |
| VRK3 | 34.8272132 | -0.5018643 | 0.43511746 | -1.1533997 | 0.24874637 | 0.99937574 |
| MYH14 | 96.9537189 | 0.35082363 | 0.25911416 | 1.35393461 | 0.17575724 | 0.99937574 |
| KCNC3 | 55.5585054 | 0.74205903 | 0.49126239 | 1.51051465 | 0.13091215 | 0.99937574 |
| NR1H2 | 54.110407 | 0.23780211 | 0.34429923 | 0.6906844 | 4.90E-01 | 0.99937574 |
| EMC10 | 115.17066 | 0.60158801 | 0.34966086 | 1.72049 | 0.08534341 | 0.99937574 |
| JOSD2 | 13.5104656 | -0.432585 | 0.57270363 | -0.7553382 | 0.45004606 | 0.99937574 |
| CLEC11A | 11.4156211 | -0.6804281 | 0.58643827 | -1.1602724 | 0.24593793 | 0.99937574 |
| ETFB | 87.570492 | 0.60917618 | 0.38034656 | 1.60163452 | 0.10923645 | 0.99937574 |
| ZNF175 | 13.5921838 | -0.4869361 | 0.5024163 | -0.9691886 | 0.3324511 | 0.99937574 |
| SPACA6 | 15.5736472 | -0.6112484 | 0.61271516 | -0.9976061 | 0.31847039 | 0.99937574 |
| ZNF577 | 27.3769065 | 0.78141138 | 0.51390828 | 1.52052694 | 0.12837859 | 0.99937574 |
| ZNF350-AS1 | 21.7950858 | 1.0596337 | 0.89687474 | 1.18147346 | 0.23741469 | 0.99937574 |
| ZNF350 | 16.4531476 | 0.25323869 | 0.69224491 | 0.36582239 | 0.71449761 | 0.99937574 |
| ZNF615 | 14.0084524 | 0.16702283 | 0.52402691 | 0.3187295 | 0.74993164 | 0.99937574 |
| ZNF432 | 27.1118752 | -0.2336971 | 0.45659802 | -0.5118224 | 0.60877532 | 0.99937574 |
| AC011468.3 | 18.5993325 | -0.1842203 | 0.6545468 | -0.2814471 | 0.77836748 | 0.99937574 |
| PPP2R1A | 125.807152 | 0.3251233 | 0.33807991 | 0.9616759 | 0.33621243 | 0.99937574 |
| ZNF766 | 35.5267377 | -0.2313301 | 0.34748993 | -0.6657176 | 0.50559165 | 0.99937574 |
| ZNF480 | 26.474456 | -0.6041048 | 0.39547165 | -1.5275554 | 0.12662297 | 0.99937574 |
| ZNF880 | 11.4605336 | 0.6502312 | 0.53780965 | 1.20903596 | 0.22664903 | 0.99937574 |
| ZNF528 | 19.5581478 | -0.1737711 | 0.53480494 | -0.3249243 | 0.74523834 | 0.99937574 |
| ZNF808 | 24.0679139 | -0.2360902 | 0.44780004 | -0.5272225 | 0.59803911 | 0.99937574 |
| ZNF701 | 36.5869203 | -0.2577206 | 0.36556538 | -0.7049918 | 0.48081536 | 0.99937574 |
| ZNF83 | 99.5534892 | -0.2017081 | 0.29478014 | -0.6842663 | 0.49380703 | 0.99937574 |
| ZNF611 | 61.456328 | -0.4168668 | 0.3417653 | -1.2197458 | 0.22256126 | 0.99937574 |
| ZNF600 | 15.6256855 | -0.4754356 | 0.58696786 | -0.8099857 | 0.41794839 | 0.99937574 |
| ZNF28 | 21.4782055 | -0.5091191 | 0.4116314 | -1.2368326 | 0.21614925 | 0.99937574 |
| AC010487.3 | 18.8666765 | -0.7067535 | 0.71044753 | -0.9948004 | 0.31983334 | 0.99937574 |
| ZNF160 | 67.9582315 | -0.5823498 | 0.42043741 | -1.3851046 | 0.1660205 | 0.99937574 |
| ZNF347 | 22.7626258 | -0.4419537 | 0.40327905 | -1.0959005 | 0.27312231 | 0.99937574 |
| ZNF677 | 20.521243 | 0.12370117 | 0.41902318 | 0.29521319 | 0.76783102 | 0.99937574 |
| ZNF525 | 20.9043913 | -0.4475499 | 0.50711974 | -0.8825329 | 0.37748868 | 0.99937574 |
| ZNF765 | 36.4311138 | 0.42188138 | 0.41013092 | 1.02865051 | 0.30364393 | 0.99937574 |
| ZNF761 | 37.870338 | -0.5652578 | 0.4910328 | -1.1511609 | 0.24966604 | 0.99937574 |
| ZNF813 | 13.9476924 | -0.7214717 | 0.63056425 | -1.1441684 | 0.25255382 | 0.99937574 |
| ZNF331 | 48.6039637 | -0.1614903 | 0.4293431 | -0.3761335 | 0.70681768 | 0.99937574 |
| NDUFA3 | 29.0961076 | 0.67350918 | 0.60495316 | 1.11332451 | 0.26556908 | 0.99937574 |
| PRPF31 | 30.1527358 | -0.0366429 | 0.46295468 | -0.0791501 | 0.9369132 | 0.99937574 |
| CNOT3 | 20.5402247 | -0.5818831 | 0.47215682 | -1.2323937 | 0.21780206 | 0.99937574 |
| LENG1 | 10.3138149 | -0.2258047 | 0.55931164 | -0.4037189 | 0.68641947 | 0.99937574 |
| TMC4 | 71.7856601 | -0.157159 | 0.31023273 | -0.5065842 | 0.61244658 | 0.99937574 |
| MBOAT7 | 35.6758636 | -0.4555199 | 0.47951572 | -0.9499581 | 0.34213352 | 0.99937574 |

| | | | | | | |
|---|---|---|---|---|---|---|
| TSEN34 | 20.4725897 | 0.24294289 | 0.63583281 | 0.38208612 | 0.70239748 | 0.99937574 |
| RPS9 | 377.0241 | -0.1351569 | 0.4882143 | -0.2768394 | 0.78190344 | 0.99937574 |
| LENG8-AS1 | 11.9572198 | -0.4822449 | 0.59651078 | -0.8084429 | 0.41883566 | 0.99937574 |
| LENG8 | 114.02577 | -0.4282682 | 0.36499631 | -1.1733494 | 0.24065573 | 0.99937574 |
| AC011476.3 | 13.9760587 | -0.1941513 | 0.86553454 | -0.2243138 | 0.82251315 | 0.99937574 |
| RDH13 | 41.342733 | -0.3039611 | 0.41343108 | -0.735216 | 0.46220796 | 0.99937574 |
| EPS8L1 | 20.439259 | 0.65875822 | 0.55922916 | 1.17797546 | 0.23880639 | 0.99937574 |
| PPP1R12C | 30.780447 | -0.5796655 | 0.47937327 | -1.2092154 | 0.22658011 | 0.99937574 |
| PPP6R1 | 40.2161944 | -0.1867755 | 0.45945593 | -0.4065145 | 0.68436463 | 0.99937574 |
| HSPBP1 | 13.185448 | -0.0436153 | 0.67442437 | -0.0646705 | 0.94843639 | 0.99937574 |
| TMEM238 | 23.0893177 | -0.5126478 | 0.62909334 | -0.8148995 | 0.41512986 | 0.99937574 |
| RPL28 | 301.121319 | -0.1094881 | 0.48673816 | -0.2249425 | 0.82202397 | 0.99937574 |
| UBE2S | 13.9146474 | -0.4765428 | 0.66666567 | -0.7148153 | 0.47472319 | 0.99937574 |
| ISOC2 | 26.7593846 | -0.6621919 | 0.65556298 | -1.0101118 | 0.31244171 | 0.99937574 |
| NAT14 | 10.8041447 | 0.8774936 | 0.64748869 | 1.35522615 | 0.17534551 | 0.99937574 |
| CCDC106 | 13.7769097 | 0.01541448 | 0.69590902 | 0.02215014 | 0.98232819 | 0.99937574 |
| U2AF2 | 25.7260205 | 0.19825339 | 0.42251357 | 0.46922373 | 0.63890973 | 0.99937574 |
| EPN1 | 85.1784548 | 0.05839299 | 0.31385708 | 0.18604961 | 0.85240585 | 0.99937574 |
| ZNF787 | 24.7442704 | -0.4026078 | 0.47529511 | -0.8470691 | 0.39695659 | 0.99937574 |
| ZNF444 | 22.4664457 | -0.526137 | 0.41438826 | -1.2696715 | 0.20420165 | 0.99937574 |
| ZSCAN5A | 74.8741231 | -0.2989693 | 0.31822079 | -0.9395027 | 0.34747269 | 0.99937574 |
| ZNF582 | 12.1839871 | 0.3328541 | 0.60345731 | 0.55157854 | 0.58123714 | 0.99937574 |
| ZNF667-AS1 | 21.5439715 | 0.72566268 | 0.42907784 | 1.69121452 | 0.09079584 | 0.99937574 |
| ZNF471 | 18.532272 | 0.42646614 | 0.49405196 | 0.863201 | 0.38802697 | 0.99937574 |
| ZNF470 | 12.9451442 | -0.2719068 | 0.69477075 | -0.3913619 | 0.69552978 | 0.99937574 |
| ZNF264 | 38.9176337 | 0.26512168 | 0.33787678 | 0.78466971 | 0.43264725 | 0.99937574 |
| ZNF460 | 15.8664694 | -0.4765606 | 0.55002407 | -0.8664359 | 0.38625117 | 0.99937574 |
| AC005261.1 | 57.5303588 | 0.35205981 | 0.45971085 | 0.7658288 | 0.44377818 | 0.99937574 |
| AC004076.1 | 24.1534015 | 0.07807603 | 0.57101253 | 0.13673261 | 0.89124215 | 0.99937574 |
| ZNF549 | 23.1493058 | 0.0693711 | 0.51375636 | 0.13502723 | 0.89259034 | 0.99937574 |
| ZNF134 | 11.4910897 | 0.20272067 | 0.58705044 | 0.3453207 | 0.72985329 | 0.99937574 |
| AC003006.1 | 14.5891152 | -0.0817772 | 0.53226657 | -0.1536396 | 0.8778939 | 0.99937574 |
| ZNF154 | 12.9711717 | -0.242694 | 0.51496765 | -0.4712802 | 0.63744068 | 0.99937574 |
| ZNF586 | 108.822062 | 0.23934898 | 0.30121388 | 0.79461469 | 0.42683767 | 0.99937574 |
| ZNF552 | 32.9544004 | 0.61699972 | 0.49564987 | 1.24482979 | 0.21319432 | 0.99937574 |
| ZNF814 | 55.8269281 | -0.0050583 | 0.35475122 | -0.0142587 | 0.98862358 | 0.99937574 |
| ZNF587 | 30.2856297 | 0.00734046 | 0.36632199 | 0.02003826 | 0.98401285 | 0.99937574 |
| ZNF606 | 20.9513223 | 0.28242911 | 0.41410585 | 0.68202155 | 0.49522533 | 0.99937574 |
| ZNF135 | 33.995441 | 0.21249906 | 0.36071923 | 0.58909822 | 0.55579539 | 0.99937574 |
| ZSCAN18 | 84.0978393 | 0.29863474 | 0.36026507 | 0.82893059 | 0.40714369 | 0.99937574 |
| ZNF329 | 21.4040039 | 0.5533485 | 0.465088 | 1.1897716 | 0.23413617 | 0.99937574 |
| ZNF274 | 48.4384576 | -0.2437006 | 0.46806936 | -0.5206506 | 0.6026102 | 0.99937574 |
| AC020915.3 | 21.0756712 | 0.56951617 | 0.46844064 | 1.21577018 | 0.22407248 | 0.99937574 |
| RPS5 | 126.179532 | -0.1449442 | 0.50042648 | -0.2896414 | 0.77209058 | 0.99937574 |
| TRIM28 | 43.1132249 | -0.079246 | 0.46392071 | -0.1708181 | 0.86436682 | 0.99937574 |
| CHMP2A | 43.9089416 | -0.5471969 | 0.40770398 | -1.3421426 | 0.17954975 | 0.99937574 |
| UBE2M | 39.2340114 | -0.3207428 | 0.43076925 | -0.7445815 | 0.45652475 | 0.99937574 |
| MZF1-AS1 | 12.6688677 | 0.61672792 | 0.74642501 | 0.8262423 | 0.40866666 | 0.99937574 |
| MZF1 | 44.5839061 | 0.36378918 | 0.40837896 | 0.89081276 | 0.37302963 | 0.99937574 |
| RPS4Y1 | 63.2189704 | -0.1308959 | 2.2337975 | -0.0585979 | 0.95327238 | 0.99937574 |
| ZFY | 34.7491431 | 0.39786898 | 2.56936615 | 0.15485102 | 0.87693877 | 0.99937574 |
| LINC00278 | 116.73689 | 0.12658886 | 2.59977205 | 0.04869229 | 0.96116452 | 0.99937574 |
| TTTY15 | 29.0828069 | 0.81104041 | 2.11040232 | 0.38430607 | 0.70075161 | 0.99937574 |
| USP9Y | 122.189568 | 1.03718281 | 2.91139916 | 0.35624892 | 0.72165416 | 0.99937574 |

| | | | | | | |
|---|---|---|---|---|---|---|
| DDX3Y | 49.0179642 | 0.65266781 | 2.69852167 | 0.24186125 | 0.80888768 | 0.99937574 |
| UTY | 344.568093 | 0.66416934 | 2.84229232 | 0.23367383 | 0.8152382 | 0.99937574 |
| TTTY14 | 335.688655 | 0.77765875 | 0.94899567 | 0.81945448 | 0.41252717 | 0.99937574 |
| KDM5D | 35.8807267 | 0.66167749 | 2.57179108 | 0.25728275 | 0.7969605 | 0.99937574 |
| TTTY10 | 12.2948063 | 0.26671089 | 1.37452595 | 0.19403845 | 0.84614577 | 0.99937574 |
| EIF1AY | 22.1556927 | 1.12051925 | 2.44980479 | 0.45739124 | 0.64738986 | 0.99937574 |
| IL17RA | 29.4987977 | 0.12432881 | 0.45538778 | 0.27301745 | 0.7848398 | 0.99937574 |
| HDHD5 | 16.5815407 | -0.575649 | 0.65687166 | -0.8763493 | 0.38084015 | 0.99937574 |
| ADA2 | 12.4535881 | 0.19328724 | 0.92268743 | 0.20948291 | 0.83407128 | 0.99937574 |
| CECR2 | 41.4526019 | 0.23846561 | 0.89518457 | 0.26638709 | 0.78994111 | 0.99937574 |
| ATP6V1E1 | 155.181285 | 0.20205715 | 0.29948073 | 0.67469164 | 0.49987169 | 0.99937574 |
| BCL2L13 | 180.795232 | 0.17979465 | 0.36574956 | 0.49157859 | 0.62301728 | 0.99937574 |
| MICAL3 | 98.1406666 | 0.17835772 | 0.35374107 | 0.50420415 | 0.61411792 | 0.99937574 |
| PEX26 | 41.8230466 | 0.04863032 | 0.41252679 | 0.11788403 | 0.90615955 | 0.99937574 |
| DGCR2 | 66.2472146 | -0.0559169 | 0.43731443 | -0.1278643 | 0.89825634 | 0.99937574 |
| SLC25A1 | 35.8214625 | -0.0557281 | 0.39309227 | -0.1417686 | 0.8872628 | 0.99937574 |
| CLTCL1 | 22.009636 | -0.0413203 | 0.47960828 | -0.0861542 | 0.93134384 | 0.99937574 |
| HIRA | 22.0330784 | -0.3583178 | 0.48542175 | -0.7381577 | 0.4604186 | 0.99937574 |
| MRPL40 | 27.1033691 | 0.19984832 | 0.45538919 | 0.4388517 | 0.660769 | 0.99937574 |
| UFD1 | 20.4107787 | -0.0891523 | 0.55703145 | -0.1600489 | 0.87284255 | 0.99937574 |
| TXNRD2 | 37.2856496 | 0.01404146 | 0.41148326 | 0.03412401 | 0.97277826 | 0.99937574 |
| COMT | 59.7943659 | 0.38026799 | 0.30466326 | 1.24815834 | 0.21197308 | 0.99937574 |
| TANGO2 | 65.4453761 | 0.07917621 | 0.63274559 | 0.12513119 | 0.90041969 | 0.99937574 |
| DGCR8 | 26.2271698 | -0.5574535 | 0.48231355 | -1.1557908 | 0.24776675 | 0.99937574 |
| RANBP1 | 19.8484379 | -0.3477497 | 0.50489829 | -0.6887519 | 0.4909794 | 0.99937574 |
| ZDHHC8 | 15.7628551 | -0.2643927 | 0.59581514 | -0.4437495 | 0.65722368 | 0.99937574 |
| DGCR6L | 11.4104637 | -0.182193 | 0.77820319 | -0.2341201 | 0.81489172 | 0.99937574 |
| KLHL22 | 62.6867181 | 0.13464341 | 0.37793962 | 0.35625642 | 0.72164854 | 0.99937574 |
| MED15 | 82.4224286 | -0.5903256 | 0.35776272 | -1.6500478 | 0.09893315 | 0.99937574 |
| PI4KA | 130.280403 | -0.1714686 | 0.3646987 | -0.4701652 | 0.638237 | 0.99937574 |
| SNAP29 | 24.6828329 | -0.0628293 | 0.49861723 | -0.126007 | 0.89972636 | 0.99937574 |
| CRKL | 34.9829326 | 0.06556991 | 0.35737552 | 0.18347621 | 0.85442438 | 0.99937574 |
| LZTR1 | 10.3636564 | 0.21393486 | 0.5828375 | 0.36705748 | 0.71357614 | 0.99937574 |
| THAP7 | 12.8977609 | 0.39813603 | 0.53415082 | 0.74536256 | 0.45605257 | 0.99937574 |
| HIC2 | 35.0123799 | 0.07668493 | 0.49928967 | 0.15358807 | 0.87793455 | 0.99937574 |
| UBE2L3 | 94.7784105 | -0.0418288 | 0.23682733 | -0.1766214 | 0.85980577 | 0.99937574 |
| SDF2L1 | 33.9491295 | -0.1206902 | 0.43835564 | -0.2753248 | 0.78306675 | 0.99937574 |
| PPIL2 | 92.3081104 | -0.2250067 | 0.33118719 | -0.6793945 | 0.49688796 | 0.99937574 |
| MAPK1 | 174.455215 | -0.0768354 | 0.22941813 | -0.3349144 | 0.73768965 | 0.99937574 |
| PPM1F | 14.9353139 | 0.13355478 | 0.59881046 | 0.22303348 | 0.82350945 | 0.99937574 |
| TOP3B | 20.7357056 | -0.4120879 | 0.48904883 | -0.8426313 | 0.39943467 | 0.99937574 |
| AC245060.5 | 21.7932783 | 0.26107845 | 0.57247011 | 0.45605603 | 0.64834969 | 0.99937574 |
| BCR | 114.30137 | 0.16238867 | 0.50290445 | 0.32290164 | 0.74676973 | 0.99937574 |
| CHCHD10 | 277.499977 | 0.15486421 | 0.42438368 | 0.36491555 | 0.71517445 | 0.99937574 |
| SMARCB1 | 48.8105379 | 0.28462133 | 0.36550406 | 0.77870908 | 0.43615111 | 0.99937574 |
| DDT | 90.73461 | 0.20098927 | 0.55902293 | 0.35953673 | 0.71919361 | 0.99937574 |
| CABIN1 | 141.883501 | 0.08360894 | 0.35066124 | 0.23843223 | 0.81154587 | 0.99937574 |
| GUCD1 | 31.6664807 | -0.2829347 | 0.41254843 | -0.6858219 | 0.49282545 | 0.99937574 |
| SNRPD3 | 52.1963786 | -0.1354739 | 0.32677977 | -0.4145726 | 0.67845484 | 0.99937574 |
| SGSM1 | 100.347651 | 0.14895135 | 0.38548926 | 0.38639559 | 0.69920371 | 0.99937574 |
| KIAA1671 | 90.6731438 | 0.30433656 | 0.4293465 | 0.70883672 | 0.47842581 | 0.99937574 |
| LRP5L | 57.9422628 | -0.1022692 | 0.65859602 | -0.1552837 | 0.87659764 | 0.99937574 |
| GRK3 | 35.9218898 | 0.06581409 | 0.53950947 | 0.12198875 | 0.90290793 | 0.99937574 |
| HPS4 | 65.1680467 | -0.3914612 | 0.53950893 | -0.7255881 | 0.46809134 | 0.99937574 |

| | | | | | | |
|---|---|---|---|---|---|---|
| SRRD | 13.0738055 | 0.17927528 | 0.52973549 | 0.33842414 | 0.73504358 | 0.99937574 |
| TFIP11 | 26.2049725 | 0.01455625 | 0.40723167 | 0.03574441 | 0.97148616 | 0.99937574 |
| TPST2 | 74.1886789 | 0.25194891 | 0.33805185 | 0.74529664 | 0.45609241 | 0.99937574 |
| MIATNB | 53.9410147 | -0.2999995 | 0.35955422 | -0.8343652 | 0.40407522 | 0.99937574 |
| PITPNB | 163.608454 | -0.0628486 | 0.33948396 | -0.1851297 | 0.85312734 | 0.99937574 |
| TTC28 | 221.655262 | -0.4399477 | 0.36655487 | -1.2002234 | 0.23005261 | 0.99937574 |
| HSCB | 35.8744592 | 0.69190091 | 0.32691176 | 2.11647605 | 0.03430434 | 0.99937574 |
| CCDC117 | 25.604731 | 0.14618909 | 0.53635527 | 0.27256019 | 0.78519132 | 0.99937574 |
| XBP1 | 64.9606537 | 0.25119647 | 0.30082316 | 0.83503034 | 0.40370063 | 0.99937574 |
| Z93930.2 | 74.01662 | 0.91549163 | 0.41908627 | 2.18449445 | 0.02892593 | 0.99049468 |
| ZNRF3 | 296.840401 | 0.33133515 | 0.38650612 | 0.8572572 | 0.39130276 | 0.99937574 |
| KREMEN1 | 108.268771 | 0.08535681 | 0.39394275 | 0.21667313 | 0.82846309 | 0.99937574 |
| RHBDD3 | 19.6468968 | -0.8247363 | 0.53878929 | -1.5307214 | 0.12583827 | 0.99937574 |
| EWSR1 | 122.198289 | -0.4045635 | 0.27624825 | -1.4644924 | 0.14305946 | 0.99937574 |
| GAS2L1 | 20.0428818 | 0.07192537 | 0.57716747 | 0.12461786 | 0.90082609 | 0.99937574 |
| AP1B1 | 85.726026 | 0.21783817 | 0.38979139 | 0.55885835 | 0.57625839 | 0.99937574 |
| THOC5 | 13.3550229 | -0.0696483 | 0.54052774 | -0.1288524 | 0.89747444 | 0.99937574 |
| NIPSNAP1 | 17.4990027 | 0.23349219 | 0.60251133 | 0.38753161 | 0.69836269 | 0.99937574 |
| NF2 | 103.614322 | 0.1008966 | 0.32854941 | 0.30709717 | 7.59E-01 | 9.99E-01 |
| UQCR10 | 202.566909 | 0.32771358 | 0.51112334 | 0.6411634 | 0.52141653 | 0.99937574 |
| ASCC2 | 37.3747267 | -0.5127263 | 0.41199072 | -1.2445094 | 0.21331215 | 0.99937574 |
| MTMR3 | 154.66944 | -0.4003684 | 0.23053906 | -1.7366617 | 0.0824469 | 0.99937574 |
| TBC1D10A | 14.7386668 | -0.8912412 | 0.56502577 | -1.5773461 | 0.11471592 | 0.99937574 |
| SF3A1 | 28.1718795 | 0.21556344 | 0.42918471 | 0.50226262 | 0.6154828 | 0.99937574 |
| RNF215 | 15.4611713 | 0.49397386 | 0.49022236 | 1.00765264 | 0.31362125 | 0.99937574 |
| PES1 | 15.1273692 | -0.4056161 | 0.55698715 | -0.7282324 | 0.46647136 | 0.99937574 |
| DUSP18 | 13.2857826 | 0.05802087 | 0.5949535 | 0.09752169 | 0.92231211 | 0.99937574 |
| OSBP2 | 39.8694218 | 0.34261597 | 0.46479574 | 0.73713234 | 0.46104187 | 0.99937574 |
| MORC2 | 21.2772882 | -0.5589365 | 0.458947 | -1.2178671 | 0.22327447 | 0.99937574 |
| SMTN | 12.6165843 | 0.0865377 | 0.7690581 | 0.11252427 | 0.91040772 | 0.99937574 |
| SELENOM | 107.025176 | -0.1667614 | 0.54486246 | -0.3060614 | 0.75955791 | 0.99937574 |
| INPP5J | 66.6264045 | 0.51662328 | 0.73816958 | 0.69987073 | 0.48400804 | 0.99937574 |
| RNF185 | 50.8206911 | -0.3149934 | 0.4035617 | -0.7805335 | 0.43507694 | 0.99937574 |
| LIMK2 | 57.7273069 | -0.0447531 | 0.33193071 | -0.1348266 | 0.89274895 | 0.99937574 |
| PIK3IP1 | 31.5116007 | 0.15808811 | 0.59565375 | 0.2654027 | 0.79069926 | 0.99937574 |
| PIK3IP1-AS1 | 20.5163257 | 0.09355834 | 0.53909055 | 0.17354848 | 0.86222033 | 0.99937574 |
| PATZ1 | 26.2315593 | -0.3043782 | 0.55916851 | -0.5443407 | 0.58620704 | 0.99937574 |
| DRG1 | 48.1285899 | -0.1407404 | 0.33315805 | -0.4224433 | 0.67270146 | 0.99937574 |
| EIF4ENIF1 | 51.4144322 | 0.08017923 | 0.34413728 | 0.23298617 | 0.81577214 | 0.99937574 |
| SFI1 | 40.601204 | -0.9981284 | 0.39730528 | -2.5122456 | 0.01199656 | 0.77519564 |
| PISD | 149.06194 | -0.8709119 | 0.26658554 | -3.2669135 | 0.00108727 | 0.32272706 |
| PRR14L | 124.168555 | -0.3674076 | 0.30827574 | -1.1918148 | 0.23333386 | 0.99937574 |
| YWHAH | 71.0548719 | -0.0544134 | 0.30600539 | -0.1778183 | 8.59E-01 | 0.99937574 |
| RTCB | 59.0364185 | 0.27407692 | 0.30231637 | 0.90658973 | 0.36462379 | 0.99937574 |
| FBXO7 | 83.3509552 | 0.10306467 | 0.36201686 | 0.28469576 | 0.77587724 | 0.99937574 |
| TIMP3 | 50.1713358 | 0.65445553 | 0.61305031 | 1.06753968 | 0.28572821 | 0.99937574 |
| LARGE1 | 123.893979 | -0.2929904 | 0.48269464 | -0.6069891 | 0.54385817 | 0.99937574 |
| HMGXB4 | 34.9399319 | 0.17519082 | 0.40344059 | 0.43424193 | 0.66411277 | 0.99937574 |
| TOM1 | 86.8395412 | -0.3138313 | 0.37792931 | -0.8303969 | 0.40631442 | 0.99937574 |
| APOL6 | 44.8219324 | -0.9332816 | 0.62024535 | -1.5046974 | 0.13240189 | 0.99937574 |
| RBFOX2 | 360.106212 | 0.28358426 | 0.2828054 | 1.00275404 | 0.31597955 | 0.99937574 |
| APOL2 | 42.8222274 | -0.7671349 | 0.48705849 | -1.5750365 | 0.11524802 | 0.99937574 |
| APOL1 | 27.7611801 | -1.1817679 | 0.52235455 | -2.2623865 | 0.02367353 | 0.96546668 |
| MYH9 | 191.935839 | -0.1840443 | 0.32200622 | -0.5715551 | 0.56762343 | 0.99937574 |

| | | | | | | |
|---|---|---|---|---|---|---|
| TXN2 | 46.1191041 | 0.37330578 | 0.35120623 | 1.06292472 | 0.28781609 | 0.99937574 |
| EIF3D | 54.2921267 | -0.2922038 | 0.37797554 | -0.773076 | 0.43947741 | 0.99937574 |
| IFT27 | 24.8234347 | -0.0904315 | 0.43468227 | -0.2080404 | 0.83519739 | 0.99937574 |
| PVALB | 15.5819905 | -0.6784732 | 0.92983268 | -0.7296723 | 0.46559049 | 0.99937574 |
| TST | 24.9107223 | -0.1190192 | 0.46600368 | -0.2554039 | 0.79841115 | 0.99937574 |
| MPST | 21.1892263 | -0.0715507 | 0.50890145 | -0.1405983 | 0.88818727 | 0.99937574 |
| CARD10 | 11.0385748 | 0.79687794 | 0.68828182 | 1.15777857 | 0.24695441 | 0.99937574 |
| GGA1 | 47.3412228 | 0.00025679 | 0.36087856 | 0.00071157 | 0.99943225 | 0.99979721 |
| LGALS1 | 17.2549514 | -0.3274481 | 0.69050205 | -0.4742174 | 0.6353449 | 0.99937574 |
| TRIOBP | 95.2271274 | -0.4977562 | 0.34342913 | -1.4493711 | 0.14723399 | 0.99937574 |
| H1F0 | 129.915094 | -1.4920908 | 0.68986171 | -2.162884 | 0.0305501 | 0.99049468 |
| GCAT | 12.9106743 | 0.15838184 | 0.76056673 | 0.20824188 | 0.83504011 | 0.99937574 |
| ANKRD54 | 16.4978043 | -0.2008444 | 0.45363698 | -0.4427427 | 0.65795188 | 0.99937574 |
| EIF3L | 88.8435944 | -0.1233216 | 0.32195366 | -0.3830414 | 0.70168908 | 0.99937574 |
| MICALL1 | 22.1431522 | -0.8701574 | 0.42242384 | -2.0599155 | 0.03940662 | 0.99937574 |
| POLR2F | 25.5700869 | -0.60492 | 0.50713091 | -1.1928282 | 0.23293667 | 0.99937574 |
| PICK1 | 23.0969846 | -0.5556508 | 0.45751743 | -1.214491 | 0.2245603 | 0.99937574 |
| AL031587.2 | 10.9644621 | 0.30728011 | 0.67552936 | 0.45487306 | 0.64920056 | 0.99937574 |
| PLA2G6 | 26.2924987 | 0.19482256 | 0.47891428 | 0.40680048 | 0.68415454 | 0.99937574 |
| MAFF | 18.6138365 | -1.5017989 | 0.71177121 | -2.1099461 | 0.034863 | 0.99937574 |
| TMEM184B | 35.6342 | -0.4985033 | 0.45081048 | -1.1057935 | 0.26881591 | 0.99937574 |
| CSNK1E | 19.697759 | 0.22709005 | 0.49031931 | 0.46314727 | 0.64325881 | 0.99937574 |
| TPTEP2-CSNI | 85.1333226 | 0.47752991 | 0.39685317 | 1.20329116 | 0.22886367 | 0.99937574 |
| DDX17 | 663.874236 | 0.12500742 | 0.25132083 | 0.49740175 | 0.61890577 | 0.99937574 |
| CBY1 | 26.1168417 | 0.38850377 | 0.42691645 | 0.91002295 | 0.3628104 | 0.99937574 |
| TOMM22 | 14.7953829 | 0.19436125 | 0.71420503 | 0.27213649 | 0.78551708 | 0.99937574 |
| JOSD1 | 35.8972162 | -0.7573871 | 0.45520306 | -1.6638445 | 0.09614351 | 0.99937574 |
| GTPBP1 | 37.7912862 | -0.5650347 | 0.43967522 | -1.2851183 | 0.19875094 | 0.99937574 |
| SUN2 | 40.4101066 | 0.24299759 | 0.42168168 | 0.57625834 | 0.56444058 | 0.99937574 |
| AL021707.1 | 13.3883967 | -0.0888467 | 0.552552 | -0.1607934 | 0.87225614 | 0.99937574 |
| CBX6 | 29.8555592 | -0.3608424 | 0.71931551 | -0.5016469 | 0.61591589 | 0.99937574 |
| CBX7 | 21.228078 | -0.0649299 | 0.40635041 | -0.159788 | 0.87304804 | 0.99937574 |
| RPL3 | 219.086279 | 0.21140287 | 0.50775386 | 0.41634912 | 0.67715455 | 0.99937574 |
| SYNGR1 | 16.4225789 | 0.39226378 | 0.58603618 | 0.6693508 | 0.50327173 | 0.99937574 |
| TAB1 | 19.5131652 | 0.50039966 | 0.515995 | 0.9697762 | 0.33215806 | 0.99937574 |
| MIEF1 | 20.2512245 | -0.1755348 | 0.44563065 | -0.3939019 | 0.69365344 | 0.99937574 |
| ATF4 | 63.7234088 | -0.9299913 | 0.5257445 | -1.7689036 | 0.07690996 | 0.99937574 |
| RPS19BP1 | 49.2040889 | -0.2621723 | 0.67852068 | -0.3863881 | 0.6992093 | 0.99937574 |
| FAM83F | 55.9357601 | 0.1029293 | 0.50461564 | 0.20397566 | 0.83837252 | 0.99937574 |
| TNRC6B | 664.969241 | -0.0889522 | 0.29623585 | -0.3002751 | 0.76396736 | 0.99937574 |
| MKL1 | 196.097663 | -0.219347 | 0.25816541 | -0.8496373 | 0.39552676 | 0.99937574 |
| SLC25A17 | 45.6889318 | -0.2066589 | 0.37784928 | -0.5469347 | 0.58442361 | 0.99937574 |
| ST13 | 130.505229 | -0.1936416 | 0.33906571 | -0.5711035 | 0.5679295 | 0.99937574 |
| XPNPEP3 | 58.0075323 | 0.26541524 | 0.28413137 | 0.93412861 | 0.35023756 | 0.99937574 |
| RBX1 | 64.9725009 | -0.0209556 | 0.30406007 | -0.0689191 | 0.945054 | 0.99937574 |
| EP300 | 171.349876 | -0.1022409 | 0.27405917 | -0.3730612 | 0.70910288 | 0.99937574 |
| L3MBTL2 | 19.0343149 | -0.1436423 | 0.44670799 | -0.3215574 | 0.74778802 | 0.99937574 |
| RANGAP1 | 23.5663499 | -0.310542 | 0.47299121 | -0.6565491 | 0.51147087 | 0.99937574 |
| ZC3H7B | 46.1198407 | 0.46788143 | 0.34040704 | 1.37447635 | 0.16929385 | 0.99937574 |
| TEF | 14.6919202 | -1.3075498 | 0.65657498 | -1.9914706 | 0.04642917 | 0.99937574 |
| TOB2 | 47.2284718 | -0.0736784 | 0.36427693 | -0.2022593 | 0.83971398 | 0.99937574 |
| PHF5A | 11.4550143 | -1.3056735 | 0.60248727 | -2.1671387 | 0.03022428 | 0.99049468 |
| ACO2 | 162.898128 | 0.34168169 | 0.29739102 | 1.14893076 | 0.25058453 | 0.99937574 |
| POLR3H | 12.4880492 | -0.1740291 | 0.63296325 | -0.2749435 | 0.78335964 | 0.99937574 |

| | | | | | | |
|---|---|---|---|---|---|---|
| PMM1 | 17.2458438 | 0.05470954 | 0.51643314 | 0.10593731 | 0.91563209 | 0.99937574 |
| DESI1 | 65.5688733 | 0.25574251 | 0.40086777 | 0.63797225 | 0.52349174 | 0.99937574 |
| XRCC6 | 131.165908 | 0.24525927 | 0.26359861 | 0.93042702 | 0.35215003 | 0.99937574 |
| SNU13 | 100.464421 | -0.0610224 | 0.41514449 | -0.1469908 | 0.88313932 | 0.99937574 |
| CCDC134 | 33.2398788 | 0.90629083 | 0.71882222 | 1.26079968 | 0.20738103 | 0.99937574 |
| SREBF2 | 508.850334 | 0.09872106 | 0.39921147 | 0.24729014 | 0.80468369 | 0.99937574 |
| WBP2NL | 24.7712457 | 0.38111443 | 0.58521343 | 0.65124006 | 0.51489154 | 0.99937574 |
| NAGA | 12.8744773 | 0.68477853 | 0.49694455 | 1.37797774 | 0.16821016 | 0.99937574 |
| SMDT1 | 64.7436667 | 0.63931648 | 0.51480194 | 1.24186883 | 0.21428496 | 0.99937574 |
| NDUFA6 | 75.4449413 | -0.1749101 | 0.55728798 | -0.3138595 | 0.75362779 | 0.99937574 |
| TCF20 | 147.381463 | -0.0725019 | 0.28111261 | -0.2579106 | 0.79647589 | 0.99937574 |
| RRP7A | 24.4730128 | -0.0129726 | 0.53902365 | -0.0240668 | 0.98079934 | 0.99937574 |
| SERHL2 | 16.8096034 | 0.03044185 | 0.50828693 | 0.05989107 | 0.95224239 | 0.99937574 |
| POLDIP3 | 37.4480134 | -0.3020137 | 0.33200767 | -0.9096587 | 0.36300251 | 0.99937574 |
| CYB5R3 | 95.9266999 | 0.48172597 | 0.36790312 | 1.30938266 | 0.19040476 | 0.99937574 |
| A4GALT | 64.0423131 | 0.74348752 | 0.39075483 | 1.90269568 | 5.71E-02 | 9.99E-01 |
| ARFGAP3 | 134.209085 | 0.06649178 | 0.28228364 | 0.23554955 | 8.14E-01 | 0.99937574 |
| PACSIN2 | 194.101379 | -0.4996193 | 0.3455569 | -1.445838 | 0.14822264 | 0.99937574 |
| MCAT | 13.6781281 | 0.58326903 | 0.55487952 | 1.05116337 | 0.29318356 | 0.99937574 |
| TSPO | 161.110777 | 0.43695474 | 0.44518241 | 0.98151842 | 0.32633716 | 0.99937574 |
| EFCAB6 | 51.3051515 | 0.52058586 | 0.46946672 | 1.10888768 | 0.26747864 | 0.99937574 |
| SAMM50 | 17.0480566 | 0.22904692 | 0.51126843 | 0.44799738 | 0.65415509 | 0.99937574 |
| PARVB | 176.124387 | 0.02828909 | 0.41373125 | 0.06837552 | 0.94548671 | 0.99937574 |
| PHF21B | 18.3212401 | 1.46560459 | 0.99341071 | 1.47532594 | 0.14012493 | 0.99937574 |
| NUP50-DT | 117.973645 | -0.1308559 | 0.37349319 | -0.3503568 | 0.72607096 | 0.99937574 |
| NUP50 | 49.3647898 | -0.2753636 | 0.34576495 | -0.7963895 | 0.4258057 | 0.99937574 |
| KIAA0930 | 15.48707 | -0.9037412 | 0.58961559 | -1.5327635 | 0.12533414 | 0.99937574 |
| FAM118A | 18.2397748 | 0.03682455 | 0.73228212 | 0.05028738 | 0.95989338 | 0.99937574 |
| ATXN10 | 187.410572 | 0.14021188 | 0.19682063 | 0.71238405 | 0.47622699 | 0.99937574 |
| MIRLET7BHG | 92.6860207 | 0.63772447 | 0.36492622 | 1.74754357 | 0.08054309 | 0.99937574 |
| PPARA | 132.233312 | -0.0295863 | 0.33689246 | -0.0878212 | 0.9300188 | 0.99937574 |
| TTC38 | 34.5172981 | 0.62822405 | 0.55428895 | 1.13338728 | 0.25705165 | 0.99937574 |
| TRMU | 34.5605321 | 0.51879523 | 0.41592711 | 1.24732249 | 0.21227927 | 0.99937574 |
| CELSR1 | 48.7583874 | 0.15743031 | 0.49145462 | 0.3203354 | 0.74871409 | 0.99937574 |
| GRAMD4 | 44.6205304 | 0.30111153 | 0.45904162 | 0.65595692 | 0.51185185 | 0.99937574 |
| CERK | 26.1017951 | 0.01455658 | 0.61825643 | 0.02354457 | 0.98121589 | 0.99937574 |
| TBC1D22A | 240.383679 | -0.3214391 | 0.4016451 | -0.8003064 | 0.42353332 | 0.99937574 |
| BRD1 | 55.4894259 | -0.4120054 | 0.36243059 | -1.1367843 | 0.25562847 | 0.99937574 |
| ZBED4 | 35.3299177 | 0.06302483 | 0.45637462 | 0.1380989 | 0.89016225 | 0.99937574 |
| ALG12 | 19.229803 | 0.49663533 | 0.52217878 | 0.95108295 | 0.34156227 | 0.99937574 |
| CRELD2 | 17.6592863 | 0.25646149 | 0.50412846 | 0.5087225 | 0.61094675 | 0.99937574 |
| PIM3 | 118.070244 | -0.2223879 | 0.53389535 | -0.4165384 | 0.67701607 | 0.99937574 |
| TRABD | 31.9421235 | 0.38944433 | 0.39738498 | 0.98001776 | 0.32707735 | 0.99937574 |
| SELENOO | 48.2880431 | -0.0308381 | 0.40839135 | -0.0755111 | 0.93980804 | 0.99937574 |
| TUBGCP6 | 15.09219 | -0.3329659 | 0.51719584 | -0.6437907 | 0.51971118 | 0.99937574 |
| PLXNB2 | 47.426049 | 0.06638117 | 0.46223114 | 0.14361034 | 0.88580818 | 0.99937574 |
| PPP6R2 | 200.777344 | -0.2908249 | 0.28549509 | -1.0186685 | 0.30836038 | 0.99937574 |
| SBF1 | 16.166574 | -0.1230614 | 0.57922274 | -0.2124596 | 0.83174852 | 0.99937574 |
| LMF2 | 17.3618539 | -0.3219684 | 0.58700218 | -0.5484961 | 0.58335129 | 0.99937574 |
| NCAPH2 | 10.4748449 | -0.464286 | 0.67841918 | -0.6843644 | 0.49374507 | 0.99937574 |
| ARSA | 13.2630439 | 0.13219334 | 0.50780754 | 0.26032174 | 0.7946156 | 0.99937574 |
| RABL2B | 14.2117558 | -0.3379895 | 0.54782163 | -0.61697 | 0.5372545 | 0.99937574 |
| SMIM34B | 143.702697 | 1.0836085 | 0.65386401 | 1.65723833 | 0.0974713 | 0.99937574 |
| FP671120.1 | 412.713352 | -1.1256839 | 0.4396332 | -2.560507 | 0.01045196 | 0.76274616 |

| | | | | | | |
|---|---|---|---|---|---|---|
| FP236383.1 | 271.648849 | -1.152614 | 0.58950801 | -1.9552135 | 0.05055787 | 0.99937574 |
| CR381653.1 | 65.8590546 | 0.39087908 | 0.96329853 | 0.4057715 | 0.6849105 | 0.99937574 |
| RBM11 | 17.0984946 | 0.54109172 | 0.56078884 | 0.96487606 | 0.33460689 | 0.99937574 |
| HSPA13 | 20.1569468 | -0.3938975 | 0.48097365 | -0.8189585 | 0.41281009 | 0.99937574 |
| NRIP1 | 39.1667608 | -0.2333004 | 0.36915428 | -0.6319861 | 0.52739598 | 0.99937574 |
| USP25 | 262.0101 | -0.3386141 | 0.33096377 | -1.0231153 | 0.30625335 | 0.99937574 |
| MIR99AHG | 543.578903 | 0.45981426 | 0.40996319 | 1.12159887 | 2.62E-01 | 0.99937574 |
| CXADR | 93.7089023 | 0.46554342 | 0.43686107 | 1.06565553 | 0.28657939 | 0.99937574 |
| BTG3 | 72.1939755 | -0.43679 | 0.41157371 | -1.061268 | 0.28856811 | 0.99937574 |
| C21orf91 | 20.8399427 | 0.26856733 | 0.43856556 | 0.6123767 | 0.54028856 | 0.99937574 |
| LINC01687 | 321.77421 | 0.38281477 | 1.23743845 | 0.30936065 | 0.7570472 | 0.99937574 |
| MRPL39 | 19.4603919 | -0.04197 | 0.51128455 | -0.0820873 | 0.9345773 | 0.99937574 |
| ATP5PF | 126.030096 | 0.40582751 | 0.49611226 | 0.81801548 | 0.41334835 | 0.99937574 |
| GABPA | 34.6990902 | -0.2301735 | 0.34415436 | -0.668809 | 0.50361732 | 0.99937574 |
| APP | 540.126293 | -0.5672784 | 0.37760662 | -1.5023 | 0.13301965 | 0.99937574 |
| ADAMTS1 | 262.874437 | 0.40218882 | 0.82329763 | 0.48850964 | 6.25E-01 | 9.99E-01 |
| ADAMTS5 | 50.3529133 | -0.3444143 | 0.88049622 | -0.3911594 | 6.96E-01 | 0.99937574 |
| LTN1 | 83.0519414 | -0.0165333 | 0.31995289 | -0.051674 | 0.95878843 | 0.99937574 |
| USP16 | 64.6370088 | -0.5870353 | 0.2710881 | -2.1654779 | 0.0303511 | 0.99049468 |
| CCT8 | 51.7960386 | 0.01531152 | 0.35291554 | 0.04338578 | 0.96539401 | 0.99937574 |
| MAP3K7CL | 77.0327597 | 0.12404416 | 0.38804844 | 0.31966153 | 0.74922493 | 0.99937574 |
| BACH1 | 417.148631 | -0.0438619 | 0.55568935 | -0.0789324 | 0.93708639 | 0.99937574 |
| GRIK1 | 33.0432068 | 0.23625224 | 0.73348258 | 0.32209659 | 7.47E-01 | 0.99937574 |
| CLDN8 | 112.787245 | 0.32360922 | 0.47958359 | 0.67477125 | 0.49982111 | 0.99937574 |
| SOD1 | 94.3265561 | 0.01778108 | 0.47252029 | 0.03763031 | 0.96998244 | 0.99937574 |
| SCAF4 | 56.4658621 | -0.1904798 | 0.33239736 | -0.5730485 | 0.56661182 | 0.99937574 |
| HUNK | 23.8590527 | 1.51802652 | 0.8468874 | 1.79247738 | 0.07305653 | 0.99937574 |
| URB1 | 18.188331 | -0.1699194 | 0.59095457 | -0.2875338 | 0.77370362 | 0.99937574 |
| SYNJ1 | 72.5874759 | -0.1788285 | 0.35809597 | -0.4993871 | 0.61750672 | 0.99937574 |
| PAXBP1 | 92.5747539 | -0.2168566 | 0.34531734 | -0.6279923 | 0.53000898 | 0.99937574 |
| C21orf62-AS | 20.2750835 | 0.15834705 | 0.60646367 | 0.26109899 | 0.79401617 | 0.99937574 |
| AP000282.1 | 142.889763 | 0.87386073 | 0.68777297 | 1.27056567 | 0.20388321 | 0.99937574 |
| OLIG1 | 16.245218 | 1.07291795 | 0.71894811 | 1.49234406 | 0.13560897 | 0.99937574 |
| IFNAR1 | 124.632037 | 0.06447466 | 0.29072187 | 0.22177438 | 0.82448952 | 0.99937574 |
| IFNGR2 | 167.152836 | -0.3674099 | 0.3811809 | -0.9638728 | 0.33510968 | 0.99937574 |
| TMEM50B | 114.941872 | -0.2488651 | 0.44284525 | -0.5619686 | 0.57413743 | 0.99937574 |
| GART | 37.4962529 | -0.0885852 | 0.36280656 | -0.2441664 | 0.80710198 | 0.99937574 |
| SON | 277.206581 | 0.14502103 | 0.28411363 | 0.51043321 | 0.609748 | 0.99937574 |
| AP000311.1 | 56.3203078 | 0.30225358 | 0.55857142 | 0.54111895 | 0.5884256 | 0.99937574 |
| ITSN1 | 195.167682 | 0.23219046 | 0.21645629 | 1.07268983 | 0.28341031 | 0.99937574 |
| LINC00649 | 15.1393111 | 0.70263151 | 0.58258444 | 1.2060595 | 0.22779455 | 0.99937574 |
| MRPS6 | 491.420922 | 0.13533373 | 0.80020595 | 0.16912363 | 0.8656994 | 0.99937574 |
| LINC00310 | 41.5941162 | 0.28907916 | 1.04625759 | 0.27629827 | 0.78231899 | 0.99937574 |
| AP000317.1 | 36.471452 | -0.4548958 | 1.05862779 | -0.4297032 | 0.66741152 | 0.99937574 |
| RCAN1 | 43.8815926 | -0.0373856 | 0.4317187 | -0.0865972 | 0.93099171 | 0.99937574 |
| CLIC6 | 20.5851618 | 0.98272742 | 0.63719015 | 1.54228283 | 0.12300488 | 0.99937574 |
| RUNX1 | 52.0218915 | -0.1317367 | 0.56780838 | -0.2320091 | 0.81653091 | 0.99937574 |
| SETD4 | 47.688416 | -0.0678238 | 0.32784136 | -0.2068801 | 0.83610353 | 0.99937574 |
| CBR1 | 114.930805 | -0.1348273 | 0.33936135 | -0.3972972 | 6.91E-01 | 0.99937574 |
| CBR3 | 53.1868709 | -0.2708925 | 0.54867204 | -0.4937238 | 0.62150122 | 0.99937574 |
| DOPEY2 | 93.5276184 | 0.29693246 | 0.68544343 | 0.43319761 | 0.66487121 | 0.99937574 |
| MORC3 | 110.596998 | -0.4471365 | 0.28082502 | -1.5922245 | 0.11133427 | 0.99937574 |
| HLCS | 165.447755 | 0.21735819 | 0.34093591 | 0.63753385 | 0.52377717 | 0.99937574 |
| PIGP | 17.0535002 | 0.81848309 | 0.60772096 | 1.34680741 | 0.17804227 | 0.99937574 |

| | | | | | | |
|---|---|---|---|---|---|---|
| TTC3 | 304.873992 | 0.03433507 | 0.25084714 | 0.13687646 | 0.89112845 | 0.99937574 |
| VPS26C | 66.9773205 | -0.4209537 | 0.30643499 | -1.373713 | 0.1695308 | 0.99937574 |
| DYRK1A | 412.161281 | -0.0362969 | 0.28581955 | -0.1269923 | 0.89894647 | 0.99937574 |
| KCNJ15 | 49.1640608 | 0.57649266 | 0.59338909 | 0.97152555 | 0.33128663 | 0.99937574 |
| ETS2 | 182.519055 | -0.0586028 | 0.41614874 | -0.1408218 | 8.88E-01 | 9.99E-01 |
| AP001043.1 | 34.2033433 | -1.1121751 | 1.09913072 | -1.0118679 | 0.31160123 | 0.99937574 |
| BRWD1 | 312.986169 | -0.005833 | 0.21806034 | -0.0267495 | 0.9786595 | 0.99937574 |
| HMGN1 | 108.054171 | 0.01805221 | 0.2572712 | 0.070168 | 0.94405994 | 0.99937574 |
| WRB | 26.3177948 | 0.4456354 | 0.55612295 | 0.80132532 | 0.42294334 | 0.99937574 |
| DSCAM | 108.77815 | 0.35773726 | 0.58994551 | 0.60639034 | 0.54425558 | 0.99937574 |
| BACE2 | 719.829641 | -0.7186528 | 0.81504331 | -0.8817358 | 0.3779197 | 0.99937574 |
| FAM3B | 53.0408345 | 0.52676671 | 0.29072648 | 1.81189792 | 0.07000197 | 0.99937574 |
| MX1 | 126.426088 | -3.202139 | 1.00166164 | -3.196827 | 0.00138948 | 0.34724223 |
| TMPRSS2 | 590.835615 | 0.31324277 | 0.30650672 | 1.02197684 | 0.30679186 | 0.99937574 |
| RIPK4 | 13.9860164 | -0.5038137 | 0.59424477 | -0.8478218 | 0.39653723 | 0.99937574 |
| PRDM15 | 16.2642783 | 0.0532599 | 0.55035947 | 0.09677292 | 0.92290673 | 0.99937574 |
| C2CD2 | 53.381983 | 0.39849414 | 0.37729747 | 1.05618027 | 0.29088586 | 0.99937574 |
| ZBTB21 | 41.6932948 | -0.0897335 | 0.40473122 | -0.2217113 | 0.8245386 | 0.99937574 |
| UMODL1 | 13.5816582 | 0.32161503 | 1.27519375 | 0.25220875 | 0.80087971 | 0.99937574 |
| ABCG1 | 42.0242426 | 0.55103918 | 0.60977244 | 0.90368003 | 0.36616509 | 0.99937574 |
| SLC37A1 | 93.0767343 | -0.2091925 | 0.50734354 | -0.4123291 | 0.6800982 | 0.99937574 |
| PDE9A | 85.4527869 | 0.19268531 | 0.45440203 | 0.42404149 | 0.67153555 | 0.99937574 |
| WDR4 | 15.8542126 | -0.4660088 | 0.6685356 | -0.697059 | 0.48576586 | 0.99937574 |
| NDUFV3 | 73.6126333 | 0.37252694 | 0.32313112 | 1.15286619 | 0.24896531 | 0.99937574 |
| PKNOX1 | 49.3855201 | -0.1021123 | 0.47291753 | -0.21592 | 0.82905012 | 0.99937574 |
| RRP1B | 25.9276805 | 0.0469232 | 0.49076813 | 0.09561175 | 0.92382893 | 0.99937574 |
| PDXK | 99.5697581 | -0.508958 | 0.3577268 | -1.4227562 | 0.15480686 | 0.99937574 |
| CSTB | 152.123517 | -0.3633504 | 0.52194862 | -0.696142 | 0.4863399 | 0.99937574 |
| AGPAT3 | 131.161213 | 0.11108998 | 0.3149759 | 0.35269359 | 0.72431816 | 0.99937574 |
| TRAPPC10 | 135.331568 | -0.1905485 | 0.32862938 | -0.5798279 | 0.56203066 | 0.99937574 |
| PFKL | 134.437496 | 0.22711586 | 0.23256083 | 0.97658688 | 3.29E-01 | 9.99E-01 |
| C21orf2 | 23.6047396 | -0.2096986 | 0.49448015 | -0.424079 | 0.67150821 | 0.99937574 |
| UBE2G2 | 83.6715719 | 0.12461209 | 0.28695134 | 0.43426209 | 0.66409813 | 0.99937574 |
| SUMO3 | 48.9765547 | -0.4423811 | 0.49059536 | -0.901723 | 0.36720404 | 0.99937574 |
| PTTG1IP | 148.259716 | -0.07294 | 0.37591049 | -0.1940356 | 0.84614803 | 0.99937574 |
| FAM207A | 13.98914 | -0.5616088 | 0.55939091 | -1.0039648 | 0.31539556 | 0.99937574 |
| ADARB1 | 47.4526222 | 1.61885647 | 0.43037927 | 3.76146476 | 0.00016892 | 0.13113484 |
| POFUT2 | 24.5326383 | 0.16759515 | 0.46106172 | 0.3634983 | 0.71623268 | 0.99937574 |
| COL18A1 | 149.870879 | -0.2129968 | 0.29244598 | -0.7283287 | 0.46641243 | 0.99937574 |
| PCBP3 | 87.8479624 | -0.4886698 | 0.45793456 | -1.0671171 | 0.28591898 | 0.99937574 |
| COL6A1 | 121.096639 | 1.20304078 | 0.67869533 | 1.77257853 | 0.07629857 | 0.99937574 |
| LSS | 45.422251 | -0.0444516 | 0.62541016 | -0.0710759 | 0.94333734 | 0.99937574 |
| MCM3AP-AS | 11.9791499 | 0.40619369 | 0.60985327 | 0.66605151 | 0.50537818 | 0.99937574 |
| MCM3AP | 79.4148322 | -0.1782613 | 0.27036732 | -0.6593301 | 0.50968379 | 0.99937574 |
| YBEY | 15.792101 | -0.9953155 | 0.53575481 | -1.8577817 | 0.06320002 | 0.99937574 |
| PCNT | 31.5644228 | -0.4746144 | 0.47197143 | -1.0056 | 0.31460805 | 0.99937574 |
| DIP2A | 129.664807 | -0.046332 | 0.23564666 | -0.1966163 | 0.8441278 | 0.99937574 |
| PRMT2 | 106.981506 | 0.24412059 | 0.3739132 | 0.65288039 | 0.51383339 | 0.99937574 |
| AL592183.1 | 74.0766296 | -0.3553086 | 0.57891166 | -0.6137527 | 0.53937876 | 0.99937574 |
| AL354822.1 | 13.7926501 | -0.5786035 | 0.84460602 | -0.6850572 | 0.49330781 | 0.99937574 |

**Collecting duct intercalated cells type B**

*This means that positive fold changes imply higher mean value in non-COVID AKI than in COVID AKI and vice versa.

| | baseMean | log2 fold cha | lfcSE | stat | pvalue | padj |
|---|---|---|---|---|---|---|
| AL669831.5 | 6.85036901 | 1.07244905 | 0.61013203 | 1.75773275 | 0.07879299 | NA |
| NOC2L | 4.32635912 | -0.5892801 | 0.79948231 | -0.7370771 | 0.46107544 | NA |
| ISG15 | 73.1567311 | -1.8666287 | 1.96332618 | -0.9507481 | 0.34173226 | 0.8003972 |
| AGRN | 12.2711978 | -0.4273642 | 0.5692586 | -0.7507383 | 0.45281017 | NA |
| C1orf159 | 7.38299122 | -1.052317 | 0.603946 | -1.7424025 | 0.08143804 | NA |
| AL390719.2 | 2.79983962 | -0.2933384 | 0.83970581 | -0.3493347 | 0.72683807 | NA |
| SDF4 | 13.2809507 | -0.2387178 | 0.42647527 | -0.5597459 | 0.57565277 | NA |
| B3GALT6 | 3.13529671 | -0.464641 | 0.99062203 | -0.4690396 | 0.63904133 | NA |
| UBE2J2 | 10.8502805 | -0.2922222 | 0.50491342 | -0.578757 | 0.56275317 | NA |
| ACAP3 | 4.77179459 | -1.6647809 | 0.88824983 | -1.874226 | 0.06089928 | NA |
| INTS11 | 7.89135668 | 0.04531365 | 0.52099463 | 0.08697527 | 0.93069117 | NA |
| CPTP | 5.45870049 | 0.0068965 | 0.67122595 | 0.01027449 | 0.99180229 | NA |
| DVL1 | 5.66247424 | -0.8011677 | 0.64931013 | -1.2338752 | 0.21724942 | NA |
| AURKAIP1 | 25.010472 | -0.3283593 | 0.49748484 | -0.6600389 | 0.50922887 | NA |
| CCNL2 | 24.2245333 | -0.2095003 | 0.34889557 | -0.6004671 | 0.54819501 | NA |
| MRPL20 | 21.1558157 | -0.3067772 | 0.33621646 | -0.9124394 | 0.36153744 | NA |
| VWA1 | 6.50836665 | 0.01830726 | 0.60213386 | 0.03040398 | 0.97574487 | NA |
| ATAD3C | 4.62484246 | -1.741488 | 0.8019512 | -2.1715635 | 0.0298886 | NA |
| ATAD3B | 8.57115054 | -1.3270162 | 0.58350398 | -2.2742196 | 0.02295279 | NA |
| ATAD3A | 5.02987711 | -2.0681381 | 0.76414421 | -2.7064762 | 0.00680015 | NA |
| SSU72 | 22.5773763 | -0.2611044 | 0.41435807 | -0.630142 | 0.52860165 | NA |
| MIB2 | 4.68663638 | -0.3054884 | 0.75510223 | -0.4045656 | 0.68579682 | NA |
| CDK11B | 6.1679364 | -0.5108628 | 0.70510099 | -0.7245242 | 0.46874395 | NA |
| SLC35E2B | 7.00258062 | -0.3858037 | 0.58329039 | -0.6614264 | 0.50833891 | NA |
| CDK11A | 4.23278121 | -0.2294252 | 0.84824102 | -0.2704717 | 0.7867974 | NA |
| SLC35E2A | 7.25853404 | 0.61425127 | 0.59943925 | 1.0247098 | 0.30550014 | NA |
| NADK | 12.2247804 | -0.4803829 | 0.48032803 | -1.0001142 | 0.31725525 | NA |
| GNB1 | 97.9028032 | -0.7654466 | 0.34859216 | -2.1958227 | 0.02810464 | 0.39934064 |
| PRKCZ | 16.0370395 | 0.10888516 | 0.45145983 | 0.24118461 | 0.80941204 | NA |
| FAAP20 | 13.1922234 | 0.0463748 | 0.44051703 | 0.10527358 | 0.91615873 | NA |
| SKI | 14.4139254 | -0.3753787 | 0.4583781 | -0.8189281 | 0.41282743 | NA |
| MORN1 | 4.86781624 | 0.43436241 | 0.6767968 | 0.64179146 | 0.5210086 | NA |
| RER1 | 22.6157582 | 0.01697957 | 0.5030044 | 0.0337563 | 0.97307148 | NA |
| PEX10 | 4.90471173 | 0.08481578 | 0.61999775 | 0.13680015 | 0.89118877 | NA |
| PANK4 | 2.87252796 | -0.262067 | 0.86080847 | -0.3044429 | 0.7607905 | NA |
| TNFRSF14-A | 4.69784715 | -0.5843953 | 0.75804369 | -0.7709257 | 0.44075099 | NA |
| TNFRSF14 | 10.5317113 | -1.0016599 | 0.60237106 | -1.662862 | 0.09634007 | NA |
| LINC00982 | 7.19576218 | -0.3527434 | 0.58251563 | -0.6055518 | 0.5448124 | NA |
| PRDM16 | 61.0618915 | -0.3417692 | 0.43414213 | -0.7872288 | 0.43114794 | 0.84223493 |
| ARHGEF16 | 5.9831691 | 0.75365631 | 0.6248292 | 1.20617972 | 0.2277482 | NA |
| TPRG1L | 8.20611174 | -0.0514598 | 0.53744683 | -0.0957487 | 0.92372017 | NA |
| WRAP73 | 3.37239962 | -0.102346 | 0.96089703 | -0.1065109 | 0.91517697 | NA |
| LRRC47 | 6.5651721 | -0.8828577 | 0.64999026 | -1.3582629 | 0.17438026 | NA |
| CEP104 | 12.1647757 | -0.7240728 | 0.54145702 | -1.3372674 | 0.18113538 | NA |
| C1orf174 | 3.27842003 | -0.9262511 | 0.87350145 | -1.0603887 | 0.2889678 | NA |
| NPHP4 | 5.47554662 | 0.74705726 | 0.62764609 | 1.1902524 | 0.23394721 | NA |
| RPL22 | 15.7182922 | -0.2246578 | 0.52096072 | -0.4312376 | 0.66629564 | NA |
| RNF207 | 7.4570565 | 0.22980151 | 0.63414515 | 0.36237999 | 0.71706809 | NA |
| ICMT | 8.33840892 | 0.01582303 | 0.51203169 | 0.03090245 | 0.97534734 | NA |
| ACOT7 | 4.44653687 | -0.8043867 | 0.79515329 | -1.0116121 | 0.31172357 | NA |
| NOL9 | 10.7585306 | -0.5135687 | 0.61765274 | -0.8314846 | 0.40569994 | NA |
| ZBTB48 | 2.54596463 | 0.36145001 | 0.90642643 | 0.39876375 | 0.69006729 | NA |

| | | | | | | |
|---|---|---|---|---|---|---|
| KLHL21 | 8.05456258 | -0.2379049 | 0.60258726 | -0.3948058 | 0.69298622 | NA |
| PHF13 | 3.21983802 | -0.1023371 | 0.81154584 | -0.1261015 | 0.89965161 | NA |
| THAP3 | 4.18308121 | -0.3939568 | 0.71867637 | -0.5481699 | 0.58357522 | NA |
| DNAJC11 | 18.8523018 | -0.4269598 | 0.44893303 | -0.9510546 | 0.34157664 | NA |
| CAMTA1 | 85.4418843 | 0.47940194 | 0.29422821 | 1.62935411 | 0.10323808 | 0.5772605 |
| VAMP3 | 10.6551454 | 1.09768654 | 0.47777928 | 2.29747624 | 0.02159162 | NA |
| PER3 | 22.5019506 | -0.0570489 | 0.72283422 | -0.0789239 | 0.93709315 | NA |
| PARK7 | 35.5177207 | 0.5622008 | 0.62206236 | 0.90376919 | 0.3661178 | 0.80952111 |
| ERRFI1 | 15.78837 | 0.46622594 | 0.69471514 | 0.67110377 | 0.50215443 | NA |
| RERE | 253.714551 | 0.19431643 | 0.26182648 | 0.74215729 | 0.45799204 | 0.85802171 |
| ENO1 | 75.7334702 | -0.3471962 | 0.32252658 | -1.0764885 | 0.28170885 | 0.76854599 |
| GPR157 | 3.78943433 | 0.61000191 | 0.77997148 | 0.78208233 | 0.43416619 | NA |
| H6PD | 10.4436061 | -0.3428634 | 0.46585849 | -0.7359818 | 0.46174178 | NA |
| SPSB1 | 4.16658612 | -1.3263868 | 0.88940819 | -1.4913139 | 0.13587909 | NA |
| SLC25A33 | 12.1107131 | -0.3203877 | 0.52947426 | -0.6051054 | 0.54510896 | NA |
| TMEM201 | 2.88818283 | -0.6560494 | 0.95413525 | -0.6875854 | 0.49171392 | NA |
| CLSTN1 | 19.3910964 | -0.0241183 | 0.35912711 | -0.0671581 | 0.94645588 | NA |
| CTNNBIP1 | 5.70345399 | -0.3779453 | 0.65574773 | -0.5763577 | 0.56437341 | NA |
| LZIC | 8.73015985 | 0.50779348 | 0.5312764 | 0.95579906 | 0.33917376 | NA |
| NMNAT1 | 12.1840167 | -0.0959489 | 0.46691094 | -0.2054973 | 0.8371836 | NA |
| UBE4B | 38.6399112 | -0.1806695 | 0.33259641 | -0.5432093 | 0.5869857 | 0.89472578 |
| KIF1B | 84.617566 | -0.3860409 | 0.29507458 | -1.3082826 | 0.19077746 | 0.68530961 |
| PGD | 9.0915598 | 0.00405696 | 0.47998024 | 0.00845235 | 0.99325608 | NA |
| DFFA | 9.88745539 | 0.50066016 | 0.50935779 | 0.98292432 | 0.32564469 | NA |
| PEX14 | 30.3576594 | -0.2563056 | 0.39367267 | -0.6510629 | 0.51500592 | 0.87158522 |
| CASZ1 | 52.9189872 | 0.54484042 | 0.39163233 | 1.39120388 | 0.16416361 | 0.64995633 |
| SRM | 4.60227055 | -1.848303 | 0.83949837 | -2.2016755 | 0.02768824 | NA |
| EXOSC10 | 9.10569245 | -0.2268917 | 0.53634947 | -0.4230296 | 0.67227368 | NA |
| MTOR | 24.6550816 | 0.21565407 | 0.36512617 | 0.59062891 | 0.55476909 | NA |
| UBIAD1 | 2.8119876 | 0.53678121 | 0.91697957 | 0.58537968 | 0.55829245 | NA |
| FBXO2 | 4.11938176 | 0.01732265 | 0.67704556 | 0.02558565 | 0.97958783 | NA |
| FBXO44 | 3.06206633 | -0.7303542 | 0.98859228 | -0.738782 | 0.46003937 | NA |
| MAD2L2 | 4.57970565 | 0.60068176 | 0.70611965 | 0.85067985 | 0.39494722 | NA |
| AGTRAP | 6.30213816 | 0.07393545 | 0.60039877 | 0.1231439 | 0.90199315 | NA |
| MTHFR | 6.17998716 | -0.8323149 | 0.6155206 | -1.3522129 | 0.17630721 | NA |
| CLCN6 | 8.52144691 | -1.1612794 | 0.74802533 | -1.5524599 | 0.12055221 | NA |
| KIAA2013 | 9.18547811 | 0.27756096 | 0.60296974 | 0.46032321 | 0.64528425 | NA |
| PLOD1 | 5.05902495 | -0.2305265 | 0.61950189 | -0.372116 | 0.70980651 | NA |
| MFN2 | 14.1338735 | -1.1478563 | 0.59502032 | -1.9291043 | 0.05371791 | NA |
| VPS13D | 89.0462139 | -0.0063137 | 0.25101289 | -0.0251529 | 0.97993303 | 0.99902646 |
| DHRS3 | 21.9980751 | -0.927655 | 0.54974471 | -1.6874287 | 0.09152095 | NA |
| PRDM2 | 47.7095684 | 0.16668641 | 0.26867735 | 0.62039621 | 0.53499697 | 0.88254911 |
| KAZN | 313.956202 | -0.0041473 | 0.3028151 | -0.0136958 | 0.98907264 | 0.99909657 |
| TMEM51-AS | 20.9511453 | 0.464799 | 0.57196906 | 0.81262961 | 0.41643045 | NA |
| TMEM51 | 143.893451 | 0.27323381 | 0.29229422 | 0.93479034 | 0.34989636 | 0.8003972 |
| C1orf195 | 5.59535445 | 1.04865019 | 0.68508074 | 1.53069576 | 0.1258446 | NA |
| EFHD2 | 4.37377097 | -1.0549676 | 0.69019285 | -1.5285114 | 0.12638561 | NA |
| CASP9 | 5.45713541 | -0.1641982 | 0.68011093 | -0.2414285 | 0.809223 | NA |
| DNAJC16 | 12.5599405 | 0.51901035 | 0.43880025 | 1.1827941 | 0.23689076 | NA |
| DDI2 | 17.671688 | 0.46060385 | 0.44002809 | 1.04676009 | 0.29521024 | NA |
| PLEKHM2 | 11.8762434 | -1.5018211 | 0.48356904 | -3.1057014 | 0.00189828 | NA |
| UQCRHL | 5.97888307 | 0.71851354 | 0.61377605 | 1.17064447 | 0.24174171 | NA |
| AL450998.2 | 2.21857323 | 1.54219578 | 0.95783961 | 1.61007727 | 0.10738099 | NA |
| SPEN | 46.1713753 | -0.5220118 | 0.31325002 | -1.6664382 | 0.09562617 | 0.56358504 |

| | | | | | | |
|---|---|---|---|---|---|---|
| ZBTB17 | 6.99493963 | -0.1784264 | 0.58489182 | -0.3050588 | 0.76032139 | NA |
| SRARP | 2.61171047 | 0.33154603 | 1.02768566 | 0.32261425 | 0.7469874 | NA |
| CLCNKA | 25.0518976 | 0.2396517 | 0.5118338 | 0.46822171 | 6.40E-01 | NA |
| CLCNKB | 54.5990382 | 0.85666824 | 0.61121923 | 1.40157278 | 0.16104286 | 0.64709575 |
| FAM131C | 4.06623725 | -0.4120082 | 0.87318986 | -0.4718426 | 0.63703913 | NA |
| EPHA2 | 5.46042377 | -0.5693746 | 0.72257127 | -0.787984 | 0.43070609 | NA |
| FBXO42 | 37.9346354 | -0.0365309 | 0.32426836 | -0.1126565 | 0.91030291 | 0.9836431 |
| SZRD1 | 18.3115678 | -0.8689842 | 0.39445438 | -2.2030032 | 0.02759453 | NA |
| NECAP2 | 6.49706839 | 0.18166177 | 0.56026244 | 0.32424406 | 0.74575327 | NA |
| NBPF1 | 6.26894141 | 0.76042474 | 0.65233888 | 1.16568974 | 0.24373991 | NA |
| CROCC | 7.09500754 | 0.58155763 | 0.56728059 | 1.02516751 | 0.30528416 | NA |
| ATP13A2 | 4.48983179 | 0.19022733 | 0.71385842 | 0.26647768 | 0.78987135 | NA |
| SDHB | 21.6753873 | 0.1543055 | 0.36285782 | 0.4252506 | 0.670654 | NA |
| RCC2 | 9.40539299 | 0.18506324 | 0.56582208 | 0.32706966 | 0.7436152 | NA |
| ARHGEF10L | 30.9529386 | 0.43524989 | 0.52724132 | 0.82552311 | 0.40907467 | 0.82839172 |
| KLHDC7A | 3.89769691 | 0.58548276 | 1.1289306 | 0.51861714 | 0.60402775 | NA |
| UBR4 | 31.2606269 | -0.3494303 | 0.31625581 | -1.1048976 | 0.26920396 | 0.75632146 |
| AL035413.1 | 5.7538753 | 0.47119451 | 0.65719346 | 0.71697991 | 0.47338649 | NA |
| EMC1 | 7.48650494 | -0.1811918 | 0.61209453 | -0.2960194 | 0.76721528 | NA |
| MRTO4 | 5.1293632 | 0.62868377 | 0.62473299 | 1.00632395 | 0.31425976 | NA |
| AKR7A2 | 6.45397624 | 1.37465197 | 0.61818693 | 2.22368331 | 0.02616976 | NA |
| PQLC2 | 2.6267441 | -0.8391211 | 0.89826632 | -0.9341563 | 0.35022327 | NA |
| CAPZB | 79.3037392 | -0.6560472 | 0.28803178 | -2.2776903 | 0.02274503 | 0.35709704 |
| NBL1 | 4.68583487 | -1.4036366 | 0.67282365 | -2.086188 | 0.0369616 | NA |
| TMCO4 | 46.4746345 | 0.18995081 | 0.45690204 | 0.4157364 | 0.6776029 | 0.9267216 |
| OTUD3 | 5.8162413 | -0.0555298 | 0.61968298 | -0.0896101 | 0.92859708 | NA |
| VWA5B1 | 24.9615817 | 1.58113877 | 0.75123389 | 2.10472238 | 0.03531548 | NA |
| CAMK2N1 | 65.2270819 | 0.59545786 | 0.32218807 | 1.84816859 | 0.06457796 | 0.52254021 |
| MUL1 | 5.75346037 | -2.0097284 | 0.69100804 | -2.9084009 | 0.00363282 | NA |
| CDA | 55.6758934 | -0.3675298 | 0.34610726 | -1.0618956 | 0.28828309 | 0.7725847 |
| PINK1 | 19.9223089 | 0.73698973 | 0.36732458 | 2.00637193 | 0.04481658 | NA |
| DDOST | 9.24342279 | 0.19837974 | 0.52483559 | 0.37798454 | 0.70544208 | NA |
| HP1BP3 | 58.9668425 | 0.40223462 | 0.28396936 | 1.41647193 | 0.15663738 | 0.63932163 |
| EIF4G3 | 135.272825 | -0.5756126 | 0.31766309 | -1.8120224 | 0.06998274 | 0.52654172 |
| ECE1 | 32.5737795 | 0.77349493 | 0.44874811 | 1.72367283 | 0.08476692 | 0.55323508 |
| RAP1GAP | 16.1827203 | 0.47909017 | 0.57937568 | 0.82690762 | 0.40828942 | NA |
| USP48 | 33.8460518 | -0.2446786 | 0.33963445 | -0.7204174 | 0.47126806 | 0.86206794 |
| HSPG2 | 14.1575107 | 0.52561809 | 0.56002248 | 0.93856605 | 0.34795359 | NA |
| CDC42 | 50.9330829 | -0.4562569 | 0.3028129 | -1.5067288 | 0.1318802 | 0.60375343 |
| ZBTB40 | 14.0736287 | -0.5378067 | 0.46192679 | -1.1642683 | 0.2443153 | NA |
| EPHB2 | 4.03958242 | -0.5043012 | 0.91097248 | -0.5535855 | 0.57986254 | NA |
| KDM1A | 27.6699488 | 0.49185571 | 0.31843489 | 1.54460372 | 0.12244215 | 0.58553957 |
| AL031428.1 | 6.25894223 | 0.73266351 | 0.7059375 | 1.03785888 | 0.29933576 | NA |
| LUZP1 | 26.5856457 | -0.2863917 | 0.4262504 | -0.6718861 | 0.50165623 | 0.87050612 |
| LINC01355 | 4.45605524 | 0.93731293 | 0.80087366 | 1.17036304 | 0.2418549 | NA |
| HNRNPR | 35.4900755 | -0.2333386 | 0.35895873 | -0.650043 | 0.51566445 | 0.87158522 |
| TCEA3 | 12.3431527 | 1.58796955 | 0.47758499 | 3.32499891 | 0.00088419 | NA |
| ASAP3 | 13.4292487 | 0.58048296 | 0.50334511 | 1.15325043 | 0.24880761 | NA |
| ID3 | 12.636544 | -1.4066914 | 0.77361819 | -1.8183277 | 0.06901406 | NA |
| RPL11 | 107.029845 | 0.12078263 | 0.44042147 | 0.27424328 | 0.78389767 | 0.96299238 |
| ELOA | 9.28260422 | -0.5996712 | 0.54847235 | -1.0933481 | 0.27424098 | NA |
| ELOA-AS1 | 4.18110488 | 1.26437289 | 0.71024195 | 1.78020025 | 0.0750432 | NA |
| PITHD1 | 11.2677958 | -0.7279771 | 0.5104714 | -1.426088 | 0.15384295 | NA |
| LYPLA2 | 10.3358579 | -0.4392575 | 0.61777934 | -0.7110265 | 0.47706782 | NA |

| | | | | | | |
|---|---|---|---|---|---|---|
| GALE | 10.283336 | -0.6087994 | 0.4476953 | -1.359852 | 0.17387676 | NA |
| HMGCL | 10.1685789 | 0.77389356 | 0.55760449 | 1.38788975 | 0.16517062 | NA |
| FUCA1 | 9.80450664 | 1.26317833 | 0.47041312 | 2.68525319 | 0.00724749 | NA |
| PNRC2 | 14.347247 | -0.158518 | 0.39407888 | -0.4022494 | 0.6875005 | NA |
| SRSF10 | 25.940578 | 0.23572591 | 0.38039537 | 0.61968661 | 0.53546413 | 0.88254911 |
| IFNLR1 | 5.99046654 | 0.38536835 | 0.60688286 | 0.63499626 | 0.52543086 | NA |
| NIPAL3 | 13.6266315 | 1.03498902 | 0.51674795 | 2.00288946 | 0.04518915 | NA |
| RCAN3 | 9.14998408 | -0.3028734 | 0.5531352 | -0.5475576 | 0.58399572 | NA |
| SRRM1 | 26.4910997 | -0.3370745 | 0.33634167 | -1.002179 | 0.31625717 | 0.780966 |
| CLIC4 | 16.0541102 | -0.7920272 | 0.62592559 | -1.2653696 | 0.20573885 | NA |
| SYF2 | 15.4665116 | -0.02995 | 0.49434508 | -0.0605853 | 0.95168949 | NA |
| RSRP1 | 74.5985639 | 0.51474833 | 0.26298216 | 1.9573508 | 0.05030624 | 0.46620608 |
| TMEM50A | 19.2350681 | 0.26759627 | 0.39683104 | 0.67433301 | 0.50009962 | NA |
| MACO1 | 47.1838774 | -0.295256 | 0.32707022 | -0.9027298 | 0.36666932 | 0.80952111 |
| LDLRAP1 | 6.01737455 | -0.5437204 | 0.64910574 | -0.8376454 | 0.40222988 | NA |
| MAN1C1 | 55.3639537 | 0.73257012 | 0.46429159 | 1.57782335 | 0.1146062 | 0.57860396 |
| AL031280.1 | 5.60664667 | 0.06015676 | 0.85018818 | 0.07075699 | 0.94359116 | NA |
| MTFR1L | 10.8288222 | -0.4326539 | 0.74587267 | -0.5800641 | 0.56187142 | NA |
| PAQR7 | 2.40908922 | -0.4805773 | 1.02789302 | -0.4675363 | 0.6401162 | NA |
| STMN1 | 6.68934662 | -1.0617673 | 0.57089268 | -1.859837 | 0.06290859 | NA |
| PAFAH2 | 5.90649871 | 0.07137628 | 0.81682607 | 0.08738248 | 0.9303675 | NA |
| ZNF593 | 7.69005033 | -0.7891347 | 0.62585241 | -1.2608959 | 0.20734637 | NA |
| CNKSR1 | 6.50859207 | -0.3369853 | 0.61548682 | -0.5475102 | 0.58402823 | NA |
| CEP85 | 5.48836983 | -0.9642764 | 0.6946284 | -1.3881904 | 0.16507909 | NA |
| SH3BGRL3 | 14.6459695 | 0.25622389 | 0.39032266 | 0.65644125 | 0.51154026 | NA |
| DHDDS | 14.3585653 | -0.850172 | 0.53194825 | -1.5982232 | 0.10999331 | NA |
| HMGN2 | 17.958651 | 0.39669594 | 0.37717083 | 1.05176729 | 0.29290633 | NA |
| RPS6KA1 | 9.76154699 | 0.18773357 | 0.48564004 | 0.38656937 | 0.69907503 | NA |
| ARID1A | 52.144073 | 0.06324349 | 0.29857563 | 0.21181731 | 0.83224956 | 0.97154892 |
| GPN2 | 7.03862649 | -0.9749856 | 0.71068502 | -1.3718956 | 0.17009595 | NA |
| GPATCH3 | 2.44773403 | 0.53808454 | 0.88852477 | 0.60559318 | 0.54478493 | NA |
| NUDC | 14.7042765 | -0.5288272 | 0.47598212 | -1.1110232 | 0.26655835 | NA |
| NR0B2 | 3.1924327 | 0.89368129 | 0.87175539 | 1.02515145 | 0.30529173 | NA |
| KDF1 | 2.56169403 | 0.09029526 | 0.96664423 | 0.09341106 | 0.925577 | NA |
| SLC9A1 | 4.67542995 | -0.8327053 | 0.72037136 | -1.1559389 | 0.24770615 | NA |
| WDTC1 | 18.7656072 | -0.3175613 | 0.46507423 | -0.6828185 | 0.49472155 | NA |
| TMEM222 | 5.472443 | -0.0078137 | 0.72587136 | -0.0107645 | 0.99141132 | NA |
| WASF2 | 35.3087054 | -0.0018805 | 0.33754197 | -0.0055712 | 0.99555481 | 0.9992989 |
| AHDC1 | 6.14897579 | 0.83083819 | 0.8373953 | 0.99216964 | 0.32111478 | NA |
| IFI6 | 48.2886808 | -2.3983414 | 1.3462873 | -1.7814484 | 0.07483922 | 0.53407991 |
| FAM76A | 7.66251816 | 0.40089824 | 0.50461361 | 0.79446576 | 0.42692433 | NA |
| STX12 | 19.0796232 | 0.38072014 | 0.39402694 | 0.96622871 | 0.33392975 | NA |
| PPP1R8 | 7.51713294 | -0.1586526 | 0.63922325 | -0.2481959 | 0.80398286 | NA |
| RPA2 | 8.02246441 | 0.9974314 | 0.51917611 | 1.92118122 | 0.05470887 | NA |
| EYA3 | 52.0547484 | -0.3275669 | 0.31592277 | -1.0368574 | 0.29980231 | 0.77521265 |
| DNAJC8 | 14.137935 | -0.4911429 | 0.46328244 | -1.0601371 | 0.28908221 | NA |
| ATP5IF1 | 36.0509816 | -0.3074978 | 0.38314958 | -0.802553 | 0.42223316 | 0.84037309 |
| SESN2 | 6.5148651 | -0.8052925 | 0.60858094 | -1.32323 | 0.18575891 | NA |
| PHACTR4 | 81.4157253 | -0.3054045 | 0.24374834 | -1.2529502 | 0.21022384 | 0.70690231 |
| RCC1 | 10.8169052 | -0.9301933 | 0.49129686 | -1.8933427 | 0.05831231 | NA |
| TRNAU1AP | 21.0974157 | -0.4312257 | 0.53317949 | -0.8087815 | 0.41864085 | NA |
| TAF12 | 8.46723695 | -0.0808723 | 0.56071198 | -0.1442315 | 0.88531769 | NA |
| GMEB1 | 8.46857843 | -1.0615811 | 0.5581756 | -1.9018765 | 0.0571873 | NA |
| YTHDF2 | 28.8453806 | -0.5843009 | 0.32277611 | -1.8102358 | 0.07025923 | 0.52674689 |

| | | | | | | |
|---|---|---|---|---|---|---|
| EPB41 | 61.0226404 | -0.2039136 | 0.33047338 | -0.6170348 | 0.53721176 | 0.88254911 |
| SRSF4 | 61.2717634 | -0.4461747 | 0.32461193 | -1.3744864 | 0.16929073 | 0.659616 |
| MECR | 6.85342826 | 0.83533971 | 0.57000529 | 1.46549467 | 0.14278601 | NA |
| PTPRU | 3.29503731 | 0.8028725 | 0.77618572 | 1.03438196 | 0.30095764 | NA |
| PUM1 | 92.3824291 | -0.3152669 | 0.27894112 | -1.1302274 | 0.25838043 | 0.75272108 |
| SNRNP40 | 11.470275 | 0.39451184 | 0.42700821 | 0.92389754 | 0.35553966 | NA |
| ZCCHC17 | 9.74272804 | 0.04783372 | 0.50647891 | 0.09444366 | 0.92475674 | NA |
| SERINC2 | 24.5885458 | -0.2042618 | 0.38688046 | -0.5279714 | 0.5975192 | NA |
| TINAGL1 | 11.5441827 | -0.9389327 | 0.55050126 | -1.7055959 | 0.08808334 | NA |
| PEF1 | 6.96155571 | -0.2691507 | 0.59799247 | -0.4500904 | 0.65264527 | NA |
| PTP4A2 | 9.23277907 | -0.4779024 | 0.59639448 | -0.8013194 | 0.42294679 | NA |
| KHDRBS1 | 43.6448539 | -0.7971055 | 0.32759732 | -2.4331868 | 0.01496658 | 0.30435125 |
| TMEM39B | 2.9139718 | -0.0699472 | 0.81209935 | -0.0861314 | 0.93136199 | NA |
| KPNA6 | 35.1021168 | -0.0001711 | 0.33586881 | -0.0005094 | 0.99959352 | 0.99960611 |
| TXLNA | 9.22204458 | -1.0116651 | 0.51227153 | -1.974861 | 0.04828392 | NA |
| EIF3I | 11.5738568 | 0.10741353 | 0.47056426 | 0.22826539 | 0.81943993 | NA |
| HDAC1 | 17.4784454 | -0.0640727 | 0.38947363 | -0.1645111 | 0.86932879 | NA |
| MARCKSL1 | 9.39307142 | -1.5784179 | 0.88322402 | -1.7871093 | 0.07391982 | NA |
| FAM229A | 3.76699897 | 0.48280687 | 0.90415937 | 0.53398426 | 0.59335242 | NA |
| BSDC1 | 15.1652612 | -0.4905695 | 0.39669194 | -1.2366511 | 0.21621665 | NA |
| ZBTB8A | 3.49237052 | -0.4389503 | 0.9056271 | -0.4846921 | 0.62789477 | NA |
| ZBTB8OS | 19.4889727 | -0.2649452 | 0.53456916 | -0.4956238 | 0.62015984 | NA |
| RBBP4 | 26.4060954 | -0.3005589 | 0.339794 | -0.8845328 | 0.37640868 | 0.81744083 |
| SYNC | 6.17316367 | -0.2840034 | 0.60120108 | -0.4723934 | 0.63664603 | NA |
| KIAA1522 | 41.2289984 | -0.2684969 | 0.31309971 | -0.8575444 | 0.39114406 | 0.8258317 |
| YARS | 14.6932101 | -0.8893976 | 0.42373756 | -2.0989351 | 0.03582262 | NA |
| S100PBP | 18.2653753 | 0.3107649 | 0.37772162 | 0.82273528 | 0.41065855 | NA |
| RNF19B | 9.96867973 | -0.3614491 | 0.64081961 | -0.5640418 | 0.57272565 | NA |
| AL020995.1 | 3.771367 | 0.49514854 | 0.70119632 | 0.70614823 | 0.48009596 | NA |
| AK2 | 22.9732257 | -0.1163733 | 0.34446561 | -0.3378372 | 0.73548589 | NA |
| ZNF362 | 4.85150279 | 2.02178803 | 0.6994158 | 2.89068112 | 0.00384408 | NA |
| PHC2 | 24.1852312 | -0.5461139 | 0.35905032 | -1.5209955 | 0.12826096 | NA |
| SMIM12 | 9.3364724 | -0.6463812 | 0.55247581 | -1.1699719 | 0.24201229 | NA |
| ZMYM1 | 14.0674153 | 0.69189929 | 0.43493955 | 1.59079416 | 0.11165591 | NA |
| SFPQ | 56.0665702 | -0.1808788 | 0.3286095 | -0.550437 | 0.58201966 | 0.89263108 |
| ZMYM4 | 115.30853 | -0.1252636 | 0.21699108 | -0.5772755 | 0.56375333 | 0.89263108 |
| KIAA0319L | 48.7196138 | 0.29195937 | 0.27057208 | 1.07904468 | 0.28056781 | 0.76854599 |
| PSMB2 | 12.5585109 | 0.20691917 | 0.42357512 | 0.48850644 | 0.62519117 | NA |
| AGO4 | 17.839783 | 0.71728163 | 0.40137674 | 1.78705332 | 0.07392887 | NA |
| AGO1 | 9.80264311 | 0.38943442 | 0.49911262 | 0.7802536 | 0.43524162 | NA |
| AGO3 | 64.2740511 | -0.2975254 | 0.26466039 | -1.124178 | 0.26093753 | 0.75479335 |
| ADPRHL2 | 4.53729227 | -0.1940947 | 0.7477927 | -0.2595568 | 0.79520564 | NA |
| TRAPPC3 | 7.19395527 | -0.3033222 | 0.59540771 | -0.5094361 | 0.6104466 | NA |
| MAP7D1 | 5.87146961 | -0.8503797 | 0.76855346 | -1.1064679 | 0.26852406 | NA |
| THRAP3 | 57.4210348 | -0.6397263 | 0.30199085 | -2.1183633 | 0.03414431 | 0.41688714 |
| EVA1B | 3.14363573 | 0.019763 | 0.76509924 | 0.02583063 | 0.97939243 | NA |
| STK40 | 16.4990237 | -0.9446187 | 0.4217732 | -2.2396367 | 0.02511452 | NA |
| LSM10 | 2.70539649 | 0.18356488 | 0.89889811 | 0.204211 | 0.83818862 | NA |
| MRPS15 | 8.07875656 | 0.34094681 | 0.60749714 | 0.56123196 | 0.57463942 | NA |
| MEAF6 | 9.55036795 | 0.25683242 | 0.48685487 | 0.52753383 | 0.59782293 | NA |
| SNIP1 | 4.86176722 | -0.6586607 | 0.73809167 | -0.8923834 | 0.37218748 | NA |
| GNL2 | 4.02720987 | -1.0408687 | 0.75702369 | -1.3749486 | 0.16914737 | NA |
| C1orf109 | 3.53357493 | 0.38623306 | 0.7540167 | 0.51223409 | 0.60848719 | NA |
| C1orf122 | 13.558538 | 0.30653622 | 0.45923236 | 0.66749699 | 0.50445473 | NA |

| | | | | | | |
|---|---|---|---|---|---|---|
| MTF1 | 7.1585961 | 0.3282042 | 0.58442877 | 0.56158119 | 0.5744014 | NA |
| INPP5B | 19.7218088 | 0.41602585 | 0.4143107 | 1.00413977 | 0.31531125 | NA |
| SF3A3 | 8.04758801 | 0.23682124 | 0.5678203 | 0.41707074 | 0.67662667 | NA |
| UTP11 | 3.19223624 | 0.28052535 | 0.74323392 | 0.37743885 | 0.7058475 | NA |
| RRAGC | 15.2947427 | -1.3216266 | 0.66114545 | -1.9989953 | 0.04560887 | NA |
| AL139260.1 | 26.0458812 | -1.1670573 | 0.75306938 | -1.5497341 | 0.12120535 | 0.58553957 |
| RHBDL2 | 3.81277201 | -0.2498754 | 0.83068544 | -0.3008063 | 0.76356222 | NA |
| AKIRIN1 | 26.4557049 | -0.9951837 | 0.37574894 | -2.6485336 | 0.00808418 | 0.27648451 |
| NDUFS5 | 33.3998731 | 0.06113515 | 0.3333328 | 0.18340574 | 0.85447967 | 0.97168963 |
| MACF1 | 258.897882 | 0.03499121 | 0.25924626 | 0.13497287 | 0.89263332 | 0.97769824 |
| PABPC4 | 9.29673258 | -0.4035911 | 0.53110511 | -0.7599082 | 0.44730945 | NA |
| PPIE | 5.83769735 | 0.72485335 | 0.73415142 | 0.98733494 | 0.32347847 | NA |
| TRIT1 | 10.7647745 | 0.0219168 | 0.52652329 | 0.0416255 | 0.96679724 | NA |
| CAP1 | 23.3950179 | -0.2185903 | 0.38971262 | -0.5609012 | 0.57486489 | NA |
| PPT1 | 6.17122388 | -0.3773083 | 0.64538317 | -0.5846268 | 0.55879868 | NA |
| RLF | 34.9546505 | -0.5772589 | 0.34127588 | -1.6914728 | 0.09074654 | 0.56211136 |
| ZMPSTE24 | 9.39744817 | -0.1262828 | 0.53438651 | -0.2363137 | 0.81318929 | NA |
| SMAP2 | 10.6429986 | -0.4588644 | 0.50661506 | -0.9057457 | 0.36507049 | NA |
| AL603839.3 | 5.05573676 | -0.3290491 | 0.74403629 | -0.4422487 | 0.65830924 | NA |
| ZNF684 | 5.41986736 | 1.02936006 | 0.69619347 | 1.47855461 | 0.13925939 | NA |
| NFYC | 32.268393 | 0.60243064 | 0.35862765 | 1.67982208 | 0.09299194 | 0.56220784 |
| CITED4 | 16.6882429 | -0.0987236 | 0.49483015 | -0.1995101 | 0.84186373 | NA |
| CTPS1 | 3.07754397 | -1.3261761 | 0.90011786 | -1.4733361 | 0.14066044 | NA |
| SCMH1 | 79.3407558 | 0.17797054 | 0.26562945 | 0.66999551 | 0.50286065 | 0.87094108 |
| HIVEP3 | 10.0725839 | -1.0113677 | 0.66591952 | -1.5187536 | 0.12882452 | NA |
| FOXJ3 | 51.4524629 | -0.2179254 | 0.28538337 | -0.7636234 | 0.44509168 | 0.84914203 |
| PPCS | 10.1239298 | 0.25946019 | 0.51874756 | 0.50016658 | 0.61695779 | NA |
| CCDC30 | 14.7803869 | 0.95325941 | 0.4131356 | 2.30737659 | 0.02103383 | NA |
| PPIH | 3.31090598 | -0.0069517 | 0.94656725 | -0.0073441 | 0.99414029 | NA |
| YBX1 | 37.7204312 | -0.4543363 | 0.36926222 | -1.2303894 | 0.21855131 | 0.72103021 |
| P3H1 | 3.04492614 | -0.3721309 | 0.86617598 | -0.429625 | 0.66746843 | NA |
| SVBP | 5.4788846 | -0.2142963 | 0.80703687 | -0.2655347 | 0.79059761 | NA |
| ERMAP | 4.60307587 | -0.0454623 | 0.78099015 | -0.0582111 | 0.95358045 | NA |
| SLC2A1 | 13.6959017 | -1.0236094 | 0.53690495 | -1.9065002 | 0.05658534 | NA |
| EBNA1BP2 | 5.02497954 | -0.3106164 | 0.75633174 | -0.410688 | 0.68130129 | NA |
| ELOVL1 | 5.01742035 | 0.16176044 | 0.78446035 | 0.20620601 | 0.83662999 | NA |
| MED8 | 4.93076083 | 0.72438731 | 0.73747978 | 0.98224701 | 0.32597817 | NA |
| SZT2 | 9.427282 | -0.3579822 | 0.54354978 | -0.6586006 | 0.51015226 | NA |
| PTPRF | 27.3911765 | 0.48450562 | 0.41425298 | 1.16958872 | 0.24216652 | 0.73323864 |
| IPO13 | 3.71804205 | -0.5239361 | 0.93406166 | -0.5609224 | 0.57485046 | NA |
| ATP6V0B | 108.586252 | 0.26846515 | 0.3986057 | 0.67351057 | 0.50062253 | 0.87036626 |
| B4GALT2 | 6.69414262 | -0.648772 | 0.70124016 | -0.925178 | 0.35487332 | NA |
| CCDC24 | 5.76970099 | 0.67462796 | 0.63598075 | 1.06076789 | 0.2887954 | NA |
| DMAP1 | 4.68069512 | -0.6294099 | 0.71390737 | -0.8816408 | 0.37797106 | NA |
| ERI3 | 19.4882505 | 0.07582158 | 0.42091298 | 0.180136 | 0.8570458 | NA |
| RNF220 | 6.40961935 | -0.5668894 | 0.62062462 | -0.9134176 | 0.36102296 | NA |
| TMEM53 | 4.93704109 | 1.17407308 | 0.7475263 | 1.57061105 | 0.11627302 | NA |
| RPS8 | 62.6047898 | 0.19717598 | 0.34533431 | 0.5709713 | 0.5680191 | 0.89263108 |
| EIF2B3 | 14.7043506 | -0.2838391 | 0.47406117 | -0.5987394 | 0.54934665 | NA |
| HECTD3 | 2.28804727 | -0.3603904 | 0.94965719 | -0.3794953 | 0.7043201 | NA |
| UROD | 4.75956948 | 0.46897014 | 0.77508647 | 0.60505525 | 0.54514229 | NA |
| ZSWIM5 | 28.7541548 | -0.3529909 | 0.58067192 | -0.6079008 | 0.54325327 | 0.88410237 |
| TESK2 | 4.96865244 | 0.53784216 | 0.68442247 | 0.78583359 | 0.43196498 | NA |
| PRDX1 | 55.6242414 | -0.2337194 | 0.40102179 | -0.5828096 | 0.56002146 | 0.89263108 |

| | | | | | | |
|---|---|---|---|---|---|---|
| AKR1A1 | 5.24263394 | -0.7463092 | 0.65964122 | -1.1313866 | 0.25789242 | NA |
| NASP | 15.0896314 | 0.19853004 | 0.39483855 | 0.5028132 | 0.61509561 | NA |
| GPBP1L1 | 46.2432361 | -0.0040883 | 0.27478251 | -0.0148784 | 0.98812923 | 0.99909657 |
| TMEM69 | 3.91722755 | -0.5010899 | 0.76654632 | -0.6536981 | 0.51330631 | NA |
| IPP | 11.3933915 | -0.3903889 | 0.57963669 | -0.6735061 | 0.50062535 | NA |
| MAST2 | 21.5446686 | 0.14000278 | 0.4368178 | 0.32050612 | 0.74858469 | NA |
| TSPAN1 | 24.4385514 | 0.75078236 | 0.65161099 | 1.15219413 | 0.24924131 | NA |
| POMGNT1 | 8.65592541 | 1.00441565 | 0.48967434 | 2.05119111 | 0.04024834 | NA |
| LURAP1 | 4.76653955 | 0.47571949 | 0.68478738 | 0.69469663 | 0.48724544 | NA |
| LRRC41 | 11.2498643 | -0.8109376 | 0.47610836 | -1.7032626 | 0.08851893 | NA |
| UQCRH | 47.2532246 | 0.22731876 | 0.32283408 | 0.70413495 | 0.48134874 | 0.86222804 |
| NSUN4 | 6.94205163 | 0.3731633 | 0.5744021 | 0.64965519 | 0.51591497 | NA |
| FAAH | 5.43970691 | -0.0677074 | 0.72667867 | -0.0931738 | 0.92576551 | NA |
| MKNK1 | 12.3972286 | 0.54517407 | 0.50656106 | 1.07622577 | 0.2818263 | NA |
| ATPAF1 | 10.0355094 | 0.28146328 | 0.53487218 | 0.52622532 | 0.59873166 | NA |
| EFCAB14 | 23.1052024 | 0.22606017 | 0.35235483 | 0.64156966 | 0.52115264 | NA |
| PDZK1IP1 | 15.5774118 | 0.90696366 | 0.65466534 | 1.38538517 | 0.16593472 | NA |
| CMPK1 | 20.0627571 | 0.03641546 | 0.36893702 | 0.09870373 | 0.9213735 | NA |
| TRABD2B | 19.8863943 | -0.1582671 | 0.71653052 | -0.2208798 | 0.82518603 | NA |
| SPATA6 | 24.1576135 | 1.04093929 | 0.46860723 | 2.22134708 | 0.02632746 | NA |
| AGBL4 | 22.8197694 | 0.68741944 | 0.60264546 | 1.14066974 | 0.25400739 | NA |
| FAF1 | 167.89764 | -0.0644597 | 0.20836161 | -0.3093647 | 0.75704409 | 0.95323728 |
| RNF11 | 29.3967569 | -0.1560058 | 0.38321287 | -0.4070996 | 0.68393484 | 0.92710517 |
| TTC39A | 7.52631926 | -0.4902183 | 0.60708356 | -0.8074972 | 0.41938008 | NA |
| EPS15 | 55.9329579 | 0.06478125 | 0.34588201 | 0.18729291 | 0.85143098 | 0.97168963 |
| OSBPL9 | 55.7311724 | 0.09619756 | 0.27906741 | 0.34471082 | 0.73031178 | 0.94309578 |
| NRDC | 52.7921187 | 0.04887041 | 0.31512487 | 0.15508267 | 0.87675614 | 0.9751578 |
| TXNDC12 | 8.96231281 | -0.3658506 | 0.49758089 | -0.7352585 | 0.46218211 | NA |
| BTF3L4 | 16.1960269 | -0.3212463 | 0.3657517 | -0.8783179 | 0.37977122 | NA |
| ZFYVE9 | 37.6530139 | -0.2555109 | 0.34440354 | -0.741894 | 0.45815154 | 0.85802171 |
| CC2D1B | 11.2708462 | -1.7608696 | 0.5600533 | -3.1441108 | 0.00166592 | NA |
| PRPF38A | 8.48545128 | 0.02129217 | 0.58821543 | 0.0361979 | 0.97112456 | NA |
| TUT4 | 52.5002935 | -0.2203383 | 0.24043155 | -0.9164285 | 0.35944221 | 0.80341474 |
| ZYG11B | 46.9006515 | -0.0021215 | 0.37118081 | -0.0057155 | 0.99543972 | 0.9992989 |
| ECHDC2 | 46.7762741 | 0.67269875 | 0.32341572 | 2.07998159 | 0.03752722 | 0.42032585 |
| SCP2 | 31.8374649 | 0.15039296 | 0.38786566 | 0.38774497 | 0.69820477 | 0.9303056 |
| CPT2 | 7.76521391 | 0.57035038 | 0.56841603 | 1.00340306 | 0.31566643 | NA |
| C1orf123 | 3.13587944 | 0.44076717 | 0.83990166 | 0.52478425 | 0.59973318 | NA |
| MAGOH | 8.59224102 | -0.3426905 | 0.58806901 | -0.5827385 | 0.56006933 | NA |
| NDC1 | 5.38692003 | 0.23054311 | 0.80041954 | 0.28802784 | 0.77332543 | NA |
| YIPF1 | 9.46835503 | 0.37061285 | 0.52098746 | 0.71136615 | 0.47685737 | NA |
| HSPB11 | 11.1165385 | 0.34736395 | 0.58974823 | 0.58900381 | 0.55585872 | NA |
| LRRC42 | 5.41645016 | 0.44191551 | 0.63398373 | 0.69704551 | 0.48577431 | NA |
| TMEM59 | 75.4172964 | 0.26094426 | 0.28115878 | 0.92810285 | 0.35335422 | 0.8003972 |
| TCEANC2 | 7.64755205 | 0.66034953 | 0.50735795 | 1.30154563 | 0.19307176 | NA |
| MRPL37 | 7.84722761 | 0.34735537 | 0.55313704 | 0.62797344 | 0.53002134 | NA |
| SSBP3 | 91.0302597 | 0.4487719 | 0.46039967 | 0.97474418 | 0.32968717 | 0.79273429 |
| ACOT11 | 68.855004 | -0.2227637 | 0.34364028 | -0.6482469 | 0.51682527 | 0.87158522 |
| MROH7-TTC4 | 3.64452128 | 0.55931287 | 1.21879146 | 0.45890777 | 0.6463004 | NA |
| DHCR24 | 9.64106386 | -1.8311697 | 0.84779134 | -2.1599298 | 0.03077811 | NA |
| TMEM61 | 6.93808135 | 0.77176448 | 0.80881741 | 0.95418875 | 0.33998811 | NA |
| BSND | 8.41197729 | 0.46197787 | 0.56132443 | 0.823014 | 0.41050003 | NA |
| USP24 | 59.1055506 | -0.4973148 | 0.28872916 | -1.7224265 | 0.08499228 | 0.55323508 |
| PLPP3 | 132.867475 | -1.1196635 | 0.32391642 | -3.4566434 | 0.00054695 | 0.09487868 |

| | | | | | | |
|---|---|---|---|---|---|---|
| PRKAA2 | 134.128712 | -0.4174708 | 0.24769523 | -1.6854212 | 0.09190734 | 0.56220784 |
| FYB2 | 89.3999666 | 0.6696686 | 0.43484528 | 1.54001582 | 0.1235565 | 0.58553957 |
| DAB1 | 22.7819941 | 0.61374331 | 0.70493241 | 0.87064136 | 0.38395001 | NA |
| TACSTD2 | 24.597894 | -0.3693922 | 0.50662678 | -0.7291209 | 0.46592769 | NA |
| MYSM1 | 28.5146126 | -0.5260255 | 0.32685117 | -1.6093731 | 0.10753479 | 0.57769936 |
| JUN | 23.0007283 | -0.4445869 | 0.68069709 | -0.6531348 | 0.51366939 | NA |
| LINC01135 | 7.96236431 | 2.20182091 | 0.71454239 | 3.08144198 | 0.00206001 | NA |
| FGGY | 33.4228056 | 0.45820416 | 0.40366785 | 1.13510194 | 0.25633261 | 0.75170774 |
| HOOK1 | 21.9322266 | -0.4426145 | 0.38917155 | -1.137325 | 0.25540247 | NA |
| NFIA | 27.0628149 | 0.62123406 | 0.55661594 | 1.11609104 | 0.26438317 | 0.75479335 |
| TM2D1 | 24.4951747 | -0.2391442 | 0.34436092 | -0.6944581 | 0.48739495 | NA |
| PATJ | 229.782002 | 0.13507212 | 0.17862378 | 0.7561822 | 0.44953997 | 0.85123446 |
| USP1 | 8.95606277 | 0.88022646 | 0.49693937 | 1.77129546 | 0.07651158 | NA |
| DOCK7 | 57.2903182 | 0.26617503 | 0.31168345 | 0.85399155 | 0.39310967 | 0.8258317 |
| ATG4C | 8.49109622 | 1.31780546 | 0.538581 | 2.44681014 | 0.01441267 | NA |
| ALG6 | 4.07100022 | 0.77434881 | 0.88545952 | 0.87451633 | 0.38183713 | NA |
| ITGB3BP | 14.9937834 | 1.19016081 | 0.60760453 | 1.9587754 | 0.0501391 | NA |
| EFCAB7 | 6.6612814 | 1.00070709 | 0.67885329 | 1.47411394 | 0.14045092 | NA |
| PGM1 | 19.6972989 | -0.0174639 | 0.40272155 | -0.0433648 | 0.96541072 | NA |
| RAVER2 | 15.8796361 | 0.36081112 | 0.60654908 | 0.59485891 | 0.55193779 | NA |
| JAK1 | 62.5995345 | 0.1182641 | 0.32844749 | 0.36007003 | 0.71879477 | 0.93714576 |
| AK4 | 7.49937502 | -0.6287338 | 0.92883092 | -0.6769088 | 0.49846384 | NA |
| DNAJC6 | 8.4923085 | -0.1893635 | 0.6851381 | -0.2763874 | 0.78225056 | NA |
| LEPR | 8.84306017 | -0.4485064 | 0.59963869 | -0.747961 | 0.45448368 | NA |
| SGIP1 | 14.8304965 | 0.23621542 | 0.85803051 | 0.27529956 | 0.7830861 | NA |
| WDR78 | 5.96539376 | 0.40073063 | 0.68203108 | 0.58755479 | 0.55683117 | NA |
| MIER1 | 40.2575967 | -0.135017 | 0.30666912 | -0.4402692 | 0.65974217 | 0.92276785 |
| SLC35D1 | 11.9163777 | 0.19572039 | 0.43974892 | 0.44507304 | 0.65626697 | NA |
| SERBP1 | 35.6841033 | 0.15254145 | 0.31631838 | 0.48224022 | 0.62963531 | 0.91751762 |
| GADD45A | 39.9071725 | -0.5634226 | 0.5517556 | -1.0211452 | 0.30718563 | 0.77729327 |
| GNG12 | 44.1988607 | 0.2548992 | 0.27882607 | 0.91418712 | 0.36061853 | 0.80341474 |
| GNG12-AS1 | 10.5410184 | 0.44440398 | 0.73145581 | 0.60756094 | 0.54347871 | NA |
| WLS | 14.7198618 | 0.68547707 | 0.58079085 | 1.18024771 | 0.23790171 | NA |
| LRRC40 | 11.1902936 | -0.5751816 | 0.52132857 | -1.1032996 | 0.26989709 | NA |
| SRSF11 | 65.6380516 | -0.0134787 | 0.22937482 | -0.058763 | 0.9531409 | 0.99488973 |
| ANKRD13C | 27.064331 | 0.29421961 | 0.39226714 | 0.75004909 | 0.45322514 | 0.85557861 |
| PTGER3 | 236.048233 | 0.18985566 | 0.25662143 | 0.73982776 | 0.45940451 | 0.85844137 |
| ZRANB2-AS1 | 4.43175958 | 1.46573874 | 0.68542272 | 2.13844492 | 0.03248065 | NA |
| ZRANB2 | 46.7867459 | -0.5977587 | 0.2851888 | -2.0960103 | 0.03608128 | 0.41688714 |
| NEGR1 | 6.26473127 | -0.2514932 | 0.74435745 | -0.3378661 | 0.73546406 | NA |
| CRYZ | 25.9344482 | -0.2404597 | 0.51459252 | -0.4672818 | 0.64029831 | NA |
| TYW3 | 7.48738391 | 0.62588395 | 0.55093237 | 1.13604497 | 0.25593775 | NA |
| ACADM | 33.9908064 | 0.129868 | 0.29522181 | 0.43989975 | 0.66000971 | 0.92276785 |
| RABGGTB | 8.88349186 | -0.0283163 | 0.53082028 | -0.0533444 | 0.95745747 | NA |
| PIGK | 10.9990447 | 0.41042402 | 0.45874087 | 0.89467507 | 0.37096081 | NA |
| AC118549.1 | 73.9329983 | -0.1136331 | 0.27688845 | -0.4103931 | 0.68151758 | 0.9267216 |
| USP33 | 46.8659719 | 0.18914766 | 0.29832267 | 0.63403715 | 0.52605658 | 0.87857739 |
| MIGA1 | 27.4872931 | 0.31519776 | 0.38586901 | 0.81685169 | 0.41401319 | 0.83459305 |
| NEXN | 12.9637261 | -0.5559026 | 0.63339452 | -0.8776562 | 0.38013033 | NA |
| FUBP1 | 30.2455406 | -0.2749072 | 0.32645106 | -0.8421086 | 0.39972719 | 0.8269015 |
| GIPC2 | 8.74157856 | 0.75131551 | 0.64967969 | 1.15643988 | 0.24750128 | NA |
| PTGFR | 41.0742135 | 1.16305737 | 0.48853696 | 2.38069473 | 0.01728003 | 0.32722884 |
| IFI44L | 17.0126481 | -2.6997274 | 2.18046116 | -1.2381451 | 0.21566225 | NA |
| IFI44 | 7.2172313 | 1.1981285 | 1.60953655 | 0.74439348 | 0.45663847 | NA |

| | | | | | | |
|---|---|---|---|---|---|---|
| ADGRL2 | 53.6847914 | 0.1810311 | 0.35135396 | 0.51523853 | 0.60638635 | 0.90821838 |
| LINC01725 | 40.6256221 | 0.4545525 | 0.49395445 | 0.92023162 | 0.35745173 | 0.80238744 |
| PRKACB | 30.0769293 | 0.36447142 | 0.30924523 | 1.17858383 | 0.23856393 | 0.73271248 |
| RPF1 | 5.68840153 | -0.7557288 | 0.61936775 | -1.2201617 | 0.22240359 | NA |
| GNG5 | 20.6430323 | -0.0163876 | 0.36388928 | -0.0450346 | 0.96407974 | NA |
| SPATA1 | 3.5941299 | 2.10091241 | 0.85512032 | 2.45686177 | 0.01401566 | NA |
| CTBS | 9.22464345 | 0.30851241 | 0.58046799 | 0.5314891 | 0.59507989 | NA |
| SSX2IP | 6.84472614 | -0.1085461 | 0.7032465 | -0.15435 | 0.87733378 | NA |
| SYDE2 | 7.40900548 | -0.4018753 | 0.58530055 | -0.6866136 | 0.49232625 | NA |
| C1orf52 | 7.68927261 | 0.31394953 | 0.56502839 | 0.55563497 | 0.57846042 | NA |
| BCL10 | 7.87357043 | -0.5379455 | 0.52915178 | -1.0166184 | 0.30933498 | NA |
| AL078459.1 | 10.5789255 | 0.49475014 | 0.66597165 | 0.74289971 | 0.4575424 | NA |
| DDAH1 | 55.8990778 | -0.4054119 | 0.3627937 | -1.1174723 | 0.26379244 | 0.75479335 |
| ZNHIT6 | 14.6329125 | -0.2991877 | 0.42872161 | -0.6978601 | 0.48526472 | NA |
| COL24A1 | 10.1004571 | 0.36547845 | 0.6220194 | 0.5875676 | 0.55682256 | NA |
| ODF2L | 21.363046 | -0.1880532 | 0.34446962 | -0.5459211 | 0.58512018 | NA |
| SH3GLB1 | 36.0972986 | -0.0242289 | 0.3799469 | -0.0637692 | 0.94915403 | 0.99344789 |
| SELENOF | 34.0056688 | -0.0652954 | 0.28700216 | -0.2275083 | 0.82002848 | 0.96845695 |
| HS2ST1 | 20.2711187 | -0.5724571 | 0.46818343 | -1.2227196 | 0.22143561 | NA |
| LMO4 | 8.19754697 | 0.54303443 | 0.5587513 | 0.97187144 | 0.33111451 | NA |
| PKN2-AS1 | 3.86404893 | 0.61826984 | 0.87415266 | 0.70727902 | 0.4793931 | NA |
| PKN2 | 97.2211809 | -0.0440562 | 0.20948258 | -0.2103095 | 0.83342609 | 0.97154892 |
| GTF2B | 11.7586814 | -0.6639271 | 0.46160345 | -1.4383063 | 0.15034717 | NA |
| KYAT3 | 15.325972 | -0.0507945 | 0.41986941 | -0.1209769 | 0.90370931 | NA |
| GBP3 | 4.17345648 | -0.4143511 | 0.740737 | -0.5593769 | 0.57590455 | NA |
| GBP2 | 13.3997314 | 0.73902977 | 0.44707645 | 1.65302774 | 9.83E-02 | NA |
| LRRC8B | 11.0657585 | -0.7450282 | 0.56656453 | -1.3149926 | 0.18851238 | NA |
| LRRC8D | 2.45160779 | 0.6577117 | 0.88855834 | 0.74020092 | 0.45917809 | NA |
| ZNF326 | 19.8085011 | -0.3789529 | 0.41039693 | -0.9233814 | 0.35580846 | NA |
| ZNF644 | 92.9605603 | -0.1557969 | 0.25531222 | -0.610221 | 0.54171542 | 0.88410237 |
| HFM1 | 9.15923891 | 0.42171918 | 0.52456148 | 0.80394614 | 0.42142808 | NA |
| TGFBR3 | 20.7587341 | 1.16342563 | 0.47631458 | 2.44255725 | 0.01458362 | NA |
| BTBD8 | 8.78816636 | 1.06378135 | 0.66495545 | 1.59977839 | 0.10964775 | NA |
| GLMN | 8.68900754 | 0.12719976 | 0.57702574 | 0.22044036 | 0.82552821 | NA |
| RPAP2 | 33.6579149 | 0.3301054 | 0.37696665 | 0.87568861 | 0.38119934 | 0.82153751 |
| EVI5 | 60.4932717 | -0.2346322 | 0.26361719 | -0.8900488 | 0.37343966 | 0.81364119 |
| RPL5 | 50.0698162 | 0.39081347 | 0.36961861 | 1.05734249 | 0.29035531 | 0.77286135 |
| FAM69A | 6.32626887 | 0.18041999 | 0.56187899 | 0.32110116 | 0.74813373 | NA |
| MTF2 | 38.0261081 | -0.121823 | 0.32037758 | -0.3802483 | 0.70376114 | 0.9303056 |
| TMED5 | 25.6423882 | 0.1644011 | 0.39061051 | 0.42088243 | 0.67384093 | NA |
| CCDC18 | 6.62138975 | 0.05514435 | 0.63472286 | 0.08687941 | 0.93076737 | NA |
| DR1 | 7.20991955 | 0.38747797 | 0.60096323 | 0.64476152 | 0.51908174 | NA |
| FNBP1L | 125.517438 | -0.2291493 | 0.24312628 | -0.9425115 | 0.34593084 | 0.8003972 |
| BCAR3 | 15.5989042 | 0.02810172 | 0.44773932 | 0.06276358 | 0.94995477 | NA |
| AL109613.1 | 2.35584139 | 1.53303961 | 1.13507119 | 1.35061097 | 0.17682008 | NA |
| DNTTIP2 | 5.81735967 | -1.1874038 | 0.64908638 | -1.8293463 | 0.06734774 | NA |
| GCLM | 14.5895356 | -0.3775368 | 0.51672301 | -0.7306366 | 0.46500113 | NA |
| ARHGAP29 | 31.694049 | 0.39097507 | 0.64169434 | 0.60928553 | 0.5423352 | 0.88410237 |
| ABCD3 | 69.614244 | 0.16038264 | 0.28553873 | 0.56168438 | 0.57433108 | 0.89263108 |
| SLC44A3 | 30.0249934 | 1.4621344 | 0.44937274 | 3.25372295 | 0.00113903 | 0.11692613 |
| CNN3 | 62.1015379 | -0.0194107 | 0.39699337 | -0.0488943 | 0.96100355 | 0.99656976 |
| AC105942.1 | 4.11739231 | 0.66349875 | 0.82793853 | 0.80138649 | 0.42290793 | NA |
| ALG14 | 9.34033901 | 1.5753227 | 0.5476699 | 2.8764091 | 0.00402228 | NA |
| PTBP2 | 61.1345907 | 0.16994956 | 0.30807616 | 0.55164789 | 0.58118962 | 0.89263108 |

| | | | | | | |
|---|---|---|---|---|---|---|
| DPYD | 33.5495378 | 0.82397691 | 0.37493159 | 2.19767266 | 0.02797244 | 0.39934064 |
| SNX7 | 16.1631899 | 0.11392748 | 0.41306307 | 0.27581135 | 0.78269296 | NA |
| FRRS1 | 14.3614561 | 0.18406745 | 0.52887772 | 0.34803403 | 0.72781462 | NA |
| AGL | 29.0857452 | 0.25016354 | 0.32099605 | 0.77933526 | 0.43578225 | 0.84282814 |
| SASS6 | 3.99497719 | 0.05746908 | 0.79697786 | 0.07210876 | 0.94251536 | NA |
| TRMT13 | 14.1676924 | -0.0409196 | 0.40475695 | -0.1010967 | 0.91947366 | NA |
| DBT | 13.8295283 | 1.02803068 | 0.4235867 | 2.42696638 | 0.01522567 | NA |
| RTCA | 13.6043605 | 0.01190047 | 0.49425002 | 0.02407783 | 0.98079052 | NA |
| CDC14A | 41.7238601 | -0.6579049 | 0.46081657 | -1.4276936 | 0.15338006 | 0.63864841 |
| EXTL2 | 11.671544 | 1.0297573 | 0.50180855 | 2.05209198 | 0.04016072 | NA |
| SLC30A7 | 22.221015 | 0.07556277 | 0.3750382 | 0.2014802 | 0.84032311 | NA |
| DPH5 | 5.46454508 | 1.47751086 | 0.78791599 | 1.8752137 | 0.06076333 | NA |
| AC093157.1 | 6.67353513 | 0.96818702 | 0.58390502 | 1.65812417 | 0.09729241 | NA |
| RNPC3 | 28.5654681 | 0.7829954 | 0.37133045 | 2.10862161 | 0.03497726 | 0.41688714 |
| AMY2B | 5.80507652 | 0.49211109 | 0.75851469 | 0.64878254 | 0.51647895 | NA |
| VAV3 | 72.8549891 | 0.55647149 | 0.36910917 | 1.5076068 | 0.13165519 | 0.60375343 |
| SLC25A24 | 14.5018186 | 0.38334898 | 0.39926525 | 0.96013611 | 0.33698672 | NA |
| FAM102B | 3.85162111 | -0.1045905 | 0.82919005 | -0.1261357 | 0.89962449 | NA |
| PRPF38B | 23.2401802 | -0.2792335 | 0.47163599 | -0.592053 | 0.55381507 | NA |
| STXBP3 | 37.1233591 | 0.26417204 | 0.30845575 | 0.85643413 | 0.39175769 | 0.8258317 |
| AL449266.1 | 9.5792412 | -1.2047436 | 0.72646526 | -1.6583637 | 0.09724408 | NA |
| GPSM2 | 81.1584985 | -1.3556248 | 0.53746688 | -2.522248 | 0.01166075 | 0.28665769 |
| WDR47 | 5.32513354 | -0.6528182 | 0.64819961 | -1.0071253 | 0.31387457 | NA |
| TAF13 | 6.89022615 | -0.9169579 | 0.55290309 | -1.6584424 | 0.0972282 | NA |
| TMEM167B | 6.30541706 | 0.33826605 | 0.56974467 | 0.59371516 | 0.55270264 | NA |
| SARS | 16.8933823 | -0.1553937 | 0.46430448 | -0.3346805 | 0.7378661 | NA |
| CELSR2 | 3.05521975 | 0.04790543 | 0.94447141 | 0.05072195 | 0.95954708 | NA |
| SORT1 | 36.1619947 | 0.13046197 | 0.27170796 | 0.48015511 | 0.6311171 | 0.9183778 |
| PSMA5 | 16.5863685 | -0.5055529 | 0.41741078 | -1.211164 | 0.22583255 | NA |
| SYPL2 | 3.50609574 | 0.18886476 | 0.79636697 | 0.23715795 | 0.81253425 | NA |
| CYB561D1 | 2.45460171 | 0.40872749 | 0.94893711 | 0.43072137 | 0.66667098 | NA |
| GNAI3 | 24.8052938 | -0.5699625 | 0.34907489 | -1.63278 | 0.10251527 | NA |
| GSTM4 | 3.96434353 | 0.79519439 | 0.78902093 | 1.0078242 | 0.31353886 | NA |
| GSTM3 | 7.48362844 | -0.7518017 | 0.75991148 | -0.989328 | 0.32250268 | NA |
| LINC01768 | 4.4641826 | -0.6640332 | 1.20588291 | -0.5506615 | 0.58186577 | NA |
| AHCYL1 | 29.5204647 | 0.21748109 | 0.33397953 | 0.6511809 | 0.51492972 | 0.87158522 |
| STRIP1 | 9.3420854 | -0.3461636 | 0.5920744 | -0.5846623 | 0.5587748 | NA |
| KCNC4 | 9.23078464 | 0.68722827 | 0.59612115 | 1.15283322 | 0.24897885 | NA |
| RBM15 | 8.09188769 | -0.7099202 | 0.58356275 | -1.2165276 | 0.223784 | NA |
| SLC16A4 | 6.84435346 | -0.674721 | 0.74799654 | -0.9020376 | 0.36703688 | NA |
| LAMTOR5 | 29.2076422 | -0.0032975 | 0.34430464 | -0.0095773 | 0.99235856 | 0.99909657 |
| LRIF1 | 5.56658965 | -0.5468296 | 0.6652965 | -0.8219337 | 0.41111462 | NA |
| DRAM2 | 13.1416122 | 1.03322682 | 0.53003888 | 1.94934156 | 0.05125465 | NA |
| CEPT1 | 20.6400686 | 0.1391606 | 0.36326328 | 0.38308468 | 0.70165698 | NA |
| DENND2D | 8.25127771 | 1.339851 | 0.64683442 | 2.07139721 | 0.03832169 | NA |
| WDR77 | 3.50352803 | -0.2375556 | 0.8478664 | -0.2801805 | 0.77933902 | NA |
| ATP5PB | 28.3279379 | 0.21913446 | 0.37280819 | 0.58779411 | 0.5566705 | 0.89234448 |
| RAP1A | 59.7319056 | -0.2757638 | 0.35926835 | -0.7675705 | 0.44274241 | 0.84768974 |
| INKA2 | 3.90313405 | 0.01467909 | 0.84044554 | 0.01746585 | 0.98606498 | NA |
| DDX20 | 4.34419992 | 0.07254661 | 0.78321542 | 0.09262663 | 0.92620018 | NA |
| CTTNBP2NL | 16.3709111 | 0.16024942 | 0.40350557 | 0.39714303 | 0.69126199 | NA |
| ST7L | 17.7789456 | -0.2502303 | 0.40411912 | -0.6191994 | 0.535785 | NA |
| CAPZA1 | 28.9279334 | -0.0741175 | 0.35220459 | -0.2104389 | 0.83332516 | 0.97154892 |
| MOV10 | 10.4091948 | 0.26091316 | 0.50231543 | 0.51942096 | 0.60346722 | NA |

| | | | | | | |
|---|---|---|---|---|---|---|
| RHOC | 12.531206 | -0.7636606 | 0.46730671 | -1.6341742 | 0.10222228 | NA |
| SLC16A1 | 8.27262134 | 0.18768184 | 0.87710569 | 0.21397859 | 0.83056375 | NA |
| SLC16A1-AS1 | 5.61710748 | -0.0423515 | 0.82588982 | -0.0512798 | 0.95910256 | NA |
| LRIG2 | 16.9929007 | 0.36860715 | 0.39633556 | 0.93003804 | 0.35235139 | NA |
| MAGI3 | 445.880806 | -0.2758791 | 0.25390652 | -1.0865379 | 0.27724106 | 0.76520827 |
| PHTF1 | 6.58338611 | -0.4339057 | 0.65428093 | -0.6631796 | 0.50721556 | NA |
| RSBN1 | 18.2644526 | 0.28260779 | 0.39725877 | 0.71139472 | 0.47683967 | NA |
| HIPK1 | 17.0460867 | 0.22808528 | 0.42408256 | 0.53783226 | 0.59069286 | NA |
| TRIM33 | 54.9414603 | -0.2707731 | 0.28730875 | -0.9424465 | 0.34596407 | 0.8003972 |
| BCAS2 | 6.18956574 | 0.08168282 | 0.74036561 | 0.11032767 | 0.91214951 | NA |
| NRAS | 7.12043502 | -0.714648 | 0.63163672 | -1.1314225 | 0.25787732 | NA |
| CSDE1 | 47.2450461 | 0.15524959 | 0.30520368 | 0.50867536 | 0.6109798 | 0.90955758 |
| SIKE1 | 6.95636153 | 0.16364841 | 0.55132025 | 0.29683003 | 0.76659627 | NA |
| VANGL1 | 2.75072902 | 1.14195641 | 0.89048638 | 1.28239627 | 0.19970367 | NA |
| SLC22A15 | 25.9527806 | 0.00819886 | 0.44731583 | 0.01832902 | 0.98537638 | 0.99902646 |
| MAB21L3 | 4.39376195 | 0.2016334 | 0.99731574 | 0.20217609 | 0.83977906 | NA |
| ATP1A1 | 160.94428 | -0.4775333 | 0.357396 | -1.3361462 | 0.18150148 | 0.67460052 |
| LINC01762 | 14.5453304 | 0.56464564 | 0.83923262 | 0.67281183 | 0.50106701 | NA |
| CD58 | 9.01037489 | 0.1999334 | 0.46422701 | 0.43068023 | 0.66670089 | NA |
| IGSF3 | 7.87457371 | 0.43084139 | 0.52038983 | 0.82792047 | 0.40771554 | NA |
| PTGFRN | 3.88597181 | -0.1472686 | 0.72918041 | -0.2019645 | 0.83994447 | NA |
| TTF2 | 5.06040552 | -0.1309038 | 0.67482621 | -0.1939814 | 0.84619042 | NA |
| MAN1A2 | 47.449435 | -0.1642726 | 0.30430409 | -0.5398305 | 0.58931392 | 0.89534694 |
| GDAP2 | 15.877594 | 0.29685506 | 0.41849383 | 0.70934154 | 0.47811255 | NA |
| WDR3 | 3.55456351 | 0.01256498 | 0.76950415 | 0.01632867 | 0.98697219 | NA |
| SPAG17 | 7.1944319 | 0.40268772 | 0.6061873 | 0.66429588 | 0.50650096 | NA |
| WARS2 | 14.7556258 | -0.0149399 | 0.53294374 | -0.0280328 | 0.97763599 | NA |
| AL359915.2 | 28.1552496 | 0.37700706 | 0.45504474 | 0.82850549 | 0.40738429 | 0.82808596 |
| HMGCS2 | 4.20145821 | 0.35447325 | 1.31320526 | 0.26992981 | 0.78721425 | NA |
| NOTCH2 | 4.23651761 | 0.40018184 | 0.74441169 | 0.53758136 | 0.5908661 | NA |
| SEC22B | 16.815891 | -0.4366853 | 0.48288225 | -0.9043309 | 0.36581999 | NA |
| LINC00623 | 21.0299182 | 0.2285153 | 0.36548093 | 0.62524548 | 0.53180996 | NA |
| AC239800.3 | 3.38071852 | 0.003388 | 0.80436831 | 0.004212 | 0.99663932 | NA |
| NBPF15 | 2.85540574 | -0.4755468 | 0.83232968 | -0.5713443 | 0.5677663 | NA |
| FP700111.1 | 5.60450312 | -0.0782286 | 0.63303164 | -0.1235778 | 0.90164959 | NA |
| SRGAP2B | 5.28998495 | 0.45896477 | 0.72069091 | 0.63683996 | 0.5242291 | NA |
| GPR89A | 6.33983934 | 1.19840973 | 0.69966574 | 1.71283181 | 0.08674349 | NA |
| PDZK1 | 13.2097149 | 0.40122747 | 0.61323262 | 0.65428266 | 0.51292972 | NA |
| RNF115 | 53.2239185 | -0.2518013 | 0.31617247 | -0.7964048 | 0.42579681 | 0.84057082 |
| POLR3C | 6.73854414 | -0.4805904 | 0.65622374 | -0.7323576 | 0.46395035 | NA |
| PEX11B | 3.63152885 | -0.3518483 | 0.7098149 | -0.4956903 | 0.62011294 | NA |
| RBM8A | 12.6342093 | 0.14174692 | 0.43191988 | 0.32817872 | 0.74277653 | NA |
| TXNIP | 131.513475 | 1.58921523 | 0.54144855 | 2.93511774 | 0.00333421 | 0.20225024 |
| NBPF12 | 8.32364658 | 0.17448484 | 0.6466467 | 0.26983024 | 0.78729086 | NA |
| PRKAB2 | 6.66260746 | 0.7111615 | 0.61526122 | 1.15586922 | 0.24773466 | NA |
| AC242426.2 | 11.1382645 | 0.40572472 | 0.47450663 | 0.85504543 | 0.392526 | NA |
| FMO5 | 5.24743552 | -0.0461719 | 0.72121775 | -0.0640194 | 0.94895477 | NA |
| CHD1L | 15.9397841 | 0.49196156 | 0.44995951 | 1.09334628 | 0.27424179 | NA |
| BCL9 | 18.8351334 | 0.87112778 | 0.44942958 | 1.93829648 | 0.05258706 | NA |
| ACP6 | 8.40793189 | 0.77040258 | 0.61655805 | 1.24952157 | 0.21147437 | NA |
| GPR89B | 4.68173911 | 0.24080462 | 0.6887065 | 0.34964766 | 0.72660314 | NA |
| NBPF11 | 10.7249121 | 0.27446293 | 0.53081742 | 0.51705712 | 0.60511628 | NA |
| AC239809.3 | 4.47934969 | 0.86147039 | 0.96560092 | 0.89215987 | 0.37230725 | NA |
| PDE4DIP | 12.7015051 | 0.27563404 | 0.55064229 | 0.50056823 | 0.61667503 | NA |

| | | | | | | |
|---|---|---|---|---|---|---|
| NBPF9 | 2.62099155 | 0.1350352 | 0.89526463 | 0.15083272 | 0.88010768 | NA |
| AC245297.3 | 25.5164622 | -0.0039927 | 0.52002781 | -0.0076779 | 0.99387402 | NA |
| NBPF19 | 3.51621651 | 0.54158321 | 0.91633616 | 0.59103114 | 0.55449956 | NA |
| SF3B4 | 4.85083205 | -1.5902874 | 0.69437435 | -2.290245 | 0.02200712 | NA |
| OTUD7B | 27.589641 | -0.4244503 | 0.32407434 | -1.3097312 | 0.19028678 | 0.68499719 |
| VPS45 | 25.7434821 | 0.13971844 | 0.35139891 | 0.39760635 | 0.69092038 | NA |
| ANP32E | 6.44006026 | 0.48143851 | 0.73344101 | 0.65641067 | 0.51155992 | NA |
| APH1A | 10.7888951 | 0.1759709 | 0.46953922 | 0.37477358 | 0.70782886 | NA |
| MRPS21 | 17.0624974 | 0.42944344 | 0.37368786 | 1.1492036 | 0.25047204 | NA |
| PRPF3 | 13.5210641 | -1.7566434 | 0.49216616 | -3.569208 | 0.00035806 | NA |
| RPRD2 | 47.4625939 | -0.0797081 | 0.30298971 | -0.2630721 | 0.79249501 | 0.96341299 |
| TARS2 | 3.18079107 | 1.15253475 | 0.9352239 | 1.23236239 | 0.21781374 | NA |
| MCL1 | 22.5026234 | -0.0968345 | 0.34646066 | -0.2794963 | 0.779864 | NA |
| ENSA | 33.1624806 | -0.2128205 | 0.32735717 | -0.6501171 | 0.51561659 | 0.87158522 |
| GOLPH3L | 13.4643177 | 0.70752877 | 0.46864174 | 1.50974341 | 0.13110891 | NA |
| ARNT | 20.2286633 | -0.2593489 | 0.38099835 | -0.6807089 | 0.49605573 | NA |
| SETDB1 | 13.6285938 | -0.4070491 | 0.46879496 | -0.8682881 | 0.38523662 | NA |
| CERS2 | 18.7856992 | 0.1183982 | 0.51257444 | 0.23098733 | 0.81732464 | NA |
| MINDY1 | 39.5490357 | -0.1092194 | 0.27972226 | -0.3904565 | 0.69619904 | 0.9303056 |
| PRUNE1 | 18.6015937 | -0.3574111 | 0.47608873 | -0.7507237 | 0.45281894 | NA |
| BNIPL | 2.4141187 | 0.00380185 | 1.02800632 | 0.00369828 | 0.99704921 | NA |
| C1orf56 | 5.69528546 | -0.4245546 | 0.75589257 | -0.5616599 | 0.57434774 | NA |
| CDC42SE1 | 10.1055133 | -0.4180947 | 0.51892184 | -0.8056988 | 0.42041652 | NA |
| GABPB2 | 18.6016177 | 0.79528365 | 0.47812199 | 1.66334883 | 0.09624263 | NA |
| SCNM1 | 4.47663032 | 0.00428081 | 0.70428888 | 0.0060782 | 0.99515033 | NA |
| VPS72 | 5.2012367 | 0.50452207 | 0.69921245 | 0.72155761 | 0.47056651 | NA |
| PIP5K1A | 62.9660544 | -0.4602126 | 0.30739003 | -1.4971616 | 0.13435121 | 0.60594834 |
| PSMD4 | 13.7138631 | 0.04291376 | 0.42930794 | 0.09996032 | 0.92037583 | NA |
| PI4KB | 16.8385287 | -0.7030757 | 0.42693543 | -1.6467963 | 0.09959992 | NA |
| SELENBP1 | 2.99604798 | -0.0258002 | 1.34909699 | -0.019124 | 0.98474215 | NA |
| PSMB4 | 9.32701413 | 0.29464856 | 0.50721692 | 0.58091232 | 0.56129955 | NA |
| POGZ | 71.0572376 | 0.17915301 | 0.28763897 | 0.62283982 | 0.53338979 | 0.88254911 |
| CGN | 9.94790389 | 1.52859013 | 0.55869794 | 2.73598669 | 0.00621935 | NA |
| TUFT1 | 10.8987798 | 0.3742173 | 0.50585582 | 0.73977068 | 0.45943915 | NA |
| SNX27 | 19.9808189 | 0.55774331 | 0.34380802 | 1.62225221 | 0.10474936 | NA |
| MRPL9 | 3.41160135 | 0.14649396 | 0.72371375 | 0.20241975 | 0.83958859 | NA |
| RORC | 7.64986997 | 0.07158836 | 0.75307651 | 0.0950612 | 0.92426621 | NA |
| THEM4 | 24.2732821 | -0.0914011 | 0.35910853 | -0.2545222 | 0.79909213 | NA |
| S100A10 | 35.9419145 | -1.0286897 | 0.51370188 | -2.0025032 | 0.04523064 | 0.44887658 |
| S100A11 | 105.722708 | -0.3349735 | 0.34604566 | -0.9680036 | 3.33E-01 | 0.79447068 |
| FLG-AS1 | 16.3034379 | 0.28263019 | 0.55618479 | 0.50815879 | 0.61134199 | NA |
| S100A6 | 239.089826 | -0.3929947 | 0.46733486 | -0.8409273 | 0.40038865 | 0.8269015 |
| S100A4 | 14.9068041 | 0.54624929 | 0.74096103 | 0.7372173 | 0.46099022 | NA |
| S100A2 | 46.8092088 | -0.2150341 | 0.54255402 | -0.3963368 | 0.69185661 | 0.92932325 |
| S100A16 | 7.65932643 | 0.17780754 | 0.76315669 | 0.23298955 | 0.81576952 | NA |
| S100A14 | 4.92898366 | -0.7140733 | 0.66508501 | -1.0736572 | 0.28297637 | NA |
| S100A13 | 29.0839781 | 0.11448136 | 0.43506501 | 0.2631362 | 0.7924456 | 0.96341299 |
| CHTOP | 14.9404301 | 3.44E-05 | 0.45161846 | 7.62E-05 | 0.9999392 | NA |
| ILF2 | 12.3498175 | -0.3185712 | 0.46161799 | -0.6901187 | 0.49011953 | NA |
| INTS3 | 10.1387958 | -0.0108933 | 0.48313578 | -0.0225471 | 0.98201154 | NA |
| GATAD2B | 64.2470282 | -0.2141972 | 0.29585094 | -0.7240038 | 0.46906338 | 0.85973853 |
| DENND4B | 7.80859874 | -1.0528106 | 0.54912344 | -1.9172568 | 0.05520532 | NA |
| CRTC2 | 5.29398269 | 0.1788336 | 0.647722 | 0.27609622 | 0.78247416 | NA |
| SLC39A1 | 11.7234454 | -0.2359801 | 0.60317052 | -0.3912329 | 0.69562512 | NA |

| | | | | | | |
|---|---|---|---|---|---|---|
| JTB | 14.7237906 | -0.0114779 | 0.46008329 | -0.0249475 | 0.98009682 | NA |
| RAB13 | 17.34601 | 0.04408556 | 0.41727717 | 0.10565054 | 0.91585963 | NA |
| RPS27 | 24.9706504 | 0.14482723 | 0.41066367 | 0.35266628 | 0.72433864 | NA |
| TPM3 | 24.7897559 | -0.382728 | 0.35948925 | -1.0646438 | 0.28703713 | NA |
| C1orf43 | 19.6766307 | -0.3775722 | 0.4233724 | -0.8918206 | 0.3724891 | NA |
| UBAP2L | 22.3872437 | -0.6505538 | 0.39219859 | -1.6587357 | 0.09716907 | NA |
| HAX1 | 5.85475079 | 0.72585796 | 0.67495302 | 1.07541998 | 0.28218674 | NA |
| IL6R | 13.3639151 | 0.29814475 | 0.46780267 | 0.63733017 | 0.52390981 | NA |
| SHE | 7.36639744 | -1.0193357 | 0.63302013 | -1.6102737 | 0.10733811 | NA |
| TDRD10 | 8.14584227 | 0.47512193 | 0.66256574 | 0.71709403 | 0.47331608 | NA |
| UBE2Q1 | 8.87860808 | -0.4543197 | 0.53514207 | -0.8489702 | 0.39589786 | NA |
| ADAR | 35.198776 | -0.7222518 | 0.39840723 | -1.8128481 | 0.06985526 | 0.52654172 |
| PMVK | 12.1062766 | 0.20127088 | 0.49251693 | 0.40865779 | 0.68279082 | NA |
| PBXIP1 | 7.4592048 | 0.29207234 | 0.57042371 | 0.512027 | 0.60863211 | NA |
| PYGO2 | 4.29827375 | 0.13895414 | 0.73042767 | 0.19023669 | 0.84912366 | NA |
| FLAD1 | 3.47016839 | 0.39885485 | 0.77310377 | 0.51591372 | 0.60591467 | NA |
| ZBTB7B | 3.12279648 | 0.46133678 | 0.84369383 | 0.54680592 | 0.58451208 | NA |
| EFNA1 | 27.0915782 | -0.7070877 | 0.45910598 | -1.5401405 | 0.12352611 | 0.58553957 |
| SLC50A1 | 3.35122054 | -0.2456795 | 0.75620669 | -0.3248841 | 0.74526879 | NA |
| DPM3 | 10.0190122 | 0.49802988 | 0.51797865 | 0.96148726 | 0.33630723 | NA |
| MUC1 | 4.02760093 | 0.12301775 | 0.70426676 | 0.17467493 | 0.86133507 | NA |
| MTX1 | 3.31210814 | 0.32393797 | 0.83660508 | 0.38720536 | 0.69860418 | NA |
| GBA | 4.22222248 | -0.9770243 | 0.78582553 | -1.2433094 | 0.21375384 | NA |
| SCAMP3 | 7.70511781 | 0.42894527 | 0.5410037 | 0.79286939 | 0.42785392 | NA |
| CLK2 | 4.76002072 | -0.5567675 | 0.66563738 | -0.8364426 | 0.40290593 | NA |
| FDPS | 16.3900668 | 0.4504532 | 0.44225076 | 1.01854704 | 0.30841806 | NA |
| RUSC1 | 3.20588292 | 0.3140754 | 0.78468926 | 0.40025449 | 0.68896909 | NA |
| ASH1L | 122.943015 | -0.1039833 | 0.18576046 | -0.5597712 | 0.57563554 | 0.89263108 |
| YY1AP1 | 19.4796845 | -0.5162133 | 0.36980896 | -1.3958919 | 0.16274706 | NA |
| DAP3 | 24.9075879 | -0.2166314 | 0.35826745 | -0.604664 | 0.54540226 | NA |
| GON4L | 25.3049105 | -0.4924717 | 0.40772581 | -1.2078501 | 0.22710493 | NA |
| RIT1 | 8.29650591 | -0.9729535 | 0.66507301 | -1.4629274 | 0.14348725 | NA |
| KHDC4 | 21.4595552 | -0.2563656 | 0.36261977 | -0.7069817 | 0.47957782 | NA |
| ARHGEF2 | 5.86822543 | 0.29456426 | 0.63804975 | 0.46166348 | 0.64432267 | NA |
| SSR2 | 15.5974131 | -0.5216824 | 0.42492623 | -1.227701 | 0.21955924 | NA |
| UBQLN4 | 4.58924327 | -0.2655024 | 0.74604738 | -0.3558787 | 0.72193141 | NA |
| LAMTOR2 | 6.85681877 | -0.2389967 | 0.61779163 | -0.3868565 | 0.69886246 | NA |
| RAB25 | 9.66027833 | -0.1205268 | 0.58241694 | -0.2069424 | 0.83605484 | NA |
| LMNA | 21.4475951 | -0.706785 | 0.39876564 | -1.7724321 | 0.07632285 | NA |
| SEMA4A | 2.49621127 | 0.63171668 | 0.88923683 | 0.71040319 | 0.47745415 | NA |
| SLC25A44 | 5.15546077 | -1.3045114 | 0.67521437 | -1.931996 | 0.05336 | NA |
| SMG5 | 17.5988977 | -0.4556684 | 0.43864251 | -1.038815 | 0.29889079 | NA |
| GLMP | 3.94927491 | -0.3422897 | 0.84690307 | -0.4041663 | 0.68609041 | NA |
| CCT3 | 19.5298208 | 0.43006933 | 0.44338 | 0.9699791 | 0.33205691 | NA |
| RHBG | 48.085235 | 1.70541911 | 0.52740395 | 3.23361081 | 0.00122236 | 0.1198504 |
| MEF2D | 9.39028637 | -0.7926667 | 0.53605594 | -1.4787015 | 0.13922011 | NA |
| NAXE | 8.99289688 | 0.46530483 | 0.60601183 | 0.76781477 | 0.44259724 | NA |
| GPATCH4 | 4.81940012 | -0.0055859 | 0.73762653 | -0.0075729 | 0.99395779 | NA |
| ISG20L2 | 2.5243468 | -0.7657949 | 0.99475657 | -0.7698314 | 0.4413999 | NA |
| RRNAD1 | 5.75196286 | 0.36027678 | 0.61391307 | 0.58685308 | 0.55730239 | NA |
| MRPL24 | 4.19755521 | 1.3500879 | 0.83583642 | 1.61525374 | 0.10625576 | NA |
| HDGF | 14.1698549 | -0.1496823 | 0.4270607 | -0.3504943 | 0.72596777 | NA |
| PRCC | 11.2798011 | 0.05141376 | 0.45698021 | 0.11250762 | 0.91042093 | NA |
| NTRK1 | 18.8134141 | -0.6858545 | 0.64246926 | -1.067529 | 0.28573305 | NA |

| | | | | | | |
|---|---|---|---|---|---|---|
| INSRR | 68.2878016 | 0.24911467 | 0.32153466 | 0.77476769 | 0.43847694 | 0.84462749 |
| ARHGEF11 | 28.626412 | -0.5975462 | 0.33487339 | -1.7843944 | 0.0743596 | 0.53207981 |
| ETV3 | 4.65682331 | -1.0894352 | 0.77376905 | -1.4079591 | 0.15914318 | NA |
| DUSP23 | 14.8859632 | 0.4569833 | 0.53061461 | 0.86123392 | 0.38910922 | NA |
| TAGLN2 | 12.9974865 | -0.3829685 | 0.51986807 | -0.7366648 | 0.46132623 | NA |
| IGSF8 | 4.23609656 | 0.28854319 | 0.76944128 | 0.37500352 | 0.70765785 | NA |
| PEA15 | 3.64434615 | -1.4172567 | 0.80504014 | -1.7604795 | 0.07832654 | NA |
| DCAF8 | 9.03499114 | 0.69051251 | 0.58718194 | 1.17597709 | 0.23960403 | NA |
| COPA | 39.0344746 | -0.657472 | 0.32845251 | -2.0017263 | 0.04531417 | 0.44887658 |
| NCSTN | 7.84862254 | 0.42992524 | 0.53004438 | 0.81111179 | 0.41730147 | NA |
| F11R | 12.2833918 | -0.8045939 | 0.42773645 | -1.8810505 | 0.05996505 | NA |
| AL591806.3 | 3.64802206 | 0.64067598 | 1.1011838 | 0.58180658 | 0.56069697 | NA |
| USF1 | 2.68908378 | -0.9125435 | 0.86710568 | -1.0524017 | 0.29261529 | NA |
| PFDN2 | 19.6255529 | -0.7603476 | 0.4531102 | -1.6780634 | 0.09333473 | NA |
| NIT1 | 8.70578389 | 0.37352473 | 0.51460631 | 0.72584561 | 0.46793342 | NA |
| DEDD | 3.85901109 | -0.6382608 | 0.8049869 | -0.7928835 | 0.42784573 | NA |
| UFC1 | 9.75643798 | 0.70993375 | 0.53350425 | 1.33069934 | 0.18328796 | NA |
| B4GALT3 | 3.78554433 | 0.21495228 | 1.00463961 | 0.21395959 | 0.83057857 | NA |
| NDUFS2 | 18.8273944 | -0.1916288 | 0.38255421 | -0.5009194 | 0.61642787 | NA |
| SDHC | 23.2293493 | 0.438136 | 0.33514839 | 1.30728958 | 0.19111438 | NA |
| AL592295.4 | 2.88899525 | 1.58177258 | 0.90795994 | 1.74211715 | 0.08148795 | NA |
| DUSP12 | 4.23964665 | -0.0948799 | 0.70719322 | -0.1341641 | 0.89327284 | NA |
| ATF6 | 41.1585002 | -0.3624576 | 0.25888591 | -1.4000669 | 0.16149329 | 0.64709575 |
| NOS1AP | 51.4778773 | -0.520619 | 0.45372355 | -1.1474366 | 0.25120122 | 0.74329941 |
| UHMK1 | 51.1894664 | 0.16370073 | 0.30013478 | 0.54542405 | 0.58546191 | 0.89393468 |
| UAP1 | 22.794493 | -1.2481758 | 0.5997226 | -2.0812552 | 0.03741055 | NA |
| DDR2 | 5.13081505 | -3.328887 | 0.82460844 | -4.0369306 | 5.42E-05 | NA |
| HSD17B7 | 4.25455204 | -0.2297241 | 0.83293112 | -0.2758021 | 0.78270007 | NA |
| PBX1 | 26.2150417 | 0.92712215 | 0.59130997 | 1.56791226 | 0.11690161 | 0.57860396 |
| MGST3 | 53.5037117 | 0.2256984 | 0.32546241 | 0.69346995 | 0.48801468 | 0.86354292 |
| ALDH9A1 | 9.60432557 | 0.60138804 | 0.54691316 | 1.09960427 | 0.27150458 | NA |
| TMCO1 | 24.4433741 | 0.33716242 | 0.35395379 | 0.95256056 | 0.34081277 | NA |
| UCK2 | 5.5834257 | -0.103868 | 0.71801817 | -0.1446593 | 0.8849799 | NA |
| FAM78B | 2.99699178 | 0.09869288 | 0.94875917 | 0.10402311 | 0.91715101 | NA |
| POGK | 9.08220879 | -0.2689999 | 0.52305814 | -0.514283 | 0.60705414 | NA |
| TADA1 | 4.32577723 | -0.214401 | 0.78692969 | -0.2724525 | 0.78527407 | NA |
| POU2F1 | 60.2027021 | 0.18060152 | 0.32746829 | 0.55150841 | 0.58128521 | 0.89263108 |
| CREG1 | 6.59851737 | 0.0818452 | 0.71773095 | 0.11403326 | 0.90921143 | NA |
| MPZL1 | 19.0860008 | -0.0207927 | 0.45509941 | -0.0456882 | 0.9635588 | NA |
| MPC2 | 14.2102308 | 0.13639248 | 0.43728482 | 0.31190765 | 0.75511071 | NA |
| DCAF6 | 71.5673557 | 0.27038127 | 0.36411011 | 0.74258106 | 0.45773536 | 0.85802171 |
| TIPRL | 8.430217 | -0.3148004 | 0.63444447 | -0.4961828 | 0.61976543 | NA |
| SFT2D2 | 13.0549361 | 0.38697822 | 0.42128236 | 0.91857208 | 0.35831944 | NA |
| ATP1B1 | 175.597651 | -0.1285498 | 0.26153959 | -0.491512 | 0.62306439 | 0.91493825 |
| NME7 | 29.5450413 | -0.05906 | 0.32302105 | -0.1828362 | 0.85492651 | 0.97168963 |
| BLZF1 | 10.3477498 | -0.0166499 | 0.53963015 | -0.0308543 | 0.9753857 | NA |
| SLC19A2 | 29.3819946 | -1.2413284 | 0.51527033 | -2.4090819 | 0.01599271 | 0.31287044 |
| SCYL3 | 9.13392536 | 0.61549797 | 0.56685031 | 1.08582101 | 0.27755819 | NA |
| KIFAP3 | 18.0405041 | 0.20340631 | 0.44302887 | 0.45912654 | 0.6461433 | NA |
| GORAB | 2.975757 | -0.286907 | 0.88572293 | -0.3239241 | 0.7459955 | NA |
| FMO4 | 6.909784 | 0.67831232 | 0.66208493 | 1.02450953 | 0.30559467 | NA |
| PRRC2C | 77.7721962 | -0.3314528 | 0.25311407 | -1.3094997 | 0.19036514 | 0.68499719 |
| VAMP4 | 5.50356578 | 0.50930478 | 0.6475386 | 0.7865242 | 0.43156045 | NA |
| METTL13 | 3.31222458 | 1.04777674 | 0.78820586 | 1.32931864 | 0.18374287 | NA |

| | | | | | | |
|---|---|---|---|---|---|---|
| DNM3 | 15.2251574 | 0.97329512 | 0.53855067 | 1.80724894 | 0.07072347 | NA |
| PIGC | 7.66482119 | 0.39578499 | 0.59664465 | 0.66335128 | 0.5071056 | NA |
| SUCO | 26.1847294 | -0.5936511 | 0.34981103 | -1.6970622 | 0.08968489 | 0.56122412 |
| AL645568.1 | 4.66906156 | 1.80374979 | 0.91047792 | 1.98110216 | 0.04757982 | NA |
| PRDX6 | 36.1499168 | 1.02841318 | 0.41724917 | 2.46474588 | 0.01371105 | 0.30435125 |
| KLHL20 | 16.2716691 | 0.22847231 | 0.46856412 | 0.48760096 | 0.62583252 | NA |
| DARS2 | 4.27012847 | 0.00123709 | 0.78689731 | 0.00157211 | 0.99874563 | NA |
| ZBTB37 | 6.84742586 | 0.51830904 | 0.59492408 | 0.87121879 | 0.3836347 | NA |
| RC3H1 | 50.0832738 | -0.2820263 | 0.27970875 | -1.0082855 | 0.31331741 | 0.78031266 |
| RABGAP1L | 156.432306 | 0.18294695 | 0.19035635 | 0.96107618 | 0.33651387 | 0.8003972 |
| CACYBP | 6.82727828 | 0.08486017 | 0.54993396 | 0.15430974 | 0.87736552 | NA |
| MRPS14 | 7.90661895 | 0.22917977 | 0.59389864 | 0.38589039 | 0.69957785 | NA |
| KIAA0040 | 27.2341806 | -0.7022233 | 0.42226654 | -1.6629859 | 0.09631526 | 0.56358504 |
| COP1 | 87.7459395 | -0.1653651 | 0.25111583 | -0.6585214 | 0.51020317 | 0.87158522 |
| PAPPA2 | 7.26792431 | 0.99820341 | 0.65634052 | 1.52086208 | 0.12829445 | NA |
| RASAL2 | 54.7131972 | -0.0953568 | 0.40979356 | -0.2326946 | 0.81599856 | 0.96845695 |
| RALGPS2 | 92.8250881 | 0.11481226 | 0.34050391 | 0.33718338 | 0.73597867 | 0.94590222 |
| ANGPTL1 | 17.9525979 | 1.28929191 | 0.45760473 | 2.81747941 | 0.00484022 | NA |
| FAM20B | 12.878876 | 0.45367326 | 0.42943676 | 1.05643789 | 0.2907682 | NA |
| TOR3A | 4.14772674 | 0.43035431 | 0.75917804 | 0.56686876 | 0.57080335 | NA |
| ABL2 | 36.7512426 | -0.065441 | 0.36888786 | -0.1774009 | 0.85919353 | 0.97168963 |
| SOAT1 | 46.9557978 | -0.2389874 | 0.3933349 | -0.6075928 | 0.54345761 | 0.88410237 |
| TOR1AIP2 | 29.5217335 | -0.3199444 | 0.35731319 | -0.8954174 | 0.370564 | 0.81199861 |
| TOR1AIP1 | 13.5082506 | -0.5331443 | 0.44155924 | -1.2074129 | 0.22727318 | NA |
| CEP350 | 53.572098 | -0.3228963 | 0.28384682 | -1.1375723 | 0.25529912 | 0.75126058 |
| QSOX1 | 11.4385706 | 0.49117758 | 0.51974385 | 0.9450378 | 0.34463958 | NA |
| ACBD6 | 65.50893 | -0.0833863 | 0.2605015 | -0.3200992 | 0.74889316 | 0.94999802 |
| XPR1 | 43.8713628 | -0.4598595 | 0.31070858 | -1.4800349 | 0.13886393 | 0.612608 |
| KIAA1614 | 2.68142692 | 1.13832079 | 0.89614144 | 1.27024679 | 0.20399674 | NA |
| STX6 | 18.4577376 | 0.27876303 | 0.44969974 | 0.61988702 | 0.53533217 | NA |
| MR1 | 12.009339 | 0.69077976 | 0.504357 | 1.3696246 | 0.17080412 | NA |
| GLUL | 24.2484431 | 0.66927525 | 0.3723889 | 1.79724812 | 0.07229624 | NA |
| RNASEL | 2.93176458 | 1.31950461 | 0.99786737 | 1.32232464 | 0.18606007 | NA |
| DHX9 | 36.1697897 | -0.2249993 | 0.31783329 | -0.707916 | 0.47899741 | 0.86206794 |
| LAMC1 | 40.2420487 | -0.3380667 | 0.33116041 | -1.0208549 | 0.30732318 | 0.77729327 |
| SMG7 | 60.8390076 | -0.1257087 | 0.27498639 | -0.457145 | 0.64756682 | 0.9218151 |
| ARPC5 | 18.489907 | -0.0524609 | 0.50470298 | -0.103944 | 0.91721376 | NA |
| RGL1 | 5.81366886 | -0.1765477 | 0.616309 | -0.2864597 | 0.77452604 | NA |
| COLGALT2 | 5.24671005 | -0.5362763 | 1.04258479 | -0.5143719 | 0.60699201 | NA |
| TSEN15 | 16.0387764 | 0.47561155 | 0.4847242 | 0.98120034 | 0.32649396 | NA |
| C1orf21 | 16.2199437 | -0.4175097 | 0.46767926 | -0.8927265 | 0.37200365 | NA |
| EDEM3 | 23.4846133 | -0.0314982 | 0.40480684 | -0.0778104 | 0.93797884 | NA |
| RNF2 | 13.7245123 | -0.0220397 | 0.40901464 | -0.0538849 | 0.95702686 | NA |
| TRMT1L | 9.96694013 | 0.35081398 | 0.51505149 | 0.68112409 | 0.49579297 | NA |
| SWT1 | 22.7272737 | 0.49336369 | 0.39673069 | 1.24357331 | 0.21365664 | NA |
| IVNS1ABP | 47.5941697 | -0.9148293 | 0.43546878 | -2.1007921 | 0.03565922 | 0.41688714 |
| TPR | 34.2447127 | -0.0369942 | 0.29263885 | -0.126416 | 0.89940263 | 0.97863828 |
| ODR4 | 21.2059168 | 0.07978071 | 0.36017949 | 0.22150264 | 0.82470108 | NA |
| PLA2G4A | 7.65468738 | -2.0030031 | 0.66664721 | -3.0045923 | 0.00265937 | NA |
| BRINP3 | 11.4737759 | 0.50896835 | 1.0063752 | 0.50574413 | 0.61303629 | NA |
| AL390957.1 | 113.762878 | -1.5081761 | 0.74493523 | -2.0245735 | 0.04291118 | 0.44186191 |
| AL136987.1 | 5.33263188 | 1.4472164 | 1.1697285 | 1.23722419 | 0.21600386 | NA |
| AL136454.1 | 23.3436948 | -0.1803868 | 0.45815907 | -0.393721 | 0.69378705 | NA |
| UCHL5 | 10.3898498 | 0.22900838 | 0.47501974 | 0.48210288 | 0.62973287 | NA |

| | | | | | | |
|---|---|---|---|---|---|---|
| TROVE2 | 27.8320877 | -0.0458732 | 0.31442228 | -0.1458967 | 0.88400297 | 0.97769824 |
| GLRX2 | 4.09997547 | -0.019348 | 0.68410609 | -0.0282821 | 0.97743715 | NA |
| CDC73 | 59.4480319 | -0.5505516 | 0.27627644 | -1.9927562 | 0.04628815 | 0.45088713 |
| ZBTB41 | 9.68163735 | 0.34544755 | 0.49241045 | 0.7015439 | 0.48296365 | NA |
| CRB1 | 5.29757166 | -3.7036382 | 1.0141145 | -3.6520907 | 0.00026011 | NA |
| DENND1B | 74.45172 | -0.3679208 | 0.313946 | -1.1719239 | 0.24122759 | 0.73323864 |
| NEK7 | 98.9272968 | -0.2451177 | 0.27773853 | -0.8825484 | 0.37748033 | 0.81843624 |
| ATP6V1G3 | 99.458827 | 1.72789286 | 0.47772049 | 3.61695362 | 0.00029809 | 0.0795604 |
| PTPRC | 27.9358387 | 0.61875529 | 0.42921455 | 1.44159907 | 0.14941551 | 0.63 |
| ZNF281 | 5.03271555 | 0.12213022 | 0.64907011 | 0.18816183 | 0.85074979 | NA |
| DDX59 | 7.9299269 | 0.2341226 | 0.55030505 | 0.42544149 | 0.67051486 | NA |
| CAMSAP2 | 49.5414123 | -0.6001928 | 0.38296819 | -1.5672132 | 0.11706487 | 0.57860396 |
| TMEM9 | 5.9211023 | 0.7958942 | 0.59115878 | 1.34632898 | 0.17819644 | NA |
| CSRP1 | 7.13877056 | 0.0107106 | 0.57729608 | 0.01855305 | 0.98519766 | NA |
| AC096677.1 | 7.30399366 | 0.04668622 | 0.6791125 | 0.06874593 | 0.94519185 | NA |
| IPO9 | 15.9275084 | -0.7079004 | 0.473383 | -1.4954072 | 0.13480819 | NA |
| TIMM17A | 10.5563237 | -0.8918707 | 0.57969383 | -1.5385203 | 0.12392144 | NA |
| RNPEP | 12.2695721 | -0.6138917 | 0.46020283 | -1.333959 | 0.1822173 | NA |
| ELF3 | 54.7295139 | 0.12178555 | 0.45686617 | 0.26656724 | 0.78980238 | 0.96325251 |
| ARL8A | 8.90533751 | -1.2936652 | 0.6554837 | -1.973604 | 0.04842679 | NA |
| PPP1R12B | 64.8841334 | 0.45012249 | 0.34625987 | 1.29995567 | 0.19361616 | 0.6871829 |
| KDM5B | 39.4958994 | 0.16304938 | 0.26298222 | 0.62000155 | 0.53525677 | 0.88254911 |
| KLHL12 | 8.44384298 | -0.0680505 | 0.56722677 | -0.1199705 | 0.90450651 | NA |
| ADIPOR1 | 6.07463232 | -0.2934738 | 0.62322266 | -0.4708973 | 0.6377141 | NA |
| CYB5R1 | 6.50890239 | 0.23130545 | 0.70851565 | 0.32646484 | 0.74407269 | NA |
| TMEM183A | 2.70028481 | -0.0009122 | 0.82525786 | -0.0011053 | 0.99911807 | NA |
| BTG2 | 10.7165707 | 0.15478749 | 0.64538184 | 0.23983863 | 0.81045536 | NA |
| ATP2B4 | 12.3319522 | 1.06826417 | 0.49554289 | 2.15574514 | 0.03110357 | NA |
| SNRPE | 10.020086 | 0.03701461 | 0.61704194 | 0.05998719 | 0.95216584 | NA |
| SOX13 | 10.0382865 | -0.5416001 | 0.51019155 | -1.0615623 | 0.28843444 | NA |
| ETNK2 | 2.64232172 | 1.38802004 | 1.04022216 | 1.33434961 | 0.1820893 | NA |
| GOLT1A | 7.3140579 | -0.9734065 | 0.56817549 | -1.7132146 | 0.08667306 | NA |
| PLEKHA6 | 43.7769611 | -0.4095496 | 0.27117613 | -1.5102715 | 0.13097416 | 0.60270696 |
| PPP1R15B | 9.98000545 | -0.9475198 | 0.56034 | -1.690973 | 0.09084196 | NA |
| MDM4 | 37.7497504 | -0.2426646 | 0.3648653 | -0.66508 | 0.50599931 | 0.87094108 |
| LRRN2 | 11.7722505 | -0.315292 | 0.46267726 | -0.6814512 | 0.49558606 | NA |
| RBBP5 | 8.13167726 | -0.4840668 | 0.5579373 | -0.8676007 | 0.38561296 | NA |
| DSTYK | 13.8483438 | 0.25270297 | 0.4981273 | 0.507306 | 0.61194013 | NA |
| NUAK2 | 13.9526046 | 1.39102006 | 0.66293469 | 2.09827616 | 0.03588076 | NA |
| CDK18 | 5.40196566 | 0.00297299 | 0.63982676 | 0.00464655 | 0.9962926 | NA |
| MFSD4A | 4.42359098 | 0.37534365 | 0.838623 | 0.44757138 | 0.65446256 | NA |
| ELK4 | 20.8906866 | 0.06323631 | 0.3466206 | 0.18243667 | 0.85524005 | NA |
| NUCKS1 | 53.8036885 | 0.06027557 | 0.29817624 | 0.20214745 | 0.83980145 | 0.97168963 |
| RAB29 | 5.92846703 | 0.29228233 | 0.68363537 | 0.42754126 | 0.66898514 | NA |
| SLC41A1 | 3.95809727 | -0.7796667 | 0.81822227 | -0.9528788 | 0.34065147 | NA |
| RHEX | 12.7891271 | -0.4087451 | 0.770806 | -0.5302827 | 0.59591592 | NA |
| SRGAP2 | 5.38624109 | 0.88111261 | 0.6496347 | 1.35632012 | 0.17499734 | NA |
| EIF2D | 4.50247387 | 0.53053606 | 0.67645751 | 0.78428586 | 0.43287239 | NA |
| MAPKAPK2 | 27.5163207 | -0.3079029 | 0.39168439 | -0.7860995 | 0.43180919 | 0.84223493 |
| PIGR | 20.7716985 | 1.36706846 | 0.78044905 | 1.75164343 | 0.07983514 | NA |
| C1orf116 | 5.99873583 | 0.25093863 | 0.64004764 | 0.39206242 | 0.69501209 | NA |
| PFKFB2 | 13.2070257 | -0.0941174 | 0.45177787 | -0.2083267 | 0.83497392 | NA |
| YOD1 | 4.69730853 | -0.5972947 | 0.72370908 | -0.8253242 | 0.40918753 | NA |
| CD55 | 12.5787465 | 0.46278605 | 0.50347838 | 0.91917761 | 0.35800268 | NA |

| | | | | | | |
|---|---|---|---|---|---|---|
| CD46 | 71.0151493 | 0.45025242 | 0.25186062 | 1.7877047 | 0.07382366 | 0.53207981 |
| MIR29B2CH( | 36.3465473 | 0.42448165 | 0.34167884 | 1.24234106 | 0.21411075 | 0.71319378 |
| PLXNA2 | 5.03613886 | 0.42186401 | 0.82317528 | 0.51248381 | 0.60831245 | NA |
| G0S2 | 8.89563516 | 2.12667042 | 0.96860318 | 2.19560545 | 0.0281202 | NA |
| TRAF3IP3 | 6.98734 | 2.79301001 | 1.02191427 | 2.73311578 | 0.00627383 | NA |
| IRF6 | 10.8204606 | 0.26400321 | 0.53157442 | 0.49664393 | 0.61944017 | NA |
| UTP25 | 2.7127952 | -0.0441236 | 1.00169614 | -0.0440489 | 0.96486546 | NA |
| HHAT | 6.23237531 | 0.61494177 | 0.60946326 | 1.00898907 | 0.31297988 | NA |
| RCOR3 | 44.5949802 | 0.5019649 | 0.3620837 | 1.38632281 | 0.16564835 | 0.64995633 |
| TRAF5 | 3.54820241 | 0.17729013 | 0.76752773 | 0.23098857 | 0.81732368 | NA |
| LINC00467 | 6.29221735 | -0.5546972 | 0.64313356 | -0.8624914 | 0.38841716 | NA |
| SLC30A1 | 4.22092544 | -0.2976406 | 0.77849609 | -0.3823276 | 0.70221837 | NA |
| LPGAT1 | 59.6200523 | -0.3846958 | 0.28155777 | -1.3663123 | 0.17184097 | 0.66182328 |
| INTS7 | 15.0044705 | -0.3011552 | 0.4495857 | -0.6698506 | 0.50295305 | NA |
| PPP2R5A | 153.443712 | 0.22858092 | 0.33317451 | 0.68606967 | 0.49266917 | 0.86619885 |
| AL360091.3 | 21.4464079 | 0.46046384 | 0.58557352 | 0.78634676 | 0.43166436 | NA |
| TMEM206 | 3.00403824 | 0.02100648 | 0.85583241 | 0.02454509 | 0.98041782 | NA |
| NENF | 12.5292487 | -0.2510479 | 0.60091665 | -0.4177749 | 0.67611171 | NA |
| ATF3 | 4.04742316 | -1.6151873 | 0.86972256 | -1.8571294 | 0.06329274 | NA |
| NSL1 | 16.8750889 | -0.09593 | 0.36840761 | -0.260391 | 0.79456218 | NA |
| TATDN3 | 7.29229494 | 0.51997089 | 0.80044717 | 0.64960051 | 0.51595031 | NA |
| FLVCR1 | 5.48830823 | -0.2572962 | 0.75571965 | -0.3404652 | 0.73350625 | NA |
| ANGEL2 | 13.4355168 | -0.2006234 | 0.45165232 | -0.4441987 | 0.65689893 | NA |
| RPS6KC1 | 40.3736909 | -0.7125498 | 0.2776134 | -2.5666981 | 0.0102672 | 0.28266532 |
| PROX1 | 5.09539062 | 0.21252904 | 0.74078803 | 0.28689589 | 0.77419203 | NA |
| SMYD2 | 14.151949 | -0.4963417 | 0.51431117 | -0.9650612 | 0.33451417 | NA |
| PTPN14 | 41.773983 | 0.05269354 | 0.3451668 | 0.15266108 | 0.87866555 | 0.97516771 |
| KCTD3 | 27.2614139 | -0.4148588 | 0.36330481 | -1.141903 | 0.25349435 | 0.74749396 |
| ESRRG | 631.253721 | 0.5815183 | 0.26279732 | 2.21280147 | 0.02691134 | 0.39506514 |
| GPATCH2 | 50.2333412 | 0.13087647 | 0.324269 | 0.40360463 | 0.68650348 | 0.92726927 |
| SPATA17 | 9.19738109 | -1.8854836 | 0.70463744 | -2.6758209 | 0.00745465 | NA |
| RRP15 | 5.89572146 | 0.21343563 | 0.59172358 | 0.36070158 | 0.71832254 | NA |
| LYPLAL1 | 43.1047843 | 0.98537228 | 0.37372868 | 2.63659797 | 0.0083742 | 0.27648451 |
| ZC3H11B | 2.69010633 | -0.3397367 | 0.99056899 | -0.3429712 | 0.7316201 | NA |
| EPRS | 18.0270097 | -0.0965281 | 0.35465354 | -0.2721757 | 0.78548694 | NA |
| BPNT1 | 5.27970728 | -0.6980763 | 0.85026699 | -0.8210084 | 0.41164148 | NA |
| IARS2 | 16.8435448 | 0.27506892 | 0.38875693 | 0.70756015 | 0.47921845 | NA |
| RAB3GAP2 | 53.4509546 | -0.0213013 | 0.2934273 | -0.0725949 | 0.94212851 | 0.99153825 |
| MARK1 | 15.79579 | 0.32656295 | 0.46100092 | 0.70837809 | 0.4787105 | NA |
| 2-Mar | 11.3433282 | 0.4999309 | 0.51189324 | 0.97663118 | 0.32875178 | NA |
| AL513314.2 | 4.71982446 | -0.0827187 | 0.70266659 | -0.1177211 | 0.90628862 | NA |
| TAF1A | 3.69441964 | 0.28149274 | 0.8977236 | 0.31356282 | 0.7538531 | NA |
| MIA3 | 12.8895489 | 0.5984875 | 0.41329962 | 1.44807174 | 0.14759699 | NA |
| AIDA | 5.80793758 | 0.2867494 | 0.62059065 | 0.46205885 | 0.64403912 | NA |
| BROX | 13.2846427 | 0.54850839 | 0.47026145 | 1.16639029 | 0.24345668 | NA |
| FAM177B | 5.28006714 | -1.8552842 | 0.73233454 | -2.5333834 | 0.01129673 | NA |
| AC093152.1 | 7.76636455 | 0.62113974 | 0.60401994 | 1.0283431 | 0.30378846 | NA |
| SUSD4 | 8.72856126 | -0.0526049 | 0.64221291 | -0.0819119 | 0.93471675 | NA |
| CAPN2 | 50.0517239 | 0.37716091 | 0.29114436 | 1.29544295 | 0.19516747 | 0.69049094 |
| TP53BP2 | 12.3414126 | -1.0440922 | 0.45052083 | -2.3175225 | 0.02047528 | NA |
| FBXO28 | 22.0804063 | 0.09184547 | 0.39541374 | 0.23227688 | 0.81632297 | NA |
| DEGS1 | 6.14334548 | -0.7680743 | 0.65790966 | -1.1674464 | 0.24303013 | NA |
| NVL | 18.3799882 | 0.16493524 | 0.43950171 | 0.37527782 | 0.70745386 | NA |
| CNIH4 | 15.5574294 | -0.0604617 | 0.47891834 | -0.1262463 | 0.89953696 | NA |

| | | | | | | |
|---|---|---|---|---|---|---|
| WDR26 | 42.2211408 | 0.15656407 | 0.29516124 | 0.53043573 | 0.59580986 | 0.89893528 |
| CNIH3 | 9.93419414 | 0.7513859 | 0.59729886 | 1.25797311 | 0.20840148 | NA |
| DNAH14 | 55.2301474 | -0.5209891 | 0.26045144 | -2.0003309 | 0.04546454 | 0.44887658 |
| LBR | 12.6108997 | -0.5237712 | 0.49799016 | -1.0517702 | 0.29290502 | NA |
| ENAH | 6.94156885 | 0.23462744 | 0.73203996 | 0.32051179 | 0.74858039 | NA |
| SRP9 | 23.9535979 | 0.07161003 | 0.35888243 | 0.19953619 | 0.84184334 | NA |
| EPHX1 | 3.93714883 | 0.3042956 | 0.66969406 | 0.45438003 | 0.64955533 | NA |
| TMEM63A | 11.5580745 | 0.71966317 | 0.48444062 | 1.48555497 | 0.13739687 | NA |
| SDE2 | 5.20507913 | 0.60667285 | 0.74291091 | 0.81661588 | 0.41414798 | NA |
| H3F3A | 6.01566596 | -0.4673038 | 0.63613711 | -0.7345961 | 0.46258551 | NA |
| ACBD3 | 24.2420935 | -0.320086 | 0.41672246 | -0.7681035 | 0.4424257 | NA |
| PARP1 | 13.4665184 | 0.35641459 | 0.46949121 | 0.75915072 | 0.4477624 | NA |
| STUM | 12.1752373 | 0.25666597 | 0.71967468 | 0.35664166 | 0.72136009 | NA |
| ITPKB | 11.6523406 | -0.9026189 | 0.67879031 | -1.3297464 | 0.18360186 | NA |
| PSEN2 | 16.7292493 | -0.3090391 | 0.469792 | -0.6578212 | 0.51065306 | NA |
| COQ8A | 21.6327562 | 0.14824782 | 0.41785622 | 0.3547819 | 0.72275299 | NA |
| CDC42BPA | 136.245692 | 0.08536528 | 0.25985089 | 0.32851641 | 0.74252124 | 0.94831173 |
| ZNF678 | 15.2043368 | 0.0616383 | 0.41805885 | 0.14743928 | 0.8827853 | NA |
| SNAP47 | 5.19270268 | -0.086288 | 0.68893607 | -0.1252483 | 0.90032701 | NA |
| JMJD4 | 2.53987284 | 0.02788336 | 0.95946744 | 0.02906129 | 0.97681571 | NA |
| ARF1 | 38.4922868 | -0.4166175 | 0.44083836 | -0.9450573 | 0.34462961 | 0.8003972 |
| C1orf35 | 3.26802404 | 0.38406025 | 0.80831962 | 0.47513414 | 0.63469137 | NA |
| MRPL55 | 5.4310157 | -0.3701704 | 0.67977749 | -0.5445465 | 0.58606548 | NA |
| GUK1 | 35.0817122 | -0.2062253 | 0.33802379 | -0.610091 | 0.5418015 | 0.88410237 |
| IBA57 | 3.43959542 | -0.5638243 | 0.84785945 | -0.6649974 | 0.50605219 | NA |
| OBSCN | 10.1357769 | 0.0013556 | 0.55182993 | 0.00245656 | 0.99803995 | NA |
| TRIM11 | 2.56775847 | -1.0491395 | 0.86960623 | -1.2064535 | 0.22764267 | NA |
| RNF187 | 6.94475981 | -0.0870605 | 0.61645485 | -0.1412278 | 0.88769001 | NA |
| RHOU | 8.01391918 | 0.23092588 | 0.58271678 | 0.39629179 | 0.69188978 | NA |
| RAB4A | 10.169556 | 1.45336875 | 0.58806044 | 2.47146153 | 0.0134562 | NA |
| NUP133 | 12.8124187 | 0.19818559 | 0.48094115 | 0.41207866 | 0.68028177 | NA |
| ABCB10 | 6.05841618 | -0.1002738 | 0.60842174 | -0.1648097 | 0.86909378 | NA |
| TAF5L | 5.65331042 | -0.1076674 | 0.78078844 | -0.1378957 | 0.89032282 | NA |
| GALNT2 | 18.2577996 | 0.20623506 | 0.42775062 | 0.48213854 | 0.62970754 | NA |
| COG2 | 9.84816921 | -0.2739611 | 0.50001361 | -0.5479073 | 0.58375553 | NA |
| C1orf198 | 4.84830499 | -1.3687248 | 0.76798388 | -1.7822311 | 0.07471155 | NA |
| TTC13 | 6.99974798 | -0.3591494 | 0.60163943 | -0.5969512 | 0.55053995 | NA |
| ARV1 | 7.48358013 | 0.14231777 | 0.5753486 | 0.2473592 | 0.80463025 | NA |
| C1orf131 | 4.7628208 | 0.75462337 | 0.6247695 | 1.20784285 | 0.22710772 | NA |
| GNPAT | 14.8011625 | 0.17483653 | 0.47003871 | 0.37196198 | 0.70992116 | NA |
| EXOC8 | 2.92284317 | -0.1618646 | 0.83929589 | -0.1928577 | 0.84707045 | NA |
| SPRTN | 3.16597043 | -0.3202266 | 0.96807852 | -0.3307858 | 0.74080628 | NA |
| EGLN1 | 75.4174783 | -0.5774036 | 0.29426092 | -1.9622163 | 0.04973731 | 0.46415693 |
| DISC1 | 18.2935781 | 1.3361891 | 0.8971717 | 1.48933487 | 0.13639921 | NA |
| SIPA1L2 | 31.5406503 | -1.7626843 | 0.55566162 | -3.1722261 | 0.00151275 | 0.13196343 |
| NTPCR | 9.69881547 | 0.14733822 | 0.49383302 | 0.29835636 | 0.76543119 | NA |
| PCNX2 | 23.0208306 | 0.25592297 | 0.40522213 | 0.63156218 | 0.52767301 | NA |
| MAP3K21 | 24.8385487 | 0.789119 | 0.36954111 | 2.13540245 | 0.03272815 | NA |
| SLC35F3 | 279.460988 | -0.3750564 | 0.43725533 | -0.8577515 | 0.39102967 | 0.8258317 |
| AL355472.3 | 20.61704 | 0.33998837 | 0.41051456 | 0.82820053 | 0.40755695 | NA |
| COA6 | 9.47314286 | 0.67273516 | 0.56685909 | 1.18677672 | 0.2353157 | NA |
| TARBP1 | 14.7860284 | 0.32015823 | 0.49552516 | 0.64609883 | 0.51821535 | NA |
| IRF2BP2 | 19.6086798 | 0.601395 | 0.50456363 | 1.19191112 | 0.2332961 | NA |
| TOMM20 | 26.7452297 | 0.50389807 | 0.39583717 | 1.27299332 | 0.20302041 | 0.700066 |

| | | | | | | |
|---|---|---|---|---|---|---|
| ARID4B | 105.6244 | -0.000529 | 0.23987409 | -0.0022052 | 0.99824047 | 0.99960611 |
| B3GALNT2 | 16.2524057 | -0.0121634 | 0.47112339 | -0.0258178 | 0.97940264 | NA |
| LYST | 48.9757001 | 0.18397421 | 0.32210654 | 0.57115951 | 0.56789152 | 0.89263108 |
| GPR137B | 9.53160357 | 0.27180868 | 0.54481151 | 0.49890407 | 0.61784697 | NA |
| ERO1B | 10.323631 | 1.2105024 | 0.64559117 | 1.87502935 | 0.06078869 | NA |
| EDARADD | 10.1204252 | -0.4441792 | 0.65385997 | -0.6793185 | 0.4969361 | NA |
| LGALS8 | 9.41801581 | -0.0696675 | 0.51270638 | -0.1358819 | 0.89191466 | NA |
| HEATR1 | 9.75750702 | 0.23508343 | 0.54750445 | 0.42937263 | 0.66765207 | NA |
| MTR | 61.927002 | 0.3930224 | 0.32969187 | 1.19209006 | 0.23322593 | 0.73156941 |
| RGS7 | 48.2927443 | -0.15913 | 0.71398051 | -0.2228772 | 0.82363107 | 0.96845695 |
| FH | 8.2938509 | 0.48978464 | 0.55892101 | 0.87630386 | 0.38086486 | NA |
| OPN3 | 10.1513102 | -0.6878833 | 0.60235975 | -1.1419809 | 0.25346198 | NA |
| CEP170 | 13.9383514 | 0.34753956 | 0.49154619 | 0.70703336 | 0.47954575 | NA |
| SDCCAG8 | 60.4891449 | 0.64457512 | 0.29620104 | 2.17614061 | 0.02954475 | 0.40646873 |
| AKT3 | 43.7551363 | 0.14230439 | 0.52156001 | 0.27284376 | 0.78497332 | 0.96299238 |
| ZBTB18 | 6.26083821 | -0.3335089 | 0.60737221 | -0.5491014 | 0.58293589 | NA |
| AL645465.1 | 3.54100162 | 1.35659347 | 0.84475982 | 1.60589251 | 0.10829753 | NA |
| ADSS | 15.2097897 | 0.33714426 | 0.47833101 | 0.70483463 | 0.48091316 | NA |
| CATSPERE | 14.0332706 | -1.1454572 | 0.64117207 | -1.7865052 | 0.0740175 | NA |
| DESI2 | 17.0623671 | -0.4868688 | 0.47188234 | -1.0317589 | 0.30218508 | NA |
| COX20 | 19.0195696 | -0.2677057 | 0.37815863 | -0.7079192 | 0.47899546 | NA |
| HNRNPU | 68.6953704 | -0.1746306 | 0.26555156 | -0.6576146 | 0.51078581 | 0.87158522 |
| EFCAB2 | 26.0472909 | -0.2629898 | 0.46259872 | -0.5685052 | 0.56969197 | 0.89263108 |
| KIF26B | 8.24819099 | 0.36759167 | 0.703989 | 0.52215542 | 0.60156212 | NA |
| SMYD3 | 120.622007 | 0.09911703 | 0.276451 | 0.35853382 | 0.71994386 | 0.93714576 |
| TFB2M | 4.00592952 | 0.0094597 | 0.78830681 | 0.01200003 | 0.99042559 | NA |
| CNST | 16.4071922 | -0.2706995 | 0.39932894 | -0.677886 | 0.49784398 | NA |
| SCCPDH | 15.7706082 | 0.02192841 | 0.50471897 | 0.04344678 | 0.96534539 | NA |
| AHCTF1 | 28.6175279 | 0.00034145 | 0.29944013 | 0.00114028 | 0.99909018 | 0.99960611 |
| ZNF124 | 8.70527925 | 0.13218449 | 0.48352372 | 0.27337747 | 0.78456307 | NA |
| AL390728.6 | 3.5327868 | -1.1524683 | 0.79955998 | -1.4413782 | 0.14947787 | NA |
| ZNF496 | 6.22548601 | 0.27409647 | 0.62523813 | 0.43838731 | 0.66110554 | NA |
| SH3BP5L | 3.10590079 | -0.1852493 | 0.88314635 | -0.2097606 | 0.83385452 | NA |
| ZNF672 | 6.90931008 | -0.6331802 | 0.60569385 | -1.04538 | 0.2958474 | NA |
| ZNF692 | 3.42244753 | -0.8384168 | 0.80947654 | -1.0357518 | 0.30031796 | NA |
| FAM110C | 7.35283958 | -0.2604015 | 0.55874335 | -0.4660485 | 0.64118075 | NA |
| SH3YL1 | 29.7798241 | 0.55064909 | 0.34707888 | 1.58652436 | 0.11262041 | 0.57860396 |
| ACP1 | 10.87694 | -0.0269263 | 0.48662955 | -0.0553321 | 0.95587385 | NA |
| TMEM18 | 2.49457768 | 0.52662246 | 0.82814345 | 0.63590729 | 0.52483685 | NA |
| LINC01250 | 3.41377804 | -0.590026 | 0.91322885 | -0.6460878 | 0.51822249 | NA |
| EIPR1 | 43.7023588 | 0.27089371 | 0.40142044 | 0.67483786 | 0.49977879 | 0.87036626 |
| TRAPPC12 | 45.1269912 | 0.01128795 | 0.30948893 | 0.03647289 | 0.9709053 | 0.99743889 |
| ADI1 | 21.7295676 | 0.53684982 | 0.45258483 | 1.18618606 | 0.23554883 | NA |
| AC108488.2 | 2.71598776 | -0.4150137 | 0.93134243 | -0.4456081 | 0.65588035 | NA |
| RPS7 | 47.1965551 | 0.01407775 | 0.41519949 | 0.03390598 | 0.97295212 | 0.99779576 |
| RNF144A | 6.44062508 | -0.662403 | 0.60291547 | -1.0986665 | 0.27191355 | NA |
| ID2 | 6.0494217 | -0.6168272 | 0.74044603 | -0.8330482 | 0.40481755 | NA |
| KIDINS220 | 62.1547309 | -0.4249659 | 0.24600405 | -1.7274752 | 0.08408236 | 0.55323508 |
| MBOAT2 | 109.533054 | -0.393489 | 0.27711307 | -1.4199583 | 0.15561982 | 0.63932163 |
| AC093904.4 | 2.8763073 | -0.3894723 | 1.25565561 | -0.3101744 | 0.75642831 | NA |
| ASAP2 | 208.58298 | -0.3391979 | 0.33282823 | -1.019138 | 0.30813745 | 0.77806893 |
| ITGB1BP1 | 15.7239926 | -0.0900426 | 0.42249058 | -0.2131234 | 0.83123069 | NA |
| CPSF3 | 8.17728874 | -0.7748263 | 0.68891753 | -1.124701 | 0.26071574 | NA |
| IAH1 | 12.4878853 | 0.04708675 | 0.4402741 | 0.10694871 | 0.91482967 | NA |

| | | | | | | |
|---|---|---|---|---|---|---|
| ADAM17 | 27.2246521 | -0.7310165 | 0.3800035 | -1.92371 | 0.05439095 | 0.49209981 |
| YWHAQ | 31.871898 | -0.780531 | 0.3672821 | -2.125154 | 0.03357376 | 0.41688714 |
| TAF1B | 22.1135625 | 0.25370993 | 0.40745811 | 0.62266507 | 0.53350464 | NA |
| KLF11 | 5.10609135 | 0.00031702 | 0.83044679 | 0.00038174 | 0.99969541 | NA |
| CYS1 | 9.83517619 | -0.1875827 | 0.50206067 | -0.3736255 | 0.70868299 | NA |
| HPCAL1 | 18.9598265 | 0.2805216 | 0.42001529 | 0.66788424 | 0.50420749 | NA |
| ODC1 | 6.71004647 | -0.8912761 | 0.62363086 | -1.4291725 | 0.15295467 | NA |
| NOL10 | 26.6182098 | -0.4201781 | 0.35156917 | -1.1951505 | 0.23202823 | 0.73156941 |
| ATP6V1C2 | 95.1638313 | -0.5518799 | 0.41465694 | -1.3309313 | 0.18321162 | 0.67478132 |
| PDIA6 | 27.7406368 | 0.11388681 | 0.40087498 | 0.28409559 | 0.77633713 | 0.96105928 |
| PQLC3 | 12.0391421 | 1.18212201 | 0.52257163 | 2.26212435 | 0.02368972 | NA |
| ROCK2 | 76.1387716 | -0.0313579 | 0.25609378 | -0.1224471 | 0.90254496 | 0.98003205 |
| E2F6 | 5.07799225 | -1.2092815 | 0.76290293 | -1.5851053 | 0.11294241 | NA |
| LPIN1 | 22.6216236 | -0.6529553 | 0.46130477 | -1.4154532 | 0.15693567 | NA |
| MIR3681HG | 5.5529001 | 1.00718309 | 0.75696093 | 1.33056153 | 0.18333333 | NA |
| TRIB2 | 12.7248104 | -0.043741 | 0.50027272 | -0.0874343 | 0.93032634 | NA |
| NBAS | 60.7531914 | -0.404725 | 0.33859062 | -1.1953227 | 0.23196099 | 0.73156941 |
| DDX1 | 19.4283609 | -0.2743287 | 0.38439901 | -0.7136562 | 0.47543979 | NA |
| SMC6 | 11.0629855 | 1.1759031 | 0.46688798 | 2.51859792 | 0.01178231 | NA |
| GEN1 | 3.94904703 | -0.40218 | 0.79524958 | -0.505728 | 0.6130476 | NA |
| RDH14 | 6.45828767 | 0.23967279 | 0.5743714 | 0.41727843 | 0.67647476 | NA |
| LINC01376 | 11.3874983 | 0.0728231 | 0.63745283 | 0.11424077 | 0.90904693 | NA |
| TTC32 | 3.39384461 | -0.4447689 | 0.83697585 | -0.5313999 | 0.59514169 | NA |
| WDR35 | 10.8193241 | -0.0193726 | 0.64214163 | -0.0301688 | 0.97593247 | NA |
| LAPTM4A | 35.9681495 | 0.30139785 | 0.32246429 | 0.93467048 | 0.34995815 | 0.8003972 |
| SDC1 | 5.49041273 | 0.00266495 | 0.65127987 | 0.00409187 | 0.99673517 | NA |
| PUM2 | 88.3926458 | -0.2855908 | 0.2219107 | -1.2869628 | 0.19810726 | 0.69343545 |
| RHOB | 7.69931787 | -0.5215947 | 0.77272495 | -0.675007 | 0.49967133 | NA |
| HS1BP3 | 3.68807098 | 0.25638377 | 0.75164856 | 0.34109527 | 0.73303186 | NA |
| LDAH | 26.272256 | 0.63648155 | 0.36822255 | 1.72852407 | 0.08389431 | 0.55323508 |
| KLHL29 | 8.63626111 | -3.2279666 | 0.93020888 | -3.4701524 | 0.00052016 | NA |
| ATAD2B | 58.0164345 | -0.3769284 | 0.30105517 | -1.2520243 | 0.21056101 | 0.70690231 |
| UBXN2A | 16.04723 | 0.73197568 | 0.45381939 | 1.61292289 | 0.10676126 | NA |
| SF3B6 | 16.4976622 | -0.3104239 | 0.42544136 | -0.7296514 | 0.46560327 | NA |
| FAM228B | 3.48202276 | 0.70677997 | 0.76105316 | 0.92868673 | 0.35305146 | NA |
| TP53I3 | 13.4194139 | 0.21483754 | 0.56426369 | 0.38073961 | 0.70339648 | NA |
| ITSN2 | 81.3087233 | 0.29278728 | 0.28679754 | 1.0208849 | 0.30730897 | 0.77729327 |
| NCOA1 | 137.102143 | 0.10893727 | 0.21022605 | 0.5181911 | 0.60432495 | 0.9071672 |
| PTRHD1 | 3.12381219 | -0.2597733 | 1.09937724 | -0.2362913 | 0.81320662 | NA |
| DNAJC27 | 9.79128685 | -0.3376116 | 0.52863883 | -0.6386431 | 0.52305511 | NA |
| DNMT3A | 28.2146975 | 0.38556094 | 0.3698607 | 1.04244909 | 0.29720352 | 0.77521265 |
| DTNB | 87.7202784 | -0.077722 | 0.27577137 | -0.2818348 | 0.77807016 | 0.96133329 |
| ASXL2 | 53.8607566 | -0.1045064 | 0.32915504 | -0.317499 | 0.75086498 | 0.9515948 |
| RAB10 | 90.1542874 | -0.4348012 | 0.26599836 | -1.6346011 | 0.1021327 | 0.5750956 |
| HADHA | 43.2169187 | 0.08049953 | 0.32480029 | 0.24784317 | 0.80425575 | 0.96743263 |
| HADHB | 36.848272 | 0.28968586 | 0.32416248 | 0.89364401 | 0.37151239 | 0.81199861 |
| SELENOI | 40.6180005 | -0.3751907 | 0.37385881 | -1.0035625 | 0.31558955 | 0.780966 |
| AC013472.2 | 20.2540332 | 1.37232466 | 0.60526823 | 2.26730002 | 0.0233719 | NA |
| MAPRE3 | 14.2321103 | 0.11268734 | 0.54917327 | 0.20519451 | 0.83742016 | NA |
| TMEM214 | 4.53767662 | -1.2194379 | 0.80491268 | -1.5149941 | 0.1297739 | NA |
| AGBL5 | 5.31848983 | 0.52557009 | 0.73287555 | 0.71713416 | 0.47329132 | NA |
| OST4 | 21.8293443 | -0.0236954 | 0.45186099 | -0.0524396 | 0.95817845 | NA |
| PREB | 2.53740238 | 0.51443433 | 0.85095168 | 0.60454 | 0.54548469 | NA |
| ATRAID | 12.0815703 | 0.1475717 | 0.47632125 | 0.30981549 | 0.75670127 | NA |

| | | | | | | |
|---|---|---|---|---|---|---|
| MPV17 | 4.13198777 | -0.0440944 | 0.70060394 | -0.0629377 | 0.94981609 | NA |
| GTF3C2 | 13.2373384 | -1.1610337 | 0.52314235 | -2.2193456 | 0.02646322 | NA |
| EIF2B4 | 3.10173247 | -0.3576648 | 0.86111923 | -0.4153488 | 0.67788658 | NA |
| SNX17 | 7.12631126 | -0.7291988 | 0.5733138 | -1.2719018 | 0.20340802 | NA |
| ZNF513 | 2.58462865 | -1.1419581 | 0.99260343 | -1.1504676 | 0.24995132 | NA |
| PPM1G | 24.9460461 | 0.49802531 | 0.3603845 | 1.38192766 | 0.16699391 | NA |
| NRBP1 | 9.50462364 | -0.5552212 | 0.52744128 | -1.0526693 | 0.29249261 | NA |
| IFT172 | 7.76549145 | 0.66249904 | 0.58874003 | 1.12528281 | 0.26046921 | NA |
| ZNF512 | 15.494838 | -0.828032 | 0.42757893 | -1.9365595 | 0.05279922 | NA |
| GPN1 | 3.97137935 | 1.04240143 | 0.74031707 | 1.40804727 | 0.15911707 | NA |
| SUPT7L | 7.16664472 | -0.3102091 | 0.57151597 | -0.5427829 | 0.58727928 | NA |
| SLC4A1AP | 6.54571062 | 0.57662528 | 0.60130759 | 0.95895226 | 0.3375828 | NA |
| MRPL33 | 23.0816776 | 0.07728382 | 0.41204542 | 0.18756141 | 0.85122048 | NA |
| RBKS | 12.4725411 | 0.92026385 | 0.54034056 | 1.70311821 | 0.08854595 | NA |
| BABAM2 | 112.866936 | 0.0529763 | 0.29294254 | 0.18084194 | 0.85649165 | 0.97168963 |
| FOSL2 | 39.5293063 | 0.2440478 | 0.34434076 | 0.70873921 | 0.47848633 | 0.86206794 |
| PPP1CB | 27.535168 | -0.210652 | 0.48029759 | -0.4385864 | 0.66096122 | 0.92276785 |
| TRMT61B | 11.2278784 | -0.1565309 | 0.48672186 | -0.3216024 | 0.74775396 | NA |
| WDR43 | 18.4523254 | -0.2506067 | 0.48760412 | -0.5139552 | 0.60728333 | NA |
| CLIP4 | 13.1773332 | 0.23708317 | 0.5537514 | 0.42814009 | 0.66854913 | NA |
| YPEL5 | 14.2870753 | 0.2783965 | 0.43740406 | 0.63647444 | 0.52446724 | NA |
| LCLAT1 | 17.3703729 | 0.10828498 | 0.40493941 | 0.26741031 | 0.78915326 | NA |
| CAPN13 | 4.52919202 | 0.56734702 | 0.75683751 | 0.74962856 | 0.45347844 | NA |
| GALNT14 | 18.6134221 | 0.73947937 | 0.54616838 | 1.35394029 | 0.17575543 | NA |
| MEMO1 | 2.84922819 | 0.47759391 | 0.92798302 | 0.51465803 | 0.60679201 | NA |
| DPY30 | 11.8274571 | 1.02779244 | 0.47390754 | 2.16876153 | 0.03010079 | NA |
| SPAST | 19.1638603 | -0.0444121 | 0.44497091 | -0.099809 | 0.92049599 | NA |
| SLC30A6 | 16.4196458 | -0.1811316 | 0.39089622 | -0.4633751 | 0.6430955 | NA |
| YIPF4 | 23.8194606 | 0.04153364 | 0.43449658 | 0.09559027 | 0.923846 | NA |
| BIRC6 | 141.286709 | -0.2362121 | 0.25382513 | -0.9306098 | 0.35205547 | 0.8003972 |
| TTC27 | 21.289371 | -0.1460908 | 0.40109114 | -0.3642334 | 0.71568371 | NA |
| LINC00486 | 23.6509981 | -0.4175249 | 0.39128297 | -1.0670663 | 0.2859419 | NA |
| LTBP1 | 4.93784485 | 0.06018913 | 0.70272024 | 0.08565162 | 0.93174336 | NA |
| RASGRP3 | 8.73149839 | 0.62652712 | 0.65113483 | 0.96220797 | 0.33594515 | NA |
| FAM98A | 8.09935676 | -0.6161635 | 0.53666779 | -1.1481284 | 0.25091558 | NA |
| LINC01320 | 50.4062656 | 0.82574674 | 0.53865727 | 1.53297243 | 0.12528265 | 0.5866305 |
| AC012593.1 | 19.0529731 | 0.11189598 | 0.88398988 | 0.12658061 | 0.89927235 | NA |
| CRIM1 | 41.1257423 | 0.41429107 | 0.34298057 | 1.20791411 | 0.2270803 | 0.73054639 |
| FEZ2 | 25.1768636 | 0.48806001 | 0.35578577 | 1.37178056 | 0.17013176 | NA |
| STRN | 62.906182 | 0.00826912 | 0.31606786 | 0.02616247 | 0.97912775 | 0.99902646 |
| HEATR5B | 21.6436436 | -0.4173529 | 0.3386305 | -1.2324729 | 0.21777247 | NA |
| GPATCH11 | 3.15933818 | -0.0308733 | 0.82864677 | -0.0372575 | 0.97027972 | NA |
| EIF2AK2 | 24.5509811 | -0.3261489 | 0.63227442 | -0.5158344 | 0.60597005 | NA |
| CEBPZOS | 11.263341 | 0.14123735 | 0.57604366 | 0.24518515 | 0.80631307 | NA |
| CEBPZ | 15.3883463 | -0.151783 | 0.43725625 | -0.347126 | 0.72849663 | NA |
| NDUFAF7 | 9.49556859 | 0.01760861 | 0.50095592 | 0.03515002 | 0.97196012 | NA |
| PRKD3 | 20.4295162 | -0.0980172 | 0.39657824 | -0.2471573 | 0.80478653 | NA |
| CDC42EP3 | 4.61349272 | -1.6188854 | 0.7305871 | -2.2158691 | 0.02670047 | NA |
| RMDN2 | 14.4103572 | 0.06236817 | 0.48358421 | 0.12897065 | 0.89738087 | NA |
| ATL2 | 138.496659 | 0.0328288 | 0.26617939 | 0.12333337 | 0.90184312 | 0.98003205 |
| HNRNPLL | 14.4982787 | -0.6234887 | 0.42031732 | -1.4833762 | 0.13797448 | NA |
| GALM | 8.66192125 | 0.41005176 | 0.55603475 | 0.73745707 | 0.46084444 | NA |
| SRSF7 | 19.4314038 | 0.19399142 | 0.56255323 | 0.34484101 | 0.7302139 | NA |
| DHX57 | 11.7052977 | 0.34658669 | 0.44956535 | 0.77093727 | 0.44074411 | NA |

| | | | | | | |
|---|---|---|---|---|---|---|
| SOS1 | 89.8391478 | 0.0790122 | 0.24795357 | 0.31865725 | 0.74998644 | 0.95093292 |
| MAP4K3 | 99.9350473 | 0.15011539 | 0.25926107 | 0.57901246 | 0.56258076 | 0.89263108 |
| AC007388.1 | 42.8448043 | 0.65815741 | 0.34722665 | 1.89546916 | 0.05803027 | 0.50751937 |
| THUMPD2 | 11.8100106 | -0.6385993 | 0.50967468 | -1.2529548 | 0.21022217 | NA |
| SLC8A1 | 24.3686941 | -0.1976805 | 1.2870907 | -0.1535871 | 0.87793534 | NA |
| PKDCC | 2.79233371 | -0.9835163 | 1.26920456 | -0.7749076 | 0.43839425 | NA |
| EML4 | 51.4470543 | -0.1183317 | 0.43399028 | -0.2726597 | 0.7851148 | 0.96299238 |
| COX7A2L | 24.2883405 | 0.34782187 | 0.43780006 | 0.79447653 | 0.42691806 | NA |
| MTA3 | 27.5177553 | 0.37554195 | 0.34893876 | 1.07624029 | 0.2818198 | 0.76854599 |
| ZFP36L2 | 26.4851215 | -0.0074614 | 0.33497309 | -0.0222746 | 0.98222892 | 0.99902646 |
| THADA | 44.5379002 | -0.0143263 | 0.28844111 | -0.049668 | 0.96038699 | 0.99656976 |
| PLEKHH2 | 21.6481872 | -0.279849 | 0.56852562 | -0.4922364 | 0.62255226 | NA |
| DYNC2LI1 | 13.3794566 | 0.00064158 | 0.45737104 | 0.00140275 | 0.99888077 | NA |
| LRPPRC | 41.8485037 | -0.1897315 | 0.30236191 | -0.6274979 | 0.53033293 | 0.88254911 |
| PPM1B | 39.8686055 | 0.21701627 | 0.27705155 | 0.78330648 | 0.43344716 | 0.84223493 |
| SLC3A1 | 14.7218188 | 1.34103004 | 0.58035723 | 2.31069758 | 0.02084957 | NA |
| PREPL | 22.4572462 | 0.41285894 | 0.36966996 | 1.11683121 | 0.2640665 | NA |
| CAMKMT | 93.7921652 | -0.2527238 | 0.33397109 | -0.7567235 | 0.44921553 | 0.85123446 |
| SRBD1 | 29.8214833 | -0.1869092 | 0.34472838 | -0.5421926 | 0.58768584 | 0.89477098 |
| PRKCE | 172.060674 | 0.19859674 | 0.36488079 | 0.5442784 | 0.58624991 | 0.89411486 |
| EPAS1 | 28.8164049 | -0.214949 | 0.43007973 | -0.4997887 | 0.61722386 | 0.91469766 |
| RHOQ | 22.2939732 | -0.1553389 | 0.33043718 | -0.4701012 | 0.6382827 | NA |
| AC018682.1 | 3.38731245 | 0.1710741 | 0.80088808 | 0.2136055 | 0.83085471 | NA |
| PIGF | 11.4906324 | 0.35238486 | 0.48779508 | 0.72240348 | 0.47004645 | NA |
| CRIPT | 3.92514704 | 1.15980198 | 0.7644475 | 1.51717675 | 0.12922207 | NA |
| SOCS5 | 13.1524715 | -0.4230942 | 0.44108923 | -0.9592031 | 0.33745643 | NA |
| MCFD2 | 17.216504 | 0.12036439 | 0.45042212 | 0.26722576 | 0.78929535 | NA |
| TTC7A | 11.2756178 | -0.054588 | 0.51339392 | -0.1063277 | 0.91532236 | NA |
| AC106869.1 | 5.38420807 | 1.55641534 | 0.82529477 | 1.88589022 | 0.05930975 | NA |
| EPCAM | 65.9074673 | 0.35306033 | 0.31199905 | 1.13160709 | 0.25779966 | 0.75272108 |
| MSH2 | 13.4437369 | 0.29582069 | 0.45835461 | 0.64539701 | 0.51866994 | NA |
| MSH6 | 9.22205187 | -0.11052 | 0.52782745 | -0.2093866 | 0.83414644 | NA |
| FBXO11 | 73.2016001 | -0.0482742 | 0.25601474 | -0.1885603 | 0.85043746 | 0.97168963 |
| FOXN2 | 8.39971628 | -0.1868603 | 0.51058951 | -0.3659697 | 0.71438769 | NA |
| PPP1R21 | 31.7322292 | 0.1485061 | 0.34072011 | 0.4358595 | 0.66293867 | 0.92276785 |
| AC010967.1 | 9.45720343 | -1.1786975 | 1.88437948 | -0.6255096 | 0.53163663 | NA |
| GPR75-ASB3 | 10.7052969 | 0.17844319 | 0.43958554 | 0.40593507 | 0.68479031 | NA |
| ERLEC1 | 10.1916881 | -0.2623172 | 0.46376013 | -0.5656312 | 0.57164454 | NA |
| PSME4 | 53.0070432 | -0.5920292 | 0.28923889 | -2.0468519 | 0.04067262 | 0.43422093 |
| ACYP2 | 49.2721692 | 0.14049213 | 0.32316225 | 0.43474176 | 0.66374988 | 0.92276785 |
| SPTBN1 | 91.9689937 | 0.0521104 | 0.30413456 | 0.17133996 | 0.86395646 | 0.97217819 |
| AC093110.1 | 4.02281741 | 0.56201432 | 0.77954302 | 0.72095356 | 0.47093809 | NA |
| EML6 | 6.07728613 | -0.1047473 | 0.73100543 | -0.1432921 | 0.88605948 | NA |
| RTN4 | 129.790736 | 0.01821844 | 0.30242983 | 0.06024022 | 0.95196431 | 0.99443943 |
| CLHC1 | 8.44024529 | -0.5344163 | 0.60019276 | -0.8904078 | 0.37324696 | NA |
| RPS27A | 99.4475817 | 0.39925404 | 0.41686044 | 0.95776428 | 0.33818164 | 0.8003972 |
| MTIF2 | 11.0255766 | 0.02627802 | 0.45577911 | 0.05765517 | 0.9540233 | NA |
| AC012358.3 | 4.59054671 | 1.30461158 | 0.87971169 | 1.48299901 | 0.13807467 | NA |
| CCDC88A | 7.71202526 | -0.3004354 | 0.6210601 | -0.4837461 | 0.62856604 | NA |
| CFAP36 | 9.7573421 | 0.66225548 | 0.46366178 | 1.42831588 | 0.15320096 | NA |
| PPP4R3B | 36.9571054 | -0.1998789 | 0.30227972 | -0.6612381 | 0.50845965 | 0.87158522 |
| PNPT1 | 13.6170181 | -0.3827309 | 0.5006325 | -0.7644947 | 0.44457247 | NA |
| EFEMP1 | 21.0334972 | -0.2616526 | 0.54997644 | -0.4757524 | 0.63425082 | NA |
| VRK2 | 27.8726792 | 0.15119898 | 0.40102621 | 0.37703016 | 0.70615119 | 0.93152073 |

| | | | | | | |
|---|---|---|---|---|---|---|
| FANCL | 16.9976126 | 0.24836 | 0.5085629 | 0.4883565 | 0.62529735 | NA |
| LINC01122 | 4.16599753 | -3.5767995 | 1.07752525 | -3.3194577 | 0.00090192 | NA |
| PAPOLG | 9.70937443 | -0.1990042 | 0.50535512 | -0.3937908 | 0.69373547 | NA |
| REL | 11.5402907 | 0.47032108 | 0.48092682 | 0.97794728 | 0.3281004 | NA |
| PUS10 | 4.10064563 | 0.35479295 | 0.93386901 | 0.37991725 | 0.70400684 | NA |
| PEX13 | 10.2034382 | -0.402301 | 0.56386516 | -0.7134702 | 0.47555487 | NA |
| KIAA1841 | 3.93177349 | -1.173589 | 0.81168196 | -1.445873 | 0.14821284 | NA |
| C2orf74 | 2.92529108 | -0.3196675 | 0.95978197 | -0.3330626 | 0.73908704 | NA |
| USP34 | 183.08853 | -0.5914558 | 0.31909572 | -1.8535371 | 0.06380541 | 0.52238232 |
| AC016727.1 | 5.7875468 | -0.2813999 | 0.59300551 | -0.4745316 | 0.63512086 | NA |
| XPO1 | 60.3594853 | -0.8373329 | 0.31519375 | -2.6565657 | 0.00789411 | 0.27648451 |
| FAM161A | 4.32641533 | -0.4388366 | 0.88087245 | -0.4981841 | 0.61835432 | NA |
| CCT4 | 16.2285138 | -0.2546509 | 0.43355084 | -0.587361 | 0.55696126 | NA |
| COMMD1 | 64.4255007 | -0.1472773 | 0.32369231 | -0.4549917 | 0.64911521 | 0.92202688 |
| AC018462.1 | 3.15494764 | 0.22939203 | 0.78754574 | 0.29127455 | 0.77084135 | NA |
| B3GNT2 | 29.2733864 | -0.1206426 | 0.38987242 | -0.3094412 | 0.75698592 | 0.95323728 |
| EHBP1 | 35.9034303 | 0.59879763 | 0.40070135 | 1.49437388 | 0.13507792 | 0.60594834 |
| WDPCP | 46.756977 | 0.31175638 | 0.37369322 | 0.83425751 | 0.40413589 | 0.82808596 |
| MDH1 | 20.7734031 | 0.2762156 | 0.39628821 | 0.69700687 | 0.48579849 | NA |
| UGP2 | 48.0802715 | 0.38753924 | 0.31683657 | 1.22315185 | 0.22127235 | 0.72409625 |
| VPS54 | 42.2709241 | 0.2588304 | 0.32540608 | 0.7954074 | 0.42637656 | 0.84057082 |
| PELI1 | 6.84647192 | -0.0900833 | 0.71292399 | -0.1263576 | 0.89944888 | NA |
| AC008074.1 | 3.48770498 | -0.8472267 | 0.8804432 | -0.962273 | 0.33591249 | NA |
| AFTPH | 40.2411484 | -0.1256618 | 0.27020993 | -0.4650526 | 0.64189377 | 0.92153493 |
| SERTAD2 | 5.84792849 | 0.27382142 | 0.72003752 | 0.38028771 | 0.70373186 | NA |
| CEP68 | 9.20619253 | 0.61435746 | 0.52438801 | 1.17157038 | 0.24136958 | NA |
| RAB1A | 59.0547247 | -0.8919583 | 0.30786771 | -2.8972129 | 0.00376494 | 0.2109414 |
| ACTR2 | 44.9485843 | -0.1190064 | 0.25445409 | -0.4676931 | 0.64000408 | 0.92153493 |
| SPRED2 | 41.2177891 | 0.02580869 | 0.33469014 | 0.07711218 | 0.9385343 | 0.99100968 |
| ETAA1 | 5.02440928 | 0.21538683 | 0.71398618 | 0.30166807 | 0.76290511 | NA |
| C1D | 9.22962212 | 0.02602676 | 0.58263499 | 0.04467078 | 0.96436972 | NA |
| PNO1 | 3.3777738 | -0.1407638 | 0.83167347 | -0.1692537 | 0.86559711 | NA |
| APLF | 8.54120192 | 1.45430695 | 0.59492092 | 2.44453828 | 0.01450377 | NA |
| GFPT1 | 31.9010949 | -0.7828265 | 0.46737731 | -1.6749346 | 0.09394706 | 0.56220784 |
| NFU1 | 8.99171051 | 0.12661875 | 0.492192 | 0.25725478 | 0.7969821 | NA |
| AAK1 | 70.4907383 | -0.1863538 | 0.25201372 | -0.739459 | 0.45962834 | 0.85844137 |
| ANXA4 | 133.62282 | 0.2829025 | 0.33919409 | 0.83404313 | 0.40425668 | 0.82808596 |
| GMCL1 | 23.5450718 | -0.7465207 | 0.36070447 | -2.0696186 | 0.03848808 | NA |
| MXD1 | 12.1259511 | -1.1887187 | 0.50765153 | -2.3416037 | 0.01920109 | NA |
| PCBP1 | 28.2025279 | -0.6589829 | 0.48695116 | -1.3532834 | 0.17596511 | 0.66719288 |
| C2orf42 | 5.74843138 | -0.5644764 | 0.6513922 | -0.8665692 | 0.38617809 | NA |
| PCYOX1 | 17.1664251 | 0.64841919 | 0.43168033 | 1.50208185 | 0.13307597 | NA |
| SNRPG | 11.6912489 | 0.07478604 | 0.49613695 | 0.15073669 | 0.88018343 | NA |
| FAM136A | 6.4274476 | 0.27171793 | 0.60579816 | 0.44852881 | 0.6537716 | NA |
| TGFA | 7.00682713 | -0.9510775 | 0.88606092 | -1.0733771 | 0.28310196 | NA |
| ATP6V1B1 | 15.3249335 | -0.2373155 | 0.7346665 | -0.3230248 | 7.47E-01 | NA |
| AC007040.2 | 33.3343524 | 2.46605434 | 0.74893708 | 3.29273903 | 0.00099217 | 0.1151343 |
| NAGK | 6.4621381 | -0.4078033 | 0.68678681 | -0.5937843 | 0.55265636 | NA |
| MCEE | 8.55095918 | 0.14103268 | 0.57745308 | 0.24423228 | 0.80705093 | NA |
| MPHOSPH10 | 6.98611653 | 0.13427269 | 0.64167275 | 0.20925415 | 0.83424984 | NA |
| PAIP2B | 2.56332095 | 2.09826329 | 0.90669158 | 2.31419738 | 0.0206569 | NA |
| ZNF638 | 106.624354 | -0.0498492 | 0.19831337 | -0.2513656 | 0.80153146 | 0.96742897 |
| EXOC6B | 137.114945 | -0.3468777 | 0.29712466 | -1.1674484 | 0.24302932 | 0.73323864 |
| SPR | 4.06458201 | 1.43214806 | 0.81401237 | 1.75936892 | 0.07851487 | NA |

| | | | | | | |
|---|---|---|---|---|---|---|
| EMX1 | 5.72504953 | 0.03343052 | 0.95118923 | 0.03514603 | 0.9719633 | NA |
| SFXN5 | 10.9593184 | 0.0988182 | 0.47081101 | 0.20988932 | 0.83375406 | NA |
| RAB11FIP5 | 4.00415281 | -0.3675574 | 0.74268437 | -0.494904 | 0.62066791 | NA |
| PRADC1 | 2.96370861 | 0.15538985 | 0.81243359 | 0.19126468 | 0.84831823 | NA |
| CCT7 | 15.4645583 | 0.20860939 | 0.43547298 | 0.47904095 | 0.6319095 | NA |
| ALMS1 | 22.1383782 | 0.05114729 | 0.35684259 | 0.14333291 | 0.88602726 | NA |
| NAT8 | 5.10509671 | 0.92527975 | 1.2499236 | 0.74026905 | 0.45913676 | NA |
| TPRKB | 2.77707687 | -0.7838818 | 0.83724857 | -0.9362593 | 0.34913969 | NA |
| DUSP11 | 4.18924783 | 1.32194275 | 0.69267265 | 1.90846678 | 0.05633091 | NA |
| STAMBP | 14.0588705 | -0.0275974 | 0.45983872 | -0.0600154 | 0.95214333 | NA |
| DGUOK | 14.5361525 | -0.3755557 | 0.39114701 | -0.9601395 | 0.33698501 | NA |
| TET3 | 25.0166859 | -0.1741083 | 0.51372833 | -0.3389112 | 0.73467665 | NA |
| BOLA3 | 3.60542658 | 1.49431577 | 0.76597426 | 1.95086943 | 0.05107258 | NA |
| MOB1A | 25.7116948 | 0.11990187 | 0.33501004 | 0.35790532 | 0.72041417 | NA |
| DCTN1 | 2.63169616 | 0.4837336 | 0.96364229 | 0.50198461 | 0.61567835 | NA |
| RTKN | 9.67086103 | -0.3282035 | 0.46998485 | -0.6983279 | 0.48497215 | NA |
| MOGS | 5.29471995 | -0.1218102 | 0.65577862 | -0.185749 | 0.85264161 | NA |
| TTC31 | 3.49078791 | -0.1242431 | 0.82521212 | -0.150559 | 0.88032363 | NA |
| AUP1 | 5.13169287 | 0.22645609 | 0.72824586 | 0.31096104 | 0.75583024 | NA |
| HTRA2 | 3.89941303 | 0.98811066 | 0.71679165 | 1.37851865 | 0.16804321 | NA |
| POLE4 | 7.67312451 | -0.4328061 | 0.55071097 | -0.7859043 | 0.43192355 | NA |
| TACR1 | 3.8037896 | 0.5962602 | 0.84900193 | 0.70230724 | 0.48248758 | NA |
| AC007681.1 | 2.63843344 | 1.25706642 | 0.91507905 | 1.3737244 | 0.16952726 | NA |
| EVA1A | 4.00564488 | 0.30059196 | 0.72887667 | 0.41240441 | 0.68004303 | NA |
| MRPL19 | 23.1816479 | 0.90890629 | 0.43877731 | 2.07145236 | 0.03831654 | NA |
| GCFC2 | 24.6916109 | 0.73118544 | 0.38951307 | 1.8771782 | 0.06049367 | NA |
| CTNNA2 | 3.21689425 | -0.8472938 | 0.88303081 | -0.9595291 | 0.33729225 | NA |
| SUCLG1 | 38.5681729 | 0.42975196 | 0.31430284 | 1.3673181 | 0.17152562 | 0.66182328 |
| DNAH6 | 4.04851555 | 1.17651239 | 0.86828157 | 1.35498947 | 0.17542091 | NA |
| TMSB10 | 72.71865 | -0.7204235 | 0.40119597 | -1.7956897 | 0.07254388 | 0.53207981 |
| KCMF1 | 32.1734175 | -0.308252 | 0.30067351 | -1.0252052 | 0.30526638 | 0.77729327 |
| TCF7L1 | 40.4847246 | -0.7636017 | 0.47822444 | -1.5967434 | 0.11032292 | 0.57860396 |
| TGOLN2 | 39.2524672 | 0.01337792 | 0.31737884 | 0.04215126 | 0.96637812 | 0.99656976 |
| RETSAT | 6.16184847 | 0.04548476 | 0.59633999 | 0.0762732 | 0.93920175 | NA |
| ELMOD3 | 12.6487761 | 0.53829324 | 0.51797693 | 1.03922242 | 0.29870131 | NA |
| CAPG | 14.5018259 | 0.63344749 | 0.51531761 | 1.22923704 | 0.21898295 | NA |
| MAT2A | 14.8452564 | -0.084706 | 0.41885076 | -0.2022344 | 0.83973347 | NA |
| GGCX | 6.01695057 | -0.2846846 | 0.68012218 | -0.4185785 | 0.67552417 | NA |
| VAMP8 | 32.0456729 | 0.54975777 | 0.36208009 | 1.51833195 | 0.12893074 | 0.59663251 |
| RNF181 | 5.49371733 | -1.3185901 | 0.68548998 | -1.9235731 | 0.05440811 | NA |
| USP39 | 14.6306593 | -0.0251474 | 0.52992236 | -0.0474549 | 0.96215068 | NA |
| C2orf68 | 6.96625037 | 0.20714739 | 0.55897949 | 0.37058137 | 0.71094936 | NA |
| ST3GAL5 | 7.45106015 | -0.4053904 | 0.64735756 | -0.6262233 | 0.53116848 | NA |
| POLR1A | 11.3004194 | -0.9732471 | 0.56601059 | -1.7194857 | 0.08552597 | NA |
| PTCD3 | 21.857349 | -0.3321081 | 0.4008112 | -0.8285899 | 0.40733648 | NA |
| IMMT | 34.9299275 | -0.0700379 | 0.28747737 | -0.2436293 | 0.80751791 | 0.96822341 |
| MRPL35 | 6.27441711 | 0.64226716 | 0.58454721 | 1.098743 | 0.27188018 | NA |
| REEP1 | 2.95636991 | -0.7988485 | 0.98327275 | -0.8124383 | 0.41654016 | NA |
| KDM3A | 32.2096116 | 0.51629166 | 0.45477696 | 1.13526343 | 0.25626496 | 0.75170774 |
| CHMP3 | 12.8857914 | 0.24723382 | 0.4125986 | 0.59921148 | 0.54903187 | NA |
| RNF103-CHN | 7.56890108 | -1.4719978 | 1.01325225 | -1.4527456 | 0.14629439 | NA |
| RMND5A | 25.0037743 | -0.3403557 | 0.39801767 | -0.855127 | 0.39248084 | NA |
| PLGLB1 | 4.18305321 | -0.0616759 | 0.83765535 | -0.0736292 | 0.94130543 | NA |
| CYTOR | 4.99676291 | -0.2462479 | 0.63086206 | -0.3903355 | 6.96E-01 | NA |

| | | | | | | |
|---|---|---|---|---|---|---|
| KRCC1 | 12.8196331 | 1.00060513 | 0.43320865 | 2.30975332 | 0.02090181 | NA |
| THNSL2 | 4.21804908 | 0.11085053 | 0.85118346 | 0.13023107 | 0.89638362 | NA |
| EIF2AK3 | 19.6462102 | -1.0467431 | 0.40479939 | -2.5858318 | 0.00971443 | NA |
| RPIA | 10.9118593 | 0.64196472 | 0.5742625 | 1.11789419 | 0.26361219 | NA |
| MAL | 130.327323 | 1.25460517 | 0.4129671 | 3.03802688 | 0.00238133 | 0.16646881 |
| MRPS5 | 42.0564297 | 0.06302721 | 0.26899016 | 0.23431048 | 0.81474395 | 0.96845695 |
| AC092835.1 | 3.21510293 | -0.3708391 | 0.95825957 | -0.3869923 | 0.6987619 | NA |
| FAHD2A | 3.5489945 | 1.04054172 | 0.80153073 | 1.29819317 | 0.19422096 | NA |
| ANKRD36C | 12.3623297 | 0.08465667 | 0.59240525 | 0.14290331 | 8.86E-01 | NA |
| STARD7 | 46.6152165 | -0.0511818 | 0.2997229 | -0.1707639 | 0.86440945 | 0.97217819 |
| TMEM127 | 16.5920843 | -0.1555383 | 0.40463568 | -0.3843909 | 0.70068871 | NA |
| CIAO1 | 10.9595338 | 0.42721257 | 0.48029241 | 0.88948431 | 0.37374285 | NA |
| SNRNP200 | 18.3498523 | -0.2297374 | 0.42553599 | -0.5398778 | 0.58928128 | NA |
| KANSL3 | 26.6720667 | -0.2688735 | 0.3671345 | -0.7323569 | 0.46395079 | 0.85894565 |
| LMAN2L | 7.83305681 | 0.11960827 | 0.55443747 | 0.21572904 | 0.82919896 | NA |
| CNNM4 | 16.2711849 | -0.8883015 | 0.6475334 | -1.3718235 | 0.17011839 | NA |
| ANKRD23 | 4.43945367 | 0.02078059 | 0.74147153 | 0.02802615 | 0.97764129 | NA |
| ANKRD36 | 17.2441867 | 0.89506592 | 0.60094518 | 1.48943023 | 0.13637411 | NA |
| AC092683.1 | 12.691125 | 0.85614928 | 0.5198561 | 1.64689665 | 0.09957929 | NA |
| ANKRD36B | 7.51010349 | 1.66291766 | 0.86952536 | 1.91244296 | 0.0558194 | NA |
| COX5B | 47.1582342 | 0.43300357 | 0.41656196 | 1.03946976 | 0.29858632 | 0.77521265 |
| ACTR1B | 5.54239347 | 0.8400423 | 0.5983314 | 1.40397495 | 0.16032631 | NA |
| TMEM131 | 53.8150993 | 0.07779848 | 0.31048068 | 0.25057432 | 0.80214324 | 0.96742897 |
| INPP4A | 84.6349178 | -1.5380286 | 0.47791056 | -3.2182353 | 0.00128982 | 0.1198504 |
| COA5 | 7.24588719 | -0.2259853 | 0.53405511 | -0.4231497 | 0.67218605 | NA |
| UNC50 | 9.6798729 | 0.24003535 | 0.45160801 | 0.53151261 | 0.59506361 | NA |
| MGAT4A | 24.131515 | -0.0938641 | 0.380898 | -0.2464284 | 0.80535061 | NA |
| TSGA10 | 29.0817249 | 0.36413115 | 0.43308406 | 0.84078631 | 0.40046766 | 0.8269015 |
| AC079447.1 | 9.37795427 | 0.71336192 | 0.61665802 | 1.15681933 | 0.24734618 | NA |
| MITD1 | 7.95486783 | 0.7107457 | 0.68178997 | 1.04247017 | 0.29719375 | NA |
| TXNDC9 | 7.18443638 | -0.328479 | 0.56908764 | -0.5772029 | 0.56380242 | NA |
| EIF5B | 31.7026288 | -0.6336336 | 0.32182583 | -1.9688713 | 0.04896787 | 0.46182067 |
| REV1 | 59.7996609 | 0.24729721 | 0.26359107 | 0.93818509 | 0.3481493 | 0.8003972 |
| LONRF2 | 13.8010721 | 0.47912274 | 0.55484044 | 0.86353248 | 0.38784477 | NA |
| CHST10 | 5.45348757 | -0.2705644 | 0.71331774 | -0.3793042 | 0.70446196 | NA |
| PDCL3 | 5.04955884 | -0.0570739 | 0.78235613 | -0.0729513 | 0.94184486 | NA |
| NPAS2 | 11.920122 | -0.9650894 | 0.51787104 | -1.863571 | 0.06238198 | NA |
| RPL31 | 29.053153 | 0.14056323 | 0.32256574 | 0.43576614 | 0.66300641 | 0.92276785 |
| TBC1D8 | 28.3654402 | -0.3641324 | 0.36847675 | -0.9882099 | 0.32304987 | 0.78622244 |
| CNOT11 | 10.3377522 | -0.0304829 | 0.45117448 | -0.0675634 | 0.94613318 | NA |
| RNF149 | 82.8173247 | -0.5868268 | 0.32100785 | -1.8280762 | 0.06753812 | 0.52400942 |
| MAP4K4 | 51.6209107 | -0.6352292 | 0.30965904 | -2.0513828 | 0.04022968 | 0.43121698 |
| IL1R1 | 19.6819678 | 0.34052138 | 0.51472778 | 0.66155626 | 0.50825564 | NA |
| SLC9A2 | 12.0061393 | 0.77220428 | 0.66512441 | 1.16099224 | 0.24564506 | NA |
| MFSD9 | 3.34196095 | 0.42911362 | 0.83073892 | 0.5165445 | 0.60547417 | NA |
| TMEM182 | 3.01021212 | 0.52357058 | 0.83148456 | 0.62968167 | 0.52890288 | NA |
| POU3F3 | 17.3593197 | -0.6079175 | 0.43443472 | -1.3993299 | 0.16171408 | NA |
| MRPS9 | 15.6782201 | 0.60611322 | 0.43943123 | 1.37931302 | 0.16779826 | NA |
| TGFBRAP1 | 7.79895801 | 0.35993609 | 0.59679058 | 0.6031196 | 0.54642913 | NA |
| C2orf49 | 6.06484502 | 0.39431778 | 0.61067792 | 0.64570499 | 0.51847043 | NA |
| NCK2 | 16.9060861 | 0.16824554 | 0.39577741 | 0.42510144 | 0.67076273 | NA |
| UXS1 | 12.3078569 | -0.9910039 | 0.45640809 | -2.171311 | 0.02990767 | NA |
| SULT1C2 | 63.9884909 | 0.17740332 | 0.38232549 | 0.46401123 | 0.6426397 | 0.92153493 |
| GCC2 | 58.0195783 | 0.19804493 | 0.25912292 | 0.7642895 | 0.44469474 | 0.84899161 |

| | | | | | | |
|---|---|---|---|---|---|---|
| GCC2-AS1 | 13.3067112 | 1.31318944 | 0.54418814 | 2.41311659 | 0.01581676 | NA |
| LIMS1 | 256.12961 | -0.9766555 | 0.2783934 | -3.5081849 | 0.00045118 | 0.09262975 |
| RANBP2 | 48.1173855 | -0.096139 | 0.2738209 | -0.3511018 | 0.72551199 | 0.93925046 |
| CCDC138 | 9.7190123 | -0.5174544 | 0.59159695 | -0.8746739 | 0.38175135 | NA |
| 10-Sep | 47.5618252 | -0.0697018 | 0.29749257 | -0.2342976 | 0.81475396 | 0.96845695 |
| SOWAHC | 10.9952325 | 0.03700866 | 0.4396361 | 0.08418022 | 0.93291314 | NA |
| NPHP1 | 3.77786877 | 0.55382225 | 0.82417931 | 0.67196815 | 0.50160397 | NA |
| SMIM37 | 11.7930766 | -0.5055712 | 0.63057844 | -0.8017577 | 0.42269311 | NA |
| LINC01106 | 3.88922758 | -0.0082673 | 0.74418533 | -0.0111092 | 0.99113629 | NA |
| AC112229.3 | 1.87199246 | 0.81928747 | 1.07288757 | 0.76362845 | 0.44508869 | NA |
| MIR4435-2H | 19.5401289 | 8.32E-05 | 0.37728231 | 0.0002206 | 1.00E+00 | NA |
| BCL2L11 | 4.32412299 | -0.25035 | 0.69231525 | -0.3616127 | 0.7176415 | NA |
| ANAPC1 | 19.1040216 | 0.0820513 | 0.41821788 | 0.1961927 | 0.84445934 | NA |
| MERTK | 5.39861479 | 0.85382859 | 0.67884662 | 1.25776362 | 0.20847726 | NA |
| TMEM87B | 18.9158681 | -0.3441982 | 0.39230807 | -0.8773672 | 0.3802872 | NA |
| ZC3H8 | 13.5485394 | -0.402869 | 0.42174294 | -0.9552478 | 0.33945241 | NA |
| ZC3H6 | 14.6924194 | 0.54926683 | 0.57427562 | 0.95645158 | 0.33884414 | NA |
| RGPD8 | 5.4498961 | -1.164749 | 0.73756328 | -1.5791851 | 0.1142936 | NA |
| TTL | 11.6152095 | -0.0416044 | 0.48438764 | -0.0858907 | 0.93155329 | NA |
| POLR1B | 5.59798217 | 0.5410717 | 0.6206203 | 0.87182404 | 0.38330438 | NA |
| CHCHD5 | 6.21575989 | 0.41770776 | 0.5995914 | 0.69665403 | 0.48601934 | NA |
| SLC20A1 | 13.4433444 | -1.3594322 | 0.49591707 | -2.7412491 | 0.00612061 | NA |
| PSD4 | 6.38935476 | 0.26659454 | 0.66759557 | 0.39933539 | 0.6896461 | NA |
| PAX8 | 265.918252 | 0.01118275 | 0.27729741 | 0.04032763 | 0.96783193 | 0.99656976 |
| CBWD2 | 8.67675286 | -0.2648958 | 0.55070837 | -0.4810092 | 0.63050999 | NA |
| RABL2A | 2.679361 | 1.87332888 | 0.96123431 | 1.9488785 | 0.05130994 | NA |
| SLC35F5 | 56.051436 | -0.1158187 | 0.32328517 | -0.3582555 | 0.7201521 | 0.93714576 |
| ACTR3 | 52.1354613 | -0.4128706 | 0.36754067 | -1.1233331 | 0.26129606 | 0.75479335 |
| DDX18 | 14.8151539 | -0.9855351 | 0.46823985 | -2.1047655 | 0.03531173 | NA |
| CCDC93 | 28.1110636 | -0.2024269 | 0.35855424 | -0.5645643 | 0.57237012 | 0.89263108 |
| INSIG2 | 25.1676925 | -0.0447349 | 0.44950264 | -0.0995208 | 0.92072475 | NA |
| C2orf76 | 3.56082327 | -0.4813084 | 0.86054427 | -0.559307 | 0.57595224 | NA |
| DBI | 60.1888562 | -0.1897991 | 0.42078078 | -0.4510641 | 0.65194338 | 0.92248109 |
| TMEM37 | 4.34819732 | 0.46756881 | 0.82597097 | 0.56608383 | 0.57133679 | NA |
| CFAP221 | 14.1657765 | 0.45921838 | 0.57003727 | 0.80559362 | 0.42047721 | NA |
| PTPN4 | 43.1628667 | 0.35439047 | 0.36971294 | 0.95855574 | 0.3377826 | 0.8003972 |
| EPB41L5 | 66.2571064 | -0.1738352 | 0.26211083 | -0.6632124 | 0.50719451 | 0.87158522 |
| RALB | 7.2664089 | 0.16689138 | 0.52547722 | 0.31759965 | 0.75078864 | NA |
| TFCP2L1 | 485.64959 | -0.7118306 | 0.21758137 | -3.2715603 | 0.00106956 | 0.11692613 |
| CLASP1 | 98.7336587 | 0.05106532 | 0.29603208 | 0.17249929 | 0.86304502 | 0.97217819 |
| NIFK-AS1 | 5.03231761 | 0.96302263 | 0.77983496 | 1.23490568 | 0.21686561 | NA |
| NIFK | 5.99481759 | 0.23161535 | 0.60189153 | 0.38481245 | 0.70037638 | NA |
| TSN | 11.3955737 | -0.1713644 | 0.51562635 | -0.3323422 | 0.73963085 | NA |
| BIN1 | 5.96665765 | -0.0835078 | 0.70417649 | -0.1185893 | 0.90560073 | NA |
| ERCC3 | 9.79335034 | -0.7345544 | 0.51677798 | -1.421412 | 0.15519702 | NA |
| MAP3K2 | 57.3852212 | 0.14083952 | 0.2753481 | 0.51149624 | 0.60900362 | 0.9088434 |
| IWS1 | 22.889875 | 0.11637233 | 0.3905781 | 0.29794894 | 0.76574213 | NA |
| WDR33 | 46.5289706 | -0.4838395 | 0.29448288 | -1.6430141 | 0.10038001 | 0.57124575 |
| SFT2D3 | 2.53778291 | -0.30627 | 0.99304658 | -0.3084145 | 0.75776695 | NA |
| POLR2D | 4.33304489 | 0.58027546 | 0.71304598 | 0.81379809 | 0.41576063 | NA |
| AMMECR1L | 8.3218622 | -0.0338081 | 0.61798654 | -0.0547068 | 0.95637205 | NA |
| SAP130 | 11.3692651 | 0.46658816 | 0.48803378 | 0.95605711 | 0.33904338 | NA |
| UGGT1 | 22.1092372 | 0.24892224 | 0.43992862 | 0.56582415 | 0.57151333 | NA |
| HS6ST1 | 21.8418128 | -0.3018596 | 0.4615958 | -0.6539479 | 0.51314538 | NA |

| | | | | | | |
|---|---|---|---|---|---|---|
| SMPD4 | 11.7284439 | -0.8587198 | 0.4491471 | -1.9118898 | 0.05589033 | NA |
| MZT2B | 17.2114239 | -0.6618307 | 0.45274582 | -1.4618151 | 0.14379189 | NA |
| CCDC115 | 5.45095597 | 0.45200348 | 0.67921197 | 0.6654822 | 0.50574212 | NA |
| IMP4 | 5.00125135 | -0.11716 | 0.63706276 | -0.1839065 | 0.85408678 | NA |
| PTPN18 | 9.43905451 | 0.33798524 | 0.53782835 | 0.62842586 | 0.529725 | NA |
| FAM168B | 38.5267415 | -0.3620493 | 0.36879876 | -0.9816988 | 0.32624826 | 0.78801502 |
| PLEKHB2 | 62.7334764 | -0.329545 | 0.31896578 | -1.0331673 | 0.30152559 | 0.77521265 |
| MZT2A | 8.46950706 | 0.17646347 | 0.47589184 | 0.37080583 | 0.71078216 | NA |
| GPR39 | 13.340715 | -0.1372454 | 0.59638906 | -0.2301273 | 0.81799288 | NA |
| NCKAP5 | 28.216703 | 1.20806038 | 0.59489217 | 2.0307216 | 0.04228324 | 0.44186191 |
| MGAT5 | 52.0394505 | -0.3473019 | 0.40277566 | -0.8622713 | 0.38853822 | 0.8258317 |
| TMEM163 | 55.4546006 | -0.9023172 | 0.52454875 | -1.720178 | 0.08540009 | 0.55323508 |
| CCNT2-AS1 | 9.71748158 | -1.0824216 | 0.87706797 | -1.2341365 | 0.21715204 | NA |
| ACMSD | 3.23238462 | 1.18301176 | 1.13139607 | 1.04562124 | 0.29573594 | NA |
| CCNT2 | 48.2576996 | -0.9443863 | 0.36324753 | -2.5998425 | 0.00932666 | 0.27655812 |
| RAB3GAP1 | 48.1346018 | 0.03115886 | 0.2715849 | 0.11472971 | 0.90865937 | 0.98306115 |
| ZRANB3 | 19.4963659 | 0.5003755 | 0.48897287 | 1.02331955 | 0.30615678 | NA |
| R3HDM1 | 42.6040939 | 0.08548501 | 0.28219209 | 0.30293197 | 0.76194171 | 0.95478238 |
| UBXN4 | 44.606394 | -0.2861551 | 0.32681929 | -0.8755758 | 0.38126067 | 0.82153751 |
| DARS | 36.4811433 | 0.33678655 | 0.31298469 | 1.07604802 | 0.28190578 | 0.76854599 |
| HNMT | 19.830209 | 0.57723018 | 0.50094165 | 1.15229025 | 0.24920182 | NA |
| AC114763.1 | 7.31172946 | 1.23447744 | 0.65068218 | 1.89720492 | 0.0578009 | NA |
| SPOPL | 31.8561359 | 0.07013346 | 0.32800036 | 0.21382129 | 0.83068642 | 0.97113537 |
| NXPH2 | 16.7566745 | -0.8856777 | 0.95138899 | -0.9309312 | 0.35188917 | NA |
| LRP1B | 56.5610513 | 0.42800378 | 0.64207581 | 0.66659384 | 5.05E-01 | 0.87094108 |
| GTDC1 | 19.9825802 | -0.0934959 | 0.44804076 | -0.2086773 | 0.83470015 | NA |
| TEX41 | 21.5587696 | 0.28908504 | 1.20311921 | 0.24027963 | 0.81011349 | NA |
| ACVR2A | 21.501587 | 0.0339917 | 0.37078318 | 0.09167542 | 0.92695593 | NA |
| ORC4 | 37.4302214 | -0.2487045 | 0.32156677 | -0.7734149 | 0.43927685 | 0.84524078 |
| MBD5 | 131.865994 | -0.1470668 | 0.20653356 | -0.712072 | 0.47642017 | 0.86206794 |
| EPC2 | 54.2461444 | -0.6023314 | 0.32149311 | -1.8735438 | 0.06099333 | 0.51625203 |
| AC105402.3 | 5.90041171 | -0.9095285 | 1.04232864 | -0.8725929 | 0.38288503 | NA |
| MMADHC | 8.31216359 | -0.151997 | 0.54886345 | -0.2769305 | 0.78183351 | NA |
| AC007364.1 | 3.72954515 | -1.3738433 | 0.7842234 | -1.751852 | 0.07979926 | NA |
| NMI | 4.59703017 | 0.56401332 | 0.73278538 | 0.7696842 | 0.44148725 | NA |
| RIF1 | 37.5986537 | -0.076363 | 0.38673401 | -0.1974562 | 0.84347059 | 0.97168963 |
| ARL5A | 9.82597061 | 0.60166896 | 0.50256252 | 1.1972022 | 0.23122775 | NA |
| CACNB4 | 6.90846237 | -0.5305712 | 0.67615805 | -0.7846852 | 0.43263817 | NA |
| STAM2 | 26.1471849 | -0.1984165 | 0.32924497 | -0.6026409 | 0.54674762 | 0.88410237 |
| FMNL2 | 66.5343282 | -0.0209894 | 0.32755376 | -0.0640793 | 0.94890711 | 0.99344789 |
| PRPF40A | 37.3458944 | 0.06206093 | 0.31570452 | 0.19657915 | 0.84415689 | 0.97168963 |
| ARL6IP6 | 14.0802476 | 0.41623389 | 0.47834525 | 0.8701537 | 0.38421642 | NA |
| KCNJ3 | 27.460158 | -1.1330139 | 0.704152 | -1.6090474 | 0.10760599 | 0.57769936 |
| LINC01876 | 12.6053055 | -0.4853362 | 0.78611537 | -0.6173855 | 0.53698048 | NA |
| GPD2 | 26.4762542 | 0.08729067 | 0.35650762 | 0.24484939 | 0.80657305 | 0.96800006 |
| CYTIP | 22.1840577 | 1.19312815 | 0.65421976 | 1.82374215 | 0.06819106 | NA |
| ACVR1 | 39.7425658 | -0.1853735 | 0.3418869 | -0.5422071 | 0.58767586 | 0.89477098 |
| UPP2 | 6.71775705 | 1.02644914 | 0.67857291 | 1.5126586 | 0.1303664 | NA |
| CCDC148 | 17.508009 | 0.32630067 | 0.48587461 | 0.67157382 | 0.50185505 | NA |
| PKP4 | 103.647004 | 0.18042982 | 0.30609805 | 0.58945106 | 0.55555873 | 0.89109751 |
| TANC1 | 57.8804184 | 0.00920238 | 0.30132751 | 0.03053947 | 0.97563682 | 0.99853281 |
| WDSUB1 | 31.4101259 | -0.0869542 | 0.36505628 | -0.2381939 | 0.81173072 | 0.96845695 |
| BAZ2B | 162.005709 | 0.27670915 | 0.1928648 | 1.4347312 | 0.15136371 | 0.63620431 |
| AC009961.1 | 3.68096952 | 0.30409975 | 0.73232311 | 0.41525352 | 0.67795632 | NA |

| | | | | | | |
|---|---|---|---|---|---|---|
| 7-Mar | 55.5476316 | -0.6431395 | 0.252347 | -2.5486316 | 0.01081465 | 0.28312744 |
| PLA2R1 | 4.01645379 | 0.68661013 | 0.68325738 | 1.00490701 | 0.31494163 | NA |
| ITGB6 | 43.9169118 | -1.1178865 | 1.1920236 | -0.9378057 | 0.34834426 | 0.8003972 |
| RBMS1 | 50.410286 | -0.2355112 | 0.37454259 | -0.6287969 | 0.52948206 | 0.88254911 |
| TANK | 30.4142671 | 0.60377433 | 0.3726135 | 1.62037697 | 0.10515132 | 0.5772605 |
| PSMD14 | 21.8056794 | -0.6221867 | 0.40315208 | -1.5433053 | 0.12275672 | NA |
| IFIH1 | 5.45809799 | -0.0357595 | 0.88410919 | -0.0404469 | 0.96773684 | NA |
| GCA | 8.06576156 | 0.17971533 | 0.61226321 | 0.29352626 | 0.76911993 | NA |
| FIGN | 8.39744152 | 0.82360953 | 0.77350735 | 1.06477272 | 0.28697879 | NA |
| GRB14 | 46.8693298 | -0.2936519 | 0.47455814 | -0.6187901 | 0.53605462 | 0.88254911 |
| COBLL1 | 336.021199 | -0.0334624 | 0.25269107 | -0.1324241 | 0.89464885 | 0.97769824 |
| AC019197.1 | 170.813349 | 1.68589814 | 0.58358308 | 2.88887425 | 0.00386624 | 0.2109414 |
| SLC38A11 | 39.7848262 | 0.96371612 | 0.72326307 | 1.33245588 | 0.18271043 | 0.67478132 |
| SCN3A | 13.0396352 | 0.56481379 | 0.56842847 | 0.99364092 | 0.32039772 | NA |
| SCN2A | 18.1980573 | 2.15393453 | 0.67802254 | 3.17678897 | 0.00148915 | NA |
| CSRNP3 | 10.6260035 | 0.99532032 | 0.73044202 | 1.3626274 | 0.17299997 | NA |
| GALNT3 | 11.5136473 | 0.19194823 | 0.44117376 | 0.43508534 | 0.66350048 | NA |
| TTC21B | 18.8953846 | 0.35206054 | 0.38845065 | 0.90631987 | 0.36476657 | NA |
| AC010127.1 | 3.66027211 | -0.3851966 | 1.09996588 | -0.3501896 | 0.72619643 | NA |
| SCN9A | 17.1228207 | 0.49204365 | 0.56065051 | 0.8776299 | 0.3801446 | NA |
| STK39 | 17.7923555 | 0.04528228 | 0.78180043 | 0.05792051 | 0.95381194 | NA |
| CERS6 | 35.1657441 | -1.141297 | 0.4671637 | -2.4430344 | 0.01456435 | 0.30435125 |
| LRP2 | 20.3578145 | 0.03188125 | 0.63325666 | 0.05034492 | 0.95984753 | NA |
| FASTKD1 | 38.2445923 | -0.7742603 | 0.33712512 | -2.2966556 | 0.02163843 | 0.35108799 |
| PPIG | 32.905227 | -0.1463997 | 0.33825698 | -0.4328062 | 0.66515555 | 0.92276785 |
| KLHL23 | 44.5679376 | -0.2837902 | 0.33551506 | -0.8458343 | 0.39764518 | 0.82656931 |
| SSB | 15.6004853 | -0.3596849 | 0.46356063 | -0.7759176 | 0.43779763 | NA |
| METTL5 | 3.90211859 | -0.3863884 | 0.6820449 | -0.5665146 | 0.57104402 | NA |
| UBR3 | 100.857516 | -0.2405981 | 0.25791042 | -0.9328745 | 0.35088476 | 0.8003972 |
| MYO3B | 19.5557502 | 0.67502923 | 0.72348803 | 0.9330206 | 0.35080933 | NA |
| GORASP2 | 14.9744299 | -0.4573967 | 0.44636864 | -1.0247062 | 0.30550185 | NA |
| TLK1 | 62.0173038 | -0.4670461 | 0.2326956 | -2.0071117 | 0.04473777 | 0.44769598 |
| METTL8 | 6.47817153 | -0.5328324 | 0.72010145 | -0.7399408 | 0.45933594 | NA |
| DCAF17 | 7.87634727 | 0.70746787 | 0.57643159 | 1.22732322 | 0.21970113 | NA |
| DYNC1I2 | 35.7945703 | 0.00476411 | 0.37225706 | 0.0127979 | 0.98978903 | 0.99909657 |
| SLC25A12 | 39.3793776 | 0.74989427 | 0.35694246 | 2.10088277 | 0.03565126 | 0.41688714 |
| HAT1 | 12.8362971 | -0.2394246 | 0.44227318 | -0.54135 | 0.58826639 | NA |
| METAP1D | 4.73247303 | 1.02639283 | 0.85582531 | 1.19930179 | 0.23041062 | NA |
| ITGA6 | 205.994785 | 0.62649104 | 0.44822469 | 1.39771648 | 0.16219822 | 0.64709575 |
| AC078883.1 | 7.66664842 | 1.23332726 | 0.63113579 | 1.95413932 | 0.05068474 | NA |
| PDK1 | 14.3576165 | -0.6499445 | 0.62797641 | -1.0349824 | 0.30067714 | NA |
| MAP3K20 | 5.48708393 | -0.9473787 | 0.63070397 | -1.5020973 | 0.13307198 | NA |
| SP3 | 28.0918094 | -0.253497 | 0.35311671 | -0.7178846 | 0.47282846 | 0.86206794 |
| OLA1 | 52.8780643 | -0.3241949 | 0.36146168 | -0.8968999 | 0.36977235 | 0.81199861 |
| CIR1 | 14.3906256 | 0.44474684 | 0.41523641 | 1.07106899 | 0.28413841 | NA |
| SCRN3 | 6.09081114 | 0.39212449 | 0.62596734 | 0.62642963 | 0.53103318 | NA |
| GPR155 | 6.21840687 | 0.51342243 | 0.87100168 | 0.58946204 | 0.55555137 | NA |
| AC010894.3 | 37.9637737 | 0.4033863 | 0.35000635 | 1.15251138 | 0.249111 | 0.74122324 |
| WIPF1 | 4.50413574 | -0.0864794 | 0.73365527 | -0.1178747 | 0.90616694 | NA |
| CHN1 | 5.40632256 | -1.0597887 | 0.99562218 | -1.0644487 | 0.28712549 | NA |
| ATF2 | 37.0285089 | -0.0425785 | 0.27550475 | -0.1545471 | 0.87717836 | 0.9751578 |
| ATP5MC3 | 48.0104235 | 0.13323288 | 0.43082181 | 0.30925286 | 0.75712919 | 0.95323728 |
| LNPK | 14.2327887 | 0.25690858 | 0.4856434 | 0.52900664 | 0.59680084 | NA |
| HOXD10 | 7.25171037 | 0.85844012 | 0.68439351 | 1.25430779 | 0.20973015 | NA |

| | | | | | | |
|---|---|---|---|---|---|---|
| HOXD-AS2 | 2.51881713 | 0.55225459 | 0.99411317 | 0.55552487 | 0.5785357 | NA |
| HOXD9 | 2.83607785 | 0.34449782 | 0.8831055 | 0.39009815 | 0.69646397 | NA |
| HOXD8 | 7.68824016 | 0.26067085 | 0.60068595 | 0.43395529 | 0.66432091 | NA |
| HOXD3 | 7.59087556 | 0.24468117 | 0.54273802 | 0.45082741 | 0.65211395 | NA |
| HAGLR | 3.43217869 | 0.19478486 | 0.7884629 | 0.24704379 | 0.80487434 | NA |
| MTX2 | 54.3169617 | 0.21027971 | 0.26232044 | 0.80161391 | 0.42277633 | 0.84057082 |
| AC074286.1 | 9.647254 | 0.08219295 | 0.71319401 | 0.11524627 | 0.90824993 | NA |
| HNRNPA3 | 54.8942856 | -0.1936825 | 0.25551914 | -0.7579962 | 0.44845324 | 0.85123446 |
| NFE2L2 | 127.020912 | 0.114941 | 0.23375136 | 0.49172336 | 0.62291492 | 0.91493825 |
| AGPS | 29.5651931 | -0.2368483 | 0.34110028 | -0.6943656 | 0.48745299 | 0.86354292 |
| RBM45 | 5.55952657 | 0.14858182 | 0.65263963 | 0.22766288 | 0.81990833 | NA |
| AC009948.1 | 8.71224867 | -0.1201577 | 0.78076395 | -0.1538976 | 0.8776905 | NA |
| PRKRA | 22.7226096 | 0.25831906 | 0.43016922 | 0.60050569 | 0.54816927 | NA |
| PJVK | 2.89157031 | 0.87923867 | 0.89684519 | 0.98036839 | 0.32690431 | NA |
| PLEKHA3 | 12.2547667 | 0.78299501 | 0.46926882 | 1.66854257 | 0.09520807 | NA |
| TTN-AS1 | 29.3521911 | 0.13274365 | 0.30614915 | 0.43359143 | 0.66458516 | 0.92276785 |
| SESTD1 | 99.7411239 | 0.06932429 | 0.29622574 | 0.23402522 | 0.8149654 | 0.96845695 |
| ZNF385B | 9.25800288 | 2.17934674 | 0.84736735 | 2.57190315 | 0.01011412 | NA |
| CWC22 | 13.6542633 | 0.52958397 | 0.4931085 | 1.07397048 | 0.28283591 | NA |
| UBE2E3 | 39.8184476 | -0.1669531 | 0.36656461 | -0.4554535 | 0.648783 | 0.92202688 |
| SSFA2 | 29.8754767 | -0.8630058 | 0.41117234 | -2.0988906 | 0.03582655 | 0.41688714 |
| PDE1A | 53.3397916 | -0.8655768 | 0.73767515 | -1.1733848 | 0.24064152 | 0.73323864 |
| DNAJC10 | 14.6987113 | 0.29764362 | 0.54275544 | 0.54839362 | 0.58342165 | NA |
| NCKAP1 | 63.3944437 | 0.05700841 | 0.23212343 | 0.24559526 | 0.80599556 | 0.96800006 |
| NUP35 | 8.38799507 | -0.8207744 | 0.67776747 | -1.2109971 | 0.22589651 | NA |
| LINC01473 | 6.20062129 | 0.29982714 | 0.71320343 | 0.42039498 | 0.67419694 | NA |
| ZC3H15 | 20.8131929 | -0.5189992 | 0.37079141 | -1.3997067 | 0.16160116 | NA |
| ITGAV | 55.3521553 | -0.7954254 | 0.40593382 | -1.9594954 | 0.0500548 | 0.46549223 |
| AC017101.1 | 4.6442429 | -0.4423916 | 0.84872374 | -0.5212434 | 0.60219724 | NA |
| AC007319.1 | 12.7616617 | -0.2166789 | 0.73489998 | -0.2948413 | 0.76811513 | NA |
| TFPI | 19.5556041 | -0.2697232 | 0.675118 | -0.3995202 | 0.68950998 | NA |
| LINC01090 | 6.89788486 | -0.1819457 | 0.68688687 | -0.2648846 | 0.79109838 | NA |
| GULP1 | 134.622286 | -0.7617443 | 0.35440857 | -2.1493394 | 0.0316075 | 0.4126665 |
| WDR75 | 7.45398245 | -0.1467153 | 0.56538127 | -0.2594981 | 0.79525098 | NA |
| SLC40A1 | 5.94282678 | 0.77485653 | 0.67341471 | 1.15063796 | 0.24988121 | NA |
| ASNSD1 | 4.10888024 | 0.14264971 | 0.84534029 | 0.16874827 | 0.86599465 | NA |
| ANKAR | 16.7919472 | 0.22544427 | 0.45091192 | 0.49997407 | 0.61709333 | NA |
| ORMDL1 | 12.9854903 | 0.45872846 | 0.53440897 | 0.85838467 | 0.39068009 | NA |
| PMS1 | 12.9158347 | 0.22644214 | 0.42993511 | 0.5266891 | 0.59840951 | NA |
| C2orf88 | 4.47993979 | -0.0591536 | 0.82367883 | -0.0718163 | 0.9427481 | NA |
| HIBCH | 27.8289447 | 0.51464914 | 0.38238783 | 1.34588264 | 0.17834037 | 0.66946617 |
| INPP1 | 5.3346801 | 0.37952254 | 0.7235297 | 0.52454314 | 0.59990082 | NA |
| MFSD6 | 13.0519591 | -0.1949714 | 0.4700227 | -0.4148128 | 0.67827897 | NA |
| NAB1 | 6.35411669 | 0.44060125 | 0.68134347 | 0.64666541 | 0.51784852 | NA |
| GLS | 157.035745 | -0.4862532 | 0.34330062 | -1.4164064 | 0.15665656 | 0.63932163 |
| STAT1 | 33.436856 | -0.6138692 | 0.74809148 | -0.8205804 | 0.41188534 | 0.83218924 |
| MYO1B | 23.6243122 | 0.00221279 | 0.46446986 | 0.00476412 | 0.9961988 | NA |
| SLC39A10 | 7.52523634 | 0.9310646 | 0.59029615 | 1.57728388 | 0.11473022 | NA |
| DNAH7 | 7.02746829 | 0.59414576 | 0.520731 | 1.14098404 | 0.25387656 | NA |
| GTF3C3 | 17.9152001 | -0.0723724 | 0.44144802 | -0.1639431 | 0.86977594 | NA |
| PGAP1 | 26.4966829 | -1.1782783 | 0.38581196 | -3.0540222 | 0.00225795 | 0.16287781 |
| ANKRD44 | 40.9130345 | -0.8971493 | 0.41424936 | -2.1657228 | 0.03033237 | 0.40822098 |
| SF3B1 | 50.2145621 | 0.1447144 | 0.26838189 | 0.53921075 | 0.58974145 | 0.89534694 |
| COQ10B | 9.27044929 | -0.7961534 | 0.48225534 | -1.6508959 | 0.09875983 | NA |

| | | | | | | |
|---|---|---|---|---|---|---|
| HSPD1 | 42.1490233 | -0.1160929 | 0.30736042 | -0.3777095 | 0.70564642 | 0.93143932 |
| MOB4 | 5.01876175 | 0.39465321 | 0.70752185 | 0.5577965 | 0.57698336 | NA |
| AC011997.1 | 2.52611161 | -0.5066386 | 0.94929193 | -0.5337015 | 0.59354805 | NA |
| PLCL1 | 351.462353 | 0.65215621 | 0.29664864 | 2.19841295 | 0.02791969 | 0.39934064 |
| FTCDNL1 | 9.12016768 | 0.78939618 | 0.50847149 | 1.55248856 | 0.12054537 | NA |
| C2orf69 | 6.35985303 | 0.44847728 | 0.60748993 | 0.73824645 | 0.4603647 | NA |
| TYW5 | 8.59667124 | 0.48375419 | 0.52863726 | 0.91509666 | 0.36014089 | NA |
| MAIP1 | 3.89355923 | 0.45015495 | 0.78179255 | 0.57579846 | 0.56475141 | NA |
| SPATS2L | 31.6094297 | -0.3950603 | 0.37117519 | -1.0643501 | 0.28717015 | 0.77250927 |
| KCTD18 | 3.8938726 | 0.50346796 | 0.7700493 | 0.65381263 | 0.51323253 | NA |
| BZW1 | 9.42668114 | -0.08032 | 0.51382042 | -0.1563193 | 0.87578135 | NA |
| CLK1 | 27.5565771 | -0.0185 | 0.34475906 | -0.0536607 | 0.95720547 | 0.9964046 |
| PPIL3 | 6.2165138 | 0.1002554 | 0.57119435 | 0.1755189 | 0.86067192 | NA |
| NIF3L1 | 4.68767676 | 0.6843704 | 0.6827164 | 1.00242267 | 0.3161395 | NA |
| ORC2 | 25.3265636 | -0.6597957 | 0.36453351 | -1.8099726 | 0.07030003 | NA |
| AC005037.1 | 6.16919092 | -0.1669961 | 0.87198663 | -0.1915122 | 0.84812431 | NA |
| FAM126B | 33.9852877 | 0.29463213 | 0.31341944 | 0.94005699 | 0.34718833 | 0.8003972 |
| NDUFB3 | 19.0482071 | 0.08128083 | 0.40921682 | 0.19862534 | 0.84255583 | NA |
| CFLAR | 45.3724916 | -0.0980566 | 0.29963906 | -0.327249 | 0.74347959 | 0.94831173 |
| CFLAR-AS1 | 5.03513677 | 0.5412612 | 0.74786102 | 0.72374571 | 0.46922186 | NA |
| CASP10 | 5.40605035 | 0.44992656 | 0.75068307 | 0.5993562 | 0.54893538 | NA |
| CASP8 | 10.3058641 | -0.3911362 | 0.59256709 | -0.6600707 | 0.50920845 | NA |
| ALS2CR12 | 3.03116838 | -0.3361837 | 1.0123672 | -0.3320768 | 0.73983126 | NA |
| TRAK2 | 24.1186757 | -0.2849577 | 0.44539947 | -0.6397801 | 0.52231558 | NA |
| STRADB | 9.04081326 | 0.38489148 | 0.6643634 | 0.57933877 | 0.56236061 | NA |
| TMEM237 | 2.79153275 | 0.27780888 | 0.95944988 | 0.28955018 | 0.77216039 | NA |
| ALS2 | 17.9684174 | 0.23788385 | 0.36999791 | 0.64293297 | 0.52026759 | NA |
| FZD7 | 5.1384523 | -3.2303591 | 1.06684935 | -3.0279431 | 0.00246224 | NA |
| SUMO1 | 21.1371907 | 0.24561523 | 0.38394135 | 0.6397207 | 0.5223542 | NA |
| NOP58 | 22.1567168 | -0.5484132 | 0.34645421 | -1.5829312 | 0.11343715 | NA |
| BMPR2 | 90.0453637 | 0.09733729 | 0.26156369 | 0.37213608 | 0.70979153 | 0.93413886 |
| FAM117B | 50.3752566 | 0.36933317 | 0.39673709 | 0.93092673 | 0.35189146 | 0.8003972 |
| ICA1L | 12.739692 | -0.4725541 | 0.50706502 | -0.9319398 | 0.35136761 | NA |
| WDR12 | 10.2280762 | 0.19501723 | 0.58497005 | 0.33337985 | 0.73884757 | NA |
| CARF | 15.155418 | 0.61766748 | 0.51855764 | 1.19112599 | 0.23360413 | NA |
| NBEAL1 | 143.375719 | 0.13218204 | 0.24501443 | 0.53948675 | 0.58955104 | 0.89534694 |
| CYP20A1 | 31.5929733 | 0.0459107 | 0.30037869 | 0.15284273 | 0.8785223 | 0.97516771 |
| ABI2 | 37.9811529 | -0.0224335 | 0.2884438 | -0.0777741 | 0.93800775 | 0.99100968 |
| RAPH1 | 48.1573421 | -0.0620913 | 0.30733301 | -0.2020327 | 0.83989117 | 0.97168963 |
| PARD3B | 213.043473 | 0.36143876 | 0.25299864 | 1.42861937 | 0.15311366 | 0.63864841 |
| NRP2 | 12.8614222 | -0.2980178 | 0.59031858 | -0.5048424 | 0.61366956 | NA |
| INO80D | 72.1354065 | -0.637115 | 0.4784457 | -1.3316348 | 0.18298022 | 0.67478132 |
| NDUFS1 | 43.5758155 | 0.26957007 | 0.28943444 | 0.93136835 | 0.35166306 | 0.8003972 |
| EEF1B2 | 21.9101746 | -0.018255 | 0.48797451 | -0.0374097 | 0.97015832 | NA |
| ZDBF2 | 11.2823088 | -0.3689327 | 0.47505417 | -0.7766118 | 0.43738781 | NA |
| FASTKD2 | 10.4779538 | 0.02679551 | 0.4930177 | 0.05434999 | 0.95665632 | NA |
| KLF7 | 25.8374549 | -0.5426459 | 0.45792325 | -1.185015 | 0.2360115 | NA |
| LINC01802 | 5.42125432 | 0.36449793 | 0.90646088 | 0.40211104 | 0.68760231 | NA |
| CREB1 | 23.5286781 | 0.00480001 | 0.35907155 | 0.01336785 | 0.98933432 | NA |
| METTL21A | 6.17221456 | -0.1003875 | 0.59260672 | -0.1693999 | 0.86548208 | NA |
| CCNYL1 | 8.35551351 | -0.9112515 | 0.75028456 | -1.2145412 | 0.22454114 | NA |
| PLEKHM3 | 14.5044593 | 0.66130672 | 0.46981005 | 1.40760445 | 0.15924823 | NA |
| IDH1 | 9.54137856 | 0.29011148 | 0.64343527 | 0.45087905 | 0.65207672 | NA |
| PIKFYVE | 28.2816229 | -0.5353779 | 0.41607749 | -1.2867265 | 0.19818965 | 0.69343545 |

| | | | | | | |
|---|---|---|---|---|---|---|
| PTH2R | 60.0523088 | -0.0686283 | 0.58387204 | -0.1175399 | 0.90643222 | 0.9814473 |
| RPE | 12.7021402 | -0.0189489 | 0.448375 | -0.0422613 | 0.96629038 | NA |
| KANSL1L | 98.1674042 | 1.00296948 | 0.35894628 | 2.79420496 | 0.00520275 | 0.22396996 |
| AC007038.1 | 10.7779615 | 0.37918763 | 0.58260738 | 0.65084592 | 0.51514596 | NA |
| ACADL | 9.02796869 | 0.72684268 | 0.65940608 | 1.10226868 | 0.27034488 | NA |
| LANCL1 | 14.8144272 | 0.20404039 | 0.42683895 | 0.47802664 | 0.63263125 | NA |
| ERBB4 | 330.897468 | 0.37543232 | 0.33577794 | 1.11809702 | 0.26352556 | 0.75479335 |
| IKZF2 | 11.6747699 | 1.88997105 | 0.66417927 | 2.84557366 | 0.00443315 | NA |
| SPAG16 | 36.7027162 | 0.57241832 | 0.47506037 | 1.20493806 | 0.22822721 | 0.73156941 |
| AC068051.1 | 6.2304035 | 1.11542501 | 0.78005821 | 1.42992536 | 0.15273844 | NA |
| BARD1 | 6.0898378 | -0.2572265 | 0.63781039 | -0.4032962 | 0.68673031 | NA |
| AC016708.1 | 6.53396127 | -0.2502974 | 0.80738658 | -0.3100094 | 0.7565538 | NA |
| ATIC | 9.00746936 | 0.41350578 | 0.52428251 | 0.78870795 | 0.43028272 | NA |
| MREG | 11.4863034 | 0.47105758 | 0.49015328 | 0.96104137 | 0.33653137 | NA |
| PECR | 17.3352496 | 0.81111493 | 0.40386605 | 2.00837613 | 0.04460334 | NA |
| XRCC5 | 67.6843046 | -0.0768789 | 0.24321058 | -0.3161002 | 0.75192648 | 0.95164077 |
| SMARCAL1 | 6.74502125 | -0.3150698 | 0.60207608 | -0.5233056 | 0.60076161 | NA |
| RPL37A | 55.7084537 | -0.4073127 | 0.38876835 | -1.0477003 | 0.2947767 | 0.77460836 |
| IGFBP2 | 2.93062488 | 0.15573521 | 1.02330623 | 0.15218827 | 0.87903844 | NA |
| IGFBP5 | 935.612562 | -0.5914514 | 0.38457204 | -1.5379469 | 0.12406159 | 0.58553957 |
| AC007563.2 | 60.619955 | 0.09134053 | 0.57555676 | 0.15869944 | 0.87390568 | 0.9751578 |
| DIRC3-AS1 | 3.43314256 | -1.8191644 | 1.23116761 | -1.4775928 | 0.1395168 | NA |
| DIRC3 | 16.7285147 | -0.1230089 | 0.72045312 | -0.1707383 | 0.86442957 | NA |
| TNS1 | 35.458503 | 0.89281282 | 0.41891706 | 2.13124003 | 0.03306937 | 0.41688714 |
| ARPC2 | 32.104906 | -0.6107755 | 0.29382126 | -2.0787313 | 0.03764206 | 0.42032585 |
| AAMP | 8.15044831 | -0.3827642 | 0.55554464 | -0.6889891 | 0.49083014 | NA |
| PNKD | 22.0980747 | 0.01577446 | 0.37273856 | 0.04232043 | 0.96624326 | NA |
| TMBIM1 | 17.9560626 | 0.11493061 | 0.42024705 | 0.27348344 | 0.78448162 | NA |
| CTDSP1 | 13.9411165 | -0.2946992 | 0.45630647 | -0.6458362 | 0.51838545 | NA |
| USP37 | 24.4411901 | -0.1471348 | 0.35867473 | -0.410218 | 0.68164601 | NA |
| CNOT9 | 11.0106413 | -0.5695444 | 0.46217578 | -1.2323112 | 0.21783287 | NA |
| PLCD4 | 6.40091424 | 0.93116016 | 0.84442656 | 1.10271301 | 0.27015181 | NA |
| ZNF142 | 4.84395179 | 0.00140479 | 0.67610952 | 0.00207776 | 0.99834219 | NA |
| STK36 | 2.55837048 | 0.65218045 | 0.89756312 | 0.72661235 | 0.46746346 | NA |
| TTLL4 | 4.79427816 | 0.50633131 | 0.63514348 | 0.79719201 | 0.42533952 | NA |
| CYP27A1 | 8.03983746 | 0.63667905 | 0.64101501 | 0.99323578 | 0.32059506 | NA |
| CNPPD1 | 10.8184771 | 0.18101579 | 0.50040317 | 0.36173989 | 0.71754642 | NA |
| RETREG2 | 13.5888533 | -0.5979413 | 0.47692874 | -1.253733 | 0.20993909 | NA |
| ZFAND2B | 5.18278666 | -0.096834 | 0.65233115 | -0.1484429 | 0.88199321 | NA |
| ANKZF1 | 5.68393971 | 0.16922436 | 0.6424816 | 0.26339176 | 0.79224864 | NA |
| STK16 | 3.43948 | 0.24786223 | 0.84090082 | 0.29475799 | 0.76817876 | NA |
| TUBA4A | 9.897005 | -0.2824705 | 0.65225237 | -0.4330693 | 0.66496439 | NA |
| DNAJB2 | 10.5840471 | -0.7598519 | 0.50758694 | -1.4969887 | 0.1343962 | NA |
| DNPEP | 5.14960525 | 0.32435297 | 0.67778866 | 0.47854587 | 0.63226173 | NA |
| GMPPA | 2.82648666 | -0.1967849 | 0.82069785 | -0.2397775 | 0.81050276 | NA |
| CHPF | 6.05571018 | -0.267827 | 0.61316358 | -0.4367954 | 0.66225976 | NA |
| OBSL1 | 5.12474772 | 1.0673942 | 0.76541858 | 1.39452351 | 0.16315958 | NA |
| SGPP2 | 63.5451785 | 0.20360096 | 0.37702306 | 0.54002255 | 0.58918148 | 0.89534694 |
| FARSB | 7.87320048 | -0.4736538 | 0.524241 | -0.903504 | 0.36625846 | NA |
| ACSL3 | 68.4536075 | -0.6806891 | 0.31909215 | -2.1332053 | 0.03290788 | 0.41688714 |
| AP1S3 | 27.9116379 | -1.555452 | 0.37345032 | -4.1650841 | 3.11E-05 | 0.04054737 |
| WDFY1 | 18.9559077 | -0.3023956 | 0.44963101 | -0.6725416 | 0.50123897 | NA |
| MRPL44 | 4.91071705 | -0.0905016 | 0.63999634 | -0.1414095 | 0.88754642 | NA |
| SERPINE2 | 5.996786 | -3.1611009 | 0.76828219 | -4.1145049 | 3.88E-05 | NA |

| | | | | | | |
|---|---|---|---|---|---|---|
| CUL3 | 66.981088 | -0.0679641 | 0.25247612 | -0.2691902 | 0.78778335 | 0.96325251 |
| DOCK10 | 32.3784091 | 1.48357781 | 0.78276296 | 1.89530916 | 0.05805146 | 0.50751937 |
| IRS1 | 33.4459992 | 1.09245309 | 0.51698353 | 2.11312937 | 0.0345897 | 0.41688714 |
| RHBDD1 | 22.8928617 | 0.05715875 | 0.40520525 | 0.14106121 | 0.88782158 | NA |
| COL4A4 | 67.8321164 | 0.45994095 | 0.5241585 | 0.87748448 | 0.38022354 | 0.82153751 |
| COL4A3 | 57.78214 | 0.17888248 | 0.47123962 | 0.37959983 | 0.70424249 | 0.9303056 |
| AC097662.1 | 9.56313206 | 0.42734798 | 0.85847898 | 0.49779667 | 0.61862736 | NA |
| MFF | 29.2194317 | -0.0935106 | 0.39895704 | -0.2343877 | 0.814684 | 0.96845695 |
| AGFG1 | 46.4773982 | -0.7131179 | 0.29615383 | -2.4079306 | 0.01604323 | 0.31287044 |
| DNER | 2.30512363 | 0.15546947 | 1.11254319 | 0.13974241 | 0.88886352 | NA |
| TRIP12 | 74.6188068 | 0.03419346 | 0.22122084 | 0.15456709 | 0.87716262 | 0.9751578 |
| FBXO36 | 4.14623237 | -0.5226242 | 0.74030205 | -0.7059608 | 0.4802125 | NA |
| SP110 | 18.2284137 | 0.26160203 | 0.78336537 | 0.33394638 | 0.73842002 | NA |
| SP140L | 11.1312793 | 0.35214064 | 0.58634648 | 0.6005675 | 0.54812809 | NA |
| SP100 | 24.7735812 | -0.4319965 | 0.44812869 | -0.964001 | 0.33504541 | NA |
| CAB39 | 76.1678946 | -0.4371355 | 0.32850642 | -1.3306756 | 0.18329579 | 0.67478132 |
| ITM2C | 20.1686352 | 0.67339068 | 0.40333226 | 1.66956811 | 0.09500484 | NA |
| C2orf72 | 9.39992975 | 0.6233759 | 0.7140847 | 0.87297194 | 0.38267837 | NA |
| PSMD1 | 36.6587654 | -0.2322229 | 0.3395516 | -0.6839106 | 0.49403164 | 0.86634063 |
| ARMC9 | 34.4381484 | 0.48570396 | 0.30668302 | 1.58373283 | 0.11325453 | 0.57860396 |
| NCL | 34.9253766 | -0.3653689 | 0.39594003 | -0.9227885 | 0.35611745 | 0.80177915 |
| PTMA | 118.95973 | -0.1230625 | 0.34061087 | -0.3612995 | 0.71787559 | 0.93714576 |
| PDE6D | 11.8464925 | -0.4359986 | 0.48938638 | -0.8909086 | 0.37297818 | NA |
| COPS7B | 6.94115656 | 0.81498984 | 0.56200239 | 1.4501537 | 0.14701566 | NA |
| DIS3L2 | 53.2271181 | 0.1250983 | 0.28150203 | 0.44439574 | 0.65675649 | 0.92276785 |
| EIF4E2 | 10.6386824 | -0.0509133 | 0.45144505 | -0.1127785 | 0.91020617 | NA |
| EFHD1 | 83.7358522 | 0.0591417 | 0.28948602 | 0.20429899 | 0.83811986 | 0.97154892 |
| GIGYF2 | 109.714234 | 0.21356329 | 0.21647265 | 0.98656018 | 0.3238583 | 0.78622244 |
| ATG16L1 | 5.48076227 | -0.3380625 | 0.64672863 | -0.522727 | 0.60116425 | NA |
| DGKD | 8.60416092 | 0.20809872 | 0.55723934 | 0.37344586 | 0.70881663 | NA |
| USP40 | 34.1637445 | 0.07022183 | 0.41853595 | 0.16777968 | 0.86675661 | 0.97217819 |
| SH3BP4 | 25.8034879 | -1.0739696 | 0.33238896 | -3.2310628 | 0.00123331 | NA |
| AGAP1 | 112.679846 | -0.0527507 | 0.26258354 | -0.2008913 | 0.8407836 | 0.97168963 |
| IQCA1 | 4.33655865 | -0.8427452 | 0.85098394 | -0.9903186 | 0.32201841 | NA |
| COPS8 | 6.09346577 | 0.4327656 | 0.62286724 | 0.69479589 | 0.48718322 | NA |
| RAB17 | 20.9758654 | -0.1539355 | 0.35378578 | -0.4351092 | 0.66348313 | NA |
| LRRFIP1 | 45.0613109 | -0.453249 | 0.34985591 | -1.2955305 | 0.1951373 | 0.69049094 |
| ILKAP | 18.9391942 | -0.333946 | 0.36673293 | -0.9105972 | 0.36250765 | NA |
| PER2 | 7.70412387 | -0.912467 | 0.6224169 | -1.4660062 | 0.1426466 | NA |
| TRAF3IP1 | 13.137449 | 0.36798279 | 0.48456694 | 0.75940549 | 0.44761003 | NA |
| ASB1 | 4.26317416 | -0.195409 | 0.77625393 | -0.2517334 | 0.80124717 | NA |
| HDAC4 | 37.7422466 | -0.0504494 | 0.42583041 | -0.118473 | 0.90569287 | 0.98144306 |
| NDUFA10 | 34.4049918 | 0.26215699 | 0.31704227 | 0.8268834 | 0.40830315 | 0.82808596 |
| COPS9 | 9.49912801 | -0.097212 | 0.51011111 | -0.1905702 | 0.84886237 | NA |
| GPC1 | 3.61551261 | -0.8883302 | 0.77386249 | -1.1479174 | 0.25100268 | NA |
| ANKMY1 | 7.04496628 | 0.50378279 | 0.70819485 | 0.71136185 | 0.47686004 | NA |
| RNPEPL1 | 8.85588051 | 0.06550219 | 0.51235812 | 0.12784454 | 0.898272 | NA |
| CAPN10 | 3.91009594 | 0.4497229 | 0.79440191 | 0.56611508 | 0.57131555 | NA |
| SNED1 | 50.1961944 | 1.13453601 | 0.72089771 | 1.57378224 | 0.1155378 | 0.57860396 |
| AC005237.1 | 11.8049438 | 0.64419591 | 0.82116927 | 0.78448613 | 0.43275492 | NA |
| MTERF4 | 11.8749602 | 0.69793558 | 0.44436663 | 1.57063003 | 0.11626861 | NA |
| PPP1R7 | 10.3147966 | 0.4780246 | 0.6332096 | 0.75492317 | 0.45029508 | NA |
| HDLBP | 53.7990361 | -0.7156572 | 0.30592701 | -2.3393069 | 0.01931955 | 0.33831016 |
| 2-Sep | 52.691216 | -0.0260457 | 0.27324238 | -0.095321 | 0.92405984 | 0.98758497 |

| | | | | | | |
|---|---|---|---|---|---|---|
| FARP2 | 18.1992505 | -0.208672 | 0.38104069 | -0.547637 | 0.58394121 | NA |
| STK25 | 10.9298211 | -0.4598215 | 0.45549291 | -1.0095032 | 0.31273339 | NA |
| BOK | 8.1220277 | -0.0652885 | 0.56997697 | -0.1145458 | 0.9088051 | NA |
| THAP4 | 15.4088201 | -0.6067121 | 0.39284939 | -1.5443884 | 0.12249426 | NA |
| ATG4B | 19.6259397 | -0.2230894 | 0.40012417 | -0.5575503 | 0.5771515 | NA |
| DTYMK | 2.42340743 | -0.0332053 | 0.98124022 | -0.0338401 | 0.97300462 | NA |
| ING5 | 13.1845402 | -0.5546084 | 0.57584142 | -0.9631269 | 0.33548385 | NA |
| D2HGDH | 13.4256018 | 0.03068613 | 0.5240085 | 0.05856037 | 0.95330227 | NA |
| CHL1 | 25.900206 | -0.72767 | 1.2688995 | -0.5734655 | 0.56632958 | NA |
| LINC01266 | 10.659606 | 0.33447865 | 0.72610349 | 0.46064872 | 0.64505065 | NA |
| CNTN4 | 92.7005374 | -0.2970044 | 0.45199638 | -0.6570945 | 0.51112013 | 0.87158522 |
| TRNT1 | 15.4612886 | -0.0773663 | 0.39247028 | -0.1971264 | 0.84372859 | NA |
| CRBN | 14.1205319 | 0.6371132 | 0.48795572 | 1.3056783 | 0.19166198 | NA |
| AC034195.1 | 15.2359178 | 1.67818891 | 0.53457985 | 3.13926708 | 0.00169371 | NA |
| SUMF1 | 83.6862511 | 0.49247926 | 0.37358149 | 1.31826461 | 0.18741508 | 0.68214045 |
| SETMAR | 6.4463612 | 0.27222109 | 0.76863101 | 0.35416356 | 0.72321631 | NA |
| ITPR1 | 51.4265556 | -0.2456613 | 0.42074668 | -0.58387 | 0.5593078 | 0.89263108 |
| BHLHE40 | 54.2208152 | -1.2107922 | 0.65585327 | -1.8461327 | 0.06487294 | 0.52254021 |
| ARL8B | 31.1371642 | -0.7909632 | 0.33207267 | -2.3818981 | 0.01722366 | 0.32722884 |
| AC026202.2 | 4.37442725 | -0.2968457 | 0.8340973 | -0.3558885 | 0.72192403 | NA |
| EDEM1 | 6.3104695 | -0.2142431 | 0.60267447 | -0.3554873 | 0.72222458 | NA |
| LMCD1-AS1 | 12.3504135 | 0.80563433 | 0.60801046 | 1.32503366 | 0.18515999 | NA |
| RAD18 | 3.89327376 | -0.6087592 | 0.86246613 | -0.7058355 | 0.48029045 | NA |
| SRGAP3 | 36.2052248 | 0.12853881 | 0.56321339 | 0.228224 | 0.8194721 | 0.96845695 |
| THUMPD3-A: | 23.129722 | 0.18988718 | 0.40948601 | 0.46372079 | 0.6428478 | NA |
| THUMPD3 | 7.29254377 | 0.28875432 | 0.62378034 | 0.46291026 | 0.6434287 | NA |
| SETD5 | 67.8133328 | -0.1346474 | 0.24078553 | -0.5592007 | 0.57602473 | 0.89263108 |
| MTMR14 | 14.257654 | -0.0847091 | 0.43040682 | -0.1968116 | 0.84397496 | NA |
| BRPF1 | 2.02946275 | -0.3572276 | 1.10067963 | -0.3245518 | 0.74552028 | NA |
| OGG1 | 6.16777298 | 0.21895964 | 0.71036285 | 0.30823633 | 0.75790251 | NA |
| CAMK1 | 11.2754002 | 0.02718639 | 0.48094837 | 0.05652663 | 0.95492228 | NA |
| TADA3 | 11.3547829 | -0.1093399 | 0.53816501 | -0.2031716 | 0.83900089 | NA |
| RPUSD3 | 3.5174179 | 0.56763438 | 0.82047438 | 0.69183681 | 0.48903982 | NA |
| JAGN1 | 4.50094583 | 0.13562227 | 0.75478279 | 0.17968384 | 0.85740078 | NA |
| IL17RC | 3.28948515 | 0.33227195 | 0.89909263 | 0.36956365 | 0.71170764 | NA |
| CRELD1 | 5.00787117 | 0.58013556 | 0.67526197 | 0.85912665 | 0.39027064 | NA |
| EMC3 | 11.5462658 | 0.54494754 | 0.48030585 | 1.13458444 | 0.25654948 | NA |
| BRK1 | 13.4711583 | -0.148528 | 0.474256 | -0.313181 | 0.75414312 | NA |
| VHL | 7.12658493 | 0.53158907 | 0.59076878 | 0.89982593 | 0.36821289 | NA |
| SEC13 | 10.5937082 | -0.7558213 | 0.54490748 | -1.3870636 | 0.16542237 | NA |
| ATP2B2 | 5.40202025 | -2.2106805 | 0.92300052 | -2.3951021 | 0.01661574 | NA |
| ATG7 | 58.8692767 | -0.1225286 | 0.43481917 | -0.281792 | 0.77810303 | 0.96133329 |
| VGLL4 | 76.2348186 | 0.10402644 | 0.29383673 | 0.35402803 | 0.72331787 | 0.93851989 |
| TAMM41 | 17.03386 | 0.05945899 | 0.39492729 | 0.1505568 | 0.88032534 | NA |
| PPARG | 93.4021919 | -0.2841112 | 0.42652377 | -0.6661088 | 0.50534158 | 0.87094108 |
| TSEN2 | 13.8194455 | 1.05802794 | 0.48270748 | 2.19186151 | 0.02838951 | NA |
| MKRN2 | 4.04293067 | 0.71471019 | 0.74745091 | 0.95619683 | 0.3389728 | NA |
| RAF1 | 40.3860775 | -0.1978098 | 0.27076573 | -0.7305569 | 0.46504982 | 0.85894565 |
| RPL32 | 75.686411 | -0.3469242 | 0.46932304 | -0.7392013 | 0.4597848 | 0.85844137 |
| IQSEC1 | 6.21639171 | 0.34839712 | 0.63633376 | 0.5475069 | 0.58403053 | NA |
| NUP210 | 10.3323162 | -0.2223256 | 0.59355289 | -0.3745675 | 0.70798217 | NA |
| HDAC11 | 4.1600137 | 0.58106449 | 0.73570838 | 0.78980273 | 0.42964298 | NA |
| CHCHD4 | 2.16937254 | -0.8118438 | 0.96915276 | -0.837684 | 0.40220819 | NA |
| TMEM43 | 3.32746846 | -1.1972011 | 0.77907574 | -1.5366942 | 0.12436822 | NA |

| | | | | | | |
|---|---|---|---|---|---|---|
| XPC | 14.904172 | 0.14470579 | 0.40567163 | 0.3567067 | 0.72131139 | NA |
| LSM3 | 11.4683372 | 0.08088989 | 0.50005429 | 0.16176222 | 0.87149311 | NA |
| SLC6A6 | 10.7763627 | 0.21352381 | 0.69216587 | 0.30848648 | 0.75771219 | NA |
| CCDC174 | 7.6495081 | 0.26246847 | 0.61750969 | 0.42504348 | 0.67080498 | NA |
| FGD5-AS1 | 16.8359183 | 0.22275252 | 0.42041421 | 0.5298406 | 0.59622245 | NA |
| NR2C2 | 27.2965179 | -0.3179773 | 0.35802984 | -0.8881306 | 0.3744705 | 0.81522167 |
| MRPS25 | 26.0425568 | -0.0101014 | 0.33018318 | -0.0305933 | 0.97559385 | 0.99853281 |
| RBSN | 7.08486891 | 0.37348592 | 0.67429463 | 0.55389129 | 0.57965325 | NA |
| CAPN7 | 33.2916344 | -0.2122222 | 0.29779604 | -0.7126427 | 0.47606687 | 0.86206794 |
| METTL6 | 8.23615536 | -0.2100444 | 0.55061062 | -0.3814754 | 0.70285055 | NA |
| EAF1 | 4.0534804 | -1.1915595 | 0.78322168 | -1.5213567 | 0.12817035 | NA |
| COLQ | 3.14866166 | -1.3484295 | 0.86497421 | -1.5589245 | 0.11901425 | NA |
| HACL1 | 12.9710071 | -0.1471987 | 0.50961238 | -0.2888444 | 0.77270046 | NA |
| BTD | 3.30375832 | -0.1134629 | 0.85296753 | -0.1330214 | 0.89417648 | NA |
| ANKRD28 | 94.3276509 | -0.1142679 | 0.2601236 | -0.4392831 | 0.66045643 | 0.92276785 |
| DPH3 | 3.57133352 | 0.52506989 | 0.72704935 | 0.72219292 | 0.47017588 | NA |
| OXNAD1 | 25.9717186 | -0.2105147 | 0.39342265 | -0.5350854 | 0.5925908 | 0.89685273 |
| RFTN1 | 38.9474913 | 0.57239887 | 0.4547057 | 1.25883371 | 0.2080904 | 0.70439682 |
| PLCL2 | 295.699916 | -0.9568018 | 0.53551952 | -1.7866796 | 0.07398929 | 0.53207981 |
| TBC1D5 | 157.292843 | 0.47406619 | 0.30153041 | 1.57220028 | 0.11590411 | 0.57860396 |
| KCNH8 | 7.19823712 | -1.2424063 | 0.69575905 | -1.7856847 | 0.07415033 | NA |
| RAB5A | 27.7548073 | -0.322338 | 0.33648543 | -0.9579554 | 0.33808526 | 0.8003972 |
| KAT2B | 22.3974708 | 0.34578766 | 0.42191125 | 0.81957441 | 0.41245877 | NA |
| ZNF385D | 8.65269947 | 1.20286184 | 0.89498225 | 1.34400636 | 0.17894633 | NA |
| UBE2E2 | 176.459581 | -0.5207639 | 0.28993276 | -1.7961541 | 0.07247001 | 0.53207981 |
| UBE2E1 | 79.5196021 | -0.254844 | 0.28091954 | -0.907178 | 0.36431268 | 0.80760012 |
| NKIRAS1 | 7.49967113 | -0.5330983 | 0.55623834 | -0.958399 | 0.33786159 | NA |
| RPL15 | 125.53268 | 0.57810702 | 0.35868695 | 1.61173141 | 0.1070204 | 0.57769936 |
| NR1D2 | 34.7102426 | 0.02943729 | 0.38889771 | 0.07569417 | 0.93966241 | 0.99100968 |
| THRB | 35.2330113 | -0.0413189 | 0.4369778 | -0.094556 | 0.92466752 | 0.98758497 |
| RARB | 21.27223 | -0.1309438 | 0.42431069 | -0.3086037 | 0.75762303 | NA |
| TOP2B | 36.8062998 | -0.0877084 | 0.31397446 | -0.2793489 | 0.77997708 | 0.96139889 |
| NGLY1 | 24.0789532 | -0.4039814 | 0.37967853 | -1.0640092 | 0.28732454 | NA |
| OXSM | 3.07260752 | 0.81117169 | 0.85028127 | 0.95400395 | 0.34008164 | NA |
| NEK10 | 8.58737629 | -0.8354357 | 0.73581765 | -1.1353841 | 0.25621441 | NA |
| SLC4A7 | 16.1756131 | -0.9756696 | 0.5023196 | -1.9423283 | 0.05209737 | NA |
| CMC1 | 41.0295386 | 0.15232638 | 0.36409664 | 0.418368 | 0.67567808 | 0.92617027 |
| AZI2 | 17.2963103 | 0.28789966 | 0.4004739 | 0.71889744 | 0.47220412 | NA |
| TGFBR2 | 12.5158387 | -0.5177494 | 0.84610691 | -0.6119195 | 0.54059099 | NA |
| STT3B | 57.5386636 | -0.339844 | 0.36601886 | -0.9284876 | 0.35315467 | 0.8003972 |
| OSBPL10 | 41.7110694 | -0.6667931 | 0.50669818 | -1.3159572 | 0.18818842 | 0.68214045 |
| GPD1L | 75.6130123 | 0.36522293 | 0.27894587 | 1.30929678 | 0.19043384 | 0.68499719 |
| CMTM8 | 9.89341497 | 0.15817423 | 0.50804036 | 0.31134186 | 0.75554075 | NA |
| CMTM7 | 7.34465468 | 0.22749495 | 0.52191123 | 0.43588821 | 0.66291784 | NA |
| CMTM6 | 21.6758518 | 0.03323647 | 0.38762333 | 0.08574426 | 0.93166972 | NA |
| DYNC1LI1 | 10.9022053 | -0.4824433 | 0.59050888 | -0.8169959 | 0.41393078 | NA |
| CNOT10 | 18.4116552 | 0.23121644 | 0.44266194 | 0.52233188 | 0.60143927 | NA |
| GLB1 | 20.2167497 | -0.2032214 | 0.35648568 | -0.5700689 | 0.56863098 | NA |
| CRTAP | 8.77732525 | -0.4091454 | 0.48801662 | -0.8383841 | 0.40181501 | NA |
| FBXL2 | 32.2832478 | -0.4373988 | 0.33946759 | -1.2884847 | 0.1975773 | 0.69343545 |
| UBP1 | 20.8298092 | -0.1752808 | 0.46445565 | -0.3773898 | 0.70588398 | NA |
| CLASP2 | 69.0669764 | -0.3543394 | 0.32230862 | -1.0993793 | 0.27160264 | 0.75979923 |
| PDCD6IP | 51.8376037 | 0.13354188 | 0.29058791 | 0.45955761 | 0.64583379 | 0.92178095 |
| STAC | 19.1564695 | -0.4104666 | 0.49400902 | -0.8308888 | 0.40603646 | NA |

| | | | | | | |
|---|---|---|---|---|---|---|
| TRANK1 | 13.3861525 | 0.49614127 | 0.61522165 | 0.80644313 | 0.41998739 | NA |
| EPM2AIP1 | 16.7218622 | -0.1177553 | 0.42765845 | -0.275349 | 0.7830481 | NA |
| MLH1 | 6.71013022 | -0.6443123 | 0.71162214 | -0.9054135 | 0.36524637 | NA |
| LRRFIP2 | 106.450803 | -0.0562082 | 0.28605637 | -0.1964936 | 0.84422383 | 0.97168963 |
| GOLGA4 | 121.254779 | -0.7145002 | 0.3007592 | -2.3756553 | 0.01751782 | 0.32926096 |
| ITGA9 | 18.0958843 | -1.9117669 | 0.55898969 | -3.4200397 | 0.00062612 | NA |
| CTDSPL | 131.287647 | -0.5703956 | 0.33759846 | -1.6895681 | 0.09111062 | 0.56211136 |
| PLCD1 | 3.3206326 | 0.24315463 | 0.76895918 | 0.31621266 | 0.7518411 | NA |
| ACAA1 | 13.2126733 | 0.15879027 | 0.48611089 | 0.32665442 | 0.74392928 | NA |
| OXSR1 | 32.7721271 | -0.3416625 | 0.28966537 | -1.1795075 | 0.23819615 | 0.73242571 |
| XYLB | 2.60698622 | -0.2336668 | 1.08429003 | -0.2155022 | 0.82937583 | NA |
| ACVR2B | 6.23455801 | 0.87445739 | 0.64010955 | 1.36610584 | 0.17190575 | NA |
| EXOG | 6.57234071 | -0.4932967 | 0.60142117 | -0.8202184 | 0.41209159 | NA |
| WDR48 | 13.7210033 | -0.2640502 | 0.50172968 | -0.5262798 | 0.59869381 | NA |
| GORASP1 | 5.9312802 | 0.29787873 | 0.5982258 | 0.49793695 | 0.61852848 | NA |
| TTC21A | 3.16772745 | -0.0898079 | 0.98794964 | -0.0909033 | 0.92756942 | NA |
| SLC25A38 | 6.18631051 | 0.24748931 | 0.6190803 | 0.39976933 | 0.68932643 | NA |
| RPSA | 53.8500153 | 0.0504054 | 0.3564699 | 0.14140155 | 0.88755273 | 0.97769824 |
| MYRIP | 166.634022 | -0.9872322 | 0.43437193 | -2.2727808 | 0.02303939 | 0.35751239 |
| EIF1B | 15.2238042 | -0.0267419 | 0.44666051 | -0.0598709 | 0.9522585 | NA |
| ENTPD3-AS1 | 4.86841452 | -0.0738995 | 0.77372399 | -0.0955114 | 0.92390861 | NA |
| RPL14 | 102.506063 | 0.34135424 | 0.45847131 | 0.74454875 | 0.45654456 | 0.85802171 |
| ZNF621 | 13.3638834 | 0.29431747 | 0.44872544 | 0.65589656 | 0.51189068 | NA |
| CTNNB1 | 33.8478319 | -0.5083603 | 0.28472473 | -1.785445 | 0.07418917 | 0.53207981 |
| ULK4 | 34.2495429 | 0.53908787 | 0.32134642 | 1.677591 | 0.09342697 | 0.56220784 |
| TRAK1 | 48.488137 | -0.1481163 | 0.26906394 | -0.5504875 | 0.58198507 | 0.89263108 |
| SEC22C | 8.39754619 | -0.0488543 | 0.65191409 | -0.0749398 | 0.94026263 | NA |
| SS18L2 | 4.20500716 | -0.2490651 | 0.98915746 | -0.2517952 | 0.80119937 | NA |
| NKTR | 72.7977809 | -0.1279943 | 0.25269101 | -0.5065251 | 0.6124881 | 0.9107135 |
| AC006059.1 | 7.33009782 | 0.90479669 | 0.71348243 | 1.26814151 | 0.20474742 | NA |
| AC099329.3 | 4.77241533 | 0.5281319 | 0.65768013 | 0.80302243 | 0.42196177 | NA |
| SNRK | 25.0590822 | -0.5121431 | 0.46066486 | -1.1117477 | 0.26624666 | NA |
| ANO10 | 38.3821448 | 0.29150571 | 0.30427136 | 0.95804517 | 0.33803998 | 0.8003972 |
| ABHD5 | 9.61596574 | -0.3237839 | 0.46950708 | -0.6896252 | 0.49042993 | NA |
| TCAIM | 46.4133115 | 0.54214854 | 0.43065916 | 1.25888077 | 0.2080734 | 0.70439682 |
| ZNF445 | 12.9898197 | -0.0565973 | 0.46611011 | -0.1214247 | 0.90335469 | NA |
| ZNF852 | 4.09305827 | -0.6265761 | 0.74967898 | -0.8357925 | 0.40327165 | NA |
| AC099669.1 | 13.1045776 | -0.1142809 | 0.49090068 | -0.2327985 | 0.81591789 | NA |
| ZNF197 | 8.05089861 | 0.59210008 | 0.5119322 | 1.15659862 | 0.24743638 | NA |
| ZNF35 | 2.97400479 | -2.4252243 | 0.97493941 | -2.4875641 | 0.01286213 | NA |
| ZNF502 | 3.38270398 | -0.2571784 | 0.85475799 | -0.3008786 | 0.76350704 | NA |
| KIAA1143 | 4.83748819 | 0.59388038 | 0.65537376 | 0.90617052 | 0.36484561 | NA |
| ZDHHC3 | 9.6150677 | -0.3182058 | 0.48482248 | -0.6563348 | 0.51160876 | NA |
| EXOSC7 | 4.03575682 | 0.08427557 | 0.6444204 | 0.13077731 | 0.89595147 | NA |
| CDCP1 | 13.2119851 | -0.3771375 | 0.76384758 | -0.493734 | 0.62149405 | NA |
| LARS2 | 18.7132722 | 0.44460206 | 0.44854847 | 0.99120182 | 0.32158704 | NA |
| LIMD1 | 14.1117582 | -0.4777048 | 0.43304639 | -1.1031261 | 0.26997243 | NA |
| SACM1L | 20.7052953 | -0.1348752 | 0.42724478 | -0.3156859 | 0.75224091 | NA |
| LZTFL1 | 4.68520545 | -1.2338556 | 1.00837627 | -1.2236063 | 0.22110077 | NA |
| FYCO1 | 11.1088113 | 0.10266211 | 0.47220229 | 0.21741128 | 0.82788784 | NA |
| LTF | 24.219742 | 1.4347372 | 0.77371249 | 1.85435445 | 0.06368847 | NA |
| ALS2CL | 7.82166931 | 0.37926371 | 0.59015056 | 0.64265585 | 0.52044744 | NA |
| PTH1R | 14.5605108 | 1.42473889 | 0.76819416 | 1.85465988 | 0.06364481 | NA |
| CCDC12 | 27.8454679 | -0.1457813 | 0.31088504 | -0.4689234 | 0.63912437 | 0.92121485 |

| | | | | | | |
|---|---|---|---|---|---|---|
| NBEAL2 | 6.52208724 | 0.20949861 | 0.64136451 | 0.32664515 | 0.74393629 | NA |
| SETD2 | 107.192405 | -0.36184 | 0.2327306 | -1.554759 | 0.12000348 | 0.58383225 |
| KIF9-AS1 | 24.2944559 | -0.00251 | 0.43361482 | -0.0057885 | 0.99538149 | NA |
| KLHL18 | 12.8028154 | 0.25494715 | 0.49276226 | 0.51738368 | 0.60488835 | NA |
| PTPN23 | 6.78198645 | -0.6456024 | 0.6392256 | -1.0099759 | 0.31250685 | NA |
| SCAP | 19.0608615 | -0.4515706 | 0.39070913 | -1.1557718 | 0.24777451 | NA |
| ELP6 | 6.12332281 | 0.94194455 | 0.57655921 | 1.63373429 | 0.10231465 | NA |
| SMARCC1 | 85.5943321 | -0.0521055 | 0.27397448 | -0.1901839 | 0.84916501 | 0.97168963 |
| DHX30 | 12.5518465 | 0.35225964 | 0.43517122 | 0.80947366 | 0.41824275 | NA |
| MAP4 | 112.233413 | -0.109702 | 0.27318341 | -0.4015689 | 0.68800135 | 0.92736198 |
| ZNF589 | 3.17501795 | 0.30976156 | 0.85613292 | 0.3618148 | 0.71749043 | NA |
| NME6 | 4.65299219 | 0.50051482 | 0.68287479 | 0.73295255 | 0.46358737 | NA |
| PLXNB1 | 16.9201649 | -0.3608609 | 0.50572452 | -0.7135523 | 0.47550404 | NA |
| TMA7 | 26.8426717 | 0.12018635 | 0.51452434 | 0.2335873 | 0.81530539 | 0.96845695 |
| SHISA5 | 12.8995092 | -0.1913026 | 0.49027907 | -0.3901912 | 0.6963952 | NA |
| UQCRC1 | 17.8059115 | 0.01226883 | 0.42650038 | 0.02876629 | 0.97705099 | NA |
| NCKIPSD | 2.34200416 | -1.0684432 | 1.09320249 | -0.9773516 | 0.3283951 | NA |
| IP6K2 | 22.9246814 | -0.8091278 | 0.4355055 | -1.8579049 | 0.06318252 | NA |
| PRKAR2A | 34.5578978 | -0.2288184 | 0.34079667 | -0.6714221 | 0.50195164 | 0.87050612 |
| SLC25A20 | 5.38487772 | 0.2059535 | 0.69553677 | 0.29610727 | 0.76714816 | NA |
| ARIH2 | 32.5206388 | -0.1320054 | 0.33518054 | -0.3938337 | 0.69370386 | 0.93004933 |
| P4HTM | 6.97135475 | -0.0192016 | 0.60865499 | -0.0315475 | 0.97483288 | NA |
| WDR6 | 19.7022966 | -0.7600129 | 0.45989054 | -1.6525952 | 0.09841329 | NA |
| DALRD3 | 4.5367909 | -0.1033387 | 0.81738868 | -0.1264254 | 0.89939517 | NA |
| NDUFAF3 | 7.93084329 | -0.2192996 | 0.51799954 | -0.4233587 | 0.67203358 | NA |
| IMPDH2 | 6.93852457 | -0.0477587 | 0.62468538 | -0.0764525 | 0.93905912 | NA |
| QRICH1 | 19.4490574 | -0.3722274 | 0.38571785 | -0.9650252 | 0.3345322 | NA |
| QARS | 7.56838828 | 0.70449472 | 0.57095854 | 1.23388068 | 0.21724737 | NA |
| USP19 | 4.33157458 | -1.6532394 | 0.82801743 | -1.996624 | 0.04586605 | NA |
| LAMB2 | 8.29157911 | 0.32696046 | 0.59941955 | 0.54546179 | 0.58543595 | NA |
| KLHDC8B | 2.9673965 | -0.3550774 | 1.02905414 | -0.3450522 | 0.73005509 | NA |
| C3orf62 | 2.32967766 | -1.8305336 | 1.06765582 | -1.7145353 | 0.08643046 | NA |
| USP4 | 25.1057186 | -0.6260993 | 0.44022621 | -1.4222218 | 0.15496187 | NA |
| RHOA | 46.0707009 | -0.4558945 | 0.33607809 | -1.3565135 | 0.17493584 | 0.6651652 |
| TCTA | 6.25569143 | 0.35827105 | 0.57959075 | 0.61814488 | 0.53647984 | NA |
| DAG1 | 18.1830813 | -0.3272417 | 0.40545762 | -0.8070923 | 0.41961333 | NA |
| APEH | 7.79831145 | -0.1431534 | 0.65925409 | -0.2171445 | 0.82809576 | NA |
| MST1 | 4.2570648 | -1.0185996 | 0.85663267 | -1.1890739 | 0.23441058 | NA |
| RNF123 | 3.89120138 | -0.9541899 | 0.75143094 | -1.2698304 | 0.20414505 | NA |
| IP6K1 | 28.566503 | -1.1516534 | 0.37006498 | -3.1120302 | 0.00185805 | 0.14585731 |
| RBM6 | 119.882389 | -0.397901 | 0.21632398 | -1.8393756 | 0.06585996 | 0.5230103 |
| RBM5 | 36.2179791 | -0.336409 | 0.29070633 | -1.1572126 | 0.24718551 | 0.73796211 |
| SEMA3F | 14.4631449 | 0.4158072 | 0.46159518 | 0.90080491 | 0.36769206 | NA |
| GNAI2 | 15.0598401 | -0.4120579 | 0.44360234 | -0.9288903 | 0.35294595 | NA |
| SEMA3B | 8.82751526 | -0.8091865 | 0.52845124 | -1.5312416 | 0.12570969 | NA |
| IFRD2 | 3.76602035 | 0.15128499 | 0.77909878 | 0.19417947 | 0.84603536 | NA |
| HYAL2 | 3.7771728 | -1.0997868 | 0.91942306 | -1.1961706 | 0.23162999 | NA |
| TUSC2 | 4.5863413 | -0.4036924 | 0.71172938 | -0.5671993 | 0.57057877 | NA |
| RASSF1 | 3.03951675 | -1.5226048 | 1.06061226 | -1.4355904 | 0.15111894 | NA |
| TMEM115 | 3.9474349 | 0.39189562 | 0.72055265 | 0.54388201 | 0.58652267 | NA |
| HEMK1 | 4.74127544 | 0.19731506 | 0.76948059 | 0.2564263 | 0.79762168 | NA |
| CISH | 6.55446415 | 0.40560027 | 0.64775119 | 0.62616677 | 0.53120556 | NA |
| MAPKAPK3 | 8.23404292 | -0.6385559 | 0.56207791 | -1.136063 | 0.25593019 | NA |
| DOCK3 | 38.2987246 | -0.623855 | 0.47717368 | -1.3073961 | 0.19107821 | 0.68546741 |

| | | | | | | |
|---|---|---|---|---|---|---|
| MANF | 9.24591285 | 0.87303121 | 0.59247025 | 1.47354439 | 0.1406043 | NA |
| RBM15B | 6.20646954 | -0.7268587 | 0.9123733 | -0.7966681 | 0.42564382 | NA |
| DCAF1 | 20.2215276 | 0.30173813 | 0.41471295 | 0.72758309 | 0.46686883 | NA |
| RAD54L2 | 25.9650953 | -0.3208704 | 0.37778283 | -0.8493514 | 0.39568577 | 0.8258317 |
| TEX264 | 19.7042625 | 0.41830956 | 0.38137125 | 1.09685656 | 0.2727041 | NA |
| PCBP4 | 3.01919942 | 0.87014592 | 0.83691092 | 1.03971152 | 0.29847395 | NA |
| RPL29 | 94.4147903 | 0.08065572 | 0.41483663 | 0.19442768 | 0.84584102 | 0.97168963 |
| ALAS1 | 6.29542137 | -0.1354977 | 0.61434035 | -0.220558 | 0.82543662 | NA |
| WDR82 | 18.5248421 | -0.383162 | 0.37597482 | -1.019116 | 0.30814789 | NA |
| BAP1 | 4.67673182 | -1.4591213 | 0.81604378 | -1.7880429 | 0.07376908 | NA |
| PHF7 | 2.89847143 | -0.7798294 | 1.06242021 | -0.7340122 | 0.46294128 | NA |
| NISCH | 14.5277432 | 0.47303175 | 0.43773439 | 1.08063647 | 0.27985885 | NA |
| SMIM4 | 11.1852911 | -0.0562912 | 0.49803353 | -0.1130269 | 0.91000919 | NA |
| PBRM1 | 52.0868428 | -0.8333878 | 0.33533947 | -2.4852064 | 0.01294763 | 0.30435125 |
| GNL3 | 9.74125452 | -1.3225398 | 0.69192888 | -1.9113811 | 0.05595562 | NA |
| GLT8D1 | 3.24176497 | 0.73914523 | 0.90440448 | 0.81727286 | 0.41377252 | NA |
| SPCS1 | 20.6902537 | 0.52849157 | 0.39001556 | 1.35505253 | 0.17540082 | NA |
| NEK4 | 12.5899499 | 0.30203732 | 0.45112584 | 0.669519 | 0.50316447 | NA |
| SFMBT1 | 13.2233819 | 0.0339879 | 0.44672533 | 0.07608231 | 0.93935361 | NA |
| RFT1 | 2.51492394 | 0.31493942 | 0.9103927 | 0.34593799 | 0.72938932 | NA |
| PRKCD | 19.9935688 | 0.62907389 | 0.51153151 | 1.22978522 | 0.21877754 | NA |
| TKT | 20.0629583 | -0.1485731 | 0.41142912 | -0.3611148 | 0.71801363 | NA |
| DCP1A | 27.5351627 | -0.3505951 | 0.34358613 | -1.0203995 | 0.30753904 | 0.77729327 |
| CACNA1D | 58.5484305 | -0.4852208 | 0.43675597 | -1.1109655 | 0.26658322 | 0.75497438 |
| CHDH | 8.70727468 | 0.65471678 | 0.71152942 | 0.9201542 | 0.35749218 | NA |
| IL17RB | 3.7973049 | 0.50375045 | 1.24986405 | 0.4030442 | 0.68691571 | NA |
| ACTR8 | 5.23238277 | -0.8077774 | 0.72571052 | -1.1130849 | 0.26567198 | NA |
| SELENOK | 10.2147877 | -0.1004683 | 0.489487 | -0.2052523 | 0.83737503 | NA |
| CACNA2D3 | 31.327614 | -0.7852939 | 0.5896224 | -1.331859 | 0.18290651 | 0.67478132 |
| ERC2 | 17.5811573 | 0.68031891 | 0.38086948 | 1.78622588 | 0.07406269 | NA |
| CCDC66 | 24.6674227 | 0.07969722 | 0.41323491 | 0.19286179 | 0.84706721 | NA |
| FAM208A | 30.4920404 | 0.23573616 | 0.38229322 | 0.61663705 | 0.53747415 | 0.88254911 |
| ARHGEF3 | 92.9681272 | 0.13130291 | 0.50938748 | 0.25776628 | 0.79658729 | 0.96525806 |
| APPL1 | 21.1931965 | 0.02967289 | 0.40116764 | 0.07396631 | 0.94103719 | NA |
| PDE12 | 7.59123618 | -0.4815623 | 0.70630105 | -0.6818089 | 0.49535981 | NA |
| ARF4 | 21.5346851 | -1.0031542 | 0.4165557 | -2.4082114 | 0.0160309 | NA |
| DENND6A | 25.5335179 | 0.08251436 | 0.36414222 | 0.22659926 | 0.82073537 | NA |
| DENND6A-AS | 5.56146305 | 1.38706796 | 0.68944616 | 2.01185827 | 0.04423488 | NA |
| SLMAP | 75.3612918 | -0.1453811 | 0.30301398 | -0.4797834 | 0.63138142 | 0.9183778 |
| FLNB | 71.596024 | 0.00105644 | 0.31136346 | 0.00339293 | 0.99729284 | 0.99960611 |
| ABHD6 | 3.74039217 | 1.2271733 | 0.87942513 | 1.3954267 | 0.16288721 | NA |
| RPP14 | 3.22095231 | 0.84015187 | 0.80825969 | 1.03945784 | 0.29859186 | NA |
| PXK | 82.9051715 | -0.7337441 | 0.4195146 | -1.7490312 | 0.08028563 | 0.54552914 |
| PDHB | 10.899904 | -0.1255801 | 0.60296677 | -0.2082704 | 0.83501786 | NA |
| KCTD6 | 3.10627809 | -1.0034549 | 0.91972221 | -1.0910413 | 0.2752547 | NA |
| FAM107A | 11.9082325 | 0.63212689 | 0.73055431 | 0.86527023 | 0.38689049 | NA |
| FHIT | 632.05105 | 0.58512467 | 0.4305441 | 1.35903538 | 0.17413538 | 0.6651652 |
| PTPRG | 179.210956 | 0.38485082 | 0.4234416 | 0.90886399 | 0.36342193 | 0.80736493 |
| THOC7 | 10.2477371 | 0.3873001 | 0.55619468 | 0.69633909 | 0.4862165 | NA |
| ATXN7 | 7.64514238 | 0.10497637 | 0.70716029 | 0.14844777 | 0.88198941 | NA |
| PSMD6-AS2 | 3.02533327 | -0.0066021 | 0.79531524 | -0.0083012 | 0.99337666 | NA |
| PSMD6 | 5.2721314 | -0.289455 | 0.64333033 | -0.4499321 | 0.65275942 | NA |
| PRICKLE2 | 18.0354656 | 0.50246592 | 0.46194091 | 1.08772769 | 0.27671534 | NA |
| ADAMTS9 | 15.3381594 | -0.6013099 | 0.67024492 | -0.8971495 | 0.36963913 | NA |

| | | | | | | |
|---|---|---|---|---|---|---|
| ADAMTS9-AS | 14.6584681 | -0.5458238 | 0.8418475 | -0.6483643 | 0.51674937 | NA |
| ADAMTS9-AS | 55.8902703 | 1.2602915 | 0.67704006 | 1.86147258 | 0.06267747 | 0.51631533 |
| MAGI1 | 2078.5482 | -0.1036607 | 0.23075674 | -0.4492207 | 0.65327249 | 0.92248109 |
| MAGI1-AS1 | 33.2969552 | 0.65986232 | 0.58304993 | 1.13174239 | 0.25774275 | 0.75272108 |
| SLC25A26 | 24.2564391 | -0.3620969 | 0.43794007 | -0.8268184 | 0.40834003 | NA |
| LRIG1 | 5.09781459 | 1.27493005 | 0.66824683 | 1.90787293 | 0.05640764 | NA |
| SUCLG2 | 52.6991243 | 0.03722623 | 0.31577196 | 0.11788959 | 0.90615515 | 0.9814473 |
| SUCLG2-AS1 | 30.6173572 | 1.32605175 | 0.5450687 | 2.4328158 | 0.01498192 | 0.30435125 |
| FAM19A1 | 7.41803797 | -0.9711096 | 0.71915852 | -1.3503415 | 0.17690646 | NA |
| EOGT | 3.97297907 | 0.31799033 | 0.75067595 | 0.42360532 | 0.67185367 | NA |
| TMF1 | 26.1712129 | -0.5295796 | 0.43187835 | -1.226224 | 0.22011438 | 0.72261413 |
| UBA3 | 17.884959 | 0.0076738 | 0.38164441 | 0.0201072 | 0.98395786 | NA |
| ARL6IP5 | 19.7497489 | -0.0545206 | 0.44033412 | -0.1238165 | 0.9014606 | NA |
| FRMD4B | 12.4767405 | -1.1579056 | 0.55492684 | -2.0865915 | 0.03692508 | NA |
| MITF | 77.4857832 | 0.3510766 | 0.37630606 | 0.93295496 | 0.35084323 | 0.8003972 |
| FOXP1 | 638.096548 | -0.1249311 | 0.27737961 | -0.4503975 | 0.65242385 | 0.92248109 |
| EIF4E3 | 6.11385733 | 0.52957868 | 0.54766932 | 0.96696794 | 0.33356006 | NA |
| RYBP | 32.4118197 | 0.0649093 | 0.41054179 | 0.15810644 | 0.87437293 | 0.9751578 |
| SHQ1 | 11.7408747 | 0.63518071 | 0.42896658 | 1.48072307 | 0.13868038 | NA |
| GXYLT2 | 4.86045927 | 0.20196159 | 0.69443035 | 0.29083059 | 0.77118089 | NA |
| PPP4R2 | 33.4480316 | -0.3024423 | 0.36277801 | -0.8336842 | 0.40445894 | 0.82808596 |
| PDZRN3 | 55.3120206 | 0.20367742 | 0.56109828 | 0.36299776 | 0.71660656 | 0.93664197 |
| CNTN3 | 14.4727327 | 0.50348516 | 0.63342986 | 0.79485542 | 0.42669761 | NA |
| ZNF717 | 10.7849994 | 0.60742795 | 0.6471182 | 0.93866616 | 0.34790217 | NA |
| ROBO2 | 5.18462022 | -0.3077801 | 1.0410521 | -0.2956433 | 0.76750247 | NA |
| GBE1 | 67.2926978 | -0.0339553 | 0.37969692 | -0.0894274 | 0.92874223 | 0.98921679 |
| CHMP2B | 13.5622987 | 0.09089149 | 0.48107079 | 0.18893579 | 0.85014314 | NA |
| CGGBP1 | 43.8360642 | 0.37313908 | 0.31818369 | 1.17271592 | 0.24090974 | 0.73323864 |
| ZNF654 | 56.9405593 | -0.1288313 | 0.30815577 | -0.4180721 | 0.67589442 | 0.92617027 |
| C3orf38 | 5.53612145 | -0.1196784 | 0.62403668 | -0.191781 | 0.84791375 | NA |
| ARL13B | 6.52389682 | -0.4448259 | 0.59175822 | -0.7517022 | 0.45223019 | NA |
| NSUN3 | 12.6636602 | 0.07872878 | 0.41346388 | 0.19041271 | 0.84898574 | NA |
| MTRNR2L12 | 115.697749 | -0.7066296 | 0.4764611 | -1.4830793 | 0.13805333 | 0.61206702 |
| ARL6 | 11.9049377 | 0.19422494 | 0.55914472 | 0.34736077 | 0.7283203 | NA |
| CRYBG3 | 47.5134353 | -0.0758565 | 0.34807012 | -0.2179345 | 0.82748011 | 0.96865983 |
| RIOX2 | 10.89346 | 1.01413015 | 0.51154418 | 1.98248792 | 0.04742466 | NA |
| CLDND1 | 8.56853531 | 0.02163643 | 0.47742645 | 0.04531888 | 0.96385314 | NA |
| CPOX | 2.26823678 | -0.3679673 | 0.95881138 | -0.3837745 | 0.70114562 | NA |
| ST3GAL6 | 41.1523703 | -0.1081456 | 0.48168303 | -0.2245162 | 0.82235567 | 0.96845695 |
| DCBLD2 | 15.1269854 | 0.18103301 | 0.43088855 | 0.42013883 | 0.67438404 | NA |
| CMSS1 | 30.9927132 | -0.2187714 | 0.35997837 | -0.6077349 | 0.54336334 | 0.88410237 |
| FILIP1L | 7.16871612 | 1.05856948 | 0.54164499 | 1.95436034 | 0.05065862 | NA |
| TBC1D23 | 17.5425026 | -0.0990919 | 0.36899741 | -0.2685436 | 0.78828094 | NA |
| NIT2 | 5.71915604 | 0.50332853 | 0.59621455 | 0.84420706 | 0.39855371 | NA |
| TOMM70 | 11.5080742 | -0.1982713 | 0.45745293 | -0.4334246 | 0.66470637 | NA |
| TFG | 37.0006305 | -0.5987547 | 0.36653158 | -1.6335692 | 0.10234935 | 0.5750956 |
| SENP7 | 29.0451829 | 0.08413648 | 0.35353848 | 0.23798394 | 0.81189355 | 0.96845695 |
| TRMT10C | 7.97462149 | -0.0905289 | 0.51824069 | -0.1746851 | 0.86132709 | NA |
| PCNP | 29.2182214 | -0.1298331 | 0.36522418 | -0.3554888 | 0.72222342 | 0.93818522 |
| ZBTB11 | 11.454951 | -0.8629698 | 0.45962066 | -1.8775697 | 0.06044005 | NA |
| RPL24 | 37.4824435 | -0.4316367 | 0.47731141 | -0.9043083 | 0.36583194 | 0.80952111 |
| CEP97 | 5.1959975 | 1.28243504 | 0.77282777 | 1.659406 | 0.09703401 | NA |
| NFKBIZ | 19.9543533 | -0.1845893 | 0.53842538 | -0.3428318 | 7.32E-01 | NA |
| ZPLD1 | 13.6173806 | -0.830471 | 1.04574269 | -0.7941447 | 0.4271112 | NA |

| | | | | | | |
|---|---|---|---|---|---|---|
| ALCAM | 25.3260367 | -1.1874279 | 0.68824534 | -1.7252975 | 0.08447388 | NA |
| CBLB | 54.2248545 | 0.7009361 | 0.37985715 | 1.84526237 | 0.06499938 | 0.52254021 |
| BBX | 102.432354 | 0.34129435 | 0.3449108 | 0.98951483 | 0.32241132 | 0.78622244 |
| CD47 | 9.77524104 | 0.38002902 | 0.51011046 | 0.74499358 | 0.4562756 | NA |
| IFT57 | 6.52727882 | 0.61758336 | 0.61305777 | 1.00738199 | 0.31375124 | NA |
| DZIP3 | 10.9108758 | 1.51019447 | 0.56442028 | 2.67565593 | 0.00745832 | NA |
| NECTIN3 | 9.64667077 | -1.0733802 | 0.56530088 | -1.8987767 | 0.05759384 | NA |
| PHLDB2 | 196.118417 | -0.1586547 | 0.2936468 | -0.5402908 | 0.5889965 | 0.89534694 |
| ABHD10 | 28.0982217 | -0.6561194 | 0.45008863 | -1.457756 | 0.14490782 | 0.62683788 |
| ATG3 | 7.74634157 | -0.6803525 | 0.5763155 | -1.180521 | 0.23779306 | NA |
| SLC35A5 | 6.78537434 | -0.334697 | 0.63813224 | -0.5244948 | 0.59993443 | NA |
| GTPBP8 | 6.4830807 | 0.22209371 | 0.62775517 | 0.35379033 | 0.72349602 | NA |
| NEPRO | 10.3072833 | -0.1874363 | 0.50946562 | -0.3679077 | 0.71294205 | NA |
| CFAP44 | 9.16917493 | 0.8966013 | 0.47462702 | 1.88906503 | 0.05888312 | NA |
| SPICE1 | 9.43532536 | 0.24803884 | 0.5687777 | 0.436091 | 0.66277071 | NA |
| USF3 | 16.9847466 | 0.71305346 | 0.42320986 | 1.68486969 | 0.09201372 | NA |
| NAA50 | 38.6497702 | -0.8591462 | 0.31606981 | -2.7182165 | 0.00656349 | 0.23699006 |
| ATP6V1A | 202.985651 | -0.419589 | 0.30924523 | -1.3568164 | 0.17483956 | 0.6651652 |
| GRAMD1C | 47.7209122 | 0.13503126 | 0.41221463 | 0.32757513 | 0.74323293 | 0.94831173 |
| ZDHHC23 | 12.4401071 | -0.0478988 | 0.49917696 | -0.0959556 | 0.92355581 | NA |
| CCDC191 | 9.45870694 | -0.0918734 | 0.71081252 | -0.1292512 | 0.89715888 | NA |
| QTRT2 | 6.26689449 | -0.4266669 | 0.6534243 | -0.6529707 | 0.51377516 | NA |
| ZBTB20 | 1513.39964 | 0.18649642 | 0.22313725 | 0.83579238 | 0.40327171 | 0.82808596 |
| ZBTB20-AS1 | 5.50773034 | -0.1257864 | 0.89916174 | -0.139893 | 0.88874455 | NA |
| ZBTB20-AS5 | 25.2965974 | 0.83971852 | 0.60875826 | 1.37939568 | 0.16777279 | NA |
| ZBTB20-AS2 | 5.66199081 | -0.0843405 | 0.64886552 | -0.1299815 | 0.89658103 | NA |
| LSAMP | 9.62315048 | -0.0007902 | 0.78593055 | -0.0010054 | 0.9991978 | NA |
| B4GALT4 | 7.07929052 | -0.3473974 | 0.53675912 | -0.6472128 | 0.51749425 | NA |
| TMEM39A | 8.55345813 | 0.12903671 | 0.70712028 | 0.18248198 | 0.8552045 | NA |
| POGLUT1 | 4.95467961 | -0.4317099 | 0.83188505 | -0.5189538 | 0.60379296 | NA |
| TIMMDC1 | 10.4297039 | -0.2148681 | 0.52511931 | -0.4091796 | 0.68240786 | NA |
| COX17 | 12.1626223 | -0.3303757 | 0.46283301 | -0.713812 | 0.47534346 | NA |
| GSK3B | 62.6297665 | -0.65742 | 0.34491943 | -1.9060104 | 0.05664886 | 0.503587 |
| LRRC58 | 8.46718071 | -0.1326546 | 0.55476818 | -0.2391172 | 0.8110147 | NA |
| NDUFB4 | 15.9819738 | -0.1458172 | 0.3795933 | -0.3841406 | 0.70087422 | NA |
| HGD | 4.38134088 | 1.27299374 | 0.92186047 | 1.38089633 | 0.16731084 | NA |
| RABL3 | 7.61477929 | -0.1245963 | 0.61068343 | -0.2040277 | 0.83833186 | NA |
| GTF2E1 | 2.50095713 | 0.4022165 | 0.94233909 | 0.42682778 | 0.66950477 | NA |
| GOLGB1 | 65.6000206 | -0.0542391 | 0.37885405 | -0.1431663 | 0.88615886 | 0.97769824 |
| IQCB1 | 29.4455518 | 0.0015613 | 0.39678484 | 0.00393488 | 0.99686043 | 0.99960611 |
| ILDR1 | 4.15389134 | 1.02534572 | 0.70878111 | 1.4466324 | 0.14799991 | NA |
| CASR | 16.1298942 | -0.6522419 | 0.76974393 | -0.8473492 | 0.3968005 | NA |
| CCDC58 | 5.50049777 | 0.67647724 | 0.73024121 | 0.92637506 | 0.3542511 | NA |
| FAM162A | 25.5378834 | -0.2033423 | 0.38697508 | -0.5254662 | 0.59925917 | NA |
| AC083798.2 | 6.13869862 | 0.12464215 | 0.67922831 | 0.18350552 | 0.85440138 | NA |
| KPNA1 | 33.5442019 | -0.3353714 | 0.36573106 | -0.9169891 | 0.35914836 | 0.80341474 |
| PARP9 | 19.9177347 | -0.5390072 | 0.73100542 | -0.7373505 | 0.46090925 | NA |
| DTX3L | 7.56143561 | -1.1754254 | 0.64339428 | -1.826913 | 0.06771285 | NA |
| PARP14 | 37.6765096 | -0.3795451 | 0.68317179 | -0.5555631 | 0.57850955 | 0.89263108 |
| HSPBAP1 | 8.97530053 | -0.3702015 | 0.54153204 | -0.6836189 | 0.49421583 | NA |
| DIRC2 | 12.9782655 | -0.0356487 | 0.65742752 | -0.0542246 | 0.95675625 | NA |
| PDIA5 | 5.21500748 | -1.0046824 | 0.68077781 | -1.4757861 | 0.14000132 | NA |
| SEC22A | 24.5401288 | -0.876791 | 0.42768044 | -2.0501079 | 0.04035391 | NA |
| HACD2 | 15.4747255 | 0.18422779 | 0.4195435 | 0.43911488 | 0.6605783 | NA |

| | | | | | | |
|---|---|---|---|---|---|---|
| MYLK | 7.66921015 | -0.0150013 | 0.57282273 | -0.0261883 | 0.97910713 | NA |
| CCDC14 | 19.5191526 | 0.38708991 | 0.46120046 | 0.83930948 | 0.40129567 | NA |
| KALRN | 11.0691095 | -0.1897419 | 0.58925142 | -0.322005 | 0.74744894 | NA |
| UMPS | 5.81115787 | 0.29634157 | 0.66645077 | 0.44465636 | 0.65656811 | NA |
| ITGB5 | 23.9378005 | 0.45491513 | 0.37963382 | 1.19829978 | 0.23080033 | NA |
| ZNF148 | 74.7240311 | -0.1037065 | 0.23122887 | -0.4485014 | 0.65379135 | 0.92248109 |
| SNX4 | 35.159155 | -0.1136157 | 0.27938133 | -0.4066688 | 0.68425123 | 0.92710517 |
| OSBPL11 | 11.5031684 | -0.5976476 | 0.43752197 | -1.3659831 | 0.17194427 | NA |
| SLC41A3 | 6.82282785 | 0.04313403 | 0.59203238 | 0.07285755 | 0.94191948 | NA |
| ALDH1L1 | 22.6311418 | 1.37301261 | 0.52217932 | 2.62938906 | 0.00855384 | NA |
| KLF15 | 5.69606855 | -0.1573627 | 0.73146311 | -0.2151341 | 0.82966273 | NA |
| ZXDC | 40.7851671 | 0.33243842 | 0.26714115 | 1.24442985 | 0.2133414 | 0.71284112 |
| TXNRD3NB | 3.12477374 | 1.08227194 | 0.88163342 | 1.2275759 | 0.21960621 | NA |
| TXNRD3 | 4.26347842 | 0.19086131 | 0.74020173 | 0.25785039 | 0.79652237 | NA |
| CHCHD6 | 21.7040392 | 0.16758859 | 0.41702276 | 0.40186917 | 0.68778031 | NA |
| PLXNA1 | 5.21594427 | -0.308008 | 0.67346511 | -0.4573481 | 0.64742088 | NA |
| TPRA1 | 5.23470491 | -0.1630149 | 0.68943548 | -0.236447 | 0.81308583 | NA |
| PODXL2 | 4.83584752 | 0.75172903 | 0.67192988 | 1.11876112 | 0.26324206 | NA |
| MGLL | 15.2026204 | -0.1178881 | 0.61823014 | -0.1906865 | 0.84877124 | NA |
| KBTBD12 | 8.99025423 | 0.13758925 | 0.5491965 | 0.25052828 | 0.80217884 | NA |
| SEC61A1 | 19.944167 | -0.8997674 | 0.4947835 | -1.8185072 | 0.06898665 | NA |
| RUVBL1 | 6.74745309 | -0.0705055 | 0.64528379 | -0.1092628 | 0.91299404 | NA |
| EEFSEC | 16.7971268 | 0.23425326 | 0.41957722 | 0.55830787 | 0.57663417 | NA |
| GATA2 | 13.3108096 | 0.0427303 | 0.55526541 | 0.07695472 | 0.93865956 | NA |
| GATA2-AS1 | 5.95832048 | -0.7205383 | 0.73221093 | -0.9840584 | 0.32508681 | NA |
| RPN1 | 16.485846 | -0.1420389 | 0.47653985 | -0.298063 | 0.76565508 | NA |
| RAB7A | 66.2238076 | -0.6063761 | 0.28781454 | -2.1068292 | 0.03513239 | 0.41688714 |
| ACAD9 | 10.9786283 | 0.0304637 | 0.46708739 | 0.06522057 | 0.94799839 | NA |
| CNBP | 30.305603 | 0.61761801 | 0.53821117 | 1.14753843 | 0.25115916 | 0.74329941 |
| COPG1 | 11.6894783 | -0.2622557 | 0.60040175 | -0.4368004 | 0.66225612 | NA |
| HMCES | 4.68956522 | -0.2332254 | 0.71254981 | -0.3273111 | 0.74343262 | NA |
| H1FX | 10.8791139 | -1.615593 | 0.63595055 | -2.540438 | 0.01107137 | NA |
| MBD4 | 16.1634398 | -0.0703856 | 0.38784131 | -0.1814805 | 0.85599046 | NA |
| IFT122 | 13.7494256 | -0.0524964 | 0.42045137 | -0.1248572 | 0.90063659 | NA |
| TMCC1 | 135.085682 | 0.17070223 | 0.30357785 | 0.56230133 | 0.57391073 | 0.89263108 |
| PIK3R4 | 9.02410594 | 0.26359797 | 0.53924429 | 0.48882849 | 0.62496313 | NA |
| ATP2C1 | 27.5987337 | 0.20105887 | 0.34334662 | 0.58558569 | 0.55815396 | 0.89263108 |
| NEK11 | 9.54228166 | 0.5034873 | 0.59266176 | 0.84953567 | 0.39558329 | NA |
| NUDT16 | 9.5663884 | 0.03095069 | 0.5024394 | 0.06160084 | 0.95088071 | NA |
| MRPL3 | 15.9271605 | -0.4700805 | 0.41588089 | -1.1303248 | 0.25833937 | NA |
| ACPP | 5.07985728 | 0.51029639 | 0.72888815 | 0.70010247 | 0.48386331 | NA |
| DNAJC13 | 26.9401187 | -0.4355953 | 0.34317293 | -1.2693171 | 0.20432799 | 0.700066 |
| UBA5 | 12.5275331 | 0.36883385 | 0.50548719 | 0.72966013 | 0.46559796 | NA |
| CDV3 | 21.3478176 | -0.5999424 | 0.43130271 | -1.3910007 | 0.16422523 | NA |
| TOPBP1 | 9.27995787 | -0.7268019 | 0.5682747 | -1.2789623 | 0.20091034 | NA |
| SRPRB | 5.54245477 | 0.01403823 | 0.61887922 | 0.02268331 | 0.98190289 | NA |
| RYK | 46.7919846 | 0.05824372 | 0.26354787 | 0.22099864 | 0.82509349 | 0.96845695 |
| AMOTL2 | 21.3716695 | -0.4648444 | 0.50579654 | -0.9190343 | 0.35807764 | NA |
| ANAPC13 | 7.36319677 | -0.1080598 | 0.59048065 | -0.183003 | 0.85479563 | NA |
| CEP63 | 12.896294 | 0.63099628 | 0.55017401 | 1.1469031 | 0.25142167 | NA |
| EPHB1 | 3.95167498 | -0.8192925 | 0.80420395 | -1.0187621 | 0.30831592 | NA |
| PPP2R3A | 103.765599 | -0.1654066 | 0.36694258 | -0.4507697 | 0.65215553 | 0.92248109 |
| MSL2 | 17.0595906 | 0.04015156 | 0.39394596 | 0.1019215 | 0.91881898 | NA |
| PCCB | 9.85604186 | 0.38866453 | 0.48151662 | 0.80716742 | 0.41957003 | NA |

| | | | | | | |
|---|---|---|---|---|---|---|
| STAG1 | 160.23565 | -0.2887009 | 0.30547159 | -0.9450989 | 0.34460841 | 0.8003972 |
| NCK1 | 13.1432943 | 0.26906983 | 0.40792352 | 0.65960853 | 0.50950508 | NA |
| DBR1 | 3.50919475 | -0.6460123 | 0.79064791 | -0.817067 | 0.41389014 | NA |
| ARMC8 | 32.8047621 | 0.23267921 | 0.34100418 | 0.68233537 | 0.49502692 | 0.86687543 |
| MRAS | 14.233544 | 0.02158146 | 0.4932574 | 0.04375294 | 0.96510134 | NA |
| ESYT3 | 2.17983611 | 0.09355267 | 1.03367085 | 0.09050528 | 0.9278857 | NA |
| CEP70 | 29.6213353 | -0.3910615 | 0.39983431 | -0.9780589 | 0.32804522 | 0.79021 |
| FAIM | 6.17445537 | 0.31707416 | 0.5836473 | 0.54326331 | 0.58694852 | NA |
| PIK3CB | 42.8214471 | -0.0208025 | 0.30051206 | -0.0692235 | 0.94481174 | 0.99240556 |
| MRPS22 | 3.8116666 | -0.1906098 | 0.80755509 | -0.2360331 | 0.81340695 | NA |
| COPB2 | 23.1484012 | -0.5035976 | 0.40496457 | -1.2435596 | 0.21366169 | NA |
| AC097103.2 | 13.0486561 | 0.64622616 | 0.50314634 | 1.28437019 | 0.19901246 | NA |
| NMNAT3 | 13.3830273 | 0.42917864 | 0.53631196 | 0.80024067 | 0.42357137 | NA |
| SLC25A36 | 45.4422256 | 0.2189329 | 0.31588185 | 0.69308476 | 0.48825636 | 0.86354292 |
| PXYLP1 | 3.01263476 | 0.14259961 | 0.84932512 | 0.16789756 | 0.86666387 | NA |
| ZBTB38 | 26.9828137 | -0.0257138 | 0.35306848 | -0.0728294 | 0.94194187 | 0.99153825 |
| RASA2 | 42.3391788 | -0.1819351 | 0.37686148 | -0.482764 | 0.62926334 | 0.91751762 |
| RNF7 | 10.9305343 | -0.7197466 | 0.60448126 | -1.1906847 | 0.23377739 | NA |
| ATP1B3 | 25.8643011 | -1.1129208 | 0.48078468 | -2.3148008 | 0.02062383 | NA |
| TFDP2 | 58.9638891 | 0.58086286 | 0.38107342 | 1.5242807 | 0.12743861 | 0.59256732 |
| GK5 | 28.5068879 | 0.6040108 | 0.35483737 | 1.70221866 | 0.08871439 | 0.56122412 |
| XRN1 | 82.597662 | -0.002856 | 0.34766675 | -0.0082148 | 0.99344563 | 0.99923383 |
| ATR | 41.3017047 | -0.5390544 | 0.33554961 | -1.606482 | 0.10816805 | 0.57769936 |
| PLS1 | 27.7197315 | 0.31582529 | 0.33195489 | 0.95141024 | 0.34139616 | 0.8003972 |
| TRPC1 | 133.871614 | -0.6205313 | 0.28471786 | -2.1794604 | 0.02929748 | 0.40645706 |
| PCOLCE2 | 6.57205261 | -0.2278739 | 0.72235709 | -0.3154589 | 0.75241328 | NA |
| U2SURP | 62.5937749 | -0.3805693 | 0.31820539 | -1.1959862 | 0.23170194 | 0.73156941 |
| CHST2 | 30.9717814 | -1.5061533 | 0.54301927 | -2.7736646 | 0.00554288 | 0.22794137 |
| SLC9A9 | 12.9469002 | 0.57958382 | 0.80450605 | 0.72042196 | 0.47126524 | NA |
| C3orf58 | 4.64178396 | -0.6868347 | 0.64113591 | -1.0712778 | 0.28404454 | NA |
| AC107021.1 | 7.37324153 | 0.77638943 | 0.63866322 | 1.21564762 | 0.22411919 | NA |
| PLOD2 | 56.2113461 | 0.18622401 | 0.45412308 | 0.41007388 | 0.68175175 | 0.9267216 |
| PLSCR4 | 5.64463782 | 0.90468338 | 0.79776149 | 1.13402739 | 0.25678306 | NA |
| PLSCR2 | 5.96942339 | 0.14706597 | 0.70901231 | 0.20742371 | 0.83567896 | NA |
| PLSCR1 | 44.8620658 | 0.13004463 | 0.46465037 | 0.27987631 | 0.7795724 | 0.96139889 |
| GYG1 | 13.9325043 | 0.16647603 | 0.52810322 | 0.31523388 | 0.75258408 | NA |
| HLTF | 9.89134249 | 0.7720653 | 0.49718723 | 1.55286633 | 0.12045507 | NA |
| HPS3 | 9.82560314 | 1.37630694 | 0.48250197 | 2.85243799 | 0.00433853 | NA |
| WWTR1 | 44.8165114 | -0.5256292 | 0.43675834 | -1.2034783 | 0.22879129 | 0.73156941 |
| COMMD2 | 8.05748054 | -0.5040182 | 0.50126874 | -1.0054851 | 0.31466332 | NA |
| ANKUB1 | 16.0529094 | 0.28946534 | 0.53488634 | 0.54117168 | 0.58838926 | NA |
| RNF13 | 36.9409974 | 0.20900175 | 0.34722098 | 0.60192719 | 0.5472226 | 0.88410237 |
| AC117386.2 | 10.767642 | 0.85882553 | 0.93103393 | 0.92244278 | 0.35629766 | NA |
| TSC22D2 | 56.7870713 | -0.4380478 | 0.30212774 | -1.4498761 | 0.14709308 | 0.63 |
| SERP1 | 30.0183286 | -0.4015947 | 0.36335147 | -1.1052515 | 0.26905065 | 0.75632146 |
| EIF2A | 13.6736315 | -0.0765117 | 0.43817259 | -0.1746154 | 0.86138182 | NA |
| SELENOT | 20.0282933 | -0.2962317 | 0.3974018 | -0.7454212 | 0.45601715 | NA |
| SIAH2 | 2.6811043 | -0.4367812 | 1.09027385 | -0.400616 | 0.68870286 | NA |
| MBNL1 | 139.194284 | 0.27579812 | 0.25107025 | 1.09848986 | 0.27199064 | 0.75979923 |
| RAP2B | 30.0978693 | -0.825496 | 0.41771979 | -1.9761956 | 0.04813262 | 0.45717429 |
| DHX36 | 43.6519354 | -0.259397 | 0.26318602 | -0.9856033 | 0.3243278 | 0.78622244 |
| MME | 4.51458217 | 0.27864058 | 0.93263041 | 0.29876849 | 0.7651167 | NA |
| PLCH1 | 30.1948234 | 1.13626712 | 0.60864845 | 1.86686932 | 0.06191984 | 0.51631533 |
| GMPS | 18.0453884 | -0.1878233 | 0.44657026 | -0.4205906 | 0.67405402 | NA |

| | | | | | | |
|---|---|---|---|---|---|---|
| SSR3 | 11.7479499 | -0.3888733 | 0.51281884 | -0.7583054 | 0.4482682 | NA |
| TIPARP | 4.20260037 | -0.1872837 | 0.67465811 | -0.2775979 | 0.78132102 | NA |
| LINC00886 | 20.9610454 | 1.49059944 | 0.53938667 | 2.76350814 | 0.00571837 | NA |
| CCNL1 | 30.1825745 | -0.3613047 | 0.35709605 | -1.0117859 | 0.31164044 | 0.778534 |
| VEPH1 | 8.98605185 | 0.97129824 | 0.61510635 | 1.57907369 | 0.11431916 | NA |
| RSRC1 | 58.472155 | 0.05383877 | 0.27978329 | 0.19243024 | 0.84740521 | 0.97168963 |
| GFM1 | 17.3756762 | -0.0729793 | 0.41001098 | -0.1779936 | 0.858728 | NA |
| RARRES1 | 3.25868072 | 0.73720755 | 0.83669684 | 0.88109278 | 0.3782676 | NA |
| MFSD1 | 9.36477911 | -0.4163654 | 0.61994014 | -0.6716219 | 0.50182441 | NA |
| IQCJ-SCHIP1 | 13.890743 | -0.2792281 | 0.66889915 | -0.4174442 | 0.67635351 | NA |
| SMC4 | 8.12253063 | -0.267708 | 0.54568686 | -0.490589 | 0.62371715 | NA |
| KPNA4 | 34.0853523 | -0.5525303 | 0.34399257 | -1.6062275 | 0.10822393 | 0.57769936 |
| PPM1L | 71.93444 | -0.2325797 | 0.38588995 | -0.6027099 | 0.54670171 | 0.88410237 |
| NMD3 | 16.4839608 | -0.4732351 | 0.41276589 | -1.1464975 | 0.25158935 | NA |
| LINC01322 | 4.47189819 | 0.72997029 | 0.82920671 | 0.88032366 | 0.378684 | NA |
| PDCD10 | 26.3078066 | -0.2966433 | 0.3121772 | -0.9502402 | 0.3419902 | 0.8003972 |
| GOLIM4 | 28.6579648 | -0.5456675 | 0.33717112 | -1.6183698 | 0.10558294 | 0.57745683 |
| MECOM | 586.717818 | 0.00868651 | 0.25030504 | 0.03470369 | 0.97231602 | 0.99779576 |
| MYNN | 5.94001291 | 0.09029462 | 0.60827518 | 0.14844371 | 0.88199261 | NA |
| SEC62 | 37.7290119 | -0.19722 | 0.32075108 | -0.6148693 | 0.53864106 | 0.88254911 |
| GPR160 | 17.4616892 | 0.16892739 | 0.62256458 | 0.27134115 | 0.78612866 | NA |
| PHC3 | 62.904835 | -0.0164134 | 0.26781691 | -0.061286 | 0.95113142 | 0.99443943 |
| PRKCI | 46.3450675 | -1.2397894 | 0.39129659 | -3.1684135 | 0.00153273 | 0.13196343 |
| SKIL | 17.5276676 | -0.1556357 | 0.42945748 | -0.3624007 | 0.71705264 | NA |
| RPL22L1 | 2.83201312 | -0.3806593 | 0.82781609 | -0.4598356 | 0.64563424 | NA |
| TNIK | 15.9943758 | 0.17610861 | 0.5544926 | 0.31760317 | 0.75078597 | NA |
| PLD1 | 14.4618027 | -0.9252461 | 0.65647198 | -1.409422 | 0.15871042 | NA |
| FNDC3B | 165.579863 | -0.1913246 | 0.27155926 | -0.7045409 | 0.48109597 | 0.86222804 |
| TNFSF10 | 15.2956454 | -0.0160173 | 0.46546916 | -0.0344111 | 0.97254933 | NA |
| NCEH1 | 24.2883264 | 0.3282739 | 0.4848382 | 0.67707928 | 0.49835565 | NA |
| ECT2 | 5.43579581 | -0.6305198 | 0.74952929 | -0.841221 | 0.40022413 | NA |
| NLGN1 | 65.527955 | -0.2040398 | 0.42846226 | -0.4762141 | 0.63392187 | 0.92072524 |
| NAALADL2 | 136.007185 | 0.29646901 | 0.43912121 | 0.67514163 | 0.49958579 | 0.87036626 |
| TBL1XR1 | 137.412078 | -0.7068512 | 0.42900246 | -1.6476625 | 0.09942196 | 0.57091823 |
| KCNMB2-AS1 | 9.55843729 | 1.58409672 | 0.87615888 | 1.80800167 | 0.07060624 | NA |
| ZMAT3 | 7.79687279 | -0.1115592 | 0.53011416 | -0.2104437 | 0.83332142 | NA |
| PIK3CA | 48.2027182 | 0.11710971 | 0.28562134 | 0.41001738 | 0.6817932 | 0.9267216 |
| KCNMB3 | 9.17550526 | -0.9321235 | 0.58281772 | -1.5993397 | 0.10974514 | NA |
| ZNF639 | 5.88560518 | -0.6836245 | 0.83869177 | -0.8151082 | 0.41501039 | NA |
| MFN1 | 29.4390766 | -0.7606748 | 0.34319864 | -2.2164272 | 0.02666226 | 0.39506514 |
| GNB4 | 5.18380336 | 0.03058156 | 0.87493677 | 0.03495288 | 0.97211731 | NA |
| ACTL6A | 6.32282448 | -0.5594828 | 0.63819345 | -0.8766664 | 0.38066784 | NA |
| MRPL47 | 6.93874511 | -1.2831582 | 0.62574939 | -2.0505945 | 0.04030645 | NA |
| NDUFB5 | 16.1349658 | 0.95530338 | 0.45481909 | 2.10040301 | 0.0356934 | NA |
| USP13 | 3.00278154 | 1.94323288 | 0.90006681 | 2.15898737 | 0.03085115 | NA |
| TTC14 | 32.9557975 | 0.20918105 | 0.43268375 | 0.48345021 | 0.62877611 | 0.91751762 |
| FXR1 | 37.0998864 | 0.34739309 | 0.27495321 | 1.26346258 | 0.20642299 | 0.70363085 |
| DNAJC19 | 6.01439492 | 0.96923662 | 0.6965811 | 1.39141965 | 0.16409821 | NA |
| ATP11B | 40.3551598 | -0.0973306 | 0.30247206 | -0.3217839 | 0.74761644 | 0.94920315 |
| DCUN1D1 | 20.161931 | 0.34635748 | 0.3607048 | 0.96022421 | 0.33694239 | NA |
| MCCC1 | 29.4158015 | 0.00777689 | 0.31005786 | 0.02508205 | 0.97998952 | 0.99902646 |
| B3GNT5 | 5.11888328 | 0.68622189 | 0.66900363 | 1.02573716 | 0.30501549 | NA |
| KLHL24 | 52.6163512 | -0.3560483 | 0.30677204 | -1.1606281 | 0.24579315 | 0.73462702 |
| YEATS2 | 21.2483201 | -1.0695545 | 0.42640694 | -2.5082953 | 0.01213152 | NA |

| | | | | | | |
|---|---|---|---|---|---|---|
| ABCC5 | 66.8100176 | -0.1503247 | 0.31334071 | -0.4797485 | 0.63140624 | 0.9183778 |
| EIF2B5 | 5.73971895 | 0.02755371 | 0.61443089 | 0.04484428 | 0.96423143 | NA |
| DVL3 | 12.1443831 | -0.9312809 | 0.46096301 | -2.0202942 | 0.04335288 | NA |
| AP2M1 | 17.3053256 | -0.5824187 | 0.4393719 | -1.3255711 | 0.18498179 | NA |
| ABCF3 | 4.2038037 | 0.36169817 | 0.72326022 | 0.50009411 | 0.61700882 | NA |
| ALG3 | 3.10795554 | -0.2489722 | 0.78574021 | -0.3168632 | 0.75134739 | NA |
| PSMD2 | 8.72670268 | -0.3341059 | 0.66443946 | -0.5028388 | 0.61507763 | NA |
| EIF4G1 | 19.7760384 | -0.2499137 | 0.47075377 | -0.53088 | 0.59550196 | NA |
| FAM131A | 3.06244019 | 0.98088575 | 0.8211923 | 1.19446536 | 0.23229598 | NA |
| POLR2H | 13.840132 | -0.7317018 | 0.47438742 | -1.5424141 | 0.12297301 | NA |
| MAGEF1 | 8.07331507 | -0.1071323 | 0.61121934 | -0.1752764 | 0.86086243 | NA |
| VPS8 | 40.7683177 | 0.03716645 | 0.27104814 | 0.13712123 | 0.89093497 | 0.97769824 |
| EHHADH | 9.62555057 | 0.11266228 | 0.64740356 | 0.17402172 | 0.86184839 | NA |
| MAP3K13 | 35.746362 | -0.1808341 | 0.38478504 | -0.4699614 | 0.63838262 | 0.92121485 |
| TMEM41A | 6.23546896 | 1.18771096 | 0.67672082 | 1.7550974 | 0.07924265 | NA |
| LIPH | 4.20529773 | -1.2734363 | 0.79621084 | -1.5993707 | 0.10973825 | NA |
| SENP2 | 15.36202 | 0.07947888 | 0.41113776 | 0.19331448 | 0.84671268 | NA |
| IGF2BP2 | 17.5059295 | 0.39164823 | 0.8916403 | 0.43924465 | 0.66048427 | NA |
| TRA2B | 26.8957772 | -0.2631396 | 0.35086779 | -0.7499681 | 0.45327394 | 0.85557861 |
| ETV5 | 4.53306148 | 0.70932097 | 0.69293949 | 1.02364055 | 0.30600508 | NA |
| AC068631.2 | 4.16206027 | 0.29296064 | 0.7867946 | 0.37234704 | 0.70963448 | NA |
| EIF4A2 | 39.618816 | 0.06162219 | 0.38475299 | 0.1601604 | 0.87275472 | 0.9751578 |
| ST6GAL1 | 137.640486 | -0.2075415 | 0.29498567 | -0.7035647 | 0.48170391 | 0.86228554 |
| BCL6 | 47.3935922 | -0.0958497 | 0.50961595 | -0.1880822 | 0.85081224 | 0.97168963 |
| LPP | 389.230029 | -0.3222118 | 0.33285497 | -0.9680245 | 0.33303215 | 0.79447068 |
| TPRG1 | 24.2533572 | -0.4100547 | 0.83940866 | -0.4885043 | 0.62519268 | NA |
| P3H2 | 69.6038528 | -1.0275613 | 0.39335687 | -2.6122876 | 0.00899386 | 0.27648451 |
| CLDN1 | 49.4286107 | -0.306434 | 0.35028902 | -0.8748032 | 0.38168097 | 0.82153751 |
| CLDN16 | 3.61012392 | 0.10625375 | 0.78121324 | 0.13601119 | 0.89181243 | NA |
| CCDC50 | 39.1320664 | 0.48170602 | 0.38223125 | 1.26024761 | 0.20758005 | 0.70439682 |
| FGF12 | 16.0629754 | -3.1718593 | 1.03406083 | -3.0673818 | 0.00215943 | NA |
| MB21D2 | 3.78921991 | -0.0307671 | 0.81859254 | -0.0375853 | 0.97001831 | NA |
| OPA1 | 35.1531546 | -0.1041946 | 0.28146438 | -0.3701874 | 0.71124284 | 0.93492235 |
| HES1 | 6.33076885 | -0.092773 | 0.91386367 | -0.1015173 | 0.91913981 | NA |
| ATP13A3 | 38.9177787 | -0.1745115 | 0.31970067 | -0.545859 | 0.58516286 | 0.89393468 |
| LSG1 | 7.93107694 | 0.23101845 | 0.59136778 | 0.39065106 | 0.69605518 | NA |
| XXYLT1 | 7.25139099 | 0.39596259 | 0.62400808 | 0.63454722 | 0.52572376 | NA |
| ACAP2 | 59.7281405 | -0.4643374 | 0.28057274 | -1.6549627 | 0.09793206 | 0.56821884 |
| PPP1R2 | 14.0775801 | -0.5093245 | 0.45791811 | -1.1122611 | 0.26602592 | NA |
| MUC20-OT1 | 229.957019 | -0.1428262 | 0.3497162 | -0.4084061 | 0.68297555 | 0.92710517 |
| TNK2 | 7.71772683 | 0.22290477 | 0.58990653 | 0.37786456 | 0.70553121 | NA |
| TFRC | 31.5658073 | -0.5484872 | 0.53275468 | -1.0295306 | 0.30323043 | 0.77521265 |
| LINC00885 | 2.20047379 | 1.50869349 | 1.25371797 | 1.20337551 | 0.22883104 | NA |
| PCYT1A | 8.38532677 | -0.5086374 | 0.54611369 | -0.9313764 | 0.35165891 | NA |
| UBXN7 | 18.7792726 | -0.2523287 | 0.42463835 | -0.5942202 | 0.55236484 | NA |
| RNF168 | 12.8250888 | -0.5869003 | 0.428869 | -1.3684839 | 0.17116066 | NA |
| WDR53 | 2.52145457 | -0.0273419 | 0.88182036 | -0.0310062 | 0.97526458 | NA |
| PIGX | 4.69095056 | 0.22702086 | 0.80764964 | 0.2810883 | 0.77864268 | NA |
| PAK2 | 41.6198325 | -0.252303 | 0.26976827 | -0.9352581 | 0.34965531 | 0.8003972 |
| SENP5 | 42.9359204 | -0.6087227 | 0.3352429 | -1.8157661 | 0.06940625 | 0.52654172 |
| NCBP2 | 9.4529888 | -0.7294164 | 0.50170842 | -1.4538652 | 0.14598369 | NA |
| NCBP2-AS2 | 5.49185739 | 0.42981774 | 0.6184455 | 0.69499697 | 0.4870572 | NA |
| DLG1 | 114.254557 | -0.5317386 | 0.25436552 | -2.0904508 | 0.03657732 | 0.41899084 |
| BDH1 | 18.8060483 | 0.56730962 | 0.55562529 | 1.02102916 | 0.30724062 | NA |

| | | | | | | |
|---|---|---|---|---|---|---|
| AC024560.1 | 5.83912502 | -1.5419377 | 1.1371762 | -1.3559356 | 0.17511965 | NA |
| RUBCN | 9.21009138 | -0.0670455 | 0.50912334 | -0.1316881 | 0.895231 | NA |
| FYTTD1 | 11.674937 | -0.1698599 | 0.48128277 | -0.3529316 | 0.72413974 | NA |
| LRCH3 | 38.8061241 | -0.2117141 | 0.30742593 | -0.688667 | 0.49103282 | 0.86506451 |
| IQCG | 4.87765517 | 0.52864746 | 0.61839964 | 0.85486378 | 0.39262656 | NA |
| RPL35A | 44.081038 | 0.26307144 | 0.3840084 | 0.6850669 | 0.4933017 | 0.86619885 |
| LMLN | 5.61350447 | 0.11601936 | 0.72969426 | 0.15899722 | 0.87367106 | NA |
| ZNF595 | 9.60542197 | 0.4490742 | 0.60290977 | 0.74484479 | 0.45636556 | NA |
| ZNF718 | 18.399809 | 0.12778472 | 0.4205742 | 0.30383394 | 0.76125441 | NA |
| ZNF141 | 21.319134 | -0.0607935 | 0.35107552 | -0.1731636 | 0.86252281 | NA |
| AC092574.2 | 3.39145815 | 0.96749371 | 0.95903549 | 1.0088195 | 0.31306121 | NA |
| ZNF721 | 45.7920945 | -0.052106 | 0.28862446 | -0.1805323 | 0.85673471 | 0.97168963 |
| PIGG | 11.7677452 | -0.0844838 | 0.43343726 | -0.1949159 | 0.84545877 | NA |
| ATP5ME | 13.7314514 | 0.12212075 | 0.4391562 | 0.27808045 | 0.7809506 | NA |
| PCGF3 | 17.4486177 | -0.2735347 | 0.44862598 | -0.6097165 | 0.54204961 | NA |
| AC139887.2 | 22.9898598 | 0.57389952 | 0.41812286 | 1.37256193 | 0.16988857 | NA |
| GAK | 28.6127301 | -0.9751015 | 0.33424088 | -2.9173617 | 0.00353006 | 0.20482034 |
| TMEM175 | 4.12592143 | 0.75086391 | 0.69955005 | 1.07335266 | 0.28311292 | NA |
| IDUA | 4.91855918 | -0.1773124 | 0.77171251 | -0.2297648 | 0.8182745 | NA |
| FGFRL1 | 4.23374271 | 0.27717064 | 0.73558079 | 0.37680516 | 0.7063184 | NA |
| RNF212 | 8.109092 | -0.4424063 | 0.59339189 | -0.7455549 | 0.45593631 | NA |
| AC092535.3 | 18.092059 | 0.72872818 | 0.56463205 | 1.2906249 | 0.19683378 | NA |
| CTBP1-AS | 3.91861502 | 0.16645202 | 0.9073678 | 0.18344493 | 0.85444893 | NA |
| CTBP1 | 12.9933394 | -0.3349019 | 0.42674675 | -0.7847791 | 0.43258309 | NA |
| CTBP1-DT | 2.17032024 | -0.3414725 | 0.98523305 | -0.3465906 | 0.72889893 | NA |
| MAEA | 16.891596 | -0.6991641 | 0.38966992 | -1.794247 | 0.07277375 | NA |
| UVSSA | 12.2089053 | 0.14208343 | 0.47939489 | 0.29638079 | 0.7669393 | NA |
| SLBP | 18.1445079 | -0.3985547 | 0.38704451 | -1.0297385 | 0.30313276 | NA |
| TMEM129 | 8.06435409 | -0.2641861 | 0.58201372 | -0.4539173 | 0.64988834 | NA |
| LETM1 | 8.13478672 | -1.3270037 | 0.57888719 | -2.2923355 | 0.02188629 | NA |
| NSD2 | 31.7326786 | -0.5222605 | 0.33309884 | -1.5678846 | 0.11690807 | 0.57860396 |
| NELFA | 12.7903209 | -0.1514404 | 0.4769753 | -0.3175015 | 0.75086311 | NA |
| NAT8L | 1.77289568 | -1.3978899 | 1.24592887 | -1.1219661 | 0.26187687 | NA |
| POLN | 4.62067978 | -0.1880279 | 0.80265311 | -0.234258 | 0.81478472 | NA |
| MXD4 | 14.7826506 | -0.035603 | 0.4815754 | -0.0739303 | 0.94106587 | NA |
| ZFYVE28 | 11.3331754 | 0.58077632 | 0.46539632 | 1.24791773 | 0.21206119 | NA |
| RNF4 | 16.450253 | -0.2687318 | 0.43101655 | -0.6234837 | 0.53296674 | NA |
| FAM193A | 47.5065356 | -0.794802 | 0.31008457 | -2.5631782 | 0.01037188 | 0.28266532 |
| TNIP2 | 7.51198765 | 0.11539103 | 0.62802625 | 0.18373599 | 0.85422057 | NA |
| SH3BP2 | 8.92967283 | -0.1665123 | 0.51940321 | -0.320584 | 0.74852568 | NA |
| ADD1 | 42.3283407 | -0.2082825 | 0.28587443 | -0.7285803 | 0.46625843 | 0.85894565 |
| MFSD10 | 14.7642757 | 0.01366743 | 0.4342405 | 0.03147433 | 0.97489127 | NA |
| NOP14 | 5.96191224 | -0.3052661 | 0.6394739 | -0.4773707 | 0.63309816 | NA |
| GRK4 | 4.36248461 | 0.89819707 | 0.78360244 | 1.14624077 | 0.25169554 | NA |
| HTT | 31.7785167 | -0.1216946 | 0.31473331 | -0.3866595 | 0.69900831 | 0.9303056 |
| RGS12 | 11.2729853 | -0.3239339 | 0.61050226 | -0.5306023 | 0.59569439 | NA |
| LRPAP1 | 18.3124186 | 0.30250061 | 0.37052835 | 0.81640341 | 0.41426945 | NA |
| TMEM128 | 4.21838349 | 0.41000747 | 0.70846791 | 0.57872412 | 0.56277534 | NA |
| ZBTB49 | 3.51824521 | -0.6391121 | 0.80089153 | -0.7980008 | 0.42487002 | NA |
| STX18 | 57.0499564 | 0.27975636 | 0.30585455 | 0.91467126 | 0.36036424 | 0.80341474 |
| STX18-AS1 | 26.1078499 | 0.56594286 | 0.45715492 | 1.23796732 | 0.21572818 | 0.71525281 |
| EVC2 | 14.0241525 | -0.1849785 | 0.48327227 | -0.3827625 | 0.70189585 | NA |
| EVC | 15.2875809 | -0.4166048 | 0.54081418 | -0.7703289 | 0.44110483 | NA |
| CRMP1 | 3.65155353 | -0.7507287 | 1.048291 | -0.7161454 | 0.47390159 | NA |

| | | | | | | |
|---|---|---|---|---|---|---|
| MAN2B2 | 8.16316013 | 0.85365599 | 0.60158224 | 1.41901793 | 0.15589379 | NA |
| MRFAP1 | 16.3774854 | 0.11759007 | 0.49522587 | 0.23744735 | 0.81230976 | NA |
| MRFAP1L1 | 4.55949657 | -0.5629247 | 0.67599698 | -0.8327326 | 0.40499554 | NA |
| BLOC1S4 | 4.8985172 | -0.5207434 | 0.6927043 | -0.7517543 | 0.45219886 | NA |
| KIAA0232 | 50.5366768 | -0.2094492 | 0.27980478 | -0.7485549 | 0.4541255 | 0.85658019 |
| TBC1D14 | 77.5679212 | 0.5567885 | 0.35656622 | 1.56152902 | 0.11839898 | 0.57982915 |
| TADA2B | 4.68644613 | -0.2657172 | 0.69317164 | -0.3833353 | 0.70147115 | NA |
| GRPEL1 | 7.95108876 | -0.265859 | 0.5602423 | -0.474543 | 0.63511276 | NA |
| SORCS2 | 6.62938111 | -0.9816765 | 0.62574 | -1.5688249 | 0.11668874 | NA |
| AFAP1 | 5.28379317 | 0.27028491 | 0.71869352 | 0.37607813 | 0.70685882 | NA |
| ABLIM2 | 3.52280451 | 1.30058832 | 0.81895667 | 1.58810394 | 0.11226284 | NA |
| SH3TC1 | 38.0110152 | -0.6559738 | 0.36664007 | -1.7891492 | 0.0735908 | 0.53207981 |
| ACOX3 | 2.88079957 | 0.14578418 | 0.9154068 | 0.15925617 | 0.87346705 | NA |
| TRMT44 | 4.69704382 | 1.26115617 | 0.90912886 | 1.38721388 | 0.16537655 | NA |
| SLC2A9 | 10.9948955 | -0.6758345 | 0.5637085 | -1.1989079 | 0.23056378 | NA |
| WDR1 | 37.788814 | 0.12309083 | 0.36473442 | 0.33748071 | 0.73575455 | 0.94590222 |
| ZNF518B | 4.28835129 | -1.0766462 | 0.75466685 | -1.426651 | 0.15368053 | NA |
| CLNK | 412.289879 | 0.96279437 | 0.34972631 | 2.75299381 | 0.0059053 | 0.22794137 |
| AC084048.1 | 4.37621725 | 1.30015494 | 1.10643935 | 1.17508017 | 0.23996264 | NA |
| HS3ST1 | 4.2956502 | 0.10652516 | 0.76264188 | 0.13967914 | 0.88891351 | NA |
| AC005699.1 | 4.29855344 | 0.99727392 | 2.30122186 | 0.43336713 | 0.66474807 | NA |
| RAB28 | 45.5465664 | -0.4273525 | 0.30824903 | -1.3863873 | 0.16562867 | 0.64995633 |
| BOD1L1 | 42.0059345 | -0.3299253 | 0.45440654 | -0.7260575 | 0.46780353 | 0.85894565 |
| LINC01182 | 15.7980242 | 0.01714109 | 0.63359739 | 0.02705359 | 0.97841699 | NA |
| CPEB2 | 51.2563569 | -1.3158392 | 0.38079971 | -3.4554628 | 0.00054935 | 0.09487868 |
| AC098829.1 | 13.3470697 | 0.00227007 | 0.54717013 | 0.00414874 | 0.99668979 | NA |
| CC2D2A | 3.73436675 | 1.69868348 | 1.15802813 | 1.46687583 | 0.14240984 | NA |
| FBXL5 | 30.0434678 | 0.1609861 | 0.29328188 | 0.54891254 | 0.58306547 | 0.89282946 |
| FAM200B | 8.34571612 | 0.33932374 | 0.67947057 | 0.49939431 | 0.61750162 | NA |
| PROM1 | 8.36096223 | -0.692472 | 0.99961775 | -0.6927368 | 0.48847477 | NA |
| TAPT1 | 20.3647518 | 0.51765127 | 0.44010299 | 1.17620486 | 0.23951302 | NA |
| TAPT1-AS1 | 31.4069151 | 1.04365761 | 0.55883848 | 1.86754787 | 0.06182512 | 0.51631533 |
| LDB2 | 5.06346087 | -0.2724426 | 0.98058273 | -0.2778374 | 0.78113717 | NA |
| QDPR | 6.87389531 | 1.35929348 | 0.64444697 | 2.10924024 | 0.03492385 | NA |
| LAP3 | 11.4374679 | -0.0020237 | 0.80851243 | -0.002503 | 0.99800287 | NA |
| MED28 | 16.7805404 | -0.7949643 | 0.38732546 | -2.0524453 | 0.0401264 | NA |
| FAM184B | 3.55205328 | -0.2474546 | 0.82188152 | -0.301083 | 0.76335119 | NA |
| DCAF16 | 15.7674198 | -0.3936464 | 0.42167081 | -0.9335397 | 0.3505414 | NA |
| LCORL | 51.6558913 | -0.02323 | 0.32424046 | -0.0716443 | 0.94288497 | 0.99182204 |
| AC024230.1 | 7.69712691 | 0.13717498 | 0.58278382 | 0.23537883 | 0.81391471 | NA |
| SLIT2 | 8.26111826 | 0.39733711 | 0.88291009 | 0.45003122 | 0.65268793 | NA |
| PACRGL | 12.1162795 | 0.88947926 | 0.5207879 | 1.70794916 | 0.08764578 | NA |
| KCNIP4 | 700.444243 | 0.72831506 | 0.60664372 | 1.20056474 | 2.30E-01 | 7.32E-01 |
| ADGRA3 | 19.8240105 | 0.04767588 | 0.4608468 | 0.10345276 | 0.91760364 | NA |
| PPARGC1A | 224.947469 | -0.4678852 | 0.22173749 | -2.1100861 | 0.03485094 | 0.41688714 |
| DHX15 | 39.2993684 | -0.5615175 | 0.30340253 | -1.8507344 | 0.06420778 | 0.52254021 |
| CCDC149 | 7.2551221 | 0.0179458 | 0.54022872 | 0.03321889 | 0.97350003 | NA |
| SEPSECS | 4.9162142 | 0.60840617 | 0.70192072 | 0.86677335 | 0.38606621 | NA |
| ZCCHC4 | 4.60543183 | -0.1884607 | 0.73727092 | -0.2556193 | 0.7982448 | NA |
| ANAPC4 | 17.0839503 | 0.25767505 | 0.3849178 | 0.66942878 | 0.50322199 | NA |
| SEL1L3 | 5.29784202 | -0.1331779 | 0.73861554 | -0.1803074 | 0.85691122 | NA |
| SMIM20 | 4.29874617 | 0.06194797 | 0.69357192 | 0.08931729 | 0.92882975 | NA |
| RBPJ | 52.8400893 | -0.3311311 | 0.25570204 | -1.2949881 | 0.19532434 | 0.69049094 |
| TBC1D19 | 19.5619905 | 0.59679565 | 0.43691674 | 1.36592534 | 0.1719624 | NA |

| | | | | | | |
|---|---|---|---|---|---|---|
| STIM2 | 19.7007103 | -0.2827955 | 0.49271906 | -0.5739488 | 0.56600244 | NA |
| ARAP2 | 19.6042296 | 0.09142121 | 0.62005158 | 0.1474413 | 0.88278371 | NA |
| C4orf19 | 62.2623588 | 0.1750401 | 0.30731987 | 0.56956975 | 0.56896955 | 0.89263108 |
| RELL1 | 5.41944131 | 0.2010416 | 0.7212616 | 0.27873604 | 0.7804474 | NA |
| PGM2 | 5.14250943 | -0.3861084 | 0.65037555 | -0.5936699 | 0.55273293 | NA |
| TBC1D1 | 235.305258 | 0.28881449 | 0.28262667 | 1.02189397 | 0.30683109 | 0.77729327 |
| KLF3 | 22.0782041 | -0.41286 | 0.46134441 | -0.8949063 | 0.37083719 | NA |
| FAM114A1 | 11.7636015 | -0.7701764 | 0.45656765 | -1.6868835 | 0.09162575 | NA |
| KLHL5 | 33.3469435 | 0.13922182 | 0.41590295 | 0.33474592 | 0.73781676 | 0.94658914 |
| WDR19 | 11.4601385 | -0.299874 | 0.51835041 | -0.5785159 | 0.56291587 | NA |
| RFC1 | 17.299784 | -0.2438239 | 0.43786469 | -0.5568477 | 0.57763151 | NA |
| RPL9 | 4.72425792 | -0.6155435 | 0.64081916 | -0.9605573 | 0.33677483 | NA |
| LIAS | 7.30764743 | 0.55702895 | 0.66559017 | 0.8368948 | 0.40265171 | NA |
| UGDH | 8.30000876 | -0.0828717 | 0.52782524 | -0.157006 | 0.87524011 | NA |
| UGDH-AS1 | 8.78931681 | 0.18310843 | 0.58154169 | 0.31486725 | 0.75286245 | NA |
| SMIM14 | 29.6100394 | 0.05127277 | 0.38474369 | 0.13326473 | 0.89398402 | 0.97769824 |
| UBE2K | 83.2069161 | -0.4666926 | 0.27409882 | -1.7026436 | 0.08863479 | 0.56122412 |
| PDS5A | 62.3080506 | -0.0488481 | 0.26406337 | -0.1849862 | 0.8532399 | 0.97168963 |
| N4BP2 | 13.0771523 | -0.3370548 | 0.44534537 | -0.7568391 | 0.44914624 | NA |
| RBM47 | 240.017838 | -0.1271343 | 0.24858407 | -0.5114337 | 0.60904739 | 0.9088434 |
| NSUN7 | 30.7170548 | 0.44301435 | 0.41535737 | 1.06658597 | 0.28615884 | 0.77103639 |
| APBB2 | 102.446014 | -0.0296847 | 0.36153533 | -0.0821074 | 0.93456132 | 0.9907482 |
| UCHL1 | 3.38040975 | -0.4234242 | 0.71035023 | -0.5960781 | 0.55112306 | NA |
| LIMCH1 | 82.4368777 | 0.81142401 | 0.37524785 | 2.16236819 | 0.03058981 | 0.40822098 |
| TMEM33 | 30.5243782 | -0.0049761 | 0.29120395 | -0.0170881 | 0.98636632 | 0.99907833 |
| SLC30A9 | 47.2515038 | -0.2947397 | 0.26629051 | -1.1068353 | 0.26836514 | 0.75632146 |
| ATP8A1 | 26.0661051 | 0.93146855 | 0.39842479 | 2.33787803 | 0.01939358 | 0.33831016 |
| KCTD8 | 14.4271157 | 0.21816117 | 0.70994703 | 0.30729218 | 0.75862098 | NA |
| GUF1 | 9.27551127 | 0.49814038 | 0.55512186 | 0.8973532 | 0.36953048 | NA |
| GNPDA2 | 5.49702031 | 0.83559524 | 0.65721786 | 1.27141286 | 0.20358181 | NA |
| COMMD8 | 4.58887782 | -0.1322497 | 0.64953612 | -0.2036064 | 0.83866112 | NA |
| ATP10D | 65.3005437 | -0.3670336 | 0.37026776 | -0.9912653 | 3.22E-01 | 0.78592775 |
| AC107068.2 | 3.23575266 | 0.25164414 | 0.91807753 | 0.274099 | 0.78400854 | NA |
| NFXL1 | 15.0991189 | 0.18302482 | 0.45342205 | 0.40365223 | 0.68646847 | NA |
| NIPAL1 | 5.98664627 | -0.6705574 | 0.70349239 | -0.9531836 | 0.34049704 | NA |
| TXK | 3.68674475 | -0.3070211 | 0.89783947 | -0.3419555 | 0.73238441 | NA |
| TEC | 13.2314193 | 0.14695146 | 0.43723935 | 0.33608929 | 0.73680354 | NA |
| SLAIN2 | 32.8547328 | -0.3244605 | 0.32903296 | -0.9861033 | 0.32408242 | 0.78622244 |
| FRYL | 70.5897294 | 0.01238506 | 0.30343454 | 0.04081626 | 0.96744238 | 0.99656976 |
| OCIAD1 | 41.1052045 | 0.24170996 | 0.32405011 | 0.74590303 | 0.455726 | 0.85802171 |
| OCIAD2 | 5.83381067 | -0.4033705 | 0.72302865 | -0.5578901 | 0.57691943 | NA |
| DCUN1D4 | 20.3158791 | 0.74046233 | 0.40799045 | 1.81490113 | 0.0695391 | NA |
| SGCB | 3.88528349 | -0.3992415 | 0.78790944 | -0.5067099 | 0.61235837 | NA |
| USP46 | 6.58690924 | -0.1920922 | 0.60219454 | -0.318987 | 0.74973636 | NA |
| SCFD2 | 50.0288746 | -0.5152989 | 0.3230033 | -1.5953363 | 0.11063706 | 0.57860396 |
| AC058822.1 | 23.2235188 | 0.49054332 | 0.589353 | 0.8323421 | 0.40521587 | NA |
| LNX1 | 49.2405344 | 0.5116177 | 0.4113817 | 1.24365692 | 0.21362586 | 0.71284112 |
| CHIC2 | 22.4716495 | 0.38388438 | 0.36130222 | 1.06250214 | 0.28800779 | NA |
| KIT | 19.4893412 | -1.5864022 | 0.4660808 | -3.4037063 | 0.00066478 | NA |
| TMEM165 | 40.9603116 | -0.1642252 | 0.29450359 | -0.5576338 | 0.57709449 | 0.89263108 |
| CLOCK | 26.9202942 | 0.15728786 | 0.35252632 | 0.44617337 | 0.65547202 | 0.92276785 |
| EXOC1 | 30.5106264 | -0.190162 | 0.40564987 | -0.4687835 | 0.63922439 | 0.92121485 |
| CEP135 | 9.3338659 | -0.4578791 | 0.51339808 | -0.8918599 | 0.37246805 | NA |
| AASDH | 13.703312 | 0.0389059 | 0.44942044 | 0.08656904 | 0.93101407 | NA |

| | | | | | | |
|---|---|---|---|---|---|---|
| PPAT | 3.55651943 | -0.0320396 | 1.01602034 | -0.0315344 | 0.97484339 | NA |
| PAICS | 7.00066912 | 0.39363329 | 0.57856498 | 0.68036142 | 0.49627564 | NA |
| SRP72 | 16.2518025 | -0.624123 | 0.41042693 | -1.5206678 | 0.12834322 | NA |
| REST | 25.7728353 | 0.07248849 | 0.38754601 | 0.18704486 | 0.85162546 | NA |
| NOA1 | 3.89116283 | -0.7224813 | 0.82028473 | -0.8807689 | 0.37844291 | NA |
| POLR2B | 22.3440372 | -0.3545909 | 0.42658764 | -0.8312264 | 0.40584575 | NA |
| IGFBP7 | 117.172223 | -0.0008067 | 0.3693888 | -0.0021838 | 0.99825758 | 0.99960611 |
| AC104806.2 | 3.58280978 | 0.36059673 | 0.99098839 | 0.36387584 | 0.71595073 | NA |
| CENPC | 20.3438581 | 0.28784405 | 0.45696674 | 0.62990152 | 0.52875902 | NA |
| UBA6 | 31.4417857 | -0.7036799 | 0.39229889 | -1.7937341 | 0.07285562 | 0.53207981 |
| UBA6-AS1 | 30.5626951 | -0.8580597 | 0.47966 | -1.7888916 | 0.07363228 | 0.53207981 |
| YTHDC1 | 44.3018824 | -0.3890207 | 0.28380863 | -1.370715 | 0.17046383 | 0.66182328 |
| UGT2B7 | 29.8184251 | 0.09206774 | 0.70552175 | 0.13049596 | 0.89617405 | 0.97769824 |
| UTP3 | 5.28155349 | -0.2138198 | 0.62312041 | -0.3431436 | 0.73149044 | NA |
| RUFY3 | 66.6905695 | -0.1492953 | 0.32408149 | -0.4606722 | 0.64503377 | 0.92153493 |
| GRSF1 | 16.0271298 | -0.1898527 | 0.43638764 | -0.4350552 | 0.66352239 | NA |
| MOB1B | 78.3955515 | -0.2773827 | 0.43083728 | -0.6438224 | 0.51969063 | 0.87214537 |
| SLC4A4 | 15.2741701 | 0.30571731 | 0.9812017 | 0.31157438 | 0.75536401 | NA |
| COX18 | 3.87190164 | -0.33111 | 0.79182917 | -0.4181583 | 0.67583135 | NA |
| ANKRD17 | 163.16303 | -0.3585479 | 0.22125077 | -1.6205499 | 0.10511421 | 0.5772605 |
| AC053527.1 | 11.9256953 | 1.32104682 | 0.44613098 | 2.96111875 | 0.00306524 | NA |
| MTHFD2L | 13.6336762 | -0.9347244 | 0.44808004 | -2.0860657 | 0.03697267 | NA |
| PARM1 | 3.30728585 | -0.144441 | 1.32870112 | -0.1087084 | 0.91343378 | NA |
| RCHY1 | 15.4452441 | -0.2927229 | 0.41885075 | -0.6988717 | 0.48463221 | NA |
| THAP6 | 5.90548583 | 0.25873165 | 0.66355752 | 0.38991593 | 0.69659871 | NA |
| CDKL2 | 3.52936308 | 0.38838488 | 0.75287409 | 0.51586963 | 0.60594547 | NA |
| G3BP2 | 20.9870139 | -0.3508084 | 0.35300619 | -0.993774 | 0.32033292 | NA |
| USO1 | 42.769858 | -0.2090785 | 0.27345499 | -0.7645812 | 0.44452099 | 0.84899161 |
| SDAD1 | 10.4463188 | -0.3500235 | 0.60456994 | -0.5789627 | 0.56261433 | NA |
| NUP54 | 22.7142924 | -0.0242706 | 0.36955224 | -0.0656758 | 0.94763597 | NA |
| SCARB2 | 38.4276686 | -0.0121039 | 0.28514803 | -0.0424478 | 0.96614172 | 0.99656976 |
| SHROOM3 | 102.244591 | -0.4713082 | 0.47933255 | -0.9832593 | 0.32547986 | 0.78758454 |
| 11-Sep | 25.3667013 | -0.7545641 | 0.47613773 | -1.5847601 | 0.11302085 | NA |
| CCNI | 87.4253375 | 0.49976445 | 0.28830018 | 1.73348644 | 0.08300925 | 0.55234224 |
| CCNG2 | 22.8488172 | 0.56021097 | 0.3626191 | 1.54490196 | 0.12236998 | NA |
| CNOT6L | 51.8051989 | 0.19095588 | 0.43876311 | 0.43521407 | 0.66340705 | 0.92276785 |
| MRPL1 | 32.4248574 | -0.6881311 | 0.4176962 | -1.6474439 | 0.09946683 | 0.57091823 |
| FRAS1 | 53.0402273 | -0.1834147 | 0.43237894 | -0.424199 | 0.67142071 | 0.92614389 |
| BMP2K | 21.5021861 | -0.5353066 | 0.44357543 | -1.2067996 | 0.22750933 | NA |
| PAQR3 | 3.62702346 | -0.1129837 | 0.71747699 | -0.1574736 | 0.8748716 | NA |
| RASGEF1B | 112.381873 | -0.7506045 | 0.48478028 | -1.5483396 | 0.12154056 | 0.58553957 |
| HNRNPD | 45.416445 | -0.5561884 | 0.29672235 | -1.8744406 | 0.06086972 | 0.51625203 |
| HNRNPDL | 46.5741964 | -0.3281491 | 0.31108828 | -1.0548423 | 0.29149746 | 0.77286135 |
| ENOPH1 | 7.98146935 | -0.2859728 | 0.56092432 | -0.5098242 | 0.61017461 | NA |
| TMEM150C | 15.7011602 | -0.4660302 | 0.46125669 | -1.010349 | 0.3123281 | NA |
| SEC31A | 65.7454507 | -0.7173308 | 0.24197532 | -2.9644791 | 0.00303196 | 0.19010422 |
| THAP9 | 3.4357489 | -0.0366178 | 0.83215726 | -0.0440035 | 0.96490162 | NA |
| LIN54 | 7.17612543 | 0.32956013 | 0.60175679 | 0.54766334 | 0.58392309 | NA |
| COPS4 | 10.3417074 | -0.371825 | 0.47143769 | -0.7887045 | 0.43028476 | NA |
| COQ2 | 4.39352055 | -0.7595249 | 0.98799148 | -0.7687566 | 0.44203783 | NA |
| HPSE | 12.5423047 | -0.7156187 | 0.95179688 | -0.7518607 | 0.45213484 | NA |
| HELQ | 9.94177868 | -0.2120119 | 0.49095832 | -0.4318328 | 0.66586293 | NA |
| MRPS18C | 3.966493 | 0.21965105 | 0.73151252 | 0.3002697 | 0.76397144 | NA |
| ABRAXAS1 | 5.64730902 | 0.30208426 | 0.75242367 | 0.4014816 | 0.68806558 | NA |

| | | | | | | |
|---|---|---|---|---|---|---|
| GPAT3 | 54.6301951 | -0.5148106 | 0.49655819 | -1.0367578 | 0.29984875 | 0.77521265 |
| CDS1 | 16.1018804 | -0.2705458 | 0.40790604 | -0.6632552 | 0.50716715 | NA |
| WDFY3 | 99.1569864 | 0.09031885 | 0.26798588 | 0.33702838 | 0.73609551 | 0.94590222 |
| ARHGAP24 | 59.9185345 | 0.17864881 | 0.49009945 | 0.36451542 | 0.71547316 | 0.93629694 |
| MAPK10 | 46.3638617 | 0.59750215 | 0.58023572 | 1.0297576 | 0.30312381 | 0.77521265 |
| PTPN13 | 37.5071552 | 0.25839951 | 0.46399224 | 0.55690481 | 0.57759247 | 0.89263108 |
| AC093827.5 | 5.22104222 | 0.30386098 | 0.80566441 | 0.37715577 | 0.70605785 | NA |
| AFF1 | 166.335894 | -0.4074727 | 0.25912068 | -1.5725211 | 0.11582975 | 0.57860396 |
| KLHL8 | 24.6946929 | 0.16707106 | 0.42890215 | 0.38953187 | 0.69688274 | NA |
| HSD17B11 | 8.88343458 | 0.20589385 | 0.5929352 | 0.34724511 | 0.72840718 | NA |
| NUDT9 | 20.8036004 | -0.8607811 | 0.37878338 | -2.2724891 | 0.02305699 | NA |
| SPP1 | 781.961217 | -0.9145651 | 0.50730996 | -1.8027737 | 7.14E-02 | 5.32E-01 |
| PKD2 | 54.800204 | -0.038297 | 0.30932776 | -0.1238073 | 0.90146786 | 0.98003205 |
| PPM1K | 9.68606026 | -0.0036266 | 0.48933095 | -0.0074113 | 0.9940867 | NA |
| HERC6 | 5.45948339 | -1.6101487 | 1.07450883 | -1.4984974 | 0.13400407 | NA |
| HERC5 | 9.04321253 | -0.2961065 | 0.8937858 | -0.3312947 | 0.7404219 | NA |
| PYURF | 6.46222323 | 0.53992666 | 0.63482751 | 0.85050925 | 0.39504202 | NA |
| HERC3 | 18.8127484 | 0.39231846 | 0.45900949 | 0.85470664 | 0.39271357 | NA |
| FAM13A-AS1 | 7.04812352 | 1.37048978 | 0.76078213 | 1.80142215 | 0.07163637 | NA |
| FAM13A | 45.5004277 | 0.15508717 | 0.44011044 | 0.35238241 | 0.72455149 | 0.93920735 |
| GPRIN3 | 38.6425232 | 0.37997976 | 0.6312053 | 0.60199076 | 0.54718029 | 0.88410237 |
| CCSER1 | 94.9106069 | 0.32428261 | 0.50141173 | 0.64673917 | 0.51780076 | 0.87186213 |
| SMARCAD1 | 25.6950817 | 0.11410811 | 0.35601977 | 0.32051061 | 0.74858129 | NA |
| PDLIM5 | 95.927584 | -0.2892451 | 0.27289083 | -1.0599298 | 0.28917652 | 0.7725847 |
| BMPR1B | 356.933607 | -0.6927685 | 0.2515499 | -2.7540004 | 0.00588717 | 0.22794137 |
| UNC5C | 3.45885602 | 2.37310068 | 1.16605937 | 2.03514567 | 0.04183622 | NA |
| STPG2 | 76.2937985 | 0.90164909 | 0.5089853 | 1.7714639 | 0.07648359 | 0.54291145 |
| RAP1GDS1 | 27.7971412 | 0.18807712 | 0.40813242 | 0.46082377 | 0.64492505 | 0.92153493 |
| TSPAN5 | 70.8912601 | -0.624827 | 0.5166719 | -1.2093302 | 0.226536 | 0.73022292 |
| EIF4E | 24.2161973 | -0.6279008 | 0.42659455 | -1.4718913 | 0.14105024 | NA |
| METAP1 | 17.765081 | 0.17806751 | 0.39219207 | 0.45403139 | 0.64980624 | NA |
| ADH5 | 15.218344 | 0.35782201 | 0.38146578 | 0.93801865 | 0.34823483 | NA |
| AP002026.1 | 6.16550051 | 0.20083516 | 0.799546 | 0.2511865 | 0.80166992 | NA |
| TRMT10A | 3.12504584 | -0.5843728 | 0.81807441 | -0.7143272 | 0.47502488 | NA |
| LAMTOR3 | 8.89819818 | -0.3823471 | 0.66226965 | -0.5773285 | 0.56371757 | NA |
| DNAJB14 | 36.2070918 | -0.0063852 | 0.29850014 | -0.0213908 | 0.98293392 | 0.99902646 |
| H2AFZ | 14.5532424 | -0.2694756 | 0.56557108 | -0.4764664 | 0.63374214 | NA |
| DDIT4L | 4.44509085 | -1.0457195 | 1.10734241 | -0.9443506 | 0.34499052 | NA |
| PPP3CA | 263.10667 | -0.3943914 | 0.27695987 | -1.4240021 | 0.15444588 | 0.63864841 |
| AP002075.1 | 2.48738383 | -0.3009003 | 1.41920692 | -0.2120201 | 0.83209138 | NA |
| NFKB1 | 28.9117247 | -0.0393985 | 0.45064711 | -0.0874265 | 0.93033249 | 0.98921679 |
| MANBA | 15.1436507 | 0.82260347 | 0.50654288 | 1.62395626 | 0.10438514 | NA |
| UBE2D3 | 74.364418 | -0.0719857 | 0.20888264 | -0.3446227 | 0.73037803 | 0.94309578 |
| CISD2 | 8.24729666 | 0.06107787 | 0.50107538 | 0.12189357 | 0.90298331 | NA |
| TET2 | 43.5501585 | -0.22378 | 0.31099187 | -0.7195686 | 0.47179067 | 0.86206794 |
| PPA2 | 42.7551884 | 0.54298831 | 0.2510473 | 2.16289244 | 0.03054945 | 0.40822098 |
| ARHGEF38 | 44.7594263 | -0.2888298 | 0.47123444 | -0.6129216 | 0.53992815 | 0.88409095 |
| INTS12 | 13.3465517 | -0.0102261 | 0.4286288 | -0.0238578 | 0.98096602 | NA |
| GSTCD | 15.6715533 | 0.08447119 | 0.44639489 | 0.18922975 | 0.84991275 | NA |
| NPNT | 46.8698526 | -0.1408027 | 0.30370288 | -0.4636198 | 0.64292016 | 0.92153493 |
| TBCK | 71.4640392 | 0.21138639 | 0.37887034 | 0.55793861 | 0.57688631 | 0.89263108 |
| AIMP1 | 11.308367 | 0.64774726 | 0.49388341 | 1.3115388 | 0.18967579 | NA |
| PAPSS1 | 20.7366088 | 0.44147617 | 0.41369689 | 1.06714888 | 0.28590461 | NA |
| SGMS2 | 42.0183237 | -0.5658874 | 0.3921996 | -1.4428556 | 0.14906114 | 0.63 |

| | | | | | | |
|---|---|---|---|---|---|---|
| AC096564.1 | 7.77939389 | 0.36009919 | 0.91452196 | 0.39375674 | 0.69376064 | NA |
| HADH | 5.75359155 | 1.60440082 | 0.67343228 | 2.38242339 | 0.01719911 | NA |
| RPL34-AS1 | 4.72674462 | 0.982588 | 0.86151375 | 1.14053664 | 0.2540628 | NA |
| RPL34 | 50.6146291 | -0.0807634 | 0.38958261 | -0.2073076 | 0.83576962 | 0.97154892 |
| OSTC | 8.04946522 | 0.42026432 | 0.5533621 | 0.75947435 | 0.44756885 | NA |
| SEC24B | 55.3628873 | -0.2117742 | 0.31379016 | -0.6748912 | 0.49974491 | 0.87036626 |
| CASP6 | 6.76139941 | 0.82966953 | 0.61313963 | 1.35314941 | 0.1760079 | NA |
| PLA2G12A | 8.79109515 | 0.4538543 | 0.59164352 | 0.7671077 | 0.44301749 | NA |
| GAR1 | 3.10136084 | -0.7728412 | 0.92251207 | -0.8377573 | 0.40216703 | NA |
| EGF | 26.4022971 | 0.0759513 | 1.17703308 | 0.06452775 | 0.94855001 | 0.99344789 |
| ELOVL6 | 22.8816109 | 0.42898233 | 0.44509505 | 0.96379937 | 0.33514652 | NA |
| ENPEP | 7.37219881 | 0.47440047 | 0.86829709 | 0.54635732 | 0.58482034 | NA |
| FAM241A | 10.6628795 | 0.09299872 | 0.42537151 | 0.21862941 | 0.82693874 | NA |
| AP1AR | 15.5149861 | -0.0575093 | 0.41904152 | -0.1372402 | 0.89084096 | NA |
| ALPK1 | 29.2744396 | 0.84131016 | 0.42454638 | 1.98166843 | 0.04751636 | 0.45661231 |
| ZGRF1 | 4.34417616 | -0.1107236 | 0.84294166 | -0.1313538 | 0.89549546 | NA |
| LARP7 | 20.3675808 | -0.2108508 | 0.3903742 | -0.5401249 | 0.5891109 | NA |
| ANK2 | 11.0250468 | -1.6918847 | 0.56117585 | -3.0148922 | 0.00257071 | NA |
| CAMK2D | 77.2633418 | -0.5746736 | 0.35976605 | -1.5973537 | 0.11018689 | 0.57860396 |
| ARSJ | 107.368185 | 0.42082719 | 0.31156904 | 1.35067075 | 0.17680093 | 0.6676327 |
| UGT8 | 59.6862161 | 0.51517437 | 0.36206882 | 1.42286311 | 0.15477584 | 0.63864841 |
| SNHG8 | 6.44554311 | 0.08540414 | 0.61087726 | 0.13980573 | 0.88881349 | NA |
| METTL14 | 11.0846245 | -0.1470185 | 0.45781431 | -0.3211313 | 0.74811089 | NA |
| SEC24D | 15.2887397 | -2.031273 | 0.63439588 | -3.2019013 | 0.00136524 | NA |
| USP53 | 70.8006424 | -0.0551606 | 0.32553139 | -0.1694478 | 0.86544446 | 0.97217819 |
| C4orf3 | 20.4527063 | 0.0752189 | 0.3750912 | 0.20053497 | 0.84106221 | NA |
| PDE5A | 20.1568417 | 0.16082019 | 0.50512458 | 0.31837728 | 0.75019877 | NA |
| AC108866.1 | 14.9512547 | -0.1830204 | 0.52584009 | -0.3480533 | 0.72780016 | NA |
| PRDM5 | 10.6424242 | -0.2850252 | 0.52360339 | -0.5443533 | 0.58619836 | NA |
| ANXA5 | 55.9478032 | 0.26464057 | 0.30983743 | 0.85412719 | 0.39303452 | 0.8258317 |
| EXOSC9 | 2.45289421 | 0.54264662 | 1.01288924 | 0.53574132 | 0.59213734 | NA |
| BBS7 | 5.15067158 | 0.83466945 | 0.74074434 | 1.12679827 | 0.25982778 | NA |
| KIAA1109 | 70.4961838 | 0.10979606 | 0.28585349 | 0.38409908 | 0.70090502 | 0.9303056 |
| SPATA5 | 38.4366996 | -0.0829703 | 0.35348647 | -0.2347198 | 0.81442624 | 0.96845695 |
| SPRY1 | 5.88542689 | 0.60962549 | 0.57787132 | 1.05495024 | 0.29144808 | NA |
| LINC01091 | 67.0690424 | 0.52507058 | 0.33779845 | 1.55439013 | 0.12009139 | 0.58383225 |
| ANKRD50 | 5.24161611 | -0.0047632 | 0.63456367 | -0.0075062 | 0.99401098 | NA |
| INTU | 8.55652271 | 0.75867947 | 0.63921719 | 1.18688839 | 0.23527165 | NA |
| HSPA4L | 19.6651072 | 0.37319237 | 0.43225476 | 0.86336208 | 0.38793842 | NA |
| MFSD8 | 14.0343585 | -0.0028761 | 0.49034221 | -0.0058655 | 0.99532002 | NA |
| ABHD18 | 43.4897215 | -0.280078 | 0.28790302 | -0.9728208 | 0.33064238 | 0.79360119 |
| LARP1B | 48.9071866 | -0.6237751 | 0.29644399 | -2.104192 | 0.03536171 | 0.41688714 |
| PGRMC2 | 25.6702686 | -1.3009295 | 0.58904873 | -2.2085262 | 0.02720761 | NA |
| AC108062.1 | 13.4263706 | -0.8395698 | 0.4389737 | -1.9125742 | 0.05580258 | NA |
| AC078850.2 | 8.68608633 | 2.16845455 | 0.82133722 | 2.64015132 | 0.0082869 | NA |
| JADE1 | 104.90353 | -0.2145721 | 0.30220627 | -0.7100188 | 0.47769248 | 0.86206794 |
| SCLT1 | 32.1464841 | -0.3861127 | 0.28594003 | -1.3503276 | 0.17691093 | 0.6676327 |
| C4orf33 | 3.55291623 | 1.05330887 | 0.86587002 | 1.21647458 | 0.22380419 | NA |
| AC096711.2 | 12.8482753 | -0.0225058 | 0.60700479 | -0.0370768 | 0.97042378 | NA |
| AC109927.1 | 6.5121095 | 0.3375307 | 0.75109273 | 0.44938619 | 0.65315309 | NA |
| NOCT | 3.03178248 | -2.0852813 | 1.08161326 | -1.9279362 | 0.05386307 | NA |
| ELF2 | 152.763006 | 0.53859434 | 0.2427071 | 2.21911247 | 0.02647907 | 0.39506514 |
| NDUFC1 | 17.4780254 | 0.02911467 | 0.49395829 | 0.05894155 | 0.95299866 | NA |
| NAA15 | 30.55138 | -0.0150527 | 0.31881222 | -0.0472149 | 0.96234199 | 0.99656976 |

| | | | | | | |
|---|---|---|---|---|---|---|
| AC097376.2 | 36.4141996 | -0.1851367 | 0.3946774 | -0.4690836 | 0.63900989 | 0.92121485 |
| RAB33B | 3.341692 | -0.9416764 | 0.92830466 | -1.0144044 | 0.3103898 | NA |
| SETD7 | 7.84411024 | 0.09496387 | 0.53415709 | 0.17778266 | 0.85889367 | NA |
| MGST2 | 21.6516334 | 0.7473633 | 0.58546539 | 1.27652859 | 0.20176873 | NA |
| MAML3 | 73.105005 | 0.43601629 | 0.45918904 | 0.9495355 | 0.34234832 | 0.8003972 |
| SCOC | 27.2259815 | 0.29281787 | 0.40151294 | 0.72928625 | 0.46582658 | 0.85894565 |
| ELMOD2 | 8.04496019 | -0.3232334 | 0.53399955 | -0.6053065 | 0.54497533 | NA |
| TBC1D9 | 48.14767 | -0.1394445 | 0.33935772 | -0.4109072 | 0.68114055 | 0.9267216 |
| RNF150 | 142.632342 | 0.03188178 | 0.39132649 | 0.08147106 | 0.93506734 | 0.9907482 |
| ZNF330 | 9.32063245 | -0.3134881 | 0.56079573 | -0.5590059 | 0.57615772 | NA |
| IL15 | 7.0299353 | -0.7990786 | 0.87204827 | -0.9163238 | 0.35949708 | NA |
| INPP4B | 181.767335 | -0.3455205 | 0.31791194 | -1.0868433 | 0.27710607 | 0.76520827 |
| AC104596.1 | 3.51715005 | -0.2350343 | 1.1232239 | -0.2092497 | 0.83425332 | NA |
| USP38 | 10.849153 | -0.4921376 | 0.48141769 | -1.0222673 | 0.3066544 | NA |
| GAB1 | 51.7909467 | 0.39320651 | 0.35920791 | 1.09464882 | 0.27367052 | 0.76165444 |
| SMARCA5 | 29.0053036 | -0.1973225 | 0.30416976 | -0.6487247 | 0.5165163 | 0.87158522 |
| ANAPC10 | 27.0702884 | 0.01974692 | 0.3324332 | 0.05940116 | 0.95263259 | 0.9947482 |
| ABCE1 | 9.14308877 | 0.00401123 | 0.54544574 | 0.00735404 | 0.99413238 | NA |
| OTUD4 | 33.3575747 | 0.41252773 | 0.36517571 | 1.12966913 | 0.25861567 | 0.75272108 |
| SMAD1 | 42.0936583 | 0.26452945 | 0.46656159 | 0.56697648 | 0.57073016 | 0.89263108 |
| MMAA | 4.33989964 | 0.43994819 | 0.71623472 | 0.61425141 | 0.5390492 | NA |
| ZNF827 | 46.0767254 | 0.13904236 | 0.27570565 | 0.50431451 | 0.61404038 | 0.91163569 |
| LSM6 | 5.61452662 | 0.00101282 | 0.60366723 | 0.00167777 | 0.99866133 | NA |
| SLC10A7 | 26.0302769 | 0.00343189 | 0.39686949 | 0.0086474 | 0.99310046 | 0.99923383 |
| TMEM184C | 13.4356935 | 0.31005246 | 0.45301116 | 0.68442567 | 0.49370641 | NA |
| PRMT9 | 8.93290087 | -0.3442141 | 0.52691911 | -0.653258 | 0.51358998 | NA |
| ARHGAP10 | 6.28072845 | -0.7249958 | 0.83455799 | -0.8687183 | 0.38500122 | NA |
| NR3C2 | 363.117911 | 0.15019433 | 0.23042071 | 0.65182651 | 0.5145131 | 0.87158522 |
| AC069272.1 | 4.18962527 | 1.39175757 | 0.77748858 | 1.79006819 | 0.07344295 | NA |
| DCLK2 | 69.5647768 | -0.8798418 | 0.36190798 | -2.4311203 | 0.01505222 | 0.30435125 |
| LRBA | 223.028941 | -0.0280463 | 0.23031152 | -0.1217754 | 0.90307691 | 0.98003205 |
| RPS3A | 32.6531325 | -0.063504 | 0.40537335 | -0.1566557 | 0.87551623 | 0.9751578 |
| SH3D19 | 62.0144245 | -0.3920192 | 0.39158345 | -1.0011127 | 0.31677231 | 0.780966 |
| FAM160A1 | 85.2833229 | -0.5738893 | 0.34720221 | -1.6528963 | 0.09835196 | 0.56941734 |
| GATB | 6.59009588 | 0.49730489 | 0.59592745 | 0.83450576 | 0.40399604 | NA |
| FBXW7 | 68.6507407 | 0.13255117 | 0.26853845 | 0.49360219 | 0.62158716 | 0.91493825 |
| ARFIP1 | 44.7639508 | 0.16290197 | 0.35224067 | 0.46247349 | 0.64374181 | 0.92153493 |
| FHDC1 | 3.51549904 | 0.71307993 | 0.77762868 | 0.91699285 | 0.35914639 | NA |
| TRIM2 | 107.176113 | -0.3809082 | 0.2780238 | -1.3700563 | 0.17066934 | 0.66182328 |
| TMEM131L | 17.3648842 | -0.3005398 | 0.41624535 | -0.7220257 | 0.4702787 | NA |
| PLRG1 | 7.08132548 | 0.02848694 | 0.58696875 | 0.0485323 | 0.96129202 | NA |
| MAP9 | 5.25389407 | 0.08346858 | 0.75261084 | 0.11090536 | 0.91169139 | NA |
| GUCY1A1 | 7.76619936 | -0.3227966 | 0.6969142 | -0.4631798 | 6.43E-01 | NA |
| CTSO | 4.04504279 | 0.30956785 | 0.72418114 | 0.42747296 | 0.66903487 | NA |
| PDGFC | 23.5347298 | 0.01875761 | 0.604308 | 0.03103982 | 0.97523778 | NA |
| ETFDH | 16.1133768 | -0.2117809 | 0.4608676 | -0.4595265 | 0.64585614 | NA |
| PPID | 9.8140036 | 0.4987155 | 0.50754118 | 0.9826109 | 0.32579898 | NA |
| FNIP2 | 140.607668 | -0.6240048 | 0.33347643 | -1.8712112 | 0.06131581 | 0.51625203 |
| C4orf45 | 5.28122915 | 0.35347487 | 0.79193056 | 0.44634579 | 0.65534749 | NA |
| RAPGEF2 | 124.110168 | 0.40245052 | 0.26773865 | 1.5031469 | 0.13280117 | 0.60574082 |
| NAF1 | 7.71593806 | 0.34016567 | 0.51824266 | 0.656383 | 0.51157773 | NA |
| TMA16 | 5.26483547 | -0.5645789 | 0.65895567 | -0.8567782 | 0.39156746 | NA |
| 1-Mar | 3.29845148 | -0.8463351 | 0.86115782 | -0.9827875 | 0.32571205 | NA |
| TMEM192 | 12.5627184 | 0.44075358 | 0.46382497 | 0.95025841 | 0.34198097 | NA |

| | | | | | | |
|---|---|---|---|---|---|---|
| KLHL2 | 24.0473476 | -0.1153342 | 0.43769148 | -0.2635058 | 0.79216076 | NA |
| MSMO1 | 17.1214875 | -0.9932307 | 0.57297627 | -1.7334587 | 0.08301417 | NA |
| DDX60 | 13.2669899 | 0.00799879 | 1.81324201 | 0.00441132 | 0.99648029 | NA |
| DDX60L | 10.6550426 | -0.8649835 | 0.91320145 | -0.947199 | 0.34353737 | NA |
| PALLD | 60.9393312 | 1.3985684 | 0.64207712 | 2.17819379 | 0.02939161 | 0.40645706 |
| CBR4 | 63.1170978 | 0.44302587 | 0.33125258 | 1.3374262 | 0.18108356 | 0.67407534 |
| SH3RF1 | 109.034634 | 0.07623111 | 0.33865061 | 0.22510253 | 0.82189951 | 0.96845695 |
| NEK1 | 35.4048111 | 0.17811749 | 0.32688173 | 0.54489888 | 0.58582307 | 0.89397471 |
| CLCN3 | 41.8463998 | 0.14719175 | 0.30383574 | 0.48444513 | 0.62807 | 0.91751762 |
| HPF1 | 6.75167448 | 0.48112705 | 0.5575646 | 0.86290818 | 0.38818795 | NA |
| MFAP3L | 4.68615995 | 0.14511855 | 0.75147793 | 0.19311086 | 0.84687215 | NA |
| AADAT | 4.58192795 | -0.1861547 | 0.95315652 | -0.1953034 | 0.84515541 | NA |
| AC105285.1 | 3.01960753 | 1.13286089 | 1.11256249 | 1.01824472 | 0.30856167 | NA |
| GALNT7 | 68.2530429 | 0.05127022 | 0.23406243 | 0.21904504 | 0.82661496 | 0.96845695 |
| HMGB2 | 9.79845234 | 0.43429467 | 0.47390218 | 0.9164226 | 0.35944528 | NA |
| FBXO8 | 8.25545509 | 1.36144871 | 0.55695901 | 2.44443248 | 0.01450802 | NA |
| CEP44 | 11.4392139 | 0.65301895 | 0.49991345 | 1.30626401 | 0.19146279 | NA |
| WDR17 | 4.94646783 | -0.0449357 | 0.70667964 | -0.0635871 | 0.949299 | NA |
| ASB5 | 3.55404234 | -2.5860123 | 1.06854984 | -2.4201139 | 0.01551565 | NA |
| SPCS3 | 26.953585 | -0.2204932 | 0.41093007 | -0.5365711 | 0.59156393 | 0.89660782 |
| AC078881.1 | 8.37066724 | 1.76971021 | 0.76996454 | 2.29843079 | 0.02153728 | NA |
| TENM3 | 5.40604372 | 1.39705635 | 0.89025261 | 1.56928083 | 0.11658252 | NA |
| DCTD | 9.10737289 | -0.2451831 | 0.51243942 | -0.4784626 | 0.63232095 | NA |
| WWC2 | 54.0032145 | -0.3100405 | 0.30160688 | -1.0279624 | 0.30396751 | 0.77561116 |
| CDKN2AIP | 4.74563705 | -0.5337995 | 0.76063754 | -0.7017791 | 0.48281695 | NA |
| RWDD4 | 9.66688323 | -0.2290985 | 0.52874551 | -0.4332869 | 0.66480634 | NA |
| TRAPPC11 | 18.6320231 | 0.18789647 | 0.39294863 | 0.47817056 | 0.63252881 | NA |
| STOX2 | 97.7193069 | 0.21386762 | 0.48527206 | 0.44071695 | 0.65941792 | 0.92276785 |
| IRF2 | 15.7103892 | 0.63294191 | 0.47190515 | 1.34124816 | 0.17983989 | NA |
| CASP3 | 7.05659983 | -0.1639954 | 0.59066337 | -0.2776462 | 0.78128395 | NA |
| PRIMPOL | 7.50630155 | 1.91810048 | 0.60902177 | 3.14947768 | 0.00163563 | NA |
| ACSL1 | 48.0402187 | -0.3204977 | 0.44706251 | -0.7168969 | 0.47343769 | 0.86206794 |
| SLC25A4 | 17.740016 | 0.82018655 | 0.49691208 | 1.65056672 | 0.09882708 | NA |
| CFAP97 | 15.2419949 | 0.65175564 | 0.39502059 | 1.64992829 | 0.0989576 | NA |
| SNX25 | 25.1110857 | -0.0258693 | 0.34219169 | -0.0755988 | 0.93973832 | NA |
| AC112722.1 | 2.75057077 | 0.79020989 | 0.98480009 | 0.80240639 | 0.42231792 | NA |
| UFSP2 | 7.35584256 | 0.10173711 | 0.56160811 | 0.18115321 | 0.85624732 | NA |
| SORBS2 | 54.7689361 | 1.81987488 | 0.39931313 | 4.55751327 | 5.18E-06 | 0.0138155 |
| TLR3 | 4.159327 | 0.80933906 | 0.77170714 | 1.04876451 | 0.29428651 | NA |
| FAM149A | 7.0117632 | -0.3018958 | 0.67260793 | -0.4488436 | 0.65354446 | NA |
| CYP4V2 | 5.90381339 | -0.1101656 | 0.69499289 | -0.1585133 | 0.87405235 | NA |
| KLKB1 | 4.47117974 | -0.3021956 | 0.93628011 | -0.322762 | 0.7468755 | NA |
| FAT1 | 9.52034184 | 0.04100044 | 0.56386329 | 0.07271345 | 0.94203415 | NA |
| FRG1-DT | 10.9167557 | 1.32251201 | 0.70669279 | 1.87141009 | 0.06128826 | NA |
| FRG1 | 5.90888731 | 0.7748323 | 0.60606783 | 1.27845805 | 0.20108797 | NA |
| CCDC127 | 2.21621376 | 0.98349825 | 0.96043495 | 1.02401339 | 0.30582895 | NA |
| SDHA | 37.066027 | 0.04089805 | 0.31144856 | 0.13131559 | 0.89552566 | 0.97769824 |
| PDCD6 | 10.9597997 | -0.7879692 | 0.54714402 | -1.4401495 | 0.1498251 | NA |
| EXOC3 | 13.153037 | -0.7432226 | 0.52096202 | -1.4266349 | 0.15368517 | NA |
| SLC9A3 | 7.57619902 | 0.43303754 | 0.7212168 | 0.60042631 | 0.54822216 | NA |
| TPPP | 3.27937555 | 2.19501558 | 0.95714247 | 2.29330078 | 0.02183069 | NA |
| ZDHHC11B | 8.66254007 | 1.31105918 | 0.75203232 | 1.74335484 | 0.08127165 | NA |
| BRD9 | 9.63066253 | -0.5686679 | 0.5093411 | -1.1164775 | 0.26421779 | NA |
| SLC12A7 | 10.1568527 | -1.1034289 | 0.49243982 | -2.2407387 | 0.02504301 | NA |

| | | | | | | |
|---|---|---|---|---|---|---|
| CLPTM1L | 11.7690245 | -0.738625 | 0.47469046 | -1.556014 | 0.11970476 | NA |
| LPCAT1 | 3.80684188 | -0.3386551 | 0.68207241 | -0.496509 | 0.61953536 | NA |
| MRPL36 | 4.91458292 | 0.90797353 | 0.66587794 | 1.36357352 | 0.17270183 | NA |
| NDUFS6 | 21.5336635 | -0.1597841 | 0.45002713 | -0.3550545 | 0.72254878 | NA |
| ADAMTS16 | 3.98683968 | -0.719968 | 0.93346493 | -0.7712856 | 0.44053767 | NA |
| ICE1 | 26.9850508 | -0.3599144 | 0.34129246 | -1.054563 | 0.29162524 | 0.77286135 |
| MED10 | 8.4711884 | -1.5194024 | 0.6686365 | -2.2723892 | 0.02306301 | NA |
| UBE2QL1 | 12.6523699 | 0.89049546 | 0.61210685 | 1.45480395 | 0.14572354 | NA |
| NSUN2 | 14.5661716 | -0.0339892 | 0.40993942 | -0.0829128 | 0.93392086 | NA |
| SRD5A1 | 4.20975509 | 0.39491942 | 0.76721491 | 0.51474419 | 0.60673179 | NA |
| TENT4A | 14.6152368 | -0.2844777 | 0.47991752 | -0.5927637 | 0.5533393 | NA |
| MTRR | 10.1209505 | -0.3019383 | 0.50430152 | -0.5987258 | 0.54935577 | NA |
| MIR4458HG | 3.64133452 | 0.10012878 | 0.80978372 | 0.12364879 | 0.90159336 | NA |
| SEMA5A | 77.6703274 | 0.22929593 | 0.59977626 | 0.38230244 | 0.70223704 | 0.9303056 |
| FAM173B | 6.18274287 | 0.8687063 | 0.59856869 | 1.45130594 | 0.14669469 | NA |
| CCT5 | 14.9361122 | -0.2030064 | 0.47168854 | -0.4303824 | 0.6669175 | NA |
| CMBL | 5.80830453 | 1.10001599 | 0.91553561 | 1.20149994 | 0.22955733 | NA |
| 6-Mar | 61.7336322 | -0.0110275 | 0.22835242 | -0.0482915 | 0.9614839 | 0.99656976 |
| ANKRD33B | 6.38394842 | -0.7950652 | 1.06675092 | -0.7453148 | 0.45608145 | NA |
| DAP | 58.4780759 | 0.23603617 | 0.25420925 | 0.92851134 | 0.35314239 | 0.8003972 |
| DNAH5 | 22.5955575 | 0.03270222 | 0.70467194 | 0.04640772 | 9.63E-01 | NA |
| TRIO | 119.192313 | -0.2429276 | 0.28900615 | -0.840562 | 0.40059335 | 0.8269015 |
| OTULIN | 10.2571835 | 0.00131741 | 0.5349201 | 0.00246282 | 0.99803496 | NA |
| ANKH | 15.735109 | 0.28173856 | 0.47879679 | 0.58843035 | 0.55624347 | NA |
| ZNF622 | 4.62885813 | -0.9841448 | 0.68769558 | -1.4310763 | 0.15240835 | NA |
| RETREG1 | 72.4539881 | 0.17180233 | 0.42827753 | 0.40114719 | 0.68831176 | 0.92736198 |
| MYO10 | 167.153898 | -0.6036465 | 0.35495166 | -1.7006443 | 0.0890098 | 0.56122412 |
| LINC02241 | 8.33965876 | 0.77180892 | 0.54966915 | 1.40413361 | 0.16027907 | NA |
| CDH12 | 63.8602918 | -1.155993 | 0.50839899 | -2.2737908 | 0.02297857 | 0.35751239 |
| C5orf17 | 13.9746167 | 0.8288485 | 1.01121441 | 0.81965653 | 0.41241194 | NA |
| CDH6 | 9.22702094 | -0.153667 | 0.80332943 | -0.1912877 | 0.84830019 | NA |
| DROSHA | 20.4553642 | 0.07133359 | 0.34870616 | 0.20456648 | 0.83791085 | NA |
| C5orf22 | 5.84406884 | -0.9253678 | 0.6464917 | -1.4313684 | 0.15232466 | NA |
| PDZD2 | 347.446794 | 0.41749071 | 0.4113795 | 1.01485539 | 0.31017476 | 0.778534 |
| GOLPH3 | 38.653279 | -0.6095046 | 0.316053 | -1.9284886 | 0.05379439 | 0.48835787 |
| MTMR12 | 18.3086567 | 0.37977562 | 0.36787286 | 1.03235565 | 0.30190554 | NA |
| ZFR | 58.4071041 | -0.1090354 | 0.27407259 | -0.3978342 | 0.69075243 | 0.92830727 |
| SUB1 | 40.5656186 | -0.1495722 | 0.30171502 | -0.4957399 | 0.62007793 | 0.91493825 |
| NPR3 | 3.81882464 | 1.3839482 | 0.94919318 | 1.45802586 | 0.14483342 | NA |
| TARS | 8.6047267 | -0.7252649 | 0.56268243 | -1.2889417 | 0.19741834 | NA |
| C1QTNF3-AN | 22.4668007 | 0.10852948 | 0.51651894 | 0.21011714 | 0.83357625 | NA |
| RAI14 | 35.1640242 | -0.2923215 | 0.40981815 | -0.7132958 | 0.47566276 | 0.86206794 |
| RAD1 | 7.36678019 | 1.07431851 | 0.65447278 | 1.64150221 | 0.10069321 | NA |
| BRIX1 | 5.31418946 | -0.434183 | 0.67777303 | -0.6406023 | 0.52178111 | NA |
| DNAJC21 | 14.9035499 | 0.61216143 | 0.40554289 | 1.50948626 | 0.13117456 | NA |
| PRLR | 41.6499222 | 0.80140467 | 0.52952069 | 1.51345299 | 0.13016463 | 0.60105434 |
| SPEF2 | 6.4004384 | 1.33502039 | 0.77858083 | 1.71468438 | 0.08640311 | NA |
| LMBRD2 | 10.3996455 | 0.53192113 | 0.49380998 | 1.07717774 | 0.28140087 | NA |
| NADK2 | 8.11670155 | 0.04661961 | 0.58070759 | 0.08028069 | 0.93601401 | NA |
| RANBP3L | 12.3253906 | 0.82083906 | 1.03255675 | 0.79495781 | 0.42663804 | NA |
| NIPBL | 137.256249 | -0.0780892 | 0.24135701 | -0.3235424 | 0.74628447 | 0.94858421 |
| CPLANE1 | 38.0441505 | 0.16249893 | 0.33285431 | 0.48819837 | 0.62540934 | 0.91664883 |
| NUP155 | 7.96498025 | 0.30034564 | 0.54555676 | 0.55053051 | 0.58195556 | NA |
| WDR70 | 68.5051484 | 0.13427362 | 0.29389114 | 0.45688214 | 0.64775576 | 0.9218151 |

| | | | | | | |
|---|---|---|---|---|---|---|
| LIFR | 31.3925027 | 0.30291287 | 0.44098351 | 0.68690294 | 0.4921439 | 0.86587479 |
| OSMR-AS1 | 19.7538403 | 0.80472478 | 0.37792602 | 2.12931826 | 0.03322794 | NA |
| OSMR | 27.0382146 | -0.0706321 | 0.44667785 | -0.1581276 | 8.74E-01 | 9.75E-01 |
| RICTOR | 89.4786795 | -0.0665368 | 0.28016068 | -0.2374953 | 0.81227258 | 0.96845695 |
| FYB1 | 17.7473964 | -1.1527881 | 0.97383336 | -1.1837632 | 2.37E-01 | NA |
| TTC33 | 14.4012397 | 0.39175588 | 0.40593748 | 0.96506458 | 0.33451247 | NA |
| PRKAA1 | 20.2877469 | 0.28977895 | 0.39516361 | 0.73331386 | 0.46336702 | NA |
| RPL37 | 31.8705117 | 0.03308157 | 0.39450815 | 0.08385523 | 0.93317154 | 0.99070598 |
| PLCXD3 | 7.34903558 | -1.568776 | 0.84844325 | -1.8490052 | 0.06445707 | NA |
| OXCT1 | 105.617592 | -0.9002704 | 0.39677269 | -2.2689828 | 0.02326937 | 0.35899398 |
| C5orf51 | 5.05057235 | 0.35562408 | 0.74169719 | 0.4794734 | 0.63160189 | NA |
| FBXO4 | 1.93308748 | -0.6017469 | 1.12995681 | -0.5325397 | 0.59435225 | NA |
| GHR | 5.28245868 | 0.36735414 | 0.70870975 | 0.51834215 | 0.60421957 | NA |
| SELENOP | 50.6548828 | 0.94665954 | 0.42693891 | 2.2173185 | 0.02660133 | 0.39506514 |
| ZNF131 | 27.3823351 | -0.5725888 | 0.37521422 | -1.5260316 | 0.12700199 | 0.59256732 |
| HMGCS1 | 21.67067 | -1.2738527 | 0.59792017 | -2.1304728 | 0.0331326 | NA |
| TMEM267 | 11.5267389 | 0.13915281 | 0.43033507 | 0.32335921 | 0.74642322 | NA |
| PAIP1 | 14.7378331 | -0.8380464 | 0.44934117 | -1.8650559 | 0.06217357 | NA |
| NNT-AS1 | 21.4867674 | 0.58504287 | 0.39669909 | 1.47477746 | 0.14027238 | NA |
| NNT | 69.771428 | 0.21812805 | 0.2971467 | 0.73407532 | 0.46290284 | 0.85894565 |
| MRPS30 | 7.56590008 | 0.4545287 | 0.56666331 | 0.80211422 | 0.42248689 | NA |
| PARP8 | 19.3322707 | -0.1902709 | 0.43667844 | -0.4357231 | 0.66303767 | NA |
| ITGA1 | 8.30232803 | 0.44229005 | 0.66341206 | 0.6666898 | 0.50497029 | NA |
| ITGA2 | 32.2488516 | -0.63069 | 0.43733268 | -1.4421287 | 0.14926608 | 0.63 |
| MOCS2 | 5.78008747 | -0.063101 | 0.72542572 | -0.0869847 | 0.93068365 | NA |
| NDUFS4 | 41.1575223 | 0.03244812 | 0.32564712 | 0.09964196 | 0.92062858 | 0.98745437 |
| ARL15 | 949.017502 | 0.11151615 | 0.26999353 | 0.41303266 | 0.67958269 | 0.9267216 |
| SNX18 | 24.8153419 | 0.90022114 | 0.44034737 | 2.04434317 | 0.04091966 | NA |
| DHX29 | 16.8785606 | -0.1235644 | 0.40455052 | -0.3054362 | 0.76003399 | NA |
| MTREX | 34.5152668 | -0.0264431 | 0.30307251 | -0.0872502 | 0.93047263 | 0.98921679 |
| PLPP1 | 80.6349793 | 0.10669503 | 0.46141974 | 0.23123204 | 0.81713454 | 0.96845695 |
| SLC38A9 | 21.353535 | 0.61021201 | 0.36963785 | 1.65083745 | 0.09877177 | NA |
| IL6ST | 63.899324 | -0.0373994 | 0.276812 | -0.1351076 | 0.89252682 | 0.97769824 |
| AC008892.1 | 3.92961671 | 0.52315833 | 0.7982212 | 0.65540521 | 0.5122069 | NA |
| MAP3K1 | 44.003413 | 0.69121709 | 0.37116758 | 1.86227768 | 0.06256396 | 0.51631533 |
| MIER3 | 8.95504611 | -0.0777291 | 0.59866645 | -0.129837 | 0.89669537 | NA |
| GPBP1 | 80.964494 | -0.1818274 | 0.25465418 | -0.714017 | 0.4752167 | 0.86206794 |
| PDE4D | 844.122062 | 0.02815715 | 0.36499306 | 0.07714435 | 0.93850871 | 0.99100968 |
| ELOVL7 | 41.4273906 | -0.6648028 | 0.3975374 | -1.6723026 | 0.09446467 | 0.56264113 |
| ERCC8 | 12.4083559 | 0.20046817 | 0.53839662 | 0.37234292 | 0.70963754 | NA |
| NDUFAF2 | 29.2762336 | -0.4695722 | 0.43078596 | -1.0900359 | 0.27569733 | 0.76410819 |
| SMIM15 | 7.65917244 | 0.36258934 | 0.53738676 | 0.67472696 | 0.49984925 | NA |
| ZSWIM6 | 181.50795 | -0.7748211 | 0.42359051 | -1.8291748 | 0.06737343 | 0.52400942 |
| KIF2A | 20.3067473 | -0.0377421 | 0.34274919 | -0.1101159 | 0.91231744 | NA |
| DIMT1 | 1.99590837 | 0.46934943 | 1.05466476 | 0.44502239 | 0.65630357 | NA |
| IPO11 | 10.1218369 | 0.29094548 | 0.55194874 | 0.5271241 | 0.59810741 | NA |
| RNF180 | 21.9409288 | 0.3182017 | 0.35964358 | 0.88476958 | 0.37628092 | NA |
| SREK1IP1 | 13.2723014 | -0.1109242 | 0.43886353 | -0.2527534 | 0.80045878 | NA |
| CWC27 | 36.0202754 | 0.29461334 | 0.31154156 | 0.94566305 | 0.34432047 | 0.8003972 |
| PPWD1 | 10.6515199 | 0.34114104 | 0.47680076 | 0.71547922 | 0.47431298 | NA |
| TRIM23 | 6.97183105 | 0.71129866 | 0.56048211 | 1.26908361 | 0.20441125 | NA |
| TRAPPC13 | 11.49998 | 0.62097367 | 0.50541543 | 1.22864013 | 0.21920676 | NA |
| NLN | 7.848645 | -0.5044122 | 0.58578189 | -0.8610922 | 0.38918728 | NA |
| ERBIN | 114.768311 | -0.0295563 | 0.22017506 | -0.1342399 | 0.89321287 | 0.97769824 |

| | | | | | | |
|---|---|---|---|---|---|---|
| SREK1 | 45.0639617 | -0.8463543 | 0.27513865 | -3.076101 | 0.00209727 | 0.15993164 |
| MAST4 | 96.3731986 | -0.6597825 | 0.36679869 | -1.798759 | 0.0720568 | 0.53207981 |
| PIK3R1 | 54.0011904 | -0.4432123 | 0.41662459 | -1.0638169 | 0.28741165 | 0.77250927 |
| SLC30A5 | 13.9990865 | -0.167311 | 0.44504352 | -0.3759429 | 0.70695933 | NA |
| MRPS36 | 13.1256889 | 0.30738511 | 0.43748793 | 0.70261393 | 0.48229638 | NA |
| CDK7 | 6.82072005 | -0.8992719 | 0.59695504 | -1.5064315 | 0.13195645 | NA |
| CCDC125 | 9.81509001 | -0.0844541 | 0.48609148 | -0.1737411 | 0.86206894 | NA |
| AK6 | 6.75454493 | 0.26639975 | 0.58624197 | 0.45441945 | 0.64952696 | NA |
| RAD17 | 15.2292702 | 0.38364266 | 0.49443975 | 0.77591386 | 0.43779985 | NA |
| MARVELD2 | 10.3074547 | 0.65370957 | 0.47487863 | 1.37658241 | 0.16864139 | NA |
| OCLN | 33.8202514 | 0.28089396 | 0.41909577 | 0.67023811 | 0.50270601 | 0.87094108 |
| GTF2H2C | 12.2798277 | 0.30860544 | 0.48190094 | 0.64039185 | 0.52191788 | NA |
| NAIP | 28.3291821 | 0.59431603 | 0.41768792 | 1.42287103 | 0.15477355 | 0.63864841 |
| GTF2H2 | 9.17521276 | -0.2809661 | 0.58990974 | -0.4762866 | 0.63387019 | NA |
| BDP1 | 51.5224537 | -0.441017 | 0.28695527 | -1.5368842 | 0.12432167 | 0.58553957 |
| MCCC2 | 21.96181 | 0.00440921 | 0.35506544 | 0.01241801 | 0.99009212 | NA |
| MRPS27 | 18.0471466 | -0.1921274 | 0.44018035 | -0.4364743 | 0.66249262 | NA |
| PTCD2 | 2.90900399 | 1.18555748 | 0.90675111 | 1.30747839 | 0.19105029 | NA |
| TNPO1 | 46.3858191 | -0.1494668 | 0.26492867 | -0.5641777 | 0.57263323 | 0.89263108 |
| FCHO2 | 41.7425721 | 0.41594664 | 0.36221206 | 1.14835116 | 0.25082363 | 0.74329941 |
| BTF3 | 23.5013068 | 0.82318429 | 0.3522978 | 2.33661486 | 0.01945922 | NA |
| ANKRA2 | 6.36407205 | 0.51706076 | 0.62223867 | 0.83096853 | 0.40599141 | NA |
| ARHGEF28 | 108.92263 | 0.2752626 | 0.38431604 | 0.71624021 | 0.47384304 | 0.86206794 |
| ENC1 | 3.73283485 | 0.51000804 | 1.00162113 | 0.50918259 | 0.61062425 | NA |
| HEXB | 9.0331481 | -0.0706465 | 0.49628447 | -0.1423508 | 0.88680293 | NA |
| GFM2 | 9.06447631 | -0.3049114 | 0.54708955 | -0.5573337 | 0.57729948 | NA |
| NSA2 | 10.9145991 | -0.3043515 | 0.52286958 | -0.5820792 | 0.56051336 | NA |
| FAM169A | 25.5919495 | -0.1491103 | 0.36631331 | -0.4070568 | 0.68396628 | NA |
| AC116337.3 | 2.99248225 | 1.09919978 | 0.93299944 | 1.17813552 | 0.23874258 | NA |
| GCNT4 | 3.02516155 | 0.00254548 | 0.9185312 | 0.00277125 | 0.99778887 | NA |
| HMGCR | 13.0530767 | -1.0947929 | 0.55697203 | -1.9656156 | 0.04934303 | NA |
| COL4A3BP | 54.1429657 | 0.13346043 | 0.33965466 | 0.39292978 | 0.69437134 | 0.9303056 |
| POLK | 33.3488179 | 0.0188729 | 0.33756467 | 0.055909 | 0.9554143 | 0.99609405 |
| ANKDD1B | 10.1450326 | 0.76980716 | 0.85321458 | 0.90224332 | 0.36692763 | NA |
| POC5 | 4.65303665 | -0.2044102 | 0.68586981 | -0.2980306 | 0.76567978 | NA |
| IQGAP2 | 235.855055 | 0.00259821 | 0.25868759 | 0.01004381 | 0.99198633 | 0.99909657 |
| F2RL1 | 7.27420143 | -0.71949 | 0.64895407 | -1.1086916 | 0.26756323 | NA |
| ZBED3 | 4.92424315 | 0.18394976 | 0.6251642 | 0.29424231 | 0.76857275 | NA |
| WDR41 | 17.5353674 | -0.5944424 | 0.43193069 | -1.3762449 | 0.16874584 | NA |
| TBCA | 60.1060705 | -0.0292658 | 0.24085658 | -0.1215074 | 0.90328919 | 0.98003205 |
| AP3B1 | 79.082809 | 0.43586315 | 0.32096556 | 1.35797483 | 0.17447167 | 0.6651652 |
| SCAMP1 | 30.7211399 | 0.73981168 | 0.43581818 | 1.69752366 | 0.0895977 | 0.56122412 |
| LHFPL2 | 2.43522919 | -0.7281492 | 1.00629498 | -0.7235942 | 0.46931489 | NA |
| ARSB | 8.66134535 | -0.2125525 | 0.68510432 | -0.3102484 | 0.75637207 | NA |
| DMGDH | 17.184892 | 0.21886068 | 0.5751933 | 0.38049936 | 0.70357477 | NA |
| BHMT2 | 12.9889179 | 0.37521574 | 0.49722533 | 0.75461913 | 0.45047754 | NA |
| BHMT | 4.36271413 | 1.1523886 | 1.08379991 | 1.06328539 | 0.28765255 | NA |
| JMY | 33.0979383 | 0.56800681 | 0.42374626 | 1.34044089 | 0.18010205 | 0.67135806 |
| HOMER1 | 20.9819967 | -0.5563339 | 0.58050935 | -0.9583547 | 0.33788392 | NA |
| TENT2 | 75.7687546 | -0.2559491 | 0.26934229 | -0.9502743 | 0.34197288 | 0.8003972 |
| CMYA5 | 11.7103813 | -0.4929726 | 0.60079698 | -0.8205311 | 0.4119134 | NA |
| MTX3 | 7.12676807 | 0.31274296 | 0.55957971 | 0.55888903 | 0.57623745 | NA |
| THBS4 | 3.8074506 | 1.40268879 | 0.77198503 | 1.81698962 | 0.0692187 | NA |
| SERINC5 | 46.1578874 | 0.54986954 | 0.52120201 | 1.05500273 | 0.29142408 | 0.77286135 |

| | | | | | | |
|---|---|---|---|---|---|---|
| AC010260.1 | 3.28745223 | 0.88164444 | 0.87580425 | 1.00666838 | 0.31409416 | NA |
| ZFYVE16 | 67.6492528 | 0.44059564 | 0.27989986 | 1.57411885 | 0.11545998 | 0.57860396 |
| FAM151B | 8.53857633 | 0.6388062 | 0.53958166 | 1.18389161 | 0.23645597 | NA |
| DHFR | 181.916753 | -1.1686143 | 0.34277974 | -3.4092281 | 0.00065147 | 0.09555626 |
| MSH3 | 28.8825424 | -0.0110473 | 0.41289769 | -0.0267557 | 0.97865462 | 0.99902646 |
| RASGRF2 | 5.56001823 | -0.0424221 | 1.07044949 | -0.0396302 | 0.96838796 | NA |
| CKMT2-AS1 | 13.5513321 | 0.56761069 | 0.41235702 | 1.37650303 | 0.16866595 | NA |
| CKMT2 | 9.33257578 | 1.65883946 | 0.59274928 | 2.7985516 | 0.00513324 | NA |
| ZCCHC9 | 3.91187017 | -0.5303896 | 0.73449856 | -0.7221111 | 0.47022615 | NA |
| SSBP2 | 105.280835 | 0.00729156 | 0.30555572 | 0.02386328 | 0.98096167 | 0.99902646 |
| ATG10 | 34.5423852 | -0.1077899 | 0.37788846 | -0.2852426 | 0.77545828 | 0.96105928 |
| RPS23 | 89.374662 | 0.05014304 | 0.37696119 | 0.1330191 | 0.89417828 | 0.97769824 |
| ATP6AP1L | 6.72330191 | -0.1631071 | 0.74920669 | -0.2177064 | 0.82765784 | NA |
| TMEM167A | 14.1693729 | -0.5211096 | 0.46679183 | -1.1163639 | 0.26426638 | NA |
| XRCC4 | 20.3667553 | -0.0480704 | 0.57149909 | -0.0841128 | 0.93296677 | NA |
| COX7C | 58.8956681 | 0.55244768 | 0.34041215 | 1.62287888 | 0.1046153 | 0.5772605 |
| RASA1 | 51.1583342 | 0.26044454 | 0.2612154 | 0.99704898 | 0.31874073 | 0.78335494 |
| CCNH | 41.0310638 | 0.71881758 | 0.39287138 | 1.82965116 | 0.06730212 | 0.52400942 |
| TMEM161B | 24.9101901 | -0.8663629 | 0.3775816 | -2.2945051 | 0.0217615 | NA |
| TMEM161B-/ | 27.4091513 | 0.13985406 | 0.36367107 | 0.38456197 | 0.70056198 | 0.9303056 |
| CETN3 | 4.48193507 | 0.85157788 | 0.83399996 | 1.02107663 | 0.30721813 | NA |
| LYSMD3 | 7.86118622 | -0.8283154 | 0.67143065 | -1.2336574 | 0.21733059 | NA |
| ADGRV1 | 21.4831415 | -0.3485239 | 0.36389147 | -0.9577687 | 0.33817938 | NA |
| ARRDC3 | 37.6757843 | -0.0379482 | 0.43596319 | -0.0870445 | 0.93063614 | 0.98921679 |
| ARRDC3-AS1 | 2.60485553 | 0.25096085 | 0.83920318 | 0.29904659 | 0.76490449 | NA |
| FAM172A | 82.9866475 | 0.03621385 | 0.21702022 | 0.16686855 | 0.86747348 | 0.97217819 |
| KIAA0825 | 16.5875331 | 0.7898875 | 0.4363228 | 1.81032828 | 0.07024489 | NA |
| SLF1 | 5.49487192 | 0.04706859 | 0.75998252 | 0.06193378 | 0.95061557 | NA |
| MCTP1 | 4.58978196 | -1.1247717 | 0.77627012 | -1.4489437 | 0.1473533 | NA |
| TTC37 | 37.6142057 | -0.5123768 | 0.3212667 | -1.5948644 | 0.11074257 | 0.57860396 |
| ARSK | 5.05263042 | -0.4550672 | 0.70054203 | -0.649593 | 0.51595515 | NA |
| RHOBTB3 | 108.885642 | -0.0530207 | 0.29617072 | -0.1790206 | 0.85792149 | 0.97168963 |
| GLRX | 19.0670689 | 0.63011173 | 0.43141988 | 1.46055331 | 0.14413807 | NA |
| ELL2 | 38.38724 | 0.0705312 | 0.31537171 | 0.22364466 | 0.82303381 | 0.96845695 |
| AC104123.1 | 8.04152098 | 0.62085274 | 0.84948451 | 0.73085822 | 0.46486575 | NA |
| CAST | 53.6909003 | 0.16415147 | 0.37040101 | 0.44317229 | 0.65764112 | 0.92276785 |
| ERAP1 | 13.2945813 | 0.57207381 | 0.58268623 | 0.98178708 | 0.32620475 | NA |
| AC009126.1 | 2.62507377 | 1.1262107 | 1.02468644 | 1.09907836 | 0.27173389 | NA |
| ERAP2 | 7.15941245 | 0.9888546 | 0.7574677 | 1.30547428 | 0.1917314 | NA |
| LNPEP | 71.2218035 | 0.38937268 | 0.31014876 | 1.25543844 | 0.20931964 | 0.70681427 |
| LIX1-AS1 | 8.61447577 | 0.46329658 | 0.59623676 | 0.77703458 | 0.43713837 | NA |
| RIOK2 | 3.24582692 | 0.06453507 | 0.93537932 | 0.06899348 | 0.94499481 | NA |
| CHD1 | 40.9106583 | 0.16108009 | 0.30021553 | 0.53654817 | 0.59157976 | 0.89660782 |
| LINC02062 | 7.99391436 | 0.35820881 | 0.83310194 | 0.42996996 | 0.66721749 | NA |
| FAM174A | 7.05212581 | 0.42338099 | 0.56510528 | 0.74920727 | 0.45373229 | NA |
| SLCO4C1 | 9.10775382 | 0.75699244 | 0.70510595 | 1.0735868 | 0.28300792 | NA |
| PAM | 38.9666487 | 0.30740114 | 0.4145995 | 0.74144117 | 0.45842599 | 0.85802171 |
| GIN1 | 4.69876642 | 1.04229592 | 0.74317308 | 1.40249418 | 0.16076773 | NA |
| PPIP5K2 | 23.8285442 | 0.06592713 | 0.36079244 | 0.1827287 | 0.8550109 | NA |
| NUDT12 | 5.22137149 | 0.91512181 | 0.61788706 | 1.48105029 | 0.13859317 | NA |
| EFNA5 | 158.161542 | 0.38440839 | 0.52956009 | 0.72590137 | 4.68E-01 | 0.85894565 |
| FBXL17 | 161.549161 | 0.41041441 | 0.21151924 | 1.94031712 | 0.05234116 | 0.48171918 |
| FER | 84.4455866 | -0.1209025 | 0.29904763 | -0.4042917 | 0.68599819 | 0.92726927 |
| AC008467.1 | 4.81437652 | -1.9439193 | 0.79634989 | -2.4410367 | 0.01464517 | NA |

| | | | | | | |
|---|---|---|---|---|---|---|
| PJA2 | 38.9164804 | -0.2057583 | 0.33332119 | -0.6172972 | 0.53703869 | 0.88254911 |
| MAN2A1 | 202.137862 | -0.9852493 | 0.36010979 | -2.7359693 | 0.00621968 | 0.23056019 |
| TMEM232 | 22.6827675 | 1.38948141 | 0.48115323 | 2.88781481 | 0.00387928 | NA |
| SLC25A46 | 18.3323007 | -0.6077995 | 0.37068143 | -1.6396816 | 0.10107138 | NA |
| WDR36 | 10.5699507 | -0.5157073 | 0.47463935 | -1.0865245 | 0.27724701 | NA |
| NREP | 6.21941396 | -0.0337648 | 0.84926902 | -0.0397575 | 0.96828647 | NA |
| EPB41L4A | 85.6242638 | -0.1159028 | 0.30518875 | -0.3797743 | 0.70411299 | 0.9303056 |
| EPB41L4A-A: | 3.29751404 | 0.15023773 | 0.7582443 | 0.19813895 | 0.84293635 | NA |
| APC | 17.0295879 | 0.04750538 | 0.39255865 | 0.12101472 | 0.90367937 | NA |
| SRP19 | 13.1505568 | -0.2087819 | 0.43778986 | -0.4768997 | 0.63343354 | NA |
| REEP5 | 64.5743173 | 0.35339001 | 0.30417097 | 1.16181372 | 0.24531114 | 0.73462702 |
| DCP2 | 26.3188968 | 0.46637167 | 0.34103617 | 1.3675138 | 0.17146432 | 0.66182328 |
| MCC | 19.7495505 | -0.5026761 | 0.50605468 | -0.9933237 | 0.32055224 | NA |
| YTHDC2 | 34.4181729 | 0.04786608 | 0.33661967 | 0.14219632 | 0.88692494 | 0.97769824 |
| PGGT1B | 13.3530821 | 0.01601198 | 0.4236532 | 0.03779502 | 0.96985112 | NA |
| CCDC112 | 3.17139343 | -0.3236719 | 0.87586034 | -0.3695474 | 0.71171979 | NA |
| FEM1C | 4.26792275 | -0.0901506 | 0.70235729 | -0.1283543 | 0.89786861 | NA |
| TMED7-TICA | 3.39919036 | -0.3538739 | 0.91057685 | -0.3886261 | 0.69755276 | NA |
| ATG12 | 11.7256696 | -0.2363671 | 0.5167884 | -0.457377 | 0.64740011 | NA |
| AP3S1 | 16.1386114 | -0.4201581 | 0.45114566 | -0.9313137 | 0.35169132 | NA |
| COMMD10 | 65.4225354 | 0.05767627 | 0.22972275 | 0.25106904 | 0.80176073 | 0.96742897 |
| SEMA6A | 6.85351651 | 0.06254145 | 0.59464323 | 0.10517474 | 0.91623715 | NA |
| SEMA6A-AS1 | 8.46069059 | -0.0709145 | 0.58160336 | -0.1219294 | 0.90295495 | NA |
| DTWD2 | 21.9131749 | 0.02107103 | 0.45525012 | 0.04628451 | 0.96308349 | NA |
| AC008629.1 | 3.96449304 | 1.23312933 | 0.85446856 | 1.44315354 | 0.14897722 | NA |
| DMXL1 | 145.465009 | -0.4109074 | 0.22553244 | -1.8219434 | 0.06846357 | 0.52508412 |
| TNFAIP8 | 70.055818 | -0.4448043 | 0.38248206 | -1.1629416 | 0.24485321 | 0.73428453 |
| HSD17B4 | 39.8955738 | 0.6546453 | 0.34806945 | 1.88078928 | 0.06000059 | 0.51625203 |
| SRFBP1 | 12.7044546 | -0.1518954 | 0.5134075 | -0.2958573 | 0.76733907 | NA |
| SNX2 | 24.0600895 | -0.2646878 | 0.37533153 | -0.7052106 | 0.48067921 | NA |
| SNX24 | 49.6879166 | -0.1253816 | 0.27457443 | -0.4566398 | 0.64792999 | 0.9218151 |
| PPIC | 9.24788418 | -0.0760328 | 0.50101751 | -0.1517567 | 0.8793788 | NA |
| CEP120 | 8.83936427 | 0.07900056 | 0.51166307 | 0.15439957 | 0.87729469 | NA |
| CSNK1G3 | 26.6821597 | -0.0748204 | 0.34594693 | -0.216277 | 0.82877184 | 0.96974662 |
| ZNF608 | 36.6435329 | -0.3522908 | 0.45473467 | -0.7747173 | 0.43850671 | 0.84462749 |
| GRAMD2B | 69.1405215 | 0.65119536 | 0.40962325 | 1.58974217 | 0.11189293 | 0.57860396 |
| ALDH7A1 | 10.7649176 | 0.64368253 | 0.48414283 | 1.32953023 | 0.1836731 | NA |
| PHAX | 6.44124723 | -0.8001922 | 0.69006156 | -1.1595954 | 0.24621357 | NA |
| 3-Mar | 6.81513933 | -1.1228572 | 0.96977942 | -1.157848 | 0.24692607 | NA |
| C5orf63 | 6.44253459 | 0.52890855 | 0.64798027 | 0.81624176 | 0.41436188 | NA |
| PRRC1 | 14.5766464 | -0.5879202 | 0.64273005 | -0.9147233 | 0.3603369 | NA |
| LINC01184 | 13.7572288 | 0.1440687 | 0.496871 | 0.28995191 | 0.77185303 | NA |
| SLC12A2 | 9.06539775 | -1.7171291 | 0.62523545 | -2.7463719 | 0.00602584 | NA |
| SLC27A6 | 3.46096958 | -0.2901999 | 2.21357946 | -0.1310999 | 0.89569632 | NA |
| ISOC1 | 4.46283812 | -0.1856395 | 0.70077701 | -0.2649053 | 0.79108241 | NA |
| HINT1 | 59.0263708 | 0.18652516 | 0.38492257 | 0.48457841 | 0.62797544 | 0.91751762 |
| LYRM7 | 7.86532406 | 0.0246776 | 0.65250483 | 0.0378198 | 0.96983136 | NA |
| CDC42SE2 | 138.049561 | -0.2867742 | 0.31049079 | -0.9236159 | 0.35568631 | 0.80177915 |
| P4HA2 | 2.78002964 | -0.815223 | 0.87290955 | -0.9339146 | 0.35034793 | NA |
| SLC22A4 | 2.57041471 | 1.42226473 | 1.04262207 | 1.36412298 | 0.17252886 | NA |
| SLC22A5 | 14.6725259 | 1.00888336 | 0.56795759 | 1.77633572 | 0.0756776 | NA |
| AC116366.3 | 19.4659664 | -0.0673215 | 0.58488501 | -0.1151021 | 0.90836419 | NA |
| IRF1 | 5.15753636 | 0.07645529 | 0.67134812 | 0.11388323 | 0.90933036 | NA |
| KIF3A | 13.9801456 | 0.30807174 | 0.42575465 | 0.72358984 | 0.46931758 | NA |

| | | | | | | |
|---|---|---|---|---|---|---|
| 8-Sep | 8.95539948 | -0.1225017 | 0.51611461 | -0.2373536 | 0.81238245 | NA |
| SOWAHA | 3.27781726 | 0.79301088 | 0.90155476 | 0.87960367 | 0.37907405 | NA |
| UQCRQ | 20.6590011 | 0.32604253 | 0.41868828 | 0.7787238 | 0.43614243 | NA |
| AFF4 | 87.9803472 | -0.6442387 | 0.29414477 | -2.1902097 | 0.02850903 | 0.40259583 |
| ZCCHC10 | 10.656666 | -0.4732797 | 0.51274403 | -0.9230331 | 0.35598994 | NA |
| HSPA4 | 22.9287933 | -0.5354215 | 0.39278166 | -1.3631531 | 0.17283426 | NA |
| FSTL4 | 14.7836927 | 1.07158922 | 0.70259735 | 1.52518255 | 0.12721357 | NA |
| C5orf15 | 14.6731254 | 0.11289913 | 0.4577322 | 0.24664888 | 0.80517997 | NA |
| VDAC1 | 43.3137207 | 0.54262479 | 0.30289075 | 1.79148683 | 0.07321521 | 0.53207981 |
| TCF7 | 4.91787231 | 0.95990785 | 0.76640219 | 1.25248578 | 0.21039291 | NA |
| SKP1 | 41.6746039 | -0.0175474 | 0.41711994 | -0.0420679 | 0.96644457 | 0.99656976 |
| CDKL3 | 5.6314746 | -0.6516619 | 0.85401925 | -0.763053 | 0.44543179 | NA |
| UBE2B | 44.1664854 | -0.0174145 | 0.30815444 | -0.0565123 | 0.95493369 | 0.99598203 |
| CDKN2AIPNL | 2.36386282 | 0.39096372 | 0.99970987 | 0.39107718 | 0.69574019 | NA |
| JADE2 | 10.7614651 | 1.19904699 | 0.65203955 | 1.83891758 | 0.06592731 | NA |
| SAR1B | 15.0501338 | -0.3847333 | 0.41262794 | -0.9323975 | 0.35113112 | NA |
| SEC24A | 19.1033258 | -0.7673938 | 0.40430399 | -1.8980615 | 0.05768798 | NA |
| CAMLG | 9.27533872 | 0.41063031 | 0.50288731 | 0.81654537 | 0.41418829 | NA |
| DDX46 | 35.0728708 | 0.09075391 | 0.30063214 | 0.30187695 | 0.76274587 | 0.95478238 |
| C5orf24 | 21.2889163 | -0.178168 | 0.41467063 | -0.4296616 | 0.66744182 | NA |
| TXNDC15 | 6.55649533 | -1.0041819 | 0.58042586 | -1.7300778 | 0.08361638 | NA |
| PCBD2 | 29.965888 | -0.1875915 | 0.37386943 | -0.5017568 | 0.61583858 | 0.91318974 |
| H2AFY | 42.9101341 | -0.756007 | 0.43031025 | -1.7568883 | 0.07893685 | 0.54552914 |
| CXCL14 | 8.21437463 | 0.82139369 | 0.77572737 | 1.05886903 | 0.28965943 | NA |
| SMAD5 | 24.2375734 | -0.2614183 | 0.33895924 | -0.7712382 | 0.44056576 | NA |
| KLHL3 | 40.590304 | 1.41214676 | 0.51291531 | 2.75317728 | 0.00590199 | 0.22794137 |
| HNRNPA0 | 22.1849809 | -0.8053169 | 0.46398827 | -1.7356407 | 0.08262739 | NA |
| AC106791.1 | 4.03014793 | 1.39924287 | 0.80135965 | 1.746086 | 0.08079601 | NA |
| FAM13B | 59.5851295 | -0.6964765 | 0.28387362 | -2.4534737 | 0.01414839 | 0.30435125 |
| BRD8 | 12.4976731 | 0.60716124 | 0.48612206 | 1.24898927 | 0.211669 | NA |
| CDC23 | 4.67158274 | 0.94023145 | 0.66021418 | 1.42413095 | 0.15440857 | NA |
| FAM53C | 4.24519429 | -0.8043508 | 0.7328198 | -1.0976106 | 0.27237455 | NA |
| KDM3B | 30.1624743 | 0.25189255 | 0.35264325 | 0.7142985 | 0.47504262 | 0.86206794 |
| EGR1 | 12.3166515 | -2.3645958 | 0.94513339 | -2.5018646 | 0.01235411 | NA |
| ETF1 | 23.3633913 | -0.9893547 | 0.34385047 | -2.8772817 | 0.00401117 | NA |
| HSPA9 | 42.1425044 | -0.4102475 | 0.33367581 | -1.2294793 | 0.21889216 | 0.72126316 |
| CTNNA1 | 96.8397133 | -0.3361925 | 0.24042871 | -1.3983044 | 0.16202166 | 0.64709575 |
| SIL1 | 35.8010302 | -0.5190882 | 0.38870002 | -1.3354468 | 0.18173015 | 0.67460052 |
| PAIP2 | 20.0873201 | 0.10213719 | 0.36799801 | 0.27754821 | 0.78135919 | NA |
| TMEM173 | 3.49541144 | -0.8169159 | 0.71862432 | -1.1367774 | 0.25563135 | NA |
| UBE2D2 | 70.5840941 | -0.4677234 | 0.29647786 | -1.5775997 | 0.1146576 | 0.57860396 |
| CXXC5 | 27.377135 | 0.20129416 | 0.49874491 | 0.40360142 | 0.68650584 | 0.92726927 |
| AC008667.1 | 2.451077 | 1.35919614 | 1.28746226 | 1.05571727 | 0.29109741 | NA |
| PURA | 28.2240056 | 0.14110872 | 0.31522535 | 0.44764396 | 0.65441017 | 0.9226734 |
| IGIP | 3.75716651 | -0.0173707 | 0.86945823 | -0.0199788 | 0.98406032 | NA |
| CYSTM1 | 94.3978818 | 0.01461934 | 0.3228068 | 0.04528821 | 0.96387759 | 0.99656976 |
| PFDN1 | 13.2013072 | -0.1225781 | 0.42947076 | -0.2854167 | 0.77532492 | NA |
| SLC4A9 | 530.304392 | 1.50352112 | 0.4613478 | 3.25897542 | 0.00111815 | 0.11692613 |
| AC008438.1 | 108.293441 | 1.6171335 | 0.63162778 | 2.56026344 | 0.01045928 | 0.28266532 |
| ANKHD1-EIF4 | 1.93661999 | 0.91973052 | 1.22640868 | 0.74993803 | 0.45329203 | NA |
| SRA1 | 7.28658795 | -0.6386024 | 0.54604835 | -1.1694979 | 0.2422031 | NA |
| SLC35A4 | 9.77911317 | -0.2909175 | 0.56793664 | -0.5122359 | 0.60848592 | NA |
| NDUFA2 | 16.8353314 | 0.43699979 | 0.4839202 | 0.90304103 | 0.36650412 | NA |
| IK | 5.83867229 | 0.32283669 | 0.68053012 | 0.47439001 | 0.63522181 | NA |

| | | | | | | |
|---|---|---|---|---|---|---|
| WDR55 | 4.43659883 | -0.7542999 | 0.72948053 | -1.0340233 | 0.30112525 | NA |
| HARS | 9.63202074 | -0.1811075 | 0.50228266 | -0.3605688 | 0.71842182 | NA |
| ZMAT2 | 7.55379363 | 0.15193889 | 0.5486592 | 0.27692762 | 0.78183568 | NA |
| TAF7 | 9.79874796 | -0.3226725 | 0.58247359 | -0.5539694 | 0.5795998 | NA |
| DIAPH1 | 90.8519927 | -0.5288315 | 0.29174263 | -1.8126645 | 0.06988359 | 0.52654172 |
| HDAC3 | 7.42994474 | -0.0911237 | 0.60697048 | -0.1501287 | 0.88066311 | NA |
| PCDH1 | 13.6250476 | 1.16678253 | 0.50530471 | 2.30906717 | 0.02093985 | NA |
| DELE1 | 10.156883 | -0.1558529 | 0.53984781 | -0.2886978 | 0.77281264 | NA |
| RNF14 | 12.5661875 | -0.4409143 | 0.46283164 | -0.9526451 | 0.34076992 | NA |
| GNPDA1 | 6.06501017 | -0.6489905 | 0.62955741 | -1.0308679 | 0.30260278 | NA |
| NDFIP1 | 60.564503 | 0.07422154 | 0.24767777 | 0.29966978 | 0.76442905 | 0.95550417 |
| ARHGAP26 | 15.8990889 | -0.1374802 | 0.60832533 | -0.2259978 | 0.82120313 | NA |
| NR3C1 | 19.0573467 | -0.7083104 | 0.45288642 | -1.5639912 | 0.11781964 | NA |
| YIPF5 | 6.90680924 | -1.3209755 | 0.58221495 | -2.2688794 | 0.02327566 | NA |
| PRELID2 | 11.6972 | -0.2540785 | 0.75860073 | -0.3349305 | 0.73767751 | NA |
| SH3RF2 | 4.32060667 | 0.13913351 | 0.98579833 | 0.14113791 | 0.887761 | NA |
| LARS | 30.7635918 | 0.15017711 | 0.36218076 | 0.41464684 | 0.67840045 | 0.9267216 |
| TCERG1 | 28.6754115 | -0.1435047 | 0.34891395 | -0.4112896 | 0.68086022 | 0.9267216 |
| STK32A | 6.22830936 | -0.0559446 | 0.82299973 | -0.0679765 | 0.94580436 | NA |
| SPINK5 | 6.98038359 | -0.2944223 | 0.7851892 | -0.3749699 | 0.70768289 | NA |
| AC091948.1 | 5.53049098 | 0.67127753 | 0.74477985 | 0.90131 | 0.36742352 | NA |
| FBXO38 | 18.553786 | 0.25439557 | 0.36461535 | 0.69770943 | 0.48535893 | NA |
| AFAP1L1 | 2.66026903 | -2.3713722 | 1.21427483 | -1.9529122 | 0.05083 | NA |
| GRPEL2 | 3.1018926 | -0.3712558 | 0.83093315 | -0.4467938 | 0.65502396 | NA |
| CSNK1A1 | 124.798138 | -0.341398 | 0.21318427 | -1.6014221 | 0.10928346 | 0.57860396 |
| ARHGEF37 | 6.8426162 | 0.97438184 | 0.60107917 | 1.62105407 | 0.10500604 | NA |
| SLC26A2 | 4.16649866 | -0.9924932 | 0.91343907 | -1.0865456 | 0.27723768 | NA |
| HMGXB3 | 13.1674758 | -0.4770792 | 0.47529689 | -1.0037498 | 0.31549921 | NA |
| TCOF1 | 4.44128821 | 0.12064421 | 0.75472525 | 0.15985182 | 0.8729978 | NA |
| CD74 | 30.1475806 | 0.03959298 | 0.72723763 | 0.05444298 | 0.95658224 | 0.99636969 |
| RPS14 | 90.7128806 | 0.12106204 | 0.37073424 | 0.32654669 | 0.74401077 | 0.94831173 |
| NDST1 | 16.2891529 | -0.5574317 | 0.50615201 | -1.1013129 | 0.27076049 | NA |
| RBM22 | 7.38368259 | -0.6616843 | 0.54328546 | -1.2179311 | 0.22325015 | NA |
| DCTN4 | 19.349337 | -0.186145 | 0.37658547 | -0.4942967 | 0.62109662 | NA |
| GPX3 | 124.529119 | -0.347545 | 0.57335745 | -0.6061576 | 0.54441008 | 0.88410237 |
| TNIP1 | 21.1007148 | 0.14707544 | 0.50626942 | 0.29050823 | 0.77142745 | NA |
| ANXA6 | 5.67331604 | 0.38789748 | 0.66872743 | 0.5800532 | 0.56187875 | NA |
| SLC36A1 | 4.382967 | -1.5668274 | 1.04125966 | -1.5047422 | 0.13239037 | NA |
| SPARC | 6.15528575 | 0.52461311 | 0.7194132 | 0.72922364 | 0.46586488 | NA |
| ATOX1 | 17.7698495 | 0.11692418 | 0.38883093 | 0.300707 | 0.76363793 | NA |
| G3BP1 | 35.5225458 | -0.1695519 | 0.32428419 | -0.5228497 | 0.60107886 | 0.90381943 |
| FAM114A2 | 8.34915309 | 0.30511303 | 0.51612039 | 0.5911664 | 0.55440893 | NA |
| MFAP3 | 12.8725604 | -0.0503816 | 0.47973195 | -0.1050203 | 0.9163597 | NA |
| GALNT10 | 42.9658385 | 0.19790287 | 0.40281421 | 0.49130062 | 0.62321384 | 0.91493825 |
| SAP30L-AS1 | 5.47566858 | 1.28169149 | 0.74158606 | 1.72831121 | 0.08393244 | NA |
| SAP30L | 4.80387412 | -0.110826 | 0.6786103 | -0.1633132 | 0.87027187 | NA |
| LARP1 | 49.9401651 | -0.2612152 | 0.30834023 | -0.8471654 | 0.39690295 | 0.82631356 |
| FAXDC2 | 8.62467008 | 1.68496145 | 0.83762001 | 2.01160601 | 0.04426148 | NA |
| CNOT8 | 9.2048558 | -0.0091926 | 0.46517214 | -0.0197617 | 0.98423344 | NA |
| GEMIN5 | 4.93709859 | -0.0297009 | 0.75558753 | -0.0393083 | 0.96864455 | NA |
| MRPL22 | 7.03620348 | -0.2178641 | 0.5394209 | -0.4038852 | 0.68629713 | NA |
| SGCD | 31.3677885 | -0.5177505 | 1.28073433 | -0.4042607 | 0.68602104 | 0.92726927 |
| HAVCR2 | 3.41103414 | 0.70797525 | 0.81962433 | 0.86378018 | 0.38770866 | NA |
| CYFIP2 | 46.1364581 | 0.31130071 | 0.30982528 | 1.00476215 | 0.31501139 | 0.780966 |

| | | | | | | |
|---|---|---|---|---|---|---|
| CLINT1 | 37.3715757 | -0.4303213 | 0.38051848 | -1.1308814 | 0.258105 | 0.75272108 |
| RNF145 | 11.7805095 | -0.0004817 | 0.46885067 | -0.0010273 | 0.9991803 | NA |
| UBLCP1 | 9.03792015 | 0.09828691 | 0.50147875 | 0.19599417 | 0.84461473 | NA |
| AC008691.1 | 4.1911962 | 0.30330326 | 0.82260236 | 0.36871188 | 0.71234249 | NA |
| TTC1 | 20.9413414 | 0.15912836 | 0.34108279 | 0.46653881 | 0.64082987 | NA |
| PWWP2A | 22.5117895 | 0.43219319 | 0.42996235 | 1.00518846 | 0.31480611 | NA |
| CCNJL | 13.6526153 | 0.0479207 | 0.53253754 | 0.08998558 | 0.92829868 | NA |
| SLU7 | 6.55004394 | -0.1285014 | 0.56863652 | -0.2259817 | 0.82121568 | NA |
| AC113414.1 | 34.3479276 | 0.92522302 | 0.7576316 | 1.22120438 | 0.22200865 | 0.72433127 |
| CCNG1 | 16.3052269 | -0.2454028 | 0.37946234 | -0.6467119 | 0.5178184 | NA |
| NUDCD2 | 3.98853752 | 0.86044965 | 0.72534707 | 1.18625922 | 0.23551995 | NA |
| MAT2B | 13.5988317 | 0.67698068 | 0.43128984 | 1.56966527 | 0.116493 | NA |
| WWC1 | 108.997042 | 0.24652545 | 0.39816111 | 0.61916003 | 0.53581094 | 0.88254911 |
| RARS | 17.5335238 | -0.3064386 | 0.39740978 | -0.7710897 | 0.44065377 | NA |
| PANK3 | 22.7066284 | -0.4827027 | 0.43044496 | -1.121404 | 0.26211594 | NA |
| FOXI1 | 99.4034142 | -0.4490705 | 0.37824178 | -1.1872579 | 0.23512591 | 0.73156941 |
| LINC01187 | 219.887426 | 0.53274146 | 0.28946532 | 1.84043275 | 0.06570473 | 0.5230103 |
| RANBP17 | 39.2022257 | -0.2447618 | 0.34995523 | -0.6994088 | 0.48429659 | 0.86354292 |
| NPM1 | 28.5740387 | -0.0074087 | 0.3658248 | -0.0202521 | 0.98384229 | 0.99902646 |
| FBXW11 | 78.2427478 | -0.5288572 | 0.31213877 | -1.6943015 | 0.090208 | 0.56122412 |
| STK10 | 4.5714208 | -0.0545638 | 0.67424787 | -0.0809254 | 0.93550129 | NA |
| UBTD2 | 22.0561968 | -0.3580749 | 0.43132142 | -0.830181 | 0.40643644 | NA |
| DUSP1 | 19.4913742 | -1.0230012 | 0.6152386 | -1.6627714 | 0.0963582 | NA |
| ERGIC1 | 25.6236978 | 0.56863855 | 0.37387963 | 1.52091343 | 0.12828156 | NA |
| RPL26L1 | 4.07278159 | 0.43810251 | 0.68841019 | 0.63639748 | 0.52451738 | NA |
| ATP6V0E1 | 121.89605 | 0.03472223 | 0.24247205 | 0.14320097 | 0.88613146 | 0.97769824 |
| CREBRF | 59.7042895 | 0.14436425 | 0.41215508 | 0.35026681 | 0.72613847 | 0.93943944 |
| BNIP1 | 3.10546781 | -0.3442936 | 0.7841157 | -0.4390852 | 0.66059978 | NA |
| BOD1 | 6.31131361 | -0.0271482 | 0.59183707 | -0.0458711 | 0.96341296 | NA |
| CPEB4 | 162.803965 | -0.067397 | 0.22771168 | -0.2959754 | 0.76724891 | 0.95690997 |
| SFXN1 | 29.8692786 | -0.0625181 | 0.28449094 | -0.2197544 | 0.82606242 | 0.96845695 |
| THOC3 | 6.92232296 | -0.3294438 | 0.75586901 | -0.4358477 | 0.66294721 | NA |
| SIMC1 | 27.1654757 | 0.00649549 | 0.3586758 | 0.01810964 | 0.98555139 | 0.99902646 |
| KIAA1191 | 16.1647258 | 0.7729698 | 0.39622366 | 1.9508421 | 0.05107583 | NA |
| ARL10 | 4.07988342 | 0.16737363 | 0.72651009 | 0.23038032 | 0.81779625 | NA |
| CLTB | 13.4258666 | -0.6255366 | 0.48062276 | -1.3015126 | 0.19308307 | NA |
| FAF2 | 20.4913006 | -0.4430916 | 0.38087262 | -1.163359 | 0.24468387 | NA |
| RNF44 | 5.74636507 | -0.479803 | 0.63399509 | -0.7567929 | 0.44917395 | NA |
| TSPAN17 | 4.69585371 | -0.741741 | 0.72283098 | -1.0261611 | 0.30481567 | NA |
| UIMC1 | 29.9800948 | -0.2562327 | 0.33242259 | -0.7708041 | 0.44082308 | 0.84705313 |
| ZNF346 | 14.5062882 | 0.24452683 | 0.40423158 | 0.60491769 | 0.54523369 | NA |
| NSD1 | 69.5156307 | -0.1624812 | 0.28184016 | -0.5765011 | 0.56427652 | 0.89263108 |
| PRELID1 | 13.8161208 | -0.7780943 | 0.45616316 | -1.7057368 | 0.0880571 | NA |
| LMAN2 | 19.5658076 | 0.17847031 | 0.43587046 | 0.40945723 | 0.68220415 | NA |
| RGS14 | 4.01939146 | 0.90920474 | 0.73555041 | 1.2360876 | 0.216426 | NA |
| DBN1 | 11.6234125 | -0.6609936 | 0.49287361 | -1.3411017 | 0.17988744 | NA |
| DDX41 | 2.84848285 | -0.0613319 | 0.83739286 | -0.0732415 | 0.94161392 | NA |
| FAM193B | 12.8677479 | -0.0996702 | 0.47220798 | -0.2110726 | 0.83283063 | NA |
| TMED9 | 17.9752078 | -0.4538461 | 0.46883222 | -0.9680351 | 0.33302683 | NA |
| B4GALT7 | 5.6016878 | 0.16140311 | 0.67264442 | 0.23995309 | 0.81036662 | NA |
| RMND5B | 11.1500598 | -0.5520758 | 0.52058096 | -1.0604994 | 0.28891747 | NA |
| NHP2 | 6.84198225 | 0.19576108 | 0.71350819 | 0.27436417 | 0.78380478 | NA |
| HNRNPAB | 13.1011327 | -0.5563309 | 0.52367 | -1.0623692 | 0.28806812 | NA |
| PHYKPL | 26.3115655 | 0.02609217 | 0.37676654 | 0.0692529 | 0.94478832 | 0.99240556 |

| | | | | | | |
|---|---|---|---|---|---|---|
| CLK4 | 12.1593816 | 0.03722499 | 0.57368746 | 0.06488724 | 0.94826378 | NA |
| ZNF354A | 5.66130367 | 1.16417486 | 0.65675468 | 1.77261754 | 0.0762921 | NA |
| ZNF354B | 4.47826031 | -0.6902175 | 0.68580708 | -1.0064311 | 0.31420826 | NA |
| ZNF354C | 3.99102605 | 0.61966805 | 0.70576202 | 0.87801274 | 0.37993681 | NA |
| RUFY1 | 21.5322352 | -0.0938386 | 0.42283042 | -0.2219297 | 0.82436863 | NA |
| HNRNPH1 | 69.7874446 | -0.3764384 | 0.26107305 | -1.441889 | 0.14933369 | 0.63 |
| CANX | 66.468962 | 0.21634183 | 0.32829361 | 0.65898886 | 0.50990292 | 0.87158522 |
| MAML1 | 9.72694096 | -0.3167416 | 0.55943504 | -0.5661811 | 0.57127066 | NA |
| MGAT4B | 14.4418072 | -0.2489839 | 0.4078294 | -0.6105099 | 0.54152408 | NA |
| SQSTM1 | 63.1738283 | -0.9678771 | 0.41363755 | -2.3399159 | 0.01928808 | 0.33831016 |
| MRNIP | 8.75397809 | 1.0091552 | 0.56373749 | 1.79011546 | 0.07343535 | NA |
| TBC1D9B | 14.1844183 | -0.3117561 | 0.40698644 | -0.766011 | 0.44366977 | NA |
| RNF130 | 53.9971632 | -1.0494619 | 0.2969032 | -3.5346938 | 0.00040825 | 0.09080115 |
| MAPK9 | 15.6514531 | -0.7342509 | 0.49154013 | -1.4937762 | 0.13523413 | NA |
| CNOT6 | 20.1060881 | -0.1868657 | 0.43388339 | -0.4306819 | 0.66669972 | NA |
| MGAT1 | 16.8444441 | -0.9460455 | 0.52347512 | -1.8072405 | 0.07072479 | NA |
| LINC00847 | 2.27685964 | 2.20679455 | 1.01065643 | 2.18352597 | 0.0289971 | NA |
| ZFP62 | 5.30697008 | -0.9128443 | 0.73033166 | -1.2499038 | 0.2113347 | NA |
| BTNL9 | 4.14577808 | 0.39691345 | 0.73907552 | 0.53704044 | 0.59123968 | NA |
| TRIM41 | 5.58803633 | 0.32365655 | 0.66516517 | 0.48658072 | 0.62655549 | NA |
| RACK1 | 54.4641596 | 0.11644533 | 0.40560913 | 0.28708754 | 0.7740453 | 0.9604383 |
| TRIM52 | 7.34440479 | -0.5301735 | 0.66649612 | -0.7954638 | 0.42634376 | NA |
| DUSP22 | 6.60592446 | 0.34207266 | 0.54958079 | 0.6224247 | 0.53366264 | NA |
| EXOC2 | 27.0908408 | -0.4749805 | 0.44087225 | -1.0773654 | 0.28131705 | 0.76854599 |
| GMDS | 70.1765849 | -0.6440172 | 0.35457649 | -1.8162999 | 0.06932438 | 0.52654172 |
| GMDS-DT | 92.3106643 | 0.31370929 | 0.38097961 | 0.82342805 | 0.41026462 | 0.82967024 |
| WRNIP1 | 12.2253293 | -0.108497 | 0.44799958 | -0.242181 | 0.80863992 | NA |
| SERPINB1 | 29.5423349 | 0.83730863 | 0.45438166 | 1.84274302 | 0.06536654 | 0.5230103 |
| SERPINB6 | 24.4779826 | 0.32535621 | 0.40742028 | 0.79857637 | 0.4245361 | NA |
| NQO2 | 9.41520579 | 0.67690439 | 0.5909331 | 1.14548395 | 0.25200874 | NA |
| RIPK1 | 13.906559 | -0.0002651 | 0.55949764 | -0.0004739 | 0.99962188 | NA |
| BPHL | 6.41358217 | 0.20044334 | 0.73473704 | 0.27280963 | 0.78499956 | NA |
| PSMG4 | 12.2257078 | 0.69950159 | 0.58325911 | 1.19929817 | 0.23041203 | NA |
| SLC22A23 | 68.1724668 | 0.41331562 | 0.47681608 | 0.86682401 | 0.38603844 | 0.8258317 |
| PRPF4B | 35.6915665 | -0.1724372 | 0.37463634 | -0.4602788 | 0.64531614 | 0.92153493 |
| ECI2 | 7.91391495 | 0.22092918 | 0.64187784 | 0.34419195 | 0.73070194 | NA |
| CDYL | 54.4406587 | -0.3655644 | 0.37031185 | -0.9871799 | 0.32355447 | 0.78622244 |
| LYRM4 | 16.7370232 | -0.2440833 | 0.4498054 | -0.5426421 | 0.58737626 | NA |
| FARS2 | 77.0014579 | 0.24675776 | 0.36936346 | 0.66806218 | 0.5040939 | 0.87094108 |
| LY86-AS1 | 22.4615122 | 1.45386116 | 1.16825222 | 1.24447541 | 0.21332464 | NA |
| AL031123.1 | 13.585481 | 2.13098837 | 0.58868473 | 3.61991446 | 0.0002947 | NA |
| RREB1 | 67.2818686 | -0.1578097 | 0.35434843 | -0.4453519 | 0.65606549 | 0.92276785 |
| SSR1 | 20.7848252 | -0.3227005 | 0.37288124 | -0.8654244 | 0.38680587 | NA |
| AL139095.4 | 2.3439713 | -0.0947077 | 0.99293113 | -0.0953819 | 0.92401147 | NA |
| RIOK1 | 4.55222468 | -0.503833 | 0.78066155 | -0.6453923 | 0.51867297 | NA |
| DSP | 76.6979019 | -0.0532454 | 0.25359181 | -0.2099652 | 0.83369487 | 0.97154892 |
| SNRNP48 | 7.43300555 | 0.12644599 | 0.57491627 | 0.2199381 | 0.82591936 | NA |
| BLOC1S5-TXI | 4.50539702 | -0.1816295 | 0.86562391 | -0.209825 | 0.83380429 | NA |
| SLC35B3 | 7.88382461 | 0.24141232 | 0.50947706 | 0.47384335 | 0.63561162 | NA |
| TFAP2A | 7.94909204 | 0.10098363 | 0.53770641 | 0.1878044 | 0.85102999 | NA |
| GCNT2 | 18.982728 | 0.20096523 | 0.60312611 | 0.333206 | 0.73897879 | NA |
| PAK1IP1 | 2.8458626 | 0.08601777 | 0.81608951 | 0.10540237 | 0.91605654 | NA |
| TMEM14C | 10.8379006 | 0.65127245 | 0.50325886 | 1.29411025 | 0.19562734 | NA |
| MAK | 8.55902224 | 1.0193877 | 0.82867258 | 1.23014532 | 0.21864269 | NA |

| | | | | | | |
|---|---|---|---|---|---|---|
| NEDD9 | 10.7793451 | -1.8250326 | 0.78852662 | -2.3144845 | 0.02064116 | NA |
| TMEM170B | 3.32906947 | -0.6496305 | 0.79367468 | -0.8185098 | 0.41306616 | NA |
| HIVEP1 | 32.7626685 | -0.3193417 | 0.37602878 | -0.8492481 | 0.39574325 | 0.8258317 |
| PHACTR1 | 13.1043432 | 0.53316024 | 0.53419606 | 0.99806097 | 0.31824979 | NA |
| TBC1D7 | 3.79712169 | -0.5348621 | 0.91982459 | -0.5814827 | 0.56091515 | NA |
| SIRT5 | 8.87598288 | 1.15154768 | 0.7237032 | 1.59118778 | 0.11156732 | NA |
| NOL7 | 12.9114249 | -0.6420217 | 0.58426967 | -1.0988448 | 0.27183577 | NA |
| RANBP9 | 61.9905086 | 0.13903071 | 0.30758703 | 0.45200447 | 0.65126576 | 0.92248109 |
| MCUR1 | 9.06129605 | 0.23101938 | 0.52057109 | 0.44378065 | 0.65720118 | NA |
| JARID2 | 45.5695547 | 0.53664976 | 0.33359703 | 1.60867667 | 0.10768706 | 0.57769936 |
| DTNBP1 | 35.1598043 | -0.006705 | 0.40300012 | -0.0166378 | 0.98672554 | 0.99907833 |
| MYLIP | 12.495244 | -0.8979886 | 0.4627703 | -1.9404628 | 0.05232347 | NA |
| GMPR | 22.7533089 | 1.09912378 | 0.60101159 | 1.82878965 | 0.06743113 | NA |
| ATXN1 | 78.4287805 | 0.55553497 | 0.47767152 | 1.16300626 | 0.24482696 | 0.73428453 |
| CAP2 | 10.1113468 | -0.0875431 | 0.57016779 | -0.1535391 | 0.87797314 | NA |
| FAM8A1 | 4.37015665 | -0.3651929 | 0.66642771 | -0.5479858 | 0.58370169 | NA |
| NUP153 | 27.5903028 | 0.2229174 | 0.35183904 | 0.63357778 | 0.5263564 | 0.87857739 |
| KIF13A | 38.0069292 | 0.15264661 | 0.32793981 | 0.46547142 | 0.6415939 | 0.92153493 |
| TPMT | 6.77711003 | 0.02507718 | 0.61338663 | 0.04088316 | 0.96738904 | NA |
| KDM1B | 11.6982602 | 0.11071234 | 0.58059768 | 0.19068685 | 0.84877094 | NA |
| DEK | 23.8156134 | 0.47096996 | 0.34967781 | 1.34686831 | 0.17802265 | NA |
| RNF144B | 10.2965377 | -0.5988132 | 0.46343057 | -1.2921314 | 1.96E-01 | NA |
| AL022068.1 | 4.86436714 | 0.39464401 | 0.72611009 | 0.54350438 | 0.58678258 | NA |
| ID4 | 4.49390759 | -0.536502 | 0.84263944 | -0.6366922 | 0.52432533 | NA |
| MBOAT1 | 36.4357075 | -0.1756846 | 0.53683412 | -0.3272605 | 0.74347086 | 0.94831173 |
| E2F3 | 8.72729628 | 0.69188928 | 0.52921485 | 1.30738824 | 0.19108089 | NA |
| CDKAL1 | 118.876794 | -0.0684543 | 0.30420383 | -0.2250279 | 0.82195758 | 0.96845695 |
| SOX4 | 40.782087 | -1.5183288 | 0.68984668 | -2.2009655 | 0.02773846 | 0.39934064 |
| CASC15 | 30.9087557 | -0.4046949 | 0.39293671 | -1.029924 | 0.30304568 | 0.77521265 |
| DCDC2 | 115.215762 | -0.1057901 | 0.42194278 | -0.2507214 | 0.80202949 | 0.96742897 |
| MRS2 | 12.7345027 | -0.469426 | 0.46409023 | -1.0114973 | 0.31177848 | NA |
| ALDH5A1 | 5.49376865 | 0.80206375 | 0.70269207 | 1.14141569 | 0.25369698 | NA |
| TDP2 | 9.54543237 | 0.15143374 | 0.52911273 | 0.28620318 | 0.77472251 | NA |
| ACOT13 | 5.11349652 | 0.09487254 | 0.81548696 | 0.11633851 | 0.90738426 | NA |
| C6orf62 | 38.119186 | -0.4029247 | 0.2905657 | -1.3866904 | 0.1655362 | 0.64995633 |
| GMNN | 15.7632433 | -0.4564333 | 0.44751317 | -1.0199326 | 0.30776044 | NA |
| RIPOR2 | 32.5840575 | -0.7440613 | 0.39749476 | -1.8718769 | 0.06122364 | 0.51625203 |
| CARMIL1 | 207.493532 | -0.1133995 | 0.26071711 | -0.4349524 | 0.66359696 | 0.92276785 |
| SLC17A1 | 7.91069181 | 1.63600297 | 1.03534547 | 1.58015175 | 0.11407212 | NA |
| TRIM38 | 16.3591293 | -0.401926 | 0.47308598 | -0.8495834 | 0.39555673 | NA |
| HIST1H1C | 7.23957367 | -0.8530968 | 0.62234546 | -1.3707769 | 0.17044453 | NA |
| HIST1H4C | 12.350524 | 1.02961284 | 0.53433653 | 1.92689957 | 0.05399215 | NA |
| HIST1H2AC | 6.95962493 | 0.29583111 | 0.57648541 | 0.51316322 | 0.60783715 | NA |
| HIST1H1E | 2.6755165 | 0.40833629 | 0.9383212 | 0.43517751 | 0.66343358 | NA |
| BTN3A2 | 5.58654907 | 1.16045646 | 0.97929051 | 1.18499714 | 0.23601858 | NA |
| BTN3A1 | 4.99923905 | 1.14526821 | 0.95642143 | 1.19745144 | 0.23113065 | NA |
| BTN2A1 | 5.63203488 | -0.007815 | 0.64079619 | -0.0121958 | 0.99026938 | NA |
| HCG11 | 3.43180387 | 0.66909089 | 0.89714461 | 0.74580048 | 0.45578795 | NA |
| HMGN4 | 7.90159194 | 0.54147628 | 0.60097546 | 0.90099566 | 0.36759063 | NA |
| ABT1 | 3.93172784 | -0.5542409 | 0.74605294 | -0.7428976 | 0.45754368 | NA |
| ZNF322 | 4.25531689 | 0.13113461 | 0.81403369 | 0.16109237 | 0.87202065 | NA |
| AL121944.1 | 4.18560094 | -0.5098619 | 0.8640846 | -0.5900602 | 0.5551503 | NA |
| ZNF165 | 4.71252991 | -1.1582277 | 0.98716932 | -1.1732817 | 0.24068287 | NA |
| ZSCAN16-AS | 6.66127291 | 0.24846614 | 0.7178434 | 0.34612861 | 0.72924606 | NA |

| | | | | | | |
|---|---|---|---|---|---|---|
| ZKSCAN8 | 6.07943716 | 0.58010137 | 0.76673773 | 0.75658383 | 0.44929923 | NA |
| ZSCAN9 | 3.76570468 | 0.3766213 | 0.73854828 | 0.50994811 | 0.61008782 | NA |
| ZSCAN31 | 4.92294115 | 2.21379451 | 0.91964108 | 2.40723752 | 0.01607371 | NA |
| ZSCAN12 | 2.72709465 | -0.2256448 | 0.90655961 | -0.2489023 | 0.80343636 | NA |
| ZSCAN23 | 1.8769922 | -0.0136135 | 1.14309802 | -0.0119093 | 0.99049798 | NA |
| TRIM27 | 16.0644136 | -0.2311005 | 0.44096548 | -0.5240783 | 0.60022405 | NA |
| GABBR1 | 3.56185135 | 0.22931097 | 0.92573957 | 0.2477057 | 0.80436212 | NA |
| HLA-F | 8.61054891 | 0.91010711 | 0.68498398 | 1.3286546 | 0.18396195 | NA |
| HLA-A | 130.577946 | -0.0536894 | 0.4156833 | -0.1291593 | 0.89723159 | 0.97769824 |
| PPP1R11 | 5.82880274 | -0.6459338 | 0.77330241 | -0.8352926 | 0.40355298 | NA |
| TRIM26 | 13.318693 | -0.4330848 | 0.47940844 | -0.9033734 | 0.36632775 | NA |
| HCG17 | 17.0109329 | -0.6315809 | 0.45914743 | -1.3755515 | 0.16896054 | NA |
| HLA-E | 63.5127499 | -0.2903712 | 0.44057196 | -0.6590778 | 0.50984581 | 0.87158522 |
| GNL1 | 10.742229 | -0.8753246 | 0.48373735 | -1.8095038 | 0.07037277 | NA |
| PRR3 | 4.07037984 | 0.11909881 | 0.78087997 | 0.15251872 | 0.87877783 | NA |
| ABCF1 | 13.5950492 | -0.7985952 | 0.52248579 | -1.5284534 | 0.12640001 | NA |
| PPP1R10 | 13.2294212 | 0.05422407 | 0.45414199 | 0.11939894 | 0.9049593 | NA |
| MRPS18B | 11.7995627 | 0.81043286 | 0.48747442 | 1.66251363 | 0.09640983 | NA |
| ATAT1 | 4.59014735 | -0.9512959 | 0.82307899 | -1.1557771 | 0.24777233 | NA |
| C6orf136 | 8.74062006 | 0.00304254 | 0.5450896 | 0.00558172 | 0.99554645 | NA |
| DHX16 | 5.71805922 | 0.14060676 | 0.62833087 | 0.22377821 | 0.82292988 | NA |
| MDC1 | 6.06606834 | 0.18182679 | 0.58003098 | 0.31347772 | 0.75391774 | NA |
| TUBB | 26.6849164 | -1.0650072 | 0.66196069 | -1.6088678 | 0.10764526 | 0.57769936 |
| FLOT1 | 27.3278021 | -0.1171975 | 0.33396206 | -0.3509305 | 0.72564049 | 0.93925046 |
| IER3 | 12.2847643 | -0.7378852 | 0.74675744 | -0.988119 | 0.32309437 | NA |
| DDR1 | 25.2684865 | 0.69181582 | 0.3379896 | 2.04685537 | 0.04067229 | NA |
| VARS2 | 2.9927628 | 0.80239161 | 0.82372764 | 0.97409819 | 0.33000779 | NA |
| CCHCR1 | 1.8388055 | 0.09822594 | 1.22369038 | 0.08027026 | 0.93602231 | NA |
| POU5F1 | 16.7208854 | -0.2844485 | 0.51025912 | -0.5574589 | 0.57721395 | NA |
| HLA-C | 115.852775 | 0.22085681 | 0.60023456 | 0.36795084 | 0.71290989 | 0.93547517 |
| HLA-B | 183.385456 | 0.4152883 | 0.54898408 | 0.75646692 | 0.4493693 | 0.85123446 |
| MICA | 5.5708125 | -0.2603426 | 0.65039025 | -0.4002868 | 0.68894532 | NA |
| NFKBIL1 | 4.06159487 | 0.08796557 | 0.75734546 | 0.11614986 | 0.90753377 | NA |
| PRRC2A | 12.8922466 | -0.8545364 | 0.50399596 | -1.6955223 | 0.08997639 | NA |
| BAG6 | 8.18200323 | -0.6661208 | 0.56429926 | -1.1804389 | 0.23782568 | NA |
| C6orf47 | 3.69613386 | -0.4416338 | 0.88147289 | -0.501018 | 0.61635845 | NA |
| GPANK1 | 3.72608578 | 1.07807941 | 0.80324348 | 1.34215768 | 0.17954487 | NA |
| DDAH2 | 7.41601762 | 0.35126256 | 0.59300956 | 0.59233877 | 0.55362376 | NA |
| CLIC1 | 31.7137656 | 0.07460107 | 0.30520491 | 0.24442945 | 0.80689823 | 0.96800006 |
| VARS | 2.98249333 | -1.3536809 | 0.94493446 | -1.432566 | 0.15198192 | NA |
| LSM2 | 4.8007908 | 0.41458447 | 0.64457843 | 0.643187 | 0.52010277 | NA |
| HSPA1A | 36.3620479 | -0.1656263 | 0.44943558 | -0.3685206 | 0.7124851 | 0.93547517 |
| HSPA1B | 15.0762923 | -0.0983152 | 0.62547414 | -0.1571851 | 0.87509898 | NA |
| C6orf48 | 10.8245326 | 0.1096872 | 0.4671088 | 0.23482153 | 0.81434726 | NA |
| NEU1 | 8.68419042 | -1.1420614 | 0.61053023 | -1.8706058 | 0.06139974 | NA |
| EHMT2 | 4.4234105 | 0.40331733 | 0.73394639 | 0.54951879 | 0.58264948 | NA |
| NELFE | 3.00214661 | -1.8360791 | 0.90512978 | -2.0285258 | 0.04250661 | NA |
| SKIV2L | 2.31128576 | -0.0719765 | 0.95555393 | -0.0753244 | 0.93995661 | NA |
| STK19 | 3.86188309 | -0.0301553 | 0.6883216 | -0.0438099 | 0.96505595 | NA |
| ATF6B | 7.82595727 | -0.8117206 | 0.72915019 | -1.113242 | 0.2656045 | NA |
| AGPAT1 | 7.15293517 | 0.17737474 | 0.55958449 | 0.3169758 | 0.75126197 | NA |
| RNF5 | 8.02917737 | 0.00056118 | 0.59138513 | 0.00094892 | 0.99924287 | NA |
| PBX2 | 5.73119203 | 0.28251626 | 0.61367596 | 0.46036717 | 0.6452527 | NA |
| HLA-DRA | 10.1668553 | 0.86478088 | 0.70996426 | 1.21806255 | 0.2232002 | NA |

| | | | | | | |
|---|---|---|---|---|---|---|
| HLA-DRB1 | 5.20186436 | -0.6364161 | 1.17191464 | -0.5430567 | 0.58709077 | NA |
| PSMB8 | 16.3956902 | -0.1261876 | 0.49991457 | -0.2524183 | 0.80071776 | NA |
| PSMB9 | 6.86911474 | 1.06719182 | 1.17533753 | 0.90798752 | 0.36388481 | NA |
| TAP1 | 8.37821905 | -0.515167 | 0.69240368 | -0.744027 | 0.45686016 | NA |
| BRD2 | 23.0987334 | -1.2561156 | 0.39668219 | -3.1665541 | 0.00154257 | NA |
| RXRB | 3.68759797 | -0.333726 | 0.82171142 | -0.4061353 | 0.68464321 | NA |
| SLC39A7 | 12.0881816 | -1.2749559 | 0.63697656 | -2.0015743 | 0.04533054 | NA |
| RING1 | 6.85483408 | 0.04339044 | 0.65715181 | 0.06602803 | 0.94735551 | NA |
| VPS52 | 6.82264866 | -0.3337454 | 0.66486742 | -0.5019729 | 0.61568659 | NA |
| RPS18 | 47.6825532 | -0.2039056 | 0.3626052 | -0.5623349 | 0.57388786 | 0.89263108 |
| WDR46 | 3.99289675 | 0.35609676 | 0.75582379 | 0.47113728 | 0.63754271 | NA |
| PFDN6 | 4.85575092 | 0.34094575 | 0.72050197 | 0.47320585 | 0.63606633 | NA |
| RGL2 | 3.69624141 | 0.37960156 | 0.69795066 | 0.54388021 | 0.58652391 | NA |
| TAPBP | 26.9007699 | 0.11199891 | 0.44591195 | 0.25116822 | 0.80168406 | 0.96742897 |
| DAXX | 3.3317432 | -1.0767956 | 0.81586149 | -1.3198264 | 0.18689297 | NA |
| PHF1 | 7.58093369 | -0.349229 | 0.56217961 | -0.6212055 | 0.53446444 | NA |
| CUTA | 18.4616746 | 0.37008757 | 0.44379418 | 0.83391714 | 0.40432768 | NA |
| SYNGAP1 | 3.00393163 | -0.1245176 | 0.90080234 | -0.1382297 | 0.8900589 | NA |
| ITPR3 | 7.32556663 | -0.0699673 | 0.58840412 | -0.1189103 | 0.90534643 | NA |
| UQCC2 | 8.04020842 | 1.04787066 | 0.52050288 | 2.01318897 | 0.04409475 | NA |
| LEMD2 | 6.83405529 | -0.2101595 | 0.602156 | -0.3490118 | 0.72708046 | NA |
| HMGA1 | 4.8943099 | -0.1857223 | 0.69619523 | -0.2667676 | 0.78964812 | NA |
| SMIM29 | 4.43027529 | 0.46048518 | 0.98070686 | 0.46954416 | 0.63868073 | NA |
| NUDT3 | 17.4629695 | 0.02448211 | 0.51014891 | 0.04799013 | 0.96172411 | NA |
| C6orf106 | 31.944803 | -0.706613 | 0.30934047 | -2.2842566 | 0.02235646 | 0.35517491 |
| SNRPC | 6.33389845 | 0.22503096 | 0.58043802 | 0.38769164 | 0.69824425 | NA |
| UHRF1BP1 | 16.267402 | 0.12322653 | 0.4474161 | 0.27541819 | 0.78299497 | NA |
| TAF11 | 10.0165999 | 0.30797881 | 0.54874624 | 0.56124085 | 0.57463336 | NA |
| ANKS1A | 77.7382694 | -0.286399 | 0.2566277 | -1.1160095 | 0.26441805 | 0.75479335 |
| ZNF76 | 10.1983814 | -0.1531085 | 0.49406654 | -0.3098945 | 0.75664117 | NA |
| PPARD | 11.8870895 | -0.424297 | 0.47986322 | -0.884204 | 0.37658612 | NA |
| RPL10A | 49.7804985 | 0.47726553 | 0.50133668 | 0.95198604 | 0.34110406 | 0.8003972 |
| TEAD3 | 6.25526819 | -0.5385434 | 0.83788651 | -0.6427402 | 0.52039268 | NA |
| FKBP5 | 19.6630342 | 0.73773018 | 1.29880032 | 0.56800893 | 0.57002891 | NA |
| SRPK1 | 20.9421272 | -0.1179153 | 0.35448752 | -0.332636 | 0.73940903 | NA |
| MAPK14 | 32.5192991 | -0.1433646 | 0.32775438 | -0.4374149 | 0.66181051 | 0.92276785 |
| MAPK13 | 7.5623654 | 0.26822078 | 0.52812625 | 0.50787246 | 0.61154279 | NA |
| BRPF3 | 10.3951999 | -0.0628319 | 0.54278361 | -0.1157587 | 0.90784375 | NA |
| KCTD20 | 26.7352277 | -0.2852147 | 0.35632918 | -0.8004248 | 0.42346471 | 0.84057082 |
| STK38 | 39.2958116 | 0.12688725 | 0.40568982 | 0.31276913 | 0.75445607 | 0.95323728 |
| SRSF3 | 31.9453412 | -0.9145474 | 0.35855909 | -2.5506183 | 0.0107532 | 0.28312744 |
| CDKN1A | 8.63738482 | -0.1376129 | 0.62117817 | -0.2215353 | 0.82467567 | NA |
| PPIL1 | 3.2857277 | -1.2845539 | 0.83820898 | -1.5324984 | 0.1253995 | NA |
| C6orf89 | 18.8828555 | -0.1141791 | 0.38238827 | -0.2985947 | 0.76524934 | NA |
| MTCH1 | 19.1920426 | -0.3062193 | 0.41689096 | -0.7345309 | 0.46262522 | NA |
| PIM1 | 16.0347067 | -0.1611986 | 0.56711374 | -0.2842438 | 0.77622355 | NA |
| TBC1D22B | 14.8611675 | -0.2158712 | 0.44334566 | -0.4869139 | 0.62631932 | NA |
| RNF8 | 9.66433871 | -0.0012936 | 0.50127927 | -0.0025806 | 0.99794101 | NA |
| CMTR1 | 5.48956436 | -0.1110139 | 0.69817086 | -0.1590068 | 0.87366351 | NA |
| ZFAND3 | 326.015955 | -0.6331752 | 0.22725605 | -2.7861753 | 0.0053334 | 0.22594999 |
| BTBD9 | 177.610914 | -0.4864032 | 0.26663706 | -1.8242146 | 0.06811963 | 0.52508412 |
| GLO1 | 18.8385051 | 0.30378166 | 0.39909689 | 0.76117271 | 0.44655392 | NA |
| SAYSD1 | 5.42941943 | 0.71437963 | 0.60316018 | 1.18439455 | 0.23625692 | NA |
| DAAM2 | 2.50936694 | 1.54728216 | 1.21513462 | 1.27334217 | 0.20289665 | NA |

| | | | | | | |
|---|---|---|---|---|---|---|
| MOCS1 | 5.76780851 | -0.1097454 | 0.61703057 | -0.1778605 | 0.85883254 | NA |
| OARD1 | 7.18352794 | 0.9388806 | 0.58226482 | 1.6124632 | 0.10686118 | NA |
| NFYA | 6.47725396 | -0.6593398 | 0.74267968 | -0.8877849 | 0.37465647 | NA |
| FOXP4 | 5.5358663 | -0.4152908 | 0.6385916 | -0.650323 | 0.51548357 | NA |
| TFEB | 27.176797 | 0.3512797 | 0.45447743 | 0.77293102 | 0.43956321 | 0.84524078 |
| USP49 | 8.33998435 | 1.32199318 | 0.53408178 | 2.47526356 | 0.01331379 | NA |
| CCND3 | 15.2312515 | -0.1037124 | 0.62986176 | -0.1646589 | 0.86921246 | NA |
| TAF8 | 8.6415666 | 0.96936931 | 0.52973302 | 1.82992049 | 0.06726183 | NA |
| MRPS10 | 5.17824144 | -0.6427864 | 0.6948383 | -0.9250877 | 0.35492028 | NA |
| TRERF1 | 5.57935609 | 0.20448893 | 0.64851701 | 0.31531776 | 0.7525204 | NA |
| UBR2 | 72.0057919 | 0.06978085 | 0.30792551 | 0.226616 | 0.82072235 | 0.96845695 |
| TBCC | 2.42566365 | 0.9912287 | 0.92208404 | 1.07498738 | 0.28238038 | NA |
| BICRAL | 31.6384219 | -0.2625357 | 0.35326784 | -0.7431632 | 0.45738287 | 0.85802171 |
| RPL7L1 | 18.3225519 | 0.27332261 | 0.39476078 | 0.6923753 | 0.48870167 | NA |
| C6orf226 | 2.98852351 | 1.9630774 | 0.85382316 | 2.29916158 | 0.02149577 | NA |
| CNPY3 | 4.4684993 | 1.26244129 | 0.7059997 | 1.78816122 | 0.07375 | NA |
| PEX6 | 7.47596012 | 0.55921573 | 0.58203506 | 0.9607939 | 0.33665581 | NA |
| PPP2R5D | 2.33044691 | 0.14424757 | 0.90110877 | 0.16007787 | 0.87281973 | NA |
| MEA1 | 5.72076986 | 0.28002159 | 0.61014767 | 0.45894069 | 0.64627676 | NA |
| KLHDC3 | 5.27912653 | 0.54216263 | 0.63996999 | 0.84716883 | 0.39690102 | NA |
| RRP36 | 8.78452415 | -0.2243652 | 0.57854461 | -0.3878097 | 0.69815684 | NA |
| CUL7 | 6.65268003 | -0.9916114 | 0.63809537 | -1.5540175 | 0.12018025 | NA |
| KLC4 | 7.56571304 | 0.70399406 | 0.58537911 | 1.20262928 | 0.22911981 | NA |
| MRPL2 | 4.68944506 | 0.0882895 | 0.67616449 | 0.130574 | 0.89611231 | NA |
| PTK7 | 10.9437675 | -0.6433208 | 0.59850415 | -1.0748811 | 0.28242798 | NA |
| SRF | 3.49243679 | -0.6334663 | 1.10286569 | -0.5743821 | 0.56570925 | NA |
| CUL9 | 4.31089141 | 0.35424415 | 0.73482277 | 0.48208108 | 0.62974835 | NA |
| DNPH1 | 6.5462222 | -0.8091613 | 0.60394916 | -1.3397838 | 0.18031565 | NA |
| ZNF318 | 5.51537626 | 0.28712653 | 0.73079931 | 0.39289382 | 0.69439791 | NA |
| ABCC10 | 6.61701127 | 0.23987438 | 0.55997688 | 0.42836479 | 0.66838555 | NA |
| TJAP1 | 5.70160671 | -0.4619245 | 0.64328084 | -0.718076 | 0.47271045 | NA |
| POLR1C | 8.26188109 | -0.5994656 | 0.61070803 | -0.9815912 | 0.32630127 | NA |
| YIPF3 | 10.6659198 | -0.8285111 | 0.55911166 | -1.4818348 | 0.13838425 | NA |
| XPO5 | 39.7536668 | 0.14193505 | 0.38561037 | 0.36807893 | 0.71281438 | 0.93547517 |
| POLH | 7.25516953 | -0.2225392 | 0.60752052 | -0.3663074 | 0.71413575 | NA |
| GTPBP2 | 5.11250372 | 0.14914889 | 0.70682577 | 0.21101224 | 0.83287772 | NA |
| MAD2L1BP | 2.43242709 | -1.1600913 | 0.92754262 | -1.2507148 | 0.21103855 | NA |
| MRPS18A | 6.05954663 | 0.7896433 | 0.5684773 | 1.38904985 | 0.16481759 | NA |
| VEGFA | 57.5528579 | -0.2262401 | 0.32551797 | -0.6950157 | 0.48704545 | 0.86354292 |
| MRPL14 | 11.8056411 | 0.62545876 | 0.45691001 | 1.3688883 | 0.17103419 | NA |
| TMEM63B | 6.7394847 | -0.2462013 | 0.54356313 | -0.4529396 | 0.65059222 | NA |
| SLC29A1 | 7.1286705 | 0.33758946 | 0.6044441 | 0.55851228 | 0.57649462 | NA |
| HSP90AB1 | 80.1163624 | -0.2642677 | 0.25332114 | -1.0432121 | 0.29685005 | 0.77521265 |
| SLC35B2 | 4.53588268 | -0.2252602 | 0.75718407 | -0.2974973 | 0.76608685 | NA |
| CDC5L | 9.78021201 | -0.0934067 | 0.51335093 | -0.1819549 | 0.85561811 | NA |
| SUPT3H | 113.065956 | -0.3727312 | 0.33182461 | -1.1232777 | 0.26131957 | 0.75479335 |
| RUNX2 | 31.9911309 | -1.3761312 | 0.49097219 | -2.80287 | 0.00506501 | 0.22356834 |
| ENPP4 | 11.7698501 | -0.2981226 | 0.4669186 | -0.6384894 | 0.52315517 | NA |
| ENPP5 | 3.1891637 | 0.61638587 | 0.88357253 | 0.69760643 | 0.48542336 | NA |
| RCAN2 | 382.001007 | 0.51672972 | 0.31948776 | 1.61736938 | 0.10579857 | 0.57745683 |
| CYP39A1 | 9.69125395 | 1.67156413 | 0.53448363 | 3.12743747 | 0.00176337 | NA |
| SLC25A27 | 27.9522072 | 0.40543919 | 0.42581704 | 0.95214411 | 0.3410239 | 0.8003972 |
| ADGRF1 | 60.2350819 | -0.4059429 | 0.6025491 | -0.6737093 | 0.50049618 | 0.87036626 |
| TNFRSF21 | 5.86556309 | -1.7359615 | 0.75761426 | -2.2913527 | 0.02194302 | NA |

| | | | | | | |
|---|---|---|---|---|---|---|
| CD2AP | 86.5471131 | -0.0444346 | 0.26058412 | -0.1705194 | 0.86460172 | 0.97217819 |
| PTCHD4 | 8.37576218 | 1.16544895 | 1.09237046 | 1.06689899 | 0.28601745 | NA |
| MUT | 12.5105051 | 0.50603241 | 0.47408824 | 1.06738023 | 0.28580017 | NA |
| PKHD1 | 81.7042543 | -0.0408596 | 0.60365035 | -0.0676875 | 0.94603443 | 0.99251804 |
| MCM3 | 4.78918393 | 0.41152593 | 0.68900979 | 0.59727152 | 0.55032612 | NA |
| EFHC1 | 8.39100263 | 0.72968493 | 0.66022195 | 1.10521157 | 0.26906792 | NA |
| TRAM2 | 4.15412244 | 0.27083484 | 0.80383226 | 0.33692955 | 0.73617001 | NA |
| TMEM14A | 2.88908243 | 0.30956679 | 0.84229363 | 0.36752835 | 0.71322495 | NA |
| GSTA4 | 4.29177377 | 0.60409991 | 0.90180275 | 0.66988031 | 0.50293409 | NA |
| ICK | 5.84396699 | 0.50043861 | 0.63105581 | 0.79301798 | 0.42776735 | NA |
| FBXO9 | 17.923693 | 0.16364937 | 0.4065969 | 0.40248553 | 0.68732674 | NA |
| ELOVL5 | 18.8248498 | -1.606083 | 0.52913244 | -3.0353138 | 0.00240286 | NA |
| GCLC | 18.1087798 | -0.1328694 | 0.39311378 | -0.3379923 | 0.735369 | NA |
| LRRC1 | 20.6071626 | -0.4288172 | 0.44269882 | -0.9686433 | 0.33272321 | NA |
| TINAG | 6.78488454 | 0.82990526 | 0.94106441 | 0.88187933 | 0.37784206 | NA |
| FAM83B | 47.0540055 | -0.461526 | 0.32005505 | -1.4420207 | 0.14929653 | 0.63 |
| DST | 72.3136447 | -0.6481796 | 0.35067051 | -1.8484007 | 0.0645444 | 0.52254021 |
| KIAA1586 | 3.02741771 | -0.2976111 | 0.81525182 | -0.3650542 | 0.71507095 | NA |
| ZNF451 | 33.0983524 | -0.3118549 | 0.29898556 | -1.0430434 | 0.2969282 | 0.77521265 |
| ZNF451-AS1 | 3.30042665 | 0.73938776 | 0.92394879 | 0.80024755 | 0.42356739 | NA |
| PRIM2 | 25.1584445 | -0.1980087 | 0.35562491 | -0.5567908 | 0.57767038 | NA |
| AL021368.2 | 3.3035651 | 0.02245768 | 0.90193626 | 0.02489941 | 0.9801352 | NA |
| PTP4A1 | 17.4183168 | -1.7643972 | 0.54941839 | -3.2113908 | 0.00132094 | NA |
| PHF3 | 61.4389528 | -0.0499696 | 0.27521622 | -0.181565 | 0.85592412 | 0.97168963 |
| EYS | 3.55604569 | 0.53566596 | 1.01129051 | 0.52968554 | 0.59632998 | NA |
| LMBRD1 | 46.9624033 | 0.03836511 | 0.29541058 | 0.12987048 | 0.8966689 | 0.97769824 |
| COL19A1 | 10.8922322 | 0.27142265 | 0.54020343 | 0.50244525 | 0.61535436 | NA |
| FAM135A | 181.511192 | -0.5998225 | 0.35481029 | -1.6905444 | 0.09092385 | 0.56211136 |
| SDHAF4 | 2.64741149 | 0.06559215 | 0.84346662 | 0.07776496 | 0.93801502 | NA |
| SMAP1 | 80.8178915 | -0.3233862 | 0.25566439 | -1.2648857 | 0.20591232 | 0.70278771 |
| OGFRL1 | 43.8664529 | 0.56802036 | 0.38490022 | 1.47576003 | 0.14000832 | 0.61443563 |
| AL136164.2 | 26.9147895 | 0.27792923 | 0.43094756 | 0.64492587 | 0.51897523 | 0.87214537 |
| LINC00472 | 11.5273823 | 0.14924298 | 0.53027962 | 0.28144204 | 0.77837137 | NA |
| OOEP | 49.3866618 | 0.06262797 | 0.34563749 | 0.18119553 | 0.85621411 | 0.97168963 |
| MTO1 | 9.71418712 | -0.1999473 | 0.50343903 | -0.3971629 | 0.69124735 | NA |
| EEF1A1 | 285.746871 | 0.10954534 | 0.40907046 | 0.26779088 | 0.78886029 | 0.96325251 |
| SLC17A5 | 8.23112582 | -0.3523685 | 0.57023327 | -0.6179375 | 0.53661656 | NA |
| COX7A2 | 31.4717751 | 0.30895427 | 0.35262022 | 0.8761672 | 0.38093914 | 0.82153751 |
| TMEM30A | 28.7558261 | 0.15406166 | 0.35131397 | 0.43852984 | 0.66100224 | 0.92276785 |
| SENP6 | 96.2410211 | 0.07298603 | 0.24616901 | 0.29648748 | 0.76685783 | 0.95686935 |
| MYO6 | 225.311865 | -0.4800945 | 0.33649189 | -1.4267641 | 0.15364791 | 0.63864841 |
| MEI4 | 4.61659412 | -0.1865806 | 0.67902246 | -0.2747783 | 0.78348657 | NA |
| IRAK1BP1 | 14.5462076 | 1.03030299 | 0.53341956 | 1.93150585 | 0.05342053 | NA |
| PHIP | 72.4008062 | 0.53587422 | 0.28218105 | 1.89904395 | 0.0575587 | 0.50751937 |
| HMGN3 | 33.7554395 | 0.26404222 | 0.41097193 | 0.64248235 | 0.52056004 | 0.87214537 |
| LCA5 | 3.17183753 | 0.4700467 | 0.91582852 | 0.5132475 | 0.6077782 | NA |
| SH3BGRL2 | 5.87386297 | 0.6705433 | 0.63784626 | 1.05126163 | 0.29313844 | NA |
| BCKDHB | 46.7983952 | 0.51846394 | 0.35685853 | 1.45285566 | 0.14626383 | 0.62964221 |
| TENT5A | 8.53086034 | 0.28876172 | 0.5828692 | 0.49541428 | 0.62030772 | NA |
| IBTK | 42.5313159 | -0.7482517 | 0.30361909 | -2.4644424 | 0.01372266 | 0.30435125 |
| UBE3D | 25.9989756 | 0.16819249 | 0.33359432 | 0.5041827 | 0.614133 | 0.91163569 |
| DOPEY1 | 38.6588889 | -0.2108894 | 0.28746726 | -0.7336118 | 0.46318538 | 0.85894565 |
| PGM3 | 9.254339 | -1.6896063 | 0.53466833 | -3.1601016 | 0.00157714 | NA |
| CYB5R4 | 14.0175899 | 0.06105157 | 0.43027068 | 0.14189107 | 0.88716605 | NA |

| | | | | | | |
|---|---|---|---|---|---|---|
| CEP162 | 13.7381017 | 0.39337152 | 0.42554504 | 0.92439457 | 0.35528092 | NA |
| SNX14 | 42.9141395 | -0.2304943 | 0.30307754 | -0.7605125 | 0.4469483 | 0.85123446 |
| SYNCRIP | 17.2942331 | -0.3722112 | 0.43213241 | -0.8613361 | 0.38905298 | NA |
| ZNF292 | 128.572173 | -0.5349606 | 0.23481356 | -2.2782357 | 0.02271254 | 0.35709704 |
| SMIM8 | 19.6404794 | 0.96117369 | 0.41305179 | 2.32700525 | 0.01996499 | NA |
| RARS2 | 25.0277255 | -0.2712892 | 0.37618501 | -0.7211589 | 0.47081174 | NA |
| ORC3 | 14.4255636 | 0.36927204 | 0.45696696 | 0.80809352 | 0.41903675 | NA |
| AKIRIN2 | 8.0565604 | 0.52510853 | 0.54724093 | 0.95955639 | 0.33727853 | NA |
| RNGTT | 58.2802115 | -0.4422426 | 0.26373944 | -1.6768162 | 0.09357843 | 0.56220784 |
| PNRC1 | 16.204682 | 0.28311656 | 0.43422625 | 0.65200241 | 0.51439962 | NA |
| PM20D2 | 12.6546913 | 0.04975322 | 0.51007794 | 0.09754042 | 0.92229723 | NA |
| UBE2J1 | 14.6553787 | -0.2160725 | 0.45390309 | -0.4760322 | 0.63405141 | NA |
| RRAGD | 29.7587191 | -0.2986841 | 0.52348502 | -0.5705685 | 0.56829219 | 0.89263108 |
| ANKRD6 | 24.4637415 | -0.4449276 | 0.35676859 | -1.2471042 | 0.21235931 | NA |
| LYRM2 | 11.2005972 | 0.30654612 | 0.53370094 | 0.57437808 | 0.56571199 | NA |
| MDN1 | 27.1283513 | 0.04118568 | 0.41486855 | 0.09927404 | 0.92092069 | 0.98745437 |
| CASP8AP2 | 15.4670073 | -0.4119712 | 0.43485597 | -0.947374 | 0.34344824 | NA |
| BACH2 | 60.4434678 | -0.1256911 | 0.45440157 | -0.276608 | 0.78208109 | 0.96199854 |
| MAP3K7 | 24.5856783 | -0.2888418 | 0.34982129 | -0.825684 | 0.4089834 | NA |
| MANEA | 17.4983552 | 0.10803803 | 0.38851337 | 0.27808059 | 0.7809505 | NA |
| UFL1 | 16.4787745 | 0.21033849 | 0.43889332 | 0.47924742 | 0.63176262 | NA |
| NDUFAF4 | 10.3373374 | -0.982124 | 0.57315236 | -1.7135479 | 0.08661179 | NA |
| KLHL32 | 12.3270608 | 0.46624746 | 0.54517367 | 0.8552274 | 0.39242527 | NA |
| MMS22L | 11.9792333 | 0.60468828 | 0.49874063 | 1.21243036 | 0.22534768 | NA |
| FBXL4 | 26.1168291 | 0.06987291 | 0.4045336 | 0.17272461 | 0.8628679 | 0.97217819 |
| COQ3 | 3.49944656 | -0.794426 | 0.99504071 | -0.7983854 | 0.42464688 | NA |
| PNISR | 79.2502254 | -0.1098946 | 0.23596575 | -0.4657226 | 0.64141404 | 0.92153493 |
| AL513550.1 | 3.20040558 | 0.74137301 | 0.89964712 | 0.8240709 | 0.40989927 | NA |
| USP45 | 16.8130141 | 0.77899794 | 0.40161712 | 1.93965321 | 0.05242185 | NA |
| CCNC | 28.046753 | -0.1320634 | 0.43517762 | -0.3034702 | 0.76153159 | 0.95478238 |
| SIM1 | 55.7951023 | 0.66537571 | 0.37226259 | 1.78738271 | 0.07387565 | 0.53207981 |
| ASCC3 | 59.5003321 | -0.0511641 | 0.33222247 | -0.1540054 | 0.87760549 | 0.9751578 |
| HACE1 | 30.449026 | -1.7606429 | 0.51362771 | -3.4278582 | 0.00060836 | 0.09551302 |
| PREP | 11.1369622 | 0.58528565 | 0.53557127 | 1.09282497 | 0.27447066 | NA |
| ATG5 | 37.3164671 | 0.237113 | 0.30166354 | 0.7860181 | 0.43185689 | 0.84223493 |
| CRYBG1 | 19.2227916 | -0.4401415 | 0.52246517 | -0.8424322 | 0.39954607 | NA |
| RTN4IP1 | 10.8190261 | 0.38200397 | 0.5568201 | 0.68604558 | 0.49268436 | NA |
| QRSL1 | 15.5334561 | -0.1112827 | 0.48417965 | -0.2298376 | 0.81821795 | NA |
| LINC02532 | 54.6253314 | -0.3157699 | 0.5221745 | -0.6047211 | 0.54536434 | 0.88410237 |
| CD24 | 92.9713945 | -0.4109426 | 0.26006481 | -1.5801547 | 1.14E-01 | 5.79E-01 |
| C6orf203 | 10.4114318 | 0.0292078 | 0.51532385 | 0.05667854 | 0.95480127 | NA |
| PDSS2 | 59.92082 | -0.071548 | 0.26849449 | -0.2664785 | 0.78987069 | 0.96325251 |
| SEC63 | 33.0527959 | -0.3367965 | 0.34297422 | -0.9819878 | 0.32610587 | 0.78801502 |
| OSTM1 | 17.0700299 | -0.2852703 | 0.39707126 | -0.718436 | 0.4724885 | NA |
| SNX3 | 30.255748 | 0.21017012 | 0.27966987 | 0.75149362 | 0.45235564 | 0.85557861 |
| AFG1L | 23.6387262 | 0.37256569 | 0.42839342 | 0.86968116 | 0.38447467 | NA |
| FOXO3 | 102.100264 | 0.3437325 | 0.28276904 | 1.21559455 | 0.22413941 | 0.72657496 |
| ARMC2 | 6.6891018 | -0.0467665 | 0.68431073 | -0.068341 | 0.94551421 | NA |
| SESN1 | 45.6379913 | 0.2396919 | 0.32933176 | 0.72781288 | 0.46672814 | 0.85894565 |
| CEP57L1 | 6.74477289 | 0.82741406 | 0.59936967 | 1.38047369 | 0.16744085 | NA |
| CD164 | 41.4001905 | -0.3856271 | 0.34300034 | -1.1242762 | 0.26089587 | 0.75479335 |
| AK9 | 10.6666772 | 1.23306815 | 0.57854533 | 2.13132503 | 0.03306237 | NA |
| FIG4 | 14.9753091 | 0.42712812 | 0.39692988 | 1.07607952 | 0.2818917 | NA |
| WASF1 | 8.08629822 | 0.29257244 | 0.59532164 | 0.49145273 | 0.62310628 | NA |

| | | | | | | |
|---|---|---|---|---|---|---|
| CDC40 | 9.18191523 | 0.01730616 | 0.56691465 | 0.03052692 | 0.97564683 | NA |
| CDK19 | 29.8218431 | 0.55764871 | 0.34711651 | 1.60651738 | 0.10816028 | 0.57769936 |
| AMD1 | 19.5361284 | -0.4867064 | 0.37044608 | -1.3138388 | 0.18890045 | NA |
| GTF3C6 | 15.9708184 | 0.29906726 | 0.42147589 | 0.70957145 | 0.47796993 | NA |
| RPF2 | 5.06587983 | -0.6354708 | 0.74300766 | -0.8552681 | 0.39240272 | NA |
| SLC16A10 | 57.9831761 | 0.40445811 | 0.36318979 | 1.11362743 | 0.26543906 | 0.75497438 |
| MFSD4B | 6.62301235 | -0.2496421 | 0.56276054 | -0.4436027 | 0.65732985 | NA |
| REV3L | 74.9016102 | 0.35987278 | 0.25970405 | 1.3857034 | 0.16583749 | 0.64995633 |
| TRAF3IP2-AS | 23.5791567 | 0.00844191 | 0.44215637 | 0.01909259 | 0.98476724 | NA |
| TRAF3IP2 | 4.35529698 | -0.2440903 | 0.71130022 | -0.3431608 | 0.73147753 | NA |
| TUBE1 | 6.42751363 | 0.61073246 | 0.63522182 | 0.96144755 | 0.33632719 | NA |
| MARCKS | 71.6427305 | -0.3950003 | 0.35608765 | -1.1092783 | 0.26731015 | 0.75497438 |
| HDAC2 | 15.2998715 | 0.07702882 | 0.41756254 | 0.18447255 | 0.85364276 | NA |
| FRK | 63.670624 | 0.24215405 | 0.37336837 | 0.64856604 | 0.51661891 | 0.87158522 |
| NT5DC1 | 57.5365063 | 0.42095527 | 0.30075402 | 1.39966629 | 0.16161327 | 0.64709575 |
| COL10A1 | 2.54939249 | 1.14262573 | 1.26532921 | 0.90302643 | 0.36651187 | NA |
| TSPYL4 | 4.03154923 | 0.09792311 | 0.75893047 | 0.12902778 | 0.89733567 | NA |
| DSE | 12.1356155 | 0.37673744 | 0.48611268 | 0.77500023 | 0.43833952 | NA |
| TSPYL1 | 14.7063138 | 0.52532777 | 0.39479583 | 1.33063152 | 0.18331028 | NA |
| RWDD1 | 10.4375094 | 0.16554138 | 0.62564064 | 0.26459499 | 0.79132148 | NA |
| ZUP1 | 8.00346757 | 0.13212225 | 0.55554208 | 0.23782582 | 0.81201619 | NA |
| KPNA5 | 16.2676763 | -0.0012949 | 0.41050966 | -0.0031544 | 0.99748318 | NA |
| GPRC6A | 20.3889357 | 0.49992644 | 0.56433929 | 0.8858615 | 0.37569217 | NA |
| DCBLD1 | 8.44770659 | -0.0176678 | 0.54749957 | -0.03227 | 0.9742567 | NA |
| GOPC | 15.2155054 | 0.95965558 | 0.40676864 | 2.35921721 | 0.01831353 | NA |
| NUS1 | 21.9366308 | 0.12020444 | 0.40446289 | 0.29719522 | 0.76631747 | NA |
| SLC35F1 | 3.79982874 | -0.2230752 | 0.71857785 | -0.3104399 | 0.75622647 | NA |
| CEP85L | 10.4714584 | 0.0716814 | 0.61215804 | 0.11709622 | 0.9067838 | NA |
| MCM9 | 32.9752322 | 0.10994133 | 0.32606048 | 0.33718078 | 0.73598063 | 0.94590222 |
| AL365275.1 | 10.7365211 | -0.70104 | 0.8535857 | -0.8212883 | 0.41148206 | NA |
| FAM184A | 11.5438479 | 0.1669966 | 0.51326794 | 0.32535949 | 0.74490901 | NA |
| MAN1A1 | 50.8037983 | 0.46288579 | 0.49360426 | 0.937767 | 0.34836416 | 0.8003972 |
| TBC1D32 | 21.0313445 | -0.3761835 | 0.41855142 | -0.8987749 | 0.36877257 | NA |
| HSF2 | 6.24018392 | -0.2886223 | 0.5747059 | -0.5022088 | 0.61552068 | NA |
| SERINC1 | 35.3060551 | -0.0134892 | 0.312011 | -0.0432332 | 0.96551567 | 0.99656976 |
| PKIB | 9.58943115 | 0.60946009 | 0.75195796 | 0.81049756 | 0.41765427 | NA |
| SMPDL3A | 7.52862617 | 0.15756393 | 0.69811896 | 0.22569782 | 0.82143645 | NA |
| NKAIN2 | 5.35253769 | 0.31181668 | 0.80552184 | 0.38709897 | 0.69868294 | NA |
| RNF217 | 65.0828329 | 0.19549822 | 0.41501385 | 0.47106433 | 0.6375948 | 0.92121485 |
| TPD52L1 | 13.0064046 | 1.47077031 | 0.47168587 | 3.11811397 | 0.00182012 | NA |
| HDDC2 | 12.7383013 | 0.3426736 | 0.55410816 | 0.61842366 | 0.5362961 | NA |
| NCOA7 | 268.46308 | -0.5291699 | 0.22528661 | -2.348874 | 0.01883028 | 0.33831016 |
| NCOA7-AS1 | 8.03839537 | 0.53010332 | 0.54313403 | 0.9760083 | 0.32906035 | NA |
| HINT3 | 9.68795396 | -0.2285863 | 0.49659763 | -0.4603049 | 0.64529735 | NA |
| TRMT11 | 13.0911018 | -0.6404832 | 0.43380305 | -1.4764378 | 0.13982641 | NA |
| AC020559.1 | 3.35286212 | 0.83450428 | 0.79961188 | 1.04363667 | 0.29665352 | NA |
| RNF146 | 11.7527304 | 0.96623034 | 0.51664875 | 1.87018811 | 0.0614577 | NA |
| ECHDC1 | 21.0551315 | -0.1846416 | 0.40608121 | -0.4546914 | 0.64933125 | NA |
| PTPRK | 122.802764 | -0.1062128 | 0.28315131 | -0.3751098 | 0.7075788 | 0.93204498 |
| LAMA2 | 112.47931 | -1.6027549 | 1.1224689 | -1.4278835 | 0.15332538 | 0.63864841 |
| AL356124.1 | 21.1479176 | 0.31076001 | 0.50944932 | 0.60999201 | 0.5418671 | NA |
| ARHGAP18 | 157.031596 | -0.3107901 | 0.3269896 | -0.9504588 | 0.34187919 | 0.8003972 |
| L3MBTL3 | 8.9793608 | 0.55480668 | 0.51937168 | 1.06822668 | 0.28541827 | NA |
| EPB41L2 | 10.9561145 | 0.3581603 | 0.61179458 | 0.58542574 | 0.55826148 | NA |

| | | | | | | |
|---|---|---|---|---|---|---|
| AKAP7 | 14.4360131 | 0.10600396 | 0.62064588 | 0.17079621 | 0.86438401 | NA |
| MED23 | 17.3367432 | 0.25123253 | 0.41382363 | 0.6071005 | 0.54378422 | NA |
| ENPP1 | 12.9922962 | -1.1975666 | 0.96211049 | -1.2447288 | 0.21323146 | NA |
| CTGF | 9.94811402 | -1.7954237 | 0.79362379 | -2.2623108 | 0.02367821 | NA |
| STX7 | 32.8121667 | 0.1507022 | 0.3126397 | 0.48203153 | 0.62978355 | 0.91751762 |
| SLC18B1 | 8.70057332 | 0.23442224 | 0.53657565 | 0.43688573 | 0.66219422 | NA |
| RPS12 | 46.1769054 | 0.2028787 | 0.33633136 | 0.60321079 | 0.54636847 | 0.88410237 |
| EYA4 | 43.1988568 | -0.1689887 | 0.43946019 | -0.3845369 | 0.70058052 | 0.9303056 |
| TBPL1 | 7.01940247 | -0.5531149 | 0.65422406 | -0.8454518 | 0.39785865 | NA |
| SGK1 | 5.15938242 | -0.6833239 | 0.67161274 | -1.0174374 | 0.30894539 | NA |
| HBS1L | 16.2380766 | -0.040225 | 0.39022778 | -0.1030807 | 0.91789892 | NA |
| AHI1 | 57.1382358 | 0.09210463 | 0.32538961 | 0.28305954 | 0.7771312 | 0.96133329 |
| LINC00271 | 14.3998121 | 0.49283654 | 0.62576773 | 0.78757104 | 0.43094766 | NA |
| PDE7B | 50.2039284 | -0.5547622 | 0.43036547 | -1.2890489 | 0.19738107 | 0.69343545 |
| AL138828.1 | 7.92041207 | -1.3648963 | 0.73854541 | -1.8480872 | 6.46E-02 | NA |
| BCLAF1 | 38.8051848 | 0.17727677 | 0.33247693 | 0.53320023 | 0.59389498 | 0.89756835 |
| MAP7 | 92.4188782 | -0.1386525 | 0.20238249 | -0.6851014 | 0.49327994 | 0.86619885 |
| MAP3K5 | 133.312829 | 0.21990232 | 0.46746682 | 0.47041268 | 0.63806021 | 0.92121485 |
| PEX7 | 14.6061377 | -0.0633176 | 0.429509 | -0.1474186 | 0.88280164 | NA |
| IFNGR1 | 11.0591441 | 0.17282529 | 0.43002886 | 0.40189229 | 0.68776329 | NA |
| AL357060.1 | 5.95905159 | -1.1540976 | 1.22155482 | -0.9447776 | 0.34477245 | NA |
| TNFAIP3 | 8.06395055 | -1.370488 | 1.14940309 | -1.1923476 | 0.23312497 | NA |
| PERP | 29.1573655 | -0.1183958 | 0.29479855 | -0.4016159 | 0.6879667 | 0.92736198 |
| ARFGEF3 | 9.04319338 | -0.6280827 | 0.55019352 | -1.1415668 | 0.25363412 | NA |
| HEBP2 | 16.4154428 | 0.28197978 | 0.49410486 | 0.57068812 | 0.56821107 | NA |
| NHSL1 | 48.087804 | -1.0491802 | 0.33413678 | -3.1399723 | 0.00168964 | 0.14092639 |
| CCDC28A | 6.57883781 | 0.10768634 | 0.58106351 | 0.18532628 | 0.85297313 | NA |
| REPS1 | 68.7813254 | 0.04861381 | 0.35881858 | 0.13548297 | 0.89223003 | 0.97769824 |
| ABRACL | 3.92779659 | -0.567854 | 0.76460801 | -0.7426734 | 0.45767944 | NA |
| HECA | 8.88692597 | -0.064791 | 0.52373385 | -0.1237098 | 0.90154507 | NA |
| CITED2 | 22.2937431 | -1.9230277 | 0.55593622 | -3.4590797 | 0.00054202 | NA |
| VTA1 | 14.5049668 | -0.1807014 | 0.39412194 | -0.4584912 | 0.64659961 | NA |
| HIVEP2 | 70.2180397 | -0.2120079 | 0.3684726 | -0.5753696 | 0.56504136 | 0.89263108 |
| AIG1 | 36.9782596 | 0.56915504 | 0.47845322 | 1.18957301 | 0.23421426 | 0.73156941 |
| ADAT2 | 4.26132928 | 0.41529294 | 0.83615465 | 0.49667001 | 0.61942177 | NA |
| PEX3 | 10.6502131 | -0.068915 | 0.4782478 | -0.144099 | 0.88542228 | NA |
| FUCA2 | 7.98069063 | -0.6521736 | 0.5861431 | -1.1126526 | 0.26585766 | NA |
| PHACTR2 | 50.7658069 | -0.0583545 | 0.39718043 | -0.1469218 | 0.88319377 | 0.97769824 |
| PLAGL1 | 19.859442 | -0.1062543 | 0.4038603 | -0.2630967 | 0.79247607 | NA |
| SF3B5 | 16.1325494 | 0.08375819 | 0.38379507 | 0.21823674 | 0.82724466 | NA |
| UTRN | 51.6910581 | 0.28790159 | 0.31035772 | 0.92764437 | 0.35359207 | 0.8003972 |
| EPM2A | 11.8828683 | 0.37736741 | 0.60407561 | 0.62470228 | 0.53216648 | NA |
| FBXO30 | 7.34163624 | -0.1214361 | 0.63610672 | -0.1909053 | 0.84859979 | NA |
| AL356599.1 | 7.62758321 | 0.82713294 | 0.6736179 | 1.22789631 | 0.2194859 | NA |
| SHPRH | 46.0443853 | 0.26999483 | 0.30228944 | 0.8931666 | 0.37176796 | 0.81199861 |
| RAB32 | 3.87958809 | -0.0219502 | 0.69694732 | -0.0314948 | 0.9748749 | NA |
| STXBP5-AS1 | 3.1843462 | 1.6537291 | 0.79010087 | 2.09306073 | 0.03634373 | NA |
| STXBP5 | 35.7161351 | 0.4659918 | 0.34435742 | 1.35322133 | 0.17598493 | 0.66719288 |
| SAMD5 | 15.8764771 | -0.4232445 | 0.56063171 | -0.7549421 | 0.45028373 | NA |
| AL033504.1 | 8.03038043 | -0.6203575 | 0.68238315 | -0.9091044 | 0.36329503 | NA |
| SASH1 | 50.1886277 | -0.5143958 | 0.40451392 | -1.2716394 | 0.20350128 | 0.700066 |
| UST | 29.1021074 | -0.7004226 | 0.6502521 | -1.0771555 | 0.28141079 | 0.76854599 |
| TAB2 | 39.9113693 | 0.00339036 | 0.29798414 | 0.01137765 | 0.99092214 | 0.99909657 |
| PPIL4 | 17.662048 | 0.28487297 | 0.39026013 | 0.72995661 | 0.4654167 | NA |

| | | | | | | |
|---|---|---|---|---|---|---|
| GINM1 | 23.1937255 | 0.40037071 | 0.32326424 | 1.23852461 | 0.2155216 | NA |
| KATNA1 | 12.2130455 | -0.931068 | 0.47487518 | -1.9606584 | 0.04991889 | NA |
| LATS1 | 29.4300106 | -0.0718505 | 0.39255901 | -0.1830312 | 0.85477356 | 0.97168963 |
| NUP43 | 8.12401012 | 0.34320431 | 0.55589393 | 0.61739171 | 0.53697639 | NA |
| PCMT1 | 19.6303127 | -0.1749461 | 0.41758741 | -0.4189449 | 0.67525643 | NA |
| LRP11 | 6.40265673 | 1.07736657 | 0.92206548 | 1.1684274 | 0.24263441 | NA |
| PLEKHG1 | 39.2806069 | -0.6471234 | 0.39334784 | -1.6451684 | 0.09993509 | 0.57114936 |
| MTHFD1L | 36.2615099 | -0.2875205 | 0.41268726 | -0.6967031 | 4.86E-01 | 0.86354292 |
| AKAP12 | 6.45418904 | -0.6648073 | 0.80582639 | -0.8250007 | 0.40937119 | NA |
| RMND1 | 10.524325 | -0.6125865 | 0.51898296 | -1.1803597 | 0.2378572 | NA |
| ARMT1 | 7.66102243 | 0.6936948 | 0.57268719 | 1.21129792 | 0.22578125 | NA |
| ESR1 | 33.2088984 | 0.22645164 | 0.67718881 | 0.33439956 | 0.73807807 | 0.94658914 |
| SYNE1 | 37.2373324 | 0.43370396 | 0.38722776 | 1.1200229 | 0.262704 | 0.75479335 |
| MTRF1L | 5.71073166 | -0.6790338 | 0.68077271 | -0.9974458 | 0.31854819 | NA |
| IPCEF1 | 17.3160793 | -0.3274865 | 0.577148 | -0.5674221 | 0.57042744 | NA |
| CNKSR3 | 111.742064 | -0.0264227 | 0.33232294 | -0.0795092 | 0.9366276 | 0.99100968 |
| SCAF8 | 76.3575128 | 0.11971867 | 0.3194684 | 0.37474339 | 0.70785132 | 0.93204498 |
| TIAM2 | 37.593853 | 0.91984648 | 0.32047482 | 2.87026133 | 0.00410133 | 0.21182958 |
| TFB1M | 17.1123065 | -0.2779269 | 0.43148948 | -0.6441105 | 0.5195038 | NA |
| ARID1B | 252.108804 | -0.0922917 | 0.23338663 | -0.3954453 | 0.69251425 | 0.9295338 |
| TMEM242 | 6.79479253 | 0.64926315 | 0.60768307 | 1.06842396 | 0.28532931 | NA |
| ZDHHC14 | 22.8452901 | 0.61305975 | 0.83721417 | 0.73226155 | 0.46400894 | NA |
| SNX9 | 41.7317521 | -0.5198088 | 0.32608958 | -1.5940675 | 0.11092091 | 0.57860396 |
| SYNJ2 | 4.75519765 | 0.14597376 | 0.66813497 | 0.21847945 | 0.82705557 | NA |
| SERAC1 | 5.53651677 | -0.4002184 | 0.70175188 | -0.5703132 | 0.56846526 | NA |
| GTF2H5 | 8.30089863 | 0.36593612 | 0.71618614 | 0.51095114 | 0.60938527 | NA |
| TULP4 | 62.7636874 | -0.4943305 | 0.32180475 | -1.5361193 | 0.12450912 | 0.58553957 |
| TMEM181 | 23.1369199 | -0.5723967 | 0.57650304 | -0.9928772 | 0.32076981 | NA |
| DYNLT1 | 21.6697884 | 0.27335362 | 0.36299869 | 0.753043 | 0.45142408 | NA |
| SYTL3 | 5.54549038 | 0.42408358 | 0.75266985 | 0.56343905 | 0.57313595 | NA |
| EZR | 100.206775 | -0.7999353 | 0.39562881 | -2.0219339 | 0.04318318 | 0.44186191 |
| RSPH3 | 7.26717509 | 0.78372714 | 0.55174797 | 1.42044409 | 0.15547843 | NA |
| AL035530.2 | 5.63078846 | 0.7340482 | 0.7471955 | 0.98240448 | 0.32590062 | NA |
| SOD2.1 | 8.16366399 | 0.1117152 | 0.49966989 | 0.223578 | 0.82308568 | NA |
| WTAP | 34.4995386 | -0.2378646 | 0.33103481 | -0.7185486 | 0.4724191 | 0.86206794 |
| ACAT2 | 7.1501408 | -0.8213702 | 0.64959631 | -1.2644318 | 0.20607509 | NA |
| TCP1 | 13.7243432 | 0.04446909 | 0.47054607 | 0.09450527 | 0.92470779 | NA |
| MRPL18 | 6.63417993 | 0.21108918 | 0.5344182 | 0.39498876 | 0.69285118 | NA |
| IGF2R | 79.487731 | -0.6247277 | 0.32182095 | -1.9412276 | 0.05223068 | 0.48171918 |
| SLC22A1 | 2.60462278 | -1.3521413 | 1.0511069 | -1.2863975 | 0.19830437 | NA |
| MAP3K4 | 26.7269925 | -0.0944267 | 0.35185572 | -0.2683678 | 0.78841624 | 0.96325251 |
| PRKN | 262.805888 | 0.67031106 | 0.32584591 | 2.05714125 | 0.03967264 | 0.43117507 |
| PACRG | 567.225742 | 0.15681302 | 0.41066927 | 0.38184746 | 0.70257451 | 0.9303056 |
| PACRG-AS2 | 3.77347173 | 1.10609802 | 1.03132438 | 1.07250254 | 0.28349437 | NA |
| PACRG-AS3 | 13.4636694 | 0.98089883 | 0.71013047 | 1.38129381 | 0.16718864 | NA |
| QKI | 162.856049 | -0.8238454 | 0.27298397 | -3.0179257 | 0.00254511 | 0.16982264 |
| PDE10A | 50.95315 | -0.9034906 | 0.54135349 | -1.6689475 | 0.09512778 | 0.56358504 |
| SFT2D1 | 16.1089853 | 0.11082448 | 0.39833759 | 0.27821749 | 0.78084541 | NA |
| MPC1 | 51.403114 | -0.1013775 | 0.34913575 | -0.2903671 | 0.77153541 | 0.95994327 |
| RPS6KA2 | 26.8485552 | -0.0878986 | 0.39769885 | -0.221018 | 0.8250784 | 0.96845695 |
| Z94721.1 | 10.7917264 | -0.992508 | 0.5926303 | -1.6747506 | 0.09398317 | NA |
| FGFR1OP | 23.4699097 | 0.08422718 | 0.39785992 | 0.21170058 | 0.83234063 | NA |
| AFDN | 114.082426 | 0.14207801 | 0.23261463 | 0.61078708 | 0.54134055 | 0.88410237 |
| WDR27 | 25.8619243 | 0.78554273 | 0.4233234 | 1.85565628 | 0.06350257 | NA |

| | | | | | | |
|---|---|---|---|---|---|---|
| C6orf120 | 2.21393596 | 0.20967343 | 0.9619585 | 0.21796515 | 0.82745626 | NA |
| PHF10 | 16.1701995 | 0.15946689 | 0.43419519 | 0.36727004 | 0.7134176 | NA |
| ERMARD | 5.7874688 | 0.20407104 | 0.67153889 | 0.30388566 | 0.76121501 | NA |
| DLL1 | 41.1534865 | -1.2385591 | 0.52018694 | -2.3809884 | 0.01726625 | 0.32722884 |
| FAM120B | 93.6887502 | -0.1846112 | 0.33530132 | -0.550583 | 0.5819196 | 0.89263108 |
| PSMB1 | 24.830695 | 0.20095017 | 0.45206951 | 0.44451167 | 0.6566727 | NA |
| TBP | 6.17792749 | 0.4764198 | 0.61457227 | 0.77520549 | 0.43821824 | NA |
| PDCD2 | 7.9238921 | 0.71792949 | 0.54110172 | 1.32679211 | 0.18457746 | NA |
| FAM20C | 24.1656503 | -0.0300114 | 0.50820621 | -0.0590535 | 0.95290946 | NA |
| PRKAR1B | 8.27318794 | -0.2996844 | 0.61687751 | -0.4858086 | 0.62710289 | NA |
| DNAAF5 | 6.99218785 | -0.1393415 | 0.77014325 | -0.1809293 | 0.85642306 | NA |
| SUN1 | 24.7742502 | 0.09187089 | 0.32430466 | 0.28328576 | 0.77695779 | NA |
| ADAP1 | 3.81760917 | -0.7162991 | 0.81567195 | -0.8781705 | 0.37985117 | NA |
| COX19 | 13.8747322 | -0.1673014 | 0.42262373 | -0.3958637 | 0.69220556 | NA |
| C7orf50 | 22.1759451 | -0.3912809 | 0.37145554 | -1.0533722 | 0.29217043 | NA |
| GPR146 | 3.70236993 | -0.3051672 | 0.99964871 | -0.3052744 | 0.76015715 | NA |
| MICALL2 | 3.53199403 | 0.01691684 | 0.73503264 | 0.02301508 | 0.98163824 | NA |
| INTS1 | 11.7149133 | 0.3523436 | 0.66718929 | 0.5281014 | 0.59742896 | NA |
| MAFK | 5.66085938 | -0.9511258 | 0.72160741 | -1.3180655 | 0.18748172 | NA |
| PSMG3 | 4.32332161 | -1.0157214 | 0.83427662 | -1.2174876 | 0.22341877 | NA |
| MAD1L1 | 15.3916276 | -0.1054866 | 0.50399317 | -0.2093016 | 0.83421277 | NA |
| MRM2 | 3.4779031 | 1.31765943 | 0.82527672 | 1.59662741 | 0.11034879 | NA |
| SNX8 | 6.25552463 | -1.6872637 | 0.72939485 | -2.3132378 | 0.02070957 | NA |
| EIF3B | 16.5139004 | -0.8690056 | 0.49826447 | -1.744065 | 0.08114776 | NA |
| CHST12 | 9.91055613 | -0.535029 | 0.53286099 | -1.0040687 | 0.3153455 | NA |
| BRAT1 | 3.31140173 | 1.43807445 | 0.84979974 | 1.69225099 | 0.09059813 | NA |
| IQCE | 5.40800487 | 1.84580575 | 0.79276936 | 2.32830106 | 0.01989612 | NA |
| GNA12 | 16.25417 | -0.1286834 | 0.38912672 | -0.330698 | 0.74087261 | NA |
| SDK1 | 71.995973 | 0.28965744 | 0.34440901 | 0.84102747 | 0.40033255 | 0.8269015 |
| FOXK1 | 37.6540284 | 0.13750421 | 0.40560059 | 0.33901383 | 0.73459931 | 0.94580104 |
| AP5Z1 | 6.44324254 | -1.0564716 | 0.67105449 | -1.5743455 | 0.11540759 | NA |
| RBAK-RBAKD | 15.7921745 | -1.011285 | 0.46247067 | -2.1867008 | 0.02876438 | NA |
| WIPI2 | 18.6559604 | 0.23648551 | 0.40318215 | 0.58654758 | 0.5575076 | NA |
| SLC29A4 | 2.76157528 | 0.76490432 | 1.09342 | 0.69955216 | 0.48420703 | NA |
| TNRC18 | 49.8825806 | -0.1212387 | 0.39493747 | -0.3069819 | 0.75885713 | 0.9535733 |
| FBXL18 | 4.52109203 | 0.12400294 | 0.83892556 | 0.14781161 | 0.88249144 | NA |
| ACTB | 224.783335 | -0.4727365 | 0.39062972 | -1.2101908 | 0.22620567 | 0.73003982 |
| RNF216 | 33.0097144 | -0.1878468 | 0.31104444 | -0.6039227 | 0.54589503 | 0.88410237 |
| CCZ1 | 7.12610826 | 0.25769598 | 0.68643697 | 0.37541099 | 0.70735483 | NA |
| PMS2 | 11.9156751 | -0.0289272 | 0.61843585 | -0.0467748 | 0.96269275 | NA |
| AIMP2 | 11.8272665 | 0.38866369 | 0.54353969 | 0.71506036 | 0.47457175 | NA |
| EIF2AK1 | 22.3353633 | 0.02677494 | 0.39800505 | 0.06727285 | 0.94636449 | NA |
| USP42 | 21.9858645 | -0.0865029 | 0.38367887 | -0.2254564 | 0.82162424 | NA |
| CYTH3 | 20.6091749 | -0.2748728 | 0.38738307 | -0.7095634 | 0.47797492 | NA |
| RAC1 | 44.936901 | -0.1160685 | 0.27364014 | -0.4241648 | 0.67144564 | 0.92614389 |
| DAGLB | 4.27217602 | -0.2197741 | 0.7266407 | -0.3024522 | 0.76230736 | NA |
| ZDHHC4 | 10.3649949 | 0.20219211 | 0.45427694 | 0.44508557 | 0.65625792 | NA |
| C7orf26 | 6.84669346 | -0.387582 | 0.58274146 | -0.6651011 | 0.50598584 | NA |
| ZNF316 | 7.21088167 | -0.3879294 | 0.72050886 | -0.5384104 | 0.59029378 | NA |
| AC073343.2 | 4.79745604 | 0.00599168 | 0.6508298 | 0.00920621 | 0.99265461 | NA |
| ZNF12 | 11.1049425 | -1.1078775 | 0.52397203 | -2.1143829 | 0.03448258 | NA |
| CCZ1B | 8.11094785 | -0.6109811 | 0.53414674 | -1.1438451 | 0.25268787 | NA |
| C1GALT1 | 31.513374 | 0.03387551 | 0.3444113 | 0.09835772 | 0.92164824 | 0.98745437 |
| COL28A1 | 13.3640923 | -0.6534421 | 0.56548001 | -1.1555529 | 0.24786408 | NA |

| | | | | | | |
|---|---|---|---|---|---|---|
| AC004982.2 | 3.64657196 | -0.6906048 | 0.97859976 | -0.7057071 | 0.48037028 | NA |
| MIOS | 13.5498676 | -0.4425041 | 0.47942064 | -0.9229976 | 0.35600845 | NA |
| RPA3 | 9.62828861 | 0.47932815 | 0.59221688 | 0.80937941 | 0.41829694 | NA |
| UMAD1 | 81.929818 | 0.09667211 | 0.24259153 | 0.39849745 | 0.69026353 | 0.92830727 |
| GLCCI1 | 100.771587 | 0.13369957 | 0.51883422 | 0.25769228 | 0.7966444 | 0.96525806 |
| ICA1 | 49.3107097 | -0.0693889 | 0.35951152 | -0.1930088 | 0.84695206 | 0.97168963 |
| NDUFA4 | 57.2141979 | 0.24192424 | 0.45272781 | 0.53437018 | 0.59308545 | 0.89685273 |
| PHF14 | 41.4762437 | -0.0577384 | 0.35758474 | -0.1614678 | 0.87172496 | 0.97488194 |
| THSD7A | 174.071389 | -0.7008335 | 0.45086018 | -1.5544365 | 0.12008034 | 0.58383225 |
| TMEM106B | 18.6332674 | 0.08171506 | 0.44729683 | 0.18268643 | 0.85504406 | NA |
| SCIN | 249.02282 | 0.4559983 | 0.29878252 | 1.52618798 | 0.12696306 | 0.59256732 |
| ARL4A | 4.93838858 | -1.4500367 | 0.84685848 | -1.7122539 | 0.0868499 | NA |
| AC011287.1 | 4.99454636 | 0.14589417 | 0.90710971 | 0.1608341 | 0.87222407 | NA |
| ETV1 | 10.2419803 | -1.0545897 | 0.84371966 | -1.249929 | 0.21132547 | NA |
| DGKB | 56.1284886 | -1.0317282 | 1.24625137 | -0.8278653 | 0.40774681 | 0.82808596 |
| AGMO | 5.31014616 | 0.73250633 | 0.94758571 | 0.77302383 | 0.43950828 | NA |
| ISPD | 34.380763 | -0.3067346 | 0.51701801 | -0.5932765 | 0.55299613 | 0.88807709 |
| ANKMY2 | 9.15289028 | 0.28607316 | 0.54073092 | 0.52904902 | 0.59677145 | NA |
| BZW2 | 8.71156139 | -0.1789118 | 0.52371687 | -0.3416193 | 0.73263737 | NA |
| AC073332.1 | 4.66132768 | 1.24925681 | 0.67157443 | 1.86019114 | 0.06285849 | NA |
| SNX13 | 48.4160473 | -0.0907935 | 0.27942155 | -0.3249339 | 0.74523112 | 0.94858421 |
| HDAC9 | 15.9334676 | -0.278403 | 0.87591582 | -0.3178421 | 7.51E-01 | NA |
| TWISTNB | 4.93748035 | 0.39216254 | 0.63876532 | 0.61393837 | 0.53925606 | NA |
| MACC1 | 32.5024725 | 0.53358295 | 0.4371748 | 1.22052542 | 0.22226576 | 0.72433127 |
| ITGB8 | 29.2920092 | 0.488276 | 0.51777941 | 0.94301934 | 0.34567101 | 0.8003972 |
| SP4 | 16.2026163 | -0.7294387 | 0.49021536 | -1.4879964 | 0.13675184 | NA |
| DNAH11 | 39.2162998 | -3.1926875 | 1.21995813 | -2.6170467 | 0.00886942 | 0.27648451 |
| RAPGEF5 | 6.85084651 | 0.09405123 | 0.59433737 | 0.15824553 | 8.74E-01 | NA |
| STEAP1B | 49.9005444 | -0.0024511 | 0.38298905 | -0.0063999 | 0.99489363 | 0.9992989 |
| TOMM7 | 24.40134 | 0.26586413 | 0.47132007 | 0.56408405 | 0.57269693 | NA |
| FAM126A | 3.65910599 | -0.3468598 | 0.79392921 | -0.4368901 | 0.66219103 | NA |
| KLHL7 | 11.0290219 | -0.0934276 | 0.44679972 | -0.209104 | 0.83436702 | NA |
| NUPL2 | 13.73743 | 0.4338974 | 0.44294233 | 0.9795799 | 0.32729353 | NA |
| MALSU1 | 9.05111196 | 0.35828786 | 0.62305246 | 0.57505248 | 0.5652558 | NA |
| TRA2A | 68.1186568 | 0.03249525 | 0.37223069 | 0.08729868 | 0.93043411 | 0.98921679 |
| CCDC126 | 13.2100434 | -0.1764458 | 0.45355156 | -0.3890313 | 0.69725296 | NA |
| FAM221A | 9.69155142 | -0.5812716 | 0.55777781 | -1.0421202 | 0.29735593 | NA |
| MPP6 | 10.6441708 | -0.8316578 | 0.83726533 | -0.9933025 | 0.32056255 | NA |
| OSBPL3 | 112.807628 | 0.31097315 | 0.43943406 | 0.70766738 | 0.47915184 | 0.86206794 |
| CYCS | 33.5761795 | -0.6199623 | 0.42371828 | -1.4631474 | 0.14342706 | 0.62448095 |
| HNRNPA2B1 | 108.139148 | -0.0294976 | 0.22884436 | -0.128898 | 0.8974384 | 0.97769824 |
| CBX3 | 18.3441745 | -0.5663266 | 0.40882286 | -1.3852615 | 0.16597253 | NA |
| SNX10 | 14.6317385 | 0.46023841 | 0.53946825 | 0.85313344 | 0.39358531 | NA |
| AC004540.1 | 10.785956 | 1.26894732 | 0.67177068 | 1.88895908 | 0.05889731 | NA |
| SKAP2 | 29.329983 | 0.77696047 | 0.36188767 | 2.14696583 | 0.031796 | 0.4126665 |
| HOXA3 | 4.29027207 | -0.289968 | 0.83019089 | -0.3492787 | 0.72688007 | NA |
| HOXA9 | 3.33200574 | 1.03716537 | 0.87560649 | 1.18451083 | 0.23621091 | NA |
| HIBADH | 53.0579708 | 0.75436409 | 0.32488994 | 2.3219066 | 0.02023797 | 0.34471691 |
| TAX1BP1 | 79.2601418 | 0.12353554 | 0.26089364 | 0.47350923 | 0.63584992 | 0.92121485 |
| JAZF1 | 11.3816969 | 0.85833071 | 0.53082496 | 1.61697503 | 0.10588367 | NA |
| CPVL | 7.89414612 | -0.1955191 | 0.76074677 | -0.2570095 | 0.79717146 | NA |
| SCRN1 | 24.4672098 | 0.13546995 | 0.35490472 | 0.38170796 | 0.702678 | NA |
| FKBP14 | 4.68784286 | -0.349534 | 0.73321835 | -0.476712 | 0.63356719 | NA |
| PLEKHA8 | 28.5616358 | -0.6432604 | 0.36217077 | -1.7761247 | 0.07571237 | 0.53887019 |

| | | | | | | |
|---|---|---|---|---|---|---|
| MTURN | 31.5753173 | -0.0079314 | 0.36913889 | -0.0214863 | 0.98285775 | 0.99902646 |
| ZNRF2 | 16.8230698 | 0.67898505 | 0.42329033 | 1.60406462 | 0.10869981 | NA |
| AC006978.2 | 15.6862882 | 0.09860493 | 0.50390079 | 0.19568323 | 0.84485811 | NA |
| AC005154.6 | 8.23804453 | 0.65091411 | 0.58676918 | 1.10931885 | 0.26729266 | NA |
| GARS | 16.9628003 | -0.9767654 | 0.42531662 | -2.2965607 | 0.02164384 | NA |
| AQP1 | 7.87406643 | -0.0418275 | 0.71224963 | -0.0587259 | 0.95317045 | NA |
| CCDC129 | 4.13460426 | 1.13606033 | 0.77004318 | 1.47532029 | 0.14012645 | NA |
| PDE1C | 538.087515 | -0.6064325 | 0.24056585 | -2.5208586 | 0.01170689 | 0.28665769 |
| LSM5 | 5.93310699 | -0.118685 | 0.63754172 | -0.1861604 | 0.85231897 | NA |
| AVL9 | 41.8931843 | 0.08553252 | 0.270489 | 0.3162144 | 0.75183978 | 0.95164077 |
| KBTBD2 | 17.9294202 | 0.30405878 | 0.60835632 | 0.49980376 | 0.61721326 | NA |
| FKBP9 | 5.25047863 | 0.26253223 | 0.841324 | 0.31204652 | 0.75500517 | NA |
| NT5C3A | 16.1005974 | 0.19309932 | 0.51738224 | 0.37322372 | 0.70898194 | NA |
| RP9 | 2.99321974 | -0.6780424 | 0.86168441 | -0.7868802 | 0.43135202 | NA |
| BBS9 | 36.6113668 | 0.36614465 | 0.36342596 | 1.00748073 | 0.31370381 | 0.78031266 |
| DPY19L1 | 20.5953931 | -0.0873004 | 0.41264943 | -0.2115608 | 0.83244972 | NA |
| HERPUD2 | 15.18731 | -0.1719076 | 0.41423777 | -0.4149975 | 0.67814375 | NA |
| SEPT7-AS1 | 3.47466409 | 1.34781039 | 0.78165831 | 1.72429612 | 0.0846544 | NA |
| 7-Sep | 48.5582275 | -0.155442 | 0.28382665 | -0.5476651 | 0.58392187 | 0.8933319 |
| EEPD1 | 16.9232199 | 1.38227566 | 0.58557653 | 2.36053803 | 0.01824845 | NA |
| KIAA0895 | 7.70777632 | -0.3206563 | 0.60655094 | -0.5286552 | 0.59704468 | NA |
| EPDR1 | 4.8466083 | 0.17969401 | 0.78435405 | 0.22909808 | 0.81879269 | NA |
| STARD3NL | 10.8184502 | -0.5400946 | 0.47172621 | -1.1449324 | 0.25223715 | NA |
| TRGC2 | 2.57162322 | -1.5778668 | 1.08442933 | -1.4550204 | 0.1456636 | NA |
| TRGC1 | 21.9607106 | -1.066186 | 0.59567461 | -1.7898799 | 0.07347322 | NA |
| VPS41 | 46.2755875 | 0.18867069 | 0.33917824 | 0.55625823 | 0.57803434 | 0.89263108 |
| YAE1D1 | 3.85571541 | 0.32198137 | 0.71670392 | 0.44925297 | 0.65324918 | NA |
| RALA | 38.0171244 | -0.540059 | 0.3626077 | -1.4893754 | 0.13638854 | 0.60594834 |
| LINC00265 | 6.09741933 | 0.20024977 | 0.64949403 | 0.30831657 | 0.75784146 | NA |
| CDK13 | 103.805609 | 0.11293422 | 0.20101842 | 0.56181029 | 0.57424528 | 0.89263108 |
| MPLKIP | 9.14599223 | 0.86837271 | 0.49134059 | 1.76735388 | 0.07716898 | NA |
| SUGCT | 19.3666333 | 0.22796319 | 0.66503909 | 0.34278164 | 0.73176274 | NA |
| MRPL32 | 9.44051035 | 0.59090511 | 0.46890801 | 1.26017279 | 0.20760704 | NA |
| STK17A | 6.73984932 | 0.1323085 | 0.55073712 | 0.24023894 | 0.81014503 | NA |
| COA1 | 41.3985491 | 0.4622776 | 0.29912581 | 1.54542868 | 0.12224261 | 0.58553957 |
| BLVRA | 5.74523249 | 0.33530535 | 0.55313579 | 0.60618994 | 0.54438863 | NA |
| URGCP | 5.65956178 | 0.56412539 | 0.82708247 | 0.68206668 | 0.49519679 | NA |
| UBE2D4 | 4.74409808 | -0.1274728 | 0.72549097 | -0.1757055 | 0.86052529 | NA |
| AC004951.1 | 2.37043292 | 0.12023761 | 1.08843121 | 0.11046873 | 0.91203765 | NA |
| DBNL | 9.50035005 | 0.6669561 | 0.49326657 | 1.35212103 | 0.1763366 | NA |
| POLM | 6.34543146 | 0.43170836 | 0.61414153 | 0.70294604 | 0.48208938 | NA |
| POLD2 | 3.06430749 | 0.63757664 | 0.82946492 | 0.76866017 | 0.44209508 | NA |
| YKT6 | 6.13244477 | -0.5976348 | 0.77510849 | -0.7710337 | 0.44068693 | NA |
| NUDCD3 | 35.6282478 | -0.2382457 | 0.3039487 | -0.7838352 | 0.43313681 | 0.84223493 |
| DDX56 | 8.35158451 | -0.3924965 | 0.56712467 | -0.6920815 | 0.48888617 | NA |
| TMED4 | 21.0465321 | 0.03009248 | 0.35244308 | 0.08538252 | 0.93195729 | NA |
| OGDH | 73.0051501 | -0.0981178 | 0.27161822 | -0.3612344 | 0.71792423 | 0.93714576 |
| ZMIZ2 | 14.1380644 | -0.2623695 | 0.43335254 | -0.6054413 | 0.54488582 | NA |
| PPIA | 83.4956581 | -0.3564918 | 0.42039883 | -0.8479848 | 0.39644645 | 0.82600747 |
| H2AFV | 29.5629062 | 0.00385348 | 0.3291356 | 0.01170789 | 0.99065867 | 0.99909657 |
| PURB | 14.5314846 | -0.5237026 | 0.46135193 | -1.1351477 | 0.25631343 | NA |
| SNHG15 | 4.11633826 | -2.1418767 | 0.89691161 | -2.3880578 | 0.01693768 | NA |
| CCM2 | 24.4435754 | -0.0437419 | 0.35814465 | -0.1221347 | 0.90279237 | NA |
| TBRG4 | 5.68613428 | 0.06787399 | 0.69462786 | 0.09771274 | 0.9221604 | NA |

| | | | | | | |
|---|---|---|---|---|---|---|
| ADCY1 | 33.162242 | 1.79653296 | 1.1409801 | 1.57455241 | 0.1153598 | 0.57860396 |
| TNS3 | 156.15424 | 0.42837896 | 0.45118886 | 0.9494449 | 0.34239438 | 0.8003972 |
| HUS1 | 7.63486686 | 0.22339455 | 0.5843192 | 0.38231594 | 0.70222703 | NA |
| UPP1 | 17.6122365 | -0.7800861 | 0.63247888 | -1.2333789 | 0.21743442 | NA |
| GRB10 | 115.598542 | -0.6050745 | 0.31698088 | -1.9088674 | 0.0562792 | 0.503587 |
| COBL | 39.2665031 | 0.13409363 | 0.41338838 | 0.32437687 | 0.74565273 | 0.94858421 |
| SEC61G | 8.83525694 | -0.0799369 | 0.55395469 | -0.1443022 | 0.88526182 | NA |
| EGFR | 225.597299 | 0.05769329 | 0.23766682 | 0.2427486 | 0.80820016 | 0.96845695 |
| LANCL2 | 9.49545436 | -0.6167457 | 0.6160157 | -1.001185 | 0.31673738 | NA |
| VOPP1 | 20.672961 | -0.1125496 | 0.42415538 | -0.26535 | 0.79073986 | NA |
| NIPSNAP2 | 51.4204773 | 0.06285032 | 0.28401206 | 0.22129453 | 0.82486311 | 0.96845695 |
| PSPH | 4.86453713 | 0.44066861 | 0.65537295 | 0.67239365 | 0.50133312 | NA |
| CCT6A | 18.5526041 | 0.12451699 | 0.56736563 | 0.21946517 | 0.82628771 | NA |
| SUMF2 | 12.6327952 | -0.0702876 | 0.4310135 | -0.1630751 | 0.87045929 | NA |
| PHKG1 | 4.13024192 | -0.4913719 | 0.69143658 | -0.7106537 | 0.47729889 | NA |
| CHCHD2 | 52.3603531 | 0.23267775 | 0.48972965 | 0.4751147 | 0.63470522 | 0.92072524 |
| NUPR2 | 5.884761 | 0.39487579 | 0.66316419 | 0.59544197 | 0.55154808 | NA |
| ZNF736 | 10.638851 | 1.00077809 | 0.55392746 | 1.80669521 | 0.07080981 | NA |
| ZNF680 | 17.2080418 | 0.33685716 | 0.42355697 | 0.79530543 | 0.42643586 | NA |
| ZNF107 | 3.56356887 | 0.1898166 | 0.83276095 | 0.22793649 | 0.81969562 | NA |
| ZNF138 | 9.33369696 | 0.70518948 | 0.48137503 | 1.46494819 | 0.14293505 | NA |
| ZNF117 | 5.67124753 | 0.49074411 | 0.58459116 | 0.8394655 | 0.40120814 | NA |
| VKORC1L1 | 28.0583089 | -0.228977 | 0.35140672 | -0.6516012 | 0.51465845 | 0.87158522 |
| GUSB | 5.12419162 | -0.8248049 | 0.70574699 | -1.1686978 | 0.24252542 | NA |
| AC068533.4 | 3.25007083 | -0.5024586 | 0.93144282 | -0.5394411 | 0.5895825 | NA |
| CRCP | 4.95202864 | -0.5770587 | 0.7016991 | -0.8223735 | 0.41086436 | NA |
| AC068533.3 | 4.2938841 | 0.88077487 | 0.74472711 | 1.18268135 | 0.23693546 | NA |
| TPST1 | 73.5395276 | -0.1756478 | 0.23970373 | -0.7327704 | 0.46369849 | 0.85894565 |
| AC027644.4 | 16.7735051 | -0.1403777 | 0.44019258 | -0.3189006 | 0.74980186 | NA |
| AC027644.3 | 6.3767741 | 1.03240035 | 0.60923239 | 1.69459203 | 0.09015284 | NA |
| TMEM248 | 21.1163996 | -0.3897547 | 0.39510657 | -0.9864547 | 0.32391004 | NA |
| SBDS | 16.4779762 | -0.1283289 | 0.44231898 | -0.2901276 | 0.77171863 | NA |
| TYW1 | 33.1059831 | -0.0866722 | 0.30920454 | -0.280307 | 0.77924197 | 0.96139889 |
| AC006480.2 | 3.13207723 | 0.28404196 | 0.81659643 | 0.34783639 | 0.72796306 | NA |
| AUTS2 | 56.5596024 | -0.9562019 | 0.37817948 | -2.5284342 | 0.01145726 | 0.28578895 |
| GALNT17 | 91.1930712 | 1.77921313 | 1.15688885 | 1.53792918 | 0.12406593 | 0.58553957 |
| CALN1 | 5.23965104 | 0.65213856 | 0.82750458 | 0.78807849 | 0.43065079 | NA |
| TYW1B | 25.1200212 | 0.28434808 | 0.39205106 | 0.72528329 | 0.46827826 | NA |
| POM121 | 25.8915893 | -0.7523755 | 0.38412213 | -1.9586883 | 0.0501493 | NA |
| NSUN5 | 7.41067699 | 0.47467487 | 0.66965947 | 0.70883022 | 0.47842984 | NA |
| BAZ1B | 32.3531145 | -0.7992463 | 0.32128872 | -2.4876264 | 0.01285988 | 0.30435125 |
| BCL7B | 8.43204388 | -0.8680357 | 0.57180922 | -1.5180512 | 0.1290015 | NA |
| TBL2 | 3.15989706 | -0.9906105 | 0.84750512 | -1.1688548 | 0.24246212 | NA |
| MLXIPL | 7.32533441 | -0.6809425 | 0.70778396 | -0.9620768 | 0.33601104 | NA |
| VPS37D | 6.03819362 | 0.36123441 | 0.60620418 | 0.59589562 | 0.55124496 | NA |
| DNAJC30 | 2.92123296 | 0.13138785 | 1.0262099 | 0.12803214 | 0.89812354 | NA |
| BUD23 | 12.5243552 | -0.4851394 | 0.47815774 | -1.0146012 | 0.31029596 | NA |
| ABHD11 | 5.18647942 | 0.37112607 | 0.78856913 | 0.47063225 | 0.63790337 | NA |
| CLDN3 | 7.07316301 | 0.35312141 | 0.60193548 | 0.5866433 | 0.5574433 | NA |
| CLDN4 | 21.1147484 | 0.76260178 | 0.44005233 | 1.73297976 | 0.08309927 | NA |
| LIMK1 | 3.09548554 | 1.33398842 | 0.91406241 | 1.45940627 | 0.14445332 | NA |
| EIF4H | 17.7963575 | -0.1400273 | 0.42285198 | -0.3311496 | 0.74053146 | NA |
| RFC2 | 3.54141843 | -0.7055673 | 1.00057593 | -0.7051611 | 0.48070996 | NA |
| CLIP2 | 5.56041059 | 0.17304263 | 0.66894232 | 0.25868094 | 0.79588143 | NA |

| | | | | | | |
|---|---|---|---|---|---|---|
| GTF2IRD1 | 41.4072697 | -0.3703412 | 0.47814326 | -0.7745403 | 0.43861135 | 0.84462749 |
| GTF2I | 87.3722714 | 0.30754325 | 0.24765662 | 1.24181315 | 0.21430551 | 0.71319378 |
| GTF2IRD2 | 10.8190396 | -0.3578212 | 0.63937477 | -0.5596423 | 0.57572344 | NA |
| CASTOR2 | 16.9261601 | 0.31030686 | 0.50898512 | 0.60965802 | 0.54208837 | NA |
| RCC1L | 13.7984331 | -0.2528992 | 0.43466897 | -0.5818203 | 0.56068774 | NA |
| GTF2IRD2B | 17.8555259 | -0.7660032 | 0.4343685 | -1.7634869 | 0.0778184 | NA |
| TRIM73 | 9.2870213 | 0.23689516 | 0.5870902 | 0.40350727 | 0.68657509 | NA |
| POM121C | 22.0503503 | -0.7217955 | 0.41358899 | -1.7452001 | 0.08095004 | NA |
| SPDYE5 | 5.63802073 | 0.11912645 | 0.62095052 | 0.19184531 | 0.84786338 | NA |
| HIP1 | 29.0725642 | -0.0204841 | 0.48631327 | -0.0421212 | 0.96640209 | 0.99656976 |
| RHBDD2 | 6.94326857 | 0.28656936 | 0.83552342 | 0.34298184 | 0.73161212 | NA |
| POR | 11.1388162 | -0.8070672 | 0.47828501 | -1.6874189 | 0.09152284 | NA |
| TMEM120A | 3.97078582 | 1.05563685 | 0.75521949 | 1.39778815 | 0.16217669 | NA |
| STYXL1 | 10.0278254 | 0.01598179 | 0.50879952 | 0.03141078 | 0.97494194 | NA |
| MDH2 | 16.8812272 | 0.36779014 | 0.56661092 | 0.64910528 | 0.51627033 | NA |
| HSPB1 | 35.8682262 | -0.1440562 | 0.44822148 | -0.3213952 | 0.74791091 | 0.94920315 |
| YWHAG | 17.5423385 | -0.7250054 | 0.38001539 | -1.9078316 | 0.05641299 | NA |
| DTX2 | 8.92985897 | -0.1910803 | 0.53773547 | -0.3553425 | 0.72233302 | NA |
| POMZP3 | 5.66888332 | -0.3028168 | 0.7223726 | -0.4191976 | 0.67507176 | NA |
| CCDC146 | 14.5817021 | 0.77842075 | 0.60112568 | 1.29493845 | 0.19534146 | NA |
| FGL2 | 7.74082743 | 0.00063431 | 0.8679877 | 0.00073078 | 0.99941692 | NA |
| GSAP | 12.3008201 | 0.83514781 | 0.55454526 | 1.50600476 | 0.13206595 | NA |
| PTPN12 | 36.5529505 | -0.0307002 | 0.31912969 | -0.0961998 | 0.92336192 | 0.98758497 |
| APTR | 3.60320836 | -0.1312278 | 0.75210124 | -0.1744815 | 0.86148706 | NA |
| RSBN1L | 17.4558178 | -0.1835144 | 0.44180724 | -0.4153721 | 0.6778695 | NA |
| PHTF2 | 12.2396176 | -0.0499693 | 0.49814502 | -0.1003108 | 0.92009756 | NA |
| MAGI2 | 203.929654 | -0.706654 | 0.43594636 | -1.6209656 | 0.10502503 | 0.5772605 |
| GNAI1 | 11.2124725 | -0.2941434 | 0.58553641 | -0.5023486 | 0.61542231 | NA |
| SEMA3C | 182.577094 | -0.8933733 | 0.44520514 | -2.0066554 | 0.04478637 | 0.44769598 |
| CACNA2D1 | 29.0626073 | -0.5129044 | 0.41509805 | -1.2356222 | 0.21659903 | 0.71673468 |
| PCLO | 55.4653029 | 0.24203866 | 0.32880256 | 0.73612158 | 0.46165671 | 0.85894565 |
| KIAA1324L | 7.31852984 | 0.36021796 | 0.77773424 | 0.4631633 | 0.64324732 | NA |
| DMTF1 | 31.9841767 | 0.21973417 | 0.37038707 | 0.59325552 | 0.55301016 | 0.88807709 |
| TMEM243 | 12.1136772 | 0.34324167 | 0.45905498 | 0.74771364 | 0.4546329 | NA |
| CROT | 5.82475123 | 0.62856888 | 0.64350203 | 0.97679394 | 0.32867118 | NA |
| ABCB1 | 6.8851692 | -0.2594622 | 0.79621427 | -0.3258698 | 0.74452286 | NA |
| SLC25A40 | 12.299083 | 0.58957929 | 0.50657812 | 1.16384674 | 0.24448613 | NA |
| DBF4 | 3.88925416 | -0.9640053 | 0.81915099 | -1.1768347 | 0.23926148 | NA |
| ADAM22 | 28.8095902 | 0.92964209 | 0.45273519 | 2.05339039 | 0.04003473 | 0.43121698 |
| SRI | 20.3682795 | 0.72069814 | 0.37549451 | 1.91933068 | 0.0549425 | NA |
| CFAP69 | 4.89164669 | 0.89402343 | 0.80295201 | 1.11342076 | 0.26552777 | NA |
| GTPBP10 | 11.3874829 | -0.030846 | 0.51074984 | -0.0603936 | 0.95184212 | NA |
| CLDN12 | 11.0736526 | -0.6199333 | 0.48661232 | -1.2739778 | 0.20267129 | NA |
| CDK14 | 53.4060999 | 0.3340187 | 0.33326566 | 1.00225958 | 0.31621824 | 0.780966 |
| AC079760.1 | 8.92010278 | 1.99967889 | 0.64995111 | 3.07666049 | 0.00209334 | NA |
| MTERF1 | 37.3647316 | -0.2762663 | 0.39338021 | -0.7022883 | 0.48249941 | 0.86255253 |
| AKAP9 | 117.397599 | 0.05085752 | 0.20626488 | 0.24656411 | 0.80524558 | 0.96800006 |
| KRIT1 | 35.162733 | -0.258194 | 0.310683 | -0.8310528 | 0.40594378 | 0.82808596 |
| ANKIB1 | 54.8111808 | -0.2591421 | 0.23949335 | -1.0820428 | 0.2792335 | 0.76753266 |
| GATAD1 | 10.6613257 | 0.5947472 | 0.50900174 | 1.16845808 | 0.24262204 | NA |
| AC007566.1 | 5.37423244 | 0.28244722 | 0.70512691 | 0.40056225 | 0.68874244 | NA |
| PEX1 | 11.6662793 | 0.09600725 | 0.4869161 | 0.19717411 | 0.84369129 | NA |
| RBM48 | 2.44791085 | -0.2895918 | 0.98498914 | -0.2940051 | 0.76875402 | NA |
| FAM133B | 17.7847987 | -0.0771142 | 0.36991832 | -0.2084629 | 0.83486756 | NA |

| | | | | | | |
|---|---|---|---|---|---|---|
| CDK6 | 31.0155683 | 0.16421562 | 0.45021177 | 0.36475195 | 0.71529658 | 0.93629694 |
| SAMD9 | 4.27367674 | 1.14555883 | 0.98486962 | 1.16315785 | 0.24476546 | NA |
| HEPACAM2 | 62.7371143 | 1.13160121 | 0.37326657 | 3.0316168 | 0.00243248 | 0.16646881 |
| VPS50 | 33.880983 | -0.0261942 | 0.30828159 | -0.0849683 | 0.93228657 | 0.99030945 |
| GNGT1 | 6.25092056 | 0.00586158 | 0.8772135 | 0.00668204 | 0.99466854 | NA |
| BET1 | 6.98900009 | 0.27371786 | 0.6226711 | 0.43958658 | 0.66023657 | NA |
| CASD1 | 23.1617969 | -0.206866 | 0.41188319 | -0.5022442 | 0.61549573 | NA |
| SGCE | 7.64329771 | 0.01560315 | 0.70299706 | 0.02219519 | 0.98229225 | NA |
| PPP1R9A | 107.730098 | 0.20010529 | 0.23269887 | 0.85993237 | 0.38982633 | 0.8258317 |
| AC002429.2 | 9.946002 | 0.462846 | 0.66942199 | 0.69141141 | 0.48930704 | NA |
| PON2 | 15.317229 | 0.13954204 | 0.49837996 | 0.27999128 | 0.77948419 | NA |
| PDK4 | 34.9243126 | -0.1819196 | 0.76822553 | -0.236805 | 0.81280811 | 0.96845695 |
| SLC25A13 | 39.0815714 | -0.4086706 | 0.34182339 | -1.1955606 | 0.23186808 | 0.73156941 |
| SEM1 | 27.1332556 | -0.1435586 | 0.33777629 | -0.4250109 | 0.67082872 | 0.92614389 |
| SDHAF3 | 5.54325341 | 0.78684033 | 0.60218198 | 1.30664875 | 0.19133203 | NA |
| TAC1 | 20.8963187 | -2.9393593 | 2.79805938 | -1.0504992 | 0.29348864 | NA |
| ASNS | 5.51595917 | -0.7389241 | 0.80051943 | -0.9230558 | 0.35597812 | NA |
| LMTK2 | 30.6768546 | -0.4262185 | 0.33229255 | -1.2826605 | 0.19961105 | 0.69642079 |
| TECPR1 | 3.82927815 | -0.2935946 | 0.82504687 | -0.3558521 | 0.72195136 | NA |
| BRI3 | 38.8639117 | -0.5259122 | 0.3601141 | -1.4604044 | 0.14417895 | 0.62469744 |
| BAIAP2L1 | 103.005559 | 0.23568172 | 0.33625337 | 0.70090516 | 0.4833622 | 0.86351655 |
| TRRAP | 35.2420744 | -0.1169737 | 0.30064829 | -0.3890714 | 0.69722333 | 0.9303056 |
| SMURF1 | 85.153288 | 0.13995653 | 0.38418917 | 0.36429066 | 0.71564097 | 0.93629694 |
| ARPC1B | 7.25222191 | 0.34175968 | 0.51490684 | 0.66373109 | 0.50686244 | NA |
| PDAP1 | 9.33025946 | 0.04005342 | 0.50569108 | 0.07920531 | 0.93686932 | NA |
| BUD31 | 8.057839 | -0.3736309 | 0.49742281 | -0.7511334 | 0.45257235 | NA |
| CPSF4 | 3.49880444 | -0.183444 | 0.72193173 | -0.2541016 | 0.79941704 | NA |
| ZNF789 | 7.40465696 | 0.8476674 | 0.60393718 | 1.40356882 | 0.16044728 | NA |
| ZNF394 | 4.18680279 | 0.43730062 | 0.80439335 | 0.54364027 | 0.58668904 | NA |
| ZKSCAN5 | 4.44786523 | -0.3820371 | 0.78984002 | -0.4836892 | 0.62860644 | NA |
| FAM200A | 5.20949179 | 1.52049372 | 0.65769415 | 2.31185533 | 0.02078566 | NA |
| ZSCAN25 | 4.12156329 | 0.59455535 | 0.80417083 | 0.73933961 | 0.4597008 | NA |
| TRIM4 | 13.3738781 | -0.1332535 | 0.45445571 | -0.2932155 | 0.76935741 | NA |
| AZGP1 | 12.5879076 | 0.31358306 | 0.76203628 | 0.41150673 | 0.68070101 | NA |
| ZKSCAN1 | 35.5089738 | 0.15885247 | 0.32743436 | 0.48514296 | 0.62757494 | 0.91751762 |
| ZNF3 | 7.52396372 | -1.19833 | 0.6468475 | -1.8525696 | 0.06394407 | NA |
| COPS6 | 7.56901865 | 0.51081026 | 0.52422468 | 0.97441094 | 0.32985254 | NA |
| MCM7 | 2.90548058 | 0.7225163 | 0.98589479 | 0.73285335 | 0.46364788 | NA |
| AP4M1 | 4.14756283 | -0.1429981 | 0.69041187 | -0.2071199 | 0.83591619 | NA |
| TAF6 | 3.92898504 | 1.14751945 | 0.73373963 | 1.56393276 | 0.11783336 | NA |
| LAMTOR4 | 26.5856004 | -0.0668885 | 0.39434977 | -0.1696171 | 0.86531127 | 0.97217819 |
| STAG3 | 2.74414849 | -0.426588 | 0.92935065 | -0.4590172 | 0.64622179 | NA |
| CASTOR3 | 18.879825 | 0.38607016 | 0.38327081 | 1.00730384 | 0.31378879 | NA |
| SPDYE3 | 5.37444715 | 0.6421601 | 0.79795691 | 0.80475536 | 0.42096086 | NA |
| PILRB | 15.3534333 | -0.5299029 | 0.54695312 | -0.9688269 | 0.33263155 | NA |
| MEPCE | 6.20301848 | -0.1509398 | 0.58333133 | -0.2587548 | 0.79582445 | NA |
| PPP1R35 | 6.40659757 | -0.0320978 | 0.54023962 | -0.0594139 | 0.9526224 | NA |
| TSC22D4 | 11.8151213 | 0.14701574 | 0.517288 | 0.28420482 | 0.77625342 | NA |
| AGFG2 | 5.16482526 | 0.22041668 | 0.66399241 | 0.33195663 | 0.73992201 | NA |
| SAP25 | 2.22366189 | -0.7947138 | 0.99402475 | -0.799491 | 0.42400579 | NA |
| LRCH4 | 9.14694088 | -0.748404 | 0.60793277 | -1.2310638 | 0.21829899 | NA |
| PCOLCE | 3.64381667 | 1.14667937 | 0.79095524 | 1.44973989 | 0.14713107 | NA |
| GNB2 | 22.9193244 | 0.65130435 | 0.40868942 | 1.59364131 | 0.1110164 | NA |
| GIGYF1 | 14.3507401 | -0.3355513 | 0.43983275 | -0.7629065 | 0.44551916 | NA |

| | | | | | | |
|---|---|---|---|---|---|---|
| POP7 | 3.68060209 | -0.208632 | 0.89904141 | -0.2320605 | 0.81649102 | NA |
| SLC12A9 | 3.55729561 | -0.1890755 | 0.8354938 | -0.2263039 | 0.82096505 | NA |
| TRIP6 | 5.00790527 | -0.6012138 | 0.63533332 | -0.9462967 | 0.34399729 | NA |
| SRRT | 6.83214086 | -0.6704221 | 0.64240697 | -1.0436096 | 0.29666606 | NA |
| TRIM56 | 21.8017653 | -0.0070613 | 0.43597192 | -0.0161967 | 0.98707748 | NA |
| AP1S1 | 4.70599522 | -0.0045944 | 0.74789669 | -0.0061431 | 0.99509852 | NA |
| PLOD3 | 3.97246637 | -0.2435402 | 0.72467292 | -0.3360692 | 0.73681871 | NA |
| ZNHIT1 | 12.4889779 | 0.05807015 | 0.4070904 | 0.14264681 | 0.88656913 | NA |
| FIS1 | 24.4828952 | 0.24328533 | 0.36482604 | 0.66685297 | 0.50486605 | NA |
| CUX1 | 129.981378 | -0.0707898 | 0.22764654 | -0.3109638 | 0.75582814 | 0.95323728 |
| PRKRIP1 | 30.0390315 | -0.4706938 | 0.37075606 | -1.2695513 | 0.20424451 | 0.700066 |
| ORAI2 | 3.28017386 | -0.6501209 | 1.26281603 | -0.5148184 | 0.60667995 | NA |
| ALKBH4 | 2.75300581 | 0.96246483 | 0.8718922 | 1.10388054 | 0.26964496 | NA |
| POLR2J | 8.26270164 | 0.30210059 | 0.64752732 | 0.46654494 | 0.64082549 | NA |
| SPDYE2 | 3.52809254 | 0.21778185 | 0.80539063 | 0.27040525 | 0.7868485 | NA |
| AC105052.3 | 8.84606616 | 0.53359324 | 0.68261484 | 0.78169006 | 0.43439674 | NA |
| FAM185A | 18.0987721 | 0.76775214 | 0.42901899 | 1.78955283 | 0.07352583 | NA |
| FBXL13 | 6.69105345 | 0.18000376 | 0.65114508 | 0.27644186 | 0.78220871 | NA |
| ARMC10 | 11.0896747 | -0.628635 | 0.59924947 | -1.0490373 | 0.29416097 | NA |
| NAPEPLD | 18.0978652 | 0.18992062 | 0.36845856 | 0.51544635 | 0.60624115 | NA |
| PMPCB | 19.8621755 | -0.3215315 | 0.37487665 | -0.8576995 | 0.39105841 | NA |
| DNAJC2 | 11.4349588 | 0.15442191 | 0.53159724 | 0.29048666 | 0.77144395 | NA |
| PSMC2 | 8.96989632 | 0.20838343 | 0.5305195 | 0.39279126 | 0.69447366 | NA |
| ORC5 | 11.0508802 | 0.77853814 | 0.46554015 | 1.67233295 | 0.09445869 | NA |
| LHFPL3 | 14.577094 | -0.2294129 | 0.48641054 | -0.4716446 | 0.63718049 | NA |
| LINC01004 | 29.6481508 | 0.49737263 | 0.39176963 | 1.26955382 | 0.20424361 | 0.700066 |
| KMT2E-AS1 | 2.72644541 | 0.53527212 | 0.8579033 | 0.6239306 | 0.53267316 | NA |
| KMT2E | 103.836912 | -0.1903745 | 0.22842973 | -0.8334051 | 0.40461628 | 0.82808596 |
| SRPK2 | 123.509001 | -0.186271 | 0.25027854 | -0.7442547 | 0.45672239 | 0.85802171 |
| PUS7 | 6.85776538 | 0.36483808 | 0.57971878 | 0.62933631 | 0.52912891 | NA |
| RINT1 | 7.71516021 | -0.3116462 | 0.54846987 | -0.5682102 | 0.56989222 | NA |
| EFCAB10 | 3.71563324 | -1.0791624 | 0.86057833 | -1.2539967 | 0.20984322 | NA |
| ATXN7L1 | 43.6809026 | 0.32931392 | 0.3904555 | 0.8434096 | 0.3989994 | 0.8269015 |
| CDHR3 | 5.80792573 | 1.02173873 | 0.88010731 | 1.16092516 | 0.24567234 | NA |
| SYPL1 | 21.871227 | 0.17246153 | 0.42944757 | 0.40158925 | 0.68798634 | NA |
| NAMPT | 43.313012 | 0.41984798 | 0.41379132 | 1.01463699 | 0.3102789 | 0.778534 |
| HBP1 | 21.3738304 | 0.299556 | 0.3899578 | 0.76817544 | 0.44238296 | NA |
| COG5 | 123.469227 | -0.2697317 | 0.2602741 | -1.0363372 | 0.30004485 | 0.77521265 |
| DUS4L | 3.3399702 | 1.26394296 | 0.93320564 | 1.3544099 | 0.17560564 | NA |
| BCAP29 | 68.0441403 | 0.6273611 | 0.48889823 | 1.2832141 | 0.19941708 | 0.69642079 |
| SLC26A4-AS1 | 2.65329429 | 1.15029817 | 1.27515263 | 0.90208666 | 0.36701084 | NA |
| SLC26A4 | 467.772246 | 0.31074772 | 0.64016601 | 0.48541741 | 0.62738029 | 0.91751762 |
| CBLL1 | 5.2074889 | 1.23726136 | 0.74094131 | 1.6698507 | 0.09494891 | NA |
| SLC26A3 | 89.6380812 | -0.9660306 | 0.47571001 | -2.0307131 | 0.0422841 | 0.44186191 |
| DLD | 24.9656221 | -0.0653629 | 0.37425176 | -0.1746494 | 0.86135511 | NA |
| LAMB1 | 65.5556871 | -0.509621 | 0.43714912 | -1.1657829 | 0.24370223 | 0.73323864 |
| NRCAM | 12.9964402 | -1.7985752 | 0.77006351 | -2.3356193 | 0.01951109 | NA |
| PNPLA8 | 63.9190496 | -1.034411 | 0.42435073 | -2.4376321 | 0.01478381 | 0.30435125 |
| DNAJB9 | 4.91276876 | -1.0223554 | 0.64082167 | -1.5953821 | 0.11062681 | NA |
| IMMP2L | 305.963235 | 0.53906243 | 0.35717242 | 1.50924989 | 0.13123494 | 0.60286754 |
| DOCK4 | 42.1420899 | -0.8218171 | 0.39566414 | -2.0770575 | 0.03779625 | 0.42032585 |
| ZNF277 | 28.1987372 | 0.29451476 | 0.40357344 | 0.72976746 | 0.46553234 | 0.85894565 |
| IFRD1 | 24.9242405 | -0.3419648 | 0.43955326 | -0.7779827 | 0.4365792 | NA |
| LSMEM1 | 3.03690664 | -0.6354671 | 1.09201407 | -0.5819221 | 0.56061914 | NA |

| | | | | | | |
|---|---|---|---|---|---|---|
| TMEM168 | 19.0367599 | -0.3837903 | 0.43581398 | -0.8806287 | 0.37851882 | NA |
| BMT2 | 22.955786 | 0.31258894 | 0.39851204 | 0.78439021 | 0.43281118 | NA |
| SMIM30 | 4.28885714 | -0.5562599 | 0.7304208 | -0.7615609 | 0.44632213 | NA |
| FOXP2 | 18.9475501 | -1.4108292 | 1.21646007 | -1.1597826 | 0.24613732 | NA |
| MDFIC | 49.1783424 | 0.41915024 | 0.39163395 | 1.07026021 | 0.2845022 | 0.76977033 |
| TES | 21.6785047 | -0.8391255 | 0.4401077 | -1.9066368 | 0.05656764 | NA |
| CAV2 | 5.03548589 | 0.8788765 | 0.75338579 | 1.16656899 | 0.24338447 | NA |
| MET | 21.8253179 | -0.7725565 | 0.34976999 | -2.2087559 | 0.02719163 | NA |
| CAPZA2 | 43.0957251 | -0.2555601 | 0.26194011 | -0.9756431 | 0.32924137 | 0.79237621 |
| ST7 | 85.7064955 | -0.6291272 | 0.28243724 | -2.2274937 | 0.02591429 | 0.39467204 |
| ST7-AS2 | 4.35359074 | -0.177681 | 0.69986699 | -0.2538783 | 0.7995896 | NA |
| CFTR | 103.731803 | -0.1224339 | 0.53466475 | -0.2289919 | 0.81887518 | 0.96845695 |
| AC000061.1 | 5.09002193 | 1.85045979 | 0.89513552 | 2.06723981 | 0.03871156 | NA |
| CTTNBP2 | 31.0331942 | 0.52100731 | 0.45079124 | 1.15576184 | 0.24777858 | 0.73890619 |
| LSM8 | 12.9756308 | -0.0973427 | 0.43361985 | -0.2244886 | 0.82237717 | NA |
| TSPAN12 | 7.81764602 | 1.07654216 | 0.58362311 | 1.84458453 | 0.065098 | NA |
| ING3 | 7.62158783 | 0.44611285 | 0.61487561 | 0.72553349 | 0.46812481 | NA |
| FAM3C | 28.9793707 | -0.5826555 | 0.35407437 | -1.6455739 | 0.09985151 | 0.57114936 |
| AASS | 14.5305359 | -0.0620528 | 0.44046851 | -0.140879 | 0.88796554 | NA |
| CADPS2 | 80.6867672 | -0.2159323 | 0.44544632 | -0.4847549 | 0.62785022 | 0.91751762 |
| AC004594.1 | 3.62894826 | 3.08464302 | 2.08502176 | 1.47942965 | 0.13902552 | NA |
| SLC13A1 | 5.65393038 | 1.74037238 | 1.15083644 | 1.51226735 | 0.13046586 | NA |
| IQUB | 4.78019645 | 0.03320582 | 1.07851402 | 0.03078849 | 0.97543822 | NA |
| NDUFA5 | 31.5188245 | 0.40049111 | 0.36052967 | 1.11084091 | 0.26663684 | 0.75497438 |
| WASL | 39.8515117 | -0.0302694 | 0.26710307 | -0.1133248 | 0.90977305 | 0.98346872 |
| POT1 | 9.86160769 | 0.29805289 | 0.49820892 | 0.59824881 | 0.54967393 | NA |
| POT1-AS1 | 6.53087178 | 1.58153549 | 0.76153381 | 2.07677646 | 0.0378222 | NA |
| GRM8 | 97.4913287 | -0.5612868 | 0.79256609 | -0.7081893 | 0.47882771 | 0.86206794 |
| ZNF800 | 21.3899559 | 0.20261517 | 0.35562918 | 0.56973721 | 0.56885595 | NA |
| ARF5 | 6.53601983 | -0.4409978 | 0.63088782 | -0.6990115 | 0.48454487 | NA |
| SND1 | 78.9381462 | -0.825881 | 0.3882215 | -2.1273449 | 0.03339144 | 0.41688714 |
| RBM28 | 11.2026964 | 0.41598503 | 0.48146976 | 0.86398993 | 0.38759342 | NA |
| METTL2B | 9.96746091 | -0.2355772 | 0.56106245 | -0.4198769 | 0.67457538 | NA |
| AC090114.2 | 2.80402362 | -0.6650692 | 0.95451942 | -0.6967581 | 0.48595417 | NA |
| FAM71F2 | 5.0020709 | -1.3004505 | 0.87977144 | -1.4781686 | 0.13936266 | NA |
| CALU | 20.2107 | -1.2613531 | 0.61262127 | -2.0589443 | 0.03949958 | NA |
| ATP6V1F | 40.3894366 | 0.27749876 | 0.53186283 | 0.52174874 | 0.60184529 | 0.90446231 |
| TNPO3 | 38.2757687 | -0.3457389 | 0.29587403 | -1.1685341 | 0.24259139 | 0.73323864 |
| TSPAN33 | 5.36011364 | -1.3566252 | 0.72371205 | -1.8745372 | 0.06085642 | NA |
| AHCYL2 | 75.5618817 | -0.3705621 | 0.31560831 | -1.1741201 | 0.24034692 | 0.73323864 |
| NRF1 | 37.0079364 | 0.25022146 | 0.27745109 | 0.90185793 | 0.36713234 | 0.80981506 |
| UBE2H | 182.206297 | -0.0623162 | 0.27345522 | -0.2278844 | 0.81973614 | 0.96845695 |
| ZC3HC1 | 6.78996351 | -1.0775537 | 0.63359816 | -1.7006894 | 0.08900132 | NA |
| KLHDC10 | 29.8334516 | 0.22974285 | 0.33591052 | 0.68394063 | 0.49401265 | 0.86634063 |
| TMEM209 | 7.2738442 | -0.2060105 | 0.56954406 | -0.3617112 | 0.71756784 | NA |
| CEP41 | 4.50036835 | 0.47322029 | 0.77778324 | 0.60842181 | 0.54290775 | NA |
| MEST | 4.03185524 | -0.8577838 | 0.91763221 | -0.9347795 | 0.34990193 | NA |
| COPG2 | 10.8532734 | 0.18604063 | 0.45481937 | 0.4090429 | 0.68250818 | NA |
| AC016831.7 | 362.59672 | -0.3294743 | 0.31136575 | -1.0581584 | 0.28998325 | 0.77286135 |
| LINC00513 | 30.642022 | 0.01306802 | 0.58358941 | 0.0223925 | 0.98213486 | 0.99902646 |
| AC058791.1 | 7.81953539 | -0.0798844 | 0.66705992 | -0.119756 | 0.90467647 | NA |
| MKLN1 | 127.435021 | -0.5212598 | 0.24892629 | -2.0940327 | 0.03625707 | 0.41711255 |
| PODXL | 3.49257549 | -0.487162 | 0.94448744 | -0.5157951 | 0.60599753 | NA |
| CHCHD3 | 149.033938 | 0.09772277 | 0.19065307 | 0.51256857 | 0.60825314 | 0.9088434 |

| | | | | | | |
|---|---|---|---|---|---|---|
| EXOC4 | 169.683379 | 0.27789409 | 0.33378595 | 0.8325518 | 0.40509755 | 0.82808596 |
| LRGUK | 4.16073375 | -0.8636006 | 0.8697795 | -0.9928961 | 0.32076061 | NA |
| SLC35B4 | 11.441287 | 0.32437221 | 0.45912939 | 0.70649411 | 0.47988092 | NA |
| AKR1B1 | 16.7110524 | 0.10891768 | 0.59597121 | 0.18275661 | 0.854989 | NA |
| BPGM | 22.2940931 | -0.6501465 | 0.53198263 | -1.2221198 | 0.22166232 | NA |
| CALD1 | 9.74370672 | -0.0720504 | 0.69079591 | -0.1043006 | 0.91693079 | NA |
| AGBL3 | 3.55638396 | -0.104357 | 0.81799947 | -0.1275759 | 0.89848459 | NA |
| CYREN | 7.79053455 | 0.07241988 | 0.58195219 | 0.124443 | 0.90096453 | NA |
| TMEM140 | 10.7132871 | -0.6297764 | 0.579922 | -1.0859674 | 0.27749341 | NA |
| WDR91 | 7.94423678 | 0.82912698 | 0.52518051 | 1.57874667 | 0.11439418 | NA |
| CNOT4 | 59.1126395 | 0.13743129 | 0.3118216 | 0.44073691 | 0.65940348 | 0.92276785 |
| NUP205 | 14.3566299 | -0.0874839 | 0.4150653 | -0.2107714 | 0.83306566 | NA |
| STMP1 | 16.8652375 | 0.13720968 | 0.48605319 | 0.28229356 | 0.77771842 | NA |
| MTPN | 27.5472161 | -0.3994806 | 0.37291839 | -1.071228 | 0.28406691 | 0.76977033 |
| PTN | 18.3396737 | -1.8335962 | 0.63811432 | -2.8734604 | 0.00406002 | NA |
| DGKI | 346.143669 | -1.0726054 | 0.37720814 | -2.8435374 | 0.00446158 | 0.21776517 |
| CREB3L2 | 26.1845678 | -0.2855276 | 0.40392963 | -0.7068746 | 0.4796444 | 0.86206794 |
| TRIM24 | 28.2319071 | -0.5059842 | 0.31317115 | -1.6156796 | 0.10616361 | 0.57769936 |
| SVOPL | 56.5317697 | -0.9213366 | 0.50625363 | -1.8199112 | 0.06877253 | 0.52594237 |
| ATP6V0A4 | 540.773191 | 0.21153211 | 0.28203764 | 0.75001373 | 0.45324643 | 0.85557861 |
| TMEM213 | 388.89685 | 0.91703268 | 0.25731355 | 3.56387248 | 0.00036542 | 0.08866504 |
| KIAA1549 | 12.0601471 | 0.30749351 | 0.61200357 | 0.50243745 | 0.61535984 | NA |
| ZC3HAV1 | 19.0186863 | -0.2190716 | 0.46823113 | -0.4678707 | 0.63987707 | NA |
| TTC26 | 2.76566751 | 0.40814996 | 0.89796484 | 0.45452778 | 0.649449 | NA |
| UBN2 | 25.2014435 | 0.37307779 | 0.39944536 | 0.93398953 | 0.35030929 | NA |
| LUC7L2 | 9.79428011 | 0.04068562 | 0.50742831 | 0.08018004 | 0.93609407 | NA |
| HIPK2 | 289.608825 | -0.113707 | 0.27900079 | -0.4075509 | 0.68360342 | 0.92710517 |
| TBXAS1 | 41.5522576 | -0.1660848 | 0.4955593 | -0.3351461 | 7.38E-01 | 0.94658914 |
| PARP12 | 13.1404283 | -0.5538302 | 0.65329856 | -0.8477444 | 0.39658031 | NA |
| KDM7A | 46.7716052 | -0.9491063 | 0.44835038 | -2.1168852 | 0.03426959 | 0.41688714 |
| SLC37A3 | 13.8031886 | -0.5203832 | 0.51154987 | -1.0172678 | 0.30902604 | NA |
| MKRN1 | 15.9721165 | 0.23306501 | 0.437891 | 0.53224434 | 0.59455678 | NA |
| DENND2A | 6.81109603 | -1.0296787 | 1.15345086 | -0.8926941 | 0.37202103 | NA |
| NDUFB2 | 35.5773002 | 0.78792478 | 0.34391212 | 2.2910643 | 0.0219597 | 0.35108799 |
| BRAF | 107.218573 | 0.05988125 | 0.22855439 | 0.262 | 0.79332144 | 0.96341299 |
| MRPS33 | 7.41418672 | 0.18995795 | 0.63400147 | 0.29961752 | 0.76446892 | NA |
| TMEM178B | 6.65043646 | 0.85065761 | 0.87206325 | 0.97545402 | 0.3293351 | NA |
| AGK | 11.2569243 | -0.2617639 | 0.52665961 | -0.4970267 | 0.61917024 | NA |
| AC004918.1 | 3.39776236 | 0.48407656 | 1.03859109 | 0.46608965 | 0.64115133 | NA |
| KIAA1147 | 8.9937742 | 0.79597754 | 0.47708333 | 1.66842456 | 0.09523148 | NA |
| SSBP1 | 13.9424985 | 0.07998716 | 0.41882241 | 0.19098108 | 0.84854041 | NA |
| GSTK1 | 19.4842181 | 0.39301977 | 0.54004242 | 0.72775723 | 0.46676221 | NA |
| TMEM139 | 2.43502501 | -1.0526829 | 0.94348091 | -1.1157437 | 0.26453186 | NA |
| CASP2 | 2.76959946 | -0.2925808 | 0.89822877 | -0.3257308 | 0.74462804 | NA |
| TCAF2 | 6.70923884 | -1.2798323 | 1.13095656 | -1.131637 | 0.25778706 | NA |
| TCAF1 | 12.4012411 | 0.70420933 | 0.52398924 | 1.34393854 | 0.17896826 | NA |
| AC004889.1 | 5.34091999 | 0.59578532 | 0.77363032 | 0.77011631 | 0.4412309 | NA |
| AC074386.1 | 13.3111383 | 0.06446199 | 0.43245761 | 0.14905968 | 0.88150654 | NA |
| ARHGEF5 | 4.00650739 | -0.3858755 | 0.69700516 | -0.5536193 | 0.57983939 | NA |
| TPK1 | 9.4960449 | 0.55291679 | 0.45631568 | 1.21169799 | 0.22562801 | NA |
| CNTNAP2 | 3.02189548 | -0.0855677 | 0.81058629 | -0.1055627 | 0.91592934 | NA |
| CUL1 | 38.7060205 | -0.1173523 | 0.30176782 | -0.3888828 | 0.69736284 | 0.9303056 |
| EZH2 | 2.98155314 | 0.36752304 | 0.84241673 | 0.43627225 | 0.66263921 | NA |
| PDIA4 | 5.65500371 | -0.1987985 | 0.71727016 | -0.2771599 | 0.78165732 | NA |

| | | | | | | |
|---|---|---|---|---|---|---|
| ZNF786 | 4.0150839 | -0.4116719 | 0.8491663 | -0.4847954 | 0.6278215 | NA |
| ZNF425 | 2.53048181 | -0.6020377 | 1.16507442 | -0.5167375 | 0.6053394 | NA |
| ZNF398 | 9.71535989 | 0.44401032 | 0.47857171 | 0.9277822 | 0.35352056 | NA |
| ZNF282 | 6.66112361 | -0.8175173 | 0.58962634 | -1.3865006 | 0.16559411 | NA |
| ZNF212 | 3.39260049 | -0.3054179 | 0.89618235 | -0.3407988 | 0.73325503 | NA |
| ZNF783 | 4.26865001 | 0.21481812 | 0.90294254 | 0.23790896 | 0.81195171 | NA |
| ZNF777 | 4.75254513 | 0.10826992 | 0.65971427 | 0.16411638 | 0.86963953 | NA |
| ZNF746 | 6.45410634 | 0.27532632 | 0.67250748 | 0.40940262 | 0.68224422 | NA |
| KRBA1 | 2.82836234 | 0.03632507 | 0.84616501 | 0.04292906 | 0.96575808 | NA |
| ZNF467 | 2.50664285 | -0.2364412 | 1.03705821 | -0.2279922 | 0.81965229 | NA |
| ZNF862 | 9.02774658 | 1.40327136 | 0.58847361 | 2.38459523 | 0.01709792 | NA |
| ATP6V0E2-A: | 2.14742242 | -0.6578263 | 1.10832639 | -0.5935312 | 0.5528257 | NA |
| ATP6V0E2 | 6.71339469 | 0.00580947 | 0.6040125 | 0.00961813 | 0.99232596 | NA |
| ACTR3C | 67.2155843 | 0.78837181 | 0.38738886 | 2.0350916 | 0.04184166 | 0.44186191 |
| LRRC61 | 9.72827446 | 0.64375867 | 0.486142 | 1.32421941 | 0.18543019 | NA |
| RARRES2 | 36.2103822 | 0.62941702 | 0.4876588 | 1.29069141 | 0.1968107 | 0.69343545 |
| TMEM176B | 8.0125566 | 0.31469492 | 0.65076185 | 0.48357924 | 0.62868451 | NA |
| TMEM176A | 5.61704119 | -0.2473514 | 0.84249667 | -0.2935933 | 0.76906867 | NA |
| AOC1 | 2.81190174 | -0.2191699 | 1.10326154 | -0.1986563 | 0.84253161 | NA |
| ABCB8 | 4.31740279 | 0.6550784 | 0.70011392 | 0.93567402 | 0.34944105 | NA |
| SLC4A2 | 8.15924364 | -1.0396095 | 0.73812954 | -1.4084378 | 0.15900148 | NA |
| FASTK | 6.20423006 | 0.15292113 | 0.62537664 | 0.24452645 | 0.80682312 | NA |
| TMUB1 | 9.29939908 | -0.7480186 | 0.50189702 | -1.4903826 | 0.13612367 | NA |
| AGAP3 | 8.17989735 | 0.16017115 | 0.59144868 | 0.27081157 | 0.78653596 | NA |
| CHPF2 | 4.30740479 | -1.4959743 | 0.73840689 | -2.0259485 | 0.04277006 | NA |
| SMARCD3 | 7.21926081 | 0.80661544 | 0.66306501 | 1.21649527 | 0.22379631 | NA |
| NUB1 | 13.5711159 | -0.3209456 | 0.49176604 | -0.6526388 | 0.51398919 | NA |
| RHEB | 47.157584 | -0.2544607 | 0.2978114 | -0.8544357 | 0.39286362 | 0.8258317 |
| PRKAG2 | 93.636748 | -0.529247 | 0.53964609 | -0.9807298 | 0.32672602 | 0.78845547 |
| GALNT11 | 35.9730697 | -0.1505675 | 0.34997187 | -0.4302274 | 0.66703025 | 0.92307876 |
| KMT2C | 156.446738 | -0.2265118 | 0.27038651 | -0.8377333 | 0.40218053 | 0.82808596 |
| ACTR3B | 12.3240096 | -0.2252514 | 0.50396286 | -0.4469604 | 0.65490368 | NA |
| DPP6 | 34.0453868 | 0.90209925 | 0.39377048 | 2.29092652 | 0.02196766 | 0.35108799 |
| PAXIP1-AS2 | 4.66151855 | 0.95516699 | 0.76623599 | 1.24657026 | 0.21255511 | NA |
| PAXIP1 | 9.44097353 | 0.63318591 | 0.51015882 | 1.24115449 | 0.21454868 | NA |
| PAXIP1-AS1 | 3.49165536 | 1.13428938 | 0.81279432 | 1.39554295 | 0.16285217 | NA |
| INSIG1 | 10.9349148 | -2.3858516 | 0.87982806 | -2.7117248 | 0.00669341 | NA |
| AC009403.1 | 2.19259584 | 0.3397586 | 0.98057132 | 0.34649046 | 0.72897416 | NA |
| RBM33 | 49.6912979 | -0.3710991 | 0.27520548 | -1.3484435 | 0.17751577 | 0.6676327 |
| LINC01006 | 12.2244708 | -0.9666965 | 0.59703133 | -1.6191721 | 0.10541023 | NA |
| LMBR1 | 48.2894846 | -0.2583136 | 0.27730502 | -0.9315145 | 0.35158751 | 0.8003972 |
| NOM1 | 5.1187153 | 0.1168604 | 0.72605185 | 0.16095324 | 0.87213023 | NA |
| UBE3C | 67.871117 | -0.4695846 | 0.2867204 | -1.6377789 | 0.10146782 | 0.5750956 |
| DNAJB6 | 44.7038185 | -0.4013562 | 0.27945805 | -1.4361947 | 0.15094696 | 0.63545337 |
| ESYT2 | 51.3612639 | 0.08702243 | 0.29176094 | 0.2982662 | 0.7655 | 0.95562184 |
| WDR60 | 22.07636 | 0.29623723 | 0.35591033 | 0.8323367 | 0.40521892 | NA |
| GTPBP6 | 13.2862409 | 0.28407004 | 0.57241881 | 0.49626258 | 0.61970916 | NA |
| LINC00685 | 3.38995491 | -1.5371926 | 0.91724149 | -1.6758865 | 0.09376043 | NA |
| PPP2R3B | 3.42061711 | -0.0647817 | 0.76310981 | -0.0848917 | 0.93234752 | NA |
| IL3RA | 5.07211493 | -1.4108955 | 1.00452731 | -1.4045367 | 0.16015909 | NA |
| SLC25A6 | 112.566787 | 0.4076602 | 0.33548608 | 1.21513296 | 0.22431539 | 0.72657496 |
| LINC00106 | 17.6716163 | -0.835588 | 0.64214506 | -1.3012449 | 0.19317464 | NA |
| ASMTL-AS1 | 8.82073203 | -1.2364737 | 0.67586632 | -1.8294649 | 0.06732999 | NA |
| ASMTL | 8.08355684 | 0.35459932 | 0.53652739 | 0.66091559 | 0.50866645 | NA |

| | | | | | | |
|---|---|---|---|---|---|---|
| AKAP17A | 7.54615934 | -0.5923859 | 0.56338261 | -1.0514807 | 0.29303787 | NA |
| DHRSX | 50.3862273 | -0.148707 | 0.29972938 | -0.4961374 | 0.61979748 | 0.91493825 |
| CD99 | 6.12449136 | -0.3720626 | 0.67533399 | -0.5509312 | 0.58168084 | NA |
| ARSD | 15.2450857 | 0.29253702 | 0.52981092 | 0.55215363 | 0.5808431 | NA |
| ARSE | 9.01340553 | 0.97666466 | 0.51931022 | 1.88069603 | 0.06001328 | NA |
| PRKX | 35.7499545 | -0.4924646 | 0.35724429 | -1.3785093 | 0.16804611 | 0.65572379 |
| BX890604.1 | 2.67984718 | 0.25365399 | 0.89689585 | 0.28281321 | 0.77732003 | NA |
| NLGN4X | 18.3942632 | -0.2719205 | 0.84506207 | -0.3217757 | 0.74762259 | NA |
| PUDP | 6.84887774 | -0.3900018 | 0.75376433 | -0.5174055 | 0.60487313 | NA |
| STS | 2.92334152 | -2.0678991 | 1.01718863 | -2.0329554 | 0.04205703 | NA |
| PNPLA4 | 3.58677891 | 0.79734189 | 0.78676698 | 1.01344097 | 0.31084958 | NA |
| ANOS1 | 4.77014746 | -2.3695815 | 1.32759703 | -1.784865 | 0.07428322 | NA |
| TBL1X | 212.35792 | -0.5575082 | 0.3968101 | -1.4049749 | 0.16002875 | 0.64519142 |
| SHROOM2 | 20.436383 | 0.46230984 | 0.51074834 | 0.90516171 | 0.36537973 | NA |
| WWC3 | 8.06306504 | 0.44049421 | 0.61117763 | 0.72073025 | 0.47107549 | NA |
| MID1 | 15.3013586 | 0.87443045 | 0.48724519 | 1.79464151 | 0.07271083 | NA |
| AC073529.1 | 11.0675595 | 1.2889236 | 0.51442229 | 2.50557492 | 0.01222525 | NA |
| HCCS | 3.95047748 | 0.33356312 | 0.79347335 | 0.42038352 | 0.6742053 | NA |
| ARHGAP6 | 126.991461 | -0.4785322 | 0.33568894 | -1.4255226 | 0.15400619 | 0.63864841 |
| MSL3 | 3.35208443 | 0.23519362 | 0.76821418 | 0.30615631 | 0.75948564 | NA |
| FRMPD4 | 21.7373837 | 0.16909453 | 0.67580713 | 0.25021123 | 0.802424 | NA |
| PRPS2 | 4.6144599 | 0.10522072 | 0.74120089 | 0.14195979 | 0.88711177 | NA |
| TMSB4X | 147.264306 | -0.5864871 | 0.41812541 | -1.4026584 | 0.16071873 | 0.64699592 |
| RAB9A | 15.2670815 | 0.14782418 | 0.45563848 | 0.32443305 | 0.7456102 | NA |
| TRAPPC2 | 5.78453862 | 0.76523622 | 0.5815194 | 1.31592552 | 0.18819904 | NA |
| OFD1 | 30.7926597 | 0.05242101 | 0.34017214 | 0.15410142 | 0.87752977 | 0.9751578 |
| GEMIN8 | 6.0430808 | -0.3414641 | 0.65012059 | -0.5252319 | 0.59942198 | NA |
| MOSPD2 | 10.1443183 | 0.4061313 | 0.48451294 | 0.83822591 | 0.40190384 | NA |
| PIGA | 2.35569479 | -0.6433045 | 0.89080895 | -0.7221577 | 0.47019753 | NA |
| PIR | 3.82782713 | 1.87495083 | 0.84667281 | 2.21449278 | 0.0267949 | NA |
| BMX | 2.69470343 | 0.22730191 | 0.84579077 | 0.26874485 | 0.78812604 | NA |
| CLTRN | 9.68135465 | -0.2600337 | 0.5153991 | -0.5045288 | 0.61388985 | NA |
| CA5B | 16.0162349 | -0.2511221 | 0.41761546 | -0.6013238 | 0.54762432 | NA |
| ZRSR2 | 8.81626739 | 0.04750637 | 0.50707749 | 0.0936866 | 0.92535811 | NA |
| AP1S2 | 13.5656722 | -0.8696176 | 0.45959561 | -1.8921364 | 0.05847281 | NA |
| CTPS2 | 13.2671672 | 0.50039584 | 0.48910007 | 1.02309501 | 0.30626293 | NA |
| SYAP1 | 17.3425095 | -0.4170512 | 0.48264958 | -0.864087 | 0.38754009 | NA |
| TXLNG | 8.74241841 | -0.0838235 | 0.62855624 | -0.1333588 | 0.89390959 | NA |
| RBBP7 | 15.1422134 | 0.21092466 | 0.37936276 | 0.55599726 | 0.57821272 | NA |
| REPS2 | 6.50228544 | -0.3804486 | 0.74775177 | -0.5087899 | 0.61089948 | NA |
| NHS | 143.721872 | -0.4743826 | 0.41564579 | -1.1413145 | 0.25373905 | 0.74749396 |
| SCML1 | 52.8558806 | 0.58730563 | 0.45548979 | 1.28939362 | 0.19726128 | 0.69343545 |
| RAI2 | 8.45885929 | 1.06434386 | 0.53784064 | 1.97892048 | 0.04782496 | NA |
| SCML2 | 3.77606735 | 0.17616377 | 0.7267041 | 0.24241472 | 0.80845883 | NA |
| CDKL5 | 14.1087641 | -0.0716722 | 0.43954255 | -0.1630609 | 0.87047048 | NA |
| PHKA2 | 10.4745948 | 0.12837853 | 0.55145655 | 0.23279899 | 0.8159175 | NA |
| ADGRG2 | 2.82517732 | -1.8146755 | 0.95580583 | -1.8985818 | 0.05761948 | NA |
| PDHA1 | 24.6827637 | 0.3772626 | 0.36330134 | 1.03842888 | 0.29907043 | NA |
| SH3KBP1 | 2.8768945 | -0.8605136 | 0.88557709 | -0.9716982 | 0.33120072 | NA |
| BCLAF3 | 20.5991859 | 0.0279288 | 0.44504376 | 0.06275517 | 0.94996147 | NA |
| EIF1AX | 34.8632528 | -0.079723 | 0.48049156 | -0.1659197 | 0.86822011 | 0.9724211 |
| RPS6KA3 | 83.0895407 | -0.4926918 | 0.30555497 | -1.6124488 | 0.1068643 | 0.57769936 |
| MBTPS2 | 7.79554973 | 0.91735838 | 0.61072982 | 1.50206909 | 0.13307927 | NA |
| SMS | 21.2117819 | -0.0926821 | 0.44802975 | -0.2068659 | 0.83611457 | NA |

| | | | | | | |
|---|---|---|---|---|---|---|
| PRDX4 | 3.88728006 | -0.0650559 | 0.78480269 | -0.0828946 | 0.93393534 | NA |
| ACOT9 | 9.77349477 | -0.4209372 | 0.52476503 | -0.8021441 | 0.42246961 | NA |
| SAT1 | 134.492299 | 0.1382727 | 0.32406295 | 0.42668469 | 0.669609 | 0.92504473 |
| APOO | 130.380266 | -0.1770571 | 0.40682931 | -0.4352123 | 0.66340837 | 0.92276785 |
| KLHL15 | 4.0357457 | 0.46478344 | 0.804381 | 0.57781504 | 0.563389 | NA |
| EIF2S3 | 17.5283079 | -0.3459978 | 0.3884649 | -0.8906797 | 0.37310101 | NA |
| ZFX | 27.4057445 | -0.5516993 | 0.39720066 | -1.3889688 | 0.16484225 | 0.64995633 |
| PDK3 | 6.0649096 | -0.4097125 | 0.89633219 | -0.4570989 | 0.64759994 | NA |
| POLA1 | 17.7838654 | 0.52337379 | 0.43220927 | 1.21092682 | 0.22592345 | NA |
| GK | 28.8318018 | -0.3118785 | 0.41872708 | -0.7448253 | 4.56E-01 | 0.85802171 |
| TAB3 | 15.7856534 | 0.11693434 | 0.43020706 | 0.27180944 | 0.78576854 | NA |
| DMD | 8.51896119 | 1.10963613 | 0.60441096 | 1.83589676 | 0.06637293 | NA |
| AF241726.2 | 11.8454733 | 0.20618315 | 0.52982944 | 0.38915005 | 0.69716515 | NA |
| DYNLT3 | 16.4349186 | 0.03887288 | 0.47024818 | 0.08266461 | 0.93411822 | NA |
| RPGR | 7.21482972 | 0.99874477 | 0.58355909 | 1.71147152 | 0.0869941 | NA |
| TSPAN7 | 8.68927003 | 0.61357549 | 0.66527107 | 0.92229396 | 0.35637526 | NA |
| MID1IP1 | 4.17619309 | 0.59008412 | 0.7087242 | 0.8326005 | 0.40507008 | NA |
| BCOR | 23.154072 | -0.0957316 | 0.41314307 | -0.2317154 | 0.81675908 | NA |
| ATP6AP2 | 141.496758 | 0.60419261 | 0.2223052 | 2.71785187 | 0.00657072 | 0.23699006 |
| CXorf38 | 16.0114171 | 0.33033417 | 0.42226289 | 0.78229506 | 0.43404119 | NA |
| MED14 | 29.1551081 | 0.17719743 | 0.40099336 | 0.44189617 | 0.65856434 | 0.92276785 |
| USP9X | 88.0812127 | -0.4545654 | 0.30733805 | -1.4790404 | 0.13912951 | 0.61276677 |
| DDX3X | 28.3216373 | -0.1431254 | 0.33956777 | -0.421493 | 0.67339513 | 0.92617027 |
| CASK | 23.6424716 | 0.01687371 | 0.36018987 | 0.04684671 | 0.9626354 | NA |
| MAOA | 12.1182545 | -0.0851191 | 0.60562235 | -0.1405482 | 0.88822686 | NA |
| FUNDC1 | 3.8006174 | -1.219344 | 1.03648052 | -1.1764273 | 0.23942415 | NA |
| KDM6A | 75.9667106 | -0.3148999 | 0.37371056 | -0.8426305 | 0.39943513 | 0.8269015 |
| MIR222HG | 23.8760701 | -1.1047089 | 0.49494213 | -2.2319962 | 0.02561522 | NA |
| KRBOX4 | 9.9587584 | 0.37487546 | 0.51715557 | 0.72487948 | 0.46852597 | NA |
| ZNF674 | 9.11155278 | -0.0189353 | 0.54012983 | -0.0350569 | 0.97203437 | NA |
| SLC9A7 | 18.2486961 | 0.07070405 | 0.47198767 | 0.14980063 | 0.88092191 | NA |
| RP2 | 2.97706362 | -0.8037068 | 0.77385859 | -1.0385706 | 0.29900449 | NA |
| JADE3 | 22.1777167 | 0.26864016 | 0.44970832 | 0.59736533 | 0.55026349 | NA |
| RGN | 4.92312931 | 1.46387188 | 0.86571674 | 1.69093633 | 0.09084896 | NA |
| NDUFB11 | 19.9173712 | 0.65529767 | 0.36711864 | 1.78497521 | 0.07426534 | NA |
| RBM10 | 18.8080453 | -0.2540249 | 0.36628656 | -0.6935142 | 0.48798693 | NA |
| UBA1 | 24.8432585 | -0.1553801 | 0.33780066 | -0.4599758 | 0.64553359 | NA |
| CDK16 | 10.2465504 | -0.4140965 | 0.48130704 | -0.8603584 | 0.38959153 | NA |
| USP11 | 8.82049126 | -0.4489301 | 0.55446904 | -0.8096577 | 0.41813694 | NA |
| ZNF41 | 6.15281322 | 0.15646832 | 0.61751553 | 0.25338362 | 0.79997179 | NA |
| ARAF | 7.68337996 | -0.5875385 | 0.52700814 | -1.1148567 | 0.26491185 | NA |
| TIMP1 | 4.30334751 | -0.9126444 | 0.69095536 | -1.3208442 | 0.18655332 | NA |
| ELK1 | 3.13265103 | 0.75944007 | 0.90888751 | 0.83557103 | 0.40339627 | NA |
| UXT | 3.80897377 | 0.23487291 | 0.83940018 | 0.27981041 | 0.77962296 | NA |
| ZNF81 | 9.23074937 | 0.124251 | 0.55665302 | 0.22321087 | 0.82337139 | NA |
| ZNF182 | 2.4810766 | 0.68219551 | 1.02669935 | 0.664455 | 0.50639914 | NA |
| EBP | 8.1427132 | 0.20978087 | 0.6268862 | 0.33463947 | 0.73789707 | NA |
| TBC1D25 | 3.92755753 | -0.4351771 | 0.7156607 | -0.6080774 | 0.54313612 | NA |
| RBM3 | 16.8631576 | -0.2636273 | 0.51459699 | -0.5122986 | 0.60844207 | NA |
| WDR13 | 9.4354174 | -0.4036364 | 0.51482223 | -0.7840306 | 0.43302217 | NA |
| HDAC6 | 5.74359192 | -0.2417025 | 0.68646063 | -0.3520995 | 0.72476361 | NA |
| PCSK1N | 18.8118458 | -0.4031933 | 0.53279145 | -0.7567562 | 0.44919591 | NA |
| TIMM17B | 5.96300996 | -0.0215867 | 0.76391731 | -0.0282579 | 0.97745649 | NA |
| PQBP1 | 4.08805413 | -0.5766459 | 0.76540284 | -0.7533888 | 0.45121632 | NA |

| | | | | | | |
|---|---|---|---|---|---|---|
| SLC35A2 | 3.34111568 | -0.8647156 | 0.79925928 | -1.0818963 | 0.27929863 | NA |
| OTUD5 | 12.8328306 | 0.33593611 | 0.44083497 | 0.76204506 | 0.44603311 | NA |
| GRIPAP1 | 9.42604259 | 0.10695401 | 0.49750511 | 0.21498072 | 0.82978235 | NA |
| TFE3 | 6.46028038 | -1.1082951 | 0.61675624 | -1.7969743 | 0.0723397 | NA |
| CCDC120 | 4.20174169 | -0.8929186 | 0.74182415 | -1.2036796 | 0.22871344 | NA |
| WDR45 | 13.0404782 | 0.10036441 | 0.49641329 | 0.20217914 | 0.83977668 | NA |
| PLP2 | 5.44677333 | 0.22776122 | 0.5725822 | 0.39777907 | 0.69079305 | NA |
| CLCN5 | 76.1547429 | 0.0270976 | 0.29093095 | 0.09314101 | 0.92579153 | 0.98758497 |
| SHROOM4 | 5.04293551 | -0.4454039 | 0.77957368 | -0.5713429 | 0.56776722 | NA |
| MAGED1 | 44.4149507 | -0.0849787 | 0.32627175 | -0.2604539 | 0.7945137 | 0.96388958 |
| FAM156A | 2.3585203 | 0.83134606 | 0.95435183 | 0.87111067 | 0.38369373 | NA |
| KANTR | 18.6927554 | 0.0183316 | 0.51976017 | 0.03526934 | 0.97186497 | NA |
| KDM5C | 9.06383717 | -1.3050655 | 0.76859642 | -1.6979853 | 0.08951053 | NA |
| IQSEC2 | 4.12392105 | -0.1030081 | 0.86570194 | -0.1189879 | 0.90528491 | NA |
| SMC1A | 12.5934322 | -0.5630658 | 0.48967986 | -1.1498651 | 0.25019944 | NA |
| HSD17B10 | 6.31937969 | 1.30802273 | 0.60198628 | 2.17284476 | 0.029792 | NA |
| HUWE1 | 50.5528209 | 0.18029798 | 0.26009428 | 0.6932024 | 0.48818254 | 0.86354292 |
| PHF8 | 21.2174717 | 0.45739152 | 0.3743373 | 1.22187001 | 0.22175679 | NA |
| FAM120C | 8.77885448 | 0.4082816 | 0.50620583 | 0.80655254 | 0.41992432 | NA |
| WNK3 | 30.3579716 | -0.2951287 | 0.36187324 | -0.8155581 | 0.41475292 | 0.83545324 |
| TSR2 | 5.8434503 | -0.2942929 | 0.78828392 | -0.3733336 | 0.70890018 | NA |
| FGD1 | 2.65127286 | 1.38844967 | 0.89791274 | 1.54630801 | 0.1220302 | NA |
| GNL3L | 3.58972673 | -0.0019383 | 0.78783609 | -0.0024603 | 0.99803693 | NA |
| MAGED2 | 15.0155696 | 0.50668857 | 0.46046424 | 1.10038638 | 0.27116381 | NA |
| RRAGB | 15.3990444 | 0.00977875 | 0.43967815 | 0.02224069 | 0.98225596 | NA |
| AL445472.1 | 8.98259797 | 1.0005157 | 0.52423043 | 1.90854183 | 0.05632122 | NA |
| KLF8 | 9.5361268 | 0.13444252 | 0.6274033 | 0.21428405 | 0.83032556 | NA |
| UBQLN2 | 6.51911326 | -0.1153218 | 0.60711355 | -0.189951 | 0.84934755 | NA |
| NBDY | 28.415572 | 0.20667591 | 0.33110595 | 0.62419871 | 0.5324971 | 0.88254911 |
| SPIN3 | 5.86768235 | 0.16114234 | 0.6046326 | 0.26651282 | 0.78984429 | NA |
| FAAH2 | 66.4970004 | 0.7573529 | 0.34923481 | 2.16860657 | 0.03011257 | 0.40822098 |
| ARHGEF9 | 7.80658042 | -0.0001831 | 0.57490057 | -0.0003185 | 0.99974586 | NA |
| MTMR8 | 2.98839975 | 0.70867497 | 0.80684158 | 0.87833223 | 0.37976345 | NA |
| ZC4H2 | 5.79052276 | 0.67407326 | 0.7029135 | 0.95897044 | 0.33757364 | NA |
| LAS1L | 4.12746455 | -0.07471 | 0.754535 | -0.0990146 | 0.92112668 | NA |
| MSN | 28.6700436 | 0.69933248 | 0.38870098 | 1.79915286 | 0.0719945 | 0.53207981 |
| HEPH | 4.01091719 | 2.20161359 | 1.28512062 | 1.71315717 | 0.08668363 | NA |
| OPHN1 | 32.3304357 | 0.69923627 | 0.441515 | 1.5837203 | 0.11325738 | 0.57860396 |
| YIPF6 | 16.1841027 | -0.5276044 | 0.42405068 | -1.2442013 | 0.21342548 | NA |
| EDA | 8.01623328 | 1.19578393 | 0.62577786 | 1.91087607 | 0.05602051 | NA |
| IGBP1 | 11.0788469 | 0.27467898 | 0.50558271 | 0.54329187 | 0.58692886 | NA |
| PDZD11 | 3.22272869 | 1.53007771 | 0.80375541 | 1.90366087 | 0.05695437 | NA |
| DLG3 | 9.74375911 | 0.07394366 | 0.47488041 | 0.15571005 | 0.87626157 | NA |
| SNX12 | 5.02551889 | -1.2081403 | 0.68090057 | -1.7743271 | 0.07600906 | NA |
| MED12 | 1.80946218 | -0.2650632 | 1.10119649 | -0.2407047 | 0.80978398 | NA |
| ZMYM3 | 2.23064149 | 0.81436905 | 1.0431921 | 0.78065109 | 0.43500773 | NA |
| NONO | 19.7123222 | -0.2853644 | 0.35528428 | -0.8032003 | 0.42185897 | NA |
| TAF1 | 22.1736055 | -0.0201194 | 0.39598359 | -0.0508086 | 0.95947801 | NA |
| OGT | 38.6259058 | -0.4121285 | 0.41699645 | -0.9883261 | 0.32299297 | 0.78622244 |
| GCNA | 3.75268855 | 0.41678509 | 0.84691454 | 0.49212178 | 0.62263326 | NA |
| NHSL2 | 5.91912451 | 0.56701056 | 0.60137464 | 0.94285745 | 0.34575382 | NA |
| PIN4 | 9.43985173 | -0.2542723 | 0.61018336 | -0.4167146 | 0.67688721 | NA |
| RPS4X | 84.3574197 | 0.19755896 | 0.39141604 | 0.50472884 | 0.6137493 | 0.91163569 |
| HDAC8 | 56.3408565 | 0.47390702 | 0.25429374 | 1.86362046 | 0.06237502 | 0.51631533 |

| | | | | | | |
|---|---|---|---|---|---|---|
| PHKA1 | 5.20946341 | 0.87032679 | 0.65764184 | 1.32340544 | 0.18570058 | NA |
| CHIC1 | 19.1998606 | -0.3743317 | 0.38115979 | -0.9820859 | 0.32605752 | NA |
| XIST | 81.6449425 | -0.3337701 | 3.82503756 | -0.0872593 | 0.9304654 | 0.98921679 |
| JPX | 61.6920453 | 0.57366212 | 0.35353746 | 1.62263461 | 0.10466754 | 0.5772605 |
| FTX | 472.67041 | 0.29264117 | 0.26129558 | 1.11996217 | 0.26272988 | 0.75479335 |
| RLIM | 11.664305 | -0.6308793 | 0.45950569 | -1.372952 | 0.16976726 | NA |
| NEXMIF | 5.17697335 | 0.47916789 | 0.62843247 | 0.76248111 | 0.44577291 | NA |
| ABCB7 | 25.0373082 | -0.2333122 | 0.33580989 | -0.6947747 | 0.48719652 | NA |
| UPRT | 5.45817182 | -0.6457864 | 0.68703277 | -0.9399644 | 0.34723581 | NA |
| ZDHHC15 | 7.64273616 | 0.51918494 | 0.53428793 | 0.97173249 | 0.33118365 | NA |
| PBDC1 | 11.3004125 | 0.18667051 | 0.50145304 | 0.37225921 | 0.70969987 | NA |
| ATRX | 101.470692 | -0.364144 | 0.28302945 | -1.2865942 | 0.19823576 | 0.69343545 |
| MAGT1 | 17.0506519 | -0.4752801 | 0.45986193 | -1.0335278 | 0.30135696 | NA |
| COX7B | 41.230003 | 0.13642963 | 0.35965457 | 0.37933519 | 0.70443897 | 0.9303056 |
| PGK1 | 39.8868417 | -0.1917187 | 0.44781668 | -0.4281188 | 0.66856463 | 0.92455906 |
| BRWD3 | 30.9463947 | 0.01668481 | 0.4270679 | 0.03906827 | 0.96883596 | 0.99684779 |
| HMGN5 | 4.95894352 | 0.0643291 | 0.72663515 | 0.08853013 | 0.92945534 | NA |
| SH3BGRL | 32.7411171 | 0.63032749 | 0.40122633 | 1.57100228 | 0.11618212 | 0.57860396 |
| POU3F4 | 3.69047672 | -0.2477636 | 0.85877018 | -0.2885098 | 0.77295652 | NA |
| RPS6KA6 | 29.4684106 | 0.21582671 | 0.46817382 | 0.46099697 | 0.64480078 | 0.92153493 |
| HDX | 9.78677406 | -1.621151 | 0.53746732 | -3.0162782 | 0.00255898 | NA |
| APOOL | 31.3858106 | -0.2109867 | 0.32347242 | -0.6522558 | 0.51423619 | 0.87158522 |
| ZNF711 | 2.71958165 | 0.5550894 | 0.93026908 | 0.59669767 | 0.55070925 | NA |
| CHM | 25.5676624 | -0.0913797 | 0.31965522 | -0.2858695 | 0.77497804 | NA |
| PCDH11X | 2.51505359 | -0.3499496 | 0.8995982 | -0.3890066 | 0.69727127 | NA |
| DIAPH2 | 85.5768082 | -0.007157 | 0.26960694 | -0.0265461 | 0.97882175 | 0.99902646 |
| DIAPH2-AS1 | 3.92668261 | 0.54086161 | 0.70325381 | 0.76908451 | 0.44184314 | NA |
| TSPAN6 | 6.44955432 | 0.29927924 | 0.58985065 | 0.50738139 | 0.61188724 | NA |
| TRMT2B | 12.5723764 | 0.69492741 | 0.4334962 | 1.60307614 | 0.10891784 | NA |
| HNRNPH2 | 10.0934893 | -0.5991219 | 0.47589377 | -1.2589403 | 0.20805189 | NA |
| ARMCX4 | 24.1271817 | 0.46819274 | 0.51187623 | 0.91466006 | 0.36037012 | NA |
| ARMCX6 | 3.09294748 | -0.1247312 | 0.76799247 | -0.162412 | 0.8709814 | NA |
| ARMCX3 | 12.2731223 | -0.5626602 | 0.44550944 | -1.2629592 | 0.20660384 | NA |
| ZMAT1 | 21.6920704 | 0.71465346 | 0.43661772 | 1.63679442 | 0.10167343 | NA |
| TCEAL2 | 2.71501245 | 0.05321813 | 0.83311257 | 0.06387868 | 0.94906683 | NA |
| BEX5 | 20.2106732 | -0.966261 | 1.21504579 | -0.7952466 | 0.42647007 | NA |
| ARMCX5 | 2.7716263 | -0.5118647 | 0.89715056 | -0.5705449 | 0.56830816 | NA |
| LINC00630 | 11.9588757 | 0.23683455 | 0.49906802 | 0.47455365 | 0.63510515 | NA |
| BEX4 | 14.1725283 | -0.0728302 | 0.59573201 | -0.1222532 | 0.90269848 | NA |
| TCEAL8 | 5.9643829 | -0.5970091 | 0.58635751 | -1.0181657 | 0.3085992 | NA |
| BEX2 | 7.49649505 | 1.49667924 | 0.72927974 | 2.05227043 | 0.04014339 | NA |
| TCEAL9 | 11.1189481 | -0.7629236 | 0.53413045 | -1.4283469 | 0.15319204 | NA |
| BEX3 | 17.468101 | 0.98782949 | 0.57401522 | 1.72091166 | 0.08526686 | NA |
| TCEAL4 | 17.328037 | -0.3100616 | 0.38571695 | -0.8038579 | 0.42147904 | NA |
| TCEAL3 | 5.55857469 | -0.1650775 | 0.63871471 | -0.2584527 | 0.79605755 | NA |
| MORF4L2 | 28.3973791 | -0.6268278 | 0.37503147 | -1.6714007 | 0.09464256 | 0.56264113 |
| SLC25A53 | 13.4815131 | -0.1935771 | 0.45319295 | -0.4271407 | 0.66927686 | NA |
| FAM199X | 20.0299038 | -0.1889919 | 0.38776449 | -0.4873883 | 0.62598321 | NA |
| IL1RAPL2 | 2.84532738 | 0.26036029 | 0.91767757 | 0.28371652 | 0.77662763 | NA |
| NRK | 16.4207101 | 0.47320262 | 0.66821527 | 0.70815894 | 0.47884656 | NA |
| RNF128 | 4.70308503 | -0.8349166 | 0.88740629 | -0.9408505 | 0.34678148 | NA |
| TBC1D8B | 6.97608714 | 0.59661478 | 0.66782184 | 0.89337416 | 0.37165683 | NA |
| MORC4 | 8.36812891 | 0.44559794 | 0.54272034 | 0.82104521 | 0.41162052 | NA |
| RBM41 | 16.2757377 | 0.04271338 | 0.4564481 | 0.09357773 | 0.9254446 | NA |

| | | | | | | |
|---|---|---|---|---|---|---|
| NUP62CL | 3.28261607 | 0.39155714 | 0.88264165 | 0.44361961 | 0.65731763 | NA |
| MID2 | 6.28415183 | -0.1761138 | 0.67456651 | -0.2610769 | 0.79403319 | NA |
| ATG4A | 2.33997936 | -0.6009787 | 0.93655109 | -0.6416935 | 0.52107221 | NA |
| COL4A5 | 14.4981072 | 0.78156874 | 0.56710564 | 1.37817136 | 0.16815039 | NA |
| ACSL4 | 168.867085 | -0.3322522 | 0.48297722 | -0.6879253 | 0.49149984 | 0.86531205 |
| TMEM164 | 13.9308669 | 0.26431742 | 0.4575958 | 0.57762205 | 0.56351932 | NA |
| AMMECR1 | 17.1363627 | -0.0603792 | 0.3903722 | -0.1546709 | 0.87708079 | NA |
| ALG13 | 15.1134416 | -0.0134286 | 0.47619311 | -0.0281998 | 0.97750278 | NA |
| PLS3 | 48.7606853 | -0.0269189 | 0.32440811 | -0.0829785 | 0.93386864 | 0.9907482 |
| DANT2 | 123.687522 | -0.5969506 | 0.41280018 | -1.4461006 | 0.14814898 | 0.63 |
| KLHL13 | 16.7027297 | 0.26886732 | 0.68182437 | 0.39433516 | 0.69333361 | NA |
| WDR44 | 17.055227 | -0.2266281 | 0.41467951 | -0.5465138 | 0.58471282 | NA |
| IL13RA1 | 14.9376195 | 0.3366649 | 0.40806834 | 0.82502088 | 0.40935974 | NA |
| PGRMC1 | 22.0989971 | 0.3568268 | 0.33210113 | 1.07445221 | 0.28262005 | NA |
| SLC25A43 | 7.38980572 | 0.33503815 | 0.58397527 | 0.57371976 | 0.56615746 | NA |
| SLC25A5 | 38.5131657 | 0.87884801 | 0.43445243 | 2.02288664 | 0.04308484 | 0.44186191 |
| CXorf56 | 3.52449732 | -0.3707187 | 0.80228741 | -0.4620771 | 0.644026 | NA |
| UBE2A | 10.1933523 | -0.3186948 | 0.50449578 | -0.6317096 | 0.52757669 | NA |
| NKRF | 3.2449761 | 0.33478409 | 0.79581912 | 0.42067862 | 0.67398978 | NA |
| 6-Sep | 12.810441 | 0.08587829 | 0.47779562 | 0.17973854 | 0.85735783 | NA |
| RPL39 | 3.43681025 | 0.36548268 | 0.8982448 | 0.40688538 | 0.68409217 | NA |
| UPF3B | 10.4164847 | -0.6085487 | 0.53916097 | -1.1286958 | 0.25902618 | NA |
| NDUFA1 | 24.3946586 | 0.21987568 | 0.34068826 | 0.6453867 | 0.51867662 | NA |
| NKAP | 4.1334398 | 0.5290974 | 0.77215515 | 0.68522161 | 0.49320409 | NA |
| RHOXF1-AS1 | 5.16472311 | 1.34628056 | 0.69273882 | 1.94341723 | 0.05196577 | NA |
| ZBTB33 | 2.31709984 | -0.3917739 | 0.89607382 | -0.4372117 | 0.66195785 | NA |
| TMEM255A | 15.5884786 | -0.6859811 | 0.58921557 | -1.1642278 | 0.2443317 | NA |
| LAMP2 | 28.3703242 | -0.0369001 | 0.35245115 | -0.1046958 | 0.91661722 | 0.98528045 |
| CUL4B | 8.81286098 | -0.5618688 | 0.54401614 | -1.0328164 | 0.30168985 | NA |
| MCTS1 | 6.76534044 | -0.1484295 | 0.60385478 | -0.2458033 | 0.80583448 | NA |
| C1GALT1C1 | 3.77772584 | 0.03356575 | 0.86000733 | 0.03902961 | 0.96886678 | NA |
| THOC2 | 39.4835376 | -0.1145063 | 0.29399915 | -0.3894782 | 0.6969224 | 0.9303056 |
| XIAP | 38.173629 | -0.4725287 | 0.3560238 | -1.3272392 | 0.18442958 | 0.67802005 |
| STAG2 | 55.9348642 | -0.2277264 | 0.29042982 | -0.7841014 | 0.43298061 | 0.84223493 |
| SMARCA1 | 13.0123122 | 0.41635495 | 0.59128978 | 0.70414703 | 0.48134121 | NA |
| OCRL | 5.00039257 | 0.18360755 | 0.6718241 | 0.27329706 | 0.78462487 | NA |
| ZDHHC9 | 19.2933905 | -0.490819 | 0.38114235 | -1.2877576 | 0.19783034 | NA |
| UTP14A | 3.87822547 | -1.3808643 | 0.77280412 | -1.7868232 | 0.07396607 | NA |
| BCORL1 | 3.95620132 | 1.22381032 | 0.72288838 | 1.69294507 | 0.09046592 | NA |
| ELF4 | 4.90966461 | 0.49244419 | 0.67914455 | 0.72509482 | 0.46839386 | NA |
| AIFM1 | 12.8568585 | 0.44867929 | 0.45594004 | 0.98407522 | 0.32507853 | NA |
| ZNF280C | 4.25398459 | -0.9867738 | 0.83934987 | -1.1756407 | 0.2397385 | NA |
| SLC25A14 | 3.34387959 | -0.1638301 | 0.90619818 | -0.1807884 | 0.85653365 | NA |
| RBMX2 | 3.13150143 | -1.0824684 | 0.79898414 | -1.3548058 | 0.17547944 | NA |
| ENOX2 | 24.4144107 | -0.2764859 | 0.35096741 | -0.7877822 | 0.43082413 | NA |
| STK26 | 4.56203222 | -1.087959 | 0.73510544 | -1.480004 | 0.13887218 | NA |
| FRMD7 | 7.65587697 | -0.5011213 | 0.72418051 | -0.691984 | 0.48894739 | NA |
| RAP2C | 13.665579 | 0.57662824 | 0.46972405 | 1.22758935 | 0.21960116 | NA |
| RAP2C-AS1 | 18.1888808 | 0.08999153 | 0.53694421 | 0.16759941 | 0.86689843 | NA |
| MBNL3 | 10.7092972 | 0.287429 | 0.50492203 | 0.56925423 | 0.56918362 | NA |
| GPC4 | 6.94679152 | -0.4688911 | 0.66605471 | -0.7039828 | 0.48144348 | NA |
| GPC3 | 20.6805142 | 0.8067652 | 0.72758499 | 1.10882606 | 0.26750523 | NA |
| PHF6 | 9.71633773 | -0.2503244 | 0.51726393 | -0.4839394 | 0.6284289 | NA |
| HPRT1 | 31.1331437 | 0.77638561 | 0.47476338 | 1.63531066 | 0.10198394 | 0.5750956 |

| | | | | | | |
|---|---|---|---|---|---|---|
| FAM122B | 5.72134357 | 0.79005521 | 0.67312525 | 1.17371203 | 0.24051039 | NA |
| MOSPD1 | 3.20349584 | -0.0131076 | 0.88775869 | -0.0147648 | 0.98821983 | NA |
| RTL8C | 9.89295346 | -0.0215951 | 0.50361669 | -0.0428801 | 0.96579712 | NA |
| RTL8A | 5.79898796 | 0.15340729 | 0.62324139 | 0.24614425 | 0.80557057 | NA |
| ZNF75D | 11.2437984 | -0.5304738 | 0.57874028 | -0.9166008 | 0.35935186 | NA |
| INTS6L | 4.70810222 | -0.2272883 | 0.88442223 | -0.2569907 | 0.79718598 | NA |
| MMGT1 | 6.355309 | 0.75347966 | 0.58039924 | 1.29820925 | 0.19421544 | NA |
| SLC9A6 | 4.57552827 | -1.0967867 | 0.66057126 | -1.6603609 | 0.09684188 | NA |
| HTATSF1 | 8.26753827 | -0.0641093 | 0.51991386 | -0.1233075 | 0.90186357 | NA |
| AL683813.1 | 4.43515397 | 0.2994581 | 0.70007808 | 0.42774958 | 0.66883345 | NA |
| RBMX | 34.5355477 | -0.1056626 | 0.32697646 | -0.3231504 | 0.74658131 | 0.94858421 |
| FGF13 | 17.3988809 | 1.07573887 | 1.24717914 | 0.86253758 | 0.38839176 | NA |
| ATP11C | 16.1697436 | 0.80335618 | 0.52501316 | 1.5301639 | 0.12597617 | NA |
| FMR1 | 20.1568489 | 0.24256914 | 0.43383084 | 0.559133 | 0.57607095 | NA |
| IDS | 12.704669 | -0.4040165 | 0.48918064 | -0.8259045 | 0.40885826 | NA |
| TMEM185A | 5.05847549 | 0.33836444 | 0.64204107 | 0.5270137 | 0.59818407 | NA |
| LINC00894 | 25.1161346 | 0.18298425 | 0.40962803 | 0.44670832 | 0.65508567 | NA |
| MTM1 | 11.3840869 | 0.32003133 | 0.57022719 | 0.56123477 | 0.5746375 | NA |
| MTMR1 | 15.6300225 | -0.4654704 | 0.44998331 | -1.034417 | 0.30094125 | NA |
| CD99L2 | 9.29904558 | 0.49601359 | 0.51053773 | 0.97155128 | 0.33127383 | NA |
| HMGB3 | 2.58595473 | 0.11975245 | 0.87790269 | 0.13640744 | 0.8914992 | NA |
| VMA21 | 11.1351598 | -0.3859803 | 0.50669528 | -0.7617603 | 0.44620311 | NA |
| CETN2 | 2.8028599 | 0.8248693 | 0.83846519 | 0.98378479 | 0.32522134 | NA |
| SLC6A8 | 7.24757897 | -0.2874709 | 0.5464951 | -0.5260265 | 0.59886978 | NA |
| BCAP31 | 15.5667054 | -0.4832274 | 0.44080544 | -1.0962373 | 0.27297493 | NA |
| IDH3G | 6.10132208 | 0.0933154 | 0.61087139 | 0.15275785 | 0.87858924 | NA |
| SSR4 | 12.8397685 | 0.46119459 | 0.41231359 | 1.11855298 | 0.26333089 | NA |
| RENBP | 2.52404057 | -0.933447 | 1.26708197 | -0.7366903 | 0.46131073 | NA |
| HCFC1 | 6.71014534 | -0.5598078 | 0.5835034 | -0.9593908 | 0.33736189 | NA |
| IRAK1 | 7.18359858 | -0.6504102 | 0.75497368 | -0.8615005 | 0.38896242 | NA |
| MECP2 | 42.5201563 | -0.1613621 | 0.38379534 | -0.420438 | 0.67416553 | 0.92617027 |
| FLNA | 6.03610242 | -2.3364932 | 0.99560906 | -2.3467979 | 0.01893552 | NA |
| EMD | 9.4954116 | -0.3166173 | 0.49098326 | -0.6448637 | 0.51901552 | NA |
| RPL10 | 102.124372 | 0.18118024 | 0.58297514 | 0.31078553 | 0.75596367 | 0.95323728 |
| DNASE1L1 | 2.60894634 | -0.2424766 | 0.85026892 | -0.2851763 | 0.77550906 | NA |
| TAZ | 6.3330182 | 0.88740446 | 0.59508956 | 1.49121162 | 0.13590594 | NA |
| AC244090.1 | 3.29242327 | -0.6886902 | 1.0235428 | -0.6728494 | 0.50104308 | NA |
| ATP6AP1 | 20.7979175 | -0.205966 | 0.51483953 | -0.4000586 | 0.68911335 | NA |
| GDI1 | 9.53748854 | -0.38481 | 0.46959119 | -0.8194575 | 0.41252546 | NA |
| FAM50A | 8.54154101 | -1.1723373 | 0.59903844 | -1.9570319 | 0.05034372 | NA |
| PLXNA3 | 5.56414218 | -0.7132396 | 0.69251455 | -1.0299272 | 0.30304418 | NA |
| UBL4A | 3.62930342 | -0.3250289 | 0.80503368 | -0.4037457 | 0.68639976 | NA |
| FAM3A | 3.97730246 | -0.1851453 | 0.73587828 | -0.2515977 | 0.80135204 | NA |
| G6PD | 2.6838592 | -0.1143336 | 0.88294663 | -0.1294909 | 0.89696923 | NA |
| IKBKG | 3.1368312 | -0.1862872 | 0.77184971 | -0.2413517 | 0.80928256 | NA |
| DKC1 | 6.77016996 | -0.1949108 | 0.5442369 | -0.3581359 | 0.72024162 | NA |
| MPP1 | 4.46929157 | 0.10887086 | 0.72229913 | 0.15072821 | 0.88019012 | NA |
| F8 | 2.59381281 | 1.52654744 | 1.02915016 | 1.48330876 | 0.1379924 | NA |
| F8A1 | 2.1187921 | -0.1539802 | 1.12850498 | -0.1364462 | 0.89146853 | NA |
| FUNDC2 | 14.9019483 | -0.0042441 | 0.55861389 | -0.0075976 | 0.99393804 | NA |
| BRCC3 | 8.53662222 | 0.66979976 | 0.59304729 | 1.12942048 | 0.2587205 | NA |
| VBP1 | 6.91843123 | -0.5737261 | 0.5860233 | -0.9790159 | 0.32757214 | NA |
| TMLHE-AS1 | 8.6021096 | 1.02549425 | 0.69079226 | 1.48451902 | 0.13767128 | NA |
| TMLHE | 24.830004 | 0.34262042 | 0.45010072 | 0.76120832 | 0.44653265 | NA |

| | | | | | | |
|---|---|---|---|---|---|---|
| VAMP7 | 8.65929868 | -0.1167247 | 0.48000191 | -0.2431755 | 0.80786944 | NA |
| ZNF596 | 4.94558798 | 0.05825379 | 0.66123257 | 0.08809879 | 0.92979816 | NA |
| FBXO25 | 22.1101285 | 0.3110252 | 0.4298556 | 0.7235574 | 0.4693375 | NA |
| TDRP | 12.7542451 | -0.1309806 | 0.49624466 | -0.2639436 | 0.79182337 | NA |
| ERICH1 | 28.8124169 | -0.2659069 | 0.44332421 | -0.5998022 | 0.54863804 | 0.88448477 |
| AC100810.1 | 2.92353916 | -0.1174681 | 1.00818251 | -0.1165147 | 0.90724465 | NA |
| AC019257.8 | 4.05792082 | -0.6972435 | 0.87529026 | -0.7965855 | 0.4256918 | NA |
| CSMD1 | 13.8381127 | -0.4969973 | 0.71200854 | -0.6980215 | 0.48516374 | NA |
| MCPH1 | 35.2787915 | 0.04041391 | 0.4135521 | 0.09772386 | 0.92215157 | 0.98745437 |
| ANGPT2 | 5.50695471 | 0.08146334 | 0.76427363 | 0.10658924 | 0.91511486 | NA |
| MCPH1-AS1 | 9.73904123 | 0.32953285 | 0.48916527 | 0.67366362 | 0.5005252 | NA |
| AGPAT5 | 18.4835767 | -0.6869596 | 0.51377502 | -1.3370825 | 0.18119572 | NA |
| AF233439.1 | 58.3722638 | -0.7113669 | 0.99868135 | -0.7123061 | 0.47627522 | 0.86206794 |
| DEFB1 | 144.827611 | -0.2498205 | 0.50685752 | -0.4928811 | 0.6220966 | 0.91493825 |
| FAM85B | 17.3464976 | 1.13194618 | 0.4322071 | 2.61899023 | 0.00881905 | NA |
| PRAG1 | 10.7682268 | -0.1552758 | 0.59343674 | -0.2616552 | 0.79358728 | NA |
| MFHAS1 | 198.092413 | -0.3614921 | 0.34936704 | -1.0347057 | 0.30080638 | 0.77521265 |
| ERI1 | 12.7129053 | 0.30737667 | 0.55154648 | 0.55729967 | 0.5773227 | NA |
| TNKS | 63.2761964 | -0.1624382 | 0.28068599 | -0.5787185 | 0.56277916 | 0.89263108 |
| MSRA | 32.4723085 | -0.1195962 | 0.32329764 | -0.3699259 | 0.71143773 | 0.93492235 |
| XKR6 | 109.247216 | -0.289105 | 0.37012895 | -0.7810926 | 0.43474801 | 0.84265972 |
| MTMR9 | 15.6151591 | 0.31056685 | 0.38570782 | 0.80518681 | 0.42071189 | NA |
| FDFT1 | 25.0115938 | -0.5803682 | 0.42489123 | -1.3659218 | 0.1719635 | NA |
| CTSB | 44.0118807 | 0.29136472 | 0.328501 | 0.88695231 | 0.37510458 | 0.81593653 |
| FAM66D | 2.9877214 | 0.63331632 | 0.94775753 | 0.6682261 | 0.50398928 | NA |
| LONRF1 | 10.59987 | 0.26126426 | 0.43429364 | 0.60158435 | 0.54745085 | NA |
| TRMT9B | 81.3232339 | 0.757528 | 0.43663203 | 1.73493455 | 0.0827524 | 0.55216537 |
| DLC1 | 15.3409465 | -0.9985465 | 0.61723725 | -1.6177678 | 0.10571265 | NA |
| TUSC3 | 18.6253338 | -0.7717502 | 0.37577591 | -2.0537511 | 0.03999979 | NA |
| MSR1 | 5.86610784 | -0.1650011 | 0.74650829 | -0.2210306 | 0.82506864 | NA |
| MICU3 | 20.0006278 | -0.3526098 | 0.38054082 | -0.9266017 | 0.35413335 | NA |
| ZDHHC2 | 17.8923311 | 0.04470265 | 0.39908741 | 0.11201218 | 0.91081375 | NA |
| CNOT7 | 8.87674706 | -0.1950056 | 0.50730127 | -0.3843981 | 0.70068342 | NA |
| VPS37A | 14.9188817 | -0.0993681 | 0.43539487 | -0.2282251 | 0.81947121 | NA |
| SLC7A2 | 5.69762287 | 0.15186993 | 1.00865857 | 0.15056624 | 0.8803179 | NA |
| MTUS1 | 61.7434644 | -0.6534625 | 0.37964308 | -1.7212549 | 0.08520457 | 0.55323508 |
| AC027117.2 | 5.73393418 | -0.7735784 | 0.65035368 | -1.1894735 | 0.23425341 | NA |
| PCM1 | 93.43895 | -0.0580147 | 0.23678064 | -0.2450147 | 0.80644501 | 0.96800006 |
| ASAH1 | 22.9777049 | 0.25210504 | 0.33801637 | 0.74583678 | 0.45576602 | NA |
| PSD3 | 250.02537 | -0.8564031 | 0.4220161 | -2.0293139 | 0.04242633 | 0.44186191 |
| SH2D4A | 42.6416567 | 1.08365523 | 0.43760794 | 2.4763153 | 0.01327463 | 0.30435125 |
| CSGALNACT1 | 31.3299902 | -0.5644774 | 0.5878436 | -0.960251 | 0.33692891 | 0.8003972 |
| INTS10 | 25.7505246 | 0.27505878 | 0.33318043 | 0.82555503 | 0.40905656 | NA |
| ATP6V1B2 | 8.09360977 | -0.4761917 | 0.52868459 | -0.9007103 | 0.36774237 | NA |
| XPO7 | 20.1647174 | -0.5615669 | 0.3501902 | -1.6036053 | 0.10880108 | NA |
| DMTN | 19.5567156 | -0.3539117 | 0.44339096 | -0.7981933 | 0.42475833 | NA |
| FAM160B2 | 6.77841627 | 0.42904018 | 0.64565867 | 0.66449999 | 0.50637036 | NA |
| SLC39A14 | 8.15276511 | -0.5516031 | 0.68700066 | -0.8029149 | 0.42202392 | NA |
| PPP3CC | 11.4932513 | -1.0735475 | 0.47187352 | -2.2750746 | 0.02290145 | NA |
| CCAR2 | 4.29692621 | -0.2732428 | 0.73941635 | -0.3695385 | 0.71172642 | NA |
| BIN3 | 9.9106287 | -1.1992751 | 0.568788 | -2.1084746 | 0.03498996 | NA |
| TNFRSF10B | 11.402443 | -0.0306941 | 0.46497437 | -0.0660125 | 0.94736784 | NA |
| TNFRSF10D | 6.18462906 | -0.5553631 | 0.57159389 | -0.9716044 | 0.33124742 | NA |
| CHMP7 | 5.33886829 | -0.948265 | 0.68488456 | -1.3845619 | 0.16618648 | NA |

| | | | | | | |
|---|---|---|---|---|---|---|
| R3HCC1 | 4.77220331 | 0.54393612 | 0.68455476 | 0.79458379 | 0.42685565 | NA |
| ENTPD4 | 11.2489311 | -0.0233531 | 0.48803099 | -0.0478517 | 0.96183447 | NA |
| SLC25A37 | 13.8650789 | 0.62698843 | 0.47857612 | 1.31011225 | 0.19015786 | NA |
| DOCK5 | 100.496633 | 0.43944928 | 0.55202392 | 0.79606927 | 0.42599178 | 0.84057082 |
| KCTD9 | 12.9646696 | 0.10858912 | 0.41326718 | 0.26275767 | 0.79273736 | NA |
| PPP2R2A | 47.8997072 | -0.1097925 | 0.30726569 | -0.3573209 | 0.72085157 | 0.93759885 |
| BNIP3L | 20.4471156 | 0.11486112 | 0.63965657 | 0.17956686 | 0.85749262 | NA |
| DPYSL2 | 3.67535659 | 2.17258274 | 0.88139686 | 2.46493134 | 0.01370395 | NA |
| TRIM35 | 2.91635366 | -1.891101 | 1.19342253 | -1.5846031 | 0.11305654 | NA |
| PTK2B | 23.7522474 | 0.85108741 | 0.42239431 | 2.01491212 | 0.04391386 | NA |
| EPHX2 | 7.80078585 | 0.75185484 | 0.69279946 | 1.08524167 | 0.27781463 | NA |
| CLU | 33.0463566 | 0.48692374 | 0.56785116 | 0.85748481 | 0.39117701 | 0.8258317 |
| CCDC25 | 6.79460724 | 0.29674575 | 0.58958587 | 0.50331219 | 0.6147448 | NA |
| ELP3 | 24.3226116 | 0.24554333 | 0.39330644 | 0.62430541 | 0.53242703 | NA |
| ZNF395 | 5.95898556 | 0.58075584 | 0.64701292 | 0.89759543 | 0.36940127 | NA |
| FZD3 | 7.5911875 | 1.13321412 | 0.61622085 | 1.83897398 | 0.06591901 | NA |
| EXTL3 | 7.92849893 | 0.30760742 | 0.71380433 | 0.43094082 | 0.6665114 | NA |
| INTS9 | 23.0940418 | 0.46595492 | 0.37612112 | 1.23884274 | 0.21540374 | NA |
| HMBOX1 | 158.640977 | 0.28517765 | 0.32033385 | 0.89025137 | 0.37333092 | 0.81364119 |
| KIF13B | 293.075221 | -0.2298834 | 0.27064497 | -0.8493909 | 0.3956638 | 0.8258317 |
| AC108449.1 | 12.674678 | -0.3654707 | 0.71143885 | -0.5137063 | 0.60745731 | NA |
| SARAF | 31.5155056 | 0.18273904 | 0.28406382 | 0.64330277 | 0.52002766 | 0.87214537 |
| LEPROTL1 | 10.7990129 | 0.40663863 | 0.43383008 | 0.93732237 | 0.34859276 | NA |
| DCTN6 | 6.00957322 | 0.41521167 | 0.64453845 | 0.6442 | 0.51944575 | NA |
| RBPMS | 266.571224 | -0.0709721 | 0.36832902 | -0.1926867 | 0.84720433 | 0.97168963 |
| GTF2E2 | 16.2593137 | -0.004892 | 0.42698009 | -0.0114571 | 0.99085876 | NA |
| GSR | 16.3794744 | -0.4885234 | 0.40433838 | -1.2082044 | 0.22696866 | NA |
| PPP2CB | 35.0454034 | -0.1943397 | 0.34366622 | -0.5654897 | 0.57174072 | 0.89263108 |
| WRN | 37.3407623 | -0.3814311 | 0.30895204 | -1.2345964 | 0.21698075 | 0.71673468 |
| NRG1 | 12.0336017 | -1.8338643 | 1.57266035 | -1.1660905 | 0.24357785 | NA |
| FUT10 | 11.3042275 | 0.46868615 | 0.51598545 | 0.90833211 | 0.36370278 | NA |
| TTI2 | 5.10309159 | 0.15105009 | 0.63030403 | 0.23964639 | 0.81060439 | NA |
| UNC5D | 24.6396927 | -0.5241636 | 0.6703135 | -0.7819679 | 0.43423347 | NA |
| ERLIN2 | 10.8653653 | -0.0721675 | 0.51440755 | -0.1402926 | 0.88842885 | NA |
| PLPBP | 7.96225556 | 0.7116311 | 0.52754293 | 1.34895391 | 0.17735177 | NA |
| RAB11FIP1 | 19.8512332 | -0.1462793 | 0.40514964 | -0.3610501 | 0.71806199 | NA |
| ASH2L | 11.1012372 | -0.7825344 | 0.51952912 | -1.5062377 | 0.13200616 | NA |
| LSM1 | 7.86900534 | -0.3311618 | 0.54796553 | -0.6043478 | 0.54561242 | NA |
| BAG4 | 16.6128024 | -0.7021248 | 0.41933661 | -1.6743704 | 0.09405783 | NA |
| DDHD2 | 16.2817623 | -0.1947884 | 0.38484422 | -0.5061486 | 0.61275233 | NA |
| PLPP5 | 4.81307136 | -0.5565115 | 0.66916755 | -0.8316474 | 0.40560799 | NA |
| NSD3 | 47.1436696 | 0.29937564 | 0.30080956 | 0.99523311 | 0.3196229 | 0.78335494 |
| FGFR1 | 7.24512401 | -0.6884892 | 0.59392401 | -1.159221 | 0.24636612 | NA |
| TACC1 | 29.2224412 | -0.3218836 | 0.42581659 | -0.7559207 | 0.44969674 | 0.85123446 |
| PLEKHA2 | 13.5326555 | -0.0086522 | 0.43083667 | -0.0200824 | 0.98397765 | NA |
| TM2D2 | 3.88084078 | 0.37524393 | 0.77810511 | 0.48225352 | 0.62962586 | NA |
| ADAM9 | 25.9215069 | 0.20400613 | 0.33642536 | 0.60639343 | 0.54425353 | NA |
| TCIM | 37.3362048 | 0.29628387 | 0.5380079 | 0.55070543 | 0.58183562 | 0.89263108 |
| GOLGA7 | 10.0690112 | -0.0472747 | 0.47375418 | -0.0997873 | 0.92051318 | NA |
| GPAT4 | 13.9160684 | -0.3153395 | 0.41308952 | -0.7633683 | 0.44524376 | NA |
| KAT6A | 68.6137931 | -0.0260218 | 0.26194858 | -0.0993392 | 0.92086897 | 0.98745437 |
| AP3M2 | 4.52461959 | -0.0635692 | 0.71170791 | -0.0893192 | 0.92882825 | NA |
| IKBKB | 22.2549784 | -0.2759217 | 0.36212983 | -0.7619415 | 0.44609492 | NA |
| POLB | 6.42844098 | -0.4111037 | 0.68565171 | -0.599581 | 0.54878551 | NA |

| | | | | | | |
|---|---|---|---|---|---|---|
| VDAC3 | 16.7952454 | 0.09306388 | 0.4025328 | 0.23119578 | 0.81716271 | NA |
| SLC20A2 | 38.5110541 | -0.1942447 | 0.29292376 | -0.6631238 | 0.50725125 | 0.87158522 |
| SMIM19 | 8.92363252 | 0.22252901 | 0.48758984 | 0.45638567 | 0.64811267 | NA |
| THAP1 | 3.35837874 | 1.40084966 | 0.79879682 | 1.75369961 | 0.07948199 | NA |
| RNF170 | 10.9271134 | 1.06498049 | 0.4984638 | 2.13652527 | 0.03263662 | NA |
| HOOK3 | 27.1509573 | 0.06101712 | 0.31068166 | 0.19639757 | 0.844299 | 0.97168963 |
| FNTA | 11.0423797 | 0.58961725 | 0.56632776 | 1.0411237 | 0.29781814 | NA |
| HGSNAT | 26.2343804 | 0.6869641 | 0.37413038 | 1.83616233 | 0.06633365 | 0.5230103 |
| SPIDR | 146.162399 | -0.2941696 | 0.24800054 | -1.1861654 | 0.235557 | 0.73189946 |
| CEBPD | 80.5605211 | -0.0554705 | 0.31209221 | -0.1777376 | 0.85892905 | 0.97168963 |
| PRKDC | 89.9382829 | 0.07172317 | 0.27488067 | 0.26092474 | 0.79415055 | 0.96388714 |
| MCM4 | 10.3976695 | -0.8801496 | 0.68439831 | -1.2860195 | 0.19843625 | NA |
| UBE2V2 | 15.1500253 | 0.27634719 | 0.41945213 | 0.65882892 | 0.51000564 | NA |
| PXDNL | 10.7999621 | 0.74589938 | 0.52137904 | 1.43062785 | 0.1525369 | NA |
| PCMTD1 | 68.4266619 | 0.45657613 | 0.37807143 | 1.20764517 | 0.22718378 | 0.73054639 |
| AC064807.1 | 3.22794889 | 0.67519056 | 1.06547151 | 0.63370119 | 0.52627585 | NA |
| RB1CC1 | 47.4445946 | 0.00507325 | 0.26185791 | 0.01937407 | 0.98454269 | 0.99902646 |
| ATP6V1H | 96.7457878 | -0.1482671 | 0.29017407 | -0.5109592 | 0.60937961 | 0.9088434 |
| TCEA1 | 20.7691703 | 0.26731876 | 0.4215316 | 0.63416065 | 0.52597598 | NA |
| LYPLA1 | 43.796725 | -0.067305 | 0.30104861 | -0.2235686 | 0.82309299 | 0.96845695 |
| MRPL15 | 17.2188439 | 0.10745159 | 0.48035826 | 0.22369053 | 0.82299812 | NA |
| TMEM68 | 8.29989415 | 0.18997288 | 0.53693498 | 0.35380984 | 0.7234814 | NA |
| TGS1 | 12.7663547 | 0.01088487 | 0.44451956 | 0.02448681 | 0.9804643 | NA |
| LYN | 15.5810894 | -0.0767551 | 0.45798054 | -0.1675947 | 0.86690215 | NA |
| RPS20 | 35.2289401 | 0.11589305 | 0.4121334 | 0.28120277 | 0.77855488 | 0.96133329 |
| PLAG1 | 18.7653305 | 1.01420796 | 0.39896895 | 2.54207241 | 0.01101973 | NA |
| CHCHD7 | 7.60464698 | -0.3468768 | 0.61359168 | -0.5653219 | 0.57185486 | NA |
| IMPAD1 | 16.4366452 | 0.18522062 | 0.41528921 | 0.44600393 | 0.6555944 | NA |
| LINC01606 | 9.63052951 | 1.39369851 | 0.75465412 | 1.84680435 | 0.0647755 | NA |
| FAM110B | 62.4678472 | 0.37785474 | 0.43005482 | 0.87861993 | 0.37960739 | 0.82153751 |
| AC027698.1 | 4.89832754 | 1.24133137 | 0.82013652 | 1.51356674 | 0.13013576 | NA |
| UBXN2B | 11.0371447 | -0.5745633 | 0.46841049 | -1.2266235 | 0.21996413 | NA |
| SDCBP | 15.9822998 | 0.14115697 | 0.42868565 | 0.32927849 | 0.7419452 | NA |
| NSMAF | 12.8639428 | -0.3840955 | 0.52712234 | -0.7286648 | 0.46620672 | NA |
| TOX | 17.4395625 | -0.5279247 | 0.64579099 | -0.8174854 | 0.41365108 | NA |
| CA8 | 408.332064 | -2.106851 | 0.54511338 | -3.8649777 | 0.0001111 | 0.04332663 |
| LINC01301 | 5.31442582 | 0.17692292 | 0.71071519 | 0.24893646 | 0.80340993 | NA |
| RAB2A | 68.4776133 | 0.11126437 | 0.23675361 | 0.46995849 | 0.63838468 | 0.92121485 |
| CHD7 | 58.2612991 | 0.27680112 | 0.27942162 | 0.9906217 | 0.32187034 | 0.78597615 |
| CLVS1 | 2.34640996 | -0.968836 | 1.05241015 | -0.9205878 | 0.35726565 | NA |
| ASPH | 19.1598784 | 0.34984655 | 0.40400446 | 0.86594723 | 0.3865191 | NA |
| YTHDF3 | 29.4789155 | -0.0387223 | 0.34762836 | -0.1113899 | 0.91130714 | 0.98375925 |
| CYP7B1 | 27.4441189 | 0.41130654 | 0.47757751 | 0.86123517 | 0.38910854 | 0.8258317 |
| ARMC1 | 12.4939701 | -0.9813944 | 0.4723876 | -2.0775193 | 0.03775365 | NA |
| MTFR1 | 16.5578229 | 0.1329839 | 0.36343207 | 0.3659113 | 0.71443127 | NA |
| PDE7A | 56.4694725 | -0.1971208 | 0.40271991 | -0.4894737 | 0.62450635 | 0.91582826 |
| ADHFE1 | 9.88605581 | 0.97150382 | 0.64981897 | 1.49503765 | 0.13490462 | NA |
| VCPIP1 | 8.70233204 | -0.3262664 | 0.5640039 | -0.5784826 | 0.56293837 | NA |
| SGK3 | 17.7333625 | 0.3026849 | 0.40650346 | 0.74460596 | 0.45650997 | NA |
| CSPP1 | 52.7085755 | 0.47778082 | 0.40180388 | 1.18908962 | 0.2344044 | 0.73156941 |
| ARFGEF1 | 136.568056 | 0.00926015 | 0.2661151 | 0.03479752 | 0.97224119 | 0.99779576 |
| NCOA2 | 149.640458 | 0.36379337 | 0.24331836 | 1.49513325 | 0.13487967 | 0.60594834 |
| TRAM1 | 28.7000134 | 0.20092571 | 0.36083262 | 0.55683908 | 0.57763738 | 0.89263108 |
| LACTB2 | 8.43543189 | 0.66761591 | 0.50187077 | 1.33025462 | 0.18343439 | NA |

| | | | | | | |
|---|---|---|---|---|---|---|
| TERF1 | 10.5903962 | -0.0799848 | 0.49710042 | -0.1609026 | 0.87217011 | NA |
| RPL7 | 78.8503582 | 0.16855971 | 0.44747934 | 0.37668713 | 0.70640613 | 0.93152073 |
| RDH10 | 26.8197055 | -0.0837232 | 0.49856809 | -0.1679274 | 0.86664042 | 0.97217819 |
| RDH10-AS1 | 6.23235856 | 0.98498866 | 0.69318987 | 1.42095074 | 0.15533108 | NA |
| STAU2 | 45.0037883 | 0.38324823 | 0.282085 | 1.35862675 | 0.17426489 | 0.6651652 |
| AC027018.1 | 5.45329067 | 0.94177098 | 0.79930945 | 1.17823076 | 0.23870462 | NA |
| ELOC | 10.5080063 | -0.2255223 | 0.52794768 | -0.4271679 | 0.66925706 | NA |
| TMEM70 | 4.81959223 | 0.10184335 | 0.69360722 | 0.14683145 | 0.88326506 | NA |
| GDAP1 | 20.635907 | -0.416798 | 0.4878597 | -0.8543398 | 0.39291672 | NA |
| MIR2052HG | 4.60132975 | 1.07626612 | 0.83961325 | 1.28185939 | 0.19989198 | NA |
| HNF4G | 38.4790907 | -0.5746725 | 0.49087979 | -1.170699 | 0.24171979 | 0.73323864 |
| PEX2 | 12.531078 | 0.381972 | 0.47633102 | 0.80190453 | 0.42260819 | NA |
| PKIA | 5.26986376 | 1.79163961 | 0.86170855 | 2.07917121 | 0.03760162 | NA |
| ZC2HC1A | 19.0772654 | 0.40069334 | 0.55564265 | 0.72113496 | 0.47082648 | NA |
| IL7 | 10.2318917 | 1.29807501 | 0.65709272 | 1.97548225 | 0.04821345 | NA |
| TPD52 | 89.5416593 | -0.5254923 | 0.31325008 | -1.6775487 | 0.09343523 | 0.56220784 |
| AC034114.2 | 8.14959101 | -2.6698429 | 1.09204412 | -2.4448123 | 0.01449276 | NA |
| ZBTB10 | 18.5509618 | -0.9847843 | 0.46204166 | -2.1313755 | 0.03305822 | NA |
| ZNF704 | 58.1869318 | 1.0408271 | 0.39869923 | 2.6105571 | 0.00903949 | 0.27648451 |
| IMPA1 | 12.3702919 | -0.364601 | 0.44790551 | -0.8140133 | 0.41563735 | NA |
| ZFAND1 | 10.3478169 | 0.41108641 | 0.47050962 | 0.87370458 | 0.38227916 | NA |
| CHMP4C | 6.85355448 | -0.3467669 | 0.55497099 | -0.6248378 | 0.5320775 | NA |
| SNX16 | 10.0904418 | -0.5901701 | 0.47179008 | -1.2509167 | 0.21096487 | NA |
| RALYL | 17.8027056 | 0.66617079 | 0.89535189 | 0.74403237 | 0.45685689 | NA |
| LRRCC1 | 2.63109077 | 0.83136509 | 0.97917077 | 0.84905016 | 0.39585338 | NA |
| E2F5 | 7.56081231 | 0.67385309 | 0.61268549 | 1.09983524 | 0.27140392 | NA |
| C8orf59 | 8.38943188 | -0.0968152 | 0.53612622 | -0.1805828 | 0.85669505 | NA |
| CA13 | 10.1823256 | 0.27094734 | 0.47391455 | 0.57172192 | 0.56751038 | NA |
| CA3 | 12.0590773 | 0.06881928 | 0.56345128 | 0.12213883 | 0.90278907 | NA |
| CA2 | 55.7920683 | -0.761584 | 0.4699465 | -1.6205759 | 0.10510862 | 0.5772605 |
| ATP6V0D2 | 374.919506 | -0.5950952 | 0.24112794 | -2.4679646 | 0.01358838 | 0.30435125 |
| AC023194.3 | 55.6402328 | 0.20308406 | 0.49268484 | 0.41219872 | 0.68019377 | 0.9267216 |
| WWP1 | 93.8630662 | 0.07683222 | 0.28154889 | 0.27289121 | 0.78493684 | 0.96299238 |
| RMDN1 | 19.7042889 | 0.47386752 | 0.38042848 | 1.24561525 | 0.21290568 | NA |
| CPNE3 | 23.0450489 | 0.15684171 | 0.37435801 | 0.4189618 | 0.67524405 | NA |
| CNGB3 | 3.77256541 | 0.48629838 | 1.1680711 | 0.41632601 | 0.67717146 | NA |
| AF117829.1 | 37.833204 | 0.13812667 | 0.38505415 | 0.35872012 | 0.71980447 | 0.93714576 |
| RIPK2 | 5.43220396 | 0.69320629 | 0.61069572 | 1.13510913 | 0.2563296 | NA |
| OSGIN2 | 10.1785072 | -1.0087991 | 0.72679187 | -1.3880165 | 0.16513202 | NA |
| NBN | 12.2636077 | 0.22382932 | 0.44081512 | 0.50776234 | 0.61162002 | NA |
| DECR1 | 26.2856384 | 0.53969008 | 0.3720054 | 1.45075871 | 0.14684706 | 0.63 |
| CALB1 | 14.3971414 | 0.81205316 | 1.91118158 | 0.42489587 | 0.67091259 | NA |
| TMEM64 | 3.35462679 | 1.43674556 | 0.80563462 | 1.78337119 | 0.0745259 | NA |
| OTUD6B-AS1 | 7.51477976 | 1.48402576 | 0.57306333 | 2.58963658 | 0.00960773 | NA |
| OTUD6B | 7.04155247 | -0.1416789 | 0.62579453 | -0.2263985 | 0.82089153 | NA |
| LRRC69 | 6.38338408 | 0.56559312 | 0.71480336 | 0.79125695 | 0.42879407 | NA |
| SLC26A7 | 4.68897019 | 0.33645397 | 1.20490222 | 0.27923757 | 0.78006251 | NA |
| TRIQK | 11.1939931 | 0.34528915 | 0.50614191 | 0.68219829 | 0.49511358 | NA |
| LINC00535 | 4.25279127 | 0.234846 | 0.8105913 | 0.28972184 | 0.77202905 | NA |
| FAM92A | 30.238866 | -0.5191044 | 0.45984112 | -1.1288777 | 0.25894945 | 0.75287154 |
| RBM12B | 14.6140196 | 0.01214605 | 0.51549864 | 0.02356174 | 0.98120219 | NA |
| TMEM67 | 14.6156938 | 0.11309769 | 0.44647144 | 0.2533145 | 0.80002519 | NA |
| PDP1 | 22.1292173 | -0.7190019 | 0.56632139 | -1.2696005 | 0.20422697 | NA |
| VIRMA | 15.4717709 | -0.2463816 | 0.49446925 | -0.4982749 | 0.61829033 | NA |

| | | | | | | |
|---|---|---|---|---|---|---|
| ESRP1 | 19.8487976 | -0.4872105 | 0.41919716 | -1.1622466 | 0.24513531 | NA |
| DPY19L4 | 17.027159 | 0.27374412 | 0.43793295 | 0.62508227 | 0.53191706 | NA |
| INTS8 | 10.7705613 | -0.5078902 | 0.5106969 | -0.9945041 | 0.3199775 | NA |
| NDUFAF6 | 11.944513 | 0.10876654 | 0.54064564 | 0.20117899 | 0.84055862 | NA |
| TP53INP1 | 7.17728125 | 0.47725281 | 0.54707848 | 0.87236627 | 0.3830086 | NA |
| PLEKHF2 | 9.03411911 | 0.17940899 | 0.64559268 | 0.27789811 | 0.78109057 | NA |
| C8orf37-AS1 | 21.8789841 | 0.47938923 | 0.75266106 | 0.63692578 | 0.52417319 | NA |
| UQCRB | 49.0683111 | 0.49047315 | 0.3647106 | 1.34482836 | 0.17868067 | 0.66980156 |
| MTERF3 | 5.54255167 | -0.1940309 | 0.73697134 | -0.2632815 | 0.79233365 | NA |
| PTDSS1 | 5.11434283 | -0.7150562 | 0.70322602 | -1.0168227 | 0.30923779 | NA |
| SDC2 | 97.2790444 | -0.2729491 | 0.35534728 | -0.7681192 | 0.44241636 | 0.84767356 |
| CPQ | 71.5295716 | 0.36846479 | 0.32927675 | 1.11901248 | 0.26313482 | 0.75479335 |
| TSPYL5 | 4.38080381 | -1.2166094 | 0.70896427 | -1.7160377 | 0.08615514 | NA |
| MTDH | 90.406439 | -0.3805131 | 0.32397315 | -1.1745204 | 0.24018665 | 0.73323864 |
| LAPTM4B | 22.9629062 | -0.5071599 | 0.34236964 | -1.4813225 | 0.13852065 | NA |
| RPL30 | 54.9045089 | 0.19194362 | 0.38874338 | 0.49375406 | 0.62147988 | 0.91493825 |
| RIDA | 5.08329795 | 0.57656771 | 0.7734974 | 0.74540354 | 0.4560278 | NA |
| POP1 | 3.76217609 | -0.6367895 | 0.74547079 | -0.8542112 | 0.39298799 | NA |
| NIPAL2 | 7.39429989 | -0.8237269 | 0.60045898 | -1.3718287 | 0.17011677 | NA |
| STK3 | 180.113093 | -0.0837758 | 0.26868639 | -0.3117977 | 0.75519428 | 0.95323728 |
| AC104986.2 | 3.89206721 | 1.09315285 | 0.79425632 | 1.3763225 | 0.16872181 | NA |
| VPS13B | 112.716103 | -0.2937243 | 0.2217957 | -1.3243012 | 0.18540304 | 0.67879383 |
| COX6C | 51.4623515 | -0.1461503 | 0.27641319 | -0.5287384 | 0.59698692 | 0.899694 |
| POLR2K | 9.90995211 | -0.3346554 | 0.69492026 | -0.4815737 | 0.63010878 | NA |
| SPAG1 | 8.77723055 | -1.2197977 | 0.52025529 | -2.3446137 | 0.0190468 | NA |
| RNF19A | 47.8480317 | -0.1191315 | 0.41185369 | -0.2892568 | 0.77238487 | 0.95994327 |
| ANKRD46 | 8.49584883 | 0.76215808 | 0.53533188 | 1.42371136 | 0.15453004 | NA |
| PABPC1 | 48.3032377 | -0.3207972 | 0.31496656 | -1.0185118 | 0.3084348 | 0.77808363 |
| YWHAZ | 68.1492721 | -0.7170231 | 0.27657878 | -2.5924733 | 0.00952886 | 0.27655812 |
| ZNF706 | 18.84545 | 0.07881439 | 0.35375378 | 0.22279449 | 0.82369546 | NA |
| GRHL2 | 48.7090705 | -0.2311859 | 0.30494271 | -0.758129 | 0.44837377 | 0.85123446 |
| RRM2B | 15.7291912 | 0.20713874 | 0.38467823 | 0.53847273 | 0.59025072 | NA |
| UBR5-AS1 | 4.46381329 | 1.83020868 | 0.72162831 | 2.53622073 | 0.01120561 | NA |
| UBR5 | 88.3771466 | -0.1695899 | 0.25590811 | -0.6626983 | 0.50752377 | 0.87158522 |
| KLF10 | 11.4540989 | 0.6390806 | 0.53644632 | 1.19132256 | 0.23352698 | NA |
| AZIN1 | 18.875176 | 0.23019869 | 0.41598073 | 0.55338787 | 0.57999784 | NA |
| AZIN1-AS1 | 16.2382282 | -0.2469368 | 0.51750883 | -0.4771644 | 0.63324506 | NA |
| ATP6V1C1 | 22.0283067 | -0.7466469 | 0.47977946 | -1.5562295 | 0.11965354 | NA |
| BAALC | 7.14040251 | 1.05886758 | 0.72736113 | 1.45576597 | 0.14545733 | NA |
| BAALC-AS1 | 6.38434569 | 0.92185935 | 0.70651844 | 1.30479164 | 0.19196381 | NA |
| FZD6 | 25.8213982 | -0.1461498 | 0.32679449 | -0.4472224 | 0.65471448 | NA |
| SLC25A32 | 11.0431511 | -0.6688315 | 0.52186696 | -1.2816131 | 0.1999784 | NA |
| DCAF13 | 7.06679897 | -1.0316377 | 0.59473839 | -1.7346075 | 0.08281035 | NA |
| DPYS | 7.51989332 | 1.32040441 | 1.01606875 | 1.29952271 | 0.19376461 | NA |
| LRP12 | 18.7635827 | -0.372956 | 0.47542302 | -0.7844718 | 0.4327633 | NA |
| ZFPM2 | 9.5964489 | 1.05153162 | 0.7784625 | 1.35078006 | 0.1767659 | NA |
| AC027031.2 | 2.56653081 | -0.2883933 | 1.04042624 | -0.2771876 | 0.78163605 | NA |
| OXR1 | 241.333849 | 0.38721471 | 0.29049452 | 1.33295012 | 0.18254817 | 0.67478132 |
| AC090579.1 | 24.1285178 | 1.15562239 | 0.69828247 | 1.65494973 | 0.0979347 | NA |
| ANGPT1 | 13.2450556 | 0.1739639 | 0.87189926 | 0.19952294 | 0.8418537 | NA |
| EIF3E | 26.6995501 | -0.2239376 | 0.38728673 | -0.5782217 | 0.56311443 | 0.89263108 |
| EMC2 | 17.7969142 | 0.94410056 | 0.45155516 | 2.09077568 | 0.03654818 | NA |
| NUDCD1 | 8.03280471 | 0.3738601 | 0.55065477 | 0.67893736 | 0.49717755 | NA |
| ENY2 | 14.4377409 | -0.3487171 | 0.47627961 | -0.7321689 | 0.46406547 | NA |

| | | | | | | |
|---|---|---|---|---|---|---|
| EBAG9 | 8.54438112 | 0.72308201 | 0.53102779 | 1.36166511 | 0.1733036 | NA |
| CSMD3 | 134.322404 | -1.7016804 | 0.42542299 | -3.9999728 | 6.33E-05 | 0.04227013 |
| TRPS1 | 83.3898233 | -0.2636037 | 0.33387939 | -0.7895176 | 0.42980954 | 0.84223493 |
| EIF3H | 43.9624356 | -0.1851961 | 0.27493889 | -0.6735901 | 0.50057193 | 0.87036626 |
| UTP23 | 4.35538545 | 0.71393624 | 0.65960957 | 1.08236185 | 0.27909177 | NA |
| RAD21 | 25.5693227 | -0.2720091 | 0.34606741 | -0.7860003 | 0.43186728 | NA |
| MED30 | 3.16972207 | 0.73417223 | 0.83712061 | 0.87702086 | 0.38047531 | NA |
| EXT1 | 179.009045 | -0.192729 | 0.31313216 | -0.6154877 | 0.53823272 | 0.88254911 |
| SAMD12 | 190.219964 | 0.51319379 | 0.30572948 | 1.67858786 | 0.09323239 | 0.56220784 |
| AC023590.1 | 19.1667879 | 1.18046856 | 0.71044202 | 1.66159733 | 0.09659354 | NA |
| SAMD12-AS1 | 16.5147657 | 0.41231315 | 0.4902101 | 0.84109476 | 0.40029485 | NA |
| MAL2 | 41.4442293 | -0.5428523 | 0.28856202 | -1.8812326 | 0.05994028 | 0.51625203 |
| TAF2 | 25.9004864 | -0.4883399 | 0.39066867 | -1.2500104 | 0.21129574 | NA |
| DEPTOR | 49.516553 | 0.4808739 | 0.46515887 | 1.03378422 | 0.30123705 | 0.77521265 |
| MRPL13 | 9.432395 | 0.08285268 | 0.64147367 | 0.1291599 | 0.89723112 | NA |
| MTBP | 6.02544231 | 0.66182825 | 0.64010479 | 1.03393735 | 0.30116545 | NA |
| SNTB1 | 402.455127 | -0.0549375 | 0.24009139 | -0.2288191 | 0.81900953 | 0.96845695 |
| AC104958.1 | 5.82413777 | 1.238271 | 0.91225171 | 1.35737866 | 0.17466092 | NA |
| ZHX2 | 69.5224441 | -0.1152273 | 0.281036 | -0.4100089 | 0.68179943 | 0.9267216 |
| DERL1 | 7.66085767 | 0.07227077 | 0.5615718 | 0.12869373 | 0.89759999 | NA |
| TBC1D31 | 3.23911158 | 1.44935469 | 0.86626672 | 1.67310443 | 0.09430674 | NA |
| C8orf76 | 3.148523 | -0.6017001 | 0.7842156 | -0.7672636 | 0.4429248 | NA |
| WDYHV1 | 6.74171511 | -0.2999828 | 0.6173814 | -0.4858954 | 0.62704136 | NA |
| FBXO32 | 13.6912163 | -0.0165109 | 0.60949113 | -0.0270897 | 0.97838818 | NA |
| FAM91A1 | 23.242862 | 0.02637891 | 0.35570197 | 0.07416015 | 0.94088295 | NA |
| TMEM65 | 47.1351628 | 0.02519042 | 0.32888689 | 0.07659297 | 0.93894735 | 0.99100968 |
| RNF139 | 2.77352728 | -0.4997408 | 0.95239627 | -0.5247194 | 0.59977826 | NA |
| TATDN1 | 16.2741564 | -0.0861867 | 0.44751001 | -0.1925917 | 0.84727873 | NA |
| NDUFB9 | 22.3854586 | -0.2649764 | 0.51493591 | -0.5145813 | 0.6068456 | NA |
| MTSS1 | 55.8047635 | -0.2979737 | 0.3488964 | -0.8540465 | 0.39307922 | 0.8258317 |
| SQLE | 8.33725996 | -1.2813995 | 0.7469386 | -1.7155352 | 0.08624714 | NA |
| WASHC5 | 16.6944056 | -0.4285957 | 0.39374257 | -1.0885176 | 0.27636665 | NA |
| NSMCE2 | 103.28128 | -0.4338283 | 0.2587457 | -1.6766588 | 0.09360922 | 0.56220784 |
| TRIB1 | 14.1175927 | -0.6156147 | 0.57437849 | -1.0717927 | 0.28381313 | NA |
| FAM84B | 9.9118566 | -0.3837494 | 0.60976558 | -0.6293392 | 0.52912699 | NA |
| PCAT1 | 58.3337865 | 0.23686855 | 0.2777742 | 0.85273777 | 0.39380474 | 0.8258317 |
| PVT1 | 12.4401336 | 0.41659236 | 0.53065354 | 0.78505527 | 0.43242117 | NA |
| FAM49B | 75.1975136 | -0.1315245 | 0.22622566 | -0.5813865 | 0.56097999 | 0.89263108 |
| ASAP1 | 33.2590863 | -0.188343 | 0.36148089 | -0.5210316 | 0.60234473 | 0.90470348 |
| EFR3A | 34.2804238 | -0.137606 | 0.31692841 | -0.4341865 | 0.66415301 | 0.92276785 |
| KCNQ3 | 9.78022493 | 3.89381286 | 1.52416181 | 2.55472406 | 0.0106272 | NA |
| LRRC6 | 6.02525161 | 1.4377706 | 0.6834687 | 2.1036378 | 0.03541005 | NA |
| PHF20L1 | 32.8420186 | -0.1749688 | 0.38867651 | -0.4501656 | 0.65259104 | 0.92248109 |
| TG | 6.36919204 | 2.08911376 | 1.01266175 | 2.06299266 | 0.03911333 | NA |
| NDRG1 | 136.102736 | -0.1611063 | 0.53337055 | -0.3020532 | 0.76261151 | 0.95478238 |
| ST3GAL1 | 68.0472203 | -0.0524114 | 0.41327095 | -0.126821 | 0.89908205 | 0.97863828 |
| ZFAT | 12.1878124 | -1.0311221 | 0.53058837 | -1.943356 | 0.05197315 | NA |
| AC083843.3 | 6.70784668 | -0.3480551 | 0.60342974 | -0.5767947 | 0.56407816 | NA |
| KHDRBS3 | 27.667053 | -0.4545384 | 0.52962342 | -0.8582294 | 0.39076581 | 0.8258317 |
| TRAPPC9 | 119.955277 | 0.15729145 | 0.30796317 | 0.51074761 | 0.6095278 | 0.9088434 |
| CHRAC1 | 6.764939 | 0.01123681 | 0.59006402 | 0.01904337 | 0.9848065 | NA |
| AGO2 | 28.3695448 | -0.4581166 | 0.38565454 | -1.1878936 | 0.23487533 | 0.73156941 |
| PTK2 | 174.475961 | 0.01237365 | 0.2593636 | 0.04770774 | 0.96194917 | 0.99656976 |
| DENND3 | 4.14989237 | 0.45148246 | 0.70931224 | 0.63650736 | 0.52444578 | NA |

| | | | | | | |
|---|---|---|---|---|---|---|
| SLC45A4 | 13.575012 | -0.2761625 | 0.53983534 | -0.511568 | 0.6089534 | NA |
| PTP4A3 | 5.13606415 | -1.0091923 | 0.71125123 | -1.4188971 | 0.15592903 | NA |
| TSNARE1 | 11.2023375 | -0.2851006 | 0.50064163 | -0.5694704 | 0.56903694 | NA |
| JRK | 3.21207204 | 0.87439097 | 0.83421986 | 1.0481541 | 0.29456762 | NA |
| THEM6 | 4.08465301 | -0.117973 | 0.83553879 | -0.1411939 | 0.88771677 | NA |
| LY6E | 8.78405015 | 0.44833023 | 0.94889533 | 0.47247596 | 0.6365871 | NA |
| GLI4 | 2.21672264 | -0.1583613 | 0.96435228 | -0.1642152 | 0.86956174 | NA |
| TOP1MT | 4.22178685 | 0.47665446 | 0.76420488 | 0.62372601 | 0.53280754 | NA |
| RHPN1 | 2.745751 | 0.14515481 | 0.95568786 | 0.15188517 | 0.8792775 | NA |
| ZC3H3 | 6.64250302 | -0.2814509 | 0.71092909 | -0.3958917 | 0.69218494 | NA |
| GSDMD | 7.67586823 | 0.45701899 | 0.60374161 | 0.75697779 | 0.44906317 | NA |
| NAPRT | 2.91034659 | 0.65988907 | 0.8881758 | 0.74297124 | 0.4574991 | NA |
| EEF1D | 51.3618834 | -0.0762868 | 0.4029623 | -0.189315 | 0.84984591 | 0.97168963 |
| TSTA3 | 3.91349562 | -0.4418867 | 0.777671 | -0.568218 | 0.56988693 | NA |
| ZNF623 | 6.24318294 | 0.34722988 | 0.63153314 | 0.54982052 | 0.58244248 | NA |
| FAM83H | 7.18030987 | -0.7697276 | 0.59344945 | -1.2970398 | 0.19461748 | NA |
| IQANK1 | 3.10022322 | 0.56747609 | 0.8255503 | 0.68739129 | 0.49183619 | NA |
| SCRIB | 7.23821124 | 0.46755711 | 0.55654182 | 0.84011136 | 0.40084595 | NA |
| PUF60 | 7.99153067 | -0.5417779 | 0.64986143 | -0.8336822 | 0.40446007 | NA |
| NRBP2 | 4.30658206 | 1.37384921 | 0.73285943 | 1.8746422 | 0.06084196 | NA |
| PLEC | 8.33311005 | 0.33932855 | 0.55825585 | 0.60783698 | 0.5432956 | NA |
| PARP10 | 5.60687037 | -0.4380831 | 0.75709376 | -0.5786378 | 0.56283358 | NA |
| GRINA | 10.562003 | -0.2516963 | 0.4715941 | -0.5337138 | 0.59353958 | NA |
| OPLAH | 1.74625789 | -0.5909347 | 1.43409852 | -0.4120601 | 0.6802954 | NA |
| EXOSC4 | 2.53326977 | 1.02886003 | 0.91481295 | 1.1246671 | 0.26073013 | NA |
| GPAA1 | 10.4239679 | 0.84332817 | 0.51922533 | 1.62420459 | 0.10433215 | NA |
| CYC1 | 17.188766 | 0.02128354 | 0.53174933 | 0.04002552 | 0.96807278 | NA |
| SHARPIN | 6.80570579 | 0.01581269 | 0.57181754 | 0.02765338 | 0.97793861 | NA |
| MAF1 | 9.48259454 | -0.1982072 | 0.53927232 | -0.3675456 | 0.71321205 | NA |
| MROH1 | 28.5521495 | -0.8273921 | 0.41061257 | -2.015019 | 0.04390266 | 0.44553691 |
| BOP1 | 1.97306213 | -1.1916456 | 1.24144357 | -0.9598871 | 0.33711206 | NA |
| HSF1 | 14.2596202 | 0.18679016 | 0.46031395 | 0.40578861 | 0.68489793 | NA |
| DGAT1 | 4.77000723 | 0.15257729 | 0.67860248 | 0.22484045 | 0.82210339 | NA |
| SLC52A2 | 5.84901457 | -0.5717154 | 0.5866347 | -0.974568 | 0.32977457 | NA |
| ADCK5 | 2.20500091 | -1.6016898 | 1.02813058 | -1.5578662 | 0.11926498 | NA |
| CPSF1 | 3.54356598 | 0.00077021 | 0.91916696 | 0.00083794 | 0.99933142 | NA |
| VPS28 | 21.5861803 | 0.50919906 | 0.35506454 | 1.43410285 | 0.15154292 | NA |
| CYHR1 | 11.292155 | 0.46767259 | 0.50117053 | 0.93316059 | 0.35073706 | NA |
| KIFC2 | 4.13832829 | 0.18916235 | 0.82927618 | 0.22810538 | 0.81956432 | NA |
| PPP1R16A | 7.1870758 | -0.5920639 | 0.58887517 | -1.005415 | 0.31469707 | NA |
| LRRC14 | 2.33003109 | 0.27804326 | 0.92078604 | 0.30196294 | 0.76268032 | NA |
| ARHGAP39 | 4.46798326 | 0.36030327 | 0.73265882 | 0.49177497 | 0.62287843 | NA |
| ZNF251 | 17.8852825 | -0.0773323 | 0.36766443 | -0.2103339 | 0.83340708 | NA |
| RPL8 | 104.014972 | 0.02545088 | 0.38486686 | 0.06612905 | 0.94727509 | 0.99342916 |
| ZNF7 | 5.60112166 | -0.0706124 | 0.73793006 | -0.0956898 | 0.92376697 | NA |
| COMMD5 | 3.66877641 | 0.55304811 | 0.7044058 | 0.78512714 | 0.43237903 | NA |
| ZNF250 | 5.42445795 | -0.6442919 | 0.70179538 | -0.9180623 | 0.35858625 | NA |
| ZNF16 | 4.07713424 | 0.51352964 | 0.66324882 | 0.77426394 | 0.43877472 | NA |
| C8orf33 | 5.97225327 | -0.7607791 | 0.62355314 | -1.2200711 | 0.22243793 | NA |
| WASHC1 | 25.524864 | 0.21384218 | 0.38812656 | 0.55095992 | 0.58166115 | NA |
| CBWD1 | 7.44346695 | 0.55922854 | 0.68791018 | 0.8129383 | 0.41625343 | NA |
| DOCK8 | 84.2992821 | 0.48549696 | 0.4551303 | 1.06672081 | 0.28609793 | 0.77103639 |
| KANK1 | 91.0244506 | 0.08916462 | 0.30321388 | 0.29406511 | 0.76870815 | 0.957884 |
| SMARCA2 | 44.9535164 | 0.53446945 | 0.42004883 | 1.27239838 | 0.20323161 | 0.700066 |

| | | | | | | |
|---|---|---|---|---|---|---|
| PUM3 | 35.6745677 | 0.19547048 | 0.4113496 | 0.47519308 | 0.63464936 | 0.92072524 |
| RFX3 | 31.7629159 | 0.37635514 | 0.38883196 | 0.96791203 | 0.33308831 | 0.79447068 |
| RFX3-AS1 | 22.6750096 | 0.31158205 | 0.37814514 | 0.82397475 | 0.4099539 | NA |
| GLIS3 | 115.712653 | 0.12254932 | 0.3750546 | 0.32675061 | 7.44E-01 | 0.94831173 |
| SLC1A1 | 16.9528258 | -0.667219 | 0.4882299 | -1.3666083 | 1.72E-01 | NA |
| SPATA6L | 16.5339466 | 0.09045519 | 0.45699614 | 0.19793424 | 0.84309651 | NA |
| CDC37L1 | 13.3453998 | 0.07131817 | 0.48449571 | 0.14720084 | 0.8829735 | NA |
| AK3 | 52.1686704 | 0.74823542 | 0.28930314 | 2.58633701 | 0.0097002 | 0.27838541 |
| RCL1 | 11.5664473 | -0.0793456 | 0.49333559 | -0.1608349 | 0.87222346 | NA |
| JAK2 | 18.0224986 | -0.643908 | 0.40339713 | -1.5962136 | 0.11044112 | NA |
| PLGRKT | 10.947386 | 0.41273902 | 0.46498793 | 0.88763383 | 0.37473775 | NA |
| RIC1 | 39.2814005 | -0.511687 | 0.36863108 | -1.3880735 | 0.16511467 | 0.64995633 |
| ERMP1 | 33.4867433 | 0.44197743 | 0.34504049 | 1.28094366 | 0.20021346 | 0.6976106 |
| KIAA2026 | 49.1224719 | -0.0727097 | 0.28425438 | -0.255791 | 0.79811223 | 0.96649797 |
| RANBP6 | 5.36096015 | -0.3274362 | 0.71215882 | -0.4597797 | 0.64567434 | NA |
| UHRF2 | 24.7206195 | -0.0355792 | 0.39092418 | -0.0910131 | 0.92748218 | NA |
| GLDC | 4.30733988 | -0.6766816 | 0.75938587 | -0.8910906 | 0.37288056 | NA |
| KDM4C | 92.723693 | 0.00785069 | 0.23686459 | 0.03314423 | 0.97355957 | 0.99779576 |
| DMAC1 | 7.11166655 | 0.66735965 | 0.69391493 | 0.96173121 | 0.33618464 | NA |
| PTPRD | 11.9946394 | -0.4164064 | 0.6138486 | -0.6783536 | 0.49754754 | NA |
| MPDZ | 17.169851 | 0.01846537 | 0.40228399 | 0.04590133 | 0.96338889 | NA |
| NFIB | 37.5883803 | 0.04289656 | 0.56268344 | 0.07623569 | 0.93923159 | 0.99100968 |
| ZDHHC21 | 58.2613401 | -0.0623862 | 0.29626117 | -0.2105785 | 0.8332162 | 0.97154892 |
| FREM1 | 111.051678 | 0.25680836 | 0.38248145 | 0.67142697 | 0.50194857 | 0.87050612 |
| TTC39B | 78.0503206 | -0.4393728 | 0.3736827 | -1.1757911 | 0.23967835 | 0.73323864 |
| SNAPC3 | 21.4429732 | 0.25633603 | 0.33924053 | 0.75561736 | 0.44987864 | NA |
| PSIP1 | 10.9357376 | -0.023103 | 0.61284963 | -0.0376976 | 0.96992878 | NA |
| CCDC171 | 21.6647377 | 0.45724876 | 0.41882878 | 1.09173196 | 0.27495093 | NA |
| BNC2 | 28.0996785 | 0.18680609 | 0.57406514 | 0.32540922 | 0.74487138 | 0.94858421 |
| CNTLN | 16.7020566 | 0.73625761 | 0.60704458 | 1.21285592 | 0.22518491 | NA |
| SH3GL2 | 37.2370406 | -1.8010244 | 1.15174604 | -1.563734 | 0.11788006 | 0.57941414 |
| ADAMTSL1 | 101.779811 | 0.36966276 | 0.63199342 | 0.58491552 | 0.55860452 | 0.89263108 |
| RRAGA | 5.30077502 | -0.0914505 | 0.71760343 | -0.1274387 | 0.89859316 | NA |
| HAUS6 | 4.06381388 | -1.2693426 | 0.96302297 | -1.3180814 | 0.1874764 | NA |
| PLIN2 | 13.3538691 | -0.0802814 | 0.64301501 | -0.1248515 | 0.9006411 | NA |
| DENND4C | 60.982452 | 0.15502158 | 0.25071577 | 0.61831603 | 0.53636704 | 0.88254911 |
| RPS6 | 103.450224 | -0.005623 | 0.43835969 | -0.0128274 | 0.98976547 | 0.99909657 |
| SLC24A2 | 11.848799 | -0.4306442 | 0.69402039 | -0.6205065 | 0.53492438 | NA |
| MLLT3 | 100.819256 | 0.70609498 | 0.4213185 | 1.67591734 | 0.09375439 | 0.56220784 |
| FOCAD | 46.9231389 | 0.27297844 | 0.48776439 | 0.55965225 | 0.57571666 | 0.89263108 |
| KLHL9 | 9.49213922 | -0.3970045 | 0.49424978 | -0.8032466 | 0.42183221 | NA |
| LINC01239 | 3.53708794 | 2.81619938 | 1.15164126 | 2.44537903 | 0.01447 | NA |
| AL391117.1 | 5.14790129 | 1.25406649 | 1.21949154 | 1.02835194 | 0.30378431 | NA |
| TUSC1 | 6.93542662 | -0.9201124 | 0.65732312 | -1.3997871 | 0.16157708 | NA |
| CAAP1 | 30.8650719 | 0.37882262 | 0.35396864 | 1.07021521 | 0.28452245 | 0.76977033 |
| PLAA | 9.79670651 | -0.7679768 | 0.63183911 | -1.2154625 | 0.22418973 | NA |
| IFT74 | 16.6013293 | 0.15549361 | 0.47422644 | 0.32788895 | 0.74299563 | NA |
| MOB3B | 48.3377141 | -0.1966283 | 0.4410209 | -0.445848 | 0.65570702 | 0.92276785 |
| C9orf72 | 6.64666759 | 0.38243767 | 0.60441161 | 0.63274375 | 0.526901 | NA |
| ACO1 | 11.421579 | -0.2405171 | 0.491557 | -0.4892965 | 0.6246318 | NA |
| DDX58 | 21.4638507 | -0.9714473 | 0.87693932 | -1.1077703 | 0.26796105 | NA |
| TOPORS | 6.88170171 | -0.3592726 | 0.64767224 | -0.5547136 | 0.57909055 | NA |
| NDUFB6 | 12.9372124 | -0.1535768 | 0.42794856 | -0.3588674 | 0.71969427 | NA |
| APTX | 15.3336383 | 0.30274212 | 0.4118265 | 0.73512054 | 0.46226609 | NA |

| | | | | | | |
|---|---|---|---|---|---|---|
| DNAJA1 | 23.9350165 | -0.0568425 | 0.36574048 | -0.1554175 | 0.8764922 | NA |
| SMU1 | 11.6488952 | -0.2959748 | 0.46674203 | -0.6341293 | 0.52599644 | NA |
| B4GALT1 | 26.3341275 | -0.0151773 | 0.33403792 | -0.0454359 | 0.96375989 | 0.99656976 |
| BAG1 | 17.2393841 | -0.2593892 | 0.43810616 | -0.5920693 | 0.55380418 | NA |
| CHMP5 | 14.0067998 | 0.38372128 | 0.56308692 | 0.68146011 | 0.4955804 | NA |
| NFX1 | 38.3183726 | -0.6945416 | 0.31949219 | -2.1738922 | 0.02971323 | 0.40669034 |
| AQP7 | 7.08061582 | 1.25263648 | 0.72205861 | 1.73481274 | 0.08277398 | NA |
| AQP3 | 16.5973842 | -0.9730462 | 0.84125834 | -1.1566556 | 0.24741309 | NA |
| UBE2R2 | 49.3625678 | -0.4089189 | 0.27989594 | -1.4609674 | 0.14402438 | 0.62469744 |
| UBAP2 | 40.0317598 | -0.3905349 | 0.4112208 | -0.9496963 | 0.34226659 | 0.8003972 |
| DCAF12 | 8.45346678 | -0.2730194 | 0.51389557 | -0.5312741 | 0.59522886 | NA |
| UBAP1 | 51.6585388 | -0.7294388 | 0.31429904 | -2.3208433 | 0.0202953 | 0.34471691 |
| NUDT2 | 4.61368506 | 0.40905362 | 0.78752121 | 0.51941917 | 0.60346846 | NA |
| FAM219A | 14.2958154 | -0.532045 | 0.43516744 | -1.2226214 | 0.22147271 | NA |
| RPP25L | 4.94037524 | 0.52350571 | 0.71752722 | 0.729597 | 0.46563656 | NA |
| DCTN3 | 5.56080655 | 1.43319146 | 0.71786828 | 1.99645464 | 0.04588446 | NA |
| VCP | 21.5236301 | -0.3189788 | 0.45592735 | -0.6996264 | 0.48416063 | NA |
| PIGO | 3.32371669 | -0.2535329 | 0.85883747 | -0.2952047 | 0.7678375 | NA |
| STOML2 | 7.43799265 | 0.69211748 | 0.54263196 | 1.27548232 | 0.20213858 | NA |
| UNC13B | 58.6911068 | -0.2442736 | 0.27349461 | -0.8931571 | 0.37177306 | 0.81199861 |
| RUSC2 | 11.424567 | -0.8091155 | 0.52707416 | -1.5351076 | 0.12475741 | NA |
| TESK1 | 3.53538573 | -0.7709814 | 0.89937026 | -0.8572458 | 0.39130905 | NA |
| CCDC107 | 5.55297263 | -0.6741029 | 0.71578331 | -0.9417696 | 0.34631062 | NA |
| TLN1 | 26.0249864 | -0.0902279 | 0.42043326 | -0.2146071 | 0.83007369 | 0.97084429 |
| CREB3 | 5.15319022 | -0.4321859 | 0.65036695 | -0.6645262 | 0.50635357 | NA |
| GBA2 | 20.0437286 | -0.5059184 | 0.43285632 | -1.1687907 | 0.24248798 | NA |
| RGP1 | 8.80204953 | 0.02507664 | 0.56266096 | 0.04456794 | 0.9644517 | NA |
| AL133410.1 | 7.03922093 | 1.21114301 | 0.59785691 | 2.0258075 | 0.04278451 | NA |
| HINT2 | 4.90705683 | 0.42607969 | 0.6527653 | 0.6527303 | 0.51393016 | NA |
| TMEM8B | 3.79323663 | 0.06522451 | 0.82253032 | 0.07929739 | 0.93679608 | NA |
| RECK | 5.17948951 | 0.68389458 | 0.61536674 | 1.11136098 | 0.266413 | NA |
| CLTA | 55.2310845 | -0.3664159 | 0.29963663 | -1.2228675 | 0.22137975 | 0.72409625 |
| GNE | 13.3816434 | 0.18300343 | 0.51240101 | 0.35714885 | 0.72098037 | NA |
| RNF38 | 79.2562824 | -0.5386328 | 0.22319475 | -2.4132862 | 0.0158094 | 0.31287044 |
| EBLN3P | 13.9295906 | 0.01402221 | 0.45080984 | 0.03110449 | 0.97518621 | NA |
| ZCCHC7 | 105.9755 | -0.3636631 | 0.30701427 | -1.1845152 | 0.23620917 | 0.7323945 |
| GRHPR | 10.986443 | 0.64221989 | 0.46136716 | 1.39199306 | 0.1639245 | NA |
| ZBTB5 | 5.85644502 | 0.23381217 | 0.69211069 | 0.33782482 | 0.7354952 | NA |
| AL138752.2 | 14.7683059 | -0.4264891 | 0.41511702 | -1.0273948 | 0.30423459 | NA |
| FRMPD1 | 2.96103464 | 2.77386828 | 1.2912434 | 2.14821488 | 0.03169669 | NA |
| TRMT10B | 12.3273625 | 0.63142049 | 0.46049716 | 1.3711713 | 0.17032156 | NA |
| DCAF10 | 36.4799551 | -0.0290446 | 0.31849795 | -0.0911923 | 0.9273398 | 0.98844645 |
| ANKRD18A | 2.89624372 | -0.2179924 | 0.84852557 | -0.2569073 | 0.79725033 | NA |
| GLIDR | 9.63245436 | 0.59060964 | 0.58986298 | 1.00126582 | 0.31669831 | NA |
| BMS1P14 | 4.46319766 | -0.0781804 | 1.03722439 | -0.0753747 | 0.93991662 | NA |
| CNTNAP3B | 20.6714684 | 0.31516803 | 0.64852642 | 0.48597562 | 0.62698445 | NA |
| FAM27C | 3.75583178 | -1.0333574 | 0.79646337 | -1.2974324 | 0.19448244 | NA |
| CBWD5 | 7.95080965 | 0.09018891 | 0.58715475 | 0.15360331 | 0.87792253 | NA |
| ZNF658 | 3.75061404 | 0.74189423 | 0.78937332 | 0.93985217 | 0.34729339 | NA |
| PGM5 | 5.25218697 | -0.1678649 | 0.76960122 | -0.2181193 | 0.82733617 | NA |
| PIP5K1B | 56.8601987 | -1.0430753 | 0.6745632 | -1.5462973 | 0.12203277 | 0.58553957 |
| FAM189A2 | 11.7096044 | -1.4436277 | 0.65890979 | -2.1909338 | 0.02845659 | NA |
| PTAR1 | 34.3161874 | 0.31785315 | 0.30325453 | 1.04813981 | 0.2945742 | 0.77460836 |
| MAMDC2-AS | 9.48677894 | -0.3823292 | 0.54461712 | -0.7020146 | 0.48267003 | NA |

| | | | | | | |
|---|---|---|---|---|---|---|
| MAMDC2 | 52.8472638 | -0.3884862 | 0.5351527 | -0.7259352 | 0.46787851 | 0.85894565 |
| SMC5-AS1 | 15.9566946 | 0.15854414 | 0.4044566 | 0.39199295 | 0.69506343 | NA |
| SMC5 | 43.9561922 | -0.6191972 | 0.28328153 | -2.1858015 | 0.02883013 | 0.40498749 |
| KLF9 | 10.772364 | 0.68804385 | 0.57522718 | 1.19612543 | 0.23164762 | NA |
| TRPM3 | 21.8266657 | 2.04474905 | 0.92767407 | 2.20416752 | 0.02751256 | NA |
| CEMIP2 | 62.7325083 | -0.5705874 | 0.41987954 | -1.3589312 | 0.17416839 | 0.6651652 |
| ABHD17B | 9.92982432 | 0.32734641 | 0.50773103 | 0.64472405 | 0.51910603 | NA |
| C9orf85 | 19.8199791 | 0.59688925 | 0.40689398 | 1.46694048 | 0.14239225 | NA |
| GDA | 8.9617016 | -0.2757822 | 0.84113831 | -0.3278679 | 0.74301156 | NA |
| ZFAND5 | 65.919427 | 0.21113286 | 0.30954222 | 0.68208098 | 0.49518774 | 0.86687543 |
| ALDH1A1 | 28.3635622 | 0.91799595 | 0.42769837 | 2.14636297 | 0.03184403 | 0.4126665 |
| C9orf41-AS1 | 3.41839295 | 1.45874362 | 0.90669582 | 1.60885667 | 0.10764769 | NA |
| CARNMT1 | 24.3700157 | 0.08861963 | 0.39456638 | 0.22460006 | 0.82229042 | NA |
| NMRK1 | 7.81150561 | 1.04846969 | 0.66697242 | 1.57198359 | 0.11595436 | NA |
| OSTF1 | 10.6484544 | 0.19413431 | 0.51703476 | 0.37547632 | 0.70730625 | NA |
| PCSK5 | 5.79638786 | 0.33736284 | 0.87833113 | 0.38409528 | 0.70090783 | NA |
| RFK | 10.3979515 | 0.0991974 | 0.48151055 | 0.20601294 | 0.83678081 | NA |
| GCNT1 | 12.0356239 | 0.7778958 | 0.60205298 | 1.292072 | 0.19633221 | NA |
| PRUNE2 | 9.44495752 | -0.414559 | 0.81174126 | -0.5107033 | 0.6095588 | NA |
| VPS13A | 45.8163534 | 0.32329892 | 0.28939873 | 1.11714008 | 0.26393444 | 0.75479335 |
| GNAQ | 289.300598 | -0.2821341 | 0.26342962 | -1.0710038 | 0.28416773 | 0.76977033 |
| CEP78 | 34.6175617 | 0.30582193 | 0.39107554 | 0.78200219 | 0.43421329 | 0.84223493 |
| TLE4 | 18.6517067 | 0.00957398 | 0.48807615 | 0.01961575 | 0.9843499 | NA |
| TLE1 | 46.6628894 | -0.8442584 | 0.35653246 | -2.3679707 | 0.01788595 | 0.33195057 |
| AL591368.1 | 2.21255684 | -1.1131591 | 1.56808524 | -0.7098843 | 0.47777587 | NA |
| RASEF | 3.52432971 | 0.80227177 | 0.75248533 | 1.06616267 | 0.28635012 | NA |
| FRMD3 | 10.4297376 | -1.1275348 | 0.50802717 | -2.2194379 | 0.02645695 | NA |
| IDNK | 2.65384454 | 1.1793307 | 1.01318431 | 1.16398436 | 0.24443035 | NA |
| UBQLN1 | 29.3640526 | -0.093965 | 0.32282491 | -0.291071 | 0.77099704 | 0.95979063 |
| GKAP1 | 15.7673358 | 0.31810048 | 0.46441567 | 0.68494777 | 0.49337688 | NA |
| KIF27 | 6.79068459 | 0.38048954 | 0.59003202 | 0.64486253 | 0.51901627 | NA |
| C9orf64 | 2.76959765 | 0.40697209 | 0.83961166 | 0.48471467 | 0.62787876 | NA |
| HNRNPK | 53.0236387 | -0.7112991 | 0.28053855 | -2.5354773 | 0.01122942 | 0.28544118 |
| RMI1 | 3.74464884 | -0.0437132 | 0.8164922 | -0.0535378 | 0.95730345 | NA |
| AGTPBP1 | 31.2312981 | 0.16730476 | 0.32983393 | 0.50723939 | 0.61198686 | 0.91047543 |
| NAA35 | 31.8244969 | -0.3582637 | 0.30561132 | -1.1722853 | 0.24108253 | 0.73323864 |
| GOLM1 | 60.440048 | 0.19010555 | 0.45306414 | 0.41959964 | 0.67477795 | 0.92617027 |
| ISCA1 | 14.1571082 | 0.4609993 | 0.42674068 | 1.0802797 | 0.28001764 | NA |
| TUT7 | 26.8390754 | -0.0540302 | 0.3042787 | -0.1775683 | 0.85906203 | 0.97168963 |
| DAPK1 | 61.2588104 | 0.27726577 | 0.37892069 | 0.73172507 | 0.46433639 | 0.85894565 |
| CTSL | 17.0151329 | -0.4744211 | 0.52193354 | -0.9089684 | 0.36336681 | NA |
| SPIN1 | 69.0910552 | 0.1693732 | 0.2440568 | 0.69399091 | 0.48768791 | 0.86354292 |
| SECISBP2 | 38.1856149 | 0.18519707 | 0.35927544 | 0.51547378 | 0.60622198 | 0.90821838 |
| SEMA4D | 32.65419 | 0.36059047 | 0.37063079 | 0.9729102 | 0.33059794 | 0.79360119 |
| GADD45G | 17.0169163 | -0.3376016 | 0.79641857 | -0.4238997 | 0.67163896 | NA |
| SYK | 46.8571545 | 0.64552793 | 0.31152301 | 2.07216773 | 0.0382498 | 0.42360465 |
| AUH | 53.4010111 | 0.68078072 | 0.4103626 | 1.65897358 | 0.09712112 | 0.5647486 |
| NFIL3 | 3.02563466 | -0.2433143 | 0.79731375 | -0.3051675 | 0.76023856 | NA |
| SPTLC1 | 21.1628432 | 0.11974368 | 0.38737154 | 0.30911842 | 0.75723145 | NA |
| IARS | 19.7861146 | -0.867376 | 0.41002902 | -2.1154015 | 0.03439574 | NA |
| NOL8 | 7.05453403 | 0.13699673 | 0.59361336 | 0.23078445 | 0.81748226 | NA |
| CENPP | 14.6217129 | -0.5325012 | 0.62547926 | -0.8513491 | 0.39457544 | NA |
| IPPK | 6.91347307 | -0.9505497 | 0.65701826 | -1.4467631 | 0.1479633 | NA |
| BICD2 | 8.40262706 | -0.3423593 | 0.57788561 | -0.5924344 | 0.55355972 | NA |

| | | | | | | |
|---|---|---|---|---|---|---|
| ZNF484 | 3.72740213 | -0.0662835 | 0.72868829 | -0.0909627 | 0.92752222 | NA |
| FGD3 | 4.61154997 | 2.26142187 | 0.90879592 | 2.48837151 | 0.01283296 | NA |
| SUSD3 | 4.68413533 | 0.26061719 | 0.74594811 | 0.34937711 | 0.72680622 | NA |
| CARD19 | 7.32057544 | 0.00046329 | 0.54008816 | 0.00085781 | 0.99931557 | NA |
| NINJ1 | 5.16753312 | 1.34249574 | 0.62731541 | 2.14006497 | 0.03234952 | NA |
| FAM120AOS | 5.43553616 | 0.47194349 | 0.60661029 | 0.77800113 | 0.43656835 | NA |
| FAM120A | 41.6164091 | 0.47933439 | 0.30715714 | 1.56055102 | 0.11862972 | 0.57989511 |
| PHF2 | 9.27842898 | 0.75319557 | 0.55412707 | 1.35924703 | 0.17406832 | NA |
| PTPDC1 | 12.6060652 | -0.3391808 | 0.42941242 | -0.789872 | 0.42960253 | NA |
| AL158152.2 | 12.952409 | 0.00586585 | 0.44072497 | 0.01330954 | 0.98938084 | NA |
| ZNF169 | 5.69235198 | 0.98681673 | 0.61388594 | 1.60749197 | 0.10794649 | NA |
| MFSD14B | 15.1020917 | -0.5461275 | 0.47642367 | -1.1463065 | 0.25166837 | NA |
| FBP1 | 5.59722221 | 1.11062548 | 1.06137919 | 1.0463984 | 0.29537713 | NA |
| C9orf3 | 84.3719345 | 0.10225284 | 0.24218788 | 0.42220463 | 0.67287566 | 0.92617027 |
| FANCC | 41.9791033 | 0.08281626 | 0.30512913 | 0.27141382 | 0.78607277 | 0.96325251 |
| PTCH1 | 11.3434952 | 0.18891916 | 0.80797193 | 0.23381896 | 0.81512553 | NA |
| LINC00476 | 15.0624441 | 0.24399086 | 0.43130623 | 0.56570214 | 0.57159628 | NA |
| ERCC6L2 | 27.3234683 | -0.3618966 | 0.35054933 | -1.0323701 | 0.3018988 | 0.77521265 |
| SLC35D2 | 3.85352667 | 1.20026365 | 0.70673229 | 1.69832858 | 0.08944576 | NA |
| HABP4 | 9.07701362 | 0.06781014 | 0.54754097 | 0.12384487 | 0.90143811 | NA |
| CDC14B | 32.5877692 | 0.00954024 | 0.42783146 | 0.02229905 | 0.9822094 | 0.99902646 |
| AAED1 | 3.57694974 | -0.0687493 | 0.73683082 | -0.093304 | 0.92566206 | NA |
| AL589843.1 | 3.59028258 | 0.89432105 | 1.01514308 | 0.8809803 | 0.37832848 | NA |
| ZNF510 | 3.46506506 | 0.01070156 | 0.89719101 | 0.01192785 | 0.99048318 | NA |
| ZNF782 | 10.8275235 | -0.3157694 | 0.50243704 | -0.6284755 | 0.52969248 | NA |
| MFSD14C | 35.2108432 | -0.2630428 | 0.32409245 | -0.8116289 | 0.4170046 | 0.8374607 |
| TDRD7 | 11.8430431 | 0.4177168 | 0.54481809 | 0.76670876 | 0.4432547 | NA |
| TSTD2 | 9.93043106 | -0.6842483 | 0.52051006 | -1.3145726 | 0.18865359 | NA |
| NCBP1 | 8.1287087 | -0.208494 | 0.57854793 | -0.3603747 | 0.71856697 | NA |
| XPA | 12.0616853 | 0.70262066 | 0.52125652 | 1.34793646 | 0.17767882 | NA |
| TRMO | 3.64718695 | 0.4796126 | 0.88712336 | 0.540638 | 0.58875712 | NA |
| ANP32B | 14.1214504 | 0.30954812 | 0.44873532 | 0.68982338 | 0.49030526 | NA |
| NANS | 3.52317855 | -0.8941522 | 0.78705963 | -1.1360667 | 0.25592865 | NA |
| TRIM14 | 5.13111016 | 0.00726302 | 0.90524227 | 0.00802329 | 0.99359841 | NA |
| TBC1D2 | 6.14064997 | 2.05759547 | 0.70934337 | 2.90070446 | 0.00372325 | NA |
| ANKS6 | 7.78222599 | 0.5620132 | 0.60182866 | 0.93384254 | 0.35038512 | NA |
| TGFBR1 | 17.3073296 | -0.5854834 | 0.45051799 | -1.2995783 | 0.19374553 | NA |
| ALG2 | 5.92837402 | -0.533654 | 0.8165262 | -0.6535663 | 0.51339125 | NA |
| SEC61B | 22.9398321 | -0.0881091 | 0.4326289 | -0.2036598 | 0.83861937 | NA |
| AL359710.1 | 20.6918124 | 0.65474949 | 0.51982997 | 1.25954549 | 0.20783337 | NA |
| STX17 | 20.3312665 | -0.0250464 | 0.38612521 | -0.0648661 | 0.94828061 | NA |
| ERP44 | 27.2416893 | -0.3639431 | 0.37827749 | -0.9621062 | 0.33599628 | 0.8003972 |
| INVS | 57.9089385 | 0.52501635 | 0.33499973 | 1.56721426 | 0.11706462 | 0.57860396 |
| TEX10 | 6.70904799 | 0.16555387 | 0.62619867 | 0.26437916 | 0.79148777 | NA |
| PLPPR1 | 8.81384181 | 0.51465012 | 0.84066754 | 0.61219221 | 0.54041059 | NA |
| MRPL50 | 5.27558956 | -0.0754599 | 0.66518855 | -0.1134414 | 0.90968066 | NA |
| ZNF189 | 9.85348889 | -0.9927171 | 0.5746893 | -1.7273979 | 0.08409622 | NA |
| ALDOB | 22.6262863 | 0.60773304 | 1.21905783 | 0.49852683 | 0.61811276 | NA |
| RNF20 | 7.34181492 | -0.0252644 | 0.56913115 | -0.0443913 | 0.96459253 | NA |
| SMC2 | 2.80611418 | 0.50712016 | 0.95621837 | 0.53033928 | 0.59587672 | NA |
| NIPSNAP3A | 4.41065721 | 1.33396171 | 0.76078048 | 1.75341211 | 0.07953129 | NA |
| ABCA1 | 20.4635177 | 0.39215935 | 0.68854327 | 0.56954932 | 0.56898341 | NA |
| SLC44A1 | 29.97825 | -0.1042233 | 0.33864495 | -0.3077657 | 0.75826064 | 0.95327256 |
| FSD1L | 19.4277001 | -0.2196262 | 0.36626206 | -0.5996422 | 0.54874474 | NA |

| | | | | | | |
|---|---|---|---|---|---|---|
| FKTN | 11.7548161 | 0.14587341 | 0.55077705 | 0.26485021 | 0.79112486 | NA |
| TMEM38B | 4.18399769 | -1.2551874 | 0.74958345 | -1.6745131 | 0.09402979 | NA |
| LINC01505 | 5.17719804 | -1.2422414 | 0.83650408 | -1.4850392 | 0.13753343 | NA |
| ZNF462 | 6.61882738 | 0.97047921 | 0.6697675 | 1.44897926 | 0.14734337 | NA |
| AL807761.4 | 3.14485604 | 2.40673773 | 1.35917251 | 1.7707375 | 0.07660436 | NA |
| RAD23B | 53.4840107 | -0.2086567 | 0.26235522 | -0.7953212 | 0.42642671 | 0.84057082 |
| ELP1 | 10.694347 | 1.01026429 | 0.52917695 | 1.90912377 | 0.05624613 | NA |
| FAM206A | 2.77591239 | 0.47537197 | 0.8421197 | 0.56449454 | 0.57241761 | NA |
| CTNNAL1 | 11.3785809 | -0.6763026 | 0.49480998 | -1.3667926 | 0.17169031 | NA |
| TMEM245 | 38.2037214 | 0.23526317 | 0.33908566 | 0.69381633 | 0.4877974 | 0.86354292 |
| EPB41L4B | 11.6611254 | 0.23655267 | 0.47333543 | 0.49975696 | 0.61724622 | NA |
| PTPN3 | 68.45147 | 0.17244143 | 0.30836807 | 0.5592065 | 0.57602079 | 0.89263108 |
| PALM2 | 8.8687688 | 0.23411616 | 0.52564272 | 0.44539029 | 0.65603773 | NA |
| TXN | 22.7948407 | -0.0159908 | 0.41253815 | -0.0387619 | 0.96908023 | NA |
| SVEP1 | 5.08383138 | -1.3345732 | 0.84732695 | -1.5750392 | 0.1152474 | NA |
| LPAR1 | 22.7894645 | -0.4741427 | 0.51702884 | -0.9170527 | 0.35911504 | NA |
| ECPAS | 62.0153158 | -0.5560107 | 0.31815479 | -1.7476107 | 0.08053147 | 0.54552914 |
| ZNF483 | 11.2292949 | -0.0872525 | 0.50676372 | -0.1721759 | 0.86329927 | NA |
| PTGR1 | 32.3754008 | -0.0741364 | 0.37741785 | -0.1964306 | 0.84427313 | 0.97168963 |
| C9orf84 | 8.04637488 | 0.86934711 | 0.58880004 | 1.47647257 | 0.13981707 | NA |
| UGCG | 42.7970335 | 0.10667559 | 0.28024592 | 0.38064992 | 0.70346303 | 0.9303056 |
| SUSD1 | 61.4812159 | -0.2297809 | 0.28616232 | -0.8029741 | 0.42198971 | 0.84037309 |
| PTBP3 | 226.566242 | -0.1297711 | 0.27151921 | -0.4779445 | 0.63268973 | 0.91974341 |
| HSDL2 | 28.6331338 | 0.57270894 | 0.35934975 | 1.59373687 | 0.11099498 | 0.57860396 |
| C9orf147 | 6.71448136 | 0.87200408 | 0.65267946 | 1.33603726 | 0.18153709 | NA |
| KIAA1958 | 71.5396801 | 0.33678788 | 0.28209439 | 1.19388366 | 0.23252348 | 0.73156941 |
| INIP | 9.58998525 | -0.4247356 | 0.53548028 | -0.7931863 | 0.42766929 | NA |
| SNX30 | 17.5964901 | -0.5795307 | 0.44817391 | -1.2930932 | 0.19597881 | NA |
| FKBP15 | 8.56424947 | 0.64713602 | 0.53027256 | 1.22038376 | 0.22231943 | NA |
| SLC31A1 | 7.39668957 | -0.3759889 | 0.55021549 | -0.6833485 | 0.49438666 | NA |
| CDC26 | 6.75581786 | 1.00279088 | 0.57426428 | 1.74621846 | 0.080773 | NA |
| PRPF4 | 4.86909904 | -0.4006887 | 0.86069594 | -0.4655404 | 0.64154452 | NA |
| BSPRY | 5.95803912 | 0.22660134 | 0.56225396 | 0.40302312 | 0.68693122 | NA |
| HDHD3 | 9.59792599 | 0.22954018 | 0.56368304 | 0.40721499 | 0.68385009 | NA |
| ALAD | 3.52784819 | 1.12938886 | 0.79419094 | 1.42206213 | 0.15500822 | NA |
| POLE3 | 2.26821318 | -0.8989323 | 1.08461026 | -0.8288067 | 0.40721378 | NA |
| RGS3 | 7.6041921 | 0.11552988 | 0.60649361 | 0.19048821 | 0.84892658 | NA |
| ZNF618 | 72.2859236 | 0.20160468 | 0.31488648 | 0.6402456 | 0.52201294 | 0.87311773 |
| KIF12 | 19.0893727 | 0.64891749 | 0.56837238 | 1.14171186 | 0.25357381 | NA |
| COL27A1 | 7.35410749 | -0.8344323 | 0.73153157 | -1.1406648 | 0.25400944 | NA |
| WHRN | 6.76783152 | -0.0131681 | 0.62194666 | -0.0211724 | 0.98310817 | NA |
| ATP6V1G1 | 33.9662897 | -0.4373634 | 0.35727492 | -1.2241648 | 0.22089007 | 0.72409625 |
| TMEM268 | 5.80494508 | 0.44563737 | 0.62543394 | 0.71252509 | 0.47613968 | NA |
| PAPPA | 15.689915 | -1.4819034 | 0.91322502 | -1.6227144 | 0.10465047 | NA |
| ASTN2 | 120.587607 | -0.6422302 | 0.27392394 | -2.3445565 | 0.01904972 | 0.33831016 |
| CDK5RAP2 | 15.5621883 | 0.66799251 | 0.46236342 | 1.44473475 | 0.1485324 | NA |
| MEGF9 | 71.6054471 | 0.41531017 | 0.28653168 | 1.44943892 | 0.14721505 | 0.63 |
| FBXW2 | 13.4403624 | -0.6363937 | 0.52839462 | -1.204391 | 0.22843851 | NA |
| PSMD5 | 7.75387873 | -0.0156394 | 0.56349036 | -0.0277545 | 0.97785794 | NA |
| CNTRL | 5.38421783 | 0.8006271 | 0.60473174 | 1.32393762 | 0.18552376 | NA |
| RAB14 | 31.9242214 | -0.0453343 | 0.30368493 | -0.1492808 | 0.88133206 | 0.97767052 |
| GSN | 19.9628258 | 0.07049383 | 0.37401174 | 0.18848027 | 0.85050018 | NA |
| STOM | 9.85352287 | -0.4804619 | 0.52320889 | -0.9182984 | 0.35846266 | NA |
| DAB2IP | 40.1566345 | 0.59892667 | 0.32416078 | 1.84762225 | 0.06465701 | 0.52254021 |

| | | | | | | |
|---|---|---|---|---|---|---|
| TTLL11 | 5.57259612 | 0.46424201 | 0.65204931 | 0.71197378 | 0.47648101 | NA |
| NDUFA8 | 12.2738694 | 0.4629466 | 0.45084192 | 1.02684905 | 0.30449155 | NA |
| RBM18 | 4.75312729 | -0.1138928 | 0.69555666 | -0.1637434 | 0.86993313 | NA |
| MRRF | 20.2899634 | 0.50246928 | 0.43361286 | 1.158797 | 0.24653894 | NA |
| PDCL | 7.33301951 | -0.1796594 | 0.56695385 | -0.3168853 | 0.75133061 | NA |
| RC3H2 | 24.5436616 | -0.3732717 | 0.3882221 | -0.9614901 | 0.3363058 | NA |
| ZBTB26 | 3.54200131 | 0.01481098 | 0.74527715 | 0.01987311 | 0.98414459 | NA |
| RABGAP1 | 64.3804228 | 0.09035198 | 0.3262539 | 0.27693763 | 0.781828 | 0.96199854 |
| STRBP | 61.9461934 | -0.1264003 | 0.28759655 | -0.4395055 | 0.6602953 | 0.92276785 |
| DENND1A | 65.5197621 | 0.22902247 | 0.2617224 | 0.87505874 | 0.38154195 | 0.82153751 |
| NEK6 | 12.3006628 | 0.31262262 | 0.49012335 | 0.63784478 | 0.52357472 | NA |
| PSMB7 | 19.8785051 | 0.53168361 | 0.40230964 | 1.32157811 | 0.18630868 | NA |
| NR6A1 | 95.7024735 | -0.587066 | 0.38225397 | -1.5358008 | 0.12458724 | 0.58553957 |
| MIR181A2HG | 6.76701736 | -0.4746669 | 0.64631574 | -0.7344196 | 0.46269307 | NA |
| RPL35 | 36.3424112 | -0.4482686 | 0.37909118 | -1.1824822 | 0.23701444 | 0.73242571 |
| ARPC5L | 11.4934083 | -0.5870486 | 0.50622881 | -1.1596508 | 0.246191 | NA |
| GOLGA1 | 14.5479991 | -0.0860414 | 0.46025215 | -0.186944 | 0.85170457 | NA |
| SCAI | 48.9667721 | -0.3547989 | 0.37804551 | -0.9385084 | 0.3479832 | 0.8003972 |
| PPP6C | 25.8835605 | -0.5137411 | 0.39640847 | -1.2959891 | 0.19497923 | NA |
| RABEPK | 5.04396202 | -0.2034263 | 0.68831614 | -0.295542 | 0.7675799 | NA |
| HSPA5 | 37.00532 | -0.5795137 | 0.58223626 | -0.995324 | 0.31957869 | 0.78335494 |
| GAPVD1 | 34.488218 | -0.0142605 | 0.33939738 | -0.0420173 | 0.96648494 | 0.99656976 |
| MAPKAP1 | 75.9367256 | 0.57589684 | 0.37445896 | 1.53794383 | 0.12406235 | 0.58553957 |
| PBX3 | 56.3433531 | 0.52824393 | 0.353561 | 1.49406732 | 0.13515802 | 0.60594834 |
| MVB12B | 179.582295 | 0.27817598 | 0.33527298 | 0.82969995 | 0.40670845 | 0.82808596 |
| ZBTB43 | 25.2146239 | -0.4330227 | 0.35900792 | -1.2061649 | 0.22775392 | NA |
| ZBTB34 | 4.45664422 | -1.3380493 | 0.86869297 | -1.5403018 | 0.12348681 | NA |
| RALGPS1 | 64.0119806 | 0.03023523 | 0.28857107 | 0.10477568 | 0.91655381 | 0.98528045 |
| GARNL3 | 8.26048436 | -0.1459188 | 0.63327988 | -0.2304176 | 0.81776728 | NA |
| SLC2A8 | 5.77723047 | 0.03508162 | 0.63307366 | 0.05541476 | 0.95580804 | NA |
| RPL12 | 16.4604193 | -0.497021 | 0.43222821 | -1.1499041 | 0.25018336 | NA |
| LRSAM1 | 13.2976699 | -0.1219168 | 0.53937553 | -0.2260333 | 0.82117556 | NA |
| FAM129B | 11.9045204 | -0.0905677 | 0.50266093 | -0.1801766 | 0.85701391 | NA |
| STXBP1 | 7.28054436 | 0.16249155 | 0.5773995 | 0.28141962 | 0.77838857 | NA |
| PTRH1 | 3.05220643 | -0.912742 | 0.80162004 | -1.1386217 | 0.25486097 | NA |
| CDK9 | 5.48218537 | 0.43662996 | 0.73731973 | 0.59218537 | 0.55372646 | NA |
| FPGS | 6.24859201 | -0.8522236 | 0.72814086 | -1.1704104 | 0.24183587 | NA |
| DPM2 | 7.79357899 | -0.5450923 | 0.552954 | -0.9857823 | 0.32423996 | NA |
| FAM102A | 17.8955519 | 0.0598737 | 0.40113665 | 0.14926012 | 0.88134839 | NA |
| SLC25A25 | 9.64678799 | -0.5228996 | 0.48099178 | -1.087128 | 0.27698026 | NA |
| SLC25A25-AS | 13.4991841 | 0.50762448 | 0.46433364 | 1.09323219 | 0.27429187 | NA |
| PTGES2 | 9.67485028 | -0.4504593 | 0.52028473 | -0.8657939 | 0.38660321 | NA |
| C9orf16 | 24.9088642 | -0.3366583 | 0.5338007 | -0.6306817 | 0.5282487 | NA |
| CIZ1 | 7.45592855 | -0.3741103 | 0.66665709 | -0.5611735 | 0.57467925 | NA |
| DNM1 | 6.85146768 | 1.5734914 | 0.67624435 | 2.32680894 | 0.01997544 | NA |
| GOLGA2 | 19.1971751 | -0.6427459 | 0.46183302 | -1.3917279 | 0.16400482 | NA |
| SWI5 | 9.88719037 | -0.240256 | 0.50308692 | -0.4775637 | 0.63296079 | NA |
| TRUB2 | 5.2469951 | -0.1954524 | 0.66370977 | -0.2944847 | 0.76838756 | NA |
| COQ4 | 13.5631271 | 0.58466949 | 0.43232451 | 1.35238571 | 0.17625196 | NA |
| SLC27A4 | 12.573024 | 0.13058308 | 0.49231203 | 0.26524455 | 0.79082108 | NA |
| URM1 | 7.59800033 | -0.548229 | 0.52263847 | -1.0489642 | 0.2941946 | NA |
| ODF2 | 5.5275861 | 0.16360416 | 0.63071262 | 0.25939574 | 0.79532992 | NA |
| GLE1 | 9.01041036 | 0.83529298 | 0.51597852 | 1.61885225 | 0.10547906 | NA |
| SPTAN1 | 30.9080833 | -0.1314837 | 0.41774283 | -0.3147479 | 0.7529531 | 0.95243214 |

| | | | | | | |
|---|---|---|---|---|---|---|
| WDR34 | 7.48767867 | 0.4938333 | 0.54496066 | 0.90618156 | 0.36483976 | NA |
| SET | 39.5258295 | 0.03900438 | 0.31497526 | 0.12383314 | 0.90144739 | 0.98003205 |
| ZDHHC12 | 5.37683363 | -0.6874285 | 0.76613181 | -0.8972718 | 0.36957389 | NA |
| ZER1 | 21.2386449 | 0.29207318 | 0.36519546 | 0.7997722 | 0.42384279 | NA |
| TBC1D13 | 6.64571494 | -0.8609516 | 0.60980641 | -1.4118442 | 0.15799584 | NA |
| ENDOG | 6.61325894 | 0.47173655 | 0.62073228 | 0.75996781 | 0.44727383 | NA |
| SPOUT1 | 3.77464839 | 0.00908795 | 0.78543698 | 0.01157056 | 0.99076823 | NA |
| KYAT1 | 7.1757405 | -1.0666807 | 0.61691561 | -1.7290545 | 0.08379934 | NA |
| LRRC8A | 4.10836925 | -1.3195127 | 0.74020273 | -1.7826369 | 0.07464543 | NA |
| PHYHD1 | 3.27451843 | 0.29883314 | 0.808237 | 0.36973454 | 0.71158029 | NA |
| NUP188 | 6.4953346 | -0.8014966 | 0.67341989 | -1.1901885 | 0.23397233 | NA |
| SH3GLB2 | 12.8706629 | -0.6114224 | 0.43300916 | -1.4120312 | 0.15794078 | NA |
| MIGA2 | 8.13299167 | -0.1840959 | 0.56585334 | -0.3253421 | 0.7449222 | NA |
| CRAT | 4.26753384 | -0.056981 | 0.80864835 | -0.0704645 | 0.94382393 | NA |
| PTPA | 11.2520858 | 0.13249827 | 0.53396553 | 0.24814011 | 0.804026 | NA |
| IER5L | 4.44202795 | -1.0868085 | 0.66339307 | -1.6382573 | 0.10136803 | NA |
| LINC01503 | 7.30128267 | -0.4799002 | 0.99483232 | -0.482393 | 0.62952676 | NA |
| NTMT1 | 5.00013345 | 0.54429142 | 0.61448253 | 0.885772 | 0.3757404 | NA |
| ASB6 | 3.19495861 | -0.3079262 | 0.78800066 | -0.390769 | 0.69596798 | NA |
| TOR1B | 5.2166384 | 1.23365679 | 0.65555192 | 1.88185979 | 0.05985506 | NA |
| TOR1A | 3.35221852 | -0.1337641 | 0.84248585 | -0.1587731 | 0.87384766 | NA |
| C9orf78 | 10.3994671 | -0.2318898 | 0.52884811 | -0.4384809 | 0.66103773 | NA |
| USP20 | 6.73043448 | -0.18359 | 0.65279593 | -0.2812364 | 0.77852909 | NA |
| FNBP1 | 32.1044952 | 0.17982387 | 0.40095725 | 0.44848639 | 0.65380221 | 0.92248109 |
| GPR107 | 21.9926151 | -0.2019251 | 0.45889065 | -0.4400288 | 0.65991624 | NA |
| FUBP3 | 26.0696244 | -0.5997344 | 0.37592191 | -1.5953697 | 0.11062958 | 0.57860396 |
| EXOSC2 | 3.04738797 | -0.1740237 | 0.9123989 | -0.190732 | 0.84873555 | NA |
| ABL1 | 43.8474872 | -0.0633823 | 0.31050261 | -0.204128 | 0.83825348 | 0.97154892 |
| AIF1L | 26.8837313 | 0.71495143 | 0.48272794 | 1.48106496 | 0.13858926 | 0.61240852 |
| NUP214 | 34.3989603 | 0.0033483 | 0.3010501 | 0.01112206 | 0.99112606 | 0.99909657 |
| AL157938.2 | 9.55205815 | 0.69178515 | 0.84945367 | 0.81438832 | 0.41542253 | NA |
| PRRC2B | 80.15218 | -0.2295486 | 0.2489947 | -0.9219015 | 0.35657994 | 0.80177915 |
| POMT1 | 4.64334053 | 0.20740815 | 0.64073387 | 0.32370404 | 0.74616211 | NA |
| UCK1 | 3.87835036 | 0.60216754 | 0.70042752 | 0.85971429 | 0.38994656 | NA |
| RAPGEF1 | 61.1261887 | -0.0123528 | 0.37712134 | -0.0327555 | 0.97386959 | 0.99779576 |
| MED27 | 29.8764643 | -0.3527493 | 0.39174625 | -0.9004535 | 0.36787897 | 0.81079189 |
| SETX | 43.4551576 | -0.3977048 | 0.34506136 | -1.1525627 | 0.24908993 | 0.74122324 |
| TTF1 | 8.20566235 | -0.0970428 | 0.55874458 | -0.1736801 | 0.86211689 | NA |
| DDX31 | 8.32021058 | -0.1744776 | 0.5820222 | -0.2997783 | 0.76434628 | NA |
| GTF3C4 | 3.42687145 | -0.3835121 | 0.81003247 | -0.4734527 | 0.63589021 | NA |
| TSC1 | 21.0906614 | 0.03748187 | 0.39729796 | 0.09434196 | 0.92483752 | NA |
| GTF3C5 | 5.94403031 | 0.30863959 | 0.6013181 | 0.51327174 | 0.60776125 | NA |
| RALGDS | 3.33633882 | -0.3627166 | 0.80092369 | -0.4528729 | 0.65064029 | NA |
| ABO | 12.2741494 | 1.24375406 | 0.51549306 | 2.41274645 | 0.01583283 | NA |
| SURF6 | 3.54322745 | -0.1453613 | 0.80489239 | -0.1805971 | 0.8566838 | NA |
| MED22 | 2.80511964 | -1.8393353 | 0.95504742 | -1.9259099 | 0.05411562 | NA |
| RPL7A | 61.2747418 | -0.0413094 | 0.41944042 | -0.0984869 | 0.9215457 | 0.98745437 |
| SURF1 | 5.34347103 | 1.09208927 | 0.65024242 | 1.67951095 | 0.09305251 | NA |
| SURF2 | 2.8861624 | -1.2235674 | 0.89222095 | -1.3713727 | 0.17025881 | NA |
| SURF4 | 19.476496 | -0.5938077 | 0.40319089 | -1.4727706 | 0.14081291 | NA |
| REXO4 | 4.70340911 | 0.13872763 | 0.70446018 | 0.19692757 | 0.84388422 | NA |
| CACFD1 | 3.82958892 | -1.0706816 | 0.78239694 | -1.3684634 | 0.17116707 | NA |
| VAV2 | 45.5155057 | -0.177417 | 0.30689209 | -0.5781088 | 0.56319069 | 0.89263108 |
| BRD3 | 12.2513381 | 0.24642146 | 0.51063699 | 0.48257661 | 0.62939639 | NA |

| | | | | | | |
|---|---|---|---|---|---|---|
| WDR5 | 4.30433623 | 0.44248888 | 0.84934232 | 0.52097825 | 0.60238192 | NA |
| RXRA | 11.8178083 | 0.13117965 | 0.51434947 | 0.25503992 | 0.79869227 | NA |
| PPP1R26-AS: | 4.42341608 | 0.15277326 | 0.87395785 | 0.17480621 | 0.86123191 | NA |
| PPP1R26 | 6.1704083 | -0.4637912 | 0.66372487 | -0.6987703 | 0.48469562 | NA |
| MRPS2 | 5.6415249 | 0.31459997 | 0.74338317 | 0.42320029 | 0.67214913 | NA |
| CAMSAP1 | 11.8932766 | 0.21306942 | 0.52294653 | 0.40744017 | 0.68368473 | NA |
| UBAC1 | 11.9194296 | -0.4339453 | 0.52048771 | -0.8337282 | 0.40443415 | NA |
| NACC2 | 12.4505666 | 0.17023343 | 0.48611052 | 0.3501949 | 0.72619243 | NA |
| TMEM250 | 3.29304072 | -1.7025933 | 1.01565396 | -1.6763518 | 0.09366931 | NA |
| QSOX2 | 2.71802208 | -0.3427643 | 0.94354987 | -0.3632709 | 0.71640249 | NA |
| GPSM1 | 4.55647441 | -0.2001777 | 0.79879867 | -0.2505985 | 0.80212458 | NA |
| SNAPC4 | 3.77598286 | -0.7114361 | 0.77361269 | -0.9196283 | 0.35776706 | NA |
| ENTR1 | 5.85718841 | -0.7924209 | 0.62314394 | -1.2716499 | 0.20349753 | NA |
| PMPCA | 8.14346099 | -0.0317603 | 0.59994199 | -0.0529389 | 0.9577806 | NA |
| INPP5E | 3.16522276 | 0.39654281 | 0.89357664 | 0.44377034 | 0.65720863 | NA |
| SEC16A | 13.8920226 | -0.4703162 | 0.47139044 | -0.9977211 | 0.31841461 | NA |
| NOTCH1 | 5.2015178 | 0.3104766 | 0.73135337 | 0.42452337 | 0.67118416 | NA |
| EGFL7 | 5.64216522 | -0.2935999 | 0.61316782 | -0.4788248 | 0.63206329 | NA |
| AGPAT2 | 19.1864538 | 0.48760375 | 0.42024938 | 1.16027239 | 0.24593792 | NA |
| FAM69B | 2.41220185 | -0.1583405 | 0.89959121 | -0.1760138 | 0.86028311 | NA |
| SNHG7 | 18.431759 | 0.57311627 | 0.40589254 | 1.41199017 | 0.15795286 | NA |
| RABL6 | 19.0884281 | -0.2884581 | 0.43906385 | -0.6569844 | 0.51119092 | NA |
| PHPT1 | 16.0793242 | 0.63907596 | 0.48161976 | 1.32693051 | 0.18453167 | NA |
| EDF1 | 27.9315301 | -0.6434675 | 0.37943023 | -1.6958784 | 0.08990891 | 0.56122412 |
| TRAF2 | 3.06769537 | 0.85924898 | 0.85127566 | 1.00936632 | 0.31279899 | NA |
| FBXW5 | 12.2257064 | -0.5681972 | 0.46686876 | -1.2170384 | 0.2235896 | NA |
| PAXX | 5.81123602 | 0.51175054 | 0.70226337 | 0.72871598 | 0.46617541 | NA |
| UAP1L1 | 2.799142 | -1.4214966 | 1.07340257 | -1.3242903 | 0.18540666 | NA |
| MAN1B1 | 6.22505668 | -1.4123326 | 0.62657687 | -2.2540452 | 0.02419332 | NA |
| DPP7 | 9.04541994 | -0.2055004 | 0.53457967 | -0.3844149 | 0.70067097 | NA |
| SSNA1 | 8.53528081 | 0.13492758 | 0.47657827 | 0.28311736 | 0.77708687 | NA |
| TPRN | 5.21654267 | 0.48498651 | 0.62649567 | 0.77412587 | 0.43885636 | NA |
| TMEM203 | 4.20028969 | -0.4035689 | 0.87149453 | -0.4630768 | 0.64330933 | NA |
| NDOR1 | 3.55630303 | 1.38428161 | 0.8658619 | 1.59873256 | 0.10988004 | NA |
| RNF208 | 2.61244614 | 0.41274008 | 0.9381562 | 0.43994814 | 0.65997467 | NA |
| TUBB4B | 26.223648 | -0.4083908 | 0.51752329 | -0.7891255 | 0.43003866 | 0.84223493 |
| NELFB | 9.95175322 | -0.1694249 | 0.50083686 | -0.3382835 | 0.73514955 | NA |
| EXD3 | 11.9223514 | -0.0747353 | 0.49401402 | -0.1512818 | 0.87975346 | NA |
| NOXA1 | 3.56566135 | 1.10365732 | 0.76532559 | 1.44207554 | 0.14928106 | NA |
| NSMF | 4.16059505 | -0.1879097 | 0.70052762 | -0.2682402 | 0.78851443 | NA |
| PNPLA7 | 4.58606954 | 0.72117222 | 0.75614421 | 0.95374958 | 0.34021042 | NA |
| MRPL41 | 29.8627457 | 0.20909903 | 0.3336451 | 0.62671091 | 0.53084875 | 0.88254911 |
| DPH7 | 5.61116401 | 0.50710452 | 0.66247803 | 0.76546618 | 0.443994 | NA |
| ZMYND19 | 2.28728889 | -1.7607415 | 1.0293413 | -1.7105517 | 0.08716391 | NA |
| ARRDC1 | 9.80476151 | -0.9357506 | 0.55654357 | -1.6813609 | 0.09269283 | NA |
| ARRDC1-AS1 | 4.01295003 | -0.163658 | 0.71354594 | -0.2293587 | 0.81859015 | NA |
| EHMT1 | 69.1865233 | -0.1844683 | 0.33005634 | -0.5588993 | 0.57623043 | 0.89263108 |
| BET1L | 6.38634397 | -0.0219196 | 0.56158726 | -0.0390315 | 0.96886523 | NA |
| RIC8A | 5.37647547 | 0.05206244 | 0.60955348 | 0.08541078 | 0.93193482 | NA |
| SIRT3 | 4.09750458 | 0.82626699 | 0.75440634 | 1.09525456 | 0.27340513 | NA |
| PSMD13 | 10.4086608 | -0.479338 | 0.48494136 | -0.9884453 | 0.3229346 | NA |
| IFITM2 | 32.9241287 | -0.0493575 | 0.58210299 | -0.0847917 | 0.93242699 | 0.99030945 |
| IFITM3 | 65.6280452 | -0.5811028 | 0.55257418 | -1.0516286 | 0.29296996 | 0.77342911 |
| PKP3 | 3.78448488 | -1.0727172 | 0.77501568 | -1.3841232 | 0.16632073 | NA |

| | | | | | | |
|---|---|---|---|---|---|---|
| SIGIRR | 8.78123957 | 0.07268607 | 0.62997015 | 0.11538018 | 0.90814379 | NA |
| ANO9 | 3.02815147 | -0.9797899 | 0.93383075 | -1.0492157 | 0.29407884 | NA |
| PTDSS2 | 10.4815031 | -0.5713414 | 0.50965398 | -1.1210378 | 0.26227178 | NA |
| RNH1 | 14.6437316 | 0.64444253 | 0.429112 | 1.50180494 | 0.13314749 | NA |
| HRAS | 12.1651457 | -0.0652795 | 0.50015701 | -0.1305181 | 0.89615654 | NA |
| RASSF7 | 11.8444842 | -0.5923873 | 0.48105148 | -1.2314427 | 0.21815735 | NA |
| PHRF1 | 9.76180125 | -0.0725305 | 0.48265046 | -0.1502753 | 0.88054739 | NA |
| IRF7 | 5.18266617 | -1.0203809 | 0.81631671 | -1.2499816 | 0.21130626 | NA |
| DEAF1 | 28.1833851 | -0.4608924 | 0.35705271 | -1.2908244 | 0.19676456 | 0.69343545 |
| AC131934.1 | 4.77195299 | -0.3939017 | 0.79299973 | -0.4967236 | 0.61938397 | NA |
| EPS8L2 | 25.3033596 | -0.0619786 | 0.50349513 | -0.1230967 | 0.9020305 | NA |
| AP006621.4 | 2.85271855 | 1.59958657 | 1.14589934 | 1.39592241 | 0.16273786 | NA |
| TALDO1 | 20.3713765 | -0.1587255 | 0.39562209 | -0.4012048 | 0.68826932 | NA |
| GATD1 | 6.54462408 | -0.7046689 | 0.65912366 | -1.0690997 | 0.28502475 | NA |
| AP006621.3 | 4.86819301 | -0.104249 | 0.81190165 | -0.1284011 | 0.89783159 | NA |
| RPLP2 | 45.1469354 | -0.2930578 | 0.31379851 | -0.9339043 | 0.35035328 | 0.8003972 |
| PNPLA2 | 6.39809301 | 0.07584679 | 0.59836382 | 0.12675699 | 0.89913274 | NA |
| CD151 | 16.7973615 | -0.1988425 | 0.38270039 | -0.5195774 | 0.60335816 | NA |
| POLR2L | 25.1821001 | -0.163205 | 0.37295112 | -0.4376042 | 0.66167324 | NA |
| TSPAN4 | 3.25499622 | -0.002747 | 0.73558414 | -0.0037344 | 0.99702039 | NA |
| CHID1 | 6.43972145 | 0.42360902 | 0.59363876 | 0.71358046 | 0.47548664 | NA |
| AP2A2 | 38.7691712 | -0.2690856 | 0.35023938 | -0.7682904 | 0.44231467 | 0.84767356 |
| TOLLIP | 11.9594417 | -1.1107195 | 0.50901276 | -2.1821054 | 0.02910175 | NA |
| MOB2 | 8.46802422 | 0.02851638 | 0.60357242 | 0.047246 | 0.96231716 | NA |
| MRPL23 | 3.56892697 | 1.24919678 | 0.82841793 | 1.50793064 | 0.13157228 | NA |
| CD81 | 39.5034382 | 0.12964019 | 0.45988851 | 0.28189482 | 0.77802416 | 0.96133329 |
| TSSC4 | 7.23696215 | -0.6326903 | 0.54969302 | -1.1509883 | 0.24973704 | NA |
| KCNQ1 | 53.4963136 | 0.3645141 | 0.3108658 | 1.17257704 | 0.24096546 | 0.73323864 |
| KCNQ1OT1 | 78.9321426 | 0.20761336 | 0.40820159 | 0.50860499 | 0.61102913 | 0.90955758 |
| CDKN1C | 38.1070166 | 0.53108909 | 0.56748892 | 0.93585807 | 0.34934627 | 0.8003972 |
| SLC22A18 | 6.6967403 | 0.48957469 | 0.61094223 | 0.80134367 | 0.42293271 | NA |
| NAP1L4 | 57.7971555 | 0.10730315 | 0.32187931 | 0.33336453 | 0.73885913 | 0.94658914 |
| CARS | 20.78288 | -0.4768036 | 0.45115831 | -1.0568431 | 0.29058319 | NA |
| OSBPL5 | 5.05550246 | 1.6560595 | 0.71058347 | 2.33056295 | 0.01977642 | NA |
| ZNF195 | 8.69059486 | -0.6359875 | 0.54372145 | -1.1696935 | 0.24212432 | NA |
| AC127526.1 | 2.76107024 | -1.1405355 | 1.12200073 | -1.0165193 | 0.30938214 | NA |
| NUP98 | 29.7731966 | -0.507789 | 0.32953717 | -1.5409157 | 0.1233373 | 0.58553957 |
| PGAP2 | 5.08147513 | 0.28629157 | 0.73441312 | 0.38982361 | 0.69666698 | NA |
| RHOG | 6.6802855 | 0.30703199 | 0.6449429 | 0.47606074 | 0.63403111 | NA |
| STIM1 | 82.7690926 | -0.1717999 | 0.30295194 | -0.5670862 | 0.5706556 | 0.89263108 |
| RRM1 | 7.16752684 | 0.63438621 | 0.63597071 | 0.99750853 | 0.31851774 | NA |
| FAM160A2 | 6.6336256 | 0.2403275 | 0.60884607 | 0.39472621 | 0.69304496 | NA |
| SMPD1 | 5.62792314 | -0.63858 | 0.70211815 | -0.9095051 | 0.36308357 | NA |
| APBB1 | 6.27279428 | 1.14489655 | 0.56909374 | 2.01178904 | 0.04424218 | NA |
| ARFIP2 | 5.61744244 | -0.3570436 | 0.58289559 | -0.6125343 | 0.5401843 | NA |
| AC084337.2 | 4.93642943 | 1.16567283 | 0.72553293 | 1.6066436 | 0.10813257 | NA |
| DNHD1 | 5.47938031 | -0.1946092 | 0.64801272 | -0.3003169 | 0.76393544 | NA |
| RRP8 | 3.55455597 | -0.1578063 | 0.76610823 | -0.2059843 | 0.83680317 | NA |
| ILK | 10.7881649 | 0.13078531 | 0.51187102 | 0.25550443 | 0.79833352 | NA |
| TAF10 | 9.86301833 | -0.4696048 | 0.60021542 | -0.7823938 | 0.43398321 | NA |
| TPP1 | 8.47473168 | 0.14173155 | 0.62056447 | 0.22839134 | 0.81934202 | NA |
| MRPL17 | 7.23009028 | -0.3257279 | 0.61271731 | -0.5316121 | 0.5949947 | NA |
| SYT9 | 3.20227166 | 0.69828129 | 0.76235658 | 0.91595103 | 0.35969258 | NA |
| AC107884.1 | 3.52173581 | 0.86633559 | 0.93037233 | 0.93117084 | 0.3517652 | NA |

| | | | | | | |
|---|---|---|---|---|---|---|
| PPFIBP2 | 26.8181688 | -0.2612851 | 0.38818988 | -0.6730858 | 0.50089268 | 0.87036626 |
| EIF3F | 16.5383864 | -0.0509532 | 0.45405731 | -0.1122175 | 0.91065094 | NA |
| TUB | 8.83683261 | 0.00954624 | 0.58775557 | 0.01624185 | 0.98704145 | NA |
| RIC3 | 20.5163245 | -0.1669601 | 0.48545884 | -0.3439224 | 0.73090467 | NA |
| STK33 | 64.4239847 | 0.36344429 | 0.34892177 | 1.04162115 | 0.29758736 | 0.77521265 |
| TRIM66 | 18.6040208 | -0.3499264 | 0.50873888 | -0.687831 | 0.49155919 | NA |
| RPL27A | 56.8142999 | -0.4095428 | 0.49356917 | -0.8297577 | 0.40667582 | 0.82808596 |
| ST5 | 23.2495545 | 0.29834793 | 0.51915586 | 0.57467892 | 0.56550847 | NA |
| AKIP1 | 3.41996237 | -0.0182561 | 0.89659827 | -0.0203616 | 0.98375495 | NA |
| TMEM9B | 18.1642523 | 0.46988646 | 0.39790402 | 1.18090402 | 0.23764085 | NA |
| TMEM9B-AS | 4.10794066 | -0.1621289 | 0.90989669 | -0.1781839 | 0.85857856 | NA |
| DENND5A | 19.5589217 | -0.4443597 | 0.43312311 | -1.0259431 | 0.30491839 | NA |
| TMEM41B | 23.9503531 | -0.6526204 | 0.39718466 | -1.6431159 | 0.10035895 | NA |
| IPO7 | 33.4868508 | -0.0130724 | 0.3183675 | -0.0410607 | 0.96724749 | 0.99656976 |
| ZNF143 | 7.13158035 | -1.1804443 | 0.60379662 | -1.9550362 | 0.0505788 | NA |
| SWAP70 | 23.6633302 | -0.2224316 | 0.40447966 | -0.5499203 | 0.58237401 | NA |
| SBF2-AS1 | 19.4296576 | -0.7766766 | 0.58214914 | -1.3341539 | 0.18215341 | NA |
| SBF2 | 185.075322 | -0.0013587 | 0.1924456 | -0.0070602 | 0.99436683 | 0.99923383 |
| AC011092.2 | 6.67231544 | 0.71153554 | 0.6938344 | 1.02551205 | 0.30512164 | NA |
| ADM | 12.4185329 | -0.0846681 | 0.45852541 | -0.184653 | 0.85350119 | NA |
| AMPD3 | 16.0428654 | 0.72910483 | 0.4668367 | 1.56179842 | 0.11833548 | NA |
| MTRNR2L8 | 70.7128147 | -1.1686284 | 0.47785524 | -2.44557 | 0.01446234 | 0.30435125 |
| RNF141 | 8.91986411 | 1.13492002 | 0.49844483 | 2.27692203 | 0.02279088 | NA |
| MRVI1 | 3.41328687 | 0.74739027 | 0.97719447 | 0.76483268 | 0.44437119 | NA |
| CTR9 | 14.0710075 | -0.3782983 | 0.4362221 | -0.8672149 | 0.38582427 | NA |
| EIF4G2 | 47.5544864 | -0.05334 | 0.2790935 | -0.1911186 | 0.84843265 | 0.97168963 |
| ZBED5 | 16.9758388 | 0.29998652 | 0.38602483 | 0.77711717 | 0.43708965 | NA |
| GALNT18 | 13.3324856 | 0.11378633 | 0.56574799 | 0.20112546 | 0.84060047 | NA |
| USP47 | 62.6335218 | -0.0915006 | 0.2398402 | -0.3815063 | 0.70282756 | 0.9303056 |
| PARVA | 34.0582712 | -0.1730314 | 0.3277755 | -0.5278959 | 0.59757157 | 0.90006689 |
| TEAD1 | 83.043864 | -0.2670255 | 0.37879289 | -0.7049381 | 0.48084878 | 0.86222804 |
| LINC00958 | 6.33361269 | -0.025282 | 0.63818752 | -0.0396153 | 0.96839986 | NA |
| ARNTL | 8.21367939 | -0.4068092 | 0.55380665 | -0.734569 | 0.46260201 | NA |
| BTBD10 | 14.4012436 | -0.3106425 | 0.42660143 | -0.7281798 | 0.46650353 | NA |
| FAR1 | 41.5415422 | -0.8389234 | 0.36238248 | -2.3150219 | 0.02061173 | 0.34599194 |
| RRAS2 | 25.8725152 | -0.8088221 | 0.46811909 | -1.7278127 | 0.08402181 | NA |
| COPB1 | 23.6050188 | -0.4741046 | 0.41944712 | -1.1303085 | 0.25834628 | NA |
| CYP2R1 | 3.53121846 | 0.16912574 | 0.95567508 | 0.17696992 | 0.85953202 | NA |
| SOX6 | 256.620929 | 0.05359093 | 0.23337575 | 0.22963368 | 0.81837643 | 0.96845695 |
| C11orf58 | 29.1773202 | -0.0376401 | 0.33156158 | -0.1135236 | 0.90961545 | 0.98346872 |
| PLEKHA7 | 137.462286 | 0.19115843 | 0.27757891 | 0.68866339 | 0.49103512 | 0.86506451 |
| RPS13 | 37.564421 | 0.37929612 | 0.3918232 | 0.96802876 | 0.33303001 | 0.79447068 |
| PIK3C2A | 56.8898431 | -0.3492226 | 0.27750351 | -1.2584441 | 0.20823121 | 0.70439682 |
| NUCB2 | 5.17210642 | 0.27579939 | 0.74496676 | 0.37021704 | 0.71122078 | NA |
| KCNC1 | 3.62446041 | 0.65640334 | 0.84982242 | 0.7724006 | 0.4398772 | NA |
| SERGEF | 41.2870855 | 0.00018363 | 0.37197672 | 0.00049367 | 0.99960611 | 0.99960611 |
| SAAL1 | 3.00360157 | -0.4430457 | 0.85259966 | -0.519641 | 0.60331381 | NA |
| HPS5 | 8.3343948 | -0.246783 | 0.56441046 | -0.4372403 | 0.66193705 | NA |
| GTF2H1 | 13.8348834 | -0.7628745 | 0.48371911 | -1.5771023 | 0.11477198 | NA |
| LDHA | 54.5752959 | 0.12861751 | 0.35209756 | 0.36528941 | 7.15E-01 | 0.93629694 |
| TSG101 | 26.6734415 | 0.10954206 | 0.30754845 | 0.35617822 | 0.7217071 | 0.93818522 |
| UEVLD | 19.2861256 | 0.40050282 | 0.39821922 | 1.00573453 | 0.31454329 | NA |
| SPTY2D1OS | 5.25075915 | 0.75510947 | 0.67134014 | 1.12477926 | 0.26068259 | NA |
| SPTY2D1 | 17.9361707 | -1.8144987 | 0.53498145 | -3.391704 | 0.00069459 | NA |

| | | | | | | |
|---|---|---|---|---|---|---|
| ZDHHC13 | 20.5794592 | -0.0685901 | 0.42470185 | -0.1615018 | 0.87169821 | NA |
| NAV2 | 69.8504115 | -0.9642051 | 0.532214 | -1.8116869 | 0.07003459 | 0.52654172 |
| HTATIP2 | 5.39427496 | 0.40878167 | 0.63470855 | 0.64404627 | 0.51954543 | NA |
| PRMT3 | 18.743943 | -0.092768 | 0.36160906 | -0.2565422 | 0.79753216 | NA |
| NELL1 | 27.406892 | 0.90727804 | 0.84850678 | 1.06926434 | 0.28495057 | 0.76977033 |
| ANO5 | 5.88195995 | 1.59935607 | 0.73095288 | 2.18804266 | 0.0286665 | NA |
| SVIP | 12.3154692 | 1.22239936 | 0.43727441 | 2.79549716 | 0.005182 | NA |
| SLC5A12 | 5.97201654 | 1.664794 | 1.05102066 | 1.58397837 | 0.11319864 | NA |
| BBOX1 | 7.03609246 | 0.67621695 | 0.81204269 | 0.83273572 | 0.4049938 | NA |
| BBOX1-AS1 | 5.62529154 | 1.10331544 | 1.18547229 | 0.93069695 | 0.35201035 | NA |
| CCDC34 | 2.58877727 | 3.32518907 | 1.21723333 | 2.73175979 | 0.00629971 | NA |
| LGR4 | 208.136383 | 0.67541664 | 0.35341799 | 1.91109865 | 0.0559919 | 0.50317304 |
| LIN7C | 12.9338651 | -0.0197846 | 0.44876791 | -0.0440865 | 0.96483545 | NA |
| BDNF-AS | 31.08712 | 0.74995483 | 0.34344561 | 2.18362039 | 0.02899015 | 0.40510325 |
| METTL15 | 50.1766014 | 0.530637 | 0.44600858 | 1.18974617 | 0.23414617 | 0.73156941 |
| ARL14EP | 7.26396187 | 0.33543286 | 0.58553747 | 0.57286317 | 0.56673734 | NA |
| MPPED2 | 14.0261618 | -0.094077 | 0.7466149 | -0.1260047 | 0.89972818 | NA |
| DCDC1 | 5.8971136 | -0.6364777 | 0.72923598 | -0.8728007 | 0.3827717 | NA |
| DNAJC24 | 11.9919263 | -0.603574 | 0.57298602 | -1.0533835 | 0.29216527 | NA |
| IMMP1L | 27.154998 | -0.1775251 | 0.35843215 | -0.4952823 | 0.62040088 | 0.91493825 |
| ELP4 | 41.2522055 | 0.58199314 | 0.33260523 | 1.74980149 | 0.08015257 | 0.54552914 |
| AC131571.1 | 4.66768067 | 1.60943891 | 0.72665956 | 2.21484584 | 0.02677065 | NA |
| RCN1 | 8.4451141 | -0.3510465 | 0.58679029 | -0.5982487 | 0.54967401 | NA |
| EIF3M | 17.6794496 | -0.8167252 | 0.39806253 | -2.0517509 | 0.04019387 | NA |
| CCDC73 | 3.22825204 | -0.9974265 | 0.82334364 | -1.2114341 | 0.22572909 | NA |
| PRRG4 | 10.78709 | 0.56869278 | 0.46103156 | 1.23352246 | 0.2173809 | NA |
| QSER1 | 22.6730379 | 0.29382639 | 0.4231313 | 0.69440949 | 0.48742545 | NA |
| TCP11L1 | 4.54241166 | 0.59192148 | 0.69846297 | 0.84746294 | 0.39673714 | NA |
| CSTF3 | 17.892358 | -0.0020417 | 0.38436866 | -0.0053118 | 0.99576178 | NA |
| HIPK3 | 98.7263287 | -0.270396 | 0.23099811 | -1.1705552 | 0.24177762 | 0.73323864 |
| FBXO3 | 13.5638108 | 0.07166046 | 0.43898232 | 0.16324226 | 0.8703277 | NA |
| CAPRIN1 | 32.7344246 | -0.3681569 | 0.31569039 | -1.1661961 | 0.24353519 | 0.73323864 |
| NAT10 | 8.18231381 | 0.07141127 | 0.54236792 | 0.13166574 | 0.89524869 | NA |
| ABTB2 | 74.4312082 | -0.2118017 | 0.46348837 | -0.4569731 | 0.64769035 | 0.9218151 |
| CAT | 26.1069764 | 0.20504926 | 0.4339592 | 0.47250814 | 0.63656413 | 0.92121485 |
| EHF | 6.43210886 | 0.87203756 | 0.83081191 | 1.04962092 | 0.29389243 | NA |
| APIP | 16.3023599 | -0.7455504 | 0.44281063 | -1.6836777 | 0.09224398 | NA |
| PDHX | 32.0391876 | -0.2593157 | 0.31216147 | -0.8307102 | 0.40613737 | 0.82808596 |
| TRIM44 | 60.9018969 | -0.2155413 | 0.2404668 | -0.8963454 | 0.37006831 | 0.81199861 |
| COMMD9 | 3.59314575 | 0.2058271 | 0.81744374 | 0.25179359 | 0.80120061 | NA |
| TRAF6 | 9.11459084 | 0.34561558 | 0.58232126 | 0.5935136 | 0.55283749 | NA |
| C11orf74 | 3.74288776 | 1.64890634 | 0.79862357 | 2.06468529 | 0.03895279 | NA |
| API5 | 22.5200883 | -0.6930705 | 0.34437771 | -2.01253 | 0.04416409 | NA |
| TTC17 | 80.7505807 | 0.05050494 | 0.24213575 | 0.20858111 | 0.83477526 | 0.97154892 |
| HSD17B12 | 92.9920943 | 0.13268877 | 0.23371623 | 0.56773451 | 0.57021526 | 0.89263108 |
| ALKBH3 | 4.78491789 | 0.45354206 | 0.65999815 | 0.68718687 | 0.49196498 | NA |
| ACCS | 5.6870328 | 0.27910918 | 0.78149096 | 0.35714959 | 0.72097982 | NA |
| EXT2 | 20.7640059 | 0.12679154 | 0.36839163 | 0.34417594 | 0.73071397 | NA |
| CD82 | 23.7304847 | 1.21339194 | 0.49336372 | 2.45942675 | 0.01391591 | NA |
| PRDM11 | 6.59644801 | -0.1820481 | 0.55947233 | -0.3253925 | 0.74488405 | NA |
| CRY2 | 11.3003572 | 0.4190506 | 0.44938948 | 0.93248866 | 0.35108404 | NA |
| PEX16 | 3.94302416 | -0.5510716 | 0.73380281 | -0.7509805 | 0.4526644 | NA |
| LARGE2 | 5.42009651 | -0.1630638 | 0.66820277 | -0.2440334 | 0.80720492 | NA |
| PHF21A | 67.4799119 | -0.055366 | 0.33934812 | -0.163154 | 0.87039718 | 0.97418878 |

| | | | | | | |
|---|---|---|---|---|---|---|
| DGKZ | 6.14445631 | -0.8889277 | 0.63669702 | -1.3961549 | 0.16266784 | NA |
| AMBRA1 | 132.61006 | 0.16962406 | 0.29958409 | 0.56619849 | 0.57125886 | 0.89263108 |
| ATG13 | 13.0478659 | 0.06704341 | 0.42122029 | 0.15916472 | 0.8735391 | NA |
| ARHGAP1 | 9.88530137 | 0.24144587 | 0.58555679 | 0.41233552 | 0.68009351 | NA |
| CKAP5 | 18.5726017 | -0.0972986 | 0.36256734 | -0.26836 | 0.78842222 | NA |
| LRP4 | 16.9364635 | -0.2737879 | 0.60050102 | -0.4559324 | 0.64843855 | NA |
| C11orf49 | 50.2337953 | 0.07367391 | 0.29674868 | 0.24827037 | 0.80392522 | 0.96743263 |
| ARFGAP2 | 9.36222091 | -0.6229696 | 0.54854975 | -1.1356666 | 0.25609611 | NA |
| DDB2 | 2.65423102 | 1.32407935 | 0.96837103 | 1.36732648 | 0.171523 | NA |
| ACP2 | 3.962772 | -0.3451458 | 0.69330753 | -0.497825 | 0.61860737 | NA |
| NR1H3 | 4.24606984 | -0.6931487 | 0.76744837 | -0.9031861 | 0.36642715 | NA |
| MADD | 7.84914213 | -0.19625 | 0.57830611 | -0.3393531 | 0.73434371 | NA |
| SLC39A13 | 8.15610521 | -1.0884543 | 0.54638051 | -1.9921178 | 0.04635813 | NA |
| PSMC3 | 6.11591173 | -0.5325435 | 0.69260664 | -0.7688975 | 0.44195415 | NA |
| CELF1 | 87.7471162 | -0.4671543 | 0.29865102 | -1.5642146 | 0.11776718 | 0.57941414 |
| NDUFS3 | 10.9975236 | -0.1513931 | 0.53747536 | -0.2816746 | 0.77819306 | NA |
| MTCH2 | 26.5693693 | -0.4939305 | 0.34748915 | -1.421427 | 0.15519266 | 0.63921176 |
| FNBP4 | 53.3469115 | 0.11807931 | 0.25786397 | 0.45791317 | 0.64701483 | 0.9218151 |
| NUP160 | 53.5414806 | -0.4810834 | 0.28322716 | -1.6985779 | 0.08939874 | 0.56122412 |
| PTPRJ | 243.907144 | -0.2630258 | 0.35715176 | -0.736454 | 0.46145446 | 0.85894565 |
| TNKS1BP1 | 5.21009095 | -0.2517423 | 0.69706484 | -0.3611462 | 0.71799015 | NA |
| SSRP1 | 6.46583469 | 0.36543221 | 0.65611961 | 0.55695974 | 0.57755493 | NA |
| TIMM10 | 7.39839565 | 0.72429222 | 0.66792254 | 1.08439553 | 0.27818946 | NA |
| UBE2L6 | 4.93554521 | -0.4325342 | 1.09330499 | -0.3956208 | 0.69238482 | NA |
| SERPING1 | 48.6537593 | -0.3255041 | 0.46549176 | -0.6992692 | 0.48438379 | 0.86354292 |
| ZDHHC5 | 19.5553017 | -0.2199448 | 0.41733365 | -0.5270238 | 0.59817704 | NA |
| MED19 | 3.72431575 | -0.934942 | 0.8532553 | -1.0957354 | 0.2731946 | NA |
| CTNND1 | 39.6584378 | -0.1495296 | 0.29803363 | -0.5017204 | 0.61586419 | 0.91318974 |
| LPXN | 8.47422194 | -0.1784674 | 0.62996923 | -0.2832954 | 0.77695038 | NA |
| GLYAT | 5.82535424 | 1.66746414 | 1.27369825 | 1.30915163 | 0.19048299 | NA |
| FAM111A | 10.9796323 | 0.43458788 | 0.57026454 | 0.7620812 | 0.44601154 | NA |
| OSBP | 32.5154325 | -0.9045825 | 0.35343296 | -2.5594174 | 0.01048478 | 0.28266532 |
| PATL1 | 14.2861773 | -0.7098732 | 0.43850572 | -1.6188458 | 0.10548044 | NA |
| STX3 | 19.6794517 | -0.4176152 | 0.47096726 | -0.8867182 | 0.37523064 | NA |
| MRPL16 | 6.46020079 | 1.48265681 | 0.61269305 | 2.41990148 | 0.01552471 | NA |
| PRPF19 | 7.91416814 | 0.09716476 | 0.65046989 | 0.14937627 | 0.88125674 | NA |
| TMEM109 | 5.62539017 | 0.79262611 | 0.63705484 | 1.2442039 | 0.21342453 | NA |
| VPS37C | 6.04273123 | -0.5488109 | 0.62014709 | -0.8849689 | 0.3761734 | NA |
| DDB1 | 30.7538494 | -0.2038698 | 0.3411993 | -0.5975093 | 0.55016738 | 0.88557663 |
| TKFC | 4.85623183 | 0.84557461 | 0.71166263 | 1.18816778 | 0.23476731 | NA |
| CYB561A3 | 3.8074361 | -0.0849584 | 0.83358772 | -0.1019189 | 0.91882104 | NA |
| TMEM138 | 3.14910345 | 0.99842059 | 0.83660065 | 1.19342555 | 0.23270275 | NA |
| CPSF7 | 18.736424 | 0.031081 | 0.41470916 | 0.07494651 | 0.94025727 | NA |
| AP003108.2 | 7.38359319 | 0.30748771 | 0.54670636 | 0.56243668 | 0.57381854 | NA |
| TMEM258 | 7.80500403 | 0.6397207 | 0.63089641 | 1.01398691 | 0.310589 | NA |
| FADS1 | 3.72442797 | -0.631856 | 0.95486392 | -0.6617236 | 0.50814836 | NA |
| FADS3 | 2.7379584 | 0.39541618 | 0.97987075 | 0.40353912 | 0.68655166 | NA |
| FTH1 | 205.661021 | 0.17368829 | 0.42280463 | 0.41080035 | 0.68121894 | 0.9267216 |
| ASRGL1 | 5.44477835 | -0.4556571 | 0.78636895 | -0.5794444 | 0.56228934 | NA |
| AHNAK | 50.0600216 | 0.20517754 | 0.35849186 | 0.57233527 | 0.56709486 | 0.89263108 |
| B3GAT3 | 7.21904142 | 0.48149902 | 0.55442294 | 0.8684688 | 0.38513775 | NA |
| GANAB | 19.1168281 | -0.0031728 | 0.38005805 | -0.0083482 | 0.99333918 | NA |
| INTS5 | 2.76823732 | 0.57579267 | 0.8566383 | 0.67215378 | 0.5014858 | NA |
| UQCC3 | 6.845737 | 0.7662195 | 0.59626889 | 1.28502342 | 0.1987841 | NA |

| | | | | | | |
|---|---|---|---|---|---|---|
| UBXN1 | 8.71668646 | 1.03450348 | 0.49549898 | 2.0878014 | 0.03681575 | NA |
| TTC9C | 7.47980222 | 0.01390722 | 0.68796331 | 0.02021506 | 0.98387181 | NA |
| POLR2G | 4.35158846 | 1.23509517 | 0.81975804 | 1.5066582 | 0.13189829 | NA |
| TAF6L | 4.51903419 | -0.2386836 | 0.7298882 | -0.327014 | 0.74365731 | NA |
| TMEM223 | 5.24940815 | 1.20937006 | 0.69516249 | 1.73969408 | 0.08191275 | NA |
| TMEM179B | 11.8711749 | 1.34702905 | 0.485617 | 2.77385069 | 0.00553971 | NA |
| NXF1 | 6.00980777 | -0.266945 | 0.67764942 | -0.3939279 | 0.69363425 | NA |
| STX5 | 12.1565256 | -0.9046086 | 0.46234687 | -1.9565583 | 0.05039943 | NA |
| SLC3A2 | 40.8528948 | -1.8959735 | 0.67712415 | -2.8000383 | 0.00510965 | 0.22356834 |
| SLC22A6 | 5.55648423 | 0.83916523 | 1.07230983 | 0.78257721 | 0.43387543 | NA |
| RARRES3 | 96.0584394 | 0.92534055 | 0.67978774 | 1.36121984 | 0.17344423 | 0.6651652 |
| PLA2G16 | 21.3861899 | 0.07419425 | 0.44843215 | 0.16545257 | 0.86858778 | NA |
| ATL3 | 20.2364226 | 0.60828356 | 0.35423691 | 1.71716597 | 0.08594885 | NA |
| RTN3 | 54.5288572 | 0.04708001 | 0.23717627 | 0.19850217 | 0.84265218 | 0.97168963 |
| C11orf95 | 2.24690133 | 1.67909442 | 1.16767865 | 1.43797647 | 0.15044073 | NA |
| MARK2 | 29.0059974 | -0.254051 | 0.36411049 | -0.6977305 | 0.48534572 | 0.86354292 |
| NAA40 | 7.5568157 | -0.2225148 | 0.54748669 | -0.4064295 | 0.68442703 | NA |
| OTUB1 | 10.5820702 | 0.48745927 | 0.50479946 | 0.96564935 | 0.33421967 | NA |
| MACROD1 | 13.8851965 | 0.89032218 | 0.58608687 | 1.51909594 | 0.12873835 | NA |
| STIP1 | 13.0675295 | 0.44365536 | 0.50717916 | 0.87475075 | 0.38170954 | NA |
| NUDT22 | 8.99983114 | 0.04447318 | 0.52164867 | 0.08525504 | 0.93205863 | NA |
| AP001453.1 | 1.93286182 | 2.48169898 | 1.13329534 | 2.1898078 | 0.02853818 | NA |
| DNAJC4 | 4.04529295 | -0.2886566 | 0.78509738 | -0.3676699 | 0.71311942 | NA |
| VEGFB | 18.2974465 | 0.43910963 | 0.5233168 | 0.83908951 | 0.40141908 | NA |
| FKBP2 | 19.2069287 | 0.34188736 | 0.44110621 | 0.77506812 | 0.43829941 | NA |
| PPP1R14B | 13.3760954 | -0.6470979 | 0.46396736 | -1.3947058 | 0.16310459 | NA |
| BAD | 8.04365508 | 0.11463501 | 0.65434957 | 0.17518925 | 0.86093093 | NA |
| GPR137 | 5.11371724 | 0.20566028 | 0.64159532 | 0.32054517 | 0.7485551 | NA |
| ESRRA | 32.3600959 | -0.1885859 | 0.40946018 | -0.4605719 | 0.64510575 | 0.92153493 |
| TRMT112 | 13.0919249 | 0.28362803 | 0.51619803 | 0.54945585 | 0.58269265 | NA |
| PRDX5 | 27.8747742 | -0.2195388 | 0.3901465 | -0.5627087 | 0.57363324 | 0.89263108 |
| SF1 | 54.2475239 | -0.2902878 | 0.28148165 | -1.0312849 | 0.30240724 | 0.77521265 |
| MAP4K2 | 2.34931867 | 1.57011286 | 1.00311638 | 1.565235 | 0.11752782 | NA |
| MEN1 | 3.16475914 | 0.46338115 | 0.81080533 | 0.57150728 | 0.56765583 | NA |
| EHD1 | 3.76898045 | -0.1555225 | 0.88516102 | -0.1756997 | 0.86052985 | NA |
| ATG2A | 8.37997029 | 0.63348723 | 0.53233071 | 1.19002572 | 0.23403628 | NA |
| PPP2R5B | 3.08520213 | 0.793883 | 0.84978777 | 0.93421325 | 0.3501939 | NA |
| ZFPL1 | 3.41100677 | -0.543388 | 0.7971513 | -0.6816623 | 0.49545253 | NA |
| VPS51 | 19.6754156 | 0.28429291 | 0.35677154 | 0.79684861 | 0.42553896 | NA |
| TM7SF2 | 3.94874893 | -0.0070493 | 0.74962844 | -0.0094037 | 0.99249702 | NA |
| FAU | 49.7207279 | -0.0364868 | 0.31393159 | -0.1162253 | 0.907474 | 0.98217685 |
| SYVN1 | 3.88774431 | 0.14224297 | 0.76857372 | 0.18507394 | 0.85317104 | NA |
| MRPL49 | 5.5917214 | -0.1063297 | 0.59380789 | -0.1790641 | 0.8578874 | NA |
| CAPN1 | 16.4243589 | -0.1053795 | 0.40284619 | -0.2615875 | 0.79363949 | NA |
| DPF2 | 8.9845358 | 0.33021226 | 0.51889249 | 0.63637895 | 0.52452946 | NA |
| SLC25A45 | 3.02802899 | 0.12054798 | 0.82791609 | 0.14560411 | 0.88423392 | NA |
| FRMD8 | 6.63109022 | -1.1854246 | 0.596188 | -1.9883403 | 0.04677407 | NA |
| NEAT1 | 1338.02 | -0.1165356 | 0.32798227 | -0.3553106 | 0.72235692 | 0.93818522 |
| MALAT1 | 18227.4228 | -0.2568451 | 0.32113648 | -0.7998005 | 0.42382641 | 0.84057082 |
| SCYL1 | 8.84354129 | -0.308091 | 0.70803705 | -0.435134 | 0.66346516 | NA |
| LTBP3 | 11.0847569 | 0.88745672 | 0.54582241 | 1.62590743 | 0.10396935 | NA |
| SSSCA1 | 3.89178113 | 0.96420769 | 0.82022995 | 1.17553339 | 0.23978139 | NA |
| FAM89B | 4.87267901 | -0.279272 | 0.70029515 | -0.3987918 | 0.69004662 | NA |
| MAP3K11 | 4.80054889 | -0.3348407 | 0.8289546 | -0.4039313 | 0.68626322 | NA |

| | | | | | | |
|---|---|---|---|---|---|---|
| PCNX3 | 5.16134791 | 0.302134 | 0.71608567 | 0.42192438 | 0.67308021 | NA |
| RELA | 8.18857051 | 0.08749373 | 0.51700152 | 0.16923303 | 0.86561335 | NA |
| KAT5 | 7.08234224 | -0.2292754 | 0.75003284 | -0.3056872 | 0.75984283 | NA |
| RNASEH2C | 6.94324238 | 0.34145331 | 0.55224927 | 0.61829562 | 0.53638049 | NA |
| AP5B1 | 2.27733748 | -1.2058147 | 1.22063401 | -0.9878593 | 0.32322156 | NA |
| CFL1 | 43.7504283 | -0.1137803 | 0.36578568 | -0.3110572 | 0.75575713 | 0.95323728 |
| MUS81 | 3.22944639 | 1.29370411 | 0.893777 | 1.44745738 | 0.14776886 | NA |
| FIBP | 3.15798683 | 1.30443071 | 0.86386318 | 1.50999687 | 0.13104422 | NA |
| CCDC85B | 6.0156036 | -0.0417172 | 0.73881263 | -0.0564652 | 0.95497122 | NA |
| C11orf68 | 3.26546937 | -0.0624624 | 0.82627541 | -0.0755951 | 0.93974125 | NA |
| DRAP1 | 6.6184788 | 0.13577109 | 0.61058677 | 0.22236167 | 0.82403235 | NA |
| SART1 | 5.79853625 | -0.5317611 | 0.65515286 | -0.8116596 | 0.41698697 | NA |
| EIF1AD | 3.62495526 | -1.4347787 | 0.82177806 | -1.7459443 | 0.08082064 | NA |
| BANF1 | 12.7323592 | -0.4300667 | 0.48483382 | -0.8870394 | 0.37505769 | NA |
| SF3B2 | 15.6331802 | -0.2300061 | 0.42246961 | -0.5444323 | 0.58614401 | NA |
| PACS1 | 106.25205 | 0.2032222 | 0.30840641 | 0.65894284 | 0.50993247 | 0.87158522 |
| KLC2 | 5.77125857 | -0.6090012 | 0.62876154 | -0.9685726 | 0.33275848 | NA |
| RAB1B | 9.84588335 | -0.2141611 | 0.58903083 | -0.3635822 | 0.71617005 | NA |
| YIF1A | 5.09561705 | 0.35535433 | 0.69249929 | 0.51314756 | 0.6078481 | NA |
| BRMS1 | 4.50189347 | -1.2365343 | 0.8331372 | -1.4841905 | 0.13775839 | NA |
| SLC29A2 | 5.89104587 | 0.06374763 | 0.65637252 | 0.09712111 | 0.92263022 | NA |
| MRPL11 | 6.01911162 | 0.98534766 | 0.60278378 | 1.63466186 | 0.10211995 | NA |
| ZDHHC24 | 4.02918058 | 0.0605008 | 0.70449818 | 0.08587786 | 0.93156351 | NA |
| CTSF | 4.33825688 | 1.75743592 | 0.69805734 | 2.5176097 | 0.01181542 | NA |
| CCS | 8.11859866 | 0.44017182 | 0.53448702 | 0.8235407 | 0.41020058 | NA |
| RBM14-RBM | 7.3228199 | -0.652281 | 0.56143938 | -1.1618013 | 0.24531617 | NA |
| RBM4 | 3.67555274 | 0.74813796 | 0.83607608 | 0.89482043 | 0.37088309 | NA |
| RBM4B | 5.85486031 | 0.56268923 | 0.63875946 | 0.88090943 | 0.37836684 | NA |
| SPTBN2 | 18.4375051 | -0.5258132 | 0.51898104 | -1.0131647 | 0.31098152 | NA |
| C11orf80 | 61.0069274 | 0.1506802 | 0.36082648 | 0.41759741 | 0.67624149 | 0.92617027 |
| RCE1 | 2.33844344 | 1.54617593 | 1.08582653 | 1.42396219 | 0.15445742 | NA |
| PC | 3.22854433 | 0.17663216 | 0.79997145 | 0.22079808 | 0.82524966 | NA |
| LRFN4 | 2.72334365 | -0.5180235 | 0.92485217 | -0.5601149 | 0.57540105 | NA |
| RHOD | 3.82329484 | -0.7841365 | 0.78306247 | -1.0013716 | 0.3166472 | NA |
| KDM2A | 90.3218058 | -0.3031036 | 0.25304098 | -1.197844 | 0.23097777 | 0.73156941 |
| GRK2 | 7.55672335 | -0.409498 | 0.63540065 | -0.644472 | 0.51926938 | NA |
| ANKRD13D | 7.85578168 | 0.63651674 | 0.66795117 | 0.95293903 | 0.34062096 | NA |
| SSH3 | 7.85805544 | 0.3950243 | 0.53013461 | 0.74513961 | 0.45618733 | NA |
| RAD9A | 25.6094874 | 0.3435981 | 0.35481805 | 0.96837831 | 0.33285547 | NA |
| PPP1CA | 15.6229234 | -0.0320772 | 0.48927669 | -0.0655605 | 0.9477277 | NA |
| RPS6KB2 | 4.90892702 | -0.3425277 | 0.65908568 | -0.5197013 | 0.60327178 | NA |
| CORO1B | 9.03956387 | 0.28818343 | 0.49020573 | 0.58788263 | 0.55661107 | NA |
| TMEM134 | 6.33473067 | 0.35160357 | 0.66382924 | 0.52965967 | 0.59634792 | NA |
| AIP | 4.6459098 | -1.2279639 | 0.7361982 | -1.6679801 | 0.09531969 | NA |
| PITPNM1 | 2.55483144 | -1.3665167 | 0.91311888 | -1.4965376 | 0.13451362 | NA |
| CDK2AP2 | 12.7968236 | -0.3255862 | 0.62069912 | -0.5245475 | 0.59989779 | NA |
| GSTP1 | 100.870143 | 0.06481458 | 0.32630515 | 0.19863179 | 0.84255078 | 0.97168963 |
| NDUFV1 | 18.0129232 | 0.23150457 | 0.45934294 | 0.50399069 | 0.61426792 | NA |
| NUDT8 | 2.91353133 | -0.963521 | 0.99873407 | -0.9647423 | 0.3346739 | NA |
| UNC93B1 | 4.5921468 | -0.0425707 | 0.85602146 | -0.0497309 | 0.96033687 | NA |
| NDUFS8 | 21.015534 | 0.17059721 | 0.43148978 | 0.3953679 | 0.6925714 | NA |
| TCIRG1 | 3.80037185 | -2.5487366 | 0.9176047 | -2.7775976 | 0.00547624 | NA |
| CHKA | 35.4358039 | -0.3287324 | 0.38325971 | -0.8577276 | 0.39104291 | 0.8258317 |
| KMT5B | 37.6085283 | -0.5260846 | 0.31476562 | -1.6713535 | 0.09465188 | 0.56264113 |

| | | | | | | |
|---|---|---|---|---|---|---|
| LRP5 | 26.0156648 | 1.21062434 | 0.49475082 | 2.4469375 | 0.01440758 | 0.30435125 |
| PPP6R3 | 122.088856 | -0.1391899 | 0.27231533 | -0.5111348 | 0.60925664 | 0.9088434 |
| TESMIN | 8.78389654 | 0.23966044 | 0.55627449 | 0.43083127 | 0.66659106 | NA |
| CPT1A | 65.819299 | -0.4232383 | 0.32014624 | -1.3220155 | 0.18616298 | 0.68006555 |
| MRPL21 | 4.38018394 | 0.04266515 | 0.67976307 | 0.06276473 | 0.94995385 | NA |
| IGHMBP2 | 7.06239923 | -0.7550241 | 0.62026085 | -1.2172687 | 0.22350202 | NA |
| TPCN2 | 6.65810474 | 0.22157652 | 0.71432587 | 0.31018969 | 0.75641671 | NA |
| CCND1 | 3.13026917 | 0.73477942 | 0.80880105 | 0.90847979 | 0.36362478 | NA |
| ORAOV1 | 5.97839687 | 0.93322698 | 0.71989732 | 1.29633347 | 0.19486062 | NA |
| PPFIA1 | 64.3055708 | -0.5020913 | 0.24661154 | -2.0359603 | 0.04175435 | 0.44186191 |
| AP002336.2 | 22.3204465 | 0.31525149 | 0.44767805 | 0.70419244 | 0.48131294 | NA |
| AP000487.1 | 2.66673573 | -0.4170715 | 0.86834075 | -0.4803085 | 0.63100803 | NA |
| CTTN | 45.3047789 | -0.4139511 | 0.27780048 | -1.4901023 | 0.13619733 | 0.60594834 |
| SHANK2 | 139.388321 | -0.0044571 | 0.31446034 | -0.0141737 | 0.9886914 | 0.99909657 |
| DHCR7 | 4.27653777 | -2.2186827 | 0.88190673 | -2.5157793 | 0.01187695 | NA |
| NADSYN1 | 12.3456448 | 0.37419885 | 0.45707241 | 0.81868616 | 0.4129655 | NA |
| KRTAP5-7 | 6.7393015 | 2.42637736 | 1.15863548 | 2.09416801 | 0.03624502 | NA |
| KRTAP5-8 | 19.9541095 | 1.01500023 | 0.78808483 | 1.28793271 | 0.19776939 | NA |
| AP002495.2 | 17.6568928 | -0.1184601 | 0.44757394 | -0.2646716 | 0.79126244 | NA |
| RNF121 | 24.2804654 | -0.7865954 | 0.40576988 | -1.9385259 | 0.05255909 | NA |
| IL18BP | 3.18512549 | 0.1129622 | 0.90843083 | 0.12434871 | 0.90103918 | NA |
| NUMA1 | 34.8417645 | 0.50515037 | 0.34572673 | 1.46112615 | 0.14398082 | 0.62469744 |
| LAMTOR1 | 21.1867318 | -0.1453327 | 0.43713497 | -0.3324665 | 0.73953707 | NA |
| ANAPC15 | 5.50668837 | -0.0230206 | 0.6981845 | -0.0329721 | 0.97369684 | NA |
| FOLR1 | 5.29423616 | -1.9962922 | 0.889429 | -2.244465 | 0.0248025 | NA |
| INPPL1 | 11.0534317 | -0.1933737 | 0.52384606 | -0.3691422 | 0.71202172 | NA |
| CLPB | 14.0404258 | -0.1795009 | 0.45670837 | -0.3930319 | 0.69429595 | NA |
| ARAP1 | 18.8286936 | -0.3590596 | 0.4485779 | -0.8004397 | 0.42345608 | NA |
| ATG16L2 | 4.48420157 | 0.29995328 | 0.83053887 | 0.36115502 | 0.71798357 | NA |
| FCHSD2 | 34.3115389 | -0.2378976 | 0.36338003 | -0.65468 | 0.51267379 | 0.87158522 |
| ARHGEF17 | 7.07592168 | 0.19868838 | 0.52398501 | 0.37918715 | 0.7045489 | NA |
| FAM168A | 60.1378044 | -0.5260952 | 0.39893712 | -1.3187421 | 0.18725535 | 0.68214045 |
| PLEKHB1 | 2.5922083 | 1.18952686 | 1.09171238 | 1.08959729 | 0.27589058 | NA |
| RAB6A | 56.7310527 | -0.6274675 | 0.24718864 | -2.5384157 | 0.01113556 | 0.28544118 |
| MRPL48 | 12.8934076 | -0.1114299 | 0.42154141 | -0.2643392 | 0.79151852 | NA |
| COA4 | 8.21078034 | 0.03877751 | 0.5341392 | 0.07259813 | 0.94212591 | NA |
| PAAF1 | 8.66516172 | 0.66427919 | 0.61164993 | 1.08604474 | 0.2774592 | NA |
| UCP2 | 6.82948508 | 0.00376319 | 0.77156011 | 0.00487738 | 0.99610843 | NA |
| C2CD3 | 16.6369307 | 0.2935236 | 0.43331917 | 0.67738431 | 0.49816215 | NA |
| PPME1 | 21.2482095 | 0.04040872 | 0.35204229 | 0.11478372 | 0.90861656 | NA |
| POLD3 | 7.39244696 | 0.46856983 | 0.61151975 | 0.76623826 | 0.44353454 | NA |
| RNF169 | 31.3763094 | 0.52871203 | 0.30281749 | 1.74597588 | 0.08081514 | 0.54606485 |
| XRRA1 | 25.3470482 | -0.2538148 | 0.73442279 | -0.3455977 | 0.72964505 | NA |
| SPCS2 | 28.8160518 | -0.0943129 | 0.33564729 | -0.280988 | 0.77871959 | 0.96133329 |
| NEU3 | 6.09722805 | -0.9380014 | 0.6824373 | -1.3744872 | 0.16929047 | NA |
| ARRB1 | 3.67130045 | -0.1265296 | 0.89104078 | -0.1420021 | 0.88707838 | NA |
| RPS3 | 97.0786625 | -0.0486073 | 0.31336257 | -0.1551151 | 0.87673056 | 0.9751578 |
| SERPINH1 | 3.01363063 | -0.0295461 | 0.91658763 | -0.0322349 | 0.97428474 | NA |
| UVRAG | 38.5988067 | -0.4602302 | 0.29105996 | -1.5812211 | 0.11382749 | 0.57860396 |
| THAP12 | 9.36841048 | -0.1937479 | 0.63240056 | -0.306369 | 0.75932371 | NA |
| EMSY | 27.362797 | 0.18861549 | 0.35965268 | 0.52443788 | 0.59997401 | 0.90266665 |
| ACER3 | 31.449391 | 0.20963548 | 0.31529156 | 0.66489406 | 0.50611825 | 0.87094108 |
| CAPN5 | 3.2229411 | 0.74932269 | 0.95362245 | 0.78576453 | 0.43200545 | NA |
| PAK1 | 15.8852221 | -0.1880804 | 0.42108209 | -0.4466598 | 0.65512071 | NA |

| | | | | | | |
|---|---|---|---|---|---|---|
| CLNS1A | 14.6058886 | 0.17754632 | 0.49359655 | 0.35969928 | 0.71907203 | NA |
| RSF1 | 58.1553673 | 0.00834266 | 0.33445424 | 0.0249441 | 0.98009955 | 0.99902646 |
| AAMDC | 11.8401912 | 0.41694482 | 0.44933506 | 0.92791517 | 0.35345158 | NA |
| AP002812.2 | 6.09073311 | -0.0944276 | 0.68540375 | -0.1377693 | 0.89042271 | NA |
| INTS4 | 17.9735565 | -0.714782 | 0.55035334 | -1.2987692 | 0.19402313 | NA |
| ALG8 | 6.66631823 | 0.46413634 | 0.61916638 | 0.7496149 | 0.45348667 | NA |
| GAB2 | 34.0868276 | 0.06082361 | 0.46819427 | 0.12991104 | 0.89663681 | 0.97769824 |
| AP003086.2 | 9.29018851 | 0.31183938 | 0.62681879 | 0.49749527 | 0.61883984 | NA |
| NARS2 | 19.9672822 | -0.0309038 | 0.37193784 | -0.0830886 | 0.93378111 | NA |
| TENM4 | 9.99126217 | -1.4036268 | 0.82545746 | -1.700423 | 0.0890514 | NA |
| MIR4300HG | 2.97760261 | -0.7447827 | 1.26178217 | -0.5902625 | 0.5550147 | NA |
| PRCP | 9.11306598 | 0.62803427 | 0.53041667 | 1.18403946 | 0.23639744 | NA |
| RAB30 | 9.04639676 | 0.61488333 | 0.65245354 | 0.94241704 | 0.34597917 | NA |
| RAB30-AS1 | 15.0869386 | 0.37331741 | 0.52604519 | 0.70966795 | 0.47791007 | NA |
| PCF11 | 15.7580478 | 0.28103238 | 0.39711453 | 0.70768596 | 0.4791403 | NA |
| ANKRD42 | 6.07640063 | 0.74307921 | 0.58103531 | 1.27888821 | 0.20093643 | NA |
| CCDC90B | 9.12408558 | 0.15204399 | 0.48993621 | 0.31033427 | 0.75630678 | NA |
| AP000446.1 | 6.19387354 | 0.51516562 | 0.75578548 | 0.68162943 | 0.4954733 | NA |
| DLG2 | 39.1578621 | 0.52693036 | 0.42062328 | 1.25273704 | 0.21030142 | 0.70690231 |
| TMEM126B | 6.85635111 | 0.19979818 | 0.53009119 | 0.37691285 | 0.70623837 | NA |
| TMEM126A | 4.85755417 | 0.17324023 | 0.69401989 | 0.24961853 | 0.80288236 | NA |
| CREBZF | 21.1125628 | 0.02596011 | 0.36835692 | 0.07047542 | 0.94381526 | NA |
| SYTL2 | 14.0347452 | -0.2570862 | 0.52123051 | -0.4932293 | 0.62185057 | NA |
| PICALM | 439.349492 | -0.4679311 | 0.32992753 | -1.4182846 | 0.15610768 | 0.63932163 |
| EED | 11.5532877 | 0.61597082 | 0.55795204 | 1.10398525 | 0.26959954 | NA |
| HIKESHI | 9.08979666 | 0.83316415 | 0.49866633 | 1.67078484 | 0.09476418 | NA |
| AP001831.1 | 7.0531376 | 0.88097312 | 0.70897819 | 1.24259552 | 0.21401692 | NA |
| ME3 | 25.319177 | 0.23172445 | 0.40406799 | 0.57347885 | 0.56632052 | NA |
| PRSS23 | 19.4301078 | 0.10772262 | 0.5160919 | 0.2087276 | 0.83466089 | NA |
| FZD4 | 5.48180337 | -1.0018167 | 0.77203493 | -1.2976313 | 0.19441406 | NA |
| TMEM135 | 44.746112 | -0.5535605 | 0.37163234 | -1.489538 | 0.13634575 | 0.60594834 |
| CTSC | 5.20279837 | -0.4763176 | 0.74698211 | -0.6376559 | 0.52369769 | NA |
| NOX4 | 15.3471516 | 0.04964902 | 0.84127388 | 0.05901648 | 0.95293899 | NA |
| CHORDC1 | 12.0104952 | -0.1049738 | 0.48589074 | -0.216044 | 0.82895342 | NA |
| SLC36A4 | 24.4794118 | -0.4990025 | 0.3753475 | -1.3294414 | 0.18370239 | NA |
| CEP295 | 11.6535525 | -0.0888322 | 0.49017955 | -0.1812238 | 0.85619193 | NA |
| TAF1D | 27.2314305 | -0.3079586 | 0.38332411 | -0.8033896 | 0.42174956 | 0.84037309 |
| PANX1 | 5.19161583 | -1.2136598 | 0.6700666 | -1.8112525 | 0.07010177 | NA |
| MRE11 | 14.4622764 | 0.63552255 | 0.47973315 | 1.32474178 | 0.18525681 | NA |
| ANKRD49 | 4.78382874 | 0.30492531 | 0.64603571 | 0.47199452 | 0.6369307 | NA |
| AMOTL1 | 30.2789922 | -0.38411 | 0.48119266 | -0.7982457 | 0.42472793 | 0.84057082 |
| CWC15 | 7.99926991 | 0.09179151 | 0.53462525 | 0.17169318 | 0.86367874 | NA |
| SRSF8 | 7.64994669 | -0.0391813 | 0.52070028 | -0.0752474 | 0.94001787 | NA |
| ENDOD1 | 4.84666546 | 0.46742049 | 0.61346444 | 0.76193575 | 0.44609835 | NA |
| AP000787.1 | 4.38422243 | 0.35447431 | 0.82878336 | 0.42770442 | 0.66886633 | NA |
| SESN3 | 5.26021544 | 0.50143765 | 0.84476015 | 0.59358582 | 0.55278917 | NA |
| FAM76B | 7.30790267 | 0.61092882 | 0.59749202 | 1.02248867 | 0.30654967 | NA |
| CEP57 | 21.558398 | 0.83406373 | 0.4080812 | 2.04386707 | 0.04096668 | NA |
| MTMR2 | 25.5327149 | -0.7699018 | 0.38075894 | -2.022019 | 0.04317439 | NA |
| MAML2 | 65.3649315 | 0.01499026 | 0.63852386 | 0.02347644 | 0.98127024 | 0.99902646 |
| CCDC82 | 13.3033303 | -0.3807128 | 0.40982903 | -0.9289551 | 0.35291237 | NA |
| JRKL | 3.58081739 | 1.34596772 | 0.80141515 | 1.67948875 | 0.09305683 | NA |
| CNTN5 | 5.28710183 | 2.92126704 | 0.85300129 | 3.42469239 | 0.0006155 | NA |
| ARHGAP42 | 29.406591 | 0.22801179 | 0.55058898 | 0.41412341 | 0.67878373 | 0.9267216 |

| | | | | | | |
|---|---|---|---|---|---|---|
| PGR | 19.9079552 | -0.7932113 | 0.56842863 | -1.3954458 | 0.16288146 | NA |
| YAP1 | 101.00714 | 0.23198906 | 0.33508029 | 0.69233872 | 0.48872464 | 0.86354292 |
| BIRC3 | 7.95815369 | 1.61857303 | 1.02277481 | 1.58253118 | 0.11352836 | NA |
| BIRC2 | 18.660606 | -0.0680482 | 0.34950781 | -0.1946972 | 0.84563003 | NA |
| TMEM123 | 45.8363199 | -0.0041649 | 0.29137415 | -0.0142941 | 0.98859533 | 0.99909657 |
| MMP7 | 13.7224126 | -0.8132967 | 0.53716296 | -1.5140596 | 1.30E-01 | NA |
| DCUN1D5 | 9.74527922 | -0.0869975 | 0.52378668 | -0.1660935 | 0.86808339 | NA |
| DYNC2H1 | 26.0091907 | 0.57733597 | 0.37657736 | 1.53311388 | 0.1252478 | 0.5866305 |
| CASP4 | 10.3586148 | 0.14592837 | 0.57078097 | 0.2556644 | 0.79820999 | NA |
| MSANTD4 | 6.01816354 | -0.0388916 | 0.59301693 | -0.0655826 | 0.94771014 | NA |
| KBTBD3 | 3.54195731 | 0.04861503 | 0.80520735 | 0.06037578 | 0.95185634 | NA |
| AASDHPPT | 9.3978223 | 0.46244688 | 0.48964875 | 0.94444616 | 0.3449417 | NA |
| CWF19L2 | 29.9489591 | -0.0486935 | 0.35506987 | -0.1371377 | 0.89092196 | 0.97769824 |
| ALKBH8 | 4.46138418 | -0.3866127 | 0.73884424 | -0.5232668 | 0.60078859 | NA |
| CUL5 | 102.945379 | -0.6461461 | 0.23590357 | -2.7390262 | 0.00616215 | 0.23056019 |
| ACAT1 | 81.0117939 | -0.2458121 | 0.26032688 | -0.9442439 | 3.45E-01 | 0.8003972 |
| NPAT | 10.2538331 | -0.4250135 | 0.48733886 | -0.8721108 | 0.38314795 | NA |
| ATM | 43.3094393 | 0.03523356 | 0.37780903 | 0.0932576 | 0.92569892 | 0.98758497 |
| C11orf65 | 7.8737181 | 1.6327814 | 0.65934796 | 2.4763577 | 0.01327305 | NA |
| KDELC2 | 4.33937091 | 0.45626557 | 0.88219865 | 0.51719142 | 0.60502254 | NA |
| EXPH5 | 17.8897171 | -0.4173171 | 0.42918615 | -0.9723453 | 0.3308788 | NA |
| DDX10 | 29.8335824 | -0.0762961 | 0.36768907 | -0.2075017 | 0.83561807 | 0.97154892 |
| RDX | 41.2127408 | 0.08093274 | 0.27526889 | 0.2940134 | 0.76874767 | 0.957884 |
| ZC3H12C | 19.635542 | -0.2341251 | 0.40994578 | -0.5711125 | 0.5679234 | NA |
| FDX1 | 17.709375 | 0.12008969 | 0.34788542 | 0.34519898 | 0.72994479 | NA |
| COLCA1 | 2.62664455 | 0.7403742 | 1.10233156 | 0.67164384 | 0.50181046 | NA |
| COLCA2 | 2.43949219 | -1.4770038 | 1.04547669 | -1.4127563 | 0.15772739 | NA |
| SIK2 | 66.1939873 | 0.04814097 | 0.27995187 | 0.17196159 | 0.86346773 | 0.97217819 |
| PPP2R1B | 19.3361703 | -0.1472433 | 0.42367791 | -0.347536 | 0.72818868 | NA |
| CRYAB | 63.0184807 | -0.0003965 | 0.68044313 | -0.0005827 | 0.99953508 | 0.99960611 |
| DIXDC1 | 4.85637076 | -0.6401866 | 0.70697539 | -0.9055288 | 0.36518532 | NA |
| DLAT | 13.0227142 | -0.3321338 | 0.44639479 | -0.744036 | 0.45685472 | NA |
| NKAPD1 | 4.41907467 | 0.28165469 | 0.80724609 | 0.34890808 | 0.72715832 | NA |
| TIMM8B | 11.5902741 | 0.15043953 | 0.49841954 | 0.30183313 | 0.76277927 | NA |
| AP002884.3 | 14.2414761 | 0.0098084 | 0.47511265 | 0.02064437 | 0.98352935 | NA |
| IL18 | 322.334784 | -0.1054726 | 0.24419558 | -0.4319186 | 0.66580055 | 0.92276785 |
| BCO2 | 94.407806 | 0.41596622 | 0.35685832 | 1.16563408 | 0.24376242 | 0.73323864 |
| PTS | 24.4884068 | 0.11802193 | 0.37553967 | 0.31427287 | 0.7533138 | NA |
| AP002884.1 | 5.66855854 | 0.58580333 | 0.79158301 | 0.74004031 | 0.45927554 | NA |
| TTC12 | 7.26571478 | 0.65109825 | 0.55349275 | 1.17634468 | 0.23945717 | NA |
| ZW10 | 5.25631621 | -0.0491945 | 0.72904524 | -0.067478 | 0.9462012 | NA |
| USP28 | 13.4205344 | 0.95490752 | 0.46754056 | 2.04240572 | 0.0411113 | NA |
| ZBTB16 | 76.0057178 | 0.63000578 | 0.52967865 | 1.18941131 | 0.23427785 | 0.73156941 |
| AP002518.2 | 4.08938828 | 1.78611109 | 1.03285899 | 1.72928842 | 0.08375749 | NA |
| NNMT | 12.3310464 | 1.16517924 | 0.51249528 | 2.2735414 | 0.02299357 | NA |
| C11orf71 | 2.27043767 | 0.68428497 | 0.89469971 | 0.76482082 | 0.44437825 | NA |
| REXO2 | 9.02182294 | 0.17838928 | 0.50343383 | 0.35434505 | 0.72308031 | NA |
| CADM1 | 24.2880999 | -0.4686013 | 0.5737169 | -0.8167814 | 0.41405335 | NA |
| BUD13 | 3.12485331 | 1.28151291 | 0.87942979 | 1.4572089 | 0.14505872 | NA |
| ZPR1 | 3.87143003 | -0.6645219 | 0.78021932 | -0.8517116 | 0.39437417 | NA |
| SIK3 | 253.772072 | -0.2424149 | 0.2418855 | -1.0021888 | 0.31625241 | 0.780966 |
| PAFAH1B2 | 43.4401821 | -0.5208253 | 0.3072113 | -1.6953325 | 0.09001235 | 0.56122412 |
| SIDT2 | 6.98757145 | -0.9641853 | 0.5988064 | -1.6101787 | 0.10735886 | NA |
| TAGLN | 4.69318801 | -1.1309481 | 0.80572426 | -1.4036417 | 0.16042558 | NA |

| | | | | | | |
|---|---|---|---|---|---|---|
| PCSK7 | 20.0037869 | -0.4062974 | 0.37204443 | -1.0920668 | 0.27480376 | NA |
| RNF214 | 11.9825406 | -0.1353856 | 0.50862672 | -0.2661787 | 0.79010162 | NA |
| BACE1 | 8.17958536 | 0.72409028 | 0.55422703 | 1.30648678 | 0.19138707 | NA |
| CEP164 | 8.72770016 | 0.70792592 | 0.63760236 | 1.11029376 | 0.26687246 | NA |
| DSCAML1 | 19.1937992 | 0.21489538 | 1.19823765 | 0.17934287 | 0.85766849 | NA |
| FXYD2 | 16.4610543 | 0.62669134 | 0.58690644 | 1.06778746 | 0.28561639 | NA |
| AP000757.1 | 14.6486581 | 1.35334825 | 0.60228892 | 2.24700838 | 0.02463949 | NA |
| MPZL3 | 3.62093794 | -1.0217177 | 0.827029 | -1.2354073 | 0.21667896 | NA |
| UBE4A | 19.2133154 | 0.33960026 | 0.43058047 | 0.78870334 | 0.43028541 | NA |
| ATP5MG | 48.7097198 | -0.1730942 | 0.38379604 | -0.4510056 | 0.65198553 | 0.92248109 |
| KMT2A | 54.6771385 | 0.1657223 | 0.28250245 | 0.58662252 | 0.55745726 | 0.89263108 |
| AP001267.3 | 2.14005143 | 3.50268059 | 1.23580178 | 2.83433851 | 0.00459207 | NA |
| IFT46 | 5.03114026 | 0.76432212 | 0.69476911 | 1.10010954 | 0.2712844 | NA |
| ARCN1 | 13.4379534 | -0.2736571 | 0.42357865 | -0.6460598 | 0.51824062 | NA |
| AP002954.1 | 6.76903377 | 0.52809106 | 0.66516864 | 0.79392057 | 0.42724168 | NA |
| DDX6 | 41.44284 | -0.212427 | 0.29358135 | -0.7235713 | 0.46932898 | 0.85973853 |
| BCL9L | 4.17272122 | 0.30137176 | 0.74977172 | 0.40195136 | 0.68771982 | NA |
| CCDC84 | 25.6302317 | -0.6072916 | 0.36631629 | -1.6578339 | 0.09735099 | NA |
| RPS25 | 60.6196218 | 0.02090187 | 0.44747626 | 0.04671056 | 0.96274392 | 0.99656976 |
| TRAPPC4 | 11.3250572 | 0.25871824 | 0.43969967 | 0.58839763 | 0.55626543 | NA |
| SLC37A4 | 4.0119761 | 1.13091596 | 0.74998979 | 1.50790847 | 0.13157795 | NA |
| HYOU1 | 7.40406646 | -0.8754957 | 0.6842014 | -1.2795878 | 0.20069016 | NA |
| VPS11 | 5.55990689 | -0.5184441 | 0.61751673 | -0.8395628 | 0.40115355 | NA |
| H2AFX | 2.87761805 | 0.05208764 | 0.9912926 | 0.05254517 | 0.9580943 | NA |
| HINFP | 3.53196077 | -0.3065633 | 0.77626209 | -0.3949224 | 0.69290012 | NA |
| NLRX1 | 4.3657275 | 0.4187065 | 0.69026855 | 0.60658492 | 0.54412641 | NA |
| CBL | 17.3876763 | -0.0403267 | 0.38637205 | -0.1043728 | 0.91687354 | NA |
| NECTIN1 | 3.85965103 | -0.9555451 | 0.85498971 | -1.11761 | 0.26373359 | NA |
| OAF | 10.4134091 | 0.40885531 | 0.57015765 | 0.71709169 | 0.47331752 | NA |
| POU2F3 | 1.86179511 | 1.33828228 | 1.53025489 | 0.8745486 | 0.38181956 | NA |
| ARHGEF12 | 163.960779 | -0.0647243 | 0.32445922 | -0.1994835 | 0.84188455 | 0.97168963 |
| SC5D | 9.6035985 | -1.3210705 | 0.54610416 | -2.4190817 | 0.01555975 | NA |
| SORL1 | 28.9249573 | 1.14313137 | 0.56249493 | 2.03225188 | 0.04212816 | 0.44186191 |
| UBASH3B | 35.5951649 | -1.184664 | 0.59572047 | -1.988624 | 0.04674272 | 0.45218426 |
| HSPA8 | 36.5162612 | 0.25393636 | 0.30414781 | 0.83491103 | 0.40376781 | 0.82808596 |
| GRAMD1B | 59.1675299 | -0.4351915 | 0.42137369 | -1.0327924 | 0.30170108 | 0.77521265 |
| ZNF202 | 2.76774294 | 0.51614652 | 0.90398128 | 0.57097037 | 0.56801973 | NA |
| TBRG1 | 13.4348277 | -0.0672127 | 0.46481397 | -0.1446013 | 0.88502565 | NA |
| SIAE | 7.41641462 | 0.79097196 | 0.58955057 | 1.34165243 | 0.17970872 | NA |
| MSANTD2 | 15.1720117 | -0.0724495 | 0.42095388 | -0.1721079 | 0.86335268 | NA |
| EI24 | 14.5412523 | 0.30939279 | 0.40768938 | 0.75889342 | 0.44791632 | NA |
| STT3A | 8.00899065 | -1.1034695 | 0.61562791 | -1.7924293 | 0.07306422 | NA |
| CDON | 5.55109954 | 0.70954588 | 0.66506877 | 1.06687596 | 0.28602785 | NA |
| RPUSD4 | 4.19988549 | -1.1449678 | 0.76709974 | -1.4925931 | 0.13554373 | NA |
| FAM118B | 5.93753412 | -0.0998581 | 0.59914743 | -0.1666669 | 0.86763214 | NA |
| SRPRA | 12.4392418 | -0.2033243 | 0.42752061 | -0.4755895 | 0.63436687 | NA |
| FOXRED1 | 7.19759636 | 0.08027468 | 0.61464662 | 0.13060298 | 0.89608939 | NA |
| TIRAP | 3.08697451 | -0.0654154 | 0.89693971 | -0.0729317 | 0.94186043 | NA |
| DCPS | 10.7586146 | -0.1446299 | 0.56676688 | -0.2551842 | 0.79858085 | NA |
| ST3GAL4 | 4.73821899 | -2.038877 | 0.72654757 | -2.8062539 | 0.00501212 | NA |
| KCNJ1 | 9.21946629 | -0.8172605 | 0.86009623 | -0.9501966 | 0.34201234 | NA |
| ARHGAP32 | 53.1944775 | -0.3568005 | 0.32387713 | -1.1016538 | 0.27061219 | 0.75947839 |
| TMEM45B | 11.7451019 | 0.35103443 | 0.41900459 | 0.83778182 | 0.40215325 | NA |
| NFRKB | 8.48078638 | 0.18482654 | 0.51394188 | 0.35962537 | 0.71912731 | NA |

| | | | | | | |
|---|---|---|---|---|---|---|
| PRDM10 | 9.2151448 | -0.4527765 | 0.49840525 | -0.9084506 | 0.36364021 | NA |
| APLP2 | 85.9874125 | -0.0246387 | 0.22994605 | -0.1071501 | 0.91466993 | 0.98477372 |
| ST14 | 43.0994283 | -0.041803 | 0.34233257 | -0.1221123 | 0.90281005 | 0.98003205 |
| ZBTB44 | 37.8814454 | 0.34924465 | 0.40078036 | 0.87141159 | 0.38352946 | 0.82352384 |
| AP002986.1 | 4.67392065 | -0.0698533 | 0.9825767 | -0.0710919 | 0.94332458 | NA |
| SNX19 | 22.2131451 | -0.3652142 | 0.41335659 | -0.8835331 | 0.37694829 | NA |
| NCAPD3 | 4.30611706 | 0.95032556 | 0.7981967 | 1.19059068 | 0.23381431 | NA |
| VPS26B | 5.25784164 | 0.54851846 | 0.66536758 | 0.82438411 | 0.40972134 | NA |
| ACAD8 | 4.54456805 | 1.68828481 | 0.7878002 | 2.14303679 | 0.03211014 | NA |
| ZMYND11 | 55.6525867 | 0.12705351 | 0.28001267 | 0.45374202 | 0.65001452 | 0.92248109 |
| DIP2C | 162.602354 | -1.115878 | 0.41789442 | -2.6702389 | 0.00757973 | 0.26973729 |
| LARP4B | 71.0489259 | -0.2662089 | 0.2649638 | -1.004699 | 0.31504182 | 0.780966 |
| GTPBP4 | 8.97151151 | -1.0605121 | 0.5176859 | -2.0485628 | 0.04050489 | NA |
| IDI1 | 13.0843932 | -1.0142109 | 0.61896303 | -1.6385646 | 0.10130397 | NA |
| WDR37 | 25.0211913 | -0.062364 | 0.36666334 | -0.1700852 | 0.86494316 | NA |
| ADARB2 | 104.704842 | -1.9014292 | 1.54976755 | -1.2269125 | 0.21985548 | 0.72261413 |
| PFKP | 17.8902644 | -0.0388607 | 0.54010862 | -0.0719498 | 0.94264183 | NA |
| PITRM1 | 8.79817906 | -0.2145585 | 0.54210612 | -0.395787 | 0.69226219 | NA |
| KLF6 | 84.733578 | -1.1592618 | 0.57660712 | -2.0104883 | 0.04437954 | 0.44769598 |
| AKR1C1 | 18.0430047 | 0.60002506 | 0.52491256 | 1.14309525 | 2.53E-01 | NA |
| AKR1C2 | 7.14156565 | -0.3221324 | 0.79153752 | -0.4069705 | 0.68402966 | NA |
| AKR1C3 | 4.8410308 | 1.70729473 | 0.71308296 | 2.39424417 | 0.01665466 | NA |
| NET1 | 4.72037538 | 0.62178697 | 0.69351633 | 0.89657149 | 0.36994761 | NA |
| ASB13 | 8.08278571 | 0.48159411 | 0.66281583 | 0.72658812 | 0.46747831 | NA |
| FAM208B | 46.5486505 | -0.6210046 | 0.32600327 | -1.9049029 | 0.05679269 | 0.503587 |
| AL365356.1 | 3.5594157 | 0.34151102 | 0.89607439 | 0.38111906 | 0.70311491 | NA |
| GDI2 | 29.9158548 | -0.4175608 | 0.32998789 | -1.2653823 | 0.20573433 | 0.70278771 |
| ANKRD16 | 3.00470537 | 0.00342519 | 1.02576159 | 0.00333917 | 0.99733573 | NA |
| FBH1 | 25.9815995 | 0.08288067 | 0.32768185 | 0.25293032 | 0.80032206 | 0.96742897 |
| RBM17 | 16.3000392 | -0.1320148 | 0.41829386 | -0.3156031 | 0.75230378 | NA |
| PFKFB3 | 17.7871155 | -0.6690058 | 0.6956808 | -0.9616563 | 0.33622228 | NA |
| PRKCQ | 15.7995437 | -0.8072011 | 0.55988119 | -1.4417365 | 0.14937673 | NA |
| SFMBT2 | 18.2868833 | -1.2133713 | 0.44519202 | -2.7255011 | 0.0064204 | NA |
| ITIH5 | 96.752174 | 0.72628699 | 0.49151953 | 1.47763607 | 0.13950521 | 0.61340925 |
| KIN | 7.91246519 | -0.2787507 | 0.56084537 | -0.4970189 | 0.61917575 | NA |
| ATP5F1C | 29.2361235 | 0.15767577 | 0.38477671 | 0.40978511 | 0.68196359 | 0.9267216 |
| TAF3 | 43.1073097 | -0.3586096 | 0.32737667 | -1.0954035 | 0.27333989 | 0.76165444 |
| GATA3 | 12.9542632 | 0.14751218 | 0.61921301 | 0.23822526 | 0.81170639 | NA |
| SFTA1P | 3.67934924 | 0.66668635 | 1.04421701 | 0.63845574 | 0.52317706 | NA |
| CELF2 | 669.189927 | -0.1939157 | 0.29148827 | -0.6652608 | 0.50588372 | 0.87094108 |
| CELF2-AS2 | 5.45327876 | 0.3921221 | 0.75234924 | 0.52119691 | 0.60222961 | NA |
| USP6NL | 37.3864288 | -0.2478545 | 0.26958325 | -0.9193986 | 0.35788712 | 0.80268969 |
| ECHDC3 | 8.16122267 | 1.22029556 | 0.65208845 | 1.87136509 | 0.06129449 | NA |
| PROSER2 | 16.375558 | -0.1619381 | 0.44599468 | -0.3630943 | 0.71653442 | NA |
| UPF2 | 30.1775828 | -0.3483327 | 0.32819907 | -1.0613458 | 0.28853276 | 0.7725847 |
| DHTKD1 | 15.8018364 | 0.67128546 | 0.43667872 | 1.5372525 | 0.12423149 | NA |
| SEC61A2 | 4.66405159 | 0.6469268 | 0.64265374 | 1.00664908 | 0.31410344 | NA |
| NUDT5 | 15.3895557 | -0.2282709 | 0.4875291 | -0.4682201 | 0.6396272 | NA |
| CDC123 | 14.0790931 | 0.06446738 | 0.49009998 | 0.13153925 | 0.89534874 | NA |
| CAMK1D | 54.2967721 | 0.97974552 | 1.11816683 | 0.87620693 | 0.38091755 | 0.82153751 |
| OPTN | 11.3411112 | -0.2564254 | 0.6023591 | -0.4257018 | 0.67032514 | NA |
| PHYH | 12.9174346 | 1.19569136 | 0.51959872 | 2.30118228 | 0.02138133 | NA |
| SEPHS1 | 9.00473687 | 0.30757778 | 0.52846081 | 0.58202571 | 0.56054936 | NA |
| BEND7 | 31.6510378 | 0.17306629 | 0.35042651 | 0.49387326 | 0.62139569 | 0.91493825 |

| | | | | | | |
|---|---|---|---|---|---|---|
| PRPF18 | 3.73090197 | -0.2373217 | 0.81075309 | -0.2927176 | 0.76973804 | NA |
| AL157392.5 | 9.23678105 | 0.61818446 | 0.52857458 | 1.16953119 | 0.24218968 | NA |
| FRMD4A | 12.3822939 | -0.1230793 | 0.5055553 | -0.2434537 | 0.80765396 | NA |
| FAM107B | 106.317833 | 0.1276388 | 0.32612421 | 0.39138093 | 0.69551569 | 0.9303056 |
| HSPA14.1 | 2.90544506 | 0.25185869 | 0.94665709 | 0.2660506 | 0.79020024 | NA |
| DCLRE1C | 8.20246427 | -0.3045571 | 0.52113386 | -0.5844124 | 0.55894288 | NA |
| RPP38 | 11.2281219 | 0.2680762 | 0.48383228 | 0.55406844 | 0.57953201 | NA |
| NMT2 | 4.50761791 | 0.36990363 | 0.71453969 | 0.51768101 | 0.60468085 | NA |
| FAM171A1 | 13.0528052 | 0.09504396 | 0.7420938 | 0.12807541 | 0.8980893 | NA |
| MINDY3 | 19.3852333 | 0.0175648 | 0.34980548 | 0.05021306 | 0.9599526 | NA |
| PTER | 28.5777585 | 0.7625485 | 0.36082928 | 2.11332211 | 0.03457321 | 0.41688714 |
| RSU1 | 22.4351582 | -0.2555871 | 0.41798533 | -0.6114738 | 0.54088598 | NA |
| CUBN | 25.4604903 | -0.2560935 | 1.34944361 | -0.1897771 | 0.84948382 | NA |
| TRDMT1 | 3.15623842 | -0.0303068 | 0.8649085 | -0.0350404 | 0.9720475 | NA |
| VIM | 5.87776385 | 0.48939596 | 0.68277142 | 0.71677863 | 0.4735107 | NA |
| STAM | 14.6541172 | -0.3253071 | 0.43750704 | -0.7435472 | 0.45715045 | NA |
| CACNB2 | 36.7805163 | 0.63684202 | 0.57343028 | 1.11058317 | 0.26674781 | 0.75497438 |
| AL450384.2 | 6.58907025 | 0.73891647 | 0.68330065 | 1.08139291 | 0.27952237 | NA |
| NSUN6 | 23.5078707 | 0.65265881 | 0.47396864 | 1.37700841 | 0.16850965 | NA |
| ARL5B | 8.53704515 | -0.7187259 | 0.55598517 | -1.292707 | 0.19611241 | NA |
| MALRD1 | 5.1486625 | 1.57703901 | 0.9159372 | 1.72177635 | 0.08511004 | NA |
| PLXDC2 | 128.454697 | -0.1649631 | 0.40571136 | -0.4066022 | 0.68430018 | 0.92710517 |
| NEBL | 356.938448 | 0.06805923 | 0.27158062 | 0.25060413 | 0.80212019 | 0.96742897 |
| MLLT10 | 78.9599012 | -0.3298295 | 0.32223076 | -1.0235817 | 0.30603289 | 0.77729327 |
| DNAJC1 | 73.5155243 | -0.8007012 | 0.31197527 | -2.5665534 | 0.01027148 | 0.28266532 |
| BMI1 | 8.19179035 | -0.6970065 | 0.57198214 | -1.2185808 | 0.22300333 | NA |
| PIP4K2A | 31.931323 | 0.03291114 | 0.48292804 | 0.06814916 | 0.9456669 | 0.99251804 |
| MSRB2 | 10.6117713 | 0.92482526 | 0.48947098 | 1.8894384 | 0.05883311 | NA |
| OTUD1 | 6.08879286 | -1.6196168 | 0.72231884 | -2.2422464 | 0.02494545 | NA |
| KIAA1217 | 172.296873 | 0.20917405 | 0.34877181 | 0.59974471 | 0.54867639 | 0.88448477 |
| ARHGAP21 | 33.5976519 | 0.14465442 | 0.31789577 | 0.45503725 | 0.64908244 | 0.92202688 |
| PDSS1 | 7.60699461 | -0.6002084 | 0.61926016 | -0.9692346 | 0.33242816 | NA |
| ABI1 | 47.9613113 | 0.23037177 | 0.27569315 | 0.83560934 | 0.40337471 | 0.82808596 |
| ANKRD26 | 29.3942043 | 0.72164952 | 0.3149799 | 2.29109702 | 0.0219578 | 0.35108799 |
| YME1L1 | 29.6584171 | -0.3299435 | 0.36351251 | -0.9076538 | 0.36406115 | 0.80760012 |
| MASTL | 7.06742132 | -0.0821647 | 0.56659606 | -0.1450145 | 0.88469938 | NA |
| ACBD5 | 20.9171247 | 0.07960724 | 0.38533888 | 0.20659022 | 0.8363299 | NA |
| RAB18 | 29.2855116 | -0.178317 | 0.4244853 | -0.4200781 | 0.67442841 | 0.92617027 |
| MPP7 | 45.9084919 | 0.62371938 | 0.30216698 | 2.06415468 | 0.03900305 | 0.43016177 |
| WAC-AS1 | 7.21495971 | -0.3659125 | 0.67013663 | -0.5460267 | 0.58504761 | NA |
| WAC | 102.642578 | -0.360349 | 0.22858068 | -1.5764629 | 0.11491916 | 0.57860396 |
| BAMBI | 4.0414672 | -0.2926026 | 0.80368944 | -0.3640742 | 0.71580258 | NA |
| SVIL | 37.2012402 | -0.1043528 | 0.45512643 | -0.229283 | 0.81864895 | 0.96845695 |
| MTPAP | 6.69283401 | -1.3546184 | 0.62700661 | -2.1604532 | 0.0307376 | NA |
| MAP3K8 | 37.6615811 | -0.6555112 | 0.37655102 | -1.7408298 | 0.08171343 | 0.54935298 |
| ZNF438 | 19.0056273 | -0.1488129 | 0.45012089 | -0.3306066 | 0.74094167 | NA |
| ZEB1 | 14.1778038 | -1.3240991 | 0.62482992 | -2.1191352 | 0.03407904 | NA |
| ARHGAP12 | 50.1960809 | -0.4302464 | 0.26883244 | -1.6004257 | 0.10950417 | 0.57860396 |
| KIF5B | 37.7802662 | -0.0684975 | 0.33246298 | -0.2060304 | 0.8367672 | 0.97154892 |
| EPC1 | 57.4747904 | -0.2841229 | 0.27789051 | -1.0224276 | 0.30657858 | 0.77729327 |
| CCDC7 | 29.5832056 | -0.1430069 | 0.47614845 | -0.300341 | 0.76391706 | 0.95543329 |
| ITGB1 | 39.9280967 | -0.0410603 | 0.27943165 | -0.1469422 | 0.88317764 | 0.97769824 |
| NRP1 | 31.3856941 | -0.0734029 | 0.58084995 | -0.1263715 | 0.89943788 | 0.97863828 |
| PARD3 | 241.611356 | 0.17978311 | 0.24635773 | 0.72976443 | 0.46553419 | 0.85894565 |

| | | | | | | |
|---|---|---|---|---|---|---|
| CUL2 | 15.1218122 | -0.3779005 | 0.43367073 | -0.8713996 | 0.38353603 | NA |
| CREM | 37.1820083 | -0.1630291 | 0.38913567 | -0.4189518 | 0.67525137 | 0.92617027 |
| CCNY | 78.4658304 | 0.18352057 | 0.30329962 | 0.60508012 | 0.54512576 | 0.88410237 |
| AL117336.3 | 4.94046425 | 0.71528222 | 0.6552194 | 1.09166827 | 0.27497894 | NA |
| ANKRD30A | 6.51290635 | 1.26401424 | 0.95285023 | 1.32656131 | 0.18465384 | NA |
| ZNF248 | 18.929814 | 0.75396061 | 0.43322147 | 1.74035837 | 0.08179611 | NA |
| ZNF33A | 52.9429352 | 0.20167243 | 0.25729447 | 0.7838195 | 0.43314603 | 0.84223493 |
| ZNF33B | 32.53421 | 0.30465499 | 0.39199434 | 0.77719233 | 0.43704531 | 0.8440477 |
| BMS1 | 12.3355659 | -0.1596492 | 0.44613663 | -0.3578482 | 0.72045694 | NA |
| CSGALNACT2 | 6.74120425 | 0.15912924 | 0.66955814 | 0.23766307 | 0.81214243 | NA |
| FXYD4 | 10.8517166 | -1.3044265 | 0.68648832 | -1.9001438 | 0.05741425 | NA |
| HNRNPF | 31.7278238 | -0.138216 | 0.33106741 | -0.4174861 | 0.67632291 | 0.92617027 |
| ZNF487 | 3.50408541 | -0.9232842 | 0.83430336 | -1.1066529 | 0.26844403 | NA |
| ZNF32 | 8.37530878 | 0.56339408 | 0.54909401 | 1.02604303 | 0.3048713 | NA |
| TMEM72-AS | 3.84293218 | 0.74623068 | 1.31589658 | 0.56708916 | 0.57065361 | NA |
| TMEM72 | 27.1775277 | 0.99928947 | 0.41042002 | 2.43479708 | 0.01490014 | 0.30435125 |
| RASSF4 | 13.4023876 | 0.65647463 | 0.65806891 | 0.99757733 | 0.31848436 | NA |
| DEPP1 | 21.0844538 | 0.41998981 | 0.76572583 | 0.54848589 | 0.58335831 | NA |
| ZNF22 | 5.59010251 | 1.08764373 | 0.69505247 | 1.56483687 | 0.11762117 | NA |
| 8-Mar | 23.1289562 | -0.138377 | 0.44710933 | -0.3094925 | 0.75694694 | NA |
| ZFAND4 | 10.7344101 | -0.1717417 | 0.47129422 | -0.3644045 | 0.71555597 | NA |
| WASHC2C | 13.3082255 | -0.6011764 | 0.50016838 | -1.201948 | 0.22938369 | NA |
| AGAP4 | 7.89616524 | 0.37383701 | 0.5560384 | 0.67232228 | 0.50137855 | NA |
| TIMM23 | 12.0249929 | 0.00500439 | 0.5129616 | 0.00975587 | 0.99221607 | NA |
| NCOA4 | 20.3109916 | -0.2342115 | 0.38546257 | -0.6076116 | 0.54344513 | NA |
| PTPN20 | 16.5732402 | 0.73780678 | 0.45971893 | 1.60490842 | 0.10851396 | NA |
| AGAP9 | 6.37254349 | 0.58605394 | 0.56875874 | 1.03040868 | 0.3028182 | NA |
| MAPK8 | 88.1455261 | -0.0462189 | 0.25021704 | -0.1847154 | 0.8534523 | 0.97168963 |
| VSTM4 | 10.0264063 | 0.5460912 | 0.55416152 | 0.98543688 | 0.32440952 | NA |
| ERCC6 | 15.093375 | 0.31342828 | 0.41333392 | 0.75829315 | 0.44827551 | NA |
| OGDHL | 65.8172002 | 0.71369404 | 0.40579645 | 1.75874884 | 0.07862018 | 0.54552914 |
| PARG | 31.3384501 | 0.06034344 | 0.32850151 | 0.18369304 | 0.85425427 | 0.97168963 |
| TIMM23B | 27.0225128 | -1.0234773 | 0.49769626 | -2.0564296 | 0.03974113 | 0.43117507 |
| LINC00843 | 6.5806337 | -0.1143859 | 0.56450145 | -0.2026318 | 0.83942285 | NA |
| AGAP6 | 7.97750156 | 0.16709677 | 0.58279291 | 0.28671724 | 0.77432884 | NA |
| WASHC2A | 11.0944208 | -1.685716 | 0.74807744 | -2.2533977 | 0.02423409 | NA |
| SGMS1 | 158.434929 | -0.5328762 | 0.29490165 | -1.8069625 | 0.07076813 | 0.52907601 |
| PRKG1 | 27.4114338 | 0.80644623 | 0.56972307 | 1.41550566 | 0.15692029 | 0.63942024 |
| IPMK | 11.2410294 | 0.19779937 | 0.44192332 | 0.44758754 | 0.6544509 | NA |
| CISD1 | 32.3660066 | -0.0822795 | 0.37050807 | -0.2220722 | 0.82425771 | 0.96845695 |
| UBE2D1 | 13.158781 | 0.64043434 | 0.43362803 | 1.47692098 | 0.13969682 | NA |
| TFAM | 9.58713133 | 0.11973535 | 0.50094647 | 0.23901825 | 0.81109143 | NA |
| BICC1 | 153.070651 | -0.0963045 | 0.49806176 | -0.1933586 | 0.84667814 | 0.97168963 |
| FAM13C | 14.4869272 | 0.7413455 | 0.94766099 | 0.78228977 | 0.4340443 | NA |
| CCDC6 | 19.5314348 | 0.18892167 | 0.4355249 | 0.43377925 | 0.66444875 | NA |
| ANK3 | 183.585199 | 0.33079381 | 0.35504158 | 0.93170442 | 0.3514893 | 0.8003972 |
| RHOBTB1 | 10.7498365 | -1.2636627 | 0.78710037 | -1.6054657 | 0.10839135 | NA |
| ARID5B | 26.5311302 | -0.3977158 | 0.49039164 | -0.8110166 | 0.41735611 | 0.83753644 |
| ADO | 3.33422024 | -0.5785488 | 0.80458444 | -0.7190654 | 0.47210061 | NA |
| NRBF2 | 4.09874387 | -0.6339846 | 0.74617954 | -0.8496408 | 0.39552484 | NA |
| JMJD1C | 215.671771 | -0.0694111 | 0.20426151 | -0.339815 | 0.73399586 | 0.94548018 |
| REEP3 | 38.8696266 | -0.2220264 | 0.31325872 | -0.7087636 | 0.47847118 | 0.86206794 |
| SIRT1 | 13.2905965 | -0.9148772 | 0.43328717 | -2.11148 | 0.03473108 | NA |
| HERC4 | 72.777215 | -0.1919319 | 0.32037503 | -0.5990849 | 0.54911625 | 0.88448477 |

| | | | | | | |
|---|---|---|---|---|---|---|
| PBLD | 7.64590178 | -0.145865 | 0.5529035 | -0.2638165 | 0.79192136 | NA |
| HNRNPH3 | 25.0345764 | -0.2994543 | 0.38196361 | -0.7839864 | 0.43304807 | NA |
| RUFY2 | 18.8821678 | -0.1953223 | 0.47117348 | -0.4145443 | 0.67847555 | NA |
| SLC25A16 | 24.5912874 | -0.3871418 | 0.34871576 | -1.1101929 | 0.2669159 | NA |
| TET1 | 4.08048504 | -0.9035731 | 0.89492967 | -1.0096582 | 0.31265906 | NA |
| CCAR1 | 47.4303556 | -0.5750052 | 0.31326655 | -1.8355141 | 0.06642956 | 0.5230103 |
| STOX1 | 4.62561527 | 0.67392745 | 0.85264598 | 0.79039539 | 0.4292969 | NA |
| DDX50 | 17.6944444 | 0.46896253 | 0.41881079 | 1.11974796 | 0.26282118 | NA |
| DDX21 | 20.8781173 | -0.9120144 | 0.39826111 | -2.2899911 | 0.02202183 | NA |
| KIF1BP | 8.19172369 | -0.0206747 | 0.48781272 | -0.0423825 | 0.96619377 | NA |
| VPS26A | 19.8766744 | -0.5048957 | 0.35919516 | -1.4056306 | 0.15983385 | NA |
| SUPV3L1 | 7.02594315 | -1.1152305 | 0.63059629 | -1.7685332 | 0.0769718 | NA |
| HKDC1 | 6.08730372 | -1.022545 | 0.75568025 | -1.353145 | 0.17600931 | NA |
| HK1 | 29.1456343 | -0.4428218 | 0.32645307 | -1.3564638 | 0.17495166 | 0.6651652 |
| TSPAN15 | 8.47925877 | 0.19442882 | 0.49873699 | 0.3898424 | 6.97E-01 | NA |
| H2AFY2 | 8.50222128 | 0.42435128 | 0.51402931 | 0.82553907 | 0.40906561 | NA |
| AIFM2 | 8.59607096 | -0.8195508 | 0.52947464 | -1.5478566 | 0.12165684 | NA |
| TYSND1 | 3.51480055 | -0.5711863 | 0.85412179 | -0.668741 | 0.5036607 | NA |
| SAR1A | 16.0908909 | -0.5943849 | 0.46655402 | -1.2739896 | 0.20266711 | NA |
| PPA1 | 29.8087676 | 0.10197338 | 0.34874649 | 0.29239974 | 0.76998101 | 0.95897308 |
| EIF4EBP2 | 25.2641935 | -0.3210806 | 0.47616391 | -0.6743068 | 0.50011629 | NA |
| SGPL1 | 25.693729 | 0.29439936 | 0.32239346 | 0.91316791 | 0.36115424 | NA |
| PCBD1 | 14.6627877 | 0.38702867 | 0.48389651 | 0.79981703 | 0.42381682 | NA |
| PSAP | 68.0583027 | 0.33103541 | 0.34491412 | 0.95976185 | 0.33717509 | 0.8003972 |
| CHST3 | 14.0931605 | -0.1339061 | 0.47343561 | -0.2828392 | 0.77730013 | NA |
| ASCC1 | 14.4662152 | -0.2952823 | 0.44862275 | -0.6581973 | 0.51041133 | NA |
| ANAPC16 | 45.4724884 | 0.40899688 | 0.32093067 | 1.27440883 | 0.20251856 | 0.700066 |
| DDIT4 | 7.68912757 | -0.8152407 | 0.73759144 | -1.105274 | 0.26904087 | NA |
| DNAJB12 | 7.39838491 | -5.05E-01 | 0.57718405 | -8.75E-01 | 0.38154959 | NA |
| MICU1 | 51.5635964 | -0.5224075 | 0.30167254 | -1.7317039 | 0.0833263 | 0.55234224 |
| MCU | 75.317111 | -0.7594724 | 0.5400286 | -1.4063558 | 0.1596185 | 0.64451101 |
| P4HA1 | 29.6342746 | -0.3350394 | 0.75806945 | -0.4419639 | 0.6585153 | 0.92276785 |
| ECD | 7.94334957 | 0.20223849 | 0.53692983 | 0.37665721 | 0.70642837 | NA |
| FAM149B1 | 14.4748063 | 0.66704159 | 0.43458381 | 1.53489748 | 0.12480901 | NA |
| DNAJC9 | 12.2675918 | 0.30509366 | 0.55702547 | 0.54771941 | 0.58388458 | NA |
| MRPS16 | 12.0623879 | -0.0110693 | 0.43663855 | -0.0253512 | 0.9797748 | NA |
| CFAP70 | 9.95806421 | 1.05313344 | 0.579866 | 1.8161669 | 0.06934477 | NA |
| ANXA7 | 27.9951149 | -0.2475588 | 0.31316193 | -0.7905138 | 0.42922777 | 0.84223493 |
| PPP3CB | 33.7443391 | -0.2548744 | 0.34516116 | -0.7384215 | 0.46025837 | 0.85844137 |
| USP54 | 14.0268908 | 0.05663377 | 0.44135721 | 0.12831731 | 0.89789787 | NA |
| AGAP5 | 3.12728134 | 0.30335839 | 0.83202962 | 0.36460047 | 0.71540966 | NA |
| SEC24C | 10.4652882 | -0.5312161 | 0.47846363 | -1.1102539 | 0.26688964 | NA |
| FUT11 | 6.93149933 | -0.5555547 | 0.67329319 | -0.8251305 | 0.4092975 | NA |
| CHCHD1 | 9.53228633 | 0.18368591 | 0.59905233 | 0.30662749 | 0.75912692 | NA |
| ZSWIM8 | 13.7370017 | -0.5164403 | 0.46883312 | -1.1015439 | 0.27066001 | NA |
| CAMK2G | 6.53027067 | 0.08683854 | 0.62248824 | 0.1395023 | 0.88905324 | NA |
| VCL | 47.1036965 | -0.8159756 | 0.30928919 | -2.6382286 | 0.00833404 | 0.27648451 |
| AP3M1 | 7.88185351 | -0.4603589 | 0.61678754 | -0.7463816 | 0.45543694 | NA |
| ADK | 118.147956 | -0.4469394 | 0.22592854 | -1.9782334 | 0.04790238 | 0.45661231 |
| KAT6B | 62.570272 | 0.00352097 | 0.28670739 | 0.0122807 | 0.99020166 | 0.99909657 |
| SAMD8 | 15.1555457 | -0.2094944 | 0.47194046 | -0.4439001 | 0.6571148 | NA |
| VDAC2 | 24.0827285 | -0.0807249 | 0.37579806 | -0.2148094 | 0.82991595 | NA |
| COMTD1 | 25.9622208 | 0.15988372 | 0.48529972 | 0.32945355 | 0.7418129 | 0.94831173 |
| AC010997.3 | 2.47456278 | 2.18375204 | 1.12475469 | 1.94153629 | 0.05219326 | NA |

| | | | | | | |
|---|---|---|---|---|---|---|
| ZNF503-AS2 | 4.08511746 | -1.4565869 | 0.84369753 | -1.7264326 | 0.08426961 | NA |
| LRMDA | 460.53504 | -0.3962454 | 0.27943583 | -1.4180194 | 0.15618509 | 0.63932163 |
| KCNMA1 | 11.4662808 | -2.2657281 | 0.98974085 | -2.2892135 | 0.02206695 | NA |
| DLG5 | 89.7904544 | -0.3395067 | 0.43335852 | -0.7834316 | 0.43337373 | 0.84223493 |
| POLR3A | 10.1135045 | 0.01149305 | 0.47821769 | 0.0240331 | 0.98082621 | NA |
| RPS24 | 107.230911 | -0.0904153 | 0.3510106 | -0.2575857 | 0.79672668 | 0.96525806 |
| ZMIZ1 | 61.2296255 | 0.24118914 | 0.43638943 | 0.55269245 | 0.58047402 | 0.89263108 |
| PPIF | 13.0090856 | -0.9360107 | 0.47388315 | -1.9751931 | 0.04824624 | NA |
| NUTM2B-AS | 38.2281278 | 0.20694069 | 0.2934187 | 0.70527438 | 0.4806395 | 0.86222804 |
| TMEM254 | 3.16617642 | -0.3316635 | 0.82927573 | -0.3999435 | 0.6891981 | NA |
| ANXA11 | 84.8577265 | 0.19243496 | 0.26944198 | 0.71419814 | 0.47510467 | 0.86206794 |
| FAM213A | 23.3337151 | 0.792592 | 0.36257013 | 2.18603778 | 0.02881284 | NA |
| TSPAN14 | 59.3547109 | 0.53079089 | 0.40066834 | 1.32476374 | 0.18524953 | 0.67879383 |
| NRG3 | 57.291499 | -0.8296755 | 1.03046723 | -0.805145 | 0.42073602 | 0.83977933 |
| GHITM | 30.4448485 | 0.32002135 | 0.39108516 | 0.8182907 | 0.41319122 | 0.83356565 |
| CCSER2 | 52.7983301 | -0.0198562 | 0.27840149 | -0.0713221 | 0.94314138 | 0.99182204 |
| WAPL | 45.6351181 | -0.192486 | 0.26426091 | -0.7283939 | 0.4663725 | 0.85894565 |
| BMPR1A | 52.0313835 | 0.13881515 | 0.2872772 | 0.48320978 | 0.62894679 | 0.91751762 |
| MMRN2 | 3.01368244 | -0.358847 | 1.03837035 | -0.3455867 | 0.72965336 | NA |
| ADIRF | 11.126579 | 0.72316432 | 0.599958 | 1.20535824 | 0.22806504 | NA |
| GLUD1 | 88.120003 | 0.48147163 | 0.31192234 | 1.54356253 | 0.12269435 | 0.58553957 |
| SHLD2 | 52.3934384 | -0.3126612 | 0.30346689 | -1.0302977 | 0.30287028 | 0.77521265 |
| NUTM2A-AS | 48.4989403 | 0.01898093 | 0.31395629 | 0.06045723 | 0.95179148 | 0.99443943 |
| MINPP1 | 6.15049743 | -0.1989288 | 0.66489424 | -0.2991886 | 0.76479618 | NA |
| ATAD1 | 29.9735318 | -0.0836628 | 0.3097718 | -0.2700788 | 0.78709961 | 0.96325251 |
| PTEN | 49.8986711 | -0.0392878 | 0.28012342 | -0.1402517 | 0.88846109 | 0.97769824 |
| RNLS | 17.5740699 | -1.0271415 | 0.46054744 | -2.2302621 | 0.02573005 | NA |
| ANKRD22 | 5.50551181 | 0.54958289 | 0.81630913 | 0.67325339 | 0.5007861 | NA |
| LIPA | 10.0320982 | 0.39650869 | 0.59664605 | 0.66456266 | 0.50633026 | NA |
| IFIT2 | 5.11703632 | 0.32432475 | 1.04095368 | 0.31156501 | 0.75537113 | NA |
| IFIT3 | 14.8773978 | -1.6645681 | 1.27388344 | -1.3066879 | 0.19131873 | NA |
| IFIT1 | 14.2978113 | -1.5140601 | 1.70511982 | -0.8879494 | 0.37456796 | NA |
| IFIT5 | 3.05344804 | -0.6433594 | 0.98425679 | -0.65365 | 0.51333735 | NA |
| SLC16A12 | 17.6409543 | 0.42580882 | 0.87249511 | 0.48803576 | 0.62552451 | NA |
| PANK1 | 6.06812054 | 0.43864517 | 0.93534154 | 0.46896791 | 0.63909257 | NA |
| KIF20B | 2.79713894 | 0.1695646 | 0.97504196 | 0.17390492 | 0.86194018 | NA |
| RPP30 | 3.36958079 | 0.24961275 | 0.86117611 | 0.28985099 | 0.77193023 | NA |
| PCGF5 | 60.2141768 | 0.04494766 | 0.26969963 | 0.16665821 | 0.86763899 | 0.97217819 |
| HECTD2 | 20.9650787 | -0.5313116 | 0.37936244 | -1.4005382 | 0.16135222 | NA |
| PPP1R3C | 6.51169929 | 0.07125198 | 0.96445893 | 0.07387768 | 0.94110772 | NA |
| TNKS2 | 48.346239 | -0.4580691 | 0.33490757 | -1.3677477 | 0.17139106 | 0.66182328 |
| BTAF1 | 38.3347452 | -0.9761594 | 0.34351343 | -2.8416921 | 0.00448748 | 0.21776517 |
| CPEB3 | 61.8499352 | 0.4140322 | 0.39306358 | 1.05334664 | 0.29218215 | 0.77287826 |
| 5-Mar | 18.42728 | 0.3561346 | 0.44379323 | 0.80247868 | 0.42227612 | NA |
| IDE | 17.6274257 | 0.36251404 | 0.42238505 | 0.8582549 | 0.39075173 | NA |
| EXOC6 | 38.5337936 | -1.1760692 | 0.34616531 | -3.3974206 | 0.00068024 | 0.09555626 |
| MYOF | 21.255875 | -0.3456547 | 0.53339551 | -0.648027 | 0.51696749 | NA |
| FRA10AC1 | 12.4730906 | -0.3673838 | 0.46883448 | -0.7836108 | 0.4332685 | NA |
| PLCE1 | 25.0278138 | 0.04542756 | 0.59381104 | 0.07650171 | 0.93901996 | NA |
| NOC3L | 4.89723988 | 0.26035422 | 0.65810824 | 0.39561003 | 0.69239273 | NA |
| TBC1D12 | 17.8000751 | 0.54748854 | 0.36839352 | 1.48615139 | 0.13723908 | NA |
| HELLS | 4.06496715 | 0.1022677 | 0.75338183 | 0.13574484 | 0.892023 | NA |
| SORBS1 | 9.5949535 | -1.0437289 | 0.66132467 | -1.5782398 | 0.11451054 | NA |
| ALDH18A1 | 9.87322716 | -0.3650238 | 0.5384841 | -0.6778729 | 0.49785229 | NA |

| | | | | | | |
|---|---|---|---|---|---|---|
| TCTN3 | 11.1115583 | -0.4252277 | 0.46884022 | -0.906978 | 0.36441845 | NA |
| ENTPD1-AS1 | 30.6962807 | 1.3436473 | 0.32954927 | 4.07722738 | 4.56E-05 | 0.04054737 |
| ZNF518A | 46.1099895 | -0.2570571 | 0.32837151 | -0.7828239 | 0.43373054 | 0.84223493 |
| BLNK | 26.4611252 | -0.1448066 | 0.40310832 | -0.359225 | 0.71942675 | 0.93714576 |
| TM9SF3 | 50.7852503 | -0.075322 | 0.24866434 | -0.3029064 | 0.76196123 | 0.95478238 |
| LCOR | 99.1122241 | -0.1982669 | 0.29365614 | -0.6751668 | 0.4995698 | 0.87036626 |
| RRP12 | 3.824478 | -0.4679848 | 0.8584677 | -0.5451397 | 0.58565742 | NA |
| PGAM1 | 23.7149268 | -0.266382 | 0.37349546 | -0.7132135 | 0.47571365 | NA |
| EXOSC1 | 6.0525978 | 0.40415701 | 0.60777077 | 0.66498264 | 0.5060616 | NA |
| ZDHHC16 | 6.71213368 | 0.77980218 | 0.60041092 | 1.2987808 | 0.19401916 | NA |
| MMS19 | 13.976146 | -0.7558542 | 0.44588938 | -1.6951608 | 0.09004493 | NA |
| UBTD1 | 6.5301397 | 0.07952031 | 0.61056299 | 0.13024095 | 0.89637579 | NA |
| ZFYVE27 | 9.34963966 | -0.4599807 | 0.60809847 | -0.7564246 | 0.44939464 | NA |
| R3HCC1L | 24.3284886 | -0.3276008 | 0.37786975 | -0.8669676 | 0.38595979 | NA |
| HPS1 | 7.83245121 | -0.3083994 | 0.56595021 | -0.5449231 | 0.58580638 | NA |
| GOT1 | 43.0569254 | -0.2343756 | 0.3312626 | -0.707522 | 0.47924212 | 0.86206794 |
| AL391684.1 | 2.72514436 | 2.94368442 | 1.1136515 | 2.64327254 | 0.00821089 | NA |
| SLC25A28 | 7.26776196 | -0.0380868 | 0.55101523 | -0.0691211 | 0.94489326 | NA |
| CUTC | 8.3706273 | 0.37633812 | 0.50951427 | 0.73862135 | 0.46013696 | NA |
| COX15 | 10.1068782 | -0.0170704 | 0.46921217 | -0.0363809 | 0.97097864 | NA |
| ABCC2 | 5.13475231 | -1.140521 | 0.88998901 | -1.2815001 | 0.20001805 | NA |
| DNMBP | 22.4672385 | 0.21641142 | 0.37488441 | 0.57727508 | 0.56375365 | NA |
| ERLIN1 | 6.22744898 | 0.45451151 | 0.56320113 | 0.80701454 | 0.4196581 | NA |
| CHUK | 12.133668 | -0.3179033 | 0.4546142 | -0.6992814 | 0.48437619 | NA |
| CWF19L1 | 9.29130902 | -0.5630757 | 0.52665313 | -1.0691585 | 0.28499824 | NA |
| BLOC1S2 | 7.6167134 | 0.11320011 | 0.57746237 | 0.19603028 | 0.84458647 | NA |
| SCD | 4.65292675 | -3.4590381 | 1.23719523 | -2.7958709 | 0.00517601 | NA |
| OLMALINC | 4.31687916 | -0.3351055 | 1.01909441 | -0.3288268 | 0.74228662 | NA |
| SEC31B | 3.68777109 | 1.40588495 | 0.88043893 | 1.59680008 | 0.11031028 | NA |
| HIF1AN | 7.7972471 | 0.35993506 | 0.53097497 | 0.67787576 | 0.49785047 | NA |
| PAX2 | 42.9513733 | 0.05923275 | 0.38814655 | 0.15260408 | 0.8787105 | 0.97516771 |
| SLF2 | 16.5343611 | 0.35567829 | 0.41938187 | 0.84810125 | 0.39638159 | NA |
| MRPL43 | 8.72396927 | 1.01677951 | 0.56965441 | 1.78490588 | 0.07427658 | NA |
| LZTS2 | 6.60882257 | -0.8761389 | 0.5961739 | -1.469603 | 0.14166931 | NA |
| BTRC | 47.1254233 | -0.1606507 | 0.34841083 | -0.4610957 | 0.64472992 | 0.92153493 |
| POLL | 3.14292293 | 1.12166192 | 0.96475437 | 1.16263989 | 0.24497564 | NA |
| FBXW4 | 41.9464652 | 0.37915808 | 0.34293078 | 1.10564028 | 0.26888225 | 0.75632146 |
| OGA | 68.9757929 | 0.11145675 | 0.27091159 | 0.41141373 | 0.68076919 | 0.9267216 |
| ARMH3 | 49.5764023 | 0.29661149 | 0.29275139 | 1.01318559 | 0.31097152 | 0.778534 |
| LDB1 | 5.80067348 | -0.0958295 | 0.65772492 | -0.1456984 | 0.8841595 | NA |
| PPRC1 | 5.97363109 | -0.7032521 | 0.71203529 | -0.9876647 | 0.32331689 | NA |
| NOLC1 | 5.67228446 | -0.0924477 | 0.56721363 | -0.1629857 | 0.87052973 | NA |
| GBF1 | 55.4157657 | -0.6387204 | 0.27474493 | -2.324776 | 0.02008394 | 0.34471691 |
| NFKB2 | 3.91742219 | 1.04339571 | 0.7684172 | 1.35785054 | 0.17451111 | NA |
| FBXL15 | 7.67500972 | 0.2530402 | 0.61766925 | 0.40966942 | 0.68204847 | NA |
| CUEDC2 | 6.91441968 | 0.72668877 | 0.57404418 | 1.26591088 | 0.20554499 | NA |
| ACTR1A | 10.0588456 | -0.1349721 | 0.51026868 | -0.2645118 | 0.79138554 | NA |
| SUFU | 14.0184975 | 0.19673566 | 0.45394845 | 0.43338767 | 0.66473315 | NA |
| TRIM8 | 31.4518437 | 0.24696254 | 0.30987928 | 0.79696372 | 0.4254721 | 0.84057082 |
| ARL3 | 8.14475508 | 1.0370453 | 0.5959557 | 1.74013825 | 0.08183475 | NA |
| SFXN2 | 6.07485285 | 0.3465431 | 0.67505351 | 0.51335648 | 0.60770198 | NA |
| WBP1L | 32.0015124 | -0.7196895 | 0.30567307 | -2.3544418 | 0.01855055 | 0.33831016 |
| CNNM2 | 70.7417885 | -0.2010507 | 0.43007496 | -0.4674782 | 0.64015778 | 0.92153493 |
| NT5C2 | 80.5966341 | 0.07192717 | 0.32586769 | 0.22072508 | 0.8253065 | 0.96845695 |

| | | | | | | |
|---|---|---|---|---|---|---|
| PCGF6 | 8.7239558 | -0.3227135 | 0.57559577 | -0.56066 | 0.57502935 | NA |
| ATP5MD | 20.078821 | 0.45149191 | 0.36244148 | 1.24569603 | 0.21287601 | NA |
| PDCD11 | 7.76143572 | -0.4967612 | 0.72122528 | -0.688774 | 0.49096553 | NA |
| NEURL1 | 58.158877 | 1.29049021 | 0.60731056 | 2.12492635 | 0.03359276 | 0.41688714 |
| SH3PXD2A | 8.72685887 | 0.56938353 | 0.57802279 | 0.98505377 | 0.32459766 | NA |
| AL121929.2 | 2.02649192 | 1.35639683 | 1.47936642 | 0.91687686 | 0.35920717 | NA |
| STN1 | 4.03787565 | -0.0522932 | 0.76284489 | -0.0685502 | 0.94534767 | NA |
| SLK | 26.299512 | 0.33260519 | 0.30288462 | 1.09812504 | 0.27214989 | 0.75979923 |
| CFAP43 | 3.62554708 | -0.0079724 | 0.94683565 | -0.00842 | 0.99328188 | NA |
| GSTO1 | 13.8931284 | 0.02576073 | 0.42355367 | 0.06082045 | 0.9515022 | NA |
| GSTO2 | 15.5437325 | 0.29454094 | 0.47269397 | 0.62311127 | 0.53321141 | NA |
| XPNPEP1 | 10.1041124 | -0.0650045 | 0.48031697 | -0.1353366 | 0.89234577 | NA |
| ADD3 | 41.9368446 | 0.50160135 | 0.34474086 | 1.45500983 | 0.14566654 | 0.628084 |
| MXI1 | 18.41326 | 0.08712738 | 0.41019048 | 0.21240711 | 0.83178944 | NA |
| SMNDC1 | 9.06090259 | -0.6181166 | 0.54077508 | -1.1430198 | 0.25303035 | NA |
| SMC3 | 12.0561418 | -0.2423205 | 0.503907 | -0.4808834 | 0.63059938 | NA |
| PDCD4 | 14.5685055 | 0.2893119 | 0.52734438 | 0.54862044 | 0.58326596 | NA |
| BBIP1 | 10.9788535 | 1.1645369 | 0.49077375 | 2.37285899 | 0.017651 | NA |
| SHOC2 | 33.1422555 | -0.1811232 | 0.38147819 | -0.474793 | 0.63493452 | 0.92072524 |
| GPAM | 3.70581682 | 0.21130319 | 0.83183513 | 0.25402052 | 0.79947971 | NA |
| ZDHHC6 | 7.81586478 | 0.0586163 | 0.54199034 | 0.10815009 | 0.91387663 | NA |
| VTI1A | 126.712968 | -0.225072 | 0.25132858 | -0.8955287 | 0.37050454 | 0.81199861 |
| TCF7L2 | 96.194562 | 0.27418506 | 0.35033341 | 0.78264035 | 0.43383834 | 0.84223493 |
| CASP7 | 3.88784222 | -0.0004498 | 0.77902174 | -0.0005774 | 0.9995393 | NA |
| NHLRC2 | 22.6779915 | 0.71945475 | 0.32739632 | 2.19750406 | 0.02798447 | NA |
| CCDC186 | 26.8533417 | -0.7423073 | 0.3215937 | -2.3082148 | 0.02098719 | 0.34990513 |
| ABLIM1 | 83.9548658 | 0.35138119 | 0.43266855 | 0.81212557 | 0.41671958 | 0.8374607 |
| FAM160B1 | 10.3878066 | 0.30719772 | 0.45319846 | 0.6778437 | 0.4978708 | NA |
| TRUB1 | 7.98403695 | 0.46247825 | 0.63493505 | 0.7283867 | 0.4663769 | NA |
| HSPA12A | 9.44178277 | -0.2021134 | 0.63408791 | -0.3187466 | 0.74991866 | NA |
| SHTN1 | 51.5722094 | -0.6346458 | 0.28872597 | -2.1980903 | 0.02794267 | 0.39934064 |
| PDZD8 | 30.9660614 | 0.06610018 | 0.42658249 | 0.15495287 | 0.87685847 | 0.9751578 |
| EMX2OS | 16.2188519 | 0.68009811 | 0.48833106 | 1.39269885 | 0.16371088 | NA |
| EMX2 | 8.33205279 | 0.35635446 | 0.5675736 | 0.62785594 | 0.53009832 | NA |
| RAB11FIP2 | 19.3218361 | -0.3276851 | 0.3750127 | -0.8737973 | 0.38222866 | NA |
| CASC2 | 8.61351734 | 0.51518563 | 0.71265447 | 0.72291082 | 0.46973467 | NA |
| FAM204A | 15.2930144 | 0.04705285 | 0.41026909 | 0.11468779 | 0.9086926 | NA |
| CACUL1 | 36.7796197 | -0.2279549 | 0.29633261 | -0.7692537 | 0.44174274 | 0.84767356 |
| EIF3A | 50.3060699 | -0.7070724 | 0.35841403 | -1.9727811 | 0.04852051 | 0.45922422 |
| FAM45A | 9.74068961 | 0.24083516 | 0.48271537 | 0.49891752 | 0.61783749 | NA |
| SFXN4 | 19.8341454 | 0.12389898 | 0.4028866 | 0.30752816 | 0.75844139 | NA |
| PRDX3 | 21.9004886 | 0.02496654 | 0.35479714 | 0.07036849 | 0.94390037 | NA |
| GRK5 | 6.29308519 | 0.53975847 | 0.95081277 | 0.56768113 | 0.57025151 | NA |
| TIAL1 | 17.8671603 | -0.0524063 | 0.39166728 | -0.1338032 | 0.89355817 | NA |
| BAG3 | 9.83302488 | -0.1453992 | 0.49497534 | -0.2937504 | 0.76894868 | NA |
| INPP5F | 13.3255873 | -0.0159028 | 0.41370001 | -0.0384405 | 0.96933651 | NA |
| MCMBP | 17.1679939 | -0.3276693 | 0.40838411 | -0.8023557 | 0.42234724 | NA |
| SEC23IP | 25.3482889 | -0.1739845 | 0.41665831 | -0.4175711 | 0.67626075 | NA |
| PLPP4 | 5.08433646 | 0.5147363 | 0.72269916 | 0.71224145 | 0.47631528 | NA |
| WDR11-AS1 | 3.3635449 | -0.2658107 | 0.92035872 | -0.2888121 | 0.7727252 | NA |
| WDR11 | 29.0712941 | 0.32718801 | 0.31109092 | 1.05174401 | 0.29291702 | 0.77342911 |
| FGFR2 | 26.7413939 | 0.24726136 | 0.63649371 | 0.38847416 | 0.69766517 | 0.9303056 |
| ATE1 | 57.259622 | 0.12892194 | 0.38877975 | 0.33160664 | 0.74018631 | 0.94705525 |
| NSMCE4A | 5.79627852 | -0.7039604 | 0.65601668 | -1.0730831 | 0.28323383 | NA |

| | | | | | | |
|---|---|---|---|---|---|---|
| TACC2 | 62.5415952 | -0.2283392 | 0.57781028 | -0.3951802 | 0.6927099 | 0.9295338 |
| PLEKHA1 | 64.33415 | -0.4651316 | 0.28854825 | -1.6119716 | 0.10696811 | 0.57769936 |
| C10orf88 | 4.83491653 | -0.1489894 | 0.76539744 | -0.1946563 | 0.84566203 | NA |
| PSTK | 2.68934523 | 0.14367532 | 0.80563381 | 0.17833825 | 0.85845734 | NA |
| IKZF5 | 6.85642228 | 0.11031904 | 0.62090715 | 0.17767398 | 0.85897902 | NA |
| ACADSB | 11.9037495 | 0.82675301 | 0.5103685 | 1.61991387 | 0.10525078 | NA |
| HMX2 | 42.6757148 | 0.01724538 | 0.8403465 | 0.02052175 | 0.98362716 | 0.99902646 |
| BUB3 | 17.2307912 | -0.313644 | 0.38676193 | -0.8109486 | 0.41739519 | NA |
| CPXM2 | 23.8600208 | -0.2769535 | 0.66125618 | -0.4188293 | 0.6753409 | NA |
| OAT | 9.09967648 | -0.5121976 | 0.55934089 | -0.9157163 | 0.35981569 | NA |
| LHPP | 6.08102381 | 1.00784281 | 0.66504858 | 1.51544239 | 0.12966041 | NA |
| ABRAXAS2 | 6.89670686 | -0.3999909 | 0.58931216 | -0.6787421 | 0.49730132 | NA |
| ZRANB1 | 24.884206 | -0.4279982 | 0.31483176 | -1.3594505 | 0.17400387 | NA |
| CTBP2 | 112.764801 | 0.01224859 | 0.27741127 | 0.04415318 | 0.9647823 | 0.99656976 |
| EDRF1 | 11.7982104 | -0.4289463 | 0.50176157 | -0.8548807 | 0.39261718 | NA |
| UROS | 6.78348073 | 0.06413511 | 0.74115473 | 0.08653403 | 0.9310419 | NA |
| BCCIP | 12.9274018 | -0.6835927 | 0.46777255 | -1.4613783 | 0.14391164 | NA |
| DHX32 | 26.2196172 | 0.20662294 | 0.33047311 | 0.62523375 | 0.53181765 | 0.88254911 |
| FANK1 | 5.40951701 | 0.08116604 | 0.72763085 | 0.11154838 | 0.9111815 | NA |
| DOCK1 | 139.797484 | -0.2619336 | 0.19912232 | -1.3154409 | 0.18836175 | 0.68214045 |
| FAM196A | 6.73541877 | 0.13420406 | 0.65019836 | 0.2064048 | 0.83647472 | NA |
| MGMT | 31.6994464 | 0.75032227 | 0.33709777 | 2.22582983 | 0.02602558 | 0.39467204 |
| AL157832.3 | 2.19703019 | 0.11100987 | 1.10053458 | 0.10086905 | 0.91965442 | NA |
| C10orf143 | 3.27792526 | 0.59069265 | 0.78109022 | 0.75624126 | 0.44950456 | NA |
| GLRX3 | 18.4298276 | -0.4852572 | 0.44646257 | -1.0868933 | 0.27708397 | NA |
| PPP2R2D | 36.7604261 | -1.0891572 | 0.32894269 | -3.3110851 | 0.00092935 | 0.11274696 |
| BNIP3 | 31.1436967 | -0.0717449 | 0.50861171 | -0.1410603 | 0.8878223 | 0.97769824 |
| STK32C | 5.19673277 | -0.3976555 | 0.70634179 | -0.5629788 | 0.57344931 | NA |
| LRRC27 | 3.22522124 | 0.06256394 | 0.83928987 | 0.0745439 | 0.94057761 | NA |
| PWWP2B | 4.4304292 | -0.3149239 | 0.81943565 | -0.384318 | 0.70074277 | NA |
| INPP5A | 21.3860279 | -0.1869571 | 0.43831107 | -0.4265399 | 0.66971451 | NA |
| ADGRA1 | 4.46942298 | 1.0663248 | 0.88969255 | 1.19853179 | 0.23071005 | NA |
| KNDC1 | 2.01309487 | 0.96102916 | 1.20206775 | 0.79948003 | 0.42401212 | NA |
| TUBGCP2 | 14.7423308 | -0.1933446 | 0.42376881 | -0.4562503 | 0.64821001 | NA |
| ECHS1 | 19.5074344 | 0.80425304 | 0.41035131 | 1.95991342 | 0.05000591 | NA |
| SCART1 | 4.18749259 | -0.9693108 | 0.91056495 | -1.0645158 | 0.2870951 | NA |
| SLC6A12 | 14.8040568 | -1.3668011 | 0.44983514 | -3.0384489 | 0.00237799 | NA |
| SLC6A13 | 4.68639177 | 0.37132764 | 1.16211124 | 0.31952848 | 0.7493258 | NA |
| KDM5A | 40.4014748 | 0.20308318 | 0.3042582 | 0.66746986 | 0.50447205 | 0.87094108 |
| CCDC77 | 3.33960452 | -2.1158198 | 0.99242381 | -2.131972 | 0.03300915 | NA |
| B4GALNT3 | 14.9695568 | 0.55778328 | 0.58270103 | 0.95723751 | 0.33844739 | NA |
| WNK1 | 53.5968911 | 0.06560369 | 0.27978193 | 0.2344815 | 0.8146112 | 0.96845695 |
| RAD52 | 9.19430652 | 0.65978651 | 0.51324491 | 1.28551984 | 0.19861069 | NA |
| ERC1 | 128.289402 | -0.2002015 | 0.2636875 | -0.7592378 | 0.4477103 | 0.85123446 |
| FBXL14 | 3.21424767 | 0.10377211 | 0.78124333 | 0.13282943 | 0.89432828 | NA |
| ADIPOR2 | 50.8156798 | -0.6085386 | 0.32644137 | -1.864159 | 0.06229937 | 0.51631533 |
| DCP1B | 9.97722813 | 1.08733364 | 0.53439692 | 2.03469295 | 0.04188178 | NA |
| CACNA1C | 6.31149943 | 0.45170239 | 1.02597399 | 0.4402669 | 0.65974381 | NA |
| FKBP4 | 11.1951953 | -0.6645574 | 0.57823986 | -1.1492763 | 0.25044207 | NA |
| ITFG2 | 17.0048273 | -0.914714 | 0.51457392 | -1.7776144 | 0.07546721 | NA |
| TULP3 | 14.6997757 | 0.2175916 | 0.44898309 | 0.48463205 | 0.62793738 | NA |
| TEAD4 | 9.74022444 | -0.2463641 | 0.63102361 | -0.3904198 | 0.69622618 | NA |
| TSPAN9 | 53.7767173 | 0.67435579 | 0.40913653 | 1.64824145 | 0.09930313 | 0.57091823 |
| PARP11 | 7.10823102 | 0.38612572 | 0.64360112 | 0.59994569 | 0.54854243 | NA |

| | | | | | | |
|---|---|---|---|---|---|---|
| TIGAR | 4.81062449 | -1.1179849 | 0.75001815 | -1.4906105 | 0.13606378 | NA |
| C12orf4 | 8.72093014 | 0.633905 | 0.53081726 | 1.1942057 | 0.23239751 | NA |
| ANO2 | 8.8521474 | 0.78386222 | 0.62286874 | 1.25847095 | 0.20822149 | NA |
| VWF | 12.4265572 | -0.3770197 | 0.57242688 | -0.6586338 | 0.51013096 | NA |
| CD9 | 276.847718 | -0.0436889 | 0.36431138 | -0.119922 | 0.90454497 | 0.98064783 |
| TNFRSF1A | 14.169121 | -0.2758402 | 0.43068328 | -0.640471 | 0.52186641 | NA |
| SCNN1A | 111.036143 | 0.28478417 | 0.28601651 | 0.99569136 | 0.31940013 | 0.78335494 |
| LTBR | 10.8849141 | -0.5319707 | 0.44134648 | -1.2053357 | 0.22807373 | NA |
| TAPBPL | 4.58747386 | 0.32580917 | 0.73280895 | 0.44460316 | 0.65660657 | NA |
| MRPL51 | 13.4221346 | 0.44059455 | 0.47145818 | 0.93453581 | 0.35002757 | NA |
| NCAPD2 | 5.61043882 | 0.35128832 | 0.75199063 | 0.46714454 | 0.64039647 | NA |
| GAPDH | 128.830084 | -0.2785285 | 0.432368 | -0.644193 | 0.51945026 | 0.87214537 |
| CHD4 | 8.33655477 | 0.19742378 | 0.56134913 | 0.35169518 | 0.72506688 | NA |
| ING4 | 4.46688172 | 0.77499887 | 0.81696425 | 0.94863254 | 0.34280754 | NA |
| ZNF384 | 5.74972997 | 0.08943749 | 0.62427012 | 0.14326729 | 0.88607909 | NA |
| COPS7A | 7.10880355 | -0.340471 | 0.59769049 | -0.5696443 | 0.56891896 | NA |
| MLF2 | 12.033203 | -0.2306952 | 0.4948668 | -0.4661764 | 0.64108925 | NA |
| PTMS | 15.8974457 | 0.12891579 | 0.61478093 | 0.20969386 | 0.83390662 | NA |
| USP5 | 5.65719972 | 0.29473722 | 0.67650064 | 0.43567914 | 0.66306954 | NA |
| TPI1 | 43.5992043 | -0.3812769 | 0.3362688 | -1.1338457 | 0.2568593 | 0.75170774 |
| LRRC23 | 5.40874054 | -0.6077465 | 0.67704427 | -0.8976466 | 0.36937398 | NA |
| ATN1 | 13.3496821 | -0.5189995 | 0.46332003 | -1.120175 | 0.26263921 | NA |
| C12orf57 | 11.459442 | 0.38572953 | 0.48859851 | 0.78946112 | 0.42984255 | NA |
| PTPN6 | 4.07580921 | -0.5910319 | 0.77829183 | -0.7593963 | 0.44761554 | NA |
| PHB2 | 17.4576868 | -0.0349249 | 0.44782019 | -0.0779888 | 0.93783698 | NA |
| LPCAT3 | 39.8535925 | -0.0371921 | 0.37454065 | -0.0993006 | 0.92089961 | 0.98745437 |
| C1S | 8.73632812 | 0.04563597 | 0.53415153 | 0.08543637 | 0.93191448 | NA |
| C1RL | 9.12321008 | 0.17180642 | 0.48903553 | 0.35131684 | 0.72535066 | NA |
| PEX5 | 4.16788523 | -0.8204009 | 0.73198959 | -1.1207821 | 0.26238061 | NA |
| CD163L1 | 3.40590637 | -0.0906774 | 0.8938379 | -0.1014472 | 0.91919546 | NA |
| FOXJ2 | 3.77905189 | -1.4622927 | 0.91749902 | -1.5937812 | 0.11098504 | NA |
| NECAP1 | 6.28034076 | -1.4445188 | 0.64898234 | -2.2258214 | 0.02602615 | NA |
| LINC00937 | 2.68662037 | 1.15745649 | 0.8587093 | 1.34790259 | 0.17768971 | NA |
| RIMKLB | 55.711983 | -1.3600537 | 0.57132056 | -2.380544 | 0.0172871 | 0.32722884 |
| A2ML1-AS1 | 7.46349036 | -0.8699289 | 0.89341629 | -0.9737106 | 0.33020024 | NA |
| PHC1 | 4.55131643 | 0.09982653 | 0.66422496 | 0.15029024 | 0.88053563 | NA |
| M6PR | 18.1875331 | 0.06472473 | 0.41878297 | 0.15455435 | 0.87717266 | NA |
| CLEC2D | 9.23465971 | 0.81038825 | 0.51040249 | 1.58774352 | 0.11234435 | NA |
| TMEM52B | 2.53134997 | 1.35576723 | 1.18445874 | 1.14463018 | 0.25236238 | NA |
| GABARAPL1 | 46.1390458 | -0.4604566 | 0.31372393 | -1.4677127 | 0.14218229 | 0.62108763 |
| KLRD1 | 7.18102817 | 0.19598548 | 0.7812191 | 0.25087133 | 0.80191359 | NA |
| MAGOHB | 2.85954936 | 0.56406553 | 0.90029836 | 0.62653178 | 0.5309662 | NA |
| YBX3 | 46.6303882 | -0.041881 | 0.38473609 | -0.1088565 | 0.9133163 | 0.98415124 |
| SMIM10L1 | 5.23072931 | 0.33280912 | 0.73633191 | 0.45198248 | 0.6512816 | NA |
| AC244131.2 | 6.83289706 | 1.19936365 | 0.69090773 | 1.73592449 | 0.08257719 | NA |
| ETV6 | 83.060516 | 0.37664535 | 0.32998311 | 1.14140798 | 0.25370018 | 0.74749396 |
| BCL2L14 | 9.20242182 | 0.40280959 | 0.49673631 | 0.81091232 | 0.41741603 | NA |
| LRP6 | 29.0689348 | 0.10175902 | 0.37238496 | 0.27326296 | 0.78465109 | 0.96299238 |
| MANSC1 | 3.34196907 | -0.862429 | 0.82412359 | -1.0464802 | 0.2953394 | NA |
| BORCS5 | 11.9927575 | 0.23452316 | 0.50739892 | 0.46220666 | 0.64393313 | NA |
| DUSP16 | 22.896749 | 0.56810482 | 0.4169392 | 1.36256035 | 0.17302111 | NA |
| CREBL2 | 16.0049531 | 0.29531504 | 0.42008219 | 0.70299347 | 0.48205982 | NA |
| CDKN1B | 6.29545996 | 0.29519093 | 0.58365259 | 0.50576479 | 0.61302178 | NA |
| APOLD1 | 2.35714234 | 1.5330586 | 1.01901746 | 1.50444782 | 0.1324661 | NA |

| | | | | | | |
|---|---|---|---|---|---|---|
| GPRC5A | 4.87064621 | -0.6509889 | 0.72738736 | -0.8949687 | 0.37080381 | NA |
| GPRC5D-AS1 | 9.83177713 | 0.3849085 | 0.52726226 | 0.73001336 | 0.46538202 | NA |
| HEBP1 | 9.23678026 | 0.73781303 | 0.5436378 | 1.35717758 | 0.17472479 | NA |
| FAM234B | 6.71546204 | 0.89677676 | 0.64116347 | 1.39867102 | 0.16191166 | NA |
| ATF7IP | 53.5845643 | 0.46472105 | 0.31505441 | 1.47505014 | 0.14019906 | 0.61443563 |
| PLBD1 | 5.60180171 | 0.07918956 | 0.73816244 | 0.10727931 | 0.9145674 | NA |
| PLBD1-AS1 | 2.86396682 | 2.50333068 | 0.96639792 | 2.59037259 | 0.00958721 | NA |
| AC010168.1 | 2.33279808 | 0.30555287 | 0.89097045 | 0.34294388 | 0.73164068 | NA |
| H2AFJ | 18.3938054 | 0.33315145 | 0.42040549 | 0.79245265 | 0.42809679 | NA |
| WBP11 | 11.5678559 | 0.15857848 | 0.53703568 | 0.29528481 | 0.76777632 | NA |
| MGP | 11.7594687 | -1.0340394 | 0.66892209 | -1.5458293 | 0.12214579 | NA |
| ERP27 | 6.39594283 | 0.88702283 | 0.7540589 | 1.17633095 | 0.23946265 | NA |
| RERG | 19.5686039 | 0.60310349 | 0.78808948 | 0.76527286 | 0.44410908 | NA |
| EPS8 | 543.218437 | -0.2795405 | 0.25586675 | -1.0925239 | 0.27460289 | 0.76265881 |
| STRAP | 17.181566 | -0.6179369 | 0.43650309 | -1.4156531 | 0.1568771 | NA |
| DERA | 49.0740503 | 0.33949185 | 0.32726602 | 1.03735747 | 0.29956929 | 0.77521265 |
| MGST1 | 17.61895 | -0.4366784 | 0.57726267 | -0.7564639 | 0.4493711 | NA |
| PIK3C2G | 18.6614766 | 1.15325424 | 0.62738906 | 1.83818036 | 0.06603583 | NA |
| PLEKHA5 | 233.714645 | 0.06888413 | 0.19488761 | 0.35345568 | 0.72374685 | 0.93862018 |
| AEBP2 | 49.9782059 | 0.33670563 | 0.29308494 | 1.14883291 | 0.25062489 | 0.74329941 |
| PYROXD1 | 9.19057245 | -0.1768782 | 0.47750207 | -0.370424 | 0.71106662 | NA |
| RECQL | 8.63078716 | -0.2438678 | 0.52658308 | -0.4631135 | 0.64328298 | NA |
| GOLT1B | 9.91597344 | -0.4720659 | 0.52966355 | -0.8912562 | 0.37279173 | NA |
| LDHB | 62.5668559 | 0.26210789 | 0.40826377 | 0.64200623 | 0.52086914 | 0.87214537 |
| CMAS | 6.72495921 | 0.90461806 | 0.61611161 | 1.46826978 | 0.14203096 | NA |
| C2CD5 | 29.0191548 | 0.54083627 | 0.36878172 | 1.46654847 | 0.14249893 | 0.62145365 |
| ETNK1 | 53.6178272 | -0.2551821 | 0.3082072 | -0.8279564 | 0.40769518 | 0.82808596 |
| SOX5 | 11.6036552 | 0.25707719 | 0.50977198 | 0.50429839 | 0.61405171 | NA |
| ETFRF1 | 7.79798436 | 0.27810767 | 0.626119 | 0.44417701 | 0.65691461 | NA |
| KRAS | 25.8584224 | -0.2680194 | 0.35111883 | -0.7633297 | 0.44526682 | NA |
| RASSF8-AS1 | 5.83636359 | -0.400295 | 0.86093391 | -0.4649544 | 0.64196412 | NA |
| RASSF8 | 36.3498732 | -0.1966795 | 0.30362956 | -0.6477614 | 0.51713931 | 0.87158522 |
| ITPR2 | 122.33643 | 0.09809437 | 0.32793777 | 0.29912494 | 0.76484472 | 0.95550417 |
| INTS13 | 5.07262218 | 0.22119204 | 0.65296675 | 0.33874932 | 0.73479858 | NA |
| FGFR1OP2 | 15.108329 | -0.1937174 | 0.40886246 | -0.473796 | 0.63564536 | NA |
| TM7SF3 | 14.3342633 | -0.0369444 | 0.47961336 | -0.0770294 | 0.93860012 | NA |
| MED21 | 8.85482902 | -0.040779 | 0.48446943 | -0.0841725 | 0.93291927 | NA |
| STK38L | 11.9579871 | 0.21691206 | 0.46469615 | 0.46678256 | 0.64065546 | NA |
| PPFIBP1 | 32.478279 | 0.34900432 | 0.38249158 | 0.91244967 | 0.36153205 | 0.80477819 |
| MRPS35 | 18.4365605 | 0.39459385 | 0.39814379 | 0.99108378 | 0.32164467 | NA |
| KLHL42 | 19.3227956 | 0.62910086 | 0.36644945 | 1.71674666 | 0.08602547 | NA |
| CCDC91 | 83.1984159 | 0.01569919 | 0.23145258 | 0.06782895 | 0.9459218 | 0.99251804 |
| ERGIC2 | 14.0221686 | -0.6090245 | 0.4762684 | -1.2787422 | 0.20098787 | NA |
| IPO8 | 18.4629844 | 0.31473896 | 0.36319327 | 0.8665881 | 0.38616774 | NA |
| CAPRIN2 | 11.5239533 | 0.44899774 | 0.51620765 | 0.86980063 | 0.38440937 | NA |
| DDX11 | 2.8904914 | -0.2951157 | 0.99410198 | -0.2968666 | 0.76656833 | NA |
| SINHCAF | 43.2758009 | 0.22348773 | 0.27071299 | 0.8255523 | 0.4090581 | 0.82839172 |
| DENND5B | 40.1242134 | 0.38163237 | 0.34070115 | 1.12013819 | 0.26265488 | 0.75479335 |
| AMN1 | 8.37545468 | 0.88131099 | 0.5263772 | 1.67429553 | 0.09407253 | NA |
| KIAA1551 | 10.4916484 | 1.36727192 | 0.46398177 | 2.94682248 | 0.00321057 | NA |
| BICD1 | 41.2479817 | 0.36371529 | 0.35009979 | 1.03889033 | 0.29885574 | 0.77521265 |
| FGD4 | 229.295293 | -0.2143004 | 0.21156035 | -1.0129515 | 0.31108333 | 0.778534 |
| DNM1L | 37.3267813 | -0.1776206 | 0.30966812 | -0.5735837 | 0.56624958 | 0.89263108 |
| YARS2 | 9.47944512 | -0.2320739 | 0.52930314 | -0.4384517 | 0.66105888 | NA |

| | | | | | | |
|---|---|---|---|---|---|---|
| PKP2 | 9.10557105 | 0.19101958 | 0.51514349 | 0.37080848 | 0.71078018 | NA |
| SYT10 | 3.73238061 | -1.1518878 | 1.16671452 | -0.9872919 | 0.32349956 | NA |
| ALG10B | 3.46017187 | -0.414237 | 0.88715594 | -0.4669269 | 0.64055217 | NA |
| CPNE8 | 39.9126055 | -0.7198227 | 0.4741532 | -1.5181226 | 0.1289835 | 0.59663251 |
| KIF21A | 53.1777564 | -0.4263058 | 0.43235414 | -0.9860106 | 0.32412791 | 0.78622244 |
| SLC2A13 | 16.1205263 | -0.5547655 | 0.63893511 | -0.8682657 | 0.38524889 | NA |
| LRRK2 | 20.7646063 | 0.23876318 | 0.60369862 | 0.39550062 | 0.69247347 | NA |
| GXYLT1 | 10.4314308 | 0.35377531 | 0.52597432 | 0.67260947 | 0.50119577 | NA |
| YAF2 | 41.3608151 | -0.2331897 | 0.29668471 | -0.7859849 | 0.43187636 | 0.84223493 |
| PPHLN1 | 35.3730421 | -0.0312774 | 0.30195926 | -0.1035816 | 0.9175014 | 0.98583383 |
| ZCRB1 | 8.85354081 | 0.97030752 | 0.61381455 | 1.58078285 | 0.1139277 | NA |
| PRICKLE1 | 14.8874162 | -1.3639583 | 0.69637601 | -1.9586521 | 0.05015354 | NA |
| PUS7L | 8.65438019 | 0.18778928 | 0.52874434 | 0.35516083 | 0.7224691 | NA |
| IRAK4 | 12.3724675 | 1.02898864 | 0.4597464 | 2.23816572 | 0.02521025 | NA |
| TWF1 | 12.5600116 | -0.4641894 | 0.46492285 | -0.9984223 | 0.31807461 | NA |
| TMEM117 | 258.605797 | -0.0236682 | 0.27199076 | -0.0870183 | 0.93065693 | 0.98921679 |
| ANO6 | 27.7129818 | 0.17327106 | 0.32269588 | 0.53694848 | 0.5913032 | 0.89660782 |
| AC008124.1 | 3.67535949 | 0.6939805 | 0.80214679 | 0.86515399 | 0.38695427 | NA |
| ARID2 | 80.0864424 | 0.36412413 | 0.30528208 | 1.19274647 | 0.23296868 | 0.73156941 |
| SCAF11 | 80.5155362 | 0.09831664 | 0.21076405 | 0.46647726 | 0.64087392 | 0.92153493 |
| SLC38A1 | 51.8017899 | -0.7402067 | 0.35906438 | -2.0614874 | 0.03925657 | 0.43110269 |
| SLC38A2 | 35.2794024 | -0.9288497 | 0.32451718 | -2.8622513 | 0.00420643 | 0.21182958 |
| AC008014.1 | 45.0547929 | 1.416623 | 0.42373703 | 3.34316543 | 0.00082829 | 0.11053464 |
| PCED1B | 7.02608559 | -1.5223436 | 0.59926795 | -2.5403387 | 0.01107452 | NA |
| RPAP3 | 10.2409943 | 0.10527548 | 0.559914 | 0.18802081 | 0.85086034 | NA |
| RAPGEF3 | 10.4821454 | 0.23755047 | 0.57426633 | 0.41365907 | 0.67912381 | NA |
| SLC48A1 | 6.02320219 | 1.15089012 | 0.6631969 | 1.73536714 | 0.0826758 | NA |
| HDAC7 | 21.6412385 | -0.7025595 | 0.39431712 | -1.7817118 | 0.07479624 | NA |
| VDR | 24.3382794 | 0.07762501 | 0.37981052 | 0.20437826 | 0.83805792 | NA |
| TMEM106C | 9.90097518 | 0.18695153 | 0.50680978 | 0.36887908 | 0.71221786 | NA |
| SENP1 | 12.5051703 | 0.16921239 | 0.46457751 | 0.36422855 | 0.71568735 | NA |
| PFKM | 6.31045703 | 0.5074848 | 0.62071026 | 0.81758727 | 0.4135929 | NA |
| ASB8 | 7.82943268 | -0.1368833 | 0.57416129 | -0.2384057 | 0.81156644 | NA |
| ZNF641 | 4.34390062 | -0.2386821 | 0.73807988 | -0.3233825 | 0.74640555 | NA |
| KANSL2 | 5.23601389 | -1.2859975 | 0.71296122 | -1.8037411 | 0.0712719 | NA |
| CCNT1 | 14.554552 | -0.5339299 | 0.45946261 | -1.1620747 | 0.24520511 | NA |
| ADCY6 | 6.48923393 | 0.53985272 | 0.74554326 | 0.72410649 | 0.46900036 | NA |
| DDX23 | 6.49794615 | -0.837334 | 0.63688191 | -1.3147398 | 0.18859736 | NA |
| CCDC65 | 4.65947358 | -0.4878122 | 0.907222 | -0.5376988 | 0.59078502 | NA |
| PRKAG1 | 10.1214465 | -0.5739932 | 0.58604258 | -0.9794394 | 0.32736292 | NA |
| KMT2D | 9.28522963 | -0.4675232 | 0.53247028 | -0.8780269 | 0.37992911 | NA |
| LMBR1L | 7.87317755 | -0.2540658 | 0.60572263 | -0.4194424 | 0.67489281 | NA |
| TUBA1B | 25.6249557 | -0.6577443 | 0.46826181 | -1.4046508 | 0.16012513 | NA |
| TUBA1A | 4.32010216 | -1.1372594 | 0.92557129 | -1.2287108 | 0.21918027 | NA |
| TUBA1C | 21.4010206 | -1.2320799 | 0.54429178 | -2.2636386 | 0.02359635 | NA |
| SPATS2 | 17.2274165 | -0.1584457 | 0.44022754 | -0.3599179 | 0.71890857 | NA |
| MCRS1 | 4.62107067 | -1.0294233 | 0.73902273 | -1.3929521 | 0.16363426 | NA |
| PRPF40B | 15.3350809 | -0.1159332 | 0.41486506 | -0.2794481 | 0.77990099 | NA |
| FMNL3 | 3.54208592 | -0.7354179 | 0.82400874 | -0.892488 | 0.37213145 | NA |
| TMBIM6 | 93.6894021 | -0.2478725 | 0.29110747 | -0.8514812 | 0.39450208 | 0.8258317 |
| NCKAP5L | 4.17466335 | -0.1719301 | 0.73776329 | -0.2330424 | 0.81572846 | NA |
| BCDIN3D | 2.84755082 | 0.85479727 | 0.9870413 | 0.86601976 | 0.38647932 | NA |
| AQP2 | 62.7490284 | -1.511971 | 1.15301768 | -1.3113164 | 0.18975089 | 0.68499719 |
| SMARCD1 | 7.56597935 | -0.8220741 | 0.57137328 | -1.4387689 | 0.15021601 | NA |

| | | | | | | |
|---|---|---|---|---|---|---|
| AC074032.1 | 16.96288 | -0.1021223 | 0.49595881 | -0.2059088 | 0.83686219 | NA |
| CERS5 | 13.3498041 | -0.0356858 | 0.51494691 | -0.0693 | 0.94475086 | NA |
| LIMA1 | 57.8876052 | 0.48912497 | 0.2976107 | 1.64350602 | 0.10027828 | 0.57124575 |
| LARP4 | 65.2029972 | 0.13363391 | 0.27656463 | 0.48319234 | 0.62895918 | 0.91751762 |
| DIP2B | 44.582847 | -0.4046295 | 0.28379543 | -1.4257788 | 0.1539322 | 0.63864841 |
| ATF1 | 14.7020516 | -0.9976432 | 0.44302701 | -2.251879 | 0.02432992 | NA |
| METTL7A | 17.4284244 | 0.58980599 | 0.40517423 | 1.45568485 | 0.14547976 | NA |
| SLC11A2 | 30.8257479 | -0.2457182 | 0.34563634 | -0.7109154 | 0.47713667 | 0.86206794 |
| LETMD1 | 17.8184185 | 0.46173013 | 0.38735921 | 1.19199473 | 0.23326331 | NA |
| CSRNP2 | 5.13367659 | -0.3682221 | 0.71321033 | -0.5162882 | 0.60565315 | NA |
| TFCP2 | 38.9634567 | -0.1471684 | 0.34764929 | -0.4233243 | 0.6720587 | 0.92617027 |
| POU6F1 | 8.75297922 | -0.1615444 | 0.56466758 | -0.2860877 | 0.77481099 | NA |
| DAZAP2 | 38.0974661 | -0.1221608 | 0.35679688 | -0.3423819 | 0.73206349 | 0.94354489 |
| ACVR1B | 31.0848524 | -0.0007004 | 0.32702856 | -0.0021416 | 0.99829125 | 0.99960611 |
| NR4A1 | 17.597824 | -1.8741588 | 0.80005591 | -2.3425348 | 0.01915325 | NA |
| ATG101 | 6.07899898 | -0.406914 | 0.63723763 | -0.6385593 | 0.52310964 | NA |
| KRT7 | 288.604698 | 0.62075466 | 0.46204689 | 1.34348843 | 0.17911387 | 0.67032316 |
| KRT8 | 49.4566776 | 0.24911242 | 0.40998339 | 0.60761588 | 0.54344226 | 0.88410237 |
| KRT18 | 42.4549261 | 0.29653109 | 0.38785266 | 0.76454571 | 0.44454211 | 0.84899161 |
| EIF4B | 40.7704343 | -0.0143341 | 0.33901183 | -0.0422821 | 0.96627382 | 0.99656976 |
| AC068888.1 | 8.54748973 | -0.4508706 | 0.58870446 | -0.7658692 | 0.44375413 | NA |
| TNS2 | 11.395631 | 0.96980044 | 0.60882231 | 1.59291213 | 0.1111799 | NA |
| SPRYD3 | 4.22213278 | -0.7535112 | 0.76432638 | -0.98585 | 0.32420672 | NA |
| CSAD | 15.1829706 | 0.61528344 | 0.50202025 | 1.22561479 | 0.22034367 | NA |
| ZNF740 | 2.33111843 | 0.69993568 | 0.9362452 | 0.74759868 | 0.45470226 | NA |
| MFSD5 | 3.92196883 | -1.5689847 | 0.81715988 | -1.9200462 | 0.05485207 | NA |
| PFDN5 | 31.7648451 | 0.42918234 | 0.42209216 | 1.01679769 | 0.30924968 | 0.778534 |
| C12orf10 | 3.53812974 | 0.66857052 | 0.78322508 | 0.85361225 | 0.39331987 | NA |
| AAAS | 2.43848267 | 0.91356003 | 1.05142321 | 0.86887946 | 0.38491307 | NA |
| SP1 | 18.7820204 | 0.04407673 | 0.38389593 | 0.11481427 | 0.90859234 | NA |
| PRR13 | 22.0063835 | 0.19671584 | 0.32140738 | 0.61204518 | 0.54050787 | NA |
| PCBP2 | 62.0835647 | -0.4002398 | 0.28771247 | -1.3911102 | 0.16419201 | 0.64995633 |
| MAP3K12 | 4.67319911 | 1.41829934 | 0.65902941 | 2.15210326 | 0.03138922 | NA |
| ATP5MC2 | 45.85568 | 0.30144084 | 0.43604535 | 0.69130617 | 0.48937316 | 0.86384719 |
| CALCOCO1 | 8.33962057 | 0.4250286 | 0.61229817 | 0.69415297 | 0.48758628 | NA |
| SMUG1 | 4.82304057 | 0.59561838 | 0.65383061 | 0.91096741 | 0.36231254 | NA |
| LINC02381 | 6.79035261 | 0.07798236 | 0.66530171 | 0.11721353 | 0.90669085 | NA |
| CBX5 | 18.1883089 | -0.1273253 | 0.54550449 | -0.2334083 | 0.81544435 | NA |
| HNRNPA1 | 22.9865549 | -0.2853048 | 0.4164201 | -0.6851369 | 0.49325753 | NA |
| COPZ1 | 14.8426096 | 0.04121821 | 0.39283965 | 0.10492375 | 0.91643631 | NA |
| PPP1R1A | 4.93529987 | -0.9796739 | 0.67646669 | -1.448222 | 0.14755497 | NA |
| CD63 | 41.6051847 | -0.0211186 | 0.42158483 | -0.0500933 | 0.96004807 | 0.99656976 |
| ORMDL2 | 6.08629587 | 0.04966922 | 0.64204233 | 0.07736129 | 0.93833614 | NA |
| PYM1 | 3.87513409 | -1.7465791 | 0.80152237 | -2.1790772 | 0.02932593 | NA |
| RAB5B | 15.6660643 | 0.12370759 | 0.40237128 | 0.30744637 | 0.75850364 | NA |
| SUOX | 4.21511228 | 0.40854134 | 0.73541986 | 0.55552123 | 0.57853819 | NA |
| IKZF4 | 2.70144531 | -1.0044612 | 0.86247899 | -1.164621 | 0.24417243 | NA |
| RPS26 | 7.78680646 | -0.1114878 | 0.5441302 | -0.2048917 | 0.8376567 | NA |
| ERBB3 | 13.2633637 | 1.25739192 | 0.51965314 | 2.41967541 | 0.01553437 | NA |
| PA2G4 | 7.05142566 | -0.5786104 | 0.62209425 | -0.9301009 | 0.35231886 | NA |
| RPL41 | 143.600885 | 0.22072618 | 0.34486065 | 0.64004456 | 0.52214363 | 0.87311773 |
| ESYT1 | 5.85040246 | -0.1992308 | 0.59637495 | -0.3340697 | 0.738327 | NA |
| MYL6B | 4.5131951 | 0.11460044 | 0.69695485 | 0.16443022 | 0.86939247 | NA |
| MYL6 | 65.6686955 | -0.4092905 | 0.36945097 | -1.1078344 | 0.26793333 | 0.75593451 |

| | | | | | | |
|---|---|---|---|---|---|---|
| SMARCC2 | 16.6023668 | 0.59783965 | 0.4157427 | 1.43800396 | 0.15043293 | NA |
| RNF41 | 4.26847555 | -0.637101 | 0.69598079 | -0.9154002 | 0.35998156 | NA |
| NABP2 | 1.78724355 | -0.4275041 | 1.15348906 | -0.3706183 | 0.71092186 | NA |
| ANKRD52 | 6.25759113 | 0.54232236 | 0.58999351 | 0.91920055 | 0.35799068 | NA |
| COQ10A | 2.50953901 | 0.20089971 | 1.30703557 | 0.15370638 | 0.87784125 | NA |
| CS | 14.2487915 | -0.3878429 | 0.43028495 | -0.9013629 | 0.3673954 | NA |
| AC073896.2 | 3.67610871 | 0.22651021 | 0.91106593 | 0.24862109 | 0.80365389 | NA |
| PAN2 | 7.3426853 | 0.80274059 | 0.70089647 | 1.14530551 | 0.25208262 | NA |
| STAT2 | 12.7901892 | -0.4775799 | 0.50139119 | -0.9525095 | 0.34083866 | NA |
| TIMELESS | 2.49587771 | -0.5131779 | 1.02818728 | -0.4991093 | 0.61770236 | NA |
| SPRYD4 | 3.50715566 | 0.65853977 | 0.76998264 | 0.85526574 | 0.39240405 | NA |
| RBMS2 | 14.1194244 | -0.0015751 | 0.47228496 | -0.0033351 | 0.99733894 | NA |
| BAZ2A | 18.9677339 | -0.4390643 | 0.37234251 | -1.1791948 | 0.23832062 | NA |
| ATP5F1B | 69.2431586 | 0.36529241 | 0.38620027 | 0.94586265 | 0.34421865 | 0.8003972 |
| PTGES3 | 54.9462399 | -0.2315487 | 0.28774423 | -0.8047033 | 0.42099091 | 0.83977933 |
| NACA | 35.6286807 | 0.04664376 | 0.35032999 | 0.13314234 | 0.89408081 | 0.97769824 |
| NEMP1 | 3.63154811 | -0.637458 | 0.7952055 | -0.8016268 | 0.42276887 | NA |
| STAT6 | 29.3288713 | 0.34614838 | 0.32315716 | 1.07114564 | 0.28410395 | 0.76977033 |
| SHMT2 | 3.04192586 | 0.26785073 | 0.773688 | 0.34619993 | 0.72919247 | NA |
| R3HDM2 | 66.7448478 | -0.1967693 | 0.24819696 | -0.7927948 | 0.42789736 | 0.84223493 |
| MARS | 8.18785923 | -1.6920379 | 0.57660316 | -2.9344929 | 0.00334093 | NA |
| DDIT3 | 12.2752084 | -0.6545311 | 0.89191617 | -0.7338482 | 0.46304123 | NA |
| MBD6 | 7.81188059 | -0.2305067 | 0.63415306 | -0.3634875 | 0.71624076 | NA |
| DCTN2 | 16.0158972 | 0.21595497 | 0.37483349 | 0.57613575 | 0.56452343 | NA |
| PIP4K2C | 9.93069417 | -0.4066646 | 0.47211688 | -0.8613643 | 0.38903745 | NA |
| DTX3 | 2.4366263 | -1.2299306 | 1.00184626 | -1.227664 | 0.21957313 | NA |
| OS9 | 22.285791 | 0.37835977 | 0.38488441 | 0.98304779 | 0.32558392 | NA |
| TSPAN31 | 5.12405667 | -0.5958927 | 0.73418689 | -0.8116363 | 0.41700037 | NA |
| CDK4 | 5.632804 | -1.1081671 | 0.66116139 | -1.6760917 | 0.09372024 | NA |
| 9-Mar | 5.50522663 | 0.04363144 | 0.68940814 | 0.06328825 | 0.94953697 | NA |
| TSFM | 6.88060453 | 0.27749111 | 0.59407292 | 0.46709941 | 0.64042875 | NA |
| AVIL | 13.6792721 | 0.72904626 | 0.60853942 | 1.19802635 | 0.23090676 | NA |
| CTDSP2 | 21.8700462 | 0.34836948 | 0.40874475 | 0.85229102 | 0.39405259 | NA |
| LRIG3 | 7.56257133 | -0.2139803 | 0.76319053 | -0.2803759 | 0.77918909 | NA |
| SLC16A7 | 24.3966873 | 1.14867052 | 0.46776984 | 2.45563184 | 0.01406372 | NA |
| FAM19A2 | 8.1431922 | 0.97767974 | 0.67626865 | 1.44569726 | 0.14826213 | NA |
| USP15 | 51.8488629 | -0.0469814 | 0.27922066 | -0.1682591 | 0.86637946 | 0.97217819 |
| MON2 | 86.3421176 | -0.0957421 | 0.25457118 | -0.3760916 | 0.70684881 | 0.93164418 |
| AC048341.1 | 5.5383813 | -0.8683936 | 0.64983437 | -1.3363306 | 0.18144124 | NA |
| PPM1H | 28.9816517 | -0.9344611 | 0.43802457 | -2.1333531 | 0.03289578 | 0.41688714 |
| AVPR1A | 52.3741926 | 1.15101786 | 0.30034493 | 3.83231992 | 0.00012694 | 0.04332663 |
| DPY19L2 | 110.009158 | 0.44141067 | 0.47965565 | 0.92026576 | 0.3574339 | 0.80238744 |
| RXYLT1 | 9.41719055 | 0.75396327 | 0.47864282 | 1.57521064 | 0.11520783 | NA |
| SRGAP1 | 29.714577 | -0.4998746 | 0.4942371 | -1.0114064 | 0.31182197 | 0.778534 |
| XPOT | 6.06685585 | -0.2182002 | 0.64595712 | -0.3377936 | 0.73551872 | NA |
| TBK1 | 25.8184514 | -0.3495883 | 0.33248286 | -1.0514477 | 0.29305303 | NA |
| RASSF3 | 4.24023614 | 0.59300258 | 0.68365938 | 0.86739479 | 0.38572573 | NA |
| GNS | 29.0987704 | -0.1912402 | 0.38800618 | -0.4928793 | 0.62209787 | 0.91493825 |
| TBC1D30 | 10.2862091 | -0.2091275 | 0.51960231 | -0.402476 | 0.68733374 | NA |
| LEMD3 | 30.5565029 | 0.42200108 | 0.3746496 | 1.12638871 | 0.26000102 | 0.75479335 |
| LLPH | 4.98770057 | -0.1051218 | 0.87882858 | -0.1196158 | 0.90478748 | NA |
| HELB | 9.63103433 | 0.58427975 | 0.61594639 | 0.94858865 | 0.34282987 | NA |
| GRIP1 | 47.5496102 | 0.3542241 | 0.57299349 | 0.61819916 | 0.53644407 | 0.88254911 |
| CAND1 | 30.3713059 | -0.0740427 | 0.35761561 | -0.2070454 | 0.83597441 | 0.97154892 |

| | | | | | | |
|---|---|---|---|---|---|---|
| DYRK2 | 2.05444739 | 1.10543497 | 0.99805263 | 1.10759186 | 0.26803812 | NA |
| MDM1 | 4.201345 | 0.07735051 | 0.69615996 | 0.11111025 | 0.91152892 | NA |
| RAP1B | 52.7178071 | -0.5319606 | 0.29708388 | -1.7906074 | 0.07335631 | 0.53207981 |
| NUP107 | 14.4974165 | 0.57553008 | 0.47285867 | 1.21712916 | 0.22355508 | NA |
| SLC35E3 | 8.76791171 | -0.7520086 | 0.55207177 | -1.3621574 | 0.17314822 | NA |
| MDM2 | 16.348 | -0.7367089 | 0.44449782 | -1.657396 | 0.09743944 | NA |
| CPM | 9.99973017 | -0.3931352 | 0.82736957 | -0.4751628 | 0.63467095 | NA |
| CPSF6 | 20.6369853 | 0.00252713 | 0.41974541 | 0.00602062 | 0.99519627 | NA |
| YEATS4 | 3.46798615 | 0.23306909 | 0.90940247 | 0.25628817 | 0.79772833 | NA |
| FRS2 | 28.5678654 | -0.1696836 | 0.32967071 | -0.5147064 | 0.60675819 | 0.90826562 |
| CCT2 | 12.8364221 | 0.19374599 | 0.47515451 | 0.40775366 | 0.68345454 | NA |
| RAB3IP | 25.1020958 | 0.25487092 | 0.39972423 | 0.63761689 | 0.5237231 | NA |
| MYRFL | 5.22761864 | 1.79678605 | 0.97055725 | 1.85129322 | 0.06412738 | NA |
| AC025159.1 | 5.21915137 | 0.89100834 | 1.05509282 | 0.84448338 | 0.39839935 | NA |
| CNOT2 | 50.900375 | -0.2055706 | 0.26014457 | -0.7902166 | 0.42940129 | 0.84223493 |
| TSPAN8 | 64.8735935 | -0.3753963 | 0.41469589 | -0.9052327 | 0.36534212 | 0.80921006 |
| ZFC3H1 | 41.4419759 | -0.378725 | 0.3910357 | -0.9685178 | 0.33278585 | 0.79447068 |
| TMEM19 | 6.06537622 | 0.58457525 | 0.60101211 | 0.97265136 | 0.33072661 | NA |
| RAB21 | 34.7349538 | -0.1738144 | 0.30845853 | -0.5634935 | 0.57309888 | 0.89263108 |
| TBC1D15 | 27.4367722 | -0.0412834 | 0.39402683 | -0.1047731 | 0.91655589 | 0.98528045 |
| TRHDE | 105.240422 | 0.80294437 | 0.28547975 | 2.81261407 | 0.00491406 | 0.22229871 |
| AC090502.1 | 6.35785004 | 1.11964027 | 0.7462487 | 1.50035808 | 0.13352167 | NA |
| ATXN7L3B | 12.1339701 | -0.1062828 | 0.44432929 | -0.2391982 | 0.81095188 | NA |
| CAPS2 | 16.1754495 | 0.74770278 | 0.4923534 | 1.51863028 | 0.12885558 | NA |
| KRR1 | 10.6610043 | 0.07973822 | 0.47017705 | 0.16959189 | 0.8653311 | NA |
| PHLDA1 | 14.313171 | -0.6697819 | 0.60301677 | -1.1107185 | 0.26668953 | NA |
| NAP1L1 | 30.5201085 | -0.2637913 | 0.43760541 | -0.6028063 | 0.54663755 | 0.88410237 |
| OSBPL8 | 52.274961 | -0.2365306 | 0.29201327 | -0.8099994 | 0.41794055 | 0.83807913 |
| ZDHHC17 | 24.8788535 | 0.15264976 | 0.39260871 | 0.3888089 | 0.69741752 | NA |
| AC027288.3 | 4.56993566 | -2.1982339 | 1.00364021 | -2.1902609 | 0.02850532 | NA |
| PAWR | 50.3815273 | 0.18860213 | 0.27153439 | 0.69457918 | 0.48731906 | 0.86354292 |
| AC073569.2 | 5.21591144 | 2.02036748 | 0.87144491 | 2.31841101 | 0.02042699 | NA |
| PPP1R12A | 58.8854594 | -0.1349641 | 0.29322224 | -0.4602791 | 0.64531591 | 0.92153493 |
| ACSS3 | 17.3709768 | 1.11862832 | 0.5467067 | 2.04612147 | 0.04074442 | NA |
| CCDC59 | 4.04254373 | 0.01106943 | 0.82132067 | 0.0134776 | 0.98924676 | NA |
| METTL25 | 29.4341588 | -0.0090784 | 0.3237784 | -0.0280388 | 0.97763122 | 0.99902646 |
| TMTC2 | 45.7036272 | 0.27475684 | 0.47857021 | 0.57412024 | 0.56588644 | 0.89263108 |
| C12orf29 | 2.23222205 | -0.3279211 | 0.97924283 | -0.3348721 | 0.73772154 | NA |
| CEP290 | 17.771431 | 0.28890093 | 0.37963636 | 0.76099382 | 0.44666076 | NA |
| TMTC3 | 22.9511305 | 0.16196917 | 0.37226421 | 0.43509198 | 0.66349566 | NA |
| KITLG | 16.9747003 | 0.2895119 | 0.90843513 | 0.31869298 | 0.74995934 | NA |
| POC1B | 13.8603399 | 0.08863271 | 0.5198013 | 0.17051267 | 0.86460697 | NA |
| POC1B-AS1 | 2.5773461 | 0.48749861 | 1.05832372 | 0.46063279 | 0.64506208 | NA |
| ATP2B1 | 44.9498507 | -1.1627319 | 0.49546446 | -2.3467514 | 0.01893789 | 0.33831016 |
| BTG1 | 99.3417196 | 0.5623775 | 0.27012313 | 2.0819302 | 0.03734885 | 0.42032585 |
| AC025164.1 | 2.82439282 | 1.38589441 | 0.90618165 | 1.52937814 | 0.12617073 | NA |
| EEA1 | 50.5374829 | -0.5542561 | 0.32423311 | -1.7094369 | 0.08737004 | 0.56122412 |
| AC138123.1 | 10.9619408 | -0.5649421 | 0.76741093 | -0.7361664 | 0.46162945 | NA |
| NUDT4 | 39.1492804 | 0.05706914 | 0.35901984 | 0.15895818 | 0.87370183 | 0.9751578 |
| UBE2N | 20.8284188 | -0.8809797 | 0.39310172 | -2.2410986 | 0.02501969 | NA |
| MRPL42 | 14.3698543 | -0.1297676 | 0.39078744 | -0.332067 | 0.7398387 | NA |
| CRADD | 86.9148188 | 0.50856825 | 0.30280787 | 1.67950803 | 0.09305308 | 0.56220784 |
| AC012085.2 | 8.34954275 | 0.81127627 | 0.7953529 | 1.02002051 | 0.30771873 | NA |
| CEP83 | 13.1563247 | 0.16207501 | 0.46263112 | 0.35033313 | 0.72608871 | NA |

| | | | | | | |
|---|---|---|---|---|---|---|
| TMCC3 | 8.63235916 | -0.0814481 | 0.68731746 | -0.1185015 | 0.90567034 | NA |
| NDUFA12 | 15.3570944 | -0.0334326 | 0.48200919 | -0.0693609 | 0.94470232 | NA |
| NR2C1 | 17.2305171 | -0.0285138 | 0.46255046 | -0.0616446 | 0.95084584 | NA |
| FGD6 | 8.93463077 | -0.7978247 | 0.6797506 | -1.173702 | 0.2405144 | NA |
| VEZT | 38.1393158 | -0.0627299 | 0.28150756 | -0.2228356 | 0.82366343 | 0.96845695 |
| METAP2 | 23.7200174 | 0.06076472 | 0.3772267 | 0.16108277 | 0.87202821 | NA |
| NTN4 | 50.9393765 | -0.3152044 | 0.51150817 | -0.6162255 | 0.53774567 | 0.88254911 |
| SNRPF | 6.15007984 | -0.0552675 | 0.61533361 | -0.0898171 | 0.92843253 | NA |
| LTA4H | 9.62488503 | 0.22962955 | 0.58881683 | 0.3899847 | 0.69654786 | NA |
| CDK17 | 22.4166329 | -0.0433805 | 0.42791259 | -0.101377 | 0.91925116 | NA |
| NEDD1 | 8.56341494 | -0.4435941 | 0.50222731 | -0.8832537 | 0.37709922 | NA |
| TMPO | 22.5788767 | 0.5706119 | 0.40959449 | 1.39311422 | 0.16358525 | NA |
| SLC25A3 | 57.1271746 | 0.38945286 | 0.36726291 | 1.06041979 | 0.28895366 | 0.7725847 |
| APAF1 | 7.92509735 | -0.4218533 | 0.55836091 | -0.7555208 | 0.44993655 | NA |
| ANKS1B | 5.1142502 | 0.77796556 | 0.87475266 | 0.88935489 | 0.37381238 | NA |
| UHRF1BP1L | 20.0986314 | -0.3685138 | 0.40934241 | -0.9002582 | 0.36798287 | NA |
| ACTR6 | 7.10132858 | 0.51937605 | 0.58386533 | 0.88954767 | 0.37370881 | NA |
| SCYL2 | 26.4739536 | -0.0458469 | 0.34652241 | -0.1323056 | 0.89474254 | 0.97769824 |
| NR1H4 | 8.37885569 | 1.0388766 | 0.80939918 | 1.28351576 | 0.19931144 | NA |
| ANO4 | 18.395731 | -0.7170872 | 1.61861291 | -0.4430258 | 0.65774709 | NA |
| UTP20 | 16.9322231 | -0.1160389 | 0.4427441 | -0.2620902 | 0.79325189 | NA |
| ARL1 | 16.8483926 | 0.05945924 | 0.3694739 | 0.16092948 | 0.87214894 | NA |
| CHPT1 | 35.5230475 | 0.34575868 | 0.3855066 | 0.89689434 | 0.3697753 | 0.81199861 |
| GNPTAB | 26.4261594 | 0.56167677 | 0.32589913 | 1.72346815 | 0.0848039 | 0.55323508 |
| DRAM1 | 21.0504964 | -0.2526473 | 0.39990245 | -0.6317724 | 0.52753563 | NA |
| WASHC3 | 18.3384959 | 0.75236135 | 0.52850071 | 1.4235768 | 0.15456902 | NA |
| NUP37 | 5.51886327 | 0.99195188 | 0.66763689 | 1.48576555 | 0.13734115 | NA |
| HSP90B1 | 49.6868655 | -0.0850261 | 0.44366804 | -0.1916436 | 0.84802139 | 0.97168963 |
| C12orf73 | 3.07358506 | 1.51183662 | 0.98977844 | 1.52744955 | 0.12664926 | NA |
| TDG | 11.3582661 | -0.7548611 | 0.5540116 | -1.3625366 | 0.17302859 | NA |
| HCFC2 | 10.9560964 | 0.1504471 | 0.46331286 | 0.32472031 | 0.74539276 | NA |
| NFYB | 5.74995073 | 0.90066786 | 0.71090475 | 1.26693184 | 0.20517966 | NA |
| TXNRD1 | 89.6685967 | -0.1440441 | 0.39990034 | -0.3602 | 0.71869758 | 0.93714576 |
| SLC41A2 | 6.87720244 | -0.8565749 | 0.62446593 | -1.371692 | 0.17015934 | NA |
| C12orf45 | 3.74242787 | -0.6189941 | 0.72341459 | -0.8556561 | 0.39218806 | NA |
| WASHC4 | 42.6748122 | -0.0427119 | 0.28843592 | -0.1480812 | 0.88227872 | 0.97769824 |
| APPL2 | 19.0450489 | 0.00750175 | 0.39062471 | 0.01920449 | 0.98467797 | NA |
| C12orf75 | 44.296052 | -0.4601297 | 0.36475935 | -1.2614611 | 0.20714278 | 0.70428546 |
| CKAP4 | 3.79545781 | -0.1478737 | 0.74049793 | -0.199695 | 0.84171912 | NA |
| TCP11L2 | 7.74483407 | -0.5328334 | 0.67631171 | -0.7878518 | 0.43078342 | NA |
| POLR3B | 9.86585615 | -0.7403149 | 0.61758416 | -1.1987272 | 0.23063404 | NA |
| RIC8B | 15.2044796 | 0.00258179 | 0.50256953 | 0.00513718 | 0.99590114 | NA |
| TMEM263 | 13.1414351 | -0.3294187 | 0.41203105 | -0.7994996 | 0.42400078 | NA |
| MTERF2 | 3.63712329 | 0.6275337 | 0.82162682 | 0.76376974 | 0.44500447 | NA |
| CRY1 | 15.0058385 | -0.7837392 | 0.53860708 | -1.4551223 | 0.14563541 | NA |
| BTBD11 | 15.1005185 | -1.4082208 | 0.63305301 | -2.224491 | 0.02611542 | NA |
| PWP1 | 8.82911556 | -0.1547152 | 0.48333547 | -0.320099 | 0.74889329 | NA |
| PRDM4 | 34.9209062 | 0.03364936 | 0.44810511 | 0.07509256 | 0.94014107 | 0.99100968 |
| AC126177.4 | 3.95021784 | -1.6899541 | 0.8869582 | -1.9053368 | 0.05673631 | NA |
| SART3 | 10.1432661 | -0.3304762 | 0.48343158 | -0.683605 | 0.49422462 | NA |
| ISCU | 31.1875274 | 0.62693808 | 0.33346901 | 1.88004899 | 0.0601014 | 0.51625203 |
| CORO1C | 6.76984503 | -0.9331451 | 0.52668733 | -1.7717249 | 0.07644024 | NA |
| SSH1 | 16.7150684 | -0.1917048 | 0.3700664 | -0.5180281 | 0.60443867 | NA |
| USP30 | 5.45164222 | -0.0725317 | 0.71728938 | -0.1011192 | 0.91945586 | NA |

| | | | | | | |
|---|---|---|---|---|---|---|
| UNG | 4.65766744 | 0.62718183 | 0.80765619 | 0.77654556 | 0.43742693 | NA |
| ACACB | 33.6884177 | -0.075964 | 0.40929146 | -0.1855987 | 0.85275945 | 0.97168963 |
| KCTD10 | 6.73837344 | 0.10175576 | 0.68275126 | 0.14903782 | 0.88152379 | NA |
| UBE3B | 8.40073812 | 0.90428742 | 0.59318365 | 1.52446452 | 0.12739271 | NA |
| MMAB | 7.6611948 | -0.5308888 | 0.59839231 | -0.8871919 | 0.3749756 | NA |
| MVK | 3.17591975 | -0.8039234 | 0.80456801 | -0.9991988 | 0.31769839 | NA |
| GLTP | 13.9251364 | 0.45348432 | 0.46993734 | 0.96498889 | 0.33455038 | NA |
| TCHP | 8.62797311 | 0.42890223 | 0.53609817 | 0.8000442 | 0.42368519 | NA |
| GIT2 | 18.1604606 | -0.5030402 | 0.36960341 | -1.361027 | 0.17350515 | NA |
| ANKRD13A | 36.5823746 | -0.4644488 | 0.3527589 | -1.3166183 | 0.1879666 | 0.68214045 |
| C12orf76 | 5.62713233 | -0.1348394 | 0.63941108 | -0.2108807 | 0.83298039 | NA |
| IFT81 | 7.17744411 | -0.276168 | 0.54201686 | -0.5095192 | 0.61038834 | NA |
| ATP2A2 | 62.6981079 | -0.0137807 | 0.33389994 | -0.041272 | 0.96707904 | 0.99656976 |
| ANAPC7 | 12.2815288 | -0.0101525 | 0.45619019 | -0.0222549 | 0.98224465 | NA |
| ARPC3 | 41.6512138 | 0.32312506 | 0.27282356 | 1.18437376 | 0.23626514 | 0.7323945 |
| GPN3 | 3.33498182 | 0.07606722 | 0.82690575 | 0.0919902 | 0.92670583 | NA |
| VPS29 | 18.7372131 | 0.1857733 | 0.36918053 | 0.5032045 | 0.6148205 | NA |
| PPTC7 | 28.612661 | -0.4614178 | 0.39117119 | -1.1795803 | 0.23816719 | 0.73242571 |
| TCTN1 | 6.10412935 | 0.90582799 | 0.71065769 | 1.27463335 | 0.20243905 | NA |
| PPP1CC | 15.1191672 | 0.02378792 | 0.46362953 | 0.05130802 | 0.95908007 | NA |
| SH2B3 | 2.72044928 | 1.07278728 | 0.82599835 | 1.29877653 | 0.19402063 | NA |
| ATXN2 | 78.0840055 | -0.3782804 | 0.26223404 | -1.4425297 | 0.149153 | 0.63 |
| BRAP | 10.51848 | 0.34364905 | 0.5246396 | 0.65501927 | 0.51245535 | NA |
| ACAD10 | 11.8776681 | -0.2336265 | 0.500713 | -0.4665876 | 0.64079496 | NA |
| ALDH2 | 18.2867752 | 0.75157445 | 0.47218974 | 1.5916789 | 0.11145687 | NA |
| MAPKAPK5-A | 2.94658439 | 0.29579693 | 0.81242049 | 0.36409339 | 0.71578827 | NA |
| MAPKAPK5 | 14.2037417 | -0.1327067 | 0.48980939 | -0.2709354 | 0.78644069 | NA |
| TMEM116 | 59.465816 | -0.1879955 | 0.35174272 | -0.5344687 | 0.59301732 | 0.89685273 |
| ERP29 | 33.9857438 | 0.49907723 | 0.36378009 | 1.37192012 | 0.17008831 | 0.66175756 |
| NAA25 | 18.6584896 | -0.4829537 | 0.38125462 | -1.2667486 | 0.20524521 | NA |
| TRAFD1 | 7.29358057 | 0.18922367 | 0.62721167 | 0.3016903 | 0.76288817 | NA |
| HECTD4 | 61.6256266 | -0.0595412 | 0.29071255 | -0.2048114 | 0.83771949 | 0.97154892 |
| RPL6 | 78.5350165 | 0.46243717 | 0.40379667 | 1.14522284 | 0.25211686 | 0.74518262 |
| PTPN11 | 24.0301818 | -0.5615638 | 0.36484107 | -1.5392012 | 0.12375518 | NA |
| OAS1 | 6.60880611 | 0.81862851 | 1.65694152 | 0.49405999 | 0.62126381 | NA |
| DDX54 | 9.89642282 | -0.3418457 | 0.50785278 | -0.6731198 | 0.50087111 | NA |
| TPCN1 | 26.6893814 | 0.52238669 | 0.44173385 | 1.18258243 | 0.23697468 | 0.73242571 |
| SLC8B1 | 11.3676816 | 0.45310566 | 0.44463266 | 1.01905618 | 0.3081763 | NA |
| PLBD2 | 4.04144561 | -1.0021643 | 0.78328921 | -1.2794308 | 0.2007454 | NA |
| RBM19 | 11.5469266 | -0.6382274 | 0.55010453 | -1.160193 | 0.24597024 | NA |
| TBX3 | 10.1799956 | 0.31606608 | 0.62282828 | 0.50746906 | 0.61182574 | NA |
| MED13L | 217.61346 | -0.1628871 | 0.22667229 | -0.7186016 | 0.47238643 | 0.86206794 |
| MAP1LC3B2 | 2.96131895 | 0.20979525 | 0.79090658 | 0.26525921 | 0.79080979 | NA |
| C12orf49 | 8.42551244 | -0.4549073 | 0.58399968 | -0.7789512 | 0.43600845 | NA |
| FBXW8 | 16.7914997 | -0.3426102 | 0.42122888 | -0.8133588 | 0.4160124 | NA |
| FBXO21 | 39.8938937 | 0.20216325 | 0.31280213 | 0.64629755 | 0.51808667 | 0.87186213 |
| NOS1 | 6.03339232 | 2.90991966 | 1.11360883 | 2.61305368 | 0.00897372 | NA |
| KSR2 | 13.497063 | -2.0295923 | 0.94582629 | -2.1458404 | 0.03188571 | NA |
| WSB2 | 10.1994777 | 0.08629504 | 0.52958951 | 0.16294703 | 0.87056014 | NA |
| VSIG10 | 10.9030231 | -0.393195 | 0.8076539 | -0.4868359 | 0.62637459 | NA |
| PEBP1 | 105.736098 | 0.28098256 | 0.36034767 | 0.77975407 | 0.43553565 | 0.84282814 |
| TAOK3 | 55.2841215 | 0.49081732 | 0.2617048 | 1.87546168 | 0.06072924 | 0.51625203 |
| SUDS3 | 11.9889427 | 0.30737609 | 0.43336451 | 0.7092784 | 0.47815173 | NA |
| AC002070.1 | 8.67590726 | 0.36650774 | 0.70184007 | 0.52220976 | 0.60152429 | NA |

| | | | | | | |
|---|---|---|---|---|---|---|
| PRKAB1 | 15.1513055 | 0.10111358 | 0.38441221 | 0.26303426 | 0.79252417 | NA |
| CIT | 14.6485617 | -0.1200657 | 0.73338463 | -0.1637146 | 0.86995585 | NA |
| BICDL1 | 44.5559129 | -0.6972776 | 0.49394801 | -1.4116417 | 0.15805548 | 0.64166827 |
| RAB35 | 9.00238236 | -0.708762 | 0.54812628 | -1.2930633 | 0.19598917 | NA |
| GCN1 | 13.2568634 | -0.4111697 | 0.43853164 | -0.9376054 | 0.34844721 | NA |
| RPLP0 | 89.7095543 | 0.17947018 | 0.43842452 | 0.40935251 | 0.68228098 | 0.9267216 |
| PXN | 27.7184429 | 0.53123125 | 0.31962142 | 1.66206397 | 0.09649995 | 0.56358504 |
| GATC | 5.95068588 | 0.76069399 | 0.62786751 | 1.21155176 | 0.22568401 | NA |
| SRSF9 | 18.2589076 | -0.3087929 | 0.48444965 | -0.6374097 | 0.52385799 | NA |
| DYNLL1 | 29.0850703 | 0.14589237 | 0.3815484 | 0.38236923 | 0.70218751 | 0.9303056 |
| COQ5 | 6.06324374 | -0.0560687 | 0.63014409 | -0.0889776 | 0.92909972 | NA |
| RNF10 | 31.980025 | 0.19892877 | 0.35014853 | 0.5681268 | 0.56994887 | 0.89263108 |
| POP5 | 8.0063257 | 0.69976785 | 0.51758957 | 1.35197439 | 0.17638351 | NA |
| MLEC | 21.2902842 | 0.24508553 | 0.33739277 | 0.72641015 | 0.46758737 | NA |
| UNC119B | 5.26991844 | 0.93200136 | 0.73141482 | 1.27424457 | 0.20257675 | NA |
| ACADS | 6.77696706 | 0.66658217 | 0.60872444 | 1.09504748 | 0.27349584 | NA |
| SPPL3 | 74.0146461 | -0.5673479 | 0.24704382 | -2.2965478 | 0.02164458 | 0.35108799 |
| C12orf43 | 3.10636606 | -1.4101078 | 0.92368232 | -1.5266156 | 0.12685663 | NA |
| P2RX4 | 9.05023799 | 0.76019719 | 0.50007187 | 1.52017587 | 0.12846678 | NA |
| CAMKK2 | 5.20170902 | 0.20323242 | 0.67917854 | 0.29923268 | 0.76476251 | NA |
| ANAPC5 | 23.7763486 | 0.25093471 | 0.38727198 | 0.64795472 | 0.51701423 | NA |
| RNF34 | 13.1717787 | -0.015806 | 0.46966368 | -0.033654 | 0.9731531 | NA |
| KDM2B | 13.6127311 | 0.30104895 | 0.44393947 | 0.67813061 | 0.49768888 | NA |
| ORAI1 | 3.88666312 | -0.3566123 | 0.7477788 | -0.4768955 | 0.63343654 | NA |
| TMEM120B | 12.9228257 | 0.28809803 | 0.46080916 | 0.62520031 | 0.5318396 | NA |
| RHOF | 3.50617723 | -0.3770524 | 0.68697689 | -0.5488575 | 0.58310323 | NA |
| SETD1B | 9.00983402 | 0.23733194 | 0.50822369 | 0.46698322 | 0.64051188 | NA |
| BCL7A | 14.1588342 | 0.88259996 | 0.50977414 | 1.73135489 | 0.08338849 | NA |
| MLXIP | 19.7407004 | -0.0508471 | 0.39619527 | -0.1283384 | 0.89788118 | NA |
| CLIP1 | 62.3159855 | -0.339785 | 0.31324099 | -1.0847399 | 0.27803688 | 0.76661203 |
| ZCCHC8 | 15.7902961 | 0.14133628 | 0.39840737 | 0.35475319 | 0.7227745 | NA |
| RSRC2 | 25.9808543 | -0.3401314 | 0.36716063 | -0.9263832 | 0.35424687 | 0.8003972 |
| KNTC1 | 5.60486179 | 0.9535337 | 0.79145851 | 1.20478039 | 0.22828809 | NA |
| DENR | 27.8809324 | -0.1037438 | 0.35897231 | -0.2890022 | 0.77257969 | 0.95994327 |
| HIP1R | 23.0252338 | 0.27430526 | 0.33806895 | 0.81138851 | 0.4171426 | NA |
| VPS37B | 28.5815213 | -0.3593856 | 0.343088 | -1.0475027 | 0.29486778 | 0.77460836 |
| ABCB9 | 3.79632213 | 1.91218502 | 0.87189807 | 2.19312909 | 0.02829808 | NA |
| ARL6IP4 | 22.5212022 | 0.23608992 | 0.36131029 | 0.65342706 | 0.513481 | NA |
| PITPNM2 | 15.7119183 | 0.32574492 | 0.45497028 | 0.71596967 | 0.47401008 | NA |
| MPHOSPH9 | 14.8046159 | -0.1281732 | 0.51303825 | -0.2498317 | 0.80271751 | NA |
| C12orf65 | 10.1845011 | 0.40437154 | 0.51456742 | 0.78584753 | 0.43195682 | NA |
| SBNO1 | 28.0177786 | -0.4280548 | 0.36289152 | -1.1795669 | 0.23817253 | 0.73242571 |
| KMT5A | 5.93069356 | 0.43010219 | 0.59200245 | 0.72652097 | 0.46751946 | NA |
| RILPL2 | 5.11520545 | 0.10183587 | 0.88669106 | 0.11484933 | 0.90856455 | NA |
| SNRNP35 | 9.72154191 | -0.0131927 | 0.49068205 | -0.0268865 | 0.97855022 | NA |
| RILPL1 | 15.4137143 | 0.80441971 | 0.4781537 | 1.68234547 | 0.09250187 | NA |
| TMED2 | 13.9161332 | -0.3558894 | 0.40002753 | -0.8896623 | 0.37364723 | NA |
| DDX55 | 6.20000217 | -1.0866177 | 0.64028439 | -1.6970861 | 0.08968039 | NA |
| EIF2B1 | 7.03288766 | -0.8049942 | 0.59952407 | -1.3427221 | 0.17936198 | NA |
| GTF2H3 | 5.64970584 | 0.32903301 | 0.63196041 | 0.52065447 | 0.6026075 | NA |
| ATP6V0A2 | 5.57310244 | 0.00154193 | 0.68310905 | 0.00225722 | 0.998199 | NA |
| CCDC92 | 20.7167971 | -0.126162 | 0.36913631 | -0.3417762 | 0.7325193 | NA |
| RFLNA | 2.79829042 | 0.05524832 | 1.00208559 | 0.05513334 | 0.95603224 | NA |
| NCOR2 | 31.3566898 | 0.20069991 | 0.32318093 | 0.62101409 | 0.53459035 | 0.88254911 |

| | | | | | | |
|---|---|---|---|---|---|---|
| UBC | 85.1137002 | -0.4374962 | 0.29139884 | -1.5013655 | 0.13326104 | 0.60591775 |
| DHX37 | 3.79749472 | 0.258666 | 0.73764988 | 0.35066229 | 0.72584172 | NA |
| BRI3BP | 8.93700438 | 0.92080952 | 0.5247817 | 1.7546525 | 0.07931877 | NA |
| AACS | 7.72443156 | -0.1048379 | 0.52079029 | -0.2013054 | 0.84045977 | NA |
| SLC15A4 | 9.44396454 | -1.2087477 | 0.5949459 | -2.0316935 | 0.04218469 | NA |
| RIMBP2 | 14.9616605 | 1.35896291 | 0.5053147 | 2.68933974 | 0.00715935 | NA |
| STX2 | 13.0350648 | -0.03914 | 0.43272714 | -0.0904497 | 0.92792985 | NA |
| RAN | 21.0228146 | 0.31222212 | 0.46967991 | 0.66475512 | 0.50620713 | NA |
| SFSWAP | 23.0818915 | -0.3543351 | 0.39674162 | -0.893113 | 0.37179666 | NA |
| ULK1 | 14.3638079 | 0.44821156 | 0.46665099 | 0.96048562 | 0.33681086 | NA |
| PUS1 | 4.36201892 | -0.8871384 | 0.6994972 | -1.2682516 | 0.20470812 | NA |
| EP400 | 25.7193198 | 0.22135585 | 0.42763722 | 0.51762531 | 0.60471972 | NA |
| DDX51 | 2.56467906 | -0.8785805 | 0.98695262 | -0.8901952 | 0.37336107 | NA |
| FBRSL1 | 19.2177915 | -0.4532172 | 0.41760912 | -1.0852665 | 0.27780363 | NA |
| POLE | 3.16323385 | 0.11992653 | 0.89872447 | 0.13344082 | 0.89384476 | NA |
| PGAM5 | 3.28178959 | -0.1710855 | 0.87667228 | -0.1951534 | 0.84527289 | NA |
| ANKLE2 | 21.058298 | -0.534652 | 0.39966003 | -1.3377669 | 0.18097243 | NA |
| GOLGA3 | 19.2608963 | -0.878372 | 0.58281349 | -1.5071237 | 0.13177897 | NA |
| CHFR | 17.1271315 | -0.3102102 | 0.45258787 | -0.6854142 | 0.49308257 | NA |
| ZNF26 | 11.3053187 | -0.68656 | 0.50438233 | -1.3611897 | 0.17345376 | NA |
| ZNF84 | 16.1848981 | 0.02623977 | 0.41324417 | 0.06349702 | 0.94937073 | NA |
| ZNF140 | 5.35643371 | -0.0761154 | 0.74700553 | -0.101894 | 0.91884081 | NA |
| ZNF891 | 4.70836059 | 0.82336106 | 0.70468489 | 1.16841025 | 0.24264132 | NA |
| ZNF268 | 5.13784159 | 0.03234585 | 0.72163883 | 0.04482277 | 0.96424858 | NA |
| MPHOSPH8 | 50.9362061 | 0.55120329 | 0.25428646 | 2.16764707 | 0.03018555 | 0.40822098 |
| PSPC1 | 48.4627769 | -0.2207321 | 0.29937039 | -0.7373211 | 0.46092712 | 0.85894565 |
| ZMYM5 | 16.7485567 | -0.233086 | 0.43172641 | -0.5398928 | 0.58927096 | NA |
| ZMYM2 | 98.3513077 | 0.1030707 | 0.23717355 | 0.43457922 | 0.66386788 | 0.92276785 |
| CRYL1 | 22.2879343 | 0.42127214 | 0.55637537 | 0.75717251 | 0.44894652 | NA |
| IFT88 | 23.7805966 | 0.0876018 | 0.40145771 | 0.21820929 | 0.82726604 | NA |
| EEF1AKMT1 | 3.01478256 | 0.34699884 | 0.98323406 | 0.35291581 | 0.72415156 | NA |
| XPO4 | 33.2937513 | -0.2562602 | 0.30077647 | -0.8519955 | 0.3942166 | 0.8258317 |
| LATS2 | 24.0427232 | -1.0287073 | 0.37536477 | -2.7405538 | 0.00613357 | NA |
| SAP18 | 21.9739409 | -0.1062062 | 0.33363693 | -0.3183288 | 0.75023554 | NA |
| MRPL57 | 16.9798977 | 0.12771957 | 0.45843111 | 0.27860146 | 0.7805507 | NA |
| ZDHHC20 | 30.489063 | 0.51807795 | 0.40748929 | 1.27139036 | 0.20358981 | 0.700066 |
| MICU2 | 60.1191666 | 0.1386615 | 0.34764596 | 0.39885837 | 0.68999756 | 0.92830727 |
| FGF9 | 65.333794 | -0.9086727 | 0.3770324 | -2.4100652 | 0.01594967 | 0.31287044 |
| AL136962.1 | 26.4458824 | -1.0748185 | 1.37711219 | -0.7804872 | 0.43510415 | 0.84273801 |
| SACS | 48.083379 | -1.825768 | 0.56780448 | -3.2154871 | 0.00130223 | 0.1198504 |
| MIPEP | 22.1505685 | -0.2216481 | 0.36296558 | -0.6106589 | 0.54142545 | NA |
| PARP4 | 39.4631345 | -0.2314799 | 0.38281242 | -0.6046824 | 0.54539007 | 0.88410237 |
| MTMR6 | 9.52415278 | -0.1013447 | 0.47068793 | -0.2153119 | 0.82952414 | NA |
| NUP58 | 25.9619988 | -0.2378609 | 0.34920821 | -0.6811436 | 0.49578063 | 0.86712877 |
| SHISA2 | 5.6880854 | -1.1850107 | 0.85404501 | -1.3875272 | 0.16528106 | NA |
| RNF6 | 5.59881508 | -0.1031954 | 0.67969573 | -0.1518259 | 0.87932424 | NA |
| CDK8 | 55.9238993 | 0.10116366 | 0.2599738 | 0.38913023 | 0.69717981 | 0.9303056 |
| WASF3 | 8.63365443 | -0.1260676 | 0.52564377 | -0.2398347 | 0.81045844 | NA |
| USP12 | 37.4261837 | 0.21030271 | 0.28709387 | 0.73252246 | 0.46384974 | 0.85894565 |
| RPL21 | 30.6704978 | 0.70140328 | 0.46855219 | 1.4969587 | 0.13440401 | 0.60594834 |
| GTF3A | 8.75048538 | 0.23553343 | 0.52240783 | 0.45086122 | 0.65208957 | NA |
| MTIF3 | 15.9401799 | 0.54264708 | 0.46203126 | 1.17448132 | 0.24020229 | NA |
| LNX2 | 16.4638026 | 0.10598799 | 0.43422964 | 0.24408281 | 0.80716669 | NA |
| POLR1D | 14.6382587 | 0.72466358 | 0.4526103 | 1.60107621 | 0.10936004 | NA |

| | | | | | | |
|---|---|---|---|---|---|---|
| PAN3 | 93.5191326 | 0.22161191 | 0.25868925 | 0.85667228 | 0.39162603 | 0.8258317 |
| POMP | 12.3616386 | -0.249744 | 0.48258926 | -0.5175085 | 0.60480126 | NA |
| SLC46A3 | 2.88822971 | 1.7198171 | 0.92840086 | 1.85245099 | 0.06396109 | NA |
| SLC7A1 | 9.25263363 | -1.4268898 | 0.49428863 | -2.8867542 | 0.00389238 | NA |
| UBL3 | 86.6771127 | 0.54622209 | 0.27781042 | 1.96616843 | 0.04927916 | 0.46265757 |
| KATNAL1 | 5.63964023 | -0.1007113 | 0.6149486 | -0.1637718 | 0.86991077 | NA |
| HMGB1 | 49.2669663 | 0.09210254 | 0.27451072 | 0.33551526 | 0.73723643 | 0.94658914 |
| USPL1 | 6.99802275 | 0.08614351 | 0.63726819 | 0.13517623 | 0.89247254 | NA |
| HSPH1 | 22.6369949 | 0.07902202 | 0.4368563 | 0.1808879 | 0.85645557 | NA |
| B3GLCT | 16.3345742 | 0.43413687 | 0.49192486 | 0.88252679 | 0.377492 | NA |
| FRY | 14.8988288 | 0.39359633 | 0.64889157 | 0.60656718 | 0.54413818 | NA |
| N4BP2L1 | 10.7177965 | 0.09121315 | 0.53398689 | 0.17081534 | 0.86436897 | NA |
| N4BP2L2 | 176.547073 | -0.036801 | 0.19793667 | -0.1859229 | 0.85250526 | 0.97168963 |
| PDS5B | 46.1400771 | -0.218602 | 0.30850999 | -0.7085736 | 0.47858912 | 0.86206794 |
| KL | 4.1611019 | -0.7924988 | 1.03395694 | -0.7664718 | 0.44339565 | NA |
| STARD13 | 17.5102494 | 0.29007628 | 0.70747959 | 0.41001363 | 0.68179595 | NA |
| AL139383.1 | 4.6404683 | -0.1267921 | 0.71580148 | -0.177133 | 0.85940391 | NA |
| RFC3 | 4.74372377 | 1.05580752 | 0.68777378 | 1.53510873 | 0.12475712 | NA |
| LINC02343 | 24.9893305 | 0.15497739 | 0.67306341 | 0.23025675 | 0.81789227 | NA |
| NBEA | 54.8902959 | -0.4970392 | 0.28641699 | -1.7353693 | 0.08267542 | 0.55216537 |
| SPART | 16.695779 | 0.01731888 | 0.43399515 | 0.0399057 | 0.96816831 | NA |
| RFXAP | 3.24462299 | 0.47414174 | 0.96499743 | 0.4913399 | 0.62318606 | NA |
| ALG5 | 8.69913657 | 0.3659294 | 0.58245942 | 0.62824876 | 0.529841 | NA |
| EXOSC8 | 8.01460644 | -0.280203 | 0.58467253 | -0.4792478 | 0.63176236 | NA |
| SUPT20H | 23.0975218 | 0.14077954 | 0.34771433 | 0.40487126 | 0.68557214 | NA |
| LINC00571 | 8.37106295 | -0.6647087 | 0.78388239 | -0.84797 | 0.3964547 | NA |
| UFM1 | 16.4081965 | -0.6376049 | 0.41658012 | -1.5305698 | 0.12587576 | NA |
| FREM2 | 11.6294886 | -0.2817271 | 0.60759841 | -0.4636731 | 0.64288195 | NA |
| PROSER1 | 7.47938256 | -0.0494127 | 0.56869492 | -0.0868878 | 0.93076069 | NA |
| NHLRC3 | 12.1231609 | 0.42534585 | 0.50855526 | 0.83638078 | 0.40294072 | NA |
| COG6 | 22.936321 | -0.2411862 | 0.39329015 | -0.6132527 | 0.53970926 | NA |
| LINC00598 | 8.31486086 | 0.77042803 | 0.60128563 | 1.28130125 | 0.20008787 | NA |
| FOXO1 | 260.00454 | 0.39204611 | 0.30862613 | 1.27029464 | 0.2039797 | 0.700066 |
| MRPS31 | 16.7885057 | 0.36581855 | 0.43980582 | 0.83177286 | 0.40553716 | NA |
| SLC25A15 | 3.15641266 | -0.2545444 | 0.9066811 | -0.280743 | 0.77890749 | NA |
| ELF1 | 86.0826473 | -0.2101454 | 0.32720083 | -0.6422519 | 0.52070964 | 0.87214537 |
| WBP4 | 12.2200658 | 0.21029643 | 0.48017946 | 0.43795382 | 0.66141976 | NA |
| MTRF1 | 6.68428515 | 0.020925 | 0.5950995 | 0.03516218 | 0.97195042 | NA |
| NAA16 | 20.593091 | 0.59293679 | 0.38111998 | 1.55577459 | 0.1197617 | NA |
| VWA8 | 83.922158 | 0.01140308 | 0.28592152 | 0.03988185 | 0.96818732 | 0.99656976 |
| DGKH | 22.9806727 | 0.78764259 | 0.4580038 | 1.7197294 | 0.08548164 | NA |
| AKAP11 | 18.4659664 | 0.66510122 | 0.41430131 | 1.60535629 | 0.10841542 | NA |
| EPSTI1 | 14.2550844 | -0.2823341 | 1.00847103 | -0.2799625 | 0.77950628 | NA |
| DNAJC15 | 18.0379266 | 0.18178277 | 0.45844643 | 0.39651909 | 0.69172212 | NA |
| ENOX1 | 4.86061124 | -0.0121211 | 0.67314103 | -0.0180068 | 0.98563339 | NA |
| CCDC122 | 7.69826662 | 0.58212137 | 0.66066087 | 0.88111979 | 0.37825299 | NA |
| SMIM2-AS1 | 7.27067553 | 0.42572197 | 1.07942322 | 0.39439764 | 0.69328749 | NA |
| TSC22D1 | 38.6390503 | -0.3399735 | 0.51127209 | -0.664956 | 0.50607861 | 0.87094108 |
| NUFIP1 | 4.40546274 | 1.04001437 | 0.65533071 | 1.58700692 | 0.11251108 | NA |
| GPALPP1 | 11.9922183 | 0.78071766 | 0.45238877 | 1.72576713 | 0.08438932 | NA |
| GTF2F2 | 32.9185944 | -0.6070398 | 0.33113901 | -1.8331871 | 0.06677475 | 0.52400942 |
| TPT1 | 196.031722 | 0.27463292 | 0.42100994 | 0.65231933 | 0.51419519 | 0.87158522 |
| SLC25A30 | 16.084154 | -0.7038194 | 0.68110181 | -1.0333542 | 0.30143817 | NA |
| COG3 | 19.4337379 | -0.2839839 | 0.42538613 | -0.6675909 | 0.50439478 | NA |

| | | | | | | |
|---|---|---|---|---|---|---|
| ZC3H13 | 32.0755573 | -0.1135953 | 0.33187802 | -0.3422803 | 0.73213993 | 0.94354489 |
| AL138686.1 | 12.7780751 | -0.5509673 | 0.54333436 | -1.0140483 | 0.31055968 | NA |
| LRCH1 | 48.6859925 | 0.78597002 | 0.30697989 | 2.56033065 | 0.01045726 | 0.28266532 |
| ESD | 14.7733514 | 0.25854332 | 0.43241622 | 0.59790383 | 0.5499041 | NA |
| SUCLA2 | 33.810675 | -0.6604187 | 0.40391809 | -1.6350314 | 0.10204247 | 0.5750956 |
| NUDT15 | 2.87864223 | -0.4375241 | 1.02232486 | -0.4279698 | 0.66867314 | NA |
| MED4 | 17.2944163 | -0.1389137 | 0.37556119 | -0.3698831 | 0.7114696 | NA |
| ITM2B | 114.984152 | 0.52394789 | 0.3797085 | 1.37986874 | 0.16762706 | 0.65504631 |
| RB1 | 54.7608869 | -0.3877275 | 0.2887358 | -1.3428452 | 0.17932212 | 0.67032316 |
| RCBTB2 | 7.40451985 | -0.0223519 | 0.76092937 | -0.0293744 | 0.97656596 | NA |
| FNDC3A | 110.215003 | -0.7674412 | 0.38385723 | -1.999288 | 0.0455772 | 0.44887658 |
| CDADC1 | 5.06616561 | 0.00511475 | 0.70192249 | 0.00728677 | 0.99418605 | NA |
| CAB39L | 30.2641916 | -0.2648419 | 0.38266655 | -0.6920957 | 0.48887724 | 0.86354292 |
| SETDB2 | 22.915262 | 0.19081598 | 0.40859001 | 0.46701088 | 0.64049209 | NA |
| PHF11 | 16.0416914 | -0.4111134 | 0.46428915 | -0.8854685 | 0.37590399 | NA |
| RCBTB1 | 5.92769848 | 0.01325773 | 0.65995787 | 0.02008875 | 0.98397258 | NA |
| EBPL | 7.42134903 | 0.58598536 | 0.74683591 | 0.78462398 | 0.43267406 | NA |
| KPNA3 | 31.4610031 | -0.0160463 | 0.30290723 | -0.0529742 | 0.9577525 | 0.99656976 |
| SPRYD7 | 4.91469075 | -0.08645 | 0.68391066 | -0.1264054 | 0.89941106 | NA |
| DLEU2 | 35.8750128 | 0.0996265 | 0.37496094 | 0.26569834 | 0.79047154 | 0.96336463 |
| TRIM13 | 8.80051474 | 0.61552884 | 0.49113967 | 1.25326638 | 0.21010878 | NA |
| RNASEH2B | 8.7137853 | 0.24698852 | 0.53501213 | 0.46165032 | 0.64433211 | NA |
| INTS6 | 49.054767 | -0.0810725 | 0.44185848 | -0.1834807 | 0.8544209 | 0.97168963 |
| INTS6-AS1 | 2.97324443 | 0.32108405 | 0.85063675 | 0.37746318 | 0.70582942 | NA |
| WDFY2 | 55.1136476 | -0.2648519 | 0.36918776 | -0.7173908 | 0.47313301 | 0.86206794 |
| ATP7B | 4.01762435 | 0.67036366 | 0.78075999 | 0.85860402 | 0.39055902 | NA |
| ALG11 | 3.90642538 | 0.26260135 | 0.7230047 | 0.36320836 | 0.71644925 | NA |
| UTP14C | 6.05696328 | -0.4502924 | 0.68177909 | -0.6604668 | 0.50895433 | NA |
| NEK3 | 7.87024933 | 0.00351573 | 0.57453416 | 0.00611928 | 0.99511755 | NA |
| VPS36 | 17.2148956 | 0.67996202 | 0.39819625 | 1.70760527 | 0.08770961 | NA |
| CKAP2 | 1.79829961 | 1.88754133 | 1.24794074 | 1.5125248 | 0.13040041 | NA |
| SUGT1 | 14.1841633 | 0.49642957 | 0.41919281 | 1.18425115 | 0.23631366 | NA |
| TDRD3 | 43.6668921 | 0.43379937 | 0.31183633 | 1.39111234 | 0.16419136 | 0.64995633 |
| PCDH9 | 39.4434836 | 1.88333811 | 0.64506286 | 2.91961952 | 0.00350459 | 0.20482034 |
| DACH1 | 304.869227 | -0.3697075 | 0.35789201 | -1.0330142 | 0.30159724 | 0.77521265 |
| MZT1 | 4.23540553 | -0.4156309 | 0.68791544 | -0.6041889 | 0.54571807 | NA |
| BORA | 2.97790719 | 0.87841596 | 0.84699445 | 1.03709766 | 0.29969035 | NA |
| DIS3 | 10.6986703 | -0.5065592 | 0.48953341 | -1.0347795 | 0.30077188 | NA |
| PIBF1 | 45.71161 | -0.239938 | 0.28395877 | -0.8449747 | 0.39812499 | 0.8269015 |
| KLF5 | 10.993026 | -0.3897214 | 0.48868497 | -0.79749 | 0.42516654 | NA |
| KLF12 | 41.1793221 | -0.0455422 | 0.47953097 | -0.0949725 | 0.92433671 | 0.98758497 |
| TBC1D4 | 255.962385 | -0.4215388 | 0.21827614 | -1.931218 | 0.05345609 | 0.48694305 |
| COMMD6 | 20.002491 | 0.80085995 | 0.44348219 | 1.80584467 | 0.07094261 | NA |
| UCHL3 | 17.8597828 | -0.3205201 | 0.40398377 | -0.7933984 | 0.42754576 | NA |
| LMO7 | 86.8327296 | 0.10171371 | 0.33599772 | 0.30272143 | 0.76210216 | 0.95478238 |
| KCTD12 | 13.6475959 | -0.6044757 | 0.47905431 | -1.2618103 | 0.20701706 | NA |
| AC001226.2 | 5.6663149 | 0.2891992 | 0.82591848 | 0.35015466 | 0.72622263 | NA |
| FBXL3 | 28.4777421 | 0.33532837 | 0.33274406 | 1.00776667 | 0.31356649 | 0.78031266 |
| MYCBP2 | 53.4764369 | -0.2099672 | 0.27737144 | -0.7569891 | 0.44905637 | 0.85123446 |
| SLAIN1 | 4.50348642 | 1.02197892 | 0.74093322 | 1.37931313 | 0.16779823 | NA |
| RNF219 | 4.89795038 | -0.7665259 | 0.64879112 | -1.1814679 | 0.23741688 | NA |
| RBM26 | 57.598031 | 0.00285432 | 0.26783514 | 0.010657 | 0.99149711 | 0.99909657 |
| NDFIP2 | 24.7287744 | 0.27136856 | 0.33997755 | 0.79819554 | 0.42475703 | NA |
| GPC5 | 41.039523 | 3.80288159 | 1.44711572 | 2.62790427 | 0.00859127 | 0.27648451 |

| | | | | | | |
|---|---|---|---|---|---|---|
| GPC6 | 604.696264 | -2.62058 | 0.93034014 | -2.8167977 | 0.00485051 | 0.22229871 |
| TGDS | 7.37018908 | -0.2226548 | 0.57130346 | -0.3897313 | 0.69673524 | NA |
| GPR180 | 7.63182556 | -0.7838629 | 0.59544612 | -1.3164295 | 0.18802991 | NA |
| ABCC4 | 59.2972216 | -0.1371537 | 0.3547573 | -0.3866127 | 0.69904297 | 0.9303056 |
| CLDN10 | 15.6350676 | -0.6764604 | 0.78142776 | -0.8656724 | 0.38666984 | NA |
| CLDN10-AS1 | 9.8576897 | 0.48864568 | 0.73547571 | 0.66439405 | 0.50643814 | NA |
| DNAJC3-DT | 3.18716046 | 1.44424099 | 0.76308311 | 1.89263918 | 0.05840587 | NA |
| DNAJC3 | 37.1103727 | -0.3619724 | 0.34837321 | -1.0390364 | 0.29878781 | 0.77521265 |
| UGGT2 | 58.1950843 | 0.00934334 | 0.30688216 | 0.03044602 | 0.97571134 | 0.99853281 |
| MBNL2 | 81.707732 | -0.3333964 | 0.27848385 | -1.197184 | 0.23123483 | 0.73156941 |
| RAP2A | 3.6352229 | -0.2716577 | 0.73174262 | -0.3712476 | 0.71045315 | NA |
| IPO5 | 21.1619174 | 0.21300659 | 0.35943346 | 0.59261759 | 0.5534371 | NA |
| FARP1 | 163.407873 | 0.10857053 | 0.27043365 | 0.40146826 | 0.6880754 | 0.92736198 |
| STK24 | 58.7926481 | -0.1714785 | 0.33237405 | -0.5159204 | 0.60591002 | 0.90821838 |
| DOCK9 | 28.4313815 | 0.20028568 | 0.31069699 | 0.64463346 | 0.51916475 | 0.87214537 |
| UBAC2 | 67.7904186 | 0.09267425 | 0.26673714 | 0.34743661 | 0.72826333 | 0.94127595 |
| TM9SF2 | 36.8493843 | -0.1950766 | 0.32031333 | -0.6090182 | 0.54251237 | 0.88410237 |
| CLYBL | 49.7168103 | 0.41485814 | 0.34256407 | 1.21103811 | 0.2258808 | 0.72987392 |
| PCCA | 162.68237 | -0.0048016 | 0.23328169 | -0.0205827 | 0.98357856 | 0.99902646 |
| GGACT | 11.8327878 | 0.30341556 | 0.59828697 | 0.50714052 | 0.61205623 | NA |
| TMTC4 | 8.81597645 | 0.27690511 | 0.52053259 | 0.53196499 | 0.59475025 | NA |
| FGF14 | 10.1778725 | -0.6922673 | 0.86791433 | -0.7976217 | 0.42509004 | NA |
| TPP2 | 27.6827247 | 0.09615111 | 0.33545448 | 0.28662939 | 0.77439611 | 0.9604383 |
| TEX30 | 3.30433666 | 0.69457084 | 0.91198355 | 0.76160457 | 0.44629605 | NA |
| EFNB2 | 27.9418122 | -1.1201097 | 0.68897867 | -1.6257538 | 0.10400204 | 0.5772605 |
| ARGLU1 | 57.2630076 | 0.15688037 | 0.29454957 | 0.53261108 | 0.59430283 | 0.89767643 |
| LIG4 | 5.02867859 | -0.748318 | 0.6951346 | -1.076508 | 0.28170011 | NA |
| ABHD13 | 7.53707691 | 0.00574165 | 0.59848287 | 0.00959368 | 0.99234547 | NA |
| TNFSF13B | 4.38531235 | 0.3784773 | 0.84808982 | 0.4462703 | 0.65540201 | NA |
| IRS2 | 19.4537903 | -0.324581 | 0.50485175 | -0.6429234 | 0.5202738 | NA |
| COL4A1 | 9.98717402 | -0.5233633 | 0.8674853 | -0.6033108 | 0.54630192 | NA |
| COL4A2 | 8.36656473 | 0.21173172 | 0.63101018 | 0.33554407 | 0.73721471 | NA |
| RAB20 | 5.60543129 | 0.12343324 | 0.69080217 | 0.17868103 | 0.85818817 | NA |
| NAXD | 7.72440645 | -0.302979 | 0.55797579 | -0.5429966 | 0.58713213 | NA |
| CARS2 | 14.6269627 | 0.07355858 | 0.42029128 | 0.17501809 | 0.86106542 | NA |
| ANKRD10 | 81.9419441 | -0.0719138 | 0.27006199 | -0.2662864 | 0.79001864 | 0.96325251 |
| ARHGEF7 | 87.7400313 | -0.0623315 | 0.35439791 | -0.1758799 | 0.86038827 | 0.97180546 |
| TUBGCP3 | 13.3998512 | 0.02213008 | 0.43836852 | 0.05048282 | 0.95973764 | NA |
| ATP11A | 38.516382 | 0.07939358 | 0.30119936 | 0.26359148 | 0.79209472 | 0.96341299 |
| MCF2L | 26.8851183 | 0.48993178 | 0.44146797 | 1.10977876 | 0.26709437 | 0.75497438 |
| PCID2 | 12.1947552 | 0.49957369 | 0.43425061 | 1.15042713 | 0.24996799 | NA |
| CUL4A | 36.202065 | -0.3993738 | 0.32332626 | -1.2352038 | 0.21675466 | 0.71673468 |
| LAMP1 | 34.3621565 | -0.8267021 | 0.49232126 | -1.6791924 | 0.09311455 | 0.56220784 |
| GRTP1 | 9.89068805 | 1.06085822 | 0.55568858 | 1.90908768 | 0.05625078 | NA |
| DCUN1D2 | 12.8050442 | -0.2516896 | 0.46922843 | -0.5363904 | 0.59168876 | NA |
| TMCO3 | 13.7702124 | -0.1669052 | 0.43810532 | -0.3809704 | 0.70322522 | NA |
| TFDP1 | 20.7941847 | -1.1763923 | 0.49572878 | -2.3730563 | 0.01764157 | NA |
| GAS6 | 25.4599288 | 0.11738184 | 0.4460033 | 0.26318604 | 0.79240719 | NA |
| CDC16 | 18.9101266 | 0.07490438 | 0.43475113 | 0.17229255 | 0.86320754 | NA |
| UPF3A | 23.0720708 | 0.21091529 | 0.39735772 | 0.53079451 | 0.59556119 | NA |
| CHAMP1 | 5.1232738 | -0.3775377 | 0.67990749 | -0.555278 | 0.57870452 | NA |
| TTC5 | 4.30286213 | 1.10004934 | 0.72753477 | 1.51202305 | 0.130528 | NA |
| CCNB1IP1 | 9.99937408 | 0.29784283 | 0.4588348 | 0.64912869 | 0.5162552 | NA |
| PARP2 | 2.53470017 | -0.1956034 | 0.89346348 | -0.2189272 | 0.82670679 | NA |

| | | | | | | |
|---|---|---|---|---|---|---|
| TEP1 | 7.40117479 | -0.3327415 | 0.60746225 | -0.5477568 | 0.58385894 | NA |
| OSGEP | 5.95471609 | -0.8533427 | 0.65712551 | -1.2985993 | 0.19408149 | NA |
| APEX1 | 8.98015176 | -0.1224836 | 0.53947554 | -0.2270421 | 0.82039104 | NA |
| PIP4P1 | 3.80905084 | -0.7676257 | 0.80339409 | -0.9554783 | 0.33933585 | NA |
| PNP | 7.4574816 | -0.5512585 | 0.59639515 | -0.9243176 | 0.35532098 | NA |
| METTL17 | 7.01733088 | 0.3399167 | 0.58977465 | 0.57635014 | 0.56437854 | NA |
| NDRG2 | 12.2612943 | 1.37713291 | 0.53252999 | 2.58601946 | 0.00970914 | NA |
| ARHGEF40 | 3.67684521 | 0.00044375 | 0.76559698 | 0.00057961 | 0.99953754 | NA |
| ZNF219 | 4.634041 | -1.2598254 | 0.95520192 | -1.31891 | 0.18719919 | NA |
| HNRNPC | 84.525718 | -0.3498684 | 0.23494407 | -1.4891563 | 0.13644622 | 0.60594834 |
| SUPT16H | 15.7500207 | 0.05630169 | 0.40316843 | 0.13964806 | 0.88893806 | NA |
| CHD8 | 21.8612604 | -0.3333805 | 0.38177686 | -0.873234 | 0.38253555 | NA |
| RAB2B | 11.1975728 | 0.09229544 | 0.44466027 | 0.20756394 | 0.83556946 | NA |
| TOX4 | 13.1826263 | 0.58276031 | 0.43891534 | 1.32772829 | 0.18426789 | NA |
| METTL3 | 5.47476902 | 0.55426782 | 0.59517264 | 0.93127234 | 0.35171271 | NA |
| DAD1 | 18.8849061 | 0.00723849 | 0.37593658 | 0.01925456 | 0.98463803 | NA |
| ABHD4 | 4.42924437 | 1.32295845 | 0.70880361 | 1.86646685 | 0.06197608 | NA |
| OXA1L | 8.66133578 | 0.42394868 | 0.52634564 | 0.80545681 | 0.42055612 | NA |
| SLC7A7 | 3.50597967 | 1.29992213 | 1.03938972 | 1.25065902 | 0.21105891 | NA |
| MRPL52 | 5.96381565 | 0.82295041 | 0.6693002 | 1.22956845 | 0.21885875 | NA |
| MMP14 | 4.32979888 | 0.43485337 | 0.7904808 | 0.5501125 | 0.58224221 | NA |
| LRP10 | 20.3863371 | 0.31053704 | 0.5049565 | 0.61497781 | 0.53856938 | NA |
| RBM23 | 13.4942125 | 0.24506894 | 0.55075448 | 0.44496949 | 0.6563418 | NA |
| PRMT5 | 2.60989922 | 0.0137283 | 0.88516868 | 0.01550925 | 0.9876259 | NA |
| C14orf93 | 3.08036855 | 0.95326091 | 0.86915189 | 1.09677138 | 0.27274135 | NA |
| PSMB5 | 11.9871578 | 0.86145561 | 0.46543111 | 1.85087673 | 0.06418729 | NA |
| ACIN1 | 29.5402588 | -0.0089268 | 0.31959099 | -0.0279319 | 0.97771648 | 0.99902646 |
| C14orf119 | 6.61834489 | 0.15687099 | 0.56218975 | 0.27903566 | 0.78021746 | NA |
| SLC7A8 | 5.89652515 | -0.3208053 | 0.92147831 | -0.348142 | 0.72773354 | NA |
| HOMEZ | 2.36381364 | 2.03130576 | 0.98814356 | 2.0556788 | 0.03981349 | NA |
| BCL2L2-PABI | 5.75335964 | 0.29205271 | 0.59627046 | 0.48979905 | 0.6242761 | NA |
| PABPN1 | 20.6992298 | -0.5430934 | 0.46694091 | -1.163088 | 0.24479379 | NA |
| NGDN | 5.66774125 | -0.5049359 | 0.6785698 | -0.7441178 | 0.4568052 | NA |
| ZFHX2 | 2.88179707 | -0.1920765 | 0.96417377 | -0.1992135 | 0.84209571 | NA |
| AP1G2 | 6.67423436 | -0.5130132 | 0.57864974 | -0.8865695 | 0.37531071 | NA |
| DHRS4-AS1 | 4.72993524 | 1.35805092 | 0.77032863 | 1.76295008 | 0.07790891 | NA |
| DHRS4L2 | 5.79780366 | 1.5773794 | 0.7560795 | 2.08626129 | 0.03695496 | NA |
| NRL | 2.54930964 | 0.86412718 | 0.89342279 | 0.96720969 | 0.33343922 | NA |
| PSME1 | 14.9331502 | 0.35360841 | 0.45647266 | 0.77465407 | 0.4385441 | NA |
| EMC9 | 3.68651531 | -0.6175283 | 0.76421051 | -0.8080605 | 0.41905575 | NA |
| PSME2 | 3.26593422 | -0.4538076 | 0.84227345 | -0.5387889 | 0.59003253 | NA |
| AL136295.1 | 3.04453303 | 0.64826508 | 0.91278404 | 0.71020641 | 0.47757614 | NA |
| GMPR2 | 4.15757155 | 0.64974292 | 0.80141084 | 0.81074885 | 0.41750991 | NA |
| TINF2 | 3.15053421 | -0.1675278 | 0.87009767 | -0.1925391 | 0.84731995 | NA |
| DHRS1 | 4.49502241 | 0.46419227 | 0.77270762 | 0.60073468 | 0.54801672 | NA |
| KHNYN | 3.16358017 | -0.4022612 | 0.91075342 | -0.4416796 | 0.65872108 | NA |
| SDR39U1 | 6.03072003 | -0.321958 | 0.5981135 | -0.5382891 | 0.59037748 | NA |
| STXBP6 | 7.2830154 | -1.2106165 | 0.81672 | -1.4822908 | 0.13826294 | NA |
| NOVA1 | 9.93221604 | -0.7630368 | 0.83845368 | -0.9100524 | 0.36279488 | NA |
| PRKD1 | 75.8367465 | -0.3218741 | 0.38805258 | -0.82946 | 0.40684418 | 0.82808596 |
| G2E3 | 10.3355275 | -0.7612406 | 0.57353894 | -1.3272693 | 0.18441963 | NA |
| SCFD1 | 39.9588728 | -0.8126837 | 0.31211425 | -2.6038019 | 0.0092196 | 0.27648451 |
| COCH | 5.41187235 | -1.3847916 | 0.67522846 | -2.050849 | 0.04028165 | NA |
| STRN3 | 79.1860755 | -0.2348364 | 0.29085658 | -0.807396 | 0.4194384 | 0.83856261 |

| | | | | | | |
|---|---|---|---|---|---|---|
| AP4S1 | 6.09365615 | 0.59361369 | 0.60832014 | 0.97582449 | 0.32915145 | NA |
| HECTD1 | 96.4898645 | -0.0703788 | 0.25888195 | -0.2718568 | 0.7857321 | 0.96325251 |
| AL136418.1 | 4.00764949 | -1.4649446 | 1.04769014 | -1.3982613 | 0.16203462 | NA |
| HEATR5A | 12.5664045 | 0.0094699 | 0.45701869 | 0.02072103 | 0.9834682 | NA |
| AL139353.1 | 2.75732144 | 2.45747295 | 0.99414663 | 2.47194215 | 0.01343813 | NA |
| NUBPL | 29.9410681 | 0.64776533 | 0.38072411 | 1.7014035 | 0.08886724 | 0.56122412 |
| ARHGAP5 | 95.4451727 | -0.4327159 | 0.20778374 | -2.0825302 | 0.03729407 | 0.42032585 |
| AKAP6 | 46.7962497 | -0.3571755 | 0.42003006 | -0.8503569 | 0.39512669 | 0.8258317 |
| NPAS3 | 26.4683177 | 1.12197804 | 0.74679307 | 1.50239481 | 0.13299517 | 0.60574082 |
| EAPP | 13.3610101 | -0.2049148 | 0.40926441 | -0.5006906 | 0.61658892 | NA |
| SNX6 | 39.2767144 | 0.03348635 | 0.27937165 | 0.1198631 | 0.90459159 | 0.98064783 |
| CFL2 | 9.15624815 | -0.1527603 | 0.58072957 | -0.2630489 | 0.79251287 | NA |
| BAZ1A | 14.9102543 | -0.4350013 | 0.41029711 | -1.0602105 | 0.28904885 | NA |
| SRP54 | 26.4202661 | -0.6171755 | 0.43746685 | -1.4107937 | 0.15830544 | 0.64166827 |
| FAM177A1 | 17.8169628 | -0.4514368 | 0.6132599 | -0.7361265 | 0.46165375 | NA |
| PPP2R3C | 17.3383032 | -0.9438087 | 0.41431406 | -2.278003 | 0.0227264 | NA |
| NFKBIA | 11.6623636 | 0.60813149 | 0.79036456 | 0.76943164 | 0.44163711 | NA |
| RALGAPA1 | 86.9970392 | -0.0459428 | 0.27373369 | -0.1678375 | 0.86671114 | 0.97217819 |
| BRMS1L | 7.39844317 | -0.0198004 | 0.66175123 | -0.0299213 | 0.97612983 | NA |
| MBIP | 7.97810011 | 1.01345458 | 0.62629767 | 1.61816756 | 0.10562649 | NA |
| SLC25A21 | 3.48630725 | -1.6376803 | 1.1750422 | -1.3937204 | 0.16340205 | NA |
| MIPOL1 | 25.346126 | 0.47715352 | 0.38713526 | 1.23252407 | 0.21775338 | NA |
| SEC23A | 31.1080066 | -0.8672776 | 0.33455565 | -2.5923269 | 0.00953291 | 0.27655812 |
| GEMIN2 | 2.53710716 | 2.04668962 | 1.00337747 | 2.03980025 | 0.04137023 | NA |
| TRAPPC6B | 16.9883781 | 0.88991116 | 0.42113122 | 2.11314459 | 0.03458839 | NA |
| PNN | 32.2410943 | -0.153274 | 0.4406976 | -0.3477986 | 0.72799145 | 0.94127595 |
| MIA2 | 50.6088332 | 0.66035957 | 0.31274042 | 2.11152616 | 0.03472711 | 0.41688714 |
| FBXO33 | 22.1681396 | -0.0514607 | 0.37061616 | -0.1388517 | 0.88956731 | NA |
| AL049875.1 | 6.46552056 | -0.0451603 | 0.67246978 | -0.0671559 | 0.94645761 | NA |
| KLHL28 | 15.1474299 | -0.0749473 | 0.53892545 | -0.139068 | 0.88939645 | NA |
| TOGARAM1 | 23.617293 | 0.07523926 | 0.32445286 | 0.23189583 | 0.81661893 | NA |
| PRPF39 | 12.4785611 | -0.01503 | 0.47040907 | -0.031951 | 0.97451114 | NA |
| FKBP3 | 12.4730758 | 0.05117698 | 0.4534952 | 0.1128501 | 0.9101494 | NA |
| FANCM | 5.68580948 | -0.7927355 | 0.66697269 | -1.1885576 | 0.23461379 | NA |
| MIS18BP1 | 10.633288 | -0.109748 | 0.68390675 | -0.1604722 | 0.87250911 | NA |
| RPS29 | 31.4203273 | 0.23927598 | 0.32851571 | 0.72835474 | 0.46639646 | 0.85894565 |
| RPL36AL | 38.2087348 | -0.1163046 | 0.43263491 | -0.2688284 | 0.78806173 | 0.96325251 |
| DNAAF2 | 3.86380626 | -0.1344765 | 0.73804372 | -0.1822067 | 0.85542051 | NA |
| POLE2 | 2.67464454 | -0.0524135 | 0.86253534 | -0.0607667 | 0.95154498 | NA |
| KLHDC1 | 9.6612635 | -0.0001555 | 0.78419457 | -0.0001983 | 0.99984175 | NA |
| KLHDC2 | 13.5840688 | 0.25874203 | 0.40831216 | 0.63368682 | 0.52628523 | NA |
| NEMF | 51.7109818 | -0.1275531 | 0.29659017 | -0.430065 | 0.66714834 | 0.92307876 |
| ARF6 | 10.3759281 | 0.0763381 | 0.47869945 | 0.1594698 | 0.87329875 | NA |
| VCPKMT | 5.60345554 | -0.3974302 | 0.634142 | -0.6267211 | 0.53084205 | NA |
| SOS2 | 63.0853042 | 0.35624657 | 0.25096483 | 1.41950791 | 0.15575099 | 0.63932163 |
| L2HGDH | 19.6255685 | 0.05667575 | 0.43786126 | 0.12943769 | 0.89701132 | NA |
| ATP5S | 15.1389704 | 0.42054101 | 0.44867495 | 0.93729549 | 0.34860658 | NA |
| CDKL1 | 15.5823194 | 0.38297166 | 0.46191191 | 0.82910106 | 0.40704722 | NA |
| MAP4K5 | 51.9110878 | -0.2490209 | 0.26995766 | -0.9224445 | 0.35629678 | 0.80177915 |
| ATL1 | 7.39228767 | 0.87365143 | 0.67156351 | 1.30092152 | 0.19328532 | NA |
| SAV1 | 16.2737743 | 0.51070743 | 0.46356071 | 1.10170561 | 0.27058968 | NA |
| NIN | 38.7473257 | -0.4265386 | 0.36467409 | -1.1696434 | 0.24214452 | 0.73323864 |
| TMX1 | 7.39983784 | -0.1267404 | 0.57189197 | -0.221616 | 0.82461282 | NA |
| FRMD6 | 4.78124699 | -0.9693267 | 1.13082427 | -0.857186 | 0.39134212 | NA |

| | | | | | | |
|---|---|---|---|---|---|---|
| RTRAF | 15.3687381 | 0.45213145 | 0.38578789 | 1.17196899 | 0.24120951 | NA |
| TXNDC16 | 5.41475883 | 1.16109468 | 0.7317047 | 1.58683507 | 0.11255 | NA |
| ERO1A | 56.0457208 | -0.5474474 | 0.51940331 | -1.0539929 | 0.29188619 | 0.77286135 |
| AL133453.1 | 6.75254957 | 0.8264695 | 0.81537944 | 1.01360111 | 0.31077313 | NA |
| PSMC6 | 23.6869441 | -0.2211911 | 0.38479785 | -0.5748242 | 0.56541017 | NA |
| STYX | 10.4320541 | 0.64066238 | 0.53320547 | 1.20153001 | 0.22954567 | NA |
| GNPNAT1 | 23.8523585 | -0.2292877 | 0.58302275 | -0.3932741 | 0.69411707 | NA |
| FERMT2 | 21.1057476 | 0.10407239 | 0.42135718 | 0.24699328 | 0.80491343 | NA |
| DDHD1 | 9.47333372 | 0.53131855 | 0.56880731 | 0.93409234 | 0.35025626 | NA |
| BMP4 | 3.127495 | 0.7493444 | 0.89698151 | 0.83540673 | 0.40348874 | NA |
| CNIH1 | 8.47008918 | 0.20737879 | 0.60415946 | 0.34325175 | 0.73140907 | NA |
| GMFB | 10.2306215 | -0.4199656 | 0.46300543 | -0.9070424 | 0.36438437 | NA |
| CGRRF1 | 4.53759538 | -0.1889953 | 0.72502 | -0.260676 | 0.79434238 | NA |
| SAMD4A | 75.6444581 | -0.5909181 | 0.37378469 | -1.5809049 | 0.11389978 | 0.57860396 |
| GCH1 | 5.42817599 | -0.2468816 | 0.88421513 | -0.2792099 | 0.78008373 | NA |
| WDHD1 | 2.5520715 | -2.080202 | 1.04952844 | -1.9820349 | 0.04747533 | NA |
| SOCS4 | 8.88410847 | -0.0696053 | 0.58467984 | -0.1190485 | 0.90523693 | NA |
| MAPK1IP1L | 30.6968238 | -0.2523361 | 0.31235834 | -0.8078417 | 0.41918173 | 0.83856261 |
| LGALS3 | 69.7178736 | 0.74291675 | 0.46645667 | 1.59268116 | 0.11123174 | 0.57860396 |
| FBXO34 | 70.7610658 | -0.6325876 | 0.32191204 | -1.9650945 | 0.0494033 | 0.46265757 |
| AL158801.3 | 3.85764975 | -0.5101037 | 0.83765932 | -0.6089632 | 0.54254881 | NA |
| ATG14 | 9.23382424 | -0.3731247 | 0.68232902 | -0.5468399 | 0.58448875 | NA |
| KTN1 | 81.9217461 | -0.0925231 | 0.228582 | -0.4047697 | 0.68564683 | 0.92726927 |
| PELI2 | 70.4698126 | -0.4312071 | 0.30932593 | -1.3940219 | 0.16331098 | 0.64995633 |
| TMEM260 | 38.2011194 | 0.04695536 | 0.32516729 | 0.1444037 | 0.8851817 | 0.97769824 |
| EXOC5 | 23.8653655 | 0.01188465 | 0.35438492 | 0.033536 | 0.97324715 | NA |
| AP5M1 | 18.5303748 | -0.1622975 | 0.4241673 | -0.3826261 | 0.70199699 | NA |
| NAA30 | 14.0921348 | -0.380577 | 0.44198255 | -0.8610679 | 0.38920064 | NA |
| CCDC198 | 24.9614706 | -0.4155838 | 0.55857756 | -0.7440038 | 0.45687419 | NA |
| ARMH4 | 13.1031111 | -0.2651197 | 0.62060695 | -0.4271942 | 0.66923788 | NA |
| ACTR10 | 19.7748884 | -0.1139066 | 0.48258141 | -0.2360361 | 0.81340469 | NA |
| PSMA3 | 15.4668318 | 0.08296965 | 0.48281839 | 0.17184442 | 0.86355985 | NA |
| ARID4A | 26.9266423 | 0.58698286 | 0.42263868 | 1.38885267 | 0.16487755 | 0.64995633 |
| TIMM9 | 9.48834027 | -0.6055924 | 0.53971153 | -1.1220669 | 0.26183401 | NA |
| KIAA0586 | 20.4410617 | 0.48560801 | 0.39355253 | 1.23390903 | 0.21723681 | NA |
| DACT1 | 2.30042976 | -1.6547709 | 1.20764457 | -1.3702466 | 0.17060992 | NA |
| DAAM1 | 29.6167773 | 0.24539226 | 0.32056821 | 0.76549156 | 0.44397889 | 0.84899161 |
| L3HYPDH | 4.33317411 | 1.64326867 | 0.74311241 | 2.21133257 | 0.02701282 | NA |
| JKAMP | 9.49740872 | 0.80211563 | 0.48508488 | 1.65355728 | 0.0982175 | NA |
| RTN1 | 51.4372205 | 0.89340039 | 0.52357224 | 1.70635552 | 0.08794191 | 0.56122412 |
| LRRC9 | 22.5775221 | 2.10812921 | 0.74801715 | 2.81828996 | 0.00482802 | NA |
| AL157911.1 | 66.6154937 | 0.50847755 | 0.41451749 | 1.22667334 | 0.21994539 | 0.72261413 |
| PCNX4 | 130.836913 | -0.2368683 | 0.31218697 | -0.7587387 | 0.44800888 | 0.85123446 |
| DHRS7 | 32.8870749 | 0.26026057 | 0.36014055 | 0.72266389 | 0.4698864 | 0.86016928 |
| PPM1A | 37.6093706 | -0.4244033 | 0.33298735 | -1.2745328 | 0.20247464 | 0.700066 |
| SIX4 | 3.80877564 | 0.53191619 | 0.82839987 | 0.64210076 | 0.52080776 | NA |
| MNAT1 | 79.6440945 | -0.361248 | 0.27478601 | -1.3146522 | 0.18862683 | 0.68217481 |
| TRMT5 | 2.62315189 | -0.9814627 | 1.12288976 | -0.8740508 | 0.3820906 | NA |
| SLC38A6 | 5.86827936 | -0.0037041 | 0.6354577 | -0.0058291 | 0.99534911 | NA |
| PRKCH | 10.5857368 | 0.35794893 | 0.64191843 | 0.55762369 | 0.57710137 | NA |
| TMEM30B | 9.74929718 | 0.06066529 | 0.46218456 | 0.13125772 | 0.89557144 | NA |
| HIF1A | 130.734445 | -0.4202029 | 0.41387528 | -1.0152886 | 3.10E-01 | 0.778534 |
| HIF1A-AS2 | 24.4405875 | -0.157103 | 0.4720415 | -0.3328161 | 7.39E-01 | NA |
| SNAPC1 | 12.6667516 | -0.9909007 | 0.52389076 | -1.8914262 | 0.05856747 | NA |

| | | | | | | |
|---|---|---|---|---|---|---|
| KCNH5 | 14.3841425 | 0.03136814 | 1.06566055 | 0.02943539 | 0.97651735 | NA |
| PPP2R5E | 80.7958975 | -0.4887572 | 0.3837231 | -1.2737238 | 0.20276132 | 0.700066 |
| WDR89 | 5.59752884 | 0.66167986 | 0.60611767 | 1.09166897 | 0.27497863 | NA |
| SGPP1 | 10.1083335 | -1.0014308 | 0.62655005 | -1.5983253 | 0.10997061 | NA |
| SYNE2 | 338.262959 | 0.20706693 | 0.37710937 | 0.54908985 | 0.58294379 | 0.89282946 |
| ESR2 | 20.4402571 | 0.72658805 | 0.58346523 | 1.24529793 | 0.21302225 | NA |
| MTHFD1 | 4.76976955 | 1.32730755 | 0.64998383 | 2.0420624 | 0.04114534 | NA |
| ZBTB25 | 6.7155197 | 0.38953864 | 0.64119952 | 0.60751549 | 0.54350887 | NA |
| ZBTB1 | 12.04464 | 0.26516445 | 0.49519935 | 0.53547011 | 0.59232482 | NA |
| PLEKHG3 | 9.166103 | 0.50934261 | 0.458949 | 1.10980222 | 0.26708426 | NA |
| CHURC1-FNT | 6.28420703 | 0.03798659 | 0.6107821 | 0.06219336 | 0.95040885 | NA |
| RAB15 | 3.20949497 | 0.60454607 | 0.7935833 | 0.76179283 | 0.44618366 | NA |
| MAX | 19.9523976 | 0.1744564 | 0.37607372 | 0.46388883 | 0.6427274 | NA |
| FUT8 | 16.0079549 | 0.50149727 | 0.41067898 | 1.22114178 | 0.22203234 | NA |
| AL359232.1 | 40.4548085 | 0.54233484 | 0.4959769 | 1.09346794 | 0.2741884 | 0.76230087 |
| GPHN | 808.001176 | -0.8652505 | 0.38465097 | -2.2494432 | 0.02448432 | 0.37556689 |
| MPP5 | 28.5228965 | -0.5874087 | 0.33279543 | -1.7650744 | 0.07755126 | 0.54469554 |
| ATP6V1D | 24.3437323 | -0.5439848 | 0.46221431 | -1.1769103 | 0.23923133 | NA |
| EIF2S1 | 13.663363 | 0.14452011 | 0.39658631 | 0.36441024 | 0.71555169 | NA |
| PLEK2 | 14.2055129 | -0.160381 | 0.4906328 | -0.3268859 | 0.74375417 | NA |
| PLEKHH1 | 15.4347306 | 0.23352637 | 0.47824719 | 0.48829639 | 0.62533992 | NA |
| PIGH | 8.49653037 | 0.53538264 | 0.56555738 | 0.94664601 | 0.34381919 | NA |
| ARG2 | 22.985297 | -0.6280333 | 0.64275211 | -0.9771003 | 0.3285195 | NA |
| VTI1B | 11.4623898 | 0.87580404 | 0.44004798 | 1.99024671 | 0.04656377 | NA |
| ZFYVE26 | 8.54456954 | -0.5556285 | 0.50957602 | -1.0903741 | 0.2755484 | NA |
| RAD51B | 68.9790562 | 0.75911171 | 0.37553754 | 2.02140035 | 0.04323834 | 0.44186191 |
| ZFP36L1 | 42.5052717 | -0.2854175 | 0.39355373 | -0.7252313 | 0.46831013 | 0.85905137 |
| ACTN1 | 11.7464739 | 0.15498032 | 0.62050137 | 0.24976629 | 0.80276809 | NA |
| DCAF5 | 44.8491947 | -0.01885 | 0.28899132 | -0.065227 | 0.94799328 | 0.99344789 |
| AL359317.2 | 3.61835443 | 1.62448326 | 0.96923506 | 1.67604673 | 0.09372904 | NA |
| EXD2 | 8.95890873 | 0.39216516 | 0.53925544 | 0.7272345 | 0.46708232 | NA |
| ERH | 17.1550033 | -0.2164225 | 0.40457118 | -0.534943 | 0.59268927 | NA |
| SLC39A9 | 22.373373 | 0.13590977 | 0.4679176 | 0.29045663 | 0.77146693 | NA |
| SUSD6 | 37.0264885 | 0.39429021 | 0.36271356 | 1.08705672 | 0.27701174 | 0.76520827 |
| SRSF5 | 55.8237104 | 0.36376562 | 0.3035521 | 1.19836308 | 0.2307757 | 0.73156941 |
| AL357153.2 | 9.62142447 | -0.4066245 | 0.49826826 | -0.8160754 | 0.414457 | NA |
| MED6 | 9.41722708 | -0.4313733 | 0.52488655 | -0.821841 | 0.41116742 | NA |
| TTC9 | 9.64821608 | -0.2256394 | 0.56355669 | -0.4003847 | 0.68887323 | NA |
| MAP3K9 | 14.3768257 | -0.9415281 | 0.49064658 | -1.9189538 | 0.05499018 | NA |
| PCNX1 | 77.1382044 | -0.1632346 | 0.28403366 | -0.5747017 | 0.56549304 | 0.89263108 |
| SIPA1L1 | 58.2537058 | 0.18283232 | 0.33014369 | 0.5537962 | 0.57971833 | 0.89263108 |
| DCAF4 | 3.48645392 | 0.16939961 | 0.75741766 | 0.22365416 | 0.82302642 | NA |
| ZFYVE1 | 23.2896752 | -0.3896642 | 0.37931588 | -1.0272816 | 0.30428788 | NA |
| RBM25 | 41.6805989 | -0.0715889 | 0.28874394 | -0.2479321 | 0.80418696 | 0.96743263 |
| PSEN1 | 39.9941398 | -0.3431146 | 0.27011767 | -1.2702413 | 0.20399869 | 0.700066 |
| PAPLN | 3.68359436 | 0.11463019 | 0.77103498 | 0.14867055 | 0.8818136 | NA |
| NUMB | 131.891843 | -0.2479474 | 0.24103728 | -1.0286684 | 0.30363552 | 0.77550546 |
| HEATR4 | 7.06199428 | 0.90948632 | 0.56988436 | 1.59591381 | 0.11050804 | NA |
| AC005225.2 | 4.09480742 | -0.4457004 | 0.74040214 | -0.6019708 | 0.54719361 | NA |
| DNAL1 | 6.83044472 | 0.2398456 | 0.61412829 | 0.39054641 | 0.69613255 | NA |
| PNMA1 | 3.51101565 | 0.25953302 | 0.80756574 | 0.32137696 | 0.74792474 | NA |
| ELMSAN1 | 13.30604 | 0.18002429 | 0.42311968 | 0.42546896 | 0.67049484 | NA |
| AC005480.1 | 5.95336861 | 0.58751094 | 0.72656592 | 0.80861341 | 0.41873755 | NA |
| COQ6 | 3.62234629 | 1.24963701 | 0.82800584 | 1.50921281 | 0.13124441 | NA |

| | | | | | | |
|---|---|---|---|---|---|---|
| ENTPD5 | 13.3782968 | 0.86042709 | 0.47619866 | 1.80686582 | 0.0707832 | NA |
| BBOF1 | 7.72680296 | 0.25246851 | 0.74273049 | 0.33991941 | 0.73391722 | NA |
| ALDH6A1 | 15.7940285 | 1.17498177 | 0.53240316 | 2.20693988 | 0.02731826 | NA |
| LIN52 | 22.4155582 | -0.9940644 | 0.44420008 | -2.2378753 | 0.02522919 | NA |
| ABCD4 | 15.9431275 | 0.3755083 | 0.41600084 | 0.90266236 | 0.36670512 | NA |
| NPC2 | 23.7490418 | 0.60632998 | 0.44378657 | 1.36626483 | 0.17185586 | NA |
| ISCA2 | 8.19076603 | 0.76864729 | 0.53183588 | 1.4452716 | 0.14838161 | NA |
| LTBP2 | 2.98189457 | 0.83259361 | 1.03666832 | 0.80314368 | 0.42189169 | NA |
| AREL1 | 21.3449058 | -0.4520012 | 0.3850144 | -1.1739852 | 0.24040095 | NA |
| FCF1 | 6.56578246 | 0.20932037 | 0.60826011 | 0.3441297 | 0.73074875 | NA |
| YLPM1 | 37.9036594 | 0.00673361 | 0.37013812 | 0.01819215 | 0.98548557 | 0.99902646 |
| DLST | 17.6868106 | -0.3405362 | 0.37563014 | -0.9065731 | 0.3646326 | NA |
| EIF2B2 | 3.35051231 | -0.2097755 | 0.91508971 | -0.2292404 | 0.81868206 | NA |
| MLH3 | 22.6496572 | 0.35823719 | 0.43675802 | 0.8202189 | 0.41209133 | NA |
| ACYP1 | 13.7561875 | -0.1215317 | 0.48945248 | -0.2483012 | 0.80390133 | NA |
| ZC2HC1C | 5.16922853 | 0.64076168 | 0.6870127 | 0.93267807 | 0.35098621 | NA |
| NEK9 | 19.0862725 | -0.0123314 | 0.3787176 | -0.032561 | 0.97402467 | NA |
| TMED10 | 28.1593209 | 0.36116723 | 0.34097254 | 1.05922674 | 0.28949653 | 0.77266624 |
| AL691403.1 | 23.7412433 | 0.65843338 | 0.58102951 | 1.13321848 | 0.25712252 | NA |
| AF111167.1 | 4.56088464 | 1.07510755 | 0.7414119 | 1.45008134 | 0.14703584 | NA |
| FOS | 10.7068889 | -1.8192543 | 0.87391714 | -2.081724 | 0.03736768 | NA |
| TTLL5 | 24.0926385 | 0.40504976 | 0.33536709 | 1.20778026 | 0.2271318 | NA |
| ERG28 | 6.96698158 | 0.32617305 | 0.54450131 | 0.5990308 | 0.54915235 | NA |
| IFT43 | 17.7140521 | 0.09537256 | 0.40726681 | 0.23417711 | 0.81484749 | NA |
| GPATCH2L | 37.8720961 | -0.2711285 | 0.32576721 | -0.8322767 | 0.40525277 | 0.82808596 |
| ESRRB | 15.1077435 | -0.4133623 | 0.71274114 | -0.5799613 | 0.56194069 | NA |
| ANGEL1 | 5.43194939 | -0.268043 | 0.72685966 | -0.3687686 | 0.71230021 | NA |
| IRF2BPL | 7.50465321 | 0.30018264 | 0.74000805 | 0.4056478 | 0.6850014 | NA |
| POMT2 | 5.16134205 | -0.4550044 | 0.6863608 | -0.6629231 | 0.50737983 | NA |
| GSTZ1 | 3.42453224 | 1.79778731 | 0.86890865 | 2.06901763 | 0.03854443 | NA |
| TMED8 | 14.1366579 | 0.25867048 | 0.45646038 | 0.5666877 | 0.57092638 | NA |
| VIPAS39 | 4.99592857 | -0.0001743 | 0.64636763 | -0.0002697 | 0.99978481 | NA |
| AHSA1 | 11.6932605 | -0.4045567 | 0.46721858 | -0.8658833 | 0.38655419 | NA |
| SPTLC2 | 130.406797 | 0.01325888 | 0.2447581 | 0.05417138 | 0.95679862 | 0.99636969 |
| ALKBH1 | 8.06344398 | -0.3161748 | 0.56352967 | -0.5610615 | 0.5747556 | NA |
| SLIRP | 12.5475086 | -0.2284296 | 0.54742979 | -0.4172764 | 0.67647621 | NA |
| SNW1 | 14.134443 | -0.3612686 | 0.39039606 | -0.92539 | 0.35476307 | NA |
| ADCK1 | 6.09649972 | -1.9275365 | 0.72325265 | -2.6650942 | 0.00769668 | NA |
| NRXN3 | 27.7574751 | -0.0128483 | 0.82409464 | -0.0155908 | 0.98756082 | 0.99909657 |
| DIO2 | 8.78218922 | -3.149267 | 1.33103586 | -2.3660271 | 0.01798013 | NA |
| CEP128 | 8.15012901 | -0.0823143 | 0.54558945 | -0.1508722 | 0.88007655 | NA |
| GTF2A1 | 13.7120114 | 0.35758873 | 0.47052982 | 0.75997039 | 0.44727228 | NA |
| STON2 | 36.9265877 | 0.78661683 | 0.39759318 | 1.97844652 | 0.04787836 | 0.45661231 |
| SEL1L | 44.3092387 | -0.1427151 | 0.39313782 | -0.3630156 | 0.71659326 | 0.93664197 |
| GALC | 4.07109085 | 0.1496422 | 0.75575643 | 0.19800321 | 0.84304256 | NA |
| SPATA7 | 7.69332625 | -0.2537018 | 0.56957282 | -0.4454247 | 0.65601285 | NA |
| PTPN21 | 44.5380258 | -0.4804836 | 0.35638843 | -1.3482021 | 0.17759339 | 0.6676327 |
| ZC3H14 | 34.0129321 | 0.16471908 | 0.29641172 | 0.55571042 | 0.57840883 | 0.89263108 |
| EML5 | 18.4953036 | -0.1063822 | 0.50904501 | -0.2089839 | 0.83446078 | NA |
| TTC8 | 14.9430328 | 0.27375372 | 0.46000081 | 0.59511575 | 0.5517661 | NA |
| FOXN3 | 303.9611 | -0.084739 | 0.40524234 | -0.2091069 | 0.83436475 | 0.97154892 |
| AL356805.1 | 5.19483122 | -0.5539529 | 0.63770716 | -0.8686635 | 0.3850312 | NA |
| EFCAB11 | 7.8907836 | 1.43698805 | 0.61401132 | 2.34032827 | 0.0192668 | NA |
| TDP1 | 5.82820334 | 0.48404525 | 0.73280374 | 0.66053873 | 0.50890817 | NA |

| | | | | | | |
|---|---|---|---|---|---|---|
| PSMC1 | 8.89015378 | -0.5380109 | 0.53133875 | -1.0125572 | 0.31127171 | NA |
| NRDE2 | 10.4086039 | -0.4583244 | 0.51622161 | -0.8878442 | 0.37462456 | NA |
| CALM1 | 30.8477197 | 0.10784894 | 0.32859159 | 0.32821575 | 0.74274854 | 0.94831173 |
| TTC7B | 22.6097209 | -0.6715291 | 0.53037481 | -1.2661406 | 0.20546275 | NA |
| RPS6KA5 | 13.1402985 | -1.2136689 | 0.54193656 | -2.2395037 | 0.02512316 | NA |
| DGLUCY | 47.9299398 | 0.19919058 | 0.38847247 | 0.51275339 | 0.60812384 | 0.9088434 |
| CCDC88C | 2.92622604 | -0.7625974 | 0.92057502 | -0.8283924 | 0.40744829 | NA |
| PPP4R3A | 37.4277807 | -0.436955 | 0.29719298 | -1.4702735 | 0.14148769 | 0.61906663 |
| CATSPERB | 3.41827753 | -0.0215254 | 0.79967085 | -0.0269178 | 0.97852526 | NA |
| TC2N | 9.71061849 | -0.3633208 | 0.49953104 | -0.7273237 | 0.46702767 | NA |
| TRIP11 | 39.8208333 | 0.19504793 | 0.27938017 | 0.69814521 | 0.48508639 | 0.86354292 |
| ATXN3 | 19.7646078 | -0.056722 | 0.37028831 | -0.1531833 | 0.87825376 | NA |
| NDUFB1 | 15.0651619 | 0.52414936 | 0.39522698 | 1.32619833 | 0.18477401 | NA |
| CPSF2 | 13.5957667 | 1.01138426 | 0.47581727 | 2.12557282 | 0.03353884 | NA |
| LGMN | 13.3846461 | 0.27873121 | 0.47682178 | 0.58456057 | 0.55884323 | NA |
| GOLGA5 | 10.8987121 | -0.4566696 | 0.51046033 | -0.8946231 | 0.37098861 | NA |
| ITPK1 | 64.0572563 | 0.19549385 | 0.30141017 | 0.6485974 | 0.51659864 | 0.87158522 |
| MOAP1 | 5.15763348 | -0.0226382 | 0.64558815 | -0.0350659 | 0.97202717 | NA |
| GON7 | 3.87591092 | -0.3656269 | 0.79986542 | -0.4571105 | 0.64759164 | NA |
| UBR7 | 8.11208443 | 1.13791402 | 0.54926404 | 2.07170675 | 0.0382928 | NA |
| BTBD7 | 54.2347225 | 0.02649183 | 0.24347178 | 0.10880861 | 0.9133543 | 0.98415124 |
| DDX24 | 32.3135926 | -0.294428 | 0.29181539 | -1.0089528 | 0.31299728 | 0.78031266 |
| IFI27L1 | 2.90897043 | -0.0719296 | 0.82202114 | -0.0875033 | 0.93027147 | NA |
| IFI27 | 16.701678 | -1.5224435 | 1.42647127 | -1.0672795 | 0.28584566 | NA |
| SERPINA1 | 16.2996176 | -1.2728719 | 0.92463898 | -1.3766151 | 0.16863129 | NA |
| DICER1 | 50.920548 | 0.33158236 | 0.26285416 | 1.26146896 | 0.20713994 | 0.70428546 |
| CLMN | 119.200904 | 0.08021984 | 0.34297393 | 0.23389487 | 0.81506659 | 0.96845695 |
| GLRX5 | 12.6865488 | -0.2436609 | 0.47575308 | -0.5121582 | 0.60854028 | NA |
| C14orf132 | 2.00717745 | 0.12145782 | 1.01416309 | 0.11976163 | 0.90467198 | NA |
| ATG2B | 30.0867264 | 0.40694421 | 0.32139319 | 1.26618804 | 0.20544577 | 0.70278771 |
| GSKIP | 6.21227491 | -0.7632142 | 0.63958915 | -1.1932882 | 0.23275651 | NA |
| PAPOLA | 63.1771968 | 0.02297525 | 0.24607622 | 0.0933664 | 0.92561248 | 0.98758497 |
| VRK1 | 4.99126115 | -0.7213568 | 0.78178058 | -0.9227101 | 0.35615833 | NA |
| SETD3 | 31.1100191 | 0.27001324 | 0.2972 | 0.90852371 | 0.36360159 | 0.80736493 |
| CCNK | 8.30040654 | -0.699155 | 0.53597276 | -1.3044599 | 0.19207682 | NA |
| CCDC85C | 11.5798022 | 0.81436852 | 0.48141067 | 1.69162957 | 0.09071662 | NA |
| EVL | 23.3291027 | -0.2640732 | 0.43316447 | -0.6096372 | 0.54210219 | NA |
| DEGS2 | 3.85192002 | 1.41791267 | 0.96987437 | 1.46195499 | 0.14375355 | NA |
| YY1 | 34.5301311 | -0.3429605 | 0.32163455 | -1.0663048 | 0.2862859 | 0.77103639 |
| SLC25A29 | 19.4308052 | -0.9606097 | 0.47787748 | -2.010159 | 0.04441436 | NA |
| WARS | 6.75005558 | -0.6027574 | 0.95794752 | -0.6292176 | 0.52920664 | NA |
| WDR25 | 5.84552773 | -0.6832992 | 0.67453276 | -1.0129963 | 0.31106192 | NA |
| PPP2R5C | 39.653392 | -0.1369957 | 0.3379992 | -0.4053137 | 0.68524697 | 0.92726927 |
| DYNC1H1 | 36.4735963 | -0.2416771 | 0.34269072 | -0.7052338 | 0.48066473 | 0.86222804 |
| HSP90AA1 | 143.680814 | -0.6369536 | 0.25625271 | -2.4856461 | 0.01293165 | 0.30435125 |
| WDR20 | 38.4781458 | -0.2820002 | 0.33237662 | -0.8484356 | 0.3961954 | 0.82600747 |
| MOK | 6.28741709 | -1.0550148 | 0.79297844 | -1.3304457 | 0.18337146 | NA |
| ZNF839 | 5.33267702 | 1.02639048 | 0.67408897 | 1.52263355 | 0.12785041 | NA |
| CINP | 4.3573248 | 0.6990228 | 0.67597191 | 1.03410038 | 0.30108924 | NA |
| TECPR2 | 15.7741717 | 0.11607552 | 0.55364505 | 0.20965692 | 0.83393545 | NA |
| ANKRD9 | 14.2678495 | 0.39257675 | 0.42042686 | 0.93375755 | 0.35042897 | NA |
| RCOR1 | 42.1720161 | -0.1677763 | 0.45833234 | -0.3660582 | 0.71432166 | 0.93629694 |
| TRAF3 | 17.3236765 | -0.0196367 | 0.4305026 | -0.0456134 | 0.96361839 | NA |
| CDC42BPB | 80.6166325 | -0.4852153 | 0.26631102 | -1.8219873 | 0.06845692 | 0.52508412 |

| | | | | | | |
|---|---|---|---|---|---|---|
| EIF5 | 26.6383403 | -0.0809503 | 0.45301311 | -0.1786931 | 0.85817865 | 0.97168963 |
| MARK3 | 88.4463563 | -0.1391078 | 0.32073676 | -0.4337132 | 0.66449669 | 0.92276785 |
| CKB | 53.6064877 | 0.34290609 | 0.39972771 | 0.85784919 | 0.39097574 | 0.8258317 |
| TRMT61A | 2.94803937 | -0.7224341 | 0.87435674 | -0.8262464 | 0.40866435 | NA |
| BAG5 | 10.9698575 | 0.24221267 | 0.46212932 | 0.52412315 | 0.60019289 | NA |
| KLC1 | 22.5916749 | -0.3526813 | 0.46436064 | -0.7594988 | 0.44755424 | NA |
| ZFYVE21 | 8.29856488 | 0.16529743 | 0.52648975 | 0.31396135 | 0.75355039 | NA |
| PPP1R13B | 70.2257078 | -0.9082027 | 0.42629792 | -2.1304414 | 0.03313519 | 0.41688714 |
| ATP5MPL | 20.7708945 | 0.20449809 | 0.3867549 | 0.52875372 | 0.59697631 | NA |
| SIVA1 | 10.894765 | -0.424953 | 0.47740666 | -0.8901279 | 0.37339719 | NA |
| AKT1 | 11.5105025 | -1.0210012 | 0.52720196 | -1.9366414 | 0.05278919 | NA |
| CEP170B | 7.35328613 | -0.2980129 | 0.64626191 | -0.4611332 | 0.64470301 | NA |
| AHNAK2 | 2.6556429 | -1.2702266 | 0.90718372 | -1.4001867 | 0.1614574 | NA |
| NUDT14 | 6.57861437 | 0.25432758 | 0.58366676 | 0.43574107 | 0.6630246 | NA |
| BRF1 | 9.54050355 | -0.4179938 | 0.52597768 | -0.7946987 | 0.4267888 | NA |
| BTBD6 | 5.31580585 | -0.0216728 | 0.64509409 | -0.0335964 | 0.97319902 | NA |
| PACS2 | 19.608448 | -0.0928439 | 0.51020074 | -0.1819753 | 0.85560214 | NA |
| MTA1 | 7.2454381 | -0.8974883 | 0.58863193 | -1.524702 | 0.12733344 | NA |
| NIPA1 | 11.6055861 | 0.11271905 | 0.42911702 | 0.26267673 | 0.79279976 | NA |
| NIPA2 | 15.9949209 | -0.0491472 | 0.46094409 | -0.1066229 | 0.91508813 | NA |
| CYFIP1 | 18.4730145 | 0.30342761 | 0.41022713 | 0.7396576 | 0.45950778 | NA |
| TUBGCP5 | 10.522401 | 0.30828725 | 0.53638352 | 0.57475153 | 0.56545935 | NA |
| PWRN3 | 6.3437326 | -1.6245394 | 1.62554083 | -0.9993839 | 0.31760874 | NA |
| PWRN1 | 48.5422807 | -0.6522083 | 0.80691825 | -0.8082706 | 0.41893483 | 0.83856261 |
| AC124312.1 | 64.9202908 | 0.20207321 | 0.28576916 | 0.70712042 | 0.47949165 | 0.86206794 |
| PWAR6 | 13.2048844 | 0.16316687 | 0.48672916 | 0.33523133 | 0.73745059 | NA |
| UBE3A | 59.4338177 | -0.2498722 | 0.28516454 | -0.8762387 | 0.38090028 | 0.82153751 |
| GABRB3 | 3.9655965 | 0.37951775 | 0.97638497 | 0.38869684 | 0.69750042 | NA |
| HERC2 | 51.557238 | -0.2374388 | 0.27795921 | -0.8542219 | 0.39298207 | 0.8258317 |
| FAM189A1 | 24.9206733 | 0.61146328 | 0.53361518 | 1.14588807 | 0.25184147 | NA |
| AC022613.1 | 4.60064699 | 0.57177469 | 0.85100163 | 0.67188437 | 0.50165731 | NA |
| TJP1 | 58.0529137 | -0.0662449 | 0.28224231 | -0.2347094 | 0.81443429 | 0.96845695 |
| FAN1 | 11.350198 | 1.04305265 | 0.48643063 | 2.1442989 | 0.03200895 | NA |
| MTMR10 | 38.0056515 | -0.0937496 | 0.45094917 | -0.207894 | 0.83531172 | 0.97154892 |
| KLF13 | 21.5874256 | -0.3805078 | 0.4391984 | -0.8663688 | 0.38628794 | NA |
| OTUD7A | 20.8342301 | 0.90694398 | 0.42311828 | 2.14347627 | 0.03207487 | NA |
| LINC02256 | 27.9358036 | 0.60913342 | 0.4891988 | 1.24516542 | 0.21307095 | 0.71284112 |
| AC123768.3 | 4.14807131 | -0.016624 | 0.82641605 | -0.0201157 | 0.98395105 | NA |
| GOLGA8N | 4.16808594 | 1.4126011 | 0.87267179 | 1.61870833 | 0.10551003 | NA |
| FMN1 | 14.5377779 | 0.68752247 | 0.56430754 | 1.21834711 | 0.22309209 | NA |
| RYR3 | 7.96134672 | 0.08763527 | 0.81076318 | 0.10808985 | 0.91392442 | NA |
| AVEN | 21.35757 | 0.00139281 | 0.41418454 | 0.00336279 | 0.99731689 | NA |
| EMC7 | 10.2865008 | 0.99462601 | 0.45589558 | 2.18169698 | 0.0291319 | NA |
| KATNBL1 | 35.2092338 | -0.154771 | 0.33813849 | -0.4577149 | 0.64715727 | 0.9218151 |
| EMC4 | 5.31571698 | 0.07489447 | 0.62336303 | 0.12014584 | 0.90436762 | NA |
| SLC12A6 | 14.0422792 | 0.53141771 | 0.5653328 | 0.94000863 | 0.34721313 | NA |
| NOP10 | 18.0780675 | 0.04035674 | 0.46377586 | 0.08701777 | 0.93065739 | NA |
| GOLGA8A | 12.6069827 | -0.2748757 | 0.60108835 | -0.4572968 | 0.64745776 | NA |
| GOLGA8B | 15.6356689 | 0.45712036 | 0.41927172 | 1.09027236 | 0.27559319 | NA |
| AQR | 21.4909354 | 0.06078071 | 0.34891271 | 0.17420033 | 0.86170803 | NA |
| ZNF770 | 12.8206753 | 0.25770657 | 0.42129022 | 0.61170793 | 0.54073101 | NA |
| AC021231.1 | 23.629322 | 1.25902989 | 0.49645378 | 2.53604654 | 0.01121118 | NA |
| DPH6 | 84.1717017 | 0.85725824 | 0.31041172 | 2.76168134 | 0.00575046 | 0.22794137 |
| DPH6-DT | 10.6137932 | 0.9963102 | 0.91932155 | 1.08374507 | 0.27847784 | NA |

| | | | | | | |
|---|---|---|---|---|---|---|
| C15orf41 | 14.58915 | 1.61340959 | 0.48979117 | 3.29407652 | 0.00098746 | NA |
| SPRED1 | 27.67775 | 0.20228201 | 0.37801445 | 0.53511715 | 0.59256885 | 0.89685273 |
| FAM98B | 10.203208 | 0.06975112 | 0.57834026 | 0.12060569 | 0.90400336 | NA |
| RASGRP1 | 4.41567155 | 0.59240346 | 0.88398478 | 0.6701512 | 0.50276141 | NA |
| C15orf53 | 90.4305776 | 2.80002051 | 0.73170399 | 3.82671212 | 0.00012987 | 0.04332663 |
| AC013652.1 | 18.6635051 | 1.82389052 | 0.67594452 | 2.69828435 | 0.00696979 | NA |
| THBS1 | 15.5289698 | -1.2258052 | 0.72499803 | -1.6907704 | 0.09088067 | NA |
| FSIP1 | 3.53402502 | -0.7524104 | 0.76494106 | -0.9836187 | 0.32530301 | NA |
| EIF2AK4 | 14.7459523 | 0.44105525 | 0.43735912 | 1.00845102 | 0.31323799 | NA |
| SRP14 | 81.4011333 | 0.07818629 | 0.31495597 | 0.24824515 | 0.80394473 | 0.96743263 |
| SRP14-AS1 | 4.72460497 | 0.96605036 | 0.92301883 | 1.04662043 | 0.29527467 | NA |
| BMF | 6.66708734 | 0.20412647 | 0.618737 | 0.3299083 | 0.74146925 | NA |
| IVD | 15.0460136 | 0.05032672 | 0.43118834 | 0.11671633 | 0.90708485 | NA |
| BAHD1 | 8.20895454 | -0.3872604 | 0.56217792 | -0.6888573 | 0.49091305 | NA |
| CCDC32 | 14.180633 | -0.5379913 | 0.48894981 | -1.1002997 | 0.27120158 | NA |
| RMDN3 | 7.68913874 | 0.11612948 | 0.53007801 | 0.21907999 | 0.82658774 | NA |
| GCHFR | 6.15704431 | 0.98902309 | 0.60181921 | 1.64338904 | 0.10030247 | NA |
| DNAJC17 | 10.0311194 | 0.22409565 | 0.52914188 | 0.42350768 | 0.6719249 | NA |
| ZFYVE19 | 2.95416185 | 0.13748632 | 0.9763303 | 0.14081948 | 0.88801256 | NA |
| SPINT1-AS1 | 5.96943586 | 0.68758071 | 0.62273075 | 1.10413805 | 0.26953326 | NA |
| SPINT1 | 16.5060469 | -0.0218098 | 0.43372161 | -0.0502853 | 0.95989506 | NA |
| VPS18 | 4.22063066 | -0.8102114 | 0.85168236 | -0.951307 | 0.34144854 | NA |
| INO80 | 32.882702 | -0.5894756 | 0.32062605 | -1.8385143 | 0.06598666 | 0.5230103 |
| CHP1 | 22.2298993 | -0.3206199 | 0.49373353 | -0.6493785 | 0.51609378 | NA |
| NDUFAF1 | 2.44314614 | 0.10349688 | 1.01728809 | 0.10173803 | 0.91896462 | NA |
| RTF1 | 21.3702161 | -0.3977254 | 0.3862797 | -1.0296306 | 0.30318343 | NA |
| RPAP1 | 4.90410305 | 0.16452346 | 0.70522529 | 0.23329205 | 0.81553463 | NA |
| MGA | 31.9015595 | -0.0709817 | 0.31877015 | -0.2226735 | 0.82378964 | 0.96845695 |
| MAPKBP1 | 21.9812657 | -0.5949747 | 0.47583715 | -1.2503747 | 0.21116272 | NA |
| EHD4 | 9.67884634 | -0.9210726 | 0.54658058 | -1.6851542 | 0.09195882 | NA |
| PLA2G4F | 8.70259573 | 1.28742142 | 0.79643765 | 1.61647483 | 0.10599169 | NA |
| VPS39 | 18.7558707 | 0.88002948 | 0.43248015 | 2.03484364 | 0.04186661 | NA |
| TMEM87A | 45.350585 | -0.2700719 | 0.28317316 | -0.9537342 | 0.34021819 | 0.8003972 |
| GANC | 10.1306928 | 0.7147678 | 0.54675388 | 1.30729351 | 0.19111305 | NA |
| ZNF106 | 20.943003 | -0.1555578 | 0.46215094 | -0.3365951 | 0.73642214 | NA |
| SNAP23 | 18.2390886 | 0.24307329 | 0.39073526 | 0.62209203 | 0.53388135 | NA |
| LRRC57 | 3.31826923 | 0.49688019 | 0.82762399 | 0.60036949 | 0.54826002 | NA |
| HAUS2 | 4.45957541 | -0.0516476 | 0.71852205 | -0.0718804 | 0.94269709 | NA |
| TTBK2 | 29.4138359 | 0.5325674 | 0.37432977 | 1.42272253 | 0.15481661 | 0.63864841 |
| UBR1 | 44.3834109 | -0.3101724 | 0.28450663 | -1.0902114 | 0.27562002 | 0.76410819 |
| AC068724.3 | 10.1230653 | 0.39840529 | 0.51497844 | 0.77363488 | 0.43914674 | NA |
| TMEM62 | 10.5018722 | -0.4655891 | 0.49947704 | -0.9321531 | 0.35125739 | NA |
| CCNDBP1 | 11.5880909 | 0.07695239 | 0.54862362 | 0.14026444 | 0.88845106 | NA |
| ADAL | 9.36698977 | 0.53399335 | 0.51041978 | 1.04618468 | 0.29547578 | NA |
| ZSCAN29 | 3.11555652 | 0.2927671 | 0.80183676 | 0.36512057 | 0.7150214 | NA |
| TUBGCP4 | 8.00869671 | 0.44110738 | 0.51827545 | 0.85110607 | 0.39471043 | NA |
| TP53BP1 | 26.9121812 | -0.2349121 | 0.32363723 | -0.7258501 | 0.46793067 | 0.85894565 |
| PPIP5K1 | 13.8907086 | 0.11255559 | 0.44699023 | 0.25180771 | 0.8011897 | NA |
| CATSPER2 | 11.3119518 | -0.3141566 | 0.59404391 | -0.5288441 | 0.59691359 | NA |
| PDIA3 | 30.6758794 | -0.0220665 | 0.3786631 | -0.0582748 | 0.95352976 | 0.99490654 |
| SERF2 | 140.531471 | -0.5489471 | 0.35013031 | -1.5678366 | 0.11691927 | 0.57860396 |
| MFAP1 | 6.78439025 | 0.28808902 | 0.56847818 | 0.50677234 | 0.61231457 | NA |
| FRMD5 | 3.77074026 | -1.093468 | 0.99616962 | -1.0976725 | 0.27234751 | NA |
| CASC4 | 59.1013123 | 0.13216512 | 0.23997661 | 0.55074168 | 0.58181077 | 0.89263108 |

| | | | | | | |
|---|---|---|---|---|---|---|
| AC025043.1 | 4.81222101 | 1.88313942 | 0.79889046 | 2.35719353 | 0.01841365 | NA |
| CTDSPL2 | 34.5848945 | -0.3254167 | 0.31819357 | -1.0227004 | 0.30644954 | 0.77729327 |
| EIF3J-DT | 6.88850355 | 0.40670222 | 0.6240227 | 0.65174268 | 0.51456719 | NA |
| EIF3J | 19.9808276 | -0.0751072 | 0.4226813 | -0.1776922 | 0.85896473 | NA |
| SPG11 | 59.4305067 | 0.09898327 | 0.2669988 | 0.37072553 | 0.71084197 | 0.93492235 |
| B2M | 366.816083 | 0.12468708 | 0.4746175 | 0.26271068 | 0.79277359 | 0.96341299 |
| TRIM69 | 5.16727282 | -0.6414205 | 0.75193479 | -0.8530268 | 0.39364446 | NA |
| SORD | 7.24370278 | -0.1727493 | 0.55104647 | -0.3134932 | 0.75390595 | NA |
| GATM | 11.3053507 | 0.54883332 | 1.00957498 | 0.54362809 | 0.58669742 | NA |
| SPATA5L1 | 4.71907534 | 0.12081997 | 0.7372016 | 0.16389 | 0.86981774 | NA |
| SQOR | 4.19937027 | 1.79932839 | 0.84705763 | 2.12421012 | 0.03365258 | NA |
| MYEF2 | 23.0990278 | -0.2062527 | 0.39332254 | -0.5243857 | 0.60001031 | NA |
| SLC12A1 | 51.9288626 | -0.479097 | 1.03211284 | -0.4641905 | 0.64251125 | 0.92153493 |
| DUT | 10.774279 | 1.30802547 | 0.4992248 | 2.62011315 | 0.00879006 | NA |
| CEP152 | 10.3217749 | 0.94619083 | 0.62230355 | 1.52046511 | 0.12839412 | NA |
| EID1 | 32.1821161 | 0.12640625 | 0.37911977 | 0.33342036 | 0.738817 | 0.94658914 |
| SECISBP2L | 48.4540718 | 0.03634487 | 0.28206736 | 0.12885173 | 0.89747497 | 0.97769824 |
| COPS2 | 11.1271936 | -0.1841374 | 0.45955118 | -0.4006895 | 0.68864871 | NA |
| GALK2 | 22.5714751 | 0.76983735 | 0.37316888 | 2.06297306 | 0.03911519 | NA |
| FAM227B | 10.776999 | 0.88691918 | 0.62942961 | 1.40908397 | 0.15881034 | NA |
| DTWD1 | 11.9353975 | 0.21811422 | 0.75435958 | 0.28913827 | 0.77247557 | NA |
| ATP8B4 | 28.6961927 | -0.5328673 | 0.54833997 | -0.9717828 | 0.33115863 | 0.79412614 |
| GABPB1 | 15.6829303 | -1.0512329 | 0.49154351 | -2.1386366 | 0.03246511 | NA |
| GABPB1-AS1 | 12.9052776 | 1.47222768 | 0.57895435 | 2.54290805 | 0.01099342 | NA |
| USP8 | 31.7287548 | -0.1315929 | 0.30318884 | -0.4340296 | 0.66426691 | 0.92276785 |
| USP50 | 4.35256161 | -0.9529849 | 0.71245409 | -1.3376088 | 0.18102399 | NA |
| TRPM7 | 64.7573531 | -0.7037421 | 0.30517023 | -2.306064 | 0.02110705 | 0.34990513 |
| SPPL2A | 45.5062731 | -0.3802722 | 0.33475537 | -1.1359705 | 0.25596892 | 0.75170774 |
| AP4E1 | 16.2266523 | 0.78058244 | 0.47525532 | 1.64244861 | 0.10049707 | NA |
| DMXL2 | 16.6064662 | -0.3348108 | 0.42501813 | -0.7877566 | 0.43083907 | NA |
| LYSMD2 | 4.13867256 | -1.1233178 | 0.91661345 | -1.2255088 | 0.22038358 | NA |
| TMOD2 | 4.43366931 | 0.6279405 | 0.64680906 | 0.97082824 | 0.33163382 | NA |
| TMOD3 | 37.8745938 | 0.07929203 | 0.32003731 | 0.24775872 | 0.8043211 | 0.96743263 |
| LEO1 | 6.61350003 | 0.46173032 | 0.68315371 | 0.67588057 | 0.49911648 | NA |
| MAPK6 | 19.6227041 | -0.4448144 | 0.37128883 | -1.1980278 | 0.23090618 | NA |
| GNB5 | 9.98397158 | 0.19695103 | 0.46236089 | 0.42596819 | 0.67013103 | NA |
| MYO5C | 18.8756452 | 0.24427777 | 0.5779741 | 0.42264485 | 0.6725544 | NA |
| MYO5A | 26.2952772 | -0.5770416 | 0.38360423 | -1.5042628 | 0.13251372 | 0.60561493 |
| ARPP19 | 33.2078957 | -0.0945975 | 0.33958075 | -0.2785714 | 0.78057377 | 0.96139889 |
| FAM214A | 33.4667221 | 0.5167194 | 0.35742688 | 1.44566464 | 0.14827129 | 0.63 |
| WDR72 | 806.917911 | -0.5912054 | 0.26722457 | -2.2123915 | 0.02693962 | 0.39506514 |
| UNC13C | 83.8270698 | -0.6877574 | 0.45978343 | -1.495829 | 0.13469823 | 0.60594834 |
| RSL24D1 | 10.7867183 | -0.4077517 | 0.42586702 | -0.9574624 | 0.3383339 | NA |
| RAB27A | 16.7840447 | -0.0955442 | 0.47565521 | -0.2008686 | 0.84080131 | NA |
| PIGB | 8.06754444 | -0.2524464 | 0.51882696 | -0.4865714 | 0.62656208 | NA |
| CCPG1 | 30.5077548 | 0.38559222 | 0.3399829 | 1.13415179 | 0.25673088 | 0.75170774 |
| DNAAF4 | 4.69737815 | 0.30398368 | 0.69091605 | 0.43997195 | 0.65995742 | NA |
| PRTG | 4.12708143 | 0.09343925 | 0.77612952 | 0.12039131 | 0.90417317 | NA |
| NEDD4 | 4.70393167 | 0.97808986 | 0.75243259 | 1.29990363 | 0.193634 | NA |
| RFX7 | 31.5911365 | 0.09181145 | 0.38524729 | 0.23831823 | 0.81163428 | 0.96845695 |
| TEX9 | 9.27926559 | 0.08619205 | 0.60428801 | 0.14263406 | 0.8865792 | NA |
| ZNF280D | 14.722977 | 0.32569362 | 0.41825974 | 0.77868748 | 0.43616383 | NA |
| AC090517.4 | 5.52814836 | 0.36623765 | 0.65281669 | 0.56101146 | 0.57478972 | NA |
| TCF12 | 183.561628 | -0.0721643 | 0.22338657 | -0.3230469 | 0.74665973 | 0.94858421 |

| | | | | | | |
|---|---|---|---|---|---|---|
| CGNL1 | 214.331801 | 0.62926104 | 0.30548648 | 2.05986539 | 0.03941141 | 0.43110269 |
| POLR2M | 5.79062923 | -0.64761 | 0.67616137 | -0.9577743 | 0.33817657 | NA |
| ALDH1A2 | 16.7765304 | -0.6297447 | 0.69042751 | -0.9121083 | 0.36171171 | NA |
| ADAM10 | 63.3265639 | 0.01774882 | 0.27332989 | 0.06493551 | 0.94822535 | 0.99344789 |
| MINDY2 | 57.8664458 | 0.34453191 | 0.28911916 | 1.19166062 | 0.23339435 | 0.73156941 |
| RNF111 | 50.7172925 | -0.0432646 | 0.24553862 | -0.176203 | 0.86013445 | 0.97180546 |
| SLTM | 59.9545856 | -0.4218895 | 0.25357213 | -1.6637849 | 0.09615543 | 0.56358504 |
| MYO1E | 48.8294021 | -0.6122338 | 0.33348357 | -1.8358738 | 0.06637632 | 0.5230103 |
| FAM81A | 3.05778304 | 0.22726407 | 0.99487531 | 0.22843472 | 0.8193083 | NA |
| GTF2A2 | 14.3249541 | 0.16359799 | 0.46054881 | 0.355224 | 0.72242179 | NA |
| BNIP2 | 15.7843272 | -0.5101678 | 0.43534403 | -1.1718728 | 0.24124812 | NA |
| ANXA2 | 29.5542285 | -0.2544449 | 0.39781744 | -0.6396022 | 5.22E-01 | 8.73E-01 |
| ICE2 | 21.4966611 | 0.44075163 | 0.35876097 | 1.22853842 | 0.21924492 | NA |
| RORA-AS1 | 12.1075006 | 0.1317228 | 0.77502544 | 0.16995933 | 0.86504212 | NA |
| RORA | 655.045979 | 0.72243253 | 0.55523562 | 1.30112785 | 0.1932147 | 0.68667113 |
| AC012404.1 | 20.8067299 | 0.79346903 | 0.82721338 | 0.9592072 | 0.33745437 | NA |
| AC009554.1 | 9.3282761 | 0.73847149 | 0.54286053 | 1.36033373 | 0.17372434 | NA |
| VPS13C | 72.7228439 | 0.30992493 | 0.28478799 | 1.08826543 | 0.27647795 | 0.76520827 |
| TLN2 | 13.5537871 | 0.81864944 | 0.582242 | 1.40602952 | 0.15971536 | NA |
| TPM1 | 48.6683799 | 0.33051001 | 0.32201284 | 1.02638765 | 0.3047089 | 0.77676031 |
| LACTB | 8.69749949 | -0.0003232 | 0.59204807 | -0.0005458 | 0.9995645 | NA |
| RPS27L | 12.1292527 | 0.2569109 | 0.48015518 | 0.53505806 | 0.59260971 | NA |
| RAB8B | 11.8505683 | -0.4701502 | 0.47742169 | -0.9847692 | 0.32473747 | NA |
| APH1B | 8.69868533 | -0.2451182 | 0.48399921 | -0.5064434 | 0.61254542 | NA |
| CA12 | 1258.06054 | -0.2947506 | 0.33938061 | -0.8684957 | 0.38512305 | 0.8256172 |
| LINC02568 | 25.9665656 | -0.6063041 | 0.75835946 | -0.7994943 | 0.42400383 | 0.84057082 |
| USP3 | 112.222001 | -0.7431241 | 0.28256571 | -2.6299161 | 0.00854059 | 0.27648451 |
| USP3-AS1 | 27.9391791 | 0.56252065 | 0.4309022 | 1.30544856 | 0.19174015 | 0.68667113 |
| HERC1 | 93.275961 | 0.2860255 | 0.24002861 | 1.19163086 | 0.23340602 | 0.73156941 |
| DAPK2 | 8.63911317 | -0.9443964 | 0.92881812 | -1.0167722 | 0.30926183 | NA |
| FAM96A | 13.715605 | 0.42173122 | 0.42169984 | 1.00007442 | 0.31727449 | NA |
| SNX1 | 23.520729 | -0.3930935 | 0.33617916 | -1.1692976 | 0.24228375 | NA |
| PPIB | 30.816745 | -0.2261955 | 0.42572823 | -0.5313143 | 0.595201 | 0.89852459 |
| CSNK1G1 | 7.01242885 | 0.26245914 | 0.60275852 | 0.43542999 | 0.66325034 | NA |
| TRIP4 | 16.3275722 | -0.0154829 | 0.41396551 | -0.0374015 | 0.9701649 | NA |
| ZNF609 | 84.8913856 | -0.2978734 | 0.24735338 | -1.2042424 | 0.2284959 | 0.73156941 |
| OAZ2 | 22.4620538 | 0.07744161 | 0.48865695 | 0.15847848 | 0.87407978 | NA |
| RBPMS2 | 5.85215496 | -0.3337681 | 0.7452391 | -0.4478671 | 0.65424912 | NA |
| SPG21 | 12.8522525 | -0.8989632 | 0.52628801 | -1.7081202 | 0.08761404 | NA |
| MTFMT | 8.85518456 | 0.14938858 | 0.51278838 | 0.29132599 | 0.77080201 | NA |
| UBAP1L | 6.18873259 | -0.6016309 | 0.66290581 | -0.9075661 | 0.36410749 | NA |
| PDCD7 | 6.43493193 | -0.0917441 | 0.60982379 | -0.1504437 | 0.88041457 | NA |
| CLPX | 20.3085327 | 0.63594122 | 0.39327458 | 1.61704125 | 0.10586938 | NA |
| PARP16 | 4.17714425 | 0.15305604 | 0.72143023 | 0.21215639 | 0.83198502 | NA |
| DPP8 | 26.4859201 | -0.3489686 | 0.32535528 | -1.0725771 | 0.28346091 | 0.76977033 |
| HACD3 | 31.0689346 | 0.35589243 | 0.29326588 | 1.21354868 | 0.22492011 | 0.72765065 |
| INTS14 | 6.89297326 | -0.5397111 | 0.56256548 | -0.9593748 | 0.33736999 | NA |
| SLC24A1 | 5.41447475 | 0.1552058 | 0.83159035 | 0.18663732 | 0.851945 | NA |
| DENND4A | 78.0402124 | 0.0058901 | 0.29479442 | 0.01998037 | 0.98405903 | 0.99902646 |
| RAB11A | 41.9601214 | 0.06213925 | 0.38530448 | 0.1612731 | 0.87187831 | 0.97488194 |
| DIS3L | 6.26178252 | 0.97390067 | 0.59271682 | 1.64311293 | 0.10035957 | NA |
| TIPIN | 5.2835358 | 0.8338135 | 0.69644942 | 1.19723482 | 0.23121504 | NA |
| MAP2K1 | 19.8919176 | -0.7236629 | 0.41296323 | -1.7523664 | 0.07971083 | NA |
| SNAPC5 | 4.31644902 | -0.0325194 | 0.69133019 | -0.0470389 | 0.96248222 | NA |

| | | | | | | |
|---|---|---|---|---|---|---|
| RPL4 | 63.9849929 | 0.05721159 | 0.34199796 | 0.16728634 | 0.86714475 | 0.97217819 |
| ZWILCH | 2.57921858 | -0.2759639 | 1.06675197 | -0.2586955 | 0.79587018 | NA |
| SMAD6 | 7.24026862 | -0.3134015 | 0.71382085 | -0.4390479 | 0.66062683 | NA |
| SMAD3 | 57.6271984 | -0.261237 | 0.31911039 | -0.8186415 | 0.41299098 | 0.83356565 |
| AAGAB | 24.0760671 | -0.4828581 | 0.39539514 | -1.221204 | 0.22200878 | NA |
| IQCH | 9.68095092 | 1.42480823 | 0.53524568 | 2.66197056 | 0.00776847 | NA |
| IQCH-AS1 | 16.7380585 | 1.27607492 | 0.59622912 | 2.14024254 | 0.03233517 | NA |
| C15orf61 | 4.0355904 | 0.58852418 | 0.76865386 | 0.76565566 | 0.44388121 | NA |
| MAP2K5 | 40.821765 | 0.49763264 | 0.33584381 | 1.48173832 | 0.13840994 | 0.61240852 |
| PIAS1 | 80.1349871 | -0.1911713 | 0.24265716 | -0.7878249 | 0.43079916 | 0.84223493 |
| FEM1B | 10.6471225 | -0.4228337 | 0.53903899 | -0.7844213 | 0.43279296 | NA |
| CORO2B | 18.951075 | -1.3884506 | 0.62382547 | -2.2257036 | 0.02603404 | NA |
| ANP32A | 20.4976043 | 0.75132998 | 0.38971884 | 1.92787696 | 0.05387043 | NA |
| SPESP1 | 3.00278681 | 1.37633295 | 0.88161743 | 1.56114534 | 0.11848946 | NA |
| GLCE | 11.321494 | 0.63286102 | 0.50802623 | 1.24572509 | 0.21286534 | NA |
| PAQR5 | 22.5999779 | -0.3919308 | 0.44928664 | -0.8723401 | 0.38302286 | NA |
| RPLP1 | 114.631332 | 0.0666845 | 0.37035931 | 0.18005353 | 0.85711054 | 0.97168963 |
| TLE3 | 11.0111744 | -0.257873 | 0.45169593 | -0.5708995 | 0.56806778 | NA |
| UACA | 47.2498648 | -0.2514256 | 0.46012424 | -0.5464299 | 0.58477048 | 0.89390172 |
| LRRC49 | 4.98990628 | 0.9765995 | 0.67851134 | 1.43932671 | 0.15005798 | NA |
| THSD4 | 15.6397554 | 0.17321705 | 0.59486243 | 0.29118842 | 0.77090722 | NA |
| THSD4-AS1 | 18.0242755 | 0.01454329 | 0.41527459 | 0.03502089 | 0.97206308 | NA |
| MYO9A | 136.146796 | 0.36857661 | 0.29368443 | 1.255009 | 0.20947549 | 0.70681427 |
| AC022872.1 | 5.08934817 | 0.52978678 | 0.97766113 | 0.54189203 | 0.58789289 | NA |
| SENP8 | 2.81594653 | 0.54736138 | 1.03980926 | 0.52640556 | 0.59860645 | NA |
| PKM | 74.4400537 | -0.5193116 | 0.30799364 | -1.6861115 | 0.09177432 | 0.56220784 |
| PARP6 | 12.5328157 | 0.03108417 | 0.47804817 | 0.0650231 | 0.94815561 | NA |
| AC009690.2 | 7.52174855 | -0.2800059 | 0.69312214 | -0.4039777 | 0.6862291 | NA |
| AC009690.1 | 29.4344183 | -0.5542685 | 0.59726952 | -0.928004 | 0.35340549 | 0.8003972 |
| ARIH1 | 111.605843 | -0.5018121 | 0.28583339 | -1.7556106 | 0.07915492 | 0.54552914 |
| BBS4 | 10.7858756 | 0.62083123 | 0.60109497 | 1.03283385 | 0.30168166 | NA |
| ADPGK | 13.6728672 | -0.1742685 | 0.4343681 | -0.4012001 | 0.68827283 | NA |
| NEO1 | 42.5491785 | 0.6671042 | 0.41770631 | 1.59706518 | 0.11025118 | 0.57860396 |
| NPTN | 49.7478566 | -0.2436557 | 0.2754303 | -0.8846366 | 0.37635268 | 0.81744083 |
| CD276 | 3.99404397 | -1.0678928 | 0.80391676 | -1.3283624 | 0.18405842 | NA |
| PML | 4.9632279 | -0.1002298 | 0.80769589 | -0.1240934 | 0.9012413 | NA |
| ARID3B | 3.49209303 | 1.03582895 | 0.82145818 | 1.26096371 | 0.20732192 | NA |
| CLK3 | 5.52011678 | -0.1981825 | 0.63255108 | -0.3133067 | 0.75404766 | NA |
| EDC3 | 11.2177361 | -0.1939437 | 0.49895473 | -0.3887001 | 0.69749802 | NA |
| CSK | 3.23886882 | 0.6862998 | 1.00914783 | 0.68007855 | 0.49645472 | NA |
| ULK3 | 3.79829667 | 1.01355514 | 0.81530292 | 1.24316389 | 0.21380745 | NA |
| SCAMP2 | 14.0912088 | -0.4711098 | 0.45954365 | -1.0251688 | 0.30528353 | NA |
| FAM219B | 4.72818068 | -0.028497 | 0.65287444 | -0.0436485 | 0.96518456 | NA |
| COX5A | 31.7910884 | -0.1100775 | 0.41910141 | -0.2626513 | 0.79281939 | 0.96341299 |
| SCAMP5 | 4.20465295 | -0.8770242 | 0.7909991 | -1.108755 | 0.2675359 | NA |
| PPCDC | 7.85925112 | -0.2473428 | 0.60162449 | -0.4111249 | 0.68098096 | NA |
| COMMD4 | 9.50758701 | 0.31146159 | 0.47567555 | 0.65477739 | 0.51261109 | NA |
| NEIL1 | 4.33628022 | 1.07669372 | 0.80603541 | 1.33578959 | 0.18161806 | NA |
| MAN2C1 | 8.93122431 | -0.3224143 | 0.56689493 | -0.5687374 | 0.56953438 | NA |
| SIN3A | 20.5634374 | 0.53189124 | 0.35346991 | 1.50477089 | 0.13238299 | NA |
| PTPN9 | 35.8107336 | -0.2648385 | 0.28968282 | -0.9142363 | 0.36059271 | 0.80341474 |
| SNUPN | 3.75507319 | -0.3325596 | 0.78414837 | -0.4241029 | 0.67149074 | NA |
| IMP3 | 6.93094789 | 0.67989669 | 0.58155989 | 1.16909144 | 0.24236679 | NA |
| SNX33 | 3.38426969 | -1.1914323 | 0.87455199 | -1.3623344 | 0.17309237 | NA |

| | | | | | | |
|---|---|---|---|---|---|---|
| UBE2Q2 | 13.3439226 | -0.3994275 | 0.49855559 | -0.8011694 | 0.42303356 | NA |
| FBXO22 | 13.9810569 | 0.04934101 | 0.43095925 | 0.11449114 | 0.90884847 | NA |
| NRG4 | 9.88729605 | 1.21249192 | 0.70937697 | 1.70923496 | 0.08740744 | NA |
| ETFA | 29.4500384 | -0.0392387 | 0.30397813 | -0.1290838 | 0.89729133 | 0.97769824 |
| SCAPER | 81.6098589 | 0.65801853 | 0.28122946 | 2.33979231 | 0.01929447 | 0.33831016 |
| RCN2 | 18.6641184 | -0.0861149 | 0.40850964 | -0.2108026 | 0.83304129 | NA |
| TSPAN3 | 21.2368939 | -0.0801174 | 0.4280207 | -0.1871812 | 0.8515186 | NA |
| PEAK1 | 36.6579046 | -0.0290157 | 0.31735235 | -0.0914306 | 0.92715046 | 0.98844645 |
| HMG20A | 23.5442234 | 0.23672778 | 0.36124804 | 0.65530537 | 0.51227116 | NA |
| LINGO1 | 165.352523 | -0.71998 | 0.4825304 | -1.4920925 | 0.13567491 | 0.60594834 |
| TBC1D2B | 5.93226592 | -0.4852407 | 0.60758578 | -0.7986374 | 0.42450072 | NA |
| IDH3A | 30.4086152 | -0.5897686 | 0.32240973 | -1.8292519 | 0.06736188 | 0.52400942 |
| DNAJA4 | 2.4893282 | -0.0698076 | 1.01624373 | -0.0686918 | 0.94523495 | NA |
| WDR61 | 8.62182731 | 0.44811934 | 0.51084568 | 0.87721077 | 0.38037217 | NA |
| IREB2 | 28.0972928 | 0.4394997 | 0.32754995 | 1.34177918 | 0.17966761 | 0.6706753 |
| PSMA4 | 15.1036874 | -0.2246044 | 0.40924727 | -0.5488232 | 0.58312675 | NA |
| MORF4L1 | 36.7784509 | -0.0496821 | 0.2726457 | -0.1822223 | 0.85540828 | 0.97168963 |
| CTSH | 14.7033489 | -0.0358615 | 0.55289164 | -0.0648618 | 0.94828404 | NA |
| TMED3 | 6.44668773 | -1.3992823 | 0.66488442 | -2.1045497 | 0.03533052 | NA |
| MTHFS | 1.96900976 | -0.037873 | 1.01050934 | -0.0374792 | 0.97010296 | NA |
| ZFAND6 | 72.4599877 | -0.2420889 | 0.29404094 | -0.823317 | 0.41032773 | 0.82967024 |
| FAH | 3.1592045 | 1.17335237 | 0.90770642 | 1.29265624 | 0.19612998 | NA |
| ARNT2 | 22.8003105 | 0.0067453 | 0.4448254 | 0.01516392 | 0.98790141 | NA |
| ABHD17C | 10.0120857 | -0.0985213 | 0.56356335 | -0.1748186 | 0.8612222 | NA |
| CEMIP | 7.08572387 | -1.0719858 | 0.84290723 | -1.271772 | 0.20345414 | NA |
| AC027808.1 | 5.0920007 | 0.57442453 | 0.64425112 | 0.89161588 | 0.37259886 | NA |
| MESD | 15.6654685 | 0.480781 | 0.47890101 | 1.00392562 | 0.31541447 | NA |
| EFL1 | 15.9200706 | -0.1244751 | 0.4119185 | -0.3021837 | 0.76251204 | NA |
| RPS17 | 5.86760742 | -0.4893047 | 0.70019567 | -0.6988114 | 0.48466988 | NA |
| WHAMM | 8.59427164 | 0.61739724 | 0.55048083 | 1.12155993 | 0.26204959 | NA |
| HOMER2 | 5.00158414 | -0.2227547 | 0.84049394 | -0.2650283 | 0.79098765 | NA |
| C15orf40 | 7.15538724 | 1.3066411 | 0.54305275 | 2.40610344 | 0.0161237 | NA |
| BTBD1 | 24.0118104 | 0.11513232 | 0.34191745 | 0.33672548 | 0.73632386 | NA |
| HDGFL3 | 19.4222329 | 0.6252601 | 0.43969244 | 1.42203969 | 0.15501473 | NA |
| ADAMTSL3 | 225.799816 | -0.9294415 | 0.58932592 | -1.5771265 | 0.11476641 | 0.57860396 |
| UBE2Q2L | 3.72126447 | 1.47349966 | 0.88165966 | 1.67127943 | 0.0946665 | NA |
| AC048382.5 | 2.28743747 | 0.57194584 | 0.90829977 | 0.62968842 | 0.52889846 | NA |
| ZSCAN2 | 2.90368861 | -0.7685249 | 0.96080901 | -0.7998727 | 0.42378454 | NA |
| WDR73 | 11.5160689 | 0.34250217 | 0.49040609 | 0.69840521 | 0.48492382 | NA |
| SEC11A | 47.2162506 | -0.1232416 | 0.28600172 | -0.4309122 | 0.66653219 | 0.92307876 |
| ZNF592 | 19.3030963 | -0.0386437 | 0.37266698 | -0.103695 | 0.91741139 | NA |
| ALPK3 | 20.7083672 | -0.6379087 | 0.5302373 | -1.2030626 | 0.22895209 | NA |
| SLC28A1 | 4.32154579 | 0.74572771 | 1.05844194 | 0.70455231 | 0.48108889 | NA |
| PDE8A | 125.902068 | -0.4735243 | 0.3954177 | -1.1975294 | 0.23110027 | 0.73156941 |
| AKAP13 | 102.459404 | -0.049075 | 0.28053969 | -0.1749306 | 0.86113415 | 0.97217819 |
| AC087286.2 | 4.3527347 | -0.0691052 | 0.96746342 | -0.0714293 | 0.94305609 | NA |
| KLHL25 | 4.87150782 | -0.4296893 | 0.64664682 | -0.6644884 | 0.50637776 | NA |
| MRPL46 | 4.7007058 | 0.34547897 | 0.85289504 | 0.40506622 | 0.68542884 | NA |
| MRPS11 | 5.30077763 | 0.65331092 | 0.66708133 | 0.97935723 | 0.3274035 | NA |
| DET1 | 5.09062156 | 0.63378179 | 0.69785543 | 0.90818494 | 0.36378052 | NA |
| ABHD2 | 23.4824825 | 0.66649566 | 0.41331483 | 1.61256168 | 0.10683977 | NA |
| POLG | 5.64396157 | 0.29738202 | 0.62117979 | 0.47873743 | 0.63212543 | NA |
| RHCG | 3.81108466 | -1.1032308 | 1.22833691 | -0.89815 | 0.36910561 | NA |
| ANPEP | 6.49042969 | -0.418496 | 0.94192453 | -0.4442988 | 0.65682654 | NA |

| | | | | | | |
|---|---|---|---|---|---|---|
| ZNF710 | 7.26123422 | 0.36529842 | 0.6223714 | 0.58694602 | 0.55723996 | NA |
| ZNF710-AS1 | 3.94429281 | 0.8666363 | 0.76690157 | 1.13004892 | 0.25845561 | NA |
| IDH2 | 42.9854153 | 0.54875832 | 0.39730672 | 1.38119569 | 0.1672188 | 0.65440905 |
| SEMA4B | 26.5230844 | -0.3178986 | 0.34715849 | -0.915716 | 0.35981586 | 0.80341474 |
| CIB1 | 28.469699 | 0.24505155 | 0.44345447 | 0.55259686 | 0.58053949 | 0.89263108 |
| AC091167.6 | 2.55554503 | -0.7075929 | 0.96283317 | -0.734907 | 0.46239613 | NA |
| IQGAP1 | 68.0905195 | -0.2512013 | 0.27081621 | -0.9275712 | 0.35363003 | 0.8003972 |
| CRTC3 | 43.7862833 | 0.17432982 | 0.31529522 | 0.55290982 | 0.58032517 | 0.89263108 |
| CRTC3-AS1 | 3.64281042 | -1.2136849 | 0.87862919 | -1.3813392 | 0.1671747 | NA |
| FURIN | 7.32149359 | 0.03982747 | 0.56066897 | 0.07103562 | 0.94336941 | NA |
| MAN2A2 | 4.06403378 | 0.86634905 | 0.75384574 | 1.14923917 | 0.25045738 | NA |
| HDDC3 | 3.47753345 | 1.14471481 | 0.8539804 | 1.34044624 | 0.18010031 | NA |
| UNC45A | 7.25855591 | -0.1990116 | 0.5991087 | -0.3321794 | 0.7397538 | NA |
| RCCD1 | 2.46295379 | 0.55675329 | 0.87042453 | 0.63963419 | 0.52241045 | NA |
| AC104035.1 | 10.568872 | 0.13371919 | 0.71552912 | 0.18688155 | 0.85175351 | NA |
| SLCO3A1 | 11.7904897 | 0.38816532 | 0.67855557 | 0.57204647 | 0.56729049 | NA |
| FAM174B | 5.56990968 | -1.757785 | 0.67182906 | -2.616417 | 0.0088858 | NA |
| AC013394.1 | 37.116916 | -0.262998 | 0.28158616 | -0.9339878 | 0.3503102 | 0.8003972 |
| CHD2 | 74.9788777 | -0.6562236 | 0.31431708 | -2.0877759 | 0.03681806 | 0.41994613 |
| MCTP2 | 5.06098951 | 0.36869378 | 0.64194634 | 0.57433739 | 0.56573952 | NA |
| NR2F2-AS1 | 20.9586555 | 0.49437908 | 0.64354402 | 0.76821331 | 0.44236047 | NA |
| NR2F2 | 9.9258957 | -0.9152992 | 0.63580054 | -1.4396011 | 0.14998029 | NA |
| ARRDC4 | 7.3159417 | 0.94604493 | 0.80422528 | 1.17634319 | 0.23945776 | NA |
| IGF1R | 91.6147817 | 0.31448791 | 0.33201856 | 0.94719979 | 0.34353698 | 0.8003972 |
| TTC23 | 11.8005879 | -0.3835068 | 0.62120814 | -0.6173563 | 0.53699973 | NA |
| LRRC28 | 22.386505 | 0.56140816 | 0.44921168 | 1.24976306 | 0.21138611 | NA |
| MEF2A | 129.992172 | -0.244215 | 0.2414269 | -1.0115483 | 0.31175406 | 0.778534 |
| AC090825.1 | 5.62299074 | -0.1332773 | 0.77399542 | -0.1721939 | 0.86328506 | NA |
| ADAMTS17 | 8.20088192 | 0.19553735 | 1.0123819 | 0.19314584 | 0.84684475 | NA |
| LINS1 | 5.91171043 | 0.17103924 | 0.60202599 | 0.28410607 | 0.7763291 | NA |
| ASB7 | 9.40699717 | -0.478286 | 0.55484458 | -0.8620179 | 0.38867767 | NA |
| ALDH1A3 | 9.52950813 | -0.6488906 | 0.59906923 | -1.0831646 | 0.27873536 | NA |
| CHSY1 | 17.657665 | 0.2292693 | 0.42502813 | 0.53942146 | 0.58959607 | NA |
| SELENOS | 19.4157857 | -0.0727686 | 0.52069115 | -0.1397539 | 0.88885446 | NA |
| SNRPA1 | 9.49873666 | 0.10888945 | 0.57231671 | 0.19026082 | 0.84910475 | NA |
| TM2D3 | 12.7708974 | -0.1571401 | 0.48597811 | -0.323348 | 0.7464317 | NA |
| TARSL2 | 17.9957888 | 0.34079979 | 0.38322994 | 0.88928279 | 0.37385112 | NA |
| SNRNP25 | 3.97947151 | -1.0042187 | 0.787742 | -1.2748067 | 0.20237768 | NA |
| MPG | 8.27594994 | 0.16887247 | 0.51140746 | 0.3302112 | 0.74124039 | NA |
| NPRL3 | 16.4391091 | 0.67908423 | 0.42086386 | 1.61354847 | 0.1066254 | NA |
| LUC7L | 33.5918226 | -0.3383197 | 0.32257958 | -1.0487944 | 0.29427275 | 0.77460836 |
| FAM234A | 5.34528079 | -0.0455738 | 0.71947607 | -0.063343 | 0.94949334 | NA |
| AXIN1 | 7.6384459 | -0.9333153 | 0.59071879 | -1.5799654 | 0.11411479 | NA |
| MRPL28 | 5.48581508 | -0.2531078 | 0.74419349 | -0.3401102 | 0.73377353 | NA |
| TMEM8A | 16.3773932 | 0.05766971 | 0.3787901 | 0.15224715 | 0.87899201 | NA |
| NME4 | 3.84881077 | -0.6565919 | 0.79011472 | -0.8310083 | 0.40596897 | NA |
| DECR2 | 2.53098619 | -0.1595929 | 0.90688295 | -0.1759796 | 0.86030999 | NA |
| RAB11FIP3 | 36.3337215 | -0.446113 | 0.35578399 | -1.2538872 | 0.20988302 | 0.70690231 |
| CAPN15 | 9.05281203 | -1.0203911 | 0.59075063 | -1.7272789 | 0.08411759 | NA |
| PIGQ | 8.21656678 | 0.8273315 | 0.61753008 | 1.33974283 | 0.18032897 | NA |
| RAB40C | 18.040226 | -0.0819699 | 0.43797154 | -0.187158 | 0.85153679 | NA |
| METTL26 | 4.16072353 | -0.0909065 | 0.73430812 | -0.1237988 | 0.9014746 | NA |
| MCRIP2 | 20.5864358 | -1.1665112 | 0.44636283 | -2.6133699 | 0.00896542 | NA |
| RHOT2 | 9.54311912 | 0.73917973 | 0.56863361 | 1.29992269 | 0.19362747 | NA |

| | | | | | | |
|---|---|---|---|---|---|---|
| STUB1 | 14.3563084 | 0.2762266 | 0.4930088 | 0.56028737 | 0.57528344 | NA |
| JMJD8 | 4.83823485 | -0.4189288 | 0.79416079 | -0.5275113 | 0.59783855 | NA |
| METRN | 9.51562253 | 0.46906287 | 0.53370946 | 0.87887307 | 0.3794701 | NA |
| FAM173A | 10.1584312 | 0.60484439 | 0.58828404 | 1.02815026 | 0.30387915 | NA |
| HAGHL | 5.45836933 | -0.6101526 | 0.74380034 | -0.8203177 | 0.41203501 | NA |
| NARFL | 4.51912796 | 0.14989398 | 0.67082646 | 0.22344673 | 0.82318784 | NA |
| LMF1 | 13.9439589 | 0.15836139 | 0.49501305 | 0.31991356 | 0.74903385 | NA |
| UBE2I | 24.5070759 | -0.3387429 | 0.358997 | -0.9435815 | 0.34538355 | NA |
| TSR3 | 8.88152243 | -0.1949691 | 0.52676698 | -0.370124 | 0.71129008 | NA |
| GNPTG | 8.6405264 | -1.3185315 | 0.67195942 | -1.9622189 | 0.04973701 | NA |
| UNKL | 4.68053366 | 0.04240988 | 0.6801789 | 0.06235107 | 0.95028326 | NA |
| CLCN7 | 9.91095469 | -0.8243302 | 0.59619182 | -1.3826593 | 0.16676936 | NA |
| TELO2 | 3.71253128 | 0.05075597 | 0.76978726 | 0.06593506 | 0.94742953 | NA |
| IFT140 | 6.38809688 | 0.12627268 | 0.57551094 | 0.2194097 | 0.82633091 | NA |
| JPT2 | 10.0656811 | 0.25922464 | 0.46050199 | 0.56291752 | 0.57349105 | NA |
| MAPK8IP3 | 19.5693426 | 0.02989306 | 0.36696344 | 0.08146061 | 0.93507565 | NA |
| NME3 | 12.5222407 | 0.66832597 | 0.50555687 | 1.32196002 | 0.18618147 | NA |
| MRPS34 | 10.4957271 | -0.2392694 | 0.45674305 | -0.5238601 | 0.60037588 | NA |
| EME2 | 3.00548455 | -0.7678626 | 1.03717508 | -0.7403404 | 0.45909347 | NA |
| SPSB3 | 12.6878763 | 0.48695984 | 0.45130096 | 1.07901352 | 0.2805817 | NA |
| NUBP2 | 4.69281092 | 1.05192544 | 0.67349486 | 1.56189082 | 0.11831371 | NA |
| HAGH | 15.9512595 | 0.47629443 | 0.43052169 | 1.10631924 | 0.26858837 | NA |
| FAHD1 | 5.13791037 | -0.4999438 | 0.70840759 | -0.7057291 | 0.48035663 | NA |
| MSRB1 | 2.67882267 | 1.62190222 | 0.99375552 | 1.6320938 | 0.10265973 | NA |
| NDUFB10 | 20.274287 | 0.52485862 | 0.3801493 | 1.38066444 | 0.16738216 | NA |
| RPS2 | 65.9958499 | -0.2372642 | 0.41878126 | -0.5665588 | 0.57101395 | 0.89263108 |
| SNHG9 | 2.96200699 | 0.5295074 | 1.14798067 | 0.46125115 | 0.64461843 | NA |
| GFER | 2.9544614 | -0.23839 | 0.86951862 | -0.2741632 | 0.78395924 | NA |
| ZNF598 | 4.49142962 | -1.0731899 | 0.78095224 | -1.3742069 | 0.16937746 | NA |
| TSC2 | 12.2169007 | -0.5381336 | 0.54600048 | -0.9855919 | 0.32433343 | NA |
| PKD1 | 7.90792417 | -0.2716556 | 0.54697194 | -0.4966536 | 0.61943332 | NA |
| SNHG19 | 3.85413649 | -0.1001014 | 0.73154735 | -0.1368352 | 0.8911611 | NA |
| TRAF7 | 6.07754668 | 0.22059739 | 0.67589044 | 0.32638039 | 0.74413657 | NA |
| MLST8 | 2.33588567 | -2.488567 | 1.12602477 | -2.2100464 | 0.02710194 | NA |
| PGP | 10.3535462 | 0.81604846 | 0.4523434 | 1.80404636 | 0.07122404 | NA |
| E4F1 | 6.11670169 | 0.212563 | 0.75028906 | 0.28330814 | 0.77694064 | NA |
| ECI1 | 34.4825545 | 0.2136923 | 0.47850041 | 0.4465875 | 0.65517292 | 0.92276785 |
| RNPS1 | 17.6423173 | -0.903626 | 0.39290997 | -2.2998297 | 0.02145787 | NA |
| ABCA3 | 4.44090674 | 0.52643459 | 0.73845272 | 0.71288869 | 0.47591464 | NA |
| AMDHD2 | 4.01356865 | -2.8815844 | 1.0710732 | -2.6903711 | 0.00713726 | NA |
| PDPK1 | 32.6652275 | -0.2615939 | 0.39712303 | -0.6587225 | 0.51007395 | 0.87158522 |
| AC093525.4 | 2.78606012 | 0.16255976 | 0.93667083 | 0.17355057 | 0.86221868 | NA |
| KCTD5 | 3.04630776 | -1.0487746 | 0.90987237 | -1.1526612 | 0.24904949 | NA |
| SRRM2 | 75.5759075 | -0.2523894 | 0.3255646 | -0.7752361 | 0.43820014 | 0.84462749 |
| ELOB | 18.433625 | -0.1030261 | 0.44949713 | -0.2292029 | 0.8187112 | NA |
| FLYWCH2 | 5.32635429 | -0.0933302 | 0.76216799 | -0.1224536 | 0.9025398 | NA |
| FLYWCH1 | 9.46879343 | -0.0698077 | 0.48499006 | -0.1439363 | 0.88555078 | NA |
| TNFRSF12A | 8.16585685 | -1.4866102 | 0.77627506 | -1.915056 | 0.05548536 | NA |
| HCFC1R1 | 6.72174578 | -0.429768 | 0.8012757 | -0.5363547 | 0.59171342 | NA |
| BICDL2 | 3.05430239 | 1.26427336 | 0.85427938 | 1.47992963 | 0.13889203 | NA |
| IL32 | 5.36377844 | 0.49711297 | 0.67422888 | 0.73730596 | 0.46093631 | NA |
| ZNF213-AS1 | 3.63482965 | 0.20404531 | 0.80152971 | 0.25456987 | 0.79905534 | NA |
| ZNF205 | 2.87385342 | -0.1083169 | 0.8758568 | -0.1236697 | 0.90157684 | NA |
| ZNF213 | 8.1203409 | 0.25154044 | 0.54811157 | 0.45892197 | 0.6462902 | NA |

| | | | | | | |
|---|---|---|---|---|---|---|
| ZNF263 | 5.61397062 | -0.191703 | 0.67373947 | -0.2845358 | 0.7759998 | NA |
| TIGD7 | 15.4007224 | -2.6017578 | 1.04490293 | -2.4899517 | 0.01277605 | NA |
| ZNF75A | 25.927475 | -0.7347485 | 0.64174349 | -1.1449255 | 0.25224002 | NA |
| AC025283.3 | 5.53190208 | -0.3684763 | 0.6685771 | -0.551135 | 0.58154111 | NA |
| ZSCAN32 | 3.83579169 | 0.03837026 | 0.74543006 | 0.05147398 | 0.95894783 | NA |
| NAA60 | 6.53728989 | -0.5209162 | 0.59443412 | -0.8763228 | 0.38085455 | NA |
| CLUAP1 | 8.80279773 | 0.55651535 | 0.57120073 | 0.97429033 | 0.3299124 | NA |
| SLX4 | 2.51921408 | -0.6027079 | 0.99669724 | -0.6047051 | 0.54537495 | NA |
| DNASE1 | 23.0135445 | 0.24577544 | 0.49358904 | 0.49793536 | 0.6185296 | NA |
| TRAP1 | 14.9533372 | 0.33870116 | 0.41477516 | 0.81658978 | 0.4141629 | NA |
| CREBBP | 58.7822851 | -0.5380945 | 0.31213044 | -1.723941 | 0.08471849 | 0.55323508 |
| ADCY9 | 23.7986715 | -0.1299417 | 0.40114084 | -0.3239305 | 0.74599067 | NA |
| GLIS2 | 9.98313983 | -0.440745 | 0.5969928 | -0.7382753 | 0.4603472 | NA |
| VASN | 11.5838651 | 0.87980897 | 0.46862574 | 1.87742349 | 0.06046007 | NA |
| DNAJA3 | 8.23597213 | 0.11019853 | 0.53123657 | 0.20743778 | 0.83566798 | NA |
| NMRAL1 | 8.23515279 | 0.93414182 | 0.5590932 | 1.67081591 | 0.09475804 | NA |
| HMOX2 | 12.1219509 | -0.1284878 | 0.47131244 | -0.272617 | 0.78514763 | NA |
| CDIP1 | 8.25865132 | -0.744985 | 0.68851108 | -1.0820233 | 0.27924219 | NA |
| UBALD1 | 4.16255738 | -2.0972973 | 0.81134062 | -2.5849775 | 0.00973853 | NA |
| MGRN1 | 20.0971769 | 0.06768383 | 0.4592406 | 0.14738206 | 0.88283046 | NA |
| NUDT16L1 | 7.47529387 | 0.15382013 | 0.65659696 | 0.23426872 | 0.81477637 | NA |
| ANKS3 | 14.4889292 | 0.54699599 | 0.42647182 | 1.28260758 | 0.19962959 | NA |
| ZNF500 | 4.23941936 | 0.28906971 | 0.83587949 | 0.34582701 | 0.72947272 | NA |
| SMIM22 | 4.76733574 | 0.00981692 | 0.62255796 | 0.01576869 | 0.98741893 | NA |
| ROGDI | 3.44942827 | 1.35180152 | 0.89911123 | 1.50348641 | 0.13271366 | NA |
| GLYR1 | 34.3010049 | -0.5403759 | 0.34842426 | -1.5509135 | 0.12092242 | 0.58553957 |
| UBN1 | 12.5637259 | -1.4783686 | 0.59959195 | -2.4656245 | 0.01367746 | NA |
| PPL | 11.0374652 | -0.5814627 | 0.49750047 | -1.1687681 | 0.24249708 | NA |
| NAGPA | 2.11785513 | -0.4327653 | 1.1045788 | -0.3917921 | 0.69521182 | NA |
| ALG1 | 2.28315435 | -0.8789507 | 1.10942282 | -0.7922595 | 0.4282094 | NA |
| RBFOX1 | 16.6657027 | 0.58697195 | 0.74283913 | 0.7901737 | 0.42942633 | NA |
| METTL22 | 3.08531502 | -0.4042849 | 0.96504217 | -0.4189298 | 0.67526745 | NA |
| ABAT | 5.57713857 | 1.27117515 | 0.74680263 | 1.70215676 | 0.08872599 | NA |
| PMM2 | 18.3718378 | -1.1944055 | 0.52620753 | -2.2698374 | 0.02321745 | NA |
| CARHSP1 | 10.7758588 | -0.4625428 | 0.48883038 | -0.9462236 | 0.34403455 | NA |
| USP7 | 26.7184363 | 0.14870783 | 0.37165129 | 0.4001273 | 0.68906276 | 0.9279054 |
| C16orf72 | 23.3484474 | -0.6919448 | 0.35639055 | -1.9415351 | 0.0521934 | NA |
| AC087190.1 | 2.66295453 | 1.02202481 | 1.06086692 | 0.96338644 | 0.33535363 | NA |
| GRIN2A | 13.3481967 | -0.7075288 | 0.85190277 | -0.8305277 | 0.4062405 | NA |
| ATF7IP2 | 35.0948975 | 1.26122874 | 0.37911756 | 3.32674842 | 0.00087866 | 0.11167307 |
| NUBP1 | 5.29935464 | -0.0001984 | 0.61403038 | -0.000323 | 0.99974225 | NA |
| DEXI | 6.8571782 | -0.1449416 | 0.6290925 | -0.2303979 | 0.81778261 | NA |
| CLEC16A | 74.151106 | -0.0656737 | 0.26878216 | -0.2443381 | 0.80696896 | 0.96800006 |
| AC007014.1 | 5.06410491 | 0.26188287 | 0.63242082 | 0.41409591 | 0.67880387 | NA |
| SOCS1 | 30.4452268 | 0.2262046 | 0.59154872 | 0.38239386 | 0.70216924 | 0.9303056 |
| AC099489.1 | 8.94343138 | -0.6486882 | 0.5848067 | -1.1092353 | 0.26732869 | NA |
| LITAF | 161.188242 | -0.0910881 | 0.31997042 | -0.2846767 | 7.76E-01 | 0.96105928 |
| SNN | 4.8560686 | -1.4813666 | 0.77628563 | -1.9082752 | 0.05635566 | NA |
| TXNDC11 | 32.6207569 | -0.408891 | 0.3945581 | -1.0363265 | 0.30004984 | 0.77521265 |
| ZC3H7A | 27.2344355 | 0.03235095 | 0.32084192 | 0.10083143 | 0.91968428 | 0.98745437 |
| RSL1D1 | 9.06078886 | -0.331664 | 0.52490674 | -0.6318531 | 0.52748286 | NA |
| AC007216.4 | 14.8462949 | -1.140119 | 0.57210864 | -1.9928366 | 0.04627935 | NA |
| GSPT1 | 42.1632297 | -0.160252 | 0.29070348 | -0.5512558 | 0.58145832 | 0.89263108 |
| SNX29 | 232.197837 | 0.63016765 | 0.33303229 | 1.89221188 | 0.05846276 | 0.50826415 |

| | | | | | | |
|---|---|---|---|---|---|---|
| AC010333.2 | 9.90721199 | -0.3314974 | 0.65427482 | -0.5066638 | 0.61239073 | NA |
| CPPED1 | 32.8171996 | -0.705692 | 0.372537 | -1.8942869 | 0.05818693 | 0.50751937 |
| ERCC4 | 3.51811447 | 0.06506526 | 0.81289967 | 0.08004095 | 0.93620468 | NA |
| MKL2 | 59.2786134 | 0.33621821 | 0.27620691 | 1.21726937 | 0.22350174 | 0.72657496 |
| MIR193BHG | 7.70923084 | -1.8357461 | 0.7079209 | -2.5931515 | 0.00951009 | NA |
| PARN | 33.714236 | 0.16042453 | 0.29303446 | 0.54745961 | 0.58406301 | 0.8933319 |
| BFAR | 15.2941177 | -0.3301295 | 0.45385612 | -0.727388 | 0.4669883 | NA |
| NOMO1 | 4.13513805 | -0.4996818 | 0.90516738 | -0.5520325 | 0.58092608 | NA |
| PDXDC1 | 58.0755244 | -0.2273621 | 0.27071101 | -0.8398703 | 0.40098111 | 0.82706227 |
| NTAN1 | 6.81293943 | 0.08130939 | 0.54369247 | 0.14955034 | 0.88111939 | NA |
| RRN3 | 8.88178124 | -0.0377251 | 0.56420388 | -0.0668643 | 0.9466897 | NA |
| C16orf45 | 16.602736 | -0.5939739 | 0.45584905 | -1.3030056 | 0.19257283 | NA |
| MARF1 | 14.2959317 | -0.3303988 | 0.4402211 | -0.7505293 | 0.45293596 | NA |
| NDE1 | 5.0176221 | 0.49753268 | 0.68863745 | 0.72248856 | 0.46999416 | NA |
| FOPNL | 5.53278312 | 0.42299149 | 0.70494924 | 0.60003113 | 0.54848549 | NA |
| ABCC1 | 16.0673145 | -0.2047612 | 0.43133638 | -0.4747136 | 0.63499113 | NA |
| RPS15A | 11.070955 | 0.37747189 | 0.53741777 | 0.70238073 | 0.48244176 | NA |
| SMG1 | 69.2138672 | -0.1052468 | 0.2643957 | -0.3980654 | 0.690582 | 0.92830727 |
| COQ7 | 5.40327845 | -0.6387073 | 0.66492511 | -0.9605702 | 0.3367683 | NA |
| GDE1 | 7.04959746 | -0.2590021 | 0.64952338 | -0.3987572 | 0.69007212 | NA |
| CCP110 | 4.18103533 | -0.3443695 | 0.70798652 | -0.4864068 | 0.62667876 | NA |
| VPS35L | 30.2037909 | 0.21508952 | 0.40661572 | 0.52897492 | 0.59682285 | 0.899694 |
| KNOP1 | 6.72842575 | -0.7866881 | 0.79580534 | -0.9885433 | 0.32288662 | NA |
| IQCK | 30.2258047 | 0.53864077 | 0.38667113 | 1.39302039 | 0.16361362 | 0.64995633 |
| GPRC5B | 38.5860943 | -0.0391886 | 0.41731581 | -0.0939063 | 0.92518356 | 0.98758497 |
| UMOD | 41.4180609 | -0.6628661 | 0.82203707 | -0.8063701 | 0.42002946 | 0.83911574 |
| ACSM2A | 8.09830714 | 1.28121422 | 0.95235262 | 1.34531495 | 0.17852355 | NA |
| ACSM2B | 10.8241269 | 0.56099488 | 1.44485666 | 0.38827027 | 0.69781604 | NA |
| ACSM3 | 9.63548862 | 0.89799626 | 0.71364848 | 1.25831733 | 0.20827702 | NA |
| THUMPD1 | 18.5784889 | 0.33653385 | 0.39162347 | 0.85933014 | 0.3901584 | NA |
| ERI2 | 4.74574314 | 0.41965589 | 0.7086472 | 0.59219297 | 0.55372138 | NA |
| LYRM1 | 3.64023288 | 0.75244249 | 0.82818015 | 0.90854929 | 0.36358808 | NA |
| METTL9 | 19.5619784 | 0.16590158 | 0.39219305 | 0.42300998 | 0.67228798 | NA |
| NPIPB4 | 7.35064674 | -1.3815244 | 0.63293441 | -2.1827291 | 0.02905577 | NA |
| UQCRC2 | 34.5015319 | -0.1673911 | 0.31878994 | -0.5250827 | 0.59952569 | 0.90250089 |
| MOSMO | 24.6060651 | -0.2540369 | 0.39980365 | -0.6354041 | 0.52516491 | NA |
| EEF2K | 4.10376265 | -0.9539595 | 1.00429485 | -0.9498799 | 0.34217326 | NA |
| POLR3E | 9.15778397 | -0.1516553 | 0.52557932 | -0.2885488 | 0.77292664 | NA |
| CDR2 | 36.3825828 | 0.04952315 | 0.33841815 | 0.14633715 | 0.88365524 | 0.97769824 |
| NPIPB5 | 17.8121579 | -0.3039277 | 0.45431523 | -0.6689797 | 0.50350844 | NA |
| USP31 | 11.4652639 | -0.5553517 | 0.56656954 | -0.9802005 | 0.32698718 | NA |
| SCNN1G | 7.71755338 | -1.5543878 | 1.10935391 | -1.4011649 | 0.16116476 | NA |
| SCNN1B | 4.64859425 | -0.2322134 | 0.98921291 | -0.2347456 | 0.81440619 | NA |
| COG7 | 14.6134975 | -0.3058057 | 0.48316685 | -0.6329195 | 0.52678621 | NA |
| GGA2 | 23.5742773 | -0.7659371 | 0.46842542 | -1.6351313 | 0.10202153 | NA |
| EARS2 | 4.94012473 | -0.6135189 | 0.66580609 | -0.9214678 | 0.35680623 | NA |
| UBFD1 | 7.12316996 | -0.5239755 | 0.61945575 | -0.8458644 | 0.39762842 | NA |
| NDUFAB1 | 16.034836 | 0.55554311 | 0.44820744 | 1.23947767 | 0.21516865 | NA |
| PALB2 | 4.09989327 | -0.0996848 | 0.75881645 | -0.1313689 | 0.89548352 | NA |
| DCTN5 | 9.77921055 | -0.5877646 | 0.75890573 | -0.7744897 | 0.43864127 | NA |
| RBBP6 | 19.1413577 | -0.3280321 | 0.50364354 | -0.6513179 | 0.5148413 | NA |
| TNRC6A | 78.8450755 | -0.4919696 | 0.28014387 | -1.7561319 | 0.07906588 | 0.54552914 |
| ARHGAP17 | 16.8179696 | -0.2053642 | 0.40283193 | -0.5098012 | 0.61019074 | NA |
| LCMT1 | 14.1731254 | 0.8956972 | 0.43584376 | 2.05508782 | 0.03987052 | NA |

| | | | | | | |
|---|---|---|---|---|---|---|
| NSMCE1 | 7.96193978 | 0.13726249 | 0.58513316 | 0.23458333 | 0.81453215 | NA |
| GTF3C1 | 10.3087737 | -0.2752606 | 0.56180595 | -0.4899567 | 0.62416453 | NA |
| KIAA0556 | 13.5038367 | 0.33149962 | 0.54067105 | 0.61312626 | 0.53979286 | NA |
| XPO6 | 21.4123408 | -0.9664335 | 0.38028071 | -2.5413687 | 0.01104194 | NA |
| NUPR1 | 7.45049633 | -0.4768004 | 1.01054193 | -0.4718265 | 0.63705064 | NA |
| SGF29 | 3.98675509 | -0.7503908 | 0.75271617 | -0.9969107 | 0.31880786 | NA |
| SULT1A1 | 3.70408641 | 1.97685001 | 0.92538303 | 2.13625056 | 0.032659 | NA |
| ATXN2L | 12.0584097 | -0.5644662 | 0.45857745 | -1.2309071 | 0.21835761 | NA |
| TUFM | 31.8992505 | 0.69578839 | 0.40832309 | 1.70401431 | 0.08837841 | 0.56122412 |
| SH2B1 | 19.758015 | 0.1234623 | 0.4315997 | 0.28605743 | 0.77483414 | NA |
| RABEP2 | 11.9292764 | -0.5062783 | 0.45863108 | -1.1038901 | 0.2696408 | NA |
| NFATC2IP | 20.9006524 | 0.20142349 | 0.52747546 | 0.38186324 | 0.70256281 | NA |
| SPNS1 | 5.99241705 | -0.2916773 | 0.93983771 | -0.3103486 | 0.75629588 | NA |
| QPRT | 2.39070576 | 2.76991286 | 1.07406192 | 2.57891357 | 0.00991116 | NA |
| KIF22 | 2.72701587 | -0.0162041 | 0.9294824 | -0.0174335 | 0.98609081 | NA |
| MAZ | 5.60010488 | -0.5933437 | 0.63774326 | -0.9303802 | 0.35217426 | NA |
| AC009133.1 | 6.14765438 | -0.3883227 | 0.64088321 | -0.605918 | 0.54456918 | NA |
| MVP | 22.9094968 | 0.36187106 | 0.39478459 | 0.91662915 | 0.359337 | NA |
| CDIPT | 7.92624097 | 0.3749998 | 0.52584247 | 0.71314095 | 0.47575854 | NA |
| KCTD13 | 9.34219564 | -0.6204193 | 0.55349477 | -1.1209127 | 0.26232501 | NA |
| TMEM219 | 11.386779 | 0.29439045 | 0.47338731 | 0.62188073 | 0.5340203 | NA |
| TAOK2 | 4.86817902 | -0.1986672 | 0.68164934 | -0.2914507 | 0.77070663 | NA |
| INO80E | 6.63242128 | 1.04002066 | 0.63092791 | 1.64839857 | 0.09927091 | NA |
| PPP4C | 13.6935247 | -0.4605279 | 0.41856054 | -1.100266 | 0.27121624 | NA |
| YPEL3 | 11.685099 | 0.62006881 | 0.48342985 | 1.28264487 | 0.19961652 | NA |
| MAPK3 | 2.81659356 | -0.0516818 | 0.94757196 | -0.0545412 | 0.95650395 | NA |
| NPIPB13 | 6.14737399 | -1.091877 | 0.79376318 | -1.3755703 | 0.16895472 | NA |
| CD2BP2 | 6.41094338 | -0.9560404 | 0.60225066 | -1.587446 | 0.11241167 | NA |
| TBC1D10B | 4.77622597 | -0.751321 | 0.66588307 | -1.1283077 | 0.25918998 | NA |
| ZNF771 | 2.63737146 | 0.26478319 | 0.84096638 | 0.31485586 | 0.7528711 | NA |
| DCTPP1 | 3.76949042 | 0.98623817 | 0.73860054 | 1.33527952 | 0.18178488 | NA |
| SEPHS2 | 16.8285881 | 0.2298911 | 0.39446567 | 0.58279114 | 0.56003391 | NA |
| ZNF768 | 7.14217234 | -0.792841 | 0.63785534 | -1.2429793 | 0.21387547 | NA |
| ZNF688 | 2.13075363 | 0.05080225 | 1.15553555 | 0.04396425 | 0.9649329 | NA |
| ZNF785 | 4.97893197 | 0.19290578 | 0.8024094 | 0.24040817 | 0.81001384 | NA |
| PRR14 | 4.71398313 | -0.3902813 | 0.6415593 | -0.6083324 | 0.54296701 | NA |
| FBRS | 5.18024182 | -0.1776831 | 0.64283935 | -0.2764036 | 0.78223812 | NA |
| PHKG2 | 4.73844464 | 0.79505656 | 0.65387139 | 1.21592193 | 0.22401466 | NA |
| RNF40 | 8.61960688 | -0.1362816 | 0.65014364 | -0.2096177 | 0.8339661 | NA |
| ZNF629 | 3.90131627 | -0.6001421 | 0.77900053 | -0.7704002 | 0.44106255 | NA |
| BCL7C | 17.4452134 | 0.45736506 | 0.39162042 | 1.16787847 | 0.24285579 | NA |
| MIR762HG | 2.70874897 | -0.8962986 | 1.00609454 | -0.8908692 | 0.37299937 | NA |
| FBXL19 | 6.35006829 | -0.3547218 | 0.69165462 | -0.5128598 | 0.60804942 | NA |
| SETD1A | 3.42075845 | -0.323514 | 0.92505819 | -0.3497228 | 0.72654672 | NA |
| STX4 | 3.60254524 | -0.1688915 | 0.74452556 | -0.2268444 | 0.82054471 | NA |
| BCKDK | 4.59341416 | 1.17526781 | 0.72998065 | 1.60999859 | 0.10739817 | NA |
| KAT8 | 11.9394433 | -0.8638142 | 0.55598558 | -1.553663 | 0.12026481 | NA |
| PRSS8 | 17.6483182 | -0.6503683 | 0.44795575 | -1.4518584 | 0.14654097 | NA |
| FUS | 44.205055 | 0.05295879 | 0.29661119 | 0.17854616 | 0.85829407 | 0.97168963 |
| AC026471.4 | 3.15912079 | -0.1006151 | 0.94918802 | -0.1060012 | 0.91558142 | NA |
| C16orf58 | 7.64519023 | 0.15027842 | 0.58251873 | 0.25798042 | 0.79642202 | NA |
| ZNF720 | 12.4646869 | 0.66307765 | 0.43558247 | 1.52227807 | 0.12793942 | NA |
| ZNF267 | 3.5298491 | -0.0794875 | 0.76659704 | -0.1036888 | 0.91741631 | NA |
| VPS35 | 16.9291007 | 0.53682932 | 0.385246 | 1.39347151 | 0.16347725 | NA |

| | | | | | | |
|---|---|---|---|---|---|---|
| C16orf87 | 5.68216476 | 0.3766916 | 0.57328403 | 0.65707674 | 0.51113157 | NA |
| GPT2 | 3.56579405 | 0.06618172 | 0.99844592 | 0.06628473 | 0.94715114 | NA |
| DNAJA2 | 10.7094756 | 0.65157583 | 0.50050246 | 1.3018434 | 0.19296992 | NA |
| NETO2 | 3.53280254 | -1.1262657 | 0.93601 | -1.2032625 | 0.22887477 | NA |
| ITFG1 | 93.8152446 | 0.01658354 | 0.23515467 | 0.07052186 | 0.9437783 | 0.99210094 |
| PHKB | 73.9036583 | -0.1766875 | 0.26036016 | -0.6786272 | 0.49737408 | 0.86934605 |
| LONP2 | 48.400599 | -0.1919767 | 0.29203621 | -0.6573728 | 0.51094125 | 0.87158522 |
| SIAH1 | 16.3703921 | -0.2455193 | 0.39092026 | -0.6280548 | 0.52996804 | NA |
| N4BP1 | 41.8875058 | 0.17303047 | 0.28719448 | 0.60248537 | 0.5468511 | 0.88410237 |
| CNEP1R1 | 2.70762626 | -0.4512648 | 0.87739821 | -0.5143216 | 0.6070272 | NA |
| HEATR3 | 4.47622035 | -0.1208491 | 0.88328799 | -0.1368173 | 0.89117518 | NA |
| TENT4B | 15.1552589 | 0.69864579 | 0.45669887 | 1.52977343 | 0.12607282 | NA |
| BRD7 | 15.9243193 | -0.0948186 | 0.40941156 | -0.2315972 | 0.81685091 | NA |
| CYLD | 12.1269588 | 0.79567497 | 0.46616737 | 1.70684397 | 0.08785106 | NA |
| SALL1 | 8.35705382 | -0.4155516 | 0.53403277 | -0.7781388 | 0.4364872 | NA |
| AC087564.1 | 18.0779647 | 0.0467515 | 0.59988259 | 0.07793442 | 0.93788022 | NA |
| HNRNPA1P4 | 4.24558919 | -0.7173003 | 0.74166467 | -0.9671491 | 0.33346951 | NA |
| TOX3 | 110.627517 | 0.03254298 | 0.41604922 | 0.07821907 | 0.9376538 | 0.99100968 |
| CHD9 | 139.942908 | 0.15673491 | 0.21222911 | 0.73851749 | 0.46020004 | 0.85844137 |
| RBL2 | 37.8752538 | 0.57418597 | 0.34460489 | 1.6662154 | 0.09567052 | 0.56358504 |
| AKTIP | 5.97610697 | -0.1498126 | 0.61158216 | -0.2449591 | 0.80648808 | NA |
| FTO | 75.5017678 | 0.05744791 | 0.23835532 | 0.24101793 | 0.80954122 | 0.96845695 |
| LPCAT2 | 4.22860869 | 0.46429443 | 0.73867068 | 0.62855403 | 0.52964107 | NA |
| AMFR | 52.2288123 | 0.5072459 | 0.31242064 | 1.62359918 | 0.10446138 | 0.5772605 |
| NUDT21 | 17.1279374 | 0.42381646 | 0.3848631 | 1.10121354 | 0.27080373 | NA |
| OGFOD1 | 7.44560153 | 0.08677345 | 0.636507 | 0.13632757 | 0.89156233 | NA |
| BBS2 | 10.0387358 | 0.39060412 | 0.50185036 | 0.77832785 | 0.43637576 | NA |
| MT2A | 84.2858745 | -1.0462789 | 0.60252446 | -1.7364919 | 0.08247689 | 0.55216537 |
| MT1E | 26.0565597 | -1.1693029 | 0.6684981 | -1.7491492 | 0.08026522 | 0.54552914 |
| MT1G | 48.0617984 | -1.8790908 | 1.15739071 | -1.6235579 | 0.1044702 | 0.5772605 |
| MT1H | 14.1358311 | -1.155141 | 1.01684593 | -1.136004 | 0.25595491 | NA |
| MT1X | 17.6845424 | -0.3224274 | 0.70336365 | -0.4584078 | 0.64665951 | NA |
| NUP93 | 11.5562196 | -0.1309701 | 0.48382575 | -0.2706969 | 0.78662414 | NA |
| SLC12A3 | 9.82869136 | 1.58226727 | 0.90109476 | 1.75593881 | 0.07909886 | NA |
| HERPUD1 | 28.379563 | -0.4442812 | 0.34082088 | -1.3035623 | 0.19238287 | 0.68667113 |
| NLRC5 | 27.8405477 | -1.4487135 | 0.65986066 | -2.1954839 | 0.0281289 | 0.39934064 |
| CPNE2 | 10.4487397 | -1.3474523 | 0.72429573 | -1.860362 | 0.06283432 | NA |
| FAM192A | 13.4402672 | 0.00882234 | 0.5291426 | 0.01667289 | 0.98669757 | NA |
| RSPRY1 | 20.5294239 | -0.3219613 | 0.37296192 | -0.8632553 | 0.38799712 | NA |
| ARL2BP | 15.782485 | -0.2092883 | 0.46874302 | -0.4464883 | 0.65524459 | NA |
| PLLP | 21.6809922 | 1.07550265 | 0.52015825 | 2.06764507 | 0.03867341 | NA |
| CX3CL1 | 3.70083477 | -0.0395216 | 0.93822833 | -0.0421236 | 0.96640017 | NA |
| CIAPIN1 | 12.7749081 | -0.2490509 | 0.52039177 | -0.4785835 | 0.63223495 | NA |
| COQ9 | 24.8851459 | 0.22856322 | 0.37819425 | 0.60435402 | 0.5456083 | NA |
| POLR2C | 5.25957687 | 0.99428299 | 0.70635536 | 1.40762433 | 0.15924234 | NA |
| ADGRG1 | 36.5548627 | 0.04298312 | 0.39369618 | 0.10917841 | 0.91306098 | 0.98415124 |
| KIFC3 | 20.7004676 | 0.10385192 | 0.50297311 | 0.20647608 | 0.83641905 | NA |
| USB1 | 2.70707914 | -0.3546395 | 0.90756157 | -0.3907609 | 0.695974 | NA |
| MMP15 | 2.56012821 | 0.74922481 | 0.96921068 | 0.77302575 | 0.43950714 | NA |
| CFAP20 | 3.39781967 | -0.1718685 | 0.90521174 | -0.1898655 | 0.84941454 | NA |
| CSNK2A2 | 11.541127 | 0.57201458 | 0.44155839 | 1.29544493 | 0.19516679 | NA |
| SETD6 | 4.40947988 | 0.02406479 | 0.78302763 | 0.030733 | 0.97548248 | NA |
| CNOT1 | 43.6388649 | -0.2599169 | 0.27714515 | -0.9378369 | 0.34832823 | 0.8003972 |
| SLC38A7 | 7.27777798 | -0.930773 | 0.56288815 | -1.6535665 | 0.09821563 | NA |

| | | | | | | |
|---|---|---|---|---|---|---|
| GOT2 | 14.6636791 | 0.08603996 | 0.40192759 | 0.21406831 | 0.83049378 | NA |
| AC138305.1 | 92.6620244 | 0.18321468 | 0.32722087 | 0.55991136 | 0.5755399 | 0.89263108 |
| AC092138.2 | 3.24507846 | -0.3473433 | 1.54571175 | -0.2247142 | 0.82220164 | NA |
| TK2 | 2.7849733 | 1.15355878 | 0.99469211 | 1.15971441 | 0.2461651 | NA |
| CMTM4 | 65.5192267 | 0.23745874 | 0.29166108 | 0.81415987 | 0.41555337 | 0.83580403 |
| DYNC1LI2 | 35.0189224 | -0.5520649 | 0.31378361 | -1.7593808 | 0.07851286 | 0.54552914 |
| AC018557.1 | 5.96162611 | 0.9660324 | 0.79495033 | 1.21521101 | 0.22428562 | NA |
| NAE1 | 10.0293499 | -0.0888511 | 0.49501236 | -0.1794927 | 0.85755082 | NA |
| PDP2 | 7.46064501 | -0.7934449 | 0.60563772 | -1.3100983 | 0.19016259 | NA |
| CDH16 | 36.0034812 | -0.0629977 | 0.57519056 | -0.1095249 | 0.9127862 | 0.98415124 |
| RRAD | 7.50785309 | 0.03127436 | 1.01933714 | 0.03068108 | 0.97552388 | NA |
| FAM96B | 15.4101354 | 0.20794704 | 0.40976068 | 0.50748413 | 0.61181517 | NA |
| CES2 | 8.07836366 | 0.31021059 | 0.70647695 | 0.43909513 | 0.66059261 | NA |
| CBFB | 49.6348472 | 0.02485957 | 0.31115208 | 0.07989523 | 0.93632058 | 0.99100968 |
| C16orf70 | 5.23929674 | -0.1405066 | 0.62695331 | -0.2241102 | 0.82267159 | NA |
| TRADD | 2.11156341 | 0.60168917 | 0.95896097 | 0.62743864 | 0.53037174 | NA |
| KIAA0895L | 3.45289354 | 0.11923313 | 0.75788839 | 0.15732281 | 0.87499044 | NA |
| E2F4 | 6.77740897 | -0.637078 | 0.63845227 | -0.9978475 | 0.31835331 | NA |
| LRRC29 | 2.46083268 | -0.5031764 | 1.22461973 | -0.4108838 | 0.68115778 | NA |
| TMEM208 | 11.1097962 | 0.49612595 | 0.43378028 | 1.14372637 | 0.25273713 | NA |
| ZDHHC1 | 3.22642605 | 0.86870801 | 0.82405366 | 1.05418863 | 0.29179656 | NA |
| HSD11B2 | 51.7790523 | 1.1333016 | 0.51086478 | 2.2183984 | 0.02652768 | 0.39506514 |
| ATP6V0D1 | 19.7896834 | -0.4344349 | 0.52030061 | -0.834969 | 0.40373514 | NA |
| AC009061.2 | 5.29194534 | -0.2009947 | 0.72089168 | -0.2788141 | 0.78038753 | NA |
| AC027682.6 | 3.41733875 | 1.93441987 | 1.03362685 | 1.87148763 | 0.06127752 | NA |
| RIPOR1 | 15.1325403 | 1.04086044 | 0.47593433 | 2.18698331 | 0.02874374 | NA |
| CTCF | 24.3800262 | -0.5447571 | 0.40360051 | -1.3497433 | 0.17709834 | NA |
| GFOD2 | 6.6002686 | 0.30317074 | 0.61556292 | 0.49250975 | 0.62235903 | NA |
| RANBP10 | 26.4558137 | -0.1685608 | 0.38959327 | -0.4326585 | 0.66526291 | 0.92276785 |
| CENPT | 2.73032145 | 0.3857878 | 0.84277309 | 0.45775999 | 0.64712489 | NA |
| NUTF2 | 5.0521768 | -0.6348715 | 0.66751041 | -0.9511035 | 0.34155183 | NA |
| PSKH1 | 6.81970023 | 1.00548786 | 0.6740242 | 1.49176819 | 0.13575993 | NA |
| DUS2 | 4.66441037 | 0.18968258 | 0.74348913 | 0.25512488 | 0.79862665 | NA |
| NFATC3 | 36.9273543 | 0.17938189 | 0.3184342 | 0.56332482 | 0.57321372 | 0.89263108 |
| AC020978.5 | 2.67615488 | 1.46603028 | 0.96443029 | 1.52009978 | 0.1284859 | NA |
| ESRP2 | 4.87834847 | 0.36632636 | 0.68725856 | 0.53302554 | 0.5940159 | NA |
| PLA2G15 | 11.2548315 | 0.92192246 | 0.52219609 | 1.76547177 | 0.07748451 | NA |
| SLC7A6 | 19.7437254 | -0.3122017 | 0.40155467 | -0.7774823 | 0.43687427 | NA |
| SLC7A6OS | 8.45198055 | -0.5131017 | 0.58495178 | -0.8771691 | 0.38039478 | NA |
| PRMT7 | 6.68729369 | 0.68411259 | 0.75618856 | 0.90468518 | 0.36563219 | NA |
| ZFP90 | 7.60951072 | 1.14128904 | 0.65080212 | 1.75366521 | 0.07948789 | NA |
| CDH1 | 64.3121406 | -0.1372896 | 0.2322357 | -0.5911651 | 0.55440981 | 0.88978941 |
| TANGO6 | 20.133682 | -0.182096 | 0.40254155 | -0.4523658 | 0.65100549 | NA |
| CHTF8 | 9.98277894 | 0.36583263 | 0.53561261 | 0.68301722 | 0.49459596 | NA |
| UTP4 | 5.98586531 | -0.7592451 | 0.65745114 | -1.1548312 | 0.24815954 | NA |
| SNTB2 | 10.998939 | -0.293025 | 0.55604447 | -0.5269812 | 0.59820667 | NA |
| VPS4A | 6.39638468 | 0.34243102 | 0.58733487 | 0.58302518 | 0.55987635 | NA |
| COG8 | 3.95341591 | 0.96905687 | 0.79347133 | 1.22128781 | 0.22197707 | NA |
| NIP7 | 4.15072205 | -0.028001 | 0.79417191 | -0.0352581 | 0.97187395 | NA |
| TERF2 | 8.79728986 | -0.4763654 | 0.49168958 | -0.9688336 | 0.3326282 | NA |
| CYB5B | 21.3784768 | -0.8245078 | 0.44062216 | -1.8712355 | 0.06131245 | NA |
| NFAT5 | 157.258464 | -0.2628624 | 0.23656685 | -1.1111548 | 0.26650171 | 0.75497438 |
| NQO1 | 5.75389665 | -0.2819661 | 0.74105632 | -0.3804922 | 0.70358012 | NA |
| NOB1 | 2.4001733 | -0.0592525 | 1.00923658 | -0.0587102 | 0.95318294 | NA |

| | | | | | | |
|---|---|---|---|---|---|---|
| WWP2 | 57.3024237 | -0.4842576 | 0.2866892 | -1.6891377 | 0.09119304 | 0.56211136 |
| PDPR | 18.0979581 | -0.8727921 | 0.45245077 | -1.9290322 | 0.05372687 | NA |
| AC009060.1 | 4.48795104 | 0.54716726 | 0.72801128 | 0.75159174 | 0.45229661 | NA |
| EXOSC6 | 9.60291918 | -0.6766336 | 0.54309193 | -1.2458914 | 0.21280427 | NA |
| AARS | 12.3600143 | 0.11916229 | 0.47579406 | 0.25044929 | 0.80223991 | NA |
| DDX19A | 2.81164423 | -1.2616848 | 0.88612692 | -1.4238195 | 0.15449872 | NA |
| ST3GAL2 | 10.1029596 | 0.88526493 | 0.5640064 | 1.56960086 | 0.116508 | NA |
| FUK | 4.80961059 | 0.55705249 | 0.61823593 | 0.90103545 | 0.36756947 | NA |
| COG4 | 10.6434091 | -0.1778248 | 0.60311795 | -0.2948425 | 0.76811417 | NA |
| SF3B3 | 16.0963495 | -0.5870807 | 0.40439304 | -1.4517578 | 0.14656896 | NA |
| MTSS1L | 3.06194556 | 0.07699754 | 0.8243994 | 0.09339835 | 0.9255871 | NA |
| VAC14 | 12.683583 | -0.702401 | 0.6331317 | -1.1094074 | 0.26725447 | NA |
| HYDIN | 8.57452499 | -0.0399723 | 0.66087686 | -0.0604838 | 0.95177031 | NA |
| CMTR2 | 13.4052447 | -1.544948 | 0.48371538 | -3.1939196 | 0.00140355 | NA |
| ZNF19 | 3.81784003 | 0.9922945 | 0.72026915 | 1.37767181 | 0.16830464 | NA |
| MARVELD3 | 4.90223399 | -0.7071249 | 0.71777496 | -0.9851624 | 0.32454431 | NA |
| PHLPP2 | 9.69606088 | -0.0005351 | 0.47457648 | -0.0011276 | 0.99910029 | NA |
| AP1G1 | 49.1026003 | -0.5785483 | 0.33508782 | -1.7265572 | 0.08424721 | 0.55323508 |
| IST1 | 32.2208747 | -0.1814687 | 0.30186735 | -0.6011538 | 0.54773757 | 0.88439902 |
| ZNF821 | 16.0453386 | 0.34404361 | 0.54587068 | 0.63026579 | 0.5285207 | NA |
| TXNL4B | 4.74924757 | 0.62783919 | 0.62312849 | 1.00755975 | 0.31366586 | NA |
| DHX38 | 5.02937594 | -0.6425879 | 0.65845444 | -0.9759033 | 0.32911236 | NA |
| LINC01572 | 7.43853253 | 0.64101924 | 0.79327254 | 0.80806937 | 0.41905065 | NA |
| AC004943.2 | 6.92800171 | 0.63243614 | 0.66847097 | 0.94609365 | 0.34410082 | NA |
| ZFHX3 | 73.3245527 | -0.5549282 | 0.27461275 | -2.0207663 | 0.04330397 | 0.44186191 |
| AC138627.1 | 11.0538143 | 0.11067187 | 0.81070123 | 0.13651375 | 0.89141515 | NA |
| PSMD7 | 13.2207043 | -0.4312562 | 0.45231853 | -0.9534348 | 0.3403698 | NA |
| NPIPB15 | 3.51565651 | 0.57382055 | 1.09473425 | 0.52416424 | 0.60016431 | NA |
| GLG1 | 83.6444728 | -0.2134846 | 0.2459905 | -0.8678572 | 0.38547249 | 0.82570312 |
| RFWD3 | 10.1439627 | 0.61093902 | 0.5111787 | 1.19515742 | 0.23202553 | NA |
| FA2H | 7.81441827 | -0.5815756 | 0.76184576 | -0.763377 | 0.44523858 | NA |
| WDR59 | 29.237481 | 0.30589044 | 0.34616811 | 0.88364708 | 0.37688676 | 0.81781362 |
| ZNRF1 | 16.1376666 | -0.1281587 | 0.52541231 | -0.2439202 | 0.80729259 | NA |
| LDHD | 3.36534549 | 1.847258 | 0.95194458 | 1.94051003 | 0.05231774 | NA |
| BCAR1 | 14.4444353 | -1.0935588 | 0.44095889 | -2.4799563 | 0.01313985 | NA |
| CFDP1 | 45.4560034 | -0.6025921 | 0.28054481 | -2.1479353 | 0.03171889 | 0.4126665 |
| GABARAPL2 | 18.1108165 | 1.05718488 | 0.47514689 | 2.22496431 | 0.02608363 | NA |
| ADAT1 | 5.82832923 | -0.546365 | 0.76918862 | -0.7103134 | 0.47750984 | NA |
| KARS | 9.18877338 | -0.0284913 | 0.49119735 | -0.0580038 | 0.9537456 | NA |
| TERF2IP | 26.5605383 | 0.02789129 | 0.36140189 | 0.07717526 | 0.93848412 | 0.99100968 |
| MON1B | 2.27229084 | -1.5512254 | 1.3470309 | -1.1515886 | 0.24949017 | NA |
| ADAMTS18 | 27.68408 | -0.0765227 | 0.46950087 | -0.1629874 | 0.87052838 | 0.97418878 |
| NUDT7 | 3.43367746 | 1.03551127 | 0.9452981 | 1.09543356 | 0.27332674 | NA |
| VAT1L | 7.5655178 | -2.1906372 | 1.03067546 | -2.1254384 | 0.03355005 | NA |
| WWOX | 722.598657 | 0.07601359 | 0.3946863 | 0.19259243 | 0.84727817 | 0.97168963 |
| AC046158.1 | 4.39306895 | 1.11498626 | 0.7618683 | 1.4634895 | 0.14333348 | NA |
| MAF | 13.5380294 | -1.5949354 | 0.63653256 | -2.5056619 | 0.01222224 | NA |
| CDYL2 | 13.7400967 | 0.62401389 | 0.42864355 | 1.45578743 | 0.14545139 | NA |
| CMC2 | 26.2682141 | 0.28356524 | 0.46591622 | 0.60861851 | 0.54277733 | 0.88410237 |
| ATMIN | 8.54666451 | -0.7566642 | 0.56713065 | -1.3341974 | 0.18213917 | NA |
| GAN | 10.3360298 | 0.80146642 | 0.54116664 | 1.48099745 | 0.13860725 | NA |
| CMIP | 81.8220987 | 0.05804943 | 0.39278736 | 0.14778845 | 0.88250973 | 0.97769824 |
| PLCG2 | 710.594706 | -0.087043 | 0.39614152 | -0.2197271 | 0.82608373 | 0.96845695 |
| MPHOSPH6 | 12.4611124 | -0.0352497 | 0.47159418 | -0.0747459 | 0.94041686 | NA |

| | | | | | | |
|---|---|---|---|---|---|---|
| HSBP1 | 21.723829 | 0.08916883 | 0.42277616 | 0.21091263 | 0.83295545 | NA |
| AC009119.2 | 3.01107794 | 0.02945946 | 0.83291537 | 0.03536909 | 0.97178543 | NA |
| MBTPS1 | 23.4378959 | -0.2462147 | 0.39435009 | -0.6243558 | 0.53239398 | NA |
| HSDL1 | 5.030834 | 0.31255695 | 0.75911349 | 0.41173942 | 0.68053043 | NA |
| TAF1C | 3.73821052 | -0.3371064 | 0.87558059 | -0.385009 | 0.70023076 | NA |
| ATP2C2 | 11.1206966 | 0.58579918 | 1.05871507 | 0.55331145 | 0.58005016 | NA |
| TLDC1 | 8.18951483 | 0.92109808 | 0.55781886 | 1.65124945 | 0.09868765 | NA |
| KLHL36 | 11.8700836 | 0.24301508 | 0.45993039 | 0.52837361 | 0.59724005 | NA |
| USP10 | 30.4854044 | -0.4802806 | 0.30302633 | -1.5849469 | 0.11297839 | 0.57860396 |
| ZDHHC7 | 18.3391467 | 0.36415986 | 0.37108864 | 0.98132851 | 0.32643077 | NA |
| KIAA0513 | 2.2421201 | -0.9731232 | 1.24466683 | -0.7818343 | 0.43431197 | NA |
| GSE1 | 91.0613178 | 0.61647166 | 0.37079566 | 1.6625644 | 0.09639966 | 0.56358504 |
| EMC8 | 6.44107517 | 0.0170311 | 0.58692756 | 0.02901737 | 0.97685074 | NA |
| COX4I1 | 90.7730149 | 0.11470327 | 0.31946361 | 0.35904958 | 0.719558 | 0.93714576 |
| MTHFSD | 3.1737344 | 0.49133427 | 0.83344446 | 0.58952251 | 0.55551081 | NA |
| FBXO31 | 8.54780415 | -1.1072094 | 0.51151162 | -2.164583 | 0.03041963 | NA |
| MAP1LC3B | 31.6438231 | -0.0128143 | 0.33398373 | -0.0383681 | 0.96939422 | 0.99703783 |
| ZCCHC14 | 69.8062649 | -0.3939188 | 0.39145037 | -1.0063058 | 0.31426848 | 0.780966 |
| KLHDC4 | 15.2460073 | -0.0443903 | 0.47117202 | -0.0942126 | 0.92494031 | NA |
| BANP | 22.914369 | -0.3079097 | 0.42063589 | -0.73201 | 0.46416245 | NA |
| ZFPM1 | 8.26430333 | -0.2328512 | 0.53326639 | -0.4366508 | 0.66236462 | NA |
| ZC3H18 | 12.1831531 | -0.9385883 | 0.46877439 | -2.0022175 | 0.04526134 | NA |
| CYBA | 11.944561 | 0.19825556 | 0.56647103 | 0.34998358 | 0.72635102 | NA |
| MVD | 7.09518544 | -0.5052523 | 0.67068013 | -0.7533432 | 0.45124371 | NA |
| SNAI3-AS1 | 3.16301188 | 0.18622757 | 0.88594847 | 0.21020135 | 0.83351052 | NA |
| PIEZO1 | 5.58966751 | 0.88099507 | 0.76975865 | 1.14450818 | 0.25241295 | NA |
| APRT | 15.9343513 | 0.79694555 | 0.39359327 | 2.02479465 | 0.04288845 | NA |
| GALNS | 6.39516135 | 0.44734735 | 0.5749968 | 0.77799972 | 0.43656918 | NA |
| TRAPPC2L | 6.03033301 | -0.0292053 | 0.60032709 | -0.048649 | 0.96119902 | NA |
| ACSF3 | 7.97237827 | -0.298506 | 0.64025026 | -0.4662333 | 0.64104851 | NA |
| ZNF778 | 3.06446366 | -1.1003071 | 0.93105824 | -1.1817812 | 0.23729254 | NA |
| ANKRD11 | 98.7614696 | -0.7122102 | 0.30412616 | -2.3418247 | 0.01918973 | 0.33831016 |
| SPG7 | 33.3220451 | -0.3119251 | 0.37290349 | -0.8364767 | 0.40288676 | 0.82808596 |
| RPL13 | 160.873152 | -0.086353 | 0.45459502 | -0.189956 | 0.84934363 | 0.97168963 |
| CHMP1A | 10.3187908 | -0.0541169 | 0.63304498 | -0.0854866 | 0.93187455 | NA |
| SPATA33 | 2.7804614 | 0.03554771 | 0.88138302 | 0.04033174 | 0.96782865 | NA |
| CDK10 | 10.2183187 | 0.7341638 | 0.45192628 | 1.62452113 | 0.10426464 | NA |
| VPS9D1 | 8.03447437 | -0.5063381 | 0.61463944 | -0.823797 | 0.41005491 | NA |
| ZNF276 | 4.27829632 | 0.1066717 | 0.69141423 | 0.15428045 | 0.87738861 | NA |
| SPIRE2 | 11.2246703 | 0.47543965 | 0.60380803 | 0.78740199 | 0.43104658 | NA |
| TCF25 | 53.4827915 | -0.7120983 | 0.27166052 | -2.62128 | 0.00876003 | 0.27648451 |
| DEF8 | 5.39269843 | 0.32378704 | 0.59698541 | 0.54237011 | 0.58756357 | NA |
| GAS8 | 5.85200033 | -0.1806601 | 0.68696923 | -0.2629813 | 0.79256496 | NA |
| FAM157C | 8.91593368 | -0.1015887 | 0.50861226 | -0.1997369 | 0.84168631 | NA |
| LINC02193 | 3.86869344 | -1.3351994 | 0.77667909 | -1.7191134 | 0.08559373 | NA |
| RPH3AL | 19.320613 | 0.04090138 | 0.38455304 | 0.10636082 | 0.91529608 | NA |
| AC129507.2 | 2.92144379 | 1.7943192 | 0.96234384 | 1.86453026 | 0.06224728 | NA |
| C17orf97 | 1.40124129 | -0.6489129 | 1.38108361 | -0.4698578 | 0.63845662 | NA |
| VPS53 | 41.9189646 | -0.8603854 | 0.33853278 | -2.541513 | 0.01103739 | 0.28544118 |
| GLOD4 | 9.46023854 | 0.9793397 | 0.53098038 | 1.84439903 | 0.06512501 | NA |
| TIMM22 | 3.37332678 | 0.04470101 | 0.7757935 | 0.05761973 | 0.95405153 | NA |
| ABR | 20.6723284 | -0.5524354 | 0.4744626 | -1.1643392 | 0.24428656 | NA |
| YWHAE | 78.2399631 | -0.251759 | 0.21018766 | -1.1977821 | 0.23100184 | 0.73156941 |
| CRK | 26.1375818 | -0.743614 | 0.47213115 | -1.5750157 | 0.11525282 | 0.57860396 |

| | | | | | | |
|---|---|---|---|---|---|---|
| MYO1C | 23.8736318 | -0.0024587 | 0.44963353 | -0.0054682 | 0.995637 | NA |
| INPP5K | 5.96418477 | 0.44539569 | 0.64315158 | 0.69252056 | 0.48861048 | NA |
| PITPNA | 27.2941073 | -0.5164108 | 0.32982485 | -1.5657123 | 0.11741598 | 0.57926663 |
| SLC43A2 | 99.2955112 | 1.19238282 | 0.60263421 | 1.97861789 | 0.04785904 | 0.45661231 |
| SCARF1 | 7.26141111 | -0.7202902 | 0.70188116 | -1.0262282 | 0.30478403 | NA |
| RILP | 5.70136009 | -0.1761075 | 0.65334356 | -0.269548 | 0.787508 | NA |
| PRPF8 | 31.1067442 | -0.1907376 | 0.34085945 | -0.5595784 | 0.57576703 | 0.89263108 |
| MIR22HG | 10.9522777 | -0.810441 | 0.68999316 | -1.1745638 | 0.24016927 | NA |
| WDR81 | 9.6291616 | -1.1788984 | 0.7628266 | -1.5454343 | 0.12224124 | NA |
| SERPINF2 | 13.7248798 | -0.2465308 | 0.49782849 | -0.4952123 | 0.62045025 | NA |
| SMYD4 | 7.00844241 | -0.7351679 | 0.68960384 | -1.0660728 | 0.28639072 | NA |
| RPA1 | 11.2184331 | -0.3353539 | 0.47664495 | -0.7035717 | 0.48169955 | NA |
| SMG6 | 59.780459 | -0.5708528 | 0.29959982 | -1.9053842 | 0.05673015 | 0.503587 |
| SRR | 3.20661673 | -0.6206562 | 0.84181767 | -0.737281 | 0.46095148 | NA |
| TSR1 | 4.80843077 | -0.0581071 | 0.85946386 | -0.0676085 | 0.94609727 | NA |
| SGSM2 | 9.62991511 | 0.17012976 | 0.50384331 | 0.33766403 | 0.73561639 | NA |
| MNT | 7.26937462 | -0.7275761 | 0.58989782 | -1.2333934 | 0.21742903 | NA |
| METTL16 | 23.2564303 | -0.380141 | 0.35861482 | -1.0600258 | 0.28913288 | NA |
| PAFAH1B1 | 79.3505965 | -0.7746736 | 0.22479825 | -3.4460836 | 0.00056877 | 0.09487868 |
| CLUH | 16.727273 | -0.5341918 | 0.39889632 | -1.3391746 | 0.18051384 | NA |
| RAP1GAP2 | 25.7030508 | 0.46872922 | 0.43933637 | 1.06690283 | 0.28601572 | NA |
| TRPV1 | 2.33024171 | -0.8299847 | 0.97661289 | -0.8498605 | 0.39540265 | NA |
| CTNS | 9.026306 | -1.1118934 | 0.85409465 | -1.3018386 | 0.19297156 | NA |
| TAX1BP3 | 6.35080572 | -1.314405 | 0.67174843 | -1.9566924 | 0.05038365 | NA |
| EMC6 | 8.99493288 | 0.35580734 | 0.65781348 | 0.54089396 | 0.58858068 | NA |
| ITGAE | 4.10730869 | 0.03079538 | 0.76083478 | 0.04047577 | 0.96771382 | NA |
| NCBP3 | 17.9254429 | -0.1553917 | 0.37985835 | -0.409078 | 0.68248241 | NA |
| ZZEF1 | 35.2197671 | -0.4680559 | 0.30696254 | -1.5247983 | 0.12730942 | 0.59256732 |
| CYB5D2 | 9.32399016 | 0.54709692 | 0.56970401 | 0.96031783 | 0.33689528 | NA |
| ANKFY1 | 22.500611 | -0.8722798 | 0.3727134 | -2.3403499 | 0.01926568 | NA |
| UBE2G1 | 36.959745 | -0.1019137 | 0.32246922 | -0.3160416 | 0.75197092 | 0.95164077 |
| MYBBP1A | 2.75616807 | -0.2727154 | 0.82842881 | -0.329196 | 0.74200755 | NA |
| GGT6 | 3.84381613 | 0.73599161 | 0.83597589 | 0.88039812 | 0.37864367 | NA |
| SMTNL2 | 4.54122 | 0.20535418 | 1.03204805 | 0.19897735 | 0.84228046 | NA |
| PELP1 | 7.35897446 | -0.8397684 | 0.58643525 | -1.4319883 | 0.15214717 | NA |
| ARRB2 | 4.03422514 | 0.7050965 | 0.86926522 | 0.81114082 | 0.41728481 | NA |
| MED11 | 4.0603918 | 1.60811309 | 0.82404421 | 1.95148886 | 0.05099892 | NA |
| PSMB6 | 10.8708268 | 0.29771983 | 0.47068365 | 0.6325264 | 0.52704296 | NA |
| MINK1 | 12.6664957 | -0.0122727 | 0.50638459 | -0.0242358 | 0.98066449 | NA |
| SLC25A11 | 11.8548676 | 0.48519036 | 0.46558053 | 1.04211909 | 0.29735647 | NA |
| RNF167 | 10.1904304 | -0.0293302 | 0.62489628 | -0.0469361 | 0.96256418 | NA |
| PFN1 | 36.1624203 | -0.5576341 | 0.45035987 | -1.2381967 | 0.21564315 | 0.71525281 |
| SPAG7 | 12.1643468 | 0.01236724 | 0.60859257 | 0.02032105 | 0.98378726 | NA |
| CAMTA2 | 6.4381682 | -0.2268122 | 0.62021577 | -0.3656988 | 0.71458981 | NA |
| KIF1C | 10.2202203 | 0.38270814 | 0.49319239 | 0.77598144 | 0.43775994 | NA |
| ZFP3 | 3.60255103 | -0.3546356 | 1.03591894 | -0.3423392 | 0.73209566 | NA |
| ZNF594 | 5.89019633 | 0.2006742 | 0.6534334 | 0.30710736 | 0.75876166 | NA |
| AC087500.1 | 6.84287488 | 0.66600062 | 0.68964952 | 0.96570882 | 0.33418991 | NA |
| RABEP1 | 52.4818093 | -0.5127857 | 0.29022355 | -1.7668647 | 0.0772509 | 0.54401752 |
| NUP88 | 29.7501166 | 0.33645368 | 0.39498043 | 0.85182367 | 0.39431196 | 0.8258317 |
| RPAIN | 14.3523214 | 0.59268823 | 0.39847473 | 1.48739226 | 0.13691124 | NA |
| C1QBP | 14.1608344 | 0.63606821 | 0.45796869 | 1.38889018 | 0.16486615 | NA |
| DERL2 | 7.65534103 | -0.0521543 | 0.56433429 | -0.0924173 | 0.92636647 | NA |
| MIS12 | 4.50234905 | 1.57774353 | 0.78334599 | 2.01410813 | 0.04399818 | NA |

| | | | | | | |
|---|---|---|---|---|---|---|
| KIAA0753 | 11.2468568 | 0.76729205 | 0.55113007 | 1.3922159 | 0.16385703 | NA |
| TXNDC17 | 6.13181713 | 0.074497 | 0.65040476 | 0.11453944 | 0.90881018 | NA |
| MED31 | 3.62899247 | -0.1948112 | 0.73648406 | -0.2645152 | 0.79138299 | NA |
| XAF1 | 7.92174431 | -2.8978716 | 1.7975108 | -1.6121581 | 0.10692754 | NA |
| ALOX12-AS1 | 5.90094265 | 0.59159023 | 0.78207898 | 0.75643284 | 0.44938973 | NA |
| ACADVL | 42.2307386 | -0.1575308 | 0.40011316 | -0.3937156 | 0.69379101 | 0.93004933 |
| DVL2 | 5.55342348 | -0.5303866 | 0.62516195 | -0.8483987 | 0.39621597 | NA |
| PHF23 | 4.58907277 | -0.0573066 | 0.64391856 | -0.0889966 | 0.9290846 | NA |
| ELP5 | 4.98243376 | 0.75302844 | 0.69415262 | 1.08481682 | 0.2780028 | NA |
| EIF5A | 13.8844163 | -0.5397313 | 0.62080925 | -0.8693996 | 0.38462858 | NA |
| GPS2 | 9.99351049 | -0.6566947 | 0.51108571 | -1.2849013 | 0.19882679 | NA |
| ZBTB4 | 23.9176281 | 0.43294273 | 0.46053242 | 0.94009175 | 0.3471705 | NA |
| POLR2A | 28.2545117 | 0.2327052 | 0.40376404 | 0.5763396 | 0.56438566 | 0.89263108 |
| EIF4A1 | 12.8457203 | -0.4525524 | 0.49088095 | -0.9219188 | 0.35657095 | NA |
| MPDU1 | 7.3836095 | -0.287738 | 0.57584901 | -0.4996762 | 0.61730311 | NA |
| FXR2 | 5.5407635 | -0.0666605 | 0.63310127 | -0.1052921 | 0.91614406 | NA |
| SAT2 | 9.46276545 | 0.01011786 | 0.49465404 | 0.02045441 | 0.98368088 | NA |
| TP53 | 5.34734846 | -0.7333801 | 0.66566488 | -1.1017258 | 0.27058092 | NA |
| KDM6B | 10.2029596 | -1.3370292 | 0.6521829 | -2.0500832 | 0.04035631 | NA |
| NAA38 | 16.2457505 | 0.38826378 | 0.36538935 | 1.06260289 | 0.28796208 | NA |
| CHD3 | 9.86444655 | -0.9241051 | 0.62522948 | -1.4780255 | 0.13940096 | NA |
| TRAPPC1 | 4.05792604 | -0.3796825 | 0.7510261 | -0.5055517 | 0.61317142 | NA |
| CNTROB | 2.66083919 | 1.17769618 | 0.95282128 | 1.23600954 | 0.21645502 | NA |
| PER1 | 6.13163604 | -0.6604052 | 0.75294908 | -0.8770915 | 0.38043694 | NA |
| BORCS6 | 2.23065914 | 0.28809935 | 0.99555445 | 0.28938583 | 0.77228614 | NA |
| CTC1 | 6.73018109 | 0.83152518 | 0.61016176 | 1.36279465 | 0.17294724 | NA |
| SLC25A35 | 4.47085332 | 1.20943722 | 0.83402845 | 1.45011507 | 0.14702643 | NA |
| RANGRF | 4.06449326 | 0.9640538 | 0.71195795 | 1.35408811 | 0.17570827 | NA |
| NDEL1 | 18.3233652 | -0.6325061 | 0.53167869 | -1.1896398 | 0.23418799 | NA |
| MYH10 | 35.6181577 | -0.1096265 | 0.30949334 | -0.3542128 | 0.7231794 | 0.93851989 |
| NTN1 | 68.4354368 | 0.3931055 | 0.37837351 | 1.03893503 | 0.29883495 | 0.77521265 |
| STX8 | 59.7044477 | -0.3202113 | 0.29720817 | -1.0773973 | 0.2813028 | 0.76854599 |
| USP43 | 7.69176823 | 0.75419535 | 0.55136338 | 1.36787348 | 0.17135168 | NA |
| GAS7 | 4.40273703 | 0.2017047 | 0.75757306 | 0.26625115 | 0.79004579 | NA |
| SCO1 | 10.4391557 | 0.67586983 | 0.45992064 | 1.46953579 | 0.14168752 | NA |
| TMEM220 | 6.50601593 | 1.00169491 | 0.63130774 | 1.58669827 | 0.112581 | NA |
| TMEM220-A! | 5.18401342 | 0.00516936 | 0.72692525 | 0.00711127 | 0.99432608 | NA |
| MAP2K4 | 57.1175922 | -0.4140441 | 0.26196761 | -1.5805165 | 0.11398862 | 0.57860396 |
| ARHGAP44 | 5.80831808 | -0.8838174 | 0.81233358 | -1.0879981 | 0.27659595 | NA |
| ELAC2 | 7.0777287 | 0.12435563 | 0.52833361 | 0.23537332 | 0.81391899 | NA |
| COX10 | 21.5776736 | -0.3082825 | 0.42583967 | -0.7239402 | 0.46910246 | NA |
| TRIM16 | 2.50880233 | 0.67541504 | 1.14363756 | 0.59058487 | 0.5547986 | NA |
| AC015922.4 | 2.58409548 | 0.03604288 | 0.85123309 | 0.04234196 | 0.9662261 | NA |
| ZSWIM7 | 13.7403851 | 0.24432599 | 0.40220908 | 0.60746016 | 0.54354557 | NA |
| TTC19 | 31.2202311 | -0.6026132 | 0.35498719 | -1.6975633 | 0.08959021 | 0.56122412 |
| NCOR1 | 94.4852824 | -0.1454677 | 0.20698801 | -0.7027833 | 0.4821908 | 0.86255253 |
| PIGL | 19.5004215 | -0.0480271 | 0.4140529 | -0.1159926 | 0.90765844 | NA |
| UBB | 61.2832299 | 0.31233003 | 0.25116009 | 1.24354959 | 0.21366538 | 0.71284112 |
| LRRC75A | 4.43104178 | -0.3592524 | 0.72972231 | -0.4923138 | 0.6224975 | NA |
| AC098850.3 | 15.2072432 | -0.6247403 | 0.7905557 | -0.7902546 | 0.4293791 | NA |
| MPRIP | 40.8357475 | -0.2886729 | 0.28953493 | -0.9970226 | 0.31875353 | 0.78335494 |
| COPS3 | 6.81715155 | -0.7581332 | 0.62603145 | -1.2110146 | 0.22588982 | NA |
| NT5M | 3.73591861 | -0.1732083 | 0.81313444 | -0.2130131 | 0.83131678 | NA |
| MED9 | 8.89735555 | 0.23691475 | 0.53571281 | 0.44224208 | 0.65831404 | NA |

| | | | | | | |
|---|---|---|---|---|---|---|
| RASD1 | 22.3893988 | -0.6754598 | 1.17819683 | -0.5732996 | 0.56644186 | NA |
| PEMT | 3.62886714 | -0.2522794 | 0.82277873 | -0.3066188 | 0.75913357 | NA |
| RAI1 | 24.152323 | -0.4897038 | 0.37368857 | -1.3104596 | 0.19004039 | NA |
| SREBF1 | 16.7289506 | -0.8370536 | 0.62991128 | -1.3288436 | 0.18389957 | NA |
| TOM1L2 | 35.317069 | 0.21403255 | 0.31660743 | 0.67601872 | 0.49902876 | 0.87036626 |
| ATPAF2 | 4.89437683 | 0.66691784 | 0.65762245 | 1.01413483 | 0.31051842 | NA |
| GID4 | 5.49025705 | -1.2845474 | 0.80641898 | -1.5929033 | 0.11118189 | NA |
| DRG2 | 5.14537848 | -0.4846552 | 0.68090983 | -0.7117759 | 0.47660356 | NA |
| ALKBH5 | 14.1319972 | -1.1005812 | 0.45835024 | -2.4011795 | 0.01634232 | NA |
| FLII | 11.3992589 | 0.1248626 | 0.4705318 | 0.26536485 | 0.79072841 | NA |
| MIEF2 | 3.31361804 | 0.1919275 | 0.83505615 | 0.22983784 | 0.81821778 | NA |
| TOP3A | 10.5728036 | 0.04754088 | 0.47121022 | 0.10089102 | 0.91963697 | NA |
| SMCR8 | 4.72054236 | -0.5458244 | 0.78946504 | -0.6913851 | 0.48932355 | NA |
| SHMT1 | 11.9202051 | 0.52802123 | 0.44590662 | 1.18415202 | 0.23635289 | NA |
| TVP23B | 6.96125571 | 0.19864545 | 0.6203339 | 0.32022343 | 0.74879897 | NA |
| PRPSAP2 | 17.6367632 | -0.3348372 | 0.50583553 | -0.6619487 | 0.50800411 | NA |
| SLC5A10 | 3.15334688 | -0.4424214 | 1.15214423 | -0.3839983 | 0.70097973 | NA |
| FAM83G | 2.24368683 | -2.5917776 | 1.15209036 | -2.2496305 | 0.02447241 | NA |
| EPN2 | 30.8496478 | 0.29595459 | 0.38054272 | 0.77771711 | 0.4367358 | 0.8440477 |
| B9D1 | 2.89107856 | -0.000397 | 0.93335191 | -0.0004254 | 0.99966062 | NA |
| MAPK7 | 1.96056533 | -0.9844128 | 1.25040945 | -0.7872724 | 0.43112244 | NA |
| AC004448.2 | 6.47598947 | -0.6411095 | 0.645143 | -0.9937478 | 0.32034565 | NA |
| SLC47A1 | 11.0646908 | 0.01978477 | 0.6269164 | 0.03155886 | 0.97482385 | NA |
| ALDH3A2 | 8.13972839 | 0.37920374 | 0.59136922 | 0.64123009 | 0.5213732 | NA |
| ULK2 | 11.2882769 | 1.02538072 | 0.54516723 | 1.88085538 | 0.05999159 | NA |
| AKAP10 | 39.7026132 | -0.1620987 | 0.34156601 | -0.4745749 | 0.63508999 | 0.92072524 |
| SPECC1 | 4.68074695 | 0.46041485 | 0.68850904 | 0.66871286 | 0.50367866 | NA |
| CCDC144NL-/ | 8.30022576 | 0.78216159 | 0.65118662 | 1.20113276 | 0.22969971 | NA |
| USP22 | 37.8949807 | -0.7388601 | 0.3068921 | -2.4075567 | 0.01605967 | 0.31287044 |
| DHRS7B | 23.575961 | 0.66137465 | 0.36283308 | 1.82280692 | 0.06833264 | NA |
| TMEM11 | 5.17589859 | 0.63916105 | 0.63771525 | 1.00226715 | 0.31621458 | NA |
| AC087294.1 | 2.94320251 | 0.06854356 | 0.88766255 | 0.07721803 | 0.9384501 | NA |
| MAP2K3 | 5.66842955 | -1.8988075 | 0.68774475 | -2.760919 | 0.0057639 | NA |
| WSB1 | 184.845056 | -0.632294 | 0.36521516 | -1.7312917 | 0.08339975 | 0.55234224 |
| KSR1 | 17.5794992 | -0.2308226 | 0.46183243 | -0.4997973 | 0.61721778 | NA |
| NLK | 56.2171006 | -0.2300969 | 0.26886183 | -0.8558183 | 0.39209827 | 0.8258317 |
| IFT20 | 4.29132982 | -0.5353982 | 0.72483809 | -0.7386452 | 0.46012245 | NA |
| TNFAIP1 | 5.68904086 | -0.4169103 | 0.70351755 | -0.5926082 | 0.55344339 | NA |
| POLDIP2 | 9.27468004 | -0.1645997 | 0.47430173 | -0.3470359 | 0.72856432 | NA |
| TMEM199 | 3.42771249 | -0.899419 | 0.79808993 | -1.1269645 | 0.25975748 | NA |
| SARM1 | 2.85432248 | 0.3258098 | 0.80221242 | 0.40613907 | 0.68464042 | NA |
| AC005726.1 | 4.08501712 | 0.27232391 | 0.8254082 | 0.32992634 | 0.74145562 | NA |
| KIAA0100 | 18.6397054 | 0.09912549 | 0.41447755 | 0.23915768 | 0.81098331 | NA |
| SDF2 | 7.90443462 | -0.7133566 | 0.56828997 | -1.2552687 | 0.20938122 | NA |
| SUPT6H | 20.6841344 | -0.7098706 | 0.40642335 | -1.7466286 | 0.08070179 | NA |
| RAB34 | 9.37586777 | 0.82722509 | 0.50630272 | 1.63385474 | 0.10228935 | NA |
| RPL23A | 63.2349854 | 0.22748666 | 0.39171835 | 0.58074037 | 0.56141545 | 0.89263108 |
| TLCD1 | 3.52425111 | -0.7589738 | 0.93328923 | -0.8132247 | 0.41608926 | NA |
| NEK8 | 6.61579401 | 0.7593078 | 0.64549576 | 1.17631725 | 0.23946812 | NA |
| TRAF4 | 13.0901122 | -0.4209751 | 0.45866463 | -0.9178278 | 0.35870903 | NA |
| FAM222B | 59.3410818 | -0.934192 | 0.3239366 | -2.883873 | 0.00392817 | 0.2109414 |
| ERAL1 | 4.04657782 | 0.45550645 | 0.85334843 | 0.53378718 | 0.59348879 | NA |
| FLOT2 | 24.0586662 | 0.00585106 | 0.38701809 | 0.01511831 | 0.98793779 | NA |
| DHRS13 | 3.37058593 | -0.1401818 | 0.88574092 | -0.158265 | 0.87424801 | NA |

| | | | | | | |
|---|---|---|---|---|---|---|
| PHF12 | 14.2700437 | -0.0988735 | 0.42459039 | -0.232868 | 0.81586392 | NA |
| MYO18A | 10.5111954 | -0.4812708 | 0.54021815 | -0.8908823 | 0.37299233 | NA |
| NUFIP2 | 43.6044813 | -0.1187516 | 0.32540101 | -0.3649391 | 0.71515686 | 0.93629694 |
| TAOK1 | 77.9837997 | -0.578484 | 0.29889537 | -1.9354063 | 0.05294046 | 0.48389756 |
| ABHD15-AS1 | 5.41715393 | 0.12017088 | 0.60062386 | 0.20007677 | 0.84142054 | NA |
| ABHD15 | 3.3833845 | 0.16154972 | 0.88480638 | 0.182582 | 0.85512601 | NA |
| GIT1 | 4.37648058 | 0.08983714 | 0.68568799 | 0.13101752 | 0.89576145 | NA |
| SSH2 | 102.244533 | -0.1647336 | 0.2830127 | -0.5820714 | 0.56051856 | 0.89263108 |
| AC023389.1 | 3.36482755 | 0.42207051 | 0.75371934 | 0.55998366 | 0.57549058 | NA |
| NSRP1 | 33.9626246 | -0.5886863 | 0.30397848 | -1.9366051 | 0.05279363 | 0.48389756 |
| AC104984.4 | 9.80475379 | 1.14235032 | 0.55549708 | 2.05644704 | 0.03973945 | NA |
| BLMH | 9.42627695 | 0.43213511 | 0.52217462 | 0.8275682 | 0.40791508 | NA |
| CPD | 29.1598552 | 0.18548851 | 0.37282774 | 0.49751799 | 0.61882382 | 0.91493825 |
| GOSR1 | 26.7467541 | -0.3853081 | 0.34555412 | -1.115044 | 0.26483156 | 0.75497438 |
| CRLF3 | 17.3190629 | 0.06975647 | 0.42692292 | 0.16339358 | 0.87020855 | NA |
| ATAD5 | 3.54151763 | 0.14783135 | 0.84306804 | 0.17534926 | 0.86080521 | NA |
| TEFM | 8.2148962 | 0.5916914 | 0.51106951 | 1.15775131 | 0.24696554 | NA |
| RNF135 | 5.62596697 | 0.42150393 | 0.7686106 | 0.54839723 | 0.58341918 | NA |
| NF1 | 171.19909 | -0.3664781 | 0.23312869 | -1.571999 | 0.11595078 | 0.57860396 |
| OMG | 4.47693287 | -0.3377883 | 0.66162249 | -0.5105454 | 0.60966943 | NA |
| RAB11FIP4 | 10.5575825 | -0.2760816 | 0.50719545 | -0.5443299 | 0.5862145 | NA |
| COPRS | 14.6936926 | -0.4308485 | 0.56202627 | -0.7665986 | 0.44332023 | NA |
| UTP6 | 8.90001298 | 0.23968946 | 0.56783758 | 0.42210919 | 0.67294532 | NA |
| AC004253.1 | 3.2434351 | 0.94305184 | 0.91278285 | 1.03316122 | 0.30152846 | NA |
| SUZ12 | 37.9344676 | -0.1669641 | 0.30352903 | -0.5500763 | 0.58226704 | 0.89263108 |
| LRRC37B | 10.3575159 | -0.651904 | 0.65056086 | -1.0020645 | 0.31631244 | NA |
| RHOT1 | 22.0712619 | 0.73848914 | 0.3685741 | 2.00363817 | 0.04510883 | NA |
| C17orf75 | 3.52754126 | 1.66869937 | 0.87197713 | 1.91369627 | 0.05565898 | NA |
| ZNF207 | 50.5782103 | -0.1346815 | 0.27033475 | -0.4982025 | 0.6183413 | 0.91493825 |
| PSMD11 | 17.1220567 | -0.7425372 | 0.41293875 | -1.7981775 | 0.07214889 | NA |
| MYO1D | 58.6034782 | -0.5971762 | 0.33914459 | -1.7608306 | 0.07826708 | 0.54552914 |
| ZNF830 | 2.68497622 | -0.7656191 | 0.87926882 | -0.8707452 | 0.38389329 | NA |
| LIG3 | 5.76825913 | -0.0403703 | 0.61812891 | -0.0653105 | 0.94792677 | NA |
| SLFN5 | 4.65476468 | 0.7431683 | 0.78958355 | 0.94121553 | 0.34659442 | NA |
| LINC02001 | 4.02286842 | -0.1330737 | 0.69762325 | -0.190753 | 0.84871912 | NA |
| AP2B1 | 34.613228 | 0.11310101 | 0.37844288 | 0.29885886 | 0.76504774 | 0.95550417 |
| TAF15 | 61.8151731 | 0.18982806 | 0.28477889 | 0.6665805 | 0.50504013 | 0.87094108 |
| ZNHIT3 | 6.30378247 | 0.49744566 | 0.64568861 | 0.77041108 | 0.44105608 | NA |
| MYO19 | 46.5404484 | -0.0584921 | 0.40702037 | -0.143708 | 0.88573108 | 0.97769824 |
| GGNBP2 | 31.2463785 | -0.2259434 | 0.43693634 | -0.5171082 | 0.60508064 | 0.90779102 |
| DHRS11 | 3.54962139 | 0.17051544 | 1.03875921 | 0.164153 | 0.8696107 | NA |
| LHX1-DT | 6.65049897 | 1.1937664 | 0.70755759 | 1.687165 | 0.09157163 | NA |
| LHX1 | 13.1177051 | 1.45594259 | 0.5684926 | 2.56105812 | 0.01043539 | NA |
| AATF | 29.6028828 | -0.6735785 | 0.35869578 | -1.8778545 | 0.06040107 | 0.51625203 |
| ACACA | 44.8908922 | -0.8155635 | 0.31243501 | -2.6103462 | 0.00904506 | 0.27648451 |
| TADA2A | 15.256667 | 0.12656321 | 0.47906834 | 0.26418612 | 0.79163651 | NA |
| SYNRG | 35.2817341 | -0.0254202 | 0.31173236 | -0.0815451 | 0.93500845 | 0.9907482 |
| DDX52 | 31.6788079 | -0.1515421 | 0.35898895 | -0.4221358 | 0.67292587 | 0.92617027 |
| AC243571.2 | 11.0127025 | 1.60481087 | 0.87373031 | 1.83673479 | 0.06624906 | NA |
| HNF1B | 41.7573834 | 0.50255282 | 0.46999649 | 1.06926931 | 0.28494834 | 0.76977033 |
| MRPL45 | 7.5095991 | 0.94746676 | 0.55056712 | 1.72089237 | 0.08527036 | NA |
| SOCS7 | 13.0529571 | -0.0686977 | 0.48749624 | -0.1409194 | 0.88793358 | NA |
| MLLT6 | 31.6866515 | 0.37826459 | 0.33731744 | 1.12139055 | 0.26212166 | 0.75479335 |
| CISD3 | 9.51260859 | 1.29508318 | 0.48340426 | 2.67908928 | 0.00738227 | NA |

| | | | | | | |
|---|---|---|---|---|---|---|
| PCGF2 | 7.94195557 | 0.03650595 | 0.62892572 | 0.05804493 | 0.95371284 | NA |
| PSMB3 | 9.74555648 | -0.2420059 | 0.54515103 | -0.4439245 | 0.65709715 | NA |
| PIP4K2B | 9.60043822 | -0.3776215 | 0.55672974 | -0.678285 | 0.49759098 | NA |
| CWC25 | 6.42185673 | -0.555571 | 0.60139153 | -0.9238091 | 0.35558571 | NA |
| RPL23 | 41.3762646 | -0.0468192 | 0.35374755 | -0.132352 | 0.8947059 | 0.97769824 |
| LASP1 | 12.3488505 | -0.31915 | 0.46188499 | -0.6909729 | 0.4895826 | NA |
| RPL19 | 77.2780581 | 0.02144511 | 0.45868784 | 0.04675317 | 0.96270996 | 0.99656976 |
| STAC2 | 52.3106345 | -1.2748009 | 0.68271294 | -1.8672575 | 0.06186564 | 0.51631533 |
| FBXL20 | 60.6214882 | -0.0414864 | 0.31229799 | -0.1328424 | 0.89431805 | 0.97769824 |
| MED1 | 13.0026874 | -0.8404457 | 0.5174144 | -1.6243183 | 0.10430789 | NA |
| CDK12 | 26.5710408 | -0.3505047 | 0.35637423 | -0.9835298 | 0.32534674 | 0.78758454 |
| STARD3 | 6.32815025 | 0.5186529 | 0.71908661 | 0.72126624 | 0.47074572 | NA |
| PGAP3 | 4.33699655 | 2.07571151 | 0.87570561 | 2.37033027 | 0.0177722 | NA |
| ERBB2 | 30.7494781 | 0.12477332 | 0.40965347 | 0.3045826 | 0.76068407 | 0.95478238 |
| MIEN1 | 11.8351749 | 0.18975486 | 0.43443898 | 0.43678138 | 0.6622699 | NA |
| GRB7 | 16.6099287 | -0.0395303 | 0.47819089 | -0.0826664 | 0.93411681 | NA |
| ORMDL3 | 3.32098758 | 0.44045175 | 0.79373929 | 0.55490733 | 0.57895805 | NA |
| PSMD3 | 8.97332616 | -0.1982698 | 0.52187693 | -0.3799168 | 0.70400716 | NA |
| MED24 | 8.95194094 | -0.034371 | 0.55299377 | -0.0621544 | 0.95043991 | NA |
| THRA | 12.1209381 | 0.78250049 | 0.60279056 | 1.29812996 | 0.19424268 | NA |
| MSL1 | 13.559805 | 0.32557629 | 0.50397892 | 0.64601171 | 0.51827177 | NA |
| CASC3 | 16.6058728 | -0.3712926 | 0.44759399 | -0.8295299 | 0.40680462 | NA |
| WIPF2 | 28.6712636 | -0.0278478 | 0.44293658 | -0.0628709 | 0.9498693 | 0.9938068 |
| RARA | 4.0287034 | -1.3767237 | 0.76455684 | -1.8006819 | 0.07175303 | NA |
| IGFBP4 | 6.64984898 | 0.66093924 | 0.56179568 | 1.1764762 | 0.23940464 | NA |
| KRT10 | 12.1681446 | -0.4969782 | 0.61009512 | -0.8145913 | 0.41530627 | NA |
| TMEM99 | 3.41984768 | 0.16556863 | 0.79335741 | 0.20869362 | 0.83468742 | NA |
| KRT19 | 25.5317759 | 0.06618917 | 0.70780899 | 0.09351275 | 0.92549622 | NA |
| EIF1 | 96.8566663 | 0.07198538 | 0.41327649 | 0.17418214 | 0.86172232 | 0.97217819 |
| JUP | 66.6312919 | -0.817381 | 0.38823689 | -2.1053665 | 0.03525942 | 0.41688714 |
| NT5C3B | 8.09140838 | -0.0811553 | 0.53043155 | -0.1529987 | 0.8783993 | NA |
| ACLY | 10.7983896 | 0.30739121 | 0.50037083 | 0.61432681 | 0.53899939 | NA |
| CNP | 8.31004658 | -0.2562594 | 0.53195418 | -0.4817322 | 0.62999623 | NA |
| DNAJC7 | 26.4018227 | -0.1142449 | 0.34443849 | -0.3316844 | 0.74012758 | 0.94705525 |
| NKIRAS2 | 3.32819008 | -0.3895716 | 0.81205575 | -0.479735 | 0.63141581 | NA |
| KAT2A | 3.9952003 | 0.51325155 | 0.74250192 | 0.69124609 | 0.4894109 | NA |
| GHDC | 2.78097692 | 0.11046629 | 1.04483916 | 0.10572564 | 0.91580003 | NA |
| STAT5B | 29.2778319 | -0.2926134 | 0.32861863 | -0.8904346 | 0.3732326 | 0.81364119 |
| STAT5A | 4.90735309 | 0.78190646 | 0.70607778 | 1.10739423 | 0.26812352 | NA |
| STAT3 | 76.7048121 | -0.222629 | 0.33805611 | -0.6585564 | 5.10E-01 | 0.87158522 |
| ATP6V0A1 | 26.3574085 | -0.4954924 | 0.3209698 | -1.5437353 | 0.12265248 | 0.58553957 |
| NAGLU | 8.17783525 | -1.1794556 | 0.61031588 | -1.932533 | 0.05329375 | NA |
| COASY | 4.13013453 | 0.76014202 | 0.80675577 | 0.94222074 | 0.34607964 | NA |
| MLX | 5.93734636 | -0.0259313 | 0.60610339 | -0.0427837 | 0.96587399 | NA |
| RETREG3 | 16.1359022 | -0.0080446 | 0.42949635 | -0.0187304 | 0.98505616 | NA |
| TUBG1 | 2.23279775 | -0.05444 | 0.94055079 | -0.057881 | 0.9538434 | NA |
| TUBG2 | 5.22978559 | -0.2145083 | 0.62644168 | -0.3424234 | 0.73203226 | NA |
| EZH1 | 16.216994 | -0.133058 | 0.43763243 | -0.3040406 | 0.76109697 | NA |
| VPS25 | 7.1285611 | -0.2485172 | 0.5414261 | -0.4590049 | 0.64623066 | NA |
| COA3 | 19.5888662 | 0.06127853 | 0.39804808 | 0.15394755 | 0.87765109 | NA |
| BECN1 | 8.56045006 | -0.78105 | 0.60381826 | -1.2935183 | 0.19583186 | NA |
| PSME3 | 8.09805429 | -0.7135387 | 0.5244479 | -1.360552 | 0.17365529 | NA |
| LINC00671 | 6.55423796 | 0.61115639 | 0.92628168 | 0.6597954 | 0.50938514 | NA |
| RUNDC1 | 7.08901089 | -0.725657 | 0.67867919 | -1.0692195 | 0.28497076 | NA |

| | | | | | | |
|---|---|---|---|---|---|---|
| RPL27 | 24.2359839 | -0.0425242 | 0.3981481 | -0.106805 | 0.91494369 | NA |
| IFI35 | 5.10839408 | 0.51889851 | 1.1845188 | 0.43806692 | 0.66133777 | NA |
| VAT1 | 6.19654192 | 0.27796319 | 0.55135485 | 0.50414573 | 0.61415897 | NA |
| BRCA1 | 6.04257769 | 0.49613369 | 0.64228832 | 0.77244701 | 0.43984972 | NA |
| NBR1 | 22.3374352 | 0.71680197 | 0.49130912 | 1.4589633 | 0.14457521 | NA |
| LINC00854 | 34.0294596 | -1.4793447 | 0.73642318 | -2.0088242 | 0.04455579 | 0.44769598 |
| ARL4D | 36.3056934 | 0.59857616 | 0.45900476 | 1.30407397 | 0.19220836 | 0.68667113 |
| DHX8 | 9.40251437 | -0.5530988 | 0.51262554 | -1.0789528 | 0.28060876 | NA |
| DUSP3 | 10.7482699 | -0.6588306 | 0.5338166 | -1.2341891 | 0.21713245 | NA |
| TMEM101 | 17.3372772 | 0.89468364 | 0.4760568 | 1.87936321 | 0.06019492 | NA |
| LSM12 | 13.7954425 | -0.8746125 | 0.42820181 | -2.042524 | 0.04109958 | NA |
| G6PC3 | 4.52541541 | -0.0520512 | 0.75282744 | -0.0691409 | 0.94487747 | NA |
| HDAC5 | 21.2179121 | -0.1649251 | 0.4639863 | -0.3554524 | 0.72225067 | NA |
| TMUB2 | 7.50555684 | -0.1414956 | 0.61652457 | -0.2295052 | 0.81847625 | NA |
| ATXN7L3 | 5.64947058 | -1.3765799 | 0.6748111 | -2.0399485 | 0.04135545 | NA |
| UBTF | 13.4319569 | 0.41958651 | 0.48296247 | 0.86877664 | 0.38496931 | NA |
| SLC25A39 | 14.5262336 | -0.4283186 | 0.50602171 | -0.846443 | 0.39730564 | NA |
| GRN | 14.2029814 | -0.3108733 | 0.56473073 | -0.5504805 | 0.58198983 | NA |
| GPATCH8 | 101.794534 | -0.0167296 | 0.22790763 | -0.0734054 | 0.94148355 | 0.99153825 |
| MEIOC | 5.49862169 | -0.0280083 | 0.66550983 | -0.0420855 | 0.96643052 | NA |
| CCDC43 | 4.66313354 | -0.6629688 | 0.67474693 | -0.9825444 | 0.32583172 | NA |
| EFTUD2 | 9.4674816 | -0.5160643 | 0.51547337 | -1.0011464 | 0.31675605 | NA |
| DCAKD | 9.19389342 | 0.13271888 | 0.62273662 | 0.21312201 | 0.8312318 | NA |
| NMT1 | 14.5565231 | 0.42241006 | 0.41127239 | 1.02708099 | 0.30438233 | NA |
| ACBD4 | 5.15179538 | 0.74576836 | 0.75824147 | 0.98354995 | 0.32533684 | NA |
| HEXIM1 | 7.65742005 | -0.4431329 | 0.84557145 | -0.5240632 | 0.60023456 | NA |
| AC138150.1 | 6.96294202 | -0.0729806 | 0.83461365 | -0.0874424 | 0.9303199 | NA |
| MAP3K14 | 7.2261802 | 0.6552548 | 0.56189273 | 1.1661564 | 0.24355122 | NA |
| ARHGAP27 | 5.1268741 | -0.8245739 | 0.64915856 | -1.2702196 | 0.20400641 | NA |
| PLEKHM1 | 9.78646623 | -0.5455111 | 0.48695767 | -1.1202434 | 0.26261003 | NA |
| LINC02210-C | 5.5129192 | 0.69177453 | 0.68430017 | 1.01092264 | 0.31205346 | NA |
| KANSL1 | 174.047416 | 0.22777714 | 0.2657294 | 0.85717702 | 0.39134706 | 0.8258317 |
| ARL17B | 36.9711174 | 1.45292269 | 1.07769517 | 1.34817593 | 0.1776018 | 0.6676327 |
| LRRC37A2 | 8.82952727 | 0.38998635 | 0.67435514 | 0.57831004 | 0.56305482 | NA |
| ARL17A | 11.7249578 | -0.4413911 | 0.6810286 | -0.6481241 | 0.51690466 | NA |
| NSF | 44.3791354 | -0.0691829 | 0.37838927 | -0.1828352 | 0.85492736 | 0.97168963 |
| AC005670.2 | 5.77858472 | 0.97285081 | 0.67921669 | 1.43231288 | 0.1520543 | NA |
| CDC27 | 48.5498191 | -0.1404541 | 0.28525311 | -0.492384 | 0.62244788 | 0.91493825 |
| AC068234.1 | 29.9858634 | 1.25281278 | 0.51151701 | 2.4492104 | 0.01431698 | 0.30435125 |
| AC068234.2 | 6.61045936 | 2.3565645 | 0.69957086 | 3.36858583 | 0.00075555 | NA |
| EFCAB13 | 15.8446867 | 1.28340183 | 0.40698283 | 3.15345448 | 0.0016135 | NA |
| NPEPPS | 89.9811991 | -0.4028053 | 0.26906936 | -1.4970316 | 0.13438503 | 0.60594834 |
| AC025682.1 | 10.4175915 | 0.54574789 | 0.58564788 | 0.93187033 | 0.35140354 | NA |
| KPNB1 | 44.6268483 | -0.5425846 | 0.33502282 | -1.6195451 | 0.10533004 | 0.5772605 |
| MRPL10 | 4.6091294 | -0.5924387 | 0.75245455 | -0.7873415 | 0.43108201 | NA |
| SCRN2 | 3.23016222 | 0.80052888 | 0.91959601 | 0.87052235 | 0.38401501 | NA |
| SP2 | 6.3704646 | -0.4768694 | 0.62850572 | -0.7587352 | 0.44801096 | NA |
| PNPO | 5.00318256 | 0.24976734 | 0.87613554 | 0.28507842 | 0.77558406 | NA |
| CDK5RAP3 | 8.28557019 | -0.2013068 | 0.53889162 | -0.3735572 | 0.70873378 | NA |
| NFE2L1 | 34.6603935 | -0.0999303 | 0.3466678 | -0.2882594 | 0.77314816 | 0.95994327 |
| CBX1 | 12.9462596 | -0.4586565 | 0.48029897 | -0.9549395 | 0.3396083 | NA |
| SNX11 | 5.0913479 | -1.3147185 | 0.83027067 | -1.5834818 | 0.11331169 | NA |
| SKAP1 | 132.881387 | 0.30632741 | 0.25964153 | 1.17980899 | 0.23807619 | 0.73242571 |
| HOXB2 | 6.76142251 | 0.05337247 | 0.67012653 | 0.07964536 | 0.93651932 | NA |

| | | | | | | |
|---|---|---|---|---|---|---|
| HOXB-AS1 | 3.35427568 | -0.6812144 | 0.91343797 | -0.7457698 | 0.4558065 | NA |
| HOXB3 | 75.0541021 | 0.09236049 | 0.29712241 | 0.31084996 | 0.75591469 | 0.95323728 |
| HOXB-AS3 | 23.7304178 | 0.77917554 | 0.42972303 | 1.81320405 | 0.06980036 | NA |
| HOXB7 | 12.9716996 | -0.8758927 | 0.48674232 | -1.7994997 | 0.07193967 | NA |
| AC103702.2 | 5.43751828 | 0.15357931 | 0.65795605 | 0.2334188 | 0.81543622 | NA |
| LINC02086 | 10.5970449 | 0.82838483 | 0.70179537 | 1.18037943 | 0.23784934 | NA |
| TTLL6 | 5.04840437 | 0.83317978 | 0.81654903 | 1.02036712 | 0.30755438 | NA |
| CALCOCO2 | 23.3679428 | 0.25543441 | 0.35642906 | 0.71664866 | 0.47359091 | NA |
| ATP5MC1 | 22.2297399 | 0.82225691 | 0.40583267 | 2.02609835 | 0.04275471 | NA |
| UBE2Z | 19.8735192 | -0.3202595 | 0.43509932 | -0.7360606 | 0.46169382 | NA |
| SNF8 | 10.5722932 | -0.5600921 | 0.4682977 | -1.1960171 | 0.23168987 | NA |
| B4GALNT2 | 47.5396968 | 1.10439707 | 0.65539773 | 1.68507917 | 0.0919733 | 0.56220784 |
| ZNF652 | 38.7209851 | 0.26618986 | 0.28870898 | 0.92200065 | 0.35652824 | 0.80177915 |
| PHB | 25.5035335 | 0.65937692 | 0.36165449 | 1.8232234 | 0.06826956 | NA |
| SPOP | 26.9321855 | 0.01770617 | 0.32673672 | 0.05419094 | 0.95678304 | 0.99636969 |
| SLC35B1 | 4.11236828 | -0.3994684 | 0.73798039 | -0.5412994 | 0.58830121 | NA |
| KAT7 | 11.8839898 | 0.04093997 | 0.49008201 | 0.08353698 | 0.93342458 | NA |
| ITGA3 | 20.930275 | -0.4388157 | 0.42385673 | -1.0352925 | 3.01E-01 | NA |
| PDK2 | 4.23936309 | 1.00232226 | 0.87161459 | 1.1499604 | 0.25016018 | NA |
| XYLT2 | 3.94936885 | 0.44191144 | 0.78426749 | 0.56347031 | 0.57311467 | NA |
| MRPL27 | 12.3137018 | 0.32449718 | 0.4757988 | 0.68200503 | 0.49523577 | NA |
| LRRC59 | 7.256738 | -1.7380332 | 0.64547407 | -2.692646 | 0.00708875 | NA |
| ACSF2 | 4.97716091 | 1.06539072 | 0.80776635 | 1.31893428 | 0.18719108 | NA |
| RSAD1 | 4.7383057 | 0.79948644 | 0.78502112 | 1.01842666 | 0.30847524 | NA |
| SPATA20 | 3.57918982 | 0.94096131 | 0.81278802 | 1.15769584 | 0.24698818 | NA |
| ABCC3 | 9.71386556 | 0.22855349 | 0.58164041 | 0.39294637 | 0.69435909 | NA |
| ANKRD40 | 9.71473348 | -0.3812117 | 0.61671091 | -0.6181367 | 0.53648521 | NA |
| LUC7L3 | 58.3817975 | 0.00036363 | 0.25135479 | 0.00144666 | 0.99884573 | 0.99960611 |
| TOB1 | 9.69889211 | -0.4405629 | 0.54329433 | -0.8109102 | 0.41741724 | NA |
| SPAG9 | 156.21643 | -0.6987063 | 0.27953711 | -2.4995117 | 0.01243646 | 0.30175371 |
| AC005920.2 | 12.0575331 | 0.01782408 | 0.55467478 | 0.03213429 | 0.97436496 | NA |
| MBTD1 | 34.0051167 | -0.0736511 | 0.35821616 | -0.2056052 | 0.83709933 | 0.97154892 |
| UTP18 | 9.85986891 | 0.50420529 | 0.52420622 | 0.96184529 | 0.33612732 | NA |
| TOM1L1 | 37.624121 | -0.1452235 | 0.32270334 | -0.4500218 | 0.6526947 | 0.92248109 |
| COX11 | 7.9104384 | 0.05322884 | 0.53043887 | 0.10034867 | 0.92006752 | NA |
| STXBP4 | 25.5778713 | -0.1152706 | 0.43614675 | -0.2642932 | 0.79155402 | NA |
| MMD | 21.9702152 | 0.05323988 | 0.34055692 | 0.15633182 | 0.87577147 | NA |
| PCTP | 2.76167877 | 0.24659128 | 1.13625549 | 0.21702098 | 0.828192 | NA |
| DGKE | 3.10232578 | 0.8205859 | 0.93428394 | 0.87830462 | 0.37977843 | NA |
| TRIM25 | 5.06549107 | -0.4252472 | 0.69631317 | -0.6107126 | 0.54138987 | NA |
| COIL | 5.58598061 | 0.41327741 | 0.62932888 | 0.65669545 | 0.51137676 | NA |
| SCPEP1 | 12.5904774 | 0.42997437 | 0.48706639 | 0.8827839 | 0.37735304 | NA |
| AKAP1 | 22.0178654 | 0.52505152 | 0.37694816 | 1.39290113 | 0.16364969 | NA |
| MSI2 | 407.266424 | 0.00977215 | 0.28967309 | 0.03373508 | 0.9730884 | 0.99779576 |
| MRPS23 | 6.3283105 | 0.30518372 | 0.55445057 | 0.55042548 | 0.58202758 | NA |
| CUEDC1 | 23.014712 | 0.16498357 | 0.35698043 | 0.4621642 | 0.64396358 | NA |
| VEZF1 | 20.3287778 | 0.22388882 | 0.41560634 | 0.53870406 | 0.59009107 | NA |
| AC015813.2 | 6.09131554 | -0.1365566 | 0.66536984 | -0.2052341 | 0.83738926 | NA |
| SRSF1 | 3.98311419 | 0.33711348 | 0.83728198 | 0.40262836 | 0.68722164 | NA |
| DYNLL2 | 18.0685138 | 0.09993047 | 0.41094022 | 0.24317519 | 0.80786969 | NA |
| TSPOAP1 | 4.53856365 | 0.19847898 | 0.95290245 | 0.20828887 | 0.83500342 | NA |
| SUPT4H1 | 7.27579756 | -0.6099966 | 0.59657579 | -1.0224963 | 0.30654605 | NA |
| RNF43 | 11.1743838 | 0.41953335 | 0.59904869 | 0.70033263 | 0.4837196 | NA |
| MTMR4 | 7.04520762 | 0.1723023 | 0.59560925 | 0.28928748 | 0.77236139 | NA |

| | | | | | | |
|---|---|---|---|---|---|---|
| RAD51C | 3.98414257 | 0.55415797 | 0.74717483 | 0.74167108 | 0.45828664 | NA |
| PPM1E | 265.620918 | -0.6778806 | 0.4848317 | -1.3981772 | 0.16205986 | 0.64709575 |
| TRIM37 | 13.1266268 | -0.4745954 | 0.42944859 | -1.1051273 | 0.26910443 | NA |
| SKA2 | 8.11841311 | -0.0287731 | 0.56416842 | -0.051001 | 0.95932474 | NA |
| YPEL2 | 48.5583995 | 0.0986118 | 0.51233224 | 0.19247627 | 0.84736915 | 0.97168963 |
| DHX40 | 9.37123055 | 0.67722915 | 0.49621325 | 1.36479458 | 0.17231762 | NA |
| CLTC | 60.487988 | -0.1760164 | 0.26972286 | -0.6525825 | 0.51402544 | 0.87158522 |
| PTRH2 | 6.29215762 | 0.21510185 | 0.64184048 | 0.33513289 | 0.73752484 | NA |
| VMP1 | 74.0123356 | -0.3978599 | 0.25379397 | -1.567649 | 0.11696306 | 0.57860396 |
| TUBD1 | 3.83406223 | 0.33893464 | 1.00170436 | 0.33835795 | 0.73509346 | NA |
| RPS6KB1 | 14.8243355 | -0.0444079 | 0.48165609 | -0.0921983 | 0.92654046 | NA |
| RNFT1 | 4.22572896 | 0.69994689 | 0.77432564 | 0.90394384 | 0.36602518 | NA |
| HEATR6 | 4.37597384 | 0.84023071 | 0.73185 | 1.14809142 | 0.25093082 | NA |
| USP32 | 50.4705377 | -0.6254386 | 0.29140582 | -2.1462804 | 0.03185062 | 0.4126665 |
| APPBP2 | 33.1134508 | 0.11144477 | 0.36135276 | 0.30840992 | 0.75777044 | 0.95323728 |
| PPM1D | 11.8799294 | 0.74593165 | 0.53089476 | 1.40504617 | 0.16000755 | NA |
| BCAS3 | 126.439611 | 0.358504 | 0.31137341 | 1.15136356 | 0.2495827 | 0.7417998 |
| AC005746.2 | 3.3505272 | 2.37543783 | 1.02035066 | 2.32806028 | 0.0199089 | NA |
| TBX2 | 15.5064623 | -0.4047315 | 0.42861935 | -0.9442679 | 0.34503278 | NA |
| NACA2 | 5.51177188 | 0.56491359 | 0.67004607 | 0.84309664 | 0.3991744 | NA |
| INTS2 | 8.93717792 | 0.00713172 | 0.53915027 | 0.0132277 | 0.98944613 | NA |
| MED13 | 97.2767581 | -0.1683217 | 0.28075698 | -0.599528 | 0.54882085 | 0.88448477 |
| METTL2A | 7.09714919 | -1.6318512 | 0.68178429 | -2.3935008 | 0.01668845 | NA |
| TLK2 | 52.7748688 | 0.08538047 | 0.27622482 | 0.30909776 | 0.75724716 | 0.95323728 |
| 10-Mar | 5.02787117 | 2.59990378 | 1.05874915 | 2.45563718 | 0.01406351 | NA |
| TANC2 | 68.0716232 | 0.24522707 | 0.32931558 | 0.74465676 | 0.45647925 | 0.85802171 |
| AC015923.1 | 5.49400913 | 0.08207617 | 0.69919993 | 0.11738583 | 0.90655431 | NA |
| CYB561 | 3.30382196 | 0.2245385 | 0.81825086 | 0.27441279 | 0.78376741 | NA |
| DCAF7 | 38.6874186 | 0.22835874 | 0.35256494 | 0.64770688 | 0.51717455 | 0.87158522 |
| TACO1 | 2.61673395 | 0.68810004 | 0.8844355 | 0.77801042 | 0.43656287 | NA |
| MAP3K3 | 37.7379001 | -0.3142768 | 0.33459932 | -0.9392632 | 0.34759565 | 0.8003972 |
| CCDC47 | 9.77354611 | -0.9266111 | 0.53937829 | -1.7179244 | 0.0858104 | NA |
| DDX42 | 23.4028957 | -0.7907718 | 0.36171924 | -2.186148 | 0.02880478 | NA |
| FTSJ3 | 4.47571474 | -0.8702495 | 0.71447096 | -1.2180334 | 0.22321129 | NA |
| PSMC5 | 7.85456089 | -0.2176256 | 0.63071519 | -0.3450458 | 0.73005991 | NA |
| SMARCD2 | 5.87092096 | -0.4084943 | 0.8097428 | -0.5044742 | 0.6139282 | NA |
| ERN1 | 7.16690985 | 0.21310133 | 0.5465449 | 0.38990635 | 0.6966058 | NA |
| SNHG25 | 3.00946578 | 0.926843 | 0.86784573 | 1.06798129 | 0.28552895 | NA |
| TEX2 | 9.88495201 | 0.59104203 | 0.56348488 | 1.04890485 | 0.29422191 | NA |
| POLG2 | 7.41962802 | 1.13356946 | 0.60847261 | 1.86297533 | 0.06246574 | NA |
| DDX5 | 89.95171 | 0.37451527 | 0.25889172 | 1.44660967 | 0.14800628 | 0.63 |
| CEP95 | 28.8807896 | -0.4046006 | 0.4075718 | -0.9927101 | 0.32085124 | 0.785589 |
| SMURF2 | 30.9090733 | -0.4973937 | 0.29884712 | -1.664375 | 0.09603751 | 0.56358504 |
| LRRC37A3 | 28.8046351 | 0.41480967 | 0.44797715 | 0.92596167 | 0.3544659 | 0.8003972 |
| GNA13 | 20.6174663 | -0.2622607 | 0.3303853 | -0.7938025 | 0.42731039 | NA |
| CEP112 | 22.6733544 | 0.49192181 | 0.68069535 | 0.72267543 | 0.46987931 | NA |
| PRKCA | 139.998106 | -0.0717596 | 0.34738639 | -0.2065701 | 0.83634565 | 0.97154892 |
| PRKCA-AS1 | 4.74848265 | -0.6347166 | 0.90845122 | -0.6986799 | 0.48475208 | NA |
| HELZ | 68.469922 | -0.0482112 | 0.25243257 | -0.1909866 | 0.8485361 | 0.97168963 |
| PSMD12 | 13.2575088 | -0.1646133 | 0.4212668 | -0.3907578 | 0.69597627 | NA |
| PITPNC1 | 15.7847951 | -0.1566679 | 0.68597696 | -0.2283866 | 0.81934574 | NA |
| NOL11 | 7.27883838 | 0.02962473 | 0.60738816 | 0.04877397 | 0.96109943 | NA |
| BPTF | 78.6091768 | -0.1733661 | 0.28196626 | -0.6148469 | 0.53865586 | 0.88254911 |
| AC005332.7 | 2.41442585 | -0.6908729 | 0.93034253 | -0.7426006 | 0.45772352 | NA |

| | | | | | | |
|---|---|---|---|---|---|---|
| AMZ2 | 4.35330262 | 0.77397183 | 0.69166561 | 1.11899712 | 0.26314137 | NA |
| ARSG | 8.43192294 | 0.2311862 | 0.6063346 | 0.38128485 | 0.70299189 | NA |
| WIPI1 | 5.62979748 | 0.01526995 | 0.70799431 | 0.0215679 | 0.98279264 | NA |
| PRKAR1A | 47.220773 | -0.1857492 | 0.33152157 | -0.5602928 | 0.57527973 | 0.89263108 |
| FAM20A | 8.24964091 | -0.9778875 | 0.96346134 | -1.0149733 | 0.31011857 | NA |
| ABCA9 | 3.32785102 | -0.8782202 | 1.11146334 | -0.7901477 | 0.42944154 | NA |
| ABCA6 | 3.5932527 | -1.4675361 | 0.81963357 | -1.7904783 | 0.07337706 | NA |
| ABCA10 | 16.8252069 | -0.5679777 | 0.58820833 | -0.9656064 | 0.33424118 | NA |
| ABCA5 | 86.8115601 | -1.0074626 | 0.32895619 | -3.0626041 | 0.0021942 | 0.16267567 |
| AC005208.1 | 8.90650897 | 0.68140395 | 0.66317571 | 1.02748628 | 0.30419154 | NA |
| KCNJ16 | 27.6565595 | -0.59396 | 0.52978074 | -1.1211431 | 0.26222696 | 0.75479335 |
| SOX9 | 12.8139145 | 0.37506084 | 0.60686609 | 0.61802899 | 0.53655623 | NA |
| SLC39A11 | 35.4425625 | 0.22220951 | 0.36125015 | 0.61511259 | 0.53848037 | 0.88254911 |
| COG1 | 5.09867576 | 0.22784151 | 0.77632922 | 0.29348568 | 0.76915094 | NA |
| FAM104A | 7.37764297 | 0.0242082 | 0.55205986 | 0.04385068 | 0.96502343 | NA |
| C17orf80 | 5.56147485 | 0.29464128 | 0.61875394 | 0.47618489 | 0.63394266 | NA |
| CDC42EP4 | 6.52919242 | 0.37092156 | 0.57710487 | 0.64272818 | 0.52040049 | NA |
| SDK2 | 14.5068132 | -0.6657475 | 0.85610038 | -0.7776512 | 0.43677469 | NA |
| RPL38 | 19.893163 | -0.1656775 | 0.5349203 | -0.3097238 | 0.75677104 | NA |
| GPRC5C | 28.9793042 | 0.54851869 | 0.33567722 | 1.63406588 | 0.10224502 | 0.5750956 |
| SLC9A3R1 | 9.66463957 | 0.65442918 | 0.58974829 | 1.10967543 | 0.26713892 | NA |
| NAT9 | 4.74863471 | -0.4579433 | 0.6165244 | -0.7427822 | 0.45761358 | NA |
| TMEM104 | 4.64714445 | -2.0615795 | 0.78526171 | -2.6253406 | 0.00865623 | NA |
| MRPL58 | 4.03958189 | 0.02034657 | 0.81777025 | 0.02488054 | 0.98015025 | NA |
| KCTD2 | 8.90640634 | -0.5603259 | 0.56985031 | -0.9832861 | 0.32546666 | NA |
| ATP5PD | 15.5135394 | 0.23385575 | 0.570621 | 0.40982674 | 0.68193305 | NA |
| SLC16A5 | 40.3507708 | 0.79855486 | 0.60237156 | 1.32568486 | 0.1849441 | 0.67879383 |
| ARMC7 | 16.5615812 | 0.80748363 | 0.50228823 | 1.6076101 | 0.1079206 | NA |
| NT5C | 13.3963929 | 0.07158773 | 0.43867529 | 0.16319071 | 0.87036828 | NA |
| JPT1 | 20.6022672 | 0.07717302 | 0.40912965 | 0.18862729 | 0.85038494 | NA |
| SUMO2 | 6.90037469 | 0.10916153 | 0.54706315 | 0.199541 | 0.84183957 | NA |
| NUP85 | 6.1937155 | -0.1987723 | 0.65364399 | -0.3040988 | 0.76105263 | NA |
| GGA3 | 8.77528914 | -0.4613684 | 0.51963922 | -0.887863 | 0.37461446 | NA |
| MRPS7 | 9.92947359 | 0.46828948 | 0.53596701 | 0.87372817 | 0.38226631 | NA |
| GRB2 | 31.9338224 | 0.06233136 | 0.35388436 | 0.17613483 | 0.86018802 | 0.97180546 |
| AC011933.4 | 3.46605735 | 0.56588567 | 0.75929079 | 0.74528189 | 0.45610133 | NA |
| TMEM94 | 10.1118276 | 0.74006548 | 0.50952705 | 1.45245571 | 0.14637494 | NA |
| CASKIN2 | 4.5309188 | -0.1960063 | 0.67877925 | -0.288763 | 0.77276278 | NA |
| TSEN54 | 7.07706194 | -0.5979479 | 0.58163899 | -1.0280395 | 0.30393124 | NA |
| LLGL2 | 69.6615272 | 0.39609616 | 0.30106624 | 1.31564459 | 0.18829336 | 0.68214045 |
| MYO15B | 11.8419252 | 0.65038868 | 0.47532738 | 1.36829626 | 0.17121936 | NA |
| RECQL5 | 8.26470771 | 0.40730185 | 0.57011686 | 0.71441817 | 0.47496864 | NA |
| SMIM5 | 13.1882368 | 0.67672279 | 0.53342109 | 1.26864649 | 0.20456718 | NA |
| SMIM6 | 8.27565671 | 0.99157507 | 0.66548921 | 1.48999421 | 0.13622576 | NA |
| SAP30BP | 15.5907535 | -0.5553155 | 0.38559506 | -1.440152 | 0.1498244 | NA |
| ITGB4 | 7.87651589 | 0.84676178 | 0.5537619 | 1.52910806 | 0.12623766 | NA |
| H3F3B | 58.9637203 | -0.1694185 | 0.27165468 | -0.623654 | 0.53285483 | 0.88254911 |
| UNK | 17.1006819 | 0.39311337 | 0.46826891 | 0.83950345 | 0.40118685 | NA |
| WBP2 | 13.1978001 | -0.0296701 | 0.4031037 | -0.0736041 | 0.94132541 | NA |
| TRIM47 | 5.49713193 | -0.5247839 | 0.60872825 | -0.8620988 | 0.38863314 | NA |
| TRIM65 | 1.43279446 | 0.65643985 | 1.27482859 | 0.51492401 | 0.60660612 | NA |
| ACOX1 | 18.1252442 | 0.73578561 | 0.39826576 | 1.84747393 | 0.06467848 | NA |
| TEN1 | 2.68404377 | -0.7419288 | 0.94299891 | -0.7867759 | 0.43141306 | NA |
| EVPL | 2.90778345 | 0.30772407 | 1.00730405 | 0.30549274 | 0.7599909 | NA |

| | | | | | | |
|---|---|---|---|---|---|---|
| SRP68 | 8.24867068 | -0.8479918 | 0.65219176 | -1.3002185 | 0.1935261 | NA |
| EXOC7 | 15.6093281 | -0.4434819 | 0.45429584 | -0.9761962 | 0.32896724 | NA |
| RNF157 | 7.88837725 | 0.36994432 | 0.63487605 | 0.58270322 | 0.56009311 | NA |
| UBALD2 | 9.13602375 | 0.41118309 | 0.52039145 | 0.7901419 | 0.4294449 | NA |
| PRPSAP1 | 8.83137413 | 0.30171173 | 0.52273505 | 0.57717906 | 0.5638185 | NA |
| UBE2O | 13.2041483 | -0.0469287 | 0.49196386 | -0.0953905 | 0.92400463 | NA |
| RHBDF2 | 4.05317633 | 0.00484163 | 0.63852598 | 0.00758251 | 0.99395009 | NA |
| MXRA7 | 14.7455005 | 0.01737156 | 0.5333241 | 0.03257224 | 0.9740157 | NA |
| JMJD6 | 7.26992269 | -0.8015479 | 0.58486189 | -1.3704908 | 0.17053374 | NA |
| METTL23 | 2.79068979 | -0.1197609 | 0.87046631 | -0.1375824 | 0.89057043 | NA |
| SRSF2 | 27.2403709 | -0.526549 | 0.36840983 | -1.4292479 | 0.15293298 | 0.63864841 |
| MFSD11 | 13.4436205 | 0.30157985 | 0.51141967 | 0.58969153 | 0.55539748 | NA |
| SEC14L1 | 36.7407164 | -0.0644245 | 0.34217551 | -0.1882791 | 0.85065789 | 0.97168963 |
| 9-Sep | 72.472422 | 0.08080878 | 0.25019954 | 0.32297733 | 0.74671241 | 0.94858421 |
| TNRC6C | 27.7536185 | -0.3148874 | 0.50350505 | -0.6253908 | 0.53171461 | 0.88254911 |
| TMC6 | 2.83039646 | -0.7778549 | 0.84040089 | -0.925576 | 0.35466637 | NA |
| SYNGR2 | 8.4475843 | -0.3308736 | 0.55303939 | -0.5982822 | 0.54965165 | NA |
| AFMID | 6.09662191 | 0.10655453 | 0.76892836 | 0.13857537 | 0.88978571 | NA |
| PGS1 | 7.35113868 | 0.34879181 | 0.55718929 | 0.62598442 | 0.53132516 | NA |
| CYTH1 | 131.109378 | -0.9470091 | 0.3307541 | -2.8631817 | 4.19E-03 | 0.21182958 |
| USP36 | 17.0276569 | -1.5202954 | 0.48793552 | -3.1157711 | 0.00183465 | NA |
| TIMP2 | 20.9885994 | 0.04244244 | 0.43128811 | 0.09840856 | 0.92160788 | NA |
| LGALS3BP | 29.1457663 | -0.0811241 | 0.59812925 | -0.1356297 | 0.89211406 | 0.97769824 |
| CANT1 | 5.01155552 | 0.62545817 | 0.70742159 | 0.88413779 | 0.37662183 | NA |
| ENGASE | 3.80314348 | 0.09020055 | 0.77401736 | 0.11653556 | 0.90722811 | NA |
| TBC1D16 | 22.6732078 | 0.04162586 | 0.42094939 | 0.09888566 | 0.92122905 | NA |
| GAA | 6.26076454 | 0.74615787 | 0.73321921 | 1.01764638 | 0.30884604 | NA |
| EIF4A3 | 5.1701853 | -0.3705906 | 0.73017482 | -0.5075369 | 0.61177817 | NA |
| AC087741.1 | 3.80418375 | 0.97748865 | 0.69591542 | 1.40460841 | 0.16013775 | NA |
| SGSH | 3.99436563 | -0.9630211 | 0.87477634 | -1.1008769 | 0.27095024 | NA |
| SLC26A11 | 8.95923707 | -0.5703074 | 0.58415359 | -0.9762969 | 0.32891735 | NA |
| RNF213 | 91.359848 | -0.0663447 | 0.59702057 | -0.1111263 | 0.91151616 | 0.98375925 |
| AC124319.2 | 21.5629887 | 0.00179573 | 0.4599926 | 0.00390383 | 0.9968852 | NA |
| ENDOV | 5.65183701 | -0.5041811 | 0.71360403 | -0.7065278 | 0.47985998 | NA |
| RPTOR | 49.3291539 | -0.0643059 | 0.34201965 | -0.1880181 | 0.85086243 | 0.97168963 |
| CHMP6 | 4.17737088 | -0.132744 | 0.78933394 | -0.1681721 | 0.86644786 | NA |
| TEPSIN | 2.34989475 | -0.6456688 | 0.99889505 | -0.6463831 | 0.51803131 | NA |
| NDUFAF8 | 9.11802014 | -0.4289257 | 0.70470821 | -0.6086572 | 0.54275166 | NA |
| SLC38A10 | 19.1964595 | 0.28905507 | 0.35701712 | 0.80963925 | 0.41814754 | NA |
| BAHCC1 | 5.07794358 | -0.2528262 | 0.82346052 | -0.3070289 | 0.75882136 | NA |
| ACTG1 | 148.061782 | -0.5460835 | 0.38719463 | -1.4103592 | 0.15843364 | 0.64166827 |
| FAAP100 | 2.6659641 | -0.7270916 | 0.93526857 | -0.7774148 | 0.43691407 | NA |
| NPLOC4 | 26.4302013 | -0.8497755 | 0.34358546 | -2.4732582 | 0.01338874 | 0.30435125 |
| OXLD1 | 5.239444 | -0.0332783 | 0.62489137 | -0.0532545 | 0.95752914 | NA |
| CCDC137 | 6.80557164 | -0.4599342 | 0.64112092 | -0.7173908 | 0.473133 | NA |
| ARL16 | 5.06037434 | -0.5174553 | 0.67743234 | -0.7638479 | 0.44495788 | NA |
| HGS | 10.757639 | -0.8508331 | 0.55599979 | -1.530276 | 0.12594843 | NA |
| GCGR | 11.0474143 | 0.96733517 | 0.61496543 | 1.57299115 | 0.11572087 | NA |
| MCRIP1 | 12.1395256 | -0.2549779 | 0.46054436 | -0.5536445 | 0.57982216 | NA |
| P4HB | 31.8359482 | -0.6820925 | 0.3593075 | -1.898353 | 0.05764959 | 0.50751937 |
| ARHGDIA | 31.4622068 | -0.7841165 | 0.44372871 | -1.7671079 | 0.07721016 | 0.54401752 |
| ALYREF | 2.96055739 | 0.41480207 | 0.98258484 | 0.42215395 | 0.67291265 | NA |
| ANAPC11 | 13.2832336 | -0.141921 | 0.45119825 | -0.3145425 | 0.75310901 | NA |
| PCYT2 | 5.5160208 | 0.20974094 | 0.72767371 | 0.28823487 | 0.77316696 | NA |

| | | | | | | |
|---|---|---|---|---|---|---|
| SIRT7 | 5.83044516 | -1.1531773 | 0.66999999 | -1.7211601 | 0.08522177 | NA |
| MAFG | 5.28320737 | -0.7109828 | 0.78405511 | -0.9068021 | 0.36451147 | NA |
| ASPSCR1 | 9.073099 | -0.4144129 | 0.55859783 | -0.7418807 | 0.45815964 | NA |
| CENPX | 5.28174399 | -0.0157765 | 0.89601467 | -0.0176075 | 0.985952 | NA |
| DCXR | 9.98613518 | 0.19653347 | 0.51193901 | 0.38390015 | 0.70105245 | NA |
| RFNG | 3.98554514 | -1.0545167 | 0.85096394 | -1.2392026 | 0.21527049 | NA |
| GPS1 | 6.58008193 | -0.1570483 | 0.75607188 | -0.2077161 | 0.83545065 | NA |
| DUS1L | 6.45944586 | -1.6812719 | 0.69192121 | -2.4298603 | 0.01510464 | NA |
| FASN | 4.13008072 | 0.08065351 | 0.73802956 | 0.10928223 | 0.91297864 | NA |
| CCDC57 | 21.0211419 | 0.29268758 | 0.3476637 | 0.84186981 | 0.39986083 | NA |
| SLC16A3 | 26.6749936 | -0.7401185 | 0.42203147 | -1.7537046 | 0.07948114 | 0.54552914 |
| CSNK1D | 21.5247251 | -0.601135 | 0.39495632 | -1.5220289 | 0.12800183 | NA |
| OGFOD3 | 5.94784252 | 0.53231868 | 0.6075456 | 0.87617897 | 0.38093275 | NA |
| HEXDC | 12.7210888 | -0.6597553 | 0.52471389 | -1.257362 | 0.20862259 | NA |
| CYBC1 | 7.09448937 | 0.55335265 | 0.65093678 | 0.85008662 | 0.39527693 | NA |
| NARF | 9.91447549 | 0.04993029 | 0.69466485 | 0.07187681 | 0.94269995 | NA |
| FOXK2 | 38.793588 | -0.5280848 | 0.30194819 | -1.7489253 | 0.08030393 | 0.54552914 |
| WDR45B | 27.3712251 | -0.6585083 | 0.42826857 | -1.537606 | 0.12414498 | 0.58553957 |
| RAB40B | 40.5394046 | -0.8964647 | 0.38444156 | -2.3318621 | 0.01970795 | 0.34156176 |
| FN3KRP | 5.53609296 | -0.2101516 | 0.68932439 | -0.3048661 | 0.76046817 | NA |
| FN3K | 9.61183043 | -0.4475775 | 0.56467268 | -0.7926317 | 0.42799243 | NA |
| TBCD | 19.1131735 | -0.9452142 | 0.42300224 | -2.234537 | 0.02544776 | NA |
| METRNL | 29.1867801 | -0.4386551 | 0.46934287 | -0.9346156 | 0.34998646 | 0.8003972 |
| AC144831.1 | 3.37342164 | -2.030794 | 1.12356664 | -1.8074531 | 0.07069166 | NA |
| USP14 | 26.2972876 | -0.4009138 | 0.36992174 | -1.08378 | 0.27846234 | 0.76699277 |
| THOC1 | 19.7316042 | -0.4991548 | 0.39788808 | -1.2545106 | 0.20965647 | NA |
| ENOSF1 | 43.3596921 | 0.35757829 | 0.33718797 | 1.06047166 | 0.28893007 | 0.7725847 |
| YES1 | 32.1977099 | -0.0662814 | 0.32200043 | -0.2058426 | 0.83691386 | 0.97154892 |
| METTL4 | 6.94472823 | -0.870556 | 0.61380169 | -1.4183017 | 0.15610269 | NA |
| SMCHD1 | 49.4554493 | -0.0839253 | 0.273862 | -0.3064511 | 0.75926119 | 0.95363205 |
| AP001011.1 | 7.87012293 | 1.39961147 | 0.72648197 | 1.92656049 | 0.05403443 | NA |
| LPIN2 | 14.6614653 | -0.267051 | 0.53364863 | -0.5004249 | 0.61677595 | NA |
| MYL12A | 27.6529759 | 0.03319002 | 0.29634842 | 0.11199662 | 0.91082608 | 0.98375925 |
| MYL12B | 67.072505 | 0.00267182 | 0.36310036 | 0.00735834 | 0.99412895 | 0.99923383 |
| TGIF1 | 5.51380422 | -1.0699773 | 0.62834282 | -1.702856 | 0.08859502 | NA |
| DLGAP1 | 5.21167831 | -0.2963793 | 0.7314763 | -0.4051796 | 0.68534547 | NA |
| LINC00667 | 4.15425248 | 0.19056715 | 0.72555737 | 0.26264931 | 0.79282089 | NA |
| EPB41L3 | 46.8855403 | -1.1000421 | 0.47960586 | -2.2936378 | 0.02181131 | 0.35108799 |
| L3MBTL4 | 93.0123602 | 0.30616567 | 0.35228501 | 0.86908513 | 0.38480057 | 0.825589 |
| AP001021.2 | 5.0055986 | 1.31155918 | 0.79859142 | 1.64234069 | 0.10051942 | NA |
| ARHGAP28 | 4.85895648 | -0.0999169 | 0.88688515 | -0.1126605 | 0.91029972 | NA |
| LAMA1 | 6.72119283 | -0.009269 | 0.72497394 | -0.0127853 | 0.98979911 | NA |
| PTPRM | 89.8469545 | 0.29571208 | 0.30520705 | 0.96889006 | 0.33260005 | 0.79447068 |
| RAB12 | 12.4817625 | -0.2777559 | 0.50222488 | -0.5530509 | 0.58022856 | NA |
| MTCL1 | 4.89816036 | -1.7302361 | 0.98671665 | -1.7535288 | 0.07951127 | NA |
| NDUFV2 | 22.4344856 | 0.58720091 | 0.40824345 | 1.43835966 | 0.15033203 | NA |
| ANKRD12 | 112.137555 | -0.7763183 | 0.26936861 | -2.8819923 | 0.00395169 | 0.2109414 |
| TWSG1 | 16.1263349 | 0.00081426 | 0.49531969 | 0.00164392 | 0.99868834 | NA |
| RALBP1 | 36.0879287 | 0.50035524 | 0.307354 | 1.62794447 | 0.10353666 | 0.5772605 |
| PPP4R1 | 26.3417508 | -0.4449988 | 0.3734214 | -1.19168 | 0.23338674 | 0.73156941 |
| RAB31 | 5.72552792 | -1.1480597 | 0.75060693 | -1.5295085 | 0.12613843 | NA |
| VAPA | 73.5539847 | -0.430876 | 0.25112394 | -1.7157904 | 0.08620041 | 0.55706756 |
| NAPG | 21.9217073 | -0.2278717 | 0.38922393 | -0.5854513 | 0.5582443 | NA |
| GNAL | 5.17138166 | 1.3530473 | 0.7490855 | 1.80626552 | 0.07087688 | NA |

| | | | | | | |
|---|---|---|---|---|---|---|
| MPPE1 | 13.2397545 | 0.35984415 | 0.43542583 | 0.82641894 | 0.40856648 | NA |
| IMPA2 | 45.4562697 | 0.57392685 | 0.28771195 | 1.99479667 | 0.04606506 | 0.45035768 |
| AFG3L2 | 36.3583111 | -0.1401111 | 0.36717407 | -0.3815932 | 0.7027631 | 0.9303056 |
| SPIRE1 | 127.828509 | -0.4655422 | 0.32135176 | -1.4486998 | 0.14742144 | 0.63 |
| PSMG2 | 21.0290639 | 0.42443211 | 0.36062191 | 1.1769449 | 0.2392175 | NA |
| CEP76 | 6.54794125 | 0.88010538 | 0.57796379 | 1.52276906 | 0.12781649 | NA |
| PTPN2 | 42.3142345 | -0.0464951 | 0.34274728 | -0.1356541 | 0.89209473 | 0.97769824 |
| SEH1L | 8.80542301 | -1.5086776 | 0.61272867 | -2.4622279 | 0.01380769 | NA |
| CEP192 | 36.6052417 | 0.0560161 | 0.39417599 | 0.14210937 | 0.88699362 | 0.97769824 |
| LDLRAD4 | 10.1387467 | 0.56810924 | 0.73027315 | 0.77794075 | 0.43660395 | NA |
| FAM210A | 12.4292368 | -1.1091504 | 0.48856126 | -2.2702381 | 0.02319314 | NA |
| RNMT | 22.2795738 | -0.7442671 | 0.34930023 | -2.1307375 | 0.03311077 | NA |
| ROCK1 | 65.4492549 | -0.183044 | 0.25718126 | -0.7117314 | 0.4766311 | 0.86206794 |
| ESCO1 | 19.8888295 | -0.5616051 | 0.39627992 | -1.4171928 | 0.15642656 | NA |
| SNRPD1 | 7.54812208 | -0.3664112 | 0.59753883 | -0.6132007 | 0.53974364 | NA |
| ABHD3 | 66.5066261 | -0.9825505 | 0.46435294 | -2.1159563 | 0.03434853 | 0.41688714 |
| MIB1 | 50.0461901 | -0.498512 | 0.29129441 | -1.7113684 | 0.08701312 | 0.56096141 |
| RBBP8 | 52.8416125 | 0.17819551 | 0.28924028 | 0.61608124 | 0.5378409 | 0.88254911 |
| CABLES1 | 8.63518894 | -0.5264787 | 0.66103629 | -0.7964445 | 0.4257737 | NA |
| TMEM241 | 4.90237087 | -0.3785537 | 0.67386805 | -0.5617624 | 0.57427792 | NA |
| RIOK3 | 29.0679767 | -0.5840325 | 0.34209676 | -1.7072142 | 0.08778224 | 0.56122412 |
| RMC1 | 9.26134853 | 0.26715981 | 0.47987583 | 0.55672695 | 0.577714 | NA |
| NPC1 | 24.7286915 | -1.1095041 | 0.44755518 | -2.4790331 | 0.01317391 | NA |
| LAMA3 | 4.59072768 | 0.80492073 | 1.22415861 | 0.65752977 | 0.51084033 | NA |
| TTC39C | 3.67633259 | 0.83326431 | 0.78579802 | 1.0604052 | 0.2889603 | NA |
| OSBPL1A | 23.0448716 | 0.59500388 | 0.53763869 | 1.10669839 | 0.26842435 | NA |
| IMPACT | 4.51528781 | 0.74909672 | 0.70508034 | 1.06242746 | 0.28804167 | NA |
| SS18 | 32.9970265 | 0.37612609 | 0.33861479 | 1.1107787 | 0.26666362 | 0.75497438 |
| TAF4B | 5.030808 | 0.5164669 | 0.74991275 | 0.68870265 | 0.49101041 | NA |
| KCTD1 | 49.7703749 | 0.55345841 | 0.31427808 | 1.76104678 | 0.07823049 | 0.54552914 |
| CDH2 | 5.59772475 | -0.8334495 | 0.85757947 | -0.9718627 | 0.33111888 | NA |
| DSC2 | 11.2830404 | -0.0606829 | 0.5044769 | -0.1202887 | 0.90425443 | NA |
| DSG2 | 39.2247087 | -0.0781621 | 0.34862055 | -0.2242038 | 0.82259873 | 0.96845695 |
| B4GALT6 | 3.43120148 | 0.38239962 | 0.76157336 | 0.5021179 | 0.61558459 | NA |
| TRAPPC8 | 34.5075005 | 0.05473739 | 0.37794485 | 0.14482905 | 0.88484585 | 0.97769824 |
| RNF138 | 14.5737093 | -0.1649532 | 0.49456828 | -0.3335298 | 0.73873443 | NA |
| GAREM1 | 200.860735 | -0.120695 | 0.31699161 | -0.3807515 | 0.70338762 | 0.9303056 |
| CCDC178 | 16.5916511 | 0.89497955 | 0.49551995 | 1.80614232 | 0.07089611 | NA |
| ASXL3 | 18.0313503 | -0.1119228 | 0.46682988 | -0.2397506 | 0.81052357 | NA |
| DTNA | 5.97088728 | 1.06535864 | 0.59395689 | 1.79366325 | 7.29E-02 | NA |
| MAPRE2 | 29.5281181 | 0.00485217 | 0.49904601 | 0.00972289 | 0.99224238 | 0.99909657 |
| ZNF397 | 13.016678 | 0.36204466 | 0.44416079 | 0.81512069 | 0.41500323 | NA |
| ZSCAN30 | 8.25604791 | 0.93654425 | 0.5638356 | 1.66102364 | 0.0967087 | NA |
| ZNF24 | 15.0657175 | 0.08049813 | 0.41829462 | 0.19244363 | 0.84739472 | NA |
| GALNT1 | 16.9158746 | -0.5652179 | 0.43806183 | -1.2902697 | 0.19695704 | NA |
| RPRD1A | 30.481365 | -0.3077577 | 0.31019194 | -0.9921525 | 0.32112312 | 0.785589 |
| SLC39A6 | 9.0855122 | -0.6754548 | 0.51266154 | -1.3175453 | 0.1876559 | NA |
| ELP2 | 19.9326834 | -0.2501918 | 0.39662751 | -0.6307979 | 0.52817269 | NA |
| MOCOS | 7.81985344 | -0.2499723 | 0.72332409 | -0.3455882 | 0.72965218 | NA |
| FHOD3 | 60.3613981 | 0.61029093 | 0.28352681 | 2.15249812 | 0.03135814 | 0.4126665 |
| TPGS2 | 10.0060734 | 0.44373016 | 0.55294873 | 0.80247975 | 0.4222755 | NA |
| KIAA1328 | 60.4787325 | 0.71912213 | 0.35051586 | 2.05161083 | 0.0402075 | 0.43121698 |
| AC016205.1 | 12.157597 | 0.83496764 | 0.51517384 | 1.62074932 | 0.10507141 | NA |
| PIK3C3 | 31.8732431 | -0.1283665 | 0.30500954 | -0.4208605 | 0.67385694 | 0.92617027 |

| | | | | | | |
|---|---|---|---|---|---|---|
| LINC00907 | 5.59235057 | 0.45420097 | 0.68207876 | 0.66590692 | 0.5054706 | NA |
| SETBP1 | 127.48312 | 0.11822045 | 0.28245415 | 0.41854741 | 0.67554693 | 0.92617027 |
| EPG5 | 20.9853488 | 0.07036867 | 0.40924708 | 0.17194667 | 0.86347946 | NA |
| ATP5F1A | 45.8521418 | 0.18918228 | 0.30331731 | 0.62371082 | 0.53281752 | 0.88254911 |
| HAUS1 | 5.47118616 | -1.1158272 | 0.75087974 | -1.4860266 | 0.13727209 | NA |
| C18orf25 | 21.62144 | -0.1805768 | 0.39635633 | -0.4555922 | 0.64868326 | NA |
| ST8SIA5 | 5.37345441 | -0.707481 | 0.80771953 | -0.8758993 | 0.3810848 | NA |
| PIAS2 | 50.8315142 | -0.798427 | 0.28907452 | -2.762011 | 0.00574465 | 0.22794137 |
| KATNAL2 | 35.6526175 | -0.057995 | 0.53609368 | -0.1081808 | 0.91385229 | 0.98429046 |
| HDHD2 | 3.29767375 | 0.76441759 | 0.8333625 | 0.91726901 | 0.35900169 | NA |
| SMAD2 | 63.5296239 | -0.3167742 | 0.29380877 | -1.0781647 | 0.28096027 | 0.76854599 |
| CTIF | 17.7961802 | -1.1476228 | 0.48971044 | -2.3434721 | 0.01910519 | NA |
| SMAD7 | 5.38879611 | -0.7966886 | 0.93731144 | -0.8499722 | 0.39534056 | NA |
| DYM | 72.2149348 | -0.0502247 | 0.26965685 | -0.1862542 | 0.85224542 | 0.97168963 |
| ACAA2 | 10.2747871 | 0.35415656 | 0.51287061 | 0.69053784 | 0.48985603 | NA |
| MYO5B | 45.1867325 | -0.1887876 | 0.38036499 | -0.4963328 | 0.61965963 | 0.91493825 |
| MBD1 | 7.75082796 | -0.9179434 | 0.57209707 | -1.6045239 | 0.10859863 | NA |
| CXXC1 | 4.64749931 | -0.7894066 | 0.77508901 | -1.0184722 | 0.3084536 | NA |
| MAPK4 | 3.14071704 | -0.7823547 | 0.89227512 | -0.8768088 | 0.3805905 | NA |
| ME2 | 8.73333971 | -0.6062054 | 0.53533712 | -1.1323807 | 0.25747442 | NA |
| SMAD4 | 17.4222701 | -0.328796 | 0.38822086 | -0.8469303 | 0.39703398 | NA |
| MEX3C | 4.34741411 | -1.5006701 | 0.72476409 | -2.0705636 | 0.0383996 | NA |
| MBD2 | 39.5894958 | -0.3851472 | 0.2958223 | -1.3019547 | 0.19293186 | 0.68667113 |
| POLI | 12.8032799 | 0.22725061 | 0.47184467 | 0.48162165 | 0.63007475 | NA |
| RAB27B | 3.46855139 | -1.1853774 | 0.94733158 | -1.2512804 | 0.21083221 | NA |
| CCDC68 | 33.5506633 | -1.5848153 | 0.52732794 | -3.0053695 | 0.00265258 | 0.17267672 |
| TCF4 | 17.7156964 | -0.1514012 | 0.6673745 | -0.2268609 | 0.8205319 | NA |
| TXNL1 | 13.1742468 | -0.330287 | 0.4564323 | -0.7236276 | 0.4692944 | NA |
| WDR7 | 38.4728547 | 0.07962782 | 0.3642248 | 0.21862273 | 0.82694395 | 0.96845695 |
| FECH | 4.9761335 | 1.02770057 | 0.63858842 | 1.60933167 | 0.10754384 | NA |
| NARS | 25.728363 | -0.1107527 | 0.35156058 | -0.3150317 | 0.75273758 | NA |
| AC027097.2 | 10.2198326 | 0.49832851 | 0.52489497 | 0.94938709 | 0.34242377 | NA |
| ATP8B1 | 26.053421 | 0.4982738 | 0.35293399 | 1.41180452 | 0.15800753 | 0.64166827 |
| NEDD4L | 738.010891 | -0.21983 | 0.30266102 | -0.7263241 | 0.4676401 | 0.85894565 |
| MALT1 | 12.4368918 | 0.0426896 | 0.47286597 | 0.09027844 | 0.92806596 | NA |
| ZNF532 | 39.9232086 | -0.7680644 | 0.38021466 | -2.0200811 | 0.04337498 | 0.44186191 |
| SEC11C | 8.05253757 | 0.88741122 | 0.59271428 | 1.49719899 | 0.13434149 | NA |
| LMAN1 | 27.8433047 | -0.3103026 | 0.41849815 | -0.7414672 | 0.45841022 | 0.85802171 |
| CCBE1 | 54.8947799 | -0.9827098 | 0.39940208 | -2.4604523 | 0.0138762 | 0.30435125 |
| AC010776.3 | 1.03712355 | -1.5968199 | 2.07056013 | -0.7712019 | 0.44058728 | NA |
| CDH20 | 7.08007421 | 1.31398955 | 0.86522416 | 1.51866951 | 0.12884571 | NA |
| AC105094.2 | 9.13760165 | 0.30475517 | 0.60040751 | 0.50758053 | 0.61174754 | NA |
| LINC01544 | 4.85638391 | -0.0977066 | 0.79246151 | -0.1232951 | 0.90187341 | NA |
| RNF152 | 655.886615 | 0.25733276 | 0.30602088 | 0.84089935 | 0.40040432 | 0.8269015 |
| PIGN | 37.3479087 | 0.24070093 | 0.30190625 | 0.79727044 | 0.42529398 | 0.84057082 |
| KIAA1468 | 37.4916883 | 0.36218271 | 0.33072058 | 1.09513204 | 0.27345879 | 0.76165444 |
| ZCCHC2 | 19.4486431 | -0.1830551 | 0.39280784 | -0.4660168 | 0.64120346 | NA |
| PHLPP1 | 39.9409794 | 0.16346596 | 0.3641925 | 0.44884494 | 0.65354352 | 0.92248109 |
| BCL2 | 49.3700149 | -0.2755914 | 0.44533729 | -0.6188374 | 0.5360235 | 0.88254911 |
| KDSR | 20.0360928 | 0.42467859 | 0.36474057 | 1.16433053 | 0.24429009 | NA |
| VPS4B | 21.7712116 | -0.0318281 | 0.35336304 | -0.090072 | 0.92822998 | NA |
| TMX3 | 18.7824226 | 0.44673276 | 0.37701588 | 1.18491762 | 0.23605002 | NA |
| DOK6 | 6.25842732 | -0.9364011 | 0.78602635 | -1.1913101 | 0.23353188 | NA |
| RTTN | 11.7809794 | 0.54501591 | 0.50484079 | 1.07957977 | 0.28032935 | NA |

| | | | | | | |
|---|---|---|---|---|---|---|
| SOCS6 | 9.1320839 | 0.11218602 | 0.51581864 | 0.21749122 | 0.82782555 | NA |
| TIMM21 | 4.45332614 | 0.70963137 | 0.70918901 | 1.00062375 | 0.31700874 | NA |
| CYB5A | 34.3404175 | 0.51172363 | 0.48809177 | 1.04841685 | 0.2944466 | 0.77460836 |
| CNDP2 | 43.8416314 | 0.19121209 | 0.43733141 | 0.43722469 | 0.66194841 | 0.92276785 |
| LINC00909 | 2.52195214 | -0.6316801 | 1.02720174 | -0.6149523 | 0.53858622 | NA |
| ZNF407 | 72.0568539 | -0.8502371 | 0.30929263 | -2.7489731 | 0.00597823 | 0.22794137 |
| ZADH2 | 7.99795133 | 0.82612247 | 0.80016891 | 1.0324351 | 0.30186834 | NA |
| TSHZ1 | 9.80343967 | 0.59971244 | 0.56745669 | 1.05684266 | 0.2905834 | NA |
| ZNF516 | 26.3386778 | -1.1852606 | 0.46509007 | -2.548454 | 0.01082016 | 0.28312744 |
| ZNF236 | 16.4897275 | -0.5122127 | 0.42163558 | -1.2148233 | 0.22443348 | NA |
| MBP | 38.6194973 | 0.12263472 | 0.3169537 | 0.38691683 | 0.69881778 | 0.9303056 |
| ATP9B | 64.8320816 | -0.4657436 | 0.34085277 | -1.3664069 | 0.1718113 | 0.66182328 |
| NFATC1 | 4.56994894 | -0.4554226 | 0.733845 | -0.6205979 | 0.53486425 | NA |
| CTDP1 | 8.24528216 | -0.5115964 | 0.51269677 | -0.9978537 | 0.3183503 | NA |
| PQLC1 | 11.7587805 | 0.12655988 | 0.48273899 | 0.26217041 | 0.79319007 | NA |
| HSBP1L1 | 9.86964251 | -0.4706884 | 0.52696306 | -0.8932094 | 0.37174505 | NA |
| TXNL4A | 17.6626166 | 0.15377937 | 0.54095613 | 0.28427328 | 0.77620096 | NA |
| ADNP2 | 18.8359137 | -0.4613138 | 0.46308134 | -0.9961831 | 0.31916118 | NA |
| PARD6G-AS1 | 3.6808848 | -0.1338803 | 0.9238389 | -0.1449174 | 0.88477609 | NA |
| C20orf96 | 5.47103459 | 1.57932212 | 0.7954793 | 1.98537174 | 0.04710313 | NA |
| ZCCHC3 | 3.43794038 | -0.0644444 | 0.75385307 | -0.0854867 | 0.9318745 | NA |
| NRSN2-AS1 | 20.3153415 | 0.10881717 | 0.38390231 | 0.28345015 | 0.77683179 | NA |
| SOX12 | 4.8518919 | 0.02709328 | 0.75243648 | 0.0360074 | 0.97127646 | NA |
| NRSN2 | 4.35329135 | -0.0859218 | 0.74324015 | -0.1156044 | 0.90796611 | NA |
| RBCK1 | 16.0643601 | 0.4563076 | 0.44755488 | 1.01955674 | 0.30793873 | NA |
| TBC1D20 | 8.62449748 | -0.0894836 | 0.50467083 | -0.1773108 | 0.85926423 | NA |
| CSNK2A1 | 53.9113409 | -0.260513 | 0.2986039 | -0.8724367 | 0.38297017 | 0.82298502 |
| SLC52A3 | 7.44640442 | 1.32345284 | 0.6332653 | 2.08988689 | 0.03662796 | NA |
| PSMF1 | 21.1335149 | -0.2147761 | 0.42259693 | -0.5082293 | 0.61129255 | NA |
| AL136531.2 | 8.33388055 | -0.6262048 | 0.70686483 | -0.8858904 | 0.37567656 | NA |
| SDCBP2-AS1 | 7.5169275 | 0.28564005 | 0.59516147 | 0.47993706 | 0.63127215 | NA |
| NSFL1C | 9.95669282 | 0.04395651 | 0.5035834 | 0.08728746 | 0.93044302 | NA |
| STK35 | 6.3771659 | -0.0962447 | 0.59597742 | -0.1614906 | 0.87170704 | NA |
| SNRPB | 12.6350831 | -0.0892384 | 0.51595235 | -0.1729586 | 0.86268401 | NA |
| NOP56 | 5.62139008 | -0.9977311 | 0.84773215 | -1.1769414 | 0.23921889 | NA |
| IDH3B | 5.23862551 | 0.06187729 | 0.61812204 | 0.1001053 | 0.92026073 | NA |
| PCED1A | 6.60939974 | 0.25197772 | 0.63589757 | 0.3962552 | 0.69191677 | NA |
| PTPRA | 45.5142268 | -0.0959481 | 0.31824349 | -0.3014926 | 0.76303889 | 0.95478238 |
| MRPS26 | 13.1700738 | 0.82210427 | 0.5129947 | 1.602559 | 0.10903205 | NA |
| UBOX5-AS1 | 3.81288673 | 0.99718137 | 0.91415661 | 1.09082116 | 0.27535158 | NA |
| UBOX5 | 4.93894793 | -0.1150763 | 0.68325413 | -0.1684238 | 0.86624988 | NA |
| LZTS3 | 5.52452282 | 0.22407118 | 0.69502416 | 0.32239337 | 0.74715471 | NA |
| DDRGK1 | 10.5316148 | 0.17054804 | 0.45221114 | 0.3771425 | 0.70606771 | NA |
| ITPA | 10.701871 | 0.62146507 | 0.45045889 | 1.37962659 | 0.16770165 | NA |
| C20orf194 | 110.664626 | 0.02011264 | 0.25315333 | 0.07944847 | 0.93667592 | 0.99100968 |
| ATRN | 39.7132559 | 0.07078451 | 0.36772233 | 0.19249445 | 0.84735492 | 0.97168963 |
| CENPB | 8.09730643 | -0.1502054 | 0.69270656 | -0.2168384 | 0.82833427 | NA |
| CDC25B | 3.11533158 | 0.1952751 | 0.78258568 | 0.24952553 | 0.8029543 | NA |
| AP5S1 | 2.10404699 | -1.9950123 | 2.66029412 | -0.7499217 | 0.45330186 | NA |
| MAVS | 29.2906318 | -0.0535769 | 0.38371735 | -0.1396258 | 0.88895563 | 0.97769824 |
| PANK2 | 15.0614686 | -0.4838332 | 0.42554327 | -1.1369778 | 0.25554758 | NA |
| RNF24 | 22.9621875 | 0.10039621 | 0.40024192 | 0.25083881 | 0.80193873 | NA |
| PRNP | 9.47099169 | -0.1014978 | 0.49841915 | -0.2036394 | 0.83863532 | NA |
| SLC23A2 | 8.10740471 | 0.02218381 | 0.69547822 | 0.0318972 | 0.97455403 | NA |

| | | | | | | |
|---|---|---|---|---|---|---|
| TMEM230 | 20.4376928 | 0.23687884 | 0.36099977 | 0.6561745 | 0.51171185 | NA |
| PCNA | 5.49095706 | 1.17522698 | 0.63061865 | 1.8636096 | 0.06237655 | NA |
| CDS2 | 20.2736341 | 0.01714222 | 0.3933052 | 0.04358504 | 0.96523518 | NA |
| GPCPD1 | 28.1121151 | -0.4344996 | 0.45457155 | -0.9558442 | 0.33915097 | 0.8003972 |
| SHLD1 | 14.3076982 | -0.2181214 | 0.45202764 | -0.48254 | 0.62942237 | NA |
| TRMT6 | 3.51136438 | 0.01022451 | 0.78753417 | 0.01298294 | 0.9896414 | NA |
| MCM8 | 3.41454311 | 1.14698632 | 0.83463474 | 1.37423745 | 0.16936798 | NA |
| CRLS1 | 19.576238 | -0.4387595 | 0.38110675 | -1.1512771 | 0.24961824 | NA |
| TMX4 | 11.7082832 | -0.1932867 | 0.54717254 | -0.3532464 | 0.72390374 | NA |
| PLCB1 | 52.1140053 | -0.695133 | 0.76292547 | -0.9111415 | 0.36222083 | 0.8056395 |
| PLCB4 | 19.3436084 | 0.98184099 | 0.53036563 | 1.85125307 | 0.06413315 | NA |
| MKKS | 6.61545296 | 0.42621727 | 0.62283724 | 0.68431564 | 0.49377587 | NA |
| SLX4IP | 65.5065306 | 0.17741165 | 0.3780497 | 0.46928129 | 0.63886859 | 0.92121485 |
| JAG1 | 21.1045233 | 0.04502715 | 0.60771107 | 0.07409303 | 0.94093636 | NA |
| AL050403.2 | 3.31998022 | -1.0733399 | 0.98369334 | -1.0911327 | 0.27521451 | NA |
| BTBD3 | 9.67630202 | 0.54698063 | 0.50606915 | 1.08084167 | 0.27976755 | NA |
| SPTLC3 | 115.454476 | 0.52825499 | 0.26468265 | 1.99580511 | 0.04595514 | 0.45035768 |
| TASP1 | 39.7177706 | 0.07833806 | 0.3325797 | 0.23554673 | 0.81378441 | 0.96845695 |
| ESF1 | 18.0752046 | -0.6690616 | 0.49278838 | -1.3577057 | 0.17455709 | NA |
| NDUFAF5 | 13.2651987 | -0.1739791 | 0.44917592 | -0.3873294 | 0.69851234 | NA |
| MACROD2 | 232.377434 | 0.66063424 | 0.42286708 | 1.56227398 | 0.11822346 | 0.57982915 |
| KIF16B | 54.7133476 | 0.37735797 | 0.30886125 | 1.22177182 | 0.22179393 | 0.72433127 |
| SNRPB2 | 10.8528236 | -0.2769899 | 0.52153869 | -0.5311013 | 0.5953486 | NA |
| DSTN | 43.3545775 | 0.09778625 | 0.36306407 | 0.26933607 | 0.78767108 | 0.96325251 |
| RRBP1 | 22.916995 | -0.1779498 | 0.50986455 | -0.3490139 | 0.72707889 | NA |
| SNX5 | 9.96214891 | -0.4294979 | 0.54142762 | -0.7932694 | 0.42762089 | NA |
| OVOL2 | 7.70783422 | 0.23863013 | 0.55734192 | 0.42815751 | 0.66853645 | NA |
| MGME1 | 7.29887779 | -0.0284689 | 0.61561893 | -0.0462443 | 0.96311552 | NA |
| KAT14 | 3.06335981 | 0.88972215 | 0.86364766 | 1.03019111 | 0.3029203 | NA |
| ZNF133 | 8.48358029 | 1.21729471 | 0.55504233 | 2.19315654 | 0.0282961 | NA |
| DZANK1 | 3.11157743 | 0.4740167 | 0.80191517 | 0.59110579 | 0.55444954 | NA |
| POLR3F | 3.56991075 | 0.0383707 | 0.81307379 | 0.04719215 | 0.96236008 | NA |
| RBBP9 | 2.42728278 | 0.18448873 | 1.04954505 | 0.17577971 | 0.86046701 | NA |
| SEC23B | 19.4567414 | -0.2753933 | 0.38023797 | -0.7242656 | 0.46890269 | NA |
| DTD1 | 3.45658812 | 0.11891822 | 0.74120421 | 0.16043921 | 0.87253511 | NA |
| AL121761.1 | 9.52374169 | 0.88091253 | 0.48667266 | 1.81007197 | 0.07028463 | NA |
| RIN2 | 56.6974831 | 0.80654194 | 0.48951541 | 1.64763338 | 0.09942793 | 0.57091823 |
| NAA20 | 15.6057306 | 0.77071849 | 0.40303801 | 1.91227245 | 0.05584126 | NA |
| CRNKL1 | 4.47802817 | -0.0312493 | 0.66444625 | -0.0470306 | 0.96248882 | NA |
| RALGAPA2 | 187.231384 | 0.24558905 | 0.37321986 | 0.65802783 | 0.51052025 | 0.87158522 |
| KIZ | 15.3671668 | 0.31104876 | 0.42188471 | 0.73728381 | 0.46094978 | NA |
| KIZ-AS1 | 5.31323365 | -0.5123548 | 0.62921184 | -0.8142803 | 0.41548439 | NA |
| XRN2 | 30.8654239 | -0.227671 | 0.32609076 | -0.698183 | 0.48506275 | 0.86354292 |
| NAPB | 14.3892085 | 0.77300294 | 0.41147114 | 1.87863222 | 0.06029473 | NA |
| CST3 | 29.9989638 | 0.56276913 | 0.3532749 | 1.59300627 | 0.11115879 | 0.57860396 |
| AL035661.1 | 2.77553371 | 1.07436228 | 0.83054539 | 1.29356239 | 0.19581662 | NA |
| APMAP | 10.1333553 | 0.36909994 | 0.49992223 | 0.73831472 | 0.46032322 | NA |
| ACSS1 | 18.9625096 | 0.25382567 | 0.42019748 | 0.60406283 | 0.54580187 | NA |
| ENTPD6 | 9.92191864 | -0.1137389 | 0.47017466 | -0.2419077 | 0.80885165 | NA |
| PYGB | 18.7455762 | 0.49557517 | 0.35851139 | 1.38231358 | 0.16687544 | NA |
| ABHD12 | 33.1197762 | -0.0812187 | 0.30321858 | -0.2678551 | 0.78881083 | 0.96325251 |
| NINL | 77.5059291 | -0.064302 | 0.34258499 | -0.1876966 | 0.85111447 | 0.97168963 |
| ZNF337-AS1 | 5.17152438 | -0.3160881 | 0.77119802 | -0.4098663 | 0.68190404 | NA |
| ZNF337 | 9.32524383 | 0.59392925 | 0.4813381 | 1.23391282 | 0.21723539 | NA |

| | | | | | | |
|---|---|---|---|---|---|---|
| FAM182B | 6.35044501 | 0.39399423 | 0.65535252 | 0.60119436 | 0.54771054 | NA |
| HM13 | 22.0436811 | -0.7520781 | 0.48290029 | -1.5574191 | 0.11937101 | NA |
| ID1 | 13.2540937 | -0.5897071 | 0.68955749 | -0.8551965 | 0.39244238 | NA |
| BCL2L1 | 44.3534666 | -0.3824903 | 0.32199955 | -1.1878596 | 0.23488872 | 0.73156941 |
| AL117381.1 | 5.6795633 | 0.9556734 | 0.59167337 | 1.61520435 | 0.10626645 | NA |
| DUSP15 | 13.9672139 | -0.0402115 | 0.47857534 | -0.0840232 | 0.93303795 | NA |
| PDRG1 | 3.59225401 | -0.3004953 | 0.7628468 | -0.393913 | 0.69364527 | NA |
| TM9SF4 | 24.6711029 | -0.1196467 | 0.40854419 | -0.2928611 | 0.76962831 | NA |
| PLAGL2 | 18.7501971 | 0.46441204 | 0.45165087 | 1.02825449 | 0.30383013 | NA |
| POFUT1 | 25.4661481 | 0.36698818 | 0.34984457 | 1.0490035 | 0.29417651 | NA |
| KIF3B | 56.2700454 | -0.0690875 | 0.2624606 | -0.26323 | 0.7923733 | 0.96341299 |
| ASXL1 | 267.445075 | -0.233089 | 0.31895748 | -0.7307839 | 0.46491114 | 0.85894565 |
| NOL4L | 7.57156997 | 0.14881823 | 0.66977332 | 0.22219193 | 0.82416447 | NA |
| MAPRE1 | 13.1708523 | -0.7750423 | 0.53611794 | -1.4456563 | 0.14827362 | NA |
| AL035071.2 | 2.7208144 | -1.2133927 | 0.93022136 | -1.3044128 | 0.19209287 | NA |
| AL035071.1 | 4.09915397 | -1.5987481 | 0.82321469 | -1.9420792 | 0.05212751 | NA |
| BPIFA2 | 9.86766566 | 1.24967298 | 0.61686869 | 2.02583305 | 0.0427819 | NA |
| CDK5RAP1 | 11.2075624 | -0.435212 | 0.46018456 | -0.9457337 | 0.34428444 | NA |
| SNTA1 | 4.82836736 | -1.3040354 | 0.64849992 | -2.0108489 | 0.04434142 | NA |
| CBFA2T2 | 68.4959896 | -0.4832696 | 0.3195981 | -1.5121166 | 0.1305042 | 0.60158154 |
| NECAB3 | 4.13179167 | 1.19185344 | 0.80707521 | 1.47675634 | 0.13974096 | NA |
| PXMP4 | 3.03349837 | 0.6573779 | 0.82311029 | 0.79865106 | 0.42449277 | NA |
| CHMP4B | 16.9619107 | -0.0152402 | 0.38094858 | -0.0400059 | 0.96808846 | NA |
| RALY | 28.9083342 | -0.8814223 | 0.35939545 | -2.4525138 | 0.0141862 | 0.30435125 |
| EIF2S2 | 21.6837585 | -0.1631809 | 0.39700221 | -0.4110327 | 0.68104856 | NA |
| AHCY | 6.53718371 | 0.53190756 | 0.58682998 | 0.90640829 | 0.36471978 | NA |
| ITCH | 69.7758098 | -0.3162542 | 0.27103254 | -1.1668494 | 0.24327117 | 0.73323864 |
| DYNLRB1 | 37.0412277 | -0.4102039 | 0.33736 | -1.2159233 | 0.22401413 | 0.72657496 |
| MAP1LC3A | 11.9364677 | -1.8846816 | 0.78410921 | -2.4035958 | 0.01623471 | NA |
| PIGU | 15.1119574 | -0.5360206 | 0.53539411 | -1.0011702 | 0.31674453 | NA |
| NCOA6 | 50.0050892 | -0.3079446 | 0.27990158 | -1.1001888 | 0.27124985 | 0.75967037 |
| TP53INP2 | 12.7260836 | -1.5567548 | 0.68058138 | -2.2873897 | 0.02217309 | NA |
| GGT7 | 15.089727 | -0.18022 | 0.42517244 | -0.4238751 | 0.67165687 | NA |
| ACSS2 | 17.8847778 | -0.0243089 | 0.43855578 | -0.0554295 | 0.9557963 | NA |
| GSS | 6.99749412 | 0.70451505 | 0.58407546 | 1.20620552 | 0.22773825 | NA |
| TRPC4AP | 37.5933261 | 0.21781634 | 0.36577995 | 0.59548464 | 0.55151956 | 0.88675043 |
| EDEM2 | 7.83042887 | 0.57516645 | 0.56427999 | 1.01929265 | 0.30806405 | NA |
| MMP24OS | 5.74569642 | -0.3351827 | 0.67842092 | -0.494063 | 0.62126169 | NA |
| MMP24 | 11.2106086 | 0.18866979 | 0.51728988 | 0.36472739 | 0.71531491 | NA |
| UQCC1 | 23.5025912 | -0.2806634 | 0.35071107 | -0.8002697 | 0.42355455 | NA |
| CEP250 | 3.8935892 | -0.7688946 | 0.72876915 | -1.0550592 | 0.29139824 | NA |
| ERGIC3 | 12.9503969 | -0.1239542 | 0.43340513 | -0.2860007 | 0.77487757 | NA |
| CPNE1 | 15.0568523 | 0.03267002 | 0.45550945 | 0.07172194 | 0.9428232 | NA |
| ROMO1 | 12.7253503 | -1.0480449 | 0.463833 | -2.2595307 | 0.02385039 | NA |
| RBM39 | 95.8597154 | -0.4489424 | 0.21337056 | -2.1040506 | 0.03537404 | 0.41688714 |
| PHF20 | 38.9070914 | -0.5893269 | 0.30812482 | -1.912624 | 0.0557962 | 0.50310834 |
| SCAND1 | 27.5312361 | -0.2251404 | 0.32156442 | -0.7001409 | 0.4838393 | 0.86354292 |
| NORAD | 76.6774963 | -0.6791714 | 0.38971743 | -1.7427279 | 0.08138116 | 0.54850081 |
| EPB41L1 | 56.0861113 | 0.25127714 | 0.40391353 | 0.62210626 | 0.533872 | 0.88254911 |
| AAR2 | 5.36706769 | 0.14660516 | 0.65308405 | 0.22448131 | 0.82238281 | NA |
| DLGAP4 | 22.9035858 | -0.8153688 | 0.4792579 | -1.7013153 | 0.08888381 | NA |
| DLGAP4-AS1 | 10.013129 | -0.1652629 | 0.52236202 | -0.3163762 | 0.75171697 | NA |
| MYL9 | 3.85715652 | -0.4071976 | 0.71939806 | -0.5660254 | 0.57137652 | NA |
| RAB5IF | 4.56141332 | -0.2780303 | 0.69327481 | -0.401039 | 0.68839138 | NA |

| | | | | | | |
|---|---|---|---|---|---|---|
| NDRG3 | 11.6611981 | -0.2660111 | 0.50304779 | -0.5287988 | 0.59694501 | NA |
| SOGA1 | 40.4390134 | -0.3455469 | 0.4082026 | -0.8465083 | 0.39726924 | 0.82643149 |
| TLDC2 | 82.0095879 | 0.82121797 | 0.39463683 | 2.08094609 | 0.03743884 | 0.42032585 |
| SAMHD1 | 41.9899525 | 1.40923804 | 0.59524798 | 2.36748059 | 0.01790966 | 0.33195057 |
| RBL1 | 5.82927667 | -0.1177439 | 0.67359976 | -0.174798 | 0.86123832 | NA |
| RPN2 | 23.8309642 | -0.0486187 | 0.36097289 | -0.134688 | 0.89285858 | NA |
| MANBAL | 7.51056525 | -0.520976 | 0.51945978 | -1.0029188 | 0.31590002 | NA |
| SRC | 10.0338506 | 0.54932481 | 0.48792645 | 1.12583529 | 0.26023524 | NA |
| BLCAP | 19.6295089 | -0.7547267 | 0.46682375 | -1.6167273 | 0.10593716 | NA |
| CTNNBL1 | 12.7867928 | 0.07344367 | 0.4404274 | 0.16675544 | 0.86756248 | NA |
| TTI1 | 8.429966 | 0.51652913 | 0.5108154 | 1.01118551 | 0.31192765 | NA |
| RPRD1B | 16.0469447 | -0.0411315 | 0.43379263 | -0.0948183 | 0.92445914 | NA |
| TGM2 | 6.19506995 | -0.1467655 | 0.73200568 | -0.2004978 | 0.84109129 | NA |
| RALGAPB | 35.2006997 | -0.4221883 | 0.33173017 | -1.2726859 | 0.20312953 | 0.700066 |
| ACTR5 | 2.86588689 | -1.4123968 | 0.9299808 | -1.5187376 | 0.12882856 | NA |
| DHX35 | 7.87849301 | -0.1226972 | 0.57191564 | -0.2145373 | 0.83012808 | NA |
| TOP1 | 52.766565 | -0.6697968 | 0.28885662 | -2.3187864 | 0.02040662 | 0.34471691 |
| PLCG1 | 7.593705 | -0.6831745 | 0.6600715 | -1.0350007 | 0.30066857 | NA |
| ZHX3 | 50.3399757 | 0.37015695 | 0.44739198 | 0.82736607 | 0.40802961 | 0.82808596 |
| LPIN3 | 8.0339528 | 0.82850627 | 0.51459743 | 1.61000856 | 0.10739599 | NA |
| CHD6 | 53.7080208 | -0.1832474 | 0.29057427 | -0.6306389 | 0.52827667 | 0.88123152 |
| PTPRT | 2.36200648 | -1.5279598 | 1.37592818 | -1.1104939 | 0.26678627 | NA |
| SRSF6 | 19.7376566 | -0.2201205 | 0.40718841 | -0.5405865 | 0.58879264 | NA |
| L3MBTL1 | 4.22335189 | -0.7492925 | 0.74452158 | -1.006408 | 0.31421934 | NA |
| SGK2 | 11.1447652 | 0.31138103 | 0.51939389 | 0.59950846 | 0.54883387 | NA |
| IFT52 | 8.56356782 | -0.4060727 | 0.56898653 | -0.7136772 | 0.47542682 | NA |
| OSER1 | 14.5528382 | -0.162819 | 0.51049798 | -0.3189415 | 0.74977085 | NA |
| FITM2 | 2.99989048 | 2.74160067 | 1.06263858 | 2.57999355 | 0.00988022 | NA |
| TTPAL | 3.31494389 | -0.4348793 | 0.84457937 | -0.5149063 | 0.60661849 | NA |
| SERINC3 | 18.0918121 | 0.2609829 | 0.35658163 | 0.73190224 | 0.46422824 | NA |
| PKIG | 13.3026854 | 0.40057763 | 0.43923463 | 0.91199009 | 0.36177394 | NA |
| YWHAB | 41.9183584 | -0.2828109 | 0.35749092 | -0.7910995 | 0.42888596 | 0.84223493 |
| PABPC1L | 6.32112684 | -0.1523789 | 0.61368056 | -0.2483034 | 0.80389969 | NA |
| TOMM34 | 4.05448615 | -0.2116822 | 0.80064408 | -0.2643899 | 0.79147947 | NA |
| STK4 | 18.0852764 | -0.3012223 | 0.42517852 | -0.7084609 | 0.47865911 | NA |
| SLPI | 43.4216019 | -0.140502 | 0.71038446 | -0.1977831 | 8.43E-01 | 9.72E-01 |
| SDC4 | 63.6883051 | 0.22318606 | 0.39805613 | 0.56068993 | 0.57500893 | 0.89263108 |
| PIGT | 6.87030352 | 1.17355831 | 0.65639559 | 1.78788268 | 0.07379494 | NA |
| WFDC2 | 1006.65492 | 0.02555246 | 0.59038638 | 0.0432809 | 0.96547761 | 0.99656976 |
| DNTTIP1 | 5.12278705 | -0.7862487 | 0.83766803 | -0.9386161 | 0.34792787 | NA |
| SNX21 | 6.52671873 | -0.5796344 | 0.83612081 | -0.6932424 | 0.48815742 | NA |
| ACOT8 | 6.98172669 | 0.47659847 | 0.57610047 | 0.8272836 | 0.40807634 | NA |
| ZSWIM3 | 5.22258449 | 0.83800953 | 0.69932761 | 1.19830752 | 0.23079732 | NA |
| CTSA | 22.3971337 | -0.7958177 | 0.56426588 | -1.4103595 | 0.15843356 | NA |
| PCIF1 | 2.18727748 | -0.4622666 | 1.01691126 | -0.4545791 | 0.64941209 | NA |
| ZNF335 | 6.60775097 | -0.7609148 | 0.64452633 | -1.1805798 | 0.23776969 | NA |
| NCOA5 | 14.7494421 | -0.7257566 | 0.51995068 | -1.3958182 | 0.16276924 | NA |
| CDH22 | 8.20186341 | 0.15555111 | 0.74759656 | 0.20806825 | 0.83517568 | NA |
| SLC35C2 | 11.0236408 | 0.70909074 | 0.49727894 | 1.42594163 | 0.15388519 | NA |
| ELMO2 | 8.06595071 | -0.3118772 | 0.56210452 | -0.5548385 | 0.57900515 | NA |
| ZNF334 | 7.47566554 | 0.98806723 | 0.71321294 | 1.38537478 | 0.1659379 | NA |
| SLC13A3 | 14.4640635 | -0.3218692 | 0.63757746 | -0.5048315 | 0.61367722 | NA |
| TP53RK | 3.03088581 | -0.0579905 | 0.81187109 | -0.0714282 | 0.94305694 | NA |
| ZMYND8 | 42.9346581 | 1.06900501 | 0.37876518 | 2.82234237 | 0.00476743 | 0.22229871 |

| | | | | | | |
|---|---|---|---|---|---|---|
| NCOA3 | 45.8071095 | -0.2910925 | 0.28184897 | -1.0327959 | 0.3016994 | 0.77521265 |
| SULF2 | 4.15911636 | 0.57310334 | 0.6744833 | 0.8496924 | 0.39549612 | NA |
| ARFGEF2 | 63.7547387 | -0.224426 | 0.22412488 | -1.0013436 | 0.3166607 | 0.780966 |
| CSE1L | 14.7606884 | 0.12674613 | 0.41603234 | 0.30465451 | 0.7606293 | NA |
| STAU1 | 44.3992562 | -0.1354981 | 0.26558745 | -0.5101826 | 0.60992357 | 0.90892575 |
| DDX27 | 7.35110095 | -0.2042783 | 0.53957905 | -0.3785883 | 0.70499365 | NA |
| ZNFX1 | 13.9900969 | -0.303631 | 0.54746693 | -0.5546106 | 0.57916105 | NA |
| ZFAS1 | 13.3974296 | -0.8276165 | 0.44928313 | -1.8420822 | 0.06546312 | NA |
| B4GALT5 | 43.0429021 | -0.190927 | 0.29484025 | -0.6475609 | 0.51726901 | 0.87158522 |
| SLC9A8 | 15.4668413 | 0.06855081 | 0.57524773 | 0.11916746 | 0.90514269 | NA |
| RNF114 | 12.6271991 | 0.17282756 | 0.42406843 | 0.4075464 | 0.68360672 | NA |
| CEBPB | 4.92618713 | -0.0415306 | 0.63563771 | -0.0653369 | 0.94790579 | NA |
| PTPN1 | 28.1619734 | -0.3903608 | 0.37664758 | -1.0364085 | 0.30001158 | 0.77521265 |
| RIPOR3 | 8.31427301 | 0.81586215 | 0.64932447 | 1.25647836 | 0.20894259 | NA |
| PARD6B | 41.0938151 | 0.23940682 | 0.35949887 | 0.66594594 | 0.50544565 | 0.87094108 |
| BCAS4 | 13.1910168 | 1.76313425 | 0.55078876 | 3.20110789 | 0.001369 | NA |
| ADNP | 59.3908192 | 0.20641691 | 0.36539366 | 0.56491651 | 0.57213055 | 0.89263108 |
| DPM1 | 15.2544191 | -0.0116313 | 0.42266504 | -0.027519 | 0.97804575 | NA |
| NFATC2 | 6.285982 | -0.0283908 | 0.65803819 | -0.0431445 | 0.96558632 | NA |
| ATP9A | 47.1559552 | -0.3310561 | 0.32098996 | -1.0313596 | 0.30237224 | 0.77521265 |
| ZFP64 | 46.4202041 | -1.1870796 | 0.42154882 | -2.8159956 | 0.00486263 | 0.22229871 |
| AL157838.1 | 3.48519523 | -1.3941531 | 1.25206579 | -1.1134823 | 0.26550135 | NA |
| ZNF217 | 27.5846282 | -0.204003 | 0.4118669 | -0.4953129 | 0.62037925 | 0.91493825 |
| PFDN4 | 4.90288314 | 0.22237391 | 0.66018058 | 0.336838 | 0.73623903 | NA |
| FAM210B | 4.23018901 | 1.16195497 | 0.86874584 | 1.33750853 | 0.1810567 | NA |
| CSTF1 | 3.0613611 | 0.29963134 | 0.77865015 | 0.38480869 | 0.70037916 | NA |
| RTF2 | 9.35805161 | -1.2206717 | 0.52921882 | -2.3065538 | 0.02107971 | NA |
| RAE1 | 6.32642414 | -0.4309107 | 0.63927899 | -0.6740574 | 0.50027485 | NA |
| RBM38 | 2.03923752 | 0.41426561 | 1.05546856 | 0.39249451 | 0.69469287 | NA |
| PCK1 | 25.2673565 | 0.7293929 | 0.88984124 | 0.81968881 | 0.41239353 | NA |
| PMEPA1 | 3.77484202 | -0.2916027 | 1.06728147 | -0.27322 | 0.78468407 | NA |
| RAB22A | 20.5249637 | -0.5372669 | 0.42391504 | -1.2673928 | 0.20501486 | NA |
| VAPB | 32.0438717 | 0.08479718 | 0.35433053 | 0.23931661 | 0.81086009 | 0.96845695 |
| NPEPL1 | 5.75298361 | 0.47084217 | 0.70609215 | 0.66682822 | 0.50488186 | NA |
| GNAS | 123.119381 | 0.22775462 | 0.27960026 | 0.8145723 | 0.41531717 | 0.83580403 |
| NELFCD | 9.87027086 | -0.3687435 | 0.47995785 | -0.768283 | 0.44231907 | NA |
| CTSZ | 13.7728793 | -0.1118072 | 0.47313181 | -0.2363129 | 0.81318984 | NA |
| ATP5F1E | 59.8538072 | -0.3465934 | 0.37796716 | -0.9169934 | 0.3591461 | 0.80341474 |
| PRELID3B | 7.33289461 | -0.8588266 | 0.68413345 | -1.2553495 | 0.20935193 | NA |
| SYCP2 | 8.33517252 | 0.41046555 | 0.54599624 | 0.75177358 | 0.45218723 | NA |
| FAM217B | 4.17710828 | 0.76705166 | 0.72684509 | 1.05531656 | 0.29128057 | NA |
| MIR646HG | 20.2721035 | 1.12504489 | 0.87629105 | 1.28387125 | 0.19918701 | NA |
| TAF4 | 15.7889501 | -0.3698508 | 0.49637323 | -0.7451063 | 0.45620743 | NA |
| LSM14B | 3.92195018 | -0.5383382 | 0.78165673 | -0.6887144 | 0.49100304 | NA |
| PSMA7 | 36.0002319 | 0.20774889 | 0.41952412 | 0.4952013 | 0.62045804 | 0.91493825 |
| SS18L1 | 10.9183382 | -1.2454039 | 0.55307885 | -2.2517655 | 0.02433709 | NA |
| MTG2 | 11.3801555 | 0.03385462 | 0.43789144 | 0.07731282 | 0.93837469 | NA |
| OSBPL2 | 14.3752948 | -0.3514933 | 0.40127439 | -0.8759425 | 0.38106132 | NA |
| ADRM1 | 5.10129521 | -0.6216012 | 0.61424402 | -1.0119776 | 0.31154875 | NA |
| LAMA5 | 14.6755672 | 0.60883097 | 0.42931861 | 1.4181332 | 0.15615188 | NA |
| RPS21 | 6.74515552 | -0.1959556 | 0.60523642 | -0.3237671 | 0.7461144 | NA |
| SLCO4A1 | 9.64357211 | -0.3955902 | 0.50213363 | -0.7878185 | 0.43080288 | NA |
| MRGBP | 3.06232577 | -0.4254419 | 0.85617281 | -0.4969112 | 0.61925164 | NA |
| OGFR | 9.527744 | -0.8500214 | 0.66605278 | -1.2762072 | 0.20188228 | NA |

| | | | | | | |
|---|---|---|---|---|---|---|
| TCFL5 | 6.94769649 | -0.6009144 | 0.6006373 | -1.0004613 | 0.31708732 | NA |
| DIDO1 | 27.9284512 | -0.1791004 | 0.30815756 | -0.5811975 | 0.56110736 | 0.89263108 |
| GID8 | 11.2705964 | -0.3107679 | 0.45733774 | -0.6795151 | 0.49681153 | NA |
| YTHDF1 | 10.7120287 | -1.1774768 | 0.49996713 | -2.3551084 | 0.0185173 | NA |
| ARFGAP1 | 7.04214073 | -1.7226328 | 0.71389021 | -2.413022 | 0.01582086 | NA |
| PPDPF | 58.9603764 | -0.325182 | 0.47688614 | -0.6818861 | 0.49531099 | 0.86687543 |
| GMEB2 | 5.2610904 | -0.7816198 | 0.63754493 | -1.2259838 | 0.22020475 | NA |
| MHENCR | 3.15665282 | -0.6038021 | 0.90485094 | -0.6672946 | 0.504584 | NA |
| ARFRP1 | 5.75601232 | 0.08101283 | 0.58786833 | 0.13780777 | 0.89039234 | NA |
| SLC2A4RG | 6.57892985 | 1.13774353 | 0.62045728 | 1.83371775 | 0.0666959 | NA |
| ZBTB46 | 10.5747194 | -0.5773344 | 0.53158346 | -1.0860654 | 0.27745005 | NA |
| TPD52L2 | 11.280739 | -0.6884383 | 0.62494804 | -1.1015929 | 0.27063869 | NA |
| DNAJC5 | 30.0422703 | -0.3963612 | 0.37309618 | -1.0623565 | 0.28807387 | 0.7725847 |
| UCKL1 | 7.75707395 | 0.28533001 | 0.52714393 | 0.54127535 | 0.58831781 | NA |
| ZNF512B | 4.38124157 | -1.1143703 | 0.79164574 | -1.4076629 | 0.15923092 | NA |
| PRPF6 | 14.5375338 | -0.2477899 | 0.4613181 | -0.5371345 | 0.59117471 | NA |
| TCEA2 | 3.94311054 | 0.01139576 | 0.7358711 | 0.01548608 | 0.98764439 | NA |
| PCMTD2 | 48.5319978 | 0.5720686 | 0.54225892 | 1.05497315 | 0.29143761 | 0.77286135 |
| MIER2 | 4.24777123 | 0.02870132 | 0.71557037 | 0.04010971 | 0.96800566 | NA |
| TPGS1 | 6.79348707 | 0.53488044 | 0.54233905 | 0.98624734 | 0.32401175 | NA |
| CDC34 | 5.49773143 | -0.6676762 | 0.63075291 | -1.0585385 | 0.28981002 | NA |
| BSG | 84.9557208 | 0.28630031 | 0.36604467 | 0.78214582 | 0.43412888 | 0.84223493 |
| HCN2 | 2.96877601 | 2.16932012 | 1.40274511 | 1.54648204 | 0.12198819 | NA |
| RNF126 | 5.06022709 | -0.3114832 | 0.67551593 | -0.4611042 | 0.64472385 | NA |
| PALM | 2.67984062 | -0.1402863 | 1.12386256 | -0.1248252 | 0.90066196 | NA |
| PTBP1 | 19.1345707 | -0.8710753 | 0.43989011 | -1.9802111 | 0.04767982 | NA |
| MED16 | 7.28445417 | -0.5221426 | 0.54511432 | -0.9578588 | 0.33813396 | NA |
| R3HDM4 | 5.75557732 | -0.0720428 | 0.68935914 | -0.1045069 | 0.91676707 | NA |
| ARID3A | 22.2076462 | -0.060666 | 0.4742166 | -0.1279289 | 0.89820523 | NA |
| WDR18 | 4.43000717 | -0.4273833 | 0.80130127 | -0.5333615 | 0.59378335 | NA |
| TMEM259 | 16.5850474 | -0.2932184 | 0.41980956 | -0.6984559 | 0.48489216 | NA |
| CNN2 | 7.20039788 | -1.253199 | 0.73453459 | -1.706113 | 0.08798704 | NA |
| ABCA7 | 3.14208683 | -0.3033864 | 0.91793748 | -0.3305088 | 0.74101557 | NA |
| POLR2E | 9.34177854 | 0.10283105 | 0.56473494 | 0.18208727 | 0.85551424 | NA |
| GPX4 | 40.8263139 | 0.15307689 | 0.56083727 | 0.27294351 | 0.78489664 | 0.96299238 |
| SBNO2 | 16.5859971 | -0.6286112 | 0.4926706 | -1.2759259 | 0.20198173 | NA |
| STK11 | 15.8459858 | -0.3872231 | 0.38594889 | -1.0033014 | 0.31571545 | NA |
| ATP5F1D | 43.1862761 | -0.0113125 | 0.3372774 | -0.0335406 | 0.97324348 | 0.99779576 |
| MIDN | 23.6810953 | -0.570002 | 0.53625248 | -1.0629358 | 0.28781105 | NA |
| CIRBP | 38.2856829 | 0.59117153 | 0.37659406 | 1.56978454 | 0.11646524 | 0.57860396 |
| C19orf24 | 7.80003716 | -0.0470585 | 0.66514448 | -0.0707493 | 0.94359732 | NA |
| MUM1 | 17.29302 | 0.29503203 | 0.37022574 | 0.79689769 | 0.42551046 | NA |
| NDUFS7 | 11.4891583 | 0.52969654 | 0.48866744 | 1.08396118 | 0.27838201 | NA |
| DAZAP1 | 21.1660044 | 0.03980047 | 0.41378732 | 0.09618581 | 0.923373 | NA |
| RPS15 | 97.349655 | -0.1721929 | 0.38409174 | -0.4483119 | 0.65392815 | 0.92248109 |
| C19orf25 | 12.8122679 | -0.2873876 | 0.45762667 | -0.6279957 | 0.53000673 | NA |
| MBD3 | 9.96273791 | -0.4841896 | 0.74065863 | -0.6537284 | 0.51328683 | NA |
| UQCR11 | 1.45908637 | 0.93363003 | 1.29468429 | 0.72112563 | 0.47083222 | NA |
| TCF3 | 22.4990053 | 0.19641994 | 0.4294013 | 0.45742744 | 0.64736385 | NA |
| REXO1 | 7.24456271 | -0.514575 | 0.57949523 | -0.887971 | 0.37455634 | NA |
| ABHD17A | 11.4045938 | 0.01489883 | 0.47833531 | 0.03114725 | 0.97515211 | NA |
| SCAMP4 | 8.82024086 | -0.4635393 | 0.50844036 | -0.9116887 | 0.36193262 | NA |
| CSNK1G2 | 11.8269449 | -0.8506679 | 0.47042415 | -1.8082997 | 0.07055986 | NA |
| BTBD2 | 12.4760457 | -0.1099299 | 0.48418638 | -0.2270405 | 0.82039225 | NA |

| | | | | | | |
|---|---|---|---|---|---|---|
| MKNK2 | 10.0740478 | -0.2375914 | 0.47488089 | -0.5003179 | 0.61685124 | NA |
| MOB3A | 7.77142962 | -0.4171408 | 0.72888853 | -0.5722971 | 0.5671207 | NA |
| AP3D1 | 20.8642999 | -0.1820519 | 0.44476592 | -0.4093207 | 0.68230432 | NA |
| DOT1L | 16.0908018 | -0.2008069 | 0.49270159 | -0.4075628 | 0.68359466 | NA |
| PLEKHJ1 | 22.573216 | -0.3626792 | 0.46938256 | -0.772673 | 0.43971593 | NA |
| SF3A2 | 2.64639712 | -0.9061419 | 1.1571773 | -0.7830623 | 0.43359052 | NA |
| LSM7 | 16.0689423 | 0.52714654 | 0.38463434 | 1.37051346 | 0.17052668 | NA |
| SPPL2B | 9.12366811 | -0.4283633 | 0.49736232 | -0.8612701 | 0.38908928 | NA |
| TIMM13 | 14.7264885 | 0.25094325 | 0.45195604 | 0.55523818 | 0.57873176 | NA |
| LMNB2 | 4.76428519 | -0.5285606 | 0.72248971 | -0.7315822 | 0.46442362 | NA |
| GADD45B | 31.2850696 | -0.9534906 | 0.76884087 | -1.2401664 | 0.21491386 | 0.71432763 |
| GNG7 | 16.3092936 | 0.9270561 | 0.49091297 | 1.8884327 | 0.05896788 | NA |
| DIRAS1 | 2.41322646 | -1.0572037 | 0.98154662 | -1.0770795 | 0.28144477 | NA |
| SGTA | 11.7162451 | -0.3832445 | 0.45211456 | -0.8476713 | 0.39662104 | NA |
| THOP1 | 6.75100085 | -0.5566001 | 0.67231045 | -0.8278915 | 0.40773193 | NA |
| ZNF554 | 2.33002206 | 0.50097533 | 0.96024243 | 0.52171755 | 0.601867 | NA |
| ZNF555 | 4.90132991 | 0.56626216 | 0.73785506 | 0.76744363 | 0.4428178 | NA |
| TLE2 | 16.00113 | 1.27116818 | 0.48715754 | 2.60935748 | 0.00907124 | NA |
| AES | 62.4611459 | -0.7192391 | 0.29069573 | -2.4741992 | 0.01335352 | 0.30435125 |
| GNA11 | 48.622099 | -0.0498515 | 0.37143539 | -0.1342131 | 0.89323406 | 0.97769824 |
| NCLN | 7.08670653 | -0.8592645 | 0.58668526 | -1.464609 | 0.14302764 | NA |
| NFIC | 55.3333517 | 0.045442 | 0.28959448 | 0.15691598 | 0.87531106 | 0.9751578 |
| SMIM24 | 34.5423673 | 0.76090052 | 0.40929402 | 1.85905603 | 0.0630192 | 0.51753303 |
| DOHH | 3.59857205 | -0.7018848 | 0.87049794 | -0.8063026 | 0.42006839 | NA |
| FZR1 | 8.03385845 | -0.4195605 | 0.54569212 | -0.7688593 | 0.44197685 | NA |
| MFSD12 | 8.77322223 | -0.8977089 | 0.62538897 | -1.435441 | 0.15116148 | NA |
| HMG20B | 6.67744584 | 0.25448319 | 0.62328117 | 0.40829597 | 0.6830564 | NA |
| PIP5K1C | 7.82566479 | -0.8694207 | 0.51789036 | -1.6787737 | 0.09319616 | NA |
| TJP3 | 8.32100622 | 0.80420996 | 0.53501866 | 1.50314375 | 0.13280198 | NA |
| MRPL54 | 11.5766283 | -0.1176985 | 0.50498707 | -0.2330724 | 0.8157052 | NA |
| DAPK3 | 6.65686015 | -0.0400396 | 0.62300456 | -0.0642685 | 0.94875641 | NA |
| EEF2 | 105.948299 | 0.20989557 | 0.37110795 | 0.56559168 | 0.57167139 | 0.89263108 |
| PIAS4 | 7.7972653 | -0.3922341 | 0.60339659 | -0.6500436 | 0.51566405 | NA |
| ZBTB7A | 21.695281 | -0.3170817 | 0.35829179 | -0.8849817 | 0.37616651 | NA |
| MAP2K2 | 18.9779687 | -0.0468112 | 0.46891234 | -0.0998293 | 0.92047988 | NA |
| SIRT6 | 3.75116721 | -0.0199126 | 0.73892971 | -0.0269479 | 0.97850132 | NA |
| YJU2 | 4.63155316 | -0.0689904 | 0.73467597 | -0.0939059 | 0.92518388 | NA |
| SH3GL1 | 10.2281622 | -0.2249982 | 0.47884899 | -0.469873 | 0.63844578 | NA |
| UBXN6 | 8.85408966 | -0.2042649 | 0.55520525 | -0.3679089 | 0.71294117 | NA |
| HDGFL2 | 5.85562947 | 0.13268112 | 0.63815943 | 0.20791219 | 0.83529753 | NA |
| PLIN5 | 16.1281001 | 0.15515395 | 0.60829172 | 0.25506503 | 0.79867287 | NA |
| MYDGF | 13.6076129 | -0.1296454 | 0.52949368 | -0.2448479 | 0.80657423 | NA |
| DPP9 | 9.80783264 | -0.8688714 | 0.54714591 | -1.5880068 | 0.11228481 | NA |
| FEM1A | 3.29412026 | -1.7956125 | 0.84361812 | -2.1284661 | 0.03329846 | NA |
| PLIN3 | 7.63476216 | 0.13497991 | 0.54579069 | 0.24731077 | 0.80466773 | NA |
| KDM4B | 38.3209083 | 0.06674663 | 0.32026953 | 0.20840769 | 0.83491065 | 0.97154892 |
| PTPRS | 10.3842517 | 0.42370692 | 0.53890805 | 0.78623231 | 0.4317314 | NA |
| TINCR | 8.04657457 | -0.7261845 | 0.51685985 | -1.4049931 | 0.16002332 | NA |
| SAFB2 | 22.4146861 | -0.8041213 | 0.35191205 | -2.2850066 | 0.02231244 | NA |
| SAFB | 25.8333279 | -0.5056525 | 0.44730002 | -1.1304549 | 0.25828459 | NA |
| RPL36 | 55.5259051 | -0.4347049 | 0.46132372 | -0.9422991 | 0.34603953 | 0.8003972 |
| C19orf70 | 20.3835455 | 0.93387214 | 0.4712244 | 1.98179919 | 0.04750172 | NA |
| LONP1 | 14.8616045 | -0.7347089 | 0.46777031 | -1.5706616 | 0.11626127 | NA |
| CAPS | 8.72470732 | 0.8943733 | 0.7563578 | 1.18247383 | 0.23701774 | NA |

| | | | | | | |
|---|---|---|---|---|---|---|
| RANBP3 | 8.87279761 | -0.0773886 | 0.62667661 | -0.1234906 | 0.90171865 | NA |
| RFX2 | 3.97233264 | 0.75659857 | 0.87710194 | 0.8626119 | 0.38835089 | NA |
| MLLT1 | 13.1747042 | 0.17044418 | 0.56630169 | 0.3009777 | 0.7634315 | NA |
| CLPP | 5.65731217 | 0.73438003 | 0.63187822 | 1.16221766 | 0.24514706 | NA |
| ALKBH7 | 13.0990017 | 0.50891832 | 0.53570508 | 0.94999718 | 0.34211368 | NA |
| GTF2F1 | 8.96974126 | -0.5290347 | 0.48763113 | -1.0849076 | 0.27796257 | NA |
| KHSRP | 13.6314027 | -0.6165153 | 0.50374192 | -1.2238713 | 0.2210008 | NA |
| SLC25A23 | 12.2484205 | 0.82238863 | 0.55160017 | 1.4909144 | 0.13598397 | NA |
| CRB3 | 5.39569288 | -1.2104874 | 0.70461361 | -1.717945 | 0.08580664 | NA |
| AC010503.4 | 5.76428299 | -1.4255313 | 0.66069865 | -2.1576119 | 0.03095802 | NA |
| TRIP10 | 5.86334053 | 0.62901977 | 0.6232302 | 1.00928962 | 0.31283576 | NA |
| SH2D3A | 10.0705846 | -0.201122 | 0.51728126 | -0.3888059 | 0.69741974 | NA |
| ZNF557 | 2.10810722 | -0.079898 | 1.04972318 | -0.0761134 | 0.93932891 | NA |
| INSR | 101.56425 | -0.0639302 | 0.23118817 | -0.2765288 | 0.78214194 | 0.96199854 |
| ZNF358 | 6.56926506 | 0.1452485 | 0.75096336 | 0.19341622 | 0.84663301 | NA |
| MCOLN1 | 2.75757542 | -1.6638042 | 0.92319083 | -1.8022322 | 0.07150888 | NA |
| PNPLA6 | 7.62189826 | -0.8331107 | 0.7631966 | -1.0916069 | 0.27500592 | NA |
| CAMSAP3 | 7.38937485 | -0.4500258 | 0.54310793 | -0.8286121 | 0.40732392 | NA |
| XAB2 | 3.36031855 | 0.53004999 | 0.86141725 | 0.61532317 | 0.53834132 | NA |
| STXBP2 | 7.83431852 | -1.0559781 | 0.5539627 | -1.9062259 | 0.0566209 | NA |
| EVI5L | 17.7641871 | -0.8683324 | 0.44992188 | -1.9299626 | 0.05361147 | NA |
| MAP2K7 | 11.0838672 | -0.0655782 | 0.49730902 | -0.131866 | 0.89509025 | NA |
| TIMM44 | 7.11621884 | -1.0866131 | 0.62615431 | -1.7353759 | 0.08267425 | NA |
| ELAVL1 | 25.1455713 | -0.496672 | 0.39131349 | -1.2692433 | 0.20435429 | NA |
| FBN3 | 19.4909392 | -1.5871 | 0.71195242 | -2.229222 | 0.02579914 | NA |
| CERS4 | 132.716065 | 0.66990968 | 0.49642343 | 1.34947231 | 0.17718531 | 0.6676327 |
| AC022146.2 | 27.1750628 | 1.01094564 | 0.578373 | 1.74791293 | 0.0804791 | 0.54552914 |
| CD320 | 4.54321239 | 0.58751976 | 0.74055934 | 0.79334596 | 0.42757629 | NA |
| RPS28 | 23.5145738 | -0.5554962 | 0.42797695 | -1.2979583 | 0.19430165 | NA |
| RAB11B | 8.98406255 | -0.3153877 | 0.48841928 | -0.6457314 | 0.5184533 | NA |
| 2-Mar | 6.24048314 | -0.503414 | 0.7112134 | -0.7078241 | 0.47905452 | NA |
| HNRNPM | 41.5493015 | -0.5938828 | 0.32571281 | -1.8233328 | 0.06825301 | 0.52508412 |
| ZNF317 | 4.42247534 | -0.1473415 | 0.77704828 | -0.189617 | 0.84960929 | NA |
| ZNF559-ZNF | 3.72141231 | 1.42103665 | 0.92221607 | 1.54089339 | 0.12334274 | NA |
| ZNF266 | 7.22109276 | -0.6275301 | 0.54732746 | -1.1465351 | 0.25157382 | NA |
| ZNF426 | 4.49007904 | 0.13532695 | 0.67030086 | 0.20188987 | 0.84000283 | NA |
| ZNF121 | 7.38317874 | 0.06967803 | 0.61350633 | 0.11357346 | 0.90957593 | NA |
| ZNF561 | 4.59089236 | 0.16188575 | 0.75241576 | 0.21515465 | 0.82964675 | NA |
| ZNF562 | 15.8492161 | -0.3235481 | 0.38959713 | -0.8304684 | 0.406274 | NA |
| ZNF846 | 8.30309944 | 1.26093317 | 0.61321885 | 2.05625311 | 0.03975813 | NA |
| FBXL12 | 4.15354353 | 0.09649329 | 0.69050366 | 0.13974334 | 0.88886278 | NA |
| UBL5 | 12.9426296 | -0.0111738 | 0.49513366 | -0.0225673 | 0.98199542 | NA |
| PIN1 | 7.65759354 | 0.27917683 | 0.71544046 | 0.39021672 | 0.6963763 | NA |
| C19orf66 | 7.11204668 | 0.47863071 | 0.80128037 | 0.59733238 | 0.55028549 | NA |
| EIF3G | 8.53604275 | 0.85854823 | 0.53240494 | 1.61258502 | 0.1068347 | NA |
| DNMT1 | 5.68717634 | -1.2070115 | 0.66818426 | -1.8064052 | 0.07085507 | NA |
| MRPL4 | 7.57871664 | 0.70746012 | 0.55689759 | 1.27035945 | 0.20395662 | NA |
| TYK2 | 8.40615533 | 0.28379102 | 0.52164268 | 0.54403336 | 0.58641851 | NA |
| CDC37 | 12.4633353 | -1.1190207 | 0.44694484 | -2.503711 | 0.01228984 | NA |
| KEAP1 | 6.93391112 | 0.24277903 | 0.57010212 | 0.42585183 | 0.67021581 | NA |
| ATG4D | 6.84083583 | -0.7572675 | 0.68422613 | -1.1067504 | 0.26840187 | NA |
| KRI1 | 3.33050135 | 0.34875456 | 0.85030771 | 0.41015101 | 0.68169518 | NA |
| AP1M2 | 10.8942587 | 0.41581502 | 0.49224667 | 0.84472895 | 0.39826219 | NA |
| SLC44A2 | 11.3773073 | -0.8888164 | 0.46799467 | -1.899202 | 0.05753792 | NA |

| | | | | | | |
|---|---|---|---|---|---|---|
| ILF3-DT | 7.48013512 | -0.0522894 | 0.68569633 | -0.0762573 | 0.93921439 | NA |
| ILF3 | 30.8715243 | -0.1669744 | 0.43307645 | -0.385554 | 0.69982697 | 0.9303056 |
| QTRT1 | 6.90957404 | -0.5848157 | 0.6743337 | -0.8672497 | 0.3858052 | NA |
| DNM2 | 46.0042119 | -0.2294266 | 0.29300705 | -0.7830071 | 0.43362297 | 0.84223493 |
| TMED1 | 4.50703852 | 0.58554327 | 0.73212423 | 0.79978676 | 0.42383435 | NA |
| CARM1 | 5.0326186 | -0.1359357 | 0.65432259 | -0.2077503 | 0.83542394 | NA |
| YIPF2 | 6.21662093 | -0.1874328 | 0.59961849 | -0.3125868 | 0.75459461 | NA |
| TIMM29 | 4.9702203 | -0.5307335 | 0.66755117 | -0.7950454 | 0.42658709 | NA |
| SMARCA4 | 35.5301338 | -0.3494502 | 0.41103916 | -0.8501628 | 0.3952346 | 0.8258317 |
| LDLR | 21.3601006 | -1.2659855 | 0.64893936 | -1.9508533 | 0.0510745 | NA |
| KANK2 | 11.2579785 | 0.64690448 | 0.4811736 | 1.34443054 | 0.1788092 | NA |
| DOCK6 | 10.8546424 | -0.0677588 | 0.56120464 | -0.1207381 | 0.90389846 | NA |
| EPOR | 8.2925095 | 1.5072895 | 0.71829021 | 2.09844081 | 0.03586622 | NA |
| RGL3 | 61.4754332 | 0.55106791 | 0.36910054 | 1.49300218 | 0.13543662 | 0.60594834 |
| PRKCSH | 13.8515326 | -0.7781456 | 0.44575913 | -1.7456637 | 0.0808694 | NA |
| ECSIT | 9.3249757 | -0.4017693 | 0.55851898 | -0.7193475 | 0.47192681 | NA |
| ELOF1 | 3.95790366 | -0.8631134 | 0.75357704 | -1.1453553 | 0.25206202 | NA |
| ZNF627 | 6.37180241 | 0.17871048 | 0.71857491 | 0.24870125 | 0.80359188 | NA |
| AC008543.1 | 3.19968268 | 1.98238351 | 0.94707879 | 2.09315586 | 0.03633524 | NA |
| ZNF823 | 8.792587 | -0.0464364 | 0.56547311 | -0.0821195 | 0.93455171 | NA |
| ZNF441 | 5.0269646 | 0.32164319 | 0.66879666 | 0.48092822 | 0.63056751 | NA |
| ZNF440 | 8.45858354 | -0.7824054 | 0.58796119 | -1.3307093 | 0.1832847 | NA |
| ZNF439 | 9.50053581 | -0.3359722 | 0.54281374 | -0.6189456 | 0.53595217 | NA |
| ZNF69 | 10.922034 | -0.3720231 | 0.46943638 | -0.7924889 | 0.42807568 | NA |
| ZNF844 | 11.4969185 | -0.0449312 | 0.45720484 | -0.0982737 | 0.92171498 | NA |
| ZNF788P | 3.29824258 | -0.2705913 | 0.86937975 | -0.3112463 | 0.75561336 | NA |
| ZNF136 | 5.0667888 | -0.295557 | 0.87635214 | -0.3372583 | 0.73592219 | NA |
| ZNF44 | 48.5225978 | -0.6134362 | 0.32657883 | -1.8783709 | 0.06033044 | 0.51625203 |
| AC008758.5 | 2.38077861 | -0.5869853 | 1.03718463 | -0.565941 | 0.57143386 | NA |
| ZNF490 | 7.15997081 | 0.55666243 | 0.59744675 | 0.93173564 | 0.35147316 | NA |
| ZNF791 | 12.1617761 | -0.3564515 | 0.45219656 | -0.7882667 | 0.43054071 | NA |
| WDR83 | 1.86478403 | -0.7566616 | 1.2899166 | -0.5865973 | 0.55747417 | NA |
| DHPS | 4.85801603 | 0.46157042 | 0.66038704 | 0.69893925 | 0.48459 | NA |
| TNPO2 | 13.0438167 | -0.1714223 | 0.44318704 | -0.3867944 | 0.69890843 | NA |
| TRIR | 24.2827477 | -0.3520439 | 0.37162017 | -0.9473218 | 0.34347483 | NA |
| ASNA1 | 4.54862481 | 0.86669658 | 0.71023566 | 1.22029438 | 0.2223533 | NA |
| HOOK2 | 21.6969217 | 0.01160979 | 0.42268616 | 0.02746668 | 0.97808751 | NA |
| JUNB | 15.8775115 | -0.4402297 | 0.5062005 | -0.8696746 | 0.38447827 | NA |
| DNASE2 | 5.70615646 | 0.88441605 | 0.65979799 | 1.34043459 | 0.1801041 | NA |
| GCDH | 6.01418336 | 0.22849755 | 0.63663779 | 0.35891295 | 0.71966021 | NA |
| FARSA | 4.7563053 | -0.0370104 | 0.66408163 | -0.0557317 | 0.95555556 | NA |
| CALR | 37.7555973 | -0.2751225 | 0.39913724 | -0.6892929 | 0.49063896 | 0.86506451 |
| AC092069.1 | 5.11848114 | -1.1713005 | 0.66056151 | -1.7731892 | 0.07619736 | NA |
| RAD23A | 11.9286954 | -0.0233855 | 0.4779628 | -0.0489274 | 0.96097718 | NA |
| GADD45GIP1 | 18.9197104 | -0.0127385 | 0.39203175 | -0.0324936 | 0.97407842 | NA |
| NFIX | 3.87810454 | 0.02760369 | 0.8217997 | 0.03358932 | 0.97320464 | NA |
| NACC1 | 4.97431553 | -0.1795371 | 0.69372275 | -0.2588025 | 0.79578766 | NA |
| STX10 | 4.07101372 | 0.46919251 | 0.72847254 | 0.64407714 | 0.51952541 | NA |
| IER2 | 11.3961412 | 0.23601307 | 0.60793114 | 0.38822336 | 0.69785075 | NA |
| CCDC130 | 10.9612662 | -0.1408534 | 0.48710032 | -0.289167 | 0.77245355 | NA |
| MRI1 | 5.19995119 | 0.11141525 | 0.70827961 | 0.15730406 | 0.87500522 | NA |
| C19orf53 | 20.7345341 | 0.05837728 | 0.50251068 | 0.11617122 | 0.90751685 | NA |
| ZSWIM4 | 4.64111865 | -1.3161076 | 0.90006603 | -1.4622345 | 0.14367697 | NA |
| CC2D1A | 8.59295607 | -0.0387354 | 0.5355735 | -0.072325 | 0.94234327 | NA |

| | | | | | | |
|---|---|---|---|---|---|---|
| DCAF15 | 3.1686954 | 0.05235859 | 0.86511959 | 0.06052179 | 0.95174006 | NA |
| RFX1 | 5.72153566 | -0.3003109 | 0.61347631 | -0.4895233 | 0.62447129 | NA |
| SAMD1 | 4.70291558 | 0.31520533 | 0.69856387 | 0.45121906 | 0.65183168 | NA |
| PRKACA | 9.73889011 | -0.7532595 | 0.55990564 | -1.345333 | 0.17851773 | NA |
| ADGRL1 | 2.50563619 | 0.58004952 | 0.94244251 | 0.6154747 | 0.53824128 | NA |
| DDX39A | 6.40642315 | -1.5360446 | 0.70028365 | -2.1934606 | 0.02827421 | NA |
| PKN1 | 10.7902924 | -1.3454787 | 0.58031127 | -2.3185466 | 0.02041963 | NA |
| GIPC1 | 17.3506897 | -0.2364805 | 0.45945825 | -0.5146941 | 0.6067668 | NA |
| DNAJB1 | 9.71058217 | -0.5975044 | 0.56000308 | -1.0669663 | 0.28598707 | NA |
| TECR | 7.76347469 | -0.0665262 | 0.57848537 | -0.1150007 | 0.90844459 | NA |
| NDUFB7 | 25.1924061 | 0.24357235 | 0.37758268 | 0.64508348 | 0.51887309 | NA |
| ZNF333 | 4.47526804 | -0.4044468 | 0.75461229 | -0.5359663 | 0.59198182 | NA |
| ILVBL | 6.65114837 | 0.28754454 | 0.53488966 | 0.5375773 | 0.59086891 | NA |
| BRD4 | 55.9749015 | -0.3360284 | 0.32327279 | -1.0394577 | 0.29859192 | 0.77521265 |
| AKAP8 | 10.6312045 | -0.2529486 | 0.52268285 | -0.4839429 | 0.62842641 | NA |
| AKAP8L | 31.6343432 | -0.3756581 | 0.32240956 | -1.1651581 | 0.24395501 | 0.73323864 |
| WIZ | 3.85281373 | -0.5057732 | 0.70524466 | -0.7171599 | 0.47327546 | NA |
| TPM4 | 7.27201732 | -0.1407074 | 0.63817228 | -0.220485 | 0.82549343 | NA |
| RAB8A | 7.90208816 | 0.16994269 | 0.60713225 | 0.2799105 | 0.77954617 | NA |
| FAM32A | 6.51837497 | 0.46830444 | 0.5976485 | 0.78357837 | 0.43328755 | NA |
| AP1M1 | 4.60952482 | -0.5432454 | 0.70841293 | -0.7668485 | 0.44317161 | NA |
| EPS15L1 | 22.5771621 | -0.605297 | 0.38029252 | -1.5916616 | 0.11146075 | NA |
| C19orf44 | 2.48646345 | 0.37117812 | 1.05345746 | 0.35234277 | 0.72458121 | NA |
| TMEM38A | 18.8276738 | -0.965501 | 0.49426626 | -1.9534027 | 0.0507719 | NA |
| SIN3B | 12.718928 | -0.5733153 | 0.46088901 | -1.2439336 | 0.21352399 | NA |
| MYO9B | 16.0939679 | -0.6032282 | 0.46825947 | -1.2882349 | 0.19766422 | NA |
| USE1 | 4.56224381 | -0.5499511 | 0.77216072 | -0.7122237 | 0.47632629 | NA |
| NR2F6 | 11.0954611 | -0.6502711 | 0.53116271 | -1.2242408 | 0.22086138 | NA |
| ABHD8 | 2.11515962 | 1.13215322 | 1.00347794 | 1.12822931 | 0.25922309 | NA |
| MRPL34 | 18.9592935 | -0.0217982 | 0.39519504 | -0.0551582 | 0.95601246 | NA |
| DDA1 | 5.76004009 | 0.24916467 | 0.5724349 | 0.43527162 | 0.66336528 | NA |
| BST2 | 8.08452515 | 0.23770891 | 1.04844175 | 0.22672591 | 0.82063688 | NA |
| MVB12A | 6.25547615 | 0.38252797 | 0.62800296 | 0.6091181 | 0.54244616 | NA |
| SLC27A1 | 4.48908201 | 0.91239237 | 0.67564406 | 1.35040389 | 0.17688646 | NA |
| AC010618.3 | 4.37505145 | -0.1242527 | 0.7758769 | -0.1601449 | 0.87276692 | NA |
| PGLS | 12.4422659 | 0.94596038 | 0.4934224 | 1.91714113 | 0.05522 | NA |
| COLGALT1 | 3.10329949 | 0.15844712 | 0.86044522 | 0.1841455 | 0.85389931 | NA |
| MAP1S | 3.2505661 | -0.9750714 | 0.86675203 | -1.1249716 | 0.26060107 | NA |
| RPL18A | 4.0902928 | 0.15432435 | 0.68697696 | 0.22464269 | 0.82225725 | NA |
| CCDC124 | 6.8032626 | -0.314256 | 0.6635785 | -0.4735778 | 0.63580103 | NA |
| ARRDC2 | 3.57964292 | 0.33514067 | 0.73248401 | 0.45753992 | 0.64728302 | NA |
| MAST3 | 3.45299648 | 0.15024414 | 0.74504425 | 0.20165802 | 0.84018409 | NA |
| JUND | 50.3273697 | -1.4652595 | 0.40064515 | -3.65725 | 0.00025494 | 0.0756026 |
| LSM4 | 16.1689957 | 0.57721225 | 0.38572848 | 1.49642115 | 0.13454394 | NA |
| PGPEP1 | 9.08565165 | -0.1056393 | 0.55357094 | -0.1908324 | 0.84865687 | NA |
| GDF15 | 21.3967955 | -1.6367348 | 0.62730653 | -2.6091468 | 0.00907683 | NA |
| SSBP4 | 5.92282472 | 0.40027409 | 0.68960673 | 0.5804382 | 0.56161915 | NA |
| ELL | 14.5386795 | -0.6237816 | 0.42078414 | -1.4824265 | 0.13822685 | NA |
| FKBP8 | 13.3815142 | -0.2463059 | 0.41225601 | -0.5974586 | 0.55020125 | NA |
| KXD1 | 5.83055259 | 0.52371981 | 0.62429038 | 0.83890418 | 0.40152308 | NA |
| UBA52 | 36.337744 | -0.0515755 | 0.53158866 | -0.0970215 | 0.92270933 | 0.98745437 |
| REX1BD | 8.19378785 | 1.14233793 | 0.63465254 | 1.79994227 | 0.07186975 | NA |
| KLHL26 | 2.50548311 | -0.7325782 | 1.03383832 | -0.7086004 | 0.4785725 | NA |
| CRTC1 | 27.0967237 | 0.06485878 | 0.43732276 | 0.14830872 | 0.88209914 | 0.97769824 |

| | | | | | | |
|---|---|---|---|---|---|---|
| UPF1 | 14.4223464 | -0.3186658 | 0.40382635 | -0.7891159 | 0.43004426 | NA |
| COPE | 12.7073124 | -0.5166335 | 0.60627948 | -0.8521376 | 0.39413775 | NA |
| DDX49 | 3.19933663 | -0.5453396 | 0.94778459 | -0.5753835 | 0.56503198 | NA |
| SUGP2 | 25.9451512 | 0.01665338 | 0.37353857 | 0.04458275 | 0.96443989 | 0.99656976 |
| ARMC6 | 3.58766331 | -1.7463395 | 0.86737881 | -2.0133527 | 0.04407754 | NA |
| SLC25A42 | 4.92489926 | -0.0207885 | 0.70562942 | -0.0294609 | 0.97649697 | NA |
| TMEM161A | 4.8448293 | 0.14689437 | 0.74240859 | 0.19786189 | 0.84315312 | NA |
| RFXANK | 3.45958407 | 0.36793265 | 0.74352532 | 0.49484885 | 0.62070682 | NA |
| SUGP1 | 5.47164862 | -0.7020209 | 0.6328901 | -1.1092304 | 0.26733081 | NA |
| MAU2 | 14.9191639 | -0.2465837 | 0.43410715 | -0.568025 | 0.57001797 | NA |
| GATAD2A | 34.755543 | -0.2711587 | 0.30177932 | -0.898533 | 0.36890146 | 0.81199861 |
| ATP13A1 | 7.16486837 | 0.08996983 | 0.66418038 | 0.13545994 | 0.89224824 | NA |
| ZNF101 | 3.48584921 | -0.4588653 | 0.80132392 | -0.572634 | 0.56689253 | NA |
| ZNF14 | 5.28585988 | 0.51884703 | 0.68284305 | 0.75983351 | 0.44735411 | NA |
| AC011477.4 | 3.15868001 | -0.0479048 | 0.86340888 | -0.0554833 | 0.95575345 | NA |
| ZNF253 | 7.24987604 | 0.46809963 | 0.58845885 | 0.79546707 | 0.42634186 | NA |
| ZNF93 | 3.91664853 | -0.0309212 | 0.69956485 | -0.0442007 | 0.96474446 | NA |
| ZNF682 | 2.07851788 | -0.9632401 | 1.05403297 | -0.9138615 | 0.36078965 | NA |
| ZNF90 | 35.2215934 | -0.112145 | 0.42094273 | -0.2664138 | 0.78992052 | 0.96325251 |
| AC011447.3 | 6.48950223 | 0.58990452 | 0.56351817 | 1.04682432 | 0.29518061 | NA |
| ZNF486 | 4.57434031 | -0.5400561 | 0.77029358 | -0.7011043 | 0.48323796 | NA |
| AC008554.1 | 6.05476775 | 1.515049 | 1.02999461 | 1.47092906 | 0.1413103 | NA |
| ZNF737 | 10.2052333 | -0.076819 | 0.60441737 | -0.127096 | 0.89886445 | NA |
| ZNF66 | 16.4570782 | 1.33435851 | 0.40884997 | 3.26368743 | 0.00109972 | NA |
| ZNF85 | 5.80257141 | 1.75056869 | 0.64969154 | 2.69446126 | 0.00705025 | NA |
| ZNF430 | 14.6796782 | -0.8591546 | 0.49375651 | -1.740037 | 0.08185251 | NA |
| ZNF714 | 9.40022255 | -0.4866147 | 0.57422484 | -0.8474289 | 0.39675609 | NA |
| ZNF431 | 57.9676973 | 0.03691113 | 0.38007975 | 0.09711418 | 0.92263572 | 0.98745437 |
| ZNF708 | 20.1918823 | -0.107341 | 0.39030184 | -0.2750206 | 0.7833004 | NA |
| ZNF738 | 4.05060623 | -0.0366596 | 0.71229252 | -0.051467 | 0.95895339 | NA |
| ZNF493 | 6.7505999 | -0.2566931 | 0.66414988 | -0.3864988 | 0.69912728 | NA |
| AC010615.4 | 10.555125 | 1.4215607 | 0.71564222 | 1.98641258 | 0.04698753 | NA |
| ZNF429 | 16.835126 | 0.94472505 | 0.37465109 | 2.52161296 | 0.01168182 | NA |
| AC123912.1 | 4.23008984 | 1.26682299 | 0.87811065 | 1.44266897 | 0.14911374 | NA |
| ZNF100 | 16.943642 | -0.66429 | 0.48881118 | -1.3589911 | 0.17414942 | NA |
| ZNF43 | 16.1680721 | -0.3081309 | 0.5009623 | -0.6150781 | 0.53850314 | NA |
| ZNF208 | 34.5459178 | -0.5842456 | 0.47929322 | -1.2189732 | 0.22285436 | 0.72536376 |
| ZNF257 | 2.31440613 | 1.55964234 | 0.90720665 | 1.71916987 | 0.08558344 | NA |
| ZNF676 | 3.50194322 | -0.2492672 | 0.83978668 | -0.296822 | 0.76660241 | NA |
| ZNF98 | 16.8120407 | -0.2503308 | 0.45477383 | -0.5504513 | 0.5820099 | NA |
| ZNF91 | 70.1561188 | 0.48445169 | 0.3722528 | 1.30140508 | 0.19311984 | 0.68667113 |
| ZNF675 | 5.95136237 | -0.2246117 | 0.61342569 | -0.3661596 | 0.71424602 | NA |
| ZNF726 | 3.5813299 | 0.66290755 | 0.9863352 | 0.67209155 | 0.50152542 | NA |
| ZNF254 | 46.9890966 | -0.3729572 | 0.33891813 | -1.1004346 | 0.27114283 | 0.75967037 |
| LINC00662 | 13.2649991 | -0.0811533 | 0.42309979 | -0.1918065 | 0.8478938 | NA |
| AC006504.5 | 7.35237943 | 0.76620292 | 0.63969127 | 1.19776985 | 0.23100663 | NA |
| LINC01532 | 3.01283519 | 0.07204244 | 1.21761578 | 0.05916681 | 0.95281925 | NA |
| UQCRFS1 | 27.6099183 | 0.41432057 | 0.45098497 | 0.91870151 | 0.35825172 | 0.80283278 |
| POP4 | 3.94124847 | -0.2955138 | 0.72765009 | -0.4061208 | 0.68465387 | NA |
| C19orf12 | 13.5262278 | 0.0742085 | 0.42378853 | 0.17510739 | 0.86099525 | NA |
| URI1 | 40.9555027 | -0.0665383 | 0.29747719 | -0.2236754 | 0.82300992 | 0.96845695 |
| ZNF507 | 12.9755298 | -0.2467832 | 0.43814636 | -0.5632437 | 0.57326893 | NA |
| DPY19L3 | 18.1553963 | 0.20058353 | 0.38519486 | 0.52073262 | 0.60255305 | NA |
| PDCD5 | 5.74919213 | 0.10031666 | 0.58887794 | 0.17035221 | 0.86473315 | NA |

| | | | | | | |
|---|---|---|---|---|---|---|
| ANKRD27 | 15.4412773 | 0.06769347 | 0.38754396 | 0.17467302 | 0.86133657 | NA |
| NUDT19 | 3.27771569 | -0.7588751 | 0.83330156 | -0.9106848 | 0.36246146 | NA |
| CEP89 | 6.92990381 | -0.3256233 | 0.74657029 | -0.436159 | 0.66272139 | NA |
| RHPN2 | 32.3290385 | 0.34440344 | 0.33965198 | 1.0139892 | 0.3105879 | 0.778534 |
| GPATCH1 | 3.74969139 | -0.5135792 | 0.73773439 | -0.6961573 | 0.48633033 | NA |
| CEBPG | 4.39120507 | 0.13398402 | 0.68257926 | 0.19629078 | 0.84438258 | NA |
| PEPD | 12.4406887 | -0.4143893 | 0.53368174 | -0.7764727 | 0.43746993 | NA |
| LSM14A | 65.0097621 | -0.1355362 | 0.33036069 | -0.4102675 | 0.68160976 | 0.9267216 |
| KIAA0355 | 30.1504891 | -0.2326139 | 0.42006308 | -0.5537594 | 0.57974349 | 0.89263108 |
| GPI | 37.5029114 | -1.3794127 | 0.44180462 | -3.1222233 | 0.00179491 | 0.14516993 |
| UBA2 | 25.889366 | -0.2546768 | 0.38639501 | -0.6591099 | 0.5098252 | NA |
| SCGB2B2 | 17.2210048 | 0.566756 | 0.49250484 | 1.15076229 | 0.24983004 | NA |
| ZNF302 | 15.8468122 | 0.14191133 | 0.39736229 | 0.35713335 | 0.72099198 | NA |
| ZNF181 | 1.82342531 | -0.4057871 | 1.13866406 | -0.3563712 | 0.72156259 | NA |
| GRAMD1A | 8.00289563 | -0.1876524 | 0.5478331 | -0.3425357 | 0.73194776 | NA |
| HPN | 27.9513319 | 0.62503169 | 0.42946758 | 1.45536408 | 0.14556849 | 0.628084 |
| LSR | 14.4626803 | -0.1109668 | 0.46189271 | -0.2402437 | 0.81014134 | NA |
| USF2 | 17.2632435 | 0.28052822 | 0.46942755 | 0.59759642 | 0.55010925 | NA |
| MAG | 2.57062726 | 1.99623808 | 1.22753348 | 1.62621884 | 0.10390311 | NA |
| CD22 | 5.66237016 | 0.66807258 | 0.96294753 | 0.6937788 | 0.48782094 | NA |
| DMKN | 13.4802892 | 0.41516528 | 0.52105341 | 0.79678066 | 0.42557843 | NA |
| TMEM147 | 6.79585894 | 0.12163077 | 0.56423337 | 0.21556819 | 0.82932436 | NA |
| RBM42 | 4.43444583 | -0.7621661 | 0.70558231 | -1.0801944 | 0.2800556 | NA |
| COX6B1 | 38.9465514 | -0.0904782 | 0.31406195 | -0.2880903 | 0.77327764 | 0.95994327 |
| KMT2B | 7.69486476 | -0.4376803 | 0.55225238 | -0.7925367 | 0.42804779 | NA |
| PROSER3 | 2.42525618 | 0.52204074 | 1.00308497 | 0.52043521 | 0.60276027 | NA |
| SDHAF1 | 2.06950427 | -0.5281405 | 1.11444144 | -0.473906 | 0.63556695 | NA |
| TBCB | 6.42339323 | -0.0017773 | 0.56813688 | -0.0031282 | 0.99750404 | NA |
| CAPNS1 | 28.3045463 | 0.35508873 | 0.41272573 | 0.86035036 | 0.38959594 | 0.8258317 |
| COX7A1 | 1.49215712 | -1.3503697 | 1.47630555 | -0.9146953 | 0.36035163 | NA |
| ZNF565 | 13.0508358 | 0.04411759 | 0.41238783 | 0.10698083 | 0.91480418 | NA |
| ZNF146 | 16.703133 | -0.5788988 | 0.42125899 | -1.3742111 | 0.16937615 | NA |
| LINC00665 | 3.88958289 | 1.3954475 | 0.79863429 | 1.74729225 | 0.08058666 | NA |
| ZFP14 | 10.9037079 | 0.94061451 | 0.49305906 | 1.90771165 | 0.05642849 | NA |
| ZFP82 | 5.93431912 | -0.7011212 | 0.69904577 | -1.0029689 | 0.31587586 | NA |
| ZNF566 | 7.52603883 | 0.31356173 | 0.65893995 | 0.47585782 | 0.63417568 | NA |
| ZNF260 | 7.93222132 | 0.10995448 | 0.59424187 | 0.18503321 | 0.85320299 | NA |
| ZNF529 | 9.78295046 | 0.09217334 | 0.49518027 | 0.18614098 | 0.85233421 | NA |
| ZNF529-AS1 | 5.37447748 | 0.54626157 | 0.6356619 | 0.85935868 | 0.39014266 | NA |
| ZNF382 | 4.6742944 | 0.08057663 | 0.6978509 | 0.11546396 | 0.90807739 | NA |
| ZNF461 | 4.32319302 | -1.2408578 | 0.81490448 | -1.5227033 | 0.12783294 | NA |
| ZNF567 | 4.53103565 | 0.24513105 | 0.71731474 | 0.3417343 | 0.73255086 | NA |
| AC020928.1 | 7.74133298 | 0.76539403 | 0.58151932 | 1.31619708 | 0.1881079 | NA |
| ZNF790 | 7.17582241 | 1.26382141 | 0.68105981 | 1.85566876 | 0.06350079 | NA |
| ZNF345 | 10.1197844 | 0.52009553 | 0.46548277 | 1.117325 | 0.26385539 | NA |
| ZNF568 | 9.52316037 | -1.0673208 | 0.52950973 | -2.0156775 | 0.04383371 | NA |
| ZNF420 | 17.0514505 | -0.1436905 | 0.43808421 | -0.3279975 | 0.74291357 | NA |
| ZNF383 | 4.85445679 | 0.64006675 | 0.77860875 | 0.82206467 | 0.41104009 | NA |
| HKR1 | 17.7925664 | -0.0054612 | 0.41816787 | -0.0130598 | 0.98958006 | NA |
| ZNF569 | 6.26067898 | 0.23091217 | 0.62483743 | 0.3695556 | 0.71171364 | NA |
| ZNF793 | 6.62388884 | 0.09675177 | 0.62124132 | 0.15573942 | 0.87623842 | NA |
| ZNF540 | 5.80145818 | 1.55567467 | 0.63162544 | 2.46297025 | 0.01377913 | NA |
| ZNF571 | 3.53647648 | 0.42297857 | 0.77708261 | 0.54431609 | 0.58622398 | NA |
| ZFP30 | 8.36316976 | 0.52230936 | 0.63043658 | 0.82848835 | 0.40739399 | NA |

| | | | | | | |
|---|---|---|---|---|---|---|
| ZNF573 | 7.58029122 | 0.07849926 | 0.61499064 | 0.12764302 | 0.89843148 | NA |
| SIPA1L3 | 96.0104671 | -0.1045506 | 0.33915058 | -0.3082719 | 0.75787543 | 0.95323728 |
| SPINT2 | 4.71625012 | -1.3384552 | 0.72134277 | -1.8555051 | 0.06352413 | NA |
| CATSPERG | 3.35462075 | -0.2872567 | 1.04173896 | -0.2757472 | 0.7827422 | NA |
| PSMD8 | 14.199317 | 0.0435857 | 0.38741791 | 0.11250305 | 0.91042455 | NA |
| EIF3K | 23.6907905 | 0.01762639 | 0.34532889 | 0.05104231 | 0.9592918 | NA |
| ACTN4 | 65.9280812 | -0.2167894 | 0.24831162 | -0.8730539 | 0.3826337 | 0.82292454 |
| ECH1 | 14.0344193 | 0.32726086 | 0.49702915 | 0.65843393 | 0.51025934 | NA |
| HNRNPL | 14.1629538 | -0.9836277 | 0.45547845 | -2.1595483 | 0.03080766 | NA |
| SIRT2 | 10.4318531 | -0.3993904 | 0.4823751 | -0.8279665 | 0.40768947 | NA |
| NFKBIB | 4.43036282 | -1.2083784 | 0.70251052 | -1.7200858 | 0.08541684 | NA |
| MRPS12 | 10.7294649 | 0.72739175 | 0.45917008 | 1.5841445 | 0.11316084 | NA |
| FBXO17 | 19.2859687 | 0.66603532 | 0.56348315 | 1.18199687 | 0.23720694 | NA |
| PAK4 | 17.849666 | -0.3766405 | 0.39874798 | -0.9445577 | 0.34488471 | NA |
| NCCRP1 | 18.9936258 | 0.79920294 | 0.59361027 | 1.34634285 | 0.17819197 | NA |
| SAMD4B | 24.1132177 | -0.1474906 | 0.34421311 | -0.4284862 | 0.6682972 | NA |
| PAF1 | 5.4572688 | -0.2853962 | 0.60675713 | -0.4703632 | 0.63809555 | NA |
| MED29 | 8.02620971 | 0.85317551 | 0.50517303 | 1.6888778 | 0.09124285 | NA |
| ZFP36 | 13.7493608 | -0.0495466 | 0.74696764 | -0.0663303 | 0.94711482 | NA |
| RPS16 | 45.6906904 | -0.1803595 | 0.6466775 | -0.2789018 | 0.78032019 | 0.96139889 |
| SUPT5H | 18.8966471 | -0.3372269 | 0.43908944 | -0.7680141 | 0.44247878 | NA |
| TIMM50 | 4.45519709 | 0.70931469 | 0.72331682 | 0.98064178 | 0.32676943 | NA |
| EID2 | 2.70484962 | 0.55925059 | 0.81518976 | 0.68603732 | 0.49268958 | NA |
| FBL | 5.09822958 | -0.2886761 | 0.67642685 | -0.4267662 | 0.66954961 | NA |
| PSMC4 | 7.50825686 | -0.3856631 | 0.59843943 | -0.6444481 | 0.51928493 | NA |
| ZNF546 | 3.28184226 | 0.25326451 | 0.74167085 | 0.34147831 | 0.73274353 | NA |
| ZNF780B | 20.908197 | 0.65434218 | 0.44750583 | 1.4621981 | 0.14368693 | NA |
| ZNF780A | 4.45592275 | 0.70578577 | 0.73372331 | 0.96192359 | 0.33608799 | NA |
| MAP3K10 | 2.84201597 | -0.3681189 | 0.98418888 | -0.3740328 | 0.70837992 | NA |
| AKT2 | 16.2819416 | -0.0955376 | 0.39175466 | -0.243871 | 0.80733075 | NA |
| PLD3 | 18.4857447 | -0.0163445 | 0.39577983 | -0.041297 | 0.96705915 | NA |
| SERTAD3 | 3.05155826 | -0.4580394 | 0.79760746 | -0.5742667 | 0.56578732 | NA |
| BLVRB | 23.3645837 | 0.56235455 | 0.50228342 | 1.11959609 | 0.26288592 | NA |
| SPTBN4 | 4.69467342 | -0.0415119 | 0.78895753 | -0.0526161 | 0.95803777 | NA |
| SHKBP1 | 3.37527993 | -0.1239891 | 0.89682314 | -0.1382537 | 0.89003991 | NA |
| COQ8B | 4.65677045 | 0.87974951 | 0.77017215 | 1.14227645 | 0.25333913 | NA |
| ITPKC | 7.10662843 | -0.4810544 | 0.61080682 | -0.7875721 | 0.43094704 | NA |
| SNRPA | 7.21748725 | -0.8834189 | 0.61985991 | -1.4251912 | 0.15410195 | NA |
| AC008537.1 | 4.02324753 | -1.6851978 | 0.97866401 | -1.721937 | 0.08508092 | NA |
| HNRNPUL1 | 13.5317127 | 0.13339975 | 0.4679696 | 0.28506071 | 0.77559763 | NA |
| TGFB1 | 10.133911 | -0.0645679 | 0.57224924 | -0.1128319 | 0.91016386 | NA |
| CCDC97 | 3.77216914 | -0.9923937 | 0.73010566 | -1.3592467 | 0.17406843 | NA |
| DMAC2 | 5.26776048 | 0.1309404 | 0.62220537 | 0.21044562 | 0.83331989 | NA |
| RPS19 | 118.40472 | 0.00555192 | 0.33088752 | 0.01677888 | 0.98661302 | 0.99907833 |
| ARHGEF1 | 4.42652697 | -0.6805382 | 0.74143782 | -0.9178628 | 0.3586907 | NA |
| RABAC1 | 20.0139781 | 0.44140142 | 0.45630648 | 0.96733541 | 0.33337639 | NA |
| GRIK5 | 4.57201718 | -0.6261027 | 0.79781795 | -0.7847689 | 0.43258905 | NA |
| DEDD2 | 3.03298371 | -0.5014117 | 0.96044398 | -0.5220624 | 0.60162688 | NA |
| GSK3A | 5.26669089 | 0.38903334 | 0.69288569 | 0.56146828 | 0.57447835 | NA |
| CIC | 6.36357296 | -1.0036713 | 0.67108634 | -1.495592 | 0.13476001 | NA |
| PAFAH1B3 | 3.52340774 | -0.272219 | 0.76824416 | -0.3543392 | 0.72308469 | NA |
| MEGF8 | 10.6943606 | 0.7007097 | 0.53898892 | 1.30004472 | 0.19358564 | NA |
| LIPE-AS1 | 7.11069622 | 0.59079453 | 0.73823573 | 0.80027897 | 0.42354918 | NA |
| ETHE1 | 3.89911553 | -0.0800865 | 0.81194974 | -0.0986348 | 0.92142823 | NA |

| | | | | | | |
|---|---|---|---|---|---|---|
| ZNF428 | 13.9131156 | -0.4367692 | 0.57016956 | -0.7660339 | 0.44365612 | NA |
| SMG9 | 4.49241381 | -0.0934484 | 0.70464249 | -0.1326181 | 0.8944954 | NA |
| ZNF283 | 4.9368458 | 0.66401034 | 0.70704217 | 0.93913824 | 0.34765977 | NA |
| ZNF45 | 2.33583098 | -0.0959529 | 0.96981229 | -0.0989396 | 0.92118621 | NA |
| ZNF155 | 5.76034227 | 0.3820414 | 0.61631023 | 0.6198849 | 0.53533357 | NA |
| ZNF224 | 7.83837587 | 0.14434587 | 0.5230963 | 0.27594511 | 0.78259022 | NA |
| ZNF234 | 3.583593 | 1.85329032 | 0.89667809 | 2.0668402 | 0.03874921 | NA |
| ZNF226 | 7.08430609 | -0.0660428 | 0.60059209 | -0.1099627 | 0.91243892 | NA |
| ZNF227 | 3.54434814 | -0.1752413 | 0.72823006 | -0.24064 | 0.80983416 | NA |
| ZNF235 | 9.85805763 | 0.69443687 | 0.51591812 | 1.34602146 | 0.17829559 | NA |
| ZNF229 | 2.42964671 | -0.2853403 | 0.93714636 | -0.3044778 | 0.76076389 | NA |
| ZNF180 | 2.88602526 | -1.1374444 | 0.9584326 | -1.1867755 | 0.23531617 | NA |
| AC243964.2 | 7.44007455 | -0.7580499 | 0.55882463 | -1.3565077 | 0.1749377 | NA |
| PVR | 4.39605263 | -0.4710702 | 0.77335256 | -0.6091273 | 0.54244005 | NA |
| CEACAM19 | 3.15397043 | 0.02137651 | 0.88757439 | 0.02408419 | 0.98078545 | NA |
| CBLC | 3.72428799 | 0.2601392 | 0.73006414 | 0.35632376 | 0.72159812 | NA |
| BCAM | 28.5183934 | -0.1041403 | 0.47419434 | -0.2196153 | 0.82617077 | 0.96845695 |
| NECTIN2 | 20.0531537 | -0.1150372 | 0.38016817 | -0.3025956 | 0.7621981 | NA |
| TOMM40 | 8.32413669 | -1.5372008 | 0.61215326 | -2.5111371 | 0.0120343 | NA |
| APOE | 31.5961269 | -0.7618358 | 0.57639544 | -1.3217243 | 0.18625999 | 0.68006555 |
| CLPTM1 | 12.304177 | -0.423688 | 0.59434067 | -0.7128705 | 0.47592587 | NA |
| RELB | 9.02963758 | 0.04437591 | 0.71996649 | 0.06163607 | 0.95085265 | NA |
| CLASRP | 15.7279323 | -0.4071065 | 0.41722871 | -0.9757393 | 0.32919365 | NA |
| GEMIN7-AS1 | 4.04611857 | -1.0603529 | 0.9674969 | -1.0959755 | 0.2730895 | NA |
| MARK4 | 11.9716419 | 0.4499525 | 0.53692919 | 0.83801087 | 0.4020246 | NA |
| TRAPPC6A | 6.51718006 | 1.1166427 | 0.65450772 | 1.70608024 | 0.08799314 | NA |
| PPP1R13L | 5.31045073 | -0.3707057 | 0.60734596 | -0.6103698 | 0.54161684 | NA |
| ERCC1 | 4.2283666 | -0.6330565 | 0.80402239 | -0.7873617 | 0.43107014 | NA |
| VASP | 7.12635176 | -0.6604252 | 0.59331501 | -1.1131105 | 0.26566096 | NA |
| OPA3 | 14.568404 | -0.8109863 | 0.53444992 | -1.5174224 | 0.12916007 | NA |
| EML2 | 9.39821625 | 1.35850742 | 0.57605605 | 2.35829034 | 0.01835933 | NA |
| SNRPD2 | 13.6586617 | -0.1079286 | 0.44692145 | -0.2414934 | 0.80917272 | NA |
| FBXO46 | 6.63379052 | -0.1261633 | 0.55615097 | -0.2268508 | 0.82053973 | NA |
| DMPK | 3.60129518 | -0.4627389 | 0.89582313 | -0.5165516 | 0.60546919 | NA |
| SYMPK | 18.367565 | -0.4536326 | 0.38902762 | -1.166068 | 0.24358697 | NA |
| IRF2BP1 | 2.92796698 | -1.154088 | 0.95883969 | -1.2036298 | 0.2287327 | NA |
| MYPOP | 3.77957073 | 0.09900545 | 0.84182734 | 0.11760779 | 0.90637843 | NA |
| PPP5C | 11.8241654 | -0.1654392 | 0.45019747 | -0.3674814 | 0.71325997 | NA |
| CCDC8 | 2.48109956 | 0.83408274 | 0.98784453 | 0.84434617 | 0.39847599 | NA |
| PPP5D1 | 3.27905094 | 0.65105294 | 0.87391324 | 0.74498579 | 0.45628031 | NA |
| CALM3 | 25.50054 | 0.18777511 | 0.40308176 | 0.46584869 | 0.64132381 | NA |
| PRKD2 | 11.8035288 | -0.2244466 | 0.52086281 | -0.430913 | 0.6665316 | NA |
| STRN4 | 5.51651536 | -1.5181386 | 0.65160376 | -2.3298493 | 0.01981412 | NA |
| FKRP | 4.32022151 | -0.54026 | 0.78279271 | -0.6901699 | 0.49008732 | NA |
| SLC1A5 | 3.13033258 | -0.6193608 | 0.86897459 | -0.7127491 | 0.47600104 | NA |
| AP2S1 | 7.37716959 | 0.24443071 | 0.56369655 | 0.4336211 | 0.66456361 | NA |
| ARHGAP35 | 30.9673781 | -0.1750747 | 0.40980762 | -0.4272119 | 0.66922501 | 0.92499304 |
| TMEM160 | 12.2795374 | 0.28100625 | 0.5781033 | 0.48608311 | 0.62690824 | NA |
| ZC3H4 | 10.2324436 | -0.3149864 | 0.50648277 | -0.6219093 | 0.53400148 | NA |
| SAE1 | 11.2926349 | -0.8237221 | 0.51324692 | -1.6049236 | 0.10851062 | NA |
| CCDC9 | 2.62587845 | 0.43627614 | 0.87846797 | 0.49663295 | 0.61944791 | NA |
| INAFM1 | 2.13647526 | -0.7982046 | 1.11982717 | -0.7127927 | 0.47597406 | NA |
| C5AR1 | 2.83256519 | 0.02191703 | 0.877714 | 0.02497058 | 0.98007843 | NA |
| DHX34 | 6.06961034 | -0.7665539 | 0.64969655 | -1.1798645 | 0.23805413 | NA |

| | | | | | | |
|---|---|---|---|---|---|---|
| NAPA-AS1 | 3.26926023 | 0.08897449 | 0.79598787 | 0.1117787 | 0.91099888 | NA |
| NAPA | 23.7575922 | -0.6834839 | 0.40389998 | -1.6922107 | 0.0906058 | NA |
| NOP53 | 12.7132376 | 0.49636448 | 0.48019302 | 1.03367699 | 0.30128719 | NA |
| SELENOW | 18.850856 | -0.153449 | 0.48038781 | -0.3194273 | 0.74940249 | NA |
| LIG1 | 1.88352878 | 0.12439907 | 1.02426691 | 0.12145181 | 0.90333318 | NA |
| ZSWIM9 | 3.64046096 | -0.1912982 | 0.77263909 | -0.2475906 | 0.80445115 | NA |
| CARD8 | 7.98880038 | 0.45544212 | 0.63246102 | 0.72011096 | 0.47145668 | NA |
| TMEM143 | 4.17713343 | 0.8356872 | 0.84958708 | 0.98363925 | 0.32529292 | NA |
| KDELR1 | 13.024377 | -0.215718 | 0.42852296 | -0.503399 | 0.61468377 | NA |
| GRWD1 | 2.22949936 | 0.51795732 | 1.02104381 | 0.50728217 | 0.61195684 | NA |
| CYTH2 | 6.15710644 | -1.5465191 | 0.6955416 | -2.2234746 | 0.02618382 | NA |
| RPL18 | 57.3893953 | 0.04750333 | 0.36751685 | 0.12925482 | 0.89715602 | 0.97769824 |
| SPHK2 | 3.7427291 | -0.0702278 | 0.930428 | -0.075479 | 0.93983357 | NA |
| BCAT2 | 8.09947168 | 0.03181368 | 0.63202686 | 0.05033596 | 0.95985467 | NA |
| PPP1R15A | 6.96840723 | -1.8747052 | 0.81304476 | -2.3057835 | 0.02112273 | NA |
| NUCB1 | 17.7562083 | -0.1101088 | 0.4186944 | -0.2629812 | 0.79256508 | NA |
| BAX | 5.80622975 | 0.53757319 | 0.62937162 | 0.85414271 | 0.39302592 | NA |
| FTL | 280.124358 | 0.43994448 | 0.54983733 | 0.8001357 | 0.42363218 | 0.84057082 |
| GYS1 | 10.6166676 | -0.3412799 | 0.64345467 | -0.5303869 | 0.59584371 | NA |
| RUVBL2 | 3.86151446 | -0.1458728 | 0.83514436 | -0.1746677 | 0.86134073 | NA |
| SNRNP70 | 46.3620512 | -0.2228919 | 0.38376277 | -0.5808065 | 0.56137084 | 0.89263108 |
| TRPM4 | 6.58028118 | -0.3220423 | 0.73999521 | -0.4351952 | 0.66342077 | NA |
| SLC6A16 | 11.3586583 | -0.424225 | 0.47221807 | -0.8983667 | 0.36899011 | NA |
| TEAD2 | 3.18331991 | -1.0422322 | 0.95164145 | -1.0951942 | 0.27343156 | NA |
| PIH1D1 | 7.10444287 | 0.20828194 | 0.57949117 | 0.35942211 | 0.71927934 | NA |
| RPL13A | 80.148994 | 0.08460117 | 0.38539405 | 0.21951862 | 0.82624608 | 0.96845695 |
| RPS11 | 66.9111292 | -0.3191114 | 0.37166703 | -0.8585949 | 0.39056404 | 0.8258317 |
| FCGRT | 13.1757434 | 0.84945542 | 0.50783282 | 1.67270682 | 0.09438503 | NA |
| NOSIP | 9.34740406 | -0.2311684 | 0.50153875 | -0.4609183 | 0.64485719 | NA |
| PRRG2 | 4.80213533 | 1.70054781 | 0.82355127 | 2.0648961 | 0.03893283 | NA |
| PRR12 | 5.74316411 | -0.7038605 | 0.67647042 | -1.0404898 | 0.29811242 | NA |
| SCAF1 | 5.85183296 | -0.1379862 | 0.61637955 | -0.2238657 | 0.8228618 | NA |
| IRF3 | 10.6007581 | 0.21814987 | 0.45916151 | 0.47510486 | 0.63471223 | NA |
| PRMT1 | 10.5529229 | 0.22639043 | 0.55271439 | 0.40959749 | 0.68210124 | NA |
| AP2A1 | 7.11625559 | -1.2892923 | 0.60018437 | -2.1481604 | 0.03170101 | NA |
| MED25 | 10.4590101 | -0.349596 | 0.67867596 | -0.5151148 | 0.60647283 | NA |
| PTOV1 | 13.5419106 | -0.537351 | 0.50554194 | -1.0629207 | 0.28781791 | NA |
| PNKP | 6.36879976 | 0.39297602 | 0.56784048 | 0.69205356 | 0.4889037 | NA |
| TBC1D17 | 6.92464892 | 0.0569617 | 0.61040505 | 0.09331788 | 0.92565103 | NA |
| ATF5 | 2.20951257 | -1.507337 | 1.02087621 | -1.476513 | 0.13980623 | NA |
| VRK3 | 9.98516325 | 0.50489373 | 0.50119127 | 1.00738732 | 0.31374868 | NA |
| MYH14 | 16.7004827 | -0.0549986 | 0.47465681 | -0.1158702 | 0.90775543 | NA |
| KCNC3 | 10.2848327 | 0.04292915 | 0.72199049 | 0.05945944 | 0.95258617 | NA |
| NR1H2 | 13.5632807 | 0.03665013 | 0.41369986 | 0.08859111 | 0.92940687 | NA |
| EMC10 | 22.9284447 | -0.2337137 | 0.40486505 | -0.5772631 | 0.56376171 | NA |
| JOSD2 | 5.72392429 | 0.64847159 | 0.68681231 | 0.94417584 | 0.3450798 | NA |
| CLEC11A | 3.54434747 | 0.13342056 | 0.9052096 | 0.14739189 | 0.88282271 | NA |
| C19orf48 | 3.38910737 | -0.9622947 | 0.88650266 | -1.0854955 | 0.27770224 | NA |
| CTU1 | 2.90232914 | -1.1483063 | 1.00007391 | -1.1482214 | 0.25087716 | NA |
| ETFB | 23.5704559 | 0.95171694 | 0.43488255 | 2.18844592 | 0.02863714 | NA |
| ZNF175 | 5.19656673 | -0.0710527 | 0.69530934 | -0.1021886 | 0.91860695 | NA |
| SPACA6 | 4.52898947 | 0.24754376 | 0.763532 | 0.32420875 | 0.74578 | NA |
| ZNF577 | 5.87072307 | 0.73059841 | 0.67561557 | 1.08138185 | 0.27952729 | NA |
| ZNF350-AS1 | 5.03883917 | 1.67701002 | 0.74813201 | 2.24159641 | 0.02498747 | NA |

| | | | | | | |
|---|---|---|---|---|---|---|
| ZNF350 | 4.86268354 | -0.774089 | 0.95983253 | -0.8064834 | 0.41996416 | NA |
| ZNF615 | 2.38836754 | 0.0019064 | 0.93780371 | 0.00203284 | 0.99837803 | NA |
| ZNF432 | 6.81850643 | -0.2264205 | 0.81535236 | -0.2776965 | 0.78124534 | NA |
| ZNF616 | 3.09477024 | -0.550809 | 0.75662409 | -0.7279823 | 0.4666244 | NA |
| AC011468.3 | 6.12604453 | 1.49107401 | 0.65931614 | 2.26154634 | 0.02372545 | NA |
| PPP2R1A | 32.1756482 | -0.6610199 | 0.33242701 | -1.9884662 | 0.04676016 | 0.45218426 |
| ZNF766 | 6.81469775 | 0.00690325 | 0.59510182 | 0.01160012 | 0.99074465 | NA |
| ZNF480 | 7.179967 | -0.4655158 | 0.67984688 | -0.6847363 | 0.49351036 | NA |
| ZNF528 | 3.370449 | -0.4521132 | 0.83456714 | -0.5417338 | 0.58800189 | NA |
| ZNF808 | 6.86011314 | 0.12103454 | 0.65992255 | 0.18340718 | 0.85447854 | NA |
| ZNF701 | 7.16104129 | -0.1666469 | 0.54039915 | -0.3083775 | 0.75779508 | NA |
| ZNF83 | 26.2728624 | -0.0722074 | 0.41948012 | -0.1721354 | 0.86333108 | 0.97217819 |
| ZNF611 | 19.208673 | 0.17521508 | 0.44307336 | 0.39545388 | 0.69250795 | NA |
| ZNF600 | 3.91250187 | -0.7632842 | 0.78294861 | -0.9748841 | 0.32961775 | NA |
| ZNF28 | 4.3333294 | 0.95719405 | 0.75871879 | 1.26159264 | 0.2070954 | NA |
| ZNF160 | 17.2637708 | 0.4321004 | 0.39478469 | 1.09452168 | 0.27372624 | NA |
| ZNF347 | 4.86119977 | 0.26450249 | 0.6597597 | 0.40090731 | 0.68848837 | NA |
| ZNF677 | 4.69858504 | 0.0154518 | 0.77367338 | 0.019972 | 0.98406571 | NA |
| ZNF525 | 3.77604883 | 0.20147073 | 0.71780545 | 0.28067595 | 0.77895895 | NA |
| ZNF765 | 8.21338072 | 1.27804191 | 0.57065855 | 2.23959128 | 0.02511747 | NA |
| ZNF761 | 9.59746079 | 0.35798474 | 0.46281009 | 0.77350244 | 0.43922508 | NA |
| ZNF813 | 4.44268555 | -0.7331021 | 0.75663541 | -0.9688974 | 0.33259638 | NA |
| ZNF331 | 10.9397617 | 0.38927932 | 0.69668844 | 0.55875668 | 0.57632779 | NA |
| NDUFA3 | 7.22741151 | 0.02012678 | 0.73740382 | 0.02729411 | 0.97822515 | NA |
| PRPF31 | 10.7427669 | 0.16267146 | 0.47535891 | 0.34220766 | 0.73219462 | NA |
| CNOT3 | 5.45739289 | -0.0961969 | 0.70458056 | -0.1365308 | 0.8914017 | NA |
| TMC4 | 22.6736865 | -0.6061407 | 0.34714641 | -1.7460665 | 0.0807994 | NA |
| MBOAT7 | 10.7865736 | -0.0051885 | 0.53068601 | -0.0097769 | 0.99219926 | NA |
| TSEN34 | 5.47035835 | 0.42988733 | 0.63609699 | 0.67582041 | 0.49915468 | NA |
| RPS9 | 127.324669 | 0.06749843 | 0.46444665 | 0.14533086 | 0.88444965 | 0.97769824 |
| LENG8-AS1 | 5.02852732 | 0.32365713 | 0.73629041 | 0.43957808 | 0.66024272 | NA |
| LENG8 | 23.8357151 | -0.8406168 | 0.39982049 | -2.1024856 | 0.03551076 | NA |
| AC011476.3 | 4.94476123 | 0.26283859 | 0.94653804 | 0.27768413 | 0.78125484 | NA |
| RDH13 | 13.5006033 | -0.0318527 | 0.48267199 | -0.0659924 | 0.94738384 | NA |
| EPS8L1 | 4.10134532 | 0.60003495 | 0.70328648 | 0.8531871 | 0.39355556 | NA |
| PPP1R12C | 8.60084417 | -0.7347319 | 0.5212691 | -1.409506 | 0.15868559 | NA |
| PPP6R1 | 7.67579567 | -0.7978452 | 0.61874017 | -1.2894672 | 0.1972357 | NA |
| HSPBP1 | 4.37863757 | 0.06563698 | 0.7607148 | 0.0862833 | 0.93124121 | NA |
| TMEM238 | 7.0807919 | 0.44258541 | 0.56857106 | 0.77841705 | 0.43632319 | NA |
| RPL28 | 81.0740617 | -0.0081417 | 0.33339964 | -0.0244202 | 0.98051744 | 0.99902646 |
| UBE2S | 5.06922181 | 0.16791022 | 0.6450467 | 0.26030708 | 0.79462691 | NA |
| ISOC2 | 9.40467511 | 0.07968802 | 0.55047491 | 0.14476232 | 0.88489853 | NA |
| FIZ1 | 2.47689964 | 0.31874844 | 1.05655693 | 0.30168601 | 0.76289144 | NA |
| ZNF524 | 3.92681307 | 0.3828188 | 0.74557138 | 0.51345693 | 0.60763173 | NA |
| CCDC106 | 3.65617083 | -0.1699118 | 0.79403274 | -0.2139858 | 0.8305581 | NA |
| U2AF2 | 9.04927686 | -0.9191089 | 0.54205921 | -1.6955876 | 0.08996402 | NA |
| EPN1 | 30.9208665 | 0.0895493 | 0.39732673 | 0.22537949 | 0.82168406 | 0.96845695 |
| ZNF787 | 7.14115612 | -1.3162166 | 0.65761712 | -2.0014938 | 0.0453392 | NA |
| ZNF444 | 6.97995707 | 0.69788664 | 0.624328 | 1.1178205 | 0.26364366 | NA |
| ZSCAN5A | 18.893461 | -0.3496914 | 0.46457785 | -0.7527078 | 0.45162549 | NA |
| ZNF582 | 3.19927565 | 0.22753502 | 1.0421604 | 0.21833013 | 0.8271719 | NA |
| ZNF667 | 2.63523137 | 0.67201374 | 0.99756419 | 0.67365464 | 0.50053091 | NA |
| ZNF667-AS1 | 7.17751635 | 0.51788852 | 0.59568489 | 0.86940013 | 0.38462831 | NA |
| ZNF471 | 4.60310455 | 2.23731821 | 0.73953611 | 3.02529949 | 0.00248387 | NA |

| | | | | | | |
|---|---|---|---|---|---|---|
| ZFP28 | 2.68953377 | 0.99730281 | 0.97435797 | 1.02354868 | 0.3060485 | NA |
| ZNF470 | 1.85069449 | -0.229477 | 1.18840039 | -0.1930974 | 0.8468827 | NA |
| ZNF264 | 11.0674776 | -0.0634588 | 0.47078783 | -0.1347928 | 0.89277573 | NA |
| ZNF460 | 2.67194612 | -0.4807964 | 0.92319065 | -0.5207986 | 0.6025071 | NA |
| AC005261.1 | 12.1800397 | -0.1409458 | 0.55531867 | -0.2538106 | 0.79964185 | NA |
| ZNF304 | 2.34572841 | 0.49863529 | 1.00499243 | 0.49615826 | 0.61978276 | NA |
| AC004076.1 | 7.52485874 | -0.336664 | 0.56460745 | -0.5962797 | 0.55098839 | NA |
| ZNF549 | 4.53213969 | 0.97309058 | 0.72149745 | 1.34870965 | 0.17743025 | NA |
| ZNF550 | 4.45576511 | 0.78760926 | 0.75203917 | 1.04729819 | 0.29496207 | NA |
| ZNF134 | 2.78754284 | 1.80883421 | 0.92264216 | 1.96049378 | 0.0499381 | NA |
| AC003006.1 | 3.96090923 | -0.9885602 | 0.7620994 | -1.2971539 | 0.19457823 | NA |
| ZNF154 | 4.41277531 | -0.4810763 | 0.72756848 | -0.6612111 | 0.50847696 | NA |
| ZNF671 | 1.78508177 | 0.50414892 | 1.05325508 | 0.47865796 | 0.63218198 | NA |
| ZNF586 | 30.0558889 | -0.1389875 | 0.34496657 | -0.4029013 | 0.68702084 | 0.92736198 |
| ZNF552 | 9.68868232 | -0.049895 | 0.47800932 | -0.1043808 | 0.91686716 | NA |
| ZNF814 | 18.4202447 | 0.25524349 | 0.34522811 | 0.73934735 | 0.4596961 | NA |
| ZNF587 | 7.84382847 | -0.7910066 | 0.62549751 | -1.264604 | 0.20601332 | NA |
| ZNF418 | 3.7242661 | -1.4652794 | 0.98598711 | -1.486104 | 0.13725162 | NA |
| C19orf18 | 2.10912287 | 0.08523133 | 1.19784145 | 0.0711541 | 0.94327511 | NA |
| ZNF606 | 5.59167269 | 1.06560708 | 0.68821951 | 1.54835349 | 0.12153721 | NA |
| ZNF135 | 8.74345561 | 0.59135954 | 0.56391114 | 1.04867505 | 0.2943277 | NA |
| ZSCAN18 | 10.9557758 | 0.26571613 | 0.63818457 | 0.41636251 | 0.67714476 | NA |
| ZNF329 | 5.36314484 | 0.78830368 | 0.69980008 | 1.12646982 | 0.25996671 | NA |
| ZNF274 | 10.5621115 | -0.9609187 | 0.60844773 | -1.5792955 | 0.1142683 | NA |
| AC020915.3 | 4.54295305 | -0.4554926 | 0.85461172 | -0.532982 | 0.59404606 | NA |
| AC010642.2 | 2.93031123 | 1.06086023 | 0.89481791 | 1.1855599 | 0.23579615 | NA |
| RPS5 | 35.1993278 | -0.0316326 | 0.52439891 | -0.0603217 | 0.95189944 | 0.99443943 |
| SLC27A5 | 2.89387419 | -0.4835979 | 0.85080929 | -0.5683976 | 0.56976505 | NA |
| TRIM28 | 11.1366301 | -0.851441 | 0.52323733 | -1.6272557 | 0.10368279 | NA |
| CHMP2A | 11.105938 | -0.4361006 | 0.49653763 | -0.878283 | 0.37979015 | NA |
| UBE2M | 10.2590559 | -0.2604034 | 0.53483544 | -0.486885 | 0.62633981 | NA |
| MZF1-AS1 | 2.86136363 | -0.1095816 | 1.00462463 | -0.1090771 | 0.91314133 | NA |
| MZF1 | 13.38624 | 0.6725942 | 0.46018842 | 1.46156264 | 0.1438611 | NA |
| RPS4Y1 | 21.2700444 | 0.35595984 | 2.23597774 | 0.1591965 | 0.87351406 | NA |
| ZFY | 12.2112332 | 0.23464111 | 2.03579956 | 0.11525747 | 0.90824105 | NA |
| LINC00278 | 45.8879744 | 0.61494663 | 2.46455993 | 0.24951579 | 0.80296183 | 0.96743263 |
| TBL1Y | 4.31208502 | -1.2366277 | 1.41889301 | -0.871544 | 0.3834572 | NA |
| TTTY15 | 8.0653839 | -0.0843785 | 1.94317476 | -0.043423 | 0.96536435 | NA |
| USP9Y | 32.8952089 | -0.0884325 | 2.36482748 | -0.0373949 | 0.97017013 | 0.99739413 |
| DDX3Y | 7.62773495 | 0.47511255 | 1.9109527 | 0.24862601 | 0.80365008 | NA |
| UTY | 105.329538 | 0.70059814 | 2.10371761 | 0.3330286 | 0.73911269 | 0.94658914 |
| NLGN4Y | 42.593778 | 0.55682608 | 2.12610599 | 0.26189949 | 0.79339893 | 0.96341299 |
| TTTY14 | 72.1921421 | 0.53428439 | 1.2373409 | 0.43180048 | 0.66588643 | 0.92276785 |
| KDM5D | 8.3741756 | 0.28688213 | 1.95622153 | 0.14665115 | 0.88340738 | NA |
| TTTY10 | 5.18919814 | 0.75581242 | 1.19895618 | 0.63039203 | 0.52843812 | NA |
| EIF1AY | 8.29005569 | 0.45722227 | 1.9864294 | 0.23017293 | 0.8179574 | NA |
| IL17RA | 7.88947949 | 0.14655562 | 0.52720167 | 0.27798777 | 0.78102175 | NA |
| HDHD5 | 8.86795384 | -0.1484879 | 0.4943505 | -0.3003698 | 0.76389513 | NA |
| CECR2 | 8.62317153 | 0.38676494 | 0.74349391 | 0.52019921 | 0.60292474 | NA |
| ATP6V1E1 | 36.9893565 | 0.12537979 | 0.35720551 | 0.35100183 | 0.72558698 | 0.93925046 |
| BCL2L13 | 38.6929427 | 0.21100962 | 0.32430455 | 0.65065265 | 0.51527073 | 0.87158522 |
| MICAL3 | 43.8727152 | 0.28244822 | 0.37276706 | 0.75770703 | 0.44862639 | 0.85123446 |
| PEX26 | 11.2865139 | 0.46393561 | 0.48798484 | 0.95071727 | 0.34174792 | NA |
| DGCR2 | 22.0972187 | 0.50960123 | 0.46645208 | 1.092505 | 0.27461119 | NA |

| | | | | | | |
|---|---|---|---|---|---|---|
| SLC25A1 | 10.190097 | -0.2408832 | 0.50668337 | -0.4754116 | 0.63449362 | NA |
| CLTCL1 | 2.67496323 | -0.2332682 | 0.89176744 | -0.2615796 | 0.79364555 | NA |
| HIRA | 8.03720751 | -0.2223528 | 0.65177216 | -0.3411511 | 0.73298982 | NA |
| MRPL40 | 7.4155935 | 0.55505928 | 0.57588134 | 0.96384314 | 0.33512457 | NA |
| UFD1 | 8.19046014 | -0.8817898 | 0.56259056 | -1.567374 | 0.1170273 | NA |
| TXNRD2 | 10.4368651 | -0.3223486 | 0.52378031 | -0.615427 | 0.53827275 | NA |
| COMT | 21.8868561 | 0.75895697 | 0.35971862 | 2.1098629 | 0.03487017 | NA |
| TANGO2 | 10.7844054 | 0.13927573 | 0.60105544 | 0.23171861 | 0.81675657 | NA |
| DGCR8 | 8.66255778 | -0.7783647 | 0.57289769 | -1.3586452 | 0.17425905 | NA |
| TRMT2A | 2.25172317 | -0.0740127 | 1.00875666 | -0.0733702 | 0.94151154 | NA |
| RANBP1 | 5.21198648 | 0.52035822 | 0.7019413 | 0.74131301 | 0.45850367 | NA |
| ZDHHC8 | 4.34105681 | -0.83358 | 0.77279189 | -1.0786604 | 0.28073913 | NA |
| DGCR6L | 4.65021557 | -0.0410493 | 0.70729671 | -0.0580369 | 0.95371921 | NA |
| KLHL22 | 15.7349618 | -0.6198693 | 0.46199699 | -1.3417171 | 0.17968772 | NA |
| MED15 | 23.62176 | -0.743867 | 0.37116943 | -2.0041172 | 0.04505751 | NA |
| PI4KA | 37.238654 | -0.3323708 | 0.36820453 | -0.90268 | 0.36669577 | 0.80952111 |
| SNAP29 | 7.17876338 | -0.9613908 | 0.56674827 | -1.6963278 | 0.08982382 | NA |
| CRKL | 6.46926618 | -0.6779395 | 0.65562242 | -1.0340396 | 0.30111766 | NA |
| LZTR1 | 3.0927816 | 0.43336241 | 0.86985867 | 0.49819864 | 0.61834404 | NA |
| THAP7 | 3.0212434 | 0.2892531 | 0.8411804 | 0.34386571 | 0.73094728 | NA |
| HIC2 | 10.4328672 | -0.8566848 | 0.460877 | -1.8588143 | 0.06305346 | NA |
| UBE2L3 | 26.8263654 | -0.310275 | 0.32956832 | -0.9414587 | 0.34646983 | 0.8003972 |
| YDJC | 2.74388133 | -0.322226 | 0.94058862 | -0.3425791 | 0.73191512 | NA |
| SDF2L1 | 8.05066687 | -0.5812476 | 0.80736716 | -0.7199297 | 0.47156825 | NA |
| PPIL2 | 19.9078101 | -0.2139654 | 0.35679443 | -0.5996881 | 0.54871413 | NA |
| YPEL1 | 5.04625801 | 0.14083055 | 0.75010395 | 0.18774805 | 0.85107416 | NA |
| MAPK1 | 47.853788 | -0.3209486 | 0.27636979 | -1.1613015 | 0.24551932 | 0.73462702 |
| PPM1F | 3.0458848 | 0.71830838 | 0.87607589 | 0.8199157 | 0.41226416 | NA |
| TOP3B | 4.78996151 | 0.72374569 | 0.68165233 | 1.06175195 | 0.28834831 | NA |
| AC245060.5 | 7.28875656 | -1.4903983 | 0.75098622 | -1.984588 | 0.04719033 | NA |
| BCR | 32.6586005 | 0.22259082 | 0.37280248 | 0.5970744 | 0.55045771 | 0.88557663 |
| CHCHD10 | 68.9714027 | 0.08290168 | 0.32917111 | 0.25184982 | 0.80115715 | 0.96742897 |
| SMARCB1 | 10.558654 | -0.1947047 | 0.48339422 | -0.4027866 | 0.68710525 | NA |
| DDT | 43.5802026 | 0.40709231 | 0.49032432 | 0.83025111 | 0.40639682 | 0.82808596 |
| CABIN1 | 35.1786385 | 0.11791877 | 0.34420912 | 0.34257886 | 0.73191531 | 0.94354489 |
| ADORA2A-AS | 6.26999079 | -0.4046902 | 0.65715824 | -0.6158185 | 0.53801428 | NA |
| GUCD1 | 6.90258762 | 0.26149016 | 0.58023628 | 0.45066152 | 0.65223352 | NA |
| SNRPD3 | 12.8573106 | -0.551239 | 0.44937357 | -1.2266832 | 0.21994169 | NA |
| SGSM1 | 16.3833355 | 1.02511505 | 0.58615516 | 1.74888002 | 0.08031176 | NA |
| KIAA1671 | 27.28812 | -0.0274476 | 0.33160973 | -0.0827709 | 0.93403368 | 0.9907482 |
| LRP5L | 10.7551286 | 0.29772704 | 0.49145519 | 0.6058071 | 0.54464286 | NA |
| GRK3 | 12.517561 | -1.073443 | 0.4726309 | -2.2712079 | 0.0231344 | NA |
| HPS4 | 12.7051335 | -0.4244072 | 0.45287043 | -0.9371493 | 0.34868175 | NA |
| SRRD | 3.23277686 | -0.8436507 | 0.90442171 | -0.9328067 | 0.35091977 | NA |
| TFIP11 | 5.01001581 | -0.0024486 | 0.59915771 | -0.0040868 | 0.99673924 | NA |
| TPST2 | 9.48534755 | 0.72449883 | 0.59522983 | 1.21717495 | 0.22353766 | NA |
| MIATNB | 9.94221196 | 0.04546885 | 0.68303879 | 0.06656848 | 0.94692524 | NA |
| PITPNB | 39.7925321 | -0.34392 | 0.32563321 | -1.0561577 | 0.29089619 | 0.77286135 |
| TTC28 | 60.3558721 | -0.2003011 | 0.30598951 | -0.6546011 | 0.51272462 | 0.87158522 |
| HSCB | 6.842409 | 0.28198981 | 0.53675288 | 0.52536245 | 0.59933125 | NA |
| CCDC117 | 6.13833916 | 0.24858645 | 0.58097241 | 0.42787995 | 0.66873853 | NA |
| XBP1 | 12.245882 | -0.5416858 | 0.44923947 | -1.205784 | 0.22790078 | NA |
| Z93930.2 | 9.64315485 | 0.98146816 | 0.52267428 | 1.87778162 | 0.06041105 | NA |
| ZNRF3 | 93.6723285 | 0.05525766 | 0.29609931 | 0.18661867 | 0.85195962 | 0.97168963 |

| | | | | | | |
|---|---|---|---|---|---|---|
| KREMEN1 | 15.3873264 | 0.84977994 | 0.50359332 | 1.68743289 | 0.09152014 | NA |
| RHBDD3 | 3.35013642 | -1.2105504 | 0.99054666 | -1.2221034 | 0.22166854 | NA |
| EWSR1 | 30.2452576 | -0.9917511 | 0.33489618 | -2.9613687 | 0.00306275 | 0.19010422 |
| GAS2L1 | 4.94447002 | -0.0968548 | 0.89587988 | -0.1081114 | 0.91390733 | NA |
| AP1B1 | 15.2736769 | -0.0191945 | 0.41055758 | -0.0467524 | 0.96271058 | NA |
| THOC5 | 3.81552848 | 0.29231291 | 0.75744237 | 0.38592099 | 0.69955519 | NA |
| NIPSNAP1 | 4.26731132 | 1.67517012 | 0.80443572 | 2.08241638 | 0.03730445 | NA |
| NF2 | 11.7826484 | 0.22131887 | 0.4838897 | 0.45737462 | 0.64740181 | NA |
| ZMAT5 | 2.72936879 | -0.3826196 | 0.93749007 | -0.4081319 | 0.68317688 | NA |
| UQCR10 | 41.7862939 | 0.29769504 | 0.38743409 | 0.76837595 | 0.44226386 | 0.84767356 |
| ASCC2 | 12.3792488 | 0.14115205 | 0.45120535 | 0.31283329 | 0.75440732 | NA |
| MTMR3 | 48.774154 | -0.5241969 | 0.35061803 | -1.4950654 | 0.13489736 | 0.60594834 |
| TBC1D10A | 5.5653991 | 0.52428667 | 0.63242958 | 0.82900403 | 0.40710213 | NA |
| SF3A1 | 8.06045294 | 0.53237478 | 0.70435608 | 0.75583188 | 0.44975 | NA |
| RNF215 | 4.51489982 | 0.81726995 | 0.74171249 | 1.10186893 | 0.27051866 | NA |
| PES1 | 3.29983212 | -0.2710296 | 0.86019433 | -0.3150795 | 0.7527013 | NA |
| DUSP18 | 3.01735334 | 0.36548036 | 0.96349816 | 0.37932648 | 0.70444544 | NA |
| OSBP2 | 16.1578088 | -1.2036881 | 0.45978282 | -2.6179492 | 0.008846 | NA |
| MORC2 | 8.26004348 | 0.50980273 | 0.50818573 | 1.00318191 | 0.3157731 | NA |
| SMTN | 3.27170194 | 0.4105487 | 1.01114671 | 0.40602288 | 0.68472579 | NA |
| SELENOM | 16.5313641 | -0.2322021 | 0.49298815 | -0.4710095 | 0.63763396 | NA |
| INPP5J | 6.35690278 | 0.83978954 | 0.69461305 | 1.20900339 | 0.22666154 | NA |
| RNF185 | 18.5687012 | -0.4860465 | 0.52960494 | -0.917753 | 0.35874822 | NA |
| LIMK2 | 11.9374013 | -0.0458456 | 0.50811775 | -0.0902264 | 0.92810731 | NA |
| PIK3IP1 | 4.75209434 | 1.92244188 | 0.73581782 | 2.61266014 | 0.00898406 | NA |
| PATZ1 | 5.3007507 | 1.11932952 | 0.65252905 | 1.71537118 | 0.0862772 | NA |
| DRG1 | 13.7044042 | -0.2847614 | 0.46273604 | -0.6153863 | 0.53829965 | NA |
| EIF4ENIF1 | 13.7925958 | -0.1777334 | 0.53652376 | -0.3312684 | 0.74044177 | NA |
| SFI1 | 10.0018492 | -0.6117503 | 0.4739512 | -1.2907453 | 0.19679202 | NA |
| PISD | 41.1229516 | -1.133065 | 0.43346029 | -2.6139995 | 0.00894892 | 0.27648451 |
| PRR14L | 31.6283921 | -0.1380205 | 0.40163599 | -0.3436456 | 0.7311128 | 0.94354489 |
| YWHAH | 10.9144235 | -0.8854967 | 0.5314111 | -1.6663119 | 0.0956513 | NA |
| RTCB | 14.6645975 | 0.2117354 | 0.3935567 | 0.53800481 | 0.59057373 | NA |
| FBXO7 | 15.5887281 | 0.2052835 | 0.41082908 | 0.49968104 | 0.61729968 | NA |
| SYN3 | 3.00575507 | 1.78180974 | 0.98712752 | 1.80504515 | 0.07106762 | NA |
| TIMP3 | 32.3885402 | 0.10503796 | 0.66281145 | 0.15847336 | 0.87408381 | 0.9751578 |
| LARGE1 | 43.344566 | 0.70928236 | 0.33331741 | 2.12794871 | 0.03334134 | 0.41688714 |
| HMGXB4 | 10.4840976 | -0.3018827 | 0.60405612 | -0.4997593 | 0.61724455 | NA |
| TOM1 | 16.5720575 | -0.4349575 | 0.38783031 | -1.1215149 | 0.26206874 | NA |
| HMOX1 | 6.9138578 | -1.0216082 | 0.80558391 | -1.2681587 | 0.20474129 | NA |
| APOL6 | 7.44394464 | -0.4241562 | 0.89547572 | -0.4736658 | 0.63573827 | NA |
| RBFOX2 | 91.2234986 | -0.0181854 | 0.24139999 | -0.075333 | 0.93994976 | 0.99100968 |
| APOL4 | 8.99967386 | 0.24533664 | 0.81728562 | 0.3001847 | 0.76403627 | NA |
| APOL2 | 19.4975976 | 0.28217809 | 0.77348426 | 0.36481426 | 0.71525006 | NA |
| APOL1 | 13.7730155 | 0.09605014 | 0.77088631 | 0.12459702 | 0.90084259 | NA |
| MYH9 | 43.9034775 | -0.3467723 | 0.29373217 | -1.1805729 | 0.23777242 | 0.73242571 |
| TXN2 | 14.2787771 | 0.51768169 | 0.42717744 | 1.21186571 | 0.22556379 | NA |
| EIF3D | 11.8523474 | 0.53501741 | 0.63200255 | 0.84654312 | 0.39724983 | NA |
| IFT27 | 3.55441358 | -0.3450107 | 0.86613225 | -0.3983349 | 0.6903833 | NA |
| PVALB | 4.31406478 | 1.28933641 | 0.94067412 | 1.37065152 | 0.17048361 | NA |
| TST | 11.7456658 | 0.02421701 | 0.4637893 | 0.05221554 | 0.95835695 | NA |
| MPST | 10.7047227 | 0.01193414 | 0.48093662 | 0.02481438 | 0.98020302 | NA |
| CARD10 | 6.97135872 | 1.14682882 | 0.79715102 | 1.43865941 | 0.15024704 | NA |
| CDC42EP1 | 5.53574518 | -0.4440662 | 0.70694321 | -0.6281498 | 0.52990581 | NA |

| | | | | | | |
|---|---|---|---|---|---|---|
| GGA1 | 14.6019163 | 0.19378915 | 0.44844951 | 0.43213148 | 0.66564586 | NA |
| TRIOBP | 20.4128702 | 0.66751391 | 0.35491321 | 1.88078068 | 0.06000176 | NA |
| H1F0 | 34.6417146 | -2.4107676 | 0.61789454 | -3.9015843 | 9.56E-05 | 0.04332663 |
| GCAT | 3.12316226 | 0.08402674 | 0.86885915 | 0.09670927 | 0.92295727 | NA |
| ANKRD54 | 5.26777559 | -0.3529869 | 0.62231957 | -0.5672116 | 0.57057042 | NA |
| EIF3L | 37.6588896 | -0.5962986 | 0.28427251 | -2.0976301 | 0.03593784 | 0.41688714 |
| MICALL1 | 6.33910586 | -0.4033169 | 0.70325246 | -0.5735023 | 0.56630464 | NA |
| POLR2F | 5.69712184 | -1.1301505 | 0.64529979 | -1.7513572 | 0.07988439 | NA |
| PICK1 | 4.28293659 | -0.5942596 | 0.7592036 | -0.7827408 | 0.43377934 | NA |
| PLA2G6 | 4.69311436 | 0.10278767 | 0.72924055 | 0.14095167 | 0.88790812 | NA |
| MAFF | 5.7791125 | -0.6197718 | 0.60385337 | -1.0263614 | 0.30472124 | NA |
| TMEM184B | 12.1429905 | -0.4637655 | 0.57594342 | -0.8052276 | 0.42068834 | NA |
| CSNK1E | 3.25867515 | -0.8740865 | 0.93364778 | -0.9362058 | 0.34916723 | NA |
| TPTEP2-CSNI | 14.7503593 | 1.04167449 | 0.50059123 | 2.08088839 | 0.03744412 | NA |
| DDX17 | 161.301472 | -0.0015199 | 0.19112585 | -0.0079524 | 0.99365497 | 0.99923383 |
| CBY1 | 5.4787528 | -0.1751373 | 0.66336027 | -0.2640154 | 0.79176807 | NA |
| TOMM22 | 5.67071748 | -0.2003065 | 0.72757974 | -0.2753052 | 0.7830818 | NA |
| JOSD1 | 7.22692088 | -0.5039146 | 0.58753424 | -0.857677 | 0.39107087 | NA |
| GTPBP1 | 6.36599696 | -0.1986378 | 0.57601864 | -0.3448461 | 0.73021006 | NA |
| SUN2 | 13.5889262 | 0.14440543 | 0.47647485 | 0.30307041 | 0.76183621 | NA |
| AL021707.1 | 3.28383556 | -0.3319715 | 0.8130177 | -0.4083201 | 0.68303867 | NA |
| CBX6 | 9.19117663 | -0.4707358 | 0.63229102 | -0.7444923 | 0.45657873 | NA |
| CBX7 | 7.20541632 | 1.72156781 | 0.60397436 | 2.85039883 | 0.00436644 | NA |
| RPL3 | 67.2982895 | 0.26896816 | 0.43207184 | 0.62250797 | 0.53360791 | 0.88254911 |
| TAB1 | 5.25790469 | 0.80083747 | 0.70477329 | 1.13630507 | 0.25582891 | NA |
| MIEF1 | 5.02224126 | -0.5402049 | 0.63517672 | -0.8504798 | 0.39505841 | NA |
| ATF4 | 20.4995115 | -1.3243178 | 0.62514558 | -2.1184151 | 0.03413993 | NA |
| RPS19BP1 | 16.247157 | -0.4536873 | 0.4359037 | -1.0407972 | 0.29796969 | NA |
| FAM83F | 10.443952 | -0.9095289 | 0.53418733 | -1.7026403 | 0.0886354 | NA |
| TNRC6B | 190.318985 | 0.38054082 | 0.32055554 | 1.18712916 | 0.23517668 | 0.73156941 |
| MKL1 | 93.1164291 | -0.2985891 | 0.26645067 | -1.1206168 | 0.262451 | 0.75479335 |
| SLC25A17 | 14.3419648 | -0.2482495 | 0.40073072 | -0.6194921 | 0.53559224 | NA |
| ST13 | 45.9152281 | 0.06713894 | 0.35044177 | 0.19158373 | 0.84806829 | 0.97168963 |
| XPNPEP3 | 16.5534585 | 1.40331141 | 0.4257805 | 3.29585647 | 0.00098122 | NA |
| RBX1 | 19.7866752 | 0.10204159 | 0.41443938 | 0.24621596 | 0.80551506 | NA |
| EP300 | 49.1329085 | -0.2314956 | 0.29479964 | -0.7852641 | 0.43229872 | 0.84223493 |
| L3MBTL2 | 3.80938888 | 0.66893116 | 0.70944203 | 0.94289756 | 0.3457333 | NA |
| RANGAP1 | 5.14467901 | 0.02136 | 0.65400136 | 0.03266048 | 0.97394534 | NA |
| ZC3H7B | 14.8632427 | -0.0347702 | 0.43071286 | -0.0807271 | 0.93565901 | NA |
| TEF | 4.1283622 | 0.05093691 | 0.91465086 | 0.05569001 | 0.95558876 | NA |
| TOB2 | 13.0151728 | -0.2645228 | 0.49644561 | -0.5328335 | 0.59414887 | NA |
| PHF5A | 3.3319089 | -1.2523694 | 0.86683242 | -1.4447652 | 0.14852384 | NA |
| ACO2 | 43.75443 | -0.2868428 | 0.33754758 | -0.8497847 | 0.39544477 | 0.8258317 |
| POLR3H | 5.47449154 | -0.6842472 | 0.71072998 | -0.9627387 | 0.33567869 | NA |
| PMM1 | 6.16190982 | 1.31615456 | 0.6509742 | 2.02182291 | 0.04319465 | NA |
| DESI1 | 10.4931573 | -0.4476295 | 0.54481653 | -0.8216151 | 0.41129597 | NA |
| XRCC6 | 30.8928392 | -0.0779678 | 0.32779568 | -0.2378548 | 0.81199372 | 0.96845695 |
| SNU13 | 29.8622719 | 0.08574865 | 0.33989106 | 0.25228274 | 0.80082253 | 0.96742897 |
| C22orf46 | 2.41999335 | 1.02894992 | 0.9749697 | 1.05536605 | 0.29125794 | NA |
| CCDC134 | 3.40947113 | -0.2643069 | 0.93400235 | -0.2829831 | 0.7771898 | NA |
| SREBF2 | 68.8944665 | -0.6215693 | 0.35157259 | -1.7679685 | 0.07706616 | 0.54401752 |
| WBP2NL | 5.14424863 | 0.29003025 | 0.85384135 | 0.33967698 | 0.7340998 | NA |
| SMDT1 | 15.6329427 | 0.65497122 | 0.42661124 | 1.53528825 | 0.12471304 | NA |
| NDUFA6 | 14.8829702 | -0.1720196 | 0.43815188 | -0.3926028 | 0.69461288 | NA |

| | | | | | | |
|---|---|---|---|---|---|---|
| TCF20 | 34.3091599 | -0.4160464 | 0.42529711 | -0.9782489 | 0.32795123 | 0.79021 |
| RRP7A | 8.75861536 | -0.3345546 | 0.57601856 | -0.5808052 | 0.56137177 | NA |
| POLDIP3 | 9.69783342 | -0.6609204 | 0.57525937 | -1.1489086 | 0.25059367 | NA |
| CYB5R3 | 27.4561014 | 0.30454489 | 0.37679233 | 0.8082566 | 0.41894288 | 0.83856261 |
| A4GALT | 9.25638749 | 0.5346869 | 0.6001629 | 0.89090295 | 0.37298124 | NA |
| ARFGAP3 | 30.3188919 | -0.0214203 | 0.29340618 | -0.0730057 | 0.9418016 | 0.99153825 |
| PACSIN2 | 41.0355056 | -0.0626123 | 0.30685915 | -0.2040424 | 0.8383204 | 0.97154892 |
| BIK | 3.17225882 | 0.08487135 | 0.8297523 | 0.10228517 | 0.91853033 | NA |
| MCAT | 3.04389691 | 0.19556939 | 0.82592358 | 0.23678873 | 0.8128207 | NA |
| TSPO | 26.5393453 | 0.54307154 | 0.4546446 | 1.19449685 | 0.23228367 | 0.73156941 |
| TTLL12 | 3.69500976 | -0.1198635 | 0.81428489 | -0.147201 | 0.88297338 | NA |
| EFCAB6 | 12.0285324 | 0.40427149 | 0.5307603 | 0.76168375 | 0.44624878 | NA |
| SAMM50 | 4.9707052 | -0.6072688 | 0.7246194 | -0.838052 | 0.40200148 | NA |
| PARVB | 60.653012 | 0.18213484 | 0.37168778 | 0.49002105 | 0.624119 | 0.91576339 |
| RTL6 | 2.29612169 | -0.7129837 | 1.1348755 | -0.6282484 | 0.52984123 | NA |
| NUP50-DT | 15.8828676 | 1.00047848 | 0.45376488 | 2.20483896 | 0.0274654 | NA |
| NUP50 | 10.9250155 | -0.3110979 | 0.4679321 | -0.6648355 | 0.5061557 | NA |
| KIAA0930 | 4.9901044 | -0.1550181 | 0.76713097 | -0.2020752 | 0.83985794 | NA |
| FAM118A | 3.49536439 | -0.6216608 | 1.15247649 | -0.539413 | 0.58960192 | NA |
| ATXN10 | 60.5641581 | -0.2219518 | 0.24843898 | -0.8933854 | 0.3716508 | 0.81199861 |
| MIRLET7BHG | 24.9471919 | -0.0717467 | 0.43352042 | -0.1654979 | 0.86855209 | NA |
| PPARA | 34.9436412 | -0.0165999 | 0.3490354 | -0.0475594 | 0.96206735 | 0.99656976 |
| TTC38 | 8.87319297 | 1.15247947 | 0.55032251 | 2.09418923 | 0.03624313 | NA |
| TRMU | 8.02783882 | -0.4297114 | 0.54418626 | -0.7896404 | 0.42973782 | NA |
| CELSR1 | 16.5252584 | 0.37324485 | 0.46120834 | 0.80927602 | 0.4183564 | NA |
| GRAMD4 | 8.43765825 | 0.04064513 | 0.52641534 | 0.07721114 | 0.93845558 | NA |
| CERK | 8.73528838 | 0.2419243 | 0.51088277 | 0.47354171 | 0.63582675 | NA |
| TBC1D22A | 83.0097518 | -0.0891164 | 0.37988452 | -0.2345882 | 0.81452837 | 0.96845695 |
| BRD1 | 18.3336149 | -1.0476541 | 0.3994114 | -2.6229949 | 0.00871606 | NA |
| ZBED4 | 9.42966675 | -1.1622792 | 0.55807277 | -2.0826659 | 0.03728168 | NA |
| ALG12 | 2.58115955 | -0.1111219 | 0.99894079 | -0.1112397 | 0.91142627 | NA |
| CRELD2 | 3.02229871 | 0.40786424 | 1.00574423 | 0.40553476 | 0.68508447 | NA |
| PIM3 | 18.4958324 | -0.6254984 | 0.37275636 | -1.6780356 | 0.09334015 | NA |
| TRABD | 10.9983598 | 0.2886793 | 0.49135012 | 0.58752259 | 0.55685278 | NA |
| SELENOO | 14.3646779 | -0.419207 | 0.43648815 | -0.9604087 | 0.33684958 | NA |
| TUBGCP6 | 3.93083163 | 0.53175872 | 0.80774067 | 0.65832852 | 0.51032705 | NA |
| PLXNB2 | 29.7627226 | 0.06275846 | 0.37547814 | 0.16714277 | 0.86725771 | 0.97217819 |
| PPP6R2 | 57.902488 | -0.4203977 | 0.27374911 | -1.5357044 | 0.12461089 | 0.58553957 |
| SBF1 | 5.141354 | 0.11375509 | 0.63898695 | 0.17802412 | 0.85870403 | NA |
| MIOX | 5.2505424 | -1.1042266 | 1.09623489 | -1.0072902 | 0.31379536 | NA |
| LMF2 | 3.47286322 | -0.1315172 | 0.82167274 | -0.1600603 | 0.87283358 | NA |
| NCAPH2 | 2.36370951 | 0.15240208 | 0.87574011 | 0.1740266 | 0.86184455 | NA |
| ARSA | 3.64064766 | 0.19837051 | 0.79347625 | 0.25000183 | 0.80258593 | NA |
| RABL2B | 3.81580557 | 0.09492905 | 0.72250521 | 0.13138875 | 0.89546779 | NA |
| FP671120.1 | 132.529757 | -1.1509785 | 0.44140015 | -2.6075625 | 0.00911894 | 0.27648451 |
| FP236383.1 | 77.8039301 | -1.0514522 | 0.42398538 | -2.4799257 | 0.01314098 | 0.30435125 |
| CR381653.1 | 7.70733801 | -0.3505062 | 0.76257001 | -0.459638 | 0.64577607 | NA |
| HSPA13 | 3.86399703 | -0.2266843 | 0.72674991 | -0.3119152 | 0.755105 | NA |
| AF127577.4 | 3.53818759 | 1.32585554 | 0.79837607 | 1.66069047 | 0.09677563 | NA |
| NRIP1 | 37.8052047 | -0.0748025 | 0.34107468 | -0.2193142 | 0.82640526 | 0.96845695 |
| AJ009632.2 | 7.19465668 | -0.2910194 | 0.74467036 | -0.390803 | 0.69594286 | NA |
| USP25 | 88.0359966 | -0.2961321 | 0.2661594 | -1.112612 | 0.2658751 | 0.75497438 |
| MIR99AHG | 117.900042 | -0.3245724 | 0.37680797 | -0.8613734 | 0.38903243 | 0.8258317 |
| CXADR | 18.9428807 | -0.2346732 | 0.4984248 | -0.4708296 | 0.63776241 | NA |

| | | | | | | |
|---|---|---|---|---|---|---|
| BTG3 | 11.4913573 | -0.5093122 | 0.6302792 | -0.8080739 | 0.41904803 | NA |
| C21orf91 | 2.95036646 | 0.94599474 | 0.85915685 | 1.10107339 | 0.27086472 | NA |
| LINC01687 | 25.3962813 | -0.1343897 | 0.48048496 | -0.2796961 | 0.77971071 | NA |
| MRPL39 | 4.45080428 | 0.1909277 | 0.6321358 | 0.3020359 | 0.7626247 | NA |
| ATP5PF | 34.2936864 | 0.35280803 | 0.37047909 | 0.95230214 | 0.34094377 | 0.8003972 |
| GABPA | 9.69262552 | 0.35254235 | 0.4944624 | 0.7129811 | 0.47585745 | NA |
| APP | 160.257265 | 0.00582503 | 0.27040642 | 0.02154176 | 0.9828135 | 0.99902646 |
| ADAMTS1 | 20.7902082 | -0.402135 | 0.51828406 | -0.775897 | 4.38E-01 | NA |
| N6AMT1 | 4.5153108 | 1.29377488 | 0.92895846 | 1.39271554 | 0.16370583 | NA |
| LTN1 | 18.6441016 | 0.08361026 | 0.46323753 | 0.18049113 | 0.85676702 | NA |
| USP16 | 15.5917362 | -1.0983332 | 0.45451527 | -2.4164934 | 0.01567081 | NA |
| CCT8 | 13.9881772 | -0.3091137 | 0.41092274 | -0.7522428 | 0.45190508 | NA |
| MAP3K7CL | 10.5329704 | 1.18014613 | 0.6719269 | 1.75636089 | 0.07902681 | NA |
| BACH1 | 212.361434 | -0.5428885 | 0.32724483 | -1.6589674 | 0.09712237 | 0.5647486 |
| GRIK1 | 8.35702443 | 0.65605441 | 0.6074531 | 1.08000833 | 0.28013847 | NA |
| CLDN8 | 16.8486654 | 0.60983724 | 0.55748537 | 1.09390716 | 0.2739957 | NA |
| TIAM1 | 56.2307435 | -0.0174856 | 0.43416946 | -0.0402737 | 0.96787491 | 0.99656976 |
| SOD1 | 28.2217336 | 0.38576897 | 0.35897045 | 1.07465382 | 0.28252975 | 0.76946112 |
| SCAF4 | 21.2164301 | -0.0647787 | 0.3655577 | -0.1772052 | 0.85934723 | NA |
| URB1 | 7.86164085 | -0.2403194 | 0.56554134 | -0.4249369 | 0.6708827 | NA |
| EVA1C | 48.6910341 | -0.604624 | 0.48414113 | -1.248859 | 0.21171665 | 0.70988913 |
| SYNJ1 | 14.2187334 | 0.12407893 | 0.40341095 | 0.30757452 | 0.75840611 | NA |
| PAXBP1 | 29.230161 | -0.3265494 | 0.36558983 | -0.8932125 | 0.37174339 | 0.81199861 |
| C21orf62-AS | 3.65077521 | 1.56777082 | 0.78465071 | 1.99804934 | 0.04571131 | NA |
| AP000282.1 | 6.05328351 | -0.4183091 | 1.08509451 | -0.3855048 | 0.69986345 | NA |
| IFNAR1 | 29.6247109 | 0.09650385 | 0.33304736 | 0.28976014 | 0.77199974 | 0.95994327 |
| IFNGR2 | 31.9442059 | 0.41505893 | 0.31854696 | 1.30297561 | 0.19258309 | 0.68667113 |
| TMEM50B | 19.6215011 | 0.53285759 | 0.44120227 | 1.2077399 | 0.22714732 | NA |
| GART | 11.0519035 | -0.1658657 | 0.49319513 | -0.3363085 | 0.73663825 | NA |
| SON | 70.859842 | -0.2497153 | 0.26330247 | -0.9483972 | 0.34292727 | 0.8003972 |
| AP000311.1 | 12.771732 | 0.50658184 | 0.60976003 | 0.83078886 | 0.40609292 | NA |
| ITSN1 | 52.5496029 | 0.13198149 | 0.34666947 | 0.38071275 | 0.7034164 | 0.9303056 |
| MRPS6 | 59.9202126 | -0.334714 | 0.41921011 | -0.7984396 | 0.42461541 | 0.84057082 |
| LINC00310 | 5.55052404 | -1.1232566 | 0.75952107 | -1.4789012 | 0.13916672 | NA |
| AP000317.1 | 5.86621751 | -1.7573478 | 0.8153197 | -2.1554094 | 0.03112981 | NA |
| RCAN1 | 11.0223765 | -0.100748 | 0.56660867 | -0.1778087 | 0.85887321 | NA |
| RUNX1 | 16.5686246 | 0.97881911 | 0.46401186 | 2.10947001 | 0.03490403 | NA |
| SETD4 | 8.96756936 | 0.00210102 | 0.49705748 | 0.00422691 | 0.99662743 | NA |
| CBR1 | 18.3231204 | 0.34849065 | 0.37459277 | 0.93031866 | 0.35220612 | NA |
| CBR3 | 16.6624925 | 0.2584485 | 0.68250791 | 0.37867472 | 0.70492943 | NA |
| DOPEY2 | 34.1870384 | -0.0168363 | 0.45508228 | -0.0369962 | 0.97048803 | 0.99739413 |
| MORC3 | 29.2309226 | -0.4157801 | 0.30550234 | -1.3609718 | 0.17352261 | 0.6651652 |
| SIM2 | 9.72613199 | -2.5206256 | 1.04100559 | -2.4213373 | 0.01546352 | NA |
| HLCS | 41.9711541 | -0.1481261 | 0.31378581 | -0.4720613 | 0.63688303 | 0.92121485 |
| PIGP | 4.31539626 | 1.64951191 | 0.77817052 | 2.11973065 | 0.03402877 | NA |
| TTC3 | 60.9961816 | -0.3295739 | 0.32978533 | -0.9993589 | 0.31762084 | 0.78203878 |
| VPS26C | 13.4838767 | 0.07997393 | 0.46030173 | 0.17374241 | 0.86206791 | NA |
| DYRK1A | 120.880668 | -0.6283908 | 0.24841922 | -2.529558 | 0.01142063 | 0.28578895 |
| KCNJ6 | 30.2507378 | -0.4182447 | 0.72890733 | -0.5737968 | 0.56610532 | 0.89263108 |
| KCNJ15 | 12.4602706 | 0.7101577 | 0.7934924 | 0.89497731 | 0.37079922 | NA |
| ETS2 | 22.0669278 | 0.32663167 | 0.42620281 | 0.76637616 | 4.43E-01 | NA |
| PSMG1 | 4.63192106 | 0.63844628 | 0.71999089 | 0.88674216 | 0.37521773 | NA |
| BRWD1 | 102.496358 | 0.33195124 | 0.30653065 | 1.08293001 | 0.27883949 | 0.7672398 |
| HMGN1 | 31.5530193 | 0.30398063 | 0.32799381 | 0.92678771 | 0.35403677 | 0.8003972 |

| | | | | | | |
|---|---|---|---|---|---|---|
| WRB | 7.11979225 | 0.61888481 | 0.57983407 | 1.06734813 | 0.28581466 | NA |
| PCP4 | 14.7932627 | -2.0867559 | 0.53535055 | -3.8979243 | 9.70E-05 | NA |
| DSCAM | 9.90327437 | -0.9819064 | 0.83817953 | -1.1714751 | 0.24140787 | NA |
| BACE2 | 174.161312 | -0.8324114 | 0.54684594 | -1.5222046 | 0.12795782 | 0.59394683 |
| FAM3B | 17.6076634 | 0.40133972 | 0.45293927 | 0.88607843 | 0.37557527 | NA |
| MX1 | 32.3357665 | -0.9851401 | 1.15178277 | -0.8553176 | 0.39237534 | 0.8258317 |
| TMPRSS2 | 68.2943933 | 0.5707782 | 0.40042393 | 1.42543477 | 0.15403156 | 0.63864841 |
| RIPK4 | 4.95584797 | -0.4432006 | 0.70186625 | -0.6314602 | 0.52773966 | NA |
| PRDM15 | 3.9710894 | -0.3755363 | 0.74178319 | -0.5062615 | 0.6126731 | NA |
| C2CD2 | 11.4479404 | 0.60283496 | 0.44844606 | 1.34427531 | 0.17885937 | NA |
| ZBTB21 | 11.0416722 | -0.3208821 | 0.48739181 | -0.6583657 | 0.51030314 | NA |
| ABCG1 | 9.15816928 | -0.4210757 | 1.0136298 | -0.4154137 | 0.6778391 | NA |
| SLC37A1 | 14.9023678 | -0.5833499 | 0.5962312 | -0.9783955 | 0.32787874 | NA |
| PDE9A | 23.4773518 | 0.47336155 | 0.36726518 | 1.28888221 | 0.19743904 | NA |
| WDR4 | 5.67631724 | -0.6800333 | 0.64853826 | -1.0485631 | 0.29437927 | NA |
| NDUFV3 | 19.0815635 | 0.43517956 | 0.36618283 | 1.18842154 | 0.23466738 | NA |
| PKNOX1 | 18.6876748 | -0.7148866 | 0.47674972 | -1.4995007 | 0.13374378 | NA |
| RRP1B | 6.09767954 | 0.2822353 | 0.61332627 | 0.46017155 | 0.64539309 | NA |
| PDXK | 37.3457849 | -0.4464536 | 0.31736325 | -1.406759 | 0.15949886 | 0.64451101 |
| CSTB | 40.1854487 | -0.6444232 | 0.35889396 | -1.7955812 | 0.07256115 | 0.53207981 |
| RRP1 | 3.06823535 | -1.5867674 | 0.94935474 | -1.6714167 | 0.0946394 | NA |
| AGPAT3 | 31.4293127 | -0.0370811 | 0.35322264 | -0.1049793 | 0.91639221 | 0.98528045 |
| TRAPPC10 | 34.9682806 | 0.11144552 | 0.29081519 | 0.38321768 | 0.70155837 | 0.9303056 |
| PFKL | 23.8723218 | 0.11079088 | 0.3850256 | 0.28774939 | 0.77353858 | NA |
| C21orf2 | 5.10525843 | 0.0907006 | 0.72467043 | 0.12516117 | 0.90039596 | NA |
| UBE2G2 | 23.8987031 | -0.3130788 | 0.39734702 | -0.7879229 | 0.43074179 | NA |
| SUMO3 | 8.44981884 | 0.45604654 | 0.5241577 | 0.87005598 | 0.38426981 | NA |
| PTTG1IP | 39.3534623 | 0.0798848 | 0.28072587 | 0.28456515 | 0.77597731 | 0.96105928 |
| FAM207A | 3.98929766 | -0.3453244 | 0.81946225 | -0.4214037 | 0.6734603 | NA |
| ADARB1 | 4.42570304 | 0.19641529 | 0.96150203 | 0.20427964 | 0.83813498 | NA |
| POFUT2 | 5.43508203 | -1.4700793 | 0.65987947 | -2.2277997 | 0.02589387 | NA |
| COL18A1 | 64.1182075 | -0.0144364 | 0.3620364 | -0.0398755 | 0.96819235 | 0.99656976 |
| COL6A1 | 53.1271296 | 0.43839681 | 0.43237364 | 1.01393047 | 0.31061593 | 0.778534 |
| LSS | 10.9902337 | -0.8129338 | 0.57898572 | -1.4040653 | 0.16029941 | NA |
| MCM3AP-AS | 3.02510639 | 0.9136988 | 0.83732348 | 1.09121365 | 0.27517888 | NA |
| MCM3AP | 20.6547386 | -0.1353002 | 0.38839657 | -0.3483559 | 0.7275729 | NA |
| YBEY | 2.78742018 | 0.55949967 | 0.81366808 | 0.68762642 | 0.49168806 | NA |
| PCNT | 10.5693713 | 0.34045572 | 0.54056906 | 0.62980986 | 0.52881899 | NA |
| DIP2A | 32.2208749 | -0.3693919 | 0.36907304 | -1.0008641 | 0.31689253 | 0.780966 |
| PRMT2 | 25.013152 | 0.20983699 | 0.33699176 | 0.62267692 | 0.53349685 | NA |
| AC011043.1 | 3.7841956 | -0.0245197 | 1.06799556 | -0.0229586 | 0.9816833 | NA |
| AL592183.1 | 17.8492647 | 0.35016334 | 0.48061792 | 0.72856904 | 0.46626532 | NA |

**Endothelial cells**

*This means that positive fold changes imply higher mean value in non-COVID AKI than in COVID AKI and vice versa.

| | baseMean | log2 fold cha | lfcSE | stat | pvalue | padj |
|---|---|---|---|---|---|---|
| AL669831.5 | 68.3277555 | -0.2633976 | 0.38227821 | -0.6890207 | 0.49081026 | 0.99989001 |
| NOC2L | 51.4037925 | 0.2167726 | 0.37243713 | 0.58203811 | 0.56054101 | 0.99989001 |
| HES4 | 120.661348 | -0.0508648 | 0.64643401 | -0.0786853 | 0.93728295 | 0.99989001 |
| ISG15 | 960.572051 | -0.4096145 | 1.04961116 | -0.3902536 | 0.69634903 | 0.99989001 |
| AGRN | 65.6699905 | 0.17269628 | 0.45303417 | 0.38119924 | 0.70305542 | 0.99989001 |
| C1orf159 | 54.8618688 | -0.1335759 | 0.40662033 | -0.3285027 | 0.74253157 | 0.99989001 |
| SDF4 | 147.114366 | -0.1195394 | 0.4454335 | -0.2683665 | 0.78841725 | 0.99989001 |
| UBE2J2 | 96.8407577 | 0.03092809 | 0.24232472 | 0.12763078 | 0.89844117 | 0.99989001 |
| ACAP3 | 38.898441 | -0.0897756 | 0.32239282 | -0.2784667 | 0.78065416 | NA |
| INTS11 | 77.5876756 | -0.0582165 | 0.21927877 | -0.2654908 | 0.79063142 | 0.99989001 |
| AURKAIP1 | 107.357535 | -0.1546662 | 0.53298847 | -0.2901868 | 0.77167333 | 0.99989001 |
| CCNL2 | 290.869238 | -0.2040657 | 0.17127628 | -1.1914419 | 0.23348016 | 0.99989001 |
| MRPL20 | 145.610027 | -0.1480168 | 0.35461199 | -0.4174051 | 0.67638213 | 0.99989001 |
| VWA1 | 217.813115 | -0.7639361 | 1.06191502 | -0.7193948 | 0.47189771 | 0.99989001 |
| ATAD3B | 51.6977469 | -0.3358359 | 0.28712267 | -1.1696599 | 0.24213784 | 0.99989001 |
| SSU72 | 127.159893 | -0.4237417 | 0.30594406 | -1.3850301 | 0.16604326 | 0.99989001 |
| CDK11B | 81.9564948 | -0.0145475 | 0.36001414 | -0.040408 | 0.96776782 | 0.99989001 |
| SLC35E2B | 56.3373981 | -0.3081661 | 0.34249341 | -0.8997723 | 0.36824142 | 0.99989001 |
| CDK11A | 42.179953 | -0.133333 | 0.52181103 | -0.2555197 | 0.79832174 | NA |
| SLC35E2A | 48.2931542 | -0.0572308 | 0.39215484 | -0.1459392 | 0.88396936 | NA |
| NADK | 72.2880691 | -0.1346064 | 0.323628 | -0.4159294 | 0.67746169 | 0.99989001 |
| GNB1 | 1049.60744 | 0.09891066 | 0.15271039 | 0.64770092 | 0.51717841 | 0.99989001 |
| FAAP20 | 68.7277317 | -0.7349236 | 0.42017257 | -1.7490994 | 0.08027383 | 0.9364867 |
| SKI | 274.635157 | 0.67750941 | 0.31580809 | 2.14531997 | 0.03192727 | 0.67281094 |
| RER1 | 121.454341 | 0.01655338 | 0.23775298 | 0.06962428 | 0.94449271 | 0.99989001 |
| TNFRSF14 | 133.466951 | 0.14220523 | 0.25648447 | 0.55443994 | 0.5792778 | 0.99989001 |
| TPRG1L | 58.0571772 | -0.0607612 | 0.37887882 | -0.1603709 | 0.87258887 | 0.99989001 |
| CEP104 | 82.771229 | 0.24947585 | 0.28316868 | 0.88101501 | 0.37830969 | 0.99989001 |
| AJAP1 | 110.030435 | 0.82798458 | 0.74351486 | 1.11360865 | 0.26544711 | 0.99989001 |
| NPHP4 | 43.7030377 | 0.04342453 | 0.30494452 | 0.14240143 | 0.88676294 | NA |
| RPL22 | 78.5907625 | -0.2445755 | 0.33589726 | -0.728126 | 0.46653644 | 0.99989001 |
| ICMT | 57.9689186 | -0.120008 | 0.35144684 | -0.3414685 | 0.73275089 | 0.99989001 |
| ACOT7 | 64.6217966 | 0.16443291 | 0.3366506 | 0.4884379 | 0.6252397 | 0.99989001 |
| PLEKHG5 | 56.2592823 | 0.14621466 | 0.3672468 | 0.39813733 | 0.69052896 | 0.99989001 |
| NOL9 | 90.3506282 | -0.1465066 | 0.31768423 | -0.4611704 | 0.64467637 | 0.99989001 |
| DNAJC11 | 117.429669 | -0.2401376 | 0.26463076 | -0.9074439 | 0.36417212 | 0.99989001 |
| CAMTA1 | 571.060946 | 0.0464229 | 0.29385673 | 0.15797799 | 0.87447414 | 0.99989001 |
| VAMP3 | 89.7243 | -0.0877872 | 0.31386218 | -0.2796999 | 0.77970777 | 0.99989001 |
| PER3 | 172.072844 | -0.7019377 | 0.57847404 | -1.2134298 | 0.22496552 | 0.99989001 |
| PARK7 | 200.02518 | -0.023304 | 0.44459813 | -0.0524158 | 0.95819741 | 0.99989001 |
| ERRFI1 | 98.4131324 | 0.18151126 | 0.6739623 | 0.2693196 | 0.78768375 | 0.99989001 |
| RERE | 1677.75841 | 0.02806412 | 0.17564889 | 0.15977398 | 0.87305913 | 0.99989001 |
| ENO1 | 386.771348 | -0.2783505 | 0.35590812 | -0.7820853 | 4.34E-01 | 0.99989001 |
| H6PD | 85.7629277 | -0.8516434 | 0.23509624 | -3.6225311 | 2.92E-04 | 0.07335471 |
| SPSB1 | 164.333763 | -0.0429231 | 0.70582376 | -0.0608127 | 0.95150835 | 0.99989001 |
| SLC25A33 | 44.9043751 | -0.3391747 | 0.30647933 | -1.1066805 | 0.26843208 | NA |
| CLSTN1 | 173.210177 | 0.38479562 | 0.28545156 | 1.34802426 | 0.17765058 | 0.99989001 |
| CTNNBIP1 | 73.0512778 | -0.9177485 | 0.3390692 | -2.7066702 | 0.00679617 | 0.39158411 |
| LZIC | 65.8904807 | -0.3270168 | 0.30238609 | -1.0814545 | 0.27949501 | 0.99989001 |
| NMNAT1 | 67.765334 | 0.29183653 | 0.3337841 | 0.87432723 | 0.38194007 | 0.99989001 |
| RBP7 | 390.061938 | -1.8297278 | 1.19513652 | -1.530978 | 0.12577482 | 0.99989001 |
| UBE4B | 368.575974 | -0.1055566 | 0.22746967 | -0.4640468 | 0.64261421 | 0.99989001 |

| | | | | | | |
|---|---|---|---|---|---|---|
| KIF1B | 486.324071 | 0.52159673 | 0.23118601 | 2.2561777 | 0.0240595 | 0.6148456 |
| DFFA | 67.355302 | -0.0867519 | 0.33130739 | -0.2618472 | 0.79343928 | 0.99989001 |
| PEX14 | 165.624798 | -0.1560094 | 0.32656801 | -0.4777242 | 0.6328465 | 0.99989001 |
| SRM | 66.6895679 | -0.4691544 | 0.58930632 | -0.796113 | 0.42596637 | 0.99989001 |
| EXOSC10 | 81.7154945 | -0.2928048 | 0.25571442 | -1.145046 | 0.25219011 | 0.99989001 |
| MTOR | 187.355071 | -0.2097923 | 0.23642502 | -0.8873523 | 0.37488927 | 0.99989001 |
| AGTRAP | 70.9489876 | 0.28797158 | 0.38485215 | 0.74826548 | 0.45430004 | 0.99989001 |
| MTHFR | 72.7298403 | 0.29265241 | 0.38705676 | 0.75609688 | 0.44959111 | 0.99989001 |
| CLCN6 | 50.3170992 | -0.0745747 | 0.38103208 | -0.1957176 | 0.84483118 | 0.99989001 |
| KIAA2013 | 59.8447368 | -0.1953755 | 0.35397394 | -0.5519489 | 0.58098334 | 0.99989001 |
| PLOD1 | 100.763717 | -0.2470393 | 0.38123345 | -0.648 | 0.51698494 | 0.99989001 |
| MFN2 | 64.1005112 | 0.01236002 | 0.26517741 | 0.0466104 | 0.96282375 | 0.99989001 |
| TNFRSF1B | 408.818399 | -0.0274846 | 0.53555057 | -0.0513203 | 0.95907028 | 0.99989001 |
| VPS13D | 596.603455 | -0.0515951 | 0.18852469 | -0.2736784 | 0.78433174 | 0.99989001 |
| DHRS3 | 460.892127 | -1.0352846 | 0.39112755 | -2.6469232 | 0.00812278 | 0.41229803 |
| PRDM2 | 267.045491 | 0.19174573 | 0.23272435 | 0.82391779 | 0.40998627 | 0.99989001 |
| KAZN | 357.529463 | -0.7668965 | 0.60748128 | -1.2624199 | 0.20679772 | 0.99989001 |
| TMEM51 | 178.965586 | 0.24983892 | 0.46394175 | 0.53851356 | 0.59022255 | 0.99989001 |
| EFHD2 | 69.2302065 | -0.7093612 | 0.3117968 | -2.2750754 | 0.02290141 | 0.60652573 |
| DNAJC16 | 90.1750601 | -0.1498964 | 0.24173665 | -0.6200812 | 0.53520432 | 0.99989001 |
| DDI2 | 127.37476 | 0.17606631 | 0.25215776 | 0.69823871 | 0.48502792 | 0.99989001 |
| PLEKHM2 | 135.9242 | -0.0891333 | 0.20905686 | -0.426359 | 0.66984625 | 0.99989001 |
| FBLIM1 | 88.574951 | -0.1650296 | 0.5575532 | -0.295989 | 0.76723851 | 0.99989001 |
| SPEN | 390.699236 | -0.096891 | 0.21622276 | -0.4481072 | 0.65407581 | 0.99989001 |
| ZBTB17 | 55.7748626 | -0.2991386 | 0.36006861 | -0.8307822 | 0.40609668 | 0.99989001 |
| EPHA2 | 82.9234653 | -0.454683 | 0.4714248 | -0.9644869 | 0.33480189 | 0.99989001 |
| FBXO42 | 263.03331 | 0.11207403 | 0.21929217 | 0.51107171 | 0.60930085 | 0.99989001 |
| SZRD1 | 99.180183 | -0.1319431 | 0.21960572 | -0.6008181 | 0.54796115 | 0.99989001 |
| NECAP2 | 107.360425 | 0.45330385 | 0.21992269 | 2.06119641 | 0.03928431 | 0.72459869 |
| NBPF1 | 66.3818709 | 0.10358374 | 0.30460783 | 0.34005605 | 0.73381432 | 0.99989001 |
| CROCC | 88.1832714 | 0.17578456 | 0.3112205 | 0.56482319 | 0.57219403 | 0.99989001 |
| ATP13A2 | 67.2358521 | -0.1995324 | 0.39842451 | -0.5008036 | 0.61650933 | 0.99989001 |
| SDHB | 94.799216 | -0.1923472 | 0.21757643 | -0.8840441 | 0.37667242 | 0.99989001 |
| RCC2 | 87.2905033 | 0.24360571 | 0.36522822 | 0.66699585 | 0.50477478 | 0.99989001 |
| ARHGEF10L | 46.1502646 | -0.4065422 | 0.50460828 | -0.805659 | 0.4204395 | NA |
| UBR4 | 324.99303 | 0.00505461 | 0.22421656 | 0.02254344 | 0.98201446 | 0.99989001 |
| AL035413.1 | 57.7325043 | 0.20515363 | 0.35727796 | 0.57421293 | 0.56582372 | 0.99989001 |
| EMC1 | 44.7002886 | -0.1104891 | 0.34225889 | -0.3228231 | 0.74682925 | NA |
| CAPZB | 528.186978 | -0.0679817 | 0.19377437 | -0.3508293 | 0.72571643 | 0.99989001 |
| TMCO4 | 88.4329438 | -0.2036012 | 0.48412332 | -0.4205565 | 0.67407894 | 0.99989001 |
| CDA | 107.324289 | -0.0641269 | 0.43007603 | -0.1491059 | 0.88147008 | 0.99989001 |
| PINK1 | 160.908226 | -0.1118213 | 0.37505212 | -0.2981487 | 0.76558966 | 0.99989001 |
| DDOST | 78.4078587 | 0.09673805 | 0.4143836 | 0.23345048 | 0.81541162 | 0.99989001 |
| HP1BP3 | 427.084027 | 0.00417168 | 0.23027772 | 0.01811588 | 0.98554641 | 0.99989001 |
| EIF4G3 | 1603.02666 | -0.0320914 | 0.25588994 | -0.1254109 | 0.90019827 | 0.99989001 |
| ECE1 | 800.433837 | 0.04804227 | 0.28899948 | 0.16623653 | 0.86797081 | 0.99989001 |
| ALPL | 87.069048 | -0.0852531 | 0.6972789 | -0.1222654 | 0.90268885 | 0.99989001 |
| USP48 | 317.630797 | -0.1086333 | 0.19153471 | -0.5671732 | 0.57059654 | 0.99989001 |
| HSPG2 | 1577.52118 | -0.1445315 | 0.58212065 | -0.2482844 | 0.80391437 | 0.99989001 |
| CDC42 | 391.527587 | -0.0347054 | 0.23787428 | -0.1458981 | 0.88400183 | 0.99989001 |
| ZBTB40 | 80.5214603 | 0.00795615 | 0.3556942 | 0.02236795 | 0.98215444 | 0.99989001 |
| KDM1A | 151.283772 | -0.0034492 | 0.16783271 | -0.0205514 | 0.9836035 | 0.99989001 |
| AL031428.1 | 75.2797537 | 0.50578509 | 0.23839924 | 2.12158855 | 0.0338723 | 0.67281094 |
| LUZP1 | 478.923421 | 0.21596569 | 0.24217091 | 0.89179042 | 0.37250528 | 0.99989001 |

| | | | | | | |
|---|---|---|---|---|---|---|
| HNRNPR | 351.117942 | 0.10924089 | 0.172012 | 0.63507714 | 0.5253781 | 0.99989001 |
| ASAP3 | 72.0756754 | -0.2965697 | 0.3910484 | -0.7583964 | 0.44821372 | 0.99989001 |
| ID3 | 711.368895 | 0.09707203 | 1.00412706 | 0.09667306 | 0.92298604 | 0.99989001 |
| RPL11 | 638.329873 | -0.1592062 | 0.58179566 | -0.2736463 | 0.78435646 | 0.99989001 |
| ELOA | 65.5832505 | 0.17130376 | 0.30896829 | 0.55443798 | 0.57927914 | 0.99989001 |
| PITHD1 | 62.7462875 | -0.1512487 | 0.31154128 | -0.4854852 | 0.62733223 | 0.99989001 |
| LYPLA2 | 46.848377 | -0.4216423 | 0.373277 | -1.1295695 | 0.25865766 | NA |
| HMGCL | 64.262043 | -0.2637222 | 0.26366358 | -1.0002223 | 0.31720292 | 0.99989001 |
| PNRC2 | 97.0790125 | -0.1774122 | 0.3100707 | -0.572167 | 0.56720886 | 0.99989001 |
| SRSF10 | 220.058775 | -0.116935 | 0.1817092 | -0.6435284 | 0.51988128 | 0.99989001 |
| NIPAL3 | 69.5680622 | 0.26925006 | 0.27561146 | 0.97691897 | 0.32860927 | 0.99989001 |
| SRRM1 | 231.94426 | -0.1464091 | 0.19804724 | -0.7392637 | 0.45974688 | 0.99989001 |
| CLIC4 | 1442.43816 | -0.0319912 | 0.37920804 | -0.0843631 | 0.93276773 | 0.99989001 |
| SYF2 | 80.7970092 | 0.1918532 | 0.3632218 | 0.52819847 | 0.59736159 | 0.99989001 |
| RSRP1 | 531.697455 | 0.4776153 | 0.29172093 | 1.63723358 | 0.10158167 | 0.99989001 |
| TMEM50A | 205.311029 | -0.2365707 | 0.32698376 | -0.7234939 | 0.46937653 | 0.99989001 |
| MACO1 | 225.352588 | 0.11015089 | 0.17594202 | 0.62606362 | 0.53127321 | 0.99989001 |
| MAN1C1 | 220.48915 | -0.2973249 | 0.58287577 | -0.5101 | 0.60998143 | 0.99989001 |
| FAM110D | 106.396757 | -0.582912 | 0.71427869 | -0.8160848 | 0.41445163 | 0.99989001 |
| SH3BGRL3 | 327.132377 | -0.5296111 | 0.63793231 | -0.8301995 | 0.40642599 | 0.99989001 |
| DHDDS | 57.8957963 | -0.1721511 | 0.41144095 | -0.4184102 | 0.6756472 | 0.99989001 |
| HMGN2 | 97.6050961 | -0.2329786 | 0.38185047 | -0.6101305 | 0.54177538 | 0.99989001 |
| ARID1A | 387.278664 | 0.07449638 | 0.21874624 | 0.34056074 | 0.73343429 | 0.99989001 |
| GPN2 | 39.1792793 | -0.3418567 | 0.28498723 | -1.1995511 | 0.23031372 | NA |
| NUDC | 97.0510762 | -0.1042084 | 0.30159379 | -0.3455258 | 0.72969913 | 0.99989001 |
| SLC9A1 | 60.2096635 | -0.4391292 | 0.34371441 | -1.2775991 | 0.20139081 | 0.99989001 |
| WDTC1 | 96.1845699 | 0.18405645 | 0.21378902 | 0.86092567 | 0.38927898 | 0.99989001 |
| TMEM222 | 50.2324614 | -0.1773974 | 0.31080344 | -0.5707705 | 0.56815524 | 0.99989001 |
| MAP3K6 | 100.895583 | 0.30539779 | 0.50408369 | 0.60584741 | 0.54461609 | 0.99989001 |
| FCN3 | 243.393506 | 1.17053869 | 1.24345581 | 0.9413593 | 0.34652076 | 0.99989001 |
| WASF2 | 555.29823 | 0.1412792 | 0.17875985 | 0.7903296 | 0.4293353 | 0.99989001 |
| AHDC1 | 111.373915 | 0.24787244 | 0.26810072 | 0.92454969 | 0.35520019 | 0.99989001 |
| IFI6 | 259.036567 | 0.12593609 | 0.96355803 | 0.13069902 | 0.89601341 | 0.99989001 |
| FAM76A | 58.3662621 | -0.3778984 | 0.35549078 | -1.063033 | 0.28776701 | 0.99989001 |
| STX12 | 491.774278 | -0.2977381 | 0.32255174 | -0.9230708 | 0.35597031 | 0.99989001 |
| PPP1R8 | 53.499651 | 0.1662191 | 0.38119972 | 0.43604204 | 0.66280623 | 0.99989001 |
| RPA2 | 51.5613554 | -0.1278236 | 0.28280529 | -0.4519843 | 0.65128028 | 0.99989001 |
| EYA3 | 242.78506 | -0.0551822 | 0.25883625 | -0.2131933 | 0.83117617 | 0.99989001 |
| DNAJC8 | 103.123744 | -0.0455274 | 0.28121661 | -0.1618945 | 0.87138891 | 0.99989001 |
| ATP5IF1 | 165.511432 | 0.00345794 | 0.49252733 | 0.00702081 | 0.99439825 | 0.99989001 |
| PHACTR4 | 462.520617 | -0.0663946 | 0.31413675 | -0.2113556 | 0.83260982 | 0.99989001 |
| RCC1 | 75.5105013 | -0.0414792 | 0.52715253 | -0.0786854 | 0.93728285 | 0.99989001 |
| TRNAU1AP | 108.435357 | -0.0839468 | 0.23301386 | -0.3602651 | 0.71864888 | 0.99989001 |
| TAF12 | 60.7401594 | 0.2731809 | 0.25240968 | 1.08229169 | 0.27912294 | 0.99989001 |
| GMEB1 | 72.7563295 | -0.1308718 | 0.3833305 | -0.3414071 | 0.73279712 | 0.99989001 |
| YTHDF2 | 167.643243 | 0.11061469 | 0.18891677 | 0.58552077 | 0.55819761 | 0.99989001 |
| EPB41 | 210.78375 | 0.3953718 | 0.44532003 | 0.88783745 | 0.37462819 | 0.99989001 |
| SRSF4 | 654.63099 | -0.0379534 | 0.30983944 | -0.1224936 | 0.9025081 | 0.99989001 |
| LAPTM5 | 49.5552613 | 0.23168957 | 0.5275981 | 0.43914027 | 0.6605599 | NA |
| PUM1 | 656.244772 | 0.04881796 | 0.18913825 | 0.25810729 | 0.79632411 | 0.99989001 |
| SNRNP40 | 95.6687341 | -0.149217 | 0.19235071 | -0.7757547 | 0.43789382 | 0.99989001 |
| ZCCHC17 | 100.362256 | -0.0546475 | 0.26462981 | -0.2065053 | 0.83639619 | 0.99989001 |
| TINAGL1 | 336.968368 | -0.6686444 | 0.51119708 | -1.3079973 | 0.19087421 | 0.99989001 |
| PEF1 | 76.6410551 | -0.1200429 | 0.27791284 | -0.4319443 | 0.66578191 | 0.99989001 |

| | | | | | | |
|---|---|---|---|---|---|---|
| PTP4A2 | 82.8850001 | -0.0537305 | 0.44785551 | -0.1199729 | 0.9045046 | 0.99989001 |
| KHDRBS1 | 378.348267 | 0.16764844 | 0.17640092 | 0.95038298 | 0.34191769 | 0.99989001 |
| TMEM39B | 62.1833161 | -0.2463556 | 0.38946402 | -0.6325503 | 0.52702738 | 0.99989001 |
| KPNA6 | 239.814002 | 0.16950275 | 0.19034236 | 0.89051509 | 0.37318937 | 0.99989001 |
| TXLNA | 92.2887611 | 0.12259418 | 0.31289269 | 0.39180901 | 0.69519933 | 0.99989001 |
| EIF3I | 119.451109 | -0.3651576 | 0.46901264 | -0.7785666 | 0.43623507 | 0.99989001 |
| FAM167B | 313.754673 | -0.7826981 | 0.74999029 | -1.0436109 | 0.29666544 | 0.99989001 |
| HDAC1 | 204.082201 | -0.1240411 | 0.25812396 | -0.4805485 | 0.63083744 | 0.99989001 |
| MARCKSL1 | 152.432752 | -0.5914974 | 0.46736256 | -1.2656072 | 0.20565376 | 0.99989001 |
| BSDC1 | 156.023642 | -0.0345679 | 0.36300126 | -0.095228 | 0.92413374 | 0.99989001 |
| ZBTB8A | 47.3028992 | -0.4549389 | 0.30804097 | -1.4768779 | 0.13970838 | NA |
| ZBTB8OS | 148.371872 | 0.15365447 | 0.21913581 | 0.70118374 | 0.48318835 | 0.99989001 |
| RBBP4 | 171.690221 | 0.11912551 | 0.155851 | 0.76435508 | 0.44465567 | 0.99989001 |
| SYNC | 49.8573453 | 0.21310007 | 0.54750342 | 0.38922145 | 0.69711234 | NA |
| KIAA1522 | 69.5605508 | -0.4077646 | 0.39512609 | -1.0319859 | 0.30207872 | 0.99989001 |
| YARS | 125.863299 | -0.1687728 | 0.46807583 | -0.3605672 | 0.71842302 | 0.99989001 |
| S100PBP | 184.028354 | 0.2210907 | 0.26096219 | 0.84721353 | 0.39687611 | 0.99989001 |
| RNF19B | 96.6806313 | -0.1045087 | 0.55862626 | -0.1870816 | 0.85159664 | 0.99989001 |
| AK2 | 141.004001 | -0.0876792 | 0.19587855 | -0.4476204 | 0.65442715 | 0.99989001 |
| ZNF362 | 90.1380803 | 0.34714245 | 0.37829617 | 0.91764727 | 0.35880356 | 0.99989001 |
| AL513327.1 | 104.885392 | 0.66529839 | 0.73497655 | 0.90519676 | 0.36536116 | 0.99989001 |
| PHC2 | 624.138549 | 0.18348829 | 0.4473432 | 0.41017341 | 0.68167875 | 0.99989001 |
| SMIM12 | 112.775607 | -0.146553 | 0.22467624 | -0.6522852 | 0.51421723 | 0.99989001 |
| GJA4 | 173.127053 | -0.6177865 | 0.68123584 | -0.9068614 | 0.36448009 | 0.99989001 |
| ZMYM1 | 89.9541781 | 0.22540744 | 0.24067652 | 0.93655767 | 0.34898614 | 0.99989001 |
| SFPQ | 423.349244 | -0.0500783 | 0.2441688 | -0.2050971 | 0.8374963 | 0.99989001 |
| ZMYM4 | 702.12204 | 0.01768477 | 0.15546203 | 0.11375622 | 0.90943104 | 0.99989001 |
| KIAA0319L | 327.204393 | 0.14566829 | 0.28121774 | 0.51799112 | 0.60446446 | 0.99989001 |
| PSMB2 | 136.636307 | 0.00821292 | 0.27458859 | 0.02990992 | 0.97613889 | 0.99989001 |
| AGO4 | 249.788254 | 0.93904979 | 0.25319917 | 3.70873956 | 0.00020829 | 0.06550832 |
| AGO1 | 66.2080328 | -0.0609342 | 0.3582006 | -0.1701119 | 0.86492212 | 0.99989001 |
| AGO3 | 495.43704 | 0.116526 | 0.1871855 | 0.62251615 | 0.53360253 | 0.99989001 |
| TRAPPC3 | 60.7038328 | 0.28907149 | 0.33864004 | 0.85362467 | 0.39331299 | 0.99989001 |
| MAP7D1 | 98.2710251 | 0.16027098 | 0.31627442 | 0.50674658 | 0.61233265 | 0.99989001 |
| THRAP3 | 344.422336 | 0.01198704 | 0.1921787 | 0.06237442 | 0.95026466 | 0.99989001 |
| EVA1B | 155.099202 | 0.37979924 | 0.44597561 | 0.85161436 | 0.39442817 | 0.99989001 |
| STK40 | 113.954065 | 0.27067444 | 0.30136279 | 0.89816808 | 0.36909595 | 0.99989001 |
| MRPS15 | 41.6175952 | -0.1146613 | 0.3889461 | -0.2948001 | 0.76814662 | NA |
| MEAF6 | 81.3328934 | -0.0697785 | 0.22895494 | -0.3047696 | 0.76054162 | 0.99989001 |
| SNIP1 | 56.564562 | -0.0791636 | 0.37872494 | -0.2090266 | 0.83442748 | 0.99989001 |
| GNL2 | 44.0875071 | -0.4741863 | 0.47013363 | -1.0086202 | 0.31315682 | NA |
| C1orf122 | 59.989181 | -0.0791697 | 0.38190893 | -0.2073001 | 0.83577552 | 0.99989001 |
| MTF1 | 61.0967562 | 0.59263356 | 0.3967715 | 1.49363946 | 0.13526988 | 0.99989001 |
| INPP5B | 145.867598 | 0.50609184 | 0.22910444 | 2.20900054 | 0.0271746 | 0.64986727 |
| SF3A3 | 75.2079814 | -0.3021055 | 0.30894092 | -0.9778747 | 0.32813631 | 0.99989001 |
| RRAGC | 57.8522203 | -0.1690362 | 0.39507857 | -0.4278548 | 0.66875686 | 0.99989001 |
| AL139260.1 | 79.7735558 | -0.2839182 | 0.70093747 | -0.4050549 | 0.68543716 | 0.99989001 |
| AKIRIN1 | 94.5089959 | 0.09204006 | 0.2878203 | 0.31978308 | 0.74913278 | 0.99989001 |
| NDUFS5 | 163.862073 | -0.1597588 | 0.41718777 | -0.3829421 | 0.70176267 | 0.99989001 |
| MACF1 | 3336.8294 | 0.06384083 | 0.34208008 | 0.1866254 | 0.85195435 | 0.99989001 |
| PABPC4 | 54.1668733 | 0.24500379 | 0.27724175 | 0.88371895 | 0.37684795 | 0.99989001 |
| PPIE | 41.6735876 | 0.05223794 | 0.303396 | 0.17217742 | 0.86329805 | NA |
| TRIT1 | 71.9858824 | -0.257297 | 0.24524042 | -1.0491623 | 0.29410343 | 0.99989001 |
| CAP1 | 237.335395 | -0.0218085 | 0.35086525 | -0.0621564 | 0.95043826 | 0.99989001 |

| | | | | | | |
|---|---|---|---|---|---|---|
| PPT1 | 69.1288421 | -0.5370471 | 0.41280194 | -1.3009801 | 0.19326526 | 0.99989001 |
| RLF | 346.437978 | -0.1749952 | 0.35074397 | -0.4989258 | 0.61783166 | 0.99989001 |
| ZMPSTE24 | 88.2805045 | 0.02472855 | 0.28080871 | 0.0880619 | 0.92982748 | 0.99989001 |
| SMAP2 | 87.5571552 | 0.59817801 | 0.27997105 | 2.1365709 | 0.03263291 | 0.67281094 |
| NFYC | 179.036492 | 0.19395546 | 0.18512103 | 1.04772245 | 0.2947665 | 0.99989001 |
| CITED4 | 53.8851667 | -0.2978272 | 0.56335116 | -0.5286705 | 0.59703402 | 0.99989001 |
| CTPS1 | 43.4996043 | -0.1228544 | 0.530361 | -0.2316429 | 0.81681539 | NA |
| SCMH1 | 418.524391 | 0.14288028 | 0.26995609 | 0.52927231 | 0.59661656 | 0.99989001 |
| HIVEP3 | 1119.35153 | 0.07826131 | 0.46214419 | 0.16934392 | 0.86552613 | 0.99989001 |
| FOXJ3 | 494.400739 | 0.2126897 | 0.1765171 | 1.20492407 | 0.22823262 | 0.99989001 |
| PPCS | 48.7561114 | 0.33383197 | 0.33198375 | 1.0055672 | 0.31462381 | NA |
| CCDC30 | 75.0328296 | 0.1592274 | 0.37917941 | 0.41992629 | 0.6745393 | 0.99989001 |
| YBX1 | 283.615245 | -0.3044193 | 0.39813638 | -0.7646106 | 0.44450343 | 0.99989001 |
| P3H1 | 45.6304081 | 0.10439677 | 0.32365348 | 0.32255723 | 0.74703059 | NA |
| SLC2A1 | 97.0239992 | -1.020469 | 0.58958163 | -1.7308357 | 0.08348106 | 0.9503998 |
| TIE1 | 439.704313 | -0.2260268 | 0.40593062 | -0.5568113 | 0.57765637 | 0.99989001 |
| ELOVL1 | 63.0288875 | -0.251423 | 0.41981107 | -0.5988956 | 0.54924251 | 0.99989001 |
| SZT2 | 91.2473195 | -0.0968049 | 0.29271082 | -0.3307187 | 0.74085697 | 0.99989001 |
| PTPRF | 44.9429354 | 0.67874598 | 0.39699064 | 1.7097279 | 0.0873162 | NA |
| ATP6V0B | 144.316349 | 0.16895696 | 0.38939278 | 0.43389854 | 0.66436212 | 0.99989001 |
| ERI3 | 162.330579 | 0.19752091 | 0.28693519 | 0.68838163 | 0.49121249 | 0.99989001 |
| RNF220 | 102.843758 | -0.0074761 | 0.42289351 | -0.0176783 | 0.98589547 | 0.99989001 |
| RPS8 | 509.547433 | -0.0766672 | 0.56570263 | -0.1355257 | 0.89219628 | 0.99989001 |
| EIF2B3 | 115.631245 | -0.0751592 | 0.39516541 | -0.1901967 | 0.849155 | 0.99989001 |
| ZSWIM5 | 65.6665274 | 0.30568293 | 0.3760013 | 0.8129837 | 0.41622741 | 0.99989001 |
| TESK2 | 99.0435628 | -0.0255048 | 0.30520323 | -0.0835665 | 0.93340109 | 0.99989001 |
| PRDX1 | 467.606121 | 0.07858922 | 0.56749047 | 0.13848553 | 0.88985671 | 0.99989001 |
| AKR1A1 | 58.9899372 | -0.123411 | 0.41600347 | -0.2966585 | 0.76672722 | 0.99989001 |
| NASP | 207.116046 | 0.121254 | 0.26685633 | 0.45437934 | 0.64955582 | 0.99989001 |
| GPBP1L1 | 411.614999 | 0.18177372 | 0.1897227 | 0.95810213 | 0.33801126 | 0.99989001 |
| IPP | 56.9915012 | -0.4721213 | 0.39099702 | -1.2074806 | 0.22724711 | 0.99989001 |
| MAST2 | 155.05868 | 0.15077791 | 0.20170856 | 0.74750376 | 0.45475953 | 0.99989001 |
| PIK3R3 | 183.541871 | -0.2267586 | 0.58242996 | -0.389332 | 0.69703057 | 0.99989001 |
| TSPAN1 | 51.0389101 | 0.45400544 | 0.34266411 | 1.32492848 | 0.18519487 | 0.99989001 |
| LRRC41 | 78.6793953 | -0.1969143 | 0.2766933 | -0.7116701 | 0.47666909 | 0.99989001 |
| UQCRH | 118.573033 | 0.11412416 | 0.42631559 | 0.26769878 | 0.78893119 | 0.99989001 |
| NSUN4 | 77.4206928 | -0.2630932 | 0.33457952 | -0.7863399 | 0.43166841 | 0.99989001 |
| MKNK1 | 73.0713959 | 0.16078514 | 0.20979957 | 0.76637499 | 0.44345321 | 0.99989001 |
| ATPAF1 | 67.7214922 | 0.00463719 | 0.32295372 | 0.01435869 | 0.98854381 | 0.99989001 |
| EFCAB14 | 212.584859 | -0.109831 | 0.29582437 | -0.371271 | 0.7104357 | 0.99989001 |
| PDZK1IP1 | 125.380197 | 1.77346251 | 0.6144589 | 2.8862183 | 0.00389902 | 0.31113256 |
| TAL1 | 103.710615 | 0.77310629 | 0.34116716 | 2.266063 | 0.02344753 | 0.60818532 |
| CMPK1 | 189.80345 | -0.100935 | 0.2521041 | -0.4003705 | 0.68888365 | 0.99989001 |
| TRABD2B | 106.251338 | 0.91960837 | 0.74389736 | 1.2362033 | 0.21638301 | 0.99989001 |
| SPATA6 | 293.36228 | 0.27024709 | 0.36625795 | 0.73786001 | 0.46059952 | 0.99989001 |
| FAF1 | 960.503442 | 0.06272304 | 0.21756946 | 0.28828971 | 0.77312498 | 0.99989001 |
| RNF11 | 175.655245 | 0.04841676 | 0.3245505 | 0.14918097 | 0.88141084 | 0.99989001 |
| EPS15 | 761.271231 | 0.03697408 | 0.19872932 | 0.18605249 | 0.8524036 | 0.99989001 |
| OSBPL9 | 336.556234 | -0.0416143 | 0.21117583 | -0.1970601 | 0.84378054 | 0.99989001 |
| NRDC | 399.495132 | 0.11271918 | 0.19360803 | 0.58220304 | 0.56042993 | 0.99989001 |
| TXNDC12 | 98.3850992 | -0.0706556 | 0.26893605 | -0.2627226 | 0.79276439 | 0.99989001 |
| BTF3L4 | 113.683238 | 0.06516655 | 0.20264673 | 0.32157712 | 0.74777308 | 0.99989001 |
| ZFYVE9 | 203.402539 | 0.14036363 | 0.16826036 | 0.83420496 | 0.4041655 | 0.99989001 |
| CC2D1B | 47.2737178 | -0.4468525 | 0.42521192 | -1.0508936 | 0.29330745 | NA |

| | | | | | | |
|---|---|---|---|---|---|---|
| PRPF38A | 74.3529408 | -0.0859921 | 0.23932691 | -0.3593081 | 0.71936461 | 0.99989001 |
| TUT4 | 402.618577 | -0.0322311 | 0.22697698 | -0.1420017 | 0.88707864 | 0.99989001 |
| ZYG11B | 170.929507 | -0.1622997 | 0.18701258 | -0.8678543 | 0.38547408 | 0.99989001 |
| ECHDC2 | 206.299288 | 0.52432002 | 0.40191275 | 1.30456181 | 0.1920421 | 0.99989001 |
| SCP2 | 219.949358 | 0.12925697 | 0.26214669 | 0.49307115 | 0.62196231 | 0.99989001 |
| C1orf123 | 40.2942303 | -0.3170918 | 0.38583607 | -0.8218304 | 0.41117341 | NA |
| MAGOH | 62.3178249 | -0.2567861 | 0.33235627 | -0.7726229 | 0.43974556 | 0.99989001 |
| NDC1 | 65.2207419 | -0.1367384 | 0.24737936 | -0.5527476 | 0.58043625 | 0.99989001 |
| YIPF1 | 52.7783384 | 0.3176114 | 0.32740345 | 0.9700918 | 0.33200074 | 0.99989001 |
| HSPB11 | 58.4211274 | 0.06476334 | 0.25739875 | 0.25160704 | 0.80134481 | 0.99989001 |
| TMEM59 | 299.495362 | 0.01733808 | 0.44847352 | 0.0386602 | 0.9691613 | 0.99989001 |
| SSBP3 | 442.753356 | 0.94846059 | 0.48697943 | 1.94763996 | 0.05145806 | 0.81256367 |
| USP24 | 412.159164 | -0.1713721 | 0.19583673 | -0.8750762 | 0.38153244 | 0.99989001 |
| AL603840.1 | 84.3580426 | -0.063064 | 1.01138387 | -0.0623542 | 0.9502808 | 0.99989001 |
| AC119674.2 | 144.804489 | 0.32509103 | 0.48529431 | 0.66988428 | 0.50293156 | 0.99989001 |
| PLPP3 | 1755.22323 | 0.2067255 | 0.46045614 | 0.44895806 | 0.65346191 | 0.99989001 |
| PRKAA2 | 49.9252799 | 0.17199112 | 0.32777566 | 0.52472207 | 0.59977641 | 0.99989001 |
| DAB1 | 62.3772183 | 0.79284139 | 0.65786082 | 1.20518106 | 0.22813341 | 0.99989001 |
| TACSTD2 | 47.5061765 | -2.0057673 | 0.7042159 | -2.8482278 | 0.00439634 | NA |
| MYSM1 | 213.420138 | 0.0168451 | 0.25703073 | 0.06553729 | 0.94774622 | 0.99989001 |
| AL136985.3 | 103.887606 | -2.4113698 | 0.64524375 | -3.7371456 | 0.00018612 | 0.06108011 |
| JUN | 476.703581 | -1.5421304 | 0.73567068 | -2.0962239 | 0.03606234 | 0.69794491 |
| FGGY | 190.22205 | -0.3177606 | 0.3567425 | -0.8907281 | 0.37307507 | 0.99989001 |
| NFIA | 1640.33663 | 0.42589714 | 0.24125162 | 1.7653649 | 0.07750246 | 0.92274869 |
| TM2D1 | 215.716125 | -0.1152054 | 0.19440523 | -0.5926043 | 0.55344602 | 0.99989001 |
| PATJ | 230.678572 | 0.03848129 | 0.36610844 | 0.105109 | 0.91628932 | 0.99989001 |
| USP1 | 57.9982478 | 0.33446006 | 0.32946677 | 1.01515566 | 0.31003163 | 0.99989001 |
| DOCK7 | 245.103941 | 0.18622623 | 0.20645546 | 0.90201648 | 0.36704811 | 0.99989001 |
| ATG4C | 133.374609 | 0.23234515 | 0.31950208 | 0.72721014 | 0.46709724 | 0.99989001 |
| ALG6 | 56.7926494 | -0.1537452 | 0.27326547 | -0.5626221 | 0.57369225 | 0.99989001 |
| ITGB3BP | 100.951626 | 0.19238457 | 0.3531829 | 0.54471654 | 0.58594849 | 0.99989001 |
| EFCAB7 | 45.7301143 | -0.2238724 | 0.24524587 | -0.9128487 | 0.36132214 | NA |
| CACHD1 | 460.566996 | -0.8936934 | 0.39395793 | -2.2684995 | 0.02329878 | 0.60818532 |
| RAVER2 | 95.7801903 | 0.64900841 | 0.51970706 | 1.2487966 | 0.21173948 | 0.99989001 |
| JAK1 | 646.152956 | 0.26774356 | 0.15057535 | 1.77813668 | 0.07538141 | 0.9206525 |
| AK4 | 117.479154 | 0.34368707 | 0.53588645 | 0.6413431 | 0.52129979 | 0.99989001 |
| LEPR | 467.323943 | -1.1004036 | 0.4922481 | -2.2354654 | 0.02538681 | 0.63387007 |
| PDE4B | 408.049202 | 1.44025863 | 1.12185567 | 1.28381811 | 0.19920561 | 0.99989001 |
| SGIP1 | 115.266807 | -1.9297972 | 0.62184442 | -3.103344 | 0.00191347 | 0.20835966 |
| TCTEX1D1 | 69.4929894 | -1.789736 | 0.71010487 | -2.5203827 | 1.17E-02 | 0.48516963 |
| WDR78 | 56.5037431 | 0.36451761 | 0.5278408 | 0.69058249 | 0.48982795 | 0.99989001 |
| MIER1 | 318.963365 | -0.0500843 | 0.21843806 | -0.2292836 | 0.81864849 | 0.99989001 |
| SLC35D1 | 72.0490302 | 0.62865987 | 0.23907431 | 2.62955841 | 0.00854958 | 0.42668269 |
| SERBP1 | 286.336029 | -0.0632522 | 0.31240012 | -0.2024719 | 0.83954782 | 0.99989001 |
| GADD45A | 79.996872 | 0.40229139 | 0.47987761 | 0.83832082 | 0.40185055 | 0.99989001 |
| GNG12 | 321.191719 | 0.1216542 | 0.20802484 | 0.58480614 | 0.55867807 | 0.99989001 |
| GNG12-AS1 | 84.9617027 | 0.09221517 | 0.36106839 | 0.2553953 | 0.7984178 | 0.99989001 |
| WLS | 114.027771 | 0.26318542 | 0.36269577 | 0.72563686 | 0.46806142 | 0.99989001 |
| LRRC40 | 110.993381 | -0.1089903 | 0.26905612 | -0.4050841 | 0.68541572 | 0.99989001 |
| SRSF11 | 496.0315 | 0.19546065 | 0.15559343 | 1.25622687 | 0.20903373 | 0.99989001 |
| ANKRD13C | 289.548309 | -0.097058 | 0.17875343 | -0.5429713 | 0.58714954 | 0.99989001 |
| PTGER3 | 49.9582433 | 0.43899678 | 0.60377091 | 0.72709165 | 0.46716982 | 0.99989001 |
| ZRANB2 | 369.57171 | 0.02165702 | 0.18931042 | 0.11439949 | 0.90892112 | 0.99989001 |
| TYW3 | 69.2370829 | -0.1071212 | 0.32820576 | -0.3263844 | 0.74413352 | 0.99989001 |

| | | | | | | |
|---|---|---|---|---|---|---|
| ACADM | 123.050632 | 0.27473363 | 0.21143657 | 1.29936665 | 0.19381813 | 0.99989001 |
| RABGGTB | 58.085089 | 0.04363001 | 0.4959855 | 0.08796631 | 0.92990345 | 0.99989001 |
| ST6GALNAC3 | 5633.99896 | -0.0139264 | 0.41318966 | -0.0337045 | 0.97311276 | 0.99989001 |
| PIGK | 100.465604 | -0.3907409 | 0.22161412 | -1.7631587 | 0.07787373 | 0.92274869 |
| AC118549.1 | 459.542341 | -0.1197033 | 0.18460148 | -0.6484417 | 0.5166993 | 0.99989001 |
| USP33 | 366.081676 | 0.44544051 | 0.17748299 | 2.50976447 | 0.01208117 | 0.49000287 |
| MIGA1 | 224.310378 | 0.08045405 | 0.21013129 | 0.38287514 | 0.70181235 | 0.99989001 |
| FUBP1 | 306.472114 | -0.1901902 | 0.21365041 | -0.8901932 | 0.37336214 | 0.99989001 |
| GIPC2 | 158.491864 | 0.34341121 | 0.36376846 | 0.94403791 | 0.34515027 | 0.99989001 |
| IFI44L | 616.117113 | -0.0700865 | 1.06530561 | -0.0657901 | 0.94754498 | 0.99989001 |
| IFI44 | 462.860121 | 0.0566111 | 0.51937609 | 0.10899828 | 0.91320386 | 0.99989001 |
| ADGRL4 | 1987.63584 | -0.1045672 | 0.27855966 | -0.3753853 | 0.70737396 | 0.99989001 |
| ADGRL2 | 1720.90676 | 0.51290033 | 0.54388401 | 0.94303256 | 0.34566425 | 0.99989001 |
| PRKACB | 145.486604 | -0.3025906 | 0.32002801 | -0.9455128 | 0.34439713 | 0.99989001 |
| GNG5 | 196.189055 | -0.4923679 | 0.48613791 | -1.0128154 | 0.31114836 | 0.99989001 |
| SPATA1 | 59.363325 | 0.13723769 | 0.59734213 | 0.22974722 | 0.8182882 | 0.99989001 |
| CTBS | 111.551888 | -0.0735676 | 0.38286029 | -0.1921525 | 0.84762275 | 0.99989001 |
| C1orf52 | 83.2068732 | -0.1052211 | 0.30774907 | -0.3419056 | 0.73242191 | 0.99989001 |
| BCL10 | 51.4567977 | -0.0766684 | 0.30706963 | -0.2496775 | 0.80283678 | 0.99989001 |
| AL078459.1 | 58.5311466 | 0.18646617 | 0.54228914 | 0.34385008 | 0.73095903 | 0.99989001 |
| DDAH1 | 224.024494 | 0.37486988 | 0.49626667 | 0.75537993 | 0.45002105 | 0.99989001 |
| CYR61 | 175.479768 | -0.9455362 | 0.94985863 | -0.9954494 | 0.31951775 | 0.99989001 |
| ZNHIT6 | 170.049107 | -0.0227392 | 0.28691478 | -0.079254 | 0.93683056 | 0.99989001 |
| ODF2L | 520.099047 | -0.045736 | 0.33527918 | -0.1364117 | 0.89149583 | 0.99989001 |
| SH3GLB1 | 348.342115 | 0.1020633 | 0.17462699 | 0.58446465 | 0.55890774 | 0.99989001 |
| SELENOF | 240.060575 | -0.0412423 | 0.27048972 | -0.1524725 | 0.87881425 | 0.99989001 |
| HS2ST1 | 191.581491 | -0.0350676 | 0.18779945 | -0.1867291 | 0.85187308 | 0.99989001 |
| LMO4 | 86.9247043 | 0.26143683 | 0.31186889 | 0.83829083 | 0.40186738 | 0.99989001 |
| PKN2 | 579.463196 | -0.0367622 | 0.14375857 | -0.2557218 | 0.79816568 | 0.99989001 |
| GTF2B | 103.372456 | 0.02100209 | 0.26926832 | 0.07799688 | 0.93783053 | 0.99989001 |
| KYAT3 | 100.892102 | 0.34890169 | 0.31692114 | 1.10091011 | 0.27093578 | 0.99989001 |
| GBP3 | 83.242309 | 0.08695657 | 0.32380447 | 0.26854652 | 0.78827867 | 0.99989001 |
| GBP1 | 119.157273 | 0.43625282 | 0.70209926 | 0.62135491 | 0.53436613 | 0.99989001 |
| GBP2 | 161.898998 | -0.1922412 | 0.36519087 | -0.526413 | 0.59860127 | 0.99989001 |
| GBP4 | 358.522258 | 0.33775926 | 0.57504375 | 0.58736272 | 0.55696012 | 0.99989001 |
| LRRC8B | 131.478281 | -0.0035204 | 0.31100019 | -0.0113195 | 0.99096857 | 0.99989001 |
| LRRC8C-DT | 46.9427583 | 0.0953588 | 0.34927042 | 0.27302284 | 0.78483566 | NA |
| ZNF326 | 123.905015 | -0.0174944 | 0.23612642 | -0.074089 | 0.94093959 | 0.99989001 |
| ZNF644 | 537.836842 | 0.11660545 | 0.23369686 | 0.49896026 | 0.61780738 | 0.99989001 |
| TGFBR3 | 933.564348 | -0.4059353 | 0.51112652 | -0.7941974 | 0.42708055 | 0.99989001 |
| BTBD8 | 55.6299258 | 0.11510771 | 0.50265114 | 0.2290012 | 0.81886798 | 0.99989001 |
| GLMN | 90.2182153 | 0.00621992 | 0.24693383 | 0.0251886 | 0.97990453 | 0.99989001 |
| RPAP2 | 228.5937 | -0.0668861 | 0.20469322 | -0.3267625 | 0.74384756 | 0.99989001 |
| EVI5 | 405.919873 | 0.43044174 | 0.20713165 | 2.07810708 | 0.0376995 | 0.70738774 |
| RPL5 | 271.664485 | 0.10083664 | 0.34348976 | 0.29356518 | 0.76909018 | 0.99989001 |
| FAM69A | 51.8741188 | 0.54159819 | 0.28799382 | 1.8805896 | 0.06002777 | 0.85885114 |
| MTF2 | 266.528466 | 0.05419416 | 0.22867257 | 0.23699457 | 0.812661 | 0.99989001 |
| TMED5 | 123.57911 | 0.10896034 | 0.29874328 | 0.364729 | 0.71531371 | 0.99989001 |
| DR1 | 85.4906739 | 0.12703749 | 0.24049382 | 0.52823599 | 0.59733555 | 0.99989001 |
| FNBP1L | 465.822106 | 0.25692936 | 0.36711475 | 0.69986118 | 0.484014 | 0.99989001 |
| BCAR3 | 325.199044 | 0.25526218 | 0.34156783 | 0.747325 | 0.4548674 | 0.99989001 |
| GCLM | 93.1202317 | 0.49143088 | 0.43892442 | 1.11962531 | 0.26287347 | 0.99989001 |
| ARHGAP29 | 1680.68869 | -0.3265749 | 0.22802757 | -1.4321728 | 0.15209437 | 0.99989001 |
| ABCD3 | 177.430144 | 0.06933601 | 0.21075022 | 0.32899613 | 0.74215862 | 0.99989001 |

| | | | | | | |
|---|---|---|---|---|---|---|
| CNN3 | 867.617674 | -0.5310258 | 0.52336015 | -1.0146469 | 0.31027417 | 0.99989001 |
| AC105942.1 | 54.337145 | -0.7904343 | 0.36728658 | -2.1520915 | 0.03139015 | 0.67281094 |
| ALG14 | 78.5428407 | -0.3387807 | 0.21564191 | -1.5710335 | 0.11617487 | 0.99989001 |
| PTBP2 | 660.336701 | -0.025516 | 0.20683238 | -0.1233654 | 0.90181773 | 0.99989001 |
| DPYD | 1482.20486 | -0.3686204 | 0.29006082 | -1.2708384 | 0.20378616 | 0.99989001 |
| DPYD-AS1 | 65.8746705 | -0.1583594 | 0.73039743 | -0.2168126 | 0.82835438 | 0.99989001 |
| SNX7 | 83.4707764 | 0.2596033 | 0.26133619 | 0.99336912 | 0.32053011 | 0.99989001 |
| PALMD | 634.550133 | -1.0035949 | 0.47914009 | -2.094575 | 0.03620879 | 0.69898709 |
| AGL | 73.530609 | -0.1700169 | 0.3383826 | -0.5024398 | 0.61535822 | 0.99989001 |
| TRMT13 | 89.7238336 | -0.0278172 | 0.19790498 | -0.1405586 | 0.88821866 | 0.99989001 |
| DBT | 79.2686249 | 0.08391684 | 0.23494886 | 0.35717066 | 0.72096405 | 0.99989001 |
| RTCA | 94.7992049 | 0.27146938 | 0.3042158 | 0.89235792 | 0.37220112 | 0.99989001 |
| CDC14A | 465.250489 | 0.13715179 | 0.22092139 | 0.62081716 | 0.53471993 | 0.99989001 |
| VCAM1 | 819.720728 | -0.3163989 | 0.75206992 | -0.4207041 | 0.67397119 | 0.99989001 |
| SLC30A7 | 371.204454 | 0.10592124 | 0.33370924 | 0.31740578 | 0.75093572 | 0.99989001 |
| AC093157.1 | 87.0461698 | 0.34310135 | 0.25468903 | 1.34713831 | 0.17793569 | 0.99989001 |
| S1PR1 | 281.661035 | -0.1353727 | 0.51406788 | -0.2633363 | 0.79229138 | 0.99989001 |
| RNPC3 | 147.560857 | 0.04641849 | 0.28991164 | 0.16011253 | 0.87279243 | 0.99989001 |
| VAV3 | 478.204337 | 1.59658125 | 0.33748255 | 4.73085566 | 2.24E-06 | 0.00222857 |
| SLC25A24 | 141.93815 | 0.3524116 | 0.32129777 | 1.09683798 | 0.27271222 | 0.99989001 |
| FAM102B | 61.9324208 | 0.70503494 | 0.47520686 | 1.48363798 | 0.13790499 | 0.99989001 |
| PRPF38B | 134.294929 | -0.0463173 | 0.3278551 | -0.1412738 | 0.88765365 | 0.99989001 |
| STXBP3 | 180.036322 | 0.04534853 | 0.23646579 | 0.19177627 | 0.84791746 | 0.99989001 |
| WDR47 | 80.2746491 | -0.0387369 | 0.26279515 | -0.1474033 | 0.88281373 | 0.99989001 |
| TAF13 | 59.6976506 | 0.03119146 | 0.34061702 | 0.0915734 | 0.92703699 | 0.99989001 |
| TMEM167B | 55.1397254 | -0.3012602 | 0.2751401 | -1.0949339 | 0.27354559 | 0.99989001 |
| SARS | 127.732941 | 0.18226112 | 0.2953665 | 0.61706768 | 0.53719009 | 0.99989001 |
| SORT1 | 111.275914 | -0.602537 | 0.48196146 | -1.2501768 | 0.21123497 | 0.99989001 |
| PSMA5 | 128.597689 | -0.497304 | 0.30217305 | -1.645759 | 0.09981338 | 0.99989001 |
| GNAI3 | 185.591667 | 0.10186427 | 0.19953837 | 0.51049968 | 0.60970144 | 0.99989001 |
| CSF1 | 72.038776 | 0.32466524 | 0.50196639 | 0.6467868 | 0.51776994 | 0.99989001 |
| AHCYL1 | 187.358235 | 0.54674214 | 0.22219897 | 2.4605971 | 0.0138706 | 0.50652625 |
| STRIP1 | 68.5051265 | -0.1144712 | 0.23536914 | -0.4863475 | 0.62672078 | 0.99989001 |
| KCNC4 | 59.7445835 | -0.1879945 | 0.40030082 | -0.4696332 | 0.63861712 | 0.99989001 |
| RBM15 | 51.1009325 | -0.6187011 | 0.28673813 | -2.1577218 | 0.03094947 | 0.67281094 |
| SLC16A4 | 52.66518 | 0.23574606 | 0.323186 | 0.72944391 | 0.46573016 | 0.99989001 |
| LAMTOR5 | 156.229774 | 0.01356487 | 0.41711601 | 0.03252061 | 0.97405688 | 0.99989001 |
| CD53 | 117.171064 | -1.4188035 | 0.65726822 | -2.158637 | 0.03087834 | 0.67281094 |
| DRAM2 | 102.334104 | 0.33322422 | 0.2606697 | 1.27833888 | 0.20112997 | 0.99989001 |
| CEPT1 | 126.880005 | 0.37035675 | 0.27467681 | 1.34833642 | 0.1775502 | 0.99989001 |
| ATP5PB | 130.790241 | 0.25712618 | 0.34866484 | 0.73745944 | 0.460843 | 0.99989001 |
| RAP1A | 602.61659 | 0.40284825 | 0.21799711 | 1.84795231 | 0.06460924 | 0.86610729 |
| AL445426.1 | 62.5634204 | 0.34407857 | 0.76232858 | 0.45135205 | 0.65173583 | 0.99989001 |
| CTTNBP2NL | 554.147194 | 0.43439606 | 0.30030588 | 1.446512 | 0.14803365 | 0.99989001 |
| WNT2B | 197.900444 | 0.75017701 | 0.50273429 | 1.49219385 | 0.13564833 | 0.99989001 |
| ST7L | 191.587959 | -0.1057246 | 0.22111352 | -0.478146 | 0.63254627 | 0.99989001 |
| CAPZA1 | 337.562117 | 0.01215471 | 0.22050482 | 0.0551222 | 0.95604111 | 0.99989001 |
| MOV10 | 117.632053 | -0.4650687 | 0.30998866 | -1.5002765 | 0.1335428 | 0.99989001 |
| RHOC | 378.368932 | -0.1762512 | 0.5006212 | -0.352065 | 0.72478953 | 0.99989001 |
| SLC16A1-AS1 | 140.590622 | 0.24179912 | 0.38404384 | 0.62961333 | 0.5289476 | 0.99989001 |
| LRIG2 | 127.634955 | -0.0946266 | 0.18377337 | -0.5149093 | 0.60661639 | 0.99989001 |
| MAGI3 | 551.887863 | 0.20152725 | 0.27641824 | 0.7290664 | 0.46596105 | 0.99989001 |
| PHTF1 | 118.271992 | 0.16560628 | 0.28447736 | 0.58214221 | 0.5604709 | 0.99989001 |
| RSBN1 | 132.923857 | 0.06948794 | 0.20248054 | 0.3431833 | 0.73146057 | 0.99989001 |

| | | | | | | |
|---|---|---|---|---|---|---|
| HIPK1 | 128.162795 | 0.05138624 | 0.18969338 | 0.27089108 | 0.7864748 | 0.99989001 |
| TRIM33 | 344.126812 | 0.01889638 | 0.21711639 | 0.08703342 | 0.93064494 | 0.99989001 |
| BCAS2 | 56.4232914 | -0.1781935 | 0.32402824 | -0.5499319 | 0.58236611 | 0.99989001 |
| DENND2C | 76.7401478 | -0.5448421 | 0.41085442 | -1.3261197 | 0.18480006 | 0.99989001 |
| NRAS | 58.1396995 | -0.2349488 | 0.30574575 | -0.768445 | 0.44222288 | 0.99989001 |
| CSDE1 | 366.63978 | -0.0621761 | 0.18431626 | -0.3373339 | 0.73586523 | 0.99989001 |
| SIKE1 | 50.6010673 | -0.1203048 | 0.46236091 | -0.2601966 | 0.7947121 | 0.99989001 |
| TSPAN2 | 55.462722 | 0.00706281 | 0.6390722 | 0.01105165 | 0.99118223 | 0.99989001 |
| VANGL1 | 63.0695275 | 0.26791274 | 0.35909933 | 0.74606861 | 0.45562597 | 0.99989001 |
| ATP1A1 | 677.745888 | -0.1972347 | 0.39422018 | -0.5003161 | 0.61685254 | 0.99989001 |
| CD58 | 193.108065 | 0.67459755 | 0.25484272 | 2.6471133 | 0.00811821 | 0.41229803 |
| TTF2 | 45.1640183 | -0.5347817 | 0.39282457 | -1.3613754 | 0.17339507 | NA |
| MAN1A2 | 353.677524 | 0.24756729 | 0.14828099 | 1.66958216 | 0.09500206 | 0.99989001 |
| GDAP2 | 117.169827 | 0.29420537 | 0.20573877 | 1.42999481 | 0.15271851 | 0.99989001 |
| WARS2 | 63.8547371 | -0.118687 | 0.38064678 | -0.3118036 | 0.75518982 | 0.99989001 |
| AL359915.2 | 149.067363 | -0.4269428 | 0.42005188 | -1.0164048 | 0.30943665 | 0.99989001 |
| SEC22B | 135.8289 | -0.0772487 | 0.29938658 | -0.2580231 | 0.79638908 | 0.99989001 |
| LINC00623 | 165.082992 | 0.14711725 | 0.37005224 | 0.39755806 | 0.69095598 | 0.99989001 |
| SRGAP2C | 156.475457 | 0.27008296 | 0.23378743 | 1.15525017 | 0.247988 | 0.99989001 |
| AC239800.3 | 37.6721239 | -0.5774043 | 0.48515208 | -1.1901512 | 0.23398696 | NA |
| NBPF15 | 187.93481 | 0.65585139 | 0.39879649 | 1.64457666 | 0.10005714 | 0.99989001 |
| FP700111.1 | 92.2593208 | -0.5276425 | 0.40498043 | -1.302884 | 0.19261438 | 0.99989001 |
| SRGAP2B | 205.230137 | 0.18949394 | 0.24768675 | 0.76505484 | 0.44423889 | 0.99989001 |
| RNF115 | 437.482971 | -0.0132843 | 0.17969536 | -0.0739266 | 0.94106877 | 0.99989001 |
| POLR3C | 52.3575218 | -0.2419144 | 0.29783451 | -0.8122444 | 0.41665138 | 0.99989001 |
| ITGA10 | 56.9782184 | 0.08159161 | 0.60822905 | 0.13414619 | 0.89328698 | 0.99989001 |
| RBM8A | 112.322663 | -0.0700511 | 0.32167334 | -0.2177709 | 0.82760762 | 0.99989001 |
| LIX1L | 102.391007 | -0.2853692 | 0.26824322 | -1.0638448 | 0.28739901 | 0.99989001 |
| TXNIP | 1544.22815 | 0.95386573 | 0.60676113 | 1.57206136 | 0.11593632 | 0.99989001 |
| NBPF12 | 57.0365272 | 0.29004057 | 0.34648636 | 0.83709087 | 0.40254149 | 0.99989001 |
| PRKAB2 | 40.4735685 | 0.20955393 | 0.2822268 | 0.74250189 | 0.45778331 | NA |
| AC242426.2 | 102.691837 | -0.4254412 | 0.30823428 | -1.3802528 | 0.16750883 | 0.99989001 |
| CHD1L | 113.243928 | 0.36383882 | 0.22605206 | 1.60953552 | 0.10749929 | 0.99989001 |
| BCL9 | 62.8538705 | 0.8230856 | 0.36409908 | 2.26060882 | 0.02378349 | 0.61268871 |
| GJA5 | 352.425029 | 0.66023296 | 0.72642483 | 0.90887995 | 0.3634135 | 0.99989001 |
| NBPF11 | 64.7846905 | -0.1621299 | 0.30761354 | -0.5270571 | 0.59815396 | 0.99989001 |
| PDE4DIP | 52.1244373 | 0.23499188 | 0.34377281 | 0.68356737 | 0.4942484 | 0.99989001 |
| NBPF9 | 50.6317407 | 0.37192443 | 0.50432828 | 0.73746495 | 0.46083965 | 0.99989001 |
| AC245297.3 | 89.6081264 | 0.37899521 | 0.40767545 | 0.92964932 | 0.35255268 | 0.99989001 |
| OTUD7B | 186.302997 | 0.04840012 | 0.29590334 | 0.16356735 | 0.87007175 | 0.99989001 |
| VPS45 | 253.816167 | -0.1192026 | 0.19633906 | -0.6071264 | 0.54376702 | 0.99989001 |
| PLEKHO1 | 161.557309 | -0.5351637 | 0.48786066 | -1.0969601 | 0.27265885 | 0.99989001 |
| ANP32E | 74.6515432 | 0.04529459 | 0.24593066 | 0.18417625 | 0.85387519 | 0.99989001 |
| APH1A | 97.4163188 | 0.06003865 | 0.41908815 | 0.1432602 | 0.88608469 | 0.99989001 |
| MRPS21 | 125.263924 | -0.1761446 | 0.30376906 | -0.5798635 | 0.5620067 | 0.99989001 |
| PRPF3 | 170.559658 | -0.2868346 | 0.30210912 | -0.9494402 | 0.34239675 | 0.99989001 |
| RPRD2 | 349.034455 | 0.07524187 | 0.21627882 | 0.34789293 | 0.72792059 | 0.99989001 |
| MCL1 | 257.797323 | -0.2461322 | 0.4729159 | -0.5204567 | 0.60274531 | 0.99989001 |
| ENSA | 284.572049 | -0.2207671 | 0.21255171 | -1.0386513 | 0.29896692 | 0.99989001 |
| GOLPH3L | 111.487486 | -0.1164432 | 0.21161552 | -0.5502582 | 0.58214228 | 0.99989001 |
| CTSS | 79.9608966 | -0.27888 | 0.39948615 | -0.6980969 | 0.48511662 | 0.99989001 |
| ARNT | 181.174013 | 0.20402301 | 0.19912425 | 1.02460149 | 0.30555126 | 0.99989001 |
| SETDB1 | 110.886481 | -0.2711804 | 0.30703516 | -0.8832227 | 0.37711594 | 0.99989001 |
| CERS2 | 81.0088689 | 0.08592693 | 0.34934657 | 0.2459647 | 0.80570956 | 0.99989001 |

| | | | | | | |
|---|---|---|---|---|---|---|
| C1orf56 | 46.8329823 | -0.4786849 | 0.66709898 | -0.717562 | 0.47302737 | NA |
| CDC42SE1 | 129.100414 | 0.41205733 | 0.2718764 | 1.51560534 | 0.12961918 | 0.99989001 |
| GABPB2 | 88.9935671 | 0.406613 | 0.4524747 | 0.89864252 | 0.3688431 | 0.99989001 |
| PIP5K1A | 273.325831 | -0.2771025 | 0.17386599 | -1.5937706 | 0.11098742 | 0.99989001 |
| PSMD4 | 122.564313 | 0.14180624 | 0.26529452 | 0.53452383 | 0.59297917 | 0.99989001 |
| PI4KB | 108.636095 | 0.04165811 | 0.21224203 | 0.19627643 | 0.84439381 | 0.99989001 |
| PSMB4 | 86.4290713 | 0.06096147 | 0.52250746 | 0.11667101 | 0.90712077 | 0.99989001 |
| POGZ | 509.644939 | 0.05523642 | 0.20750003 | 0.26619959 | 0.7900855 | 0.99989001 |
| SNX27 | 133.44988 | -0.119714 | 0.20073205 | -0.5963872 | 0.55091657 | 0.99989001 |
| THEM4 | 65.1376925 | 0.23008082 | 0.44431096 | 0.51783738 | 0.60457173 | 0.99989001 |
| S100A10 | 465.625157 | -0.0548962 | 0.56171518 | -0.0977296 | 0.92214704 | 0.99989001 |
| S100A11 | 698.126896 | -0.567181 | 0.51834258 | -1.0942203 | 0.27385838 | 0.99989001 |
| S100A6 | 1663.94738 | -1.1002421 | 0.71823617 | -1.5318667 | 0.12555533 | 0.99989001 |
| S100A4 | 440.027894 | -2.4285378 | 1.25209387 | -1.9395812 | 0.0524306 | 0.81270649 |
| S100A16 | 250.029191 | -0.367856 | 0.46733454 | -0.7871363 | 0.43120209 | 0.99989001 |
| S100A13 | 148.500977 | 0.29499556 | 0.47582606 | 0.61996512 | 0.53528075 | 0.99989001 |
| CHTOP | 105.18355 | -0.0134698 | 0.18303791 | -0.0735902 | 0.94133651 | 0.99989001 |
| ILF2 | 101.874054 | 0.50570923 | 0.3335103 | 1.5163227 | 0.12943777 | 0.99989001 |
| NPR1 | 242.72297 | 0.39827272 | 0.35951912 | 1.10779287 | 0.26795128 | 0.99989001 |
| INTS3 | 175.024589 | -0.0546023 | 0.29772058 | -0.1834011 | 0.85448333 | 0.99989001 |
| SLC27A3 | 61.5571514 | 0.42600306 | 0.32159258 | 1.32466697 | 0.18528163 | 0.99989001 |
| GATAD2B | 538.608514 | 0.19778457 | 0.23849146 | 0.8293151 | 0.40692613 | 0.99989001 |
| DENND4B | 108.827907 | -0.3595061 | 0.22907444 | -1.5693853 | 0.11655818 | 0.99989001 |
| CRTC2 | 41.7636596 | 0.02460154 | 0.31716768 | 0.07756635 | 0.93817301 | NA |
| SLC39A1 | 96.8353502 | -0.1675289 | 0.47929917 | -0.3495289 | 0.72669231 | 0.99989001 |
| JTB | 94.4747553 | -0.019092 | 0.46767544 | -0.0408232 | 0.96743683 | 0.99989001 |
| RAB13 | 143.478227 | 0.13819335 | 0.44716723 | 0.30904178 | 0.75728975 | 0.99989001 |
| RPS27 | 109.176373 | -0.0006366 | 0.54679653 | -0.0011641 | 0.99907115 | 0.99989001 |
| TPM3 | 356.378003 | -0.0207171 | 0.27213769 | -0.0761273 | 0.93931782 | 0.99989001 |
| C1orf43 | 139.23385 | -0.203892 | 0.23633705 | -0.862717 | 0.38829311 | 0.99989001 |
| UBAP2L | 157.574199 | -0.0951358 | 0.23903478 | -0.3979999 | 0.69063027 | 0.99989001 |
| HAX1 | 42.0020727 | -0.1475233 | 0.41550148 | -0.3550487 | 0.72255307 | NA |
| ATP8B2 | 66.9665953 | 0.4728129 | 0.31875262 | 1.48332243 | 0.13798877 | 0.99989001 |
| SHE | 426.476289 | 0.03903239 | 0.39802984 | 0.09806398 | 0.92188149 | 0.99989001 |
| TDRD10 | 140.72775 | 0.51188402 | 0.26566956 | 1.92676958 | 0.05400835 | 0.81854801 |
| UBE2Q1 | 74.6065343 | -0.302338 | 0.32382125 | -0.9336571 | 0.35048078 | 0.99989001 |
| ADAR | 364.87772 | -0.2362765 | 0.35731052 | -0.6612638 | 0.50844315 | 0.99989001 |
| KCNN3 | 304.33765 | -0.2488674 | 0.42558872 | -0.5847603 | 0.55870892 | 0.99989001 |
| ADAM15 | 352.890148 | -0.6417724 | 0.42023251 | -1.527184 | 0.12671526 | 0.99989001 |
| EFNA1 | 249.889383 | -0.1166577 | 0.34062326 | -0.342483 | 0.73198745 | 0.99989001 |
| SLC50A1 | 40.5237579 | -0.1620041 | 0.26138642 | -0.6197877 | 0.53539757 | NA |
| DPM3 | 57.6538605 | -0.264465 | 0.49512779 | -0.5341349 | 0.59324819 | 0.99989001 |
| SCAMP3 | 77.0445875 | -0.1799203 | 0.32139138 | -0.5598168 | 0.57560443 | 0.99989001 |
| CLK2 | 47.8997308 | 0.2548223 | 0.25623672 | 0.99448005 | 0.31998921 | NA |
| FDPS | 268.306868 | 0.13175066 | 0.33035134 | 0.39881981 | 0.69002598 | 0.99989001 |
| ASH1L | 931.037478 | 0.05816389 | 0.18092779 | 0.32147569 | 0.74784994 | 0.99989001 |
| YY1AP1 | 179.324468 | 0.0561214 | 0.20711585 | 0.27096622 | 0.78641701 | 0.99989001 |
| DAP3 | 186.457901 | 0.00324955 | 0.29581937 | 0.01098492 | 0.99123547 | 0.99989001 |
| GON4L | 258.720456 | -0.0268159 | 0.25444134 | -0.1053914 | 0.91606525 | 0.99989001 |
| SYT11 | 82.4981839 | 0.06165383 | 0.44448082 | 0.13870977 | 0.8896795 | 0.99989001 |
| RIT1 | 87.8924695 | -0.3205079 | 0.35185977 | -0.9108967 | 0.36234978 | 0.99989001 |
| KHDC4 | 157.580588 | 0.05018395 | 0.1874078 | 0.26777941 | 0.78886912 | 0.99989001 |
| ARHGEF2 | 217.005756 | -0.1285971 | 0.18479807 | -0.6958792 | 0.48650447 | 0.99989001 |
| SSR2 | 151.940214 | -0.0011429 | 0.48432248 | -0.0023597 | 0.99811719 | 0.99989001 |

| | | | | | | |
|---|---|---|---|---|---|---|
| UBQLN4 | 42.6957704 | 0.48692039 | 0.32407887 | 1.50247498 | 0.13297449 | NA |
| LAMTOR2 | 49.9646935 | -0.2354507 | 0.42647526 | -0.5520851 | 0.58089003 | 0.99989001 |
| LMNA | 286.802005 | -0.0475147 | 0.45628886 | -0.1041329 | 0.91706391 | 0.99989001 |
| SMG5 | 85.6847905 | -0.1239445 | 0.25175546 | -0.4923211 | 0.62249236 | 0.99989001 |
| CCT3 | 153.904987 | 0.14634628 | 0.36509876 | 0.40084026 | 0.68853774 | 0.99989001 |
| MEF2D | 131.593723 | 0.19599416 | 0.27733877 | 0.70669586 | 0.4797555 | 0.99989001 |
| NES | 203.039853 | -0.4998661 | 0.455455 | -1.0975092 | 0.27241886 | 0.99989001 |
| HDGF | 121.505487 | -0.0533053 | 0.36072364 | -0.1477733 | 0.88252172 | 0.99989001 |
| PRCC | 86.729746 | -0.1141641 | 0.26674431 | -0.4279909 | 0.66865772 | 0.99989001 |
| PEAR1 | 87.1655187 | 0.07591747 | 0.23613058 | 0.3215063 | 0.74782674 | 0.99989001 |
| ARHGEF11 | 185.508512 | 0.13884714 | 0.28928605 | 0.47996487 | 0.63125238 | 0.99989001 |
| KIRREL1 | 145.005467 | 0.14617819 | 0.44110179 | 0.33139334 | 0.7403474 | 0.99989001 |
| IFI16 | 743.04559 | -0.1765912 | 0.2112658 | -0.8358724 | 0.4032267 | 0.99989001 |
| ACKR1 | 751.083754 | -0.7714123 | 1.32455126 | -0.5823952 | 0.56030051 | 0.99989001 |
| DUSP23 | 162.313452 | 0.18414216 | 0.43232843 | 0.42593119 | 6.70E-01 | 0.99989001 |
| TAGLN2 | 548.023441 | -0.0721686 | 0.52866837 | -0.1365101 | 0.89141801 | 0.99989001 |
| PEA15 | 202.493271 | -0.2558957 | 0.41155367 | -0.6217797 | 0.53408672 | 0.99989001 |
| DCAF8 | 49.9514586 | -0.2106576 | 0.30029867 | -0.7014937 | 0.48299494 | 0.99989001 |
| COPA | 336.376257 | -0.0768229 | 0.23757447 | -0.3233634 | 0.74642005 | 0.99989001 |
| NCSTN | 68.5932408 | 0.14967978 | 0.29409231 | 0.50895509 | 0.61078371 | 0.99989001 |
| F11R | 136.363063 | -0.4681674 | 0.3192691 | -1.4663724 | 0.14254685 | 0.99989001 |
| USF1 | 40.9490123 | -0.7080158 | 0.44333152 | -1.5970346 | 0.110258 | NA |
| PFDN2 | 129.989425 | 0.0187088 | 0.43359592 | 0.04314801 | 0.96558355 | 0.99989001 |
| NIT1 | 101.281215 | -0.2791149 | 0.31990926 | -0.8724814 | 0.38294584 | 0.99989001 |
| DEDD | 63.7019262 | -0.311584 | 0.2927347 | -1.0643903 | 0.28715191 | 0.99989001 |
| UFC1 | 76.1570252 | -0.1333014 | 0.58133612 | -0.2293017 | 0.81863441 | 0.99989001 |
| B4GALT3 | 38.9893113 | -0.0138011 | 0.50102651 | -0.0275457 | 0.9780245 | NA |
| ADAMTS4 | 178.195367 | -1.1093965 | 1.36464773 | -0.8129545 | 0.41624413 | 0.99989001 |
| NDUFS2 | 131.24935 | -0.2982538 | 0.46944309 | -0.6353355 | 0.52520965 | 0.99989001 |
| SDHC | 131.873871 | 0.23820118 | 0.19402452 | 1.22768598 | 0.21956487 | 0.99989001 |
| AL592295.4 | 45.7847901 | -0.0796398 | 0.37012354 | -0.215171 | 0.829634 | NA |
| ATF6 | 543.90447 | 0.10089469 | 0.3179028 | 0.31737592 | 0.75095838 | 0.99989001 |
| NOS1AP | 190.107604 | -0.4669278 | 0.31051196 | -1.5037353 | 0.13264954 | 0.99989001 |
| UHMK1 | 172.951727 | -0.3794669 | 0.19591588 | -1.9368869 | 0.05275917 | 0.81270649 |
| UAP1 | 58.6507985 | -0.4232588 | 0.33744566 | -1.2543022 | 0.20973217 | 0.99989001 |
| PBX1 | 4179.25664 | 0.42728289 | 0.32202517 | 1.32686176 | 0.18455441 | 0.99989001 |
| MGST3 | 332.166991 | -0.0778335 | 0.40110018 | -0.1940501 | 0.84613668 | 0.99989001 |
| ALDH9A1 | 79.9833063 | 0.6682957 | 0.27925381 | 2.39314801 | 0.0167045 | 0.55059202 |
| TMCO1 | 140.660215 | 0.03185785 | 0.23489199 | 0.13562767 | 0.89211563 | 0.99989001 |
| UCK2 | 133.763691 | 0.39670437 | 0.50226071 | 0.78983755 | 0.42962264 | 0.99989001 |
| POGK | 61.0749392 | -0.1831228 | 0.28901902 | -0.6336013 | 0.52634107 | 0.99989001 |
| POU2F1 | 510.165475 | 0.09911777 | 0.20178976 | 0.49119326 | 0.62328977 | 0.99989001 |
| CREG1 | 50.4064424 | 0.20755191 | 0.34799453 | 0.59642291 | 0.55089274 | 0.99989001 |
| RCSD1 | 136.457903 | -0.1711421 | 0.70919989 | -0.2413171 | 0.80930933 | 0.99989001 |
| MPZL1 | 232.742953 | 0.31672909 | 0.33880361 | 0.93484566 | 0.34986784 | 0.99989001 |
| MPC2 | 65.5931199 | 0.39770619 | 0.36744659 | 1.08235102 | 0.27909658 | 0.99989001 |
| DCAF6 | 379.501202 | -0.0155484 | 0.22731701 | -0.0683997 | 0.94546745 | 0.99989001 |
| TIPRL | 106.889206 | 0.15697637 | 0.23365546 | 0.67182836 | 0.50169297 | 0.99989001 |
| SFT2D2 | 209.292021 | -0.132721 | 0.2131316 | -0.6227186 | 0.53346943 | 0.99989001 |
| TBX19 | 63.3666631 | -0.3402965 | 0.44904673 | -0.7578199 | 0.44855881 | 0.99989001 |
| ATP1B1 | 354.377483 | -0.2700958 | 0.5461835 | -0.4945148 | 0.62094268 | 0.99989001 |
| NME7 | 207.988489 | -0.1222074 | 0.25109031 | -0.4867068 | 0.62646613 | 0.99989001 |
| BLZF1 | 97.3755309 | 0.19053248 | 0.37875037 | 0.50305556 | 0.61492521 | 0.99989001 |
| SELP | 176.049028 | -0.3091373 | 1.11030717 | -0.278425 | 0.78068614 | 0.99989001 |

| | | | | | | |
|---|---|---|---|---|---|---|
| SELE | 353.568081 | 1.80617045 | 1.93641789 | 0.93273795 | 0.35095529 | 0.99989001 |
| SCYL3 | 63.8193327 | 0.3485828 | 0.24915787 | 1.39904391 | 0.16179982 | 0.99989001 |
| KIFAP3 | 195.357856 | 0.11775052 | 0.25672773 | 0.45865913 | 0.64647897 | 0.99989001 |
| FMO2 | 169.146105 | -1.7433273 | 1.05160115 | -1.6577838 | 0.09736112 | 0.99989001 |
| PRRC2C | 796.59864 | 0.07622691 | 0.26729897 | 0.28517472 | 0.77551029 | 0.99989001 |
| VAMP4 | 95.1548253 | 0.084439 | 0.26265939 | 0.32147719 | 0.7478488 | 0.99989001 |
| DNM3 | 1578.53473 | -0.1556723 | 0.38796915 | -0.4012493 | 0.68823662 | 0.99989001 |
| DNM3OS | 65.744699 | 0.32046831 | 0.34605276 | 0.92606776 | 0.35441077 | 0.99989001 |
| PIGC | 82.4187556 | -0.0512999 | 0.39318415 | -0.1304729 | 0.89619226 | 0.99989001 |
| SUCO | 184.445474 | -0.1801391 | 0.21947611 | -0.8207685 | 0.41177812 | 0.99989001 |
| AL645568.1 | 65.6745457 | -0.2043867 | 0.31712959 | -0.6444896 | 0.51925799 | 0.99989001 |
| PRDX6 | 143.353931 | 0.21934515 | 0.32025935 | 0.6848985 | 0.49340797 | 0.99989001 |
| KLHL20 | 177.909005 | -0.0529987 | 0.24243285 | -0.218612 | 0.82695231 | 0.99989001 |
| ZBTB37 | 88.315447 | 0.18377234 | 0.31104186 | 0.59082832 | 0.55463546 | 0.99989001 |
| RC3H1 | 358.76675 | 0.18693347 | 0.19613526 | 0.95308448 | 0.34054727 | 0.99989001 |
| RABGAP1L | 1723.03644 | -0.1088095 | 0.28760629 | -0.3783281 | 0.70518688 | 0.99989001 |
| CACYBP | 53.8256371 | -0.3315039 | 0.34771259 | -0.9533848 | 0.34039513 | 0.99989001 |
| MRPS14 | 45.1100521 | 0.18132444 | 0.36955967 | 0.49064997 | 0.62367403 | NA |
| COP1 | 774.531355 | 0.05504072 | 0.13873034 | 0.39674613 | 0.69155468 | 0.99989001 |
| SEC16B | 65.1000826 | -0.30415 | 0.77248078 | -0.3937314 | 0.69377933 | 0.99989001 |
| RASAL2 | 2517.28146 | 0.17072907 | 0.25452801 | 0.67076732 | 5.02E-01 | 0.99989001 |
| RALGPS2 | 52.062635 | 0.16349447 | 0.38348843 | 0.42633481 | 0.6698639 | 0.99989001 |
| FAM20B | 110.528108 | -0.2577902 | 0.23257351 | -1.1084248 | 0.2676784 | 0.99989001 |
| ABL2 | 415.020505 | -0.0429462 | 0.20318694 | -0.2113628 | 0.83260415 | 0.99989001 |
| SOAT1 | 130.993272 | 0.42749403 | 0.18623787 | 2.29541948 | 0.0217091 | 0.60140868 |
| TOR1AIP2 | 187.868076 | -0.0911805 | 0.27008106 | -0.3376041 | 0.73566158 | 0.99989001 |
| TOR1AIP1 | 113.390172 | -0.0241761 | 0.23181226 | -0.1042917 | 0.9169379 | 0.99989001 |
| CEP350 | 529.373088 | 0.18467127 | 0.16497569 | 1.11938478 | 0.26297602 | 0.99989001 |
| QSOX1 | 153.768296 | -0.8201677 | 0.46427627 | -1.766551 | 0.07730346 | 0.92274869 |
| ACBD6 | 388.984732 | -0.2269997 | 0.23029554 | -0.9856885 | 0.32428601 | 0.99989001 |
| XPR1 | 690.896643 | -0.185546 | 0.2841745 | -0.6529297 | 0.51380158 | 0.99989001 |
| STX6 | 99.3640087 | 0.30198088 | 0.30404341 | 0.99321632 | 0.32060455 | 0.99989001 |
| IER5 | 78.0965961 | 0.66969623 | 0.37106391 | 1.80480025 | 0.07110595 | 0.91166457 |
| GLUL | 215.853662 | -0.300168 | 0.40731906 | -0.7369359 | 0.46116134 | 0.99989001 |
| DHX9 | 212.722485 | 0.08909465 | 0.17227484 | 0.51716579 | 0.60504043 | 0.99989001 |
| LAMC1 | 488.914642 | -0.445628 | 0.36160965 | -1.2323453 | 0.21782012 | 0.99989001 |
| SMG7 | 290.591309 | 0.17857418 | 0.24703909 | 0.722858 | 0.46976713 | 0.99989001 |
| ARPC5 | 195.085274 | 0.09729916 | 0.24876186 | 0.39113377 | 0.69569836 | 0.99989001 |
| RGL1 | 403.012238 | 0.57041147 | 0.43815726 | 1.30184188 | 0.19297044 | 0.99989001 |
| TSEN15 | 92.0083577 | 0.04043427 | 0.23035599 | 0.17552948 | 0.86066361 | 0.99989001 |
| C1orf21 | 891.599665 | -0.7643746 | 0.48756968 | -1.5677238 | 0.1169456 | 0.99989001 |
| EDEM3 | 243.01998 | -0.1099764 | 0.23133373 | -0.4754015 | 0.63450085 | 0.99989001 |
| FAM129A | 102.948347 | 0.15900185 | 0.47046107 | 0.33797025 | 0.73538561 | 0.99989001 |
| RNF2 | 132.935524 | -0.039137 | 0.22663181 | -0.1726898 | 0.86289528 | 0.99989001 |
| TRMT1L | 75.730625 | 0.20808775 | 0.28328109 | 0.7345628 | 0.4626058 | 0.99989001 |
| SWT1 | 166.523015 | -0.0948085 | 0.32600959 | -0.2908152 | 0.77119267 | 0.99989001 |
| IVNS1ABP | 366.270579 | -1.1761255 | 0.73182651 | -1.6071098 | 0.10803029 | 0.99989001 |
| HMCN1 | 210.0906 | -0.3561849 | 0.70653969 | -0.5041258 | 0.61417296 | 0.99989001 |
| TPR | 291.625629 | 0.13425577 | 0.22291446 | 0.60227482 | 0.54699122 | 0.99989001 |
| ODR4 | 160.120188 | -0.1662778 | 0.18700646 | -0.8891552 | 0.37391966 | 0.99989001 |
| AL136454.1 | 143.082758 | -0.4647243 | 0.51531725 | -0.9018217 | 0.36715158 | 0.99989001 |
| UCHL5 | 89.8443625 | -0.0721082 | 0.23515699 | -0.3066388 | 0.75911834 | 0.99989001 |
| TROVE2 | 225.530225 | 0.17598219 | 0.19576037 | 0.89896739 | 0.36867003 | 0.99989001 |
| CDC73 | 402.485216 | -0.0736762 | 0.23359329 | -0.3154036 | 0.75245521 | 0.99989001 |

| | | | | | | |
|---|---|---|---|---|---|---|
| KCNT2 | 77.5990179 | -0.3288865 | 0.58010404 | -0.5669441 | 0.57075218 | 0.99989001 |
| CFH | 259.422703 | -1.2431735 | 0.62950365 | -1.9748472 | 0.04828549 | 0.78123673 |
| ZBTB41 | 59.5209602 | 0.10906895 | 0.24838398 | 0.43911426 | 0.66057875 | 0.99989001 |
| DENND1B | 165.198386 | -0.0540124 | 0.26294598 | -0.2054124 | 8.37E-01 | 0.99989001 |
| NEK7 | 578.218057 | 0.1812732 | 0.17725195 | 1.02268667 | 0.30645602 | 0.99989001 |
| MIR181A1HG | 202.965614 | 0.14016204 | 0.61044672 | 0.22960568 | 0.81839819 | 0.99989001 |
| NR5A2 | 382.482339 | -1.7701695 | 0.54903524 | -3.2241454 | 0.00126349 | 0.1693992 |
| ZNF281 | 47.4519389 | 0.14436229 | 0.30944399 | 0.46652156 | 0.64084222 | NA |
| DDX59 | 52.3939097 | 0.09893389 | 0.38354736 | 0.2579444 | 0.79644981 | 0.99989001 |
| CAMSAP2 | 512.350725 | 0.19532899 | 0.14691842 | 1.32950647 | 0.18368093 | 0.99989001 |
| CSRP1 | 125.453949 | -0.2281634 | 0.51989468 | -0.4388646 | 0.66075966 | 0.99989001 |
| NAV1 | 992.37255 | -0.0965981 | 0.28241223 | -0.3420464 | 0.73231599 | 0.99989001 |
| IPO9-AS1 | 180.481769 | 0.10290712 | 0.39429226 | 0.26099199 | 0.79409869 | 0.99989001 |
| IPO9 | 199.011814 | -0.0532579 | 0.23813641 | -0.2236446 | 0.82303385 | 0.99989001 |
| TIMM17A | 48.2904727 | -0.0478245 | 0.39879672 | -0.1199221 | 0.90454485 | NA |
| RNPEP | 88.500461 | -0.0624514 | 0.32688248 | -0.1910516 | 0.8484852 | 0.99989001 |
| ARL8A | 71.0107798 | 0.08138365 | 0.26164199 | 0.31104966 | 0.75576287 | 0.99989001 |
| PPP1R12B | 338.359111 | 0.00512143 | 0.1923835 | 0.02662095 | 0.97876206 | 0.99989001 |
| KDM5B | 233.244688 | 0.28764315 | 0.30454362 | 0.94450561 | 0.34491134 | 0.99989001 |
| KLHL12 | 51.5918975 | -0.1345183 | 0.31806525 | -0.4229267 | 0.67234874 | 0.99989001 |
| ADIPOR1 | 54.2809838 | 0.15866409 | 0.35982532 | 0.44094754 | 0.65925098 | 0.99989001 |
| CYB5R1 | 39.5017969 | -0.165494 | 0.41314637 | -0.4005699 | 0.68873679 | NA |
| BTG2 | 87.3634562 | -0.2452176 | 0.54277168 | -0.4517878 | 0.65142187 | 0.99989001 |
| ATP2B4 | 403.657436 | 0.25651645 | 0.24379046 | 1.05220055 | 0.29270755 | 0.99989001 |
| SOX13 | 237.357404 | 0.26700256 | 0.38204569 | 0.698876 | 0.48462953 | 0.99989001 |
| PLEKHA6 | 101.377496 | 0.40911248 | 0.48273826 | 0.84748303 | 0.39672594 | 0.99989001 |
| AL606489.1 | 79.5719786 | 0.10765763 | 0.57729092 | 0.18648765 | 0.85206236 | 0.99989001 |
| PPP1R15B | 89.2689464 | 0.05991844 | 0.37086285 | 0.16156497 | 0.87164845 | 0.99989001 |
| PIK3C2B | 785.538875 | -0.0856118 | 0.41167816 | -0.207958 | 0.83526173 | 0.99989001 |
| MDM4 | 293.509522 | -0.1261477 | 0.22039106 | -0.5723812 | 0.56706378 | 0.99989001 |
| NFASC | 86.2217294 | 0.93799039 | 0.48267617 | 1.94331199 | 0.05197847 | 0.81270649 |
| RBBP5 | 67.5430637 | 0.34075225 | 0.24585264 | 1.38600199 | 0.1657463 | 0.99989001 |
| DSTYK | 165.271126 | 0.04978812 | 0.36042403 | 0.13813763 | 0.89013165 | 0.99989001 |
| ELK4 | 167.17872 | 0.11867665 | 0.2032856 | 0.5837927 | 0.55935979 | 0.99989001 |
| NUCKS1 | 449.300054 | -0.0732788 | 0.19586584 | -0.3741275 | 0.70830948 | 0.99989001 |
| RAB29 | 65.9959634 | 0.18574213 | 0.38355281 | 0.48426743 | 0.62819609 | 0.99989001 |
| SLC41A1 | 56.2298144 | 0.09252468 | 0.40353172 | 0.22928727 | 0.81864565 | 0.99989001 |
| SRGAP2 | 377.45377 | 0.26765173 | 0.22435989 | 1.1929571 | 0.23288617 | 0.99989001 |
| MAPKAPK2 | 125.339668 | 0.03725808 | 0.25227764 | 0.14768681 | 0.88258994 | 0.99989001 |
| CD55 | 293.696858 | -0.6221902 | 0.45252209 | -1.3749388 | 0.16915042 | 0.99989001 |
| CD46 | 499.88695 | -0.3609216 | 0.37470989 | -0.9632028 | 0.33544577 | 0.99989001 |
| MIR29B2CHG | 390.591022 | 0.14692849 | 0.32129282 | 0.45730399 | 0.64745257 | 0.99989001 |
| CD34 | 644.59922 | -0.4404359 | 0.49176488 | -0.895623 | 0.37045416 | 0.99989001 |
| PLXNA2 | 708.651851 | -0.2301044 | 0.38514769 | -0.5974446 | 0.55021061 | 0.99989001 |
| HHAT | 64.4778712 | -0.1602225 | 0.58227055 | -0.2751684 | 0.78318687 | 0.99989001 |
| RCOR3 | 255.361333 | -0.0046526 | 0.22813604 | -0.0203939 | 0.98372916 | 0.99989001 |
| TRAF5 | 63.9672448 | 0.07128772 | 0.33475089 | 0.21295752 | 0.8313601 | 0.99989001 |
| SLC30A1 | 72.8026944 | -0.3406706 | 0.48776477 | -0.6984321 | 0.48490701 | 0.99989001 |
| LPGAT1 | 291.668287 | -0.0963044 | 0.14749469 | -0.6529346 | 0.51379847 | 0.99989001 |
| INTS7 | 74.280414 | 0.01526924 | 0.37494811 | 0.0407236 | 0.96751625 | 0.99989001 |
| PPP2R5A | 479.675061 | -0.0591045 | 0.17719971 | -0.3335472 | 0.73872127 | 0.99989001 |
| AL360091.3 | 67.8255893 | 0.10110402 | 0.54824688 | 0.18441331 | 0.85368923 | 0.99989001 |
| NENF | 77.8444689 | -0.2702298 | 0.40091338 | -0.6740354 | 0.50028881 | 0.99989001 |
| NSL1 | 140.607175 | 0.23658143 | 0.26379043 | 0.89685373 | 0.36979697 | 0.99989001 |

| | | | | | | |
|---|---|---|---|---|---|---|
| TATDN3 | 46.162127 | -0.2228861 | 0.49040004 | -0.4544986 | 0.64946999 | NA |
| FLVCR1 | 63.4304306 | 0.2618125 | 0.30576766 | 0.85624655 | 0.39186142 | 0.99989001 |
| ANGEL2 | 112.445502 | -0.31546 | 0.23304716 | -1.3536314 | 0.175854 | 0.99989001 |
| RPS6KC1 | 440.715674 | 0.17380571 | 0.38336398 | 0.45336995 | 0.65028237 | 0.99989001 |
| PTPN14 | 772.063702 | -0.3826768 | 0.33906564 | -1.1286214 | 0.25905756 | 0.99989001 |
| KCTD3 | 221.173725 | 0.03982928 | 0.18489704 | 0.21541327 | 0.82944513 | 0.99989001 |
| ESRRG | 152.40916 | -1.2831745 | 0.603576 | -2.1259535 | 0.03350713 | 0.67281094 |
| GPATCH2 | 269.86839 | 0.08436465 | 0.33257862 | 0.25366829 | 0.79975184 | 0.99989001 |
| RRP15 | 59.9171148 | -0.1759126 | 0.32716733 | -0.5376838 | 0.59079536 | 0.99989001 |
| TGFB2 | 65.4640216 | 1.39939595 | 0.42920053 | 3.26047113 | 0.00111227 | 0.16145069 |
| LYPLAL1 | 222.210092 | 0.10365769 | 0.35422378 | 0.29263334 | 0.76980243 | 0.99989001 |
| EPRS | 148.3855 | 0.16396599 | 0.24203027 | 0.67746069 | 0.49811371 | 0.99989001 |
| IARS2 | 131.461863 | -0.3012815 | 0.19829336 | -1.5193726 | 0.12866873 | 0.99989001 |
| RAB3GAP2 | 362.712325 | -0.254725 | 0.19759469 | -1.2891287 | 0.19735334 | 0.99989001 |
| MARK1 | 56.6807065 | 0.30464642 | 0.57747871 | 0.52754572 | 0.59781468 | 0.99989001 |
| C1orf115 | 49.6898421 | -0.1484133 | 0.49426985 | -0.3002677 | 0.76397295 | NA |
| 2-Mar | 109.72462 | 0.11824262 | 0.3780739 | 0.31275002 | 0.75447059 | 0.99989001 |
| HLX | 66.5977791 | -0.0052987 | 0.33096414 | -0.0160099 | 0.98722653 | 0.99989001 |
| MIA3 | 147.842999 | -0.0015938 | 0.16432492 | -0.0096988 | 0.99226158 | 0.99989001 |
| AIDA | 86.546337 | 0.31014859 | 0.24396736 | 1.27127085 | 0.20363231 | 0.99989001 |
| BROX | 101.861136 | -0.0577386 | 0.22677623 | -0.2546059 | 0.79902754 | 0.99989001 |
| AC093152.1 | 59.3907052 | 0.43827296 | 0.32791107 | 1.33656045 | 0.18136616 | 0.99989001 |
| CAPN2 | 265.756008 | -0.1814423 | 0.29074737 | -0.6240548 | 0.53259159 | 0.99989001 |
| TP53BP2 | 103.581737 | -0.1855729 | 0.30548776 | -0.6074641 | 0.54354294 | 0.99989001 |
| FBXO28 | 185.917839 | 0.08543821 | 0.26458937 | 0.3229087 | 0.74676438 | 0.99989001 |
| DEGS1 | 106.171247 | 0.13644398 | 0.43901163 | 0.3107981 | 0.75595411 | 0.99989001 |
| NVL | 149.099893 | -0.1714517 | 0.20511387 | -0.8358855 | 0.40321929 | 0.99989001 |
| CNIH4 | 87.2950821 | -0.1267475 | 0.35253934 | -0.3595273 | 0.71920065 | 0.99989001 |
| WDR26 | 315.584259 | -0.0406061 | 0.23392823 | -0.1735834 | 0.8621929 | 0.99989001 |
| CNIH3 | 100.717702 | -0.1084886 | 0.42021602 | -0.2581735 | 0.79627299 | 0.99989001 |
| LBR | 145.761318 | 0.11136534 | 0.21421731 | 0.51987086 | 0.60315359 | 0.99989001 |
| ENAH | 192.342224 | -0.0502513 | 0.35849056 | -0.1401747 | 0.88852194 | 0.99989001 |
| SRP9 | 174.519124 | 0.01735084 | 0.33543463 | 0.05172644 | 0.95874667 | 0.99989001 |
| EPHX1 | 112.833764 | 0.24548928 | 0.31262307 | 0.78525644 | 0.43230323 | 0.99989001 |
| TMEM63A | 91.0443688 | 0.11265978 | 0.35735198 | 0.31526279 | 0.75256213 | 0.99989001 |
| SDE2 | 57.9727471 | -0.4896863 | 0.44648694 | -1.096754 | 0.27274895 | 0.99989001 |
| H3F3A | 74.8843972 | -0.1785015 | 0.42750897 | -0.4175387 | 0.67628441 | 0.99989001 |
| ACBD3 | 169.71765 | 0.03366507 | 0.24931168 | 0.13503208 | 0.89258651 | 0.99989001 |
| PARP1 | 85.7409282 | 0.12326981 | 0.3344467 | 0.36857835 | 0.71244203 | 0.99989001 |
| ITPKB | 447.452084 | -0.7739612 | 0.41568689 | -1.8618851 | 0.06261929 | 0.85940342 |
| COQ8A | 77.4132279 | -0.1946791 | 0.41086605 | -0.4738262 | 0.63562385 | 0.99989001 |
| CDC42BPA | 1632.15781 | -0.2594685 | 0.18764483 | -1.3827641 | 0.16673721 | 0.99989001 |
| ZNF678 | 112.912753 | 0.0160964 | 0.23282143 | 0.06913626 | 0.94488116 | 0.99989001 |
| ARF1 | 334.21819 | 0.03451623 | 0.30234513 | 0.1141617 | 0.90910962 | 0.99989001 |
| GUK1 | 300.087341 | -0.2025416 | 0.42488517 | -0.4766972 | 0.63357775 | 0.99989001 |
| RNF187 | 45.6428121 | 0.04285779 | 0.41025172 | 0.10446707 | 0.9167987 | NA |
| RHOU | 111.494189 | 1.39101965 | 0.42451109 | 3.27675695 | 0.00105007 | 0.15851814 |
| RAB4A | 50.4206239 | 0.13249683 | 0.34918145 | 0.3794498 | 0.70435387 | 0.99989001 |
| NUP133 | 124.169216 | -0.1381501 | 0.19454812 | -0.7101077 | 0.47763736 | 0.99989001 |
| ABCB10 | 70.3092646 | 0.34521221 | 0.27292506 | 1.2648608 | 0.20592123 | 0.99989001 |
| GALNT2 | 193.415345 | 0.00213251 | 0.23783846 | 0.00896623 | 0.99284608 | 0.99989001 |
| COG2 | 55.5842004 | -0.2232428 | 0.30113188 | -0.7413456 | 0.45848395 | 0.99989001 |
| TTC13 | 109.513015 | -0.2172063 | 0.37633622 | -0.5771603 | 0.56383115 | 0.99989001 |
| FAM89A | 58.8011502 | 0.03442824 | 0.45409031 | 0.07581805 | 9.40E-01 | 0.99989001 |

| | | | | | | |
|---|---|---|---|---|---|---|
| GNPAT | 127.055553 | -0.0693689 | 0.21470062 | -0.3230961 | 0.74662245 | 0.99989001 |
| SPRTN | 46.7267736 | 0.2502595 | 0.3781092 | 0.66187098 | 0.50805391 | NA |
| EGLN1 | 377.034336 | -0.043631 | 0.48955369 | -0.089124 | 0.92898339 | 0.99989001 |
| DISC1 | 544.046043 | 0.38336385 | 0.34837907 | 1.1004216 | 0.27114847 | 0.99989001 |
| SIPA1L2 | 636.720115 | -0.0136057 | 0.44713001 | -0.030429 | 0.97572492 | 0.99989001 |
| NTPCR | 77.4494875 | -0.2977933 | 0.33673016 | -0.8843677 | 0.37649778 | 0.99989001 |
| PCNX2 | 84.9705452 | -0.7712619 | 0.50336447 | -1.5322137 | 0.12546972 | 0.99989001 |
| TARBP1 | 169.205296 | -0.5183266 | 0.42586101 | -1.2171262 | 0.22355621 | 0.99989001 |
| IRF2BP2 | 199.775837 | -0.0550853 | 0.58436707 | -0.0942649 | 0.92489873 | 0.99989001 |
| TOMM20 | 145.782432 | -0.1731283 | 0.40922976 | -0.423059 | 0.67225221 | 0.99989001 |
| ARID4B | 961.978253 | 0.12757324 | 0.27831832 | 0.4583717 | 0.64668542 | 0.99989001 |
| B3GALNT2 | 85.7768476 | -0.038363 | 0.33151892 | -0.1157189 | 0.90787533 | 0.99989001 |
| LYST | 402.982848 | 0.47247667 | 0.32131886 | 1.4704293 | 0.14144552 | 0.99989001 |
| NID1 | 309.236901 | 0.77583225 | 0.41190371 | 1.88352817 | 0.05962883 | 0.85885114 |
| GPR137B | 69.9482048 | 0.2291875 | 0.44691959 | 0.51281597 | 0.60808006 | 0.99989001 |
| ERO1B | 135.094921 | 0.43426427 | 0.32259806 | 1.34614657 | 0.17825525 | 0.99989001 |
| LGALS8 | 132.553751 | 0.11105094 | 0.24296208 | 0.45707108 | 0.64761996 | 0.99989001 |
| HEATR1 | 114.440685 | -0.1714256 | 0.31764747 | -0.5396726 | 0.58942282 | 0.99989001 |
| MTR | 261.475065 | -0.1595573 | 0.19768156 | -0.8071431 | 0.41958407 | 0.99989001 |
| RYR2 | 105.934276 | -3.1297849 | 0.68896117 | -4.5427595 | 5.55E-06 | 0.00411251 |
| CHRM3 | 1359.03014 | 0.36123884 | 0.35748328 | 1.01050556 | 3.12E-01 | 0.99989001 |
| CHRM3-AS2 | 55.0967873 | 1.69072745 | 0.4473361 | 3.77954619 | 0.00015711 | 0.05647142 |
| CEP170 | 454.577953 | -0.147763 | 0.15894461 | -0.9296511 | 0.35255174 | 0.99989001 |
| SDCCAG8 | 657.573522 | 0.10804885 | 0.24558835 | 0.4399592 | 0.65996666 | 0.99989001 |
| AKT3 | 2496.79977 | 0.02916123 | 0.26530915 | 0.10991416 | 0.91247745 | 0.99989001 |
| ZBTB18 | 57.8012919 | -0.4717054 | 0.39265541 | -1.2013216 | 0.22962649 | 0.99989001 |
| ADSS | 101.285891 | 0.27828729 | 0.30528104 | 0.91157739 | 0.36199123 | 0.99989001 |
| CATSPERE | 60.0050796 | -0.179917 | 0.40193815 | -0.4476236 | 0.65442483 | 0.99989001 |
| DESI2 | 148.079995 | 0.00067115 | 0.1553522 | 0.00432016 | 0.99655302 | 0.99989001 |
| COX20 | 80.5266725 | -0.5906736 | 0.31807244 | -1.8570412 | 0.06330529 | 0.85940342 |
| HNRNPU | 627.391721 | -0.1016448 | 0.24584505 | -0.4134505 | 0.67927658 | 0.99989001 |
| EFCAB2 | 191.07933 | -0.5366634 | 0.28499142 | -1.8830862 | 0.05968869 | 0.85885114 |
| KIF26B | 569.059566 | 1.22605661 | 0.63689529 | 1.9250521 | 0.05422284 | 0.81854801 |
| SMYD3 | 1686.13397 | -0.1018801 | 0.19002425 | -0.5361425 | 0.59186008 | 0.99989001 |
| CNST | 388.542075 | 0.00220628 | 0.39101559 | 0.00564244 | 0.99549801 | 0.99989001 |
| AHCTF1 | 273.306218 | -0.3980676 | 0.21843544 | -1.8223579 | 0.0684007 | 0.88709358 |
| ZNF124 | 66.3653947 | -0.0898807 | 0.3084798 | -0.2913665 | 0.770771 | 0.99989001 |
| AL390728.6 | 100.975135 | -0.1571027 | 0.356208 | -0.441042 | 0.65918261 | 0.99989001 |
| ZNF496 | 72.9404322 | -0.0913381 | 0.3318859 | -0.2752093 | 0.78315547 | 0.99989001 |
| AL390860.1 | 80.0105144 | -0.5300761 | 0.62742851 | -0.8448391 | 0.39820068 | 0.99989001 |
| SH3YL1 | 91.7493249 | 0.17220433 | 0.23974928 | 0.71826839 | 0.47259182 | 0.99989001 |
| ACP1 | 71.2415074 | 0.02602039 | 0.23476725 | 0.11083483 | 0.91174732 | 0.99989001 |
| PXDN | 222.806516 | 0.481219 | 0.59816856 | 0.80448729 | 0.42111561 | 0.99989001 |
| EIPR1 | 112.72943 | -0.007785 | 0.38850357 | -0.0200383 | 0.98401282 | 0.99989001 |
| TRAPPC12 | 196.600038 | 0.03908469 | 0.2600142 | 0.15031754 | 0.8805141 | 0.99989001 |
| ADI1 | 66.3158706 | 0.45408676 | 0.40340226 | 1.12564258 | 0.26031684 | 0.99989001 |
| RPS7 | 399.91366 | -0.0555571 | 0.60968302 | -0.0911245 | 0.92739364 | 0.99989001 |
| NRIR | 86.397777 | -1.3647986 | 0.79447781 | -1.7178561 | 0.08582287 | 0.95514629 |
| CMPK2 | 88.9105365 | -0.3703565 | 0.72179057 | -0.5131079 | 0.60787583 | 0.99989001 |
| RSAD2 | 313.84666 | -0.2967601 | 1.30996169 | -0.226541 | 0.82078066 | 0.99989001 |
| RNF144A | 517.709111 | 0.93525925 | 0.37848697 | 2.4710474 | 0.0134718 | 0.50339166 |
| ID2 | 274.584914 | 2.23742636 | 0.52706212 | 4.24509039 | 2.19E-05 | 0.01099519 |
| KIDINS220 | 369.803163 | 0.06863009 | 0.18336725 | 0.37427668 | 0.70819848 | 0.99989001 |
| MBOAT2 | 160.521756 | 1.34462844 | 0.5080859 | 2.64645887 | 0.00813394 | 0.41229803 |

| | | | | | | |
|---|---|---|---|---|---|---|
| ASAP2 | 717.633764 | 0.28223828 | 0.32225918 | 0.87581146 | 0.38113254 | 0.99989001 |
| ITGB1BP1 | 138.259823 | -0.0228515 | 0.20649926 | -0.1106616 | 0.91188469 | 0.99989001 |
| CPSF3 | 50.6038548 | -0.2009222 | 0.26237265 | -0.7657895 | 0.44380154 | 0.99989001 |
| IAH1 | 66.517293 | 0.09880479 | 0.36111606 | 0.27360951 | 0.78438472 | 0.99989001 |
| ADAM17 | 344.562812 | -0.0485622 | 0.21916074 | -0.2215827 | 0.82463872 | 0.99989001 |
| YWHAQ | 266.406611 | 0.06370997 | 0.17624297 | 0.3614894 | 0.71773363 | 0.99989001 |
| TAF1B | 107.73577 | -0.0220966 | 0.37005292 | -0.059712 | 0.95238504 | 0.99989001 |
| KLF11 | 73.2085368 | 0.31813136 | 0.39703677 | 0.80126422 | 0.4229787 | 0.99989001 |
| HPCAL1 | 652.569173 | 0.13420904 | 0.22190899 | 0.60479318 | 0.54531642 | 0.99989001 |
| ODC1 | 44.3537388 | 0.39074696 | 0.41293669 | 0.9462636 | 0.34401416 | NA |
| NOL10 | 246.652038 | -0.1033557 | 0.23547164 | -0.4389308 | 0.66071171 | 0.99989001 |
| PDIA6 | 216.32464 | 0.20768896 | 0.36520953 | 0.5686844 | 0.56957034 | 0.99989001 |
| PQLC3 | 49.7426192 | 0.11350483 | 0.35121031 | 0.32318193 | 0.74655746 | NA |
| ROCK2 | 835.22668 | -0.0987096 | 0.28086744 | -0.3514454 | 0.72525424 | 0.99989001 |
| E2F6 | 50.812126 | 0.3689801 | 0.32498541 | 1.13537436 | 0.2562185 | 0.99989001 |
| LPIN1 | 77.0705235 | -0.3180198 | 0.31453042 | -1.0110939 | 0.31197147 | 0.99989001 |
| MIR3681HG | 55.3924527 | -0.0700229 | 0.48746507 | -0.1436471 | 0.88577915 | 0.99989001 |
| TRIB2 | 109.211905 | 0.43626948 | 0.39837979 | 1.09510948 | 0.27346868 | 0.99989001 |
| NBAS | 503.675492 | -0.1735479 | 0.19873104 | -0.8732803 | 0.38251032 | 0.99989001 |
| DDX1 | 115.340368 | -0.1126519 | 0.20496012 | -0.5496283 | 0.58257437 | 0.99989001 |
| SMC6 | 172.875156 | 0.02990142 | 0.19944812 | 0.14992077 | 0.88082712 | 0.99989001 |
| LINC01376 | 80.58354 | 0.48128294 | 0.56838718 | 0.84675192 | 0.39713341 | 0.99989001 |
| WDR35 | 61.6817685 | -0.7402683 | 0.35430279 | -2.0893663 | 0.03667476 | 0.70081289 |
| LAPTM4A | 317.993903 | -0.0491801 | 0.42792663 | -0.1149264 | 0.90850345 | 0.99989001 |
| PUM2 | 650.908007 | -0.0367847 | 0.1429378 | -0.2573475 | 0.79691056 | 0.99989001 |
| RHOB | 336.826212 | 0.44107493 | 0.464167 | 0.95025052 | 0.34198497 | 0.99989001 |
| LDAH | 173.829802 | 0.28155088 | 0.26335289 | 1.06910119 | 0.28502408 | 0.99989001 |
| KLHL29 | 70.1834512 | -1.3391509 | 0.48922351 | -2.7372987 | 0.0061946 | 0.3832529 |
| ATAD2B | 451.917369 | 0.04807257 | 0.19740938 | 0.24351718 | 0.80760478 | 0.99989001 |
| UBXN2A | 145.687895 | 0.24757803 | 0.32920709 | 0.75204342 | 0.45202495 | 0.99989001 |
| SF3B6 | 98.334981 | -0.1338236 | 0.49727641 | -0.269113 | 0.78784271 | 0.99989001 |
| FAM228B | 84.3911959 | -0.2152493 | 0.36345758 | -0.5922269 | 0.55369867 | 0.99989001 |
| ITSN2 | 777.820278 | 0.13880119 | 0.16254273 | 0.85393663 | 0.3931401 | 0.99989001 |
| NCOA1 | 1106.48976 | 0.01750246 | 0.19095061 | 0.0916596 | 0.9269685 | 0.99989001 |
| EFR3B | 47.8583748 | -1.4708267 | 0.57149613 | -2.5736425 | 0.01006342 | NA |
| DNMT3A | 167.037862 | 0.09481769 | 0.40894709 | 0.23185808 | 0.81664825 | 0.99989001 |
| DTNB | 153.99701 | 0.28894665 | 0.38114983 | 0.75809201 | 0.4483959 | 0.99989001 |
| ASXL2 | 292.775341 | -0.0072031 | 0.23463575 | -0.0306992 | 0.97550945 | 0.99989001 |
| KIF3C | 45.3201761 | 0.39377193 | 0.44244888 | 0.88998288 | 0.37347508 | NA |
| RAB10 | 572.663407 | 0.11500237 | 0.26357098 | 0.43632408 | 0.66260161 | 0.99989001 |
| HADHA | 192.188702 | 0.20643302 | 0.20370783 | 1.01337791 | 0.31087969 | 0.99989001 |
| HADHB | 239.54803 | 0.1172071 | 0.19899648 | 0.58899082 | 0.55586744 | 0.99989001 |
| SELENOI | 114.140539 | -0.2528692 | 0.27405229 | -0.9227042 | 0.35616138 | 0.99989001 |
| TMEM214 | 39.7719252 | 0.28065187 | 0.35068454 | 0.80029723 | 0.42353861 | NA |
| OST4 | 107.327298 | -0.269185 | 0.61813127 | -0.435482 | 0.66321263 | 0.99989001 |
| ATRAID | 85.0206338 | -0.3647003 | 0.54998456 | -0.6631101 | 0.50726007 | 0.99989001 |
| MPV17 | 53.2034229 | -0.1624625 | 0.24284722 | -0.6689907 | 0.50350142 | 0.99989001 |
| GTF3C2 | 106.767436 | -0.0296381 | 0.26313237 | -0.1126358 | 0.91031931 | 0.99989001 |
| SNX17 | 54.4185298 | 0.05068913 | 0.42003994 | 0.12067694 | 0.90394693 | 0.99989001 |
| PPM1G | 191.007349 | 0.25009735 | 0.36883639 | 0.67807124 | 0.49772652 | 0.99989001 |
| NRBP1 | 91.5071058 | 0.14018777 | 0.2588001 | 0.54168361 | 0.58803648 | 0.99989001 |
| IFT172 | 65.8563875 | -0.4157971 | 0.33574509 | -1.2384308 | 0.21555636 | 0.99989001 |
| ZNF512 | 148.428458 | 0.24534745 | 0.31247528 | 0.78517395 | 0.43235159 | 0.99989001 |
| SUPT7L | 47.8961857 | 0.09318052 | 0.34299419 | 0.27166792 | 0.78587737 | NA |

| | | | | | | |
|---|---|---|---|---|---|---|
| SLC4A1AP | 54.7941276 | -0.1836931 | 0.23616335 | -0.7778222 | 0.43667381 | 0.99989001 |
| MRPL33 | 234.669448 | -0.1533402 | 0.47102149 | -0.3255483 | 0.74476615 | 0.99989001 |
| RBKS | 71.4856814 | -0.2257358 | 0.45820174 | -0.492656 | 0.62225567 | 0.99989001 |
| BABAM2 | 790.913551 | 0.10249072 | 0.26994714 | 0.37966959 | 0.7041907 | 0.99989001 |
| FOSL2 | 505.923953 | -1.0053341 | 0.52384884 | -1.9191301 | 0.05496787 | 0.82320925 |
| PPP1CB | 218.369942 | -0.1676399 | 0.43365603 | -0.3865734 | 0.69907204 | 0.99989001 |
| TRMT61B | 62.5662915 | 0.06173716 | 0.27247342 | 0.22658047 | 0.82074998 | 0.99989001 |
| WDR43 | 156.91954 | -0.2637004 | 0.35723912 | -0.7381623 | 0.46041585 | 0.99989001 |
| YPEL5 | 104.32426 | -0.1003103 | 0.30427079 | -0.3296744 | 0.74164602 | 0.99989001 |
| LCLAT1 | 208.708767 | 0.20134015 | 0.30178298 | 0.66716867 | 0.5046644 | 0.99989001 |
| EHD3 | 164.475001 | 1.55671361 | 0.58291334 | 2.6705747 | 0.00757215 | 0.4053518 |
| DPY30 | 75.7395274 | 0.09895117 | 0.30405778 | 0.3254354 | 0.74485156 | 0.99989001 |
| SPAST | 159.163419 | -0.0595737 | 0.24039041 | -0.2478207 | 0.80427317 | 0.99989001 |
| SLC30A6 | 124.195397 | -0.1992871 | 0.20845676 | -0.9560118 | 0.33906628 | 0.99989001 |
| YIPF4 | 233.970292 | -0.0696209 | 0.13854429 | -0.5025171 | 0.61530379 | 0.99989001 |
| BIRC6 | 1244.77436 | -0.0413145 | 0.18002586 | -0.229492 | 0.81848653 | 0.99989001 |
| TTC27 | 183.424016 | -0.1417603 | 0.29220418 | -0.4851413 | 0.62757609 | 0.99989001 |
| LINC00486 | 207.829946 | -0.1765261 | 0.22876312 | -0.7716547 | 0.44031899 | 0.99989001 |
| LTBP1 | 384.12475 | -0.2257532 | 0.6193232 | -0.364516 | 0.71547275 | 0.99989001 |
| RASGRP3 | 494.365833 | 0.2829078 | 0.28907213 | 0.97867545 | 3.28E-01 | 0.99989001 |
| FAM98A | 55.8953296 | -0.2858354 | 0.30716461 | -0.9305608 | 0.3520808 | 0.99989001 |
| LINC01320 | 56.9947194 | 0.73308833 | 0.40040332 | 1.83087475 | 0.06711924 | 0.87873839 |
| CRIM1 | 1530.91326 | -0.6360866 | 0.50679175 | -1.2551242 | 0.20943367 | 0.99989001 |
| FEZ2 | 523.368079 | -0.6133408 | 0.42520562 | -1.4424569 | 0.14917353 | 0.99989001 |
| STRN | 314.874856 | -0.1305229 | 0.16614568 | -0.7855933 | 0.43210576 | 0.99989001 |
| HEATR5B | 241.558572 | -0.0680111 | 0.19252679 | -0.3532551 | 0.72389724 | 0.99989001 |
| EIF2AK2 | 334.136178 | -0.3681094 | 0.54486245 | -0.6756005 | 0.49929431 | 0.99989001 |
| CEBPZOS | 96.5904241 | -0.3807214 | 0.28628133 | -1.3298854 | 0.18355603 | 0.99989001 |
| CEBPZ | 119.413463 | 0.25961976 | 0.27587633 | 0.94107297 | 0.34666746 | 0.99989001 |
| NDUFAF7 | 75.8357826 | -0.1849569 | 0.249376 | -0.7416789 | 0.4582819 | 0.99989001 |
| PRKD3 | 328.154469 | 0.27104399 | 0.19089867 | 1.41983172 | 0.15565668 | 0.99989001 |
| CDC42EP3 | 196.080727 | 0.17042784 | 0.35437897 | 0.48091972 | 6.31E-01 | 1.00E+00 |
| ATL2 | 173.210851 | 0.09504119 | 0.16811737 | 0.56532641 | 0.57185177 | 0.99989001 |
| HNRNPLL | 132.168776 | 0.25098054 | 0.2807723 | 0.89389355 | 0.37137885 | 0.99989001 |
| GALM | 61.6642394 | 0.33972947 | 0.37717536 | 0.90072022 | 0.3677371 | 0.99989001 |
| SRSF7 | 205.750407 | 0.47689953 | 0.33317743 | 1.43136806 | 0.15232476 | 0.99989001 |
| DHX57 | 84.3448924 | -0.265897 | 0.3382112 | -0.7861864 | 0.43175832 | 0.99989001 |
| SOS1 | 828.045344 | 0.15300226 | 0.17245363 | 0.8872081 | 0.37496687 | 0.99989001 |
| MAP4K3 | 610.980682 | -0.3450287 | 0.20056993 | -1.7202415 | 0.08538855 | 0.95514629 |
| AC007388.1 | 195.565425 | -1.0960471 | 0.36913332 | -2.9692447 | 0.00298533 | 0.27818834 |
| THUMPD2 | 87.0658756 | -1.0186774 | 0.23014972 | -4.426151 | 9.59E-06 | 0.00556981 |
| SLC8A1 | 483.024864 | 0.60220721 | 0.4128106 | 1.45879783 | 0.14462076 | 0.99989001 |
| EML4 | 341.820224 | -0.0846799 | 0.2825475 | -0.2997015 | 0.76440489 | 0.99989001 |
| COX7A2L | 130.354017 | -0.1917532 | 0.30818504 | -0.6222015 | 0.53380936 | 0.99989001 |
| MTA3 | 217.083728 | 0.36294362 | 0.19703108 | 1.8420628 | 0.06546596 | 0.8666545 |
| ZFP36L2 | 347.213761 | 0.21111472 | 0.28453655 | 0.74195993 | 0.45811161 | 0.99989001 |
| THADA | 478.623064 | 0.00039978 | 0.25838979 | 0.00154719 | 0.99876552 | 0.99989001 |
| PLEKHH2 | 69.2787018 | -0.5183844 | 0.55044934 | -0.9417476 | 0.34632187 | 0.99989001 |
| DYNC2LI1 | 76.6662067 | -0.1283933 | 0.29866834 | -0.429886 | 0.66727859 | 0.99989001 |
| LRPPRC | 261.147466 | -0.2907773 | 0.22496659 | -1.2925355 | 0.19617178 | 0.99989001 |
| PPM1B | 196.432946 | -0.0560069 | 0.20076172 | -0.2789719 | 0.78026643 | 0.99989001 |
| SLC3A1 | 69.9274741 | 1.71866347 | 0.53873603 | 3.19017733 | 0.00142186 | 0.18190106 |
| PREPL | 156.789459 | 0.05766462 | 0.36785367 | 0.15675967 | 0.87543426 | 0.99989001 |
| CAMKMT | 579.525201 | 0.18195658 | 0.23641742 | 0.76964116 | 0.44151278 | 0.99989001 |

| | | | | | | |
|---|---|---|---|---|---|---|
| SRBD1 | 365.95101 | 0.01356895 | 0.25090467 | 0.0540801 | 0.95687135 | 0.99989001 |
| PRKCE | 1608.41537 | -0.0442224 | 0.42462265 | -0.1041452 | 0.91705409 | 0.99989001 |
| EPAS1 | 3405.33722 | 0.60686536 | 0.38435436 | 1.57892148 | 0.11435407 | 0.99989001 |
| RHOQ | 128.584934 | -0.6028856 | 0.46566315 | -1.2946817 | 0.19543005 | 0.99989001 |
| PIGF | 109.599287 | 0.08571649 | 0.27502119 | 0.3116723 | 0.75528958 | 0.99989001 |
| SOCS5 | 196.496385 | -0.1744068 | 0.25277613 | -0.6899654 | 0.49021593 | 0.99989001 |
| MCFD2 | 128.924348 | -0.11862 | 0.33300922 | -0.3562065 | 0.72168594 | 0.99989001 |
| TTC7A | 73.163466 | 0.13906341 | 0.32925334 | 0.42235989 | 0.67276235 | 0.99989001 |
| AC106869.1 | 104.811114 | 0.11452217 | 0.44560063 | 0.25700629 | 0.79717391 | 0.99989001 |
| EPCAM | 60.8189839 | 0.12431937 | 0.47379781 | 0.26238908 | 0.79302149 | 0.99989001 |
| MSH2 | 101.264379 | -0.0376095 | 0.27219843 | -0.1381695 | 0.89010643 | 0.99989001 |
| MSH6 | 51.4547482 | 0.30388236 | 0.30848453 | 0.98508136 | 0.32458411 | 0.99989001 |
| FBXO11 | 780.238624 | 0.07096126 | 0.21584635 | 0.32875819 | 0.74233846 | 0.99989001 |
| FOXN2 | 206.637803 | 0.11832356 | 0.2424246 | 0.48808397 | 0.62549036 | 0.99989001 |
| PPP1R21 | 244.938777 | 0.06735242 | 0.24828679 | 0.27126866 | 0.78618441 | 0.99989001 |
| GPR75-ASB3 | 81.5318513 | 0.44107418 | 0.36181826 | 1.21904899 | 0.2228256 | 0.99989001 |
| ERLEC1 | 83.4516472 | 0.09197779 | 0.24411805 | 0.37677589 | 0.70634016 | 0.99989001 |
| PSME4 | 335.353536 | -0.0009529 | 0.27010112 | -0.0035281 | 0.99718499 | 0.99989001 |
| ACYP2 | 429.516785 | -0.315357 | 0.22088783 | -1.4276792 | 0.15338422 | 0.99989001 |
| SPTBN1 | 3064.20807 | 0.13974651 | 0.1788749 | 0.78125274 | 0.43465386 | 0.99989001 |
| AC092839.1 | 120.254601 | 0.34228551 | 0.54347278 | 0.6298117 | 0.52881779 | 0.99989001 |
| AC093110.1 | 84.918413 | -0.0760368 | 0.72624159 | -0.104699 | 0.91661462 | 0.99989001 |
| EML6 | 50.3236883 | -0.1341958 | 0.29510871 | -0.4547335 | 0.649301 | 0.99989001 |
| RTN4 | 1270.01949 | 0.28956513 | 0.38608612 | 0.75000139 | 0.45325387 | 0.99989001 |
| CLHC1 | 56.2795497 | -0.4464259 | 0.64909649 | -0.6877652 | 0.49160068 | 0.99989001 |
| RPS27A | 618.078962 | -0.113574 | 0.59626987 | -0.1904741 | 0.84893761 | 0.99989001 |
| MTIF2 | 43.6630161 | -0.0300705 | 0.35007528 | -0.0858971 | 0.93154819 | NA |
| AC012358.3 | 64.2523359 | -0.0235374 | 0.36692095 | -0.0641485 | 0.94885202 | 0.99989001 |
| CCDC88A | 267.598268 | 0.47239848 | 0.23453887 | 2.0141586 | 0.04399288 | 0.77097498 |
| CFAP36 | 91.3755888 | -0.1476297 | 0.25776972 | -0.5727194 | 0.5668347 | 0.99989001 |
| PPP4R3B | 325.863265 | -0.223612 | 0.16409637 | -1.3626868 | 0.17298123 | 0.99989001 |
| PNPT1 | 173.149778 | -0.4326415 | 0.48372223 | -0.8944008 | 0.37110749 | 0.99989001 |
| EFEMP1 | 115.893482 | 0.57612245 | 0.79034053 | 0.72895471 | 0.46602937 | 0.99989001 |
| CCDC85A | 388.770739 | -0.4045906 | 0.66715158 | -0.6064447 | 0.54421946 | 0.99989001 |
| VRK2 | 246.954616 | -0.4003876 | 0.24446474 | -1.6378133 | 0.10146066 | 0.99989001 |
| FANCL | 241.51499 | -0.3017907 | 0.29919236 | -1.0086846 | 0.31312591 | 0.99989001 |
| AC007100.1 | 371.650795 | 0.17752578 | 0.57026789 | 0.31130242 | 0.75557073 | 0.99989001 |
| PAPOLG | 94.8377942 | 0.1823719 | 0.31422921 | 0.58037857 | 0.56165935 | 0.99989001 |
| REL | 212.081341 | 0.51122284 | 0.39453657 | 1.29575529 | 0.19505981 | 0.99989001 |
| PUS10 | 87.2363653 | 0.08272733 | 0.27581091 | 0.29994221 | 0.76422123 | 0.99989001 |
| PEX13 | 74.3934425 | 0.32004925 | 0.29875096 | 1.07129112 | 0.28403855 | 0.99989001 |
| KIAA1841 | 160.956428 | 0.40290916 | 0.32524016 | 1.23880509 | 0.21541769 | 0.99989001 |
| USP34 | 1263.52106 | -0.1046679 | 0.21767634 | -0.4808417 | 0.63062901 | 0.99989001 |
| XPO1 | 423.945971 | 0.16781521 | 0.42621319 | 0.39373536 | 0.69377643 | 0.99989001 |
| CCT4 | 146.361479 | 0.11463053 | 0.30071618 | 0.38119175 | 0.70306097 | 0.99989001 |
| COMMD1 | 494.016504 | 0.08240985 | 0.2674875 | 0.30808861 | 0.75801491 | 0.99989001 |
| B3GNT2 | 86.6372707 | 0.16019296 | 0.30668777 | 0.52233239 | 0.60143892 | 0.99989001 |
| EHBP1 | 736.355967 | -0.2097186 | 0.23635956 | -0.8872863 | 0.37492477 | 0.99989001 |
| WDPCP | 516.997112 | 0.02231344 | 0.24746208 | 0.09016913 | 0.92815282 | 0.99989001 |
| MDH1 | 91.736481 | 0.38537596 | 0.27855087 | 1.38350298 | 0.16651069 | 0.99989001 |
| UGP2 | 194.017001 | 0.55946023 | 0.24646943 | 2.26989704 | 0.02321383 | 0.60818532 |
| VPS54 | 459.392244 | -0.0180339 | 0.18212153 | -0.099021 | 0.92112158 | 0.99989001 |
| PELI1 | 947.109268 | 0.04549184 | 0.61651501 | 0.0737887 | 0.94117852 | 0.99989001 |
| AFTPH | 251.798519 | 0.06932661 | 0.19232124 | 0.360473 | 0.71849345 | 0.99989001 |

| | | | | | | |
|---|---|---|---|---|---|---|
| SERTAD2 | 115.589641 | 0.37030352 | 0.34553557 | 1.07167987 | 0.28386385 | 0.99989001 |
| CEP68 | 186.005449 | -0.2030918 | 0.35779317 | -0.5676234 | 0.57029068 | 0.99989001 |
| RAB1A | 597.026626 | 0.01501941 | 0.23623576 | 0.06357806 | 0.9493062 | 0.99989001 |
| ACTR2 | 432.599943 | 0.05168679 | 0.24570438 | 0.2103617 | 0.83338538 | 0.99989001 |
| SPRED2 | 382.952136 | 0.36740102 | 0.2604248 | 1.41077584 | 0.15831072 | 0.99989001 |
| MEIS1 | 206.885367 | 0.07973965 | 0.30532796 | 0.26116064 | 0.79396863 | 0.99989001 |
| ETAA1 | 43.2453069 | -0.1148009 | 0.36059684 | -0.3183636 | 0.75020918 | NA |
| C1D | 92.2502853 | -0.1472962 | 0.25381961 | -0.5803186 | 0.56169982 | 0.99989001 |
| CNRIP1 | 150.453577 | 0.04468975 | 0.19390161 | 0.23047644 | 0.81772157 | 0.99989001 |
| APLF | 100.602037 | 0.29150658 | 0.34368114 | 0.84818906 | 0.39633269 | 0.99989001 |
| ARHGAP25 | 106.296594 | -0.1686413 | 0.45425373 | -0.371249 | 0.71045204 | 0.99989001 |
| GFPT1 | 160.583215 | 0.17595911 | 0.27568665 | 0.63825763 | 0.52330599 | 0.99989001 |
| NFU1 | 79.2024769 | 0.20840234 | 0.25079976 | 0.8309511 | 0.40600126 | 0.99989001 |
| AAK1 | 692.854455 | 0.16662098 | 0.22048328 | 0.75570802 | 0.44982427 | 0.99989001 |
| ANXA4 | 211.027128 | 0.12426243 | 0.25897708 | 0.47982017 | 0.63135527 | 0.99989001 |
| GMCL1 | 83.6692072 | -0.1596142 | 0.22680595 | -0.7037479 | 0.48158977 | 0.99989001 |
| MXD1 | 50.2671714 | -0.6182807 | 0.45746725 | -1.3515301 | 0.1765257 | 0.99989001 |
| PCBP1 | 213.184483 | 0.00193076 | 0.47864603 | 0.0040338 | 0.9967815 | 0.99989001 |
| C2orf42 | 48.1175826 | -0.2129871 | 0.31387697 | -0.6785689 | 0.49741107 | NA |
| PCYOX1 | 93.0622716 | 0.1222543 | 0.47803151 | 0.25574527 | 0.79814753 | 0.99989001 |
| SNRPG | 81.9675461 | -0.4080199 | 0.38143192 | -1.0697057 | 0.28475181 | 0.99989001 |
| NAGK | 51.1723155 | -0.1786737 | 0.29599844 | -0.6036306 | 0.54608927 | 0.99989001 |
| MCEE | 47.0175657 | 0.06639267 | 0.33133526 | 0.20037912 | 0.84118409 | NA |
| MPHOSPH10 | 67.4295053 | 0.16706726 | 0.41345474 | 0.4040763 | 0.68615661 | 0.99989001 |
| ZNF638 | 1022.6906 | -0.0880262 | 0.2219388 | -0.396624 | 0.69164478 | 0.99989001 |
| DYSF | 971.490795 | -0.6355825 | 0.34853018 | -1.8236082 | 0.06821132 | 0.88709358 |
| CYP26B1 | 234.83894 | -2.5732053 | 0.93059559 | -2.7651165 | 0.00569025 | 0.36709383 |
| EXOC6B | 1191.56673 | -0.3466467 | 0.23526097 | -1.4734562 | 0.14062806 | 0.99989001 |
| SFXN5 | 96.7236827 | -0.0061779 | 0.35217394 | -0.0175421 | 0.98600414 | 0.99989001 |
| CCT7 | 122.622877 | 0.17985445 | 0.41193046 | 0.43661363 | 0.66239157 | 0.99989001 |
| ALMS1 | 194.998084 | 0.20024736 | 0.2441121 | 0.82030904 | 0.41203996 | 0.99989001 |
| TPRKB | 89.3054654 | -0.448415 | 0.46265724 | -0.9692165 | 0.33243718 | 0.99989001 |
| DUSP11 | 157.815756 | -0.5292801 | 0.43060531 | -1.2291536 | 0.21901424 | 0.99989001 |
| STAMBP | 85.4435207 | 0.04116198 | 0.29416058 | 0.13993031 | 0.88871505 | 0.99989001 |
| DGUOK | 99.7882239 | -0.0985439 | 0.22281612 | -0.4422657 | 0.65829693 | 0.99989001 |
| TET3 | 117.715816 | -0.196583 | 0.38702014 | -0.5079401 | 0.61149535 | 0.99989001 |
| MOB1A | 165.911088 | 0.03833454 | 0.29424719 | 0.13028006 | 0.89634485 | 0.99989001 |
| MOGS | 46.6070602 | 0.06421167 | 0.4186928 | 0.15336226 | 0.87811261 | NA |
| AUP1 | 58.1257117 | -0.239024 | 0.33907945 | -0.7049205 | 0.48085973 | 0.99989001 |
| POLE4 | 107.332603 | -0.698681 | 0.49531634 | -1.4105753 | 0.15836988 | 0.99989001 |
| TACR1 | 420.35239 | 0.28186892 | 0.42304768 | 0.66628168 | 5.05E-01 | 0.99989001 |
| AC007681.1 | 252.690079 | 0.46415395 | 0.59028104 | 0.78632705 | 0.43167591 | 0.99989001 |
| MRPL19 | 125.929439 | 0.09777226 | 0.29264896 | 0.33409399 | 0.73830864 | 0.99989001 |
| GCFC2 | 154.367464 | 0.39555651 | 0.22811013 | 1.73405941 | 0.08290754 | 0.94799019 |
| SUCLG1 | 115.793363 | 0.26219265 | 0.22322349 | 1.17457461 | 0.24016495 | 0.99989001 |
| TMSB10 | 2791.17544 | -0.5323939 | 0.66985336 | -0.7947917 | 0.42673468 | 0.99989001 |
| KCMF1 | 328.147294 | 0.1359248 | 0.21972725 | 0.61860693 | 0.53617534 | 0.99989001 |
| TCF7L1 | 1069.44558 | 0.225177 | 0.28037087 | 0.80313976 | 0.42189396 | 0.99989001 |
| TGOLN2 | 289.62822 | -0.0941943 | 0.28018968 | -0.3361805 | 0.73673479 | 0.99989001 |
| ELMOD3 | 93.7222073 | -0.2748531 | 0.27060936 | -1.0156822 | 0.30978074 | 0.99989001 |
| MAT2A | 130.965735 | -1.1469353 | 0.41837002 | -2.7414375 | 0.0061171 | 0.38158569 |
| GGCX | 73.4833637 | -0.3286923 | 0.4009211 | -0.8198428 | 0.41230572 | 0.99989001 |
| VAMP8 | 132.588614 | 0.05326277 | 0.52413131 | 0.10162105 | 0.91905747 | 0.99989001 |
| VAMP5 | 459.658441 | -0.4268439 | 0.58997996 | -0.7234889 | 0.46937958 | 0.99989001 |

| | | | | | | |
|---|---|---|---|---|---|---|
| RNF181 | 40.1048662 | -0.4097716 | 0.41221433 | -0.9940741 | 0.32018679 | NA |
| USP39 | 128.275198 | -0.2281528 | 0.27771573 | -0.8215335 | 0.41134245 | 0.99989001 |
| C2orf68 | 46.0095091 | -0.4561547 | 0.36794149 | -1.2397478 | 0.21506869 | NA |
| POLR1A | 66.758874 | 0.1305559 | 0.53794818 | 0.24269234 | 0.80824374 | 0.99989001 |
| PTCD3 | 120.972627 | 0.24755788 | 0.24357617 | 1.01634688 | 0.30946423 | 0.99989001 |
| IMMT | 195.469983 | 0.24352726 | 0.19230906 | 1.26633275 | 0.20539398 | 0.99989001 |
| KDM3A | 213.488651 | 0.13345821 | 0.40980002 | 0.32566667 | 0.74467657 | 0.99989001 |
| CHMP3 | 74.6331751 | 0.11423193 | 0.51761765 | 0.22068786 | 0.82533549 | 0.99989001 |
| RMND5A | 139.732716 | 0.37656434 | 0.21870587 | 1.72178435 | 0.08510859 | 0.95514629 |
| CYTOR | 110.314145 | -1.2010905 | 0.44630478 | -2.6911889 | 0.00711979 | 0.39807515 |
| KRCC1 | 109.318789 | 0.65420664 | 0.31390311 | 2.0841037 | 0.03715074 | 0.70556534 |
| EIF2AK3 | 149.297518 | -0.0608825 | 0.3274296 | -0.1859406 | 0.85249135 | 0.99989001 |
| RPIA | 70.9242828 | -0.3362344 | 0.29550208 | -1.137841 | 0.25518688 | 0.99989001 |
| MRPS5 | 128.267325 | 0.03880293 | 0.22171668 | 0.17501132 | 0.86107074 | 0.99989001 |
| ANKRD36C | 194.816318 | 0.10336267 | 0.34709507 | 0.29779355 | 0.76586074 | 0.99989001 |
| STARD7 | 135.737282 | -0.0983143 | 0.33500379 | -0.2934723 | 0.76916115 | 0.99989001 |
| TMEM127 | 83.9812981 | 0.1873163 | 0.28925987 | 0.647571 | 0.51726246 | 0.99989001 |
| CIAO1 | 85.4785125 | 0.13759024 | 0.30569824 | 0.45008516 | 0.65264903 | 0.99989001 |
| SNRNP200 | 145.963027 | -0.4169589 | 0.23495272 | -1.7746501 | 0.07595568 | 0.92274869 |
| ARID5A | 110.626513 | -0.8919803 | 0.43996138 | -2.0274059 | 0.04262092 | 0.76066939 |
| KANSL3 | 146.10949 | -0.1146605 | 0.21894243 | -0.5237015 | 0.60048617 | 0.99989001 |
| LMAN2L | 40.1766175 | -0.04428 | 0.29929065 | -0.1479498 | 0.88238241 | NA |
| ANKRD23 | 45.7476668 | -0.1829395 | 0.44784741 | -0.4084863 | 0.68291668 | NA |
| SEMA4C | 128.474618 | 0.61703239 | 0.33403524 | 1.84720746 | 0.06471708 | 0.86610729 |
| ANKRD36 | 268.792125 | -0.0702161 | 0.30072283 | -0.2334912 | 0.81538002 | 0.99989001 |
| AC092683.1 | 161.00787 | -0.0970536 | 0.41162674 | -0.2357805 | 0.81360301 | 0.99989001 |
| ANKRD36B | 80.538795 | 0.09386636 | 0.46326722 | 0.20261818 | 0.83943348 | 0.99989001 |
| COX5B | 166.293517 | 0.04802276 | 0.453869 | 0.10580753 | 0.91573506 | 0.99989001 |
| ACTR1B | 77.7450085 | 0.02413668 | 0.27483344 | 0.08782295 | 0.93001739 | 0.99989001 |
| TMEM131 | 420.863176 | -0.1375352 | 0.17496966 | -0.7860519 | 0.43183709 | 0.99989001 |
| INPP4A | 194.414205 | 0.48913932 | 0.2510246 | 1.94857125 | 0.05134665 | 0.8125042 |
| COA5 | 47.1701322 | -0.3530391 | 0.36175202 | -0.9759146 | 0.3291068 | NA |
| UNC50 | 78.3945559 | 0.11799193 | 0.28369855 | 0.41590601 | 0.67747878 | 0.99989001 |
| MGAT4A | 488.303578 | -0.6743653 | 0.5016183 | -1.3443794 | 0.17882574 | 0.99989001 |
| KIAA1211L | 146.236649 | -1.1339099 | 0.36089233 | -3.1419617 | 0.0016782 | 0.20106436 |
| AC079447.1 | 122.1045 | 0.11055492 | 0.35955305 | 0.30747875 | 0.75847899 | 0.99989001 |
| MITD1 | 80.0939403 | -0.0388565 | 0.22454823 | -0.173043 | 0.86261764 | 0.99989001 |
| TXNDC9 | 68.6487518 | -0.0280869 | 0.33841129 | -0.0829964 | 0.93385439 | 0.99989001 |
| EIF5B | 230.076902 | 0.05413806 | 0.36951191 | 0.14651235 | 0.88351694 | 0.99989001 |
| REV1 | 272.000108 | 0.13466931 | 0.21135611 | 0.63716778 | 0.52401556 | 0.99989001 |
| AFF3 | 210.926945 | -2.4005189 | 0.66100939 | -3.6315958 | 0.00028167 | 0.07335471 |
| NPAS2 | 551.41786 | -2.1687585 | 0.72873105 | -2.9760753 | 0.00291963 | 0.27546724 |
| RPL31 | 174.524561 | 0.02662201 | 0.43133302 | 0.06172033 | 0.95078555 | 0.99989001 |
| TBC1D8 | 724.548159 | 0.57355409 | 0.490847 | 1.16849871 | 0.24260566 | 0.99989001 |
| CNOT11 | 63.4996633 | 0.09711223 | 0.28412069 | 0.34179922 | 0.732502 | 0.99989001 |
| RNF149 | 388.648174 | 0.08235455 | 0.4411839 | 0.18666716 | 0.85192161 | 0.99989001 |
| RFX8 | 50.6174472 | -0.2398068 | 1.07294828 | -0.2235026 | 0.82314434 | 0.99989001 |
| MAP4K4 | 1167.39824 | 0.1953237 | 0.44262689 | 0.44128296 | 0.65900816 | 0.99989001 |
| IL1R1 | 467.638066 | -0.2465559 | 0.60492999 | -0.4075775 | 0.68358386 | 0.99989001 |
| IL18R1 | 209.863003 | 1.45640536 | 0.58406976 | 2.49354695 | 0.01264739 | 0.49000287 |
| MRPS9 | 95.1592123 | -0.375724 | 0.25572486 | -1.469251 | 0.14176473 | 0.99989001 |
| TGFBRAP1 | 46.754389 | 0.03831791 | 0.400715 | 0.09562385 | 0.92381932 | NA |
| C2orf49 | 41.0576696 | 0.35386907 | 0.26854192 | 1.31774238 | 0.1875899 | NA |
| NCK2 | 159.789042 | 0.82669959 | 0.28731846 | 2.87729365 | 0.00401102 | 0.31536656 |

| | | | | | | |
|---|---|---|---|---|---|---|
| UXS1 | 79.4929367 | -0.1782733 | 0.29456852 | -0.6052014 | 0.5450452 | 0.99989001 |
| GCC2 | 305.144259 | 0.51430385 | 0.25041604 | 2.05379753 | 0.03999529 | 0.73029084 |
| LIMS1 | 457.266496 | 0.30872582 | 0.36293815 | 0.85062928 | 0.39497532 | 0.99989001 |
| RANBP2 | 281.67591 | 0.07327311 | 0.20091256 | 0.36470151 | 0.71533423 | 0.99989001 |
| CCDC138 | 58.6042194 | -0.1397734 | 0.38103683 | -0.3668239 | 0.71375041 | 0.99989001 |
| SH3RF3 | 2133.14451 | 0.42792809 | 0.72598895 | 0.5894416 | 0.55556508 | 0.99989001 |
| 10-Sep | 229.036231 | 0.16607356 | 0.19490744 | 0.85206374 | 0.39417872 | 0.99989001 |
| MALL | 95.2001042 | 0.03238723 | 0.58444104 | 0.05541574 | 0.95580726 | 0.99989001 |
| SMIM37 | 73.0254334 | -0.2512547 | 0.53091683 | -0.4732468 | 0.63603712 | 0.99989001 |
| MIR4435-2H | 421.49763 | -0.6915065 | 0.35534732 | -1.9460017 | 0.05165453 | 0.81270649 |
| BCL2L11 | 69.1903001 | 0.39761625 | 0.3739805 | 1.0632005 | 0.28769104 | 0.99989001 |
| ANAPC1 | 191.203034 | 0.00630929 | 0.20524972 | 0.0307396 | 0.97547721 | 0.99989001 |
| MERTK | 126.467169 | -1.1650939 | 0.52998044 | -2.1983714 | 0.02792265 | 0.65805899 |
| TMEM87B | 328.829091 | -0.3442809 | 0.26042336 | -1.3220046 | 0.18616661 | 0.99989001 |
| ZC3H8 | 73.3993167 | -0.1334945 | 0.2922308 | -0.4568118 | 0.64780632 | 0.99989001 |
| ZC3H6 | 174.863531 | -0.4328134 | 0.43947307 | -0.9848463 | 0.32469959 | 0.99989001 |
| TTL | 140.925486 | 0.1371143 | 0.27149769 | 0.50502935 | 0.61353822 | 0.99989001 |
| SLC20A1 | 165.976868 | 0.2234517 | 0.43468274 | 0.51405698 | 0.60721215 | 0.99989001 |
| PAX8 | 108.376037 | -0.3909195 | 0.41960318 | -0.931641 | 0.35152211 | 0.99989001 |
| CBWD2 | 48.5542504 | -0.4118392 | 0.42865951 | -0.9607606 | 0.33667257 | NA |
| SLC35F5 | 241.757299 | -0.0454011 | 0.20910844 | -0.2171176 | 0.8281167 | 0.99989001 |
| ACTR3 | 514.994027 | 0.02507859 | 0.26157319 | 0.09587598 | 0.92361907 | 0.99989001 |
| DDX18 | 134.995617 | 0.02437388 | 0.22610373 | 0.10779954 | 9.14E-01 | 0.99989001 |
| CCDC93 | 186.241284 | -0.0385187 | 0.20555387 | -0.1873898 | 0.85135502 | 0.99989001 |
| INSIG2 | 60.8600393 | 0.08032737 | 0.44186873 | 0.18179011 | 0.85574744 | 0.99989001 |
| DBI | 150.229512 | 0.50264011 | 0.51372859 | 0.97841569 | 0.32786877 | 0.99989001 |
| PTPN4 | 297.112669 | 0.03445447 | 0.2026182 | 0.17004629 | 0.86497373 | 0.99989001 |
| EPB41L5 | 153.141831 | -0.0396733 | 0.37332533 | -0.10627 | 0.91536817 | 0.99989001 |
| RALB | 173.274376 | 0.150536 | 0.28909787 | 0.52070948 | 0.60256917 | 0.99989001 |
| INHBB | 123.708098 | -0.673593 | 0.57647119 | -1.1684765 | 2.43E-01 | 1.00E+00 |
| CLASP1 | 835.43189 | 0.27879521 | 0.25394739 | 1.09784634 | 0.27227159 | 0.99989001 |
| NIFK-AS1 | 57.6717058 | -0.542135 | 0.42262416 | -1.2827828 | 0.19956817 | 0.99989001 |
| TSN | 71.3173364 | 0.01185636 | 0.26660399 | 0.04447179 | 0.96452834 | 0.99989001 |
| GYPC | 341.023682 | 0.21371328 | 0.4623128 | 0.46226987 | 0.6438878 | 0.99989001 |
| ERCC3 | 84.5804753 | -0.1580674 | 0.24754892 | -0.6385298 | 0.52312885 | 0.99989001 |
| MAP3K2 | 393.275347 | 0.14472413 | 0.16092196 | 0.89934357 | 0.36846969 | 0.99989001 |
| IWS1 | 197.131441 | 0.15790062 | 0.20296098 | 0.77798508 | 0.43657781 | 0.99989001 |
| AC010976.2 | 77.894767 | -2.1314951 | 0.68748114 | -3.1004416 | 0.00193232 | 0.20835966 |
| LIMS2 | 740.808199 | -0.0425532 | 0.42870224 | -0.0992606 | 0.9209314 | 0.99989001 |
| WDR33 | 334.061521 | 0.11418374 | 0.20200626 | 0.56524849 | 0.57190476 | 0.99989001 |
| AMMECR1L | 52.7558496 | 0.30135726 | 0.27096044 | 1.11218177 | 0.26606001 | 0.99989001 |
| SAP130 | 107.753893 | 0.00212228 | 0.38765838 | 0.00547462 | 0.99563191 | 0.99989001 |
| UGGT1 | 173.952816 | 0.10718408 | 0.21841707 | 0.49073126 | 0.62361653 | 0.99989001 |
| HS6ST1 | 141.850284 | 0.13269076 | 0.39197528 | 0.33851818 | 0.73497273 | 0.99989001 |
| SMPD4 | 87.2719587 | -0.3801047 | 0.22693596 | -1.6749425 | 0.09394551 | 0.99989001 |
| MZT2B | 124.803565 | 0.04270246 | 0.50679105 | 0.08426049 | 0.93284933 | 0.99989001 |
| PTPN18 | 74.3806936 | -0.1452963 | 0.31734879 | -0.4578443 | 0.64706432 | 0.99989001 |
| FAM168B | 161.118013 | -0.4511448 | 0.18193269 | -2.4797349 | 0.01314801 | 0.49620582 |
| PLEKHB2 | 169.251561 | -0.0389033 | 0.26626318 | -0.1461083 | 0.88383586 | 0.99989001 |
| MZT2A | 40.9507622 | -0.118633 | 0.40748243 | -0.2911365 | 0.77094693 | NA |
| NCKAP5 | 141.370476 | 1.57663117 | 0.73938591 | 2.1323522 | 0.0329779 | 0.67281094 |
| MGAT5 | 349.176981 | 0.06653132 | 0.31470074 | 0.2114114 | 0.83256626 | 0.99989001 |
| CCNT2 | 234.639632 | 0.06246781 | 0.27379527 | 0.22815519 | 0.81952559 | 0.99989001 |
| RAB3GAP1 | 502.362704 | 0.02679994 | 0.19401272 | 0.13813498 | 0.89013374 | 0.99989001 |

| | | | | | | |
|---|---|---|---|---|---|---|
| ZRANB3 | 142.518132 | -0.2149306 | 0.25662352 | -0.8375326 | 0.40229326 | 0.99989001 |
| R3HDM1 | 425.756485 | 0.16227813 | 0.16149538 | 1.00484693 | 0.31497056 | 0.99989001 |
| UBXN4 | 335.437618 | -0.1152351 | 0.16355276 | -0.7045745 | 0.48107506 | 0.99989001 |
| DARS | 267.480978 | 0.13328742 | 0.2633482 | 0.5061262 | 0.61276806 | 0.99989001 |
| AC114763.1 | 95.6672293 | -0.5531383 | 0.3518962 | -1.5718791 | 0.11597859 | 0.99989001 |
| SPOPL | 150.027615 | 0.19846134 | 0.16388492 | 1.21097985 | 0.22590313 | 0.99989001 |
| GTDC1 | 284.885776 | 0.12021529 | 0.14923944 | 0.80551959 | 0.4205199 | 0.99989001 |
| ZEB2 | 209.874932 | 0.31512852 | 0.34213006 | 0.92107814 | 0.35700963 | 0.99989001 |
| ACVR2A | 152.741491 | 0.18559934 | 0.23395123 | 0.79332491 | 0.42758855 | 0.99989001 |
| ORC4 | 181.581109 | 0.04059566 | 0.25667983 | 0.15815679 | 0.87433325 | 0.99989001 |
| MBD5 | 1421.20884 | -0.0158266 | 0.24464107 | -0.064693 | 0.94841842 | 0.99989001 |
| EPC2 | 285.015376 | -0.022592 | 0.22348425 | -0.1010897 | 0.91947922 | 0.99989001 |
| LYPD6B | 378.14079 | -1.2133562 | 0.62736571 | -1.9340494 | 0.05310706 | 0.81308738 |
| MMADHC | 75.9497523 | 0.09731913 | 0.29636929 | 0.32837118 | 0.74263103 | 0.99989001 |
| AC007364.1 | 93.1378505 | 0.31555931 | 0.24689326 | 1.2781204 | 0.20120698 | 0.99989001 |
| RND3 | 67.8978231 | -0.5734417 | 0.51814998 | -1.10671 | 0.26841935 | 0.99989001 |
| RBM43 | 41.452781 | -0.4348973 | 0.4324681 | -1.0056171 | 0.31459978 | NA |
| NMI | 85.9946635 | 0.20410182 | 0.45051206 | 0.45304409 | 0.650517 | 0.99989001 |
| RIF1 | 310.31806 | 0.02789518 | 0.228735 | 0.12195414 | 0.90293534 | 0.99989001 |
| ARL5A | 103.419739 | 0.34011962 | 0.37090923 | 0.91698883 | 0.35914849 | 0.99989001 |
| STAM2 | 174.265526 | 0.13706981 | 0.21990939 | 0.62330133 | 0.53308653 | 0.99989001 |
| FMNL2 | 2176.29721 | -0.8170828 | 0.50077682 | -1.6316306 | 0.10275732 | 0.99989001 |
| PRPF40A | 259.77922 | 0.0839621 | 0.22444003 | 0.37409591 | 0.70833296 | 0.99989001 |
| ARL6IP6 | 123.089121 | 0.37460355 | 0.27243038 | 1.37504323 | 0.16911804 | 0.99989001 |
| GPD2 | 230.286308 | 0.06062081 | 0.3173172 | 0.19104168 | 0.84849294 | 0.99989001 |
| ACVR1 | 145.602654 | 0.06332225 | 0.24253271 | 0.26108747 | 0.79402505 | 0.99989001 |
| CCDC148 | 63.2797291 | -0.2960432 | 0.5171136 | -0.5724915 | 0.56698903 | 0.99989001 |
| PKP4 | 2188.83945 | -0.0375403 | 0.36922293 | -0.1016737 | 0.91901569 | 0.99989001 |
| PKP4-AS1 | 51.8647002 | -0.6648079 | 0.65820341 | -1.0100342 | 0.31247891 | 0.99989001 |
| TANC1 | 1474.41764 | -0.5582364 | 0.24489913 | -2.2794545 | 0.02264006 | 0.60384164 |
| WDSUB1 | 110.776565 | 0.06978315 | 0.29844848 | 0.23381976 | 0.81512491 | 0.99989001 |
| BAZ2B | 1483.72157 | -0.056612 | 0.27242082 | -0.207811 | 0.83537653 | 0.99989001 |
| 7-Mar | 504.236326 | 0.12454011 | 0.15857425 | 0.78537408 | 0.43223427 | 0.99989001 |
| PLA2R1 | 75.4443659 | 1.17456671 | 0.5004523 | 2.34701031 | 0.01892473 | 0.56657733 |
| ITGB6 | 56.5787001 | -1.4197509 | 0.71748035 | -1.978801 | 0.04783841 | 0.77819898 |
| RBMS1 | 1944.70934 | 0.39888909 | 0.33890745 | 1.17698533 | 0.23920136 | 0.99989001 |
| TANK | 369.777987 | -0.0903238 | 0.28524111 | -0.3166576 | 0.7515034 | 0.99989001 |
| PSMD14 | 182.980359 | 0.08198323 | 0.40831457 | 0.20078448 | 0.8408671 | 0.99989001 |
| IFIH1 | 238.351563 | 0.1543185 | 0.62169839 | 0.24822085 | 0.80396354 | 0.99989001 |
| GCA | 132.497485 | -0.0267627 | 0.45296089 | -0.0590839 | 0.95288528 | 0.99989001 |
| COBLL1 | 500.538384 | -1.532582 | 0.54448064 | -2.8147594 | 0.00488138 | 0.34759088 |
| AC019197.1 | 171.24113 | -0.5234424 | 0.46248689 | -1.1317994 | 0.25771878 | 0.99989001 |
| CSRNP3 | 56.0328654 | -1.0621101 | 0.6144923 | -1.7284352 | 0.08391022 | 0.95384695 |
| TTC21B | 234.778476 | -0.0864841 | 0.23437001 | -0.3690068 | 0.71212263 | 0.99989001 |
| STK39 | 144.851268 | 0.00687692 | 0.32155194 | 0.02138665 | 0.98293722 | 0.99989001 |
| CERS6 | 245.628708 | 0.03925747 | 0.21920573 | 0.17908962 | 0.85786733 | 0.99989001 |
| NOSTRIN | 1119.65668 | -0.0551731 | 0.31706303 | -0.1740129 | 0.86185534 | 0.99989001 |
| SPC25 | 200.838898 | -0.1980704 | 0.64279409 | -0.3081397 | 0.75797601 | 0.99989001 |
| LRP2 | 100.484321 | 3.46597839 | 0.78039885 | 4.44129097 | 8.94E-06 | 0.00556981 |
| FASTKD1 | 104.57177 | -0.126624 | 0.2697289 | -0.4694491 | 0.63874864 | 0.99989001 |
| PPIG | 294.695417 | -0.0172129 | 0.15725491 | -0.1094586 | 0.91283872 | 0.99989001 |
| SSB | 113.013228 | 0.08712421 | 0.29935321 | 0.2910415 | 0.77101958 | 0.99989001 |
| METTL5 | 43.4650576 | -0.1781104 | 0.36271924 | -0.491042 | 0.62339673 | NA |
| UBR3 | 690.020824 | -0.0683497 | 0.17384432 | -0.3931662 | 0.69419674 | 0.99989001 |

| | | | | | | |
|---|---|---|---|---|---|---|
| GORASP2 | 106.522347 | 0.12087539 | 0.25017259 | 0.48316799 | 0.62897646 | 0.99989001 |
| TLK1 | 388.055345 | -0.0324123 | 0.17188478 | -0.1885698 | 0.85042999 | 0.99989001 |
| METTL8 | 82.9831764 | -0.1286716 | 0.34857393 | -0.3691372 | 0.71202549 | 0.99989001 |
| DCAF17 | 73.7629414 | 0.07346576 | 0.23924086 | 0.30707864 | 0.75878351 | 0.99989001 |
| CYBRD1 | 92.5795093 | 0.79565801 | 0.42317368 | 1.88021621 | 0.06007862 | 0.85885114 |
| DYNC1I2 | 308.081771 | 0.11866725 | 0.15740885 | 0.75387917 | 0.45092178 | 0.99989001 |
| SLC25A12 | 153.804825 | 0.06707283 | 0.22627426 | 0.29642271 | 0.76690729 | 0.99989001 |
| HAT1 | 121.951407 | -0.2014736 | 0.22675061 | -0.8885252 | 0.37425829 | 0.99989001 |
| ITGA6 | 455.272714 | 0.12448678 | 0.41629371 | 0.29903593 | 0.76491263 | 0.99989001 |
| AC078883.1 | 103.575659 | 0.11475786 | 0.550465 | 0.2084744 | 0.83485857 | 0.99989001 |
| PDK1 | 146.262506 | -0.4473164 | 0.57360523 | -0.7798332 | 0.43548909 | 0.99989001 |
| RAPGEF4 | 1583.82166 | 0.08829646 | 0.32578564 | 0.27102624 | 0.78637085 | 0.99989001 |
| MAP3K20 | 149.537534 | 0.41517072 | 0.31303822 | 1.32626208 | 0.1847529 | 0.99989001 |
| SP3 | 262.424297 | 0.0335699 | 0.14797942 | 0.22685518 | 0.82053636 | 0.99989001 |
| OLA1 | 246.607904 | -0.3386086 | 0.27543051 | -1.2293795 | 0.21892954 | 0.99989001 |
| CIR1 | 131.560936 | 0.02352541 | 0.24262619 | 0.09696155 | 0.92275693 | 0.99989001 |
| GPR155 | 60.3660439 | 0.14115039 | 0.29683347 | 0.47552045 | 0.63441606 | 0.99989001 |
| AC010894.3 | 314.466849 | 0.03636677 | 0.42479933 | 0.0856093 | 0.931777 | 0.99989001 |
| WIPF1 | 310.521962 | 0.21172786 | 0.32771015 | 0.64608269 | 0.5182258 | 0.99989001 |
| CHN1 | 677.515946 | 0.346229 | 0.25368398 | 1.36480436 | 0.17231455 | 0.99989001 |
| ATF2 | 325.618766 | -0.1449754 | 0.23539568 | -0.6158797 | 0.53797392 | 0.99989001 |
| ATP5MC3 | 152.857559 | 0.28334544 | 0.52957775 | 0.53504032 | 0.59262198 | 0.99989001 |
| LNPK | 86.4027477 | -0.0705135 | 0.21499884 | -0.3279714 | 0.7429333 | 0.99989001 |
| HOXD8 | 48.5065846 | -0.2215593 | 0.40718292 | -0.5441272 | 0.58635392 | NA |
| HOXD3 | 139.602241 | -0.5418994 | 0.46342321 | -1.1693403 | 0.24226656 | 0.99989001 |
| MTX2 | 243.63877 | -0.0446095 | 0.27179469 | -0.1641294 | 0.86962927 | 0.99989001 |
| AC074286.1 | 228.076338 | -3.2905576 | 1.00679009 | -3.2683651 | 0.00108171 | 0.16009268 |
| HNRNPA3 | 499.860813 | -0.1359071 | 0.19905568 | -0.6827593 | 0.49475894 | 0.99989001 |
| NFE2L2 | 747.42371 | -0.066027 | 0.22076251 | -0.299086 | 0.76487441 | 0.99989001 |
| AGPS | 196.250751 | 0.01757428 | 0.22111085 | 0.07948175 | 0.93664945 | 0.99989001 |
| RBM45 | 60.3327958 | -0.0037402 | 0.26112567 | -0.0143232 | 0.98857214 | 0.99989001 |
| AC009948.1 | 95.499836 | 0.01190884 | 0.35551944 | 0.03349702 | 0.97327824 | 0.99989001 |
| PRKRA | 107.174586 | 0.1521126 | 0.22333285 | 0.68110264 | 0.49580655 | 0.99989001 |
| PLEKHA3 | 99.2246276 | 0.27306364 | 0.24825942 | 1.09991249 | 0.27137025 | 0.99989001 |
| TTN-AS1 | 281.853031 | 0.33666824 | 0.23690886 | 1.4210876 | 0.1552913 | 0.99989001 |
| SESTD1 | 302.304867 | -0.2465198 | 0.30184441 | -0.8167116 | 0.41409327 | 0.99989001 |
| CWC22 | 109.339154 | 0.07465132 | 0.2011729 | 0.3710804 | 0.71057765 | 0.99989001 |
| UBE2E3 | 294.980682 | 0.31699151 | 0.21138584 | 1.49958728 | 0.13372134 | 0.99989001 |
| SSFA2 | 329.216641 | -0.2065754 | 0.29685047 | -0.6958906 | 0.48649737 | 0.99989001 |
| PDE1A | 88.3392831 | -0.3122913 | 0.53300664 | -0.5859051 | 0.55793927 | 0.99989001 |
| DNAJC10 | 182.412954 | -0.111112 | 0.27812579 | -0.3995025 | 0.68952295 | 0.99989001 |
| NCKAP1 | 840.028195 | -0.0353227 | 0.24607914 | -0.1435422 | 0.88586202 | 0.99989001 |
| NUP35 | 59.1604517 | -0.1797397 | 0.28657761 | -0.6271938 | 0.53053222 | 0.99989001 |
| ZC3H15 | 142.76821 | -0.2775549 | 0.27237862 | -1.0190041 | 0.30820103 | 0.99989001 |
| ITGAV | 516.299137 | 0.31638093 | 0.37657028 | 0.84016434 | 0.40081625 | 0.99989001 |
| AC017101.1 | 51.6461231 | 0.49356176 | 0.68536923 | 0.72013995 | 0.47143883 | 0.99989001 |
| FAM171B | 65.2427875 | -0.254613 | 0.52872743 | -0.4815581 | 0.63011986 | 0.99989001 |
| AC007319.1 | 1150.39847 | 0.19598382 | 0.56489198 | 0.34694034 | 0.72863614 | 0.99989001 |
| CALCRL | 1685.33027 | -0.1188247 | 0.52225217 | -0.2275236 | 0.82001664 | 0.99989001 |
| TFPI | 1415.08759 | 0.41200621 | 0.44246567 | 0.93115972 | 0.35177095 | 0.99989001 |
| GULP1 | 362.620739 | -0.2094911 | 0.517969 | -0.4044471 | 0.68588395 | 0.99989001 |
| COL3A1 | 225.838865 | -1.9264858 | 1.19185118 | -1.6163812 | 0.10601193 | 0.99989001 |
| COL5A2 | 91.126137 | -0.7570486 | 0.53715253 | -1.4093736 | 0.15872472 | 0.99989001 |
| WDR75 | 105.30173 | -0.2939796 | 0.20841515 | -1.4105479 | 0.15837796 | 0.99989001 |

| | | | | | | |
|---|---|---|---|---|---|---|
| SLC40A1 | 128.687403 | 0.06861303 | 0.57413001 | 0.11950783 | 0.90487304 | 0.99989001 |
| ASNSD1 | 52.1453223 | 0.57479414 | 0.3231152 | 1.77891394 | 0.07525387 | 0.9206525 |
| ANKAR | 188.526158 | 0.00250435 | 0.34318289 | 0.00729743 | 0.99417755 | 0.99989001 |
| ORMDL1 | 128.578792 | 0.16796926 | 0.25148393 | 0.66791251 | 0.50418944 | 0.99989001 |
| PMS1 | 155.795731 | -0.156742 | 0.18623027 | -0.8416568 | 0.3999801 | 0.99989001 |
| C2orf88 | 50.3273599 | 0.15499538 | 0.31679085 | 0.48926724 | 0.62465251 | 0.99989001 |
| HIBCH | 391.331957 | 0.03657685 | 0.26424314 | 0.1384212 | 0.88990755 | 0.99989001 |
| INPP1 | 147.449835 | -0.1727972 | 0.45989362 | -0.375733 | 0.7071154 | 0.99989001 |
| MFSD6 | 89.8102882 | 0.61394258 | 0.40399321 | 1.51968541 | 0.12859006 | 0.99989001 |
| NAB1 | 147.277704 | -0.1447007 | 0.19152387 | -0.7555231 | 0.44993516 | 0.99989001 |
| GLS | 835.606866 | -0.237216 | 0.31179491 | -0.7608077 | 0.44677194 | 0.99989001 |
| STAT1 | 254.955013 | 0.04691058 | 0.6324368 | 0.07417433 | 0.94087167 | 0.99989001 |
| MYO1B | 348.561089 | -0.7919498 | 0.58808565 | -1.3466572 | 0.17809067 | 0.99989001 |
| CAVIN2 | 562.234093 | -0.5219881 | 0.54770328 | -0.9530492 | 0.34056517 | 0.99989001 |
| AC098617.1 | 200.63323 | -0.2282092 | 0.43159522 | -0.5287575 | 0.59697367 | 0.99989001 |
| SLC39A10 | 187.255786 | -0.1023026 | 0.64303125 | -0.1590943 | 0.87359456 | 0.99989001 |
| HECW2 | 545.485185 | 0.72778289 | 0.35517719 | 2.04906992 | 0.04045528 | 0.73402991 |
| GTF3C3 | 86.8941041 | -0.2317995 | 0.20878659 | -1.110222 | 0.26690336 | 0.99989001 |
| PGAP1 | 100.236617 | 0.20530123 | 0.2488575 | 0.82497506 | 0.40938575 | 0.99989001 |
| ANKRD44 | 594.684799 | 0.65011069 | 0.28100514 | 2.31351881 | 0.02069413 | 0.58699182 |
| SF3B1 | 325.682137 | 0.02170619 | 0.20981282 | 0.103455 | 0.91760186 | 0.99989001 |
| COQ10B | 73.3381909 | -0.1853122 | 0.2759461 | -0.671552 | 0.50186894 | 0.99989001 |
| HSPD1 | 186.696717 | -0.0037811 | 0.34924689 | -0.0108264 | 0.99136198 | 0.99989001 |
| MOB4 | 57.7118382 | 0.18473062 | 0.47326181 | 0.39033494 | 0.69628889 | 0.99989001 |
| RFTN2 | 106.69319 | 0.25229203 | 0.49587343 | 0.50878313 | 0.61090425 | 0.99989001 |
| PLCL1 | 534.714327 | 0.206818 | 0.29297534 | 0.70592291 | 0.48023608 | 0.99989001 |
| C2orf69 | 46.5530841 | -0.1234059 | 0.39493012 | -0.3124753 | 0.75467934 | NA |
| TYW5 | 51.0471531 | -0.2177868 | 0.23932791 | -0.9099934 | 0.36282599 | 0.99989001 |
| SPATS2L | 397.435382 | 0.35606301 | 0.38010978 | 0.93673731 | 0.3488937 | 0.99989001 |
| BZW1 | 83.9424757 | 0.04439638 | 0.29645594 | 0.14975711 | 0.88095625 | 0.99989001 |
| CLK1 | 276.475034 | -0.093544 | 0.39026192 | -0.2396954 | 0.81056638 | 0.99989001 |
| ORC2 | 228.733603 | -0.3632355 | 0.20533663 | -1.7689756 | 0.07689795 | 0.92274869 |
| FAM126B | 254.238572 | 0.07630298 | 0.24309366 | 0.31388306 | 0.75360986 | 0.99989001 |
| NDUFB3 | 96.9093087 | 0.05504745 | 0.59334201 | 0.09277524 | 0.92608212 | 0.99989001 |
| CFLAR | 1236.04714 | 0.01998357 | 0.23710365 | 0.08428199 | 9.33E-01 | 0.99989001 |
| CFLAR-AS1 | 105.4567 | 0.15840988 | 0.48400396 | 0.32729046 | 0.74344821 | 0.99989001 |
| CASP10 | 208.065764 | 0.11654433 | 0.26656473 | 0.43720835 | 0.66196026 | 0.99989001 |
| CASP8 | 60.3270909 | 0.39164484 | 0.42316791 | 0.92550696 | 0.35470227 | 0.99989001 |
| TRAK2 | 173.912281 | 0.02179561 | 0.27210156 | 0.08010102 | 0.93615691 | 0.99989001 |
| STRADB | 53.871626 | 0.34632983 | 0.4817987 | 0.71882682 | 0.47224763 | 0.99989001 |
| ALS2 | 139.800373 | -0.2336725 | 0.22553187 | -1.0360954 | 0.30015762 | 0.99989001 |
| SUMO1 | 153.307935 | -0.0631499 | 0.31287902 | -0.2018349 | 0.8400458 | 0.99989001 |
| NOP58 | 138.712398 | -0.188479 | 0.21049752 | -0.8953979 | 0.37057444 | 0.99989001 |
| BMPR2 | 2496.13784 | 0.25036391 | 0.35215887 | 0.71094025 | 0.47712127 | 0.99989001 |
| FAM117B | 88.3683229 | 0.00485691 | 0.45780802 | 0.01060905 | 0.99153536 | 0.99989001 |
| ICA1L | 52.9008986 | 0.23312974 | 0.40435502 | 0.57654718 | 0.56424539 | 0.99989001 |
| WDR12 | 66.8980659 | -0.1419893 | 0.40799592 | -0.3480166 | 0.72782775 | 0.99989001 |
| CARF | 88.4535082 | 0.16412276 | 0.49718251 | 0.33010566 | 0.74132013 | 0.99989001 |
| NBEAL1 | 858.05394 | -0.1460215 | 0.19138889 | -0.7629568 | 0.44548915 | 0.99989001 |
| CYP20A1 | 207.958933 | -0.0122981 | 0.35709391 | -0.0344394 | 0.97252674 | 0.99989001 |
| ABI2 | 308.964176 | 0.31525643 | 0.19158521 | 1.64551549 | 0.09986355 | 0.99989001 |
| RAPH1 | 327.843295 | 0.76648621 | 0.303317 | 2.52701366 | 0.0115037 | 0.48516963 |
| PARD3B | 600.046576 | 0.10806626 | 0.24232299 | 0.44595959 | 0.65562644 | 0.99989001 |
| NRP2 | 224.922994 | 1.1661276 | 0.45678506 | 2.55290224 | 0.01068295 | 0.47431649 |

| | | | | | | |
|---|---|---|---|---|---|---|
| INO80D | 514.722094 | -0.027433 | 0.61115442 | -0.0448871 | 0.9641973 | 0.99989001 |
| NDUFS1 | 227.905525 | 0.2320396 | 0.15242875 | 1.52228235 | 0.12793835 | 0.99989001 |
| EEF1B2 | 134.213598 | -0.0945515 | 0.51873201 | -0.1822743 | 0.85536745 | 0.99989001 |
| ZDBF2 | 69.9513975 | 0.37518636 | 0.37780277 | 0.99307466 | 0.32067357 | 0.99989001 |
| FASTKD2 | 72.14534 | -0.0344792 | 0.26820188 | -0.1285568 | 0.89770838 | 0.99989001 |
| KLF7 | 861.954828 | 0.24385914 | 0.27723301 | 0.87961797 | 0.3790663 | 0.99989001 |
| CREB1 | 326.444872 | 0.2876064 | 0.1680357 | 1.71157915 | 0.08697425 | 0.95976851 |
| METTL21A | 79.6042999 | 0.18854673 | 0.35326675 | 0.53372339 | 0.59353292 | 0.99989001 |
| CCNYL1 | 145.472033 | -0.3989745 | 0.22416411 | -1.7798323 | 0.07510341 | 0.9206525 |
| PLEKHM3 | 174.339301 | 0.32848452 | 0.28295328 | 1.16091435 | 0.24567673 | 0.99989001 |
| IDH1 | 44.8012064 | 0.06122382 | 0.46981312 | 0.13031527 | 0.896317 | NA |
| PIKFYVE | 296.670542 | -0.1028273 | 0.19383185 | -0.5304975 | 0.59576704 | 0.99989001 |
| PTH2R | 94.5957084 | -0.2082755 | 0.47295294 | -0.4403725 | 0.65966735 | 0.99989001 |
| MAP2 | 362.675282 | -0.4215508 | 0.4961878 | -0.8495792 | 0.39555908 | 0.99989001 |
| RPE | 77.0879456 | 0.01544792 | 0.2475348 | 0.06240708 | 0.95023866 | 0.99989001 |
| KANSL1L | 377.006956 | 0.32029804 | 0.28855769 | 1.10999658 | 0.2670005 | 0.99989001 |
| AC007038.1 | 72.1516381 | -0.2983268 | 0.27949726 | -1.0673693 | 0.28580512 | 0.99989001 |
| LANCL1 | 103.710795 | 0.13229203 | 0.20353061 | 0.64998591 | 0.51570132 | 0.99989001 |
| ERBB4 | 199.576786 | -0.5205762 | 0.40434041 | -1.2874702 | 0.19793045 | 0.99989001 |
| IKZF2 | 49.3527476 | 0.10821203 | 0.40375806 | 0.26801207 | 0.78869003 | NA |
| SPAG16 | 162.426255 | 0.04100302 | 0.26988505 | 0.15192773 | 0.87924394 | 0.99989001 |
| BARD1 | 78.6025998 | 0.12689766 | 0.32246986 | 0.39351791 | 0.693937 | 0.99989001 |
| FN1 | 1116.45958 | -0.3458392 | 0.5373186 | -0.643639 | 0.51980953 | 0.99989001 |
| XRCC5 | 525.32262 | 0.06549036 | 0.19815504 | 0.33050061 | 0.74102173 | 0.99989001 |
| SMARCAL1 | 79.4240976 | -0.2702334 | 0.2822429 | -0.9574498 | 0.33834025 | 0.99989001 |
| RPL37A | 346.59733 | -0.1130838 | 0.45021753 | -0.2511759 | 0.80167813 | 0.99989001 |
| IGFBP2 | 128.919575 | -1.1195756 | 0.77500492 | -1.4446045 | 0.148569 | 0.99989001 |
| IGFBP5 | 4716.48657 | -0.2508499 | 0.67255982 | -0.3729778 | 0.70916495 | 0.99989001 |
| AC007563.2 | 219.08366 | 0.02435913 | 0.6221237 | 0.0391548 | 0.96876697 | 0.99989001 |
| DIRC3 | 139.33102 | 0.80849423 | 0.4665641 | 1.7328685 | 0.08311904 | 0.94799019 |
| TNS1 | 856.171305 | 0.29702585 | 0.40433068 | 0.7346112 | 0.46257631 | 0.99989001 |
| ARPC2 | 431.978881 | -0.1412115 | 0.22670911 | -0.6228755 | 0.53336635 | 0.99989001 |
| AAMP | 57.8602611 | -0.0074476 | 0.26125871 | -0.0285065 | 0.97725819 | 0.99989001 |
| PNKD | 90.3317058 | -0.106996 | 0.34614656 | -0.309106 | 0.7572409 | 0.99989001 |
| TMBIM1 | 142.840098 | 0.00139577 | 0.25897729 | 0.00538953 | 0.99569979 | 0.99989001 |
| CTDSP1 | 119.505788 | 0.08244458 | 0.29826219 | 0.27641648 | 0.7822282 | 0.99989001 |
| USP37 | 210.601297 | -0.2185279 | 0.28987064 | -0.7538807 | 0.45092087 | 0.99989001 |
| CNOT9 | 123.199992 | -0.1532935 | 0.22371597 | -0.6852149 | 0.49320831 | 0.99989001 |
| CYP27A1 | 61.3660482 | 0.30433847 | 0.39596731 | 0.76859495 | 0.44213381 | 0.99989001 |
| RETREG2 | 53.8249619 | -0.383949 | 0.37420181 | -1.0260479 | 0.304869 | 0.99989001 |
| ZFAND2B | 39.6621557 | -0.40498 | 0.34902641 | -1.1603134 | 0.24592124 | NA |
| ANKZF1 | 45.9201137 | -0.1313452 | 0.34120881 | -0.3849407 | 0.70028134 | NA |
| TUBA4A | 57.8945982 | 0.69923361 | 0.60331976 | 1.15897681 | 0.24646564 | 0.99989001 |
| DNAJB2 | 48.9714259 | -0.0444492 | 0.35911196 | -0.1237754 | 0.90149311 | NA |
| DNPEP | 66.2770494 | -0.1921165 | 0.37542587 | -0.5117295 | 0.60884036 | 0.99989001 |
| EPHA4 | 395.721643 | -0.2687761 | 0.33251805 | -0.8083053 | 0.41891487 | 0.99989001 |
| FARSB | 91.6003582 | -0.3645159 | 0.43433197 | -0.8392565 | 0.4013254 | 0.99989001 |
| ACSL3 | 299.612158 | 0.43689997 | 0.16824974 | 2.59673489 | 0.00941145 | 0.44677767 |
| KCNE4 | 86.4073886 | -1.7597047 | 1.12991647 | -1.5573759 | 0.11938127 | 0.99989001 |
| WDFY1 | 296.880685 | 0.29957432 | 0.20974151 | 1.4283025 | 0.15320481 | 0.99989001 |
| SERPINE2 | 269.337982 | 0.25360833 | 0.47781953 | 0.53076176 | 0.59558388 | 0.99989001 |
| CUL3 | 502.314125 | 0.05690272 | 0.15627859 | 0.3641108 | 0.71577527 | 0.99989001 |
| RHBDD1 | 185.192868 | 0.28671932 | 0.31489029 | 0.91053717 | 0.36253929 | 0.99989001 |
| COL4A3 | 44.0452953 | -0.02729 | 0.46906501 | -0.0581795 | 0.95360567 | NA |

| | | | | | | |
|---|---|---|---|---|---|---|
| MFF | 142.669491 | 0.23703387 | 0.22325438 | 1.06172103 | 0.28836235 | 0.99989001 |
| AGFG1 | 391.028122 | -0.0267178 | 0.20331949 | -0.131408 | 0.89545255 | 0.99989001 |
| TRIP12 | 712.970249 | -0.0815258 | 0.22130793 | -0.3683818 | 0.71258855 | 0.99989001 |
| SLC16A14 | 62.548911 | -1.4593929 | 0.57257354 | -2.5488304 | 0.01080848 | 0.47431649 |
| SP110 | 208.69762 | -0.0422425 | 0.57986344 | -0.072849 | 0.94192628 | 0.99989001 |
| SP140L | 158.835071 | -0.2806168 | 0.26281826 | -1.0677218 | 0.28564604 | 0.99989001 |
| SP100 | 913.297889 | -0.0575391 | 0.34320525 | -0.1676522 | 0.86685694 | 0.99989001 |
| CAB39 | 380.301844 | 0.13983796 | 0.18529937 | 0.75465969 | 0.4504532 | 0.99989001 |
| PSMD1 | 341.06318 | 0.08515959 | 0.3004822 | 0.28340977 | 0.77686274 | 0.99989001 |
| ARMC9 | 214.699369 | -0.2906122 | 0.3628142 | -0.8009947 | 0.42313474 | 0.99989001 |
| NCL | 241.543911 | 0.08890255 | 0.45091898 | 0.19715858 | 0.84370344 | 0.99989001 |
| PTMA | 1252.78861 | -0.1087489 | 0.4883261 | -0.2226973 | 0.82377111 | 0.99989001 |
| PDE6D | 113.783219 | 0.24759889 | 0.2900696 | 0.8535844 | 0.3933353 | 0.99989001 |
| COPS7B | 60.6293025 | -0.3178437 | 0.2999212 | -1.0597572 | 0.28925506 | 0.99989001 |
| DIS3L2 | 351.616491 | 0.21553229 | 0.28283677 | 0.76203773 | 0.44603748 | 0.99989001 |
| EIF4E2 | 75.6924111 | 0.36096897 | 0.29224469 | 1.23516007 | 0.21677094 | 0.99989001 |
| GIGYF2 | 444.658013 | -0.1605748 | 0.21275758 | -0.7547313 | 0.4504102 | 0.99989001 |
| INPP5D | 729.706665 | 0.70625922 | 0.38341164 | 1.84203905 | 0.06546944 | 0.8666545 |
| ATG16L1 | 56.0462142 | -0.1868951 | 0.26683398 | -0.7004172 | 0.48366679 | 0.99989001 |
| DGKD | 106.925813 | -0.232451 | 0.24390919 | -0.9530226 | 0.34057861 | 0.99989001 |
| USP40 | 205.542149 | 0.21354083 | 0.32939014 | 0.6482915 | 0.51679643 | 0.99989001 |
| SH3BP4 | 118.102195 | 0.30251631 | 0.37384493 | 0.80920265 | 0.41839859 | 0.99989001 |
| AGAP1 | 652.011257 | -0.0534865 | 0.3115743 | -0.1716654 | 0.86370057 | 0.99989001 |
| ACKR3 | 78.2625044 | 0.70708486 | 0.5893044 | 1.19986353 | 0.23019234 | 0.99989001 |
| COPS8 | 61.0927975 | 0.00500977 | 0.29387172 | 0.01704747 | 0.98639875 | 0.99989001 |
| LRRFIP1 | 1062.80966 | -0.3118437 | 0.24728296 | -1.2610804 | 0.20727986 | 0.99989001 |
| ILKAP | 153.652551 | 0.23723666 | 0.29476766 | 0.80482592 | 0.42092014 | 0.99989001 |
| PER2 | 49.8908281 | -0.4315185 | 0.43087967 | -1.0014826 | 0.31659357 | 0.99989001 |
| TRAF3IP1 | 67.3072097 | 0.14778795 | 0.37036684 | 0.39903127 | 0.68987016 | 0.99989001 |
| HDAC4 | 376.565026 | 0.09102081 | 0.38798979 | 0.23459588 | 0.81452241 | 0.99989001 |
| NDUFA10 | 205.879698 | -0.153517 | 0.20803357 | -0.7379433 | 0.46054893 | 0.99989001 |
| COPS9 | 78.2256524 | -0.164997 | 0.50269074 | -0.3282277 | 0.74273947 | 0.99989001 |
| ANKMY1 | 82.4056336 | -0.2891901 | 0.40136803 | -0.7205111 | 0.47121038 | 0.99989001 |
| RNPEPL1 | 60.323703 | -0.2407601 | 0.42204564 | -0.5704598 | 0.56836589 | 0.99989001 |
| SNED1 | 764.532895 | 0.50374397 | 0.38084392 | 1.32270452 | 0.18593366 | 0.99989001 |
| AC005237.1 | 78.2034641 | 0.53785568 | 0.51357134 | 1.04728523 | 0.29496805 | 0.99989001 |
| MTERF4 | 133.759756 | 0.35317953 | 0.23988611 | 1.47228004 | 0.14094527 | 0.99989001 |
| PPP1R7 | 66.7312209 | 0.09995547 | 0.25743204 | 0.38827907 | 0.69780952 | 0.99989001 |
| HDLBP | 315.808103 | -0.1242614 | 0.18337877 | -0.6776218 | 0.49801155 | 0.99989001 |
| 2-Sep | 526.964191 | 0.13920606 | 0.17369124 | 0.80145701 | 0.42286712 | 0.99989001 |
| FARP2 | 154.506939 | 0.25886098 | 0.25864399 | 1.00083898 | 0.31690466 | 0.99989001 |
| STK25 | 100.932036 | -0.0202662 | 0.26039665 | -0.0778282 | 0.93796471 | 0.99989001 |
| THAP4 | 61.2412233 | -0.2616048 | 0.26296446 | -0.9948296 | 0.31981916 | 0.99989001 |
| ATG4B | 145.009917 | -0.0963649 | 0.20764301 | -0.4640893 | 0.64258378 | 0.99989001 |
| ING5 | 66.7171288 | -0.3205692 | 0.39055344 | -0.8208074 | 0.41175597 | 0.99989001 |
| D2HGDH | 87.6332108 | 0.11406407 | 0.38670333 | 0.29496533 | 0.76802037 | 0.99989001 |
| CNTN4 | 146.592634 | -0.8678488 | 0.77072483 | -1.1260164 | 0.26015859 | 0.99989001 |
| TRNT1 | 123.054892 | -0.1333805 | 0.25676068 | -0.519474 | 0.60343023 | 0.99989001 |
| CRBN | 258.591365 | -0.1760586 | 0.29146702 | -0.6040429 | 0.54581514 | 0.99989001 |
| AC034195.1 | 96.3242156 | 0.75600748 | 0.4194754 | 1.80226894 | 0.0715031 | 0.91166457 |
| SUMF1 | 532.053804 | -0.2387396 | 0.37223663 | -0.6413651 | 0.52128547 | 0.99989001 |
| ITPR1 | 350.115468 | -0.1523669 | 0.30247043 | -0.5037415 | 0.61444304 | 0.99989001 |
| AC018816.1 | 279.910331 | -0.4764534 | 0.44448704 | -1.0719174 | 0.28375713 | 0.99989001 |
| BHLHE40 | 470.989684 | -1.4893562 | 0.61649899 | -2.4158291 | 0.01569943 | 0.52666344 |

| | | | | | | |
|---|---|---|---|---|---|---|
| ARL8B | 260.62919 | -0.2959608 | 0.22093633 | -1.3395752 | 0.1803835 | 0.99989001 |
| EDEM1 | 72.7370712 | -0.0216226 | 0.29805983 | -0.0725446 | 0.94216849 | 0.99989001 |
| LMCD1-AS1 | 226.152412 | -0.5539454 | 0.40986233 | -1.3515401 | 0.17652248 | 0.99989001 |
| LMCD1 | 391.992905 | -1.0077263 | 0.5467332 | -1.8431775 | 0.0653031 | 0.8666545 |
| RAD18 | 69.1523632 | -0.3649273 | 0.22122584 | -1.6495689 | 0.09903115 | 0.99989001 |
| THUMPD3-A: | 168.373651 | 0.09105862 | 0.28409803 | 0.32051829 | 0.74857547 | 0.99989001 |
| SETD5 | 719.456912 | 0.23139059 | 0.18019526 | 1.28411032 | 0.19910336 | 0.99989001 |
| MTMR14 | 105.680415 | 0.12349235 | 0.27167821 | 0.45455374 | 0.64943032 | 0.99989001 |
| OGG1 | 65.4826535 | -0.120233 | 0.27756249 | -0.4331746 | 0.66488792 | 0.99989001 |
| CAMK1 | 58.9057696 | 0.71927895 | 0.387913 | 1.8542275 | 0.06370662 | 0.86329904 |
| TADA3 | 42.0512169 | -0.6952811 | 0.29396144 | -2.3652117 | 0.01801977 | NA |
| CRELD1 | 54.9616244 | -0.0732309 | 0.42792261 | -0.1711312 | 0.86412062 | 0.99989001 |
| EMC3 | 116.285439 | -0.176912 | 0.20994049 | -0.8426767 | 0.39940927 | 0.99989001 |
| BRK1 | 144.940456 | 0.016584 | 0.41540677 | 0.03992231 | 0.96815506 | 0.99989001 |
| VHL | 65.2959043 | -0.0798709 | 0.31352814 | -0.2547489 | 0.79891705 | 0.99989001 |
| IRAK2 | 242.752468 | 0.2055002 | 0.39690623 | 0.51775503 | 0.6046292 | 0.99989001 |
| SEC13 | 127.841002 | 0.18875043 | 0.31529145 | 0.59865382 | 0.54940376 | 0.99989001 |
| HRH1 | 164.571938 | 0.33395342 | 0.40753412 | 0.81944899 | 0.41253029 | 0.99989001 |
| ATG7 | 732.783279 | -0.0743888 | 0.42821381 | -0.1737187 | 0.86208651 | 0.99989001 |
| VGLL4 | 848.00126 | 0.52511019 | 0.24750407 | 2.12162243 | 0.03386946 | 0.67281094 |
| TAMM41 | 140.6153 | -0.0084998 | 0.31253987 | -0.0271958 | 0.9783036 | 0.99989001 |
| PPARG | 956.135045 | -0.2052349 | 0.5201084 | -0.3946003 | 0.69313788 | 0.99989001 |
| TSEN2 | 99.7266299 | -0.2139424 | 0.36300849 | -0.5893593 | 0.55562031 | 0.99989001 |
| RAF1 | 295.930546 | -0.05208 | 0.18383169 | -0.2833028 | 0.77694477 | 0.99989001 |
| RPL32 | 571.829328 | -0.2495927 | 0.56302305 | -0.4433081 | 0.65754292 | 0.99989001 |
| IQSEC1 | 249.296047 | 1.26073399 | 0.2630609 | 4.79255556 | 1.65E-06 | 0.00207155 |
| FBLN2 | 240.109929 | -0.835409 | 0.57303625 | -1.4578641 | 0.14487801 | 0.99989001 |
| XPC | 138.5166 | 0.28721841 | 0.20264353 | 1.41735791 | 0.15637831 | 0.99989001 |
| LSM3 | 71.5076616 | -0.0600038 | 0.53131428 | -0.1129347 | 0.91008234 | 0.99989001 |
| SLC6A6 | 263.352927 | 0.08974793 | 0.43504558 | 0.20629546 | 0.83656012 | 0.99989001 |
| GRIP2 | 50.1489841 | 0.96861143 | 0.59489 | 1.62821938 | 1.03E-01 | 1.00E+00 |
| CCDC174 | 59.8428568 | -0.4164716 | 0.28062829 | -1.4840686 | 0.13779073 | 0.99989001 |
| FGD5 | 961.150971 | 0.28978423 | 0.30872297 | 0.93865459 | 0.34790811 | 0.99989001 |
| FGD5-AS1 | 142.604336 | -0.0863469 | 0.31388387 | -0.2750918 | 0.78324574 | 0.99989001 |
| NR2C2 | 315.932268 | 0.04916874 | 0.17262219 | 0.2848344 | 0.77577102 | 0.99989001 |
| MRPS25 | 84.2275359 | -0.4317771 | 0.26057894 | -1.6569916 | 0.09752117 | 0.99989001 |
| RBSN | 77.5261359 | -0.2404779 | 0.23936644 | -1.0046435 | 0.31506854 | 0.99989001 |
| CAPN7 | 288.708641 | -0.0448992 | 0.24072163 | -0.1865193 | 0.85203758 | 0.99989001 |
| SH3BP5 | 1839.70889 | 0.12545787 | 0.47942925 | 0.26168172 | 0.79356684 | 0.99989001 |
| METTL6 | 47.24576 | 0.08799041 | 0.32105132 | 0.27406962 | 0.78403112 | NA |
| HACL1 | 101.264374 | -0.1724724 | 0.2471298 | -0.6979021 | 0.4852384 | 0.99989001 |
| BTD | 40.7943309 | -0.3329516 | 0.30768373 | -1.082123 | 0.27919787 | NA |
| ANKRD28 | 611.660966 | -1.1264633 | 0.44292639 | -2.5432291 | 0.01098332 | 0.4764488 |
| GALNT15 | 767.562782 | 0.0465597 | 0.5168223 | 0.09008841 | 9.28E-01 | 0.99989001 |
| OXNAD1 | 100.929221 | -0.044352 | 0.26806245 | -0.1654538 | 0.86858681 | 0.99989001 |
| RFTN1 | 338.45093 | 0.60281711 | 0.21761636 | 2.77009099 | 0.00560406 | 0.36465062 |
| PLCL2 | 504.683619 | 0.45805498 | 0.25882967 | 1.76971589 | 0.07677448 | 0.92274869 |
| TBC1D5 | 1635.21311 | 0.03588426 | 0.26184392 | 0.13704448 | 0.89099564 | 0.99989001 |
| RAB5A | 254.103993 | -0.2517519 | 0.17721985 | -1.4205627 | 0.15544394 | 0.99989001 |
| KAT2B | 316.392121 | -0.4763945 | 0.41141635 | -1.1579376 | 0.24688949 | 0.99989001 |
| ZNF385D | 1011.71814 | -3.0831095 | 0.84540801 | -3.6468894 | 0.00026543 | 0.07335471 |
| UBE2E2 | 816.954991 | 0.15878008 | 0.51858519 | 0.30617935 | 0.75946809 | 0.99989001 |
| UBE2E1 | 680.036747 | 0.05038022 | 0.29613443 | 0.17012619 | 0.86491089 | 0.99989001 |
| RPL15 | 704.093049 | 0.1517609 | 0.56180134 | 0.27013267 | 0.78705819 | 0.99989001 |

| | | | | | | |
|---|---|---|---|---|---|---|
| NR1D2 | 175.611885 | -0.1463924 | 0.33646842 | -0.4350851 | 0.66350064 | 0.99989001 |
| THRB | 337.175687 | -0.4414329 | 0.67802295 | -0.6510589 | 0.51500849 | 0.99989001 |
| AC092422.1 | 106.04396 | -0.4731513 | 0.3834935 | -1.2337922 | 0.21728036 | 0.99989001 |
| RARB | 911.333422 | -0.788522 | 0.23414642 | -3.3676448 | 0.00075813 | 0.1300541 |
| TOP2B | 300.356498 | -0.0601122 | 0.21664408 | -0.2774697 | 0.78141946 | 0.99989001 |
| NGLY1 | 207.551767 | -0.3230949 | 0.37995342 | -0.850354 | 0.39512827 | 0.99989001 |
| SLC4A7 | 317.305866 | -0.6242766 | 0.47809972 | -1.3057457 | 0.19163907 | 0.99989001 |
| CMC1 | 177.32008 | -0.1225777 | 0.26784006 | -0.4576525 | 0.64720209 | 0.99989001 |
| AZI2 | 133.629467 | 0.25236099 | 0.38572028 | 0.65425906 | 0.51294492 | 0.99989001 |
| RBMS3 | 1256.06314 | -0.2141788 | 0.30035951 | -0.7130748 | 0.47579944 | 0.99989001 |
| RBMS3-AS3 | 540.490308 | -0.397918 | 0.48013643 | -0.8287603 | 0.40724006 | 0.99989001 |
| RBMS3-AS2 | 141.742942 | -0.1037583 | 0.59938595 | -0.1731076 | 0.86256681 | 0.99989001 |
| TGFBR2 | 2538.35157 | 0.70194089 | 0.30642819 | 2.29071905 | 0.02197967 | 0.60384164 |
| STT3B | 426.705455 | 0.05450499 | 0.28398694 | 0.19192779 | 0.84779877 | 0.99989001 |
| OSBPL10 | 484.425164 | 0.16381218 | 0.37645436 | 0.43514487 | 0.66345728 | 0.99989001 |
| GPD1L | 43.0349745 | 0.22969308 | 0.43401624 | 0.52922694 | 0.59664803 | NA |
| CMTM8 | 597.658535 | 0.62474212 | 0.42552112 | 1.46818125 | 0.142055 | 0.99989001 |
| CMTM7 | 130.33951 | 0.07844509 | 0.48534844 | 0.16162633 | 0.87160012 | 0.99989001 |
| CMTM6 | 165.785081 | 0.27516947 | 0.30862097 | 0.89160974 | 0.37260215 | 0.99989001 |
| DYNC1LI1 | 123.483513 | 0.37531783 | 0.22038944 | 1.70297559 | 0.08857264 | 0.97313867 |
| CNOT10 | 171.671414 | 0.06346158 | 0.24763302 | 0.25627269 | 0.79774028 | 0.99989001 |
| GLB1 | 130.607357 | 0.02424091 | 0.20313521 | 0.11933387 | 0.90501085 | 0.99989001 |
| CRTAP | 91.0934482 | -0.0050368 | 0.38101446 | -0.0132195 | 0.98945268 | 0.99989001 |
| FBXL2 | 66.4425813 | -0.6457954 | 0.34960229 | -1.8472286 | 0.06471401 | 0.86610729 |
| UBP1 | 175.994876 | -0.02741 | 0.2587177 | -0.1059457 | 0.91562543 | 0.99989001 |
| CLASP2 | 541.213947 | -0.1145154 | 0.24731866 | -0.4630275 | 0.64334463 | 0.99989001 |
| PDCD6IP | 403.377702 | 0.07331382 | 0.19765884 | 0.37091094 | 0.71070387 | 0.99989001 |
| TRANK1 | 110.192161 | 0.7761417 | 0.56196347 | 1.38112482 | 0.16724059 | 0.99989001 |
| EPM2AIP1 | 81.0882921 | 0.38851328 | 0.23264228 | 1.67000288 | 0.09491879 | 0.99989001 |
| MLH1 | 67.5980288 | -0.2475001 | 0.27069573 | -0.9143111 | 0.36055338 | 0.99989001 |
| LRRFIP2 | 448.480805 | 0.11030608 | 0.24722145 | 0.4461833 | 0.65546485 | 0.99989001 |
| GOLGA4 | 697.15413 | -0.2803889 | 0.21187056 | -1.3233972 | 0.18570332 | 0.99989001 |
| ITGA9 | 877.798296 | 0.0739663 | 0.3081502 | 0.24003325 | 0.81030448 | 0.99989001 |
| CTDSPL | 188.885893 | 0.0928192 | 0.44523983 | 0.20847011 | 0.83486192 | 0.99989001 |
| ACAA1 | 62.3185248 | 0.63251273 | 0.34429581 | 1.83712006 | 0.06619217 | 0.8719346 |
| OXSR1 | 223.771027 | -0.2224995 | 0.28514742 | -0.7802965 | 0.4352164 | 0.99989001 |
| ACVR2B | 47.9414102 | 0.53245515 | 0.35298074 | 1.50845382 | 0.13143842 | NA |
| EXOG | 77.8092686 | -0.1341352 | 0.2770349 | -0.4841815 | 0.62825709 | 0.99989001 |
| WDR48 | 133.229678 | -0.0298555 | 0.30475513 | -0.0979655 | 0.92195965 | 0.99989001 |
| GORASP1 | 60.2226183 | -0.2001498 | 0.35418825 | -0.5650944 | 0.57200953 | 0.99989001 |
| CSRNP1 | 63.3887466 | -1.0026341 | 0.71548067 | -1.4013434 | 0.1611114 | 0.99989001 |
| RPSA | 297.788297 | -0.100695 | 0.47841036 | -0.2104784 | 0.83329431 | 0.99989001 |
| MYRIP | 388.148182 | -0.826645 | 0.65154789 | -1.2687402 | 0.20453373 | 0.99989001 |
| EIF1B | 87.7886119 | -0.0818526 | 0.41822304 | -0.1957152 | 0.84483305 | 0.99989001 |
| ENTPD3-AS1 | 47.0598637 | 0.0370382 | 0.41553864 | 0.08913299 | 0.92897622 | NA |
| RPL14 | 540.764389 | -0.157188 | 0.60543138 | -0.2596298 | 0.79514934 | 0.99989001 |
| ZNF621 | 105.37693 | -0.2961461 | 0.25086996 | -1.1804765 | 0.23781075 | 0.99989001 |
| CTNNB1 | 1002.91291 | -0.3785441 | 0.28211906 | -1.3417884 | 0.17966461 | 0.99989001 |
| ULK4 | 375.892127 | -0.4886109 | 0.32126271 | -1.5209077 | 0.128283 | 0.99989001 |
| TRAK1 | 466.189819 | -0.202231 | 0.20809403 | -0.9718253 | 3.31E-01 | 1.00E+00 |
| SEC22C | 85.3054542 | 0.02742108 | 0.27149972 | 0.10099855 | 0.91955161 | 0.99989001 |
| NKTR | 958.421539 | 0.11670067 | 0.28386206 | 0.41111753 | 0.68098636 | 0.99989001 |
| AC006059.1 | 157.810766 | 0.22163151 | 0.55715445 | 0.39779186 | 0.69078362 | 0.99989001 |
| SNRK | 1443.5095 | 0.04684531 | 0.38779659 | 0.12079867 | 0.90385051 | 0.99989001 |

| | | | | | | |
|---|---|---|---|---|---|---|
| ANO10 | 431.803609 | 0.12809107 | 0.20961077 | 0.6110901 | 0.54113993 | 0.99989001 |
| ABHD5 | 86.8563224 | -0.1211493 | 0.34519726 | -0.3509566 | 0.7256209 | 0.99989001 |
| TCAIM | 171.389009 | 0.38179079 | 0.29102971 | 1.31186188 | 0.18956674 | 0.99989001 |
| ZNF445 | 72.4011707 | 0.26599367 | 0.25651297 | 1.03695995 | 0.29975452 | 0.99989001 |
| AC099669.1 | 92.9476584 | -0.1113924 | 0.3145635 | -0.3541175 | 0.72325084 | 0.99989001 |
| ZNF197 | 70.4841565 | 0.32967603 | 0.34719397 | 0.94954422 | 0.34234389 | 0.99989001 |
| KIAA1143 | 56.8593026 | -0.1030692 | 0.28526828 | -0.3613063 | 0.71787049 | 0.99989001 |
| ZDHHC3 | 106.91669 | 0.50952461 | 0.2386319 | 2.13519065 | 0.03274544 | 0.67281094 |
| LARS2 | 132.640085 | -0.3014762 | 0.33857943 | -0.8904151 | 0.37324304 | 0.99989001 |
| LIMD1 | 127.357772 | 0.3234952 | 0.35542474 | 0.91016511 | 0.36273544 | 0.99989001 |
| SACM1L | 131.378872 | 0.05104835 | 0.2164754 | 0.23581595 | 0.81357549 | 0.99989001 |
| FYCO1 | 68.6673849 | -0.1155738 | 0.27031679 | -0.4275495 | 0.66897917 | 0.99989001 |
| LTF | 100.312237 | 1.29276035 | 0.48239141 | 2.67989918 | 0.00736443 | 0.4053518 |
| ALS2CL | 67.8099421 | -0.3011462 | 0.45479911 | -0.6621521 | 0.50787377 | 0.99989001 |
| CCDC12 | 138.583485 | -0.2228678 | 0.24306596 | -0.9169027 | 0.35919363 | 0.99989001 |
| NBEAL2 | 86.1616363 | -0.4703752 | 0.31935571 | -1.4728879 | 0.14078125 | 0.99989001 |
| SETD2 | 655.711764 | 0.12755818 | 0.21094362 | 0.6047027 | 0.54537655 | 0.99989001 |
| KIF9-AS1 | 118.393777 | 0.20535886 | 0.35955454 | 0.57114802 | 0.56789931 | 0.99989001 |
| KLHL18 | 77.1239143 | 0.14491204 | 0.29852774 | 0.48542236 | 0.62737677 | 0.99989001 |
| PTPN23 | 48.5681989 | -0.3614658 | 0.34602525 | -1.0446227 | 0.29619737 | NA |
| SCAP | 95.4970793 | 0.01348985 | 0.284744 | 0.04737535 | 0.96221407 | 0.99989001 |
| ELP6 | 49.0437328 | -0.3393298 | 0.38723733 | -0.8762839 | 0.38087571 | NA |
| SMARCC1 | 403.41159 | -0.1500739 | 0.2824527 | -0.5313239 | 0.59519436 | 0.99989001 |
| DHX30 | 96.8697609 | -0.1809518 | 0.20740425 | -0.8724597 | 0.38295767 | 0.99989001 |
| MAP4 | 1089.74354 | -0.262497 | 0.21005712 | -1.2496459 | 0.21142894 | 0.99989001 |
| ZNF589 | 78.7420262 | -0.3068144 | 0.33783431 | -0.90818 | 0.36378314 | 0.99989001 |
| TMA7 | 156.592464 | -0.2126701 | 0.51472208 | -0.4131747 | 0.67947865 | 0.99989001 |
| SHISA5 | 255.251988 | -0.250275 | 0.30914988 | -0.8095589 | 0.41819372 | 0.99989001 |
| UQCRC1 | 47.3141505 | 0.23006302 | 0.31657651 | 0.72672172 | 0.46739644 | NA |
| NCKIPSD | 53.7227738 | 0.22522139 | 0.37293744 | 0.60391199 | 0.54590216 | 0.99989001 |
| IP6K2 | 156.522363 | -0.2261028 | 0.30905139 | -0.7316027 | 0.46441109 | 0.99989001 |
| PRKAR2A | 326.177913 | 0.091977 | 0.1925008 | 0.47780059 | 0.63279214 | 0.99989001 |
| ARIH2 | 278.617253 | 0.11485493 | 0.19050475 | 0.60289799 | 0.54657655 | 0.99989001 |
| WDR6 | 83.261437 | -0.5018769 | 0.31627563 | -1.5868339 | 0.11255027 | 0.99989001 |
| NDUFAF3 | 61.1207682 | -0.2932837 | 0.52207821 | -0.561762 | 0.57427816 | 0.99989001 |
| QRICH1 | 158.381936 | -0.1523693 | 0.20921315 | -0.7282969 | 0.46643183 | 0.99989001 |
| QARS | 39.4079787 | -0.370308 | 0.3281542 | -1.1284574 | 0.25912681 | NA |
| LAMB2 | 216.213275 | -0.0849949 | 0.50752109 | -0.1674706 | 0.86699976 | 0.99989001 |
| USP4 | 175.369712 | 0.01533335 | 0.24084195 | 0.0636656 | 0.9492365 | 0.99989001 |
| RHOA | 620.505149 | 0.245706 | 0.25849499 | 0.95052522 | 0.34184544 | 0.99989001 |
| DAG1 | 93.8838773 | -0.0754535 | 0.32135004 | -0.2348015 | 0.81436278 | 0.99989001 |
| IP6K1 | 195.930524 | 0.06297772 | 0.26679454 | 0.23605327 | 0.81339133 | 0.99989001 |
| UBA7 | 50.9906829 | -0.6556969 | 0.57260073 | -1.1451206 | 0.2521592 | 0.99989001 |
| RBM6 | 890.691448 | 0.05007039 | 0.2141094 | 0.23385424 | 0.81509814 | 0.99989001 |
| RBM5 | 332.595616 | -0.0450766 | 0.16114728 | -0.2797231 | 0.77968995 | 0.99989001 |
| SEMA3F | 389.914485 | -0.4670983 | 0.3591074 | -1.3007204 | 0.19335416 | 0.99989001 |
| GNAI2 | 688.904152 | 0.24954436 | 0.30592227 | 0.81571163 | 0.41466509 | 0.99989001 |
| IFRD2 | 41.6697804 | -0.1318159 | 0.48885175 | -0.2696439 | 0.78743422 | NA |
| HYAL1 | 176.954544 | -0.6595783 | 0.49326093 | -1.3371794 | 0.18116409 | 0.99989001 |
| HYAL2 | 716.527226 | -0.3712144 | 0.49519581 | -0.7496316 | 0.45347661 | 0.99989001 |
| RASSF1 | 134.536632 | -0.034355 | 0.3904779 | -0.087982 | 0.92989099 | 0.99989001 |
| MANF | 74.5930604 | 0.46622269 | 0.49945082 | 0.93347065 | 0.35057702 | 0.99989001 |
| DCAF1 | 125.603878 | -0.6126164 | 0.21986166 | -2.7863721 | 0.00533017 | 0.35652986 |
| RAD54L2 | 135.855006 | -0.1852132 | 0.266169 | -0.6958481 | 0.48652395 | 0.99989001 |

| | | | | | | |
|---|---|---|---|---|---|---|
| TEX264 | 75.8330116 | -0.7579217 | 0.36617201 | -2.0698514 | 0.03846626 | 0.71513134 |
| RPL29 | 743.555554 | -0.0596708 | 0.59773097 | -0.0998289 | 0.92048014 | 0.99989001 |
| DUSP7 | 42.5661835 | 0.76702765 | 0.52868838 | 1.4508124 | 0.14683211 | NA |
| ALAS1 | 42.852444 | 0.03648733 | 0.395885 | 0.09216648 | 0.92656577 | NA |
| WDR82 | 157.330145 | -0.0579218 | 0.23141547 | -0.2502935 | 0.80236037 | 0.99989001 |
| DNAH1 | 50.4448673 | -0.0236974 | 0.40098202 | -0.0590984 | 0.9528737 | 0.99989001 |
| BAP1 | 39.4183134 | -0.2946539 | 0.3755719 | -0.7845474 | 0.43271899 | NA |
| SEMA3G | 231.676377 | -0.0210503 | 0.68076027 | -0.0309218 | 0.97533189 | 0.99989001 |
| NISCH | 134.292539 | 0.1594641 | 0.24020988 | 0.6638532 | 0.50678427 | 0.99989001 |
| STAB1 | 170.532138 | -0.7511769 | 0.53157232 | -1.4131227 | 0.15761965 | 0.99989001 |
| SMIM4 | 53.6134447 | -0.0807194 | 0.31581751 | -0.2555887 | 0.79826843 | 0.99989001 |
| PBRM1 | 434.539099 | 0.10845773 | 0.26104997 | 0.41546733 | 0.67779983 | 0.99989001 |
| GNL3 | 69.8068649 | -0.4131057 | 0.42111278 | -0.980986 | 0.32659965 | 0.99989001 |
| GLT8D1 | 47.7942396 | -0.1764388 | 0.29177768 | -0.604703 | 0.54537634 | NA |
| SPCS1 | 148.156864 | 0.06996732 | 0.45018171 | 0.15542018 | 0.87649007 | 0.99989001 |
| NEK4 | 79.5389254 | -0.0406565 | 0.24554866 | -0.1655742 | 0.86849203 | 0.99989001 |
| SFMBT1 | 48.0222824 | 0.31885572 | 0.40376136 | 0.78971332 | 0.42969521 | NA |
| TKT | 90.2607067 | 0.98443626 | 0.55677047 | 1.76811867 | 0.07704107 | 0.92274869 |
| DCP1A | 237.691348 | 0.19723807 | 0.31396608 | 0.62821457 | 0.52986338 | 0.99989001 |
| SELENOK | 103.109303 | -0.1344984 | 0.53640932 | -0.2507384 | 0.80201639 | 0.99989001 |
| CACNA2D3 | 53.5717157 | -1.2962176 | 0.65134875 | -1.9900516 | 0.04658525 | 0.77516398 |
| CCDC66 | 189.300702 | 0.02966216 | 0.2326004 | 0.12752411 | 0.89852559 | 0.99989001 |
| FAM208A | 266.220472 | 0.02930879 | 0.22873487 | 0.12813432 | 0.89804267 | 0.99989001 |
| ARHGEF3 | 1255.79887 | 0.09045104 | 0.35918768 | 0.25182111 | 0.80117934 | 0.99989001 |
| APPL1 | 183.321988 | -0.0354375 | 0.20814699 | -0.1702523 | 0.86481174 | 0.99989001 |
| PDE12 | 46.093282 | -0.1288872 | 0.27443598 | -0.4696439 | 0.63860946 | NA |
| ARF4 | 162.861005 | -0.0232839 | 0.35388388 | -0.0657953 | 0.94754076 | 0.99989001 |
| DENND6A | 299.616059 | 0.08607623 | 0.18636846 | 0.46186051 | 0.64418136 | 0.99989001 |
| DENND6A-AS | 69.2583368 | 0.0036669 | 0.41461646 | 0.00884407 | 0.99294355 | 0.99989001 |
| SLMAP | 454.236261 | -0.025128 | 0.16787517 | -0.1496828 | 0.88101487 | 0.99989001 |
| FLNB | 876.683453 | -0.6591081 | 0.30307938 | -2.1747046 | 0.02965226 | 0.6652708 |
| DNASE1L3 | 403.741967 | -0.9581149 | 0.40734801 | -2.3520795 | 0.01866878 | 0.56643441 |
| ABHD6 | 111.720873 | 0.09357935 | 0.39083084 | 0.23943695 | 0.81076678 | 0.99989001 |
| PXK | 123.924153 | 0.56442076 | 0.27247159 | 2.0714848 | 0.03831351 | 0.71513134 |
| PDHB | 49.4147041 | 0.05814225 | 0.33062053 | 0.17585796 | 0.86040554 | NA |
| FAM107A | 234.948823 | -0.6356787 | 0.55750219 | -1.1402264 | 0.25419199 | 0.99989001 |
| FHIT | 707.459226 | -0.2585727 | 0.34367515 | -0.7523753 | 0.45182537 | 0.99989001 |
| PTPRG | 5333.57268 | -0.0009793 | 0.2446104 | -0.0040033 | 0.99680583 | 0.99989001 |
| THOC7 | 67.0031036 | 0.09936917 | 0.29556961 | 0.33619548 | 0.73672346 | 0.99989001 |
| ATXN7 | 47.1328593 | 0.43492938 | 0.36711095 | 1.18473552 | 0.23612203 | NA |
| PSMD6 | 47.8786136 | -0.0027998 | 0.36136106 | -0.0077479 | 0.99381816 | NA |
| PRICKLE2 | 111.619625 | 0.82975611 | 0.40891941 | 2.02914338 | 0.04244369 | 0.76066939 |
| ADAMTS9 | 3568.57614 | -0.9948323 | 0.73970896 | -1.3448969 | 0.17865852 | 0.99989001 |
| ADAMTS9-AS | 157.843999 | -0.4472599 | 0.59864507 | -0.7471203 | 0.45499097 | 0.99989001 |
| ADAMTS9-AS | 563.903189 | 0.51166249 | 0.61300956 | 0.83467293 | 0.40390188 | 0.99989001 |
| MAGI1 | 2513.0468 | 0.04113037 | 0.2744554 | 0.14986175 | 0.88087369 | 0.99989001 |
| SLC25A26 | 167.270923 | -0.3679324 | 0.36203775 | -1.016282 | 0.30949512 | 0.99989001 |
| LRIG1 | 68.3251925 | -0.2334561 | 0.48347202 | -0.4828741 | 0.62918516 | 0.99989001 |
| SUCLG2 | 454.620632 | -0.1213361 | 0.12719186 | -0.9539614 | 0.34010317 | 0.99989001 |
| SUCLG2-AS1 | 119.168927 | -0.3925967 | 0.52947288 | -0.741486 | 0.45839881 | 0.99989001 |
| EOGT | 118.412282 | -0.6382543 | 0.3193055 | -1.9988828 | 0.04562104 | 0.77228819 |
| TMF1 | 175.973774 | -0.1532167 | 0.18704422 | -0.8191471 | 0.41270249 | 0.99989001 |
| UBA3 | 143.399769 | 0.18411709 | 0.16541654 | 1.11305124 | 0.26568642 | 0.99989001 |
| ARL6IP5 | 188.233623 | -0.0316675 | 0.52185385 | -0.0606826 | 0.95161198 | 0.99989001 |

| | | | | | | |
|---|---|---|---|---|---|---|
| FRMD4B | 1208.60449 | -0.20298 | 0.20817967 | -0.9750231 | 0.32954882 | 0.99989001 |
| MITF | 78.0870776 | 0.34414659 | 0.36374838 | 0.94611165 | 0.34409164 | 0.99989001 |
| FOXP1 | 1501.92215 | 0.26354891 | 0.25699243 | 1.02551232 | 0.30512152 | 0.99989001 |
| EIF4E3 | 58.5959002 | 0.55746199 | 0.44324508 | 1.25768341 | 0.20850627 | 0.99989001 |
| RYBP | 346.17432 | 0.30534965 | 0.28363399 | 1.07656227 | 0.28167587 | 0.99989001 |
| SHQ1 | 116.76865 | 0.0418491 | 0.28378977 | 0.14746516 | 0.88276488 | 0.99989001 |
| GXYLT2 | 47.4510558 | 0.36507638 | 0.42668961 | 0.85560176 | 0.3922181 | NA |
| PPP4R2 | 273.52514 | 0.14416786 | 0.21557756 | 0.66875172 | 0.50365387 | 0.99989001 |
| PDZRN3 | 148.596394 | -1.4811216 | 0.68699284 | -2.1559491 | 0.03108764 | 0.67281094 |
| ZNF717 | 66.0505315 | 0.38545122 | 0.43617883 | 0.88369996 | 0.3768582 | 0.99989001 |
| ROBO1 | 825.858802 | 0.7388906 | 0.67589101 | 1.0932097 | 0.27430174 | 0.99989001 |
| GBE1 | 717.790416 | -0.891972 | 0.46878599 | -1.9027275 | 0.05707611 | 0.83978648 |
| CHMP2B | 86.4651385 | -0.2256898 | 0.29139837 | -0.774506 | 0.43863163 | 0.99989001 |
| CGGBP1 | 249.50701 | 0.02833582 | 0.30890606 | 0.09172959 | 0.92691289 | 0.99989001 |
| ZNF654 | 310.053998 | 0.02879255 | 0.28338442 | 0.10160245 | 0.91907223 | 0.99989001 |
| C3orf38 | 58.6146505 | 0.00931883 | 0.27538375 | 0.03383944 | 0.97300519 | 0.99989001 |
| EPHA3 | 154.539081 | -1.787772 | 0.96956734 | -1.8438864 | 0.0651997 | 0.8666545 |
| PROS1 | 280.412143 | 0.21369002 | 0.3591723 | 0.59495128 | 0.55187604 | 0.99989001 |
| ARL13B | 176.465088 | -0.0220263 | 0.21414527 | -0.1028569 | 0.91807658 | 0.99989001 |
| NSUN3 | 97.5721217 | -0.1673662 | 0.19894009 | -0.8412896 | 0.40018574 | 0.99989001 |
| MTRNR2L12 | 258.707358 | 0.04698996 | 0.4697779 | 0.10002591 | 0.92032375 | 0.99989001 |
| ARL6 | 48.7897966 | -0.2443377 | 0.3460141 | -0.7061497 | 0.48009507 | NA |
| CRYBG3 | 469.190968 | -0.1915448 | 0.27199819 | -0.7042134 | 0.48129987 | 0.99989001 |
| RIOX2 | 48.6813724 | -0.4417884 | 0.2903995 | -1.5213127 | 0.12818139 | NA |
| CLDND1 | 122.010146 | -0.0501317 | 0.29030983 | -0.1726834 | 0.86290033 | 0.99989001 |
| ST3GAL6 | 126.708972 | -0.6266418 | 0.61220495 | -1.0235817 | 0.30603291 | 0.99989001 |
| DCBLD2 | 58.5252181 | -0.1390316 | 0.44547812 | -0.3120953 | 0.75496812 | 0.99989001 |
| COL8A1 | 355.286198 | 2.30388291 | 0.74704218 | 3.08400646 | 0.00204233 | 0.21410443 |
| CMSS1 | 453.500551 | 0.15736637 | 0.34225377 | 0.45979442 | 0.64566379 | 0.99989001 |
| FILIP1L | 191.686384 | 0.23452271 | 0.44857321 | 0.52281925 | 0.60110004 | 0.99989001 |
| TBC1D23 | 164.392275 | -0.2191506 | 0.25967879 | -0.8439294 | 0.39870887 | 0.99989001 |
| TOMM70 | 79.7054794 | 0.16923066 | 0.26196363 | 0.64600824 | 0.51827402 | 0.99989001 |
| TFG | 248.080243 | 0.05491765 | 0.22852887 | 0.24030944 | 0.81009037 | 0.99989001 |
| ABI3BP | 322.65871 | 0.60320393 | 0.58388795 | 1.03308164 | 0.30156569 | 0.99989001 |
| SENP7 | 576.572206 | -0.2186946 | 0.27711534 | -0.7891828 | 0.4300052 | 0.99989001 |
| TRMT10C | 50.0793229 | -0.2113325 | 0.38550632 | -0.5481945 | 0.58355834 | 0.99989001 |
| PCNP | 230.735992 | -0.1498815 | 0.21346804 | -0.702126 | 0.48260056 | 0.99989001 |
| ZBTB11 | 96.4327812 | 0.07364367 | 0.19438422 | 0.37885621 | 0.70479464 | 0.99989001 |
| RPL24 | 237.916318 | -0.2284713 | 0.49178214 | -0.4645783 | 0.64223348 | 0.99989001 |
| CEP97 | 55.9744051 | 0.02033786 | 0.27806647 | 0.07314028 | 0.94169448 | 0.99989001 |
| NXPE3 | 131.282885 | -0.058187 | 0.3739873 | -0.1555855 | 0.87635977 | 0.99989001 |
| NFKBIZ | 208.119588 | 0.26192176 | 0.53315304 | 0.49126938 | 0.62323593 | 0.99989001 |
| ALCAM | 113.876131 | 0.34200777 | 0.41869799 | 0.81683642 | 0.41402191 | 0.99989001 |
| CBLB | 705.626363 | 0.17146599 | 0.39529823 | 0.43376362 | 0.6644601 | 0.99989001 |
| LINC00882 | 89.0089064 | -0.9528419 | 1.00364893 | -0.9493777 | 0.34242856 | 0.99989001 |
| BBX | 613.231437 | 0.23127241 | 0.24412303 | 0.94736006 | 0.34345534 | 0.99989001 |
| CD47 | 219.941122 | 0.2198496 | 0.26005598 | 0.84539338 | 0.39789124 | 0.99989001 |
| IFT57 | 66.5093109 | -0.233732 | 0.25216701 | -0.9268938 | 0.35398166 | 0.99989001 |
| DZIP3 | 76.8892357 | -0.2717182 | 0.54778372 | -0.496032 | 0.61987186 | 0.99989001 |
| NECTIN3 | 193.245279 | 0.04859893 | 0.37801636 | 0.12856304 | 0.89770342 | 0.99989001 |
| PHLDB2 | 413.146313 | 0.0121126 | 0.30838447 | 0.03927759 | 0.96866907 | 0.99989001 |
| CD200 | 159.291272 | 0.81936349 | 0.59216214 | 1.38368098 | 0.16645616 | 0.99989001 |
| ATG3 | 78.871332 | 0.03203201 | 0.25070946 | 0.12776547 | 0.89833458 | 0.99989001 |
| SLC35A5 | 53.0489751 | -0.0682787 | 0.26534949 | -0.257316 | 0.79693483 | 0.99989001 |

| | | | | | | |
|---|---|---|---|---|---|---|
| NEPRO | 70.6107504 | 0.10441098 | 0.23847901 | 0.43782045 | 0.66151645 | 0.99989001 |
| AC078785.1 | 76.9990589 | -0.3958766 | 0.38928667 | -1.0169283 | 0.30918755 | 0.99989001 |
| BOC | 377.646215 | 0.09205164 | 0.32277613 | 0.28518726 | 0.77550068 | 0.99989001 |
| CFAP44 | 128.093888 | 0.19827769 | 0.30532138 | 0.64940651 | 0.51607566 | 0.99989001 |
| SPICE1 | 91.1803952 | 0.09750593 | 0.28528384 | 0.34178567 | 0.73251219 | 0.99989001 |
| SIDT1 | 81.4361054 | 0.0191972 | 0.51498172 | 0.03727743 | 0.9702638 | 0.99989001 |
| USF3 | 111.599509 | 0.07555418 | 0.22049966 | 0.34264988 | 0.73186187 | 0.99989001 |
| NAA50 | 231.850557 | -0.1438635 | 0.18243361 | -0.7885798 | 0.43035767 | 0.99989001 |
| ATP6V1A | 234.590576 | 0.03854065 | 0.20280506 | 0.19003792 | 0.84927941 | 0.99989001 |
| GRAMD1C | 115.257887 | -0.1238543 | 0.32283232 | -0.3836489 | 0.70123872 | 0.99989001 |
| CCDC191 | 80.5766398 | -0.195466 | 0.47611085 | -0.4105473 | 0.68140452 | 0.99989001 |
| QTRT2 | 82.8101572 | -0.207167 | 0.21676313 | -0.9557297 | 0.33920884 | 0.99989001 |
| ZBTB20 | 4960.95318 | -0.2903491 | 0.32174349 | -0.9024242 | 0.36683159 | 0.99989001 |
| ZBTB20-AS5 | 104.794809 | -0.6559942 | 1.02463852 | -0.6402202 | 0.52202948 | 0.99989001 |
| LSAMP | 76.0604823 | -0.1443539 | 0.4475333 | -0.3225545 | 0.74703262 | 0.99989001 |
| ARHGAP31 | 1256.13055 | 0.44763588 | 0.20170637 | 2.21924519 | 0.02647005 | 0.64658875 |
| TMEM39A | 78.3532629 | 0.04301532 | 0.28451043 | 0.15119065 | 0.87982533 | 0.99989001 |
| POGLUT1 | 83.723222 | -0.152853 | 0.31958597 | -0.4782845 | 0.63244771 | 0.99989001 |
| TIMMDC1 | 142.331245 | -0.082521 | 0.24160022 | -0.3415602 | 0.73268193 | 0.99989001 |
| PLA1A | 104.535638 | -0.3011564 | 0.72563289 | -0.4150258 | 0.67812303 | 0.99989001 |
| COX17 | 110.410045 | -0.166087 | 0.38098318 | -0.4359432 | 0.66287791 | 0.99989001 |
| GSK3B | 1045.51952 | -0.0519774 | 0.23454754 | -0.221607 | 0.82461984 | 0.99989001 |
| LRRC58 | 98.5055686 | -0.1241442 | 0.268139 | -0.4629845 | 0.64337547 | 0.99989001 |
| FSTL1 | 175.677548 | 0.09150658 | 0.36914185 | 0.24789002 | 0.80421951 | 0.99989001 |
| NDUFB4 | 110.688789 | -0.2677972 | 0.3487725 | -0.7678276 | 0.44258959 | 0.99989001 |
| RABL3 | 57.221203 | -0.2007399 | 0.27202631 | -0.7379429 | 0.46054917 | 0.99989001 |
| HCLS1 | 90.9671555 | -0.1612929 | 0.40971035 | -0.3936754 | 0.69382067 | 0.99989001 |
| GOLGB1 | 326.101235 | -0.018384 | 0.25196623 | -0.072962 | 0.94183634 | 0.99989001 |
| IQCB1 | 173.662864 | 0.08575304 | 0.26263128 | 0.32651497 | 0.74403477 | 0.99989001 |
| FAM162A | 90.5290976 | -0.7825976 | 0.51015792 | -1.53403 | 0.12502226 | 0.99989001 |
| AC083798.2 | 49.7971051 | 0.32224468 | 0.42456869 | 0.75899305 | 0.44785671 | NA |
| KPNA1 | 314.398842 | 0.02458898 | 0.24342004 | 0.10101461 | 0.91953886 | 0.99989001 |
| PARP9 | 339.251536 | -0.1312078 | 0.51139467 | -0.2565686 | 0.79751184 | 0.99989001 |
| DTX3L | 131.426075 | -0.0466363 | 0.4139585 | -0.1126594 | 0.91030056 | 0.99989001 |
| PARP14 | 1217.17298 | -0.4043473 | 0.59183694 | -0.6832073 | 0.49447586 | 0.99989001 |
| HSPBAP1 | 77.1176714 | -0.2016787 | 0.2118015 | -0.9522061 | 0.34099246 | 0.99989001 |
| DIRC2 | 86.769184 | -0.1479762 | 0.30467449 | -0.4856864 | 0.62718956 | 0.99989001 |
| SEMA5B | 166.378354 | -0.1578452 | 0.60282592 | -0.2618421 | 0.79344316 | 0.99989001 |
| PDIA5 | 523.421756 | 0.10972123 | 0.36951205 | 0.29693545 | 0.76651579 | 0.99989001 |
| SEC22A | 243.265316 | 0.38792664 | 0.23283598 | 1.66609405 | 0.09569468 | 0.99989001 |
| HACD2 | 92.7363518 | 0.02497214 | 0.27515527 | 0.09075653 | 0.92768605 | 0.99989001 |
| MYLK | 152.104276 | 0.11902609 | 0.49965492 | 0.23821659 | 0.81171311 | 0.99989001 |
| CCDC14 | 162.197808 | -0.3343631 | 0.30550736 | -1.0944519 | 0.27375682 | 0.99989001 |
| KALRN | 2016.82762 | -0.0668473 | 0.37737986 | -0.1771352 | 0.85940217 | 0.99989001 |
| UMPS | 42.9271833 | -0.143988 | 0.289632 | -0.4971411 | 0.61908953 | NA |
| ITGB5 | 312.593036 | -0.0423125 | 0.33797329 | -0.1251948 | 0.90036936 | 0.99989001 |
| HEG1 | 1105.7464 | -0.072823 | 0.49027385 | -0.1485354 | 0.88192024 | 0.99989001 |
| ZNF148 | 674.022846 | -0.0020941 | 0.1626148 | -0.0128778 | 0.98972529 | 0.99989001 |
| SNX4 | 195.769099 | 0.03279698 | 0.15669912 | 0.20929905 | 0.8342148 | 0.99989001 |
| OSBPL11 | 72.811726 | 0.01116709 | 0.27564666 | 0.04051234 | 0.96768467 | 0.99989001 |
| SLC41A3 | 48.9428034 | -0.0279086 | 0.42488426 | -0.0656851 | 0.94762854 | NA |
| ZXDC | 176.261123 | 0.03471196 | 0.31532342 | 0.11008368 | 0.91234301 | 0.99989001 |
| CHCHD6 | 69.200603 | 0.16426711 | 0.45736505 | 0.35915972 | 0.71947561 | 0.99989001 |
| TPRA1 | 43.4278416 | 0.15599012 | 0.34645086 | 0.45025179 | 0.6525289 | NA |

| | | | | | | |
|---|---|---|---|---|---|---|
| MGLL | 230.031322 | 0.01177244 | 0.53010368 | 0.02220781 | 0.98228219 | 0.99989001 |
| SEC61A1 | 182.721674 | -0.1480632 | 0.41233624 | -0.3590837 | 0.71953247 | 0.99989001 |
| RUVBL1 | 108.560296 | 0.19579801 | 0.30506637 | 0.64182103 | 0.52098939 | 0.99989001 |
| EEFSEC | 176.345095 | 0.30276988 | 0.30623589 | 0.9886819 | 0.3228188 | 0.99989001 |
| GATA2 | 140.15823 | 0.36634758 | 0.48981117 | 0.74793635 | 0.45449855 | 0.99989001 |
| GATA2-AS1 | 136.434266 | 0.17242849 | 0.38397379 | 0.44906317 | 0.65338609 | 0.99989001 |
| RPN1 | 141.951705 | 0.07604328 | 0.30728348 | 0.24746948 | 0.80454491 | 0.99989001 |
| RAB7A | 541.042594 | -0.3130944 | 0.21209958 | -1.4761668 | 0.13989912 | 0.99989001 |
| ACAD9 | 78.3818797 | -0.1718718 | 0.24025183 | -0.7153817 | 0.47437321 | 0.99989001 |
| EFCC1 | 74.9668105 | 0.1580412 | 0.35074918 | 0.45058181 | 0.65229098 | 0.99989001 |
| CNBP | 329.774329 | 0.19305829 | 0.41690512 | 0.46307489 | 0.64331069 | 0.99989001 |
| COPG1 | 71.2839022 | -0.1935466 | 0.26764121 | -0.7231568 | 0.46958357 | 0.99989001 |
| HMCES | 83.4702096 | -0.0930738 | 0.25407344 | -0.3663262 | 0.71412165 | 0.99989001 |
| H1FX | 196.33975 | -0.1377198 | 0.6771305 | -0.2033874 | 0.83883226 | 0.99989001 |
| MBD4 | 82.9322731 | 0.06361826 | 0.27706251 | 0.229617 | 0.81838939 | 0.99989001 |
| IFT122 | 101.330311 | 0.56363951 | 0.25135233 | 2.24242805 | 0.02493372 | 0.62943051 |
| PLXND1 | 488.049686 | 0.08343118 | 0.31861001 | 0.26185989 | 0.79342946 | 0.99989001 |
| TMCC1 | 617.498178 | 0.15108864 | 0.23622322 | 0.63960111 | 0.52243196 | 0.99989001 |
| PIK3R4 | 107.712021 | -0.2615304 | 0.19776806 | -1.3224095 | 0.18603184 | 0.99989001 |
| ATP2C1 | 301.777303 | 0.15540392 | 0.17569795 | 0.88449481 | 0.37642917 | 0.99989001 |
| NEK11 | 62.845171 | -0.7730618 | 0.43091199 | -1.7940133 | 0.07281105 | 0.9155703 |
| MRPL3 | 89.7613001 | -0.1145089 | 0.33362035 | -0.3432313 | 0.7314245 | 0.99989001 |
| DNAJC13 | 254.049339 | -0.124728 | 0.21769251 | -0.5729551 | 0.56667509 | 0.99989001 |
| UBA5 | 93.1414788 | 0.00084208 | 0.32915983 | 0.00255828 | 0.99795879 | 0.99989001 |
| CDV3 | 171.179793 | -0.1201271 | 0.28954101 | -0.414888 | 0.67822391 | 0.99989001 |
| TOPBP1 | 93.7349816 | -0.2061548 | 0.18848195 | -1.093764 | 0.27405851 | 0.99989001 |
| SRPRB | 44.6209842 | -0.3356113 | 0.45642707 | -0.735301 | 0.4621562 | NA |
| SLCO2A1 | 2271.29416 | 1.09408606 | 0.43017972 | 2.54332319 | 0.01098036 | 0.4764488 |
| RYK | 379.971482 | 0.04790788 | 0.21684931 | 0.22092707 | 0.82514922 | 0.99989001 |
| AMOTL2 | 58.9508649 | -0.1630979 | 0.3371532 | -0.48375 | 6.29E-01 | 1.00E+00 |
| CEP63 | 108.930744 | -0.0055687 | 0.20830563 | -0.0267332 | 0.97867254 | 0.99989001 |
| EPHB1 | 83.5150044 | -0.3502511 | 0.50186244 | -0.6979027 | 0.48523805 | 0.99989001 |
| PPP2R3A | 150.685206 | -0.0165896 | 0.21692629 | -0.0764758 | 0.93904056 | 0.99989001 |
| MSL2 | 127.150611 | 0.41034492 | 0.28020377 | 1.46445179 | 0.14307055 | 0.99989001 |
| PCCB | 60.6816388 | -0.0495622 | 0.29655019 | -0.1671291 | 0.86726846 | 0.99989001 |
| STAG1 | 1598.30444 | 0.05661453 | 0.19842447 | 0.28532031 | 0.77539876 | 0.99989001 |
| SLC35G2 | 159.059918 | 0.43878754 | 0.22262674 | 1.97095612 | 0.04872889 | 0.78323466 |
| NCK1 | 266.510922 | 0.44674391 | 0.30121981 | 1.48311598 | 0.1380436 | 0.99989001 |
| ARMC8 | 295.366878 | -0.0282862 | 0.19582559 | -0.1444461 | 0.88514821 | 0.99989001 |
| MRAS | 51.4477258 | 0.27238029 | 0.41737161 | 0.65260858 | 0.51400865 | 0.99989001 |
| CEP70 | 105.601504 | 0.00671828 | 0.26019577 | 0.02582009 | 0.97940083 | 0.99989001 |
| PIK3CB | 253.33075 | 0.2433541 | 0.25624621 | 0.94968859 | 0.34227051 | 0.99989001 |
| COPB2 | 151.56733 | 0.06976446 | 0.19945759 | 0.34977092 | 0.72651062 | 0.99989001 |
| AC097103.2 | 97.6735872 | -0.34009 | 0.4043111 | -0.8411591 | 0.40025882 | 0.99989001 |
| RBP1 | 51.3171888 | -0.2801621 | 0.80864244 | -0.3464598 | 0.72899718 | 0.99989001 |
| NMNAT3 | 62.8467172 | -0.2897597 | 0.37468911 | -0.7733337 | 0.43932491 | 0.99989001 |
| SLC25A36 | 211.582344 | -0.1201256 | 0.26403191 | -0.4549662 | 0.64913356 | 0.99989001 |
| ZBTB38 | 460.990089 | -0.2145553 | 0.24452066 | -0.8774526 | 0.38024083 | 0.99989001 |
| RASA2 | 505.405678 | -0.150526 | 0.20527238 | -0.7332987 | 0.46337627 | 0.99989001 |
| RNF7 | 132.567191 | -0.1960882 | 0.55677289 | -0.352187 | 0.72469805 | 0.99989001 |
| ATP1B3 | 532.804569 | -0.3442329 | 0.46011683 | -0.7481423 | 0.45437431 | 0.99989001 |
| TFDP2 | 325.621147 | -0.0396639 | 0.28553431 | -0.1389112 | 0.88952031 | 0.99989001 |
| GK5 | 92.6830448 | -0.0872463 | 0.20923607 | -0.4169755 | 0.67669634 | 0.99989001 |
| XRN1 | 610.201408 | 0.04133534 | 0.40488478 | 0.10209161 | 0.91868396 | 0.99989001 |

| | | | | | | |
|---|---|---|---|---|---|---|
| ATR | 307.188855 | -0.3423425 | 0.29893152 | -1.1452206 | 0.2521178 | 0.99989001 |
| TRPC1 | 215.08291 | -0.0481877 | 0.34277928 | -0.1405794 | 0.88820219 | 0.99989001 |
| U2SURP | 307.059508 | -0.1759322 | 0.20383523 | -0.86311 | 0.388077 | 0.99989001 |
| SLC9A9 | 233.627249 | 0.31670468 | 0.42645366 | 0.74264736 | 0.45769521 | 0.99989001 |
| C3orf58 | 80.2726732 | -0.0292373 | 0.40178176 | -0.0727691 | 0.94198986 | 0.99989001 |
| PLOD2 | 420.105145 | -0.5816327 | 0.61526407 | -0.9453382 | 0.34448622 | 0.99989001 |
| PLSCR4 | 578.843799 | -0.2039689 | 0.38716795 | -0.5268229 | 0.59831657 | 0.99989001 |
| PLSCR1 | 845.612338 | -0.2034477 | 0.4111724 | -0.4947991 | 0.62074194 | 0.99989001 |
| GYG1 | 58.2264057 | -0.1214278 | 0.23652766 | -0.5133767 | 0.60768787 | 0.99989001 |
| HLTF | 88.9028848 | 0.10034774 | 0.39973883 | 0.25103326 | 0.80178839 | 0.99989001 |
| HPS3 | 183.599427 | 0.70300076 | 0.35989995 | 1.95332272 | 0.05078137 | 0.80694265 |
| CP | 224.591414 | -0.0683383 | 1.0702172 | -0.0638546 | 0.94908597 | 0.99989001 |
| TM4SF18 | 348.234581 | -0.6131617 | 0.72417518 | -0.8467035 | 0.39716038 | 0.99989001 |
| TM4SF1 | 1279.95455 | -0.2456909 | 0.612661 | -0.4010226 | 0.68840346 | 0.99989001 |
| WWTR1 | 2594.82142 | 0.1745887 | 0.36746682 | 0.47511418 | 0.63470559 | 0.99989001 |
| COMMD2 | 86.4095987 | 0.27360281 | 0.30933657 | 0.88448258 | 0.37643576 | 0.99989001 |
| ANKUB1 | 112.941057 | 0.28949827 | 0.45788909 | 0.6322454 | 0.52722654 | 0.99989001 |
| RNF13 | 368.84275 | 0.03377912 | 0.20988517 | 0.16094095 | 0.87213991 | 0.99989001 |
| TSC22D2 | 330.917027 | 0.00079091 | 0.34170004 | 0.00231464 | 0.99815319 | 0.99989001 |
| SERP1 | 186.852393 | -0.0053007 | 0.32230019 | -0.0164464 | 0.98687823 | 0.99989001 |
| EIF2A | 123.490885 | -0.0902677 | 0.32431515 | -0.2783332 | 0.78075656 | 0.99989001 |
| SELENOT | 120.451128 | 0.19257994 | 0.31973263 | 0.60231558 | 0.54696409 | 0.99989001 |
| MBNL1 | 2190.24357 | -0.0159044 | 0.3235785 | -0.0491516 | 0.96079847 | 0.99989001 |
| DHX36 | 291.400808 | -0.0817059 | 0.23783066 | -0.3435465 | 0.73118737 | 0.99989001 |
| GMPS | 129.660409 | 0.10386786 | 0.19581667 | 0.53043421 | 0.59581091 | 0.99989001 |
| SSR3 | 125.600522 | -0.2917053 | 0.4215614 | -0.6919639 | 0.48896 | 0.99989001 |
| TIPARP | 69.6135093 | -0.6828492 | 0.5817531 | -1.1737784 | 0.24048381 | 0.99989001 |
| CCNL1 | 351.547848 | -0.4648496 | 0.45073909 | -1.0313052 | 0.3023977 | 0.99989001 |
| RSRC1 | 409.440981 | -0.3022955 | 0.16254536 | -1.8597609 | 0.06291937 | 0.85940342 |
| GFM1 | 136.62832 | -0.2180876 | 0.27749927 | -0.7859032 | 0.43192418 | 0.99989001 |
| RARRES1 | 52.8343316 | 0.15267778 | 0.34292574 | 0.4452211 | 6.56E-01 | 0.99989001 |
| MFSD1 | 64.9327481 | -0.1703097 | 0.31459055 | -0.5413693 | 0.58825309 | 0.99989001 |
| IQCJ-SCHIP1 | 399.060531 | -1.041156 | 0.46352656 | -2.2461625 | 0.0246936 | 0.62546071 |
| SMC4 | 145.76439 | 0.23735423 | 0.29419585 | 0.80678986 | 0.41978756 | 0.99989001 |
| KPNA4 | 314.117864 | -0.0606359 | 0.3104499 | -0.1953163 | 0.84514532 | 0.99989001 |
| PPM1L | 83.1597237 | 0.78560338 | 0.44386223 | 1.76992617 | 0.07673944 | 0.92274869 |
| NMD3 | 133.744818 | 0.24059642 | 0.1992164 | 1.20771396 | 0.2271573 | 0.99989001 |
| PDCD10 | 183.176797 | -0.2540972 | 0.22799641 | -1.114479 | 0.26507376 | 0.99989001 |
| GOLIM4 | 203.537584 | 0.13589814 | 0.29400794 | 0.46222609 | 0.64391919 | 0.99989001 |
| MECOM | 3184.68456 | 0.09904686 | 0.43701626 | 0.22664343 | 0.82070102 | 0.99989001 |
| MYNN | 54.620793 | -0.1041492 | 0.31397046 | -0.3317165 | 0.74010331 | 0.99989001 |
| SEC62 | 291.415251 | -0.0181529 | 0.23541512 | -0.0771102 | 0.93853589 | 0.99989001 |
| GPR160 | 50.3411189 | 0.9259479 | 0.43342325 | 2.13635956 | 0.03265012 | 0.67281094 |
| PHC3 | 511.925518 | 0.07318997 | 0.20767627 | 0.35242337 | 0.72452077 | 0.99989001 |
| PRKCI | 174.745935 | 0.05417375 | 0.20747037 | 0.26111562 | 0.79400335 | 0.99989001 |
| SKIL | 441.2756 | 0.40275682 | 0.29129328 | 1.38265058 | 0.16677203 | 0.99989001 |
| RPL22L1 | 50.3081748 | -0.1120008 | 0.67858199 | -0.1650513 | 0.86890362 | 0.99989001 |
| PLD1 | 248.454185 | 0.51057522 | 0.35251792 | 1.4483667 | 0.14751452 | 0.99989001 |
| FNDC3B | 2135.61964 | 0.26711246 | 0.3131387 | 0.85301646 | 0.39365018 | 0.99989001 |
| TNFSF10 | 573.295944 | 0.57724101 | 0.5117141 | 1.12805374 | 0.25929723 | 0.99989001 |
| NCEH1 | 99.3021189 | -0.5812371 | 0.35529796 | -1.6359146 | 0.10185747 | 0.99989001 |
| ECT2 | 75.7883691 | -0.6123287 | 0.46965036 | -1.3037968 | 0.19230286 | 0.99989001 |
| NLGN1 | 150.442032 | 0.65580955 | 0.48513473 | 1.35180912 | 0.17643639 | 0.99989001 |
| NAALADL2 | 338.64549 | -0.0733343 | 0.43359253 | -0.1691317 | 0.86569303 | 0.99989001 |

| | | | | | | |
|---|---|---|---|---|---|---|
| TBL1XR1 | 897.609404 | -0.1422871 | 0.26879102 | -0.5293595 | 0.59655612 | 0.99989001 |
| ZMAT3 | 73.1500866 | -0.1425201 | 0.57761127 | -0.2467405 | 0.80510907 | 0.99989001 |
| PIK3CA | 334.380233 | 0.11771241 | 0.24980034 | 0.47122597 | 0.63747937 | 0.99989001 |
| KCNMB3 | 46.2930397 | -0.9140532 | 0.35853565 | -2.5494067 | 0.01079064 | NA |
| ZNF639 | 41.4927647 | 0.14404482 | 0.32780572 | 0.43942133 | 0.66035628 | NA |
| MFN1 | 155.196664 | -0.0986597 | 0.23295429 | -0.4235152 | 0.67191941 | 0.99989001 |
| GNB4 | 400.976455 | 0.57120955 | 0.28256948 | 2.02148357 | 0.04322973 | 0.76595778 |
| ACTL6A | 58.4800109 | -0.2337264 | 0.29632945 | -0.7887382 | 0.43026501 | 0.99989001 |
| NDUFB5 | 70.6657924 | 0.35357599 | 0.38689653 | 0.91387738 | 0.36078128 | 0.99989001 |
| USP13 | 224.177945 | 0.38860975 | 0.49961336 | 0.77782097 | 0.43667456 | 0.99989001 |
| TTC14 | 269.289229 | -0.1218958 | 0.22244063 | -0.5479923 | 0.58369719 | 0.99989001 |
| FXR1 | 296.415 | 0.01080377 | 0.19240352 | 0.05615163 | 0.95522101 | 0.99989001 |
| ATP11B | 284.637864 | -0.0559754 | 0.25800561 | -0.2169543 | 0.82824398 | 0.99989001 |
| DCUN1D1 | 132.233621 | 0.30099025 | 0.22632482 | 1.3299039 | 0.18354994 | 0.99989001 |
| MCCC1 | 88.497386 | 0.08656092 | 0.32852438 | 0.263484 | 0.79217755 | 0.99989001 |
| MCF2L2 | 106.201988 | -0.6564042 | 0.41123863 | -1.5961637 | 0.11045225 | 0.99989001 |
| B3GNT5 | 106.061445 | -0.2407362 | 0.42137651 | -0.571309 | 0.56779021 | 0.99989001 |
| KLHL24 | 222.867738 | -0.1424936 | 0.38985396 | -0.3655051 | 0.71473438 | 0.99989001 |
| YEATS2 | 323.586428 | 0.01450179 | 0.2975861 | 0.04873139 | 0.96113336 | 0.99989001 |
| ABCC5 | 75.1823166 | -0.2619948 | 0.3025246 | -0.8660281 | 0.38647475 | 0.99989001 |
| EIF2B5 | 47.9851417 | -0.3367294 | 0.29317628 | -1.1485562 | 0.25073903 | NA |
| DVL3 | 75.3690731 | -0.3463089 | 0.23116284 | -1.4981164 | 0.13410301 | 0.99989001 |
| AP2M1 | 190.17172 | -0.0188756 | 0.41203071 | -0.0458111 | 0.96346082 | 0.99989001 |
| PSMD2 | 94.9045432 | -0.0465295 | 0.35362749 | -0.1315777 | 0.89531835 | 0.99989001 |
| EIF4G1 | 168.47808 | -0.1972859 | 0.25579694 | -0.7712598 | 0.44055296 | 0.99989001 |
| POLR2H | 91.3268683 | -0.2014245 | 0.39844048 | -0.5055323 | 0.613185 | 0.99989001 |
| VPS8 | 307.951731 | -0.0447297 | 0.18815692 | -0.2377255 | 0.81209404 | 0.99989001 |
| EHHADH | 60.5414637 | 0.70134126 | 0.40920689 | 1.71390386 | 0.08654638 | 0.95784767 |
| MAP3K13 | 225.946914 | 0.15626485 | 0.26946999 | 0.57989702 | 0.56198407 | 0.99989001 |
| TMEM41A | 47.7542389 | 0.21255875 | 0.31175367 | 0.68181636 | 0.49535508 | NA |
| SENP2 | 128.017661 | -0.1045983 | 0.27867869 | -0.3753366 | 0.70741012 | 0.99989001 |
| IGF2BP2 | 273.732751 | 0.85039119 | 0.51891185 | 1.63879703 | 0.10125554 | 0.99989001 |
| TRA2B | 237.633049 | 0.04479319 | 0.24717146 | 0.18122316 | 0.85619242 | 0.99989001 |
| DGKG | 245.730653 | -0.8800748 | 0.66659577 | -1.3202525 | 0.18675072 | 0.99989001 |
| EIF4A2 | 255.685067 | 0.05076553 | 0.37150408 | 0.13664864 | 0.89130853 | 0.99989001 |
| ST6GAL1 | 2076.47745 | 0.55521513 | 0.47711495 | 1.16369258 | 0.24454862 | 0.99989001 |
| BCL6 | 128.996619 | -0.5039622 | 0.44912261 | -1.1221039 | 2.62E-01 | 0.99989001 |
| LPP | 2606.57129 | -0.0702252 | 0.21912421 | -0.3204811 | 0.74860365 | 0.99989001 |
| P3H2 | 147.277164 | -0.6721344 | 0.459121 | -1.4639593 | 0.14320508 | 0.99989001 |
| IL1RAP | 148.429247 | 0.08527303 | 0.57034192 | 0.14951211 | 0.88114955 | 0.99989001 |
| CCDC50 | 463.888385 | 0.07109662 | 0.38846814 | 0.18301789 | 0.85478398 | 0.99989001 |
| FGF12 | 179.334954 | -1.1951109 | 0.67152237 | -1.7797038 | 0.07512444 | 0.9206525 |
| MB21D2 | 150.816945 | 1.19028375 | 0.73863581 | 1.61146229 | 0.107079 | 0.99989001 |
| OPA1 | 283.636624 | -0.0122149 | 0.19065642 | -0.0640674 | 0.94891657 | 0.99989001 |
| HES1 | 509.750424 | -1.6045102 | 0.74474041 | -2.1544557 | 0.03120446 | 0.67281094 |
| ATP13A3 | 761.624626 | 0.26313913 | 0.34698435 | 0.75836021 | 0.44823537 | 0.99989001 |
| TMEM44 | 240.926283 | -0.2998772 | 0.31108813 | -0.9639621 | 0.33506492 | 0.99989001 |
| LSG1 | 86.5122906 | 0.0152855 | 0.27310802 | 0.05596869 | 0.95536675 | 0.99989001 |
| FAM43A | 115.848602 | -0.8340802 | 0.49966854 | -1.6692669 | 0.09506449 | 0.99989001 |
| XXYLT1 | 165.369456 | -0.2290584 | 0.29124504 | -0.7864801 | 0.43158627 | 0.99989001 |
| ACAP2 | 744.207988 | -0.0501892 | 0.17946431 | -0.279661 | 0.77973759 | 0.99989001 |
| PPP1R2 | 170.118019 | 0.46308704 | 0.28963759 | 1.59884992 | 0.10985395 | 0.99989001 |
| MUC20-OT1 | 431.602816 | 0.08926137 | 0.3876295 | 0.23027496 | 0.81787812 | 0.99989001 |
| TNK2 | 74.7317438 | 0.29435937 | 0.38120523 | 0.77218081 | 0.44000735 | 0.99989001 |

| | | | | | | |
|---|---|---|---|---|---|---|
| PCYT1A | 86.4620222 | -0.1136435 | 0.27676856 | -0.4106084 | 0.68135968 | 0.99989001 |
| UBXN7 | 166.820016 | 0.35384894 | 0.18955256 | 1.86675895 | 0.06193526 | 0.85940342 |
| RNF168 | 215.344661 | 0.05864623 | 0.18160155 | 0.32293903 | 0.74674142 | 0.99989001 |
| PIGX | 87.0567391 | -0.1934553 | 0.42470548 | -0.4555045 | 0.64874634 | 0.99989001 |
| PAK2 | 406.736449 | -0.0407175 | 0.13865613 | -0.2936582 | 0.76901909 | 0.99989001 |
| SENP5 | 341.968878 | -0.1186867 | 0.30487493 | -0.3892962 | 0.69705704 | 0.99989001 |
| NCBP2 | 57.8866085 | -0.1817603 | 0.32710229 | -0.555668 | 0.57843784 | 0.99989001 |
| DLG1 | 840.08528 | -0.0530381 | 0.2485422 | -0.2133967 | 0.83101757 | 0.99989001 |
| RUBCN | 94.6160413 | 0.14982058 | 0.29946612 | 0.50029228 | 0.61686929 | 0.99989001 |
| FYTTD1 | 126.953659 | -0.0970913 | 0.22164066 | -0.4380573 | 0.66134474 | 0.99989001 |
| LRCH3 | 453.976368 | 0.27666239 | 0.21237168 | 1.30272736 | 0.19266786 | 0.99989001 |
| IQCG | 59.048037 | 0.21715589 | 0.48316231 | 0.44944708 | 0.65310917 | 0.99989001 |
| RPL35A | 308.267676 | -0.0420976 | 0.60168711 | -0.069966 | 0.94422076 | 0.99989001 |
| ZNF595 | 83.6982854 | 0.09469051 | 0.24617045 | 0.38465427 | 0.70049358 | 0.99989001 |
| ZNF718 | 196.357759 | -0.0857593 | 0.21065034 | -0.407117 | 0.68392203 | 0.99989001 |
| ZNF141 | 135.170793 | -0.0561421 | 0.20990972 | -0.2674585 | 0.78911617 | 0.99989001 |
| ZNF721 | 329.891618 | 0.42030794 | 0.29776254 | 1.41155414 | 0.15808128 | 0.99989001 |
| PIGG | 95.4119555 | -0.3154389 | 0.24696735 | -1.2772494 | 0.2015142 | 0.99989001 |
| ATP5ME | 66.7304566 | -0.0756235 | 0.46051292 | -0.1642157 | 0.86956131 | 0.99989001 |
| PCGF3 | 188.367889 | 0.23638431 | 0.20626759 | 1.14600801 | 0.25179184 | 0.99989001 |
| AC139887.2 | 220.262135 | 0.56910964 | 0.31612173 | 1.80028637 | 0.07181543 | 0.91256377 |
| CPLX1 | 100.437238 | -0.9392923 | 0.53546732 | -1.7541542 | 0.07940408 | 0.93210268 |
| GAK | 405.810252 | -0.1171724 | 0.4540223 | -0.2580764 | 0.79634795 | 0.99989001 |
| CTBP1 | 85.7453221 | 0.02364712 | 0.29436809 | 0.08033182 | 0.93597335 | 0.99989001 |
| MAEA | 80.4226361 | -0.0674285 | 0.32814789 | -0.205482 | 0.83719555 | 0.99989001 |
| UVSSA | 96.7248066 | -0.6314948 | 0.29484897 | -2.1417568 | 0.03221306 | 0.67281094 |
| SLBP | 96.2281006 | -0.3338302 | 0.22895863 | -1.4580374 | 0.14483024 | 0.99989001 |
| LETM1 | 55.6785319 | -0.1541943 | 0.39600635 | -0.3893733 | 0.69700005 | 0.99989001 |
| NSD2 | 163.982744 | 0.15771421 | 0.30189671 | 0.52241114 | 0.6013841 | 0.99989001 |
| NELFA | 84.8369585 | -0.0544017 | 0.21193713 | -0.2566881 | 0.79741957 | 0.99989001 |
| POLN | 51.1372668 | 0.11044593 | 0.43133784 | 0.25605436 | 0.79790887 | 0.99989001 |
| MXD4 | 255.472986 | -0.2089958 | 0.38266436 | -0.5461595 | 0.5849563 | 0.99989001 |
| ZFYVE28 | 300.282442 | 0.3054649 | 0.3277625 | 0.93197024 | 0.35135191 | 0.99989001 |
| RNF4 | 191.181063 | 0.00976993 | 0.33736692 | 0.02895937 | 0.976897 | 0.99989001 |
| FAM193A | 360.522727 | -0.0027435 | 0.28076424 | -0.0097714 | 0.99220364 | 0.99989001 |
| TNIP2 | 91.6965389 | 0.23001666 | 0.43901249 | 0.52394105 | 0.60031954 | 0.99989001 |
| SH3BP2 | 148.348187 | -0.2317704 | 0.33935304 | -0.6829771 | 0.49462129 | 0.99989001 |
| ADD1 | 607.725973 | -0.2070445 | 0.27337102 | -0.7573756 | 0.44882486 | 0.99989001 |
| MFSD10 | 66.6687238 | -0.2928707 | 0.41315198 | -0.7088692 | 0.47840564 | 0.99989001 |
| NOP14 | 47.1867523 | -0.3757265 | 0.41408423 | -0.9073673 | 0.36421261 | NA |
| HTT | 286.766593 | -0.3062655 | 0.17810571 | -1.7195717 | 0.08551033 | 0.95514629 |
| RGS12 | 313.264118 | 0.00526585 | 0.31705956 | 0.01660839 | 0.98674903 | 0.99989001 |
| LRPAP1 | 106.06307 | 0.19044806 | 0.39629958 | 0.4805659 | 0.63082505 | 0.99989001 |
| TMEM128 | 42.6410163 | 0.29746892 | 0.29246509 | 1.01710917 | 0.3091015 | NA |
| STX18 | 216.994352 | 0.00239976 | 0.27870822 | 0.0086103 | 0.99313006 | 0.99989001 |
| STX18-AS1 | 83.4592182 | -0.5668465 | 0.53034931 | -1.0688173 | 0.28515198 | 0.99989001 |
| WFS1 | 64.9915426 | 0.17477735 | 0.31378848 | 0.55699097 | 0.5775336 | 0.99989001 |
| MRFAP1 | 188.870391 | 0.1691271 | 0.4506429 | 0.37530182 | 0.70743601 | 0.99989001 |
| MRFAP1L1 | 46.0903747 | -0.0972011 | 0.27363554 | -0.355221 | 0.72242406 | NA |
| KIAA0232 | 256.086801 | -0.0424534 | 0.24288998 | -0.1747845 | 0.86124898 | 0.99989001 |
| TBC1D14 | 92.2819173 | -0.1562351 | 0.25858788 | -0.6041857 | 0.54572021 | 0.99989001 |
| GRPEL1 | 51.3161609 | -0.2502278 | 0.4281847 | -0.5843922 | 0.55895647 | 0.99989001 |
| AFAP1 | 578.212089 | 0.14075359 | 0.37585863 | 0.37448546 | 0.70804317 | 0.99989001 |
| ABLIM2 | 156.761097 | 0.37175146 | 0.59412158 | 0.62571614 | 0.53150114 | 0.99989001 |

| | | | | | | |
|---|---|---|---|---|---|---|
| SH3TC1 | 131.514723 | 0.06019746 | 0.30269383 | 0.19887245 | 8.42E-01 | 0.99989001 |
| ACOX3 | 57.2646807 | -0.0581653 | 0.25325586 | -0.2296702 | 0.81834802 | 0.99989001 |
| SLC2A9 | 166.356266 | 0.6319247 | 0.5194695 | 1.21648084 | 0.22380181 | 0.99989001 |
| WDR1 | 193.820979 | 0.1053947 | 0.29521841 | 0.35700587 | 0.72108741 | 0.99989001 |
| HS3ST1 | 113.8745 | -1.2388931 | 0.53793844 | -2.3030388 | 0.02127666 | 0.59480081 |
| AC005699.1 | 130.571136 | -1.9281183 | 0.69214594 | -2.7857106 | 0.00534105 | 0.35652986 |
| LINC02360 | 173.153909 | -3.560771 | 1.46535161 | -2.4299772 | 0.01509977 | 0.51942334 |
| RAB28 | 246.326634 | 0.03386973 | 0.24930392 | 0.13585721 | 0.89193417 | 0.99989001 |
| BOD1L1 | 264.12473 | 0.02704702 | 0.28773633 | 0.09399931 | 0.92510971 | 0.99989001 |
| CPEB2 | 450.382501 | 0.14174178 | 0.27825318 | 0.5093986 | 0.61047286 | 0.99989001 |
| AC098829.1 | 132.557563 | -0.0930853 | 0.26637609 | -0.3494505 | 0.72675111 | 0.99989001 |
| CC2D2A | 85.9574631 | 0.27231524 | 0.57970107 | 0.46975115 | 0.63853282 | 0.99989001 |
| FBXL5 | 290.600613 | 0.13786509 | 0.17824586 | 0.77345467 | 0.43925335 | 0.99989001 |
| FAM200B | 46.7699114 | -0.027089 | 0.34802507 | -0.0778362 | 0.93795832 | NA |
| CD38 | 108.248951 | 0.4745473 | 0.99084323 | 0.47893278 | 0.63198645 | 0.99989001 |
| TAPT1 | 167.928524 | 0.18724663 | 0.26664988 | 0.70221907 | 0.48254256 | 0.99989001 |
| TAPT1-AS1 | 100.155685 | -0.1095369 | 0.52249464 | -0.2096421 | 0.83394701 | 0.99989001 |
| AC097515.1 | 44.5760007 | 0.31867199 | 0.68306743 | 0.46653079 | 0.64083561 | NA |
| LDB2 | 11411.3621 | 0.29426612 | 0.28709624 | 1.02497378 | 0.30537556 | 0.99989001 |
| QDPR | 46.5924996 | -0.0004365 | 0.28139667 | -0.0015513 | 0.99876223 | NA |
| LAP3 | 253.093239 | -0.1805988 | 0.44884079 | -0.4023673 | 0.68741375 | 0.99989001 |
| MED28 | 99.7413838 | -0.2893814 | 0.22128375 | -1.307739 | 0.19096184 | 0.99989001 |
| DCAF16 | 70.497042 | -0.1884334 | 0.25684192 | -0.7336552 | 0.4631589 | 0.99989001 |
| LCORL | 393.149775 | 0.02678895 | 0.22499726 | 0.11906343 | 0.9052251 | 0.99989001 |
| AC024230.1 | 51.1460227 | 0.50345055 | 0.64483587 | 0.78074217 | 0.43495415 | 0.99989001 |
| PACRGL | 105.399164 | -0.3382904 | 0.27458127 | -1.2320228 | 0.21794056 | 0.99989001 |
| KCNIP4 | 4946.8846 | -0.1685046 | 0.7220748 | -0.2333617 | 0.81548056 | 0.99989001 |
| ADGRA3 | 46.6564154 | -0.3015996 | 0.46159763 | -0.653382 | 0.51351002 | NA |
| DHX15 | 303.389254 | -0.1709709 | 0.18110274 | -0.9440545 | 0.34514181 | 0.99989001 |
| CCDC149 | 159.359591 | -0.3242677 | 0.22675671 | -1.4300245 | 0.15270999 | 0.99989001 |
| SEPSECS | 51.1536159 | -0.605559 | 0.24641447 | -2.4574816 | 0.0139915 | 0.50734455 |
| ZCCHC4 | 49.9493104 | -0.5137743 | 0.3078198 | -1.6690748 | 0.09510256 | 0.99989001 |
| ANAPC4 | 105.323321 | -0.1430846 | 0.30944423 | -0.4623921 | 0.64380019 | 0.99989001 |
| SEL1L3 | 76.7781838 | -0.1856321 | 0.30989112 | -0.5990237 | 0.54915705 | 0.99989001 |
| RBPJ | 330.729054 | 0.0938461 | 0.183371 | 0.51178268 | 0.60880311 | 0.99989001 |
| TBC1D19 | 186.547695 | 0.18734566 | 0.28573132 | 0.65567072 | 0.51203601 | 0.99989001 |
| STIM2 | 502.366445 | 0.0293971 | 0.19233414 | 0.1528439 | 0.87852137 | 0.99989001 |
| PCDH7 | 123.774494 | 0.31845838 | 0.69058578 | 0.4611424 | 0.64469645 | 0.99989001 |
| ARAP2 | 206.03463 | -0.5978237 | 0.37472327 | -1.5953737 | 0.11062869 | 0.99989001 |
| C4orf19 | 190.146557 | -0.849988 | 0.5367138 | -1.5836895 | 1.13E-01 | 0.99989001 |
| RELL1 | 470.598726 | -1.1602743 | 0.42773894 | -2.7125757 | 6.68E-03 | 0.390188 |
| PGM2 | 67.4133571 | -0.2381087 | 0.27420875 | -0.8683482 | 0.38520376 | 0.99989001 |
| TBC1D1 | 786.186102 | -0.0120492 | 0.19847366 | -0.0607091 | 9.52E-01 | 0.99989001 |
| KLF3 | 364.845071 | 0.38466243 | 0.25062313 | 1.53482416 | 0.12482703 | 0.99989001 |
| FAM114A1 | 128.424244 | -0.0485691 | 0.31796867 | -0.152748 | 0.87859701 | 0.99989001 |
| KLHL5 | 754.457775 | 0.22984218 | 0.3340943 | 0.68795601 | 0.49148048 | 0.99989001 |
| WDR19 | 94.1259208 | -0.50689 | 0.3963222 | -1.2789847 | 0.20090245 | 0.99989001 |
| RFC1 | 230.728478 | -0.1649472 | 0.34019993 | -0.4848538 | 0.62778008 | 0.99989001 |
| UGDH | 46.5775314 | 0.03486389 | 0.37535114 | 0.0928834 | 0.92599619 | NA |
| UGDH-AS1 | 74.5110308 | 0.2745041 | 0.32286527 | 0.85021255 | 0.39520693 | 0.99989001 |
| SMIM14 | 264.117003 | -0.1477942 | 0.23699981 | -0.6236047 | 0.53288722 | 0.99989001 |
| UBE2K | 680.846386 | -0.0079769 | 0.2364002 | -0.0337431 | 0.973082 | 0.99989001 |
| PDS5A | 589.322852 | -0.1088423 | 0.20673411 | -0.5264842 | 0.59855181 | 0.99989001 |
| N4BP2 | 58.6827719 | 0.06898061 | 0.2769111 | 0.2491074 | 0.8032777 | 0.99989001 |

| | | | | | | |
|---|---|---|---|---|---|---|
| RBM47 | 79.8895885 | -0.1879388 | 0.48138955 | -0.390409 | 0.69623416 | 0.99989001 |
| APBB2 | 2013.92466 | -0.203344 | 0.24074308 | -0.8446515 | 0.39830547 | 0.99989001 |
| LIMCH1 | 2438.96826 | 0.20696919 | 0.23105809 | 0.89574528 | 0.37038882 | 0.99989001 |
| TMEM33 | 115.38249 | -0.1040448 | 0.23599147 | -0.4408835 | 0.65929731 | 0.99989001 |
| SLC30A9 | 268.14969 | 0.1863514 | 0.16944736 | 1.09975987 | 0.27143676 | 0.99989001 |
| SHISA3 | 52.7527836 | -0.6220908 | 1.05569645 | -0.5892705 | 0.55567983 | 0.99989001 |
| ATP8A1 | 279.966583 | -0.4171424 | 0.89656636 | -0.4652666 | 0.64174056 | 0.99989001 |
| GUF1 | 50.766997 | -0.1678175 | 0.26514971 | -0.6329161 | 0.52678843 | 0.99989001 |
| GNPDA2 | 58.1121718 | -0.0162538 | 0.3292127 | -0.0493717 | 0.96062306 | 0.99989001 |
| ATP10D | 202.131653 | 0.23183754 | 0.24775527 | 0.93575219 | 0.34940079 | 0.99989001 |
| NFXL1 | 83.8166459 | -0.0519762 | 0.292253 | -0.1778465 | 0.85884351 | 0.99989001 |
| SLAIN2 | 222.482786 | 0.46564494 | 0.16911369 | 2.75344319 | 0.0058972 | 0.37352207 |
| FRYL | 1303.26837 | 0.27274078 | 0.20746924 | 1.31460832 | 0.18864157 | 0.99989001 |
| OCIAD1 | 300.384024 | -0.1613564 | 0.19897505 | -0.8109376 | 0.41740149 | 0.99989001 |
| OCIAD2 | 62.9601964 | -0.2809315 | 0.43570956 | -0.6447678 | 0.51907766 | 0.99989001 |
| DCUN1D4 | 220.352472 | 0.01049485 | 0.18191941 | 0.05768957 | 0.9539959 | 0.99989001 |
| SGCB | 48.4986948 | 0.40775903 | 0.28130847 | 1.44950855 | 0.14719561 | NA |
| USP46 | 59.5454454 | 0.01638604 | 0.27573111 | 0.05942761 | 0.95261152 | 0.99989001 |
| SCFD2 | 408.936376 | -0.2441651 | 0.45669973 | -0.5346293 | 0.59290618 | 0.99989001 |
| AC058822.1 | 174.55827 | 0.25714094 | 0.52659266 | 0.48831091 | 0.62532964 | 0.99989001 |
| LNX1 | 239.825304 | 1.09950063 | 0.69924373 | 1.57241401 | 0.11585457 | 0.99989001 |
| CHIC2 | 371.406772 | 0.26350928 | 0.34608643 | 0.76139732 | 0.44641979 | 0.99989001 |
| KDR | 644.564389 | 0.36902347 | 0.34426393 | 1.07192022 | 0.28375587 | 0.99989001 |
| TMEM165 | 565.166437 | 0.08817472 | 0.27224951 | 0.32387467 | 0.74603292 | 0.99989001 |
| CLOCK | 339.44132 | 0.20856302 | 0.1896088 | 1.09996486 | 0.27134743 | 0.99989001 |
| EXOC1 | 315.752474 | 0.13075105 | 0.20560242 | 0.6359412 | 0.52481475 | 0.99989001 |
| CEP135 | 148.25602 | 0.04832817 | 0.21383742 | 0.22600429 | 0.82119808 | 0.99989001 |
| AASDH | 66.5014325 | 0.08945362 | 0.22836514 | 0.3917131 | 0.69527021 | 0.99989001 |
| PAICS | 51.314506 | -0.6051831 | 0.45047436 | -1.3434353 | 0.17913107 | 0.99989001 |
| SRP72 | 153.091873 | 0.08550063 | 0.21344181 | 0.40058052 | 0.68872899 | 0.99989001 |
| REST | 136.163201 | 0.51028148 | 0.26585946 | 1.91936548 | 0.0549381 | 0.82320925 |
| POLR2B | 186.861944 | -0.2119977 | 0.18805326 | -1.1273278 | 0.25960389 | 0.99989001 |
| IGFBP7 | 2567.15011 | -0.7879611 | 0.5145792 | -1.5312727 | 0.125702 | 0.99989001 |
| AC116049.2 | 54.6616669 | -0.2333795 | 0.55704622 | -0.418959 | 0.67524612 | 0.99989001 |
| AC104806.2 | 73.0221609 | 0.6132053 | 0.29304779 | 2.09250957 | 0.03639296 | 0.70003115 |
| CENPC | 198.977114 | -0.5319613 | 0.22058263 | -2.4116193 | 0.01588185 | 0.53042584 |
| UBA6 | 283.132143 | -0.2869364 | 0.27842491 | -1.0305701 | 0.30274246 | 0.99989001 |
| UBA6-AS1 | 123.448348 | -0.838395 | 0.51012881 | -1.6434966 | 0.10028023 | 0.99989001 |
| YTHDC1 | 322.266079 | -0.0538189 | 0.25394933 | -0.2119278 | 0.83216337 | 0.99989001 |
| UGT2B7 | 91.5914092 | 1.26586627 | 0.60581421 | 2.08952883 | 0.03666015 | 0.70081289 |
| RUFY3 | 364.688543 | 0.02332129 | 0.24109835 | 0.09672935 | 0.92294133 | 0.99989001 |
| GRSF1 | 77.0237702 | 0.15240022 | 0.22367618 | 0.68134312 | 0.49565441 | 0.99989001 |
| MOB1B | 167.303613 | 0.07344328 | 0.19574779 | 0.37519341 | 0.70751663 | 0.99989001 |
| SLC4A4 | 95.0028059 | 2.52874254 | 0.52542398 | 4.81276573 | 1.49E-06 | 0.00207155 |
| ANKRD17 | 1233.7247 | -0.0436383 | 0.21026755 | -0.2075369 | 0.83559056 | 0.99989001 |
| AC053527.1 | 82.7208482 | 0.78324397 | 0.55402332 | 1.41373828 | 0.15743876 | 0.99989001 |
| CXCL2 | 166.432677 | 1.75543899 | 1.09145147 | 1.60835276 | 0.10775794 | 0.99989001 |
| MTHFD2L | 119.943905 | -0.0144948 | 0.25852783 | -0.0560667 | 0.95528868 | 0.99989001 |
| RCHY1 | 96.2237114 | 0.3838036 | 0.30969665 | 1.23928884 | 0.21523855 | 0.99989001 |
| THAP6 | 53.1419584 | -0.0211054 | 0.38935496 | -0.0542061 | 0.95677097 | 0.99989001 |
| G3BP2 | 177.393358 | 0.01005608 | 0.24399428 | 0.0412144 | 0.96712497 | 0.99989001 |
| USO1 | 241.822268 | 0.20412755 | 0.18460074 | 1.10577861 | 0.26882236 | 0.99989001 |
| SDAD1 | 93.0531455 | -0.2726367 | 0.32813544 | -0.8308664 | 0.40604912 | 0.99989001 |
| NUP54 | 139.741638 | -0.1713071 | 0.21779017 | -0.7865695 | 0.43153394 | 0.99989001 |

| | | | | | | |
|---|---|---|---|---|---|---|
| SCARB2 | 341.29398 | 0.2639467 | 0.22959635 | 1.14961191 | 0.25030375 | 0.99989001 |
| 11-Sep | 173.522804 | 0.50211861 | 0.21667769 | 2.3173526 | 0.02048453 | 0.58699182 |
| CCNI | 446.345997 | 0.15095133 | 0.28292319 | 0.53354174 | 0.59365862 | 0.99989001 |
| CCNG2 | 98.7036493 | 0.10053202 | 0.50749423 | 0.1980949 | 0.84297081 | 0.99989001 |
| CNOT6L | 468.033003 | -0.2556165 | 0.32435601 | -0.788074 | 0.4306534 | 0.99989001 |
| MRPL1 | 146.114665 | 0.01652825 | 0.27532541 | 0.0600317 | 0.95213038 | 0.99989001 |
| ANXA3 | 125.962683 | -1.7477576 | 1.05464083 | -1.6572064 | 0.09747775 | 0.99989001 |
| BMP2K | 267.386509 | 0.05421569 | 0.23179797 | 0.23389197 | 0.81506884 | 0.99989001 |
| ANTXR2 | 176.773094 | 0.93662884 | 0.33415663 | 2.80296348 | 0.00506354 | 0.34893315 |
| RASGEF1B | 1007.20406 | -0.3065843 | 0.66421088 | -0.4615767 | 0.64438489 | 0.99989001 |
| HNRNPD | 382.884629 | -0.0074571 | 0.18713232 | -0.0398491 | 0.9682134 | 0.99989001 |
| HNRNPDL | 325.045903 | -0.0497628 | 0.20988182 | -0.2370989 | 0.81258004 | 0.99989001 |
| ENOPH1 | 73.2180311 | 0.01098589 | 0.25710342 | 0.04272944 | 0.96591721 | 0.99989001 |
| TMEM150C | 110.230946 | 0.42030379 | 0.32820984 | 1.28059473 | 0.20033605 | 0.99989001 |
| SEC31A | 439.609203 | 0.01868091 | 0.14969809 | 0.12479054 | 0.90068938 | 0.99989001 |
| LIN54 | 88.8292328 | 0.34140331 | 0.3083301 | 1.10726559 | 0.26817912 | 0.99989001 |
| COPS4 | 67.8837648 | 0.33598205 | 0.32152751 | 1.04495583 | 0.29604339 | 0.99989001 |
| HELQ | 77.6945548 | 0.00187375 | 0.24737425 | 0.00757454 | 0.99395645 | 0.99989001 |
| MRPS18C | 46.8844484 | 0.01452158 | 0.31479978 | 0.04612957 | 0.96320698 | NA |
| WDFY3 | 940.198343 | -0.2161202 | 0.19206651 | -1.1252361 | 0.26048899 | 0.99989001 |
| ARHGAP24 | 118.714081 | -1.6524521 | 0.66982466 | -2.466992 | 0.01362534 | 0.50652625 |
| MAPK10 | 62.1430269 | -0.3738946 | 0.4549843 | -0.8217747 | 0.41120516 | 0.99989001 |
| PTPN13 | 63.0318757 | 0.63751078 | 0.47063135 | 1.3545863 | 0.1755494 | 0.99989001 |
| AC093827.5 | 53.0813661 | -0.0362014 | 0.27107425 | -0.133548 | 0.89376001 | 0.99989001 |
| AFF1 | 1740.94851 | 0.16355173 | 0.23617567 | 0.69250034 | 0.48862318 | 0.99989001 |
| KLHL8 | 115.592609 | 0.47675814 | 0.35164186 | 1.35580601 | 0.1751609 | 0.99989001 |
| HSD17B11 | 145.914755 | 0.29767865 | 0.26591205 | 1.11946281 | 0.26294275 | 0.99989001 |
| NUDT9 | 100.775058 | 0.0202203 | 0.26208108 | 0.07715284 | 0.93850196 | 0.99989001 |
| SPARCL1 | 985.254881 | 0.10060737 | 0.69065563 | 0.14566937 | 0.8841824 | 0.99989001 |
| SPP1 | 2239.94144 | 0.70114255 | 0.69263158 | 1.01228788 | 3.11E-01 | 0.99989001 |
| PKD2 | 339.346664 | -0.4906859 | 0.29407732 | -1.6685609 | 0.09520443 | 0.99989001 |
| ABCG2 | 99.9350478 | 0.90819121 | 0.63364531 | 1.43328011 | 0.1517778 | 0.99989001 |
| PPM1K | 97.8321566 | 0.39355217 | 0.52234061 | 0.75343973 | 0.45118572 | 0.99989001 |
| HERC6 | 250.220842 | -0.3523904 | 1.10085964 | -0.3201047 | 0.74888893 | 0.99989001 |
| HERC5 | 160.546743 | -0.1695717 | 0.72844604 | -0.2327855 | 0.81592794 | 0.99989001 |
| PYURF | 69.8434151 | -0.1079974 | 0.52799285 | -0.2045433 | 0.83792892 | 0.99989001 |
| HERC3 | 105.198854 | 0.03172808 | 0.32584211 | 0.09737255 | 0.92243054 | 0.99989001 |
| FAM13A-AS1 | 45.7256267 | -0.3450974 | 0.31298927 | -1.1025855 | 0.27020722 | NA |
| FAM13A | 359.892046 | -1.3081767 | 0.58724841 | -2.2276377 | 0.02590469 | 0.63690091 |
| SNCA | 92.4696867 | -3.2755673 | 1.46798303 | -2.2313387 | 0.0256587 | 0.6349898 |
| CCSER1 | 315.571711 | -1.5295877 | 0.64865522 | -2.3580905 | 0.01836921 | 0.56643441 |
| SMARCAD1 | 218.850345 | -0.0932904 | 0.1831845 | -0.5092699 | 0.61056306 | 0.99989001 |
| PDLIM5 | 1537.56472 | -0.0345492 | 0.33189069 | -0.1040982 | 0.9170914 | 0.99989001 |
| STPG2 | 114.264379 | -0.6935422 | 0.44241061 | -1.5676436 | 0.11696433 | 0.99989001 |
| RAP1GDS1 | 373.547369 | 0.42927018 | 0.17217504 | 2.49321949 | 0.01265906 | 0.49000287 |
| TSPAN5 | 119.558819 | 0.70499491 | 0.45270574 | 1.55729174 | 0.11940123 | 0.99989001 |
| EIF4E | 171.528767 | -0.0741834 | 0.1773877 | -0.4181994 | 0.67580131 | 0.99989001 |
| METAP1 | 82.4539835 | 0.18459161 | 0.25146973 | 0.734051 | 0.46291766 | 0.99989001 |
| ADH5 | 80.3924818 | 0.07210738 | 0.34227343 | 0.21067186 | 0.83314333 | 0.99989001 |
| LAMTOR3 | 45.9046465 | -0.2391923 | 0.2713832 | -0.8813822 | 0.37811097 | NA |
| DNAJB14 | 331.773559 | 0.14308375 | 0.20035688 | 0.71414442 | 0.47513788 | 0.99989001 |
| H2AFZ | 162.105547 | -0.3353069 | 0.94710445 | -0.3540337 | 0.72331359 | 0.99989001 |
| EMCN | 3739.13588 | 0.37718471 | 0.53699484 | 0.70239913 | 4.82E-01 | 0.99989001 |
| PPP3CA | 1105.69932 | 0.25391159 | 0.35985508 | 0.70559401 | 0.48044065 | 0.99989001 |

| | | | | | | |
|---|---|---|---|---|---|---|
| SLC39A8 | 57.6970028 | -0.2641691 | 0.46423026 | -0.5690477 | 0.5693238 | 0.99989001 |
| NFKB1 | 588.744088 | 0.86167619 | 0.51505553 | 1.67297725 | 0.09433177 | 0.99989001 |
| MANBA | 114.129372 | -0.0403507 | 0.38350431 | -0.1052159 | 0.91620452 | 0.99989001 |
| UBE2D3 | 779.223721 | -0.0533764 | 0.23867492 | -0.2236363 | 0.82304029 | 0.99989001 |
| CISD2 | 53.3306302 | -0.0302544 | 0.27005546 | -0.1120304 | 0.91079933 | 0.99989001 |
| SLC9B2 | 76.5798588 | 0.0863576 | 0.28229452 | 0.30591311 | 0.7596708 | 0.99989001 |
| TET2 | 255.614909 | 0.00844249 | 0.25453289 | 0.03316856 | 0.97354017 | 0.99989001 |
| PPA2 | 313.804882 | -0.1127553 | 0.22821298 | -0.4940793 | 0.62125017 | 0.99989001 |
| INTS12 | 101.853566 | -0.280915 | 0.36888055 | -0.7615339 | 0.44633823 | 0.99989001 |
| GSTCD | 92.9915775 | -0.2233686 | 0.286125 | -0.7806679 | 0.43499787 | 0.99989001 |
| NPNT | 169.566775 | -1.0372669 | 0.65844207 | -1.5753351 | 0.11517912 | 0.99989001 |
| TBCK | 456.854295 | 0.20335305 | 0.25726433 | 0.79044402 | 0.4292685 | 0.99989001 |
| AIMP1 | 78.5125208 | 0.04093852 | 0.28335611 | 0.14447728 | 0.88512359 | 0.99989001 |
| PAPSS1 | 115.652835 | -0.0621605 | 0.30206188 | -0.2057872 | 0.8369571 | 0.99989001 |
| LEF1 | 109.277917 | -2.1005205 | 0.67916431 | -3.0928016 | 0.00198277 | 0.21078759 |
| RPL34 | 296.553753 | 0.06391878 | 0.56186362 | 0.1137621 | 0.90942638 | 0.99989001 |
| OSTC | 94.6448538 | 0.10586315 | 0.43839746 | 0.24147756 | 0.80918501 | 0.99989001 |
| SEC24B | 489.438352 | 0.02446505 | 0.20811323 | 0.11755644 | 0.90641912 | 0.99989001 |
| MCUB | 87.9922875 | 0.10922644 | 0.4427653 | 0.24669151 | 0.80514698 | 0.99989001 |
| EGF | 55.4061104 | -0.323594 | 0.85824565 | -0.3770412 | 7.06E-01 | 0.99989001 |
| FAM241A | 445.396929 | 0.05462104 | 0.35079602 | 0.15570599 | 0.87626478 | 0.99989001 |
| AP1AR | 89.0962102 | -0.3386778 | 0.288251 | -1.1749406 | 0.24001849 | 0.99989001 |
| ALPK1 | 285.046118 | -0.8075259 | 0.24987049 | -3.2317778 | 0.00123023 | 0.16883184 |
| LARP7 | 109.596184 | 0.23726944 | 0.24355488 | 0.97419291 | 0.32996076 | 0.99989001 |
| ANK2 | 63.3683167 | -0.060043 | 0.57705953 | -0.1040499 | 0.91712979 | 0.99989001 |
| CAMK2D | 900.503733 | -0.1219255 | 0.2517396 | -0.4843319 | 0.62815032 | 0.99989001 |
| ARSJ | 66.6930342 | 0.59124835 | 0.58602824 | 1.00890761 | 0.31301895 | 0.99989001 |
| SNHG8 | 47.7721465 | -0.1180147 | 0.72621082 | -0.1625075 | 0.87090625 | NA |
| METTL14 | 95.1657552 | -0.0576487 | 0.25587032 | -0.2253044 | 0.82174248 | 0.99989001 |
| SEC24D | 245.291815 | 0.0675257 | 0.25193159 | 0.26803187 | 0.78867479 | 0.99989001 |
| SYNPO2 | 70.1083832 | 0.04224857 | 0.484806 | 0.08714531 | 0.93055601 | 0.99989001 |
| USP53 | 303.600309 | -0.6644763 | 0.37360832 | -1.7785372 | 0.07531568 | 0.9206525 |
| C4orf3 | 229.071003 | -0.1984423 | 0.39210577 | -0.5060939 | 0.61279074 | 0.99989001 |
| PDE5A | 91.7717497 | -0.5046185 | 0.45657108 | -1.1052353 | 0.26905763 | 0.99989001 |
| AC108866.1 | 84.8820067 | -0.2465906 | 0.39152995 | -0.6298128 | 0.52881708 | 0.99989001 |
| PRDM5 | 133.331056 | -0.4434676 | 0.29437636 | -1.5064648 | 0.13194791 | 0.99989001 |
| ANXA5 | 328.382289 | -0.2957521 | 0.38386587 | -0.7704569 | 0.44102892 | 0.99989001 |
| BBS7 | 78.5999744 | -0.069881 | 0.26749303 | -0.2612441 | 0.7939043 | 0.99989001 |
| KIAA1109 | 812.116064 | -0.262348 | 0.28992826 | -0.9048721 | 0.36553317 | 0.99989001 |
| FGF2 | 220.911302 | 0.1917602 | 0.48809328 | 0.39287614 | 0.69441097 | 0.99989001 |
| SPATA5 | 430.00608 | -0.2472647 | 0.19877296 | -1.2439554 | 0.21351599 | 0.99989001 |
| SPRY1 | 770.039111 | -0.8989692 | 0.59700593 | -1.5057961 | 1.32E-01 | 0.99989001 |
| ANKRD50 | 201.502854 | 0.70794105 | 0.38084504 | 1.85886901 | 0.06304571 | 0.85940342 |
| FAT4 | 541.099523 | 0.15648935 | 0.40847441 | 0.38310688 | 0.70164052 | 0.99989001 |
| INTU | 62.1503334 | -0.2299328 | 0.31782499 | -0.7234574 | 0.46939894 | 0.99989001 |
| MFSD8 | 90.4866975 | -0.1887834 | 0.25524707 | -0.7396104 | 0.45953643 | 0.99989001 |
| ABHD18 | 240.119092 | -0.0174017 | 0.20370193 | -0.0854271 | 0.93192181 | 0.99989001 |
| LARP1B | 142.235415 | 0.55942645 | 0.22330297 | 2.50523514 | 0.012237 | 0.49000287 |
| PGRMC2 | 151.66218 | 0.29013206 | 0.27512561 | 1.05454403 | 0.29163392 | 0.99989001 |
| AC108062.1 | 76.7786348 | 0.33150716 | 0.46184845 | 0.71778341 | 0.47289084 | 0.99989001 |
| JADE1 | 95.4987628 | -0.0385201 | 0.27818046 | -0.1384717 | 0.88986767 | 0.99989001 |
| SCLT1 | 395.438833 | 0.21685875 | 0.38998307 | 0.55607222 | 0.57816148 | 0.99989001 |
| AC096711.2 | 100.83612 | -0.5585183 | 0.40653105 | -1.3738639 | 0.16948393 | 0.99989001 |
| AC079380.1 | 87.3187676 | -0.4204972 | 0.63175547 | -0.6656013 | 0.505666 | 0.99989001 |

| | | | | | | |
|---|---|---|---|---|---|---|
| AC109927.1 | 81.7704522 | 0.4386321 | 0.41487621 | 1.0572602 | 0.29039285 | 0.99989001 |
| ELF2 | 1096.75824 | 0.20283689 | 0.19753764 | 1.02682653 | 0.30450216 | 0.99989001 |
| NDUFC1 | 87.6588017 | 0.05420403 | 0.29998971 | 0.18068631 | 0.8566138 | 0.99989001 |
| NAA15 | 261.586866 | -0.0254985 | 0.24702913 | -0.1032206 | 0.91778791 | 0.99989001 |
| AC097376.2 | 135.434818 | -0.2223954 | 0.26732067 | -0.8319426 | 0.40544135 | 0.99989001 |
| SETD7 | 119.462731 | 0.2263267 | 0.20490679 | 1.10453489 | 0.26936118 | 0.99989001 |
| MGST2 | 272.33229 | -0.1970121 | 0.3788021 | -0.5200925 | 0.60299912 | 0.99989001 |
| MAML3 | 1208.76068 | 0.11092139 | 0.34548933 | 0.3210559 | 0.74816803 | 0.99989001 |
| SCOC | 85.0716602 | 0.01342039 | 0.38211989 | 0.0351209 | 0.97198334 | 0.99989001 |
| ELMOD2 | 62.1754756 | -0.1287126 | 0.25823896 | -0.4984245 | 0.61818486 | 0.99989001 |
| TBC1D9 | 391.965132 | -0.0934159 | 0.29844884 | -0.3130047 | 0.75427712 | 0.99989001 |
| RNF150 | 63.6399047 | 0.93752093 | 0.68227083 | 1.37411844 | 0.16940492 | 0.99989001 |
| ZNF330 | 63.738764 | -0.5295485 | 0.4264594 | -1.2417325 | 0.21433527 | 0.99989001 |
| IL15 | 298.058513 | 0.01960492 | 0.39035966 | 0.05022271 | 0.95994492 | 0.99989001 |
| INPP4B | 353.007714 | -1.4582895 | 0.72912826 | -2.0000452 | 0.04549539 | 0.77228819 |
| USP38 | 83.7407988 | -0.2024116 | 0.24472467 | -0.8270994 | 0.40818072 | 0.99989001 |
| GAB1 | 636.870432 | -0.1436926 | 0.30804076 | -0.4664727 | 0.64087721 | 0.99989001 |
| SMARCA5 | 234.607986 | 0.20798379 | 0.2300707 | 0.90399945 | 0.36599569 | 0.99989001 |
| ANAPC10 | 202.815061 | 0.10128929 | 0.1698866 | 0.59621705 | 0.55103024 | 0.99989001 |
| ABCE1 | 118.236754 | -0.216556 | 0.31730819 | -0.6824784 | 0.49493649 | 0.99989001 |
| OTUD4 | 222.42534 | -0.179774 | 0.20788121 | -0.864792 | 0.38715295 | 0.99989001 |
| SMAD1 | 429.041194 | -0.3541914 | 0.33292228 | -1.0638861 | 0.28738031 | 0.99989001 |
| ZNF827 | 127.195926 | 0.43036257 | 0.37696711 | 1.14164488 | 0.25360166 | 0.99989001 |
| LSM6 | 69.9733949 | -0.1723906 | 0.33843189 | -0.5093805 | 0.61048555 | 0.99989001 |
| SLC10A7 | 273.95567 | 0.22546325 | 0.18047979 | 1.24924379 | 0.21157592 | 0.99989001 |
| TMEM184C | 69.4453116 | -0.1245139 | 0.27016322 | -0.4608841 | 0.64488173 | 0.99989001 |
| PRMT9 | 81.2948987 | 0.0746499 | 0.34341343 | 0.21737618 | 0.82791519 | 0.99989001 |
| ARHGAP10 | 174.731301 | 0.05691326 | 0.31355755 | 0.18150819 | 0.8559687 | 0.99989001 |
| NR3C2 | 901.057742 | 0.09782488 | 0.36482555 | 0.26814153 | 0.78859038 | 0.99989001 |
| DCLK2 | 433.787816 | 0.0544935 | 0.38049045 | 0.14321911 | 0.88611714 | 0.99989001 |
| LRBA | 936.785494 | -0.1277052 | 0.31518244 | -0.4051787 | 0.68534615 | 0.99989001 |
| RPS3A | 194.754555 | 0.06680698 | 0.54152168 | 0.12336898 | 0.90181492 | 0.99989001 |
| SH3D19 | 735.649765 | 0.2948106 | 0.27755086 | 1.06218585 | 0.28815132 | 0.99989001 |
| FBXW7 | 662.059269 | 0.20032387 | 0.23522966 | 0.85160973 | 0.39443074 | 0.99989001 |
| ARFIP1 | 359.995088 | 0.44789845 | 0.2400244 | 1.86605378 | 0.06203384 | 0.85940342 |
| TRIM2 | 169.375962 | 0.30138301 | 0.34607995 | 0.87084792 | 0.3838372 | 0.99989001 |
| TMEM131L | 311.351082 | 0.11983636 | 0.29404037 | 0.40755068 | 0.68360358 | 0.99989001 |
| PLRG1 | 51.2995263 | 0.13826601 | 0.26965095 | 0.5127592 | 0.60811977 | 0.99989001 |
| GUCY1A1 | 70.7934209 | -0.4818579 | 0.7047346 | -0.6837438 | 0.49413695 | 0.99989001 |
| CTSO | 65.7768404 | -0.2452683 | 0.30279991 | -0.8100012 | 0.41793947 | 0.99989001 |
| PDGFC | 113.229863 | -0.4093192 | 0.61284619 | -0.6678987 | 0.50419828 | 0.99989001 |
| FAM198B | 364.429804 | 0.41879878 | 0.65252327 | 0.64181433 | 0.52099375 | 0.99989001 |
| FAM198B-AS | 48.3123027 | 0.11820365 | 0.70300407 | 0.16814078 | 0.86647253 | NA |
| RXFP1 | 162.658697 | 1.58774801 | 0.52191394 | 3.04216442 | 2.35E-03 | 2.36E-01 |
| ETFDH | 135.088896 | -0.2630325 | 0.40439644 | -0.6504323 | 0.51541299 | 0.99989001 |
| PPID | 62.9578385 | 0.14893625 | 0.30657894 | 0.48580064 | 0.62710852 | 0.99989001 |
| FNIP2 | 472.880038 | -0.0708267 | 0.28940005 | -0.2447364 | 0.80666055 | 0.99989001 |
| RAPGEF2 | 1186.46228 | 0.10515252 | 0.27459856 | 0.3829318 | 0.70177033 | 0.99989001 |
| NAF1 | 86.5145113 | -0.1914177 | 0.21697234 | -0.8822215 | 0.37765706 | 0.99989001 |
| NPY1R | 92.5541595 | -0.8306213 | 0.70827989 | -1.1727303 | 0.24090399 | 0.99989001 |
| TMA16 | 73.1039774 | -0.1020577 | 0.30292443 | -0.3369081 | 0.73618619 | 0.99989001 |
| TMEM192 | 71.2970309 | -0.309369 | 0.44065503 | -0.7020661 | 0.48263794 | 0.99989001 |
| KLHL2 | 534.863148 | 0.19002218 | 0.21588993 | 0.88018082 | 0.37876136 | 0.99989001 |
| MSMO1 | 53.2138818 | 0.02723954 | 0.46717499 | 0.05830694 | 0.95350414 | 0.99989001 |

| | | | | | | |
|---|---|---|---|---|---|---|
| CPE | 148.988479 | 2.32812397 | 0.61933371 | 3.75907839 | 0.00017054 | 0.05851086 |
| TLL1 | 1773.08578 | -0.639352 | 0.49509696 | -1.2913672 | 0.19657639 | 0.99989001 |
| DDX60 | 294.33605 | 0.24042168 | 0.63283954 | 0.37990939 | 0.70401268 | 0.99989001 |
| DDX60L | 442.831125 | 0.0485385 | 0.71530478 | 0.06785709 | 0.9458994 | 0.99989001 |
| PALLD | 72.5202677 | 0.5973088 | 0.31365027 | 1.90437841 | 0.05686092 | 0.83825432 |
| CBR4 | 292.338727 | -0.1163525 | 0.2795794 | -0.4161696 | 0.67728587 | 0.99989001 |
| SH3RF1 | 203.299954 | 0.26437801 | 0.27899275 | 0.94761605 | 0.34332495 | 0.99989001 |
| NEK1 | 257.828633 | -0.0232384 | 0.21534779 | -0.1079109 | 0.91406636 | 0.99989001 |
| CLCN3 | 244.006691 | 0.08155538 | 0.20064544 | 0.40646516 | 0.68440085 | 0.99989001 |
| HPF1 | 57.149785 | -0.2263412 | 0.36379909 | -0.6221599 | 0.53383676 | 0.99989001 |
| GALNT7 | 98.5831562 | 0.2757544 | 0.25686617 | 1.07353335 | 0.28303188 | 0.99989001 |
| HMGB2 | 89.0899583 | 0.11738644 | 0.63362255 | 0.18526242 | 0.85302321 | 0.99989001 |
| FBXO8 | 53.5108569 | 0.11707078 | 0.35400286 | 0.33070574 | 0.74086676 | 0.99989001 |
| CEP44 | 76.0723372 | -0.0020348 | 0.29155056 | -0.0069793 | 0.99443133 | 0.99989001 |
| GPM6A | 2209.73796 | -0.9210922 | 0.67610857 | -1.3623436 | 0.17308946 | 0.99989001 |
| SPCS3 | 293.02936 | -0.0015225 | 0.43987889 | -0.0034613 | 0.99723833 | 0.99989001 |
| VEGFC | 1255.64297 | -0.8354618 | 0.31901005 | -2.6189198 | 0.00882087 | 0.42668269 |
| AC078881.1 | 89.4527126 | -0.2830639 | 0.45802555 | -0.6180089 | 0.53656947 | 0.99989001 |
| DCTD | 51.1511668 | -0.0836164 | 0.37321358 | -0.2240442 | 0.82272287 | 0.99989001 |
| WWC2 | 606.807334 | 0.09031089 | 0.20328857 | 0.44424974 | 0.65686203 | 0.99989001 |
| RWDD4 | 49.1101957 | 0.31209981 | 0.27231947 | 1.14607968 | 0.25176219 | NA |
| TRAPPC11 | 150.236273 | -0.0304894 | 0.24309659 | -0.125421 | 0.90019029 | 0.99989001 |
| STOX2 | 1338.91277 | 0.12850098 | 0.28941585 | 0.44400119 | 0.65704172 | 0.99989001 |
| IRF2 | 294.506156 | 0.54109169 | 0.32298843 | 1.67526645 | 0.09388196 | 0.99989001 |
| CASP3 | 78.6130421 | 0.27039393 | 0.22903483 | 1.18057995 | 0.23776963 | 0.99989001 |
| PRIMPOL | 101.91879 | 0.185289 | 0.33549271 | 0.55228921 | 0.58075022 | 0.99989001 |
| ACSL1 | 56.6667547 | 0.7091972 | 0.39604124 | 1.79071551 | 0.07333896 | 0.9158596 |
| CFAP97 | 121.06424 | 0.17478024 | 0.31783819 | 0.54990319 | 0.58238577 | 0.99989001 |
| SNX25 | 574.826629 | -0.5615704 | 0.56901769 | -0.9869121 | 0.32368573 | 0.99989001 |
| AC112722.1 | 46.6899163 | 0.06756061 | 0.49024473 | 0.13780996 | 0.89039061 | NA |
| ANKRD37 | 200.681445 | -3.1558775 | 1.45964611 | -2.162084 | 0.0306117 | 0.67281094 |
| UFSP2 | 88.003707 | -0.4376324 | 0.34012349 | -1.2866869 | 0.19820345 | 0.99989001 |
| PDLIM3 | 82.3363897 | 1.37252429 | 0.71841008 | 1.91050255 | 0.05606854 | 0.83144465 |
| SORBS2 | 938.166774 | -0.1231522 | 0.38869561 | -0.3168345 | 0.75136919 | 0.99989001 |
| TLR3 | 42.9521595 | -0.2445018 | 0.54090193 | -0.4520261 | 0.6512502 | NA |
| FAM149A | 47.4329826 | -1.1614493 | 0.37078233 | -3.1324288 | 0.00173366 | NA |
| CYP4V2 | 50.9627255 | -0.6379866 | 0.49482479 | -1.2893183 | 0.19728746 | 0.99989001 |
| FRG1-DT | 77.2946163 | -0.4781446 | 0.48535912 | -0.9851356 | 0.32455745 | 0.99989001 |
| FRG1 | 50.7696009 | 0.01733831 | 0.28098963 | 0.06170445 | 0.9507982 | 0.99989001 |
| SDHA | 153.218346 | 0.16419219 | 0.16657459 | 0.98569771 | 0.32428147 | 0.99989001 |
| PDCD6 | 92.2943948 | 0.18175707 | 0.38571488 | 0.47122131 | 0.6374827 | 0.99989001 |
| EXOC3 | 77.3535554 | -0.3250147 | 0.19283649 | -1.6854421 | 0.09190331 | 0.99175102 |
| BRD9 | 69.7268442 | 0.00804834 | 0.33592244 | 0.02395893 | 0.98088537 | 0.99989001 |
| SLC12A7 | 151.511778 | -0.0990323 | 0.39588722 | -0.2501527 | 0.80246927 | 0.99989001 |
| CLPTM1L | 92.3430973 | 0.01376137 | 0.32180206 | 0.04276345 | 0.9658901 | 0.99989001 |
| LPCAT1 | 80.620693 | -0.4245411 | 0.34980378 | -1.213655 | 0.2248795 | 0.99989001 |
| MRPL36 | 48.8851496 | 0.23798847 | 0.5460338 | 0.43584934 | 0.66294604 | NA |
| NDUFS6 | 112.835485 | -0.2818589 | 0.38795107 | -0.726532 | 0.46751268 | 0.99989001 |
| ICE1 | 209.617183 | 0.13388392 | 0.2438627 | 0.5490135 | 0.58299618 | 0.99989001 |
| MED10 | 94.2119784 | 0.00040466 | 0.30765715 | 0.0013153 | 0.99895054 | 0.99989001 |
| NSUN2 | 95.3576423 | -0.2297134 | 0.2887151 | -0.7956404 | 0.42624105 | 0.99989001 |
| TENT4A | 200.98029 | 0.08180482 | 0.19455083 | 0.42048044 | 0.67413452 | 0.99989001 |
| MTRR | 59.3940084 | 0.15272033 | 0.27811867 | 0.54911931 | 0.58292358 | 0.99989001 |
| CCT5 | 138.622878 | 0.34052363 | 0.36778029 | 0.92588875 | 0.3545038 | 0.99989001 |

| | | | | | | |
|---|---|---|---|---|---|---|
| 6-Mar | 498.358665 | -0.000179 | 0.22834745 | -0.0007841 | 0.9993744 | 0.99989001 |
| ANKRD33B | 128.669146 | 0.83328257 | 0.47416418 | 1.75737138 | 0.07885453 | 0.92927888 |
| DAP | 167.18944 | -0.2909583 | 0.31867257 | -0.9130322 | 0.3612256 | 0.99989001 |
| TRIO | 1540.17038 | -0.1603056 | 0.22659129 | -0.707466 | 0.47927691 | 0.99989001 |
| OTULIN | 109.828897 | 0.29035203 | 0.27990534 | 1.03732223 | 0.29958571 | 0.99989001 |
| ANKH | 351.375562 | 0.11120701 | 0.19180446 | 0.57979364 | 0.56205378 | 0.99989001 |
| FBXL7 | 3031.5822 | -0.2101359 | 0.24709571 | -0.8504231 | 0.39508989 | 0.99989001 |
| RETREG1 | 55.7748835 | 1.16191766 | 0.43477432 | 2.67246159 | 0.0075297 | 0.4053518 |
| MYO10 | 589.791863 | -0.2490989 | 0.37188443 | -0.6698287 | 0.50296698 | 0.99989001 |
| BASP1 | 62.1757946 | -1.6108715 | 0.73694117 | -2.1858889 | 0.02882374 | 0.65805899 |
| LINC02100 | 58.7946713 | 0.85185075 | 0.59410126 | 1.43384774 | 0.15161572 | 0.99989001 |
| LINC02241 | 39.6641037 | -0.261094 | 0.40450229 | -0.6454697 | 0.51862285 | NA |
| DROSHA | 158.146539 | 0.00253835 | 0.23085107 | 0.01099561 | 0.99122695 | 0.99989001 |
| C5orf22 | 55.7927966 | 0.01755719 | 0.26544004 | 0.0661437 | 0.94726342 | 0.99989001 |
| PDZD2 | 857.710914 | 0.2438926 | 0.37872669 | 0.6439805 | 0.51958808 | 0.99989001 |
| GOLPH3 | 624.530285 | 0.03544288 | 0.34944475 | 0.10142628 | 0.91921208 | 0.99989001 |
| MTMR12 | 267.034598 | -0.0728325 | 0.31912117 | -0.2282284 | 0.81946868 | 0.99989001 |
| ZFR | 491.725285 | 0.08498402 | 0.15747471 | 0.53966771 | 0.58942621 | 0.99989001 |
| SUB1 | 288.068997 | -0.1629598 | 0.3956496 | -0.4118791 | 0.68042802 | 0.99989001 |
| NPR3 | 91.3825635 | 1.14497022 | 0.61138596 | 1.87274536 | 0.06110356 | 0.85923309 |
| TARS | 87.2073532 | 0.13612593 | 0.37299517 | 0.36495359 | 0.71514605 | 0.99989001 |
| C1QTNF3-AN | 150.675285 | -1.0184707 | 0.38898974 | -2.6182458 | 0.00883831 | 0.42668269 |
| RAI14 | 870.586544 | 0.31805657 | 0.42029474 | 0.75674648 | 0.44920176 | 0.99989001 |
| BRIX1 | 46.3985421 | -0.0739834 | 0.32520879 | -0.2274952 | 0.82003873 | NA |
| DNAJC21 | 98.9858385 | -0.0808677 | 0.27960933 | -0.2892167 | 0.77241553 | 0.99989001 |
| SPEF2 | 51.1197213 | 0.48265393 | 0.62363495 | 0.77393663 | 0.43896827 | 0.99989001 |
| LMBRD2 | 75.1128849 | -0.1835321 | 0.29439423 | -0.6234228 | 0.53300669 | 0.99989001 |
| NADK2 | 115.151924 | -0.783222 | 0.51424216 | -1.5230607 | 0.12774352 | 0.99989001 |
| RANBP3L | 378.72121 | -2.9664914 | 1.50802958 | -1.9671308 | 0.04916814 | 0.78717121 |
| NIPBL | 1091.53239 | 0.0883365 | 0.19459551 | 0.45394931 | 0.64986531 | 0.99989001 |
| CPLANE1 | 275.548586 | -0.0202825 | 0.17723082 | -0.114441 | 0.90888824 | 0.99989001 |
| NUP155 | 88.2850378 | 0.08614626 | 0.22379023 | 0.384942 | 0.70028039 | 0.99989001 |
| WDR70 | 564.20571 | -0.0263823 | 0.17371369 | -0.1518722 | 0.87928775 | 0.99989001 |
| LIFR | 950.982591 | 0.21124751 | 0.43722338 | 0.48315693 | 0.62898431 | 0.99989001 |
| OSMR-AS1 | 241.891131 | 0.84791744 | 0.3128387 | 2.71039812 | 0.00672025 | 0.390188 |
| OSMR | 733.930969 | -0.0231912 | 0.23039173 | -0.1006597 | 0.91982061 | 0.99989001 |
| RICTOR | 640.96636 | -0.1370505 | 0.20992871 | -0.6528428 | 0.51385762 | 0.99989001 |
| TTC33 | 132.8515 | -0.1345922 | 0.29079891 | -0.4628361 | 0.64348186 | 0.99989001 |
| PRKAA1 | 263.062855 | -0.3288311 | 0.27717586 | -1.1863628 | 0.23547905 | 0.99989001 |
| RPL37 | 234.647185 | -0.0833755 | 0.4284135 | -0.1946145 | 0.84569472 | 0.99989001 |
| CARD6 | 76.9236448 | 0.36077774 | 0.29626659 | 1.21774698 | 0.22332014 | 0.99989001 |
| PLCXD3 | 154.213224 | 1.62228485 | 1.09016991 | 1.48810276 | 0.1367238 | 0.99989001 |
| OXCT1 | 114.644077 | -0.1599036 | 0.22726313 | -0.7036056 | 0.4816784 | 0.99989001 |
| GHR | 240.873027 | 0.10321026 | 0.26652252 | 0.3872478 | 0.69857277 | 0.99989001 |
| SELENOP | 179.544046 | 0.24712317 | 0.48622426 | 0.50824937 | 0.61127847 | 0.99989001 |
| ZNF131 | 275.073579 | -0.0772733 | 0.17228031 | -0.4485325 | 0.65376897 | 0.99989001 |
| HMGCS1 | 75.452341 | -0.199256 | 0.27409757 | -0.7269528 | 0.46725484 | 0.99989001 |
| TMEM267 | 89.4526707 | 0.13051048 | 0.25123684 | 0.51947191 | 0.6034317 | 0.99989001 |
| PAIP1 | 97.1008935 | -0.4594477 | 0.20574273 | -2.2331174 | 0.0255412 | 0.63416113 |
| NNT-AS1 | 62.6301814 | -0.3752935 | 0.33055142 | -1.1353558 | 0.25622628 | 0.99989001 |
| NNT | 180.799847 | -0.0459147 | 0.25248944 | -0.1818481 | 0.85570193 | 0.99989001 |
| EMB | 59.8253068 | 2.47207645 | 0.74553814 | 3.31582826 | 0.00091372 | 0.14992947 |
| PARP8 | 137.828596 | 0.25254026 | 0.43310359 | 0.58309435 | 0.55982979 | 0.99989001 |
| ITGA1 | 1000.45828 | 0.71379715 | 0.24680788 | 2.89211655 | 0.00382656 | 0.31056852 |

| | | | | | | |
|---|---|---|---|---|---|---|
| ITGA2 | 143.332296 | -1.027142 | 0.49981245 | -2.0550549 | 0.0398737 | 0.73029084 |
| NDUFS4 | 228.964086 | -0.1622178 | 0.17061923 | -0.9507592 | 0.34172665 | 0.99989001 |
| ARL15 | 7198.91035 | -0.6270307 | 0.56372512 | -1.1122987 | 0.26600974 | 0.99989001 |
| SNX18 | 77.0628217 | 0.48146564 | 0.28494431 | 1.6896833 | 0.09108856 | 0.99068653 |
| ESM1 | 276.138749 | 1.11305428 | 0.72007789 | 1.54574151 | 0.12216701 | 0.99989001 |
| DHX29 | 135.738618 | 0.05063795 | 0.21866221 | 0.2315807 | 0.8168637 | 0.99989001 |
| MTREX | 257.237186 | 0.03438757 | 0.22774025 | 0.15099469 | 0.8799799 | 0.99989001 |
| PLPP1 | 5242.1184 | 0.08254325 | 0.46635624 | 0.17699614 | 0.85951142 | 0.99989001 |
| SLC38A9 | 176.309304 | 0.07677377 | 0.17250956 | 0.44504067 | 0.65629036 | 0.99989001 |
| IL6ST | 1188.80111 | 0.21928617 | 0.39926944 | 0.54921852 | 0.58285549 | 0.99989001 |
| MAP3K1 | 318.885441 | 0.17653113 | 0.16429981 | 1.07444517 | 2.83E-01 | 0.99989001 |
| MIER3 | 51.2140606 | -0.007564 | 0.32495611 | -0.0232771 | 0.98142924 | 0.99989001 |
| GPBP1 | 561.340634 | 0.13508341 | 0.29250365 | 0.46181788 | 0.64421193 | 0.99989001 |
| PLK2 | 177.958013 | 0.27860781 | 0.40503431 | 0.68786224 | 0.49153954 | 0.99989001 |
| RAB3C | 150.960344 | -0.8014199 | 0.72299433 | -1.1084733 | 2.68E-01 | 0.99989001 |
| PDE4D | 3656.47386 | -0.0500303 | 0.34489958 | -0.1450575 | 0.88466546 | 0.99989001 |
| ELOVL7 | 176.798077 | -0.8531655 | 0.51939625 | -1.64261 | 0.10046365 | 0.99989001 |
| ERCC8 | 103.838154 | -0.179084 | 0.24751195 | -0.7235367 | 0.46935019 | 0.99989001 |
| NDUFAF2 | 222.37951 | -0.0488774 | 0.30360633 | -0.1609894 | 0.87210178 | 0.99989001 |
| ZSWIM6 | 1624.84091 | 0.21363201 | 0.3089564 | 0.69146329 | 0.48927444 | 0.99989001 |
| KIF2A | 186.107357 | 0.13763001 | 0.19856038 | 0.69313933 | 0.48822212 | 0.99989001 |
| DIMT1 | 50.7093211 | -0.3623057 | 0.31538321 | -1.1487793 | 0.25064699 | 0.99989001 |
| IPO11 | 226.869251 | -0.1670197 | 0.24872828 | -0.6714945 | 0.50190557 | 0.99989001 |
| RNF180 | 290.17249 | 0.02687755 | 0.18434564 | 0.14579978 | 0.88407945 | 0.99989001 |
| RGS7BP | 110.59646 | -0.1498154 | 0.72430517 | -0.2068402 | 0.83613464 | 0.99989001 |
| SREK1IP1 | 117.461437 | 0.00796953 | 0.20644834 | 0.03860303 | 0.96920689 | 0.99989001 |
| CWC27 | 283.255106 | 0.00437388 | 0.18912157 | 0.02312735 | 0.98154869 | 0.99989001 |
| ADAMTS6 | 993.893412 | -0.9257059 | 0.51609777 | -1.7936638 | 0.07286684 | 0.9155703 |
| PPWD1 | 132.296617 | -0.0129914 | 0.21865212 | -0.0594156 | 0.95262107 | 0.99989001 |
| TRIM23 | 80.1841656 | -0.2316608 | 0.25530242 | -0.9073977 | 0.36419655 | 0.99989001 |
| TRAPPC13 | 102.101909 | -0.0327905 | 0.22936398 | -0.1429626 | 0.88631971 | 0.99989001 |
| ERBIN | 812.380315 | -0.128138 | 0.1915582 | -0.6689246 | 0.50354355 | 0.99989001 |
| SREK1 | 282.268499 | -0.1897772 | 0.20162425 | -0.9412418 | 0.34658098 | 0.99989001 |
| MAST4 | 954.230215 | 0.23146774 | 0.24917445 | 0.92893851 | 0.35292096 | 0.99989001 |
| PIK3R1 | 221.825892 | 0.43562715 | 0.28502861 | 1.52836291 | 0.12642246 | 0.99989001 |
| SLC30A5 | 121.659133 | 0.02380064 | 0.25241314 | 0.09429241 | 0.92487688 | 0.99989001 |
| MRPS36 | 58.4801405 | -0.0491272 | 0.24703485 | -0.1988676 | 0.84236634 | 0.99989001 |
| CDK7 | 53.7491313 | -0.3184324 | 0.31840003 | -1.0001016 | 0.31726135 | 0.99989001 |
| CCDC125 | 48.1968939 | -0.0519325 | 0.48931885 | -0.1061322 | 0.91547743 | NA |
| AK6 | 51.2827551 | -0.309243 | 0.2994091 | -1.0328442 | 0.30167681 | 0.99989001 |
| RAD17 | 88.0361014 | -0.2889354 | 0.23073863 | -1.2522196 | 0.21048986 | 0.99989001 |
| OCLN | 67.3471222 | 0.24404376 | 0.45556416 | 0.5356957 | 0.59216888 | 0.99989001 |
| GTF2H2C | 57.5908703 | -0.0061821 | 0.23684379 | -0.0261021 | 0.97917592 | 0.99989001 |
| NAIP | 332.464137 | -0.4511008 | 0.49166538 | -0.9174955 | 0.35888303 | 0.99989001 |
| GTF2H2 | 56.3400103 | -0.2647516 | 0.51920262 | -0.5099197 | 0.61010774 | 0.99989001 |
| BDP1 | 356.280039 | -0.0299444 | 0.24602948 | -0.1217104 | 0.90312835 | 0.99989001 |
| MCCC2 | 118.622421 | -0.4378562 | 0.24225504 | -1.8074183 | 0.07069708 | 0.90906573 |
| MAP1B | 158.128915 | -0.2184254 | 0.48115106 | -0.4539643 | 0.64985452 | 0.99989001 |
| MRPS27 | 169.886216 | -0.1916596 | 0.24079753 | -0.7959369 | 0.42606874 | 0.99989001 |
| ZNF366 | 357.561438 | 0.01079348 | 0.34194337 | 0.0315651 | 0.97481888 | 0.99989001 |
| TNPO1 | 423.114076 | 0.12287877 | 0.18423104 | 0.6669819 | 0.5047837 | 0.99989001 |
| FCHO2 | 639.513886 | 0.18543093 | 0.2359259 | 0.78597107 | 0.43188444 | 0.99989001 |
| BTF3 | 203.619681 | -0.2315665 | 0.51667211 | -0.4481886 | 0.65401712 | 0.99989001 |
| ANKRA2 | 51.0957822 | 0.28368246 | 0.3390103 | 0.83679599 | 0.40270725 | 0.99989001 |

| | | | | | | |
|---|---|---|---|---|---|---|
| ARHGEF28 | 289.296614 | -1.1250111 | 0.43898283 | -2.5627679 | 0.01038414 | 0.46378417 |
| HEXB | 105.210479 | -0.2409944 | 0.28557674 | -0.8438868 | 0.39873266 | 0.99989001 |
| GFM2 | 62.5697125 | -0.3760839 | 0.23872588 | -1.5753795 | 0.11516887 | 0.99989001 |
| NSA2 | 59.2576727 | -0.1954971 | 0.36504501 | -0.5355425 | 0.59227478 | 0.99989001 |
| HMGCR | 54.1247823 | 0.31188235 | 0.35539902 | 0.87755547 | 0.380185 | 0.99989001 |
| COL4A3BP | 389.116527 | 0.24859756 | 0.23951866 | 1.03790477 | 0.29931439 | 0.99989001 |
| POLK | 370.386403 | 0.26401728 | 0.24407226 | 1.08171771 | 0.27937798 | 0.99989001 |
| POC5 | 57.4530559 | 0.00900579 | 0.2847835 | 0.03162327 | 0.97477249 | 0.99989001 |
| IQGAP2 | 149.452118 | 1.22034526 | 0.60795339 | 2.00730068 | 0.04471766 | 0.77228819 |
| F2R | 162.981254 | -0.1223973 | 0.35808185 | -0.3418137 | 0.7324911 | 0.99989001 |
| CRHBP | 390.780424 | 2.41398993 | 1.00972897 | 2.39073059 | 0.01681489 | 0.5510556 |
| WDR41 | 131.355205 | 0.13499061 | 0.26588164 | 0.50770942 | 0.61165714 | 0.99989001 |
| TBCA | 333.920169 | -0.1190976 | 0.24324094 | -0.4896282 | 0.62439699 | 0.99989001 |
| AP3B1 | 562.808009 | -0.0221424 | 0.17734062 | -0.1248578 | 0.9006361 | 0.99989001 |
| SCAMP1 | 227.410773 | 0.24184851 | 0.20931908 | 1.15540595 | 0.24792423 | 0.99989001 |
| LHFPL2 | 116.816861 | -0.5200096 | 0.23794429 | -2.1854257 | 0.02885765 | 0.65805899 |
| ARSB | 199.31061 | -0.4478826 | 0.38398354 | -1.1664109 | 0.24344837 | 0.99989001 |
| DMGDH | 79.0546642 | -0.3193933 | 0.39055295 | -0.8177976 | 0.41347277 | 0.99989001 |
| BHMT2 | 56.4382248 | 1.13821991 | 0.57066339 | 1.99455567 | 0.04609136 | 0.77310579 |
| JMY | 126.017051 | 0.01208945 | 0.36857035 | 0.03280094 | 0.97383333 | 0.99989001 |
| TENT2 | 558.178966 | -0.2205492 | 0.20266257 | -1.0882581 | 0.27648118 | 0.99989001 |
| SERINC5 | 206.695396 | 0.19997003 | 0.37553487 | 0.5324939 | 0.59438397 | 0.99989001 |
| ZFYVE16 | 280.580807 | 0.15777181 | 0.20656884 | 0.7637735 | 0.44500223 | 0.99989001 |
| DHFR | 351.455056 | 0.04043279 | 0.31967354 | 0.12648152 | 0.89935078 | 0.99989001 |
| MSH3 | 246.986224 | -0.1860657 | 0.19332403 | -0.9624552 | 0.33582098 | 0.99989001 |
| RASGRF2 | 1109.52061 | -0.3267563 | 0.32050179 | -1.0195149 | 0.30795858 | 0.99989001 |
| CKMT2-AS1 | 140.845292 | -0.0292865 | 0.29896837 | -0.0979586 | 0.92196514 | 0.99989001 |
| ZCCHC9 | 43.6259968 | 0.3945867 | 0.27021108 | 1.46029059 | 0.14421023 | NA |
| SSBP2 | 930.315733 | 0.32686049 | 0.42070379 | 0.77693736 | 0.43719573 | 0.99989001 |
| ATG10 | 254.181041 | -0.1324024 | 0.24863905 | -0.5325083 | 0.59437402 | 0.99989001 |
| RPS23 | 596.788555 | -0.3938991 | 0.64439131 | -0.6112731 | 0.54101882 | 0.99989001 |
| TMEM167A | 98.2637018 | -0.1368608 | 0.31222776 | -0.4383366 | 0.66114232 | 0.99989001 |
| XRCC4 | 131.956545 | -0.3890406 | 0.16938345 | -2.2968041 | 0.02162995 | 0.60140868 |
| VCAN | 224.117247 | -0.1036741 | 0.58612395 | -0.1768808 | 8.60E-01 | 1.00E+00 |
| EDIL3 | 428.976495 | -2.1922526 | 0.73456373 | -2.984428 | 0.00284109 | 0.27392168 |
| AC113383.1 | 83.8082118 | -3.8297476 | 1.18017575 | -3.2450655 | 0.00117424 | 0.16624537 |
| COX7C | 176.853007 | 0.11895991 | 0.47398378 | 0.25097887 | 0.80183045 | 0.99989001 |
| RASA1 | 553.143272 | 0.04085778 | 0.18996197 | 0.21508399 | 0.82970184 | 0.99989001 |
| CCNH | 780.53809 | 0.32001882 | 0.3145823 | 1.01728172 | 0.30901943 | 0.99989001 |
| TMEM161B | 107.488275 | -0.1666953 | 0.23157701 | -0.7198265 | 0.47163185 | 0.99989001 |
| TMEM161B-/ | 184.724997 | -0.41809 | 0.34075324 | -1.2269584 | 0.21983824 | 0.99989001 |
| MEF2C | 2130.29459 | 0.06308436 | 0.29253154 | 0.21564977 | 0.82926076 | 0.99989001 |
| MEF2C-AS1 | 363.156248 | 0.01199111 | 0.41324504 | 0.02901694 | 0.97685108 | 0.99989001 |
| LYSMD3 | 55.6252068 | 0.32456517 | 0.29630941 | 1.09535897 | 0.2733594 | 0.99989001 |
| ARRDC3 | 910.124185 | -1.0187781 | 0.69822238 | -1.4591027 | 0.14453685 | 0.99989001 |
| NR2F1-AS1 | 598.010625 | -0.2732047 | 0.33094009 | -0.8255412 | 0.40906443 | 0.99989001 |
| NR2F1 | 183.150034 | -0.5988182 | 0.53373154 | -1.1219464 | 0.26188524 | 0.99989001 |
| FAM172A | 1015.89187 | -0.2003705 | 0.174003 | -1.1515346 | 0.24951237 | 0.99989001 |
| KIAA0825 | 222.596702 | -0.0464238 | 0.36378828 | -0.1276121 | 0.89845595 | 0.99989001 |
| SLF1 | 132.533041 | -0.3486288 | 0.39630782 | -0.879692 | 0.37902617 | 0.99989001 |
| MCTP1 | 941.33812 | -1.3281297 | 0.61086612 | -2.1741747 | 0.02969202 | 0.6652708 |
| TTC37 | 321.35491 | -0.2288582 | 0.16597822 | -1.3788451 | 0.16794252 | 0.99989001 |
| RHOBTB3 | 69.3499742 | 0.27914363 | 0.40057685 | 0.69685411 | 0.4858941 | 0.99989001 |
| GLRX | 141.666415 | 1.18301449 | 0.51147076 | 2.31296603 | 0.02072451 | 0.58699182 |

| | | | | | | |
|---|---|---|---|---|---|---|
| ELL2 | 276.932947 | -0.5699654 | 0.67222873 | -0.8478742 | 0.39650801 | 0.99989001 |
| CAST | 564.540747 | 0.04627845 | 0.14241054 | 0.32496506 | 0.74520752 | 0.99989001 |
| ERAP1 | 152.07462 | -0.0635484 | 0.21831536 | -0.2910852 | 0.77098613 | 0.99989001 |
| ERAP2 | 165.778159 | 0.66873829 | 0.66422512 | 1.00679465 | 0.31403347 | 0.99989001 |
| LNPEP | 258.919918 | 0.15847172 | 0.17126787 | 0.92528577 | 0.35481728 | 0.99989001 |
| LIX1-AS1 | 61.9621141 | -0.367191 | 0.48297714 | -0.7602659 | 0.44709567 | 0.99989001 |
| CHD1 | 417.455376 | -0.0266759 | 0.281521 | -0.0947563 | 0.92450839 | 0.99989001 |
| LINC02062 | 66.3245222 | 0.28802588 | 0.47844676 | 0.60200194 | 0.54717284 | 0.99989001 |
| FAM174A | 64.1982183 | -0.1657958 | 0.25701038 | -0.6450938 | 0.51886639 | 0.99989001 |
| ST8SIA4 | 67.3528352 | -0.2760195 | 0.32290636 | -0.8547975 | 0.39266326 | 0.99989001 |
| PAM | 1079.69375 | -0.2464846 | 0.31063458 | -0.7934872 | 0.42749405 | 0.99989001 |
| PPIP5K2 | 253.916808 | -0.2015427 | 0.18460691 | -1.0917397 | 0.27494751 | 0.99989001 |
| EFNA5 | 185.131029 | 1.59639147 | 0.68142616 | 2.34272112 | 0.01914369 | 0.56665312 |
| FBXL17 | 778.452679 | -0.1946155 | 0.19269061 | -1.0099896 | 0.31250026 | 0.99989001 |
| FER | 892.024475 | -0.1737841 | 0.21869389 | -0.7946453 | 0.42681987 | 0.99989001 |
| PJA2 | 376.401213 | 0.19641516 | 0.23536715 | 0.83450541 | 0.40399624 | 0.99989001 |
| MAN2A1 | 480.589288 | 0.34463801 | 0.23615965 | 1.45934334 | 0.14447063 | 0.99989001 |
| SLC25A46 | 129.567763 | -0.4477017 | 0.19574116 | -2.2872129 | 0.0221834 | 0.60384164 |
| WDR36 | 126.158585 | -0.4188105 | 0.26157702 | -1.6010983 | 0.10935516 | 0.99989001 |
| STARD4 | 158.85902 | -0.4764506 | 0.55563442 | -0.8574893 | 0.39117453 | 0.99989001 |
| NREP | 201.308465 | -0.7825323 | 0.40951931 | -1.9108557 | 0.05602313 | 0.83144465 |
| EPB41L4A | 1721.33537 | 0.26833124 | 0.29591696 | 0.90677885 | 0.36452375 | 0.99989001 |
| APC | 242.911719 | 0.14667742 | 0.19245985 | 0.76211958 | 0.44598864 | 0.99989001 |
| SRP19 | 94.3544003 | 0.0450254 | 0.28882713 | 0.15589049 | 0.87611934 | 0.99989001 |
| REEP5 | 232.970703 | 0.01053574 | 0.33544194 | 0.03140853 | 0.97494374 | 0.99989001 |
| DCP2 | 208.723315 | 0.11837073 | 0.17408001 | 0.67997889 | 0.49651783 | 0.99989001 |
| MCC | 751.430913 | 0.4886029 | 0.31996746 | 1.52703931 | 0.12675124 | 0.99989001 |
| YTHDC2 | 293.068594 | -0.2586687 | 0.21628067 | -1.1959862 | 0.23170193 | 0.99989001 |
| PGGT1B | 124.530053 | 0.09394244 | 0.2272614 | 0.41336734 | 0.6793375 | 0.99989001 |
| FEM1C | 69.5787296 | -0.5324857 | 0.35251454 | -1.5105353 | 0.13090689 | 0.99989001 |
| ATG12 | 102.578716 | -0.0559573 | 0.29763017 | -0.1880094 | 0.85086929 | 0.99989001 |
| AP3S1 | 158.786325 | 0.19841807 | 0.18698633 | 1.06113675 | 0.28862776 | 0.99989001 |
| COMMD10 | 757.456616 | 0.06271024 | 0.21265376 | 0.29489365 | 0.76807512 | 0.99989001 |
| SEMA6A | 877.676689 | 0.74732171 | 0.35517658 | 2.104085 | 0.03537103 | 0.69174161 |
| SEMA6A-AS1 | 153.536115 | 0.05502704 | 0.26623825 | 0.20668347 | 0.83625707 | 0.99989001 |
| LINC02147 | 1495.65786 | -0.4567093 | 0.64340897 | -0.7098273 | 4.78E-01 | 0.99989001 |
| DTWD2 | 63.1147836 | -0.3283176 | 0.29237366 | -1.1229384 | 0.26146366 | 0.99989001 |
| DMXL1 | 361.973704 | -0.1379253 | 0.17731567 | -0.7778519 | 0.4366563 | 0.99989001 |
| TNFAIP8 | 83.7962487 | 0.95200296 | 0.31772562 | 2.99630534 | 0.00273273 | 0.26787818 |
| HSD17B4 | 194.364548 | 0.20272733 | 0.34602125 | 0.58588114 | 0.55795539 | 0.99989001 |
| SRFBP1 | 129.393901 | -0.3245289 | 0.27953678 | -1.1609523 | 0.24566131 | 0.99989001 |
| SNCAIP | 350.098214 | -0.8577209 | 0.50170647 | -1.709607 | 0.08733856 | 0.96238172 |
| SNX2 | 140.423706 | 0.0380904 | 0.22909546 | 0.16626431 | 0.86794895 | 0.99989001 |
| SNX24 | 144.124269 | 0.29133077 | 0.28178038 | 1.03389301 | 0.30118618 | 0.99989001 |
| PPIC | 70.8760082 | -0.7743347 | 0.50338732 | -1.5382484 | 0.1239879 | 0.99989001 |
| CEP120 | 109.788644 | -0.2921994 | 0.32967456 | -0.8863268 | 0.37544146 | 0.99989001 |
| CSNK1G3 | 285.314122 | -0.0845654 | 0.23264571 | -0.3634944 | 0.71623557 | 0.99989001 |
| ZNF608 | 263.002301 | -0.3500254 | 0.31284621 | -1.1188418 | 0.26320761 | 0.99989001 |
| GRAMD2B | 486.51016 | 0.82885353 | 0.49592353 | 1.67133334 | 0.09465586 | 0.99989001 |
| ALDH7A1 | 128.069223 | 0.90624523 | 0.38461516 | 2.35623896 | 0.01846104 | 0.56643441 |
| PHAX | 57.9359233 | -0.0165388 | 0.33671372 | -0.0491183 | 0.96082504 | 0.99989001 |
| 3-Mar | 789.302097 | 0.79323525 | 0.70951186 | 1.1180014 | 0.2635664 | 0.99989001 |
| PRRC1 | 89.7625533 | -0.1456512 | 0.30867781 | -0.471855 | 0.63703027 | 0.99989001 |
| LINC01184 | 229.79671 | -0.3615828 | 0.35827433 | -1.0092344 | 0.31286222 | 0.99989001 |

| | | | | | | |
|---|---|---|---|---|---|---|
| SLC12A2 | 205.362781 | -0.0915465 | 0.33962972 | -0.269548 | 0.78750801 | 0.99989001 |
| CHSY3 | 219.148067 | 0.80992909 | 0.45751257 | 1.77028816 | 0.07667915 | 0.92274869 |
| HINT1 | 371.722247 | 0.30733541 | 0.50031341 | 0.61428578 | 0.5390265 | 0.99989001 |
| LYRM7 | 44.3111216 | 0.07114475 | 0.36293931 | 0.19602381 | 0.84459153 | NA |
| CDC42SE2 | 383.118161 | 0.34391921 | 0.29506773 | 1.16556023 | 0.24379229 | 0.99989001 |
| AC116366.3 | 552.864321 | -0.3288814 | 0.48748822 | -0.6746448 | 0.49990143 | 0.99989001 |
| IRF1 | 140.609053 | 0.08310243 | 0.38030846 | 0.21851323 | 0.82702925 | 0.99989001 |
| KIF3A | 69.2973826 | -0.3158725 | 0.26157701 | -1.2075696 | 0.22721286 | 0.99989001 |
| 8-Sep | 58.7622155 | 0.00283649 | 0.32995694 | 0.00859656 | 0.99314102 | 0.99989001 |
| SHROOM1 | 80.0442383 | -1.3563977 | 0.6246941 | -2.1712991 | 0.02990857 | 0.6659289 |
| UQCRQ | 99.4395359 | -0.0194276 | 0.49182215 | -0.0395014 | 0.96849067 | 0.99989001 |
| AFF4 | 584.072782 | -0.3160202 | 0.2402885 | -1.3151699 | 0.18845282 | 0.99989001 |
| ZCCHC10 | 92.6410745 | -0.2372415 | 0.25926229 | -0.9150635 | 0.36015829 | 0.99989001 |
| HSPA4 | 143.512805 | 0.00285364 | 0.26796093 | 0.01064945 | 0.99150313 | 0.99989001 |
| C5orf15 | 87.298807 | -0.2449502 | 0.28199223 | -0.8686417 | 0.38504316 | 0.99989001 |
| VDAC1 | 137.539421 | 0.41632992 | 0.28772894 | 1.44695184 | 0.14791041 | 0.99989001 |
| SKP1 | 248.715235 | 0.1317475 | 0.3865985 | 0.34078637 | 0.73326441 | 0.99989001 |
| CDKL3 | 66.6666302 | -0.3767088 | 0.39328803 | -0.9578446 | 0.33814114 | 0.99989001 |
| UBE2B | 334.814092 | 0.00107512 | 0.30807933 | 0.00348976 | 0.99721558 | 0.99989001 |
| JADE2 | 114.128579 | 0.04698234 | 0.25269445 | 0.18592547 | 0.85250321 | 0.99989001 |
| SAR1B | 123.070805 | 0.12772817 | 0.21045614 | 0.60691114 | 0.54390988 | 0.99989001 |
| SEC24A | 148.119412 | 0.06821463 | 0.26452283 | 0.25787805 | 0.79650103 | 0.99989001 |
| CAMLG | 67.9825396 | 0.03221734 | 0.24377565 | 0.13215981 | 0.89485789 | 0.99989001 |
| DDX46 | 266.89766 | -0.0481607 | 0.20997964 | -0.229359 | 0.81858992 | 0.99989001 |
| C5orf24 | 155.583042 | 0.1639677 | 0.35552023 | 0.46120497 | 0.64465155 | 0.99989001 |
| TXNDC15 | 89.9995006 | -0.2408174 | 0.24832196 | -0.9697789 | 0.33215671 | 0.99989001 |
| PCBD2 | 87.6584172 | -0.5092962 | 0.55245338 | -0.9218809 | 0.35659068 | 0.99989001 |
| H2AFY | 215.060712 | 0.07124552 | 0.23475904 | 0.30348362 | 0.76152133 | 0.99989001 |
| CXCL14 | 77.0959592 | 3.02889042 | 0.65206915 | 4.64504478 | 3.40E-06 | 0.00285149 |
| SMAD5 | 220.471096 | 0.0091707 | 0.18527145 | 0.04949871 | 0.96052187 | 0.99989001 |
| KLHL3 | 144.577561 | -0.3151177 | 0.45990423 | -0.6851811 | 0.49322962 | 0.99989001 |
| HNRNPA0 | 137.202179 | -0.0427564 | 0.43933032 | -0.0973218 | 0.92247088 | 0.99989001 |
| FAM13B | 353.504158 | 0.13427034 | 0.14182643 | 0.94672293 | 0.34377998 | 0.99989001 |
| BRD8 | 74.7818658 | 0.11474949 | 0.2471716 | 0.46425031 | 0.64246842 | 0.99989001 |
| CDC23 | 45.0706874 | 0.05273389 | 0.27122261 | 0.1944303 | 0.84583897 | NA |
| FAM53C | 41.8632469 | -0.4636738 | 0.33318595 | -1.3916368 | 0.16403241 | NA |
| KDM3B | 213.938181 | 0.25006671 | 0.24320658 | 1.02820699 | 0.30385247 | 0.99989001 |
| EGR1 | 91.789625 | -2.8193421 | 1.27482412 | -2.2115538 | 0.02699751 | 0.64986727 |
| ETF1 | 136.617864 | -0.2578731 | 0.24166711 | -1.0670593 | 0.28594505 | 0.99989001 |
| HSPA9 | 195.539434 | -0.2464783 | 0.28422307 | -0.8672003 | 0.38583225 | 0.99989001 |
| CTNNA1 | 974.592587 | 0.06439082 | 0.20737374 | 0.31050612 | 0.7561761 | 0.99989001 |
| SIL1 | 217.396153 | 0.17476507 | 0.24050725 | 0.72665198 | 0.46743918 | 0.99989001 |
| PAIP2 | 184.080407 | 0.33484223 | 0.29292463 | 1.14310029 | 0.25299695 | 0.99989001 |
| DNAJC18 | 126.404931 | -0.404735 | 0.44692855 | -0.9055922 | 0.36515175 | 0.99989001 |
| AC142391.1 | 45.6061154 | -0.4417909 | 0.53933409 | -0.8191414 | 0.41270577 | NA |
| TMEM173 | 127.303591 | -0.168575 | 0.42163663 | -0.3998111 | 0.68929564 | 0.99989001 |
| UBE2D2 | 400.546896 | 0.16639212 | 0.21920069 | 0.75908577 | 0.44780125 | 0.99989001 |
| CXXC5 | 66.572191 | 0.33344982 | 0.39023597 | 0.85448253 | 0.39283769 | 0.99989001 |
| NRG2 | 184.42925 | 0.2624048 | 0.68965917 | 0.38048476 | 0.70358561 | 0.99989001 |
| PURA | 214.355957 | 0.00699509 | 0.19181478 | 0.03646796 | 0.97090923 | 0.99989001 |
| CYSTM1 | 366.890139 | -0.3881658 | 0.44596151 | -0.870402 | 0.38408073 | 0.99989001 |
| PFDN1 | 125.048356 | 0.03082508 | 0.1998609 | 0.15423266 | 0.87742629 | 0.99989001 |
| SRA1 | 64.6908539 | 0.22139829 | 0.31019191 | 0.71374618 | 0.47538415 | 0.99989001 |
| SLC35A4 | 50.4997032 | -0.1814641 | 0.41215132 | -0.4402852 | 0.65973057 | 0.99989001 |

| | | | | | | |
|---|---|---|---|---|---|---|
| CD14 | 57.6657928 | -0.3769415 | 1.05933692 | -0.3558278 | 0.72196955 | 0.99989001 |
| NDUFA2 | 71.6242469 | -0.0072132 | 0.47682723 | -0.0151275 | 0.98793046 | 0.99989001 |
| IK | 55.5238585 | 0.15550471 | 0.33689063 | 0.46158811 | 0.64437672 | 0.99989001 |
| HARS | 88.5511722 | -0.0966938 | 0.23538974 | -0.4107818 | 0.68123254 | 0.99989001 |
| ZMAT2 | 53.3980083 | 0.18302738 | 0.31415181 | 0.58260808 | 0.56015717 | 0.99989001 |
| TAF7 | 100.009524 | 0.25679515 | 0.22646394 | 1.13393394 | 0.25682226 | 0.99989001 |
| DIAPH1 | 259.632653 | -0.0138487 | 0.37984291 | -0.036459 | 0.97091635 | 0.99989001 |
| HDAC3 | 69.9409603 | 0.02898594 | 0.29895881 | 0.0969563 | 0.9227611 | 0.99989001 |
| ARAP3 | 360.811701 | -0.0170662 | 0.23587371 | -0.0723533 | 0.94232074 | 0.99989001 |
| PCDH1 | 106.157792 | -0.5074203 | 0.44320002 | -1.1449013 | 0.25225004 | 0.99989001 |
| DELE1 | 181.80844 | -0.4082238 | 0.16348477 | -2.4970141 | 0.0125244 | 0.49000287 |
| PCDH12 | 127.709604 | 0.15972594 | 0.42175332 | 0.37871887 | 0.70489664 | 0.99989001 |
| RNF14 | 86.9657734 | -0.1900393 | 0.21351655 | -0.8900446 | 0.37344194 | 0.99989001 |
| NDFIP1 | 303.197759 | 0.11576821 | 0.17799087 | 0.65041658 | 0.51542317 | 0.99989001 |
| SPRY4 | 85.1663658 | 0.53636725 | 0.50809458 | 1.05564449 | 0.29113067 | 0.99989001 |
| SPRY4-AS1 | 57.9973427 | 0.30883011 | 0.34466542 | 0.89602871 | 0.37023743 | 0.99989001 |
| ARHGAP26 | 4081.74788 | 0.41537933 | 0.5589303 | 0.74316839 | 0.45737974 | 0.99989001 |
| ARHGAP26-A | 67.7704188 | -0.1200033 | 0.71349239 | -0.1681914 | 0.86643271 | 0.99989001 |
| NR3C1 | 594.286321 | 0.49554688 | 0.19194791 | 2.58167374 | 9.83E-03 | 4.50E-01 |
| YIPF5 | 68.7092673 | 0.32064714 | 0.34145882 | 0.93905068 | 0.34770472 | 0.99989001 |
| KCTD16 | 173.605669 | -2.712095 | 0.70828813 | -3.8290843 | 0.00012862 | 0.04854155 |
| PRELID2 | 308.286974 | 0.06679444 | 0.45050825 | 0.14826464 | 0.88213392 | 0.99989001 |
| LARS | 216.682166 | 0.06826319 | 0.2762737 | 0.24708539 | 0.80484214 | 0.99989001 |
| TCERG1 | 241.682262 | -0.4007994 | 0.23977592 | -1.6715583 | 0.09461145 | 0.99989001 |
| DPYSL3 | 165.03932 | -0.0661116 | 0.69872017 | -0.0946181 | 0.92461818 | 0.99989001 |
| AC091948.1 | 65.9574906 | 0.45153124 | 0.46042312 | 0.98068757 | 0.32674683 | 0.99989001 |
| FBXO38 | 141.699979 | 0.07117289 | 0.18566735 | 0.38333549 | 0.70147103 | 0.99989001 |
| ABLIM3 | 137.752486 | 0.73461635 | 0.32014035 | 2.29466966 | 0.02175206 | 0.60140868 |
| AFAP1L1 | 534.376695 | -0.1557455 | 0.38426838 | -0.4053041 | 0.68525403 | 0.99989001 |
| GRPEL2 | 93.347235 | -0.410785 | 0.43215021 | -0.9505607 | 0.3418274 | 0.99989001 |
| CSNK1A1 | 1070.97777 | -0.1161994 | 0.22814711 | -0.5093177 | 0.61052955 | 0.99989001 |
| ARHGEF37 | 47.2584743 | 0.02822745 | 0.54295111 | 0.05198893 | 0.95853752 | NA |
| SLC26A2 | 91.9946529 | -0.2389641 | 0.37362209 | -0.6395878 | 0.52244063 | 0.99989001 |
| HMGXB3 | 113.392811 | -0.5217501 | 0.27875256 | -1.871732 | 0.06124369 | 0.85923309 |
| TCOF1 | 46.0391689 | -0.3594074 | 0.38381696 | -0.936403 | 0.34906571 | NA |
| CD74 | 1919.09379 | 0.03031123 | 0.4526344 | 0.06696624 | 0.94660858 | 0.99989001 |
| RPS14 | 507.674723 | -0.1635688 | 0.53291163 | -0.3069343 | 0.7588934 | 0.99989001 |
| NDST1 | 219.701342 | -0.4172295 | 0.29688044 | -1.4053789 | 0.15990862 | 0.99989001 |
| SYNPO | 446.878724 | -0.1662523 | 0.35689199 | -0.4658337 | 0.64133452 | 0.99989001 |
| RBM22 | 67.7168969 | 0.0818437 | 0.24662735 | 0.33185169 | 0.74000125 | 0.99989001 |
| DCTN4 | 208.693437 | -0.1085584 | 0.22503524 | -0.4824061 | 0.62951746 | 0.99989001 |
| SMIM3 | 155.325242 | 0.02503925 | 0.62241311 | 0.04022932 | 0.96791031 | 0.99989001 |
| GPX3 | 1024.89623 | 1.00901516 | 0.58830972 | 1.71510877 | 0.08632529 | 0.95680362 |
| TNIP1 | 277.896056 | 1.0839396 | 0.60508273 | 1.79139073 | 0.07323061 | 0.9158596 |
| ANXA6 | 102.66604 | 0.09828104 | 0.41794667 | 0.23515211 | 0.81409068 | 0.99989001 |
| CCDC69 | 220.616187 | -0.0673241 | 0.27406235 | -0.2456526 | 0.80595114 | 0.99989001 |
| SPARC | 1167.51441 | -0.1904482 | 0.58382836 | -0.3262058 | 0.74426869 | 0.99989001 |
| ATOX1 | 146.834078 | -0.1960977 | 0.36995199 | -0.5300627 | 0.59606848 | 0.99989001 |
| G3BP1 | 330.215293 | 0.05136453 | 0.22467159 | 0.22862051 | 0.81916388 | 0.99989001 |
| FAM114A2 | 108.613561 | -0.0808078 | 0.23892443 | -0.3382147 | 0.73520138 | 0.99989001 |
| MFAP3 | 239.768545 | -0.0450784 | 0.21589835 | -0.2087948 | 8.35E-01 | 1.00E+00 |
| GALNT10 | 261.968502 | 0.33073454 | 0.27305926 | 1.21121891 | 0.22581152 | 0.99989001 |
| SAP30L-AS1 | 67.3727633 | 0.33085785 | 0.35653402 | 0.92798397 | 0.35341589 | 0.99989001 |
| SAP30L | 51.2654799 | 0.17381357 | 0.33171859 | 0.52397899 | 0.60029315 | 0.99989001 |

| | | | | | | |
|---|---|---|---|---|---|---|
| LARP1 | 264.460423 | -0.0939658 | 0.34063899 | -0.2758514 | 0.78266218 | 0.99989001 |
| FAXDC2 | 116.458077 | 0.17034295 | 0.60100676 | 0.28342934 | 0.77684774 | 0.99989001 |
| CNOT8 | 127.993584 | 0.08530386 | 0.32243895 | 0.26455818 | 0.79134984 | 0.99989001 |
| GEMIN5 | 68.710464 | -0.0597934 | 0.27230282 | -0.2195842 | 0.826195 | 0.99989001 |
| MRPL22 | 72.6926524 | -0.158313 | 0.30499431 | -0.5190688 | 0.60371275 | 0.99989001 |
| ADAM19 | 144.945774 | 0.28169736 | 0.59167763 | 0.4760994 | 0.63400357 | 0.99989001 |
| CLINT1 | 254.16006 | -0.0430823 | 0.18823341 | -0.2288768 | 0.81896465 | 0.99989001 |
| EBF1 | 2112.78274 | -0.2367746 | 0.23783929 | -0.9955233 | 0.31948181 | 0.99989001 |
| RNF145 | 460.914869 | 0.47861276 | 0.38399512 | 1.24640323 | 0.2126164 | 0.99989001 |
| UBLCP1 | 90.1366941 | 0.0091782 | 0.29922547 | 0.0306732 | 0.97553016 | 0.99989001 |
| TTC1 | 140.766653 | -0.1886593 | 0.27953125 | -0.6749132 | 0.49973089 | 0.99989001 |
| PWWP2A | 177.268098 | 0.32349398 | 0.26639577 | 1.21433602 | 0.22461944 | 0.99989001 |
| SLU7 | 48.939007 | 0.19142902 | 0.27881267 | 0.68658651 | 0.49234333 | NA |
| CCNG1 | 49.7790872 | 0.33376354 | 0.32165978 | 1.03762906 | 0.29944278 | NA |
| NUDCD2 | 44.487795 | -0.1991348 | 0.39635246 | -0.5024184 | 0.61537325 | NA |
| MAT2B | 92.5043655 | 0.51100129 | 0.3194031 | 1.59986328 | 0.10962892 | 0.99989001 |
| RARS | 130.127157 | 0.07260344 | 0.24469368 | 0.29671155 | 0.76668674 | 0.99989001 |
| PANK3 | 130.211808 | 0.16590778 | 0.19604576 | 0.84627065 | 0.39740178 | 0.99989001 |
| SLIT3 | 221.402367 | -3.1305122 | 0.65213762 | -4.8003859 | 1.58E-06 | 0.00207155 |
| FAM196B | 60.7669118 | 0.21410014 | 1.49181976 | 0.14351609 | 0.8858826 | 0.99989001 |
| LCP2 | 84.9217695 | -0.2633466 | 0.55154283 | -0.4774727 | 0.63302557 | 0.99989001 |
| RANBP17 | 409.239103 | 0.44245974 | 0.3118236 | 1.41894245 | 0.15591579 | 0.99989001 |
| NPM1 | 127.873736 | -0.1319815 | 0.3887724 | -0.3394827 | 0.73424612 | 0.99989001 |
| FBXW11 | 453.865488 | 0.00415387 | 0.21807565 | 0.01904786 | 0.98480293 | 0.99989001 |
| STK10 | 339.684635 | -0.0722343 | 0.22650875 | -0.318903 | 0.74980008 | 0.99989001 |
| UBTD2 | 122.577326 | 0.42634814 | 0.26636642 | 1.6006077 | 0.10946384 | 0.99989001 |
| SH3PXD2B | 110.814033 | 0.32859308 | 0.2784035 | 1.18027642 | 2.38E-01 | 0.99989001 |
| NEURL1B | 353.862004 | 0.52419987 | 0.44701772 | 1.17266016 | 0.24093212 | 0.99989001 |
| DUSP1 | 734.2757 | -0.689008 | 0.77653572 | -0.8872844 | 0.37492579 | 0.99989001 |
| ERGIC1 | 209.580534 | 0.11532774 | 0.26144795 | 0.44111167 | 0.65913215 | 0.99989001 |
| ATP6V0E1 | 363.336201 | -0.0156485 | 0.33550274 | -0.046642 | 0.96279855 | 0.99989001 |
| CREBRF | 273.917279 | -0.3453127 | 0.37331843 | -0.9249817 | 0.35497545 | 0.99989001 |
| STC2 | 63.7775825 | 1.57612789 | 1.30075093 | 1.21170614 | 0.22562489 | 0.99989001 |
| CPEB4 | 282.337452 | 0.26550296 | 0.24076538 | 1.1027456 | 0.27013766 | 0.99989001 |
| SFXN1 | 79.3144585 | -0.7030858 | 0.35903536 | -1.9582633 | 0.05019912 | 0.79937335 |
| THOC3 | 62.8451228 | -0.1886288 | 0.39309225 | -0.479859 | 0.63132769 | 0.99989001 |
| SIMC1 | 144.997251 | 0.4028853 | 0.27091636 | 1.48712061 | 0.13698296 | 0.99989001 |
| KIAA1191 | 66.9651857 | 0.40169048 | 0.27289918 | 1.47193729 | 0.14103781 | 0.99989001 |
| CLTB | 106.400865 | -0.4213488 | 0.35736804 | -1.1790333 | 0.23838491 | 0.99989001 |
| FAF2 | 179.302055 | -0.1447646 | 0.19517582 | -0.7417137 | 0.45826083 | 0.99989001 |
| RNF44 | 46.9073183 | -0.3508688 | 0.32678948 | -1.0736844 | 0.28296414 | NA |
| UIMC1 | 283.611784 | 0.04641679 | 0.2529348 | 0.18351288 | 0.85439561 | 0.99989001 |
| ZNF346 | 133.81714 | -0.1194109 | 0.28119732 | -0.4246518 | 0.67109054 | 0.99989001 |
| NSD1 | 536.518429 | 0.12977819 | 0.2090713 | 0.62073654 | 0.53477298 | 0.99989001 |
| PRELID1 | 111.690658 | -0.0092311 | 0.55533658 | -0.0166226 | 0.98673771 | 0.99989001 |
| LMAN2 | 129.392206 | -0.1107795 | 0.36324495 | -0.3049718 | 0.76038766 | 0.99989001 |
| DBN1 | 56.4640315 | -0.1416444 | 0.39916288 | -0.3548536 | 0.72269924 | 0.99989001 |
| PDLIM7 | 87.1778446 | -0.2276286 | 0.39659725 | -0.573954 | 0.56599896 | 0.99989001 |
| FAM193B | 183.778535 | 0.11995189 | 0.21289263 | 0.5634384 | 0.57313639 | 0.99989001 |
| TMED9 | 145.974129 | -0.1686161 | 0.51119457 | -0.3298472 | 0.74151543 | 0.99989001 |
| RMND5B | 64.9790397 | -0.1840417 | 0.30899055 | -0.5956224 | 0.55142751 | 0.99989001 |
| NHP2 | 71.567667 | -0.5651193 | 0.60714913 | -0.9307751 | 0.35196993 | 0.99989001 |
| HNRNPAB | 116.357171 | -0.1189337 | 0.4630386 | -0.2568547 | 0.79729091 | 0.99989001 |
| PHYKPL | 277.449326 | 0.4810892 | 0.34134705 | 1.40938438 | 0.15872154 | 0.99989001 |

| | | | | | | |
|---|---|---|---|---|---|---|
| CLK4 | 100.274072 | 0.3087767 | 0.27041118 | 1.14187845 | 0.25350455 | 0.99989001 |
| ZNF354A | 49.5669646 | -0.2810322 | 0.41927836 | -0.670276 | 0.50268186 | NA |
| ZNF354B | 44.9890196 | -0.1801717 | 0.31519256 | -0.5716242 | 0.56757659 | NA |
| ZNF354C | 44.3613981 | -0.0518217 | 0.32430963 | -0.1597909 | 0.87304582 | NA |
| RUFY1 | 142.064555 | -0.0002185 | 0.28889319 | -0.0007563 | 0.99939659 | 0.99989001 |
| HNRNPH1 | 548.007646 | 0.04071824 | 0.29873547 | 0.136302 | 0.89158254 | 0.99989001 |
| CANX | 391.905026 | 0.12927386 | 0.40271981 | 0.32100198 | 0.74820889 | 0.99989001 |
| MAML1 | 93.9480183 | 0.01731614 | 0.23966195 | 0.07225236 | 0.94240108 | 0.99989001 |
| LTC4S | 58.0850815 | 1.01556362 | 0.7858057 | 1.29238516 | 0.19622379 | 0.99989001 |
| MGAT4B | 70.8767139 | -0.0260357 | 0.40234052 | -0.0647107 | 0.94840432 | 0.99989001 |
| SQSTM1 | 469.965054 | -0.1371087 | 0.38865488 | -0.3527775 | 0.72425526 | 0.99989001 |
| MRNIP | 61.8414363 | 0.01240021 | 0.33806809 | 0.03667962 | 0.97074046 | 0.99989001 |
| TBC1D9B | 111.387514 | -0.0798708 | 0.28672147 | -0.2785658 | 0.78057807 | 0.99989001 |
| RNF130 | 286.542155 | 0.00497604 | 0.18606861 | 0.02674301 | 0.9786647 | 0.99989001 |
| MAPK9 | 105.053883 | 0.19121946 | 0.23881419 | 0.80070393 | 0.42330306 | 0.99989001 |
| CNOT6 | 135.787257 | 0.20251623 | 0.24431364 | 0.82891904 | 0.40715022 | 0.99989001 |
| FLT4 | 314.091292 | -0.123981 | 0.37132747 | -0.3338859 | 0.73846563 | 0.99989001 |
| MGAT1 | 269.988928 | -0.1826518 | 0.31216223 | -0.5851181 | 0.55846832 | 0.99989001 |
| BTNL9 | 753.119362 | 0.86595615 | 0.60437312 | 1.4328171 | 0.15191011 | 0.99989001 |
| RACK1 | 335.320712 | 0.04241286 | 0.51022407 | 0.08312596 | 0.93375139 | 0.99989001 |
| TRIM52 | 52.6129634 | 0.38008757 | 0.25542092 | 1.48808314 | 0.13672897 | 0.99989001 |
| DUSP22 | 122.64481 | 0.35615046 | 0.24887419 | 1.43104622 | 0.15241697 | 0.99989001 |
| EXOC2 | 344.692476 | 0.01885809 | 0.17957183 | 0.10501695 | 0.91636236 | 0.99989001 |
| FOXC1 | 69.78504 | 0.48223023 | 0.29786632 | 1.61894847 | 0.10545835 | 0.99989001 |
| GMDS | 931.537063 | 0.22597088 | 0.22936997 | 0.98518077 | 0.32453529 | 0.99989001 |
| GMDS-DT | 648.686054 | -0.036112 | 0.43434695 | -0.0831409 | 0.93373952 | 0.99989001 |
| WRNIP1 | 104.807995 | 0.12910449 | 0.19939947 | 0.64746657 | 0.51733002 | 0.99989001 |
| SERPINB1 | 131.956905 | -0.0379146 | 0.38706369 | -0.0979543 | 0.92196859 | 0.99989001 |
| SERPINB9 | 109.193842 | 0.3404722 | 0.43622052 | 0.78050478 | 0.43509381 | 0.99989001 |
| SERPINB6 | 912.127472 | -0.3206938 | 0.46212749 | -0.693951 | 0.48771294 | 0.99989001 |
| NQO2 | 91.4227596 | -0.0906018 | 0.26394619 | -0.3432586 | 0.73140394 | 0.99989001 |
| RIPK1 | 185.825264 | 0.21138564 | 0.33150076 | 0.6376626 | 0.52369334 | 0.99989001 |
| PSMG4 | 101.309827 | -0.0035784 | 0.2217622 | -0.0161361 | 0.98712582 | 0.99989001 |
| SLC22A23 | 597.275613 | -0.1141477 | 0.64447751 | -0.1771166 | 0.85941681 | 0.99989001 |
| PXDC1 | 100.963831 | -0.3172625 | 0.20018988 | -1.5848078 | 0.11301002 | 0.99989001 |
| PRPF4B | 307.650639 | 0.07413786 | 0.20496884 | 0.3617031 | 0.71757391 | 0.99989001 |
| ECI2 | 85.5264818 | 0.35519101 | 0.30853392 | 1.15122194 | 0.24964094 | 0.99989001 |
| CDYL | 348.677849 | 0.03044393 | 0.2408252 | 0.12641505 | 0.89940339 | 0.99989001 |
| LYRM4 | 173.001733 | -0.1448254 | 0.38287525 | -0.3782575 | 0.70523929 | 0.99989001 |
| FARS2 | 542.267933 | 0.11885786 | 0.29354718 | 0.40490207 | 0.6855495 | 0.99989001 |
| NRN1 | 84.2499098 | -0.328532 | 0.60586766 | -0.5422505 | 0.58764598 | 0.99989001 |
| RREB1 | 597.52388 | -0.1165209 | 0.33798382 | -0.3447529 | 0.73028015 | 0.99989001 |
| SSR1 | 166.64369 | 0.02268429 | 0.33061972 | 0.06861142 | 0.94529893 | 0.99989001 |
| RIOK1 | 54.7407627 | -0.2609331 | 0.35414327 | -0.7368009 | 0.46124341 | 0.99989001 |
| DSP | 58.0300106 | 0.43923935 | 0.55726271 | 0.78820877 | 0.4305746 | 0.99989001 |
| SNRNP48 | 68.2047789 | -0.0703373 | 0.25285296 | -0.2781749 | 0.7808781 | 0.99989001 |
| BMP6 | 283.722844 | 0.98821158 | 0.49713264 | 1.98782277 | 0.04683129 | 0.77516398 |
| SLC35B3 | 84.8361599 | 0.34087316 | 0.31854806 | 1.07008393 | 0.28458153 | 0.99989001 |
| GCNT2 | 263.258875 | -0.6142532 | 0.35768351 | -1.7173093 | 0.08592267 | 0.95514629 |
| TMEM14C | 79.8892061 | -0.0785345 | 0.47851676 | -0.1641207 | 0.86963615 | 0.99989001 |
| MAK | 79.1671982 | 0.27002655 | 0.4806679 | 0.56177363 | 0.57427026 | 0.99989001 |
| SYCP2L | 80.7769041 | 0.37472349 | 0.45243421 | 0.82823864 | 0.40753537 | 0.99989001 |
| SMIM13 | 69.3256321 | 0.03946708 | 0.23205955 | 0.17007305 | 0.86495268 | 0.99989001 |
| AL139807.1 | 303.798019 | -0.0606572 | 0.74855157 | -0.0810327 | 0.93541592 | 0.99989001 |

| | | | | | | |
|---|---|---|---|---|---|---|
| NEDD9 | 1055.70098 | 0.08826879 | 0.568778 | 0.15519024 | 0.87667134 | 0.99989001 |
| HIVEP1 | 253.179538 | 0.13123097 | 0.40178213 | 0.3266222 | 0.74395365 | 0.99989001 |
| EDN1 | 189.221354 | 0.7915569 | 0.61941381 | 1.27791292 | 0.20128013 | 0.99989001 |
| PHACTR1 | 923.909746 | -0.2056605 | 0.42445221 | -0.4845316 | 0.62800865 | 0.99989001 |
| GFOD1 | 745.120865 | 0.21691917 | 0.33752741 | 0.64267127 | 0.52043743 | 0.99989001 |
| SIRT5 | 75.8320748 | -0.046389 | 0.36012031 | -0.1288152 | 0.89750387 | 0.99989001 |
| NOL7 | 107.483796 | -0.3308732 | 0.3444239 | -0.9606568 | 0.33672475 | 0.99989001 |
| RANBP9 | 318.661826 | -0.074142 | 0.22740798 | -0.3260309 | 0.744401 | 0.99989001 |
| MCUR1 | 52.4942334 | 0.2065465 | 0.30370338 | 0.68009284 | 0.49644568 | 0.99989001 |
| JARID2 | 525.625037 | -0.0491235 | 0.28839685 | -0.170333 | 0.86474822 | 0.99989001 |
| DTNBP1 | 229.472315 | -0.3611159 | 0.31058038 | -1.1627132 | 0.24494589 | 0.99989001 |
| MYLIP | 133.139932 | 0.90597695 | 0.22556199 | 4.01653209 | 5.91E-05 | 0.02622301 |
| GMPR | 68.9949244 | -0.564438 | 0.55924851 | -1.0092794 | 0.31284064 | 0.99989001 |
| ATXN1 | 770.771419 | -0.2549991 | 0.40992962 | -0.6220557 | 0.53390521 | 0.99989001 |
| NUP153 | 232.785222 | -0.2082355 | 0.18195937 | -1.1444066 | 0.25245507 | 0.99989001 |
| KIF13A | 782.482785 | 0.01689571 | 0.24722077 | 0.06834259 | 9.46E-01 | 0.99989001 |
| KDM1B | 66.6534884 | 0.28524687 | 0.26250758 | 1.08662338 | 0.27720329 | 0.99989001 |
| DEK | 202.52171 | 0.06288912 | 0.21745277 | 0.28920818 | 0.77242207 | 0.99989001 |
| RNF144B | 712.559001 | -0.6011568 | 0.38949232 | -1.543437 | 0.12272479 | 0.99989001 |
| E2F3 | 124.946668 | -0.0136276 | 0.39300895 | -0.034675 | 0.97233886 | 0.99989001 |
| CDKAL1 | 1036.36706 | -0.1077477 | 0.19145927 | -0.562771 | 0.57359085 | 0.99989001 |
| SOX4 | 177.514368 | 0.13452666 | 0.50123745 | 0.26838908 | 0.78839984 | 0.99989001 |
| CASC15 | 641.788558 | -0.0980644 | 0.46024011 | -0.2130723 | 0.83127057 | 0.99989001 |
| DCDC2 | 45.2967621 | -0.0551279 | 0.55183704 | -0.0998988 | 0.92042463 | NA |
| MRS2 | 48.3907155 | -0.2035188 | 0.29553865 | -0.6886369 | 0.49105182 | NA |
| TDP2 | 73.7047698 | 0.40686157 | 0.31638381 | 1.2859747 | 0.19845189 | 0.99989001 |
| ACOT13 | 54.9872215 | 0.14340407 | 0.36679439 | 0.39096584 | 0.69582249 | 0.99989001 |
| C6orf62 | 196.736969 | -0.0297673 | 0.24654733 | -0.1207368 | 0.90389949 | 0.99989001 |
| RIPOR2 | 136.0193 | 2.01613707 | 0.33002676 | 6.10901096 | 1.00E-09 | 4.47E-06 |
| CARMIL1 | 609.688834 | -0.0106971 | 0.36757726 | -0.0291016 | 0.97678357 | 0.99989001 |
| TRIM38 | 178.562294 | 0.35947673 | 0.26214114 | 1.37130986 | 0.17027838 | 0.99989001 |
| HIST1H1C | 47.7994802 | 0.02960141 | 0.4273932 | 0.06926036 | 0.94478238 | NA |
| HIST1H4C | 58.7492086 | 0.90160047 | 0.73534718 | 1.22608816 | 0.2201655 | 0.99989001 |
| HIST1H2AC | 290.776374 | 1.54312235 | 0.89584829 | 1.72252641 | 0.0849742 | 0.95514629 |
| HIST1H1E | 49.0153124 | -0.028849 | 0.49367415 | -0.0584374 | 0.95340023 | NA |
| BTN3A2 | 116.727861 | 1.30214598 | 0.55511863 | 2.3457076 | 0.018991 | 0.56657733 |
| BTN3A1 | 78.3530889 | 0.61237071 | 0.69783637 | 0.87752765 | 0.38020011 | 0.99989001 |
| BTN3A3 | 62.4229301 | 0.79194412 | 0.57675216 | 1.37310993 | 0.16971817 | 0.99989001 |
| BTN2A1 | 76.6257598 | 0.74706416 | 0.30593492 | 2.4419055 | 0.01460997 | 0.5186081 |
| ZKSCAN8 | 55.6276255 | 0.05255482 | 0.29068879 | 0.18079411 | 0.85652918 | 0.99989001 |
| TRIM27 | 158.563699 | 0.082968 | 0.23965392 | 0.34619922 | 0.729193 | 0.99989001 |
| GABBR1 | 119.37029 | 0.62590319 | 0.34452666 | 1.81670464 | 0.06926235 | 0.89213691 |
| HLA-F | 160.822395 | 0.36587334 | 0.37441166 | 0.97719537 | 0.32847244 | 0.99989001 |
| HLA-A | 1680.91238 | 0.07609368 | 0.50642151 | 0.1502576 | 0.88056139 | 0.99989001 |
| PPP1R11 | 44.1490717 | 0.41809531 | 0.3519514 | 1.18793478 | 0.2348591 | NA |
| TRIM26 | 93.9572956 | -0.0225803 | 0.40605885 | -0.0556084 | 0.95565375 | 0.99989001 |
| HCG17 | 123.265514 | 0.14574396 | 0.30440983 | 0.47877546 | 0.63209838 | 0.99989001 |
| HLA-E | 1881.44902 | 0.38904607 | 0.39597613 | 0.9824988 | 0.32585417 | 0.99989001 |
| GNL1 | 111.496419 | 0.15458971 | 0.2472992 | 0.62511205 | 0.53189752 | 0.99989001 |
| ABCF1 | 92.548502 | -0.1585108 | 0.31898844 | -0.496917 | 0.6192476 | 0.99989001 |
| PPP1R10 | 110.34372 | -0.0555402 | 0.30391836 | -0.1827471 | 0.85499643 | 0.99989001 |
| MRPS18B | 67.0891131 | -0.0328793 | 0.24905046 | -0.1320188 | 0.89496942 | 0.99989001 |
| DHX16 | 42.0506848 | -0.0501271 | 0.27922599 | -0.1795217 | 0.85752811 | NA |
| PPP1R18 | 84.8861254 | -0.231929 | 0.33454059 | -0.6932762 | 0.48813625 | 0.99989001 |

| | | | | | | |
|---|---|---|---|---|---|---|
| MDC1 | 47.2267225 | -0.057797 | 0.3281661 | -0.1761212 | 0.86019869 | NA |
| TUBB | 286.346366 | 0.36445928 | 0.69656715 | 0.52322203 | 0.60081975 | 0.99989001 |
| FLOT1 | 375.475821 | 0.09858271 | 0.30517032 | 0.3230416 | 0.74666374 | 0.99989001 |
| IER3 | 124.812381 | 0.45832822 | 0.65549217 | 0.69921235 | 4.84E-01 | 0.99989001 |
| HLA-C | 1104.23925 | 0.41188051 | 0.6302611 | 0.65350774 | 0.513429 | 0.99989001 |
| HLA-B | 1953.62613 | 0.52401236 | 0.49468282 | 1.05928958 | 0.28946792 | 0.99989001 |
| MICA | 68.3465709 | 0.44130634 | 0.31880933 | 1.38423283 | 0.16628717 | 0.99989001 |
| PRRC2A | 103.32707 | -0.1152558 | 0.29116378 | -0.3958453 | 0.69221913 | 0.99989001 |
| BAG6 | 75.3954218 | -0.1414888 | 0.33176158 | -0.4264774 | 0.66976003 | 0.99989001 |
| DDAH2 | 77.4900918 | 0.08168072 | 0.34607653 | 0.23601923 | 0.81341775 | 0.99989001 |
| CLIC1 | 408.404902 | 0.19563773 | 0.49893791 | 0.39210837 | 0.69497815 | 0.99989001 |
| LSM2 | 72.5304492 | -0.1731368 | 0.47018902 | -0.3682281 | 0.71270312 | 0.99989001 |
| HSPA1A | 581.701739 | 0.93489606 | 1.09524024 | 0.85359908 | 0.39332717 | 0.99989001 |
| HSPA1B | 205.92033 | 0.98879103 | 1.32278658 | 0.74750609 | 0.45475813 | 0.99989001 |
| C6orf48 | 51.60008 | -0.4902519 | 0.46966546 | -1.0438322 | 0.29656303 | 0.99989001 |
| NEU1 | 42.5595169 | -0.1404666 | 0.32048178 | -0.4382981 | 0.66117018 | NA |
| EHMT2 | 46.97197 | -0.2548198 | 0.27814956 | -0.916125 | 0.35960131 | NA |
| STK19 | 46.4500925 | -0.1013078 | 0.26816874 | -0.3777764 | 0.70559669 | NA |
| TNXB | 295.896677 | -1.21531 | 0.53547244 | -2.2696033 | 0.02323166 | 0.60818532 |
| ATF6B | 99.9081818 | -0.3112953 | 0.27271577 | -1.1414642 | 0.2536768 | 0.99989001 |
| AGPAT1 | 69.3676191 | -0.0676846 | 0.51561066 | -0.1312708 | 0.89556106 | 0.99989001 |
| RNF5 | 43.7988172 | 0.10137728 | 0.48987725 | 0.20694424 | 0.83605341 | NA |
| PBX2 | 81.1254374 | 0.0345758 | 0.30331 | 0.11399493 | 0.90924181 | 0.99989001 |
| NOTCH4 | 625.095328 | -0.0846901 | 0.27224975 | -0.3110749 | 0.75574367 | 0.99989001 |
| HLA-DRA | 871.286699 | 0.19747091 | 0.5366064 | 0.36799953 | 0.71287358 | 0.99989001 |
| HLA-DRB5 | 356.684833 | 0.70899345 | 0.72767806 | 0.97432298 | 0.3298962 | 0.99989001 |
| HLA-DRB1 | 957.234881 | -0.7841042 | 0.73926588 | -1.0606525 | 0.28884785 | 0.99989001 |
| HLA-DQA1 | 162.750447 | 0.35742709 | 0.51702098 | 0.69132028 | 0.48936429 | 0.99989001 |
| HLA-DQB1 | 191.704866 | 0.13869621 | 0.67968971 | 0.20405813 | 0.83830807 | 0.99989001 |
| PSMB8 | 166.506011 | 0.13293725 | 0.40937953 | 0.32472862 | 0.74538647 | 0.99989001 |
| PSMB9 | 166.125745 | 0.83900092 | 0.42329541 | 1.98206949 | 0.04747146 | 0.77516398 |
| TAP1 | 145.677901 | 0.0532591 | 0.46972432 | 0.11338374 | 0.90972633 | 0.99989001 |
| HLA-DMB | 64.5646833 | -0.3807275 | 0.50093421 | -0.7600348 | 0.44723376 | 0.99989001 |
| BRD2 | 188.798211 | 0.21603298 | 0.27314913 | 0.79089755 | 0.42900378 | 0.99989001 |
| HLA-DOA | 54.1721672 | 0.29718935 | 0.48344795 | 0.61472875 | 0.53873387 | 0.99989001 |
| HLA-DPA1 | 400.076567 | -0.0648513 | 0.52487137 | -0.1235565 | 0.90166642 | 0.99989001 |
| HLA-DPB1 | 296.162511 | -0.223358 | 0.55070973 | -0.4055821 | 0.68504967 | 0.99989001 |
| RXRB | 39.5827242 | -0.0082898 | 0.30804971 | -0.0269107 | 0.97853095 | NA |
| SLC39A7 | 60.0145712 | -0.2205123 | 0.49023084 | -0.4498131 | 0.65284521 | 0.99989001 |
| RING1 | 53.6590691 | -0.215161 | 0.28956074 | -0.74306 | 0.45744538 | 0.99989001 |
| VPS52 | 57.494857 | -0.0362278 | 0.23035019 | -0.1572729 | 0.87502981 | 0.99989001 |
| RPS18 | 318.232028 | -0.1829945 | 0.58100222 | -0.3149636 | 0.7527893 | 0.99989001 |
| RGL2 | 132.780448 | -0.0626594 | 0.29093982 | -0.2153688 | 0.82947978 | 0.99989001 |
| TAPBP | 355.774515 | -0.0718695 | 0.30802969 | -0.2333202 | 0.81551281 | 0.99989001 |
| PHF1 | 57.0250833 | -0.3934902 | 0.29039698 | -1.3550078 | 0.17541508 | 0.99989001 |
| CUTA | 58.0054911 | -0.2130948 | 0.54807881 | -0.3888032 | 0.6974217 | 0.99989001 |
| ITPR3 | 193.363829 | -0.5667862 | 0.28216335 | -2.0087166 | 0.04456719 | 0.77228819 |
| UQCC2 | 42.768047 | 0.13280365 | 0.41769592 | 0.31794339 | 0.75052788 | NA |
| LEMD2 | 77.9371156 | -0.1527813 | 0.24168061 | -0.6321621 | 0.52728097 | 0.99989001 |
| NUDT3 | 75.9583185 | -0.0521246 | 0.46851488 | -0.1112549 | 0.91141421 | 0.99989001 |
| C6orf106 | 260.747416 | 0.16242382 | 0.21583088 | 0.75255137 | 0.45171955 | 0.99989001 |
| SNRPC | 52.1017679 | -0.4552975 | 0.35738034 | -1.273986 | 0.20266838 | 0.99989001 |
| UHRF1BP1 | 102.016882 | 0.543605 | 0.2738077 | 1.98535321 | 0.04710519 | 0.77516398 |
| TAF11 | 95.7134411 | -0.2502036 | 0.26051434 | -0.9604215 | 0.33684314 | 0.99989001 |

| | | | | | | |
|---|---|---|---|---|---|---|
| ANKS1A | 756.360073 | -0.0495956 | 0.39878271 | -0.1243674 | 0.9010244 | 0.99989001 |
| ZNF76 | 78.6440819 | 0.02640388 | 0.23167376 | 0.1139701 | 0.9092615 | 0.99989001 |
| PPARD | 119.960015 | -0.3733851 | 0.2973068 | -1.2558917 | 0.20915525 | 0.99989001 |
| RPL10A | 286.493859 | -0.1047828 | 0.60847599 | -0.1722053 | 0.86327614 | 0.99989001 |
| FKBP5 | 839.03194 | 0.5726311 | 0.64356525 | 0.88977939 | 0.37358436 | 0.99989001 |
| SRPK1 | 154.722729 | 0.24691588 | 0.28557541 | 0.86462582 | 0.38724419 | 0.99989001 |
| MAPK14 | 317.706241 | 0.09498822 | 0.2330178 | 0.40764363 | 0.68353533 | 0.99989001 |
| KCTD20 | 184.56088 | 0.33637392 | 0.21804099 | 1.54270957 | 0.12290126 | 0.99989001 |
| STK38 | 276.235091 | 0.40278737 | 0.260118 | 1.54847946 | 0.1215069 | 0.99989001 |
| SRSF3 | 257.444374 | -0.0016804 | 0.28353381 | -0.0059265 | 0.99527134 | 0.99989001 |
| CDKN1A | 187.67161 | -1.8975818 | 0.93669287 | -2.0258313 | 0.04278207 | 0.76160158 |
| CPNE5 | 178.569661 | -0.3079188 | 0.61319646 | -0.5021536 | 0.61555951 | 0.99989001 |
| C6orf89 | 149.554538 | 0.00787012 | 0.22816972 | 0.0344924 | 0.97248451 | 0.99989001 |
| MTCH1 | 180.882578 | 0.06626057 | 0.24796549 | 0.26721688 | 0.78930218 | 0.99989001 |
| TBC1D22B | 101.868141 | 0.02789964 | 0.39245663 | 0.07108973 | 0.94332634 | 0.99989001 |
| RNF8 | 70.1500016 | 0.17128049 | 0.26767487 | 0.63988259 | 0.52224894 | 0.99989001 |
| CMTR1 | 77.9588528 | -0.0613993 | 0.41761905 | -0.1470224 | 0.88311436 | 0.99989001 |
| ZFAND3 | 2296.80808 | -0.0639848 | 0.2996922 | -0.2135016 | 0.83093573 | 0.99989001 |
| BTBD9 | 665.867841 | -0.1323158 | 0.33942221 | -0.3898267 | 0.69666471 | 0.99989001 |
| GLO1 | 109.403268 | 0.14126558 | 0.20037401 | 0.70500953 | 0.48080431 | 0.99989001 |
| OARD1 | 62.9618333 | -0.6331549 | 0.27912477 | -2.2683581 | 0.02330739 | 0.60818532 |
| NFYA | 62.9611009 | 0.26251459 | 0.34851207 | 0.75324388 | 0.45130338 | 0.99989001 |
| TFEB | 66.3088482 | 0.03112574 | 0.424812 | 0.07326944 | 0.94159171 | 0.99989001 |
| USP49 | 65.0362314 | 0.21948693 | 0.37241263 | 0.58936489 | 0.55561653 | 0.99989001 |
| CCND3 | 325.014001 | 0.43328407 | 0.50790564 | 0.85307984 | 0.39361503 | 0.99989001 |
| TAF8 | 78.7852043 | 0.24366535 | 0.33728093 | 0.72244034 | 0.47002379 | 0.99989001 |
| TRERF1 | 85.3856855 | 0.49711215 | 0.4511689 | 1.1018316 | 0.27053489 | 0.99989001 |
| UBR2 | 534.069688 | 0.07944518 | 0.21157133 | 0.37550069 | 0.70728814 | 0.99989001 |
| BICRAL | 263.418461 | 0.41296016 | 0.29553035 | 1.3973528 | 0.16230751 | 0.99989001 |
| RPL7L1 | 136.714123 | -0.0715663 | 0.3626586 | -0.197338 | 0.84356303 | 0.99989001 |
| CNPY3 | 39.4306956 | -0.2181416 | 0.47613605 | -0.4581498 | 0.64684481 | NA |
| PEX6 | 50.8413946 | 0.77909092 | 0.30264658 | 2.5742598 | 0.01004548 | 0.45403168 |
| KLHDC3 | 48.9669719 | 0.22971067 | 0.30206019 | 0.76047979 | 0.44696785 | NA |
| RRP36 | 83.6889923 | 0.06601245 | 0.31327445 | 0.21071764 | 0.83310761 | 0.99989001 |
| PTK7 | 44.0251475 | 0.96878474 | 0.37208939 | 2.60363441 | 0.00922411 | NA |
| CUL9 | 53.6591258 | -0.3591554 | 0.24570238 | -1.4617497 | 0.14380982 | 0.99989001 |
| ZNF318 | 53.7932158 | -0.0042988 | 0.29828102 | -0.014412 | 0.98850125 | 0.99989001 |
| ABCC10 | 39.6716717 | 0.02762416 | 0.28710107 | 0.09621755 | 0.92334779 | NA |
| TJAP1 | 49.0124082 | -0.0039687 | 0.26617222 | -0.0149101 | 0.98810388 | NA |
| POLR1C | 70.1659684 | -0.6370351 | 0.36968866 | -1.7231663 | 0.08485845 | 0.95514629 |
| YIPF3 | 88.5474003 | -0.1486949 | 0.40840913 | -0.3640833 | 0.71579582 | 0.99989001 |
| XPO5 | 237.815921 | -0.0852892 | 0.39713346 | -0.2147621 | 0.82995283 | 0.99989001 |
| GTPBP2 | 71.528117 | -0.0744302 | 0.39201224 | -0.1898669 | 0.84941341 | 0.99989001 |
| MRPS18A | 46.8211899 | 0.65134767 | 0.28211618 | 2.30879233 | 0.02095511 | NA |
| VEGFA | 85.5074517 | 0.24715514 | 0.58311839 | 0.42385071 | 0.6716747 | 0.99989001 |
| MRPL14 | 56.0095771 | -0.2458468 | 0.43985602 | -0.5589255 | 0.57621256 | 0.99989001 |
| SLC29A1 | 38.7642291 | 0.10521775 | 0.58378008 | 0.18023525 | 0.85696789 | NA |
| HSP90AB1 | 564.492855 | 0.02944058 | 0.36328094 | 0.0810408 | 0.93540951 | 0.99989001 |
| CDC5L | 99.1904992 | -0.0385022 | 0.22411568 | -0.1717962 | 0.86359778 | 0.99989001 |
| SUPT3H | 285.783061 | -0.8641125 | 0.34301344 | -2.5191798 | 0.01176286 | 0.48516963 |
| CLIC5 | 52.0741758 | 1.20459333 | 0.65806882 | 1.83049751 | 0.06717558 | 0.87873839 |
| ENPP4 | 64.9000025 | 0.0626114 | 0.55411789 | 0.11299291 | 0.91003617 | 0.99989001 |
| RCAN2 | 327.873528 | 0.57719523 | 0.43161954 | 1.33727781 | 1.81E-01 | 1.00E+00 |
| SLC25A27 | 93.71106 | -0.2493162 | 0.56231401 | -0.4433754 | 0.65749423 | 0.99989001 |

| | | | | | | |
|---|---|---|---|---|---|---|
| ADGRF5 | 1329.26281 | -0.2429377 | 0.32471846 | -0.7481486 | 0.4543705 | 0.99989001 |
| CD2AP | 408.756593 | -0.1620199 | 0.18487853 | -0.8763586 | 0.38083514 | 0.99989001 |
| PTCHD4 | 114.033258 | -1.0435946 | 1.00742331 | -1.0359047 | 0.30024658 | 0.99989001 |
| MUT | 50.6408295 | -0.0041496 | 0.29653363 | -0.0139938 | 0.98883491 | 0.99989001 |
| PKHD1 | 124.508754 | -0.5383238 | 0.60720884 | -0.8865547 | 0.37531869 | 0.99989001 |
| MCM3 | 59.8238118 | -0.0914587 | 0.38369241 | -0.2383645 | 0.81159839 | 0.99989001 |
| EFHC1 | 82.0144118 | 0.33325427 | 0.328324 | 1.01501648 | 0.31009797 | 0.99989001 |
| TRAM2 | 167.995836 | -0.0272077 | 0.33353965 | -0.0815725 | 0.9349867 | 0.99989001 |
| GSTA4 | 48.5140807 | -0.2553742 | 0.44327781 | -0.5761042 | 0.56454472 | NA |
| ICK | 54.426921 | 0.31790151 | 0.24830108 | 1.28030659 | 0.20043733 | 0.99989001 |
| FBXO9 | 125.075018 | 0.18487151 | 0.19895336 | 0.92922034 | 0.35277491 | 0.99989001 |
| ELOVL5 | 342.381894 | -0.0149804 | 0.31701777 | -0.0472541 | 0.96231071 | 0.99989001 |
| GCLC | 87.4961219 | -0.0431486 | 0.36133435 | -0.1194147 | 0.90494682 | 0.99989001 |
| LRRC1 | 93.8129762 | 0.67805504 | 0.49695219 | 1.36442712 | 0.17243318 | 0.99989001 |
| DST | 662.084481 | -0.1786709 | 0.30807787 | -0.5799536 | 0.56194593 | 0.99989001 |
| ZNF451 | 321.977723 | -0.0711648 | 0.17575692 | -0.4049044 | 0.68554775 | 0.99989001 |
| PRIM2 | 253.893756 | -0.4238858 | 0.21510581 | -1.9705919 | 0.04877057 | 0.78323466 |
| AL445250.1 | 82.0203182 | -0.1445172 | 0.60144089 | -0.2402849 | 0.8101094 | 0.99989001 |
| PTP4A1 | 79.0226239 | -0.2346735 | 0.44704704 | -0.5249414 | 0.5996239 | 0.99989001 |
| PHF3 | 375.230143 | 0.31294394 | 0.22708172 | 1.37811152 | 0.16816886 | 0.99989001 |
| ADGRB3 | 1621.41912 | -0.9767263 | 0.53570845 | -1.8232423 | 0.0682667 | 0.88709358 |
| LMBRD1 | 270.586065 | 0.11994381 | 0.2703877 | 0.44359936 | 0.65733227 | 0.99989001 |
| COL19A1 | 69.354787 | -0.0786136 | 0.35835984 | -0.2193705 | 0.82636147 | 0.99989001 |
| FAM135A | 332.254822 | -0.2545557 | 0.25102479 | -1.0140661 | 0.31055119 | 0.99989001 |
| SMAP1 | 352.326923 | 0.18921148 | 0.26380453 | 0.7172412 | 0.47322528 | 0.99989001 |
| OGFRL1 | 51.1360272 | -0.0233518 | 0.31751908 | -0.0735446 | 0.94137272 | 0.99989001 |
| AL136164.2 | 120.516911 | -0.3071325 | 0.46177619 | -0.6651112 | 0.5059794 | 0.99989001 |
| OOEP | 334.54519 | -0.2761008 | 0.57873678 | -0.4770749 | 0.63330876 | 0.99989001 |
| CGAS | 57.5475957 | 0.59414695 | 0.40890938 | 1.45300396 | 0.14622265 | 0.99989001 |
| MTO1 | 67.9618377 | 0.14592141 | 0.25842484 | 0.56465704 | 0.57230706 | 0.99989001 |
| EEF1A1 | 1316.99409 | -0.0264101 | 0.54639621 | -0.048335 | 0.96144923 | 0.99989001 |
| SLC17A5 | 116.690277 | -0.0797219 | 0.44196576 | -0.1803802 | 0.8568541 | 0.99989001 |
| CD109 | 402.566464 | 0.03359976 | 0.39483385 | 0.08509847 | 0.93218311 | 0.99989001 |
| COX7A2 | 134.644331 | 0.12962038 | 0.5560386 | 0.233114 | 0.81567288 | 0.99989001 |
| TMEM30A | 266.947052 | -0.1033439 | 0.44813027 | -0.2306113 | 0.81761679 | 0.99989001 |
| FILIP1 | 117.007989 | -0.2094113 | 0.66323977 | -0.3157399 | 0.75219993 | 0.99989001 |
| SENP6 | 680.975 | -0.2125774 | 0.17424338 | -1.2200028 | 0.22246381 | 0.99989001 |
| MYO6 | 646.930535 | 0.37271051 | 0.22249358 | 1.67515178 | 9.39E-02 | 0.99989001 |
| IRAK1BP1 | 69.9169083 | -0.0009676 | 0.40407514 | -0.0023945 | 0.99808946 | 0.99989001 |
| PHIP | 661.550424 | -0.1003345 | 0.22954858 | -0.4370948 | 0.66204262 | 0.99989001 |
| HMGN3 | 217.672115 | 0.2548081 | 0.38493852 | 0.66194492 | 0.50800652 | 0.99989001 |
| LCA5 | 64.5602151 | -0.1778028 | 0.26795537 | -0.6635538 | 0.50697596 | 0.99989001 |
| SH3BGRL2 | 228.58605 | 0.06541051 | 0.39937402 | 0.16378258 | 0.86990231 | 0.99989001 |
| BCKDHB | 181.535667 | -0.1385136 | 0.25165204 | -0.5504173 | 0.58203318 | 0.99989001 |
| IBTK | 237.299812 | -0.1366241 | 0.20397686 | -0.669802 | 0.502984 | 0.99989001 |
| UBE3D | 158.944503 | -0.0060362 | 0.31392025 | -0.0192286 | 0.98465874 | 0.99989001 |
| DOPEY1 | 270.359565 | -0.1207549 | 0.22531913 | -0.5359284 | 0.59200805 | 0.99989001 |
| PGM3 | 55.2834336 | 0.24490918 | 0.34704876 | 0.70569098 | 0.48038033 | 0.99989001 |
| CYB5R4 | 111.777389 | 0.14152178 | 0.27618645 | 0.51241391 | 0.60836136 | 0.99989001 |
| CEP162 | 115.985426 | 0.09919435 | 0.28764852 | 0.34484567 | 0.7302104 | 0.99989001 |
| SNX14 | 262.273768 | -0.0793147 | 0.22874867 | -0.3467331 | 0.72879182 | 0.99989001 |
| SYNCRIP | 121.781414 | -0.2326762 | 0.38261886 | -0.6081149 | 0.54311126 | 0.99989001 |
| ZNF292 | 730.8216 | -0.4422226 | 0.38367809 | -1.1525877 | 0.24907966 | 0.99989001 |
| SMIM8 | 93.6962922 | -0.2011725 | 0.33069823 | -0.6083265 | 0.54297098 | 0.99989001 |

| | | | | | | |
|---|---|---|---|---|---|---|
| RARS2 | 257.555414 | 0.10725198 | 0.21640972 | 0.49559687 | 0.62017886 | 0.99989001 |
| ORC3 | 128.735983 | -0.0367268 | 0.35880792 | -0.1023579 | 0.9184726 | 0.99989001 |
| AKIRIN2 | 71.3168071 | 0.02540477 | 0.28846755 | 0.08806804 | 0.9298226 | 0.99989001 |
| RNGTT | 373.993339 | -0.0468635 | 0.2650569 | -0.1768054 | 0.85966127 | 0.99989001 |
| PNRC1 | 213.168383 | -0.0798105 | 0.49156808 | -0.162359 | 0.87102313 | 0.99989001 |
| UBE2J1 | 318.891132 | -0.4552504 | 0.40687482 | -1.1188955 | 0.26318471 | 0.99989001 |
| ANKRD6 | 154.778147 | 0.10832335 | 0.26843064 | 0.40354317 | 0.68654869 | 0.99989001 |
| LYRM2 | 65.737608 | -0.3555139 | 0.27254895 | -1.3044039 | 0.1920959 | 0.99989001 |
| MDN1 | 200.25603 | -0.0122078 | 0.26253131 | -0.0465003 | 0.96291151 | 0.99989001 |
| CASP8AP2 | 134.018842 | -0.2407787 | 0.19325664 | -1.2459014 | 0.21280059 | 0.99989001 |
| BACH2 | 165.811835 | 0.32785981 | 0.43066195 | 0.76129273 | 0.44648224 | 0.99989001 |
| MAP3K7 | 229.427217 | 0.15314844 | 0.2370401 | 0.64608667 | 0.51822323 | 0.99989001 |
| MANEA | 54.0178309 | 0.06579814 | 0.38320024 | 0.17170693 | 0.86366794 | 0.99989001 |
| UFL1 | 127.60141 | 0.05130749 | 0.18934325 | 0.27097608 | 0.78640943 | 0.99989001 |
| MMS22L | 96.4828582 | -0.711526 | 0.22720601 | -3.1316336 | 0.00173837 | 0.20186448 |
| FBXL4 | 228.843442 | -0.1058126 | 0.28723058 | -0.3683891 | 0.71258315 | 0.99989001 |
| PNISR | 577.832908 | -0.0729534 | 0.23059377 | -0.3163718 | 0.75172031 | 0.99989001 |
| USP45 | 139.204999 | -0.0130525 | 0.24558397 | -0.0531488 | 0.95761333 | 0.99989001 |
| CCNC | 166.731881 | -0.2045324 | 0.23869399 | -0.8568812 | 0.39151057 | 0.99989001 |
| ASCC3 | 593.5081 | 0.25076861 | 0.26090132 | 0.9611627 | 0.33647037 | 0.99989001 |
| HACE1 | 190.503652 | -0.0900131 | 0.32858782 | -0.2739393 | 0.78413127 | 0.99989001 |
| PREP | 144.973695 | 0.30889668 | 0.22818538 | 1.35370932 | 0.17582913 | 0.99989001 |
| ATG5 | 287.919094 | 0.19188827 | 0.15747048 | 1.21856657 | 0.22300874 | 0.99989001 |
| PRDM1 | 85.9599724 | 1.22908549 | 0.70043173 | 1.75475416 | 0.07930137 | 0.93210268 |
| CRYBG1 | 101.337101 | -0.8561807 | 0.41553926 | -2.0604086 | 0.03935949 | 0.72459869 |
| QRSL1 | 63.743756 | -0.2275702 | 0.28765619 | -0.7911187 | 0.42887475 | 0.99989001 |
| CD24 | 141.633315 | -0.0823196 | 0.32058287 | -0.2567812 | 0.79734772 | 0.99989001 |
| PDSS2 | 433.383834 | -0.1127343 | 0.27551664 | -0.4091744 | 0.6824117 | 0.99989001 |
| SEC63 | 285.007546 | -0.0573376 | 0.17968957 | -0.3190925 | 0.74965637 | 0.99989001 |
| OSTM1 | 107.781777 | -0.0415496 | 0.24818821 | -0.1674116 | 0.86704623 | 0.99989001 |
| SNX3 | 280.719821 | 0.00575167 | 0.25346421 | 0.02269225 | 0.98189576 | 0.99989001 |
| AFG1L | 108.864502 | -0.2774329 | 0.31914516 | -0.8693001 | 0.384683 | 0.99989001 |
| FOXO3 | 514.133691 | 0.27802601 | 0.30861786 | 0.90087466 | 0.36765497 | 0.99989001 |
| ARMC2 | 67.9644804 | 0.3158798 | 0.50509502 | 0.62538688 | 0.53171717 | 0.99989001 |
| SESN1 | 325.157409 | -0.0651266 | 0.26986989 | -0.2413258 | 0.80930265 | 0.99989001 |
| CEP57L1 | 101.260898 | 0.09210237 | 0.29917105 | 0.30785856 | 0.75818996 | 0.99989001 |
| CD164 | 213.790338 | 0.04030204 | 0.27090997 | 0.14876542 | 0.88173874 | 0.99989001 |
| AK9 | 109.785423 | -0.2097101 | 0.36967569 | -0.5672814 | 0.57052302 | 0.99989001 |
| FIG4 | 87.1342186 | -0.097681 | 0.27802146 | -0.3513435 | 0.72533066 | 0.99989001 |
| CDC40 | 87.7345289 | -0.1181755 | 0.25406057 | -0.465147 | 0.64182616 | 0.99989001 |
| CDK19 | 288.476471 | -0.0960679 | 0.3642373 | -0.263751 | 0.79197183 | 0.99989001 |
| AMD1 | 97.4704678 | -0.292029 | 0.30122038 | -0.9694863 | 0.33230262 | 0.99989001 |
| GTF3C6 | 72.9530663 | -0.3661062 | 0.42721386 | -0.8569624 | 0.39146569 | 0.99989001 |
| RPF2 | 62.0827024 | -0.3638808 | 0.28845166 | -1.2614967 | 0.20712993 | 0.99989001 |
| REV3L | 753.640153 | -0.2696042 | 0.37042235 | -0.7278292 | 0.46671814 | 0.99989001 |
| TRAF3IP2-AS | 275.772117 | 0.11993127 | 0.2783943 | 0.43079645 | 0.66661638 | 0.99989001 |
| TRAF3IP2 | 57.144213 | -0.5491447 | 0.27454065 | -2.0002308 | 0.04547534 | 0.77228819 |
| FYN | 1166.70846 | 0.76087526 | 0.33308914 | 2.28429921 | 0.02235395 | 0.60384164 |
| TUBE1 | 80.3589762 | 0.15738646 | 0.38270207 | 0.4112506 | 0.68088879 | 0.99989001 |
| LAMA4 | 174.245084 | -0.0308939 | 0.506098 | -0.0610433 | 0.9513247 | 0.99989001 |
| MARCKS | 695.247144 | -0.549473 | 0.54300626 | -1.0119091 | 0.31158153 | 0.99989001 |
| HDAC2 | 134.635965 | 0.08852588 | 0.2706366 | 0.32710241 | 0.74359043 | 0.99989001 |
| NT5DC1 | 199.276338 | 0.10547878 | 0.20387553 | 0.51736853 | 0.60489893 | 0.99989001 |
| DSE | 111.798596 | 0.06898218 | 0.39644096 | 0.17400366 | 0.86186259 | 0.99989001 |

| | | | | | | |
|---|---|---|---|---|---|---|
| TSPYL1 | 54.6735146 | 0.19610449 | 0.39441837 | 0.49719919 | 0.61904859 | 0.99989001 |
| RWDD1 | 75.6943902 | 0.11648789 | 0.45787553 | 0.25440951 | 0.7991792 | 0.99989001 |
| ZUP1 | 52.8261299 | 0.21560882 | 0.30285941 | 0.71191059 | 0.47652014 | 0.99989001 |
| KPNA5 | 139.534897 | 0.19874447 | 0.23508779 | 0.84540534 | 0.39788457 | 0.99989001 |
| DCBLD1 | 116.064599 | 0.91745132 | 0.260871 | 3.51687742 | 0.00043666 | 0.09416785 |
| GOPC | 103.063353 | 0.19512297 | 0.25918496 | 0.75283291 | 0.45155033 | 0.99989001 |
| NUS1 | 128.965532 | 0.24307856 | 0.23270993 | 1.044556 | 0.29622822 | 0.99989001 |
| CEP85L | 258.489875 | 0.46641549 | 0.26053081 | 1.79025084 | 0.0734136 | 0.9158596 |
| MCM9 | 166.938419 | -0.1068362 | 0.26562928 | -0.4022005 | 0.68753646 | 0.99989001 |
| MAN1A1 | 930.602041 | 0.09265268 | 0.27228508 | 0.3402782 | 0.73364703 | 0.99989001 |
| TBC1D32 | 155.947317 | -0.5863413 | 0.33458511 | -1.7524427 | 0.07969771 | 0.93383766 |
| GJA1 | 298.532243 | 0.67047605 | 0.55071543 | 1.21746372 | 0.22342783 | 0.99989001 |
| HSF2 | 51.0211105 | -0.4661771 | 0.34237436 | -1.3616003 | 0.17332405 | 0.99989001 |
| SERINC1 | 307.647098 | 0.26541396 | 0.30662323 | 0.86560293 | 0.38670795 | 0.99989001 |
| NKAIN2 | 239.492504 | 0.32116758 | 1.23756194 | 0.25951637 | 0.79523685 | 0.99989001 |
| RNF217 | 312.894773 | 0.4953881 | 0.32408144 | 1.5285914 | 0.12636577 | 0.99989001 |
| TPD52L1 | 249.050773 | -1.3002538 | 0.97013991 | -1.3402745 | 0.18015611 | 0.99989001 |
| NCOA7 | 1047.89782 | 0.5200443 | 0.36096387 | 1.44071012 | 0.14966659 | 0.99989001 |
| HINT3 | 54.3922482 | 0.22837736 | 0.31297993 | 0.72968693 | 0.46558157 | 0.99989001 |
| TRMT11 | 91.7534824 | -0.2083908 | 0.30941121 | -0.6735075 | 0.50062447 | 0.99989001 |
| RSPO3 | 65.6554853 | -0.7665596 | 0.66913867 | -1.1455915 | 0.2519642 | 0.99989001 |
| RNF146 | 82.8271175 | -0.0922664 | 0.29116133 | -0.3168908 | 0.75132643 | 0.99989001 |
| ECHDC1 | 137.539632 | 0.15579069 | 0.24203412 | 0.64367242 | 0.51978788 | 0.99989001 |
| PTPRK | 932.746094 | -0.1303179 | 0.30122684 | -0.4326236 | 0.66528821 | 0.99989001 |
| ARHGAP18 | 320.207291 | -0.0502765 | 0.3518369 | -0.1428972 | 8.86E-01 | 0.99989001 |
| L3MBTL3 | 78.1706436 | 0.0206207 | 0.31641036 | 0.06517076 | 0.94803804 | 0.99989001 |
| EPB41L2 | 554.934854 | 0.38270554 | 0.15761669 | 2.42807747 | 0.0151791 | 0.51942334 |
| AKAP7 | 105.740751 | -0.0985878 | 0.45025043 | -0.2189621 | 0.82667954 | 0.99989001 |
| MED23 | 150.379591 | -0.0918713 | 0.18906724 | -0.4859186 | 0.62702492 | 0.99989001 |
| CTGF | 244.156067 | -0.1256081 | 1.09597314 | -0.1146087 | 0.90875529 | 0.99989001 |
| STX7 | 263.198936 | 0.16937312 | 0.23484913 | 0.72119967 | 0.47078667 | 0.99989001 |
| SLC18B1 | 85.4412578 | -0.3616658 | 0.34720669 | -1.041644 | 0.29757675 | 0.99989001 |
| RPS12 | 319.660301 | 0.04235374 | 0.58195156 | 0.0727788 | 0.94198214 | 0.99989001 |
| TBPL1 | 68.6394713 | 0.303675 | 0.22139375 | 1.37165122 | 0.17017204 | 0.99989001 |
| SGK1 | 512.78734 | 0.68306522 | 0.63861305 | 1.06960736 | 0.28479608 | 0.99989001 |
| AL078590.2 | 146.832232 | -0.6101673 | 1.45179059 | -0.420286 | 0.67427652 | 0.99989001 |
| AL078590.3 | 355.304864 | -0.9797519 | 1.50920292 | -0.649185 | 0.51621882 | 0.99989001 |
| HBS1L | 166.099201 | -0.2776633 | 0.19830956 | -1.4001507 | 0.16146819 | 0.99989001 |
| AHI1 | 436.92457 | -0.2767026 | 0.14263197 | -1.9399763 | 0.05238257 | 0.81270649 |
| PDE7B | 823.274393 | 0.50041106 | 0.50050129 | 0.99981973 | 0.31739775 | 0.99989001 |
| AL138828.1 | 157.426576 | -0.0506391 | 0.69448657 | -0.0729159 | 0.94187305 | 0.99989001 |
| BCLAF1 | 346.698645 | 0.05012415 | 0.21795297 | 0.2299769 | 0.81810972 | 0.99989001 |
| MAP3K5 | 244.362671 | 1.50681518 | 0.5103537 | 2.95249192 | 0.0031522 | 0.28843642 |
| IFNGR1 | 245.580441 | 0.31183084 | 0.41063547 | 0.75938603 | 0.44762167 | 0.99989001 |
| TNFAIP3 | 368.302923 | 1.75285285 | 1.00699567 | 1.74067565 | 0.08174045 | 0.9437794 |
| ARFGEF3 | 103.23318 | -0.2399057 | 0.47224222 | -0.508014 | 0.61144352 | 0.99989001 |
| HEBP2 | 86.4032729 | 0.35139041 | 0.31021256 | 1.13274075 | 0.25732314 | 0.99989001 |
| NHSL1 | 744.325179 | 0.53257624 | 0.45447253 | 1.17185575 | 0.24125497 | 0.99989001 |
| CCDC28A | 45.3227351 | -0.1828872 | 0.46472318 | -0.39354 | 0.69392068 | NA |
| REPS1 | 251.461865 | -0.0856911 | 0.23254514 | -0.3684923 | 0.71250619 | 0.99989001 |
| ABRACL | 42.2129939 | 0.16722953 | 0.53476236 | 0.31271747 | 0.75449532 | NA |
| HECA | 111.312823 | 0.18129831 | 0.23486628 | 0.77192142 | 0.44016097 | 0.99989001 |
| VTA1 | 130.724334 | 0.03704156 | 0.19216699 | 0.19275715 | 0.84714917 | 0.99989001 |
| ADGRG6 | 110.45235 | 0.40330409 | 0.65755466 | 0.61333926 | 0.53965204 | 0.99989001 |

| | | | | | | |
|---|---|---|---|---|---|---|
| HIVEP2 | 761.12527 | -0.068387 | 0.53637122 | -0.1274994 | 0.89854512 | 0.99989001 |
| AIG1 | 177.118165 | -0.2259198 | 0.32946445 | -0.6857183 | 0.49289077 | 0.99989001 |
| PEX3 | 119.467521 | -0.3571554 | 0.23757778 | -1.5033198 | 0.13275659 | 0.99989001 |
| FUCA2 | 41.7438146 | -0.1661106 | 0.35629912 | -0.4662112 | 0.64106436 | NA |
| PHACTR2 | 466.728456 | -0.0026495 | 0.17683649 | -0.0149825 | 0.98804614 | 0.99989001 |
| PLAGL1 | 62.0245517 | -0.3016643 | 0.31731347 | -0.9506823 | 0.34176569 | 0.99989001 |
| SF3B5 | 108.264023 | -0.2226356 | 0.48164493 | -0.4622401 | 0.64390917 | 0.99989001 |
| STX11 | 54.181729 | 0.54659192 | 0.53257075 | 1.02632733 | 0.30473732 | 0.99989001 |
| UTRN | 4112.57986 | -0.2027091 | 0.33532924 | -0.6045077 | 0.54550617 | 0.99989001 |
| AL024474.2 | 56.7901913 | 0.49467241 | 0.55418691 | 0.89260933 | 0.37206642 | 0.99989001 |
| AL356599.1 | 68.2950386 | 0.31042558 | 0.42215506 | 0.73533545 | 0.46213523 | 0.99989001 |
| SHPRH | 332.965081 | -0.0621957 | 0.16002287 | -0.3886677 | 0.69752198 | 0.99989001 |
| RAB32 | 43.3970962 | 0.60980755 | 0.41225871 | 1.47918657 | 0.13909046 | NA |
| STXBP5-AS1 | 103.333794 | 0.03616008 | 0.31664203 | 0.11419862 | 0.90908035 | 0.99989001 |
| STXBP5 | 388.254204 | 0.31461979 | 0.19252369 | 1.6341874 | 0.10221951 | 0.99989001 |
| SAMD5 | 103.858019 | 0.7984084 | 0.6208405 | 1.28601212 | 0.19843883 | 0.99989001 |
| SASH1 | 2473.19739 | -0.1408328 | 0.3105778 | -0.4534543 | 0.65022167 | 0.99989001 |
| TAB2 | 305.057508 | 0.17252034 | 0.24495712 | 0.7042879 | 0.4812535 | 0.99989001 |
| PPIL4 | 127.155242 | 0.2023011 | 0.26387231 | 0.76666287 | 0.44328198 | 0.99989001 |
| GINM1 | 168.54616 | 0.04730996 | 0.1792507 | 0.26393182 | 0.79183246 | 0.99989001 |
| KATNA1 | 55.2012664 | 0.18666816 | 0.24784087 | 0.7531775 | 0.45134326 | 0.99989001 |
| LATS1 | 211.363438 | 0.39856137 | 0.24331774 | 1.63802841 | 0.10141577 | 0.99989001 |
| NUP43 | 75.0899041 | -0.436155 | 0.20521707 | -2.1253351 | 0.03355866 | 0.67281094 |
| PCMT1 | 152.843437 | 0.08634796 | 0.23013375 | 0.37520773 | 0.70750598 | 0.99989001 |
| LRP11 | 64.965101 | 0.12590877 | 0.49461996 | 0.25455659 | 0.79906559 | 0.99989001 |
| PLEKHG1 | 2813.10499 | -0.6895246 | 0.39486823 | -1.7462143 | 0.08077371 | 0.93972046 |
| MTHFD1L | 405.714599 | -0.0650116 | 0.50909306 | -0.1277008 | 0.89838576 | 0.99989001 |
| AKAP12 | 972.166167 | -1.157348 | 0.78592912 | -1.4725857 | 0.14086278 | 0.99989001 |
| RMND1 | 59.0365067 | 0.15118013 | 0.31880278 | 0.47421209 | 0.63534867 | 0.99989001 |
| SYNE1 | 3348.84709 | 0.37684917 | 0.39781744 | 0.94729172 | 3.43E-01 | 0.99989001 |
| MYCT1 | 219.22543 | 0.01504136 | 0.30934714 | 0.04862293 | 0.9612198 | 0.99989001 |
| MTRF1L | 55.0109833 | -0.011267 | 0.31419752 | -0.0358595 | 0.9713944 | 0.99989001 |
| CNKSR3 | 465.040053 | 0.52742445 | 0.34001698 | 1.5511709 | 0.12086073 | 0.99989001 |
| SCAF8 | 525.092746 | -0.1138585 | 0.19867564 | -0.5730876 | 0.56658541 | 0.99989001 |
| TIAM2 | 414.034114 | -0.1579289 | 0.26072587 | -0.6057279 | 0.54469543 | 0.99989001 |
| TFB1M | 84.4585672 | -0.2640715 | 0.28122645 | -0.9389997 | 0.34773092 | 0.99989001 |
| ARID1B | 1745.18553 | 0.15436356 | 0.19229073 | 0.80276133 | 0.4221127 | 0.99989001 |
| TMEM242 | 77.866129 | -0.0633167 | 0.24243332 | -0.2611716 | 0.79396016 | 0.99989001 |
| ZDHHC14 | 697.822609 | 0.77645263 | 0.38148341 | 2.03535098 | 0.04181557 | 0.75508116 |
| SNX9 | 586.556492 | -0.3043452 | 0.29925712 | -1.0170024 | 0.30915231 | 0.99989001 |
| SYNJ2 | 438.962213 | -0.8012383 | 0.37138968 | -2.1574058 | 0.03097406 | 0.67281094 |
| GTF2H5 | 63.0104633 | -0.5329062 | 0.41071394 | -1.2975118 | 0.19445514 | 0.99989001 |
| TULP4 | 443.755574 | 0.23393394 | 0.21320862 | 1.09720674 | 0.27255103 | 0.99989001 |
| TMEM181 | 231.924343 | 0.02624436 | 0.22941154 | 0.11439861 | 0.90892182 | 0.99989001 |
| DYNLT1 | 183.854183 | 0.08748385 | 0.30707719 | 0.28489203 | 0.77572686 | 0.99989001 |
| SYTL3 | 121.030922 | -0.6816244 | 0.44894083 | -1.5182946 | 0.12894014 | 0.99989001 |
| EZR | 92.4461269 | 0.62883471 | 0.32046443 | 1.96226053 | 0.04973216 | 0.79361172 |
| SOD2.1 | 63.4301297 | -0.0114963 | 0.54893879 | -0.0209427 | 0.98329135 | 0.99989001 |
| WTAP | 395.451408 | 0.34350835 | 0.52121415 | 0.65905414 | 0.509861 | 0.99989001 |
| TCP1 | 107.627215 | 0.29045062 | 0.31331321 | 0.92702959 | 0.35391117 | 0.99989001 |
| MRPL18 | 55.0807458 | 0.02380229 | 0.40675629 | 0.05851732 | 0.95333657 | 0.99989001 |
| IGF2R | 490.974549 | -0.1432289 | 0.33076584 | -0.433022 | 0.66499881 | 0.99989001 |
| MAP3K4 | 235.028541 | -0.1031192 | 0.18949228 | -0.5441866 | 0.58631306 | 0.99989001 |
| AGPAT4 | 248.195412 | 0.08598534 | 0.19628022 | 0.43807439 | 0.66133236 | 0.99989001 |

| | | | | | | |
|---|---|---|---|---|---|---|
| PRKN | 868.278646 | -0.3908914 | 0.28305573 | -1.3809696 | 0.1672883 | 0.99989001 |
| QKI | 1766.68303 | 0.30231298 | 0.19199648 | 1.57457564 | 0.11535443 | 0.99989001 |
| PDE10A | 1599.35131 | 0.39094734 | 0.47698387 | 0.81962382 | 0.41243059 | 0.99989001 |
| SFT2D1 | 84.4251848 | 0.07792552 | 0.26271912 | 0.29661152 | 0.76676311 | 0.99989001 |
| MPC1 | 86.7054751 | 0.30573431 | 0.2543079 | 1.20222103 | 0.2292779 | 0.99989001 |
| RPS6KA2 | 1050.51125 | 0.0622501 | 0.27832365 | 0.22366085 | 0.82302121 | 0.99989001 |
| FGFR1OP | 77.0015442 | -0.0534525 | 0.32399143 | -0.1649812 | 0.86895877 | 0.99989001 |
| AFDN | 839.677309 | 0.119133 | 0.21801176 | 0.54645216 | 0.58475516 | 0.99989001 |
| WDR27 | 208.567325 | 0.43071519 | 0.34317923 | 1.25507356 | 0.20945206 | 0.99989001 |
| PHF10 | 185.715255 | -0.6282888 | 0.41785998 | -1.5035869 | 0.13268777 | 0.99989001 |
| ERMARD | 46.6259813 | -0.3958304 | 0.34835244 | -1.1362928 | 0.25583403 | NA |
| DLL1 | 89.8038507 | -0.331754 | 0.48155256 | -0.6889259 | 0.49086991 | 0.99989001 |
| FAM120B | 470.651966 | -0.0627069 | 0.27722875 | -0.2261918 | 0.82105226 | 0.99989001 |
| PSMB1 | 221.14963 | 0.08958406 | 0.45457657 | 0.19707143 | 0.84377164 | 0.99989001 |
| TBP | 56.0366521 | -0.167487 | 0.25592906 | -0.6544273 | 0.51283656 | 0.99989001 |
| PDCD2 | 73.1194337 | 0.14040561 | 0.35045416 | 0.40063901 | 0.68868592 | 0.99989001 |
| PRKAR1B | 128.349244 | 0.12515024 | 0.38415586 | 0.32577985 | 0.74459093 | 0.99989001 |
| DNAAF5 | 53.3665578 | -0.4536696 | 0.27735965 | -1.6356728 | 0.1019081 | 0.99989001 |
| SUN1 | 440.633273 | -0.1128551 | 0.28407647 | -0.3972702 | 0.69116824 | 0.99989001 |
| COX19 | 60.2783998 | 0.33441547 | 0.37122676 | 0.90083881 | 0.36767403 | 0.99989001 |
| C7orf50 | 100.520244 | 0.10430914 | 0.34798935 | 0.29974807 | 0.76436933 | 0.99989001 |
| GPR146 | 52.2178123 | -0.7503597 | 0.62058823 | -1.2091105 | 0.2266204 | 0.99989001 |
| MICALL2 | 67.9804892 | 0.05828465 | 0.28230066 | 0.20646304 | 0.83642924 | 0.99989001 |
| INTS1 | 73.6313096 | -0.3087108 | 0.29500491 | -1.0464599 | 0.29534876 | 0.99989001 |
| MAFK | 55.5603107 | -0.754108 | 0.35888381 | -2.1012593 | 0.03561821 | 0.6933146 |
| MAD1L1 | 166.803663 | 0.01795159 | 0.35619806 | 0.05039777 | 0.95980541 | 0.99989001 |
| SNX8 | 73.8921126 | -0.5996314 | 0.44672789 | -1.3422743 | 0.17950709 | 0.99989001 |
| EIF3B | 173.431398 | -0.3338273 | 0.28690474 | -1.1635477 | 0.24460736 | 0.99989001 |
| CHST12 | 161.457025 | -0.3665302 | 0.71864844 | -0.5100272 | 0.61003244 | 0.99989001 |
| BRAT1 | 43.8093784 | -0.193204 | 0.33171126 | -0.5824464 | 0.56026604 | NA |
| IQCE | 38.800425 | 0.02175849 | 0.40536532 | 0.05367624 | 0.95719311 | NA |
| TTYH3 | 80.4249055 | -0.2626726 | 0.38477488 | -0.6826656 | 0.49481818 | 0.99989001 |
| GNA12 | 221.035848 | 0.16555862 | 0.24905791 | 0.66473945 | 0.50621716 | 0.99989001 |
| SDK1 | 146.597785 | -0.0950997 | 0.42585279 | -0.223316 | 0.82328959 | 0.99989001 |
| FOXK1 | 314.488644 | 0.139918 | 0.29991803 | 0.46652081 | 0.64084275 | 0.99989001 |
| AP5Z1 | 48.6812767 | -0.491142 | 0.38284881 | -1.2828614 | 0.19954064 | NA |
| RBAK-RBAKI | 122.069208 | -0.0339414 | 0.2841779 | -0.1194373 | 0.90492894 | 0.99989001 |
| WIPI2 | 116.817844 | 0.10095542 | 0.24822814 | 0.40670416 | 0.68422529 | 0.99989001 |
| TNRC18 | 382.949742 | 0.23100693 | 0.31861242 | 0.72504056 | 0.46842715 | 0.99989001 |
| FBXL18 | 61.8142459 | -0.2574126 | 0.40570169 | -0.6344875 | 0.52576272 | 0.99989001 |
| ACTB | 1707.76835 | -0.3462118 | 0.50894832 | -0.6802494 | 0.49634654 | 0.99989001 |
| FSCN1 | 109.891502 | 0.1856801 | 0.50265547 | 0.36939835 | 0.71183083 | 0.99989001 |
| RNF216 | 372.927707 | -0.1186133 | 0.21794809 | -0.5442271 | 0.58628521 | 0.99989001 |
| CCZ1 | 55.5511114 | 0.14558846 | 0.28338411 | 0.51374954 | 0.60742711 | 0.99989001 |
| PMS2 | 51.5708215 | -0.1005052 | 0.27348982 | -0.3674916 | 0.71325237 | 0.99989001 |
| EIF2AK1 | 98.9506515 | -0.1032475 | 0.1948332 | -0.5299276 | 0.59616216 | 0.99989001 |
| USP42 | 143.208948 | -0.0151374 | 0.30763792 | -0.0492053 | 0.96075567 | 0.99989001 |
| CYTH3 | 385.238616 | -0.1496953 | 0.3188124 | -0.4695403 | 0.63868345 | 0.99989001 |
| RAC1 | 604.134285 | 0.09482916 | 0.28884061 | 0.32830966 | 0.74267754 | 0.99989001 |
| DAGLB | 53.8426885 | 0.00390595 | 0.30276252 | 0.01290103 | 0.98970676 | 0.99989001 |
| ZNF12 | 67.4929349 | 0.28822853 | 0.25324658 | 1.13813397 | 0.25506454 | 0.99989001 |
| CCZ1B | 71.5913792 | -0.4255001 | 0.31690305 | -1.3426824 | 0.17937484 | 0.99989001 |
| C1GALT1 | 99.158889 | -0.0363111 | 0.33543667 | -0.1082501 | 0.91379729 | 0.99989001 |
| MIOS | 57.1399211 | 0.0963079 | 0.28141095 | 0.34223223 | 0.73217613 | 0.99989001 |

| | | | | | | |
|---|---|---|---|---|---|---|
| UMAD1 | 376.284021 | 0.05583866 | 0.23392568 | 0.23870258 | 0.81133621 | 0.99989001 |
| GLCCI1 | 106.198209 | -0.1365336 | 0.38443802 | -0.3551512 | 0.7224763 | 0.99989001 |
| ICA1 | 454.660486 | 0.00732999 | 0.33654693 | 0.02178 | 0.98262345 | 0.99989001 |
| NDUFA4 | 215.518463 | -0.0200607 | 0.57536306 | -0.0348662 | 0.97218642 | 0.99989001 |
| PHF14 | 619.692992 | -0.0752769 | 0.17109169 | -0.4399797 | 0.6599518 | 0.99989001 |
| THSD7A | 2718.26802 | -1.1535778 | 0.46247542 | -2.4943548 | 0.01261863 | 0.49000287 |
| TMEM106B | 140.780175 | -0.1067003 | 0.2790682 | -0.3823448 | 0.70220561 | 0.99989001 |
| SCIN | 330.739059 | -2.7538598 | 0.90396073 | -3.0464374 | 0.00231571 | 0.23620208 |
| ARL4A | 76.7634358 | 0.234982 | 0.66081597 | 0.35559371 | 0.72214486 | 0.99989001 |
| ISPD | 174.041316 | 0.20092696 | 0.48213224 | 0.41674657 | 0.67686379 | 0.99989001 |
| ANKMY2 | 45.3370204 | 0.06187645 | 0.34209156 | 0.18087688 | 0.85646422 | NA |
| BZW2 | 63.0874101 | -0.1506115 | 0.3457878 | -0.4355606 | 0.66315559 | 0.99989001 |
| TSPAN13 | 110.095937 | -0.2655487 | 0.57591299 | -0.4610917 | 0.64473282 | 0.99989001 |
| SNX13 | 414.000147 | -0.1564555 | 0.28107585 | -0.5566309 | 0.57777962 | 0.99989001 |
| HDAC9 | 396.445761 | -0.4163777 | 0.63558323 | -0.6551112 | 0.51239615 | 0.99989001 |
| TWISTNB | 43.5633374 | 0.07662277 | 0.48821283 | 0.15694542 | 0.87528786 | NA |
| ITGB8 | 54.109278 | 0.99036008 | 0.50196521 | 1.97296557 | 0.04849948 | 0.78220964 |
| SP4 | 89.2865109 | -0.292137 | 0.26037767 | -1.1219743 | 0.26187337 | 0.99989001 |
| RAPGEF5 | 3195.35857 | -0.1064804 | 0.45070789 | -0.2362516 | 0.81323748 | 0.99989001 |
| STEAP1B | 305.457674 | -0.1990166 | 0.38509461 | -0.5167993 | 6.05E-01 | 0.99989001 |
| TOMM7 | 153.949504 | 0.01220987 | 0.55518976 | 0.02199224 | 0.98245414 | 0.99989001 |
| FAM126A | 250.63472 | 0.01847758 | 0.33677462 | 0.05486631 | 0.95624497 | 0.99989001 |
| KLHL7 | 99.8930183 | 0.03075784 | 0.22444719 | 0.13703821 | 0.89100059 | 0.99989001 |
| NUPL2 | 86.2123722 | 0.07822396 | 0.32246348 | 0.24258239 | 0.80832892 | 0.99989001 |
| MALSU1 | 60.5944723 | 0.01021315 | 0.2739134 | 0.03728607 | 0.97025691 | 0.99989001 |
| TRA2A | 441.10149 | 0.37266197 | 0.28707896 | 1.29811661 | 0.19424727 | 0.99989001 |
| CCDC126 | 72.9860304 | 0.16107148 | 0.27528468 | 0.58510878 | 0.55847458 | 0.99989001 |
| OSBPL3 | 236.906016 | -0.620731 | 0.30986521 | -2.0032291 | 0.0451527 | 0.77228819 |
| CYCS | 81.5283354 | 0.15144329 | 0.48497805 | 0.31226834 | 0.7548366 | 0.99989001 |
| NFE2L3 | 46.7412394 | 0.22929747 | 0.33617912 | 0.68206934 | 0.49519511 | NA |
| HNRNPA2B1 | 899.582211 | -0.0521477 | 0.26312862 | -0.1981834 | 0.84290155 | 0.99989001 |
| CBX3 | 156.644721 | -0.2007154 | 0.2925801 | -0.6860185 | 0.49270147 | 0.99989001 |
| SKAP2 | 564.869303 | 0.68501767 | 0.27343697 | 2.50521234 | 0.01223779 | 0.49000287 |
| HOTAIRM1 | 88.6515222 | 0.26541954 | 0.32021176 | 0.82888757 | 0.40716803 | 0.99989001 |
| HIBADH | 218.575484 | 0.21584932 | 0.23872465 | 0.90417693 | 0.36590159 | 0.99989001 |
| TAX1BP1 | 419.424072 | 0.08890423 | 0.15725658 | 0.56534502 | 0.57183911 | 0.99989001 |
| JAZF1 | 372.004062 | 0.63279372 | 0.35549543 | 1.78003334 | 0.0750705 | 0.9206525 |
| CREB5 | 227.335585 | -0.2495917 | 0.6439153 | -0.3876158 | 0.69830037 | 0.99989001 |
| AC005162.3 | 86.6622214 | -1.0323123 | 0.7529901 | -1.3709507 | 0.17039034 | 0.99989001 |
| CPVL | 58.0217057 | -1.042776 | 0.78232302 | -1.3329225 | 0.18255724 | 0.99989001 |
| PLEKHA8 | 81.4403119 | -0.4381958 | 0.22051699 | -1.9871296 | 0.04690803 | 0.77516398 |
| MTURN | 90.1537986 | 0.45044041 | 0.35731785 | 1.26061547 | 0.20744742 | 0.99989001 |
| ZNRF2 | 118.019898 | 0.11273096 | 0.23864954 | 0.47237031 | 0.63666249 | 0.99989001 |
| AC006978.2 | 108.906686 | 0.09884197 | 0.20827075 | 0.47458402 | 0.6350835 | 0.99989001 |
| NOD1 | 268.26122 | 0.44056169 | 0.27726821 | 1.58893689 | 0.11207464 | 0.99989001 |
| AC005154.6 | 91.4621511 | -0.056608 | 0.34922191 | -0.1620974 | 0.87122918 | 0.99989001 |
| GARS | 100.152389 | 0.07203918 | 0.37735098 | 0.19090762 | 0.84859797 | 0.99989001 |
| AQP1 | 1056.83819 | -0.0489389 | 0.6266704 | -0.0780936 | 0.9377536 | 0.99989001 |
| LSM5 | 41.6536896 | -0.7989729 | 0.36077419 | -2.2146067 | 0.02678707 | NA |
| AVL9 | 362.873884 | -0.0876977 | 0.21916808 | -0.4001389 | 0.68905425 | 0.99989001 |
| KBTBD2 | 233.349611 | 0.12768913 | 0.25259029 | 0.50551876 | 0.61319452 | 0.99989001 |
| FKBP9 | 203.655309 | -0.4641527 | 0.43271039 | -1.0726637 | 0.28342205 | 0.99989001 |
| NT5C3A | 231.165572 | 0.1026602 | 0.33887677 | 0.30294257 | 0.76193363 | 0.99989001 |
| BBS9 | 507.132853 | 0.17628336 | 0.28379186 | 0.62117131 | 0.53448692 | 0.99989001 |

| | | | | | | |
|---|---|---|---|---|---|---|
| DPY19L1 | 149.664547 | 0.15480697 | 0.41546805 | 0.37260862 | 0.70943976 | 0.99989001 |
| HERPUD2 | 145.952787 | 0.29411004 | 0.20137028 | 1.46054339 | 0.14414079 | 0.99989001 |
| 7-Sep | 527.468844 | 0.01635647 | 0.22989781 | 0.0711467 | 0.943281 | 0.99989001 |
| EEPD1 | 260.679785 | 0.17391021 | 0.55206077 | 0.31502005 | 0.75274643 | 0.99989001 |
| ELMO1 | 2027.6526 | 0.16045517 | 0.15056106 | 1.06571492 | 0.28655253 | 0.99989001 |
| ELMO1-AS1 | 66.8424475 | 0.64187077 | 0.46689416 | 1.37476719 | 0.16920363 | 0.99989001 |
| STARD3NL | 100.269128 | 0.23953537 | 0.17592135 | 1.36160486 | 0.17332262 | 0.99989001 |
| VPS41 | 228.1419 | -0.4019753 | 0.25780429 | -1.5592264 | 0.1189428 | 0.99989001 |
| RALA | 280.885201 | -0.0268788 | 0.34807641 | -0.0772209 | 0.9384478 | 0.99989001 |
| CDK13 | 648.564341 | -0.0402324 | 0.14959939 | -0.2689341 | 0.78798041 | 0.99989001 |
| MPLKIP | 68.5674787 | 0.02052613 | 0.25398067 | 0.08081769 | 0.93558694 | 0.99989001 |
| SUGCT | 88.0918504 | -0.367598 | 0.42750397 | -0.8598703 | 0.38986054 | 0.99989001 |
| GLI3 | 134.764174 | 0.33347003 | 0.35903793 | 0.92878776 | 0.35299909 | 0.99989001 |
| MRPL32 | 59.8192931 | 0.00834761 | 0.3917278 | 0.02130972 | 0.98299859 | 0.99989001 |
| STK17A | 82.6964026 | -0.4906909 | 0.31499692 | -1.5577641 | 0.11928918 | 0.99989001 |
| COA1 | 335.105605 | -0.1595836 | 0.19867286 | -0.8032482 | 0.42183127 | 0.99989001 |
| BLVRA | 45.8538303 | 0.51567085 | 0.296721 | 1.73789805 | 0.08222878 | NA |
| DBNL | 93.9372638 | -0.0439559 | 0.20924768 | -0.2100664 | 0.83361584 | 0.99989001 |
| YKT6 | 37.9596942 | -0.2502637 | 0.44565853 | -0.5615594 | 0.57441624 | NA |
| NUDCD3 | 301.76005 | -0.1816476 | 0.18302402 | -0.9924796 | 0.32096362 | 0.99989001 |
| DDX56 | 41.0378844 | -0.112908 | 0.34924994 | -0.323287 | 0.74647793 | NA |
| TMED4 | 67.705628 | -0.1115721 | 0.30119463 | -0.3704319 | 0.71106069 | 0.99989001 |
| OGDH | 216.717595 | 0.06032421 | 0.22083473 | 0.27316449 | 0.78472677 | 0.99989001 |
| ZMIZ2 | 101.675698 | 0.324912 | 0.29728103 | 1.0929456 | 0.27441769 | 0.99989001 |
| PPIA | 446.68518 | -0.1356487 | 0.51914041 | -0.2612947 | 0.79386522 | 0.99989001 |
| H2AFV | 164.065425 | -0.1047059 | 0.26486478 | -0.3953183 | 0.69260803 | 0.99989001 |
| PURB | 124.530496 | 0.19315992 | 0.25280242 | 0.76407463 | 0.44482277 | 0.99989001 |
| CCM2 | 258.067276 | 0.19976861 | 0.29682258 | 0.67302363 | 0.50093226 | 0.99989001 |
| RAMP3 | 1024.14282 | -0.1946196 | 0.32733243 | -0.5945627 | 0.55213582 | 0.99989001 |
| ADCY1 | 53.3114398 | 0.35992607 | 0.52372827 | 0.6872382 | 4.92E-01 | 1.00E+00 |
| IGFBP3 | 883.470416 | -0.0115505 | 1.30021944 | -0.0088835 | 0.99291211 | 0.99989001 |
| TNS3 | 303.459473 | -0.1681465 | 0.41380472 | -0.4063426 | 0.6844909 | 0.99989001 |
| HUS1 | 99.37126 | -0.2916301 | 0.32489165 | -0.8976226 | 0.36938678 | 0.99989001 |
| UPP1 | 202.596382 | -0.5440615 | 0.41768757 | -1.3025562 | 0.19272633 | 0.99989001 |
| GRB10 | 1825.99345 | 0.64873942 | 0.37406649 | 1.73428906 | 0.08286681 | 0.94799019 |
| SEC61G | 84.9679887 | -0.1520931 | 0.533527 | -0.285071 | 0.77558975 | 0.99989001 |
| EGFR | 139.171437 | 0.5719025 | 0.43086089 | 1.32734835 | 0.18439348 | 0.99989001 |
| LANCL2 | 59.0602709 | -0.8871028 | 0.4428476 | -2.0031784 | 0.04515814 | 0.77228819 |
| VOPP1 | 498.515981 | -0.0530468 | 0.25792347 | -0.2056689 | 0.83704958 | 0.99989001 |
| NIPSNAP2 | 135.026608 | -0.1224069 | 0.23361009 | -0.5239793 | 0.60029292 | 0.99989001 |
| CCT6A | 127.444209 | 0.05435477 | 0.32761255 | 0.16591175 | 0.8682264 | 0.99989001 |
| SUMF2 | 100.533703 | -0.1791275 | 0.21944736 | -0.8162662 | 0.4143479 | 0.99989001 |
| CHCHD2 | 239.176952 | -0.0849316 | 0.51323908 | -0.1654815 | 0.86856501 | 0.99989001 |
| ZNF736 | 99.9793697 | 0.05296742 | 0.24704408 | 0.21440474 | 0.83023145 | 0.99989001 |
| ZNF680 | 130.838835 | 0.14304386 | 0.33795953 | 0.42325738 | 0.67210749 | 0.99989001 |
| ZNF107 | 73.7344385 | -0.2116591 | 0.36403274 | -0.5814289 | 0.56095146 | 0.99989001 |
| ZNF138 | 72.6640781 | -0.3284051 | 0.33366988 | -0.9842217 | 0.3250065 | 0.99989001 |
| ZNF273 | 51.0184429 | 0.28580364 | 0.49059459 | 0.58256582 | 0.56018562 | 0.99989001 |
| ZNF117 | 64.5208119 | 0.09070222 | 0.23623955 | 0.38394174 | 0.70102163 | 0.99989001 |
| VKORC1L1 | 169.781874 | 0.06068532 | 0.32717617 | 0.18548209 | 0.85285093 | 0.99989001 |
| GUSB | 78.5953507 | -0.329798 | 0.22952006 | -1.4369028 | 0.15074562 | 0.99989001 |
| CRCP | 42.1915874 | -0.2232511 | 0.30433375 | -0.7335734 | 0.4632088 | NA |
| TPST1 | 333.017204 | 0.19514636 | 0.25889382 | 0.75376986 | 0.45098743 | 0.99989001 |
| AC027644.4 | 74.4858279 | -0.1658816 | 0.47346868 | -0.350354 | 0.72607308 | 0.99989001 |

| | | | | | | |
|---|---|---|---|---|---|---|
| TMEM248 | 169.747584 | 0.06911873 | 0.2429095 | 0.28454518 | 0.77599261 | 0.99989001 |
| SBDS | 147.623807 | 0.03765371 | 0.41638517 | 0.09043 | 0.92794552 | 0.99989001 |
| TYW1 | 257.451388 | -0.0312267 | 0.17790126 | -0.1755281 | 0.86066472 | 0.99989001 |
| AUTS2 | 1214.6212 | -0.3435779 | 0.20397757 | -1.6843907 | 0.09210618 | 0.99175102 |
| TYW1B | 175.319378 | 0.03065569 | 0.18567344 | 0.16510541 | 0.86886102 | 0.99989001 |
| POM121 | 196.101534 | -0.1148835 | 0.19828587 | -0.5793834 | 0.56233051 | 0.99989001 |
| BAZ1B | 238.540765 | 0.1512064 | 0.22203891 | 0.68099054 | 0.49587748 | 0.99989001 |
| BCL7B | 60.4899231 | -0.1403167 | 0.25294819 | -0.554725 | 0.57908274 | 0.99989001 |
| BUD23 | 73.5865409 | -0.0869562 | 0.38060003 | -0.2284714 | 0.81927981 | 0.99989001 |
| ELN | 141.312659 | 0.07224114 | 0.55692607 | 0.12971406 | 0.89679266 | 0.99989001 |
| EIF4H | 168.428848 | -0.1491161 | 0.32859867 | -0.453794 | 0.64997712 | 0.99989001 |
| CLIP2 | 161.363668 | -0.2846735 | 0.3269529 | -0.8706865 | 0.38392537 | 0.99989001 |
| GTF2IRD1 | 126.561904 | -0.0336723 | 0.37218369 | -0.0904722 | 0.92791196 | 0.99989001 |
| GTF2I | 731.441899 | 0.07460313 | 0.16303386 | 0.45759287 | 0.64724497 | 0.99989001 |
| CASTOR2 | 46.6361593 | 0.56944346 | 0.42635722 | 1.33560179 | 0.18167946 | NA |
| RCC1L | 79.1013689 | -0.0971657 | 0.32456123 | -0.2993755 | 0.76465355 | 0.99989001 |
| GTF2IRD2B | 44.849088 | -0.2279631 | 0.45408846 | -0.5020235 | 0.61565098 | NA |
| POM121C | 156.802194 | -0.1103092 | 0.23601285 | -0.4673864 | 0.64022342 | 0.99989001 |
| HIP1 | 1009.78377 | 0.32728759 | 0.33705842 | 0.97101146 | 0.33154257 | 0.99989001 |
| RHBDD2 | 47.9122158 | -0.0129561 | 0.30102419 | -0.04304 | 0.96566962 | NA |
| POR | 99.3617284 | 0.15424268 | 0.2330995 | 0.66170315 | 0.50816149 | 0.99989001 |
| STYXL1 | 59.5764743 | 0.37216505 | 0.43868235 | 0.84837023 | 0.39623182 | 0.99989001 |
| MDH2 | 74.2357661 | 0.30205947 | 0.31314206 | 0.96460843 | 0.33474097 | 0.99989001 |
| HSPB1 | 659.238772 | 0.04705345 | 0.52069901 | 0.09036592 | 0.92799644 | 0.99989001 |
| YWHAG | 219.165395 | 0.07616675 | 0.27117925 | 0.28087235 | 0.77880831 | 0.99989001 |
| DTX2 | 74.5873983 | 0.26802952 | 0.38711651 | 0.69237429 | 0.48870231 | 0.99989001 |
| CCDC146 | 141.832542 | 0.10850715 | 0.31935627 | 0.33976836 | 0.73403098 | 0.99989001 |
| FGL2 | 265.662999 | 0.74067107 | 0.53841649 | 1.37564709 | 0.16893092 | 0.99989001 |
| GSAP | 98.9509422 | 0.01474531 | 0.48779387 | 0.03022856 | 0.97588477 | 0.99989001 |
| PTPN12 | 1223.81379 | 0.15451172 | 0.14403491 | 1.072738 | 0.28338869 | 0.99989001 |
| RSBN1L | 241.828856 | 0.1581667 | 0.20212834 | 0.78250632 | 0.43391707 | 0.99989001 |
| PHTF2 | 174.343436 | -0.3164978 | 0.24572871 | -1.287997 | 0.19774702 | 0.99989001 |
| MAGI2 | 935.654146 | -0.5823652 | 0.31586254 | -1.8437298 | 0.06522252 | 0.8666545 |
| GNAI1 | 196.76143 | 0.18901919 | 0.42815332 | 0.44147548 | 0.65886881 | 0.99989001 |
| CD36 | 231.213079 | -0.1301654 | 0.97541371 | -0.1334464 | 0.89384038 | 0.99989001 |
| CACNA2D1 | 847.072667 | -0.6397173 | 0.48026616 | -1.3320057 | 0.18285831 | 0.99989001 |
| SEMA3A | 371.808046 | 1.62819604 | 0.65256239 | 2.49508102 | 0.01259284 | 0.49000287 |
| SEMA3D | 436.802513 | -1.100154 | 0.63888936 | -1.7219789 | 0.08507333 | 0.95514629 |
| KIAA1324L | 141.937445 | -0.0093498 | 0.34744993 | -0.0269098 | 0.97853171 | 0.99989001 |
| DMTF1 | 294.431018 | 0.30764204 | 0.35581346 | 0.86461608 | 0.38724955 | 0.99989001 |
| TMEM243 | 85.2662385 | 0.22239973 | 0.32334151 | 0.68781682 | 0.49156814 | 0.99989001 |
| CROT | 64.0572224 | -0.0597883 | 0.3247199 | -0.1841226 | 0.85391726 | 0.99989001 |
| ABCB1 | 136.84454 | -2.6715885 | 1.00600898 | -2.6556309 | 0.00791602 | 0.41229803 |
| SLC25A40 | 54.4886839 | -0.1882957 | 0.31815444 | -0.5918373 | 0.55395953 | 0.99989001 |
| ADAM22 | 46.9289998 | 0.35018341 | 0.39718233 | 0.88166916 | 0.37795574 | NA |
| SRI | 87.6426084 | 0.22328461 | 0.37850735 | 0.58990825 | 0.55525216 | 0.99989001 |
| GTPBP10 | 70.9550051 | 0.08060834 | 0.28969426 | 0.27825314 | 0.78081805 | 0.99989001 |
| CDK14 | 299.493257 | 0.43608094 | 0.36254718 | 1.20282535 | 0.22904391 | 0.99989001 |
| MTERF1 | 153.614956 | 0.33045061 | 0.49790106 | 0.6636873 | 0.50689047 | 0.99989001 |
| AKAP9 | 504.969348 | -0.0080028 | 0.19611492 | -0.0408069 | 0.96744984 | 0.99989001 |
| KRIT1 | 248.971912 | 0.10402621 | 0.25909639 | 0.40149617 | 0.68805485 | 0.99989001 |
| ANKIB1 | 509.503924 | -0.0146379 | 0.20343758 | -0.0719526 | 0.94263962 | 0.99989001 |
| GATAD1 | 92.5293582 | 0.13388668 | 0.3134168 | 0.42718411 | 0.66924523 | 0.99989001 |
| AC007566.1 | 64.235024 | 0.28654207 | 0.46826154 | 0.61192741 | 0.54058579 | 0.99989001 |

| | | | | | | |
|---|---|---|---|---|---|---|
| PEX1 | 76.5397308 | -0.1903135 | 0.22987035 | -0.8279165 | 0.40771776 | 0.99989001 |
| FAM133B | 139.895865 | -0.0800047 | 0.2919429 | -0.2740424 | 0.78405206 | 0.99989001 |
| CDK6 | 433.428267 | -0.1920401 | 0.32973864 | -0.582401 | 0.56029663 | 0.99989001 |
| SAMD9 | 99.4080994 | 0.53868939 | 0.40640479 | 1.32549959 | 0.1850055 | 0.99989001 |
| SAMD9L | 204.312101 | -0.0121929 | 0.68975008 | -0.0176773 | 0.98589628 | 0.99989001 |
| VPS50 | 219.455625 | -0.2310024 | 0.23116462 | -0.9992981 | 0.3176503 | 0.99989001 |
| TFPI2 | 58.0821127 | -0.9995868 | 0.60643121 | -1.6483103 | 0.09928901 | 0.99989001 |
| GNG11 | 569.487606 | -0.3759796 | 0.58345519 | -0.6444018 | 0.51931492 | 0.99989001 |
| BET1 | 87.6003846 | -0.0463439 | 0.38109943 | -0.1216059 | 0.90321112 | 0.99989001 |
| CASD1 | 198.517809 | 0.10311955 | 0.25020823 | 0.41213491 | 0.68024054 | 0.99989001 |
| SGCE | 94.8924611 | 0.65992351 | 0.48513229 | 1.36029599 | 0.17373628 | 0.99989001 |
| PPP1R9A | 147.514402 | -0.4129988 | 0.44861297 | -0.9206127 | 0.35725268 | 0.99989001 |
| PON2 | 172.563768 | 0.18571499 | 0.28979646 | 0.64084631 | 0.52162254 | 0.99989001 |
| PDK4 | 405.522222 | 1.11908624 | 0.95944226 | 1.16639249 | 0.24345579 | 0.99989001 |
| DYNC1I1 | 94.9822906 | 1.32758419 | 0.55743479 | 2.38159551 | 0.01723782 | 0.55366411 |
| SLC25A13 | 449.671914 | -0.4927143 | 0.19481899 | -2.5290874 | 0.01143595 | 0.48516963 |
| SEM1 | 296.693662 | 0.16459598 | 0.30551535 | 0.53874864 | 0.59006031 | 0.99989001 |
| LMTK2 | 169.017566 | 0.01111291 | 0.2811028 | 0.03953326 | 0.96846524 | 0.99989001 |
| TECPR1 | 199.56256 | 0.08239634 | 0.32369523 | 0.25454914 | 0.79907135 | 0.99989001 |
| BRI3 | 403.32007 | -0.3565594 | 0.3412459 | -1.0448753 | 0.2960806 | 0.99989001 |
| BAIAP2L1 | 504.798824 | -0.1489956 | 0.56523774 | -0.263598 | 0.79208967 | 0.99989001 |
| TRRAP | 273.544632 | -0.0644544 | 0.20752123 | -0.3105919 | 0.75611086 | 0.99989001 |
| SMURF1 | 448.065168 | 0.32923498 | 0.26872167 | 1.22518951 | 0.22050383 | 0.99989001 |
| ARPC1B | 112.180641 | -0.3699873 | 0.59911355 | -0.6175579 | 0.5368668 | 0.99989001 |
| PDAP1 | 83.0793582 | -0.1454354 | 0.3889291 | -0.373938 | 0.70845042 | 0.99989001 |
| BUD31 | 56.8451886 | -0.2657563 | 0.32998025 | -0.8053704 | 0.42060598 | 0.99989001 |
| ZNF789 | 67.1696774 | 0.21819547 | 0.28007071 | 0.77907279 | 0.43593684 | 0.99989001 |
| ZNF394 | 64.7101764 | 0.42624491 | 0.31281613 | 1.36260529 | 0.17300694 | 0.99989001 |
| ZKSCAN5 | 41.2219395 | 0.18769341 | 0.33199374 | 0.56535225 | 0.5718342 | NA |
| TRIM4 | 121.060168 | 0.234228 | 0.32818186 | 0.71371404 | 0.47540403 | 0.99989001 |
| ZKSCAN1 | 255.64591 | 0.22495769 | 0.1932808 | 1.16389051 | 0.24446839 | 0.99989001 |
| ZNF3 | 44.4393 | -0.3405142 | 0.44241864 | -0.7696652 | 0.44149854 | NA |
| COPS6 | 88.0225621 | -0.1020587 | 0.33876378 | -0.3012679 | 0.76321018 | 0.99989001 |
| LAMTOR4 | 180.025017 | -0.1983228 | 0.47722217 | -0.4155774 | 0.67771926 | 0.99989001 |
| CASTOR3 | 62.3594847 | 0.27300803 | 0.41417356 | 0.65916334 | 0.50979088 | 0.99989001 |
| SPDYE3 | 64.7808704 | 0.17655471 | 0.39122141 | 0.45129102 | 0.65177982 | 0.99989001 |
| PILRB | 133.89853 | -0.1930123 | 0.38867665 | -0.4965883 | 0.6194794 | 0.99989001 |
| MEPCE | 51.9395713 | 0.03582202 | 0.42780441 | 0.08373457 | 0.93326747 | 0.99989001 |
| PPP1R35 | 53.5993519 | -0.229544 | 0.4247618 | -0.5404064 | 0.5889168 | 0.99989001 |
| TSC22D4 | 72.1261554 | 0.07884732 | 0.27795136 | 0.28367307 | 0.77666093 | 0.99989001 |
| AGFG2 | 72.1150499 | -0.055428 | 0.38630083 | -0.1434842 | 0.88590782 | 0.99989001 |
| GNB2 | 212.262923 | -0.0264176 | 0.41684999 | -0.0633742 | 0.9494685 | 0.99989001 |
| GIGYF1 | 83.1082968 | -0.1187675 | 0.28003398 | -0.424118 | 0.67147974 | 0.99989001 |
| EPHB4 | 295.256337 | -0.111657 | 0.31982703 | -0.3491169 | 0.72700155 | 0.99989001 |
| SLC12A9 | 94.2555202 | 0.0082453 | 0.35224941 | 0.02340756 | 0.98132518 | 0.99989001 |
| TRIP6 | 49.7338501 | -0.2619184 | 0.38479327 | -0.6806731 | 0.49607835 | NA |
| SRRT | 46.6601789 | -0.0133565 | 0.31787075 | -0.0420187 | 0.96648377 | NA |
| TRIM56 | 369.408203 | -0.0819494 | 0.39640677 | -0.2067306 | 0.83622027 | 0.99989001 |
| SERPINE1 | 816.722209 | 1.38713219 | 1.191709 | 1.16398566 | 2.44E-01 | 0.99989001 |
| AP1S1 | 45.3464292 | 0.12500591 | 0.4631342 | 0.26991292 | 0.78722724 | NA |
| PLOD3 | 94.6638278 | -0.0723159 | 0.34250767 | -0.2111365 | 0.83278079 | 0.99989001 |
| ZNHIT1 | 106.027742 | 0.11557758 | 0.4682957 | 0.24680469 | 0.80505938 | 0.99989001 |
| CLDN15 | 140.951373 | -1.8611212 | 0.96060029 | -1.9374564 | 0.05268957 | 0.81270649 |
| FIS1 | 197.534537 | -0.2657622 | 0.45768859 | -0.5806616 | 0.56146851 | 0.99989001 |

| | | | | | | |
|---|---|---|---|---|---|---|
| CUX1 | 849.704621 | 0.43837399 | 0.31290571 | 1.40097789 | 0.16122069 | 0.99989001 |
| SH2B2 | 84.5655132 | 0.87246044 | 0.62597344 | 1.3937659 | 0.16338831 | 0.99989001 |
| PRKRIP1 | 358.497702 | 0.10489815 | 0.27783714 | 0.37755265 | 0.70576294 | 0.99989001 |
| ORAI2 | 98.2358262 | -0.0374657 | 0.40915886 | -0.0915675 | 0.92704166 | 0.99989001 |
| POLR2J | 53.5075714 | -0.332712 | 0.5394265 | -0.6167884 | 0.53737433 | 0.99989001 |
| AC105052.3 | 73.0894038 | 0.71895101 | 0.46519337 | 1.54548853 | 0.12222814 | 0.99989001 |
| FAM185A | 102.976207 | 0.06420959 | 0.26362628 | 0.24356294 | 0.80756933 | 0.99989001 |
| FBXL13 | 49.650395 | -0.3165269 | 0.2996194 | -1.0564299 | 0.29077186 | NA |
| ARMC10 | 69.0627553 | 0.05375012 | 0.30398914 | 0.17681592 | 0.85965299 | 0.99989001 |
| NAPEPLD | 68.2902368 | 0.47971914 | 0.24111882 | 1.98955489 | 0.04663999 | 0.77516398 |
| PMPCB | 84.6529048 | -0.0007395 | 0.26924424 | -0.0027465 | 0.9978086 | 0.99989001 |
| DNAJC2 | 93.8948813 | -0.0724561 | 0.25656032 | -0.2824134 | 0.77762656 | 0.99989001 |
| PSMC2 | 88.0911088 | 0.31295661 | 0.23193835 | 1.34930945 | 0.17723759 | 0.99989001 |
| SLC26A5 | 98.8361665 | 0.35343355 | 0.43000035 | 0.82193783 | 0.41111228 | 0.99989001 |
| RELN | 166.168205 | -0.8351305 | 1.18507137 | -0.704709 | 0.48099135 | 0.99989001 |
| ORC5 | 124.137038 | -0.2469156 | 0.20326087 | -1.2147718 | 0.22445314 | 0.99989001 |
| LHFPL3 | 53.4464407 | -0.1385727 | 0.35191788 | -0.3937642 | 0.69375513 | 0.99989001 |
| LINC01004 | 260.436521 | -0.1561583 | 0.28326792 | -0.5512741 | 0.5814458 | 0.99989001 |
| KMT2E | 1214.4405 | -0.1376304 | 0.41374473 | -0.3326456 | 0.73940178 | 0.99989001 |
| SRPK2 | 910.888433 | -0.1630827 | 0.19574243 | -0.8331497 | 0.40476033 | 0.99989001 |
| PUS7 | 88.5035169 | -0.2087882 | 0.46656426 | -0.4475015 | 0.65451297 | 0.99989001 |
| RINT1 | 56.3821841 | -0.1527843 | 0.24565272 | -0.6219523 | 0.53397325 | 0.99989001 |
| ATXN7L1 | 299.540216 | 0.20386451 | 0.28437372 | 0.7168894 | 0.47344234 | 0.99989001 |
| CDHR3 | 55.8342526 | -0.3749415 | 0.62963966 | -0.5954859 | 0.55151873 | 0.99989001 |
| SYPL1 | 227.951011 | 0.32591241 | 0.32255512 | 1.01040843 | 0.31229965 | 0.99989001 |
| NAMPT | 1457.72557 | -0.1671108 | 0.82882928 | -0.2016227 | 8.40E-01 | 0.99989001 |
| PRKAR2B | 51.85413 | -2.0414533 | 0.65111259 | -3.1353308 | 0.0017166 | 0.20186448 |
| HBP1 | 195.523873 | -0.4287137 | 0.41999117 | -1.0207684 | 0.30736417 | 0.99989001 |
| COG5 | 541.809116 | -0.3028324 | 0.16175385 | -1.8721805 | 0.06118164 | 0.85923309 |
| BCAP29 | 220.597175 | -0.0166059 | 0.38699079 | -0.0429103 | 0.96577304 | 0.99989001 |
| CBLL1 | 47.8996799 | 0.13473745 | 0.30032447 | 0.44863959 | 0.65369167 | NA |
| SLC26A3 | 767.431721 | -0.4034223 | 0.73902453 | -0.5458848 | 0.58514512 | 0.99989001 |
| DLD | 102.207307 | -0.0185196 | 0.22812722 | -0.081181 | 0.93529799 | 0.99989001 |
| LAMB1 | 172.794531 | -0.1582706 | 0.4193879 | -0.3773846 | 0.70588777 | 0.99989001 |
| NRCAM | 126.249065 | -0.70646 | 0.66269438 | -1.0660419 | 0.2864047 | 0.99989001 |
| PNPLA8 | 212.863574 | -0.118792 | 0.36531742 | -0.3251749 | 0.74504874 | 0.99989001 |
| IMMP2L | 1570.03626 | 0.22322976 | 0.25780294 | 0.86589298 | 0.38654885 | 0.99989001 |
| DOCK4 | 5089.02163 | 0.05499356 | 0.26499014 | 0.20753059 | 0.83559551 | 0.99989001 |
| ZNF277 | 182.671772 | 0.31044697 | 0.21319386 | 1.4561722 | 0.14534502 | 0.99989001 |
| IFRD1 | 167.530426 | -0.2553668 | 0.47827945 | -0.533928 | 0.59339135 | 0.99989001 |
| TMEM168 | 178.860901 | -0.0603995 | 0.22938941 | -0.2633054 | 0.79231519 | 0.99989001 |
| BMT2 | 147.121888 | 0.0816213 | 0.24935103 | 0.32733494 | 0.74341457 | 0.99989001 |
| MDFIC | 389.508805 | 0.48488357 | 0.25739164 | 1.88383576 | 0.0595872 | 0.85885114 |
| TES | 162.420042 | 0.01195065 | 0.37632039 | 0.03175659 | 0.97466616 | 0.99989001 |
| CAV2 | 153.706473 | -0.2471291 | 0.35792782 | -0.6904439 | 0.48991511 | 0.99989001 |
| CAV1 | 244.664818 | 0.00190108 | 0.58554338 | 0.0032467 | 0.99740952 | 0.99989001 |
| MET | 145.171873 | -0.3324873 | 0.40155904 | -0.8279912 | 0.40767551 | 0.99989001 |
| CAPZA2 | 519.466042 | 0.38920245 | 0.32880777 | 1.18367778 | 0.23654064 | 0.99989001 |
| ST7 | 335.755584 | 0.04090923 | 0.31350858 | 0.1304884 | 0.89618003 | 0.99989001 |
| LSM8 | 110.492946 | -0.0532198 | 0.24891895 | -0.2138036 | 0.83070025 | 0.99989001 |
| TSPAN12 | 130.070727 | 1.68417529 | 0.39528618 | 4.26064804 | 2.04E-05 | 0.01098962 |
| ING3 | 63.1436837 | 0.23683343 | 0.32720789 | 0.72380108 | 0.46918787 | 0.99989001 |
| CPED1 | 130.979166 | 1.51833643 | 0.46006178 | 3.30028813 | 0.00096586 | 0.1530645 |
| FAM3C | 169.935305 | -0.1737139 | 0.38001433 | -0.4571245 | 0.64758156 | 0.99989001 |

| | | | | | | |
|---|---|---|---|---|---|---|
| AASS | 115.506272 | 0.15464034 | 0.32948853 | 0.46933452 | 0.63883054 | 0.99989001 |
| CADPS2 | 1522.44795 | 0.18758799 | 0.22758632 | 0.82424982 | 0.40979762 | 0.99989001 |
| AC004594.1 | 75.4850261 | 0.35370891 | 0.58612554 | 0.60346953 | 0.54619638 | 0.99989001 |
| NDUFA5 | 98.4217641 | 0.26498899 | 0.27684963 | 0.95715856 | 0.33848723 | 0.99989001 |
| WASL | 209.059157 | 0.39893505 | 0.18527467 | 2.15320877 | 0.03130227 | 0.67281094 |
| POT1 | 135.760229 | -0.3515406 | 0.24312159 | -1.4459457 | 0.14819244 | 0.99989001 |
| ZNF800 | 259.669815 | 0.1765912 | 0.27845887 | 0.63417337 | 0.52596769 | 0.99989001 |
| ARF5 | 46.5166283 | -0.0944404 | 0.41916711 | -0.2253049 | 0.82174212 | NA |
| SND1 | 439.843072 | -0.003532 | 0.33877942 | -0.0104255 | 0.99168177 | 0.99989001 |
| RBM28 | 81.0233142 | -0.3932222 | 0.30904589 | -1.2723747 | 0.20324 | 0.99989001 |
| HILPDA | 102.502663 | -1.0538983 | 1.30245358 | -0.8091638 | 0.41842091 | 0.99989001 |
| METTL2B | 79.744842 | -0.1669404 | 0.27586924 | -0.6051431 | 0.54508395 | 0.99989001 |
| CALU | 225.448325 | -0.1163439 | 0.34021709 | -0.3419695 | 0.73237386 | 0.99989001 |
| ATP6V1F | 108.362428 | -0.2027485 | 0.47293281 | -0.4287047 | 0.66813816 | 0.99989001 |
| TNPO3 | 241.239517 | 0.03445492 | 0.18200663 | 0.18930584 | 0.84985312 | 0.99989001 |
| AHCYL2 | 342.025614 | 0.04049873 | 0.31740946 | 0.12759143 | 0.89847231 | 0.99989001 |
| NRF1 | 315.464792 | 0.12342942 | 0.21369467 | 0.57759708 | 0.56353618 | 0.99989001 |
| UBE2H | 680.198307 | -0.4197295 | 0.25453665 | -1.6489944 | 0.09914878 | 0.99989001 |
| ZC3HC1 | 65.3088411 | -0.087105 | 0.29406617 | -0.2962089 | 0.76707052 | 0.99989001 |
| KLHDC10 | 287.84354 | 0.18469883 | 0.22151369 | 0.83380321 | 0.40439189 | 0.99989001 |
| TMEM209 | 68.1240289 | -0.3098264 | 0.21702121 | -1.4276319 | 0.15339783 | 0.99989001 |
| COPG2 | 75.3618851 | -0.2529721 | 0.441006 | -0.5736251 | 0.56622151 | 0.99989001 |
| AC016831.7 | 2071.59504 | 0.30562547 | 0.36106875 | 0.84644675 | 0.39730357 | 0.99989001 |
| LINC00513 | 151.983835 | 0.39357404 | 0.57633974 | 0.68288548 | 0.49467921 | 0.99989001 |
| MKLN1 | 1486.18067 | 0.08349675 | 0.28364871 | 0.29436675 | 0.76847767 | 0.99989001 |
| PODXL | 1334.45606 | -0.4167032 | 0.34516706 | -1.2072506 | 0.22733563 | 0.99989001 |
| CHCHD3 | 696.830379 | -0.037072 | 0.21119809 | -0.175532 | 0.86066167 | 0.99989001 |
| EXOC4 | 1511.85881 | 0.0581849 | 0.15752865 | 0.36936071 | 0.71185888 | 0.99989001 |
| SLC35B4 | 74.8099467 | -0.2897128 | 0.28443646 | -1.0185501 | 0.30841661 | 0.99989001 |
| AKR1B1 | 248.063564 | -1.0821509 | 1.19707568 | -0.9039954 | 0.36599783 | 0.99989001 |
| CALD1 | 851.900355 | -0.4048021 | 0.31941547 | -1.2673216 | 0.20504032 | 0.99989001 |
| AGBL3 | 92.0178345 | 0.37527654 | 0.4123825 | 0.91002054 | 0.36281168 | 0.99989001 |
| CYREN | 65.5087607 | 0.01797588 | 0.25012681 | 0.07186707 | 0.9427077 | 0.99989001 |
| TMEM140 | 126.089314 | -0.1753537 | 0.39130399 | -0.4481265 | 0.65406192 | 0.99989001 |
| CNOT4 | 449.120026 | 0.11501295 | 0.18698111 | 0.61510466 | 0.5384856 | 0.99989001 |
| NUP205 | 110.500657 | -0.2477953 | 0.25255 | -0.9811734 | 0.32650722 | 0.99989001 |
| STMP1 | 115.979454 | 0.26632426 | 0.33119554 | 0.80412998 | 0.42132191 | 0.99989001 |
| MTPN | 202.085282 | 0.07985843 | 0.2807196 | 0.28447757 | 0.77604442 | 0.99989001 |
| DGKI | 122.435482 | -1.5943235 | 0.56804593 | -2.8066807 | 0.00500548 | 0.34893315 |
| CREB3L2 | 195.928278 | 0.61593053 | 0.23194443 | 2.65550904 | 0.00791888 | 0.41229803 |
| TRIM24 | 241.213171 | 0.36157345 | 0.31348244 | 1.15340893 | 0.24874258 | 0.99989001 |
| KIAA1549 | 118.482256 | -0.9311938 | 0.50373314 | -1.8485855 | 0.06451768 | 0.86610729 |
| ZC3HAV1 | 552.213886 | 0.19198499 | 0.63182741 | 0.30385669 | 0.76123707 | 0.99989001 |
| UBN2 | 209.82747 | 0.31339609 | 0.22513343 | 1.39204597 | 0.16390848 | 0.99989001 |
| LUC7L2 | 77.2822979 | 0.12842718 | 0.30481706 | 0.42132543 | 0.67351746 | 0.99989001 |
| HIPK2 | 2178.16404 | 0.14278795 | 0.44485969 | 0.320973 | 0.74823085 | 0.99989001 |
| TBXAS1 | 110.395914 | 0.99714814 | 0.54786565 | 1.82005959 | 0.06874993 | 0.88856931 |
| PARP12 | 211.210222 | 0.09162364 | 0.53887621 | 0.17002726 | 0.8649887 | 0.99989001 |
| KDM7A | 176.803118 | 0.40799357 | 0.26457295 | 1.54208343 | 0.12305332 | 0.99989001 |
| SLC37A3 | 93.6900928 | 0.10019415 | 0.22078045 | 0.45381803 | 0.64995981 | 0.99989001 |
| MKRN1 | 73.4662082 | 0.10247479 | 0.25455822 | 0.40255936 | 0.68727241 | 0.99989001 |
| NDUFB2 | 124.246269 | 0.22954352 | 0.34355144 | 0.6681489 | 0.50403855 | 0.99989001 |
| BRAF | 871.989744 | 0.05318696 | 0.23102319 | 0.23022349 | 0.81791811 | 0.99989001 |
| AGK | 113.897755 | -0.1756302 | 0.26091453 | -0.6731332 | 0.50086258 | 0.99989001 |

| | | | | | | |
|---|---|---|---|---|---|---|
| AC004918.1 | 71.453831 | -0.4455372 | 0.44098556 | -1.0103215 | 0.31234128 | 0.99989001 |
| KIAA1147 | 188.215676 | 0.15748441 | 0.35799654 | 0.43990483 | 0.66000604 | 0.99989001 |
| SSBP1 | 157.102994 | -0.1553848 | 0.24111764 | -0.6444357 | 0.51929292 | 0.99989001 |
| GSTK1 | 114.56148 | -0.0782477 | 0.33856068 | -0.2311186 | 0.81722266 | 0.99989001 |
| CASP2 | 46.1706056 | 0.1321136 | 0.35673331 | 0.37034276 | 0.71112712 | NA |
| ZYX | 71.5424238 | -0.2846654 | 0.32798576 | -0.8679199 | 0.3854382 | 0.99989001 |
| TCAF2 | 391.356867 | 0.86349859 | 0.48088426 | 1.79564744 | 0.07255061 | 0.9155703 |
| TCAF1 | 148.378434 | 0.62834177 | 0.25583623 | 2.45603126 | 0.01404809 | 0.50734455 |
| AC004889.1 | 84.0633022 | -0.8543138 | 0.37689857 | -2.2666942 | 0.02340891 | 0.60818532 |
| AC074386.1 | 49.3084707 | -0.3330611 | 0.36261885 | -0.9184881 | 0.3583634 | NA |
| TPK1 | 342.898291 | 0.03891625 | 0.32453969 | 0.11991215 | 0.90455274 | 0.99989001 |
| CUL1 | 325.276611 | 0.18464102 | 0.21482168 | 0.85950829 | 0.39006015 | 0.99989001 |
| EZH2 | 49.1088005 | -0.035336 | 0.3515894 | -0.1005037 | 0.91994445 | NA |
| PDIA4 | 80.9731501 | -0.0822276 | 0.47187499 | -0.1742571 | 0.86166343 | 0.99989001 |
| ZNF398 | 69.1418122 | 0.22772825 | 0.38475588 | 0.59187723 | 0.5539328 | 0.99989001 |
| ZNF282 | 44.7507236 | 0.12957416 | 0.36044652 | 0.35948235 | 0.71923428 | NA |
| ZNF783 | 42.2114538 | -0.0682098 | 0.2875403 | -0.2372183 | 0.81248744 | NA |
| ZNF467 | 57.8821637 | 0.64323863 | 0.41306102 | 1.55724845 | 0.11941151 | 0.99989001 |
| ZNF862 | 79.835764 | 0.12015346 | 0.25638921 | 0.46863696 | 0.63932915 | 0.99989001 |
| GIMAP8 | 390.167849 | 0.49047386 | 0.30021277 | 1.63375413 | 0.10231049 | 0.99989001 |
| GIMAP7 | 410.851642 | 0.42029963 | 0.36284487 | 1.15834527 | 0.24672316 | 0.99989001 |
| GIMAP4 | 287.334611 | 0.65735099 | 0.45990332 | 1.4293243 | 0.15291104 | 0.99989001 |
| GIMAP6 | 114.361817 | 0.06098464 | 0.45162908 | 0.13503258 | 0.89258611 | 0.99989001 |
| GIMAP2 | 62.8748613 | 0.59391167 | 0.28283814 | 2.09982883 | 0.0357439 | 0.69356032 |
| NOS3 | 288.208319 | -0.6714268 | 0.36072724 | -1.8613144 | 0.0626998 | 0.85940342 |
| SLC4A2 | 45.5592694 | -0.4088917 | 0.35260444 | -1.1596329 | 0.2461983 | NA |
| FASTK | 36.8511564 | -0.123634 | 0.38156416 | -0.3240188 | 0.74592379 | NA |
| AGAP3 | 148.888945 | 0.20678147 | 0.26267371 | 0.78721798 | 0.43115427 | 0.99989001 |
| NUB1 | 137.547325 | -0.0231704 | 0.32559868 | -0.0711626 | 0.94326835 | 0.99989001 |
| RHEB | 253.028735 | 0.04518729 | 0.23013137 | 0.19635432 | 0.84433284 | 0.99989001 |
| PRKAG2 | 175.735971 | 0.50388448 | 0.45812154 | 1.09989257 | 0.27137893 | 0.99989001 |
| GALNT11 | 173.449519 | 0.21370498 | 0.25683708 | 0.83206435 | 0.40537262 | 0.99989001 |
| KMT2C | 1120.12748 | -0.029005 | 0.217695 | -0.1332366 | 0.89400623 | 0.99989001 |
| ACTR3B | 76.8761715 | 0.11302725 | 0.34659082 | 0.32611149 | 0.74434 | 0.99989001 |
| PAXIP1-AS2 | 59.6002695 | -0.3645475 | 0.4069993 | -0.8956957 | 0.37041529 | 0.99989001 |
| PAXIP1 | 61.2576782 | 0.28954397 | 0.3091938 | 0.93644818 | 0.34904248 | 0.99989001 |
| INSIG1 | 69.8451997 | -1.0333851 | 0.57311249 | -1.8031104 | 0.07137088 | 0.91166457 |
| RBM33 | 428.282302 | 0.07786197 | 0.22816554 | 0.34125209 | 0.73291382 | 0.99989001 |
| LINC01006 | 54.1378974 | -0.4019945 | 0.662417 | -0.6068601 | 0.54394373 | 0.99989001 |
| LMBR1 | 609.325252 | -0.333998 | 0.31452967 | -1.0618966 | 0.28828262 | 0.99989001 |
| NOM1 | 50.8622065 | -0.6986689 | 0.42146006 | -1.6577347 | 0.09737103 | 0.99989001 |
| UBE3C | 477.523602 | -0.1772277 | 0.19104244 | -0.9276878 | 0.35356956 | 0.99989001 |
| DNAJB6 | 320.055969 | 0.1451277 | 0.2531091 | 0.57338002 | 0.56638742 | 0.99989001 |
| PTPRN2 | 83.7457006 | 0.04812733 | 0.30337471 | 0.1586399 | 0.87395259 | 0.99989001 |
| ESYT2 | 673.202467 | -0.5999814 | 0.54861724 | -1.0936248 | 0.27411955 | 0.99989001 |
| WDR60 | 310.558711 | -0.2492928 | 0.27229952 | -0.9155096 | 0.35992419 | 0.99989001 |
| GTPBP6 | 82.7926059 | 0.20817791 | 0.39077474 | 0.53273124 | 0.59421964 | 0.99989001 |
| IL3RA | 160.323689 | -0.4369584 | 0.56570881 | -0.7724086 | 0.43987245 | 0.99989001 |
| SLC25A6 | 188.344535 | 0.31608332 | 0.52867484 | 0.59787849 | 0.54992101 | 0.99989001 |
| ASMTL | 59.2833524 | -0.6765579 | 0.35141486 | -1.92524 | 0.05419935 | 0.81854801 |
| P2RY8 | 240.383486 | -0.3307258 | 0.50808902 | -0.650921 | 0.51509751 | 0.99989001 |
| AKAP17A | 74.3613513 | -0.0665682 | 0.3120979 | -0.2132926 | 0.83109878 | 0.99989001 |
| DHRSX | 234.718279 | 0.22844176 | 0.30373372 | 0.75211195 | 0.45198374 | 0.99989001 |
| CD99 | 324.489273 | 0.38072578 | 0.43123516 | 0.88287278 | 0.37730501 | 0.99989001 |

| | | | | | | |
|---|---|---|---|---|---|---|
| PRKX | 257.420177 | -0.0898009 | 0.26681081 | -0.3365716 | 0.73643988 | 0.99989001 |
| PUDP | 79.3430676 | 0.40448714 | 0.35572853 | 1.1370669 | 0.25551032 | 0.99989001 |
| TBL1X | 340.933035 | 0.24704217 | 0.30512484 | 0.80964292 | 0.41814543 | 0.99989001 |
| SHROOM2 | 282.980439 | 0.62697084 | 0.42865972 | 1.46263064 | 0.14356847 | 0.99989001 |
| WWC3 | 313.592764 | 0.2572904 | 0.19868924 | 1.29493877 | 0.19534135 | 0.99989001 |
| MID1 | 550.563727 | 0.49847494 | 0.22133353 | 2.25214382 | 0.02431318 | 0.61789872 |
| ARHGAP6 | 200.574746 | 0.18491968 | 0.41975956 | 0.44053713 | 0.65954812 | 0.99989001 |
| MSL3 | 55.7559804 | -0.0734104 | 0.40610254 | -0.1807682 | 0.85654954 | 0.99989001 |
| TMSB4X | 2749.07695 | -0.4719572 | 0.63408726 | -0.7443095 | 0.45668926 | 0.99989001 |
| RAB9A | 58.3393271 | -0.0366619 | 0.37049916 | -0.0989528 | 0.92117574 | 0.99989001 |
| OFD1 | 102.309974 | -0.3366971 | 0.33895981 | -0.9933246 | 0.3205518 | 0.99989001 |
| MOSPD2 | 105.375817 | -0.0515775 | 0.32338996 | -0.1594901 | 0.87328278 | 0.99989001 |
| PIR | 150.859241 | 0.35322499 | 0.28275104 | 1.24924384 | 0.2115759 | 0.99989001 |
| BMX | 56.9960576 | 0.63668426 | 0.43704163 | 1.4568046 | 0.14517032 | 0.99989001 |
| CA5B | 125.98088 | -0.0609038 | 0.26827123 | -0.2270232 | 0.82040567 | 0.99989001 |
| ZRSR2 | 57.5332335 | -0.2227149 | 0.35471664 | -0.6278672 | 0.53009093 | 0.99989001 |
| AP1S2 | 266.448425 | -0.2895023 | 0.25061877 | -1.15515 | 0.248029 | 0.99989001 |
| CTPS2 | 147.746157 | 0.11095734 | 0.33434503 | 0.33186477 | 0.73999137 | 0.99989001 |
| SYAP1 | 103.636984 | 0.19251065 | 0.23533339 | 0.81803374 | 0.41333792 | 0.99989001 |
| TXLNG | 86.039767 | -0.0943156 | 0.36969246 | -0.255119 | 0.7986312 | 0.99989001 |
| RBBP7 | 92.2113113 | -0.0326008 | 0.2490459 | -0.1309029 | 0.89585212 | 0.99989001 |
| REPS2 | 53.2234262 | 0.48980432 | 0.44417383 | 1.10273116 | 0.27014393 | 0.99989001 |
| NHS | 118.854948 | -0.6534198 | 0.52689995 | -1.2401212 | 0.21493058 | 0.99989001 |
| SCML1 | 133.678813 | -0.3441675 | 0.36971055 | -0.9309106 | 0.35189983 | 0.99989001 |
| RAI2 | 73.552167 | 1.01690937 | 0.38397828 | 2.64835126 | 0.00808854 | 0.41229803 |
| CDKL5 | 150.040466 | 0.5205111 | 0.27793458 | 1.87278279 | 0.06109839 | 0.85923309 |
| PHKA2 | 65.942127 | -0.1957438 | 0.24316172 | -0.8049943 | 0.42082298 | 0.99989001 |
| PDHA1 | 61.9705459 | 0.37397871 | 0.29449846 | 1.26988341 | 0.20412616 | 0.99989001 |
| SH3KBP1 | 252.825781 | 1.30179304 | 0.39573879 | 3.28952597 | 0.00100356 | 0.15458966 |
| BCLAF3 | 89.8554367 | 0.06783435 | 0.27480766 | 0.24684301 | 0.80502973 | 0.99989001 |
| EIF1AX | 187.413779 | 0.3183312 | 0.27424545 | 1.16075293 | 0.24574239 | 0.99989001 |
| RPS6KA3 | 531.618525 | -0.1016073 | 0.21414775 | -0.4744727 | 0.63516285 | 0.99989001 |
| MBTPS2 | 60.3066668 | -0.5318073 | 0.23565324 | -2.2567366 | 0.02402454 | 0.6148456 |
| SMS | 77.7427035 | -0.0783498 | 0.29861151 | -0.2623804 | 0.79302821 | 0.99989001 |
| ACOT9 | 120.844324 | 0.33287876 | 0.24506175 | 1.35834646 | 0.17435377 | 0.99989001 |
| SAT1 | 602.266421 | -0.4465345 | 0.65802084 | -0.6786023 | 0.49738987 | 0.99989001 |
| APOO | 733.842256 | -0.0921344 | 0.58020789 | -0.1587955 | 0.87382997 | 0.99989001 |
| EIF2S3 | 123.633115 | -0.2171759 | 0.19182666 | -1.1321468 | 0.25757273 | 0.99989001 |
| ZFX | 190.113103 | 0.0916125 | 0.25904451 | 0.35365546 | 0.72359711 | 0.99989001 |
| PDK3 | 65.965079 | 0.24662466 | 0.54614566 | 0.45157305 | 0.65157659 | 0.99989001 |
| POLA1 | 215.580559 | -0.0872406 | 0.26455747 | -0.3297603 | 0.74158106 | 0.99989001 |
| TAB3 | 85.8287071 | 0.09529685 | 0.24466403 | 0.38950084 | 0.69690569 | 0.99989001 |
| DMD | 80.4097844 | -0.1114083 | 0.51292865 | -0.2172003 | 0.82805223 | 0.99989001 |
| TMEM47 | 196.92524 | 0.06788024 | 0.55761177 | 0.12173388 | 0.90310978 | 0.99989001 |
| AF241726.2 | 113.305427 | 0.42191274 | 0.52343496 | 0.80604617 | 0.42021623 | 0.99989001 |
| DYNLT3 | 68.4892216 | 0.13548575 | 0.45109017 | 0.30035181 | 0.76390882 | 0.99989001 |
| RPGR | 535.857267 | 0.44404015 | 0.44683595 | 0.9937431 | 0.32034795 | 0.99989001 |
| TSPAN7 | 138.61189 | 1.0864356 | 0.77940834 | 1.3939235 | 0.16334071 | 0.99989001 |
| BCOR | 158.035445 | 0.717447 | 0.31099911 | 2.30691013 | 0.02105983 | 0.59254339 |
| ATP6AP2 | 151.450652 | -0.0189391 | 0.32053281 | -0.0590862 | 0.95288346 | 0.99989001 |
| CXorf38 | 70.0638136 | 0.23333254 | 0.22587673 | 1.03300828 | 0.30160002 | 0.99989001 |
| MED14 | 154.684585 | 0.02580564 | 0.20878799 | 0.12359734 | 0.9016341 | 0.99989001 |
| USP9X | 454.504629 | 0.19868605 | 0.27195401 | 0.73058694 | 0.46503149 | 0.99989001 |
| DDX3X | 365.866047 | -0.2175669 | 0.3326789 | -0.6539847 | 0.51312165 | 0.99989001 |

| | | | | | | |
|---|---|---|---|---|---|---|
| CASK | 391.015068 | 0.08312592 | 0.2092261 | 0.39730187 | 0.69114487 | 0.99989001 |
| MAOA | 63.0974276 | 0.77381367 | 0.55029331 | 1.40618404 | 0.15966948 | 0.99989001 |
| KDM6A | 686.703128 | -0.0883045 | 0.37869158 | -0.2331833 | 0.81561909 | 0.99989001 |
| CXorf36 | 433.821494 | -0.6923929 | 0.24714283 | -2.8015903 | 0.00508514 | 0.34893315 |
| MIR222HG | 185.363205 | -0.9693779 | 0.67633338 | -1.4332842 | 0.15177665 | 0.99989001 |
| KRBOX4 | 129.145218 | 0.35438401 | 0.28086366 | 1.26176528 | 0.20703326 | 0.99989001 |
| RP2 | 49.8147278 | -0.268568 | 0.2585126 | -1.0388973 | 0.29885248 | NA |
| JADE3 | 179.32816 | -0.1507769 | 0.20759978 | -0.7262863 | 0.46766329 | 0.99989001 |
| NDUFB11 | 95.9174655 | 0.19274151 | 0.5332336 | 0.36145793 | 0.71775715 | 0.99989001 |
| RBM10 | 88.454117 | 0.10356928 | 0.27225189 | 0.38041711 | 0.70363582 | 0.99989001 |
| UBA1 | 130.248016 | -0.0705837 | 0.22928788 | -0.3078389 | 0.75820489 | 0.99989001 |
| CDK16 | 44.6365892 | -0.0531173 | 0.31651287 | -0.1678202 | 0.86672473 | NA |
| USP11 | 39.6020757 | 0.50297935 | 0.32397658 | 1.55251764 | 0.12053841 | NA |
| ZNF41 | 50.7654059 | 0.11581373 | 0.27294363 | 0.42431375 | 0.67133701 | 0.99989001 |
| ARAF | 40.6275187 | 0.0770453 | 0.30773744 | 0.2503605 | 0.80230857 | NA |
| TIMP1 | 579.744294 | -0.4459752 | 0.76291505 | -0.5845673 | 0.55883867 | 0.99989001 |
| ZNF81 | 93.9193463 | 0.24549879 | 0.2723346 | 0.90146016 | 0.36734371 | 0.99989001 |
| RBM3 | 152.45696 | -0.3129959 | 0.46634808 | -0.6711636 | 0.5021163 | 0.99989001 |
| WDR13 | 89.2190027 | -0.3169381 | 0.24416917 | -1.2980266 | 0.1942782 | 0.99989001 |
| HDAC6 | 47.566527 | -0.0913892 | 0.24542763 | -0.3723674 | 0.70961934 | NA |
| PCSK1N | 51.372933 | 0.6630962 | 0.62247576 | 1.06525626 | 0.28675998 | 0.99989001 |
| OTUD5 | 92.2648426 | 0.09302771 | 0.22392137 | 0.41544811 | 0.6778139 | 0.99989001 |
| GRIPAP1 | 97.1414637 | -0.0038341 | 0.26320655 | -0.0145669 | 0.98837772 | 0.99989001 |
| TFE3 | 63.1839509 | 0.13763839 | 0.26748165 | 0.51457132 | 0.60685261 | 0.99989001 |
| WDR45 | 52.1460375 | 0.11573584 | 0.24014863 | 0.48193419 | 0.62985269 | 0.99989001 |
| CLCN5 | 140.31215 | 0.2969072 | 0.32811551 | 0.90488622 | 0.36552567 | 0.99989001 |
| SHROOM4 | 545.776688 | 0.20585505 | 0.23118893 | 0.89041917 | 0.37324086 | 0.99989001 |
| MAGED1 | 89.7206409 | 0.03883347 | 0.32535738 | 0.11935634 | 0.90499305 | 0.99989001 |
| GPR173 | 79.9869285 | 1.16228766 | 0.36089219 | 3.22059526 | 0.00127925 | 0.1693992 |
| TSPYL2 | 43.123247 | -0.5331406 | 0.41966449 | -1.2703972 | 0.20394319 | NA |
| KANTR | 179.729185 | -0.1666539 | 0.31408124 | -0.5306077 | 0.59569067 | 0.99989001 |
| KDM5C | 88.6743422 | -0.1210398 | 0.28448329 | -0.4254726 | 0.67049218 | 0.99989001 |
| IQSEC2 | 58.7040636 | 0.17886118 | 0.352649 | 0.50719322 | 0.61201925 | 0.99989001 |
| SMC1A | 114.834707 | -0.5483806 | 0.24947081 | -2.1981756 | 0.0279366 | 0.65805899 |
| HSD17B10 | 41.8663368 | 0.2637316 | 0.54392516 | 0.48486744 | 0.62777038 | NA |
| HUWE1 | 412.810845 | 0.07009297 | 0.18231828 | 0.38445386 | 0.70064208 | 0.99989001 |
| PHF8 | 126.390428 | -0.0381895 | 0.16900703 | -0.2259641 | 0.82122931 | 0.99989001 |
| FAM120C | 71.5434432 | -0.0784784 | 0.26208931 | -0.2994339 | 0.76460897 | 0.99989001 |
| MAGED2 | 84.1734341 | 0.03850696 | 0.51497096 | 0.074775 | 0.94039373 | 0.99989001 |
| RRAGB | 115.359746 | -0.5665764 | 0.24402523 | -2.3217942 | 0.02024402 | 0.58699182 |
| AL445472.1 | 114.134718 | -0.1242138 | 0.38735794 | -0.3206693 | 0.74846103 | 0.99989001 |
| KLF8 | 47.9164278 | 0.17650676 | 0.48311134 | 0.36535421 | 0.71484701 | NA |
| UBQLN2 | 38.5725472 | 0.28086819 | 0.38196581 | 0.73532285 | 0.46214289 | NA |
| NBDY | 327.422251 | -0.0021414 | 0.21158723 | -0.0101205 | 0.99192515 | 0.99989001 |
| ARHGEF9 | 73.9761462 | 0.06198149 | 0.35213565 | 0.17601594 | 0.86028142 | 0.99989001 |
| MTMR8 | 59.66408 | -0.1364914 | 0.30744438 | -0.4439549 | 0.65707517 | 0.99989001 |
| ZC4H2 | 53.9030066 | -0.3959221 | 0.35264825 | -1.1227112 | 0.26156017 | 0.99989001 |
| MSN | 1054.37579 | 0.00085163 | 0.28392864 | 0.00299945 | 0.99760679 | 0.99989001 |
| HEPH | 51.2263272 | 0.16332777 | 0.62701491 | 0.26048467 | 0.79448994 | 0.99989001 |
| AR | 218.680276 | 0.18239707 | 0.31108301 | 0.58632926 | 0.55765427 | 0.99989001 |
| OPHN1 | 389.849434 | 0.00103125 | 0.19278715 | 0.00534917 | 0.995732 | 0.99989001 |
| YIPF6 | 115.055737 | -0.1552707 | 0.26305783 | -0.5902531 | 0.55502099 | 0.99989001 |
| STARD8 | 130.746667 | 0.04210496 | 0.24362016 | 0.17283038 | 0.86278476 | 0.99989001 |
| EFNB1 | 225.446188 | -0.2466363 | 0.33148553 | -0.7440333 | 0.45685632 | 0.99989001 |

| | | | | | | |
|---|---|---|---|---|---|---|
| EDA | 82.6011732 | 0.6409624 | 0.54938202 | 1.16669709 | 0.24333272 | 0.99989001 |
| IGBP1 | 79.6265504 | 0.04584709 | 0.2383691 | 0.19233656 | 0.84747858 | 0.99989001 |
| MED12 | 42.4094566 | -0.4911378 | 0.30455368 | -1.6126478 | 0.10682104 | NA |
| NONO | 140.890205 | -0.2576126 | 0.31529994 | -0.8170398 | 0.41390569 | 0.99989001 |
| TAF1 | 336.964254 | -0.1545132 | 0.24557902 | -0.6291792 | 0.52923172 | 0.99989001 |
| OGT | 368.687621 | -0.1533699 | 0.26523358 | -0.5782447 | 0.5630989 | 0.99989001 |
| NHSL2 | 251.740624 | -0.0424398 | 0.36731276 | -0.1155414 | 0.90801598 | 0.99989001 |
| PIN4 | 88.395537 | 0.15063841 | 0.29726439 | 0.50674893 | 0.61233099 | 0.99989001 |
| RPS4X | 412.253237 | -0.0113419 | 0.53832175 | -0.021069 | 0.98319064 | 0.99989001 |
| HDAC8 | 613.955948 | -0.0186187 | 0.21007449 | -0.088629 | 0.92937678 | 0.99989001 |
| PHKA1 | 62.1669104 | -0.7458266 | 0.52470003 | -1.4214342 | 0.15519056 | 0.99989001 |
| CHIC1 | 170.228472 | -0.6436247 | 0.32442905 | -1.9838688 | 0.04727047 | 0.77516398 |
| XIST | 404.008731 | -1.1195365 | 4.21521922 | -0.2655939 | 0.79055199 | 0.99989001 |
| JPX | 472.231469 | -0.1820902 | 0.24356102 | -0.7476163 | 0.45469164 | 0.99989001 |
| FTX | 3212.3435 | 0.13359352 | 0.29052533 | 0.45983432 | 0.64563515 | 0.99989001 |
| SLC16A2 | 51.7958228 | 0.34671833 | 0.32265913 | 1.07456538 | 0.28256936 | 0.99989001 |
| RLIM | 96.8742106 | -0.3895371 | 0.25181022 | -1.546947 | 0.12187602 | 0.99989001 |
| ABCB7 | 152.190161 | -0.1209823 | 0.19657369 | -0.6154554 | 0.53825403 | 0.99989001 |
| ATRX | 947.43061 | 0.03926443 | 0.19200586 | 0.20449598 | 0.83796593 | 0.99989001 |
| MAGT1 | 167.363731 | 0.30145857 | 0.16908394 | 1.78289301 | 0.07460372 | 0.9206525 |
| COX7B | 132.602786 | 0.08346863 | 0.42224289 | 0.19767919 | 0.84329607 | 0.99989001 |
| PGK1 | 261.087455 | -0.0724666 | 0.3665837 | -0.1976809 | 0.84329472 | 0.99989001 |
| ITM2A | 105.425708 | -0.3084191 | 1.01810432 | -0.3029347 | 0.76193965 | 0.99989001 |
| BRWD3 | 274.989657 | -0.1361881 | 0.23839023 | -0.5712822 | 0.56780834 | 0.99989001 |
| HMGN5 | 49.1505859 | -0.1435715 | 0.34449549 | -0.4167588 | 0.67685483 | NA |
| SH3BGRL | 324.088311 | 0.03550254 | 0.21704734 | 0.16357051 | 0.87006927 | 0.99989001 |
| HDX | 116.136136 | 0.13763386 | 0.3556475 | 0.38699514 | 0.6987598 | 0.99989001 |
| APOOL | 102.895223 | -0.05854 | 0.23506177 | -0.249041 | 0.80332909 | 0.99989001 |
| CHM | 223.920988 | -0.0668579 | 0.21329517 | -0.3134525 | 0.7539369 | 0.99989001 |
| KLHL4 | 94.4088766 | -0.8025105 | 0.54182278 | -1.481131 | 0.13857168 | 0.99989001 |
| DIAPH2 | 770.111799 | -0.021871 | 0.28616415 | -0.0764283 | 0.93907837 | 0.99989001 |
| SYTL4 | 115.81495 | -0.4209754 | 0.39539373 | -1.0646991 | 0.28701212 | 0.99989001 |
| TRMT2B | 53.618092 | -0.3147681 | 0.39735042 | -0.7921674 | 0.42826305 | 0.99989001 |
| HNRNPH2 | 62.3583596 | -0.1122527 | 0.46542624 | -0.2411825 | 0.80941365 | 0.99989001 |
| ARMCX4 | 248.088244 | 0.14923247 | 0.42132498 | 0.354198 | 0.7231905 | 0.99989001 |
| ARMCX3 | 97.4938225 | 0.10639023 | 0.33750727 | 0.31522351 | 0.75259196 | 0.99989001 |
| ZMAT1 | 91.7121132 | -0.1165897 | 0.38774683 | -0.3006851 | 0.7636546 | 0.99989001 |
| BHLHB9 | 44.0491977 | 0.10327848 | 0.46050307 | 0.22427316 | 0.82254476 | NA |
| LINC00630 | 130.615093 | 0.14371312 | 0.28492443 | 0.50439029 | 0.61398714 | 0.99989001 |
| BEX4 | 49.3795984 | 0.1968665 | 0.46374418 | 0.4245153 | 0.67119005 | NA |
| TCEAL8 | 56.5065389 | -0.0629463 | 0.49594948 | -0.1269209 | 0.89900302 | 0.99989001 |
| TCEAL9 | 164.789252 | 0.32392696 | 0.48656362 | 0.6657443 | 0.50557455 | 0.99989001 |
| BEX3 | 170.668996 | -0.0274557 | 0.42235339 | -0.0650066 | 0.94816877 | 0.99989001 |
| TCEAL4 | 129.443012 | -0.0703069 | 0.3279715 | -0.2143689 | 0.83025938 | 0.99989001 |
| TCEAL3 | 45.8759621 | -0.1764061 | 0.34749153 | -0.5076558 | 0.61169473 | NA |
| MORF4L2 | 175.579637 | -0.3372373 | 0.31872514 | -1.058082 | 0.29001805 | 0.99989001 |
| SLC25A53 | 83.0344356 | 0.01785828 | 0.25150388 | 0.07100599 | 0.94339299 | 0.99989001 |
| FAM199X | 100.00381 | 0.35828114 | 0.23412363 | 1.53030748 | 0.12594064 | 0.99989001 |
| TBC1D8B | 56.9739201 | 0.24956142 | 0.38880958 | 0.64186026 | 0.52096392 | 0.99989001 |
| MORC4 | 47.6905925 | 0.40094851 | 0.41805517 | 0.95908039 | 0.33751825 | NA |
| RBM41 | 99.0714925 | 0.25875092 | 0.27385341 | 0.94485193 | 0.34473447 | 0.99989001 |
| TSC22D3 | 262.212531 | -0.0198859 | 0.58589476 | -0.033941 | 0.97292417 | 0.99989001 |
| MID2 | 115.881935 | 0.34125709 | 0.33737747 | 1.01149935 | 0.31177749 | 0.99989001 |
| ATG4A | 56.2484801 | 0.08789453 | 0.25600405 | 0.34333256 | 0.73134829 | 0.99989001 |

| | | | | | | |
|---|---|---|---|---|---|---|
| ACSL4 | 335.135759 | 0.07365537 | 0.28142638 | 0.26172164 | 0.79353606 | 0.99989001 |
| TMEM164 | 128.293006 | -0.055716 | 0.22832695 | -0.2440185 | 0.8072165 | 0.99989001 |
| AMMECR1 | 133.748398 | -0.0606818 | 0.23375589 | -0.2595947 | 0.79517643 | 0.99989001 |
| ALG13 | 358.907763 | 0.01792465 | 0.2819814 | 0.06356678 | 0.94931519 | 0.99989001 |
| LRCH2 | 89.588818 | -0.3051394 | 0.46291552 | -0.6591687 | 0.50978742 | 0.99989001 |
| PLS3 | 416.722956 | -0.2956862 | 0.35153027 | -0.84114 | 0.40026953 | 0.99989001 |
| DANT2 | 90.0864259 | 0.13704999 | 1.13537999 | 0.12070847 | 0.90392195 | 0.99989001 |
| WDR44 | 182.766808 | -0.0441161 | 0.17121013 | -0.2576723 | 0.79665983 | 0.99989001 |
| IL13RA1 | 259.15006 | -0.2472962 | 0.4538401 | -0.5448971 | 0.58582429 | 0.99989001 |
| PGRMC1 | 105.694493 | 0.2123886 | 0.43009411 | 0.49381891 | 0.62143408 | 0.99989001 |
| SLC25A43 | 73.6671223 | 0.52380265 | 0.41993556 | 1.24734053 | 0.21227266 | 0.99989001 |
| SLC25A5 | 116.358655 | 0.49565667 | 0.56249608 | 0.88117355 | 0.37822389 | 0.99989001 |
| UBE2A | 70.2793229 | -0.0633921 | 0.28385504 | -0.2233257 | 0.82328199 | 0.99989001 |
| 6-Sep | 194.866048 | 0.30788738 | 0.31124698 | 0.98920598 | 0.32256237 | 0.99989001 |
| UPF3B | 109.07656 | -0.5185293 | 0.2939916 | -1.7637555 | 0.07777316 | 0.92274869 |
| NDUFA1 | 110.132395 | 0.08405198 | 0.43713105 | 0.19228097 | 0.84752213 | 0.99989001 |
| NKAP | 49.4583866 | -0.1310598 | 0.29751394 | -0.4405165 | 0.65956304 | NA |
| LAMP2 | 223.464462 | -0.2091465 | 0.29778846 | -0.7023324 | 0.48247191 | 0.99989001 |
| CUL4B | 91.4644076 | 0.13738651 | 0.23937486 | 0.57393876 | 0.56600925 | 0.99989001 |
| MCTS1 | 64.4966697 | -0.1209222 | 0.21761039 | -0.5556821 | 0.57842823 | 0.99989001 |
| THOC2 | 409.938171 | -0.0983912 | 0.21735691 | -0.4526711 | 0.65078563 | 0.99989001 |
| XIAP | 309.515785 | 0.16149574 | 0.18358139 | 0.8796956 | 0.37902424 | 0.99989001 |
| STAG2 | 439.709484 | 0.01837501 | 0.22228216 | 0.08266526 | 0.93411771 | 0.99989001 |
| SMARCA1 | 100.165658 | -0.0830471 | 0.41726149 | -0.199029 | 0.84224009 | 0.99989001 |
| OCRL | 61.7016865 | -0.2777817 | 0.23571982 | -1.1784401 | 0.2386212 | 0.99989001 |
| ZDHHC9 | 66.9420124 | -0.3124335 | 0.34192287 | -0.9137543 | 0.36084596 | 0.99989001 |
| BCORL1 | 43.6633215 | 0.44571071 | 0.439515 | 1.01409669 | 0.31053662 | NA |
| ELF4 | 53.2976415 | -0.0896513 | 0.24861839 | -0.3605981 | 0.71839989 | 0.99989001 |
| AIFM1 | 51.5660217 | 0.4752371 | 0.34440581 | 1.37987539 | 0.16762502 | 0.99989001 |
| ZNF280C | 47.1308762 | 0.14001846 | 0.28189191 | 0.49670975 | 0.61939375 | NA |
| ENOX2 | 185.411663 | -0.0982869 | 0.29298324 | -0.3354692 | 0.73727117 | 0.99989001 |
| RAP2C | 47.7571995 | 0.07526279 | 0.32270212 | 0.23322682 | 0.81558528 | NA |
| RAP2C-AS1 | 92.4215933 | 0.3925753 | 0.44011049 | 0.89199261 | 0.37239689 | 0.99989001 |
| PHF6 | 80.4545274 | 0.11706797 | 0.22503703 | 0.52021648 | 0.6029127 | 0.99989001 |
| HPRT1 | 124.961282 | -0.0103369 | 0.27949561 | -0.036984 | 0.97049774 | 0.99989001 |
| FAM122B | 54.8275513 | -0.224316 | 0.28464421 | -0.7880573 | 0.43066317 | 0.99989001 |
| FAM122C | 93.5520218 | 0.14570145 | 0.37938258 | 0.38404887 | 0.70094223 | 0.99989001 |
| RTL8C | 64.5265979 | 0.06119799 | 0.424598 | 0.14413161 | 0.88539654 | 0.99989001 |
| RTL8A | 49.8844225 | -0.1366696 | 0.55497964 | -0.2462606 | 0.80548051 | NA |
| ZNF75D | 131.855452 | 0.46836845 | 0.28701191 | 1.63187808 | 0.10270517 | 0.99989001 |
| INTS6L | 76.0738404 | -0.1617019 | 0.33980077 | -0.4758727 | 0.63416507 | 0.99989001 |
| SLC9A6 | 52.527702 | 0.09699153 | 0.27643367 | 0.35086727 | 0.72568792 | 0.99989001 |
| FHL1 | 109.42409 | 0.03036797 | 0.3646796 | 0.08327302 | 0.93363446 | 0.99989001 |
| MAP7D3 | 62.608798 | 0.04686422 | 0.23339245 | 0.2007958 | 0.84085825 | 0.99989001 |
| HTATSF1 | 40.6951221 | 0.12743745 | 0.44364413 | 0.28725153 | 0.77391974 | NA |
| ARHGEF6 | 181.95316 | 0.60723674 | 0.25087801 | 2.4204463 | 0.01550147 | 0.52234412 |
| AL683813.1 | 59.7991812 | 0.83935315 | 0.42345712 | 1.98214437 | 0.04746308 | 0.77516398 |
| RBMX | 277.307821 | 0.09613145 | 0.1788034 | 0.53763773 | 0.59082718 | 0.99989001 |
| FGF13 | 117.058575 | 0.68625395 | 0.70645951 | 0.97139884 | 0.33134971 | 0.99989001 |
| ATP11C | 650.580376 | -0.2669268 | 0.38166897 | -0.6993673 | 0.48432252 | 0.99989001 |
| FMR1 | 121.504236 | 0.03085076 | 0.28467895 | 0.10837036 | 0.91370191 | 0.99989001 |
| IDS | 75.8853459 | 0.30568209 | 0.37441313 | 0.81642994 | 0.41425428 | 0.99989001 |
| LINC00894 | 146.989799 | 0.03948504 | 0.40785639 | 0.09681113 | 0.92287638 | 0.99989001 |
| MAMLD1 | 69.8208809 | 0.43617471 | 0.41564956 | 1.04938089 | 0.29400285 | 0.99989001 |

| | | | | | | |
|---|---|---|---|---|---|---|
| MTM1 | 178.613157 | 0.34788985 | 0.20446373 | 1.70147461 | 0.0888539 | 0.97339514 |
| MTMR1 | 95.6415955 | -0.1847131 | 0.22955709 | -0.8046501 | 0.42102162 | 0.99989001 |
| CD99L2 | 117.807955 | -0.488864 | 0.19756526 | -2.474443 | 0.01334441 | 0.50111255 |
| VMA21 | 62.8623672 | -0.2880938 | 0.45456958 | -0.6337726 | 0.52622921 | 0.99989001 |
| GABRE | 172.287942 | 0.53714255 | 0.55768793 | 0.96315973 | 0.33546737 | 0.99989001 |
| BGN | 81.4203561 | 0.93550528 | 1.0826406 | 0.86409588 | 0.38753522 | 0.99989001 |
| BCAP31 | 151.560237 | -0.3622639 | 0.45970275 | -0.7880395 | 0.43067357 | 0.99989001 |
| IDH3G | 39.8778289 | -0.2245673 | 0.2906715 | -0.7725811 | 0.43977034 | NA |
| SSR4 | 75.2891996 | -0.2418744 | 0.72420731 | -0.333985 | 0.73839087 | 0.99989001 |
| ARHGAP4 | 44.4887243 | -0.5805149 | 0.79950229 | -0.7260954 | 0.46778029 | NA |
| HCFC1 | 51.7674669 | 0.02713618 | 0.28709563 | 0.09451965 | 0.92469637 | 0.99989001 |
| IRAK1 | 45.4321909 | -0.9302487 | 0.45620047 | -2.0391227 | 0.04143778 | NA |
| MECP2 | 451.805692 | -0.7685628 | 0.2761569 | -2.7830656 | 0.00538479 | 0.35652986 |
| FLNA | 320.159772 | -0.7251072 | 0.49384541 | -1.4682879 | 0.14202604 | 0.99989001 |
| EMD | 55.8392023 | -0.2775164 | 0.39762808 | -0.6979295 | 0.48522131 | 0.99989001 |
| RPL10 | 814.053377 | -0.0562221 | 0.63435828 | -0.0886284 | 0.92937727 | 0.99989001 |
| DNASE1L1 | 49.8649413 | -0.4791494 | 0.3598492 | -1.3315284 | 0.18301522 | NA |
| TAZ | 80.8456656 | -0.1041276 | 0.30760044 | -0.3385157 | 0.7349746 | 0.99989001 |
| ATP6AP1 | 103.59893 | 0.11982228 | 0.42918858 | 0.2791833 | 0.78010416 | 0.99989001 |
| GDI1 | 87.460198 | 0.25546924 | 0.30424223 | 0.83969026 | 0.40108208 | 0.99989001 |
| FAM50A | 51.540734 | -0.607995 | 0.31276404 | -1.9439416 | 0.05190249 | 0.81270649 |
| GAB3 | 67.7184432 | 0.183027 | 0.27631969 | 0.66237406 | 0.50773153 | 0.99989001 |
| DKC1 | 44.3502287 | 0.02070367 | 0.35629947 | 0.05810749 | 0.95366301 | NA |
| F8 | 727.46561 | 0.78510381 | 0.58780134 | 1.33566183 | 0.18165983 | 0.99989001 |
| FUNDC2 | 89.6535334 | 0.08065321 | 0.35018136 | 0.23031839 | 0.81784437 | 0.99989001 |
| BRCC3 | 66.4882516 | -0.0207364 | 0.28582834 | -0.0725483 | 0.94216559 | 0.99989001 |
| VBP1 | 44.589107 | -0.1013358 | 0.2720656 | -0.3724684 | 0.70954417 | NA |
| CLIC2 | 206.180728 | 0.20735169 | 0.24082608 | 0.86100181 | 0.38923704 | 0.99989001 |
| TMLHE-AS1 | 70.7222913 | -0.1193552 | 0.4557982 | -0.2618598 | 0.7934295 | 0.99989001 |
| TMLHE | 147.019709 | 0.01542117 | 0.4048475 | 0.03809132 | 0.96961487 | 0.99989001 |
| VAMP7 | 71.4258962 | 0.07866104 | 0.26506191 | 0.29676477 | 0.7666461 | 0.99989001 |
| FBXO25 | 135.591346 | -0.3416354 | 0.29913234 | -1.1420878 | 0.25341752 | 0.99989001 |
| TDRP | 225.528783 | -0.1475459 | 0.22473806 | -0.6565237 | 5.11E-01 | 0.99989001 |
| ERICH1 | 324.178715 | 0.00422151 | 0.45023562 | 0.00937622 | 0.99251897 | 0.99989001 |
| DLGAP2 | 109.797816 | 0.50713377 | 0.73382401 | 0.69108365 | 0.48951297 | 0.99989001 |
| CSMD1 | 154.407632 | 1.33079085 | 1.35698894 | 0.98069396 | 0.32674369 | 0.99989001 |
| MCPH1 | 458.274281 | -0.0666493 | 0.19050968 | -0.3498474 | 0.7264532 | 0.99989001 |
| ANGPT2 | 261.644542 | -1.019283 | 0.9309101 | -1.0949317 | 0.27354656 | 0.99989001 |
| MCPH1-AS1 | 48.9455467 | 0.28411391 | 0.34126125 | 0.83254078 | 0.40510377 | NA |
| AGPAT5 | 122.410294 | -0.4067566 | 0.30942063 | -1.314575 | 0.18865279 | 0.99989001 |
| DEFB1 | 130.27819 | 0.81469932 | 0.53143379 | 1.53302129 | 0.12527061 | 0.99989001 |
| FAM85B | 134.282249 | 0.31131396 | 0.51697918 | 0.6021789 | 0.54705506 | 0.99989001 |
| PRAG1 | 114.615898 | 1.24869428 | 0.56418024 | 2.21328964 | 0.02687768 | 0.64986727 |
| MFHAS1 | 263.035335 | 0.2685483 | 0.28206528 | 0.95207851 | 0.34105717 | 0.99989001 |
| ERI1 | 73.2860503 | 0.15248233 | 0.3076956 | 0.49556227 | 0.62020328 | 0.99989001 |
| PPP1R3B | 88.1147362 | -0.7355266 | 0.56184021 | -1.3091385 | 0.19048745 | 0.99989001 |
| TNKS | 554.944082 | -0.0540626 | 0.22234615 | -0.2431459 | 0.80789235 | 0.99989001 |
| MSRA | 361.487451 | -0.2719476 | 0.29084157 | -0.935037 | 0.34976923 | 0.99989001 |
| SOX7 | 142.51075 | -0.0050008 | 0.57871157 | -0.0086413 | 0.99310533 | 0.99989001 |
| AC105001.1 | 71.0729182 | 0.26632512 | 0.61622177 | 0.43219037 | 0.66560306 | 0.99989001 |
| XKR6 | 331.827554 | -0.9806076 | 0.40361349 | -2.4295708 | 0.01511671 | 0.51942334 |
| MTMR9 | 141.911211 | 0.60516121 | 0.23990452 | 2.52250859 | 0.01165211 | 0.48516963 |
| FDFT1 | 89.4811992 | 0.11139495 | 0.34914204 | 0.3190534 | 0.74968602 | 0.99989001 |
| CTSB | 239.312659 | 0.15533011 | 0.43768171 | 0.35489285 | 0.72266987 | 0.99989001 |

| | | | | | | |
|---|---|---|---|---|---|---|
| FAM66D | 54.9570487 | 0.04061263 | 0.70971639 | 0.05722375 | 0.95436696 | 0.99989001 |
| LONRF1 | 146.308865 | -0.3969124 | 0.44336319 | -0.8952308 | 0.3706637 | 0.99989001 |
| TRMT9B | 88.5186865 | -0.5091853 | 0.56551485 | -0.9003924 | 0.36791145 | 0.99989001 |
| DLC1 | 1223.5663 | 0.04032722 | 0.35916616 | 0.11228012 | 0.91060131 | 0.99989001 |
| TUSC3 | 65.4991027 | -0.6736574 | 0.80729884 | -0.8344585 | 0.40402266 | 0.99989001 |
| MICU3 | 163.362562 | -0.0441743 | 0.20836638 | -0.212003 | 0.83210471 | 0.99989001 |
| ZDHHC2 | 124.672918 | -0.3269171 | 0.52818841 | -0.6189403 | 0.53595567 | 0.99989001 |
| CNOT7 | 81.9385562 | -0.2226396 | 0.30168192 | -0.7379946 | 0.46051775 | 0.99989001 |
| VPS37A | 214.034205 | 0.36204337 | 0.4063443 | 0.89097686 | 0.37294158 | 0.99989001 |
| MTUS1 | 728.250922 | -1.3488539 | 0.5644574 | -2.3896469 | 0.01686458 | 0.5510556 |
| PCM1 | 739.920246 | 0.07525357 | 0.17719268 | 0.42469908 | 0.67105606 | 0.99989001 |
| ASAH1 | 176.827466 | -0.0568215 | 0.40049379 | -0.1418785 | 0.88717596 | 0.99989001 |
| PSD3 | 185.949355 | -0.5734394 | 0.3858602 | -1.4861326 | 0.13724405 | 0.99989001 |
| CSGALNACT1 | 951.12489 | -1.5038607 | 0.45597663 | -3.2981091 | 0.00097338 | 0.1530645 |
| INTS10 | 202.149963 | -0.0449238 | 0.25651995 | -0.1751278 | 0.86097919 | 0.99989001 |
| ATP6V1B2 | 85.5997369 | 0.01221236 | 0.35471899 | 0.03442828 | 0.97253564 | 0.99989001 |
| XPO7 | 193.877049 | -0.054579 | 0.17637576 | -0.3094471 | 0.75698144 | 0.99989001 |
| FAM160B2 | 73.7961033 | -0.0351973 | 0.28918315 | -0.121713 | 0.90312634 | 0.99989001 |
| BMP1 | 86.4833408 | 0.18922631 | 0.51817088 | 0.36518128 | 0.71497609 | 0.99989001 |
| SLC39A14 | 273.495805 | -0.193493 | 0.60565671 | -0.3194763 | 7.49E-01 | 0.99989001 |
| PPP3CC | 442.165273 | 0.22492329 | 0.21373803 | 1.0523316 | 0.29264744 | 0.99989001 |
| SORBS3 | 87.7907876 | 0.06082487 | 0.33978453 | 0.17901013 | 0.85792975 | 0.99989001 |
| CCAR2 | 65.9740953 | -0.113724 | 0.26482897 | -0.4294244 | 0.66761442 | 0.99989001 |
| BIN3 | 180.821692 | -0.3021378 | 0.24802945 | -1.218153 | 0.22316583 | 0.99989001 |
| TNFRSF10B | 188.893405 | 0.19634752 | 0.49643066 | 0.39551852 | 0.69246026 | 0.99989001 |
| TNFRSF10D | 206.175994 | 0.2377235 | 0.43113529 | 0.55138956 | 0.58136665 | 0.99989001 |
| TNFRSF10A | 121.855122 | -0.332034 | 0.45025758 | -0.7374311 | 0.4608602 | 0.99989001 |
| LOXL2 | 91.106408 | -0.0352865 | 0.2898656 | -0.1217342 | 0.90310955 | 0.99989001 |
| ENTPD4 | 119.680729 | -0.1916074 | 0.26752156 | -0.7162314 | 0.47384845 | 0.99989001 |
| SLC25A37 | 436.234249 | 0.54381184 | 0.52122238 | 1.0433394 | 0.29679113 | 0.99989001 |
| STC1 | 346.606761 | -0.7447347 | 1.07297637 | -0.694083 | 0.48763016 | 0.99989001 |
| DOCK5 | 123.098071 | 1.12129377 | 0.46028972 | 2.43606088 | 0.01484819 | 0.51942334 |
| KCTD9 | 137.463928 | 0.11834226 | 0.31394385 | 0.37695358 | 0.7062081 | 0.99989001 |
| PPP2R2A | 585.11975 | 0.21857049 | 0.23214699 | 0.94151765 | 0.34643965 | 0.99989001 |
| BNIP3L | 262.131941 | 0.06408832 | 0.48237586 | 0.13285971 | 0.89430433 | 0.99989001 |
| DPYSL2 | 340.292253 | 0.77249662 | 0.38934961 | 1.98406933 | 0.04724811 | 0.77516398 |
| TRIM35 | 45.7014257 | 0.74286077 | 0.35865728 | 2.07122737 | 0.03833755 | NA |
| PTK2B | 90.5211377 | 0.76922479 | 0.58238202 | 1.3208251 | 0.18655969 | 0.99989001 |
| EPHX2 | 39.4945335 | 0.32773042 | 0.50154492 | 0.65344181 | 0.51347149 | NA |
| CLU | 212.077691 | 1.15855539 | 0.38856134 | 2.98165382 | 0.00286696 | 0.27392168 |
| CCDC25 | 68.4684193 | -0.1484847 | 0.26172716 | -0.5673262 | 0.57049255 | 0.99989001 |
| ELP3 | 141.447074 | -0.068121 | 0.1964979 | -0.3466756 | 0.72883502 | 0.99989001 |
| EXTL3 | 86.8877971 | -0.1525429 | 0.25137779 | -0.6068271 | 0.54396567 | 0.99989001 |
| INTS9 | 159.894243 | -0.0829536 | 0.24351737 | -0.3406477 | 0.73336883 | 0.99989001 |
| HMBOX1 | 959.631146 | -0.0882621 | 0.26272436 | -0.3359493 | 0.73690913 | 0.99989001 |
| KIF13B | 334.267176 | -0.118329 | 0.23034221 | -0.5137097 | 0.60745499 | 0.99989001 |
| SARAF | 266.873963 | 0.34605368 | 0.39293643 | 0.88068617 | 0.3784877 | 0.99989001 |
| LEPROTL1 | 201.696843 | 0.00086788 | 0.54938787 | 0.00157972 | 0.99873956 | 0.99989001 |
| DCTN6 | 67.2045345 | -0.0076403 | 0.4439118 | -0.0172113 | 0.98626805 | 0.99989001 |
| RBPMS | 1215.52272 | -0.188109 | 0.27584369 | -0.6819406 | 0.49527652 | 0.99989001 |
| GTF2E2 | 134.372505 | -0.0552556 | 0.29042439 | -0.190258 | 0.84910697 | 0.99989001 |
| GSR | 73.2719876 | 0.04381015 | 0.35823767 | 0.12229353 | 0.90266655 | 0.99989001 |
| PPP2CB | 250.31966 | 0.06532187 | 0.21003438 | 0.3110056 | 0.75579636 | 0.99989001 |
| WRN | 265.855282 | -0.2523818 | 0.21930575 | -1.1508215 | 0.24980567 | 0.99989001 |

| | | | | | | |
|---|---|---|---|---|---|---|
| FUT10 | 97.8599611 | 1.05833709 | 0.38915544 | 2.71957421 | 0.0065366 | 0.390188 |
| TTI2 | 43.5916049 | -0.1385415 | 0.38606421 | -0.3588561 | 0.71970274 | NA |
| RNF122 | 45.8543651 | -0.5978949 | 0.52559637 | -1.1375553 | 0.25530623 | NA |
| PLPBP | 53.9289053 | -0.0325569 | 0.29576963 | -0.110075 | 0.91234986 | 0.99989001 |
| ADGRA2 | 47.6670963 | 0.10038387 | 0.53625915 | 0.18719284 | 0.85150944 | NA |
| EIF4EBP1 | 50.6910173 | 0.2077133 | 0.62592535 | 0.33184995 | 0.74000256 | 0.99989001 |
| ASH2L | 91.9847534 | -0.0192469 | 0.3186131 | -0.0604083 | 0.95183047 | 0.99989001 |
| LSM1 | 45.2106292 | -0.2796847 | 0.36101467 | -0.7747183 | 0.43850616 | NA |
| BAG4 | 53.3386898 | -0.1022777 | 0.25721875 | -0.3976294 | 0.69090338 | 0.99989001 |
| DDHD2 | 128.596959 | -0.0740406 | 0.24300584 | -0.3046865 | 0.76060491 | 0.99989001 |
| PLPP5 | 48.1475335 | -0.3275025 | 0.27571342 | -1.1878366 | 0.2348978 | NA |
| NSD3 | 429.350277 | 0.05732107 | 0.24182955 | 0.23703086 | 0.81263285 | 0.99989001 |
| FGFR1 | 456.878323 | -1.0563109 | 0.66999775 | -1.5765887 | 0.1148902 | 0.99989001 |
| TACC1 | 2329.05286 | -0.1852302 | 0.50295472 | -0.3682841 | 0.71266142 | 0.99989001 |
| PLEKHA2 | 158.851024 | 0.41238872 | 0.30587095 | 1.34824415 | 0.17757987 | 0.99989001 |
| ADAM9 | 230.722048 | 0.11576883 | 0.32966662 | 0.35116941 | 0.72546126 | 0.99989001 |
| TCIM | 693.446665 | -0.1538108 | 0.71991449 | -0.2136514 | 0.8308189 | 0.99989001 |
| SFRP1 | 85.7427241 | -0.7632456 | 0.31025436 | -2.4600641 | 0.01389122 | 0.50652625 |
| GOLGA7 | 71.5709351 | -0.0664216 | 0.37340268 | -0.177882 | 0.85881565 | 0.99989001 |
| GPAT4 | 135.56916 | 0.48951345 | 0.43510431 | 1.12504851 | 0.26056848 | 0.99989001 |
| KAT6A | 1003.27042 | 0.10260265 | 0.23578566 | 0.43515222 | 0.66345194 | 0.99989001 |
| AP3M2 | 134.105504 | 0.2789993 | 0.34882972 | 0.79981515 | 0.4238179 | 0.99989001 |
| PLAT | 1445.30589 | -0.6065749 | 0.44471682 | -1.3639577 | 0.17258088 | 0.99989001 |
| IKBKB | 439.715124 | -0.0658297 | 0.24000255 | -0.2742874 | 0.78386373 | 0.99989001 |
| POLB | 109.205686 | -0.1576667 | 0.41768599 | -0.3774765 | 0.70581951 | 0.99989001 |
| VDAC3 | 123.055927 | -0.1764587 | 0.25114913 | -0.7026054 | 0.48230168 | 0.99989001 |
| SLC20A2 | 274.999522 | -0.2577983 | 0.24514556 | -1.0516132 | 0.29297707 | 0.99989001 |
| SMIM19 | 55.8163607 | -0.3047598 | 0.37328347 | -0.8164301 | 0.4142542 | 0.99989001 |
| RNF170 | 103.605802 | 0.22513438 | 0.29731799 | 0.75721748 | 0.44891958 | 0.99989001 |
| HOOK3 | 352.373489 | 0.0699117 | 0.23915249 | 0.29233104 | 0.77003353 | 0.99989001 |
| FNTA | 89.6322821 | 0.21913288 | 0.22178617 | 0.98803674 | 0.32313465 | 0.99989001 |
| HGSNAT | 263.073286 | 0.03530867 | 0.20361285 | 0.1734108 | 0.86232853 | 0.99989001 |
| SPIDR | 1202.54425 | -0.0936202 | 0.23205538 | -0.4034392 | 0.68662513 | 0.99989001 |
| CEBPD | 687.773079 | -0.8391802 | 0.69869419 | -1.2010694 | 2.30E-01 | 0.99989001 |
| PRKDC | 375.18346 | -0.0011357 | 0.2474158 | -0.0045903 | 0.99633751 | 0.99989001 |
| UBE2V2 | 126.046989 | -0.0591694 | 0.21972332 | -0.2692906 | 0.78770609 | 0.99989001 |
| PXDNL | 60.3748932 | 0.54014973 | 0.35384718 | 1.52650569 | 0.12688397 | 0.99989001 |
| PCMTD1 | 506.993003 | 0.07757298 | 0.23783866 | 0.32615798 | 0.74430483 | 0.99989001 |
| RB1CC1 | 392.984859 | 0.00249198 | 0.18188323 | 0.01370101 | 0.98906852 | 0.99989001 |
| ATP6V1H | 169.853105 | -0.0348894 | 0.21445262 | -0.1626905 | 0.87076214 | 0.99989001 |
| TCEA1 | 139.637508 | -0.1298752 | 0.21845967 | -0.5945043 | 0.55217487 | 0.99989001 |
| LYPLA1 | 210.687398 | 0.21139769 | 0.22372684 | 0.94489196 | 0.34471404 | 0.99989001 |
| SOX17 | 109.893424 | -0.8461793 | 0.63523811 | -1.3320664 | 0.18283839 | 0.99989001 |
| TMEM68 | 75.1134018 | -0.0264542 | 0.30807423 | -0.0858696 | 0.93157004 | 0.99989001 |
| TGS1 | 89.0459715 | 0.11408611 | 0.32062995 | 0.35581862 | 0.7219764 | 0.99989001 |
| LYN | 223.533035 | 0.24030184 | 0.34858047 | 0.68937262 | 0.49058881 | 0.99989001 |
| RPS20 | 242.44601 | -0.0297754 | 0.50277431 | -0.0592222 | 0.95277511 | 0.99989001 |
| PLAG1 | 68.7652788 | 0.19745963 | 0.54492949 | 0.36235813 | 0.71708443 | 0.99989001 |
| IMPAD1 | 103.796058 | 0.10415774 | 0.27633242 | 0.37692913 | 0.70622627 | 0.99989001 |
| FAM110B | 59.9372568 | -1.2991412 | 0.61213688 | -2.1223051 | 0.03381213 | 0.67281094 |
| SDCBP | 704.725832 | -0.5205896 | 0.38631588 | -1.347575 | 0.17779512 | 0.99989001 |
| NSMAF | 118.15449 | -0.3567926 | 0.33690643 | -1.0590257 | 0.28958808 | 0.99989001 |
| CA8 | 308.113364 | -0.5859451 | 0.70289927 | -0.8336117 | 0.40449983 | 0.99989001 |
| RAB2A | 564.780103 | 0.201713 | 0.19657026 | 1.02616237 | 0.30481505 | 0.99989001 |

| | | | | | | |
|---|---|---|---|---|---|---|
| CHD7 | 408.932145 | 0.47188408 | 0.35890188 | 1.31479968 | 0.18857724 | 0.99989001 |
| ASPH | 335.136302 | -0.7540989 | 0.3098209 | -2.4339833 | 0.01493368 | 0.51942334 |
| YTHDF3 | 206.493758 | 0.12410817 | 0.1995627 | 0.62190064 | 0.5340072 | 0.99989001 |
| CYP7B1 | 64.2156653 | 0.18127516 | 0.49014186 | 0.36984224 | 0.71150004 | 0.99989001 |
| ARMC1 | 82.5499475 | -0.1322423 | 0.20184567 | -0.6551653 | 0.51236131 | 0.99989001 |
| MTFR1 | 160.645976 | 0.55324051 | 0.31722264 | 1.74401335 | 0.08115676 | 0.94203683 |
| PDE7A | 156.237992 | 0.35786163 | 0.24773045 | 1.44456053 | 0.14858136 | 0.99989001 |
| VCPIP1 | 72.5237292 | 0.20713421 | 0.21540089 | 0.9616219 | 0.33623957 | 0.99989001 |
| SGK3 | 226.858049 | 0.19417451 | 0.23245331 | 0.83532694 | 0.40353366 | 0.99989001 |
| CSPP1 | 255.931331 | -0.0327564 | 0.1961956 | -0.1669581 | 0.86740301 | 0.99989001 |
| ARFGEF1 | 672.956184 | 0.04150419 | 0.15487288 | 0.26798875 | 0.78870798 | 0.99989001 |
| PREX2 | 1073.09498 | 0.2337327 | 0.35561856 | 0.65725674 | 0.51101585 | 0.99989001 |
| SULF1 | 364.96234 | 2.06748376 | 0.70512881 | 2.93206538 | 0.00336716 | 0.29068264 |
| NCOA2 | 900.202911 | 0.44960387 | 0.24713639 | 1.81925405 | 0.06887268 | 0.88863418 |
| TRAM1 | 246.800081 | -0.0999492 | 0.26434618 | -0.3780996 | 0.70535661 | 0.99989001 |
| LACTB2 | 54.1908293 | 0.06437937 | 0.39818545 | 0.16168186 | 0.87155639 | 0.99989001 |
| TERF1 | 104.312284 | 0.19982661 | 0.24410627 | 0.81860498 | 0.41301182 | 0.99989001 |
| RPL7 | 351.809539 | 0.10783869 | 0.57501756 | 0.18753982 | 0.8512374 | 0.99989001 |
| RDH10 | 85.2522641 | -0.8263039 | 0.6862675 | -1.2040551 | 0.22856828 | 0.99989001 |
| STAU2 | 592.658271 | -0.0876073 | 0.30653938 | -0.2857947 | 0.77503536 | 0.99989001 |
| AC027018.1 | 76.6103858 | -0.2461749 | 0.55893264 | -0.4404376 | 0.6596202 | 0.99989001 |
| ELOC | 128.552443 | -0.3561149 | 0.32208557 | -1.1056531 | 0.26887671 | 0.99989001 |
| TMEM70 | 53.6077866 | -0.8361451 | 0.53629293 | -1.5591201 | 0.11896796 | 0.99989001 |
| LY96 | 78.0569905 | -0.489852 | 0.3498774 | -1.4000675 | 0.16149309 | 0.99989001 |
| GDAP1 | 103.19477 | -0.6953619 | 0.29327269 | -2.371042 | 0.01773801 | 0.56254844 |
| PEX2 | 50.9473179 | 0.17539803 | 0.35458433 | 0.49465815 | 0.62084145 | 0.99989001 |
| ZC2HC1A | 96.5662361 | 0.50546448 | 0.35237132 | 1.43446543 | 0.15143949 | 0.99989001 |
| IL7 | 82.423345 | 0.98712654 | 0.43768289 | 2.25534643 | 0.02411159 | 0.6148456 |
| HEY1 | 50.4826666 | -0.0002124 | 0.82828321 | -0.0002564 | 0.99979541 | 0.99992789 |
| ZBTB10 | 105.192103 | 0.30860991 | 0.39634756 | 0.77863456 | 0.43619502 | 0.99989001 |
| ZNF704 | 356.167623 | 0.35050616 | 0.44194081 | 0.79310658 | 4.28E-01 | 0.99989001 |
| PAG1 | 66.5374795 | 0.72880961 | 0.46616044 | 1.56343083 | 0.11795129 | 0.99989001 |
| FABP5 | 143.780762 | -3.2460138 | 1.74574868 | -1.8593821 | 0.062973 | 0.85940342 |
| IMPA1 | 69.342194 | -0.0918517 | 0.32165838 | -0.2855566 | 0.77521774 | 0.99989001 |
| ZFAND1 | 71.5620198 | -0.1891728 | 0.33887168 | -0.5582432 | 0.57667835 | 0.99989001 |
| SNX16 | 67.0819248 | -0.388568 | 0.3161401 | -1.2291008 | 0.21903403 | 0.99989001 |
| C8orf59 | 61.0047063 | -0.3553278 | 0.26239434 | -1.3541746 | 0.17568069 | 0.99989001 |
| CA2 | 115.264143 | 0.41065308 | 0.77855342 | 0.52745653 | 0.5978766 | 0.99989001 |
| WWP1 | 629.075335 | 0.311225 | 0.27235076 | 1.14273595 | 0.25314824 | 0.99989001 |
| RMDN1 | 131.547009 | -0.0055543 | 0.33473435 | -0.0165931 | 0.98676126 | 0.99989001 |
| CPNE3 | 110.689495 | -0.128481 | 0.31784213 | -0.4042291 | 0.68604427 | 0.99989001 |
| AF117829.1 | 242.235059 | 0.25548205 | 0.46900508 | 0.54473194 | 0.5859379 | 0.99989001 |
| RIPK2 | 71.2762344 | 0.31431079 | 0.44173849 | 0.71153137 | 0.47675502 | 0.99989001 |
| NBN | 112.0046 | 0.21485202 | 0.33030557 | 0.65046444 | 0.51539227 | 0.99989001 |
| DECR1 | 201.20896 | 0.28343662 | 0.31008729 | 0.91405428 | 0.36068832 | 0.99989001 |
| PIP4P2 | 62.2339315 | 0.00995583 | 0.40976181 | 0.02429662 | 0.98061601 | 0.99989001 |
| OTUD6B-AS1 | 61.1374536 | -0.121468 | 0.37209703 | -0.3264417 | 0.74409021 | 0.99989001 |
| OTUD6B | 59.1766555 | 0.17377083 | 0.2750251 | 0.63183627 | 0.52749387 | 0.99989001 |
| LRRC69 | 86.4744343 | -0.3027536 | 0.23351465 | -1.296508 | 0.19480051 | 0.99989001 |
| RUNX1T1 | 690.840909 | 0.43093265 | 0.46247469 | 0.93179728 | 0.3514413 | 0.99989001 |
| TRIQK | 123.149313 | 0.15985071 | 0.47020452 | 0.33995995 | 0.73388669 | 0.99989001 |
| RBM12B | 88.449764 | 0.04248057 | 0.22438447 | 0.18932044 | 0.84984167 | 0.99989001 |
| TMEM67 | 55.275178 | -0.326102 | 0.41630164 | -0.783331 | 0.43343276 | 0.99989001 |
| VIRMA | 138.287244 | -0.0381475 | 0.21665385 | -0.1760757 | 0.86023447 | 0.99989001 |

| | | | | | | |
|---|---|---|---|---|---|---|
| DPY19L4 | 154.589033 | 0.00164241 | 0.16167612 | 0.01015867 | 0.9918947 | 0.99989001 |
| INTS8 | 116.35609 | 0.23362767 | 0.2283805 | 1.02297557 | 0.3063194 | 0.99989001 |
| NDUFAF6 | 156.570942 | 0.24911695 | 0.25944813 | 0.96018017 | 0.33696455 | 0.99989001 |
| TP53INP1 | 74.9214721 | 0.13158446 | 0.57460754 | 0.22899884 | 0.81886982 | 0.99989001 |
| PLEKHF2 | 58.3774121 | 0.03609962 | 0.27438581 | 0.13156518 | 0.89532823 | 0.99989001 |
| UQCRB | 158.728232 | -0.1320203 | 0.54583012 | -0.2418707 | 0.80888034 | 0.99989001 |
| PTDSS1 | 68.3388334 | -0.062449 | 0.34739921 | -0.1797615 | 0.85733979 | 0.99989001 |
| SDC2 | 75.8324747 | -0.6341044 | 0.37724345 | -1.6808891 | 0.09278447 | 0.99479708 |
| CPQ | 521.157604 | -0.06379 | 0.2600307 | -0.2453174 | 0.80621072 | 0.99989001 |
| MTDH | 525.653578 | 0.01259509 | 0.19307161 | 0.06523535 | 0.94798662 | 0.99989001 |
| LAPTM4B | 155.792242 | -0.4464517 | 0.493113 | -0.9053739 | 0.36526733 | 0.99989001 |
| MATN2 | 139.191707 | 0.40710762 | 0.69564069 | 0.58522687 | 0.55839518 | 0.99989001 |
| RPL30 | 456.868074 | -0.286121 | 0.57084292 | -0.5012254 | 0.6162125 | 0.99989001 |
| NIPAL2 | 99.8177704 | -0.5896977 | 0.29366917 | -2.0080339 | 0.04463969 | 0.77228819 |
| STK3 | 1068.97098 | -0.1102293 | 0.27917696 | -0.3948365 | 0.69296353 | 0.99989001 |
| VPS13B | 950.632075 | -0.0046633 | 0.16280579 | -0.0286433 | 0.97714908 | 0.99989001 |
| COX6C | 201.024501 | -0.2232277 | 0.41193233 | -0.5419037 | 0.58788484 | 0.99989001 |
| POLR2K | 62.3106741 | -0.2426677 | 0.5999628 | -0.4044712 | 0.68586627 | 0.99989001 |
| SPAG1 | 58.8442186 | -0.2206953 | 0.38683861 | -0.5705099 | 0.56833189 | 0.99989001 |
| RNF19A | 505.788631 | -0.3154083 | 0.25355812 | -1.2439292 | 0.21352562 | 0.99989001 |
| ANKRD46 | 62.2158278 | -0.1978658 | 0.23581994 | -0.8390546 | 0.40143867 | 0.99989001 |
| PABPC1 | 422.59526 | -0.0985417 | 0.37430016 | -0.2632691 | 0.79234317 | 0.99989001 |
| YWHAZ | 448.351313 | 0.34960614 | 0.22977052 | 1.52154481 | 0.12812318 | 0.99989001 |
| ZNF706 | 185.814997 | 0.11137985 | 0.2720522 | 0.40940617 | 0.68224161 | 0.99989001 |
| NCALD | 340.724983 | 0.51551975 | 0.46066995 | 1.11906529 | 2.63E-01 | 0.99989001 |
| RRM2B | 71.909201 | -0.0948757 | 0.39908812 | -0.2377313 | 0.81208954 | 0.99989001 |
| UBR5 | 871.822559 | -0.1362702 | 0.2643557 | -0.5154804 | 0.60621736 | 0.99989001 |
| KLF10 | 159.295396 | 0.72740756 | 0.46130839 | 1.57683576 | 0.11483332 | 0.99989001 |
| AZIN1 | 121.562387 | 0.21864059 | 0.2880323 | 0.75908357 | 0.44780257 | 0.99989001 |
| AZIN1-AS1 | 67.0122208 | -0.9819174 | 0.57155487 | -1.7179757 | 0.08580105 | 0.95514629 |
| ATP6V1C1 | 153.225244 | -0.3994065 | 0.25076459 | -1.5927547 | 0.11121524 | 0.99989001 |
| BAALC | 94.4988586 | -0.7345116 | 0.50583729 | -1.4520708 | 0.14648191 | 0.99989001 |
| BAALC-AS1 | 68.4974982 | -0.5349798 | 0.45985109 | -1.1633762 | 0.24467691 | 0.99989001 |
| FZD6 | 159.54772 | -0.3517751 | 0.38211761 | -0.9205937 | 0.35726259 | 0.99989001 |
| SLC25A32 | 117.609794 | -0.010232 | 0.31067746 | -0.0329346 | 0.97372674 | 0.99989001 |
| DCAF13 | 58.4331541 | -0.2889274 | 0.31504962 | -0.9170853 | 0.35909794 | 0.99989001 |
| LRP12 | 74.8080802 | 0.03804812 | 0.22683122 | 0.16773757 | 0.86678974 | 0.99989001 |
| ZFPM2 | 514.602625 | 0.86418078 | 0.40045008 | 2.15802376 | 0.03092598 | 0.67281094 |
| OXR1 | 423.244293 | 0.0946368 | 0.24057247 | 0.39338167 | 0.6940376 | 0.99989001 |
| EIF3E | 238.716447 | -0.0544143 | 0.18667575 | -0.2914909 | 0.77067588 | 0.99989001 |
| EMC2 | 145.743404 | -0.1086171 | 0.26789702 | -0.4054436 | 0.68515145 | 0.99989001 |
| NUDCD1 | 205.19097 | 0.08488714 | 0.21490827 | 0.3949924 | 0.69284849 | 0.99989001 |
| ENY2 | 159.281708 | -0.0012266 | 0.38348905 | -0.0031985 | 0.99744798 | 0.99989001 |
| PKHD1L1 | 324.474603 | 0.79150766 | 1.29184674 | 0.61269471 | 0.54007822 | 0.99989001 |
| EBAG9 | 54.5184767 | -0.0839859 | 0.24761169 | -0.339184 | 0.73447114 | 0.99989001 |
| SYBU | 71.2518797 | -0.1761153 | 0.41773617 | -0.4215945 | 0.67332105 | 0.99989001 |
| TRPS1 | 509.812047 | 0.76512051 | 0.32973887 | 2.3203831 | 0.02032016 | 0.58699182 |
| EIF3H | 330.194594 | -0.060896 | 0.24469713 | -0.2488627 | 0.80346701 | 0.99989001 |
| UTP23 | 49.2645867 | 0.32354832 | 0.30646692 | 1.05573652 | 0.29108861 | NA |
| RAD21 | 210.385399 | 0.26236761 | 0.27656909 | 0.94865125 | 0.34279801 | 0.99989001 |
| EXT1 | 1215.80812 | 0.12011995 | 0.30935736 | 0.38828864 | 0.69780244 | 0.99989001 |
| SAMD12 | 261.439661 | -0.5044126 | 0.5537793 | -0.910855 | 0.3623718 | 0.99989001 |
| ENPP2 | 1904.30382 | -1.9061193 | 0.85700759 | -2.2241568 | 0.0261379 | 0.64054821 |
| TAF2 | 240.586731 | -0.0922854 | 0.18723556 | -0.4928838 | 0.62209469 | 0.99989001 |

| | | | | | | |
|---|---|---|---|---|---|---|
| DEPTOR | 128.744289 | -0.4783877 | 0.54214841 | -0.8823926 | 0.37756455 | 0.99989001 |
| MRPL13 | 55.7263026 | 0.15243375 | 0.30955186 | 0.49243366 | 0.62241281 | 0.99989001 |
| SNTB1 | 72.6034174 | 0.63756955 | 0.56674682 | 1.12496361 | 2.61E-01 | 0.99989001 |
| ZHX2 | 447.958971 | 0.11735037 | 0.32712013 | 0.35873784 | 0.71979121 | 0.99989001 |
| DERL1 | 93.7734508 | 0.07533074 | 0.24162823 | 0.31176298 | 0.75522066 | 0.99989001 |
| WDYHV1 | 48.5077136 | 0.03736038 | 0.37726136 | 0.0990305 | 0.92111405 | NA |
| FAM91A1 | 203.855672 | -0.1821481 | 0.2376855 | -0.766341 | 0.44347341 | 0.99989001 |
| TMEM65 | 377.651718 | 0.08495946 | 0.2993161 | 0.28384527 | 0.77652896 | 0.99989001 |
| TATDN1 | 80.1806406 | 0.11514556 | 0.2985745 | 0.38565101 | 0.69975515 | 0.99989001 |
| NDUFB9 | 110.789529 | -0.0869596 | 0.33150014 | -0.2623215 | 0.79307362 | 0.99989001 |
| MTSS1 | 1026.9273 | 0.06273013 | 0.37647001 | 0.16662718 | 0.86766341 | 0.99989001 |
| SQLE | 108.756228 | 0.66323234 | 0.58355659 | 1.13653475 | 0.25573283 | 0.99989001 |
| WASHC5 | 112.44774 | 0.03217069 | 0.31886605 | 0.10089091 | 0.91963706 | 0.99989001 |
| NSMCE2 | 763.967534 | 0.02734327 | 0.27491792 | 0.09945974 | 0.92077325 | 0.99989001 |
| FAM84B | 229.172117 | -0.1392893 | 0.57685174 | -0.2414646 | 0.80919506 | 0.99989001 |
| PCAT1 | 962.44474 | 0.10165648 | 0.36748334 | 0.27662881 | 0.78206514 | 0.99989001 |
| PVT1 | 122.661639 | 0.02067042 | 0.44362939 | 0.0465939 | 9.63E-01 | 1.00E+00 |
| FAM49B | 240.399378 | -0.1862365 | 0.29577219 | -0.6296619 | 0.5289158 | 0.99989001 |
| ASAP1 | 2494.60559 | -0.0810061 | 0.27026515 | -0.2997281 | 0.76438456 | 0.99989001 |
| EFR3A | 307.399497 | -0.0276794 | 0.21424139 | -0.1291973 | 0.89720155 | 0.99989001 |
| LRRC6 | 44.5508472 | -0.1172183 | 0.38905619 | -0.3012889 | 0.76319419 | NA |
| PHF20L1 | 295.994405 | 0.18777145 | 0.20603535 | 0.9113555 | 0.36210809 | 0.99989001 |
| NDRG1 | 734.566898 | -0.4199372 | 0.54590388 | -0.7692512 | 0.44174421 | 0.99989001 |
| ST3GAL1 | 415.895818 | 0.41722455 | 0.3827811 | 1.08998211 | 0.27572102 | 0.99989001 |
| ZFAT | 89.4147576 | -0.1634223 | 0.299927 | -0.5448735 | 0.58584055 | 0.99989001 |
| KHDRBS3 | 210.204217 | 1.0833088 | 0.3027353 | 3.57840267 | 0.0003457 | 0.07907115 |
| TRAPPC9 | 578.924513 | 0.04167701 | 0.29285909 | 0.14231079 | 0.88683452 | 0.99989001 |
| AGO2 | 244.52944 | 0.14313134 | 0.21596862 | 0.66274138 | 0.50749621 | 0.99989001 |
| PTK2 | 2178.33604 | 0.16204658 | 0.19847516 | 0.8164577 | 0.41423841 | 0.99989001 |
| DENND3 | 380.04836 | 0.86657849 | 0.36513482 | 2.373311 | 0.01762941 | 0.56214542 |
| SLC45A4 | 166.724891 | -0.8646201 | 0.54029311 | -1.6002798 | 0.10953653 | 0.99989001 |
| PTP4A3 | 45.194029 | -1.6181513 | 0.72320574 | -2.2374703 | 0.02525562 | NA |
| TSNARE1 | 50.4461745 | -0.1002299 | 0.32416645 | -0.3091926 | 0.757175 | 0.99989001 |
| THEM6 | 39.082282 | -0.3148719 | 0.73799134 | -0.4266607 | 0.66962648 | NA |
| LY6E | 316.270573 | 0.03330247 | 0.45432225 | 0.07330144 | 0.94156625 | 0.99989001 |
| GPIHBP1 | 79.0049406 | -0.3588093 | 0.80627076 | -0.4450234 | 0.65630287 | 0.99989001 |
| ZC3H3 | 66.624691 | 0.32844544 | 0.3198868 | 1.02675521 | 0.30453574 | 0.99989001 |
| GSDMD | 162.521375 | 0.22270604 | 0.44400877 | 0.50158027 | 0.6159628 | 0.99989001 |
| NAPRT | 37.703263 | 0.1896142 | 0.44802385 | 0.42322345 | 0.67213224 | NA |
| EEF1D | 284.254727 | -0.1721145 | 0.33620279 | -0.5119364 | 0.6086955 | 0.99989001 |
| PUF60 | 70.3337233 | -0.0717918 | 0.51334127 | -0.139852 | 0.8887769 | 0.99989001 |
| PLEC | 228.528403 | -0.108191 | 0.44289938 | -0.244279 | 0.80701474 | 0.99989001 |
| PARP10 | 56.334731 | -0.1342154 | 0.61313295 | -0.2189009 | 0.82672725 | 0.99989001 |
| GRINA | 80.8938594 | -0.0152159 | 0.41043397 | -0.0370728 | 0.97042698 | 0.99989001 |
| GPAA1 | 54.4320038 | 0.10574323 | 0.52414094 | 0.20174579 | 0.84011546 | 0.99989001 |
| CYC1 | 58.581386 | 0.48621558 | 0.40432279 | 1.2025431 | 0.22915318 | 0.99989001 |
| SHARPIN | 46.1729474 | -0.2011033 | 0.2623807 | -0.7664561 | 0.44340494 | NA |
| MAF1 | 68.009361 | 0.01897971 | 0.47006555 | 0.04037673 | 0.96779278 | 0.99989001 |
| MROH1 | 115.377917 | -0.4698429 | 0.40042701 | -1.1733545 | 0.24065366 | 0.99989001 |
| HSF1 | 153.197248 | -0.1667763 | 0.29336003 | -0.5685039 | 0.56969286 | 0.99989001 |
| DGAT1 | 44.5283059 | -0.072966 | 0.35397432 | -0.2061336 | 0.83668657 | NA |
| VPS28 | 164.161401 | -0.0361708 | 0.39065022 | -0.0925913 | 0.92622824 | 0.99989001 |
| CYHR1 | 59.382898 | -0.2769604 | 0.43537853 | -0.636137 | 0.5246871 | 0.99989001 |
| ZNF251 | 133.379344 | 0.28401763 | 0.30086324 | 0.94400906 | 0.34516502 | 0.99989001 |

| | | | | | | |
|---|---|---|---|---|---|---|
| RPL8 | 548.761386 | -0.1160485 | 0.58493212 | -0.1983965 | 0.84273488 | 0.99989001 |
| ZNF7 | 48.0847486 | 0.09979206 | 0.35539557 | 0.28079152 | 0.77887031 | NA |
| C8orf33 | 45.6566331 | -0.498987 | 0.40762878 | -1.224121 | 0.22090659 | NA |
| WASHC1 | 148.029164 | -0.1784419 | 0.33761423 | -0.528538 | 0.597126 | 0.99989001 |
| KANK1 | 140.40738 | -0.5181887 | 0.41982972 | -1.234283 | 0.21709746 | 0.99989001 |
| SMARCA2 | 638.721786 | 0.23742625 | 0.30913364 | 0.76803756 | 0.44246487 | 0.99989001 |
| PUM3 | 246.78991 | -0.1896023 | 0.32178248 | -0.5892251 | 0.55571027 | 0.99989001 |
| RFX3 | 1176.68128 | -0.1151262 | 0.32476616 | -0.3544896 | 0.72297201 | 0.99989001 |
| RFX3-AS1 | 165.155551 | -0.7240607 | 0.41787299 | -1.7327292 | 0.08314381 | 0.94799019 |
| GLIS3 | 804.437142 | -0.0849072 | 0.3462597 | -0.2452124 | 0.80629195 | 0.99989001 |
| SLC1A1 | 154.059531 | -0.1524275 | 0.56888912 | -0.2679388 | 0.78874645 | 0.99989001 |
| SPATA6L | 105.950357 | 0.18792855 | 0.3725514 | 0.50443658 | 0.61395462 | 0.99989001 |
| CDC37L1 | 72.404625 | 0.21835333 | 0.30224472 | 0.72243885 | 0.47002471 | 0.99989001 |
| AK3 | 215.298366 | 0.03503119 | 0.29678656 | 0.11803496 | 0.90603996 | 0.99989001 |
| RCL1 | 121.790685 | -0.0636101 | 0.2936883 | -0.2165904 | 0.8285276 | 0.99989001 |
| JAK2 | 363.176184 | 0.22441131 | 0.42110161 | 0.53291487 | 0.59409251 | 0.99989001 |
| PLGRKT | 60.4914484 | -0.1892549 | 0.40437347 | -0.4680201 | 0.63977021 | 0.99989001 |
| PDCD1LG2 | 49.1877116 | -0.1443987 | 0.65912149 | -0.2190775 | 0.82658967 | NA |
| RIC1 | 348.898305 | 0.10453824 | 0.24949948 | 0.4189918 | 0.67522213 | 0.99989001 |
| ERMP1 | 120.095006 | 0.02174978 | 0.36656867 | 0.05933344 | 0.95268653 | 0.99989001 |
| KIAA2026 | 460.411677 | 0.13592256 | 0.16951492 | 0.80183245 | 0.42264988 | 0.99989001 |
| IL33 | 134.81745 | 1.29658758 | 0.61217385 | 2.1180055 | 0.0341746 | 0.67703384 |
| UHRF2 | 269.538179 | 0.10228721 | 0.2079341 | 0.49192129 | 0.62277499 | 0.99989001 |
| KDM4C | 630.734799 | -0.0060173 | 0.18557452 | -0.0324253 | 0.97413292 | 0.99989001 |
| DMAC1 | 46.0990163 | -0.2610819 | 0.29000516 | -0.9002665 | 0.36797846 | NA |
| MPDZ | 656.210823 | -0.4043685 | 0.33278429 | -1.2151068 | 0.22432535 | 0.99989001 |
| NFIB | 2027.10349 | -0.2188258 | 0.24968577 | -0.8764047 | 0.38081008 | 0.99989001 |
| ZDHHC21 | 266.732533 | -0.4831008 | 0.39391264 | -1.226416 | 0.22004218 | 0.99989001 |
| TTC39B | 149.352226 | 0.04202333 | 0.2478082 | 0.16958007 | 0.86534039 | 0.99989001 |
| SNAPC3 | 173.449328 | 0.0221345 | 0.18290809 | 0.12101435 | 0.90367967 | 0.99989001 |
| PSIP1 | 197.005644 | 0.11062029 | 0.29371376 | 0.37662618 | 0.70645143 | 0.99989001 |
| CCDC171 | 80.1086831 | -0.8456115 | 0.40247705 | -2.101018 | 0.03563938 | 0.6933146 |
| BNC2 | 530.01422 | -1.3320858 | 0.4806324 | -2.7715272 | 0.0055794 | 0.36465062 |
| CNTLN | 361.770466 | -0.0210255 | 0.17470801 | -0.1203466 | 0.9042086 | 0.99989001 |
| ADAMTSL1 | 404.27455 | 0.2120177 | 0.70535866 | 0.30058141 | 0.76373371 | 0.99989001 |
| RRAGA | 79.7891811 | -0.1204696 | 0.6104988 | -0.1973298 | 0.84356943 | 0.99989001 |
| HAUS6 | 69.6912553 | 0.07107475 | 0.31806897 | 0.22345704 | 0.82317981 | 0.99989001 |
| PLIN2 | 442.37024 | 0.09402315 | 0.66598631 | 0.1411788 | 0.88772869 | 0.99989001 |
| DENND4C | 720.664888 | -0.0694067 | 0.31596725 | -0.2196642 | 0.8261327 | 0.99989001 |
| RPS6 | 698.142419 | -0.2058418 | 0.63665792 | -0.3233162 | 0.74645576 | 0.99989001 |
| ACER2 | 395.488956 | 0.0057497 | 0.90474455 | 0.00635505 | 0.99492944 | 0.99989001 |
| SLC24A2 | 78.8754534 | -4.0086996 | 1.37301835 | -2.9196257 | 0.00350452 | 0.29068264 |
| MLLT3 | 699.363055 | -0.2565456 | 0.35521158 | -0.7222331 | 0.47015119 | 0.99989001 |
| FOCAD | 340.324812 | -0.2254155 | 0.31868945 | -0.7073203 | 0.47936746 | 0.99989001 |
| KLHL9 | 64.9019616 | 0.21275003 | 0.38238242 | 0.55638025 | 0.57795094 | 0.99989001 |
| CAAP1 | 102.313263 | -0.2269073 | 0.21489004 | -1.055923 | 0.2910034 | 0.99989001 |
| PLAA | 121.099555 | -0.0005041 | 0.33305164 | -0.0015136 | 0.99879236 | 0.99989001 |
| IFT74 | 105.525896 | 0.1217762 | 0.25882638 | 0.47049376 | 0.63800229 | 0.99989001 |
| TEK | 1585.43649 | -0.3424989 | 0.31781469 | -1.0776685 | 0.28118172 | 0.99989001 |
| MOB3B | 80.0922951 | 0.77823476 | 0.37903984 | 2.05317408 | 0.0400557 | 0.73029084 |
| C9orf72 | 66.2529902 | -0.1919439 | 0.303958 | -0.6314816 | 0.5277257 | 0.99989001 |
| ACO1 | 153.856035 | 0.10121516 | 0.22662881 | 0.44661206 | 0.65515518 | 0.99989001 |
| DDX58 | 514.271008 | 0.25598759 | 0.72612801 | 0.35253782 | 0.72443496 | 0.99989001 |
| TOPORS | 64.6565657 | -0.0321113 | 0.29027274 | -0.1106246 | 0.91191406 | 0.99989001 |

| | | | | | | |
|---|---|---|---|---|---|---|
| NDUFB6 | 79.4351774 | -0.0221624 | 0.36645715 | -0.0604775 | 0.9517753 | 0.99989001 |
| APTX | 106.702439 | -0.2093438 | 0.29416531 | -0.7116535 | 0.47667934 | 0.99989001 |
| DNAJA1 | 249.900598 | 0.12631317 | 0.25650997 | 0.49242984 | 0.62241551 | 0.99989001 |
| SMU1 | 102.283881 | 0.08829431 | 0.19561099 | 0.45137704 | 0.65171782 | 0.99989001 |
| B4GALT1 | 397.941456 | 0.23531042 | 0.37986038 | 0.61946556 | 0.5356097 | 0.99989001 |
| BAG1 | 69.9181245 | 0.09392307 | 0.37727904 | 0.24894855 | 0.80340058 | 0.99989001 |
| CHMP5 | 134.977481 | 0.17728263 | 0.35145556 | 0.50442402 | 0.61396344 | 0.99989001 |
| NFX1 | 236.63115 | -0.1763418 | 0.25899219 | -0.6808769 | 0.49594936 | 0.99989001 |
| UBE2R2 | 437.931976 | -0.0791036 | 0.2238474 | -0.3533816 | 0.72380235 | 0.99989001 |
| UBAP2 | 253.777373 | -0.1271482 | 0.20913248 | -0.6079792 | 0.54320125 | 0.99989001 |
| DCAF12 | 71.3250543 | -0.1178597 | 0.24323187 | -0.4845571 | 0.62799055 | 0.99989001 |
| UBAP1 | 240.54701 | -0.3225732 | 0.31452183 | -1.0255988 | 0.30508072 | 0.99989001 |
| FAM219A | 319.247231 | 0.09879715 | 0.28397932 | 0.34790261 | 0.72791333 | 0.99989001 |
| DCTN3 | 54.662337 | 0.02168716 | 0.51899909 | 0.04178652 | 0.96666888 | 0.99989001 |
| VCP | 256.224487 | 0.12264986 | 0.35734125 | 0.34322895 | 0.73142623 | 0.99989001 |
| STOML2 | 50.9159543 | 0.13285218 | 0.51552486 | 0.25770277 | 0.79663631 | 0.99989001 |
| UNC13B | 355.314267 | 0.3502682 | 0.26017141 | 1.34629781 | 0.17820649 | 0.99989001 |
| RUSC2 | 237.250961 | 0.05227551 | 0.28973679 | 0.18042416 | 0.85681959 | 0.99989001 |
| CCDC107 | 73.2568225 | -0.8144822 | 0.42964502 | -1.8957097 | 0.05799844 | 0.8467548 |
| TLN1 | 188.20731 | 0.09277854 | 0.28644695 | 0.3238943 | 0.74601806 | 0.99989001 |
| GBA2 | 89.1981472 | 0.0656643 | 0.33823376 | 0.1941388 | 0.8460672 | 0.99989001 |
| SPAAR | 243.362553 | -0.4466216 | 0.68480646 | -0.6521866 | 0.51428079 | 0.99989001 |
| RECK | 205.939347 | 0.41285974 | 0.2715895 | 1.52016088 | 0.12847055 | 0.99989001 |
| CLTA | 288.938608 | -0.4095162 | 0.30146423 | -1.3584237 | 0.17432927 | 0.99989001 |
| GNE | 99.3477998 | -0.3649655 | 0.26867979 | -1.3583661 | 0.17434753 | 0.99989001 |
| RNF38 | 570.392814 | 0.17339044 | 0.17758674 | 0.97637041 | 0.32888094 | 0.99989001 |
| EBLN3P | 65.163833 | 0.10708275 | 0.30417238 | 0.35204626 | 0.72480357 | 0.99989001 |
| ZCCHC7 | 844.830149 | 0.1463817 | 0.19286481 | 0.75898607 | 0.44786089 | 0.99989001 |
| GRHPR | 66.2806878 | -0.0633309 | 0.32187797 | -0.1967544 | 0.84401971 | 0.99989001 |
| AL138752.2 | 92.9287312 | 0.18463986 | 0.22234349 | 0.8304262 | 0.40629786 | 0.99989001 |
| TRMT10B | 86.7680988 | 0.56269235 | 0.36485806 | 1.54222263 | 0.1230195 | 0.99989001 |
| DCAF10 | 291.519018 | 0.27570962 | 0.20353234 | 1.3546231 | 0.17553767 | 0.99989001 |
| CNTNAP3 | 114.080993 | -0.6488645 | 0.57630066 | -1.1259132 | 0.26020226 | 0.99989001 |
| GLIDR | 51.5780073 | -0.4208417 | 0.40982676 | -1.026877 | 0.30447839 | 0.99989001 |
| BMS1P14 | 245.655847 | -1.1288982 | 0.62702758 | -1.8003964 | 0.07179806 | 0.91256377 |
| CNTNAP3B | 1514.45411 | 0.30267088 | 0.51585513 | 0.5867362 | 0.55738089 | 0.99989001 |
| CBWD5 | 80.1132574 | -0.4461965 | 0.27165781 | -1.6424947 | 0.10048753 | 0.99989001 |
| PGM5 | 611.55479 | -0.34875 | 0.42318376 | -0.8241101 | 0.409877 | 0.99989001 |
| APBA1 | 79.6681181 | 0.48276412 | 0.34961234 | 1.38085551 | 0.16732339 | 0.99989001 |
| PTAR1 | 262.186514 | -0.0587418 | 0.20981747 | -0.2799664 | 0.77950332 | 0.99989001 |
| MAMDC2 | 256.837914 | -0.0429037 | 0.60061434 | -0.0714331 | 0.94305306 | 0.99989001 |
| SMC5 | 428.137278 | 0.00548251 | 0.1756847 | 0.03120651 | 0.97510484 | 0.99989001 |
| KLF9 | 491.52115 | -0.4035828 | 0.63172507 | -0.6388583 | 0.52291515 | 0.99989001 |
| TRPM3 | 120.623315 | -0.5108465 | 0.40610598 | -1.2579142 | 0.20842279 | 0.99989001 |
| CEMIP2 | 1019.97911 | -0.7285452 | 0.5897109 | -1.2354277 | 0.21667136 | 0.99989001 |
| ABHD17B | 112.618312 | 0.47811954 | 0.39391474 | 1.21376403 | 0.22483785 | 0.99989001 |
| C9orf85 | 149.188373 | -0.1399508 | 0.22106333 | -0.6330802 | 0.52668125 | 0.99989001 |
| ZFAND5 | 229.696896 | -0.1657994 | 0.32776127 | -0.5058542 | 0.612959 | 0.99989001 |
| ALDH1A1 | 105.556315 | 0.24023822 | 0.38623148 | 0.6220058 | 0.53393805 | 0.99989001 |
| ANXA1 | 224.252154 | -0.5568546 | 0.49830047 | -1.1175078 | 0.26377729 | 0.99989001 |
| CARNMT1 | 85.4502742 | -0.1697343 | 0.26658167 | -0.6367065 | 0.52431607 | 0.99989001 |
| NMRK1 | 117.004109 | 0.20786461 | 0.45035619 | 0.46155602 | 0.64439974 | 0.99989001 |
| OSTF1 | 246.143822 | 0.56587972 | 0.16292991 | 3.47314825 | 0.00051439 | 0.09952814 |
| PCSK5 | 793.797725 | 0.44898991 | 0.33190471 | 1.35276752 | 0.17612991 | 0.99989001 |

| | | | | | | |
|---|---|---|---|---|---|---|
| RFK | 151.952187 | -0.3987636 | 0.50456457 | -0.7903123 | 0.42934541 | 0.99989001 |
| GCNT1 | 53.7389894 | 0.01062949 | 0.51413975 | 0.02067432 | 0.98350546 | 0.99989001 |
| PRUNE2 | 82.1919454 | 0.21978862 | 0.58941386 | 0.37289354 | 0.70922768 | 0.99989001 |
| VPS13A | 455.064428 | 0.09893494 | 0.27304037 | 0.36234546 | 0.71709389 | 0.99989001 |
| GNA14 | 2765.57026 | 0.08433034 | 0.38836559 | 0.21714163 | 0.82809797 | 0.99989001 |
| GNA14-AS1 | 433.952319 | -0.3084571 | 0.58155665 | -0.5303991 | 0.59583528 | 0.99989001 |
| GNAQ | 2409.68103 | 0.27150868 | 0.2316579 | 1.17202429 | 0.24118731 | 0.99989001 |
| CEP78 | 54.2884564 | -0.3858316 | 0.28391423 | -1.3589725 | 0.17415529 | 0.99989001 |
| TLE4 | 189.000339 | 0.07894075 | 0.31683815 | 0.24915166 | 0.80324347 | 0.99989001 |
| TLE1 | 150.467348 | 0.30789657 | 0.38570672 | 0.79826604 | 0.42471612 | 0.99989001 |
| FRMD3 | 527.891439 | -0.8377809 | 0.40329163 | -2.0773574 | 0.03776858 | 0.70738774 |
| UBQLN1 | 237.409736 | 0.06524192 | 0.17177962 | 0.37980011 | 0.7040938 | 0.99989001 |
| GKAP1 | 71.8270658 | 0.07128797 | 0.36692262 | 0.19428611 | 0.84595186 | 0.99989001 |
| KIF27 | 86.8606502 | -0.0505866 | 0.33110493 | -0.1527812 | 0.87857085 | 0.99989001 |
| HNRNPK | 463.267286 | -0.1907813 | 0.36302438 | -0.5255329 | 0.59921276 | 0.99989001 |
| AGTPBP1 | 537.302031 | -0.5907847 | 0.48896526 | -1.2082345 | 0.22695708 | 0.99989001 |
| NAA35 | 195.206344 | -0.166087 | 0.19579629 | -0.848264 | 0.39629094 | 0.99989001 |
| GOLM1 | 145.386339 | -0.0451798 | 0.5896664 | -0.0766193 | 0.93892642 | 0.99989001 |
| ISCA1 | 55.3783497 | 0.1161421 | 0.30292773 | 0.38339869 | 0.70142417 | 0.99989001 |
| TUT7 | 196.149429 | 0.21640787 | 0.1894494 | 1.14229906 | 0.25332973 | 0.99989001 |
| DAPK1 | 369.129537 | 0.3307386 | 0.40026549 | 0.82629806 | 0.40863503 | 0.99989001 |
| CTSL | 183.739264 | -0.5186729 | 0.46620311 | -1.112547 | 0.26590303 | 0.99989001 |
| SPIN1 | 211.113033 | -0.0280958 | 0.17551559 | -0.1600759 | 0.87282127 | 0.99989001 |
| SECISBP2 | 301.378014 | -0.0584604 | 0.36255841 | -0.1612441 | 0.87190114 | 0.99989001 |
| AUH | 289.749116 | -0.106989 | 0.24888869 | -0.4298667 | 0.66729262 | 0.99989001 |
| NFIL3 | 50.9641045 | -0.6752349 | 0.69795798 | -0.9674435 | 0.33332236 | 0.99989001 |
| SPTLC1 | 192.054258 | 0.01753546 | 0.18128895 | 0.09672656 | 0.92294355 | 0.99989001 |
| IARS | 157.087891 | -0.2103619 | 0.35860443 | -0.5866128 | 0.55746381 | 0.99989001 |
| NOL8 | 82.9512095 | -0.351359 | 0.36028614 | -0.9752222 | 0.32945004 | 0.99989001 |
| CENPP | 177.120005 | -0.587938 | 0.43124844 | -1.3633395 | 0.17277553 | 0.99989001 |
| IPPK | 79.5966794 | 0.12482486 | 0.39743716 | 0.31407444 | 0.7534645 | 0.99989001 |
| BICD2 | 90.2200894 | 0.28456859 | 0.23655702 | 1.20295985 | 0.22899186 | 0.99989001 |
| ZNF484 | 53.4267326 | -0.175024 | 0.32275352 | -0.5422839 | 0.58762293 | 0.99989001 |
| CARD19 | 42.1627666 | 0.68454955 | 0.34185718 | 2.00244306 | 0.0452371 | NA |
| NINJ1 | 54.3152825 | 0.72449875 | 0.34046685 | 2.12795681 | 0.03334066 | 0.67281094 |
| FAM120A | 396.688824 | 2.80E-01 | 0.17353144 | 1.6163433 | 0.10602011 | 0.99989001 |
| PHF2 | 115.064184 | 0.14636744 | 0.24577514 | 0.59553393 | 0.55148662 | 0.99989001 |
| PTPDC1 | 89.3957337 | 0.11145228 | 0.26339523 | 0.42313706 | 0.67219527 | 0.99989001 |
| AL158152.2 | 124.899398 | 0.0452386 | 0.33982882 | 0.13312172 | 0.89409712 | 0.99989001 |
| ZNF169 | 50.5812221 | -0.0767574 | 0.31835648 | -0.2411053 | 0.80947351 | 0.99989001 |
| MFSD14B | 145.480226 | -0.1896415 | 0.25007514 | -0.758338 | 0.44824865 | 0.99989001 |
| C9orf3 | 1081.62143 | -0.0471529 | 0.22358465 | -0.2108951 | 0.83296909 | 0.99989001 |
| FANCC | 261.193992 | -0.5066947 | 0.24993235 | -2.0273275 | 0.04262893 | 0.76066939 |
| LINC00476 | 100.164594 | -0.3304469 | 0.46991268 | -0.7032091 | 0.48192545 | 0.99989001 |
| ERCC6L2 | 249.108556 | -0.2429868 | 0.19887538 | -1.2218043 | 0.22178165 | 0.99989001 |
| SLC35D2 | 47.6793142 | 0.15606504 | 0.33489865 | 0.46600678 | 0.64121065 | NA |
| HABP4 | 114.665377 | -0.0192838 | 0.38837637 | -0.0496525 | 0.96039933 | 0.99989001 |
| CDC14B | 213.244371 | 0.53221842 | 0.26670784 | 1.99551097 | 0.04598718 | 0.77310579 |
| AAED1 | 49.6944491 | -0.2369787 | 0.48874377 | -0.484873 | 0.62776644 | NA |
| ZNF782 | 122.570601 | 0.03900792 | 0.22360626 | 0.17444916 | 0.86151248 | 0.99989001 |
| MFSD14C | 302.169312 | 0.02743262 | 0.30014401 | 0.09139818 | 0.9271762 | 0.99989001 |
| TDRD7 | 114.105673 | 0.37372078 | 0.37648649 | 0.99265389 | 0.32087866 | 0.99989001 |
| TMOD1 | 110.706635 | 0.94217224 | 0.47907616 | 1.96664395 | 0.04922427 | 0.78717121 |
| TSTD2 | 77.8805987 | -0.1092921 | 0.30509987 | -0.3582176 | 0.72018051 | 0.99989001 |

| | | | | | | |
|---|---|---|---|---|---|---|
| NCBP1 | 74.2238161 | -0.0619525 | 0.23729348 | -0.2610797 | 0.79403106 | 0.99989001 |
| XPA | 76.162648 | -0.0963758 | 0.26921122 | -0.3579931 | 0.72034848 | 0.99989001 |
| ANP32B | 127.651944 | -0.2522642 | 0.24615355 | -1.0248245 | 0.30544599 | 0.99989001 |
| NANS | 52.7517287 | -0.1359383 | 0.34463228 | -0.3944446 | 0.69325286 | 0.99989001 |
| TBC1D2 | 85.7890014 | -0.9140078 | 0.47130986 | -1.9392928 | 0.05246569 | 0.81270649 |
| COL15A1 | 98.3708374 | 0.40140828 | 0.98339879 | 0.40818464 | 0.68313812 | 0.99989001 |
| TGFBR1 | 126.392281 | 0.22302639 | 0.28479301 | 0.78311748 | 0.43355813 | 0.99989001 |
| SEC61B | 168.248864 | 0.02821059 | 0.45526282 | 0.06196551 | 0.9505903 | 0.99989001 |
| STX17 | 131.200202 | -0.1229942 | 0.32502406 | -0.3784156 | 0.7051219 | 0.99989001 |
| ERP44 | 213.644692 | 0.12095955 | 0.15950171 | 0.75835895 | 0.44823613 | 0.99989001 |
| INVS | 461.270083 | 0.27431766 | 0.22818837 | 1.20215447 | 0.22930369 | 0.99989001 |
| TEX10 | 55.8703382 | 0.04223722 | 0.31672847 | 0.13335466 | 0.8939129 | 0.99989001 |
| ALDOB | 169.8956 | 2.31386176 | 1.03548347 | 2.23457141 | 0.0254455 | 0.63387007 |
| RNF20 | 49.3540687 | 0.06132147 | 0.39707936 | 0.15443127 | 0.8772697 | NA |
| NIPSNAP3A | 44.6339706 | 0.14049734 | 0.29851912 | 0.4706477 | 0.63789234 | NA |
| NIPSNAP3B | 50.8719741 | 0.02004523 | 0.42806444 | 0.0468276 | 0.96265063 | 0.99989001 |
| ABCA1 | 147.505196 | 0.23283508 | 0.63453684 | 0.36693706 | 7.14E-01 | 0.99989001 |
| SLC44A1 | 398.070043 | 0.42068914 | 0.32258737 | 1.30410913 | 0.19219638 | 0.99989001 |
| FSD1L | 87.4680133 | 0.16155387 | 0.25239513 | 0.64008313 | 0.52211856 | 0.99989001 |
| FKTN | 99.3158209 | 0.01778442 | 0.30465193 | 0.0583762 | 0.95344897 | 0.99989001 |
| ZNF462 | 286.609855 | 0.49524107 | 0.32068602 | 1.54431761 | 0.12251141 | 0.99989001 |
| RAD23B | 302.551081 | -0.0077835 | 0.18208638 | -0.0427464 | 0.96590369 | 0.99989001 |
| KLF4 | 135.834894 | -0.3207156 | 0.56139233 | -0.5712861 | 0.56780571 | 0.99989001 |
| ELP1 | 101.267789 | 0.02438507 | 0.2281885 | 0.10686373 | 0.91489709 | 0.99989001 |
| CTNNAL1 | 247.347193 | 0.71843529 | 0.44469583 | 1.61556561 | 0.10618826 | 0.99989001 |
| TMEM245 | 323.047782 | 0.00146437 | 0.22590452 | 0.00648225 | 0.99482795 | 0.99989001 |
| TXN | 134.72473 | -0.0090482 | 0.42846961 | -0.0211175 | 0.98315191 | 0.99989001 |
| ECPAS | 456.462208 | -0.1170319 | 0.30846951 | -0.3793955 | 0.70439419 | 0.99989001 |
| PTGR1 | 134.730478 | 0.17170288 | 0.30089938 | 0.57063221 | 0.56824898 | 0.99989001 |
| C9orf84 | 172.585924 | 0.48573033 | 0.68375709 | 0.71038433 | 0.47746584 | 0.99989001 |
| UGCG | 824.646172 | -0.1115759 | 0.69633057 | -0.1602341 | 0.87269669 | 0.99989001 |
| SUSD1 | 83.6915365 | 0.0028713 | 0.31741412 | 0.0090459 | 0.99278251 | 0.99989001 |
| PTBP3 | 471.854453 | -0.0357595 | 0.24600339 | -0.1453619 | 0.88442512 | 0.99989001 |
| HSDL2 | 167.207141 | 0.51759943 | 0.31055128 | 1.66671164 | 0.09557176 | 0.99989001 |
| KIAA1958 | 68.8897463 | 0.32140544 | 0.39573551 | 0.81217234 | 0.41669275 | 0.99989001 |
| INIP | 54.9375648 | -0.2913828 | 0.32229239 | -0.9040947 | 0.36594518 | 0.99989001 |
| SNX30 | 54.6966262 | 0.75208627 | 0.38083817 | 1.97481851 | 0.04828875 | 0.78123673 |
| FKBP15 | 124.826068 | 0.1393802 | 0.30960951 | 0.45018062 | 0.65258021 | 0.99989001 |
| SLC31A1 | 54.4707611 | -0.3597579 | 0.33236179 | -1.0824285 | 0.27906217 | 0.99989001 |
| CDC26 | 48.1087247 | -0.157368 | 0.43166517 | -0.3645602 | 0.7154397 | NA |
| RGS3 | 437.819516 | -0.2098506 | 0.39505687 | -0.5311908 | 0.59528658 | 0.99989001 |
| ZNF618 | 152.68803 | 0.0695838 | 0.3254692 | 0.21379534 | 0.83070666 | 0.99989001 |
| ATP6V1G1 | 159.229487 | 0.17430136 | 0.43031679 | 0.4050536 | 0.68543811 | 0.99989001 |
| ASTN2 | 49.8713018 | -0.1057153 | 0.49894708 | -0.2118768 | 0.83220313 | NA |
| CDK5RAP2 | 157.738396 | -0.2195375 | 0.23805926 | -0.9221968 | 0.35642595 | 0.99989001 |
| MEGF9 | 106.201706 | 0.25587997 | 0.26793003 | 0.95502536 | 0.33956487 | 0.99989001 |
| FBXW2 | 139.492478 | -0.1505574 | 0.21394005 | -0.7037362 | 0.48159706 | 0.99989001 |
| PSMD5 | 112.692728 | -0.089011 | 0.20832857 | -0.4272627 | 0.66918798 | 0.99989001 |
| CNTRL | 96.7963086 | -0.0021877 | 0.37351943 | -0.005857 | 0.9953268 | 0.99989001 |
| RAB14 | 229.165487 | 0.14863611 | 0.22129774 | 0.67165672 | 0.50180226 | 0.99989001 |
| GSN | 769.386208 | -0.4880714 | 0.6628265 | -0.7363486 | 0.46151857 | 0.99989001 |
| STOM | 1181.27495 | -0.5894606 | 0.49432623 | -1.1924526 | 0.23308384 | 0.99989001 |
| DAB2IP | 437.099646 | 0.12614777 | 0.3094297 | 0.40767827 | 0.68350989 | 0.99989001 |
| TTLL11 | 71.5944163 | 0.38311765 | 0.37629749 | 1.01812437 | 0.30861885 | 0.99989001 |

| | | | | | | |
|---|---|---|---|---|---|---|
| NDUFA8 | 75.1414576 | -0.1072751 | 0.41823146 | -0.2564969 | 0.79756719 | 0.99989001 |
| LHX6 | 46.4937382 | -0.2488236 | 0.7551461 | -0.3295039 | 0.74177486 | NA |
| MRRF | 139.076257 | -0.2287884 | 0.31512874 | -0.7260156 | 0.46782921 | 0.99989001 |
| PDCL | 76.475167 | 0.15286475 | 0.36284266 | 0.42129761 | 0.67353777 | 0.99989001 |
| RC3H2 | 258.859437 | 0.15784532 | 0.18847185 | 0.83750078 | 0.40231114 | 0.99989001 |
| RABGAP1 | 447.441637 | 0.11170182 | 0.21000496 | 0.53190088 | 0.59479465 | 0.99989001 |
| STRBP | 131.187333 | -0.528957 | 0.21836273 | -2.4223777 | 0.01541931 | 0.52205731 |
| DENND1A | 577.602567 | 0.24648523 | 0.34625305 | 0.71186442 | 0.47654874 | 0.99989001 |
| NEK6 | 107.019769 | 0.09176545 | 0.51900699 | 0.17680967 | 0.8596579 | 0.99989001 |
| PSMB7 | 182.814239 | -0.2467622 | 0.24336572 | -1.0139563 | 0.31060359 | 0.99989001 |
| NR6A1 | 141.70028 | -0.0258863 | 0.4751661 | -0.0544784 | 0.95655404 | 0.99989001 |
| OLFML2A | 58.1543438 | 0.42617518 | 0.53543841 | 0.79593689 | 0.42606872 | 0.99989001 |
| RPL35 | 247.436924 | -0.0609617 | 0.5078251 | -0.1200446 | 0.9044478 | 0.99989001 |
| ARPC5L | 127.250941 | -0.5304442 | 0.41899653 | -1.2659872 | 0.20551768 | 0.99989001 |
| GOLGA1 | 84.6806497 | -0.2082566 | 0.35635441 | -0.5844086 | 0.55894542 | 0.99989001 |
| SCAI | 152.974859 | -0.0573155 | 0.28261357 | -0.2028053 | 0.83928724 | 0.99989001 |
| PPP6C | 181.877714 | -0.1665266 | 0.1529694 | -1.0886269 | 0.27631846 | 0.99989001 |
| RABEPK | 59.9392372 | -0.2888785 | 0.39432066 | -0.7325979 | 0.46380374 | 0.99989001 |
| HSPA5 | 365.316514 | 0.04816964 | 0.66890982 | 0.07201215 | 0.94259224 | 0.99989001 |
| GAPVD1 | 272.418551 | 0.02905655 | 0.1571974 | 0.18484118 | 0.85335361 | 0.99989001 |
| MAPKAP1 | 433.275476 | 0.12060689 | 0.26132834 | 0.46151475 | 0.64442934 | 0.99989001 |
| PBX3 | 430.807713 | 0.22791945 | 0.25197348 | 0.90453747 | 0.36571048 | 0.99989001 |
| MVB12B | 135.682213 | -0.5245198 | 0.33963216 | -1.5443762 | 0.12249723 | 0.99989001 |
| ZBTB43 | 81.8893712 | 0.18584301 | 0.37103589 | 0.5008761 | 0.61645832 | 0.99989001 |
| RALGPS1 | 107.126541 | -0.5560687 | 0.4794911 | -1.159706 | 0.24616852 | 0.99989001 |
| ANGPTL2 | 91.4560696 | -1.8563336 | 0.716149 | -2.5921053 | 0.00953906 | 0.44953399 |
| GARNL3 | 43.5034361 | 0.15620677 | 0.44919418 | 0.34774888 | 0.72802879 | NA |
| RPL12 | 92.7415098 | -0.1005723 | 0.4424086 | -0.2273291 | 0.82016787 | 0.99989001 |
| LRSAM1 | 93.6346108 | -0.1507359 | 0.39820532 | -0.378538 | 0.70503094 | 0.99989001 |
| FAM129B | 168.031102 | -0.2187971 | 0.4505715 | -0.485599 | 0.62725149 | 0.99989001 |
| STXBP1 | 222.447312 | 0.39371516 | 0.26224978 | 1.50129836 | 0.13327841 | 0.99989001 |
| SH2D3C | 572.916438 | -0.1836648 | 0.32672825 | -0.5621332 | 0.57402527 | 0.99989001 |
| CDK9 | 92.7900017 | -0.1592343 | 0.36300031 | -0.4386615 | 0.66090683 | 0.99989001 |
| FPGS | 86.4453476 | 0.00292183 | 0.42298824 | 0.00690759 | 0.99448858 | 0.99989001 |
| ENG | 2129.27885 | 0.3322827 | 0.62670179 | 0.53020863 | 0.59596729 | 0.99989001 |
| DPM2 | 41.3746151 | -0.5223167 | 0.43208819 | -1.2088197 | 0.22673211 | NA |
| FAM102A | 84.6381491 | -0.0698072 | 0.48475649 | -0.1440047 | 0.88549675 | 0.99989001 |
| SLC25A25 | 52.5024893 | 0.16981 | 0.38712171 | 0.43864758 | 0.66091691 | 0.99989001 |
| C9orf16 | 106.457229 | -0.4400247 | 0.52114317 | -0.8443452 | 0.39847656 | 0.99989001 |
| CIZ1 | 56.1808012 | -0.1906966 | 0.32373365 | -0.5890541 | 0.55582498 | 0.99989001 |
| GOLGA2 | 91.3612906 | -0.1989529 | 0.29416251 | -0.6763368 | 0.49882685 | 0.99989001 |
| SWI5 | 47.549534 | -0.5390427 | 0.44680038 | -1.2064509 | 0.22764367 | NA |
| COQ4 | 65.8450334 | -0.047714 | 0.34920464 | -0.1366362 | 0.89131834 | 0.99989001 |
| SLC27A4 | 79.7579401 | -0.4546954 | 0.35863174 | -1.2678615 | 0.20484741 | 0.99989001 |
| URM1 | 69.217903 | -0.0651978 | 0.32058312 | -0.2033725 | 0.83884387 | 0.99989001 |
| ODF2 | 95.4164635 | -0.2451723 | 0.31252784 | -0.7844815 | 0.43275766 | 0.99989001 |
| GLE1 | 64.2348653 | 0.17663573 | 0.291638 | 0.60566776 | 0.5447354 | 0.99989001 |
| SPTAN1 | 436.095633 | 0.29178913 | 0.27985505 | 1.04264379 | 2.97E-01 | 0.99989001 |
| SET | 234.331622 | 0.00149662 | 0.38786214 | 0.00385864 | 0.99692126 | 0.99989001 |
| PKN3 | 158.570509 | -0.5735253 | 0.29567158 | -1.9397376 | 0.05241159 | 0.81270649 |
| ZER1 | 149.823774 | 0.2406804 | 0.27016448 | 0.89086618 | 0.37300097 | 0.99989001 |
| TBC1D13 | 52.0617732 | -0.2511548 | 0.26576136 | -0.9450387 | 0.34463913 | 0.99989001 |
| KYAT1 | 48.9012352 | 0.36504137 | 0.40390418 | 0.90378212 | 0.36611094 | NA |
| LRRC8A | 137.810538 | 0.35747236 | 0.31327445 | 1.14108367 | 0.25383511 | 0.99989001 |

| | | | | | | |
|---|---|---|---|---|---|---|
| NUP188 | 68.838043 | 0.1193341 | 0.29417347 | 0.40565897 | 0.68499319 | 0.99989001 |
| SH3GLB2 | 40.9237827 | -0.1516182 | 0.37337311 | -0.4060769 | 0.68468611 | NA |
| MIGA2 | 41.6906099 | 0.06521212 | 0.27890706 | 0.23381307 | 0.8151301 | NA |
| PTPA | 84.4603833 | -0.0996504 | 0.35363504 | -0.2817887 | 0.77810552 | 0.99989001 |
| IER5L | 94.3823567 | -0.4629116 | 0.41461995 | -1.1164721 | 0.26422009 | 0.99989001 |
| NTMT1 | 42.5392311 | 0.11049833 | 0.34189069 | 0.32319782 | 0.74654543 | NA |
| C9orf78 | 64.2500389 | 0.08888143 | 0.38931093 | 0.22830448 | 0.81940954 | 0.99989001 |
| USP20 | 47.0675628 | -0.3347636 | 0.35376595 | -0.9462856 | 0.34400295 | NA |
| FNBP1 | 592.896309 | 0.52563385 | 0.2047878 | 2.56672441 | 1.03E-02 | 0.46125545 |
| GPR107 | 206.026467 | 0.2097498 | 0.19086141 | 1.09896392 | 0.2717838 | 0.99989001 |
| HMCN2 | 71.2004672 | -3.6399656 | 1.04417569 | -3.4859705 | 0.00049036 | 0.09952814 |
| ASS1 | 86.3399518 | 0.2476532 | 0.44836253 | 0.55235034 | 0.58070834 | 0.99989001 |
| FUBP3 | 199.770441 | 0.11872834 | 0.22003498 | 0.53958847 | 0.58948087 | 0.99989001 |
| ABL1 | 648.19369 | 0.23410078 | 0.31249351 | 0.74913804 | 0.45377401 | 0.99989001 |
| AIF1L | 176.735851 | -0.0985614 | 0.40775572 | -0.2417167 | 0.80899965 | 0.99989001 |
| NUP214 | 259.516957 | -0.1968591 | 0.17294824 | -1.1382546 | 0.25501417 | 0.99989001 |
| AL157938.2 | 62.1174102 | -0.1146911 | 0.54789974 | -0.2093286 | 0.83419171 | 0.99989001 |
| PRRC2B | 453.705761 | 0.068104 | 0.20173881 | 0.33758503 | 0.73567593 | 0.99989001 |
| POMT1 | 43.9023584 | -0.5128843 | 0.30349058 | -1.6899513 | 0.09103727 | NA |
| RAPGEF1 | 1026.04298 | 0.0357901 | 0.35176887 | 0.10174321 | 0.9189605 | 0.99989001 |
| MED27 | 254.546068 | -0.0015516 | 0.31240651 | -0.0049666 | 0.99603728 | 0.99989001 |
| SETX | 694.586934 | 0.26484486 | 0.30644565 | 0.86424741 | 0.387452 | 0.99989001 |
| TTF1 | 90.0549598 | 0.11906882 | 0.26812763 | 0.44407516 | 0.65698825 | 0.99989001 |
| DDX31 | 60.4840663 | -0.369217 | 0.47521992 | -0.7769392 | 0.43719462 | 0.99989001 |
| TSC1 | 101.409844 | -0.2545631 | 0.20745308 | -1.2270878 | 0.21978961 | 0.99989001 |
| GTF3C5 | 43.2838042 | -0.4057162 | 0.36106192 | -1.1236749 | 0.26115097 | NA |
| RALGDS | 96.1469046 | 0.0359048 | 0.48858089 | 0.07348794 | 0.94141784 | 0.99989001 |
| ABO | 126.932427 | 0.34731559 | 0.44640687 | 0.77802475 | 0.43655442 | 0.99989001 |
| RPL7A | 295.158124 | -0.176475 | 0.55269653 | -0.3192981 | 0.74950044 | 0.99989001 |
| SURF4 | 142.891379 | 0.20567512 | 0.36647205 | 0.56123003 | 0.57464073 | 0.99989001 |
| VAV2 | 114.620251 | 0.80456176 | 0.4834178 | 1.66431967 | 0.09604856 | 0.99989001 |
| BRD3 | 60.1694096 | 0.25126673 | 0.33034628 | 0.76061617 | 0.44688636 | 0.99989001 |
| WDR5 | 42.141735 | 0.09899894 | 0.37806396 | 0.26185766 | 0.79343118 | NA |
| RXRA | 141.056807 | 0.23319499 | 0.43360881 | 0.5378004 | 0.59071485 | 0.99989001 |
| CAMSAP1 | 100.968601 | 0.12729735 | 0.35760519 | 0.35597177 | 0.72186171 | 0.99989001 |
| UBAC1 | 60.3852555 | 0.15213914 | 0.27071076 | 0.56199887 | 0.57411679 | 0.99989001 |
| NACC2 | 214.950562 | -0.3652737 | 0.24699957 | -1.4788434 | 0.13918219 | 0.99989001 |
| ENTR1 | 40.5649159 | -0.2983871 | 0.32166781 | -0.9276251 | 0.3536021 | NA |
| PMPCA | 47.3671689 | -0.2287433 | 0.28958308 | -0.7899056 | 0.4295829 | NA |
| SEC16A | 115.744434 | -0.1083853 | 0.23117537 | -0.4688445 | 0.63918082 | 0.99989001 |
| NOTCH1 | 378.235626 | 0.16904046 | 0.32064192 | 0.52719388 | 0.59805896 | 0.99989001 |
| AL590226.1 | 81.9308293 | 1.38985977 | 0.48177818 | 2.88485415 | 0.00391595 | 0.31113256 |
| EGFL7 | 1657.70085 | -0.2565819 | 0.34762083 | -0.7381085 | 0.46044851 | 0.99989001 |
| AGPAT2 | 41.6450062 | 0.39924966 | 0.31347797 | 1.27361313 | 0.20280056 | NA |
| FAM69B | 63.3446499 | 0.38640984 | 0.63086033 | 0.61251251 | 0.54019873 | 0.99989001 |
| SNHG7 | 151.035424 | -0.1230207 | 0.44202912 | -0.278309 | 0.78077518 | 0.99989001 |
| RABL6 | 124.052328 | 0.03634473 | 0.29655383 | 0.12255696 | 0.90245794 | 0.99989001 |
| PHPT1 | 95.9389883 | 0.18140786 | 0.43719241 | 0.41493825 | 0.67818711 | 0.99989001 |
| EDF1 | 180.575748 | -0.1535342 | 0.49603286 | -0.3095242 | 0.75692279 | 0.99989001 |
| FBXW5 | 104.0247 | 0.16203546 | 0.35619554 | 0.45490591 | 0.64917693 | 0.99989001 |
| PAXX | 61.1376287 | 0.26252309 | 0.54452915 | 0.48211026 | 0.62972762 | 0.99989001 |
| AL807752.6 | 40.3232357 | -1.0525046 | 0.64636896 | -1.6283341 | 0.10345406 | NA |
| NPDC1 | 78.2403903 | -0.2002884 | 0.48382154 | -0.4139716 | 0.67889491 | 0.99989001 |
| MAN1B1 | 57.4899923 | -0.1417062 | 0.31906986 | -0.4441227 | 0.6569539 | 0.99989001 |

| | | | | | | |
|---|---|---|---|---|---|---|
| DPP7 | 68.8329727 | -0.2724442 | 0.44671767 | -0.6098801 | 0.54194126 | 0.99989001 |
| SSNA1 | 75.1981448 | -0.1608882 | 0.41348425 | -0.3891036 | 0.69719951 | 0.99989001 |
| TUBB4B | 159.140755 | 0.59218642 | 0.65162038 | 0.90879052 | 0.36346071 | 0.99989001 |
| NELFB | 87.8944693 | -0.0579078 | 0.35697614 | -0.1622175 | 0.87113459 | 0.99989001 |
| EXD3 | 174.046856 | -0.252073 | 0.26516503 | -0.9506268 | 0.34179383 | 0.99989001 |
| MRPL41 | 120.021081 | -0.2702342 | 0.43762152 | -0.6175066 | 0.53690061 | 0.99989001 |
| DPH7 | 51.0213333 | -0.2747026 | 0.26864192 | -1.0225605 | 0.3065157 | 0.99989001 |
| ARRDC1 | 58.9385397 | -0.143533 | 0.32411425 | -0.442847 | 0.65787638 | 0.99989001 |
| EHMT1 | 550.062965 | 0.10937676 | 0.24363662 | 0.448934 | 0.65347927 | 0.99989001 |
| RIC8A | 41.2740958 | -0.1125824 | 0.39126373 | -0.2877406 | 0.77354534 | NA |
| PSMD13 | 130.836303 | -0.0015024 | 0.25425104 | -0.005909 | 0.99528536 | 0.99989001 |
| PGGHG | 52.2074564 | -0.0341414 | 0.44335491 | -0.0770069 | 0.93861804 | 0.99989001 |
| IFITM2 | 1388.06726 | -0.4872215 | 0.60850218 | -0.8006899 | 0.4233112 | 0.99989001 |
| IFITM3 | 2508.56448 | -0.2619632 | 0.53624581 | -0.4885132 | 0.62518637 | 0.99989001 |
| SIGIRR | 99.2094257 | -0.5772779 | 0.38121401 | -1.5143145 | 0.12994609 | 0.99989001 |
| PTDSS2 | 56.2183097 | -0.1766729 | 0.30966543 | -0.5705282 | 0.56831952 | 0.99989001 |
| RNH1 | 107.830592 | -0.0877921 | 0.43173174 | -0.2033487 | 0.83886251 | 0.99989001 |
| HRAS | 53.8187455 | -0.1460068 | 0.40730677 | -0.3584689 | 0.71999245 | 0.99989001 |
| PHRF1 | 73.6264854 | -0.1427343 | 0.25898917 | -0.5511206 | 0.58155097 | 0.99989001 |
| DEAF1 | 107.956006 | -0.4577013 | 0.1904898 | -2.40276 | 0.01627186 | 0.54105741 |
| TALDO1 | 93.5756392 | 0.32341419 | 0.28937195 | 1.11764183 | 0.26372 | 0.99989001 |
| RPLP2 | 327.266416 | -0.3605128 | 0.56621114 | -0.636711 | 0.52431314 | 0.99989001 |
| PNPLA2 | 60.6435585 | 0.05830035 | 0.2754236 | 0.21167522 | 0.83236042 | 0.99989001 |
| CD151 | 281.730513 | -0.0913606 | 0.56631928 | -0.1613235 | 0.87183861 | 0.99989001 |
| POLR2L | 182.267209 | -0.200838 | 0.47506727 | -0.422757 | 0.67247255 | 0.99989001 |
| TSPAN4 | 115.207414 | -0.2048532 | 0.36396664 | -0.562835 | 0.57354726 | 0.99989001 |
| AP2A2 | 292.883698 | 0.10449678 | 0.22653134 | 0.46129061 | 0.64459012 | 0.99989001 |
| TOLLIP | 59.8450496 | -0.3068875 | 0.27926055 | -1.0989288 | 0.27179914 | 0.99989001 |
| MOB2 | 112.417685 | -0.1708023 | 0.20338938 | -0.8397798 | 0.40103185 | 0.99989001 |
| CD81 | 767.372064 | -0.2058893 | 0.46363205 | -0.444079 | 0.65698544 | 0.99989001 |
| TSSC4 | 48.2311758 | 0.08745078 | 0.54458852 | 0.1605814 | 0.87242311 | NA |
| KCNQ1 | 161.40556 | 0.37661761 | 0.25828542 | 1.45814507 | 0.14480056 | 0.99989001 |
| KCNQ1OT1 | 216.420238 | 0.25154375 | 0.44263348 | 0.56828902 | 0.56983873 | 0.99989001 |
| CDKN1C | 98.8521906 | 0.05951924 | 0.53553471 | 0.11113983 | 0.91150547 | 0.99989001 |
| NAP1L4 | 244.454196 | 0.12663499 | 0.187027 | 0.67709472 | 0.49834586 | 0.99989001 |
| CARS | 108.587061 | 0.17961948 | 0.2745256 | 0.65429045 | 0.5129247 | 0.99989001 |
| OSBPL5 | 221.302658 | 0.34951786 | 0.31164755 | 1.12151644 | 0.2620681 | 0.99989001 |
| ZNF195 | 73.7048556 | -0.1030381 | 0.39014382 | -0.2641027 | 0.79170076 | 0.99989001 |
| NUP98 | 296.15327 | -0.1954238 | 0.25394993 | -0.7695366 | 0.44157485 | 0.99989001 |
| RHOG | 53.9654526 | 0.10487845 | 0.38158312 | 0.27485086 | 0.78343082 | 0.99989001 |
| STIM1 | 383.91639 | 0.47540645 | 0.35618574 | 1.33471498 | 0.18196965 | 0.99989001 |
| RRM1 | 50.0569755 | -0.2369558 | 0.36317799 | -0.652451 | 0.51411028 | 0.99989001 |
| AC104389.4 | 128.995528 | 0.36207164 | 0.47440345 | 0.76321461 | 0.44533542 | 0.99989001 |
| TRIM5 | 86.5012964 | -0.1644078 | 0.46284813 | -0.355209 | 0.72243303 | 0.99989001 |
| TRIM22 | 225.086017 | 0.04827678 | 0.53828911 | 0.08968559 | 0.92853707 | 0.99989001 |
| FAM160A2 | 60.3422207 | -0.5146223 | 0.26465225 | -1.9445225 | 0.05183247 | 0.81270649 |
| CAVIN3 | 105.783601 | -1.1917132 | 0.58674867 | -2.0310455 | 0.04225038 | 0.76066939 |
| SMPD1 | 46.8489004 | -0.2696238 | 0.37666122 | -0.7158257 | 0.474099 | NA |
| APBB1 | 42.7103889 | 0.30670123 | 0.36400512 | 0.84257396 | 0.39946676 | NA |
| AC084337.2 | 71.0276395 | -0.2452603 | 0.32441143 | -0.7560162 | 0.44963948 | 0.99989001 |
| DNHD1 | 94.019106 | -0.141585 | 0.30768974 | -0.4601552 | 0.64540481 | 0.99989001 |
| ILK | 162.587629 | 0.26023479 | 0.24561856 | 1.05950785 | 0.28936855 | 0.99989001 |
| TAF10 | 43.8420014 | -0.3909284 | 0.50594375 | -0.7726717 | 0.43971668 | NA |
| TPP1 | 102.351021 | -0.1118728 | 0.46628465 | -0.2399239 | 0.81038924 | 0.99989001 |

| | | | | | | |
|---|---|---|---|---|---|---|
| DCHS1 | 94.5621529 | 0.11719971 | 0.40432678 | 0.28986383 | 0.77192042 | 0.99989001 |
| MRPL17 | 99.2078798 | 0.07623347 | 0.53010886 | 0.14380719 | 0.88565272 | 0.99989001 |
| EIF3F | 101.491243 | 0.03184951 | 0.38656537 | 0.082391 | 0.93433579 | 0.99989001 |
| TRIM66 | 102.089919 | -0.2309595 | 0.3629213 | -0.6363901 | 0.52452219 | 0.99989001 |
| RPL27A | 351.905559 | 0.00025668 | 0.54623087 | 0.0004699 | 0.99962507 | 0.99989001 |
| ST5 | 139.440103 | -0.0721771 | 0.42465519 | -0.1699664 | 0.86503657 | 0.99989001 |
| TMEM9B | 81.2541482 | -0.0473115 | 0.34789373 | -0.1359942 | 0.89182589 | 0.99989001 |
| DENND5A | 608.072677 | -0.1032528 | 0.35952946 | -0.2871887 | 0.77396782 | 0.99989001 |
| TMEM41B | 128.830982 | 0.10572985 | 0.32304277 | 0.32729366 | 0.74344579 | 0.99989001 |
| IPO7 | 238.124662 | -0.0995323 | 0.31207161 | -0.3189407 | 0.74977151 | 0.99989001 |
| ZNF143 | 80.2401246 | -0.0067802 | 0.30184794 | -0.0224622 | 0.98207923 | 0.99989001 |
| SWAP70 | 778.021838 | -0.4659557 | 0.32357726 | -1.4400138 | 0.1498635 | 0.99989001 |
| SBF2-AS1 | 47.6931067 | 0.2564341 | 0.34908117 | 0.73459734 | 0.46258476 | NA |
| SBF2 | 1869.23905 | 0.12343606 | 0.24070879 | 0.51280247 | 0.6080895 | 0.99989001 |
| AC011092.2 | 72.5097095 | 0.12969159 | 0.57224908 | 0.22663486 | 0.82070768 | 0.99989001 |
| ADM | 125.953362 | 0.13985044 | 1.11128868 | 0.12584529 | 0.89985439 | 0.99989001 |
| MTRNR2L8 | 97.0804134 | 0.33992997 | 0.71286607 | 0.4768497 | 0.63346916 | 0.99989001 |
| RNF141 | 50.3094114 | -0.4116915 | 0.30687944 | -1.3415416 | 0.17974468 | 0.99989001 |
| LYVE1 | 122.94912 | -0.0914932 | 0.55032602 | -0.1662528 | 8.68E-01 | 0.99989001 |
| CTR9 | 86.0178714 | 0.53917689 | 0.27998516 | 1.9257338 | 0.05413762 | 0.81854801 |
| EIF4G2 | 388.901436 | 0.15171239 | 0.33351815 | 0.45488495 | 0.64919201 | 0.99989001 |
| ZBED5 | 97.7785077 | 0.01145794 | 0.23722801 | 0.04829928 | 0.96147773 | 0.99989001 |
| GALNT18 | 1600.77967 | 1.06739372 | 0.59379033 | 1.79759365 | 0.07224143 | 0.91336395 |
| USP47 | 433.799718 | 0.07824786 | 0.17913199 | 0.43681676 | 0.66224424 | 0.99989001 |
| PARVA | 218.403429 | 0.14530967 | 0.35460561 | 0.40977826 | 0.68196861 | 0.99989001 |
| TEAD1 | 962.856767 | -0.2105787 | 0.303652 | -0.693487 | 0.48800398 | 0.99989001 |
| ARNTL | 73.5124189 | 0.12312674 | 0.35610708 | 0.34575761 | 0.72952488 | 0.99989001 |
| BTBD10 | 221.511935 | 0.02206371 | 0.21177996 | 0.10418222 | 0.91702474 | 0.99989001 |
| FAR1 | 419.958858 | 0.22507532 | 0.20444334 | 1.10091785 | 0.27093241 | 0.99989001 |
| RRAS2 | 132.854902 | -0.1373552 | 0.28535259 | -0.4813525 | 0.630266 | 0.99989001 |
| COPB1 | 198.280441 | 0.18144924 | 0.22346399 | 0.81198424 | 0.41680067 | 0.99989001 |
| PDE3B | 68.5972073 | 1.97500237 | 0.53722117 | 3.67633009 | 0.00023661 | 0.0714383 |
| SOX6 | 211.221994 | -1.2540996 | 0.5089073 | -2.4642988 | 0.01372816 | 0.50652625 |
| C11orf58 | 253.393013 | -0.3053746 | 0.21869911 | -1.396323 | 0.16261725 | 0.99989001 |
| PLEKHA7 | 570.251289 | -0.2686205 | 0.48370365 | -0.555341 | 0.57866145 | 0.99989001 |
| RPS13 | 307.0742 | -0.3575727 | 0.59420975 | -0.6017617 | 0.54733279 | 0.99989001 |
| PIK3C2A | 1434.51472 | -0.0600441 | 0.22516536 | -0.2666668 | 0.78972573 | 0.99989001 |
| NUCB2 | 205.507193 | -0.1304056 | 0.27630241 | -0.471967 | 0.63695031 | 0.99989001 |
| SERGEF | 129.001874 | -0.0151126 | 0.21026513 | -0.0718742 | 0.94270199 | 0.99989001 |
| HPS5 | 65.513085 | 0.24834407 | 0.2864659 | 0.86692367 | 0.38598383 | 0.99989001 |
| GTF2H1 | 118.303706 | -0.4255761 | 0.22609032 | -1.882328 | 0.05979149 | 0.85885114 |
| LDHA | 384.991555 | -0.0018516 | 0.48574774 | -0.0038119 | 0.99695852 | 0.99989001 |
| TSG101 | 180.418599 | 0.13437606 | 0.26701795 | 0.5032473 | 0.61479041 | 0.99989001 |
| UEVLD | 134.738941 | 0.11281757 | 0.26705609 | 0.42244895 | 0.67269735 | 0.99989001 |
| SPTY2D1 | 47.0552841 | -0.1937983 | 0.30861326 | -0.6279651 | 0.53002682 | NA |
| ZDHHC13 | 145.981641 | -0.6527874 | 0.3090987 | -2.1119059 | 0.03469452 | 0.6855346 |
| NAV2 | 730.68557 | -0.9008892 | 0.54163137 | -1.6632885 | 0.09625469 | 0.99989001 |
| HTATIP2 | 41.714333 | 0.32322681 | 0.38757553 | 0.83397114 | 0.40429725 | NA |
| PRMT3 | 111.066533 | -0.2556832 | 0.24461679 | -1.0452399 | 0.29591211 | 0.99989001 |
| SVIP | 92.4573053 | 0.36138814 | 0.27900327 | 1.29528284 | 0.19522268 | 0.99989001 |
| LGR4 | 408.766185 | -0.2401056 | 0.46371419 | -0.5177879 | 0.60460623 | 0.99989001 |
| LIN7C | 77.1663208 | 0.07930103 | 0.35634782 | 0.22253827 | 0.82389489 | 0.99989001 |
| BDNF-AS | 228.861409 | -0.2877796 | 0.43568531 | -0.6605217 | 0.50891909 | 0.99989001 |
| METTL15 | 430.344739 | -0.0700814 | 0.28726156 | -0.2439638 | 0.80725889 | 0.99989001 |

| | | | | | | |
|---|---|---|---|---|---|---|
| ARL14EP | 60.5327178 | 0.3624131 | 0.29308508 | 1.23654573 | 0.21625578 | 0.99989001 |
| MPPED2 | 174.180805 | -0.0998795 | 0.47786605 | -0.2090116 | 0.83443919 | 0.99989001 |
| DNAJC24 | 79.4895655 | -0.1896873 | 0.30011311 | -0.6320526 | 0.52735251 | 0.99989001 |
| IMMP1L | 156.410583 | 0.02547941 | 0.24367781 | 0.10456186 | 0.91672348 | 0.99989001 |
| ELP4 | 352.178093 | -0.0297616 | 0.22644438 | -0.1314301 | 0.89543504 | 0.99989001 |
| RCN1 | 89.136996 | -0.3775329 | 0.57882109 | -0.6522445 | 0.51424347 | 0.99989001 |
| EIF3M | 107.053378 | -0.0142699 | 0.29797025 | -0.0478905 | 0.96180354 | 0.99989001 |
| PRRG4 | 56.6544436 | -0.4677215 | 0.59759257 | -0.7826763 | 0.43381723 | 0.99989001 |
| QSER1 | 221.935741 | 0.00792251 | 0.15878806 | 0.04989359 | 0.96020718 | 0.99989001 |
| TCP11L1 | 72.5407945 | -0.072199 | 0.30541554 | -0.2363959 | 0.81312548 | 0.99989001 |
| CSTF3 | 133.229109 | 0.05991463 | 0.24641038 | 0.2431498 | 0.80788936 | 0.99989001 |
| HIPK3 | 884.92984 | 0.1156225 | 0.355975 | 0.32480512 | 0.74532857 | 0.99989001 |
| KIAA1549L | 81.7151849 | 0.02301621 | 0.5621833 | 0.04094076 | 0.96734313 | 0.99989001 |
| CD59 | 98.0791046 | -0.8433262 | 0.5536013 | -1.5233457 | 0.12767224 | 0.99989001 |
| FBXO3 | 130.319646 | 0.22443766 | 0.25826932 | 0.86900629 | 0.38484369 | 0.99989001 |
| LMO2 | 537.494804 | -0.1373317 | 0.55350181 | -0.2481142 | 0.80404603 | 0.99989001 |
| CAPRIN1 | 248.873154 | -0.0194746 | 0.21633689 | -0.0900197 | 0.92827156 | 0.99989001 |
| NAT10 | 74.2553519 | -0.2004407 | 0.38799297 | -0.516609 | 0.60542913 | 0.99989001 |
| CAT | 105.693689 | 0.09360367 | 0.4825598 | 0.1939732 | 0.84619687 | 0.99989001 |
| APIP | 70.9093037 | -0.6917811 | 0.32186271 | -2.1493049 | 0.03161024 | 0.67281094 |
| PDHX | 120.210705 | -0.1088821 | 0.19916271 | -0.5466993 | 0.58458531 | 0.99989001 |
| TRIM44 | 417.366631 | 0.05440592 | 0.24146594 | 0.22531508 | 0.82173417 | 0.99989001 |
| LDLRAD3 | 408.819856 | 0.89838476 | 0.32857289 | 2.73420233 | 0.00625316 | 0.38373055 |
| PRR5L | 176.780927 | -0.4197337 | 0.32301619 | -1.2994199 | 0.19379985 | 0.99989001 |
| TRAF6 | 80.710691 | 0.19965415 | 0.25394975 | 0.78619551 | 0.43175295 | 0.99989001 |
| C11orf74 | 42.1445723 | 0.06533561 | 0.35099346 | 0.1861448 | 0.85233121 | NA |
| API5 | 134.982383 | -0.1526543 | 0.17441992 | -0.8752114 | 0.38145889 | 0.99989001 |
| TTC17 | 747.44759 | 0.00791639 | 0.22295714 | 0.03550635 | 0.97167599 | 0.99989001 |
| HSD17B12 | 563.58488 | 0.00749654 | 0.2192297 | 0.0341949 | 0.97272173 | 0.99989001 |
| ALKBH3 | 64.9113155 | -0.1443084 | 0.30681566 | -0.4703425 | 0.63811035 | 0.99989001 |
| C11orf96 | 208.678329 | -0.3807239 | 0.92346066 | -0.4122795 | 0.6801346 | 0.99989001 |
| ACCS | 50.8994887 | -0.1377518 | 0.61138335 | -0.2253117 | 0.82173681 | 0.99989001 |
| EXT2 | 231.761719 | 0.07303683 | 0.16778158 | 0.43530901 | 0.66333815 | 0.99989001 |
| TSPAN18 | 199.674416 | 1.50120551 | 0.53667213 | 2.79724888 | 0.00515398 | 0.35047074 |
| TP53I11 | 97.5299543 | 0.27078235 | 0.52179127 | 0.51894764 | 0.60379725 | 0.99989001 |
| PRDM11 | 65.3865093 | -0.0951314 | 0.26794229 | -0.3550442 | 0.72255644 | 0.99989001 |
| CRY2 | 73.8855553 | 0.22433416 | 0.31317545 | 0.71632103 | 0.47379314 | 0.99989001 |
| PHF21A | 770.782247 | 0.47005878 | 0.30081319 | 1.56262687 | 0.11814038 | 0.99989001 |
| DGKZ | 164.896952 | 0.00956727 | 0.30284231 | 0.03159158 | 0.97479776 | 0.99989001 |
| MDK | 52.3518193 | -0.8200787 | 0.78898993 | -1.0394033 | 0.29861723 | 0.99989001 |
| AMBRA1 | 875.107408 | 0.04205412 | 0.30156759 | 0.13945173 | 0.8890932 | 0.99989001 |
| ATG13 | 81.9928612 | -0.3472876 | 0.30931648 | -1.1227582 | 0.26154019 | 0.99989001 |
| ARHGAP1 | 64.7655682 | -0.1878348 | 0.26902112 | -0.6982156 | 0.48504234 | 0.99989001 |
| CKAP5 | 116.779595 | -0.0296741 | 0.20557759 | -0.1443451 | 0.88522796 | 0.99989001 |
| LRP4 | 138.169673 | 0.68492968 | 0.42150376 | 1.62496695 | 0.1041696 | 0.99989001 |
| C11orf49 | 248.813477 | -0.1608936 | 0.24864129 | -0.6470914 | 0.51757279 | 0.99989001 |
| ARFGAP2 | 67.1834829 | 0.0955486 | 0.31643076 | 0.30195738 | 0.76268455 | 0.99989001 |
| DDB2 | 113.286044 | -0.1421579 | 0.56990598 | -0.249441 | 0.80301965 | 0.99989001 |
| MADD | 111.569988 | 0.12757364 | 0.24086311 | 0.52965206 | 0.59635319 | 0.99989001 |
| PSMC3 | 43.4232602 | -0.1290885 | 0.47910564 | -0.2694363 | 0.78759395 | NA |
| CELF1 | 578.60055 | -0.0474813 | 0.2381516 | -0.1993745 | 0.84196983 | 0.99989001 |
| NDUFS3 | 50.6885759 | -0.1182984 | 0.46524425 | -0.2542716 | 0.79928571 | 0.99989001 |
| MTCH2 | 97.4032941 | -0.4008278 | 0.31652143 | -1.2663526 | 0.20538686 | 0.99989001 |
| FNBP4 | 412.570265 | 0.06953425 | 0.23664039 | 0.2938393 | 0.7688807 | 0.99989001 |

| | | | | | | |
|---|---|---|---|---|---|---|
| NUP160 | 225.218018 | 0.11603221 | 0.21053167 | 0.55113896 | 0.58153842 | 0.99989001 |
| PTPRJ | 172.385652 | 0.99007301 | 0.42158 | 2.34848193 | 0.01885011 | 0.56657733 |
| APLNR | 38.963228 | 1.01208155 | 1.25035351 | 0.80943632 | 0.41826422 | NA |
| TNKS1BP1 | 106.625638 | -0.0778353 | 0.23730585 | -0.3279957 | 0.74291493 | 0.99989001 |
| SSRP1 | 66.6484308 | -0.0658086 | 0.43307764 | -0.1519556 | 0.87922197 | 0.99989001 |
| TIMM10 | 42.3867225 | 0.1284141 | 0.38473773 | 0.33377048 | 0.73855276 | NA |
| UBE2L6 | 79.7808429 | 0.01078576 | 0.53900073 | 0.02001065 | 0.98403488 | 0.99989001 |
| SERPING1 | 274.855438 | 0.05933332 | 0.37893426 | 0.15657944 | 0.8755763 | 0.99989001 |
| ZDHHC5 | 106.173202 | 0.01596484 | 0.20151415 | 0.07922441 | 0.93685413 | 0.99989001 |
| CTNND1 | 432.348917 | 0.05491319 | 0.17339069 | 0.31670208 | 0.75146967 | 0.99989001 |
| LPXN | 62.1508071 | -0.0042142 | 0.34615131 | -0.0121745 | 0.99028637 | 0.99989001 |
| FAM111A | 130.017847 | 0.11582935 | 0.30625146 | 0.37821649 | 0.70526978 | 0.99989001 |
| OSBP | 121.222326 | -0.0122002 | 0.1945939 | -0.0626957 | 0.95000882 | 0.99989001 |
| PATL1 | 113.494195 | -0.1009555 | 0.45669829 | -0.2210552 | 0.82504946 | 0.99989001 |
| STX3 | 91.5316479 | -0.3137751 | 0.3943714 | -0.7956336 | 0.42624501 | 0.99989001 |
| MRPL16 | 53.1755287 | -0.4647973 | 0.35832945 | -1.2971228 | 0.19458895 | 0.99989001 |
| PRPF19 | 65.5219216 | 0.32487538 | 0.36696381 | 0.88530631 | 0.37599145 | 0.99989001 |
| TMEM109 | 123.904448 | 0.04136074 | 0.47223099 | 0.08758583 | 0.93020586 | 0.99989001 |
| TMEM132A | 45.6900448 | 0.54956901 | 0.53293978 | 1.03120284 | 0.30244571 | NA |
| SLC15A3 | 57.476126 | -0.0584868 | 0.46311151 | -0.126291 | 0.8995016 | 0.99989001 |
| VPS37C | 53.1293252 | 0.05413877 | 0.37166135 | 0.14566695 | 0.88418432 | 0.99989001 |
| DDB1 | 129.895457 | 0.15792069 | 0.24355262 | 0.64840483 | 0.51672315 | 0.99989001 |
| CPSF7 | 116.442872 | 0.03013092 | 0.24799502 | 0.12149809 | 0.90329653 | 0.99989001 |
| AP003108.2 | 105.13406 | 0.24137608 | 0.38272289 | 0.63068106 | 0.52824908 | 0.99989001 |
| TMEM258 | 63.9387509 | -0.206133 | 0.53084568 | -0.3883106 | 0.69778618 | 0.99989001 |
| FADS1 | 39.4740381 | -0.0407567 | 0.53398854 | -0.0763251 | 0.93916043 | NA |
| FADS3 | 52.4670324 | -1.504031 | 0.4507879 | -3.3364495 | 0.00084856 | 0.14233155 |
| FTH1 | 1831.3912 | 0.80360869 | 0.47474146 | 1.69272912 | 0.09050704 | 0.98720685 |
| ASRGL1 | 491.817264 | -0.4343585 | 0.39603666 | -1.0967634 | 0.27274483 | 0.99989001 |
| AHNAK | 553.38012 | -0.3830746 | 0.34793066 | -1.1010085 | 0.27089295 | 0.99989001 |
| EML3 | 87.8034089 | -0.3369931 | 0.4244072 | -0.7940326 | 0.42717648 | 0.99989001 |
| B3GAT3 | 56.6530377 | -0.1737307 | 0.3388921 | -0.5126432 | 0.60820094 | 0.99989001 |
| GANAB | 166.235889 | -0.0085209 | 0.37799175 | -0.0225427 | 0.98201508 | 0.99989001 |
| UQCC3 | 50.0040677 | -0.4778153 | 0.41729313 | -1.1450351 | 0.25219463 | 0.99989001 |
| UBXN1 | 85.2490275 | 0.00299004 | 0.45437179 | 0.0065806 | 0.99474947 | 0.99989001 |
| TTC9C | 65.5625267 | 0.04814133 | 0.26884258 | 0.17906885 | 0.85788364 | 0.99989001 |
| TMEM223 | 60.069487 | -0.1751003 | 0.36708721 | -0.4769991 | 0.63336273 | 0.99989001 |
| TMEM179B | 84.7867894 | 0.00299217 | 0.35152474 | 0.00851198 | 0.9932085 | 0.99989001 |
| NXF1 | 47.4542674 | 0.12845801 | 0.35717135 | 0.35965374 | 0.71910609 | NA |
| STX5 | 140.724638 | 0.18633521 | 0.18898269 | 0.98599086 | 0.3241376 | 0.99989001 |
| SLC3A2 | 270.932652 | -0.4430373 | 0.24301056 | -1.8231195 | 0.0682853 | 0.88709358 |
| RARRES3 | 84.7562281 | 0.94571395 | 0.43514918 | 2.17330973 | 0.02975701 | 0.6652708 |
| PLA2G16 | 128.96872 | -0.0278346 | 0.43584611 | -0.0638635 | 0.94907894 | 0.99989001 |
| ATL3 | 215.623801 | -0.0147744 | 0.22052151 | -0.0669975 | 0.94658371 | 0.99989001 |
| RTN3 | 199.361131 | 0.09292682 | 0.20008058 | 0.46444697 | 0.64232755 | 0.99989001 |
| MARK2 | 183.81425 | 0.19170287 | 0.31978597 | 0.59947242 | 0.5488579 | 0.99989001 |
| OTUB1 | 53.7061077 | -0.0706532 | 0.25724726 | -0.2746508 | 0.78358452 | 0.99989001 |
| MACROD1 | 56.6823099 | 0.26152537 | 0.33063382 | 0.79098191 | 0.42895455 | 0.99989001 |
| STIP1 | 84.8636239 | 0.09584827 | 0.36379313 | 0.26346917 | 0.79218898 | 0.99989001 |
| VEGFB | 54.0313355 | -0.3829937 | 0.3489788 | -1.0974699 | 0.27243603 | 0.99989001 |
| FKBP2 | 79.1521031 | 0.01111671 | 0.41055132 | 0.02707752 | 0.97839791 | 0.99989001 |
| PPP1R14B | 51.1212162 | -0.4819253 | 0.40756078 | -1.1824623 | 0.23702232 | 0.99989001 |
| BAD | 55.7773458 | -0.3178049 | 0.44869671 | -0.7082845 | 0.47876862 | 0.99989001 |
| ESRRA | 58.4628991 | 0.08385471 | 0.36679633 | 0.22861381 | 0.81916909 | 0.99989001 |

| | | | | | | |
|---|---|---|---|---|---|---|
| TRMT112 | 116.8739 | -0.163627 | 0.52835861 | -0.3096893 | 0.75679723 | 0.99989001 |
| PRDX5 | 80.3059374 | -0.2717205 | 0.47581567 | -0.5710624 | 0.56795732 | 0.99989001 |
| SF1 | 327.141106 | -0.0444859 | 0.29273776 | -0.151965 | 0.87921454 | 0.99989001 |
| MAP4K2 | 69.5733899 | 0.71149388 | 0.35909082 | 1.98137583 | 0.04754914 | 0.77516398 |
| PPP2R5B | 59.5707053 | 0.20677669 | 0.27645461 | 0.74795892 | 0.45448494 | 0.99989001 |
| VPS51 | 138.637338 | 0.27095368 | 0.18594353 | 1.45718262 | 0.14506598 | 0.99989001 |
| FAU | 352.830067 | -0.3309541 | 0.56823075 | -0.5824291 | 0.56027771 | 0.99989001 |
| CAPN1 | 89.4945008 | -0.109376 | 0.2125812 | -0.514514 | 0.60689268 | 0.99989001 |
| POLA2 | 54.7204652 | -0.2657538 | 0.27811151 | -0.9555655 | 0.33929182 | 0.99989001 |
| CDC42EP2 | 49.6281604 | 0.41183495 | 0.47506307 | 0.86690584 | 0.3859936 | NA |
| DPF2 | 67.6052751 | 0.14024201 | 0.23720481 | 0.59122749 | 0.55436801 | 0.99989001 |
| FRMD8 | 149.668124 | 0.27411688 | 0.33381278 | 0.82116952 | 0.41154972 | 0.99989001 |
| NEAT1 | 11801.758 | -0.0641633 | 0.47087193 | -0.1362648 | 0.89161192 | 0.99989001 |
| MALAT1 | 73401.3909 | -0.3184344 | 0.30365053 | -1.0486872 | 0.29432212 | 0.99989001 |
| SCYL1 | 54.9796843 | 0.12682295 | 0.28561934 | 0.44402788 | 0.65702242 | 0.99989001 |
| LTBP3 | 164.947702 | -0.2969011 | 0.48481394 | -0.6124022 | 0.54027168 | 0.99989001 |
| FAM89B | 59.7769148 | -0.3469597 | 0.61563506 | -0.5635801 | 0.57303994 | 0.99989001 |
| EHBP1L1 | 178.549753 | -0.4250159 | 0.30261304 | -1.4044863 | 0.16017408 | 0.99989001 |
| MAP3K11 | 200.028519 | 0.26842958 | 0.37782612 | 0.71045797 | 0.47742018 | 0.99989001 |
| SIPA1 | 97.5425386 | -0.0423668 | 0.30430391 | -0.1392253 | 0.8892721 | 0.99989001 |
| RELA | 100.105878 | 0.09415842 | 0.30223241 | 0.31154311 | 0.75538778 | 0.99989001 |
| KAT5 | 52.0958606 | -0.1202338 | 0.38763073 | -0.3101761 | 0.75642704 | 0.99989001 |
| RNASEH2C | 68.6731453 | 0.46552356 | 0.46878775 | 0.99303696 | 0.32069194 | 0.99989001 |
| CFL1 | 449.631846 | -0.3762705 | 0.53176098 | -0.7075934 | 0.47919782 | 0.99989001 |
| MUS81 | 43.6998393 | -0.0580173 | 0.30757611 | -0.1886275 | 0.85038477 | NA |
| CCDC85B | 432.270754 | -0.4063434 | 0.59298298 | -0.685253 | 0.49318428 | 0.99989001 |
| DRAP1 | 57.803211 | 0.02744144 | 0.47072911 | 0.05829561 | 0.95351316 | 0.99989001 |
| SART1 | 42.1203423 | 0.02059631 | 0.37642881 | 0.05471501 | 0.95636551 | NA |
| BANF1 | 86.0262607 | -0.5730731 | 0.51758863 | -1.107198 | 0.26820831 | 0.99989001 |
| SF3B2 | 120.58959 | -0.1194456 | 0.21895204 | -0.5455331 | 0.5853869 | 0.99989001 |
| PACS1 | 550.233093 | 0.19487933 | 0.23230758 | 0.83888494 | 0.40153388 | 0.99989001 |
| RAB1B | 94.0604802 | -0.1686384 | 0.44889905 | -0.3756711 | 0.70716141 | 0.99989001 |
| BRMS1 | 38.5605143 | -0.0485363 | 0.49397441 | -0.0982567 | 0.92172845 | NA |
| ZDHHC24 | 53.5597107 | -0.2315142 | 0.35833581 | -0.6460817 | 0.51822642 | 0.99989001 |
| CTSF | 35.9098581 | 0.75984407 | 0.49073917 | 1.54836646 | 0.12153409 | NA |
| CCS | 62.6518853 | -0.0399314 | 0.33130875 | -0.1205263 | 0.90406621 | 0.99989001 |
| RBM14-RBM | 65.8876856 | 0.19664766 | 0.34572458 | 0.56879862 | 0.56949281 | 0.99989001 |
| RBM4 | 57.5773633 | -0.284598 | 0.36931857 | -0.7706029 | 0.44094232 | 0.99989001 |
| RBM4B | 46.6743818 | 0.31434003 | 0.37167057 | 0.84574906 | 0.39769275 | NA |
| C11orf80 | 62.3552169 | 0.63141029 | 0.39469756 | 1.59973191 | 0.10965807 | 0.99989001 |
| KDM2A | 724.22366 | 0.12297157 | 0.22292606 | 0.55162493 | 0.58120535 | 0.99989001 |
| GRK2 | 82.086579 | 0.45926213 | 0.28603813 | 1.60559758 | 0.10836236 | 0.99989001 |
| ANKRD13D | 86.904259 | 0.43410655 | 0.30530159 | 1.42189415 | 0.15505698 | 0.99989001 |
| RAD9A | 122.274636 | 0.02901874 | 0.28203346 | 0.10289114 | 0.91804937 | 0.99989001 |
| PPP1CA | 82.4931743 | 0.23254093 | 0.47414733 | 0.49044025 | 0.6238224 | 0.99989001 |
| CORO1B | 59.1776708 | -0.3923323 | 0.46328498 | -0.8468488 | 0.39707938 | 0.99989001 |
| TMEM134 | 42.2692017 | 0.12653939 | 0.43870212 | 0.28844034 | 0.77300969 | NA |
| AIP | 49.6057268 | -0.0717442 | 0.47429429 | -0.1512652 | 0.87976654 | NA |
| CDK2AP2 | 46.5984512 | 0.2066118 | 0.52415932 | 0.39417749 | 0.69345001 | NA |
| GSTP1 | 329.648656 | -0.0643546 | 0.47539301 | -0.1353714 | 0.89231822 | 0.99989001 |
| NDUFV1 | 90.3408424 | 0.22531757 | 0.26759347 | 0.84201444 | 0.39977987 | 0.99989001 |
| NDUFS8 | 81.9369011 | -0.1501791 | 0.40511056 | -0.3707115 | 0.71085242 | 0.99989001 |
| TCIRG1 | 51.3032172 | -0.0009056 | 0.34592991 | -0.0026178 | 0.99791133 | 0.99989001 |
| CHKA | 139.410039 | -0.2215473 | 0.24868868 | -0.8908622 | 0.37300311 | 0.99989001 |

| | | | | | | |
|---|---|---|---|---|---|---|
| KMT5B | 283.828149 | 0.40393898 | 0.30356449 | 1.33065293 | 0.18330324 | 0.99989001 |
| LRP5 | 250.400807 | 0.4670363 | 0.47114811 | 0.99127277 | 0.32155241 | 0.99989001 |
| PPP6R3 | 957.630158 | 0.07421 | 0.24997981 | 0.29686397 | 0.76657036 | 0.99989001 |
| CPT1A | 247.608147 | 0.34507718 | 0.34891065 | 0.98901304 | 0.32265676 | 0.99989001 |
| IGHMBP2 | 39.3850142 | -0.2406099 | 0.40255887 | -0.5977011 | 0.55003942 | NA |
| TPCN2 | 51.4044344 | 0.435948 | 0.47635052 | 0.91518321 | 0.36009546 | 0.99989001 |
| CCND1 | 100.859795 | -0.2948139 | 0.62472935 | -0.4719066 | 0.63699347 | 0.99989001 |
| ORAOV1 | 50.0156032 | -0.1993465 | 0.3498661 | -0.5697793 | 0.56882739 | 0.99989001 |
| PPFIA1 | 293.901982 | -0.3429113 | 0.26870328 | -1.2761709 | 0.20189511 | 0.99989001 |
| AP002336.2 | 112.97797 | 0.0038329 | 0.37874868 | 0.01011991 | 0.99192562 | 0.99989001 |
| CTTN | 290.371679 | -0.2872496 | 0.20363812 | -1.4105885 | 0.15836598 | 0.99989001 |
| NADSYN1 | 73.2007444 | 0.08422255 | 0.31473693 | 0.26759667 | 0.7890098 | 0.99989001 |
| AP002495.2 | 88.8276549 | 0.09502947 | 0.30933942 | 0.30720129 | 0.75869016 | 0.99989001 |
| RNF121 | 80.873112 | 0.23038978 | 0.31265624 | 0.73687888 | 0.46119601 | 0.99989001 |
| IL18BP | 68.5706773 | -0.4566524 | 0.44916961 | -1.0166592 | 0.30931557 | 0.99989001 |
| NUMA1 | 214.492841 | -0.1178277 | 0.24710402 | -0.4768343 | 0.6334801 | 0.99989001 |
| LAMTOR1 | 121.579483 | -0.0784684 | 0.34983552 | -0.2243008 | 0.82252329 | 0.99989001 |
| INPPL1 | 51.2058694 | -0.4229625 | 0.29568658 | -1.430442 | 0.15259019 | 0.99989001 |
| CLPB | 61.2150808 | -0.1363206 | 0.3414317 | -0.3992618 | 0.68970028 | 0.99989001 |
| PDE2A | 375.678913 | 0.38498189 | 0.33208787 | 1.15927719 | 0.24634322 | 0.99989001 |
| ARAP1 | 54.1178416 | 0.07323258 | 0.27147562 | 0.26975747 | 0.78734684 | 0.99989001 |
| ATG16L2 | 111.255255 | 0.76423114 | 0.22241441 | 3.43606842 | 0.00059022 | 0.10865841 |
| FCHSD2 | 1887.76667 | -0.0007754 | 0.29993439 | -0.0025852 | 0.99793732 | 0.99989001 |
| FAM168A | 381.311666 | -0.0048248 | 0.17372835 | -0.0277723 | 0.97784377 | 0.99989001 |
| RAB6A | 477.969169 | -0.0465263 | 0.16777922 | -0.2773065 | 0.78154476 | 0.99989001 |
| MRPL48 | 54.9187019 | -0.0291797 | 0.28094657 | -0.1038621 | 0.91727876 | 0.99989001 |
| COA4 | 59.8286291 | -0.3602841 | 0.3967631 | -0.9080584 | 0.36384739 | 0.99989001 |
| PAAF1 | 59.3729267 | -0.0481816 | 0.30292138 | -0.1590565 | 0.87362436 | 0.99989001 |
| C2CD3 | 124.183773 | 0.18342927 | 0.24642062 | 0.74437468 | 0.45664983 | 0.99989001 |
| PPME1 | 153.837592 | 0.08857423 | 0.25307688 | 0.34998942 | 0.72634664 | 0.99989001 |
| PGM2L1 | 90.7162442 | 0.6864688 | 0.36881934 | 1.86126029 | 0.06270743 | 0.85940342 |
| POLD3 | 88.4860598 | 0.02426996 | 0.24900668 | 0.09746709 | 0.92235547 | 0.99989001 |
| RNF169 | 202.783879 | -0.3975841 | 0.28617171 | -1.3893201 | 0.16473544 | 0.99989001 |
| XRRA1 | 210.374531 | -0.404797 | 0.61332443 | -0.6600047 | 0.50925079 | 0.99989001 |
| SPCS2 | 191.377497 | 0.31071514 | 0.35979432 | 0.863591 | 0.38781261 | 0.99989001 |
| ARRB1 | 443.60974 | -0.1524496 | 0.28654586 | -0.532025 | 0.59470868 | 0.99989001 |
| RPS3 | 596.641595 | -0.1427931 | 0.57035935 | -0.2503563 | 0.8023118 | 0.99989001 |
| GDPD5 | 89.2699727 | -0.2155513 | 0.45080711 | -0.4781454 | 0.63254671 | 0.99989001 |
| SERPINH1 | 255.858293 | -0.1697732 | 0.57369628 | -0.2959287 | 0.76728456 | 0.99989001 |
| UVRAG | 839.360393 | -0.1934118 | 0.39685549 | -0.4873607 | 0.62600272 | 0.99989001 |
| THAP12 | 61.4442223 | 0.29107654 | 0.2402568 | 1.21152259 | 0.22569518 | 0.99989001 |
| EMSY | 266.389117 | -0.1783451 | 0.26371222 | -0.6762869 | 0.49885852 | 0.99989001 |
| LRRC32 | 757.703761 | -0.4352896 | 0.47714389 | -0.9122816 | 0.36162052 | 0.99989001 |
| ACER3 | 523.622915 | 0.07702137 | 0.39149042 | 0.19673885 | 0.84403191 | 0.99989001 |
| CLNS1A | 137.928362 | 0.00126835 | 0.29252844 | 0.00433582 | 0.99654053 | 0.99989001 |
| RSF1 | 523.386792 | 0.2088868 | 0.18489669 | 1.12974873 | 0.25858212 | 0.99989001 |
| AAMDC | 161.968011 | 0.21676191 | 0.28923456 | 0.74943295 | 0.4535963 | 0.99989001 |
| AP002812.2 | 57.0756119 | 0.2622858 | 0.34010866 | 0.77118234 | 0.44059886 | 0.99989001 |
| INTS4 | 117.727002 | 0.20281939 | 0.23341268 | 0.86893045 | 0.38488518 | 0.99989001 |
| ALG8 | 59.7350831 | -0.358044 | 0.36436505 | -0.982652 | 0.32577876 | 0.99989001 |
| GAB2 | 377.489872 | 0.0801936 | 0.27845762 | 0.2879921 | 0.77335279 | 0.99989001 |
| AP003086.2 | 106.887442 | 0.38716447 | 0.45955484 | 0.8424772 | 0.3995209 | 0.99989001 |
| NARS2 | 147.307806 | -0.259655 | 0.31966779 | -0.8122652 | 0.41663948 | 0.99989001 |
| PRCP | 467.078495 | 0.5071603 | 0.45781961 | 1.10777322 | 0.26795977 | 0.99989001 |

| | | | | | | |
|---|---|---|---|---|---|---|
| RAB30 | 100.217112 | -0.0497459 | 0.36560193 | -0.1360657 | 0.89176935 | 0.99989001 |
| RAB30-AS1 | 99.7408069 | 0.07264006 | 0.35162652 | 0.20658301 | 0.83633553 | 0.99989001 |
| PCF11 | 176.259478 | -0.1507694 | 0.20880726 | -0.7220506 | 0.47026335 | 0.99989001 |
| CCDC90B | 62.4473838 | 0.2715625 | 0.31962845 | 0.84961928 | 0.39553679 | 0.99989001 |
| DLG2 | 676.813043 | -0.2385634 | 0.43616463 | -0.5469572 | 0.58440815 | 0.99989001 |
| TMEM126B | 54.7623043 | -0.0092936 | 0.31456015 | -0.0295448 | 0.97643009 | 0.99989001 |
| CREBZF | 144.710133 | -0.0879712 | 0.25514558 | -0.3447884 | 0.73025343 | 0.99989001 |
| SYTL2 | 71.518848 | -0.9686481 | 0.57672068 | -1.6795792 | 0.09303921 | 0.99611342 |
| PICALM | 2017.2628 | -0.3401471 | 0.40262728 | -0.8448188 | 0.39821204 | 0.99989001 |
| EED | 77.1820366 | -0.2800474 | 0.27107307 | -1.0331068 | 0.30155392 | 0.99989001 |
| HIKESHI | 91.2878193 | 0.34446655 | 0.24160833 | 1.42572296 | 0.15394833 | 0.99989001 |
| AP001831.1 | 47.7039238 | -0.1596192 | 0.4934571 | -0.3234712 | 0.74633841 | NA |
| ME3 | 175.452767 | -0.1785613 | 0.25249264 | -0.707194 | 0.47944594 | 0.99989001 |
| PRSS23 | 617.19901 | 0.00371518 | 0.50698782 | 0.00732795 | 0.9941532 | 0.99989001 |
| AP001528.2 | 210.774012 | 0.97101787 | 0.56449965 | 1.72013902 | 0.08540718 | 0.95514629 |
| FZD4 | 322.896196 | 0.62818939 | 0.29597063 | 2.12247204 | 0.03379812 | 0.67281094 |
| TMEM135 | 568.06786 | 0.45894837 | 0.22739754 | 2.0182644 | 0.04356373 | 0.76826882 |
| CTSC | 111.300113 | 0.00319777 | 0.33742041 | 0.0094771 | 0.99243848 | 0.99989001 |
| NOX4 | 86.7440439 | 1.64922848 | 0.6749651 | 2.4434278 | 0.01454848 | 0.5186081 |
| CHORDC1 | 95.8475413 | -0.1121091 | 0.4358599 | -0.2572136 | 0.79701389 | 0.99989001 |
| SLC36A4 | 115.445407 | -0.3230416 | 0.27772858 | -1.1631557 | 0.24476634 | 0.99989001 |
| SMCO4 | 55.7185999 | -0.3742017 | 0.44511243 | -0.8406902 | 0.40052152 | 0.99989001 |
| CEP295 | 97.367481 | -0.2708054 | 0.27548783 | -0.9830032 | 0.32560586 | 0.99989001 |
| TAF1D | 222.876682 | 0.11752822 | 0.2894211 | 0.40608034 | 0.68468357 | 0.99989001 |
| MRE11 | 122.123838 | -0.2926343 | 0.20092127 | -1.4564627 | 0.14526474 | 0.99989001 |
| AMOTL1 | 403.678279 | 0.47676729 | 0.2671539 | 1.78461659 | 0.07432353 | 0.9206525 |
| CWC15 | 74.6714714 | -0.2370494 | 0.26258772 | -0.9027437 | 0.36666192 | 0.99989001 |
| SRSF8 | 73.1782603 | -0.1153219 | 0.34649009 | -0.332829 | 0.73926338 | 0.99989001 |
| ENDOD1 | 165.312253 | -0.3832292 | 0.32598936 | -1.1755882 | 0.23975947 | 0.99989001 |
| AP000787.1 | 252.708949 | -1.719309 | 0.6432734 | -2.67275 | 0.00752323 | 0.4053518 |
| SESN3 | 540.211695 | -3.2946709 | 1.04630849 | -3.1488523 | 0.00163913 | 0.20106436 |
| FAM76B | 90.6812767 | -0.3615461 | 0.25754553 | -1.4038142 | 0.16037418 | 0.99989001 |
| CEP57 | 148.282479 | -0.0944704 | 0.21895997 | -0.4314507 | 0.66614068 | 0.99989001 |
| MTMR2 | 240.039321 | 0.20881002 | 0.25804414 | 0.80920273 | 0.41839855 | 0.99989001 |
| MAML2 | 1381.9023 | 0.91813387 | 0.39022311 | 2.35284338 | 0.01863048 | 0.56643441 |
| CCDC82 | 154.033925 | 0.06123021 | 0.18807772 | 0.32555801 | 0.74475878 | 0.99989001 |
| YAP1 | 328.255093 | 0.2163006 | 0.32118503 | 0.67344547 | 0.50066393 | 0.99989001 |
| BIRC3 | 102.20359 | 1.32613943 | 0.6532487 | 2.03006821 | 0.04234961 | 0.76066939 |
| BIRC2 | 134.929977 | 0.48895608 | 0.23448726 | 2.08521383 | 0.0370499 | 0.70556534 |
| TMEM123 | 300.248381 | 0.16728763 | 0.3512217 | 0.47630209 | 0.63385918 | 0.99989001 |
| DCUN1D5 | 75.3975902 | -0.1602311 | 0.22459475 | -0.7134232 | 0.47558391 | 0.99989001 |
| DYNC2H1 | 238.880615 | -0.0173224 | 0.30058817 | -0.0576282 | 0.95404477 | 0.99989001 |
| AP002989.1 | 123.063258 | 0.89992884 | 0.57727694 | 1.55892046 | 0.11901521 | 0.99989001 |
| PDGFD | 545.264998 | 0.69645189 | 0.37387869 | 1.86277506 | 0.06249392 | 0.85940342 |
| CASP4 | 172.836672 | -0.1102134 | 0.33093286 | -0.3330387 | 0.73910511 | 0.99989001 |
| CASP1 | 41.602884 | 0.11065295 | 0.45300971 | 0.24426176 | 0.8070281 | NA |
| AASDHPPT | 79.9320087 | 0.03403952 | 0.26655685 | 0.12770078 | 0.89838577 | 0.99989001 |
| CWF19L2 | 255.49615 | -0.2244347 | 0.19827526 | -1.1319348 | 0.25766184 | 0.99989001 |
| ALKBH8 | 44.9352349 | -0.2622518 | 0.32453678 | -0.8080804 | 0.4190443 | NA |
| CUL5 | 344.115757 | -0.0685275 | 0.19975623 | -0.3430556 | 0.7315566 | 0.99989001 |
| ACAT1 | 133.37603 | 0.17090222 | 0.26183389 | 0.65271239 | 0.51394171 | 0.99989001 |
| NPAT | 145.94225 | 0.03652742 | 0.2621279 | 0.13934961 | 0.88917389 | 0.99989001 |
| ATM | 261.197198 | -0.077144 | 0.24506582 | -0.314789 | 0.75292187 | 0.99989001 |
| KDELC2 | 48.3414511 | -0.3330149 | 0.4013116 | -0.8298162 | 0.4066427 | NA |

| | | | | | | |
|---|---|---|---|---|---|---|
| DDX10 | 307.688325 | -0.1472862 | 0.34584794 | -0.4258699 | 0.67020267 | 0.99989001 |
| RDX | 827.976512 | 0.39014565 | 0.31514866 | 1.23797338 | 0.21572593 | 0.99989001 |
| ZC3H12C | 174.625204 | 0.25891522 | 0.34180971 | 0.75748352 | 0.44876023 | 0.99989001 |
| FDX1 | 81.0245059 | 0.18964909 | 0.22566678 | 0.84039438 | 0.4006873 | 0.99989001 |
| LAYN | 62.8146034 | -0.2011432 | 0.30157797 | -0.6669693 | 0.50479174 | 0.99989001 |
| SIK2 | 695.765679 | 0.07161047 | 0.43539526 | 0.16447233 | 0.86935933 | 0.99989001 |
| PPP2R1B | 169.355388 | -0.3216105 | 0.30757555 | -1.045631 | 0.29573143 | 0.99989001 |
| CRYAB | 614.454502 | 2.01323624 | 1.11651318 | 1.80314597 | 0.07136529 | 0.91166457 |
| DIXDC1 | 114.921299 | -0.1975247 | 0.47064069 | -0.4196932 | 0.67470962 | 0.99989001 |
| DLAT | 45.4532847 | 0.13921632 | 0.39775437 | 0.35000576 | 0.72633437 | NA |
| TIMM8B | 63.2356492 | 0.15530806 | 0.50668648 | 0.30651707 | 0.75921098 | 0.99989001 |
| PTS | 170.797708 | -0.3716257 | 0.17447601 | -2.1299531 | 0.03317549 | 0.67281094 |
| AP002884.1 | 66.0055491 | -0.5100716 | 0.39403467 | -1.2944842 | 0.19549823 | 0.99989001 |
| ZW10 | 43.8186401 | 0.27715415 | 0.2797737 | 0.99063689 | 0.32186292 | NA |
| USP28 | 64.4776793 | 0.26001548 | 0.2636873 | 0.98607507 | 0.32409628 | 0.99989001 |
| ZBTB16 | 1081.45135 | 0.51114797 | 0.82798737 | 0.61733788 | 0.53701188 | 0.99989001 |
| AP002518.2 | 128.224557 | 0.15069617 | 1.12504668 | 0.1339466 | 0.8934448 | 0.99989001 |
| NNMT | 454.473825 | 0.22943223 | 0.52319458 | 0.43852181 | 6.61E-01 | 1.00E+00 |
| REXO2 | 80.4509542 | 0.380619 | 0.41662299 | 0.91358138 | 0.36093686 | 0.99989001 |
| CADM1 | 148.514582 | 0.27159995 | 0.45068097 | 0.6026435 | 0.54674588 | 0.99989001 |
| SIK3 | 1219.74067 | -0.0379393 | 0.35649177 | -0.106424 | 0.91524592 | 0.99989001 |
| PAFAH1B2 | 180.866544 | 0.10404598 | 0.17095319 | 0.60862263 | 0.5427746 | 0.99989001 |
| SIDT2 | 43.7643705 | 0.64688228 | 0.2649838 | 2.44121445 | 0.01463796 | NA |
| TAGLN | 96.1316319 | -0.7660696 | 0.74747289 | -1.0248794 | 0.30542008 | 0.99989001 |
| PCSK7 | 182.743153 | 0.05317097 | 0.22670894 | 0.23453406 | 0.81457039 | 0.99989001 |
| RNF214 | 92.4303094 | -0.0776345 | 0.23483411 | -0.3305931 | 0.74095182 | 0.99989001 |
| BACE1 | 64.4621222 | 0.19524016 | 0.27284073 | 0.71558291 | 0.47424893 | 0.99989001 |
| CEP164 | 82.9886717 | -0.0683372 | 0.38492691 | -0.1775328 | 0.8590899 | 0.99989001 |
| DSCAML1 | 181.803875 | 1.2086998 | 0.71907036 | 1.68092007 | 0.09277844 | 0.99479708 |
| FXYD2 | 46.8527699 | 0.54850221 | 0.400259 | 1.37036821 | 0.17057199 | NA |
| MPZL2 | 289.379527 | -0.4970053 | 0.51930155 | -0.9570648 | 0.33853453 | 0.99989001 |
| UBE4A | 154.207203 | -0.1059675 | 0.17716261 | -0.5981371 | 0.54974849 | 0.99989001 |
| ATP5MG | 179.350918 | -0.0092077 | 0.38211135 | -0.0240969 | 0.98077532 | 0.99989001 |
| KMT2A | 527.423165 | 0.0988895 | 0.22334495 | 0.44276579 | 0.65793515 | 0.99989001 |
| ARCN1 | 111.66005 | -0.0922098 | 0.27811558 | -0.3315519 | 0.74022764 | 0.99989001 |
| PHLDB1 | 186.235509 | 0.12359034 | 0.26813201 | 0.46093093 | 0.64484816 | 0.99989001 |
| AP002954.1 | 46.1898534 | 0.10196741 | 0.66434996 | 0.15348449 | 0.87801622 | NA |
| DDX6 | 280.143344 | 0.11560027 | 0.15628301 | 0.73968549 | 0.45949086 | 0.99989001 |
| BCL9L | 59.6602211 | 0.21590322 | 0.34903995 | 0.61856304 | 0.53620425 | 0.99989001 |
| CCDC84 | 161.452073 | -0.4074776 | 0.27984464 | -1.4560852 | 0.14536907 | 0.99989001 |
| RPS25 | 379.772847 | -0.4462456 | 0.53354927 | -0.8363718 | 0.40294578 | 0.99989001 |
| TRAPPC4 | 72.4109307 | 0.28771584 | 0.35171047 | 0.81804742 | 0.41333011 | 0.99989001 |
| HYOU1 | 40.6632613 | 0.03984081 | 0.48274717 | 0.08252934 | 0.93422579 | NA |
| CBL | 277.344849 | -0.0418617 | 0.18740634 | -0.2233741 | 0.82324433 | 0.99989001 |
| MCAM | 388.074353 | -1.1949299 | 0.45649628 | -2.6176114 | 0.00885476 | 0.42668269 |
| OAF | 82.2615427 | -0.550893 | 0.56395693 | -0.9768353 | 0.32865072 | 0.99989001 |
| ARHGEF12 | 1163.6534 | 0.08549632 | 0.23857173 | 0.35836737 | 0.72006841 | 0.99989001 |
| UBASH3B | 127.225488 | 0.26304364 | 0.57405742 | 0.45821834 | 0.64679558 | 0.99989001 |
| JHY | 50.6474158 | 0.63616556 | 0.34718484 | 1.83235409 | 0.06689868 | 0.8781761 |
| HSPA8 | 288.970448 | 0.25635502 | 0.45549144 | 0.56280974 | 0.57356445 | 0.99989001 |
| GRAMD1B | 117.300221 | -2.1715646 | 0.55885804 | -3.8857178 | 1.02E-04 | 0.04053195 |
| TBRG1 | 128.516887 | 0.15941675 | 0.2605462 | 0.61185598 | 0.54063305 | 0.99989001 |
| SIAE | 53.9537037 | -0.4461392 | 0.44365007 | -1.0056106 | 0.31460294 | 0.99989001 |
| ESAM | 398.014711 | -0.4511236 | 0.39463336 | -1.1431461 | 0.25297795 | 0.99989001 |

| | | | | | | |
|---|---|---|---|---|---|---|
| MSANTD2 | 146.145736 | 0.03648191 | 0.25216393 | 0.14467536 | 0.88496719 | 0.99989001 |
| ROBO4 | 370.511426 | -0.5640785 | 0.38010771 | -1.4839964 | 0.13780988 | 0.99989001 |
| EI24 | 95.361619 | 0.01692818 | 0.32432751 | 0.0521947 | 0.95837356 | 0.99989001 |
| STT3A | 60.7993922 | 0.20702863 | 0.26026908 | 0.79544075 | 0.42635716 | 0.99989001 |
| FAM118B | 64.8151116 | 0.12486663 | 0.30510167 | 0.40926235 | 0.68234714 | 0.99989001 |
| SRPRA | 87.912911 | -0.0744334 | 0.32275443 | -0.2306192 | 0.81761067 | 0.99989001 |
| FOXRED1 | 48.1689891 | 0.1266937 | 0.39718063 | 0.31898258 | 0.74973973 | NA |
| ETS1 | 1392.4882 | 0.41568068 | 0.29693646 | 1.39989775 | 0.16154394 | 0.99989001 |
| FLI1 | 1151.04673 | -0.0824351 | 0.37202616 | -0.2215843 | 0.82463752 | 0.99989001 |
| ARHGAP32 | 164.002725 | 0.59128909 | 0.2516911 | 2.34926498 | 0.01881051 | 0.56657733 |
| NFRKB | 55.5044219 | 0.09649769 | 0.28311727 | 0.34084 | 0.73322403 | 0.99989001 |
| PRDM10 | 63.7194275 | -0.3437735 | 0.39734542 | -0.8651755 | 0.38694248 | 0.99989001 |
| APLP2 | 347.577869 | 0.097569 | 0.2973462 | 0.32813267 | 0.74281135 | 0.99989001 |
| ZBTB44 | 182.541374 | 0.22197155 | 0.24809536 | 0.89470253 | 0.37094612 | 0.99989001 |
| SNX19 | 133.37765 | -0.5839124 | 0.3079164 | -1.8963343 | 0.05791585 | 0.8467548 |
| OPCML | 196.504566 | -2.5073253 | 0.95784465 | -2.6176743 | 0.00885313 | 0.42668269 |
| JAM3 | 296.234524 | 0.35439966 | 0.19311759 | 1.83514954 | 0.06648353 | 0.8742469 |
| NCAPD3 | 47.6469689 | -0.1303844 | 0.25574067 | -0.5098306 | 0.61017017 | NA |
| VPS26B | 57.0172996 | 0.61371387 | 0.32629403 | 1.88086149 | 0.05999076 | 0.85885114 |
| ZMYND11 | 408.758345 | 0.39162838 | 0.17880798 | 2.19021756 | 0.02850846 | 0.65805899 |
| DIP2C | 468.702588 | -0.2026498 | 0.45186972 | -0.4484696 | 0.65381436 | 0.99989001 |
| LARP4B | 463.625957 | -0.2329839 | 0.2907258 | -0.8013869 | 0.42290768 | 0.99989001 |
| GTPBP4 | 69.9061644 | -0.2660326 | 0.37838817 | -0.703068 | 0.48201338 | 0.99989001 |
| IDI1 | 66.8044944 | 0.21307645 | 0.40116921 | 0.5311386 | 0.59532274 | 0.99989001 |
| WDR37 | 224.508729 | -0.1212712 | 0.23256186 | -0.5214578 | 0.60204792 | 0.99989001 |
| PFKP | 55.8340364 | 0.3397297 | 0.5712272 | 0.59473657 | 0.55201958 | 0.99989001 |
| PITRM1 | 66.2618288 | -0.1410081 | 0.31980283 | -0.4409219 | 0.65926956 | 0.99989001 |
| KLF6 | 402.535068 | 0.03487586 | 0.49073612 | 0.07106846 | 0.94334327 | 0.99989001 |
| AKR1E2 | 47.1328208 | 0.01455432 | 0.40299722 | 0.03611518 | 0.97119052 | NA |
| NET1 | 100.601218 | 0.12258686 | 0.33719927 | 0.36354426 | 0.71619836 | 0.99989001 |
| ASB13 | 37.9325011 | 0.37796229 | 0.50785557 | 0.74423185 | 0.45673622 | NA |
| FAM208B | 550.273549 | 0.09266029 | 0.25322612 | 0.36591917 | 0.71442539 | 0.99989001 |
| GDI2 | 312.468052 | -0.1227524 | 0.2447027 | -0.5016391 | 0.61592142 | 0.99989001 |
| ANKRD16 | 42.3179295 | -0.2077584 | 0.45339409 | -0.4582292 | 0.64678775 | NA |
| FBH1 | 209.046539 | 0.39226433 | 0.21260126 | 1.8450706 | 0.06502727 | 0.8666545 |
| IL15RA | 323.953129 | 0.05960475 | 0.36842436 | 0.16178287 | 0.87147685 | 0.99989001 |
| RBM17 | 302.783225 | 0.04571388 | 0.27708326 | 0.16498245 | 0.86895779 | 0.99989001 |
| PFKFB3 | 222.32861 | -0.2104769 | 0.4243907 | -0.4959507 | 0.61992917 | 0.99989001 |
| SFMBT2 | 656.543164 | 0.18179333 | 0.41344982 | 0.43969865 | 0.66015538 | 0.99989001 |
| ITIH5 | 184.280933 | -3.3725766 | 1.05659886 | -3.1919177 | 0.00141332 | 0.18190106 |
| KIN | 64.0810436 | 0.14072245 | 0.27957298 | 0.50334782 | 0.61471975 | 0.99989001 |
| ATP5F1C | 149.138192 | 0.25377967 | 0.18330955 | 1.3844324 | 0.16622609 | 0.99989001 |
| TAF3 | 373.209549 | 0.25598966 | 0.24133241 | 1.06073469 | 0.28881049 | 0.99989001 |
| CELF2 | 212.045181 | -2.0946249 | 0.59714632 | -3.5077248 | 0.00045196 | 0.09476018 |
| USP6NL | 252.170191 | 0.08117318 | 0.20201589 | 0.40181581 | 0.68781959 | 0.99989001 |
| PROSER2 | 47.4545205 | -0.0940665 | 0.42958857 | -0.2189688 | 0.82667437 | NA |
| UPF2 | 237.932059 | -0.0654477 | 0.17831718 | -0.3670296 | 0.71359698 | 0.99989001 |
| DHTKD1 | 64.0529646 | 0.24189645 | 0.34983201 | 0.69146459 | 0.48927362 | 0.99989001 |
| NUDT5 | 100.970615 | -0.03353 | 0.26227293 | -0.127844 | 0.89827245 | 0.99989001 |
| CDC123 | 125.875605 | -0.1332687 | 0.24678499 | -0.5400194 | 0.58918363 | 0.99989001 |
| CAMK1D | 605.416285 | 0.16278854 | 0.29050276 | 0.56036831 | 0.57522824 | 0.99989001 |
| CCDC3 | 386.475478 | -0.0003869 | 0.48426358 | -0.000799 | 0.99936252 | 0.99989001 |
| OPTN | 223.043506 | 0.29966479 | 0.37559871 | 0.79783232 | 0.4249678 | 0.99989001 |
| SEPHS1 | 80.1844192 | 0.15511549 | 0.26654357 | 0.58195174 | 0.56059919 | 0.99989001 |

| | | | | | | |
|---|---|---|---|---|---|---|
| BEND7 | 214.455156 | 0.34564191 | 0.21664512 | 1.59542903 | 0.11061633 | 0.99989001 |
| AL157392.5 | 78.8216 | 0.01752226 | 0.26798991 | 0.06538403 | 0.94786824 | 0.99989001 |
| FRMD4A | 477.677658 | 0.60761542 | 0.55187997 | 1.10099198 | 0.27090015 | 0.99989001 |
| FAM107B | 179.15354 | 0.62120523 | 0.53680063 | 1.15723642 | 0.24717578 | 0.99989001 |
| DCLRE1C | 60.2524282 | 0.05456914 | 0.28671792 | 0.19032343 | 0.84905569 | 0.99989001 |
| RPP38 | 83.2417675 | 0.28668711 | 0.24094054 | 1.18986662 | 0.23409882 | 0.99989001 |
| NMT2 | 112.833083 | 0.09770444 | 0.35489754 | 0.27530322 | 0.78308329 | 0.99989001 |
| FAM171A1 | 384.946202 | -0.4725499 | 0.34963191 | -1.3515639 | 0.17651486 | 0.99989001 |
| ITGA8 | 568.744048 | 1.410063 | 0.59937591 | 2.35255199 | 0.01864508 | 0.56643441 |
| MINDY3 | 247.032634 | -0.071711 | 0.21715693 | -0.3302268 | 0.74122857 | 0.99989001 |
| PTER | 60.984438 | -0.154467 | 0.48527806 | -0.3183062 | 0.75025268 | 0.99989001 |
| RSU1 | 462.981072 | -0.1445974 | 0.1607483 | -0.899527 | 0.368372 | 0.99989001 |
| CUBN | 154.444748 | 0.19198729 | 0.54247954 | 0.35390697 | 0.72340861 | 0.99989001 |
| TRDMT1 | 50.1825943 | -0.1796936 | 0.44392906 | -0.40478 | 0.68563921 | 0.99989001 |
| VIM | 1530.06691 | -0.1325714 | 0.65651923 | -0.2019307 | 0.83997087 | 0.99989001 |
| ST8SIA6 | 283.080077 | -0.7213265 | 1.33245924 | -0.5413498 | 0.58826652 | 0.99989001 |
| STAM | 119.447559 | -0.2519651 | 0.25307262 | -0.9956236 | 0.31943308 | 0.99989001 |
| MRC1 | 95.8055956 | 1.30097187 | 0.62557478 | 2.07964245 | 0.03755834 | 0.70738774 |
| NSUN6 | 159.870979 | -0.3584867 | 0.4020382 | -0.8916731 | 0.37256817 | 0.99989001 |
| ARL5B | 103.102367 | -0.9415602 | 0.53358639 | -1.7645881 | 0.07763301 | 0.92274869 |
| PLXDC2 | 927.069486 | 0.274202 | 0.31203265 | 0.87876061 | 0.37953109 | 0.99989001 |
| NEBL | 124.857008 | -0.5290876 | 0.43388022 | -1.2194324 | 0.22268012 | 0.99989001 |
| MLLT10 | 702.392281 | -0.001882 | 0.19516192 | -0.0096435 | 0.99230571 | 0.99989001 |
| DNAJC1 | 969.581261 | 0.6984649 | 0.42338849 | 1.64970212 | 0.09900388 | 0.99989001 |
| BMI1 | 45.9391853 | 0.44658826 | 0.48645344 | 0.91804934 | 0.35859303 | NA |
| PIP4K2A | 1210.85235 | 0.5080744 | 0.35480923 | 1.43196501 | 0.15215384 | 0.99989001 |
| MSRB2 | 56.4218495 | 0.16951641 | 0.40496301 | 0.41859728 | 0.67551048 | 0.99989001 |
| KIAA1217 | 1818.99632 | 0.54437217 | 0.50846104 | 1.07062709 | 0.28433714 | 0.99989001 |
| ARHGAP21 | 314.11939 | 5.43E-01 | 0.23791269 | 2.28042145 | 0.0225827 | 0.60384164 |
| ABI1 | 418.83976 | 0.20267879 | 0.22651154 | 0.89478348 | 0.37090284 | 0.99989001 |
| ANKRD26 | 214.121413 | -0.0394857 | 0.30249571 | -0.1305329 | 0.89614479 | 0.99989001 |
| YME1L1 | 257.424936 | 0.05000487 | 0.16478644 | 0.30345257 | 0.76154499 | 0.99989001 |
| ACBD5 | 81.599597 | 0.02068794 | 0.31926053 | 0.06479956 | 0.94833359 | 0.99989001 |
| RAB18 | 197.054046 | 0.18072034 | 0.16330774 | 1.10662447 | 0.26845632 | 0.99989001 |
| WAC-AS1 | 52.621927 | -0.4455471 | 0.5185521 | -0.8592137 | 0.39022262 | 0.99989001 |
| WAC | 961.76554 | -0.0733711 | 0.23367005 | -0.3139943 | 0.7535254 | 0.99989001 |
| SVIL | 1077.61306 | 0.48333168 | 0.43829276 | 1.10275991 | 0.27013144 | 0.99989001 |
| JCAD | 336.174754 | 0.30916828 | 0.30269393 | 1.0213891 | 0.30707013 | 0.99989001 |
| MTPAP | 60.055513 | -0.0245941 | 0.23319782 | -0.1054645 | 0.91600723 | 0.99989001 |
| MAP3K8 | 118.723233 | 0.53072631 | 0.39415994 | 1.34647449 | 0.17814954 | 0.99989001 |
| ZNF438 | 559.026093 | -0.0377493 | 0.32932657 | -0.1146257 | 0.90874179 | 0.99989001 |
| ZEB1-AS1 | 112.027931 | -0.0597324 | 0.38428316 | -0.1554384 | 0.8764757 | 0.99989001 |
| ZEB1 | 3385.21289 | -0.1013281 | 0.22159961 | -0.4572576 | 0.64748593 | 0.99989001 |
| ARHGAP12 | 261.545337 | 0.42976266 | 0.22050142 | 1.94902446 | 0.0512925 | 0.8125042 |
| KIF5B | 331.352719 | 0.31595368 | 0.21711334 | 1.45524767 | 0.14560071 | 0.99989001 |
| EPC1 | 483.697022 | 0.20988429 | 0.22340048 | 0.93949791 | 0.34747516 | 0.99989001 |
| CCDC7 | 208.360268 | -0.2148942 | 0.419606 | -0.5121332 | 0.60855781 | 0.99989001 |
| ITGB1 | 907.344324 | -0.3116075 | 0.31016393 | -1.0046543 | 0.31506334 | 0.99989001 |
| NRP1 | 1012.65222 | 1.28368643 | 0.39602585 | 3.2414208 | 0.00118935 | 0.16624537 |
| PARD3 | 835.913775 | 0.29082911 | 0.39201532 | 0.741882 | 0.45815883 | 0.99989001 |
| CUL2 | 162.989063 | -0.0360756 | 0.18562018 | -0.1943519 | 0.84590036 | 0.99989001 |
| CREM | 328.060365 | -0.4098345 | 0.52514839 | -0.7804165 | 0.43514577 | 0.99989001 |
| CCNY | 2142.44503 | -0.0096265 | 0.45284688 | -0.0212576 | 0.98304014 | 0.99989001 |
| AL117336.3 | 63.8014582 | 0.30715411 | 0.35526454 | 0.86457857 | 0.38727014 | 0.99989001 |

| | | | | | | |
|---|---|---|---|---|---|---|
| ZNF248 | 136.262095 | -0.2151286 | 0.23886089 | -0.900644 | 0.36777761 | 0.99989001 |
| ZNF25 | 62.8009513 | 0.5003377 | 0.2505431 | 1.99701251 | 0.04582382 | 0.77310579 |
| ZNF33A | 294.989129 | -0.0153418 | 0.30529386 | -0.0502527 | 0.95992105 | 0.99989001 |
| ZNF33B | 119.124954 | 0.32743709 | 0.25646362 | 1.27673897 | 0.20169442 | 0.99989001 |
| BMS1 | 119.248194 | -0.0238334 | 0.26068563 | -0.0914257 | 0.92715432 | 0.99989001 |
| CSGALNACT2 | 102.054922 | 0.14245653 | 0.26628673 | 0.53497419 | 0.5926677 | 0.99989001 |
| HNRNPF | 223.604771 | -0.215892 | 0.24783034 | -0.8711283 | 0.3836841 | 0.99989001 |
| ZNF32 | 44.0499513 | 0.19796648 | 0.31620178 | 0.62607644 | 0.53126481 | NA |
| CXCL12 | 105.5558 | -0.2947072 | 1.11448831 | -0.2644327 | 0.79144651 | 0.99989001 |
| RASSF4 | 42.6720991 | 0.41472754 | 0.40437364 | 1.02560478 | 0.30507791 | NA |
| DEPP1 | 671.265787 | -0.4248989 | 0.54459378 | -0.7802125 | 4.35E-01 | 1.00E+00 |
| ZNF22 | 93.7044932 | 0.02988966 | 0.41719699 | 0.07164401 | 0.94288522 | 0.99989001 |
| 8-Mar | 122.895214 | -0.3500075 | 0.25471494 | -1.3741146 | 0.1694061 | 0.99989001 |
| ZFAND4 | 79.2234889 | -0.1073948 | 0.22283361 | -0.4819505 | 0.62984111 | 0.99989001 |
| WASHC2C | 70.95733 | -0.2509061 | 0.29659389 | -0.8459583 | 0.39757603 | 0.99989001 |
| AGAP4 | 71.3686171 | 0.00633783 | 0.25975578 | 0.02439919 | 0.98053419 | 0.99989001 |
| TIMM23 | 76.6943589 | -0.1022925 | 0.26937877 | -0.3797347 | 0.70414234 | 0.99989001 |
| NCOA4 | 133.160575 | 0.29568032 | 0.29738825 | 0.9942569 | 0.32009781 | 0.99989001 |
| AL356056.2 | 51.4334619 | 0.09251019 | 0.45774734 | 0.2020988 | 0.83983948 | 0.99989001 |
| MAPK8 | 472.646691 | 0.29622268 | 0.20674898 | 1.43276488 | 0.15192504 | 0.99989001 |
| TMEM273 | 50.9550445 | -0.4163338 | 0.63398487 | -0.6566936 | 0.51137794 | 0.99989001 |
| ERCC6 | 68.1568001 | 0.13523734 | 0.27830102 | 0.48593908 | 0.62701036 | 0.99989001 |
| PARG | 247.447319 | -0.0941812 | 0.16504114 | -0.5706528 | 0.56823499 | 0.99989001 |
| TIMM23B | 202.221677 | -0.2041215 | 0.43139091 | -0.4731706 | 0.63609146 | 0.99989001 |
| LINC00843 | 53.686164 | 0.45371034 | 0.27139989 | 1.67174104 | 0.0945754 | 0.99989001 |
| AGAP6 | 82.4758269 | 0.34543031 | 0.33593885 | 1.02825354 | 0.30383058 | 0.99989001 |
| WASHC2A | 61.6069484 | -0.620168 | 0.31966456 | -1.9400585 | 0.05237258 | 0.81270649 |
| SGMS1 | 408.899164 | 0.25265649 | 0.23633365 | 1.06906696 | 0.2850395 | 0.99989001 |
| PRKG1 | 783.578889 | 1.64204764 | 0.57531489 | 2.85417197 | 0.00431492 | 0.32568991 |
| IPMK | 127.242155 | -0.2558491 | 0.40483439 | -0.6319845 | 0.527397 | 0.99989001 |
| CISD1 | 112.348942 | -0.1314445 | 0.3776783 | -0.348033 | 0.72781537 | 0.99989001 |
| UBE2D1 | 67.1851223 | 0.36369923 | 0.24891781 | 1.4611218 | 0.14398202 | 0.99989001 |
| TFAM | 48.0523258 | 0.25962402 | 0.28129791 | 0.92295039 | 0.35603306 | NA |
| BICC1 | 105.941078 | -0.3785043 | 0.50865952 | -0.7441211 | 0.45680319 | 0.99989001 |
| CCDC6 | 179.408575 | 0.00481565 | 0.20846005 | 0.02310107 | 0.98156965 | 0.99989001 |
| ANK3 | 337.50651 | 0.26695555 | 0.40721901 | 0.65555767 | 0.51210877 | 0.99989001 |
| RHOBTB1 | 129.413708 | 0.28178761 | 0.26102945 | 1.07952423 | 2.80E-01 | 1.00E+00 |
| ARID5B | 399.518937 | 0.56751944 | 0.38197899 | 1.4857347 | 0.13734931 | 0.99989001 |
| JMJD1C | 3157.81751 | -0.1724849 | 0.40678575 | -0.4240189 | 0.671552 | 0.99989001 |
| REEP3 | 624.344901 | -0.1535735 | 0.39622162 | -0.387595 | 0.69831577 | 0.99989001 |
| LINC01515 | 113.001998 | -1.6011014 | 0.65021409 | -2.4624218 | 0.01380023 | 0.50652625 |
| SIRT1 | 100.475791 | 0.12950956 | 0.22081664 | 0.58650273 | 0.55753773 | 0.99989001 |
| HERC4 | 921.745834 | -0.2242231 | 0.20175833 | -1.111345 | 0.26641987 | 0.99989001 |
| HNRNPH3 | 226.065285 | 0.15904505 | 0.2786838 | 0.57070075 | 0.56820251 | 0.99989001 |
| RUFY2 | 212.288938 | -0.076617 | 0.24482746 | -0.3129428 | 0.75432411 | 0.99989001 |
| SLC25A16 | 168.945083 | -0.2227072 | 0.21438522 | -1.038818 | 0.29888938 | 0.99989001 |
| TET1 | 58.4852618 | 0.35850351 | 0.51088448 | 0.70173106 | 0.4828469 | 0.99989001 |
| CCAR1 | 326.277732 | -0.1864007 | 0.23401854 | -0.796521 | 0.4257293 | 0.99989001 |
| DDX50 | 136.690677 | -0.041717 | 0.30115992 | -0.1385211 | 0.88982862 | 0.99989001 |
| DDX21 | 217.401849 | -0.035633 | 0.50444218 | -0.0706384 | 0.94368555 | 0.99989001 |
| KIF1BP | 64.486807 | 0.32223443 | 0.27230621 | 1.18335324 | 0.23666918 | 0.99989001 |
| SRGN | 683.293498 | -1.0413485 | 0.7123671 | -1.4618144 | 0.14379207 | 0.99989001 |
| VPS26A | 164.322918 | -0.2963717 | 0.22348734 | -1.3261232 | 0.18479891 | 0.99989001 |
| SUPV3L1 | 51.209416 | -0.5823096 | 0.30168015 | -1.9302219 | 0.05357935 | 0.81700389 |

| | | | | | | |
|---|---|---|---|---|---|---|
| HK1 | 326.553521 | 0.01271566 | 0.18506293 | 0.06870994 | 0.94522051 | 0.99989001 |
| TSPAN15 | 140.589294 | 0.03797217 | 0.34512866 | 0.11002322 | 0.91239096 | 0.99989001 |
| COL13A1 | 56.5807835 | -0.3023605 | 0.54519207 | -0.5545945 | 5.79E-01 | 0.99989001 |
| SAR1A | 118.796968 | -0.0221841 | 0.28801452 | -0.0770244 | 0.93860416 | 0.99989001 |
| PPA1 | 225.26773 | 0.20101512 | 0.49245425 | 0.40819045 | 0.68313386 | 0.99989001 |
| EIF4EBP2 | 109.291219 | 0.1554153 | 0.37449517 | 0.41499947 | 0.67814229 | 0.99989001 |
| PALD1 | 275.897411 | -0.0048026 | 0.34134887 | -0.0140695 | 0.98877449 | 0.99989001 |
| SGPL1 | 70.6342587 | 0.1901018 | 0.24557177 | 0.77411908 | 0.43886037 | 0.99989001 |
| UNC5B | 147.059191 | 0.7269187 | 0.46676389 | 1.55735846 | 0.1193854 | 0.99989001 |
| VSIR | 77.9046486 | -0.5970131 | 0.53453463 | -1.1168839 | 0.26404396 | 0.99989001 |
| PSAP | 450.625836 | -0.1562142 | 0.39564763 | -0.3948317 | 0.69296713 | 0.99989001 |
| CHST3 | 67.4845821 | 0.7022893 | 0.29862591 | 2.351736 | 0.01868603 | 0.56643441 |
| ASCC1 | 250.502838 | -0.1675645 | 0.16783864 | -0.9983666 | 0.31810163 | 0.99989001 |
| ANAPC16 | 324.528639 | 0.03856637 | 0.27056533 | 0.14253996 | 0.88665352 | 0.99989001 |
| DDIT4 | 628.81694 | -0.5125067 | 0.6704458 | -0.7644267 | 0.44461299 | 0.99989001 |
| DNAJB12 | 62.1558829 | -0.2991243 | 0.30030168 | -0.9960793 | 0.31921161 | 0.99989001 |
| MICU1 | 376.598011 | -0.1629712 | 0.28047566 | -0.5810528 | 0.5612049 | 0.99989001 |
| MCU | 442.043366 | 0.13323873 | 0.31358859 | 0.42488385 | 6.71E-01 | 0.99989001 |
| P4HA1 | 396.700147 | -0.8427377 | 0.66499242 | -1.2672891 | 0.20505193 | 0.99989001 |
| ECD | 97.5974767 | -0.2620805 | 0.30654188 | -0.8549581 | 0.39257432 | 0.99989001 |
| FAM149B1 | 83.2759681 | -0.0544336 | 0.35015727 | -0.1554546 | 0.87646293 | 0.99989001 |
| DNAJC9 | 75.719809 | -0.3844404 | 0.29805444 | -1.2898329 | 0.19710869 | 0.99989001 |
| MRPS16 | 63.6052249 | -0.344308 | 0.28962581 | -1.188803 | 0.2345172 | 0.99989001 |
| ANXA7 | 206.36615 | 0.17828245 | 0.19917372 | 0.89511029 | 0.37072813 | 0.99989001 |
| PPP3CB | 285.770298 | 0.10580579 | 0.20390487 | 0.51889781 | 0.603832 | 0.99989001 |
| USP54 | 96.9916478 | -0.7912811 | 0.28761658 | -2.7511664 | 0.00593835 | 0.37352207 |
| SEC24C | 79.5960847 | 0.01900703 | 0.2272193 | 0.0836506 | 0.93333424 | 0.99989001 |
| FUT11 | 141.70178 | 0.05068276 | 0.31893614 | 0.15891195 | 0.87373825 | 0.99989001 |
| CHCHD1 | 56.617565 | -0.1201528 | 0.48526426 | -0.2476029 | 0.80444168 | 0.99989001 |
| ZSWIM8 | 100.157431 | 0.15081159 | 0.2796289 | 0.53932764 | 0.5896608 | 0.99989001 |
| CAMK2G | 123.698846 | -0.145554 | 0.42709838 | -0.3407974 | 0.73325611 | 0.99989001 |
| VCL | 364.731316 | -0.2134511 | 0.34702182 | -0.6150942 | 0.53849252 | 0.99989001 |
| AP3M1 | 64.5201161 | -0.0713451 | 0.26488584 | -0.269343 | 0.78766578 | 0.99989001 |
| ADK | 537.261358 | -0.0797953 | 0.25635025 | -0.3112746 | 0.75559187 | 0.99989001 |
| KAT6B | 349.158569 | 0.32561491 | 0.28730962 | 1.13332404 | 0.2570782 | 0.99989001 |
| SAMD8 | 177.92242 | 0.06140202 | 0.14979055 | 0.40991917 | 0.68186524 | 0.99989001 |
| VDAC2 | 94.9005991 | 0.30696561 | 0.40892233 | 0.75066973 | 0.45285144 | 0.99989001 |
| ZNF503-AS1 | 185.860867 | 0.66351702 | 0.37614429 | 1.76399599 | 0.07773265 | 0.92274869 |
| AC010997.3 | 81.69052 | 0.54417514 | 0.51007407 | 1.06685513 | 0.28603726 | 0.99989001 |
| LRMDA | 3962.6869 | -0.5924329 | 0.31802935 | -1.8628247 | 0.06248693 | 0.85940342 |
| DLG5 | 93.5611831 | 0.67657071 | 0.30512344 | 2.2173672 | 0.02659801 | 0.64761862 |
| POLR3A | 89.8110259 | -0.124828 | 0.21809078 | -0.5723671 | 0.56707327 | 0.99989001 |
| RPS24 | 567.340486 | -0.1584503 | 0.56471344 | -0.2805854 | 0.77902844 | 0.99989001 |
| ZMIZ1 | 630.647829 | 0.16785519 | 0.41990086 | 0.39974956 | 0.68934099 | 0.99989001 |
| ZCCHC24 | 144.340096 | 0.09443359 | 0.41618455 | 0.22690316 | 0.82049905 | 0.99989001 |
| NUTM2B-AS | 440.64105 | 0.08234226 | 0.27273243 | 0.30191591 | 0.76271617 | 0.99989001 |
| ANXA11 | 200.379417 | 0.30097197 | 0.29043118 | 1.03629359 | 0.3000652 | 0.99989001 |
| FAM213A | 204.322113 | 0.00422405 | 0.55021547 | 0.00767708 | 0.99387463 | 0.99989001 |
| TSPAN14 | 613.648609 | -0.2633768 | 0.2158725 | -1.220057 | 0.22244327 | 0.99989001 |
| NRG3 | 1928.38055 | 1.71462433 | 0.70785782 | 2.42227222 | 0.01542379 | 0.52205731 |
| GHITM | 103.327518 | 0.10986286 | 0.35057005 | 0.31338348 | 0.75398933 | 0.99989001 |
| CCSER2 | 458.959387 | 0.34259777 | 0.20131428 | 1.7018056 | 0.08879182 | 0.97339514 |
| WAPL | 328.274912 | 0.15264087 | 0.14833205 | 1.02904846 | 3.03E-01 | 0.99989001 |
| MMRN2 | 548.468215 | -0.3479491 | 0.54100943 | -0.643148 | 0.52012807 | 0.99989001 |

| | | | | | | |
|---|---|---|---|---|---|---|
| SNCG | 90.8100838 | -0.7482367 | 0.46611232 | -1.6052713 | 0.10843412 | 0.99989001 |
| ADIRF | 345.899625 | -0.3655027 | 0.95633735 | -0.3821902 | 0.70232031 | 0.99989001 |
| GLUD1 | 227.39366 | 0.30825848 | 0.22979117 | 1.34147227 | 0.17976717 | 0.99989001 |
| SHLD2 | 209.883634 | -0.1334235 | 0.32864174 | -0.4059845 | 0.68475398 | 0.99989001 |
| NUTM2A-AS | 579.5234 | -0.0961657 | 0.21046073 | -0.4569293 | 0.64772187 | 0.99989001 |
| MINPP1 | 47.9214508 | 0.45192103 | 0.26121087 | 1.73010038 | 0.08361234 | NA |
| PAPSS2 | 227.12745 | 2.4457508 | 0.60033299 | 4.07399031 | 4.62E-05 | 0.02180165 |
| ATAD1 | 113.806884 | -0.1388069 | 0.21513201 | -0.6452172 | 0.51878644 | 0.99989001 |
| PTEN | 479.298993 | 0.12162083 | 0.17704903 | 0.68693306 | 0.49212491 | 0.99989001 |
| RNLS | 88.4889199 | -1.0215026 | 0.42657113 | -2.394683 | 0.01663474 | 0.55059202 |
| LIPA | 173.877955 | 0.02222282 | 0.27525484 | 0.08073543 | 0.93565236 | 0.99989001 |
| IFIT2 | 220.614786 | 0.87955523 | 1.12917224 | 0.77893805 | 0.43601621 | 0.99989001 |
| IFIT3 | 207.259333 | -0.1541193 | 0.95014545 | -0.162206 | 0.87114361 | 0.99989001 |
| IFIT1 | 262.668838 | -0.0494201 | 0.98563428 | -0.0501404 | 0.96001054 | 0.99989001 |
| IFIT5 | 53.0566477 | 0.6112077 | 0.34331586 | 1.78030718 | 0.0750257 | 0.9206525 |
| RPP30 | 41.2633394 | -0.5922761 | 0.32745427 | -1.8087291 | 0.07049309 | NA |
| PCGF5 | 410.410079 | -0.1623228 | 0.40074162 | -0.4050561 | 0.68543629 | 0.99989001 |
| HECTD2 | 154.814402 | 0.12372691 | 0.26836137 | 0.4610459 | 0.64476568 | 0.99989001 |
| TNKS2 | 295.724957 | -0.068565 | 0.20837943 | -0.3290394 | 0.74212592 | 0.99989001 |
| BTAF1 | 300.00824 | -0.1990837 | 0.34132452 | -0.5832681 | 0.55971286 | 0.99989001 |
| CPEB3 | 493.32588 | 0.00267272 | 0.27624142 | 0.0096753 | 0.99228035 | 0.99989001 |
| 5-Mar | 193.817468 | 0.00575593 | 0.23529688 | 0.02446241 | 0.98048377 | 0.99989001 |
| IDE | 205.471964 | 0.25816977 | 0.26619163 | 0.96986435 | 0.33211411 | 0.99989001 |
| HHEX | 86.0120558 | 0.27083414 | 0.41642477 | 0.65037951 | 0.51544711 | 0.99989001 |
| EXOC6 | 1434.6294 | -0.7758618 | 0.35881054 | -2.1623162 | 0.03059381 | 0.67281094 |
| MYOF | 324.741469 | -1.3932885 | 0.49424989 | -2.8189961 | 0.00481741 | 0.34759088 |
| FRA10AC1 | 139.566736 | -0.152321 | 0.2478924 | -0.6144644 | 0.53890851 | 0.99989001 |
| SLC35G1 | 45.1739447 | 0.84385626 | 0.46515724 | 1.81413119 | 0.06965753 | NA |
| NOC3L | 57.2839574 | -0.0135081 | 0.37061785 | -0.0364475 | 0.97092551 | 0.99989001 |
| TBC1D12 | 132.382619 | 0.36930841 | 0.19772125 | 1.86782362 | 0.06178666 | 0.85940342 |
| PDLIM1 | 567.656673 | 0.21262323 | 0.66671807 | 0.31891025 | 0.74979457 | 0.99989001 |
| SORBS1 | 743.88191 | -0.0369068 | 0.59227305 | -0.0623138 | 0.9503129 | 0.99989001 |
| ALDH18A1 | 45.4204154 | -0.3834596 | 0.30048602 | -1.2761311 | 0.20190917 | NA |
| TCTN3 | 78.2692485 | -0.3204941 | 0.32108871 | -0.9981481 | 0.31820753 | 0.99989001 |
| ENTPD1 | 445.089293 | -0.0354817 | 0.43072844 | -0.0823761 | 0.93434761 | 0.99989001 |
| ENTPD1-AS1 | 390.07211 | -0.3207134 | 0.54707271 | -0.5862354 | 0.55771735 | 0.99989001 |
| AL365273.2 | 228.91926 | -0.0163281 | 0.49725456 | -0.0328365 | 0.973805 | 0.99989001 |
| CC2D2B | 175.659956 | -0.3272646 | 0.70597747 | -0.4635624 | 0.64296129 | 0.99989001 |
| ZNF518A | 238.607481 | -0.0023765 | 0.28794998 | -0.0082533 | 0.99341488 | 0.99989001 |
| TM9SF3 | 407.010784 | 0.12868484 | 0.23422126 | 0.54941569 | 0.58272021 | 0.99989001 |
| LCOR | 482.195067 | 0.06283926 | 0.21226991 | 0.29603468 | 0.76720359 | 0.99989001 |
| RRP12 | 48.428625 | -0.2793194 | 0.43256622 | -0.6457263 | 0.51845659 | NA |
| PGAM1 | 166.568454 | 0.23831691 | 0.39786073 | 0.5989958 | 0.54917568 | 0.99989001 |
| EXOSC1 | 65.0445449 | 0.20449405 | 0.28363994 | 0.72096352 | 0.47093196 | 0.99989001 |
| MMS19 | 128.457778 | 0.0163008 | 0.21900925 | 0.07442975 | 0.94066844 | 0.99989001 |
| UBTD1 | 97.3638035 | -0.4044302 | 0.40039361 | -1.0100815 | 0.31245623 | 0.99989001 |
| ZFYVE27 | 65.9472691 | -0.031293 | 0.27819581 | -0.1124855 | 0.91043844 | 0.99989001 |
| R3HCC1L | 205.306617 | 0.02361603 | 0.21090349 | 0.11197552 | 0.91084281 | 0.99989001 |
| HPS1 | 58.1554851 | -0.352283 | 0.29925899 | -1.1771843 | 0.23912194 | 0.99989001 |
| GOT1 | 53.4102282 | 0.18498484 | 0.35080658 | 0.5273129 | 0.59797631 | 0.99989001 |
| SLC25A28 | 101.010005 | 0.14144317 | 0.46252239 | 0.30580827 | 0.75975063 | 0.99989001 |
| CUTC | 87.3304408 | 0.24925182 | 0.27157501 | 0.91780102 | 0.35872305 | 0.99989001 |
| COX15 | 59.9010715 | -0.1011094 | 0.23459254 | -0.431 | 0.66646838 | 0.99989001 |
| DNMBP | 246.522345 | 0.29722445 | 0.35371222 | 0.84030022 | 0.40074008 | 0.99989001 |

| | | | | | | |
|---|---|---|---|---|---|---|
| ERLIN1 | 60.621418 | 0.01237428 | 0.23169122 | 0.0534085 | 0.95740644 | 0.99989001 |
| CHUK | 67.6861912 | -0.0175858 | 0.37249596 | -0.0472107 | 0.96234527 | 0.99989001 |
| CWF19L1 | 85.1857004 | -0.1958042 | 0.19921198 | -0.9828938 | 0.32565969 | 0.99989001 |
| HIF1AN | 48.2315301 | 0.26294777 | 0.29512484 | 0.89097131 | 0.37294456 | NA |
| SLF2 | 184.34409 | -0.0612614 | 0.21904114 | -0.2796799 | 0.77972308 | 0.99989001 |
| MRPL43 | 41.1457683 | 0.40254533 | 0.41546965 | 0.96889227 | 0.33259894 | NA |
| LZTS2 | 69.5184841 | -0.3050603 | 0.27423681 | -1.1123973 | 0.26596738 | 0.99989001 |
| SFXN3 | 41.9598001 | 0.24143626 | 0.35728059 | 0.67576092 | 0.49919245 | NA |
| BTRC | 321.771211 | 0.09551193 | 0.19547954 | 0.48860319 | 0.62512266 | 0.99989001 |
| FBXW4 | 220.098054 | 0.11048231 | 0.2284926 | 0.48352689 | 0.62872167 | 0.99989001 |
| OGA | 528.199939 | -0.0324462 | 0.3683003 | -0.0880971 | 0.9297995 | 0.99989001 |
| ARMH3 | 335.848406 | 0.00357787 | 0.20944857 | 0.01708233 | 0.98637094 | 0.99989001 |
| PPRC1 | 47.1912055 | -0.7057163 | 0.52040988 | -1.3560777 | 0.17507443 | NA |
| NOLC1 | 41.7989372 | 0.01436543 | 0.48008015 | 0.02992298 | 0.97612848 | NA |
| GBF1 | 382.163862 | 0.01264891 | 0.23853355 | 0.0530278 | 0.95770976 | 0.99989001 |
| NFKB2 | 73.9977103 | 0.53401548 | 0.66443961 | 0.80370808 | 0.42156558 | 0.99989001 |
| FBXL15 | 46.3465224 | -0.039532 | 0.45010388 | -0.0878286 | 0.93001289 | NA |
| CUEDC2 | 46.7828147 | 0.06451949 | 0.3585215 | 0.17995991 | 0.85718405 | NA |
| ACTR1A | 89.8693761 | 0.11907418 | 0.2942285 | 0.40469968 | 0.68569828 | 0.99989001 |
| SUFU | 175.484183 | 0.48948301 | 0.34502895 | 1.41867227 | 0.15599458 | 0.99989001 |
| TRIM8 | 136.247857 | 0.51630079 | 0.38956204 | 1.32533649 | 0.18505957 | 0.99989001 |
| ARL3 | 84.9024381 | 0.22077103 | 0.28698862 | 0.76926755 | 0.4417345 | 0.99989001 |
| WBP1L | 214.16102 | 0.03363489 | 0.20131791 | 0.1670735 | 0.86731221 | 0.99989001 |
| CNNM2 | 140.807083 | 0.14389244 | 0.34675204 | 0.41497214 | 0.6781623 | 0.99989001 |
| NT5C2 | 494.957687 | -0.1129587 | 0.15752733 | -0.7170736 | 0.47332869 | 0.99989001 |
| PCGF6 | 70.0514742 | 0.11292463 | 0.250336 | 0.45109226 | 0.65192305 | 0.99989001 |
| ATP5MD | 58.3038523 | -0.2327754 | 0.42046484 | -0.5536144 | 0.57984278 | 0.99989001 |
| PDCD11 | 57.1825175 | -0.1594381 | 0.53139017 | -0.3000395 | 0.764147 | 0.99989001 |
| SH3PXD2A | 162.323988 | 0.61695034 | 0.39901864 | 1.54616923 | 0.1220637 | 0.99989001 |
| STN1 | 115.535436 | -0.0880254 | 0.27667812 | -0.3181509 | 0.75037046 | 0.99989001 |
| SLK | 449.395518 | 0.20576322 | 0.26480036 | 0.77705039 | 0.43712904 | 0.99989001 |
| GSTO1 | 228.622543 | -0.2358881 | 0.33455156 | -0.7050875 | 0.48075579 | 0.99989001 |
| ITPRIP | 556.567078 | -0.6094151 | 0.3512297 | -1.7350898 | 0.0827249 | 0.94799019 |
| XPNPEP1 | 107.906666 | 0.34292915 | 0.25463771 | 1.34673356 | 0.17806606 | 0.99989001 |
| ADD3 | 479.83761 | 0.79994168 | 0.33843695 | 2.36363578 | 0.01809659 | 0.56443426 |
| MXI1 | 100.114863 | -0.1959723 | 0.43764074 | -0.4477927 | 0.65430283 | 0.99989001 |
| SMNDC1 | 78.7008415 | -0.0555918 | 0.35942222 | -0.1546699 | 0.87708159 | 0.99989001 |
| DUSP5 | 192.757846 | -0.4663317 | 0.47576194 | -0.9801786 | 0.32699797 | 0.99989001 |
| SMC3 | 147.515814 | -0.0870513 | 0.22917704 | -0.379843 | 0.70406197 | 0.99989001 |
| PDCD4 | 262.962137 | -1.2862119 | 0.40603215 | -3.1677588 | 0.00153619 | 0.19325256 |
| BBIP1 | 77.5027725 | 0.13362452 | 0.26711348 | 0.50025375 | 0.61689642 | 0.99989001 |
| SHOC2 | 294.159793 | 0.08659372 | 0.16785317 | 0.51588969 | 0.60593146 | 0.99989001 |
| ACSL5 | 139.397465 | -0.1709793 | 0.36107908 | -0.4735231 | 0.63584004 | 0.99989001 |
| ZDHHC6 | 79.1561108 | 0.12208435 | 0.23431821 | 0.52101948 | 0.6023532 | 0.99989001 |
| VTI1A | 982.430186 | 0.03603981 | 0.24903514 | 0.14471777 | 0.88493371 | 0.99989001 |
| TCF7L2 | 943.734871 | -0.063921 | 0.3098558 | -0.2062928 | 0.83656222 | 0.99989001 |
| CASP7 | 106.179119 | -0.182147 | 0.22720246 | -0.8016945 | 0.42272972 | 0.99989001 |
| NHLRC2 | 233.584954 | 0.11109847 | 0.14441826 | 0.76928269 | 0.44172551 | 0.99989001 |
| CCDC186 | 113.264543 | -0.1697299 | 0.28360354 | -0.5984758 | 0.54952252 | 0.99989001 |
| AFAP1L2 | 94.6744339 | -0.6061966 | 0.54020012 | -1.1221704 | 0.26179 | 0.99989001 |
| ABLIM1 | 607.72115 | -0.1097353 | 0.3800578 | -0.2887331 | 0.77278564 | 0.99989001 |
| FAM160B1 | 153.552684 | 0.22463395 | 0.16547541 | 1.35750651 | 0.17462033 | 0.99989001 |
| TRUB1 | 42.0418129 | 0.00449125 | 0.27921336 | 0.01608539 | 0.98716627 | NA |
| GFRA1 | 99.1617139 | -0.862568 | 1.39483243 | -0.6184026 | 0.53631 | 0.99989001 |

| | | | | | | |
|---|---|---|---|---|---|---|
| PDZD8 | 212.809314 | 0.15693476 | 0.18214325 | 0.86160075 | 0.38890726 | 0.99989001 |
| RAB11FIP2 | 112.783585 | 0.02596916 | 0.20131284 | 0.12899904 | 0.89735841 | 0.99989001 |
| FAM204A | 132.357333 | -0.2329124 | 0.19475885 | -1.1959017 | 0.23173494 | 0.99989001 |
| CACUL1 | 234.032813 | -0.0194358 | 0.1648677 | -0.1178872 | 0.90615701 | 0.99989001 |
| EIF3A | 307.201987 | -0.0246532 | 0.32319116 | -0.0762805 | 0.93919596 | 0.99989001 |
| FAM45A | 49.621871 | -0.0848395 | 0.25439245 | -0.3334986 | 0.73875791 | NA |
| PRDX3 | 93.445617 | 0.06824055 | 0.34305636 | 0.19891936 | 0.84232582 | 0.99989001 |
| GRK5 | 537.709559 | 0.83491749 | 0.45083274 | 1.85194513 | 0.0640337 | 0.86610729 |
| TIAL1 | 210.602053 | -0.1553028 | 0.21264632 | -0.7303339 | 0.46518609 | 0.99989001 |
| BAG3 | 110.957199 | -0.1002693 | 0.406399 | -0.2467263 | 0.80512005 | 0.99989001 |
| INPP5F | 153.497558 | 0.06269676 | 0.34900248 | 0.1796456 | 0.8574308 | 0.99989001 |
| MCMBP | 176.894251 | 0.1182563 | 0.26970961 | 0.43845788 | 0.66105439 | 0.99989001 |
| SEC23IP | 125.857459 | -0.0448877 | 0.21379077 | -0.2099608 | 0.8336983 | 0.99989001 |
| WDR11 | 206.108056 | 0.16969353 | 0.17877965 | 0.94917697 | 0.34253061 | 0.99989001 |
| ATE1 | 447.383317 | 0.21258417 | 0.22595281 | 0.94083438 | 0.34678974 | 0.99989001 |
| NSMCE4A | 62.6635462 | -0.1072484 | 0.35416942 | -0.3028167 | 0.76202956 | 0.99989001 |
| TACC2 | 133.019375 | -0.2573794 | 0.33905244 | -0.7591137 | 0.44778455 | 0.99989001 |
| PLEKHA1 | 331.480142 | -0.0617054 | 0.16664064 | -0.3702901 | 0.71116636 | 0.99989001 |
| HTRA1 | 195.794953 | -0.6134368 | 0.70543405 | -0.8695877 | 0.38452577 | 0.99989001 |
| IKZF5 | 52.8904801 | 0.17050852 | 0.32473066 | 0.52507675 | 0.59952984 | 0.99989001 |
| ACADSB | 56.5057859 | 0.54052213 | 0.3201251 | 1.68847156 | 0.09132074 | 0.99169284 |
| BUB3 | 84.2068092 | 0.27441765 | 0.30267992 | 0.90662656 | 0.36460431 | 0.99989001 |
| CPXM2 | 65.6871403 | 0.86280337 | 1.07207116 | 0.80480046 | 0.42093483 | 0.99989001 |
| CHST15 | 331.636769 | 1.54181472 | 0.6172358 | 2.49793468 | 0.01249192 | 0.49000287 |
| OAT | 48.2191762 | 0.17970269 | 0.54050552 | 0.33247152 | 0.73953324 | NA |
| ABRAXAS2 | 59.3835341 | -0.074085 | 0.30080345 | -0.2462902 | 0.80545757 | 0.99989001 |
| ZRANB1 | 160.692807 | 0.07466221 | 0.18454199 | 0.40458114 | 0.68578543 | 0.99989001 |
| CTBP2 | 971.402198 | 0.1125006 | 0.29846362 | 0.37693236 | 0.70622388 | 0.99989001 |
| EDRF1 | 110.408397 | -0.0130183 | 0.19815799 | -0.0656964 | 0.94761956 | 0.99989001 |
| UROS | 40.972422 | -0.1828889 | 0.42166163 | -0.4337337 | 0.6644818 | NA |
| BCCIP | 76.8253086 | -0.16378 | 0.20977734 | -0.7807327 | 0.43495973 | 0.99989001 |
| DHX32 | 130.543252 | -0.3209377 | 0.21061937 | -1.5237805 | 0.12756355 | 0.99989001 |
| DOCK1 | 1455.06427 | -0.0525159 | 0.23490981 | -0.2235575 | 0.82310163 | 0.99989001 |
| FAM196A | 56.6873513 | 0.02853329 | 0.55504349 | 0.05140731 | 0.95900096 | 0.99989001 |
| PTPRE | 657.294116 | 0.37454213 | 0.36387704 | 1.02930958 | 0.30333422 | 0.99989001 |
| MGMT | 271.29686 | -0.4010202 | 0.38499357 | -1.0416284 | 0.297584 | 0.99989001 |
| GLRX3 | 150.348165 | 0.25426145 | 0.30263884 | 0.84014813 | 0.40082534 | 0.99989001 |
| PPP2R2D | 206.475449 | 0.10963192 | 0.28266749 | 0.38784763 | 0.6981288 | 0.99989001 |
| BNIP3 | 85.8041833 | 0.09192851 | 0.58666241 | 0.15669747 | 0.87548328 | 0.99989001 |
| INPP5A | 210.857697 | 0.06069479 | 0.32505061 | 0.18672412 | 0.85187695 | 0.99989001 |
| TUBGCP2 | 109.863258 | -0.2163084 | 0.22013801 | -0.9826037 | 0.32580252 | 0.99989001 |
| ECHS1 | 92.7990508 | 0.59049598 | 0.34402962 | 1.71641032 | 0.08608697 | 0.95556536 |
| KDM5A | 351.57819 | 0.268506 | 0.20103761 | 1.33560083 | 0.18167978 | 0.99989001 |
| WNK1 | 666.783089 | 0.45807561 | 0.21470112 | 2.13355013 | 0.03287962 | 0.67281094 |
| RAD52 | 70.7757061 | -0.2122253 | 0.30905855 | -0.686683 | 0.49228249 | 0.99989001 |
| ERC1 | 1274.26005 | 0.12453865 | 0.12768206 | 0.97538098 | 0.32937131 | 0.99989001 |
| WNT5B | 79.8205049 | 0.19008453 | 0.59972359 | 0.31695356 | 0.75127884 | 0.99989001 |
| FBXL14 | 49.1961835 | 0.0528039 | 0.39211248 | 0.13466519 | 0.8928766 | NA |
| ADIPOR2 | 580.624475 | -0.1814026 | 0.42606403 | -0.4257638 | 0.67027999 | 0.99989001 |
| DCP1B | 68.5165733 | 0.34579064 | 0.30010618 | 1.15222766 | 0.24922754 | 0.99989001 |
| CACNA1C | 1632.14218 | 0.2270299 | 0.47385639 | 0.4791112 | 0.63185952 | 0.99989001 |
| FKBP4 | 54.2273389 | 0.11678218 | 0.40409559 | 0.28899644 | 0.7725841 | 0.99989001 |
| ITFG2 | 86.8172763 | -0.1517677 | 0.45220875 | -0.3356143 | 0.73716172 | 0.99989001 |
| TULP3 | 123.922707 | 0.44255604 | 0.2602372 | 1.70058714 | 0.08902054 | 0.9738073 |

| | | | | | | |
|---|---|---|---|---|---|---|
| TEAD4 | 277.160337 | -0.1354921 | 0.33897235 | -0.3997144 | 0.68936685 | 0.99989001 |
| TSPAN9 | 855.461164 | 0.36333022 | 0.34858397 | 1.04230329 | 0.29727109 | 0.99989001 |
| PARP11 | 112.223365 | -0.0874943 | 0.21633866 | -0.4044323 | 0.68589484 | 0.99989001 |
| TIGAR | 49.0758774 | 0.14471033 | 0.34039944 | 0.42511918 | 0.6707498 | NA |
| C12orf4 | 84.1180575 | 0.23918007 | 0.21797879 | 1.09726308 | 0.27252641 | 0.99989001 |
| ANO2 | 625.098504 | 0.21700379 | 0.7310229 | 0.29684951 | 0.7665814 | 0.99989001 |
| VWF | 1223.01858 | 0.20454628 | 0.61776987 | 0.33110433 | 0.74056568 | 0.99989001 |
| CD9 | 1374.34902 | 0.04276542 | 0.50758636 | 0.0842525 | 0.93285568 | 0.99989001 |
| TNFRSF1A | 532.051957 | 0.00362949 | 0.41597267 | 0.00872532 | 0.99303829 | 0.99989001 |
| LTBR | 133.076637 | -0.0763328 | 0.26432108 | -0.288788 | 0.77274363 | 0.99989001 |
| MRPL51 | 93.6062243 | -0.0410609 | 0.47809157 | -0.0858851 | 0.93155778 | 0.99989001 |
| NCAPD2 | 45.6540007 | -0.2408888 | 0.28458473 | -0.8464572 | 0.39729776 | NA |
| GAPDH | 902.312833 | -0.1792231 | 0.57141305 | -0.3136489 | 0.75378768 | 0.99989001 |
| CHD4 | 55.4289825 | 0.43126369 | 0.36325308 | 1.18722655 | 0.23513827 | 0.99989001 |
| ING4 | 50.0474145 | 0.12454673 | 0.29812167 | 0.41777147 | 0.67611421 | 0.99989001 |
| ZNF384 | 53.6135615 | 0.22739889 | 0.34825445 | 0.65296764 | 0.51377714 | 0.99989001 |
| COPS7A | 47.9649241 | 0.3581833 | 0.31647426 | 1.13179282 | 0.25772155 | NA |
| MLF2 | 89.6868604 | -0.1677354 | 0.38554883 | -0.4350562 | 0.66352161 | 0.99989001 |
| PTMS | 385.056695 | -0.2964216 | 0.54437902 | -0.5445133 | 0.58608828 | 0.99989001 |
| USP5 | 39.4464204 | 0.35328389 | 0.41134016 | 0.85886068 | 0.39041739 | NA |
| TPI1 | 224.457701 | 0.23596341 | 0.44133061 | 0.5346636 | 0.5928825 | 0.99989001 |
| LRRC23 | 46.6180471 | -0.2949808 | 0.37321245 | -0.790383 | 0.42930412 | NA |
| ATN1 | 178.169765 | -0.3126268 | 0.24890782 | -1.2559944 | 0.20911801 | 0.99989001 |
| C12orf57 | 90.8245984 | -0.3071971 | 0.68939776 | -0.4456021 | 0.65588469 | 0.99989001 |
| PHB2 | 85.5149295 | 0.04829904 | 0.41467782 | 0.11647366 | 0.90727716 | 0.99989001 |
| LPCAT3 | 140.181513 | 0.15669697 | 0.26140679 | 0.59943726 | 0.54888134 | 0.99989001 |
| C1S | 98.5673339 | 0.37591927 | 0.36850192 | 1.02012839 | 0.30766758 | 0.99989001 |
| C1R | 136.324947 | 0.95905046 | 0.61905172 | 1.54922509 | 0.12132762 | 0.99989001 |
| C1RL | 137.093971 | -0.0897915 | 0.30935005 | -0.2902586 | 7.72E-01 | 0.99989001 |
| RBP5 | 124.598538 | 0.06239615 | 0.41145946 | 0.15164593 | 0.8794662 | 0.99989001 |
| CLSTN3 | 90.5002219 | -0.4329946 | 0.31217494 | -1.3870254 | 0.16543402 | 0.99989001 |
| PEX5 | 40.7904783 | -0.4611923 | 0.30445476 | -1.5148138 | 0.12981956 | NA |
| SLC2A3 | 1230.14677 | -0.7907488 | 0.66626452 | -1.1868392 | 0.23529106 | 0.99989001 |
| FOXJ2 | 48.5109073 | 0.0067656 | 0.32674305 | 0.02070619 | 0.98348003 | NA |
| NECAP1 | 48.866091 | 0.05125215 | 0.29320034 | 0.17480249 | 0.86123483 | NA |
| RIMKLB | 464.113241 | -0.3229852 | 0.41577061 | -0.7768351 | 0.43725608 | 0.99989001 |
| A2ML1-AS1 | 61.8202601 | -0.6395498 | 0.55528975 | -1.1517406 | 0.24942768 | 0.99989001 |
| PHC1 | 61.102123 | 0.1066415 | 0.28391151 | 0.37561527 | 0.70720294 | 0.99989001 |
| M6PR | 94.6238783 | -0.149827 | 0.30481815 | -0.4915293 | 0.62305215 | 0.99989001 |
| A2M | 947.953196 | -0.8140656 | 0.63025481 | -1.2916452 | 0.19648004 | 0.99989001 |
| CLEC2D | 68.763056 | -0.1369138 | 0.52326973 | -0.2616506 | 0.79359083 | 0.99989001 |
| CLEC2B | 274.758415 | -0.2349618 | 0.66168662 | -0.3550953 | 0.72251818 | 0.99989001 |
| CLEC1A | 298.066703 | 0.17563883 | 0.37107153 | 0.47332876 | 0.63597865 | 0.99989001 |
| GABARAPL1 | 211.928182 | -0.6857489 | 0.57793511 | -1.18655 | 0.23540517 | 0.99989001 |
| KLRD1 | 88.2153527 | 0.28645645 | 0.78455098 | 0.36512152 | 0.7150207 | 0.99989001 |
| YBX3 | 660.325936 | -0.413347 | 0.45925677 | -0.9000346 | 0.36810182 | 0.99989001 |
| AC244131.2 | 74.4366439 | -0.0986609 | 0.61928614 | -0.159314 | 0.87342153 | 0.99989001 |
| ETV6 | 1704.06892 | -0.1613445 | 0.38117924 | -0.4232772 | 0.67209303 | 0.99989001 |
| BCL2L14 | 123.816658 | 0.23580399 | 0.32755328 | 0.71989505 | 0.47158961 | 0.99989001 |
| LRP6 | 413.742371 | 0.24596312 | 0.2216417 | 1.10973306 | 0.26711407 | 0.99989001 |
| MANSC1 | 80.1878561 | -0.0928068 | 0.43514775 | -0.2132766 | 0.83111124 | 0.99989001 |
| BORCS5 | 99.1369368 | -0.071192 | 0.25374379 | -0.2805663 | 0.77904306 | 0.99989001 |
| DUSP16 | 296.080465 | -0.1996448 | 0.46542958 | -0.4289474 | 0.66796147 | 0.99989001 |
| CREBL2 | 70.9692946 | 0.03487308 | 0.20925978 | 0.16664969 | 0.8676457 | 0.99989001 |

| | | | | | | |
|---|---|---|---|---|---|---|
| CDKN1B | 93.0518556 | -0.0555544 | 0.48202657 | -0.1152517 | 0.90824559 | 0.99989001 |
| APOLD1 | 457.162817 | -1.455328 | 0.76105803 | -1.9122432 | 0.05584501 | 0.83144465 |
| HEBP1 | 140.842886 | -0.661849 | 0.56912412 | -1.1629257 | 0.24485967 | 0.99989001 |
| EMP1 | 519.535302 | -0.4764604 | 0.55677565 | -0.8557493 | 0.39213648 | 0.99989001 |
| ATF7IP | 469.749177 | 0.28846766 | 0.29928474 | 0.96385691 | 0.33511767 | 0.99989001 |
| H2AFJ | 165.226393 | -0.8474718 | 0.61029166 | -1.388634 | 0.16494406 | 0.99989001 |
| WBP11 | 78.5981063 | -0.0768028 | 0.21736227 | -0.35334 | 0.72383359 | 0.99989001 |
| MGP | 975.042843 | 2.28865065 | 1.18514437 | 1.93111549 | 0.05346877 | 0.8169682 |
| ARHGDIB | 107.542712 | 0.3521975 | 0.46513556 | 0.75719323 | 0.4489341 | 0.99989001 |
| EPS8 | 221.293325 | -0.5928477 | 0.36314355 | -1.6325436 | 0.10256502 | 0.99989001 |
| STRAP | 134.447391 | 0.20084741 | 0.26698118 | 0.7522905 | 0.45187638 | 0.99989001 |
| DERA | 206.556438 | -0.1774662 | 0.21927413 | -0.8093347 | 0.41832268 | 0.99989001 |
| MGST1 | 83.2744738 | 1.64480713 | 0.71134546 | 2.31224802 | 0.02076402 | 0.58699182 |
| PLEKHA5 | 382.362753 | 0.37429549 | 0.36111551 | 1.03649796 | 0.29996989 | 0.99989001 |
| AEBP2 | 332.336449 | 0.29741809 | 0.28281702 | 1.05162727 | 0.29297059 | 0.99989001 |
| PDE3A | 745.11158 | 1.47178492 | 0.51753397 | 2.84384215 | 0.00445731 | 0.33310687 |
| PYROXD1 | 56.4733313 | -0.0949019 | 0.22475864 | -0.4222394 | 0.67285032 | 0.99989001 |
| RECQL | 124.080325 | 0.01147883 | 0.19020222 | 0.06035065 | 0.95187636 | 0.99989001 |
| GOLT1B | 64.1228923 | -0.237793 | 0.22091401 | -1.0764053 | 0.28174604 | 0.99989001 |
| LDHB | 169.440129 | 0.62697754 | 0.38431891 | 1.63139914 | 0.10280613 | 0.99989001 |
| CMAS | 69.4331485 | 0.37588706 | 0.22143635 | 1.69749488 | 0.08960314 | 0.97876192 |
| C2CD5 | 252.165199 | 0.05488502 | 0.2754142 | 0.19928173 | 0.84204237 | 0.99989001 |
| ETNK1 | 384.640183 | -0.0106305 | 0.24320268 | -0.0437104 | 0.96513523 | 0.99989001 |
| SOX5 | 1208.11224 | -1.0926889 | 0.50288054 | -2.1728597 | 0.02979088 | 0.6652708 |
| LRMP | 83.9681101 | 0.26980098 | 0.58162763 | 0.46387237 | 0.64273919 | 0.99989001 |
| ETFRF1 | 58.1466005 | 0.2741902 | 0.27148832 | 1.0099521 | 0.31251824 | 0.99989001 |
| KRAS | 208.349876 | 0.49619058 | 0.24869668 | 1.99516362 | 0.04602504 | 0.77310579 |
| RASSF8 | 258.622626 | 0.10701316 | 0.36745334 | 0.29122925 | 0.770876 | 0.99989001 |
| ITPR2 | 678.75888 | 0.67877232 | 0.30974095 | 2.19141942 | 0.02842145 | 0.65805899 |
| INTS13 | 44.7872917 | 0.02181734 | 0.30651071 | 0.07117971 | 0.94325473 | NA |
| FGFR1OP2 | 214.201941 | 0.3120274 | 0.34295146 | 0.90982965 | 0.36291236 | 0.99989001 |
| TM7SF3 | 159.550719 | 0.00419383 | 0.35607715 | 0.01177787 | 0.99060284 | 0.99989001 |
| MED21 | 59.6601033 | 0.07606023 | 0.31877697 | 0.23860014 | 0.81141566 | 0.99989001 |
| STK38L | 341.002232 | -0.4698849 | 0.5640101 | -0.8331143 | 0.4047803 | 0.99989001 |
| PPFIBP1 | 1464.72539 | 0.32257434 | 0.29711972 | 1.08567125 | 0.27762446 | 0.99989001 |
| MRPS35 | 70.9834505 | 0.14181596 | 0.36464875 | 0.38891115 | 0.69734188 | 0.99989001 |
| KLHL42 | 78.1060139 | -0.1633142 | 0.32733097 | -0.4989269 | 0.6178309 | 0.99989001 |
| CCDC91 | 338.278831 | -0.602744 | 0.19181659 | -3.1422933 | 0.0016763 | 0.20106436 |
| FAR2 | 602.11578 | 0.07587024 | 0.23182921 | 0.32726782 | 0.74346533 | 0.99989001 |
| ERGIC2 | 123.470049 | -0.0996985 | 0.24092499 | -0.4138154 | 0.67900929 | 0.99989001 |
| OVCH1-AS1 | 144.756857 | 0.13100023 | 0.94227441 | 0.13902556 | 0.88942995 | 0.99989001 |
| OVCH1 | 83.4711303 | 0.69621939 | 0.76479839 | 0.91033062 | 0.36264818 | 0.99989001 |
| TMTC1 | 2853.81838 | 0.14409268 | 0.38308326 | 0.37613933 | 0.70681331 | 0.99989001 |
| IPO8 | 160.273856 | 0.03553622 | 0.20063213 | 0.17712127 | 0.85941313 | 0.99989001 |
| CAPRIN2 | 89.4084728 | 0.47972067 | 0.33728243 | 1.42231148 | 0.15493585 | 0.99989001 |
| SINHCAF | 164.890039 | -0.3352377 | 0.28883394 | -1.160659 | 2.46E-01 | 0.99989001 |
| DENND5B | 420.398821 | 0.02827841 | 0.26945515 | 0.10494665 | 0.91641814 | 0.99989001 |
| AMN1 | 99.0323648 | -0.3258792 | 0.38683987 | -0.8424137 | 0.39955644 | 0.99989001 |
| KIAA1551 | 389.124607 | 0.5419541 | 0.2878237 | 1.8829377 | 0.05970881 | 0.85885114 |
| BICD1 | 710.62788 | 0.23107311 | 0.24051811 | 0.96073063 | 0.33668762 | 0.99989001 |
| FGD4 | 278.292136 | -0.4176314 | 0.48193803 | -0.8665667 | 0.38617949 | 0.99989001 |
| DNM1L | 235.551031 | 0.00865696 | 0.20544365 | 0.04213787 | 0.96638879 | 0.99989001 |
| YARS2 | 79.4598431 | -0.1073345 | 0.25949504 | -0.4136284 | 0.67914627 | 0.99989001 |
| ALG10B | 41.7630164 | -0.4513268 | 0.46742317 | -0.9655635 | 0.33426265 | NA |

| | | | | | | |
|---|---|---|---|---|---|---|
| CPNE8 | 998.63114 | -0.1830309 | 0.18831045 | -0.9719637 | 0.33106859 | 0.99989001 |
| SLC2A13 | 212.028858 | 0.34914831 | 0.40488743 | 0.86233427 | 0.3885036 | 0.99989001 |
| LRRK2 | 113.695999 | 0.60410058 | 0.32806438 | 1.84140862 | 0.0655617 | 0.8666545 |
| AC090630.1 | 87.1771942 | -0.8939608 | 1.42598953 | -0.6269056 | 0.53072113 | 0.99989001 |
| GXYLT1 | 119.582708 | 0.43349526 | 0.22475636 | 1.92873413 | 0.05376388 | 0.8181648 |
| YAF2 | 296.361456 | 0.0879411 | 0.17081147 | 0.51484308 | 0.60666268 | 0.99989001 |
| PPHLN1 | 269.763512 | -0.1022161 | 0.17451467 | -0.5857162 | 0.55806628 | 0.99989001 |
| ZCRB1 | 45.1879774 | -0.0941687 | 0.2799457 | -0.3363821 | 0.73658272 | NA |
| PRICKLE1 | 97.2449005 | 1.39813117 | 0.4496756 | 3.10919954 | 0.00187595 | 0.20823042 |
| PUS7L | 75.5380584 | 0.14843557 | 0.23954048 | 0.61966802 | 0.53547638 | 0.99989001 |
| IRAK4 | 100.751939 | -0.1154456 | 0.30608797 | -0.3771648 | 0.70605114 | 0.99989001 |
| TWF1 | 51.7197904 | 0.04612844 | 0.39289194 | 0.11740746 | 0.90653718 | 0.99989001 |
| TMEM117 | 259.087458 | 0.13371905 | 0.25873946 | 0.51680963 | 0.60528906 | 0.99989001 |
| ANO6 | 689.482527 | 0.19922581 | 0.21181015 | 0.9405867 | 0.3469167 | 0.99989001 |
| ARID2 | 640.500387 | 0.17250574 | 0.19953673 | 0.86453128 | 0.3872961 | 0.99989001 |
| SCAF11 | 696.301516 | 0.07027639 | 0.12125138 | 0.57959244 | 0.56218949 | 0.99989001 |
| SLC38A2 | 624.631147 | 0.04715202 | 0.64433293 | 0.07317959 | 0.94166321 | 0.99989001 |
| AC008014.1 | 387.620436 | -0.3035426 | 0.72854844 | -0.4166403 | 0.67694152 | 0.99989001 |
| PCED1B | 779.206788 | 0.46073406 | 0.31368649 | 1.46877239 | 0.14189453 | 0.99989001 |
| RPAP3 | 45.3568163 | -0.5084843 | 0.29617187 | -1.7168556 | 0.08600556 | NA |
| RAPGEF3 | 186.97409 | -0.4043974 | 0.39978959 | -1.0115256 | 0.31176495 | 0.99989001 |
| SLC48A1 | 72.0695387 | -0.3075379 | 0.29496108 | -1.0426388 | 0.29711561 | 0.99989001 |
| HDAC7 | 408.655713 | -0.2147231 | 0.23565032 | -0.9111936 | 0.36219336 | 0.99989001 |
| SENP1 | 105.253005 | -0.1151227 | 0.22314833 | -0.5159023 | 0.60592263 | 0.99989001 |
| ASB8 | 45.6447716 | 0.19218208 | 0.32806397 | 0.58580672 | 0.55800541 | NA |
| ZNF641 | 38.8426731 | 0.07427 | 0.4047448 | 0.18349835 | 0.85440701 | NA |
| KANSL2 | 44.394597 | 0.20645475 | 0.30016504 | 0.68780413 | 0.49157613 | NA |
| CCNT1 | 118.643674 | -0.3327976 | 0.34217865 | -0.9725843 | 0.33075995 | 0.99989001 |
| ADCY6 | 55.5769732 | 0.33131076 | 0.27480729 | 1.20561121 | 0.22796743 | 0.99989001 |
| DDX23 | 63.4336944 | 0.06809101 | 0.25131316 | 0.27094087 | 0.78643651 | 0.99989001 |
| CCDC65 | 46.9120618 | 0.13399288 | 0.46132254 | 0.2904538 | 0.77146909 | NA |
| PRKAG1 | 73.3468302 | 0.22323239 | 0.39004715 | 0.57232157 | 0.56710414 | 0.99989001 |
| KMT2D | 101.182021 | 0.0216199 | 0.3068814 | 0.07045033 | 0.94383523 | 0.99989001 |
| LMBR1L | 108.67281 | 0.10114346 | 0.2278957 | 0.4438147 | 0.65717656 | 0.99989001 |
| TUBA1B | 354.186085 | 0.476442 | 0.63940463 | 0.7451338 | 0.45619084 | 0.99989001 |
| TUBA1A | 110.188208 | 1.28348863 | 0.50334295 | 2.54992869 | 0.0107745 | 0.47431649 |
| TUBA1C | 112.267186 | 0.09748346 | 0.3782428 | 0.25772721 | 0.79661744 | 0.99989001 |
| SPATS2 | 359.040216 | 0.15456385 | 0.19360552 | 0.79834426 | 0.42467074 | 0.99989001 |
| PRPF40B | 106.757245 | -0.138797 | 0.31516469 | -0.4403952 | 0.65965088 | 0.99989001 |
| FMNL3 | 441.186854 | -0.0030503 | 0.2524695 | -0.012082 | 0.99036016 | 0.99989001 |
| TMBIM6 | 441.958732 | 0.05267051 | 0.3127876 | 0.16839066 | 0.86627595 | 0.99989001 |
| NCKAP5L | 56.659824 | -0.1442906 | 0.26377276 | -0.547026 | 0.58436084 | 0.99989001 |
| AQP2 | 72.1553787 | -1.639201 | 0.56100389 | -2.9219066 | 0.00347896 | 0.29068264 |
| SMARCD1 | 64.2428248 | -0.0793337 | 0.28411174 | -0.279234 | 0.78006524 | 0.99989001 |
| AC074032.1 | 155.191623 | -0.1209831 | 0.19762206 | -0.6121941 | 0.54040934 | 0.99989001 |
| CERS5 | 131.897885 | -0.04824 | 0.21778313 | -0.2215049 | 0.82469931 | 0.99989001 |
| LIMA1 | 720.548558 | -0.0492423 | 0.21640015 | -0.2275519 | 0.8199946 | 0.99989001 |
| LARP4 | 275.511626 | -0.1890747 | 0.17453381 | -1.083313 | 0.27866953 | 0.99989001 |
| DIP2B | 525.631899 | 0.04020997 | 0.27600543 | 0.14568542 | 0.88416973 | 0.99989001 |
| ATF1 | 128.844384 | -0.0831832 | 0.2787521 | -0.298413 | 0.76538799 | 0.99989001 |
| METTL7A | 72.9588835 | -0.1632825 | 0.63693542 | -0.2563564 | 0.79767565 | 0.99989001 |
| SLC11A2 | 83.8446766 | 0.0264167 | 0.2690144 | 0.09819808 | 0.92177501 | 0.99989001 |
| LETMD1 | 84.567369 | -0.1755354 | 0.26322847 | -0.6668557 | 0.50486434 | 0.99989001 |
| CSRNP2 | 54.4386205 | -0.0811818 | 0.24045495 | -0.3376176 | 0.73565136 | 0.99989001 |

| | | | | | | |
|---|---|---|---|---|---|---|
| TFCP2 | 338.67077 | 0.22618239 | 0.24830101 | 0.91092011 | 0.36233747 | 0.99989001 |
| POU6F1 | 74.6735602 | -0.3699129 | 0.38464864 | -0.9616903 | 0.33620519 | 0.99989001 |
| DAZAP2 | 233.124638 | 0.03846535 | 0.3253908 | 0.11821278 | 0.90589906 | 0.99989001 |
| SMAGP | 73.1393788 | 0.03056839 | 0.36620937 | 0.08347245 | 0.93347588 | 0.99989001 |
| SCN8A | 51.6146145 | 0.03997986 | 0.53298664 | 0.075011 | 0.94020596 | 0.99989001 |
| ACVRL1 | 261.819113 | 0.26887488 | 0.28085256 | 0.95735242 | 0.33838941 | 0.99989001 |
| ACVR1B | 147.73126 | 0.89716007 | 0.30708367 | 2.92154922 | 0.00348295 | 0.29068264 |
| GRASP | 356.624151 | 0.05284209 | 0.32374903 | 0.16321929 | 0.87034578 | 0.99989001 |
| NR4A1 | 192.426338 | -0.8071156 | 0.70323697 | -1.1477149 | 2.51E-01 | 0.99989001 |
| EIF4B | 218.833228 | -0.1943806 | 0.21594122 | -0.9001552 | 0.36803764 | 0.99989001 |
| AC068888.1 | 84.4629244 | -0.3514397 | 0.31721213 | -1.1079013 | 0.26790444 | 0.99989001 |
| TNS2 | 341.186065 | -0.25585 | 0.32362039 | -0.7905869 | 0.42918512 | 0.99989001 |
| CSAD | 118.84912 | -0.200785 | 0.31000475 | -0.6476836 | 0.51718963 | 0.99989001 |
| RARG | 46.5682674 | -0.0997966 | 0.5421068 | -0.1840904 | 0.85394255 | NA |
| PFDN5 | 184.967275 | -0.3339492 | 0.44353337 | -0.7529292 | 0.45149248 | 0.99989001 |
| SP1 | 129.34774 | -0.0641258 | 0.23795411 | -0.269488 | 0.78755416 | 0.99989001 |
| PRR13 | 87.1089401 | 0.30996166 | 0.2741614 | 1.13058096 | 0.2582315 | 0.99989001 |
| PCBP2 | 331.322043 | -0.2087702 | 0.1996832 | -1.0455069 | 0.29578873 | 0.99989001 |
| MAP3K12 | 45.8054364 | 0.505228 | 0.35421363 | 1.42633698 | 0.1537711 | NA |
| ATP5MC2 | 218.375523 | -0.5066151 | 0.32875277 | -1.5410217 | 0.12331151 | 0.99989001 |
| CALCOCO1 | 87.3079068 | 0.273509 | 0.26822617 | 1.01969545 | 0.30787292 | 0.99989001 |
| CBX5 | 130.922026 | -0.1767301 | 0.26717941 | -0.661466 | 0.50831349 | 0.99989001 |
| HNRNPA1 | 131.013615 | -0.5796751 | 0.36799667 | -1.5752184 | 0.11520603 | 0.99989001 |
| COPZ1 | 79.1677142 | -0.114688 | 0.28873461 | -0.3972089 | 0.69121343 | 0.99989001 |
| ITGA5 | 456.81561 | -0.1588624 | 0.299555 | -0.5303279 | 0.59588459 | 0.99989001 |
| CD63 | 309.317018 | -0.4320772 | 0.49155786 | -0.8789956 | 0.37940365 | 0.99989001 |
| RAB5B | 79.1734093 | 0.12751399 | 0.24709485 | 0.51605282 | 0.60581752 | 0.99989001 |
| RPS26 | 65.0653586 | -0.8853144 | 0.49464267 | -1.7898059 | 0.07348512 | 0.9158596 |
| PA2G4 | 62.5306927 | 0.10956357 | 0.4558806 | 0.24033391 | 0.81007141 | 0.99989001 |
| RPL41 | 800.407015 | -0.0242918 | 0.55653305 | -0.0436484 | 0.96518467 | 0.99989001 |
| ESYT1 | 70.9508426 | -0.5354281 | 0.32385818 | -1.6532796 | 0.09827397 | 0.99989001 |
| MYL6 | 623.118813 | -0.3625035 | 0.51890134 | -0.698598 | 0.48480327 | 0.99989001 |
| SMARCC2 | 152.21048 | 0.07001462 | 0.28905272 | 0.24222095 | 0.80860896 | 0.99989001 |
| RNF41 | 52.5394662 | 0.36363648 | 0.29145719 | 1.2476497 | 0.21215937 | 0.99989001 |
| ANKRD52 | 50.7140112 | 0.33611497 | 0.26201894 | 1.2827888 | 0.19956608 | 0.99989001 |
| CS | 74.9005004 | -0.0298041 | 0.37557401 | -0.0793561 | 0.93674935 | 0.99989001 |
| PAN2 | 55.6089773 | -0.1625939 | 0.24712729 | -0.6579359 | 0.51057931 | 0.99989001 |
| STAT2 | 177.282655 | -0.3323825 | 0.35122519 | -0.9463515 | 0.34396935 | 0.99989001 |
| RBMS2 | 893.598475 | 0.15053884 | 0.36892845 | 0.40804347 | 0.68324176 | 0.99989001 |
| BAZ2A | 228.179591 | 0.09467601 | 0.20038589 | 0.47246842 | 0.63659247 | 0.99989001 |
| ATP5F1B | 206.38784 | 0.32548727 | 0.34324929 | 0.94825329 | 0.34300052 | 0.99989001 |
| PTGES3 | 391.376967 | 0.10281829 | 0.25621033 | 0.40130424 | 0.68819614 | 0.99989001 |
| NACA | 261.082048 | 0.18050092 | 0.48046118 | 0.37568263 | 0.70715285 | 0.99989001 |
| NEMP1 | 42.438199 | -0.1722128 | 0.37208632 | -0.4628303 | 0.64348604 | NA |
| STAT6 | 183.343341 | 0.14057887 | 0.26490666 | 0.53067322 | 0.59564524 | 0.99989001 |
| R3HDM2 | 426.62668 | 0.20986763 | 0.26492109 | 0.79218924 | 0.42825034 | 0.99989001 |
| MARS | 70.5521297 | 0.12233965 | 0.372219 | 0.32867654 | 0.74240019 | 0.99989001 |
| MBD6 | 49.7304211 | -0.2331744 | 0.34940142 | -0.6673538 | 0.50454619 | NA |
| DCTN2 | 119.120688 | 0.192697 | 0.27288321 | 0.70615191 | 0.48009367 | 0.99989001 |
| OS9 | 167.115383 | -0.016103 | 0.28752233 | -0.056006 | 0.95533703 | 0.99989001 |
| TSPAN31 | 49.1473601 | -0.0812955 | 0.48001548 | -0.1693602 | 0.86551335 | NA |
| CDK4 | 41.4669136 | -0.4194247 | 0.35194763 | -1.1917247 | 0.23336922 | NA |
| TSFM | 40.2806907 | 0.29751535 | 0.37159343 | 0.8006475 | 0.42333575 | NA |
| AVIL | 58.4325552 | 0.87189491 | 0.43819212 | 1.98975486 | 0.04661795 | 0.77516398 |

| | | | | | | |
|---|---|---|---|---|---|---|
| CTDSP2 | 163.339967 | 0.05151616 | 0.24872222 | 0.20712326 | 0.83591359 | 0.99989001 |
| LRIG3 | 73.1564139 | 0.22511449 | 0.47557553 | 0.47335171 | 0.63596228 | 0.99989001 |
| SLC16A7 | 264.636808 | -1.0358484 | 0.51323384 | -2.0182778 | 0.04356234 | 0.76826882 |
| USP15 | 586.973137 | 0.2118031 | 0.2897278 | 0.7310417 | 0.46475368 | 0.99989001 |
| MON2 | 612.515856 | -0.1034377 | 0.16928075 | -0.6110423 | 0.54117157 | 0.99989001 |
| AC048341.1 | 82.0633214 | -0.1232089 | 0.39047698 | -0.3155344 | 0.75235595 | 0.99989001 |
| PPM1H | 75.4605106 | -1.5517135 | 0.60221711 | -2.5766679 | 0.00997577 | 0.45359728 |
| RXYLT1 | 45.0029145 | -0.133531 | 0.33922742 | -0.3936328 | 0.69385217 | NA |
| SRGAP1 | 385.923427 | 0.32131054 | 0.33548087 | 0.95776111 | 0.33818323 | 0.99989001 |
| XPOT | 103.155916 | -0.0712738 | 0.29602252 | -0.2407715 | 0.80973218 | 0.99989001 |
| TBK1 | 289.034473 | 0.03888513 | 0.24760608 | 0.15704431 | 0.87520993 | 0.99989001 |
| RASSF3 | 418.414813 | 0.00907085 | 0.26819917 | 0.03382133 | 0.97301963 | 0.99989001 |
| GNS | 218.187101 | -0.1754905 | 0.24088431 | -0.7285263 | 0.4662915 | 0.99989001 |
| TBC1D30 | 123.075032 | 0.54577753 | 0.39355518 | 1.38678782 | 0.16550647 | 0.99989001 |
| LEMD3 | 212.164771 | 0.15417029 | 0.1766559 | 0.87271521 | 0.38281833 | 0.99989001 |
| MSRB3 | 267.552429 | 0.26862255 | 0.43197872 | 0.62184209 | 0.53404571 | 0.99989001 |
| IRAK3 | 653.954692 | 0.58511398 | 0.46612929 | 1.25526113 | 0.20938398 | 0.99989001 |
| HELB | 67.8911589 | 0.19808512 | 0.44104559 | 0.44912619 | 0.65334063 | 0.99989001 |
| CAND1 | 193.450363 | -0.0315015 | 0.17576407 | -0.1792262 | 0.85776007 | 0.99989001 |
| MDM1 | 46.3068506 | -0.3430876 | 0.37022896 | -0.9266903 | 0.35408734 | NA |
| LINC02384 | 186.265114 | 0.21882492 | 0.70339689 | 0.31109737 | 0.7557266 | 0.99989001 |
| RAP1B | 1235.50726 | 0.01833854 | 0.33771179 | 0.05430233 | 0.95669429 | 0.99989001 |
| NUP107 | 146.35444 | -0.3797348 | 0.17286338 | -2.1967338 | 0.02803946 | 0.65805899 |
| SLC35E3 | 113.860694 | 0.07853766 | 0.33431124 | 0.23492379 | 0.81426788 | 0.99989001 |
| MDM2 | 240.144291 | 0.37981599 | 0.33506623 | 1.13355498 | 0.25698127 | 0.99989001 |
| CPM | 70.1727759 | -0.3190088 | 0.55194841 | -0.5779685 | 0.56328537 | 0.99989001 |
| CPSF6 | 355.272587 | -0.0631767 | 0.25160963 | -0.25109 | 0.80174455 | 0.99989001 |
| FRS2 | 248.899498 | 0.17665843 | 0.2599939 | 0.67947143 | 0.4968392 | 0.99989001 |
| CCT2 | 92.1484172 | 0.12460831 | 0.35065577 | 0.35535794 | 0.72232145 | 0.99989001 |
| AC025159.1 | 144.380656 | -0.1266971 | 0.3781553 | -0.3350397 | 0.73759511 | 0.99989001 |
| CNOT2 | 672.601538 | -0.3324407 | 0.45311639 | -0.7336762 | 0.4631461 | 0.99989001 |
| KCNMB4 | 337.751244 | 0.22480089 | 0.48902386 | 0.45969309 | 0.64573653 | 0.99989001 |
| AC025569.1 | 122.335109 | 0.07703529 | 0.65360582 | 0.11786201 | 0.906177 | 0.99989001 |
| PTPRB | 3001.72607 | 0.2352323 | 0.35916633 | 0.65493973 | 0.51250656 | 0.99989001 |
| ZFC3H1 | 348.533255 | 0.15402899 | 0.23823587 | 0.64653987 | 0.51792978 | 0.99989001 |
| RAB21 | 259.591166 | 0.1638538 | 0.29732864 | 0.55108649 | 0.58157438 | 0.99989001 |
| TBC1D15 | 464.615024 | -0.0410946 | 0.34463988 | -0.1192393 | 0.90508577 | 0.99989001 |
| ATXN7L3B | 74.4994904 | -0.0701613 | 0.283529 | -0.2474573 | 0.80455435 | 0.99989001 |
| CAPS2 | 92.4469742 | -0.2494456 | 0.62844682 | -0.396924 | 0.69142353 | 0.99989001 |
| KRR1 | 81.3088976 | -0.3319463 | 0.28795375 | -1.1527764 | 0.24900218 | 0.99989001 |
| NAP1L1 | 305.44424 | -0.109704 | 0.29619495 | -0.3703777 | 0.71110108 | 0.99989001 |
| OSBPL8 | 427.791278 | 0.12691257 | 0.16800451 | 0.75541167 | 0.45000201 | 0.99989001 |
| ZDHHC17 | 302.286705 | -0.2079169 | 0.22723141 | -0.9150006 | 0.3601913 | 0.99989001 |
| CSRP2 | 52.2750396 | 0.11202668 | 0.50574781 | 0.221507 | 0.82469769 | 0.99989001 |
| NAV3 | 412.37026 | 1.63988699 | 0.61970158 | 2.64625272 | 0.0081389 | 0.41229803 |
| SYT1 | 93.9707949 | -3.7956731 | 1.06349953 | -3.5690407 | 0.00035829 | 0.07954055 |
| PAWR | 199.621465 | 0.3317022 | 0.23301812 | 1.42350386 | 0.15459014 | 0.99989001 |
| PPP1R12A | 800.740995 | 0.35131679 | 0.19631884 | 1.78952156 | 0.07353086 | 0.9158596 |
| ACSS3 | 59.7011213 | -2.3800145 | 0.79310407 | -3.0008855 | 0.00269196 | 0.26735394 |
| CCDC59 | 59.555952 | -0.0467265 | 0.24006918 | -0.1946378 | 0.84567654 | 0.99989001 |
| METTL25 | 118.777255 | 0.14968527 | 0.33621236 | 0.4452105 | 0.65616764 | 0.99989001 |
| TMTC2 | 1386.6744 | -0.316783 | 0.46023859 | -0.6883016 | 0.49126285 | 0.99989001 |
| RASSF9 | 95.7457512 | 0.52806542 | 0.41892251 | 1.26053244 | 0.20747735 | 0.99989001 |
| CEP290 | 186.273877 | -0.100359 | 0.20020591 | -0.501279 | 0.61617476 | 0.99989001 |

| | | | | | | |
|---|---|---|---|---|---|---|
| TMTC3 | 104.604653 | 0.12957578 | 0.21607676 | 0.59967475 | 0.54872302 | 0.99989001 |
| KITLG | 112.752316 | -0.3956922 | 0.41362386 | -0.9566474 | 0.33874524 | 0.99989001 |
| DUSP6 | 449.605175 | 0.04263107 | 0.49456384 | 0.08619934 | 0.93130796 | 0.99989001 |
| POC1B | 133.120965 | 0.29055643 | 0.25620017 | 1.13409929 | 0.2567529 | 0.99989001 |
| POC1B-AS1 | 48.7008564 | -0.5008145 | 0.30735106 | -1.6294544 | 0.10321687 | NA |
| ATP2B1 | 231.523061 | 0.29641683 | 0.33118624 | 0.89501553 | 0.37077878 | 0.99989001 |
| BTG1 | 427.984023 | -0.7375395 | 0.55746471 | -1.3230245 | 0.18582724 | 0.99989001 |
| EEA1 | 565.238878 | -0.6019273 | 0.24272094 | -2.479915 | 0.01314137 | 0.49620582 |
| NUDT4 | 134.652029 | -1.2997409 | 0.56776662 | -2.2892169 | 0.02206676 | 0.60384164 |
| UBE2N | 135.961953 | 0.18856754 | 0.19773 | 0.95366174 | 0.3402549 | 0.99989001 |
| MRPL42 | 123.180082 | -0.2894136 | 0.19080978 | -1.516765 | 0.12932604 | 0.99989001 |
| SOCS2 | 166.137012 | -0.5324194 | 0.46497306 | -1.1450542 | 0.2521867 | 0.99989001 |
| CRADD | 586.942565 | -0.2775801 | 0.28673798 | -0.9680619 | 0.33301345 | 0.99989001 |
| CEP83 | 147.828997 | -0.0540224 | 0.2748794 | -0.1965313 | 0.8441943 | 0.99989001 |
| TMCC3 | 253.718556 | -0.3095036 | 0.38337134 | -0.8073208 | 0.41948171 | 0.99989001 |
| NDUFA12 | 149.836088 | -0.2669437 | 0.51851434 | -0.5148241 | 0.60667592 | 0.99989001 |
| NR2C1 | 177.581595 | -0.3610254 | 0.1527224 | -2.3639322 | 0.01808212 | 0.56443426 |
| FGD6 | 283.708111 | 0.08342361 | 0.18943204 | 0.44038804 | 6.60E-01 | 0.99989001 |
| VEZT | 355.794923 | 0.15126213 | 0.19930483 | 0.75894864 | 0.44788328 | 0.99989001 |
| METAP2 | 170.09802 | -0.1094763 | 0.24414666 | -0.448404 | 0.65386167 | 0.99989001 |
| NTN4 | 798.164602 | 0.32029466 | 0.46737678 | 0.68530289 | 0.49315281 | 0.99989001 |
| SNRPF | 52.2074568 | -0.6496588 | 0.32305397 | -2.0109917 | 0.04432634 | 0.77228819 |
| LTA4H | 101.073098 | 0.27439279 | 0.31603094 | 0.8682466 | 0.38525935 | 0.99989001 |
| ELK3 | 633.19233 | -0.0007151 | 0.28987669 | -0.0024668 | 0.99803174 | 0.99989001 |
| CDK17 | 1644.13792 | -0.3841717 | 0.41090623 | -0.9349376 | 0.34982044 | 0.99989001 |
| CFAP54 | 106.461373 | 0.58103432 | 0.4763928 | 1.21965385 | 0.22259612 | 0.99989001 |
| NEDD1 | 100.921197 | 0.05580105 | 0.26736838 | 0.20870477 | 0.83467872 | 0.99989001 |
| TMPO | 166.437852 | 0.00142974 | 0.28414355 | 0.00503175 | 0.99598526 | 0.99989001 |
| SLC25A3 | 221.823208 | 0.45939544 | 0.41207908 | 1.11482349 | 0.26492607 | 0.99989001 |
| IKBIP | 103.486626 | 0.0510991 | 0.26949763 | 0.18960871 | 0.84961576 | 0.99989001 |
| APAF1 | 99.9375028 | 0.47334668 | 0.27976784 | 1.69192672 | 0.09065995 | 0.98744773 |
| UHRF1BP1L | 128.465383 | 0.29180473 | 0.23617232 | 1.23555857 | 0.21662269 | 0.99989001 |
| ACTR6 | 53.3866408 | -0.0529001 | 0.25874656 | -0.2044474 | 0.83800389 | 0.99989001 |
| SCYL2 | 218.652614 | -0.2285568 | 0.25415722 | -0.8992731 | 0.36850719 | 0.99989001 |
| UTP20 | 112.686804 | -0.2655124 | 0.41742331 | -0.6360748 | 0.52472769 | 0.99989001 |
| ARL1 | 81.8293389 | -0.0196822 | 0.37018508 | -0.0531686 | 0.9575976 | 0.99989001 |
| CHPT1 | 134.247438 | 1.08797136 | 0.40985396 | 2.65453424 | 0.0079418 | 0.41229803 |
| GNPTAB | 135.890027 | 0.09215576 | 0.28494876 | 0.3234117 | 0.74638347 | 0.99989001 |
| DRAM1 | 197.464141 | -0.0136984 | 0.19464928 | -0.0703747 | 0.94389546 | 0.99989001 |
| WASHC3 | 116.628186 | -0.2942545 | 0.24721683 | -1.190269 | 0.23394067 | 0.99989001 |
| NUP37 | 45.2626658 | -0.2753864 | 0.28344739 | -0.9715607 | 0.33126912 | NA |
| HSP90B1 | 492.609231 | -0.0833612 | 0.5555106 | -0.1500623 | 0.88071543 | 0.99989001 |
| TDG | 88.4756185 | -0.1545307 | 0.24112786 | -0.6408661 | 0.52160965 | 0.99989001 |
| HCFC2 | 119.440876 | 0.05439563 | 0.23568634 | 0.2307967 | 0.81747275 | 0.99989001 |
| NFYB | 52.8766018 | 0.31811485 | 0.31156453 | 1.02102394 | 0.3072431 | 0.99989001 |
| TXNRD1 | 783.829832 | -0.118847 | 0.43297743 | -0.2744876 | 0.7837099 | 0.99989001 |
| CHST11 | 681.450298 | 1.04812032 | 0.33566218 | 3.12254521 | 0.00179295 | 0.20504779 |
| WASHC4 | 410.87844 | 0.18522879 | 0.1884497 | 0.98290839 | 0.32565253 | 0.99989001 |
| APPL2 | 208.562922 | 0.21664076 | 0.25395644 | 0.85306267 | 0.39362455 | 0.99989001 |
| NUAK1 | 430.747251 | -0.5428073 | 0.49577814 | -1.0948593 | 0.27357827 | 0.99989001 |
| CKAP4 | 61.1743265 | 0.45849614 | 0.45314396 | 1.01181121 | 0.31162833 | 0.99989001 |
| TCP11L2 | 74.6413914 | 0.17955963 | 0.47866691 | 0.37512437 | 0.70756797 | 0.99989001 |
| POLR3B | 45.298236 | -0.2016332 | 0.40991984 | -0.4918845 | 0.62280102 | NA |
| RIC8B | 163.246351 | 0.11869274 | 0.20093483 | 0.59070266 | 0.55471967 | 0.99989001 |

| | | | | | | |
|---|---|---|---|---|---|---|
| TMEM263 | 86.5358947 | 0.20667893 | 0.28278467 | 0.73087034 | 0.46485835 | 0.99989001 |
| CRY1 | 178.182076 | -0.4122625 | 0.38260892 | -1.0775036 | 0.28125536 | 0.99989001 |
| PWP1 | 66.1763027 | 0.1032214 | 0.27299458 | 0.37810785 | 0.70535048 | 0.99989001 |
| PRDM4 | 66.9410448 | 0.00579735 | 0.36280883 | 0.01597909 | 0.98725107 | 0.99989001 |
| CMKLR1 | 96.9166242 | 2.04076123 | 0.58872323 | 3.46641875 | 0.00052744 | 0.09952814 |
| SART3 | 78.6223528 | 0.04045949 | 0.20871633 | 0.19384919 | 0.84629397 | 0.99989001 |
| ISCU | 120.697468 | 0.23780967 | 0.51796561 | 0.4591225 | 0.6461462 | 0.99989001 |
| CORO1C | 174.956972 | -0.5480441 | 0.32984924 | -1.6614989 | 0.0966133 | 0.99989001 |
| SSH1 | 379.784853 | 0.17695079 | 0.41823866 | 0.42308569 | 0.67223275 | 0.99989001 |
| ACACB | 484.752334 | 0.07999948 | 0.53440813 | 0.14969736 | 0.88100339 | 0.99989001 |
| KCTD10 | 89.7431036 | 0.11519333 | 0.34581007 | 0.33311154 | 0.73905009 | 0.99989001 |
| UBE3B | 87.372168 | 0.16884118 | 0.25959468 | 0.65040309 | 0.51543188 | 0.99989001 |
| GLTP | 86.9115231 | 0.11427831 | 0.26263026 | 0.43513004 | 0.66346804 | 0.99989001 |
| TCHP | 108.936988 | -0.1524282 | 0.28542907 | -0.534032 | 0.59331942 | 0.99989001 |
| GIT2 | 232.939479 | 0.09032657 | 0.14948971 | 0.60423269 | 0.54568895 | 0.99989001 |
| ANKRD13A | 110.25153 | 0.37576778 | 0.29783749 | 1.26165373 | 0.20707341 | 0.99989001 |
| IFT81 | 59.9686292 | 0.1856523 | 0.36902668 | 0.50308639 | 0.61490353 | 0.99989001 |
| ATP2A2 | 263.017463 | -0.0064081 | 0.31060345 | -0.0206312 | 0.98353987 | 0.99989001 |
| ANAPC7 | 99.3171212 | -0.1978738 | 0.33902971 | -0.5836475 | 0.55945749 | 0.99989001 |
| ARPC3 | 316.49293 | -0.2745097 | 0.28014753 | -0.9798754 | 0.3271476 | 0.99989001 |
| VPS29 | 150.863361 | 0.20514173 | 0.21118007 | 0.97140666 | 0.33134581 | 0.99989001 |
| PPTC7 | 99.9559215 | 0.30682816 | 0.22584982 | 1.35854952 | 0.17428938 | 0.99989001 |
| PPP1CC | 96.7307566 | 0.04702483 | 0.28461514 | 0.16522253 | 0.86876883 | 0.99989001 |
| SH2B3 | 349.273651 | 0.34145467 | 0.26340609 | 1.29630517 | 0.19487037 | 0.99989001 |
| ATXN2 | 697.076721 | 0.06040925 | 0.21169711 | 0.28535701 | 0.77537064 | 0.99989001 |
| BRAP | 85.3335656 | -0.111992 | 0.24491981 | -0.4572597 | 0.64748437 | 0.99989001 |
| ACAD10 | 52.6763356 | 0.26613587 | 0.40781736 | 0.65258591 | 0.51402327 | 0.99989001 |
| ALDH2 | 73.5174129 | 1.12004683 | 0.44117496 | 2.53878152 | 0.01112393 | 0.47956379 |
| MAPKAPK5 | 92.5152796 | -0.1435646 | 0.20832611 | -0.6891341 | 0.49073891 | 0.99989001 |
| ERP29 | 96.7837366 | -0.1661946 | 0.46278846 | -0.3591157 | 0.71950855 | 0.99989001 |
| NAA25 | 184.073048 | -0.3307579 | 0.2871329 | -1.1519332 | 0.24934853 | 0.99989001 |
| TRAFD1 | 59.8512084 | -0.1532313 | 0.37926301 | -0.4040238 | 0.68619522 | 0.99989001 |
| HECTD4 | 388.833222 | -0.0048455 | 0.18995465 | -0.025509 | 0.979649 | 0.99989001 |
| RPL6 | 341.08485 | 0.10870729 | 0.50863743 | 0.21372254 | 0.83076343 | 0.99989001 |
| PTPN11 | 193.083228 | -0.0644274 | 0.21958027 | -0.2934115 | 0.76920763 | 0.99989001 |
| OAS1 | 98.552085 | -0.4386364 | 0.96237508 | -0.4557853 | 0.64854438 | 0.99989001 |
| OAS3 | 96.6986103 | -0.3970858 | 1.1394217 | -0.3484977 | 0.72746644 | 0.99989001 |
| OAS2 | 125.693327 | -0.2115986 | 0.73044585 | -0.2896842 | 0.77205783 | 0.99989001 |
| DDX54 | 53.5176562 | -0.1462845 | 0.43856479 | -0.3335527 | 0.73871711 | 0.99989001 |
| TPCN1 | 167.144105 | 0.607681 | 0.46196364 | 1.31543036 | 0.18836531 | 0.99989001 |
| SLC8B1 | 112.91572 | 0.03480809 | 0.36401064 | 0.09562384 | 0.92381933 | 0.99989001 |
| PLBD2 | 41.6775566 | 0.12617605 | 0.26385647 | 0.47819956 | 0.63250818 | NA |
| RBM19 | 71.8891299 | -0.3615064 | 0.41114853 | -0.8792599 | 0.3792604 | 0.99989001 |
| TBX3 | 138.221931 | 1.47577189 | 0.50199196 | 2.93983173 | 0.00328391 | 0.29068264 |
| MED13L | 1713.05664 | 0.5838084 | 0.20429408 | 2.85768627 | 0.00426742 | 0.32568991 |
| FBXW8 | 132.031898 | -0.3740242 | 0.41826807 | -0.8942213 | 0.37120352 | 0.99989001 |
| FBXO21 | 154.956327 | 0.24226288 | 0.23054436 | 1.05082978 | 0.29333677 | 0.99989001 |
| WSB2 | 66.3990431 | 0.42163611 | 0.29457028 | 1.43135999 | 0.15232707 | 0.99989001 |
| VSIG10 | 74.7309158 | 0.19084967 | 0.46572417 | 0.4097912 | 0.68195912 | 0.99989001 |
| PEBP1 | 259.892325 | 0.42477997 | 0.36689029 | 1.1577847 | 0.24695191 | 0.99989001 |
| TAOK3 | 405.500475 | 0.29271534 | 0.25584408 | 1.14411615 | 0.25257547 | 0.99989001 |
| SUDS3 | 66.5202399 | -0.1623467 | 0.25223778 | -0.6436258 | 0.51981812 | 0.99989001 |
| HSPB8 | 47.0554497 | 0.62031324 | 0.34337248 | 1.80653163 | 0.07083534 | NA |
| CCDC60 | 52.6466134 | 0.25721733 | 0.57050503 | 0.45085901 | 0.65209117 | 0.99989001 |

| | | | | | | |
|---|---|---|---|---|---|---|
| AC002070.1 | 430.945071 | 0.2447131 | 0.36176456 | 0.67644297 | 0.49875944 | 0.99989001 |
| TMEM233 | 171.581265 | 0.21517648 | 0.55592826 | 0.387058 | 0.69871327 | 0.99989001 |
| PRKAB1 | 76.7970201 | -0.2219709 | 0.35467629 | -0.6258407 | 0.53141944 | 0.99989001 |
| RAB35 | 74.7822748 | -0.0400083 | 0.32922993 | -0.1215207 | 0.9032786 | 0.99989001 |
| GCN1 | 85.9480137 | -0.0836035 | 0.25580016 | -0.3268312 | 0.74379558 | 0.99989001 |
| RPLP0 | 501.744343 | -0.2521761 | 0.58535899 | -0.4308059 | 0.66660948 | 0.99989001 |
| PXN | 797.082779 | -0.1481746 | 0.29412624 | -0.503779 | 0.61441668 | 0.99989001 |
| SRSF9 | 128.047024 | -0.0983326 | 0.38812251 | -0.2533545 | 0.79999431 | 0.99989001 |
| DYNLL1 | 339.479081 | 0.50708987 | 0.5916812 | 0.85703225 | 0.39142706 | 0.99989001 |
| RNF10 | 195.809434 | 0.08627576 | 0.14994759 | 0.57537275 | 0.56503923 | 0.99989001 |
| MLEC | 106.080637 | -0.0630042 | 0.35003272 | -0.1799953 | 0.85715625 | 0.99989001 |
| SPPL3 | 465.339574 | -0.1099028 | 0.16843379 | -0.6524985 | 0.51407965 | 0.99989001 |
| CAMKK2 | 100.748833 | 0.25541304 | 0.40587068 | 0.6292966 | 0.5291549 | 0.99989001 |
| ANAPC5 | 250.131022 | -0.0794555 | 0.18977138 | -0.4186907 | 0.67544217 | 0.99989001 |
| RNF34 | 86.3670083 | 0.4019412 | 0.39430156 | 1.01937513 | 0.30802491 | 0.99989001 |
| KDM2B | 87.6819972 | 0.0919881 | 0.33053895 | 0.27829732 | 0.78078414 | 0.99989001 |
| ORAI1 | 63.9676525 | -0.0738755 | 0.60429449 | -0.1222508 | 0.90270042 | 0.99989001 |
| TMEM120B | 58.4597061 | -0.1424299 | 0.34321055 | -0.4149928 | 0.67814717 | 0.99989001 |
| SETD1B | 51.8115286 | 0.18449004 | 0.34349998 | 0.53708894 | 0.59120618 | 0.99989001 |
| MLXIP | 210.064147 | 0.25032615 | 0.28885226 | 0.86662348 | 0.38614834 | 0.99989001 |
| CLIP1 | 360.892004 | -0.0192562 | 0.23496764 | -0.0819525 | 0.93468445 | 0.99989001 |
| ZCCHC8 | 133.725312 | 0.12292292 | 0.25757491 | 0.47723174 | 0.63319711 | 0.99989001 |
| RSRC2 | 200.025136 | -0.0982757 | 0.25048345 | -0.3923439 | 0.69480413 | 0.99989001 |
| DENR | 126.289455 | 0.05040088 | 0.34025496 | 0.14812681 | 0.88224269 | 0.99989001 |
| HIP1R | 94.2227733 | -0.1050675 | 0.32674208 | -0.321561 | 0.74778531 | 0.99989001 |
| VPS37B | 222.089859 | -0.1473512 | 0.27577672 | -0.5343135 | 0.59312462 | 0.99989001 |
| ARL6IP4 | 166.70962 | -0.0537842 | 0.48298731 | -0.1113575 | 0.91133289 | 0.99989001 |
| PITPNM2 | 655.598353 | -0.4940552 | 0.36121681 | -1.3677524 | 0.17138959 | 0.99989001 |
| MPHOSPH9 | 104.883368 | -0.0719288 | 0.230068 | -0.3126413 | 0.75455321 | 0.99989001 |
| C12orf65 | 52.3323411 | 0.59362971 | 0.3292847 | 1.80278557 | 0.0714219 | 0.91166457 |
| SBNO1 | 194.08528 | -0.194449 | 0.20812339 | -0.9342967 | 0.35015086 | 0.99989001 |
| KMT5A | 65.4064609 | -0.4505904 | 0.2438373 | -1.847914 | 0.06461478 | 0.86610729 |
| RILPL2 | 186.691695 | 0.00712459 | 0.36411803 | 0.0195667 | 0.98438903 | 0.99989001 |
| SNRNP35 | 68.0404561 | -0.1074864 | 0.26320695 | -0.4083723 | 0.68300038 | 0.99989001 |
| RILPL1 | 200.601587 | 0.03485952 | 0.3747412 | 0.09302291 | 0.92588536 | 0.99989001 |
| TMED2 | 87.5366358 | 0.26257449 | 0.36711338 | 0.71524086 | 0.47446022 | 0.99989001 |
| DDX55 | 53.2718419 | -0.2096169 | 0.34319687 | -0.6107776 | 0.54134685 | 0.99989001 |
| EIF2B1 | 63.5278936 | -0.0364393 | 0.26518139 | -0.1374127 | 0.89070462 | 0.99989001 |
| GTF2H3 | 43.3255725 | 0.11329103 | 0.23377938 | 0.4846066 | 0.62795543 | NA |
| ATP6V0A2 | 86.941396 | 0.02056733 | 0.38789883 | 0.05302241 | 0.95771406 | 0.99989001 |
| CCDC92 | 126.101132 | 0.34076278 | 0.21989236 | 1.54967996 | 0.12121835 | 0.99989001 |
| NCOR2 | 453.939242 | -0.154585 | 0.40359359 | -0.3830215 | 0.70170381 | 0.99989001 |
| SCARB1 | 46.7698023 | -0.1389547 | 0.60029667 | -0.2314767 | 0.81694452 | NA |
| UBC | 1252.94254 | -0.9736242 | 0.42054343 | -2.3151573 | 0.02060432 | 0.58699182 |
| SLC15A4 | 54.1533016 | -0.0530857 | 0.31168983 | -0.1703158 | 0.86476179 | 0.99989001 |
| STX2 | 99.2932339 | 0.31164243 | 0.36162366 | 0.86178661 | 0.38880495 | 0.99989001 |
| RAN | 142.998314 | 0.29465766 | 0.62579471 | 0.47085354 | 0.63774533 | 0.99989001 |
| SFSWAP | 249.168741 | -0.1890645 | 0.25148331 | -0.7517974 | 0.45217293 | 0.99989001 |
| ULK1 | 89.8217615 | -0.3025091 | 0.28573319 | -1.0587119 | 0.28973101 | 0.99989001 |
| EP400 | 226.482148 | -0.0580369 | 0.30106917 | -0.1927692 | 0.84713969 | 0.99989001 |
| FBRSL1 | 91.220821 | -0.1059176 | 0.2246556 | -0.4714664 | 0.63730772 | 0.99989001 |
| ANKLE2 | 159.314177 | -0.2190516 | 0.26320979 | -0.8322321 | 0.40527796 | 0.99989001 |
| GOLGA3 | 131.047392 | -0.3562016 | 0.22055739 | -1.6150067 | 0.10630925 | 0.99989001 |
| CHFR | 185.331799 | -0.1522218 | 0.30797134 | -0.4942726 | 0.62111369 | 0.99989001 |

| | | | | | | |
|---|---|---|---|---|---|---|
| ZNF26 | 79.4059124 | 0.01612548 | 0.30229327 | 0.05334384 | 0.95745795 | 0.99989001 |
| ZNF84 | 83.2046438 | 0.33433912 | 0.31481264 | 1.06202571 | 0.28822402 | 0.99989001 |
| ZNF140 | 49.2773812 | 0.41474867 | 0.32680212 | 1.26911255 | 0.20440092 | NA |
| ZNF891 | 50.1782729 | -0.0010577 | 0.28541602 | -0.0037058 | 0.99704317 | 0.99989001 |
| MPHOSPH8 | 289.957209 | 0.30947257 | 0.25521408 | 1.21259991 | 0.22528282 | 0.99989001 |
| PSPC1 | 443.322504 | -0.0247047 | 0.23131762 | -0.1067998 | 0.91494778 | 0.99989001 |
| ZMYM5 | 119.301727 | 0.11466082 | 0.28608953 | 0.40078649 | 0.68857733 | 0.99989001 |
| ZMYM2 | 662.562819 | -0.1136612 | 0.26367448 | -0.4310664 | 0.6664201 | 0.99989001 |
| CRYL1 | 128.969361 | 0.27181111 | 0.30127492 | 0.9022029 | 0.36694909 | 0.99989001 |
| IFT88 | 207.418173 | -0.226945 | 0.30701544 | -0.7391973 | 0.45978718 | 0.99989001 |
| XPO4 | 243.264851 | -0.0145585 | 0.20874975 | -0.0697414 | 0.94439948 | 0.99989001 |
| LATS2 | 222.059357 | -0.081345 | 0.25845369 | -0.3147371 | 0.75296127 | 0.99989001 |
| SAP18 | 147.997437 | -0.0778804 | 0.46548206 | -0.1673112 | 0.86712519 | 0.99989001 |
| MRPL57 | 99.3337395 | -0.0700402 | 0.43393698 | -0.1614064 | 0.87177334 | 0.99989001 |
| ZDHHC20 | 316.514652 | -0.008698 | 0.257314 | -0.0338029 | 0.97303432 | 0.99989001 |
| MICU2 | 250.111661 | -0.0335987 | 0.21826189 | -0.1539377 | 0.87765886 | 0.99989001 |
| SACS | 64.3692473 | -0.2056286 | 0.2334455 | -0.880842 | 0.37840336 | 0.99989001 |
| MIPEP | 100.448641 | -0.3361051 | 0.3571029 | -0.9411997 | 0.34660255 | 0.99989001 |
| PARP4 | 275.273061 | -0.4204326 | 0.3408272 | -1.2335653 | 0.21736492 | 0.99989001 |
| MTMR6 | 134.934309 | 0.51679587 | 0.20803958 | 2.48412284 | 0.0129871 | 0.495084 |
| NUP58 | 176.384333 | -0.385679 | 0.3246998 | -1.1878019 | 0.23491148 | 0.99989001 |
| RNF6 | 63.1815952 | -0.0108562 | 0.36658097 | -0.0296148 | 0.97637428 | 0.99989001 |
| CDK8 | 319.848805 | 0.10099287 | 0.24206037 | 0.41722181 | 0.67651617 | 0.99989001 |
| WASF3 | 63.828765 | -0.2101026 | 0.24462388 | -0.8588802 | 0.3904066 | 0.99989001 |
| USP12 | 253.943717 | 0.26882602 | 0.27401175 | 0.98107477 | 0.32655587 | 0.99989001 |
| RPL21 | 185.453589 | -0.0627767 | 0.54806649 | -0.1145421 | 0.90880807 | 0.99989001 |
| GTF3A | 64.6954385 | -0.2858799 | 0.26863484 | -1.0641952 | 0.2872403 | 0.99989001 |
| MTIF3 | 102.868581 | 0.17108963 | 0.26235889 | 0.65212056 | 0.5143234 | 0.99989001 |
| LNX2 | 119.209722 | 0.58632761 | 0.30296927 | 1.93527088 | 0.05295706 | 0.81308738 |
| POLR1D | 110.226289 | -0.3013222 | 0.30083377 | -1.0016237 | 0.31652539 | 0.99989001 |
| PAN3 | 934.112805 | 0.0334039 | 0.18955431 | 0.17622337 | 0.86011846 | 0.99989001 |
| FLT1 | 7180.33784 | 0.06999283 | 0.64094279 | 0.10920292 | 0.91304154 | 0.99989001 |
| POMP | 250.777009 | -0.177054 | 0.37047561 | -0.4779101 | 0.63271419 | 0.99989001 |
| SLC46A3 | 57.3906982 | 0.24267184 | 0.45043129 | 0.53875439 | 0.59005634 | 0.99989001 |
| SLC7A1 | 86.2326319 | -0.8658111 | 0.60498361 | -1.4311315 | 0.15239253 | 0.99989001 |
| UBL3 | 275.09688 | 0.17707404 | 0.22819253 | 0.77598526 | 0.43775769 | 0.99989001 |
| KATNAL1 | 130.388582 | 0.23016948 | 0.24110956 | 0.95462608 | 0.33976682 | 0.99989001 |
| HMGB1 | 510.883121 | 0.00552044 | 0.26281873 | 0.02100476 | 0.98324186 | 0.99989001 |
| USPL1 | 131.493066 | -0.304048 | 0.2575675 | -1.1804594 | 0.23781757 | 0.99989001 |
| MEDAG | 55.6959898 | -1.3014806 | 0.44768145 | -2.9071578 | 0.00364729 | 0.29923647 |
| HSPH1 | 172.984849 | -0.4875202 | 0.60890573 | -0.8006497 | 0.42333446 | 0.99989001 |
| B3GLCT | 110.592999 | -0.0711806 | 0.29984517 | -0.2373911 | 0.8123534 | 0.99989001 |
| FRY | 361.356572 | -0.0607275 | 0.34361913 | -0.1767291 | 0.85972118 | 0.99989001 |
| N4BP2L1 | 154.352852 | 0.28446484 | 0.39929271 | 0.71242183 | 0.4762036 | 0.99989001 |
| N4BP2L2 | 1266.16522 | -0.2009328 | 0.23279799 | -0.8631207 | 0.38807109 | 0.99989001 |
| PDS5B | 733.280441 | -0.003483 | 0.26001259 | -0.0133954 | 0.98931235 | 0.99989001 |
| KL | 70.9147365 | 0.07147131 | 0.29716785 | 0.24050823 | 0.80993628 | 0.99989001 |
| STARD13 | 849.964504 | 0.31657968 | 0.43325449 | 0.73070144 | 0.46496154 | 0.99989001 |
| AL139383.1 | 61.9706073 | 0.07992991 | 0.71516877 | 0.1117637 | 0.91101077 | 0.99989001 |
| RFC3 | 204.142206 | -0.5406779 | 0.60854481 | -0.8884768 | 0.37428435 | 0.99989001 |
| NBEA | 609.348315 | -0.1341502 | 0.31374922 | -0.4275716 | 0.66896308 | 0.99989001 |
| SPART | 214.973324 | 0.0482857 | 0.27242301 | 0.17724531 | 0.85931571 | 0.99989001 |
| RFXAP | 44.3320219 | -0.1242469 | 0.30670746 | -0.4050991 | 0.68540465 | NA |
| SMAD9 | 397.993618 | 0.87616595 | 0.46648223 | 1.87824078 | 0.06034823 | 0.85923309 |

| | | | | | | |
|---|---|---|---|---|---|---|
| ALG5 | 82.4622826 | 0.01835836 | 0.2850735 | 0.06439868 | 0.94865278 | 0.99989001 |
| EXOSC8 | 50.8873545 | -0.2993398 | 0.2916999 | -1.0261909 | 0.30480162 | 0.99989001 |
| SUPT20H | 174.889537 | 0.20321772 | 0.22470887 | 0.90436005 | 0.36580451 | 0.99989001 |
| UFM1 | 145.336954 | -0.0760038 | 0.32463494 | -0.2341209 | 0.81489116 | 0.99989001 |
| PROSER1 | 67.019788 | -0.3248013 | 0.22514176 | -1.4426524 | 0.14911841 | 0.99989001 |
| NHLRC3 | 78.6188096 | 0.04134884 | 0.43685961 | 0.09465019 | 0.92459268 | 0.99989001 |
| LHFPL6 | 982.81425 | 0.20414996 | 0.19508379 | 1.04647317 | 0.29534262 | 0.99989001 |
| COG6 | 167.626308 | -0.2634963 | 0.32541971 | -0.8097122 | 0.4181056 | 0.99989001 |
| FOXO1 | 1243.86108 | 0.24851711 | 0.32917735 | 0.75496419 | 0.45027047 | 0.99989001 |
| MRPS31 | 114.92346 | -0.015115 | 0.20434973 | -0.0739663 | 0.94103719 | 0.99989001 |
| ELF1 | 639.127419 | -0.0118108 | 0.31008821 | -0.0380885 | 0.9696171 | 0.99989001 |
| WBP4 | 87.9559674 | -0.2845707 | 0.23819275 | -1.1947077 | 0.23220125 | 0.99989001 |
| NAA16 | 184.963241 | 0.07846866 | 0.21008363 | 0.37351153 | 0.70876777 | 0.99989001 |
| RGCC | 500.155789 | -0.9484009 | 1.16004634 | -0.8175543 | 0.41361173 | 0.99989001 |
| VWA8 | 322.545279 | -0.3830531 | 0.27411903 | -1.3973969 | 0.16229426 | 0.99989001 |
| DGKH | 969.50615 | -0.0048261 | 0.21129402 | -0.0228407 | 0.98177735 | 0.99989001 |
| AKAP11 | 189.873627 | 0.15079924 | 0.18269143 | 0.82543138 | 0.40912672 | 0.99989001 |
| EPSTI1 | 366.295239 | 0.24902983 | 0.96583617 | 0.25783858 | 0.79653149 | 0.99989001 |
| DNAJC15 | 155.479508 | 0.11476607 | 0.1911939 | 0.60026012 | 0.54833289 | 0.99989001 |
| CCDC122 | 70.7683725 | 1.00939746 | 0.44238371 | 2.28172385 | 0.02250565 | 0.60384164 |
| TSC22D1 | 741.641946 | 0.12243021 | 0.27740663 | 0.44133842 | 0.65896801 | 0.99989001 |
| GPALPP1 | 111.327428 | -0.0648783 | 0.21097245 | -0.3075204 | 0.75844732 | 0.99989001 |
| GTF2F2 | 180.716299 | -0.1149547 | 0.27730534 | -0.4145421 | 0.67847713 | 0.99989001 |
| TPT1 | 1240.24161 | 0.12461783 | 0.56566937 | 0.22030153 | 0.82563633 | 0.99989001 |
| COG3 | 167.757307 | 0.02012751 | 0.21509436 | 0.09357528 | 0.92544655 | 0.99989001 |
| ZC3H13 | 246.376376 | -0.0527621 | 0.20254871 | -0.260491 | 0.79448505 | 0.99989001 |
| AL138686.1 | 69.1675697 | -0.2966606 | 0.73196079 | -0.4052958 | 0.68526009 | 0.99989001 |
| LRCH1 | 1219.99739 | 0.19008081 | 0.26116886 | 0.72780808 | 0.46673108 | 0.99989001 |
| ESD | 100.301415 | -0.1727851 | 0.27358438 | -0.6315604 | 0.52767417 | 0.99989001 |
| SUCLA2 | 162.488196 | -0.2012602 | 0.30917012 | -0.650969 | 0.51506648 | 0.99989001 |
| MED4 | 181.709626 | -0.0207087 | 0.16199964 | -0.1278317 | 0.89828213 | 0.99989001 |
| ITM2B | 1392.07084 | 0.10987235 | 0.42647665 | 0.25762806 | 0.79669397 | 0.99989001 |
| RB1 | 846.951771 | 0.20615286 | 0.20586216 | 1.0014121 | 0.31662761 | 0.99989001 |
| LPAR6 | 243.367486 | -0.1074223 | 0.42695449 | -0.2516012 | 0.8013493 | 0.99989001 |
| RCBTB2 | 96.280811 | 0.12930276 | 0.24966442 | 0.51790622 | 0.6045237 | 0.99989001 |
| FNDC3A | 727.081622 | -0.1673234 | 0.37956652 | -0.4408276 | 0.6593378 | 0.99989001 |
| CDADC1 | 56.7424858 | 0.15561831 | 0.36626467 | 0.4248794 | 0.67092459 | 0.99989001 |
| CAB39L | 63.8885174 | 0.29395337 | 0.26211521 | 1.12146627 | 0.26208944 | 0.99989001 |
| SETDB2 | 80.3128312 | 0.12342056 | 0.25080716 | 0.49209344 | 0.62265329 | 0.99989001 |
| PHF11 | 149.52019 | -0.1320002 | 0.37584515 | -0.3512089 | 0.72543163 | 0.99989001 |
| RCBTB1 | 72.9036068 | 0.01301698 | 0.27867908 | 0.04670958 | 0.96274469 | 0.99989001 |
| EBPL | 71.2111735 | -0.6824032 | 0.47319297 | -1.4421246 | 0.14926722 | 0.99989001 |
| KPNA3 | 297.488705 | 0.03935412 | 0.20758323 | 0.18958235 | 0.84963642 | 0.99989001 |
| SPRYD7 | 47.811867 | 0.18556929 | 0.26004623 | 0.71360118 | 0.47547383 | NA |
| DLEU2 | 898.289854 | 0.49000925 | 0.3186579 | 1.53772821 | 0.12411508 | 0.99989001 |
| TRIM13 | 134.399339 | 0.11594398 | 0.30258284 | 0.38318094 | 0.70158561 | 0.99989001 |
| RNASEH2B | 88.461311 | -0.5774819 | 0.24218489 | -2.3844669 | 0.01710389 | 0.55366411 |
| FAM124A | 62.4167412 | 0.4126004 | 0.31985417 | 1.28996411 | 0.19706312 | 0.99989001 |
| INTS6 | 416.145322 | -0.0300505 | 0.30318905 | -0.0991148 | 0.9210471 | 0.99989001 |
| WDFY2 | 208.941916 | 0.54955515 | 0.23027702 | 2.38649584 | 0.0170098 | 0.55340504 |
| UTP14C | 41.4037743 | 0.14601485 | 0.32178215 | 0.45376927 | 0.6499949 | NA |
| NEK3 | 121.58023 | 0.00368654 | 0.39200855 | 0.00940423 | 0.99249662 | 0.99989001 |
| THSD1 | 93.9979308 | 0.11186558 | 0.31732734 | 0.35252423 | 0.72444515 | 0.99989001 |
| VPS36 | 246.997535 | -0.0533793 | 0.2034389 | -0.2623848 | 0.79302479 | 0.99989001 |

| | | | | | | |
|---|---|---|---|---|---|---|
| SUGT1 | 124.269247 | 0.32335725 | 0.22270919 | 1.45192593 | 0.1465222 | 0.99989001 |
| PCDH17 | 714.099028 | -0.3823394 | 0.57417268 | -0.6658962 | 0.50547746 | 0.99989001 |
| TDRD3 | 328.024731 | -0.0394137 | 0.18794189 | -0.2097119 | 0.8338925 | 0.99989001 |
| PCDH9 | 264.979688 | -1.8830968 | 0.75111868 | -2.5070563 | 0.01217413 | 0.49000287 |
| DACH1 | 394.841205 | -1.2233401 | 0.50099068 | -2.441842 | 0.01461254 | 0.5186081 |
| DIS3 | 104.786066 | 0.04955518 | 0.20357108 | 0.24342939 | 0.80767278 | 0.99989001 |
| PIBF1 | 517.273951 | 0.08490078 | 0.2196751 | 0.38648338 | 0.69913871 | 0.99989001 |
| KLF12 | 517.576516 | -0.3629665 | 0.50360886 | -0.7207309 | 0.47107509 | 0.99989001 |
| TBC1D4 | 343.384461 | -1.1043257 | 0.55211134 | -2.0001866 | 0.04548012 | 0.77228819 |
| COMMD6 | 97.9974606 | -0.3980244 | 0.53390974 | -0.74549 | 0.45597554 | 0.99989001 |
| UCHL3 | 60.2393887 | -0.2253609 | 0.42329233 | -0.5324001 | 0.59444895 | 0.99989001 |
| LMO7 | 137.702938 | -1.2399967 | 0.48031006 | -2.5816589 | 0.00983267 | 0.44980004 |
| KCTD12 | 601.090419 | 0.92901865 | 0.56125984 | 1.65523805 | 0.09787622 | 0.99989001 |
| AC001226.2 | 65.7513783 | -0.0210119 | 0.5105341 | -0.0411568 | 0.96717092 | 0.99989001 |
| FBXL3 | 145.025923 | 0.37322058 | 0.24159885 | 1.54479456 | 0.12239596 | 0.99989001 |
| MYCBP2 | 1044.93959 | -0.0224969 | 0.24963402 | -0.0901195 | 0.92819225 | 0.99989001 |
| EDNRB | 151.515871 | 0.73631198 | 0.49366897 | 1.49150953 | 0.13582777 | 0.99989001 |
| RNF219 | 63.4867007 | 0.04457726 | 0.30969718 | 0.14393823 | 0.88554925 | 0.99989001 |
| RBM26 | 508.177845 | -0.0332742 | 0.11845601 | -0.280899 | 0.77878789 | 0.99989001 |
| NDFIP2 | 107.243131 | 0.31762175 | 0.29928429 | 1.06127104 | 0.28856674 | 0.99989001 |
| GPC5 | 1379.73663 | -1.4118552 | 0.74290715 | -1.9004464 | 0.05737456 | 0.84253542 |
| GPR180 | 60.237894 | 0.11919162 | 0.29923068 | 0.39832686 | 0.69038927 | 0.99989001 |
| ABCC4 | 326.835239 | 0.08346363 | 0.41069527 | 0.2032252 | 0.83895902 | 0.99989001 |
| CLDN10 | 48.0930117 | 0.43703087 | 0.4939058 | 0.88484661 | 0.37623937 | NA |
| DNAJC3 | 284.876967 | 0.31219008 | 0.25405906 | 1.22880907 | 0.2191434 | 0.99989001 |
| UGGT2 | 480.14659 | -0.0565054 | 0.22381475 | -0.2524649 | 0.80068177 | 0.99989001 |
| MBNL2 | 788.753557 | -0.222278 | 0.31029365 | -0.7163473 | 0.47377693 | 0.99989001 |
| RAP2A | 119.962508 | 0.00121629 | 0.29058121 | 0.00418572 | 0.99666029 | 0.99989001 |
| IPO5 | 113.144892 | -0.0767696 | 0.22952748 | -0.3344679 | 0.73802654 | 0.99989001 |
| FARP1 | 404.662601 | 0.4071443 | 0.47071788 | 0.86494334 | 0.38706989 | 0.99989001 |
| STK24 | 378.436306 | 0.18723855 | 0.21245831 | 0.8812955 | 0.3781579 | 0.99989001 |
| DOCK9 | 1921.62386 | 0.24079377 | 0.28320065 | 0.85025854 | 0.39518136 | 0.99989001 |
| UBAC2 | 380.556924 | -0.2614263 | 0.17517913 | -1.492337 | 0.13561083 | 0.99989001 |
| TM9SF2 | 252.523807 | -0.0932457 | 0.19473391 | -0.4788363 | 0.63205511 | 0.99989001 |
| CLYBL | 130.911224 | -0.2905849 | 0.41527802 | -0.6997357 | 0.48409236 | 0.99989001 |
| PCCA | 786.318186 | -0.0790399 | 0.19831855 | -0.3985503 | 0.69022459 | 0.99989001 |
| GGACT | 86.2159056 | -0.6307215 | 0.35063236 | -1.7988114 | 0.07204851 | 0.91336395 |
| TMTC4 | 61.2160581 | -0.158669 | 0.42755163 | -0.3711107 | 0.7105551 | 0.99989001 |
| TPP2 | 234.039992 | 0.23804445 | 0.20152017 | 1.1812438 | 0.23750589 | 0.99989001 |
| EFNB2 | 1105.56024 | 0.49472391 | 0.41479886 | 1.19268386 | 0.23299321 | 0.99989001 |
| ARGLU1 | 1053.60212 | -0.0406158 | 0.28393526 | -0.1430461 | 0.88625379 | 0.99989001 |
| LINC00551 | 75.9389994 | 0.89055505 | 0.57883462 | 1.53853108 | 0.12391881 | 0.99989001 |
| FAM155A | 384.999634 | 0.41357018 | 0.63368498 | 0.65264319 | 0.51398634 | 0.99989001 |
| ABHD13 | 72.2072947 | 0.25381404 | 0.31549964 | 0.80448282 | 0.42111819 | 0.99989001 |
| COL4A1 | 1108.89194 | -0.2915592 | 0.51142976 | -0.5700865 | 0.56861905 | 0.99989001 |
| COL4A2 | 1239.93122 | -0.2790858 | 0.42215443 | -0.6610989 | 0.50854892 | 0.99989001 |
| RAB20 | 96.7344347 | -0.1177585 | 0.44896423 | -0.2622894 | 0.79309836 | 0.99989001 |
| NAXD | 42.085863 | -0.0047243 | 0.2503183 | -0.0188731 | 0.98494237 | NA |
| CARS2 | 104.622779 | -0.0344087 | 0.2637207 | -0.1304739 | 0.89619154 | 0.99989001 |
| ANKRD10 | 608.344396 | -0.1643418 | 0.22147241 | -0.7420418 | 0.45806201 | 0.99989001 |
| ARHGEF7 | 725.347842 | 0.24760109 | 0.31333176 | 0.7902202 | 0.42939918 | 0.99989001 |
| TUBGCP3 | 196.632374 | 0.0994229 | 0.27891508 | 0.35646296 | 0.72149389 | 0.99989001 |
| ATP11A | 661.763937 | 0.2473461 | 0.42521498 | 0.58169659 | 0.56077107 | 0.99989001 |
| MCF2L | 328.870057 | 0.49412517 | 0.47781841 | 1.03412752 | 0.30107655 | 0.99989001 |

| | | | | | | |
|---|---|---|---|---|---|---|
| PCID2 | 99.7756341 | 0.17254096 | 0.21028141 | 0.82052409 | 0.4119174 | 0.99989001 |
| CUL4A | 249.263906 | -0.1186774 | 0.2168416 | -0.5472998 | 0.58417279 | 0.99989001 |
| LAMP1 | 270.0627 | 0.10603529 | 0.43718436 | 0.24254136 | 0.80836071 | 0.99989001 |
| DCUN1D2 | 136.803488 | 0.07002708 | 0.27284646 | 0.25665378 | 0.79744606 | 0.99989001 |
| TMCO3 | 187.634442 | 0.21344452 | 0.17951512 | 1.1890058 | 0.23443738 | 0.99989001 |
| TFDP1 | 72.1089781 | -0.8857984 | 0.30199646 | -2.9331418 | 0.00335551 | 0.29068264 |
| TMEM255B | 450.725092 | 0.07124362 | 0.49975064 | 0.14255833 | 0.88663901 | 0.99989001 |
| GAS6 | 357.662361 | 0.31181719 | 0.55402471 | 0.56282182 | 0.57355622 | 0.99989001 |
| RASA3 | 72.2328792 | 0.19361159 | 0.56329548 | 0.34371231 | 0.73106265 | 0.99989001 |
| CDC16 | 143.153412 | -0.0054152 | 0.1898087 | -0.0285298 | 0.97723959 | 0.99989001 |
| UPF3A | 332.483185 | -0.0667773 | 0.22299732 | -0.2994535 | 0.76459403 | 0.99989001 |
| TTC5 | 49.6690182 | -0.0818795 | 0.28688628 | -0.2854075 | 0.77533194 | NA |
| CCNB1IP1 | 46.50234 | -0.3372155 | 0.29876871 | -1.128684 | 0.25903116 | NA |
| TEP1 | 84.443497 | -0.0144181 | 0.2081206 | -0.0692776 | 0.9447687 | 0.99989001 |
| OSGEP | 50.9268784 | -0.0211594 | 0.28447195 | -0.0743813 | 0.94070702 | 0.99989001 |
| APEX1 | 48.5769863 | -0.3999228 | 0.46198132 | -0.8656688 | 0.38667184 | NA |
| PNP | 116.530243 | -0.6728017 | 0.69993162 | -0.9612391 | 0.33643194 | 0.99989001 |
| RNASE1 | 792.200657 | -0.1447899 | 0.73062896 | -0.1981715 | 0.84291085 | 0.99989001 |
| NDRG2 | 53.6729271 | 0.50213832 | 0.37770691 | 1.32943907 | 0.18370316 | 0.99989001 |
| ARHGEF40 | 45.330712 | 0.46549478 | 0.35633396 | 1.30634414 | 0.19143555 | NA |
| HNRNPC | 641.140617 | 0.0671829 | 0.25557234 | 0.26287233 | 0.79264898 | 0.99989001 |
| SUPT16H | 132.540305 | -0.066795 | 0.2201235 | -0.3034432 | 0.76155213 | 0.99989001 |
| CHD8 | 187.787193 | 0.07844255 | 0.26208697 | 0.29929969 | 0.76471139 | 0.99989001 |
| RAB2B | 84.5917685 | -0.4267202 | 0.25593541 | -1.6672963 | 0.0954555 | 0.99989001 |
| TOX4 | 102.988468 | -0.0708855 | 0.24334223 | -0.2912996 | 0.77082218 | 0.99989001 |
| METTL3 | 47.0003869 | 0.28877788 | 0.28167131 | 1.02523001 | 0.30525467 | NA |
| DAD1 | 188.121361 | -0.2802666 | 0.47840275 | -0.5858383 | 0.5579842 | 0.99989001 |
| OXA1L | 50.7167811 | -0.0146393 | 0.33678883 | -0.0434673 | 0.96532906 | 0.99989001 |
| MRPL52 | 51.4964121 | 0.07043992 | 0.39884585 | 0.17660939 | 0.85981522 | 0.99989001 |
| LRP10 | 116.216659 | 0.18688084 | 0.38344155 | 0.48737765 | 0.62599073 | 0.99989001 |
| RBM23 | 95.3458981 | -0.0266066 | 0.24120619 | -0.1103065 | 0.91216632 | 0.99989001 |
| PSMB5 | 129.240016 | -0.0814116 | 0.3727507 | -0.2184077 | 0.82711147 | 0.99989001 |
| ACIN1 | 242.299985 | -0.1307956 | 0.25703069 | -0.5088715 | 0.61084227 | 0.99989001 |
| C14orf119 | 47.5435731 | 0.05433659 | 0.3306563 | 0.16432952 | 0.86947174 | NA |
| PABPN1 | 97.1058413 | -0.3420196 | 0.29529479 | -1.1582311 | 0.24676973 | 0.99989001 |
| NGDN | 55.0187711 | 0.11341841 | 0.3281685 | 0.34561029 | 0.72963562 | 0.99989001 |
| DHRS4L2 | 48.1737598 | 0.16627454 | 0.43959272 | 0.3782468 | 0.70524726 | NA |
| PSME1 | 135.060128 | -0.1249984 | 0.4469233 | -0.2796866 | 0.779718 | 0.99989001 |
| TINF2 | 41.8312254 | 0.17621469 | 0.37246999 | 0.4730977 | 0.63614348 | NA |
| ADCY4 | 215.813716 | -0.4310407 | 0.24604866 | -1.7518516 | 0.07979932 | 0.93383766 |
| KHNYN | 45.8755882 | -0.0295526 | 0.27888483 | -0.105967 | 0.91560857 | NA |
| SDR39U1 | 43.7949804 | -0.1129205 | 0.3006684 | -0.375565 | 0.70724031 | NA |
| NOVA1 | 69.5962848 | -0.223328 | 0.37087613 | -0.6021634 | 0.54706539 | 0.99989001 |
| PRKD1 | 1104.65689 | -0.3969921 | 0.49123531 | -0.8081507 | 0.41900386 | 0.99989001 |
| G2E3 | 96.9693203 | 0.16494248 | 0.23528428 | 0.70103484 | 0.48328127 | 0.99989001 |
| SCFD1 | 336.932759 | -0.0617701 | 0.23939311 | -0.2580279 | 0.79638538 | 0.99989001 |
| STRN3 | 614.907903 | -0.0258405 | 0.2391916 | -0.1080327 | 0.91396979 | 0.99989001 |
| AP4S1 | 69.2546044 | 0.33676199 | 0.35329048 | 0.95321557 | 0.34048086 | 0.99989001 |
| HECTD1 | 523.03613 | 0.08101081 | 0.20193261 | 0.40117748 | 0.68828946 | 0.99989001 |
| HEATR5A | 73.1528606 | -0.1094053 | 0.27459603 | -0.3984227 | 0.69031861 | 0.99989001 |
| NUBPL | 287.887153 | -0.0353797 | 0.37874916 | -0.0934121 | 0.9255762 | 0.99989001 |
| ARHGAP5 | 497.451798 | -0.0507317 | 0.33559172 | -0.1511708 | 0.87984096 | 0.99989001 |
| NPAS3 | 952.216602 | 0.36183189 | 0.57715772 | 0.6269203 | 0.53071148 | 0.99989001 |
| EGLN3 | 161.83769 | -1.2534478 | 0.97379471 | -1.2871787 | 0.19803201 | 0.99989001 |

| | | | | | | |
|---|---|---|---|---|---|---|
| EAPP | 104.971036 | -0.0081182 | 0.32942974 | -0.0246432 | 0.98033959 | 0.99989001 |
| SNX6 | 314.18796 | 0.0251988 | 0.16506971 | 0.15265548 | 0.87866997 | 0.99989001 |
| CFL2 | 55.2399067 | -0.2210371 | 0.32966811 | -0.6704838 | 0.50254941 | 0.99989001 |
| BAZ1A | 354.738672 | 0.23856907 | 0.36426437 | 0.65493384 | 0.51251035 | 0.99989001 |
| SRP54 | 196.423377 | -0.183163 | 0.20573611 | -0.8902815 | 0.37331477 | 0.99989001 |
| FAM177A1 | 206.290917 | 0.19110481 | 0.17864721 | 1.06973296 | 0.28473953 | 0.99989001 |
| PPP2R3C | 189.202579 | 0.11218473 | 0.19497039 | 0.57539371 | 0.56502506 | 0.99989001 |
| NFKBIA | 968.669028 | 0.69626547 | 0.58687779 | 1.18638919 | 0.23546864 | 0.99989001 |
| RALGAPA1 | 754.680222 | -0.2886418 | 0.23251624 | -1.2413833 | 0.21446419 | 0.99989001 |
| BRMS1L | 57.3377248 | -0.3115136 | 0.33234586 | -0.9373175 | 0.34859528 | 0.99989001 |
| MBIP | 47.0364759 | 0.17781775 | 0.24996776 | 0.71136275 | 0.47685948 | NA |
| MIPOL1 | 264.288992 | 0.11117068 | 0.23114485 | 0.48095678 | 0.63054722 | 0.99989001 |
| CLEC14A | 414.467592 | 0.30409782 | 0.49115892 | 0.61914344 | 0.53582187 | 0.99989001 |
| LINC00639 | 138.500597 | -2.5314363 | 1.44979677 | -1.7460629 | 0.08080002 | 0.93972046 |
| SEC23A | 291.263243 | 0.06002083 | 0.19413653 | 0.30916813 | 0.75719363 | 0.99989001 |
| TRAPPC6B | 87.3263696 | 0.14416359 | 0.22518515 | 0.64020022 | 0.52204244 | 0.99989001 |
| PNN | 209.765021 | 0.00558269 | 0.24339143 | 0.02293709 | 0.98170046 | 0.99989001 |
| MIA2 | 235.103079 | 0.23185567 | 0.25724024 | 0.90131958 | 0.36741842 | 0.99989001 |
| FBXO33 | 143.983567 | 0.32027604 | 0.18177001 | 1.7619851 | 0.07807182 | 0.92364586 |
| KLHL28 | 117.981942 | 0.34144852 | 0.20493392 | 1.66613956 | 0.09568562 | 0.99989001 |
| TOGARAM1 | 191.339867 | 0.02164784 | 0.29049963 | 0.07451935 | 0.94059715 | 0.99989001 |
| PRPF39 | 90.1490408 | -0.2902157 | 0.27476945 | -1.0562153 | 0.29086985 | 0.99989001 |
| FKBP3 | 68.0162931 | -0.2120982 | 0.30672685 | -0.6914888 | 0.48925843 | 0.99989001 |
| FANCM | 65.2645633 | -0.3869242 | 0.29557808 | -1.3090424 | 0.19052 | 0.99989001 |
| MIS18BP1 | 278.325331 | -0.0807101 | 0.34228263 | -0.2357996 | 0.81358821 | 0.99989001 |
| RPS29 | 194.838843 | 0.05162581 | 0.44573463 | 0.11582185 | 0.90779373 | 0.99989001 |
| RPL36AL | 258.959952 | 0.09919928 | 0.4318824 | 0.22969049 | 0.81833228 | 0.99989001 |
| KLHDC1 | 127.59068 | -0.0868199 | 0.37520862 | -0.2313909 | 0.81701113 | 0.99989001 |
| KLHDC2 | 86.7826724 | 0.2991175 | 0.29174287 | 1.02527782 | 0.30523212 | 0.99989001 |
| NEMF | 402.83197 | -0.1270013 | 0.16704349 | -0.7602889 | 0.44708194 | 0.99989001 |
| ARF6 | 88.5082811 | 0.10354886 | 0.32123959 | 0.32234154 | 0.74719396 | 0.99989001 |
| VCPKMT | 51.1652677 | 0.13879674 | 0.31501896 | 0.44059804 | 0.65950402 | 0.99989001 |
| SOS2 | 491.98762 | 0.29820386 | 0.17358214 | 1.71794091 | 0.08580739 | 0.95514629 |
| L2HGDH | 43.3510231 | -0.3233803 | 0.36889411 | -0.8766208 | 0.38069263 | NA |
| ATP5S | 47.3365673 | -0.0338352 | 0.30660106 | -0.1103559 | 0.91212716 | NA |
| CDKL1 | 104.953015 | 0.51657497 | 0.3224574 | 1.60199446 | 0.10915683 | 0.99989001 |
| MAP4K5 | 508.055294 | 0.28701944 | 0.19140235 | 1.49956075 | 0.13372822 | 0.99989001 |
| SAV1 | 150.208403 | 0.06280118 | 0.2876372 | 0.21833471 | 0.82716833 | 0.99989001 |
| NIN | 317.443983 | 0.42446926 | 0.27453116 | 1.54616058 | 0.12206579 | 0.99989001 |
| PYGL | 82.8778283 | 0.50554013 | 0.66335291 | 0.76209832 | 0.44600133 | 0.99989001 |
| TMX1 | 87.4597774 | -0.0610603 | 0.24758223 | -0.2466265 | 0.80519733 | 0.99989001 |
| FRMD6 | 130.163367 | 0.0271931 | 0.2611549 | 0.10412633 | 0.9170691 | 0.99989001 |
| GNG2 | 148.45175 | 0.43723853 | 0.22773248 | 1.91996558 | 0.05486225 | 0.82320925 |
| RTRAF | 115.318561 | 0.09626087 | 0.47492809 | 0.20268515 | 0.83938113 | 0.99989001 |
| TXNDC16 | 68.0211249 | -0.3848024 | 0.51215264 | -0.7513431 | 0.45244618 | 0.99989001 |
| GPR137C | 55.9554219 | -0.0047101 | 0.80494798 | -0.0058514 | 0.99533126 | 0.99989001 |
| ERO1A | 354.474153 | -0.9627679 | 0.63544671 | -1.515104 | 0.12974607 | 0.99989001 |
| PSMC6 | 175.35225 | -0.1343524 | 0.24544104 | -0.5473916 | 0.58410973 | 0.99989001 |
| STYX | 76.1281048 | -0.0981683 | 0.21679974 | -0.4528062 | 0.65068829 | 0.99989001 |
| GNPNAT1 | 60.6211048 | -0.3982478 | 0.34241437 | -1.1630582 | 0.24480589 | 0.99989001 |
| FERMT2 | 466.08291 | 0.08330173 | 0.21588622 | 0.38585941 | 0.6996008 | 0.99989001 |
| DDHD1 | 195.73947 | 0.00689105 | 0.25480377 | 0.02704452 | 0.97842423 | 0.99989001 |
| CNIH1 | 88.4093135 | 0.07912982 | 0.39133894 | 0.20220278 | 0.8397582 | 0.99989001 |
| CGRRF1 | 52.943034 | 0.32547258 | 0.27234952 | 1.19505473 | 0.23206565 | 0.99989001 |

| | | | | | | |
|---|---|---|---|---|---|---|
| SAMD4A | 550.440301 | 0.05539685 | 0.66278324 | 0.08358216 | 0.93338865 | 0.99989001 |
| GCH1 | 209.344456 | 1.12318213 | 0.64667811 | 1.73684884 | 0.08241386 | 0.94799019 |
| SOCS4 | 74.3315358 | 0.1242684 | 0.28310658 | 0.43894565 | 0.66070092 | 0.99989001 |
| MAPK1IP1L | 214.157692 | 0.06300426 | 0.27443992 | 0.22957397 | 0.81842283 | 0.99989001 |
| LGALS3 | 170.187784 | -0.0573926 | 0.52577475 | -0.1091581 | 0.91307712 | 0.99989001 |
| FBXO34 | 235.432179 | -0.0308908 | 0.25709664 | -0.1201526 | 0.90436225 | 0.99989001 |
| ATG14 | 66.1941956 | -0.3208097 | 0.27815654 | -1.1533422 | 0.24876994 | 0.99989001 |
| KTN1 | 618.258898 | 0.05321756 | 0.24704258 | 0.21541855 | 0.82944101 | 0.99989001 |
| PELI2 | 333.649706 | 0.45223693 | 0.30963524 | 1.46054737 | 0.1441397 | 0.99989001 |
| TMEM260 | 158.841893 | -0.0296637 | 0.22092802 | -0.1342687 | 0.89319013 | 0.99989001 |
| EXOC5 | 264.454207 | 0.09339295 | 0.16352657 | 0.5711179 | 0.56791972 | 0.99989001 |
| AP5M1 | 97.9206363 | -0.0077614 | 0.25944906 | -0.0299148 | 0.97613497 | 0.99989001 |
| NAA30 | 60.1186923 | 0.1223239 | 0.26017324 | 0.47016328 | 0.63823836 | 0.99989001 |
| ARMH4 | 118.330172 | 0.27385749 | 0.28132715 | 0.97344846 | 0.33033046 | 0.99989001 |
| ACTR10 | 160.456208 | 0.1868217 | 0.15716685 | 1.18868391 | 0.23456407 | 0.99989001 |
| PSMA3 | 143.551601 | -0.1198406 | 0.24354245 | -0.4920726 | 0.62266806 | 0.99989001 |
| ARID4A | 248.111245 | 0.27001163 | 0.31827392 | 0.84836239 | 0.39623618 | 0.99989001 |
| TIMM9 | 51.9080219 | 0.01041123 | 0.29837834 | 0.0348927 | 0.9721653 | 0.99989001 |
| KIAA0586 | 173.104985 | -0.0032243 | 0.2222974 | -0.0145045 | 0.9884275 | 0.99989001 |
| DAAM1 | 280.284319 | 0.18242519 | 0.34462343 | 0.52934645 | 0.59656514 | 0.99989001 |
| JKAMP | 72.3696206 | 0.12917763 | 0.31958363 | 0.40420603 | 0.68606122 | 0.99989001 |
| PCNX4 | 313.539117 | -0.0776201 | 0.27843133 | -0.2787766 | 0.78041631 | 0.99989001 |
| DHRS7 | 69.5829426 | 0.05032811 | 0.27905866 | 0.18034955 | 0.85687816 | 0.99989001 |
| PPM1A | 123.397536 | 0.36105837 | 0.22930114 | 1.57460347 | 0.115348 | 0.99989001 |
| MNAT1 | 387.692188 | -0.1277855 | 0.23913572 | -0.5343641 | 0.59308967 | 0.99989001 |
| SLC38A6 | 83.1987714 | -0.3696967 | 0.3269459 | -1.1307581 | 0.25815694 | 0.99989001 |
| PRKCH | 1779.3818 | 0.04952569 | 0.21142937 | 0.23424225 | 0.81479692 | 0.99989001 |
| AL359220.1 | 106.254601 | -0.1115688 | 0.28722668 | -0.3884346 | 0.69769441 | 0.99989001 |
| HIF1A | 1403.97161 | -1.3669588 | 0.54686405 | -2.4996318 | 0.01243224 | 0.49000287 |
| HIF1A-AS2 | 422.542105 | -1.1688241 | 0.94979958 | -1.2306007 | 0.21847223 | 0.99989001 |
| SNAPC1 | 96.6591308 | -0.79533 | 0.42383472 | -1.8765097 | 0.06058533 | 0.85923309 |
| RHOJ | 815.87291 | -0.1860066 | 0.44202347 | -0.420807 | 0.673896 | 0.99989001 |
| PPP2R5E | 519.779829 | 0.12645311 | 0.23104828 | 0.54730165 | 0.58417151 | 0.99989001 |
| WDR89 | 55.6334109 | -0.3911086 | 0.27466334 | -1.4239564 | 0.15445908 | 0.99989001 |
| SGPP1 | 88.5462616 | -0.2122673 | 0.28162275 | -0.7537291 | 0.45101188 | 0.99989001 |
| SYNE2 | 2220.61987 | -0.0526132 | 0.1935217 | -0.2718723 | 7.86E-01 | 0.99989001 |
| ESR2 | 269.287939 | -2.0539457 | 0.37238727 | -5.5156175 | 3.48E-08 | 8.74E-05 |
| MTHFD1 | 43.2593029 | 0.00418151 | 0.29976983 | 0.01394907 | 0.98887062 | NA |
| ZBTB25 | 83.3481838 | -0.3300876 | 0.45233486 | -0.7297417 | 0.46554806 | 0.99989001 |
| ZBTB1 | 229.500641 | 0.12312781 | 0.33671004 | 0.36567906 | 0.71460457 | 0.99989001 |
| HSPA2 | 62.7131071 | -2.3452875 | 0.83255421 | -2.8169787 | 0.00484777 | 0.34759088 |
| CHURC1-FNT | 43.5655948 | -0.1544019 | 0.37301144 | -0.4139334 | 0.67892286 | NA |
| MAX | 308.416828 | -0.0288019 | 0.30131752 | -0.0955865 | 0.923849 | 0.99989001 |
| FUT8 | 500.411105 | 0.33599191 | 0.36233292 | 0.92730163 | 0.35376995 | 0.99989001 |
| AL359232.1 | 100.068655 | -0.6914103 | 0.4560637 | -1.516039 | 0.1295095 | 0.99989001 |
| GPHN | 1145.45058 | 0.10670374 | 0.17315647 | 0.6162273 | 0.53774451 | 0.99989001 |
| MPP5 | 150.635493 | -0.060826 | 0.19376522 | -0.3139158 | 0.75358501 | 0.99989001 |
| ATP6V1D | 110.98138 | -0.4883246 | 0.40226935 | -1.2139245 | 0.22477654 | 0.99989001 |
| EIF2S1 | 114.471118 | -0.0536139 | 0.31229767 | -0.1716756 | 0.86369256 | 0.99989001 |
| VTI1B | 53.2748092 | -0.2103729 | 0.39863318 | -0.5277356 | 0.59768286 | 0.99989001 |
| ZFYVE26 | 67.0837778 | -0.3069789 | 0.37297809 | -0.8230482 | 0.41048059 | 0.99989001 |
| RAD51B | 824.05254 | -0.195554 | 0.37667206 | -0.5191626 | 0.60364738 | 0.99989001 |
| ZFP36L1 | 585.322092 | 0.11222768 | 0.37922821 | 0.29593706 | 0.76727815 | 0.99989001 |
| ACTN1 | 406.339089 | -0.8161552 | 0.45875391 | -1.7790698 | 0.07522832 | 0.9206525 |

| | | | | | | |
|---|---|---|---|---|---|---|
| DCAF5 | 379.562295 | 0.30421177 | 0.17155902 | 1.77321936 | 0.07619237 | 0.92274869 |
| EXD2 | 42.3655055 | 0.05553457 | 0.29695259 | 0.18701494 | 0.85164892 | NA |
| ERH | 131.48003 | -0.2180424 | 0.35333096 | -0.6171051 | 0.53716537 | 0.99989001 |
| SLC39A9 | 129.574733 | -0.1509874 | 0.19540244 | -0.7726997 | 0.43970015 | 0.99989001 |
| SUSD6 | 574.249561 | 0.03314598 | 0.4448325 | 0.07451339 | 0.94060188 | 0.99989001 |
| SRSF5 | 382.856574 | -0.1891452 | 0.28837533 | -0.6558995 | 0.51188879 | 0.99989001 |
| AL357153.2 | 104.854658 | 0.53182876 | 0.39648215 | 1.34136872 | 0.17980077 | 0.99989001 |
| MED6 | 77.1571753 | 0.06931211 | 0.20623884 | 0.33607692 | 0.73681286 | 0.99989001 |
| PCNX1 | 745.638163 | 0.44325845 | 0.27593588 | 1.60638208 | 0.10818999 | 0.99989001 |
| SIPA1L1 | 484.950995 | 0.12090927 | 0.3346338 | 0.36131815 | 0.71786163 | 0.99989001 |
| DPF3 | 51.1336789 | -0.2952335 | 0.44746174 | -0.6597962 | 0.50938465 | 0.99989001 |
| ZFYVE1 | 91.6577262 | 0.1315007 | 0.33628984 | 0.39103382 | 0.69577224 | 0.99989001 |
| RBM25 | 346.054529 | -0.1350982 | 0.25608078 | -0.5275609 | 0.59780416 | 0.99989001 |
| PSEN1 | 229.649867 | 0.23041124 | 0.35603596 | 0.64715721 | 0.5175302 | 0.99989001 |
| NUMB | 948.688635 | 0.23375295 | 0.17414975 | 1.34225259 | 0.17951411 | 0.99989001 |
| DNAL1 | 50.956484 | -0.1644564 | 0.31548693 | -0.5212781 | 0.60217308 | 0.99989001 |
| ELMSAN1 | 266.782908 | -0.0883571 | 0.34844489 | -0.2535755 | 0.79982353 | 0.99989001 |
| AC005480.1 | 50.1854722 | 0.27760239 | 0.43592058 | 0.63681872 | 0.52424293 | 0.99989001 |
| ENTPD5 | 54.9584389 | 0.33660149 | 0.35208781 | 0.95601575 | 0.33906428 | 0.99989001 |
| ALDH6A1 | 54.9667329 | 0.63290273 | 0.41421476 | 1.52795793 | 0.12652298 | 0.99989001 |
| LIN52 | 138.241443 | -0.0657426 | 0.31757932 | -0.2070117 | 0.83600071 | 0.99989001 |
| ABCD4 | 201.064801 | 0.00602301 | 0.26343588 | 0.02286329 | 0.98175932 | 0.99989001 |
| NPC2 | 140.760942 | -0.0110875 | 0.4405518 | -0.0251674 | 0.97992147 | 0.99989001 |
| AREL1 | 97.3900857 | 0.27246292 | 0.24216875 | 1.1250953 | 0.26054865 | 0.99989001 |
| FCF1 | 51.1135936 | 0.06505201 | 0.25787908 | 0.25225779 | 0.80084181 | 0.99989001 |
| YLPM1 | 299.954942 | 0.11115064 | 0.16227773 | 0.68494078 | 0.49338129 | 0.99989001 |
| DLST | 103.86005 | 0.14732376 | 0.30063374 | 0.490044 | 0.62410276 | 0.99989001 |
| MLH3 | 123.858482 | -0.3277933 | 0.27872479 | -1.1760464 | 0.23957634 | 0.99989001 |
| ACYP1 | 63.0050182 | -1.0333535 | 0.45414638 | -2.2753753 | 0.02288343 | 0.60652573 |
| NEK9 | 159.878568 | -0.1683165 | 0.18980008 | -0.8868096 | 0.37518144 | 0.99989001 |
| TMED10 | 219.817376 | -0.0423751 | 0.36044805 | -0.1175622 | 0.90641458 | 0.99989001 |
| FOS | 76.0469642 | -1.6286082 | 1.04237483 | -1.5624017 | 0.11819338 | 0.99989001 |
| JDP2 | 69.4614367 | 0.78569964 | 0.42117156 | 1.86550974 | 0.06210999 | 0.85940342 |
| TTLL5 | 337.043482 | -0.2940399 | 0.21896975 | -1.3428335 | 0.1793259 | 0.99989001 |
| IFT43 | 269.525364 | 0.22260593 | 0.29275896 | 0.76037272 | 4.47E-01 | 0.99989001 |
| GPATCH2L | 302.277917 | 0.30980688 | 0.27724762 | 1.1174375 | 0.26380731 | 0.99989001 |
| AC008050.1 | 93.9767144 | -0.6363261 | 0.57555989 | -1.1055776 | 0.2689094 | 0.99989001 |
| AC007376.2 | 74.6596532 | -0.3080135 | 0.39505841 | -0.7796656 | 0.43558773 | 0.99989001 |
| VASH1 | 118.418637 | -0.3074247 | 0.46725161 | -0.6579426 | 0.51057501 | 0.99989001 |
| ANGEL1 | 126.718925 | -0.717303 | 0.28854361 | -2.485943 | 0.01292087 | 0.49505934 |
| IRF2BPL | 112.798002 | 0.14879403 | 0.57176362 | 0.26023697 | 0.79468099 | 0.99989001 |
| TMED8 | 77.2104732 | -0.1575933 | 0.27490522 | -0.5732643 | 0.56646578 | 0.99989001 |
| VIPAS39 | 43.6384161 | 0.25615538 | 0.3171575 | 0.80765984 | 0.41928643 | NA |
| AHSA1 | 66.3382249 | 0.28789522 | 0.29073696 | 0.99022576 | 0.32206379 | 0.99989001 |
| SPTLC2 | 657.659305 | 0.60872016 | 0.42546881 | 1.43070454 | 0.15251491 | 0.99989001 |
| ALKBH1 | 108.730445 | 0.18397241 | 0.43961275 | 0.41848744 | 0.67559077 | 0.99989001 |
| SLIRP | 106.491764 | 0.10708567 | 0.21952318 | 0.4878103 | 0.62568421 | 0.99989001 |
| SNW1 | 128.329343 | -0.0315453 | 0.23959473 | -0.131661 | 0.89525243 | 0.99989001 |
| CEP128 | 114.395106 | -0.8832483 | 0.31447467 | -2.8086469 | 0.00497502 | 0.34893315 |
| GTF2A1 | 130.454566 | 0.3470581 | 0.20851104 | 1.66445908 | 0.09602072 | 0.99989001 |
| STON2 | 212.565178 | 0.23780342 | 0.3913939 | 0.60758081 | 0.54346553 | 0.99989001 |
| SEL1L | 226.618692 | 0.09978588 | 0.22970858 | 0.43440207 | 0.66399649 | 0.99989001 |
| FLRT2 | 465.151183 | -0.2556803 | 0.44873492 | -0.5697803 | 0.56882673 | 0.99989001 |
| LINC02328 | 165.883565 | 0.46865266 | 0.52720944 | 0.88893072 | 0.37404032 | 0.99989001 |

| | | | | | | |
|---|---|---|---|---|---|---|
| GALC | 57.0933516 | -0.0949517 | 0.29880384 | -0.3177727 | 0.75065738 | 0.99989001 |
| SPATA7 | 75.6653386 | -0.2084356 | 0.28462857 | -0.7323074 | 0.46398096 | 0.99989001 |
| PTPN21 | 101.070108 | -0.0627133 | 0.27836622 | -0.2252907 | 0.82175316 | 0.99989001 |
| ZC3H14 | 181.42702 | -0.1000356 | 0.16747331 | -0.5973227 | 0.55029196 | 0.99989001 |
| FOXN3 | 896.020746 | 0.57255661 | 0.24178904 | 2.36800067 | 0.0178845 | 0.56443426 |
| EFCAB11 | 132.689476 | 0.04367483 | 0.30461289 | 0.14337815 | 0.88599154 | 0.99989001 |
| PSMC1 | 102.86357 | -0.246066 | 0.33710724 | -0.729934 | 0.46543054 | 0.99989001 |
| NRDE2 | 95.9238714 | 0.36698839 | 0.36460757 | 1.00652983 | 0.31416077 | 0.99989001 |
| CALM1 | 595.745451 | 0.12471665 | 0.36231301 | 0.34422349 | 0.73067822 | 0.99989001 |
| TTC7B | 271.081945 | 0.15193692 | 0.32940943 | 0.46124033 | 0.64462619 | 0.99989001 |
| RPS6KA5 | 250.36315 | -0.2633263 | 0.27349236 | -0.9628287 | 0.3356335 | 0.99989001 |
| DGLUCY | 282.186566 | -0.3192723 | 0.45873863 | -0.6959787 | 0.48644219 | 0.99989001 |
| CCDC88C | 172.388361 | -0.5863137 | 0.48371274 | -1.2121114 | 0.22546973 | 0.99989001 |
| PPP4R3A | 306.669645 | -0.1028484 | 0.17280764 | -0.5951613 | 0.55173566 | 0.99989001 |
| TC2N | 75.8475879 | -0.195774 | 0.39456835 | -0.4961727 | 0.6197726 | 0.99989001 |
| FBLN5 | 238.627208 | -0.7876984 | 0.9667908 | -0.8147558 | 0.41521209 | 0.99989001 |
| TRIP11 | 311.247171 | 0.2478175 | 0.17771465 | 1.39446855 | 0.16317616 | 0.99989001 |
| ATXN3 | 356.204384 | 0.18559803 | 0.22347755 | 0.83049967 | 0.40625633 | 0.99989001 |
| NDUFB1 | 71.3636443 | 0.03625575 | 0.3273779 | 0.11074586 | 0.91181788 | 0.99989001 |
| CPSF2 | 143.629982 | 0.0740467 | 0.24868459 | 0.29775348 | 0.76589133 | 0.99989001 |
| RIN3 | 404.075572 | 0.08247952 | 0.32825737 | 0.25126479 | 0.8016094 | 0.99989001 |
| LGMN | 102.693766 | 0.39401313 | 0.40710053 | 0.96785218 | 0.3331182 | 0.99989001 |
| GOLGA5 | 78.7224483 | -0.1484543 | 0.26940347 | -0.5510481 | 0.58160069 | 0.99989001 |
| ITPK1 | 119.909817 | -0.2741544 | 0.31071894 | -0.8823228 | 0.37760228 | 0.99989001 |
| BTBD7 | 542.551446 | 0.3752924 | 0.1802485 | 2.08208333 | 0.03733486 | 0.70627448 |
| DDX24 | 241.639554 | -0.0324666 | 0.24685709 | -0.1315198 | 0.89536409 | 0.99989001 |
| IFI27 | 1397.00415 | -0.4597608 | 0.57194243 | -0.8038585 | 0.42147869 | 0.99989001 |
| IFI27L2 | 65.0999771 | -0.4585863 | 0.63633853 | -0.720664 | 0.47111627 | 0.99989001 |
| SERPINA1 | 95.3381615 | -0.2160518 | 0.95598859 | -0.2259983 | 0.82120271 | 0.99989001 |
| DICER1 | 455.251207 | -0.0051858 | 0.21474484 | -0.0241484 | 0.98073421 | 0.99989001 |
| CLMN | 62.6756395 | -1.2850453 | 0.55137776 | -2.3306077 | 0.01977405 | 0.57850605 |
| SYNE3 | 47.6050112 | -0.1346631 | 0.39391323 | -0.3418599 | 0.73245636 | NA |
| GLRX5 | 60.9109272 | -0.0864641 | 0.50959618 | -0.1696718 | 0.86526825 | 0.99989001 |
| ATG2B | 204.249223 | -0.035662 | 0.24363602 | -0.1463739 | 0.88362622 | 0.99989001 |
| PAPOLA | 423.910855 | 0.07552652 | 0.22250198 | 0.339442 | 0.73427678 | 0.99989001 |
| VRK1 | 66.9828342 | -0.1560662 | 0.32398011 | -0.4817154 | 0.63000815 | 0.99989001 |
| SETD3 | 248.622258 | 0.2235864 | 0.23294203 | 0.9598371 | 0.33713721 | 0.99989001 |
| CCNK | 88.8490061 | 0.03273907 | 0.32453109 | 0.10088115 | 0.91964481 | 0.99989001 |
| CCDC85C | 63.3184663 | -0.1012093 | 0.4789223 | -0.2113272 | 8.33E-01 | 1.00E+00 |
| EML1 | 619.965077 | 1.05543682 | 0.48247375 | 2.18755284 | 0.02870219 | 0.65805899 |
| EVL | 220.022234 | -0.0755038 | 0.32138323 | -0.2349339 | 0.81426005 | 0.99989001 |
| YY1 | 310.812739 | 0.11112948 | 0.20433792 | 0.54385149 | 0.58654367 | 0.99989001 |
| SLC25A29 | 108.577132 | -0.0816235 | 0.42790422 | -0.1907519 | 0.84871999 | 0.99989001 |
| WARS | 673.598339 | -0.3037101 | 0.45506817 | -0.6673947 | 0.50452005 | 0.99989001 |
| AL157871.4 | 115.345118 | -0.2086756 | 0.660054 | -0.3161493 | 0.7518892 | 0.99989001 |
| WDR25 | 90.3582833 | -0.1370256 | 0.3725586 | -0.3677961 | 0.71302525 | 0.99989001 |
| MEG3 | 51.385686 | 0.97668961 | 0.71524936 | 1.36552321 | 0.17208866 | 0.99989001 |
| MEG8 | 79.1822934 | 0.71507517 | 0.72032846 | 0.99270709 | 0.32085272 | 0.99989001 |
| PPP2R5C | 436.938131 | 0.13882304 | 0.30539682 | 0.45456611 | 0.64942142 | 0.99989001 |
| DYNC1H1 | 423.470011 | 0.08457834 | 0.28487622 | 0.29689507 | 0.76654662 | 0.99989001 |
| HSP90AA1 | 842.785408 | 0.19814289 | 0.49790787 | 0.39795091 | 0.69066637 | 0.99989001 |
| WDR20 | 247.433724 | 0.33169962 | 0.27794891 | 1.19338339 | 0.23271926 | 0.99989001 |
| CINP | 52.0164975 | 0.03193729 | 0.2542428 | 0.12561727 | 0.90003489 | 0.99989001 |
| TECPR2 | 160.067995 | -0.0285597 | 0.27404876 | -0.1042139 | 0.91699961 | 0.99989001 |

| | | | | | | |
|---|---|---|---|---|---|---|
| RCOR1 | 244.778133 | 0.00572293 | 0.25004069 | 0.02288799 | 0.98173962 | 0.99989001 |
| TRAF3 | 167.289409 | 0.4727916 | 0.62598198 | 0.75527989 | 0.45008106 | 0.99989001 |
| CDC42BPB | 553.945059 | -0.0458884 | 0.25114291 | -0.1827181 | 0.8550192 | 0.99989001 |
| TNFAIP2 | 101.167901 | 0.96664511 | 0.47231217 | 2.04662335 | 0.04069508 | 0.73661023 |
| EIF5 | 188.854181 | 0.01344093 | 0.41388405 | 0.03247511 | 0.97409316 | 0.99989001 |
| MARK3 | 718.652721 | 0.01777782 | 0.26137987 | 0.06801526 | 0.94577349 | 0.99989001 |
| CKB | 47.3504445 | 1.45955304 | 0.36346723 | 4.01563859 | 5.93E-05 | NA |
| BAG5 | 72.9920853 | 0.09512655 | 0.30447199 | 0.3124312 | 0.75471285 | 0.99989001 |
| KLC1 | 293.795599 | -0.0058482 | 0.21835563 | -0.0267831 | 0.97863273 | 0.99989001 |
| ZFYVE21 | 72.1987507 | -0.2440159 | 0.39223918 | -0.6221101 | 0.5338695 | 0.99989001 |
| PPP1R13B | 884.623413 | 0.04395394 | 0.30767787 | 0.14285702 | 0.88640311 | 0.99989001 |
| ATP5MPL | 87.5426684 | 0.15398865 | 0.39095434 | 0.39387886 | 0.69367048 | 0.99989001 |
| INF2 | 131.547053 | -0.031514 | 0.22222743 | -0.1418095 | 0.88723051 | 0.99989001 |
| SIVA1 | 90.4160905 | 0.20565748 | 0.37092468 | 0.5544454 | 0.57927407 | 0.99989001 |
| AKT1 | 83.7294252 | -0.0499271 | 0.27279587 | -0.1830198 | 0.85478245 | 0.99989001 |
| JAG2 | 142.95988 | -0.6320867 | 0.68792034 | -0.918837 | 0.35818084 | 0.99989001 |
| NUDT14 | 51.7175712 | -0.5080547 | 0.50700066 | -1.0020789 | 0.31630549 | 0.99989001 |
| BRF1 | 91.9189137 | 0.1914309 | 0.31577598 | 0.60622377 | 0.54436617 | 0.99989001 |
| PACS2 | 227.108393 | -0.193032 | 0.28110499 | -0.6866901 | 0.49227803 | 0.99989001 |
| MTA1 | 80.2707091 | -0.2771779 | 0.30462086 | -0.9099111 | 0.36286937 | 0.99989001 |
| CRIP2 | 446.765554 | -0.3697285 | 0.5715864 | -0.6468463 | 0.51773143 | 0.99989001 |
| NIPA1 | 59.8487734 | -0.1058327 | 0.33643053 | -0.3145753 | 0.75308416 | 0.99989001 |
| NIPA2 | 96.4030618 | 0.14627175 | 0.25318373 | 0.57772967 | 0.56344665 | 0.99989001 |
| CYFIP1 | 417.305429 | -0.1421641 | 0.27284309 | -0.5210471 | 0.60233397 | 0.99989001 |
| TUBGCP5 | 73.5996349 | -0.0698951 | 0.38829537 | -0.1800049 | 0.85714873 | 0.99989001 |
| AC124312.1 | 221.779922 | 0.29078541 | 0.21373358 | 1.36050408 | 0.17367046 | 0.99989001 |
| UBE3A | 378.270876 | 0.00860236 | 0.19048412 | 0.04516051 | 0.96397937 | 0.99989001 |
| HERC2 | 520.759004 | -0.2149804 | 0.17119924 | -1.2557321 | 0.20921312 | 0.99989001 |
| FAM189A1 | 75.4222786 | 1.09137869 | 0.38224835 | 2.85515608 | 0.00430157 | 0.32568991 |
| TJP1 | 1134.7461 | 0.00177138 | 0.32080428 | 0.00552168 | 0.99559436 | 0.99989001 |
| FAN1 | 65.9633036 | -0.2661202 | 0.27938184 | -0.9525321 | 0.34082719 | 0.99989001 |
| MTMR10 | 213.374456 | 0.02082204 | 0.30709296 | 0.06780369 | 0.9459419 | 0.99989001 |
| KLF13 | 126.356342 | 0.07593341 | 0.34421523 | 0.22059863 | 0.82540497 | 0.99989001 |
| OTUD7A | 116.895772 | 0.17211588 | 0.38627435 | 0.44557935 | 0.65590113 | 0.99989001 |
| LINC02256 | 188.338914 | -0.0169024 | 0.29716298 | -0.0568794 | 0.9546413 | 0.99989001 |
| AC123768.3 | 42.0123068 | -0.757134 | 0.50591982 | -1.4965493 | 0.13451057 | NA |
| RYR3 | 236.05233 | 1.71450948 | 1.08134227 | 1.58553821 | 0.1128441 | 0.99989001 |
| AVEN | 166.914736 | 0.0867789 | 0.41080019 | 0.21124358 | 0.83269721 | 0.99989001 |
| EMC7 | 84.427017 | -0.2872009 | 0.31339681 | -0.916413 | 0.35945035 | 0.99989001 |
| KATNBL1 | 276.030899 | -0.1239628 | 0.19452988 | -0.6372429 | 0.52396662 | 0.99989001 |
| EMC4 | 49.9567436 | -0.422339 | 0.40474178 | -1.0434776 | 0.29672714 | 0.99989001 |
| SLC12A6 | 196.511666 | 0.32387388 | 0.24770317 | 1.30750799 | 0.19104024 | 0.99989001 |
| NOP10 | 145.212003 | -0.303662 | 0.5748951 | -0.5282041 | 0.59735768 | 0.99989001 |
| GOLGA8A | 109.769238 | -0.1223995 | 0.29039508 | -0.421493 | 0.6733951 | 0.99989001 |
| GOLGA8B | 49.4807152 | -0.2939704 | 0.37969608 | -0.7742254 | 0.4387975 | NA |
| AQR | 225.704912 | -0.0394944 | 0.17008415 | -0.2322051 | 0.81637869 | 0.99989001 |
| ZNF770 | 81.3065546 | -0.1253311 | 0.28129849 | -0.4455449 | 0.655926 | 0.99989001 |
| AC021231.1 | 95.9264746 | 0.77777642 | 0.35117503 | 2.21478279 | 0.02677498 | 0.64983128 |
| DPH6 | 349.743987 | 0.16221874 | 0.3252315 | 0.49877931 | 0.61793487 | 0.99989001 |
| DPH6-DT | 366.731806 | -0.5122707 | 0.95339685 | -0.5373111 | 0.59105276 | 0.99989001 |
| C15orf41 | 370.244431 | -0.1849376 | 0.22393018 | -0.8258717 | 0.40887685 | 0.99989001 |
| MEIS2 | 6554.28123 | -0.0337838 | 0.21764838 | -0.1552217 | 8.77E-01 | 0.99989001 |
| SPRED1 | 408.651879 | 0.2976777 | 0.33498468 | 0.88863081 | 0.37420153 | 0.99989001 |
| FAM98B | 58.4104673 | -0.261437 | 0.2502908 | -1.0445331 | 0.2962388 | 0.99989001 |

| | | | | | | |
|---|---|---|---|---|---|---|
| THBS1 | 136.735181 | -0.5463166 | 0.5921154 | -0.9226522 | 0.3561885 | 0.99989001 |
| EIF2AK4 | 169.788398 | -0.3034408 | 0.26137718 | -1.1609307 | 0.2456701 | 0.99989001 |
| SRP14 | 1020.35443 | -0.3262642 | 0.73312491 | -0.4450323 | 0.65629644 | 0.99989001 |
| IVD | 92.1490275 | 0.09958463 | 0.32132881 | 0.30991503 | 0.75662558 | 0.99989001 |
| CCDC32 | 96.2144993 | -0.1469641 | 0.45815166 | -0.3207761 | 0.7483801 | 0.99989001 |
| RMDN3 | 59.9691164 | -0.0519143 | 0.32487498 | -0.1597977 | 0.87304043 | 0.99989001 |
| DNAJC17 | 77.0009932 | -0.2538918 | 0.20514665 | -1.237611 | 0.21586032 | 0.99989001 |
| DLL4 | 260.359999 | -0.571265 | 0.32151866 | -1.7767708 | 0.07560596 | 0.9206525 |
| INO80 | 289.514038 | -0.1084294 | 0.29380025 | -0.3690582 | 0.71208434 | 0.99989001 |
| CHP1 | 115.226895 | 0.16606874 | 0.21570839 | 0.76987612 | 0.44137338 | 0.99989001 |
| RTF1 | 164.780975 | 0.13546766 | 0.34410831 | 0.39367739 | 0.69381924 | 0.99989001 |
| MGA | 211.934438 | 0.05031063 | 0.25623119 | 0.19634859 | 0.84433733 | 0.99989001 |
| MAPKBP1 | 132.689513 | 0.21251684 | 0.21615622 | 0.98316318 | 0.32552713 | 0.99989001 |
| EHD4 | 653.373675 | -0.0487231 | 0.3448194 | -0.1413004 | 0.88763266 | 0.99989001 |
| VPS39 | 144.823029 | -0.0931557 | 0.21395557 | -0.4353975 | 0.66327394 | 0.99989001 |
| TMEM87A | 268.074547 | -0.068146 | 0.28197598 | -0.2416729 | 0.80903362 | 0.99989001 |
| GANC | 117.426149 | 0.1290143 | 0.39907081 | 0.32328674 | 0.7464781 | 0.99989001 |
| ZNF106 | 204.448262 | 0.17061537 | 0.21342302 | 0.79942347 | 0.42404491 | 0.99989001 |
| SNAP23 | 268.363109 | 0.04653043 | 0.34279596 | 0.13573798 | 0.89202842 | 0.99989001 |
| LRRC57 | 48.1688583 | -0.1299986 | 0.27751977 | -0.46843 | 0.63947709 | NA |
| HAUS2 | 48.9302021 | 0.14114229 | 0.25999823 | 0.54285865 | 0.58722713 | NA |
| STARD9 | 311.382218 | -0.0669795 | 0.38308091 | -0.1748444 | 0.86120192 | 0.99989001 |
| TTBK2 | 294.123017 | 0.05988029 | 0.20724061 | 0.28894093 | 0.77262658 | 0.99989001 |
| UBR1 | 565.828635 | 0.00160823 | 0.25518948 | 0.00630209 | 0.9949717 | 0.99989001 |
| AC068724.3 | 63.8347429 | -0.0658522 | 0.49496186 | -0.133045 | 0.89415776 | 0.99989001 |
| TMEM62 | 93.3348031 | -0.37062 | 0.50891074 | -0.7282613 | 0.46645363 | 0.99989001 |
| TP53BP1 | 150.69778 | 0.30787094 | 0.20312761 | 1.51565279 | 0.12960717 | 0.99989001 |
| PPIP5K1 | 62.2095541 | -0.0355218 | 0.5161898 | -0.0688155 | 0.94513651 | 0.99989001 |
| CATSPER2 | 73.0038661 | -0.3350872 | 0.59572421 | -0.562487 | 0.57378423 | 0.99989001 |
| PDIA3 | 346.742987 | -0.0256917 | 0.38585832 | -0.0665834 | 0.94691339 | 0.99989001 |
| SERF2 | 840.622404 | -0.227487 | 0.61783028 | -0.3682031 | 0.7127218 | 0.99989001 |
| MFAP1 | 52.1711373 | 0.19650502 | 0.25730157 | 0.7637148 | 0.44503722 | 0.99989001 |
| CASC4 | 292.87081 | 0.06395165 | 0.18619662 | 0.34346301 | 0.73125016 | 0.99989001 |
| CTDSPL2 | 299.496105 | 0.00394726 | 0.19211555 | 0.0205463 | 0.98360758 | 0.99989001 |
| EIF3J-DT | 64.4418307 | 0.46366738 | 0.3545191 | 1.30787698 | 0.19091504 | 0.99989001 |
| EIF3J | 162.894774 | 0.14844192 | 0.26543808 | 0.55923369 | 0.57600224 | 0.99989001 |
| SPG11 | 367.119898 | 5.69E-06 | 0.28862858 | 1.97E-05 | 0.99998426 | 0.99998426 |
| B2M | 5671.49537 | 0.15721633 | 0.57053652 | 0.27555876 | 0.78288699 | 0.99989001 |
| GATM | 49.3721595 | 2.19489269 | 0.64302436 | 3.41338962 | 0.0006416 | NA |
| SLC30A4 | 48.0215553 | 0.19205953 | 0.31887305 | 0.60230719 | 0.54696968 | NA |
| SQOR | 68.1794498 | 0.32423122 | 0.30928536 | 1.04832387 | 0.29448942 | 0.99989001 |
| SEMA6D | 83.2242671 | 0.49707736 | 0.35906515 | 1.38436538 | 0.1662466 | 0.99989001 |
| MYEF2 | 64.6948359 | -1.4431815 | 0.61588308 | -2.3432718 | 0.01911545 | 0.56665312 |
| SLC12A1 | 137.049839 | -0.0527791 | 0.64388686 | -0.0819696 | 0.93467092 | 0.99989001 |
| DUT | 104.306521 | 0.09946974 | 0.3342537 | 0.29758754 | 0.76601799 | 0.99989001 |
| FBN1 | 475.03788 | 0.07980289 | 0.36646889 | 0.21776172 | 0.82761478 | 0.99989001 |
| CEP152 | 67.4905981 | -0.5137033 | 0.27638716 | -1.8586365 | 0.06307868 | 0.85940342 |
| EID1 | 261.06319 | -0.0750552 | 0.57396743 | -0.1307657 | 0.89596066 | 0.99989001 |
| SECISBP2L | 233.950995 | 0.23124588 | 0.19273286 | 1.19982594 | 0.23020695 | 0.99989001 |
| COPS2 | 113.656262 | 0.04391098 | 0.24221959 | 0.18128584 | 0.85614322 | 0.99989001 |
| GALK2 | 274.818138 | 0.48679876 | 0.28860831 | 1.68671084 | 0.09165897 | 0.99175102 |
| FAM227B | 62.3169282 | 0.77843372 | 0.45316089 | 1.71778664 | 0.08583554 | 0.95514629 |
| DTWD1 | 109.806236 | 0.09678553 | 0.29054062 | 0.33312219 | 0.73904205 | 0.99989001 |
| GABPB1 | 171.278242 | 0.05438739 | 0.1831661 | 0.29692932 | 0.76652047 | 0.99989001 |

| | | | | | | |
|---|---|---|---|---|---|---|
| GABPB1-AS1 | 167.901536 | 0.23839217 | 0.41194021 | 0.57870575 | 0.56278774 | 0.99989001 |
| USP8 | 231.499806 | -0.0294683 | 0.18422864 | -0.1599549 | 0.87291659 | 0.99989001 |
| TRPM7 | 415.449774 | -0.1354495 | 0.20492148 | -0.6609826 | 0.50862349 | 0.99989001 |
| SPPL2A | 315.438879 | 0.15986552 | 0.20095198 | 0.79554089 | 0.42629893 | 0.99989001 |
| AP4E1 | 107.706479 | -0.1423069 | 0.20430701 | -0.6965347 | 0.48609404 | 0.99989001 |
| DMXL2 | 124.924486 | 0.26312311 | 0.31384627 | 0.83838215 | 0.40181611 | 0.99989001 |
| LYSMD2 | 50.3924468 | 0.37389483 | 0.38996455 | 0.95879184 | 0.33766362 | 0.99989001 |
| TMOD2 | 70.675021 | 1.28979424 | 0.37813471 | 3.41093847 | 0.0006474 | 0.11634649 |
| TMOD3 | 533.028115 | 0.42609882 | 0.16952546 | 2.51347976 | 0.01195466 | 0.49000287 |
| LEO1 | 41.857276 | 0.22654381 | 0.27169801 | 0.8338074 | 0.40438953 | NA |
| MAPK6 | 174.889615 | -0.1008531 | 0.25507039 | -0.3953934 | 0.69255257 | 0.99989001 |
| MYO5C | 175.175072 | -1.8102992 | 0.69887044 | -2.5903217 | 0.00958863 | 0.44953399 |
| MYO5A | 121.04468 | -0.7028291 | 0.49546174 | -1.4185335 | 0.15603505 | 0.99989001 |
| ARPP19 | 189.033215 | -0.1438502 | 0.21251564 | -0.6768925 | 0.49847416 | 0.99989001 |
| FAM214A | 335.569222 | 0.26048712 | 0.2473396 | 1.05315576 | 0.29226961 | 0.99989001 |
| WDR72 | 63.2656665 | 1.05099596 | 0.62269063 | 1.68783006 | 0.09144386 | 0.99169284 |
| RSL24D1 | 69.9326544 | 0.13320847 | 0.33296994 | 0.40006156 | 0.68911118 | 0.99989001 |
| RAB27A | 186.184284 | 0.2363779 | 0.19288926 | 1.22545906 | 0.22040231 | 0.99989001 |
| PIGB | 51.5208798 | -0.0619336 | 0.40457806 | -0.153082 | 0.87833363 | 0.99989001 |
| CCPG1 | 241.201636 | 0.35697194 | 0.28944405 | 1.23330203 | 0.2174631 | 0.99989001 |
| PRTG | 139.933327 | 0.20399399 | 0.42322688 | 0.48199678 | 0.62980823 | 0.99989001 |
| NEDD4 | 702.949769 | -0.0353304 | 0.30792207 | -0.114738 | 0.90865276 | 0.99989001 |
| RFX7 | 364.674693 | 0.00480455 | 0.16689994 | 0.028787 | 0.97703447 | 0.99989001 |
| TEX9 | 50.173493 | -1.096927 | 0.52647792 | -2.0835195 | 0.0372039 | 0.70556534 |
| ZNF280D | 229.044824 | -0.0304619 | 0.31638152 | -0.0962822 | 0.92329642 | 0.99989001 |
| AC090517.4 | 210.378085 | 0.25573912 | 0.3384959 | 0.75551616 | 0.44993934 | 0.99989001 |
| TCF12 | 2026.66556 | 0.08442424 | 0.21137211 | 0.39941049 | 0.68959077 | 0.99989001 |
| CGNL1 | 341.459524 | -0.9246484 | 0.42486682 | -2.1763253 | 0.02953095 | 0.6652708 |
| POLR2M | 41.2809911 | -0.4188369 | 0.43110186 | -0.9715497 | 0.33127462 | NA |
| ADAM10 | 795.622115 | -0.2834835 | 0.20225709 | -1.4015996 | 0.16103484 | 0.99989001 |
| MINDY2 | 242.892317 | -0.2323361 | 0.38421763 | -0.6046992 | 0.54537888 | 0.99989001 |
| RNF111 | 372.643389 | -0.0023809 | 0.19979485 | -0.0119167 | 0.99049205 | 0.99989001 |
| SLTM | 390.107962 | -0.0623419 | 0.21792808 | -0.2860664 | 0.77482724 | 0.99989001 |
| MYO1E | 1156.93752 | -0.0367441 | 0.35137171 | -0.1045733 | 0.91671436 | 0.99989001 |
| GTF2A2 | 102.572274 | -0.1772953 | 0.34400013 | -0.5153932 | 0.60627827 | 0.99989001 |
| BNIP2 | 251.402976 | -0.0581033 | 0.25636363 | -0.226644 | 0.82070061 | 0.99989001 |
| ANXA2 | 567.5164 | -0.2732498 | 0.35243729 | -0.7753147 | 0.4381537 | 0.99989001 |
| ICE2 | 167.239893 | 0.07004483 | 0.21378948 | 0.3276346 | 0.74318795 | 0.99989001 |
| RORA-AS1 | 88.5208405 | -0.4467381 | 0.54567709 | -0.8186858 | 0.41296568 | 0.99989001 |
| RORA | 1355.25856 | 0.01850869 | 0.4519985 | 0.04094857 | 0.9673369 | 0.99989001 |
| AC009554.1 | 102.381637 | -0.2244134 | 0.58692467 | -0.3823547 | 0.70219829 | 0.99989001 |
| VPS13C | 749.910061 | -0.3646964 | 0.22889086 | -1.59332 | 0.11108842 | 0.99989001 |
| TLN2 | 95.4736699 | -0.0841849 | 0.5037556 | -0.1671146 | 0.86727988 | 0.99989001 |
| TPM1 | 197.023578 | -0.1447253 | 0.33645812 | -0.4301435 | 0.66709128 | 0.99989001 |
| LACTB | 73.0987264 | 0.03413263 | 0.36666306 | 0.09308991 | 0.92583214 | 0.99989001 |
| RPS27L | 146.994991 | -0.0146364 | 0.37532128 | -0.038997 | 0.96889275 | 0.99989001 |
| RAB8B | 427.231107 | 0.37708279 | 0.39686179 | 0.9501615 | 0.3420302 | 0.99989001 |
| APH1B | 62.2977772 | -0.460846 | 0.49582588 | -0.9294512 | 0.35265531 | 0.99989001 |
| CA12 | 88.5233502 | 1.27683536 | 0.63867288 | 1.99920083 | 0.04558663 | 0.77228819 |
| USP3 | 372.588512 | -0.0884461 | 0.1914165 | -0.4620608 | 0.64403771 | 0.99989001 |
| USP3-AS1 | 67.4425274 | 0.16979057 | 0.50672931 | 0.33507154 | 0.73757112 | 0.99989001 |
| HERC1 | 1066.58738 | 0.07530964 | 0.35215521 | 0.21385353 | 0.83066128 | 0.99989001 |
| DAPK2 | 97.2523881 | -0.6601375 | 0.47164112 | -1.3996607 | 0.16161496 | 0.99989001 |
| FAM96A | 44.0958198 | 0.03757046 | 0.26871891 | 0.13981323 | 0.88880756 | NA |

| | | | | | | |
|---|---|---|---|---|---|---|
| SNX1 | 211.570126 | 0.3262053 | 0.16521698 | 1.97440545 | 0.04833566 | 0.78123673 |
| PPIB | 258.957925 | -0.0633028 | 0.52478294 | -0.1206267 | 0.9039867 | 0.99989001 |
| CSNK1G1 | 46.5694188 | 0.17263984 | 0.27871185 | 0.61942051 | 0.53563937 | NA |
| TRIP4 | 126.315491 | 0.04676629 | 0.20305789 | 0.23031015 | 0.81785078 | 0.99989001 |
| ZNF609 | 810.356927 | -0.0952685 | 0.18265152 | -0.5215859 | 0.60195865 | 0.99989001 |
| OAZ2 | 127.659831 | 0.00762196 | 0.42223535 | 0.01805145 | 0.98559781 | 0.99989001 |
| AC069368.1 | 84.7341718 | -0.4141858 | 0.53483391 | -0.7744194 | 0.43868282 | 0.99989001 |
| ANKDD1A | 78.4381951 | -0.0237476 | 0.35996197 | -0.0659726 | 0.94739967 | 0.99989001 |
| SPG21 | 97.8265936 | -0.1435048 | 0.21173577 | -0.6777541 | 0.49792761 | 0.99989001 |
| MTFMT | 69.8319566 | 0.08247177 | 0.28999053 | 0.28439469 | 0.77610793 | 0.99989001 |
| PDCD7 | 60.0394188 | 0.00999515 | 0.30550105 | 0.03271724 | 0.97390008 | 0.99989001 |
| CLPX | 106.894067 | 0.1247788 | 0.21997087 | 0.56725146 | 0.57054335 | 0.99989001 |
| PARP16 | 43.6031787 | -0.4948757 | 0.32260358 | -1.5340055 | 0.1250283 | NA |
| DPP8 | 310.105343 | 0.01313739 | 0.15253729 | 0.08612576 | 0.93136645 | 0.99989001 |
| HACD3 | 130.603687 | -0.1822722 | 0.26814811 | -0.6797444 | 0.49666635 | 0.99989001 |
| INTS14 | 58.4868982 | -0.4267552 | 0.24477125 | -1.7434857 | 0.0812488 | 0.94203683 |
| DENND4A | 570.174284 | -0.1308885 | 0.28647315 | -0.4568963 | 0.64774556 | 0.99989001 |
| RAB11A | 446.788673 | -0.1025669 | 0.38335468 | -0.2675509 | 0.78904503 | 0.99989001 |
| DIS3L | 54.4692899 | -0.0857705 | 0.28464851 | -0.3013206 | 0.76317003 | 0.99989001 |
| MAP2K1 | 295.733118 | 0.35456361 | 0.36855624 | 0.96203393 | 0.33603256 | 0.99989001 |
| RPL4 | 309.252862 | -0.074786 | 0.51574106 | -0.1450069 | 0.88470546 | 0.99989001 |
| SMAD6 | 585.816401 | 1.94354556 | 0.49559299 | 3.92165667 | 8.79E-05 | 0.03687711 |
| SMAD3 | 440.477132 | 0.08603641 | 0.55923939 | 0.1538454 | 0.87773164 | 0.99989001 |
| AAGAB | 153.244915 | -0.1389435 | 0.29525737 | -0.4705845 | 0.63793748 | 0.99989001 |
| IQCH | 53.1073333 | -0.2513025 | 0.38911658 | -0.6458282 | 0.51839061 | 0.99989001 |
| MAP2K5 | 433.258049 | -0.104933 | 0.20653371 | -0.508067 | 0.61140639 | 0.99989001 |
| PIAS1 | 1046.47094 | -0.0348639 | 0.28654716 | -0.121669 | 0.90316115 | 0.99989001 |
| FEM1B | 87.1105929 | -0.1144873 | 0.28985459 | -0.394982 | 0.69285617 | 0.99989001 |
| ANP32A | 151.855888 | 0.32917233 | 0.2654342 | 1.24012777 | 0.21492814 | 0.99989001 |
| GLCE | 77.7976906 | 0.61292589 | 0.26815311 | 2.28573107 | 0.02227 | 0.60384164 |
| RPLP1 | 723.174537 | -0.3051566 | 0.58562756 | -0.5210763 | 0.60231364 | 0.99989001 |
| TLE3 | 100.953556 | 0.27648399 | 0.48134138 | 0.57440313 | 0.56569504 | 0.99989001 |
| UACA | 1078.58525 | -0.2215282 | 0.38022689 | -0.5826211 | 0.56014843 | 0.99989001 |
| LRRC49 | 113.102283 | 0.00059664 | 0.29449242 | 0.002026 | 0.99838349 | 0.99989001 |
| THSD4 | 1338.47669 | -1.0294137 | 0.40576775 | -2.536953 | 0.0111822 | 0.47956379 |
| THSD4-AS1 | 218.037236 | -0.010237 | 0.45555489 | -0.0224715 | 0.98207186 | 0.99989001 |
| MYO9A | 651.593403 | -0.2753732 | 0.1923067 | -1.4319479 | 0.15215872 | 0.99989001 |
| PKM | 307.428195 | -0.4198582 | 0.25759499 | -1.6299159 | 1.03E-01 | 0.99989001 |
| PARP6 | 83.8299004 | -0.4155097 | 0.32847794 | -1.2649546 | 0.2058876 | 0.99989001 |
| AC009690.1 | 139.501898 | 0.23754171 | 0.50319115 | 0.47207052 | 0.63687645 | 0.99989001 |
| ARIH1 | 758.7744 | 0.08236903 | 0.18553446 | 0.44395545 | 0.65707479 | 0.99989001 |
| BBS4 | 68.3136194 | -0.1309893 | 0.29848483 | -0.4388473 | 0.66077219 | 0.99989001 |
| ADPGK | 91.5504039 | 0.12420793 | 0.33055123 | 0.37576001 | 0.70709532 | 0.99989001 |
| NEO1 | 91.0856955 | 1.00694014 | 0.47857179 | 2.10405244 | 0.03537387 | 0.69174161 |
| NPTN | 364.154955 | 0.26323365 | 0.1823664 | 1.44343285 | 0.14889857 | 0.99989001 |
| PML | 221.792103 | 0.24288611 | 0.31134814 | 0.78011102 | 0.43532553 | 0.99989001 |
| ARID3B | 49.7108665 | -0.1529593 | 0.26016442 | -0.5879333 | 0.55657707 | NA |
| CLK3 | 53.2617766 | 0.22175479 | 0.24585128 | 0.90198753 | 0.36706349 | 0.99989001 |
| EDC3 | 67.5567795 | 0.61030825 | 0.29477714 | 2.07040562 | 0.03841438 | 0.71513134 |
| CSK | 40.7346388 | -0.3591409 | 0.37669163 | -0.9534081 | 0.34038333 | NA |
| SCAMP2 | 68.0614604 | -0.0476439 | 0.26781273 | -0.1778999 | 0.85880156 | 0.99989001 |
| COX5A | 77.804489 | 0.29613596 | 0.32344568 | 0.91556628 | 0.35989442 | 0.99989001 |
| PPCDC | 192.884361 | -0.2159798 | 0.37904889 | -0.5697941 | 0.56881734 | 0.99989001 |
| COMMD4 | 59.6546251 | -0.2716951 | 0.30635245 | -0.886871 | 0.37514837 | 0.99989001 |

| | | | | | | |
|---|---|---|---|---|---|---|
| NEIL1 | 46.4023406 | -0.5789682 | 0.32697951 | -1.770656 | 0.07661793 | NA |
| MAN2C1 | 145.629496 | -0.0950605 | 0.17680635 | -0.537653 | 0.59081662 | 0.99989001 |
| SIN3A | 168.983449 | -0.1131413 | 0.20913625 | -0.540993 | 0.58851239 | 0.99989001 |
| PTPN9 | 226.261984 | 0.40136002 | 0.18333611 | 2.18920332 | 0.02858207 | 0.65805899 |
| IMP3 | 43.8381473 | 0.10890761 | 0.39936109 | 0.27270461 | 0.78508029 | NA |
| UBE2Q2 | 209.210526 | 0.42295826 | 0.23935438 | 1.76707965 | 0.07721489 | 0.92274869 |
| FBXO22 | 125.408344 | 0.27395708 | 0.23125927 | 1.1846318 | 0.23616306 | 0.99989001 |
| ETFA | 233.068318 | 0.26463943 | 0.19070278 | 1.3877062 | 0.16522653 | 0.99989001 |
| SCAPER | 638.937664 | 0.26593942 | 0.20872474 | 1.27411546 | 0.2026225 | 0.99989001 |
| RCN2 | 104.321951 | -0.2263461 | 0.36549787 | -0.6192816 | 0.53573087 | 0.99989001 |
| TSPAN3 | 132.123766 | 0.04596968 | 0.28846202 | 0.1593613 | 0.87338423 | 0.99989001 |
| PEAK1 | 1296.50928 | 0.56433257 | 0.3844528 | 1.4678852 | 0.14213541 | 0.99989001 |
| HMG20A | 157.639559 | 0.04902002 | 0.17828821 | 0.27494819 | 0.78335604 | 0.99989001 |
| LINGO1 | 1321.5447 | -0.5344297 | 0.66222168 | -0.8070254 | 0.41965187 | 0.99989001 |
| TBC1D2B | 167.715424 | 0.25624464 | 0.30434116 | 0.8419651 | 0.39980749 | 0.99989001 |
| IDH3A | 57.6522784 | 0.00869129 | 0.36975079 | 0.02350581 | 0.9812468 | 0.99989001 |
| WDR61 | 77.626616 | 0.1699246 | 0.21965359 | 0.77360265 | 0.4391658 | 0.99989001 |
| IREB2 | 199.32878 | 0.10012809 | 0.23135527 | 0.43278932 | 0.66516783 | 0.99989001 |
| PSMA4 | 141.830813 | 0.26477231 | 0.33593226 | 0.78817174 | 0.43059625 | 0.99989001 |
| MORF4L1 | 402.165171 | 0.19282685 | 0.21224783 | 0.90849858 | 0.36361486 | 0.99989001 |
| CTSH | 76.3909183 | 0.98299214 | 0.40357426 | 2.43571567 | 0.01486236 | 0.51942334 |
| ZFAND6 | 382.768002 | 0.01893298 | 0.17804119 | 0.10634045 | 0.91531224 | 0.99989001 |
| AC027808.1 | 49.6760044 | 0.2982793 | 0.52201529 | 0.57139955 | 0.56772883 | NA |
| MESD | 125.654269 | 0.2211994 | 0.21544924 | 1.02668919 | 0.30456684 | 0.99989001 |
| TLNRD1 | 103.549018 | 0.30656567 | 0.39472569 | 0.77665499 | 0.43736235 | 0.99989001 |
| CFAP161 | 158.033125 | 0.20814218 | 0.50273149 | 0.41402257 | 0.67885758 | 0.99989001 |
| EFL1 | 92.3925299 | 0.42140789 | 0.38372008 | 1.09821691 | 0.27210978 | 0.99989001 |
| WHAMM | 92.7490967 | -0.188677 | 0.26357977 | -0.7158251 | 0.47409934 | 0.99989001 |
| C15orf40 | 52.816251 | -0.5506384 | 0.25148956 | -2.189508 | 0.02855993 | 0.65805899 |
| BTBD1 | 168.06332 | -0.1510445 | 0.18491815 | -0.816818 | 0.41403242 | 0.99989001 |
| TM6SF1 | 100.912621 | -0.4926429 | 0.67122077 | -0.7339506 | 0.46297882 | 0.99989001 |
| HDGFL3 | 340.199041 | 0.23407892 | 0.22509326 | 1.0399197 | 0.29837721 | 0.99989001 |
| WDR73 | 80.5587094 | 0.14066775 | 0.26480244 | 0.53121773 | 0.59526791 | 0.99989001 |
| SEC11A | 224.259228 | -0.0334549 | 0.2009065 | -0.1665195 | 0.86774811 | 0.99989001 |
| ZNF592 | 126.911064 | 0.43100845 | 0.24763724 | 1.74048319 | 0.08177421 | 0.9437794 |
| ALPK3 | 84.8068981 | -0.0681926 | 0.45498206 | -0.1498797 | 0.88085955 | 0.99989001 |
| PDE8A | 886.86226 | 0.34056097 | 0.32223356 | 1.05687618 | 0.2905681 | 0.99989001 |
| AKAP13 | 1400.31788 | -0.1010014 | 0.28111458 | -0.3592892 | 0.71937874 | 0.99989001 |
| AC087286.2 | 68.6285517 | -0.4317216 | 0.5071062 | -0.8513435 | 0.39457858 | 0.99989001 |
| MRPS11 | 59.3698096 | -0.1853568 | 0.30870851 | -0.6004266 | 0.54822195 | 0.99989001 |
| DET1 | 45.954803 | 0.06694775 | 0.63736225 | 0.10503878 | 0.91634503 | NA |
| ISG20 | 153.964935 | -0.0324049 | 0.77571909 | -0.041774 | 0.9666789 | 0.99989001 |
| MFGE8 | 76.6215465 | -0.279413 | 0.73247914 | -0.3814621 | 0.70286038 | 0.99989001 |
| ABHD2 | 210.940615 | -0.3933873 | 0.28071582 | -1.4013719 | 0.1611029 | 0.99989001 |
| POLG | 60.9814633 | 0.34343408 | 0.2665089 | 1.28864017 | 0.19752321 | 0.99989001 |
| ANPEP | 46.5480041 | 0.97572203 | 0.70509998 | 1.38380664 | 0.16641767 | NA |
| ZNF710 | 238.28516 | 0.42282696 | 0.52101907 | 0.81153836 | 0.41705657 | 0.99989001 |
| IDH2 | 86.462748 | 0.63739951 | 0.28604023 | 2.22835615 | 0.02585678 | 0.63690091 |
| SEMA4B | 42.2309678 | -0.2229888 | 0.4881649 | -0.45679 | 0.647822 | NA |
| CIB1 | 97.0660583 | -0.0029326 | 0.42068294 | -0.006971 | 0.99443798 | 0.99989001 |
| IQGAP1 | 625.487768 | -0.1250926 | 0.25028789 | -0.499795 | 0.61721941 | 0.99989001 |
| CRTC3 | 335.156475 | 0.02684794 | 0.18038106 | 0.14884013 | 0.88167979 | 0.99989001 |
| FURIN | 124.662947 | -0.2224303 | 0.59779204 | -0.3720863 | 0.70982857 | 0.99989001 |
| FES | 144.141757 | -0.5675666 | 0.39443571 | -1.4389331 | 0.15016949 | 0.99989001 |

| | | | | | | |
|---|---|---|---|---|---|---|
| UNC45A | 75.9524147 | -0.0146398 | 0.28775323 | -0.0508763 | 0.95942407 | 0.99989001 |
| SV2B | 137.291325 | -1.4694122 | 0.55667243 | -2.6396353 | 0.00829953 | 0.41763227 |
| SLCO3A1 | 729.218357 | 0.77584036 | 0.28776045 | 2.69613272 | 0.00701497 | 0.3951418 |
| FAM174B | 122.654193 | -0.905076 | 0.3458675 | -2.6168286 | 0.00887509 | 0.42668269 |
| AC013394.1 | 243.279377 | 0.11580131 | 0.27996001 | 0.41363518 | 0.67914131 | 0.99989001 |
| CHD2 | 329.09777 | -0.188103 | 0.25225072 | -0.7456985 | 0.45584957 | 0.99989001 |
| MCTP2 | 193.358837 | -0.7312812 | 0.64175655 | -1.1394993 | 0.25449495 | 0.99989001 |
| LINC01197 | 103.580258 | 0.22675465 | 0.38538107 | 0.58839075 | 0.55627005 | 0.99989001 |
| AC012409.2 | 75.0556226 | -0.212408 | 0.78713078 | -0.269851 | 0.78727492 | 0.99989001 |
| NR2F2-AS1 | 544.236282 | 0.03919595 | 0.51277753 | 0.07643851 | 0.93907024 | 0.99989001 |
| NR2F2 | 666.836821 | 0.027073 | 0.52234268 | 0.05182995 | 0.95866419 | 0.99989001 |
| IGF1R | 986.241489 | 0.2480584 | 0.33259651 | 0.74582381 | 0.45577385 | 0.99989001 |
| TTC23 | 176.477874 | -0.0271196 | 0.26214195 | -0.103454 | 0.91760267 | 0.99989001 |
| LRRC28 | 176.189532 | 0.19165917 | 0.21867571 | 0.87645386 | 0.38078335 | 0.99989001 |
| MEF2A | 2128.9908 | -0.0503302 | 0.38813567 | -0.1296717 | 0.89682616 | 0.99989001 |
| AC090825.1 | 137.219033 | -0.0897545 | 0.21569339 | -0.416121 | 0.67732148 | 0.99989001 |
| ASB7 | 64.1588739 | 0.0768137 | 0.22763514 | 0.3374422 | 0.73578358 | 0.99989001 |
| LRRK1 | 118.921286 | -0.1050446 | 0.26967462 | -0.3895234 | 0.69688902 | 0.99989001 |
| CHSY1 | 639.023292 | -0.2036356 | 0.37038483 | -0.5497947 | 0.58246022 | 0.99989001 |
| SELENOS | 148.440125 | 0.44647882 | 0.36542134 | 1.22181924 | 0.22177599 | 0.99989001 |
| SNRPA1 | 98.1113872 | 0.02599948 | 0.28034092 | 0.09274235 | 0.92610825 | 0.99989001 |
| PCSK6 | 149.580197 | 0.21804952 | 0.68516613 | 0.31824328 | 0.7503004 | 0.99989001 |
| TM2D3 | 114.266562 | -0.0005093 | 0.25379948 | -0.0020069 | 0.99839875 | 0.99989001 |
| TARSL2 | 84.163672 | -0.2016064 | 0.26066685 | -0.7734255 | 0.43927063 | 0.99989001 |
| MPG | 83.7364354 | -0.1258088 | 0.34836843 | -0.3611372 | 0.71799687 | 0.99989001 |
| NPRL3 | 96.1783483 | -0.1122022 | 0.35992285 | -0.3117397 | 0.75523835 | 0.99989001 |
| LUC7L | 261.470318 | 0.09679876 | 0.21662078 | 0.44685815 | 0.65497748 | 0.99989001 |
| AXIN1 | 58.2579645 | -0.1053008 | 0.30349666 | -0.3469587 | 0.72862232 | 0.99989001 |
| MRPL28 | 57.9906709 | -0.0611017 | 0.50601264 | -0.1207512 | 0.90388807 | 0.99989001 |
| TMEM8A | 61.8522868 | 0.1225103 | 0.32761734 | 0.37394326 | 0.70844653 | 0.99989001 |
| RAB11FIP3 | 214.818926 | 0.396852 | 0.28510237 | 1.39196319 | 0.16393355 | 0.99989001 |
| CAPN15 | 43.9172542 | -0.1410432 | 0.4332034 | -0.3255819 | 0.7447407 | NA |
| PIGQ | 38.9289905 | 0.0183758 | 0.33012179 | 0.0556637 | 0.95560971 | NA |
| RAB40C | 76.5907746 | 0.29447303 | 0.29009538 | 1.01509039 | 0.31006274 | 0.99989001 |
| RHOT2 | 51.6792747 | 0.06798142 | 0.3184932 | 0.21344702 | 0.83097831 | 0.99989001 |
| STUB1 | 86.0614737 | -0.0360065 | 0.53278233 | -0.067582 | 0.94611838 | 0.99989001 |
| JMJD8 | 40.3716037 | -0.1956705 | 0.44734567 | -0.4374033 | 0.66181891 | NA |
| METRN | 50.3628669 | 0.35175996 | 0.50331131 | 0.69889143 | 0.48461988 | 0.99989001 |
| LMF1 | 124.576255 | -0.1719081 | 0.36868272 | -0.4662764 | 0.64101767 | 0.99989001 |
| UBE2I | 160.416478 | 0.13046389 | 0.25971978 | 0.50232558 | 0.61543852 | 0.99989001 |
| TSR3 | 45.5849333 | 0.20604178 | 0.44583688 | 0.4621461 | 0.64397655 | NA |
| GNPTG | 50.786666 | -0.0815757 | 0.25331341 | -0.3220347 | 0.74742642 | 0.99989001 |
| UNKL | 58.5670122 | 0.39472146 | 0.36904752 | 1.0695681 | 0.28481376 | 0.99989001 |
| CLCN7 | 70.7576068 | -0.3518507 | 0.28187517 | -1.24825 | 0.21193952 | 0.99989001 |
| IFT140 | 106.084143 | 0.37313142 | 0.35589302 | 1.04843702 | 0.29443731 | 0.99989001 |
| TMEM204 | 262.383283 | 0.16677136 | 0.47992536 | 0.34749436 | 0.72821995 | 0.99989001 |
| JPT2 | 52.175621 | 0.076364 | 0.2696014 | 0.2832478 | 0.77698689 | 0.99989001 |
| MAPK8IP3 | 142.852828 | -0.0018091 | 0.21482201 | -0.0084212 | 0.99328092 | 0.99989001 |
| NME3 | 153.849841 | 0.0291751 | 0.48084345 | 0.06067484 | 0.95161817 | 0.99989001 |
| MRPS34 | 72.1966724 | -0.2570321 | 0.54114882 | -0.4749749 | 0.63480486 | 0.99989001 |
| SPSB3 | 81.810722 | 0.02567262 | 0.33187962 | 0.07735523 | 0.93834096 | 0.99989001 |
| HAGH | 93.0983963 | -0.188291 | 0.27783694 | -0.6777031 | 0.49795994 | 0.99989001 |
| NDUFB10 | 98.6633856 | -0.2090621 | 0.54137107 | -0.3861715 | 0.69936963 | 0.99989001 |
| RPS2 | 389.168688 | -0.2918324 | 0.62212817 | -0.4690872 | 0.6390073 | 0.99989001 |

| | | | | | | |
|---|---|---|---|---|---|---|
| ZNF598 | 38.0623199 | -0.2542954 | 0.44027761 | -0.5775796 | 0.56354799 | NA |
| SLC9A3R2 | 787.70678 | 0.01714702 | 0.64071064 | 0.0267625 | 0.97864916 | 0.99989001 |
| TSC2 | 134.234216 | -0.3755131 | 0.30548916 | -1.2292192 | 0.21898963 | 0.99989001 |
| PKD1 | 82.5958104 | 0.02853173 | 0.26900021 | 0.10606583 | 0.91553012 | 0.99989001 |
| TRAF7 | 52.7466519 | -0.116017 | 0.46995511 | -0.2468682 | 0.80501025 | 0.99989001 |
| PGP | 83.7213385 | 0.17665608 | 0.34106658 | 0.51795188 | 0.60449184 | 0.99989001 |
| ECI1 | 77.1361958 | 0.26701472 | 0.23541419 | 1.13423377 | 0.2566965 | 0.99989001 |
| RNPS1 | 138.076909 | 0.07923938 | 0.23486553 | 0.33738188 | 0.73582904 | 0.99989001 |
| ABCA3 | 58.7616878 | 0.2114681 | 0.32560443 | 0.6494632 | 0.51603902 | 0.99989001 |
| PDPK1 | 169.726134 | 0.10256923 | 0.27768183 | 0.36937683 | 0.71184687 | 0.99989001 |
| SRRM2 | 638.568779 | -0.0682764 | 0.1478205 | -0.4618871 | 0.64416228 | 0.99989001 |
| ELOB | 158.440401 | -0.2015848 | 0.55754662 | -0.3615569 | 0.71768319 | 0.99989001 |
| FLYWCH2 | 45.837287 | 0.18020973 | 0.43266127 | 0.41651459 | 0.67703349 | NA |
| FLYWCH1 | 80.34078 | 0.0289041 | 0.29848031 | 0.09683754 | 0.92285541 | 0.99989001 |
| HCFC1R1 | 52.2696342 | -0.6113314 | 0.48858175 | -1.2512367 | 0.21084814 | 0.99989001 |
| IL32 | 215.380197 | -0.5107894 | 0.6402033 | -0.797855 | 0.42495465 | 0.99989001 |
| AC108134.3 | 70.2325662 | 0.07342971 | 0.36033814 | 0.20378 | 0.83852542 | 0.99989001 |
| ZNF213 | 51.2949082 | 0.15035493 | 0.388648 | 0.38686659 | 0.69885498 | 0.99989001 |
| ZNF263 | 53.9020168 | 0.27110235 | 0.30193786 | 0.89787466 | 0.36925237 | 0.99989001 |
| ZNF75A | 73.8237913 | 0.49808863 | 0.4842141 | 1.02865371 | 0.30364243 | 0.99989001 |
| NAA60 | 42.1524927 | 0.18739019 | 0.36723715 | 0.51027024 | 0.60986215 | NA |
| CLUAP1 | 52.5242399 | 0.03212218 | 0.34603292 | 0.09282984 | 0.92603875 | 0.99989001 |
| DNASE1 | 114.648301 | 0.91242301 | 0.33981487 | 2.68505908 | 0.0072517 | 0.40246927 |
| TRAP1 | 98.209272 | -0.2935323 | 0.28664403 | -1.0240307 | 0.30582079 | 0.99989001 |
| CREBBP | 460.460684 | 0.10829394 | 0.23739948 | 0.45616756 | 0.64826949 | 0.99989001 |
| ADCY9 | 240.905961 | 0.68913768 | 0.3842527 | 1.79344916 | 0.07290113 | 0.9155703 |
| DNAJA3 | 63.5777189 | 0.25316598 | 0.24624597 | 1.02810201 | 0.30390185 | 0.99989001 |
| NMRAL1 | 48.8090431 | 0.17868674 | 0.32088167 | 0.55686178 | 0.57762187 | NA |
| HMOX2 | 79.5945796 | 0.41765695 | 0.54539217 | 0.76579197 | 0.44380009 | 0.99989001 |
| CDIP1 | 51.5442686 | -0.2418174 | 0.30673435 | -0.7883609 | 0.43048563 | 0.99989001 |
| MGRN1 | 110.596059 | 0.11089203 | 0.28309135 | 0.39171818 | 0.69526646 | 0.99989001 |
| ANKS3 | 59.5553673 | -0.6116052 | 0.33426038 | -1.829727 | 0.06729078 | 0.87873839 |
| GLYR1 | 212.533553 | -0.0403639 | 0.23545081 | -0.1714324 | 0.86388379 | 0.99989001 |
| UBN1 | 133.768827 | 0.01656074 | 0.36119718 | 0.0458496 | 0.96343013 | 0.99989001 |
| PMM2 | 165.836452 | -0.1121951 | 0.37092081 | -0.3024771 | 0.76228838 | 0.99989001 |
| CARHSP1 | 407.594526 | -0.3299224 | 0.34098235 | -0.9675644 | 0.33326198 | 0.99989001 |
| USP7 | 178.48552 | 0.07550741 | 0.1992413 | 0.37897469 | 0.70470666 | 0.99989001 |
| C16orf72 | 160.8639 | 0.05820596 | 0.19578613 | 0.29729356 | 0.7662424 | 0.99989001 |
| ATF7IP2 | 194.592857 | 0.09183366 | 0.39280806 | 0.23378761 | 0.81514987 | 0.99989001 |
| EMP2 | 316.443757 | 0.08326712 | 0.37907702 | 0.21965753 | 0.82613788 | 0.99989001 |
| NUBP1 | 45.0418609 | -0.2762901 | 0.28568734 | -0.9671065 | 0.33349078 | NA |
| CIITA | 274.262638 | 0.42604678 | 0.47384456 | 0.89912772 | 0.36858463 | 0.99989001 |
| DEXI | 68.5727872 | 0.22602058 | 0.31490258 | 0.71774762 | 0.47291292 | 0.99989001 |
| CLEC16A | 335.931413 | 0.06173796 | 0.26896117 | 0.22954229 | 0.81844745 | 0.99989001 |
| LITAF | 155.343468 | 0.29582601 | 0.57630179 | 0.51331788 | 6.08E-01 | 0.99989001 |
| TXNDC11 | 147.852183 | 0.14539644 | 0.21593847 | 0.67332345 | 0.50074154 | 0.99989001 |
| ZC3H7A | 246.392549 | 0.25039317 | 0.23937019 | 1.04604992 | 0.29553798 | 0.99989001 |
| RSL1D1 | 58.2852711 | -0.2004766 | 0.3842312 | -0.5217602 | 0.60183728 | 0.99989001 |
| AC007216.4 | 80.5865169 | 0.20833225 | 0.34398287 | 0.60564717 | 0.54474907 | 0.99989001 |
| GSPT1 | 221.261053 | -0.0565487 | 0.24990691 | -0.226279 | 0.82098442 | 0.99989001 |
| SNX29 | 468.520898 | -0.0776898 | 0.35542663 | -0.2185818 | 0.82697581 | 0.99989001 |
| CPPED1 | 105.31093 | -0.8319633 | 0.3216037 | -2.5869206 | 0.00968379 | 0.44980004 |
| ERCC4 | 56.6612736 | -0.2886664 | 0.25105267 | -1.149824 | 0.25021639 | 0.99989001 |
| MKL2 | 550.099492 | -0.9549876 | 0.4340856 | -2.1999983 | 0.02780702 | 0.65805899 |

| | | | | | | |
|---|---|---|---|---|---|---|
| PARN | 218.537519 | -0.2076265 | 0.28297151 | -0.7337365 | 0.46310935 | 0.99989001 |
| BFAR | 105.394882 | -0.0100034 | 0.26826094 | -0.03729 | 0.97025379 | 0.99989001 |
| PDXDC1 | 398.493687 | -0.103736 | 0.22740134 | -0.4561801 | 0.64826045 | 0.99989001 |
| NTAN1 | 74.2330558 | -0.1358823 | 0.30296668 | -0.4485058 | 0.65378818 | 0.99989001 |
| RRN3 | 94.0271855 | 0.09131574 | 0.29095586 | 0.31384741 | 0.75363693 | 0.99989001 |
| C16orf45 | 366.312228 | -0.1437953 | 0.39917605 | -0.3602303 | 0.71867489 | 0.99989001 |
| MARF1 | 153.215842 | 0.00351825 | 0.26492099 | 0.01328037 | 0.98940411 | 0.99989001 |
| NDE1 | 128.718597 | 0.06871984 | 0.27678907 | 0.24827513 | 0.80392154 | 0.99989001 |
| MYH11 | 61.9235687 | -0.3371455 | 0.38577993 | -0.8739322 | 0.38215517 | 0.99989001 |
| FOPNL | 43.065216 | -0.2001663 | 0.25752224 | -0.7772778 | 0.43699491 | NA |
| ABCC1 | 445.888355 | -0.0673227 | 0.33481479 | -0.2010745 | 0.84064034 | 0.99989001 |
| XYLT1 | 182.936706 | 0.08083638 | 0.48611642 | 0.16629016 | 0.86792861 | 0.99989001 |
| RPS15A | 92.3771034 | 0.26145719 | 0.64737241 | 0.40387448 | 0.68630502 | 0.99989001 |
| SMG1 | 491.116252 | 0.07463691 | 0.24312063 | 0.30699539 | 0.75884688 | 0.99989001 |
| TMC7 | 83.6173885 | 0.08598856 | 0.44257823 | 0.19429008 | 0.84594875 | 0.99989001 |
| CCP110 | 48.5252882 | -0.2128351 | 0.28211526 | -0.7544261 | 0.4505934 | NA |
| VPS35L | 261.116979 | -0.1500309 | 0.23894808 | -0.6278809 | 0.53008195 | 0.99989001 |
| KNOP1 | 81.7520817 | -0.5917641 | 0.38198028 | -1.5492006 | 0.12133351 | 0.99989001 |
| IQCK | 321.746785 | -0.1536855 | 0.33059171 | -0.4648801 | 0.64201729 | 0.99989001 |
| GPRC5B | 307.291378 | -0.5788856 | 0.25104771 | -2.3058788 | 0.0211174 | 0.59254339 |
| UMOD | 248.554682 | -0.3313498 | 0.91364363 | -0.3626685 | 0.7168525 | 0.99989001 |
| ACSM3 | 196.794682 | 0.54188727 | 0.54160433 | 1.00052241 | 0.31705776 | 0.99989001 |
| THUMPD1 | 102.273343 | -0.0606136 | 0.20290263 | -0.2987327 | 0.76514402 | 0.99989001 |
| LYRM1 | 67.4582008 | -0.2214085 | 0.42099542 | -0.5259167 | 0.59894612 | 0.99989001 |
| METTL9 | 190.25733 | 0.08292626 | 0.21047252 | 0.39400039 | 0.69358075 | 0.99989001 |
| NPIPB4 | 68.0629491 | -0.4655177 | 0.25151677 | -1.8508417 | 0.06419233 | 0.86610729 |
| UQCRC2 | 154.330566 | 0.14534131 | 0.20916537 | 0.69486317 | 0.48714106 | 0.99989001 |
| MOSMO | 299.330228 | -0.0457519 | 0.19026562 | -0.2404635 | 0.80997097 | 0.99989001 |
| EEF2K | 155.330999 | -0.5471384 | 0.4475306 | -1.2225721 | 0.22149135 | 0.99989001 |
| POLR3E | 45.6530497 | 0.25254595 | 0.28704514 | 0.87981267 | 0.3789608 | NA |
| CDR2 | 477.31327 | 0.10743672 | 0.2830229 | 0.37960435 | 0.70423913 | 0.99989001 |
| NPIPB5 | 195.430685 | 0.18821247 | 0.28584548 | 0.65844131 | 0.5102546 | 0.99989001 |
| USP31 | 273.706961 | 0.3403607 | 0.39496423 | 0.8617507 | 0.38882472 | 0.99989001 |
| COG7 | 109.168495 | -0.0752039 | 0.26415855 | -0.2846923 | 0.77587989 | 0.99989001 |
| GGA2 | 117.107128 | -0.5717878 | 0.2103049 | -2.7188518 | 0.0065509 | 0.390188 |
| UBFD1 | 46.3261322 | 0.25704302 | 0.28599159 | 0.89877824 | 0.36877079 | NA |
| NDUFAB1 | 75.3267069 | -0.1886551 | 0.38155497 | -0.4944375 | 0.62099725 | 0.99989001 |
| DCTN5 | 101.163578 | 0.28952383 | 0.22384859 | 1.29339137 | 0.19587573 | 0.99989001 |
| RBBP6 | 188.11853 | -0.0287053 | 0.21414642 | -0.1340451 | 0.89336688 | 0.99989001 |
| TNRC6A | 724.479065 | -0.0187417 | 0.20269598 | -0.0924623 | 0.92633071 | 0.99989001 |
| ARHGAP17 | 347.516111 | 0.20199317 | 0.14510109 | 1.39208579 | 0.16389642 | 0.99989001 |
| LCMT1 | 101.883886 | 0.2745924 | 0.2432514 | 1.12884203 | 0.25896448 | 0.99989001 |
| NSMCE1 | 63.6881133 | 0.4956311 | 0.32676062 | 1.51680182 | 0.12931673 | 0.99989001 |
| IL4R | 845.792644 | -0.0379314 | 0.44025841 | -0.0861572 | 0.93134143 | 0.99989001 |
| GTF3C1 | 163.25864 | -0.1995653 | 0.27398218 | -0.7283877 | 0.46637628 | 0.99989001 |
| KIAA0556 | 143.070153 | -0.0176829 | 0.33477787 | -0.0528199 | 0.95787544 | 0.99989001 |
| XPO6 | 167.792077 | -0.135461 | 0.22536013 | -0.6010869 | 0.54778208 | 0.99989001 |
| NUPR1 | 96.1535209 | -0.1508754 | 0.6028995 | -0.2502496 | 0.80239432 | 0.99989001 |
| ATXN2L | 126.83564 | -0.1224024 | 0.22608205 | -0.5414069 | 0.58822712 | 0.99989001 |
| TUFM | 121.099309 | 0.12576143 | 0.42466552 | 0.29614231 | 0.76712141 | 0.99989001 |
| SH2B1 | 66.206367 | -0.1347456 | 0.25120986 | -0.5363866 | 0.59169142 | 0.99989001 |
| RABEP2 | 74.4541975 | -0.4453153 | 0.40698414 | -1.0941835 | 0.2738745 | 0.99989001 |
| NFATC2IP | 231.355364 | 0.029704 | 0.27970555 | 0.1061974 | 0.91542573 | 0.99989001 |
| SPNS1 | 45.2249696 | -0.461418 | 0.3453095 | -1.3362449 | 0.18146925 | NA |

| | | | | | | |
|---|---|---|---|---|---|---|
| AC009093.2 | 75.7831112 | -0.0656529 | 0.64555588 | -0.1016998 | 0.91899497 | 0.99989001 |
| MAZ | 53.8998813 | -0.1296237 | 0.47249042 | -0.2743415 | 0.78382222 | 0.99989001 |
| MVP | 181.457081 | 0.3339511 | 0.35928972 | 0.9294758 | 0.35264256 | 0.99989001 |
| CDIPT | 64.8886156 | 0.21229621 | 0.47071004 | 0.45101271 | 0.65198039 | 0.99989001 |
| KCTD13 | 64.6524602 | -0.1607534 | 0.34743946 | -0.4626803 | 0.64359355 | 0.99989001 |
| TMEM219 | 131.956964 | -0.1635617 | 0.28767478 | -0.5685645 | 0.56965173 | 0.99989001 |
| TAOK2 | 157.217564 | -0.2407005 | 0.26574085 | -0.9057716 | 0.36505677 | 0.99989001 |
| INO80E | 52.4944442 | -0.0651334 | 0.28076243 | -0.2319875 | 0.81654771 | 0.99989001 |
| PPP4C | 98.0367198 | -0.0936865 | 0.2859436 | -0.3276396 | 0.74318415 | 0.99989001 |
| YPEL3 | 101.95971 | -0.316891 | 0.53548776 | -0.5917801 | 0.55399785 | 0.99989001 |
| MAPK3 | 132.017688 | -0.2950368 | 0.42396611 | -0.6958972 | 0.48649321 | 0.99989001 |
| NPIPB13 | 54.7728705 | -0.5731965 | 0.36918953 | -1.5525805 | 0.12052339 | 0.99989001 |
| CD2BP2 | 50.0266194 | -0.2425678 | 0.32619028 | -0.7436391 | 0.45709486 | 0.99989001 |
| SEPHS2 | 71.3275022 | 0.40448652 | 0.41543314 | 0.9736501 | 0.3302303 | 0.99989001 |
| ZNF785 | 59.5594189 | 0.06211729 | 0.28772295 | 0.2158927 | 0.82907138 | 0.99989001 |
| PRR14 | 43.4269192 | 0.04208482 | 0.26143748 | 0.16097468 | 0.87211334 | NA |
| FBRS | 57.2280423 | 0.05051919 | 0.30931371 | 0.1633267 | 0.87026121 | 0.99989001 |
| RNF40 | 53.6447819 | 0.20679944 | 0.31503331 | 0.65643674 | 0.51154316 | 0.99989001 |
| BCL7C | 94.0122317 | 0.21360852 | 0.2722824 | 0.78451095 | 0.43274036 | 0.99989001 |
| SETD1A | 42.6947231 | -0.4461547 | 0.38837473 | -1.1487739 | 0.25064923 | NA |
| STX4 | 62.8489138 | -0.5133719 | 0.23398144 | -2.1940709 | 0.02823031 | 0.65805899 |
| KAT8 | 65.3553961 | -0.5794249 | 0.25331153 | -2.2874006 | 0.02217245 | 0.60384164 |
| FUS | 398.153249 | -0.0868323 | 0.32690998 | -0.2656154 | 0.79053541 | 0.99989001 |
| AC026471.4 | 42.5420268 | -0.1970658 | 0.35015814 | -0.5627907 | 0.57357739 | NA |
| TGFB1I1 | 48.8545469 | -0.0123459 | 0.39409699 | -0.0313269 | 0.97500881 | NA |
| C16orf58 | 42.4089728 | -0.248196 | 0.26604045 | -0.9329257 | 0.35085833 | NA |
| ZNF720 | 117.643121 | 0.61114538 | 0.20916224 | 2.92187244 | 0.00347934 | 0.29068264 |
| ZNF267 | 78.7042931 | 0.26212366 | 0.35782134 | 0.73255458 | 0.46383014 | 0.99989001 |
| VPS35 | 111.135018 | 0.13554103 | 0.21620954 | 0.62689665 | 0.53072698 | 0.99989001 |
| DNAJA2 | 58.8325322 | 0.23327071 | 0.29820089 | 0.78226027 | 0.43406163 | 0.99989001 |
| NETO2 | 96.0723869 | 0.40392772 | 0.54020629 | 0.74772864 | 0.45462385 | 0.99989001 |
| ITFG1 | 669.899555 | -0.278824 | 0.24402139 | -1.1426209 | 0.25319602 | 0.99989001 |
| PHKB | 587.680224 | 0.16199004 | 0.21614115 | 0.74946412 | 0.45357752 | 0.99989001 |
| LONP2 | 425.601076 | 0.01828341 | 0.18295157 | 0.09993577 | 0.92039531 | 0.99989001 |
| SIAH1 | 118.494049 | 0.09497253 | 0.23054525 | 0.41194746 | 0.68037793 | 0.99989001 |
| N4BP1 | 263.167249 | 0.26657145 | 0.42765461 | 0.62333351 | 0.53306538 | 0.99989001 |
| ZNF423 | 317.294896 | 0.55193796 | 0.35459775 | 1.55651849 | 0.11958485 | 0.99989001 |
| HEATR3 | 57.2372363 | -0.2323708 | 0.29174894 | -0.7964752 | 0.42575588 | 0.99989001 |
| TENT4B | 171.843084 | 0.36411831 | 0.27414637 | 1.32818945 | 0.18411553 | 0.99989001 |
| BRD7 | 127.172982 | 0.0498811 | 0.25706555 | 0.19404039 | 0.84614426 | 0.99989001 |
| CYLD | 167.262739 | 0.04836142 | 0.27793596 | 0.17400204 | 0.86186386 | 0.99989001 |
| CHD9 | 1080.6236 | 0.0125894 | 0.21449564 | 0.05869301 | 0.95319662 | 0.99989001 |
| RBL2 | 224.834107 | 0.13669079 | 0.20410384 | 0.66971198 | 0.50304142 | 0.99989001 |
| AKTIP | 48.6591711 | -0.2900389 | 0.27560891 | -1.0523566 | 0.29263596 | NA |
| FTO | 803.984334 | 0.14282345 | 0.21791146 | 0.65541964 | 0.51219761 | 0.99989001 |
| IRX3 | 190.51673 | -0.097394 | 0.47126133 | -0.2066666 | 0.83627021 | 0.99989001 |
| CRNDE | 139.504849 | 0.10921765 | 0.30961348 | 0.35275482 | 0.72427226 | 0.99989001 |
| IRX5 | 38.6478709 | 0.51881822 | 0.43672121 | 1.18798495 | 0.23483934 | NA |
| LPCAT2 | 106.964926 | -0.9415679 | 0.69587273 | -1.3530749 | 0.1760317 | 0.99989001 |
| AMFR | 113.05041 | -0.0741303 | 0.20606549 | -0.3597417 | 0.71904031 | 0.99989001 |
| NUDT21 | 111.664769 | 0.05555162 | 0.23241407 | 0.23902003 | 0.81109005 | 0.99989001 |
| OGFOD1 | 47.4509244 | 0.15233953 | 0.37634405 | 0.40478794 | 0.6856334 | NA |
| BBS2 | 69.3544194 | -0.3365422 | 0.25403517 | -1.324786 | 0.18524213 | 0.99989001 |
| MT2A | 1264.97669 | -0.5495009 | 0.71453207 | -0.769036 | 0.44187195 | 0.99989001 |

| | | | | | | |
|---|---|---|---|---|---|---|
| MT1E | 229.896686 | -1.3814461 | 0.63951384 | -2.1601505 | 0.03076102 | 0.67281094 |
| MT1M | 75.450367 | -0.7291077 | 0.64894644 | -1.1235252 | 0.26121451 | 0.99989001 |
| MT1G | 421.873243 | -0.2380889 | 1.1278102 | -0.2111073 | 0.83280356 | 0.99989001 |
| MT1H | 139.518253 | -0.397868 | 1.41915864 | -0.2803548 | 0.77920529 | 0.99989001 |
| MT1X | 194.397269 | -0.1493779 | 0.66504524 | -0.2246131 | 0.82228027 | 0.99989001 |
| NUP93 | 125.238952 | -0.2447873 | 0.31764147 | -0.7706403 | 0.4409202 | 0.99989001 |
| HERPUD1 | 296.303392 | 0.10178114 | 0.50515412 | 0.20148532 | 0.84031911 | 0.99989001 |
| NLRC5 | 235.596495 | -0.0818727 | 0.56977713 | -0.1436925 | 0.88574331 | 0.99989001 |
| CPNE2 | 145.616142 | -0.2383966 | 0.30791239 | -0.7742352 | 0.43879169 | 0.99989001 |
| FAM192A | 114.008616 | -0.4437715 | 0.21351338 | -2.0784247 | 0.03767026 | 0.70738774 |
| RSPRY1 | 140.915565 | 0.11753061 | 0.19854043 | 0.59197315 | 0.55386857 | 0.99989001 |
| ARL2BP | 112.311147 | 0.01155376 | 0.30751411 | 0.03757148 | 0.97002935 | 0.99989001 |
| PLLP | 74.7935901 | -0.0142515 | 0.66426128 | -0.0214547 | 0.98288293 | 0.99989001 |
| CX3CL1 | 174.897266 | 1.73877933 | 0.55805465 | 3.11578683 | 0.00183455 | 0.2066742 |
| CIAPIN1 | 50.2540822 | -0.1083291 | 0.35334544 | -0.3065814 | 0.759162 | 0.99989001 |
| COQ9 | 43.8967463 | 0.15123715 | 0.28539577 | 0.52992077 | 0.59616687 | NA |
| ADGRG1 | 259.023199 | -0.5619354 | 0.35451239 | -1.5850939 | 0.11294499 | 0.99989001 |
| KIFC3 | 301.534357 | -0.3210733 | 0.28648304 | -1.1207412 | 0.26239805 | 0.99989001 |
| CSNK2A2 | 113.898759 | 0.17930656 | 0.28704223 | 0.62466964 | 0.5321879 | 0.99989001 |
| CNOT1 | 285.748739 | -0.0033544 | 0.19158284 | -0.017509 | 0.98603053 | 0.99989001 |
| GOT2 | 79.0355198 | 0.0916971 | 0.28059229 | 0.32679835 | 0.7438204 | 0.99989001 |
| CDH5 | 511.197786 | -0.1005781 | 0.40649766 | -0.2474259 | 0.80457862 | 0.99989001 |
| CMTM3 | 49.2607278 | 0.03605105 | 0.31243776 | 0.11538636 | 0.90813889 | NA |
| CMTM4 | 91.8415858 | 0.07354996 | 0.23366063 | 0.31477258 | 0.75293433 | 0.99989001 |
| DYNC1LI2 | 262.892148 | -0.0649897 | 0.19482135 | -0.3335861 | 0.73869188 | 0.99989001 |
| NAE1 | 84.2809808 | -0.204329 | 0.22014099 | -0.9281735 | 0.35331757 | 0.99989001 |
| CDH16 | 61.4651861 | 0.32239632 | 0.327003 | 0.98591242 | 0.32417609 | 0.99989001 |
| FAM96B | 96.7540465 | -0.132971 | 0.47306769 | -0.2810824 | 0.7786472 | 0.99989001 |
| CES2 | 48.0309753 | 0.43980503 | 0.36606203 | 1.20144948 | 0.22957689 | NA |
| CBFB | 362.156634 | 0.52456058 | 0.25506722 | 2.05655821 | 0.03972874 | 0.72961688 |
| C16orf70 | 42.5539902 | 0.40577081 | 0.27960517 | 1.45122784 | 0.14671643 | NA |
| EXOC3L1 | 68.9310602 | 0.64105051 | 0.34341247 | 1.86670716 | 0.06194249 | 0.85940342 |
| E2F4 | 65.9272313 | 0.24634276 | 0.39853321 | 0.61812356 | 0.53649389 | 0.99989001 |
| TMEM208 | 55.5589607 | -0.1032105 | 0.54712361 | -0.1886421 | 0.85037337 | 0.99989001 |
| FHOD1 | 65.7201006 | 0.32091921 | 0.41022901 | 0.78229282 | 0.43404251 | 0.99989001 |
| ATP6V0D1 | 85.6811148 | 0.21780882 | 0.35836559 | 0.60778387 | 0.54333083 | 0.99989001 |
| AC009061.2 | 54.9143159 | 0.06230874 | 0.46104316 | 0.1351473 | 0.89249541 | 0.99989001 |
| RIPOR1 | 192.509788 | -0.1468064 | 0.28590935 | -0.5134717 | 0.60762141 | 0.99989001 |
| CTCF | 191.656159 | 0.0782975 | 0.25602208 | 0.30582322 | 0.75973925 | 0.99989001 |
| GFOD2 | 275.601091 | 0.24005957 | 0.24471413 | 0.98097959 | 0.32660281 | 0.99989001 |
| RANBP10 | 184.629335 | -0.1079106 | 0.28129169 | -0.3836254 | 0.7012561 | 0.99989001 |
| PSKH1 | 42.7318934 | 0.03737973 | 0.3064139 | 0.12199099 | 0.90290615 | NA |
| LCAT | 41.831058 | -0.5878857 | 0.43043684 | -1.3657886 | 1.72E-01 | NA |
| SLC12A4 | 53.3290964 | -0.0244473 | 0.29168382 | -0.0838144 | 0.93320403 | 0.99989001 |
| DUS2 | 55.3947556 | -0.378454 | 0.31967026 | -1.1838886 | 0.23645715 | 0.99989001 |
| NFATC3 | 340.306901 | 0.60506099 | 0.24926511 | 2.42737943 | 0.01520834 | 0.51942334 |
| SLC7A6 | 156.031547 | -0.2860179 | 0.3129542 | -0.9139288 | 0.36075425 | 0.99989001 |
| SLC7A6OS | 74.0690066 | -0.4014839 | 0.2838455 | -1.4144453 | 0.1572312 | 0.99989001 |
| PRMT7 | 62.0687516 | -0.0014116 | 0.46968596 | -0.0030055 | 0.99760195 | 0.99989001 |
| ZFP90 | 65.2532009 | 0.51201922 | 0.29675236 | 1.72540911 | 0.08445377 | 0.95514629 |
| TANGO6 | 234.577962 | 0.0734012 | 0.28012825 | 0.26202714 | 0.79330052 | 0.99989001 |
| CHTF8 | 78.677854 | 0.37003933 | 0.27978463 | 1.32258637 | 0.18597297 | 0.99989001 |
| UTP4 | 47.089142 | -0.2921248 | 0.36239291 | -0.8060996 | 0.42018541 | NA |
| SNTB2 | 196.123766 | -0.2488879 | 0.31927608 | -0.7795383 | 0.4356627 | 0.99989001 |

| | | | | | | |
|---|---|---|---|---|---|---|
| VPS4A | 61.3005885 | -0.0691569 | 0.30943315 | -0.2234954 | 0.82314999 | 0.99989001 |
| TERF2 | 80.175699 | -0.2450515 | 0.31479736 | -0.7784419 | 0.43630854 | 0.99989001 |
| CYB5B | 290.486231 | -0.1070598 | 0.15815875 | -0.6769136 | 0.49846077 | 0.99989001 |
| NFAT5 | 2155.91909 | -0.308145 | 0.1912942 | -1.6108433 | 0.10721388 | 0.99989001 |
| NQO1 | 112.174367 | 1.47852573 | 0.56818366 | 2.60219684 | 0.00926287 | 0.44250709 |
| WWP2 | 256.35105 | -0.3115553 | 0.27566354 | -1.1302015 | 0.25839134 | 0.99989001 |
| PDPR | 152.533882 | -0.2534175 | 0.37403479 | -0.677524 | 0.49807356 | 0.99989001 |
| EXOSC6 | 76.5275061 | -0.4979024 | 0.34326236 | -1.450501 | 0.14691887 | 0.99989001 |
| AARS | 86.4874961 | -0.0539515 | 0.35359901 | -0.1525783 | 0.87873084 | 0.99989001 |
| ST3GAL2 | 194.753684 | 0.59155416 | 0.31033562 | 1.90617552 | 0.05662744 | 0.83644598 |
| COG4 | 87.0935675 | 0.01895111 | 0.29156865 | 0.06499706 | 0.94817634 | 0.99989001 |
| SF3B3 | 173.619079 | -0.0070858 | 0.22770317 | -0.0311187 | 0.97517484 | 0.99989001 |
| MTSS1L | 39.7911271 | 0.77332565 | 0.43361653 | 1.78343214 | 0.07451599 | NA |
| VAC14 | 78.6266558 | -0.5743092 | 0.40945596 | -1.4026153 | 0.1607316 | 0.99989001 |
| CMTR2 | 43.7129315 | 0.12439126 | 0.2591446 | 0.48000713 | 0.63122232 | NA |
| PHLPP2 | 79.2693758 | 0.40851705 | 0.32241101 | 1.26706918 | 0.20513056 | 0.99989001 |
| AP1G1 | 266.647837 | 0.00948975 | 0.1883078 | 0.05039488 | 0.95980772 | 0.99989001 |
| IST1 | 190.552159 | -0.0252254 | 0.17440203 | -0.1446395 | 0.88499553 | 0.99989001 |
| TXNL4B | 46.2441272 | 0.02645172 | 0.36063293 | 0.07334805 | 0.94152916 | NA |
| DHX38 | 66.6289114 | -0.4055464 | 0.2515554 | -1.6121552 | 0.10692817 | 0.99989001 |
| ZFHX3 | 820.810115 | 0.32038786 | 0.30914006 | 1.03638416 | 0.30002296 | 0.99989001 |
| PSMD7 | 119.519226 | 0.04154473 | 0.35327149 | 0.11760001 | 0.9063846 | 0.99989001 |
| GLG1 | 645.403265 | -0.0221737 | 0.15521413 | -0.1428587 | 0.88640174 | 0.99989001 |
| RFWD3 | 69.8234085 | 0.13328056 | 0.33792788 | 0.39440533 | 0.69328181 | 0.99989001 |
| MLKL | 201.389509 | -0.0808721 | 0.41331476 | -0.195667 | 0.8448708 | 0.99989001 |
| WDR59 | 184.024291 | -0.1808149 | 0.17905208 | -1.0098452 | 0.31256947 | 0.99989001 |
| ZNRF1 | 329.24681 | -0.2794538 | 0.31545187 | -0.8858842 | 0.3756799 | 0.99989001 |
| BCAR1 | 191.732132 | 0.1114338 | 0.32535478 | 0.34249934 | 0.73197514 | 0.99989001 |
| CFDP1 | 389.929909 | -0.3766318 | 0.19601439 | -1.9214498 | 0.05467503 | 0.82320925 |
| GABARAPL2 | 211.667766 | 0.31530882 | 0.63579435 | 0.49592894 | 0.61994455 | 0.99989001 |
| ADAT1 | 48.0292559 | -0.1578449 | 0.30315266 | -0.5206781 | 0.60259104 | NA |
| KARS | 69.996014 | -0.062236 | 0.37323264 | -0.1667484 | 0.86756798 | 0.99989001 |
| TERF2IP | 277.492512 | 0.11005571 | 0.35723497 | 0.30807653 | 0.7580241 | 0.99989001 |
| WWOX | 1990.0631 | -0.4273602 | 0.29362462 | -1.4554645 | 0.14554071 | 0.99989001 |
| MAF | 233.262381 | 0.30860235 | 0.61512481 | 0.50169063 | 0.61588516 | 0.99989001 |
| CDYL2 | 211.231137 | 0.19435274 | 0.31459967 | 0.61777796 | 0.53672172 | 0.99989001 |
| CMC2 | 95.1222532 | -0.2813651 | 0.26561365 | -1.0593023 | 0.28946215 | 0.99989001 |
| ATMIN | 51.1559196 | -0.0113355 | 0.3893706 | -0.0291124 | 0.97677495 | 0.99989001 |
| GAN | 125.304256 | 0.23635473 | 0.25765172 | 0.91734195 | 0.35896348 | 0.99989001 |
| CMIP | 1449.38957 | 0.19060532 | 0.4698437 | 0.40567814 | 0.68497911 | 0.99989001 |
| PLCG2 | 706.084901 | -0.0440419 | 0.37130811 | -0.1186129 | 0.90558205 | 0.99989001 |
| CDH13 | 1416.59136 | -1.021769 | 0.34629374 | -2.9505847 | 0.00317173 | 0.28843642 |
| HSBP1 | 127.446642 | -0.1740496 | 0.44296673 | -0.392918 | 0.69438008 | 0.99989001 |
| MBTPS1 | 166.811055 | -0.0559525 | 0.19307469 | -0.2897974 | 0.77197123 | 0.99989001 |
| TAF1C | 75.8707729 | -0.1201037 | 0.35812381 | -0.3353692 | 0.73734659 | 0.99989001 |
| TLDC1 | 49.7601966 | 0.33016096 | 0.40829034 | 0.8086426 | 0.41872076 | NA |
| COTL1 | 204.648579 | 0.17471377 | 0.40112532 | 0.43555907 | 0.66315667 | 0.99989001 |
| KLHL36 | 65.289079 | 0.43170624 | 0.24863119 | 1.73633181 | 0.08250518 | 0.94799019 |
| USP10 | 191.239802 | -0.0628011 | 0.3244059 | -0.1935879 | 0.84649857 | 0.99989001 |
| ZDHHC7 | 81.7855489 | -0.3021228 | 0.22149672 | -1.3640059 | 0.17256571 | 0.99989001 |
| GSE1 | 929.957834 | 1.08869848 | 0.51307428 | 2.12191203 | 0.03384513 | 0.67281094 |
| EMC8 | 50.9718142 | -0.2071704 | 0.28722342 | -0.7212867 | 0.47073312 | 0.99989001 |
| COX4I1 | 293.681921 | -0.0672892 | 0.45864505 | -0.1467131 | 0.88335852 | 0.99989001 |
| FBXO31 | 50.6368489 | -0.3151107 | 0.32858194 | -0.9590018 | 0.33755784 | 0.99989001 |

| | | | | | | |
|---|---|---|---|---|---|---|
| MAP1LC3B | 147.695658 | -0.3693316 | 0.42947151 | -0.8599677 | 0.38980683 | 0.99989001 |
| ZCCHC14 | 233.713408 | -0.2031882 | 0.21278537 | -0.9548974 | 0.33962956 | 0.99989001 |
| KLHDC4 | 82.777149 | -0.4462127 | 0.33516496 | -1.3313227 | 0.18308286 | 0.99989001 |
| BANP | 148.413414 | -0.2316558 | 0.32558826 | -0.7114993 | 0.4767749 | 0.99989001 |
| ZC3H18 | 110.006102 | -0.1140883 | 0.29832936 | -0.382424 | 0.70214687 | 0.99989001 |
| CYBA | 130.594058 | 0.8426053 | 0.51813717 | 1.62622053 | 0.10390275 | 0.99989001 |
| MVD | 39.5376755 | 0.50264062 | 0.34167832 | 1.4710931 | 0.14126594 | NA |
| PIEZO1 | 251.544162 | -0.2861432 | 0.15376921 | -1.8608617 | 0.06276371 | 0.85940342 |
| APRT | 119.508959 | -0.1990082 | 0.49498747 | -0.402047 | 0.68764943 | 0.99989001 |
| ACSF3 | 63.3423358 | -0.1009104 | 0.31487613 | -0.3204764 | 0.74860725 | 0.99989001 |
| ANKRD11 | 1029.47316 | 0.08469114 | 0.30866072 | 0.27438264 | 0.78379058 | 0.99989001 |
| SPG7 | 332.988739 | -0.5916483 | 0.29416984 | -2.0112474 | 0.04429933 | 0.77228819 |
| RPL13 | 926.990598 | -0.1709897 | 0.55356095 | -0.3088905 | 0.75740481 | 0.99989001 |
| CHMP1A | 74.6335429 | -0.0470944 | 0.30509834 | -0.154358 | 0.87732747 | 0.99989001 |
| CDK10 | 82.3094561 | -0.070907 | 0.21552603 | -0.328995 | 0.74215947 | 0.99989001 |
| ZNF276 | 70.395713 | 0.06937311 | 0.22391585 | 0.30981777 | 0.75669953 | 0.99989001 |
| TCF25 | 445.308254 | -0.0685524 | 0.2311099 | -0.2966226 | 0.76675469 | 0.99989001 |
| DEF8 | 51.3739187 | -0.1396576 | 0.26584463 | -0.5253355 | 0.59934997 | 0.99989001 |
| FAM157C | 60.7035852 | -0.2947538 | 0.34968001 | -0.8429244 | 0.39927071 | 0.99989001 |
| DOC2B | 163.425397 | -0.4328976 | 0.38991161 | -1.1102454 | 0.26689328 | 0.99989001 |
| RPH3AL | 365.933922 | 0.43189177 | 0.44932503 | 0.96120124 | 0.336451 | 0.99989001 |
| RFLNB | 93.4018229 | -0.1701105 | 0.41058932 | -0.4143081 | 0.67864846 | 0.99989001 |
| VPS53 | 419.079299 | -0.4210674 | 0.23130008 | -1.8204375 | 0.0686924 | 0.88856931 |
| GLOD4 | 50.8384966 | -0.1339473 | 0.24575868 | -0.5450358 | 0.58572891 | 0.99989001 |
| NXN | 1192.36664 | 0.28909526 | 0.45695894 | 0.63265041 | 0.52696196 | 0.99989001 |
| ABR | 550.840741 | 0.17138851 | 0.29779267 | 0.57552963 | 0.56493316 | 0.99989001 |
| YWHAE | 503.7195 | -0.0417937 | 0.18048878 | -0.2315585 | 0.81688095 | 0.99989001 |
| CRK | 325.285742 | 0.4290812 | 0.31606958 | 1.35755295 | 0.17460558 | 0.99989001 |
| MYO1C | 361.481426 | 0.07374634 | 0.33145703 | 0.22249141 | 0.82393136 | 0.99989001 |
| INPP5K | 73.7771282 | 0.17141891 | 0.36553936 | 0.46894789 | 0.63910689 | 0.99989001 |
| PITPNA | 257.216398 | 0.32647519 | 0.29397568 | 1.11055172 | 0.26676136 | 0.99989001 |
| SCARF1 | 180.407297 | -0.0628272 | 0.2027312 | -0.3099037 | 0.75663415 | 0.99989001 |
| PRPF8 | 186.384335 | -0.0011521 | 0.26033146 | -0.0044254 | 0.99646904 | 0.99989001 |
| SMYD4 | 46.5799993 | -0.1369079 | 0.27166105 | -0.5039659 | 0.61428536 | NA |
| RPA1 | 111.783375 | -0.3102129 | 0.19723387 | -1.5728177 | 0.11576104 | 0.99989001 |
| HIC1 | 87.1078451 | -0.4361832 | 0.60253899 | -0.7239087 | 0.46912178 | 0.99989001 |
| SMG6 | 532.186673 | -0.0300255 | 0.3221096 | -0.0932153 | 0.92573253 | 0.99989001 |
| TSR1 | 40.5664335 | -0.4016805 | 0.50425421 | -0.7965833 | 0.42569305 | NA |
| SGSM2 | 67.4507072 | 0.1744048 | 0.24898812 | 0.7004543 | 0.48364364 | 0.99989001 |
| METTL16 | 162.717528 | -0.0871497 | 0.2882368 | -0.3023546 | 0.76238173 | 0.99989001 |
| PAFAH1B1 | 718.541233 | 0.00385594 | 0.20504596 | 0.01880527 | 0.98499645 | 0.99989001 |
| CLUH | 40.8767271 | -0.3434066 | 0.43664525 | -0.7864659 | 0.43159461 | NA |
| TAX1BP3 | 44.7034223 | -0.6458516 | 0.4861735 | -1.3284385 | 0.18403328 | NA |
| EMC6 | 49.5188093 | -0.1560366 | 0.63355934 | -0.2462856 | 0.80546113 | NA |
| ITGAE | 41.8767661 | -0.0098427 | 0.39090095 | -0.0251796 | 0.9799117 | NA |
| NCBP3 | 146.217304 | 0.22052013 | 0.20003555 | 1.1024047 | 0.27028577 | 0.99989001 |
| CAMKK1 | 69.0810762 | 0.1363655 | 0.44464077 | 0.30668689 | 0.75908171 | 0.99989001 |
| ATP2A3 | 124.938254 | 1.34258722 | 0.62834385 | 2.13670781 | 0.03262176 | 0.67281094 |
| ZZEF1 | 246.528029 | 0.00393863 | 0.196311 | 0.02006321 | 0.98399295 | 0.99989001 |
| CYB5D2 | 70.6392137 | -0.2112717 | 0.29012474 | -0.7282099 | 0.46648513 | 0.99989001 |
| ANKFY1 | 235.182393 | -0.0516537 | 0.37239303 | -0.1387075 | 0.88968128 | 0.99989001 |
| UBE2G1 | 254.539781 | 0.11630211 | 0.28809858 | 0.40368859 | 0.68644173 | 0.99989001 |
| SPNS2 | 180.167729 | 0.75259662 | 0.4955247 | 1.5187873 | 0.12881604 | 0.99989001 |
| PELP1 | 42.5633586 | -0.5894123 | 0.4094733 | -1.4394401 | 0.15002587 | NA |

| | | | | | | |
|---|---|---|---|---|---|---|
| PSMB6 | 96.9580682 | 0.10261777 | 0.50541225 | 0.20303776 | 0.83910552 | 0.99989001 |
| PLD2 | 45.4994761 | 0.37766069 | 0.3369168 | 1.12093162 | 0.26231697 | NA |
| MINK1 | 157.824848 | 0.12904144 | 0.25117821 | 0.51374454 | 0.60743061 | 0.99989001 |
| SLC25A11 | 46.9276428 | 0.14997497 | 0.4273127 | 0.35097242 | 0.72560904 | NA |
| RNF167 | 41.3705635 | -0.2140658 | 0.4673617 | -0.4580303 | 0.64693068 | NA |
| PFN1 | 356.6825 | -0.3558611 | 0.51471293 | -0.6913778 | 0.48932814 | 0.99989001 |
| SPAG7 | 60.2072317 | -0.2070901 | 0.33778765 | -0.6130778 | 0.53982489 | 0.99989001 |
| CAMTA2 | 67.4088537 | -0.2196474 | 0.28371863 | -0.7741734 | 0.43882825 | 0.99989001 |
| KIF1C | 73.9152038 | 0.07500565 | 0.3290132 | 0.22797157 | 0.81966834 | 0.99989001 |
| AC087500.1 | 43.298803 | -0.034991 | 0.47763215 | -0.0732593 | 0.94159977 | NA |
| RABEP1 | 338.688905 | -0.0893261 | 0.26494734 | -0.3371464 | 0.73600651 | 0.99989001 |
| NUP88 | 106.772705 | -0.0456341 | 0.2409327 | -0.1894061 | 0.84977455 | 0.99989001 |
| RPAIN | 82.6100714 | -0.0238293 | 0.24457581 | -0.0974313 | 0.92238386 | 0.99989001 |
| C1QBP | 75.276777 | -0.2246992 | 0.46926415 | -0.4788332 | 0.63205733 | 0.99989001 |
| DERL2 | 56.8607755 | 0.04588792 | 0.36275562 | 0.12649818 | 0.89933759 | 0.99989001 |
| NLRP1 | 405.28305 | 0.2854122 | 0.33932395 | 0.84112011 | 0.40028065 | 0.99989001 |
| KIAA0753 | 73.1467821 | 0.00795975 | 0.34112077 | 0.0233341 | 0.98138377 | 0.99989001 |
| XAF1 | 823.508598 | 0.40538463 | 0.74938499 | 0.54095642 | 0.58853762 | 0.99989001 |
| BCL6B | 109.600007 | -0.3155372 | 0.42519549 | -0.742099 | 0.45802735 | 0.99989001 |
| ACADVL | 162.033078 | 0.29933296 | 0.28990115 | 1.03253457 | 0.30182176 | 0.99989001 |
| EIF5A | 106.62398 | -0.4592747 | 0.56187855 | -0.8173915 | 0.41370475 | 0.99989001 |
| GPS2 | 75.5445241 | 0.10361325 | 0.38703091 | 0.26771312 | 0.78892015 | 0.99989001 |
| ZBTB4 | 126.97473 | -0.0589227 | 0.32827771 | -0.1794905 | 0.8575526 | 0.99989001 |
| POLR2A | 187.266644 | 0.13934342 | 0.27968593 | 0.49821392 | 0.61833327 | 0.99989001 |
| EIF4A1 | 136.798694 | 0.19851748 | 0.5742417 | 0.34570371 | 0.7295654 | 0.99989001 |
| MPDU1 | 44.9367987 | -0.5479982 | 0.35375988 | -1.5490683 | 0.12136529 | NA |
| FXR2 | 38.8953403 | -0.1875587 | 0.34503545 | -0.5435927 | 0.58672181 | NA |
| SAT2 | 88.689356 | -0.171846 | 0.42505521 | -0.404291 | 0.68599871 | 0.99989001 |
| TP53 | 63.7259677 | 0.01408929 | 0.36744463 | 0.03834399 | 0.96941342 | 0.99989001 |
| KDM6B | 70.2657526 | -0.5091448 | 0.44518399 | -1.1436727 | 0.2527594 | 0.99989001 |
| TMEM88 | 143.284709 | -1.1049537 | 1.04123692 | -1.0611933 | 0.28860205 | 0.99989001 |
| NAA38 | 227.437391 | -0.3995062 | 0.42050496 | -0.950063 | 0.34208025 | 0.99989001 |
| CHD3 | 112.405144 | 0.09569072 | 0.22694383 | 0.42164935 | 0.67328098 | 0.99989001 |
| PER1 | 131.102616 | -1.1893254 | 0.68969338 | -1.7244262 | 0.08463093 | 0.95514629 |
| CTC1 | 42.0012859 | -0.4719023 | 0.30158574 | -1.5647368 | 0.11764465 | NA |
| ARHGEF15 | 251.429016 | -0.1111923 | 0.33399112 | -0.33292 | 0.73919466 | 0.99989001 |
| NDEL1 | 170.891315 | 0.01592249 | 0.26606815 | 0.05984365 | 0.95228016 | 0.99989001 |
| MYH10 | 98.6425929 | 0.77983955 | 0.36484165 | 2.13747404 | 0.03255945 | 0.67281094 |
| STX8 | 526.732476 | -0.1073606 | 0.23416478 | -0.458483 | 0.64660546 | 0.99989001 |
| SCO1 | 67.6892803 | -0.1045017 | 0.30063899 | -0.3475985 | 0.72814173 | 0.99989001 |
| TMEM220 | 48.7203453 | 0.0164272 | 0.37554998 | 0.04374171 | 0.96511029 | NA |
| MAP2K4 | 349.43765 | 0.02857285 | 0.18471504 | 0.15468613 | 0.87706876 | 0.99989001 |
| ARHGAP44 | 46.2554417 | 0.50829858 | 0.48954455 | 1.03830914 | 0.29912616 | NA |
| ELAC2 | 49.7628358 | -0.2135872 | 0.33510729 | -0.6373697 | 0.52388403 | NA |
| HS3ST3A1 | 127.88443 | -1.0634189 | 0.57782427 | -1.8403847 | 0.06571178 | 0.86711984 |
| COX10 | 83.3389203 | 0.04692233 | 0.27349317 | 0.17156673 | 0.86377817 | 0.99989001 |
| PMP22 | 468.361553 | -1.0196797 | 0.21604795 | -4.7196917 | 2.36E-06 | 0.00222857 |
| ZSWIM7 | 83.327683 | -0.0374761 | 0.25164793 | -0.1489228 | 0.88161455 | 0.99989001 |
| TTC19 | 329.752254 | -0.0380772 | 0.18321982 | -0.2078224 | 0.83536765 | 0.99989001 |
| NCOR1 | 692.05851 | -0.0116901 | 0.16598976 | -0.0704268 | 0.94385399 | 0.99989001 |
| PIGL | 185.296182 | -0.4800198 | 0.39754459 | -1.2074616 | 0.22725442 | 0.99989001 |
| UBB | 425.050317 | 0.1208569 | 0.50373415 | 0.23992199 | 0.81039073 | 0.99989001 |
| AC098850.3 | 143.925245 | -0.2896226 | 0.60858834 | -0.4758925 | 0.63415099 | 0.99989001 |
| MPRIP | 473.347053 | -0.0816032 | 0.21170809 | -0.3854514 | 0.69990304 | 0.99989001 |

| | | | | | | |
|---|---|---|---|---|---|---|
| COPS3 | 48.1947064 | 0.31017026 | 0.35500367 | 0.87371003 | 0.38227619 | NA |
| RAI1 | 166.76864 | 0.34999008 | 0.36766077 | 0.95193751 | 0.34112867 | 0.99989001 |
| SREBF1 | 61.8450479 | -0.1850602 | 0.35499916 | -0.5212975 | 0.60215951 | 0.99989001 |
| TOM1L2 | 229.031641 | 0.13334001 | 0.23607236 | 0.56482688 | 0.57219152 | 0.99989001 |
| DRG2 | 37.8446341 | -0.3072057 | 0.32281289 | -0.9516525 | 0.34127325 | NA |
| ALKBH5 | 119.554328 | 0.02429917 | 0.28104104 | 0.08646128 | 0.93109973 | 0.99989001 |
| FLII | 95.7246739 | -0.0291492 | 0.32715575 | -0.0890989 | 0.92900335 | 0.99989001 |
| TOP3A | 67.1066696 | -0.0685757 | 0.25753941 | -0.2662726 | 0.79002929 | 0.99989001 |
| SHMT1 | 51.7171157 | 0.45391547 | 0.2815254 | 1.61234286 | 0.10688735 | 0.99989001 |
| PRPSAP2 | 115.344944 | -0.0834927 | 0.24071266 | -0.3468562 | 0.72869939 | 0.99989001 |
| GRAP | 71.681761 | 0.09877097 | 0.37842786 | 0.26100342 | 0.79408987 | 0.99989001 |
| GRAPL | 535.362413 | 0.5604134 | 0.50816447 | 1.10281893 | 0.27010581 | 0.99989001 |
| EPN2 | 555.099307 | 0.43208944 | 0.22993845 | 1.87915262 | 0.06022366 | 0.85923309 |
| AC004448.2 | 40.5201207 | -0.7519094 | 0.40817844 | -1.8421097 | 0.06545911 | NA |
| SLC47A1 | 49.4247825 | 0.19389082 | 0.40647404 | 0.47700666 | 0.63335738 | NA |
| ALDH3A2 | 65.6857477 | 0.49421764 | 0.36614155 | 1.3497994 | 0.17708034 | 0.99989001 |
| ULK2 | 57.0168226 | 0.94145134 | 0.38563648 | 2.44129222 | 0.01463481 | 0.5186081 |
| AKAP10 | 275.747452 | -0.0906426 | 0.18833752 | -0.4812773 | 0.63031944 | 0.99989001 |
| SPECC1 | 195.470619 | -2.4091352 | 0.95414399 | -2.5249178 | 0.01157253 | 0.48516963 |
| CCDC144NL-/ | 53.0420215 | 0.16972071 | 0.50942038 | 0.33316436 | 0.73901022 | 0.99989001 |
| USP22 | 240.537159 | -0.0552041 | 0.18500176 | -0.298398 | 0.76539945 | 0.99989001 |
| DHRS7B | 179.145039 | -0.0145361 | 0.33978121 | -0.0427808 | 0.96587625 | 0.99989001 |
| MAP2K3 | 126.7616 | -0.1323804 | 0.35786622 | -0.3699158 | 0.71144523 | 0.99989001 |
| WSB1 | 1179.11897 | 0.17583786 | 0.37963723 | 0.46317338 | 0.64324009 | 0.99989001 |
| KSR1 | 263.60841 | 0.10945201 | 0.30265752 | 0.36163653 | 0.71762366 | 0.99989001 |
| LGALS9 | 81.1013884 | -0.7874763 | 0.60606229 | -1.2993322 | 0.19382994 | 0.99989001 |
| NLK | 929.516676 | -0.0542312 | 0.30157387 | -0.1798272 | 0.85728825 | 0.99989001 |
| TNFAIP1 | 160.029802 | 0.29733881 | 0.29610052 | 1.004182 | 0.31529089 | 0.99989001 |
| POLDIP2 | 46.4275165 | -0.088518 | 0.3316371 | -0.2669124 | 0.78953663 | NA |
| KIAA0100 | 111.152445 | 0.1645886 | 0.27673634 | 0.59474878 | 0.55201141 | 0.99989001 |
| SDF2 | 66.3207718 | -0.2514239 | 0.37179173 | -0.6762492 | 0.49888244 | 0.99989001 |
| SUPT6H | 152.474012 | -0.2003595 | 0.26955012 | -0.7433107 | 0.45729357 | 0.99989001 |
| RAB34 | 42.0611335 | -0.7145023 | 0.57361982 | -1.2456026 | 0.21291033 | NA |
| RPL23A | 357.478368 | 0.02665617 | 0.54433159 | 0.04897047 | 0.96094283 | 0.99989001 |
| FAM222B | 400.751036 | 0.03365743 | 0.30701541 | 0.10962782 | 0.91270455 | 0.99989001 |
| FLOT2 | 155.547395 | -0.219832 | 0.282287 | -0.7787535 | 0.43612491 | 0.99989001 |
| PHF12 | 139.551414 | 0.13265286 | 0.22201711 | 0.59748937 | 0.5501807 | 0.99989001 |
| MYO18A | 120.057708 | 0.34176986 | 0.41400375 | 0.82552357 | 0.40907441 | 0.99989001 |
| NUFIP2 | 344.805756 | -0.1569074 | 0.18653888 | -0.8411513 | 0.40026321 | 0.99989001 |
| TAOK1 | 597.253265 | 0.00178103 | 0.17752385 | 0.01003262 | 0.99199526 | 0.99989001 |
| ABHD15-AS1 | 65.8145584 | -0.085859 | 0.46800782 | -0.1834564 | 0.85443991 | 0.99989001 |
| GIT1 | 153.045598 | 0.13930767 | 0.30676893 | 0.45411272 | 0.6497477 | 0.99989001 |
| SSH2 | 764.87939 | 0.04815643 | 0.14044356 | 0.34288809 | 0.73168265 | 0.99989001 |
| NSRP1 | 413.739033 | 0.06257403 | 0.20924168 | 0.29905147 | 0.76490078 | 0.99989001 |
| BLMH | 106.661414 | 0.46067451 | 0.26312306 | 1.75079485 | 0.07998125 | 0.9345178 |
| CPD | 319.295672 | 0.75040329 | 0.34301498 | 2.18766917 | 0.02869371 | 0.65805899 |
| GOSR1 | 185.849455 | 0.0676768 | 0.26405766 | 0.25629552 | 0.79772265 | 0.99989001 |
| CRLF3 | 168.624885 | 0.04046954 | 0.24843629 | 0.16289705 | 0.87059949 | 0.99989001 |
| TEFM | 70.8362912 | 0.3213432 | 0.31769381 | 1.01148713 | 0.31178334 | 0.99989001 |
| RNF135 | 45.7038893 | 0.00845255 | 0.28249049 | 0.02992152 | 0.97612964 | NA |
| NF1 | 1016.80594 | -0.0377678 | 0.2681104 | -0.1408667 | 0.88797526 | 0.99989001 |
| UTP6 | 80.6707271 | -0.0021789 | 0.33481239 | -0.0065079 | 0.99480747 | 0.99989001 |
| SUZ12 | 290.666563 | -0.0277107 | 0.24746205 | -0.1119795 | 0.91083966 | 0.99989001 |
| LRRC37B | 97.6335543 | 0.33357384 | 0.42518853 | 0.7845316 | 0.43272825 | 0.99989001 |

| | | | | | | |
|---|---|---|---|---|---|---|
| RHOT1 | 214.271086 | 0.00219513 | 0.28306824 | 0.00775477 | 0.99381265 | 0.99989001 |
| ZNF207 | 496.549427 | 0.07461901 | 0.20171852 | 0.36991654 | 0.71144468 | 0.99989001 |
| PSMD11 | 154.24538 | -0.0800447 | 0.27635664 | -0.2896429 | 0.77208943 | 0.99989001 |
| MYO1D | 547.708708 | -0.4007053 | 0.37723114 | -1.0622275 | 0.28813241 | 0.99989001 |
| ASIC2 | 93.4233053 | 1.89216255 | 0.7607981 | 2.48707581 | 0.01287979 | 0.49505934 |
| CCL2 | 234.615547 | -0.1609853 | 1.18048852 | -0.1363717 | 0.89152742 | 0.99989001 |
| SLFN5 | 298.053974 | 0.408845 | 0.28497602 | 1.43466454 | 0.15138271 | 0.99989001 |
| SLFN11 | 117.320877 | 0.70956891 | 0.30170689 | 2.35184852 | 0.01868038 | 0.56643441 |
| SLFN12 | 57.6803743 | 0.52733385 | 0.36967157 | 1.42649284 | 0.15372613 | 0.99989001 |
| AP2B1 | 308.268038 | -0.149988 | 0.25506586 | -0.5880365 | 0.5565078 | 0.99989001 |
| TAF15 | 521.758142 | -0.0573437 | 0.16884624 | -0.3396207 | 0.7341422 | 0.99989001 |
| MMP28 | 46.0591985 | 0.16886064 | 0.78490705 | 0.21513457 | 0.8296624 | NA |
| MYO19 | 105.205851 | -0.5009119 | 0.33179495 | -1.5097031 | 0.1311192 | 0.99989001 |
| GGNBP2 | 226.55222 | -0.0492616 | 0.19279107 | -0.2555183 | 0.79832281 | 0.99989001 |
| AATF | 158.385137 | 0.06065215 | 0.2435707 | 0.2490125 | 0.80335111 | 0.99989001 |
| ACACA | 285.786472 | -0.211344 | 0.32091638 | -0.6585642 | 0.51017564 | 0.99989001 |
| TADA2A | 110.756213 | -0.1464348 | 0.27612635 | -0.5303181 | 0.59589137 | 0.99989001 |
| SYNRG | 225.230209 | 0.18524067 | 0.17301481 | 1.07066367 | 0.28432068 | 0.99989001 |
| DDX52 | 109.780376 | 0.06166229 | 0.25604755 | 0.24082357 | 0.80969186 | 0.99989001 |
| MRPL45 | 49.8037873 | -0.1549646 | 0.24522984 | -0.6319159 | 0.52744184 | NA |
| SOCS7 | 47.9379599 | 0.14370679 | 0.32594545 | 0.44089216 | 0.65929107 | NA |
| ARHGAP23 | 217.246152 | -0.5886376 | 0.39015603 | -1.5087237 | 0.13136941 | 0.99989001 |
| MLLT6 | 205.149205 | -0.3072663 | 0.25692901 | -1.1959191 | 0.23172812 | 0.99989001 |
| CISD3 | 48.2620734 | 0.14138546 | 0.37753565 | 0.37449566 | 0.70803559 | NA |
| PCGF2 | 145.72719 | -0.2390036 | 0.33862042 | -0.7058156 | 0.48030281 | 0.99989001 |
| PSMB3 | 117.55927 | -0.2965978 | 0.39917625 | -0.7430247 | 0.45746675 | 0.99989001 |
| PIP4K2B | 94.7076015 | 0.08665268 | 0.28695172 | 0.3019765 | 0.76266998 | 0.99989001 |
| CWC25 | 67.1855535 | -0.1977368 | 0.26329102 | -0.7510199 | 0.45264067 | 0.99989001 |
| RPL23 | 265.115297 | 0.12472549 | 0.54455585 | 0.22904076 | 0.81883723 | 0.99989001 |
| LASP1 | 158.53868 | 0.04191569 | 0.24217562 | 0.17307974 | 0.86258875 | 0.99989001 |
| RPL19 | 520.468171 | -0.1431236 | 0.59974592 | -0.2386404 | 0.81138447 | 0.99989001 |
| FBXL20 | 331.949679 | -0.0124518 | 0.26689975 | -0.0466534 | 0.96278947 | 0.99989001 |
| MED1 | 106.670848 | 0.15247118 | 0.27583313 | 0.55276599 | 0.58042366 | 0.99989001 |
| CDK12 | 219.562454 | 0.01778127 | 0.2520846 | 0.07053693 | 0.94376631 | 0.99989001 |
| STARD3 | 77.0689574 | 0.22686898 | 0.24427209 | 0.92875523 | 0.35301595 | 0.99989001 |
| ERBB2 | 46.4092373 | 0.72566255 | 0.36301508 | 1.99898735 | 0.04560972 | NA |
| PSMD3 | 83.6261416 | -0.0766422 | 0.31183047 | -0.2457817 | 0.80585123 | 0.99989001 |
| MED24 | 71.4978789 | 0.27795728 | 0.25405869 | 1.09406721 | 0.2739255 | 0.99989001 |
| THRA | 68.1410707 | 0.41636527 | 0.50286721 | 0.82798254 | 0.40768039 | 0.99989001 |
| MSL1 | 135.184018 | 0.16651983 | 0.2465356 | 0.6754393 | 0.49939671 | 0.99989001 |
| CASC3 | 143.902913 | -0.0256886 | 0.3102454 | -0.082801 | 0.93400975 | 0.99989001 |
| WIPF2 | 262.162209 | -0.007774 | 0.26467184 | -0.0293722 | 0.97656777 | 0.99989001 |
| RARA | 65.0201322 | -0.0557127 | 0.36891555 | -0.1510175 | 0.87996193 | 0.99989001 |
| IGFBP4 | 1367.72856 | -0.0489533 | 0.59162869 | -0.0827432 | 0.9340557 | 0.99989001 |
| KRT10 | 104.754885 | -0.75144 | 0.55040445 | -1.3652506 | 0.17217431 | 0.99989001 |
| EIF1 | 741.471209 | 0.03785751 | 0.38905362 | 0.09730666 | 0.92248287 | 0.99989001 |
| JUP | 234.001948 | 0.54249808 | 0.39087954 | 1.38789069 | 0.16517033 | 0.99989001 |
| FKBP10 | 59.078969 | -0.30281 | 0.41132209 | -0.7361871 | 0.46161685 | 0.99989001 |
| ACLY | 124.685887 | -0.0428074 | 0.30639988 | -0.1397108 | 0.88888853 | 0.99989001 |
| CNP | 62.0613753 | 0.10265543 | 0.31925175 | 0.32155009 | 0.74779356 | 0.99989001 |
| DNAJC7 | 208.037558 | -0.1124687 | 0.20054154 | -0.5608249 | 0.57491691 | 0.99989001 |
| NKIRAS2 | 45.6148188 | 0.29464946 | 0.3903308 | 0.75487115 | 0.4503263 | NA |
| KAT2A | 51.3850634 | -1.0225358 | 0.47972883 | -2.1314872 | 0.03304903 | 0.67281094 |
| STAT5B | 396.435027 | -0.1015877 | 0.20391692 | -0.4981816 | 0.61835602 | 0.99989001 |

| | | | | | | |
|---|---|---|---|---|---|---|
| STAT5A | 101.275196 | -0.4437655 | 0.32193295 | -1.3784406 | 0.16806728 | 0.99989001 |
| STAT3 | 990.212112 | -0.0262618 | 0.30955113 | -0.0848382 | 0.93239003 | 0.99989001 |
| CAVIN1 | 349.193092 | -0.1499024 | 0.46737883 | -0.32073 | 0.74841502 | 0.99989001 |
| ATP6V0A1 | 180.641727 | -0.2579898 | 0.28215641 | -0.9143504 | 0.36053278 | 0.99989001 |
| NAGLU | 52.4767783 | -0.826309 | 0.35488022 | -2.3284166 | 0.01988999 | 0.57965116 |
| MLX | 40.3679914 | 0.13989525 | 0.34268953 | 0.4082274 | 0.68310674 | NA |
| RETREG3 | 114.329773 | 0.22128972 | 0.26996081 | 0.81971051 | 0.41238116 | 0.99989001 |
| TUBG2 | 42.1816957 | -0.0985496 | 0.32021732 | -0.3077585 | 0.75826611 | NA |
| EZH1 | 158.518021 | 0.07976238 | 0.2275737 | 0.35049035 | 0.72597073 | 0.99989001 |
| RAMP2-AS1 | 41.5505791 | 0.16304706 | 0.44000399 | 0.37055813 | 0.71096667 | NA |
| RAMP2 | 457.145431 | -0.23978 | 0.74306536 | -0.3226903 | 0.74692976 | 0.99989001 |
| VPS25 | 40.1501357 | -0.0785833 | 0.33857214 | -0.232102 | 0.81645882 | NA |
| COA3 | 63.5921457 | 0.33646381 | 0.44569909 | 0.75491249 | 0.45030149 | 0.99989001 |
| BECN1 | 74.855608 | 0.1448196 | 0.26684445 | 0.54271166 | 0.58732835 | 0.99989001 |
| RUNDC1 | 39.6135274 | -0.4256454 | 0.28005776 | -1.5198487 | 0.128549 | NA |
| RPL27 | 205.455092 | -0.0912609 | 0.58890924 | -0.154966 | 0.87684812 | 0.99989001 |
| IFI35 | 81.2870009 | -0.2895349 | 0.49722615 | -0.5823001 | 0.56036453 | 0.99989001 |
| VAT1 | 122.571008 | -0.2070448 | 0.49003885 | -0.4225069 | 0.67265505 | 0.99989001 |
| BRCA1 | 58.6552024 | 0.13652872 | 0.39302752 | 0.34737701 | 0.7283081 | 0.99989001 |
| NBR1 | 180.770458 | 0.34168327 | 0.22136626 | 1.54352014 | 0.12270463 | 0.99989001 |
| LINC00854 | 179.193655 | -1.4312086 | 0.66923079 | -2.1385875 | 0.03246909 | 0.67281094 |
| DHX8 | 82.3862079 | -0.0068137 | 0.26888368 | -0.0253407 | 0.97978325 | 0.99989001 |
| SOST | 67.0762511 | 4.76234799 | 1.05207491 | 4.52662445 | 5.99E-06 | 0.00411251 |
| DUSP3 | 98.4754328 | -0.5021719 | 0.29429278 | -1.7063685 | 0.08793949 | 0.96759079 |
| MPP3 | 51.1124551 | -1.0619564 | 0.36057287 | -2.9451922 | 0.00322754 | 0.29001765 |
| TMEM101 | 46.456959 | -0.258288 | 0.36227509 | -0.7129609 | 0.47586998 | NA |
| LSM12 | 97.3302507 | -0.1774883 | 0.26137877 | -0.6790462 | 0.49710856 | 0.99989001 |
| G6PC3 | 40.2333144 | -0.7090218 | 0.43868094 | -1.6162585 | 0.10603844 | NA |
| HDAC5 | 93.4734136 | 0.14874821 | 0.23609836 | 0.63002647 | 5.29E-01 | 0.99989001 |
| TMUB2 | 48.8109972 | 0.11184927 | 0.40759555 | 0.2744124 | 0.78376771 | NA |
| ATXN7L3 | 41.6572172 | -0.2766429 | 0.28892901 | -0.957477 | 0.33832655 | NA |
| UBTF | 92.4775722 | -0.1241193 | 0.31278075 | -0.3968251 | 0.69149644 | 0.99989001 |
| SLC25A39 | 49.6048759 | -0.079742 | 0.4506461 | -0.1769504 | 0.85954736 | NA |
| GRN | 227.295597 | -0.0163632 | 0.46903391 | -0.0348869 | 0.9721699 | 0.99989001 |
| GPATCH8 | 596.736485 | 0.12640782 | 0.25948768 | 0.48714383 | 0.62615641 | 0.99989001 |
| GJC1 | 64.6554974 | 0.44257521 | 0.42851098 | 1.03282118 | 0.30168759 | 0.99989001 |
| EFTUD2 | 96.1812124 | -0.2233813 | 0.25324593 | -0.8820726 | 0.37773754 | 0.99989001 |
| DCAKD | 56.1349386 | -0.0752633 | 0.32272851 | -0.2332095 | 0.81559874 | 0.99989001 |
| NMT1 | 149.463341 | -0.1072596 | 0.20929427 | -0.5124825 | 0.60831338 | 0.99989001 |
| AC138150.1 | 61.2730509 | -0.0103669 | 0.40668939 | -0.025491 | 0.9796633 | 0.99989001 |
| MAP3K14 | 132.207425 | 0.42158363 | 0.34254004 | 1.23075724 | 0.21841367 | 0.99989001 |
| ARHGAP27 | 119.29763 | -0.582055 | 0.37603506 | -1.5478742 | 0.12165259 | 0.99989001 |
| PLEKHM1 | 128.72195 | -0.1023552 | 0.30630294 | -0.3341632 | 0.73825645 | 0.99989001 |
| KANSL1 | 1310.54369 | 0.27757068 | 0.30801152 | 0.90116981 | 0.36749804 | 0.99989001 |
| ARL17B | 281.518741 | 0.84159285 | 1.23914827 | 0.67917042 | 0.49702989 | 0.99989001 |
| LRRC37A2 | 61.114021 | 0.07849271 | 0.34866738 | 0.22512202 | 0.82188435 | 0.99989001 |
| ARL17A | 78.4422585 | -0.3711742 | 0.50297629 | -0.7379557 | 0.46054136 | 0.99989001 |
| NSF | 318.528672 | 0.2232886 | 0.3468696 | 0.6437249 | 0.51975384 | 0.99989001 |
| CDC27 | 563.57707 | -0.1659173 | 0.18194178 | -0.911925 | 0.36180818 | 0.99989001 |
| AC068234.1 | 95.1880609 | -0.7939093 | 0.74397468 | -1.0671187 | 0.28591823 | 0.99989001 |
| EFCAB13 | 109.562675 | 0.39984691 | 0.32905312 | 1.21514396 | 0.22431119 | 0.99989001 |
| NPEPPS | 431.819842 | -0.0599349 | 0.17684034 | -0.338921 | 0.73466925 | 0.99989001 |
| AC025682.1 | 59.9103156 | 0.61251053 | 0.41217262 | 1.48605343 | 0.13726499 | 0.99989001 |
| KPNB1 | 332.236139 | 0.07610584 | 0.29591235 | 0.25719047 | 0.79703174 | 0.99989001 |

| | | | | | | |
|---|---|---|---|---|---|---|
| TBKBP1 | 44.3037016 | -0.1916244 | 0.28814287 | -0.6650327 | 0.50602957 | NA |
| SCRN2 | 39.54423 | 0.2229058 | 0.33309163 | 0.66920264 | 0.50336622 | NA |
| SP2 | 39.2848371 | -0.1388298 | 0.4507585 | -0.3079915 | 0.75808882 | NA |
| CDK5RAP3 | 55.7686969 | 0.00928197 | 0.26868808 | 0.03454552 | 0.97244214 | 0.99989001 |
| NFE2L1 | 173.485165 | 0.2014065 | 0.27800701 | 0.72446556 | 0.46877997 | 0.99989001 |
| CBX1 | 65.8616476 | -0.271352 | 0.30505357 | -0.8895224 | 0.37372238 | 0.99989001 |
| HOXB2 | 61.7130853 | -0.2529571 | 0.55776581 | -0.4535184 | 0.65017549 | 0.99989001 |
| HOXB3 | 292.20229 | 0.20066586 | 0.36369505 | 0.55174206 | 0.58112509 | 0.99989001 |
| HOXB-AS3 | 53.2847976 | 0.24809532 | 0.46202863 | 0.53696957 | 0.59128863 | 0.99989001 |
| HOXB7 | 60.9974139 | -0.0966894 | 0.44936606 | -0.2151684 | 0.82963604 | 0.99989001 |
| CALCOCO2 | 301.013977 | -0.06224 | 0.33619227 | -0.1851321 | 0.85312539 | 0.99989001 |
| ATP5MC1 | 73.5334736 | 0.10573499 | 0.42123689 | 0.25101076 | 0.80180579 | 0.99989001 |
| UBE2Z | 190.381705 | 0.03603145 | 0.20332664 | 0.1772097 | 0.85934368 | 0.99989001 |
| SNF8 | 79.9328757 | -0.157143 | 0.29788738 | -0.5275248 | 0.5978292 | 0.99989001 |
| ABI3 | 59.0892577 | 0.24159811 | 0.45562152 | 0.53026053 | 0.59593131 | 0.99989001 |
| ZNF652 | 253.080363 | 0.48404082 | 0.22545015 | 2.1469971 | 0.03179351 | 0.67281094 |
| PHB | 98.9920712 | -0.0856109 | 0.39140798 | -0.2187254 | 0.82686395 | 0.99989001 |
| SPOP | 257.813818 | 0.21702567 | 0.19143463 | 1.13368031 | 0.25692867 | 0.99989001 |
| KAT7 | 83.8086257 | 0.11961082 | 0.2516379 | 0.47532909 | 0.63455243 | 0.99989001 |
| ITGA3 | 201.781966 | -0.2830715 | 0.34851757 | -0.8122158 | 0.41666781 | 0.99989001 |
| PPP1R9B | 48.789618 | 0.24922492 | 0.35602947 | 0.70001205 | 0.48391978 | NA |
| MRPL27 | 57.496361 | 0.04510921 | 0.3632691 | 0.12417575 | 0.90117612 | 0.99989001 |
| LRRC59 | 48.854484 | -0.1398787 | 0.49633098 | -0.2818255 | 0.7780773 | NA |
| ACSF2 | 44.5711334 | 1.0717379 | 0.40418292 | 2.65161597 | 0.00801076 | NA |
| RSAD1 | 46.2147311 | -0.6350613 | 0.34886142 | -1.8203827 | 0.06870075 | NA |
| ABCC3 | 55.4102743 | 0.51279166 | 0.4365757 | 1.17457673 | 2.40E-01 | 0.99989001 |
| ANKRD40 | 80.4796711 | -0.2093359 | 0.34719407 | -0.6029363 | 0.54655109 | 0.99989001 |
| LUC7L3 | 513.574737 | 0.0211437 | 0.18648468 | 0.11338035 | 0.90972902 | 0.99989001 |
| SPAG9 | 970.657809 | -0.0411522 | 0.30512723 | -0.134869 | 0.89271549 | 0.99989001 |
| AC005920.2 | 59.7384232 | -0.135479 | 0.7021762 | -0.1929416 | 0.84700467 | 0.99989001 |
| MBTD1 | 256.992929 | 0.00972819 | 0.23421304 | 0.04153565 | 0.96686888 | 0.99989001 |
| UTP18 | 68.6449997 | -0.4668955 | 0.37080576 | -1.2591378 | 0.20798058 | 0.99989001 |
| TOM1L1 | 57.9257972 | 0.39687882 | 0.35097041 | 1.13080422 | 0.2581375 | 0.99989001 |
| COX11 | 61.7230328 | 0.20196158 | 0.28649528 | 0.70493858 | 0.48084846 | 0.99989001 |
| STXBP4 | 119.890546 | -0.1610005 | 0.2678391 | -0.601109 | 0.54776736 | 0.99989001 |
| PCTP | 54.6277156 | 1.05048158 | 0.43151443 | 2.43440658 | 0.01491623 | 0.51942334 |
| C17orf67 | 79.6977613 | 1.06811685 | 0.34720139 | 3.07636111 | 0.00209544 | 0.21666265 |
| DGKE | 221.785979 | 0.69752474 | 0.32229504 | 2.16424285 | 0.03044572 | 0.67281094 |
| TRIM25 | 102.758 | -0.0424846 | 0.44324252 | -0.0958495 | 0.92364009 | 0.99989001 |
| COIL | 44.9507882 | -0.1147684 | 0.31456895 | -0.3648432 | 0.71522844 | NA |
| SCPEP1 | 57.8144255 | 0.19256398 | 0.38504547 | 0.50010711 | 0.61699966 | 0.99989001 |
| AKAP1 | 50.2537692 | -0.0041724 | 0.34254222 | -0.0121808 | 0.99028138 | 0.99989001 |
| MSI2 | 1110.8156 | 0.56641433 | 0.41061371 | 1.37943354 | 0.16776112 | 0.99989001 |
| MRPS23 | 47.6932318 | -0.0188363 | 0.34546646 | -0.0545242 | 0.9565175 | NA |
| CUEDC1 | 170.65928 | -0.1154826 | 0.3765843 | -0.3066581 | 0.75910365 | 0.99989001 |
| VEZF1 | 123.722254 | 0.4209278 | 0.21059352 | 1.99876903 | 0.04563335 | 0.77228819 |
| AC015813.2 | 50.5061175 | 0.04544436 | 0.2817693 | 0.16128214 | 0.87187119 | 0.99989001 |
| DYNLL2 | 58.1869423 | 0.24775312 | 0.25638075 | 0.96634838 | 0.33386988 | 0.99989001 |
| SUPT4H1 | 44.4595018 | -0.0956125 | 0.45514023 | -0.2100727 | 0.83361097 | NA |
| MTMR4 | 47.3338437 | 0.27261737 | 0.26763372 | 1.01862116 | 0.30838285 | NA |
| RAD51C | 53.0406047 | -0.1241178 | 0.27024512 | -0.4592787 | 0.64603405 | 0.99989001 |
| TRIM37 | 117.861442 | -0.3952136 | 0.20433531 | -1.9341425 | 0.05309561 | 0.81308738 |
| SKA2 | 72.325301 | -0.0193913 | 0.22128259 | -0.0876316 | 0.93016949 | 0.99989001 |
| YPEL2 | 176.303941 | 0.07379261 | 0.23727609 | 0.31099895 | 0.75580142 | 0.99989001 |

| | | | | | | |
|---|---|---|---|---|---|---|
| DHX40 | 85.1211575 | -0.1419044 | 0.34995458 | -0.4054939 | 0.6851145 | 0.99989001 |
| CLTC | 454.222579 | -0.0004162 | 0.19925845 | -0.0020889 | 0.99833327 | 0.99989001 |
| PTRH2 | 81.0964768 | -0.0674288 | 0.29262354 | -0.2304285 | 0.81775884 | 0.99989001 |
| VMP1 | 588.955929 | -0.494728 | 0.38405714 | -1.2881625 | 0.1976894 | 0.99989001 |
| TUBD1 | 51.804914 | 0.13349707 | 0.29567131 | 0.45150497 | 0.65162565 | 0.99989001 |
| RPS6KB1 | 110.124877 | -0.1779979 | 0.31985153 | -0.5565017 | 0.57786794 | 0.99989001 |
| CA4 | 74.701573 | 1.66855846 | 0.82747924 | 2.01643545 | 0.04375446 | 0.76983375 |
| USP32 | 390.785754 | -0.2528455 | 0.24039754 | -1.0517806 | 0.29290023 | 0.99989001 |
| APPBP2 | 363.359439 | 0.12785875 | 0.15785501 | 0.80997586 | 0.41795405 | 0.99989001 |
| PPM1D | 245.267311 | 0.04339804 | 0.17404091 | 0.24935538 | 0.8030859 | 0.99989001 |
| BCAS3 | 1033.24004 | -0.1119121 | 0.18841035 | -0.5939809 | 0.5525249 | 0.99989001 |
| INTS2 | 66.5236805 | -0.645395 | 0.29516415 | -2.1865629 | 0.02877445 | 0.65805899 |
| MED13 | 628.230343 | -0.0228756 | 0.18034435 | -0.1268438 | 0.89906406 | 0.99989001 |
| TLK2 | 412.479505 | 0.11808 | 0.19485404 | 0.60599205 | 0.54452003 | 0.99989001 |
| MRC2 | 44.846211 | 0.66625955 | 0.51793757 | 1.28637037 | 0.19831383 | NA |
| TANC2 | 1019.62892 | 0.49459174 | 0.39026212 | 1.26733215 | 0.20503655 | 0.99989001 |
| AC015923.1 | 60.4662523 | 0.48300106 | 0.56289309 | 0.85806891 | 0.39085441 | 0.99989001 |
| DCAF7 | 198.960257 | 0.21312058 | 0.24591286 | 0.86665083 | 0.38613336 | 0.99989001 |
| MAP3K3 | 463.595373 | 0.10517073 | 0.26665893 | 0.39440168 | 0.6932845 | 0.99989001 |
| CCDC47 | 67.5191707 | -0.249783 | 0.33621324 | -0.7429303 | 0.4575239 | 0.99989001 |
| DDX42 | 180.170813 | -0.1122886 | 0.17071316 | -0.657762 | 0.51069108 | 0.99989001 |
| PSMC5 | 51.1512513 | 0.07195536 | 0.48104066 | 0.14958271 | 0.88109385 | 0.99989001 |
| SMARCD2 | 71.2783091 | -0.0890313 | 0.3461928 | -0.2571726 | 0.79704552 | 0.99989001 |
| ICAM2 | 423.389059 | -0.9019313 | 0.40838428 | -2.2085357 | 0.02720695 | 0.64986727 |
| ERN1 | 82.9085533 | 0.16954993 | 0.31377269 | 0.54035911 | 0.5889494 | 0.99989001 |
| SNHG25 | 45.2379926 | -0.1936479 | 0.46397285 | -0.417369 | 0.67640854 | NA |
| TEX2 | 99.9540028 | 0.00712871 | 0.25179865 | 0.02831116 | 0.97741398 | 0.99989001 |
| PECAM1 | 3023.74571 | 0.19511269 | 0.34126734 | 0.57172974 | 0.56750509 | 0.99989001 |
| POLG2 | 50.4036057 | -0.2492382 | 0.27069837 | -0.9207228 | 0.35719516 | 0.99989001 |
| DDX5 | 1142.39375 | 0.30624097 | 0.25021264 | 1.22392288 | 0.22098132 | 0.99989001 |
| CEP95 | 138.627644 | -0.0937386 | 0.28070432 | -0.3339406 | 0.73842441 | 0.99989001 |
| SMURF2 | 663.249446 | 0.02467418 | 0.31805983 | 0.07757718 | 0.93816439 | 0.99989001 |
| LRRC37A3 | 247.953031 | 0.22663067 | 0.36775238 | 0.61625888 | 0.53772367 | 0.99989001 |
| GNA13 | 290.099208 | 0.26792207 | 0.4556662 | 0.58797881 | 0.55654651 | 0.99989001 |
| RGS9 | 46.3300523 | 1.35590211 | 0.54614652 | 2.48267098 | 0.01304015 | NA |
| CEP112 | 935.612158 | -0.199145 | 0.25658916 | -0.7761241 | 0.43767574 | 0.99989001 |
| PRKCA | 882.676923 | 0.3328676 | 0.43951649 | 0.75734953 | 0.44884049 | 0.99989001 |
| HELZ | 480.544405 | -0.1715298 | 0.2242183 | -0.7650127 | 0.44426399 | 0.99989001 |
| PSMD12 | 135.107944 | -0.0447982 | 0.24057956 | -0.1862096 | 0.85228036 | 0.99989001 |
| PITPNC1 | 2856.64086 | -0.0593588 | 0.42281517 | -0.1403895 | 0.88835227 | 0.99989001 |
| NOL11 | 61.4704656 | -0.4809239 | 0.28586971 | -1.6823184 | 0.09250712 | 0.99464924 |
| BPTF | 683.463878 | -0.154459 | 0.25623736 | -0.6027964 | 0.54664413 | 0.99989001 |
| KPNA2 | 55.6482256 | 0.00712817 | 0.50276215 | 0.01417801 | 0.98868797 | 0.99989001 |
| AC005332.7 | 58.1628357 | -0.4963695 | 0.39569755 | -1.2544163 | 0.20969073 | 0.99989001 |
| ARSG | 76.9543628 | 0.01737197 | 0.46380963 | 0.03745496 | 0.97012225 | 0.99989001 |
| WIPI1 | 137.701539 | 0.10564856 | 0.38997168 | 0.27091342 | 0.78645762 | 0.99989001 |
| PRKAR1A | 343.252879 | 0.16985843 | 0.33649366 | 0.50478939 | 0.61370677 | 0.99989001 |
| ABCA5 | 90.7875215 | -0.4703766 | 0.47668478 | -0.9867666 | 0.32375705 | 0.99989001 |
| MAP2K6 | 91.9987767 | -0.406545 | 0.69880303 | -0.5817733 | 0.56071937 | 0.99989001 |
| KCNJ16 | 56.8370893 | 0.00532481 | 0.4459551 | 0.01194024 | 0.9904733 | 0.99989001 |
| KCNJ2 | 53.1677709 | -0.9707346 | 0.59603025 | -1.6286667 | 0.1033836 | 0.99989001 |
| SLC39A11 | 244.137143 | -0.1537991 | 0.41658007 | -0.3691946 | 0.71198265 | 0.99989001 |
| FAM104A | 64.030651 | -0.00256 | 0.23036325 | -0.0111129 | 0.99113337 | 0.99989001 |
| CDC42EP4 | 121.132081 | -0.0177616 | 0.23546066 | -0.0754336 | 0.93986972 | 0.99989001 |

| | | | | | | |
|---|---|---|---|---|---|---|
| RPL38 | 168.263064 | 0.00036143 | 0.51093285 | 0.00070739 | 0.99943559 | 0.99989001 |
| HID1 | 73.8466463 | 0.57147067 | 0.46815506 | 1.22068674 | 0.22220465 | 0.99989001 |
| CDR2L | 48.5630252 | 0.34744783 | 0.29929976 | 1.16086907 | 0.24569515 | NA |
| KCTD2 | 45.3249422 | -0.056122 | 0.26551256 | -0.2113723 | 0.83259679 | NA |
| ATP5PD | 53.7365594 | -0.0517154 | 0.38979551 | -0.132673 | 0.89445198 | 0.99989001 |
| JPT1 | 59.9288862 | 0.15941427 | 0.31070637 | 0.5130705 | 0.60790201 | 0.99989001 |
| SUMO2 | 67.817173 | -0.1501925 | 0.2904613 | -0.5170827 | 0.60509844 | 0.99989001 |
| NUP85 | 81.466805 | -0.124093 | 0.26438838 | -0.4693587 | 0.63881324 | 0.99989001 |
| GGA3 | 83.6274932 | -0.0931454 | 0.26656814 | -0.3494245 | 0.72677062 | 0.99989001 |
| MRPS7 | 49.6864571 | -0.0390146 | 0.49716722 | -0.0784738 | 0.93745119 | NA |
| GRB2 | 308.056245 | -0.0845625 | 0.20627626 | -0.4099478 | 0.68184427 | 0.99989001 |
| TMEM94 | 98.693859 | 0.08132219 | 0.30682105 | 0.26504762 | 0.79097277 | 0.99989001 |
| CASKIN2 | 237.196235 | -0.2175339 | 0.41848535 | -0.5198125 | 0.60319426 | 0.99989001 |
| LLGL2 | 70.0790418 | 0.11514466 | 0.30835829 | 0.37341192 | 0.70884189 | 0.99989001 |
| MYO15B | 233.32393 | 0.07079872 | 0.35077266 | 0.20183649 | 0.84004455 | 0.99989001 |
| SAP30BP | 136.699457 | 0.03687555 | 0.22595495 | 0.1631987 | 0.87036199 | 0.99989001 |
| ITGB4 | 102.21261 | 0.18270177 | 0.60042024 | 0.30428982 | 0.7609071 | 0.99989001 |
| H3F3B | 902.974797 | 0.17009774 | 0.38745018 | 0.43901837 | 0.66064823 | 0.99989001 |
| UNK | 188.300298 | 0.0369107 | 0.29028665 | 0.12715259 | 0.89881963 | 0.99989001 |
| WBP2 | 133.496668 | -0.0991661 | 0.28536997 | -0.3475003 | 0.72821553 | 0.99989001 |
| TRIM47 | 60.2178054 | -0.1030732 | 0.46648435 | -0.2209574 | 0.82512558 | 0.99989001 |
| TRIM65 | 43.6784544 | 0.2190865 | 0.30678793 | 0.71413011 | 0.47514673 | NA |
| ACOX1 | 116.033018 | 0.2016221 | 0.24840135 | 0.81167878 | 0.41697597 | 0.99989001 |
| SRP68 | 60.9618494 | -0.1608058 | 0.25792231 | -0.6234659 | 0.53297838 | 0.99989001 |
| EXOC7 | 93.654139 | 0.18560823 | 0.21916062 | 0.84690499 | 0.39704808 | 0.99989001 |
| UBALD2 | 80.3845137 | -0.1156092 | 0.33093992 | -0.349336 | 0.72683711 | 0.99989001 |
| PRPSAP1 | 103.298417 | 0.24080646 | 0.25320253 | 0.95104288 | 0.34158261 | 0.99989001 |
| UBE2O | 105.206602 | 0.04064354 | 0.41771319 | 0.09730011 | 0.92248807 | 0.99989001 |
| RHBDF2 | 100.697045 | 0.92683473 | 0.43115395 | 2.1496608 | 0.03158206 | 0.67281094 |
| MXRA7 | 146.832938 | -0.6354613 | 0.31763946 | -2.000574 | 0.04543832 | 0.77228819 |
| JMJD6 | 46.1234189 | -0.1681813 | 0.29702007 | -0.5662287 | 0.5712383 | NA |
| SRSF2 | 198.774722 | 0.01078932 | 0.30290088 | 0.03561997 | 0.97158539 | 0.99989001 |
| MFSD11 | 167.585671 | -0.1657524 | 0.28420077 | -0.583223 | 0.55974316 | 0.99989001 |
| SEC14L1 | 2227.1736 | -0.4672969 | 0.3576732 | -1.3064911 | 0.1913856 | 0.99989001 |
| 9-Sep | 427.820085 | -0.2658906 | 0.32920942 | -0.807664 | 0.41928404 | 0.99989001 |
| TNRC6C | 229.391404 | 0.27701746 | 0.29240609 | 0.94737242 | 0.34344904 | 0.99989001 |
| TMC6 | 85.4638282 | -0.1283202 | 0.34748964 | -0.3692777 | 0.71192076 | 0.99989001 |
| SYNGR2 | 97.3479739 | -0.0388553 | 0.52561922 | -0.073923 | 0.94107167 | 0.99989001 |
| AFMID | 96.8151541 | -0.2166352 | 0.55771435 | -0.388434 | 0.69769486 | 0.99989001 |
| SOCS3 | 379.092797 | -0.8653873 | 0.90567631 | -0.955515 | 0.33931734 | 0.99989001 |
| PGS1 | 223.149668 | -0.4329711 | 0.31454679 | -1.3764919 | 0.1686694 | 0.99989001 |
| CYTH1 | 706.77421 | 0.21777234 | 0.34859733 | 0.6247103 | 0.53216121 | 0.99989001 |
| USP36 | 198.036483 | -0.6450617 | 0.4084559 | -1.5792689 | 0.1142744 | 0.99989001 |
| TIMP2 | 469.402071 | -0.042231 | 0.49784539 | -0.0848276 | 0.93239845 | 0.99989001 |
| LGALS3BP | 92.820483 | 0.3192053 | 0.66115526 | 0.4827993 | 0.62923825 | 0.99989001 |
| CANT1 | 42.3422059 | 0.54139515 | 0.33069883 | 1.63712447 | 0.10160446 | NA |
| C1QTNF1 | 95.0971696 | -0.6614629 | 0.74091231 | -0.8927682 | 0.37198133 | 0.99989001 |
| TBC1D16 | 138.510979 | 0.22752835 | 0.31051843 | 0.73273704 | 0.46371883 | 0.99989001 |
| GAA | 50.6713202 | -0.2131895 | 0.43225204 | -0.4932064 | 0.62186673 | 0.99989001 |
| EIF4A3 | 49.1750394 | 0.29224092 | 0.37216121 | 0.78525358 | 0.43230491 | NA |
| SLC26A11 | 53.7076915 | 0.04241263 | 0.30823399 | 0.1375988 | 0.8905575 | 0.99989001 |
| RNF213 | 922.060738 | 0.06679993 | 0.46783012 | 0.14278673 | 0.88645862 | 0.99989001 |
| AC124319.2 | 132.043513 | 0.2450266 | 0.43024275 | 0.5695078 | 0.56901158 | 0.99989001 |
| ENDOV | 44.4348438 | -0.0575208 | 0.33311272 | -0.1726767 | 0.86290557 | NA |

| | | | | | | |
|---|---|---|---|---|---|---|
| RPTOR | 274.295763 | -0.1252394 | 0.28522504 | -0.4390899 | 0.66059643 | 0.99989001 |
| BAIAP2 | 88.1274932 | 0.38235146 | 0.46709883 | 0.81856651 | 0.41303378 | 0.99989001 |
| AATK | 42.3847927 | 0.15007868 | 0.34581253 | 0.43398855 | 0.66429676 | NA |
| NDUFAF8 | 46.7446976 | -0.2588252 | 0.42830588 | -0.6042998 | 0.54564432 | NA |
| SLC38A10 | 142.858967 | -0.038663 | 0.23378427 | -0.1653791 | 0.86864558 | 0.99989001 |
| BAHCC1 | 44.7359173 | -0.1019756 | 0.38234215 | -0.2667128 | 0.78969027 | NA |
| ACTG1 | 1124.98306 | -0.3119773 | 0.50561623 | -0.6170239 | 0.53721894 | 0.99989001 |
| NPLOC4 | 264.656473 | -0.2672596 | 0.2413609 | -1.1073028 | 0.26816302 | 0.99989001 |
| CCDC137 | 47.2671664 | -0.2396717 | 0.31358014 | -0.7643078 | 0.44468386 | NA |
| ARL16 | 43.2277523 | 0.44425578 | 0.43611531 | 1.01866588 | 0.30836162 | NA |
| HGS | 86.1430997 | -0.5309038 | 0.30473747 | -1.7421676 | 0.08147911 | 0.94325821 |
| MCRIP1 | 87.7730122 | -0.1278626 | 0.39767535 | -0.3215251 | 0.74781247 | 0.99989001 |
| P4HB | 253.016847 | -0.2821224 | 0.40693798 | -0.693281 | 0.48813325 | 0.99989001 |
| ARHGDIA | 221.689102 | -0.132415 | 0.50511255 | -0.2621494 | 0.79320626 | 0.99989001 |
| ANAPC11 | 105.177926 | -0.0282538 | 0.53607416 | -0.0527051 | 0.95796686 | 0.99989001 |
| ASPSCR1 | 44.2474199 | -0.734192 | 0.44174342 | -1.6620326 | 0.09650624 | NA |
| CENPX | 41.6268147 | -0.7216207 | 0.4879051 | -1.4790186 | 0.13913535 | NA |
| GPS1 | 47.672117 | 0.0059975 | 0.4367283 | 0.0137328 | 0.98904316 | NA |
| DUS1L | 53.1552044 | -0.265321 | 0.32101871 | -0.8264971 | 0.40852215 | 0.99989001 |
| CCDC57 | 109.224913 | -0.3598473 | 0.31482811 | -1.1429962 | 0.25304017 | 0.99989001 |
| SLC16A3 | 101.722115 | -0.2527474 | 0.45514462 | -0.5553123 | 0.5786811 | 0.99989001 |
| CSNK1D | 188.599066 | -0.0384083 | 0.23547281 | -0.1631115 | 0.87043065 | 0.99989001 |
| OGFOD3 | 40.3543128 | -0.4613723 | 0.37520494 | -1.2296541 | 0.21882666 | NA |
| HEXDC | 74.0110484 | -0.3578629 | 0.36288656 | -0.9861564 | 0.32405637 | 0.99989001 |
| CYBC1 | 74.6545956 | -0.5673267 | 0.36300099 | -1.5628793 | 0.11808098 | 0.99989001 |
| NARF | 193.927267 | -0.3095379 | 0.23270277 | -1.3301858 | 0.18345708 | 0.99989001 |
| FOXK2 | 313.394375 | -0.1396968 | 0.20888393 | -0.6687773 | 0.50363757 | 0.99989001 |
| WDR45B | 201.424724 | -0.5386478 | 0.28852746 | -1.8668856 | 0.06191757 | 0.85940342 |
| TBCD | 668.946918 | -0.2372892 | 0.2609839 | -0.9092102 | 0.3632392 | 0.99989001 |
| METRNL | 93.6002571 | 0.60443013 | 0.60102221 | 1.00567021 | 0.31457424 | 0.99989001 |
| USP14 | 186.378365 | -0.1282492 | 0.27645819 | -0.4639008 | 0.64271884 | 0.99989001 |
| THOC1 | 165.844767 | -0.3340819 | 0.23269313 | -1.435719 | 0.15108233 | 0.99989001 |
| ENOSF1 | 318.406646 | -0.3707227 | 0.35063353 | -1.0572938 | 0.29037753 | 0.99989001 |
| YES1 | 845.45638 | -0.3060455 | 0.31960996 | -0.9575594 | 0.338285 | 0.99989001 |
| METTL4 | 100.173655 | -0.0594332 | 0.36634967 | -0.1622309 | 0.87112406 | 0.99989001 |
| SMCHD1 | 1260.80356 | 0.16940987 | 0.50801508 | 0.33347411 | 0.73877643 | 0.99989001 |
| AP001011.1 | 201.084428 | 0.46904901 | 0.61711351 | 0.76006927 | 0.44721318 | 0.99989001 |
| LPIN2 | 308.896293 | -0.5674401 | 0.3882099 | -1.4616837 | 0.14382792 | 0.99989001 |
| MYL12A | 665.419418 | -0.3145158 | 0.56454364 | -0.5571151 | 0.5774488 | 0.99989001 |
| MYL12B | 760.796133 | 0.0866288 | 0.61023032 | 0.14196083 | 0.88711095 | 0.99989001 |
| TGIF1 | 44.5388251 | -0.2276109 | 0.31221839 | -0.729012 | 0.46599433 | NA |
| DLGAP1 | 305.52261 | -0.7648072 | 0.53473638 | -1.4302509 | 0.15264503 | 0.99989001 |
| LINC00667 | 49.7955857 | -0.4042161 | 0.27027848 | -1.4955542 | 0.13476985 | NA |
| EPB41L3 | 467.83959 | 0.25627776 | 0.51457668 | 0.4980361 | 0.6184586 | 0.99989001 |
| L3MBTL4 | 391.264476 | -0.1906894 | 0.43460849 | -0.4387614 | 0.66083445 | 0.99989001 |
| ARHGAP28 | 148.253192 | 0.4456916 | 0.39187946 | 1.13731807 | 0.25540534 | 0.99989001 |
| PTPRM | 6370.50185 | -0.1469987 | 0.25791171 | -0.5699575 | 0.56870655 | 0.99989001 |
| RAB12 | 192.325407 | 0.08516518 | 0.18240169 | 0.46691006 | 0.64056422 | 0.99989001 |
| NDUFV2 | 150.05091 | 0.16933469 | 0.19459896 | 0.87017263 | 0.38420607 | 0.99989001 |
| ANKRD12 | 931.591466 | -0.03296 | 0.24430046 | -0.134916 | 0.8926783 | 0.99989001 |
| TWSG1 | 52.8772635 | 0.06751489 | 0.30795873 | 0.21923357 | 0.82646811 | 0.99989001 |
| RALBP1 | 135.116503 | 0.16072934 | 0.27237389 | 0.59010552 | 0.55511991 | 0.99989001 |
| PPP4R1 | 277.874472 | 0.07578841 | 0.18875438 | 0.40151866 | 0.68803831 | 0.99989001 |
| RAB31 | 149.78218 | 0.56495905 | 0.41330552 | 1.36692839 | 0.17164775 | 0.99989001 |

| | | | | | | |
|---|---|---|---|---|---|---|
| VAPA | 425.615294 | 0.05374936 | 0.25802143 | 0.20831357 | 0.83498414 | 0.99989001 |
| APCDD1 | 71.1381637 | -4.1046264 | 0.67486051 | -6.0821848 | 1.19E-09 | 4.47E-06 |
| NAPG | 128.257104 | -0.0164498 | 0.21089462 | -0.0780002 | 0.93782792 | 0.99989001 |
| PIEZO2 | 194.629204 | 1.51879477 | 0.42022935 | 3.61420441 | 0.00030127 | 0.07335471 |
| GNAL | 85.988373 | -0.1210634 | 0.50264632 | -0.240852 | 0.80966985 | 0.99989001 |
| MPPE1 | 73.5234392 | 0.04720067 | 0.28227787 | 0.16721352 | 0.86720205 | 0.99989001 |
| TUBB6 | 335.23906 | -0.4613131 | 0.35843825 | -1.2870085 | 0.19809131 | 0.99989001 |
| AFG3L2 | 103.638846 | -0.0212522 | 0.33398414 | -0.0636325 | 0.94926288 | 0.99989001 |
| SPIRE1 | 375.995966 | -0.1014633 | 0.31395314 | -0.3231797 | 0.74655913 | 0.99989001 |
| PSMG2 | 115.195645 | -0.3705689 | 0.23187581 | -1.5981354 | 0.11001284 | 0.99989001 |
| PTPN2 | 162.532584 | 0.14856109 | 0.30231771 | 0.49140717 | 0.6231385 | 0.99989001 |
| SEH1L | 76.0494952 | -0.4083505 | 0.26039058 | -1.5682229 | 0.11682912 | 0.99989001 |
| CEP192 | 281.672801 | -0.052479 | 0.25896644 | -0.2026481 | 0.83941011 | 0.99989001 |
| LDLRAD4 | 993.705193 | 0.21743531 | 0.49562997 | 0.43870493 | 0.66087536 | 0.99989001 |
| FAM210A | 103.83855 | -0.8243257 | 0.46397349 | -1.7766655 | 0.07562328 | 0.9206525 |
| RNMT | 179.738326 | -0.6477894 | 0.41215087 | -1.5717287 | 0.11601348 | 0.99989001 |
| ZNF519 | 47.6596456 | -0.6380106 | 0.32862131 | -1.9414767 | 0.05220049 | NA |
| ROCK1 | 657.981699 | 0.2001007 | 0.15276539 | 1.30985618 | 0.19024449 | 0.99989001 |
| ESCO1 | 133.212644 | 0.12494181 | 0.26287852 | 0.47528346 | 0.63458495 | 0.99989001 |
| SNRPD1 | 69.1390501 | -0.1487884 | 0.43619047 | -0.3411088 | 0.73302166 | 0.99989001 |
| ABHD3 | 98.0352216 | 0.60232743 | 0.26638942 | 2.26107869 | 0.02375438 | 0.61268871 |
| MIB1 | 410.334679 | 0.16046557 | 0.13791612 | 1.16350121 | 0.24462621 | 0.99989001 |
| GATA6 | 100.620982 | 0.49681639 | 0.38266422 | 1.29830896 | 0.19418119 | 0.99989001 |
| RBBP8 | 111.933533 | 0.05590896 | 0.27877697 | 0.20055084 | 0.8410498 | 0.99989001 |
| CABLES1 | 271.253968 | -0.6501639 | 0.60448488 | -1.0755669 | 0.28212098 | 0.99989001 |
| TMEM241 | 181.77657 | -0.1263398 | 0.32587663 | -0.3876922 | 0.6982438 | 0.99989001 |
| RIOK3 | 189.717342 | 0.21507588 | 0.21348228 | 1.00746478 | 0.31371148 | 0.99989001 |
| RMC1 | 56.2609469 | -0.4106068 | 0.27877081 | -1.4729189 | 0.14077291 | 0.99989001 |
| NPC1 | 155.212405 | -0.9088665 | 0.3217682 | -2.8246001 | 0.00473397 | 0.34759088 |
| ANKRD29 | 91.312726 | -2.5698417 | 0.75535117 | -3.4021814 | 0.0006685 | 0.1173455 |
| LAMA3 | 122.810869 | -2.9047335 | 0.79286624 | -3.6635858 | 0.00024871 | 0.07220208 |
| OSBPL1A | 291.758874 | -0.7379739 | 0.38925612 | -1.895857 | 0.05797895 | 0.8467548 |
| IMPACT | 43.8670881 | -0.5620092 | 0.33701366 | -1.6676154 | 0.0953921 | NA |
| ZNF521 | 925.687906 | 0.53081217 | 0.32797529 | 1.61845172 | 0.10556528 | 0.99989001 |
| SS18 | 235.240289 | 0.30878604 | 0.21758634 | 1.41914256 | 0.15585746 | 0.99989001 |
| TAF4B | 47.697237 | -0.1147267 | 0.40699792 | -0.2818852 | 0.77803154 | NA |
| KCTD1 | 79.4845831 | 0.76610716 | 0.364113 | 2.10403684 | 0.03537523 | 0.69174161 |
| CDH2 | 99.9576244 | -2.7988874 | 0.77947084 | -3.5907532 | 0.00032972 | 0.07777358 |
| TRAPPC8 | 275.427523 | -0.0586624 | 0.15276406 | -0.3840066 | 0.70097359 | 0.99989001 |
| RNF125 | 89.6909755 | -0.024789 | 0.4589329 | -0.0540145 | 0.95692359 | 0.99989001 |
| RNF138 | 144.393935 | -0.0343996 | 0.19997805 | -0.1720168 | 0.86342436 | 0.99989001 |
| GAREM1 | 96.2334769 | -0.4338082 | 0.30828457 | -1.4071682 | 0.15937752 | 0.99989001 |
| KLHL14 | 99.0680487 | -0.5424138 | 0.52212493 | -1.0388583 | 0.29887066 | 0.99989001 |
| CCDC178 | 145.098139 | -2.2615145 | 1.06392473 | -2.1256339 | 0.03353375 | 0.67281094 |
| ASXL3 | 52.9044321 | -0.9697565 | 0.63211112 | -1.5341552 | 0.12499148 | 0.99989001 |
| DTNA | 158.35734 | -2.4926816 | 1.11331661 | -2.2389692 | 0.02515792 | 0.63297322 |
| MAPRE2 | 221.077885 | 0.01143532 | 0.3815189 | 0.02997315 | 0.97608847 | 0.99989001 |
| ZNF397 | 85.3575691 | 0.49667328 | 0.31014469 | 1.60142444 | 0.10928294 | 0.99989001 |
| ZSCAN30 | 92.0117984 | 0.18277975 | 0.31438402 | 0.58139007 | 0.56097759 | 0.99989001 |
| ZNF24 | 115.727728 | -0.0223063 | 0.30480407 | -0.0731826 | 0.94166084 | 0.99989001 |
| GALNT1 | 424.063921 | -0.2734309 | 0.21278012 | -1.2850396 | 1.99E-01 | 1.00E+00 |
| RPRD1A | 312.288215 | 0.01761512 | 0.1886093 | 0.09339475 | 0.92558996 | 0.99989001 |
| ELP2 | 156.07749 | -0.0366077 | 0.22102471 | -0.1656272 | 0.86845033 | 0.99989001 |
| TPGS2 | 80.2535799 | 0.19817556 | 0.2967713 | 0.66777199 | 0.50427915 | 0.99989001 |

| | | | | | | |
|---|---|---|---|---|---|---|
| KIAA1328 | 362.086564 | -0.3147953 | 0.27890593 | -1.1286791 | 0.25903322 | 0.99989001 |
| AC016205.1 | 70.25763 | -0.2865171 | 0.52869013 | -0.5419378 | 0.58786138 | 0.99989001 |
| PIK3C3 | 287.087217 | -0.1643844 | 0.22294905 | -0.7373182 | 0.46092888 | 0.99989001 |
| SETBP1 | 766.952425 | -0.2492613 | 0.22942469 | -1.0864624 | 0.27727445 | 0.99989001 |
| SLC14A1 | 492.121333 | -2.7829303 | 1.02364185 | -2.7186562 | 0.00655477 | 0.390188 |
| EPG5 | 176.207015 | -0.4530924 | 0.30058031 | -1.5073921 | 0.13171017 | 0.99989001 |
| ATP5F1A | 133.014938 | 0.22668558 | 0.24338793 | 0.93137561 | 0.35165931 | 0.99989001 |
| HAUS1 | 54.032466 | 0.1639715 | 0.36047764 | 0.45487288 | 0.64920069 | 0.99989001 |
| C18orf25 | 144.608951 | 0.45292834 | 0.19153496 | 2.36472942 | 0.01804325 | 0.56443426 |
| PIAS2 | 276.271412 | -0.2920219 | 0.17867809 | -1.6343463 | 0.10218616 | 0.99989001 |
| KATNAL2 | 83.847604 | -0.465038 | 0.45275949 | -1.0271193 | 0.30436429 | 0.99989001 |
| SMAD2 | 580.004513 | -0.2911508 | 0.28407028 | -1.0249251 | 0.30539851 | 0.99989001 |
| ZBTB7C | 207.817278 | -0.3206492 | 0.53026368 | -0.6046977 | 0.54537988 | 0.99989001 |
| CTIF | 303.458314 | 0.01633247 | 0.27279892 | 0.05986999 | 0.95225918 | 0.99989001 |
| SMAD7 | 182.11167 | 0.77427135 | 0.57780107 | 1.340031 | 0.18023527 | 0.99989001 |
| DYM | 775.001908 | 0.17139606 | 0.17532601 | 0.97758491 | 0.32827966 | 0.99989001 |
| ACAA2 | 49.6632275 | 1.06764338 | 0.45671991 | 2.33763266 | 0.01940631 | NA |
| MBD1 | 52.3275514 | 0.05159489 | 0.25544569 | 0.20197986 | 0.83993247 | 0.99989001 |
| ME2 | 105.543968 | -0.0354398 | 0.30969716 | -0.1144337 | 0.90889396 | 0.99989001 |
| SMAD4 | 121.615644 | -0.0093139 | 0.217053 | -0.0429107 | 0.96577274 | 0.99989001 |
| MEX3C | 46.5341417 | 0.40577118 | 0.28336263 | 1.43198553 | 0.15214796 | NA |
| MBD2 | 327.576978 | -0.1038062 | 0.16157436 | -0.6424669 | 0.52057009 | 0.99989001 |
| POLI | 65.0816623 | 0.3502861 | 0.32115517 | 1.09070671 | 0.27540196 | 0.99989001 |
| CCDC68 | 64.9120131 | 0.58454917 | 0.55607683 | 1.05120216 | 0.29316575 | 0.99989001 |
| TCF4 | 4500.04923 | -0.0522484 | 0.35704937 | -0.1463337 | 0.88365793 | 0.99989001 |
| TCF4-AS1 | 62.9493874 | 0.02779377 | 0.55274495 | 0.05028317 | 0.95989674 | 0.99989001 |
| TXNL1 | 141.818328 | 0.14528673 | 0.2786574 | 0.52138119 | 0.60210125 | 0.99989001 |
| WDR7 | 385.605583 | 0.05594333 | 0.17736698 | 0.31541007 | 0.75245032 | 0.99989001 |
| NARS | 106.738422 | 0.22439179 | 0.3644112 | 0.61576535 | 0.5380494 | 0.99989001 |
| AC027097.2 | 176.751258 | -0.2017682 | 0.25104201 | -0.8037229 | 0.42155701 | 0.99989001 |
| ATP8B1 | 806.257254 | 0.43065395 | 0.35816651 | 1.20238475 | 0.2292145 | 0.99989001 |
| NEDD4L | 443.269228 | -0.5259147 | 0.40214314 | -1.3077798 | 0.19094801 | 0.99989001 |
| MALT1 | 262.204765 | -0.3564373 | 0.24520014 | -1.4536585 | 0.146041 | 0.99989001 |
| ZNF532 | 341.186658 | -0.2037588 | 0.33658775 | -0.605366 | 0.54493581 | 0.99989001 |
| SEC11C | 46.0346536 | -0.1047441 | 0.50922891 | -0.2056915 | 0.8370319 | NA |
| LMAN1 | 180.353588 | -0.0747532 | 0.35818034 | -0.2087027 | 0.83468031 | 0.99989001 |
| RNF152 | 197.55271 | 0.20499492 | 0.38520539 | 0.53217044 | 0.59460796 | 0.99989001 |
| PIGN | 208.522555 | -0.433411 | 0.16225637 | -2.6711491 | 0.00755921 | 0.4053518 |
| KIAA1468 | 302.897651 | -0.2366311 | 0.19307784 | -1.2255737 | 0.22035912 | 0.99989001 |
| ZCCHC2 | 248.095322 | 0.13548209 | 0.45693908 | 0.29649923 | 0.76684885 | 0.99989001 |
| PHLPP1 | 526.682952 | 0.30004909 | 0.34454037 | 0.87086773 | 3.84E-01 | 0.99989001 |
| KDSR | 120.858829 | -0.134449 | 0.25857201 | -0.5199673 | 0.6030864 | 0.99989001 |
| VPS4B | 152.990858 | 0.12952569 | 0.18776303 | 0.68983596 | 0.49029735 | 0.99989001 |
| TMX3 | 166.457592 | -0.0997906 | 0.25886557 | -0.3854919 | 0.69987303 | 0.99989001 |
| CCDC102B | 62.6168833 | -0.459867 | 0.32601207 | -1.4105829 | 0.15836764 | 0.99989001 |
| DOK6 | 268.056812 | 0.93411928 | 0.48200837 | 1.93797316 | 0.0526265 | 0.81270649 |
| RTTN | 154.854349 | -0.2470679 | 0.3331447 | -0.7416233 | 0.45831558 | 0.99989001 |
| SOCS6 | 66.9272635 | 0.61920483 | 0.30747073 | 2.01386594 | 0.04402361 | 0.77097498 |
| CYB5A | 126.696912 | 0.93774084 | 0.32736501 | 2.86451149 | 0.00417653 | 0.32499431 |
| CNDP2 | 207.765556 | 0.28449341 | 0.27257886 | 1.04371045 | 0.29661937 | 0.99989001 |
| ZNF407 | 721.639279 | 0.10890379 | 0.28502158 | 0.38208962 | 0.70239489 | 0.99989001 |
| ZADH2 | 135.672405 | 0.44426186 | 0.32441704 | 1.36941592 | 0.1708693 | 0.99989001 |
| TSHZ1 | 230.501129 | 0.2590872 | 0.2937236 | 0.88207823 | 0.3777345 | 0.99989001 |
| ZNF516 | 165.768552 | 0.16288865 | 0.24518369 | 0.66435353 | 0.50646407 | 0.99989001 |

| | | | | | | |
|---|---|---|---|---|---|---|
| ZNF236 | 141.920717 | -0.1909012 | 0.18230522 | -1.0471517 | 0.2950296 | 0.99989001 |
| MBP | 87.30989 | -0.2394105 | 0.47284644 | -0.5063176 | 0.61263371 | 0.99989001 |
| ATP9B | 444.242955 | -0.0861871 | 0.28196281 | -0.3056682 | 0.75985731 | 0.99989001 |
| NFATC1 | 87.4029259 | 0.09758339 | 0.3178939 | 0.30696842 | 0.75886741 | 0.99989001 |
| CTDP1 | 54.4948312 | -0.1434467 | 0.30658878 | -0.4678797 | 0.63987061 | 0.99989001 |
| PQLC1 | 134.187435 | 0.15708829 | 0.21435067 | 0.73285655 | 0.46364593 | 0.99989001 |
| TXNL4A | 105.351558 | -0.2858245 | 0.30535884 | -0.9360282 | 0.34925867 | 0.99989001 |
| ADNP2 | 102.944621 | -0.0246173 | 0.23516767 | -0.1046798 | 0.91662989 | 0.99989001 |
| RBCK1 | 139.88601 | -0.0698647 | 0.3061428 | -0.2282094 | 0.81948344 | 0.99989001 |
| TBC1D20 | 92.0880607 | 0.09010566 | 0.32711738 | 0.27545361 | 0.78296776 | 0.99989001 |
| CSNK2A1 | 275.442925 | 0.19210099 | 0.21373549 | 0.89877913 | 0.36877032 | 0.99989001 |
| TCF15 | 42.6046651 | -0.839054 | 0.75726739 | -1.1080023 | 0.26786081 | NA |
| PSMF1 | 189.397715 | 0.05846712 | 0.33418341 | 0.1749552 | 0.86111484 | 0.99989001 |
| SDCBP2 | 55.5907625 | -0.1853382 | 0.35501691 | -0.5220546 | 6.02E-01 | 1.00E+00 |
| AL136531.2 | 442.833091 | -0.194416 | 0.43190606 | -0.4501348 | 0.65261324 | 0.99989001 |
| SDCBP2-AS1 | 229.762139 | -0.0994156 | 0.47096108 | -0.2110908 | 0.8328164 | 0.99989001 |
| NSFL1C | 97.1842906 | 0.01756479 | 0.28403921 | 0.06183931 | 0.9506908 | 0.99989001 |
| SIRPA | 41.6443348 | 0.72487375 | 0.3446424 | 2.10326345 | 0.03544275 | NA |
| STK35 | 47.0964756 | 0.02997333 | 0.28675877 | 0.10452455 | 0.91675309 | NA |
| SNRPB | 107.12276 | -0.2231972 | 0.45840382 | -0.4869008 | 0.62632864 | 0.99989001 |
| NOP56 | 72.9232823 | -0.095019 | 0.47218684 | -0.2012319 | 0.84051729 | 0.99989001 |
| IDH3B | 41.3554044 | 0.08521475 | 0.38213102 | 0.22299877 | 0.82353647 | NA |
| EBF4 | 105.110206 | 0.54783931 | 0.23458776 | 2.33532779 | 0.0195263 | 0.5734807 |
| PCED1A | 44.6971366 | -0.1642254 | 0.34676613 | -0.4735913 | 0.63579141 | NA |
| PTPRA | 464.935411 | 0.01150029 | 0.25771129 | 0.04462469 | 0.96440647 | 0.99989001 |
| MRPS26 | 48.6057416 | -0.18714 | 0.52339243 | -0.3575519 | 0.72067866 | NA |
| DDRGK1 | 53.2345753 | -0.0684651 | 0.33150443 | -0.2065285 | 0.83637811 | 0.99989001 |
| ITPA | 49.2921779 | -0.3247023 | 0.39119701 | -0.8300225 | 0.40652605 | NA |
| C20orf194 | 523.977842 | 0.38004218 | 0.28878833 | 1.31598871 | 0.18817783 | 0.99989001 |
| ATRN | 241.904733 | 0.05571969 | 0.22563611 | 0.24694493 | 0.80495084 | 0.99989001 |
| HSPA12B | 140.834361 | -0.3992951 | 0.4356729 | -0.9165021 | 0.35940359 | 0.99989001 |
| CENPB | 52.4759762 | -0.047712 | 0.60055577 | -0.0794463 | 0.93667761 | 0.99989001 |
| CDC25B | 65.7585842 | -0.042959 | 0.2475595 | -0.1735301 | 0.86223477 | 0.99989001 |
| MAVS | 139.75548 | 0.23979722 | 0.2678258 | 0.89534772 | 0.37060124 | 0.99989001 |
| PANK2 | 206.008623 | -0.1520094 | 0.19914939 | -0.7632936 | 0.44528834 | 0.99989001 |
| RNF24 | 205.28728 | -0.0740599 | 0.46205746 | -0.1602828 | 0.87265833 | 0.99989001 |
| PRNP | 88.5164585 | -0.4188668 | 0.45223418 | -0.9262165 | 0.35433348 | 0.99989001 |
| RASSF2 | 97.2579401 | -0.2692922 | 0.65564461 | -0.4107289 | 0.68127131 | 0.99989001 |
| SLC23A2 | 255.028413 | -0.2522216 | 0.31121851 | -0.8104326 | 0.41769161 | 0.99989001 |
| TMEM230 | 131.245848 | 0.00639773 | 0.2354697 | 0.02717009 | 0.97832407 | 0.99989001 |
| PCNA | 43.5552263 | 0.21655415 | 0.52592777 | 0.41175645 | 0.68051794 | NA |
| CDS2 | 237.429341 | -0.0217116 | 0.26158963 | -0.0829986 | 0.93385269 | 0.99989001 |
| GPCPD1 | 295.984044 | 0.13248975 | 0.51718671 | 0.25617394 | 0.79781653 | 0.99989001 |
| SHLD1 | 124.287136 | -0.2294337 | 0.28880817 | -0.7944155 | 0.42695356 | 0.99989001 |
| CRLS1 | 107.213862 | -0.124315 | 0.30681679 | -0.4051767 | 0.6853476 | 0.99989001 |
| TMX4 | 100.259512 | 0.16444659 | 0.27907417 | 0.58925764 | 0.55568845 | 0.99989001 |
| PLCB1 | 3111.64614 | -0.0321154 | 0.25137611 | -0.1277583 | 0.89834023 | 0.99989001 |
| PLCB4 | 814.769682 | -0.2694741 | 0.55416489 | -0.4862707 | 0.62677527 | 0.99989001 |
| SLX4IP | 168.526161 | -0.1797284 | 0.33683336 | -0.5335826 | 0.59363038 | 0.99989001 |
| JAG1 | 288.978344 | 0.1391443 | 0.4630682 | 0.30048338 | 0.76380847 | 0.99989001 |
| AL050403.2 | 117.736721 | -0.2567708 | 0.59262047 | -0.4332803 | 0.66481116 | 0.99989001 |
| BTBD3 | 260.090691 | 0.25839647 | 0.18545441 | 1.39331535 | 1.64E-01 | 0.99989001 |
| SPTLC3 | 67.6166646 | 0.88768026 | 0.35213589 | 2.52084573 | 0.01170732 | 0.48516963 |
| TASP1 | 217.52131 | -0.0211811 | 0.32691069 | -0.0647918 | 0.94833979 | 0.99989001 |

| | | | | | | |
|---|---|---|---|---|---|---|
| ESF1 | 128.827496 | -0.163427 | 0.31788839 | -0.5141019 | 0.60718075 | 0.99989001 |
| NDUFAF5 | 52.5582596 | 0.05807523 | 0.36369757 | 0.15968001 | 0.87313315 | 0.99989001 |
| MACROD2 | 692.831319 | -0.5697429 | 0.48314567 | -1.1792363 | 0.23830409 | 0.99989001 |
| KIF16B | 204.806205 | -0.3444519 | 0.19968943 | -1.7249383 | 0.0845386 | 0.95514629 |
| SNRPB2 | 77.1865884 | 0.25530224 | 0.32541386 | 0.7845463 | 0.43271962 | 0.99989001 |
| DSTN | 430.177344 | 0.13470052 | 0.43246973 | 0.31146809 | 0.7554448 | 0.99989001 |
| RRBP1 | 309.79897 | 0.11770849 | 0.32532422 | 0.36181903 | 0.71748727 | 0.99989001 |
| SNX5 | 84.6130701 | 0.17578491 | 0.22351555 | 0.78645495 | 0.431601 | 0.99989001 |
| ZNF133 | 83.7427898 | 0.33713371 | 0.2467243 | 1.36643901 | 0.17180121 | 0.99989001 |
| SEC23B | 93.0308286 | 0.09940083 | 0.24338864 | 0.40840373 | 0.6829773 | 0.99989001 |
| DTD1 | 43.6643196 | 0.22752532 | 0.36997046 | 0.61498238 | 0.53856636 | NA |
| AL121761.1 | 102.349568 | 0.66503242 | 0.44321048 | 1.50048893 | 0.1334878 | 0.99989001 |
| RIN2 | 1023.3354 | -0.1019638 | 0.23578854 | -0.4324375 | 0.66542344 | 0.99989001 |
| NAA20 | 91.1795668 | -0.1438179 | 0.32866519 | -0.4375817 | 0.66168951 | 0.99989001 |
| RALGAPA2 | 2623.2909 | -0.1360318 | 0.27402202 | -0.4964266 | 0.61959345 | 0.99989001 |
| KIZ | 100.283095 | 0.09676214 | 0.31669782 | 0.30553458 | 0.75995904 | 0.99989001 |
| KIZ-AS1 | 42.9362634 | 0.17285991 | 0.52618508 | 0.32851542 | 0.74252199 | NA |
| XRN2 | 352.313247 | -0.0824848 | 0.18883131 | -0.4368175 | 0.66224374 | 0.99989001 |
| THBD | 214.055356 | 0.06668054 | 0.54096004 | 0.12326333 | 0.90189858 | 0.99989001 |
| CD93 | 814.207435 | -0.2438965 | 0.49044414 | -0.4972973 | 0.61897945 | 0.99989001 |
| NAPB | 109.128278 | -0.2680374 | 0.27703362 | -0.9675265 | 0.3332809 | 0.99989001 |
| CST3 | 343.214878 | -0.5332216 | 0.43313464 | -1.2310759 | 0.21829448 | 0.99989001 |
| APMAP | 60.7143348 | 0.16182679 | 0.34713707 | 0.46617549 | 0.64108989 | 0.99989001 |
| ACSS1 | 64.0123159 | 0.27367576 | 0.33353986 | 0.82051889 | 0.41192037 | 0.99989001 |
| PYGB | 52.4232618 | 0.22317467 | 0.39885729 | 0.55953515 | 0.57579655 | 0.99989001 |
| ABHD12 | 102.572296 | -0.103619 | 0.33366296 | -0.3105499 | 0.75614282 | 0.99989001 |
| NINL | 48.8267906 | -0.0373461 | 0.45897867 | -0.0813677 | 0.93514952 | NA |
| ZNF337 | 82.5667843 | 0.14475829 | 0.24972538 | 0.57966991 | 0.56213724 | 0.99989001 |
| FAM182B | 65.558284 | 0.13870714 | 0.46820726 | 0.29625159 | 0.76703796 | 0.99989001 |
| HM13 | 175.140312 | 0.10068006 | 0.27878137 | 0.36114341 | 0.71799224 | 0.99989001 |
| ID1 | 1606.52163 | 0.7092891 | 0.82851524 | 0.85609662 | 0.39194434 | 0.99989001 |
| BCL2L1 | 371.001458 | -0.5045837 | 0.41062366 | -1.2288229 | 0.21913823 | 0.99989001 |
| AL117381.1 | 64.3381725 | 0.34490995 | 0.64374961 | 0.53578278 | 0.59210869 | 0.99989001 |
| CCM2L | 204.312952 | -0.0321638 | 0.3784032 | -0.0849988 | 0.93226236 | 0.99989001 |
| TM9SF4 | 144.183635 | -0.1196526 | 0.16222344 | -0.7375788 | 0.46077044 | 0.99989001 |
| POFUT1 | 54.6122334 | -0.2051812 | 0.29471015 | -0.6962136 | 0.48629508 | 0.99989001 |
| KIF3B | 113.420738 | 0.15056712 | 0.28891841 | 0.52114063 | 0.60226881 | 0.99989001 |
| ASXL1 | 655.957115 | -0.2449584 | 0.26133172 | -0.9373468 | 0.34858022 | 0.99989001 |
| NOL4L | 155.410371 | 0.11069442 | 0.3904615 | 0.28349639 | 0.77679635 | 0.99989001 |
| MAPRE1 | 75.5519673 | -0.1305268 | 0.33117804 | -0.3941288 | 0.69348593 | 0.99989001 |
| CDK5RAP1 | 78.018198 | -0.0971956 | 0.31638044 | -0.307211 | 0.75868275 | 0.99989001 |
| CBFA2T2 | 315.247376 | -0.0402921 | 0.22845261 | -0.1763697 | 0.86000348 | 0.99989001 |
| CHMP4B | 134.800717 | -0.0182648 | 0.24844589 | -0.0735164 | 0.94139521 | 0.99989001 |
| RALY | 284.511127 | 0.06796693 | 0.22698164 | 0.29943802 | 0.76460586 | 0.99989001 |
| EIF2S2 | 229.034801 | -0.0569387 | 0.4099609 | -0.1388881 | 0.88953855 | 0.99989001 |
| ITCH | 546.615118 | 0.08130016 | 0.19564711 | 0.41554488 | 0.67774307 | 0.99989001 |
| DYNLRB1 | 219.906933 | -0.0159183 | 0.42989467 | -0.0370285 | 0.97046231 | 0.99989001 |
| MAP1LC3A | 51.7994877 | -0.0424553 | 0.43407863 | -0.0978055 | 0.92208677 | 0.99989001 |
| PIGU | 103.272253 | -0.2073931 | 0.2475353 | -0.8378322 | 0.40212493 | 0.99989001 |
| NCOA6 | 285.356063 | 0.07390464 | 0.20150445 | 0.3667643 | 0.71379484 | 0.99989001 |
| ACSS2 | 109.725043 | 0.40227983 | 0.34301176 | 1.17278729 | 0.24088112 | 0.99989001 |
| TRPC4AP | 385.661983 | -0.0488098 | 0.18716488 | -0.2607849 | 0.79425836 | 0.99989001 |
| EDEM2 | 69.2596579 | -0.1745267 | 0.22356453 | -0.7806547 | 0.4350056 | 0.99989001 |
| PROCR | 194.603667 | -0.2969163 | 0.50130052 | -0.5922921 | 0.55365503 | 0.99989001 |

| | | | | | | |
|---|---|---|---|---|---|---|
| MMP24OS | 103.140408 | 0.07061787 | 0.29735716 | 0.23748503 | 0.81228053 | 0.99989001 |
| MMP24 | 52.2816654 | 0.05044369 | 0.3786748 | 0.13321111 | 0.89402643 | 0.99989001 |
| UQCC1 | 165.943086 | -0.0531696 | 0.16721621 | -0.3179695 | 0.75050811 | 0.99989001 |
| ERGIC3 | 70.7793573 | -0.3253577 | 0.28618489 | -1.1368795 | 0.25558865 | 0.99989001 |
| CPNE1 | 88.0482871 | 0.08495367 | 0.26546197 | 0.32002199 | 0.74895166 | 0.99989001 |
| ROMO1 | 58.9069107 | -0.396856 | 0.55880386 | -0.7101884 | 0.47758733 | 0.99989001 |
| RBM39 | 683.401126 | 0.05447452 | 0.25142692 | 0.21666143 | 0.82847221 | 0.99989001 |
| PHF20 | 383.686274 | 0.10143827 | 0.20763026 | 0.48855242 | 0.6251586 | 0.99989001 |
| SCAND1 | 136.584777 | -0.0885752 | 0.45196326 | -0.1959788 | 0.84462672 | 0.99989001 |
| NORAD | 289.070504 | -0.1686415 | 0.34518769 | -0.4885503 | 0.62516011 | 0.99989001 |
| EPB41L1 | 48.7286468 | 0.26861344 | 0.56788465 | 0.47300706 | 0.63620815 | NA |
| AAR2 | 42.7698254 | 0.15995301 | 0.26128207 | 0.6121852 | 0.54041523 | NA |
| DLGAP4 | 224.595541 | -0.1407614 | 0.3075383 | -0.4577038 | 0.64716526 | 0.99989001 |
| DLGAP4-AS1 | 53.2905905 | -0.0605451 | 0.3621723 | -0.1671721 | 0.86723462 | 0.99989001 |
| MYL9 | 85.1890031 | -0.2662161 | 0.52908664 | -0.5031617 | 0.61485058 | 0.99989001 |
| RAB5IF | 44.2726197 | 0.21010727 | 0.48458709 | 0.43357999 | 0.66459347 | NA |
| NDRG3 | 87.2903048 | 0.30791393 | 0.31032462 | 0.99223171 | 0.32108451 | 0.99989001 |
| SOGA1 | 144.002239 | 0.50484388 | 0.35744367 | 1.41237327 | 0.15784008 | 0.99989001 |
| SAMHD1 | 213.167742 | 0.19408424 | 0.51952852 | 0.37357764 | 0.70871857 | 0.99989001 |
| RBL1 | 51.8939178 | -0.1619446 | 0.29664554 | -0.5459197 | 0.58512114 | 0.99989001 |
| RPN2 | 167.971147 | 0.04729382 | 0.33182048 | 0.14252833 | 0.88666271 | 0.99989001 |
| BLCAP | 155.49296 | 0.48542997 | 0.22773603 | 2.1315466 | 0.03304414 | 0.67281094 |
| CTNNBL1 | 170.060081 | 0.04547406 | 0.25557338 | 0.17792954 | 0.85877831 | 0.99989001 |
| TTI1 | 64.7980722 | 0.04364977 | 0.27669109 | 0.15775632 | 0.87464882 | 0.99989001 |
| RPRD1B | 135.809942 | -0.1830993 | 0.18991142 | -0.9641299 | 0.33498079 | 0.99989001 |
| TGM2 | 1051.11662 | -0.2455968 | 0.55974874 | -0.4387626 | 0.66083354 | 0.99989001 |
| RALGAPB | 315.172065 | 0.04900603 | 0.18631359 | 0.2630298 | 0.79252761 | 0.99989001 |
| PPP1R16B | 770.255518 | 0.00527291 | 0.33876351 | 0.01556516 | 0.9875813 | 0.99989001 |
| DHX35 | 75.0820058 | 0.00447368 | 0.26894875 | 0.01663396 | 0.98672864 | 0.99989001 |
| TOP1 | 509.27685 | 0.13736586 | 0.29335238 | 0.46826229 | 0.63959703 | 0.99989001 |
| PLCG1 | 155.250564 | 0.11512285 | 0.23744718 | 0.48483562 | 0.62779295 | 0.99989001 |
| ZHX3 | 346.611304 | -0.0654727 | 0.27689252 | -0.2364553 | 0.81307943 | 0.99989001 |
| CHD6 | 377.06313 | 0.21782821 | 0.16753499 | 1.30019528 | 0.19353405 | 0.99989001 |
| SRSF6 | 100.175237 | 0.2374871 | 0.38958013 | 0.60959756 | 0.54212843 | 0.99989001 |
| IFT52 | 71.9219466 | -0.0691767 | 0.22643711 | -0.3055007 | 0.75998483 | 0.99989001 |
| TOX2 | 232.101042 | 0.66471863 | 0.28437666 | 2.33745847 | 0.01941536 | 0.57244969 |
| OSER1 | 43.0439158 | -0.0783915 | 0.44004669 | -0.1781436 | 0.85861023 | NA |
| SERINC3 | 254.45145 | -0.0076297 | 0.34942884 | -0.0218347 | 0.98257979 | 0.99989001 |
| PKIG | 387.728781 | -0.0774889 | 0.27223947 | -0.2846351 | 0.7759237 | 0.99989001 |
| YWHAB | 439.833705 | 0.36564005 | 0.38571962 | 0.94794256 | 0.3431587 | 0.99989001 |
| STK4 | 154.403134 | 0.10670535 | 0.21305501 | 0.50083476 | 0.61648742 | 0.99989001 |
| SLPI | 167.258874 | 0.16398578 | 0.94588923 | 0.17336679 | 8.62E-01 | 0.99989001 |
| SDC4 | 71.2827935 | 0.04613312 | 0.37515204 | 0.1229718 | 0.90212943 | 0.99989001 |
| PIGT | 72.9401692 | -0.4264175 | 0.42160878 | -1.0114056 | 0.31182233 | 0.99989001 |
| WFDC2 | 237.065251 | 0.95247857 | 0.58591719 | 1.62561977 | 1.04E-01 | 0.99989001 |
| DNTTIP1 | 42.8036755 | -0.0360165 | 0.27801202 | -0.1295502 | 0.89692233 | NA |
| CTSA | 93.4673781 | -0.1271527 | 0.41738694 | -0.3046398 | 0.76064053 | 0.99989001 |
| ZNF335 | 50.0765841 | -0.4988051 | 0.39090994 | -1.2760103 | 0.20195189 | 0.99989001 |
| NCOA5 | 94.3048934 | 0.14219684 | 0.30737665 | 0.46261433 | 0.64364084 | 0.99989001 |
| CD40 | 78.1351261 | 0.56839381 | 0.3235019 | 1.757003 | 0.0789173 | 0.92927888 |
| SLC35C2 | 60.4270323 | 0.32392596 | 0.359727 | 0.9004772 | 0.36786635 | 0.99989001 |
| ELMO2 | 54.8508512 | 0.10081786 | 0.25144137 | 0.40095972 | 0.68844978 | 0.99989001 |
| SLC13A3 | 73.7242385 | 0.72419181 | 0.54919079 | 1.3186525 | 0.18728531 | 0.99989001 |
| ZMYND8 | 299.821175 | -0.2105392 | 0.30219305 | -0.6967043 | 0.48598788 | 0.99989001 |

| | | | | | | |
|---|---|---|---|---|---|---|
| NCOA3 | 843.209656 | 0.14830721 | 0.19605772 | 0.75644667 | 4.49E-01 | 0.99989001 |
| SULF2 | 525.478785 | -0.1102011 | 0.47313898 | -0.2329148 | 0.81582753 | 0.99989001 |
| PREX1 | 411.302105 | 0.25657353 | 0.35299773 | 0.72684187 | 0.46732283 | 0.99989001 |
| ARFGEF2 | 355.199423 | -0.2386907 | 0.16709602 | -1.4284642 | 0.1531583 | 0.99989001 |
| CSE1L | 122.642538 | -0.1355746 | 0.21408056 | -0.6332878 | 0.52654573 | 0.99989001 |
| STAU1 | 289.805837 | 0.08199978 | 0.1687712 | 0.48586359 | 0.62706388 | 0.99989001 |
| DDX27 | 45.4670905 | -0.1421002 | 0.25491746 | -0.557436 | 0.57722959 | NA |
| ZNFX1 | 226.078445 | 0.14386485 | 0.45613266 | 0.31540133 | 0.75245695 | 0.99989001 |
| ZFAS1 | 103.807814 | -0.1820657 | 0.25315032 | -0.7192001 | 0.47201763 | 0.99989001 |
| PTGIS | 138.189328 | 0.28793731 | 0.64325454 | 0.44762577 | 0.6544233 | 0.99989001 |
| B4GALT5 | 598.498546 | -0.000945 | 0.37892438 | -0.0024939 | 0.99801018 | 0.99989001 |
| SLC9A8 | 146.270477 | -0.0083219 | 0.38012124 | -0.0218927 | 0.98253353 | 0.99989001 |
| RNF114 | 144.17113 | 0.39090078 | 0.36901476 | 1.05930933 | 0.28945893 | 0.99989001 |
| SNAI1 | 45.0757757 | -0.4632352 | 0.74976385 | -0.6178415 | 0.53667983 | NA |
| CEBPB | 158.802311 | 1.08906146 | 0.61887637 | 1.75973994 | 0.07845191 | 0.92669018 |
| PTPN1 | 336.187722 | -0.2544408 | 0.31156906 | -0.8166434 | 0.41413222 | 0.99989001 |
| ADNP | 380.533309 | 0.05738262 | 0.15224302 | 0.37691465 | 0.70623704 | 0.99989001 |
| DPM1 | 102.980183 | -0.133241 | 0.21039593 | -0.633287 | 0.52654627 | 0.99989001 |
| NFATC2 | 140.699112 | -0.8772795 | 0.96768984 | -0.9065709 | 0.36463375 | 0.99989001 |
| ATP9A | 53.5396031 | 0.28597387 | 0.41271862 | 0.69290274 | 0.48837059 | 0.99989001 |
| ZFP64 | 74.4549233 | -0.2514126 | 0.43588331 | -0.5767887 | 0.56408222 | 0.99989001 |
| TSHZ2 | 376.384218 | 0.50342634 | 0.52991894 | 0.9500063 | 0.34210905 | 0.99989001 |
| ZNF217 | 93.5873189 | 0.52866676 | 0.24780435 | 2.13340391 | 0.03289161 | 0.67281094 |
| DOK5 | 145.599861 | 0.5082284 | 0.38893933 | 1.30670354 | 0.19131342 | 0.99989001 |
| FAM210B | 50.6445659 | 0.07875897 | 0.48278726 | 0.1631339 | 0.87041301 | 0.99989001 |
| RTF2 | 85.0342974 | -0.1525298 | 0.30359452 | -0.5024128 | 0.6153772 | 0.99989001 |
| RAE1 | 45.2079944 | -0.2771899 | 0.24584272 | -1.127509 | 0.25952732 | NA |
| PCK1 | 131.124146 | 1.67695086 | 1.2554941 | 1.33568996 | 0.18165063 | 0.99989001 |
| PMEPA1 | 59.6654878 | 0.96044669 | 0.53940228 | 1.78057588 | 0.07498176 | 0.9206525 |
| RAB22A | 135.29652 | 0.12195152 | 0.25579617 | 0.47675271 | 0.63353822 | 0.99989001 |
| VAPB | 209.026522 | 0.18570315 | 0.20340905 | 0.91295421 | 0.36126662 | 0.99989001 |
| GNAS | 753.082372 | 0.1221616 | 0.41641759 | 0.2933632 | 0.76924455 | 0.99989001 |
| NELFCD | 67.873302 | -0.0494825 | 0.2493815 | -0.1984207 | 0.8427159 | 0.99989001 |
| CTSZ | 84.3315324 | -0.1000751 | 0.38621363 | -0.2591186 | 0.79554373 | 0.99989001 |
| ATP5F1E | 347.71966 | -0.0021252 | 0.52107899 | -0.0040784 | 0.99674592 | 0.99989001 |
| TAF4 | 94.6350242 | 0.40874911 | 0.29745216 | 1.37416757 | 0.16938967 | 0.99989001 |
| PSMA7 | 242.380982 | -0.0363042 | 0.4415382 | -0.082222 | 0.93447015 | 0.99989001 |
| SS18L1 | 89.7892293 | 0.02300243 | 0.26454254 | 0.08695171 | 0.93070989 | 0.99989001 |
| MTG2 | 78.8461526 | -0.0773724 | 0.26252415 | -0.294725 | 0.76820396 | 0.99989001 |
| OSBPL2 | 71.6517573 | -0.0629696 | 0.2181545 | -0.2886466 | 0.7728518 | 0.99989001 |
| ADRM1 | 48.2161846 | 0.01693995 | 0.58265638 | 0.02907365 | 0.97680585 | NA |
| LAMA5 | 306.630112 | -0.8706706 | 0.32058117 | -2.7159131 | 0.00660933 | 0.390188 |
| SLCO4A1 | 291.803523 | -0.9722699 | 0.61702221 | -1.5757454 | 0.11508448 | 0.99989001 |
| OGFR | 113.37376 | 0.08224522 | 0.39096779 | 0.21036316 | 0.83338424 | 0.99989001 |
| DIDO1 | 211.786508 | 0.2628079 | 0.21184331 | 1.24057683 | 0.21476212 | 0.99989001 |
| GID8 | 63.3354472 | -0.016158 | 0.27684205 | -0.0583655 | 0.9534575 | 0.99989001 |
| YTHDF1 | 92.942231 | -0.0213765 | 0.25867511 | -0.0826385 | 0.93413903 | 0.99989001 |
| ARFGAP1 | 56.3968938 | -0.220111 | 0.26171489 | -0.8410336 | 0.40032909 | 0.99989001 |
| PPDPF | 150.938971 | 0.15836141 | 0.45635801 | 0.34701136 | 0.72858278 | 0.99989001 |
| HELZ2 | 77.8303892 | -0.2949168 | 0.77834673 | -0.3789015 | 0.70476099 | 0.99989001 |
| GMEB2 | 52.0338194 | -0.0539682 | 0.32044827 | -0.1684147 | 0.86625707 | 0.99989001 |
| ARFRP1 | 68.9888792 | 0.05229139 | 0.3189338 | 0.16395688 | 0.86976509 | 0.99989001 |
| ZBTB46 | 663.107946 | 0.25439388 | 0.36698424 | 0.6932011 | 0.48818336 | 0.99989001 |
| TPD52L2 | 104.279406 | -0.3670568 | 0.23557796 | -1.558112 | 0.11920671 | 0.99989001 |

| | | | | | | |
|---|---|---|---|---|---|---|
| DNAJC5 | 287.919818 | -0.025129 | 0.36945236 | -0.0680168 | 0.94577229 | 0.99989001 |
| PRPF6 | 129.393675 | 0.13170261 | 0.22264102 | 0.59154693 | 0.55415402 | 0.99989001 |
| SOX18 | 600.199318 | 0.15339801 | 0.78922263 | 0.19436596 | 0.84588934 | 0.99989001 |
| TCEA2 | 99.0212211 | -0.0471908 | 0.30271777 | -0.1558906 | 0.87611927 | 0.99989001 |
| PCMTD2 | 164.609393 | 0.37770009 | 0.36163946 | 1.04441062 | 0.29629545 | 0.99989001 |
| MIER2 | 122.891 | -0.1871975 | 0.36829561 | -0.5082806 | 0.61125657 | 0.99989001 |
| SHC2 | 146.077042 | -0.1171422 | 0.29868406 | -0.3921944 | 0.69491458 | 0.99989001 |
| TPGS1 | 48.5724262 | -0.3331062 | 0.34946809 | -0.9531807 | 0.34049854 | NA |
| CDC34 | 41.2932068 | -0.1591754 | 0.33351622 | -0.4772643 | 0.63317396 | NA |
| BSG | 175.972646 | -0.3906388 | 0.40774544 | -0.9580459 | 0.33803964 | 0.99989001 |
| RNF126 | 45.949055 | -0.363981 | 0.3320668 | -1.0961079 | 0.27303156 | NA |
| PALM | 227.91651 | -0.1301292 | 0.3562263 | -0.3652992 | 0.71488808 | 0.99989001 |
| PTBP1 | 234.485949 | -0.4310841 | 0.35951956 | -1.1990559 | 0.23050621 | 0.99989001 |
| MED16 | 41.290721 | -0.1966746 | 0.36753723 | -0.5351148 | 0.59257046 | NA |
| ARID3A | 54.4097235 | 0.28952081 | 0.35771475 | 0.80936224 | 0.41830681 | 0.99989001 |
| TMEM259 | 145.051244 | -0.2507608 | 0.29296651 | -0.8559368 | 0.39203277 | 0.99989001 |
| POLR2E | 68.9298101 | -0.1388146 | 0.29644507 | -0.4682642 | 0.63959565 | 0.99989001 |
| GPX4 | 184.351451 | -0.229042 | 0.4776739 | -0.4794945 | 0.6315869 | 0.99989001 |
| SBNO2 | 238.704314 | -0.158566 | 0.49875566 | -0.3179231 | 0.75054325 | 0.99989001 |
| STK11 | 72.1029723 | -0.268871 | 0.23698934 | -1.1345277 | 0.25657326 | 0.99989001 |
| ATP5F1D | 135.94203 | -0.2543357 | 0.32843234 | -0.7743929 | 0.43869847 | 0.99989001 |
| MIDN | 291.939663 | -0.3361646 | 0.40123373 | -0.8378274 | 0.40212766 | 0.99989001 |
| CIRBP | 213.095243 | -0.1531891 | 0.31908206 | -0.4800931 | 0.63116118 | 0.99989001 |
| C19orf24 | 62.440175 | -0.3795584 | 0.43778731 | -0.8669927 | 0.385946 | 0.99989001 |
| MUM1 | 87.7035648 | 0.1242889 | 0.29065733 | 0.42761318 | 0.66893277 | 0.99989001 |
| NDUFS7 | 51.2284683 | -0.2879247 | 0.28084485 | -1.0252091 | 0.30526452 | 0.99989001 |
| DAZAP1 | 134.984489 | -0.0139633 | 0.25609994 | -0.054523 | 0.95651848 | 0.99989001 |
| RPS15 | 638.607307 | -0.2707813 | 0.57024145 | -0.4748538 | 0.6348912 | 0.99989001 |
| C19orf25 | 79.1462391 | 0.21998753 | 0.36001253 | 0.6110552 | 0.54116303 | 0.99989001 |
| MBD3 | 53.0180106 | -0.1831368 | 0.38274288 | -0.4784852 | 0.63230493 | 0.99989001 |
| TCF3 | 77.1560883 | -0.2986209 | 0.35363437 | -0.844434 | 0.39842694 | 0.99989001 |
| REXO1 | 55.8664747 | -0.3015108 | 0.23623409 | -1.2763219 | 0.20184174 | 0.99989001 |
| ABHD17A | 171.823423 | -0.1063024 | 0.44583034 | -0.2384369 | 0.81154222 | 0.99989001 |
| SCAMP4 | 39.723239 | -0.1121171 | 0.292208 | -0.3836894 | 0.70120869 | NA |
| CSNK1G2 | 88.8664611 | 0.17105447 | 0.3010458 | 0.56820082 | 0.56989861 | 0.99989001 |
| BTBD2 | 59.2732189 | -0.1817991 | 0.36997454 | -0.4913829 | 0.62315569 | 0.99989001 |
| MKNK2 | 82.7729284 | 0.16743216 | 0.30984711 | 0.54037025 | 0.58894172 | 0.99989001 |
| MOB3A | 79.9672157 | 0.48983545 | 0.30140357 | 1.62518134 | 0.10412393 | 0.99989001 |
| AP3D1 | 125.109392 | -0.0274089 | 0.25510721 | -0.1074409 | 0.91443921 | 0.99989001 |
| DOT1L | 85.7113661 | -0.2122944 | 0.27565376 | -0.7701488 | 0.44121161 | 0.99989001 |
| PLEKHJ1 | 43.0682668 | 0.49333757 | 0.2983798 | 1.65338799 | 0.09825193 | NA |
| LSM7 | 118.305286 | -0.3328833 | 0.50224345 | -0.6627927 | 0.50746332 | 0.99989001 |
| SPPL2B | 73.7467397 | -0.2174402 | 0.26667069 | -0.8153885 | 0.41484994 | 0.99989001 |
| TIMM13 | 59.7860011 | -0.1827803 | 0.48751828 | -0.3749198 | 0.70772011 | 0.99989001 |
| GADD45B | 124.453426 | -1.5658538 | 0.59824348 | -2.6174189 | 0.00885975 | 0.42668269 |
| GNG7 | 205.930376 | 0.1108786 | 0.45891363 | 0.24161103 | 0.80908157 | 0.99989001 |
| SGTA | 64.5353092 | -0.0519557 | 0.25000795 | -0.2078163 | 0.83537242 | 0.99989001 |
| TLE2 | 147.169497 | 0.7851244 | 0.3040612 | 2.58212622 | 0.00981937 | 0.44980004 |
| AES | 322.750837 | -0.2170288 | 0.32440601 | -0.6690037 | 0.5034931 | 0.99989001 |
| GNA11 | 147.537692 | 0.1988794 | 0.39607066 | 0.50213113 | 0.61557529 | 0.99989001 |
| NCLN | 56.9369671 | -0.1091158 | 0.35229962 | -0.3097245 | 0.75677048 | 0.99989001 |
| NFIC | 371.785021 | -0.0393775 | 0.28114242 | -0.1400626 | 0.88861051 | 0.99989001 |
| FZR1 | 37.8884743 | 0.01472975 | 0.33644834 | 0.04378014 | 0.96507966 | NA |
| MFSD12 | 78.5580509 | -0.2114917 | 0.27107228 | -0.7802041 | 0.43527077 | 0.99989001 |

| | | | | | | |
|---|---|---|---|---|---|---|
| HMG20B | 51.1562076 | -0.2000383 | 0.33512444 | -0.5969074 | 0.5505692 | 0.99989001 |
| PIP5K1C | 163.388039 | -0.1604186 | 0.28937609 | -0.5543604 | 0.57933222 | 0.99989001 |
| MRPL54 | 50.3107117 | -0.0178397 | 0.44818914 | -0.0398039 | 0.96824944 | 0.99989001 |
| DAPK3 | 102.096721 | -0.6391727 | 0.23704013 | -2.6964747 | 0.00700777 | 0.3951418 |
| EEF2 | 417.893284 | 0.07953585 | 0.46610583 | 0.17063903 | 0.8645076 | 0.99989001 |
| PIAS4 | 46.5410432 | -0.1326467 | 0.33211186 | -0.3994036 | 0.68959587 | NA |
| ZBTB7A | 170.999605 | 0.08487516 | 0.24284983 | 0.34949647 | 0.72671662 | 0.99989001 |
| MAP2K2 | 137.005909 | -0.2327704 | 0.23212465 | -1.0027817 | 0.31596618 | 0.99989001 |
| SH3GL1 | 263.192914 | -0.0268444 | 0.35037562 | -0.0766159 | 0.93892909 | 0.99989001 |
| UBXN6 | 57.7941132 | 0.27237405 | 0.39996886 | 0.68098814 | 0.49587899 | 0.99989001 |
| HDGFL2 | 46.6992206 | 0.00124779 | 0.28255185 | 0.00441613 | 0.99647645 | NA |
| SEMA6B | 252.502286 | -1.1116222 | 0.41216602 | -2.6970252 | 0.0069962 | 0.3951418 |
| TNFAIP8L1 | 134.677814 | 0.08667151 | 0.38485711 | 0.22520439 | 0.82182027 | 0.99989001 |
| MYDGF | 183.446535 | 0.05433092 | 0.39647567 | 0.13703468 | 0.89100339 | 0.99989001 |
| DPP9 | 93.5109058 | 0.23099482 | 0.31420806 | 0.73516517 | 0.46223891 | 0.99989001 |
| PLIN3 | 49.7999512 | -0.2426245 | 0.35795077 | -0.6778153 | 0.49788879 | NA |
| KDM4B | 242.354098 | 0.00701818 | 0.3350289 | 0.020948 | 0.98328714 | 0.99989001 |
| SAFB2 | 126.25143 | -0.2341647 | 0.27693821 | -0.8455486 | 0.39780459 | 0.99989001 |
| SAFB | 171.187507 | -0.0600303 | 0.29193632 | -0.2056281 | 0.83708143 | 0.99989001 |
| RPL36 | 383.648126 | -0.2187006 | 0.5378381 | -0.406629 | 0.68428051 | 0.99989001 |
| C19orf70 | 68.5216568 | -0.3607421 | 0.47579595 | -0.7581866 | 0.44833928 | 0.99989001 |
| LONP1 | 71.2202476 | -0.4043672 | 0.27991352 | -1.4446146 | 0.14856616 | 0.99989001 |
| RANBP3 | 73.6987382 | 0.06884819 | 0.32154062 | 0.21411974 | 0.83045368 | 0.99989001 |
| RFX2 | 200.293948 | -0.3575114 | 0.72667386 | -0.4919833 | 0.62273116 | 0.99989001 |
| MLLT1 | 104.283587 | 0.37639298 | 0.44489102 | 0.84603412 | 0.39753371 | 0.99989001 |
| CLPP | 46.7443502 | -0.1917041 | 0.37024089 | -0.517782 | 0.60461035 | NA |
| GTF2F1 | 47.7712454 | -0.0395607 | 0.31163965 | -0.1269436 | 0.89898504 | NA |
| KHSRP | 80.7979782 | -0.1802256 | 0.40811467 | -0.4416054 | 0.65877481 | 0.99989001 |
| TRIP10 | 101.998417 | 0.20505775 | 0.25929256 | 0.79083545 | 0.42904002 | 0.99989001 |
| INSR | 2902.70023 | -0.1742775 | 0.45548642 | -0.3826185 | 0.70200267 | 0.99989001 |
| ZNF358 | 50.7714107 | -0.0793251 | 0.59974341 | -0.1322651 | 0.89477464 | 0.99989001 |
| PNPLA6 | 79.9520427 | -0.6351352 | 0.30981393 | -2.0500538 | 0.04035918 | 0.73402991 |
| MAP2K7 | 57.4870556 | -0.3121108 | 0.40414194 | -0.7722802 | 0.43994848 | 0.99989001 |
| ELAVL1 | 137.461866 | -0.2773015 | 0.18097034 | -1.5323035 | 0.12544756 | 0.99989001 |
| CERS4 | 67.3197346 | 0.98386397 | 0.43111334 | 2.28214689 | 0.02248067 | 0.60384164 |
| CD320 | 75.1424368 | -0.5410397 | 0.57726891 | -0.9372404 | 0.34863493 | 0.99989001 |
| RPS28 | 160.488686 | -0.3116026 | 0.51699368 | -0.6027203 | 0.54669478 | 0.99989001 |
| KANK3 | 487.127252 | 0.38312216 | 0.39262479 | 0.97579716 | 0.32916499 | 0.99989001 |
| ANGPTL4 | 123.454957 | 0.48526931 | 0.7371969 | 0.65826282 | 5.10E-01 | 1.00E+00 |
| RAB11B | 79.647931 | -0.1441378 | 0.36683329 | -0.3929246 | 0.69437521 | 0.99989001 |
| 2-Mar | 141.212721 | -0.1549335 | 0.41706191 | -0.3714881 | 0.71027401 | 0.99989001 |
| HNRNPM | 423.423378 | -0.1206877 | 0.30103789 | -0.4009053 | 0.68848985 | 0.99989001 |
| ADAMTS10 | 68.9865891 | 0.46456887 | 0.3810256 | 1.21925895 | 0.22274592 | 0.99989001 |
| ZNF266 | 78.3158438 | 0.55846407 | 0.30066857 | 1.85740757 | 0.06325319 | 0.85940342 |
| ZNF426 | 40.9135789 | 0.36945693 | 0.31552006 | 1.17094594 | 0.24162051 | NA |
| ZNF121 | 67.4202034 | 0.11678556 | 0.27954044 | 0.41777699 | 0.67611018 | 0.99989001 |
| ZNF562 | 131.72197 | 0.04331248 | 0.26340181 | 0.16443501 | 0.8693887 | 0.99989001 |
| ZNF846 | 91.714831 | 0.18368143 | 0.35623098 | 0.51562453 | 0.60611667 | 0.99989001 |
| UBL5 | 72.2243996 | -0.2718238 | 0.5561278 | -0.4887793 | 0.62499792 | 0.99989001 |
| PIN1 | 60.1515915 | -0.1081074 | 0.29678062 | -0.364267 | 0.71565866 | 0.99989001 |
| C19orf66 | 142.429688 | -0.2704249 | 0.4250494 | -0.6362199 | 0.52463313 | 0.99989001 |
| EIF3G | 61.2652133 | 0.33270372 | 0.51218549 | 0.64957662 | 0.51596574 | 0.99989001 |
| DNMT1 | 120.837256 | -0.2240759 | 0.24339332 | -0.9206328 | 0.35724218 | 0.99989001 |
| MRPL4 | 39.3466278 | -0.0960469 | 0.48003235 | -0.2000841 | 0.84141479 | NA |

| | | | | | | |
|---|---|---|---|---|---|---|
| ICAM1 | 300.74427 | 0.42786516 | 0.52450538 | 0.81574981 | 0.41464325 | 0.99989001 |
| TYK2 | 74.2761461 | 0.03511516 | 0.28141093 | 0.12478251 | 0.90069573 | 0.99989001 |
| CDC37 | 378.72677 | -0.5242279 | 0.27958864 | -1.8749971 | 0.06079312 | 0.85923309 |
| PDE4A | 41.7120577 | 0.10256575 | 0.41365684 | 0.24794889 | 0.80417395 | NA |
| KEAP1 | 64.2053815 | 0.37470515 | 0.52950506 | 0.70765168 | 0.47916159 | 0.99989001 |
| KRI1 | 45.8768392 | 0.11695392 | 0.30636024 | 0.38175293 | 0.70264463 | NA |
| CDKN2D | 45.7543022 | -0.3870054 | 0.70177566 | -0.5514659 | 0.58131432 | NA |
| SLC44A2 | 349.851173 | -0.0191642 | 0.32234529 | -0.0594523 | 0.95259184 | 0.99989001 |
| ILF3-DT | 50.853278 | -0.0424065 | 0.37950023 | -0.1117431 | 0.91102709 | 0.99989001 |
| ILF3 | 176.0705 | -0.2561026 | 0.27081166 | -0.9456851 | 0.34430923 | 0.99989001 |
| DNM2 | 302.417625 | 0.03767299 | 0.24392068 | 0.15444771 | 0.87725674 | 0.99989001 |
| CARM1 | 56.3169362 | 0.0166864 | 0.39258409 | 0.042504 | 0.96609692 | 0.99989001 |
| SMARCA4 | 231.006179 | 0.10899593 | 0.35988992 | 0.30285908 | 0.76199726 | 0.99989001 |
| LDLR | 132.226871 | -0.2860832 | 0.48674138 | -0.587752 | 0.55669877 | 0.99989001 |
| KANK2 | 131.463544 | 0.11096939 | 0.3839661 | 0.2890083 | 0.77257503 | 0.99989001 |
| DOCK6 | 724.476768 | 0.06180288 | 0.24354692 | 0.25376168 | 0.79967968 | 0.99989001 |
| RAB3D | 48.0260568 | 0.2583336 | 0.35781135 | 0.7219827 | 0.47030512 | NA |
| RGL3 | 128.617499 | 1.12463205 | 0.50893667 | 2.20976817 | 0.02712126 | 0.64986727 |
| PRKCSH | 85.8545238 | -0.2332592 | 0.35483487 | -0.657374 | 0.51094044 | 0.99989001 |
| ECSIT | 56.0090202 | -0.1680643 | 0.32791963 | -0.5125167 | 0.60828941 | 0.99989001 |
| ZNF440 | 46.905833 | 0.05725098 | 0.38665887 | 0.14806587 | 0.88229079 | NA |
| ZNF439 | 45.2087359 | 0.38799956 | 0.26793915 | 1.44808836 | 0.14759234 | NA |
| ZNF69 | 49.4349161 | 0.30874471 | 0.2779139 | 1.11093655 | 0.26659566 | NA |
| ZNF136 | 47.8785104 | 0.08426791 | 0.27945745 | 0.3015411 | 0.76300191 | NA |
| ZNF44 | 219.563734 | 0.51238013 | 0.27347133 | 1.87361549 | 0.06098344 | 0.85923309 |
| ZNF490 | 58.3143895 | 0.47694608 | 0.29257677 | 1.63015703 | 0.10306831 | 0.99989001 |
| ZNF791 | 132.043938 | 0.0219296 | 0.22017675 | 0.09959996 | 0.92066193 | 0.99989001 |
| TNPO2 | 86.4294928 | -0.1090447 | 0.26324169 | -0.414238 | 0.67869981 | 0.99989001 |
| TRIR | 169.335786 | 0.15777292 | 0.36100058 | 0.4370434 | 0.66207987 | 0.99989001 |
| HOOK2 | 83.4474436 | -0.2894313 | 0.29408182 | -0.9841862 | 0.32502399 | 0.99989001 |
| JUNB | 363.607595 | -0.8721246 | 0.65033174 | -1.3410457 | 0.17990563 | 0.99989001 |
| FARSA | 41.7243151 | 0.28879058 | 0.46817071 | 0.61684888 | 0.53733441 | NA |
| CALR | 320.305175 | 0.14084424 | 0.58638854 | 0.24018928 | 0.81018353 | 0.99989001 |
| AC092069.1 | 61.9033791 | -0.2543491 | 0.76790379 | -0.3312252 | 0.74047436 | 0.99989001 |
| RAD23A | 112.229879 | -0.308924 | 0.34110478 | -0.9056571 | 0.36511737 | 0.99989001 |
| GADD45GIP1 | 164.183288 | -0.1279229 | 0.52880223 | -0.2419107 | 0.80884936 | 0.99989001 |
| NFIX | 101.303985 | 0.12685484 | 0.43796514 | 0.28964598 | 0.77208709 | 0.99989001 |
| LYL1 | 67.8176223 | 0.11871727 | 0.60907426 | 0.19491428 | 0.84546005 | 0.99989001 |
| IER2 | 355.188378 | -0.0151126 | 0.48173087 | -0.0313714 | 0.97497333 | 0.99989001 |
| CCDC130 | 80.128763 | -0.0546076 | 0.26465879 | -0.206332 | 0.83653156 | 0.99989001 |
| MRI1 | 46.4860717 | 0.42000319 | 0.34725656 | 1.20948958 | 0.22647481 | NA |
| C19orf53 | 130.743732 | -0.3133383 | 0.51681557 | -0.6062864 | 0.54432458 | 0.99989001 |
| ZSWIM4 | 74.6503418 | 0.30052295 | 0.29701888 | 1.01179748 | 0.31163489 | 0.99989001 |
| AC020916.1 | 75.7568758 | -0.2040359 | 0.5190459 | -0.3930981 | 0.69424706 | 0.99989001 |
| CC2D1A | 67.3983535 | -0.4033744 | 0.29694647 | -1.3584076 | 0.17433437 | 0.99989001 |
| RFX1 | 53.6140873 | -0.0254771 | 0.32653616 | -0.0780222 | 0.93781043 | 0.99989001 |
| IL27RA | 40.548032 | 0.00518397 | 0.46388007 | 0.01117525 | 0.99108363 | NA |
| PRKACA | 82.3115442 | -0.1033864 | 0.36275085 | -0.2850067 | 0.775639 | 0.99989001 |
| ADGRL1 | 64.9902764 | 0.36307709 | 0.30537475 | 1.18895582 | 0.23445705 | 0.99989001 |
| ADGRE5 | 67.9903422 | 0.58909237 | 0.4515838 | 1.3045029 | 0.19206217 | 0.99989001 |
| DDX39A | 40.668158 | -0.4796144 | 0.37322529 | -1.2850534 | 0.19877362 | NA |
| PKN1 | 231.404526 | -0.1762729 | 0.33003571 | -0.5341024 | 0.59327066 | 0.99989001 |
| GIPC1 | 98.6888803 | -0.0416008 | 0.29526439 | -0.1408936 | 0.88795404 | 0.99989001 |
| DNAJB1 | 78.628017 | -0.6405008 | 0.7518864 | -0.8518584 | 0.39429266 | 0.99989001 |

| | | | | | | |
|---|---|---|---|---|---|---|
| TECR | 53.4426301 | -0.4175133 | 0.41066339 | -1.0166801 | 0.30930564 | 0.99989001 |
| NDUFB7 | 118.091852 | -0.3532504 | 0.40386152 | -0.8746821 | 0.38174692 | 0.99989001 |
| ADGRE3 | 43.3556215 | -0.0829706 | 0.42626057 | -0.1946477 | 0.84566878 | NA |
| ZNF333 | 64.9529178 | -0.2093527 | 0.35692421 | -0.5865467 | 0.55750819 | 0.99989001 |
| BRD4 | 354.009714 | -0.0374564 | 0.2360884 | -0.158654 | 0.87394144 | 0.99989001 |
| AKAP8 | 96.4087995 | -0.2680531 | 0.25395219 | -1.0555258 | 0.2911849 | 0.99989001 |
| AKAP8L | 154.886669 | -0.0257718 | 0.21813128 | -0.118148 | 0.9059504 | 0.99989001 |
| TPM4 | 501.42088 | -0.1953462 | 0.3293968 | -0.5930421 | 0.55315296 | 0.99989001 |
| RAB8A | 38.5860703 | -0.1049026 | 0.3228564 | -0.3249202 | 0.74524144 | NA |
| FAM32A | 49.2967348 | 0.23337519 | 0.43430585 | 0.53735217 | 0.59102438 | NA |
| AP1M1 | 65.1907183 | -0.2192358 | 0.23761854 | -0.9226376 | 0.35619611 | 0.99989001 |
| KLF2 | 661.087889 | 0.3671647 | 0.54963251 | 0.66801853 | 5.04E-01 | 0.99989001 |
| EPS15L1 | 197.086727 | 0.23960099 | 0.26445241 | 0.90602687 | 0.36492163 | 0.99989001 |
| SIN3B | 210.301507 | 0.16758257 | 0.33213965 | 0.50455455 | 0.61387174 | 0.99989001 |
| F2RL3 | 188.363881 | 0.27855385 | 0.72757591 | 0.38285194 | 0.70182955 | 0.99989001 |
| CPAMD8 | 176.159379 | 1.56934361 | 0.55648187 | 2.82011631 | 0.00480062 | 0.34759088 |
| MYO9B | 256.117244 | -0.3213049 | 0.2479406 | -1.2958946 | 0.19501179 | 0.99989001 |
| NR2F6 | 57.2151534 | 0.43312632 | 0.44269285 | 0.97839013 | 0.32788141 | 0.99989001 |
| USHBP1 | 63.3446427 | 0.2039564 | 0.33876509 | 0.60205849 | 0.5471352 | 0.99989001 |
| MRPL34 | 59.2428427 | -0.2272256 | 0.55328511 | -0.4106845 | 0.68130392 | 0.99989001 |
| PLVAP | 1376.68457 | -0.440275 | 0.51635504 | -0.8526595 | 0.39384816 | 0.99989001 |
| BST2 | 676.94845 | -0.2666749 | 0.50789397 | -0.5250603 | 0.5995413 | 0.99989001 |
| MVB12A | 68.6163949 | -0.1972138 | 0.28071066 | -0.7025517 | 0.48233515 | 0.99989001 |
| SLC27A1 | 81.5206138 | -0.2754309 | 0.30084706 | -0.9155181 | 0.35991972 | 0.99989001 |
| AC010618.3 | 56.1783346 | -0.1409951 | 0.34659269 | -0.4068033 | 0.68415244 | 0.99989001 |
| PGLS | 79.7302172 | -0.1375707 | 0.46664028 | -0.294811 | 0.76813828 | 0.99989001 |
| COLGALT1 | 93.6342128 | -0.3647273 | 0.35844174 | -1.0175359 | 0.30889858 | 0.99989001 |
| MAP1S | 43.6202919 | -0.2548782 | 0.46620372 | -0.5467099 | 0.58457808 | NA |
| JAK3 | 74.9082777 | 0.88129202 | 0.64983648 | 1.35617506 | 0.17504347 | 0.99989001 |
| RPL18A | 44.1876479 | -0.4560055 | 0.58176934 | -0.7838253 | 0.43314265 | NA |
| CCDC124 | 45.1065186 | -0.211482 | 0.46935672 | -0.4505783 | 0.65229349 | NA |
| ARRDC2 | 92.3984116 | 0.51828912 | 0.31739243 | 1.63295993 | 0.10247742 | 0.99989001 |
| MAST3 | 51.155341 | -0.3239344 | 0.42021212 | -0.7708831 | 0.44077623 | 0.99989001 |
| JUND | 344.335174 | -1.1411287 | 0.53211346 | -2.1445215 | 0.03199113 | 0.67281094 |
| LSM4 | 76.2297401 | -0.0875356 | 0.35865367 | -0.2440673 | 0.80717872 | 0.99989001 |
| PGPEP1 | 56.4284163 | -0.0966769 | 0.49004968 | -0.1972798 | 0.84360857 | 0.99989001 |
| GDF15 | 74.9650315 | -1.6931344 | 0.77620777 | -2.1812902 | 0.02916196 | 0.66299531 |
| SSBP4 | 112.846093 | 0.07072007 | 0.31921443 | 0.2215441 | 0.82466881 | 0.99989001 |
| ELL | 145.327413 | 0.26641809 | 0.41893493 | 0.63594145 | 0.52481459 | 0.99989001 |
| FKBP8 | 79.096581 | 0.02453207 | 0.45846111 | 0.0535096 | 0.95732588 | 0.99989001 |
| KXD1 | 61.0822595 | -0.019654 | 0.25863823 | -0.0759903 | 0.93942683 | 0.99989001 |
| UBA52 | 234.6403 | 0.04603822 | 0.60153715 | 0.07653429 | 0.93899403 | 0.99989001 |
| REX1BD | 73.2042922 | -0.5405058 | 0.50600781 | -1.0681769 | 0.28544073 | 0.99989001 |
| CRTC1 | 73.2544088 | -0.1621025 | 0.34581893 | -0.4687497 | 0.63924853 | 0.99989001 |
| UPF1 | 82.8738419 | 0.16842324 | 0.28144351 | 0.59842644 | 0.54955543 | 0.99989001 |
| COPE | 73.8869884 | -0.0712504 | 0.44715548 | -0.1593415 | 0.87339985 | 0.99989001 |
| SUGP2 | 160.014097 | 0.23384543 | 0.18546431 | 1.26086488 | 0.20735753 | 0.99989001 |
| SUGP1 | 41.9836967 | 0.17715462 | 0.34229224 | 0.51755371 | 0.60476969 | NA |
| MAU2 | 133.686298 | 0.03227612 | 0.24143276 | 0.13368576 | 0.89365107 | 0.99989001 |
| GATAD2A | 401.064217 | 0.18643407 | 0.28721063 | 0.64911966 | 0.51626103 | 0.99989001 |
| ATP13A1 | 41.3489367 | -0.2563546 | 0.28537457 | -0.8983091 | 0.36902078 | NA |
| ZNF253 | 44.4935771 | 0.58640795 | 0.37773397 | 1.55243635 | 0.12055785 | NA |
| ZNF682 | 47.3214682 | -0.2152717 | 0.26631495 | -0.8083349 | 0.41889783 | NA |
| ZNF90 | 240.399255 | -0.0862001 | 0.50258226 | -0.1715144 | 0.86381934 | 0.99989001 |

| | | | | | | |
|---|---|---|---|---|---|---|
| AC011447.3 | 52.9762001 | -0.0327551 | 0.28499343 | -0.1149329 | 0.90849827 | 0.99989001 |
| ZNF737 | 50.7934033 | 0.45959859 | 0.30521824 | 1.50580318 | 0.13211771 | 0.99989001 |
| ZNF430 | 97.9229073 | 0.25554143 | 0.22571893 | 1.13212227 | 0.25758303 | 0.99989001 |
| ZNF431 | 157.069559 | 0.04071379 | 0.22250786 | 0.18297684 | 0.85481619 | 0.99989001 |
| ZNF708 | 126.458488 | 0.35746956 | 0.3217196 | 1.11112146 | 0.26651607 | 0.99989001 |
| ZNF493 | 39.9873475 | 0.10258943 | 0.29679295 | 0.34565993 | 0.7295983 | NA |
| AC010615.4 | 48.2669244 | 0.4639381 | 0.53892553 | 0.86085752 | 0.38931652 | NA |
| ZNF429 | 72.1878729 | 0.41163138 | 0.31872462 | 1.29149539 | 0.19653195 | 0.99989001 |
| ZNF100 | 88.1795072 | -0.1282965 | 0.41203908 | -0.3113696 | 0.75551964 | 0.99989001 |
| ZNF43 | 176.479387 | -0.0815649 | 0.25328875 | -0.3220234 | 0.747435 | 0.99989001 |
| ZNF208 | 162.415593 | 0.24896957 | 0.29027272 | 0.85770917 | 0.39105308 | 0.99989001 |
| ZNF91 | 290.02003 | 0.1592214 | 0.22948456 | 0.69382184 | 0.48779395 | 0.99989001 |
| ZNF675 | 45.9118817 | 0.12338382 | 0.3364384 | 0.36673524 | 0.71381652 | NA |
| ZNF254 | 214.317274 | 0.12866343 | 0.16150671 | 0.79664449 | 0.42565753 | 0.99989001 |
| LINC00662 | 90.5485965 | -0.1417189 | 0.34243275 | -0.4138591 | 0.6789773 | 0.99989001 |
| UQCRFS1 | 73.8942768 | 0.17514896 | 0.50628894 | 0.34594664 | 0.72938282 | 0.99989001 |
| C19orf12 | 42.5489306 | 0.00135806 | 0.30521636 | 0.00444949 | 0.99644983 | NA |
| URI1 | 236.1886 | 0.18835866 | 0.28172964 | 0.66857951 | 0.50376375 | 0.99989001 |
| TSHZ3 | 215.709635 | 0.60658188 | 0.29377254 | 2.06480112 | 0.03894182 | 0.72042371 |
| ZNF507 | 91.1357687 | -0.0752997 | 0.22770462 | -0.3306901 | 0.74087859 | 0.99989001 |
| DPY19L3 | 114.980166 | 0.05595827 | 0.2888379 | 0.19373591 | 0.84638267 | 0.99989001 |
| PDCD5 | 56.0483247 | -0.1626577 | 0.27888345 | -0.5832463 | 0.55972752 | 0.99989001 |
| ANKRD27 | 114.876144 | 0.29401249 | 0.30420347 | 0.96649946 | 0.33379432 | 0.99989001 |
| NUDT19 | 52.7888272 | 0.06851219 | 0.26416287 | 0.25935588 | 0.79536067 | 0.99989001 |
| CEP89 | 80.1278742 | -0.0001676 | 0.28719153 | -0.0005838 | 0.99953423 | 0.99989001 |
| CEBPG | 85.5274561 | 0.01491545 | 0.31794576 | 0.04691193 | 0.96258342 | 0.99989001 |
| PEPD | 142.683745 | -0.2570409 | 0.2762136 | -0.9305876 | 0.35206695 | 0.99989001 |
| KCTD15 | 87.2362716 | 0.28974452 | 0.50035853 | 0.57907381 | 0.56253937 | 0.99989001 |
| LSM14A | 506.862013 | 0.1634647 | 0.15861977 | 1.0305443 | 0.30275457 | 0.99989001 |
| KIAA0355 | 805.30374 | 0.16431833 | 0.2972409 | 0.55281197 | 0.58039217 | 0.99989001 |
| GPI | 120.361147 | -0.4461507 | 0.47832082 | -0.9327436 | 0.35095237 | 0.99989001 |
| UBA2 | 234.502817 | -0.1710464 | 0.26675754 | -0.6412056 | 0.52138913 | 0.99989001 |
| WTIP | 39.7460856 | -0.5489578 | 0.47487008 | -1.1560168 | 0.24767428 | NA |
| SCGB2B2 | 104.709855 | -0.1852588 | 0.4040155 | -0.4585438 | 0.64656181 | 0.99989001 |
| ZNF302 | 115.929981 | -0.0299527 | 0.31338714 | -0.0955774 | 0.92385624 | 0.99989001 |
| GRAMD1A | 256.524374 | -0.084892 | 0.37839937 | -0.2243449 | 8.22E-01 | 0.99989001 |
| FXYD5 | 113.62507 | -0.2387561 | 0.44154374 | -0.5407304 | 0.58869345 | 0.99989001 |
| USF2 | 72.8840364 | -0.5198821 | 0.36354362 | -1.4300405 | 0.1527054 | 0.99989001 |
| TMEM147 | 48.6317901 | 0.0414846 | 0.57791372 | 0.07178338 | 0.9427743 | NA |
| RBM42 | 44.150644 | -0.313959 | 0.43689826 | -0.718609 | 0.47238184 | NA |
| COX6B1 | 145.730262 | 0.02316793 | 0.36526808 | 0.06342718 | 0.94942634 | 0.99989001 |
| KMT2B | 56.4585751 | -0.3072974 | 0.26082505 | -1.1781744 | 0.23872708 | 0.99989001 |
| TBCB | 66.0067355 | -0.3067713 | 0.3096832 | -0.9905971 | 0.32188236 | 0.99989001 |
| CAPNS1 | 257.145008 | -0.09958 | 0.41620818 | -0.2392554 | 0.81090757 | 0.99989001 |
| COX7A1 | 57.3593379 | -0.7229608 | 0.61628113 | -1.1731023 | 0.24075476 | 0.99989001 |
| ZNF565 | 95.4031523 | 0.17805518 | 0.28485643 | 0.6250699 | 0.53192518 | 0.99989001 |
| ZNF146 | 120.661114 | 0.18966357 | 0.23125593 | 0.82014574 | 0.41213303 | 0.99989001 |
| ZFP14 | 93.1857832 | 0.44472305 | 0.26555676 | 1.67468174 | 0.09399669 | 0.99989001 |
| ZFP82 | 44.7699217 | 0.44881119 | 0.30964978 | 1.44941551 | 0.14722158 | NA |
| ZNF566 | 73.1771798 | -0.3699333 | 0.26506892 | -1.3956116 | 0.16283149 | 0.99989001 |
| ZNF260 | 48.7108757 | 0.172154 | 0.24575329 | 0.70051554 | 0.4836054 | NA |
| ZNF529 | 69.3574455 | 0.28542861 | 0.24339765 | 1.17268431 | 0.24092242 | 0.99989001 |
| ZNF529-AS1 | 44.5242499 | 0.71421012 | 0.33103573 | 2.15750162 | 0.0309666 | NA |
| ZNF567 | 44.1317969 | 0.32359224 | 0.33637208 | 0.96200684 | 0.33604617 | NA |

| | | | | | | |
|---|---|---|---|---|---|---|
| AC020928.1 | 53.490705 | 0.46726625 | 0.24667195 | 1.89428207 | 0.05818758 | 0.84787616 |
| ZNF790 | 89.2447805 | 0.38633123 | 0.30701209 | 1.25835833 | 0.2082622 | 0.99989001 |
| ZNF345 | 99.5249333 | 0.49159776 | 0.3428791 | 1.433735 | 0.1516479 | 0.99989001 |
| ZNF568 | 59.8703872 | 0.38215058 | 0.27011492 | 1.41477038 | 0.15713583 | 0.99989001 |
| ZNF420 | 110.072273 | 0.73841356 | 0.38628229 | 1.91159049 | 0.05592874 | 0.83144465 |
| HKR1 | 125.962022 | 0.32760508 | 0.23953431 | 1.36767496 | 0.17141384 | 0.99989001 |
| ZNF569 | 45.2699491 | 0.01237396 | 0.28265997 | 0.04377685 | 0.96508228 | NA |
| ZFP30 | 58.4389598 | 0.25697356 | 0.30897697 | 0.83169165 | 0.40558301 | 0.99989001 |
| ZNF573 | 60.1646166 | 0.46100635 | 0.36031844 | 1.27944146 | 0.20074164 | 0.99989001 |
| SIPA1L3 | 448.924136 | -0.0138893 | 0.34616785 | -0.040123 | 0.96799509 | 0.99989001 |
| PPP1R14A | 66.8632264 | -1.4569705 | 0.77712354 | -1.8748248 | 0.06081683 | 0.85923309 |
| YIF1B | 42.5618462 | -0.7421775 | 0.36349667 | -2.0417725 | 0.0411741 | NA |
| KCNK6 | 89.9969205 | -0.6115769 | 0.35727082 | -1.711802 | 0.08693316 | 0.95976851 |
| PSMD8 | 112.042666 | 0.23606705 | 0.38695064 | 0.61007021 | 0.5418153 | 0.99989001 |
| EIF3K | 118.692085 | -0.3105503 | 0.37473791 | -0.8287134 | 0.40726661 | 0.99989001 |
| ACTN4 | 748.735859 | -0.1800673 | 0.3324278 | -0.5416733 | 0.58804358 | 0.99989001 |
| ECH1 | 73.9232448 | 0.09241208 | 0.58089209 | 0.15908648 | 0.87360074 | 0.99989001 |
| HNRNPL | 121.78486 | -0.3438343 | 0.31046854 | -1.1074692 | 0.26809114 | 0.99989001 |
| SIRT2 | 118.090804 | 0.17270143 | 0.28279562 | 0.61069343 | 0.54140256 | 0.99989001 |
| MRPS12 | 47.1091112 | -0.0272825 | 0.52537735 | -0.0519293 | 0.95858503 | NA |
| PAK4 | 56.0549545 | -0.0940316 | 0.35309289 | -0.2663083 | 0.79000181 | 0.99989001 |
| GMFG | 94.3032129 | 0.24726964 | 0.30659807 | 0.80649442 | 0.41995783 | 0.99989001 |
| SAMD4B | 195.477145 | 0.09282173 | 0.34476638 | 0.2692308 | 0.78775208 | 0.99989001 |
| MED29 | 48.9274751 | 0.01843489 | 0.33254424 | 0.05543589 | 0.9557912 | NA |
| ZFP36 | 238.330419 | -1.9617133 | 0.98769642 | -1.98615 | 0.04701667 | 0.77516398 |
| PLEKHG2 | 65.6586025 | -0.1153042 | 0.38707438 | -0.2978864 | 0.76578986 | 0.99989001 |
| RPS16 | 286.259077 | -0.041439 | 0.52823223 | -0.0784484 | 0.93747137 | 0.99989001 |
| SUPT5H | 110.650045 | -0.0121707 | 0.22197172 | -0.05483 | 0.9562739 | 0.99989001 |
| FBL | 41.6506941 | -0.38241 | 0.39219699 | -0.9750456 | 0.32953762 | NA |
| PSMC4 | 50.4702381 | -0.0009129 | 0.41146297 | -0.0022187 | 0.99822972 | 0.99989001 |
| ZNF780B | 114.621391 | 0.63592154 | 0.29237365 | 2.17503032 | 0.02962784 | 0.6652708 |
| AKT2 | 93.6489764 | 0.11794822 | 0.26206734 | 0.45006836 | 0.65266115 | 0.99989001 |
| PLD3 | 177.176062 | -0.0138846 | 0.3222288 | -0.0430892 | 0.96563043 | 0.99989001 |
| PRX | 44.9511814 | 1.80820001 | 0.43751521 | 4.13288487 | 3.58E-05 | NA |
| SERTAD1 | 78.6096734 | 0.2000875 | 0.57425253 | 0.3484312 | 0.72751638 | 0.99989001 |
| BLVRB | 81.6733408 | -0.1499023 | 0.37574403 | -0.398948 | 0.68993151 | 0.99989001 |
| SHKBP1 | 42.1900028 | -0.1637355 | 0.35503499 | -0.4611813 | 0.64466853 | NA |
| LTBP4 | 89.0535623 | 2.04866209 | 0.58987124 | 3.47306655 | 0.00051455 | 0.09952814 |
| ITPKC | 147.667367 | 0.17444203 | 0.42723659 | 0.40830312 | 6.83E-01 | 0.99989001 |
| SNRPA | 68.1585224 | -0.0740219 | 0.3301393 | -0.2242142 | 0.8225906 | 0.99989001 |
| AC008537.1 | 195.498793 | -0.157044 | 0.48235905 | -0.3255748 | 0.74474606 | 0.99989001 |
| HNRNPUL1 | 137.015087 | 0.04243869 | 0.2342679 | 0.18115454 | 0.85624628 | 0.99989001 |
| TGFB1 | 232.578224 | -0.0267397 | 0.30902698 | -0.0865287 | 0.93104612 | 0.99989001 |
| CCDC97 | 41.2543341 | -0.1419433 | 0.3545664 | -0.4003294 | 0.68891395 | NA |
| DMAC2 | 41.5408379 | -0.3067262 | 0.30313975 | -1.0118311 | 0.31161882 | NA |
| PCAT19 | 399.104965 | -0.0248903 | 0.27476903 | -0.0905861 | 0.92782146 | 0.99989001 |
| RPS19 | 829.442401 | -0.2527177 | 0.54819143 | -0.4610027 | 0.64479667 | 0.99989001 |
| ARHGEF1 | 109.627722 | 0.09204938 | 0.22184512 | 0.41492632 | 0.67819584 | 0.99989001 |
| RABAC1 | 202.44941 | 0.04912672 | 0.58962839 | 0.0833181 | 0.93359861 | 0.99989001 |
| CIC | 56.4881056 | -0.3918827 | 0.41016329 | -0.9554308 | 0.33935986 | 0.99989001 |
| MEGF8 | 46.6129211 | -0.0438492 | 0.32225247 | -0.1360709 | 0.89176527 | NA |
| LIPE-AS1 | 410.290013 | -0.2687002 | 0.36878843 | -0.7286026 | 0.46624477 | 0.99989001 |
| LIPE | 75.3046525 | -0.2769553 | 0.34232855 | -0.8090337 | 0.41849579 | 0.99989001 |
| CEACAM1 | 1436.24294 | -0.0142379 | 0.37996224 | -0.037472 | 0.97010867 | 0.99989001 |

| | | | | | | |
|---|---|---|---|---|---|---|
| ETHE1 | 45.3975324 | 0.02551043 | 0.30783081 | 0.0828716 | 0.93395364 | NA |
| ZNF428 | 69.8283677 | -0.2502924 | 0.38297643 | -0.6535453 | 0.51340482 | 0.99989001 |
| PLAUR | 69.5955571 | -0.9656583 | 0.64488512 | -1.4974113 | 0.13428627 | 0.99989001 |
| SMG9 | 52.5292058 | -0.4019216 | 0.29682621 | -1.3540638 | 0.17571603 | 0.99989001 |
| ZNF155 | 46.1558681 | 1.09484475 | 0.33722664 | 3.24661406 | 0.00116787 | NA |
| ZNF224 | 67.4203231 | 0.48463561 | 0.27356593 | 1.77154955 | 0.07646936 | 0.92274869 |
| ZNF235 | 107.090821 | 0.68850943 | 0.3610239 | 1.90710209 | 0.05650737 | 0.83630905 |
| AC243964.2 | 94.8048776 | -0.2602921 | 0.25703807 | -1.0126599 | 3.11E-01 | 0.99989001 |
| CEACAM19 | 87.9658651 | -0.3984082 | 0.50195406 | -0.7937145 | 0.42736164 | 0.99989001 |
| BCL3 | 117.043173 | -0.5529671 | 0.51750941 | -1.0685161 | 0.28528776 | 0.99989001 |
| BCAM | 174.77387 | -0.5316009 | 0.48454981 | -1.0971027 | 0.27259652 | 0.99989001 |
| NECTIN2 | 275.640814 | -0.4792219 | 0.27128849 | -1.7664661 | 0.07731768 | 0.92274869 |
| APOE | 186.468491 | 1.0092115 | 0.73681362 | 1.36969714 | 0.17078146 | 0.99989001 |
| CLPTM1 | 101.352839 | -0.1262216 | 0.24991703 | -0.5050541 | 0.61352085 | 0.99989001 |
| RELB | 178.159439 | 0.67432283 | 0.65694758 | 1.02644846 | 0.30468025 | 0.99989001 |
| CLASRP | 102.06866 | -0.2195912 | 0.27814616 | -0.7894812 | 0.4298308 | 0.99989001 |
| MARK4 | 512.968828 | 0.32247342 | 0.36339453 | 0.88739206 | 0.37486786 | 0.99989001 |
| EXOC3L2 | 458.709972 | 0.04350282 | 0.46151594 | 0.09426071 | 0.92490206 | 0.99989001 |
| PPP1R13L | 71.12242 | -0.1968633 | 0.24895395 | -0.790762 | 0.42908288 | 0.99989001 |
| ERCC1 | 42.8970063 | -0.359704 | 0.30146389 | -1.1931909 | 0.23279464 | NA |
| VASP | 153.346535 | -0.3165193 | 0.2939465 | -1.0767921 | 0.28157314 | 0.99989001 |
| OPA3 | 93.422825 | -0.2013722 | 0.35458741 | -0.5679057 | 0.57009901 | 0.99989001 |
| EML2 | 55.8369845 | 0.01925265 | 0.2874372 | 0.06698037 | 0.94659733 | 0.99989001 |
| SNRPD2 | 109.765026 | -0.1352056 | 0.49676261 | -0.2721734 | 0.78548868 | 0.99989001 |
| FBXO46 | 42.9390234 | -0.1198182 | 0.32747802 | -0.3658817 | 0.71445337 | NA |
| DMPK | 50.4420646 | -0.0927582 | 0.36083134 | -0.257068 | 0.79712629 | 0.99989001 |
| SYMPK | 157.294186 | -0.1634872 | 0.28158519 | -0.580596 | 0.56151277 | 0.99989001 |
| NOVA2 | 268.470316 | -0.1063184 | 0.30434806 | -0.3493317 | 0.72684028 | 0.99989001 |
| HIF3A | 415.636877 | 0.3214154 | 0.64384424 | 0.49921298 | 0.61762935 | 0.99989001 |
| PPP5C | 93.8199005 | 0.13683964 | 0.26362436 | 0.51907055 | 0.60371154 | 0.99989001 |
| CALM3 | 89.8665697 | -0.1714694 | 0.45829405 | -0.3741472 | 0.70829482 | 0.99989001 |
| PRKD2 | 339.228589 | 0.00149515 | 0.29382385 | 0.00508858 | 0.99593991 | 0.99989001 |
| STRN4 | 61.0890084 | -0.0366937 | 0.31764437 | -0.1155181 | 0.90803444 | 0.99989001 |
| SLC1A5 | 45.3012869 | 0.18409441 | 0.52944515 | 0.34771195 | 0.72805652 | NA |
| AP2S1 | 149.574564 | -0.1267666 | 0.45504116 | -0.2785828 | 0.78056501 | 0.99989001 |
| ARHGAP35 | 207.323199 | 0.30243776 | 0.29147039 | 1.03762771 | 0.29944341 | 0.99989001 |
| TMEM160 | 77.9066207 | -0.2824265 | 0.45199395 | -0.6248458 | 0.53207225 | 0.99989001 |
| ZC3H4 | 67.9811371 | -0.1220684 | 0.26991517 | -0.4522473 | 0.65109087 | 0.99989001 |
| SAE1 | 126.601123 | -0.2087687 | 0.2783053 | -0.7501428 | 0.45316871 | 0.99989001 |
| NAPA | 93.8480464 | 0.08983267 | 0.349689 | 0.25689304 | 0.79726134 | 0.99989001 |
| EHD2 | 82.2435843 | -0.6933881 | 0.39160536 | -1.7706298 | 0.07662228 | 0.92274869 |
| NOP53 | 53.0544227 | 0.0671169 | 0.44191903 | 0.151876 | 0.87928473 | 0.99989001 |
| SELENOW | 339.704092 | -0.410981 | 0.55465857 | -0.7409622 | 0.45871635 | 0.99989001 |
| PLA2G4C | 164.518446 | -0.2242866 | 0.56818258 | -0.3947438 | 0.69303195 | 0.99989001 |
| CARD8 | 448.090885 | 0.22565746 | 0.13665247 | 1.65132374 | 0.09867249 | 0.99989001 |
| EMP3 | 164.105098 | -0.1983426 | 0.4068733 | -0.4874801 | 0.62591812 | 0.99989001 |
| KDELR1 | 97.444306 | -0.1256091 | 0.32428487 | -0.3873417 | 0.69850325 | 0.99989001 |
| CYTH2 | 61.5118098 | 0.01600987 | 0.35735596 | 0.04480092 | 0.96426599 | 0.99989001 |
| RPL18 | 360.734384 | 0.00919226 | 0.52195651 | 0.01761115 | 0.98594906 | 0.99989001 |
| RASIP1 | 447.168199 | 0.21695861 | 0.28784014 | 0.7537469 | 0.45100122 | 0.99989001 |
| BCAT2 | 45.5738955 | -0.2939078 | 0.27346901 | -1.0747388 | 0.28249167 | NA |
| PPP1R15A | 76.808165 | -0.3257484 | 0.58132447 | -0.5603556 | 0.57523693 | 0.99989001 |
| NUCB1 | 232.4808 | -0.2269405 | 0.35698179 | -0.63572 | 0.52495894 | 0.99989001 |
| BAX | 90.5313542 | -0.1942991 | 0.46874805 | -0.4145064 | 0.67850328 | 0.99989001 |

| | | | | | | |
|---|---|---|---|---|---|---|
| FTL | 2011.864 | 0.91232069 | 0.63015987 | 1.44776069 | 0.14768399 | 0.99989001 |
| GYS1 | 61.9141881 | 0.13711695 | 0.34623465 | 0.39602318 | 0.69208793 | 0.99989001 |
| SNRNP70 | 289.179143 | 0.01481989 | 0.28139544 | 0.0526657 | 0.95799827 | 0.99989001 |
| TRPM4 | 104.625906 | -0.1023623 | 0.28245145 | -0.3624066 | 0.71704824 | 0.99989001 |
| SLC6A16 | 47.2354912 | 0.07635059 | 0.57186186 | 0.1335123 | 0.89378824 | NA |
| TEAD2 | 117.214144 | 0.00981869 | 0.3884902 | 0.02527396 | 0.97983644 | 0.99989001 |
| PIH1D1 | 43.0807116 | 0.25744229 | 0.35354165 | 0.72818093 | 0.46650284 | NA |
| RPL13A | 425.978762 | -0.0014895 | 0.53258646 | -0.0027968 | 0.99776851 | 0.99989001 |
| RPS11 | 405.162516 | -0.0918233 | 0.558467 | -0.1644203 | 0.86940029 | 0.99989001 |
| FCGRT | 201.456486 | -0.3390169 | 0.4148945 | -0.8171159 | 0.41386221 | 0.99989001 |
| NOSIP | 85.9731406 | 0.34463783 | 0.29767237 | 1.15777571 | 0.24695558 | 0.99989001 |
| PRR12 | 71.9611214 | -0.1217062 | 0.37803113 | -0.3219476 | 0.74749241 | 0.99989001 |
| RRAS | 136.304162 | 0.07182298 | 0.47389018 | 0.15156039 | 0.87953368 | 0.99989001 |
| SCAF1 | 45.0254147 | 0.41716571 | 0.34943589 | 1.19382618 | 0.23254597 | NA |
| IRF3 | 92.07753 | -0.0159923 | 0.27495597 | -0.0581631 | 0.9536187 | 0.99989001 |
| PRMT1 | 138.322614 | 0.15283009 | 0.47204691 | 0.32376039 | 0.74611944 | 0.99989001 |
| AP2A1 | 98.1727959 | 0.06550203 | 0.25192999 | 0.2600009 | 0.79486308 | 0.99989001 |
| MED25 | 46.4256913 | 0.07862178 | 0.30429864 | 0.25837047 | 0.796121 | NA |
| PTOV1 | 65.5112592 | 0.04802699 | 0.22597087 | 0.2125362 | 0.83168874 | 0.99989001 |
| PNKP | 69.2550881 | 0.29667855 | 0.51222755 | 0.57919288 | 0.56245903 | 0.99989001 |
| TBC1D17 | 46.93567 | -0.2607879 | 0.31184565 | -0.8362724 | 0.40300167 | NA |
| VRK3 | 71.2663552 | 0.17624844 | 0.35451801 | 0.49714947 | 0.61908365 | 0.99989001 |
| NR1H2 | 72.8894972 | -0.3479729 | 0.31021395 | -1.1217191 | 0.26198191 | 0.99989001 |
| EMC10 | 72.7102396 | -0.2008999 | 0.37326346 | -0.5382254 | 0.59042144 | 0.99989001 |
| ETFB | 88.9827846 | 0.30519522 | 0.39509302 | 0.77246421 | 0.43983953 | 0.99989001 |
| SPACA6 | 66.2596036 | 0.3270785 | 0.45185032 | 0.72386471 | 0.4691488 | 0.99989001 |
| ZNF577 | 58.6558269 | 0.82565257 | 0.35605671 | 2.31887936 | 0.02040158 | 0.58699182 |
| ZNF432 | 77.3050942 | -0.1945887 | 0.30287637 | -0.6424692 | 0.52056861 | 0.99989001 |
| PPP2R1A | 169.489172 | 0.08469521 | 0.24283577 | 0.34877568 | 0.72725772 | 0.99989001 |
| ZNF766 | 53.3870076 | -0.0391589 | 0.45703718 | -0.0856799 | 0.93172088 | 0.99989001 |
| ZNF480 | 44.2384508 | -0.1924074 | 0.33250026 | -0.5786684 | 0.56281291 | NA |
| ZNF701 | 52.0322803 | 0.27562534 | 0.29323116 | 0.93995925 | 0.34723846 | 0.99989001 |
| ZNF83 | 383.528375 | 0.45270804 | 0.26090247 | 1.73516193 | 0.08271213 | 0.94799019 |
| ZNF611 | 124.027329 | -0.0340742 | 0.34318562 | -0.0992879 | 0.9209097 | 0.99989001 |
| ZNF160 | 233.970597 | -0.1077489 | 0.37695021 | -0.2858438 | 0.77499777 | 0.99989001 |
| ZNF347 | 116.726091 | -0.0116941 | 0.37115261 | -0.0315075 | 0.97486482 | 0.99989001 |
| ZNF677 | 42.4596604 | 0.4889481 | 0.25984141 | 1.88171736 | 0.0598744 | NA |
| ZNF765 | 63.5458076 | 0.65684109 | 0.34880423 | 1.88312249 | 0.05968377 | 0.85885114 |
| ZNF761 | 71.6398835 | 0.24690751 | 0.28468076 | 0.86731365 | 0.38577018 | 0.99989001 |
| ZNF331 | 63.5879627 | -0.2455907 | 0.31042003 | -0.7911562 | 0.42885285 | 0.99989001 |
| MYADM | 54.3906425 | -0.4033165 | 0.56322375 | -0.7160857 | 0.47393845 | 0.99989001 |
| PRPF31 | 66.2063173 | 0.00578203 | 0.33484773 | 0.01726765 | 0.98622309 | 0.99989001 |
| CNOT3 | 40.5076621 | -0.4102234 | 0.29507741 | -1.390223 | 0.16446117 | NA |
| MBOAT7 | 64.723431 | -0.2218819 | 0.42625136 | -0.5205425 | 0.60268553 | 0.99989001 |
| RPS9 | 667.789753 | -0.1578834 | 0.56689149 | -0.2785072 | 0.78062301 | 0.99989001 |
| LENG8 | 196.49548 | -0.0944019 | 0.30000484 | -0.3146679 | 0.75301384 | 0.99989001 |
| CDC42EP5 | 130.792411 | -0.1919347 | 0.43232297 | -0.4439614 | 0.6570705 | 0.99989001 |
| AC011476.3 | 73.4669015 | 0.16191558 | 0.52350083 | 0.30929383 | 0.75709802 | 0.99989001 |
| RDH13 | 48.0250329 | 0.31020588 | 0.29732299 | 1.04332962 | 0.29679566 | NA |
| PPP1R12C | 91.7356398 | -0.0529567 | 0.27925781 | -0.1896339 | 0.84959601 | 0.99989001 |
| PPP6R1 | 71.5502365 | -0.4491135 | 0.32522431 | -1.3809345 | 0.16729909 | 0.99989001 |
| RPL28 | 515.471739 | -0.1985409 | 0.52036954 | -0.3815382 | 0.70280391 | 0.99989001 |
| ISOC2 | 43.2117194 | 0.29562904 | 0.50734952 | 0.58269304 | 0.56009996 | NA |
| U2AF2 | 60.5671954 | 0.01016852 | 0.2360675 | 0.04307463 | 0.96564204 | 0.99989001 |

| | | | | | | |
|---|---|---|---|---|---|---|
| EPN1 | 98.8442883 | -0.3777056 | 0.27925443 | -1.3525502 | 0.17619937 | 0.99989001 |
| ZNF787 | 53.5886473 | -0.0889212 | 0.32432215 | -0.2741757 | 0.78394964 | 0.99989001 |
| ZNF444 | 50.9937021 | 0.23105524 | 0.25566848 | 0.90372983 | 0.36613868 | 0.99989001 |
| ZSCAN5A | 121.634202 | 0.113799 | 0.46071422 | 0.24700562 | 0.80490387 | 0.99989001 |
| ZNF667-AS1 | 42.8933265 | 0.3431104 | 0.28286527 | 1.2129817 | 0.22513682 | NA |
| ZNF264 | 86.6862242 | 0.41816584 | 0.2754148 | 1.51831289 | 0.12893554 | 0.99989001 |
| AC005261.1 | 113.61839 | 0.58161687 | 0.24511191 | 2.37286257 | 0.01765083 | 0.56214542 |
| ZNF586 | 182.376279 | 0.59948645 | 0.37504989 | 1.59841789 | 0.10995001 | 0.99989001 |
| ZNF814 | 108.678833 | 0.75754581 | 0.4010209 | 1.88904319 | 0.05888604 | 0.85640047 |
| ZNF587 | 53.7235419 | 0.44306569 | 0.33056127 | 1.34034365 | 0.18013365 | 0.99989001 |
| ZSCAN18 | 81.135726 | 0.63939237 | 0.33394351 | 1.91467227 | 0.05553431 | 0.83004552 |
| ZNF329 | 41.2656666 | 0.47825029 | 0.33170653 | 1.44178738 | 0.14936236 | NA |
| ZNF274 | 77.1166826 | 0.52305262 | 0.51150682 | 1.02257214 | 0.30651019 | 0.99989001 |
| RPS5 | 203.166232 | 0.04319868 | 0.55298711 | 0.07811878 | 0.93773357 | 0.99989001 |
| TRIM28 | 83.2445248 | 0.08966967 | 0.4076974 | 0.21994173 | 0.82591654 | 0.99989001 |
| CHMP2A | 93.346665 | -0.2606904 | 0.4945144 | -0.5271644 | 0.59807944 | 0.99989001 |
| UBE2M | 64.4405863 | -0.1897776 | 0.47136369 | -0.4026138 | 0.68723232 | 0.99989001 |
| MZF1 | 68.7103467 | -0.3793438 | 0.35517759 | -1.0680397 | 0.28550261 | 0.99989001 |
| RPS4Y1 | 146.633925 | 0.24068771 | 2.55645847 | 0.09414888 | 0.92499089 | 0.99989001 |
| ZFY | 67.4852667 | 0.21227616 | 2.28827664 | 0.09276683 | 0.92608881 | 0.99989001 |
| LINC00278 | 108.169962 | 0.24358417 | 2.34500039 | 0.10387383 | 0.91726948 | 0.99989001 |
| AC010737.1 | 699.831968 | -0.0490711 | 1.59343734 | -0.0307958 | 0.97543242 | 0.99989001 |
| PCDH11Y | 285.58056 | 0.38170583 | 1.76495917 | 0.21626893 | 0.82877813 | 0.99989001 |
| TTTY15 | 44.2431374 | 0.07181225 | 2.08960537 | 0.03436642 | 0.97258496 | NA |
| USP9Y | 192.760579 | 0.47717148 | 2.57373548 | 0.18540036 | 0.85291503 | 0.99989001 |
| DDX3Y | 103.524944 | 0.30467317 | 2.7390456 | 0.11123333 | 0.91143132 | 0.99989001 |
| UTY | 438.290434 | 0.24864333 | 2.24918227 | 0.11054833 | 0.91197453 | 0.99989001 |
| NLGN4Y | 159.484895 | -0.5921532 | 2.05319815 | -0.2884053 | 0.77303654 | 0.99989001 |
| TTTY14 | 398.89677 | 0.44636697 | 1.64191716 | 0.27185718 | 0.78573184 | 0.99989001 |
| KDM5D | 71.4796708 | 0.15933201 | 2.62252019 | 0.06075531 | 0.95155409 | 0.99989001 |
| TTTY10 | 50.6347737 | 0.10084662 | 1.72040075 | 0.0586181 | 0.95325629 | 0.99989001 |
| IL17RA | 102.573921 | -0.0752147 | 0.23463209 | -0.3205643 | 0.74854063 | 0.99989001 |
| HDHD5 | 58.4906411 | -0.0817402 | 0.26744191 | -0.3056373 | 0.75988083 | 0.99989001 |
| ATP6V1E1 | 148.823644 | -0.0572312 | 0.29620384 | -0.1932156 | 0.84679014 | 0.99989001 |
| BCL2L13 | 168.275717 | 0.1663024 | 0.38572187 | 0.4311459 | 0.66636229 | 0.99989001 |
| BID | 52.9459687 | 0.05387493 | 0.4416101 | 0.12199659 | 0.90290172 | 0.99989001 |
| MICAL3 | 320.868058 | -0.1453829 | 0.26168868 | -0.5555566 | 0.578514 | 0.99989001 |
| PEX26 | 80.7788059 | 0.0487826 | 0.31552277 | 0.1546088 | 0.87712973 | 0.99989001 |
| DGCR2 | 137.557277 | 0.12051304 | 0.31245257 | 0.38570026 | 0.69971867 | 0.99989001 |
| SLC25A1 | 42.9876607 | 0.39763191 | 0.41999222 | 0.94676018 | 0.343761 | NA |
| CLTCL1 | 52.3270649 | 0.07771962 | 0.36011324 | 0.21581995 | 0.8291281 | 0.99989001 |
| HIRA | 62.3783354 | -0.3205912 | 0.34167161 | -0.938302 | 0.34808922 | 0.99989001 |
| MRPL40 | 50.8965781 | -0.2746731 | 0.54151893 | -0.5072271 | 0.61199548 | 0.99989001 |
| UFD1 | 66.6374295 | -0.0175344 | 0.27199135 | -0.0644666 | 0.94859867 | 0.99989001 |
| CLDN5 | 1031.62809 | -0.7288048 | 0.60532061 | -1.203998 | 0.22859036 | 0.99989001 |
| TXNRD2 | 43.8913333 | -0.0202405 | 0.44415429 | -0.0455708 | 0.96365236 | NA |
| COMT | 167.749829 | -0.1143054 | 0.31192169 | -0.3664554 | 0.71402525 | 0.99989001 |
| TANGO2 | 52.024928 | -0.2504505 | 0.3212196 | -0.7796863 | 0.43557553 | 0.99989001 |
| DGCR8 | 71.9892605 | -0.1069504 | 0.43241592 | -0.2473322 | 0.80465113 | 0.99989001 |
| RANBP1 | 59.7161123 | -0.3386155 | 0.39970979 | -0.8471534 | 0.39690963 | 0.99989001 |
| ZDHHC8 | 56.4848887 | -0.6285321 | 0.29858388 | -2.1050437 | 0.03528751 | 0.69174161 |
| KLHL22 | 65.6502918 | 0.30745066 | 0.49061398 | 0.6266651 | 0.53087879 | 0.99989001 |
| MED15 | 233.840666 | -0.0579885 | 0.27098942 | -0.2139879 | 0.8305565 | 0.99989001 |
| PI4KA | 312.986253 | 0.36660321 | 0.27986378 | 1.30993445 | 0.19021801 | 0.99989001 |

| | | | | | | |
|---|---|---|---|---|---|---|
| SNAP29 | 66.8195061 | 0.01218248 | 0.36170889 | 0.03368034 | 0.97313206 | 0.99989001 |
| CRKL | 57.721499 | 0.2941031 | 0.29270336 | 1.00478213 | 0.31500177 | 0.99989001 |
| LZTR1 | 44.0423214 | 0.06281096 | 0.27412625 | 0.22913151 | 0.81876671 | NA |
| HIC2 | 46.7097096 | -0.1683632 | 0.32281908 | -0.5215403 | 0.60199043 | NA |
| UBE2L3 | 191.495925 | -0.0132065 | 0.22585249 | -0.058474 | 0.95337109 | 0.99989001 |
| SDF2L1 | 63.1741628 | 0.29069543 | 0.5967794 | 0.48710702 | 0.62618249 | 0.99989001 |
| PPIL2 | 92.3567316 | -0.1956226 | 0.32906949 | -0.5944721 | 0.55219638 | 0.99989001 |
| YPEL1 | 45.5071934 | 0.17841757 | 0.50452146 | 0.35363722 | 0.72361077 | NA |
| MAPK1 | 447.021315 | 0.12767561 | 0.18440763 | 0.69235534 | 0.4887142 | 0.99989001 |
| PPM1F | 380.448925 | -0.0215438 | 0.35929067 | -0.0599619 | 0.95218594 | 0.99989001 |
| BCR | 401.185654 | -0.0316637 | 0.35072524 | -0.0902806 | 0.92806425 | 0.99989001 |
| CHCHD10 | 70.1074333 | 0.18460887 | 0.31290024 | 0.58999276 | 0.5551955 | 0.99989001 |
| SMARCB1 | 80.2486159 | -0.1451874 | 0.26904054 | -0.5396489 | 0.58943919 | 0.99989001 |
| DDT | 90.6344664 | 0.18928162 | 0.43963473 | 0.43054291 | 0.66680076 | 0.99989001 |
| CABIN1 | 198.284972 | -0.0992041 | 0.17555685 | -0.5650823 | 0.5720178 | 0.99989001 |
| GGT5 | 121.083032 | 0.11772449 | 0.59722603 | 0.19711882 | 0.84373455 | 0.99989001 |
| UPB1 | 53.1874246 | -0.3748892 | 0.59412464 | -0.6309942 | 0.52804431 | 0.99989001 |
| GUCD1 | 48.1234175 | -0.2432002 | 0.33927447 | -0.7168243 | 0.47348253 | NA |
| SNRPD3 | 121.404929 | -0.1375949 | 0.30974794 | -0.4442156 | 0.65688672 | 0.99989001 |
| SGSM1 | 64.2484759 | 1.21226418 | 0.54950149 | 2.20611629 | 0.02737586 | 0.65183905 |
| KIAA1671 | 732.93016 | 0.14673827 | 0.40257102 | 0.36450282 | 0.71548257 | 0.99989001 |
| AL022323.4 | 58.5739421 | 0.42328062 | 0.38379012 | 1.10289607 | 0.2700723 | 0.99989001 |
| LRP5L | 52.5601261 | 0.38941685 | 0.54364345 | 0.71630929 | 0.47380039 | 0.99989001 |
| GRK3 | 67.7142685 | -0.0126288 | 0.35672626 | -0.035402 | 0.97175918 | 0.99989001 |
| HPS4 | 64.3486758 | -0.3853657 | 0.34447782 | -1.1186953 | 0.26327013 | 0.99989001 |
| TPST2 | 265.572567 | 0.15094366 | 0.20323923 | 0.74268958 | 0.45766964 | 0.99989001 |
| MIATNB | 111.159858 | 0.05749467 | 0.4243942 | 0.13547468 | 0.89223658 | 0.99989001 |
| MN1 | 123.736553 | 0.5145597 | 0.48323537 | 1.06482209 | 0.28695645 | 0.99989001 |
| PITPNB | 443.446118 | 0.06070609 | 0.29340436 | 0.20690247 | 0.83608603 | 0.99989001 |
| TTC28 | 2410.62059 | -0.069724 | 0.20125657 | -0.3464436 | 0.72900938 | 0.99989001 |
| HSCB | 60.5393233 | -0.0939619 | 0.30148346 | -0.3116653 | 0.75529489 | 0.99989001 |
| XBP1 | 92.3163638 | 0.85957214 | 0.43178744 | 1.99072985 | 0.0465106 | 0.77516398 |
| Z93930.2 | 112.715068 | 0.04111333 | 0.47223319 | 0.08706151 | 0.93062262 | 0.99989001 |
| ZNRF3 | 117.887088 | -0.3651189 | 0.30502864 | -1.1969986 | 0.23130709 | 0.99989001 |
| EMID1 | 186.495564 | 1.08172174 | 0.45390148 | 2.38316418 | 0.01716454 | 0.55366411 |
| EWSR1 | 282.241076 | -0.1141084 | 0.21393217 | -0.5333858 | 0.59376657 | 0.99989001 |
| GAS2L1 | 58.6369017 | 0.16669926 | 0.52378808 | 0.31825708 | 0.75028994 | 0.99989001 |
| AP1B1 | 105.831999 | 0.33297264 | 0.27503361 | 1.21066165 | 0.22602511 | 0.99989001 |
| THOC5 | 50.7857009 | -0.2194409 | 0.27352983 | -0.8022558 | 0.42240499 | 0.99989001 |
| NF2 | 89.5260906 | 0.17467211 | 0.31001657 | 0.56342828 | 0.57314328 | 0.99989001 |
| UQCR10 | 130.736303 | 0.04610409 | 0.5100879 | 0.09038459 | 0.9279816 | 0.99989001 |
| ASCC2 | 89.2236599 | 0.02250648 | 0.35680229 | 0.06307828 | 0.94970417 | 0.99989001 |
| MTMR3 | 354.351178 | 0.15683299 | 0.2173253 | 0.72165086 | 0.47050916 | 0.99989001 |
| TBC1D10A | 47.6478073 | 0.33460654 | 0.31563435 | 1.06010812 | 0.28909541 | NA |
| SF3A1 | 62.4320708 | 0.11613203 | 0.3392492 | 0.34232072 | 0.73210954 | 0.99989001 |
| RNF215 | 44.8892007 | -0.4070931 | 0.29617533 | -1.3745004 | 0.1692864 | NA |
| PES1 | 87.5857398 | 0.01909363 | 0.27808135 | 0.06866203 | 0.94525864 | 0.99989001 |
| TCN2 | 164.324295 | 0.35478026 | 0.24931797 | 1.42300318 | 0.15473523 | 0.99989001 |
| SLC35E4 | 62.9508213 | 0.01890398 | 0.38906166 | 0.04858864 | 0.96124712 | 0.99989001 |
| DUSP18 | 69.1769848 | -0.0990605 | 0.24594433 | -0.4027763 | 0.68711281 | 0.99989001 |
| OSBP2 | 75.7535472 | -0.1357423 | 0.45460266 | -0.2985955 | 0.76524867 | 0.99989001 |
| MORC2 | 58.5258534 | 0.22281248 | 0.25739225 | 0.86565343 | 0.38668025 | 0.99989001 |
| SMTN | 142.211521 | 0.2449372 | 0.24462694 | 1.0012683 | 0.31669711 | 0.99989001 |
| SELENOM | 232.025059 | -0.1561681 | 0.60594672 | -0.2577258 | 0.79661854 | 0.99989001 |

| | | | | | | |
|---|---|---|---|---|---|---|
| RNF185 | 110.798085 | -0.159883 | 0.27996483 | -0.5710823 | 0.56794384 | 0.99989001 |
| LIMK2 | 666.48807 | 0.13329561 | 0.4775997 | 0.27909485 | 0.78017204 | 0.99989001 |
| PIK3IP1 | 37.6267728 | 1.48807382 | 0.71167219 | 2.090954 | 0.03653219 | NA |
| DRG1 | 99.2765421 | -0.0200262 | 0.21770624 | -0.0919873 | 0.9267081 | 0.99989001 |
| EIF4ENIF1 | 103.048458 | 0.12918687 | 0.265135 | 0.4872494 | 0.6260816 | 0.99989001 |
| SFI1 | 90.7608134 | -0.224032 | 0.41154136 | -0.5443729 | 0.58618487 | 0.99989001 |
| PISD | 86.7149531 | -0.512494 | 0.30393789 | -1.6861801 | 0.09176111 | 0.99175102 |
| PRR14L | 218.019879 | -0.0382082 | 0.25231433 | -0.1514309 | 0.87963578 | 0.99989001 |
| YWHAH | 126.032213 | 0.0280459 | 0.52321504 | 0.053603 | 0.95725146 | 0.99989001 |
| SLC5A4-AS1 | 73.4165853 | 0.53200641 | 0.5603101 | 0.94948567 | 0.34237366 | 0.99989001 |
| SLC5A4 | 132.445778 | 0.6097893 | 0.59898999 | 1.01802918 | 0.30866409 | 0.99989001 |
| AL021937.3 | 60.6032057 | 0.65608246 | 0.70154819 | 0.93519228 | 0.34968921 | 0.99989001 |
| RTCB | 107.140881 | 0.13503042 | 0.23808006 | 0.56716389 | 0.57060284 | 0.99989001 |
| FBXO7 | 170.854796 | 0.0545756 | 0.24452607 | 0.22318929 | 0.82338818 | 0.99989001 |
| SYN3 | 517.8246 | 0.02720266 | 0.6664472 | 0.04081743 | 0.96744145 | 0.99989001 |
| TIMP3 | 6055.86241 | -0.3632028 | 0.39538421 | -0.9186072 | 0.35830108 | 0.99989001 |
| LARGE1 | 601.588273 | 0.43157311 | 0.27138323 | 1.59027186 | 0.11177354 | 0.99989001 |
| HMGXB4 | 103.551355 | -0.1650001 | 0.27752317 | -0.5945453 | 0.55214748 | 0.99989001 |
| TOM1 | 57.9278377 | -0.3002365 | 0.33789291 | -0.8885552 | 0.37424216 | 0.99989001 |
| APOL6 | 178.923032 | 0.28583081 | 0.55611988 | 0.51397337 | 0.60727061 | 0.99989001 |
| RBFOX2 | 1302.05997 | 0.14641336 | 0.18912904 | 0.77414533 | 0.43884485 | 0.99989001 |
| APOL3 | 255.097372 | 0.83935986 | 0.37528113 | 2.23661621 | 0.02531143 | 0.63387007 |
| APOL2 | 99.242933 | 0.20818308 | 0.53919783 | 0.38609777 | 0.69942426 | 0.99989001 |
| APOL1 | 184.01114 | -0.3267444 | 0.45105974 | -0.7243928 | 0.46882462 | 0.99989001 |
| MYH9 | 1318.42092 | -0.6592253 | 0.32584181 | -2.0231453 | 0.04305817 | 0.76471303 |
| TXN2 | 59.1621142 | -0.0170632 | 0.27883076 | -0.0611957 | 0.95120337 | 0.99989001 |
| EIF3D | 116.153267 | 0.1070286 | 0.36455933 | 0.2935835 | 0.76907619 | 0.99989001 |
| MPST | 45.5865247 | 0.75966809 | 0.35026713 | 2.16882498 | 0.03009597 | NA |
| MFNG | 75.662845 | 0.19933105 | 0.39714997 | 0.50190374 | 0.61573524 | 0.99989001 |
| CARD10 | 94.3865911 | 0.01438554 | 0.33884658 | 0.04245444 | 0.96613643 | 0.99989001 |
| CDC42EP1 | 74.8402053 | -0.3945472 | 0.38991237 | -1.0118869 | 0.31159216 | 0.99989001 |
| GGA1 | 85.4730076 | -0.0493874 | 0.27117934 | -0.1821206 | 0.85548804 | 0.99989001 |
| LGALS1 | 255.376636 | -0.0386817 | 0.62004967 | -0.0623849 | 0.95025633 | 0.99989001 |
| TRIOBP | 283.237824 | 0.01295356 | 0.26969608 | 0.04803023 | 0.96169215 | 0.99989001 |
| H1F0 | 94.3751894 | -0.0805816 | 0.74077954 | -0.1087795 | 0.91337741 | 0.99989001 |
| EIF3L | 142.702274 | -0.1697555 | 0.30455876 | -0.5573816 | 0.57726671 | 0.99989001 |
| MICALL1 | 78.1324589 | -0.2092095 | 0.3485331 | -0.6002573 | 0.54833477 | 0.99989001 |
| POLR2F | 65.6159621 | -0.4610486 | 0.31292347 | -1.4733589 | 0.14065428 | 0.99989001 |
| MAFF | 71.6302435 | -0.9115862 | 0.7676248 | -1.1875413 | 0.23501416 | 0.99989001 |
| TMEM184B | 317.469142 | 0.12574894 | 0.27384653 | 0.45919494 | 0.64609419 | 0.99989001 |
| CSNK1E | 76.5595677 | -0.360045 | 0.50491584 | -0.7130793 | 0.47579667 | 0.99989001 |
| TPTEP2-CSNI | 423.851546 | 0.44435354 | 0.26372031 | 1.68494245 | 0.09199968 | 0.99175102 |
| DDX17 | 1403.88874 | 0.04585655 | 0.26914548 | 0.17037832 | 0.86471262 | 0.99989001 |
| JOSD1 | 103.568216 | -0.4410803 | 0.2938779 | -1.5008966 | 0.13338231 | 0.99989001 |
| GTPBP1 | 81.1116085 | 0.10396625 | 0.50190946 | 0.20714144 | 0.8358994 | 0.99989001 |
| SUN2 | 137.137532 | -0.0008612 | 0.42816671 | -0.0020114 | 0.99839512 | 0.99989001 |
| AL021707.1 | 54.4255409 | 0.194752 | 0.54763041 | 0.3556267 | 0.72212015 | 0.99989001 |
| CBX6 | 74.1790438 | -0.3603102 | 0.52426685 | -0.6872649 | 0.49191584 | 0.99989001 |
| CBX7 | 73.206843 | -0.0631539 | 0.41272968 | -0.1530151 | 0.87838634 | 0.99989001 |
| PDGFB | 185.105672 | 0.11278174 | 0.2840913 | 0.39699118 | 0.69137397 | 0.99989001 |
| RPL3 | 340.154832 | 0.20646558 | 0.58073308 | 0.35552578 | 0.72219573 | 0.99989001 |
| TAB1 | 60.7664693 | 0.19315453 | 0.3039491 | 0.63548315 | 0.52511335 | 0.99989001 |
| MIEF1 | 39.5320449 | -0.0502849 | 0.36994367 | -0.1359259 | 0.89187987 | NA |
| ATF4 | 165.091728 | -0.1587187 | 0.29600663 | -0.5361999 | 0.59182039 | 0.99989001 |

| | | | | | | |
|---|---|---|---|---|---|---|
| RPS19BP1 | 127.467042 | -0.0665338 | 0.52711499 | -0.1262225 | 0.89955576 | 0.99989001 |
| TNRC6B | 1341.25308 | 0.12613184 | 0.22266033 | 0.56647648 | 0.57106992 | 0.99989001 |
| MKL1 | 711.577869 | 0.28888593 | 0.36650894 | 0.78820978 | 0.43057401 | 0.99989001 |
| SLC25A17 | 115.700736 | 0.14321904 | 0.27558508 | 0.51969083 | 0.60327908 | 0.99989001 |
| ST13 | 173.156136 | 0.30637948 | 0.20966924 | 1.46125144 | 0.14394645 | 0.99989001 |
| XPNPEP3 | 126.603652 | -0.0515825 | 0.29893912 | -0.1725518 | 0.86300376 | 0.99989001 |
| RBX1 | 139.652447 | -0.3884382 | 0.24374515 | -1.5936243 | 0.11102021 | 0.99989001 |
| EP300 | 314.37141 | 0.08049521 | 0.2153437 | 0.37379877 | 0.70855404 | 0.99989001 |
| L3MBTL2 | 39.7347698 | -0.2642376 | 0.26793255 | -0.9862092 | 0.32403046 | NA |
| RANGAP1 | 51.1948648 | -0.2411515 | 0.31828982 | -0.7576475 | 0.44866203 | 0.99989001 |
| ZC3H7B | 122.610693 | -0.2433026 | 0.33043518 | -0.7363097 | 0.46154223 | 0.99989001 |
| TOB2 | 79.0776801 | 0.19209863 | 0.21690782 | 0.88562333 | 0.37582054 | 0.99989001 |
| ACO2 | 111.178312 | 0.13581647 | 0.26575282 | 0.51106313 | 0.60930685 | 0.99989001 |
| DESI1 | 67.2100817 | 0.23638537 | 0.29115557 | 0.81188683 | 0.41685657 | 0.99989001 |
| XRCC6 | 226.202518 | -0.0718612 | 0.26441638 | -0.271773 | 0.78579655 | 0.99989001 |
| SNU13 | 225.539435 | -0.1132676 | 0.35864192 | -0.3158237 | 0.75213635 | 0.99989001 |
| SREBF2 | 277.917078 | 0.0567367 | 0.32502558 | 0.17456073 | 0.86142481 | 0.99989001 |
| SMDT1 | 70.2526887 | -0.3070223 | 0.40209831 | -0.7635504 | 0.44513521 | 0.99989001 |
| NDUFA6 | 55.4397701 | -0.2448656 | 0.54536741 | -0.4489921 | 0.65343738 | 0.99989001 |
| TCF20 | 180.867485 | 0.4690958 | 0.22424497 | 2.09188994 | 0.03644836 | 0.70003115 |
| RRP7A | 71.2828678 | 0.11759039 | 0.4640737 | 0.25338732 | 0.79996893 | 0.99989001 |
| POLDIP3 | 72.1024814 | -0.0146561 | 0.24988727 | -0.0586509 | 0.95323015 | 0.99989001 |
| CYB5R3 | 272.80596 | -0.3508161 | 0.4212175 | -0.8328621 | 0.40492251 | 0.99989001 |
| A4GALT | 144.104605 | 0.08312254 | 0.27489011 | 0.30238463 | 0.76235887 | 0.99989001 |
| ARFGAP3 | 193.720625 | -0.2142376 | 0.1480653 | -1.4469129 | 0.14792132 | 0.99989001 |
| PACSIN2 | 288.773126 | 0.22781472 | 0.26359287 | 0.8642674 | 0.38744102 | 0.99989001 |
| TSPO | 212.377666 | -0.2881796 | 0.55131576 | -0.5227124 | 0.60117442 | 0.99989001 |
| SCUBE1 | 77.6532964 | -0.4640368 | 0.54315881 | -0.8543298 | 0.39292228 | 0.99989001 |
| SAMM50 | 45.6428591 | -0.5338765 | 0.24331719 | -2.1941585 | 0.02822402 | NA |
| PARVB | 345.16568 | 0.27442422 | 0.25080067 | 1.09419252 | 0.27387055 | 0.99989001 |
| NUP50-DT | 96.9873597 | 0.04253589 | 0.27229879 | 0.15621037 | 0.8758672 | 0.99989001 |
| NUP50 | 85.9978529 | 0.08842719 | 0.2462038 | 0.35916257 | 0.71947348 | 0.99989001 |
| FAM118A | 94.8601791 | 0.86675394 | 0.48213335 | 1.79774733 | 0.07221706 | 0.91336395 |
| ATXN10 | 426.761225 | 0.13686003 | 0.20105805 | 0.68069909 | 0.49606191 | 0.99989001 |
| MIRLET7BHG | 281.355478 | 0.45203312 | 0.36217151 | 1.24811894 | 0.2119875 | 0.99989001 |
| PPARA | 107.927486 | 0.55298177 | 0.38258113 | 1.44539739 | 0.1483463 | 0.99989001 |
| TTC38 | 60.2014229 | 0.61908211 | 0.28809359 | 2.14889232 | 0.03164294 | 0.67281094 |
| TRMU | 57.4683946 | -0.2280889 | 0.23809326 | -0.9579813 | 0.33807217 | 0.99989001 |
| CELSR1 | 112.250402 | -0.0625691 | 0.36200903 | -0.1728384 | 0.86277844 | 0.99989001 |
| GRAMD4 | 49.6868308 | 0.56779813 | 0.36831259 | 1.54162022 | 0.12316591 | NA |
| CERK | 133.733693 | 0.22949364 | 0.33244428 | 0.69032213 | 0.48999164 | 0.99989001 |
| TBC1D22A | 933.345219 | -0.0755363 | 0.29709832 | -0.254247 | 0.79930477 | 0.99989001 |
| C22orf34 | 464.829102 | 0.76689448 | 0.59941404 | 1.27940693 | 0.2007538 | 0.99989001 |
| BRD1 | 122.457055 | 0.23877426 | 0.34483341 | 0.69243366 | 0.48866503 | 0.99989001 |
| ZBED4 | 69.9028956 | 0.22326047 | 0.25861031 | 0.86330845 | 0.3879679 | 0.99989001 |
| PIM3 | 258.29277 | 0.60653594 | 0.41267914 | 1.46975188 | 0.14162897 | 0.99989001 |
| TRABD | 43.3015592 | -0.3868567 | 0.27844869 | -1.3893284 | 0.16473291 | NA |
| SELENOO | 87.453018 | -0.1969555 | 0.26624229 | -0.7397604 | 0.45944541 | 0.99989001 |
| MAPK12 | 56.9249268 | 0.04702109 | 0.41146632 | 0.11427688 | 0.90901831 | 0.99989001 |
| MAPK11 | 73.1934321 | -0.1196461 | 0.33723885 | -0.3547814 | 0.72275337 | 0.99989001 |
| PLXNB2 | 102.802215 | 0.09755681 | 0.44277425 | 0.2203308 | 0.82561353 | 0.99989001 |
| PPP6R2 | 383.147183 | -0.0226459 | 0.26488372 | -0.0854936 | 0.93186895 | 0.99989001 |
| LMF2 | 55.8326071 | -0.1990076 | 0.31477973 | -0.6322124 | 0.52724812 | 0.99989001 |
| TYMP | 57.1523127 | 0.68530769 | 1.08980668 | 0.62883418 | 0.52945762 | 0.99989001 |

| | | | | | | |
|---|---|---|---|---|---|---|
| SHANK3 | 1157.62723 | -0.2375971 | 0.22459567 | -1.0578881 | 0.29010645 | 0.99989001 |
| FP671120.1 | 1265.30278 | -0.6819453 | 0.31810213 | -2.1437937 | 0.03204942 | 0.67281094 |
| FP236383.1 | 715.169669 | -0.7366673 | 0.4130805 | -1.7833504 | 0.07452928 | 0.9206525 |
| AP001347.1 | 82.4811549 | -0.45196 | 0.49321924 | -0.916347 | 3.59E-01 | 0.99989001 |
| HSPA13 | 48.468505 | 0.14561273 | 0.31888464 | 0.45663137 | 0.64793603 | NA |
| NRIP1 | 343.895573 | -0.306108 | 0.40481388 | -0.7561698 | 0.44954738 | 0.99989001 |
| USP25 | 607.075053 | -0.1540336 | 0.23069994 | -0.6676793 | 0.50433834 | 0.99989001 |
| MIR99AHG | 2887.36984 | -0.4495522 | 0.52649888 | -0.8538521 | 0.39318694 | 0.99989001 |
| BTG3 | 58.9489609 | 0.32381591 | 0.2796301 | 1.15801522 | 0.24685783 | 0.99989001 |
| MRPL39 | 52.5501056 | -0.2581429 | 0.40442343 | -0.6382985 | 0.52327938 | 0.99989001 |
| JAM2 | 398.512603 | -0.4958574 | 0.49637422 | -0.9989587 | 0.31781468 | 0.99989001 |
| ATP5PF | 148.153829 | 0.17980735 | 0.4782467 | 0.37597197 | 0.70693773 | 0.99989001 |
| GABPA | 146.454673 | -0.2164974 | 0.25976994 | -0.8334197 | 0.40460808 | 0.99989001 |
| APP | 4121.71781 | 0.02435777 | 0.25039152 | 0.09727873 | 0.92250505 | 0.99989001 |
| AP001439.1 | 69.4760556 | -0.1376817 | 0.46864557 | -0.2937864 | 0.7689211 | 0.99989001 |
| CYYR1-AS1 | 60.0859846 | -0.3670473 | 0.69238289 | -0.5301218 | 0.59602747 | 0.99989001 |
| CYYR1 | 826.034787 | -0.7759753 | 0.38795794 | -2.0001532 | 0.04548372 | 0.77228819 |
| ADAMTS1 | 645.579713 | -0.3088904 | 0.51004691 | -0.6056117 | 0.5447726 | 0.99989001 |
| LTN1 | 167.024448 | 0.10471704 | 0.23474443 | 0.44608955 | 0.65553256 | 0.99989001 |
| USP16 | 122.001204 | 0.00372999 | 0.24010787 | 0.01553466 | 0.98760564 | 0.99989001 |
| CCT8 | 85.2683638 | 0.19834195 | 0.27526887 | 0.72053896 | 0.47119322 | 0.99989001 |
| MAP3K7CL | 50.6245147 | 0.19645665 | 0.4304573 | 0.45639058 | 0.64810914 | 0.99989001 |
| BACH1 | 549.099952 | 0.53421354 | 0.39838084 | 1.34096193 | 0.17993281 | 0.99989001 |
| TIAM1 | 211.825301 | -0.561676 | 0.89670322 | -0.626379 | 0.53106641 | 0.99989001 |
| SOD1 | 164.071936 | 0.15201607 | 0.46081262 | 0.32988695 | 0.74148538 | 0.99989001 |
| SCAF4 | 162.890803 | 0.05043659 | 0.20133275 | 0.25051362 | 0.80219018 | 0.99989001 |
| URB1 | 75.0612661 | -0.1692687 | 0.49015409 | -0.3453377 | 0.72984052 | 0.99989001 |
| EVA1C | 967.528153 | -0.2068882 | 0.34578373 | -0.5983168 | 0.54962855 | 0.99989001 |
| SYNJ1 | 180.062139 | 0.25122198 | 0.22434779 | 1.11978805 | 0.26280409 | 0.99989001 |
| PAXBP1 | 210.855913 | -0.1747457 | 0.20312118 | -0.8603029 | 0.38962211 | 0.99989001 |
| IFNAR1 | 222.243256 | 0.2656389 | 0.26230539 | 1.01270848 | 0.31119943 | 0.99989001 |
| IFNGR2 | 201.559479 | 0.35479058 | 0.24391568 | 1.45456242 | 0.14579044 | 0.99989001 |
| TMEM50B | 182.823087 | 0.01253668 | 0.27996941 | 0.04477874 | 0.96428367 | 0.99989001 |
| GART | 90.0621104 | -0.1818461 | 0.35470725 | -0.5126654 | 0.60818541 | 0.99989001 |
| SON | 659.073416 | 0.10221353 | 0.24209355 | 0.42220676 | 0.67287411 | 0.99989001 |
| AP000311.1 | 55.466181 | 0.50569383 | 0.46095254 | 1.09706267 | 0.272614 | 0.99989001 |
| ITSN1 | 192.220159 | 0.31136666 | 0.33283249 | 0.93550561 | 0.34952779 | 0.99989001 |
| MRPS6 | 121.556842 | -1.7844666 | 0.64631425 | -2.7609892 | 0.00576266 | 0.3686148 |
| RCAN1 | 140.0604 | 0.35972294 | 0.3583902 | 1.0037187 | 0.31551422 | 0.99989001 |
| SETD4 | 45.1622539 | -0.0831037 | 0.2587845 | -0.3211309 | 0.74811119 | NA |
| CBR1 | 59.146264 | 0.45102651 | 0.41936805 | 1.07549087 | 0.28215502 | 0.99989001 |
| DOPEY2 | 93.7357746 | -0.0489579 | 0.44567767 | -0.1098505 | 0.91252796 | 0.99989001 |
| MORC3 | 216.971352 | -0.0122542 | 0.26383409 | -0.0464468 | 0.96295416 | 0.99989001 |
| AP000695.2 | 63.4659322 | -1.4934172 | 0.55093312 | -2.7107051 | 0.00671403 | 0.390188 |
| HLCS | 188.569573 | 0.0791085 | 0.3491852 | 0.2265517 | 0.82077236 | 0.99989001 |
| TTC3 | 327.664467 | -0.0552815 | 0.13651291 | -0.4049542 | 0.68551118 | 0.99989001 |
| VPS26C | 136.947826 | 0.26023418 | 0.20982727 | 1.2402305 | 0.21489015 | 0.99989001 |
| DYRK1A | 899.729193 | -0.0720551 | 0.23241287 | -0.3100305 | 0.7565378 | 0.99989001 |
| KCNJ15 | 45.2033719 | 0.94380774 | 0.53994412 | 1.74797299 | 0.0804687 | NA |
| ERG | 1489.57578 | -0.2144784 | 0.19307503 | -1.110855 | 0.26663077 | 0.99989001 |
| ETS2 | 963.219016 | -0.1789962 | 0.51115149 | -0.3501823 | 0.72620189 | 0.99989001 |
| BRWD1 | 590.065605 | -0.003457 | 0.21124592 | -0.0163648 | 0.98694338 | 0.99989001 |
| HMGN1 | 170.389107 | -0.1878831 | 0.2984727 | -0.6294817 | 0.52903376 | 0.99989001 |
| WRB | 87.6481284 | -0.0042625 | 0.27917616 | -0.015268 | 0.9878184 | 0.99989001 |

| | | | | | | |
|---|---|---|---|---|---|---|
| BACE2 | 403.559849 | -0.0265759 | 0.26112511 | -0.1017745 | 9.19E-01 | 0.99989001 |
| MX2 | 268.419675 | 0.23895246 | 1.28484516 | 0.18597763 | 0.8524623 | 0.99989001 |
| MX1 | 387.794449 | -0.3543654 | 1.09995023 | -0.3221649 | 0.74732776 | 0.99989001 |
| C2CD2 | 192.205807 | -0.8940997 | 0.24623273 | -3.6311166 | 0.0002822 | 0.07335471 |
| ZBTB21 | 55.9459814 | 0.38672036 | 0.42147857 | 0.91753269 | 0.35886357 | 0.99989001 |
| ABCG1 | 486.355302 | -0.0452976 | 0.4195282 | -0.1079727 | 9.14E-01 | 0.99989001 |
| SLC37A1 | 77.2278731 | -0.4202987 | 0.37807694 | -1.111675 | 0.26627793 | 0.99989001 |
| PDE9A | 76.4655284 | -0.3939378 | 0.57978761 | -0.6794519 | 0.4968516 | 0.99989001 |
| NDUFV3 | 86.3409064 | -0.0875935 | 0.21875512 | -0.4004181 | 0.6888486 | 0.99989001 |
| PKNOX1 | 91.5879085 | 0.1181017 | 0.32360854 | 0.36495235 | 0.71514698 | 0.99989001 |
| RRP1B | 75.9468419 | -0.0390279 | 0.25772609 | -0.1514315 | 0.87963534 | 0.99989001 |
| PDXK | 71.4948328 | -0.0609014 | 0.32349268 | -0.1882621 | 0.8506712 | 0.99989001 |
| CSTB | 188.57644 | -0.2627803 | 0.40064859 | -0.6558871 | 0.51189674 | 0.99989001 |
| AGPAT3 | 210.988504 | 0.01260122 | 0.2561414 | 0.04919635 | 0.96076282 | 0.99989001 |
| TRAPPC10 | 213.144165 | 0.19704359 | 0.27204529 | 0.72430437 | 0.46887889 | 0.99989001 |
| PFKL | 164.311423 | 0.15158929 | 0.25555146 | 0.59318499 | 0.55305736 | 0.99989001 |
| UBE2G2 | 203.301587 | -0.1008585 | 0.28548598 | -0.353287 | 0.72387331 | 0.99989001 |
| SUMO3 | 110.34774 | -0.3220044 | 0.455367 | -0.7071315 | 0.47948475 | 0.99989001 |
| PTTG1IP | 564.126082 | -0.3873354 | 0.39002235 | -0.9931107 | 0.32065601 | 0.99989001 |
| ADARB1 | 366.493526 | -0.1727344 | 0.25679294 | -0.6726603 | 5.01E-01 | 0.99989001 |
| POFUT2 | 43.378118 | -0.034899 | 0.26770788 | -0.1303622 | 0.8962799 | NA |
| COL18A1 | 292.592272 | -0.8627452 | 0.4172874 | -2.0675085 | 0.03868626 | 0.71745436 |
| PCBP3 | 103.4763 | 0.20061104 | 0.49469858 | 0.40552176 | 0.68509403 | 0.99989001 |
| COL6A1 | 67.4105233 | -0.7628734 | 0.62003347 | -1.2303745 | 0.21855688 | 0.99989001 |
| COL6A2 | 68.9295198 | -0.8076606 | 0.71307206 | -1.1326494 | 0.25736153 | 0.99989001 |
| LSS | 67.3513384 | 0.60864887 | 0.36705793 | 1.65818203 | 0.09728073 | 0.99989001 |
| MCM3AP | 156.94381 | 0.06074395 | 0.18356963 | 0.33090412 | 0.74071691 | 0.99989001 |
| PCNT | 86.6055892 | -0.0836338 | 0.29614587 | -0.2824076 | 0.77763098 | 0.99989001 |
| DIP2A | 338.030896 | -0.0521228 | 0.16705572 | -0.3120086 | 0.75503402 | 0.99989001 |
| PRMT2 | 247.812119 | -0.0288714 | 0.21157974 | -0.1364565 | 0.89146044 | 0.99989001 |
| AL592183.1 | 123.271275 | 0.26081046 | 0.60235961 | 0.43298132 | 0.66502834 | 0.99989001 |

**Interstitial cells**

*This means that positive fold changes imply higher mean value in non-COVID AKI than in COVID AKI and vice versa.

| | baseMean | log2 fold cha | lfcSE | stat | pvalue | padj |
|---|---|---|---|---|---|---|
| AL669831.5 | 32.6000421 | -0.3123431 | 0.34746822 | -0.8989114 | 0.36869985 | 0.97217293 |
| SAMD11 | 31.8417224 | 0.10913814 | 0.41082501 | 0.26565603 | 0.79050413 | 0.99592423 |
| NOC2L | 22.7672471 | 0.33126095 | 0.38111348 | 0.86919243 | 0.38474189 | 0.97254921 |
| HES4 | 66.582799 | -1.0630123 | 0.51142624 | -2.0785252 | 0.03766101 | 0.52201457 |
| ISG15 | 264.626103 | -3.1808848 | 1.25438409 | -2.535814 | 0.01121863 | 0.28321649 |
| AGRN | 19.6123781 | 0.84339463 | 0.43321378 | 1.94683244 | 0.05155483 | 0.57655256 |
| C1orf159 | 25.08242 | 0.47026387 | 0.34420025 | 1.36625081 | 0.17186025 | 0.87233975 |
| SDF4 | 42.2143163 | 0.25993211 | 0.32671116 | 0.79560219 | 0.42626329 | 0.98015476 |
| UBE2J2 | 30.6710557 | 0.00200534 | 0.33315619 | 0.00601921 | 0.99519739 | 0.99935941 |
| ACAP3 | 21.6881338 | -0.3825035 | 0.43081067 | -0.8878691 | 0.37461118 | 0.97217293 |
| INTS11 | 33.02019 | 0.12372356 | 0.29179394 | 0.42401003 | 0.6715585 | 0.9958918 |
| MXRA8 | 39.7021008 | -0.7911735 | 0.39228933 | -2.0168112 | 0.04371522 | 0.55581858 |
| AURKAIP1 | 30.4957617 | -0.4734309 | 0.46777288 | -1.0120957 | 0.31149231 | 0.9599143 |
| CCNL2 | 88.0269672 | -0.1270581 | 0.23211195 | -0.5474001 | 0.5841039 | 0.9958918 |
| MRPL20 | 44.0789191 | 0.10250932 | 0.38427351 | 0.26676135 | 0.78965291 | 0.99592423 |
| ATAD3B | 17.1170556 | 0.54678237 | 0.4028807 | 1.35718184 | 0.17472344 | 0.8740623 |
| SSU72 | 44.2019559 | 0.22556704 | 0.29888397 | 0.7546977 | 0.45043039 | 0.98092839 |
| CDK11B | 18.66467 | 0.00478423 | 0.50201842 | 0.00952999 | 0.99239628 | 0.99935941 |
| SLC35E2B | 23.7810081 | 0.06330417 | 0.39732357 | 0.1593265 | 0.87341165 | 0.9971731 |
| SLC35E2A | 18.68708 | 0.58796805 | 0.37538426 | 1.56630979 | 0.11727611 | 0.7849382 |
| NADK | 28.388152 | -0.0777457 | 0.35078143 | -0.2216359 | 0.82459735 | 0.99592423 |
| GNB1 | 238.014468 | 0.07350303 | 0.16015833 | 0.45893977 | 0.64627742 | 0.9958918 |
| FAAP20 | 26.778249 | -0.1314197 | 0.31327712 | -0.4194998 | 0.67485091 | 0.9958918 |
| SKI | 110.373669 | 0.55421146 | 0.2716888 | 2.03987595 | 0.04136268 | 0.54347495 |
| RER1 | 39.9552718 | -0.2900288 | 0.29538774 | -0.9818578 | 0.3261699 | 0.95992644 |
| TNFRSF14 | 34.0568336 | -0.437503 | 0.52425175 | -0.8345284 | 0.40398328 | 0.97575999 |
| PRDM16 | 45.0406513 | 2.00116834 | 0.64318398 | 3.11134668 | 0.00186236 | 0.09345751 |
| CEP104 | 27.9483235 | 0.19884233 | 0.3339395 | 0.59544417 | 0.55154661 | 0.9958918 |
| RPL22 | 26.8630813 | -0.1224115 | 0.41244967 | -0.2967914 | 0.76662581 | 0.9958918 |
| ICMT | 18.3536637 | -0.2150305 | 0.36992109 | -0.5812876 | 0.56104667 | 0.9958918 |
| NOL9 | 24.3118447 | 0.60520259 | 0.54490672 | 1.11065356 | 0.2667175 | 0.93533685 |
| DNAJC11 | 39.985956 | 0.24262739 | 0.35579619 | 0.681928 | 0.49528448 | 0.99176652 |
| CAMTA1 | 166.744809 | 0.25190999 | 0.19688902 | 1.27945173 | 0.20073803 | 0.89191279 |
| VAMP3 | 19.770221 | 0.66209889 | 0.34876068 | 1.89843329 | 0.05763903 | 0.60512745 |
| PER3 | 68.042059 | -0.4819212 | 0.59934455 | -0.8040804 | 0.42135052 | 0.97812179 |
| PARK7 | 62.2451629 | 0.3065385 | 0.24777196 | 1.23717991 | 0.2160203 | 0.90187565 |
| ERRFI1 | 119.071231 | -1.2399663 | 0.90264451 | -1.3737039 | 0.16953362 | 0.86883023 |
| RERE | 547.845735 | 0.01925589 | 0.18638239 | 0.10331388 | 0.91771386 | 0.99935941 |
| ENO1 | 149.14189 | 0.17248846 | 0.38550299 | 0.44743742 | 6.55E-01 | 0.9958918 |
| MIR34AHG | 36.4421647 | 0.41948062 | 0.44591311 | 0.94072277 | 0.34684695 | 0.96887708 |
| H6PD | 44.4016171 | 0.06357985 | 0.39672029 | 0.16026366 | 0.87267338 | 0.9971731 |
| SPSB1 | 70.7875287 | -0.0374897 | 0.49570964 | -0.0756284 | 0.93971473 | 0.99935941 |
| SLC25A33 | 23.0465022 | 0.00076756 | 0.37583072 | 0.00204229 | 0.99837049 | 0.99959465 |
| CLSTN1 | 55.8615058 | 0.12209718 | 0.25476805 | 0.47924839 | 0.63176193 | 0.9958918 |
| LZIC | 19.0966657 | 0.13440241 | 0.40715543 | 0.330101 | 0.74132365 | 0.9958918 |
| NMNAT1 | 19.5359129 | -0.0693757 | 0.34416226 | -0.2015784 | 0.84024632 | 0.99692952 |
| UBE4B | 147.717568 | -0.3865415 | 0.21862742 | -1.7680374 | 0.07705466 | 0.67905476 |
| KIF1B | 163.403151 | -0.1637536 | 0.27963891 | -0.5855896 | 0.55815133 | 0.9958918 |
| DFFA | 19.8069651 | 0.08027561 | 0.41393767 | 0.19393165 | 0.8462294 | 0.99692952 |
| PEX14 | 53.9474258 | 0.16305343 | 0.24327983 | 0.67022994 | 0.50271122 | 0.99339198 |
| SRM | 28.3859772 | -0.6175181 | 0.58709659 | -1.0518169 | 0.29288358 | 0.95186143 |
| EXOSC10 | 26.3302225 | -0.2804437 | 0.32757346 | -0.8561246 | 0.39192888 | 0.97297628 |

| | | | | | | |
|---|---|---|---|---|---|---|
| MTOR | 63.9463686 | -0.1685763 | 0.31158767 | -0.5410236 | 0.58849129 | 0.9958918 |
| MTHFR | 46.8408965 | 0.16969393 | 0.44831763 | 0.37851273 | 0.70504974 | 0.9958918 |
| CLCN6 | 25.6862523 | -0.28741 | 0.39520012 | -0.7272518 | 0.46707171 | 0.98834148 |
| PLOD1 | 34.8281116 | -0.0784014 | 0.26771889 | -0.2928497 | 0.76963701 | 0.9958918 |
| MFN2 | 28.1685491 | 0.16899394 | 0.3560857 | 0.47458783 | 0.63508078 | 0.9958918 |
| TNFRSF1B | 26.5236968 | -0.0426324 | 0.56311937 | -0.0757075 | 0.9396518 | 0.99935941 |
| VPS13D | 338.231813 | -0.6122351 | 0.21203934 | -2.8873656 | 0.00388483 | 0.15300235 |
| DHRS3 | 195.65804 | -1.3224639 | 0.45664894 | -2.8960188 | 0.0037793 | 0.15284924 |
| PRDM2 | 110.481676 | -0.1761331 | 0.23026932 | -0.7649005 | 0.4443308 | 0.98092839 |
| KAZN | 1241.12909 | -0.8604817 | 0.46400945 | -1.8544486 | 0.063675 | 0.62395844 |
| TMEM51 | 64.6096464 | 0.11694847 | 0.4253228 | 0.27496404 | 0.78334387 | 0.9958918 |
| DNAJC16 | 34.4718815 | 0.187728 | 0.33008178 | 0.56873179 | 0.56953817 | 0.9958918 |
| DDI2 | 70.2727118 | -0.1367771 | 0.26257864 | -0.5208996 | 0.60243675 | 0.9958918 |
| PLEKHM2 | 61.711746 | 0.13901627 | 0.30387873 | 0.45747285 | 0.64733121 | 0.9958918 |
| FBLIM1 | 123.394396 | -0.2415082 | 0.34153407 | -0.7071278 | 0.47948708 | 0.99002719 |
| SPEN | 144.831286 | 0.07674585 | 0.21861711 | 0.35105145 | 0.72554975 | 0.9958918 |
| EPHA2 | 25.3594236 | -0.7434804 | 0.45555073 | -1.6320474 | 0.10266949 | 0.75678848 |
| FBXO42 | 152.070985 | -0.2912978 | 0.22925591 | -1.2706228 | 0.20386288 | 0.89396747 |
| SZRD1 | 38.1730454 | 0.26149461 | 0.30678457 | 0.85237209 | 0.3940076 | 0.97500135 |
| NECAP2 | 18.1716372 | 0.338957 | 0.35188951 | 0.96324838 | 0.33542289 | 0.96640422 |
| CROCC | 34.3924373 | -0.1134694 | 0.35637779 | -0.3183965 | 0.7501842 | 0.9958918 |
| SDHB | 36.6131617 | -0.0395802 | 0.28053727 | -0.1410872 | 0.88780102 | 0.99935941 |
| RCC2 | 20.1523215 | -0.4922807 | 0.4240492 | -1.1609047 | 0.24568065 | 0.92559078 |
| ARHGEF10L | 82.5587764 | 0.63372827 | 0.32606051 | 1.94359101 | 0.05194479 | 0.57839737 |
| UBR4 | 135.170468 | -0.0826792 | 0.23346514 | -0.3541392 | 0.72323454 | 0.9958918 |
| AL035413.1 | 22.0230377 | -0.1603423 | 0.3220108 | -0.4979409 | 0.61852573 | 0.9958918 |
| EMC1 | 20.922311 | -0.0296209 | 0.37584158 | -0.0788123 | 0.93718196 | 0.99935941 |
| CAPZB | 296.48487 | -0.206279 | 0.31074795 | -0.6638144 | 0.50680911 | 0.99397937 |
| NBL1 | 31.6518983 | -0.437869 | 0.37945721 | -1.1539351 | 0.24852677 | 0.92624282 |
| TMCO4 | 29.0865984 | 0.44561995 | 0.36271223 | 1.22857714 | 0.21923039 | 0.90437211 |
| CAMK2N1 | 31.6096945 | 0.00805768 | 0.31601935 | 0.02549742 | 0.97965821 | 0.99935941 |
| PINK1 | 22.3895564 | 0.07650508 | 0.4337611 | 0.17637608 | 0.85999849 | 0.9971731 |
| DDOST | 19.730064 | 0.43173397 | 0.46254389 | 0.93339029 | 0.35061849 | 0.9693503 |
| HP1BP3 | 152.369288 | 6.28E-05 | 0.19308712 | 0.00032505 | 0.99974065 | 0.99978196 |
| EIF4G3 | 387.67391 | -0.2358579 | 0.24563128 | -0.9602112 | 0.33694894 | 0.96728036 |
| ECE1 | 147.54535 | 0.18356234 | 0.40225595 | 0.45633218 | 0.64815113 | 0.9958918 |
| USP48 | 104.586303 | -0.1036663 | 0.277701 | -0.3733017 | 0.70892392 | 0.9958918 |
| HSPG2 | 115.821127 | 1.00495972 | 0.38718638 | 2.59554512 | 0.0094441 | 0.25610587 |
| CDC42 | 84.5156528 | -0.1611167 | 0.24052841 | -0.6698446 | 0.50295683 | 0.99354798 |
| ZBTB40 | 32.39219 | 0.12239118 | 0.29093168 | 0.420687 | 0.67398365 | 0.9958918 |
| KDM1A | 57.7492203 | 0.00356008 | 0.25281579 | 0.01408173 | 0.98876477 | 0.99935941 |
| LUZP1 | 73.1888086 | 0.1783514 | 0.21271214 | 0.83846364 | 0.40177036 | 0.97500135 |
| HNRNPR | 124.851477 | 0.06402563 | 0.21484349 | 0.29801055 | 0.76569511 | 0.9958918 |
| ASAP3 | 25.9649012 | 0.29836895 | 0.37851508 | 0.78826174 | 0.43054362 | 0.98092839 |
| ID3 | 153.825374 | -1.4651714 | 0.58664571 | -2.4975405 | 0.01250582 | 0.29742804 |
| RPL11 | 148.173363 | -0.2785786 | 0.30195621 | -0.9225795 | 0.35622637 | 0.97217293 |
| ELOA | 20.1570403 | 0.13406242 | 0.39641354 | 0.33818829 | 0.7352213 | 0.9958918 |
| HMGCL | 23.9143118 | 0.10562936 | 0.35096979 | 0.30096424 | 0.76344176 | 0.9958918 |
| PNRC2 | 36.1733058 | 0.0633334 | 0.39318097 | 0.16107953 | 0.87203077 | 0.9971731 |
| SRSF10 | 97.7338424 | -0.2072257 | 0.20692338 | -1.0014611 | 0.31660392 | 0.95992644 |
| SRRM1 | 78.4159457 | -0.0488662 | 0.26324103 | -0.185633 | 0.85273259 | 0.9971731 |
| CLIC4 | 361.020701 | -0.1757726 | 0.37186932 | -0.4726731 | 0.63644645 | 0.9958918 |
| SYF2 | 21.3270064 | -0.003909 | 0.38137762 | -0.0102497 | 0.99182209 | 0.99935941 |
| RSRP1 | 156.721126 | 0.5951478 | 0.35346214 | 1.68376676 | 0.09222675 | 0.72957419 |

| | | | | | | |
|---|---|---|---|---|---|---|
| TMEM50A | 50.6782146 | 0.06175472 | 0.27301922 | 0.22619185 | 0.82105221 | 0.99592423 |
| MACO1 | 59.9884446 | -0.079324 | 0.21466277 | -0.3695286 | 0.71173377 | 0.9958918 |
| LDLRAP1 | 22.905408 | -0.1302479 | 0.59316777 | -0.2195801 | 0.82619818 | 0.99592423 |
| MAN1C1 | 95.6716979 | -0.1072403 | 0.50752981 | -0.2112986 | 0.83265428 | 0.99592423 |
| SH3BGRL3 | 40.2576353 | -0.2324995 | 0.44451532 | -0.5230405 | 0.60094605 | 0.9958918 |
| DHDDS | 25.3608345 | -0.1462517 | 0.37554053 | -0.3894433 | 0.69694825 | 0.9958918 |
| HMGN2 | 49.3524602 | 0.01247327 | 0.35442931 | 0.03519255 | 0.9719262 | 0.99935941 |
| ARID1A | 145.632986 | 0.2537392 | 0.1912493 | 1.32674579 | 0.18459279 | 0.87866798 |
| NUDC | 36.4626797 | 0.11722161 | 0.31230976 | 0.37533766 | 0.70740936 | 0.9958918 |
| SLC9A1 | 20.8444031 | 0.29102574 | 0.34505825 | 0.84341048 | 0.39899891 | 0.97500135 |
| WDTC1 | 36.4009807 | 0.14597648 | 0.35108645 | 0.41578499 | 0.67756734 | 0.9958918 |
| WASF2 | 139.9855 | 0.01193939 | 0.18274098 | 0.06533505 | 0.94790724 | 0.99935941 |
| AHDC1 | 51.2022639 | 0.54284325 | 0.35198573 | 1.54223086 | 0.1230175 | 0.79184923 |
| IFI6 | 91.3345261 | -2.3300058 | 1.24404998 | -1.8729197 | 0.06107947 | 0.61696136 |
| FAM76A | 23.1815665 | 0.12305211 | 0.3291625 | 0.37383393 | 0.70852788 | 0.9958918 |
| STX12 | 126.751825 | -0.431835 | 0.35871885 | -1.2038258 | 0.2286569 | 0.91375723 |
| EYA3 | 70.5066503 | 0.08578617 | 0.19626686 | 0.43708941 | 0.6620465 | 0.9958918 |
| DNAJC8 | 28.2487419 | -0.2528811 | 0.34536768 | -0.7322084 | 0.4640414 | 0.98590351 |
| ATP5IF1 | 46.7770456 | -0.0273316 | 0.31786801 | -0.0859842 | 0.93147894 | 0.99935941 |
| PHACTR4 | 124.084357 | 0.39881242 | 0.2177731 | 1.83132084 | 0.06705267 | 0.63913109 |
| RCC1 | 42.0722144 | -0.1701992 | 0.50718979 | -0.3355731 | 0.73719282 | 0.9958918 |
| TRNAU1AP | 43.2067143 | 0.20216622 | 0.32560274 | 0.62089839 | 0.53466648 | 0.9958918 |
| TAF12 | 20.4872007 | 0.19943891 | 0.3810801 | 0.52335167 | 0.60072955 | 0.9958918 |
| GMEB1 | 24.893633 | -0.0809642 | 0.34340783 | -0.235767 | 0.8136135 | 0.99592423 |
| YTHDF2 | 46.5851848 | -0.1576575 | 0.23839677 | -0.6613239 | 0.50840462 | 0.99397937 |
| EPB41 | 54.6790032 | 0.14338056 | 0.33211839 | 0.43171521 | 0.66594841 | 0.9958918 |
| SRSF4 | 183.31689 | 0.07205283 | 0.24134015 | 0.29855304 | 0.7652811 | 0.9958918 |
| MECR | 17.4119907 | -0.4725353 | 0.43441153 | -1.0877595 | 0.27670131 | 0.94273432 |
| PUM1 | 270.352801 | -0.2083205 | 0.21912614 | -0.9506877 | 0.34176294 | 0.96887708 |
| SNRNP40 | 32.6137768 | 0.30667155 | 0.32812682 | 0.93461288 | 0.34998784 | 0.9693503 |
| ZCCHC17 | 33.2869644 | 0.2036773 | 0.29540919 | 0.68947516 | 0.4905243 | 0.99046213 |
| TINAGL1 | 205.856498 | -0.1545364 | 0.36976104 | -0.4179358 | 0.67599409 | 0.9958918 |
| PEF1 | 19.5491921 | 0.32856937 | 0.31893132 | 1.03021984 | 0.30290682 | 0.95650251 |
| COL16A1 | 69.3432524 | -0.5057995 | 0.387339 | -1.3058314 | 0.19160988 | 0.88158763 |
| PTP4A2 | 38.0495619 | 0.08710941 | 0.3942713 | 0.22093774 | 0.82514091 | 0.99592423 |
| KHDRBS1 | 117.532864 | -0.136959 | 0.21682195 | -0.6316656 | 0.5276054 | 0.9958918 |
| KPNA6 | 96.9628467 | 0.07452015 | 0.24953085 | 0.29864105 | 0.76521394 | 0.9958918 |
| TXLNA | 34.9844518 | -0.0309317 | 0.31682897 | -0.097629 | 0.92222687 | 0.99935941 |
| EIF3I | 27.2392025 | 0.19628851 | 0.46459957 | 0.42248965 | 0.67266766 | 0.9958918 |
| HDAC1 | 31.2652758 | -0.2555826 | 0.29207235 | -0.875066 | 0.381538 | 0.97217293 |
| BSDC1 | 44.023378 | 0.08037635 | 0.41707695 | 0.19271349 | 0.84718336 | 0.99694964 |
| ZBTB8OS | 65.0214792 | 0.32059884 | 0.20805546 | 1.54092971 | 0.12333389 | 0.79184923 |
| RBBP4 | 59.4478798 | 0.11868393 | 0.21653938 | 0.54809398 | 0.58362737 | 0.9958918 |
| YARS | 51.0206075 | -0.5142314 | 0.38367144 | -1.340291 | 0.18015075 | 0.87815793 |
| S100PBP | 50.4433691 | 0.00593189 | 0.29362604 | 0.0202022 | 0.98388207 | 0.99935941 |
| RNF19B | 33.4789965 | -0.7918498 | 0.4867806 | -1.6267077 | 0.10379918 | 0.75827058 |
| AK2 | 45.0850024 | 0.12633032 | 0.34619671 | 0.36490908 | 0.71517927 | 0.9958918 |
| ZNF362 | 38.0406905 | 0.21945475 | 0.39436776 | 0.55647235 | 0.57788799 | 0.9958918 |
| AL513327.1 | 39.4975729 | -0.3069752 | 0.45945631 | -0.6681272 | 5.04E-01 | 0.99387859 |
| PHC2 | 238.736882 | -0.5112608 | 0.25155896 | -2.0323696 | 0.04211626 | 0.5458772 |
| SMIM12 | 35.9937653 | 0.20024726 | 0.29474745 | 0.67938587 | 0.4968934 | 0.99176652 |
| GJA4 | 51.819821 | -0.9276829 | 0.59845571 | -1.5501279 | 0.12111083 | 0.79021701 |
| ZMYM1 | 27.525694 | 0.13816765 | 0.34480963 | 0.40070705 | 0.68863582 | 0.9958918 |
| SFPQ | 165.821248 | 0.06708875 | 0.22734995 | 0.29509022 | 0.76792496 | 0.9958918 |

| | | | | | | |
|---|---|---|---|---|---|---|
| ZMYM4 | 245.684926 | -0.0090321 | 0.17082138 | -0.0528747 | 0.95783174 | 0.99935941 |
| KIAA0319L | 113.016759 | 0.1966073 | 0.24725766 | 0.79515152 | 0.42652537 | 0.98015476 |
| PSMB2 | 35.8803106 | -0.3746187 | 0.33951473 | -1.1033947 | 0.26985581 | 0.93634106 |
| AGO4 | 67.3255856 | 0.187215 | 0.37600347 | 0.49790763 | 0.61854915 | 0.9958918 |
| AGO1 | 23.8647889 | -0.0381252 | 0.33789611 | -0.112831 | 0.91016453 | 0.99935941 |
| AGO3 | 212.194937 | -0.1354799 | 0.14795265 | -0.915698 | 0.35982531 | 0.97217293 |
| MAP7D1 | 25.1649145 | 0.55466769 | 0.42069011 | 1.31847094 | 0.18734605 | 0.8797483 |
| THRAP3 | 140.287982 | -0.0508245 | 0.1729955 | -0.2937909 | 0.76891772 | 0.9958918 |
| EVA1B | 31.1346893 | -0.0318979 | 0.39853916 | -0.0800369 | 0.93620787 | 0.99935941 |
| STK40 | 41.9420224 | 0.36442993 | 0.38793549 | 0.9394086 | 0.347521 | 0.96887708 |
| MEAF6 | 21.4216602 | 0.01259635 | 0.34206203 | 0.03682476 | 0.97062473 | 0.99935941 |
| C1orf122 | 22.9887052 | -0.1160651 | 0.39858188 | -0.2911951 | 0.77090214 | 0.9958918 |
| MTF1 | 20.3391702 | 0.52607374 | 0.38041047 | 1.38291078 | 0.16669222 | 0.86269095 |
| INPP5B | 88.8273434 | 0.12632486 | 0.21747823 | 0.58086212 | 0.56133338 | 0.9958918 |
| SF3A3 | 32.6798968 | 0.06095096 | 0.40751289 | 0.14956817 | 0.88110532 | 0.9977723 |
| RRAGC | 24.4114062 | -0.4651466 | 0.37604792 | -1.2369343 | 0.21611147 | 0.90187565 |
| AL139260.1 | 35.5870887 | -0.192165 | 0.51114048 | -0.3759533 | 0.70695161 | 0.9958918 |
| AKIRIN1 | 35.2008459 | 0.32533174 | 0.28103257 | 1.15763002 | 0.24701505 | 0.92605444 |
| NDUFS5 | 59.3777688 | -0.2396822 | 0.43508695 | -0.5508834 | 0.58171359 | 0.9958918 |
| MACF1 | 788.882492 | -0.2694182 | 0.18987671 | -1.4189111 | 0.15592494 | 0.84427321 |
| PABPC4 | 33.9585265 | 0.32204242 | 0.42043189 | 0.76598 | 0.4436882 | 0.98092839 |
| HEYL | 56.280813 | -0.724014 | 0.3765404 | -1.9228056 | 0.05450446 | 0.59304667 |
| TRIT1 | 31.2642882 | -0.3364738 | 0.31438813 | -1.0702498 | 0.28450689 | 0.94505166 |
| CAP1 | 77.8477672 | 0.34815135 | 0.28616608 | 1.2166059 | 0.2237542 | 0.91093521 |
| PPT1 | 20.1778875 | 0.09196787 | 0.38385122 | 0.2395925 | 0.81064618 | 0.99592423 |
| RLF | 144.492902 | -0.028864 | 0.33992762 | -0.0849123 | 0.93233113 | 0.99935941 |
| ZMPSTE24 | 28.0217018 | -0.2945612 | 0.33093884 | -0.8900775 | 0.3734243 | 0.97217293 |
| SMAP2 | 36.140541 | 0.62908132 | 0.30416564 | 2.06821955 | 0.03861938 | 0.5265563 |
| NFYC | 46.2986407 | 0.65208459 | 0.2865261 | 2.27582967 | 0.02285621 | 0.40968356 |
| CTPS1 | 25.0227708 | -0.4953429 | 0.49051396 | -1.0098446 | 0.31256973 | 0.9599143 |
| SCMH1 | 299.741715 | -0.2043436 | 0.22258734 | -0.9180379 | 0.35859901 | 0.97217293 |
| HIVEP3 | 148.354437 | -0.3111838 | 0.27291086 | -1.1402397 | 0.25418646 | 0.92646125 |
| FOXJ3 | 141.621518 | -0.1848635 | 0.21959695 | -0.8418308 | 0.39988267 | 0.97500135 |
| CCDC30 | 24.5815705 | 0.25751824 | 0.46834174 | 0.54985115 | 0.58242147 | 0.9958918 |
| YBX1 | 77.8358103 | -0.1872434 | 0.37771541 | -0.4957262 | 0.62008763 | 0.9958918 |
| P3H1 | 19.8364384 | -0.2314668 | 0.40658886 | -0.5692896 | 0.56915966 | 0.9958918 |
| SLC2A1 | 29.9548629 | -1.0285667 | 0.64300852 | -1.599616 | 0.1096838 | 0.77303254 |
| SZT2 | 35.7151012 | 0.02762683 | 0.34036323 | 0.08116868 | 0.93530781 | 0.99935941 |
| PTPRF | 29.6574001 | 0.07767962 | 0.353448 | 0.21977665 | 0.82604511 | 0.99592423 |
| ATP6V0B | 44.2370286 | -0.1726588 | 0.38538369 | -0.4480179 | 0.65414027 | 0.9958918 |
| ERI3 | 76.2904291 | 0.21507298 | 0.24094534 | 0.89262146 | 0.37205993 | 0.97217293 |
| RPS8 | 148.539981 | -0.2372791 | 0.37817361 | -0.6274342 | 0.53037466 | 0.9958918 |
| EIF2B3 | 61.5005504 | 0.40341343 | 0.31077465 | 1.29808989 | 0.19425645 | 0.88391893 |
| ZSWIM5 | 63.1197777 | 0.34845239 | 0.29555874 | 1.17896155 | 0.23841349 | 0.92291633 |
| TESK2 | 35.583243 | 0.25975026 | 0.36175949 | 0.71801921 | 0.47274544 | 0.98910524 |
| PRDX1 | 105.391768 | 0.31657324 | 0.40229331 | 0.78692147 | 0.43132784 | 0.98092839 |
| AKR1A1 | 20.3501741 | 0.30264286 | 0.38933391 | 0.77733497 | 0.43696117 | 0.98092839 |
| NASP | 45.0896396 | 0.07489246 | 0.29479024 | 0.25405339 | 0.79945431 | 0.99592423 |
| GPBP1L1 | 115.364902 | 0.19886874 | 0.17627901 | 1.12814756 | 0.25925761 | 0.92976724 |
| IPP | 19.896797 | 0.31945529 | 0.47137484 | 0.67770967 | 0.49795579 | 0.99176652 |
| MAST2 | 71.7216081 | 0.10863979 | 0.26984986 | 0.40259346 | 0.68724732 | 0.9958918 |
| TSPAN1 | 22.0394791 | 0.40884632 | 0.37541536 | 1.08905059 | 2.76E-01 | 0.94222064 |
| LRRC41 | 38.0491916 | 0.07089398 | 0.2578373 | 0.27495627 | 0.78334983 | 0.9958918 |
| UQCRH | 47.6161909 | 0.01524892 | 0.2892073 | 0.0527266 | 0.95794974 | 0.99935941 |

| | | | | | | |
|---|---|---|---|---|---|---|
| NSUN4 | 26.7290634 | -0.3728189 | 0.35353353 | -1.0545505 | 0.29163097 | 0.95186143 |
| MKNK1 | 29.3293824 | -0.033137 | 0.32245107 | -0.1027661 | 0.91814864 | 0.99935941 |
| ATPAF1 | 32.5551877 | 0.46608879 | 0.28505131 | 1.63510489 | 0.10202706 | 0.75449741 |
| EFCAB14 | 55.6398663 | 0.2049187 | 0.26272064 | 0.77998706 | 0.43539849 | 0.98092839 |
| CYP4X1 | 22.4474949 | 0.38829691 | 0.55759443 | 0.69637874 | 0.48619167 | 0.99046213 |
| PDZK1IP1 | 53.6532816 | 0.9698281 | 0.52557747 | 1.84526195 | 0.06499944 | 0.62770158 |
| CMPK1 | 58.3203187 | -0.0213911 | 0.34387965 | -0.0622053 | 0.95039933 | 0.99935941 |
| TRABD2B | 57.8873153 | 1.23760026 | 0.59129929 | 2.09301832 | 0.03634752 | 0.51270234 |
| SPATA6 | 95.4475466 | -0.0954828 | 0.34999275 | -0.2728136 | 0.78499649 | 0.9958918 |
| FAF1 | 344.751015 | -0.1301139 | 0.15934027 | -0.8165786 | 0.41416929 | 0.97812179 |
| RNF11 | 50.0425844 | 0.21027906 | 0.23259898 | 0.90404117 | 0.36597357 | 0.97217293 |
| EPS15 | 230.704871 | -0.1732628 | 0.2248591 | -0.7705396 | 0.4409799 | 0.98092839 |
| OSBPL9 | 225.156993 | -0.5331619 | 0.24374459 | -2.1873794 | 0.02871484 | 0.45975021 |
| NRDC | 167.235292 | -0.1116535 | 0.15864719 | -0.7037852 | 0.48156657 | 0.99002719 |
| TXNDC12 | 23.8167528 | 0.5369212 | 0.40082077 | 1.33955435 | 0.18039027 | 0.87815793 |
| BTF3L4 | 40.1581619 | 0.14904189 | 0.28822922 | 0.51709501 | 0.60508984 | 0.9958918 |
| ZFYVE9 | 102.027238 | -0.0581306 | 0.21307504 | -0.2728173 | 0.78499367 | 0.9958918 |
| PRPF38A | 19.6137118 | -0.1869655 | 0.36908352 | -0.5065669 | 0.61245872 | 0.9958918 |
| TUT4 | 154.423809 | 0.0312776 | 0.24781722 | 0.12621239 | 0.89956381 | 0.99935941 |
| ZYG11B | 73.1384436 | -0.1418448 | 0.2566202 | -0.5527421 | 0.58043999 | 0.9958918 |
| ECHDC2 | 113.649552 | 0.45762242 | 0.4775801 | 0.95821082 | 0.33795646 | 0.96750939 |
| SCP2 | 78.7045295 | 0.05564197 | 0.2699832 | 0.2060942 | 0.83671733 | 0.99692952 |
| PODN | 38.49505 | -1.0653195 | 0.5462181 | -1.9503555 | 0.05113376 | 0.57655256 |
| MAGOH | 18.2038538 | -0.2906945 | 0.40970886 | -0.7095149 | 0.47800503 | 0.99002719 |
| NDC1 | 18.2756406 | 0.06594959 | 0.35446352 | 0.18605466 | 0.85240189 | 0.9971731 |
| YIPF1 | 17.7438897 | 0.18213923 | 0.33625923 | 0.54166314 | 0.58805059 | 0.9958918 |
| HSPB11 | 28.0747156 | 0.37597875 | 0.37660797 | 0.99832925 | 0.31811973 | 0.95992644 |
| TMEM59 | 101.97026 | 0.1571862 | 0.23521922 | 0.66825409 | 0.50397142 | 0.99387859 |
| SSBP3 | 463.129323 | 0.0140964 | 0.39867141 | 0.03535844 | 0.97179393 | 0.99935941 |
| USP24 | 144.959186 | -0.2389474 | 0.20251344 | -1.1799088 | 0.2380365 | 0.92291633 |
| PLPP3 | 178.925179 | -1.1395486 | 0.48690319 | -2.3404008 | 0.01926306 | 0.37253738 |
| PRKAA2 | 43.3256303 | -0.4800289 | 0.53289055 | -0.9008021 | 0.36769357 | 0.97217293 |
| DAB1 | 64.0262935 | 0.87569909 | 0.56795502 | 1.54184584 | 0.12311106 | 0.79184923 |
| MYSM1 | 58.972532 | -0.3504081 | 0.23800604 | -1.4722654 | 0.14094921 | 0.81630561 |
| JUN | 154.517971 | -1.023801 | 0.4848313 | -2.1116644 | 0.03471524 | 0.50468149 |
| FGGY | 62.6997716 | -0.2755544 | 0.38371561 | -0.7181214 | 0.47268242 | 0.98910524 |
| NFIA | 776.987714 | -0.4724842 | 0.27228911 | -1.73523 | 0.08270007 | 0.70018756 |
| TM2D1 | 79.1757979 | -0.1862518 | 0.19834381 | -0.9390353 | 0.34771262 | 0.96887708 |
| PATJ | 138.622627 | 0.21858616 | 0.34508838 | 0.6334208 | 0.52645889 | 0.9958918 |
| DOCK7 | 124.021747 | 0.40640098 | 0.25224453 | 1.61113893 | 0.10714945 | 0.77179565 |
| ATG4C | 37.4434088 | -0.0254066 | 0.31526972 | -0.0805869 | 0.93577049 | 0.99935941 |
| ITGB3BP | 30.8693919 | 0.59542809 | 0.43373214 | 1.3728014 | 0.1698141 | 0.86966118 |
| EFCAB7 | 20.3925158 | -0.1271355 | 0.33127823 | -0.3837726 | 0.70114703 | 0.9958918 |
| PGM1 | 25.2573524 | 0.46762877 | 0.42546438 | 1.09910204 | 0.27172356 | 0.93795041 |
| ROR1 | 228.855181 | 0.52813445 | 0.38773165 | 1.36211332 | 0.17316212 | 0.87233975 |
| CACHD1 | 109.65643 | -0.5847857 | 0.29803824 | -1.9621161 | 0.04974896 | 0.57344369 |
| JAK1 | 205.264745 | -0.0155128 | 0.22135137 | -0.0700825 | 0.94412803 | 0.99935941 |
| AK4 | 46.9310833 | 0.38708818 | 0.60626546 | 0.63847968 | 0.52316148 | 0.9958918 |
| LEPR | 54.7195525 | 0.38940581 | 0.34480378 | 1.12935482 | 0.25874818 | 0.92939413 |
| PDE4B | 100.24819 | 0.06486972 | 0.87338591 | 0.07427384 | 0.9407925 | 0.99935941 |
| SGIP1 | 688.673977 | -0.2860257 | 0.41678661 | -0.6862641 | 0.49254657 | 0.99046213 |
| WDR78 | 35.7059221 | -0.3518224 | 0.48620597 | -0.7236076 | 0.46930665 | 0.98910524 |
| MIER1 | 119.430353 | -0.2190784 | 0.22909958 | -0.9562585 | 0.33894164 | 0.96887708 |
| SLC35D1 | 40.223295 | 0.29390803 | 0.28059938 | 1.04742939 | 0.29490158 | 0.95304825 |

| | | | | | | |
|---|---|---|---|---|---|---|
| SERBP1 | 78.9745405 | 0.107684 | 0.36213888 | 0.29735554 | 0.76619508 | 0.9958918 |
| GADD45A | 29.1187005 | 0.62419728 | 0.49419365 | 1.2630621 | 0.20656686 | 0.89589437 |
| GNG12 | 152.509993 | 0.13232495 | 0.21842528 | 0.60581332 | 0.54463872 | 0.9958918 |
| GNG12-AS1 | 28.086249 | 0.08222147 | 0.42785555 | 0.1921711 | 0.84760818 | 0.99713023 |
| WLS | 45.8196038 | 0.02715827 | 0.29050603 | 0.09348607 | 0.92551741 | 0.99935941 |
| LRRC40 | 32.4492765 | -0.3808535 | 0.30181423 | -1.2618804 | 0.20699181 | 0.89620104 |
| SRSF11 | 217.497324 | -0.0799953 | 0.16964966 | -0.4715323 | 0.63726067 | 0.9958918 |
| ANKRD13C | 113.679903 | -0.3867513 | 0.2078619 | -1.8606166 | 0.06279834 | 0.6220697 |
| PTGER3 | 37.2962687 | -0.7234491 | 0.47038604 | -1.5379901 | 0.12405103 | 0.79195137 |
| ZRANB2 | 178.576773 | -0.0080481 | 0.22941709 | -0.0350805 | 0.97201554 | 0.99935941 |
| NEGR1 | 1257.19346 | -0.3209269 | 0.53924503 | -0.5951411 | 0.55174916 | 0.9958918 |
| CRYZ | 24.9303192 | 0.43960803 | 0.56269462 | 0.78125508 | 0.43465249 | 0.98092839 |
| TYW3 | 23.9433655 | -0.0078258 | 0.36069041 | -0.0216966 | 0.98268994 | 0.99935941 |
| ACADM | 37.3529401 | 0.55225656 | 0.3326039 | 1.66040317 | 0.09683337 | 0.74205262 |
| RABGGTB | 23.0625704 | -0.1914375 | 0.40894638 | -0.4681237 | 0.63969611 | 0.9958918 |
| ST6GALNAC3 | 28.9217063 | 0.42517088 | 0.51765805 | 0.8213354 | 0.41145525 | 0.97812179 |
| PIGK | 37.1217808 | 0.24256131 | 0.32773795 | 0.7401075 | 0.45923477 | 0.98508941 |
| AC118549.1 | 167.843375 | -0.2619065 | 0.16317436 | -1.6050714 | 0.10847808 | 0.77248589 |
| USP33 | 124.40064 | 0.38439436 | 0.23685755 | 1.62289256 | 0.10461237 | 0.76269478 |
| MIGA1 | 75.6082639 | 0.01170181 | 0.20928369 | 0.05591365 | 0.9554106 | 0.99935941 |
| NEXN | 116.431109 | 0.02876716 | 0.61117511 | 0.04706861 | 0.96245855 | 0.99935941 |
| FUBP1 | 119.327626 | -0.2530434 | 0.26623462 | -0.9504528 | 0.34188224 | 0.96887708 |
| IFI44L | 134.219307 | -3.6208387 | 1.41811277 | -2.5532798 | 0.01067137 | 0.27809901 |
| IFI44 | 54.3645479 | -2.1729878 | 1.22907175 | -1.767991 | 0.07706241 | 0.67905476 |
| ADGRL2 | 229.393435 | 0.02757045 | 0.32837411 | 0.08396047 | 0.93308786 | 0.99935941 |
| TTLL7 | 64.3576563 | 0.73736729 | 0.46359626 | 1.59053762 | 0.11171367 | 0.77793929 |
| PRKACB | 82.1385562 | 0.44366582 | 0.28562057 | 1.55333986 | 0.12034196 | 0.79021701 |
| GNG5 | 43.1539955 | -0.2248224 | 0.33209868 | -0.6769748 | 0.49842196 | 0.99176652 |
| CTBS | 35.9362944 | -0.1531038 | 0.34555965 | -0.4430604 | 0.65772207 | 0.9958918 |
| SYDE2 | 22.6109793 | -0.2079004 | 0.36830609 | -0.5644773 | 0.57242934 | 0.9958918 |
| C1orf52 | 34.5665815 | -0.0074943 | 0.31632852 | -0.0236915 | 0.98109867 | 0.99935941 |
| AL078459.1 | 25.3471627 | -0.4385365 | 0.53781431 | -0.8154051 | 0.41484049 | 0.97812179 |
| DDAH1 | 210.907859 | -0.7252791 | 0.46950709 | -1.5447672 | 0.12240258 | 0.79184923 |
| CYR61 | 524.671522 | -1.9857371 | 0.71388843 | -2.7815791 | 0.00540951 | 0.18576881 |
| ZNHIT6 | 64.471913 | -0.0485542 | 0.31504174 | -0.15412 | 0.87751508 | 0.99722158 |
| ODF2L | 76.4804992 | -0.5125806 | 0.25987167 | -1.9724374 | 0.04855969 | 0.57189932 |
| SH3GLB1 | 88.0893731 | 0.23299704 | 0.24688226 | 0.94375774 | 0.34529346 | 0.96887708 |
| SELENOF | 84.3146028 | -0.0806206 | 0.21233173 | -0.3796915 | 0.70417443 | 0.9958918 |
| HS2ST1 | 59.6894497 | -0.3695898 | 0.33460843 | -1.1045441 | 0.26935719 | 0.93634106 |
| LMO4 | 101.52057 | -0.3152927 | 0.31907929 | -0.9881329 | 0.32308758 | 0.95992644 |
| PKN2 | 191.559343 | -0.0044229 | 0.20013287 | -0.0221001 | 0.98236814 | 0.99935941 |
| GTF2B | 37.3806072 | -0.1294469 | 0.31876424 | -0.4060899 | 0.68467654 | 0.9958918 |
| KYAT3 | 38.378239 | 0.43678208 | 0.36698572 | 1.19018822 | 0.23397242 | 0.91930824 |
| GBP3 | 20.1379819 | 0.5289984 | 0.47226069 | 1.12014065 | 0.26265383 | 0.93248454 |
| GBP1 | 39.490997 | -0.9403608 | 1.00890182 | -0.9320638 | 0.35130357 | 0.97021043 |
| GBP2 | 85.4073811 | 0.01161021 | 0.48983917 | 0.02370208 | 0.98109025 | 0.99935941 |
| GBP4 | 24.612446 | 0.42234418 | 0.91038855 | 0.4639164 | 0.64270764 | 0.9958918 |
| LRRC8B | 33.9464188 | -0.9134764 | 0.44837991 | -2.0372822 | 0.04162178 | 0.54347495 |
| ZNF326 | 46.7282604 | 0.31471947 | 0.24594092 | 1.27965474 | 0.20066659 | 0.89191279 |
| ZNF644 | 180.124466 | -0.0900918 | 0.2166905 | -0.4157626 | 0.67758372 | 0.9958918 |
| CDC7 | 20.8804783 | -0.4496059 | 0.58586095 | -0.7674276 | 0.44282731 | 0.98092839 |
| TGFBR3 | 282.899061 | -0.5523787 | 0.4117938 | -1.3413962 | 0.17979185 | 0.87815793 |
| BTBD8 | 28.6368671 | -0.2147355 | 0.6276239 | -0.3421404 | 0.73224521 | 0.9958918 |
| GLMN | 39.6670401 | -0.2156228 | 0.24842411 | -0.8679627 | 0.38541477 | 0.97254921 |

| | | | | | | |
|---|---|---|---|---|---|---|
| RPAP2 | 87.6755431 | 0.21750803 | 0.25423566 | 0.85553705 | 0.3922539 | 0.97297628 |
| EVI5 | 216.928586 | -0.0034067 | 0.254352 | -0.0133938 | 0.98931365 | 0.99935941 |
| RPL5 | 98.4696983 | 0.24391613 | 0.31830395 | 0.76629942 | 0.44349817 | 0.98092839 |
| FAM69A | 77.4843334 | 0.22415319 | 0.44847131 | 0.49981611 | 0.61720456 | 0.9958918 |
| MTF2 | 60.9129375 | -0.0570601 | 0.23588979 | -0.241893 | 0.80886304 | 0.99592423 |
| TMED5 | 51.3433054 | 0.22286882 | 0.30276324 | 0.73611586 | 0.46166019 | 0.98590351 |
| CCDC18 | 21.3603672 | -0.6997657 | 0.35180375 | -1.9890797 | 0.04669241 | 0.56864121 |
| DR1 | 27.1780104 | 0.39393723 | 0.31658603 | 1.24432915 | 0.21337845 | 0.89738044 |
| FNBP1L | 104.36012 | 0.82724971 | 0.31856806 | 2.59677542 | 0.00941034 | 0.25610587 |
| BCAR3 | 83.5681373 | 1.12783028 | 0.39092823 | 2.88500599 | 0.00391406 | 0.15300235 |
| GCLM | 32.4585801 | 0.08492273 | 0.50544819 | 0.16801471 | 0.86657171 | 0.9971731 |
| ARHGAP29 | 176.167666 | -1.0677803 | 0.3548979 | -3.008697 | 0.00262371 | 0.11615431 |
| ABCD3 | 72.6807753 | -0.0105632 | 0.26798309 | -0.0394173 | 0.96855772 | 0.99935941 |
| F3 | 38.182227 | -1.0769147 | 0.39078352 | -2.7557833 | 0.00585518 | 0.19136125 |
| CNN3 | 254.300775 | -0.2384482 | 0.3888256 | -0.6132523 | 0.53970951 | 0.9958918 |
| ALG14 | 44.7542986 | 0.00269543 | 0.31303217 | 0.00861073 | 0.99312972 | 0.99935941 |
| PTBP2 | 226.016209 | -0.4754878 | 0.21861873 | -2.1749639 | 0.02963282 | 0.46634874 |
| DPYD | 508.224284 | -0.294804 | 0.22221321 | -1.3266717 | 0.18461732 | 0.87866798 |
| DPYD-AS1 | 21.3401806 | -0.4583839 | 0.39174656 | -1.1701032 | 0.24195943 | 0.92516122 |
| SNX7 | 24.2791473 | 0.46934723 | 0.49895442 | 0.94066153 | 0.34687834 | 0.96887708 |
| AGL | 29.7648872 | 0.7307036 | 0.30543978 | 2.39230003 | 0.01674315 | 0.34578631 |
| TRMT13 | 33.986606 | 0.00232178 | 0.27361827 | 0.00848546 | 0.99322966 | 0.99935941 |
| DBT | 27.9233829 | 0.10204932 | 0.40614194 | 0.25126516 | 0.80160911 | 0.99592423 |
| RTCA | 23.9726779 | 0.56416582 | 0.37367977 | 1.50975748 | 0.13110532 | 0.80565822 |
| CDC14A | 47.4445666 | 0.08994416 | 0.31349618 | 0.28690674 | 0.77418373 | 0.9958918 |
| VCAM1 | 53.6529407 | 1.54652183 | 0.67144315 | 2.30328039 | 0.02126307 | 0.3906558 |
| EXTL2 | 23.7054995 | -0.004448 | 0.55003526 | -0.0080868 | 0.99354771 | 0.99935941 |
| SLC30A7 | 98.2009065 | 0.09577306 | 0.26138479 | 0.36640638 | 0.71406186 | 0.9958918 |
| RNPC3 | 56.2183728 | -0.2283762 | 0.23905912 | -0.9553127 | 0.33941959 | 0.96887708 |
| VAV3 | 37.3980927 | 0.25139568 | 0.32293827 | 0.77846356 | 0.43629578 | 0.98092839 |
| SLC25A24 | 39.7987226 | -0.0547167 | 0.27230962 | -0.2009357 | 0.84074888 | 0.99692952 |
| FAM102B | 73.6326258 | 0.34735093 | 0.31422704 | 1.1054139 | 0.26898028 | 0.93598159 |
| PRPF38B | 55.3120336 | 0.22643102 | 0.32925878 | 0.68769926 | 0.49164219 | 0.99046213 |
| STXBP3 | 65.9243377 | -0.0690468 | 0.29345892 | -0.2352861 | 0.81398668 | 0.99592423 |
| SARS | 46.845973 | -0.1846846 | 0.25351246 | -0.7285029 | 0.46630579 | 0.98791879 |
| SORT1 | 104.505962 | 0.69718402 | 0.36267359 | 1.92234571 | 0.05456227 | 0.59304667 |
| PSMA5 | 38.6361235 | -0.1917555 | 0.3850874 | -0.4979532 | 0.61851701 | 0.9958918 |
| GNAI3 | 49.1347928 | -0.003953 | 0.32417367 | -0.0121941 | 0.99027075 | 0.99935941 |
| GSTM3 | 19.0587045 | 0.2698605 | 0.39487582 | 0.68340599 | 0.49435034 | 0.99176652 |
| CSF1 | 45.0400437 | -0.197441 | 0.48463127 | -0.4074045 | 0.68371091 | 0.9958918 |
| AHCYL1 | 100.546744 | 0.01881391 | 0.27417699 | 0.06861957 | 0.94529244 | 0.99935941 |
| STRIP1 | 21.5995997 | 0.45391835 | 0.36450026 | 1.24531696 | 0.21301526 | 0.89738044 |
| RBM15 | 18.5029464 | -0.209045 | 0.41222468 | -0.5071141 | 0.61207476 | 0.9958918 |
| SLC16A4 | 53.5397636 | 0.26806453 | 0.37637445 | 0.7122283 | 0.47632342 | 0.98968074 |
| LAMTOR5 | 48.340709 | 0.14909967 | 0.38409981 | 0.3881795 | 0.6978832 | 0.9958918 |
| DRAM2 | 27.5864785 | 0.03685219 | 0.32886253 | 0.11205956 | 0.91077619 | 0.99935941 |
| CEPT1 | 37.1685841 | -0.0166052 | 0.27380233 | -0.0606465 | 0.95164073 | 0.99935941 |
| ATP5PB | 40.4605903 | 0.28536291 | 0.28639422 | 0.99639899 | 0.31905632 | 0.95992644 |
| RAP1A | 216.310934 | -0.1047762 | 0.27515656 | -0.3807877 | 0.70336076 | 0.9958918 |
| KCND3 | 28.3048329 | 0.53629476 | 0.60187914 | 0.89103397 | 0.37291095 | 0.97217293 |
| AL445426.1 | 57.889849 | -0.2845075 | 0.59511212 | -0.4780738 | 0.63259771 | 0.9958918 |
| CTTNBP2NL | 41.7648779 | -0.1189866 | 0.27350114 | -0.4350496 | 0.6635264 | 0.9958918 |
| ST7L | 72.3751489 | 0.04257888 | 0.24813835 | 0.17159331 | 0.86375727 | 0.9971731 |
| CAPZA1 | 98.8782789 | -0.2497067 | 0.20544064 | -1.2154689 | 0.22418732 | 0.91093521 |

| | | | | | | |
|---|---|---|---|---|---|---|
| MOV10 | 24.6014718 | -0.7825145 | 0.47404425 | -1.6507203 | 0.09879569 | 0.74528964 |
| RHOC | 50.4685958 | -0.0303433 | 0.44100067 | -0.0688056 | 0.94514433 | 0.99935941 |
| SLC16A1-AS1 | 90.3950583 | 0.08405351 | 0.35651247 | 0.23576599 | 0.81361426 | 0.99592423 |
| LRIG2 | 41.9602394 | -0.0918216 | 0.32071857 | -0.2862996 | 0.77464869 | 0.9958918 |
| MAGI3 | 133.274924 | -0.181977 | 0.27247169 | -0.6678748 | 0.50421353 | 0.99387859 |
| PHTF1 | 27.1006739 | -0.0788852 | 0.28985883 | -0.2721503 | 0.78550644 | 0.9958918 |
| RSBN1 | 49.9805277 | 0.34715417 | 0.25168578 | 1.3793158 | 0.16779741 | 0.86560961 |
| HIPK1 | 53.5220756 | 0.01233055 | 0.2467103 | 0.04997989 | 0.96013842 | 0.99935941 |
| TRIM33 | 133.561527 | -0.1175576 | 0.18321561 | -0.6416354 | 0.52110997 | 0.9958918 |
| BCAS2 | 22.7014968 | -0.2429542 | 0.44358638 | -0.5477043 | 0.58389495 | 0.9958918 |
| CSDE1 | 132.581906 | 0.1198441 | 0.20384664 | 0.58791306 | 0.55659065 | 0.9958918 |
| NGF | 125.193383 | -0.1501754 | 0.50625556 | -0.2966395 | 0.76674178 | 0.9958918 |
| SLC22A15 | 22.8714071 | 0.74439103 | 0.40670705 | 1.83028799 | 0.06720689 | 0.63977125 |
| ATP1A1 | 163.441945 | 0.01857009 | 0.27339009 | 0.06792524 | 0.94584514 | 0.99935941 |
| CD58 | 21.6064515 | 0.95079993 | 0.32139579 | 2.95834598 | 0.00309295 | 0.1329244 |
| MAN1A2 | 126.666355 | 0.24080816 | 0.22252137 | 1.08217993 | 0.27917259 | 0.94419503 |
| GDAP2 | 41.2654388 | 0.24211622 | 0.29890368 | 0.81001416 | 0.41793204 | 0.97812179 |
| WARS2 | 43.3863117 | -0.0500354 | 0.30427995 | -0.1644388 | 0.86938569 | 0.9971731 |
| AL359915.2 | 87.1027323 | -0.6615832 | 0.32848386 | -2.0140507 | 0.04400421 | 0.55583613 |
| ZNF697 | 20.118781 | -0.2491255 | 0.42636813 | -0.5842968 | 0.55902066 | 0.9958918 |
| NOTCH2 | 148.876803 | -0.5566342 | 0.28570914 | -1.9482547 | 0.05138449 | 0.57655256 |
| SEC22B | 79.6979368 | -0.5765238 | 0.30347529 | -1.8997388 | 0.0574674 | 0.60512745 |
| NBPF26 | 24.3457872 | -0.0440075 | 0.51898472 | -0.0847953 | 0.93242414 | 0.99935941 |
| LINC00623 | 63.0376444 | -0.0198209 | 0.33850561 | -0.0585541 | 0.9533073 | 0.99935941 |
| SRGAP2C | 34.6979088 | -0.1125019 | 0.33560139 | -0.3352247 | 0.73745557 | 0.9958918 |
| NBPF15 | 24.8093407 | 0.47131682 | 0.33173853 | 1.42074793 | 0.15539005 | 0.84277602 |
| FP700111.1 | 24.9660625 | -0.7073983 | 0.40526317 | -1.7455282 | 0.08089297 | 0.69693132 |
| SRGAP2B | 38.9631284 | 0.23777157 | 0.28826786 | 0.8248286 | 0.40946891 | 0.97812179 |
| RNF115 | 139.558104 | 0.02415645 | 0.19043169 | 0.126851 | 0.89905833 | 0.99935941 |
| RBM8A | 24.5058091 | 0.0374422 | 0.39464457 | 0.09487575 | 0.92441352 | 0.99935941 |
| LIX1L | 33.8790141 | 0.37530243 | 0.28384347 | 1.32221616 | 0.18609618 | 0.8789337 |
| TXNIP | 354.753646 | 0.04652797 | 0.46717346 | 0.09959463 | 0.92066616 | 0.99935941 |
| NBPF12 | 23.9420246 | 0.25642851 | 0.34066933 | 0.75271967 | 0.45161839 | 0.98092839 |
| PRKAB2 | 26.4320573 | 0.4150133 | 0.40313313 | 1.02946959 | 0.30325906 | 0.95650251 |
| AC242426.2 | 27.5682247 | -0.3864058 | 0.36502255 | -1.0585806 | 0.28979083 | 0.95186143 |
| CHD1L | 57.238305 | -0.0927398 | 0.29385515 | -0.315597 | 0.7523084 | 0.9958918 |
| BCL9 | 23.1716123 | -0.4191827 | 0.43959625 | -0.9535629 | 0.34030494 | 0.96887708 |
| GJA5 | 41.1170111 | 1.19767447 | 0.56223145 | 2.13021607 | 0.03315378 | 0.49122404 |
| GPR89B | 18.9090478 | 0.20084785 | 0.35336653 | 0.5683839 | 0.56977432 | 0.9958918 |
| NBPF11 | 22.1278325 | 0.68493749 | 0.3850688 | 1.77874057 | 0.07528231 | 0.67167936 |
| NBPF14 | 28.2552577 | -0.142099 | 0.30881566 | -0.4601418 | 0.64541441 | 0.9958918 |
| PDE4DIP | 126.602725 | -0.1527046 | 0.29526599 | -0.5171765 | 0.60503296 | 0.9958918 |
| NBPF9 | 20.6301187 | 0.15588382 | 0.33448504 | 0.46604122 | 0.641186 | 0.9958918 |
| AC245297.3 | 160.605767 | -0.0407386 | 0.28022011 | -0.1453809 | 0.88441018 | 0.99862036 |
| NBPF19 | 42.2204391 | -0.1949239 | 0.86514152 | -0.2253087 | 0.82173911 | 0.99592423 |
| OTUD7B | 81.7033961 | -0.0745202 | 0.2961715 | -0.2516115 | 0.80134136 | 0.99592423 |
| VPS45 | 108.870309 | 0.0479455 | 0.22818047 | 0.21012096 | 0.83357327 | 0.99592423 |
| PLEKHO1 | 34.8871389 | 0.07226809 | 0.32808118 | 0.22027501 | 0.82565698 | 0.99592423 |
| ANP32E | 24.0247273 | 0.06536456 | 0.3213583 | 0.20340088 | 0.83882171 | 0.99692952 |
| C1orf54 | 21.4462199 | -0.104275 | 0.36611514 | -0.2848149 | 0.77578596 | 0.9958918 |
| MRPS21 | 38.6730047 | -0.1560027 | 0.25914414 | -0.6019921 | 0.54717942 | 0.9958918 |
| PRPF3 | 49.9332683 | -0.2758086 | 0.27375335 | -1.0075076 | 0.3136909 | 0.95992644 |
| RPRD2 | 143.545197 | -0.0290198 | 0.16868459 | -0.1720356 | 0.86340953 | 0.9971731 |
| MCL1 | 75.9922865 | -0.3375826 | 0.37596654 | -0.8979059 | 0.36923573 | 0.97217293 |

| | | | | | | |
|---|---|---|---|---|---|---|
| ENSA | 89.1663307 | -0.2268373 | 0.26297087 | -0.8625947 | 0.38836036 | 0.97297628 |
| GOLPH3L | 27.2616467 | 0.09972734 | 0.33761308 | 0.29538944 | 0.7676964 | 0.9958918 |
| ARNT | 109.757441 | -0.3274769 | 0.21709833 | -1.5084266 | 0.13144539 | 0.80565822 |
| SETDB1 | 37.3317207 | -0.2363614 | 0.29271202 | -0.807488 | 0.41938539 | 0.97812179 |
| CERS2 | 27.2957275 | -0.3602899 | 0.3349914 | -1.0755199 | 0.28214201 | 0.94419503 |
| CDC42SE1 | 43.2624096 | 0.01467926 | 0.35556179 | 0.04128469 | 0.96706894 | 0.99935941 |
| GABPB2 | 67.6834807 | 0.50580167 | 0.32049541 | 1.57818693 | 0.11452267 | 0.77853537 |
| PIP5K1A | 149.821808 | -0.3673017 | 0.26071018 | -1.4088505 | 0.15887937 | 0.84665568 |
| PSMD4 | 36.4377443 | 0.01912356 | 0.29646471 | 0.06450534 | 0.94856785 | 0.99935941 |
| PI4KB | 46.1010669 | 0.30817857 | 0.24501519 | 1.25779374 | 0.20846636 | 0.89634177 |
| PSMB4 | 26.509431 | -0.0839986 | 0.41772493 | -0.2010859 | 0.84063142 | 0.99692952 |
| POGZ | 195.677364 | -0.1666869 | 0.28247636 | -0.5900916 | 0.55512925 | 0.9958918 |
| SNX27 | 62.4112716 | -0.1837708 | 0.30560948 | -0.6013255 | 0.54762318 | 0.9958918 |
| THEM4 | 24.3103924 | 0.9832819 | 0.36169594 | 2.71853176 | 0.00655724 | 0.20509206 |
| S100A10 | 85.9386723 | -0.3247025 | 0.49648287 | -0.6540054 | 0.51310833 | 0.9958918 |
| S100A11 | 164.759033 | -0.3302359 | 0.32784764 | -1.0072847 | 0.31379799 | 0.95992644 |
| FLG-AS1 | 44.9050369 | -0.427894 | 0.40898321 | -1.0462385 | 0.29545095 | 0.95393362 |
| S100A6 | 272.807837 | -0.7305039 | 0.51583186 | -1.4161666 | 0.15672673 | 0.84427321 |
| S100A4 | 63.0682373 | -0.7916637 | 0.88553697 | -0.8939928 | 0.37132575 | 0.97217293 |
| S100A16 | 21.8742518 | -0.1896192 | 0.467666 | -0.4054585 | 0.68514049 | 0.9958918 |
| S100A13 | 39.984713 | -0.2986474 | 0.29212932 | -1.0223125 | 0.30663303 | 0.95793395 |
| CHTOP | 29.218578 | -0.1380054 | 0.34467686 | -0.4003908 | 0.68886872 | 0.9958918 |
| ILF2 | 25.3256024 | 0.78973569 | 0.33165485 | 2.38119747 | 0.01725646 | 0.34765232 |
| NPR1 | 31.4296147 | 0.48084855 | 0.36674265 | 1.31113342 | 0.18981269 | 0.88139754 |
| INTS3 | 44.8347924 | 0.16607023 | 0.2608429 | 0.63666762 | 0.52434137 | 0.9958918 |
| GATAD2B | 168.710392 | 0.10900669 | 0.19833601 | 0.54960613 | 0.58258955 | 0.9958918 |
| DENND4B | 22.386551 | -0.4071186 | 0.36330039 | -1.1206115 | 0.26245325 | 0.93222277 |
| SLC39A1 | 24.6314905 | -0.1115691 | 0.44106295 | -0.252955 | 0.80030301 | 0.99592423 |
| JTB | 28.972905 | 0.26847627 | 0.52554726 | 0.51085086 | 0.60945549 | 0.9958918 |
| RAB13 | 33.1584102 | 0.28017624 | 0.34642078 | 0.80877436 | 0.41864495 | 0.97812179 |
| RPS27 | 41.323994 | 0.08290771 | 0.38850619 | 0.21340126 | 0.831014 | 0.99592423 |
| TPM3 | 49.0970788 | 0.38846774 | 0.33917531 | 1.14533021 | 0.25207239 | 0.92631316 |
| C1orf43 | 44.479968 | 0.00486955 | 0.24856334 | 0.0195908 | 0.98436981 | 0.99935941 |
| UBAP2L | 67.7794572 | -0.506466 | 0.25181774 | -2.0112405 | 0.04430006 | 0.55651483 |
| ATP8B2 | 43.81097 | 0.29762491 | 0.39703817 | 0.74961284 | 0.45348792 | 0.98092839 |
| IL6R | 38.0427331 | -0.4666 | 0.52881715 | -0.8823466 | 0.37758938 | 0.97217293 |
| ADAR | 91.1707383 | -0.2217311 | 0.29401633 | -0.7541457 | 0.45076175 | 0.98092839 |
| PBXIP1 | 18.6054984 | 0.57751236 | 0.45019718 | 1.28279871 | 0.19956261 | 0.88991844 |
| SCAMP3 | 17.6257289 | -0.0771433 | 0.39776361 | -0.1939425 | 0.8462209 | 0.99692952 |
| CLK2 | 19.7085045 | 0.00613601 | 0.32705363 | 0.01876148 | 0.98503138 | 0.99935941 |
| FDPS | 43.9566295 | 0.47635364 | 0.46804757 | 1.0177462 | 0.30879859 | 0.95793395 |
| ASH1L | 415.028322 | 0.09266721 | 0.14261238 | 0.64978377 | 0.5158319 | 0.9958918 |
| YY1AP1 | 76.3933211 | -0.0439397 | 0.24459871 | -0.1796399 | 0.85743524 | 0.9971731 |
| DAP3 | 64.4911743 | -0.0936564 | 0.32770325 | -0.2857965 | 0.77503399 | 0.9958918 |
| GON4L | 93.1605053 | 0.05950877 | 0.18762854 | 0.31716268 | 0.75112017 | 0.9958918 |
| SYT11 | 19.6605404 | 0.02379458 | 0.36477309 | 0.06523118 | 0.94798994 | 0.99935941 |
| KHDC4 | 58.0613308 | -0.0296636 | 0.28914728 | -0.10259 | 0.91828836 | 0.99935941 |
| ARHGEF2 | 56.291566 | 0.71911512 | 0.30545362 | 2.35425307 | 0.01855997 | 0.36469845 |
| SSR2 | 39.5353276 | 0.18140661 | 0.37137696 | 0.48847028 | 0.62521677 | 0.9958918 |
| LMNA | 145.612504 | -0.0407464 | 0.4488789 | -0.0907738 | 0.92767234 | 0.99935941 |
| SMG5 | 29.9751158 | 0.19542484 | 0.35738043 | 0.54682581 | 0.58449841 | 0.9958918 |
| CCT3 | 45.8043901 | 0.1684624 | 0.38668684 | 0.43565588 | 0.66308642 | 0.9958918 |
| MEF2D | 76.94973 | 0.10413537 | 0.24154754 | 0.4311175 | 0.66638294 | 0.9958918 |
| NES | 33.806822 | -0.0017378 | 0.50142999 | -0.0034658 | 0.99723472 | 0.99947501 |

| | | | | | | |
|---|---|---|---|---|---|---|
| HDGF | 40.0664499 | -0.0138001 | 0.41977407 | -0.0328752 | 0.97377413 | 0.99935941 |
| PRCC | 28.4823107 | 0.39081128 | 0.29138027 | 1.3412414 | 0.17984209 | 0.87815793 |
| PEAR1 | 36.7442826 | -0.6086018 | 0.41062387 | -1.4821393 | 0.13830324 | 0.81520215 |
| ARHGEF11 | 75.35921 | -0.1152254 | 0.2737632 | -0.4208943 | 0.6738323 | 0.9958918 |
| KIRREL1 | 169.126335 | 0.07133493 | 0.26367482 | 0.2705413 | 0.78674385 | 0.99592423 |
| IFI16 | 161.797229 | -0.6115071 | 0.40602427 | -1.5060852 | 0.13204532 | 0.80565822 |
| DUSP23 | 29.4886113 | -0.1738751 | 0.43000456 | -0.4043565 | 0.68595057 | 0.9958918 |
| TAGLN2 | 71.584298 | -0.0997612 | 0.4495964 | -0.2218906 | 0.82439902 | 0.99592423 |
| PEA15 | 25.4970137 | 0.1281256 | 0.38723492 | 0.33087304 | 0.74074039 | 0.9958918 |
| DCAF8 | 19.668492 | -0.6412116 | 0.37445552 | -1.712384 | 0.08682593 | 0.71453945 |
| COPA | 157.036304 | -0.2221438 | 0.2137495 | -1.0392714 | 0.29867851 | 0.95447177 |
| NCSTN | 18.9636176 | 0.30745144 | 0.34755955 | 0.88460076 | 0.37637199 | 0.97217293 |
| PFDN2 | 32.6915578 | 0.07538093 | 0.3723537 | 0.20244444 | 0.83956929 | 0.99692952 |
| NIT1 | 25.4107044 | -0.0462445 | 0.39779707 | -0.1162515 | 0.90745326 | 0.99935941 |
| UFC1 | 21.8335823 | 0.01582921 | 0.41846325 | 0.037827 | 0.96982562 | 0.99935941 |
| ADAMTS4 | 139.298902 | -1.7468986 | 0.83029985 | -2.1039371 | 3.54E-02 | 0.50689386 |
| NDUFS2 | 54.882127 | -0.508016 | 0.39142788 | -1.2978534 | 0.19433771 | 0.88391893 |
| SDHC | 44.0910014 | -0.088248 | 0.25626621 | -0.3443605 | 0.73057519 | 0.9958918 |
| ATF6 | 148.1238 | -0.4685418 | 0.27138824 | -1.7264632 | 0.08426411 | 0.70370105 |
| NOS1AP | 35.9370446 | 1.10805104 | 0.36074228 | 3.0715863 | 0.00212925 | 0.09909091 |
| UHMK1 | 47.6286852 | -0.2058513 | 0.25594082 | -0.8042927 | 0.42122796 | 0.97812179 |
| UAP1 | 72.4419782 | -0.5432073 | 0.56261967 | -0.9654965 | 0.3342962 | 0.96640422 |
| DDR2 | 205.222271 | -1.0885304 | 0.28748802 | -3.7863506 | 0.00015288 | 0.01904212 |
| RGS5 | 278.98234 | -0.0338787 | 0.42878125 | -0.0790117 | 0.93702334 | 0.99935941 |
| PBX1 | 946.409462 | -0.5391049 | 0.33401312 | -1.6140232 | 0.10652239 | 0.77126411 |
| MGST3 | 106.224318 | 0.44900202 | 0.25797303 | 1.74049984 | 0.08177129 | 0.69958018 |
| ALDH9A1 | 24.4709884 | 0.98146243 | 0.37601146 | 2.61019285 | 0.00904912 | 0.25292422 |
| TMCO1 | 40.6169738 | 0.00685251 | 0.25417375 | 0.02695995 | 0.97849168 | 0.99935941 |
| UCK2 | 25.029153 | 0.27570031 | 0.50305871 | 0.54804797 | 0.58365896 | 0.9958918 |
| POGK | 18.4211831 | 0.01466073 | 0.39534231 | 0.03708364 | 0.97041832 | 0.99935941 |
| POU2F1 | 211.808495 | -0.1199794 | 0.18699105 | -0.6416319 | 0.52111221 | 0.9958918 |
| CREG1 | 19.5928679 | 0.20897355 | 0.36478602 | 0.57286612 | 0.56673535 | 0.9958918 |
| RCSD1 | 52.5635281 | 0.10967627 | 0.43048127 | 0.25477593 | 0.79889616 | 0.99592423 |
| MPZL1 | 137.979508 | 0.03502639 | 0.33751292 | 0.10377791 | 0.9173456 | 0.99935941 |
| MPC2 | 25.7684142 | 0.64285341 | 0.40741403 | 1.57788728 | 0.11459151 | 0.77853537 |
| DCAF6 | 196.815142 | 0.10559681 | 0.21330905 | 0.49504139 | 0.6205709 | 0.9958918 |
| TIPRL | 34.6602517 | 0.27316858 | 0.27025587 | 1.01077762 | 0.31212288 | 0.9599143 |
| SFT2D2 | 57.2619051 | 0.26510361 | 0.28003467 | 0.94668141 | 0.34380115 | 0.96887708 |
| TBX19 | 18.0935064 | 0.19002968 | 0.4544869 | 0.41811916 | 0.67585999 | 0.9958918 |
| ATP1B1 | 136.198689 | -0.0640633 | 0.37648456 | -0.1701619 | 0.86488283 | 0.9971731 |
| NME7 | 81.6329533 | -0.2012836 | 0.23872434 | -0.8431635 | 0.39913703 | 0.97500135 |
| BLZF1 | 37.4288481 | 0.00441711 | 0.32785487 | 0.01347277 | 0.98925061 | 0.99935941 |
| SLC19A2 | 42.9883919 | -0.959348 | 0.41151994 | -2.3312308 | 0.01974119 | 0.37879384 |
| SCYL3 | 25.3017065 | 0.0617824 | 0.35487024 | 0.17409856 | 0.861788 | 0.9971731 |
| KIFAP3 | 92.6277969 | -0.2478548 | 0.27312375 | -0.9074817 | 0.36415214 | 0.97217293 |
| PRRX1 | 124.930532 | 0.464365 | 0.60337213 | 0.76961626 | 0.44152756 | 0.98092839 |
| FMO3 | 33.6119326 | 1.17725408 | 1.07041484 | 1.09981106 | 0.27141445 | 0.93776436 |
| FMO2 | 55.9903287 | 0.21272802 | 1.04682274 | 0.20321303 | 0.83896853 | 0.99692952 |
| PRRC2C | 259.377601 | -0.3395977 | 0.20961696 | -1.6200871 | 0.10521358 | 0.76555899 |
| VAMP4 | 28.9840696 | -0.3156003 | 0.39255513 | -0.8039643 | 0.42141759 | 0.97812179 |
| DNM3 | 66.1792902 | -0.032831 | 0.36354577 | -0.0903078 | 0.92804263 | 0.99935941 |
| DNM3OS | 42.4604492 | -0.6304906 | 0.33848081 | -1.8627071 | 0.06250349 | 0.62156719 |
| SUCO | 65.2294966 | 0.19593773 | 0.27477881 | 0.71307438 | 0.47579973 | 0.98943187 |
| AL645568.1 | 34.2733386 | 0.3109224 | 0.32311178 | 0.96227504 | 0.33591146 | 0.96640422 |

| | | | | | | |
|---|---|---|---|---|---|---|
| PRDX6 | 82.861276 | 0.23164286 | 0.35542711 | 0.65173097 | 0.51457474 | 0.9958918 |
| KLHL20 | 49.1004578 | 0.22101562 | 0.31533389 | 0.70089395 | 0.4833692 | 0.99015594 |
| ZBTB37 | 29.9393701 | 0.13530047 | 0.30989758 | 0.43659736 | 0.66240337 | 0.9958918 |
| RC3H1 | 120.127329 | 0.13268558 | 0.17473519 | 0.75935236 | 0.4476418 | 0.98092839 |
| RABGAP1L | 355.836108 | -0.1367853 | 0.18604218 | -0.735238 | 0.46219456 | 0.98590351 |
| KIAA0040 | 27.1023514 | -0.1226908 | 0.35644498 | -0.3442067 | 0.73069084 | 0.9958918 |
| COP1 | 231.908837 | 0.1181261 | 0.18634439 | 0.63391279 | 0.52613773 | 0.9958918 |
| RASAL2 | 436.053704 | -0.4783671 | 0.35256901 | -1.3568041 | 0.17484345 | 0.8740623 |
| RALGPS2 | 47.0800335 | 0.33782065 | 0.44797378 | 0.75410809 | 0.45078432 | 0.98092839 |
| FAM20B | 35.6906293 | 0.15338644 | 0.30851213 | 0.49718122 | 0.61906127 | 0.9958918 |
| ABL2 | 123.351992 | -0.0595268 | 0.25919887 | -0.2296569 | 0.81835839 | 0.99592423 |
| SOAT1 | 50.9126015 | -0.1379259 | 0.25346411 | -0.5441635 | 0.58632895 | 0.9958918 |
| TOR1AIP2 | 52.2173938 | 0.09740672 | 0.21919623 | 0.44438134 | 0.6567669 | 0.9958918 |
| TOR1AIP1 | 51.0988791 | 0.17880647 | 0.30221476 | 0.59165366 | 0.55408253 | 0.9958918 |
| CEP350 | 230.347699 | -0.0059952 | 0.22934089 | -0.0261411 | 0.97914478 | 0.99935941 |
| QSOX1 | 38.4448128 | -0.0186491 | 0.28808714 | -0.0647342 | 0.9483856 | 0.99935941 |
| ACBD6 | 128.650908 | -0.0169229 | 0.20245781 | -0.0835874 | 0.93338452 | 0.99935941 |
| XPR1 | 111.780144 | -0.1201827 | 0.33340648 | -0.3604689 | 0.71849649 | 0.9958918 |
| STX6 | 28.8808594 | 0.49461622 | 0.33283084 | 1.4860889 | 0.13725561 | 0.81404904 |
| MR1 | 20.5487137 | 0.53628714 | 0.31190919 | 1.7193695 | 0.08554711 | 0.708721 |
| GLUL | 103.484097 | -1.0474798 | 0.37894179 | -2.7642234 | 0.00570585 | 0.1903825 |
| RGS16 | 62.6315709 | 0.22894357 | 0.93640196 | 0.24449284 | 0.80684915 | 0.99592423 |
| DHX9 | 90.7659946 | -0.2432205 | 0.24439627 | -0.9951893 | 0.31964422 | 0.95992644 |
| LAMC1 | 259.421115 | -0.4716343 | 0.2297944 | -2.0524184 | 0.04012901 | 0.53815352 |
| SMG7 | 96.8673237 | 0.17950823 | 0.2490408 | 0.72079847 | 0.47103351 | 0.98910524 |
| ARPC5 | 68.3385505 | -0.0309585 | 0.25193668 | -0.122882 | 0.90220053 | 0.99935941 |
| RGL1 | 115.541762 | -0.2600231 | 0.28771937 | -0.9037388 | 0.36613394 | 0.97217293 |
| TSEN15 | 30.6520105 | -0.0280763 | 0.37690043 | -0.0744926 | 0.94061843 | 0.99935941 |
| C1orf21 | 288.780821 | -0.6285335 | 0.5345445 | -1.1758301 | 0.23966277 | 0.92426606 |
| EDEM3 | 89.4892009 | -0.0670459 | 0.30785453 | -0.2177844 | 0.8275971 | 0.99592423 |
| FAM129A | 115.787429 | 0.34501723 | 0.37501705 | 0.92000411 | 0.35757061 | 0.97217293 |
| TRMT1L | 23.7468918 | -0.1916567 | 0.3992438 | -0.4800494 | 0.63119226 | 0.9958918 |
| SWT1 | 36.7943448 | -0.0530739 | 0.38476642 | -0.137938 | 0.89028945 | 0.99935941 |
| IVNS1ABP | 41.0708732 | -0.3156083 | 0.39650652 | -0.7959725 | 0.42604802 | 0.98015476 |
| HMCN1 | 78.9049077 | -0.9627385 | 0.51041447 | -1.8861897 | 0.05926939 | 0.61175668 |
| TPR | 101.054122 | 0.19203925 | 0.25423071 | 0.75537393 | 0.45002465 | 0.98092839 |
| ODR4 | 56.5575218 | 0.20073149 | 0.23072924 | 0.86998721 | 0.38430739 | 0.9725044 |
| PTGS2 | 72.5119386 | -1.1137409 | 0.90649543 | -1.2286228 | 0.21921327 | 0.90437211 |
| PLA2G4A | 225.288005 | -0.531814 | 0.43802203 | -1.2141262 | 0.22469954 | 0.91093521 |
| AL390957.1 | 59.8898126 | -0.004852 | 0.54613578 | -0.0088842 | 0.9929115 | 0.99935941 |
| AL136454.1 | 35.0345406 | 0.00646757 | 0.39401743 | 0.01641442 | 0.98690377 | 0.99935941 |
| RGS2 | 65.6466538 | 0.47548791 | 0.48659497 | 0.97717391 | 0.32848306 | 0.9620084 |
| UCHL5 | 35.1214567 | 0.01955647 | 0.32395009 | 0.06036876 | 0.95186194 | 0.99935941 |
| TROVE2 | 70.9871919 | 0.02118528 | 0.27433159 | 0.07722508 | 0.93844449 | 0.99935941 |
| CDC73 | 162.103081 | -0.3004217 | 0.28833247 | -1.0419279 | 0.29744509 | 0.95417903 |
| KCNT2 | 243.059656 | 0.09811952 | 0.37342518 | 0.26275551 | 0.79273903 | 0.99592423 |
| CFH | 209.938145 | -0.0384433 | 0.51276708 | -0.0749723 | 0.94023675 | 0.99935941 |
| ZBTB41 | 30.4500179 | 0.00751935 | 0.3684659 | 0.02040719 | 0.98371855 | 0.99935941 |
| DENND1B | 63.4896326 | 0.3131002 | 0.35489623 | 0.88223027 | 0.37765229 | 0.97217293 |
| NEK7 | 235.929256 | 0.07187672 | 0.21408532 | 0.33573867 | 0.73706794 | 0.9958918 |
| MIR181A1H( | 35.7295493 | -0.5766575 | 0.43291624 | -1.3320302 | 0.18285027 | 0.87866798 |
| DDX59 | 18.0007021 | 0.52063462 | 0.45090469 | 1.15464451 | 0.24823603 | 0.92624282 |
| CAMSAP2 | 187.672103 | 0.10344112 | 0.18241146 | 0.56707579 | 0.57066269 | 0.9958918 |
| PHLDA3 | 21.8597629 | 0.35433982 | 0.46542417 | 0.76132664 | 0.44646199 | 0.98092839 |

| | | | | | | |
|---|---|---|---|---|---|---|
| CSRP1 | 207.15568 | 0.14153219 | 0.30274079 | 0.46750287 | 6.40E-01 | 0.9958918 |
| NAV1 | 155.469138 | -0.7256628 | 0.40757985 | -1.7804186 | 0.07500748 | 0.67167936 |
| IPO9-AS1 | 42.3942778 | -1.1901485 | 0.34996433 | -3.4007708 | 0.00067196 | 0.05624153 |
| IPO9 | 75.1693086 | 0.06016014 | 0.25942599 | 0.23189712 | 0.81661792 | 0.99592423 |
| LMOD1 | 172.40129 | 0.52029311 | 0.49342645 | 1.05444917 | 0.29167732 | 0.95186143 |
| TIMM17A | 19.2925427 | 0.48892234 | 0.40158834 | 1.21747144 | 0.2234249 | 0.91093521 |
| RNPEP | 28.7130618 | 0.34412851 | 0.35107083 | 0.98022529 | 0.32697492 | 0.95992644 |
| ARL8A | 24.438711 | 0.23504478 | 0.33033064 | 0.71154398 | 0.47674721 | 0.98993168 |
| PPP1R12B | 444.124969 | -0.3436651 | 0.36704662 | -0.9362984 | 0.34911959 | 0.9692706 |
| KDM5B | 115.869913 | -0.1419759 | 0.30548757 | -0.4647519 | 0.64210915 | 0.9958918 |
| BTG2 | 39.4347518 | 0.31651591 | 0.48246213 | 0.65604302 | 0.51179644 | 0.9958918 |
| FMOD | 29.7161115 | -1.1753935 | 0.36889073 | -3.1862917 | 0.00144109 | 0.08339044 |
| PRELP | 67.7236297 | -0.1580213 | 0.35480365 | -0.4453769 | 0.65604743 | 0.9958918 |
| ATP2B4 | 143.155824 | -0.087235 | 0.35134979 | -0.2482854 | 0.80391362 | 0.99592423 |
| SOX13 | 85.632665 | -0.0206978 | 0.28163694 | -0.0734912 | 0.94141526 | 0.99935941 |
| REN | 643.974268 | 2.87939288 | 1.39522509 | 2.06374792 | 0.03904162 | 0.52881167 |
| PLEKHA6 | 56.9249995 | -0.0390385 | 0.29274139 | -0.133355 | 0.89391264 | 0.99935941 |
| PPP1R15B | 30.9480427 | 0.05480126 | 0.4514162 | 0.12139852 | 0.90337539 | 0.99935941 |
| PIK3C2B | 48.5619853 | -0.3681054 | 0.48987396 | -0.7514288 | 0.45239463 | 0.98092839 |
| MDM4 | 111.729673 | -0.054688 | 0.18923878 | -0.2889892 | 0.77258967 | 0.9958918 |
| NFASC | 414.323804 | -0.518726 | 0.55095878 | -0.9414969 | 0.34645029 | 0.96887708 |
| RBBP5 | 24.8926256 | -0.039311 | 0.3120646 | -0.1259708 | 0.89975508 | 0.99935941 |
| DSTYK | 60.4537912 | -0.0980849 | 0.2716845 | -0.3610249 | 0.71808083 | 0.9958918 |
| CDK18 | 44.6688315 | 0.65484313 | 0.35072833 | 1.86709506 | 0.06188831 | 0.619642 |
| ELK4 | 40.3945459 | -0.3192871 | 0.29596754 | -1.0787909 | 0.28068095 | 0.94419503 |
| NUCKS1 | 174.532959 | -0.1054652 | 0.2397197 | -0.4399523 | 0.65997168 | 0.9958918 |
| SRGAP2 | 69.1167494 | 0.42426374 | 0.21726934 | 1.95270875 | 0.05085412 | 0.57655256 |
| RASSF5 | 37.2583792 | 0.48075533 | 0.52941203 | 0.90809295 | 0.36382912 | 0.97217293 |
| MAPKAPK2 | 122.987886 | -0.568024 | 0.30192315 | -1.8813531 | 0.0599239 | 0.61332974 |
| PIGR | 25.8589822 | 1.49108708 | 0.60930291 | 2.44720164 | 1.44E-02 | 0.32858308 |
| CD55 | 54.1098278 | -0.2770698 | 0.35989317 | -0.7698669 | 0.44137884 | 0.98092839 |
| CD46 | 238.474385 | -0.0947676 | 0.2546466 | -0.3721536 | 0.7097785 | 0.9958918 |
| MIR29B2CH( | 95.3839272 | 0.2819724 | 0.31751904 | 0.88804878 | 0.37451451 | 0.97217293 |
| CD34 | 27.7844397 | 1.09865648 | 0.48906913 | 2.24642369 | 0.02467688 | 0.42470821 |
| PLXNA2 | 49.9618306 | 0.66533323 | 0.32143393 | 2.06989107 | 0.03846255 | 0.5265563 |
| G0S2 | 148.17116 | 1.18996132 | 1.03722325 | 1.14725669 | 0.25127555 | 0.92631316 |
| HHAT | 31.8364735 | -0.0437199 | 0.60171219 | -0.0726591 | 0.94207737 | 0.99935941 |
| RCOR3 | 105.341706 | -0.2228532 | 0.27705988 | -0.8043502 | 0.42119478 | 0.97812179 |
| TRAF5 | 45.8890009 | -0.2084619 | 0.38393208 | -0.5429656 | 0.5871535 | 0.9958918 |
| LPGAT1 | 104.917048 | -0.0375809 | 0.20541022 | -0.1829554 | 0.85483303 | 0.9971731 |
| INTS7 | 29.0195935 | -0.2921911 | 0.40526982 | -0.7209792 | 0.47092232 | 0.98910524 |
| PPP2R5A | 57.9831634 | 0.38582193 | 0.26095417 | 1.47850452 | 0.13927279 | 0.8153985 |
| NENF | 29.1065907 | 0.11631327 | 0.29619386 | 0.39269306 | 0.6945462 | 0.9958918 |
| ATF3 | 62.8524529 | -1.6667076 | 0.50748686 | -3.284238 | 0.00102259 | 0.07036431 |
| NSL1 | 34.5975676 | -0.0875609 | 0.34254489 | -0.2556187 | 0.7982453 | 0.99592423 |
| ANGEL2 | 35.3230077 | -0.1047949 | 0.26729783 | -0.392053 | 0.69501903 | 0.9958918 |
| RPS6KC1 | 133.3339 | -0.2037775 | 0.28528042 | -0.7143061 | 0.47503792 | 0.98910524 |
| PTPN14 | 145.965807 | 0.27802056 | 0.26430958 | 1.05187471 | 0.29285703 | 0.95186143 |
| KCTD3 | 72.5697388 | -0.1780294 | 0.2891136 | -0.6157765 | 0.53804205 | 0.9958918 |
| ESRRG | 75.2202625 | -0.4999233 | 0.4354093 | -1.1481687 | 0.25089894 | 0.92624282 |
| GPATCH2 | 95.2175074 | 0.01126721 | 0.32524244 | 0.03464251 | 0.9723648 | 0.99935941 |
| RRP15 | 23.0511268 | -0.2081034 | 0.36171514 | -0.5753239 | 0.56507228 | 0.9958918 |
| TGFB2 | 29.02693 | -0.801232 | 0.57186943 | -1.401075 | 0.16119165 | 0.85161569 |
| LYPLAL1 | 80.1065559 | -0.0740744 | 0.39732251 | -0.186434 | 0.85210441 | 0.9971731 |

| | | | | | | |
|---|---|---|---|---|---|---|
| EPRS | 49.3228349 | -0.0394576 | 0.27961161 | -0.1411156 | 0.8877786 | 0.99935941 |
| IARS2 | 37.5802947 | 0.12729755 | 0.26564279 | 0.47920575 | 0.63179226 | 0.9958918 |
| RAB3GAP2 | 153.722614 | -0.4525102 | 0.22024919 | -2.0545374 | 0.03992371 | 0.53637904 |
| MARK1 | 81.7116897 | -0.1631269 | 0.24591034 | -0.6633594 | 0.50710038 | 0.99397937 |
| 2-Mar | 51.5680931 | 0.77181758 | 0.32213078 | 2.39597588 | 0.01657618 | 0.34578631 |
| MIA3 | 59.3890253 | -0.1122434 | 0.24798772 | -0.4526168 | 0.65082471 | 0.9958918 |
| AIDA | 23.7505 | 0.19713386 | 0.34229704 | 0.57591459 | 0.56467291 | 0.9958918 |
| BROX | 23.7532586 | -0.2736891 | 0.30532907 | -0.8963742 | 0.37005295 | 0.97217293 |
| AC093152.1 | 22.63705 | -0.0374193 | 0.509351 | -0.0734647 | 0.94143635 | 0.99935941 |
| CAPN2 | 61.0833534 | 0.00185042 | 0.33690136 | 0.00549248 | 0.99561766 | 0.99935941 |
| TP53BP2 | 42.6118391 | 0.36548223 | 0.34273404 | 1.0663727 | 0.2862552 | 0.94614505 |
| FBXO28 | 68.6827041 | 0.31707295 | 0.33435866 | 0.94830187 | 0.3429758 | 0.96887708 |
| DEGS1 | 34.4531655 | -0.2037869 | 0.34168406 | -0.5964192 | 0.55089521 | 0.9958918 |
| NVL | 48.5116032 | -0.0168929 | 0.28660834 | -0.0589408 | 0.95299924 | 0.99935941 |
| CNIH4 | 27.2510417 | 0.1334823 | 0.33228793 | 0.40170675 | 0.68789985 | 0.9958918 |
| WDR26 | 138.421822 | -0.058206 | 0.23012193 | -0.2529355 | 0.80031807 | 0.99592423 |
| CNIH3 | 42.9266426 | 0.17635785 | 0.44781212 | 0.39382107 | 0.69371315 | 0.9958918 |
| LBR | 40.2358376 | -0.2580592 | 0.46441935 | -0.5556598 | 0.57844341 | 0.9958918 |
| ENAH | 234.660698 | -0.1244315 | 0.43346419 | -0.2870629 | 0.77406417 | 0.9958918 |
| SRP9 | 47.4672641 | 0.1410672 | 0.25893879 | 0.54478976 | 0.58589812 | 0.9958918 |
| EPHX1 | 35.0650699 | 0.34916848 | 0.27487502 | 1.27028089 | 0.20398459 | 0.89396747 |
| AL591895.1 | 17.6310029 | 0.52309231 | 0.47699137 | 1.09664942 | 0.27279468 | 0.93819216 |
| TMEM63A | 48.7890799 | 0.67272168 | 0.3359531 | 2.00242734 | 0.04523879 | 0.55894748 |
| ACBD3 | 82.7317232 | -0.0880681 | 0.30456642 | -0.2891591 | 0.77245964 | 0.9958918 |
| PARP1 | 20.1116806 | 0.06467886 | 0.35584792 | 0.18175984 | 0.8557712 | 0.9971731 |
| ITPKB | 54.9136808 | -0.3375279 | 0.45238729 | -0.7461039 | 0.45560463 | 0.98275269 |
| COQ8A | 22.7641738 | 0.43969993 | 0.33645388 | 1.3068654 | 0.19125843 | 0.88158763 |
| CDC42BPA | 539.905342 | -0.1705075 | 0.20793908 | -0.8199877 | 0.41222312 | 0.97812179 |
| ZNF678 | 35.8608398 | -0.1798822 | 0.27197596 | -0.6613901 | 0.50836219 | 0.99397937 |
| ARF1 | 91.8043285 | 0.14740052 | 0.35984845 | 0.40961832 | 0.68208595 | 0.9958918 |
| GUK1 | 58.4010303 | 0.18419774 | 0.25687777 | 0.71706377 | 0.47333475 | 0.98910524 |
| OBSCN | 26.8817967 | 0.28652642 | 0.54052472 | 0.53008939 | 0.59604995 | 0.9958918 |
| NUP133 | 43.9873052 | 0.16116806 | 0.2909174 | 0.55399939 | 0.57957926 | 0.9958918 |
| ABCB10 | 25.0781278 | -0.0044971 | 0.39595223 | -0.0113576 | 0.99093812 | 0.99935941 |
| GALNT2 | 114.925618 | -0.0703458 | 0.45957816 | -0.1530661 | 0.87834613 | 0.99722158 |
| COG2 | 25.1020353 | 0.42931815 | 0.32315125 | 1.32853622 | 0.18400102 | 0.87866798 |
| C1orf198 | 43.0825218 | -0.3622108 | 0.44976932 | -0.8053257 | 0.42063176 | 0.97812179 |
| TTC13 | 24.3040232 | -0.1373673 | 0.42062831 | -0.3265764 | 0.74398829 | 0.9958918 |
| FAM89A | 26.3830291 | 0.45418689 | 0.60543379 | 0.75018425 | 0.45314374 | 0.98092839 |
| GNPAT | 32.52922 | 0.11762091 | 0.27884308 | 0.42181758 | 0.67315817 | 0.9958918 |
| EGLN1 | 101.228634 | 0.12821217 | 0.35774092 | 0.35839391 | 0.72004855 | 0.9958918 |
| DISC1 | 54.4581612 | -0.2884687 | 0.35592372 | -0.8104791 | 0.41766487 | 0.97812179 |
| SIPA1L2 | 68.5182818 | 0.1053224 | 0.48697768 | 0.21627767 | 0.82877132 | 0.99592423 |
| PCNX2 | 68.8793094 | -0.3632637 | 0.23756835 | -1.5290912 | 0.12624183 | 0.79566999 |
| TARBP1 | 37.3200822 | 0.07155043 | 0.44228356 | 0.16177501 | 0.87148303 | 0.9971731 |
| IRF2BP2 | 80.6566956 | 0.02011969 | 0.36318975 | 0.05539719 | 0.95582204 | 0.99935941 |
| TOMM20 | 51.8798029 | 0.06993636 | 0.28304158 | 0.24708864 | 0.80483963 | 0.99592423 |
| ARID4B | 324.128672 | -0.2075077 | 0.26956348 | -0.7697914 | 0.44142363 | 0.98092839 |
| B3GALNT2 | 67.6612407 | -0.4026564 | 0.36328653 | -1.1083715 | 0.2677014 | 0.93549101 |
| LYST | 179.03788 | -0.3650306 | 0.22665461 | -1.610515 | 0.10728548 | 0.77179565 |
| NID1 | 436.871949 | -1.2854683 | 0.44618326 | -2.8810322 | 0.00396375 | 0.15331374 |
| GPR137B | 69.2344069 | -0.1708624 | 0.45972162 | -0.371665 | 0.71014231 | 0.9958918 |
| ERO1B | 48.8423101 | 0.34982747 | 0.32279673 | 1.08373921 | 0.27848044 | 0.94419503 |
| LGALS8 | 37.5454067 | -0.3685198 | 0.2898994 | -1.2711988 | 0.20365794 | 0.89396747 |

| | | | | | | |
|---|---|---|---|---|---|---|
| HEATR1 | 43.0991896 | 0.02723354 | 0.33865892 | 0.08041582 | 0.93590654 | 0.99935941 |
| MTR | 118.709829 | -0.0549999 | 0.23436764 | -0.2346735 | 0.81446212 | 0.99592423 |
| RYR2 | 43.9317272 | 0.3786859 | 0.60874769 | 0.62207366 | 0.53389344 | 0.9958918 |
| PLD5 | 57.9419811 | 1.46368886 | 1.09688057 | 1.33441041 | 0.18206939 | 0.87866798 |
| CEP170 | 105.372317 | -0.1529298 | 0.18598051 | -0.8222894 | 0.41091221 | 0.97812179 |
| SDCCAG8 | 207.730959 | 0.23590521 | 0.19202224 | 1.22853069 | 0.21924782 | 0.90437211 |
| AKT3 | 732.633933 | -0.3921473 | 0.28433392 | -1.3791788 | 0.16783963 | 0.86560961 |
| ADSS | 33.6851169 | -0.2048092 | 0.30062223 | -0.6812842 | 0.49569165 | 0.99176652 |
| DESI2 | 38.6668807 | -0.090039 | 0.26847923 | -0.3353666 | 0.73734854 | 0.9958918 |
| COX20 | 33.8196413 | -0.0545626 | 0.29922479 | -0.1823465 | 0.85531083 | 0.9971731 |
| HNRNPU | 254.799385 | -0.3099547 | 0.24207941 | -1.2803845 | 0.20040995 | 0.89191279 |
| EFCAB2 | 122.574876 | -0.3134974 | 0.383731 | -0.8169719 | 0.41394451 | 0.97812179 |
| KIF26B | 187.467562 | -1.9989001 | 0.44391921 | -4.5028466 | 6.70E-06 | 0.0025934 |
| SMYD3 | 409.93312 | -0.5849091 | 0.27737734 | -2.1087128 | 0.03496938 | 0.50468149 |
| CNST | 75.9816692 | -0.1925525 | 0.275264 | -0.6995194 | 0.48422749 | 0.99046213 |
| SCCPDH | 20.6091082 | -0.2374376 | 0.45307256 | -0.5240608 | 0.60023627 | 0.9958918 |
| AHCTF1 | 93.7326004 | -0.092989 | 0.2468103 | -0.3767629 | 0.70634979 | 0.9958918 |
| ZNF496 | 25.1365183 | -0.1054663 | 0.38950342 | -0.2707712 | 0.78656701 | 0.99592423 |
| SH3YL1 | 48.0447431 | 0.11830882 | 0.30906972 | 0.38279005 | 0.70187544 | 0.9958918 |
| ACP1 | 24.2302124 | 0.44890476 | 0.31879039 | 1.40815024 | 0.15908659 | 0.84665568 |
| PXDN | 119.415837 | -0.1437682 | 0.36063584 | -0.3986521 | 0.69014956 | 0.9958918 |
| EIPR1 | 26.9339809 | -0.0189519 | 0.35792369 | -0.0529495 | 0.95777217 | 0.99935941 |
| TRAPPC12 | 74.3526324 | 0.09753537 | 0.26605956 | 0.36659225 | 0.71392319 | 0.9958918 |
| ADI1 | 35.9287341 | 0.34047448 | 0.3455438 | 0.98532945 | 0.32446227 | 0.95992644 |
| RPS7 | 113.5642 | -0.4356199 | 0.44116374 | -0.9874337 | 0.32343008 | 0.95992644 |
| RNF144A | 217.531259 | 0.05611283 | 0.52017218 | 0.10787356 | 0.91409599 | 0.99935941 |
| ID2 | 82.3387517 | -0.4874321 | 0.37956022 | -1.2842024 | 0.19907116 | 0.88864367 |
| KIDINS220 | 154.694688 | -0.2083491 | 0.20096108 | -1.0367636 | 0.29984606 | 0.95587816 |
| MBOAT2 | 57.3697391 | -0.0231396 | 0.38312292 | -0.0603973 | 0.95183918 | 0.99935941 |
| ASAP2 | 244.614983 | -0.337772 | 0.34664932 | -0.9743909 | 0.32986247 | 0.96348071 |
| ITGB1BP1 | 51.989534 | 0.17528183 | 0.27442784 | 0.63871737 | 0.52300681 | 0.9958918 |
| IAH1 | 21.1822132 | -0.0503857 | 0.36436671 | -0.1382829 | 0.89001683 | 0.99935941 |
| ADAM17 | 109.292516 | -0.0725329 | 0.29947106 | -0.2422035 | 0.80862249 | 0.99592423 |
| YWHAQ | 97.8040849 | 0.04065843 | 0.22749719 | 0.17872057 | 0.85815712 | 0.9971731 |
| TAF1B | 49.1867842 | -0.0213012 | 0.32836751 | -0.06487 | 0.94827753 | 0.99935941 |
| HPCAL1 | 23.2106163 | -0.2292233 | 0.34621006 | -0.662093 | 0.50791163 | 0.99397937 |
| NOL10 | 83.5425235 | -0.1447198 | 0.27717845 | -0.5221178 | 0.60158828 | 0.9958918 |
| PDIA6 | 59.2548081 | -0.0430193 | 0.38210032 | -0.1125864 | 0.9103585 | 0.99935941 |
| ROCK2 | 485.254925 | -0.5610858 | 0.39334161 | -1.4264593 | 0.15373581 | 0.84125425 |
| LPIN1 | 54.0895403 | -0.0863269 | 0.26143025 | -0.3302102 | 0.74124118 | 0.9958918 |
| TRIB2 | 82.530608 | -0.7493533 | 0.47689309 | -1.5713235 | 0.11610753 | 0.78417352 |
| NBAS | 200.88543 | -0.398833 | 0.24767519 | -1.6103065 | 0.10733095 | 0.77179565 |
| DDX1 | 47.4913476 | 0.00963385 | 0.24620596 | 0.03912922 | 0.96878737 | 0.99935941 |
| FAM49A | 41.5945774 | -0.1658399 | 0.36487713 | -0.454509 | 0.64946252 | 0.9958918 |
| SMC6 | 36.6801451 | -0.1539391 | 0.33018597 | -0.4662193 | 0.64105856 | 0.9958918 |
| LINC01376 | 52.3229318 | 0.19285066 | 0.47257118 | 0.40808807 | 0.68320902 | 0.9958918 |
| WDR35 | 40.223963 | 0.05765026 | 0.33027189 | 0.17455393 | 0.86143015 | 0.9971731 |
| LAPTM4A | 122.898609 | 0.10020756 | 0.24442745 | 0.40996853 | 0.68182904 | 0.9958918 |
| PUM2 | 219.386781 | -0.0159405 | 0.15275098 | -0.1043561 | 0.91688677 | 0.99935941 |
| RHOB | 111.224117 | -0.6306183 | 0.54078572 | -1.166115 | 0.24356797 | 0.92534155 |
| HS1BP3 | 19.5808291 | -0.0840937 | 0.40215098 | -0.2091099 | 0.83436247 | 0.99638118 |
| LDAH | 70.9831372 | 0.06623617 | 0.22996491 | 0.2880273 | 0.77332584 | 0.9958918 |
| KLHL29 | 201.864851 | -0.1925699 | 0.25587267 | -0.7526004 | 0.45169007 | 0.98092839 |
| ATAD2B | 168.880204 | -0.0797324 | 0.2442291 | -0.3264655 | 0.7440722 | 0.9958918 |

| | | | | | | |
|---|---|---|---|---|---|---|
| UBXN2A | 52.3130682 | 0.32224502 | 0.26199993 | 1.22994317 | 0.21871839 | 0.90437211 |
| SF3B6 | 25.0178679 | -0.5413751 | 0.359877 | -1.5043338 | 0.13249543 | 0.80565822 |
| FAM228B | 36.7177454 | 0.1561237 | 0.41502174 | 0.37618198 | 0.70678161 | 0.9958918 |
| ITSN2 | 167.727523 | 0.19291253 | 0.26176944 | 0.73695587 | 0.46114919 | 0.98590351 |
| NCOA1 | 449.844235 | -0.287042 | 0.2541291 | -1.1295127 | 0.25868161 | 0.92939413 |
| ADCY3 | 230.050975 | -0.7423746 | 0.31397957 | -2.3644042 | 0.0180591 | 0.35723014 |
| DNAJC27 | 26.3687404 | -0.1504804 | 0.46869827 | -0.3210603 | 0.74816469 | 0.9958918 |
| DNMT3A | 58.8307149 | 0.50272662 | 0.28868439 | 1.74144025 | 0.08160644 | 0.69958018 |
| DTNB | 46.2801778 | 0.25857113 | 0.37097956 | 0.69699563 | 0.48580553 | 0.99046213 |
| ASXL2 | 75.6491342 | 0.10980064 | 0.30786284 | 0.35665442 | 0.72135053 | 0.9958918 |
| RAB10 | 178.929032 | 0.2689963 | 0.24362191 | 1.10415479 | 0.269526 | 0.93634106 |
| HADHA | 63.5911905 | 0.1934064 | 0.23349736 | 0.82830229 | 0.40749933 | 0.97733978 |
| HADHB | 70.9051032 | 0.16316641 | 0.28273432 | 0.57710153 | 0.56387087 | 0.9958918 |
| SELENOI | 35.4060882 | 0.11986555 | 0.34894398 | 0.34350944 | 0.73121524 | 0.9958918 |
| KCNK3 | 20.220417 | 0.82946994 | 0.67681996 | 1.22554001 | 0.22037183 | 0.90731076 |
| MAPRE3 | 22.2954258 | 0.12282614 | 0.32293295 | 0.38034564 | 0.70368886 | 0.9958918 |
| TMEM214 | 24.6479044 | -0.4487274 | 0.51029648 | -0.8793465 | 0.37921343 | 0.97217293 |
| OST4 | 36.2206847 | -0.1969563 | 0.56507143 | -0.3485512 | 0.7274263 | 0.9958918 |
| EMILIN1 | 113.39151 | -0.2045506 | 0.42149071 | -0.4853028 | 0.62746155 | 0.9958918 |
| ATRAID | 24.2520832 | -0.0489776 | 0.37575322 | -0.1303451 | 0.89629338 | 0.99935941 |
| GTF3C2 | 30.9844639 | 0.3208154 | 0.41132654 | 0.77995309 | 0.43541849 | 0.98092839 |
| PPM1G | 51.0672103 | 0.77361567 | 0.43607634 | 1.77403724 | 0.07605699 | 0.67423742 |
| NRBP1 | 29.8192532 | 0.18492264 | 0.30492538 | 0.60645211 | 0.54421457 | 0.9958918 |
| IFT172 | 28.9610554 | -0.1981845 | 0.36001887 | -0.5504836 | 0.58198769 | 0.9958918 |
| ZNF512 | 48.9564643 | -0.1864753 | 0.27502778 | -0.6780234 | 0.49775688 | 0.99176652 |
| MRPL33 | 51.1885993 | 0.12636942 | 0.30174537 | 0.4187949 | 0.67536603 | 0.9958918 |
| BABAM2 | 253.065125 | 0.0096712 | 0.25375974 | 0.03811163 | 0.96959868 | 0.99935941 |
| FOSL2 | 128.841184 | -0.8252298 | 0.37995978 | -2.1718873 | 0.02986417 | 0.46665355 |
| PPP1CB | 111.808459 | -0.7282076 | 0.48141544 | -1.5126386 | 0.13037148 | 0.80565822 |
| TRMT61B | 22.1669079 | -0.3307388 | 0.33080467 | -0.9998008 | 0.3174069 | 0.95992644 |
| WDR43 | 77.1041694 | -0.3487543 | 0.37241499 | -0.9364668 | 0.34903291 | 0.9692706 |
| CLIP4 | 101.959286 | -0.5642434 | 0.35205928 | -1.6026944 | 0.10900215 | 0.77303254 |
| YPEL5 | 38.3531569 | 0.32882629 | 0.33030922 | 0.99551046 | 0.31948806 | 0.95992644 |
| LBH | 81.6420524 | -0.3958036 | 0.40225359 | -0.9839654 | 0.32513253 | 0.95992644 |
| LCLAT1 | 105.225845 | -0.1396363 | 0.21145024 | -0.6603741 | 0.5090138 | 0.99461377 |
| EHD3 | 23.6863983 | -0.3178892 | 0.46146263 | -0.6888732 | 4.91E-01 | 0.99046213 |
| DPY30 | 23.1297271 | 0.35233844 | 0.35666865 | 0.98785928 | 0.32322157 | 0.95992644 |
| SPAST | 37.9564769 | -0.3433921 | 0.34309571 | -1.000864 | 0.31689257 | 0.95992644 |
| SLC30A6 | 43.0775109 | 0.03065758 | 0.33666372 | 0.09106292 | 0.9274426 | 0.99935941 |
| YIPF4 | 76.4779152 | 0.01367678 | 0.22408912 | 0.06103275 | 0.95133312 | 0.99935941 |
| BIRC6 | 450.220203 | -0.185039 | 0.14103428 | -1.3120144 | 0.18951528 | 0.88139754 |
| TTC27 | 66.4569563 | 0.21095495 | 0.22922636 | 0.92029098 | 0.35742072 | 0.97217293 |
| LINC00486 | 72.5289974 | -0.2157167 | 0.2272124 | -0.9494054 | 3.42E-01 | 0.96887708 |
| LTBP1 | 585.59405 | -0.875872 | 0.53431077 | -1.6392558 | 0.10115999 | 0.75167369 |
| FAM98A | 18.2486682 | 0.12810746 | 0.36041739 | 0.35544196 | 0.72225852 | 0.9958918 |
| LINC01320 | 32.6322505 | 0.43436882 | 0.34581908 | 1.25605799 | 0.20909495 | 0.89634177 |
| CRIM1 | 678.198702 | -0.4433929 | 0.46997901 | -0.9434312 | 0.34546039 | 0.96887708 |
| FEZ2 | 86.6076488 | -0.2537953 | 0.30101779 | -0.8431238 | 0.3991592 | 0.97500135 |
| STRN | 70.5034666 | -0.2192047 | 0.26504755 | -0.8270393 | 0.40821481 | 0.97782616 |
| HEATR5B | 69.6387386 | -0.136703 | 0.27100833 | -0.5044237 | 0.61396365 | 0.9958918 |
| EIF2AK2 | 92.9484311 | -0.6906746 | 0.53137078 | -1.2997978 | 0.19367028 | 0.88391893 |
| CEBPZOS | 35.3736082 | 0.20443184 | 0.28371638 | 0.72055 | 0.47118642 | 0.98910524 |
| CEBPZ | 42.2850025 | 0.12836845 | 0.29063426 | 0.44168381 | 0.65871803 | 0.9958918 |
| NDUFAF7 | 27.0687839 | 0.32809196 | 0.29494092 | 1.11239891 | 0.26596667 | 0.93533685 |

| | | | | | | |
|---|---|---|---|---|---|---|
| PRKD3 | 96.7092142 | 0.33207773 | 0.32655177 | 1.01692214 | 0.30919047 | 0.95793395 |
| CDC42EP3 | 56.6193879 | -0.7622304 | 0.48797696 | -1.5620214 | 0.11828295 | 0.78595066 |
| RMDN2 | 37.0974838 | -0.4578883 | 0.45917671 | -0.997194 | 0.31867034 | 0.95992644 |
| CYP1B1 | 120.193468 | -0.2249534 | 0.48274382 | -0.4659891 | 0.64122331 | 0.9958918 |
| CYP1B1-AS1 | 72.3100721 | 0.33336533 | 0.39950767 | 0.83444036 | 0.40403288 | 0.97575999 |
| ATL2 | 75.6797012 | -0.2323606 | 0.20348225 | -1.1419208 | 0.25348695 | 0.92646125 |
| HNRNPLL | 88.8242673 | -0.6630988 | 0.41463535 | -1.5992336 | 0.10976872 | 0.77303254 |
| GALM | 27.6235256 | 0.71365185 | 0.3605579 | 1.97929887 | 0.04778237 | 0.56981199 |
| SRSF7 | 66.2597529 | -0.0003707 | 0.46637663 | -0.0007948 | 0.99936583 | 0.99968761 |
| DHX57 | 34.6148419 | 0.22238761 | 0.33741731 | 0.65908774 | 0.50983943 | 0.99530567 |
| SOS1 | 212.651922 | 0.51887993 | 0.20714141 | 2.50495503 | 0.0122467 | 0.2958939 |
| MAP4K3 | 331.886255 | -0.4344388 | 0.28978318 | -1.4991856 | 0.1338255 | 0.80812129 |
| AC007388.1 | 99.5640304 | -0.4621622 | 0.34997051 | -1.3205747 | 0.18664322 | 0.87955894 |
| THUMPD2 | 45.9603969 | 0.09985376 | 0.24538319 | 0.40692992 | 0.68405946 | 0.9958918 |
| SLC8A1 | 414.553064 | 0.0813282 | 0.44803909 | 0.18152032 | 8.56E-01 | 0.9971731 |
| PKDCC | 67.3916755 | -0.0805705 | 0.34698579 | -0.232201 | 0.81638188 | 0.99592423 |
| EML4 | 275.348628 | -0.5883714 | 0.29741468 | -1.9782862 | 0.04789643 | 0.56981199 |
| COX7A2L | 44.3861392 | -0.1640457 | 0.31119525 | -0.5271473 | 0.59809128 | 0.9958918 |
| MTA3 | 106.851276 | 0.00416259 | 0.2219325 | 0.01875609 | 0.98503568 | 0.99935941 |
| HAAO | 17.7419156 | 0.18345439 | 0.36758178 | 0.49908455 | 0.61771982 | 0.9958918 |
| ZFP36L2 | 129.29479 | -0.6764537 | 0.29891073 | -2.2630625 | 0.02363183 | 0.41547931 |
| THADA | 169.855422 | -0.1946344 | 0.25201356 | -0.7723172 | 0.43992657 | 0.98092839 |
| PLEKHH2 | 121.641977 | -0.201388 | 0.40277602 | -0.5 | 0.61707511 | 0.9958918 |
| DYNC2LI1 | 64.7756702 | -0.1021072 | 0.36779856 | -0.2776171 | 0.7813063 | 0.9958918 |
| LRPPRC | 114.176834 | -0.2751271 | 0.25541096 | -1.0771939 | 0.28139363 | 0.94419503 |
| PPM1B | 105.362766 | 0.28456475 | 0.24224745 | 1.17468624 | 0.24012026 | 0.92426606 |
| SLC3A1 | 29.4579683 | 1.62762365 | 0.60424724 | 2.69363854 | 0.00706768 | 0.21552014 |
| PREPL | 82.4194102 | -0.3060401 | 0.31341316 | -0.9764751 | 0.3288291 | 0.9620084 |
| CAMKMT | 216.327379 | -0.0533398 | 0.27084115 | -0.1969413 | 0.8438735 | 0.99692952 |
| SRBD1 | 105.653219 | -0.2275722 | 0.23276405 | -0.977695 | 0.3282252 | 0.9620084 |
| PRKCE | 294.29896 | -0.0886899 | 0.17114372 | -0.518219 | 0.60430547 | 0.9958918 |
| EPAS1 | 635.057486 | -0.0239832 | 0.43626383 | -0.0549741 | 0.95615906 | 0.99935941 |
| RHOQ | 188.54925 | -0.5738936 | 0.32841556 | -1.7474617 | 0.08055727 | 0.69567028 |
| AC018682.1 | 34.5431817 | -0.7429199 | 0.51361619 | -1.4464496 | 1.48E-01 | 0.8301633 |
| PIGF | 32.8946467 | -0.0125083 | 0.33539222 | -0.0372946 | 0.97025013 | 0.99935941 |
| SOCS5 | 141.934374 | -0.4583382 | 0.40545167 | -1.1304386 | 0.25829144 | 0.92891801 |
| MCFD2 | 43.8893564 | -0.4936063 | 0.4201133 | -1.1749362 | 0.24002023 | 0.92426606 |
| TTC7A | 46.6484175 | 0.00877684 | 0.38600324 | 0.02273774 | 0.98185947 | 0.99935941 |
| EPCAM | 25.5064504 | 0.22975662 | 0.44310531 | 0.51851472 | 0.6040992 | 0.9958918 |
| FBXO11 | 250.392794 | 0.289959 | 0.15649238 | 1.85286338 | 0.06390194 | 0.62395844 |
| FOXN2 | 66.4404376 | 0.47360608 | 0.29290225 | 1.61694243 | 0.10589071 | 0.76820416 |
| PPP1R21 | 99.8697532 | 0.02300264 | 0.36890839 | 0.06235325 | 0.95028152 | 0.99935941 |
| STON1-GTF2 | 31.3665601 | 0.28597538 | 0.56447712 | 0.50661996 | 0.6124215 | 0.9958918 |
| AC009975.1 | 46.3693717 | -1.0156434 | 1.25534689 | -0.809054 | 0.4184841 | 0.97812179 |
| GPR75-ASB3 | 29.976582 | 0.19257554 | 0.3859324 | 0.49898775 | 0.61778801 | 0.9958918 |
| ERLEC1 | 33.675846 | 0.21049829 | 0.30175648 | 0.69757671 | 0.48544195 | 0.99046213 |
| PSME4 | 159.513291 | -0.2485547 | 0.29633196 | -0.8387711 | 0.4015978 | 0.97500135 |
| ACYP2 | 226.657448 | -0.000907 | 0.23020435 | -0.0039399 | 0.99685639 | 0.99935941 |
| SPTBN1 | 544.946578 | 0.20683166 | 0.30077605 | 0.68765999 | 0.49166692 | 0.99046213 |
| RTN4 | 342.266722 | -0.0620972 | 0.3248878 | -0.1911343 | 0.84842035 | 0.99713289 |
| CLHC1 | 27.5669432 | -1.0458435 | 0.47145217 | -2.2183447 | 0.02653134 | 0.44313361 |
| RPS27A | 163.029993 | -0.3377483 | 0.43403616 | -0.778157 | 0.43647645 | 0.98092839 |
| CCDC88A | 90.2562044 | 0.03368033 | 0.21963914 | 0.15334395 | 0.87812705 | 0.99722158 |
| CFAP36 | 32.6201759 | 0.03489058 | 0.29566471 | 0.11800727 | 0.9060619 | 0.99935941 |

| | | | | | | |
|---|---|---|---|---|---|---|
| PPP4R3B | 109.82569 | -0.2050447 | 0.1993896 | -1.0283621 | 0.30377954 | 0.9573224 |
| PNPT1 | 59.7795441 | -0.9466614 | 0.56888994 | -1.6640502 | 0.09610239 | 0.73903832 |
| EFEMP1 | 120.829967 | 0.27493938 | 0.56552593 | 0.48616581 | 0.62684961 | 0.9958918 |
| VRK2 | 62.9969975 | -0.162408 | 0.2807496 | -0.5784798 | 0.56294021 | 0.9958918 |
| FANCL | 37.9094023 | -0.968906 | 0.38700184 | -2.5036212 | 0.01229296 | 0.2958939 |
| PAPOLG | 33.9899521 | 0.07213892 | 0.28250784 | 0.25535194 | 0.79845129 | 0.99592423 |
| REL | 53.8590476 | 0.4318848 | 0.52423859 | 0.82383252 | 0.41003472 | 0.97812179 |
| PEX13 | 19.7904336 | -0.3183639 | 0.33749208 | -0.9433226 | 0.3455159 | 0.96887708 |
| KIAA1841 | 30.1831044 | 0.02996051 | 0.45113516 | 0.06641138 | 0.94705031 | 0.99935941 |
| USP34 | 459.668615 | -0.2601597 | 0.19460657 | -1.3368498 | 0.18127168 | 0.87815793 |
| XPO1 | 179.265246 | -0.467428 | 0.3943385 | -1.1853471 | 0.23588022 | 0.91962002 |
| CCT4 | 40.9872968 | 0.55657069 | 0.2742942 | 2.02910118 | 0.04244799 | 0.54728114 |
| COMMD1 | 161.933407 | 0.11197568 | 0.21052661 | 0.53188371 | 0.59480654 | 0.9958918 |
| B3GNT2 | 23.271296 | -0.8092008 | 0.35712271 | -2.2658901 | 0.02345811 | 0.41385091 |
| EHBP1 | 355.36564 | -0.2766906 | 0.25111502 | -1.1018482 | 0.27052767 | 0.93726628 |
| WDPCP | 168.138059 | -0.1831095 | 0.22641583 | -0.808731 | 0.4186699 | 0.97812179 |
| MDH1 | 36.9196727 | 0.75386446 | 0.31478595 | 2.39484787 | 0.01662726 | 0.34578631 |
| UGP2 | 114.211677 | -0.05508 | 0.40294475 | -0.1366937 | 0.89127291 | 0.99935941 |
| VPS54 | 142.39841 | 0.02703883 | 0.15621051 | 0.17309228 | 0.8625789 | 0.9971731 |
| PELI1 | 85.1463632 | 0.28020888 | 0.37945661 | 0.73844774 | 0.46024242 | 0.98590351 |
| AFTPH | 76.4379728 | 0.04751376 | 0.23894365 | 0.19884924 | 0.84238068 | 0.99692952 |
| SERTAD2 | 53.7880529 | 0.37781169 | 0.27700132 | 1.36393459 | 0.17258815 | 0.87233975 |
| CEP68 | 26.7981227 | 0.80973491 | 0.36607938 | 2.21191073 | 0.02697283 | 0.446449 |
| RAB1A | 276.107646 | -0.4486019 | 0.30060095 | -1.4923502 | 0.13560735 | 0.81154597 |
| ACTR2 | 103.046567 | 0.27290481 | 0.23379838 | 1.16726563 | 0.24310311 | 0.92534155 |
| SPRED2 | 194.255477 | 0.33401048 | 0.25198076 | 1.32553963 | 0.18499224 | 0.87866798 |
| MEIS1 | 745.299484 | -0.0198037 | 0.27323932 | -0.0724776 | 0.9422218 | 0.99935941 |
| LINC01798 | 64.5115407 | -0.4623948 | 0.43550271 | -1.0617495 | 0.28834941 | 0.95026001 |
| LINC01828 | 29.0470714 | 0.53005685 | 0.40788881 | 1.2995131 | 0.1937679 | 0.88391893 |
| C1D | 31.8653524 | -0.1886254 | 0.34482395 | -0.5470194 | 0.58436541 | 0.9958918 |
| CNRIP1 | 22.0366896 | -0.2528836 | 0.31529795 | -0.8020464 | 0.4225261 | 0.97828057 |
| APLF | 37.2236332 | 0.00429801 | 0.38742395 | 0.0110938 | 0.99114861 | 0.99935941 |
| ARHGAP25 | 31.615214 | 0.79615998 | 0.89606249 | 0.88850944 | 0.37426678 | 0.97217293 |
| ANTXR1 | 231.076496 | 0.09371044 | 0.25116949 | 0.37309643 | 0.70907668 | 0.9958918 |
| GFPT1 | 131.205433 | -0.0197133 | 0.19021216 | -0.1036383 | 0.91745641 | 0.99935941 |
| NFU1 | 34.2009299 | 0.04113862 | 0.39119498 | 0.10516142 | 0.91624772 | 0.99935941 |
| AAK1 | 216.47765 | 0.26157976 | 0.20315231 | 1.28760417 | 0.19788379 | 0.88565649 |
| ANXA4 | 73.159653 | 0.32502022 | 0.32252028 | 1.00775126 | 0.31357389 | 0.95992644 |
| GMCL1 | 49.425463 | -0.0336961 | 0.24323216 | -0.1385346 | 0.88981795 | 0.99935941 |
| MXD1 | 23.2699427 | -0.3831405 | 0.3667628 | -1.0446549 | 0.2961825 | 0.95393362 |
| PCBP1 | 55.558212 | 0.16227795 | 0.33241758 | 0.48817498 | 0.6254259 | 0.9958918 |
| C2orf42 | 18.4400393 | 0.59014595 | 0.37902303 | 1.55701872 | 0.11946604 | 0.79021701 |
| PCYOX1 | 45.4104673 | 0.91521628 | 0.41367241 | 2.21241799 | 0.0269378 | 0.446449 |
| SNRPG | 22.0970431 | -0.267778 | 0.43486957 | -0.6157662 | 0.53804881 | 0.9958918 |
| MCEE | 33.9774749 | 0.05094789 | 0.33299774 | 0.15299771 | 0.87840008 | 0.99722158 |
| MPHOSPH10 | 22.0544046 | -0.0564466 | 0.40995177 | -0.1376909 | 0.8904847 | 0.99935941 |
| ZNF638 | 440.892587 | -0.1463355 | 0.20582951 | -0.7109548 | 0.47711222 | 0.98993168 |
| EXOC6B | 361.720947 | -0.1892951 | 0.29226295 | -0.6476875 | 0.51718711 | 0.9958918 |
| SFXN5 | 49.1467708 | 0.48835493 | 0.28867632 | 1.69170414 | 0.0907024 | 0.72611321 |
| CCT7 | 38.8453927 | 0.07239052 | 0.33323204 | 0.21723759 | 0.82802319 | 0.99592423 |
| ALMS1 | 84.9544108 | 0.1411064 | 0.20252101 | 0.69674944 | 0.48595961 | 0.99046213 |
| DUSP11 | 23.8167786 | -0.2752293 | 0.37076017 | -0.7423378 | 0.45788269 | 0.98420003 |
| STAMBP | 32.3369857 | -0.1207854 | 0.31399308 | -0.3846753 | 0.70047801 | 0.9958918 |
| DGUOK | 42.0885881 | -0.2790824 | 0.36187529 | -0.7712115 | 0.44058159 | 0.98092839 |

| | | | | | | |
|---|---|---|---|---|---|---|
| TET3 | 34.5935616 | -0.1143181 | 0.34083581 | -0.3354052 | 0.73731943 | 0.9958918 |
| MOB1A | 45.933736 | -0.1386531 | 0.27697044 | -0.5006062 | 0.61664827 | 0.9958918 |
| MTHFD2 | 45.2829929 | 0.34068156 | 0.37857664 | 0.89990117 | 0.36817285 | 0.97217293 |
| AC006030.1 | 21.5129654 | 0.1931945 | 0.40165206 | 0.48099965 | 0.63051675 | 0.9958918 |
| LOXL3 | 21.5263931 | -0.1887742 | 0.56120316 | -0.3363741 | 0.73658881 | 0.9958918 |
| MRPL19 | 44.5321533 | 0.42027376 | 0.38188399 | 1.10052729 | 0.27110245 | 0.93776436 |
| GCFC2 | 59.2811614 | 0.30969916 | 0.28353652 | 1.09227253 | 0.27471333 | 0.93944545 |
| SUCLG1 | 41.0412019 | 0.52496475 | 0.24899905 | 2.10830022 | 0.03500503 | 0.50468149 |
| TMSB10 | 211.105956 | -0.3030564 | 0.40692216 | -0.7447528 | 0.4564212 | 0.98330915 |
| KCMF1 | 105.124114 | -0.0942584 | 0.21329892 | -0.4419074 | 0.65855621 | 0.9958918 |
| TCF7L1 | 232.170481 | 0.1010445 | 0.21773384 | 0.46407349 | 0.64259509 | 0.9958918 |
| TGOLN2 | 64.1027436 | 0.2924412 | 0.28136388 | 1.03937009 | 0.29863265 | 0.95447177 |
| ELMOD3 | 34.4406524 | 0.70375075 | 0.32032224 | 2.19700872 | 0.02801982 | 0.45356318 |
| MAT2A | 78.8849775 | -0.5928657 | 0.29263304 | -2.0259698 | 0.04276788 | 0.54851863 |
| GGCX | 22.1817772 | -0.1742396 | 0.32031456 | -0.5439641 | 0.58646615 | 0.9958918 |
| VAMP5 | 24.6455013 | -0.1502563 | 0.30027182 | -0.500401 | 0.61679276 | 0.9958918 |
| USP39 | 33.9939483 | -0.3631148 | 0.29185937 | -1.2441432 | 0.21344686 | 0.89738044 |
| ATOH8 | 29.3027126 | -0.5504084 | 0.45442911 | -1.2112084 | 0.22581555 | 0.91093521 |
| POLR1A | 28.9216605 | 0.11937543 | 0.34572697 | 0.34528817 | 0.72987774 | 0.9958918 |
| PTCD3 | 51.3071445 | 0.22316238 | 0.2394954 | 0.93180235 | 0.35143868 | 0.97021895 |
| IMMT | 66.7616396 | 0.3323872 | 0.23559137 | 1.41086318 | 0.15828496 | 0.84598994 |
| KDM3A | 70.5135515 | -0.08541 | 0.47005442 | -0.1817024 | 0.85581628 | 0.9971731 |
| CHMP3 | 19.4833722 | 0.71181162 | 0.38019627 | 1.87222147 | 0.06117597 | 0.61696136 |
| RMND5A | 33.4489385 | 0.00924244 | 0.37716096 | 0.02450528 | 0.98044957 | 0.99935941 |
| CYTOR | 98.2731695 | -1.3418825 | 0.48413165 | -2.7717306 | 0.00557592 | 0.18713567 |
| KRCC1 | 46.5986929 | 0.40107337 | 0.31079547 | 1.29047369 | 0.19688624 | 0.88391893 |
| EIF2AK3 | 97.2879383 | -0.5139262 | 0.32976263 | -1.558473 | 0.11912116 | 0.79009152 |
| MRPS5 | 43.5220734 | 0.53009151 | 0.31723793 | 1.6709588 | 0.09472982 | 0.73435592 |
| ANKRD36C | 67.1841462 | -0.0240178 | 0.35356476 | -0.0679304 | 0.94584103 | 0.99935941 |
| STARD7 | 31.4567489 | 0.21173701 | 0.30926324 | 0.68464978 | 0.49356494 | 0.99116088 |
| TMEM127 | 22.7291992 | 0.1665708 | 0.3690675 | 0.45132882 | 0.65175258 | 0.9958918 |
| CIAO1 | 28.5935725 | 0.59404723 | 0.3949054 | 1.50427728 | 0.13250999 | 0.80565822 |
| SNRNP200 | 45.5145575 | 0.20719896 | 0.39221869 | 0.52827407 | 0.59730913 | 0.9958918 |
| ARID5A | 27.8251575 | 0.64127416 | 0.45730258 | 1.40229726 | 0.1608265 | 0.85092986 |
| KANSL3 | 65.1551046 | -0.1387856 | 0.24920374 | -0.5569164 | 0.57758457 | 0.9958918 |
| LMAN2L | 19.7309153 | 0.09273277 | 0.43261752 | 0.21435277 | 0.83027197 | 0.99592423 |
| ANKRD23 | 18.4838349 | 0.23997808 | 0.37474647 | 0.64037448 | 0.52192917 | 0.9958918 |
| SEMA4C | 21.9613699 | 0.15393822 | 0.39098288 | 0.39372113 | 0.69378694 | 0.9958918 |
| ANKRD36 | 78.1287155 | 0.0155676 | 0.39221295 | 0.03969171 | 0.96833891 | 0.99935941 |
| AC092683.1 | 41.7676151 | 0.25657371 | 0.36568582 | 0.70162336 | 0.48291408 | 0.99002719 |
| ANKRD36B | 20.937241 | -0.1993004 | 0.44612569 | -0.446736 | 0.65506571 | 0.9958918 |
| COX5B | 47.6063001 | 0.01865963 | 0.45687974 | 0.04084145 | 0.9674223 | 0.99935941 |
| ACTR1B | 21.2911265 | -0.3807756 | 0.33837404 | -1.1253098 | 0.26045778 | 0.93055907 |
| TMEM131 | 186.372708 | -0.0352244 | 0.22271439 | -0.1581596 | 0.87433103 | 0.9971731 |
| INPP4A | 89.2712542 | 0.84460902 | 0.27161792 | 3.10954818 | 0.00187374 | 0.09345751 |
| UNC50 | 23.6161245 | 0.12864964 | 0.30216706 | 0.42575664 | 0.67028518 | 0.9958918 |
| MGAT4A | 21.5383866 | 0.22964938 | 0.37303795 | 0.61561934 | 0.53814578 | 0.9958918 |
| AC079447.1 | 48.0617018 | -0.0792503 | 0.48830223 | -0.1622977 | 0.87107141 | 0.9971731 |
| MITD1 | 23.5724493 | 0.12116347 | 0.38359452 | 0.31586341 | 0.75210619 | 0.9958918 |
| EIF5B | 87.2015099 | -0.0732495 | 0.37343691 | -0.1961496 | 0.84449307 | 0.99692952 |
| REV1 | 106.686145 | -0.2256726 | 0.23542902 | -0.9585591 | 0.33778089 | 0.96750939 |
| AFF3 | 395.421203 | -0.7944985 | 0.50402309 | -1.5763136 | 0.11495355 | 0.77932991 |
| NPAS2 | 284.069244 | -0.9975096 | 0.46971809 | -2.1236346 | 0.03370071 | 0.49530613 |
| RPL31 | 50.471434 | -0.2983191 | 0.29764485 | -1.0022653 | 0.31621546 | 0.95992644 |

| | | | | | | |
|---|---|---|---|---|---|---|
| TBC1D8 | 64.9695973 | -0.0507023 | 0.33007126 | -0.1536103 | 0.87791704 | 0.99722158 |
| CNOT11 | 19.0928866 | -0.1244837 | 0.35479397 | -0.3508619 | 0.72569196 | 0.9958918 |
| RNF149 | 114.108421 | -0.3133142 | 0.2589048 | -1.210152 | 0.22622059 | 0.91093521 |
| MAP4K4 | 261.444639 | -0.2172484 | 0.33917238 | -0.640525 | 0.52183132 | 0.9958918 |
| IL1R1 | 226.047078 | -1.0000946 | 0.34871716 | -2.8679247 | 0.00413174 | 0.15814658 |
| IL18R1 | 115.600793 | 1.31938739 | 0.78063654 | 1.69014302 | 0.0910006 | 0.72693717 |
| MRPS9 | 31.0533017 | 0.34893985 | 0.3302477 | 1.0566004 | 0.290694 | 0.95186143 |
| FHL2 | 141.815786 | 0.31809388 | 0.49789761 | 0.63887407 | 0.52290486 | 0.9958918 |
| NCK2 | 104.962322 | 0.05227096 | 0.27790447 | 0.18808968 | 0.85080635 | 0.9971731 |
| UXS1 | 39.2003654 | -0.1598113 | 0.34000921 | -0.4700206 | 0.63834027 | 0.9958918 |
| GCC2 | 141.487998 | 0.40097407 | 0.23775354 | 1.68651143 | 0.09169734 | 0.72711043 |
| LIMS1 | 247.092241 | -0.349756 | 0.18183215 | -1.9235102 | 0.05441601 | 0.59304667 |
| RANBP2 | 116.150736 | -0.247723 | 0.24019748 | -1.0313305 | 0.30238588 | 0.95650251 |
| SH3RF3 | 412.333294 | -0.0045351 | 0.34449852 | -0.0131643 | 9.89E-01 | 0.99935941 |
| 10-Sep | 80.5645382 | -0.0831524 | 0.27824087 | -0.2988503 | 0.76505427 | 0.9958918 |
| MIR4435-2H | 383.084085 | -0.7891419 | 0.38622395 | -2.0432236 | 0.0410303 | 0.54347495 |
| BCL2L11 | 22.515085 | -0.2278832 | 0.54165188 | -0.420719 | 0.67396029 | 0.9958918 |
| AC017002.5 | 35.630185 | -0.8920188 | 0.89151123 | -1.0005693 | 0.31703507 | 0.95992644 |
| ANAPC1 | 65.4693884 | -0.2511144 | 0.20512428 | -1.2242059 | 0.22087455 | 0.90885053 |
| MERTK | 41.3086283 | 0.09612576 | 0.42819212 | 0.22449213 | 0.82237439 | 0.99592423 |
| TMEM87B | 81.0399888 | -0.4467831 | 0.26573165 | -1.6813321 | 0.09269843 | 0.73016158 |
| ZC3H8 | 29.0921181 | -0.2916202 | 0.28208131 | -1.0338161 | 0.30122215 | 0.95597956 |
| ZC3H6 | 63.3390666 | -0.3065604 | 0.3958591 | -0.7744181 | 0.4386836 | 0.98092839 |
| TTL | 28.1337314 | 0.29264725 | 0.38314207 | 0.76380871 | 0.44498124 | 0.98092839 |
| SLC20A1 | 20.5462213 | -0.2313561 | 0.41505914 | -0.5574052 | 0.57725062 | 0.9958918 |
| PAX8 | 52.2528454 | -0.2845522 | 0.35553575 | -0.8003475 | 0.42350947 | 0.97828057 |
| CBWD2 | 19.5241645 | -0.464336 | 0.36519603 | -1.2714705 | 0.20356131 | 0.89396747 |
| SLC35F5 | 109.56279 | -0.0555704 | 0.3366618 | -0.1650631 | 0.86889429 | 0.9971731 |
| ACTR3 | 180.228323 | -0.2404236 | 0.26772857 | -0.8980126 | 0.36917883 | 0.97217293 |
| DDX18 | 41.0207986 | -0.107288 | 0.32073689 | -0.3345049 | 0.73799863 | 0.9958918 |
| CCDC93 | 53.8385956 | -0.2874669 | 0.26275478 | -1.0940503 | 0.27393294 | 0.9394201 |
| INSIG2 | 37.7646735 | -0.3338114 | 0.38894868 | -0.8582402 | 0.39075982 | 0.97297628 |
| DBI | 53.6737317 | 0.31878435 | 0.305246 | 1.04435227 | 0.29632244 | 0.95393362 |
| PTPN4 | 73.7355755 | 0.01376025 | 0.22389595 | 0.06145822 | 0.95099429 | 0.99935941 |
| EPB41L5 | 68.9256691 | 0.43690902 | 0.36531764 | 1.19597022 | 0.23170818 | 0.91549646 |
| RALB | 20.5228647 | 1.12756868 | 0.3536413 | 3.18845303 | 0.00143036 | 0.08339044 |
| GLI2 | 128.981197 | -0.491071 | 0.30619789 | -1.6037699 | 0.10876479 | 0.77302943 |
| CLASP1 | 308.568461 | -0.2278909 | 0.29833445 | -0.7638772 | 0.44494044 | 0.98092839 |
| GYPC | 51.7370376 | 0.40616754 | 0.41822854 | 0.9711617 | 0.33146777 | 0.9654289 |
| BIN1 | 23.2089841 | -0.025123 | 0.32752703 | -0.0767051 | 0.93885811 | 0.99935941 |
| ERCC3 | 30.2752437 | -0.041418 | 0.3121861 | -0.1326709 | 0.89445366 | 0.99935941 |
| MAP3K2 | 144.541619 | 0.21849599 | 0.18331211 | 1.19193431 | 0.233287 | 0.91861399 |
| IWS1 | 75.4490004 | -0.038437 | 0.21465058 | -0.179068 | 0.85788432 | 0.9971731 |
| LIMS2 | 30.8274955 | 0.71606074 | 0.51670495 | 1.38582132 | 0.16580147 | 0.85938344 |
| WDR33 | 109.942108 | 0.24755225 | 0.22180346 | 1.11608831 | 0.26438434 | 0.93499423 |
| AMMECR1L | 20.1979563 | 0.04333935 | 0.33872142 | 0.12794984 | 0.89818867 | 0.99935941 |
| SAP130 | 36.3866018 | 0.13223781 | 0.28527114 | 0.46355132 | 0.64296924 | 0.9958918 |
| UGGT1 | 57.762467 | 0.07125312 | 0.30032605 | 0.23725256 | 0.81246086 | 0.99592423 |
| HS6ST1 | 28.0300322 | 0.20268407 | 0.45638977 | 0.44410301 | 0.65696811 | 0.9958918 |
| SMPD4 | 30.6180179 | -0.1554974 | 0.27995135 | -0.5554445 | 0.57859067 | 0.9958918 |
| MZT2B | 37.9385065 | 0.22482364 | 0.39132961 | 0.57451221 | 0.56562124 | 0.9958918 |
| FAM168B | 60.6893219 | -0.2038273 | 0.23271561 | -0.8758643 | 0.38110383 | 0.97217293 |
| PLEKHB2 | 42.4475396 | 0.48864472 | 0.28671125 | 1.70430957 | 0.08832327 | 0.71818729 |
| NCKAP5 | 135.82669 | 1.99444762 | 0.87868973 | 2.26979736 | 0.02321988 | 0.41205662 |

| | | | | | | |
|---|---|---|---|---|---|---|
| MGAT5 | 158.500036 | 0.06598741 | 0.28384761 | 0.23247477 | 0.81616928 | 0.99592423 |
| CCNT2 | 83.09207 | 0.15496393 | 0.26683911 | 0.58073917 | 0.56141626 | 0.9958918 |
| RAB3GAP1 | 160.285361 | -0.2692211 | 0.24042297 | -1.1197812 | 0.262807 | 0.93257782 |
| ZRANB3 | 72.8112048 | 0.14821985 | 0.24945123 | 0.5941837 | 0.55238925 | 0.9958918 |
| R3HDM1 | 140.758014 | 0.2759951 | 0.26738857 | 1.03218732 | 0.30198437 | 0.95650251 |
| UBXN4 | 110.036318 | -0.1206842 | 0.20921801 | -0.5768345 | 0.56405127 | 0.9958918 |
| DARS | 99.911166 | 0.04639554 | 0.30894116 | 0.15017597 | 0.88062578 | 0.99749293 |
| HNMT | 44.5523517 | -0.3103105 | 0.53946343 | -0.5752207 | 0.56514208 | 0.9958918 |
| AC114763.1 | 27.1604612 | -0.2698209 | 0.33291939 | -0.8104693 | 0.41767052 | 0.97812179 |
| SPOPL | 56.0549881 | 0.34972587 | 0.25143105 | 1.39094146 | 0.16424318 | 0.85697309 |
| LRP1B | 39.7662427 | -0.8145915 | 0.69247856 | -1.1763418 | 0.23945831 | 0.92426606 |
| ARHGAP15 | 145.485999 | 0.43447839 | 0.46439591 | 0.93557757 | 0.34949073 | 0.9692706 |
| GTDC1 | 229.811062 | 0.02609197 | 0.27738612 | 0.09406372 | 0.92505854 | 0.99935941 |
| ZEB2 | 700.675032 | -0.1956722 | 0.24283369 | -0.8057867 | 0.42036586 | 0.97812179 |
| TEX41 | 100.895678 | 0.93138354 | 0.5148865 | 1.8089104 | 0.07046492 | 0.65633299 |
| ACVR2A | 88.5088188 | 0.06110083 | 0.30458844 | 0.20060128 | 0.84101036 | 0.99692952 |
| ORC4 | 65.2515376 | -0.2957986 | 0.33937734 | -0.871592 | 0.38343102 | 0.9725044 |
| MBD5 | 502.308227 | -0.3024767 | 0.25488039 | -1.1867398 | 0.23533028 | 0.91962002 |
| EPC2 | 130.121408 | 0.04302827 | 0.18763962 | 0.22931333 | 0.81862539 | 0.99592423 |
| MMADHC | 23.5855842 | -0.0520462 | 0.35048162 | -0.1484992 | 0.88194883 | 0.99818549 |
| AC007364.1 | 38.8542692 | 0.24261025 | 0.45364591 | 0.53480092 | 0.59278753 | 0.9958918 |
| RND3 | 99.8693433 | -0.9985807 | 0.49003127 | -2.0377898 | 0.04157096 | 0.54347495 |
| RIF1 | 128.211458 | -0.1348954 | 0.21310663 | -0.6329947 | 0.52673711 | 0.9958918 |
| ARL5A | 35.1884394 | 0.64507026 | 0.37097456 | 1.73885309 | 0.08206061 | 0.69958018 |
| STAM2 | 70.5877979 | -0.0548877 | 0.25491413 | -0.2153183 | 0.82951915 | 0.99592423 |
| FMNL2 | 291.983122 | -0.2876084 | 0.25568171 | -1.124869 | 0.26064454 | 0.93055907 |
| PRPF40A | 84.0978227 | -0.1012627 | 0.21446411 | -0.4721663 | 0.63680812 | 0.9958918 |
| ARL6IP6 | 27.0391206 | 0.15882075 | 0.45225683 | 0.3511738 | 0.72545797 | 0.9958918 |
| NR4A2 | 57.0623848 | 0.09860534 | 0.5148823 | 0.19151045 | 0.84812569 | 0.99713289 |
| GPD2 | 91.7405266 | -0.1371387 | 0.28750906 | -0.4769891 | 0.63336992 | 0.9958918 |
| ACVR1 | 65.452067 | -0.1882108 | 0.3350357 | -0.5617634 | 0.57427726 | 0.9958918 |
| PKP4 | 127.088366 | -0.1745061 | 0.17167447 | -1.016494 | 0.30939423 | 0.95793395 |
| TANC1 | 405.815603 | -0.5209553 | 0.38912535 | -1.3387854 | 0.18064054 | 0.87815793 |
| WDSUB1 | 39.5684882 | -0.024111 | 0.4159169 | -0.0579707 | 0.953772 | 0.99935941 |
| BAZ2B | 373.961996 | -0.2776151 | 0.31547084 | -0.8800026 | 0.37885792 | 0.97217293 |
| 7-Mar | 179.526206 | 0.00342166 | 0.22542964 | 0.01517841 | 0.98788984 | 0.99935941 |
| CD302 | 22.3416862 | 0.37676506 | 0.33244494 | 1.13331567 | 0.25708171 | 0.9266065 |
| PLA2R1 | 159.484179 | -0.0848864 | 0.45991015 | -0.1845717 | 0.85356502 | 0.9971731 |
| ITGB6 | 31.4803223 | -1.0614011 | 0.62561778 | -1.6965649 | 8.98E-02 | 0.72305504 |
| RBMS1 | 819.048664 | 0.13632432 | 0.2946974 | 0.46259086 | 0.64365766 | 0.9958918 |
| TANK | 208.561798 | -0.1274851 | 0.27560478 | -0.4625649 | 0.64367626 | 0.9958918 |
| PSMD14 | 63.9875219 | -0.2361556 | 0.34413483 | -0.68623 | 0.49256811 | 0.99046213 |
| IFIH1 | 33.9126122 | -1.4155329 | 1.13141029 | -1.2511225 | 0.21088979 | 0.89738044 |
| FIGN | 283.213402 | 0.40762305 | 0.39817009 | 1.023741 | 0.30595762 | 0.95793395 |
| COBLL1 | 177.704626 | -0.7444562 | 0.34714792 | -2.1444928 | 0.03199342 | 0.48478277 |
| AC019197.1 | 57.1763676 | -0.1344681 | 0.41192275 | -0.3264401 | 0.74409137 | 0.9958918 |
| SLC38A11 | 88.7326077 | -2.3091571 | 0.80666911 | -2.8625828 | 0.00420203 | 0.16000389 |
| CSRNP3 | 47.6370931 | -0.0264403 | 0.4255059 | -0.0621386 | 0.95045245 | 0.99935941 |
| TTC21B | 61.9429521 | -0.1115707 | 0.21044054 | -0.530177 | 0.59598922 | 0.9958918 |
| STK39 | 176.688666 | -0.5365496 | 0.26414633 | -2.031259 | 0.04222872 | 0.54637128 |
| CERS6 | 296.584292 | -1.3109662 | 0.41206902 | -3.1814239 | 0.00146553 | 0.08388666 |
| LRP2 | 45.5569191 | 1.95977185 | 0.63137786 | 3.10396039 | 0.00190949 | 0.09345751 |
| PPIG | 110.261259 | 0.02136115 | 0.21441414 | 0.09962564 | 0.92064154 | 0.99935941 |
| KLHL23 | 54.2572752 | -0.6073849 | 0.39454845 | -1.5394432 | 0.12369614 | 0.79184923 |

| | | | | | | |
|---|---|---|---|---|---|---|
| SSB | 46.0124024 | -0.2849011 | 0.25577208 | -1.1138866 | 0.26532782 | 0.93520104 |
| UBR3 | 329.679362 | -0.2847548 | 0.23147229 | -1.2301896 | 0.2186261 | 0.90437211 |
| GORASP2 | 42.0340759 | 0.47094293 | 0.27867307 | 1.68994776 | 0.09103795 | 0.72693717 |
| TLK1 | 109.579606 | -0.0652861 | 0.21422545 | -0.3047542 | 0.76055336 | 0.9958918 |
| METTL8 | 41.1776934 | 0.25006373 | 0.31248352 | 0.80024615 | 0.4235682 | 0.97828057 |
| DCAF17 | 26.0774914 | 0.18087537 | 0.33372504 | 0.54198922 | 0.58782594 | 0.9958918 |
| CYBRD1 | 95.851131 | 0.04774938 | 0.3636259 | 0.13131459 | 0.89552645 | 0.99935941 |
| DYNC1I2 | 93.1021474 | -0.0277224 | 0.21317263 | -0.1300465 | 0.8965296 | 0.99935941 |
| SLC25A12 | 74.1015646 | 0.34136194 | 0.20771879 | 1.64338501 | 0.1003033 | 0.75035372 |
| HAT1 | 45.3685067 | -0.0402756 | 0.26585705 | -0.1514936 | 0.87958635 | 0.99749293 |
| METAP1D | 20.7889151 | -0.1672348 | 0.3809233 | -0.4390248 | 0.6606436 | 0.9958918 |
| ITGA6 | 24.8414339 | 0.03126826 | 0.55505483 | 0.05633364 | 0.95507602 | 0.99935941 |
| PDK1 | 80.8494645 | -0.8270923 | 0.80798428 | -1.023649 | 0.30600109 | 0.95793395 |
| RAPGEF4 | 80.227213 | 0.1915269 | 0.43212774 | 0.44321826 | 0.65760787 | 0.9958918 |
| MAP3K20 | 172.678697 | 0.09638975 | 0.19148889 | 0.50336993 | 0.61470421 | 0.9958918 |
| SP3 | 111.715369 | 0.20729887 | 0.21671372 | 0.95655627 | 0.33879127 | 0.96887708 |
| OLA1 | 106.006285 | -0.056604 | 0.26551213 | -0.2131878 | 0.83118046 | 0.99592423 |
| CIR1 | 47.3820628 | 0.03036575 | 0.32476576 | 0.09350048 | 0.92550597 | 0.99935941 |
| GPR155 | 27.7933524 | 0.01549687 | 0.36983813 | 0.04190176 | 0.96657701 | 0.99935941 |
| AC010894.3 | 55.3239014 | 0.68969159 | 0.35229589 | 1.95770549 | 0.05026458 | 0.57538069 |
| WIPF1 | 79.2974348 | 0.17360409 | 0.19493155 | 0.89059003 | 0.37314915 | 0.97217293 |
| CHN1 | 38.1964811 | -0.1689361 | 0.33896622 | -0.4983863 | 0.61821182 | 0.9958918 |
| ATF2 | 108.000488 | -0.2176643 | 0.25809268 | -0.8433573 | 0.39902866 | 0.97500135 |
| ATP5MC3 | 55.1842968 | 0.11400649 | 0.40784948 | 0.27953079 | 0.77983751 | 0.9958918 |
| LNPK | 49.8303892 | 0.07053264 | 0.28318427 | 0.24906975 | 0.80330682 | 0.99592423 |
| HOXD10 | 73.0168046 | 0.027178 | 0.23860471 | 0.11390386 | 0.909314 | 0.99935941 |
| HOXD9 | 16.6774526 | -0.4208309 | 0.45880141 | -0.9172398 | 0.35901701 | 0.97217293 |
| HOXD3 | 30.8099391 | 0.10473615 | 0.31077818 | 0.33701257 | 0.73610743 | 0.9958918 |
| MTX2 | 79.6313413 | -0.1123461 | 0.23889842 | -0.4702674 | 0.63816398 | 0.9958918 |
| HNRNPA3 | 151.964103 | -0.0190682 | 0.19817575 | -0.0962186 | 0.92334696 | 0.99935941 |
| NFE2L2 | 212.575888 | 0.18093815 | 0.19983336 | 0.90544517 | 0.36522959 | 0.97217293 |
| AGPS | 80.9441478 | -0.7077699 | 0.25764984 | -2.7470225 | 0.0060139 | 0.19299633 |
| PRKRA | 44.8378561 | -0.1276877 | 0.24026472 | -0.5314461 | 0.59510972 | 0.9958918 |
| PLEKHA3 | 36.0680153 | -0.0702862 | 0.25636208 | -0.2741676 | 0.78395582 | 0.9958918 |
| TTN-AS1 | 122.804886 | 0.23405547 | 0.27226732 | 0.85965319 | 0.38998024 | 0.97297628 |
| TTN | 54.0086895 | -0.2344309 | 1.07283444 | -0.2185154 | 0.82702755 | 0.99592423 |
| SESTD1 | 124.499765 | 0.19856491 | 0.31558706 | 0.62919219 | 0.52922324 | 0.9958918 |
| CWC22 | 40.4278192 | 0.03287408 | 0.26654252 | 0.12333524 | 0.90184164 | 0.99935941 |
| UBE2E3 | 96.5694173 | 0.16605572 | 0.21965547 | 0.75598264 | 0.44965961 | 0.98092839 |
| SSFA2 | 93.4376056 | -0.5422325 | 0.33087542 | -1.6387815 | 0.10125876 | 0.75167369 |
| PDE1A | 481.590514 | 0.49981328 | 0.24825696 | 2.01329008 | 0.04408412 | 0.55583613 |
| DNAJC10 | 66.6337878 | -0.1630941 | 0.32280906 | -0.505234 | 0.61339451 | 0.9958918 |
| FRZB | 48.527021 | 0.67490958 | 0.27757307 | 2.43146632 | 1.50E-02 | 3.34E-01 |
| NCKAP1 | 246.166563 | -0.2193348 | 0.2876155 | -0.7625974 | 0.44570353 | 0.98092839 |
| NUP35 | 19.8833011 | 0.09928688 | 0.343321 | 0.28919549 | 0.77243179 | 0.9958918 |
| ZC3H15 | 54.9136727 | 0.01728775 | 0.28288615 | 0.06111205 | 0.95126997 | 0.99935941 |
| ITGAV | 165.822647 | -0.1087519 | 0.33217282 | -0.3273954 | 0.74336884 | 0.9958918 |
| AC007319.1 | 123.683361 | -0.5631015 | 0.37385708 | -1.5061945 | 0.13201724 | 0.80565822 |
| CALCRL | 66.9816643 | -1.9085985 | 0.51433085 | -3.7108381 | 0.00020657 | 0.02333707 |
| TFPI | 324.532528 | -0.1292427 | 0.25890349 | -0.4991926 | 0.61764371 | 0.9958918 |
| GULP1 | 281.737067 | -0.1039402 | 0.28472658 | -0.3650526 | 0.71507212 | 0.9958918 |
| COL3A1 | 407.890953 | -0.3649313 | 0.50525766 | -0.7222676 | 0.47012995 | 0.98910524 |
| COL5A2 | 218.191906 | -0.7361994 | 0.35046249 | -2.1006512 | 0.0356716 | 0.50762504 |
| WDR75 | 47.009406 | -0.1679929 | 0.25389179 | -0.6616713 | 0.50818188 | 0.99397937 |

| | | | | | | |
|---|---|---|---|---|---|---|
| SLC40A1 | 81.0671507 | -0.3805151 | 0.44257083 | -0.8597836 | 0.38990834 | 0.97297628 |
| ANKAR | 82.4351048 | 0.20320404 | 0.35860141 | 0.5666571 | 0.57094717 | 0.9958918 |
| ORMDL1 | 38.3903357 | 0.08353763 | 0.31249929 | 0.26732104 | 0.78922199 | 0.99592423 |
| PMS1 | 45.5416027 | 0.14377766 | 0.24050336 | 0.59781976 | 0.54996021 | 0.9958918 |
| HIBCH | 98.0558397 | 0.06571744 | 0.23629339 | 0.27811797 | 0.7809218 | 0.9958918 |
| INPP1 | 27.6383135 | -0.4267126 | 0.3501062 | -1.2188091 | 0.22291667 | 0.91093521 |
| MFSD6 | 26.2594906 | -0.2591115 | 0.42338345 | -0.6120019 | 0.54053648 | 0.9958918 |
| NAB1 | 52.1176353 | 0.05601522 | 0.24406456 | 0.22950986 | 0.81847266 | 0.99592423 |
| GLS | 364.797816 | -0.2070537 | 0.34389546 | -0.6020833 | 0.54711871 | 0.9958918 |
| STAT1 | 93.9601904 | -0.6190019 | 0.78754568 | -0.7859885 | 0.4318742 | 0.98092839 |
| STAT4 | 54.221357 | 0.2120715 | 0.41472519 | 0.51135427 | 0.60910301 | 0.9958918 |
| MYO1B | 357.690648 | -0.3316583 | 0.35978971 | -0.9218115 | 0.3566269 | 0.97217293 |
| NABP1 | 98.9160198 | 0.04492012 | 0.83975089 | 0.05349219 | 0.95733975 | 0.99935941 |
| SLC39A10 | 32.1342492 | 0.69293918 | 0.35461845 | 1.95404152 | 0.05069631 | 0.57655256 |
| STK17B | 100.912387 | 0.68980811 | 0.41721369 | 1.65336883 | 0.09825582 | 0.74364783 |
| HECW2 | 49.5959611 | 1.20909882 | 0.45848764 | 2.63714596 | 0.00836068 | 0.24190028 |
| GTF3C3 | 28.9297767 | -0.2655138 | 0.28102262 | -0.9448129 | 0.34475438 | 0.96887708 |
| PGAP1 | 79.9332709 | -0.9140708 | 0.46907461 | -1.9486683 | 0.05133505 | 0.57655256 |
| ANKRD44 | 189.537988 | 0.18924543 | 0.24978181 | 0.75764296 | 0.44866476 | 0.98092839 |
| SF3B1 | 122.331702 | -0.1181844 | 0.20698195 | -0.5709891 | 0.56800703 | 0.9958918 |
| COQ10B | 24.715806 | -0.6163245 | 0.31264612 | -1.9713167 | 0.04868766 | 0.57248898 |
| HSPD1 | 76.9492298 | -0.0686174 | 0.35388054 | -0.1938999 | 0.84625427 | 0.99692952 |
| RFTN2 | 48.2796601 | -0.3025493 | 0.43125511 | -0.7015553 | 0.48295652 | 0.99002719 |
| PLCL1 | 725.936511 | 0.74366175 | 0.42196823 | 1.76236434 | 0.07800776 | 0.68056484 |
| SPATS2L | 145.610979 | -0.0271418 | 0.48042374 | -0.0564956 | 0.95494699 | 0.99935941 |
| AOX1 | 52.0672541 | -0.1030453 | 0.47534394 | -0.2167805 | 0.82837938 | 0.99592423 |
| BZW1 | 24.3161848 | 0.0342759 | 0.40069945 | 0.08554016 | 0.93183197 | 0.99935941 |
| CLK1 | 97.5103832 | -0.3635478 | 0.26546019 | -1.3695004 | 0.1708429 | 0.87217428 |
| ORC2 | 62.7077968 | -0.1296439 | 0.26904488 | -0.4818671 | 0.62990035 | 0.9958918 |
| FAM126B | 77.3563561 | -0.1880044 | 0.20700873 | -0.9081956 | 0.36377489 | 0.97217293 |
| NDUFB3 | 23.5054362 | -0.2583033 | 0.5092469 | -0.5072261 | 0.61199616 | 0.9958918 |
| CFLAR | 235.296956 | -0.0419948 | 0.30647492 | -0.1370252 | 0.89101088 | 0.99935941 |
| TRAK2 | 68.827658 | 0.31899404 | 0.33985478 | 0.93861867 | 0.34792656 | 0.96887708 |
| STRADB | 30.1521615 | 0.35319645 | 0.55462967 | 0.63681493 | 0.5242454 | 0.9958918 |
| ALS2 | 56.9401964 | -0.2735373 | 0.30844528 | -0.886826 | 0.3751726 | 0.97217293 |
| SUMO1 | 36.6804353 | -0.1925732 | 0.2741186 | -0.7025177 | 0.48235636 | 0.99002719 |
| NOP58 | 80.3477944 | -0.2327456 | 0.24784052 | -0.939094 | 0.34768247 | 0.96887708 |
| BMPR2 | 477.512319 | 0.48268045 | 0.31029937 | 1.55553153 | 0.11981953 | 0.79021701 |
| FAM117B | 37.723965 | 0.10394297 | 0.3813589 | 0.27255943 | 0.7851919 | 0.9958918 |
| ICA1L | 27.448439 | -0.1448259 | 0.50701188 | -0.285646 | 0.77514927 | 0.9958918 |
| WDR12 | 25.9055797 | 0.01145061 | 0.3630109 | 0.03154344 | 0.97483615 | 0.99935941 |
| CARF | 29.0017538 | 0.03518243 | 0.44298959 | 0.07942045 | 0.93669821 | 0.99935941 |
| NBEAL1 | 752.720386 | 0.19287245 | 0.2544545 | 0.75798403 | 0.44846055 | 0.98092839 |
| CYP20A1 | 93.4855299 | -0.0841714 | 0.2737247 | -0.3075038 | 0.7584599 | 0.9958918 |
| ABI2 | 141.933579 | 0.12027215 | 0.27023174 | 0.44507041 | 0.65626887 | 0.9958918 |
| RAPH1 | 258.445582 | -0.0808234 | 0.26762308 | -0.3020047 | 0.76264845 | 0.9958918 |
| PARD3B | 460.340479 | 0.07978883 | 0.31914771 | 0.25000596 | 0.80258274 | 0.99592423 |
| NRP2 | 58.705038 | -0.662179 | 0.84949702 | -0.7794954 | 0.43568794 | 0.98092839 |
| INO80D | 181.991863 | -0.4346884 | 0.58435666 | -0.7438751 | 0.45695204 | 0.98335849 |
| NDUFS1 | 89.2134446 | 0.18137026 | 0.24040823 | 0.75442618 | 0.45059336 | 0.98092839 |
| EEF1B2 | 42.7556771 | 0.22456068 | 0.40853968 | 0.54966674 | 0.58254797 | 0.9958918 |
| ZDBF2 | 46.3613945 | -0.2552858 | 0.5089387 | -0.5016042 | 0.61594599 | 0.9958918 |
| FASTKD2 | 25.08684 | -0.337506 | 0.29918471 | -1.1280856 | 0.25928378 | 0.92976724 |
| KLF7 | 133.279108 | -0.0995607 | 0.33940733 | -0.2933369 | 0.76926465 | 0.9958918 |

| | | | | | | |
|---|---|---|---|---|---|---|
| CREB1 | 86.9615394 | 0.27922053 | 0.21016367 | 1.32858609 | 0.18398456 | 0.87866798 |
| CCNYL1 | 18.9890096 | 0.06665506 | 0.47009388 | 0.14179096 | 0.88724513 | 0.99935941 |
| PLEKHM3 | 57.0504244 | 0.06971066 | 0.24656536 | 0.28272689 | 0.7773862 | 0.9958918 |
| PIKFYVE | 109.028264 | -0.0155074 | 0.24835464 | -0.0624407 | 0.95021185 | 0.99935941 |
| PTH2R | 46.8256124 | 0.15484342 | 0.39210263 | 0.39490533 | 0.69291275 | 0.9958918 |
| MAP2 | 118.875053 | -0.921413 | 0.40671765 | -2.2654856 | 0.0234829 | 0.41385091 |
| RPE | 28.1261878 | -0.6161863 | 0.29225695 | -2.1083719 | 0.03499884 | 0.50468149 |
| KANSL1L | 148.6562 | 0.07112921 | 0.27743375 | 0.25638268 | 0.79765536 | 0.99592423 |
| AC007038.1 | 22.6913979 | -0.4765888 | 0.37242624 | -1.2796865 | 0.20065541 | 0.89191279 |
| LANCL1 | 36.928318 | 0.1649939 | 0.25387893 | 0.64989205 | 0.51576195 | 0.9958918 |
| ERBB4 | 159.426579 | 0.24086591 | 0.38470824 | 0.62610021 | 0.53124922 | 0.9958918 |
| IKZF2 | 33.1527519 | 0.15646771 | 0.3802093 | 0.41153048 | 0.6806836 | 0.9958918 |
| SPAG16 | 83.187237 | -0.1126184 | 0.25865879 | -0.4353938 | 0.66327661 | 0.9958918 |
| BARD1 | 24.2745888 | 0.01892809 | 0.510158 | 0.03710241 | 0.97040335 | 0.99935941 |
| ATIC | 21.6955951 | -0.007794 | 0.46577946 | -0.0167333 | 0.98664937 | 0.99935941 |
| FN1 | 366.301619 | 1.3686527 | 0.61151553 | 2.23813236 | 0.02521242 | 0.42757124 |
| XRCC5 | 165.555102 | 0.05959389 | 0.18387311 | 0.32410335 | 0.74585979 | 0.9958918 |
| SMARCAL1 | 20.5385207 | 0.22432801 | 0.33630184 | 0.66704366 | 0.50474424 | 0.99387859 |
| RPL37A | 113.14986 | -0.3132084 | 0.33772173 | -0.9274155 | 0.35371087 | 0.97217293 |
| IGFBP2 | 109.185534 | -1.0992311 | 0.81059536 | -1.3560787 | 0.17507412 | 0.8740623 |
| IGFBP5 | 522.195913 | -0.5169809 | 0.52273166 | -0.9889986 | 0.32266383 | 0.95992644 |
| AC007563.2 | 27.6781918 | -1.1797844 | 0.48944136 | -2.4104714 | 0.01593192 | 0.34135182 |
| DIRC3 | 115.062643 | 1.27924954 | 0.36457418 | 3.50888679 | 0.00044999 | 0.04186013 |
| TNS1 | 537.885748 | 0.39930174 | 0.28028948 | 1.4246048 | 0.15427147 | 0.84218718 |
| ARPC2 | 106.698403 | 0.21813357 | 0.24883035 | 0.87663571 | 0.38068454 | 0.97217293 |
| AAMP | 17.9762566 | -0.1471396 | 0.41335137 | -0.3559674 | 0.72186499 | 0.9958918 |
| PNKD | 39.8962042 | 0.6174842 | 0.28415973 | 2.17301799 | 0.02977896 | 0.46665355 |
| TMBIM1 | 41.3170213 | 0.03314836 | 0.28491017 | 0.1163467 | 0.90737778 | 0.99935941 |
| CTDSP1 | 37.5612481 | 0.30450912 | 0.33062915 | 0.92099901 | 0.35705094 | 0.97217293 |
| USP37 | 92.0235201 | -0.1799071 | 0.3779828 | -0.4759663 | 0.6340984 | 0.9958918 |
| CNOT9 | 36.4688838 | -0.3853926 | 0.3390394 | -1.1367193 | 0.25565566 | 0.9266065 |
| CYP27A1 | 20.7976 | 0.40537028 | 0.3918972 | 1.03437911 | 0.30095897 | 0.95597956 |
| DNAJB2 | 19.734913 | -0.2363677 | 0.36671563 | -0.6445531 | 0.51921682 | 0.9958918 |
| DNPEP | 20.452259 | 0.00741387 | 0.45921974 | 0.01614448 | 0.98711912 | 0.99935941 |
| DES | 48.0079167 | 3.85183771 | 1.20159271 | 3.20561009 | 0.00134776 | 0.08118626 |
| SPEG | 26.0820488 | 1.67189628 | 0.54486845 | 3.06844026 | 0.00215179 | 0.09945616 |
| OBSL1 | 19.8539484 | -0.0296984 | 0.34029769 | -0.0872719 | 0.93045539 | 0.99935941 |
| FARSB | 43.2882101 | 0.024509 | 0.36554883 | 0.06704714 | 0.94654417 | 0.99935941 |
| ACSL3 | 131.228247 | 0.03473605 | 0.20927144 | 0.16598561 | 0.86816828 | 0.9971731 |
| KCNE4 | 106.937198 | -0.5935549 | 0.41579193 | -1.4275288 | 0.15342754 | 0.84093985 |
| WDFY1 | 79.7442972 | 0.14074191 | 0.29297873 | 0.48038269 | 0.6309553 | 0.9958918 |
| SERPINE2 | 72.9602554 | -0.7145832 | 0.79582735 | -0.8979124 | 0.36923227 | 0.97217293 |
| CUL3 | 180.146935 | 0.26045715 | 0.20417305 | 1.27566858 | 0.2020727 | 0.89319989 |
| DOCK10 | 125.32322 | 0.08028362 | 0.25089716 | 0.31998618 | 7.49E-01 | 0.9958918 |
| IRS1 | 76.0430253 | -0.1281674 | 0.41671456 | -0.3075664 | 0.75841229 | 0.9958918 |
| RHBDD1 | 77.0410176 | -0.050566 | 0.21734523 | -0.2326531 | 0.81603076 | 0.99592423 |
| COL4A4 | 356.37759 | -0.8636657 | 0.31273659 | -2.7616396 | 0.00575119 | 0.19038519 |
| COL4A3 | 216.752636 | -0.8031593 | 0.42497525 | -1.8898967 | 0.05877177 | 0.6110796 |
| AC097662.1 | 35.0221287 | -0.3758318 | 0.68292957 | -0.550323 | 0.58209787 | 0.9958918 |
| MFF | 48.1726038 | 0.17929679 | 0.2735277 | 0.65549777 | 0.51214732 | 0.9958918 |
| AGFG1 | 113.777713 | -0.0291142 | 0.30208423 | -0.0963776 | 0.92322067 | 0.99935941 |
| PID1 | 386.984853 | -0.1686225 | 0.59025598 | -0.2856769 | 0.77512556 | 0.9958918 |
| TRIP12 | 236.018896 | -0.2907242 | 0.23337546 | -1.2457357 | 0.21286143 | 0.89738044 |
| SP110 | 37.9170958 | -0.4844055 | 0.62961381 | -0.7693692 | 0.44167414 | 0.98092839 |

| | | | | | | |
|---|---|---|---|---|---|---|
| SP140L | 29.0589661 | -0.0289823 | 0.40239682 | -0.0720243 | 0.94258258 | 0.99935941 |
| SP100 | 187.536218 | 0.0283665 | 0.41329818 | 0.06863446 | 0.94528059 | 0.99935941 |
| CAB39 | 105.640176 | 0.26506428 | 0.23942786 | 1.10707369 | 0.26826207 | 0.93598159 |
| ITM2C | 35.2605438 | -0.3050946 | 0.46729788 | -0.6528911 | 0.51382651 | 0.9958918 |
| PSMD1 | 115.531142 | -0.0788401 | 0.24694607 | -0.3192605 | 0.74952899 | 0.9958918 |
| ARMC9 | 118.775367 | -0.1979029 | 0.33582098 | -0.5893107 | 0.55565284 | 0.9958918 |
| NCL | 71.0029433 | 0.29482858 | 0.47015054 | 0.62709398 | 0.53059763 | 0.9958918 |
| PTMA | 402.6873 | -0.0413914 | 0.44774992 | -0.0924431 | 0.92634597 | 0.99935941 |
| PDE6D | 21.9702323 | 0.43693438 | 0.4085961 | 1.06935524 | 0.28490963 | 0.94505166 |
| COPS7B | 19.5857242 | -0.0110687 | 0.32616422 | -0.0339359 | 0.97292823 | 0.99935941 |
| DIS3L2 | 105.094766 | 0.0937521 | 0.27079451 | 0.34621123 | 0.72918397 | 0.9958918 |
| EIF4E2 | 26.4481715 | 0.08377728 | 0.31232096 | 0.26824099 | 0.78851383 | 0.99592423 |
| EFHD1 | 58.3593072 | 0.62755285 | 0.37938972 | 1.65411137 | 0.09810489 | 0.74364783 |
| GIGYF2 | 194.082028 | -0.1884131 | 0.16954963 | -1.1112562 | 0.2664581 | 0.93533685 |
| DGKD | 97.4125272 | -0.6292627 | 0.37418336 | -1.6816961 | 0.09262779 | 0.73016158 |
| USP40 | 82.541617 | -0.1394255 | 0.35402018 | -0.3938348 | 0.693703 | 0.9958918 |
| SH3BP4 | 22.7353111 | -0.4123594 | 0.44958892 | -0.9171922 | 0.35904195 | 0.97217293 |
| AGAP1 | 532.213146 | -0.0667276 | 0.26920466 | -0.2478694 | 0.80423543 | 0.99592423 |
| COPS8 | 19.5838215 | -0.2130444 | 0.34568227 | -0.6163013 | 0.53769566 | 0.9958918 |
| COL6A3 | 288.861425 | -0.3401658 | 0.52755191 | -0.6448006 | 0.51905638 | 0.9958918 |
| LRRFIP1 | 264.681453 | -0.2844657 | 0.3019124 | -0.9422126 | 0.34608383 | 0.96887708 |
| RAMP1 | 94.6776831 | 0.14379989 | 0.51317918 | 0.28021381 | 0.77931347 | 0.9958918 |
| ILKAP | 61.1823002 | 0.30496689 | 0.22374062 | 1.36303763 | 0.17287065 | 0.87233975 |
| PER2 | 28.9735618 | -0.6864665 | 0.43916346 | -1.563123 | 0.11802367 | 0.78539225 |
| TRAF3IP1 | 31.6979049 | 0.19128546 | 0.32330744 | 0.5916519 | 0.55408371 | 0.9958918 |
| TWIST2 | 52.5556971 | -0.8867788 | 0.4478638 | -1.980019 | 0.04770139 | 0.56981199 |
| HDAC4 | 143.241604 | 0.16999727 | 0.34740365 | 0.48933645 | 0.62460352 | 0.9958918 |
| NDUFA10 | 69.1014797 | -0.0666843 | 0.24180799 | -0.2757739 | 0.78272173 | 0.9958918 |
| COPS9 | 21.7090816 | -0.0652174 | 0.3832127 | -0.1701859 | 0.86486392 | 0.9971731 |
| GPC1 | 36.1172286 | 0.41900603 | 0.46198429 | 0.9069703 | 0.36442251 | 0.97217293 |
| ANKMY1 | 24.3123391 | 0.55318675 | 0.32500092 | 1.70210827 | 0.08873507 | 0.71818729 |
| SNED1 | 217.430956 | 1.04647689 | 0.33765247 | 3.09927211 | 0.00193997 | 0.09345751 |
| AC005237.1 | 27.0567925 | -0.004083 | 0.45337681 | -0.0090057 | 0.99281458 | 0.99935941 |
| MTERF4 | 36.7213754 | 0.74391561 | 0.2680164 | 2.77563468 | 0.00550941 | 0.18713567 |
| PPP1R7 | 22.8092299 | 0.4080947 | 0.3477058 | 1.17367813 | 0.24052397 | 0.92426606 |
| HDLBP | 149.136439 | -0.210827 | 0.21641839 | -0.9741639 | 0.32997515 | 0.96348071 |
| 2-Sep | 200.326788 | -0.1017919 | 0.20803914 | -0.489292 | 0.62463495 | 0.9958918 |
| FARP2 | 91.6017609 | 0.21464534 | 0.18863479 | 1.1378884 | 0.25516708 | 0.9266065 |
| STK25 | 25.2455432 | -0.1358717 | 0.33134429 | -0.4100619 | 0.68176057 | 0.9958918 |
| THAP4 | 23.9708391 | -0.0889971 | 0.34712939 | -0.2563802 | 0.79765726 | 0.99592423 |
| ATG4B | 47.2386885 | -0.4726945 | 0.25981378 | -1.8193589 | 0.06885669 | 0.64792295 |
| ING5 | 37.6204791 | -0.1419853 | 0.28184439 | -0.5037721 | 0.61442156 | 0.9958918 |
| D2HGDH | 47.1292886 | -0.1182009 | 0.3746668 | -0.3154829 | 0.75239507 | 0.9958918 |
| LINC01266 | 75.0498984 | -0.2825432 | 0.46119626 | -0.6126311 | 0.54012029 | 0.9958918 |
| CNTN4 | 307.44155 | -1.1227673 | 0.40941663 | -2.7423589 | 0.00609996 | 0.19357111 |
| TRNT1 | 32.678445 | 0.4451698 | 0.33735336 | 1.31959498 | 0.18697028 | 0.87955894 |
| CRBN | 45.1574896 | 0.13086615 | 0.34794116 | 0.37611574 | 0.70683085 | 0.9958918 |
| AC034195.1 | 34.7709645 | 0.3227649 | 0.2869193 | 1.12493269 | 0.26061756 | 0.93055907 |
| SUMF1 | 183.549337 | 0.04113899 | 0.27149343 | 0.15152848 | 0.87955884 | 0.99749293 |
| ITPR1 | 313.327295 | -0.0209898 | 0.32495689 | -0.0645927 | 0.94849831 | 0.99935941 |
| AC018816.1 | 25.8357124 | 0.50746248 | 0.44213032 | 1.14776676 | 0.25106486 | 0.92624282 |
| BHLHE40 | 99.5856254 | -0.5163371 | 0.53681966 | -0.9618447 | 0.33612762 | 0.96640422 |
| ARL8B | 101.882221 | -0.270216 | 0.22925973 | -1.1786455 | 0.23853936 | 0.92291633 |
| EDEM1 | 28.8852572 | 0.21293968 | 0.43772093 | 0.4864736 | 0.62663142 | 0.9958918 |

| | | | | | | |
|---|---|---|---|---|---|---|
| LMCD1-AS1 | 123.988601 | -0.5764377 | 0.40737263 | -1.4150132 | 0.15706462 | 0.84427321 |
| LMCD1 | 244.597057 | -0.563222 | 0.54961913 | -1.0247497 | 0.30548132 | 0.95793395 |
| RAD18 | 24.0377588 | -0.4999758 | 0.32612535 | -1.5330787 | 0.12525647 | 0.79233388 |
| THUMPD3-A! | 67.7286399 | -0.2507023 | 0.25506485 | -0.9828961 | 0.32565857 | 0.95992644 |
| SETD5 | 306.451143 | -0.1691307 | 0.20040253 | -0.843955 | 0.39869456 | 0.97500135 |
| MTMR14 | 29.3001114 | 0.49697876 | 0.37871822 | 1.31226523 | 0.18943066 | 0.88139754 |
| OGG1 | 19.6121498 | 0.14701781 | 0.45196805 | 0.32528364 | 0.74496641 | 0.9958918 |
| TADA3 | 18.7814344 | -0.2507311 | 0.38182919 | -0.6566578 | 0.51140097 | 0.9958918 |
| EMC3 | 34.2861064 | 0.04775239 | 0.29947152 | 0.15945555 | 0.87330998 | 0.9971731 |
| BRK1 | 35.7342726 | 0.49032116 | 0.40229122 | 1.21882142 | 0.22291198 | 0.91093521 |
| VHL | 18.0992148 | 0.38932808 | 0.33855786 | 1.14996022 | 0.25016025 | 0.92624282 |
| IRAK2 | 25.2018737 | 1.711719 | 0.54637496 | 3.132865 | 0.00173109 | 0.09218684 |
| SEC13 | 44.4412599 | 0.25610779 | 0.33327898 | 0.76844867 | 0.44222068 | 0.98092839 |
| ATG7 | 197.90911 | 0.22171626 | 0.28749151 | 0.77120976 | 0.44058261 | 0.98092839 |
| VGLL4 | 182.053498 | 0.44507063 | 0.18499642 | 2.40583375 | 0.01613561 | 0.34471095 |
| TAMM41 | 41.219748 | 0.00972903 | 0.29981695 | 0.03244991 | 0.97411326 | 0.99935941 |
| PPARG | 229.40748 | 0.4879168 | 0.23902029 | 2.04131963 | 0.04121907 | 0.54347495 |
| TSEN2 | 39.2152292 | -0.1709218 | 0.37581608 | -0.4548016 | 0.64925196 | 0.9958918 |
| RAF1 | 130.128882 | 0.29599493 | 0.17991768 | 1.64516873 | 0.09993502 | 0.7498958 |
| RPL32 | 158.009742 | -0.2794139 | 0.36954523 | -0.756102 | 0.44958806 | 0.98092839 |
| IQSEC1 | 40.4878658 | 0.09836347 | 0.35493714 | 0.27712928 | 0.78168084 | 0.9958918 |
| XPC | 51.1121155 | 0.0878915 | 0.25223169 | 0.34845542 | 0.72749818 | 0.9958918 |
| SLC6A6 | 25.4717339 | -0.9889442 | 0.41345903 | -2.3918796 | 0.01676234 | 0.34578631 |
| GRIP2 | 22.6314501 | -0.5644167 | 0.51731626 | -1.0910477 | 0.27525188 | 0.94084933 |
| CCDC174 | 23.3756011 | -0.2693871 | 0.38030969 | -0.7083362 | 0.4787365 | 0.99002719 |
| FGD5 | 43.681471 | 1.67389481 | 0.50237691 | 3.33195013 | 0.0008624 | 0.0644401 |
| FGD5-AS1 | 43.5976904 | -0.0141466 | 0.37138596 | -0.0380913 | 0.96961489 | 0.99935941 |
| NR2C2 | 83.8772651 | -0.1099243 | 0.23319073 | -0.4713921 | 0.63736078 | 0.9958918 |
| MRPS25 | 40.3965261 | -0.1686324 | 0.35944802 | -0.4691426 | 0.6389677 | 0.9958918 |
| RBSN | 32.4027888 | -0.0054125 | 0.26335676 | -0.0205519 | 0.98360311 | 0.99935941 |
| CAPN7 | 120.57078 | 0.23625349 | 0.18707517 | 1.26287995 | 0.20663232 | 0.89589437 |
| SH3BP5 | 205.882824 | 0.29101176 | 0.50813079 | 0.57271034 | 0.56684084 | 0.9958918 |
| METTL6 | 21.4611982 | 0.37094668 | 0.43945131 | 0.84411327 | 0.39860611 | 0.97500135 |
| COLQ | 23.0372593 | 0.75919035 | 0.58833931 | 1.29039542 | 0.1969134 | 0.88391893 |
| HACL1 | 30.3551948 | -0.1723547 | 0.30984757 | -0.5562563 | 0.57803563 | 0.9958918 |
| ANKRD28 | 105.110509 | -0.6148511 | 0.4147571 | -1.4824365 | 0.13822418 | 0.81520215 |
| OXNAD1 | 40.9394403 | 0.01541047 | 0.28921189 | 0.05328437 | 0.95750533 | 0.99935941 |
| RFTN1 | 277.413664 | 0.33797594 | 0.33909332 | 0.9967048 | 0.31890782 | 0.95992644 |
| PLCL2 | 143.906885 | -0.2065713 | 0.30025823 | -0.6879788 | 0.49146611 | 0.99046213 |
| TBC1D5 | 371.804628 | -0.2292725 | 0.2791254 | -0.8213962 | 0.41142061 | 0.97812179 |
| RAB5A | 71.8586019 | -0.1843556 | 0.23260938 | -0.7925546 | 0.4280374 | 0.98092839 |
| KAT2B | 41.6562098 | 0.67372173 | 0.40288603 | 1.67223897 | 0.09447721 | 0.7340733 |
| ZNF385D | 41.6043777 | 0.85161941 | 0.52171033 | 1.63236064 | 0.10260354 | 0.75678848 |
| UBE2E2 | 583.657888 | -0.2877257 | 0.23733987 | -1.212294 | 0.22539985 | 0.91093521 |
| UBE2E1 | 189.240978 | -0.2983718 | 0.19005366 | -1.5699343 | 0.11643039 | 0.78417352 |
| RPL15 | 190.78595 | 0.23500851 | 0.38596231 | 0.60888978 | 0.5425975 | 0.9958918 |
| NR1D2 | 52.1098554 | 0.44910773 | 0.48884133 | 0.91871881 | 0.35824267 | 0.97217293 |
| THRB | 228.285467 | 0.31565225 | 0.39145799 | 0.80635026 | 0.42004092 | 0.97812179 |
| RARB | 253.217727 | -1.2438342 | 0.40023827 | -3.1077344 | 0.00188527 | 0.09345751 |
| TOP2B | 112.845103 | -0.0790331 | 0.18770764 | -0.4210435 | 0.67372332 | 0.9958918 |
| NGLY1 | 63.0082121 | -0.0980287 | 0.40357412 | -0.2429015 | 0.80808174 | 0.99592423 |
| SLC4A7 | 122.413459 | 0.11946671 | 0.44748819 | 0.26697177 | 0.7894909 | 0.99592423 |
| CMC1 | 76.4457561 | -0.533516 | 0.36640766 | -1.4560722 | 0.14537265 | 0.82817333 |
| AZI2 | 108.74391 | -0.4286789 | 0.5109982 | -0.8389049 | 0.40152269 | 0.97500135 |

| | | | | | | |
|---|---|---|---|---|---|---|
| RBMS3 | 825.268541 | -0.1017299 | 0.27983968 | -0.3635292 | 0.71620958 | 0.9958918 |
| RBMS3-AS3 | 310.651479 | -0.8013997 | 0.49249319 | -1.6272301 | 0.10368824 | 0.75827058 |
| RBMS3-AS2 | 89.89444 | -0.4998231 | 0.52338596 | -0.95498 | 0.33958782 | 0.96887708 |
| TGFBR2 | 288.570395 | -0.3270577 | 0.3193758 | -1.0240529 | 0.30581027 | 0.95793395 |
| STT3B | 162.846124 | -0.1316957 | 0.27367462 | -0.4812127 | 0.63036532 | 0.9958918 |
| OSBPL10 | 219.278038 | -0.3060484 | 0.43832228 | -0.6982269 | 0.4850353 | 0.99046213 |
| CMTM8 | 48.238028 | 0.36147779 | 0.59715247 | 0.60533584 | 0.54495587 | 0.9958918 |
| CMTM7 | 30.6493812 | 0.68705938 | 0.35093254 | 1.95781037 | 0.05025227 | 0.57538069 |
| CMTM6 | 39.9859304 | 0.13838385 | 0.29039587 | 0.47653519 | 0.63369314 | 0.9958918 |
| DYNC1LI1 | 37.8680542 | -0.0032843 | 0.28822573 | -0.0113949 | 0.99090841 | 0.99935941 |
| CNOT10 | 57.6427918 | -0.2525776 | 0.26529373 | -0.9520675 | 0.34106273 | 0.96887708 |
| GLB1 | 43.6223403 | 0.12767488 | 0.33106069 | 0.38565401 | 0.69975293 | 0.9958918 |
| CRTAP | 42.9092639 | 0.1768984 | 0.32645556 | 0.54187591 | 0.58790399 | 0.9958918 |
| FBXL2 | 37.4639699 | -0.476479 | 0.48597645 | -0.9804571 | 0.32686055 | 0.95992644 |
| UBP1 | 51.7032288 | 0.73866236 | 0.28309444 | 2.60924358 | 0.00907426 | 0.25292422 |
| CLASP2 | 230.415796 | 0.24165067 | 0.32966152 | 0.73302662 | 0.46354219 | 0.98590351 |
| PDCD6IP | 105.609228 | 0.0668652 | 0.24034539 | 0.27820465 | 0.78085526 | 0.9958918 |
| STAC | 54.5655034 | -0.0439378 | 0.48656595 | -0.0903018 | 9.28E-01 | 0.99935941 |
| TRANK1 | 70.4973724 | -0.1966821 | 0.86528374 | -0.2273036 | 0.82018768 | 0.99592423 |
| EPM2AIP1 | 37.0036524 | -0.0360499 | 0.24397618 | -0.1477597 | 0.8825324 | 0.99840747 |
| MLH1 | 29.6664825 | -0.1665307 | 0.2883377 | -0.5775545 | 0.56356494 | 0.9958918 |
| LRRFIP2 | 175.520089 | -0.1222029 | 0.32244569 | -0.3789875 | 0.70469712 | 0.9958918 |
| GOLGA4 | 366.431203 | -0.4648617 | 0.1933419 | -2.4043507 | 0.01620122 | 0.34510952 |
| ITGA9 | 105.362263 | 0.67951542 | 0.46925903 | 1.4480604 | 0.14760016 | 0.8301633 |
| CTDSPL | 136.821109 | -0.4033403 | 0.33706523 | -1.1966239 | 0.2314532 | 0.91549646 |
| ACAA1 | 20.7260597 | 0.57855336 | 0.38545107 | 1.50097745 | 0.13336139 | 0.80664435 |
| OXSR1 | 68.8536437 | -0.2523007 | 0.31286227 | -0.8064275 | 0.41999641 | 0.97812179 |
| EXOG | 28.8988346 | -0.0944786 | 0.29071091 | -0.3249916 | 0.74518742 | 0.9958918 |
| WDR48 | 36.3002466 | -0.0570583 | 0.3795189 | -0.1503437 | 0.88049344 | 0.99749293 |
| RPSA | 90.5994957 | -0.1556849 | 0.36133031 | -0.4308658 | 0.66656599 | 0.9958918 |
| EIF1B | 28.4581746 | -0.1237705 | 0.37371089 | -0.3311933 | 0.74049852 | 0.9958918 |
| RPL14 | 157.074004 | -0.2558052 | 0.43600588 | -0.5867012 | 0.55740439 | 0.9958918 |
| ZNF621 | 36.9257151 | -0.4790394 | 0.30053755 | -1.5939419 | 0.11094905 | 0.77653769 |
| CTNNB1 | 166.931152 | 0.4149664 | 0.24845999 | 1.6701538 | 0.09488894 | 0.73481434 |
| ULK4 | 109.57767 | -0.3045752 | 0.37768116 | -0.8064348 | 0.41999217 | 0.97812179 |
| TRAK1 | 134.857193 | -0.3483731 | 0.32579799 | -1.0692919 | 0.28493816 | 0.94505166 |
| SEC22C | 24.4749889 | 0.04323643 | 0.39177689 | 0.11035984 | 0.912124 | 0.99935941 |
| NKTR | 202.169218 | -0.1187379 | 0.21395572 | -0.5549649 | 0.57891864 | 0.9958918 |
| AC006059.1 | 37.1143707 | 0.09619035 | 0.45579521 | 0.21103853 | 0.8328572 | 0.99592423 |
| SNRK | 131.691433 | 0.15173416 | 0.27594046 | 0.54988009 | 5.82E-01 | 0.9958918 |
| ANO10 | 110.13766 | 0.29893352 | 0.29989974 | 0.99677819 | 0.31887219 | 0.95992644 |
| ABHD5 | 27.3411875 | -0.2344699 | 0.45293541 | -0.5176674 | 0.60469035 | 0.9958918 |
| TCAIM | 56.6542428 | 0.84770826 | 0.26011071 | 3.25902868 | 0.00111794 | 0.07280105 |
| ZNF445 | 28.4327939 | 0.40347808 | 0.35697522 | 1.13026916 | 0.25836283 | 0.92891801 |
| AC099669.1 | 36.6162314 | 0.03025921 | 0.32728762 | 0.09245449 | 0.92633695 | 0.99935941 |
| ZNF197 | 34.592158 | 0.61229186 | 0.40515431 | 1.51125595 | 0.13072325 | 0.80565822 |
| ZDHHC3 | 33.3964887 | 0.00767601 | 0.28538888 | 0.02689668 | 0.97854214 | 0.99935941 |
| LARS2 | 42.3883385 | -0.1734816 | 0.32339849 | -0.5364329 | 0.59165939 | 0.9958918 |
| LIMD1 | 51.6824679 | 0.20137834 | 0.30791576 | 0.65400467 | 0.5131088 | 0.9958918 |
| SACM1L | 44.820671 | -0.1824501 | 0.32289177 | -0.5650505 | 0.57203945 | 0.9958918 |
| FYCO1 | 29.0108657 | 0.3040113 | 0.30203704 | 1.00653647 | 0.31415758 | 0.95992644 |
| LTF | 29.6227315 | 0.32121009 | 1.23550543 | 0.25998275 | 0.79487708 | 0.99592423 |
| PTH1R | 101.95796 | -0.0845141 | 0.37460346 | -0.2256094 | 0.8215052 | 0.99592423 |
| CCDC12 | 34.3791158 | 0.09063005 | 0.29595613 | 0.30622798 | 0.75943107 | 0.9958918 |

| | | | | | | |
|---|---|---|---|---|---|---|
| SETD2 | 238.193715 | 0.00280651 | 0.17421159 | 0.01610976 | 0.98714683 | 0.99935941 |
| KIF9-AS1 | 46.4231824 | 0.23082638 | 0.29441795 | 0.7840092 | 0.43303471 | 0.98092839 |
| KLHL18 | 23.7447594 | 0.15476831 | 0.36629434 | 0.42252444 | 0.67264226 | 0.9958918 |
| PTPN23 | 19.8374331 | -0.0389431 | 0.37253851 | -0.1045344 | 0.91674527 | 0.99935941 |
| SCAP | 32.8131283 | -0.0042738 | 0.30666232 | -0.0139364 | 0.98888072 | 0.99935941 |
| SMARCC1 | 131.096829 | -0.0984672 | 0.23756765 | -0.4144806 | 0.67852214 | 0.9958918 |
| DHX30 | 34.5880694 | -0.0478364 | 0.26424936 | -0.1810276 | 0.85634588 | 0.9971731 |
| MAP4 | 373.349329 | -0.3516174 | 0.23401968 | -1.5025121 | 0.1329649 | 0.80664435 |
| PLXNB1 | 25.2405977 | 0.41693793 | 0.3472126 | 1.20081451 | 0.22982316 | 0.91455844 |
| TMA7 | 42.4870045 | 0.04960659 | 0.42955994 | 0.11548234 | 0.90806281 | 0.99935941 |
| SHISA5 | 54.8668707 | -0.0669094 | 0.2991571 | -0.2236596 | 0.82302218 | 0.99592423 |
| UQCRC1 | 19.062994 | 1.03666114 | 0.33814066 | 3.06576894 | 0.00217111 | 0.09972181 |
| NCKIPSD | 19.6879982 | -0.3217486 | 0.36129336 | -0.8905467 | 0.37317239 | 0.97217293 |
| IP6K2 | 52.4687692 | -0.5504025 | 0.27616312 | -1.9930339 | 0.04625774 | 0.56563746 |
| PRKAR2A | 113.265236 | 0.35737419 | 0.2491686 | 1.43426657 | 0.15149621 | 0.83773184 |
| ARIH2 | 95.8184646 | 0.03151405 | 0.23075183 | 0.13657117 | 0.89136977 | 0.99935941 |
| P4HTM | 18.629156 | -0.1762656 | 0.33516452 | -0.5259076 | 0.59895241 | 0.9958918 |
| WDR6 | 31.5342132 | -0.6140129 | 0.294402 | -2.0856276 | 0.03701237 | 0.51855699 |
| QRICH1 | 51.4255816 | 0.26548752 | 0.27184206 | 0.97662415 | 0.32875526 | 0.9620084 |
| LAMB2 | 63.6222168 | 0.24203818 | 0.32369925 | 0.74772548 | 0.45462576 | 0.98173285 |
| USP4 | 54.14094 | -0.0541696 | 0.30787074 | -0.1759492 | 0.86033387 | 0.9971731 |
| RHOA | 138.424027 | 0.15943746 | 0.30688423 | 0.51953619 | 0.60338688 | 0.9958918 |
| DAG1 | 36.3672421 | 0.22677504 | 0.36360336 | 0.62368797 | 0.53283253 | 0.9958918 |
| IP6K1 | 68.9981601 | -0.1192882 | 0.27518423 | -0.433485 | 0.66466244 | 0.9958918 |
| RBM6 | 306.095854 | 0.0634499 | 0.16218574 | 0.39121751 | 0.69563646 | 0.9958918 |
| RBM5 | 110.87633 | -0.0352189 | 0.25320426 | -0.1390928 | 0.88937679 | 0.99935941 |
| GNAI2 | 64.6716324 | 0.53394465 | 0.35312446 | 1.51205794 | 0.13051912 | 0.80565822 |
| RASSF1 | 32.4479833 | 0.24812803 | 0.3235171 | 0.76697037 | 0.44309913 | 0.98092839 |
| DOCK3 | 28.1324805 | 0.68075779 | 0.35282195 | 1.92946552 | 0.0536731 | 0.58872178 |
| MANF | 21.6869736 | 0.14318809 | 0.51976607 | 0.27548563 | 0.78294316 | 0.9958918 |
| DCAF1 | 40.6531751 | 0.00821325 | 0.42139604 | 0.01949057 | 0.98444976 | 0.99935941 |
| RAD54L2 | 49.3869978 | -0.2630036 | 0.29134786 | -0.9027133 | 0.36667807 | 0.97217293 |
| RPL29 | 200.757957 | -0.089087 | 0.39442523 | -0.2258653 | 0.82130617 | 0.99592423 |
| WDR82 | 52.2171572 | 0.08563933 | 0.28459809 | 0.30091325 | 0.76348065 | 0.9958918 |
| NISCH | 57.4217652 | 0.30429964 | 0.22417191 | 1.35743874 | 0.17464184 | 0.8740623 |
| NT5DC2 | 29.5752206 | -0.1842227 | 0.59233287 | -0.3110121 | 0.75579144 | 0.9958918 |
| SMIM4 | 24.8709781 | -0.1983163 | 0.29896849 | -0.6633353 | 0.50711586 | 0.99397937 |
| PBRM1 | 149.083909 | 0.16338488 | 0.22868227 | 0.71446237 | 0.47494132 | 0.98910524 |
| GNL3 | 31.3857947 | -0.3526291 | 0.37512883 | -0.9400214 | 0.34720659 | 0.96887708 |
| SPCS1 | 40.1640617 | -0.1659758 | 0.33737622 | -0.4919605 | 0.62274724 | 0.9958918 |
| NEK4 | 30.1551435 | 0.53496741 | 0.30827707 | 1.7353461 | 0.08267952 | 0.70018756 |
| TKT | 40.6378797 | 0.75259395 | 0.47565467 | 1.5822276 | 0.11359762 | 0.77853537 |
| DCP1A | 86.1601582 | 0.28280404 | 0.21139778 | 1.33778154 | 0.18096766 | 0.87815793 |
| CACNA1D | 49.086269 | -0.728288 | 0.64788711 | -1.124097 | 0.26097187 | 0.93055907 |
| SELENOK | 26.1200888 | 0.07770923 | 0.36918677 | 0.21048758 | 0.83328715 | 0.99592423 |
| CACNA2D3 | 70.0457348 | -0.2338432 | 0.42998692 | -0.5438379 | 0.58655304 | 0.9958918 |
| CCDC66 | 68.6814894 | 0.09227669 | 0.23249643 | 0.39689509 | 0.69144483 | 0.9958918 |
| FAM208A | 92.1050076 | 0.16229683 | 0.2776885 | 0.58445645 | 0.55891326 | 0.9958918 |
| ARHGEF3 | 70.2104573 | -0.1030431 | 0.34994146 | -0.2944582 | 0.76840779 | 0.9958918 |
| APPL1 | 54.7742846 | 0.15792506 | 0.25971723 | 0.6080654 | 0.5431441 | 0.9958918 |
| ARF4 | 76.6355566 | -0.3397411 | 0.30415606 | -1.1169959 | 0.26399607 | 0.93499423 |
| DENND6A | 98.8500601 | -0.1969397 | 0.22238721 | -0.8855712 | 0.37584865 | 0.97217293 |
| DENND6A-A! | 20.9852435 | -0.4397276 | 0.43622742 | -1.0080238 | 0.31344301 | 0.95992644 |
| SLMAP | 177.347417 | 0.04753347 | 0.1957059 | 0.24288213 | 0.80809671 | 0.99592423 |

| | | | | | | |
|---|---|---|---|---|---|---|
| FLNB | 147.115963 | -0.1809146 | 0.52160299 | -0.3468436 | 0.72870884 | 0.9958918 |
| PXK | 73.4491776 | -0.0707976 | 0.3995578 | -0.1771898 | 0.85935932 | 0.9971731 |
| FHIT | 344.323492 | -0.5352486 | 0.43253882 | -1.237458 | 0.21591708 | 0.90187565 |
| PTPRG | 1491.83183 | 0.38551774 | 0.2861874 | 1.34708147 | 0.177954 | 0.87815793 |
| THOC7 | 22.9214642 | 0.05515408 | 0.33352588 | 0.16536673 | 0.86865534 | 0.9971731 |
| PRICKLE2 | 258.782296 | 0.23910655 | 0.38489997 | 0.62121736 | 0.53445662 | 0.9958918 |
| ADAMTS9 | 1316.19094 | 0.07910508 | 0.6373189 | 0.12412165 | 9.01E-01 | 9.99E-01 |
| ADAMTS9-AS | 79.7142397 | 0.04636222 | 0.4432982 | 0.10458472 | 0.91670534 | 0.99935941 |
| ADAMTS9-AS | 756.577248 | 0.92208255 | 0.48937769 | 1.88419409 | 0.05953873 | 0.61332974 |
| MAGI1 | 513.733744 | 0.51323848 | 0.29787115 | 1.72302178 | 0.08488459 | 0.70647432 |
| SLC25A26 | 57.9526315 | -0.1664342 | 0.30413802 | -0.5472326 | 0.58421896 | 0.9958918 |
| LRIG1 | 41.5959721 | 0.97943448 | 0.4232608 | 2.31402126 | 0.02066656 | 0.38830162 |
| SUCLG2 | 174.262848 | -0.0366184 | 0.20686758 | -0.1770136 | 0.85949767 | 0.9971731 |
| SUCLG2-AS1 | 49.4902192 | -0.4930859 | 0.43989617 | -1.1209142 | 0.26232439 | 0.93221565 |
| EOGT | 34.3213009 | 0.58878421 | 0.42280617 | 1.39256297 | 0.16375199 | 0.85697309 |
| TMF1 | 82.8841953 | -0.01652 | 0.21609586 | -0.0764474 | 0.93906317 | 0.99935941 |
| UBA3 | 44.0484164 | 0.11375129 | 0.29328318 | 0.38785481 | 0.69812348 | 0.9958918 |
| ARL6IP5 | 55.0150912 | 0.41876917 | 0.34938152 | 1.19860138 | 0.23068298 | 0.91488895 |
| FRMD4B | 41.5574447 | -0.1947767 | 0.46797778 | -0.4162093 | 0.67725687 | 0.9958918 |
| MITF | 196.081991 | 0.23572434 | 0.53363389 | 0.44173419 | 0.65868157 | 0.9958918 |
| FOXP1 | 397.791151 | 0.45148727 | 0.18613631 | 2.42557335 | 0.01528423 | 0.33629875 |
| RYBP | 93.4800751 | -0.1065205 | 0.36330965 | -0.2931949 | 0.76937315 | 0.9958918 |
| SHQ1 | 50.3524302 | -0.1777767 | 0.29070562 | -0.6115353 | 0.54084527 | 0.9958918 |
| GXYLT2 | 68.1875166 | -0.0669975 | 0.40514195 | -0.1653679 | 0.86865441 | 0.9971731 |
| PPP4R2 | 104.207341 | 0.00476382 | 0.20811767 | 0.02289002 | 0.981738 | 0.99935941 |
| PDZRN3 | 370.349675 | -0.2877346 | 0.33565212 | -0.8572406 | 0.39131193 | 0.97297628 |
| CNTN3 | 59.9761564 | 0.92379848 | 0.29418648 | 3.14017993 | 1.69E-03 | 9.06E-02 |
| ZNF717 | 31.0029861 | -0.0388363 | 0.36493514 | -0.1064197 | 0.91524937 | 0.99935941 |
| ROBO2 | 312.871939 | 1.03597448 | 0.45420314 | 2.28086156 | 0.02255664 | 0.40828078 |
| ROBO1 | 686.543007 | 1.71008915 | 0.42782298 | 3.99718861 | 6.41E-05 | 0.0120786 |
| GBE1 | 408.134616 | -0.7075179 | 0.41806154 | -1.6923773 | 9.06E-02 | 0.72589216 |
| VGLL3 | 66.6346405 | -1.4972376 | 0.91363087 | -1.6387774 | 0.10125962 | 0.75167369 |
| CHMP2B | 27.9409648 | -0.4735659 | 0.29996116 | -1.5787574 | 0.11439172 | 0.77853537 |
| CGGBP1 | 73.3820228 | -0.094457 | 0.23628405 | -0.3997604 | 0.689333 | 0.9958918 |
| ZNF654 | 111.506685 | -0.0738867 | 0.33383112 | -0.2213295 | 0.8248359 | 0.99592423 |
| C3orf38 | 20.6522797 | 0.17560981 | 0.31572438 | 0.55621239 | 0.57806567 | 0.9958918 |
| EPHA3 | 68.224183 | -0.2506413 | 0.49486119 | -0.5064881 | 0.61251405 | 0.9958918 |
| PROS1 | 88.4284069 | 0.3656191 | 0.41492694 | 0.881165 | 0.37822852 | 0.97217293 |
| ARL13B | 63.3436273 | -0.2922549 | 0.28933129 | -1.0101047 | 0.31244514 | 0.9599143 |
| NSUN3 | 34.1541194 | -0.2039435 | 0.31254177 | -0.6525319 | 0.51405807 | 0.9958918 |
| MTRNR2L12 | 110.390965 | -0.0060699 | 0.39324083 | -0.0154355 | 0.98768477 | 0.99935941 |
| ARL6 | 24.7798865 | -0.0679698 | 0.4038725 | -0.1682952 | 0.86635109 | 0.9971731 |
| CRYBG3 | 110.951366 | -0.3737226 | 0.29359539 | -1.272917 | 0.20304748 | 0.89396747 |
| CLDND1 | 36.7691346 | -0.0276405 | 0.3409733 | -0.0810635 | 0.93539147 | 0.99935941 |
| DCBLD2 | 102.716321 | -0.5515601 | 0.27734858 | -1.9886892 | 0.04673551 | 0.56864121 |
| COL8A1 | 131.552142 | 0.45537523 | 0.43295886 | 1.05177482 | 0.29290287 | 0.95186143 |
| CMSS1 | 194.651627 | 0.04556627 | 0.35719282 | 0.12756772 | 0.89849108 | 0.99935941 |
| FILIP1L | 204.640719 | -0.6286947 | 0.45611439 | -1.3783706 | 0.1680889 | 0.86560961 |
| TBC1D23 | 73.8916348 | -0.5726151 | 0.25872236 | -2.2132417 | 0.02688099 | 0.446449 |
| TOMM70 | 36.4238489 | 0.03950279 | 0.30789195 | 0.12830082 | 0.89791092 | 0.99935941 |
| TMEM45A | 50.6221228 | -0.0818618 | 0.57942546 | -0.141281 | 0.88764798 | 0.99935941 |
| TFG | 106.458059 | -0.1717033 | 0.25530039 | -0.6725542 | 0.50123098 | 0.99213111 |
| ABI3BP | 116.018436 | 0.60952282 | 0.99796175 | 0.61076772 | 0.54135337 | 0.9958918 |
| SENP7 | 120.711069 | 0.18555775 | 0.26470995 | 0.70098516 | 0.48331227 | 0.99015594 |

| | | | | | | |
|---|---|---|---|---|---|---|
| PCNP | 77.1348311 | -0.0373414 | 0.23936649 | -0.1560008 | 0.87603238 | 0.9971731 |
| ZBTB11 | 29.2102302 | 0.32898827 | 0.29647759 | 1.10965646 | 0.26714709 | 0.93533685 |
| RPL24 | 72.0436646 | 0.00189056 | 0.37338422 | 0.00506331 | 0.99596008 | 0.99935941 |
| CEP97 | 18.5158428 | -0.2469616 | 0.34522978 | -0.7153542 | 0.47439023 | 0.98910524 |
| NFKBIZ | 64.8238659 | 0.38282279 | 0.46228947 | 0.8281019 | 0.40761279 | 0.97733978 |
| ALCAM | 68.8607792 | 0.28633936 | 0.38643813 | 0.74097077 | 0.45871116 | 0.98482861 |
| CBLB | 806.634651 | -0.77338 | 0.31688926 | -2.4405373 | 0.01466543 | 0.33025022 |
| LINC00882 | 152.385195 | -0.3431517 | 0.51498664 | -0.6663312 | 0.50519943 | 0.99397937 |
| DUBR | 35.770605 | -0.2489608 | 0.49003491 | -0.508047 | 0.61142036 | 0.9958918 |
| BBX | 282.250827 | 0.01569364 | 0.25740339 | 0.06096903 | 0.95138387 | 0.99935941 |
| CD47 | 68.4070305 | 0.36346694 | 0.36036275 | 1.00861407 | 0.31315976 | 0.95992644 |
| IFT57 | 36.5148899 | 0.23873725 | 0.38268227 | 0.62385241 | 0.53272452 | 0.9958918 |
| DZIP3 | 42.586098 | -0.5413704 | 0.39730596 | -1.3626033 | 0.17300756 | 0.87233975 |
| NECTIN3 | 49.8863241 | -0.1145301 | 0.33405024 | -0.3428531 | 0.731709 | 0.9958918 |
| PHLDB2 | 322.957093 | -0.3308811 | 0.38575547 | -0.8577484 | 0.39103143 | 0.97297628 |
| ATG3 | 19.8104439 | 0.01120331 | 0.33720547 | 0.03322398 | 0.97349598 | 0.99935941 |
| CCDC80 | 116.402825 | -0.0200064 | 0.47360685 | -0.0422427 | 0.96630524 | 0.99935941 |
| NEPRO | 31.5433754 | 0.27728194 | 0.32225464 | 0.86044358 | 0.38954457 | 0.97297628 |
| BOC | 52.2937027 | 0.27028744 | 0.86069419 | 0.31403423 | 0.75349504 | 0.9958918 |
| CFAP44 | 35.5900521 | -0.0810499 | 0.31145 | -0.260234 | 0.79468328 | 0.99592423 |
| SPICE1 | 27.680948 | -0.3546262 | 0.33085986 | -1.071832 | 0.28379548 | 0.94505166 |
| USF3 | 35.9469164 | 0.45973298 | 0.25755906 | 1.78496144 | 0.07426757 | 0.66886321 |
| NAA50 | 77.4257713 | -0.3037632 | 0.23920231 | -1.2699009 | 0.20411994 | 0.89396747 |
| ATP6V1A | 67.9679582 | 0.14279937 | 0.24876419 | 0.57403509 | 0.56594406 | 0.9958918 |
| ZBTB20 | 2127.7386 | -0.5174021 | 0.34320572 | -1.5075566 | 0.13166805 | 0.80565822 |
| ZBTB20-AS5 | 35.2611028 | -0.9522281 | 0.6308168 | -1.509516 | 0.13116697 | 0.80565822 |
| LSAMP | 319.511104 | -2.4598973 | 0.40447834 | -6.081654 | 1.19E-09 | 4.37E-06 |
| AC092691.1 | 109.8068 | -2.9354562 | 0.56313634 | -5.2126919 | 1.86E-07 | 0.00045593 |
| ARHGAP31 | 51.6843608 | 0.48187284 | 0.33371007 | 1.44398649 | 0.14874278 | 0.8301633 |
| TMEM39A | 38.9526364 | -0.2587398 | 0.28541112 | -0.9065511 | 0.36464421 | 0.97217293 |
| POGLUT1 | 25.2632748 | 0.33669541 | 0.3675519 | 0.91604862 | 0.35964139 | 0.97217293 |
| TIMMDC1 | 34.1352029 | 0.21116399 | 0.29363472 | 0.71913837 | 0.47205567 | 0.98910524 |
| COX17 | 19.0081812 | 0.03148 | 0.3488611 | 0.09023649 | 0.92809929 | 0.99935941 |
| GSK3B | 241.803181 | -0.0271722 | 0.17109068 | -0.1588177 | 0.87381254 | 0.9971731 |
| LRRC58 | 45.1844261 | -0.1683829 | 0.26206177 | -0.6425313 | 0.5205283 | 0.9958918 |
| FSTL1 | 270.467506 | -0.3964089 | 0.33104878 | -1.1974336 | 0.23113761 | 0.91521027 |
| NDUFB4 | 36.3787228 | 0.1957703 | 0.41595174 | 0.47065628 | 0.63788621 | 0.9958918 |
| RABL3 | 20.8477679 | 0.50903999 | 0.36703299 | 1.38690525 | 0.16547066 | 0.85938344 |
| HCLS1 | 25.5704498 | 0.07472385 | 0.67453905 | 0.11077765 | 0.91179267 | 0.99935941 |
| GOLGB1 | 151.277046 | -0.160755 | 0.22145624 | -0.7258996 | 0.4679003 | 0.98910524 |
| IQCB1 | 60.7162769 | 0.21767928 | 0.27602319 | 0.78862681 | 0.43033016 | 0.98092839 |
| CASR | 41.0997923 | -2.3235799 | 0.67108185 | -3.4624389 | 0.0005353 | 0.04787234 |
| FAM162A | 62.8510844 | -1.1823638 | 0.50330941 | -2.3491788 | 0.01881487 | 0.36546347 |
| AC083798.2 | 21.5212947 | 0.49674696 | 0.40410975 | 1.22923777 | 0.21898267 | 0.90437211 |
| KPNA1 | 103.242722 | 0.0008655 | 0.21992604 | 0.0039354 | 0.99686001 | 0.99935941 |
| PARP9 | 71.5027086 | -1.1304537 | 0.78726808 | -1.4359196 | 0.15102522 | 0.83575626 |
| DTX3L | 26.5719767 | -0.2989417 | 0.55034592 | -0.5431887 | 0.58699987 | 0.9958918 |
| PARP14 | 211.886132 | -1.0955328 | 0.91361843 | -1.1991142 | 0.23048355 | 0.91459157 |
| HSPBAP1 | 29.0765411 | -0.1376486 | 0.31205222 | -0.4411074 | 0.65913522 | 0.9958918 |
| DIRC2 | 42.557835 | 0.21696754 | 0.39711101 | 0.54636495 | 0.5848151 | 0.9958918 |
| PDIA5 | 43.81716 | -0.0091509 | 0.3914385 | -0.0233777 | 0.98134903 | 0.99935941 |
| SEC22A | 86.2294124 | 0.45021248 | 0.21999301 | 2.04648542 | 0.04070864 | 0.54295419 |
| ADCY5 | 88.9281506 | -0.07282 | 0.31947818 | -0.2279342 | 0.81969738 | 0.99592423 |
| HACD2 | 30.6936405 | 0.54628126 | 0.31742877 | 1.72095698 | 0.08525863 | 0.70747633 |

| | | | | | | |
|---|---|---|---|---|---|---|
| MYLK | 329.259989 | -0.2678171 | 0.42780739 | -0.6260226 | 0.53130011 | 0.9958918 |
| CCDC14 | 90.6089524 | -0.6983953 | 0.28626035 | -2.4397206 | 0.01469863 | 0.33025022 |
| KALRN | 532.822928 | 0.28756573 | 0.37560414 | 0.76560852 | 0.44390927 | 0.98092839 |
| ITGB5 | 95.7785251 | -0.1381078 | 0.26629544 | -0.5186263 | 0.60402137 | 0.9958918 |
| HEG1 | 41.2359645 | -0.289363 | 0.39398416 | -0.7344533 | 0.46267249 | 0.98590351 |
| ZNF148 | 213.867894 | 0.15580555 | 0.14761643 | 1.05547567 | 0.29120783 | 0.95186143 |
| SNX4 | 75.3950356 | -0.3453678 | 0.24020414 | -1.4378096 | 0.15048809 | 0.83401707 |
| OSBPL11 | 23.6205461 | -0.5057899 | 0.32041868 | -1.5785281 | 0.11444433 | 0.77853537 |
| ZXDC | 60.9158965 | -0.0301657 | 0.30504206 | -0.0988902 | 0.92122546 | 0.99935941 |
| CHCHD6 | 19.1677202 | 0.00100348 | 0.39617147 | 0.00253295 | 0.997979 | 0.99948864 |
| ABTB1 | 22.0763026 | 0.73558169 | 0.34109423 | 2.15653516 | 0.0310419 | 0.47969946 |
| MGLL | 105.444182 | -0.2012081 | 0.37120751 | -0.5420367 | 0.58779325 | 0.9958918 |
| SEC61A1 | 82.6705005 | -0.4220097 | 0.38913449 | -1.0844829 | 0.27815076 | 0.94419503 |
| RUVBL1 | 30.8799848 | 0.02619712 | 0.35929357 | 0.07291285 | 0.94187547 | 0.99935941 |
| EEFSEC | 52.789724 | 0.15112008 | 0.29353274 | 0.51483211 | 0.60667034 | 0.9958918 |
| RPN1 | 42.1298167 | 0.04146576 | 0.30679841 | 0.13515638 | 0.89248824 | 0.99935941 |
| RAB7A | 170.422187 | -0.3808666 | 0.23155024 | -1.644855 | 0.09999971 | 0.7498958 |
| ACAD9 | 26.3285024 | -0.2188887 | 0.29518061 | -0.7415416 | 0.45836511 | 0.98457237 |
| CNBP | 109.89874 | 0.66931942 | 0.44421429 | 1.50674893 | 0.13187502 | 0.80565822 |
| COPG1 | 34.9055499 | -0.0542687 | 0.38375793 | -0.141414 | 0.88754289 | 0.99935941 |
| HMCES | 19.9414319 | -0.1837927 | 0.42312629 | -0.4343684 | 0.66402093 | 0.9958918 |
| H1FX | 44.2939159 | -0.233795 | 0.39011175 | -0.5993026 | 0.54897109 | 0.9958918 |
| MBD4 | 32.6157489 | 0.02323051 | 0.27640043 | 0.08404658 | 0.9330194 | 0.99935941 |
| IFT122 | 36.6214163 | 0.067883 | 0.28692703 | 0.23658629 | 0.81297775 | 0.99592423 |
| PLXND1 | 84.7857748 | -0.4800051 | 0.24610412 | -1.9504147 | 0.05112671 | 0.57655256 |
| TMCC1 | 222.503038 | -0.1211062 | 0.24640454 | -0.4914936 | 0.62307741 | 0.9958918 |
| PIK3R4 | 28.2585305 | 0.00133564 | 0.29461947 | 0.00453342 | 0.99638286 | 0.99935941 |
| ATP2C1 | 94.1292434 | -0.0751119 | 0.22228662 | -0.3379058 | 0.73543416 | 0.9958918 |
| NEK11 | 26.0631006 | -0.3286691 | 0.43640038 | -0.7531366 | 0.45136784 | 0.98092839 |
| MRPL3 | 41.8721209 | -0.1669139 | 0.32269133 | -0.5172555 | 0.6049778 | 0.9958918 |
| DNAJC13 | 79.3133251 | -0.4158575 | 0.23228465 | -1.7902925 | 0.0734069 | 0.66848492 |
| UBA5 | 37.3612069 | -0.0920426 | 0.32531041 | -0.2829378 | 0.77722449 | 0.9958918 |
| CDV3 | 45.5863016 | 0.14790439 | 0.42440465 | 0.34849851 | 0.72746583 | 0.9958918 |
| TOPBP1 | 25.9542248 | -0.2442759 | 0.29838855 | -0.8186503 | 0.41298596 | 0.97812179 |
| SLCO2A1 | 69.3233463 | 1.71268835 | 0.59307218 | 2.88782448 | 0.00387916 | 0.15300235 |
| RYK | 122.095053 | -0.1614034 | 0.20690045 | -0.7801017 | 0.43533103 | 0.98092839 |
| AMOTL2 | 41.8550409 | -1.248687 | 0.37490147 | -3.3307072 | 0.00086626 | 0.0644401 |
| CEP63 | 46.061762 | -0.0981011 | 0.28696042 | -0.3418628 | 0.73245418 | 0.9958918 |
| EPHB1 | 37.8499518 | 0.27396639 | 0.49074188 | 0.55826983 | 0.57666014 | 0.9958918 |
| PPP2R3A | 85.2492397 | 0.07086346 | 0.31866796 | 0.22237397 | 0.82402277 | 0.99592423 |
| MSL2 | 45.1284549 | 0.03708734 | 0.25355599 | 0.14626882 | 0.88370918 | 0.99862036 |
| PCCB | 23.2952432 | -0.0510884 | 0.34645644 | -0.1474599 | 0.88276901 | 0.99840747 |
| STAG1 | 570.566365 | 0.0052951 | 0.20563294 | 0.02575027 | 0.97945653 | 0.99935941 |
| NCK1 | 50.4587143 | 0.07631004 | 0.26112386 | 0.29223695 | 0.77010547 | 0.9958918 |
| DZIP1L | 18.5338501 | 0.22267852 | 0.40365909 | 0.55164995 | 0.58118821 | 0.9958918 |
| ARMC8 | 105.990169 | -0.1959612 | 0.26912013 | -0.7281551 | 0.46651867 | 0.98791879 |
| MRAS | 24.1207713 | 0.06052965 | 0.42344597 | 0.14294539 | 0.88633331 | 0.99935941 |
| CEP70 | 40.9334787 | 0.02681682 | 0.29922384 | 0.08962128 | 0.92858817 | 0.99935941 |
| PIK3CB | 59.63002 | 0.27547433 | 0.22632877 | 1.21714234 | 0.22355006 | 0.91093521 |
| COPB2 | 71.085866 | -0.1066675 | 0.22571738 | -0.472571 | 0.63651928 | 0.9958918 |
| AC097103.2 | 43.7705935 | -0.3868438 | 0.48984601 | -0.7897253 | 0.42968822 | 0.98092839 |
| CLSTN2 | 118.49006 | -0.2100274 | 0.63025263 | -0.3332432 | 0.7389507 | 0.9958918 |
| SLC25A36 | 114.115749 | -0.2857697 | 0.34398003 | -0.8307741 | 0.40610123 | 0.97637555 |
| ZBTB38 | 207.67144 | -0.3331905 | 0.24968955 | -1.3344191 | 0.18206654 | 0.87866798 |

| | | | | | | |
|---|---|---|---|---|---|---|
| RASA2 | 141.773754 | -0.0414272 | 0.19551248 | -0.2118905 | 0.83219244 | 0.99592423 |
| RNF7 | 26.073893 | 0.20607628 | 0.32825619 | 0.62779099 | 0.53014087 | 0.9958918 |
| ATP1B3 | 214.521659 | -1.0818975 | 0.62439824 | -1.7327043 | 0.08314823 | 0.7007527 |
| TFDP2 | 136.792736 | 0.18930758 | 0.31338708 | 0.6040695 | 0.54579744 | 0.9958918 |
| GK5 | 51.9295408 | 0.38496247 | 0.34731029 | 1.10841077 | 0.26768446 | 0.93549101 |
| XRN1 | 203.101176 | -0.3729753 | 0.30330186 | -1.2297166 | 0.21880326 | 0.90437211 |
| ATR | 105.003357 | -0.0706525 | 0.30194212 | -0.2339937 | 0.81498989 | 0.99592423 |
| TRPC1 | 107.644582 | 0.24087101 | 0.39586498 | 0.6084676 | 0.54287739 | 0.9958918 |
| PCOLCE2 | 31.5552329 | -0.3342781 | 0.39615249 | -0.8438118 | 0.39877461 | 0.97500135 |
| U2SURP | 107.361652 | -0.0580383 | 0.21928643 | -0.2646688 | 0.79126459 | 0.99592423 |
| SLC9A9 | 122.694792 | 0.32693759 | 0.43119304 | 0.75821629 | 0.44832151 | 0.98092839 |
| C3orf58 | 25.6937028 | 0.0534472 | 0.35906231 | 0.14885216 | 0.88167029 | 0.99805838 |
| PLOD2 | 279.448347 | -0.2738926 | 0.58265314 | -0.4700783 | 0.63829911 | 0.9958918 |
| PLSCR4 | 245.228992 | -0.3652629 | 0.42367665 | -0.8621266 | 0.38861788 | 0.97297628 |
| PLSCR1 | 205.819477 | -0.7459414 | 0.56125178 | -1.3290673 | 0.18382577 | 0.87866798 |
| AGTR1 | 120.447836 | 0.43072697 | 0.51420802 | 0.83765121 | 0.40222662 | 0.97512144 |
| GYG1 | 24.7655048 | 0.13444947 | 0.30429864 | 0.44183396 | 0.65860936 | 0.9958918 |
| HLTF | 43.6326546 | -0.3262096 | 0.40436509 | -0.8067204 | 0.4198276 | 0.97812179 |
| HPS3 | 30.6665858 | -0.2280643 | 0.45190297 | -0.5046754 | 0.61378684 | 0.9958918 |
| CP | 73.0169974 | 1.26417206 | 1.03855755 | 1.21723832 | 0.22351355 | 0.91093521 |
| TM4SF1 | 68.3841092 | -0.0967114 | 0.61216783 | -0.1579818 | 0.87447113 | 0.9971731 |
| WWTR1 | 232.681601 | -0.1987447 | 0.23891512 | -0.831863 | 0.40548627 | 0.97637555 |
| COMMD2 | 20.0151911 | 0.10941605 | 0.33833854 | 0.32339222 | 0.74639822 | 0.9958918 |
| ANKUB1 | 49.8824803 | -0.3171981 | 0.47176514 | -0.6723644 | 0.50135176 | 0.99213111 |
| RNF13 | 96.1663109 | -0.2653792 | 0.20017777 | -1.3257177 | 0.18493324 | 0.87866798 |
| TSC22D2 | 158.079306 | 0.04790711 | 0.28758661 | 0.16658325 | 0.86769797 | 0.9971731 |
| SERP1 | 66.7765305 | 0.01097026 | 0.2628107 | 0.04174205 | 0.96670433 | 0.99935941 |
| EIF2A | 42.2151575 | -0.0821055 | 0.30710372 | -0.2673544 | 0.78919633 | 0.99592423 |
| SELENOT | 40.5746884 | 0.18745707 | 0.31017617 | 0.60435676 | 0.54560648 | 0.9958918 |
| MBNL1 | 796.902545 | 0.15480061 | 0.38900424 | 0.39794067 | 0.69067392 | 0.9958918 |
| ARHGEF26 | 45.2697408 | -0.1181851 | 0.60439305 | -0.1955435 | 0.84496748 | 0.99692952 |
| DHX36 | 110.869202 | -0.2222891 | 0.21730185 | -1.022951 | 0.30633104 | 0.95793395 |
| GMPS | 48.8280832 | 0.12642876 | 0.22575109 | 0.56003609 | 0.57545482 | 0.9958918 |
| KCNAB1 | 152.826832 | 1.52300286 | 0.40694096 | 3.74256464 | 1.82E-04 | 0.0219448 |
| SSR3 | 57.7230776 | -0.1895423 | 0.35049134 | -0.5407903 | 0.58865211 | 0.9958918 |
| TIPARP | 80.5972137 | -2.0274482 | 0.62530738 | -3.2423224 | 0.0011856 | 0.07383864 |
| CCNL1 | 152.613014 | -0.5149504 | 0.34516976 | -1.4918758 | 0.13573169 | 0.81162915 |
| RSRC1 | 131.42908 | -0.3610743 | 0.18053611 | -2.0000114 | 0.04549903 | 0.55915113 |
| GFM1 | 40.6141522 | -0.5378317 | 0.26358572 | -2.0404433 | 0.04130619 | 0.54347495 |
| LXN | 39.7228794 | -1.4802534 | 0.44450459 | -3.3301196 | 0.00086809 | 0.0644401 |
| RARRES1 | 338.404614 | -1.6526491 | 0.49006902 | -3.3722783 | 0.00074549 | 0.06155743 |
| MFSD1 | 32.1032835 | -0.2168627 | 0.37415273 | -0.5796102 | 0.56217753 | 0.9958918 |
| IQCJ-SCHIP1 | 85.824679 | -0.1651596 | 0.50146984 | -0.3293511 | 0.74189036 | 0.9958918 |
| KPNA4 | 90.5356132 | -0.0492981 | 0.28458534 | -0.1732279 | 0.86247231 | 0.9971731 |
| PPM1L | 130.00728 | -0.7662701 | 0.45838332 | -1.6716796 | 0.09458752 | 0.7340733 |
| NMD3 | 104.336954 | -0.2632108 | 0.36830835 | -0.714648 | 0.4748266 | 0.98910524 |
| PDCD10 | 45.1665711 | -0.1349236 | 0.25968074 | -0.5195747 | 0.60336001 | 0.9958918 |
| GOLIM4 | 65.4988767 | -0.0260862 | 0.24814683 | -0.1051241 | 0.91627734 | 0.99935941 |
| MECOM | 1139.01894 | 1.27429372 | 0.29509101 | 4.31830747 | 1.57E-05 | 0.0042512 |
| SEC62 | 80.4411604 | -0.1779825 | 0.22443103 | -0.7930386 | 0.42775536 | 0.98092839 |
| PHC3 | 229.263262 | -0.0414406 | 0.23132462 | -0.1791448 | 0.85782398 | 0.9971731 |
| PRKCI | 78.2111292 | -0.2361927 | 0.28348518 | -0.8331747 | 0.40474622 | 0.97582411 |
| SKIL | 98.7488882 | 0.03640928 | 0.33087301 | 0.11004005 | 0.91237761 | 0.99935941 |
| TNIK | 35.4074747 | 0.19668919 | 0.40450953 | 0.48624119 | 0.62679617 | 0.9958918 |

| | | | | | | |
|---|---|---|---|---|---|---|
| PLD1 | 152.618004 | 0.4278061 | 0.41210504 | 1.03809963 | 0.29922367 | 0.95483924 |
| FNDC3B | 1224.92905 | -0.2199216 | 0.31595244 | -0.6960592 | 0.48639173 | 0.99046213 |
| TNFSF10 | 48.5626123 | -0.0531904 | 0.59842779 | -0.0888836 | 0.9291744 | 0.99935941 |
| NCEH1 | 27.5918745 | -1.0002994 | 0.48150464 | -2.0774449 | 0.03776051 | 0.52201457 |
| NLGN1 | 177.365652 | -0.260316 | 0.34512821 | -0.7542589 | 0.45069379 | 0.98092839 |
| NAALADL2 | 416.893086 | 0.26735494 | 0.32930936 | 0.81186559 | 0.41686876 | 0.97812179 |
| TBL1XR1 | 335.291439 | -0.3503303 | 0.24372086 | -1.4374244 | 0.15059745 | 0.83401707 |
| KCNMB2-AS1 | 33.2930441 | 0.49906724 | 0.49345437 | 1.01137465 | 0.31183715 | 0.9599143 |
| ZMAT3 | 50.9844177 | 0.11453072 | 0.42543038 | 0.26921142 | 0.787767 | 0.99592423 |
| PIK3CA | 138.080077 | -0.3803487 | 0.3247552 | -1.1711857 | 0.24152412 | 0.92516122 |
| MFN1 | 56.990237 | -0.2593404 | 0.29185154 | -0.888604 | 0.37421595 | 0.97217293 |
| GNB4 | 59.0724959 | -0.068372 | 0.28295348 | -0.2416368 | 0.80906159 | 0.99592423 |
| NDUFB5 | 20.1139191 | 0.23274374 | 0.33669208 | 0.69126586 | 0.48939848 | 0.99046213 |
| USP13 | 26.8593736 | -0.0131391 | 0.34127977 | -0.0384994 | 0.96928948 | 0.99935941 |
| TTC14 | 91.4385397 | 0.17053757 | 0.19387214 | 0.87963939 | 0.37905469 | 0.97217293 |
| FXR1 | 105.234102 | -0.1605438 | 0.2420921 | -0.6631517 | 0.50723337 | 0.99397937 |
| ATP11B | 130.777933 | -0.2379263 | 0.2582966 | -0.9211359 | 0.35697945 | 0.97217293 |
| DCUN1D1 | 42.1140086 | -0.0266741 | 0.2772028 | -0.096226 | 0.92334112 | 0.99935941 |
| MCCC1 | 53.7162875 | -0.1948943 | 0.32786175 | -0.5944404 | 0.55221761 | 0.9958918 |
| MCF2L2 | 27.8994428 | 1.22520341 | 0.37034029 | 3.30831788 | 0.00093858 | 0.06829346 |
| KLHL24 | 123.386834 | 0.10349971 | 0.43942778 | 0.23553292 | 0.81379513 | 0.99592423 |
| YEATS2 | 111.764001 | -0.2111754 | 0.40875715 | -0.516628 | 0.60541588 | 0.9958918 |
| ABCC5 | 40.2137489 | -0.1792893 | 0.34237827 | -0.5236585 | 0.60051608 | 0.9958918 |
| DVL3 | 34.341355 | -0.4995386 | 0.33885244 | -1.4742069 | 0.14042588 | 0.8153985 |
| AP2M1 | 60.9084465 | 0.02694989 | 0.35765345 | 0.07535197 | 0.93993467 | 0.99935941 |
| PSMD2 | 28.379542 | -0.0232626 | 0.42247566 | -0.0550625 | 0.95608864 | 0.99935941 |
| EIF4G1 | 57.986276 | -0.1217335 | 0.39103543 | -0.3113106 | 0.75556453 | 0.9958918 |
| POLR2H | 27.3434932 | -0.4913044 | 0.35280871 | -1.3925518 | 0.16375538 | 0.85697309 |
| VPS8 | 98.9945056 | -0.1870088 | 0.31170735 | -0.5999499 | 0.54853962 | 0.9958918 |
| C3orf70 | 20.4149062 | 0.46658101 | 0.53245109 | 0.87628896 | 0.38087297 | 0.97217293 |
| EHHADH | 19.1405554 | 0.88850757 | 0.39158729 | 2.26898978 | 0.02326895 | 0.41205662 |
| MAP3K13 | 102.478382 | 0.40882848 | 0.27810112 | 1.47007131 | 0.14154244 | 0.81652619 |
| SENP2 | 50.514242 | -0.1718919 | 0.28695477 | -0.5990208 | 0.54915902 | 0.9958918 |
| IGF2BP2 | 48.1486067 | -0.498865 | 0.3461722 | -1.4410892 | 0.14955948 | 0.83171145 |
| TRA2B | 93.3674822 | 0.0211213 | 0.26554849 | 0.07953841 | 0.93660438 | 0.99935941 |
| ETV5 | 22.0077675 | 0.03713746 | 0.39297323 | 0.0945038 | 0.92470896 | 0.99935941 |
| DGKG | 110.227516 | 0.87189575 | 0.50053723 | 1.74191987 | 0.08152247 | 0.69958018 |
| EIF4A2 | 88.753796 | -0.1001827 | 0.37085445 | -0.2701402 | 0.78705242 | 0.99592423 |
| ST6GAL1 | 46.9617305 | 1.29132313 | 0.33064001 | 3.90552586 | 9.40E-05 | 0.01502083 |
| BCL6 | 223.12291 | -0.0674905 | 0.40307105 | -0.1674407 | 8.67E-01 | 0.9971731 |
| LPP | 1731.69614 | 0.01542165 | 0.43825327 | 0.0351889 | 0.97192911 | 0.99935941 |
| TPRG1 | 75.5158059 | -0.2274895 | 0.33006662 | -0.6892229 | 0.49068298 | 0.99046213 |
| P3H2 | 148.110038 | -1.611475 | 0.50464204 | -3.1933031 | 0.00140655 | 0.0833609 |
| CCDC50 | 103.69033 | 0.03107382 | 0.23702639 | 0.13109856 | 0.89569734 | 0.99935941 |
| MB21D2 | 60.0470073 | 0.69993233 | 0.58325036 | 1.2000547 | 0.2301181 | 0.91455844 |
| OPA1 | 90.0458834 | 0.11845192 | 0.22215973 | 0.53318357 | 0.59390651 | 0.9958918 |
| HES1 | 39.2049071 | -0.3530921 | 0.50282506 | -0.7022167 | 0.48254404 | 0.99002719 |
| ATP13A3 | 272.287472 | -0.5116184 | 0.46206886 | -1.1072342 | 0.26819271 | 0.93598159 |
| LSG1 | 22.6916306 | -0.045216 | 0.41131671 | -0.1099299 | 0.91246497 | 0.99935941 |
| XXYLT1 | 37.2893401 | -0.2076198 | 0.34549201 | -0.6009396 | 0.54788021 | 0.9958918 |
| ACAP2 | 186.712173 | 0.00315127 | 0.17135237 | 0.01839056 | 0.98532728 | 0.99935941 |
| PPP1R2 | 40.8409017 | 0.08922342 | 0.25195072 | 0.35413045 | 0.72324112 | 0.9958918 |
| MUC20-OT1 | 175.698902 | 0.12019239 | 0.41881447 | 0.28698241 | 0.77412579 | 0.9958918 |
| TNK2 | 17.4211182 | 0.48601694 | 0.44905033 | 1.08232174 | 0.27910959 | 0.94419503 |

| | | | | | | |
|---|---|---|---|---|---|---|
| TFRC | 29.9869327 | 0.15837571 | 0.40517988 | 0.39087753 | 0.69588776 | 0.9958918 |
| PCYT1A | 24.9692577 | -0.0007371 | 0.29299224 | -0.0025159 | 0.9979926 | 0.99948864 |
| UBXN7 | 63.346678 | -0.1569303 | 0.25861776 | -0.6068039 | 0.54398107 | 0.9958918 |
| RNF168 | 57.5181792 | 0.00231228 | 0.30876693 | 0.00748874 | 0.9940249 | 0.99935941 |
| PAK2 | 92.1882795 | 0.1438355 | 0.22726923 | 0.63288592 | 0.52680815 | 0.9958918 |
| SENP5 | 104.190754 | 0.16939638 | 0.30631954 | 0.55300544 | 0.58025968 | 0.9958918 |
| DLG1 | 270.948793 | -0.39014 | 0.20297953 | -1.9220656 | 0.0545975 | 0.59304667 |
| RUBCN | 32.8166225 | 0.36450316 | 0.28771581 | 1.2668861 | 0.20519602 | 0.89589437 |
| FYTTD1 | 40.1276681 | -0.1672073 | 0.26139887 | -0.6396633 | 0.52239153 | 0.9958918 |
| LRCH3 | 191.8798 | 0.03714087 | 0.19499079 | 0.19047499 | 0.84893694 | 0.9971731 |
| IQCG | 20.4093909 | 0.24339922 | 0.34513734 | 0.70522423 | 0.4806707 | 0.99002719 |
| RPL35A | 89.3407791 | -0.3659574 | 0.34077288 | -1.0739043 | 0.28286559 | 0.94471801 |
| ZNF595 | 27.0579881 | 0.02996236 | 0.33140788 | 0.09040933 | 0.92796194 | 0.99935941 |
| ZNF718 | 54.8664166 | 0.03194841 | 0.30740318 | 0.10393 | 0.9172249 | 0.99935941 |
| ZNF141 | 57.2864108 | 0.06635525 | 0.24239347 | 0.27375016 | 0.78427663 | 0.9958918 |
| ZNF721 | 115.030432 | 0.04229534 | 0.27347216 | 0.15466049 | 0.87708898 | 0.9971731 |
| PIGG | 32.2780757 | -0.1269426 | 0.35530958 | -0.3572731 | 0.72088737 | 0.9958918 |
| ATP5ME | 22.9808834 | 0.70413577 | 0.32474516 | 2.16827181 | 0.03013801 | 0.46924629 |
| PCGF3 | 46.4064891 | -0.1996446 | 0.2451367 | -0.8144213 | 0.41540363 | 0.97812179 |
| AC139887.2 | 84.4979917 | 0.73414171 | 0.35219205 | 2.08449255 | 0.03711539 | 0.51855699 |
| GAK | 57.7562184 | 0.07436122 | 0.272831 | 0.27255416 | 0.78519595 | 0.9958918 |
| AC092535.3 | 20.7267101 | -0.1053199 | 0.40985657 | -0.2569677 | 0.79720369 | 0.99592423 |
| SPON2 | 42.9732104 | -0.9292212 | 0.53488041 | -1.7372504 | 0.08234297 | 0.69989185 |
| CTBP1 | 35.420034 | -0.482286 | 0.33175194 | -1.4537548 | 0.1460143 | 0.8301633 |
| MAEA | 28.8492788 | 0.13076794 | 0.33786284 | 0.38704447 | 0.69872328 | 0.9958918 |
| UVSSA | 37.8652433 | -0.5078791 | 0.28813812 | -1.7626237 | 0.07796397 | 0.68056484 |
| SLBP | 33.03908 | -0.335572 | 0.32472119 | -1.0334158 | 0.30140934 | 0.95597956 |
| LETM1 | 18.0684816 | -0.0994325 | 0.38394097 | -0.2589787 | 0.79565169 | 0.99592423 |
| NSD2 | 40.0369533 | 0.19834941 | 0.3633392 | 0.54590699 | 0.58512987 | 0.9958918 |
| NELFA | 22.9131096 | -0.0660764 | 0.34022744 | -0.1942123 | 0.84600965 | 0.99692952 |
| MXD4 | 52.4892763 | 0.02455915 | 0.37165483 | 0.06608054 | 0.9473137 | 0.99935941 |
| ZFYVE28 | 26.4689549 | 0.59315139 | 0.46190844 | 1.28413196 | 0.19909579 | 0.88864367 |
| RNF4 | 39.1386864 | 0.25898161 | 0.33551186 | 0.77190002 | 0.44017365 | 0.98092839 |
| FAM193A | 128.617021 | -0.0913712 | 0.24127195 | -0.3787064 | 0.70490592 | 0.9958918 |
| TNIP2 | 18.3934092 | 0.08420318 | 0.43992608 | 0.19140303 | 0.84820985 | 0.99713289 |
| SH3BP2 | 21.0770187 | 0.60702915 | 0.32412611 | 1.87281779 | 0.06109355 | 0.61696136 |
| ADD1 | 186.873363 | -0.2360687 | 0.16537761 | -1.4274525 | 0.1534495 | 0.84093985 |
| MFSD10 | 21.7041059 | -0.1704463 | 0.40385672 | -0.4220464 | 0.67299118 | 0.9958918 |
| GRK4 | 20.4396996 | 0.14185516 | 0.51137752 | 0.27739811 | 0.78147443 | 0.9958918 |
| HTT | 101.908144 | -0.1217189 | 0.19866574 | -0.6126821 | 0.54008655 | 0.9958918 |
| RGS12 | 38.8369788 | 0.12441402 | 0.29549225 | 0.42103986 | 0.67372598 | 0.9958918 |
| LRPAP1 | 44.2482753 | -0.1300409 | 0.24638552 | -0.5277945 | 0.59764201 | 0.9958918 |
| ADRA2C | 32.04177 | -0.4360951 | 0.3634222 | -1.1999682 | 0.2301517 | 0.91455844 |
| STX18 | 125.925283 | 0.58639411 | 0.30030789 | 1.952643 | 0.05086191 | 0.57655256 |
| STX18-AS1 | 44.0123122 | 0.94556048 | 0.50726387 | 1.86404066 | 0.06231599 | 0.62054234 |
| EVC2 | 34.9621342 | -0.148113 | 0.34734961 | -0.4264091 | 0.6698098 | 0.9958918 |
| EVC | 24.7055028 | 0.37855692 | 0.34226342 | 1.10603968 | 0.26870935 | 0.93598159 |
| MAN2B2 | 18.4801521 | 0.51167432 | 0.3653922 | 1.40034275 | 0.1614107 | 0.85202157 |
| MRFAP1 | 49.283759 | 0.43579321 | 0.42531515 | 1.02463598 | 0.30553498 | 0.95793395 |
| KIAA0232 | 99.8734904 | 0.20231191 | 0.23087126 | 0.87629751 | 0.38086832 | 0.97217293 |
| TBC1D14 | 38.7808103 | 0.16056899 | 0.35009044 | 0.45865003 | 0.6464855 | 0.9958918 |
| SORCS2 | 27.1637764 | -0.258109 | 0.42005803 | -0.6144602 | 0.53891124 | 0.9958918 |
| AFAP1 | 78.804706 | 0.10600406 | 0.37086393 | 0.28583005 | 0.77500829 | 0.9958918 |
| ABLIM2 | 61.6307868 | -0.2524972 | 0.49371737 | -0.5114204 | 0.60905668 | 0.9958918 |

| | | | | | | |
|---|---|---|---|---|---|---|
| ACOX3 | 25.6581294 | -0.0625538 | 0.43080559 | -0.145202 | 0.88455138 | 0.99862036 |
| SLC2A9 | 24.6165106 | -0.0594316 | 0.44049143 | -0.1349212 | 0.89267417 | 0.99935941 |
| WDR1 | 69.0395453 | 0.14875569 | 0.3372311 | 0.44110905 | 0.65913406 | 0.9958918 |
| RAB28 | 104.338756 | -0.0016552 | 0.20374697 | -0.0081239 | 0.99351816 | 0.99935941 |
| BOD1L1 | 83.0544731 | -0.1371353 | 0.27851583 | -0.4923788 | 0.62245155 | 0.9958918 |
| CPEB2 | 60.8939569 | -0.125459 | 0.35196728 | -0.3564507 | 0.72150309 | 0.9958918 |
| AC098829.1 | 137.704391 | -0.0935829 | 0.43104514 | -0.217107 | 0.82812493 | 0.99592423 |
| C1QTNF7 | 26.2736403 | 0.1664424 | 0.54313075 | 0.30644997 | 0.75926207 | 0.9958918 |
| CC2D2A | 48.1359715 | -0.0372559 | 0.33526837 | -0.1111226 | 0.91151914 | 0.99935941 |
| FBXL5 | 101.980674 | 0.06735933 | 0.24315646 | 0.27702054 | 0.78176434 | 0.9958918 |
| TAPT1 | 64.0312078 | 0.03265711 | 0.22711671 | 0.14378999 | 0.88566631 | 0.99919584 |
| TAPT1-AS1 | 56.2180917 | 0.15636496 | 0.46369506 | 0.33721507 | 0.73595478 | 0.9958918 |
| LDB2 | 544.219951 | -0.7014041 | 0.4474852 | -1.5674352 | 0.117013 | 0.78460631 |
| LAP3 | 35.070534 | 0.57837464 | 0.60868736 | 0.95019984 | 0.34201072 | 0.96887708 |
| MED28 | 28.8397004 | -0.0731551 | 0.28286784 | -0.2586195 | 0.79592886 | 0.99592423 |
| DCAF16 | 28.2322303 | 0.24354924 | 0.35344879 | 0.68906515 | 0.49078227 | 0.99046213 |
| LCORL | 81.8109811 | -0.0691415 | 0.25402662 | -0.2721822 | 0.7854819 | 0.9958918 |
| AC024230.1 | 18.8563091 | -0.2473416 | 0.43077897 | -0.574173 | 0.56585077 | 0.9958918 |
| SLIT2 | 161.844208 | 0.06067163 | 0.36226376 | 0.16747916 | 0.86699305 | 0.9971731 |
| PACRGL | 45.0777085 | -0.5074434 | 0.30465237 | -1.6656474 | 0.09578366 | 0.73786716 |
| KCNIP4 | 172.683599 | 0.39570706 | 0.37076211 | 1.0672802 | 0.28584533 | 0.94549382 |
| ADGRA3 | 47.4414129 | -0.0576874 | 0.3841389 | -0.1501733 | 0.88062786 | 0.99749293 |
| PPARGC1A | 64.2212764 | 0.53798436 | 0.4156631 | 1.29427983 | 0.19556878 | 0.88391893 |
| DHX15 | 115.955117 | -0.0591327 | 0.24334053 | -0.2430038 | 0.80800243 | 0.99592423 |
| SOD3 | 104.338094 | 0.60567124 | 0.40236285 | 1.50528617 | 0.13225052 | 0.80565822 |
| CCDC149 | 33.2826357 | 0.42231318 | 0.49363385 | 0.85551908 | 0.39226385 | 0.97297628 |
| ANAPC4 | 37.8043278 | 0.35385879 | 0.3234756 | 1.09392729 | 0.27398687 | 0.9394201 |
| RBPJ | 160.335632 | 0.34206328 | 0.29164452 | 1.17287747 | 0.24084495 | 0.92426606 |
| TBC1D19 | 62.6920189 | -0.2464747 | 0.2765108 | -0.8913744 | 0.37272835 | 0.97217293 |
| STIM2 | 52.2730298 | 0.1624535 | 0.2794652 | 0.58130137 | 0.56103736 | 0.9958918 |
| PCDH7 | 144.48129 | 1.22067978 | 0.86095653 | 1.41781813 | 0.15624387 | 0.84427321 |
| ARAP2 | 23.2484016 | -0.6452453 | 0.3766316 | -1.7132001 | 0.08667573 | 0.71410305 |
| C4orf19 | 27.2729965 | 0.03694056 | 0.33924836 | 0.10888943 | 0.91329019 | 0.99935941 |
| RELL1 | 59.7549911 | 0.54209519 | 0.26229825 | 2.06671293 | 0.03876121 | 0.52751138 |
| PGM2 | 32.6361422 | -0.0534114 | 0.25893937 | -0.2062698 | 8.37E-01 | 0.99692952 |
| TBC1D1 | 429.811431 | 0.63867727 | 0.23542059 | 2.71292023 | 0.00666932 | 0.20675326 |
| KLF3 | 41.913896 | 0.37359659 | 0.37613262 | 0.99325761 | 0.32058443 | 0.95992644 |
| FAM114A1 | 70.4508436 | -0.2759488 | 0.33471849 | -0.8244206 | 0.40970062 | 0.97812179 |
| KLHL5 | 68.9701944 | 0.1575953 | 0.45548489 | 0.34599458 | 0.72934679 | 0.9958918 |
| WDR19 | 46.2808002 | -0.4615745 | 0.24059107 | -1.9185021 | 0.05504738 | 0.59491644 |
| RFC1 | 61.1177021 | -0.3644069 | 0.27590649 | -1.3207623 | 0.18658065 | 0.87955894 |
| UGDH | 52.7967233 | -0.5203865 | 0.63266046 | -0.8225368 | 0.41077147 | 0.97812179 |
| UGDH-AS1 | 34.3031521 | 0.21392203 | 0.43876354 | 0.48755652 | 0.625864 | 0.9958918 |
| SMIM14 | 107.839835 | -0.0761673 | 0.27459665 | -0.2773787 | 0.78148931 | 0.9958918 |
| UBE2K | 218.595799 | -0.2510501 | 0.21469216 | -1.1693493 | 0.24226291 | 0.92534155 |
| PDS5A | 198.953825 | -0.1217083 | 0.14577645 | -0.8348968 | 0.40377579 | 0.97575999 |
| RBM47 | 44.6487042 | -0.0011219 | 0.37409725 | -0.0029988 | 0.99760729 | 0.99948864 |
| APBB2 | 637.98181 | -0.5005399 | 0.23419171 | -2.1373082 | 0.03257293 | 0.48564518 |
| LIMCH1 | 220.768193 | 1.64535508 | 0.36153745 | 4.55099485 | 5.34E-06 | 0.00230814 |
| TMEM33 | 31.7537274 | -0.1254391 | 0.30810983 | -0.4071246 | 0.68391648 | 0.9958918 |
| SLC30A9 | 96.8995101 | -0.1721058 | 0.21924135 | -0.7850061 | 0.43244996 | 0.98092839 |
| ATP8A1 | 28.9895993 | 1.06163855 | 0.47190814 | 2.24967203 | 0.02446977 | 0.42312555 |
| GUF1 | 20.1122834 | -0.5962356 | 0.34566583 | -1.7248903 | 0.08454724 | 0.7052641 |
| GNPDA2 | 20.4179042 | 0.15218062 | 0.3614105 | 0.42107415 | 0.67370094 | 0.9958918 |

| | | | | | | |
|---|---|---|---|---|---|---|
| ATP10D | 221.37537 | -1.25641 | 0.49904384 | -2.5176345 | 0.01181459 | 0.29332902 |
| NFXL1 | 42.306242 | -0.4764721 | 0.30674658 | -1.5533086 | 0.12034943 | 0.79021701 |
| SLAIN2 | 87.0264252 | 0.1709851 | 0.23891341 | 0.71567814 | 0.47419011 | 0.98910524 |
| FRYL | 253.566736 | 0.0461803 | 0.23481836 | 0.19666394 | 0.84409053 | 0.99692952 |
| OCIAD1 | 91.8141286 | -0.0727519 | 0.18146932 | -0.4009045 | 0.68849044 | 0.9958918 |
| DCUN1D4 | 89.7988576 | -0.2671227 | 0.30641807 | -0.8717591 | 0.3833398 | 0.9725044 |
| SGCB | 22.9637407 | 0.43147886 | 0.3073068 | 1.40406546 | 0.16029936 | 0.84995669 |
| USP46 | 34.0775796 | 0.35693687 | 0.30262138 | 1.17948334 | 0.23820577 | 0.92291633 |
| SCFD2 | 242.132908 | -0.2192298 | 0.38832227 | -0.5645562 | 0.57237564 | 0.9958918 |
| AC058822.1 | 70.0058609 | -0.4464903 | 0.49030281 | -0.9106419 | 0.3624841 | 0.97217293 |
| CHIC2 | 198.309748 | -0.4634675 | 0.30292354 | -1.5299818 | 0.12602122 | 0.79527093 |
| PDGFRA | 115.735557 | -0.6557894 | 0.42595802 | -1.5395634 | 0.12366682 | 0.79184923 |
| TMEM165 | 228.290094 | 0.00660928 | 0.2919419 | 0.02263903 | 0.98193821 | 0.99935941 |
| CLOCK | 104.828176 | -0.1678635 | 0.24893578 | -0.6743245 | 0.50010506 | 0.99213111 |
| EXOC1 | 85.1098453 | -0.1700187 | 0.29079726 | -0.584664 | 0.55877363 | 0.9958918 |
| CEP135 | 24.15904 | -0.6414087 | 0.39421501 | -1.627053 | 0.10372584 | 0.75827058 |
| KIAA1211 | 60.2532575 | 0.10277985 | 0.4718634 | 0.21781696 | 0.82757173 | 0.99592423 |
| AASDH | 24.2052622 | -0.1535555 | 0.3757969 | -0.4086131 | 0.6828236 | 0.9958918 |
| PAICS | 20.9888511 | -0.3096142 | 0.41073141 | -0.7538117 | 0.45096227 | 0.98092839 |
| SRP72 | 52.3154852 | 0.40149334 | 0.24252369 | 1.65548093 | 0.09782698 | 0.74364783 |
| REST | 44.3016171 | 0.53269029 | 0.27047864 | 1.96943572 | 0.04890308 | 0.57273715 |
| POLR2B | 63.0960418 | -0.0770239 | 0.22134584 | -0.3479798 | 0.72785534 | 0.9958918 |
| IGFBP7 | 1910.15949 | -0.1424451 | 0.31816963 | -0.4477017 | 0.65436848 | 0.9958918 |
| ADGRL3 | 239.982964 | 0.40019396 | 0.46169143 | 0.86679964 | 3.86E-01 | 0.97254921 |
| AC104806.2 | 18.6212461 | 0.81974921 | 0.38292066 | 2.14078084 | 0.03229172 | 0.48564518 |
| CENPC | 82.2272387 | -0.4607283 | 0.37863162 | -1.2168246 | 0.22367097 | 0.91093521 |
| UBA6 | 66.8477131 | -0.1183153 | 0.24399803 | -0.4849027 | 0.62774538 | 0.9958918 |
| UBA6-AS1 | 64.4347932 | -0.9091234 | 0.42752659 | -2.1264722 | 0.03346396 | 0.492839 |
| YTHDC1 | 109.586401 | -0.3410721 | 0.28816022 | -1.1836197 | 0.23656365 | 0.92093871 |
| UGT2B7 | 40.7284427 | 1.03724246 | 0.54858737 | 1.89075161 | 0.05865751 | 0.6110796 |
| RUFY3 | 190.81554 | -0.617388 | 0.23218399 | -2.6590466 | 0.00783621 | 0.23127843 |
| GRSF1 | 32.4994095 | 0.2667328 | 0.28511247 | 0.93553537 | 0.34951246 | 0.9692706 |
| MOB1B | 63.6436572 | 0.27771253 | 0.25162292 | 1.10368536 | 0.26972965 | 0.93634106 |
| SLC4A4 | 40.4025934 | 0.63659418 | 0.43849305 | 1.45177712 | 0.14656359 | 0.8301633 |
| ADAMTS3 | 86.1280166 | -1.2022021 | 0.61182856 | -1.9649329 | 0.049422 | 0.57273715 |
| ANKRD17 | 442.108863 | -0.1797939 | 0.24553109 | -0.7322652 | 0.46400673 | 0.98590351 |
| AC053527.1 | 44.2361411 | 0.76029113 | 0.45977229 | 1.65362538 | 0.09820365 | 0.74364783 |
| MTHFD2L | 30.7985141 | 0.24574936 | 0.38495095 | 0.63839138 | 0.52321894 | 0.9958918 |
| PARM1 | 152.008614 | -0.4839838 | 0.33801957 | -1.431822 | 0.15219478 | 0.83906933 |
| RCHY1 | 35.019607 | -0.0278393 | 0.34200511 | -0.0814003 | 0.93512364 | 0.99935941 |
| THAP6 | 22.1804245 | -0.3265241 | 0.37794161 | -0.8639538 | 0.38761328 | 0.97297628 |
| G3BP2 | 61.4676322 | -0.3116937 | 0.35967427 | -0.8666 | 0.38616119 | 0.97254921 |
| USO1 | 146.196261 | -0.0817781 | 0.24958843 | -0.327652 | 0.74317482 | 0.9958918 |
| SDAD1 | 36.653819 | 0.13878354 | 0.34598907 | 0.40112117 | 0.68833091 | 0.9958918 |
| NUP54 | 48.2818514 | -0.1153263 | 0.23839347 | -0.4837645 | 0.62855299 | 0.9958918 |
| SCARB2 | 106.951043 | -0.190948 | 0.27651552 | -0.6905506 | 0.48984798 | 0.99046213 |
| SHROOM3 | 43.212386 | -0.3050341 | 0.40819731 | -0.7472712 | 0.45489985 | 0.98173285 |
| 11-Sep | 237.575939 | -1.1200432 | 0.38035696 | -2.9447159 | 0.00323251 | 0.13652725 |
| CCNI | 164.680126 | 0.28960893 | 0.19203079 | 1.50813794 | 0.13151923 | 0.80565822 |
| CCNG2 | 34.1558955 | -0.5689155 | 0.42262342 | -1.3461524 | 0.17825336 | 0.87815793 |
| CNOT6L | 91.2786195 | 0.12025424 | 0.27598701 | 0.43572427 | 0.6630368 | 0.9958918 |
| MRPL1 | 46.5731475 | 0.09078143 | 0.26323325 | 0.34487069 | 0.73019159 | 0.9958918 |
| FRAS1 | 103.928583 | 0.66470625 | 0.51347745 | 1.29451888 | 0.19548625 | 0.88391893 |
| BMP2K | 77.2267029 | 0.02874964 | 0.32379674 | 0.08878917 | 0.92924946 | 0.99935941 |

| | | | | | | |
|---|---|---|---|---|---|---|
| LINC01088 | 72.8583436 | -1.8021961 | 0.84428162 | -2.1345911 | 0.03279442 | 0.48688124 |
| LINC00989 | 48.5119955 | 0.5145774 | 0.58486055 | 0.87982922 | 0.37895183 | 0.97217293 |
| ANTXR2 | 210.575861 | -0.3944649 | 0.36757134 | -1.0731655 | 0.28319687 | 0.94471801 |
| RASGEF1B | 422.560747 | -0.7930184 | 0.58278136 | -1.3607477 | 0.17359343 | 0.87233975 |
| HNRNPD | 120.899062 | -0.0690715 | 0.22153035 | -0.3117926 | 0.75519815 | 0.9958918 |
| HNRNPDL | 124.65825 | -0.3437666 | 0.23074862 | -1.4897883 | 0.13627991 | 0.81177777 |
| ENOPH1 | 18.32872 | 0.25649523 | 0.37451636 | 0.68487055 | 0.49342561 | 0.99116088 |
| SEC31A | 249.840564 | -0.3248117 | 0.19082128 | -1.7021773 | 0.08872213 | 0.71818729 |
| LIN54 | 26.6978516 | -0.0676481 | 0.34746778 | -0.1946888 | 0.84563655 | 0.99692952 |
| COPS4 | 30.331578 | 0.38571441 | 0.3253978 | 1.18536268 | 0.23587408 | 0.91962002 |
| HELQ | 30.1001768 | 0.29431152 | 0.35506912 | 0.82888514 | 0.40716941 | 0.97691413 |
| WDFY3 | 370.382976 | -0.2359076 | 0.23933414 | -0.9856831 | 0.32428864 | 0.95992644 |
| ARHGAP24 | 418.117375 | -1.5393502 | 0.35478276 | -4.338853 | 1.43E-05 | 0.0040484 |
| MAPK10 | 487.012754 | -0.4992901 | 0.32536415 | -1.5345579 | 0.12489247 | 0.79195137 |
| PTPN13 | 133.644719 | 0.31845231 | 0.22358788 | 1.42428256 | 0.15436469 | 0.84218718 |
| AFF1 | 455.888629 | 0.3464797 | 0.2716558 | 1.2754364 | 0.20215482 | 0.89319989 |
| KLHL8 | 34.0368171 | 0.17929689 | 0.29200346 | 0.61402316 | 0.53920002 | 0.9958918 |
| HSD17B11 | 34.8691408 | 0.25836511 | 0.31473183 | 0.82090554 | 0.41170007 | 0.97812179 |
| NUDT9 | 32.4622591 | -0.0550608 | 0.29102886 | -0.1891935 | 0.84994118 | 0.9971731 |
| SPARCL1 | 317.002507 | 0.85264822 | 0.41106953 | 2.07421895 | 0.03805898 | 0.52280126 |
| SPP1 | 1102.4676 | 0.23513679 | 0.5562546 | 0.42271432 | 6.73E-01 | 9.96E-01 |
| PKD2 | 229.514804 | -0.0287713 | 0.27765446 | -0.1036226 | 0.91746883 | 0.99935941 |
| PPM1K | 53.2177319 | -0.280399 | 0.49034172 | -0.571844 | 0.56742765 | 0.9958918 |
| HERC3 | 33.770327 | -0.1139216 | 0.3706382 | -0.3073659 | 0.75856486 | 0.9958918 |
| FAM13A | 393.092802 | -0.8068894 | 0.57695306 | -1.3985356 | 0.16195229 | 0.85327904 |
| GPRIN3 | 107.161157 | 0.52315155 | 0.45693783 | 1.14490751 | 0.25224747 | 0.92631316 |
| CCSER1 | 79.7050674 | -0.1909122 | 0.35627283 | -0.5358595 | 0.59205565 | 0.9958918 |
| GRID2 | 142.894688 | -0.4435987 | 0.55351716 | -0.8014181 | 0.42288962 | 0.97828057 |
| SMARCAD1 | 76.8174723 | -0.5123035 | 0.22217993 | -2.3058045 | 0.02112156 | 0.39000584 |
| PDLIM5 | 1083.22498 | -0.6342437 | 0.52246231 | -1.213951 | 0.22476643 | 0.91093521 |
| BMPR1B | 234.591233 | -2.6473635 | 0.59717566 | -4.4331403 | 9.29E-06 | 0.00341252 |
| UNC5C | 363.087398 | 0.13900498 | 0.22177819 | 0.62677479 | 0.53080687 | 0.9958918 |
| STPG2 | 25.533253 | -1.5330708 | 0.52315476 | -2.9304347 | 0.00338488 | 0.14115602 |
| RAP1GDS1 | 90.238988 | 0.47378708 | 0.24151667 | 1.96171585 | 0.04979558 | 0.57344369 |
| TSPAN5 | 33.5701642 | 0.16098574 | 0.40247376 | 0.39999065 | 0.68916341 | 0.9958918 |
| EIF4E | 54.618297 | -0.0648724 | 0.35183027 | -0.1843854 | 0.85371111 | 0.9971731 |
| METAP1 | 33.4406981 | -0.1379307 | 0.31469137 | -0.4383047 | 0.66116544 | 0.9958918 |
| ADH5 | 53.4681603 | -0.458891 | 0.24400225 | -1.8806833 | 0.06001501 | 0.61332974 |
| AP002026.1 | 39.8856383 | -0.6013219 | 0.50432118 | -1.1923391 | 0.23312829 | 0.91861399 |
| ADH1B | 223.696121 | -0.878236 | 0.62729201 | -1.4000433 | 0.16150035 | 0.85202157 |
| DNAJB14 | 94.4744474 | -0.1219086 | 0.25992186 | -0.46902 | 0.63905531 | 0.9958918 |
| H2AFZ | 26.3820904 | 0.0316837 | 0.45027774 | 0.07036479 | 0.94390331 | 0.99935941 |
| EMCN | 246.548398 | 0.28582537 | 0.4824649 | 0.59242727 | 0.55356451 | 0.9958918 |
| PPP3CA | 238.317636 | -0.0802803 | 0.28811281 | -0.2786418 | 0.78051975 | 0.9958918 |
| NFKB1 | 199.786231 | 0.54528026 | 0.46824665 | 1.16451501 | 0.24421536 | 0.92546331 |
| MANBA | 41.7913206 | 0.05157233 | 0.33067136 | 0.15596251 | 0.87606257 | 0.9971731 |
| UBE2D3 | 254.014758 | -0.0874247 | 0.21435878 | -0.4078427 | 0.6833892 | 0.9958918 |
| SLC9B2 | 20.0773288 | 0.00778634 | 0.46326308 | 0.0168076 | 0.98659011 | 0.99935941 |
| TET2 | 121.046445 | 0.16017808 | 0.31775082 | 0.50409965 | 0.61419135 | 0.9958918 |
| PPA2 | 109.648075 | 0.07062063 | 0.22238974 | 0.31755346 | 0.75082369 | 0.9958918 |
| INTS12 | 24.6312189 | 0.36921357 | 0.36109699 | 1.02247756 | 0.30655493 | 0.95793395 |
| GSTCD | 30.7233556 | -0.1702381 | 0.43587701 | -0.3905645 | 0.6961192 | 0.9958918 |
| NPNT | 46.4843662 | 0.0838567 | 0.43252762 | 0.19387595 | 0.84627301 | 0.99692952 |
| TBCK | 193.257244 | 0.14900495 | 0.33404913 | 0.44605697 | 0.65555609 | 0.9958918 |

| | | | | | | |
|---|---|---|---|---|---|---|
| AIMP1 | 27.0203592 | -0.2738975 | 0.29111231 | -0.9408655 | 0.34677379 | 0.96887708 |
| PAPSS1 | 41.17637 | -0.1959116 | 0.2834638 | -0.6911346 | 0.48948098 | 0.99046213 |
| SGMS2 | 29.7355727 | -1.2250563 | 0.42308733 | -2.8955163 | 0.00378535 | 0.15284924 |
| RPL34 | 71.1541163 | -0.2433003 | 0.38445734 | -0.6328408 | 0.52683764 | 0.9958918 |
| OSTC | 35.3564487 | 0.2212711 | 0.36433421 | 0.60733001 | 0.54363193 | 0.9958918 |
| COL25A1 | 405.458717 | -1.8261512 | 0.4633384 | -3.9412904 | 8.10E-05 | 0.01398756 |
| SEC24B | 147.676733 | -0.0735929 | 0.23979784 | -0.3068955 | 0.75892295 | 0.9958918 |
| EGF | 22.6648697 | -0.4068735 | 0.61045976 | -0.6665034 | 5.05E-01 | 0.99397937 |
| ELOVL6 | 30.0483557 | -0.109741 | 0.3817616 | -0.2874596 | 0.77376042 | 0.9958918 |
| ENPEP | 58.7111477 | 0.01351498 | 0.53011897 | 0.02549423 | 0.97966075 | 0.99935941 |
| FAM241A | 58.2010666 | 0.61159148 | 0.26488747 | 2.30887278 | 0.02095064 | 0.39000584 |
| AP1AR | 24.4391352 | -0.239142 | 0.40578599 | -0.5893303 | 0.5556397 | 0.9958918 |
| ALPK1 | 110.410296 | 0.03653517 | 0.39562336 | 0.09234835 | 0.92642127 | 0.99935941 |
| LARP7 | 37.294761 | -0.1501111 | 0.3723977 | -0.4030934 | 0.6868795 | 0.9958918 |
| ANK2 | 143.195184 | -0.6584712 | 0.28530275 | -2.3079737 | 0.0210006 | 0.39000584 |
| CAMK2D | 460.480854 | 0.08801433 | 0.25913589 | 0.33964547 | 0.73412353 | 0.9958918 |
| ARSJ | 38.6331253 | 0.40741895 | 0.37152815 | 1.09660319 | 0.27281489 | 0.93819216 |
| METTL14 | 36.5994459 | 0.02447432 | 0.31770292 | 0.07703525 | 0.9385955 | 0.99935941 |
| SEC24D | 233.097771 | -0.1951794 | 0.21233997 | -0.9191836 | 0.35799956 | 0.97217293 |
| SYNPO2 | 519.259071 | -0.4138834 | 0.54951135 | -0.7531845 | 0.45133904 | 0.98092839 |
| USP53 | 809.638375 | -0.9178021 | 0.45808336 | -2.00357 | 0.04511614 | 0.55894748 |
| C4orf3 | 59.8393454 | 0.1345459 | 0.33951378 | 0.39628996 | 0.69189113 | 0.9958918 |
| PDE5A | 211.042852 | 0.37107292 | 0.42801299 | 0.86696648 | 0.38596037 | 0.97254921 |
| AC108866.1 | 45.5451736 | -0.2248179 | 0.4072823 | -0.5519952 | 0.58095165 | 0.9958918 |
| PRDM5 | 53.4974371 | -0.5376394 | 0.28214378 | -1.905551 | 0.05670848 | 0.60440467 |
| ANXA5 | 131.057731 | -0.2979284 | 0.35012803 | -0.8509128 | 0.39481779 | 0.97500135 |
| BBS7 | 27.9739699 | -0.2645787 | 0.37350095 | -0.708375 | 0.47871244 | 0.99002719 |
| KIAA1109 | 223.661663 | -0.0026356 | 0.24708841 | -0.0106667 | 0.99148939 | 0.99935941 |
| FGF2 | 41.2346712 | -0.5264269 | 0.31572014 | -1.6673845 | 0.09543797 | 0.73751173 |
| SPATA5 | 120.548999 | -0.1175026 | 0.23834636 | -0.4929911 | 0.62201884 | 0.9958918 |
| SPRY1 | 56.9612654 | -0.4489487 | 0.39391261 | -1.1397166 | 0.25440438 | 0.92647067 |
| ANKRD50 | 31.5540194 | 0.35821975 | 0.31603017 | 1.13349857 | 0.25700494 | 0.9266065 |
| FAT4 | 101.205724 | -1.6263544 | 0.50022958 | -3.251216 | 0.00114913 | 0.07280105 |
| INTU | 40.4784075 | -0.8887354 | 0.27325913 | -3.252354 | 0.00114453 | 0.07280105 |
| HSPA4L | 21.7559057 | -0.3042841 | 0.39880402 | -0.7629915 | 0.44546847 | 0.98092839 |
| MFSD8 | 28.8269794 | 0.06964165 | 0.32499263 | 0.21428688 | 0.83032335 | 0.99592423 |
| ABHD18 | 62.1316605 | -0.0204311 | 0.291175 | -0.0701677 | 0.94406017 | 0.99935941 |
| LARP1B | 88.9166991 | 0.41382462 | 0.25209075 | 1.64157003 | 0.10067914 | 0.75064053 |
| PGRMC2 | 42.2165553 | -0.5370488 | 0.24343592 | -2.2061199 | 0.02737561 | 0.44841843 |
| JADE1 | 65.5040745 | -0.5733213 | 0.44306489 | -1.2939894 | 0.1956691 | 0.88391893 |
| SCLT1 | 117.035393 | -0.2364236 | 0.29437227 | -0.8031449 | 0.421891 | 0.97812179 |
| AC096711.2 | 35.0525699 | -0.004417 | 0.30704582 | -0.0143856 | 0.98852236 | 0.99935941 |
| PCDH10 | 24.2304989 | 0.53109428 | 0.59162476 | 0.89768772 | 0.36935206 | 0.97217293 |
| PCDH18 | 41.0646489 | -1.3459546 | 0.42622523 | -3.1578483 | 0.00158938 | 0.08716696 |
| AC109927.1 | 33.1206058 | -0.2507316 | 0.38407427 | -0.6528205 | 0.513872 | 0.9958918 |
| ELF2 | 299.627939 | 0.05003167 | 0.22553706 | 0.22183347 | 0.82444352 | 0.99592423 |
| NDUFC1 | 20.6173504 | 0.2063453 | 0.35317557 | 0.584257 | 0.55904742 | 0.9958918 |
| NAA15 | 106.027231 | -0.3939918 | 0.26333749 | -1.4961477 | 0.13461517 | 0.80890178 |
| AC097376.2 | 51.9638552 | 0.28938423 | 0.37536024 | 0.77095067 | 0.44073617 | 0.98092839 |
| SETD7 | 44.0523882 | -0.0156811 | 0.276517 | -0.0567093 | 0.95477679 | 0.99935941 |
| MGST2 | 22.506842 | 0.86478648 | 0.36590826 | 2.36339701 | 0.01810826 | 0.35723014 |
| MAML3 | 407.456605 | 0.52713141 | 0.31374099 | 1.68014836 | 0.09292845 | 0.73019523 |
| SCOC | 26.5871006 | 0.51637176 | 0.32552454 | 1.58627598 | 0.11267672 | 0.77793929 |
| ELMOD2 | 20.2677222 | 0.11526048 | 0.3637168 | 0.31689623 | 0.75132234 | 0.9958918 |

| | | | | | | |
|---|---|---|---|---|---|---|
| TBC1D9 | 33.1963129 | 0.23362672 | 0.28864227 | 0.80939883 | 0.41828578 | 0.97812179 |
| RNF150 | 179.480995 | 0.00172227 | 0.43218187 | 0.00398506 | 0.99682039 | 0.99935941 |
| ZNF330 | 21.1524762 | -0.9076029 | 0.37877097 | -2.3961787 | 0.01656701 | 0.34578631 |
| IL15 | 24.6853322 | 0.11475844 | 0.46737966 | 0.24553581 | 0.80604158 | 0.99592423 |
| INPP4B | 996.410237 | -0.2887738 | 0.44769181 | -0.645028 | 0.51890903 | 0.9958918 |
| AC139720.1 | 37.613256 | -0.766832 | 0.56539989 | -1.3562648 | 0.17501492 | 0.8740623 |
| AC104596.1 | 29.6407729 | -0.3834655 | 0.67602662 | -0.5672343 | 0.57055498 | 0.9958918 |
| USP38 | 27.9081145 | 0.03976441 | 0.36122691 | 0.11008154 | 0.91234471 | 0.99935941 |
| GAB1 | 214.148124 | -0.1617762 | 0.38359109 | -0.4217412 | 0.67321395 | 0.9958918 |
| SMARCA5 | 77.5582624 | 0.09785966 | 0.21893468 | 0.44698109 | 0.65488872 | 0.9958918 |
| ANAPC10 | 63.0269305 | 0.07076007 | 0.25065885 | 0.28229631 | 0.77771631 | 0.9958918 |
| ABCE1 | 43.9595174 | -0.2281065 | 0.40332677 | -0.5655626 | 0.57169116 | 0.9958918 |
| OTUD4 | 62.2447407 | -0.3554805 | 0.25430344 | -1.3978596 | 0.16215523 | 0.85363812 |
| SMAD1 | 39.5182871 | -0.3105103 | 0.37267529 | -0.8331927 | 0.40473607 | 0.97582411 |
| ZNF827 | 41.4280617 | 0.20231377 | 0.38777629 | 0.52172806 | 0.60185969 | 0.9958918 |
| SLC10A7 | 100.075611 | 0.09684217 | 0.28729049 | 0.33708798 | 0.73605058 | 0.9958918 |
| EDNRA | 178.228225 | 0.1254224 | 0.31337892 | 0.40022604 | 0.68899004 | 0.9958918 |
| TMEM184C | 26.1816857 | 0.24705678 | 0.38511415 | 0.6415157 | 0.52118768 | 0.9958918 |
| PRMT9 | 32.0977589 | 0.1013003 | 0.27880993 | 0.36333102 | 0.71635762 | 0.9958918 |
| ARHGAP10 | 412.0294 | -0.7616394 | 0.42177516 | -1.8057947 | 0.07095041 | 0.65918401 |
| NR3C2 | 189.967598 | 0.61995144 | 0.3963699 | 1.56407293 | 0.11780044 | 0.78539225 |
| DCLK2 | 66.0586197 | 0.20650553 | 0.38878527 | 0.53115575 | 0.59531085 | 0.9958918 |
| LRBA | 246.834237 | -0.2723494 | 0.30383042 | -0.8963864 | 0.37004645 | 0.97217293 |
| RPS3A | 58.6750758 | 0.12618554 | 0.42723691 | 0.29535262 | 0.76772452 | 0.9958918 |
| SH3D19 | 324.040125 | -0.3504806 | 0.33638654 | -1.0418985 | 0.29745872 | 0.95417903 |
| FBXW7 | 193.807535 | -0.0070278 | 0.21129628 | -0.0332606 | 0.9734668 | 0.99935941 |
| ARFIP1 | 99.0360953 | -0.0736698 | 0.19062767 | -0.386459 | 0.69915679 | 0.9958918 |
| TRIM2 | 76.1992566 | -0.0844086 | 0.33784899 | -0.2498411 | 0.80271023 | 0.99592423 |
| TMEM131L | 244.533803 | 0.19919466 | 0.24186782 | 0.82356829 | 0.4101849 | 0.97812179 |
| MAP9 | 19.9655022 | -0.196464 | 0.41515804 | -0.473227 | 0.63605123 | 0.9958918 |
| GUCY1A1 | 180.78896 | 0.18460123 | 0.30830617 | 0.59875944 | 0.54933331 | 0.9958918 |
| GUCY1B1 | 76.7048906 | 0.06094582 | 0.2177353 | 0.27990783 | 0.77954822 | 0.9958918 |
| CTSO | 19.9662865 | 0.04078081 | 0.44088647 | 0.09249731 | 0.92630293 | 0.99935941 |
| PDGFC | 150.877233 | 0.58704099 | 0.21030095 | 2.79143294 | 0.00524752 | 0.18276799 |
| FAM198B | 39.1040887 | -0.2107871 | 0.53760809 | -0.3920831 | 0.69499679 | 0.9958918 |
| RXFP1 | 89.1633287 | -0.5826331 | 0.93387677 | -0.6238865 | 0.53270211 | 0.9958918 |
| ETFDH | 59.8232865 | -0.1323205 | 0.2862709 | -0.4622213 | 0.6439226 | 0.9958918 |
| PPID | 21.0894811 | 0.31777949 | 0.38732428 | 0.82044815 | 0.41196067 | 0.97812179 |
| FNIP2 | 170.319061 | -0.2396778 | 0.30504055 | -0.7857244 | 0.43202897 | 0.98092839 |
| RAPGEF2 | 328.850853 | 0.16898044 | 0.25170917 | 0.67133205 | 0.50200902 | 0.99280525 |
| NAF1 | 64.4424582 | -0.1125582 | 0.26482828 | -0.4250234 | 0.67081963 | 0.9958918 |
| NPY1R | 183.411781 | -0.340357 | 0.43790595 | -0.7772376 | 0.43701862 | 0.98092839 |
| TMA16 | 47.7885449 | -0.4383129 | 0.31879962 | -1.3748852 | 0.16916703 | 0.86755653 |
| TMEM192 | 28.4239099 | 0.13374489 | 0.45621691 | 0.29316075 | 0.76939928 | 0.9958918 |
| KLHL2 | 56.5580764 | 0.4525606 | 0.34577672 | 1.30882323 | 0.19059424 | 0.88158763 |
| MSMO1 | 21.4230523 | -0.1145223 | 0.47787863 | -0.2396473 | 0.81060368 | 0.99592423 |
| CPE | 177.920055 | -1.2002932 | 0.4145125 | -2.8956743 | 0.00378345 | 0.15284924 |
| DDX60 | 62.8195146 | -0.7735603 | 1.01609089 | -0.7613102 | 0.44647182 | 0.98092839 |
| DDX60L | 49.9782251 | -2.3114028 | 1.14333575 | -2.0216308 | 0.04321451 | 0.55231899 |
| PALLD | 770.71159 | -1.0659644 | 0.5736188 | -1.858315 | 0.06312429 | 0.62352207 |
| CBR4 | 138.652211 | -0.268812 | 0.39038726 | -0.6885779 | 0.49108895 | 0.99046213 |
| SH3RF1 | 178.439034 | 0.05321905 | 0.18212643 | 0.29220938 | 0.77012654 | 0.9958918 |
| NEK1 | 118.538298 | -0.1056256 | 0.31744777 | -0.3327337 | 0.73933531 | 0.9958918 |
| CLCN3 | 67.0782847 | 0.21376298 | 0.23924989 | 0.8934716 | 0.37160468 | 0.97217293 |

| | | | | | | |
|---|---|---|---|---|---|---|
| GALNT7 | 39.9332896 | 0.32676628 | 0.28151439 | 1.16074451 | 0.24574582 | 0.92559078 |
| CEP44 | 31.7697381 | 0.04317448 | 0.34667335 | 0.12453936 | 0.90088824 | 0.99935941 |
| SPCS3 | 62.170157 | 0.03156434 | 0.32284522 | 0.09776926 | 0.92211552 | 0.99935941 |
| TENM3 | 338.450474 | -0.1577463 | 0.55247367 | -0.2855273 | 0.77524023 | 0.9958918 |
| DCTD | 22.6252681 | 0.29106557 | 0.37840377 | 0.7691931 | 0.44177869 | 0.98092839 |
| WWC2 | 187.295672 | -0.0572132 | 0.21195223 | -0.2699343 | 0.7872108 | 0.99592423 |
| TRAPPC11 | 47.4112564 | 0.00386074 | 0.25219086 | 0.01530881 | 0.98778581 | 0.99935941 |
| STOX2 | 26.7692407 | 0.95705059 | 0.49350694 | 1.93928495 | 0.05246665 | 0.583162 |
| IRF2 | 81.5886875 | 0.40130197 | 0.31143967 | 1.2885384 | 0.19755861 | 0.88473993 |
| PRIMPOL | 28.1345289 | -0.1453099 | 0.34416462 | -0.4222105 | 0.67287135 | 0.9958918 |
| ACSL1 | 60.7584557 | -0.0467281 | 0.46891704 | -0.0996511 | 0.92062136 | 0.99935941 |
| SLC25A4 | 30.632015 | 0.3010707 | 0.49603613 | 0.60695317 | 0.54388199 | 0.9958918 |
| CFAP97 | 42.4405856 | 0.335056 | 0.2784101 | 1.2034621 | 0.22879755 | 0.91382239 |
| SNX25 | 134.165087 | -0.1555139 | 0.35568206 | -0.4372274 | 0.66194643 | 0.9958918 |
| ANKRD37 | 65.1129243 | -3.1360599 | 1.29159101 | -2.4280596 | 0.01517985 | 0.33500515 |
| UFSP2 | 34.506376 | -0.1817494 | 0.40629492 | -0.4473337 | 0.6546341 | 0.9958918 |
| PDLIM3 | 158.444204 | -0.3123465 | 0.61181132 | -0.5105275 | 0.60968199 | 0.9958918 |
| SORBS2 | 310.775924 | 0.18141813 | 0.37494162 | 0.48385701 | 0.62848735 | 0.9958918 |
| CYP4V2 | 21.6202113 | 0.39522139 | 0.50397398 | 0.78420992 | 0.43291695 | 0.98092839 |
| FAT1 | 76.5103005 | 0.49245318 | 0.37831289 | 1.30170869 | 0.19301599 | 0.88158763 |
| SDHA | 58.6444658 | 0.08819789 | 0.30041633 | 0.29358553 | 0.76907463 | 0.9958918 |
| PDCD6 | 28.2800477 | 0.15034233 | 0.39845544 | 0.37731278 | 0.70594118 | 0.9958918 |
| EXOC3 | 27.190887 | 0.07916313 | 0.39004926 | 0.20295675 | 0.83916884 | 0.99692952 |
| BRD9 | 19.5285195 | -0.2125291 | 0.46682251 | -0.4552676 | 0.64891676 | 0.9958918 |
| SLC12A7 | 30.2834026 | -0.3488094 | 0.38093314 | -0.915671 | 0.35983948 | 0.97217293 |
| CLPTM1L | 26.9432596 | 0.03643432 | 0.42568633 | 0.08558959 | 0.93179268 | 0.99935941 |
| LPCAT1 | 32.2177733 | -0.1192321 | 0.30401922 | -0.3921862 | 0.69492063 | 0.9958918 |
| NDUFS6 | 30.8767898 | 0.275172 | 0.34969211 | 0.78689794 | 0.43134162 | 0.98092839 |
| ICE1 | 67.9144661 | -0.0239987 | 0.22303805 | -0.1075992 | 0.91431365 | 0.99935941 |
| MED10 | 20.3091265 | 0.28202227 | 0.30947053 | 0.91130573 | 0.36213431 | 0.97217293 |
| NSUN2 | 32.5874183 | -0.2434456 | 0.36659198 | -0.6640778 | 0.50664051 | 0.99397937 |
| TENT4A | 62.4195111 | -0.1052949 | 0.25511013 | -0.4127428 | 0.67979503 | 0.9958918 |
| ADCY2 | 34.8886628 | -0.8100489 | 0.42627462 | -1.9002981 | 0.05739401 | 0.60512745 |
| MTRR | 25.2407753 | -0.5235194 | 0.34989894 | -1.4962019 | 0.13460106 | 0.80890178 |
| SEMA5A | 199.169606 | -0.489455 | 0.25577818 | -1.9135916 | 0.05567236 | 0.59815227 |
| CCT5 | 35.1888124 | 0.1297833 | 0.42553406 | 0.30498921 | 0.76037437 | 0.9958918 |
| CMBL | 28.5756847 | 2.52166646 | 0.74163129 | 3.40016193 | 0.00067346 | 0.05624153 |
| 6-Mar | 164.963235 | -0.2480995 | 0.24349401 | -1.0189143 | 0.30824366 | 0.95793395 |
| DAP | 48.5026282 | -0.2080527 | 0.30832435 | -0.6747852 | 0.49981225 | 0.99213111 |
| TRIO | 432.826315 | -0.2519402 | 0.26025901 | -0.9680363 | 0.33302626 | 0.9654289 |
| OTULIN | 27.4825798 | 0.36500355 | 0.37968166 | 0.96134102 | 0.33638073 | 0.96640422 |
| ANKH | 122.880523 | -0.1932091 | 0.39095692 | -0.4941953 | 0.62116823 | 0.9958918 |
| FBXL7 | 649.338445 | -0.5262987 | 0.28169543 | -1.8683251 | 0.06171677 | 0.61962951 |
| RETREG1 | 28.5628923 | 0.49651847 | 0.42428216 | 1.17025535 | 2.42E-01 | 0.92516122 |
| MYO10 | 217.943973 | -0.3155318 | 0.34908225 | -0.9038896 | 0.36605394 | 0.97217293 |
| BASP1 | 79.5269008 | -0.1395447 | 0.39503553 | -0.3532459 | 0.72390413 | 0.9958918 |
| CDH6 | 45.1466468 | -0.5060548 | 0.37701259 | -1.3422755 | 0.17950668 | 0.87815793 |
| DROSHA | 49.7677034 | 0.00818022 | 0.22934475 | 0.03566781 | 0.97154724 | 0.99935941 |
| PDZD2 | 168.857476 | -1.6051545 | 0.42801441 | -3.7502348 | 0.00017667 | 0.02163901 |
| GOLPH3 | 137.740805 | -0.0575308 | 0.2948849 | -0.1950958 | 0.84531798 | 0.99692952 |
| MTMR12 | 44.1742263 | 0.13589623 | 0.28503575 | 0.47676907 | 0.63352657 | 0.9958918 |
| ZFR | 181.507839 | 0.12770333 | 0.25292831 | 0.5048993 | 0.61362957 | 0.9958918 |
| SUB1 | 75.3661983 | -0.0232927 | 0.31847465 | -0.0731383 | 0.94169604 | 0.99935941 |
| NPR3 | 35.5745951 | 1.80443273 | 0.65052385 | 2.77381487 | 0.00554032 | 0.18713567 |

| | | | | | | |
|---|---|---|---|---|---|---|
| TARS | 30.8908305 | -0.3917674 | 0.373931 | -1.0476996 | 0.29477704 | 0.95304825 |
| ADAMTS12 | 172.445968 | -0.4655277 | 0.53968622 | -0.8625895 | 0.38836322 | 0.97297628 |
| C1QTNF3-AM | 45.675961 | -0.2185367 | 0.46118639 | -0.4738577 | 0.63560141 | 0.9958918 |
| RAI14 | 172.313259 | -0.0582517 | 0.36344691 | -0.1602757 | 0.8726639 | 0.9971731 |
| DNAJC21 | 31.2315542 | 0.21142264 | 0.34451545 | 0.61368116 | 0.53942604 | 0.9958918 |
| SPEF2 | 29.1958621 | -0.212623 | 0.43677834 | -0.4867983 | 0.62640125 | 0.9958918 |
| LMBRD2 | 27.2007209 | 0.00964915 | 0.36342786 | 0.02655037 | 0.97881836 | 0.99935941 |
| NADK2 | 42.9604945 | -0.4368044 | 0.39396499 | -1.1087391 | 0.26754273 | 0.93549101 |
| RANBP3L | 95.3120839 | -2.0392181 | 0.97372935 | -2.094235 | 0.03623905 | 0.51215534 |
| NIPBL | 402.202906 | -0.098958 | 0.21569436 | -0.4587881 | 0.64638631 | 0.9958918 |
| CPLANE1 | 117.34332 | -0.2411624 | 0.249393 | -0.9669976 | 0.33354524 | 0.9654289 |
| NUP155 | 25.496579 | 0.46988573 | 0.34587599 | 1.35853815 | 0.17429299 | 0.8740623 |
| WDR70 | 201.623114 | -0.0865861 | 0.1667847 | -0.5191491 | 0.60365675 | 0.9958918 |
| LIFR | 210.760003 | -0.6180991 | 0.43844791 | -1.4097435 | 0.15861544 | 0.84652496 |
| OSMR-AS1 | 157.118352 | 0.46258308 | 0.28627886 | 1.61584787 | 0.10612721 | 0.76916061 |
| OSMR | 374.725256 | 0.18446247 | 0.37760898 | 0.48850128 | 0.62519482 | 0.9958918 |
| RICTOR | 186.040004 | -0.6558417 | 0.21557688 | -3.0422638 | 0.00234806 | 0.10666545 |
| TTC33 | 48.5836013 | -0.172071 | 0.27942866 | -0.6157957 | 0.53802938 | 0.9958918 |
| PTGER4 | 34.8551115 | 0.02527171 | 0.38127662 | 0.06628181 | 0.94715346 | 0.99935941 |
| PRKAA1 | 79.514238 | 0.0884526 | 0.28868604 | 0.30639721 | 0.75930223 | 0.9958918 |
| RPL37 | 74.9359412 | -0.185726 | 0.29994917 | -0.6191917 | 0.53579007 | 0.9958918 |
| C7 | 1142.63976 | -0.2285026 | 0.30691538 | -0.7445135 | 0.45656588 | 0.98330915 |
| PLCXD3 | 215.145273 | -0.4714591 | 0.41618511 | -1.1328111 | 0.25729359 | 0.9266065 |
| OXCT1 | 50.9939856 | -0.4671491 | 0.28040164 | -1.6659997 | 0.09571347 | 0.73786716 |
| GHR | 370.461266 | -0.9546938 | 0.4361328 | -2.1889978 | 0.028597 | 0.45975021 |
| SELENOP | 120.071175 | -0.3309028 | 0.4160497 | -0.7953444 | 0.42641319 | 0.98015476 |
| ZNF131 | 95.7654353 | 0.32415896 | 0.26860264 | 1.2068346 | 0.22749585 | 0.9120167 |
| HMGCS1 | 42.510418 | 0.17926289 | 0.45083789 | 0.39762161 | 0.69090913 | 0.9958918 |
| TMEM267 | 31.8491969 | 0.1123167 | 0.3514964 | 0.31953868 | 0.74931807 | 0.9958918 |
| PAIP1 | 29.4293843 | -0.0590831 | 0.2695789 | -0.2191681 | 0.8265191 | 0.99592423 |
| NNT-AS1 | 22.840583 | 0.14763041 | 0.38954841 | 0.37897833 | 0.70470396 | 0.9958918 |
| NNT | 72.4000268 | 0.14801282 | 0.2529996 | 0.5850318 | 0.55852633 | 0.9958918 |
| PARP8 | 97.8999299 | -0.0423028 | 0.35074329 | -0.1206089 | 0.90400083 | 0.99935941 |
| ITGA1 | 807.964304 | -0.6013829 | 0.39142303 | -1.5364013 | 0.12443998 | 0.79195137 |
| ITGA2 | 75.4378721 | 0.47948775 | 0.61624619 | 0.77807823 | 0.43652289 | 0.98092839 |
| NDUFS4 | 86.0015165 | -0.2873463 | 0.25109247 | -1.1443843 | 0.25246429 | 0.92631316 |
| ARL15 | 275.347312 | -0.2198297 | 0.27847152 | -0.7894155 | 0.42986922 | 0.98092839 |
| SNX18 | 35.5622765 | 0.60645965 | 0.3711536 | 1.63398563 | 0.10226187 | 0.75529896 |
| DHX29 | 57.3028351 | 0.00320829 | 0.2552181 | 0.0125708 | 0.98997022 | 0.99935941 |
| MTREX | 126.424838 | -0.1401537 | 0.22867017 | -0.6129078 | 0.53993728 | 0.9958918 |
| PLPP1 | 611.632503 | -0.2594567 | 0.46902507 | -0.553183 | 0.5801381 | 0.9958918 |
| SLC38A9 | 62.8942697 | 0.22663973 | 0.25244151 | 0.89779106 | 0.36929695 | 0.97217293 |
| IL6ST | 333.240787 | 0.05039133 | 0.30581162 | 0.164779 | 0.86911794 | 0.9971731 |
| MAP3K1 | 63.8547877 | 0.41815065 | 0.2700895 | 1.54819292 | 0.12157586 | 0.79021701 |
| MIER3 | 19.6648097 | -0.1920587 | 0.3478832 | -0.5520781 | 0.58089488 | 0.9958918 |
| GPBP1 | 191.838934 | -0.176649 | 0.29604028 | -0.5967058 | 0.55070381 | 0.9958918 |
| PDE4D | 1525.99087 | 0.54470591 | 0.38034493 | 1.43213663 | 0.15210472 | 0.83906933 |
| ERCC8 | 28.6474519 | -0.1004284 | 0.36809824 | -0.2728303 | 0.78498365 | 0.9958918 |
| NDUFAF2 | 112.339046 | -0.022667 | 0.27741595 | -0.0817076 | 0.93487927 | 0.99935941 |
| ZSWIM6 | 411.421608 | -0.0022471 | 0.22740255 | -0.0098816 | 0.99211578 | 0.99935941 |
| KIF2A | 64.3088337 | 0.03073868 | 0.23071169 | 0.13323418 | 0.89400818 | 0.99935941 |
| DIMT1 | 21.1526992 | -0.4972492 | 0.31968559 | -1.5554319 | 0.11984324 | 0.79021701 |
| IPO11 | 38.0972314 | -0.1656482 | 0.26050249 | -0.6358794 | 0.52485505 | 0.9958918 |
| RNF180 | 132.737443 | -0.3182129 | 0.2379764 | -1.3371615 | 0.18116992 | 0.87815793 |

| | | | | | | |
|---|---|---|---|---|---|---|
| RGS7BP | 50.0343413 | -1.0683227 | 0.51099407 | -2.0906753 | 0.03655718 | 0.51467183 |
| SREK1IP1 | 38.786184 | 0.08195317 | 0.30345139 | 0.27007018 | 0.78710627 | 0.99592423 |
| CWC27 | 97.2242176 | -0.1734623 | 0.19836299 | -0.8744689 | 0.38186297 | 0.97217293 |
| ADAMTS6 | 47.9546295 | 0.64602649 | 0.61437359 | 1.0515206 | 0.29301955 | 0.95186143 |
| PPWD1 | 41.8408747 | 0.14652677 | 0.32183825 | 0.45528077 | 0.64890726 | 0.9958918 |
| TRIM23 | 32.8241524 | 0.22544729 | 0.28140542 | 0.80114764 | 0.42304618 | 0.97828057 |
| TRAPPC13 | 32.7097692 | 0.32826332 | 0.28748849 | 1.14183114 | 0.25352421 | 0.92646125 |
| ERBIN | 268.337605 | -0.4487334 | 0.27338249 | -1.6414122 | 0.10071187 | 0.75064053 |
| SREK1 | 99.1300512 | -0.3203851 | 0.18188563 | -1.7614643 | 0.07815985 | 0.68056484 |
| MAST4 | 165.641716 | 0.15199229 | 0.27699968 | 0.54870925 | 0.58320499 | 0.9958918 |
| PIK3R1 | 193.872133 | -0.4783792 | 0.43572768 | -1.0978856 | 0.27225443 | 0.93813668 |
| SLC30A5 | 47.2818381 | 0.14576485 | 0.26612833 | 0.54772391 | 0.58388149 | 0.9958918 |
| CDK7 | 20.3438971 | -0.1357881 | 0.34112729 | -0.3980569 | 0.69058827 | 0.9958918 |
| RAD17 | 24.3881008 | 0.04114791 | 0.42346208 | 0.09717022 | 0.92259121 | 0.99935941 |
| GTF2H2C | 17.4712883 | 0.04844867 | 0.413609 | 0.11713639 | 0.90675197 | 0.99935941 |
| NAIP | 60.5791217 | 0.09109875 | 0.34324703 | 0.26540288 | 0.79069912 | 0.99592423 |
| GTF2H2 | 19.0074615 | -0.5400416 | 0.4254626 | -1.2693045 | 0.20433249 | 0.8943654 |
| BDP1 | 121.962379 | 0.01872544 | 0.19127297 | 0.09789905 | 0.92201245 | 0.99935941 |
| MCCC2 | 36.0273738 | -0.0729028 | 0.26567842 | -0.2744024 | 0.78377543 | 0.9958918 |
| MAP1B | 210.818218 | -0.5629435 | 0.26548979 | -2.1203962 | 0.03397265 | 0.4973406 |
| MRPS27 | 68.1041265 | 0.08800436 | 0.22473714 | 0.39158797 | 0.69536267 | 0.9958918 |
| TNPO1 | 152.573269 | -0.1627737 | 0.19744807 | -0.8243873 | 0.40971952 | 0.97812179 |
| FCHO2 | 112.547244 | 0.00583589 | 0.25443634 | 0.02293653 | 0.9817009 | 0.99935941 |
| BTF3 | 54.2534783 | 0.02221859 | 0.36587783 | 0.06072681 | 0.95157678 | 0.99935941 |
| ANKRA2 | 22.3928964 | 0.22983069 | 0.36695082 | 0.62632558 | 0.53110141 | 0.9958918 |
| ARHGEF28 | 68.9192565 | -0.391973 | 0.37512559 | -1.0449113 | 0.29606398 | 0.95393362 |
| HEXB | 27.3042725 | 0.2428647 | 0.34863597 | 0.69661399 | 0.4860444 | 0.99046213 |
| GFM2 | 22.2027625 | -0.7986663 | 0.36601883 | -2.182036 | 0.02910688 | 0.4640053 |
| NSA2 | 21.7750352 | 0.39799924 | 0.42967718 | 0.92627504 | 0.35430306 | 0.97217293 |
| HMGCR | 19.2002321 | -0.4663529 | 0.49283305 | -0.9462696 | 0.34401111 | 0.96887708 |
| COL4A3BP | 148.490124 | -0.1931413 | 0.31082262 | -0.6213876 | 0.5343446 | 0.9958918 |
| POLK | 117.888633 | -0.1977506 | 0.296181 | -0.6676682 | 0.5043454 | 0.99387859 |
| POC5 | 19.2575493 | -0.1754257 | 0.36145297 | -0.4853347 | 0.62743897 | 0.9958918 |
| IQGAP2 | 25.4177294 | 1.66676424 | 0.43993046 | 3.78869934 | 0.00015144 | 0.01904212 |
| F2R | 42.8213142 | 0.62209892 | 0.31087252 | 2.00113837 | 0.04537748 | 0.55894748 |
| AC022414.1 | 28.2908173 | -0.7699051 | 0.53432542 | -1.4408918 | 0.14961526 | 0.83171145 |
| PDE8B | 23.5989342 | -1.0771932 | 0.63088503 | -1.7074319 | 0.0877418 | 0.71805622 |
| WDR41 | 111.306945 | -0.7689701 | 0.42629855 | -1.80383 | 0.07125796 | 0.66044085 |
| TBCA | 124.750301 | -0.0970046 | 0.18970607 | -0.5113417 | 0.60911182 | 0.9958918 |
| AP3B1 | 176.736849 | -0.2065364 | 0.2019895 | -1.0225106 | 0.30653931 | 0.95793395 |
| SCAMP1 | 55.9504783 | -0.544552 | 0.28676198 | -1.8989684 | 0.05756863 | 0.60512745 |
| LHFPL2 | 80.1561965 | -1.125568 | 0.45252735 | -2.4872927 | 0.01287195 | 0.30408773 |
| ARSB | 34.9218905 | -0.1691108 | 0.45320717 | -0.3731423 | 0.70904253 | 0.9958918 |
| DMGDH | 18.499284 | -0.1039179 | 0.38708552 | -0.2684624 | 0.7883434 | 0.99592423 |
| BHMT2 | 27.1620164 | 0.79486238 | 0.58422793 | 1.36053471 | 0.17366078 | 0.87233975 |
| JMY | 52.2633132 | -0.0864935 | 0.32783331 | -0.2638338 | 0.79190802 | 0.99592423 |
| HOMER1 | 36.7943172 | 0.02470532 | 0.517608 | 0.04772979 | 0.96193159 | 0.99935941 |
| TENT2 | 196.402934 | -0.2556545 | 0.24234564 | -1.0549168 | 0.29146338 | 0.95186143 |
| SERINC5 | 82.9309608 | -0.3251875 | 0.28489712 | -1.1414209 | 0.25369483 | 0.92646125 |
| ZFYVE16 | 110.678935 | -0.2248561 | 0.28812393 | -0.7804144 | 0.435147 | 0.98092839 |
| DHFR | 158.434461 | -0.1555196 | 0.26907293 | -0.577983 | 0.56327558 | 0.9958918 |
| MSH3 | 75.6684709 | -0.1471504 | 0.27320485 | -0.5386082 | 0.59015726 | 0.9958918 |
| RASGRF2 | 28.2279742 | -0.0458618 | 0.56933483 | -0.0805532 | 0.93579727 | 0.99935941 |
| CKMT2-AS1 | 45.6635857 | -0.4075521 | 0.34377723 | -1.1855123 | 0.23581496 | 0.91962002 |

| | | | | | | |
|---|---|---|---|---|---|---|
| SSBP2 | 584.531835 | 0.00599426 | 0.3257422 | 0.01840185 | 0.98531828 | 0.99935941 |
| ATG10 | 70.9130764 | -0.3429656 | 0.33855006 | -1.0130424 | 0.31103991 | 0.95957532 |
| RPS23 | 143.637837 | -0.4186655 | 0.38081627 | -1.0993898 | 0.27159809 | 0.93795041 |
| ATP6AP1L | 22.818009 | 0.34062 | 0.53911386 | 0.63181459 | 0.52750804 | 0.9958918 |
| TMEM167A | 31.7096138 | -0.1251934 | 0.30360745 | -0.412353 | 0.68008071 | 0.9958918 |
| XRCC4 | 40.3602263 | -0.1055632 | 0.30022013 | -0.3516195 | 0.72512367 | 0.9958918 |
| VCAN | 213.350928 | -0.0903044 | 0.5388017 | -0.1676023 | 0.86689614 | 0.9971731 |
| EDIL3 | 182.379889 | 0.05121397 | 0.48875985 | 0.10478351 | 0.9165476 | 0.99935941 |
| AC113383.1 | 30.5880961 | 0.07739062 | 0.47279065 | 0.16368897 | 0.869976 | 0.9971731 |
| COX7C | 54.8064927 | 0.01509685 | 0.45356038 | 0.03328521 | 0.97344715 | 0.99935941 |
| RASA1 | 145.501655 | 0.08175259 | 0.28364061 | 0.28822599 | 0.77317376 | 0.9958918 |
| CCNH | 346.834973 | 0.20805326 | 0.32749315 | 0.63529041 | 0.52523903 | 0.9958918 |
| TMEM161B | 39.1013307 | -0.0057747 | 0.26144045 | -0.0220878 | 0.9823779 | 0.99935941 |
| TMEM161B-. | 78.1799691 | -0.1582149 | 0.28878849 | -0.5478572 | 0.58378994 | 0.9958918 |
| MEF2C | 262.214664 | 0.5901634 | 0.29159038 | 2.02394673 | 0.04297563 | 0.55022287 |
| MEF2C-AS1 | 81.3347102 | 0.0991756 | 0.51286824 | 0.19337441 | 0.84666575 | 0.99692952 |
| LYSMD3 | 21.5024637 | 0.23794428 | 0.45422569 | 0.52384593 | 0.6003857 | 0.9958918 |
| ARRDC3 | 121.818795 | 0.01102272 | 0.49775378 | 0.02214492 | 0.98233236 | 0.99935941 |
| AC124854.1 | 63.0512394 | 0.15552765 | 0.52117407 | 0.29841785 | 0.76538426 | 0.9958918 |
| NR2F1-AS1 | 368.352041 | -0.3567434 | 0.24982845 | -1.4279536 | 0.15330523 | 0.84093985 |
| NR2F1 | 277.236476 | -0.8416647 | 0.3345032 | -2.5161635 | 0.01186401 | 0.29356444 |
| FAM172A | 300.760703 | -0.252644 | 0.21828511 | -1.1574037 | 0.24710747 | 0.92605444 |
| KIAA0825 | 79.9242113 | -0.1137495 | 0.24682901 | -0.4608433 | 0.64491101 | 0.9958918 |
| TTC37 | 119.272472 | -0.5027982 | 0.22597545 | -2.2250123 | 0.02608041 | 0.43845312 |
| RHOBTB3 | 174.291027 | -0.5579661 | 0.41913126 | -1.3312443 | 0.18310864 | 0.87866798 |
| GLRX | 62.5518686 | -0.2550757 | 0.54309322 | -0.469672 | 0.6385894 | 0.9958918 |
| ELL2 | 546.038592 | -1.1557227 | 0.64739351 | -1.7851935 | 0.07422994 | 0.66886321 |
| AC104123.1 | 81.5942484 | 0.17698488 | 0.39015594 | 0.45362601 | 0.65009803 | 0.9958918 |
| CAST | 188.864276 | -0.490374 | 0.2834129 | -1.730246 | 0.08358633 | 0.70213814 |
| ERAP1 | 43.6572104 | -0.4481507 | 0.355898 | -1.2592112 | 0.20795407 | 0.89633695 |
| ERAP2 | 18.5757801 | 0.60118042 | 0.57243629 | 1.05021367 | 0.29361989 | 0.95225574 |
| LNPEP | 97.1816681 | 0.0018967 | 0.26390341 | 0.00718711 | 0.99426556 | 0.99935941 |
| LIX1-AS1 | 26.8902174 | -0.579115 | 0.45677824 | -1.2678252 | 0.20486036 | 0.89560905 |
| CHD1 | 129.190364 | -0.2216052 | 0.30846665 | -0.718409 | 0.47250516 | 0.98910524 |
| LINC02062 | 29.9987604 | -0.0228629 | 0.33590552 | -0.0680634 | 0.94573513 | 0.99935941 |
| FAM174A | 20.2866212 | 0.42975407 | 0.36092243 | 1.19071033 | 0.23376732 | 0.91918462 |
| PAM | 320.579679 | 0.05334009 | 0.31423678 | 0.16974489 | 0.86521077 | 0.9971731 |
| PPIP5K2 | 86.0005331 | -0.0366326 | 0.23523178 | -0.1557296 | 0.87624615 | 0.9971731 |
| EFNA5 | 140.440806 | -1.2876387 | 0.38964021 | -3.3046863 | 0.00095083 | 0.06850619 |
| FBXL17 | 301.164034 | -0.1450867 | 0.19534851 | -0.7427067 | 0.45765926 | 0.98400758 |
| FER | 382.632174 | -0.2573018 | 0.34011925 | -0.7565046 | 0.44934671 | 0.98092839 |
| PJA2 | 157.552715 | -0.0695356 | 0.23433264 | -0.2967388 | 0.76666593 | 0.9958918 |
| MAN2A1 | 124.502693 | 0.10733023 | 0.33667708 | 0.3187928 | 0.74988364 | 0.9958918 |
| TMEM232 | 28.278575 | -0.908931 | 0.47416131 | -1.9169236 | 0.05524764 | 0.59593882 |
| SLC25A46 | 46.8617001 | 0.09756183 | 0.2698946 | 0.36148123 | 0.71773974 | 0.9958918 |
| WDR36 | 35.442015 | 0.25553597 | 0.29608633 | 0.86304551 | 0.38811244 | 0.97297628 |
| CAMK4 | 43.2616398 | -2.1938494 | 0.68802601 | -3.1886141 | 0.00142957 | 0.08339044 |
| STARD4 | 24.8684081 | -0.9377273 | 0.42468856 | -2.2080352 | 0.02724182 | 0.44841843 |
| NREP | 23.6397967 | -1.7353097 | 0.51538029 | -3.3670472 | 0.00075978 | 0.06204001 |
| EPB41L4A | 82.8927191 | -0.0561243 | 0.33070778 | -0.1697097 | 0.86523842 | 0.9971731 |
| APC | 69.1836861 | -0.3298791 | 0.28675212 | -1.1503982 | 0.24997992 | 0.92624282 |
| SRP19 | 29.6510142 | -0.420971 | 0.38537093 | -1.0923788 | 0.27466665 | 0.93944545 |
| REEP5 | 77.8898954 | -0.0815482 | 0.20775264 | -0.3925253 | 0.69467015 | 0.9958918 |
| DCP2 | 57.2143448 | -0.1287325 | 0.25489977 | -0.5050318 | 0.61353649 | 0.9958918 |

| | | | | | | |
|---|---|---|---|---|---|---|
| MCC | 189.54446 | 0.47654051 | 0.450687 | 1.05736466 | 0.2903452 | 0.95186143 |
| YTHDC2 | 106.050182 | -0.1270118 | 0.2710789 | -0.4685418 | 0.63939716 | 0.9958918 |
| PGGT1B | 32.1366379 | 0.05397259 | 0.27932441 | 0.19322547 | 0.84678239 | 0.99692952 |
| ATG12 | 41.2278589 | -0.1154227 | 0.2708552 | -0.4261418 | 0.67000455 | 0.9958918 |
| AP3S1 | 122.586933 | 0.28227428 | 0.28771689 | 0.98108346 | 0.32655159 | 0.95992644 |
| COMMD10 | 183.263684 | -0.0137966 | 0.2184301 | -0.0631627 | 0.94963699 | 0.99935941 |
| SEMA6A | 24.7978111 | -0.6593971 | 0.30994735 | -2.1274486 | 0.03338283 | 0.492839 |
| SEMA6A-AS1 | 24.2413511 | 0.04881902 | 0.404759 | 0.12061257 | 0.90399791 | 0.99935941 |
| DTWD2 | 24.0311876 | -0.0205426 | 0.36674921 | -0.0560126 | 0.9553318 | 0.99935941 |
| DMXL1 | 161.442933 | -0.0243119 | 0.19411431 | -0.1252454 | 0.90032927 | 0.99935941 |
| TNFAIP8 | 78.1351058 | 1.01814884 | 0.39114747 | 2.60297948 | 0.00924175 | 0.25437307 |
| HSD17B4 | 99.6026758 | 0.29474857 | 0.27109299 | 1.08726002 | 0.27692191 | 0.94304045 |
| PRR16 | 338.148116 | 0.0585677 | 0.36950502 | 0.15850311 | 0.87406037 | 0.9971731 |
| SRFBP1 | 46.3572905 | -0.5230043 | 0.32426045 | -1.6129143 | 0.10676314 | 0.77179565 |
| LOX | 35.594011 | -0.7377035 | 0.69912902 | -1.055175 | 0.29134529 | 0.95186143 |
| SNCAIP | 49.2393521 | -0.5278817 | 0.45371415 | -1.1634677 | 0.24463981 | 0.92559078 |
| SNX2 | 47.7544098 | 0.18720121 | 0.2871916 | 0.65183386 | 0.51450836 | 0.9958918 |
| SNX24 | 49.3021606 | 0.20905691 | 0.35023401 | 0.59690637 | 0.5505699 | 0.9958918 |
| CEP120 | 36.4053641 | 0.58449265 | 0.36075701 | 1.62018376 | 0.10519281 | 0.76555899 |
| CSNK1G3 | 71.6156506 | 0.35011277 | 0.2463522 | 1.42118792 | 0.15526213 | 0.84277602 |
| ZNF608 | 97.9452817 | -0.2904515 | 0.81654965 | -0.3557059 | 0.72206084 | 0.9958918 |
| GRAMD2B | 304.639513 | -0.4937576 | 0.43011234 | -1.1479736 | 0.25097946 | 0.92624282 |
| AC093535.1 | 32.057697 | -0.7411126 | 0.69330602 | -1.0689546 | 0.28509015 | 0.94505166 |
| ALDH7A1 | 91.9263841 | -0.5393192 | 0.54618955 | -0.9874213 | 0.32343613 | 0.95992644 |
| PHAX | 19.6978016 | 0.12358484 | 0.39159059 | 0.31559706 | 0.75230837 | 0.9958918 |
| 3-Mar | 138.812313 | -0.2147502 | 0.88224677 | -0.2434128 | 0.80768561 | 0.99592423 |
| PRRC1 | 41.5349482 | -0.456031 | 0.3147867 | -1.4486985 | 0.14742179 | 0.8301633 |
| LINC01184 | 75.0890709 | -0.2204587 | 0.27375952 | -0.8053005 | 0.42064629 | 0.97812179 |
| SLC12A2 | 60.4256622 | -0.2837003 | 0.37812994 | -0.750272 | 0.4530909 | 0.98092839 |
| CHSY3 | 97.9409997 | -0.1010672 | 0.33979709 | -0.2974338 | 0.76613532 | 0.9958918 |
| HINT1 | 88.6232062 | 0.45218816 | 0.31643288 | 1.42901764 | 0.15299916 | 0.84093985 |
| CDC42SE2 | 72.79987 | 0.49250314 | 0.29675042 | 1.65965439 | 0.096984 | 0.74243272 |
| P4HA2 | 23.6714224 | 0.06006175 | 0.40482258 | 0.14836562 | 0.88205423 | 0.99818549 |
| AC116366.3 | 119.179412 | -0.3431419 | 0.4819739 | -0.7119512 | 0.47649499 | 0.9897574 |
| IRF1 | 29.3446078 | 0.59233954 | 0.59352268 | 0.99800659 | 0.31827616 | 0.95992644 |
| KIF3A | 29.4138521 | -0.0942646 | 0.42687609 | -0.2208243 | 0.82522922 | 0.99592423 |
| 8-Sep | 22.3763902 | -0.1227592 | 0.42328769 | -0.2900137 | 0.77180574 | 0.9958918 |
| UQCRQ | 31.9275542 | 0.2607364 | 0.39036014 | 0.66793806 | 0.50417313 | 0.99387859 |
| AFF4 | 261.203993 | -0.1871108 | 0.20866043 | -0.8967239 | 0.36986626 | 0.97217293 |
| ZCCHC10 | 20.6639887 | 0.14559063 | 0.33903313 | 0.42942893 | 0.66761111 | 0.9958918 |
| HSPA4 | 50.4986289 | 0.00613801 | 0.29795656 | 0.02060036 | 0.98356446 | 0.99935941 |
| C5orf15 | 32.6341503 | 0.17282271 | 0.3085963 | 0.56002846 | 0.57546003 | 0.9958918 |
| VDAC1 | 50.4820289 | 0.20907744 | 0.28929116 | 0.72272323 | 0.46984994 | 0.98910524 |
| SKP1 | 88.947697 | 0.33393008 | 0.25215908 | 1.32428342 | 0.18540893 | 0.87866798 |
| CDKL3 | 29.1767887 | -0.2557846 | 0.37760708 | -0.677383 | 0.49816298 | 0.99176652 |
| UBE2B | 115.341775 | -0.1615781 | 0.20078929 | -0.8047146 | 0.42098438 | 0.97812179 |
| JADE2 | 24.8184154 | -0.1126973 | 0.40876554 | -0.2757015 | 0.78277734 | 0.9958918 |
| SAR1B | 39.7532437 | 0.0817174 | 0.26959994 | 0.30310615 | 0.76180897 | 0.9958918 |
| SEC24A | 77.5956685 | 0.06148699 | 0.30890775 | 0.19904645 | 0.84222641 | 0.99692952 |
| CAMLG | 23.9910809 | -0.0499573 | 0.33139422 | -0.150749 | 0.88017373 | 0.99749293 |
| DDX46 | 92.2239788 | -0.1183775 | 0.31375721 | -0.3772901 | 0.70595805 | 0.9958918 |
| C5orf24 | 38.2836569 | 0.42837545 | 0.30260911 | 1.41560662 | 0.15689071 | 0.84427321 |
| TXNDC15 | 36.977064 | 0.36289798 | 0.29167357 | 1.24419218 | 0.21342884 | 0.89738044 |
| PCBD2 | 27.2877755 | -0.3743111 | 0.49382988 | -0.7579757 | 0.44846552 | 0.98092839 |

| | | | | | | |
|---|---|---|---|---|---|---|
| H2AFY | 60.4958346 | 0.21736461 | 0.35821681 | 0.60679624 | 0.54398614 | 0.9958918 |
| CXCL14 | 40.5553856 | 1.49566507 | 0.63013683 | 2.373556 | 0.01761772 | 0.35016405 |
| TGFBI | 94.6639737 | -0.210241 | 0.39641745 | -0.5303526 | 0.59586747 | 0.9958918 |
| SMAD5 | 99.880822 | 0.05120135 | 0.23594827 | 0.21700242 | 0.82820646 | 0.99592423 |
| SPOCK1 | 277.745249 | 0.43271702 | 0.47548189 | 0.91005994 | 0.3627909 | 0.97217293 |
| HNRNPA0 | 35.12181 | -0.2479573 | 0.42907145 | -0.5778928 | 0.56333649 | 0.9958918 |
| FAM13B | 142.518988 | 0.13003741 | 0.19258904 | 0.6752067 | 0.49954445 | 0.99213111 |
| BRD8 | 29.6090935 | 0.00769409 | 0.31006602 | 0.02481435 | 0.98020305 | 0.99935941 |
| KDM3B | 75.1742908 | 0.36983824 | 0.26381273 | 1.40189686 | 0.16094605 | 0.85092986 |
| EGR1 | 123.736033 | -1.4405541 | 0.90816199 | -1.5862304 | 0.11268706 | 0.77793929 |
| ETF1 | 63.3921223 | -0.2988967 | 0.32137236 | -0.9300636 | 0.35233818 | 0.97087862 |
| HSPA9 | 92.0551735 | -0.293006 | 0.25716166 | -1.1393846 | 0.25454276 | 0.92651547 |
| CTNNA1 | 305.245665 | 0.02007863 | 0.23567284 | 0.08519705 | 0.93210473 | 0.99935941 |
| SIL1 | 122.61461 | 0.06099688 | 0.25977831 | 0.23480359 | 0.81436118 | 0.99592423 |
| PAIP2 | 63.3694825 | 0.08553859 | 0.25908482 | 0.3301567 | 0.74128156 | 0.9958918 |
| TMEM173 | 35.6551942 | 0.11892693 | 0.34090514 | 0.34885636 | 0.72719715 | 0.9958918 |
| UBE2D2 | 129.948105 | -0.0143754 | 0.22560593 | -0.0637189 | 0.94919403 | 0.99935941 |
| CXXC5 | 35.1143044 | 0.45762473 | 0.40756213 | 1.12283428 | 0.26150788 | 0.93155278 |
| PURA | 73.156909 | 0.24855357 | 0.22159629 | 1.12165042 | 0.2620111 | 0.93155278 |
| CYSTM1 | 116.449038 | -0.2120086 | 0.32718202 | -0.6479837 | 0.51699548 | 0.9958918 |
| PFDN1 | 29.6165985 | -0.1997621 | 0.3656598 | -0.5463059 | 0.58485566 | 0.9958918 |
| SRA1 | 24.6179551 | 0.02989331 | 0.34974285 | 0.08547225 | 0.93188595 | 0.99935941 |
| NDUFA2 | 20.5060789 | -0.1943876 | 0.35647089 | -0.5453114 | 0.58553938 | 0.9958918 |
| HARS | 30.8024785 | 0.35313881 | 0.26921579 | 1.3117314 | 0.18961078 | 0.88139754 |
| TAF7 | 33.1177602 | 0.59274305 | 0.32117984 | 1.84551758 | 0.06496228 | 0.62770158 |
| DIAPH1 | 80.0326311 | 0.29886076 | 0.26707012 | 1.11903482 | 0.26312528 | 0.93280642 |
| HDAC3 | 27.9583152 | -0.171888 | 0.43267049 | -0.3972723 | 0.69116669 | 0.9958918 |
| ARAP3 | 20.3828236 | 0.76494919 | 0.41337991 | 1.850475 | 0.06424512 | 0.62609028 |
| DELE1 | 32.6972932 | -0.297978 | 0.3468018 | -0.8592171 | 0.39022077 | 0.97297628 |
| RNF14 | 26.2362446 | -0.2000208 | 0.28703446 | -0.6968529 | 0.48589485 | 0.99046213 |
| NDFIP1 | 125.045346 | 0.0873816 | 0.20487502 | 0.42651172 | 0.66973501 | 0.9958918 |
| SPRY4 | 54.454943 | -0.5693282 | 0.44482144 | -1.2799027 | 0.20057935 | 0.89191279 |
| SPRY4-AS1 | 117.376754 | -0.0577187 | 0.39103812 | -0.1476039 | 0.88265541 | 0.99840747 |
| FGF1 | 42.7483249 | -0.7469427 | 0.39418966 | -1.8948815 | 0.0581081 | 0.60892588 |
| ARHGAP26 | 148.925483 | 0.53654234 | 0.26791116 | 2.00268754 | 0.04521084 | 0.55894748 |
| NR3C1 | 288.960526 | 0.20072448 | 0.3622432 | 0.55411523 | 0.57949999 | 0.9958918 |
| YIPF5 | 24.6501715 | 0.29149321 | 0.45714896 | 0.6376329 | 0.52371267 | 0.9958918 |
| PRELID2 | 41.1547904 | 0.38900357 | 0.46570282 | 0.8353043 | 0.40354639 | 0.97575999 |
| LARS | 79.3680272 | -0.3057972 | 0.22930102 | -1.3336059 | 0.18233305 | 0.87866798 |
| TCERG1 | 90.5769046 | -0.4774755 | 0.30700315 | -1.555279 | 0.11987964 | 0.79021701 |
| DPYSL3 | 55.2781461 | 0.99691258 | 0.58785717 | 1.69584148 | 0.0899159 | 0.72305504 |
| FBXO38 | 44.648251 | 0.09005171 | 0.26494796 | 0.33988452 | 0.7339435 | 0.9958918 |
| ABLIM3 | 23.4200859 | 0.6324476 | 0.48066428 | 1.31577823 | 0.18824849 | 0.88139754 |
| CARMN | 625.763473 | 0.48193232 | 0.4421348 | 1.0900122 | 0.27570777 | 0.94196951 |
| AC131025.2 | 53.6497949 | 0.2792274 | 0.39783425 | 0.70186866 | 0.48276107 | 0.99002719 |
| CSNK1A1 | 410.483261 | -0.22803 | 0.25911581 | -0.8800314 | 0.37884232 | 0.97217293 |
| PPARGC1B | 24.1404545 | -0.0964081 | 0.63250037 | -0.1524238 | 0.87885271 | 0.99732684 |
| SLC26A2 | 18.959539 | 0.02936358 | 0.5063558 | 0.05799001 | 0.95375659 | 0.99935941 |
| HMGXB3 | 38.1440545 | -0.0413827 | 0.32330067 | -0.1280005 | 0.89814855 | 0.99935941 |
| CSF1R | 19.3312701 | 0.75762343 | 0.59980092 | 1.26312483 | 0.20654432 | 0.89589437 |
| PDGFRB | 476.661082 | 0.24592868 | 0.27171266 | 0.90510572 | 0.36540938 | 0.97217293 |
| CD74 | 81.8253919 | 0.34870563 | 0.42426098 | 0.82191305 | 0.41112639 | 0.97812179 |
| RPS14 | 136.150278 | -0.0996752 | 0.32824317 | -0.3036626 | 0.76138498 | 0.9958918 |
| NDST1 | 37.5841566 | -0.0074112 | 0.34984433 | -0.0211841 | 0.98309877 | 0.99935941 |

| | | | | | | |
|---|---|---|---|---|---|---|
| SYNPO | 48.7746195 | 0.63854825 | 0.41154837 | 1.5515752 | 0.1207639 | 0.79021701 |
| DCTN4 | 81.0609133 | -0.3716791 | 0.26426061 | -1.4064868 | 0.15957962 | 0.84762439 |
| SMIM3 | 59.5776651 | -0.2903925 | 0.29629109 | -0.980092 | 3.27E-01 | 0.95992644 |
| GPX3 | 638.555217 | -0.1890602 | 0.48875456 | -0.3868203 | 0.69888926 | 0.9958918 |
| TNIP1 | 62.440561 | 0.8598266 | 0.62047157 | 1.38576309 | 0.16581926 | 0.85938344 |
| ANXA6 | 91.9415693 | 0.29715351 | 0.32138956 | 0.92458983 | 0.35517931 | 0.97217293 |
| SLC36A1 | 18.3784233 | 0.45015596 | 0.42793064 | 1.05193674 | 0.29282857 | 0.95186143 |
| SPARC | 300.551982 | -0.4736472 | 0.38420992 | -1.2327822 | 0.21765703 | 0.90370709 |
| ATOX1 | 32.1452716 | -0.0597118 | 0.34602319 | -0.1725658 | 0.86299273 | 0.9971731 |
| G3BP1 | 92.7534916 | -0.112105 | 0.25124235 | -0.4462028 | 0.65545079 | 0.9958918 |
| FAM114A2 | 20.1324343 | 0.05224609 | 0.33753879 | 0.15478544 | 0.87699047 | 0.9971731 |
| MFAP3 | 24.2066113 | 0.7368897 | 0.41763041 | 1.76445414 | 0.07765555 | 0.68020338 |
| GALNT10 | 54.4513226 | 1.18097383 | 0.43734873 | 2.70030239 | 0.00692765 | 0.21213035 |
| LARP1 | 109.862999 | -0.0694566 | 0.27498115 | -0.2525866 | 0.80058766 | 0.99592423 |
| FAXDC2 | 35.8700905 | 0.15831546 | 0.56911946 | 0.27817614 | 0.78087715 | 0.9958918 |
| CNOT8 | 30.543138 | -0.1942462 | 0.29788123 | -0.6520928 | 0.51434131 | 0.9958918 |
| GEMIN5 | 20.0916951 | -0.0628071 | 0.35035368 | -0.1792676 | 0.85772762 | 0.9971731 |
| SGCD | 60.8841686 | 1.19832459 | 0.54722284 | 2.18982927 | 0.02853662 | 0.45975021 |
| CYFIP2 | 23.4224785 | 0.91171604 | 0.36289645 | 2.51233109 | 0.01199365 | 0.2957763 |
| ADAM19 | 56.7957665 | 0.04093076 | 0.43019393 | 0.09514491 | 0.92419973 | 0.99935941 |
| CLINT1 | 105.053373 | -0.0175425 | 0.21964248 | -0.0798686 | 0.93634175 | 0.99935941 |
| EBF1 | 1348.24024 | 0.29052574 | 0.20341014 | 1.42827565 | 0.15321253 | 0.84093985 |
| RNF145 | 108.609362 | 0.17302955 | 0.28717806 | 0.60251662 | 0.54683031 | 0.9958918 |
| UBLCP1 | 25.8179523 | 0.46526018 | 0.32045457 | 1.45187565 | 0.14653618 | 0.8301633 |
| ADRA1B | 27.7713191 | 0.69458928 | 0.64916937 | 1.06996618 | 0.28463453 | 0.94505166 |
| TTC1 | 46.0245054 | 0.02095709 | 0.31634229 | 0.06624815 | 0.94718026 | 0.99935941 |
| PWWP2A | 83.3962682 | 0.23543372 | 0.21207786 | 1.11012871 | 0.26694357 | 0.93533685 |
| MAT2B | 22.1886742 | -0.0518046 | 0.31541322 | -0.1642435 | 0.86953943 | 0.9971731 |
| WWC1 | 22.4943272 | -0.5715189 | 0.55440898 | -1.0308615 | 0.30260578 | 0.95650251 |
| RARS | 47.4959595 | 0.18188806 | 0.2880612 | 0.63142158 | 0.52776491 | 0.9958918 |
| PANK3 | 48.9106256 | -0.19583 | 0.25376646 | -0.7716937 | 0.44029585 | 0.98092839 |
| SLIT3 | 1449.2016 | -0.5075312 | 0.31551715 | -1.6085694 | 0.10771053 | 0.77225821 |
| KCNIP1 | 71.2576059 | -0.4782507 | 1.33713896 | -0.3576671 | 0.72059242 | 0.9958918 |
| NPM1 | 51.3898115 | -0.2795166 | 0.31319786 | -0.8924601 | 0.37214639 | 0.97217293 |
| FBXW11 | 169.407166 | 0.28079228 | 0.21473349 | 1.30763155 | 0.19099831 | 0.88158763 |
| STK10 | 61.6391815 | 0.09109757 | 0.25819139 | 0.35282963 | 0.72421617 | 0.9958918 |
| UBTD2 | 52.8396853 | -0.0048608 | 0.27412423 | -0.0177321 | 0.98585258 | 0.99935941 |
| SH3PXD2B | 78.7462299 | -0.2342851 | 0.38490035 | -0.6086902 | 0.54272984 | 0.9958918 |
| NEURL1B | 114.895203 | -1.3498244 | 0.37029285 | -3.645289 | 0.00026709 | 0.02764585 |
| DUSP1 | 215.326483 | -1.0976312 | 0.6027222 | -1.8211229 | 0.06858818 | 0.64792295 |
| ERGIC1 | 81.0415035 | 0.23153305 | 0.26213477 | 0.8832596 | 0.37709603 | 0.97217293 |
| ATP6V0E1 | 107.777853 | -0.2360819 | 0.18726342 | -1.2606943 | 2.07E-01 | 0.89620104 |
| CREBRF | 99.5073928 | -0.1049076 | 0.38490316 | -0.2725558 | 0.78519469 | 0.9958918 |
| CPEB4 | 142.481213 | 0.38405972 | 0.27098968 | 1.41724847 | 0.15641029 | 0.84427321 |
| SFXN1 | 53.5550237 | 0.185171 | 0.32968326 | 0.56166334 | 0.57434541 | 0.9958918 |
| HRH2 | 43.0422432 | 1.39830223 | 0.60985333 | 2.29285004 | 0.02185664 | 0.39660357 |
| SIMC1 | 49.2997085 | 0.16194098 | 0.34614631 | 0.46783968 | 0.63989924 | 0.9958918 |
| KIAA1191 | 24.1009798 | 0.12259102 | 0.30894037 | 0.39681126 | 0.69150665 | 0.9958918 |
| CLTB | 36.8989622 | -0.1928746 | 0.41936506 | -0.4599205 | 0.64557331 | 0.9958918 |
| FAF2 | 64.5125235 | -0.2949613 | 0.25276765 | -1.1669265 | 0.24324004 | 0.92534155 |
| UIMC1 | 88.8894512 | 0.03794559 | 0.20964745 | 0.18099717 | 0.8563698 | 0.9971731 |
| ZNF346 | 40.0603703 | -0.0259072 | 0.2671763 | -0.0969668 | 0.92275277 | 0.99935941 |
| NSD1 | 198.493794 | 0.17425011 | 0.1757703 | 0.99135131 | 0.32151407 | 0.95992644 |
| PRELID1 | 27.4421442 | 0.13416124 | 0.35301028 | 0.3800491 | 0.70390897 | 0.9958918 |

| | | | | | | |
|---|---|---|---|---|---|---|
| LMAN2 | 32.319779 | 0.07351268 | 0.32727385 | 0.2246213 | 0.82227389 | 0.99592423 |
| PDLIM7 | 79.3149192 | -0.094463 | 0.42466706 | -0.2224401 | 0.82397128 | 0.99592423 |
| FAM193B | 73.8838281 | 0.33657257 | 0.23038349 | 1.46092311 | 0.14403654 | 0.82247441 |
| TMED9 | 45.8281454 | -0.4169281 | 0.47302398 | -0.8814101 | 0.3780959 | 0.97217293 |
| RMND5B | 23.9526675 | 0.36608133 | 0.39167892 | 0.93464649 | 0.34997051 | 0.9693503 |
| NHP2 | 21.388639 | -0.1394649 | 0.43987495 | -0.3170558 | 0.75120129 | 0.9958918 |
| HNRNPAB | 37.898849 | 0.03313753 | 0.46589654 | 0.07112636 | 0.94329719 | 0.99935941 |
| PHYKPL | 91.0777018 | 0.57676156 | 0.36771071 | 1.5685199 | 0.11675985 | 0.78417352 |
| COL23A1 | 101.62614 | 1.619834 | 0.51523689 | 3.14386265 | 0.00166734 | 0.09009749 |
| CLK4 | 41.1723152 | 0.48288535 | 0.33430629 | 1.4444399 | 0.14861527 | 0.8301633 |
| ADAMTS2 | 182.101135 | -0.8205801 | 0.29758874 | -2.7574299 | 0.00582577 | 0.19136125 |
| RUFY1 | 55.8777187 | 0.15835348 | 0.30468962 | 0.51972062 | 0.60325832 | 0.9958918 |
| HNRNPH1 | 175.040995 | -0.2158951 | 0.26044288 | -0.8289537 | 0.40713059 | 0.97691413 |
| CANX | 126.188991 | -0.075824 | 0.27756689 | -0.2731737 | 0.78471971 | 0.9958918 |
| MAML1 | 41.8483945 | 0.31359981 | 0.26453373 | 1.18548139 | 0.23582717 | 0.91962002 |
| SQSTM1 | 197.028794 | 0.27952125 | 0.3613345 | 0.77358029 | 0.43917903 | 0.98092839 |
| MRNIP | 24.4228587 | -0.579338 | 0.37669326 | -1.537957 | 0.12405912 | 0.79195137 |
| TBC1D9B | 32.6436646 | 0.31842475 | 0.30074302 | 1.05879351 | 0.28969383 | 0.95186143 |
| RNF130 | 72.2233375 | 0.18132745 | 0.24803878 | 0.73104478 | 0.4647518 | 0.98627231 |
| MAPK9 | 26.5992783 | -0.3302101 | 0.34444796 | -0.9586646 | 0.33772775 | 0.96750939 |
| GFPT2 | 43.54058 | -0.0346558 | 0.87131019 | -0.0397744 | 0.96827301 | 0.99935941 |
| CNOT6 | 44.9134198 | 0.05107416 | 0.26552832 | 0.19234921 | 0.84746868 | 0.99712573 |
| MGAT1 | 38.0147909 | -0.2255483 | 0.33373243 | -0.6758359 | 0.49914485 | 0.99194579 |
| RACK1 | 99.4697531 | -0.074199 | 0.39541131 | -0.1876502 | 0.85115086 | 0.9971731 |
| DUSP22 | 25.5235876 | 0.71981181 | 0.31378632 | 2.29395534 | 0.02179306 | 0.39642879 |
| EXOC2 | 89.7446886 | 0.07852891 | 0.22308642 | 0.35201118 | 0.72482988 | 0.9958918 |
| GMDS | 172.398498 | 0.58956111 | 0.27588414 | 2.13698801 | 0.03259897 | 0.48564518 |
| GMDS-DT | 169.150879 | 0.36526132 | 0.37758043 | 0.96737353 | 0.33335734 | 0.9654289 |
| WRNIP1 | 32.1302546 | 0.08331117 | 0.26362498 | 0.31602152 | 0.75198618 | 0.9958918 |
| SERPINB1 | 18.6749711 | -0.115776 | 0.41008269 | -0.2823236 | 0.7776954 | 0.9958918 |
| SERPINB9 | 22.3731314 | 0.79052234 | 0.34756635 | 2.27445017 | 0.02293893 | 0.410166 |
| SERPINB6 | 85.0827006 | -0.1190028 | 0.31783077 | -0.3744217 | 0.70809057 | 0.9958918 |
| NQO2 | 31.2403006 | 0.20677682 | 0.31278112 | 0.66109113 | 0.50855387 | 0.99397937 |
| RIPK1 | 49.275446 | 0.36646691 | 0.262826 | 1.39433279 | 0.16321713 | 0.85650717 |
| PSMG4 | 19.0721533 | -0.0999789 | 0.48781395 | -0.2049529 | 0.83760894 | 0.99692952 |
| PXDC1 | 31.6004888 | 0.00100767 | 0.37500617 | 0.00268706 | 0.99785604 | 0.99948864 |
| PRPF4B | 97.2751199 | -0.034342 | 0.23688761 | -0.1449716 | 0.88473333 | 0.99862036 |
| ECI2 | 32.943574 | 0.79557644 | 0.45124478 | 1.76307068 | 0.07788857 | 0.68056484 |
| CDYL | 90.9770013 | -0.0971892 | 0.22791627 | -0.4264252 | 0.66979803 | 0.9958918 |
| LYRM4 | 66.1513776 | 0.18904953 | 0.44213373 | 0.42758451 | 0.66895365 | 0.9958918 |
| FARS2 | 169.871465 | 0.07276492 | 0.22835045 | 0.31865458 | 0.74998846 | 0.9958918 |
| RREB1 | 139.626568 | -0.2272032 | 0.25858433 | -0.8786424 | 0.37959518 | 0.97217293 |
| SSR1 | 60.6630718 | 0.16760312 | 0.3690407 | 0.45415892 | 0.64971445 | 0.9958918 |
| SNRNP48 | 17.8120955 | 0.1860134 | 0.38284811 | 0.48586735 | 0.62706122 | 0.9958918 |
| SLC35B3 | 31.1605664 | -0.2663412 | 0.31618687 | -0.8423538 | 0.39958994 | 0.97500135 |
| TMEM14C | 25.9347686 | -0.1339927 | 0.39090171 | -0.3427784 | 0.73176514 | 0.9958918 |
| MAK | 32.5516565 | 0.05848955 | 0.37073044 | 0.15776839 | 0.87463931 | 0.9971731 |
| AL139807.1 | 34.4231147 | -0.6193491 | 0.60028999 | -1.0317498 | 0.30218933 | 0.95650251 |
| NEDD9 | 74.8642342 | -0.3393338 | 0.48354489 | -0.7017629 | 0.48282706 | 0.99002719 |
| HIVEP1 | 105.22611 | 0.07483114 | 0.42303273 | 0.17689209 | 0.85959316 | 0.9971731 |
| PHACTR1 | 65.9340842 | 1.12818278 | 0.57721065 | 1.95454256 | 0.05063709 | 0.57655256 |
| GFOD1 | 24.2367974 | 1.01278255 | 0.44020878 | 2.30068685 | 2.14E-02 | 0.39236216 |
| NOL7 | 25.3944514 | -0.2187528 | 0.34665234 | -0.6310438 | 0.52801188 | 0.9958918 |
| RANBP9 | 126.102612 | -0.1168412 | 0.27338108 | -0.427393 | 0.66909312 | 0.9958918 |

| | | | | | | |
|---|---|---|---|---|---|---|
| JARID2 | 174.55573 | 0.04311049 | 0.28225914 | 0.15273373 | 0.87860826 | 0.99722158 |
| DTNBP1 | 55.1158298 | -0.0104149 | 0.32832632 | -0.0317213 | 0.97469428 | 0.99935941 |
| MYLIP | 33.769105 | 0.64348039 | 0.47514196 | 1.3542908 | 0.17564362 | 0.87571571 |
| ATXN1 | 387.678019 | -0.1222009 | 0.30954479 | -0.3947761 | 0.69300811 | 0.9958918 |
| CAP2 | 44.8209399 | 0.59611436 | 0.38541225 | 1.54669284 | 0.12193733 | 0.79162318 |
| NUP153 | 112.346563 | -0.4565015 | 0.34752526 | -1.3135779 | 0.18898829 | 0.88139754 |
| KIF13A | 258.639164 | 0.12130267 | 0.27019856 | 0.44893897 | 0.65347568 | 0.9958918 |
| DEK | 62.6578028 | -0.1061236 | 0.26946371 | -0.3938326 | 0.69370466 | 0.9958918 |
| AL022068.1 | 23.378313 | 1.170191 | 0.52281083 | 2.23826847 | 0.02520355 | 0.42757124 |
| ID4 | 55.4018387 | 0.59098423 | 0.51573664 | 1.14590313 | 0.25183524 | 0.92631316 |
| MBOAT1 | 31.1852534 | 0.19962501 | 0.43298727 | 0.46104129 | 0.64476898 | 0.9958918 |
| E2F3 | 45.1120063 | 0.47670826 | 0.31742898 | 1.50177925 | 0.13315413 | 0.80664435 |
| CDKAL1 | 318.91281 | -0.0006579 | 0.16116322 | -0.0040821 | 0.99674299 | 0.99935941 |
| SOX4 | 40.59966 | -0.1124713 | 0.52593216 | -0.2138513 | 0.83066304 | 0.99592423 |
| CASC15 | 205.795787 | -2.0106036 | 0.39706463 | -5.0636684 | 4.11E-07 | 0.00058389 |
| DCDC2 | 23.1257163 | 0.25066227 | 0.42111514 | 0.59523452 | 0.55168672 | 0.9958918 |
| MRS2 | 18.9699192 | 0.26551084 | 0.36132589 | 0.73482373 | 0.46244685 | 0.98590351 |
| TDP2 | 22.4641066 | 0.68377707 | 0.37022332 | 1.84693137 | 0.06475709 | 0.62770158 |
| C6orf62 | 68.6538445 | -0.1006593 | 0.26446691 | -0.3806121 | 0.70349109 | 0.9958918 |
| RIPOR2 | 29.7133015 | 0.25736378 | 0.47471229 | 0.54214687 | 0.58771733 | 0.9958918 |
| CARMIL1 | 201.206266 | -0.1643587 | 0.40252554 | -0.4083187 | 0.68303968 | 0.9958918 |
| TRIM38 | 75.9348702 | 0.17187681 | 0.40645391 | 0.42286912 | 0.67239075 | 0.9958918 |
| HIST1H4C | 19.1391896 | 1.53911332 | 0.7076486 | 2.17496837 | 0.02963249 | 0.46634874 |
| HIST1H2AC | 88.1887982 | -0.0957117 | 0.89242949 | -0.1072484 | 0.91459188 | 0.99935941 |
| BTN2A1 | 20.2824203 | 0.00683139 | 0.4315615 | 0.01582947 | 0.98737044 | 0.99935941 |
| TRIM27 | 40.3420194 | 0.14332516 | 0.3323593 | 0.43123558 | 0.66629709 | 0.9958918 |
| GABBR1 | 66.5973335 | 0.21453494 | 0.26145138 | 0.82055389 | 0.41190042 | 0.97812179 |
| HLA-A | 189.504929 | 0.07717155 | 0.33563482 | 0.22992713 | 0.8181484 | 0.99592423 |
| TRIM26 | 25.7793962 | 0.1349726 | 0.35804515 | 0.37697089 | 0.70619524 | 0.9958918 |
| HCG17 | 76.1323642 | -0.0804718 | 0.37798603 | -0.2128961 | 0.83140801 | 0.99592423 |
| HLA-E | 112.917264 | 0.24004946 | 0.35694956 | 0.67250246 | 0.50126388 | 0.99213111 |
| GNL1 | 34.3020135 | 0.06326511 | 0.31153678 | 0.20307427 | 0.83907698 | 0.99692952 |
| ABCF1 | 31.2062387 | 0.30128007 | 0.41458252 | 0.72670714 | 0.46740538 | 0.98876285 |
| PPP1R10 | 45.7776104 | -0.3361354 | 0.31466733 | -1.0682245 | 0.28541924 | 0.94505166 |
| MRPS18B | 20.9299323 | 0.55817675 | 0.36810922 | 1.51633461 | 0.12943476 | 0.80543275 |
| PPP1R18 | 20.9004329 | 0.4036734 | 0.53480209 | 0.75480894 | 0.45036363 | 0.98092839 |
| MDC1 | 17.3878336 | 0.42112884 | 0.37937458 | 1.11006077 | 0.26697284 | 0.93533685 |
| TUBB | 81.7035834 | 0.30990161 | 0.5431395 | 0.57057461 | 0.56828803 | 0.9958918 |
| FLOT1 | 81.8143295 | 0.13455381 | 0.26500221 | 0.50774599 | 0.61163149 | 0.9958918 |
| IER3 | 123.184096 | 0.82973196 | 0.93644067 | 0.88604862 | 0.37559133 | 0.97217293 |
| HLA-C | 112.501365 | -0.0540153 | 0.49850841 | -0.1083539 | 0.913715 | 0.99935941 |
| HLA-B | 162.864091 | 0.23771486 | 0.50374515 | 0.47189508 | 0.63700168 | 0.9958918 |
| MICA | 22.28303 | 0.50606436 | 0.5495472 | 0.92087515 | 0.35711561 | 0.97217293 |
| PRRC2A | 38.358893 | 0.09095234 | 0.36198416 | 0.25126055 | 0.80161267 | 0.99592423 |
| BAG6 | 21.5202666 | 0.08847899 | 0.39431482 | 0.22438666 | 0.82245645 | 0.99592423 |
| DDAH2 | 19.0043853 | -0.2578661 | 0.4532212 | -0.568963 | 0.56938124 | 0.9958918 |
| CLIC1 | 75.5566361 | 0.52359126 | 0.39378753 | 1.32962884 | 0.18364059 | 0.87866798 |
| HSPA1A | 165.196969 | 0.03604426 | 0.86468661 | 0.04168477 | 0.96675 | 0.99935941 |
| HSPA1B | 76.2231952 | 0.30921519 | 1.00960857 | 0.30627235 | 0.75939729 | 0.9958918 |
| C6orf48 | 25.5620314 | -0.3588903 | 0.36467675 | -0.9841328 | 0.32505025 | 0.95992644 |
| TNXB | 111.010449 | 0.033085 | 0.5477509 | 0.06040155 | 0.95183582 | 0.99935941 |
| ATF6B | 26.0195114 | -0.4148096 | 0.32779624 | -1.2654497 | 0.20571017 | 0.89589437 |
| PBX2 | 22.5027992 | -0.3011783 | 0.32645623 | -0.9225688 | 0.35623194 | 0.97217293 |
| AL662796.1 | 62.389912 | 0.43504237 | 0.35781068 | 1.21584512 | 2.24E-01 | 0.91093521 |

| | | | | | | |
|---|---|---|---|---|---|---|
| HLA-DRA | 31.9761164 | 0.19738753 | 0.53437228 | 0.36938205 | 0.71184297 | 0.9958918 |
| HLA-DRB1 | 26.1174855 | -0.5831377 | 0.73528161 | -0.7930807 | 0.4277308 | 0.98092839 |
| BRD2 | 60.5239771 | 0.15582581 | 0.25134571 | 0.61996606 | 0.53528013 | 0.9958918 |
| SLC39A7 | 26.7514446 | -0.3299521 | 0.47291551 | -0.6976978 | 0.48536623 | 0.99046213 |
| RING1 | 21.9319679 | -0.2166377 | 0.32028318 | -0.6763943 | 0.49879031 | 0.99176652 |
| VPS52 | 17.0914018 | -0.0431194 | 0.35716943 | -0.1207254 | 0.90390853 | 0.99935941 |
| RPS18 | 94.368971 | -0.5131117 | 0.35143033 | -1.4600666 | 0.14427177 | 0.82317797 |
| RGL2 | 17.4690587 | 0.67280804 | 0.43651916 | 1.54130242 | 0.1232432 | 0.79184923 |
| TAPBP | 74.7438559 | 0.41802104 | 0.33090846 | 1.2632528 | 0.20649834 | 0.89589437 |
| PHF1 | 25.0629431 | 0.03148495 | 0.32671448 | 0.09636839 | 0.923228 | 0.99935941 |
| CUTA | 18.7793244 | -0.5726399 | 0.43567641 | -1.3143699 | 0.18872176 | 0.88139754 |
| SYNGAP1 | 34.6850902 | -0.2507472 | 0.36911026 | -0.6793286 | 0.49692969 | 0.99176652 |
| LEMD2 | 20.6036258 | 0.45898136 | 0.35747315 | 1.28396037 | 0.19915582 | 0.88864367 |
| NUDT3 | 19.703557 | 0.35870463 | 0.57715396 | 0.62150597 | 0.53426677 | 0.9958918 |
| C6orf106 | 90.1275409 | 0.28057258 | 0.2048987 | 1.36932338 | 0.17089821 | 0.87217428 |
| UHRF1BP1 | 33.1298046 | 0.28036298 | 0.38035892 | 0.73710109 | 0.46106088 | 0.98590351 |
| TAF11 | 27.6608548 | 0.08229101 | 0.29370696 | 0.28018066 | 0.7793389 | 0.9958918 |
| ANKS1A | 129.755682 | 0.05848069 | 0.25667727 | 0.22783741 | 0.81977264 | 0.99592423 |
| ZNF76 | 22.6664596 | -0.0475584 | 0.36816101 | -0.1291782 | 0.89721661 | 0.99935941 |
| PPARD | 59.0506956 | -0.0488168 | 0.30926664 | -0.1578471 | 0.87457728 | 0.9971731 |
| RPL10A | 102.267011 | -0.1024378 | 0.40210401 | -0.2547545 | 0.79891269 | 0.99592423 |
| TEAD3 | 64.3515458 | -0.3102373 | 0.31504117 | -0.9847516 | 0.3247461 | 0.95992644 |
| FKBP5 | 467.03695 | -1.0320984 | 0.64650593 | -1.5964253 | 0.11039387 | 0.77370222 |
| SRPK1 | 31.9173848 | 0.28344373 | 0.28503552 | 0.99441548 | 0.32002063 | 0.95992644 |
| MAPK14 | 100.223617 | -0.005872 | 0.29561468 | -0.0198637 | 0.98415211 | 0.99935941 |
| KCTD20 | 60.8941821 | 0.32899558 | 0.32960168 | 0.99816112 | 0.31820124 | 0.95992644 |
| STK38 | 72.6795719 | 0.08085517 | 0.25580541 | 0.31608078 | 0.75194119 | 0.9958918 |
| SRSF3 | 85.3467772 | -0.3630475 | 0.25782748 | -1.4081024 | 0.15910076 | 0.84665568 |
| CDKN1A | 48.3420511 | -0.2276173 | 0.53407352 | -0.4261909 | 6.70E-01 | 0.9958918 |
| C6orf89 | 46.8148448 | 0.35842347 | 0.26535647 | 1.35072444 | 0.17678372 | 0.87771577 |
| MTCH1 | 67.0971276 | -0.1612142 | 0.27641106 | -0.5832409 | 0.55973117 | 0.9958918 |
| PIM1 | 24.9801898 | -0.8277745 | 0.66472441 | -1.2452898 | 0.21302524 | 0.89738044 |
| TBC1D22B | 36.9560957 | -0.2070713 | 0.26075317 | -0.7941275 | 0.42712121 | 0.98091054 |
| RNF8 | 20.5924836 | 0.24158777 | 0.35599264 | 0.67863136 | 0.49737146 | 0.99176652 |
| CMTR1 | 21.3111349 | -0.3882584 | 0.37909067 | -1.0241835 | 0.30574863 | 0.95793395 |
| ZFAND3 | 716.338192 | -0.0345332 | 0.26284205 | -0.1313837 | 0.89547179 | 0.99935941 |
| BTBD9 | 377.152914 | 0.15661688 | 0.30175286 | 0.51902369 | 0.60374422 | 0.9958918 |
| GLO1 | 37.4105003 | 0.30969925 | 0.34538355 | 0.89668212 | 0.36988856 | 0.97217293 |
| DAAM2 | 191.765915 | -0.0480481 | 0.35618847 | -0.1348953 | 0.89269467 | 0.99935941 |
| MOCS1 | 30.3913941 | -0.0280447 | 0.41440446 | -0.0676747 | 0.94604456 | 0.99935941 |
| NFYA | 23.6182611 | -0.1165973 | 0.31925875 | -0.3652125 | 0.71495279 | 0.9958918 |
| TFEB | 19.5269445 | 1.03468627 | 0.38656933 | 2.67658657 | 0.00743763 | 0.22219179 |
| USP49 | 28.8116571 | 0.09234414 | 0.28773954 | 0.32092963 | 0.74826372 | 0.9958918 |
| CCND3 | 142.181049 | -0.2507297 | 0.48246734 | -0.5196822 | 0.60328509 | 0.9958918 |
| TAF8 | 22.6844123 | 0.09813228 | 0.42699413 | 0.22982116 | 0.81823075 | 0.99592423 |
| TRERF1 | 302.342862 | -0.6545195 | 0.3125157 | -2.0943571 | 0.03622818 | 0.51215534 |
| UBR2 | 220.994387 | -0.1816313 | 0.21602301 | -0.8407962 | 0.40046209 | 0.97500135 |
| BICRAL | 82.6133148 | 0.13848297 | 0.38073221 | 0.36372801 | 0.71606112 | 0.9958918 |
| RPL7L1 | 39.4683311 | 0.1460341 | 0.43919989 | 0.3325003 | 0.73951151 | 0.9958918 |
| PEX6 | 20.3652265 | 0.77529303 | 0.36289332 | 2.13642137 | 0.03264508 | 0.48564518 |
| KLHDC3 | 18.9450413 | 0.09759197 | 0.33080127 | 0.29501693 | 0.76798095 | 0.9958918 |
| RRP36 | 26.8306335 | 0.08448883 | 0.36414061 | 0.23202254 | 0.81652051 | 0.99592423 |
| PTK7 | 50.3812651 | -1.7835258 | 0.44933443 | -3.9692613 | 7.21E-05 | 0.01292273 |
| ZNF318 | 18.0211777 | 0.00329971 | 0.35502759 | 0.00929422 | 0.99258439 | 0.99935941 |

| | | | | | | |
|---|---|---|---|---|---|---|
| TJAP1 | 18.2581049 | 0.22554872 | 0.33240657 | 0.67853268 | 0.49743401 | 0.99176652 |
| POLR1C | 22.6559743 | -0.2671591 | 0.41325009 | -0.6464829 | 0.51796667 | 0.9958918 |
| YIPF3 | 21.9482728 | 0.26545224 | 0.3448575 | 0.76974473 | 0.44145133 | 0.98092839 |
| XPO5 | 75.9392643 | -0.1856982 | 0.36038224 | -0.5152814 | 0.60635643 | 0.9958918 |
| GTPBP2 | 18.200862 | -0.1963686 | 0.40370717 | -0.4864136 | 0.62667396 | 0.9958918 |
| VEGFA | 129.857716 | -0.9755577 | 0.80720016 | -1.2085697 | 0.2268282 | 0.91140536 |
| MRPL14 | 19.5619763 | 0.14230955 | 0.36965326 | 0.38498119 | 7.00E-01 | 0.9958918 |
| TMEM63B | 21.4058786 | 0.29593572 | 0.41701732 | 0.70964851 | 0.47792213 | 0.99002719 |
| HSP90AB1 | 156.884634 | 0.0231148 | 0.33058865 | 0.06992012 | 0.94425724 | 0.99935941 |
| CDC5L | 32.5023413 | 0.09752419 | 0.26686654 | 0.3654418 | 0.71478164 | 0.9958918 |
| SUPT3H | 175.46817 | -0.526753 | 0.20990003 | -2.5095422 | 0.01208878 | 0.2958939 |
| RUNX2 | 91.5552742 | -0.4226085 | 0.25039639 | -1.6877578 | 0.09145772 | 0.72711043 |
| CLIC5 | 28.7309377 | 0.62993871 | 1.02696981 | 0.61339555 | 0.53961483 | 0.9958918 |
| RCAN2 | 350.611497 | 0.03866502 | 0.30824576 | 0.12543568 | 0.90017864 | 0.99935941 |
| SLC25A27 | 61.2356356 | -0.4117094 | 0.41608472 | -0.9894846 | 0.32242611 | 0.95992644 |
| ADGRF5 | 191.056098 | -0.0397511 | 0.28721163 | -0.1384035 | 0.8899215 | 0.99935941 |
| TNFRSF21 | 32.2964496 | 0.31298298 | 0.51091566 | 0.61259227 | 0.54014597 | 0.9958918 |
| CD2AP | 117.352656 | -0.0984445 | 0.24921733 | -0.3950148 | 0.69283196 | 0.9958918 |
| MUT | 27.7854408 | -0.3880994 | 0.36312974 | -1.0687625 | 0.2851767 | 0.94505166 |
| PKHD1 | 58.2315038 | -0.0344109 | 0.49754315 | -0.0691616 | 0.94486096 | 0.99935941 |
| EFHC1 | 46.2254278 | -0.8056239 | 0.33678007 | -2.3921366 | 0.01675061 | 0.34578631 |
| TRAM2 | 121.352775 | -0.9583484 | 0.41282109 | -2.3214618 | 0.02026193 | 0.38543207 |
| GSTA4 | 31.1075955 | -0.1315484 | 0.41329287 | -0.3182934 | 0.75026236 | 0.9958918 |
| ICK | 21.6354649 | 0.6766428 | 0.43646185 | 1.55029083 | 0.12107173 | 0.79021701 |
| FBXO9 | 59.2034173 | 0.21307608 | 0.21947363 | 0.97085051 | 0.33162273 | 0.9654289 |
| ELOVL5 | 165.216217 | -0.3024748 | 0.29212028 | -1.035446 | 0.30046067 | 0.95597956 |
| GCLC | 24.3541948 | -0.3051772 | 0.47428999 | -0.6434401 | 0.5199386 | 0.9958918 |
| COL21A1 | 57.4535954 | -0.5105107 | 0.3449595 | -1.4799148 | 1.39E-01 | 0.8153985 |
| DST | 604.393977 | -0.4727927 | 0.30426179 | -1.553901 | 0.12020804 | 0.79021701 |
| ZNF451 | 126.946856 | -0.1736328 | 0.31724668 | -0.5473115 | 0.58416473 | 0.9958918 |
| BAG2 | 30.8996336 | -1.1133782 | 0.5281463 | -2.1080868 | 0.03502348 | 0.50468149 |
| RAB23 | 37.2561051 | -0.9091754 | 0.33829046 | -2.6875586 | 0.00719765 | 0.21767701 |
| PRIM2 | 102.834979 | -0.7925909 | 0.25046738 | -3.1644474 | 0.00155378 | 0.0868147 |
| AL445250.1 | 39.4888833 | -1.2080159 | 0.58011706 | -2.0823658 | 0.03730907 | 0.52013314 |
| PTP4A1 | 50.6079626 | -0.8618328 | 0.46893796 | -1.8378397 | 0.06608603 | 0.63381211 |
| PHF3 | 139.818726 | -0.1678172 | 0.24121253 | -0.6957233 | 0.48660213 | 0.99046213 |
| ADGRB3 | 529.575509 | -1.3210104 | 0.44147581 | -2.99226 | 0.0027692 | 0.12186152 |
| LMBRD1 | 92.347838 | 0.09551307 | 0.28063345 | 0.34034813 | 0.73359437 | 0.9958918 |
| COL19A1 | 31.2076139 | 0.18273542 | 0.33915841 | 0.53879077 | 0.59003123 | 0.9958918 |
| FAM135A | 121.441145 | -0.1815802 | 0.20406351 | -0.889822 | 0.37356147 | 0.97217293 |
| SMAP1 | 108.269059 | 0.07343949 | 0.23592689 | 0.31128072 | 0.75558722 | 0.9958918 |
| OGFRL1 | 38.8095854 | -0.4208753 | 0.26868523 | -1.566425 | 0.11724915 | 0.7849382 |
| AL136164.2 | 31.8077916 | 0.31837011 | 0.43918048 | 0.72491864 | 0.46850194 | 0.98910524 |
| KCNQ5 | 96.3734611 | 1.47511654 | 0.42676278 | 3.45652579 | 0.00054719 | 0.04787234 |
| OOEP | 86.2643101 | -0.1812628 | 0.34203065 | -0.5299607 | 0.59613917 | 0.9958918 |
| MTO1 | 23.5069324 | 0.3426374 | 0.32810053 | 1.04430612 | 0.29634378 | 0.95393362 |
| EEF1A1 | 407.76805 | 0.27750557 | 0.36441474 | 0.76151029 | 0.44635234 | 0.98092839 |
| SLC17A5 | 41.1501072 | 0.22048737 | 0.42299067 | 0.52125823 | 0.60218689 | 0.9958918 |
| COL12A1 | 470.956735 | -0.8655559 | 0.40961322 | -2.1131055 | 0.03459174 | 0.50439422 |
| COX7A2 | 40.6870085 | -0.1445082 | 0.42974132 | -0.3362678 | 0.73666892 | 0.9958918 |
| TMEM30A | 79.5746174 | -0.112521 | 0.33801968 | -0.3328829 | 0.73922264 | 0.9958918 |
| FILIP1 | 203.875154 | -0.1982099 | 0.44027709 | -0.4501935 | 0.65257092 | 0.9958918 |
| SENP6 | 320.782221 | -0.017802 | 0.18038247 | -0.0986903 | 0.92138414 | 0.99935941 |
| MYO6 | 118.463271 | 0.24625224 | 0.2559373 | 0.96215844 | 0.33597002 | 0.96640422 |

| | | | | | | |
|---|---|---|---|---|---|---|
| PHIP | 241.578494 | 0.15553905 | 0.2121858 | 0.73303232 | 0.46353872 | 0.98590351 |
| HMGN3 | 64.8910693 | 0.09110292 | 0.26437325 | 0.34459962 | 0.73039539 | 0.9958918 |
| LCA5 | 18.2297321 | 0.54980208 | 0.35815187 | 1.53510875 | 0.12475712 | 0.79195137 |
| BCKDHB | 147.42832 | 0.99835104 | 0.32155437 | 3.10476589 | 0.0019043 | 0.09345751 |
| TENT5A | 177.133774 | -0.3446944 | 0.50527035 | -0.682198 | 0.49511376 | 0.99176652 |
| IBTK | 96.3335728 | -0.1548887 | 0.22040536 | -0.7027445 | 0.48221499 | 0.99002719 |
| UBE3D | 53.6739962 | -0.1837437 | 0.24028638 | -0.7646861 | 0.44445847 | 0.98092839 |
| DOPEY1 | 93.4022223 | -0.3383418 | 0.27957153 | -1.2102157 | 0.22619614 | 0.91093521 |
| PGM3 | 38.7348742 | -0.1193663 | 0.30451068 | -0.3919939 | 0.69506276 | 0.9958918 |
| ME1 | 31.1042038 | -0.511728 | 0.62072561 | -0.8244029 | 0.40971069 | 0.97812179 |
| CYB5R4 | 34.9413827 | 0.40483384 | 0.31323024 | 1.29244814 | 0.19620199 | 0.88391893 |
| CEP162 | 36.5668552 | -0.1394759 | 0.31788353 | -0.4387643 | 0.66083236 | 0.9958918 |
| SNX14 | 73.5969246 | -0.2573789 | 0.25619699 | -1.0046135 | 0.31508301 | 0.95992644 |
| SYNCRIP | 38.3413235 | -0.1343686 | 0.27807856 | -0.4832038 | 0.62895102 | 0.9958918 |
| ZNF292 | 266.824513 | -0.2989971 | 0.33350797 | -0.8965217 | 0.36997421 | 0.97217293 |
| SMIM8 | 29.4423747 | -0.1552709 | 0.35966819 | -0.4317058 | 0.66595523 | 0.9958918 |
| RARS2 | 96.9827052 | -0.3824119 | 0.24914208 | -1.5349149 | 0.12480473 | 0.79195137 |
| ORC3 | 56.8136054 | -0.1242558 | 0.30342531 | -0.4095104 | 0.68216516 | 0.9958918 |
| AKIRIN2 | 22.9918258 | 0.35539488 | 0.38048798 | 0.93405023 | 0.35027798 | 0.9693503 |
| RNGTT | 139.161706 | -0.2643417 | 0.31473063 | -0.8398983 | 0.40096543 | 0.97500135 |
| PNRC1 | 81.8008805 | -0.1699225 | 0.34435861 | -0.4934465 | 0.62169716 | 0.9958918 |
| PM20D2 | 19.5952863 | 0.81829392 | 0.40051433 | 2.04310771 | 0.04104178 | 0.54347495 |
| UBE2J1 | 50.2865662 | 0.1371779 | 0.26740545 | 0.5129959 | 0.60795419 | 0.9958918 |
| ANKRD6 | 53.6884307 | 0.7788688 | 0.42726842 | 1.82290281 | 0.06831811 | 0.647832 |
| LYRM2 | 23.6575876 | -0.2855047 | 0.3510657 | -0.8132515 | 0.41607385 | 0.97812179 |
| MDN1 | 90.3892162 | -0.6058033 | 0.27853796 | -2.1749398 | 0.02963463 | 0.46634874 |
| CASP8AP2 | 38.1097758 | -0.2773134 | 0.24941506 | -1.111855 | 0.26620048 | 0.93533685 |
| BACH2 | 87.9437961 | -0.0129071 | 0.4368199 | -0.0295479 | 0.97642765 | 0.99935941 |
| MAP3K7 | 76.4844128 | -0.1259681 | 0.30357722 | -0.4149458 | 0.67818156 | 0.9958918 |
| UFL1 | 54.8962043 | -0.0153911 | 0.21907544 | -0.0702546 | 0.94399104 | 0.99935941 |
| FHL5 | 117.411079 | -0.0778421 | 0.535728 | -0.1453016 | 0.88447277 | 0.99862036 |
| MMS22L | 30.274453 | -0.2664864 | 0.44213132 | -0.6027314 | 0.54668737 | 0.9958918 |
| FBXL4 | 74.7954917 | -0.133094 | 0.32085323 | -0.4148127 | 0.678279 | 0.9958918 |
| PNISR | 287.120284 | 0.06758144 | 0.21504472 | 0.31426691 | 0.75331833 | 0.9958918 |
| USP45 | 46.7168339 | 0.20949426 | 0.31934899 | 0.65600415 | 0.51182146 | 0.9958918 |
| CCNC | 49.6355756 | -0.1562265 | 0.24061488 | -0.6492804 | 0.51615716 | 0.9958918 |
| ASCC3 | 228.375348 | 0.09165612 | 0.25699468 | 0.35664598 | 0.72135685 | 0.9958918 |
| GRIK2 | 90.2055528 | 0.77388682 | 0.48236928 | 1.60434516 | 1.09E-01 | 7.73E-01 |
| HACE1 | 54.9107731 | -0.0968043 | 0.35413137 | -0.2733572 | 0.78457868 | 0.9958918 |
| PREP | 43.2715538 | 0.37726755 | 0.33839188 | 1.11488356 | 0.26490032 | 0.93499423 |
| ATG5 | 71.2096471 | 0.11987785 | 0.2244991 | 0.5339792 | 0.59335593 | 0.9958918 |
| QRSL1 | 23.1078071 | -0.0288328 | 0.34932342 | -0.0825389 | 0.93421817 | 0.99935941 |
| LINC02532 | 20.5739295 | -0.5790998 | 0.49363097 | -1.1731431 | 0.24073841 | 0.92426606 |
| CD24 | 66.3487146 | 0.08004662 | 0.29647245 | 0.26999682 | 0.7871627 | 0.99592423 |
| PDSS2 | 132.752983 | -0.0687316 | 0.20950102 | -0.3280728 | 0.74285662 | 0.9958918 |
| SOBP | 165.721525 | 0.51411071 | 0.30446303 | 1.68858171 | 0.09129962 | 0.72711043 |
| SEC63 | 136.813187 | -0.0544496 | 0.20852984 | -0.2611119 | 0.79400619 | 0.99592423 |
| OSTM1 | 44.6722984 | 0.12929713 | 0.366318 | 0.35296417 | 0.7241153 | 0.9958918 |
| SNX3 | 66.9233888 | 0.01295121 | 0.24601261 | 0.0526445 | 0.95801516 | 0.99935941 |
| AFG1L | 34.2248073 | -0.4234894 | 0.28583703 | -1.4815764 | 0.13845305 | 0.81520215 |
| FOXO3 | 444.963348 | -0.2522954 | 0.37288329 | -0.6766068 | 0.49865548 | 0.99176652 |
| ARMC2 | 51.7607175 | -0.0179843 | 0.43656937 | -0.0411945 | 0.96714081 | 0.99935941 |
| SESN1 | 141.644855 | 0.14578832 | 0.26137984 | 0.55776422 | 0.5770054 | 0.9958918 |
| CEP57L1 | 36.5381302 | 0.20815781 | 0.312714 | 0.66564916 | 0.50563538 | 0.99397937 |

| | | | | | | |
|---|---|---|---|---|---|---|
| CD164 | 70.3535483 | 0.1438358 | 0.2618546 | 0.54929646 | 0.58280202 | 0.9958918 |
| FIG4 | 32.2307417 | 0.0953975 | 0.27110735 | 0.3518809 | 0.72492759 | 0.9958918 |
| WASF1 | 34.9639334 | -0.81354 | 0.36575963 | -2.2242477 | 0.02613178 | 0.43845312 |
| CDC40 | 33.5379992 | -0.0217741 | 0.32776137 | -0.0664329 | 0.94703319 | 0.99935941 |
| METTL24 | 32.6431879 | -0.7233921 | 0.49058596 | -1.474547 | 0.14033437 | 0.8153985 |
| CDK19 | 97.2420713 | -0.2878782 | 0.34698623 | -0.8296532 | 0.40673488 | 0.97671046 |
| AMD1 | 44.7320403 | -0.4616175 | 0.36543658 | -1.2631946 | 0.20651927 | 0.89589437 |
| GTF3C6 | 24.3875753 | 0.46364354 | 0.36304193 | 1.27710739 | 0.20156434 | 0.89319989 |
| RPF2 | 28.5649642 | -0.1461363 | 0.30073543 | -0.4859299 | 0.62701687 | 0.9958918 |
| REV3L | 1025.62617 | -0.6322867 | 0.4751831 | -1.330617 | 0.18331505 | 0.87866798 |
| TRAF3IP2-AS | 231.174001 | 0.27158736 | 0.3635181 | 0.74710821 | 0.45499824 | 0.98173285 |
| TRAF3IP2 | 33.8513423 | 0.35290362 | 0.2725152 | 1.29498689 | 1.95E-01 | 0.88391893 |
| FYN | 193.950824 | -0.1082467 | 0.38380343 | -0.2820368 | 0.77791529 | 0.9958918 |
| TUBE1 | 29.6920989 | -0.6097042 | 0.39749923 | -1.5338499 | 0.12506658 | 0.79233388 |
| LAMA4 | 160.400474 | -1.3371602 | 0.41435328 | -3.2271017 | 0.00125051 | 0.07658331 |
| MARCKS | 85.3391794 | -0.1103596 | 0.4556193 | -0.2422188 | 0.80861059 | 0.99592423 |
| HDAC2 | 37.3421543 | 0.01782826 | 0.27355961 | 0.06517138 | 0.94803755 | 0.99935941 |
| NT5DC1 | 126.032436 | 1.16186716 | 0.23556594 | 4.93223743 | 8.13E-07 | 0.00074678 |
| DSE | 54.8389847 | -0.7328733 | 0.30051078 | -2.4387589 | 0.0147378 | 0.33025022 |
| TSPYL1 | 28.6022321 | 0.29928101 | 0.31979397 | 0.9358557 | 0.34934749 | 0.9692706 |
| RWDD1 | 22.4422402 | -0.111557 | 0.36344598 | -0.3069423 | 0.75888725 | 0.9958918 |
| KPNA5 | 52.3185713 | 0.03556165 | 0.27278642 | 0.13036443 | 0.8962781 | 0.99935941 |
| FAM162B | 26.2325484 | 0.12051488 | 0.65957086 | 0.1827171 | 0.85502 | 0.9971731 |
| DCBLD1 | 40.9242711 | -0.7287533 | 0.30368757 | -2.399681 | 0.01640936 | 0.34578631 |
| GOPC | 37.7759477 | 0.44435471 | 0.2855246 | 1.55627471 | 0.11964278 | 0.79021701 |
| NUS1 | 45.4832912 | 0.42023466 | 0.30321741 | 1.38591861 | 0.16577176 | 0.85938344 |
| SLC35F1 | 157.488362 | -1.0771374 | 0.4534701 | -2.3753218 | 0.01753366 | 0.35014904 |
| CEP85L | 213.869729 | -0.0915959 | 0.25713498 | -0.3562172 | 0.7216779 | 0.9958918 |
| PLN | 27.6812305 | 0.93956982 | 0.37192631 | 2.52622573 | 0.01152954 | 0.28819922 |
| MCM9 | 47.53764 | 0.2500439 | 0.29891668 | 0.8365003 | 0.40287351 | 0.97512144 |
| MAN1A1 | 126.685222 | 0.35339778 | 0.33489456 | 1.05525087 | 0.2913106 | 0.95186143 |
| TBC1D32 | 63.2419514 | -0.8271622 | 0.32286573 | -2.5619386 | 0.01040897 | 0.27417754 |
| GJA1 | 25.551719 | -0.3019318 | 0.36631917 | -0.8242314 | 0.40980807 | 0.97812179 |
| SERINC1 | 79.8245907 | -0.1728366 | 0.22754854 | -0.7595594 | 0.44751801 | 0.98092839 |
| RNF217 | 117.80118 | -0.6438693 | 0.41173213 | -1.5638064 | 0.11786305 | 0.78539225 |
| HDDC2 | 21.6177784 | 0.08930287 | 0.38615824 | 0.23125978 | 0.81711299 | 0.99592423 |
| NCOA7 | 148.342065 | 0.02304034 | 0.39388142 | 0.05849562 | 0.95335385 | 0.99935941 |
| TRMT11 | 58.1003139 | -0.4107755 | 0.25904328 | -1.5857408 | 0.11279813 | 0.77793929 |
| RSPO3 | 60.0454945 | -1.2879008 | 0.87947748 | -1.4643931 | 0.14308656 | 0.81959713 |
| RNF146 | 35.4414863 | 0.11621357 | 0.34860652 | 0.33336602 | 0.73885801 | 0.9958918 |
| ECHDC1 | 62.9365582 | -0.0992207 | 0.22353384 | -0.4438731 | 0.65713433 | 0.9958918 |
| PTPRK | 381.159716 | -0.3963909 | 0.23823422 | -1.6638704 | 0.09613834 | 0.73903832 |
| LAMA2 | 819.048325 | -0.2052046 | 0.35567421 | -0.5769454 | 0.56397636 | 0.9958918 |
| ARHGAP18 | 64.0075619 | 0.1366002 | 0.38087312 | 0.35865015 | 0.71985682 | 0.9958918 |
| L3MBTL3 | 33.679803 | 0.33634682 | 0.31148316 | 1.07982344 | 0.28022081 | 0.94419503 |
| SAMD3 | 105.471838 | 0.84133134 | 0.39803664 | 2.11370326 | 0.03454062 | 0.50439422 |
| TMEM200A | 77.3094445 | 0.71250593 | 0.51933024 | 1.37197081 | 0.17007253 | 0.87006748 |
| EPB41L2 | 448.364172 | 0.14364204 | 0.23285283 | 0.61687908 | 0.53731448 | 0.9958918 |
| AKAP7 | 33.5394088 | -0.2177199 | 0.30484418 | -0.7142005 | 0.47510321 | 0.98910524 |
| MED23 | 49.3681049 | -0.3996975 | 0.25712117 | -1.5545103 | 0.12006274 | 0.79021701 |
| ENPP1 | 54.1210628 | -0.3067128 | 0.60892825 | -0.5036929 | 0.61447719 | 0.9958918 |
| CTGF | 229.463067 | -1.8287653 | 0.86146633 | -2.1228517 | 0.03376628 | 0.49530613 |
| AL133346.1 | 50.4762772 | 1.63534067 | 0.87721088 | 1.86425033 | 0.06228656 | 0.62054234 |
| MOXD1 | 359.225319 | -0.6574476 | 0.44811289 | -1.4671473 | 0.142336 | 0.81652619 |

| | | | | | | |
|---|---|---|---|---|---|---|
| STX7 | 72.3874792 | -0.1582316 | 0.35141848 | -0.4502654 | 0.65251911 | 0.9958918 |
| RPS12 | 90.1487346 | -0.2969251 | 0.41531513 | -0.7149392 | 0.47464662 | 0.98910524 |
| EYA4 | 238.9966 | -0.7646567 | 0.21576929 | -3.5438624 | 0.00039431 | 0.03863721 |
| TARID | 76.5586671 | -1.8360652 | 0.44885178 | -4.0905823 | 4.30E-05 | 0.00854652 |
| TCF21 | 43.9961853 | -1.7252095 | 0.54574324 | -3.161211 | 0.00157115 | 0.0868147 |
| TBPL1 | 26.3532436 | -0.1211359 | 0.3405102 | -0.3557483 | 0.72202904 | 0.9958918 |
| SGK1 | 44.4288822 | -1.0933142 | 0.37861034 | -2.8877029 | 0.00388066 | 0.15300235 |
| HBS1L | 55.4297205 | -0.1167658 | 0.23542388 | -0.4959812 | 0.61990767 | 0.9958918 |
| AHI1 | 295.188952 | -0.1623303 | 0.20303492 | -0.7995191 | 4.24E-01 | 0.97828057 |
| PDE7B | 540.410609 | 0.52279951 | 0.37929914 | 1.37833031 | 0.16810133 | 0.86560961 |
| AL138828.1 | 92.9706623 | 0.21476252 | 0.57710563 | 0.37213728 | 0.70979064 | 0.9958918 |
| BCLAF1 | 135.207305 | -0.1628902 | 0.21219473 | -0.7676448 | 0.44269823 | 0.98092839 |
| MAP7 | 31.9704802 | 0.26834919 | 0.35533283 | 0.75520518 | 0.45012588 | 0.98092839 |
| MAP3K5 | 746.555468 | -0.2306249 | 0.30318167 | -0.7606821 | 0.44684695 | 0.98092839 |
| PEX7 | 23.4986348 | -0.3599586 | 0.31964497 | -1.1261202 | 0.26011466 | 0.93055907 |
| IFNGR1 | 55.5754551 | 0.07933353 | 0.27979786 | 0.28353873 | 0.77676389 | 0.9958918 |
| TNFAIP3 | 66.8628567 | 1.50536641 | 1.09924318 | 1.36945712 | 0.17085643 | 0.87217428 |
| HEBP2 | 24.6785178 | 0.35002343 | 0.30063167 | 1.16429326 | 0.24430519 | 0.92546331 |
| NHSL1 | 146.52433 | -0.6250025 | 0.41281123 | -1.5140152 | 0.13002199 | 0.80565822 |
| REPS1 | 120.594328 | 0.83905302 | 0.23663152 | 3.545821 | 0.00039139 | 0.03863721 |
| HECA | 25.9281948 | 0.19017936 | 0.32005974 | 0.59419956 | 0.55237865 | 0.9958918 |
| CITED2 | 24.0139161 | -1.3306292 | 0.41416518 | -3.2127983 | 0.00131449 | 0.07983599 |
| VTA1 | 34.1105433 | 0.10497478 | 0.26731518 | 0.39270042 | 0.69454076 | 0.9958918 |
| HIVEP2 | 373.794511 | 0.16431654 | 0.32489506 | 0.50575267 | 0.61303029 | 0.9958918 |
| AIG1 | 70.97379 | 0.11938077 | 0.26862293 | 0.44441764 | 0.65674066 | 0.9958918 |
| PEX3 | 35.6304944 | 0.06266 | 0.31570539 | 0.19847617 | 0.84267253 | 0.99692952 |
| PHACTR2 | 127.217973 | -0.6624304 | 0.27701935 | -2.3912785 | 0.01678981 | 0.34578631 |
| PLAGL1 | 114.698624 | -0.6354364 | 0.31700565 | -2.0044955 | 0.04501701 | 0.55894748 |
| SF3B5 | 27.2830205 | 0.62176622 | 0.37587805 | 1.65417007 | 0.09809296 | 0.74364783 |
| UTRN | 987.517783 | -0.8055042 | 0.30406208 | -2.6491439 | 0.0080696 | 0.23533118 |
| EPM2A | 26.9931233 | -0.1012922 | 0.41221432 | -0.245727 | 0.80589357 | 0.99592423 |
| SHPRH | 125.315071 | 0.10020421 | 0.1763837 | 0.56810359 | 0.56996463 | 0.9958918 |
| STXBP5 | 134.17422 | 0.37177021 | 0.18824781 | 1.97489791 | 0.04827973 | 0.57134906 |
| SAMD5 | 37.29667 | -0.0398005 | 0.47374662 | -0.0840121 | 0.93304679 | 0.99935941 |
| SASH1 | 129.605713 | -0.7412321 | 0.3446784 | -2.1505035 | 0.03151541 | 0.48041788 |
| UST | 152.601706 | -0.9296199 | 0.42163492 | -2.2047982 | 0.02746826 | 0.44841843 |
| TAB2 | 101.671967 | -0.0930725 | 0.26845784 | -0.346693 | 0.72882197 | 0.9958918 |
| PPIL4 | 46.0667198 | -0.2739704 | 0.30241166 | -0.9059519 | 0.36496131 | 0.97217293 |
| GINM1 | 49.2255094 | 0.32103039 | 0.2369716 | 1.35472096 | 0.17550648 | 0.875626 |
| KATNA1 | 22.4704013 | 0.27019462 | 0.35133723 | 0.76904638 | 0.44186578 | 0.98092839 |
| LATS1 | 75.4640243 | 0.0385557 | 0.21560217 | 0.17882797 | 0.85807278 | 0.9971731 |
| NUP43 | 23.0231405 | -0.2620046 | 0.32370748 | -0.8093869 | 0.41829264 | 0.97812179 |
| PCMT1 | 50.3181362 | 0.1303999 | 0.24259427 | 0.5375226 | 0.59090668 | 0.9958918 |
| PLEKHG1 | 119.188205 | -0.214541 | 0.54647875 | -0.3925881 | 0.69462374 | 0.9958918 |
| MTHFD1L | 367.034001 | 0.15026179 | 0.52044333 | 0.28871883 | 0.77279655 | 0.9958918 |
| AKAP12 | 1148.88038 | -1.3660586 | 0.56619821 | -2.4126862 | 0.01583545 | 0.34027688 |
| RMND1 | 24.2229344 | 0.06493935 | 0.30052477 | 0.21608649 | 0.82892033 | 0.99592423 |
| CCDC170 | 21.3065438 | -0.0660711 | 0.51060797 | -0.1293968 | 0.89704365 | 0.99935941 |
| ESR1 | 161.603763 | -0.3067103 | 0.45039388 | -0.6809824 | 0.49588265 | 0.99176652 |
| SYNE1 | 827.131443 | 0.83895704 | 0.3510162 | 2.39008067 | 0.01684467 | 0.34578631 |
| CNKSR3 | 111.827431 | 0.19690039 | 0.35988408 | 0.54712171 | 0.58429512 | 0.9958918 |
| SCAF8 | 203.176506 | -0.2834361 | 0.18714863 | -1.5144971 | 0.1298998 | 0.80565822 |
| TIAM2 | 124.106035 | 0.14027659 | 0.18724694 | 0.74915291 | 0.45376505 | 0.98092839 |
| TFB1M | 29.8343069 | 0.28771686 | 0.31962056 | 0.90018258 | 0.36802309 | 0.97217293 |

| | | | | | | |
|---|---|---|---|---|---|---|
| ARID1B | 706.432638 | -0.0332505 | 0.17573742 | -0.1892059 | 0.84993148 | 0.9971731 |
| TMEM242 | 27.2303967 | 0.08706685 | 0.36879654 | 0.23608368 | 0.81336773 | 0.99592423 |
| ZDHHC14 | 226.066169 | 0.11647395 | 0.26792747 | 0.43472195 | 0.66376427 | 0.9958918 |
| SNX9 | 201.391456 | 0.22315763 | 0.29334959 | 0.76072248 | 0.44682284 | 0.98092839 |
| SYNJ2 | 111.72544 | -0.8275609 | 0.32664205 | -2.5335407 | 0.01129166 | 0.28321649 |
| TULP4 | 156.516462 | 0.35920774 | 0.28135124 | 1.27672351 | 0.20169988 | 0.89319989 |
| TMEM181 | 59.9162524 | 0.01957776 | 0.33371714 | 0.05866574 | 0.95321835 | 0.99935941 |
| DYNLT1 | 47.1739919 | 0.22312565 | 0.28858486 | 0.77317169 | 0.43942078 | 0.98092839 |
| EZR | 40.7811933 | 0.37367022 | 0.28693766 | 1.30226969 | 0.19282421 | 0.88158763 |
| AL035530.2 | 21.1511768 | 0.91327326 | 0.42482671 | 2.14975482 | 0.03157461 | 0.48041788 |
| WTAP | 136.029356 | 0.46761727 | 0.42486647 | 1.10062172 | 0.27106133 | 0.93776436 |
| TCP1 | 29.0460046 | 0.25984381 | 0.32465498 | 0.8003691 | 0.42349698 | 0.97828057 |
| IGF2R | 192.450631 | -0.2126132 | 0.17689076 | -1.2019465 | 0.22938424 | 0.91455844 |
| SLC22A3 | 70.0297558 | 0.71957756 | 0.82242462 | 0.87494652 | 0.381603 | 0.97217293 |
| MAP3K4 | 100.005297 | -0.4735084 | 0.25988623 | -1.8219832 | 0.06845753 | 0.64792295 |
| AGPAT4 | 40.711212 | -0.6860949 | 0.3152178 | -2.1765741 | 0.02951236 | 0.46634874 |
| PRKN | 311.280239 | -0.1074621 | 0.242822 | -0.4425552 | 0.65808751 | 0.9958918 |
| QKI | 411.80493 | 0.12944213 | 0.24033569 | 0.53858885 | 0.59017058 | 0.9958918 |
| PDE10A | 369.22426 | 1.40162566 | 0.40759115 | 3.43880294 | 0.00058429 | 0.05051723 |
| SFT2D1 | 33.1288915 | 0.1332592 | 0.27740081 | 0.48038505 | 0.63095362 | 0.9958918 |
| MPC1 | 37.2737195 | 0.24072411 | 0.31094222 | 0.77417635 | 0.43882651 | 0.98092839 |
| RPS6KA2 | 76.4943703 | -1.3318862 | 0.28291474 | -4.7077298 | 2.50E-06 | 0.00141604 |
| FGFR1OP | 37.2750429 | -0.3042024 | 0.30241682 | -1.0059044 | 0.31446154 | 0.95992644 |
| AFDN | 122.534095 | -0.4702597 | 0.32016084 | -1.4688232 | 0.14188075 | 0.81652619 |
| SMOC2 | 87.0188173 | -0.7410571 | 0.44302647 | -1.6727152 | 0.09438338 | 0.7340733 |
| BX322234.1 | 26.5848615 | 0.04748819 | 0.53486576 | 0.08878525 | 0.92925258 | 0.99935941 |
| THBS2 | 198.420781 | -0.089371 | 0.48419215 | -0.1845776 | 0.85356033 | 0.9971731 |
| WDR27 | 59.665715 | 0.02286584 | 0.29239104 | 0.07820294 | 0.93766663 | 0.99935941 |
| PHF10 | 35.7219881 | -0.4308593 | 0.36860311 | -1.1688975 | 0.24244491 | 0.92534155 |
| ERMARD | 17.1355512 | -0.6737527 | 0.37960071 | -1.7748983 | 0.07591468 | 0.67378863 |
| FAM120B | 134.956964 | 0.04298114 | 0.24429191 | 0.17594171 | 0.86033974 | 0.9971731 |
| PSMB1 | 63.7691204 | -0.0835151 | 0.33034346 | -0.2528129 | 0.80041284 | 0.99592423 |
| PDCD2 | 17.9147865 | 0.05329862 | 0.38084293 | 0.1399491 | 0.88870021 | 0.99935941 |
| FAM20C | 61.2670621 | 0.93826369 | 0.32980025 | 2.84494535 | 0.0044419 | 0.16654875 |
| PDGFA | 44.1886082 | 0.26485352 | 0.31879435 | 0.83079743 | 0.40608808 | 0.97637555 |
| PRKAR1B | 19.4863415 | -0.0205649 | 0.43800631 | -0.0469512 | 0.9625521 | 0.99935941 |
| DNAAF5 | 17.7292661 | -0.008981 | 0.37779882 | -0.023772 | 0.98103446 | 0.99935941 |
| SUN1 | 110.326816 | -0.2557632 | 0.39509249 | -0.6473502 | 0.51740532 | 0.9958918 |
| C7orf50 | 38.2968845 | -0.3858037 | 0.26572619 | -1.4518843 | 0.14653378 | 0.8301633 |
| MICALL2 | 35.3563276 | 0.88778296 | 0.38628254 | 2.2982736 | 0.02154622 | 0.39388852 |
| INTS1 | 22.8690345 | 0.09835375 | 0.38749154 | 0.25382168 | 0.79963333 | 0.99592423 |
| MAFK | 23.3886845 | -0.3712086 | 0.40524916 | -0.9160008 | 0.35966645 | 0.97217293 |
| MAD1L1 | 48.1689179 | -0.0841607 | 0.35669391 | -0.2359466 | 0.81347414 | 0.99592423 |
| SNX8 | 19.6947595 | -0.9735989 | 0.49981997 | -1.9478991 | 0.05142704 | 0.57655256 |
| EIF3B | 60.9295351 | 0.15835267 | 0.33958758 | 0.46630878 | 0.64099449 | 0.9958918 |
| CHST12 | 19.8223148 | 0.15037736 | 0.40340552 | 0.37276972 | 0.70931984 | 0.9958918 |
| TTYH3 | 18.566433 | 0.06527923 | 0.36165116 | 0.18050332 | 0.85675745 | 0.9971731 |
| GNA12 | 111.128378 | 0.07543452 | 0.29916953 | 0.25214639 | 0.80092792 | 0.99592423 |
| SDK1 | 592.86647 | 1.23217577 | 0.43398167 | 2.83923457 | 0.00452219 | 0.16781827 |
| FOXK1 | 90.9530498 | 0.48074704 | 0.36546359 | 1.31544443 | 0.18836058 | 0.88139754 |
| RBAK-RBAKI | 48.5479682 | -0.2182931 | 0.33573341 | -0.6501979 | 0.51556441 | 0.9958918 |
| WIPI2 | 40.3900307 | -0.0788081 | 0.31207181 | -0.2525321 | 0.80062984 | 0.99592423 |
| TNRC18 | 153.645012 | 0.18724904 | 0.32020143 | 0.58478516 | 0.55869219 | 0.9958918 |
| FBXL18 | 18.1351456 | -0.0026002 | 0.47604094 | -0.0054622 | 0.99564181 | 0.99935941 |

| | | | | | | |
|---|---|---|---|---|---|---|
| ACTB | 921.92965 | -0.0270375 | 0.35846927 | -0.0754247 | 0.93987677 | 0.99935941 |
| RNF216 | 129.027503 | 0.00157663 | 0.27338605 | 0.00576705 | 0.99539859 | 0.99935941 |
| PMS2 | 17.9823525 | -0.0492462 | 0.38675012 | -0.1273333 | 0.89867659 | 0.99935941 |
| EIF2AK1 | 36.4971657 | -0.2858093 | 0.27319572 | -1.0461705 | 0.2954823 | 0.95393362 |
| USP42 | 52.7625326 | -0.0506495 | 0.27472691 | -0.1843632 | 0.85372852 | 0.9971731 |
| CYTH3 | 158.628109 | -0.8799001 | 0.30157914 | -2.9176426 | 0.00352688 | 0.14479926 |
| RAC1 | 117.359016 | 0.21229422 | 0.28792797 | 0.73731712 | 0.46092952 | 0.98590351 |
| ZNF12 | 26.7635611 | 0.20193251 | 0.29927388 | 0.67474149 | 0.49984002 | 0.99213111 |
| CCZ1B | 23.2218294 | -0.0799287 | 0.37987096 | -0.2104101 | 0.8333476 | 0.99592423 |
| C1GALT1 | 50.6889194 | 0.44785223 | 0.36281294 | 1.2343888 | 0.21705806 | 0.90308192 |
| MIOS | 20.4417171 | 0.1866031 | 0.33557266 | 0.5560736 | 0.57816054 | 0.9958918 |
| UMAD1 | 154.916502 | 0.01395524 | 0.21203143 | 0.06581686 | 0.94752363 | 0.99935941 |
| GLCCI1 | 79.1690645 | 0.1824412 | 0.33998419 | 0.53661671 | 0.5915324 | 0.9958918 |
| NDUFA4 | 84.453849 | -0.1863925 | 0.36221112 | -0.5145963 | 0.60683512 | 0.9958918 |
| PHF14 | 204.312129 | -0.2762714 | 0.17370133 | -1.5904969 | 0.11172283 | 0.77793929 |
| THSD7A | 132.397392 | -1.0935839 | 0.43163882 | -2.5335625 | 0.01129096 | 0.28321649 |
| TMEM106B | 41.3292798 | -0.3097401 | 0.26392993 | -1.1735694 | 0.24056756 | 0.92426606 |
| ETV1 | 25.2063249 | 0.26410301 | 0.44887295 | 0.58836918 | 0.55628452 | 0.9958918 |
| ISPD | 46.4202254 | 0.30295366 | 0.34812142 | 0.87025285 | 0.38416224 | 0.9725044 |
| BZW2 | 23.0090684 | 0.90029046 | 0.42780995 | 2.10441684 | 0.0353421 | 0.50689386 |
| SNX13 | 143.544191 | -0.2524107 | 0.20057394 | -1.2584419 | 0.20823197 | 0.89634177 |
| HDAC9 | 163.94054 | 1.72287099 | 0.35617877 | 4.8370963 | 1.32E-06 | 0.00096823 |
| ITGB8 | 87.8667271 | 0.98541312 | 0.52368303 | 1.88169766 | 0.05987708 | 0.61332974 |
| SP4 | 32.9460874 | 0.37536091 | 0.35750969 | 1.04993212 | 0.29374932 | 0.95225574 |
| RAPGEF5 | 520.704392 | -0.1844175 | 0.62397849 | -0.2955511 | 0.7675729 | 0.9958918 |
| STEAP1B | 72.8704074 | -0.084452 | 0.33160621 | -0.2546755 | 0.79897377 | 0.99592423 |
| TOMM7 | 40.0628632 | 0.25429414 | 0.37715735 | 0.67423885 | 0.50015947 | 0.99213111 |
| FAM126A | 88.4071176 | -0.2927574 | 0.24364514 | -1.2015728 | 0.2295291 | 0.91455844 |
| KLHL7 | 36.1759997 | 0.03600986 | 0.31726778 | 0.11349991 | 0.90963424 | 0.99935941 |
| NUPL2 | 28.6153612 | 0.41308011 | 0.39549173 | 1.04447218 | 0.29626698 | 0.95393362 |
| MALSU1 | 22.0667212 | 0.25048036 | 0.31319995 | 0.79974585 | 0.42385806 | 0.97828057 |
| TRA2A | 161.866096 | 0.16107917 | 0.30477179 | 0.52852386 | 0.59713579 | 0.9958918 |
| OSBPL3 | 33.3799867 | 1.3050973 | 0.5915846 | 2.20610425 | 0.0273767 | 0.44841843 |
| CYCS | 31.9473845 | 0.39052884 | 0.35419379 | 1.10258523 | 0.27020733 | 0.93698328 |
| HNRNPA2B1 | 283.669385 | -0.2051943 | 0.20558398 | -0.9981046 | 0.31822862 | 0.95992644 |
| CBX3 | 38.9157198 | -0.5034111 | 0.26033395 | -1.9337127 | 0.05314846 | 0.58707106 |
| SKAP2 | 206.612637 | -0.0321166 | 0.39211363 | -0.0819063 | 0.93472122 | 0.99935941 |
| HOTAIRM1 | 38.1441129 | 0.42218734 | 0.41109527 | 1.02698173 | 0.30442907 | 0.95793395 |
| HOXA3 | 52.3272954 | 0.33283322 | 0.28643348 | 1.16199133 | 0.24523899 | 0.92559078 |
| HIBADH | 113.468506 | 0.26661179 | 0.25742236 | 1.03569788 | 0.30034311 | 0.95597956 |
| TAX1BP1 | 173.63389 | 0.24170002 | 0.18455711 | 1.3096218 | 0.19032381 | 0.88158763 |
| JAZF1 | 125.704889 | -0.0114154 | 0.34917792 | -0.0326921 | 0.9739201 | 0.99935941 |
| CREB5 | 81.1357711 | -0.3182576 | 0.49558475 | -0.642186 | 0.5207524 | 0.9958918 |
| SCRN1 | 24.2151859 | 0.42362493 | 0.32118361 | 1.31894938 | 0.18718604 | 0.87955894 |
| FKBP14 | 19.8284409 | 0.07985241 | 0.44069232 | 0.18119765 | 0.85621244 | 0.9971731 |
| PLEKHA8 | 36.8633169 | -0.3359706 | 0.33765289 | -0.9950177 | 0.31972764 | 0.95992644 |
| MTURN | 31.5342935 | 0.14334534 | 0.27700296 | 0.51748667 | 0.60481647 | 0.9958918 |
| ZNRF2 | 59.0867764 | 0.58701377 | 0.32447289 | 1.80913039 | 0.07043075 | 0.65633299 |
| AC006978.2 | 40.0875101 | 0.61460483 | 0.37129311 | 1.65530902 | 0.09786183 | 0.74364783 |
| NOD1 | 62.9147884 | 0.13918522 | 0.42559826 | 0.32703428 | 0.74364196 | 0.9958918 |
| AC005154.6 | 29.985193 | -0.0705697 | 0.34326013 | -0.2055867 | 0.83711377 | 0.99692952 |
| GARS | 41.3624585 | -0.2767712 | 0.38775514 | -0.7137783 | 0.47536428 | 0.98910524 |
| AQP1 | 77.1352833 | 2.29759382 | 0.47404354 | 4.8467991 | 1.25E-06 | 0.00096823 |
| PDE1C | 187.79549 | -0.7062008 | 0.56339514 | -1.2534734 | 0.21003349 | 0.89735318 |

| | | | | | | |
|---|---|---|---|---|---|---|
| AVL9 | 131.002095 | 0.02195977 | 0.21399372 | 0.10261878 | 0.91826554 | 0.99935941 |
| KBTBD2 | 32.4634386 | 0.21613111 | 0.3295528 | 0.65583151 | 0.51193254 | 0.9958918 |
| FKBP9 | 36.4639562 | -0.1164502 | 0.29778975 | -0.3910485 | 0.69576136 | 0.9958918 |
| NT5C3A | 58.1819304 | -0.4887209 | 0.29744257 | -1.6430766 | 0.10036709 | 0.75035372 |
| BBS9 | 141.233536 | -0.0046993 | 0.31041165 | -0.0151389 | 0.98792135 | 0.99935941 |
| DPY19L1 | 34.465459 | -0.2633343 | 0.34218329 | -0.7695708 | 0.44155453 | 0.98092839 |
| HERPUD2 | 53.1241714 | -0.1713319 | 0.25678551 | -0.6672179 | 0.50463298 | 0.99387859 |
| 7-Sep | 258.294499 | -0.2678885 | 0.21999322 | -1.2177124 | 0.22333329 | 0.91093521 |
| EEPD1 | 146.480692 | -0.3695551 | 0.54619482 | -0.6765995 | 0.49866009 | 0.99176652 |
| ELMO1 | 85.0284792 | 0.97181771 | 0.35312533 | 2.75204756 | 0.00592239 | 0.19136125 |
| STARD3NL | 26.4290209 | 0.07130086 | 0.33861944 | 0.21056339 | 0.83322798 | 0.99592423 |
| AMPH | 41.4143166 | -0.1144443 | 0.47373675 | -0.2415779 | 0.80910724 | 0.99592423 |
| VPS41 | 94.1989279 | -0.313787 | 0.33131925 | -0.9470835 | 0.34359623 | 0.96887708 |
| RALA | 59.507477 | 0.23607979 | 0.281189 | 0.8395769 | 0.40114566 | 0.97500135 |
| CDK13 | 249.555755 | 0.09489442 | 0.17081947 | 0.55552461 | 0.57853588 | 0.9958918 |
| MPLKIP | 18.3890802 | 0.56747056 | 0.45445714 | 1.24867784 | 0.21178293 | 0.89738044 |
| SUGCT | 141.204224 | -0.0592632 | 0.47716815 | -0.1241978 | 0.90115866 | 0.99935941 |
| INHBA | 79.3019805 | -0.1703318 | 0.93983589 | -0.1812357 | 0.85618258 | 0.9971731 |
| GLI3 | 104.051561 | -0.256076 | 0.25060083 | -1.0218482 | 0.30685275 | 0.95793395 |
| MRPL32 | 25.6075823 | -0.1717463 | 0.32174772 | -0.533792 | 0.59348548 | 0.9958918 |
| STK17A | 24.1679222 | -0.1551835 | 0.50508361 | -0.3072431 | 0.75865832 | 0.9958918 |
| COA1 | 107.792397 | -0.069162 | 0.26542539 | -0.2605705 | 0.79442372 | 0.99592423 |
| DBNL | 21.6956928 | 0.06566987 | 0.31901873 | 0.20584959 | 0.83690841 | 0.99692952 |
| AEBP1 | 221.220647 | -0.0009409 | 0.44816804 | -0.0020995 | 0.99832486 | 0.99959465 |
| NUDCD3 | 85.8211189 | -0.023091 | 0.20898905 | -0.110489 | 0.91202161 | 0.99935941 |
| TMED4 | 25.3410245 | -0.0778132 | 0.39955396 | -0.1947501 | 0.84558857 | 0.99692952 |
| OGDH | 69.1252024 | 0.11452708 | 0.26947332 | 0.42500342 | 0.67083418 | 0.9958918 |
| ZMIZ2 | 30.5496693 | -0.0382815 | 0.35125129 | -0.108986 | 0.91321363 | 0.99935941 |
| PPIA | 124.870727 | 0.12669264 | 0.4156306 | 0.3048203 | 0.76050302 | 0.9958918 |
| H2AFV | 55.7790944 | 0.1931157 | 0.32632185 | 0.59179517 | 0.55398776 | 0.9958918 |
| PURB | 36.4311188 | 0.05385389 | 0.31840032 | 0.16913894 | 0.86568735 | 0.9971731 |
| CCM2 | 66.4111365 | 0.44826336 | 0.22524402 | 1.99012319 | 0.04657737 | 0.56859978 |
| ADCY1 | 33.4672027 | 0.42953149 | 0.89915595 | 0.47770523 | 0.63286002 | 0.9958918 |
| IGFBP3 | 73.1679937 | -1.6972149 | 0.53790737 | -3.1552179 | 0.00160378 | 0.08730519 |
| TNS3 | 382.846394 | -0.0422536 | 0.35194812 | -0.1200563 | 0.90443854 | 0.99935941 |
| HUS1 | 26.7568497 | -0.4593348 | 0.28772639 | -1.5964291 | 0.11039303 | 0.77370222 |
| UPP1 | 25.8475455 | -0.4545093 | 0.72388073 | -0.6278788 | 0.53008335 | 0.9958918 |
| GRB10 | 90.19987 | -0.1858558 | 0.32312195 | -0.5751878 | 0.56516432 | 0.9958918 |
| SEC61G | 29.1739551 | -0.1209602 | 0.535155 | -0.2260283 | 0.82117937 | 0.99592423 |
| EGFR | 304.632182 | -0.0401835 | 0.43956983 | -0.0914155 | 0.92716247 | 0.99935941 |
| LANCL2 | 26.9525006 | -0.0818604 | 0.36996754 | -0.2212637 | 0.82488715 | 0.99592423 |
| VOPP1 | 103.598785 | -0.2256563 | 0.32053303 | -0.7040032 | 0.48143079 | 0.99002719 |
| NIPSNAP2 | 53.1039978 | 0.45031292 | 0.2969507 | 1.51645681 | 0.12940388 | 0.80543275 |
| CCT6A | 40.633905 | -0.4614456 | 0.38116159 | -1.2106299 | 0.22603726 | 0.91093521 |
| SUMF2 | 39.2601361 | -0.1428124 | 0.31964982 | -0.4467776 | 0.65503562 | 0.9958918 |
| CHCHD2 | 75.1336834 | 0.23398414 | 0.37004968 | 0.63230467 | 0.52718781 | 0.9958918 |
| ZNF736 | 33.9083999 | -0.1322251 | 0.28706666 | -0.4606078 | 0.64508001 | 0.9958918 |
| ZNF680 | 45.0118349 | -0.2582365 | 0.3728758 | -0.6925536 | 0.48858976 | 0.99046213 |
| ZNF138 | 19.4492601 | -0.0723582 | 0.39137232 | -0.1848834 | 0.8533205 | 0.9971731 |
| ZNF117 | 18.3288192 | -0.386065 | 0.42944279 | -0.8989906 | 0.36865765 | 0.97217293 |
| VKORC1L1 | 73.1411872 | -0.1721299 | 0.31084017 | -0.5537571 | 0.5797451 | 0.9958918 |
| TPST1 | 212.189703 | -0.5919148 | 0.30132669 | -1.9643622 | 0.04948811 | 0.57273715 |
| AC027644.4 | 50.1364629 | 0.01804631 | 0.31937189 | 0.05650564 | 0.95493901 | 0.99935941 |
| TMEM248 | 52.1446543 | 0.11973885 | 0.28184025 | 0.42484654 | 0.67094855 | 0.9958918 |

| | | | | | | |
|---|---|---|---|---|---|---|
| SBDS | 52.4009779 | -0.0335493 | 0.40461325 | -0.0829169 | 0.93391766 | 0.99935941 |
| TYW1 | 101.932221 | -0.1498766 | 0.20358012 | -0.7362047 | 0.46160613 | 0.98590351 |
| AUTS2 | 685.930026 | -0.7878031 | 0.30902819 | -2.549292 | 0.01079419 | 0.28030558 |
| GALNT17 | 48.5260061 | 0.9353512 | 0.48031605 | 1.9473661 | 0.05149086 | 0.57655256 |
| TYW1B | 60.426583 | 0.1122336 | 0.25846729 | 0.4342275 | 0.66412325 | 0.9958918 |
| POM121 | 81.7096788 | -0.2792973 | 0.28113771 | -0.9934537 | 0.32048891 | 0.95992644 |
| BAZ1B | 97.3750023 | 0.17859555 | 0.20801395 | 0.8585749 | 0.39057509 | 0.97297628 |
| BUD23 | 21.1306643 | 0.23519752 | 0.40066338 | 0.58702026 | 0.5571901 | 0.9958918 |
| ELN | 72.7881302 | 0.2581267 | 0.53406299 | 0.48332632 | 6.29E-01 | 0.9958918 |
| EIF4H | 49.4369093 | -0.3879443 | 0.33797468 | -1.1478502 | 0.25103042 | 0.92624282 |
| CLIP2 | 104.601583 | -0.2609419 | 0.32258044 | -0.8089205 | 0.41856087 | 0.97812179 |
| GTF2IRD1 | 56.3298501 | -0.4289941 | 0.38130828 | -1.1250585 | 0.26056423 | 0.93055907 |
| GTF2I | 231.995421 | 0.10026915 | 0.16579714 | 0.6047701 | 0.54533176 | 0.9958918 |
| GTF2IRD2 | 25.7869882 | 0.1894515 | 0.55796705 | 0.33953887 | 0.73420382 | 0.9958918 |
| CASTOR2 | 23.6618936 | 0.17226358 | 0.43473864 | 0.39624631 | 0.69192333 | 0.9958918 |
| RCC1L | 27.3743719 | -0.1854048 | 0.33858924 | -0.5475803 | 0.58398015 | 0.9958918 |
| GTF2IRD2B | 41.2733249 | 0.16519761 | 0.51888674 | 0.31836931 | 0.75020481 | 0.9958918 |
| POM121C | 66.5724845 | -0.1570982 | 0.29131933 | -0.5392646 | 0.58970427 | 0.9958918 |
| HIP1 | 404.881107 | 0.1341248 | 0.31630083 | 0.42404187 | 0.67153528 | 0.9958918 |
| RHBDD2 | 23.3394498 | -0.1635666 | 0.35477536 | -0.4610427 | 0.644768 | 0.9958918 |
| POR | 59.1752781 | -0.3290209 | 0.2482203 | -1.3255197 | 0.18499884 | 0.87866798 |
| STYXL1 | 22.3446713 | 0.29520441 | 0.45020842 | 0.65570609 | 0.51201325 | 0.9958918 |
| MDH2 | 24.6749692 | 0.38013987 | 0.36521509 | 1.04086572 | 0.29793788 | 0.95447177 |
| HSPB1 | 216.26129 | 0.19032069 | 0.47990111 | 0.39658313 | 0.69167489 | 0.9958918 |
| YWHAG | 63.4374791 | 0.32275147 | 0.27389919 | 1.17835862 | 0.23865367 | 0.92291633 |
| DTX2 | 28.9417018 | 0.31327854 | 0.43881616 | 0.71391752 | 0.47527818 | 0.98910524 |
| POMZP3 | 18.2961279 | 0.3729473 | 0.40269529 | 0.92612779 | 0.35437958 | 0.97217293 |
| CCDC146 | 60.6381354 | 0.33615154 | 0.31719704 | 1.05975624 | 0.28925551 | 0.95181201 |
| GSAP | 28.9451596 | 0.31562424 | 0.38459453 | 0.82066752 | 0.41183568 | 0.97812179 |
| PTPN12 | 283.729921 | 0.15390099 | 0.22650623 | 0.67945587 | 0.49684906 | 0.99176652 |
| RSBN1L | 110.066693 | -0.0636335 | 0.24787105 | -0.2567201 | 0.79739484 | 0.99592423 |
| PHTF2 | 119.206301 | 0.68050715 | 0.33335689 | 2.04137716 | 0.04121335 | 0.54347495 |
| MAGI2 | 572.448411 | 0.10559209 | 0.25294147 | 0.41745661 | 0.67634446 | 0.9958918 |
| GNAI1 | 211.932952 | -0.2209363 | 0.3285169 | -0.6725264 | 0.50124865 | 0.99213111 |
| SEMA3C | 153.637054 | -1.5022734 | 0.39604827 | -3.7931574 | 0.00014874 | 0.01904212 |
| HGF | 158.240475 | -0.6195683 | 0.32328469 | -1.9164789 | 0.05530416 | 0.59593882 |
| CACNA2D1 | 364.982107 | -0.1376791 | 0.26527504 | -0.5190051 | 0.60375721 | 0.9958918 |
| KIAA1324L | 27.035525 | 0.02055444 | 0.39451283 | 0.0521008 | 0.95844837 | 0.99935941 |
| DMTF1 | 125.919258 | -0.3250936 | 0.34387489 | -0.9453834 | 0.34446318 | 0.96887708 |
| TMEM243 | 25.5484963 | 0.08500223 | 0.35974503 | 0.23628465 | 0.8132118 | 0.99592423 |
| ADAM22 | 57.9101865 | -1.1155545 | 0.58893704 | -1.8941829 | 0.05820073 | 0.60892588 |
| SRI | 25.5527099 | -0.3840122 | 0.41651753 | -0.9219592 | 0.35654983 | 0.97217293 |
| STEAP4 | 117.29805 | -0.7029781 | 0.4041998 | -1.7391847 | 8.20E-02 | 7.00E-01 |
| STEAP2 | 39.0984566 | -0.1474257 | 0.45685797 | -0.3226949 | 0.74692631 | 0.9958918 |
| CFAP69 | 42.2616081 | 0.12916341 | 0.49819178 | 0.25926442 | 0.79543123 | 0.99592423 |
| GTPBP10 | 29.0645214 | 0.21292502 | 0.35923906 | 0.59271121 | 0.55337444 | 0.9958918 |
| CLDN12 | 22.9302803 | -0.4218944 | 0.37914286 | -1.1127585 | 0.26581215 | 0.93533685 |
| CDK14 | 271.16698 | 0.61500194 | 0.38640347 | 1.59160564 | 0.11147334 | 0.77793929 |
| FZD1 | 28.2793741 | 0.49820919 | 0.45193904 | 1.10238139 | 0.2702959 | 0.93698328 |
| MTERF1 | 54.3920094 | -0.0552794 | 0.34664067 | -0.1594717 | 0.87329725 | 0.9971731 |
| AKAP9 | 248.409121 | 0.15446417 | 0.2297727 | 0.6722477 | 0.50142602 | 0.99213111 |
| KRIT1 | 89.9752964 | 0.10552532 | 0.24584025 | 0.42924348 | 0.66774605 | 0.9958918 |
| ANKIB1 | 173.918091 | -0.1905168 | 0.21671701 | -0.8791042 | 0.37934479 | 0.97217293 |
| GATAD1 | 31.75209 | 0.41718164 | 0.42459248 | 0.982546 | 0.32583093 | 0.95992644 |

| | | | | | | |
|---|---|---|---|---|---|---|
| AC007566.1 | 18.9080463 | 0.24974366 | 0.48318265 | 0.51687215 | 0.60524541 | 0.9958918 |
| PEX1 | 28.5780713 | 0.23966847 | 0.32481225 | 0.73786772 | 0.46059484 | 0.98590351 |
| FAM133B | 51.17839 | 0.07559229 | 0.25195321 | 0.30002511 | 0.764158 | 0.9958918 |
| CDK6 | 110.782311 | 0.1580632 | 0.20326135 | 0.77763529 | 4.37E-01 | 0.98092839 |
| SAMD9L | 29.1055958 | -1.4340198 | 1.06316613 | -1.34882 | 0.17739478 | 0.87771577 |
| VPS50 | 60.5032602 | 0.10036645 | 0.29154131 | 0.34426148 | 0.73064965 | 0.9958918 |
| GNG11 | 19.8749298 | 0.42339547 | 0.40200064 | 1.0532209 | 0.29223976 | 0.95186143 |
| BET1 | 23.7372711 | -0.0966503 | 0.40458655 | -0.2388866 | 0.81119348 | 0.99592423 |
| AC002074.1 | 30.6275981 | -0.2097228 | 0.47045134 | -0.4457906 | 0.65574851 | 0.9958918 |
| COL1A2 | 448.936815 | -0.6675342 | 0.36238156 | -1.8420755 | 0.0654641 | 0.63034109 |
| CASD1 | 50.9854509 | 0.31012738 | 0.26717855 | 1.16074955 | 0.24574377 | 0.92559078 |
| SGCE | 50.2873569 | 0.36749857 | 0.36031379 | 1.01994034 | 0.30775676 | 0.95793395 |
| PPP1R9A | 31.3438307 | -0.5278661 | 0.41892681 | -1.2600436 | 0.20765363 | 0.89620104 |
| PON2 | 34.4692961 | -0.126449 | 0.30078471 | -0.420397 | 0.67419549 | 0.9958918 |
| PDK4 | 281.214078 | -0.3209349 | 0.65717909 | -0.4883522 | 0.62530037 | 0.9958918 |
| DYNC1I1 | 87.3921502 | -0.4381167 | 0.37752798 | -1.1604879 | 0.24585021 | 0.92559078 |
| SLC25A13 | 185.686358 | 0.42862615 | 0.21464931 | 1.99686714 | 0.04583962 | 0.56239624 |
| SEM1 | 109.361159 | 0.32054846 | 0.3306016 | 0.96959136 | 0.33225022 | 0.9654289 |
| LMTK2 | 49.5922571 | 0.15133002 | 0.26059243 | 0.58071536 | 0.56143231 | 0.9958918 |
| TECPR1 | 17.4045191 | -0.0393961 | 0.57982919 | -0.0679444 | 0.94582991 | 0.99935941 |
| BRI3 | 84.2537201 | -0.0710853 | 0.32079732 | -0.2215895 | 0.82463344 | 0.99592423 |
| BAIAP2L1 | 113.570164 | 0.10371616 | 0.40406661 | 0.25668086 | 0.79742515 | 0.99592423 |
| TRRAP | 102.99757 | 0.0088716 | 0.19951801 | 0.04446517 | 0.96453361 | 0.99935941 |
| SMURF1 | 291.198424 | -0.262969 | 0.32285007 | -0.8145237 | 0.41534499 | 0.97812179 |
| PDAP1 | 26.1844617 | 0.07034535 | 0.30754983 | 0.2287283 | 0.8190801 | 0.99592423 |
| ZNF789 | 23.6280789 | 0.27610937 | 0.40300344 | 0.68512906 | 0.49326248 | 0.99116088 |
| TRIM4 | 40.920448 | 0.30200731 | 0.33977122 | 0.8888549 | 0.37408107 | 0.97217293 |
| ZKSCAN1 | 96.1401609 | 0.22879446 | 0.27059529 | 0.84552272 | 0.39781905 | 0.97500135 |
| COPS6 | 27.8857153 | 0.00766073 | 0.35521382 | 0.02156652 | 0.98279374 | 0.99935941 |
| LAMTOR4 | 44.5742822 | 0.16142808 | 0.28589465 | 0.56464184 | 0.5723174 | 0.9958918 |
| CASTOR3 | 47.7096205 | 0.20247534 | 0.43603999 | 0.46435038 | 0.64239673 | 0.9958918 |
| SPDYE3 | 21.9488959 | -0.4361923 | 0.41592314 | -1.0487329 | 0.29430108 | 0.95304825 |
| PILRB | 48.1015981 | -0.0172057 | 0.4276021 | -0.0402377 | 0.96790365 | 0.99935941 |
| TSC22D4 | 26.2248041 | 0.10005917 | 0.34982178 | 0.286029 | 0.77485591 | 0.9958918 |
| PCOLCE | 40.7270197 | -0.9079653 | 0.37596224 | -2.4150439 | 0.01573331 | 0.34027688 |
| GNB2 | 52.4084418 | 0.05490612 | 0.33174205 | 0.16550846 | 0.86854379 | 0.9971731 |
| GIGYF1 | 33.7719383 | -0.0157142 | 0.30110723 | -0.052188 | 0.95837893 | 0.99935941 |
| TRIP6 | 24.7359672 | -0.2575664 | 0.32164694 | -0.8007738 | 0.42326263 | 0.97828057 |
| SRRT | 19.921979 | -0.0652489 | 0.41591842 | -0.1568791 | 0.87534017 | 0.9971731 |
| TRIM56 | 86.012971 | 0.11427645 | 0.36576551 | 0.31243091 | 0.75471306 | 0.9958918 |
| SERPINE1 | 850.987379 | -1.7228342 | 1.04425606 | -1.6498197 | 0.09897982 | 0.74528964 |
| PLOD3 | 23.5015779 | 0.04864331 | 0.35895555 | 0.13551347 | 0.89220591 | 0.99935941 |
| ZNHIT1 | 22.3925308 | 0.39331311 | 0.40302885 | 0.97589318 | 0.3291174 | 0.96246868 |
| FIS1 | 41.4807119 | 0.39134687 | 0.33845594 | 1.15627124 | 0.24757023 | 0.92624282 |
| CUX1 | 488.683048 | 0.01859242 | 0.29048969 | 0.06400371 | 0.94896727 | 0.99935941 |
| PRKRIP1 | 110.178893 | -0.0724991 | 0.27211761 | -0.2664258 | 0.78991131 | 0.99592423 |
| AC105052.3 | 27.1813912 | 0.24533484 | 0.35213289 | 0.69671095 | 0.48598371 | 0.99046213 |
| FAM185A | 51.7743195 | -0.1052045 | 0.31191429 | -0.3372865 | 0.73590091 | 0.9958918 |
| FBXL13 | 22.9076808 | -0.5293985 | 0.39532455 | -1.3391491 | 0.18052215 | 0.87815793 |
| ARMC10 | 30.4537684 | 0.09600617 | 0.29129804 | 0.32958055 | 0.74171692 | 0.9958918 |
| NAPEPLD | 41.017654 | 0.48387582 | 0.30111386 | 1.60695299 | 0.10806468 | 0.77248589 |
| PMPCB | 27.837913 | -0.4183476 | 0.34971723 | -1.1962454 | 0.23160082 | 0.91549646 |
| DNAJC2 | 24.9670496 | 0.06365095 | 0.35954418 | 0.17703234 | 0.85948299 | 0.9971731 |
| PSMC2 | 28.8511847 | -0.0944215 | 0.27504147 | -0.3432993 | 0.73137331 | 0.9958918 |

| | | | | | | |
|---|---|---|---|---|---|---|
| ORC5 | 44.3470972 | -0.171518 | 0.3050719 | -0.5622217 | 0.57396501 | 0.9958918 |
| LINC01004 | 112.82728 | 0.00371391 | 0.32328189 | 0.01148816 | 0.99083398 | 0.99935941 |
| KMT2E | 406.317797 | -0.0457379 | 0.37080428 | -0.1233478 | 0.90183172 | 0.99935941 |
| SRPK2 | 263.11988 | -0.0608722 | 0.21048269 | -0.289203 | 0.77242605 | 0.9958918 |
| PUS7 | 41.0581248 | 0.22360269 | 0.50944559 | 0.43891379 | 0.660724 | 0.9958918 |
| RINT1 | 23.9704899 | -0.4858519 | 0.3474159 | -1.3984735 | 0.16197092 | 0.85327904 |
| ATXN7L1 | 131.291373 | -0.1289641 | 0.50220224 | -0.2567972 | 0.79733534 | 0.99592423 |
| CDHR3 | 27.734044 | 0.84159084 | 0.45337548 | 1.85627778 | 0.06341397 | 0.62395844 |
| SYPL1 | 51.4997765 | 0.17435288 | 0.24845697 | 0.70174278 | 0.48283959 | 0.99002719 |
| NAMPT | 883.324841 | -0.128288 | 0.61003795 | -0.2102951 | 8.33E-01 | 9.96E-01 |
| CCDC71L | 46.1253188 | -0.5049676 | 0.51085647 | -0.9884726 | 0.32292124 | 0.95992644 |
| PRKAR2B | 62.2455831 | -2.1969416 | 0.53354278 | -4.1176485 | 3.83E-05 | 0.00803682 |
| HBP1 | 82.344437 | -0.3716226 | 0.38550649 | -0.9639852 | 0.33505335 | 0.96640422 |
| COG5 | 221.118865 | -0.4576801 | 0.23057438 | -1.9849566 | 0.0471493 | 0.56885457 |
| BCAP29 | 61.5189962 | -0.0354716 | 0.27630904 | -0.1283764 | 0.89785113 | 0.99935941 |
| SLC26A3 | 262.906993 | -0.9605899 | 0.63656109 | -1.5090301 | 0.13129108 | 0.80565822 |
| DLD | 39.0709034 | 0.18761162 | 0.29251164 | 0.64138171 | 0.52127471 | 0.9958918 |
| LAMB1 | 172.763789 | -0.9719498 | 0.38294096 | -2.5381193 | 0.011145 | 0.28321649 |
| NRCAM | 33.7604452 | 2.20947603 | 0.51413227 | 4.29748556 | 1.73E-05 | 0.00437176 |
| PNPLA8 | 74.6822036 | -0.2881675 | 0.25261227 | -1.1407501 | 0.25397395 | 0.92646125 |
| IMMP2L | 989.62786 | 0.07991593 | 0.28288961 | 0.28249866 | 0.77756118 | 0.9958918 |
| DOCK4 | 586.61038 | -0.3649038 | 0.321073 | -1.1365134 | 0.25574175 | 0.9266065 |
| ZNF277 | 66.4475185 | 0.03847424 | 0.27203797 | 0.14142966 | 0.88753053 | 0.99935941 |
| IFRD1 | 80.8946488 | -0.3164774 | 0.42110436 | -0.7515415 | 0.45232682 | 0.98092839 |
| TMEM168 | 65.9679179 | 0.19949828 | 0.22610597 | 0.88232204 | 0.37760268 | 0.97217293 |
| BMT2 | 54.9613184 | 0.14926768 | 0.23993463 | 0.62211812 | 0.5338642 | 0.9958918 |
| FOXP2 | 75.4167264 | -0.8258561 | 0.58226204 | -1.4183582 | 0.15608621 | 0.84427321 |
| MDFIC | 66.7711259 | 0.52472747 | 0.40608339 | 1.29216678 | 0.19629939 | 0.88391893 |
| TES | 35.2017642 | 0.81155346 | 0.49934978 | 1.62522041 | 0.1041156 | 0.75982678 |
| CAV2 | 34.5065096 | -0.1379966 | 0.4067298 | -0.3392832 | 0.73439643 | 0.9958918 |
| CAV1 | 44.4354666 | -0.3628916 | 0.4491062 | -0.8080307 | 0.4190729 | 0.97812179 |
| CAPZA2 | 86.3962803 | -0.0546381 | 0.21001775 | -0.2601595 | 0.79474074 | 0.99592423 |
| ST7 | 121.475806 | -0.1140364 | 0.21351986 | -0.5340784 | 0.59328726 | 0.9958918 |
| CTTNBP2 | 36.322919 | -1.2608844 | 0.42442963 | -2.9707739 | 0.0029705 | 0.129173 |
| LSM8 | 26.1035485 | -0.1261516 | 0.3594577 | -0.3509499 | 0.72562596 | 0.9958918 |
| ING3 | 18.6893125 | 0.0238375 | 0.34818591 | 0.06846199 | 0.94541787 | 0.99935941 |
| CPED1 | 406.209996 | 0.33261963 | 0.41140594 | 0.80849496 | 0.41880571 | 0.97812179 |
| FAM3C | 76.1123571 | -0.1339602 | 0.21246137 | -0.6305158 | 0.52835719 | 0.9958918 |
| AASS | 120.890074 | -0.6743569 | 0.45418425 | -1.4847651 | 0.13760607 | 0.81404904 |
| CADPS2 | 177.590365 | 0.12037057 | 0.2303296 | 0.52260139 | 0.60125168 | 0.9958918 |
| NDUFA5 | 33.5986744 | 0.23405911 | 0.31317091 | 0.74738457 | 0.45483145 | 0.98173285 |
| WASL | 88.6809102 | -0.0052219 | 0.23198823 | -0.0225093 | 0.98204167 | 0.99935941 |
| POT1 | 48.5596388 | -0.3142371 | 0.33363649 | -0.9418548 | 0.34626697 | 0.96887708 |
| ZNF800 | 58.038414 | -0.0530779 | 0.26843273 | -0.1977328 | 0.84325415 | 0.99692952 |
| SND1 | 277.989039 | 0.06453977 | 0.37616195 | 0.17157443 | 0.86377211 | 0.9971731 |
| RBM28 | 31.6598708 | 0.00705693 | 0.34389363 | 0.02052069 | 0.98362801 | 0.99935941 |
| HILPDA | 92.554275 | -0.8364049 | 0.92218614 | -0.9069805 | 3.64E-01 | 0.97217293 |
| METTL2B | 25.3068771 | 0.39375627 | 0.49780954 | 0.79097776 | 0.42895697 | 0.98092839 |
| CALU | 127.122618 | -0.3331063 | 0.33313325 | -0.999919 | 0.31734973 | 0.95992644 |
| ATP6V1F | 29.4748163 | 0.17589863 | 0.40362228 | 0.43580009 | 0.66298178 | 0.9958918 |
| TNPO3 | 72.0977096 | 0.24776204 | 0.20379462 | 1.21574377 | 0.22408255 | 0.91093521 |
| AHCYL2 | 132.217635 | -0.0550243 | 0.24327039 | -0.2261857 | 0.82105699 | 0.99592423 |
| NRF1 | 107.872179 | 0.33071591 | 0.17850969 | 1.85264956 | 0.0639326 | 0.62395844 |
| UBE2H | 200.254261 | -0.3765758 | 0.29200579 | -1.2896178 | 0.19718341 | 0.88391893 |

| | | | | | | |
|---|---|---|---|---|---|---|
| KLHDC10 | 82.5272235 | -0.0138018 | 0.23616029 | -0.0584424 | 0.95339627 | 0.99935941 |
| COPG2 | 27.2970213 | -0.3429734 | 0.36763347 | -0.932922 | 0.35086022 | 0.9693503 |
| AC016831.7 | 1071.00628 | -0.0984803 | 0.31069825 | -0.3169646 | 0.75127047 | 0.9958918 |
| LINC00513 | 80.1266899 | -0.424223 | 0.4409374 | -0.9620935 | 0.33600265 | 0.96640422 |
| MKLN1 | 712.023888 | -0.4800694 | 0.30778703 | -1.5597453 | 0.11882008 | 0.78880646 |
| PODXL | 24.8712065 | 1.31081373 | 0.56690484 | 2.31222886 | 0.02076507 | 0.38830162 |
| PLXNA4 | 73.3943233 | -0.8956535 | 0.45723904 | -1.9588299 | 0.05013271 | 0.57538069 |
| CHCHD3 | 223.127719 | 0.01569307 | 0.28513865 | 0.05503664 | 0.95610927 | 0.99935941 |
| EXOC4 | 483.184292 | -0.31803 | 0.2033678 | -1.563817 | 0.11786056 | 0.78539225 |
| SLC35B4 | 19.9023541 | 0.10759874 | 0.47125989 | 0.22832145 | 0.81939635 | 0.99592423 |
| AKR1B1 | 26.3077125 | -0.2680054 | 0.57730235 | -0.4642375 | 0.64247757 | 0.9958918 |
| CALD1 | 2372.97465 | -0.4103524 | 0.2518217 | -1.6295353 | 0.10319975 | 0.75827058 |
| AGBL3 | 29.2267055 | 0.28931489 | 0.30587223 | 0.94586843 | 0.3442157 | 0.96887708 |
| CYREN | 21.6114894 | 0.29907601 | 0.40422828 | 0.7398691 | 0.45937943 | 0.98511217 |
| WDR91 | 18.708931 | 0.59101719 | 0.37134905 | 1.59154088 | 0.1114879 | 0.77793929 |
| CNOT4 | 162.991043 | -0.2269948 | 0.21028063 | -1.0794853 | 0.28037145 | 0.94419503 |
| NUP205 | 35.3203392 | 0.08326417 | 0.31289363 | 0.26611014 | 0.79015439 | 0.99592423 |
| STMP1 | 47.1850639 | 0.00047763 | 0.27325683 | 0.00174792 | 0.99860537 | 0.99968761 |
| MTPN | 48.2058149 | -0.0099744 | 0.25537861 | -0.0390573 | 0.9688447 | 0.99935941 |
| PTN | 56.6351878 | -0.3374361 | 0.51653664 | -0.6532666 | 0.51358442 | 0.9958918 |
| CREB3L2 | 273.004328 | 0.03502238 | 0.41407319 | 0.08458017 | 0.93259516 | 0.99935941 |
| TRIM24 | 43.9626864 | 0.3059738 | 0.28409576 | 1.07700938 | 0.28147607 | 0.94419503 |
| KIAA1549 | 32.9578986 | 1.43307945 | 0.47119145 | 3.04139531 | 0.00235484 | 0.10666545 |
| ZC3HAV1 | 67.7494383 | -0.3750785 | 0.56480298 | -0.6640874 | 0.5066344 | 0.99397937 |
| UBN2 | 89.5224737 | 0.15996915 | 0.22864784 | 0.69963112 | 0.4841577 | 0.99046213 |
| LUC7L2 | 25.5980652 | -0.5556475 | 0.42965446 | -1.2932426 | 0.19592715 | 0.88391893 |
| HIPK2 | 452.212321 | 0.40777031 | 0.32571406 | 1.25192727 | 2.11E-01 | 0.89738044 |
| PARP12 | 33.6671842 | -1.1523927 | 0.71533611 | -1.6109807 | 0.10718392 | 0.77179565 |
| KDM7A | 86.4537401 | 0.28862861 | 0.31424771 | 0.9184748 | 0.35837034 | 0.97217293 |
| SLC37A3 | 53.7698718 | 0.05656076 | 0.2618966 | 0.21596601 | 0.82901424 | 0.99592423 |
| MKRN1 | 32.540517 | 0.12840837 | 0.30125351 | 0.42624688 | 0.66992796 | 0.9958918 |
| DENND2A | 119.593341 | -0.4844839 | 0.31724115 | -1.5271786 | 0.12671661 | 0.797978 |
| NDUFB2 | 39.5323738 | 0.17041003 | 0.30225876 | 0.56378854 | 0.57289805 | 0.9958918 |
| BRAF | 309.101843 | -0.0058025 | 0.18498217 | -0.0313681 | 0.97497596 | 0.99935941 |
| AGK | 33.723707 | -0.3599143 | 0.28703767 | -1.2538923 | 0.20988116 | 0.89727554 |
| KIAA1147 | 26.2190909 | 0.43261267 | 0.37263754 | 1.16094764 | 0.24566319 | 0.92559078 |
| SSBP1 | 39.7114714 | 0.08410507 | 0.27626133 | 0.30444027 | 0.76079249 | 0.9958918 |
| GSTK1 | 25.4480403 | 0.21445586 | 0.31791946 | 0.67456035 | 0.49995513 | 0.99213111 |
| ZYX | 27.1755507 | -0.2276327 | 0.3642763 | -0.6248901 | 0.53204317 | 0.9958918 |
| TCAF2 | 38.0023518 | -0.6745615 | 0.58670621 | -1.1497433 | 0.25024963 | 0.92624282 |
| TCAF1 | 49.6850386 | 0.2027249 | 0.29224337 | 0.69368519 | 0.48787966 | 0.99046213 |
| AC004889.1 | 26.4479432 | 0.47634004 | 0.3490853 | 1.36453768 | 1.72E-01 | 0.87233975 |
| TPK1 | 41.8129221 | 0.33782453 | 0.26859586 | 1.25774287 | 0.20848476 | 0.89634177 |
| CUL1 | 144.28309 | 0.09083844 | 0.27278617 | 0.33300235 | 0.73913251 | 0.9958918 |
| PDIA4 | 28.9407717 | -0.3122406 | 0.34795746 | -0.8973528 | 0.36953069 | 0.97217293 |
| ZNF398 | 28.8847676 | -0.0663006 | 0.37848036 | -0.1751758 | 0.86094148 | 0.9971731 |
| ZNF783 | 22.5677507 | 0.18627105 | 0.31768478 | 0.58633923 | 0.55764757 | 0.9958918 |
| ZNF862 | 25.0128914 | 0.33365314 | 0.32783474 | 1.01774796 | 0.30879775 | 0.95793395 |
| RARRES2 | 52.3240696 | -0.6290041 | 0.4737269 | -1.3277779 | 0.18425149 | 0.87866798 |
| TMEM176B | 31.4907128 | 0.63375604 | 0.48683584 | 1.30178592 | 0.19298958 | 0.88158763 |
| AGAP3 | 30.2519769 | 0.17897605 | 0.28695876 | 0.62369956 | 0.53282491 | 0.9958918 |
| CHPF2 | 17.7899255 | -0.465541 | 0.50177877 | -0.9277814 | 0.35352098 | 0.97217293 |
| SMARCD3 | 42.0603267 | 0.06898851 | 0.32614163 | 0.21152929 | 0.83247428 | 0.99592423 |
| NUB1 | 37.2256572 | 0.27211874 | 0.32534926 | 0.83638959 | 0.40293576 | 0.97512144 |

| | | | | | | |
|---|---|---|---|---|---|---|
| WDR86 | 31.1818347 | -0.3308456 | 0.4497061 | -0.7356929 | 0.46191758 | 0.98590351 |
| RHEB | 116.547685 | -0.1557994 | 0.25386181 | -0.6137172 | 0.53940222 | 0.9958918 |
| PRKAG2 | 89.024174 | -0.365521 | 0.37684634 | -0.969947 | 0.3320729 | 0.9654289 |
| GALNT11 | 70.6953063 | -0.4152792 | 0.30888322 | -1.3444537 | 0.17880173 | 0.87815793 |
| KMT2C | 407.989221 | -0.2732664 | 0.2168825 | -1.2599743 | 0.20767865 | 0.89620104 |
| ACTR3B | 35.6778593 | 0.15220674 | 0.34765191 | 0.43781362 | 0.6615214 | 0.9958918 |
| DPP6 | 68.7204495 | -1.5058701 | 0.52184411 | -2.8856703 | 0.00390581 | 0.15300235 |
| PAXIP1 | 19.7172814 | 0.25862834 | 0.34687103 | 0.74560376 | 0.45590681 | 0.98311595 |
| INSIG1 | 23.0404198 | -1.2449655 | 0.63355768 | -1.9650389 | 0.04940973 | 0.57273715 |
| RBM33 | 149.440924 | 0.11758961 | 0.1832654 | 0.64163564 | 0.52110979 | 0.9958918 |
| LMBR1 | 85.1626604 | 0.08921675 | 0.28898905 | 0.30872017 | 0.7575344 | 0.9958918 |
| UBE3C | 157.551773 | -0.0424608 | 0.22120885 | -0.1919491 | 0.84778209 | 0.99713289 |
| DNAJB6 | 88.9181105 | 0.29217743 | 0.22653454 | 1.2897699 | 0.19713056 | 0.88391893 |
| ESYT2 | 350.784303 | -0.7381751 | 0.47209078 | -1.5636295 | 0.11790461 | 0.78539225 |
| WDR60 | 106.970763 | 0.01389174 | 0.24566187 | 0.0565482 | 0.9549051 | 0.99935941 |
| VIPR2 | 49.7385308 | 1.26898174 | 0.46123899 | 2.75124557 | 0.00593691 | 0.19136125 |
| GTPBP6 | 26.3217349 | 0.23489415 | 0.32888501 | 0.7142136 | 0.47509511 | 0.98910524 |
| SLC25A6 | 78.6980928 | 0.31889424 | 0.41926304 | 0.7606066 | 0.44689207 | 0.98092839 |
| ASMTL | 23.5950616 | -0.0468758 | 0.33969466 | -0.1379938 | 0.89024529 | 0.99935941 |
| AKAP17A | 20.4493856 | 0.33528071 | 0.31899827 | 1.0510424 | 0.29323912 | 0.95186143 |
| DHRSX | 107.301666 | 0.03859055 | 0.19484071 | 0.19806207 | 0.8429965 | 0.99692952 |
| CD99 | 87.2314183 | 0.50900355 | 0.36590464 | 1.39108251 | 0.16420041 | 0.85697309 |
| PRKX | 51.2274732 | 0.58596349 | 0.2697406 | 2.17232217 | 0.02983137 | 0.46665355 |
| TBL1X | 283.273283 | -0.0382023 | 0.20639191 | -0.1850959 | 0.85315379 | 0.9971731 |
| SHROOM2 | 25.7351616 | -1.023997 | 0.43083759 | -2.3767588 | 0.0174655 | 0.34973839 |
| WWC3 | 56.7677752 | 0.19430016 | 0.24707683 | 0.78639572 | 0.43163569 | 0.98092839 |
| MID1 | 159.480343 | 0.76127388 | 0.40002717 | 1.90305544 | 0.05703331 | 0.60440467 |
| ARHGAP6 | 348.017354 | -0.1192988 | 0.41378981 | -0.2883076 | 0.77311129 | 0.9958918 |
| TMSB4X | 661.913199 | -0.4889502 | 0.43587538 | -1.121766 | 0.26196194 | 0.93155278 |
| RAB9A | 28.0103667 | -0.4883039 | 0.41007425 | -1.1907695 | 0.23374407 | 0.91918462 |
| OFD1 | 40.5798779 | -0.0489945 | 0.31590574 | -0.1550923 | 0.87674858 | 0.9971731 |
| GPM6B | 86.1467244 | -0.7058589 | 0.54105866 | -1.3045886 | 0.19203298 | 0.88158763 |
| MOSPD2 | 24.9171518 | -0.2010966 | 0.44089449 | -0.4561104 | 0.64831059 | 0.9958918 |
| PIR | 36.7513864 | 0.36812359 | 0.31605846 | 1.16473259 | 0.24412725 | 0.92546331 |
| CLTRN | 22.6428248 | 1.15068838 | 0.71151643 | 1.61723375 | 0.10582783 | 0.76820416 |
| CA5B | 46.075494 | 0.11543842 | 0.35391626 | 0.32617439 | 0.74429242 | 0.9958918 |
| ZRSR2 | 19.2033554 | -0.1639718 | 0.47271026 | -0.3468759 | 0.72868457 | 0.9958918 |
| CTPS2 | 36.1211561 | 0.13469779 | 0.37447905 | 0.35969379 | 0.71907614 | 0.9958918 |
| SYAP1 | 38.933283 | 0.35610681 | 0.2825881 | 1.26016209 | 0.20761089 | 0.89620104 |
| TXLNG | 44.318255 | -0.0607141 | 0.40206974 | -0.1510038 | 0.8799727 | 0.99749293 |
| RBBP7 | 43.962873 | -0.1063339 | 0.28304188 | -0.3756825 | 0.70715295 | 0.9958918 |
| NHS | 520.271286 | 0.11233395 | 0.30552788 | 0.3676717 | 0.71311805 | 0.9958918 |
| SCML1 | 66.1298522 | 0.00443751 | 0.48222985 | 0.00920207 | 0.99265792 | 0.99935941 |
| CDKL5 | 40.4428496 | -0.1231184 | 0.41277465 | -0.2982702 | 0.76549694 | 0.9958918 |
| PHKA2 | 47.3190258 | -0.3181657 | 0.44381461 | -0.7168887 | 0.47344275 | 0.98910524 |
| ADGRG2 | 24.3010886 | -0.4728242 | 0.53340235 | -0.8864306 | 0.37538552 | 0.97217293 |
| PDHA1 | 24.1400763 | 0.42888044 | 0.33226305 | 1.29078583 | 0.19677795 | 0.88391893 |
| SH3KBP1 | 94.7557291 | 0.03972278 | 0.35979868 | 0.11040279 | 0.91208994 | 0.99935941 |
| BCLAF3 | 31.1498631 | 0.23360845 | 0.30244332 | 0.77240406 | 0.43987515 | 0.98092839 |
| EIF1AX | 64.6064882 | -0.1153542 | 0.30926638 | -0.3729931 | 0.70915355 | 0.9958918 |
| RPS6KA3 | 158.018581 | -0.0079147 | 0.30834784 | -0.0256682 | 0.97952198 | 0.99935941 |
| MBTPS2 | 29.5826522 | -0.0322257 | 0.33663197 | -0.0957298 | 0.9237352 | 0.99935941 |
| SMS | 32.4883903 | 0.24378294 | 0.4857167 | 0.50190357 | 0.61573535 | 0.9958918 |
| ACOT9 | 33.5069074 | -0.0396238 | 0.35995188 | -0.1100809 | 0.91234519 | 0.99935941 |

| | | | | | | |
|---|---|---|---|---|---|---|
| SAT1 | 143.254969 | -0.0565835 | 0.45440188 | -0.124523 | 0.90090121 | 0.99935941 |
| APOO | 222.571818 | -0.0753768 | 0.35046669 | -0.2150755 | 0.82970843 | 0.99592423 |
| EIF2S3 | 52.3163895 | -0.2203194 | 0.36611597 | -0.6017749 | 0.54732399 | 0.9958918 |
| ZFX | 90.8048111 | -0.3675133 | 0.33057434 | -1.1117416 | 0.26624925 | 0.93533685 |
| PDK3 | 53.4140681 | -0.1604521 | 0.54809343 | -0.2927459 | 0.76971637 | 0.9958918 |
| POLA1 | 61.8641579 | -0.3018164 | 0.25899981 | -1.1653153 | 0.2438914 | 0.92546331 |
| IL1RAPL1 | 96.1891964 | 0.25580719 | 0.40488257 | 0.63180589 | 5.28E-01 | 0.9958918 |
| TAB3 | 27.2669128 | -0.2818371 | 0.30299561 | -0.930169 | 0.35228357 | 0.97087862 |
| DMD | 719.001968 | 0.71455292 | 0.45330057 | 1.57633359 | 0.11494895 | 0.77932991 |
| TMEM47 | 55.3468478 | 0.42076894 | 0.31312988 | 1.34375213 | 0.17902855 | 0.87815793 |
| DYNLT3 | 27.2170931 | -0.1251395 | 0.33530982 | -0.3732057 | 0.70899535 | 0.9958918 |
| RPGR | 24.6545172 | -0.1220288 | 0.40553437 | -0.3009087 | 0.76348413 | 0.9958918 |
| BCOR | 48.5773857 | 0.53072612 | 0.33535116 | 1.58259813 | 0.11351309 | 0.77853537 |
| ATP6AP2 | 63.2203637 | 0.33306145 | 0.31122332 | 1.07016865 | 0.2845434 | 0.94505166 |
| CXorf38 | 19.105537 | 0.00964067 | 0.33214721 | 0.02902529 | 0.97684442 | 0.99935941 |
| MED14 | 66.7896774 | -0.3264276 | 0.22146726 | -1.4739314 | 0.14050006 | 0.8153985 |
| USP9X | 198.747099 | -0.2393885 | 0.25937477 | -0.9229444 | 0.35603619 | 0.97217293 |
| DDX3X | 108.440001 | -0.1398685 | 0.30275917 | -0.4619794 | 0.64409612 | 0.9958918 |
| CASK | 128.235904 | 0.09000546 | 0.26047145 | 0.34554825 | 0.72968224 | 0.9958918 |
| MAOA | 52.6484196 | 0.4895261 | 0.49842664 | 0.98214274 | 0.32602953 | 0.95992644 |
| MAOB | 34.651914 | -0.3390111 | 0.66925624 | -0.5065491 | 0.61247123 | 0.9958918 |
| KDM6A | 230.501507 | -0.5164052 | 0.35685198 | -1.4471131 | 0.14786525 | 0.8301633 |
| MIR222HG | 44.7754974 | -1.1567397 | 0.62196875 | -1.8598035 | 0.06291333 | 0.62227462 |
| KRBOX4 | 29.8062009 | -0.3218656 | 0.28382091 | -1.1340449 | 0.25677572 | 0.9266065 |
| JADE3 | 24.6094578 | -0.3184476 | 0.32446743 | -0.9814472 | 0.32637227 | 0.95992644 |
| NDUFB11 | 31.1789465 | -0.0961666 | 0.38569365 | -0.249334 | 0.8031024 | 0.99592423 |
| RBM10 | 31.6490224 | 0.10967043 | 0.28940102 | 0.37895663 | 0.70472007 | 0.9958918 |
| UBA1 | 44.3722203 | -0.2569889 | 0.2666544 | -0.9637528 | 0.33516986 | 0.96640422 |
| USP11 | 33.3088502 | 0.1220334 | 0.32302035 | 0.37778859 | 0.70558765 | 0.9958918 |
| ARAF | 17.3413495 | 0.36942371 | 0.38169707 | 0.9678453 | 0.33312164 | 0.9654289 |
| TIMP1 | 588.196824 | -0.7965399 | 0.72595923 | -1.0972241 | 0.27254345 | 0.93813668 |
| ZNF81 | 37.0608121 | -0.3852761 | 0.28199867 | -1.3662337 | 0.17186561 | 0.87233975 |
| RBM3 | 52.8860367 | -0.0700617 | 0.41009689 | -0.1708417 | 0.86434823 | 0.9971731 |
| WDR13 | 26.6675572 | -0.324114 | 0.45747061 | -0.7084913 | 0.47864021 | 0.99002719 |
| HDAC6 | 19.4782137 | 0.11263819 | 0.33867801 | 0.33258194 | 0.73944988 | 0.9958918 |
| PCSK1N | 32.2281008 | -0.1299627 | 0.45861554 | -0.2833806 | 0.77688513 | 0.9958918 |
| OTUD5 | 40.8559361 | 0.02200153 | 0.28927461 | 0.0760576 | 0.93937327 | 0.99935941 |
| GRIPAP1 | 31.2714134 | 0.22626518 | 0.28962015 | 0.78124807 | 0.43465661 | 0.98092839 |
| TFE3 | 28.2359439 | 0.10133691 | 0.30393443 | 0.33341703 | 0.73881951 | 0.9958918 |
| WDR45 | 22.1125752 | 0.06679372 | 0.30156851 | 0.22148773 | 0.82471269 | 0.99592423 |
| CLCN5 | 43.9777027 | -0.2360608 | 0.33371401 | -0.7073745 | 0.47933375 | 0.99002719 |
| SHROOM4 | 35.1135477 | 0.13586332 | 0.39306583 | 0.34565031 | 0.72960554 | 0.9958918 |
| MAGED1 | 96.8962176 | 0.00023638 | 0.21251084 | 0.00111233 | 0.99911249 | 0.99968761 |
| GPR173 | 50.9321979 | -0.177778 | 0.31447049 | -0.5653248 | 0.57185285 | 0.9958918 |
| TSPYL2 | 39.4810221 | -0.0269924 | 0.31762471 | -0.0849821 | 0.93227559 | 0.99935941 |
| KANTR | 80.3745422 | 0.06905191 | 0.31232586 | 0.2210893 | 0.8250229 | 0.99592423 |
| KDM5C | 37.233118 | -0.1017709 | 0.307536 | -0.3309234 | 0.74070235 | 0.9958918 |
| SMC1A | 37.4749976 | 0.04789246 | 0.30893401 | 0.15502489 | 0.87680169 | 0.9971731 |
| HUWE1 | 161.702007 | -0.1364344 | 0.21407115 | -0.6373318 | 0.52390873 | 0.9958918 |
| PHF8 | 49.1898005 | 0.113471 | 0.30014284 | 0.37805666 | 0.7053885 | 0.9958918 |
| FAM120C | 27.7896927 | -0.2827021 | 0.31974851 | -0.884139 | 0.37662117 | 0.97217293 |
| MAGED2 | 29.1674708 | 0.27803499 | 0.35729398 | 0.77816869 | 0.43646957 | 0.98092839 |
| RRAGB | 45.6864855 | -0.0112482 | 0.30186904 | -0.0372618 | 0.97027624 | 0.99935941 |
| AL445472.1 | 39.2300243 | -0.2802638 | 0.3712944 | -0.7548291 | 0.45035153 | 0.98092839 |

| | | | | | | |
|---|---|---|---|---|---|---|
| UBQLN2 | 17.5245592 | 0.42597165 | 0.354336 | 1.2021687 | 0.22929817 | 0.91455844 |
| NBDY | 71.2660345 | 0.40538886 | 0.24293718 | 1.6686983 | 0.09517719 | 0.73627067 |
| ARHGEF9 | 38.9889228 | 0.63661695 | 0.3129877 | 2.03399989 | 0.04195161 | 0.54566789 |
| ZC4H2 | 37.0824113 | 0.40053777 | 0.3726976 | 1.07469908 | 0.28250947 | 0.94419503 |
| ZC3H12B | 26.8193786 | -0.1099055 | 0.39513388 | -0.2781476 | 0.78089904 | 0.9958918 |
| MSN | 174.352349 | 0.41060599 | 0.22710426 | 1.80800653 | 0.07060548 | 0.65680975 |
| HEPH | 41.3514975 | -0.703277 | 0.38989013 | -1.8037824 | 0.07126542 | 0.66044085 |
| AR | 70.5504825 | -0.8735283 | 0.28146494 | -3.1035067 | 0.00191242 | 0.09345751 |
| OPHN1 | 86.693549 | -0.261594 | 0.24314157 | -1.0758918 | 0.28197567 | 0.94419503 |
| YIPF6 | 37.1016202 | -0.0809542 | 0.29343839 | -0.2758813 | 0.78263925 | 0.9958918 |
| STARD8 | 23.8015599 | -0.9576922 | 0.46005263 | -2.0817014 | 0.03736975 | 0.52013314 |
| EFNB1 | 24.2686647 | -0.5588342 | 0.38245799 | -1.4611649 | 0.14397019 | 0.82247441 |
| EDA | 71.476365 | -1.0280084 | 0.36516059 | -2.8152227 | 0.00487434 | 0.1747393 |
| IGBP1 | 25.4418817 | 0.46058953 | 0.33835056 | 1.36127903 | 0.17342553 | 0.87233975 |
| NONO | 47.7503259 | -0.0725601 | 0.32143114 | -0.2257406 | 0.82140319 | 0.99592423 |
| TAF1 | 93.1672762 | -0.0126086 | 0.30298805 | -0.0416141 | 0.96680633 | 0.99935941 |
| OGT | 144.826167 | -0.1304661 | 0.24886203 | -0.5242507 | 0.60010418 | 0.9958918 |
| NHSL2 | 49.6505572 | 0.1873635 | 0.34148877 | 0.54866665 | 0.58323424 | 0.9958918 |
| PIN4 | 28.6021175 | 0.27299185 | 0.28986984 | 0.94177392 | 0.3463084 | 0.96887708 |
| RPS4X | 129.699652 | 0.17903297 | 0.27208048 | 0.65801476 | 0.51052865 | 0.9958918 |
| HDAC8 | 180.988368 | 0.22977068 | 0.21162588 | 1.08573998 | 0.27759405 | 0.94358865 |
| CHIC1 | 63.2046342 | -0.269868 | 0.2973947 | -0.9074404 | 0.36417398 | 0.97217293 |
| XIST | 300.946339 | -1.4327855 | 3.58180871 | -0.4000173 | 0.68914376 | 0.9958918 |
| JPX | 189.195704 | -0.2010675 | 0.32844896 | -0.6121727 | 0.54042353 | 0.9958918 |
| FTX | 1476.39826 | 0.06596861 | 0.26045489 | 0.25328227 | 0.8000501 | 0.99592423 |
| SLC16A2 | 25.3282072 | 0.58906132 | 0.35976186 | 1.63736459 | 0.10155431 | 0.75234138 |
| RLIM | 26.7972979 | -0.0026315 | 0.39289752 | -0.0066977 | 0.99465604 | 0.99935941 |
| NEXMIF | 28.1116529 | -0.0122851 | 0.37702828 | -0.032584 | 0.97400631 | 0.99935941 |
| ABCB7 | 51.5263246 | -0.0178096 | 0.24886975 | -0.0715618 | 0.94295061 | 0.99935941 |
| ATRX | 314.539848 | -0.1224522 | 0.17806512 | -0.6876821 | 0.49165298 | 0.99046213 |
| MAGT1 | 51.7189158 | 0.23808969 | 0.29929427 | 0.79550366 | 0.42632059 | 0.98015476 |
| COX7B | 47.6012716 | -0.1096893 | 0.28851183 | -0.3801901 | 0.70380433 | 0.9958918 |
| PGK1 | 115.327632 | -0.2802581 | 0.48979244 | -0.5721976 | 0.56718809 | 0.9958918 |
| BRWD3 | 111.065277 | -0.2152722 | 0.33584271 | -0.6409911 | 0.52152848 | 0.9958918 |
| SH3BGRL | 137.200347 | 0.2018358 | 0.27551782 | 0.73256895 | 0.46382137 | 0.98590351 |
| HDX | 41.2204436 | 0.12131669 | 0.34834516 | 0.34826577 | 0.7276406 | 0.9958918 |
| APOOL | 33.0356943 | -0.2196609 | 0.27972807 | -0.785266 | 0.43229764 | 0.98092839 |
| CHM | 104.546941 | -0.3109397 | 0.28780961 | -1.080366 | 0.27997925 | 0.94419503 |
| DACH2 | 143.023526 | -4.8278128 | 1.11102736 | -4.3453591 | 1.39E-05 | 0.0040484 |
| DIAPH2 | 449.310607 | 0.66604219 | 0.2835993 | 2.34853256 | 0.01884755 | 0.36546347 |
| SRPX2 | 115.927409 | -1.321518 | 0.3937484 | -3.3562498 | 0.00079007 | 0.06338861 |
| SYTL4 | 78.4934703 | -0.7010515 | 0.54544958 | -1.2852728 | 0.19869699 | 0.88821421 |
| ARMCX4 | 103.489369 | 0.27573861 | 0.31478526 | 0.87595781 | 0.38105297 | 0.97217293 |
| ARMCX3 | 34.6838038 | -0.3432486 | 0.35668273 | -0.9623359 | 0.33588091 | 0.96640422 |
| ZMAT1 | 20.2286502 | -0.1950183 | 0.44583556 | -0.437422 | 0.66180535 | 0.9958918 |
| LINC00630 | 31.9172589 | -0.5248114 | 0.35647205 | -1.4722372 | 0.14095684 | 0.81630561 |
| BEX4 | 18.6918401 | -0.075721 | 0.35701925 | -0.2120922 | 0.83203508 | 0.99592423 |
| TCEAL9 | 49.1009857 | -0.4679053 | 0.30804172 | -1.5189673 | 0.12877072 | 0.80402383 |
| BEX3 | 58.9076039 | 0.22444739 | 0.28996489 | 0.77405023 | 0.43890109 | 0.98092839 |
| TCEAL4 | 75.4810413 | -0.4764682 | 0.30383324 | -1.5681899 | 0.11683681 | 0.78417352 |
| TCEAL3 | 27.5922795 | -0.2621575 | 0.31346386 | -0.8363245 | 0.40297235 | 0.97512144 |
| MORF4L2 | 149.556251 | -0.6070036 | 0.372342 | -1.6302314 | 0.1030526 | 0.75809165 |
| SLC25A53 | 23.7647238 | -0.0937398 | 0.3606931 | -0.2598879 | 0.79495021 | 0.99592423 |
| FAM199X | 31.8138168 | 0.20341657 | 0.32494379 | 0.62600541 | 0.5313114 | 0.9958918 |

| | | | | | | |
|---|---|---|---|---|---|---|
| TBC1D8B | 27.1978198 | -0.3670762 | 0.40654114 | -0.9029251 | 0.36656566 | 0.97217293 |
| MORC4 | 27.8491314 | -0.2991402 | 0.38331999 | -0.7803929 | 0.43515966 | 0.98092839 |
| RBM41 | 34.0046892 | -0.166568 | 0.40323038 | -0.4130839 | 0.67954512 | 0.9958918 |
| TSC22D3 | 80.450469 | -1.0440612 | 0.60859444 | -1.7155286 | 0.08624836 | 0.71137956 |
| MID2 | 29.7548078 | -0.0752462 | 0.34343709 | -0.2190975 | 0.82657413 | 0.99592423 |
| COL4A5 | 97.1008131 | 0.19261676 | 0.45122994 | 0.42687052 | 0.66947364 | 0.9958918 |
| ACSL4 | 252.662116 | 0.8420268 | 0.50223725 | 1.67655187 | 0.09363015 | 0.73159032 |
| TMEM164 | 43.9698123 | 0.70334286 | 0.26259862 | 2.67839507 | 0.00739759 | 0.22219179 |
| AMMECR1 | 42.4760714 | -0.479667 | 0.38334968 | -1.2512519 | 0.2108426 | 0.89738044 |
| CHRDL1 | 26.2893976 | -0.1355982 | 0.44791739 | -0.3027303 | 0.7620954 | 0.9958918 |
| PAK3 | 58.9683383 | -0.5504819 | 0.63112013 | -0.87223 | 0.38308294 | 0.9725044 |
| ALG13 | 115.454404 | -0.0776458 | 0.30349171 | -0.2558416 | 0.79807318 | 0.99592423 |
| LRCH2 | 55.9955123 | -0.6903481 | 0.47800643 | -1.4442235 | 0.1486761 | 0.8301633 |
| PLS3 | 197.87298 | -0.2120778 | 0.31991531 | -0.6629186 | 0.50738271 | 0.99397937 |
| DANT2 | 65.6908016 | 0.20334259 | 0.40899075 | 0.4971814 | 0.61906114 | 0.9958918 |
| KLHL13 | 143.06901 | -0.9935117 | 0.35079679 | -2.8321573 | 0.00462351 | 0.16989087 |
| WDR44 | 75.6864911 | -0.170389 | 0.24249379 | -0.7026531 | 0.48227196 | 0.99002719 |
| DOCK11 | 50.9733291 | -0.0685176 | 0.35404041 | -0.1935303 | 0.84654367 | 0.99692952 |
| IL13RA1 | 53.7040416 | -0.2577993 | 0.32262116 | -0.7990775 | 0.4242455 | 0.97828057 |
| PGRMC1 | 39.264894 | -0.0259044 | 0.35041422 | -0.0739249 | 0.94107011 | 0.99935941 |
| SLC25A43 | 51.6452465 | 0.1370431 | 0.30153155 | 0.45449007 | 0.64947613 | 0.9958918 |
| SLC25A5 | 38.0119242 | 0.81466981 | 0.44403761 | 1.8346865 | 0.06655215 | 0.63518413 |
| 6-Sep | 21.8651753 | -0.0672004 | 0.35656234 | -0.1884674 | 0.85051028 | 0.9971731 |
| UPF3B | 25.3806526 | -0.1383491 | 0.38776362 | -0.3567873 | 0.72125104 | 0.9958918 |
| NDUFA1 | 32.7618076 | -0.090374 | 0.38966021 | -0.2319302 | 0.8165922 | 0.99592423 |
| LAMP2 | 71.1857268 | -0.0609981 | 0.27207356 | -0.2241971 | 0.82260393 | 0.99592423 |
| CUL4B | 39.3145214 | -0.7474393 | 0.30867706 | -2.4214281 | 0.01545966 | 0.33877878 |
| MCTS1 | 21.9506108 | -0.200978 | 0.33388222 | -0.6019428 | 0.54721224 | 0.9958918 |
| THOC2 | 169.477695 | -0.5443448 | 0.22748897 | -2.3928403 | 0.01671851 | 0.34578631 |
| XIAP | 88.3923898 | 3.92E-02 | 0.17818499 | 0.22002703 | 0.8258501 | 0.99592423 |
| STAG2 | 170.332243 | -0.2241217 | 0.26638915 | -0.8413319 | 0.40016204 | 0.97500135 |
| SMARCA1 | 90.3693812 | -0.1536637 | 0.35926835 | -0.4277129 | 0.66886018 | 0.9958918 |
| OCRL | 24.4087613 | -0.1655919 | 0.37486099 | -0.441742 | 0.65867589 | 0.9958918 |
| ZDHHC9 | 26.388269 | 0.35961736 | 0.51840136 | 0.69370452 | 4.88E-01 | 0.99046213 |
| BCORL1 | 22.0480295 | -0.1272325 | 0.32193837 | -0.3952076 | 0.69268971 | 0.9958918 |
| AIFM1 | 20.2089982 | 1.10517427 | 0.35486865 | 3.11431925 | 0.0018437 | 0.09345751 |
| ENOX2 | 48.3306703 | 0.24065973 | 0.39974893 | 0.60202721 | 0.54715603 | 0.9958918 |
| RAP2C-AS1 | 43.391215 | 0.03033824 | 0.45841588 | 0.0661806 | 0.94723404 | 0.99935941 |
| GPC4 | 47.148748 | 0.14957182 | 0.42331787 | 0.35333218 | 0.72383943 | 0.9958918 |
| GPC3 | 110.090859 | -2.0465377 | 0.52029503 | -3.9334179 | 8.37E-05 | 0.01398756 |
| PHF6 | 28.674924 | -0.5035766 | 0.35920254 | -1.4019294 | 0.16093632 | 0.85092986 |
| HPRT1 | 42.2008629 | 0.34948785 | 0.29140047 | 1.19933868 | 0.23039628 | 0.91459157 |
| FAM122C | 20.2648802 | -0.7952656 | 0.59290123 | -1.3413121 | 0.17981913 | 0.87815793 |
| RTL8C | 35.2033723 | 0.22288943 | 0.32766585 | 0.68023393 | 0.49635635 | 0.99176652 |
| ZNF75D | 44.0726476 | -0.0001052 | 0.38511072 | -0.0002733 | 0.99978196 | 0.99978196 |
| INTS6L | 50.1276885 | 0.09188809 | 0.34818053 | 0.26390932 | 0.7918498 | 0.99592423 |
| FHL1 | 115.805997 | -0.3930708 | 0.59246844 | -0.663446 | 0.50704495 | 0.99397937 |
| MAP7D3 | 26.6487257 | -0.1241126 | 0.37701979 | -0.329194 | 0.74200908 | 0.9958918 |
| ARHGEF6 | 58.4407106 | -0.1053587 | 0.34735131 | -0.3033204 | 0.76164571 | 0.9958918 |
| AL683813.1 | 28.3851211 | 0.62995083 | 0.31890375 | 1.97536355 | 0.04822691 | 0.57134906 |
| RBMX | 118.012608 | 0.24846833 | 0.18595584 | 1.33616841 | 0.18149423 | 0.87865686 |
| FGF13 | 52.8810614 | 0.57221117 | 0.5005274 | 1.14321647 | 0.25294872 | 0.92646125 |
| ATP11C | 271.557453 | 0.92702487 | 0.23924326 | 3.87482132 | 0.0001067 | 0.01600327 |
| FMR1 | 49.845121 | -0.2015544 | 0.38674707 | -0.521153 | 0.6022602 | 0.9958918 |

| | | | | | | |
|---|---|---|---|---|---|---|
| IDS | 42.2285831 | 0.40217396 | 0.31758202 | 1.26636248 | 0.20538334 | 0.89589437 |
| LINC00894 | 96.7130469 | 0.14790237 | 0.3579036 | 0.41324638 | 0.67942611 | 0.9958918 |
| MAMLD1 | 150.191174 | -0.4667108 | 0.46977393 | -0.9934795 | 0.32047631 | 0.95992644 |
| MTM1 | 44.7732969 | 0.0419503 | 0.26433704 | 0.15870004 | 0.87390521 | 0.9971731 |
| MTMR1 | 40.5913841 | -0.0239079 | 0.28546745 | -0.08375 | 0.9332552 | 0.99935941 |
| CD99L2 | 37.2116947 | 0.34698744 | 0.31363413 | 1.10634464 | 0.26857738 | 0.93598159 |
| GABRE | 25.1614004 | -0.3396615 | 0.64070667 | -0.5301357 | 0.59601783 | 0.9958918 |
| BGN | 331.143221 | -0.2730284 | 0.55544852 | -0.4915458 | 0.6230405 | 0.9958918 |
| BCAP31 | 31.0586073 | -0.3869152 | 0.37578307 | -1.0296237 | 0.30318666 | 0.95650251 |
| SSR4 | 26.6403411 | 0.11194151 | 0.43650688 | 0.25644844 | 0.79760459 | 0.99592423 |
| MECP2 | 107.369691 | -0.6233218 | 0.23863029 | -2.6120815 | 0.00899928 | 0.25292422 |
| FLNA | 474.827414 | 0.13069454 | 0.29504287 | 0.44296797 | 0.6577889 | 0.9958918 |
| EMD | 27.4931705 | -0.6382662 | 0.36119553 | -1.7670933 | 0.0772126 | 0.67956337 |
| RPL10 | 238.013183 | 0.20216567 | 0.39282642 | 0.51464377 | 0.60680197 | 0.9958918 |
| TAZ | 20.2781457 | -0.4202037 | 0.3412466 | -1.2313784 | 0.21818136 | 0.90437211 |
| ATP6AP1 | 26.3186984 | 0.58293998 | 0.35330183 | 1.64997726 | 0.09894759 | 0.74528964 |
| GDI1 | 34.609411 | -0.5223172 | 0.33740596 | -1.5480378 | 0.1216132 | 0.79021701 |
| FAM50A | 20.5571249 | -0.2580469 | 0.4218901 | -0.6116449 | 0.54077275 | 0.9958918 |
| FUNDC2 | 26.982786 | -0.3683815 | 0.32014429 | -1.1506733 | 0.24986668 | 0.92624282 |
| BRCC3 | 23.9719457 | 0.35442008 | 0.45544876 | 0.77817773 | 0.43646425 | 0.98092839 |
| TMLHE | 46.5925567 | 0.25194057 | 0.40487025 | 0.62227484 | 0.53376116 | 0.9958918 |
| VAMP7 | 23.606895 | 0.18323482 | 0.35062973 | 0.52258781 | 0.60126113 | 0.9958918 |
| FBXO25 | 34.7605266 | -0.3146854 | 0.30945379 | -1.016906 | 0.30919817 | 0.95793395 |
| TDRP | 24.6396077 | -0.8794223 | 0.3411457 | -2.5778495 | 0.00994173 | 0.26559396 |
| ERICH1 | 61.0908974 | -0.0156312 | 0.34801261 | -0.0449158 | 0.96417446 | 0.99935941 |
| MYOM2 | 45.4211598 | 0.53957661 | 0.54899945 | 0.98283635 | 0.32568799 | 0.95992644 |
| CSMD1 | 98.110486 | 0.93357706 | 0.5304337 | 1.76002593 | 0.07840341 | 0.68187771 |
| MCPH1 | 136.061965 | 0.40197722 | 0.23791086 | 1.68961271 | 0.09110207 | 0.72693717 |
| ANGPT2 | 72.1263568 | 0.43663689 | 0.32918331 | 1.32642473 | 0.18469905 | 0.87866798 |
| AGPAT5 | 40.956739 | 0.47941905 | 0.32366482 | 1.48122075 | 1.39E-01 | 0.81520215 |
| DEFB1 | 64.1927138 | 0.44027889 | 0.47291801 | 0.93098356 | 0.35186207 | 0.97065853 |
| FAM85B | 106.789926 | 0.1635592 | 0.49623359 | 0.32960123 | 0.74170129 | 0.9958918 |
| PRAG1 | 24.3138696 | -0.0042979 | 0.44629476 | -0.0096302 | 0.99231635 | 0.99935941 |
| MFHAS1 | 95.5242573 | 0.30067339 | 0.43743731 | 0.68735196 | 0.49186097 | 0.99046213 |
| ERI1 | 29.9699768 | -0.1630453 | 0.35096575 | -0.4645618 | 0.64224527 | 0.9958918 |
| PPP1R3B | 30.196538 | -0.6670407 | 0.54629853 | -1.2210186 | 0.22207898 | 0.91093521 |
| TNKS | 223.77139 | -0.2015239 | 0.16827503 | -1.1975867 | 0.23107795 | 0.91521027 |
| MSRA | 76.6899388 | -0.2387701 | 0.33873508 | -0.7048875 | 0.48088024 | 0.99002719 |
| XKR6 | 56.1180661 | -0.2325185 | 0.41854243 | -0.5555434 | 0.57852305 | 0.9958918 |
| MTMR9 | 35.4172017 | 0.18949316 | 0.29678728 | 0.63848141 | 0.52316035 | 0.9958918 |
| FDFT1 | 48.890471 | 0.05666875 | 0.39786983 | 0.14243037 | 0.88674007 | 0.99935941 |
| CTSB | 89.8207116 | 0.83293122 | 0.33547572 | 2.48283609 | 0.0130341 | 0.30408773 |
| FAM66D | 31.0299519 | 0.85796088 | 0.64373686 | 1.33278198 | 0.18260336 | 0.87866798 |
| LONRF1 | 110.200189 | -0.6084618 | 0.5986556 | -1.0163804 | 0.30944825 | 0.95793395 |
| DLC1 | 1023.22359 | -0.6200708 | 0.20990467 | -2.9540591 | 0.00313624 | 0.13394152 |
| TUSC3 | 73.3733058 | -0.6355042 | 0.37287167 | -1.7043511 | 0.08831551 | 0.71818729 |
| MICU3 | 118.23666 | 0.0697694 | 0.27319958 | 0.25537888 | 0.79843048 | 0.99592423 |
| VPS37A | 49.0085519 | -0.1501517 | 0.29158548 | -0.5149492 | 0.60658853 | 0.9958918 |
| SLC7A2 | 388.294279 | 0.8288496 | 0.41414773 | 2.00133805 | 0.04535597 | 0.55894748 |
| MTUS1 | 142.433438 | -1.1721448 | 0.35242089 | -3.3259801 | 0.00088108 | 0.06475074 |
| AC027117.2 | 25.421728 | -0.6123911 | 0.62213505 | -0.9843378 | 0.32494943 | 0.95992644 |
| PCM1 | 235.379327 | -0.2027101 | 0.1944026 | -1.0427337 | 0.29707167 | 0.95417903 |
| ASAH1 | 53.2556075 | -0.0242371 | 0.38923478 | -0.0622686 | 0.95034891 | 0.99935941 |
| PSD3 | 300.16761 | -0.8212212 | 0.26472296 | -3.1021911 | 0.00192094 | 0.09345751 |

| | | | | | | |
|---|---|---|---|---|---|---|
| CSGALNACT1 | 102.88835 | 1.20512893 | 0.38641047 | 3.11877917 | 0.00181602 | 0.09345751 |
| INTS10 | 60.3867888 | 0.30888123 | 0.27668236 | 1.11637488 | 0.2642617 | 0.93499423 |
| ATP6V1B2 | 41.7430518 | -0.3991015 | 0.27161316 | -1.4693745 | 0.14173124 | 0.81652619 |
| LZTS1 | 54.5099364 | 0.58661282 | 0.31925141 | 1.83746355 | 0.06614149 | 0.63381211 |
| XPO7 | 66.7498427 | 0.0770743 | 0.24226843 | 0.31813595 | 0.75038181 | 0.9958918 |
| FAM160B2 | 24.2032328 | 0.00046266 | 0.33986586 | 0.00136131 | 0.99891383 | 0.99968761 |
| BMP1 | 40.2050635 | 0.04558335 | 0.26266176 | 0.17354393 | 0.8622239 | 0.9971731 |
| SLC39A14 | 100.202667 | 0.56894418 | 0.5612998 | 1.01361907 | 0.31076456 | 0.95956408 |
| PPP3CC | 146.94804 | -0.0216595 | 0.20375667 | -0.1063009 | 0.91534364 | 0.99935941 |
| SORBS3 | 61.5438939 | -0.2826526 | 0.31896643 | -0.8861514 | 0.37553592 | 0.97217293 |
| BIN3 | 31.7562159 | 0.80236833 | 0.32926902 | 2.43681691 | 0.01481718 | 0.33097706 |
| RHOBTB2 | 18.8990552 | 0.44456099 | 0.5188749 | 0.85677875 | 0.39156717 | 0.97297628 |
| TNFRSF10B | 32.1174011 | 0.60932235 | 0.29640951 | 2.05567744 | 0.03981362 | 0.53587963 |
| TNFRSF10D | 23.0598575 | -0.1399066 | 0.38126933 | -0.3669496 | 7.14E-01 | 0.9958918 |
| LOXL2 | 58.4244107 | -0.416831 | 0.55218026 | -0.7548821 | 0.45031971 | 0.98092839 |
| ENTPD4 | 42.2241618 | 0.04123634 | 0.25991906 | 0.15865071 | 0.87394407 | 0.9971731 |
| SLC25A37 | 59.5509319 | 0.43090262 | 0.27193674 | 1.58456936 | 0.11306421 | 0.77793929 |
| DOCK5 | 78.3108282 | -0.4098994 | 0.4147517 | -0.9883006 | 0.32300543 | 0.95992644 |
| KCTD9 | 46.8632468 | -0.4573094 | 0.39712473 | -1.151551 | 0.24950561 | 0.92624282 |
| PPP2R2A | 148.18971 | -0.4328464 | 0.27578283 | -1.5695189 | 0.11652707 | 0.78417352 |
| BNIP3L | 204.894418 | -0.3787808 | 0.48600358 | -0.7793786 | 0.43575671 | 0.98092839 |
| DPYSL2 | 90.0813495 | -0.0823276 | 0.32238589 | -0.2553697 | 0.79843758 | 0.99592423 |
| ADRA1A | 38.5151298 | 1.52840368 | 0.55143415 | 2.77168847 | 0.00557664 | 0.18713567 |
| CLU | 107.079571 | 0.55711492 | 0.46173748 | 1.20656206 | 0.22760085 | 0.9120167 |
| CCDC25 | 20.8312613 | 0.31937118 | 0.36657589 | 0.87122799 | 0.38362968 | 0.9725044 |
| ELP3 | 52.9034228 | -0.1034187 | 0.23673371 | -0.4368568 | 0.66221519 | 0.9958918 |
| ZNF395 | 20.3449994 | -0.2619692 | 0.5259853 | -0.4980543 | 0.61844576 | 0.9958918 |
| FZD3 | 23.9992512 | 0.29007819 | 0.36446042 | 0.79591137 | 0.42608355 | 0.98015476 |
| EXTL3 | 28.9336508 | 0.17077466 | 0.32372887 | 0.52752372 | 0.59782995 | 0.9958918 |
| INTS9 | 46.856473 | 0.11956626 | 0.28811189 | 0.41499941 | 0.67814234 | 0.9958918 |
| HMBOX1 | 175.956754 | -0.2331611 | 0.22180358 | -1.0512051 | 0.2931644 | 0.95186143 |
| KIF13B | 76.9371888 | -0.2417008 | 0.28531295 | -0.8471427 | 0.39691558 | 0.97500135 |
| SARAF | 71.166481 | 0.25753738 | 0.22722816 | 1.13338672 | 0.25705189 | 0.9266065 |
| LEPROTL1 | 30.440947 | 0.25715066 | 0.453467 | 0.5670769 | 0.57066193 | 0.9958918 |
| RBPMS | 1819.04818 | -0.0462438 | 0.27061264 | -0.1708855 | 0.8643138 | 0.9971731 |
| GTF2E2 | 52.8460618 | -0.1407311 | 0.25062723 | -0.5615156 | 0.57444611 | 0.9958918 |
| GSR | 26.7527883 | -0.2373568 | 0.33205673 | -0.7148081 | 0.47472765 | 0.98910524 |
| PPP2CB | 129.227313 | -0.04294 | 0.18393928 | -0.2334464 | 0.81541475 | 0.99592423 |
| WRN | 106.787531 | -0.4924694 | 0.19334253 | -2.5471343 | 0.01086116 | 0.28105167 |
| FUT10 | 29.5970352 | 0.0061574 | 0.30235415 | 0.02036485 | 0.98375233 | 0.99935941 |
| RNF122 | 19.567358 | 0.10623591 | 0.64139107 | 0.1656336 | 0.8684453 | 0.9971731 |
| ADGRA2 | 66.3800672 | -0.2903659 | 0.33174868 | -0.8752585 | 0.38143326 | 0.97217293 |
| RAB11FIP1 | 22.9736926 | 1.33619762 | 0.51474433 | 2.59584718 | 0.0094358 | 0.25610587 |
| ASH2L | 30.343557 | 0.4793945 | 0.30870069 | 1.55294274 | 0.12043682 | 0.79021701 |
| BAG4 | 23.6230095 | 0.49301752 | 0.34687383 | 1.42131656 | 0.15522475 | 0.84277602 |
| DDHD2 | 40.818763 | -0.068196 | 0.29414705 | -0.2318434 | 0.81665967 | 0.99592423 |
| PLPP5 | 20.5916461 | -0.2615357 | 0.4991981 | -0.5239116 | 0.60034004 | 0.9958918 |
| NSD3 | 165.970281 | -0.0712694 | 0.20159237 | -0.3535324 | 0.72368936 | 0.9958918 |
| FGFR1 | 134.097546 | -0.5243889 | 0.47888228 | -1.0950268 | 0.27350489 | 0.93924645 |
| TACC1 | 503.589838 | -0.6973467 | 0.48418927 | -1.4402358 | 0.14980071 | 0.83211294 |
| PLEKHA2 | 150.003639 | -0.0507022 | 0.35972669 | -0.1409465 | 0.88791217 | 0.99935941 |
| ADAM9 | 122.361337 | -0.2420195 | 0.34461298 | -0.7022935 | 0.48249612 | 0.99002719 |
| TCIM | 60.113573 | -1.5997594 | 0.85767011 | -1.8652386 | 0.06214796 | 0.62054234 |
| SFRP1 | 113.381868 | -1.1224456 | 0.35501454 | -3.161689 | 0.00156857 | 0.0868147 |

| | | | | | | |
|---|---|---|---|---|---|---|
| GOLGA7 | 20.1309792 | 0.54488494 | 0.31419074 | 1.73424889 | 0.08287393 | 0.7007527 |
| GPAT4 | 24.4655269 | -0.0340728 | 0.31034004 | -0.109792 | 0.91257436 | 0.99935941 |
| KAT6A | 215.488981 | 0.12437898 | 0.17247317 | 0.72114973 | 0.47081739 | 0.98910524 |
| AP3M2 | 29.2941577 | 0.75095788 | 0.33893965 | 2.21560946 | 0.02671826 | 0.44524376 |
| PLAT | 41.6666115 | -0.8368026 | 0.3124865 | -2.6778841 | 0.00740889 | 0.22219179 |
| IKBKB | 57.7236148 | 0.05958342 | 0.26749459 | 0.22274625 | 0.823733 | 0.99592423 |
| VDAC3 | 45.2907199 | 0.07665338 | 0.23315634 | 0.32876386 | 0.74233418 | 0.9958918 |
| SLC20A2 | 165.467329 | -0.0515207 | 0.33371964 | -0.1543832 | 0.87730762 | 0.99722158 |
| SMIM19 | 20.5645389 | 0.12363499 | 0.34071135 | 0.36287312 | 0.71669967 | 0.9958918 |
| RNF170 | 30.2819163 | -0.3388118 | 0.36240096 | -0.9349086 | 0.34983541 | 0.9693503 |
| HOOK3 | 115.841436 | -0.1266304 | 0.21638779 | -0.5852012 | 0.55841246 | 0.9958918 |
| FNTA | 25.8633513 | 0.14646698 | 0.30068128 | 0.48711707 | 0.62617537 | 0.9958918 |
| HGSNAT | 96.6295854 | 0.14849938 | 0.2675284 | 0.55507894 | 0.57884067 | 0.9958918 |
| SPIDR | 470.528582 | -0.5748729 | 0.31581623 | -1.8202767 | 0.06871688 | 0.64792295 |
| CEBPD | 323.396338 | -1.335591 | 0.48706007 | -2.7421484 | 6.10E-03 | 0.19357111 |
| PRKDC | 130.526872 | -0.157842 | 0.309843 | -0.5094257 | 0.61045386 | 0.9958918 |
| UBE2V2 | 47.7442667 | -0.3990649 | 0.2845901 | -1.4022443 | 0.1608423 | 0.85092986 |
| PCMTD1 | 130.930939 | 0.11229581 | 0.18477748 | 0.60773538 | 0.543363 | 0.9958918 |
| RB1CC1 | 148.110493 | -0.0646922 | 0.18522902 | -0.3492552 | 0.72689774 | 0.9958918 |
| ATP6V1H | 70.5079257 | -0.2259491 | 0.29526238 | -0.7652485 | 0.44412361 | 0.98092839 |
| TCEA1 | 55.5082946 | -0.0560018 | 0.25552321 | -0.2191653 | 0.82652129 | 0.99592423 |
| LYPLA1 | 41.9753701 | 0.20373125 | 0.29299695 | 0.69533573 | 0.48684493 | 0.99046213 |
| TMEM68 | 19.4496662 | -0.2067431 | 0.38000369 | -0.5440554 | 0.58640336 | 0.9958918 |
| TGS1 | 34.0761602 | 0.30524931 | 0.3094174 | 0.98652921 | 0.32387349 | 0.95992644 |
| LYN | 45.98036 | 0.34380773 | 0.29918045 | 1.14916508 | 0.25048792 | 0.92624282 |
| RPS20 | 69.4244136 | -0.2308101 | 0.33880155 | -0.6812546 | 0.49571041 | 0.99176652 |
| IMPAD1 | 44.5170561 | -0.3276424 | 0.29960202 | -1.0935921 | 0.27413393 | 0.9394201 |
| FAM110B | 108.157688 | 0.84806192 | 0.40759287 | 2.08065934 | 0.0374651 | 0.5204745 |
| SDCBP | 71.6468117 | 0.18666398 | 0.35197229 | 0.53033716 | 0.59587819 | 0.9958918 |
| NSMAF | 54.0038498 | -0.0359978 | 0.2709144 | -0.1328753 | 0.89429199 | 0.99935941 |
| TOX | 49.4332746 | 0.97180058 | 0.33775898 | 2.87720134 | 0.0040122 | 0.15437498 |
| CA8 | 82.1863226 | -5.1775125 | 1.04458365 | -4.9565322 | 7.18E-07 | 0.00074678 |
| RAB2A | 211.971665 | -0.0197542 | 0.17764641 | -0.1111997 | 0.91145797 | 0.99935941 |
| CHD7 | 33.8355669 | -0.6434233 | 0.34449886 | -1.8677082 | 0.06180275 | 0.61962951 |
| ASPH | 145.451442 | -0.1524966 | 0.19662272 | -0.7755797 | 0.43799717 | 0.98092839 |
| YTHDF3 | 76.2817952 | -0.176187 | 0.23565822 | -0.7476378 | 0.45467867 | 0.98173285 |
| CYP7B1 | 322.080069 | 0.52432244 | 0.51822667 | 1.01176275 | 0.31165151 | 0.9599143 |
| ARMC1 | 28.5845023 | 0.25174012 | 0.33028728 | 0.76218532 | 0.44594941 | 0.98092839 |
| MTFR1 | 40.7182546 | 0.39016163 | 0.35000142 | 1.11474298 | 0.26496058 | 0.93499423 |
| PDE7A | 54.6453192 | 0.09298456 | 0.30501261 | 0.30485481 | 0.76047674 | 0.9958918 |
| VCPIP1 | 20.7125998 | 0.07087619 | 0.36366687 | 0.19489318 | 0.84547657 | 0.99692952 |
| SGK3 | 44.1059828 | 0.30095364 | 0.35752643 | 0.84176613 | 0.39991887 | 0.97500135 |
| CSPP1 | 90.2547182 | -0.4642852 | 0.30707391 | -1.5119655 | 0.13054265 | 0.80565822 |
| ARFGEF1 | 186.932385 | -0.0656968 | 0.17513952 | -0.375111 | 0.70757795 | 0.9958918 |
| PREX2 | 77.1669115 | -0.3333986 | 0.56309874 | -0.5920785 | 0.55379804 | 0.9958918 |
| SULF1 | 212.05484 | -0.3850931 | 0.5480948 | -0.7026031 | 0.48230314 | 0.99002719 |
| NCOA2 | 306.691567 | -0.1161442 | 0.30857986 | -0.3763831 | 0.70663213 | 0.9958918 |
| TRAM1 | 74.1691374 | -0.1173813 | 0.25381151 | -0.4624745 | 0.64374111 | 0.9958918 |
| MSC-AS1 | 194.827161 | 1.3235889 | 0.53826335 | 2.45899874 | 0.01393251 | 0.31996881 |
| TERF1 | 47.3300984 | 0.03407136 | 0.29094053 | 0.11710763 | 0.90677476 | 0.99935941 |
| SBSPON | 33.032123 | -0.3473553 | 0.41654749 | -0.8338912 | 0.40434231 | 0.97582411 |
| RPL7 | 111.074092 | -0.0800885 | 0.44217557 | -0.1811238 | 0.85627038 | 0.9971731 |
| RDH10 | 23.6135631 | -0.7104414 | 0.46741227 | -1.519946 | 0.12852455 | 0.80316917 |
| STAU2 | 72.8682415 | 0.62026853 | 0.25060353 | 2.47509896 | 0.01331993 | 0.30782006 |

| | | | | | | |
|---|---|---|---|---|---|---|
| ELOC | 35.2581831 | 0.28402841 | 0.28646378 | 0.9914985 | 0.32144222 | 0.95992644 |
| GDAP1 | 22.2907749 | -0.1558207 | 0.43465827 | -0.3584901 | 0.7199766 | 0.9958918 |
| PI15 | 114.33977 | 0.80762104 | 0.94109184 | 0.85817452 | 3.91E-01 | 9.73E-01 |
| ZFHX4 | 55.2793368 | -0.910683 | 0.55924108 | -1.6284265 | 0.10343448 | 0.75827058 |
| PEX2 | 21.6599821 | -0.4985746 | 0.3505478 | -1.4222728 | 0.15494706 | 0.84277602 |
| ZC2HC1A | 36.8548595 | 0.3348804 | 0.34292858 | 0.97653101 | 0.32880139 | 0.9620084 |
| ZBTB10 | 42.660846 | -0.0410673 | 0.26824882 | -0.1530942 | 0.87832397 | 0.99722158 |
| ZNF704 | 150.655782 | 0.09970526 | 0.42712319 | 0.23343444 | 0.81542407 | 0.99592423 |
| PAG1 | 264.613835 | -1.3217568 | 0.55692668 | -2.3733049 | 0.0176297 | 0.35016405 |
| ZFAND1 | 25.6850784 | 0.07746096 | 0.34282183 | 0.22595107 | 0.82123947 | 0.99592423 |
| C8orf59 | 18.8412457 | -0.1800159 | 0.32602284 | -0.5521574 | 0.58084055 | 0.9958918 |
| WWP1 | 96.2976958 | 0.07963376 | 0.2444372 | 0.32578411 | 0.7445877 | 0.9958918 |
| RMDN1 | 42.741376 | -0.0805864 | 0.27417717 | -0.2939209 | 0.76881838 | 0.9958918 |
| CPNE3 | 36.4557524 | -0.066196 | 0.31653204 | -0.2091289 | 0.83434763 | 0.99638118 |
| AF117829.1 | 103.091393 | -0.0940574 | 0.43444488 | -0.2165003 | 0.8285978 | 0.99592423 |
| RIPK2 | 23.3658991 | 0.56941983 | 0.36308211 | 1.56829493 | 0.11681232 | 0.78417352 |
| NBN | 36.9436176 | -0.2835656 | 0.40041453 | -0.7081801 | 0.47883344 | 0.99002719 |
| DECR1 | 69.0601245 | 0.51006228 | 0.35289657 | 1.44535911 | 0.14835704 | 0.8301633 |
| PIP4P2 | 45.3430561 | 0.05467088 | 0.34647788 | 0.15779039 | 0.87462197 | 0.9971731 |
| OTUD6B-AS1 | 19.6877456 | 0.15316088 | 0.46727412 | 0.32777523 | 0.74308162 | 0.9958918 |
| OTUD6B | 18.3278095 | -0.5496213 | 0.41057793 | -1.3386528 | 0.18068374 | 0.87815793 |
| LRRC69 | 23.4954124 | -0.1612497 | 0.38072825 | -0.4235297 | 0.67190885 | 0.9958918 |
| RUNX1T1 | 280.833369 | -0.1179057 | 0.3379771 | -0.3488571 | 0.72719658 | 0.9958918 |
| TRIQK | 46.7562833 | -0.6102626 | 0.27238244 | -2.2404624 | 0.02506092 | 0.4274706 |
| FAM92A | 23.768385 | 0.145909 | 0.34000486 | 0.42913797 | 0.66782282 | 0.9958918 |
| RBM12B | 33.4636476 | -0.1457557 | 0.2799925 | -0.5205701 | 0.60266626 | 0.9958918 |
| TMEM67 | 32.9735906 | -0.28607 | 0.32612283 | -0.8771849 | 0.38038623 | 0.97217293 |
| GEM | 362.829244 | -0.3605812 | 0.5110613 | -0.7055537 | 0.48046575 | 0.99002719 |
| VIRMA | 49.1803877 | 0.08614931 | 0.27290346 | 0.31567688 | 0.75224778 | 0.9958918 |
| DPY19L4 | 49.9044469 | 0.09090752 | 0.34692161 | 0.26204051 | 0.79329021 | 0.99592423 |
| INTS8 | 39.4577972 | -0.2079867 | 0.32563075 | -0.6387193 | 0.52300553 | 0.9958918 |
| NDUFAF6 | 33.7334039 | 0.37409016 | 0.42783437 | 0.87438081 | 0.3819109 | 0.97217293 |
| TP53INP1 | 25.1785563 | -0.2149043 | 0.56543 | -0.3800724 | 0.70389166 | 0.9958918 |
| UQCRB | 51.3835769 | -0.1751062 | 0.41353223 | -0.4234403 | 0.67197405 | 0.9958918 |
| SDC2 | 124.241961 | -1.1684625 | 0.33239708 | -3.515261 | 0.00043932 | 0.04172562 |
| CPQ | 175.999149 | -0.3490585 | 0.35498001 | -0.9833187 | 0.32545063 | 0.95992644 |
| MTDH | 207.729158 | 0.21550958 | 0.20856142 | 1.03331472 | 0.30145664 | 0.95597956 |
| LAPTM4B | 51.06983 | -0.0068891 | 0.32212217 | -0.0213866 | 0.98293729 | 0.99935941 |
| MATN2 | 90.431467 | 0.62393504 | 0.49911338 | 1.25008679 | 0.21126784 | 0.89738044 |
| RPL30 | 127.122988 | -0.4932642 | 0.32430264 | -1.5209998 | 0.1282599 | 0.80288072 |
| NIPAL2 | 64.8644651 | 0.68695163 | 0.4512994 | 1.52216384 | 0.12796803 | 0.80197876 |
| STK3 | 474.95153 | -0.3968632 | 0.23533059 | -1.686407 | 0.09171743 | 0.72711043 |
| AC104986.2 | 20.1938349 | 0.21664389 | 0.39281926 | 0.55151036 | 0.58128386 | 0.9958918 |
| VPS13B | 421.579546 | -0.1531411 | 0.21877395 | -0.699997 | 0.48392919 | 0.99046213 |
| COX6C | 59.3300611 | -0.052593 | 0.26450267 | -0.1988375 | 0.84238987 | 0.99692952 |
| RNF19A | 110.794065 | 0.32437764 | 0.27709341 | 1.17064362 | 0.24174206 | 0.92516122 |
| PABPC1 | 143.029458 | -0.1733154 | 0.31677385 | -0.5471267 | 0.58429166 | 0.9958918 |
| YWHAZ | 146.465537 | -0.1546167 | 0.19728096 | -0.7837387 | 0.43319346 | 0.98092839 |
| ZNF706 | 57.0921716 | 0.33944115 | 0.33832452 | 1.00330049 | 0.3157159 | 0.95992644 |
| NCALD | 230.3602 | -0.4569866 | 0.55230383 | -0.8274189 | 0.40799967 | 0.97762947 |
| RRM2B | 27.6727231 | 0.17601568 | 0.35926624 | 0.48993103 | 0.6241827 | 0.9958918 |
| UBR5 | 346.711567 | -0.3227052 | 0.28455614 | -1.1340653 | 0.25676717 | 0.9266065 |
| KLF10 | 30.931747 | -0.006313 | 0.528719 | -0.0119402 | 0.99047335 | 0.99935941 |
| AZIN1 | 42.2310424 | -0.0572138 | 0.27466741 | -0.2083021 | 0.83499313 | 0.99679581 |

| | | | | | | |
|---|---|---|---|---|---|---|
| AZIN1-AS1 | 40.7759831 | -0.2548589 | 0.44659179 | -0.5706754 | 0.5682197 | 0.9958918 |
| ATP6V1C1 | 60.4676932 | -0.1120191 | 0.3059542 | -0.3661304 | 0.71426775 | 0.9958918 |
| SLC25A32 | 37.1177261 | 0.18522605 | 0.26919937 | 0.68806272 | 0.49141329 | 0.99046213 |
| LRP12 | 29.6876252 | -0.4407632 | 0.30434541 | -1.4482334 | 0.14755178 | 0.8301633 |
| ZFPM2 | 535.220392 | 0.13035336 | 0.34999381 | 0.37244476 | 0.70956174 | 0.9958918 |
| OXR1 | 296.924032 | -0.5291752 | 0.25512081 | -2.0742142 | 0.03805942 | 0.52280126 |
| ANGPT1 | 184.449242 | -1.6249466 | 0.38774186 | -4.190795 | 2.78E-05 | 0.00658989 |
| EIF3E | 98.521886 | 0.0226662 | 0.26818383 | 0.08451738 | 0.93264508 | 0.99935941 |
| EMC2 | 51.8497592 | 0.01682277 | 0.29134351 | 0.05774206 | 0.95395409 | 0.99935941 |
| NUDCD1 | 43.6456928 | -0.2488866 | 0.33215191 | -0.7493157 | 0.45366695 | 0.98092839 |
| ENY2 | 34.8709339 | -0.6157625 | 0.30585207 | -2.013269 | 0.04408634 | 0.55583613 |
| TRPS1 | 160.108191 | -0.0150606 | 0.27253047 | -0.0552621 | 0.95592969 | 0.99935941 |
| EIF3H | 101.351597 | -0.0593964 | 0.29366748 | -0.2022573 | 0.83971555 | 0.99692952 |
| UTP23 | 19.2999867 | 0.34529626 | 0.31963719 | 1.08027561 | 0.28001947 | 0.94419503 |
| RAD21 | 63.3723716 | 0.05710279 | 0.2333606 | 0.24469764 | 0.80669056 | 0.99592423 |
| EXT1 | 1057.82444 | -0.6720686 | 0.40568837 | -1.656613 | 0.09759775 | 0.74364783 |
| SAMD12 | 143.030216 | 1.21372043 | 0.25488665 | 4.76180456 | 1.92E-06 | 0.00123166 |
| ENPP2 | 24.5365493 | 1.53956432 | 0.35317273 | 4.35923898 | 1.31E-05 | 0.00399958 |
| TAF2 | 85.1744946 | -0.2738389 | 0.2181695 | -1.2551658 | 0.20941858 | 0.89634177 |
| DEPTOR | 54.5582648 | -0.2724815 | 0.30479708 | -0.8939766 | 0.37133443 | 0.97217293 |
| COL14A1 | 173.405591 | -0.8084721 | 0.52749421 | -1.5326653 | 0.12535835 | 0.79233388 |
| MRPL13 | 22.1513304 | -0.2296381 | 0.31758588 | -0.723074 | 0.46963445 | 0.98910524 |
| SNTB1 | 89.6798035 | 0.37312456 | 0.24789092 | 1.50519655 | 0.13227355 | 0.80565822 |
| ZHX2 | 272.971284 | 0.2185808 | 0.39965089 | 0.54692935 | 0.58442727 | 0.9958918 |
| DERL1 | 30.8149733 | -0.1041597 | 0.26531834 | -0.3925837 | 0.69462698 | 0.9958918 |
| ATAD2 | 24.4449123 | -0.0085774 | 0.42065026 | -0.0203907 | 0.98373167 | 0.99935941 |
| FBXO32 | 71.3698911 | 0.44232576 | 0.50307881 | 0.87923751 | 0.37927251 | 0.97217293 |
| FAM91A1 | 70.833024 | -0.0192013 | 0.32642357 | -0.0588233 | 0.95309288 | 0.99935941 |
| TMEM65 | 45.0388157 | 0.40564964 | 0.36724918 | 1.10456241 | 0.26934925 | 0.93634106 |
| TATDN1 | 40.3204492 | 0.09301189 | 0.32768247 | 0.28384761 | 0.77652717 | 0.9958918 |
| NDUFB9 | 34.8482497 | -0.2429469 | 0.38199131 | -0.636001 | 0.52477577 | 0.9958918 |
| MTSS1 | 311.429832 | -0.1907671 | 0.44473085 | -0.4289496 | 0.66795991 | 0.9958918 |
| SQLE | 21.5109529 | -0.5402999 | 0.50686884 | -1.065956 | 0.28644352 | 0.94614505 |
| WASHC5 | 32.8138921 | 0.10204205 | 0.35205831 | 0.28984417 | 0.77193546 | 0.9958918 |
| NSMCE2 | 268.299086 | -0.0090032 | 0.23211515 | -0.0387878 | 0.96905957 | 0.99935941 |
| PVT1 | 101.050548 | -0.8972923 | 0.39718565 | -2.2591257 | 0.02387557 | 0.41876263 |
| FAM49B | 38.0736507 | 0.11112393 | 0.36194621 | 0.3070178 | 0.75882982 | 0.9958918 |
| ASAP1 | 601.6166 | -0.7746816 | 0.30952244 | -2.5028285 | 0.01232052 | 0.2958939 |
| EFR3A | 112.784508 | -0.1988878 | 0.2639025 | -0.7536414 | 0.45106459 | 0.98092839 |
| KCNQ3 | 49.5223534 | -2.6201558 | 1.03302727 | -2.5363859 | 0.01120032 | 0.28321649 |
| PHF20L1 | 116.878809 | -0.1011209 | 0.23718052 | -0.4263456 | 0.66985603 | 0.9958918 |
| TG | 28.8077111 | 0.77171767 | 0.47308707 | 1.63123814 | 0.10284008 | 0.75728633 |
| SLA | 55.1893663 | 1.64544198 | 0.43013261 | 3.8254295 | 0.00013054 | 0.01776611 |
| WISP1 | 279.279597 | 0.10949338 | 0.63682233 | 0.17193709 | 0.86348698 | 0.9971731 |
| NDRG1 | 161.488789 | 0.20136827 | 0.49554359 | 0.40635834 | 0.68447932 | 0.9958918 |
| ST3GAL1 | 163.875054 | 0.83198443 | 0.34409317 | 2.41790449 | 0.01561017 | 0.34027688 |
| ZFAT | 34.3865048 | 0.0170567 | 0.26684021 | 0.06392102 | 0.94903311 | 0.99935941 |
| KHDRBS3 | 63.7705796 | -0.5721138 | 0.36120904 | -1.5838855 | 0.11321978 | 0.77793929 |
| TRAPPC9 | 168.169634 | 0.14646134 | 0.24784975 | 0.59092793 | 0.55456871 | 0.9958918 |
| AGO2 | 148.203275 | -0.0721202 | 0.23735699 | -0.3038469 | 0.76124453 | 0.9958918 |
| PTK2 | 500.898106 | -0.1642533 | 0.14442089 | -1.1373236 | 0.25540305 | 0.9266065 |
| DENND3 | 52.9163219 | -0.489839 | 0.53922436 | -0.9084142 | 0.36365945 | 0.97217293 |
| PTP4A3 | 100.183836 | -0.3173536 | 0.48530003 | -0.6539328 | 0.51315513 | 0.9958918 |
| LY6E | 84.2709318 | -1.012301 | 0.55956913 | -1.8090723 | 0.07043977 | 0.65633299 |

| | | | | | | |
|---|---|---|---|---|---|---|
| ZC3H3 | 26.0118633 | 0.46800182 | 0.37258249 | 1.25610256 | 0.20907879 | 0.89634177 |
| GSDMD | 21.6967663 | 0.48059252 | 0.40527444 | 1.18584464 | 0.23568366 | 0.91962002 |
| EEF1D | 93.9877665 | 0.05923804 | 0.33219194 | 0.17832473 | 0.85846796 | 0.9971731 |
| PUF60 | 18.6143901 | 0.78210341 | 0.46324013 | 1.6883326 | 0.0913474 | 0.72711043 |
| NRBP2 | 19.7096556 | 0.61775652 | 0.43475605 | 1.42092679 | 0.15533804 | 0.84277602 |
| PLEC | 91.6837211 | -0.4665095 | 0.33852964 | -1.3780462 | 0.16818903 | 0.86560961 |
| GRINA | 32.3504997 | 0.21869642 | 0.28799993 | 0.75936276 | 0.44763558 | 0.98092839 |
| GPAA1 | 18.8337998 | -0.1210803 | 0.36775063 | -0.3292458 | 0.74196991 | 0.9958918 |
| CYC1 | 18.3494127 | -0.3055714 | 0.48030334 | -0.6362051 | 0.52464276 | 0.9958918 |
| MAF1 | 20.0071307 | 0.44135232 | 0.36454557 | 1.21069174 | 0.22601357 | 0.91093521 |
| MROH1 | 32.506932 | 0.23110134 | 0.40620984 | 0.56892108 | 0.5694097 | 0.9958918 |
| HSF1 | 35.0374365 | 0.54500724 | 0.27435331 | 1.98651597 | 0.04697606 | 0.56885457 |
| VPS28 | 41.2141423 | 0.36384248 | 0.30035064 | 1.21139241 | 0.22574505 | 0.91093521 |
| CYHR1 | 21.4032141 | -0.0332127 | 0.32155825 | -0.1032868 | 0.91773534 | 0.99935941 |
| ZNF251 | 37.0477443 | 0.36015774 | 0.3324277 | 1.08341676 | 0.27862348 | 0.94419503 |
| RPL8 | 154.84274 | -0.0764699 | 0.42172807 | -0.1813251 | 0.85611244 | 0.9971731 |
| WASHC1 | 51.4178888 | 0.19383059 | 0.32475932 | 0.59684381 | 0.55061167 | 0.9958918 |
| DOCK8 | 172.733027 | 0.44939023 | 0.34067851 | 1.31910354 | 0.1871345 | 0.87955894 |
| KANK1 | 127.318096 | -0.1457993 | 0.37906139 | -0.3846323 | 0.70050987 | 0.9958918 |
| SMARCA2 | 142.700757 | 0.26877051 | 0.30429767 | 0.88324866 | 0.37710193 | 0.97217293 |
| VLDLR-AS1 | 41.7519952 | -0.2890835 | 0.62868227 | -0.4598245 | 6.46E-01 | 0.9958918 |
| PUM3 | 124.07803 | -0.2243039 | 0.29371602 | -0.7636761 | 0.4450603 | 0.98092839 |
| RFX3 | 214.141462 | -0.2652525 | 0.37473966 | -0.7078314 | 0.47904999 | 0.99002719 |
| RFX3-AS1 | 62.6373042 | -0.4461551 | 0.33294744 | -1.3400165 | 0.18023997 | 0.87815793 |
| GLIS3 | 639.81472 | -0.3676097 | 0.36353468 | -1.0112095 | 3.12E-01 | 0.9599143 |
| SLC1A1 | 49.8022736 | -0.538818 | 0.429216 | -1.2553539 | 0.20935032 | 0.89634177 |
| CDC37L1 | 49.334149 | -0.2114088 | 0.38466676 | -0.5495895 | 0.58260099 | 0.9958918 |
| AK3 | 76.7102906 | -0.0286798 | 0.25968148 | -0.1104424 | 0.91205852 | 0.99935941 |
| RCL1 | 56.7136035 | 0.26444214 | 0.39629248 | 0.66729033 | 0.5045867 | 0.99387859 |
| JAK2 | 190.212695 | 0.65144765 | 0.37703749 | 1.72780606 | 0.084023 | 0.70248581 |
| PLGRKT | 25.7163924 | 0.25730408 | 0.33921606 | 0.75852565 | 0.44813636 | 0.98092839 |
| RIC1 | 165.238556 | 0.03687614 | 0.2643875 | 0.13947762 | 0.88907274 | 0.99935941 |
| ERMP1 | 34.0630821 | -0.2271442 | 0.39842661 | -0.5701029 | 0.56860793 | 0.9958918 |
| KIAA2026 | 136.065888 | -0.0392208 | 0.26840132 | -0.1461274 | 0.88382082 | 0.99862036 |
| IL33 | 43.2777804 | 1.55069035 | 0.91934523 | 1.68673344 | 0.09165462 | 0.72711043 |
| UHRF2 | 72.2546553 | -0.1365003 | 0.24272096 | -0.5623756 | 0.57386014 | 0.9958918 |
| KDM4C | 230.32411 | -0.0887517 | 0.25234674 | -0.3517054 | 0.72505924 | 0.9958918 |
| PTPRD | 176.808918 | 0.96918678 | 0.40110553 | 2.41628877 | 0.01567962 | 0.34027688 |
| LURAP1L | 51.0814357 | -0.9372666 | 0.38420558 | -2.4394923 | 0.01470792 | 0.33025022 |
| MPDZ | 183.268457 | 0.02447016 | 0.32147499 | 0.07611841 | 0.9393249 | 0.99935941 |
| NFIB | 319.915908 | 0.30769849 | 0.30268262 | 1.01657136 | 0.30935738 | 0.95793395 |
| ZDHHC21 | 110.7639 | -0.2490257 | 0.32525984 | -0.7656209 | 0.44390193 | 0.98092839 |
| FREM1 | 91.9312964 | -0.559888 | 0.62538207 | -0.8952735 | 0.37064088 | 0.97217293 |
| TTC39B | 124.225167 | -0.3108556 | 0.27438057 | -1.1329358 | 0.25724119 | 0.9266065 |
| SNAPC3 | 66.0606463 | 0.14998349 | 0.25465329 | 0.58897135 | 0.55588049 | 0.9958918 |
| PSIP1 | 48.6087395 | 0.13819113 | 0.28665771 | 0.48207716 | 0.62975113 | 0.9958918 |
| CCDC171 | 53.2368871 | 0.05785662 | 0.43089183 | 0.1342718 | 0.89318765 | 0.99935941 |
| BNC2 | 277.411398 | -0.7570533 | 0.42902351 | -1.7645963 | 0.07763164 | 0.68020338 |
| CNTLN | 95.8431103 | -0.0752762 | 0.23893233 | -0.3150526 | 0.75272174 | 0.9958918 |
| ADAMTSL1 | 524.820242 | -0.4624319 | 0.4284786 | -1.0792416 | 0.28048004 | 0.94419503 |
| PLIN2 | 162.088209 | -0.4604316 | 0.62565309 | -0.7359216 | 4.62E-01 | 9.86E-01 |
| DENND4C | 191.795717 | 0.02107196 | 0.29114428 | 0.07237634 | 0.94230242 | 0.99935941 |
| RPS6 | 169.303829 | -0.1771794 | 0.37813975 | -0.4685552 | 0.63938757 | 0.9958918 |
| ACER2 | 77.8715749 | 1.84731161 | 0.48649992 | 3.79714676 | 0.00014637 | 0.01904212 |

| | | | | | | |
|---|---|---|---|---|---|---|
| MLLT3 | 189.298786 | -0.0797481 | 0.32760028 | -0.2434311 | 0.80767146 | 0.99592423 |
| FOCAD | 131.04176 | -0.3027156 | 0.26030625 | -1.1629211 | 0.24486152 | 0.92559078 |
| CAAP1 | 31.6962405 | 0.11029881 | 0.30188552 | 0.36536636 | 0.71483795 | 0.9958918 |
| PLAA | 37.4075109 | 0.27939834 | 0.25489349 | 1.09613764 | 0.27301855 | 0.93824549 |
| IFT74 | 42.0351052 | -0.1797214 | 0.30782915 | -0.5838348 | 0.55933145 | 0.9958918 |
| MOB3B | 57.8717122 | 0.47323645 | 0.25041945 | 1.88977517 | 0.05878804 | 0.6110796 |
| ACO1 | 94.7097135 | -0.3035517 | 0.31370384 | -0.9676379 | 0.33322522 | 0.9654289 |
| DDX58 | 132.31238 | -2.3347973 | 1.07529591 | -2.1713068 | 0.02990799 | 0.46665355 |
| NDUFB6 | 28.2569197 | 0.1206122 | 0.34758511 | 0.34700047 | 0.72859097 | 0.9958918 |
| APTX | 43.1955904 | 0.30436288 | 0.32018694 | 0.9505787 | 0.34181828 | 0.96887708 |
| DNAJA1 | 52.3072336 | 0.17947731 | 0.39007882 | 0.46010523 | 0.64544069 | 0.9958918 |
| SMU1 | 33.5718852 | -0.0573472 | 0.29531887 | -0.1941875 | 0.84602909 | 0.99692952 |
| B4GALT1 | 155.4248 | 0.41032679 | 0.45249039 | 0.90681879 | 0.36450263 | 0.97217293 |
| BAG1 | 23.5999473 | 0.23878704 | 0.42228635 | 0.56546237 | 0.57175931 | 0.9958918 |
| CHMP5 | 29.0180116 | 0.05909885 | 0.44936905 | 0.13151517 | 0.89536778 | 0.99935941 |
| NFX1 | 105.986602 | -0.2061828 | 0.25846612 | -0.797717 | 0.42503472 | 0.97917873 |
| UBE2R2 | 163.576088 | -0.0258111 | 0.20379711 | -0.1266509 | 0.8992167 | 0.99935941 |
| UBAP2 | 75.6373833 | -0.3625602 | 0.23116714 | -1.56839 | 0.11679014 | 0.78417352 |
| DCAF12 | 17.5077167 | 0.23481939 | 0.44912047 | 0.52284276 | 0.60108368 | 0.9958918 |
| UBAP1 | 118.766021 | -0.1701146 | 0.28318638 | -0.600716 | 0.54802919 | 0.9958918 |
| FAM219A | 65.3163869 | -0.0706233 | 0.26810745 | -0.263414 | 7.92E-01 | 0.99592423 |
| VCP | 70.5079783 | 0.33490492 | 0.31381149 | 1.06721691 | 0.2858739 | 0.94549382 |
| UNC13B | 107.714247 | 0.07436846 | 0.26676815 | 0.27877562 | 0.78041702 | 0.9958918 |
| RUSC2 | 105.604003 | 0.13363887 | 0.27677849 | 0.48283691 | 0.62921154 | 0.9958918 |
| CCDC107 | 37.4078327 | 0.64451498 | 0.4556712 | 1.41442994 | 0.15723571 | 0.84427321 |
| TPM2 | 140.455267 | 0.49939315 | 0.35331531 | 1.41344895 | 0.15752376 | 0.84427321 |
| TLN1 | 95.8201645 | 0.18576969 | 0.24161302 | 0.76887286 | 0.44196879 | 0.98092839 |
| GBA2 | 20.5910766 | 0.44224658 | 0.44254993 | 0.99931453 | 0.31764235 | 0.95992644 |
| RECK | 55.0523581 | -0.5072592 | 0.30333724 | -1.6722614 | 0.09447279 | 0.7340733 |
| CLTA | 76.8892054 | -0.0281146 | 0.28778815 | -0.0976919 | 0.92217694 | 0.99935941 |
| GNE | 28.0829633 | -0.0405192 | 0.33726491 | -0.1201406 | 0.90437174 | 0.99935941 |
| RNF38 | 142.157644 | -0.0012235 | 0.2277105 | -0.0053732 | 0.99571285 | 0.99935941 |
| EBLN3P | 26.5676984 | 0.09253594 | 0.37147129 | 0.24910657 | 0.80327834 | 0.99592423 |
| ZCCHC7 | 383.04135 | -0.3477552 | 0.20341208 | -1.7096092 | 0.08733817 | 0.71795104 |
| GRHPR | 24.6492528 | -0.0107008 | 0.36070693 | -0.0296661 | 0.97633332 | 0.99935941 |
| AL138752.2 | 27.8598548 | -0.0223321 | 0.34206776 | -0.0652857 | 0.94794651 | 0.99935941 |
| TRMT10B | 31.387798 | 0.58252579 | 0.31064093 | 1.87523837 | 0.06075994 | 0.61696136 |
| DCAF10 | 102.943964 | 0.27710533 | 0.20123908 | 1.37699565 | 0.16851359 | 0.86601846 |
| GLIDR | 28.2385124 | 0.23279625 | 0.49836706 | 0.46711805 | 0.64041542 | 0.9958918 |
| CBWD5 | 28.8569937 | -0.2030377 | 0.35817537 | -0.5668666 | 0.57080479 | 0.9958918 |
| PGM5 | 63.2721431 | 1.41372566 | 0.40241214 | 3.51312877 | 0.00044286 | 0.04172562 |
| PIP5K1B | 539.679486 | 1.17372554 | 0.36003779 | 3.26000652 | 0.0011141 | 0.07280105 |
| APBA1 | 80.4542805 | -0.671603 | 0.49845004 | -1.3473827 | 0.17785701 | 0.87815793 |
| PTAR1 | 72.0844883 | 0.12058236 | 0.21393433 | 0.56364193 | 0.57299784 | 0.9958918 |
| MAMDC2 | 112.675535 | -0.6807654 | 0.42925716 | -1.585915 | 0.1127586 | 0.77793929 |
| SMC5-AS1 | 23.9851294 | -0.6313105 | 0.39709416 | -1.5898258 | 0.11187408 | 0.77793929 |
| SMC5 | 210.228244 | -0.3110421 | 0.19227511 | -1.6176931 | 0.10572876 | 0.76820416 |
| KLF9 | 288.058135 | -0.8531618 | 0.45760744 | -1.8643968 | 0.06226601 | 0.62054234 |
| TRPM3 | 312.589597 | 1.46080592 | 0.39682711 | 3.68121505 | 0.00023213 | 0.02508658 |
| CEMIP2 | 62.0996908 | 0.73805943 | 0.39103825 | 1.88743536 | 0.05910179 | 0.61175668 |
| ABHD17B | 21.3808211 | 0.95151459 | 0.39017092 | 2.4387122 | 0.0147397 | 0.33025022 |
| C9orf85 | 43.5675875 | 0.72102533 | 0.2538248 | 2.84064179 | 0.00450229 | 0.16781827 |
| ZFAND5 | 131.39756 | -0.6736819 | 0.39450488 | -1.7076644 | 0.08769864 | 0.71805622 |
| ALDH1A1 | 65.8839581 | 0.53375467 | 0.39536985 | 1.35001357 | 0.17701163 | 0.87771577 |

| | | | | | | |
|---|---|---|---|---|---|---|
| ANXA1 | 48.7653785 | 0.25888432 | 0.51719748 | 0.50055217 | 0.61668633 | 0.9958918 |
| CARNMT1 | 26.4062467 | 0.29394543 | 0.36695192 | 0.80104617 | 0.42310492 | 0.97828057 |
| NMRK1 | 19.2753307 | 0.29547601 | 0.42858826 | 0.68941695 | 0.49056092 | 0.99046213 |
| OSTF1 | 48.7119066 | 0.10500681 | 0.29483083 | 0.35615955 | 0.72172108 | 0.9958918 |
| PCSK5 | 168.994669 | -0.0362573 | 0.41531004 | -0.0873017 | 0.9304317 | 0.99935941 |
| PRUNE2 | 62.1740062 | 2.02253249 | 0.60506983 | 3.3426431 | 0.00082985 | 0.06434787 |
| VPS13A | 123.080126 | 0.09917908 | 0.37443697 | 0.26487524 | 0.79110557 | 0.99592423 |
| GNA14 | 358.626402 | -0.2807109 | 0.52233334 | -0.5374172 | 0.59097945 | 0.9958918 |
| GNA14-AS1 | 52.2060954 | -1.2164152 | 0.61817124 | -1.9677641 | 0.04909518 | 0.57273715 |
| GNAQ | 335.024524 | 0.00480028 | 0.21480955 | 0.02234666 | 0.98217143 | 0.99935941 |
| TLE4 | 118.13737 | 0.05334426 | 0.27850757 | 0.1915361 | 0.8481056 | 0.99713289 |
| TLE1 | 113.747377 | -0.4802001 | 0.38391796 | -1.2507882 | 0.21101175 | 0.89738044 |
| FRMD3 | 122.984022 | -1.0622694 | 0.31766433 | -3.3439996 | 0.0008258 | 0.06434787 |
| UBQLN1 | 84.1438985 | -0.0848645 | 0.24505212 | -0.3463122 | 0.72910808 | 0.9958918 |
| KIF27 | 28.8140837 | -0.3135386 | 0.3007819 | -1.0424119 | 0.29722073 | 0.95417903 |
| HNRNPK | 130.696534 | -0.1432837 | 0.26318621 | -0.5444193 | 0.58615297 | 0.9958918 |
| NTRK2 | 78.97381 | 0.56288576 | 0.33957958 | 1.65759601 | 0.09739904 | 0.74364783 |
| AGTPBP1 | 63.8220623 | 0.01590217 | 0.31319887 | 0.05077338 | 0.9595061 | 0.99935941 |
| NAA35 | 87.3765198 | 0.19319664 | 0.21118236 | 0.91483323 | 0.36027919 | 0.97217293 |
| GOLM1 | 21.108006 | -0.9302749 | 0.32175529 | -2.8912497 | 0.00383713 | 0.15300235 |
| ISCA1 | 23.0031694 | -0.0239055 | 0.35893917 | -0.0666003 | 0.9468999 | 0.99935941 |
| TUT7 | 73.9075147 | 0.03477472 | 0.30947402 | 0.11236715 | 0.9105323 | 0.99935941 |
| GAS1 | 31.7068046 | -0.7304928 | 0.4440044 | -1.6452378 | 0.09992078 | 0.7498958 |
| DAPK1 | 51.6282029 | 0.24450822 | 0.44633465 | 0.54781366 | 0.58381986 | 0.9958918 |
| CTSL | 48.8600312 | -0.5473637 | 0.43954889 | -1.2452851 | 0.21302699 | 0.89738044 |
| SPIN1 | 123.324584 | -0.5173287 | 0.25348816 | -2.0408395 | 0.04126678 | 0.54347495 |
| S1PR3 | 120.212314 | 0.17551593 | 0.42191927 | 0.41599411 | 0.67741432 | 0.9958918 |
| SHC3 | 44.6101301 | -2.4867401 | 0.48798011 | -5.0959866 | 3.47E-07 | 0.00058389 |
| SECISBP2 | 97.852518 | -0.025873 | 0.24487655 | -0.1056573 | 0.91585425 | 0.99935941 |
| SEMA4D | 20.2355632 | -0.1184124 | 0.44670907 | -0.2650771 | 0.79095004 | 0.99592423 |
| AUH | 130.685262 | -0.1612952 | 0.35876018 | -0.4495906 | 0.65300568 | 0.9958918 |
| NFIL3 | 57.0373194 | -0.9075911 | 0.44552023 | -2.0371491 | 0.04163511 | 0.54347495 |
| ROR2 | 126.820496 | 0.18689896 | 0.85294345 | 0.21912234 | 0.82655475 | 0.99592423 |
| SPTLC1 | 63.6657672 | 0.01460286 | 0.29066471 | 0.05023955 | 0.9599315 | 0.99935941 |
| IARS | 77.7190758 | -0.751646 | 0.36553023 | -2.056317 | 0.03975197 | 0.53587963 |
| NOL8 | 30.4014356 | -0.3171514 | 0.29106402 | -1.0896275 | 0.27587728 | 0.94211066 |
| CENPP | 129.05069 | -1.1102282 | 0.30195614 | -3.6767864 | 0.00023619 | 0.02515602 |
| ASPN | 24.217348 | -0.6838858 | 0.60814471 | -1.1245445 | 0.2607821 | 0.93055907 |
| ECM2 | 48.1592786 | -0.3229602 | 0.57009384 | -0.5665036 | 0.57105147 | 0.9958918 |
| IPPK | 21.618305 | 0.1737329 | 0.31234934 | 0.55621344 | 0.57806495 | 0.9958918 |
| BICD2 | 33.1699215 | 0.43579632 | 0.33464432 | 1.30226718 | 0.19282507 | 0.88158763 |
| NINJ1 | 18.1182739 | 1.47617313 | 0.47898107 | 3.08190283 | 0.00205682 | 0.09689466 |
| FAM120A | 126.508357 | 0.10075741 | 0.27649081 | 0.36441505 | 0.7155481 | 0.9958918 |
| PHF2 | 39.2349147 | 0.08190473 | 0.27180483 | 0.30133655 | 0.76315787 | 0.9958918 |
| PTPDC1 | 40.9132477 | -0.8692218 | 0.39886545 | -2.1792357 | 0.02931416 | 0.46529106 |
| AL158152.2 | 54.2951193 | -0.0932688 | 0.35198206 | -0.2649818 | 0.79102351 | 0.99592423 |
| MFSD14B | 44.1829988 | -0.2082976 | 0.29948072 | -0.6955294 | 0.4867236 | 0.99046213 |
| C9orf3 | 667.041947 | -0.8097349 | 0.23419152 | -3.4575758 | 0.00054506 | 0.04787234 |
| FANCC | 94.0953009 | -0.4442471 | 0.24787546 | -1.7922188 | 0.07309792 | 0.66743052 |
| PTCH1 | 52.0946548 | -0.6174045 | 0.46776826 | -1.3198939 | 0.18687044 | 0.87955894 |
| LINC00476 | 49.4837547 | -0.5849349 | 0.47301718 | -1.2366039 | 0.21623417 | 0.90187565 |
| ERCC6L2 | 126.954981 | -0.035021 | 0.2237287 | -0.1565333 | 0.87561269 | 0.9971731 |
| HABP4 | 32.9886591 | 0.34663596 | 0.32756426 | 1.05822277 | 0.28995389 | 0.95186143 |
| CDC14B | 60.1145669 | 0.67215599 | 0.45258418 | 1.4851513 | 0.13750375 | 0.81404904 |

| | | | | | | |
|---|---|---|---|---|---|---|
| ZNF782 | 38.6068029 | -0.0584528 | 0.3059438 | -0.1910572 | 0.84848082 | 0.99713289 |
| MFSD14C | 88.5315519 | 0.04052513 | 0.236955 | 0.17102457 | 0.86420445 | 0.9971731 |
| TDRD7 | 21.6177628 | -0.1541613 | 0.53942296 | -0.2857893 | 0.77503953 | 0.9958918 |
| TMOD1 | 52.9277442 | 0.57556752 | 0.63817057 | 0.90190234 | 0.36710875 | 0.97217293 |
| TSTD2 | 37.8856158 | -0.1683546 | 0.25743006 | -0.6539817 | 0.51312356 | 0.9958918 |
| NCBP1 | 22.3656947 | -0.0305713 | 0.30535768 | -0.1001165 | 0.92025185 | 0.99935941 |
| XPA | 26.5177436 | 0.07877241 | 0.30150658 | 0.26126264 | 0.79388997 | 0.99592423 |
| ANP32B | 38.4952546 | 0.3257786 | 0.3583185 | 0.9091872 | 0.36325132 | 0.97217293 |
| COL15A1 | 160.900491 | 0.15752349 | 0.90692416 | 0.17368981 | 0.86210924 | 0.9971731 |
| TGFBR1 | 69.6276882 | -0.096716 | 0.34238682 | -0.2824757 | 0.77757876 | 0.9958918 |
| SEC61B | 57.5020145 | -0.1142391 | 0.34211446 | -0.3339207 | 0.73843941 | 0.9958918 |
| AL359710.1 | 32.5131852 | 0.12644473 | 0.59197244 | 0.21359902 | 0.83085976 | 0.99592423 |
| NR4A3 | 129.850553 | -2.0504109 | 1.04353132 | -1.9648772 | 0.04942846 | 0.57273715 |
| STX17 | 53.411643 | -0.0176136 | 0.33733072 | -0.0522146 | 0.95835771 | 0.99935941 |
| ERP44 | 77.3162655 | -0.0649891 | 0.25425262 | -0.2556085 | 0.79825318 | 0.99592423 |
| INVS | 180.37488 | -0.0497623 | 0.19507155 | -0.2550979 | 0.79864751 | 0.99592423 |
| TEX10 | 29.8309964 | -0.3243844 | 0.34108984 | -0.9510232 | 0.34159258 | 0.96887708 |
| ALDOB | 74.0105096 | 1.60309836 | 0.86153241 | 1.86075224 | 0.06277917 | 0.6220697 |
| ABCA1 | 242.214814 | -0.1786482 | 0.60064243 | -0.2974286 | 0.76613934 | 0.9958918 |
| SLC44A1 | 91.6546319 | -0.2681291 | 0.32816453 | -0.817057 | 0.41389588 | 0.97812179 |
| FSD1L | 31.3097945 | -0.1330462 | 0.28908913 | -0.4602256 | 0.64535432 | 0.9958918 |
| FKTN | 37.1782009 | 0.02011083 | 0.30902391 | 0.06507854 | 0.94811147 | 0.99935941 |
| TMEM38B | 31.4526966 | -0.1204763 | 0.37519744 | -0.3211012 | 0.74813372 | 0.9958918 |
| ZNF462 | 78.5979518 | 0.14374115 | 0.26937263 | 0.53361452 | 0.59360826 | 0.9958918 |
| RAD23B | 101.274697 | -0.1196997 | 0.23988145 | -0.4989954 | 0.61778261 | 0.9958918 |
| ELP1 | 33.681054 | 0.19384181 | 0.29858098 | 0.64921016 | 0.51620255 | 0.9958918 |
| CTNNAL1 | 70.5419771 | 0.08549468 | 0.36245445 | 0.23587703 | 0.81352809 | 0.99592423 |
| TMEM245 | 97.095132 | 0.3463206 | 0.20331862 | 1.70333934 | 0.08850458 | 0.71818729 |
| PTPN3 | 25.6315546 | 0.39586701 | 0.37183965 | 1.06461753 | 0.28704904 | 0.9476745 |
| PALM2-AKAP | 80.0549074 | -0.1974966 | 0.39171701 | -0.5041818 | 0.61413364 | 0.9958918 |
| TXN | 47.4679041 | -0.2225209 | 0.49764879 | -0.4471445 | 0.65477073 | 0.9958918 |
| SVEP1 | 512.599941 | -0.628895 | 0.47039578 | -1.3369486 | 0.18123941 | 0.87815793 |
| LPAR1 | 516.140551 | -0.8320353 | 0.41231139 | -2.0179781 | 0.04359354 | 0.55523214 |
| ECPAS | 182.407378 | -0.075107 | 0.26519568 | -0.2832135 | 0.77701318 | 0.9958918 |
| ZNF483 | 30.5860445 | -0.4265452 | 0.46782525 | -0.9117617 | 0.36189416 | 0.97217293 |
| PTGR1 | 77.9043366 | -0.4421936 | 0.25291566 | -1.7483835 | 0.08039765 | 0.69510861 |
| C9orf84 | 28.8480282 | 0.27852521 | 0.49237376 | 0.56567842 | 0.57161241 | 0.9958918 |
| UGCG | 205.30521 | -0.4449408 | 0.46252762 | -0.9619767 | 3.36E-01 | 0.96640422 |
| SUSD1 | 47.2026047 | -0.3576221 | 0.39440128 | -0.9067467 | 0.36454075 | 0.97217293 |
| PTBP3 | 142.417345 | -0.2820726 | 0.26243716 | -1.0748197 | 0.28245544 | 0.94419503 |
| HSDL2 | 70.524526 | 0.46366512 | 0.34568955 | 1.34127607 | 0.17983084 | 0.87815793 |
| KIAA1958 | 77.6527683 | 0.24157846 | 0.26378353 | 0.91582085 | 0.35976087 | 0.97217293 |
| FKBP15 | 36.1624594 | 0.14702701 | 0.33845393 | 0.43440777 | 0.66399236 | 0.9958918 |
| SLC31A1 | 25.8755849 | 0.08313224 | 0.36122007 | 0.23014291 | 0.81798073 | 0.99592423 |
| RGS3 | 30.2690939 | 0.3020217 | 0.33417405 | 0.9037856 | 0.3661091 | 0.97217293 |
| ZNF618 | 92.059807 | 0.00310417 | 0.31197231 | 0.00995016 | 0.99206105 | 0.99935941 |
| COL27A1 | 181.097626 | -0.0390565 | 0.48691382 | -0.0802123 | 0.93606839 | 0.99935941 |
| AKNA | 28.3813368 | 0.29679616 | 0.32859422 | 0.90323 | 0.36640384 | 0.97217293 |
| ATP6V1G1 | 56.8293207 | 0.26849358 | 0.34920182 | 0.76887796 | 0.44196576 | 0.98092839 |
| TNC | 720.737063 | -1.5751822 | 1.24548696 | -1.2647119 | 0.20597462 | 0.89589437 |
| PAPPA | 190.289931 | 0.23601359 | 0.83005763 | 0.28433398 | 7.76E-01 | 0.9958918 |
| ASTN2 | 43.1666167 | -0.5093626 | 0.30264072 | -1.6830604 | 0.09236341 | 0.72986954 |
| CDK5RAP2 | 60.6044993 | -0.1099746 | 0.27131667 | -0.4053366 | 0.68523011 | 0.9958918 |
| MEGF9 | 33.9719663 | 0.8868855 | 0.35404997 | 2.50497268 | 0.01224609 | 0.2958939 |

| | | | | | | |
|---|---|---|---|---|---|---|
| FBXW2 | 39.1379452 | -0.0142925 | 0.35180571 | -0.0406262 | 0.96759393 | 0.99935941 |
| PSMD5 | 42.2930886 | 0.27073995 | 0.240596 | 1.12528869 | 0.26046672 | 0.93055907 |
| CNTRL | 48.1907915 | 0.18572347 | 0.3064532 | 0.60604185 | 0.54448697 | 0.9958918 |
| RAB14 | 70.4733417 | 0.28407208 | 0.24535321 | 1.15780869 | 0.24694212 | 0.92605444 |
| GSN | 206.060234 | -0.2335819 | 0.31045276 | -0.7523912 | 0.45181584 | 0.98092839 |
| STOM | 127.470548 | -0.5974938 | 0.31119828 | -1.9199779 | 0.0548607 | 0.5937721 |
| DAB2IP | 118.84623 | -0.0092429 | 0.37137501 | -0.0248884 | 0.98014397 | 0.99935941 |
| TTLL11 | 28.4730076 | 0.8342579 | 0.34462991 | 2.42073563 | 0.01548914 | 0.33877878 |
| MRRF | 30.5843056 | 0.25778422 | 0.29832253 | 0.86411246 | 0.38752612 | 0.97297628 |
| RC3H2 | 79.5051894 | 0.20221109 | 0.22572117 | 0.89584462 | 0.37033576 | 0.97217293 |
| RABGAP1 | 224.709424 | -0.0526434 | 0.25624559 | -0.2054413 | 0.83722735 | 0.99692952 |
| STRBP | 23.8310969 | 0.21328104 | 0.50896988 | 0.41904451 | 0.6751836 | 0.9958918 |
| DENND1A | 245.079238 | -0.0094791 | 0.25886599 | -0.0366178 | 0.97078974 | 0.99935941 |
| NEK6 | 50.0853881 | 0.53042805 | 0.29573397 | 1.79359864 | 0.07287725 | 0.66701558 |
| PSMB7 | 57.4342466 | 0.10746998 | 0.22962827 | 0.46801719 | 0.6397723 | 0.9958918 |
| NR6A1 | 35.2558096 | -1.5421113 | 0.35377071 | -4.35907 | 1.31E-05 | 0.00399958 |
| RPL35 | 81.5081022 | -0.5011515 | 0.32854969 | -1.5253447 | 0.12717313 | 0.79948277 |
| GOLGA1 | 28.2427345 | -0.4437713 | 0.32484775 | -1.3660901 | 0.17191068 | 0.87233975 |
| SCAI | 37.7006626 | -0.273537 | 0.33943349 | -0.8058633 | 0.42032169 | 0.97812179 |
| PPP6C | 62.6798698 | 0.02766817 | 0.23191101 | 0.11930512 | 0.90503363 | 0.99935941 |
| RABEPK | 20.8495826 | 0.20078574 | 0.40238598 | 0.4989879 | 0.61778791 | 0.9958918 |
| HSPA5 | 145.870363 | -0.6487956 | 0.56227384 | -1.1538783 | 0.24855006 | 0.92624282 |
| GAPVD1 | 106.918653 | -0.0430468 | 0.20684521 | -0.2081113 | 0.83514204 | 0.99679581 |
| MAPKAP1 | 133.554862 | 0.03614569 | 0.21862459 | 0.16533225 | 0.86868248 | 0.9971731 |
| PBX3 | 160.267253 | -0.2242649 | 0.26738178 | -0.8387442 | 0.40161288 | 0.97500135 |
| MVB12B | 34.9471352 | 0.52096469 | 0.38843745 | 1.34118038 | 0.1798619 | 0.87815793 |
| ZBTB43 | 26.161996 | -0.5384119 | 0.42685851 | -1.2613358 | 0.2071879 | 0.89620104 |
| RALGPS1 | 21.3481169 | 0.47509493 | 0.63039383 | 0.75364781 | 0.45106073 | 0.98092839 |
| ANGPTL2 | 23.8988012 | -0.7467488 | 0.3288299 | -2.2709272 | 0.02315138 | 0.41196007 |
| GARNL3 | 82.1314215 | -0.152698 | 0.41230417 | -0.3703529 | 0.71111957 | 0.9958918 |
| RPL12 | 38.4054488 | -0.3272804 | 0.45825868 | -0.7141827 | 0.47511422 | 0.98910524 |
| LRSAM1 | 34.5059303 | 0.26672533 | 0.30297096 | 0.88036598 | 0.37866108 | 0.97217293 |
| FAM129B | 79.1925857 | -0.4587889 | 0.42691473 | -1.0746618 | 0.28252618 | 0.94419503 |
| STXBP1 | 72.1317273 | 0.01030562 | 0.39403635 | 0.02615398 | 0.97913452 | 0.99935941 |
| CDK9 | 20.6724229 | 0.28764547 | 0.32506763 | 0.88487886 | 0.37622197 | 0.97217293 |
| FPGS | 34.8870187 | 0.19921879 | 0.3658869 | 0.54448189 | 0.58610991 | 0.9958918 |
| ENG | 368.243558 | 0.63856423 | 0.43016626 | 1.48445911 | 0.13768716 | 0.81404904 |
| SLC25A25 | 42.1846291 | -0.2764039 | 0.32499541 | -0.8504856 | 0.39505515 | 0.97500135 |
| PTGES2 | 19.4385731 | 0.21151058 | 0.448831 | 0.4712477 | 0.63746386 | 0.9958918 |
| C9orf16 | 23.9862834 | -0.0494827 | 0.44603912 | -0.1109379 | 0.91166556 | 0.99935941 |
| CIZ1 | 22.1734794 | 0.42722679 | 0.36167547 | 1.18124348 | 0.23750601 | 0.92291633 |
| GOLGA2 | 49.4685116 | -0.3830461 | 0.32675 | -1.172291 | 0.24108023 | 0.92468614 |
| COQ4 | 24.8785986 | -0.32418 | 0.3703932 | -0.875232 | 0.3814477 | 0.97217293 |
| SLC27A4 | 25.7277915 | -0.1706909 | 0.31960862 | -0.5340623 | 0.5932984 | 0.9958918 |
| ODF2 | 27.6614776 | 0.10263795 | 0.40887179 | 0.25102721 | 0.80179307 | 0.99592423 |
| GLE1 | 20.5769369 | 0.19939143 | 0.43962218 | 0.45355181 | 0.65015144 | 0.9958918 |
| SPTAN1 | 113.17866 | -0.1315824 | 0.31653643 | -0.4156945 | 0.67763359 | 0.9958918 |
| SET | 56.2768214 | 0.17547415 | 0.37591121 | 0.4667968 | 0.64064526 | 0.9958918 |
| ZER1 | 55.7394859 | 0.56521482 | 0.37028347 | 1.52643817 | 0.12690078 | 0.7984536 |
| KYAT1 | 18.9790724 | -0.1110502 | 0.37181803 | -0.2986682 | 0.7651932 | 0.9958918 |
| LRRC8A | 23.830006 | 0.39356905 | 0.48429468 | 0.81266441 | 0.41641049 | 0.97812179 |
| NUP188 | 23.8699942 | 0.1468269 | 0.37972693 | 0.38666445 | 0.69900464 | 0.9958918 |
| SH3GLB2 | 16.6183425 | -0.2336837 | 0.37880377 | -0.616899 | 0.53730132 | 0.9958918 |
| PTPA | 34.333073 | -0.3362201 | 0.39845432 | -0.8438108 | 0.39877515 | 0.97500135 |

| | | | | | | |
|---|---|---|---|---|---|---|
| IER5L | 68.1645582 | -0.5649481 | 0.42718004 | -1.3225059 | 0.18599975 | 0.8789337 |
| NTMT1 | 22.8705686 | -0.2574296 | 0.4417682 | -0.5827254 | 0.56007816 | 0.9958918 |
| C9orf78 | 19.9938935 | 0.18784628 | 0.41954792 | 0.44773497 | 0.65434448 | 0.9958918 |
| USP20 | 24.7406975 | 0.68480424 | 0.37994303 | 1.80238664 | 0.0714846 | 0.66163765 |
| FNBP1 | 200.2273 | 0.17137903 | 0.3234659 | 0.529821 | 0.59623604 | 0.9958918 |
| GPR107 | 81.3413419 | 0.12343893 | 0.22580567 | 0.54666001 | 0.58461233 | 0.9958918 |
| NCS1 | 18.8050543 | -0.1046805 | 0.42660381 | -0.245381 | 0.80616141 | 0.99592423 |
| ASS1 | 18.5634948 | 0.04288193 | 0.45808164 | 0.09361197 | 0.9254174 | 0.99935941 |
| FUBP3 | 67.8151761 | -0.030971 | 0.23488834 | -0.1318541 | 0.89509972 | 0.99935941 |
| ABL1 | 244.947718 | 0.01363424 | 0.29368032 | 0.04642543 | 0.96297117 | 0.99935941 |
| LAMC3 | 129.728445 | -1.1306522 | 0.52592357 | -2.1498412 | 0.03156778 | 0.48041788 |
| AIF1L | 21.7067779 | -0.0059514 | 0.47727422 | -0.0124696 | 0.99005096 | 0.99935941 |
| NUP214 | 80.2166579 | -0.3460256 | 0.28298565 | -1.2227673 | 0.22141762 | 0.9105753 |
| PRRC2B | 159.575677 | -0.0222651 | 0.22984146 | -0.0968716 | 0.92282838 | 0.99935941 |
| RAPGEF1 | 82.993956 | 0.10959986 | 0.34234727 | 0.32014236 | 0.74886042 | 0.9958918 |
| MED27 | 114.248394 | 0.25714161 | 0.33833479 | 0.76002118 | 0.44724193 | 0.98092839 |
| SETX | 175.714469 | -0.274706 | 0.18199392 | -1.5094243 | 0.13119038 | 0.80565822 |
| TTF1 | 20.8580899 | -0.1189595 | 0.35194509 | -0.3380059 | 0.73535871 | 0.9958918 |
| DDX31 | 23.188216 | -0.2411574 | 0.39601885 | -0.6089542 | 0.54255479 | 0.9958918 |
| TSC1 | 43.5079227 | -0.0870823 | 0.29941028 | -0.290846 | 0.77116913 | 0.9958918 |
| RPL7A | 91.0885153 | -0.4035161 | 0.37429079 | -1.0780819 | 0.28099721 | 0.94419503 |
| SURF4 | 63.4986233 | 0.08266505 | 0.44479123 | 0.18585135 | 0.85256134 | 0.9971731 |
| ADAMTSL2 | 95.7698873 | -1.2676014 | 0.57667974 | -2.1981029 | 0.02794177 | 0.45329824 |
| VAV2 | 68.7951591 | -0.3618289 | 0.30523114 | -1.1854259 | 0.23584909 | 0.91962002 |
| BRD3 | 22.499394 | 0.47277831 | 0.33509349 | 1.41088477 | 0.15827859 | 0.84598994 |
| WDR5 | 21.9122938 | 0.0144226 | 0.3421497 | 0.04215288 | 0.96637683 | 0.99935941 |
| RXRA | 45.7995549 | -0.2496876 | 0.41154942 | -0.6067013 | 0.54404917 | 0.9958918 |
| COL5A1 | 141.121206 | -0.9707307 | 0.41511353 | -2.3384704 | 0.01936286 | 0.37348469 |
| CAMSAP1 | 25.83964 | 0.00179363 | 0.30336852 | 0.00591237 | 0.99528264 | 0.99935941 |
| NACC2 | 44.4546123 | 0.37813066 | 0.31502202 | 1.20033087 | 0.23001087 | 0.91455844 |
| ENTR1 | 18.1674696 | -0.2974669 | 0.39443709 | -0.7541554 | 0.45075589 | 0.98092839 |
| SEC16A | 45.6888703 | -0.0500708 | 0.30772937 | -0.1627105 | 0.87074635 | 0.9971731 |
| NOTCH1 | 19.7600756 | 0.6974927 | 0.35134264 | 1.98522077 | 0.04711991 | 0.56885457 |
| RABL6 | 45.1203304 | 0.30158049 | 0.34221194 | 0.88126818 | 0.37817269 | 0.97217293 |
| PHPT1 | 38.1187906 | 0.56391758 | 0.36390734 | 1.54961858 | 0.12123309 | 0.79021701 |
| EDF1 | 54.1695715 | -0.0020172 | 0.38275236 | -0.0052704 | 0.99579489 | 0.99935941 |
| FBXW5 | 38.9243823 | 0.63743443 | 0.33914665 | 1.87952448 | 0.06017291 | 0.61332974 |
| MAN1B1 | 22.3849002 | 0.00263331 | 0.32359764 | 0.0081376 | 0.99350721 | 0.99935941 |
| DPP7 | 19.547975 | 0.08644703 | 0.34554195 | 0.2501781 | 0.80244962 | 0.99592423 |
| TUBB4B | 43.2764115 | 0.93122638 | 0.52150227 | 1.78566122 | 0.07415413 | 0.66886321 |
| NELFB | 25.9562174 | 0.01440485 | 0.34292339 | 0.04200602 | 0.9664939 | 0.99935941 |
| EXD3 | 32.3854086 | 0.3589828 | 0.38910726 | 0.92258058 | 0.35622582 | 0.97217293 |
| MRPL41 | 38.4780985 | -0.095714 | 0.30205784 | -0.3168732 | 0.75133985 | 0.9958918 |
| EHMT1 | 164.147173 | -0.04256 | 0.21620089 | -0.1968538 | 0.84394197 | 0.99692952 |
| PSMD13 | 30.6375605 | 0.02609073 | 0.40543862 | 0.06435187 | 0.94869005 | 0.99935941 |
| PGGHG | 18.6020019 | -0.3777497 | 0.45218388 | -0.8353895 | 0.40349842 | 0.97575999 |
| IFITM2 | 262.540443 | -0.2622037 | 0.3582458 | -0.7319102 | 0.46422341 | 0.98600515 |
| IFITM3 | 443.186394 | -0.3709395 | 0.34256637 | -1.082825 | 0.27888608 | 0.94419503 |
| PTDSS2 | 23.2227923 | -0.2796364 | 0.3262275 | -0.8571821 | 0.39134426 | 0.97297628 |
| RNH1 | 37.8370052 | 0.24074823 | 0.27264957 | 0.88299511 | 0.37723891 | 0.97217293 |
| PHRF1 | 25.8318049 | -0.1390079 | 0.30697328 | -0.4528337 | 0.65066846 | 0.9958918 |
| DEAF1 | 36.4036689 | 0.22638954 | 0.2998878 | 0.75491412 | 0.45030051 | 0.98092839 |
| TALDO1 | 29.1476806 | 0.17224158 | 0.40797232 | 0.42218939 | 0.67288679 | 0.9958918 |
| RPLP2 | 102.634619 | -0.4132153 | 0.3824373 | -1.0804785 | 0.27992917 | 0.94419503 |

| | | | | | | |
|---|---|---|---|---|---|---|
| PNPLA2 | 23.576198 | 0.25620885 | 0.39029117 | 0.65645566 | 0.51153099 | 0.9958918 |
| CD151 | 88.9556374 | 0.66916817 | 0.34358177 | 1.94762423 | 0.05145994 | 0.57655256 |
| POLR2L | 56.1647728 | 0.21938767 | 0.3312723 | 0.66225781 | 0.50780601 | 0.99397937 |
| TSPAN4 | 36.5339965 | -0.5093526 | 0.40078455 | -1.2708887 | 0.20376824 | 0.89396747 |
| AP2A2 | 80.1281907 | -0.0033218 | 0.31874259 | -0.0104214 | 0.99168505 | 0.99935941 |
| TOLLIP | 21.5137927 | -0.1622637 | 0.34473072 | -0.4706969 | 0.63785718 | 0.9958918 |
| MOB2 | 29.2925951 | 0.81286746 | 0.3024039 | 2.68801908 | 0.00718773 | 0.21767701 |
| CD81 | 228.215926 | -0.0659152 | 0.25268216 | -0.2608621 | 0.79419883 | 0.99592423 |
| KCNQ1 | 34.1792978 | -0.4677558 | 0.40778791 | -1.1470565 | 0.25135826 | 0.92631316 |
| KCNQ1OT1 | 161.758675 | -0.7667658 | 0.30698478 | -2.4977323 | 0.01249905 | 0.29742804 |
| CDKN1C | 48.5544662 | -1.1728958 | 0.67221095 | -1.7448329 | 0.08101396 | 0.69715641 |
| NAP1L4 | 91.1738373 | -0.2135844 | 0.22705207 | -0.9406845 | 0.34686658 | 0.96887708 |
| CARS | 31.2106155 | 0.23103563 | 0.348315 | 0.6632951 | 0.50714157 | 0.99397937 |
| OSBPL5 | 82.2233176 | 0.23635796 | 0.26490203 | 0.8922467 | 0.37226071 | 0.97217293 |
| ZNF195 | 19.3532691 | -0.1043347 | 0.45447202 | -0.2295733 | 0.81842333 | 0.99592423 |
| NUP98 | 132.840464 | -0.2469684 | 0.26828827 | -0.9205337 | 0.35729392 | 0.97217293 |
| STIM1 | 109.051313 | 0.48201636 | 0.22545693 | 2.13795314 | 0.03252055 | 0.48564518 |
| AC104389.4 | 59.9896043 | -0.9475911 | 0.85302971 | -1.1108536 | 0.26663139 | 0.93533685 |
| TRIM22 | 66.6362856 | -1.4615325 | 0.99147098 | -1.4741052 | 0.14045327 | 0.8153985 |
| CAVIN3 | 36.9364383 | -0.1007699 | 0.63242436 | -0.159339 | 0.87340181 | 0.9971731 |
| APBB1 | 18.4731814 | 0.01111756 | 0.37367143 | 0.02975223 | 0.97626465 | 0.99935941 |
| AC084337.2 | 24.3764229 | -0.099531 | 0.39823238 | -0.2499321 | 0.80263988 | 0.99592423 |
| DNHD1 | 36.9884364 | -0.2450231 | 0.35641947 | -0.6874572 | 0.49179465 | 0.99046213 |
| ILK | 72.3240265 | 0.54167592 | 0.31747915 | 1.70617795 | 0.08797495 | 0.71818729 |
| TPP1 | 27.7356424 | 0.36959724 | 0.31699828 | 1.1659282 | 0.24364348 | 0.92534155 |
| PPFIBP2 | 25.6310507 | 0.3138644 | 0.48982842 | 0.64076397 | 0.52167604 | 0.9958918 |
| EIF3F | 28.8131487 | 0.05160314 | 0.31895484 | 0.16178822 | 0.87147263 | 0.9971731 |
| STK33 | 28.9794566 | -0.7148704 | 0.39901676 | -1.7915798 | 0.0732003 | 0.66743052 |
| TRIM66 | 36.5607399 | 0.25086209 | 0.35085508 | 0.71500201 | 0.4746078 | 0.98910524 |
| RPL27A | 115.754755 | -0.1169253 | 0.30548226 | -0.3827563 | 0.70190045 | 0.9958918 |
| ST5 | 222.736012 | -0.7126509 | 0.29814133 | -2.3903124 | 0.01683405 | 0.34578631 |
| TMEM9B | 17.925252 | 0.09353099 | 0.34797469 | 0.26878676 | 0.78809378 | 0.99592423 |
| DENND5A | 168.046214 | -0.3502431 | 0.34538247 | -1.0140733 | 0.3105478 | 0.95956408 |
| TMEM41B | 56.3761958 | -0.3798938 | 0.35233391 | -1.0782208 | 0.28093523 | 0.94419503 |
| IPO7 | 97.24457 | -0.137516 | 0.38374376 | -0.3583536 | 0.7200787 | 0.9958918 |
| ZNF143 | 29.931045 | 0.14091135 | 0.30526354 | 0.46160555 | 0.64436421 | 0.9958918 |
| WEE1 | 21.6355163 | -0.436694 | 0.6222349 | -0.7018154 | 0.48279429 | 0.99002719 |
| SWAP70 | 71.1395906 | -0.3120052 | 0.28117347 | -1.1096539 | 0.26714819 | 0.93533685 |
| SBF2 | 649.062663 | -0.0974098 | 0.25002515 | -0.3896001 | 0.6968323 | 0.9958918 |
| AC011092.2 | 23.1991831 | 0.40789 | 0.44800425 | 0.91046012 | 0.3625799 | 0.97217293 |
| ADM | 74.7984808 | -1.7892982 | 0.5921237 | -3.0218317 | 0.0025125 | 0.11258768 |
| MTRNR2L8 | 47.2138654 | 0.98174823 | 0.57115667 | 1.71887729 | 0.08563672 | 0.708721 |
| MRVI1 | 118.376325 | 0.3957657 | 0.43403537 | 0.91182824 | 0.36185914 | 0.97217293 |
| CTR9 | 29.677346 | -0.0828215 | 0.30999209 | -0.2671729 | 0.78933605 | 0.99592423 |
| EIF4G2 | 108.791519 | 0.27863728 | 0.25837307 | 1.07843006 | 0.28084188 | 0.94419503 |
| ZBED5 | 40.2087112 | 0.31807269 | 0.313458 | 1.01472188 | 0.31023842 | 0.95956408 |
| GALNT18 | 55.0420228 | 0.26474005 | 0.37448487 | 0.70694459 | 0.47960091 | 0.99002719 |
| USP47 | 231.42809 | -0.0747747 | 0.18188773 | -0.4111035 | 0.68099667 | 0.9958918 |
| DKK3 | 52.5921136 | -0.275146 | 0.42527264 | -0.6469873 | 0.51764017 | 0.9958918 |
| MICAL2 | 163.3092 | 0.04502681 | 0.33540061 | 0.13424785 | 0.89320659 | 0.99935941 |
| PARVA | 224.099395 | -0.3117628 | 0.22800139 | -1.3673724 | 0.17150861 | 0.87233975 |
| TEAD1 | 319.850806 | -0.3761557 | 0.30054655 | -1.2515723 | 0.21072576 | 0.89738044 |
| ARNTL | 35.7064221 | -0.1812034 | 0.4700067 | -0.3855337 | 0.69984207 | 0.9958918 |
| BTBD10 | 45.2667881 | 0.26414345 | 0.34653789 | 0.76223542 | 0.44591951 | 0.98092839 |

| | | | | | | |
|---|---|---|---|---|---|---|
| FAR1 | 100.46439 | 0.22256378 | 0.23505556 | 0.94685605 | 0.34371214 | 0.96887708 |
| SPON1 | 187.536915 | -1.8035186 | 0.46082783 | -3.9136494 | 9.09E-05 | 0.01484687 |
| RRAS2 | 68.4647881 | -0.0863874 | 0.43919853 | -0.1966933 | 0.84406756 | 0.99692952 |
| COPB1 | 81.5064855 | -0.2157305 | 0.2176829 | -0.9910312 | 0.32167036 | 0.95992644 |
| SOX6 | 152.184618 | -0.2664756 | 0.37934817 | -0.7024566 | 0.48239448 | 0.99002719 |
| C11orf58 | 89.9412027 | 0.00856014 | 0.2134945 | 0.04009534 | 0.96801711 | 0.99935941 |
| PLEKHA7 | 133.893803 | -0.1210949 | 0.30179598 | -0.4012474 | 0.68823797 | 0.9958918 |
| RPS13 | 90.7646821 | -0.2894288 | 0.39948233 | -0.7245096 | 0.46875293 | 0.98910524 |
| PIK3C2A | 156.635325 | 0.00988385 | 0.18874472 | 0.05236623 | 0.95823688 | 0.99935941 |
| NUCB2 | 28.2081557 | 0.13839678 | 0.34612732 | 0.39984356 | 0.68927175 | 0.9958918 |
| SERGEF | 56.7433338 | 0.15246484 | 0.31604062 | 0.48242165 | 0.62950644 | 0.9958918 |
| HPS5 | 27.4594134 | -0.1117692 | 0.30261178 | -0.3693484 | 0.71186803 | 0.9958918 |
| GTF2H1 | 40.7779498 | -0.3285673 | 0.25379177 | -1.2946335 | 0.1954467 | 0.88391893 |
| LDHA | 137.369756 | 0.08592986 | 0.37329642 | 0.23019203 | 8.18E-01 | 9.96E-01 |
| TSG101 | 53.4050035 | 0.11649726 | 0.2406335 | 0.48412736 | 0.62829549 | 0.9958918 |
| UEVLD | 34.6585641 | 0.34388412 | 0.29303777 | 1.17351467 | 0.24058948 | 0.92426606 |
| NAV2 | 799.761807 | -1.1148019 | 0.41112213 | -2.7116077 | 0.00669578 | 0.20675326 |
| PRMT3 | 52.9950426 | -0.0068084 | 0.25850593 | -0.0263376 | 0.97898803 | 0.99935941 |
| GAS2 | 49.0201183 | 1.09365456 | 0.59325097 | 1.84349395 | 6.53E-02 | 6.29E-01 |
| AC099687.1 | 23.9311805 | -0.2997552 | 0.65749289 | -0.4559064 | 0.64845731 | 0.9958918 |
| ANO3 | 136.450691 | 0.36866524 | 0.44035821 | 0.83719396 | 4.02E-01 | 9.75E-01 |
| LGR4 | 52.3341644 | 0.24180393 | 0.28675074 | 0.84325477 | 0.39908598 | 0.97500135 |
| LIN7C | 23.0974302 | -0.0945958 | 0.32121226 | -0.2944963 | 0.76837868 | 0.9958918 |
| BDNF-AS | 65.5563661 | -0.484173 | 0.26378324 | -1.8354955 | 0.0664323 | 0.63518413 |
| METTL15 | 182.61973 | -0.239738 | 0.24328584 | -0.985417 | 0.32441927 | 0.95992644 |
| ARL14EP | 21.536278 | 0.56245779 | 0.37538887 | 1.4983337 | 0.13404657 | 0.80879167 |
| DNAJC24 | 24.3621944 | -0.468138 | 0.34372389 | -1.3619595 | 0.17321067 | 0.87233975 |
| IMMP1L | 43.5471901 | 0.08192503 | 0.29605973 | 0.27671791 | 0.78199672 | 0.9958918 |
| ELP4 | 100.375936 | 0.03319031 | 0.19408411 | 0.17100993 | 0.86421596 | 0.9971731 |
| RCN1 | 31.054509 | -0.4657664 | 0.45808903 | -1.0167597 | 0.30926778 | 0.95793395 |
| WT1 | 57.822986 | 0.03888522 | 0.54680644 | 0.07111332 | 0.94330757 | 0.99935941 |
| EIF3M | 39.3652693 | 0.09791686 | 0.34606252 | 0.28294558 | 0.77721855 | 0.9958918 |
| QSER1 | 60.3004973 | -0.0669654 | 0.24966682 | -0.2682192 | 0.78853057 | 0.99592423 |
| TCP11L1 | 28.0587866 | 0.06382015 | 0.34494296 | 0.18501652 | 0.85321608 | 0.9971731 |
| CSTF3 | 46.7246791 | 0.01395312 | 0.25862011 | 0.0539522 | 0.95697325 | 0.99935941 |
| HIPK3 | 270.978195 | 0.67464247 | 0.29678868 | 2.27314086 | 0.02301769 | 0.4105753 |
| FBXO3 | 40.5834694 | 0.0220003 | 0.28775864 | 0.07645399 | 0.93905792 | 0.99935941 |
| CAPRIN1 | 83.5411486 | 0.11824529 | 0.20807614 | 0.56827894 | 0.56984558 | 0.9958918 |
| NAT10 | 28.4183056 | -0.0758466 | 0.43746375 | -0.1733781 | 0.86235426 | 0.9971731 |
| ABTB2 | 77.8874928 | 1.0320747 | 0.85956978 | 1.20068751 | 0.22987244 | 0.91455844 |
| CAT | 41.30266 | 0.26539684 | 0.42957279 | 0.61781577 | 0.53669679 | 0.9958918 |
| APIP | 28.1199792 | -1.1701812 | 0.30549929 | -3.8303892 | 0.00012794 | 0.01774031 |
| PDHX | 67.7941972 | -0.0318002 | 0.32767979 | -0.0970467 | 0.92268929 | 0.99935941 |
| CD44 | 226.208383 | -0.10434 | 0.39979032 | -0.2609867 | 0.79410273 | 0.99592423 |
| PAMR1 | 69.0483828 | -1.9526847 | 0.55223108 | -3.535992 | 0.00040625 | 0.03928301 |
| TRIM44 | 169.181266 | 0.061211 | 0.16363802 | 0.37406346 | 0.70835711 | 0.9958918 |
| LDLRAD3 | 345.378348 | -0.0981805 | 0.35304689 | -0.2780948 | 0.78093959 | 0.9958918 |
| PRR5L | 37.7621068 | -0.8473781 | 0.34123543 | -2.4832652 | 0.01301841 | 0.30408773 |
| TRAF6 | 18.5000124 | -0.4352302 | 0.33198208 | -1.311005 | 0.18985608 | 0.88139754 |
| API5 | 46.5842915 | -0.2435838 | 0.26230915 | -0.9286133 | 0.3530895 | 0.97185572 |
| TTC17 | 242.370115 | -0.0964053 | 0.16005924 | -0.6023098 | 0.54696793 | 0.9958918 |
| HSD17B12 | 167.347439 | 0.41900614 | 0.2030437 | 2.06362544 | 0.03905324 | 0.52881167 |
| ALKBH3 | 19.4996566 | -0.2345751 | 0.39946828 | -0.5872184 | 0.55705701 | 0.9958918 |
| C11orf96 | 486.599633 | -0.2702121 | 0.53317021 | -0.5068027 | 0.61229326 | 0.9958918 |

| | | | | | | |
|---|---|---|---|---|---|---|
| EXT2 | 74.4590722 | 0.21777943 | 0.21823029 | 0.99793401 | 0.31831136 | 0.95992644 |
| TP53I11 | 28.4201817 | 0.07874286 | 0.46984586 | 0.16759297 | 0.8669035 | 0.9971731 |
| PRDM11 | 31.2378035 | 0.0957009 | 0.29229573 | 0.32741122 | 0.74335688 | 0.9958918 |
| CRY2 | 34.5823998 | 0.02591894 | 0.33797646 | 0.07668859 | 0.93887128 | 0.99935941 |
| PHF21A | 358.42952 | 0.24268815 | 0.24846566 | 0.97674724 | 3.29E-01 | 0.9620084 |
| DGKZ | 18.5213623 | 0.0553751 | 0.40646769 | 0.13623493 | 0.89163556 | 0.99935941 |
| AMBRA1 | 287.191062 | -0.0240879 | 0.21104162 | -0.1141383 | 0.90912819 | 0.99935941 |
| ATG13 | 46.7079534 | -0.0338529 | 0.29447946 | -0.1149584 | 0.90847806 | 0.99935941 |
| ARHGAP1 | 62.4533204 | 0.09443247 | 0.4317138 | 0.2187386 | 0.82685368 | 0.99592423 |
| CKAP5 | 49.5812228 | -0.1455276 | 0.23688999 | -0.6143255 | 0.53900027 | 0.9958918 |
| LRP4 | 46.3019659 | 0.33562614 | 0.46530206 | 0.7213081 | 0.47071998 | 0.98910524 |
| C11orf49 | 98.1041519 | -0.3634237 | 0.29205275 | -1.2443769 | 0.21336089 | 0.89738044 |
| ARFGAP2 | 21.5290526 | 0.01801442 | 0.37808886 | 0.04764599 | 0.96199838 | 0.99935941 |
| DDB2 | 30.5381103 | 0.39677441 | 0.44750917 | 0.88662855 | 0.37527892 | 0.97217293 |
| MADD | 31.304696 | -0.0291081 | 0.30191179 | -0.0964126 | 0.92319285 | 0.99935941 |
| CELF1 | 198.273303 | 0.01419173 | 0.16798069 | 0.08448427 | 0.93267141 | 0.99935941 |
| MTCH2 | 36.3173746 | 0.07973579 | 0.29326924 | 0.27188596 | 0.78570971 | 0.9958918 |
| FNBP4 | 170.917192 | -0.0654107 | 0.23716537 | -0.2758019 | 0.78270021 | 0.9958918 |
| NUP160 | 78.6463542 | 0.20451027 | 0.21568422 | 0.948193 | 0.34303121 | 0.96887708 |
| PTPRJ | 63.0267405 | -0.087403 | 0.31633609 | -0.2762979 | 7.82E-01 | 0.9958918 |
| TNKS1BP1 | 45.3714988 | -0.1598895 | 0.26184523 | -0.610626 | 0.5414472 | 0.9958918 |
| SSRP1 | 20.0731437 | 0.28138412 | 0.52491964 | 0.53605182 | 0.59192274 | 0.9958918 |
| SERPING1 | 227.936212 | -0.4195757 | 0.30157357 | -1.3912881 | 0.16413807 | 0.85697309 |
| ZDHHC5 | 32.1828129 | 0.19721349 | 0.2892271 | 0.68186378 | 0.49532509 | 0.99176652 |
| CTNND1 | 94.6178081 | -0.1858857 | 0.31343843 | -0.5930534 | 0.55314544 | 0.9958918 |
| LPXN | 21.4114223 | -0.2754374 | 0.36746998 | -0.7495507 | 0.45352537 | 0.98092839 |
| FAM111A | 27.7465522 | 0.09230716 | 0.55793638 | 0.16544387 | 0.86859462 | 0.9971731 |
| DTX4 | 18.7485424 | 0.17226615 | 0.37362787 | 0.46106343 | 0.6447531 | 0.9958918 |
| OSBP | 51.5242353 | 0.04208213 | 0.26903936 | 0.15641627 | 0.87570491 | 0.9971731 |
| PATL1 | 39.491602 | -0.2941035 | 0.38388602 | -0.766122 | 0.4436037 | 0.98092839 |
| STX3 | 20.4719629 | -0.489573 | 0.52189105 | -0.9380751 | 0.34820582 | 0.96887708 |
| PRPF19 | 17.5579246 | -0.287254 | 0.51560706 | -0.557118 | 0.57744678 | 0.9958918 |
| TMEM109 | 21.8224943 | 0.3511234 | 0.36307385 | 0.96708536 | 0.33350137 | 0.9654289 |
| VPS37C | 18.7127736 | 0.41841972 | 0.33683839 | 1.24219726 | 0.21416379 | 0.89936554 |
| DDB1 | 56.8994844 | -0.3046718 | 0.21560151 | -1.4131245 | 0.15761911 | 0.84427321 |
| CPSF7 | 49.525414 | 0.24654915 | 0.31519537 | 0.78221057 | 0.43409083 | 0.98092839 |
| AP003108.2 | 38.4791063 | 0.6001405 | 0.33308357 | 1.80177154 | 0.07158136 | 0.66169987 |
| FADS2 | 54.0156798 | -0.4896505 | 0.37460005 | -1.3071288 | 0.19116896 | 0.88158763 |
| FADS1 | 44.9225822 | -0.0389251 | 0.55179998 | -0.0705421 | 0.94376221 | 0.99935941 |
| FADS3 | 34.601254 | 0.24066111 | 0.41320375 | 0.58242722 | 0.56027895 | 0.9958918 |
| FTH1 | 701.537203 | 0.4604564 | 0.46311594 | 0.99425728 | 0.32009763 | 0.95992644 |
| AHNAK | 171.783147 | -0.055427 | 0.27924364 | -0.1984898 | 0.84266186 | 0.99692952 |
| EML3 | 19.8396663 | -0.1429698 | 0.4127241 | -0.3464053 | 0.72903816 | 0.9958918 |
| B3GAT3 | 21.7717786 | 0.16847775 | 0.33705148 | 0.49985761 | 0.61717534 | 0.9958918 |
| GANAB | 68.0097467 | 0.0167026 | 0.27601334 | 0.06051374 | 0.95174647 | 0.99935941 |
| UBXN1 | 23.153187 | 0.68076307 | 0.40462785 | 1.68244246 | 0.09248307 | 0.7300302 |
| TTC9C | 20.1411342 | 0.06948208 | 0.3661059 | 0.18978682 | 0.84947618 | 0.9971731 |
| NXF1 | 20.2297512 | 0.82663645 | 0.37004368 | 2.23388885 | 0.02549039 | 0.43064105 |
| STX5 | 53.4988777 | 0.02588211 | 0.23917341 | 0.10821484 | 0.91382528 | 0.99935941 |
| SLC3A2 | 40.8388522 | -0.3791566 | 0.30938256 | -1.2255266 | 0.22037688 | 0.90731076 |
| RARRES3 | 29.5979883 | 0.28054293 | 0.66207993 | 0.4237297 | 0.67176296 | 0.9958918 |
| PLA2G16 | 20.5170034 | 0.72892851 | 0.33141709 | 2.19942948 | 0.0278474 | 0.45276666 |
| ATL3 | 88.0723668 | -0.0645115 | 0.30920434 | -0.2086372 | 0.83473144 | 0.99665985 |
| RTN3 | 79.4078448 | 0.06641246 | 0.23861322 | 0.27832684 | 0.78076148 | 0.9958918 |

| | | | | | | |
|---|---|---|---|---|---|---|
| MARK2 | 52.615802 | 0.32087803 | 0.26060664 | 1.23127339 | 0.21822063 | 0.90437211 |
| MACROD1 | 21.8746895 | 0.33697821 | 0.35804159 | 0.94117058 | 0.34661745 | 0.96887708 |
| STIP1 | 26.1530756 | -0.121 | 0.32834133 | -0.368519 | 0.71248627 | 0.9958918 |
| VEGFB | 32.0151297 | 0.14451241 | 0.3703518 | 0.39020309 | 0.69638638 | 0.9958918 |
| FKBP2 | 28.3538336 | -0.2051984 | 0.3933394 | -0.5216828 | 0.60189121 | 0.9958918 |
| ESRRA | 22.5832126 | 0.09722251 | 0.45542173 | 0.21347797 | 0.83095417 | 0.99592423 |
| TRMT112 | 30.3414529 | -0.1535356 | 0.35264721 | -0.43538 | 0.6632866 | 0.9958918 |
| PRDX5 | 28.0763995 | 0.08024091 | 0.38871509 | 0.20642603 | 0.83645814 | 0.99692952 |
| NRXN2 | 18.3987275 | 1.69817087 | 0.49939435 | 3.40046068 | 0.00067272 | 0.05624153 |
| RASGRP2 | 29.510636 | -0.1711512 | 0.34144247 | -0.5012593 | 0.61618864 | 0.9958918 |
| SF1 | 134.149543 | 0.01245161 | 0.19555768 | 0.0636723 | 0.94923116 | 0.99935941 |
| EHD1 | 23.3529729 | 0.38839269 | 0.53032582 | 0.73236617 | 0.4639451 | 0.98590351 |
| ATG2A | 18.2745709 | -0.1596442 | 0.43351022 | -0.3682594 | 0.71267979 | 0.9958918 |
| VPS51 | 42.6155241 | 0.12335645 | 0.24648405 | 0.50046423 | 0.61674823 | 0.9958918 |
| FAU | 97.4210229 | -0.0148532 | 0.42110045 | -0.0352722 | 0.97186266 | 0.99935941 |
| SYVN1 | 19.7770914 | -0.6013661 | 0.3802108 | -1.581665 | 0.11372607 | 0.77853537 |
| CAPN1 | 31.7455249 | 0.32447248 | 0.35883563 | 0.90423707 | 0.36586971 | 0.97217293 |
| DPF2 | 27.1812106 | 0.37122664 | 0.2941974 | 1.26182843 | 0.20701053 | 0.89620104 |
| FRMD8 | 18.3309919 | 0.00205132 | 0.35243662 | 0.00582039 | 0.99535602 | 0.99935941 |
| NEAT1 | 4726.0414 | -0.1108587 | 0.45040256 | -0.2461324 | 0.80557972 | 0.99592423 |
| MALAT1 | 41344.0507 | -0.0980638 | 0.28219382 | -0.347505 | 0.72821196 | 0.9958918 |
| SCYL1 | 27.8999077 | 0.29966567 | 0.35335727 | 0.84805294 | 0.39640849 | 0.97500135 |
| LTBP3 | 84.7873672 | 0.09197889 | 0.28399952 | 0.32386989 | 0.74603654 | 0.9958918 |
| EHBP1L1 | 61.298144 | 0.84848494 | 0.32929058 | 2.57670578 | 0.00997468 | 0.26559396 |
| RELA | 22.8502932 | 0.28890915 | 0.40723111 | 0.70944765 | 0.47804673 | 0.99002719 |
| KAT5 | 24.3144146 | -0.0173376 | 0.36030111 | -0.0481198 | 0.96162079 | 0.99935941 |
| RNASEH2C | 23.0976848 | 0.34014595 | 0.38560536 | 0.88210896 | 0.37771789 | 0.97217293 |
| CFL1 | 101.623977 | 0.0779899 | 0.42825818 | 0.18210953 | 0.85549677 | 0.9971731 |
| EFEMP2 | 26.8679337 | -0.5314253 | 0.39383512 | -1.3493598 | 0.17722144 | 0.87771577 |
| CCDC85B | 43.2696517 | -0.1218072 | 0.42872439 | -0.2841154 | 0.77632197 | 0.9958918 |
| DRAP1 | 19.6046424 | -0.3152142 | 0.42722965 | -0.7378098 | 0.46063004 | 0.98590351 |
| BANF1 | 21.8175946 | -0.291667 | 0.47221311 | -0.6176597 | 0.53679968 | 0.9958918 |
| SF3B2 | 43.2541162 | -0.1573577 | 0.3048392 | -0.516199 | 0.6057154 | 0.9958918 |
| PACS1 | 178.937108 | 0.36981405 | 0.18643112 | 1.98364983 | 0.04729489 | 0.56885457 |
| RAB1B | 22.5169568 | -0.0868254 | 0.35857374 | -0.2421409 | 0.80867102 | 0.99592423 |
| CD248 | 40.1931355 | 0.69408273 | 0.61511044 | 1.12838717 | 0.25915644 | 0.92976724 |
| CCS | 31.4252221 | 0.41882092 | 0.36257055 | 1.15514322 | 0.24803178 | 0.92624282 |
| RBM14-RBM | 27.3598441 | 0.17723798 | 0.32818595 | 0.54005354 | 0.58916011 | 0.9958918 |
| RBM4 | 22.5040051 | 0.02120888 | 0.38744254 | 0.05474071 | 0.95634504 | 0.99935941 |
| C11orf80 | 21.678035 | 0.43004462 | 0.43274496 | 0.99375998 | 0.32033974 | 0.95992644 |
| KDM2A | 287.217717 | 0.04714474 | 0.18352801 | 0.25688037 | 0.79727112 | 0.99592423 |
| GRK2 | 22.5463541 | -0.0959483 | 0.32518617 | -0.2950565 | 0.76795075 | 0.9958918 |
| ANKRD13D | 30.0538293 | 0.15539324 | 0.35133336 | 0.4422957 | 0.65827524 | 0.9958918 |
| RAD9A | 49.8082876 | -0.2319767 | 0.28751746 | -0.8068263 | 0.41976655 | 0.97812179 |
| PPP1CA | 25.3268216 | 0.43021044 | 0.43776542 | 0.98274194 | 0.32573446 | 0.95992644 |
| CDK2AP2 | 19.3584383 | 0.15399708 | 0.44616485 | 0.34515735 | 0.72997608 | 0.9958918 |
| GSTP1 | 120.97753 | -0.0190549 | 0.32590187 | -0.0584681 | 0.95337579 | 0.99935941 |
| NDUFV1 | 31.4339257 | 0.17207397 | 0.32562836 | 0.52843668 | 0.59719628 | 0.9958918 |
| NDUFS8 | 27.1344952 | 0.11254047 | 0.35771277 | 0.31461127 | 0.75305682 | 0.9958918 |
| CHKA | 37.4701265 | 0.06302457 | 0.28653235 | 0.2199562 | 0.82590527 | 0.99592423 |
| KMT5B | 97.0305201 | 0.48100401 | 0.31216587 | 1.54086033 | 0.12335078 | 0.79184923 |
| LRP5 | 42.7770459 | -0.2271626 | 0.47015902 | -0.4831611 | 0.62898135 | 0.9958918 |
| PPP6R3 | 339.426685 | -0.206154 | 0.24071116 | -0.8564374 | 0.39175591 | 0.97297628 |
| CPT1A | 66.9810829 | 0.37466479 | 0.42982449 | 0.87166925 | 0.38338884 | 0.9725044 |

| | | | | | | |
|---|---|---|---|---|---|---|
| MRGPRF | 23.0440787 | -0.6886471 | 0.4622066 | -1.4899118 | 0.13624743 | 0.81177777 |
| TPCN2 | 26.5605142 | 0.17865327 | 0.35681414 | 0.50068998 | 0.61658933 | 0.9958918 |
| CCND1 | 20.5015555 | 0.26456743 | 0.49577699 | 0.533642 | 0.59358924 | 0.9958918 |
| ORAOV1 | 17.2643674 | 0.07811212 | 0.39722648 | 0.19664378 | 0.84410631 | 0.99692952 |
| ANO1 | 70.5136957 | -0.2025822 | 0.45123982 | -0.4489457 | 0.65347083 | 0.9958918 |
| PPFIA1 | 117.236635 | 0.08362191 | 0.25323344 | 0.33021668 | 0.74123624 | 0.9958918 |
| AP002336.2 | 38.8736255 | -0.1842703 | 0.32749253 | -0.5626703 | 0.57365941 | 0.9958918 |
| CTTN | 92.5062281 | -0.149973 | 0.27192923 | -0.5515147 | 0.58128092 | 0.9958918 |
| AP002495.2 | 42.0982303 | 0.22611048 | 0.30916866 | 0.73134993 | 0.46456544 | 0.98616158 |
| RNF121 | 33.449295 | 0.05042378 | 0.30395117 | 0.16589433 | 0.86824011 | 0.9971731 |
| NUMA1 | 78.4426636 | 0.04748217 | 0.28694223 | 0.16547642 | 0.86856901 | 0.9971731 |
| LAMTOR1 | 33.2761052 | -0.0412289 | 0.31964787 | -0.1289822 | 0.89737171 | 0.99935941 |
| INPPL1 | 24.2660922 | -0.1376206 | 0.32631733 | -0.4217385 | 0.67321589 | 0.9958918 |
| CLPB | 23.9380256 | 0.11367239 | 0.43134252 | 0.26353161 | 0.79214086 | 0.99592423 |
| ARAP1 | 33.4197741 | -0.1215974 | 0.34706244 | -0.3503617 | 0.72606727 | 0.9958918 |
| ATG16L2 | 23.7650057 | 0.40540467 | 0.36642529 | 1.10637744 | 0.26856319 | 0.93598159 |
| FCHSD2 | 465.988037 | 0.22022623 | 0.33212634 | 0.66307967 | 0.50727953 | 0.99397937 |
| ARHGEF17 | 180.532353 | -0.2076811 | 0.30658076 | -0.6774106 | 0.49814544 | 0.99176652 |
| FAM168A | 135.261337 | -0.0686399 | 0.2692253 | -0.2549534 | 0.79875911 | 0.99592423 |
| RAB6A | 145.652924 | -0.326924 | 0.16179624 | -2.0205906 | 0.04332216 | 0.55273362 |
| COA4 | 18.6943809 | -0.035141 | 0.53311492 | -0.0659163 | 0.94744447 | 0.99935941 |
| PAAF1 | 25.8633633 | 0.00542661 | 0.41855862 | 0.012965 | 0.98965571 | 0.99935941 |
| C2CD3 | 45.5115083 | 0.01455859 | 0.27106147 | 0.05370955 | 0.95716657 | 0.99935941 |
| PPME1 | 81.4605976 | -0.1624366 | 0.39706262 | -0.4090958 | 0.68246936 | 0.9958918 |
| PGM2L1 | 25.2758438 | 0.24404105 | 0.45170747 | 0.54026349 | 0.58901534 | 0.9958918 |
| POLD3 | 23.5633976 | 0.09477175 | 0.39364232 | 0.24075601 | 0.80974423 | 0.99592423 |
| RNF169 | 76.2329925 | 0.04447823 | 0.22078785 | 0.20145235 | 0.84034489 | 0.99692952 |
| XRRA1 | 85.5210346 | -0.148869 | 0.55142566 | -0.2699711 | 0.78718245 | 0.99592423 |
| SPCS2 | 62.7229534 | 0.51243725 | 0.27642762 | 1.85378458 | 0.06376999 | 0.62395844 |
| ARRB1 | 58.2381667 | 0.05827055 | 0.35502142 | 0.16413248 | 0.86962685 | 0.9971731 |
| RPS3 | 159.154318 | -0.2802435 | 0.35659885 | -0.785879 | 0.43193839 | 0.98092839 |
| SERPINH1 | 81.7769222 | -0.1899801 | 0.48659848 | -0.3904248 | 0.69622247 | 0.9958918 |
| UVRAG | 224.999229 | -0.1682893 | 0.2156813 | -0.7802686 | 0.43523281 | 0.98092839 |
| THAP12 | 18.5773386 | -0.2310602 | 0.33593863 | -0.6878049 | 0.49157565 | 0.99046213 |
| EMSY | 115.785813 | -0.0872987 | 0.22128733 | -0.3945038 | 0.69320915 | 0.9958918 |
| LRRC32 | 74.2776323 | -0.7764128 | 0.43891943 | -1.7689187 | 0.07690744 | 0.67905476 |
| TSKU | 23.5048876 | -1.2024013 | 0.49021814 | -2.4527883 | 0.01417538 | 0.32453219 |
| ACER3 | 106.717077 | -0.0782139 | 0.31100907 | -0.2514844 | 0.80143961 | 0.99592423 |
| PAK1 | 32.0441026 | -0.0264007 | 0.31936818 | -0.0826655 | 0.9341175 | 0.99935941 |
| CLNS1A | 47.8344069 | 0.27004657 | 0.30935355 | 0.87293832 | 0.3826967 | 0.9725044 |
| RSF1 | 176.85402 | 0.16572516 | 0.17762665 | 0.93299717 | 0.35082143 | 0.9693503 |
| AAMDC | 48.3575073 | 0.14625072 | 0.32479488 | 0.4502864 | 0.65250394 | 0.9958918 |
| INTS4 | 38.6130023 | -0.1911647 | 0.29626423 | -0.6452505 | 0.51876485 | 0.9958918 |
| ALG8 | 23.2749936 | -0.0342755 | 0.40353123 | -0.084939 | 0.93230992 | 0.99935941 |
| GAB2 | 278.674716 | -0.2297236 | 0.29442441 | -0.7802464 | 0.43524586 | 0.98092839 |
| AP003086.2 | 70.8965166 | -0.8489345 | 0.4969017 | -1.7084557 | 0.08755183 | 0.71805622 |
| NARS2 | 73.0950092 | -0.7529109 | 0.28899076 | -2.6053113 | 0.00917908 | 0.25431499 |
| TENM4 | 34.0042978 | -0.4836527 | 0.56389998 | -0.8576924 | 0.39106236 | 0.97297628 |
| PRCP | 30.8212245 | 0.07705756 | 0.26048575 | 0.29582255 | 0.7673656 | 0.9958918 |
| RAB30 | 40.6068522 | -0.3127769 | 0.31090556 | -1.006019 | 0.31440645 | 0.95992644 |
| RAB30-AS1 | 47.6394374 | 0.19353522 | 0.36268243 | 0.53362172 | 0.59360328 | 0.9958918 |
| PCF11 | 60.0704299 | 0.0674498 | 0.26489383 | 0.25462958 | 0.79900921 | 0.99592423 |
| ANKRD42 | 23.2979305 | -0.0341114 | 0.3488638 | -0.0977785 | 0.92210817 | 0.99935941 |
| DLG2 | 789.360324 | -0.2445438 | 0.42336444 | -0.5776202 | 0.56352058 | 0.9958918 |

| | | | | | | |
|---|---|---|---|---|---|---|
| CREBZF | 64.847651 | -0.3129636 | 0.28072144 | -1.1148546 | 0.26491272 | 0.93499423 |
| SYTL2 | 178.54137 | -0.042314 | 0.57633869 | -0.0734186 | 0.94147301 | 0.99935941 |
| PICALM | 587.083034 | -0.0959573 | 0.34153313 | -0.2809605 | 0.77874073 | 0.9958918 |
| EED | 27.2854387 | -0.1903733 | 0.36565695 | -0.5206335 | 0.6026221 | 0.9958918 |
| HIKESHI | 32.9314507 | 0.20705959 | 0.29958323 | 0.69115882 | 0.48946574 | 0.99046213 |
| ME3 | 58.1689695 | -0.6438104 | 0.29858996 | -2.1561689 | 0.03107048 | 0.47969946 |
| PRSS23 | 177.192132 | -0.9978436 | 0.33980579 | -2.9365113 | 0.00331927 | 0.13939025 |
| AP001528.2 | 39.1590925 | 0.08830735 | 0.63397788 | 0.13929091 | 0.88922027 | 0.99935941 |
| FZD4 | 30.8322357 | 0.07187758 | 0.45196128 | 0.15903482 | 0.87364144 | 0.9971731 |
| TMEM135 | 180.278192 | 0.00734644 | 0.33560704 | 0.02189 | 0.9825357 | 0.99935941 |
| CTSC | 58.2738987 | -0.7820189 | 0.56789413 | -1.3770506 | 0.16849662 | 0.86601846 |
| NOX4 | 88.9287538 | -0.2643785 | 0.50144585 | -0.5272325 | 0.59803216 | 0.9958918 |
| NAALAD2 | 31.4369327 | -0.0199289 | 0.88891442 | -0.0224193 | 0.98211346 | 0.99935941 |
| CHORDC1 | 34.7380399 | -0.0269937 | 0.30417109 | -0.0887451 | 0.92928446 | 0.99935941 |
| FAT3 | 44.7629716 | -0.0939967 | 0.58853361 | -0.1597134 | 0.87310686 | 0.9971731 |
| SLC36A4 | 40.9805941 | -0.4642734 | 0.32136126 | -1.4447088 | 0.14853971 | 0.8301633 |
| CEP295 | 31.6912006 | -0.4515777 | 0.34684341 | -1.3019641 | 0.19292864 | 0.88158763 |
| TAF1D | 98.3771573 | -0.0226896 | 0.28712818 | -0.0790224 | 0.93701478 | 0.99935941 |
| PANX1 | 21.7082876 | 1.07961606 | 0.41383524 | 2.60880647 | 0.00908586 | 0.25292422 |
| MRE11 | 35.8637994 | -0.3293797 | 0.27295816 | -1.2067039 | 0.22754619 | 0.9120167 |
| AMOTL1 | 99.3416723 | 0.24058262 | 0.25020729 | 0.9615332 | 0.33628414 | 0.96640422 |
| CWC15 | 20.9512492 | 0.22399115 | 0.31784448 | 0.70471933 | 0.48098492 | 0.99002719 |
| SRSF8 | 19.260919 | 0.5209919 | 0.3684954 | 1.41383557 | 0.15741019 | 0.84427321 |
| ENDOD1 | 48.904123 | -0.1332509 | 0.34505087 | -0.3861774 | 0.69936527 | 0.9958918 |
| SESN3 | 71.4050086 | -0.8417222 | 0.38392535 | -2.1924112 | 0.02834983 | 0.45789648 |
| FAM76B | 25.4065743 | -0.4135248 | 0.29606115 | -1.3967547 | 0.16248736 | 0.85477422 |
| CEP57 | 46.8705386 | 0.33360686 | 0.25726046 | 1.29676695 | 0.19471139 | 0.88391893 |
| MTMR2 | 84.1086725 | 0.17366278 | 0.20475402 | 0.84815321 | 0.39635265 | 0.97500135 |
| MAML2 | 1277.90411 | -0.1492082 | 0.43549353 | -0.3426186 | 0.73188544 | 0.9958918 |
| CCDC82 | 51.4109112 | 0.22713498 | 0.2649687 | 0.85721436 | 0.39132643 | 0.97297628 |
| CNTN5 | 142.031976 | 1.36145168 | 0.57094345 | 2.38456483 | 0.01709934 | 0.34765232 |
| ARHGAP42 | 483.909382 | -0.7673227 | 0.25818369 | -2.9720028 | 0.00295864 | 0.129173 |
| PGR | 51.5028643 | 0.03492175 | 0.33510741 | 0.10421062 | 0.9170022 | 0.99935941 |
| TRPC6 | 43.4211499 | -0.8580055 | 0.41307149 | -2.0771355 | 0.03778905 | 0.52201457 |
| CEP126 | 49.6364524 | -0.1000079 | 0.34997393 | -0.2857581 | 0.77506342 | 0.9958918 |
| YAP1 | 420.44393 | -0.1224099 | 0.30875984 | -0.3964566 | 0.69176822 | 0.9958918 |
| BIRC3 | 27.2794122 | 0.51906416 | 0.92012872 | 0.56412124 | 0.57267162 | 0.9958918 |
| BIRC2 | 50.3794644 | 0.37332316 | 0.27434011 | 1.36080412 | 0.1735756 | 0.87233975 |
| TMEM123 | 73.4507625 | -0.128957 | 0.27530214 | -0.4684197 | 0.63948447 | 0.9958918 |
| DCUN1D5 | 26.4805202 | -0.151379 | 0.31589688 | -0.4792037 | 0.63179372 | 0.9958918 |
| DYNC2H1 | 178.532014 | -0.4858685 | 0.32635747 | -1.4887616 | 0.13655016 | 0.8125564 |
| PDGFD | 75.5724256 | 0.34258731 | 0.55446264 | 0.61787267 | 0.53665927 | 0.9958918 |
| CASP4 | 43.5972219 | -0.0820349 | 0.41486121 | -0.1977406 | 0.84324802 | 0.99692952 |
| AASDHPPT | 35.0171923 | -0.1944437 | 0.33091769 | -0.5875892 | 0.55680803 | 0.9958918 |
| GUCY1A2 | 354.452111 | -0.7567909 | 0.46287125 | -1.6349922 | 0.10205068 | 0.75449741 |
| CWF19L2 | 79.9534237 | -0.1860724 | 0.23452552 | -0.7933996 | 0.42754506 | 0.98092839 |
| CUL5 | 120.930673 | -0.0550391 | 0.17433185 | -0.3157148 | 0.75221902 | 0.9958918 |
| ACAT1 | 43.9247205 | 0.34317236 | 0.28904782 | 1.18725114 | 0.23512857 | 0.91962002 |
| NPAT | 45.2772571 | -0.0861953 | 0.32705739 | -0.2635481 | 0.79212817 | 0.99592423 |
| ATM | 131.466482 | -0.2687218 | 0.24263181 | -1.1075293 | 0.26806517 | 0.93598159 |
| KDELC2 | 29.1663752 | -0.2908656 | 0.34845068 | -0.8347395 | 0.40386438 | 0.97575999 |
| DDX10 | 120.887226 | -0.3751649 | 0.34524112 | -1.0866751 | 0.27718043 | 0.94305507 |
| RDX | 86.147956 | 0.03465496 | 0.26053895 | 0.13301258 | 0.89418343 | 0.99935941 |
| ZC3H12C | 36.5836976 | 0.56614729 | 0.38366997 | 1.47561011 | 0.14004859 | 0.8153985 |

| | | | | | | |
|---|---|---|---|---|---|---|
| FDX1 | 30.0935612 | 0.94572351 | 0.34788684 | 2.71848023 | 0.00655826 | 0.20509206 |
| ARHGAP20 | 41.2047227 | -1.7004646 | 0.5475811 | -3.1054114 | 0.00190015 | 0.09345751 |
| SIK2 | 326.348444 | -0.4380559 | 0.43874655 | -0.9984258 | 0.31807293 | 0.95992644 |
| PPP2R1B | 75.689688 | -0.100815 | 0.37340445 | -0.2699887 | 0.78716898 | 0.99592423 |
| CRYAB | 305.426719 | 0.62110804 | 0.64325877 | 0.96556482 | 0.33426198 | 0.96640422 |
| DIXDC1 | 47.5620707 | -0.6990786 | 0.39512498 | -1.7692594 | 0.0768506 | 0.67905476 |
| PTS | 57.0164666 | -0.2305562 | 0.28636224 | -0.8051209 | 0.4207499 | 0.97812179 |
| AP002884.1 | 24.9892118 | -0.1749113 | 0.42747493 | -0.4091733 | 0.68241248 | 0.9958918 |
| NCAM1 | 71.3185023 | -0.9600467 | 0.83904332 | -1.1442159 | 0.25253409 | 0.92631316 |
| USP28 | 27.6770757 | 0.45219025 | 0.35971017 | 1.25709608 | 0.20871885 | 0.89634177 |
| ZBTB16 | 726.139574 | -0.5282201 | 0.78098797 | -0.6763485 | 0.49881938 | 0.99176652 |
| AP002518.2 | 88.78272 | -0.2287074 | 1.00072043 | -0.2285427 | 0.81922435 | 0.99592423 |
| NNMT | 258.528705 | 0.32843818 | 0.5239194 | 0.62688685 | 5.31E-01 | 0.9958918 |
| REXO2 | 36.6741009 | -0.0412045 | 0.56345678 | -0.073128 | 0.94170426 | 0.99935941 |
| CADM1 | 29.2923111 | -1.0215974 | 0.37260447 | -2.7417743 | 0.00611083 | 0.19357111 |
| SIK3 | 605.827356 | -0.0644938 | 0.34040673 | -0.1894609 | 0.8497316 | 0.9971731 |
| PAFAH1B2 | 55.042108 | -0.1207473 | 0.25072868 | -0.4815853 | 0.63010055 | 0.9958918 |
| SIDT2 | 41.4128882 | 0.13087003 | 0.4062048 | 0.32217747 | 0.74731825 | 0.9958918 |
| TAGLN | 922.468189 | 0.71669293 | 0.39384953 | 1.81971253 | 0.06880279 | 0.64792295 |
| PCSK7 | 83.294581 | 0.20501667 | 0.26517692 | 0.77313163 | 0.43944448 | 0.98092839 |
| RNF214 | 31.7867671 | -0.1310473 | 0.33155564 | -0.3952497 | 0.69265862 | 0.9958918 |
| BACE1 | 31.9134593 | -0.03988 | 0.31569834 | -0.126323 | 0.89947627 | 0.99935941 |
| CEP164 | 36.6850025 | -0.0337428 | 0.25876111 | -0.1304014 | 0.89624889 | 0.99935941 |
| DSCAML1 | 50.1004907 | -0.3882419 | 1.00986008 | -0.3844511 | 0.70064411 | 0.9958918 |
| FXYD2 | 22.1896403 | -0.4774478 | 0.48386918 | -0.9867291 | 0.32377544 | 0.95992644 |
| UBE4A | 50.6648196 | -0.1384914 | 0.25160935 | -0.5504224 | 0.58202968 | 0.9958918 |
| ATP5MG | 56.3544929 | 0.02084513 | 0.33066657 | 0.06303974 | 0.94973486 | 0.99935941 |
| KMT2A | 187.098235 | 0.09038263 | 0.18117607 | 0.4988663 | 0.61787358 | 0.9958918 |
| ARCN1 | 43.619663 | 0.06449759 | 0.2549315 | 0.25299968 | 0.80026846 | 0.99592423 |
| PHLDB1 | 94.7801516 | -0.4992869 | 0.35025722 | -1.4254864 | 0.15401664 | 0.84216389 |
| DDX6 | 110.284056 | 0.0012797 | 0.2037864 | 0.00627963 | 0.99498962 | 0.99935941 |
| BCL9L | 39.3179923 | -0.1086427 | 0.38983996 | -0.2786853 | 0.78048638 | 0.9958918 |
| CCDC84 | 62.1751298 | -0.0243757 | 0.33018508 | -0.0738243 | 0.94115022 | 0.99935941 |
| RPS25 | 102.894484 | -0.3431344 | 0.35650589 | -0.9624928 | 0.3358021 | 0.96640422 |
| TRAPPC4 | 20.7159207 | -0.0275995 | 0.37140826 | -0.0743104 | 0.94076341 | 0.99935941 |
| HYOU1 | 19.2087095 | -0.0306455 | 0.53288764 | -0.0575084 | 0.95414025 | 0.99935941 |
| CBL | 76.2354118 | -0.0363044 | 0.27571404 | -0.1316741 | 0.8952421 | 0.99935941 |
| MCAM | 187.43115 | 0.88833944 | 0.41499725 | 2.14059114 | 0.03230702 | 0.48564518 |
| THY1 | 22.3719653 | -1.7532215 | 0.94873687 | -1.8479534 | 0.06460909 | 0.62722878 |
| OAF | 42.141301 | -1.3857923 | 0.33289206 | -4.1628877 | 3.14E-05 | 0.0072169 |
| ARHGEF12 | 353.807194 | -0.1613994 | 0.25297974 | -0.6379934 | 0.52347801 | 0.9958918 |
| SORL1 | 32.3175468 | 0.16272882 | 0.4705357 | 0.34583736 | 0.72946495 | 0.9958918 |
| UBASH3B | 28.0266075 | 0.17241236 | 0.6084312 | 0.28337199 | 0.7768917 | 0.9958918 |
| HSPA8 | 81.2215549 | 0.30103678 | 0.37359891 | 0.80577532 | 0.42037241 | 0.97812179 |
| CLMP | 205.520654 | -1.8886095 | 0.49514864 | -3.8142274 | 0.00013661 | 0.01825353 |
| TBRG1 | 43.3511662 | 0.05846891 | 0.29327426 | 0.19936598 | 0.84197647 | 0.99692952 |
| SIAE | 20.4171373 | 0.26638876 | 0.34731872 | 0.76698648 | 0.44308956 | 0.98092839 |
| NRGN | 39.5830232 | 1.43246104 | 0.41414565 | 3.458834 | 5.43E-04 | 0.04787234 |
| ESAM | 21.6919576 | 0.3995094 | 0.35333 | 1.13069765 | 0.25818237 | 0.92891801 |
| MSANTD2 | 43.8398798 | 0.09729885 | 0.2895214 | 0.33606791 | 0.73681966 | 0.9958918 |
| PKNOX2 | 78.2402783 | -0.1859137 | 0.41717709 | -0.4456469 | 0.65585233 | 0.9958918 |
| EI24 | 26.6773162 | 0.23621176 | 0.40662076 | 0.58091417 | 0.5612983 | 0.9958918 |
| STT3A | 29.3427542 | -0.1682628 | 0.29145592 | -0.577318 | 0.56372465 | 0.9958918 |
| FAM118B | 19.9289088 | -0.0603178 | 0.35303889 | -0.1708532 | 0.86433917 | 0.9971731 |

| | | | | | | |
|---|---|---|---|---|---|---|
| SRPRA | 27.6511075 | 0.01385063 | 0.37673984 | 0.03676444 | 0.97067282 | 0.99935941 |
| ST3GAL4 | 38.4613454 | 0.52034669 | 0.45406944 | 1.1459628 | 0.25181054 | 0.92631316 |
| ETS1 | 207.199097 | 0.06946581 | 0.29893162 | 0.23238026 | 0.81624268 | 0.99592423 |
| FLI1 | 167.09341 | 0.20420315 | 0.355656 | 0.57415915 | 0.56586011 | 0.9958918 |
| ARHGAP32 | 156.920152 | -0.290663 | 0.27958095 | -1.0396382 | 0.29850803 | 0.95447177 |
| NFRKB | 20.8260188 | 0.36650847 | 0.31638935 | 1.15840963 | 0.24669691 | 0.92600668 |
| PRDM10 | 28.0277945 | -0.3412613 | 0.32179018 | -1.0605088 | 0.2889132 | 0.95126484 |
| APLP2 | 127.156547 | 0.12286971 | 0.20597267 | 0.59653401 | 0.55081854 | 0.9958918 |
| ZBTB44 | 71.3831959 | 0.0317556 | 0.29028692 | 0.10939382 | 0.91289013 | 0.99935941 |
| SNX19 | 56.4947794 | -0.2939941 | 0.31256499 | -0.9405856 | 0.34691726 | 0.96887708 |
| NTM | 72.2353422 | -0.1649859 | 0.50408662 | -0.3272966 | 0.74344354 | 0.9958918 |
| JAM3 | 68.0997959 | 0.33517939 | 0.25590951 | 1.30975749 | 0.19027789 | 0.88158763 |
| VPS26B | 21.7179588 | 0.56638663 | 0.35677441 | 1.58752033 | 0.11239485 | 0.77793929 |
| ZMYND11 | 134.992885 | 0.13760636 | 0.20181434 | 0.68184629 | 0.49533615 | 0.99176652 |
| DIP2C | 304.218118 | -0.3204254 | 0.29951884 | -1.0698004 | 0.28470916 | 0.94505166 |
| LARP4B | 146.824383 | -0.1730965 | 0.22534761 | -0.7681309 | 0.44240945 | 0.98092839 |
| GTPBP4 | 26.9594026 | -0.1427379 | 0.46511732 | -0.3068857 | 0.75893038 | 0.9958918 |
| IDI1 | 20.5859142 | -0.0571199 | 0.43711558 | -0.1306745 | 0.89603281 | 0.99935941 |
| WDR37 | 52.2677647 | -0.0665908 | 0.23038972 | -0.2890356 | 0.77255411 | 0.9958918 |
| PITRM1 | 36.9498656 | 0.04097875 | 0.37923629 | 0.10805597 | 0.91395129 | 0.99935941 |
| KLF6 | 108.286999 | -0.2227983 | 0.41769553 | -0.5333987 | 0.5937576 | 0.9958918 |
| NET1 | 36.079053 | 1.18421055 | 0.36071456 | 3.28295749 | 0.00102724 | 0.07036431 |
| FAM208B | 169.686192 | 0.18322439 | 0.29360794 | 0.62404441 | 0.53259842 | 0.9958918 |
| GDI2 | 92.5104489 | 0.46446782 | 0.24769294 | 1.87517586 | 0.06076853 | 0.61696136 |
| ANKRD16 | 25.5276795 | 0.42848444 | 0.42405927 | 1.01043525 | 0.31228681 | 0.9599143 |
| FBH1 | 85.1573137 | 0.61482664 | 0.2225119 | 2.76311807 | 0.00572521 | 0.1903825 |
| IL15RA | 29.5659737 | 1.1529687 | 0.5859252 | 1.96777457 | 0.04909398 | 0.57273715 |
| RBM17 | 63.8397162 | 0.39239973 | 0.34855617 | 1.12578619 | 0.26025603 | 0.93055907 |
| PFKFB3 | 94.0232933 | -0.3635958 | 0.35311516 | -1.0296806 | 0.30315995 | 0.95650251 |
| SFMBT2 | 194.625592 | -0.4397359 | 0.55737996 | -0.7889337 | 0.43015074 | 0.98092839 |
| ITIH5 | 64.3000219 | 0.42356151 | 0.5464084 | 0.77517386 | 0.43823693 | 0.98092839 |
| KIN | 17.6222782 | 0.14047445 | 0.38057598 | 0.36911012 | 0.71204565 | 0.9958918 |
| ATP5F1C | 50.2962989 | 0.36763143 | 0.25873992 | 1.42085318 | 0.15535945 | 0.84277602 |
| TAF3 | 124.835425 | 0.28232033 | 0.19550219 | 1.44407755 | 0.14871716 | 0.8301633 |
| GATA3 | 85.0348857 | 2.26748368 | 0.34908936 | 6.49542475 | 8.28E-11 | 6.08E-07 |
| CELF2 | 106.657705 | 0.28389321 | 0.4493193 | 0.63182953 | 0.52749827 | 0.9958918 |
| USP6NL | 80.7872175 | -0.089936 | 0.29550223 | -0.3043497 | 0.76086152 | 0.9958918 |
| PROSER2 | 31.7851345 | -1.2640121 | 0.60104518 | -2.1030234 | 0.03546372 | 0.50704843 |
| UPF2 | 75.3846713 | 0.14738636 | 0.20731021 | 0.71094599 | 0.4771177 | 0.98993168 |
| DHTKD1 | 28.2235571 | 0.22229572 | 0.34546777 | 0.64346296 | 0.51992374 | 0.9958918 |
| SEC61A2 | 17.6009044 | -0.1891737 | 0.39715539 | -0.4763215 | 0.63384535 | 0.9958918 |
| NUDT5 | 31.0662358 | 0.12708424 | 0.30647315 | 0.41466681 | 0.67838583 | 0.9958918 |
| CDC123 | 32.7052637 | -0.1862091 | 0.31302176 | -0.594876 | 0.55192634 | 0.9958918 |
| CAMK1D | 238.388078 | -0.0448125 | 0.40081301 | -0.1118039 | 0.9109789 | 0.99935941 |
| CCDC3 | 72.9625333 | 0.34449524 | 0.55339769 | 0.62250936 | 0.53360699 | 0.9958918 |
| OPTN | 70.7572532 | -0.1565773 | 0.31222135 | -0.5014944 | 0.61602323 | 0.9958918 |
| BEND7 | 35.7525127 | 0.25318364 | 0.38382187 | 0.65963839 | 0.50948591 | 0.99507991 |
| AL157392.5 | 22.3722729 | 0.20059594 | 0.34596488 | 0.5798159 | 0.56203877 | 0.9958918 |
| FRMD4A | 367.718117 | 0.61966047 | 0.3882038 | 1.59622464 | 0.11043865 | 0.77370222 |
| FAM107B | 89.8367456 | 0.51083805 | 0.33135919 | 1.54164443 | 0.12316002 | 0.79184923 |
| DCLRE1C | 22.2154861 | 0.02748248 | 0.33334681 | 0.08244411 | 0.93429356 | 0.99935941 |
| RPP38 | 28.4115295 | 0.30957913 | 0.35844461 | 0.86367354 | 0.38776725 | 0.97297628 |
| NMT2 | 34.5795742 | 0.38833093 | 0.34191398 | 1.13575623 | 0.2560586 | 0.9266065 |
| FAM171A1 | 34.2534267 | 0.1815205 | 0.36521411 | 0.49702488 | 0.6191715 | 0.9958918 |

| | | | | | | |
|---|---|---|---|---|---|---|
| ITGA8 | 345.971714 | 0.43154583 | 0.55543769 | 0.77694733 | 0.43718985 | 0.98092839 |
| MINDY3 | 81.6954387 | 0.01447441 | 0.22552343 | 0.06418138 | 0.9488258 | 0.99935941 |
| RSU1 | 167.921863 | 0.07422385 | 0.34098057 | 0.21767766 | 0.82768027 | 0.99592423 |
| CUBN | 47.8603239 | 1.19698103 | 0.53081287 | 2.25499625 | 0.02413356 | 0.42027858 |
| VIM | 332.103477 | 0.36964322 | 0.36995965 | 0.99914467 | 0.31772462 | 0.95992644 |
| STAM | 44.0021957 | -0.1026595 | 0.24707273 | -0.4155033 | 0.67777349 | 0.9958918 |
| CACNB2 | 430.344468 | -0.0081915 | 0.38085905 | -0.0215081 | 0.98284036 | 0.99935941 |
| AL450384.2 | 24.6249509 | -0.3316316 | 0.35360426 | -0.937861 | 0.34831586 | 0.96887708 |
| NSUN6 | 48.8215928 | -0.3016447 | 0.23707642 | -1.2723522 | 0.20324799 | 0.89396747 |
| ARL5B | 28.1686121 | -0.8884693 | 0.46403439 | -1.9146627 | 0.05553554 | 0.59755589 |
| MALRD1 | 46.9735871 | -0.9196225 | 0.93126814 | -0.9874949 | 0.32340008 | 0.95992644 |
| PLXDC2 | 641.887814 | -0.1883273 | 0.42756106 | -0.4404688 | 0.65959759 | 0.9958918 |
| NEBL | 47.8149825 | -1.5706299 | 0.67728862 | -2.3189964 | 0.02039523 | 0.38630036 |
| MLLT10 | 237.765875 | -0.4502017 | 0.21165747 | -2.1270297 | 0.03341761 | 0.492839 |
| DNAJC1 | 257.837377 | -0.2043285 | 0.28326791 | -0.721326 | 0.47070898 | 0.98910524 |
| PIP4K2A | 259.545264 | 0.24247286 | 0.30922469 | 0.78413163 | 0.43296288 | 0.98092839 |
| MSRB2 | 29.4577208 | -0.2422798 | 0.32720144 | -0.7404606 | 0.45902056 | 0.98508941 |
| KIAA1217 | 997.648772 | -0.1773733 | 0.44397781 | -0.3995093 | 0.68951794 | 0.9958918 |
| ARHGAP21 | 195.359834 | 0.1725993 | 0.20741717 | 0.83213604 | 0.40533216 | 0.97637555 |
| ABI1 | 127.613186 | -0.0471933 | 0.30903181 | -0.1527134 | 0.87862427 | 0.99722158 |
| ANKRD26 | 67.6749833 | -0.0783274 | 0.26725033 | -0.2930862 | 0.76945624 | 0.9958918 |
| YME1L1 | 91.4907974 | -0.1811672 | 0.21641324 | -0.8371355 | 0.40251641 | 0.97512144 |
| ACBD5 | 22.4059401 | 0.21854945 | 0.34065786 | 0.64155117 | 0.52116464 | 0.9958918 |
| RAB18 | 72.3738421 | -0.1361865 | 0.22926882 | -0.5940034 | 0.55250985 | 0.9958918 |
| MPP7 | 35.4892082 | -0.476154 | 0.56245203 | -0.8465681 | 0.3972359 | 0.97500135 |
| WAC-AS1 | 21.3256372 | -0.2556395 | 0.45764824 | -0.5585938 | 0.57643895 | 0.9958918 |
| WAC | 314.626183 | -0.206264 | 0.20337536 | -1.0142035 | 0.31048568 | 0.95956408 |
| SVIL | 244.773624 | 0.6708516 | 0.4541689 | 1.47709717 | 0.13964959 | 0.8153985 |
| JCAD | 19.4751754 | 1.01076109 | 0.35380167 | 2.85685785 | 0.00427857 | 0.16207856 |
| MTPAP | 19.893444 | -0.0242764 | 0.32973946 | -0.0736231 | 0.94131028 | 0.99935941 |
| MAP3K8 | 98.6666914 | 0.36866039 | 0.40437308 | 0.91168382 | 0.36193518 | 0.97217293 |
| ZNF438 | 78.4738645 | -0.0967305 | 0.31530449 | -0.3067844 | 0.75900745 | 0.9958918 |
| ZEB1-AS1 | 26.3774254 | 0.04907423 | 0.36287606 | 0.13523688 | 0.89242459 | 0.99935941 |
| ZEB1 | 400.716102 | -0.1034106 | 0.26756067 | -0.386494 | 0.69913085 | 0.9958918 |
| ARHGAP12 | 90.6107693 | 0.04490274 | 0.3662702 | 0.12259457 | 0.90242815 | 0.99935941 |
| KIF5B | 114.816492 | 0.07633985 | 0.23087052 | 0.3306609 | 0.74090064 | 0.9958918 |
| EPC1 | 209.242693 | 0.30260597 | 0.20151088 | 1.50168553 | 0.13317834 | 0.80664435 |
| CCDC7 | 89.8115108 | -0.2030329 | 0.29414963 | -0.6902368 | 0.4900453 | 0.99046213 |
| ITGB1 | 338.444287 | -0.0603671 | 0.24164236 | -0.2498198 | 0.80272668 | 0.99592423 |
| NRP1 | 979.043201 | 0.27521415 | 0.38759874 | 0.71004914 | 0.47767366 | 0.99002719 |
| PARD3 | 871.855193 | -0.6181962 | 0.30511242 | -2.0261261 | 0.04275186 | 0.54851863 |
| CUL2 | 42.1816427 | -0.0203593 | 0.25055553 | -0.0812568 | 0.93523775 | 0.99935941 |
| CREM | 95.8783439 | -0.0953057 | 0.33218434 | -0.2869061 | 0.77418425 | 0.9958918 |
| CCNY | 247.768935 | 0.06830594 | 0.18097735 | 0.37742812 | 0.70585547 | 0.9958918 |
| AL117336.3 | 19.6518317 | 0.61911456 | 0.49639036 | 1.24723324 | 0.21231199 | 0.89738044 |
| ZNF248 | 61.6188829 | 0.19392437 | 0.27574921 | 0.70326354 | 0.48189153 | 0.99002719 |
| ZNF25 | 19.937468 | -0.0182219 | 0.34006432 | -0.0535838 | 0.9572668 | 0.99935941 |
| ZNF33A | 77.7603401 | 0.23406579 | 0.32657805 | 0.71672236 | 0.47354543 | 0.98910524 |
| ZNF33B | 29.18472 | 0.39782366 | 0.46615554 | 0.85341398 | 0.39342977 | 0.97500135 |
| BMS1 | 38.8713449 | -0.5297478 | 0.28046186 | -1.8888409 | 0.05891315 | 0.61151513 |
| CSGALNACT2 | 58.195527 | 0.16714594 | 0.23076005 | 0.72432792 | 0.46886444 | 0.98910524 |
| HNRNPF | 66.2934612 | 0.04565632 | 0.32928118 | 0.13865451 | 0.88972317 | 0.99935941 |
| CXCL12 | 89.6951069 | -0.2703183 | 0.33617799 | -0.8040928 | 0.4213434 | 0.97812179 |
| RASSF4 | 23.1087743 | 0.25047443 | 0.34101833 | 0.73448964 | 0.46265037 | 0.98590351 |

| | | | | | | |
|---|---|---|---|---|---|---|
| DEPP1 | 103.61596 | -0.017422 | 0.31618783 | -0.0551002 | 9.56E-01 | 9.99E-01 |
| 8-Mar | 41.1044156 | 0.28456299 | 0.31783548 | 0.89531538 | 0.37061852 | 0.97217293 |
| ZFAND4 | 27.9937868 | 0.26782073 | 0.33571184 | 0.7977697 | 0.42500415 | 0.97917873 |
| WASHC2C | 23.6683547 | -0.2076711 | 0.39131858 | -0.5306957 | 0.59562969 | 0.9958918 |
| AGAP4 | 26.376086 | -0.0309547 | 0.33441684 | -0.0925632 | 0.92625057 | 0.99935941 |
| TIMM23 | 26.7917022 | -0.136774 | 0.32212663 | -0.4245971 | 0.67113044 | 0.9958918 |
| NCOA4 | 30.9681132 | 0.35341807 | 0.30940686 | 1.14224379 | 0.2533527 | 0.92646125 |
| AGAP9 | 18.5363737 | 1.18833413 | 0.38498958 | 3.08666574 | 0.00202415 | 0.09597086 |
| MAPK8 | 157.44417 | 0.1142194 | 0.19548391 | 0.58429053 | 0.55902486 | 0.9958918 |
| VSTM4 | 50.8702594 | -0.5596253 | 0.43864067 | -1.2758173 | 0.20202012 | 0.89319989 |
| ERCC6 | 41.1821226 | -0.0488559 | 0.39593654 | -0.1233932 | 0.90179571 | 0.99935941 |
| PARG | 87.5632666 | -0.2002076 | 0.20184897 | -0.9918686 | 0.32126165 | 0.95992644 |
| TIMM23B | 81.3266794 | -0.0686183 | 0.35076565 | -0.1956244 | 0.84490417 | 0.99692952 |
| LINC00843 | 20.2008134 | 0.15865401 | 0.44324179 | 0.3579401 | 0.72038814 | 0.9958918 |
| AGAP6 | 28.5867986 | 0.36862648 | 0.35030813 | 1.05229213 | 0.29266554 | 0.95186143 |
| WASHC2A | 20.0400757 | -0.7634402 | 0.4249539 | -1.7965248 | 0.07241109 | 0.66435596 |
| SGMS1 | 165.919283 | 0.12592904 | 0.20314472 | 0.6198982 | 0.53532481 | 0.9958918 |
| PRKG1 | 3030.40969 | 0.12800036 | 0.33749422 | 0.37926681 | 0.70448974 | 0.9958918 |
| IPMK | 34.5868035 | -0.2281124 | 0.37272569 | -0.6120114 | 0.54053021 | 0.9958918 |
| CISD1 | 33.0900144 | -0.2379943 | 0.33960041 | -0.7008069 | 0.48342353 | 0.99015594 |
| UBE2D1 | 25.1857936 | 0.27845533 | 0.32944124 | 0.84523519 | 0.39797954 | 0.97500135 |
| BICC1 | 413.89479 | -0.5100952 | 0.28512712 | -1.7890096 | 0.07361327 | 0.66886321 |
| FAM13C | 50.5987518 | -0.4561757 | 1.04692832 | -0.4357277 | 0.66303428 | 0.9958918 |
| CCDC6 | 78.8057108 | -0.4369372 | 0.31970534 | -1.3666873 | 0.17172335 | 0.87233975 |
| ANK3 | 283.11314 | 1.22608469 | 0.35001768 | 3.50292219 | 0.00046018 | 0.04227364 |
| RHOBTB1 | 71.5064597 | 0.11995171 | 0.31499831 | 0.38080111 | 0.70335084 | 0.9958918 |
| TMEM26 | 24.7588383 | -0.7949549 | 0.45721029 | -1.7387074 | 0.08208625 | 0.69958018 |
| ARID5B | 485.778531 | 0.38876536 | 0.33025006 | 1.17718483 | 0.23912174 | 0.92392517 |
| JMJD1C | 818.101857 | -0.4398231 | 0.23107966 | -1.9033396 | 0.05699624 | 0.60440467 |
| REEP3 | 124.530132 | -0.2268032 | 0.26375444 | -0.8599027 | 0.38984266 | 0.97297628 |
| CTNNA3 | 83.9379934 | -0.9584498 | 0.24768422 | -3.869644 | 0.00010899 | 0.01602 |
| SIRT1 | 39.5610612 | -0.0269216 | 0.33096514 | -0.0813428 | 0.93516937 | 0.99935941 |
| HERC4 | 301.426679 | 0.09707597 | 0.15468891 | 0.62755609 | 0.53029478 | 0.9958918 |
| HNRNPH3 | 86.1079894 | -0.1184382 | 0.29024685 | -0.4080603 | 0.68322937 | 0.9958918 |
| RUFY2 | 76.2147324 | -0.2338443 | 0.26577202 | -0.879868 | 0.3789308 | 0.97217293 |
| SLC25A16 | 53.8109547 | -0.0534283 | 0.21797186 | -0.2451156 | 0.80636691 | 0.99592423 |
| TET1 | 39.6936322 | -0.7601397 | 0.41275497 | -1.8416246 | 0.06553008 | 0.63034109 |
| CCAR1 | 119.749094 | -0.1861281 | 0.17919927 | -1.0386654 | 0.29896037 | 0.9544725 |
| DDX50 | 56.0528244 | 0.28096561 | 0.28210032 | 0.99597763 | 0.31926102 | 0.95992644 |
| DDX21 | 72.9433048 | -0.037883 | 0.50948593 | -0.0743553 | 0.94072768 | 0.99935941 |
| SRGN | 152.053365 | 0.16428154 | 0.59871837 | 0.27438868 | 0.78378594 | 0.9958918 |
| VPS26A | 54.2490597 | -0.6126579 | 0.2342071 | -2.6158808 | 0.00889976 | 0.25292422 |
| SUPV3L1 | 19.8391098 | -0.0824311 | 0.34931108 | -0.2359818 | 0.81344679 | 0.99592423 |
| HK1 | 107.830351 | 0.59133127 | 0.31878844 | 1.8549332 | 0.06360577 | 0.62395844 |
| SAR1A | 49.2718103 | -0.4876388 | 0.27852964 | -1.7507611 | 0.07998707 | 0.69237339 |
| PPA1 | 52.8240683 | 0.23293603 | 0.39064156 | 0.59629096 | 0.55098087 | 0.9958918 |
| EIF4EBP2 | 43.2095269 | 0.23377145 | 0.24793806 | 0.94286231 | 0.34575133 | 0.96887708 |
| PALD1 | 22.3473355 | 0.80638288 | 0.42210657 | 1.91037749 | 0.05608463 | 0.60041061 |
| SGPL1 | 51.4711619 | -0.0989069 | 0.28486789 | -0.3472026 | 0.72843911 | 0.9958918 |
| UNC5B | 43.058491 | 0.41460102 | 0.46425733 | 0.89304141 | 0.371835 | 0.97217293 |
| VSIR | 20.013918 | -0.3365702 | 0.41487974 | -0.8112476 | 0.41722347 | 0.97812179 |
| PSAP | 189.781856 | 0.11251193 | 0.23475109 | 0.47928183 | 0.63173815 | 0.9958918 |
| CHST3 | 27.0116883 | 0.49889639 | 0.46944718 | 1.06273169 | 0.28790365 | 0.94955775 |
| ASCC1 | 74.2529822 | -0.0014989 | 0.28313514 | -0.0052939 | 0.99577613 | 0.99935941 |

| | | | | | | |
|---|---|---|---|---|---|---|
| ANAPC16 | 106.449787 | 0.54937338 | 0.23041512 | 2.38427667 | 0.01711273 | 0.34765232 |
| DDIT4 | 131.735072 | -1.1782692 | 0.92139059 | -1.2787944 | 0.20096945 | 0.89240151 |
| DNAJB12 | 22.3371148 | -0.100214 | 0.38010248 | -0.2636498 | 0.79204977 | 0.99592423 |
| MICU1 | 173.037346 | -0.1243289 | 0.30812816 | -0.4034974 | 0.68658237 | 0.9958918 |
| MCU | 170.685464 | 0.027976 | 0.46093149 | 0.06069448 | 0.95160253 | 0.99935941 |
| P4HA1 | 228.310755 | -0.741003 | 0.55379355 | -1.3380492 | 0.1808804 | 0.87815793 |
| ECD | 31.3178043 | -0.3246045 | 0.36945882 | -0.8785945 | 0.37962115 | 0.97217293 |
| FAM149B1 | 29.9601085 | 0.21322371 | 0.41909134 | 0.50877622 | 0.61090909 | 0.9958918 |
| DNAJC9 | 28.4702454 | -0.0974013 | 0.38498642 | -0.2529994 | 0.80026871 | 0.99592423 |
| MRPS16 | 22.7653671 | -0.2017735 | 0.36131355 | -0.5584444 | 0.57654095 | 0.9958918 |
| ANXA7 | 67.0038179 | 0.21201467 | 0.27731951 | 0.76451408 | 0.44456095 | 0.98092839 |
| PPP3CB | 110.515705 | -0.037474 | 0.26547272 | -0.1411595 | 0.88774391 | 0.99935941 |
| USP54 | 68.4689074 | -0.0245085 | 0.32029298 | -0.0765191 | 0.93900615 | 0.99935941 |
| SEC24C | 34.5334323 | 0.13167835 | 0.283367 | 0.4646919 | 0.64215211 | 0.9958918 |
| FUT11 | 25.1641029 | 0.40195488 | 0.394342 | 1.01930527 | 0.30805806 | 0.95793395 |
| ZSWIM8 | 44.2803798 | 0.14807261 | 0.27424941 | 0.53991953 | 0.58925253 | 0.9958918 |
| CAMK2G | 49.8063229 | 0.34106563 | 0.51136191 | 0.66697504 | 0.50478808 | 0.99387859 |
| VCL | 430.872233 | -0.3072121 | 0.37450751 | -0.8203095 | 0.4120397 | 0.97812179 |
| AP3M1 | 19.0262141 | 0.05060639 | 0.38307302 | 0.13210639 | 0.89490014 | 0.99935941 |
| ADK | 223.693941 | -0.0412433 | 0.26559721 | -0.1552852 | 0.8765965 | 0.9971731 |
| KAT6B | 187.497769 | -0.331206 | 0.24673232 | -1.3423697 | 0.17947615 | 0.87815793 |
| SAMD8 | 81.675011 | -0.1348018 | 0.26656388 | -0.5057017 | 0.61306605 | 0.9958918 |
| VDAC2 | 31.8125158 | 0.19981621 | 0.30156438 | 0.66259885 | 0.50758751 | 0.99397937 |
| LRMDA | 149.618925 | 0.08685593 | 0.3271571 | 0.26548693 | 0.79063438 | 0.99592423 |
| KCNMA1 | 102.922044 | 1.60298381 | 0.88598706 | 1.80926322 | 0.07041012 | 0.65633299 |
| DLG5 | 88.6975766 | 0.08976542 | 0.29270506 | 0.30667534 | 0.7590905 | 0.9958918 |
| POLR3A | 30.5989436 | -0.3114121 | 0.32165578 | -0.9681534 | 0.33296778 | 0.9654289 |
| RPS24 | 168.1264 | -0.3029833 | 0.32646136 | -0.9280831 | 0.35336445 | 0.97217293 |
| ZMIZ1 | 156.483621 | 0.36336583 | 0.30559735 | 1.18903461 | 0.23442605 | 0.91930824 |
| ZCCHC24 | 79.7014214 | -0.2847752 | 0.23096126 | -1.2329999 | 0.21757582 | 0.90370709 |
| NUTM2B-AS | 153.020064 | -0.1010857 | 0.28731567 | -0.3518279 | 0.72496736 | 0.9958918 |
| PLAC9 | 33.4401513 | 0.0433855 | 0.49060153 | 0.08843328 | 0.92953231 | 0.99935941 |
| ANXA11 | 83.1553642 | 0.17709731 | 0.25609945 | 0.69151776 | 0.48924023 | 0.99046213 |
| FAM213A | 42.8264915 | -1.1471973 | 0.45730473 | -2.5086059 | 0.01212086 | 0.2958939 |
| TSPAN14 | 59.8613348 | -0.2631358 | 0.2790826 | -0.9428599 | 0.34575259 | 0.96887708 |
| NRG3 | 98.8155093 | 2.16077616 | 0.77938002 | 2.7724295 | 0.00556396 | 0.18713567 |
| GHITM | 34.2595541 | 0.57429194 | 0.29640279 | 1.93753891 | 0.0526795 | 0.58392407 |
| CCSER2 | 150.23127 | -0.0420353 | 0.19617322 | -0.2142767 | 0.83033131 | 0.99592423 |
| GRID1 | 68.8267014 | 0.22386518 | 0.36270562 | 0.61720901 | 0.53709687 | 0.9958918 |
| WAPL | 102.88207 | -0.1263147 | 0.20956873 | -0.6027362 | 0.54668417 | 0.9958918 |
| LDB3 | 43.2426795 | 0.49287265 | 0.4963112 | 0.99307178 | 0.32067497 | 0.95992644 |
| BMPR1A | 236.415365 | 0.29166689 | 0.26513705 | 1.10006085 | 0.27130561 | 0.93776436 |
| ADIRF | 344.598615 | 1.02986914 | 0.48897351 | 2.10618597 | 0.0351882 | 0.5060628 |
| GLUD1 | 74.853303 | 0.24704932 | 0.4031874 | 0.61274067 | 0.54004783 | 0.9958918 |
| SHLD2 | 88.3771621 | 0.06692488 | 0.30206376 | 0.22155877 | 0.82465738 | 0.99592423 |
| NUTM2A-AS | 134.671767 | -0.1102239 | 0.20894603 | -0.5275235 | 0.59783011 | 0.9958918 |
| PAPSS2 | 90.6873064 | 0.10508174 | 0.45374096 | 0.2315897 | 0.8168567 | 0.99592423 |
| ATAD1 | 58.6703049 | -0.1576432 | 0.26025739 | -0.6057205 | 0.54470035 | 0.9958918 |
| PTEN | 688.053289 | -0.2291013 | 0.35351409 | -0.6480683 | 0.51694079 | 0.9958918 |
| RNLS | 70.7489436 | -0.4775965 | 0.2780713 | -1.7175324 | 0.08588194 | 0.70916334 |
| STAMBPL1 | 37.9415529 | -0.5806952 | 0.37436614 | -1.5511425 | 0.12086754 | 0.79021701 |
| ACTA2 | 605.625497 | 0.38981984 | 0.46392336 | 0.84026775 | 0.40075828 | 0.97500135 |
| FAS | 50.0611634 | -0.2160601 | 0.46107869 | -0.4685971 | 0.63935764 | 0.9958918 |
| LIPA | 39.7309704 | -0.3701947 | 0.26662944 | -1.3884238 | 0.16500803 | 0.85881304 |

| | | | | | | |
|---|---|---|---|---|---|---|
| IFIT3 | 47.7059503 | -2.0240845 | 1.1935326 | -1.695877 | 0.08990917 | 0.72305504 |
| RPP30 | 17.7330153 | -0.0285837 | 0.44033 | -0.0649143 | 0.9482422 | 0.99935941 |
| PCGF5 | 269.950027 | -0.5716135 | 0.25397706 | -2.25065 | 0.02440771 | 0.42312555 |
| HECTD2 | 86.7901352 | -0.183731 | 0.32672936 | -0.5623341 | 0.57388843 | 0.9958918 |
| TNKS2 | 109.745923 | -0.2172441 | 0.21101394 | -1.029525 | 0.30323304 | 0.95650251 |
| BTAF1 | 146.332279 | -0.2773892 | 0.28947935 | -0.9582348 | 0.33794437 | 0.96750939 |
| CPEB3 | 184.606839 | -0.0715038 | 0.285322 | -0.2506072 | 0.8021178 | 0.99592423 |
| 5-Mar | 64.6307438 | -0.0001535 | 0.20952878 | -0.0007325 | 0.99941555 | 0.99968761 |
| IDE | 60.918439 | -0.2100635 | 0.31499597 | -0.6668768 | 0.50485084 | 0.99387859 |
| EXOC6 | 45.2163312 | 0.08986165 | 0.35994159 | 0.24965621 | 0.80285323 | 0.99592423 |
| MYOF | 245.033052 | -0.491875 | 0.5606687 | -0.8773006 | 0.3803234 | 0.97217293 |
| FRA10AC1 | 43.2105915 | -0.2405491 | 0.34739713 | -0.6924325 | 0.48866574 | 0.99046213 |
| PLCE1 | 147.915278 | 0.11562679 | 0.36326281 | 0.31830065 | 0.75025689 | 0.9958918 |
| NOC3L | 23.3431493 | -0.1683434 | 0.42884384 | -0.3925519 | 0.6946505 | 0.9958918 |
| TBC1D12 | 70.3989279 | 0.21903889 | 0.23688071 | 0.92468017 | 0.3551323 | 0.97217293 |
| PDLIM1 | 115.259391 | -0.1055462 | 0.43006599 | -0.2454187 | 0.80613227 | 0.99592423 |
| SORBS1 | 924.392706 | 0.34730293 | 0.48250826 | 0.7197865 | 0.47165646 | 0.98910524 |
| ALDH18A1 | 29.1443382 | -0.0934577 | 0.30345674 | -0.3079771 | 0.75809974 | 0.9958918 |
| TCTN3 | 32.9904018 | 0.56167628 | 0.35466514 | 1.58368053 | 0.11326644 | 0.77793929 |
| ENTPD1 | 31.8783791 | -0.2921352 | 0.53543097 | -0.5456076 | 0.58533573 | 0.9958918 |
| ENTPD1-AS1 | 110.822269 | 0.03943057 | 0.28507144 | 0.1383182 | 0.88998894 | 0.99935941 |
| AL365273.2 | 21.5107052 | 0.08270718 | 0.40954306 | 0.2019499 | 0.83995589 | 0.99692952 |
| ZNF518A | 80.6301969 | -0.308019 | 0.28641444 | -1.0754312 | 0.28218173 | 0.94419503 |
| TM9SF3 | 142.445326 | -0.117755 | 0.22075243 | -0.5334255 | 0.59373907 | 0.9958918 |
| LCOR | 176.116975 | -0.1317821 | 0.2082917 | -0.6326807 | 0.52694219 | 0.9958918 |
| PGAM1 | 51.3209215 | 0.20816945 | 0.32820506 | 0.63426642 | 0.52590697 | 0.9958918 |
| EXOSC1 | 22.5172664 | 0.38731085 | 0.31040132 | 1.24777451 | 0.21211365 | 0.89738044 |
| MMS19 | 40.4744884 | -0.0163035 | 0.25634757 | -0.0635991 | 0.94928943 | 0.99935941 |
| UBTD1 | 20.6640377 | -0.1561549 | 0.34322436 | -0.4549646 | 0.64913471 | 0.9958918 |
| MARVELD1 | 21.9186568 | -0.0532521 | 0.44376442 | -0.1200009 | 0.90448246 | 0.99935941 |
| ZFYVE27 | 22.5390951 | -0.0076367 | 0.29030127 | -0.0263062 | 0.97901313 | 0.99935941 |
| R3HCC1L | 52.1641084 | -0.2179294 | 0.2244981 | -0.9707406 | 0.33167747 | 0.9654289 |
| HPS1 | 25.9610372 | 0.09964576 | 0.27968204 | 0.35628229 | 0.72162917 | 0.9958918 |
| HPSE2 | 338.546166 | -0.0176744 | 0.64919923 | -0.0272249 | 0.97828032 | 0.99935941 |
| GOT1 | 21.7879365 | 0.07806113 | 0.36685039 | 0.21278738 | 0.83149281 | 0.99592423 |
| SLC25A28 | 31.3869096 | -0.2870242 | 0.44336554 | -0.6473759 | 0.5173887 | 0.9958918 |
| CUTC | 28.0730527 | 0.08514468 | 0.30884949 | 0.27568339 | 0.78279125 | 0.9958918 |
| COX15 | 22.9842925 | -0.3951678 | 0.30607537 | -1.29108 | 0.19667593 | 0.88391893 |
| DNMBP | 64.0676234 | -0.2436443 | 0.36843483 | -0.6612955 | 0.50842282 | 0.99397937 |
| ERLIN1 | 20.6637482 | -0.052979 | 0.31148422 | -0.1700858 | 0.8649427 | 0.9971731 |
| CHUK | 27.1493273 | -0.0744698 | 0.38180103 | -0.1950487 | 0.84535481 | 0.99692952 |
| CWF19L1 | 28.8792813 | -0.1509553 | 0.3043861 | -0.4959337 | 0.61994119 | 0.9958918 |
| HIF1AN | 19.2743581 | -0.1619259 | 0.35808558 | -0.4521988 | 0.65112579 | 0.9958918 |
| SLF2 | 93.0094384 | -0.1382218 | 0.34098615 | -0.405359 | 0.68521367 | 0.9958918 |
| LZTS2 | 38.4971853 | -0.4786933 | 0.27700418 | -1.7281085 | 0.08396878 | 0.70248581 |
| BTRC | 131.599279 | 0.17461282 | 0.2506335 | 0.69668587 | 0.4859994 | 0.99046213 |
| FBXW4 | 107.54095 | 0.0930796 | 0.25959373 | 0.35855874 | 0.71992522 | 0.9958918 |
| OGA | 200.698187 | -0.1126394 | 0.29736127 | -0.3787964 | 0.70483905 | 0.9958918 |
| ARMH3 | 109.637453 | -0.0679172 | 0.18683066 | -0.3635228 | 0.71621436 | 0.9958918 |
| LDB1 | 20.0587597 | -0.114816 | 0.44608185 | -0.2573878 | 0.79687943 | 0.99592423 |
| PPRC1 | 25.2448589 | -0.7480046 | 0.52534027 | -1.4238479 | 0.1544905 | 0.84224829 |
| NOLC1 | 19.7124653 | 0.02432225 | 0.47238423 | 0.05148828 | 0.95893644 | 0.99935941 |
| GBF1 | 180.904564 | -0.094889 | 0.19886375 | -0.4771557 | 0.63325123 | 0.9958918 |
| NFKB2 | 23.3175615 | 0.83676763 | 0.59494727 | 1.4064568 | 0.15958853 | 0.84762439 |

| | | | | | | |
|---|---|---|---|---|---|---|
| ACTR1A | 26.4847552 | -0.1953942 | 0.35904768 | -0.5442012 | 0.586303 | 0.9958918 |
| SUFU | 42.7365837 | 0.68525038 | 0.2451287 | 2.79547185 | 0.0051824 | 0.18276799 |
| TRIM8 | 35.2301008 | 0.11369299 | 0.34194327 | 0.33249081 | 0.73951868 | 0.9958918 |
| ARL3 | 32.1941141 | 0.18679523 | 0.30987178 | 0.60281461 | 0.54663203 | 0.9958918 |
| WBP1L | 123.49861 | 0.24785535 | 0.25861386 | 0.9583993 | 3.38E-01 | 0.96750939 |
| CNNM2 | 122.783358 | 0.15209587 | 0.31893203 | 0.47689116 | 0.63343963 | 0.9958918 |
| NT5C2 | 188.513942 | -0.0609234 | 0.1885265 | -0.3231555 | 0.74657746 | 0.9958918 |
| PCGF6 | 27.1539157 | -0.1753483 | 0.31986022 | -0.5482028 | 0.58355265 | 0.9958918 |
| ATP5MD | 20.6809828 | -0.6500869 | 0.34726089 | -1.8720418 | 0.06120082 | 0.61696136 |
| PDCD11 | 23.2304409 | -0.2144736 | 0.42633994 | -0.5030577 | 0.61492373 | 0.9958918 |
| SH3PXD2A | 140.103302 | -0.0298279 | 0.34823742 | -0.0856539 | 0.93174156 | 0.99935941 |
| SLK | 78.8669148 | -0.094948 | 0.23510906 | -0.4038468 | 0.68632541 | 0.9958918 |
| GSTO1 | 36.9363534 | 0.04799376 | 0.3613974 | 0.1328005 | 0.89435116 | 0.99935941 |
| ITPRIP | 39.4817962 | -0.2263777 | 0.42221501 | -0.5361669 | 0.5918432 | 0.9958918 |
| XPNPEP1 | 38.8587238 | -0.278908 | 0.32023163 | -0.8709569 | 0.38377768 | 0.9725044 |
| ADD3 | 172.077384 | -0.504974 | 0.29070887 | -1.7370438 | 0.08237943 | 0.69989185 |
| MXI1 | 83.9405128 | -0.4724013 | 0.47397206 | -0.996686 | 0.31891693 | 0.95992644 |
| SMNDC1 | 21.5118322 | -0.5235646 | 0.35385543 | -1.4796002 | 0.13897997 | 0.8153985 |
| SMC3 | 39.1618455 | 0.16659569 | 0.27631558 | 0.6029182 | 0.54656311 | 0.9958918 |
| RBM20 | 39.2372838 | 0.89886265 | 0.50249731 | 1.78879099 | 0.07364848 | 0.66886321 |
| PDCD4 | 53.5499892 | -0.1337656 | 0.35990682 | -0.3716674 | 0.71014052 | 0.9958918 |
| BBIP1 | 33.0962814 | -0.0184861 | 0.40805564 | -0.0453028 | 0.96386596 | 0.99935941 |
| SHOC2 | 103.049447 | -0.0213992 | 0.27886329 | -0.0767373 | 0.93883253 | 0.99935941 |
| ZDHHC6 | 21.6159006 | 0.14976705 | 0.33622443 | 0.44543772 | 0.65600346 | 0.9958918 |
| VTI1A | 259.551995 | 0.27264083 | 0.19649025 | 1.38755398 | 0.1652729 | 0.85897493 |
| TCF7L2 | 359.874788 | -0.1682986 | 0.30700191 | -0.5482006 | 0.58355418 | 0.9958918 |
| CASP7 | 22.7665852 | -0.0553165 | 0.41053133 | -0.1347437 | 0.89281453 | 0.99935941 |
| NHLRC2 | 66.5166678 | 0.28855265 | 0.25840171 | 1.11668241 | 0.26413014 | 0.93499423 |
| CCDC186 | 36.9262952 | 0.25773148 | 0.28966249 | 0.8897648 | 0.37359219 | 0.97217293 |
| AFAP1L2 | 45.2747305 | 0.08217528 | 0.33227583 | 0.24731044 | 8.05E-01 | 9.96E-01 |
| ABLIM1 | 54.6352851 | 0.04086668 | 0.41730789 | 0.09792933 | 0.92198841 | 0.99935941 |
| FAM160B1 | 56.1776192 | -0.62625 | 0.27045938 | -2.3155048 | 0.02058532 | 0.38830162 |
| TRUB1 | 19.4041811 | -0.1064246 | 0.42464068 | -0.2506228 | 0.80210575 | 0.99592423 |
| ATRNL1 | 479.326459 | -1.0729795 | 0.54385038 | -1.9729314 | 0.04850338 | 0.57189932 |
| GFRA1 | 72.4276693 | 0.53493354 | 0.56279727 | 0.95049064 | 0.34186301 | 0.96887708 |
| HSPA12A | 66.1621219 | -0.0661387 | 0.33270475 | -0.198791 | 0.84242627 | 0.99692952 |
| PDZD8 | 70.5614075 | -0.007024 | 0.25272716 | -0.0277928 | 0.97782744 | 0.99935941 |
| RAB11FIP2 | 49.6048582 | 0.00153047 | 0.33368376 | 0.00458659 | 0.99634044 | 0.99935941 |
| FAM204A | 45.570127 | -0.246301 | 0.25987015 | -0.9477848 | 0.343239 | 0.96887708 |
| CACUL1 | 82.9463748 | 0.17292685 | 0.23209486 | 0.74506972 | 0.45622957 | 0.98330915 |
| EIF3A | 110.532426 | 0.05018665 | 0.38681126 | 0.12974453 | 0.89676855 | 0.99935941 |
| PRDX3 | 34.6667356 | 0.0429799 | 0.276411 | 0.15549271 | 0.8764329 | 0.9971731 |
| GRK5 | 574.017592 | -0.3475289 | 0.40047544 | -0.8677908 | 0.38550885 | 0.97254921 |
| TIAL1 | 74.7524904 | -0.2567084 | 0.27370308 | -0.9379084 | 0.34829148 | 0.96887708 |
| BAG3 | 26.921218 | 0.32741997 | 0.3469783 | 0.9436324 | 0.34535753 | 0.96887708 |
| INPP5F | 34.1353367 | 0.33609953 | 0.42692324 | 0.78725984 | 0.43112977 | 0.98092839 |
| MCMBP | 42.9127808 | 0.25503778 | 0.27160138 | 0.93901503 | 0.34772303 | 0.96887708 |
| SEC23IP | 53.9020634 | -0.0987324 | 0.29981824 | -0.3293074 | 0.74192339 | 0.9958918 |
| WDR11 | 65.0906798 | 0.21712416 | 0.22906909 | 0.94785445 | 0.34320355 | 0.96887708 |
| ATE1 | 124.871415 | 0.40680105 | 0.18143083 | 2.24218256 | 0.02494958 | 0.42739961 |
| TACC2 | 49.2193313 | 0.58288122 | 0.31578101 | 1.84583998 | 0.06491544 | 0.62770158 |
| PLEKHA1 | 78.3057163 | -0.1138793 | 0.28905381 | -0.3939726 | 0.69360126 | 0.9958918 |
| HTRA1 | 35.5837307 | 0.36051445 | 0.40217083 | 0.89642116 | 0.37002787 | 0.97217293 |
| IKZF5 | 18.6879355 | 0.02014708 | 0.40195827 | 0.05012232 | 0.96002492 | 0.99935941 |

| | | | | | | |
|---|---|---|---|---|---|---|
| ACADSB | 27.2771818 | 0.32188863 | 0.42697268 | 0.75388576 | 0.45091782 | 0.98092839 |
| BUB3 | 22.9886273 | 0.34400006 | 0.31694037 | 1.08537788 | 0.27775433 | 0.94369696 |
| CPXM2 | 29.1160315 | 0.45510742 | 0.58261486 | 0.78114626 | 0.43471648 | 0.98092839 |
| CHST15 | 118.3348 | -0.0255308 | 0.45233929 | -0.0564416 | 0.95499 | 0.99935941 |
| OAT | 39.2857067 | 0.32422747 | 0.34258883 | 0.94640409 | 0.34394252 | 0.96887708 |
| ZRANB1 | 71.075946 | -0.1322177 | 0.25412493 | -0.5202862 | 0.60286414 | 0.9958918 |
| CTBP2 | 320.286011 | 0.27301809 | 0.20927718 | 1.30457648 | 0.1920371 | 0.88158763 |
| EDRF1 | 40.3949273 | 0.17180923 | 0.32260665 | 0.53256568 | 0.59433427 | 0.9958918 |
| BCCIP | 29.7622627 | -0.0142023 | 0.28768135 | -0.0493683 | 0.9606258 | 0.99935941 |
| DHX32 | 55.2081212 | 0.3369276 | 0.34633813 | 0.9728285 | 0.33063855 | 0.9649971 |
| ADAM12 | 136.335584 | -0.2735643 | 0.4446517 | -0.6152327 | 0.53840104 | 0.9958918 |
| DOCK1 | 376.085987 | 0.09960313 | 0.20402024 | 0.48820222 | 0.62540661 | 0.9958918 |
| PTPRE | 94.9956612 | 0.84905396 | 0.94094197 | 0.90234465 | 0.36687381 | 0.97217293 |
| MGMT | 76.5763377 | 0.32625335 | 0.23414508 | 1.39338121 | 0.16350455 | 0.85697309 |
| GLRX3 | 52.6574883 | 0.15957459 | 0.2785522 | 0.5728714 | 0.56673177 | 0.9958918 |
| PPP2R2D | 79.1028039 | -0.0155381 | 0.19783743 | -0.0785397 | 0.93739876 | 0.99935941 |
| BNIP3 | 58.7477461 | -0.2248885 | 0.57303733 | -0.39245 | 0.69472574 | 0.9958918 |
| INPP5A | 118.941417 | 0.26056189 | 0.36429262 | 0.71525435 | 0.47445189 | 0.98910524 |
| TUBGCP2 | 34.4615103 | 0.35419014 | 0.40331819 | 0.87819035 | 0.37984043 | 0.97217293 |
| ECHS1 | 36.2026536 | 1.05740942 | 0.32168285 | 3.28711776 | 0.00101218 | 0.07036431 |
| KDM5A | 120.084287 | -0.2810293 | 0.19463585 | -1.4438723 | 0.14877489 | 0.8301633 |
| WNK1 | 208.928785 | 0.56956682 | 0.20383009 | 2.7943216 | 0.00520087 | 0.18276799 |
| RAD52 | 26.8361843 | -0.4553532 | 0.34423631 | -1.3227924 | 0.18590441 | 0.8789337 |
| ERC1 | 330.076468 | -0.1213246 | 0.22508436 | -0.5390184 | 0.58987415 | 0.9958918 |
| WNT5B | 33.6485955 | -0.0828046 | 0.40727735 | -0.2033125 | 0.83889075 | 0.99692952 |
| ADIPOR2 | 76.1908278 | 0.06638356 | 0.22203714 | 0.29897504 | 0.76495909 | 0.9958918 |
| DCP1B | 44.2727934 | 0.25181788 | 0.40379634 | 0.62362594 | 0.53287327 | 0.9958918 |
| CACNA1C | 1269.47262 | 0.04239909 | 0.26303933 | 0.16118917 | 0.87194441 | 0.9971731 |
| FKBP4 | 19.290997 | -0.1461423 | 0.44942142 | -0.3251788 | 0.74504575 | 0.9958918 |
| ITFG2 | 44.9227429 | 0.46253059 | 0.46640573 | 0.9916915 | 0.32134804 | 0.95992644 |
| NRIP2 | 21.0899029 | 0.62958448 | 0.48806965 | 1.28994801 | 0.19706871 | 0.88391893 |
| TULP3 | 79.2212848 | 0.12759416 | 0.22914413 | 0.55682927 | 0.57764408 | 0.9958918 |
| TEAD4 | 32.1096806 | 0.16147046 | 0.50998792 | 0.31661624 | 0.75153481 | 0.9958918 |
| TSPAN9 | 210.95879 | -0.0090151 | 0.26219365 | -0.0343832 | 0.97257154 | 0.99935941 |
| PARP11 | 39.6217179 | -0.2371387 | 0.26486667 | -0.8953136 | 0.37061948 | 0.97217293 |
| CCND2 | 48.7405553 | -1.3783927 | 0.37145089 | -3.7108344 | 0.00020658 | 0.02333707 |
| C12orf4 | 29.8054446 | 0.2900863 | 0.28641889 | 1.01280435 | 0.31115362 | 0.95957532 |
| NTF3 | 56.7482001 | -1.0664066 | 0.46805598 | -2.2783739 | 0.02270431 | 0.40828078 |
| CD9 | 87.8582188 | -1.043332 | 0.41068353 | -2.5404769 | 0.01107014 | 0.28321649 |
| TNFRSF1A | 117.942724 | -0.1678371 | 0.2898217 | -0.5791045 | 0.56251865 | 0.9958918 |
| LTBR | 43.7875352 | -0.105868 | 0.36750726 | -0.2880705 | 0.7732928 | 0.9958918 |
| MRPL51 | 22.9392139 | -0.2325666 | 0.35909799 | -0.647641 | 0.51721715 | 0.9958918 |
| GAPDH | 265.988751 | 0.00254738 | 0.44409999 | 0.00573606 | 0.99542331 | 0.99935941 |
| CHD4 | 22.2507886 | 0.49813405 | 0.46638135 | 1.06808313 | 0.28548302 | 0.94505166 |
| COPS7A | 18.8744082 | -0.2139783 | 0.36641388 | -0.5839797 | 0.55923395 | 0.9958918 |
| MLF2 | 31.7139963 | -0.0491488 | 0.41939043 | -0.117191 | 0.90670872 | 0.99935941 |
| PTMS | 87.4355404 | -0.2858101 | 0.31207169 | -0.9158475 | 0.35974689 | 0.97217293 |
| P3H3 | 22.3478614 | -0.1464898 | 0.39974436 | -0.3664588 | 0.71402277 | 0.9958918 |
| TPI1 | 80.8821609 | 0.21555868 | 0.36841423 | 0.5850987 | 0.55848135 | 0.9958918 |
| LRRC23 | 18.1457293 | -0.4121049 | 0.49925188 | -0.8254449 | 0.40911903 | 0.97812179 |
| ATN1 | 59.9123207 | -0.0133164 | 0.25195993 | -0.0528515 | 0.95785027 | 0.99935941 |
| C12orf57 | 39.0714217 | 0.427347 | 0.402906 | 1.0606618 | 0.28884362 | 0.95126484 |
| PHB2 | 28.2063411 | 0.46560177 | 0.35767423 | 1.30174819 | 0.19300248 | 0.88158763 |
| LPCAT3 | 67.7637448 | 0.35046979 | 0.31243748 | 1.12172774 | 0.26197822 | 0.93155278 |

| | | | | | | |
|---|---|---|---|---|---|---|
| C1S | 276.285755 | -0.2443164 | 0.36272382 | -0.6735603 | 0.50059088 | 0.99213111 |
| C1R | 319.675383 | -0.3432344 | 0.33482472 | -1.0251165 | 0.3053082 | 0.95793395 |
| C1RL | 92.2905944 | 0.30291471 | 0.36056377 | 0.84011412 | 0.40084441 | 0.97500135 |
| C1RL-AS1 | 18.761896 | 0.7798988 | 0.64480046 | 1.20951962 | 0.22646328 | 0.91093521 |
| CLSTN3 | 22.9420792 | 0.54192475 | 0.46926171 | 1.15484543 | 0.24815373 | 0.92624282 |
| SLC2A3 | 478.130565 | -1.8278491 | 0.55721442 | -3.2803335 | 0.00103684 | 0.07036431 |
| RIMKLB | 55.1375202 | 0.09340267 | 0.346923 | 0.26923171 | 0.78775138 | 0.99592423 |
| PHC1 | 24.0676796 | -0.2493555 | 0.34757058 | -0.7174242 | 0.47311239 | 0.98910524 |
| M6PR | 29.663056 | -0.0524167 | 0.29605946 | -0.177048 | 0.85947067 | 0.9971731 |
| A2M | 814.343216 | -0.1399523 | 0.48373911 | -0.2893136 | 0.77234144 | 0.9958918 |
| CLEC2D | 32.3414846 | 0.16908134 | 0.31938718 | 0.52939303 | 0.59653283 | 0.9958918 |
| GABARAPL1 | 68.2764915 | -0.6184512 | 0.41194449 | -1.5012974 | 0.13327866 | 0.80664435 |
| YBX3 | 152.899445 | 0.01460117 | 0.32194537 | 0.04535294 | 9.64E-01 | 0.99935941 |
| AC244131.2 | 25.550105 | -0.1495337 | 0.52730478 | -0.2835812 | 0.77673138 | 0.9958918 |
| ETV6 | 511.930993 | 0.15414624 | 0.41178759 | 0.37433435 | 0.70815558 | 0.9958918 |
| BCL2L14 | 29.730868 | -0.2174062 | 0.3500604 | -0.6210534 | 0.53456448 | 0.9958918 |
| LRP6 | 92.8338026 | 0.06024069 | 0.26225172 | 0.22970563 | 0.81832052 | 0.99592423 |
| BORCS5 | 34.4515748 | 0.25332405 | 0.28350819 | 0.89353344 | 0.37157157 | 0.97217293 |
| DUSP16 | 89.3335125 | -0.3214421 | 0.34167353 | -0.9407872 | 0.34681393 | 0.96887708 |
| CREBL2 | 21.4437177 | 0.60030739 | 0.40570314 | 1.47967154 | 0.13896092 | 0.8153985 |
| CDKN1B | 23.4491379 | -0.1487679 | 0.46796279 | -0.3179053 | 0.75055673 | 0.9958918 |
| APOLD1 | 45.4485675 | -1.4307752 | 0.68125671 | -2.1001998 | 0.03571127 | 0.50762504 |
| HEBP1 | 19.1955888 | 0.01161184 | 0.37483098 | 0.03097887 | 0.97528639 | 0.99935941 |
| EMP1 | 52.8705844 | -1.2058617 | 0.88404466 | -1.364028 | 0.17255875 | 0.87233975 |
| ATF7IP | 161.191868 | -0.2073979 | 0.30225235 | -0.6861747 | 0.49260295 | 0.99046213 |
| H2AFJ | 95.8226224 | -0.1548326 | 0.45427384 | -0.3408354 | 0.73322746 | 0.9958918 |
| WBP11 | 30.9713417 | -0.4432845 | 0.2970037 | -1.4925218 | 0.13556242 | 0.81154597 |
| C12orf60 | 33.1137453 | -0.4796549 | 0.5164273 | -0.9287946 | 0.35299553 | 0.97185572 |
| ART4 | 44.337248 | 1.45332187 | 0.5685812 | 2.5560498 | 0.0105868 | 0.27687675 |
| MGP | 2198.25889 | -0.7467768 | 0.51466239 | -1.4510033 | 0.14677895 | 0.8301633 |
| ARHGDIB | 32.5123818 | -0.4226127 | 0.3884494 | -1.0879478 | 0.27661815 | 0.94273432 |
| RERG | 553.290243 | -0.4968988 | 0.36947464 | -1.3448793 | 0.17866421 | 0.87815793 |
| EPS8 | 829.674955 | -0.3224211 | 0.42979172 | -0.75018 | 0.45314633 | 0.98092839 |
| STRAP | 49.693859 | 0.09921583 | 0.29844909 | 0.33243805 | 0.73955851 | 0.9958918 |
| DERA | 57.0388117 | -0.2468677 | 0.31745401 | -0.7776487 | 0.43677613 | 0.98092839 |
| MGST1 | 84.3516763 | 0.22289734 | 0.23321819 | 0.95574594 | 0.33920061 | 0.96887708 |
| LMO3 | 41.920949 | 0.37525893 | 0.56945557 | 0.65897842 | 0.50990963 | 0.99530567 |
| RERGL | 38.0023247 | 0.13246786 | 0.49112147 | 0.26972525 | 0.78737163 | 0.99592423 |
| PLEKHA5 | 333.544917 | -0.3331421 | 0.32301052 | -1.0313659 | 0.30236925 | 0.95650251 |
| AEBP2 | 286.716993 | 0.29910678 | 0.33821506 | 0.88436858 | 0.37649728 | 0.97217293 |
| AC024901.1 | 31.1331148 | 0.20441291 | 0.39360043 | 0.51934117 | 0.60352285 | 0.9958918 |
| PDE3A | 1118.33669 | 0.5434801 | 0.26519634 | 2.04934991 | 0.04042791 | 0.54117435 |
| PYROXD1 | 19.3368258 | 0.6427074 | 0.32648543 | 1.96856383 | 0.0490032 | 0.57273715 |
| RECQL | 31.5018009 | 0.11613128 | 0.33334208 | 0.3483847 | 0.72755129 | 0.9958918 |
| GOLT1B | 30.3156392 | -0.0895143 | 0.27095277 | -0.3303685 | 0.74112151 | 0.9958918 |
| LDHB | 91.4273053 | 0.47286103 | 0.29846755 | 1.58429628 | 0.11312631 | 0.77793929 |
| KCNJ8 | 28.4727212 | -1.1569324 | 0.42829695 | -2.701239 | 0.00690817 | 0.21213035 |
| ABCC9 | 243.079282 | -0.9576589 | 0.36820023 | -2.6009187 | 0.00929745 | 0.25495129 |
| CMAS | 20.9871104 | -0.2421181 | 0.35199119 | -0.6878528 | 0.49154549 | 0.99046213 |
| ST8SIA1 | 36.0733049 | -0.2908096 | 0.65466664 | -0.4442102 | 0.65689065 | 0.9958918 |
| C2CD5 | 88.4150345 | -0.3818751 | 0.28855609 | -1.3233999 | 0.18570241 | 0.8789337 |
| ETNK1 | 196.053022 | -0.5233098 | 0.34489544 | -1.5172999 | 0.12919098 | 0.80527948 |
| SOX5 | 1046.0803 | -0.7303621 | 0.46280307 | -1.5781271 | 0.11453641 | 0.77853537 |
| KRAS | 35.6333065 | 0.26195431 | 0.34261104 | 0.76458222 | 0.44452036 | 0.98092839 |

| | | | | | | |
|---|---|---|---|---|---|---|
| RASSF8-AS1 | 26.8072157 | -0.2488245 | 0.47683155 | -0.521829 | 0.6017894 | 0.9958918 |
| RASSF8 | 141.92263 | 0.00526055 | 0.27756357 | 0.0189526 | 0.98487892 | 0.99935941 |
| SSPN | 30.2275455 | 0.16682166 | 0.34434771 | 0.48445701 | 0.62806157 | 0.9958918 |
| ITPR2 | 348.541335 | -0.9516372 | 0.20596534 | -4.6203755 | 3.83E-06 | 0.00175938 |
| FGFR1OP2 | 70.3193004 | -0.3392015 | 0.37492006 | -0.9047303 | 0.36560829 | 0.97217293 |
| TM7SF3 | 40.6177151 | 0.62037888 | 0.35797794 | 1.73300868 | 0.08309413 | 0.7007527 |
| STK38L | 88.5561885 | 0.05565711 | 0.50985812 | 0.10916197 | 0.91307403 | 0.99935941 |
| PPFIBP1 | 246.810671 | -0.24931 | 0.22451272 | -1.1104493 | 0.26680546 | 0.93533685 |
| MRPS35 | 30.2042452 | 0.21689989 | 0.34554906 | 0.62769636 | 0.53020287 | 0.9958918 |
| KLHL42 | 49.5740599 | 0.19697965 | 0.2444259 | 0.80588697 | 0.42030802 | 0.97812179 |
| CCDC91 | 207.585268 | -0.2884236 | 0.17189269 | -1.6779284 | 0.09336109 | 0.73146123 |
| ERGIC2 | 44.6965652 | 0.20736689 | 0.23587078 | 0.87915462 | 0.37931745 | 0.97217293 |
| TMTC1 | 1053.93339 | 0.34515922 | 0.40697863 | 0.84810157 | 0.39638141 | 0.97500135 |
| IPO8 | 48.8377398 | 0.2945328 | 0.33862258 | 0.8697967 | 0.38441151 | 0.9725044 |
| CAPRIN2 | 27.5102834 | 0.43064034 | 0.32346761 | 1.33132445 | 0.18308227 | 0.87866798 |
| SINHCAF | 28.7946131 | -0.1707335 | 0.35870851 | -0.4759673 | 0.63409768 | 0.9958918 |
| DENND5B | 212.320936 | 0.16216409 | 0.42262607 | 0.38370583 | 0.70119649 | 0.9958918 |
| AMN1 | 38.3024855 | 0.29861421 | 0.38445281 | 0.77672526 | 0.43732088 | 0.98092839 |
| KIAA1551 | 76.2223076 | 0.35649665 | 0.33518561 | 1.06357983 | 0.28751908 | 0.94880006 |
| BICD1 | 151.610058 | -0.1133351 | 0.43972216 | -0.2577425 | 0.79660566 | 0.99592423 |
| FGD4 | 214.345567 | -0.2701403 | 0.35316881 | -0.7649042 | 0.44432862 | 0.98092839 |
| DNM1L | 86.0295456 | 0.00703511 | 0.22177013 | 0.03172255 | 0.97469331 | 0.99935941 |
| YARS2 | 21.7441042 | 0.34279446 | 0.3286862 | 1.04292318 | 0.29698387 | 0.95417903 |
| CPNE8 | 109.000917 | 0.13104239 | 0.27517478 | 0.47621511 | 0.63392113 | 0.9958918 |
| KIF21A | 25.1906555 | 0.73011484 | 0.49760831 | 1.46724809 | 0.14230858 | 0.81652619 |
| SLC2A13 | 108.373621 | -0.4383842 | 0.29503478 | -1.4858728 | 0.13731276 | 0.81404904 |
| LRRK2 | 50.3122167 | 0.35241574 | 0.43791071 | 0.80476622 | 0.4209546 | 0.97812179 |
| CNTN1 | 167.05156 | -0.0861303 | 0.56151664 | -0.1533886 | 0.87809183 | 0.99722158 |
| PDZRN4 | 182.357103 | 0.81902268 | 0.60514941 | 1.35342224 | 0.17592078 | 0.87624983 |
| GXYLT1 | 27.6194983 | 0.06374291 | 0.31076963 | 0.20511307 | 0.83748379 | 0.99692952 |
| YAF2 | 128.046286 | -0.3275384 | 0.18427436 | -1.77745 | 0.07549424 | 0.67167936 |
| PPHLN1 | 104.290943 | -0.004229 | 0.20259442 | -0.0208744 | 0.98334583 | 0.99935941 |
| PRICKLE1 | 132.662556 | -1.5189169 | 0.32435973 | -4.6828159 | 2.83E-06 | 0.00148534 |
| PUS7L | 19.2233686 | 0.0196751 | 0.32258668 | 0.06099167 | 0.95136584 | 0.99935941 |
| IRAK4 | 25.0332661 | 0.31503165 | 0.30202295 | 1.0430719 | 0.29691499 | 0.95417903 |
| TMEM117 | 99.0371896 | 0.25889165 | 0.4135776 | 0.62598082 | 0.53132752 | 0.9958918 |
| ANO6 | 256.521789 | -0.5814131 | 0.29582609 | -1.9653882 | 0.04936933 | 0.57273715 |
| ARID2 | 211.010463 | 0.15451747 | 0.21071112 | 0.73331426 | 0.46336678 | 0.98590351 |
| SCAF11 | 232.800715 | 0.06418235 | 0.21889619 | 0.29320909 | 0.76936234 | 0.9958918 |
| SLC38A1 | 42.9888132 | 0.32955378 | 0.45429077 | 0.72542477 | 0.46819148 | 0.98910524 |
| SLC38A2 | 377.644102 | -1.0033644 | 0.4663218 | -2.1516566 | 0.03142441 | 0.48041788 |
| AC008014.1 | 189.746402 | -0.6791939 | 0.51868109 | -1.3094635 | 0.19037741 | 0.88158763 |
| PCED1B | 49.3220817 | 0.48044171 | 0.43454756 | 1.10561364 | 0.26889379 | 0.93598159 |
| RAPGEF3 | 26.4870152 | 0.6654977 | 0.38598438 | 1.72415709 | 0.08467949 | 0.7055664 |
| HDAC7 | 83.5370461 | 0.03228949 | 0.32354433 | 0.09979929 | 0.92050367 | 0.99935941 |
| SENP1 | 29.2077028 | -0.145833 | 0.38256672 | -0.3811962 | 0.70305769 | 0.9958918 |
| ZNF641 | 20.5016126 | 0.23757694 | 0.40782004 | 0.58255337 | 0.56019401 | 0.9958918 |
| CCNT1 | 40.6869275 | -0.0650196 | 0.30556099 | -0.2127875 | 0.83149273 | 0.99592423 |
| ADCY6 | 64.1608457 | 0.72658279 | 0.26108756 | 2.78290846 | 0.0053874 | 0.18576881 |
| DDX23 | 18.4240158 | 0.16741366 | 0.34292754 | 0.48818962 | 0.62541554 | 0.9958918 |
| CCDC65 | 20.5798212 | -0.0418203 | 0.43778244 | -0.0955277 | 0.92389571 | 0.99935941 |
| PRKAG1 | 26.4426263 | 0.32303553 | 0.39527314 | 0.81724635 | 0.41378766 | 0.97812179 |
| KMT2D | 33.7219337 | 0.3006136 | 0.36580673 | 0.82178257 | 0.41120065 | 0.97812179 |
| LMBR1L | 38.0244563 | 0.2681989 | 0.32332404 | 0.82950499 | 0.40681871 | 0.97671046 |

| | | | | | | |
|---|---|---|---|---|---|---|
| TUBA1B | 94.7731134 | 0.53569562 | 0.58117438 | 0.92174679 | 0.35666067 | 0.97217293 |
| TUBA1A | 46.3232838 | 0.48465506 | 0.55726091 | 0.86970941 | 0.38445923 | 0.9725044 |
| TUBA1C | 34.7894225 | 0.36684696 | 0.43739344 | 0.83871162 | 0.40163116 | 0.97500135 |
| SPATS2 | 92.7542197 | 0.10620759 | 0.19790341 | 0.53666379 | 0.59149987 | 0.9958918 |
| PRPF40B | 40.1950843 | 0.37382711 | 0.40247439 | 0.92882211 | 0.35298128 | 0.97185572 |
| FMNL3 | 75.3796728 | 0.43453492 | 0.33554389 | 1.29501663 | 0.19531449 | 0.88391893 |
| TMBIM6 | 122.790876 | 0.30321292 | 0.27339772 | 1.10905429 | 0.26740676 | 0.93549101 |
| NCKAP5L | 30.3356456 | -0.1086928 | 0.31830271 | -0.3414761 | 0.73274522 | 0.9958918 |
| AQP2 | 32.2329427 | -1.4799127 | 0.56666741 | -2.6116073 | 0.00901177 | 0.25292422 |
| AC074032.1 | 59.2293963 | 0.25828587 | 0.29495913 | 0.87566664 | 0.38121129 | 0.97217293 |
| CERS5 | 54.007939 | 0.15597478 | 0.25684839 | 0.60726402 | 0.54367571 | 0.9958918 |
| LIMA1 | 298.034228 | -0.4948667 | 0.32195071 | -1.5370883 | 0.12427169 | 0.79195137 |
| LARP4 | 133.268426 | -0.385939 | 0.24117363 | -1.6002536 | 0.10954233 | 0.77303254 |
| DIP2B | 160.582441 | -0.1718305 | 0.29536922 | -0.5817482 | 0.56073631 | 0.9958918 |
| ATF1 | 49.7727938 | 0.27251641 | 0.32825805 | 0.83018957 | 0.40643161 | 0.97671046 |
| METTL7A | 45.7469959 | -0.0594739 | 0.38296827 | -0.1552972 | 0.87658706 | 0.9971731 |
| SLC11A2 | 46.3559448 | 0.67554322 | 0.35682752 | 1.89319257 | 0.05833227 | 0.60892588 |
| LETMD1 | 35.2029709 | -0.0224949 | 0.3137995 | -0.0716857 | 0.94285203 | 0.99935941 |
| CSRNP2 | 19.0376888 | 0.36328627 | 0.34387297 | 1.05645486 | 0.29076045 | 0.95186143 |
| TFCP2 | 108.994665 | 0.05532636 | 0.21880422 | 0.25285783 | 0.80037808 | 0.99592423 |
| POU6F1 | 27.2160403 | 0.44768046 | 0.3414732 | 1.31102661 | 0.18984877 | 0.88139754 |
| DAZAP2 | 61.2303825 | 0.36700837 | 0.30872194 | 1.18879912 | 0.23451872 | 0.91930824 |
| ACVR1B | 71.9975019 | 0.0923898 | 0.26858808 | 0.34398323 | 0.7308589 | 0.9958918 |
| NR4A1 | 271.276196 | -1.2642608 | 0.38807146 | -3.2578041 | 1.12E-03 | 0.07280105 |
| KRT8 | 19.8671893 | -0.1372257 | 0.35554709 | -0.3859565 | 0.69952891 | 0.9958918 |
| KRT18 | 22.0542195 | -0.0017122 | 0.53260244 | -0.0032148 | 0.99743499 | 0.99947501 |
| EIF4B | 74.0238977 | -0.0860878 | 0.26072389 | -0.3301874 | 0.74125835 | 0.9958918 |
| AC068888.1 | 40.2096389 | -0.2009706 | 0.30986815 | -0.6485682 | 0.51661751 | 0.9958918 |
| TNS2 | 105.948016 | -0.3930802 | 0.24695533 | -1.5917058 | 0.11145082 | 0.77793929 |
| IGFBP6 | 145.565589 | -1.898725 | 0.81426967 | -2.3318136 | 0.0197105 | 0.37879384 |
| CSAD | 54.6896729 | 0.08280325 | 0.32062273 | 0.25825757 | 0.79620813 | 0.99592423 |
| PFDN5 | 58.6447847 | -0.1364172 | 0.32716333 | -0.4169698 | 0.67670053 | 0.9958918 |
| SP1 | 44.773414 | 0.15057947 | 0.32120928 | 0.46878929 | 0.63922026 | 0.9958918 |
| PRR13 | 31.6939871 | 0.44645314 | 0.32784879 | 1.36176541 | 0.17327193 | 0.87233975 |
| PCBP2 | 121.135319 | -0.0742023 | 0.23474806 | -0.3160933 | 0.75193171 | 0.9958918 |
| MAP3K12 | 17.862668 | -0.1375706 | 0.3387293 | -0.4061372 | 0.68464176 | 0.9958918 |
| ATP5MC2 | 68.890944 | -0.0261732 | 0.25054938 | -0.1044631 | 0.91680187 | 0.99935941 |
| CALCOCO1 | 43.5136663 | 0.0597504 | 0.37961127 | 0.1573989 | 0.87493048 | 0.9971731 |
| HOXC4 | 33.6631416 | 0.43209572 | 0.34204501 | 1.26327151 | 0.20649162 | 0.89589437 |
| LINC02381 | 17.8090933 | 0.52094115 | 0.33812461 | 1.54067803 | 0.12339517 | 0.79184923 |
| CBX5 | 60.0722608 | 0.21168356 | 0.24820013 | 0.8528745 | 0.3937289 | 0.97500135 |
| HNRNPA1 | 53.4633262 | -0.0782261 | 0.27501389 | -0.2844442 | 0.77607 | 0.9958918 |
| COPZ1 | 31.8180066 | -0.2793038 | 0.28463934 | -0.9812552 | 0.3264669 | 0.95992644 |
| ITGA5 | 55.0744732 | 0.68908151 | 0.51013319 | 1.35078744 | 0.17676353 | 0.87771577 |
| ITGA7 | 71.1772401 | 0.48995928 | 0.33298605 | 1.47141085 | 0.14118004 | 0.81652619 |
| CD63 | 126.910875 | -0.2703151 | 0.3907648 | -0.6917591 | 0.48908861 | 0.99046213 |
| GDF11 | 19.7801714 | 0.10493459 | 0.39485592 | 0.26575412 | 0.79042858 | 0.99592423 |
| RAB5B | 34.2500078 | -0.1244617 | 0.26488827 | -0.4698647 | 0.63845168 | 0.9958918 |
| RPS26 | 20.7631374 | 0.12890234 | 0.42516021 | 0.30318534 | 0.76174862 | 0.9958918 |
| PA2G4 | 18.7390646 | 0.22750692 | 0.47141269 | 0.48260668 | 0.62937503 | 0.9958918 |
| RPL41 | 246.150428 | -0.0601357 | 0.37642302 | -0.1597556 | 0.87307363 | 0.9971731 |
| ESYT1 | 21.3211367 | -0.0688994 | 0.34436296 | -0.2000777 | 0.84141978 | 0.99692952 |
| MYL6 | 227.669683 | 0.0918595 | 0.4088586 | 0.22467302 | 0.82223365 | 0.99592423 |
| SMARCC2 | 51.4707416 | 0.42892114 | 0.2851569 | 1.50415839 | 0.13254059 | 0.80565822 |

| | | | | | | |
|---|---|---|---|---|---|---|
| RNF41 | 18.49108 | -0.1449017 | 0.3587056 | -0.403957 | 0.68624431 | 0.9958918 |
| ANKRD52 | 18.7283284 | 0.55184112 | 0.40798909 | 1.35258792 | 0.17618731 | 0.87664222 |
| CS | 26.6800093 | 0.26312384 | 0.36659546 | 0.71774987 | 0.47291153 | 0.98910524 |
| STAT2 | 107.584587 | -1.1907304 | 0.39168838 | -3.0399941 | 0.00236583 | 0.10666545 |
| RBMS2 | 111.882781 | -0.4123686 | 0.38338022 | -1.0756126 | 0.28210057 | 0.94419503 |
| BAZ2A | 81.3477236 | -0.2043997 | 0.21056762 | -0.9707083 | 0.33169357 | 0.9654289 |
| ATP5F1B | 75.5685849 | 0.34790834 | 0.22185644 | 1.56816878 | 0.11684175 | 0.78417352 |
| PTGES3 | 121.152935 | 0.0904178 | 0.20091273 | 0.45003521 | 0.65268505 | 0.9958918 |
| NACA | 73.3392622 | -0.0668496 | 0.32420797 | -0.2061935 | 0.83663979 | 0.99692952 |
| NAB2 | 18.9751213 | -0.313996 | 0.44058711 | -0.7126764 | 0.47604603 | 0.989664 |
| STAT6 | 89.17701 | 0.36193366 | 0.23616578 | 1.53254065 | 0.12538907 | 0.79233388 |
| LRP1 | 166.815586 | -0.154403 | 0.22788266 | -0.6775547 | 0.49805408 | 0.99176652 |
| NDUFA4L2 | 43.9467813 | 0.32312137 | 0.54220147 | 0.59594336 | 0.55121306 | 0.9958918 |
| R3HDM2 | 162.875792 | 0.19682157 | 0.22281243 | 0.88335092 | 0.3770467 | 0.97217293 |
| MARS | 28.1964434 | 0.17850208 | 0.35577647 | 0.50172537 | 0.61586072 | 0.9958918 |
| DDIT3 | 23.2732757 | -1.2451241 | 0.49681955 | -2.5061898 | 0.01220401 | 0.2958939 |
| DCTN2 | 31.7194732 | -0.2036047 | 0.38333093 | -0.5311459 | 0.59531766 | 0.9958918 |
| DTX3 | 24.9329601 | -0.1390436 | 0.42280955 | -0.3288564 | 0.74226425 | 0.9958918 |
| OS9 | 54.0430685 | 0.08663657 | 0.31055611 | 0.27897236 | 0.78026604 | 0.9958918 |
| AVIL | 24.9155712 | -0.1799013 | 0.49328924 | -0.3646975 | 0.71533725 | 0.9958918 |
| CTDSP2 | 72.1632851 | -0.1590252 | 0.23230445 | -0.6845549 | 0.49362482 | 0.99116088 |
| LRIG3 | 31.9691916 | 0.23657262 | 0.49331155 | 0.47956027 | 0.63154011 | 0.9958918 |
| SLC16A7 | 88.5629398 | -0.1222078 | 0.41472199 | -0.294674 | 0.76824294 | 0.9958918 |
| USP15 | 186.121578 | 0.13343072 | 0.24777353 | 0.53851886 | 0.59021889 | 0.9958918 |
| MON2 | 306.287882 | -0.3641572 | 0.19300979 | -1.8867294 | 0.05919673 | 0.61175668 |
| AC048341.1 | 24.9440782 | 0.03607269 | 0.41860586 | 0.08617339 | 0.93132858 | 0.99935941 |
| DPY19L2 | 37.4138716 | 0.55883626 | 0.51691898 | 1.08109062 | 0.2796568 | 0.94419503 |
| SRGAP1 | 249.966575 | -0.6014644 | 0.45719145 | -1.3155635 | 0.1883206 | 0.88139754 |
| XPOT | 36.6598919 | -0.2386459 | 0.37577511 | -0.6350765 | 0.52537855 | 0.9958918 |
| TBK1 | 82.9328665 | 0.07243905 | 0.32226722 | 0.22477946 | 0.82215085 | 0.99592423 |
| RASSF3 | 107.621104 | 0.14309705 | 0.39748158 | 0.36000928 | 0.7188402 | 0.9958918 |
| GNS | 54.1087314 | -0.1521076 | 0.2669842 | -0.5697249 | 0.56886429 | 0.9958918 |
| LEMD3 | 93.0641463 | -0.1149701 | 0.24492052 | -0.469418 | 0.63877087 | 0.9958918 |
| MSRB3 | 359.420411 | -0.1205525 | 0.36327592 | -0.3318484 | 0.74000376 | 0.9958918 |
| IRAK3 | 196.007152 | 0.30654157 | 0.5191115 | 0.590512 | 0.55484744 | 0.9958918 |
| HELB | 23.6166248 | 0.47733735 | 0.39627705 | 1.20455462 | 0.22837528 | 0.91371352 |
| GRIP1 | 39.306359 | 0.00051938 | 0.35210434 | 0.00147506 | 0.99882307 | 0.99968761 |
| CAND1 | 80.403646 | 0.05222044 | 0.20132107 | 0.25938885 | 0.79533524 | 0.99592423 |
| MDM1 | 18.6798959 | 0.60367882 | 0.36722211 | 1.64390652 | 0.10019551 | 0.75035372 |
| RAP1B | 336.926082 | 0.22704806 | 0.26428705 | 0.85909641 | 0.39028733 | 0.97297628 |
| NUP107 | 44.825791 | 0.08987955 | 0.39011687 | 0.23039134 | 0.81778769 | 0.99592423 |
| SLC35E3 | 37.0094099 | 0.33495197 | 0.39643454 | 0.84491118 | 0.39816043 | 0.97500135 |
| MDM2 | 96.7019158 | 0.07480566 | 0.32458082 | 0.23046851 | 0.81772773 | 0.99592423 |
| CPM | 221.599211 | 0.42591684 | 0.4249193 | 1.00234759 | 0.31617574 | 0.95992644 |
| CPSF6 | 175.4166 | -0.0940613 | 0.30869323 | -0.304708 | 0.76058857 | 0.9958918 |
| FRS2 | 105.000153 | -0.0635902 | 0.2105002 | -0.3020911 | 0.76258265 | 0.9958918 |
| CCT2 | 28.9983067 | -0.2518049 | 0.38000947 | -0.6626279 | 0.50756891 | 0.99397937 |
| AC025159.1 | 51.0729806 | -0.2182121 | 0.50444512 | -0.4325785 | 0.66532102 | 0.9958918 |
| CNOT2 | 157.937342 | -0.3542223 | 0.27422012 | -1.2917445 | 0.19644562 | 0.88391893 |
| ZFC3H1 | 133.271796 | -0.1529813 | 0.26157158 | -0.5848545 | 0.55864558 | 0.9958918 |
| RAB21 | 96.0267355 | 0.04133543 | 0.25513933 | 0.16201122 | 0.87129702 | 0.9971731 |
| TBC1D15 | 108.644584 | -0.3991642 | 0.27199954 | -1.4675178 | 0.14223524 | 0.81652619 |
| ATXN7L3B | 24.9315251 | 0.07221494 | 0.36833678 | 0.19605683 | 0.84456568 | 0.99692952 |
| CAPS2 | 77.5479835 | -0.1644923 | 0.4456389 | -0.3691157 | 0.71204147 | 0.9958918 |

| | | | | | | |
|---|---|---|---|---|---|---|
| GLIPR1 | 25.1168138 | -0.6949395 | 0.60315909 | -1.1521661 | 0.24925283 | 0.92624282 |
| KRR1 | 25.8342454 | -0.1080716 | 0.37387657 | -0.2890568 | 0.77253794 | 0.9958918 |
| PHLDA1 | 64.1518452 | -0.7808128 | 0.54033504 | -1.4450531 | 0.14844296 | 0.8301633 |
| NAP1L1 | 177.688841 | -0.3893524 | 0.40143515 | -0.9699012 | 0.33209574 | 0.9654289 |
| OSBPL8 | 262.579129 | 0.22148952 | 0.26138578 | 0.84736637 | 0.39679094 | 0.97500135 |
| ZDHHC17 | 100.572678 | -0.1880045 | 0.25824189 | -0.7280171 | 0.46660309 | 0.98791879 |
| CSRP2 | 57.5070843 | 0.84039896 | 0.30514599 | 2.75408819 | 0.00588559 | 0.19136125 |
| NAV3 | 162.862703 | -0.1405279 | 0.45548642 | -0.3085227 | 0.75768462 | 0.9958918 |
| SYT1 | 237.166497 | -2.2766805 | 0.45212999 | -5.0354556 | 4.77E-07 | 0.00058389 |
| AC027288.3 | 36.7777697 | -0.0668988 | 0.5223057 | -0.1280837 | 0.89808277 | 0.99935941 |
| PAWR | 245.061473 | -0.1308395 | 0.2360899 | -0.5541936 | 5.79E-01 | 0.9958918 |
| PPP1R12A | 360.890465 | 0.12914567 | 0.27239895 | 0.47410486 | 0.63542513 | 0.9958918 |
| LIN7A | 71.8374228 | -0.6401418 | 0.50406608 | -1.2699562 | 0.20410024 | 0.89396747 |
| ACSS3 | 166.484104 | -0.5541862 | 0.2840242 | -1.9511935 | 0.05103403 | 0.57655256 |
| PPFIA2 | 77.1458176 | 1.12317521 | 0.34101681 | 3.29360659 | 0.00098911 | 0.07012172 |
| METTL25 | 35.9454826 | 0.22227022 | 0.36995144 | 0.60080917 | 0.54796709 | 0.9958918 |
| TMTC2 | 113.289834 | -0.3874194 | 0.29006718 | -1.3356194 | 0.1816737 | 0.87866798 |
| ALX1 | 27.2800002 | -1.371057 | 0.62205499 | -2.2040768 | 0.02751894 | 0.44841843 |
| CEP290 | 62.4517176 | -0.312802 | 0.22437325 | -1.3941147 | 0.16328297 | 0.85650717 |
| TMTC3 | 43.6069143 | 0.22814657 | 0.27351887 | 0.83411637 | 0.40421541 | 0.97582411 |
| KITLG | 61.4722215 | 0.0744604 | 0.28769685 | 0.25881548 | 0.79577762 | 0.99592423 |
| DUSP6 | 25.6985985 | -0.610283 | 0.52387146 | -1.164948 | 0.24404006 | 0.92546331 |
| POC1B | 38.6954368 | 0.11041495 | 0.26814404 | 0.41177475 | 0.68050453 | 0.9958918 |
| ATP2B1 | 112.318709 | 0.14436049 | 0.26457447 | 0.54563271 | 0.58531844 | 0.9958918 |
| LUM | 141.219111 | 0.06766946 | 0.52927954 | 0.12785202 | 0.89826608 | 0.99935941 |
| DCN | 402.118081 | -0.5497625 | 0.36893248 | -1.4901439 | 0.13618639 | 0.81177777 |
| BTG1 | 205.324036 | 0.47798602 | 0.4467144 | 1.07000361 | 0.28461768 | 0.94505166 |
| EEA1 | 170.394544 | -0.2485985 | 0.2263781 | -1.0981562 | 0.27213628 | 0.93813668 |
| AC138123.1 | 40.9000788 | -0.7139668 | 0.50819306 | -1.4049126 | 0.16004727 | 0.84923277 |
| NUDT4 | 80.255837 | 0.39239449 | 0.34520772 | 1.13669094 | 0.25566751 | 0.9266065 |
| UBE2N | 49.0364983 | 0.28479641 | 0.33336224 | 0.85431516 | 0.39293039 | 0.97423935 |
| MRPL42 | 37.5375235 | -0.3301878 | 0.33700782 | -0.979763 | 0.32720312 | 0.95992644 |
| SOCS2 | 26.8316436 | -0.4035372 | 0.44483259 | -0.9071665 | 0.36431876 | 0.97217293 |
| CRADD | 182.300341 | 0.06263546 | 0.28567312 | 0.21925571 | 0.82645086 | 0.99592423 |
| CEP83 | 63.0393666 | 0.14763842 | 0.20549719 | 0.71844499 | 0.47248296 | 0.98910524 |
| NDUFA12 | 19.7728064 | -0.1033234 | 0.35924797 | -0.2876101 | 0.77364518 | 0.9958918 |
| NR2C1 | 54.6390409 | -0.1686313 | 0.24199215 | -0.6968461 | 0.48589909 | 0.99046213 |
| FGD6 | 34.7433176 | 0.34599559 | 0.49078834 | 0.70497924 | 0.48082315 | 0.99002719 |
| VEZT | 128.088882 | 0.08999402 | 0.32568762 | 0.27632007 | 0.78230224 | 0.9958918 |
| METAP2 | 58.4749341 | 0.0870069 | 0.23140739 | 0.37599015 | 0.70692422 | 0.9958918 |
| NTN4 | 95.2908951 | 0.99241164 | 0.38705192 | 2.56402715 | 0.01034654 | 0.27351352 |
| LTA4H | 21.3589234 | -0.0569912 | 0.33194602 | -0.1716882 | 0.86368268 | 0.9971731 |
| ELK3 | 37.9123871 | -0.4390122 | 0.45358625 | -0.9678694 | 0.3331096 | 0.9654289 |
| CDK17 | 143.337942 | -0.4091469 | 0.40199144 | -1.0178001 | 0.30877297 | 0.95793395 |
| NEDD1 | 21.4591591 | 0.14686569 | 0.35646192 | 0.41200948 | 0.68033248 | 0.9958918 |
| TMPO | 31.0182613 | 0.46208846 | 0.29550879 | 1.56370464 | 0.11788695 | 0.78539225 |
| SLC25A3 | 76.4721841 | 0.37861583 | 0.36453541 | 1.03862566 | 0.29897887 | 0.9544725 |
| IKBIP | 26.9749932 | 0.10784923 | 0.38418825 | 0.28071975 | 0.77892536 | 0.9958918 |
| APAF1 | 26.5513287 | -0.0076189 | 0.32131982 | -0.0237113 | 0.98108286 | 0.99935941 |
| ANKS1B | 81.6644492 | -1.1163106 | 0.43937966 | -2.5406515 | 0.01106461 | 0.28321649 |
| UHRF1BP1L | 76.9262344 | -0.2022497 | 0.31172064 | -0.6488171 | 0.5164566 | 0.9958918 |
| ACTR6 | 20.0301068 | 0.13955958 | 0.33050473 | 0.42226198 | 0.67283381 | 0.9958918 |
| SCYL2 | 71.724482 | -0.1170617 | 0.23024034 | -0.5084326 | 0.61114999 | 0.9958918 |
| UTP20 | 35.1794834 | 0.10675697 | 0.38053201 | 0.28054662 | 0.77905816 | 0.9958918 |

| | | | | | | |
|---|---|---|---|---|---|---|
| ARL1 | 33.2328094 | 0.12403455 | 0.36297992 | 0.34171189 | 0.73256772 | 0.9958918 |
| CHPT1 | 103.706903 | 0.45809556 | 0.3936258 | 1.1637844 | 0.2445114 | 0.92559078 |
| GNPTAB | 56.5428806 | 0.60451738 | 0.25716663 | 2.35068364 | 0.01873896 | 0.36528543 |
| DRAM1 | 104.710783 | 0.19566983 | 0.3058974 | 0.63965836 | 0.52239473 | 0.9958918 |
| WASHC3 | 38.0524408 | 0.26683911 | 0.31314983 | 0.85211322 | 0.39415125 | 0.97500135 |
| IGF1 | 112.223089 | -0.8970367 | 0.81743723 | -1.0973768 | 0.27247669 | 0.93813668 |
| HSP90B1 | 184.430979 | -0.5338027 | 0.39583131 | -1.3485611 | 0.17747798 | 0.87771577 |
| TDG | 31.5530867 | 0.24266698 | 0.29013187 | 0.83640237 | 0.40292858 | 0.97512144 |
| HCFC2 | 40.3050341 | 0.01575064 | 0.26033934 | 0.06050043 | 0.95175707 | 0.99935941 |
| NFYB | 22.6670355 | 0.26300437 | 0.31019076 | 0.84787943 | 0.39650512 | 0.97500135 |
| TXNRD1 | 300.145884 | -0.1783735 | 0.29668829 | -0.6012153 | 0.54769661 | 0.9958918 |
| CHST11 | 190.341385 | -0.0551078 | 0.34241395 | -0.1609392 | 8.72E-01 | 0.9971731 |
| SLC41A2 | 37.7928027 | 0.50974321 | 0.34527862 | 1.47632431 | 0.13985685 | 0.8153985 |
| WASHC4 | 153.259139 | -0.3873938 | 0.31599931 | -1.2259324 | 0.22022413 | 0.90731076 |
| APPL2 | 58.645468 | -0.0562641 | 0.35779002 | -0.1572545 | 0.87504425 | 0.9971731 |
| C12orf75 | 32.5072201 | 0.91248935 | 0.38017564 | 2.40017835 | 0.01638709 | 0.34578631 |
| NUAK1 | 123.35508 | 0.18713041 | 0.36296466 | 0.51556096 | 0.60616109 | 0.9958918 |
| CKAP4 | 39.6167195 | -0.1145221 | 0.4700661 | -0.2436298 | 0.80751755 | 0.99592423 |
| TCP11L2 | 23.5967505 | -0.2334378 | 0.39853764 | -0.5857358 | 0.55805308 | 0.9958918 |
| POLR3B | 30.772329 | -0.4972055 | 0.30866777 | -1.610811 | 0.10722091 | 0.77179565 |
| RIC8B | 66.4082541 | -0.1141778 | 0.21541502 | -0.5300364 | 0.59608669 | 0.9958918 |
| TMEM263 | 51.648781 | -0.1826692 | 0.2676433 | -0.68251 | 0.4949165 | 0.99176652 |
| CRY1 | 74.0360508 | -0.4704442 | 0.42705266 | -1.1016069 | 0.27063261 | 0.93726628 |
| PWP1 | 22.5425949 | 0.10082385 | 0.35361858 | 0.28512033 | 0.77555195 | 0.9958918 |
| PRDM4 | 19.1001857 | 0.01163848 | 0.40614021 | 0.0286563 | 0.97713871 | 0.99935941 |
| SART3 | 30.4053545 | -0.0886657 | 0.30257021 | -0.2930416 | 0.76949034 | 0.9958918 |
| ISCU | 41.7687201 | 0.40124686 | 0.35282155 | 1.13725157 | 0.25543313 | 0.9266065 |
| CORO1C | 110.865206 | -0.8278174 | 0.32858592 | -2.5193332 | 0.01175773 | 0.29290706 |
| SSH1 | 121.998068 | -0.1510215 | 0.39123966 | -0.3860077 | 0.69949095 | 0.9958918 |
| ACACB | 144.333071 | 0.44501964 | 0.51318026 | 0.86717997 | 0.38584341 | 0.97254921 |
| KCTD10 | 38.2557881 | -0.1466747 | 0.29740706 | -0.4931782 | 0.6218867 | 0.9958918 |
| UBE3B | 32.0179029 | 0.27770309 | 0.2891946 | 0.96026376 | 0.33692248 | 0.96728036 |
| MMAB | 19.27229 | -0.4376477 | 0.54555286 | -0.8022095 | 0.42243178 | 0.97828057 |
| GLTP | 28.0727855 | 0.47772982 | 0.41781934 | 1.14338846 | 0.25287734 | 0.92646125 |
| TCHP | 37.1860029 | -0.3652648 | 0.28228974 | -1.2939359 | 0.19568757 | 0.88391893 |
| GIT2 | 84.6988812 | 0.00137608 | 0.28188091 | 0.00488176 | 0.99610493 | 0.99935941 |
| ANKRD13A | 55.3632833 | -0.919539 | 0.51675073 | -1.7794634 | 0.07516382 | 0.67167936 |
| IFT81 | 54.7121938 | -0.2656161 | 0.29811725 | -0.8909786 | 0.37294063 | 0.97217293 |
| ATP2A2 | 202.943035 | -0.3386905 | 0.22630539 | -1.4966081 | 0.13449526 | 0.80890178 |
| ANAPC7 | 39.0717363 | 0.22452356 | 0.42960455 | 0.52262846 | 0.60123283 | 0.9958918 |
| ARPC3 | 72.1216778 | 0.03248553 | 0.2479881 | 0.13099634 | 0.8957782 | 0.99935941 |
| VPS29 | 40.1135908 | 0.07903933 | 0.28922942 | 0.27327554 | 0.78464142 | 0.9958918 |
| PPTC7 | 37.1645674 | 0.06086155 | 0.32169092 | 0.18919262 | 0.84994185 | 0.9971731 |
| PPP1CC | 36.1449159 | 0.13104302 | 0.27443268 | 0.47750516 | 0.63300245 | 0.9958918 |
| SH2B3 | 50.9841016 | 0.69473489 | 0.38626082 | 1.798616 | 0.07207944 | 0.66213979 |
| ATXN2 | 295.029493 | -0.1097233 | 0.1754056 | -0.6255406 | 0.53161629 | 0.9958918 |
| BRAP | 26.7536884 | 0.40072305 | 0.34223254 | 1.17090868 | 0.24163549 | 0.92516122 |
| ALDH2 | 41.8293928 | 0.4727307 | 0.35835039 | 1.31918567 | 0.18710705 | 0.87955894 |
| MAPKAPK5 | 28.3845197 | -0.1666154 | 0.35370799 | -0.4710536 | 0.63760246 | 0.9958918 |
| ERP29 | 29.326391 | 0.26699697 | 0.34395892 | 0.77624668 | 0.43760335 | 0.98092839 |
| NAA25 | 78.2805911 | -0.352993 | 0.29145683 | -1.211133 | 0.22584443 | 0.91093521 |
| TRAFD1 | 25.9868804 | -0.1211387 | 0.36623277 | -0.3307696 | 0.74081852 | 0.9958918 |
| HECTD4 | 164.378419 | -0.0143173 | 0.18995727 | -0.075371 | 0.93991954 | 0.99935941 |
| RPL6 | 112.472547 | 0.13351029 | 0.38079656 | 0.35060792 | 0.72588251 | 0.9958918 |

| | | | | | | |
|---|---|---|---|---|---|---|
| PTPN11 | 80.7862488 | -0.2493191 | 0.24919128 | -1.0005129 | 0.31706233 | 0.95992644 |
| DDX54 | 18.3637899 | -0.2113708 | 0.42314764 | -0.4995201 | 0.617413 | 0.9958918 |
| TPCN1 | 114.954679 | 0.45103382 | 0.28205354 | 1.59910707 | 0.10979681 | 0.77303254 |
| SLC8B1 | 32.7817338 | 0.47559224 | 0.38353376 | 1.24002706 | 0.21496538 | 0.9006731 |
| PLBD2 | 22.6558298 | -0.0518225 | 0.41679673 | -0.1243351 | 0.90104995 | 0.99935941 |
| RBM19 | 30.1272592 | 0.37567079 | 0.38780292 | 0.96871573 | 0.33268704 | 0.9654289 |
| TBX3 | 115.697919 | -0.0711318 | 0.48922397 | -0.1453971 | 0.88439733 | 0.99862036 |
| MED13L | 758.824881 | -0.2549811 | 0.28366691 | -0.898875 | 0.36871926 | 0.97217293 |
| FBXW8 | 55.1632393 | -0.157835 | 0.32646592 | -0.4834654 | 0.62876535 | 0.9958918 |
| FBXO21 | 52.2920251 | 0.22943624 | 0.28361174 | 0.80898007 | 0.41852661 | 0.97812179 |
| WSB2 | 25.3791534 | 0.02922133 | 0.30802116 | 0.09486795 | 0.92441972 | 0.99935941 |
| VSIG10 | 29.7188143 | 0.28879058 | 0.54140837 | 0.53340621 | 0.59375242 | 0.9958918 |
| PEBP1 | 126.576535 | 0.59964139 | 0.39063863 | 1.5350284 | 0.12477685 | 0.79195137 |
| TAOK3 | 151.740312 | -0.2770533 | 0.24207451 | -1.144496 | 0.252418 | 0.92631316 |
| SUDS3 | 23.6111079 | 0.02462793 | 0.30845235 | 0.07984355 | 0.93636169 | 0.99935941 |
| HSPB8 | 24.4466441 | 1.18465465 | 0.37414448 | 3.16630264 | 0.0015439 | 0.0868147 |
| RAB35 | 18.4021876 | -0.0483403 | 0.41455516 | -0.1166078 | 0.90717089 | 0.99935941 |
| GCN1 | 36.1018339 | -0.1793564 | 0.3288291 | -0.5454396 | 0.58545123 | 0.9958918 |
| RPLP0 | 171.140673 | -0.0612014 | 0.37841907 | -0.1617292 | 0.87151908 | 0.9971731 |
| PXN | 36.6126304 | 0.05984587 | 0.41172459 | 0.14535414 | 0.88443127 | 0.99862036 |
| SRSF9 | 35.4773382 | -0.0135877 | 0.34325047 | -0.0395855 | 0.96842358 | 0.99935941 |
| DYNLL1 | 73.1275426 | 0.20511582 | 0.41901433 | 0.48951982 | 0.62447372 | 0.9958918 |
| RNF10 | 74.2167228 | 0.28624221 | 0.20297468 | 1.41023603 | 0.15847 | 0.8463634 |
| MLEC | 46.3283071 | 0.37475505 | 0.36282708 | 1.03287511 | 0.30166235 | 0.95597956 |
| SPPL3 | 162.006928 | -0.1999357 | 0.20506537 | -0.9749854 | 0.32956752 | 0.96340164 |
| P2RX4 | 20.2877045 | 0.06984349 | 0.54651925 | 0.12779695 | 0.89830966 | 0.99935941 |
| CAMKK2 | 37.5256278 | 0.18050838 | 0.32142349 | 0.5615905 | 0.57439506 | 0.9958918 |
| ANAPC5 | 89.3064497 | -0.3083182 | 0.24211238 | -1.2734509 | 0.20285807 | 0.89396747 |
| RNF34 | 22.566097 | -0.0093781 | 0.40152795 | -0.0233562 | 0.98136618 | 0.99935941 |
| KDM2B | 37.7536325 | 0.01116482 | 0.4428813 | 0.02520951 | 0.97988786 | 0.99935941 |
| TMEM120B | 42.6030208 | 0.33756841 | 0.4077739 | 0.82783232 | 0.40776547 | 0.97738697 |
| SETD1B | 27.051615 | 0.01750722 | 0.4707589 | 0.03718936 | 0.97033402 | 0.99935941 |
| WDR66 | 39.6204897 | -0.490901 | 0.5657148 | -0.8677535 | 0.38552931 | 0.97254921 |
| MLXIP | 127.777252 | 0.07027403 | 0.32760633 | 0.21450754 | 0.83015129 | 0.99592423 |
| CLIP1 | 154.054869 | -0.0805753 | 0.29277667 | -0.2752108 | 0.78315427 | 0.9958918 |
| ZCCHC8 | 48.787321 | 0.14347357 | 0.25847632 | 0.55507433 | 0.57884382 | 0.9958918 |
| RSRC2 | 67.6913535 | -0.2090048 | 0.23598694 | -0.8856627 | 0.37579929 | 0.97217293 |
| DENR | 41.9933119 | 0.46789494 | 0.29039805 | 1.61121927 | 0.10713194 | 0.77179565 |
| VPS37B | 48.5244429 | -0.1959321 | 0.31860869 | -0.6149616 | 0.53858005 | 0.9958918 |
| ARL6IP4 | 54.8690328 | -0.1385652 | 0.3860602 | -0.3589212 | 0.719654 | 0.9958918 |
| PITPNM2 | 34.422788 | -0.2979585 | 0.33935135 | -0.8780235 | 0.37993095 | 0.97217293 |
| MPHOSPH9 | 41.4770915 | -0.3431433 | 0.28252511 | -1.2145586 | 0.22453449 | 0.91093521 |
| C12orf65 | 19.5089045 | 0.03443406 | 0.34681368 | 0.09928691 | 0.92091047 | 0.99935941 |
| SBNO1 | 81.1355187 | -0.2175461 | 0.20813003 | -1.0452414 | 0.29591142 | 0.95393362 |
| KMT5A | 19.0982615 | 0.14343795 | 0.40868027 | 0.35097839 | 0.72560456 | 0.9958918 |
| RILPL2 | 29.7382504 | 0.62383717 | 0.41678755 | 1.49677497 | 0.13445182 | 0.80890178 |
| SNRNP35 | 21.5656328 | 0.2494194 | 0.38540536 | 0.6471612 | 0.51752762 | 0.9958918 |
| RILPL1 | 57.9217985 | 0.06449405 | 0.30351337 | 0.21249163 | 0.83172351 | 0.99592423 |
| TMED2 | 30.9881541 | 0.30663372 | 0.32842677 | 0.93364412 | 0.3504875 | 0.9693503 |
| DDX55 | 19.5793914 | 0.05552307 | 0.4137485 | 0.13419522 | 0.8932482 | 0.99935941 |
| EIF2B1 | 19.6018654 | -0.035664 | 0.34089813 | -0.1046177 | 0.91667918 | 0.99935941 |
| ATP6V0A2 | 30.1181087 | -0.1664127 | 0.46247383 | -0.3598315 | 0.71897313 | 0.9958918 |
| CCDC92 | 59.5813 | 0.28830008 | 0.28784954 | 1.00156522 | 0.31655363 | 0.95992644 |
| NCOR2 | 168.408714 | -0.1113502 | 0.26573524 | -0.4190268 | 0.67519656 | 0.9958918 |

| | | | | | | |
|---|---|---|---|---|---|---|
| SCARB1 | 21.5691097 | -0.3846845 | 0.37806269 | -1.0175152 | 0.30890841 | 0.95793395 |
| UBC | 321.282803 | -0.5774517 | 0.31471744 | -1.8348258 | 0.0665315 | 0.63518413 |
| TMEM132C | 56.4337513 | 0.96230862 | 0.59130708 | 1.62742617 | 0.10364662 | 0.75827058 |
| SLC15A4 | 26.9561206 | 0.36664656 | 0.30952455 | 1.1845476 | 0.23619637 | 0.92036432 |
| STX2 | 26.989765 | -0.2174661 | 0.37763455 | -0.575864 | 0.56470713 | 0.9958918 |
| RAN | 40.7107443 | 0.25937766 | 0.46612128 | 0.5564596 | 0.57789671 | 0.9958918 |
| ADGRD1 | 53.7109599 | -1.5412564 | 0.5204837 | -2.9612001 | 0.00306443 | 0.13247341 |
| SFSWAP | 90.5405037 | 0.00148865 | 0.22172299 | 0.00671403 | 0.99464302 | 0.99935941 |
| ULK1 | 38.9767743 | -0.0529993 | 0.36978806 | -0.1433234 | 0.88603476 | 0.99935941 |
| EP400 | 70.6602665 | 0.2330018 | 0.24310858 | 0.95842688 | 0.33784754 | 0.96750939 |
| FBRSL1 | 36.0495512 | 0.26788169 | 0.37346956 | 0.71727851 | 0.47320226 | 0.98910524 |
| ANKLE2 | 54.4006821 | -0.1238898 | 0.32579008 | -0.3802748 | 0.70374144 | 0.9958918 |
| GOLGA3 | 63.3544961 | -0.5048219 | 0.342111 | -1.4756085 | 0.14004903 | 0.8153985 |
| CHFR | 56.6273185 | -0.2874482 | 0.29796706 | -0.9646978 | 0.3346962 | 0.96640422 |
| ZNF26 | 28.8432334 | -0.0651758 | 0.3309035 | -0.1969632 | 0.84385636 | 0.99692952 |
| ZNF84 | 30.9032249 | -0.1858406 | 0.4345839 | -0.4276287 | 0.66892146 | 0.9958918 |
| ZNF140 | 18.6284087 | 0.03405492 | 0.37755239 | 0.09019918 | 0.92812894 | 0.99935941 |
| MPHOSPH8 | 104.646672 | 0.55474577 | 0.29043624 | 1.91004322 | 0.05612765 | 0.60041061 |
| PSPC1 | 160.197585 | -0.102794 | 0.19558953 | -0.5255599 | 0.59919405 | 0.9958918 |
| ZMYM5 | 43.1722748 | 0.23849273 | 0.27976256 | 0.85248267 | 0.39394625 | 0.97500135 |
| ZMYM2 | 259.90511 | -0.2746953 | 0.30856238 | -0.8902422 | 0.37333585 | 0.97217293 |
| CRYL1 | 26.6104057 | 0.56498662 | 0.36755446 | 1.53715077 | 0.12425639 | 0.79195137 |
| IFT88 | 81.173764 | -0.2465451 | 0.26675445 | -0.9242399 | 0.35536144 | 0.97217293 |
| XPO4 | 88.3776059 | 0.17323494 | 0.23718162 | 0.73038941 | 0.46515219 | 0.98655222 |
| LATS2 | 93.8962773 | -0.4570761 | 0.3207958 | -1.4248194 | 0.15420941 | 0.84218718 |
| SAP18 | 39.5002475 | -0.0725535 | 0.36135291 | -0.2007828 | 0.84086841 | 0.99692952 |
| MRPL57 | 24.4674015 | -0.0822904 | 0.33730604 | -0.2439636 | 0.80725898 | 0.99592423 |
| ZDHHC20 | 55.4512092 | 0.32791692 | 0.31555213 | 1.03918462 | 0.29871888 | 0.95447177 |
| MICU2 | 106.661868 | 0.25569201 | 0.23544652 | 1.08598764 | 0.27748446 | 0.94358865 |
| SACS | 23.6615569 | 0.21253864 | 0.37664287 | 0.56429752 | 0.57255167 | 0.9958918 |
| MIPEP | 34.0422246 | -0.4116353 | 0.44614257 | -0.9226541 | 0.35618748 | 0.97217293 |
| PARP4 | 51.5269855 | -0.1756749 | 0.32997892 | -0.5323823 | 0.59446122 | 0.9958918 |
| MTMR6 | 34.3607845 | 0.08101374 | 0.30895932 | 0.26221491 | 0.79315576 | 0.99592423 |
| NUP58 | 85.4194942 | -0.3545289 | 0.32616105 | -1.0869751 | 0.27704781 | 0.94304045 |
| CDK8 | 149.367097 | -0.2840725 | 0.26417396 | -1.0753236 | 0.28222988 | 0.94419503 |
| WASF3 | 80.8580432 | -1.3539267 | 0.41589183 | -3.2554781 | 0.00113202 | 0.07280105 |
| USP12 | 111.476826 | -0.2104145 | 0.24255921 | -0.8674768 | 0.38568083 | 0.97254921 |
| RPL21 | 64.2917779 | 0.36471749 | 0.38572793 | 0.94553043 | 0.34438814 | 0.96887708 |
| GTF3A | 19.5745557 | -0.371262 | 0.41588332 | -0.892707 | 0.37201409 | 0.97217293 |
| MTIF3 | 27.6882071 | 0.1468744 | 0.31135685 | 0.47172368 | 0.63712404 | 0.9958918 |
| POLR1D | 26.6278886 | -0.1223045 | 0.30549043 | -0.4003545 | 0.68889543 | 0.9958918 |
| PAN3 | 306.486358 | -0.0754313 | 0.18635952 | -0.4047625 | 0.68565211 | 0.9958918 |
| FLT1 | 29.6681207 | -0.3481889 | 0.62664871 | -0.5556365 | 0.5784594 | 0.9958918 |
| POMP | 50.710421 | -0.2753138 | 0.3241809 | -0.8492597 | 0.39573681 | 0.97500135 |
| UBL3 | 80.6871305 | 0.09226599 | 0.22461651 | 0.41077118 | 0.68124032 | 0.9958918 |
| KATNAL1 | 65.2041138 | -0.4969433 | 0.37355324 | -1.3303145 | 0.18341466 | 0.87866798 |
| HMGB1 | 153.544561 | -0.0027003 | 0.2254722 | -0.0119763 | 0.99044454 | 0.99935941 |
| USPL1 | 36.2481529 | -0.1728603 | 0.24730288 | -0.6989822 | 0.48456315 | 0.99046213 |
| HSPH1 | 42.7055603 | -0.2387609 | 0.40789931 | -0.5853426 | 0.55831735 | 0.9958918 |
| B3GLCT | 29.342177 | 0.2357913 | 0.41282329 | 0.57116763 | 0.56788602 | 0.9958918 |
| FRY | 323.388005 | -0.2641369 | 0.38802612 | -0.6807194 | 0.49604903 | 0.99176652 |
| N4BP2L1 | 35.4789109 | 0.45799779 | 0.43284184 | 1.05811811 | 0.2900016 | 0.95186143 |
| N4BP2L2 | 441.715611 | 0.03187211 | 0.16705984 | 0.19078262 | 0.84869591 | 0.99713289 |
| PDS5B | 298.072553 | -0.4284191 | 0.28219591 | -1.518162 | 0.12897357 | 0.80460676 |

| | | | | | | |
|---|---|---|---|---|---|---|
| STARD13 | 513.12434 | -0.4674529 | 0.33013881 | -1.4159283 | 0.1567965 | 0.84427321 |
| AL139383.1 | 90.4286467 | -1.3719943 | 0.53060336 | -2.5857249 | 0.00971745 | 0.26158795 |
| RFC3 | 25.7123377 | -0.150825 | 0.37576242 | -0.401384 | 0.68813744 | 0.9958918 |
| NBEA | 324.036046 | 0.2090022 | 0.27794036 | 0.75196779 | 0.45207043 | 0.98092839 |
| DCLK1 | 156.945953 | -0.6137415 | 0.52511516 | -1.168775 | 0.2424943 | 0.92534155 |
| SPART | 128.072627 | -0.4596942 | 0.32461889 | -1.4161044 | 0.15674494 | 0.84427321 |
| SMAD9 | 28.4486746 | 0.88971165 | 0.34570349 | 2.57362643 | 0.01006389 | 0.26700182 |
| ALG5 | 28.0097732 | 0.0099557 | 0.31253267 | 0.03185492 | 0.97458775 | 0.99935941 |
| SUPT20H | 71.9314914 | 0.16852849 | 0.28451838 | 0.59232901 | 0.5536303 | 0.9958918 |
| POSTN | 178.237435 | 2.05369188 | 0.88490193 | 2.32081297 | 0.02029694 | 0.38543207 |
| TRPC4 | 323.42944 | -0.8050617 | 0.4282258 | -1.8799934 | 6.01E-02 | 0.61332974 |
| UFM1 | 70.6864624 | -0.2942665 | 0.26153434 | -1.1251542 | 0.26052368 | 0.93055907 |
| NHLRC3 | 33.5642818 | 0.23117352 | 0.45666753 | 0.50621843 | 0.61270332 | 0.9958918 |
| LHFPL6 | 913.98732 | 0.36719126 | 0.41850987 | 0.87737778 | 0.38028148 | 0.97217293 |
| COG6 | 65.6888735 | 0.02591723 | 0.35494832 | 0.07301691 | 0.94179266 | 0.99935941 |
| FOXO1 | 210.024358 | -0.2291494 | 0.37341169 | -0.6136642 | 0.53943723 | 0.9958918 |
| MRPS31 | 33.1842374 | 0.29056605 | 0.27653583 | 1.05073562 | 0.29338003 | 0.95189839 |
| ELF1 | 176.39257 | -0.2103041 | 0.2992739 | -0.7027146 | 0.48223365 | 0.99002719 |
| WBP4 | 37.248869 | 0.2885878 | 0.29624931 | 0.97413829 | 0.32998788 | 0.96348071 |
| NAA16 | 68.0344903 | -0.2055133 | 0.26902325 | -0.7639238 | 0.44491266 | 0.98092839 |
| RGCC | 44.1031945 | -0.891794 | 1.04589398 | -0.852662 | 0.39384679 | 0.97500135 |
| VWA8 | 210.52942 | -0.5290036 | 0.19950605 | -2.651567 | 0.00801192 | 0.23533118 |
| DGKH | 301.146149 | 0.00502385 | 0.2798055 | 0.01795478 | 0.98567493 | 0.99935941 |
| AKAP11 | 78.2845541 | 0.21626199 | 0.22684396 | 0.95335133 | 0.34041209 | 0.96887708 |
| EPSTI1 | 84.7667195 | -1.3722351 | 1.08170759 | -1.2685823 | 0.20459008 | 0.89495984 |
| DNAJC15 | 54.1227077 | 0.4396096 | 0.27906805 | 1.57527745 | 0.11519242 | 0.78022956 |
| ENOX1 | 102.175805 | 0.042095 | 0.35026943 | 0.12017892 | 0.90434142 | 0.99935941 |
| CCDC122 | 25.6341148 | 1.33480959 | 0.46793461 | 2.85255584 | 0.00433692 | 0.16344625 |
| TSC22D1 | 292.654659 | 0.08304929 | 0.2507648 | 0.33118399 | 0.74050551 | 0.9958918 |
| GPALPP1 | 38.3181223 | 0.05895718 | 0.36264507 | 0.16257544 | 0.87085273 | 0.9971731 |
| GTF2F2 | 64.3040419 | -0.1959169 | 0.30722749 | -0.6376934 | 0.5236733 | 0.9958918 |
| TPT1 | 317.063674 | 0.12746344 | 0.40571463 | 0.31417019 | 0.75339178 | 0.9958918 |
| COG3 | 69.0438445 | -0.0759102 | 0.23895764 | -0.3176721 | 0.75073368 | 0.9958918 |
| ZC3H13 | 95.7824798 | -0.1091109 | 0.22237946 | -0.4906517 | 0.6236728 | 0.9958918 |
| LRCH1 | 226.882675 | 0.21380383 | 0.25148418 | 0.85016812 | 0.39523162 | 0.97500135 |
| ESD | 42.2427402 | 0.12503209 | 0.27036036 | 0.46246459 | 0.64374819 | 0.9958918 |
| SUCLA2 | 59.5433758 | -0.3438616 | 0.38771077 | -0.8869025 | 0.37513141 | 0.97217293 |
| MED4 | 50.6741505 | 0.09070914 | 0.22084223 | 0.41074182 | 0.68126186 | 0.9958918 |
| ITM2B | 353.205117 | -0.0622687 | 0.2228492 | -0.2794206 | 0.77992206 | 0.9958918 |
| RB1 | 227.557592 | -0.0815617 | 0.13655214 | -0.5972936 | 0.55031139 | 0.9958918 |
| LPAR6 | 38.6640392 | -0.6858805 | 0.39934221 | -1.7175257 | 0.08588316 | 0.70916334 |
| RCBTB2 | 20.0106076 | 0.11887009 | 0.35873791 | 0.33135636 | 0.74037532 | 0.9958918 |
| FNDC3A | 415.328546 | 0.14001052 | 0.33226749 | 0.42137894 | 0.6734784 | 0.9958918 |
| CDADC1 | 20.5192604 | -0.3484785 | 0.36635863 | -0.951195 | 0.3415054 | 0.96887708 |
| CAB39L | 36.5354807 | 0.00284027 | 0.29515739 | 0.00962291 | 0.99232215 | 0.99935941 |
| SETDB2 | 25.9320086 | -0.5875498 | 0.45008102 | -1.3054311 | 0.1917461 | 0.88158763 |
| PHF11 | 40.3611226 | -0.1713434 | 0.39926028 | -0.4291522 | 0.6678125 | 0.9958918 |
| RCBTB1 | 23.5244154 | -0.2271311 | 0.45155526 | -0.5029974 | 0.61496609 | 0.9958918 |
| KPNA3 | 71.4919712 | 0.01671091 | 0.19323898 | 0.08647792 | 0.9310865 | 0.99935941 |
| DLEU2 | 330.829375 | 0.55826721 | 0.35147729 | 1.58834505 | 0.11220834 | 0.77793929 |
| TRIM13 | 42.9296242 | 0.03540331 | 0.33064685 | 0.10707289 | 0.91473115 | 0.99935941 |
| RNASEH2B | 49.7502616 | 0.04796467 | 0.31935475 | 0.15019244 | 0.88061279 | 0.99749293 |
| INTS6 | 156.70399 | 0.15592464 | 0.30199365 | 0.51631762 | 0.60563259 | 0.9958918 |
| WDFY2 | 130.402445 | 0.44194977 | 0.31425952 | 1.40632104 | 0.15962881 | 0.84762439 |

| | | | | | | |
|---|---|---|---|---|---|---|
| VPS36 | 61.6171354 | 0.07201865 | 0.22106272 | 0.3257838 | 0.74458794 | 0.9958918 |
| SUGT1 | 38.816004 | 0.49870392 | 0.24855063 | 2.00644799 | 0.04480847 | 0.55894748 |
| PCDH17 | 46.5186068 | 1.04661341 | 0.63892851 | 1.63807591 | 0.10140586 | 0.75199966 |
| TDRD3 | 120.394365 | 0.39619158 | 0.22828023 | 1.73554926 | 0.08264356 | 0.70018756 |
| PCDH9 | 383.101567 | 1.2074814 | 0.42551709 | 2.83768016 | 0.00454427 | 0.16781827 |
| DACH1 | 612.039034 | -1.3274914 | 0.42765751 | -3.1040993 | 0.00190859 | 0.09345751 |
| DIS3 | 34.8134297 | -0.1229469 | 0.26598369 | -0.4622346 | 0.64391307 | 0.9958918 |
| PIBF1 | 168.592978 | 0.0721744 | 0.2138582 | 0.33748716 | 0.73574969 | 0.9958918 |
| KLF12 | 296.643195 | -0.1075627 | 0.33173724 | -0.3242406 | 0.74575586 | 0.9958918 |
| TBC1D4 | 178.083378 | -0.5240271 | 0.32779647 | -1.5986357 | 0.10990156 | 0.77303254 |
| COMMD6 | 29.4547534 | 0.02703207 | 0.41544339 | 0.065068 | 0.94811986 | 0.99935941 |
| UCHL3 | 25.6828884 | -0.2111413 | 0.35371516 | -0.5969247 | 0.55055765 | 0.9958918 |
| LMO7 | 166.275294 | -0.6234337 | 0.42503919 | -1.4667675 | 0.14243932 | 0.81652619 |
| AC001226.2 | 26.750668 | -0.2850214 | 0.43338816 | -0.6576584 | 0.51075766 | 0.9958918 |
| FBXL3 | 75.3797095 | 0.43454493 | 0.31570157 | 1.3764421 | 0.1686848 | 0.86629252 |
| MYCBP2 | 383.595681 | -0.2342803 | 0.27947014 | -0.8383016 | 0.40186135 | 0.97500135 |
| EDNRB | 180.121589 | -0.9543393 | 0.45557047 | -2.0948225 | 0.03618677 | 0.51215534 |
| RBM26 | 186.773684 | -0.3197761 | 0.20638472 | -1.5494176 | 0.12128137 | 0.79021701 |
| NDFIP2 | 43.3251454 | -0.0986464 | 0.35836068 | -0.2752714 | 0.78310773 | 0.9958918 |
| SPRY2 | 34.5463371 | -2.0337438 | 0.47167556 | -4.311743 | 1.62E-05 | 0.0042512 |
| GPC5 | 36.3246148 | -0.4936389 | 0.53582261 | -0.9212731 | 0.35690787 | 0.97217293 |
| GPC6 | 760.92312 | 0.04193769 | 0.36734299 | 0.11416494 | 0.90910704 | 0.99935941 |
| GPR180 | 23.2257064 | -0.5292271 | 0.31847095 | -1.6617752 | 0.09655786 | 0.74071367 |
| ABCC4 | 88.3601462 | -2.0031975 | 0.48353833 | -4.1427894 | 3.43E-05 | 0.00741616 |
| DZIP1 | 56.2853081 | -0.04265 | 0.25283922 | -0.1686842 | 0.86604507 | 0.9971731 |
| DNAJC3 | 109.020245 | 0.0248837 | 0.27962305 | 0.08899016 | 0.92908973 | 0.99935941 |
| UGGT2 | 272.376998 | -0.1980476 | 0.27478948 | -0.7207248 | 0.47107888 | 0.98910524 |
| HS6ST3 | 252.821319 | -1.8925195 | 0.51536311 | -3.6722061 | 0.00024047 | 0.02524546 |
| MBNL2 | 418.23701 | -0.2899706 | 0.32275204 | -0.8984314 | 0.36895562 | 0.97217293 |
| RAP2A | 32.7821362 | 0.22322213 | 0.40511119 | 0.55101448 | 0.58162375 | 0.9958918 |
| IPO5 | 55.4368133 | -0.1643367 | 0.32275291 | -0.5091719 | 0.61063176 | 0.9958918 |
| FARP1 | 246.783432 | -0.038239 | 0.19822347 | -0.1929085 | 0.8470306 | 0.99692952 |
| STK24 | 112.30639 | 0.06746824 | 0.25330346 | 0.26635339 | 0.78996706 | 0.99592423 |
| DOCK9 | 93.1376422 | 0.08637439 | 0.27799615 | 0.31070355 | 0.756026 | 0.9958918 |
| UBAC2 | 114.335688 | 0.00210191 | 0.21821946 | 0.00963208 | 0.99231483 | 0.99935941 |
| TM9SF2 | 79.1502012 | 0.10460586 | 0.21393836 | 0.48895328 | 0.62487477 | 0.9958918 |
| CLYBL | 34.1766069 | -0.129236 | 0.42745045 | -0.3023416 | 0.7623917 | 0.9958918 |
| PCCA | 253.546987 | 0.05171926 | 0.21341853 | 0.24233727 | 0.80851884 | 0.99592423 |
| GGACT | 21.8734531 | 0.05452135 | 0.34617295 | 0.15749742 | 0.87485284 | 0.9971731 |
| ITGBL1 | 54.2849481 | -0.6284126 | 1.03362354 | -0.6079705 | 0.54320703 | 0.9958918 |
| FGF14 | 160.158174 | -0.6310019 | 0.6127055 | -1.0298617 | 0.30307491 | 0.95650251 |
| TPP2 | 88.1800758 | -0.1489754 | 0.22432501 | -0.6641054 | 0.50662288 | 0.99397937 |
| EFNB2 | 23.5367933 | 1.33687812 | 0.50838798 | 2.62964147 | 0.0085475 | 0.24537324 |
| ARGLU1 | 183.284131 | 0.13004232 | 0.16523986 | 0.78699121 | 0.43128701 | 0.98092839 |
| FAM155A | 55.3563654 | 1.71710603 | 0.44281731 | 3.87768497 | 0.00010546 | 0.01600327 |
| ABHD13 | 19.7037874 | 0.19388182 | 0.44202473 | 0.4386221 | 0.66093538 | 0.9958918 |
| IRS2 | 55.0978245 | 0.43824655 | 0.24640678 | 1.77854907 | 0.07531372 | 0.67167936 |
| COL4A1 | 1460.23409 | -0.9934505 | 0.55594609 | -1.7869548 | 0.0739448 | 0.66886321 |
| COL4A2 | 1161.32251 | -0.4905228 | 0.42036126 | -1.1669077 | 0.24324766 | 0.92534155 |
| RAB20 | 47.4644916 | 0.37270437 | 0.50103519 | 0.74386865 | 0.45695595 | 0.98335849 |
| NAXD | 27.9039305 | -0.0504424 | 0.31051727 | -0.1624465 | 0.87095426 | 0.9971731 |
| CARS2 | 54.4148119 | 0.38326266 | 0.27007821 | 1.41908028 | 0.15587561 | 0.84427321 |
| ANKRD10 | 259.45902 | 0.04733791 | 0.27930442 | 0.16948499 | 0.86541518 | 0.9971731 |
| ARHGEF7 | 244.888979 | -0.4853346 | 0.22556324 | -2.1516564 | 0.03142443 | 0.48041788 |

| | | | | | | |
|---|---|---|---|---|---|---|
| TUBGCP3 | 47.2279548 | -0.0072079 | 0.29695343 | -0.0242728 | 0.98063502 | 0.99935941 |
| ATP11A | 112.108125 | 0.16657582 | 0.39345397 | 0.423368 | 0.67202679 | 0.9958918 |
| PCID2 | 34.4785212 | 0.17123058 | 0.2762874 | 0.6197553 | 0.5354189 | 0.9958918 |
| CUL4A | 80.2610244 | -0.2334594 | 0.23756901 | -0.9827014 | 0.32575443 | 0.95992644 |
| LAMP1 | 71.2295847 | -0.1555877 | 0.2214529 | -0.702577 | 0.48231941 | 0.99002719 |
| DCUN1D2 | 27.7418241 | -0.3765075 | 0.29756083 | -1.2653126 | 0.2057593 | 0.89589437 |
| TMCO3 | 56.0294604 | 0.38845565 | 0.28605751 | 1.35796346 | 0.17447528 | 0.8740623 |
| TFDP1 | 26.1701619 | -0.6890635 | 0.34834588 | -1.9781016 | 0.04791724 | 0.56981199 |
| GAS6 | 69.310391 | 0.13805178 | 0.32207094 | 0.42863778 | 0.66818685 | 0.9958918 |
| RASA3 | 99.9597006 | -0.0039622 | 0.43740674 | -0.0090583 | 0.99277263 | 0.99935941 |
| CDC16 | 49.1093253 | 0.01376175 | 0.22064442 | 0.06237073 | 0.95026761 | 0.99935941 |
| UPF3A | 64.7063517 | -0.0916628 | 0.23582093 | -0.3886966 | 0.69750059 | 0.9958918 |
| CCNB1IP1 | 30.3180297 | -0.6342155 | 0.35527844 | -1.7851224 | 0.07424146 | 0.66886321 |
| TEP1 | 25.0791539 | -0.0992326 | 0.30345669 | -0.3270074 | 0.74366228 | 0.9958918 |
| NDRG2 | 49.2051495 | -0.4665289 | 0.40885427 | -1.141064 | 0.25384328 | 0.92646125 |
| ARHGEF40 | 26.3613567 | 0.38896376 | 0.39551239 | 0.98344267 | 0.32538962 | 0.95992644 |
| HNRNPC | 245.751711 | -0.0436135 | 0.2453921 | -0.1777299 | 0.85893513 | 0.9971731 |
| SUPT16H | 54.161246 | -0.0993782 | 0.26741037 | -0.3716318 | 0.71016703 | 0.9958918 |
| CHD8 | 65.5298268 | -0.1031204 | 0.23102006 | -0.4463701 | 0.65532995 | 0.9958918 |
| RAB2B | 21.6698726 | -0.4133411 | 0.3641084 | -1.1352143 | 0.25628555 | 0.9266065 |
| TOX4 | 33.9989607 | 0.11849022 | 0.26834202 | 0.44156414 | 0.65880463 | 0.9958918 |
| DAD1 | 55.6405892 | -0.2708608 | 0.41395841 | -0.6543188 | 0.51290641 | 0.9958918 |
| OXA1L | 19.0042946 | 0.3189194 | 0.42979461 | 0.74202744 | 0.45807071 | 0.98431627 |
| MMP14 | 42.4999422 | -0.0863119 | 0.38579053 | -0.2237274 | 0.82296945 | 0.99592423 |
| LRP10 | 37.9757972 | 0.09780438 | 0.3493919 | 0.27992743 | 0.77953318 | 0.9958918 |
| RBM23 | 34.5598358 | 0.42766345 | 0.32331355 | 1.32275141 | 0.18591806 | 0.8789337 |
| PSMB5 | 35.5569327 | 0.14770361 | 0.4035785 | 0.36598483 | 0.7143764 | 0.9958918 |
| ACIN1 | 100.156512 | -0.2785933 | 0.2017921 | -1.3805955 | 0.16740336 | 0.86560961 |
| SLC7A8 | 28.5333478 | 0.48125411 | 0.41017517 | 1.17328922 | 0.24067984 | 0.92426606 |
| PABPN1 | 38.8301851 | -0.3466091 | 0.33690067 | -1.0288169 | 0.30356574 | 0.95705905 |
| PSME1 | 28.0244708 | -0.2660017 | 0.43982072 | -0.6047959 | 0.54531465 | 0.9958918 |
| NFATC4 | 66.0674838 | -0.1679783 | 0.38274948 | -0.4388728 | 0.66075369 | 0.9958918 |
| KHNYN | 17.7499116 | 0.06991859 | 0.37207131 | 0.18791717 | 0.85094158 | 0.9971731 |
| NOVA1 | 88.7903822 | 0.06720094 | 0.30166352 | 0.22276786 | 0.82371619 | 0.99592423 |
| PRKD1 | 308.935288 | -0.1079 | 0.24709632 | -0.4366716 | 0.6623495 | 0.9958918 |
| G2E3 | 40.868469 | -0.0386706 | 0.26341249 | -0.1468064 | 0.88328484 | 0.99862036 |
| SCFD1 | 173.333219 | -0.3471456 | 0.1523858 | -2.2780703 | 0.02272239 | 0.40828078 |
| STRN3 | 266.167395 | -0.2196812 | 0.23984391 | -0.915934 | 0.35970151 | 0.97217293 |
| AP4S1 | 19.3817729 | -0.0665597 | 0.39442126 | -0.1687529 | 0.86599101 | 0.9971731 |
| HECTD1 | 183.134016 | -0.103381 | 0.16790481 | -0.6157118 | 0.53808475 | 0.9958918 |
| HEATR5A | 41.4120523 | 0.28178304 | 0.27275079 | 1.03311538 | 0.30154991 | 0.95597956 |
| NUBPL | 111.6647 | -0.5711692 | 0.28098215 | -2.0327598 | 0.04207679 | 0.5458772 |
| ARHGAP5 | 272.70483 | -0.3130568 | 0.18009035 | -1.7383318 | 0.08215236 | 0.69958018 |
| AKAP6 | 125.286427 | 1.39420241 | 0.44873274 | 3.10697723 | 0.00189011 | 0.09345751 |
| NPAS3 | 202.332171 | -0.6066996 | 0.43619959 | -1.3908761 | 0.16426301 | 0.85697309 |
| EAPP | 42.483959 | -0.3287498 | 0.33307341 | -0.9870192 | 0.32363324 | 0.95992644 |
| SNX6 | 89.7597421 | -0.1759192 | 0.26016225 | -0.6761905 | 0.49891969 | 0.99176652 |
| CFL2 | 19.746169 | 0.37967439 | 0.37705174 | 1.00695568 | 0.31395607 | 0.95992644 |
| BAZ1A | 81.0440376 | 0.06786084 | 0.36858857 | 0.18411 | 0.85392716 | 0.9971731 |
| SRP54 | 85.649456 | -0.3296166 | 0.22430895 | -1.4694757 | 0.1417038 | 0.81652619 |
| FAM177A1 | 45.413697 | 0.4101437 | 0.30096136 | 1.36277859 | 0.1729523 | 0.87233975 |
| PPP2R3C | 44.503053 | 0.28179493 | 0.27795906 | 1.01380013 | 0.31067813 | 0.95956408 |
| NFKBIA | 126.599633 | 0.51852833 | 0.61320997 | 0.84559671 | 0.39777776 | 0.97500135 |
| RALGAPA1 | 288.359343 | 0.06524546 | 0.21222587 | 0.30743406 | 0.75851301 | 0.9958918 |

| | | | | | | |
|---|---|---|---|---|---|---|
| BRMS1L | 26.8079981 | 0.18737234 | 0.35115928 | 0.5335822 | 0.59363063 | 0.9958918 |
| MBIP | 19.7858132 | -0.1479675 | 0.33865373 | -0.4369286 | 0.66216316 | 0.9958918 |
| MIPOL1 | 119.316271 | -0.2412467 | 0.26714466 | -0.9030566 | 0.36649587 | 0.97217293 |
| SEC23A | 118.795818 | -0.0939281 | 0.24500726 | -0.3833685 | 0.70144653 | 0.9958918 |
| TRAPPC6B | 32.6674839 | -0.0736328 | 0.31279173 | -0.2354053 | 0.81389415 | 0.99592423 |
| PNN | 67.089794 | 0.17161043 | 0.2999837 | 0.57206583 | 0.56727738 | 0.9958918 |
| MIA2 | 71.7369746 | -0.001277 | 0.21522701 | -0.0059333 | 0.99526592 | 0.99935941 |
| FBXO33 | 48.7825802 | 0.327227 | 0.26384811 | 1.24020973 | 0.21489783 | 0.9006731 |
| LRFN5 | 66.8110234 | 1.17756453 | 0.53996781 | 2.18080505 | 0.02919784 | 0.46444788 |
| KLHL28 | 38.4399304 | 0.4924284 | 0.34458802 | 1.42903515 | 0.15299413 | 0.84093985 |
| TOGARAM1 | 66.6811215 | 0.15999485 | 0.25485696 | 0.62778293 | 0.53014615 | 0.9958918 |
| PRPF39 | 40.5982926 | -0.4725422 | 0.3614736 | -1.307266 | 0.19112238 | 0.88158763 |
| FKBP3 | 20.4519132 | -0.0828657 | 0.34996812 | -0.2367807 | 0.81282695 | 0.99592423 |
| MIS18BP1 | 32.2076583 | -0.0047947 | 0.34139011 | -0.0140446 | 0.98879444 | 0.99935941 |
| MDGA2 | 62.3433449 | -1.6549566 | 0.4651286 | -3.5580624 | 0.0003736 | 0.03761083 |
| RPS29 | 61.0803461 | -0.1343942 | 0.27042677 | -0.4969709 | 0.6192096 | 0.9958918 |
| RPL36AL | 71.1810776 | 0.1694072 | 0.33033699 | 0.51283145 | 0.60806923 | 0.9958918 |
| KLHDC1 | 62.3261951 | 0.31443159 | 0.3833767 | 0.82016351 | 0.4121229 | 0.97812179 |
| KLHDC2 | 43.023896 | 0.05662773 | 0.35903886 | 0.15772034 | 0.87467717 | 0.9971731 |
| NEMF | 138.149362 | -0.1879885 | 0.17492504 | -1.0746803 | 0.2825179 | 0.94419503 |
| SOS2 | 202.983132 | 0.07018054 | 0.20650995 | 0.33984098 | 0.73397628 | 0.9958918 |
| L2HGDH | 18.9838957 | 0.01755047 | 0.40699583 | 0.043122 | 0.96560428 | 0.99935941 |
| ATP5S | 18.5225747 | -0.0659801 | 0.38812156 | -0.1699984 | 0.86501137 | 0.9971731 |
| CDKL1 | 35.9568558 | -0.6556416 | 0.45398361 | -1.4441966 | 0.14868368 | 0.8301633 |
| MAP4K5 | 176.985244 | -0.3011595 | 0.25210256 | -1.1945911 | 0.23224684 | 0.91565561 |
| ATL1 | 21.7406234 | -0.1722422 | 0.47423429 | -0.3632007 | 0.71645493 | 0.9958918 |
| SAV1 | 55.5506349 | 0.2843553 | 0.3636032 | 0.78204841 | 0.43418612 | 0.98092839 |
| NIN | 78.7850329 | 0.54601696 | 0.3173109 | 1.7207633 | 0.08529378 | 0.70747633 |
| PYGL | 45.1466982 | 0.7366523 | 0.37319031 | 1.97393204 | 0.04838947 | 0.57172704 |
| TRIM9 | 52.1144628 | 1.0320475 | 0.47896759 | 2.15473348 | 0.0311827 | 0.48041788 |
| TMX1 | 29.1539305 | -0.0211333 | 0.34293844 | -0.0616242 | 0.95086208 | 0.99935941 |
| FRMD6 | 98.0350331 | -0.2361355 | 0.28074521 | -0.8411026 | 0.40029044 | 0.97500135 |
| GNG2 | 30.8780503 | -0.4923434 | 0.41898291 | -1.1750918 | 0.23995798 | 0.92426606 |
| RTRAF | 35.6803611 | 0.05727414 | 0.30565275 | 0.18738304 | 0.85136032 | 0.9971731 |
| NID2 | 80.5808941 | -0.6914203 | 0.35811164 | -1.9307396 | 0.05351526 | 0.58786795 |
| TXNDC16 | 33.259586 | -0.2521008 | 0.50321876 | -0.5009765 | 0.61638767 | 0.9958918 |
| ERO1A | 119.691666 | -1.2068598 | 0.62008829 | -1.9462709 | 0.05162221 | 0.57655256 |
| PSMC6 | 55.8578862 | -0.4028571 | 0.22828456 | -1.7647148 | 0.0776117 | 0.68020338 |
| FERMT2 | 257.593362 | -0.251354 | 0.33364616 | -0.753355 | 0.45123659 | 0.98092839 |
| DDHD1 | 63.159521 | -0.0127141 | 0.24799346 | -0.051268 | 0.959112 | 0.99935941 |
| CNIH1 | 26.8761713 | -0.3517939 | 0.2822325 | -1.2464685 | 0.21259244 | 0.89738044 |
| GMFB | 20.7357876 | 0.14749208 | 0.32253885 | 0.4572847 | 0.64746643 | 0.9958918 |
| SAMD4A | 444.860911 | -1.0380716 | 0.47031797 | -2.2071698 | 0.0273022 | 0.44841843 |
| MAPK1IP1L | 76.6116447 | -0.1003991 | 0.25135627 | -0.3994296 | 0.6895767 | 0.9958918 |
| LGALS3 | 46.9104934 | -0.1649884 | 0.31900472 | -0.5171974 | 0.60501837 | 0.9958918 |
| FBXO34 | 77.4699312 | -0.3297852 | 0.3112432 | -1.0595741 | 0.28933842 | 0.95181201 |
| ATG14 | 26.7855374 | 0.62609102 | 0.54308452 | 1.15284271 | 0.24897495 | 0.92624282 |
| KTN1 | 179.307399 | 0.06474997 | 0.20659999 | 0.31340741 | 0.75397116 | 0.9958918 |
| PELI2 | 309.620047 | -0.4652537 | 0.50311252 | -0.9247508 | 0.35509557 | 0.97217293 |
| TMEM260 | 57.4070484 | -0.0747786 | 0.25607381 | -0.2920199 | 0.77027144 | 0.9958918 |
| EXOC5 | 63.0301604 | 0.00117049 | 0.22661832 | 0.00516501 | 0.99587894 | 0.99935941 |
| AP5M1 | 28.8218377 | 0.43651801 | 0.28824531 | 1.51439764 | 0.12992502 | 0.80565822 |
| NAA30 | 20.237099 | 0.04228076 | 0.33155028 | 0.12752443 | 0.89852534 | 0.99935941 |
| ARMH4 | 47.8415224 | 0.00714054 | 0.26907779 | 0.02653707 | 0.97882897 | 0.99935941 |

| | | | | | | |
|---|---|---|---|---|---|---|
| ACTR10 | 50.563633 | -0.008036 | 0.21770474 | -0.0369122 | 0.97055501 | 0.99935941 |
| PSMA3 | 53.6807534 | -0.3170135 | 0.31428296 | -1.008688 | 0.31312428 | 0.95992644 |
| ARID4A | 79.203461 | 0.00410174 | 0.35019106 | 0.01171286 | 0.99065471 | 0.99935941 |
| TIMM9 | 22.1732225 | -0.1281707 | 0.30751665 | -0.4167926 | 0.67683008 | 0.9958918 |
| KIAA0586 | 57.7538803 | 0.26519134 | 0.28187036 | 0.94082731 | 0.34679337 | 0.96887708 |
| DACT1 | 25.7397868 | -0.5655834 | 0.49730163 | -1.1373045 | 0.25541102 | 0.9266065 |
| DAAM1 | 278.628797 | -0.5840379 | 0.50016285 | -1.1676954 | 0.24292964 | 0.92534155 |
| GPR135 | 25.5273917 | -0.873144 | 0.42844775 | -2.0379242 | 0.04155752 | 0.54347495 |
| L3HYPDH | 21.7547062 | -0.8525648 | 0.36080894 | -2.362926 | 0.01813129 | 0.35723014 |
| JKAMP | 27.0850538 | -0.0250345 | 0.34065948 | -0.0734884 | 0.9414175 | 0.99935941 |
| PCNX4 | 118.749945 | -0.0061816 | 0.26648246 | -0.023197 | 0.98149315 | 0.99935941 |
| DHRS7 | 21.6637126 | -0.0759917 | 0.41156933 | -0.1846389 | 0.85351229 | 0.9971731 |
| PPM1A | 52.8355976 | -0.1786376 | 0.26272137 | -0.6799509 | 0.49653552 | 0.99176652 |
| MNAT1 | 157.747977 | -0.1849313 | 0.21817643 | -0.8476227 | 0.39664813 | 0.97500135 |
| SLC38A6 | 42.4909995 | -0.5728719 | 0.25671791 | -2.231523 | 0.02564651 | 0.4322848 |
| PRKCH | 74.4584745 | 0.28495428 | 0.3779448 | 0.7539574 | 0.4508748 | 0.98092839 |
| HIF1A | 755.382023 | -0.1728834 | 0.31614135 | -0.5468549 | 5.84E-01 | 0.9958918 |
| HIF1A-AS2 | 268.023071 | -0.2978154 | 0.8117721 | -0.3668707 | 7.14E-01 | 0.9958918 |
| SNAPC1 | 40.3327377 | -0.3538078 | 0.34021968 | -1.0399393 | 2.98E-01 | 0.95447177 |
| RHOJ | 65.1941559 | -0.275803 | 0.42779119 | -0.644714 | 0.51911253 | 0.9958918 |
| PPP2R5E | 171.292193 | 0.26365817 | 0.21975614 | 1.19977611 | 0.2302263 | 0.91455844 |
| SGPP1 | 42.7653716 | -0.0125626 | 0.26432869 | -0.0475263 | 0.96209378 | 0.99935941 |
| SYNE2 | 580.656115 | -0.1656661 | 0.37338697 | -0.4436848 | 0.65727047 | 0.9958918 |
| ESR2 | 93.9217994 | 0.8765676 | 0.48283939 | 1.81544342 | 0.06945579 | 0.65272459 |
| ZBTB25 | 39.841213 | -0.2071338 | 0.33959514 | -0.6099433 | 0.54189937 | 0.9958918 |
| ZBTB1 | 101.319144 | 0.0962336 | 0.34046648 | 0.2826522 | 0.77744346 | 0.9958918 |
| PLEKHG3 | 42.8166798 | -0.2451854 | 0.3754464 | -0.6530502 | 0.51372391 | 0.9958918 |
| MAX | 73.6717721 | -0.4086849 | 0.32980172 | -1.2391836 | 0.21527752 | 0.90146693 |
| FUT8 | 98.8103069 | -0.0627842 | 0.2625604 | -0.2391229 | 0.81101026 | 0.99592423 |
| AL359232.1 | 31.4424041 | -0.6172486 | 0.52923941 | -1.1662937 | 0.24349572 | 0.92534155 |
| GPHN | 502.420801 | -0.0287869 | 0.24371001 | -0.1181197 | 0.90597283 | 0.99935941 |
| MPP5 | 54.7275968 | -0.4836662 | 0.30202686 | -1.6014014 | 0.10928805 | 0.77303254 |
| ATP6V1D | 36.7973685 | -0.4555382 | 0.32059514 | -1.4209142 | 0.15534171 | 0.84277602 |
| EIF2S1 | 32.2124492 | -0.0404932 | 0.31830404 | -0.1272155 | 0.8987698 | 0.99935941 |
| ZFYVE26 | 25.344908 | -0.5485696 | 0.3658371 | -1.4994914 | 0.1337462 | 0.80812129 |
| RAD51B | 205.308615 | -0.4733659 | 0.4218858 | -1.1220239 | 0.26185229 | 0.93155278 |
| ZFP36L1 | 434.67109 | -0.7297675 | 0.4099281 | -1.7802328 | 0.07503786 | 0.67167936 |
| ACTN1 | 403.415247 | -0.185139 | 0.23868813 | -0.7756524 | 0.43795423 | 0.98092839 |
| DCAF5 | 138.212435 | 0.06795093 | 0.2687005 | 0.25288724 | 0.80035535 | 0.99592423 |
| GALNT16 | 22.3314469 | 0.72552373 | 0.62804698 | 1.15520614 | 0.24800602 | 0.92624282 |
| ERH | 39.7470223 | -0.2869833 | 0.32099101 | -0.8940542 | 0.37129292 | 0.97217293 |
| SLC39A9 | 52.0637175 | -0.2262946 | 0.25622659 | -0.8831817 | 0.37713809 | 0.97217293 |
| SUSD6 | 115.031066 | 0.68733424 | 0.30570014 | 2.24839363 | 0.0245511 | 0.42353531 |
| SRSF5 | 148.437796 | 0.09228185 | 0.25465941 | 0.36237362 | 0.71707285 | 0.9958918 |
| AL357153.2 | 38.255303 | 0.24035001 | 0.40488221 | 0.59362945 | 0.55275998 | 0.9958918 |
| MED6 | 29.5140061 | -0.4281651 | 0.33540487 | -1.2765619 | 0.20175697 | 0.89319989 |
| PCNX1 | 268.251844 | 0.02959188 | 0.29414362 | 0.10060351 | 0.91986521 | 0.99935941 |
| SIPA1L1 | 203.087753 | -0.041107 | 0.26149504 | -0.1571998 | 0.87508741 | 0.9971731 |
| RGS6 | 134.318042 | 0.50165149 | 0.37579188 | 1.33491841 | 0.18190305 | 0.87866798 |
| ZFYVE1 | 31.7440793 | 0.04634826 | 0.3414333 | 0.13574616 | 0.89202196 | 0.99935941 |
| RBM25 | 129.579352 | 0.03119587 | 0.21277079 | 0.14661724 | 0.88343415 | 0.99862036 |
| PSEN1 | 66.7654181 | -0.0721883 | 0.21446487 | -0.3365972 | 0.73642056 | 0.9958918 |
| PAPLN | 23.3455704 | 0.52375726 | 0.4893522 | 1.07030737 | 0.28448098 | 0.94505166 |
| NUMB | 197.074709 | 0.07341051 | 0.1565087 | 0.46905068 | 0.63903341 | 0.9958918 |

| | | | | | | |
|---|---|---|---|---|---|---|
| DNAL1 | 18.8631741 | 0.39317539 | 0.33982573 | 1.15699122 | 0.24727595 | 0.92621352 |
| ELMSAN1 | 59.9566279 | -0.394038 | 0.3656531 | -1.077628 | 0.2811998 | 0.94419503 |
| AC005480.1 | 22.6526202 | 0.24276863 | 0.48958073 | 0.49587047 | 0.6199858 | 0.9958918 |
| ENTPD5 | 21.7274785 | 0.07863995 | 0.42339002 | 0.18573879 | 0.8526496 | 0.9971731 |
| BBOF1 | 22.4762494 | -0.3385592 | 0.37836467 | -0.8947959 | 0.3708962 | 0.97217293 |
| ALDH6A1 | 25.5970798 | 0.69407265 | 0.43296997 | 1.60305033 | 0.10892354 | 0.77303254 |
| LIN52 | 50.5565969 | 0.22295203 | 0.28240618 | 0.78947293 | 0.42983565 | 0.98092839 |
| ABCD4 | 34.9024468 | 0.00596309 | 0.30635665 | 0.01946455 | 0.98447052 | 0.99935941 |
| NPC2 | 48.7812709 | -0.129488 | 0.41051923 | -0.315425 | 0.752439 | 0.9958918 |
| LTBP2 | 27.1928399 | 0.76895477 | 0.646265 | 1.18984436 | 0.23410757 | 0.91930824 |
| AREL1 | 34.1197724 | 0.23335791 | 0.30397329 | 0.76769215 | 0.4426701 | 0.98092839 |
| YLPM1 | 100.466149 | -0.0551309 | 0.18541948 | -0.2973304 | 0.76621425 | 0.9958918 |
| DLST | 37.1614661 | 0.26353328 | 0.26361359 | 0.99969537 | 0.31745796 | 0.95992644 |
| PGF | 25.1737775 | 0.84499192 | 0.40208104 | 2.10154629 | 0.03559304 | 0.50762504 |
| MLH3 | 58.2843617 | -0.0507263 | 0.26038189 | -0.1948151 | 0.84553772 | 0.99692952 |
| NEK9 | 53.9257111 | -0.2402201 | 0.25495903 | -0.9421911 | 0.34609479 | 0.96887708 |
| TMED10 | 93.3927383 | 0.00681961 | 0.25470476 | 0.02677457 | 0.97863953 | 0.99935941 |
| FOS | 115.112711 | -1.7877308 | 0.61033982 | -2.9290744 | 0.00339973 | 0.14115602 |
| JDP2 | 20.2933682 | -0.1240128 | 0.37357683 | -0.3319608 | 7.40E-01 | 0.9958918 |
| FLVCR2 | 46.308704 | -1.0239456 | 0.63759944 | -1.6059386 | 0.10828739 | 0.77248589 |
| TTLL5 | 106.02892 | -0.733715 | 0.29590197 | -2.479588 | 0.01315343 | 0.30493547 |
| IFT43 | 112.417164 | -0.1553077 | 0.2775983 | -0.5594692 | 0.57584155 | 0.9958918 |
| TGFB3 | 65.7736541 | -0.7317348 | 0.54581288 | -1.340633 | 0.18003963 | 0.87815793 |
| GPATCH2L | 99.6571204 | 0.01380662 | 0.20879681 | 0.06612466 | 0.94727857 | 0.99935941 |
| IRF2BPL | 29.6350599 | -0.4027667 | 0.41881547 | -0.9616805 | 0.33621013 | 0.96640422 |
| TMED8 | 32.493129 | 0.04715698 | 0.32551434 | 0.14486913 | 0.8848142 | 0.99862036 |
| AHSA1 | 24.1619678 | -0.0185102 | 0.447834 | -0.0413327 | 0.96703067 | 0.99935941 |
| SPTLC2 | 54.0558519 | 0.09158916 | 0.29857003 | 0.30675939 | 0.75902652 | 0.9958918 |
| ALKBH1 | 23.9640874 | 0.78762643 | 0.3777156 | 2.08523671 | 0.03704782 | 0.51855699 |
| SLIRP | 29.9730312 | 0.42789312 | 0.34612071 | 1.23625402 | 0.21636416 | 0.90190595 |
| SNW1 | 40.4418554 | -0.0399114 | 0.25549541 | -0.156212 | 0.87586592 | 0.9971731 |
| NRXN3 | 237.016404 | 0.88357378 | 0.51896428 | 1.70257147 | 0.08864829 | 0.71818729 |
| CEP128 | 35.5106078 | -0.5213494 | 0.4945166 | -1.0542607 | 0.29176359 | 0.95186143 |
| GTF2A1 | 44.5721471 | 0.04809866 | 0.26250687 | 0.18322819 | 0.85461898 | 0.9971731 |
| SEL1L | 74.6564402 | -0.0837159 | 0.26182138 | -0.3197442 | 0.74916227 | 0.9958918 |
| FLRT2 | 158.478121 | 0.73177668 | 0.51224861 | 1.42855766 | 0.15313141 | 0.84093985 |
| GALC | 27.2592156 | 0.00217463 | 0.3651194 | 0.00595595 | 0.99524787 | 0.99935941 |
| SPATA7 | 31.3071198 | 0.13613363 | 0.42171321 | 0.32281092 | 0.74683844 | 0.9958918 |
| PTPN21 | 55.7638163 | -0.0742476 | 0.43318449 | -0.1713996 | 0.86390959 | 0.9971731 |
| ZC3H14 | 73.9909436 | -0.1438649 | 0.20146681 | -0.7140872 | 0.47517327 | 0.98910524 |
| TTC8 | 24.2230885 | -0.3558902 | 0.33154856 | -1.0734179 | 0.28308367 | 0.94471801 |
| FOXN3 | 399.615014 | 0.10766949 | 0.33471065 | 0.3216793 | 0.74769566 | 0.9958918 |
| PSMC1 | 31.9627891 | 0.00351016 | 0.36196586 | 0.00969749 | 0.99226264 | 0.99935941 |
| NRDE2 | 32.524343 | 0.31838871 | 0.44198482 | 0.72036119 | 0.47130264 | 0.98910524 |
| CALM1 | 97.81632 | 0.56416552 | 0.21599663 | 2.61191818 | 0.00900358 | 0.25292422 |
| TTC7B | 90.5796384 | -0.0550153 | 0.23773861 | -0.2314107 | 0.81699576 | 0.99592423 |
| RPS6KA5 | 55.939268 | 0.30772058 | 0.4307286 | 0.71441875 | 0.47496828 | 0.98910524 |
| DGLUCY | 49.8135101 | -0.1651897 | 0.41918626 | -0.3940722 | 0.69352773 | 0.9958918 |
| PPP4R3A | 93.5926411 | -0.1863214 | 0.19968844 | -0.9330603 | 0.35078885 | 0.9693503 |
| CATSPERB | 27.2399013 | -0.4211954 | 0.40056765 | -1.0514963 | 0.29303072 | 0.95186143 |
| TC2N | 58.322557 | -0.836954 | 0.47510863 | -1.7616056 | 0.07813595 | 0.68056484 |
| FBLN5 | 255.863291 | -0.6813212 | 0.42411924 | -1.6064379 | 0.10817773 | 0.77248589 |
| TRIP11 | 114.948471 | 0.17279297 | 0.18299475 | 0.94425099 | 0.3450414 | 0.96887708 |
| ATXN3 | 102.270503 | 0.1080229 | 0.27915492 | 0.38696398 | 0.69878287 | 0.9958918 |

| | | | | | | |
|---|---|---|---|---|---|---|
| NDUFB1 | 22.0619828 | 0.40949017 | 0.32518364 | 1.25925819 | 0.20793709 | 0.89633695 |
| CPSF2 | 40.1588612 | 0.04106648 | 0.27333183 | 0.15024403 | 0.88057209 | 0.99749293 |
| RIN3 | 29.6164838 | 0.62102502 | 0.31846755 | 1.95004179 | 0.05117114 | 0.57655256 |
| LGMN | 36.9297372 | 0.44510947 | 0.35771154 | 1.24432515 | 0.21337992 | 0.89738044 |
| GOLGA5 | 31.554357 | -0.2349022 | 0.30222369 | -0.7772461 | 0.43701357 | 0.98092839 |
| ITPK1 | 30.1215077 | 0.65529005 | 0.35079213 | 1.86802951 | 0.06175796 | 0.61962951 |
| BTBD7 | 115.996961 | 0.12239204 | 0.23112427 | 0.52955081 | 0.59642341 | 0.9958918 |
| DDX24 | 82.8307572 | -0.0274929 | 0.21934846 | -0.1253388 | 0.90025531 | 0.99935941 |
| IFI27 | 74.1577879 | -1.7275825 | 0.99828045 | -1.7305582 | 0.08353058 | 0.70213814 |
| SERPINA1 | 44.5349063 | -0.6473868 | 0.90028697 | -0.7190894 | 0.47208585 | 0.98910524 |
| DICER1 | 193.364502 | 0.0096181 | 0.21607598 | 0.04451257 | 0.96449583 | 0.99935941 |
| CLMN | 327.80131 | -0.0858173 | 0.28705806 | -0.2989544 | 0.76497486 | 0.9958918 |
| GLRX5 | 17.8263626 | 0.4573256 | 0.40101614 | 1.14041696 | 0.25411263 | 0.92646125 |
| C14orf132 | 25.361384 | -0.0821958 | 0.54024072 | -0.1521466 | 0.87907135 | 0.99742093 |
| ATG2B | 78.6765319 | -0.3245915 | 0.21905121 | -1.4818067 | 0.13839173 | 0.81520215 |
| PAPOLA | 132.779255 | 0.02054585 | 0.24125926 | 0.08516086 | 0.93213351 | 0.99935941 |
| SETD3 | 76.1612792 | 0.2275121 | 0.25601663 | 0.88866144 | 0.37418506 | 0.97217293 |
| CCNK | 28.4808879 | -0.0064356 | 0.27418104 | -0.0234721 | 0.9812737 | 0.99935941 |
| EML1 | 40.3660934 | 0.22565916 | 0.53224872 | 0.42397314 | 6.72E-01 | 0.9958918 |
| EVL | 131.281093 | 0.36365145 | 0.46034264 | 0.78995822 | 0.42955217 | 0.98092839 |
| YY1 | 92.6073593 | 0.31926816 | 0.23410069 | 1.36380703 | 0.17262831 | 0.87233975 |
| SLC25A29 | 18.2464537 | -0.3339573 | 0.40970083 | -0.8151247 | 0.41500093 | 0.97812179 |
| WARS | 48.4868137 | -0.487536 | 0.54109508 | -0.9010172 | 0.36757915 | 0.97217293 |
| WDR25 | 33.5784357 | 0.14798979 | 0.37023436 | 0.39971922 | 0.68936333 | 0.9958918 |
| MEG3 | 73.4014973 | -0.1159064 | 0.4012261 | -0.2888806 | 0.77267276 | 0.9958918 |
| MEG8 | 98.0752905 | -0.7296851 | 0.41048756 | -1.7776058 | 0.07546862 | 0.67167936 |
| PPP2R5C | 114.911189 | -0.0187735 | 0.22650911 | -0.082882 | 0.93394533 | 0.99935941 |
| DYNC1H1 | 127.492838 | -0.1083428 | 0.24767577 | -0.437438 | 0.66179374 | 0.9958918 |
| HSP90AA1 | 249.543745 | -0.1092903 | 0.38717786 | -0.2822742 | 0.7777333 | 0.9958918 |
| WDR20 | 81.2327565 | 0.42669462 | 0.18504917 | 2.30584448 | 0.02111932 | 0.39000584 |
| MOK | 31.0561639 | -0.3423321 | 0.5496349 | -0.6228355 | 0.53339261 | 0.9958918 |
| TECPR2 | 49.7985204 | 0.16821634 | 0.30128785 | 0.55832436 | 0.57662291 | 0.9958918 |
| RCOR1 | 86.9973063 | 0.25032784 | 0.22425124 | 1.11628296 | 0.26430103 | 0.93499423 |
| TRAF3 | 95.4875126 | 0.87473266 | 0.44121562 | 1.98255145 | 0.04741756 | 0.56939806 |
| CDC42BPB | 122.001481 | -0.0175099 | 0.20349368 | -0.0860464 | 0.93142951 | 0.99935941 |
| TNFAIP2 | 53.7784776 | 1.40434396 | 0.95271907 | 1.47403784 | 0.1404714 | 0.8153985 |
| EIF5 | 55.545548 | 0.45606612 | 0.27639058 | 1.6500784 | 0.0989269 | 0.74528964 |
| MARK3 | 301.584994 | -0.1911608 | 0.24182621 | -0.7904885 | 0.42924255 | 0.98092839 |
| CKB | 40.749023 | 0.68926518 | 0.36796414 | 1.87318578 | 0.06104274 | 0.61696136 |
| KLC1 | 97.1476039 | 0.2302547 | 0.22482892 | 1.02413293 | 0.30577249 | 0.95793395 |
| ZFYVE21 | 25.397651 | -0.1535167 | 0.33948898 | -0.4521993 | 0.65112542 | 0.9958918 |
| PPP1R13B | 46.6512821 | 0.24850412 | 0.34889537 | 0.71225973 | 0.47630395 | 0.98968074 |
| ATP5MPL | 31.6636839 | -0.0651693 | 0.28667264 | -0.2273301 | 0.82016705 | 0.99592423 |
| INF2 | 36.8075324 | 0.11225424 | 0.26654195 | 0.42115037 | 0.67364528 | 0.9958918 |
| SIVA1 | 23.8264454 | -0.3725571 | 0.31478144 | -1.183542 | 0.23659441 | 0.92093871 |
| AKT1 | 21.9444711 | 0.16477858 | 0.45020598 | 0.36600709 | 0.71435979 | 0.9958918 |
| BRF1 | 29.9731306 | 0.09884648 | 0.29369532 | 0.33656131 | 0.73644762 | 0.9958918 |
| PACS2 | 43.9153443 | -0.1313265 | 0.31286658 | -0.4197523 | 0.67466639 | 0.9958918 |
| MTA1 | 23.9865202 | -0.4537562 | 0.35622903 | -1.2737766 | 0.20274262 | 0.89396747 |
| CRIP2 | 49.6826334 | 0.71460257 | 0.48490495 | 1.47369617 | 0.14056341 | 0.8153985 |
| NIPA2 | 22.6996352 | 0.1733185 | 0.30725337 | 0.56408981 | 0.57269301 | 0.9958918 |
| CYFIP1 | 102.862208 | 0.02779046 | 0.25707261 | 0.10810355 | 0.91391356 | 0.99935941 |
| TUBGCP5 | 27.7064013 | 0.09180429 | 0.33387148 | 0.27496895 | 0.78334009 | 0.9958918 |
| PWRN1 | 29.9935827 | -0.3430206 | 0.72273014 | -0.4746179 | 0.63505936 | 0.9958918 |

| | | | | | | |
|---|---|---|---|---|---|---|
| AC124312.1 | 116.564322 | 0.04559107 | 0.23600861 | 0.19317548 | 0.84682154 | 0.99692952 |
| PWAR6 | 16.8399266 | 0.79479965 | 0.38102562 | 2.08594805 | 0.03698333 | 0.51855699 |
| UBE3A | 142.595342 | 0.01799706 | 0.22735179 | 0.07915954 | 0.93690573 | 0.99935941 |
| ATP10A | 144.484495 | -0.5459227 | 0.3013778 | -1.8114233 | 0.07007536 | 0.65603033 |
| HERC2 | 149.59477 | 0.00613931 | 0.23672813 | 0.02593403 | 0.97930996 | 0.99935941 |
| TJP1 | 339.754049 | -0.2572833 | 0.27985033 | -0.9193603 | 0.35790713 | 0.97217293 |
| FAN1 | 20.7776038 | -0.336491 | 0.32483217 | -1.0358918 | 0.30025261 | 0.95597956 |
| MTMR10 | 61.4628039 | 0.02767099 | 0.26161513 | 0.10576982 | 0.91576498 | 0.99935941 |
| KLF13 | 50.6615938 | 0.37751465 | 0.2879291 | 1.31113754 | 0.1898113 | 0.88139754 |
| OTUD7A | 57.4958561 | 0.04350409 | 0.25375117 | 0.1714439 | 0.86387474 | 0.9971731 |
| LINC02256 | 96.964481 | 0.15913866 | 0.29156713 | 0.54580451 | 0.58520032 | 0.9958918 |
| AVEN | 51.7178212 | -0.0003874 | 0.3618254 | -0.0010706 | 0.99914575 | 0.99968761 |
| EMC7 | 27.2046455 | 0.32593139 | 0.28658768 | 1.13728335 | 0.25541985 | 0.9266065 |
| KATNBL1 | 88.3010117 | -0.3183716 | 0.20068712 | -1.5864075 | 0.1126469 | 0.77793929 |
| SLC12A6 | 32.7719705 | 0.0675635 | 0.41989841 | 0.16090441 | 0.87216869 | 0.9971731 |
| NOP10 | 28.7819129 | -0.2863485 | 0.4959671 | -0.5773538 | 0.56370048 | 0.9958918 |
| GOLGA8A | 41.0491136 | -0.4230359 | 0.36743706 | -1.1513154 | 0.24960251 | 0.92624282 |
| GOLGA8B | 20.8322769 | -0.0185999 | 0.43298163 | -0.0429578 | 0.96573521 | 0.99935941 |
| AQR | 63.1354534 | 0.1681917 | 0.2411037 | 0.69759069 | 0.48543321 | 0.99046213 |
| ZNF770 | 25.2597095 | 0.06892623 | 0.32855604 | 0.2097853 | 0.83383525 | 0.9960753 |
| AC021231.1 | 51.9301923 | 0.33450017 | 0.40641778 | 0.82304511 | 0.41048234 | 0.97812179 |
| DPH6 | 155.046571 | 0.12871594 | 0.35390894 | 0.36369791 | 0.7160836 | 0.9958918 |
| DPH6-DT | 109.50324 | -0.0877231 | 0.54299817 | -0.1615532 | 0.87165772 | 0.9971731 |
| C15orf41 | 77.0150173 | -0.0946722 | 0.22344744 | -0.4236889 | 0.67179272 | 0.9958918 |
| MEIS2 | 1085.25955 | -0.198522 | 0.29452977 | -0.6740303 | 0.50029203 | 0.99213111 |
| SPRED1 | 176.296497 | 0.01395619 | 0.25137149 | 0.05552017 | 0.95572406 | 0.99935941 |
| FAM98B | 18.4663228 | 0.56828458 | 0.33900792 | 1.67631651 | 0.09367621 | 0.73159032 |
| THBS1 | 597.908846 | -1.6472163 | 0.46144553 | -3.5696875 | 0.00035741 | 0.03648037 |
| FSIP1 | 47.1893232 | -1.252658 | 0.64387919 | -1.9454861 | 0.0517165 | 0.57672918 |
| GPR176 | 71.0158013 | 0.08993358 | 0.83888629 | 0.10720593 | 0.91462561 | 0.99935941 |
| EIF2AK4 | 50.5368061 | 0.0881459 | 0.23624063 | 0.37311914 | 0.70905977 | 0.9958918 |
| SRP14 | 102.608503 | 0.14545076 | 0.3350287 | 0.43414418 | 0.66418375 | 0.9958918 |
| IVD | 28.9944173 | 0.18187909 | 0.44941792 | 0.40469925 | 0.68569859 | 0.9958918 |
| CCDC32 | 28.0561632 | -0.0434835 | 0.4072496 | -0.1067735 | 0.91496867 | 0.99935941 |
| DNAJC17 | 29.0441206 | 0.23552701 | 0.35271897 | 0.66774693 | 0.50429515 | 0.99387859 |
| INO80 | 104.365524 | -0.118089 | 0.26333811 | -0.4484312 | 0.65384206 | 0.9958918 |
| CHP1 | 33.8191954 | 0.20609162 | 0.27670771 | 0.74479898 | 0.45639325 | 0.98330915 |
| RTF1 | 52.5186983 | 0.38541431 | 0.29315792 | 1.31469864 | 0.1886112 | 0.88139754 |
| MGA | 80.593102 | 0.05373368 | 0.23577469 | 0.22790267 | 0.81972191 | 0.99592423 |
| MAPKBP1 | 38.6133785 | 0.14253107 | 0.38878942 | 0.36660222 | 0.71391575 | 0.9958918 |
| VPS39 | 51.1694394 | -0.0724737 | 0.25471458 | -0.2845289 | 0.77600509 | 0.9958918 |
| TMEM87A | 105.283926 | 0.25151045 | 0.24700099 | 1.01825686 | 0.3085559 | 0.95793395 |
| GANC | 45.3046333 | -0.2210721 | 0.35654127 | -0.6200462 | 0.53522738 | 0.9958918 |
| ZNF106 | 94.2281399 | -0.0029399 | 0.29152749 | -0.0100843 | 0.99195399 | 0.99935941 |
| SNAP23 | 64.1157455 | -0.0644834 | 0.32158381 | -0.2005181 | 0.84107543 | 0.99692952 |
| STARD9 | 88.8177286 | 0.07566189 | 0.39106953 | 0.19347426 | 0.84658757 | 0.99692952 |
| TTBK2 | 96.2317708 | -0.0816037 | 0.24467345 | -0.3335208 | 0.73874121 | 0.9958918 |
| UBR1 | 214.093771 | -0.1984712 | 0.24216041 | -0.8195858 | 0.41245227 | 0.97812179 |
| AC068724.3 | 20.5419099 | -0.6643389 | 0.4561793 | -1.456311 | 0.14530666 | 0.82817333 |
| TMEM62 | 24.8911756 | -0.4335083 | 0.51440497 | -0.8427374 | 0.39937532 | 0.97500135 |
| TP53BP1 | 65.2314923 | -0.1644659 | 0.31634796 | -0.5198893 | 0.60314075 | 0.9958918 |
| CATSPER2 | 18.7227837 | 0.23641978 | 0.51038374 | 0.46321966 | 0.64320693 | 0.9958918 |
| PDIA3 | 110.56459 | 0.33207181 | 0.29868815 | 1.11176762 | 0.26623808 | 0.93533685 |
| SERF2 | 229.654433 | -0.1166938 | 0.35402253 | -0.3296224 | 0.74168527 | 0.9958918 |

| | | | | | | |
|---|---|---|---|---|---|---|
| CASC4 | 139.936935 | 0.10571007 | 0.18639351 | 0.56713388 | 0.57062322 | 0.9958918 |
| CTDSPL2 | 106.201204 | -0.0740866 | 0.19098665 | -0.3879151 | 0.69807883 | 0.9958918 |
| EIF3J-DT | 23.8677188 | 0.60045278 | 0.39417844 | 1.52330192 | 0.12768319 | 0.80131832 |
| EIF3J | 52.7719966 | -0.0472002 | 0.3381188 | -0.1395964 | 0.88897891 | 0.99935941 |
| SPG11 | 110.428021 | -0.0704122 | 0.24304205 | -0.2897121 | 0.7720365 | 0.9958918 |
| B2M | 692.142505 | -0.1496914 | 0.4348405 | -0.3442443 | 0.73066257 | 0.9958918 |
| GATM | 23.3933739 | 1.8463845 | 0.48004721 | 3.84625604 | 0.00011994 | 0.01695025 |
| SEMA6D | 22.7811579 | -0.8053814 | 0.5622332 | -1.4324687 | 0.15200974 | 0.83906933 |
| MYEF2 | 22.3108708 | 0.15927238 | 0.42438036 | 0.37530572 | 0.70743311 | 0.9958918 |
| SLC12A1 | 63.3974416 | -0.1178432 | 0.58223939 | -0.2023964 | 0.83960684 | 0.99692952 |
| DUT | 27.5657189 | -0.008235 | 0.30186402 | -0.0272804 | 0.97823608 | 0.99935941 |
| FBN1 | 305.834172 | -0.4752869 | 0.37705112 | -1.2605369 | 0.20747573 | 0.89620104 |
| EID1 | 71.7017611 | 0.3638931 | 0.41375935 | 0.87948006 | 0.37914104 | 0.97217293 |
| SECISBP2L | 96.3736539 | -0.0412444 | 0.21613601 | -0.1908263 | 0.84866169 | 0.99713289 |
| COPS2 | 43.5533151 | -0.2794032 | 0.31153116 | -0.8968709 | 0.36978779 | 0.97217293 |
| GALK2 | 90.0817577 | 0.00325614 | 0.2698397 | 0.01206695 | 0.9903722 | 0.99935941 |
| FAM227B | 45.8396641 | 0.4337028 | 0.36016568 | 1.20417582 | 0.22852163 | 0.91371352 |
| FGF7 | 261.393099 | 0.03914176 | 0.53570895 | 0.07306534 | 9.42E-01 | 0.99935941 |
| DTWD1 | 99.6943983 | -0.3133916 | 0.38406619 | -0.8159833 | 0.41450966 | 0.97812179 |
| GABPB1 | 53.1327218 | 0.00119566 | 0.37076091 | 0.00322488 | 0.99742692 | 0.99947501 |
| GABPB1-AS1 | 76.7984651 | 0.24236705 | 0.37447006 | 0.64722678 | 0.51748518 | 0.9958918 |
| USP8 | 82.285386 | -0.1663196 | 0.20700002 | -0.8034765 | 0.42169938 | 0.97812179 |
| TRPM7 | 182.432793 | -0.3270696 | 0.2226368 | -1.4690726 | 0.1418131 | 0.81652619 |
| SPPL2A | 102.279507 | 0.57535201 | 0.23001671 | 2.50134876 | 0.01237213 | 0.29616536 |
| AP4E1 | 38.8920667 | -0.0579118 | 0.24653834 | -0.2348999 | 0.81428644 | 0.99592423 |
| DMXL2 | 98.7203292 | 0.20522826 | 0.39454399 | 0.52016572 | 0.60294807 | 0.9958918 |
| TMOD3 | 129.997114 | 0.11218732 | 0.23595277 | 0.47546513 | 0.63445548 | 0.9958918 |
| MAPK6 | 57.4561702 | -0.3795819 | 0.27746179 | -1.3680512 | 0.17129604 | 0.87233975 |
| GNB5 | 33.1201863 | 0.3880456 | 0.33618316 | 1.1542684 | 0.24839015 | 0.92624282 |
| MYO5A | 61.555794 | 0.36588286 | 0.32065649 | 1.14104303 | 0.25385202 | 0.92646125 |
| ARPP19 | 50.9270075 | 0.45803211 | 0.24059531 | 1.90374497 | 0.05694341 | 0.60440467 |
| FAM214A | 130.358408 | 0.04867073 | 0.39291197 | 0.12387184 | 0.90141675 | 0.99935941 |
| WDR72 | 27.9573932 | 1.05783611 | 0.53782233 | 1.96688766 | 0.04919616 | 0.57273715 |
| RSL24D1 | 22.4504357 | 0.72649407 | 0.37701889 | 1.92694341 | 0.05398668 | 0.59127888 |
| RAB27A | 60.7618655 | 0.72610986 | 0.27663062 | 2.62483549 | 0.00866908 | 0.24789533 |
| PIGB | 22.1612762 | 0.42342662 | 0.4502536 | 0.94041807 | 0.34700316 | 0.96887708 |
| CCPG1 | 75.3623699 | 0.2681545 | 0.31152909 | 0.86076874 | 0.38936543 | 0.97297628 |
| PYGO1 | 20.8304027 | 0.11003869 | 0.38560463 | 0.28536662 | 0.77536328 | 0.9958918 |
| NEDD4 | 99.8491991 | -0.0258354 | 0.22645314 | -0.1140871 | 0.90916878 | 0.99935941 |
| RFX7 | 124.037788 | 0.1438888 | 0.20320129 | 0.70810968 | 0.47887715 | 0.99002719 |
| TEX9 | 22.7985808 | 0.24983094 | 0.45168135 | 0.55311325 | 0.58018587 | 0.9958918 |
| ZNF280D | 73.9816893 | 0.12570908 | 0.30750499 | 0.40880338 | 0.68268396 | 0.9958918 |
| AC090517.4 | 60.6220062 | -0.1267091 | 0.42856708 | -0.2956575 | 0.76749164 | 0.9958918 |
| TCF12 | 666.979451 | -0.1634448 | 0.20612981 | -0.7929218 | 0.42782341 | 0.98092839 |
| CGNL1 | 79.5744812 | -0.1179851 | 0.35343214 | -0.3338268 | 0.73851024 | 0.9958918 |
| POLR2M | 20.5141153 | -0.1927148 | 0.40215785 | -0.4792018 | 0.63179504 | 0.9958918 |
| ALDH1A2 | 123.312537 | -0.0729604 | 0.3199866 | -0.2280107 | 0.81963792 | 0.99592423 |
| ADAM10 | 199.188008 | -0.0722355 | 0.25428977 | -0.2840675 | 0.77635868 | 0.9958918 |
| MINDY2 | 100.170653 | 0.13368676 | 0.38559725 | 0.34670049 | 0.72881634 | 0.9958918 |
| RNF111 | 112.184378 | -0.1109438 | 0.26693587 | -0.4156198 | 0.67768824 | 0.9958918 |
| SLTM | 138.181892 | 0.20214048 | 0.18318007 | 1.10350697 | 0.26980707 | 0.93634106 |
| MYO1E | 395.251581 | 0.32289698 | 0.33925483 | 0.95178298 | 0.34120706 | 0.96887708 |
| GTF2A2 | 29.6852179 | 0.08989562 | 0.32920318 | 0.27307033 | 0.78479915 | 0.9958918 |
| BNIP2 | 49.3390162 | -0.1885924 | 0.28351106 | -0.6652031 | 0.50592063 | 0.99397937 |

| | | | | | | |
|---|---|---|---|---|---|---|
| ANXA2 | 111.849131 | -0.2933703 | 0.47086023 | -0.6230519 | 0.53325043 | 0.9958918 |
| ICE2 | 56.1241277 | 0.32286162 | 0.24500934 | 1.31775227 | 0.18758659 | 0.87975358 |
| RORA-AS1 | 36.3726261 | -0.9715472 | 0.64493203 | -1.5064334 | 0.13195595 | 0.80565822 |
| RORA | 1352.51862 | -0.8280227 | 0.4869853 | -1.7003032 | 0.0890739 | 0.72013653 |
| AC012404.1 | 41.1624381 | -1.1723659 | 0.58234272 | -2.0131888 | 0.04409477 | 0.55583613 |
| AC009554.1 | 25.4701677 | -0.2404271 | 0.65831257 | -0.3652173 | 0.71494922 | 0.9958918 |
| VPS13C | 224.075547 | -0.1593842 | 0.19596532 | -0.8133285 | 0.41602972 | 0.97812179 |
| TLN2 | 179.980775 | -1.0930354 | 0.25477132 | -4.2902609 | 1.78E-05 | 0.00437176 |
| TPM1 | 509.42666 | 0.11819561 | 0.36755794 | 0.32157 | 0.74777848 | 0.9958918 |
| LACTB | 32.5587751 | 0.1311444 | 0.34253675 | 0.38286228 | 0.70182188 | 0.9958918 |
| RPS27L | 58.9816878 | -0.1192708 | 0.3045179 | -0.391671 | 0.69530135 | 0.9958918 |
| RAB8B | 144.97734 | -0.4346611 | 0.40198105 | -1.0812975 | 0.27956478 | 0.94419503 |
| APH1B | 32.2947198 | -0.3883871 | 0.50452508 | -0.7698072 | 0.44141425 | 0.98092839 |
| CA12 | 122.392481 | 2.29820702 | 0.48359879 | 4.75230099 | 2.01E-06 | 0.00123166 |
| USP3 | 132.194579 | -0.1091159 | 0.22547792 | -0.4839316 | 0.62843444 | 0.9958918 |
| USP3-AS1 | 29.9478336 | -0.4137375 | 0.33130062 | -1.2488281 | 0.21172794 | 0.89738044 |
| HERC1 | 343.0487 | -0.0854672 | 0.22032606 | -0.3879125 | 0.6980808 | 0.9958918 |
| DAPK2 | 122.366557 | 0.4903921 | 0.47911907 | 1.02352866 | 0.30605795 | 0.95793395 |
| FAM96A | 18.1690682 | 0.01546784 | 0.40351134 | 0.0383331 | 0.9694221 | 0.99935941 |
| SNX1 | 89.8462121 | 0.0635179 | 0.22063779 | 0.28788315 | 0.77343619 | 0.9958918 |
| PPIB | 107.96863 | -0.4235575 | 0.39319244 | -1.077227 | 0.28137888 | 0.94419503 |
| TRIP4 | 36.8246063 | 0.24883658 | 0.29923021 | 0.83158908 | 0.40564093 | 0.97637555 |
| ZNF609 | 298.029636 | -0.385114 | 0.19933713 | -1.931973 | 0.05336283 | 0.58707106 |
| OAZ2 | 42.4595694 | 0.41038632 | 0.32116594 | 1.27780152 | 0.20131942 | 0.89319989 |
| RBPMS2 | 25.3465733 | 1.2529716 | 0.38749212 | 3.23354085 | 0.00122266 | 0.07550687 |
| ANKDD1A | 26.559388 | 0.13631491 | 0.39810981 | 0.3424053 | 0.7320459 | 0.9958918 |
| SPG21 | 32.9210252 | 0.06535859 | 0.30366286 | 0.21523405 | 0.82958485 | 0.99592423 |
| MTFMT | 25.3616783 | 0.37790505 | 0.46419195 | 0.81411375 | 0.41557979 | 0.97812179 |
| RASL12 | 26.5403267 | 0.07848918 | 0.38172867 | 0.2056151 | 0.83709158 | 0.99692952 |
| PDCD7 | 21.851523 | 0.47307836 | 0.4516412 | 1.04746502 | 0.29488516 | 0.95304825 |
| CLPX | 43.4757974 | 0.1353025 | 0.29479363 | 0.45897361 | 0.64625312 | 0.9958918 |
| DPP8 | 69.114827 | 0.02043511 | 0.26883949 | 0.07601231 | 0.9394093 | 0.99935941 |
| HACD3 | 36.4976781 | 0.41525925 | 0.29370569 | 1.4138618 | 0.15740248 | 0.84427321 |
| DENND4A | 186.244754 | -0.1349916 | 0.21604643 | -0.6248266 | 0.53208484 | 0.9958918 |
| RAB11A | 86.844813 | -0.2057753 | 0.26838694 | -0.7667111 | 0.44325327 | 0.98092839 |
| DIS3L | 18.3615034 | 0.90937951 | 0.4210898 | 2.15958568 | 0.03080476 | 0.47861346 |
| MAP2K1 | 82.2495475 | 0.51684086 | 0.38888477 | 1.32903341 | 0.18383695 | 0.87866798 |
| RPL4 | 105.643281 | 0.21458557 | 0.41193844 | 0.5209166 | 0.60242487 | 0.9958918 |
| SMAD6 | 29.6084131 | 1.22506395 | 0.46312377 | 2.64521934 | 0.0081638 | 0.23713736 |
| SMAD3 | 269.347936 | 0.18458732 | 0.42199637 | 0.43741447 | 0.66181079 | 0.9958918 |
| AAGAB | 45.7542287 | 0.20561389 | 0.36391688 | 0.56500233 | 0.57207218 | 0.9958918 |
| MAP2K5 | 138.818085 | -0.3775893 | 0.21713681 | -1.7389466 | 0.08204415 | 0.69958018 |
| PIAS1 | 220.61237 | -0.3149083 | 0.25501378 | -1.2348677 | 0.21687974 | 0.90303069 |
| FEM1B | 43.0221841 | -0.0940899 | 0.3237776 | -0.2906005 | 0.77135687 | 0.9958918 |
| ITGA11 | 199.449719 | -0.2963532 | 0.29665123 | -0.9989952 | 0.31779702 | 0.95992644 |
| CORO2B | 56.4802023 | -0.9164178 | 0.4521993 | -2.0265795 | 0.04270544 | 0.54851863 |
| ANP32A | 41.2361222 | 0.38165399 | 0.2992926 | 1.27518686 | 0.20224311 | 0.89319989 |
| GLCE | 35.4135838 | -1.1245826 | 0.52353993 | -2.148036 | 0.0317109 | 0.48149458 |
| RPLP1 | 253.010021 | -0.2566475 | 0.38910721 | -0.6595804 | 0.50952316 | 0.99507991 |
| TLE3 | 27.1168789 | 0.23413055 | 0.41396202 | 0.56558462 | 0.57167618 | 0.9958918 |
| UACA | 333.356268 | -0.3389011 | 0.36168579 | -0.9370042 | 0.34875637 | 0.96900211 |
| LRRC49 | 37.1363852 | 0.01887685 | 0.32766466 | 0.05761028 | 0.95405906 | 0.99935941 |
| THSD4 | 776.509008 | -0.5350947 | 0.44075943 | -1.2140289 | 0.22473669 | 0.91093521 |
| THSD4-AS1 | 53.0635322 | -0.4412253 | 0.33231961 | -1.3277137 | 0.18427272 | 0.87866798 |

| | | | | | | |
|---|---|---|---|---|---|---|
| MYO9A | 306.616165 | -0.1516279 | 0.22289002 | -0.6802814 | 0.4963263 | 0.99176652 |
| PKM | 129.524227 | 0.08500779 | 0.33312098 | 0.25518594 | 7.99E-01 | 0.99592423 |
| PARP6 | 38.2482888 | 0.02570677 | 0.31277817 | 0.0821885 | 0.93449682 | 0.99935941 |
| AC009690.1 | 41.1578844 | 0.52643088 | 0.45430334 | 1.15876515 | 0.24655192 | 0.92600668 |
| ARIH1 | 292.693616 | -0.1636593 | 0.22139452 | -0.7392203 | 0.45977325 | 0.98566907 |
| BBS4 | 38.9195387 | -0.1426573 | 0.2883885 | -0.4946705 | 0.6208327 | 0.9958918 |
| ADPGK | 36.9608351 | -0.0839746 | 0.37729224 | -0.2225718 | 0.8238688 | 0.99592423 |
| NEO1 | 166.074558 | 0.08122217 | 0.28189793 | 0.28812616 | 0.77325017 | 0.9958918 |
| NPTN | 159.835198 | -0.1304369 | 0.25721335 | -0.5071156 | 0.61207369 | 0.9958918 |
| LOXL1 | 30.1301107 | -0.2582534 | 0.53113531 | -0.4862291 | 0.62680475 | 0.9958918 |
| PML | 49.7658515 | 0.21307284 | 0.34131967 | 0.62426182 | 0.53245565 | 0.9958918 |
| ISLR | 42.7036903 | -0.1319415 | 0.42176321 | -0.3128332 | 0.75440741 | 0.9958918 |
| EDC3 | 21.3056738 | -0.2049546 | 0.38454832 | -0.5329749 | 0.59405094 | 0.9958918 |
| SCAMP2 | 19.0707798 | 0.02479835 | 0.36206167 | 0.06849207 | 0.94539393 | 0.99935941 |
| COX5A | 22.0663094 | 0.30410331 | 0.32274884 | 0.94222899 | 0.34607541 | 0.96887708 |
| PPCDC | 57.5645527 | 0.22048981 | 0.49482633 | 0.4455903 | 0.65589322 | 0.9958918 |
| MAN2C1 | 37.6759649 | 0.05364867 | 0.29002477 | 0.18497961 | 0.85324503 | 0.9971731 |
| SIN3A | 63.5168346 | -0.1892642 | 0.23133997 | -0.8181217 | 0.41328772 | 0.97812179 |
| PTPN9 | 103.931247 | 0.41612538 | 0.20166768 | 2.0634212 | 0.03907263 | 0.52881167 |
| SNX33 | 17.3290218 | 0.20157484 | 0.36603788 | 0.55069392 | 0.58184351 | 0.9958918 |
| CSPG4 | 53.7599067 | 0.7094698 | 0.37694877 | 1.88213849 | 0.05981721 | 0.61332974 |
| UBE2Q2 | 50.231392 | -0.3269324 | 0.2774242 | -1.1784567 | 0.23861458 | 0.92291633 |
| FBXO22 | 40.530048 | -0.1932073 | 0.30619493 | -0.6309944 | 0.52804421 | 0.9958918 |
| ETFA | 84.4209528 | 0.22203548 | 0.22660393 | 0.97983952 | 0.32716534 | 0.95992644 |
| SCAPER | 139.282428 | 0.53401585 | 0.30464293 | 1.75292383 | 0.07961508 | 0.68996609 |
| RCN2 | 41.3207231 | -0.0541547 | 0.2982225 | -0.1815917 | 0.85590319 | 0.9971731 |
| TSPAN3 | 52.3031707 | -0.0493024 | 0.23887138 | -0.2063972 | 0.83648064 | 0.99692952 |
| PEAK1 | 305.683372 | -0.4926761 | 0.28832003 | -1.708782 | 0.08749134 | 0.71805622 |
| HMG20A | 58.6064465 | 0.12099302 | 0.31701035 | 0.38166898 | 0.70270691 | 0.9958918 |
| LINGO1 | 478.091138 | -0.9570364 | 0.59554973 | -1.6069798 | 0.10805879 | 0.77248589 |
| TBC1D2B | 140.200665 | -0.2565482 | 0.23462086 | -1.0934586 | 0.27419251 | 0.9394201 |
| IDH3A | 30.9427817 | 0.03574902 | 0.38220127 | 0.09353453 | 0.92547892 | 0.99935941 |
| WDR61 | 29.0403741 | 0.61487881 | 0.27446007 | 2.24032156 | 0.02507005 | 0.4274706 |
| IREB2 | 85.7611645 | -0.1385764 | 0.19579006 | -0.7077803 | 0.47908169 | 0.99002719 |
| PSMA4 | 34.8348771 | -0.2921176 | 0.29239351 | -0.9990564 | 0.31776737 | 0.95992644 |
| MORF4L1 | 132.907919 | 0.01595475 | 0.21880442 | 0.07291785 | 0.94187149 | 0.99935941 |
| CTSH | 27.5545945 | 0.05261831 | 0.319707 | 0.16458293 | 0.86927227 | 0.9971731 |
| TMED3 | 23.7184411 | -0.7331381 | 0.42977523 | -1.7058641 | 0.08803339 | 0.71818729 |
| ZFAND6 | 117.0981 | 0.08835817 | 0.22671932 | 0.38972494 | 0.69673995 | 0.9958918 |
| ARNT2 | 24.0443209 | 0.58072831 | 0.43213553 | 1.34385688 | 0.17899467 | 0.87815793 |
| MESD | 32.7954138 | 0.26683814 | 0.33372987 | 0.79956327 | 0.42396388 | 0.97828057 |
| EFL1 | 27.3007516 | -0.1465323 | 0.41471254 | -0.3533347 | 0.72383756 | 0.9958918 |
| WHAMM | 31.1834627 | 0.23358148 | 0.30755656 | 0.75947488 | 0.44756854 | 0.98092839 |
| BTBD1 | 45.8721 | 0.32783505 | 0.26563099 | 1.23417471 | 0.21713781 | 0.90308192 |
| HDGFL3 | 69.7249706 | 0.15379626 | 0.22488292 | 0.6838948 | 0.4940416 | 0.99172677 |
| ADAMTSL3 | 127.257351 | 0.04786743 | 0.87693585 | 0.05458488 | 0.95646919 | 0.99935941 |
| WDR73 | 36.0105489 | 0.33703919 | 0.33045269 | 1.01993176 | 0.30776083 | 0.95793395 |
| SEC11A | 62.2315404 | 0.18565585 | 0.31018992 | 0.59852314 | 0.54949092 | 0.9958918 |
| ZNF592 | 46.5355571 | 0.20697296 | 0.26727255 | 0.77438914 | 0.4387007 | 0.98092839 |
| ALPK3 | 23.3143874 | 0.61754462 | 0.46957052 | 1.31512647 | 0.1884674 | 0.88139754 |
| PDE8A | 426.654465 | 0.09185578 | 0.27362321 | 0.33570171 | 0.73709581 | 0.9958918 |
| AKAP13 | 618.743334 | -0.7310687 | 0.27742572 | -2.635187 | 0.00840909 | 0.24234676 |
| AC087286.2 | 43.9313704 | -1.4863506 | 0.52580722 | -2.8267976 | 0.0047016 | 0.17190092 |
| KLHL25 | 23.7048223 | -0.1018422 | 0.50388386 | -0.2021144 | 0.83982727 | 0.99692952 |

| | | | | | | |
|---|---|---|---|---|---|---|
| NTRK3 | 402.844626 | 0.58434942 | 0.38430464 | 1.52053699 | 0.12837607 | 0.80292402 |
| DET1 | 21.4148827 | 0.91477051 | 0.63070072 | 1.45040348 | 0.14694604 | 0.8301633 |
| MFGE8 | 178.285329 | -0.2373206 | 0.28737821 | -0.8258127 | 0.40891037 | 0.97812179 |
| ABHD2 | 133.538851 | 0.08695241 | 0.35109117 | 0.24766332 | 0.80439491 | 0.99592423 |
| POLG | 17.5392362 | -0.0935957 | 0.43403067 | -0.215643 | 0.82926603 | 0.99592423 |
| KIF7 | 17.4397748 | 0.29690622 | 0.3686051 | 0.80548593 | 0.42053932 | 0.97812179 |
| ANPEP | 24.0964238 | 0.94644903 | 0.50667899 | 1.86794606 | 0.06176959 | 0.61962951 |
| ZNF710 | 30.1463955 | 0.67065705 | 0.42305636 | 1.58526646 | 0.11290581 | 0.77793929 |
| IDH2 | 25.7063788 | 0.60912224 | 0.30670563 | 1.98601585 | 0.04703156 | 0.56885457 |
| SEMA4B | 63.202739 | -0.1335865 | 0.28679454 | -0.4657918 | 6.41E-01 | 9.96E-01 |
| CIB1 | 31.269333 | -0.0998054 | 0.31469042 | -0.3171542 | 0.7511266 | 0.9958918 |
| IQGAP1 | 194.694543 | -0.0189757 | 0.21777034 | -0.0871364 | 0.93056309 | 0.99935941 |
| CRTC3 | 146.400885 | 0.03421525 | 0.2194745 | 0.15589624 | 0.87611481 | 0.9971731 |
| UNC45A | 25.45902 | 0.20975088 | 0.30796093 | 0.68109575 | 0.49581091 | 0.99176652 |
| SLCO3A1 | 496.022478 | 1.46672415 | 0.36397216 | 4.02977016 | 5.58E-05 | 0.0107975 |
| FAM174B | 23.7015357 | 0.04447895 | 0.32383642 | 0.13735005 | 0.89075411 | 0.99935941 |
| AC013394.1 | 104.17932 | -0.3026548 | 0.3345864 | -0.904564 | 0.36569641 | 0.97217293 |
| CHD2 | 166.442682 | -0.4679689 | 0.20375637 | -2.2967083 | 0.02163542 | 0.39453769 |
| MCTP2 | 39.129626 | -1.8151484 | 0.48877067 | -3.7137016 | 0.00020425 | 0.02333707 |
| LINC01197 | 115.091851 | 0.05830083 | 0.48448798 | 0.12033495 | 0.90421782 | 0.99935941 |
| LINC00924 | 131.125643 | -0.5659737 | 0.46742326 | -1.2108377 | 0.2259576 | 0.91093521 |
| AC015574.1 | 43.6738588 | 0.39265674 | 0.92006283 | 0.42677166 | 0.66954565 | 0.9958918 |
| AC012409.2 | 239.278875 | -0.0907662 | 0.49782637 | -0.182325 | 0.85532764 | 0.9971731 |
| NR2F2-AS1 | 1333.3954 | 0.22371694 | 0.45607693 | 0.49052456 | 0.62376275 | 0.9958918 |
| NR2F2 | 484.062794 | -0.3246134 | 0.23871657 | -1.3598276 | 0.17388449 | 0.8728669 |
| AC087477.2 | 86.9898869 | -0.3963588 | 0.45093654 | -0.8789679 | 0.37941867 | 0.97217293 |
| LINC00923 | 28.9871672 | -0.8906793 | 0.41287431 | -2.1572651 | 0.03098502 | 0.47969946 |
| IGF1R | 431.923116 | -0.5841258 | 0.23520916 | -2.4834313 | 0.01301234 | 0.30408773 |
| SYNM | 30.0246493 | 1.43325143 | 0.57493961 | 2.492873 | 0.01267142 | 0.30039431 |
| TTC23 | 57.3970136 | -0.0079033 | 0.25040042 | -0.0315627 | 0.9748208 | 0.99935941 |
| LRRC28 | 49.9339525 | 0.09963462 | 0.29328323 | 0.3397215 | 0.73406627 | 0.9958918 |
| MEF2A | 336.58218 | -0.6683154 | 0.35102997 | -1.90387 | 0.05692712 | 0.60440467 |
| AC090825.1 | 48.3884395 | -0.3053803 | 0.47207127 | -0.6468945 | 0.51770022 | 0.9958918 |
| ASB7 | 19.2653002 | 0.25430529 | 0.35639658 | 0.71354582 | 0.47550807 | 0.98910524 |
| LRRK1 | 52.9973145 | 0.3249233 | 0.41496153 | 0.78302028 | 0.4336152 | 0.98092839 |
| CHSY1 | 140.814466 | 0.48191131 | 0.45815559 | 1.05185076 | 0.29286802 | 0.95186143 |
| SELENOS | 48.0761591 | 0.14604625 | 0.39289824 | 0.37171521 | 0.7101049 | 0.9958918 |
| SNRPA1 | 35.2502148 | -0.0498566 | 0.29473858 | -0.1691553 | 0.86567448 | 0.9971731 |
| TM2D3 | 34.4117271 | -0.0420144 | 0.35437344 | -0.1185597 | 0.90562421 | 0.99935941 |
| TARSL2 | 45.8858204 | 0.13489342 | 0.28918591 | 0.46645916 | 0.64088687 | 0.9958918 |
| RHBDF1 | 18.4862102 | -0.182144 | 0.38689876 | -0.4707795 | 0.63779822 | 0.9958918 |
| MPG | 31.2639613 | -0.2416587 | 0.33417004 | -0.7231607 | 0.46958114 | 0.98910524 |
| NPRL3 | 44.5432536 | 0.21176104 | 0.39582787 | 0.53498264 | 0.59266186 | 0.9958918 |
| LUC7L | 105.932945 | -0.0585355 | 0.21675244 | -0.2700571 | 0.78711632 | 0.99592423 |
| AXIN1 | 22.2685443 | 0.50009206 | 0.48976964 | 1.02107607 | 0.3072184 | 0.95793395 |
| RAB11FIP3 | 75.8045778 | 0.79173147 | 0.32809364 | 2.41312651 | 0.01581633 | 0.34027688 |
| RAB40C | 27.1975619 | 0.49412316 | 0.32900286 | 1.50188103 | 0.13312784 | 0.80664435 |
| RHOT2 | 21.5010654 | 0.06258948 | 0.39597391 | 0.15806466 | 0.87440585 | 0.9971731 |
| STUB1 | 28.3034152 | 0.71269321 | 0.33283876 | 2.14125669 | 0.03225334 | 0.48564518 |
| JMJD8 | 18.1641304 | 0.15981641 | 0.41178893 | 0.38810274 | 0.69794 | 0.9958918 |
| LMF1 | 35.0091897 | 0.01987782 | 0.45606356 | 0.04358564 | 0.9652347 | 0.99935941 |
| CACNA1H | 27.6109877 | -0.2645732 | 0.45872499 | -0.5767578 | 0.5641031 | 0.9958918 |
| UBE2I | 57.5208078 | 0.12269948 | 0.32120034 | 0.38200295 | 0.70245918 | 0.9958918 |
| GNPTG | 19.2239823 | -0.1375226 | 0.4066545 | -0.3381805 | 0.73522718 | 0.9958918 |

| | | | | | | |
|---|---|---|---|---|---|---|
| UNKL | 23.9854117 | 0.41850947 | 0.49722881 | 0.84168387 | 0.39996493 | 0.97500135 |
| CLCN7 | 24.8574376 | -0.1823487 | 0.36700363 | -0.496858 | 0.61928922 | 0.9958918 |
| IFT140 | 32.7148703 | -0.3296635 | 0.3180351 | -1.0365632 | 0.29993945 | 0.95587816 |
| TMEM204 | 36.4833793 | -0.1872293 | 0.39249726 | -0.4770207 | 0.63334735 | 0.9958918 |
| MAPK8IP3 | 56.4775337 | -0.1372043 | 0.29789915 | -0.460573 | 0.64510497 | 0.9958918 |
| NME3 | 44.280763 | -0.0697194 | 0.44759803 | -0.1557634 | 0.87621953 | 0.9971731 |
| MRPS34 | 21.7327553 | -0.1111866 | 0.460166 | -0.2416229 | 0.80907241 | 0.99592423 |
| SPSB3 | 23.9301605 | -0.0795547 | 0.35077621 | -0.2267961 | 0.82058227 | 0.99592423 |
| HAGH | 26.7265376 | 0.3033773 | 0.34129843 | 0.88889158 | 0.37406135 | 0.97217293 |
| NDUFB10 | 29.2424976 | 0.1425604 | 0.42173368 | 0.33803417 | 0.73533743 | 0.9958918 |
| RPS2 | 109.98948 | -0.0965151 | 0.37982221 | -0.2541059 | 0.79941375 | 0.99592423 |
| SLC9A3R2 | 25.0507371 | 0.02026625 | 0.47766763 | 0.04242751 | 0.9661579 | 0.99935941 |
| TSC2 | 35.5851517 | 0.19100683 | 0.32444306 | 0.5887222 | 0.55604764 | 0.9958918 |
| PKD1 | 60.4316762 | 0.40236646 | 0.22794079 | 1.76522363 | 0.07752619 | 0.68020338 |
| PGP | 20.4314175 | 0.52072662 | 0.50946346 | 1.02210789 | 0.30672984 | 0.95793395 |
| ECI1 | 25.517463 | 0.70645803 | 0.35440887 | 1.99334183 | 0.04622403 | 0.56563746 |
| RNPS1 | 38.4655141 | 0.15359297 | 0.3010211 | 0.5102399 | 0.6098834 | 0.9958918 |
| PDPK1 | 70.8123687 | 0.03225915 | 0.24720324 | 0.13049647 | 0.89617365 | 0.99935941 |
| SRRM2 | 250.340163 | -0.2597815 | 0.17989548 | -1.4440691 | 0.14871953 | 0.8301633 |
| ELOB | 44.3117236 | -0.2161544 | 0.44541537 | -0.4852872 | 0.62747266 | 0.9958918 |
| FLYWCH1 | 31.2043236 | -0.029474 | 0.34713545 | -0.0849064 | 0.9323358 | 0.99935941 |
| TNFRSF12A | 31.1403486 | -0.4442917 | 0.65639669 | -0.6768646 | 0.49849185 | 0.99176652 |
| HCFC1R1 | 32.5756917 | 0.36885327 | 0.4687372 | 0.78690846 | 0.43133545 | 0.98092839 |
| IL32 | 26.0585247 | 0.58395546 | 0.50304857 | 1.16083317 | 0.24570975 | 0.92559078 |
| ZNF213 | 20.8480051 | 0.32765315 | 0.34203148 | 0.95796198 | 0.33808193 | 0.96750939 |
| ZNF263 | 19.3611243 | -0.0451348 | 0.40229043 | -0.1121945 | 0.91066919 | 0.99935941 |
| ZNF75A | 24.9389796 | 0.33195348 | 0.51664023 | 0.64252348 | 0.52053335 | 0.9958918 |
| CLUAP1 | 22.0388623 | 0.07160473 | 0.34730087 | 0.20617494 | 0.83665426 | 0.99692952 |
| DNASE1 | 39.910084 | 0.40259793 | 0.36124717 | 1.11446667 | 0.26507904 | 0.93499423 |
| TRAP1 | 33.7985895 | -0.2796356 | 0.32847387 | -0.8513177 | 0.39459291 | 0.97500135 |
| CREBBP | 192.767102 | -0.0431384 | 0.24166324 | -0.1785063 | 0.85832541 | 0.9971731 |
| ADCY9 | 121.205892 | 0.06175922 | 0.25824548 | 0.23914928 | 0.81098983 | 0.99592423 |
| VASN | 48.2197785 | -0.6665631 | 0.33988343 | -1.9611522 | 0.04986128 | 0.57344369 |
| DNAJA3 | 24.3539783 | -0.1834472 | 0.32423446 | -0.5657856 | 0.57153953 | 0.9958918 |
| NMRAL1 | 23.1504905 | -0.339955 | 0.39525822 | -0.8600833 | 0.38974312 | 0.97297628 |
| HMOX2 | 53.6734115 | -0.0442473 | 0.42880744 | -0.1031868 | 0.91781473 | 0.99935941 |
| CDIP1 | 19.2188896 | 0.51371306 | 0.38587999 | 1.33127676 | 0.18309796 | 0.87866798 |
| MGRN1 | 55.705786 | 0.47413283 | 0.28716009 | 1.65110979 | 0.09871616 | 0.74528964 |
| ANKS3 | 30.2812707 | -0.4248073 | 0.29341276 | -1.4478145 | 0.14766893 | 0.8301633 |
| GLYR1 | 71.2015913 | 0.08638155 | 0.25891195 | 0.33363294 | 0.73865656 | 0.9958918 |
| UBN1 | 33.432354 | 0.34560343 | 0.27202272 | 1.27049471 | 0.20390846 | 0.89396747 |
| RBFOX1 | 21.9552775 | -0.2189012 | 0.40691357 | -0.5379551 | 0.59060808 | 0.9958918 |
| PMM2 | 66.6298613 | -0.0801085 | 0.31365082 | -0.2554065 | 7.98E-01 | 0.99592423 |
| CARHSP1 | 55.2805383 | -0.0176637 | 0.29602168 | -0.0596704 | 0.95241818 | 0.99935941 |
| USP7 | 62.6527397 | 0.45152868 | 0.29213066 | 1.54563945 | 0.12219167 | 0.79184923 |
| C16orf72 | 51.7848187 | -0.1266784 | 0.26931531 | -0.470372 | 0.63808924 | 0.9958918 |
| GRIN2A | 147.328575 | 2.34447688 | 1.15510321 | 2.02966875 | 0.04239022 | 0.54728114 |
| ATF7IP2 | 61.527501 | 1.15032396 | 0.31061618 | 3.70336141 | 0.00021276 | 0.02333707 |
| EMP2 | 47.6469871 | 0.63394906 | 0.44234128 | 1.43316731 | 0.15181003 | 0.83883602 |
| CLEC16A | 122.59454 | 0.47686372 | 0.23131228 | 2.06155821 | 0.03924982 | 0.53023327 |
| AC099489.1 | 24.0402376 | -0.0255789 | 0.74433063 | -0.034365 | 0.97258612 | 0.99935941 |
| LITAF | 184.281185 | -0.3359731 | 0.38780518 | -0.866345 | 0.38630098 | 0.97256797 |
| TXNDC11 | 68.3223391 | 0.05199197 | 0.20179385 | 0.25764891 | 0.79667788 | 0.99592423 |
| ZC3H7A | 98.8413171 | -0.1312313 | 0.21465994 | -0.6113453 | 0.54097103 | 0.9958918 |

| | | | | | | |
|---|---|---|---|---|---|---|
| RSL1D1 | 23.9104122 | 0.1974894 | 0.41274808 | 0.47847441 | 0.63231258 | 0.9958918 |
| AC007216.4 | 34.749939 | -0.1377573 | 0.31415696 | -0.4384982 | 0.66102515 | 0.9958918 |
| GSPT1 | 84.2046867 | -0.0029122 | 0.25126814 | -0.01159 | 0.99075276 | 0.99935941 |
| SNX29 | 158.021923 | -0.0413029 | 0.2572291 | -0.1605684 | 0.87243331 | 0.9971731 |
| MKL2 | 175.908697 | -0.0862912 | 0.22518274 | -0.3832051 | 0.7015677 | 0.9958918 |
| PARN | 80.3902617 | -0.1842706 | 0.2403495 | -0.7666778 | 0.44327308 | 0.98092839 |
| BFAR | 20.9077251 | 0.32957888 | 0.38525861 | 0.8554744 | 0.39228857 | 0.97297628 |
| PDXDC1 | 197.140132 | -0.1300857 | 0.24920611 | -0.5220006 | 0.6016699 | 0.9958918 |
| RRN3 | 33.5965114 | -0.1052017 | 0.29383472 | -0.3580301 | 0.72032082 | 0.9958918 |
| C16orf45 | 95.6836441 | -0.1440144 | 0.34301604 | -0.4198475 | 0.67459683 | 0.9958918 |
| MARF1 | 47.4458743 | -0.189351 | 0.36719267 | -0.5156722 | 0.60608338 | 0.9958918 |
| NDE1 | 33.2146406 | 0.70813641 | 0.35312421 | 2.00534653 | 0.04492601 | 0.55894748 |
| MYH11 | 436.75616 | 1.28408308 | 0.5044545 | 2.5454884 | 0.0109125 | 0.28138932 |
| ABCC1 | 126.807577 | 0.33151025 | 0.2897651 | 1.14406547 | 0.25259649 | 0.92631316 |
| XYLT1 | 125.257259 | -0.3863177 | 0.37928473 | -1.0185427 | 0.30842011 | 0.95793395 |
| RPS15A | 27.1939933 | -0.1868391 | 0.47551793 | -0.392917 | 0.69438075 | 0.9958918 |
| SMG1 | 173.318863 | -0.1551384 | 0.17760143 | -0.8735198 | 0.38237984 | 0.9725044 |
| CCP110 | 30.8544937 | 1.05151561 | 0.40375451 | 2.60434394 | 0.00920503 | 0.25431499 |
| VPS35L | 87.4221731 | 0.06249192 | 0.25929788 | 0.24100434 | 0.80955175 | 0.99592423 |
| IQCK | 34.234004 | -0.2669855 | 0.41701322 | -0.6402327 | 0.52202136 | 0.9958918 |
| UMOD | 143.712794 | -0.6526157 | 0.77762711 | -0.8392399 | 0.40133467 | 0.97500135 |
| THUMPD1 | 29.9643644 | -0.1847146 | 0.31570068 | -0.5850941 | 0.55848445 | 0.9958918 |
| METTL9 | 41.9652628 | 0.25306643 | 0.2839189 | 0.89133353 | 0.37275026 | 0.97217293 |
| NPIPB4 | 20.0418059 | -0.7737779 | 0.42959543 | -1.8011782 | 0.07167479 | 0.66173122 |
| UQCRC2 | 57.2156223 | 0.05470148 | 0.23284944 | 0.2349221 | 0.8142692 | 0.99592423 |
| MOSMO | 106.790195 | -0.3431264 | 0.27397045 | -1.252421 | 0.21041648 | 0.89738044 |
| EEF2K | 42.3090186 | -0.1868629 | 0.39389275 | -0.4744004 | 0.63521439 | 0.9958918 |
| POLR3E | 20.7013558 | -0.2850247 | 0.38352423 | -0.7431726 | 0.45737722 | 0.9838379 |
| CDR2 | 129.929182 | 0.20196024 | 0.29003603 | 0.69632812 | 0.48622337 | 0.99046213 |
| NPIPB5 | 72.6524928 | 0.20117853 | 0.26841505 | 0.74950539 | 0.45355265 | 0.98092839 |
| USP31 | 62.2046652 | -0.5623804 | 0.42383971 | -1.3268704 | 0.18455155 | 0.87866798 |
| COG7 | 46.9250237 | -0.1781303 | 0.25644009 | -0.6946273 | 0.48728889 | 0.99046213 |
| GGA2 | 43.9286268 | -0.1361284 | 0.36350806 | -0.3744852 | 0.70804333 | 0.9958918 |
| UBFD1 | 18.5627718 | -0.3926105 | 0.33027187 | -1.1887496 | 0.23453822 | 0.91930824 |
| NDUFAB1 | 24.5224714 | -0.3913318 | 0.37197196 | -1.0520466 | 0.29277817 | 0.95186143 |
| DCTN5 | 25.6007796 | 0.43036058 | 0.32965951 | 1.30546992 | 0.19173288 | 0.88158763 |
| PRKCB | 65.9659423 | -2.7404074 | 0.92819419 | -2.9524074 | 0.00315307 | 0.13394152 |
| RBBP6 | 73.9701623 | 0.0217979 | 0.25237839 | 0.08636991 | 0.93117236 | 0.99935941 |
| TNRC6A | 410.340894 | -0.2384223 | 0.29109846 | -0.8190436 | 0.41276155 | 0.97812179 |
| ARHGAP17 | 145.605795 | -0.2409008 | 0.2713957 | -0.8876366 | 0.37473623 | 0.97217293 |
| LCMT1 | 39.9677966 | -0.0723385 | 0.30387189 | -0.2380558 | 0.81183783 | 0.99592423 |
| NSMCE1 | 25.4553899 | 0.82053728 | 0.33760232 | 2.43048469 | 0.01507864 | 0.33377394 |
| IL4R | 149.735179 | 0.1319998 | 0.35065049 | 0.37644265 | 0.70658785 | 0.9958918 |
| GTF3C1 | 42.3616537 | -0.0827747 | 0.27159706 | -0.3047702 | 0.76054117 | 0.9958918 |
| KIAA0556 | 37.7038744 | 0.24281145 | 0.38595787 | 0.62911387 | 0.52927451 | 0.9958918 |
| XPO6 | 79.7536663 | -0.2642077 | 0.24444707 | -1.0808381 | 0.27976912 | 0.94419503 |
| NUPR1 | 73.7936096 | -0.1680342 | 0.35057612 | -0.4793087 | 0.63171906 | 0.9958918 |
| ATXN2L | 41.5113586 | -0.3346059 | 0.32891767 | -1.0172937 | 0.30901373 | 0.95793395 |
| TUFM | 34.8692635 | 0.22373128 | 0.39027321 | 0.57326834 | 0.56646302 | 0.9958918 |
| SH2B1 | 24.9535749 | 0.13851011 | 0.33815474 | 0.40960569 | 0.68209522 | 0.9958918 |
| NFATC2IP | 56.3677062 | 0.18386149 | 0.26560625 | 0.69223331 | 0.48879082 | 0.99046213 |
| AC009093.2 | 29.2846951 | 0.44805108 | 0.44791956 | 1.00029363 | 0.31716843 | 0.95992644 |
| MAZ | 17.5179986 | 0.17746876 | 0.39655885 | 0.44752187 | 0.6544983 | 0.9958918 |
| MVP | 67.0261395 | 0.29596675 | 0.33959019 | 0.87154092 | 0.38345887 | 0.9725044 |

| | | | | | | |
|---|---|---|---|---|---|---|
| CDIPT | 21.6377721 | -0.132436 | 0.43598373 | -0.3037636 | 0.76130797 | 0.9958918 |
| KCTD13 | 18.666036 | 0.13410815 | 0.41474178 | 0.32335336 | 0.74642765 | 0.9958918 |
| TMEM219 | 28.0735232 | -0.1415334 | 0.34194346 | -0.4139088 | 0.67894089 | 0.9958918 |
| INO80E | 15.8961053 | 0.60812395 | 0.3927818 | 1.54824881 | 0.1215624 | 0.79021701 |
| PPP4C | 24.3792774 | -0.0805873 | 0.51830911 | -0.1554811 | 0.87644208 | 0.9971731 |
| YPEL3 | 36.1747434 | -0.2040881 | 0.3103876 | -0.6575265 | 0.51084247 | 0.9958918 |
| MAPK3 | 20.6624779 | 0.18153875 | 0.31859936 | 0.56980262 | 0.56881158 | 0.9958918 |
| SEPHS2 | 26.1699662 | 0.98637117 | 0.40864691 | 2.41374922 | 0.01578933 | 0.34027688 |
| ZNF785 | 17.6786132 | 0.54906128 | 0.50420113 | 1.08897274 | 0.27616591 | 0.94222064 |
| FBRS | 20.9381456 | -0.0039146 | 0.41762113 | -0.0093736 | 0.99252105 | 0.99935941 |
| RNF40 | 18.7061181 | 0.51554141 | 0.40353579 | 1.27756058 | 0.20140441 | 0.89319989 |
| BCL7C | 37.9063157 | 0.32578866 | 0.3262239 | 0.99866584 | 0.3179566 | 0.95992644 |
| KAT8 | 25.3310239 | -0.2235093 | 0.29838872 | -0.7490542 | 0.45382454 | 0.98092839 |
| FUS | 152.957088 | -0.0873965 | 0.22021888 | -0.3968619 | 0.69146929 | 0.9958918 |
| TGFB1I1 | 37.2596961 | 0.25876947 | 0.40525065 | 0.63854178 | 0.52312107 | 0.9958918 |
| C16orf58 | 20.9054722 | -0.3982414 | 0.34334582 | -1.1598841 | 0.24609601 | 0.92600668 |
| ZNF720 | 35.9211389 | 0.58879008 | 0.27535884 | 2.13826465 | 0.03249527 | 0.48564518 |
| VPS35 | 33.2170163 | 0.19006604 | 0.27366651 | 0.694517 | 0.48735804 | 0.99046213 |
| DNAJA2 | 21.0789031 | 0.19045608 | 0.31194667 | 0.6105405 | 0.54150382 | 0.9958918 |
| ITFG1 | 231.061086 | -0.0902548 | 0.17701316 | -0.5098762 | 0.61013819 | 0.9958918 |
| PHKB | 166.957962 | -0.0793999 | 0.28693769 | -0.2767149 | 0.78199901 | 0.9958918 |
| LONP2 | 148.340507 | 0.18733377 | 0.18323798 | 1.02235234 | 0.30661417 | 0.95793395 |
| SIAH1 | 35.3954547 | -0.0945714 | 0.29231007 | -0.3235312 | 0.74629301 | 0.9958918 |
| N4BP1 | 67.5161617 | -0.4312518 | 0.28528254 | -1.5116657 | 0.13061893 | 0.80565822 |
| ZNF423 | 74.0629078 | 0.27839957 | 0.50754487 | 0.54852208 | 0.58333347 | 0.9958918 |
| TENT4B | 65.4579093 | 0.45145094 | 0.31318383 | 1.44148867 | 0.14944667 | 0.83171145 |
| BRD7 | 36.9578437 | -0.0414767 | 0.25754426 | -0.161047 | 0.8720564 | 0.9971731 |
| NKD1 | 60.0199433 | -0.2130716 | 0.56516777 | -0.3770059 | 0.70616924 | 0.9958918 |
| CYLD | 49.7234367 | 0.19711284 | 0.27027925 | 0.72929327 | 0.46582229 | 0.9876884 |
| CHD9 | 402.866918 | -0.432377 | 0.24013553 | -1.8005542 | 0.07177317 | 0.6618081 |
| RBL2 | 70.6804557 | 0.37937997 | 0.23321487 | 1.62674007 | 0.10379232 | 0.75827058 |
| FTO | 432.528647 | -0.2625871 | 0.21520283 | -1.2201844 | 0.222395 | 0.91093521 |
| MMP2 | 80.0285817 | -0.3900261 | 0.36386977 | -1.0718838 | 0.28377223 | 0.94505166 |
| LPCAT2 | 22.0800808 | -0.3263601 | 0.38132725 | -0.8558531 | 0.39207904 | 0.97297628 |
| GNAO1 | 27.1214438 | 0.39799907 | 0.49332226 | 0.80677298 | 0.41979729 | 0.97812179 |
| AMFR | 40.5233504 | 0.24781942 | 0.28928875 | 0.85665073 | 0.39163794 | 0.97297628 |
| NUDT21 | 33.5196499 | -0.2120693 | 0.26936791 | -0.7872848 | 0.43111515 | 0.98092839 |
| OGFOD1 | 20.9107483 | 0.15126818 | 0.4189471 | 0.36106749 | 0.71804899 | 0.9958918 |
| BBS2 | 28.6602426 | -0.3483156 | 0.30060854 | -1.1587016 | 0.24657784 | 0.92600668 |
| MT2A | 638.234056 | -1.0907744 | 0.58626494 | -1.8605485 | 6.28E-02 | 0.6220697 |
| MT1E | 136.781208 | -1.610215 | 0.41668876 | -3.8643111 | 0.0001114 | 0.01605298 |
| MT1M | 46.5703747 | -1.4446792 | 0.46626668 | -3.0983969 | 1.95E-03 | 0.09345751 |
| MT1G | 181.906412 | -1.3270999 | 1.00398543 | -1.3218318 | 0.18622415 | 0.87897321 |
| MT1H | 54.7122132 | -1.9900031 | 1.23691121 | -1.6088488 | 0.10764941 | 0.77225821 |
| MT1X | 106.817076 | -1.8016356 | 0.46291429 | -3.8919421 | 9.94E-05 | 0.01554938 |
| NUP93 | 29.1664876 | -0.0593064 | 0.35301127 | -0.1680014 | 0.86658218 | 0.9971731 |
| HERPUD1 | 82.5070661 | -0.6137379 | 0.52397968 | -1.171301 | 0.24147782 | 0.92516122 |
| NLRC5 | 29.3481036 | -0.2415869 | 0.4587286 | -0.5266444 | 0.59844052 | 0.9958918 |
| FAM192A | 36.427105 | -0.0047966 | 0.2709006 | -0.0177062 | 0.98587327 | 0.99935941 |
| RSPRY1 | 53.2203386 | -0.0730303 | 0.31049674 | -0.2352046 | 0.81404993 | 0.99592423 |
| ARL2BP | 23.4341823 | 0.5918958 | 0.35938308 | 1.64697734 | 0.0995627 | 0.74891126 |
| COQ9 | 20.0557678 | 0.24157893 | 0.37777154 | 0.6394842 | 0.52250799 | 0.9958918 |
| ADGRG1 | 23.6895898 | 0.51050154 | 0.57312095 | 0.89073962 | 0.37306888 | 0.97217293 |
| KIFC3 | 53.8147988 | -0.4263963 | 0.32000867 | -1.3324524 | 0.18271159 | 0.87866798 |

| | | | | | | |
|---|---|---|---|---|---|---|
| CSNK2A2 | 28.9849127 | -0.2796644 | 0.29462432 | -0.9492239 | 0.34250674 | 0.96887708 |
| CNOT1 | 103.984229 | -0.2254622 | 0.19859093 | -1.1353094 | 0.2562457 | 0.9266065 |
| GOT2 | 20.7158965 | 0.50042104 | 0.35982391 | 1.39073869 | 0.16430468 | 0.85697309 |
| CDH11 | 378.028723 | -0.5732317 | 0.33131453 | -1.7301737 | 0.08359925 | 0.70213814 |
| CMTM4 | 39.9397316 | 0.2863063 | 0.33749112 | 0.84833729 | 0.39625016 | 0.97500135 |
| DYNC1LI2 | 73.5355189 | -0.0883225 | 0.24696122 | -0.3576369 | 0.72061504 | 0.9958918 |
| NAE1 | 32.6427771 | 0.3352112 | 0.32376918 | 1.03534006 | 0.30051013 | 0.95597956 |
| CDH16 | 29.6223421 | 0.51172932 | 0.41021479 | 1.24746679 | 0.21222639 | 0.89738044 |
| RRAD | 29.5491315 | -0.4607453 | 0.46917544 | -0.982032 | 0.32608409 | 0.95992644 |
| FAM96B | 27.0927425 | -0.1995994 | 0.31406842 | -0.6355283 | 0.5250839 | 0.9958918 |
| CES2 | 18.3378704 | 0.77325246 | 0.37375935 | 2.06885114 | 0.03856006 | 0.5265563 |
| CBFB | 106.538301 | -0.0924674 | 0.23484461 | -0.3937385 | 0.69377408 | 0.9958918 |
| KIAA0895L | 25.0790562 | -0.303278 | 0.47227086 | -0.6421696 | 0.52076304 | 0.9958918 |
| FHOD1 | 22.9545488 | -0.7378561 | 0.50757887 | -1.4536778 | 0.14603567 | 0.8301633 |
| ATP6V0D1 | 31.8362073 | 0.03908641 | 0.38033701 | 0.10276782 | 0.91814725 | 0.99935941 |
| RIPOR1 | 32.5628657 | 0.12206469 | 0.35808872 | 0.34087834 | 0.73319517 | 0.9958918 |
| CTCF | 58.5184522 | 0.37736085 | 0.27727491 | 1.36096287 | 0.17352542 | 0.87233975 |
| GFOD2 | 22.7639589 | 0.78945773 | 0.4816749 | 1.63898459 | 0.10121647 | 0.75167369 |
| RANBP10 | 49.6418874 | 0.18137918 | 0.33379989 | 0.543377 | 0.58687026 | 0.9958918 |
| LCAT | 19.2629018 | -0.6915845 | 0.53531676 | -1.2919165 | 0.19638607 | 0.88391893 |
| SLC12A4 | 22.6121047 | 0.27416823 | 0.39913188 | 0.68691138 | 0.49213858 | 0.99046213 |
| NFATC3 | 119.069052 | 0.2591401 | 0.20936957 | 1.23771618 | 0.21582132 | 0.90187565 |
| SLC7A6 | 92.0713881 | -0.8785331 | 0.44352059 | -1.980817 | 0.0476118 | 0.56981199 |
| SLC7A6OS | 27.2671776 | -0.5989532 | 0.32376807 | -1.8499454 | 0.06432143 | 0.62609028 |
| PRMT7 | 28.4328252 | -0.4787076 | 0.55588669 | -0.8611604 | 0.38914971 | 0.97297628 |
| ZFP90 | 25.7097333 | 0.12888422 | 0.35682641 | 0.36119585 | 0.71795304 | 0.9958918 |
| TANGO6 | 81.6823354 | 0.04128579 | 0.22513476 | 0.18338258 | 0.85449785 | 0.9971731 |
| CHTF8 | 21.8025875 | 0.91469267 | 0.40650968 | 2.25011289 | 0.02444178 | 0.42312555 |
| UTP4 | 19.8203697 | -0.0498964 | 0.41851124 | -0.1192235 | 0.90509825 | 0.99935941 |
| SNTB2 | 44.8464487 | -0.0013753 | 0.35531586 | -0.0038707 | 0.99691167 | 0.99935941 |
| VPS4A | 19.1977091 | -0.1502786 | 0.33717049 | -0.4457052 | 0.6558102 | 0.9958918 |
| TERF2 | 25.0094378 | -0.2900857 | 0.32164825 | -0.9018723 | 0.3671247 | 0.97217293 |
| CYB5B | 59.7748892 | -0.3410593 | 0.2853249 | -1.1953366 | 0.23195555 | 0.91565561 |
| NFAT5 | 529.827069 | -0.2411355 | 0.19519571 | -1.2353526 | 0.21669931 | 0.90279094 |
| NQO1 | 23.5234102 | 1.84448406 | 0.56521992 | 3.26330334 | 0.00110122 | 0.07280105 |
| WWP2 | 78.015203 | 0.09615902 | 0.30986341 | 0.3103271 | 0.75631222 | 0.9958918 |
| PDPR | 45.1547705 | -0.5056628 | 0.29649697 | -1.705457 | 0.08810923 | 0.71818729 |
| EXOSC6 | 17.6580083 | -0.5216954 | 0.40068399 | -1.3020121 | 0.19291224 | 0.88158763 |
| AARS | 31.4499545 | -0.1305363 | 0.31878655 | -0.4094789 | 0.68218828 | 0.9958918 |
| ST3GAL2 | 47.924349 | 0.6703997 | 0.33920387 | 1.97639166 | 0.04811044 | 0.57118512 |
| COG4 | 25.990288 | 0.26029409 | 0.3018044 | 0.86245955 | 0.38843468 | 0.97297628 |
| SF3B3 | 71.6366983 | -0.4033117 | 0.23667252 | -1.7040918 | 0.08836394 | 0.71818729 |
| IL34 | 34.6998644 | 0.43800699 | 0.60142559 | 0.72828126 | 0.46644143 | 0.98791879 |
| MTSS1L | 25.2297238 | -0.4909896 | 0.39074802 | -1.2565377 | 0.20892109 | 0.89634177 |
| VAC14 | 27.5658923 | 0.37458415 | 0.483971 | 0.77398057 | 0.43894228 | 0.98092839 |
| HYDIN | 27.7043307 | 0.13860869 | 0.47037579 | 0.29467649 | 0.76824103 | 0.9958918 |
| PHLPP2 | 39.1091408 | 0.41387871 | 0.34361523 | 1.20448302 | 0.22840294 | 0.91371352 |
| AP1G1 | 94.4626822 | -0.1828743 | 0.23026739 | -0.7941822 | 0.42708935 | 0.98091054 |
| IST1 | 67.5554698 | -0.0936063 | 0.26180936 | -0.3575361 | 0.7206905 | 0.9958918 |
| DHX38 | 18.6881351 | -0.2051922 | 0.3925658 | -0.5226951 | 0.60118642 | 0.9958918 |
| AC004943.2 | 21.377185 | 0.26353982 | 0.4749649 | 0.55486168 | 0.57898927 | 0.9958918 |
| ZFHX3 | 475.717866 | 0.20072545 | 0.33041538 | 0.60749429 | 0.54352293 | 0.9958918 |
| PSMD7 | 38.6900317 | -0.0821628 | 0.36898779 | -0.2226708 | 0.8237917 | 0.99592423 |
| GLG1 | 215.348987 | 0.15621451 | 0.2024543 | 0.77160382 | 0.44034911 | 0.98092839 |

| | | | | | | |
|---|---|---|---|---|---|---|
| RFWD3 | 21.5130539 | 0.53863585 | 0.44397508 | 1.21321191 | 0.22504881 | 0.91093521 |
| WDR59 | 70.4830041 | -0.1880256 | 0.24849809 | -0.756648 | 0.44926077 | 0.98092839 |
| ZNRF1 | 97.3531749 | 0.0829934 | 0.19778514 | 0.41961395 | 0.67476749 | 0.9958918 |
| BCAR1 | 58.8724171 | 0.13667528 | 0.24714911 | 0.55300738 | 0.58025836 | 0.9958918 |
| CFDP1 | 137.063438 | -0.2417 | 0.21526584 | -1.1227978 | 0.26152337 | 0.93155278 |
| GABARAPL2 | 40.318899 | 0.28959454 | 0.35913642 | 0.80636363 | 0.42003321 | 0.97812179 |
| KARS | 22.9467035 | 0.01710554 | 0.34201779 | 0.05001359 | 0.96011156 | 0.99935941 |
| TERF2IP | 96.4428455 | 0.43236425 | 0.25832953 | 1.67369271 | 0.09419101 | 0.7340733 |
| WWOX | 674.499796 | -0.1954578 | 0.28199961 | -0.6931137 | 0.48823818 | 0.99046213 |
| MAF | 35.9002558 | -0.0791712 | 0.35384534 | -0.2237452 | 0.82295558 | 0.99592423 |
| CDYL2 | 36.8145458 | 0.2798691 | 0.38138445 | 0.7338241 | 0.46305595 | 0.98590351 |
| CMC2 | 34.2390566 | -0.0171995 | 0.35684872 | -0.0481982 | 0.96155831 | 0.99935941 |
| GAN | 36.6094809 | 0.20123179 | 0.27944345 | 0.72011632 | 0.47145338 | 0.98910524 |
| CMIP | 170.566716 | 0.42306368 | 0.31560087 | 1.34050229 | 0.1800821 | 0.87815793 |
| PLCG2 | 87.4574432 | 0.76538992 | 0.442774 | 1.72862436 | 0.08387635 | 0.70248581 |
| CDH13 | 177.835706 | 0.13562005 | 0.78447347 | 0.17288036 | 0.86274547 | 0.9971731 |
| HSBP1 | 32.4725444 | -0.2585231 | 0.31483424 | -0.8211403 | 0.41156634 | 0.97812179 |
| MBTPS1 | 62.2823541 | 0.0301634 | 0.2457252 | 0.12275256 | 0.90230304 | 0.99935941 |
| WFDC1 | 86.0541134 | 1.39343087 | 0.50572834 | 2.75529521 | 0.00586392 | 0.19136125 |
| COTL1 | 23.0679406 | -0.1023565 | 0.41751466 | -0.2451566 | 0.80633515 | 0.99592423 |
| KLHL36 | 28.6295331 | 0.23021554 | 0.30652605 | 0.75104722 | 0.45262424 | 0.98092839 |
| USP10 | 71.0070644 | 0.09129567 | 0.29513679 | 0.30933342 | 0.75706791 | 0.9958918 |
| CRISPLD2 | 579.897013 | -1.6356915 | 0.49676429 | -3.2926913 | 0.00099233 | 0.07012172 |
| ZDHHC7 | 32.5190148 | -0.0362152 | 0.26084606 | -0.1388376 | 0.88957849 | 0.99935941 |
| GSE1 | 287.269811 | 0.17474123 | 0.41936529 | 0.41668023 | 0.67691231 | 0.9958918 |
| COX4I1 | 85.1628876 | 0.16093933 | 0.2980602 | 0.53995578 | 0.58922753 | 0.9958918 |
| FOXC2 | 22.252355 | -0.5148128 | 0.34099702 | -1.509728 | 0.13111283 | 0.80565822 |
| FBXO31 | 25.9662817 | -0.1091813 | 0.30188378 | -0.3616666 | 0.71760116 | 0.9958918 |
| MAP1LC3B | 37.0601208 | -0.5331628 | 0.42483559 | -1.2549861 | 0.20948382 | 0.89634177 |
| ZCCHC14 | 87.0924962 | 0.26745117 | 0.38798445 | 0.68933477 | 0.49061262 | 0.99046213 |
| KLHDC4 | 28.6195706 | -0.1467066 | 0.40606072 | -0.3612923 | 0.71788093 | 0.9958918 |
| BANP | 52.2289281 | 0.21589291 | 0.28179166 | 0.76614372 | 0.44359079 | 0.98092839 |
| ZFPM1 | 17.6103637 | 0.04904207 | 0.42693985 | 0.11486879 | 0.90854912 | 0.99935941 |
| ZC3H18 | 29.2205298 | -0.1796081 | 0.41348548 | -0.4343759 | 0.66401547 | 0.9958918 |
| CYBA | 41.5413258 | 0.37065288 | 0.40773875 | 0.90904502 | 0.36332636 | 0.97217293 |
| PIEZO1 | 36.8904568 | -0.0306463 | 0.30354524 | -0.1009612 | 0.91958127 | 0.99935941 |
| APRT | 26.6740469 | 0.32982066 | 0.37050833 | 0.8901842 | 0.37336699 | 0.97217293 |
| GALNS | 22.8302125 | -0.1577062 | 0.41287314 | -0.3819726 | 0.7024817 | 0.9958918 |
| ACSF3 | 23.4181729 | 0.10784629 | 0.38094342 | 0.28310317 | 0.77709775 | 0.9958918 |
| ANKRD11 | 326.077538 | 0.01067387 | 0.21570778 | 0.049483 | 0.96053439 | 0.99935941 |
| SPG7 | 72.9019193 | -0.1329 | 0.40001035 | -0.3322413 | 0.73970703 | 0.9958918 |
| RPL13 | 298.118216 | -0.2303422 | 0.32777706 | -0.7027405 | 0.4822175 | 0.99002719 |
| CHMP1A | 20.87801 | 0.26422688 | 0.31243616 | 0.84569878 | 0.3977208 | 0.97500135 |
| TCF25 | 166.100713 | 0.0503142 | 0.30958458 | 0.16252166 | 0.87089508 | 0.9971731 |
| FAM157C | 23.430229 | 0.00657592 | 0.38810922 | 0.01694348 | 0.9864817 | 0.99935941 |
| VPS53 | 115.882013 | -0.0801397 | 0.25439405 | -0.315022 | 0.75274493 | 0.9958918 |
| NXN | 262.515288 | -0.4938934 | 0.56799817 | -0.8695334 | 0.38455542 | 0.9725044 |
| ABR | 83.8989429 | 0.45608178 | 0.38643017 | 1.18024372 | 0.23790329 | 0.92291633 |
| YWHAE | 139.083829 | 0.01351252 | 0.17360452 | 0.07783508 | 0.93795924 | 0.99935941 |
| CRK | 64.3753979 | 0.46081154 | 0.31380823 | 1.46844949 | 0.14198217 | 0.81652619 |
| MYO1C | 75.250093 | 0.60018676 | 0.25156954 | 2.38576876 | 0.01704346 | 0.34765232 |
| PITPNA | 63.5352179 | 0.44451209 | 0.25278008 | 1.75849338 | 0.0786636 | 0.68254797 |
| SLC43A2 | 24.8062643 | 0.96603779 | 0.64502183 | 1.49768233 | 0.1342158 | 0.80890178 |
| PRPF8 | 67.8802115 | 0.10066608 | 0.28354565 | 0.355026 | 0.72257012 | 0.9958918 |

| | | | | | | |
|---|---|---|---|---|---|---|
| MIR22HG | 23.2086684 | -0.2373548 | 0.62590072 | -0.3792212 | 0.7045236 | 0.9958918 |
| SERPINF1 | 54.8858158 | -0.8324615 | 0.4627216 | -1.7990547 | 0.07201003 | 0.66213979 |
| SMYD4 | 22.2048239 | -0.3046491 | 0.36365024 | -0.837753 | 0.40216942 | 0.97512144 |
| RPA1 | 37.385082 | 0.19348714 | 0.29538651 | 0.65503039 | 0.51244819 | 0.9958918 |
| HIC1 | 40.9920398 | -0.9525882 | 0.476087 | -2.00087 | 0.0454064 | 0.55894748 |
| SMG6 | 334.389172 | -0.8691097 | 0.35630698 | -2.4392161 | 0.01471916 | 0.33025022 |
| SGSM2 | 24.7059984 | 0.0659069 | 0.3127242 | 0.21075088 | 0.83308167 | 0.99592423 |
| MNT | 18.3275781 | 0.49183765 | 0.40991024 | 1.19986672 | 0.23019111 | 0.91455844 |
| METTL16 | 70.9627975 | -0.1839748 | 0.24095653 | -0.7635186 | 0.44515419 | 0.98092839 |
| PAFAH1B1 | 263.71106 | -0.0476266 | 0.1620956 | -0.293818 | 0.76889699 | 0.9958918 |
| ASPA | 37.3440299 | -0.5410265 | 0.62353211 | -0.8676803 | 0.38556938 | 0.97254921 |
| NCBP3 | 43.8496611 | -0.0645076 | 0.2450455 | -0.2632474 | 0.79235987 | 0.99592423 |
| ATP2A3 | 20.9030665 | 1.74344681 | 0.5482451 | 3.18004994 | 0.0014725 | 0.08388666 |
| ZZEF1 | 95.32143 | 0.18542597 | 0.22720068 | 0.816133 | 0.41442407 | 0.97812179 |
| ANKFY1 | 63.9418512 | -0.0486634 | 0.32274438 | -0.15078 | 0.88014927 | 0.99749293 |
| UBE2G1 | 82.4898418 | -0.0781732 | 0.21581871 | -0.3622169 | 0.71718995 | 0.9958918 |
| PSMB6 | 21.291832 | 0.39995238 | 0.5022119 | 0.79638172 | 0.4258102 | 0.98015476 |
| MINK1 | 29.7626331 | 0.6344172 | 0.33011798 | 1.92178929 | 0.05463228 | 0.59304667 |
| PFN1 | 109.103911 | -0.0231848 | 0.46146554 | -0.0502416 | 0.95992984 | 0.99935941 |
| SPAG7 | 18.1163589 | -0.0207013 | 0.46101389 | -0.0449038 | 0.96418396 | 0.99935941 |
| CAMTA2 | 19.7665293 | -0.4705788 | 0.37206433 | -1.2647781 | 0.20595089 | 0.89589437 |
| KIF1C | 27.5315933 | 0.51028863 | 0.45793724 | 1.11432002 | 0.26514192 | 0.93499423 |
| AC087500.1 | 24.782603 | 0.32168775 | 0.35670839 | 0.90182276 | 0.36715103 | 0.97217293 |
| RABEP1 | 121.031543 | 0.1202162 | 0.24686246 | 0.48697642 | 0.62627504 | 0.9958918 |
| NUP88 | 44.0299844 | -0.070953 | 0.2733119 | -0.2596046 | 0.79516878 | 0.99592423 |
| RPAIN | 26.8778889 | 0.07355991 | 0.38955744 | 0.18882942 | 0.85022651 | 0.9971731 |
| C1QBP | 25.5387741 | -0.1077637 | 0.36869894 | -0.292281 | 0.77007181 | 0.9958918 |
| DERL2 | 21.2143266 | 0.44889793 | 0.37486575 | 1.19748984 | 0.23111568 | 0.91521027 |
| KIAA0753 | 33.779287 | -0.0981807 | 0.30514239 | -0.3217536 | 0.74763933 | 0.9958918 |
| XAF1 | 116.795375 | -0.8455637 | 1.28375308 | -0.6586653 | 0.51011069 | 0.99543375 |
| DLG4 | 18.7955586 | 0.30780359 | 0.45873338 | 0.67098581 | 0.50222957 | 0.9929742 |
| ACADVL | 81.7386586 | 0.2103861 | 0.33450074 | 0.62895555 | 0.52937815 | 0.9958918 |
| EIF5A | 32.135019 | -0.4373517 | 0.51786557 | -0.8445276 | 0.39837466 | 0.97500135 |
| GPS2 | 25.8401056 | -0.1395128 | 0.40319419 | -0.3460188 | 0.72932857 | 0.9958918 |
| ZBTB4 | 54.0436978 | 0.05344138 | 0.30664414 | 0.17427818 | 0.86164685 | 0.9971731 |
| POLR2A | 64.2238169 | 0.20003402 | 0.36652768 | 0.54575419 | 0.58523492 | 0.9958918 |
| EIF4A1 | 48.8698287 | -0.0809528 | 0.54974064 | -0.1472564 | 0.88292967 | 0.99840747 |
| FXR2 | 18.3524063 | -0.2695374 | 0.35610374 | -0.7569071 | 0.44910553 | 0.98092839 |
| SAT2 | 35.6687555 | 0.14794252 | 0.35778924 | 0.4134907 | 0.67924713 | 0.9958918 |
| TP53 | 18.7389787 | 0.6518151 | 0.40803812 | 1.59743678 | 0.11016838 | 0.77370222 |
| KDM6B | 32.3176896 | -0.2572499 | 0.51882077 | -0.4958357 | 0.62001031 | 0.9958918 |
| NAA38 | 61.5071936 | 0.06456165 | 0.30544526 | 0.21136897 | 0.83259937 | 0.99592423 |
| CHD3 | 55.0860505 | -0.0358259 | 0.36911203 | -0.0970597 | 0.92267895 | 0.99935941 |
| PER1 | 73.7396401 | -0.6521577 | 0.4621787 | -1.411051 | 0.15822958 | 0.84598994 |
| NDEL1 | 63.4672857 | -0.3512009 | 0.30505456 | -1.1512724 | 0.2496202 | 0.92624282 |
| MYH10 | 283.37979 | -0.2395801 | 0.29771405 | -0.8047323 | 4.21E-01 | 0.97812179 |
| STX8 | 143.649225 | -0.2035627 | 0.20944433 | -0.9719178 | 0.33109143 | 0.9654289 |
| GAS7 | 65.8226522 | -0.4865227 | 0.2940119 | -1.6547721 | 9.80E-02 | 7.44E-01 |
| SCO1 | 24.4429226 | 0.3999151 | 0.34428387 | 1.16158533 | 0.24540394 | 0.92559078 |
| TMEM220 | 28.2929321 | 0.07408685 | 0.42209093 | 0.17552343 | 0.86066836 | 0.9971731 |
| MAP2K4 | 142.202095 | 0.10310085 | 0.17977334 | 0.57350465 | 0.56630305 | 0.9958918 |
| MYOCD | 71.1259112 | 0.64987462 | 0.41977533 | 1.54814867 | 0.12158651 | 0.79021701 |
| ARHGAP44 | 105.217751 | 0.05116358 | 0.34219336 | 0.14951657 | 0.88114603 | 0.9977723 |
| HS3ST3A1 | 49.3109327 | 0.47488608 | 0.46862583 | 1.01335875 | 3.11E-01 | 0.95956408 |

| | | | | | | |
|---|---|---|---|---|---|---|
| COX10 | 26.3793921 | -0.0116706 | 0.33410123 | -0.0349313 | 0.97213448 | 0.99935941 |
| HS3ST3B1 | 27.0447372 | 1.38750683 | 0.59575383 | 2.32899354 | 0.01985941 | 0.38006976 |
| PMP22 | 54.6342726 | -0.1638226 | 0.31598383 | -0.5184525 | 0.60414257 | 0.9958918 |
| ZSWIM7 | 32.6644658 | 0.01216465 | 0.33191123 | 0.03665031 | 0.97076383 | 0.99935941 |
| TTC19 | 77.7512194 | -0.0299901 | 0.23578942 | -0.1271901 | 0.89878993 | 0.99935941 |
| NCOR1 | 266.116744 | -0.2616448 | 0.1629763 | -1.6054165 | 0.10840218 | 0.77248589 |
| PIGL | 80.9047777 | -0.1204345 | 0.32761034 | -0.3676152 | 0.71316016 | 0.9958918 |
| UBB | 100.154249 | 0.28948604 | 0.37817591 | 0.76547986 | 0.44398586 | 0.98092839 |
| LRRC75A | 30.3274519 | 0.37948497 | 0.33056293 | 1.14799615 | 0.25097015 | 0.92624282 |
| AC098850.3 | 64.4266703 | -0.2870063 | 0.6257871 | -0.4586326 | 0.64649806 | 0.9958918 |
| MPRIP | 280.699791 | 0.07914993 | 0.34215738 | 0.2313261 | 0.81706147 | 0.99592423 |
| RAI1 | 58.5802829 | 0.30101593 | 0.30438429 | 0.98893385 | 0.32269551 | 0.95992644 |
| SREBF1 | 45.1661032 | 0.00629066 | 0.36130589 | 0.0174109 | 0.98610881 | 0.99935941 |
| TOM1L2 | 110.236938 | 0.21938195 | 0.19024239 | 1.15317074 | 0.24884031 | 0.92624282 |
| ALKBH5 | 43.7744919 | 0.3139278 | 0.30208933 | 1.03918863 | 0.29871702 | 0.95447177 |
| FLII | 21.6509765 | 0.1536255 | 0.36561496 | 0.42018385 | 0.67435115 | 0.9958918 |
| TOP3A | 24.8584269 | 0.15716102 | 0.35243192 | 0.44593299 | 0.65564565 | 0.9958918 |
| PRPSAP2 | 38.7671754 | -0.0544536 | 0.2922237 | -0.186342 | 0.85217656 | 0.9971731 |
| EPN2 | 152.01228 | 0.10506433 | 0.1977409 | 0.53132324 | 0.59519481 | 0.9958918 |
| MFAP4 | 33.174492 | 0.39145104 | 0.39032958 | 1.00287309 | 0.3159221 | 0.95992644 |
| SLC47A1 | 23.313476 | -0.0379166 | 0.39486449 | -0.0960243 | 0.9235013 | 0.99935941 |
| ALDH3A2 | 40.490301 | -0.0393902 | 0.38026383 | -0.1035866 | 0.91749741 | 0.99935941 |
| ULK2 | 67.09787 | -0.563349 | 0.3657381 | -1.5403072 | 0.12348549 | 0.79184923 |
| AKAP10 | 110.564878 | -0.3018821 | 0.26136992 | -1.1549995 | 0.24809065 | 0.92624282 |
| SPECC1 | 134.833856 | -1.1896913 | 0.91019778 | -1.307069 | 0.19118927 | 0.88158763 |
| USP22 | 100.724512 | 0.12245489 | 0.18412292 | 0.66507143 | 0.50600481 | 0.99397937 |
| DHRS7B | 47.5786741 | 0.35689533 | 0.36714094 | 0.97209351 | 0.33100403 | 0.9654289 |
| MAP2K3 | 30.6014602 | -0.5283592 | 0.4322953 | -1.2222183 | 0.22162508 | 0.91091875 |
| WSB1 | 378.259259 | -0.3606826 | 0.45636994 | -0.7903294 | 0.4293354 | 0.98092839 |
| KSR1 | 120.846437 | 0.36283246 | 0.34688557 | 1.04597161 | 0.29557414 | 0.95393362 |
| NLK | 127.149085 | 0.11294876 | 0.17607405 | 0.64148443 | 0.521208 | 0.9958918 |
| SARM1 | 21.8471883 | 0.15853325 | 0.47954503 | 0.33059096 | 0.74095348 | 0.9958918 |
| KIAA0100 | 35.0783266 | 0.17603786 | 0.3480677 | 0.50575753 | 0.61302688 | 0.9958918 |
| SUPT6H | 54.9492294 | 0.08393483 | 0.28594643 | 0.29353341 | 0.76911447 | 0.9958918 |
| RAB34 | 29.4864889 | -0.1983931 | 0.43505501 | -0.4560184 | 0.64837673 | 0.9958918 |
| RPL23A | 115.191498 | -0.0584964 | 0.3134603 | -0.1866151 | 0.85196241 | 0.9971731 |
| FAM222B | 101.823112 | 0.05222983 | 0.236755 | 0.22060709 | 0.82539838 | 0.99592423 |
| FLOT2 | 36.0543043 | 0.62686486 | 0.32333402 | 1.93875319 | 0.0525314 | 0.583162 |
| PHF12 | 47.3734827 | 0.01577597 | 0.28538475 | 0.05527966 | 0.95591567 | 0.99935941 |
| MYO18A | 25.6964632 | 0.98808618 | 0.44159315 | 2.23754871 | 0.0252505 | 0.42757124 |
| NUFIP2 | 112.746692 | 0.1986084 | 0.19362156 | 1.02575559 | 0.3050068 | 0.95793395 |
| TAOK1 | 224.161838 | 0.08712596 | 0.20303988 | 0.42910763 | 0.6678449 | 0.9958918 |
| SSH2 | 134.248576 | -0.0137159 | 0.23985965 | -0.0571829 | 0.95439948 | 0.99935941 |
| NSRP1 | 134.310801 | -0.113197 | 0.20301028 | -0.5575926 | 0.57712261 | 0.9958918 |
| BLMH | 25.973538 | 0.45509384 | 0.2936937 | 1.54955258 | 0.12124894 | 0.79021701 |
| CPD | 79.9128695 | 0.22059107 | 0.29686918 | 0.74305816 | 0.45744647 | 0.9838379 |
| GOSR1 | 61.68892 | 0.04575357 | 0.22959718 | 0.19927759 | 0.84204561 | 0.99692952 |
| CRLF3 | 49.4899039 | 0.02027052 | 0.30995515 | 0.06539824 | 0.94785693 | 0.99935941 |
| TEFM | 22.8389097 | 0.02288752 | 0.32531581 | 0.07035478 | 0.94391128 | 0.99935941 |
| NF1 | 400.732662 | -0.1928237 | 0.21881491 | -0.8812182 | 0.37819973 | 0.97217293 |
| UTP6 | 29.4550508 | -0.0993683 | 0.31622105 | -0.3142369 | 0.75334108 | 0.9958918 |
| SUZ12 | 83.5201704 | 0.05466007 | 0.23563395 | 0.23197028 | 0.81656109 | 0.99592423 |
| LRRC37B | 33.4582278 | -0.4271659 | 0.37019531 | -1.1538933 | 0.24854391 | 0.92624282 |
| RHOT1 | 74.0952508 | 0.43997076 | 0.23405579 | 1.87976875 | 0.0601396 | 0.61332974 |

| | | | | | | |
|---|---|---|---|---|---|---|
| ZNF207 | 171.163064 | 0.15645086 | 0.17838728 | 0.87702922 | 0.38047077 | 0.97217293 |
| PSMD11 | 51.8710976 | -0.3688004 | 0.24711569 | -1.4924202 | 0.13558903 | 0.81154597 |
| MYO1D | 428.420846 | 0.40038415 | 0.33513819 | 1.19468376 | 0.23221061 | 0.91565561 |
| TMEM98 | 19.3351457 | 0.4798192 | 0.36045317 | 1.33115545 | 0.18313787 | 0.87866798 |
| CCL2 | 111.340584 | -1.083783 | 1.14302402 | -0.9481717 | 0.34304206 | 0.96887708 |
| SLFN5 | 46.227259 | 0.48257443 | 0.46040585 | 1.04815009 | 0.29456946 | 0.95304825 |
| SLFN11 | 32.2555688 | 1.02828752 | 0.4706263 | 2.18493425 | 0.02889367 | 0.46160775 |
| AP2B1 | 105.092826 | 0.0621101 | 0.25872145 | 0.24006553 | 0.81027946 | 0.99592423 |
| TAF15 | 173.872288 | 0.05966169 | 0.24550664 | 0.24301457 | 0.80799411 | 0.99592423 |
| MYO19 | 30.1394454 | -0.3562269 | 0.38784131 | -0.9184863 | 0.35836432 | 0.97217293 |
| GGNBP2 | 90.4296668 | -0.3020156 | 0.20231526 | -1.4927967 | 0.1354904 | 0.81154597 |
| AATF | 58.5807154 | -0.0274578 | 0.23554447 | -0.1165717 | 0.90719943 | 0.99935941 |
| ACACA | 123.049205 | -0.3766134 | 0.34363037 | -1.0959841 | 0.27308574 | 0.93824549 |
| TADA2A | 32.5980738 | 0.07063701 | 0.29329717 | 0.24083768 | 0.80968093 | 0.99592423 |
| SYNRG | 68.9147147 | 0.31408297 | 0.25732108 | 1.22058779 | 0.22224213 | 0.91093521 |
| DDX52 | 34.1102156 | -0.0201729 | 0.29481281 | -0.0684262 | 0.94544637 | 0.99935941 |
| SOCS7 | 27.9634573 | 0.15342592 | 0.29441833 | 0.52111537 | 0.60228641 | 0.9958918 |
| ARHGAP23 | 33.8159207 | 0.07836879 | 0.40043503 | 0.19570913 | 0.84483784 | 0.99692952 |
| MLLT6 | 82.8634432 | 0.35178388 | 0.32180381 | 1.09316256 | 0.27432243 | 0.9394201 |
| PCGF2 | 19.856201 | -0.2306585 | 0.42423041 | -0.5437104 | 0.58664081 | 0.9958918 |
| PSMB3 | 29.0405899 | -0.2645258 | 0.43646589 | -0.606063 | 0.54447294 | 0.9958918 |
| PIP4K2B | 29.9188951 | -0.1858943 | 0.31087453 | -0.5979721 | 0.54985858 | 0.9958918 |
| CWC25 | 23.7553598 | -0.0353092 | 0.32671508 | -0.1080735 | 0.91393742 | 0.99935941 |
| RPL23 | 75.2468194 | 0.09733599 | 0.31017765 | 0.31380724 | 0.75366744 | 0.9958918 |
| LASP1 | 70.5184381 | -0.0869422 | 0.36192774 | -0.2402197 | 0.81015993 | 0.99592423 |
| PLXDC1 | 23.7097705 | -1.404464 | 0.60732594 | -2.3125375 | 0.02074808 | 0.38830162 |
| RPL19 | 132.346692 | -0.2179631 | 0.38377414 | -0.5679462 | 0.57007149 | 0.9958918 |
| FBXL20 | 122.70647 | 0.20285014 | 0.28172564 | 0.72002725 | 0.47150822 | 0.98910524 |
| MED1 | 27.6640286 | 0.10909696 | 0.33963004 | 0.32122294 | 0.74804145 | 0.9958918 |
| CDK12 | 77.627155 | -0.0930692 | 0.23345733 | -0.398656 | 0.69014667 | 0.9958918 |
| ERBB2 | 24.1849205 | 0.34333315 | 0.35503941 | 0.96702827 | 0.3335299 | 0.9654289 |
| PSMD3 | 24.5635738 | -0.253332 | 0.43100535 | -0.5877701 | 0.55668664 | 0.9958918 |
| MED24 | 22.4313574 | -0.0517831 | 0.31865477 | -0.1625053 | 0.87090799 | 0.9971731 |
| THRA | 27.3308196 | -0.1145456 | 0.45656309 | -0.2508867 | 0.80190173 | 0.99592423 |
| MSL1 | 49.2690135 | -0.0709951 | 0.22947783 | -0.3093765 | 0.75703513 | 0.9958918 |
| CASC3 | 55.7824643 | -0.3582913 | 0.3130602 | -1.1444804 | 0.25242445 | 0.92631316 |
| WIPF2 | 66.5751428 | 0.0148575 | 0.24585522 | 0.06043192 | 0.95181163 | 0.99935941 |
| RARA | 21.88809 | -0.0184141 | 0.38787052 | -0.0474749 | 0.96213472 | 0.99935941 |
| IGFBP4 | 484.783236 | -0.4360031 | 0.39090903 | -1.115357 | 0.26469748 | 0.93499423 |
| KRT10 | 30.6456879 | -0.2855077 | 0.61176193 | -0.4666973 | 0.64071643 | 0.9958918 |
| EIF1 | 156.858453 | 0.1445712 | 0.35592861 | 0.40618033 | 0.68461011 | 0.9958918 |
| JUP | 37.4404371 | -0.1996879 | 0.46507455 | -0.4293674 | 0.66765586 | 0.9958918 |
| FKBP10 | 57.7232208 | 0.02697111 | 0.39505208 | 0.06827228 | 0.94556889 | 0.99935941 |
| ACLY | 37.4750517 | 0.35779575 | 0.31587958 | 1.13269666 | 0.25734166 | 0.9266065 |
| DNAJC7 | 74.41771 | 0.01063426 | 0.25324074 | 0.04199269 | 0.96650453 | 0.99935941 |
| KAT2A | 18.7004186 | -0.7497139 | 0.41028076 | -1.8273193 | 0.06765178 | 0.64317326 |
| STAT5B | 119.59916 | 0.27409771 | 0.29221759 | 0.93799181 | 0.34824862 | 0.96887708 |
| STAT5A | 18.6627306 | 0.55985414 | 0.36751163 | 1.52336441 | 0.12766756 | 0.80131832 |
| STAT3 | 484.51711 | -0.0733043 | 0.3352754 | -0.2186392 | 0.82693114 | 0.99592423 |
| CAVIN1 | 104.753981 | 0.28941829 | 0.42121977 | 0.6870957 | 0.49202242 | 0.99046213 |
| ATP6V0A1 | 99.0192627 | -0.5410862 | 0.21575261 | -2.5079012 | 0.01214506 | 0.2958939 |
| NAGLU | 28.7275149 | -0.2429762 | 0.48985807 | -0.4960134 | 0.61988492 | 0.9958918 |
| RETREG3 | 44.9432935 | 0.38490696 | 0.28535429 | 1.34887394 | 0.17737746 | 0.87771577 |
| TUBG2 | 22.2503524 | 0.13122404 | 0.42392331 | 0.30954666 | 0.75690572 | 0.9958918 |

| | | | | | | |
|---|---|---|---|---|---|---|
| PLEKHH3 | 31.5367897 | 0.18864471 | 0.34910831 | 0.54036155 | 0.58894772 | 0.9958918 |
| AC100793.2 | 21.5183124 | 1.69790943 | 0.56315616 | 3.01498864 | 0.00256989 | 0.11446129 |
| EZH1 | 49.1774877 | 0.19780446 | 0.31157612 | 0.63485115 | 0.5255255 | 0.9958918 |
| COA3 | 21.8681331 | 0.60043176 | 0.39740268 | 1.51089007 | 0.13081646 | 0.80565822 |
| BECN1 | 17.2843405 | 0.19986313 | 0.39912759 | 0.50074997 | 0.6165471 | 0.9958918 |
| AOC3 | 26.9715362 | 0.64001668 | 0.33648347 | 1.90207464 | 0.0571614 | 0.60443037 |
| RPL27 | 66.3141705 | -0.2712725 | 0.38142893 | -0.7112007 | 0.47695989 | 0.98993168 |
| NBR1 | 63.4219789 | -0.0137298 | 0.31403372 | -0.0437207 | 0.96512706 | 0.99935941 |
| LINC00854 | 97.8208005 | -2.410323 | 0.53262253 | -4.5253869 | 6.03E-06 | 0.00246131 |
| DHX8 | 23.1682942 | -0.1470707 | 0.33363377 | -0.4408146 | 0.65934719 | 0.9958918 |
| DUSP3 | 27.9256204 | 0.198876 | 0.34581231 | 0.57509809 | 0.56522496 | 0.9958918 |
| LSM12 | 29.8475899 | -0.2168389 | 0.33279753 | -0.651564 | 0.51468248 | 0.9958918 |
| HDAC5 | 44.8852637 | 0.22125196 | 0.44718621 | 0.49476473 | 0.6207662 | 0.9958918 |
| UBTF | 22.2785446 | 0.10448456 | 0.38784624 | 0.26939686 | 0.7876243 | 0.99592423 |
| SLC25A39 | 19.6685382 | -0.1482213 | 0.47100647 | -0.3146906 | 0.75299661 | 0.9958918 |
| GRN | 34.3828998 | -0.1096965 | 0.36235739 | -0.30273 | 0.7620956 | 0.9958918 |
| GPATCH8 | 247.594516 | 0.27389287 | 0.2113828 | 1.29571971 | 0.19507207 | 0.88391893 |
| GJC1 | 38.5733051 | -1.8880916 | 0.50839743 | -3.7138103 | 0.00020416 | 0.02333707 |
| EFTUD2 | 31.1471373 | -0.2806684 | 0.37857123 | -0.7413886 | 0.45845784 | 0.98457237 |
| DCAKD | 25.7383526 | -0.0267785 | 0.36569055 | -0.0732273 | 0.94162521 | 0.99935941 |
| NMT1 | 59.0471075 | 0.09302892 | 0.21986459 | 0.42311913 | 0.67220834 | 0.9958918 |
| AC138150.1 | 30.2290532 | 0.28872026 | 0.45487207 | 0.6347285 | 0.5256055 | 0.9958918 |
| MAP3K14 | 33.7370234 | 0.72529386 | 0.43746088 | 1.6579628 | 0.09732498 | 0.74364783 |
| PLEKHM1 | 36.9019827 | 0.00867481 | 0.26683606 | 0.03250989 | 0.97406543 | 0.99935941 |
| MAPT | 24.5034756 | 1.54656173 | 0.55018657 | 2.81097689 | 0.00493913 | 0.17620238 |
| KANSL1 | 462.661761 | 0.19574531 | 0.25908948 | 0.75551236 | 0.44994162 | 0.98092839 |
| ARL17B | 106.700831 | 0.66657322 | 1.09886084 | 0.60660385 | 0.54411384 | 0.9958918 |
| LRRC37A2 | 23.8362556 | -0.0806234 | 0.38346199 | -0.2102514 | 0.83347143 | 0.99592423 |
| ARL17A | 26.4697708 | -0.2911722 | 0.44683113 | -0.6516382 | 0.51463462 | 0.9958918 |
| NSF | 145.110685 | -0.0898459 | 0.2784903 | -0.3226178 | 0.74698471 | 0.9958918 |
| CDC27 | 168.115172 | -0.0487394 | 0.22503808 | -0.216583 | 0.82853332 | 0.99592423 |
| AC068234.1 | 26.0047855 | 0.52857038 | 0.65771996 | 0.80364048 | 0.42160464 | 0.97812179 |
| EFCAB13 | 42.1325229 | 0.12194646 | 0.38382133 | 0.31771674 | 0.75069981 | 0.9958918 |
| NPEPPS | 185.369574 | -0.0817096 | 0.2195804 | -0.3721168 | 0.70980588 | 0.9958918 |
| AC025682.1 | 18.3607486 | -0.4652267 | 0.37822797 | -1.2300165 | 0.21869091 | 0.90437211 |
| KPNB1 | 98.2739099 | 0.04204055 | 0.2491231 | 0.1687541 | 0.86599006 | 0.9971731 |
| CDK5RAP3 | 19.4026282 | -0.1244684 | 0.37108409 | -0.3354184 | 0.73730952 | 0.9958918 |
| NFE2L1 | 51.7697471 | 0.09052552 | 0.31307507 | 0.28914958 | 0.77246692 | 0.9958918 |
| CBX1 | 18.957123 | 0.10476238 | 0.36793514 | 0.28473057 | 0.77585057 | 0.9958918 |
| HOXB3 | 42.5952433 | 0.47462599 | 0.39228764 | 1.20989281 | 0.22632003 | 0.91093521 |
| CALCOCO2 | 80.5651593 | -0.3927816 | 0.31643706 | -1.2412631 | 0.21450859 | 0.89978517 |
| ATP5MC1 | 28.3918969 | -0.0838459 | 0.37192985 | -0.2254346 | 0.82164118 | 0.99592423 |
| UBE2Z | 49.7193801 | 0.15751101 | 0.28667731 | 0.5494366 | 0.58270586 | 0.9958918 |
| SNF8 | 20.9843114 | -0.0449981 | 0.36169432 | -0.1244091 | 0.90099139 | 0.99935941 |
| ZNF652 | 83.540644 | 0.13990979 | 0.2144608 | 0.65237931 | 0.51415651 | 0.9958918 |
| PHB | 24.0805956 | 0.05518886 | 0.3811908 | 0.14478016 | 0.88488445 | 0.99862036 |
| SPOP | 92.8091691 | 0.19993386 | 0.22018689 | 0.90801887 | 0.36386825 | 0.97217293 |
| KAT7 | 30.6050946 | 0.122851 | 0.29281856 | 0.4195465 | 0.67481677 | 0.9958918 |
| ITGA3 | 78.7132742 | 0.23690766 | 0.4630942 | 0.51157553 | 0.60894811 | 0.9958918 |
| COL1A1 | 178.161407 | -0.4347851 | 0.53208548 | -0.8171339 | 0.41385194 | 0.97812179 |
| MRPL27 | 19.4373115 | 0.25159665 | 0.34141962 | 0.73691328 | 0.46117509 | 0.98590351 |
| LRRC59 | 20.3499641 | -0.2384045 | 0.46052861 | -0.5176758 | 0.60468448 | 0.9958918 |
| ABCC3 | 26.179924 | 0.71417151 | 0.37177548 | 1.92097528 | 0.05473482 | 0.59328351 |
| ANKRD40 | 19.8748596 | 0.19361205 | 0.39026167 | 0.49610828 | 0.61981802 | 0.9958918 |

| | | | | | | |
|---|---|---|---|---|---|---|
| LUC7L3 | 216.673281 | -0.0901735 | 0.17347189 | -0.5198163 | 0.60319164 | 0.9958918 |
| TOB1 | 61.1534596 | -0.0894069 | 0.32869981 | -0.2720018 | 0.78562063 | 0.9958918 |
| SPAG9 | 312.585683 | -0.08606 | 0.26365869 | -0.3264069 | 0.7441165 | 0.9958918 |
| MBTD1 | 91.4137292 | 0.08347012 | 0.27048549 | 0.30859372 | 0.7576306 | 0.9958918 |
| UTP18 | 24.6700843 | -0.5660576 | 0.31688194 | -1.7863358 | 0.0740449 | 0.66886321 |
| TOM1L1 | 31.4462111 | 0.15649768 | 0.31675146 | 0.49407092 | 0.6212561 | 0.9958918 |
| COX11 | 21.9649771 | -0.061777 | 0.36701479 | -0.1683229 | 0.86632929 | 0.9971731 |
| STXBP4 | 128.889464 | -0.385283 | 0.32457319 | -1.1870452 | 0.2352098 | 0.91962002 |
| DGKE | 25.5513072 | 0.90736367 | 0.36578765 | 2.48057493 | 0.01311707 | 0.30493547 |
| TRIM25 | 26.1975573 | 0.15259974 | 0.47021822 | 0.32452963 | 0.7455371 | 0.9958918 |
| SCPEP1 | 33.9569972 | -0.1171966 | 0.26556606 | -0.4413088 | 0.65898947 | 0.9958918 |
| AKAP1 | 21.1730995 | 0.3424099 | 0.4020958 | 0.851563 | 0.39445668 | 0.97500135 |
| MSI2 | 353.166199 | 0.3580182 | 0.36103635 | 0.99164031 | 0.32137302 | 0.95992644 |
| CUEDC1 | 70.9433345 | 0.29895231 | 0.3537233 | 0.84515865 | 0.39802227 | 0.97500135 |
| VEZF1 | 39.1562781 | 0.09991863 | 0.26527152 | 0.37666551 | 0.7064222 | 0.9958918 |
| AC015813.2 | 23.0875603 | -0.2934227 | 0.3902495 | -0.7518848 | 0.45212033 | 0.98092839 |
| DYNLL2 | 21.1874735 | 0.28939843 | 0.33979065 | 0.85169626 | 0.3943827 | 0.97500135 |
| MTMR4 | 21.3670287 | 0.22573913 | 0.368925 | 0.61188354 | 0.54061482 | 0.9958918 |
| 4-Sep | 21.7663595 | 0.3412129 | 0.41380798 | 0.82456821 | 0.40961678 | 0.97812179 |
| TRIM37 | 58.9244155 | -0.3164205 | 0.32393594 | -0.9767995 | 0.32866843 | 0.9620084 |
| SKA2 | 29.9352461 | 0.08617555 | 0.31373738 | 0.27467415 | 0.78356659 | 0.9958918 |
| YPEL2 | 124.775338 | 0.58664886 | 0.37998189 | 1.54388636 | 0.12261587 | 0.79184923 |
| DHX40 | 33.9780578 | 0.170521 | 0.36107725 | 0.47225628 | 0.63674388 | 0.9958918 |
| CLTC | 196.39397 | 0.00657821 | 0.22188971 | 0.02964632 | 0.97634912 | 0.99935941 |
| PTRH2 | 33.1164403 | -0.1119632 | 0.34312025 | -0.3263089 | 0.74419069 | 0.9958918 |
| VMP1 | 737.943011 | -0.693411 | 0.55496255 | -1.2494735 | 0.21149194 | 0.89738044 |
| TUBD1 | 25.0797078 | 0.37516772 | 0.47536633 | 0.78921811 | 0.42998454 | 0.98092839 |
| RPS6KB1 | 50.3442157 | -0.4484019 | 0.30071258 | -1.4911313 | 0.13592703 | 0.81177777 |
| USP32 | 206.128755 | -0.5188402 | 0.31149606 | -1.6656398 | 0.09578518 | 0.73786716 |
| APPBP2 | 114.348556 | 0.2291571 | 0.22690097 | 1.00994326 | 0.31252248 | 0.9599143 |
| PPM1D | 39.7569958 | 0.17404157 | 0.3481182 | 0.49994966 | 0.61711053 | 0.9958918 |
| BCAS3 | 345.152808 | -0.345284 | 0.20558702 | -1.679503 | 0.09305406 | 0.73019523 |
| TBX2 | 113.971792 | -0.3389015 | 0.34205934 | -0.990768 | 0.32179887 | 0.95992644 |
| INTS2 | 20.7391031 | -0.2907549 | 0.33398685 | -0.870558 | 0.38399552 | 0.9725044 |
| MED13 | 235.742183 | 0.10097418 | 0.19694284 | 0.51270805 | 0.60815556 | 0.9958918 |
| TLK2 | 140.553178 | -0.3049978 | 0.23346738 | -1.3063828 | 0.19142241 | 0.88158763 |
| MRC2 | 116.283098 | -0.0147774 | 0.3954306 | -0.0373703 | 0.97018976 | 0.99935941 |
| TANC2 | 148.336085 | -1.4207116 | 0.32329907 | -4.3944192 | 1.11E-05 | 0.00371022 |
| DCAF7 | 63.1915267 | 0.52733154 | 0.31311163 | 1.68416466 | 0.09214985 | 0.72957419 |
| MAP3K3 | 74.8885905 | 0.23033418 | 0.26329557 | 0.87481219 | 0.38167611 | 0.97217293 |
| CCDC47 | 25.5352051 | -0.7916178 | 0.30653589 | -2.5824637 | 0.00980977 | 0.26310947 |
| DDX42 | 77.9904571 | -0.1739182 | 0.19033531 | -0.9137465 | 0.36085004 | 0.97217293 |
| SMARCD2 | 23.154995 | 0.07521181 | 0.3857809 | 0.1949599 | 0.84542434 | 0.99692952 |
| ERN1 | 55.9222258 | 1.1029651 | 0.39358785 | 2.8023352 | 0.00507341 | 0.18011842 |
| SNHG25 | 23.4120661 | -0.2990454 | 0.48106752 | -0.6216287 | 0.53418608 | 0.9958918 |
| TEX2 | 38.4768716 | 0.09146901 | 0.24817936 | 0.3685601 | 0.71245564 | 0.9958918 |
| PECAM1 | 41.9119623 | 1.18029428 | 0.49571814 | 2.38097857 | 0.01726672 | 0.34765232 |
| POLG2 | 22.5184802 | 0.19728745 | 0.36080347 | 0.54680032 | 0.58451592 | 0.9958918 |
| DDX5 | 325.981219 | 0.34207486 | 0.25354788 | 1.34915289 | 0.17728786 | 0.87771577 |
| CEP95 | 71.4128283 | -0.1659915 | 0.25713804 | -0.6455344 | 0.51858092 | 0.9958918 |
| SMURF2 | 131.027978 | 0.06924021 | 0.22761463 | 0.3041993 | 0.76097606 | 0.9958918 |
| LRRC37A3 | 85.3806611 | -0.032688 | 0.39199622 | -0.0833885 | 0.93354262 | 0.99935941 |
| GNA13 | 58.8889465 | 0.34059532 | 0.23107162 | 1.47398161 | 0.14048654 | 0.8153985 |
| CEP112 | 176.07957 | -0.3056714 | 0.23372523 | -1.3078238 | 0.1909331 | 0.88158763 |

| | | | | | | |
|---|---|---|---|---|---|---|
| PRKCA | 730.667021 | -0.1252003 | 0.35550807 | -0.3521729 | 0.7247086 | 0.9958918 |
| PRKCA-AS1 | 31.247789 | -0.8144427 | 0.65599308 | -1.2415415 | 0.21440577 | 0.89978517 |
| HELZ | 191.851258 | -0.1485727 | 0.26135471 | -0.5684713 | 0.56971497 | 0.9958918 |
| PSMD12 | 48.1702999 | -0.2107442 | 0.28534893 | -0.7385491 | 0.46018085 | 0.98590351 |
| PITPNC1 | 552.968031 | 0.69888556 | 0.50307106 | 1.38923825 | 0.16476031 | 0.8581315 |
| NOL11 | 22.1368301 | 0.08047009 | 0.35650065 | 0.22572214 | 0.82141754 | 0.99592423 |
| BPTF | 246.353071 | 0.03147521 | 0.25851578 | 0.12175354 | 0.90309421 | 0.99935941 |
| ARSG | 20.3550342 | -0.2040806 | 0.40744473 | -0.5008791 | 0.61645618 | 0.9958918 |
| WIPI1 | 39.3672186 | -0.2331667 | 0.31693086 | -0.7357022 | 0.46191196 | 0.98590351 |
| PRKAR1A | 133.84813 | 0.36217073 | 0.28900347 | 1.25317086 | 0.21014353 | 0.89735318 |
| FAM20A | 31.4520339 | -0.6875033 | 0.56595052 | -1.2147764 | 0.22445138 | 0.91093521 |
| LINC01482 | 29.4686476 | -1.6186924 | 0.68834542 | -2.3515699 | 0.01869438 | 0.36528543 |
| ABCA8 | 206.090172 | 0.1869103 | 0.50316903 | 0.37146622 | 0.71029031 | 0.9958918 |
| ABCA9 | 158.170365 | -0.1697042 | 0.50557198 | -0.3356677 | 0.73712146 | 0.9958918 |
| ABCA6 | 172.809613 | -0.573042 | 0.55829957 | -1.026406 | 0.30470027 | 0.95793395 |
| ABCA10 | 126.256106 | -0.0764628 | 0.50743368 | -0.1506854 | 0.88022392 | 0.99749293 |
| ABCA5 | 80.0871991 | -0.2783341 | 0.31795416 | -0.8753908 | 0.38136132 | 0.97217293 |
| KCNJ16 | 31.1999463 | 0.20615386 | 0.42536868 | 0.48464748 | 0.62792643 | 0.9958918 |
| SLC39A11 | 123.679995 | -0.2095929 | 0.24352251 | -0.8606715 | 0.38941902 | 0.97297628 |
| SSTR2 | 41.7891651 | -1.5462625 | 0.5697298 | -2.7140277 | 0.00664706 | 0.20675326 |
| FAM104A | 18.8281094 | -0.001995 | 0.32536689 | -0.0061315 | 0.99510783 | 0.99935941 |
| CDC42EP4 | 43.8599794 | 0.00261665 | 0.28295058 | 0.00924774 | 0.99262147 | 0.99935941 |
| RPL38 | 46.5360703 | 0.13203057 | 0.42359851 | 0.31168799 | 0.75527766 | 0.9958918 |
| GPRC5C | 32.3609391 | 0.88240831 | 0.31291869 | 2.81992843 | 0.00480344 | 0.1747393 |
| ATP5PD | 20.0885603 | -0.1701349 | 0.42491192 | -0.4004004 | 0.6888616 | 0.9958918 |
| NT5C | 22.8599476 | 0.29649153 | 0.36415801 | 0.81418373 | 0.4155397 | 0.97812179 |
| SUMO2 | 19.1253918 | -0.2288127 | 0.37629731 | -0.6080637 | 0.54314523 | 0.9958918 |
| NUP85 | 22.5311805 | -0.009341 | 0.3488838 | -0.0267738 | 0.97864014 | 0.99935941 |
| GGA3 | 27.7948574 | 0.00929301 | 0.32170705 | 0.02888656 | 0.97695507 | 0.99935941 |
| GRB2 | 79.698774 | 0.23427651 | 0.25832278 | 0.90691386 | 0.36445235 | 0.97217293 |
| TMEM94 | 28.2867915 | 0.18954406 | 0.36169263 | 0.52404733 | 0.60024562 | 0.9958918 |
| MYO15B | 39.7727959 | 0.33435032 | 0.50488822 | 0.66222643 | 0.50782612 | 0.99397937 |
| SAP30BP | 31.3185216 | 0.25563773 | 0.2973283 | 0.85978268 | 0.38990885 | 0.97297628 |
| H3F3B | 262.922791 | -0.2851143 | 0.30266006 | -0.9420281 | 0.34617824 | 0.96887708 |
| UNK | 70.4360951 | 0.10531893 | 0.29230257 | 0.36030792 | 0.71861687 | 0.9958918 |
| WBP2 | 34.7179923 | 0.29144543 | 0.27571767 | 1.05704299 | 0.29049197 | 0.95186143 |
| ACOX1 | 36.1248162 | 0.1756552 | 0.30413159 | 0.57756314 | 0.5635591 | 0.9958918 |
| SRP68 | 25.1262579 | 0.03681089 | 0.3799763 | 0.0968768 | 0.92282423 | 0.99935941 |
| EXOC7 | 29.6420073 | 0.01235339 | 0.45493566 | 0.02715414 | 0.97833679 | 0.99935941 |
| PRPSAP1 | 40.594761 | 0.36858846 | 0.27492525 | 1.34068609 | 0.18002239 | 0.87815793 |
| UBE2O | 41.7354555 | 0.27008795 | 0.31843263 | 0.84817925 | 0.39633815 | 0.97500135 |
| MXRA7 | 57.0102623 | -0.1535767 | 0.29818101 | -0.5150453 | 0.60652136 | 0.9958918 |
| SRSF2 | 57.7324479 | -0.1494598 | 0.25270838 | -0.5914319 | 0.55423106 | 0.9958918 |
| MFSD11 | 43.4215331 | -0.0990091 | 0.35181268 | -0.2814257 | 0.77838388 | 0.9958918 |
| SEC14L1 | 106.144643 | -0.4149168 | 0.24092831 | -1.7221589 | 0.08504074 | 0.70697328 |
| 9-Sep | 187.92281 | -0.4183227 | 0.36430639 | -1.1482716 | 0.25085645 | 0.92624282 |
| TNRC6C | 145.897183 | 0.86402389 | 0.23315836 | 3.70573836 | 0.00021078 | 0.02333707 |
| SYNGR2 | 21.3652273 | 0.30503728 | 0.38893752 | 0.7842835 | 0.43287378 | 0.98092839 |
| SOCS3 | 52.4466586 | -0.773898 | 0.5287053 | -1.4637606 | 0.14325936 | 0.81994785 |
| PGS1 | 31.4886242 | 0.27019502 | 0.40176387 | 0.67252195 | 0.50125147 | 0.99213111 |
| CYTH1 | 68.8636857 | 0.28611106 | 0.290316 | 0.98551601 | 0.32437067 | 0.95992644 |
| USP36 | 60.3754952 | -0.0996333 | 0.29190304 | -0.3413234 | 0.73286017 | 0.9958918 |
| TIMP2 | 205.278867 | 0.07543448 | 0.35029887 | 0.21534321 | 0.82949975 | 0.99592423 |
| LGALS3BP | 84.9244991 | -1.0053667 | 0.54716567 | -1.8374081 | 0.06614967 | 0.63381211 |

| | | | | | | |
|---|---|---|---|---|---|---|
| C1QTNF1 | 64.0197769 | 1.51416141 | 0.46184717 | 3.27849018 | 0.00104364 | 0.07036431 |
| TBC1D16 | 87.3881413 | -0.1296312 | 0.30453781 | -0.4256653 | 0.67035172 | 0.9958918 |
| EIF4A3 | 21.9465398 | -0.6801078 | 0.5025893 | -1.3532079 | 0.17598922 | 0.87624983 |
| RNF213 | 195.177722 | -0.3991395 | 0.50532481 | -0.7898672 | 0.42960532 | 0.98092839 |
| AC124319.2 | 36.5559943 | -0.442022 | 0.40975188 | -1.0787553 | 0.28069685 | 0.94419503 |
| RPTOR | 105.720081 | 0.17158979 | 0.26904446 | 0.63777483 | 0.52362027 | 0.9958918 |
| SLC38A10 | 47.4406621 | 0.06297027 | 0.29880964 | 0.21073709 | 0.83309243 | 0.99592423 |
| BAHCC1 | 24.0352378 | -0.0854031 | 0.50101107 | -0.1704614 | 0.86464726 | 0.9971731 |
| ACTG1 | 356.395968 | -0.4760693 | 0.30174058 | -1.5777438 | 0.11462448 | 0.77853537 |
| NPLOC4 | 85.0819528 | -0.3203103 | 0.21827097 | -1.4674895 | 0.14224295 | 0.81652619 |
| HGS | 29.0436637 | -0.2332261 | 0.30914189 | -0.7544305 | 0.45059077 | 0.98092839 |
| MCRIP1 | 27.7887586 | -0.1016988 | 0.41718216 | -0.2437756 | 0.80740463 | 0.99592423 |
| P4HB | 134.51655 | -0.194672 | 0.4210847 | -0.4623108 | 0.64385843 | 0.9958918 |
| ARHGDIA | 63.8641798 | -0.0288971 | 0.42206616 | -0.0684659 | 0.94541477 | 0.99935941 |
| ANAPC11 | 32.1251189 | 0.33565864 | 0.40407123 | 0.83069176 | 0.40614778 | 0.97637555 |
| ASPSCR1 | 16.2934574 | -0.5632775 | 0.52708343 | -1.0686687 | 0.285219 | 0.94505166 |
| DUS1L | 20.7411889 | 0.16539989 | 0.38289411 | 0.43197294 | 0.66576108 | 0.9958918 |
| CCDC57 | 65.5400018 | 0.01655462 | 0.28408124 | 0.05827425 | 0.95353018 | 0.99935941 |
| SLC16A3 | 23.4641302 | -0.2971586 | 0.48868966 | -0.6080723 | 0.54313955 | 0.9958918 |
| CSNK1D | 64.1023531 | -0.142056 | 0.21857087 | -0.649931 | 0.51573679 | 0.9958918 |
| HEXDC | 26.4878527 | -0.2663532 | 0.37073481 | -0.7184466 | 0.47248198 | 0.98910524 |
| NARF | 29.708268 | 0.20718256 | 0.29386111 | 0.70503567 | 0.48078804 | 0.99002719 |
| FOXK2 | 124.380617 | -0.3948486 | 0.25660655 | -1.5387315 | 0.12386986 | 0.79195137 |
| WDR45B | 71.484497 | -0.6634381 | 0.33441803 | -1.9838587 | 0.04727159 | 0.56885457 |
| TBCD | 55.2951167 | -0.2002209 | 0.33317059 | -0.600956 | 0.54786927 | 0.9958918 |
| USP14 | 70.1148148 | -0.2842181 | 0.22779152 | -1.2477117 | 0.21213667 | 0.89738044 |
| THOC1 | 60.3277748 | -0.0137029 | 0.28704507 | -0.0477379 | 0.96192514 | 0.99935941 |
| COLEC12 | 76.1168307 | -2.0930273 | 0.53208181 | -3.933657 | 8.37E-05 | 0.01398756 |
| ENOSF1 | 104.349974 | -0.0873279 | 0.31781995 | -0.2747717 | 0.78349164 | 0.9958918 |
| YES1 | 60.3877938 | -0.2476133 | 0.24336911 | -1.0174394 | 0.30894446 | 0.95793395 |
| METTL4 | 19.7052074 | -0.3604621 | 0.34055962 | -1.0584406 | 0.2898546 | 0.95186143 |
| SMCHD1 | 288.492539 | -0.4724453 | 0.37864498 | -1.2477263 | 0.21213132 | 0.89738044 |
| AP001011.1 | 40.3223074 | -0.4490277 | 0.49050657 | -0.9154367 | 0.35996242 | 0.97217293 |
| LPIN2 | 175.887609 | -0.7377283 | 0.38171977 | -1.9326436 | 0.05328012 | 0.58707106 |
| MYOM1 | 87.3547534 | 1.04774161 | 0.57556526 | 1.82036979 | 0.06870271 | 0.64792295 |
| MYL12A | 106.989438 | -0.0079035 | 0.38904168 | -0.0203152 | 0.98379192 | 0.99935941 |
| MYL12B | 123.222416 | 0.16968014 | 0.35048107 | 0.48413497 | 0.62829009 | 0.9958918 |
| DLGAP1 | 132.359281 | -1.7323247 | 0.7855423 | -2.2052596 | 0.02743588 | 0.44841843 |
| LINC00667 | 19.6001396 | -0.0043686 | 0.44576212 | -0.0098003 | 0.99218064 | 0.99935941 |
| EPB41L3 | 38.6631533 | -0.033725 | 0.45264122 | -0.074507 | 0.94060695 | 0.99935941 |
| L3MBTL4 | 172.852884 | 0.63942666 | 0.46171602 | 1.38489164 | 0.1660856 | 0.86015722 |
| ARHGAP28 | 30.2440379 | 0.24767542 | 0.51050759 | 0.48515521 | 6.28E-01 | 0.9958918 |
| PTPRM | 399.565544 | -0.2467599 | 0.25073108 | -0.9841614 | 0.32503616 | 0.95992644 |
| RAB12 | 46.9746534 | 0.17233342 | 0.27740617 | 0.62123138 | 0.5344474 | 0.9958918 |
| NDUFV2 | 53.0442543 | -0.4265214 | 0.27586742 | -1.5461103 | 0.12207794 | 0.79183653 |
| ANKRD12 | 329.998215 | 0.05521229 | 0.18900414 | 0.29212211 | 0.77019326 | 0.9958918 |
| TWSG1 | 50.3620793 | 0.23760112 | 0.2377355 | 0.99943479 | 0.31758411 | 0.95992644 |
| RALBP1 | 42.8468761 | 0.30423559 | 0.25873412 | 1.17586188 | 0.23965007 | 0.92426606 |
| PPP4R1 | 70.4546894 | -0.1171501 | 0.25862939 | -0.4529653 | 0.65057375 | 0.9958918 |
| RAB31 | 142.584947 | -0.0279931 | 0.40023801 | -0.0699412 | 0.94424049 | 0.99935941 |
| VAPA | 164.311193 | -0.2856016 | 0.27836149 | -1.0260096 | 0.30488704 | 0.95793395 |
| APCDD1 | 28.9598909 | -0.3936661 | 0.42661702 | -0.9227624 | 0.35613107 | 0.97217293 |
| NAPG | 44.9811867 | 0.405532 | 0.32895402 | 1.23279238 | 0.21765324 | 0.90370709 |
| PIEZO2 | 185.429657 | 0.27879898 | 0.35462188 | 0.78618665 | 0.43175815 | 0.98092839 |

| | | | | | | |
|---|---|---|---|---|---|---|
| MPPE1 | 20.6815376 | -0.2289638 | 0.33363814 | -0.6862639 | 0.4925467 | 0.99046213 |
| IMPA2 | 24.5206264 | 0.60982565 | 0.46051429 | 1.3242274 | 0.18542753 | 0.87866798 |
| TUBB6 | 94.0713153 | -0.6971867 | 0.30226569 | -2.3065361 | 0.02108069 | 0.39000584 |
| AFG3L2 | 46.3758847 | 0.0938236 | 0.34346137 | 0.27317076 | 0.78472196 | 0.9958918 |
| SPIRE1 | 139.115416 | 0.18568291 | 0.26828871 | 0.69210112 | 0.48887383 | 0.99046213 |
| PSMG2 | 34.4045012 | 0.19399194 | 0.27319437 | 0.71008762 | 0.4776498 | 0.99002719 |
| PTPN2 | 80.1910762 | -0.0556672 | 0.28212032 | -0.1973171 | 0.84357942 | 0.99692952 |
| SEH1L | 38.8155098 | -0.3254762 | 0.39073726 | -0.8329798 | 0.40485613 | 0.97582411 |
| CEP192 | 79.7681695 | -0.0413983 | 0.25839117 | -0.1602157 | 0.87271118 | 0.9971731 |
| LDLRAD4 | 432.237642 | 0.1292231 | 0.39909208 | 0.32379269 | 0.74609499 | 0.9958918 |
| FAM210A | 33.2039971 | -0.4876154 | 0.52399618 | -0.9305706 | 0.35207573 | 0.97087862 |
| RNMT | 79.9896306 | -0.6269774 | 0.41194051 | -1.5220096 | 0.12800668 | 0.80197876 |
| ROCK1 | 354.931808 | -0.0834165 | 0.2659676 | -0.313634 | 0.75379905 | 0.9958918 |
| GREB1L | 39.6054292 | -1.382885 | 0.58804141 | -2.3516796 | 0.01868886 | 0.36528543 |
| ESCO1 | 50.5927732 | 0.06375636 | 0.24471086 | 0.26053752 | 0.79444918 | 0.99592423 |
| ABHD3 | 45.6424741 | 0.30561041 | 0.35370462 | 0.86402718 | 0.38757296 | 0.97297628 |
| MIB1 | 159.174306 | -0.2859024 | 0.19870396 | -1.4388362 | 0.15019694 | 0.83354431 |
| GATA6 | 62.7064543 | -0.7480471 | 0.47147741 | -1.5866021 | 0.11260278 | 0.77793929 |
| RBBP8 | 29.7174979 | -0.5037519 | 0.36556329 | -1.3780156 | 0.16819847 | 0.86560961 |
| CABLES1 | 97.2165059 | 0.40469589 | 0.44949635 | 0.90033186 | 0.36794367 | 0.97217293 |
| TMEM241 | 61.6172728 | -0.0465578 | 0.2691507 | -0.1729802 | 0.86266697 | 0.9971731 |
| RIOK3 | 75.0398319 | 0.00404536 | 0.27345975 | 0.01479325 | 0.98819712 | 0.99935941 |
| NPC1 | 57.7485541 | -0.3657333 | 0.35078987 | -1.0425993 | 0.29713393 | 0.95417903 |
| TTC39C | 34.7092848 | 0.09154831 | 0.42580085 | 0.21500266 | 0.82976525 | 0.99592423 |
| OSBPL1A | 186.881792 | -0.0590045 | 0.35441239 | -0.1664853 | 0.86777504 | 0.9971731 |
| ZNF521 | 29.8738123 | -0.1532802 | 0.40582716 | -0.3776983 | 0.70565472 | 0.9958918 |
| SS18 | 109.18652 | -0.0971889 | 0.26071871 | -0.3727731 | 0.70931734 | 0.9958918 |
| TAF4B | 21.6233353 | -0.6136146 | 0.31884302 | -1.9245037 | 0.05429148 | 0.59304667 |
| KCTD1 | 46.6743336 | 0.58360735 | 0.31160651 | 1.87289843 | 0.06108241 | 0.61696136 |
| TRAPPC8 | 91.9731447 | 0.01232866 | 0.25398171 | 0.04854154 | 0.96128466 | 0.99935941 |
| RNF138 | 32.8149789 | -0.6128026 | 0.38169422 | -1.6054804 | 0.10838813 | 0.77248589 |
| GAREM1 | 162.999944 | -0.6251688 | 0.23443579 | -2.6666951 | 0.00766011 | 0.22699265 |
| DTNA | 60.7410823 | 0.93741487 | 0.56652905 | 1.65466339 | 0.0979928 | 0.74364783 |
| MAPRE2 | 114.271258 | -0.5463682 | 0.40474359 | -1.3499118 | 0.17704426 | 0.87771577 |
| ZNF397 | 33.3909813 | 0.33543651 | 0.3034537 | 1.10539599 | 0.26898804 | 0.93598159 |
| ZSCAN30 | 23.7339506 | -0.1759134 | 0.33925444 | -0.5185293 | 0.60408899 | 0.9958918 |
| ZNF24 | 37.7896743 | 0.11649589 | 0.3109928 | 0.37459353 | 0.70796278 | 0.9958918 |
| GALNT1 | 73.6264924 | 0.26450874 | 0.30266269 | 0.87393905 | 0.38215144 | 0.9724484 |
| RPRD1A | 95.0177224 | 0.03403154 | 0.20227297 | 0.16824562 | 0.86639006 | 0.9971731 |
| SLC39A6 | 23.6143246 | 0.45758062 | 0.31277349 | 1.46297762 | 0.1434735 | 0.82053445 |
| ELP2 | 75.7924254 | -0.3744959 | 0.28412468 | -1.3180688 | 0.18748061 | 0.87975358 |
| FHOD3 | 59.4185458 | 1.70767474 | 0.51151592 | 3.33845862 | 0.00084245 | 0.0644401 |
| TPGS2 | 29.9166926 | 0.03917394 | 0.28438391 | 0.13775018 | 0.89043786 | 0.99935941 |
| KIAA1328 | 160.597053 | -0.2796271 | 0.32187768 | -0.8687371 | 0.38499092 | 0.97254921 |
| AC016205.1 | 19.2175555 | 0.09339324 | 0.3948773 | 0.23651206 | 0.81303535 | 0.99592423 |
| PIK3C3 | 119.648352 | -0.1575344 | 0.29415128 | -0.5355559 | 0.59226553 | 0.9958918 |
| SETBP1 | 1049.35055 | -0.6292499 | 0.37069808 | -1.6974728 | 0.0896073 | 0.72305504 |
| SLC14A2-AS1 | 30.7865761 | -0.7907934 | 0.96138593 | -0.8225556 | 0.41076077 | 0.97812179 |
| EPG5 | 73.1280512 | -0.1881657 | 0.22200334 | -0.8475804 | 0.39667172 | 0.97500135 |
| PSTPIP2 | 24.9911607 | -0.1682502 | 0.37591217 | -0.4475784 | 0.65445751 | 0.9958918 |
| ATP5F1A | 52.9980221 | 0.27605344 | 0.23812526 | 1.15927827 | 0.24634278 | 0.92600668 |
| C18orf25 | 45.4223834 | 0.46883219 | 0.25250477 | 1.85672604 | 0.06335014 | 0.62395844 |
| PIAS2 | 81.323852 | -0.2381918 | 0.2257074 | -1.0553125 | 0.29128242 | 0.95186143 |
| KATNAL2 | 21.9430676 | -0.7291467 | 0.45314685 | -1.6090737 | 0.10760024 | 0.77225821 |

| | | | | | | |
|---|---|---|---|---|---|---|
| SMAD2 | 127.110777 | -0.2715677 | 0.21360522 | -1.2713535 | 0.20360293 | 0.89396747 |
| ZBTB7C | 81.2529354 | -0.8297083 | 0.24730227 | -3.355037 | 0.00079354 | 0.06338861 |
| CTIF | 103.318965 | -0.5207563 | 0.28566713 | -1.822948 | 0.06831126 | 0.647832 |
| SMAD7 | 44.3693499 | -0.0207963 | 0.30619559 | -0.0679183 | 0.94585065 | 0.99935941 |
| DYM | 268.558943 | -0.0794716 | 0.20132754 | -0.3947378 | 0.69303641 | 0.9958918 |
| SNHG22 | 23.1282243 | 0.68330077 | 0.46293927 | 1.47600519 | 0.1399425 | 0.8153985 |
| ME2 | 28.8227522 | 0.34734446 | 0.34693375 | 1.00118382 | 0.31673795 | 0.95992644 |
| SMAD4 | 56.3336637 | -0.0924641 | 0.27910366 | -0.3312896 | 0.74042575 | 0.9958918 |
| DCC | 45.1068352 | 1.87798064 | 0.58712807 | 3.19858771 | 0.00138103 | 0.08251345 |
| MBD2 | 98.7852924 | 0.0134546 | 0.17372096 | 0.07744948 | 0.93826598 | 0.99935941 |
| POLI | 25.6910233 | -0.1344785 | 0.45605589 | -0.2948729 | 0.76809097 | 0.9958918 |
| TCF4 | 406.005163 | 0.22436843 | 0.29807214 | 0.75273198 | 0.45161099 | 0.98092839 |
| TXNL1 | 45.9872722 | -0.1536985 | 0.24232377 | -0.6342692 | 0.52590515 | 0.9958918 |
| WDR7 | 140.168971 | 0.04184743 | 0.19789002 | 0.21146814 | 0.83252199 | 0.99592423 |
| NARS | 37.5268308 | 0.28535846 | 0.34306369 | 0.83179441 | 0.405525 | 0.97637555 |
| AC027097.2 | 54.9422066 | -0.0767716 | 0.32321904 | -0.2375219 | 0.81225196 | 0.99592423 |
| ATP8B1 | 137.344333 | 0.78693464 | 0.41833049 | 1.88113145 | 0.05995404 | 0.61332974 |
| NEDD4L | 50.9163331 | -0.2269645 | 0.30257091 | -0.75012 | 0.45318242 | 0.98092839 |
| MALT1 | 50.6052479 | -0.0832513 | 0.38957743 | -0.2136965 | 0.83078372 | 0.99592423 |
| ZNF532 | 138.29005 | -0.204811 | 0.34917924 | -0.5865498 | 0.55750612 | 0.9958918 |
| LMAN1 | 83.0261525 | -0.3478134 | 0.35106527 | -0.9907373 | 0.32181389 | 0.95992644 |
| CCBE1 | 284.405971 | -0.0511702 | 0.55556902 | -0.0921041 | 0.92661532 | 0.99935941 |
| RNF152 | 281.169263 | -1.173022 | 0.59096599 | -1.984923 | 0.04715304 | 0.56885457 |
| PIGN | 100.87372 | -0.6909534 | 0.28395049 | -2.4333587 | 0.01495947 | 0.33314297 |
| KIAA1468 | 96.0822561 | -0.2848974 | 0.22704079 | -1.2548292 | 0.20954078 | 0.89634177 |
| ZCCHC2 | 59.0902472 | 0.05313062 | 0.45208116 | 0.11752452 | 0.90644441 | 0.99935941 |
| PHLPP1 | 105.807385 | 0.33030465 | 0.29284675 | 1.12790955 | 0.25935812 | 0.92976724 |
| BCL2 | 142.823466 | -0.4348283 | 0.38307368 | -1.1351036 | 0.25633192 | 0.9266065 |
| KDSR | 53.0976155 | -0.3393712 | 0.22184264 | -1.5297836 | 0.1260703 | 0.79527093 |
| VPS4B | 49.3723514 | 0.14247163 | 0.28831965 | 0.4941447 | 0.62120399 | 0.9958918 |
| TMX3 | 63.8617125 | -0.234781 | 0.25924716 | -0.905626 | 0.36513383 | 0.97217293 |
| CCDC102B | 303.280862 | -1.2097718 | 0.30458576 | -3.9718594 | 7.13E-05 | 0.01292273 |
| DOK6 | 157.685907 | -0.3959751 | 0.31261024 | -1.2666734 | 0.20527208 | 0.89589437 |
| RTTN | 41.9629335 | -0.2223886 | 0.2937959 | -0.7569492 | 0.44908031 | 0.98092839 |
| CYB5A | 87.416954 | 0.10624856 | 0.3653626 | 0.29080306 | 0.77120194 | 0.9958918 |
| CNDP2 | 40.6997328 | 0.66015944 | 0.32784796 | 2.01361463 | 0.04405001 | 0.55583613 |
| ZNF407 | 213.044085 | -0.1369682 | 0.15983886 | -0.8569141 | 0.39149236 | 0.97297628 |
| ZADH2 | 23.1848023 | 0.34458326 | 0.43665773 | 0.78913812 | 0.43003128 | 0.98092839 |
| TSHZ1 | 43.1338018 | -0.1430725 | 0.29452233 | -0.4857782 | 0.62712442 | 0.9958918 |
| ZNF516 | 130.328872 | -0.9213873 | 0.39980739 | -2.304578 | 0.02119022 | 0.39029308 |
| ZNF236 | 45.5930853 | -0.1326497 | 0.26942874 | -0.4923369 | 0.62248118 | 0.9958918 |
| MBP | 82.8188425 | 0.01249919 | 0.50249924 | 0.02487404 | 0.98015543 | 0.99935941 |
| ATP9B | 173.085552 | 0.02079113 | 0.21098613 | 0.09854265 | 0.92150141 | 0.99935941 |
| NFATC1 | 37.1733577 | -0.2590463 | 0.6010259 | -0.4310069 | 0.66646332 | 0.9958918 |
| PQLC1 | 36.8237817 | 0.27092684 | 0.30774889 | 0.88035036 | 0.37866954 | 0.97217293 |
| TXNL4A | 41.8979332 | 0.02862318 | 0.29430154 | 0.097258 | 0.92252151 | 0.99935941 |
| ADNP2 | 36.2959674 | -0.0032773 | 0.29910317 | -0.0109569 | 0.99125779 | 0.99935941 |
| PARD6G | 23.9875894 | -0.3523084 | 0.45445872 | -0.7752264 | 0.43820592 | 0.98092839 |
| NRSN2-AS1 | 19.9352686 | 0.14459581 | 0.5101876 | 0.28341694 | 0.77685725 | 0.9958918 |
| RBCK1 | 35.8801248 | -0.126338 | 0.40627309 | -0.3109681 | 0.7558249 | 0.9958918 |
| TBC1D20 | 26.4934861 | -0.1415955 | 0.31778317 | -0.4455728 | 0.65590587 | 0.9958918 |
| CSNK2A1 | 101.269651 | 0.14554255 | 0.20395147 | 0.71361362 | 0.47546613 | 0.98910524 |
| PSMF1 | 67.3792567 | 0.22128968 | 0.25437667 | 0.86992913 | 0.38433913 | 0.9725044 |
| AL136531.2 | 38.4380779 | 0.10206169 | 0.35177201 | 0.2901359 | 0.77171227 | 0.9958918 |

| | | | | | | |
|---|---|---|---|---|---|---|
| SDCBP2-AS1 | 29.438062 | 0.20801148 | 0.28430832 | 0.73164049 | 0.46438803 | 0.98606981 |
| NSFL1C | 29.3542261 | -0.1546187 | 0.29279316 | -0.5280816 | 0.59744268 | 0.9958918 |
| STK35 | 18.8907612 | 0.55167525 | 0.40215854 | 1.37178552 | 0.17013022 | 0.87006748 |
| SNRPB | 27.8395324 | 0.04940159 | 0.46198134 | 0.10693418 | 0.9148412 | 0.99935941 |
| NOP56 | 24.9941286 | 0.00462333 | 0.51908511 | 0.00890669 | 0.99289358 | 0.99935941 |
| EBF4 | 41.0842066 | 0.32872021 | 0.28724021 | 1.14440873 | 0.25245417 | 0.92631316 |
| PCED1A | 27.5021856 | -0.4077395 | 0.49418753 | -0.8250705 | 0.40933158 | 0.97812179 |
| PTPRA | 159.212658 | 0.10684556 | 0.225086 | 0.47468771 | 0.63500958 | 0.9958918 |
| ITPA | 25.8815742 | -0.5350579 | 0.40371235 | -1.3253445 | 0.1850569 | 0.87866798 |
| C20orf194 | 165.58244 | 0.3445534 | 0.29309297 | 1.17557713 | 0.2397639 | 0.92426606 |
| ATRN | 87.7167394 | 0.02053451 | 0.22106453 | 0.09288923 | 0.92599156 | 0.99935941 |
| MAVS | 47.441752 | 0.31063264 | 0.38667988 | 0.80333281 | 0.4217824 | 0.97812179 |
| PANK2 | 66.8647296 | -0.2161194 | 0.21141535 | -1.0222504 | 0.3066624 | 0.95793395 |
| RNF24 | 134.024312 | 0.05644329 | 0.29244784 | 0.19300292 | 0.84695668 | 0.99692952 |
| PRNP | 53.5425602 | -0.4500569 | 0.33969839 | -1.3248721 | 0.18521357 | 0.87866798 |
| SLC23A2 | 78.5030411 | -0.4598567 | 0.29006603 | -1.5853518 | 0.11288642 | 0.77793929 |
| TMEM230 | 43.6398434 | 0.0955158 | 0.29391688 | 0.32497554 | 0.74519958 | 0.9958918 |
| CDS2 | 79.118098 | 0.0391139 | 0.3195933 | 0.12238649 | 0.90259293 | 0.99935941 |
| GPCPD1 | 131.564288 | -0.8336226 | 0.43901072 | -1.8988661 | 0.05758209 | 0.60512745 |
| SHLD1 | 25.534982 | -0.5279154 | 0.43733574 | -1.2071171 | 0.22738705 | 0.9120167 |
| CRLS1 | 44.7110065 | -0.3739946 | 0.32988426 | -1.1337145 | 0.25691432 | 0.9266065 |
| TMX4 | 34.7913117 | 0.29988773 | 0.31641989 | 0.94775246 | 0.34325549 | 0.96887708 |
| PLCB1 | 522.179066 | -1.1143364 | 0.54779941 | -2.0342051 | 0.04193092 | 0.54566789 |
| PLCB4 | 96.6157382 | -0.8731299 | 0.56898571 | -1.5345374 | 0.12489749 | 0.79195137 |
| SLX4IP | 44.8737878 | -0.4291179 | 0.33101788 | -1.2963588 | 0.1948519 | 0.88391893 |
| JAG1 | 251.106845 | -0.4776422 | 0.31007673 | -1.5403999 | 0.1234629 | 0.79184923 |
| AL050403.2 | 40.1415911 | -0.1656537 | 0.50462023 | -0.328274 | 0.74270451 | 0.9958918 |
| BTBD3 | 29.7103501 | 0.0259896 | 0.30636224 | 0.0848329 | 0.93239424 | 0.99935941 |
| SPTLC3 | 21.3777805 | 0.58762284 | 0.43663087 | 1.34581149 | 0.17836332 | 0.87815793 |
| TASP1 | 115.036777 | -0.1473204 | 0.24778344 | -0.5945531 | 0.55214223 | 0.9958918 |
| ESF1 | 51.8047136 | -0.206808 | 0.28644424 | -0.7219836 | 0.47030457 | 0.98910524 |
| NDUFAF5 | 23.8461069 | 0.50093377 | 0.34703168 | 1.44348138 | 0.14888491 | 0.8301633 |
| MACROD2 | 327.088928 | 0.18061908 | 0.57852698 | 0.3122051 | 0.75488465 | 0.9958918 |
| KIF16B | 70.4239722 | -0.455456 | 0.25900579 | -1.7584779 | 0.07866623 | 0.68254797 |
| SNRPB2 | 19.0145213 | -0.4013152 | 0.40243892 | -0.9972076 | 0.31866375 | 0.95992644 |
| DSTN | 290.845679 | 0.64220421 | 0.22789812 | 2.81794431 | 0.00483322 | 0.1747393 |
| RRBP1 | 137.91506 | -0.1819022 | 0.416918 | -0.4363021 | 0.66261754 | 0.9958918 |
| SNX5 | 34.2926703 | 0.11252671 | 0.33060166 | 0.34036946 | 0.73357831 | 0.9958918 |
| ZNF133 | 29.0527257 | 0.3876212 | 0.3648187 | 1.06250364 | 0.28800711 | 0.94955775 |
| SEC23B | 35.1670185 | 0.81864175 | 0.26483948 | 3.09108651 | 0.00199426 | 0.09516741 |
| SLC24A3 | 220.030683 | 0.30001082 | 0.5658619 | 0.53018381 | 0.5959845 | 0.9958918 |
| AL121761.1 | 42.699266 | 0.49567006 | 0.42475233 | 1.16696253 | 0.2432255 | 0.92534155 |
| RIN2 | 107.120885 | 0.08455992 | 0.34647263 | 0.24405945 | 0.80718478 | 0.99592423 |
| NAA20 | 23.8585634 | 0.0806293 | 0.31619514 | 0.25499854 | 0.79872422 | 0.99592423 |
| RALGAPA2 | 107.221031 | 0.12311834 | 0.22811237 | 0.53972671 | 0.58938552 | 0.9958918 |
| KIZ | 76.3492995 | -0.077884 | 0.32749352 | -0.2378184 | 0.81202193 | 0.99592423 |
| XRN2 | 116.565886 | -0.1752543 | 0.23915261 | -0.7328137 | 0.46367207 | 0.98590351 |
| NAPB | 43.41524 | 0.13201934 | 0.32974269 | 0.40037077 | 0.68888345 | 0.9958918 |
| CST3 | 228.517565 | -0.33885 | 0.34559747 | -0.9804759 | 0.32685126 | 0.95992644 |
| SYNDIG1 | 42.6953697 | -0.6758309 | 0.56491993 | -1.1963304 | 0.23156763 | 0.91549646 |
| APMAP | 21.2676645 | 0.36367265 | 0.34117997 | 1.06592614 | 0.28645703 | 0.94614505 |
| ACSS1 | 34.5852936 | -0.1853884 | 0.32552706 | -0.5695021 | 0.56901543 | 0.9958918 |
| PYGB | 20.0482807 | 0.56454773 | 0.47883187 | 1.17901035 | 0.23839405 | 0.92291633 |
| ABHD12 | 34.9016039 | -0.0164143 | 0.33059879 | -0.0496501 | 0.96040124 | 0.99935941 |

| | | | | | | |
|---|---|---|---|---|---|---|
| NINL | 33.5621079 | -0.4353767 | 0.31586725 | -1.3783535 | 0.16809416 | 0.86560961 |
| ZNF337 | 20.6545275 | 0.1155749 | 0.32798138 | 0.3523825 | 0.72455142 | 0.9958918 |
| FAM182B | 40.6727983 | 0.06421066 | 0.37909724 | 0.16937781 | 0.86549947 | 0.9971731 |
| HM13 | 102.902779 | 0.33301177 | 0.33983466 | 0.97992291 | 0.32712417 | 0.95992644 |
| ID1 | 94.8203083 | 0.9395216 | 0.58698466 | 1.60058968 | 0.10946783 | 0.77303254 |
| BCL2L1 | 86.0838349 | 0.06139515 | 0.30146488 | 0.20365606 | 0.83862228 | 0.99692952 |
| TM9SF4 | 57.1732002 | -0.0161324 | 0.23397851 | -0.068948 | 0.945031 | 0.99935941 |
| POFUT1 | 19.6446467 | -0.0254963 | 0.44300923 | -0.0575525 | 0.95410513 | 0.99935941 |
| KIF3B | 29.0799916 | 0.35572425 | 0.33951742 | 1.04773489 | 0.29476077 | 0.95304825 |
| ASXL1 | 196.873798 | -0.1532658 | 0.23118335 | -0.6629621 | 0.50735481 | 0.99397937 |
| MAPRE1 | 27.6936539 | 0.21873983 | 0.31589324 | 0.69244857 | 0.48865567 | 0.99046213 |
| CDK5RAP1 | 27.2721296 | 0.09373052 | 0.31000729 | 0.30234941 | 0.76238571 | 0.9958918 |
| CBFA2T2 | 130.695081 | 0.17454236 | 0.25707434 | 0.67895675 | 0.49716527 | 0.99176652 |
| CHMP4B | 37.3603382 | 0.42603615 | 0.32833346 | 1.29757156 | 0.1944346 | 0.88391893 |
| RALY | 80.7290056 | 0.00524373 | 0.24276001 | 0.02160047 | 0.98276666 | 0.99935941 |
| EIF2S2 | 55.2094079 | 0.43062814 | 0.34783883 | 1.23801054 | 0.21571215 | 0.90187565 |
| ITCH | 216.152879 | 0.04051128 | 0.17762225 | 0.2280755 | 0.81958755 | 0.99592423 |
| DYNLRB1 | 63.2953095 | 0.0675857 | 0.3161211 | 0.21379687 | 0.83070546 | 0.99592423 |
| MAP1LC3A | 23.1686143 | 0.2775725 | 0.45449087 | 0.61073285 | 0.54137645 | 0.9958918 |
| PIGU | 36.7048114 | 0.22750971 | 0.39275224 | 0.5792703 | 0.5624068 | 0.9958918 |
| NCOA6 | 104.976924 | -0.0076284 | 0.21260013 | -0.0358814 | 0.9713769 | 0.99935941 |
| GGT7 | 31.1407954 | 0.50329785 | 0.29969469 | 1.67936858 | 0.09308023 | 0.73019523 |
| ACSS2 | 53.6698366 | 0.75084389 | 0.4862076 | 1.5442866 | 0.12251892 | 0.79184923 |
| TRPC4AP | 145.498498 | -0.1119747 | 0.16698166 | -0.6705807 | 0.50248768 | 0.99321731 |
| EDEM2 | 21.3090868 | 0.19438892 | 0.32155724 | 0.60452354 | 0.54549562 | 0.9958918 |
| PROCR | 53.9061426 | -0.5641967 | 0.46602949 | -1.210646 | 0.22603112 | 0.91093521 |
| MMP24OS | 38.4429964 | 0.08837711 | 0.30276321 | 0.29190175 | 0.77036175 | 0.9958918 |
| UQCC1 | 64.7233545 | -0.2848923 | 0.24433166 | -1.1660067 | 0.24361176 | 0.92534155 |
| CEP250 | 17.8429012 | 0.24619371 | 0.33495525 | 0.73500481 | 0.46233657 | 0.98590351 |
| ERGIC3 | 24.7316365 | -0.3683102 | 0.33688008 | -1.0932977 | 0.27426312 | 0.9394201 |
| CPNE1 | 33.1088427 | 0.01697296 | 0.42779508 | 0.03967545 | 0.96835187 | 0.99935941 |
| ROMO1 | 22.2552589 | -0.4205643 | 0.49803768 | -0.8444428 | 0.39842201 | 0.97500135 |
| RBM39 | 258.681224 | -0.0818945 | 0.19857725 | -0.4124062 | 0.68004173 | 0.9958918 |
| PHF20 | 129.48831 | -0.2606202 | 0.20082524 | -1.2977463 | 0.19437454 | 0.88391893 |
| SCAND1 | 36.3519972 | -0.4104593 | 0.43110172 | -0.9521169 | 0.34103768 | 0.96887708 |
| NORAD | 92.8111858 | -0.1778991 | 0.28411471 | -0.6261524 | 0.53121497 | 0.9958918 |
| EPB41L1 | 32.7239274 | -0.091929 | 0.39402074 | -0.23331 | 0.81552068 | 0.99592423 |
| DLGAP4 | 93.0669525 | 0.36762083 | 0.29518247 | 1.24540196 | 0.21298403 | 0.89738044 |
| DLGAP4-AS1 | 28.5820291 | -0.1582029 | 0.33172186 | -0.4769144 | 0.63342304 | 0.9958918 |
| MYL9 | 393.974504 | 1.06046924 | 0.3267618 | 3.24538925 | 0.0011729 | 0.07367227 |
| NDRG3 | 25.5148292 | 0.05735324 | 0.4549049 | 0.12607743 | 0.89967063 | 0.99935941 |
| SOGA1 | 51.1364025 | 0.51021407 | 0.32267942 | 1.58117945 | 0.11383701 | 0.77853537 |
| SAMHD1 | 100.268331 | -0.0603732 | 0.80919472 | -0.0746089 | 0.94052586 | 0.99935941 |
| RPN2 | 62.544401 | -0.0080619 | 0.29930619 | -0.0269352 | 0.97851144 | 0.99935941 |
| MANBAL | 16.7579628 | 0.28410374 | 0.41287635 | 0.68810853 | 0.49138444 | 0.99046213 |
| SRC | 18.0287196 | 0.14935273 | 0.39954522 | 0.37380681 | 0.70854805 | 0.9958918 |
| BLCAP | 45.7307814 | 0.2145323 | 0.26688113 | 0.80384961 | 0.42148384 | 0.97812179 |
| CTNNBL1 | 53.4982643 | -0.2580048 | 0.28213732 | -0.9144652 | 0.36047247 | 0.97217293 |
| TTI1 | 20.1614845 | 0.15582534 | 0.34864783 | 0.44694194 | 0.65491698 | 0.9958918 |
| RPRD1B | 55.6427178 | -0.1802933 | 0.2385199 | -0.7558836 | 0.44971901 | 0.98092839 |
| TGM2 | 210.495571 | 0.80834729 | 0.54376259 | 1.48658127 | 0.13712544 | 0.81404904 |
| RALGAPB | 81.060991 | 0.13053455 | 0.18283406 | 0.71395094 | 0.47525752 | 0.98910524 |
| DHX35 | 27.247241 | 0.28203623 | 0.33369173 | 0.84519995 | 0.39799921 | 0.97500135 |
| TOP1 | 208.89054 | -0.4111995 | 0.27759809 | -1.4812766 | 0.13853289 | 0.81520215 |

| | | | | | | |
|---|---|---|---|---|---|---|
| PLCG1 | 26.3151467 | -0.1455292 | 0.34418064 | -0.4228278 | 0.67242087 | 0.9958918 |
| ZHX3 | 111.565568 | -0.1492791 | 0.32707281 | -0.4564095 | 0.64809553 | 0.9958918 |
| CHD6 | 154.493409 | -0.0671272 | 0.20927833 | -0.3207558 | 0.74839547 | 0.9958918 |
| SRSF6 | 40.7486176 | 0.38464963 | 0.40008031 | 0.96143105 | 0.33633548 | 0.96640422 |
| IFT52 | 27.5816713 | 0.28097918 | 0.32405988 | 0.86705944 | 0.38590944 | 0.97254921 |
| JPH2 | 28.4576295 | 1.19156492 | 0.52788985 | 2.25722263 | 0.02399417 | 0.41984079 |
| SERINC3 | 62.5686516 | -0.2388498 | 0.3259308 | -0.7328237 | 0.46366598 | 0.98590351 |
| PKIG | 98.3145534 | -0.1329023 | 0.26248187 | -0.5063295 | 0.61262537 | 0.9958918 |
| YWHAB | 111.529319 | 0.84966447 | 0.35738248 | 2.37746541 | 0.01743208 | 0.34973839 |
| STK4 | 58.4945096 | -0.1936395 | 0.23483112 | -0.8245903 | 0.40960424 | 0.97812179 |
| SLPI | 93.0040468 | -0.1645287 | 0.99146519 | -0.165945 | 0.86820024 | 0.9971731 |
| SDC4 | 111.697116 | -0.6503672 | 0.38427072 | -1.6924714 | 0.09055613 | 0.72589216 |
| PIGT | 25.8927103 | -0.2883934 | 0.49412324 | -0.5836468 | 0.559458 | 0.9958918 |
| WFDC2 | 115.961357 | 0.50236783 | 0.53441617 | 0.94003112 | 3.47E-01 | 0.96887708 |
| SNX21 | 22.3912403 | 0.41068469 | 0.51162643 | 0.8027042 | 0.42214572 | 0.97828057 |
| CTSA | 34.0729097 | -0.3226953 | 0.31127497 | -1.036689 | 0.29988081 | 0.95587816 |
| PLTP | 29.2130344 | -0.4527041 | 0.47681706 | -0.9494293 | 0.3424023 | 0.96887708 |
| ZNF335 | 19.1018009 | 0.24952303 | 0.40440553 | 0.6170119 | 0.53722688 | 0.9958918 |
| NCOA5 | 35.0708484 | 0.09779636 | 0.32783061 | 0.29831369 | 0.76546376 | 0.9958918 |
| SLC35C2 | 26.5030946 | 0.05340149 | 0.32633351 | 0.16364084 | 0.87001389 | 0.9971731 |
| ELMO2 | 31.312533 | 0.70311519 | 0.34334647 | 2.04782997 | 0.04057666 | 0.54217799 |
| SLC13A3 | 42.5159588 | 0.22615796 | 0.29896473 | 0.75647036 | 0.44936723 | 0.98092839 |
| EYA2 | 60.5143027 | -0.3045798 | 0.49820659 | -0.6113524 | 0.5409663 | 0.9958918 |
| ZMYND8 | 112.920568 | -0.4590837 | 0.24756222 | -1.8544176 | 0.06367945 | 0.62395844 |
| NCOA3 | 176.82996 | -0.3848617 | 0.33224676 | -1.1583611 | 0.24671671 | 0.92600668 |
| SULF2 | 32.4422078 | -0.4510574 | 0.3452958 | -1.3062929 | 0.19145299 | 0.88158763 |
| PREX1 | 23.5490826 | 1.30025287 | 0.46564181 | 2.79238859 | 0.00523205 | 0.18276799 |
| ARFGEF2 | 100.484289 | -0.0265933 | 0.22515227 | -0.1181123 | 0.90597865 | 0.99935941 |
| CSE1L | 33.0555411 | 0.13699525 | 0.34874318 | 0.39282559 | 0.6944483 | 0.9958918 |
| STAU1 | 96.9224284 | 0.21889473 | 0.18425005 | 1.18803082 | 0.23482127 | 0.91962002 |
| DDX27 | 19.5768948 | -0.1222937 | 0.3698723 | -0.3306376 | 0.7409182 | 0.9958918 |
| ZNFX1 | 67.9261324 | -0.0373314 | 0.48303243 | -0.0772854 | 0.93839647 | 0.99935941 |
| ZFAS1 | 41.4952474 | -0.6420319 | 0.35423538 | -1.8124444 | 0.06991756 | 0.65538799 |
| PTGIS | 197.980668 | -0.2914982 | 0.82555352 | -0.3530942 | 0.7240178 | 0.9958918 |
| B4GALT5 | 126.231503 | -0.4167104 | 0.21832452 | -1.9086745 | 0.0563041 | 0.60142269 |
| SLC9A8 | 38.5388501 | -0.0236142 | 0.39621742 | -0.0595991 | 0.95247491 | 0.99935941 |
| RNF114 | 48.947119 | 0.08281864 | 0.4124229 | 0.20080999 | 0.84084715 | 0.99692952 |
| CEBPB | 145.59583 | -0.6478228 | 0.44094145 | -1.469181 | 1.42E-01 | 0.81652619 |
| PTPN1 | 203.284008 | -0.348605 | 0.37191229 | -0.9373312 | 0.34858822 | 0.96890122 |
| RIPOR3 | 312.610192 | -0.6718045 | 0.46584959 | -1.442106 | 0.14927247 | 0.83169325 |
| PARD6B | 27.2888107 | -0.8927968 | 0.50086842 | -1.7824977 | 0.0746681 | 0.67164733 |
| ADNP | 147.759134 | 0.13070073 | 0.16274425 | 0.80310506 | 0.42191401 | 0.97812179 |
| DPM1 | 36.172615 | -0.1508553 | 0.26808845 | -0.5627072 | 0.57363428 | 0.9958918 |
| NFATC2 | 52.0275891 | -0.3421509 | 1.10860106 | -0.3086331 | 0.75760066 | 0.9958918 |
| ATP9A | 63.3433522 | -0.5032923 | 0.38007742 | -1.3241835 | 0.18544211 | 0.87866798 |
| ZFP64 | 25.2804778 | -0.1671882 | 0.38471446 | -0.4345773 | 0.66386928 | 0.9958918 |
| TSHZ2 | 896.691264 | -0.0745238 | 0.3832434 | -0.1944555 | 0.84581921 | 0.99692952 |
| AL109930.1 | 72.8236388 | -0.343983 | 0.39778877 | -0.8647378 | 0.38718274 | 0.97297628 |
| ZNF217 | 47.6716164 | 0.05257446 | 0.23624083 | 0.22254603 | 0.82388884 | 0.99592423 |
| DOK5 | 30.8974292 | 0.1191985 | 0.70209253 | 0.16977606 | 0.86518626 | 0.9971731 |
| FAM210B | 19.2386163 | -0.0830662 | 0.4259376 | -0.1950196 | 0.84537758 | 0.99692952 |
| RTF2 | 26.5389465 | 0.19710979 | 0.33347369 | 0.59108048 | 0.5544665 | 0.9958918 |
| PCK1 | 93.6364928 | 0.21338021 | 1.24757029 | 0.17103662 | 0.86419497 | 0.9971731 |
| PMEPA1 | 53.7300759 | 0.04708934 | 0.36395461 | 0.12938246 | 0.89705503 | 0.99935941 |

| | | | | | | |
|---|---|---|---|---|---|---|
| RAB22A | 58.4715074 | 0.16836299 | 0.26535423 | 0.6344839 | 0.52576507 | 0.9958918 |
| VAPB | 68.6167888 | 0.16922442 | 0.20919117 | 0.80894629 | 0.41854604 | 0.97812179 |
| GNAS | 279.279726 | 0.55246182 | 0.27454148 | 2.01230729 | 0.04418755 | 0.55605195 |
| NELFCD | 26.2406106 | 0.31965311 | 0.30928815 | 1.03351229 | 0.30136422 | 0.95597956 |
| CTSZ | 31.8389564 | 0.60756159 | 0.32854602 | 1.84924349 | 0.06442266 | 0.62624621 |
| ATP5F1E | 97.5488199 | 0.21160696 | 0.38242315 | 0.55333197 | 0.58003611 | 0.9958918 |
| PHACTR3 | 43.6808534 | -0.4116547 | 0.97786972 | -0.4209709 | 0.6737763 | 0.9958918 |
| MIR646HG | 81.8728676 | -0.7029359 | 0.53566261 | -1.3122736 | 0.18942785 | 0.88139754 |
| TAF4 | 27.4387008 | 0.23879308 | 0.37057936 | 0.6443777 | 0.51933054 | 0.9958918 |
| PSMA7 | 59.1168339 | 0.13198699 | 0.42042899 | 0.31393409 | 0.7535711 | 0.9958918 |
| SS18L1 | 25.6445462 | 0.0743229 | 0.30757009 | 0.24164541 | 0.80905493 | 0.99592423 |
| MTG2 | 26.4104504 | 0.50533235 | 0.33834708 | 1.49353248 | 0.13529786 | 0.81154597 |
| OSBPL2 | 23.4819626 | 0.01557376 | 0.3639848 | 0.04278684 | 0.96587145 | 0.99935941 |
| LAMA5 | 41.4653673 | 0.76302236 | 0.42862562 | 1.78016039 | 0.07504972 | 0.67167936 |
| OGFR | 27.7989599 | 0.20130271 | 0.32150064 | 0.62613472 | 0.53122658 | 0.9958918 |
| DIDO1 | 71.4350836 | -0.0591064 | 0.23067537 | -0.2562317 | 0.7977719 | 0.99592423 |
| GID8 | 21.3662909 | -0.0726136 | 0.33006119 | -0.2200003 | 0.82587091 | 0.99592423 |
| YTHDF1 | 24.2726749 | 0.03499443 | 0.32166451 | 0.10879171 | 0.9133677 | 0.99935941 |
| ARFGAP1 | 27.3675339 | -0.796101 | 0.41192843 | -1.9326197 | 0.05328307 | 0.58707106 |
| PPDPF | 65.1494186 | 0.11881652 | 0.32692226 | 0.36343968 | 0.71627647 | 0.9958918 |
| ARFRP1 | 17.2580165 | 0.18662696 | 0.38748136 | 0.48164113 | 0.63006091 | 0.9958918 |
| ZBTB46 | 221.591106 | 0.84489254 | 0.25281171 | 3.3419834 | 0.00083182 | 0.06434787 |
| TPD52L2 | 31.75515 | -0.0823355 | 0.33766527 | -0.2438377 | 0.8073565 | 0.99592423 |
| DNAJC5 | 58.4357259 | 0.01941087 | 0.36155578 | 0.05368708 | 0.95718448 | 0.99935941 |
| PRPF6 | 41.8484616 | 0.15251466 | 0.33042805 | 0.46156691 | 0.64439193 | 0.9958918 |
| PCMTD2 | 53.3022061 | 0.21641063 | 0.42707629 | 0.50672593 | 0.61234713 | 0.9958918 |
| SHC2 | 21.5476737 | -0.7777655 | 0.3994 | -1.9473347 | 0.05149462 | 0.57655256 |
| BSG | 69.4832364 | 0.24155351 | 0.25594294 | 0.94377876 | 0.34528272 | 0.96887708 |
| FSTL3 | 38.5734945 | -1.0600546 | 0.44272579 | -2.394382 | 0.0166484 | 0.34578631 |
| PALM | 27.3957107 | -0.3863181 | 0.31898701 | -1.2110778 | 0.2258656 | 0.91093521 |
| PTBP1 | 56.7931028 | -0.300888 | 0.28024351 | -1.0736664 | 0.28297225 | 0.94471801 |
| ARID3A | 42.0926742 | -0.3816011 | 0.31317017 | -1.2185105 | 0.22303005 | 0.91093521 |
| TMEM259 | 50.2059723 | -0.2139351 | 0.38089243 | -0.561668 | 0.57434224 | 0.9958918 |
| CNN2 | 40.2249186 | -0.1025066 | 0.49289061 | -0.2079702 | 0.83525222 | 0.99679581 |
| POLR2E | 19.4612343 | 0.76988069 | 0.40289732 | 1.91086078 | 0.05602247 | 0.60041061 |
| GPX4 | 61.9338342 | -0.0261029 | 0.34160454 | -0.0764126 | 0.93909082 | 0.99935941 |
| SBNO2 | 67.3132816 | 0.31472045 | 0.38915183 | 0.80873434 | 0.41866797 | 0.97812179 |
| STK11 | 30.6369536 | 0.2909129 | 0.31234459 | 0.93138448 | 0.35165472 | 0.97045081 |
| ATP5F1D | 46.1926316 | 0.21332809 | 0.30244953 | 0.70533451 | 0.48060209 | 0.99002719 |
| MIDN | 62.1530334 | 0.31341887 | 0.42901763 | 0.7305501 | 0.465054 | 0.98655222 |
| CIRBP | 92.6371401 | 0.28855412 | 0.33628932 | 0.85805318 | 0.3908631 | 0.97297628 |
| MUM1 | 37.855555 | -0.1198906 | 0.34030381 | -0.3523046 | 0.72460981 | 0.9958918 |
| DAZAP1 | 59.2565912 | 0.1395514 | 0.2832341 | 0.49270692 | 0.62221969 | 0.9958918 |
| RPS15 | 172.015872 | -0.2470262 | 0.3337105 | -0.740241 | 0.45915379 | 0.98508941 |
| C19orf25 | 22.4946008 | 0.31113321 | 0.38129914 | 0.81598195 | 0.41451046 | 0.97812179 |
| MBD3 | 21.2813819 | -0.1466271 | 0.49460028 | -0.2964557 | 0.76688209 | 0.9958918 |
| TCF3 | 20.7413526 | 0.01815044 | 0.40327138 | 0.045008 | 0.96410093 | 0.99935941 |
| ABHD17A | 24.5355943 | 0.20077367 | 0.40413199 | 0.49680222 | 0.61932853 | 0.9958918 |
| CSNK1G2 | 29.1013403 | 0.23162195 | 0.50894588 | 0.45510134 | 0.64903634 | 0.9958918 |
| BTBD2 | 28.1595578 | -0.097379 | 0.40690468 | -0.2393165 | 0.81086016 | 0.99592423 |
| MOB3A | 27.9704163 | 0.21899181 | 0.41238059 | 0.53104297 | 0.595389 | 0.9958918 |
| AP3D1 | 46.6478125 | -0.1540885 | 0.28824664 | -0.5345716 | 0.59294614 | 0.9958918 |
| DOT1L | 43.5489607 | -0.2209117 | 0.37740799 | -0.5853391 | 0.55831971 | 0.9958918 |
| LSM7 | 24.9142854 | 0.01009397 | 0.38020957 | 0.02654843 | 0.9788199 | 0.99935941 |

| | | | | | | |
|---|---|---|---|---|---|---|
| SPPL2B | 26.2312374 | -0.3004106 | 0.3620496 | -0.8297499 | 0.4066802 | 0.97671046 |
| GADD45B | 130.510638 | -1.2574535 | 0.46244246 | -2.7191566 | 6.54E-03 | 0.20509206 |
| GNG7 | 36.141443 | -0.1241576 | 0.3433269 | -0.361631 | 0.71762779 | 0.9958918 |
| SGTA | 23.9743589 | -0.1727138 | 0.29952065 | -0.5766342 | 0.56418662 | 0.9958918 |
| TLE2 | 117.29342 | 0.15496434 | 0.33747788 | 0.45918369 | 0.64610226 | 0.9958918 |
| AES | 115.588009 | 0.23011237 | 0.19520444 | 1.17882753 | 0.23846686 | 0.92291633 |
| GNA11 | 46.381104 | 0.14583149 | 0.31932613 | 0.4566851 | 0.6478974 | 0.9958918 |
| NCLN | 19.1254547 | -0.2840111 | 0.53540077 | -0.5304645 | 0.59578991 | 0.9958918 |
| NFIC | 187.448696 | -0.0060982 | 0.20261128 | -0.0300979 | 0.97598897 | 0.99935941 |
| MFSD12 | 23.9163524 | 0.02570566 | 0.41485536 | 0.06196295 | 0.95059234 | 0.99935941 |
| HMG20B | 16.9287191 | -0.0767898 | 0.42723113 | -0.1797382 | 0.85735811 | 0.9971731 |
| PIP5K1C | 39.346124 | 0.2267895 | 0.27294908 | 0.83088576 | 0.40603817 | 0.97637555 |
| DAPK3 | 44.3640586 | -0.3334214 | 0.37477964 | -0.8896464 | 0.37365578 | 0.97217293 |
| EEF2 | 155.482515 | 0.12366464 | 0.40837117 | 0.30282411 | 0.76202391 | 0.9958918 |
| ZBTB7A | 50.5363728 | 0.21531561 | 0.39534626 | 0.54462538 | 0.58601119 | 0.9958918 |
| MAP2K2 | 43.6239417 | 0.16883145 | 0.28794987 | 0.58632237 | 0.5576589 | 0.9958918 |
| SH3GL1 | 42.3172026 | -0.0860745 | 0.32757593 | -0.2627619 | 0.79273411 | 0.99592423 |
| UBXN6 | 19.3828691 | 0.04647903 | 0.38165533 | 0.12178275 | 0.90307108 | 0.99935941 |
| MYDGF | 47.0582456 | -0.0340453 | 0.45592681 | -0.0746726 | 0.94047519 | 0.99935941 |
| DPP9 | 24.3182776 | 0.5042484 | 0.36873151 | 1.36752185 | 0.17146179 | 0.87233975 |
| PLIN3 | 24.7960492 | 0.27514811 | 0.42215918 | 0.65176389 | 0.5145535 | 0.9958918 |
| KDM4B | 106.797212 | 0.0717715 | 0.34658938 | 0.20707933 | 0.8359479 | 0.99692952 |
| PTPRS | 119.863418 | -0.5825873 | 0.37395749 | -1.5578972 | 0.11925762 | 0.79021701 |
| SAFB2 | 52.2357175 | -0.1830525 | 0.245923 | -0.7443489 | 0.45666542 | 0.98330915 |
| SAFB | 67.1465724 | 0.03090421 | 0.24519606 | 0.12603876 | 0.89970124 | 0.99935941 |
| RPL36 | 96.4223757 | -0.1949982 | 0.29973679 | -0.6505648 | 0.51532745 | 0.9958918 |
| C19orf70 | 22.1537118 | -0.0226162 | 0.36159688 | -0.0625453 | 0.95012857 | 0.99935941 |
| LONP1 | 34.2739093 | -0.7024705 | 0.33835759 | -2.0761187 | 0.03788298 | 0.52233019 |
| RANBP3 | 23.4044593 | 0.17712211 | 0.32834671 | 0.53943624 | 0.58958588 | 0.9958918 |
| RFX2 | 69.7962345 | -0.4968446 | 0.62043948 | -0.8007946 | 0.42325056 | 0.97828057 |
| MLLT1 | 47.0661211 | -0.3648296 | 0.33760638 | -1.0806359 | 0.27985911 | 0.94419503 |
| KHSRP | 29.9005102 | 0.09092331 | 0.39238051 | 0.23172229 | 0.81675371 | 0.99592423 |
| C3 | 103.49143 | 0.02051101 | 0.80678874 | 0.02542303 | 0.97971754 | 0.99935941 |
| TRIP10 | 36.8020759 | -0.3248917 | 0.34553367 | -0.9402605 | 0.34708397 | 0.96887708 |
| INSR | 445.731882 | 0.90487595 | 0.62637709 | 1.44461853 | 0.14856507 | 0.8301633 |
| ZNF358 | 17.5203745 | -0.272689 | 0.45246939 | -0.6026685 | 0.54672924 | 0.9958918 |
| PNPLA6 | 19.0685658 | -0.3304897 | 0.38430558 | -0.8599659 | 0.38980785 | 0.97297628 |
| MAP2K7 | 28.8768847 | -0.088799 | 0.45137619 | -0.1967295 | 0.8440392 | 0.99692952 |
| ELAVL1 | 42.7569364 | 0.39837129 | 0.28536699 | 1.39599637 | 0.16271559 | 0.85536256 |
| CERS4 | 45.252249 | 0.56389931 | 0.37985975 | 1.48449345 | 0.13767806 | 0.81404904 |
| RPS28 | 40.50804 | -0.3895443 | 0.3171132 | -1.2284078 | 0.21929393 | 0.90437211 |
| ANGPTL4 | 65.6036501 | 0.20609453 | 0.90042355 | 0.2288862 | 0.81895737 | 0.99592423 |
| RAB11B | 26.514262 | 0.22407275 | 0.37506251 | 0.59742774 | 0.55022184 | 0.9958918 |
| HNRNPM | 115.102631 | 0.10137215 | 0.28101035 | 0.36074172 | 0.71829254 | 0.9958918 |
| ADAMTS10 | 39.0281754 | 0.20963475 | 0.31336947 | 0.66896993 | 0.50351466 | 0.99387859 |
| ZNF266 | 22.1086197 | 0.14979134 | 0.38655458 | 0.38750373 | 0.69838332 | 0.9958918 |
| ZNF121 | 22.7030733 | -0.4228895 | 0.40015272 | -1.0568202 | 0.29059365 | 0.95186143 |
| ZNF562 | 33.7813344 | 0.19362575 | 0.29779382 | 0.65020069 | 0.51556259 | 0.9958918 |
| ZNF846 | 38.9324852 | 0.46343025 | 0.35323539 | 1.31195871 | 1.90E-01 | 0.88139754 |
| COL5A3 | 36.60783 | -0.5269079 | 0.50488485 | -1.04362 | 0.29666125 | 0.95417903 |
| C19orf66 | 34.9828421 | -0.2045723 | 0.4847074 | -0.4220532 | 0.67298619 | 0.9958918 |
| EIF3G | 18.9078554 | 0.38425117 | 0.4234078 | 0.90752029 | 0.36413173 | 0.97217293 |
| DNMT1 | 38.1636139 | -0.367243 | 0.36323252 | -1.011041 | 0.31199682 | 0.9599143 |
| ICAM1 | 45.7641514 | 0.7403016 | 0.9719229 | 0.76168758 | 0.44624649 | 0.98092839 |

| | | | | | | |
|---|---|---|---|---|---|---|
| TYK2 | 23.8366957 | 0.26495018 | 0.33151683 | 0.79920583 | 0.42417107 | 0.97828057 |
| CDC37 | 48.5949579 | -0.1680269 | 0.32812371 | -0.5120839 | 0.60859229 | 0.9958918 |
| PDE4A | 19.6156734 | 0.05224511 | 0.44208484 | 0.11817893 | 0.90592589 | 0.99935941 |
| SLC44A2 | 57.2624699 | 0.25816937 | 0.23194123 | 1.11308096 | 0.26567366 | 0.93533685 |
| ILF3 | 66.9598055 | -0.0064133 | 0.31178835 | -0.0205694 | 0.98358914 | 0.99935941 |
| QTRT1 | 16.5694545 | -0.3942148 | 0.46070412 | -0.8556789 | 0.39217543 | 0.97297628 |
| DNM2 | 88.5467367 | 0.50114737 | 0.24994699 | 2.00501458 | 0.04496149 | 0.55894748 |
| CARM1 | 28.0475421 | -0.0748533 | 0.28695379 | -0.2608548 | 0.79420448 | 0.99592423 |
| SMARCA4 | 76.5787239 | 0.06458567 | 0.35184614 | 0.18356225 | 0.85435688 | 0.9971731 |
| LDLR | 69.3907192 | -0.222951 | 0.55396598 | -0.4024634 | 0.68734299 | 0.9958918 |
| KANK2 | 122.512 | -0.1895677 | 0.31166032 | -0.608251 | 0.54302102 | 0.9958918 |
| DOCK6 | 92.5537349 | 0.45504225 | 0.23512186 | 1.93534639 | 0.0529478 | 0.58601416 |
| RGL3 | 94.0536863 | 0.33640564 | 0.34961966 | 0.96220459 | 0.33594684 | 0.96640422 |
| PRKCSH | 26.1900079 | 0.11795659 | 0.35412953 | 0.33308884 | 0.73906723 | 0.9958918 |
| ECSIT | 18.5386964 | -0.2277968 | 0.4331807 | -0.5258701 | 0.59897846 | 0.9958918 |
| CNN1 | 68.6842252 | 0.33204488 | 0.4320589 | 0.76851763 | 4.42E-01 | 0.98092839 |
| ZNF439 | 28.3348626 | 0.72742065 | 0.28096031 | 2.5890513 | 0.00962408 | 0.26002696 |
| ZNF69 | 20.2332374 | -0.4199186 | 0.4057844 | -1.0348319 | 0.30074744 | 0.95597956 |
| ZNF44 | 68.3064271 | 0.16950025 | 0.26345077 | 0.64338491 | 0.51997437 | 0.9958918 |
| ZNF791 | 40.7636621 | -0.3488305 | 0.32275638 | -1.0807856 | 0.27979249 | 0.94419503 |
| TNPO2 | 35.2912876 | 0.30610731 | 0.31778078 | 0.96326567 | 0.33541422 | 0.96640422 |
| TRIR | 40.6208618 | 0.16553696 | 0.34287271 | 0.48279422 | 0.62924186 | 0.9958918 |
| HOOK2 | 31.7171296 | -0.098943 | 0.3254223 | -0.3040448 | 0.76109373 | 0.9958918 |
| JUNB | 132.188023 | -0.9503747 | 0.53524782 | -1.7755789 | 0.07580236 | 0.67360523 |
| CALR | 102.767915 | -0.135725 | 0.40465751 | -0.335407 | 0.73731809 | 0.9958918 |
| RAD23A | 32.6922726 | 0.06269577 | 0.39978927 | 0.15682204 | 0.8753851 | 0.9971731 |
| GADD45GIP1 | 42.2470431 | 0.08661329 | 0.40698234 | 0.21281829 | 0.8314687 | 0.99592423 |
| NFIX | 88.4967565 | 0.20471746 | 0.38571894 | 0.53074255 | 0.5955972 | 0.9958918 |
| IER2 | 61.6948945 | -0.874588 | 0.48762525 | -1.7935658 | 0.0728825 | 0.66701558 |
| CCDC130 | 29.0425185 | 0.0659077 | 0.3276144 | 0.20117462 | 0.84056204 | 0.99692952 |
| MRI1 | 16.9975558 | 0.66837324 | 0.42766792 | 1.5628323 | 0.11809204 | 0.78539225 |
| C19orf53 | 38.9644086 | -0.1297672 | 0.405751 | -0.3198197 | 0.74910501 | 0.9958918 |
| ZSWIM4 | 23.8658179 | 0.47328206 | 0.33477684 | 1.41372401 | 0.15744295 | 0.84427321 |
| AC020916.1 | 25.3241226 | 0.11872014 | 0.63342216 | 0.18742657 | 0.85132619 | 0.9971731 |
| RFX1 | 21.0723656 | -0.0326509 | 0.35044349 | -0.0931701 | 0.9257684 | 0.99935941 |
| PRKACA | 30.4351661 | 0.33605174 | 0.30551891 | 1.09993763 | 0.2713593 | 0.93776436 |
| ADGRL1 | 33.3526059 | 0.1757886 | 0.34167744 | 0.51448699 | 0.60691155 | 0.9958918 |
| ADGRE5 | 98.7494151 | 0.55034256 | 0.37065646 | 1.484778 | 0.13760264 | 0.81404904 |
| PKN1 | 34.4306123 | -0.0586264 | 0.46091539 | -0.1271955 | 0.89878567 | 0.99935941 |
| GIPC1 | 33.2434643 | -0.1380976 | 0.33904605 | -0.4073121 | 0.68377879 | 0.9958918 |
| DNAJB1 | 21.5673242 | -0.0354516 | 0.45480482 | -0.0779489 | 0.93786867 | 0.99935941 |
| TECR | 18.393056 | 0.14563015 | 0.41962752 | 0.34704623 | 0.72855659 | 0.9958918 |
| NDUFB7 | 35.651743 | 0.02255051 | 0.28343691 | 0.07956097 | 0.93658644 | 0.99935941 |
| ZNF333 | 24.6965186 | 0.23215311 | 0.40034731 | 0.5798793 | 0.56199602 | 0.9958918 |
| ADGRE2 | 55.7003149 | 1.30741938 | 0.90054985 | 1.45180123 | 0.14655688 | 0.8301633 |
| SYDE1 | 24.0274971 | -0.4115196 | 0.31973736 | -1.2870552 | 0.19807504 | 0.88597291 |
| NOTCH3 | 149.268236 | -0.1967039 | 0.29040041 | -0.677354 | 0.49818138 | 0.99176652 |
| BRD4 | 179.671871 | 0.02814108 | 0.18314788 | 0.15365223 | 0.87788395 | 0.99722158 |
| AKAP8 | 37.2489507 | 0.03017042 | 0.32346519 | 0.09327253 | 0.92568705 | 0.99935941 |
| AKAP8L | 78.5218052 | -0.1847838 | 0.26401739 | -0.6998927 | 0.48399433 | 0.99046213 |
| TPM4 | 292.482858 | -0.2964648 | 0.30906447 | -0.9592328 | 0.3374415 | 0.96750939 |
| AP1M1 | 24.0328956 | -0.0420757 | 0.32126388 | -0.1309691 | 0.89579972 | 0.99935941 |
| KLF2 | 52.6258238 | 0.12390349 | 0.56645701 | 0.21873415 | 0.82685715 | 0.99592423 |
| EPS15L1 | 57.6358479 | 0.36914722 | 0.25571531 | 1.4435867 | 0.14885527 | 0.8301633 |

| | | | | | | |
|---|---|---|---|---|---|---|
| SIN3B | 33.2315665 | -0.3042864 | 0.27186502 | -1.1192554 | 0.26303118 | 0.93280642 |
| MYO9B | 112.666151 | 0.00734191 | 0.29478768 | 0.02490575 | 0.98013014 | 0.99935941 |
| MRPL34 | 19.5222336 | 0.46316995 | 0.36663228 | 1.26330925 | 0.20647806 | 0.89589437 |
| BST2 | 88.2569021 | 0.52206073 | 0.59831686 | 0.87254892 | 0.382909 | 0.9725044 |
| SLC27A1 | 33.5545547 | 0.18398181 | 0.33241325 | 0.55347316 | 0.57993946 | 0.9958918 |
| AC010618.3 | 20.0937757 | 0.53776727 | 0.41281324 | 1.30268899 | 0.19268096 | 0.88158763 |
| PGLS | 21.7442053 | 0.42854152 | 0.33471562 | 1.28031529 | 0.20043427 | 0.89191279 |
| COLGALT1 | 21.3536644 | -0.1790223 | 0.40213327 | -0.4451816 | 0.65618854 | 0.9958918 |
| JAK3 | 28.1586671 | 0.71414076 | 0.56073723 | 1.27357473 | 0.20281417 | 0.89396747 |
| ARRDC2 | 25.3241978 | -0.2756577 | 0.45019349 | -0.6123094 | 0.54033305 | 0.9958918 |
| JUND | 136.066096 | -1.5433409 | 0.34915607 | -4.4202034 | 9.86E-06 | 0.00345081 |
| LSM4 | 18.8830606 | 0.17162723 | 0.47954729 | 0.35789427 | 0.72042244 | 0.9958918 |
| PGPEP1 | 28.5446384 | 0.32438512 | 0.38438375 | 0.84390957 | 0.39871994 | 0.97500135 |
| GDF15 | 39.670255 | -1.4890105 | 0.5619128 | -2.6498961 | 8.05E-03 | 0.23533118 |
| SSBP4 | 44.1425953 | -0.4374696 | 0.38071921 | -1.149061 | 0.25053081 | 0.92624282 |
| ISYNA1 | 43.3073089 | -0.080503 | 0.43010508 | -0.1871705 | 0.85152692 | 0.9971731 |
| ELL | 89.2071823 | 0.07689241 | 0.3827701 | 0.20088406 | 0.84078924 | 0.99692952 |
| FKBP8 | 36.7023367 | -0.0291771 | 0.26531581 | -0.1099712 | 0.91243225 | 0.99935941 |
| KXD1 | 22.136643 | -0.5216448 | 0.43648044 | -1.1951161 | 0.23204165 | 0.91565561 |
| UBA52 | 68.3134033 | -0.0108741 | 0.43790416 | -0.0248321 | 0.98018891 | 0.99935941 |
| CRLF1 | 26.3766041 | 0.74503863 | 0.59354844 | 1.25522803 | 0.20939599 | 0.89634177 |
| REX1BD | 16.7344057 | -0.0941813 | 0.36823136 | -0.2557667 | 0.79813096 | 0.99592423 |
| CRTC1 | 47.7186534 | 0.23842163 | 0.35057471 | 0.68008793 | 0.49644879 | 0.99176652 |
| UPF1 | 30.4935394 | 0.129287 | 0.29571222 | 0.43720548 | 0.66196233 | 0.9958918 |
| COPE | 23.1738605 | 0.16910612 | 0.37159104 | 0.45508664 | 0.64904691 | 0.9958918 |
| SUGP2 | 74.0016789 | 0.12797438 | 0.19020175 | 0.67283492 | 0.50105232 | 0.99213111 |
| SLC25A42 | 25.5338779 | 0.14027368 | 0.39290809 | 0.35701397 | 0.72108135 | 0.9958918 |
| MAU2 | 54.1854112 | 0.4173392 | 0.29256095 | 1.42650343 | 0.15372308 | 0.84125425 |
| GATAD2A | 128.221892 | 0.08061015 | 0.20965526 | 0.38448907 | 0.700616 | 0.9958918 |
| ZNF90 | 75.7441527 | -0.020118 | 0.34284476 | -0.0586797 | 0.95320727 | 0.99935941 |
| ZNF430 | 23.4738079 | -0.0813983 | 0.29858804 | -0.2726107 | 0.7851525 | 0.9958918 |
| ZNF431 | 44.5779967 | -0.2518562 | 0.25592361 | -0.984107 | 0.3250629 | 0.95992644 |
| ZNF708 | 42.7149756 | 0.34232006 | 0.2905562 | 1.1781544 | 0.23873506 | 0.92291633 |
| AC010615.4 | 21.0802658 | 1.36316493 | 0.61409386 | 2.219799 | 0.02643241 | 0.44248706 |
| ZNF429 | 26.5007439 | 0.43938087 | 0.31947593 | 1.37531762 | 0.169033 | 0.8674745 |
| ZNF100 | 21.6719764 | 0.36837276 | 0.3510244 | 1.04942208 | 0.2939839 | 0.95259599 |
| ZNF43 | 38.1121447 | 0.19574052 | 0.35837648 | 0.54618686 | 0.5849375 | 0.9958918 |
| ZNF208 | 23.4288758 | -0.7158794 | 0.52490704 | -1.3638213 | 0.17262382 | 0.87233975 |
| ZNF91 | 104.033 | 0.13992404 | 0.19193718 | 0.72900959 | 0.4659958 | 0.98777131 |
| ZNF254 | 89.572921 | -0.1188391 | 0.23357592 | -0.5087816 | 0.61090531 | 0.9958918 |
| LINC00662 | 29.9179157 | -0.088569 | 0.31931013 | -0.2773762 | 0.78149125 | 0.9958918 |
| UQCRFS1 | 27.5547271 | 0.30801243 | 0.49623212 | 0.62070232 | 0.5347955 | 0.9958918 |
| C19orf12 | 21.8324884 | 0.40504728 | 0.37908128 | 1.06849718 | 0.2852963 | 0.94505166 |
| URI1 | 112.321522 | -0.1615448 | 0.18315701 | -0.8820016 | 0.37777594 | 0.97217293 |
| TSHZ3 | 82.7806587 | -0.2004447 | 0.31555927 | -0.6352046 | 0.52529496 | 0.9958918 |
| ZNF507 | 34.8480304 | 0.05213504 | 0.29841843 | 0.17470451 | 0.86131183 | 0.9971731 |
| DPY19L3 | 47.9646826 | -0.2980014 | 0.27269453 | -1.092803 | 0.27448029 | 0.93944545 |
| ANKRD27 | 29.0604577 | -0.009596 | 0.32345121 | -0.0296675 | 0.97633226 | 0.99935941 |
| CEP89 | 29.0905755 | 0.02115271 | 0.31903844 | 0.06630143 | 0.94713784 | 0.99935941 |
| PEPD | 38.2262347 | 0.34791447 | 0.35970981 | 0.96720874 | 0.3334397 | 0.9654289 |
| LSM14A | 190.875179 | 0.0552656 | 0.15651688 | 0.35309678 | 0.72401589 | 0.9958918 |
| KIAA0355 | 76.1194488 | -0.0081678 | 0.2465039 | -0.0331346 | 0.97356723 | 0.99935941 |
| GPI | 88.2553311 | 0.2936092 | 0.47139368 | 0.62285349 | 0.5333808 | 0.9958918 |
| UBA2 | 163.947537 | 0.00617486 | 0.24983589 | 0.02471568 | 0.98028175 | 0.99935941 |

| | | | | | | |
|---|---|---|---|---|---|---|
| WTIP | 48.1072512 | 0.04376558 | 0.4325429 | 0.10118205 | 0.91940594 | 0.99935941 |
| SCGB2B2 | 29.056896 | -0.2180615 | 0.50323779 | -0.433317 | 0.66478448 | 0.9958918 |
| ZNF302 | 41.9882781 | -0.1262638 | 0.32289251 | -0.3910398 | 0.6957678 | 0.9958918 |
| GRAMD1A | 36.4357018 | 0.28055725 | 0.31553568 | 0.88914588 | 0.37392468 | 0.97217293 |
| USF2 | 21.515444 | 0.28424154 | 0.39628295 | 0.71726916 | 0.47320803 | 0.98910524 |
| COX6B1 | 48.6292217 | -0.1618728 | 0.3530294 | -0.4585251 | 0.64657524 | 0.9958918 |
| KMT2B | 20.3974832 | -0.0369736 | 0.3765058 | -0.098202 | 0.92177193 | 0.99935941 |
| HSPB6 | 46.4300131 | 0.20581136 | 0.47515211 | 0.43314835 | 0.664907 | 0.9958918 |
| CAPNS1 | 60.853476 | 0.22732574 | 0.36277171 | 0.62663579 | 0.530898 | 0.9958918 |
| ZNF565 | 44.5456623 | -0.278489 | 0.32381782 | -0.8600175 | 0.38977942 | 0.97297628 |
| ZNF146 | 50.5501461 | 0.05989647 | 0.23310477 | 0.25695085 | 0.79721671 | 0.99592423 |
| ZFP14 | 22.2959807 | 0.1653537 | 0.37396817 | 0.44215982 | 0.65837356 | 0.9958918 |
| ZNF566 | 27.9592797 | -0.0922731 | 0.28241963 | -0.3267233 | 0.74387718 | 0.9958918 |
| ZNF529 | 25.8096246 | -0.0072338 | 0.30649876 | -0.0236014 | 0.98117053 | 0.99935941 |
| AC020928.1 | 19.8826357 | 0.07461831 | 0.33571238 | 0.22226858 | 0.82410481 | 0.99592423 |
| ZNF345 | 28.2702059 | 0.43957182 | 0.48602332 | 0.90442537 | 0.36576989 | 0.97217293 |
| ZNF568 | 20.1915984 | 0.50053303 | 0.36069907 | 1.38767488 | 0.16523607 | 0.85897493 |
| ZNF420 | 20.8786272 | 0.32929726 | 0.3556761 | 0.92583466 | 0.35453191 | 0.97217293 |
| HKR1 | 42.1556329 | -0.0356749 | 0.24761718 | -0.1440727 | 0.88544306 | 0.99909735 |
| ZFP30 | 19.2637652 | 0.33474683 | 0.41519615 | 0.8062378 | 0.42010575 | 0.97812179 |
| ZNF573 | 20.6098112 | -0.1341908 | 0.3566453 | -0.3762585 | 0.70672472 | 0.9958918 |
| SIPA1L3 | 79.6899891 | 0.04835896 | 0.32411981 | 0.14920088 | 0.88139512 | 0.9979006 |
| PPP1R14A | 56.6440476 | -0.7457132 | 0.62732378 | -1.1887214 | 0.23454932 | 0.91930824 |
| KCNK6 | 22.8511388 | -1.0702503 | 0.5405745 | -1.9798386 | 0.04772167 | 0.56981199 |
| PSMD8 | 41.0373633 | 0.33095019 | 0.38419357 | 0.86141523 | 0.38900939 | 0.97297628 |
| EIF3K | 33.9450178 | 0.21118579 | 0.32482344 | 0.65015563 | 0.5155917 | 0.9958918 |
| ACTN4 | 420.25279 | -0.0904377 | 0.35787516 | -0.2527074 | 0.8004943 | 0.99592423 |
| ECH1 | 21.8152136 | 0.19539659 | 0.47632497 | 0.41021698 | 0.68164679 | 0.9958918 |
| HNRNPL | 46.0531545 | -0.3250277 | 0.31688599 | -1.0256927 | 0.30503644 | 0.95793395 |
| SIRT2 | 23.0880999 | 0.63689602 | 0.34338696 | 1.85474726 | 0.06363233 | 0.62395844 |
| FBXO17 | 31.7333957 | 0.21847273 | 0.40532214 | 0.53901012 | 0.58987987 | 0.9958918 |
| SAMD4B | 92.7048374 | 0.14024264 | 0.36448053 | 0.38477402 | 0.70040485 | 0.9958918 |
| ZFP36 | 162.202101 | -1.6728595 | 0.58041732 | -2.8821668 | 0.00394951 | 0.15331374 |
| PLEKHG2 | 51.2894992 | -0.1755518 | 0.28482542 | -0.6163488 | 0.53766435 | 0.9958918 |
| RPS16 | 85.4312839 | -0.348458 | 0.34714461 | -1.0037836 | 0.31548295 | 0.95992644 |
| SUPT5H | 41.6224208 | -0.1261227 | 0.24929127 | -0.5059249 | 0.6129094 | 0.9958918 |
| ZNF780B | 42.1693932 | 0.1597192 | 0.26305458 | 0.60717133 | 0.54373722 | 0.9958918 |
| AKT2 | 43.45475 | 0.14173955 | 0.28394254 | 0.49918391 | 0.61764983 | 0.9958918 |
| PLD3 | 73.8555969 | 0.19914218 | 0.26329904 | 0.7563346 | 0.44944861 | 0.98092839 |
| BLVRB | 32.3817619 | 0.18474334 | 0.37076887 | 0.4982709 | 0.61829312 | 0.9958918 |
| LTBP4 | 116.36491 | 0.16841092 | 0.39126718 | 0.43042435 | 0.66688698 | 0.9958918 |
| NUMBL | 27.5366147 | -0.4882581 | 0.32235979 | -1.5146371 | 0.12986434 | 0.80565822 |
| ITPKC | 31.7102091 | -0.1172149 | 0.33530244 | -0.3495795 | 0.72665428 | 0.9958918 |
| SNRPA | 20.9484703 | 0.29057519 | 0.35603949 | 0.81613192 | 0.41442469 | 0.97812179 |
| AC008537.1 | 32.0579335 | -0.1565505 | 0.41937502 | -0.3732947 | 0.70892913 | 0.9958918 |
| AXL | 168.398504 | -0.2895632 | 0.29422223 | -0.9841651 | 0.32503436 | 0.95992644 |
| HNRNPUL1 | 53.1473765 | 0.27037542 | 0.30593237 | 0.8837751 | 0.37681763 | 0.97217293 |
| TGFB1 | 50.5582967 | -0.4070131 | 0.30382586 | -1.3396261 | 0.18036693 | 0.87815793 |
| RPS19 | 218.55756 | -0.3255762 | 0.32411079 | -1.0045213 | 0.31512743 | 0.95992644 |
| ARHGEF1 | 21.4105839 | 0.14661398 | 0.35359209 | 0.41464157 | 0.67840431 | 0.9958918 |
| RABAC1 | 55.0760855 | -0.1512127 | 0.37566877 | -0.4025161 | 0.68730425 | 0.9958918 |
| ETHE1 | 23.1822781 | -0.0946602 | 0.43493082 | -0.2176444 | 0.82770622 | 0.99592423 |
| ZNF428 | 21.9060528 | 0.25317665 | 0.36932524 | 0.68551137 | 0.49302129 | 0.99103212 |
| PLAUR | 22.7764137 | -0.0614258 | 0.60259123 | -0.1019361 | 0.91880743 | 0.99935941 |

| | | | | | | |
|---|---|---|---|---|---|---|
| ZNF235 | 33.3871721 | -0.0334217 | 0.46059082 | -0.0725626 | 0.94215417 | 0.99935941 |
| BCL3 | 23.4156155 | 0.27144718 | 0.58470297 | 0.464248 | 0.64247007 | 0.9958918 |
| BCAM | 70.37452 | 0.43862028 | 0.3049098 | 1.43852469 | 0.15028524 | 0.83354431 |
| NECTIN2 | 66.6324355 | -0.0619422 | 0.24838352 | -0.2493813 | 0.80306587 | 0.99592423 |
| APOE | 127.481444 | 1.03079635 | 0.4026031 | 2.56032887 | 0.01045732 | 0.27446718 |
| CLPTM1 | 32.9864335 | 0.04823842 | 0.38972093 | 0.12377683 | 0.90149198 | 0.99935941 |
| RELB | 49.2950617 | 0.92300335 | 0.58406489 | 1.5803096 | 0.11403598 | 0.77853537 |
| CLASRP | 49.3987821 | 0.02303534 | 0.26226443 | 0.08783251 | 0.9300098 | 0.99935941 |
| MARK4 | 56.5672799 | 0.7258596 | 0.31381538 | 2.31301472 | 0.02072183 | 0.38830162 |
| ERCC1 | 38.4560966 | -0.2909529 | 0.5045887 | -0.5766141 | 0.56420019 | 0.9958918 |
| VASP | 43.061537 | 0.06906929 | 0.34548317 | 0.19992086 | 0.84154248 | 0.99692952 |
| OPA3 | 32.2207975 | -0.13281 | 0.41599684 | -0.3192571 | 0.74953154 | 0.9958918 |
| EML2 | 25.216037 | 0.36774879 | 0.30385799 | 1.21026533 | 0.2261771 | 0.91093521 |
| SNRPD2 | 32.3822859 | -0.0172973 | 0.36184571 | -0.0478031 | 0.96187318 | 0.99935941 |
| DMPK | 36.2507896 | 0.16024794 | 0.48797982 | 0.32839051 | 0.74261642 | 0.9958918 |
| SYMPK | 68.8937225 | 0.16428094 | 0.23907956 | 0.68713922 | 0.491995 | 0.99046213 |
| HIF3A | 79.8399385 | -1.4466864 | 0.85310784 | -1.6957838 | 0.08992683 | 0.72305504 |
| PPP5C | 33.3858444 | -0.0137919 | 0.2973717 | -0.0463793 | 0.96300795 | 0.99935941 |
| CALM3 | 28.1096513 | 0.47611146 | 0.3196069 | 1.48967826 | 0.13630885 | 0.81177777 |
| PTGIR | 61.3140456 | 0.41493103 | 0.45120943 | 0.91959742 | 0.35778317 | 0.97217293 |
| DACT3 | 34.5011311 | 0.50401781 | 0.43400436 | 1.16131968 | 0.24551192 | 0.92559078 |
| PRKD2 | 34.5593417 | 0.02345537 | 0.44088268 | 0.05320093 | 0.95757181 | 0.99935941 |
| SLC1A5 | 19.8196295 | -0.677224 | 0.40016378 | -1.692367 | 0.09057601 | 0.72589216 |
| AP2S1 | 40.2225186 | 0.49906329 | 0.38532865 | 1.29516268 | 0.19526412 | 0.88391893 |
| ARHGAP35 | 87.47999 | -0.0284198 | 0.24482147 | -0.1160838 | 0.9075861 | 0.99935941 |
| ZC3H4 | 23.6951906 | 0.11118988 | 0.40365405 | 0.27545836 | 0.78296411 | 0.9958918 |
| SAE1 | 50.2974106 | -0.3171128 | 0.33535769 | -0.9455957 | 0.34435484 | 0.96887708 |
| INAFM1 | 21.658101 | -0.9863991 | 0.4329014 | -2.2785768 | 0.02269223 | 0.40828078 |
| MEIS3 | 22.7630328 | -0.268236 | 0.49369255 | -0.5433261 | 0.58690531 | 0.9958918 |
| NAPA | 44.1944155 | -0.0634969 | 0.31704316 | -0.2002783 | 0.84126295 | 0.99692952 |
| EHD2 | 53.2742056 | -0.3009412 | 0.29100001 | -1.0341623 | 0.30106031 | 0.95597956 |
| NOP53 | 16.8638073 | 0.25234163 | 0.4662467 | 0.54121912 | 0.58835656 | 0.9958918 |
| SELENOW | 80.6122159 | 0.34504935 | 0.31440788 | 1.09745771 | 0.27244136 | 0.93813668 |
| PLA2G4C | 54.7509655 | 0.1103745 | 0.65496083 | 0.16852076 | 0.86617361 | 0.9971731 |
| CARD8 | 28.5015026 | 0.08684008 | 0.32838374 | 0.26444695 | 0.79143554 | 0.99592423 |
| EMP3 | 30.5003963 | -0.3531669 | 0.55828244 | -0.6325954 | 0.5269979 | 0.9958918 |
| KDELR1 | 28.6875282 | -0.0844726 | 0.36230494 | -0.2331534 | 0.81564225 | 0.99592423 |
| RPL18 | 105.776254 | -0.1432629 | 0.34021747 | -0.4210922 | 0.67368774 | 0.9958918 |
| PLEKHA4 | 51.9817847 | -0.4537647 | 0.48192994 | -0.9415574 | 0.34641931 | 0.96887708 |
| PPP1R15A | 35.1726306 | -0.2410589 | 0.43876459 | -0.5494037 | 0.58272841 | 0.9958918 |
| NUCB1 | 60.2739059 | 0.44951985 | 0.25087852 | 1.79178293 | 0.07316774 | 0.66743052 |
| FTL | 912.380886 | 0.65523603 | 0.54363863 | 1.20527865 | 0.22809575 | 0.91350172 |
| GYS1 | 61.0196917 | 0.14785943 | 0.41906732 | 0.35282978 | 0.72421605 | 0.9958918 |
| SNRNP70 | 136.85693 | 0.26483329 | 0.32800249 | 0.80741243 | 0.41942891 | 0.97812179 |
| SLC6A16 | 47.0238529 | -0.2101039 | 0.37507522 | -0.5601646 | 0.57536717 | 0.9958918 |
| TEAD2 | 25.3669706 | -0.2785606 | 0.34568942 | -0.8058118 | 0.42035135 | 0.97812179 |
| RPL13A | 146.960357 | -0.0738091 | 0.37854241 | -0.1949823 | 0.84540678 | 0.99692952 |
| RPS11 | 126.382282 | 0.00723451 | 0.3348602 | 0.02160458 | 0.98276338 | 0.99935941 |
| FCGRT | 45.6078635 | 0.13169047 | 0.34034083 | 0.38693703 | 0.69880282 | 0.9958918 |
| RCN3 | 27.3056265 | -0.4034174 | 0.47865787 | -0.8428095 | 0.39933502 | 0.97500135 |
| NOSIP | 20.3153513 | 0.24378146 | 0.37348765 | 0.65271624 | 0.51393923 | 0.9958918 |
| PRR12 | 20.7339704 | 0.23133099 | 0.46126498 | 0.50151431 | 0.61600921 | 0.9958918 |
| RRAS | 38.9736343 | 0.30602574 | 0.41746116 | 0.73306399 | 0.4635194 | 0.98590351 |
| IRF3 | 30.9408107 | 0.52197566 | 0.30119841 | 1.73299605 | 0.08309637 | 0.7007527 |

| | | | | | | |
|---|---|---|---|---|---|---|
| PRMT1 | 28.3055033 | 0.1599449 | 0.45234552 | 0.3535901 | 0.72364609 | 0.9958918 |
| AP2A1 | 50.8093116 | 0.35970063 | 0.30825525 | 1.16689216 | 0.24325392 | 0.92534155 |
| MED25 | 19.1078554 | -0.4607846 | 0.37048597 | -1.2437302 | 0.21359886 | 0.89750604 |
| PTOV1 | 23.9258592 | 0.14515428 | 0.35250075 | 0.41178431 | 0.68049752 | 0.9958918 |
| PNKP | 17.7632285 | 0.29677482 | 0.67760992 | 0.43797296 | 0.66140589 | 0.9958918 |
| TBC1D17 | 19.0666403 | -0.288792 | 0.40223277 | -0.7179724 | 0.47277428 | 0.98910524 |
| VRK3 | 20.9993003 | 0.37814869 | 0.32815574 | 1.15234519 | 0.24917925 | 0.92624282 |
| NR1H2 | 20.1463812 | 0.62612124 | 0.35324915 | 1.77246353 | 0.07631764 | 0.67573297 |
| EMC10 | 25.6529917 | 0.70409433 | 0.44835546 | 1.57039312 | 0.11632368 | 0.78417352 |
| CLEC11A | 27.3989986 | -0.5439837 | 0.35045932 | -1.552202 | 0.12061389 | 0.79021701 |
| ETFB | 30.3015395 | 0.57764827 | 0.39087578 | 1.47783084 | 0.13945306 | 0.8153985 |
| SPACA6 | 27.0048319 | 0.29374144 | 0.43706358 | 0.67207941 | 0.50153314 | 0.99213111 |
| ZNF577 | 22.1111301 | 0.63260215 | 0.38031826 | 1.66334939 | 0.09624252 | 0.73906611 |
| ZNF350-AS1 | 23.027795 | 0.17583038 | 0.47372234 | 0.3711676 | 0.7105127 | 0.9958918 |
| ZNF432 | 43.8156403 | 0.10473266 | 0.26604877 | 0.39365961 | 0.69383236 | 0.9958918 |
| PPP2R1A | 58.0205579 | 0.14895496 | 0.25689535 | 0.57982741 | 0.56203101 | 0.9958918 |
| ZNF83 | 92.0591592 | 0.62122311 | 0.19864921 | 3.12723674 | 0.00176458 | 0.09329411 |
| ZNF611 | 36.3755572 | 0.05018369 | 0.32180461 | 0.15594459 | 0.8760767 | 0.9971731 |
| ZNF160 | 42.7817635 | 0.21836208 | 0.34381071 | 0.63512296 | 0.52534822 | 0.9958918 |
| ZNF761 | 24.3369325 | -0.1145601 | 0.36811551 | -0.3112069 | 0.75564331 | 0.9958918 |
| ZNF331 | 110.329653 | -1.6379526 | 0.61343303 | -2.6701408 | 0.00758194 | 0.22558584 |
| MYADM | 47.5218265 | -0.2231496 | 0.46749236 | -0.4773331 | 0.63312498 | 0.9958918 |
| PRPF31 | 20.8639866 | 0.09259001 | 0.3345215 | 0.27678342 | 0.78194641 | 0.9958918 |
| MBOAT7 | 20.2814943 | 0.50360408 | 0.37135262 | 1.35613447 | 0.17505639 | 0.8740623 |
| RPS9 | 198.58791 | -0.226421 | 0.34486624 | -0.6565472 | 0.5114721 | 0.9958918 |
| LENG8 | 90.218287 | -0.3203997 | 0.34178513 | -0.9374303 | 0.34853729 | 0.96890122 |
| CDC42EP5 | 20.0208216 | -0.075035 | 0.3506584 | -0.2139832 | 0.83056012 | 0.99592423 |
| PPP1R12C | 29.5271412 | 0.40164221 | 0.35726754 | 1.12420572 | 0.26092577 | 0.93055907 |
| PPP6R1 | 21.676278 | 0.04941055 | 0.39587691 | 0.12481292 | 0.90067166 | 0.99935941 |
| RPL28 | 142.235734 | -0.3187411 | 0.27959124 | -1.1400254 | 0.25427573 | 0.92646125 |
| SSC5D | 22.7003821 | 1.28209262 | 0.41742619 | 3.07142352 | 0.00213041 | 0.09909091 |
| U2AF2 | 21.031852 | 0.01101097 | 0.35645419 | 0.03089027 | 0.97535705 | 0.99935941 |
| EPN1 | 22.8839916 | 0.22849763 | 0.32011411 | 0.71380053 | 0.47535053 | 0.98910524 |
| ZSCAN5A | 51.7396227 | -0.0736145 | 0.43949004 | -0.1674997 | 0.86697685 | 0.9971731 |
| ZNF264 | 26.500807 | 0.03082239 | 0.33473111 | 0.09208105 | 0.92663364 | 0.99935941 |
| AC005261.1 | 42.5961626 | 0.56143609 | 0.32157884 | 1.74587388 | 0.08083287 | 0.69693132 |
| ZNF586 | 45.1219996 | 0.66677585 | 0.39706378 | 1.67926638 | 0.09310014 | 0.73019523 |
| ZNF814 | 41.1043903 | 0.84617467 | 0.30044078 | 2.81644409 | 0.00485585 | 0.1747393 |
| ZNF587 | 22.2791194 | -0.1310915 | 0.33618409 | -0.3899395 | 0.6965813 | 0.9958918 |
| ZSCAN18 | 30.2717192 | 0.66612781 | 0.35009188 | 1.90272281 | 0.05707672 | 0.60440467 |
| ZNF274 | 24.17247 | 0.25536136 | 0.32722335 | 0.78038859 | 0.43516218 | 0.98092839 |
| RPS5 | 64.533332 | -0.0326993 | 0.38147155 | -0.0857189 | 0.93168984 | 0.99935941 |
| TRIM28 | 30.5851155 | 0.00182342 | 0.40693378 | 0.00448088 | 0.99642478 | 0.99935941 |
| CHMP2A | 21.5767655 | -0.26577 | 0.42703258 | -0.6223646 | 0.53370214 | 0.9958918 |
| MZF1 | 30.6533325 | 0.26488138 | 0.31204504 | 0.84885623 | 0.3959613 | 0.97500135 |
| RPS4Y1 | 48.8144023 | 0.45877046 | 2.12691946 | 0.21569715 | 0.82922383 | 0.99592423 |
| ZFY | 23.861884 | 0.53987548 | 1.9129306 | 0.28222429 | 0.77777153 | 0.9958918 |
| LINC00278 | 45.0509816 | 0.71764475 | 1.75676587 | 0.40850335 | 0.68290417 | 0.9958918 |
| TTTY15 | 22.1415396 | 0.82393247 | 1.90951157 | 0.43148859 | 0.66611314 | 0.9958918 |
| USP9Y | 93.2000972 | 0.72980146 | 1.9518863 | 0.37389548 | 0.70848208 | 0.9958918 |
| DDX3Y | 34.0553234 | 0.73214625 | 1.93984164 | 0.37742578 | 0.70585721 | 0.9958918 |
| UTY | 165.6505 | 0.7579689 | 1.59885911 | 0.4740686 | 0.63545099 | 0.9958918 |
| NLGN4Y | 51.7611925 | 0.16461669 | 1.54354871 | 0.1066482 | 0.91506808 | 0.99935941 |
| TTTY14 | 181.744516 | 0.77752222 | 1.49653864 | 0.51954704 | 0.60337932 | 0.9958918 |

| | | | | | | |
|---|---|---|---|---|---|---|
| KDM5D | 37.5730949 | 1.02654232 | 2.26443792 | 0.45333206 | 0.65030965 | 0.9958918 |
| TTTY10 | 20.0893119 | 0.66363295 | 1.79777779 | 0.3691407 | 0.71202285 | 0.9958918 |
| IL17RA | 26.1121838 | 0.40070352 | 0.3315006 | 1.20875656 | 0.22675639 | 0.91140536 |
| HDHD5 | 21.7477796 | 0.37780073 | 0.34723872 | 1.0880144 | 0.27658875 | 0.94273432 |
| CECR2 | 32.13235 | -2.7205205 | 0.66353749 | -4.1000254 | 4.13E-05 | 0.00843308 |
| ATP6V1E1 | 47.9460584 | 0.29559027 | 0.24249602 | 1.21894895 | 0.22286357 | 0.91093521 |
| BCL2L13 | 80.8016436 | -0.1162605 | 0.2323463 | -0.5003761 | 0.61681031 | 0.9958918 |
| MICAL3 | 101.374252 | 0.2405347 | 0.32632504 | 0.73710156 | 0.46106059 | 0.98590351 |
| PEX26 | 28.1102449 | 0.15097756 | 0.36433161 | 0.41439599 | 0.67858412 | 0.9958918 |
| DGCR2 | 51.896213 | 0.73576369 | 0.2640251 | 2.78671869 | 0.00532447 | 0.18457318 |
| CLTCL1 | 52.3162801 | 0.15452603 | 0.37224665 | 0.41511731 | 0.67805603 | 0.9958918 |
| HIRA | 21.8362626 | -0.5576735 | 0.45870311 | -1.2157614 | 0.22407583 | 0.91093521 |
| UFD1 | 16.9643541 | -0.343222 | 0.42916506 | -0.7997435 | 0.42385942 | 0.97828057 |
| TXNRD2 | 22.9061587 | -0.0735988 | 0.4229384 | -0.1740178 | 0.86185147 | 0.9971731 |
| COMT | 62.4666426 | 0.26662159 | 0.22572129 | 1.18119825 | 0.23752398 | 0.92291633 |
| ARVCF | 16.4069886 | -0.1639873 | 0.57912312 | -0.2831649 | 7.77E-01 | 0.9958918 |
| TANGO2 | 20.2695616 | 0.00936296 | 0.36681499 | 0.02552502 | 0.97963619 | 0.99935941 |
| DGCR8 | 24.2590661 | 0.21572059 | 0.37853317 | 0.56988556 | 0.56875532 | 0.9958918 |
| KLHL22 | 44.015301 | 0.34752973 | 0.3340275 | 1.04042251 | 0.29814365 | 0.95447177 |
| MED15 | 86.2495142 | -0.2250085 | 0.22307046 | -1.0086881 | 0.31312422 | 0.95992644 |
| PI4KA | 141.483951 | -0.0472656 | 0.18659519 | -0.2533054 | 0.80003222 | 0.99592423 |
| SNAP29 | 22.1744029 | 0.20746757 | 0.30980014 | 0.66968196 | 0.50306056 | 0.99354798 |
| CRKL | 18.946609 | 0.16312794 | 0.41735315 | 0.39086308 | 0.69589844 | 0.9958918 |
| LZTR1 | 17.3147651 | 0.48125728 | 0.38319878 | 1.25589461 | 0.20915418 | 0.89634177 |
| HIC2 | 23.2321769 | 0.10590585 | 0.33757854 | 0.31372211 | 0.75373211 | 0.9958918 |
| UBE2L3 | 56.3265615 | 0.17789517 | 0.28995356 | 0.61352988 | 0.53952603 | 0.9958918 |
| PPIL2 | 32.7296149 | -0.3620845 | 0.30383429 | -1.1917171 | 0.2333722 | 0.91861399 |
| MAPK1 | 155.404951 | -0.0202274 | 0.21123791 | -0.0957563 | 0.92371415 | 0.99935941 |
| PPM1F | 17.9520281 | 0.59030782 | 0.44054949 | 1.33993533 | 0.18026637 | 0.87815793 |
| BCR | 59.0464546 | 0.66841551 | 0.36832399 | 1.81474876 | 0.06956253 | 0.65289275 |
| CHCHD10 | 43.6094408 | 0.17823202 | 0.25344203 | 0.70324571 | 0.48190264 | 0.99002719 |
| SMARCB1 | 31.423221 | 0.07854405 | 0.34418053 | 0.22820598 | 0.81948611 | 0.99592423 |
| DDT | 30.5074803 | 0.07018207 | 0.34568761 | 0.20302165 | 0.83911811 | 0.99692952 |
| CABIN1 | 73.5810166 | -0.0657212 | 0.24649135 | -0.266627 | 0.78975638 | 0.99592423 |
| GGT5 | 139.07665 | -0.5463708 | 0.21516323 | -2.5393315 | 0.01110645 | 0.28321649 |
| SNRPD3 | 36.545221 | -0.5231864 | 0.27631322 | -1.8934542 | 0.05829749 | 0.60892588 |
| KIAA1671 | 45.9392346 | -0.1648668 | 0.42702689 | -0.3860806 | 0.69943696 | 0.9958918 |
| GRK3 | 95.6412348 | -0.2685943 | 0.46159389 | -0.5818845 | 0.5606445 | 0.9958918 |
| HPS4 | 20.0285347 | 0.02458473 | 0.41186655 | 0.05969102 | 0.95240172 | 0.99935941 |
| MIATNB | 29.5578681 | 0.19975265 | 0.46629028 | 0.42838691 | 0.66836945 | 0.9958918 |
| PITPNB | 194.908059 | 0.03207206 | 0.33212588 | 0.09656597 | 0.92307108 | 0.99935941 |
| TTC28 | 673.291848 | -0.1110903 | 0.19797545 | -0.5611318 | 0.57470768 | 0.9958918 |
| HSCB | 30.7022416 | -0.0978379 | 0.43181544 | -0.2265733 | 0.82075552 | 0.99592423 |
| XBP1 | 40.2073736 | 0.39671128 | 0.42874868 | 0.92527696 | 0.35482186 | 0.97217293 |
| Z93930.2 | 52.4671755 | 0.16398893 | 0.38663771 | 0.42414106 | 0.67146294 | 0.9958918 |
| ZNRF3 | 66.9138135 | 0.14709782 | 0.2373174 | 0.61983581 | 0.53536589 | 0.9958918 |
| KREMEN1 | 28.7477338 | 0.10697167 | 0.45744709 | 0.23384491 | 0.81510539 | 0.99592423 |
| EMID1 | 55.581295 | 0.73829682 | 0.56642005 | 1.30344403 | 0.19242321 | 0.88158763 |
| EWSR1 | 114.666606 | 0.01554506 | 0.21312877 | 0.07293741 | 0.94185593 | 0.99935941 |
| AP1B1 | 22.4695405 | 0.18797212 | 0.36823453 | 0.51046847 | 0.6097233 | 0.9958918 |
| THOC5 | 19.5557208 | 0.00591719 | 0.32430929 | 0.0182455 | 0.985443 | 0.99935941 |
| NF2 | 34.1612734 | 0.32108948 | 0.31655514 | 1.01432401 | 0.31042817 | 0.95956408 |
| UQCR10 | 44.5459293 | 0.06832444 | 0.41961323 | 0.16282719 | 0.8706545 | 0.9971731 |
| ASCC2 | 28.7932578 | -0.5767931 | 0.38176545 | -1.5108573 | 0.13082482 | 0.80565822 |

| | | | | | | |
|---|---|---|---|---|---|---|
| MTMR3 | 142.614055 | 0.16336853 | 0.1791166 | 0.91207925 | 0.361727 | 0.97217293 |
| RNF215 | 24.0344576 | 0.4041338 | 0.48156859 | 0.83920299 | 0.40135541 | 0.97500135 |
| MORC2 | 26.4597902 | -0.3204673 | 0.31804098 | -1.007629 | 0.31363259 | 0.95992644 |
| SMTN | 92.9620832 | -0.3151625 | 0.36238216 | -0.8696965 | 0.38446631 | 0.9725044 |
| SELENOM | 113.497827 | 0.07331836 | 0.43057874 | 0.17027862 | 0.86479102 | 0.9971731 |
| RNF185 | 36.9895838 | 0.08280267 | 0.33155247 | 0.24974227 | 0.80278666 | 0.99592423 |
| LIMK2 | 29.1464377 | 0.72052921 | 0.35858406 | 2.00937324 | 0.04449757 | 0.55709134 |
| DRG1 | 36.7689673 | 0.05491365 | 0.25490759 | 0.21542572 | 0.82943542 | 0.99592423 |
| EIF4ENIF1 | 32.6547145 | 0.07115466 | 0.30066512 | 0.23665753 | 0.81292248 | 0.99592423 |
| SFI1 | 25.3207975 | -0.3070638 | 0.42218597 | -0.7273189 | 0.46703062 | 0.98834148 |
| PISD | 33.1022625 | 0.13145826 | 0.29841273 | 0.44052498 | 0.65955693 | 0.9958918 |
| PRR14L | 69.9711154 | 0.07582913 | 0.22303254 | 0.33999133 | 0.73386305 | 0.9958918 |
| YWHAH | 20.0052779 | 0.47531657 | 0.43918093 | 1.08227964 | 0.27912829 | 0.94419503 |
| RTCB | 33.0391419 | 0.44531899 | 0.27122163 | 1.64190072 | 0.10061058 | 0.75064053 |
| FBXO7 | 43.3854613 | -0.1867262 | 0.26839967 | -0.695702 | 0.4866155 | 0.99046213 |
| SYN3 | 106.191654 | -1.7363503 | 0.85092364 | -2.0405478 | 0.0412958 | 0.54347495 |
| TIMP3 | 1163.80541 | -0.8167207 | 0.37336774 | -2.1874431 | 0.02871019 | 0.45975021 |
| LARGE1 | 507.346778 | -0.4806974 | 0.3973389 | -1.2097919 | 0.22635877 | 0.91093521 |
| HMGXB4 | 35.246914 | -0.4217755 | 0.34863338 | -1.2097965 | 0.22635698 | 0.91093521 |
| TOM1 | 31.3497824 | 0.3899396 | 0.38178079 | 1.0213704 | 0.30707898 | 0.95793395 |
| HMOX1 | 34.4595589 | -0.8605408 | 0.97410657 | -0.8834155 | 0.37701182 | 0.97217293 |
| APOL6 | 49.7614566 | -0.0426575 | 0.51021917 | -0.0836063 | 0.93336949 | 0.99935941 |
| RBFOX2 | 397.238526 | -0.1050436 | 0.17477273 | -0.6010297 | 0.54782021 | 0.9958918 |
| APOL3 | 20.3443889 | 0.26014067 | 0.54206715 | 0.47990489 | 0.63129503 | 0.9958918 |
| APOL2 | 22.0426207 | -0.6656997 | 0.4651744 | -1.4310754 | 0.1524086 | 0.83961827 |
| APOL1 | 26.7005703 | -0.7679465 | 0.4070556 | -1.8865888 | 0.05921565 | 0.61175668 |
| MYH9 | 397.079777 | -0.4061609 | 0.19250323 | -2.1098915 | 0.0348677 | 0.50468149 |
| TXN2 | 18.0968866 | 0.16208923 | 0.37367141 | 0.43377475 | 0.66445202 | 0.9958918 |
| EIF3D | 32.177933 | 0.48923228 | 0.34731165 | 1.40862618 | 0.15894573 | 0.84665568 |
| CDC42EP1 | 20.2053231 | -0.2926496 | 0.42577323 | -0.6873369 | 0.49187047 | 0.99046213 |
| GGA1 | 26.64178 | 0.27554946 | 0.35302018 | 0.78054874 | 0.43506795 | 0.98092839 |
| LGALS1 | 150.103061 | 0.1448384 | 0.60458234 | 0.2395677 | 0.81066541 | 0.99592423 |
| TRIOBP | 44.7550064 | -0.1596468 | 0.28038961 | -0.5693748 | 0.5691018 | 0.9958918 |
| H1F0 | 30.6093908 | -1.4496433 | 0.60861711 | -2.3818642 | 0.01722525 | 0.34765232 |
| EIF3L | 51.1146304 | 0.19946559 | 0.35539241 | 0.5612545 | 0.57462406 | 0.9958918 |
| POLR2F | 19.7426001 | -0.6301974 | 0.39380743 | -1.6002679 | 0.10953916 | 0.77303254 |
| MAFF | 24.861179 | -0.6207086 | 0.30883073 | -2.0098667 | 0.0444453 | 0.55709134 |
| TMEM184B | 66.387325 | -0.2578235 | 0.3318214 | -0.7769948 | 0.43716182 | 0.98092839 |
| CSNK1E | 30.1910503 | -0.4248091 | 0.35175885 | -1.2076714 | 0.2271737 | 0.9120167 |
| TPTEP2-CSNI | 57.2810166 | -0.0461719 | 0.29822003 | -0.1548248 | 0.87695942 | 0.9971731 |
| DDX17 | 496.035391 | 0.08234981 | 0.2727824 | 0.30188829 | 0.76273722 | 0.9958918 |
| CBY1 | 17.6360344 | 0.15854678 | 0.51152612 | 0.30994854 | 0.75660009 | 0.9958918 |
| JOSD1 | 28.8667985 | -0.0945581 | 0.3228009 | -0.29293 | 0.76957564 | 0.9958918 |
| GTPBP1 | 25.1514286 | -0.2712691 | 0.41553006 | -0.6528267 | 0.51386804 | 0.9958918 |
| SUN2 | 39.9821995 | 0.49856846 | 0.35873424 | 1.38979894 | 0.16458994 | 0.8578521 |
| CBX6 | 47.2783913 | 0.25753032 | 0.48530655 | 0.53065495 | 0.59565791 | 0.9958918 |
| CBX7 | 34.0566647 | 0.87709593 | 0.44672924 | 1.96337255 | 0.0496029 | 0.57316309 |
| RPL3 | 110.627996 | 0.28951214 | 0.35884309 | 0.80679314 | 0.41978567 | 0.97812179 |
| TAB1 | 18.4600494 | 0.10458642 | 0.35788069 | 0.29223823 | 0.77010449 | 0.9958918 |
| ATF4 | 57.5686396 | -0.1117679 | 0.31039424 | -0.3600837 | 0.71878455 | 0.9958918 |
| RPS19BP1 | 34.8222378 | -0.1025487 | 0.36061322 | -0.2843732 | 0.77612442 | 0.9958918 |
| TNRC6B | 402.294679 | 0.03836225 | 0.26033752 | 0.14735583 | 0.88285117 | 0.99840747 |
| MKL1 | 350.531635 | -0.3219111 | 0.34505597 | -0.9329242 | 0.35085912 | 0.9693503 |
| SLC25A17 | 41.2279041 | -0.0301177 | 0.24928713 | -0.1208153 | 0.90383732 | 0.99935941 |

| | | | | | | |
|---|---|---|---|---|---|---|
| ST13 | 73.097429 | -0.1854256 | 0.23574507 | -0.7865512 | 0.43154463 | 0.98092839 |
| XPNPEP3 | 43.7424311 | -0.2866521 | 0.33188379 | -0.8637122 | 0.38774599 | 0.97297628 |
| RBX1 | 47.2478615 | -0.4346173 | 0.24333712 | -1.7860708 | 0.0740878 | 0.66886321 |
| EP300 | 129.742149 | 0.15442468 | 0.16252203 | 0.95017691 | 0.34202237 | 0.96887708 |
| ZC3H7B | 36.2285414 | 0.46872527 | 0.3626387 | 1.2925407 | 0.19616996 | 0.88391893 |
| TOB2 | 37.8116666 | 0.22522246 | 0.32102685 | 0.70156891 | 0.48294805 | 0.99002719 |
| ACO2 | 42.6886808 | 0.10679793 | 0.36062501 | 0.29614676 | 0.767118 | 0.9958918 |
| CSDC2 | 20.4091139 | 0.17224554 | 0.5068144 | 0.33985922 | 0.73396255 | 0.9958918 |
| XRCC6 | 72.9064754 | -0.0932767 | 0.24785312 | -0.3763385 | 0.70666523 | 0.9958918 |
| SNU13 | 55.3856585 | -0.1507629 | 0.26428642 | -0.5704529 | 0.56837059 | 0.9958918 |
| SREBF2 | 147.936109 | 0.28199164 | 0.37577732 | 0.75042217 | 0.45300048 | 0.98092839 |
| SMDT1 | 23.6164822 | -0.1506435 | 0.38185366 | -0.394506 | 0.69320753 | 0.9958918 |
| TCF20 | 74.5800509 | 0.1228124 | 0.26955832 | 0.45560605 | 0.64867329 | 0.9958918 |
| RRP7A | 20.9325492 | 0.44005133 | 0.48518916 | 0.9069686 | 0.3644234 | 0.97217293 |
| POLDIP3 | 22.6136456 | 0.00824604 | 0.34938811 | 0.02360137 | 0.98117058 | 0.99935941 |
| CYB5R3 | 70.2985496 | 0.19105001 | 0.2609002 | 0.73227241 | 0.46400232 | 0.98590351 |
| A4GALT | 30.2419163 | 0.17037626 | 0.42786368 | 0.3982022 | 0.69048114 | 0.9958918 |
| ARFGAP3 | 105.127332 | -0.1557934 | 0.3754345 | -0.4149683 | 0.6781651 | 0.9958918 |
| PACSIN2 | 119.652396 | -0.0114235 | 0.35803131 | -0.0319065 | 0.97454661 | 0.99935941 |
| TSPO | 42.2967321 | -0.1680206 | 0.4190278 | -0.4009771 | 0.68843697 | 0.9958918 |
| EFCAB6 | 21.2087093 | 0.22435177 | 0.45755214 | 0.49033051 | 0.62390004 | 0.9958918 |
| NUP50-DT | 35.308108 | 0.52289289 | 0.47647953 | 1.09740892 | 0.27246267 | 0.93813668 |
| NUP50 | 24.1731727 | -0.4492487 | 0.32202073 | -1.3950924 | 0.16298796 | 0.85618195 |
| FAM118A | 49.5568373 | 0.31336533 | 0.52959426 | 0.59170831 | 0.55404593 | 0.9958918 |
| FBLN1 | 209.980147 | -0.4523549 | 0.36777797 | -1.2299675 | 0.21870929 | 0.90437211 |
| ATXN10 | 209.982924 | 0.08524856 | 0.18429578 | 0.46256382 | 0.64367704 | 0.9958918 |
| MIRLET7BHG | 97.1989147 | 0.2632834 | 0.38163958 | 0.68987447 | 0.49027313 | 0.99046213 |
| PPARA | 62.1124009 | 0.24777134 | 0.30333627 | 0.81682067 | 0.41403092 | 0.97812179 |
| TTC38 | 24.3706538 | 0.79815506 | 0.45914124 | 1.73836499 | 0.08214652 | 0.69958018 |
| TRMU | 22.6132797 | -0.0551367 | 0.30032454 | -0.1835904 | 0.85433482 | 0.9971731 |
| GRAMD4 | 22.9413229 | -0.1381002 | 0.32092232 | -0.4303229 | 0.66696077 | 0.9958918 |
| CERK | 37.4952434 | -0.1962831 | 0.3112121 | -0.6307051 | 0.52823337 | 0.9958918 |
| TBC1D22A | 197.13545 | 0.17883277 | 0.32505925 | 0.55015438 | 0.58221349 | 0.9958918 |
| BRD1 | 44.1339009 | 0.3750431 | 0.36071621 | 1.03971789 | 0.29847098 | 0.95447177 |
| ZBED4 | 22.90145 | 0.18802576 | 0.37391072 | 0.50286271 | 0.6150608 | 0.9958918 |
| PIM3 | 39.2738671 | -0.8441241 | 0.39260521 | -2.1500584 | 0.0315506 | 0.48041788 |
| SELENOO | 29.8158591 | 0.23830508 | 0.2986445 | 0.79795567 | 0.42489621 | 0.97917873 |
| PLXNB2 | 65.7001911 | 0.02260438 | 0.32799001 | 0.0689179 | 0.94505497 | 0.99935941 |
| PPP6R2 | 144.636631 | -0.1213238 | 0.20139124 | -0.6024284 | 0.54688904 | 0.9958918 |
| FP671120.1 | 548.58525 | -0.4259648 | 0.43957129 | -0.969046 | 0.33252222 | 0.9654289 |
| FP236383.1 | 317.79829 | -0.3558768 | 0.49293568 | -0.7219538 | 0.47032286 | 0.98910524 |
| HSPA13 | 22.4821157 | 0.01746038 | 0.42453038 | 0.0411287 | 0.9671933 | 0.99935941 |
| NRIP1 | 102.336833 | -0.4733607 | 0.22883395 | -2.0685772 | 0.03858578 | 0.5265563 |
| USP25 | 130.209607 | -0.3120656 | 0.27358386 | -1.140658 | 0.25401226 | 0.92646125 |
| MIR99AHG | 583.656173 | -0.4051145 | 0.45882667 | -0.8829359 | 0.37727092 | 0.97217293 |
| BTG3 | 28.4042948 | 0.30290984 | 0.40503189 | 0.74786664 | 0.4545406 | 0.98173285 |
| ATP5PF | 35.0721023 | 0.25283154 | 0.47808487 | 0.52884237 | 0.5969148 | 0.9958918 |
| GABPA | 32.7347132 | -0.0751358 | 0.30567762 | -0.2458008 | 0.80583647 | 0.99592423 |
| APP | 379.285851 | -0.2046703 | 0.20732711 | -0.9871852 | 0.32355184 | 0.95992644 |
| ADAMTS1 | 327.127658 | -1.1858996 | 0.40566469 | -2.9233494 | 0.00346288 | 0.14297014 |
| ADAMTS5 | 62.7399947 | 0.91439999 | 0.5452051 | 1.67716698 | 0.09350983 | 0.73159032 |
| LTN1 | 53.596071 | 0.07616102 | 0.24640748 | 0.30908569 | 0.75725635 | 0.9958918 |
| USP16 | 47.0954423 | -0.0251847 | 0.25081524 | -0.1004113 | 0.92001782 | 0.99935941 |
| CCT8 | 29.8272329 | -0.2054602 | 0.37287889 | -0.5510106 | 0.58162642 | 0.9958918 |

| | | | | | | |
|---|---|---|---|---|---|---|
| MAP3K7CL | 22.3126828 | 0.08052869 | 0.37195095 | 0.21650351 | 0.82859529 | 0.99592423 |
| BACH1 | 263.854884 | -0.1488302 | 0.25320809 | -0.5877782 | 0.55668118 | 0.9958918 |
| SOD1 | 53.9595304 | 0.51589133 | 0.22202678 | 2.32355451 | 0.02014938 | 0.3846177 |
| SCAF4 | 67.8345525 | 0.14666957 | 0.21790072 | 0.67310272 | 0.50088195 | 0.99213111 |
| URB1 | 51.6105322 | -0.2123107 | 0.3745714 | -0.5668096 | 0.57084356 | 0.9958918 |
| EVA1C | 159.164525 | 0.15301491 | 0.31520115 | 0.48545162 | 0.62735602 | 0.9958918 |
| SYNJ1 | 42.2956114 | -0.0985981 | 0.29394091 | -0.335435 | 0.73729694 | 0.9958918 |
| PAXBP1 | 91.3934892 | -0.4973268 | 0.22051916 | -2.2552543 | 0.02411737 | 0.42027858 |
| IFNAR1 | 48.614879 | 0.42051053 | 0.26624167 | 1.57943166 | 0.11423708 | 0.77853537 |
| IFNGR2 | 63.8304418 | 0.35051277 | 0.28159738 | 1.24473023 | 0.21323093 | 0.89738044 |
| TMEM50B | 57.3019686 | -0.2913287 | 0.29226676 | -0.9967904 | 0.31886624 | 0.95992644 |
| GART | 33.9892476 | -0.3480605 | 0.29496382 | -1.1800109 | 0.23799589 | 0.92291633 |
| SON | 241.103463 | 0.20629192 | 0.23451948 | 0.87963661 | 0.3790562 | 0.97217293 |
| AP000311.1 | 79.2084385 | -1.0027187 | 0.62727062 | -1.5985425 | 0.1099223 | 0.77303254 |
| ITSN1 | 292.453622 | -0.4807091 | 0.33969053 | -1.4151383 | 0.15702795 | 0.84427321 |
| MRPS6 | 128.699377 | -1.4506615 | 0.60893737 | -2.3822837 | 0.01720564 | 0.34765232 |
| RCAN1 | 54.8611121 | 0.52701809 | 0.35762992 | 1.47364093 | 0.1405783 | 0.8153985 |
| RUNX1 | 184.228062 | 1.0338847 | 0.61848177 | 1.67164944 | 0.09459347 | 0.7340733 |
| CBR1 | 25.1972431 | 0.05623463 | 0.42973512 | 0.13085881 | 0.895887 | 0.99935941 |
| DOPEY2 | 20.6206944 | 0.10219513 | 0.4255248 | 0.24016256 | 0.81020424 | 0.99592423 |
| MORC3 | 92.8786881 | -0.0844034 | 0.25001831 | -0.337589 | 0.73567295 | 0.9958918 |
| HLCS | 100.975646 | 0.44993742 | 0.18120028 | 2.48309455 | 0.01302465 | 0.30408773 |
| TTC3 | 221.969832 | -0.1457313 | 0.1766755 | -0.8248532 | 0.40945495 | 0.97812179 |
| VPS26C | 35.8445206 | -0.0546243 | 0.40522745 | -0.1347991 | 0.89277072 | 0.99935941 |
| DYRK1A | 311.862844 | -0.2665699 | 0.21526943 | -1.2383084 | 0.21560173 | 0.90187565 |
| KCNJ15 | 17.4794062 | 1.18673748 | 0.47968711 | 2.47398242 | 0.01336163 | 0.30782006 |
| ERG | 237.733887 | 0.4028 | 0.48339987 | 0.8332646 | 0.40469552 | 0.97582411 |
| ETS2 | 107.79257 | -0.7483069 | 0.34991337 | -2.138549 | 0.03247221 | 0.48564518 |
| BRWD1 | 197.862117 | 0.02057467 | 0.24412488 | 0.08427928 | 0.93283439 | 0.99935941 |
| HMGN1 | 46.5562567 | -0.374037 | 0.32890743 | -1.1372106 | 0.25545025 | 0.9266065 |
| WRB | 21.6464994 | -0.2487812 | 0.36077814 | -0.6895682 | 0.49046574 | 0.99046213 |
| BACE2 | 86.8813186 | 0.25376261 | 0.35310333 | 0.71866388 | 0.47234804 | 0.98910524 |
| MX2 | 59.7642201 | -1.5238591 | 1.42356894 | -1.0704498 | 0.2844169 | 0.94505166 |
| MX1 | 85.544602 | -2.4551889 | 1.41950663 | -1.7296072 | 0.08370048 | 0.70218588 |
| C2CD2 | 196.528546 | -1.3965862 | 0.29999825 | -4.6553145 | 3.23E-06 | 0.00158487 |
| ZBTB21 | 26.6050102 | -0.4188405 | 0.3248307 | -1.2894117 | 0.19725501 | 0.88391893 |
| SLC37A1 | 17.4494313 | 0.709689 | 0.49079239 | 1.44600654 | 0.14817537 | 0.8301633 |
| PDE9A | 32.4230015 | -0.2989584 | 0.44154514 | -0.6770733 | 0.49835943 | 0.99176652 |
| NDUFV3 | 38.8228562 | 0.06102866 | 0.28448822 | 0.21452087 | 0.83014089 | 0.99592423 |
| PKNOX1 | 51.5705853 | -0.0701417 | 0.25025967 | -0.2802759 | 0.77926587 | 0.9958918 |
| RRP1B | 26.5348209 | -0.0548091 | 0.31179906 | -0.1757834 | 0.86046412 | 0.9971731 |
| PDXK | 56.8418277 | 0.07597303 | 0.26626664 | 0.28532689 | 0.77539372 | 0.9958918 |
| CSTB | 47.0329719 | -0.3196567 | 0.27759001 | -1.1515426 | 0.2495091 | 0.92624282 |
| AGPAT3 | 64.2239155 | 0.14897346 | 0.34040833 | 0.43763166 | 0.66165332 | 0.9958918 |
| TRAPPC10 | 60.1676375 | 0.12161178 | 0.21070563 | 0.57716435 | 0.56382843 | 0.9958918 |
| PFKL | 53.7572231 | 0.22880416 | 0.28690719 | 0.79748494 | 0.42516945 | 0.97918216 |
| UBE2G2 | 78.0642992 | 0.24192208 | 0.27468949 | 0.88071109 | 0.37847421 | 0.97217293 |
| SUMO3 | 24.0035343 | -0.0845638 | 0.42503545 | -0.1989572 | 0.84229626 | 0.99692952 |
| PTTG1IP | 77.1117162 | 0.01110755 | 0.24276836 | 0.04575369 | 0.96350657 | 0.99935941 |
| ADARB1 | 158.812606 | -0.0738019 | 0.42143282 | -0.1751215 | 0.86098418 | 0.9971731 |
| POFUT2 | 18.9692464 | -0.768153 | 0.34243925 | -2.2431803 | 0.02488519 | 0.42729271 |
| COL18A1 | 483.736321 | -0.5502328 | 0.30584539 | -1.7990554 | 0.07200992 | 0.66213979 |
| PCBP3 | 61.6507199 | -2.2129377 | 0.53312135 | -4.1509081 | 3.31E-05 | 0.0073748 |
| COL6A1 | 136.834909 | -0.4647228 | 0.26101042 | -1.7804762 | 0.07499806 | 0.67167936 |

| | | | | | | |
|---|---|---|---|---|---|---|
| COL6A2 | 243.052385 | -0.2185303 | 0.33973398 | -0.6432394 | 0.52006878 | 0.9958918 |
| LSS | 30.0309164 | 0.04988094 | 0.4718972 | 0.10570299 | 0.91581801 | 0.99935941 |
| MCM3AP | 48.7587858 | -0.0346024 | 0.23715157 | -0.1459085 | 0.88399361 | 0.99862036 |
| PCNT | 33.4832188 | 0.13192203 | 0.41405756 | 0.31860795 | 0.75002383 | 0.9958918 |
| DIP2A | 71.9836819 | 0.09023653 | 0.22195811 | 0.40654754 | 0.68434033 | 0.9958918 |
| PRMT2 | 64.3083631 | 0.69922166 | 0.44495558 | 1.57144148 | 0.11608014 | 0.78417352 |
| AL592183.1 | 34.6911875 | 0.38820858 | 0.5683478 | 0.68304756 | 0.49457679 | 0.99176652 |

**Leukocytes**

*This means that positive fold changes imply higher mean value in non-COVID AKI than in COVID AKI and vice versa.

| | baseMean | log2 fold cha | lfcSE | stat | pvalue | padj |
|---|---|---|---|---|---|---|
| AL669831.5 | 11.4426808 | -0.3020664 | 0.5723185 | -0.5277941 | 0.59764222 | NA |
| NOC2L | 9.32746858 | 0.06649092 | 0.65877403 | 0.1009313 | 0.919605 | NA |
| ISG15 | 79.0322025 | 1.24934288 | 1.06578569 | 1.17222711 | 0.24110589 | 0.83493679 |
| C1orf159 | 8.14888496 | 0.78209387 | 0.70096105 | 1.11574511 | 0.26453125 | NA |
| SDF4 | 16.5523511 | -0.2005665 | 0.51043831 | -0.39293 | 0.69437115 | 0.9685713 |
| UBE2J2 | 18.5593821 | -0.468234 | 0.48602622 | -0.9633924 | 0.33535063 | 0.88047896 |
| INTS11 | 8.08083264 | -0.5765983 | 0.60514282 | -0.9528301 | 0.34067618 | NA |
| AURKAIP1 | 12.0702703 | -0.2167495 | 0.68714494 | -0.3154349 | 0.75243149 | NA |
| CCNL2 | 26.7462897 | -0.5197712 | 0.36409015 | -1.4275892 | 0.15341014 | 0.74419469 |
| MRPL20 | 18.4340156 | 0.31802791 | 0.43561291 | 0.73006998 | 0.46534741 | 0.9314919 |
| SSU72 | 19.2629869 | -0.4716583 | 0.45845578 | -1.0287979 | 0.30357466 | 0.86514059 |
| CDK11B | 12.4634196 | 0.14568658 | 0.51815274 | 0.28116532 | 0.7785836 | NA |
| SLC35E2B | 14.5438172 | -0.8443815 | 0.56919352 | -1.4834699 | 0.1379496 | 0.73044824 |
| SLC35E2A | 9.09848858 | 0.4403896 | 0.65549431 | 0.67184351 | 0.50168332 | NA |
| NADK | 25.0722065 | 0.23470966 | 0.51463966 | 0.45606602 | 0.6483425 | 0.96049083 |
| GNB1 | 165.529027 | -0.3004706 | 0.25979177 | -1.1565823 | 0.24744306 | 0.83817091 |
| PRKCZ | 8.47675968 | -0.828356 | 0.67209702 | -1.2324946 | 0.21776437 | NA |
| FAAP20 | 8.28089249 | -0.0901757 | 0.61227567 | -0.1472796 | 0.88291132 | NA |
| SKI | 41.6208551 | -0.0255422 | 0.42683512 | -0.0598408 | 0.95228239 | 0.99514836 |
| RER1 | 18.6174104 | -0.1521507 | 0.48130269 | -0.3161226 | 0.75190946 | 0.97859589 |
| TNFRSF14 | 19.1694129 | 0.35463889 | 0.43022329 | 0.82431357 | 0.40976141 | 0.91773508 |
| CEP104 | 17.5953909 | -0.4780308 | 0.47352488 | -1.0095158 | 0.31272734 | 0.87058449 |
| KCNAB2 | 59.5555324 | 0.59318838 | 0.41084805 | 1.44381451 | 0.14879116 | 0.7383468 |
| RPL22 | 9.59351648 | -0.2632449 | 0.53648557 | -0.490684 | 0.62364995 | NA |
| NOL9 | 13.0176432 | -1.1041524 | 0.49160602 | -2.2460106 | 0.02470333 | 0.39492781 |
| DNAJC11 | 15.7121 | 0.14585063 | 0.48116381 | 0.30312053 | 0.76179801 | 0.98163473 |
| CAMTA1 | 54.3191471 | -0.5347007 | 0.3362796 | -1.590048 | 0.11182399 | 0.69385269 |
| VAMP3 | 7.39451092 | 0.4618852 | 0.64715117 | 0.71372072 | 0.47539989 | NA |
| PARK7 | 24.5277181 | -0.1646733 | 0.41572678 | -0.3961094 | 0.69202432 | 0.9685713 |
| RERE | 194.975072 | -0.5228214 | 0.3238864 | -1.6142122 | 0.10648141 | 0.67654204 |
| ENO1 | 72.9643657 | 0.52381447 | 0.45047075 | 1.16281571 | 0.24490428 | 0.83763924 |
| MIR34AHG | 12.7678359 | 1.11379931 | 0.89577639 | 1.24338989 | 0.21372419 | NA |
| H6PD | 12.2948002 | 0.08446251 | 0.53170791 | 0.15885133 | 0.87378601 | NA |
| PIK3CD | 80.1269624 | -0.0550464 | 0.41910063 | -0.1313442 | 0.89550305 | 0.99115168 |
| CLSTN1 | 29.2897297 | -0.4956836 | 0.42783289 | -1.1585917 | 0.24662267 | 0.83817091 |
| CTNNBIP1 | 11.0994501 | -0.4224749 | 0.55495714 | -0.7612749 | 0.44649289 | NA |
| LZIC | 9.19056564 | -1.1201696 | 0.59581551 | -1.8800612 | 0.06009974 | NA |
| UBE4B | 54.3482911 | -0.1218986 | 0.32429084 | -0.3758928 | 0.70699658 | 0.97184211 |
| KIF1B | 56.344207 | 0.25369534 | 0.44078679 | 0.57555115 | 0.56491861 | 0.95415437 |
| PGD | 21.3976891 | 0.56976298 | 0.53894161 | 1.0571887 | 0.29042547 | 0.85822878 |
| DFFA | 7.69328501 | -0.2585901 | 0.62130281 | -0.4162062 | 0.67725909 | NA |
| PEX14 | 38.1458986 | -0.1622475 | 0.46853765 | -0.3462849 | 0.72912862 | 0.97605445 |
| SRM | 10.3698894 | -1.0684622 | 0.52969827 | -2.0171147 | 0.04368355 | NA |
| EXOSC10 | 12.1929609 | -0.2090997 | 0.49394405 | -0.4233266 | 0.672057 | NA |
| MTOR | 29.2796481 | -0.2527486 | 0.37348642 | -0.6767278 | 0.49857869 | 0.93916044 |
| AGTRAP | 12.2219895 | 1.22972588 | 0.64419058 | 1.90894731 | 0.05626889 | NA |
| MTHFR | 19.2484408 | -0.7548797 | 0.5481871 | -1.3770475 | 0.16849757 | 0.7567539 |
| CLCN6 | 12.2077155 | -0.0560082 | 0.57147783 | -0.098006 | 0.92192754 | NA |
| KIAA2013 | 9.11884155 | 0.49035832 | 0.64668613 | 0.75826323 | 0.44829341 | NA |
| MFN2 | 12.0313549 | -0.1390694 | 0.51077583 | -0.2722709 | 0.78541371 | NA |
| TNFRSF1B | 152.007501 | 0.11023885 | 0.61930107 | 0.17800527 | 0.85871884 | 0.98751288 |
| VPS13D | 77.2872679 | -0.1784977 | 0.24841484 | -0.7185468 | 0.47242017 | 0.9314919 |

| | | | | | | |
|---|---|---|---|---|---|---|
| DHRS3 | 21.5546907 | -0.6443461 | 0.6469901 | -0.9959133 | 0.31929226 | 0.87300562 |
| PRDM2 | 89.1391901 | -0.1173521 | 0.30815322 | -0.3808237 | 0.70333406 | 0.9714177 |
| KAZN | 29.0286656 | -1.0942803 | 0.53004589 | -2.0645012 | 0.03897022 | 0.48138665 |
| TMEM51 | 25.3421033 | 0.70616897 | 0.73157034 | 0.96527829 | 0.33440544 | 0.88047896 |
| EFHD2 | 35.3320172 | 0.23143792 | 0.7239937 | 0.31966842 | 0.74921971 | 0.97818861 |
| DNAJC16 | 14.7800157 | 0.05917402 | 0.44732477 | 0.13228424 | 0.89475947 | 0.99115168 |
| DDI2 | 27.16173 | 0.04478255 | 0.42906782 | 0.10437172 | 0.91687436 | 0.99498068 |
| PLEKHM2 | 27.1594918 | -0.7529665 | 0.4880264 | -1.5428807 | 0.12285973 | 0.71297276 |
| SPEN | 54.5565015 | -0.12133 | 0.3727873 | -0.325467 | 0.74482767 | 0.97805001 |
| FBXO42 | 56.7566239 | 0.23090718 | 0.34652671 | 0.66634742 | 0.50518906 | 0.94166217 |
| SZRD1 | 15.3024133 | -0.0002447 | 0.50466869 | -0.0004848 | 0.99961317 | 0.99961317 |
| NECAP2 | 16.6162389 | 0.31468023 | 0.5692288 | 0.55281853 | 0.58038768 | 0.95415437 |
| NBPF1 | 8.82126879 | 0.17242646 | 0.59471437 | 0.28993155 | 0.7718686 | NA |
| CROCC | 10.6168775 | 0.24137498 | 0.67539881 | 0.35738142 | 0.72080629 | NA |
| ATP13A2 | 9.15146132 | -0.2911568 | 0.65674218 | -0.4433351 | 0.6575234 | NA |
| SDHB | 18.8385209 | 0.55204519 | 0.46716589 | 1.18168985 | 0.23732879 | 0.83171006 |
| RCC2 | 20.3706556 | -0.3974881 | 0.44494982 | -0.8933324 | 0.37167918 | 0.90230329 |
| ARHGEF10L | 20.6704898 | 0.5451323 | 0.60274505 | 0.90441605 | 0.36577483 | 0.89868152 |
| IGSF21 | 17.568416 | 0.72218556 | 0.70955618 | 1.01779898 | 0.3087735 | 0.8673137 |
| UBR4 | 53.4780318 | 0.11597908 | 0.31758266 | 0.36519336 | 0.71496707 | 0.97348719 |
| CAPZB | 146.432898 | 0.07529853 | 0.33022296 | 0.22802331 | 0.81962812 | 0.98320463 |
| TMCO4 | 17.5854487 | -0.1745133 | 0.49741669 | -0.3508393 | 0.72570893 | 0.9754739 |
| DDOST | 11.747545 | 0.17138884 | 0.57014282 | 0.30060685 | 0.76371431 | NA |
| HP1BP3 | 67.1086686 | -0.1986019 | 0.3241585 | -0.612669 | 0.54009524 | 0.95415437 |
| EIF4G3 | 247.165416 | -0.2414725 | 0.29780364 | -0.8108446 | 0.41745491 | 0.91954684 |
| ECE1 | 31.6549194 | -0.1688325 | 0.47361979 | -0.3564726 | 0.7214867 | 0.97417403 |
| USP48 | 63.5701566 | -0.1139572 | 0.45484485 | -0.2505409 | 0.80216907 | 0.98320463 |
| CDC42 | 69.1786181 | 0.15385183 | 0.30938934 | 0.4972758 | 0.61899457 | 0.95688177 |
| ZBTB40 | 15.9394907 | -0.662012 | 0.47756352 | -1.3862281 | 0.16567725 | 0.75337545 |
| C1QA | 88.0504087 | 2.24878135 | 0.86707564 | 2.59352384 | 0.0094998 | 0.26141219 |
| C1QC | 56.3120088 | 1.35768715 | 0.98768347 | 1.37461767 | 0.16925001 | 0.75745073 |
| C1QB | 104.417503 | 1.91162762 | 0.86225291 | 2.217015 | 0.02662207 | 0.40712515 |
| KDM1A | 19.5796636 | 0.07922645 | 0.52333559 | 0.15138747 | 0.87967008 | 0.99019008 |
| LUZP1 | 28.4821868 | 1.03824401 | 0.51050015 | 2.03377805 | 0.04197398 | 0.50341641 |
| HNRNPR | 49.3046147 | -0.0599534 | 0.30503343 | -0.1965471 | 0.84418196 | 0.9842451 |
| RPL11 | 126.496084 | -0.3713486 | 0.58336472 | -0.6365633 | 0.52440935 | 0.95007724 |
| PITHD1 | 9.7083548 | 0.19145053 | 0.62081431 | 0.30838615 | 0.75778852 | NA |
| HMGCL | 8.49034038 | 0.40143098 | 0.55978513 | 0.7171162 | 0.4733024 | NA |
| PNRC2 | 14.4876644 | 0.06766372 | 0.45902339 | 0.147408 | 0.88280999 | 0.99019008 |
| SRSF10 | 31.3416759 | 0.00978558 | 0.40388884 | 0.0242284 | 0.98067043 | 0.99758514 |
| NIPAL3 | 19.1939655 | 0.38003006 | 0.45469377 | 0.83579342 | 0.40327113 | 0.91528253 |
| RCAN3 | 14.6884008 | 0.78957506 | 0.65860055 | 1.19886791 | 0.23057931 | 0.82393202 |
| SRRM1 | 39.3332312 | 0.17274679 | 0.30902579 | 0.55900443 | 0.57615869 | 0.95415437 |
| CLIC4 | 44.097297 | -0.888857 | 0.61618045 | -1.4425271 | 0.14915374 | 0.73895865 |
| RUNX3 | 55.8993265 | 0.45250727 | 0.32816818 | 1.37888831 | 0.16792919 | 0.7567539 |
| SYF2 | 11.9427526 | 0.38009054 | 0.52109255 | 0.72941082 | 0.4657504 | NA |
| RSRP1 | 102.7066 | 0.58972445 | 0.31505588 | 1.87180906 | 0.06123302 | 0.57337319 |
| TMEM50A | 24.0996974 | 0.10733299 | 0.40916856 | 0.26231973 | 0.79307495 | 0.98320463 |
| MACO1 | 30.1121445 | -0.5295377 | 0.35030252 | -1.5116582 | 0.13062084 | 0.72931666 |
| MAN1C1 | 22.2685189 | 0.21766962 | 0.40629308 | 0.53574533 | 0.59213457 | 0.9542025 |
| SH3BGRL3 | 97.068866 | -0.4245559 | 0.7838979 | -0.5415959 | 0.58809694 | 0.95415437 |
| UBXN11 | 9.06085099 | 0.78211959 | 0.67497527 | 1.15873814 | 0.24656294 | NA |
| CD52 | 42.9218707 | -0.6193816 | 0.73464113 | -0.8431077 | 0.39916821 | 0.91507741 |
| DHDDS | 14.344464 | -0.6122849 | 0.6645918 | -0.9212947 | 0.35689661 | 0.8957272 |

| | | | | | | |
|---|---|---|---|---|---|---|
| HMGN2 | 25.0925107 | -0.5571321 | 0.54528079 | -1.0217343 | 0.30690668 | 0.8673137 |
| RPS6KA1 | 31.6052088 | 0.50411127 | 0.4053447 | 1.24366071 | 0.21362446 | 0.80634897 |
| ARID1A | 94.233715 | -0.2914542 | 0.30459071 | -0.9568714 | 0.33863215 | 0.88437484 |
| ZDHHC18 | 9.98572724 | 0.48145799 | 0.61521661 | 0.78258288 | 0.4338721 | NA |
| NUDC | 11.7456177 | 0.51541633 | 0.53938581 | 0.95556153 | 0.33929381 | NA |
| SLC9A1 | 9.00055935 | 0.27450509 | 0.66557877 | 0.41243065 | 0.6800238 | NA |
| WDTC1 | 21.6027267 | 0.91367612 | 0.40948951 | 2.23125649 | 0.02566414 | 0.40120162 |
| SYTL1 | 11.1472321 | 0.28014994 | 0.56938099 | 0.49202545 | 0.62270135 | NA |
| WASF2 | 94.0285596 | -0.1281821 | 0.24300074 | -0.5274968 | 0.59784866 | 0.95538391 |
| AHDC1 | 10.2340879 | 0.02467448 | 0.65846562 | 0.03747269 | 0.97010811 | NA |
| FGR | 52.1709259 | -0.8563494 | 0.74308999 | -1.1524168 | 0.24914984 | 0.83865855 |
| IFI6 | 32.9386249 | 0.30994083 | 1.07591328 | 0.28807232 | 0.77329138 | 0.98264092 |
| FAM76A | 9.86488353 | 0.05707714 | 0.57136212 | 0.09989661 | 0.92042641 | NA |
| STX12 | 61.0765832 | -0.5769714 | 0.41402597 | -1.3935634 | 0.16344948 | 0.75321592 |
| THEMIS2 | 43.4914606 | -0.6611414 | 0.55911128 | -1.1824863 | 0.2370128 | 0.83171006 |
| RPA2 | 15.975956 | 0.071169 | 0.50308757 | 0.14146444 | 0.88750305 | 0.99019008 |
| EYA3 | 46.4308372 | -0.0385794 | 0.3142457 | -0.1227684 | 0.90229049 | 0.99262388 |
| PTAFR | 17.7361701 | 0.20627339 | 0.7771391 | 0.2654266 | 0.79068085 | 0.98320463 |
| DNAJC8 | 15.0078233 | -0.2531738 | 0.53989353 | -0.4689329 | 0.63911761 | 0.95933464 |
| ATP5IF1 | 24.9676514 | 0.58989689 | 0.53282725 | 1.10710721 | 0.26824758 | 0.84994492 |
| PHACTR4 | 56.6181023 | -0.0560442 | 0.38190893 | -0.1467475 | 0.88333136 | 0.99019008 |
| RCC1 | 15.975387 | -0.3186645 | 0.57984834 | -0.5495652 | 0.58261761 | 0.95415437 |
| TRNAU1AP | 19.743156 | -0.4836183 | 0.42096823 | -1.1488237 | 0.25062866 | 0.83984169 |
| TAF12 | 9.60163256 | -0.8063977 | 0.54202599 | -1.4877473 | 0.13681754 | NA |
| GMEB1 | 14.4550946 | -0.1747684 | 0.54164144 | -0.3226643 | 0.7469495 | 0.97813973 |
| YTHDF2 | 28.1648464 | -0.1893813 | 0.41668939 | -0.4544904 | 0.64947593 | 0.96084839 |
| EPB41 | 110.016164 | 0.762382 | 0.44664084 | 1.70692407 | 0.08783617 | 0.64676224 |
| SRSF4 | 66.9291157 | -0.4309541 | 0.35011697 | -1.230886 | 0.21836552 | 0.80919927 |
| LAPTM5 | 154.067318 | -0.9290494 | 0.59503104 | -1.5613461 | 0.1184421 | 0.70431251 |
| SDC3 | 10.5372664 | -1.8841792 | 0.81848829 | -2.3020234 | 0.02133385 | NA |
| PUM1 | 120.781076 | -0.2552791 | 0.21046369 | -1.2129366 | 0.22515406 | 0.81875277 |
| SNRNP40 | 12.334022 | 0.00909467 | 0.52376192 | 0.01736413 | 0.98614612 | NA |
| ZCCHC17 | 15.9695042 | 0.01097947 | 0.48401196 | 0.02268429 | 0.9819021 | 0.99758514 |
| PTP4A2 | 36.9837784 | 0.82593389 | 0.51126834 | 1.61546067 | 0.10621097 | 0.67559944 |
| KHDRBS1 | 53.7081518 | -0.1895263 | 0.28369699 | -0.6680587 | 0.5040961 | 0.94166217 |
| KPNA6 | 38.1880509 | -0.4897604 | 0.35762038 | -1.3694981 | 0.17084363 | 0.75751515 |
| TXLNA | 11.5454802 | -0.1993713 | 0.54337191 | -0.3669151 | 0.71368237 | NA |
| EIF3I | 8.96206963 | -0.5661573 | 0.70998807 | -0.7974181 | 0.42520826 | NA |
| LCK | 35.5289067 | -0.2132394 | 0.41238071 | -0.5170936 | 0.60509084 | 0.95688177 |
| HDAC1 | 21.3605272 | -0.2416702 | 0.42532325 | -0.5682035 | 0.56989679 | 0.95415437 |
| BSDC1 | 16.7130589 | 0.38300209 | 0.47889751 | 0.79975793 | 0.42385106 | 0.91985469 |
| ZBTB8OS | 33.0059037 | -0.9352004 | 0.37033458 | -2.5252851 | 0.01156044 | 0.29101961 |
| RBBP4 | 30.7224452 | 0.02322398 | 0.40737481 | 0.05700887 | 0.95453813 | 0.99514836 |
| YARS | 27.020604 | -0.7175787 | 0.43176771 | -1.6619554 | 0.09652171 | 0.66178928 |
| S100PBP | 39.4503877 | -0.4461976 | 0.44026461 | -1.0134759 | 0.31083291 | 0.8687623 |
| RNF19B | 19.0216133 | 0.23887292 | 0.65953069 | 0.3621862 | 0.71721289 | 0.97348719 |
| AK2 | 15.8893459 | 0.28615709 | 0.49709177 | 0.57566251 | 0.56484332 | 0.95415437 |
| ZNF362 | 8.84029961 | 0.39957062 | 0.55917178 | 0.71457579 | 0.47487121 | NA |
| PHC2 | 55.0683115 | 0.4422127 | 0.50170715 | 0.88141598 | 0.37809271 | 0.9051267 |
| SMIM12 | 10.8978917 | 0.35992043 | 0.60943374 | 0.59058173 | 0.55480071 | NA |
| ZMYM1 | 12.9121974 | -0.0672323 | 0.50068858 | -0.1342798 | 0.89318135 | NA |
| SFPQ | 62.9158796 | -0.3157755 | 0.38740261 | -0.8151095 | 0.41500965 | 0.91894074 |
| ZMYM4 | 94.4810469 | 0.18585838 | 0.2326533 | 0.79886412 | 0.42436921 | 0.91985469 |
| KIAA0319L | 60.0302011 | 0.16312314 | 0.38386148 | 0.42495313 | 0.67087084 | 0.96412305 |

| | | | | | | |
|---|---|---|---|---|---|---|
| PSMB2 | 17.6096466 | -0.1194403 | 0.45763543 | -0.2609944 | 0.7940968 | 0.98320463 |
| AGO4 | 53.7112547 | 0.54113155 | 0.37053283 | 1.46041458 | 0.14417617 | 0.73132073 |
| AGO1 | 20.4784145 | 0.20805032 | 0.50354435 | 0.41317179 | 0.67948075 | 0.9673431 |
| AGO3 | 87.8336192 | 0.29867692 | 0.34030564 | 0.87767255 | 0.38012145 | 0.90672073 |
| TRAPPC3 | 8.7109115 | 0.08424296 | 0.5725479 | 0.14713697 | 0.88302391 | NA |
| MAP7D1 | 15.200763 | 0.93739203 | 0.59201075 | 1.58340373 | 0.11332947 | 0.69801324 |
| THRAP3 | 61.6064652 | -0.2775159 | 0.25597861 | -1.0841371 | 0.27830401 | 0.84994492 |
| STK40 | 29.8645943 | 0.20748321 | 0.4546204 | 0.45638781 | 0.64811113 | 0.96049083 |
| CSF3R | 48.6934842 | 1.32036292 | 0.636102 | 2.07570942 | 0.03792083 | 0.47371083 |
| MEAF6 | 12.2387433 | 0.14241963 | 0.53827448 | 0.26458552 | 0.79132878 | NA |
| C1orf122 | 9.61796298 | 1.12023473 | 0.66198489 | 1.6922361 | 0.09060097 | NA |
| MTF1 | 16.253026 | 1.22012454 | 0.65796767 | 1.85438372 | 0.06368428 | 0.58463007 |
| INPP5B | 29.053023 | -0.5029653 | 0.35386282 | -1.4213567 | 0.15521308 | 0.74486619 |
| SF3A3 | 14.5216194 | -0.7903746 | 0.45391044 | -1.7412566 | 0.0816386 | 0.63740875 |
| RRAGC | 12.5459324 | -1.4735117 | 0.54607054 | -2.6983908 | 0.00696756 | NA |
| AKIRIN1 | 22.9775539 | -0.1425738 | 0.40220095 | -0.3544841 | 0.72297614 | 0.97441549 |
| NDUFS5 | 25.4190064 | -0.3964507 | 0.55407381 | -0.7155196 | 0.47428801 | 0.9314919 |
| MACF1 | 270.927133 | -0.3319765 | 0.26872901 | -1.2353578 | 0.21669737 | 0.80640512 |
| PABPC4 | 16.0170343 | -0.3743248 | 0.45185161 | -0.8284242 | 0.4074303 | 0.9169561 |
| TRIT1 | 12.7258617 | -0.4797788 | 0.56578786 | -0.8479836 | 0.39644708 | NA |
| CAP1 | 65.0872025 | 0.21262262 | 0.49569114 | 0.42894174 | 0.66796562 | 0.96412305 |
| PPT1 | 21.0471197 | 0.73312613 | 0.62046361 | 1.18157797 | 0.2373732 | 0.83171006 |
| RLF | 71.374952 | 0.08960585 | 0.32868366 | 0.27262033 | 0.78514508 | 0.98264092 |
| ZMPSTE24 | 11.4310536 | -0.6181904 | 0.53017442 | -1.1660133 | 0.24360908 | NA |
| SMAP2 | 202.071812 | 0.44845514 | 0.44220736 | 1.01412861 | 0.31052139 | 0.86858542 |
| NFYC | 32.5676681 | -0.0263996 | 0.41799733 | -0.0631573 | 0.94964127 | 0.99514836 |
| SCMH1 | 54.2654392 | 0.59318175 | 0.54974937 | 1.07900396 | 0.28058596 | 0.85036293 |
| HIVEP3 | 76.8733243 | 0.42416107 | 0.47986269 | 0.88392175 | 0.37673846 | 0.90433844 |
| FOXJ3 | 66.2788647 | -0.1595416 | 0.26523901 | -0.6015014 | 0.54750611 | 0.95415437 |
| YBX1 | 46.7405708 | -0.5674145 | 0.48695645 | -1.1652263 | 0.2439274 | 0.83666131 |
| SZT2 | 16.5081555 | -0.3020658 | 0.57334725 | -0.5268461 | 0.59830047 | 0.95567595 |
| ATP6V0B | 39.3740093 | 0.41620334 | 0.46509328 | 0.89488145 | 0.37085047 | 0.90226557 |
| ERI3 | 23.0845996 | -0.098181 | 0.40782988 | -0.2407402 | 0.8097565 | 0.98320463 |
| RNF220 | 38.3162665 | 3.49881036 | 1.47735342 | 2.36829612 | 0.01787023 | 0.34827529 |
| RPS8 | 108.483554 | -0.5991472 | 0.69880097 | -0.8573932 | 0.39122762 | 0.90910959 |
| EIF2B3 | 23.2740635 | -0.4192546 | 0.44655388 | -0.9388669 | 0.3477991 | 0.88939849 |
| ZSWIM5 | 12.9518311 | 0.05937325 | 0.72908641 | 0.08143513 | 0.93509591 | 0.99514836 |
| TESK2 | 22.2006587 | 0.7493908 | 0.51052258 | 1.46788963 | 0.14213421 | 0.73044824 |
| PRDX1 | 43.3536367 | 0.61389301 | 0.69444038 | 0.88401112 | 0.37669021 | 0.90433844 |
| AKR1A1 | 17.7271582 | 0.01113605 | 0.57075539 | 0.01951107 | 0.98443341 | 0.99811561 |
| NASP | 23.3103105 | 0.04494137 | 0.40440232 | 0.11113035 | 0.91151299 | 0.9947373 |
| GPBP1L1 | 75.8613177 | 0.29502204 | 0.34238655 | 0.86166363 | 0.38887264 | 0.90910959 |
| MAST2 | 25.5723648 | -0.1194042 | 0.49228316 | -0.2425519 | 0.80835258 | 0.98320463 |
| LRRC41 | 10.3705604 | 0.05675186 | 0.51333625 | 0.11055494 | 0.91196928 | NA |
| UQCRH | 26.1384858 | -0.1342926 | 0.49198759 | -0.2729594 | 0.78488446 | 0.98264092 |
| NSUN4 | 9.78609336 | -0.3977202 | 0.62539461 | -0.6359509 | 0.52480845 | NA |
| MKNK1 | 40.3666471 | 0.52208118 | 0.44517083 | 1.17276592 | 0.24088969 | 0.83493679 |
| ATPAF1 | 9.89067872 | 0.0148539 | 0.57834871 | 0.0256833 | 0.97950994 | NA |
| EFCAB14 | 29.8891152 | -0.2008501 | 0.44737331 | -0.4489541 | 0.6534648 | 0.96178254 |
| PDZK1IP1 | 21.1861592 | 1.86888238 | 0.75850509 | 2.46390223 | 0.01374336 | 0.31861834 |
| CMPK1 | 23.8468564 | -0.0300464 | 0.41351084 | -0.0726616 | 0.94207537 | 0.99514836 |
| SPATA6 | 21.1272686 | 0.80045213 | 0.72337181 | 1.10655697 | 0.26848552 | 0.84994492 |
| FAF1 | 160.369988 | -0.169743 | 0.2423108 | -0.7005177 | 0.48360407 | 0.93399425 |
| RNF11 | 16.7574524 | 0.20643253 | 0.52468635 | 0.39343988 | 0.69399462 | 0.9685713 |

| | | | | | | |
|---|---|---|---|---|---|---|
| EPS15 | 161.547108 | 0.17760729 | 0.28924782 | 0.61403157 | 0.53919447 | 0.95415437 |
| OSBPL9 | 63.983745 | 0.29730351 | 0.38901287 | 0.76425109 | 0.44471763 | 0.92914597 |
| NRDC | 82.3545639 | 0.14432504 | 0.3220281 | 0.44817529 | 0.65402669 | 0.96178254 |
| TXNDC12 | 17.222034 | 0.13342202 | 0.53263334 | 0.25049505 | 0.80220453 | 0.98320463 |
| BTF3L4 | 20.6247081 | -0.2893912 | 0.40358932 | -0.7170436 | 0.47334718 | 0.9314919 |
| ZFYVE9 | 11.5658061 | -1.9917452 | 0.73180566 | -2.721686 | 0.00649498 | NA |
| PRPF38A | 10.1739405 | 0.2014972 | 0.68443948 | 0.29439739 | 0.76845426 | NA |
| TUT4 | 98.2931239 | -0.9581634 | 0.30620436 | -3.1291632 | 0.00175305 | 0.0941881 |
| SHISAL2A | 12.3753773 | -0.2359297 | 0.52898824 | -0.4460017 | 0.65559601 | NA |
| ZYG11B | 22.6471897 | -0.3039691 | 0.40315888 | -0.7539684 | 0.45086819 | 0.92914597 |
| ECHDC2 | 19.3728374 | 0.32549668 | 0.57622158 | 0.5648811 | 0.57215464 | 0.95415437 |
| SCP2 | 30.1049437 | 0.48941181 | 0.37376126 | 1.30942361 | 0.1903909 | 0.78046165 |
| NDC1 | 13.3974733 | 0.14824631 | 0.52262308 | 0.28365818 | 0.77667235 | 0.98264092 |
| HSPB11 | 15.1002323 | 0.30292387 | 0.47277599 | 0.64073447 | 0.52169522 | 0.94895469 |
| TMEM59 | 37.2692598 | 0.01780499 | 0.40764139 | 0.04367807 | 0.96516102 | 0.99653147 |
| SSBP3 | 49.2810884 | 0.97767243 | 0.41604998 | 2.34989177 | 0.01877887 | 0.3571212 |
| USP24 | 115.332536 | -0.8663619 | 0.30111114 | -2.8772164 | 0.004012 | 0.15732988 |
| DAB1 | 14.8214986 | -0.6817854 | 0.79168982 | -0.8611774 | 0.38914036 | 0.90910959 |
| MYSM1 | 33.0288468 | -0.3238684 | 0.33052175 | -0.9798703 | 0.32715015 | 0.87715548 |
| JUN | 23.6337813 | -0.6701929 | 0.55205044 | -1.2140066 | 0.22474519 | 0.81875277 |
| FGGY | 26.3514743 | -0.1849381 | 0.55607788 | -0.3325758 | 0.7394545 | 0.97605445 |
| NFIA | 69.5725418 | 0.75744531 | 0.63999501 | 1.18351753 | 0.2366041 | 0.83171006 |
| TM2D1 | 35.0164132 | 0.16704165 | 0.34162706 | 0.48895907 | 0.62487067 | 0.95876809 |
| PATJ | 48.8296581 | 0.23816738 | 0.50794428 | 0.46888487 | 0.63915193 | 0.95933464 |
| USP1 | 9.11886308 | 0.26324666 | 0.58822804 | 0.44752484 | 0.65449615 | NA |
| DOCK7 | 28.7975813 | 0.13289758 | 0.39206112 | 0.33897158 | 0.73463114 | 0.97605445 |
| ATG4C | 28.6396536 | -0.3916674 | 0.47439869 | -0.825608 | 0.40902649 | 0.91716068 |
| ALG6 | 12.7873534 | 0.32565662 | 0.56115055 | 0.58033734 | 0.56168715 | NA |
| ITGB3BP | 16.7490141 | 0.52732552 | 0.56496819 | 0.93337206 | 0.3506279 | 0.89048867 |
| PGM1 | 10.2203339 | 1.28516748 | 0.57147074 | 2.24887714 | 0.02452031 | NA |
| JAK1 | 140.998808 | 0.41114922 | 0.27173846 | 1.51303288 | 0.13027131 | 0.72931666 |
| AK4 | 11.3179759 | 2.19329229 | 0.87441205 | 2.50830519 | 0.01213118 | NA |
| LEPR | 14.8298424 | 1.51523248 | 0.77322263 | 1.95963287 | 0.05003872 | 0.54084816 |
| PDE4B | 218.767956 | 1.37874539 | 0.67041887 | 2.05654324 | 0.03973018 | 0.48300729 |
| MIER1 | 58.1054909 | -0.2617255 | 0.3060333 | -0.8552189 | 0.39242997 | 0.90910959 |
| SLC35D1 | 15.6988624 | 0.95142735 | 0.82596732 | 1.15189467 | 0.24936436 | 0.83865855 |
| IL12RB2 | 18.3556597 | -0.0498165 | 0.72616779 | -0.0686019 | 0.94530648 | 0.99514836 |
| SERBP1 | 40.7280313 | -0.3947043 | 0.36112575 | -1.092983 | 0.27440126 | 0.84994492 |
| WLS | 15.3656549 | 0.43742476 | 0.87678217 | 0.49889788 | 0.61785133 | 0.95688177 |
| LRRC40 | 13.8669278 | -0.2568668 | 0.61514903 | -0.4175683 | 0.67626276 | 0.96653877 |
| SRSF11 | 82.1349094 | -0.236287 | 0.32999575 | -0.7160303 | 0.47397262 | 0.9314919 |
| ANKRD13C | 42.5993696 | -0.0660598 | 0.3635385 | -0.1817134 | 0.85580768 | 0.98751288 |
| ZRANB2 | 56.5046915 | 0.13073017 | 0.33634474 | 0.38867909 | 0.69751355 | 0.96861929 |
| NEGR1 | 27.5805707 | 2.47915832 | 0.8921135 | 2.77897187 | 0.00545312 | 0.18513861 |
| ACADM | 21.0631293 | -0.305809 | 0.45965362 | -0.6653031 | 0.50585664 | 0.94166217 |
| RABGGTB | 12.6918137 | -0.6420004 | 0.61046533 | -1.0516575 | 0.29295672 | NA |
| ST6GALNAC3 | 58.3419551 | 0.1309468 | 0.6539489 | 0.2002401 | 0.8412928 | 0.98380297 |
| PIGK | 13.8978346 | -0.160127 | 0.6063584 | -0.2640799 | 0.79171838 | 0.98320463 |
| AC118549.1 | 65.3176953 | -0.1864195 | 0.37460073 | -0.4976486 | 0.61873171 | 0.95688177 |
| USP33 | 66.9158925 | 0.24732592 | 0.27791346 | 0.8899386 | 0.37349885 | 0.90259505 |
| MIGA1 | 33.2430318 | 0.20557188 | 0.38950727 | 0.52777419 | 0.59765607 | 0.95538391 |
| FUBP1 | 62.4329902 | -0.1480211 | 0.29919446 | -0.4947321 | 0.62078927 | 0.95688177 |
| IFI44L | 48.1091392 | 1.50205365 | 1.63897953 | 0.91645663 | 0.35942745 | 0.8957272 |
| IFI44 | 34.5728539 | 1.04961642 | 0.96541653 | 1.08721613 | 0.2769413 | 0.84994492 |

| | | | | | | |
|---|---|---|---|---|---|---|
| PRKACB | 57.7552097 | 0.15658974 | 0.40455827 | 0.3870635 | 0.6987092 | 0.96884407 |
| GNG5 | 24.2454869 | 0.14429204 | 0.4642728 | 0.3107915 | 0.75595914 | 0.97989151 |
| CTBS | 15.3223628 | -0.1447267 | 0.4608547 | -0.3140398 | 0.75349084 | 0.97883059 |
| MCOLN2 | 24.4334572 | 0.20466272 | 0.75001937 | 0.27287658 | 0.78494809 | 0.98264092 |
| C1orf52 | 12.2030921 | -0.3637357 | 0.51813312 | -0.702012 | 0.48267169 | NA |
| ZNHIT6 | 19.8364293 | -0.1314417 | 0.51868308 | -0.2534143 | 0.79994804 | 0.98320463 |
| ODF2L | 34.3665906 | 0.2465818 | 0.44518943 | 0.55388062 | 0.57966055 | 0.95415437 |
| SH3GLB1 | 45.5765722 | -0.2859256 | 0.38284103 | -0.7468522 | 0.45515281 | 0.9314919 |
| SELENOF | 44.83374 | -0.1249347 | 0.36696238 | -0.3404564 | 0.73351283 | 0.97605445 |
| HS2ST1 | 34.7151394 | -0.3473741 | 0.37846006 | -0.9178621 | 0.35869108 | 0.8957272 |
| LMO4 | 25.8326515 | 0.23181473 | 0.45949087 | 0.50450344 | 0.61390764 | 0.95688177 |
| PKN2 | 102.425869 | 0.1024845 | 0.28190123 | 0.36354755 | 0.7161959 | 0.97348719 |
| GTF2B | 23.271507 | -0.071846 | 0.46074723 | -0.1559337 | 0.87608527 | 0.98939075 |
| KYAT3 | 23.1104154 | -0.4009987 | 0.36748989 | -1.0911831 | 0.27519234 | 0.84994492 |
| GBP3 | 11.1871085 | 0.34883506 | 0.59718323 | 0.58413406 | 0.55913012 | NA |
| GBP1 | 16.5178085 | 1.00140614 | 0.88045987 | 1.13736715 | 0.25538483 | 0.8406233 |
| GBP2 | 41.0588466 | -0.2623787 | 0.47777905 | -0.5491632 | 0.58289349 | 0.95415437 |
| GBP4 | 16.4582521 | 0.74945507 | 0.77738695 | 0.96406953 | 0.33501107 | 0.88047896 |
| GBP5 | 21.0561769 | -0.100097 | 0.49528774 | -0.2020986 | 0.83983962 | 0.98342625 |
| LRRC8B | 21.6785474 | 0.18138877 | 0.5093284 | 0.35613323 | 0.72174079 | 0.97417403 |
| ZNF326 | 16.9876734 | 0.07856701 | 0.58546919 | 0.13419494 | 0.89324842 | 0.99080204 |
| ZNF644 | 112.025102 | 0.05724654 | 0.27599211 | 0.20742092 | 0.83568114 | 0.98320463 |
| TGFBR3 | 51.2016277 | 0.21759971 | 0.62880833 | 0.34605094 | 0.72930444 | 0.97605445 |
| GLMN | 24.3283306 | -0.832064 | 0.47864943 | -1.7383579 | 0.08214777 | 0.63740875 |
| RPAP2 | 39.4746673 | 0.55809389 | 0.46023974 | 1.2126156 | 0.22527682 | 0.81875277 |
| EVI5 | 79.0148529 | -0.1141654 | 0.25499876 | -0.4477096 | 0.65436277 | 0.96178254 |
| RPL5 | 57.7807791 | -0.3231328 | 0.55984702 | -0.5771806 | 0.56381748 | 0.95415437 |
| FAM69A | 59.7773056 | -0.3733769 | 0.48024336 | -0.7774743 | 0.436879 | 0.92489992 |
| MTF2 | 47.2964386 | -0.0828012 | 0.30752356 | -0.2692514 | 0.7877362 | 0.98320463 |
| TMED5 | 28.8681498 | -0.5766948 | 0.39401354 | -1.4636421 | 0.14329177 | 0.73044824 |
| CCDC18 | 29.9801799 | 0.02482944 | 0.45328657 | 0.05477646 | 0.95631655 | 0.99514836 |
| DR1 | 15.9355719 | 0.0617778 | 0.42882532 | 0.14406284 | 0.88545084 | 0.99019008 |
| FNBP1L | 15.1181926 | -0.437274 | 0.5560433 | -0.7864028 | 0.43163153 | 0.92284736 |
| BCAR3 | 21.6812947 | -2.5025083 | 0.69374857 | -3.6072265 | 0.00030949 | 0.03358242 |
| GCLM | 15.8537864 | 0.21507899 | 0.50789843 | 0.42346851 | 0.67195347 | 0.96412305 |
| ARHGAP29 | 10.4238406 | -0.0436019 | 0.56354184 | -0.0773712 | 0.93832827 | NA |
| ABCD3 | 24.1333437 | 0.03048811 | 0.41059166 | 0.07425408 | 0.94080821 | 0.99514836 |
| CNN3 | 10.0885385 | -0.3327972 | 0.54633072 | -0.6091498 | 0.54242515 | NA |
| PTBP2 | 89.8559427 | 0.40990266 | 0.41380127 | 0.99057856 | 0.32189141 | 0.87363761 |
| DPYD | 644.195236 | 0.47394329 | 0.18601432 | 2.54788608 | 0.01083779 | 0.28424789 |
| DPYD-AS1 | 27.5368739 | 1.01403199 | 0.70179879 | 1.44490417 | 0.1484848 | 0.7383468 |
| AGL | 17.3004535 | 0.31659877 | 0.52054793 | 0.60820292 | 0.5430529 | 0.95415437 |
| TRMT13 | 14.9721974 | -0.4564346 | 0.44811984 | -1.0185548 | 0.30841436 | 0.8673137 |
| DBT | 13.6296456 | -0.0378532 | 0.52295782 | -0.0723828 | 0.94229725 | 0.99514836 |
| RTCA | 12.9541162 | 0.60396006 | 0.5104831 | 1.1831147 | 0.23676369 | 0.83171006 |
| CDC14A | 89.878033 | -0.0403047 | 0.31578454 | -0.1276334 | 0.8984391 | 0.99141995 |
| VCAM1 | 9.54889251 | 1.37923263 | 0.78274149 | 1.76205381 | 0.07806021 | NA |
| SLC30A7 | 49.5462595 | 0.64466073 | 0.43878608 | 1.46919139 | 0.14178088 | 0.73044824 |
| AC093157.1 | 13.2878907 | -0.2577596 | 0.69221704 | -0.3723681 | 0.70961879 | 0.97211298 |
| RNPC3 | 31.3223069 | -0.3465392 | 0.36706763 | -0.9440747 | 0.3451315 | 0.88711459 |
| VAV3 | 219.629163 | -0.7846548 | 0.34351271 | -2.2842088 | 0.02235926 | 0.38788767 |
| SLC25A24 | 24.9319009 | -0.0495499 | 0.47574719 | -0.1041518 | 0.91704891 | 0.99498068 |
| FAM102B | 52.1193145 | 1.11945378 | 0.47159503 | 2.37376076 | 0.01760795 | 0.34827529 |
| PRPF38B | 42.0573975 | -0.2962383 | 0.36071221 | -0.8212594 | 0.41149855 | 0.91823911 |

| | | | | | | |
|---|---|---|---|---|---|---|
| STXBP3 | 37.0313125 | 0.51319913 | 0.39498362 | 1.29929219 | 0.19384367 | 0.78135737 |
| WDR47 | 12.2678618 | 0.08341242 | 0.551496 | 0.15124755 | 0.87978045 | NA |
| SARS | 16.6922299 | -0.4777169 | 0.42984752 | -1.1113637 | 0.26641184 | 0.84994492 |
| SORT1 | 22.1129891 | 0.9675098 | 0.51327749 | 1.8849644 | 0.05943464 | 0.57281427 |
| PSMA5 | 29.1331804 | -0.5463653 | 0.38275938 | -1.4274381 | 0.15345365 | 0.74419469 |
| GNAI3 | 52.3851225 | 0.12020699 | 0.51453078 | 0.23362448 | 0.81527652 | 0.98320463 |
| AMPD2 | 9.46492365 | 0.9676528 | 0.94652139 | 1.02232534 | 0.30662695 | NA |
| AHCYL1 | 22.3395774 | 0.22254301 | 0.44015961 | 0.50559617 | 0.61314017 | 0.95688177 |
| STRIP1 | 9.58815944 | -0.5096426 | 0.62065241 | -0.8211402 | 0.41156644 | NA |
| RBM15 | 11.6688004 | -0.5184254 | 0.58472949 | -0.8866073 | 0.37529038 | NA |
| LAMTOR5 | 22.6076928 | 0.07610689 | 0.44178973 | 0.17226949 | 0.86322567 | 0.98751288 |
| CD53 | 124.609341 | -0.4881541 | 0.26329804 | -1.8539981 | 0.06373943 | 0.58463007 |
| LRIF1 | 8.85643186 | 0.21346706 | 0.63670207 | 0.33526993 | 0.73742147 | NA |
| DRAM2 | 33.4489707 | -0.0467407 | 0.37411784 | -0.1249357 | 0.90057448 | 0.99194856 |
| CEPT1 | 41.3114498 | 0.04128159 | 0.39810677 | 0.10369476 | 0.91741158 | 0.99498068 |
| DENND2D | 19.6020236 | -0.0055205 | 0.48471547 | -0.0113891 | 0.99091304 | 0.99827179 |
| ATP5PB | 18.9895213 | 0.62787386 | 0.44044078 | 1.42555794 | 0.15399598 | 0.74480157 |
| C1orf162 | 27.3834325 | 0.76703654 | 0.78903897 | 0.9721149 | 0.33099339 | 0.88037558 |
| RAP1A | 177.7917 | 0.14416253 | 0.26102563 | 0.55229262 | 0.58074788 | 0.95415437 |
| INKA2 | 14.5838321 | -0.1177058 | 0.75047249 | -0.1568423 | 0.87536915 | 0.98939075 |
| CTTNBP2NL | 27.7339451 | 0.91100163 | 0.51737447 | 1.76081675 | 0.07826942 | 0.62756088 |
| ST7L | 17.325787 | 0.33647106 | 0.5548368 | 0.60643248 | 0.54422761 | 0.95415437 |
| CAPZA1 | 100.002221 | -0.074349 | 0.28946703 | -0.256848 | 0.79729609 | 0.98320463 |
| RHOC | 16.6232085 | 0.2704607 | 0.68594648 | 0.39428835 | 0.69336816 | 0.9685713 |
| SLC16A1-AS1 | 26.04322 | -0.385833 | 0.45102109 | -0.8554655 | 0.39229351 | 0.90910959 |
| LRIG2 | 23.4718945 | -0.250756 | 0.39165879 | -0.6402411 | 0.52201589 | 0.94895469 |
| MAGI3 | 26.8498069 | 0.64689124 | 0.66811299 | 0.96823628 | 0.33292638 | 0.88047896 |
| PHTF1 | 20.1704053 | -0.7294282 | 0.50108336 | -1.4557024 | 0.14547491 | 0.73251288 |
| RSBN1 | 28.1922591 | 0.25876149 | 0.36503625 | 0.70886521 | 0.47840813 | 0.9318939 |
| PTPN22 | 79.5270741 | -0.5789787 | 0.39605831 | -1.4618521 | 0.14378174 | 0.73044824 |
| HIPK1 | 42.8619748 | 0.27234306 | 0.34685834 | 0.78517084 | 0.43235341 | 0.92284736 |
| TRIM33 | 76.7976247 | 0.20260208 | 0.27049373 | 0.74900843 | 0.45385213 | 0.9314919 |
| CSDE1 | 60.2162431 | -0.2896287 | 0.28158467 | -1.028567 | 0.3036832 | 0.86514059 |
| SLC22A15 | 21.9449292 | 0.83231215 | 0.47162189 | 1.76478692 | 0.07759958 | 0.62691401 |
| ATP1A1 | 77.6144127 | 0.17044166 | 0.42693098 | 0.39922534 | 0.68972718 | 0.9673431 |
| CD58 | 50.2377351 | 0.30047961 | 0.45447009 | 0.66116476 | 0.50850666 | 0.94305491 |
| CD2 | 44.6038332 | 0.89244035 | 0.47909831 | 1.86274995 | 0.06249745 | 0.58030355 |
| TTF2 | 11.6249283 | 0.18878012 | 0.55074137 | 0.34277454 | 0.73176808 | NA |
| MAN1A2 | 71.3540585 | -0.4371202 | 0.33591186 | -1.3012942 | 0.19315776 | 0.78135737 |
| TENT5C | 25.309316 | 0.32637656 | 1.11076047 | 0.29383163 | 0.76888656 | 0.98264092 |
| GDAP2 | 26.9045058 | -0.3095266 | 0.37216639 | -0.8316888 | 0.40558463 | 0.91575086 |
| AL359915.2 | 16.5355015 | 0.39783378 | 0.75781318 | 0.52497606 | 0.59959983 | 0.95624942 |
| NOTCH2 | 75.2832814 | -0.1968377 | 0.37216997 | -0.528892 | 0.5968804 | 0.95538391 |
| SEC22B | 23.712585 | 0.41102327 | 0.46486085 | 0.88418561 | 0.37659602 | 0.90433844 |
| NBPF26 | 13.9685371 | 0.50127816 | 0.78449358 | 0.63898312 | 0.52283391 | 0.94895469 |
| LINC00623 | 69.6192033 | 0.72662664 | 0.41025169 | 1.77117283 | 0.07653197 | 0.62467241 |
| SRGAP2C | 30.3148762 | -0.2362546 | 0.43740112 | -0.5401326 | 0.58910556 | 0.95415437 |
| NBPF15 | 8.41450811 | -0.2422708 | 0.59380167 | -0.4079995 | 0.68327407 | NA |
| SRGAP2B | 39.3215351 | 0.31736747 | 0.56625071 | 0.56047165 | 0.57515777 | 0.95415437 |
| RNF115 | 75.7546089 | 0.02641407 | 0.34242446 | 0.07713839 | 0.93851346 | 0.99514836 |
| POLR3C | 11.2410383 | -0.256502 | 0.56022617 | -0.4578544 | 0.64705704 | NA |
| RBM8A | 12.9617225 | 0.03865309 | 0.57624349 | 0.06707771 | 0.94651984 | 0.99514836 |
| LIX1L | 9.91034744 | 0.23642627 | 0.6511017 | 0.36311726 | 0.71651729 | NA |
| TXNIP | 199.797611 | 1.22867748 | 0.36978775 | 3.32265595 | 0.00089165 | 0.06269654 |

| | | | | | | |
|---|---|---|---|---|---|---|
| NBPF12 | 15.0990964 | 0.14838441 | 0.51084538 | 0.29046834 | 0.77145797 | 0.98264092 |
| AC242426.2 | 13.1591287 | -0.292354 | 0.59435627 | -0.4918835 | 0.62280172 | 0.95783378 |
| CHD1L | 27.4851698 | 0.01398396 | 0.34717559 | 0.04027923 | 0.96787051 | 0.99653147 |
| GPR89B | 10.9994514 | -0.1773785 | 0.52623408 | -0.3370715 | 0.73606301 | NA |
| NBPF11 | 11.9038419 | -0.2421699 | 0.64364012 | -0.3762505 | 0.70673071 | NA |
| NBPF14 | 17.3913997 | 0.38505875 | 0.55292525 | 0.6964029 | 0.48617655 | 0.93459582 |
| PDE4DIP | 69.3844882 | -1.6221788 | 0.49840688 | -3.2547279 | 0.00113501 | 0.07001544 |
| NBPF9 | 9.46016948 | -0.7165435 | 0.5617545 | -1.2755457 | 0.20211617 | NA |
| AC245297.3 | 87.2727329 | 0.81400893 | 0.36851072 | 2.20891521 | 0.02718054 | 0.40712515 |
| NBPF19 | 34.423779 | 0.99811892 | 0.88467947 | 1.12822661 | 0.25922423 | 0.84484505 |
| OTUD7B | 9.91737349 | -0.4858168 | 0.60062948 | -0.8088461 | 0.41860369 | NA |
| VPS45 | 39.0830203 | -0.0690959 | 0.30929291 | -0.2233996 | 0.82322452 | 0.98320463 |
| PLEKHO1 | 41.0059221 | -0.3951688 | 0.55158482 | -0.7164243 | 0.47372941 | 0.9314919 |
| ANP32E | 17.0464884 | -1.1908683 | 0.52762354 | -2.2570417 | 0.02400547 | 0.39492781 |
| APH1A | 12.1574911 | -0.3533455 | 0.55888903 | -0.6322283 | 0.52723769 | NA |
| MRPS21 | 16.5924927 | -0.415312 | 0.45965114 | -0.9035374 | 0.36624072 | 0.89868152 |
| PRPF3 | 22.5845281 | -0.0144101 | 0.43099637 | -0.0334343 | 0.97332827 | 0.9973754 |
| RPRD2 | 62.9831565 | -0.0502793 | 0.29840912 | -0.1684912 | 0.86619685 | 0.98751288 |
| MCL1 | 73.4687545 | 0.42971721 | 0.62611943 | 0.68631828 | 0.49251243 | 0.93678212 |
| ENSA | 38.838521 | 0.47221723 | 0.33227578 | 1.42116055 | 0.15527009 | 0.74486619 |
| GOLPH3L | 18.0959273 | -0.0598038 | 0.58180648 | -0.1027898 | 0.91812981 | 0.99498068 |
| CTSS | 149.764976 | 0.66229455 | 0.65054209 | 1.01806564 | 0.30864676 | 0.8673137 |
| ARNT | 34.3444922 | 0.14740452 | 0.3771733 | 0.39081377 | 0.69593489 | 0.9685713 |
| SETDB1 | 22.7668626 | 0.14871037 | 0.44378774 | 0.33509346 | 0.73755458 | 0.97605445 |
| CERS2 | 12.092786 | 0.10466822 | 0.52538081 | 0.19922353 | 0.84208789 | NA |
| C1orf56 | 10.6391025 | 0.66418934 | 0.90150856 | 0.73675322 | 0.46127243 | NA |
| CDC42SE1 | 34.7661616 | 0.19762529 | 0.39447176 | 0.50098717 | 0.61638015 | 0.95688177 |
| GABPB2 | 16.6294441 | -0.3763139 | 0.52172423 | -0.7212888 | 0.47073184 | 0.9314919 |
| PIP5K1A | 48.1725397 | -0.3164163 | 0.34981045 | -0.9045365 | 0.36571101 | 0.89868152 |
| PSMD4 | 17.6177014 | -0.599186 | 0.46517747 | -1.2880805 | 0.19771793 | 0.78142991 |
| PI4KB | 20.9803003 | -0.2682938 | 0.43110876 | -0.6223344 | 0.53372198 | 0.95293563 |
| PSMB4 | 9.36658087 | 0.06571045 | 0.61340401 | 0.10712425 | 0.91469041 | NA |
| POGZ | 59.7932947 | -0.2409771 | 0.31978912 | -0.7535501 | 0.4511194 | 0.92914597 |
| SNX27 | 45.3788171 | 0.02791301 | 0.47338362 | 0.05896489 | 0.95298007 | 0.99514836 |
| THEM4 | 11.1804261 | 0.31909036 | 0.61457193 | 0.51920751 | 0.60361604 | NA |
| S100A10 | 79.2212199 | -0.6574884 | 0.49201899 | -1.3363069 | 0.18144898 | 0.77059034 |
| S100A11 | 131.448577 | -0.1452825 | 0.59639125 | -0.2436027 | 0.80753855 | 0.98320463 |
| S100A9 | 462.948593 | 1.41608944 | 1.62655562 | 0.87060622 | 0.3839692 | 0.90910959 |
| S100A8 | 325.902216 | 0.60579154 | 1.51738078 | 0.39923501 | 0.68972005 | 0.9673431 |
| S100A6 | 146.641257 | -0.5241563 | 0.61314052 | -0.8548714 | 0.39262232 | 0.90910959 |
| S100A4 | 154.746597 | -0.0563366 | 1.00649521 | -0.0559731 | 0.95536325 | 0.99514836 |
| CHTOP | 13.7091203 | -0.5313353 | 0.47204666 | -1.1255992 | 0.26033523 | 0.84597484 |
| ILF2 | 11.3335401 | 0.5252154 | 0.59156606 | 0.88783896 | 0.37462738 | NA |
| INTS3 | 13.3843672 | -0.1284698 | 0.53758291 | -0.2389768 | 0.81112358 | 0.98320463 |
| GATAD2B | 101.722809 | -0.2799235 | 0.23571269 | -1.1875623 | 0.23500591 | 0.82903329 |
| DENND4B | 33.9727385 | -0.8567792 | 0.37497008 | -2.2849267 | 0.02231713 | 0.38788767 |
| CRTC2 | 10.2047546 | -0.798254 | 0.58187802 | -1.371858 | 0.17010764 | NA |
| JTB | 16.499584 | -0.1518404 | 0.58838024 | -0.2580651 | 0.79635668 | 0.98320463 |
| RPS27 | 32.8387773 | -0.047167 | 0.57867415 | -0.0815087 | 0.93503742 | 0.99514836 |
| TPM3 | 69.2238753 | 0.10402981 | 0.4032325 | 0.25798965 | 0.79641489 | 0.98320463 |
| C1orf43 | 19.3114407 | 0.37681565 | 0.4457125 | 0.84542312 | 0.39787464 | 0.91429865 |
| UBAP2L | 28.6676992 | -0.0601253 | 0.41128454 | -0.1461892 | 0.88377207 | 0.99019008 |
| ATP8B2 | 12.8276511 | 0.65415005 | 0.53932336 | 1.21290879 | 0.22516469 | NA |
| IL6R | 30.1495788 | 0.43453956 | 0.46744083 | 0.92961405 | 0.35257095 | 0.89109867 |

| | | | | | | |
|---|---|---|---|---|---|---|
| ADAR | 53.7690686 | -0.2202335 | 0.3924722 | -0.5611443 | 0.5746992 | 0.95415437 |
| PBXIP1 | 18.4619639 | -0.6525741 | 0.51697848 | -1.2622849 | 0.20684629 | 0.79234878 |
| ZBTB7B | 8.58476604 | 1.80558481 | 0.794712 | 2.27199891 | 0.02308658 | NA |
| SCAMP3 | 10.8032038 | -0.1554771 | 0.56297865 | -0.2761688 | 0.78241844 | NA |
| FDPS | 21.8819231 | 0.11742415 | 0.4391464 | 0.26739181 | 0.7891675 | 0.98320463 |
| ASH1L | 191.864082 | -0.007109 | 0.20453223 | -0.0347576 | 0.97227305 | 0.99734828 |
| YY1AP1 | 32.3960039 | -0.24853 | 0.36684415 | -0.6774812 | 0.49810071 | 0.93916044 |
| DAP3 | 35.2872422 | 0.44764379 | 0.34817473 | 1.28568716 | 0.19855226 | 0.78142991 |
| GON4L | 42.298198 | -0.7290783 | 0.38257572 | -1.9057098 | 0.05668787 | 0.56510781 |
| RIT1 | 10.7190144 | 0.3690828 | 0.61904824 | 0.59621009 | 0.55103489 | NA |
| KHDC4 | 23.000616 | -0.1380007 | 0.41772135 | -0.3303653 | 0.74112392 | 0.97605445 |
| ARHGEF2 | 42.304744 | -0.1951143 | 0.5096112 | -0.382869 | 0.70181692 | 0.97078484 |
| SSR2 | 26.2145649 | -0.1633451 | 0.5333422 | -0.306267 | 0.75940138 | 0.9808103 |
| LMNA | 26.1217897 | -0.7746326 | 1.01057259 | -0.7665284 | 0.44336196 | 0.92899684 |
| SEMA4A | 12.9832295 | 0.24369152 | 0.61881816 | 0.39380149 | 0.6937276 | 0.9685713 |
| SMG5 | 16.6987225 | 0.09185116 | 0.50383314 | 0.18230471 | 0.8553436 | 0.98751288 |
| CCT3 | 16.0504409 | 0.54130346 | 0.53983539 | 1.00271948 | 0.31599623 | 0.87263466 |
| MEF2D | 29.3775111 | -0.3354607 | 0.35610591 | -0.942025 | 0.34617984 | 0.88774757 |
| HDGF | 12.9938265 | 0.24346713 | 0.67426284 | 0.36108639 | 0.71803487 | 0.97348719 |
| PRCC | 12.7093114 | 0.13129535 | 0.54822509 | 0.23949168 | 0.81072435 | NA |
| SH2D2A | 13.9872094 | 0.02276175 | 0.75888062 | 0.02999385 | 0.97607196 | 0.99752211 |
| ARHGEF11 | 21.6334036 | 0.29481218 | 0.4419768 | 0.66703089 | 0.5047524 | 0.94166217 |
| FCRL5 | 35.358338 | -3.4036583 | 0.92325713 | -3.6865768 | 0.00022729 | 0.02990804 |
| MNDA | 51.7258839 | 0.60666021 | 0.44725671 | 1.35640272 | 0.17497107 | 0.76214163 |
| PYHIN1 | 49.7107203 | -0.3434479 | 0.31687129 | -1.0838719 | 0.2784216 | 0.84994492 |
| IFI16 | 126.562046 | -0.3794804 | 0.35395981 | -1.0721003 | 0.28367496 | 0.85176121 |
| DUSP23 | 13.1012391 | 0.14035301 | 0.65761643 | 0.21342685 | 0.83099404 | 0.98320463 |
| TAGLN2 | 34.8991108 | -0.9575333 | 0.60945456 | -1.5711315 | 0.1161521 | 0.70122655 |
| COPA | 70.7097448 | -0.1700097 | 0.31202943 | -0.5448513 | 0.58585577 | 0.95415437 |
| NCSTN | 9.216499 | 0.52285748 | 0.66185638 | 0.78998631 | 0.42953576 | NA |
| SLAMF6 | 35.2712495 | -0.7015679 | 0.46071057 | -1.5227953 | 0.12780993 | 0.72782081 |
| CD84 | 35.4957016 | -0.0211273 | 0.49684699 | -0.0425227 | 0.96608203 | 0.99653147 |
| SLAMF1 | 14.3538544 | 0.56353194 | 0.63384312 | 0.88907164 | 0.37396458 | 0.90262385 |
| CD48 | 40.6779828 | 0.39785406 | 0.60523074 | 0.6573593 | 0.51094991 | 0.94382295 |
| SLAMF7 | 20.7104603 | -0.2362907 | 0.50229593 | -0.4704214 | 0.638054 | 0.95933464 |
| LY9 | 15.3439708 | 0.37745078 | 0.78415341 | 0.48134814 | 0.63026909 | 0.95877428 |
| CD244 | 13.6369205 | 1.78141859 | 0.83843338 | 2.12469904 | 0.03361173 | 0.45147411 |
| ARHGAP30 | 53.6729412 | -0.0868154 | 0.3765003 | -0.2305851 | 0.81763715 | 0.98320463 |
| PFDN2 | 14.3145589 | -0.2902916 | 0.52926936 | -0.5484762 | 0.58336497 | 0.95415437 |
| NIT1 | 11.7850405 | -0.0178337 | 0.49477378 | -0.0360442 | 0.9712471 | NA |
| UFC1 | 11.3066063 | -0.5975497 | 0.55074803 | -1.0849784 | 0.27793121 | NA |
| NDUFS2 | 11.2320292 | -0.0750052 | 0.50690145 | -0.1479679 | 0.88236807 | NA |
| FCER1G | 54.4077935 | 1.92561088 | 0.88097207 | 2.18577971 | 0.02883173 | 0.42210257 |
| SDHC | 20.3215381 | -0.4446639 | 0.38995876 | -1.1402845 | 0.25416779 | 0.84026145 |
| FCGR2A | 60.3551218 | 1.4433097 | 0.6479733 | 2.22742157 | 0.02591911 | 0.40178254 |
| HSPA6 | 9.83682219 | 1.32332701 | 0.87635649 | 1.51003276 | 0.13103506 | NA |
| FCGR3A | 68.5003436 | 2.40249282 | 0.87978768 | 2.73076432 | 0.00631876 | 0.19738118 |
| FCGR2B | 15.717372 | 0.21169535 | 0.58630217 | 0.36106869 | 0.7180481 | 0.97348719 |
| ATF6 | 93.7361119 | 0.40092794 | 0.36326999 | 1.10366382 | 0.26973899 | 0.84994492 |
| OLFML2B | 13.582608 | 1.91747937 | 0.82386689 | 2.32741403 | 0.01994324 | 0.36765113 |
| UHMK1 | 32.1512869 | 0.1533565 | 0.4032598 | 0.38029206 | 0.70372863 | 0.9714177 |
| MGST3 | 26.9367609 | 0.53002676 | 0.43946396 | 1.20607561 | 0.22778834 | 0.82176054 |
| ALDH9A1 | 11.8325544 | 1.09681441 | 0.67139275 | 1.63364053 | 0.10233435 | NA |
| TMCO1 | 20.1976856 | 0.7297939 | 0.44077641 | 1.65570089 | 0.09778241 | 0.66447623 |

| | | | | | | |
|---|---|---|---|---|---|---|
| UCK2 | 10.318952 | 1.03381549 | 0.7010008 | 1.47477076 | 0.14027418 | NA |
| POGK | 8.14302277 | 0.48092224 | 0.61849939 | 0.777563 | 0.43682668 | NA |
| POU2F1 | 82.383964 | 0.76076235 | 0.40300213 | 1.8877378 | 0.05906116 | 0.57281427 |
| CD247 | 150.637198 | 0.90193158 | 0.40841947 | 2.20834619 | 0.02722015 | 0.40712515 |
| CREG1 | 19.46721 | 0.4468856 | 0.50426932 | 0.88620424 | 0.37550748 | 0.90341205 |
| RCSD1 | 121.668491 | -0.342035 | 0.38805483 | -0.881409 | 0.37809648 | 0.9051267 |
| MPZL1 | 23.2757647 | -0.0606796 | 0.38375142 | -0.1581221 | 0.87436056 | 0.98899412 |
| MPC2 | 9.45497328 | 1.11308037 | 0.69101078 | 1.61080029 | 0.10722326 | NA |
| DCAF6 | 70.4345034 | 0.02606105 | 0.31360978 | 0.08310024 | 0.93377183 | 0.99514836 |
| TIPRL | 13.2787907 | 0.45935453 | 0.47014185 | 0.97705518 | 0.32854183 | 0.87918303 |
| SFT2D2 | 39.5446155 | -0.4361559 | 0.3746186 | -1.1642666 | 0.24431599 | 0.83666131 |
| TBX19 | 10.1690919 | -0.7027106 | 0.56861055 | -1.2358381 | 0.21651875 | NA |
| ATP1B1 | 39.9468215 | 0.75934327 | 0.50230569 | 1.51171546 | 0.13060626 | 0.72931666 |
| NME7 | 21.3899561 | 0.27584283 | 0.47568404 | 0.57988667 | 0.56199105 | 0.95415437 |
| BLZF1 | 14.5501142 | 1.93494109 | 0.61438782 | 3.14938062 | 0.00163617 | 0.08877021 |
| C1orf112 | 11.6168412 | -0.8387156 | 0.51900377 | -1.6160106 | 0.10609202 | NA |
| SELL | 20.0182313 | -0.3186084 | 0.63195321 | -0.5041645 | 0.61414577 | 0.95688177 |
| SCYL3 | 10.548279 | 0.04024379 | 0.62915913 | 0.06396441 | 0.94899857 | NA |
| KIFAP3 | 29.610281 | -0.3360217 | 0.41247722 | -0.814643 | 0.41527671 | 0.91894074 |
| PRRC2C | 129.244309 | -0.3214286 | 0.24269717 | -1.324402 | 0.18536955 | 0.77694741 |
| VAMP4 | 14.4938964 | 0.03045214 | 0.58205532 | 0.0523183 | 0.95827507 | 0.99514836 |
| SUCO | 47.0208633 | -0.1540079 | 0.38358696 | -0.4014941 | 0.68805641 | 0.9673431 |
| AL645568.1 | 28.7032377 | 0.76613862 | 0.42578092 | 1.79937281 | 0.07195973 | 0.61092445 |
| PRDX6 | 22.2907805 | 0.53181025 | 0.44241276 | 1.20206805 | 0.22933716 | 0.82393202 |
| KLHL20 | 23.353551 | 0.34051698 | 0.46606824 | 0.73061614 | 0.46501365 | 0.9314919 |
| ZBTB37 | 15.4879349 | -0.2203729 | 0.4666793 | -0.4722149 | 0.63677339 | 0.95933464 |
| RC3H1 | 80.6892404 | -0.0321518 | 0.30392293 | -0.1057894 | 0.91574947 | 0.99498068 |
| RABGAP1L | 456.730322 | 0.02798281 | 0.25859036 | 0.10821288 | 0.91382683 | 0.99498068 |
| CACYBP | 10.6377599 | 0.17642722 | 0.58591912 | 0.30111191 | 0.76332916 | NA |
| KIAA0040 | 11.1451201 | -0.8023805 | 0.61133453 | -1.3125064 | 0.18934933 | NA |
| COP1 | 210.885155 | 0.08650104 | 0.25340829 | 0.34135049 | 0.73283975 | 0.97605445 |
| RASAL2 | 50.7754584 | -0.8791894 | 0.57456611 | -1.5301797 | 0.12597225 | 0.72092082 |
| RALGPS2 | 92.1859446 | -1.7396788 | 0.65743848 | -2.6461469 | 0.00814144 | 0.23588876 |
| FAM20B | 13.2680424 | 0.37655167 | 0.60474204 | 0.62266495 | 0.53350472 | 0.95293563 |
| TOR3A | 8.64443457 | 0.12870081 | 0.64030603 | 0.2009989 | 0.84069943 | NA |
| ABL2 | 41.8357339 | 0.3697196 | 0.36811988 | 1.00434563 | 0.31521205 | 0.87263466 |
| SOAT1 | 47.5916331 | 0.77471021 | 0.39174175 | 1.97760443 | 0.04797335 | 0.53331354 |
| TOR1AIP2 | 24.0167721 | -0.3506835 | 0.39868734 | -0.8795952 | 0.37907866 | 0.90579504 |
| TOR1AIP1 | 20.8912269 | -0.2581285 | 0.41448614 | -0.6227675 | 0.53343735 | 0.95293563 |
| CEP350 | 107.097683 | 0.19516207 | 0.26449941 | 0.73785446 | 0.46060289 | 0.9314919 |
| QSOX1 | 15.7462381 | 1.06535669 | 0.48901562 | 2.17857394 | 0.02936333 | 0.42316846 |
| ACBD6 | 57.5197268 | -0.1945064 | 0.30855266 | -0.6303832 | 0.5284439 | 0.95182065 |
| XPR1 | 67.678346 | -0.0823176 | 0.36406114 | -0.2261092 | 0.82111649 | 0.98320463 |
| STX6 | 20.7638784 | 0.38963643 | 0.49698314 | 0.78400331 | 0.43303817 | 0.92284736 |
| MR1 | 16.7694558 | 0.80576792 | 0.54570319 | 1.4765681 | 0.13979145 | 0.73044824 |
| IER5 | 13.2484561 | -0.3982934 | 0.81106687 | -0.4910734 | 0.62337451 | 0.95799904 |
| GLUL | 73.2220782 | 0.67684724 | 0.6237873 | 1.08506096 | 0.27789466 | 0.84994492 |
| NPL | 51.1514535 | 0.75289714 | 0.51198182 | 1.47055444 | 0.14141165 | 0.73044824 |
| DHX9 | 46.3631762 | -0.2838394 | 0.33583907 | -0.8451648 | 0.39801883 | 0.91429865 |
| LAMC1 | 13.1046455 | -0.6707935 | 0.79393409 | -0.8448983 | 0.39816764 | 0.91429865 |
| SMG7 | 64.6215456 | 0.01873627 | 0.28881703 | 0.06487244 | 0.94827556 | 0.99514836 |
| NCF2 | 35.8603452 | 0.89513653 | 0.65880811 | 1.35872117 | 0.17423496 | 0.76214163 |
| ARPC5 | 62.3212024 | 0.09342192 | 0.51448813 | 0.18158227 | 0.85591056 | 0.98751288 |
| RGL1 | 93.0529153 | 0.65300587 | 0.75250472 | 0.86777645 | 0.38551673 | 0.90910959 |

| | | | | | | |
|---|---|---|---|---|---|---|
| TSEN15 | 13.3904689 | -0.6221111 | 0.48192833 | -1.2908789 | 0.19674568 | 0.78135737 |
| C1orf21 | 41.2763618 | 0.21844702 | 0.65362031 | 0.33421089 | 0.73822043 | 0.97605445 |
| EDEM3 | 44.5751925 | 0.09177039 | 0.38060461 | 0.2411174 | 0.80946413 | 0.98320463 |
| FAM129A | 105.450823 | 0.95501259 | 0.42672677 | 2.23799551 | 0.02522135 | 0.39878554 |
| RNF2 | 18.0914179 | 0.4975246 | 0.43868908 | 1.13411667 | 0.25674561 | 0.84072794 |
| TRMT1L | 14.4230697 | 1.24020104 | 0.54454489 | 2.27750009 | 0.02275638 | 0.38988793 |
| SWT1 | 13.1165504 | 0.44798757 | 0.55227272 | 0.81117093 | 0.41726752 | 0.91954684 |
| IVNS1ABP | 59.9052599 | -0.2044651 | 0.45769167 | -0.446731 | 0.65506928 | 0.96178254 |
| TPR | 45.6607506 | -0.1286827 | 0.30348407 | -0.4240179 | 0.67155276 | 0.96412305 |
| ODR4 | 21.5661089 | 0.22094448 | 0.41679927 | 0.53009805 | 0.59604395 | 0.9553365 |
| PLA2G4A | 25.3067183 | 1.16053204 | 0.50560656 | 2.29532633 | 0.02171443 | 0.38269953 |
| RGS18 | 16.0449719 | 0.64275273 | 0.83417825 | 0.77052204 | 0.44099029 | 0.92898373 |
| AL390957.1 | 20.9867331 | -2.3747575 | 0.48027318 | -4.9445974 | 7.63E-07 | 0.00070375 |
| RGS1 | 67.0903263 | -0.2454174 | 0.59103742 | -0.4152316 | 0.67797238 | 0.96718429 |
| AL136454.1 | 31.1904517 | -0.2878524 | 0.60176928 | -0.4783435 | 0.63240576 | 0.95877428 |
| RGS2 | 27.284564 | 0.93999731 | 0.921392 | 1.02019261 | 0.30763712 | 0.8673137 |
| UCHL5 | 17.397712 | -0.4630226 | 0.50190605 | -0.9225284 | 0.356253 | 0.89503727 |
| TROVE2 | 30.0201701 | -0.1316012 | 0.37932811 | -0.3469324 | 0.72864212 | 0.97605445 |
| CDC73 | 74.3544382 | -0.250023 | 0.27422633 | -0.9117395 | 0.36190589 | 0.89853007 |
| ZBTB41 | 13.9401568 | 0.532853 | 0.66276057 | 0.80399019 | 0.42140264 | 0.91985469 |
| DENND1B | 108.270864 | 0.46816849 | 0.43521463 | 1.07571864 | 0.28205311 | 0.85171619 |
| NEK7 | 125.321133 | 0.2260076 | 0.32946242 | 0.68598902 | 0.49272003 | 0.93678212 |
| PTPRC | 704.641993 | 0.3181153 | 0.35968353 | 0.88443111 | 0.37646354 | 0.90433844 |
| AL157402.2 | 12.259415 | -0.0917315 | 0.54797107 | -0.167402 | 0.86705371 | NA |
| MIR181A1HG | 90.6101158 | 2.34252876 | 0.73606073 | 3.18252104 | 0.00145999 | 0.0807958 |
| CAMSAP2 | 23.9261732 | -0.2730171 | 0.54540013 | -0.5005812 | 0.61666588 | 0.95688177 |
| KIF21B | 11.6336881 | -1.5436073 | 0.50517518 | -3.0555881 | 0.0022462 | NA |
| NAV1 | 29.0145351 | 0.27670291 | 0.74273678 | 0.37254505 | 0.70948708 | 0.97211298 |
| IPO9 | 28.691699 | -0.5650062 | 0.39423894 | -1.4331567 | 0.15181305 | 0.74282354 |
| RNPEP | 18.8981826 | -0.3825668 | 0.63058854 | -0.6066822 | 0.54406186 | 0.95415437 |
| ARL8A | 14.246451 | -0.1202624 | 0.52447396 | -0.2293009 | 0.81863505 | 0.98320463 |
| PTPN7 | 21.705755 | 0.06754961 | 0.47240015 | 0.14299236 | 0.88629621 | 0.99019008 |
| PPP1R12B | 58.1510543 | 0.44041771 | 0.29506258 | 1.49262477 | 0.13553544 | 0.73044824 |
| KDM5B | 39.6755908 | 0.88485424 | 0.41223039 | 2.14650414 | 0.03183278 | 0.43969912 |
| KLHL12 | 9.2706419 | -0.313728 | 0.61038281 | -0.5139856 | 0.60726203 | NA |
| ADIPOR1 | 11.5994636 | 0.59237746 | 0.65653581 | 0.90227746 | 0.3669095 | NA |
| BTG2 | 37.6635481 | -1.020823 | 0.63851252 | -1.5987517 | 0.10987579 | 0.68862129 |
| ATP2B4 | 54.3876905 | 0.7927003 | 0.34798756 | 2.27795588 | 0.02272921 | 0.38988793 |
| PLEKHA6 | 10.5839847 | 0.14835983 | 0.71378658 | 0.207849 | 0.83534687 | NA |
| PPP1R15B | 11.0166404 | -0.5050791 | 0.60306644 | -0.8375182 | 0.40230136 | NA |
| PIK3C2B | 12.4247239 | -0.998961 | 0.48872155 | -2.044029 | 0.04095068 | NA |
| MDM4 | 63.9194448 | -0.3882054 | 0.28507781 | -1.3617523 | 0.17327607 | 0.76214163 |
| RBBP5 | 12.0194348 | 0.08834819 | 0.54143937 | 0.16317283 | 0.87038236 | NA |
| DSTYK | 18.08037 | 1.13149139 | 0.46215511 | 2.44829358 | 0.01435346 | 0.32121732 |
| ELK4 | 28.2159733 | -0.4333322 | 0.36634899 | -1.1828399 | 0.23687262 | 0.83171006 |
| NUCKS1 | 54.9860585 | -0.4536021 | 0.36674593 | -1.2368292 | 0.21615051 | 0.80634897 |
| RAB29 | 23.0775934 | -0.8425827 | 0.55683853 | -1.5131545 | 0.13024043 | 0.72931666 |
| RHEX | 38.9167613 | -1.0132816 | 1.16831849 | -0.8672991 | 0.38577816 | 0.90910959 |
| SRGAP2 | 71.9567099 | 0.13894788 | 0.38555198 | 0.3603869 | 0.71855782 | 0.97348719 |
| IKBKE | 12.4560469 | 1.29263665 | 0.551778 | 2.34267521 | 0.01914604 | NA |
| RASSF5 | 35.2497902 | 0.1645005 | 0.40241801 | 0.40878015 | 0.68270101 | 0.9673431 |
| MAPKAPK2 | 38.0372607 | -0.1328633 | 0.31955428 | -0.415777 | 0.6775732 | 0.96718429 |
| FCMR | 22.1290258 | -2.1170744 | 0.7926388 | -2.6709195 | 0.00756438 | 0.22148821 |
| PIGR | 12.7329455 | -0.2533794 | 0.74253294 | -0.3412366 | 0.73292546 | NA |

| | | | | | | |
|---|---|---|---|---|---|---|
| CD55 | 82.0651248 | 0.31276292 | 0.50778352 | 0.61593753 | 0.53793575 | 0.95415437 |
| CR1 | 54.1133399 | 0.40512826 | 0.66284445 | 0.61119657 | 0.54106945 | 0.95415437 |
| CD46 | 49.8354741 | 0.40884396 | 0.31025514 | 1.31776694 | 0.18758168 | 0.77798634 |
| MIR29B2CHG | 45.4254945 | 0.28163348 | 0.42974944 | 0.65534344 | 0.51224666 | 0.94507477 |
| TRAF3IP3 | 59.692954 | 0.74168269 | 0.39106826 | 1.8965556 | 0.05788662 | 0.57000806 |
| HHAT | 11.4749832 | 1.54777176 | 0.85819566 | 1.8035185 | 0.07130682 | NA |
| RCOR3 | 49.0217658 | 0.10778662 | 0.32619276 | 0.33043842 | 0.74106872 | 0.97605445 |
| TRAF5 | 25.5375126 | 1.23753396 | 0.55532006 | 2.22850576 | 0.02584681 | 0.40178254 |
| LPGAT1 | 61.6441412 | -0.4801338 | 0.32840245 | -1.4620287 | 0.14373334 | 0.73044824 |
| INTS7 | 19.0580137 | -0.089429 | 0.53266381 | -0.1678901 | 0.86666971 | 0.98751288 |
| PPP2R5A | 68.0988617 | 0.15625989 | 0.31596432 | 0.49454916 | 0.6209184 | 0.95688177 |
| AL360091.3 | 11.4981419 | 0.42730663 | 0.86312468 | 0.49506942 | 0.62055112 | NA |
| TMEM206 | 13.8970572 | 0.2770883 | 0.57368574 | 0.48299667 | 0.6290981 | 0.95877428 |
| NSL1 | 25.4188818 | -0.0265054 | 0.40769837 | -0.0650124 | 0.94816414 | 0.99514836 |
| TATDN3 | 16.121956 | -0.5727419 | 0.49998739 | -1.1455127 | 0.25199685 | 0.84008912 |
| FLVCR1 | 10.8211534 | 0.20775168 | 0.56891532 | 0.36517154 | 0.71498336 | NA |
| ANGEL2 | 17.6425832 | 0.1266822 | 0.45291964 | 0.27970126 | 0.77970671 | 0.98264092 |
| RPS6KC1 | 85.1358507 | 0.05431224 | 0.25477667 | 0.21317589 | 0.83118977 | 0.98320463 |
| SMYD2 | 11.0905172 | 0.48554326 | 0.56804404 | 0.85476341 | 0.39268214 | NA |
| ESRRG | 25.2876016 | -1.2261935 | 0.83617444 | -1.4664326 | 0.14253046 | 0.73044824 |
| GPATCH2 | 32.7404196 | -0.0944443 | 0.34708027 | -0.2721109 | 0.78553677 | 0.98264092 |
| LYPLAL1 | 27.3733465 | -0.4580271 | 0.47578367 | -0.9626794 | 0.33570846 | 0.88047896 |
| EPRS | 23.3123637 | 0.15929137 | 0.38995866 | 0.4084827 | 0.68291933 | 0.9673431 |
| IARS2 | 15.7064958 | -0.206807 | 0.47560177 | -0.4348322 | 0.66368424 | 0.96379377 |
| RAB3GAP2 | 58.6877926 | -0.2072029 | 0.31926218 | -0.6490054 | 0.51633489 | 0.94596835 |
| DUSP10 | 11.9707222 | 1.56675869 | 0.69058098 | 2.26875447 | 0.02328326 | NA |
| MIA3 | 23.0475799 | 0.48464847 | 0.38205268 | 1.26853834 | 0.20460577 | 0.7890511 |
| AIDA | 15.2208956 | -1.1860644 | 0.50738665 | -2.3375949 | 0.01940827 | 0.36408606 |
| BROX | 19.9231246 | 0.02918594 | 0.46853806 | 0.0622915 | 0.9503307 | 0.99514836 |
| AC093152.1 | 15.7882669 | 0.01983628 | 0.4933667 | 0.04020596 | 0.96792893 | 0.99653147 |
| TLR5 | 9.91835778 | -0.1065671 | 0.93361614 | -0.1141445 | 0.90912328 | NA |
| CAPN2 | 32.6396792 | 0.12333252 | 0.36063995 | 0.34198241 | 0.73236413 | 0.97605445 |
| TP53BP2 | 21.3779995 | -0.1326539 | 0.43838318 | -0.302598 | 0.76219626 | 0.98163473 |
| FBXO28 | 34.2572699 | 0.21892719 | 0.40127481 | 0.54557922 | 0.58535522 | 0.95415437 |
| DEGS1 | 14.4610183 | -0.07496 | 0.46049137 | -0.1627826 | 0.87068958 | 0.98751288 |
| NVL | 26.2042065 | -0.730823 | 0.42139704 | -1.734286 | 0.08286735 | 0.63740875 |
| CNIH4 | 19.5062192 | 0.89570156 | 0.47642414 | 1.88005076 | 0.06010116 | 0.57281427 |
| WDR26 | 74.0547456 | -0.1522178 | 0.38832584 | -0.3919846 | 0.69506957 | 0.9685713 |
| LBR | 38.6565489 | 0.07937419 | 0.45163641 | 0.17574798 | 0.86049194 | 0.98751288 |
| SRP9 | 18.9522016 | 0.24529259 | 0.44923046 | 0.54602839 | 0.58504642 | 0.95415437 |
| TMEM63A | 25.965832 | 0.16983793 | 0.51685188 | 0.32860078 | 0.74245746 | 0.97664835 |
| H3F3A | 24.3493317 | 0.08572145 | 0.41092052 | 0.20860835 | 0.83475399 | 0.98320463 |
| ACBD3 | 30.6536401 | 0.02682924 | 0.39712069 | 0.06755941 | 0.94613637 | 0.99514836 |
| PARP1 | 27.9723527 | -1.8677409 | 0.48066625 | -3.8857334 | 0.00010202 | 0.01821247 |
| ITPKB | 50.5907384 | -0.2933158 | 0.35478097 | -0.8267517 | 0.40837781 | 0.9169561 |
| COQ8A | 10.4894399 | 0.35950963 | 0.57334775 | 0.62703591 | 0.5306357 | NA |
| CDC42BPA | 20.71633 | 0.38916417 | 0.52348232 | 0.74341416 | 0.45723097 | 0.9314919 |
| ZNF678 | 20.4096515 | -0.7357884 | 0.48295985 | -1.5234981 | 0.12763413 | 0.72782081 |
| ARF1 | 42.0287256 | 0.11231979 | 0.37445115 | 0.29995846 | 0.76420884 | 0.98222198 |
| GUK1 | 36.4460455 | 0.06648778 | 0.39321336 | 0.1690883 | 0.86572719 | 0.98751288 |
| RAB4A | 13.4228376 | 0.2752766 | 0.49904968 | 0.5516016 | 0.58122134 | 0.95415437 |
| NUP133 | 20.1931654 | -0.0399027 | 0.40826069 | -0.0977383 | 0.92214012 | 0.99514836 |
| ABCB10 | 16.0745391 | -0.0088243 | 0.4568922 | -0.0193136 | 0.9845909 | 0.99811561 |
| GALNT2 | 50.6513628 | 0.01868407 | 0.37812204 | 0.04941281 | 0.96059032 | 0.99614757 |

| | | | | | | |
|---|---|---|---|---|---|---|
| COG2 | 14.1344357 | -0.2772316 | 0.46804922 | -0.5923129 | 0.5536411 | 0.95415437 |
| TTC13 | 43.9430184 | 0.01870785 | 0.32359155 | 0.05781315 | 0.95389746 | 0.99514836 |
| GNPAT | 21.833306 | -0.2131264 | 0.41565772 | -0.5127449 | 0.60812977 | 0.95688177 |
| EGLN1 | 32.6293951 | -0.1609135 | 0.44788379 | -0.3592752 | 0.71938924 | 0.97348719 |
| DISC1 | 100.9011 | 0.7487913 | 0.47806793 | 1.56628639 | 0.11728158 | 0.70122655 |
| SIPA1L2 | 26.6595349 | 0.55571751 | 0.57410831 | 0.96796633 | 0.33306119 | 0.88047896 |
| PCNX2 | 69.7478942 | 0.70218111 | 0.32601172 | 2.15385234 | 0.03125175 | 0.43563527 |
| TARBP1 | 17.7802788 | 0.59296627 | 0.55920888 | 1.06036632 | 0.28897798 | 0.85703857 |
| IRF2BP2 | 27.4395412 | 0.65617283 | 0.37275652 | 1.76032554 | 0.07835263 | 0.62756088 |
| TOMM20 | 17.4479246 | -0.6410808 | 0.56336921 | -1.1379408 | 0.25514518 | 0.8406233 |
| ARID4B | 188.119833 | -0.0910422 | 0.27901045 | -0.326304 | 0.74419436 | 0.97805001 |
| B3GALNT2 | 13.5008116 | 1.01697773 | 0.61276531 | 1.65965291 | 0.0969843 | 0.66178928 |
| LYST | 245.689259 | 0.44711341 | 0.40306455 | 1.10928488 | 0.26730731 | 0.84994492 |
| GPR137B | 35.7435053 | -0.0378421 | 0.46213991 | -0.0818846 | 0.9347385 | 0.99514836 |
| ERO1B | 28.7183054 | 0.47656198 | 0.48055985 | 0.99168081 | 0.32135325 | 0.87363761 |
| LGALS8 | 22.056132 | -0.0287252 | 0.43991916 | -0.0652965 | 0.94793793 | 0.99514836 |
| HEATR1 | 16.6569832 | -0.453749 | 0.53883186 | -0.8420975 | 0.39973337 | 0.91508218 |
| MTR | 42.1674077 | 0.27876993 | 0.39704334 | 0.70211461 | 0.48260769 | 0.93382901 |
| KMO | 22.7551496 | -1.0469208 | 0.57397134 | -1.8239949 | 0.06815285 | 0.59676875 |
| CEP170 | 112.607101 | -0.2470251 | 0.40886443 | -0.6041735 | 0.54572828 | 0.95415437 |
| SDCCAG8 | 111.546104 | 0.16555059 | 0.46002479 | 0.35987319 | 0.71894196 | 0.97348719 |
| AKT3 | 180.890269 | -0.8879134 | 0.4712146 | -1.8843079 | 0.05952335 | 0.57281427 |
| ZBTB18 | 11.8980102 | -0.0937069 | 0.59989795 | -0.1562047 | 0.87587168 | NA |
| ADSS | 26.184058 | -0.3239057 | 0.46455666 | -0.6972362 | 0.48565499 | 0.93417265 |
| DESI2 | 20.233335 | -0.9321724 | 0.42159793 | -2.2110459 | 0.02703266 | 0.40712515 |
| COX20 | 15.840534 | -0.0983504 | 0.50069012 | -0.1964297 | 0.84427385 | 0.9842451 |
| HNRNPU | 111.162154 | 0.11829569 | 0.26491673 | 0.44653915 | 0.65520784 | 0.96178254 |
| EFCAB2 | 15.8255486 | 1.34068525 | 1.00829122 | 1.32966074 | 0.18363008 | 0.77688373 |
| SMYD3 | 207.020099 | 0.45844766 | 0.3849245 | 1.19100672 | 0.23365095 | 0.82817204 |
| CNST | 47.667736 | -0.0315056 | 0.51793951 | -0.0608288 | 0.95149555 | 0.99514836 |
| AHCTF1 | 48.8092753 | -1.3273281 | 0.44323486 | -2.9946383 | 0.0027477 | 0.12544207 |
| ZNF124 | 22.9034358 | -0.4353371 | 0.41093066 | -1.0593932 | 0.28942076 | 0.85760168 |
| AL390728.6 | 14.4796506 | -0.0117525 | 0.55712956 | -0.0210946 | 0.98317015 | 0.99763261 |
| NLRP3 | 12.0358788 | 1.95622431 | 1.09196216 | 1.79147628 | 0.0732169 | NA |
| SH3YL1 | 11.7755066 | 0.10104583 | 0.54279239 | 0.18615925 | 0.85231988 | NA |
| ACP1 | 9.48838957 | -0.4185068 | 0.54498631 | -0.7679216 | 0.44253376 | NA |
| EIPR1 | 22.2980055 | -0.3692256 | 0.44557977 | -0.8286409 | 0.40730766 | 0.9169561 |
| TRAPPC12 | 40.4864839 | -0.0509017 | 0.39162128 | -0.1299769 | 0.89658467 | 0.99118739 |
| ADI1 | 10.6977195 | 0.45877836 | 0.77083618 | 0.59516972 | 0.55173003 | NA |
| RPS7 | 75.011404 | -0.2920692 | 0.60271797 | -0.4845868 | 0.6279695 | 0.95877428 |
| RNF144A | 18.3890443 | 1.03809006 | 0.7164084 | 1.44901994 | 0.14733201 | 0.73693374 |
| LINC00299 | 13.2599224 | 2.38855186 | 1.03444663 | 2.3090141 | 0.0209428 | 0.37507265 |
| ID2 | 40.356566 | 1.3881181 | 0.65875483 | 2.10718469 | 0.03510157 | 0.46140643 |
| KIDINS220 | 55.4340182 | 0.11211512 | 0.31984452 | 0.35053007 | 0.72594093 | 0.9754739 |
| ITGB1BP1 | 16.288185 | -0.105295 | 0.48244718 | -0.218252 | 0.82723281 | 0.98320463 |
| CPSF3 | 8.54398977 | 0.06998199 | 0.58013377 | 0.12063077 | 0.90398349 | NA |
| IAH1 | 11.2584455 | -0.0384245 | 0.50923151 | -0.0754559 | 0.93985195 | NA |
| ADAM17 | 65.3944995 | 0.27747493 | 0.38605124 | 0.71875155 | 0.47229401 | 0.9314919 |
| YWHAQ | 35.0506888 | 0.40027809 | 0.4103548 | 0.97544392 | 0.3293401 | 0.88004256 |
| TAF1B | 26.1125193 | -0.1643006 | 0.50906984 | -0.3227467 | 0.74688707 | 0.97813973 |
| HPCAL1 | 56.0370067 | 0.56825622 | 0.33780569 | 1.68219847 | 0.09253036 | 0.65902576 |
| ODC1 | 10.3060976 | -0.6272473 | 0.61241978 | -1.0242113 | 0.30573548 | NA |
| NOL10 | 28.0390817 | -0.1842542 | 0.3829191 | -0.4811832 | 0.6303863 | 0.95877428 |
| PDIA6 | 32.677512 | -0.3977553 | 0.46007072 | -0.8645525 | 0.38728444 | 0.90910959 |

| | | | | | | |
|---|---|---|---|---|---|---|
| PQLC3 | 8.66648534 | 1.1760246 | 0.62475887 | 1.88236558 | 0.0597864 | NA |
| ROCK2 | 61.1000213 | 0.39259295 | 0.32410355 | 1.21131951 | 0.22577297 | 0.81875277 |
| LPIN1 | 32.6393329 | 0.38464961 | 0.32736953 | 1.17497069 | 0.24000644 | 0.83493679 |
| TRIB2 | 7.70341432 | -0.9473349 | 0.69007406 | -1.3728018 | 0.16981398 | NA |
| NBAS | 75.7751279 | -0.099817 | 0.28768688 | -0.3469642 | 0.72861822 | 0.97605445 |
| DDX1 | 18.305314 | -0.1006995 | 0.44060671 | -0.2285474 | 0.81922074 | 0.98320463 |
| FAM49A | 125.585437 | -0.6538922 | 0.61533377 | -1.0626627 | 0.28793494 | 0.85677513 |
| SMC6 | 33.3136004 | -0.9320734 | 0.37012049 | -2.5182973 | 0.01179237 | 0.29101961 |
| GEN1 | 14.1551861 | -1.8755811 | 0.65020112 | -2.8846168 | 0.0039189 | 0.15732988 |
| LAPTM4A | 33.4814654 | 0.93621939 | 0.44423723 | 2.10747621 | 0.03507632 | 0.46140643 |
| PUM2 | 147.508267 | 0.01340704 | 0.23935744 | 0.05601264 | 0.95533174 | 0.99514836 |
| RHOB | 18.0527503 | -0.0169283 | 0.53607895 | -0.0315779 | 0.97480866 | 0.9973754 |
| HS1BP3 | 9.08928691 | 0.81167953 | 0.57964578 | 1.40030266 | 0.16142271 | NA |
| LDAH | 28.8321353 | -0.1333666 | 0.46366843 | -0.2876336 | 0.77362723 | 0.98264092 |
| ATAD2B | 96.2328509 | -0.1182924 | 0.2703805 | -0.4375033 | 0.66174641 | 0.96298563 |
| UBXN2A | 19.7277429 | 0.16237856 | 0.54488926 | 0.29800286 | 0.76570098 | 0.98264092 |
| SF3B6 | 12.0165246 | 0.10868165 | 0.6387737 | 0.17014109 | 0.86489918 | NA |
| FAM228B | 14.597938 | 0.67406917 | 0.60329341 | 1.11731566 | 0.26385938 | 0.84928231 |
| ITSN2 | 143.905064 | -0.5360522 | 0.28586105 | -1.8752194 | 0.06076255 | 0.57281427 |
| NCOA1 | 179.688199 | 0.47862534 | 0.26956587 | 1.77554128 | 0.07580856 | 0.62329636 |
| ADCY3 | 12.6984351 | 0.56890658 | 0.50603885 | 1.12423497 | 0.26091336 | NA |
| DNMT3A | 35.2123816 | -0.182645 | 0.51728083 | -0.3530868 | 0.72402339 | 0.97487724 |
| DTNB | 57.6049565 | 0.07277514 | 0.45095831 | 0.16137886 | 0.87179502 | 0.98751288 |
| ASXL2 | 69.8327263 | -0.0910928 | 0.26569185 | -0.3428511 | 0.73171045 | 0.97605445 |
| RAB10 | 163.257672 | 0.14926757 | 0.2995353 | 0.4983305 | 0.61825112 | 0.95688177 |
| HADHA | 33.1183611 | 0.15646856 | 0.41198434 | 0.3797925 | 0.70409945 | 0.9714177 |
| HADHB | 43.4489666 | 0.75360614 | 0.33049059 | 2.28026505 | 0.02259197 | 0.38988793 |
| SELENOI | 22.2198545 | 0.05569789 | 0.42482585 | 0.13110758 | 0.8956902 | 0.99115168 |
| OST4 | 18.4227099 | -0.2640207 | 0.61330142 | -0.430491 | 0.66683852 | 0.96412305 |
| ATRAID | 8.8757926 | 0.48007351 | 0.64209929 | 0.74766241 | 0.45466381 | NA |
| GTF3C2 | 19.9132244 | -0.0631731 | 0.465127 | -0.135819 | 0.89196436 | 0.99080204 |
| PPM1G | 32.3057937 | 0.34357669 | 0.49583495 | 0.69292552 | 0.48835629 | 0.93596573 |
| NRBP1 | 15.2907519 | -0.5168033 | 0.57859455 | -0.8932045 | 0.37174765 | 0.90230329 |
| ZNF512 | 20.4988459 | -0.1396815 | 0.41395064 | -0.3374351 | 0.73578892 | 0.97605445 |
| SLC4A1AP | 7.95614281 | -0.1551871 | 0.58362025 | -0.2659042 | 0.790313 | NA |
| MRPL33 | 15.9903556 | -0.7949687 | 0.51029127 | -1.5578725 | 0.11926348 | 0.70431251 |
| RBKS | 10.6405778 | 0.75605148 | 0.78952454 | 0.95760352 | 0.33826272 | NA |
| BABAM2 | 126.99163 | 0.15824176 | 0.24750363 | 0.63935125 | 0.52259445 | 0.94895469 |
| FOSL2 | 53.5517967 | 0.28378234 | 0.73276444 | 0.38727636 | 0.69855163 | 0.96884407 |
| PLB1 | 10.730892 | 1.29857852 | 0.82876372 | 1.56688629 | 0.11714127 | NA |
| PPP1CB | 55.4158902 | 0.1373008 | 0.5728732 | 0.23967048 | 0.81058572 | 0.98320463 |
| TRMT61B | 8.96940521 | -0.6985497 | 0.68390155 | -1.0214185 | 0.3070562 | NA |
| WDR43 | 25.0631843 | -0.5485094 | 0.39506601 | -1.3883993 | 0.16501548 | 0.75321592 |
| CLIP4 | 48.4360394 | 0.91552059 | 0.60445792 | 1.51461425 | 0.12987012 | 0.72931666 |
| YPEL5 | 21.0473566 | -1.2056868 | 0.44403873 | -2.7152739 | 0.0066221 | 0.20175229 |
| LBH | 24.5765958 | -1.8454684 | 0.48523825 | -3.8032211 | 0.00014283 | 0.02258295 |
| LCLAT1 | 31.8035446 | -1.2783444 | 0.51986686 | -2.4589842 | 0.01393308 | 0.31861834 |
| DPY30 | 8.9021375 | -0.5983701 | 0.607835 | -0.9844285 | 0.32490486 | NA |
| SPAST | 30.4570665 | -0.0676826 | 0.3964418 | -0.1707251 | 0.86443992 | 0.98751288 |
| SLC30A6 | 17.7025544 | -0.1120807 | 0.47334564 | -0.2367841 | 0.8128243 | 0.98320463 |
| NLRC4 | 16.2840611 | 0.83791577 | 0.65003441 | 1.28903296 | 0.19738663 | 0.78135737 |
| YIPF4 | 34.9439313 | -0.2585397 | 0.35812738 | -0.7219211 | 0.47034297 | 0.9314919 |
| BIRC6 | 243.553963 | -0.0995564 | 0.21068415 | -0.4725388 | 0.63654225 | 0.95933464 |
| TTC27 | 40.4101679 | 1.89468862 | 0.99457198 | 1.90502915 | 0.05677628 | 0.56510781 |

| | | | | | | |
|---|---|---|---|---|---|---|
| LINC00486 | 29.3342793 | -0.1259827 | 0.44990279 | -0.2800221 | 0.77946058 | 0.98264092 |
| RASGRP3 | 24.649573 | 0.80612275 | 1.1333696 | 0.71126202 | 0.47692188 | 0.9316222 |
| LINC01320 | 9.68296443 | 0.09440094 | 0.6586817 | 0.14331799 | 0.88603905 | NA |
| CRIM1 | 25.9808108 | -0.2720196 | 0.52951642 | -0.5137133 | 0.60745248 | 0.95688177 |
| FEZ2 | 42.7977759 | 0.31092311 | 0.41898484 | 0.74208677 | 0.45803476 | 0.9314919 |
| STRN | 47.4846661 | 0.12025647 | 0.31842606 | 0.37765902 | 0.70568392 | 0.97184211 |
| HEATR5B | 59.7691708 | 0.05875944 | 0.46587407 | 0.12612731 | 0.89963115 | 0.99182061 |
| EIF2AK2 | 37.8424917 | -0.0019087 | 0.71617416 | -0.0026651 | 0.99787354 | 0.99892869 |
| CEBPZOS | 11.0106891 | 0.70233761 | 0.54740269 | 1.28303647 | 0.1994793 | NA |
| CEBPZ | 14.6367625 | 0.16632579 | 0.49047403 | 0.33911233 | 0.73452511 | 0.97605445 |
| NDUFAF7 | 11.4738944 | -0.1416349 | 0.53709379 | -0.2637062 | 0.79200634 | NA |
| PRKD3 | 44.1512213 | -1.4593988 | 0.55842568 | -2.6134164 | 0.0089642 | 0.2531015 |
| AC006369.1 | 18.2339704 | -0.2234173 | 0.70139927 | -0.3185309 | 0.75008228 | 0.97818861 |
| CDC42EP3 | 24.6766734 | 0.69413333 | 0.70485162 | 0.98479356 | 0.32472548 | 0.87685391 |
| CYP1B1 | 17.3699679 | 2.12489208 | 1.08979169 | 1.9498149 | 0.05119819 | 0.54802854 |
| ATL2 | 28.8066596 | 0.02001953 | 0.44717934 | 0.04476845 | 0.96429187 | 0.99653147 |
| HNRNPLL | 37.6624757 | 1.73675326 | 0.75450116 | 2.30185631 | 0.02134328 | 0.37758452 |
| GALM | 36.3673384 | 0.42385818 | 0.59213707 | 0.71581091 | 0.47410812 | 0.9314919 |
| SRSF7 | 47.4176779 | 0.19653533 | 0.31780288 | 0.61841899 | 0.53629919 | 0.95415437 |
| DHX57 | 16.6558464 | 0.79515805 | 0.50622714 | 1.57075351 | 0.11623991 | 0.70122655 |
| SOS1 | 162.101147 | 0.13090286 | 0.2223786 | 0.58864862 | 0.55609701 | 0.95415437 |
| MAP4K3 | 83.2396692 | -1.1275569 | 0.45682598 | -2.4682416 | 0.01357786 | 0.31838937 |
| AC007388.1 | 22.3211832 | -1.1706099 | 0.70263746 | -1.6660226 | 0.09570892 | 0.66124053 |
| THUMPD2 | 14.7877301 | -0.0736323 | 0.54950638 | -0.1339972 | 0.8934048 | 0.99080204 |
| SLC8A1-AS1 | 15.0149565 | 1.32045999 | 0.64207664 | 2.05654576 | 0.03972994 | 0.48300729 |
| SLC8A1 | 400.040273 | 1.72794595 | 0.60964608 | 2.83434276 | 0.004592 | 0.17055138 |
| EML4 | 201.787796 | -0.453488 | 0.41264126 | -1.0989885 | 0.27177309 | 0.84994492 |
| COX7A2L | 22.5148993 | 0.48853068 | 0.46199779 | 1.05743077 | 0.29031503 | 0.85822878 |
| MTA3 | 23.0682427 | -0.2649627 | 0.47132048 | -0.5621709 | 0.57399959 | 0.95415437 |
| ZFP36L2 | 119.551777 | 0.55443647 | 0.41633321 | 1.33171328 | 0.18295442 | 0.77523613 |
| THADA | 73.2344808 | 0.05862182 | 0.36028249 | 0.16271072 | 0.8707462 | 0.98751288 |
| LRPPRC | 39.919065 | -0.3411876 | 0.41515283 | -0.821836 | 0.41117022 | 0.91823911 |
| PPM1B | 70.6600406 | 0.25519351 | 0.29772794 | 0.85713659 | 0.3913694 | 0.90910959 |
| SLC3A1 | 11.9328999 | 1.81061086 | 0.7370807 | 2.45646217 | 0.01403126 | NA |
| PREPL | 20.6483868 | 0.24426824 | 0.47645189 | 0.51268186 | 0.60817388 | 0.95688177 |
| CAMKMT | 112.490248 | 0.04647841 | 0.34195607 | 0.13591923 | 0.89188513 | 0.99080204 |
| SRBD1 | 77.7385937 | 0.12305421 | 0.36355179 | 0.3384778 | 0.73500315 | 0.97605445 |
| PRKCE | 133.77145 | -1.1109434 | 0.46987354 | -2.3643455 | 0.01806196 | 0.34868204 |
| EPAS1 | 20.1222434 | -0.1273068 | 0.5183006 | -0.2456235 | 0.80597369 | 0.98320463 |
| RHOQ | 56.5981797 | -0.1757446 | 0.36802861 | -0.4775298 | 0.6329849 | 0.95877428 |
| AC018682.1 | 11.8213327 | 0.60820072 | 0.57289785 | 1.06162159 | 0.28840751 | NA |
| PIGF | 19.278127 | 0.30100497 | 0.46314289 | 0.64991816 | 0.51574509 | 0.94596835 |
| SOCS5 | 14.2745699 | 0.32139461 | 0.53558105 | 0.60008585 | 0.54844902 | 0.95415437 |
| MCFD2 | 12.8493974 | 0.12355711 | 0.53676456 | 0.23018865 | 0.81794518 | NA |
| TTC7A | 45.7318095 | 0.78352433 | 0.40902541 | 1.9155884 | 0.05541751 | 0.55760088 |
| EPCAM | 9.24726217 | 0.21635776 | 0.74736274 | 0.28949498 | 0.77220262 | NA |
| MSH2 | 10.2701355 | 0.66298121 | 0.79663909 | 0.8322228 | 0.4052832 | NA |
| FBXO11 | 142.638455 | 0.18477404 | 0.23370787 | 0.79061967 | 0.42916597 | 0.92204028 |
| FOXN2 | 59.2177453 | -0.0163917 | 0.32824088 | -0.049938 | 0.96017178 | 0.99614757 |
| PPP1R21 | 57.9419532 | -0.0490403 | 0.34597382 | -0.1417456 | 0.88728098 | 0.99019008 |
| GPR75-ASB3 | 12.9477232 | 0.39095983 | 0.66348936 | 0.58924808 | 0.55569487 | NA |
| ERLEC1 | 12.1242895 | -0.3366722 | 0.56809242 | -0.5926363 | 0.55342457 | NA |
| PSME4 | 84.4375578 | 0.01602934 | 0.27110128 | 0.05912677 | 0.95285114 | 0.99514836 |
| ACYP2 | 83.0853832 | -0.5815706 | 0.39305987 | -1.4795981 | 0.13898055 | 0.73044824 |

| | | | | | | |
|---|---|---|---|---|---|---|
| SPTBN1 | 35.5046876 | 0.5647763 | 0.44129966 | 1.27980225 | 0.20061469 | 0.78404075 |
| RTN4 | 118.819507 | 0.28533968 | 0.26420903 | 1.07997701 | 0.28015242 | 0.84994492 |
| RPS27A | 133.397218 | -0.0914424 | 0.6418631 | -0.1424641 | 0.88671344 | 0.99019008 |
| AC012358.3 | 10.9569722 | 0.16360666 | 0.8021904 | 0.20394991 | 0.83839264 | NA |
| CCDC88A | 139.48766 | -0.7553086 | 0.37231545 | -2.0286791 | 0.04249099 | 0.50435669 |
| PPP4R3B | 54.504997 | -0.2329971 | 0.31534136 | -0.7388726 | 0.45998437 | 0.9314919 |
| PNPT1 | 20.2674813 | -0.6890127 | 0.75059005 | -0.9179613 | 0.3586391 | 0.8957272 |
| VRK2 | 54.8396234 | 0.22308985 | 0.41169966 | 0.54187524 | 0.58790445 | 0.95415437 |
| FANCL | 15.9183748 | 0.00956661 | 0.65045636 | 0.01470754 | 0.98826551 | 0.99811561 |
| BCL11A | 32.9577677 | -1.6841166 | 0.66015771 | -2.5510823 | 0.0107389 | 0.28299549 |
| PAPOLG | 22.5620057 | 0.00191629 | 0.38850356 | 0.00493249 | 0.99606446 | 0.99859588 |
| REL | 90.9557035 | 0.05789091 | 0.39814819 | 0.1454004 | 0.88439475 | 0.99019008 |
| PUS10 | 13.0695219 | -0.3022487 | 0.51758608 | -0.5839583 | 0.55924838 | 0.95415437 |
| PEX13 | 12.133221 | 0.49763335 | 0.54729385 | 0.90926172 | 0.36321199 | NA |
| KIAA1841 | 12.5359617 | -0.5721662 | 0.58244057 | -0.9823598 | 0.32592264 | NA |
| USP34 | 205.795181 | -0.237347 | 0.22797268 | -1.0411203 | 0.29781972 | 0.8608685 |
| XPO1 | 97.6796802 | 0.38102322 | 0.47065596 | 0.80955784 | 0.41819435 | 0.91982811 |
| CCT4 | 20.3899727 | 0.52111004 | 0.40897882 | 1.27417364 | 0.20260188 | 0.78735873 |
| COMMD1 | 57.3867612 | 0.13681464 | 0.31014185 | 0.4411357 | 0.65911476 | 0.96280444 |
| B3GNT2 | 9.84672967 | 0.09382157 | 0.59427392 | 0.15787596 | 0.87455454 | NA |
| EHBP1 | 48.9628522 | 0.90853509 | 0.46836945 | 1.93978296 | 0.05240607 | 0.54907299 |
| WDPCP | 55.6375833 | 0.51553371 | 0.38923488 | 1.32447972 | 0.18534377 | 0.77694741 |
| MDH1 | 17.4338002 | 0.79606118 | 0.5229866 | 1.52214452 | 0.12797287 | 0.72782081 |
| UGP2 | 50.9621378 | 1.24186401 | 0.53899733 | 2.30402628 | 0.02122117 | 0.37758452 |
| VPS54 | 55.511662 | -0.0020324 | 0.36194885 | -0.0056151 | 0.9955198 | 0.99859588 |
| PELI1 | 120.336463 | 0.94354827 | 0.65732035 | 1.43544661 | 0.15115987 | 0.74119263 |
| AFTPH | 70.5101743 | -0.0935203 | 0.30872161 | -0.3029275 | 0.76194511 | 0.98163473 |
| SERTAD2 | 25.961536 | 0.62563467 | 0.42996185 | 1.45509344 | 0.14564339 | 0.73251288 |
| CEP68 | 13.6191672 | 1.43464647 | 0.87428904 | 1.64092926 | 0.1008121 | 0.6683775 |
| RAB1A | 118.011862 | 0.15879381 | 0.28897379 | 0.5495094 | 0.58265592 | 0.95415437 |
| ACTR2 | 143.881971 | -0.0380774 | 0.35455582 | -0.1073946 | 0.91447594 | 0.99498068 |
| SPRED2 | 27.7767351 | -0.7517988 | 0.55624149 | -1.3515691 | 0.1765132 | 0.76409729 |
| C1D | 10.7938114 | -0.5549465 | 0.52049192 | -1.0661961 | 0.286335 | NA |
| PLEK | 47.2204075 | 0.41932276 | 0.60071556 | 0.69803879 | 0.48515294 | 0.93417265 |
| APLF | 14.8945854 | -0.2445813 | 0.52304379 | -0.4676115 | 0.64006242 | 0.95963232 |
| ARHGAP25 | 102.444597 | -0.7639108 | 0.37077458 | -2.0603105 | 0.03936887 | 0.48300729 |
| GFPT1 | 37.3490906 | 0.42946989 | 0.45003789 | 0.95429719 | 0.33993323 | 0.8844375 |
| NFU1 | 8.97113363 | -0.4187283 | 0.57998341 | -0.7219659 | 0.47031544 | NA |
| AAK1 | 170.875464 | 1.22095013 | 0.48357849 | 2.52482308 | 0.01157565 | 0.29101961 |
| ANXA4 | 40.2028161 | 0.17187632 | 0.45693791 | 0.37614809 | 0.70680681 | 0.97184211 |
| GMCL1 | 25.4887959 | 0.11168352 | 0.41184681 | 0.27117732 | 0.78625465 | 0.98300067 |
| MXD1 | 52.5431422 | 0.74772576 | 0.64057141 | 1.16727932 | 0.24309759 | 0.83507265 |
| PCBP1 | 36.5120492 | 0.22235545 | 0.55400309 | 0.40136139 | 0.68815407 | 0.9673431 |
| SNRPG | 14.2139377 | -0.5241061 | 0.51592412 | -1.0158588 | 0.30969662 | 0.86794644 |
| NAGK | 27.7623164 | 0.18168733 | 0.45515161 | 0.39917979 | 0.68976073 | 0.9673431 |
| MCEE | 11.079867 | 0.42619862 | 0.57905698 | 0.7360219 | 0.46171737 | NA |
| MPHOSPH10 | 11.018364 | -0.1483272 | 0.51138227 | -0.2900516 | 0.77177678 | NA |
| ZNF638 | 148.226919 | 0.05310724 | 0.25890698 | 0.20512092 | 0.83747765 | 0.98320463 |
| DYSF | 32.7799639 | -0.28721 | 0.61605855 | -0.4662057 | 0.6410683 | 0.95992767 |
| EXOC6B | 94.1211266 | 0.15726693 | 0.46542579 | 0.33789906 | 0.73543926 | 0.97605445 |
| SFXN5 | 16.4599916 | -0.2910702 | 0.54142592 | -0.5375994 | 0.59085365 | 0.95415437 |
| CCT7 | 13.7129469 | -0.6119888 | 0.45774183 | -1.3369737 | 0.18123122 | 0.77030229 |
| ALMS1 | 22.5970558 | -0.4745882 | 0.49406915 | -0.9605703 | 0.33676825 | 0.88143379 |
| DUSP11 | 8.35068979 | 0.47685338 | 0.61830924 | 0.7712215 | 0.44057565 | NA |

| | | | | | | |
|---|---|---|---|---|---|---|
| STAMBP | 17.0499946 | -0.3134992 | 0.47120128 | -0.6653191 | 0.50584639 | 0.94166217 |
| DGUOK | 19.5320453 | -0.4236698 | 0.43236682 | -0.979885 | 0.32714287 | 0.87715548 |
| TET3 | 77.9238442 | 0.36918152 | 0.3216814 | 1.147662 | 0.25110813 | 0.84008912 |
| MOB1A | 48.8485158 | -0.380669 | 0.37543643 | -1.0139373 | 0.31061267 | 0.86858542 |
| MTHFD2 | 11.613223 | 0.00487861 | 0.63537931 | 0.00767826 | 0.99387369 | NA |
| MOGS | 8.56873691 | -0.2970912 | 0.64047527 | -0.4638605 | 0.64274771 | NA |
| AUP1 | 9.5655338 | -0.775664 | 0.6635186 | -1.1690162 | 0.2423971 | NA |
| HK2 | 19.1340205 | 1.71437809 | 1.14771726 | 1.49372859 | 0.13524657 | 0.73044824 |
| POLE4 | 10.1039452 | -0.6087097 | 0.5976145 | -1.0185658 | 0.30840914 | NA |
| MRPL19 | 9.98315181 | 0.14250413 | 0.58562763 | 0.24333574 | 0.80774532 | NA |
| GCFC2 | 11.6811873 | 0.84693037 | 0.54203097 | 1.56251287 | 0.11816722 | NA |
| SUCLG1 | 19.2373438 | 0.16088823 | 0.44297763 | 0.36319719 | 0.71645759 | 0.97348719 |
| TMSB10 | 266.670412 | 0.14197367 | 0.61659454 | 0.23025451 | 0.81789401 | 0.98320463 |
| KCMF1 | 56.5043073 | -0.0842032 | 0.32619361 | -0.2581387 | 0.79629989 | 0.98320463 |
| TGOLN2 | 50.2763743 | 0.13481633 | 0.40541081 | 0.33254251 | 0.73947965 | 0.97605445 |
| ELMOD3 | 16.9988017 | 0.41275288 | 0.4586436 | 0.89994251 | 0.36815084 | 0.89868152 |
| CAPG | 11.2277707 | 0.61480894 | 0.61937991 | 0.99262009 | 0.32089513 | NA |
| MAT2A | 36.028713 | 0.01710585 | 0.69540774 | 0.02459831 | 0.98037537 | 0.99758514 |
| VAMP8 | 32.5938516 | -0.0951436 | 0.42811234 | -0.2222398 | 0.82412721 | 0.98320463 |
| VAMP5 | 17.4834989 | 0.05049539 | 0.51214381 | 0.09859612 | 0.92145895 | 0.99514836 |
| USP39 | 24.2771606 | -0.2246621 | 0.40159905 | -0.5594189 | 0.57587586 | 0.95415437 |
| C2orf68 | 8.74059422 | 0.33767483 | 0.64038026 | 0.52730363 | 0.59798276 | NA |
| GNLY | 65.0321973 | 1.61973407 | 1.42868863 | 1.13372084 | 0.25691166 | 0.84077419 |
| ST3GAL5 | 33.487516 | -0.3976096 | 0.3804214 | -1.0451819 | 0.29593889 | 0.8608685 |
| POLR1A | 9.82163306 | -0.4233557 | 0.55480644 | -0.7630693 | 0.44542209 | NA |
| PTCD3 | 22.944222 | 0.35397127 | 0.50895982 | 0.69547979 | 0.48675467 | 0.93497601 |
| IMMT | 34.0764012 | -0.6108779 | 0.3884442 | -1.5726272 | 0.11580517 | 0.70122655 |
| KDM3A | 37.0168729 | -0.2442777 | 0.50053812 | -0.4880303 | 0.62552841 | 0.95876809 |
| RMND5A | 29.0421804 | 0.15879246 | 0.38627404 | 0.41108757 | 0.68100832 | 0.9673431 |
| CD8A | 16.0164298 | -0.4409505 | 0.62098146 | -0.7100864 | 0.47765057 | 0.9318939 |
| CYTOR | 33.2367961 | 0.03334435 | 0.46518436 | 0.07167985 | 0.94285669 | 0.99514836 |
| KRCC1 | 21.9868238 | 0.60077583 | 0.5138138 | 1.16924815 | 0.24230366 | 0.83493679 |
| EIF2AK3 | 32.4775961 | 0.22224151 | 0.49532337 | 0.44867963 | 0.65366278 | 0.96178254 |
| IGKC | 439.7482 | 0.84820275 | 1.90529012 | 0.44518299 | 0.65618752 | 0.96219972 |
| MRPS5 | 19.3477967 | -0.2035002 | 0.40907391 | -0.4974656 | 0.61886074 | 0.95688177 |
| ANKRD36C | 34.7501906 | -0.032632 | 0.47657132 | -0.0684724 | 0.94540956 | 0.99514836 |
| DUSP2 | 16.2174816 | 0.62599262 | 0.87247037 | 0.71749441 | 0.47306908 | 0.9314919 |
| STARD7 | 17.4926176 | 0.32803148 | 0.4598359 | 0.7133664 | 0.47561906 | 0.9314919 |
| TMEM127 | 15.0919075 | 0.80592037 | 0.62756392 | 1.28420443 | 0.19907044 | 0.78142991 |
| CIAO1 | 13.3939925 | -0.12197 | 0.53996332 | -0.2258857 | 0.8212903 | 0.98320463 |
| SNRNP200 | 30.9649671 | -0.2793717 | 0.3331391 | -0.8386037 | 0.40169174 | 0.91528253 |
| ARID5A | 17.1134836 | -0.859508 | 0.42757571 | -2.010189 | 0.04441119 | 0.5181889 |
| KANSL3 | 24.5101051 | 0.43262521 | 0.49710045 | 0.87029736 | 0.38413792 | 0.90910959 |
| ANKRD23 | 9.49980809 | 0.49420084 | 0.75396341 | 0.65547058 | 0.51216482 | NA |
| ANKRD36 | 48.7582231 | 0.211693 | 0.47405669 | 0.4465563 | 0.65519545 | 0.96178254 |
| AC092683.1 | 25.4244674 | -0.10317 | 0.45936054 | -0.2245948 | 0.82229451 | 0.98320463 |
| ANKRD36B | 14.1240734 | 0.71136186 | 0.58846023 | 1.2088529 | 0.22671937 | 0.82057879 |
| COX5B | 21.7021284 | -0.3027751 | 0.439801 | -0.6884366 | 0.4911779 | 0.93633431 |
| ZAP70 | 21.8458785 | 0.33201078 | 0.5189704 | 0.63974897 | 0.52233581 | 0.94895469 |
| TMEM131 | 104.946175 | -0.49015 | 0.27633835 | -1.7737313 | 0.07610761 | 0.62396969 |
| INPP4A | 134.270313 | 0.52841703 | 0.37713024 | 1.40115264 | 0.16116843 | 0.74887162 |
| UNC50 | 9.42682177 | 1.11467502 | 0.64811559 | 1.71987071 | 0.08545595 | NA |
| MGAT4A | 99.956817 | 0.79674325 | 0.42115548 | 1.89180311 | 0.05851722 | 0.57281427 |
| TSGA10 | 10.4870479 | -0.1935208 | 0.84400302 | -0.2292892 | 0.81864414 | NA |

| | | | | | | |
|---|---|---|---|---|---|---|
| AC079447.1 | 14.471729 | -0.9601224 | 0.65948541 | -1.455866 | 0.14542968 | 0.73251288 |
| MITD1 | 16.14673 | 0.31098258 | 0.45978164 | 0.67637016 | 0.49880566 | 0.9392278 |
| EIF5B | 39.2816737 | -0.6560912 | 0.40257623 | -1.6297317 | 0.10315822 | 0.67250824 |
| REV1 | 39.7577079 | 0.10219025 | 0.36237197 | 0.28200373 | 0.77794065 | 0.98264092 |
| AFF3 | 153.70899 | -2.7148336 | 0.51896513 | -5.2312447 | 1.68E-07 | 0.00027433 |
| RPL31 | 27.3900527 | -0.2516849 | 0.50254668 | -0.5008189 | 0.61649857 | 0.95688177 |
| TBC1D8 | 71.9519445 | 1.35401645 | 0.62407392 | 2.16964113 | 0.03003404 | 0.42727094 |
| CNOT11 | 14.2223547 | 0.13382993 | 0.50586126 | 0.26455856 | 0.79134955 | 0.98320463 |
| RNF149 | 263.477877 | 0.52513322 | 0.47275307 | 1.11079812 | 0.26665526 | 0.84994492 |
| MAP4K4 | 113.957867 | -0.1890438 | 0.36223588 | -0.5218803 | 0.60175367 | 0.95688177 |
| IL1R2 | 86.2571899 | -1.195572 | 1.65334219 | -0.7231243 | 0.4696035 | 0.9314919 |
| IL1R1 | 19.6465468 | -1.6653297 | 0.54666247 | -3.0463582 | 0.00231632 | 0.10955981 |
| IL18R1 | 38.4518155 | 0.23770253 | 0.4464973 | 0.5323717 | 0.59446858 | 0.9553365 |
| IL18RAP | 16.40735 | 0.15885471 | 0.71675361 | 0.22163084 | 0.82460127 | 0.98320463 |
| MRPS9 | 12.95744 | -0.1544815 | 0.55392192 | -0.2788868 | 0.78033169 | 0.98264092 |
| TGFBRAP1 | 9.85729945 | 0.96271903 | 0.66180924 | 1.45467753 | 0.14575856 | NA |
| NCK2 | 64.4025792 | 0.20733042 | 0.36912437 | 0.56168175 | 0.57433287 | 0.95415437 |
| UXS1 | 25.5106022 | -0.6324068 | 0.45343125 | -1.3947138 | 0.16310217 | 0.75321592 |
| GCC2 | 70.0790467 | -0.0510013 | 0.4672006 | -0.1091637 | 0.91307267 | 0.99498068 |
| LIMS1 | 98.8427005 | -0.0346985 | 0.32868474 | -0.1055678 | 0.91592524 | 0.99498068 |
| RANBP2 | 60.4502204 | -0.035848 | 0.48099587 | -0.0745286 | 0.94058977 | 0.99514836 |
| CCDC138 | 11.7361656 | 0.45749961 | 0.6200923 | 0.73779277 | 0.46064039 | NA |
| SH3RF3 | 35.2351645 | 0.36508803 | 0.61415169 | 0.59445906 | 0.55220512 | 0.95415437 |
| 10-Sep | 11.025887 | -0.2042563 | 0.51253308 | -0.3985231 | 0.69024464 | NA |
| MIR4435-2H | 89.0346721 | -0.0009802 | 0.47411127 | -0.0020674 | 0.99835045 | 0.99907258 |
| BCL2L11 | 31.6963154 | -0.1921519 | 0.74870807 | -0.2566446 | 0.79745316 | 0.98320463 |
| ANAPC1 | 31.3683208 | -0.5653823 | 0.45645606 | -1.2386347 | 0.21548082 | 0.80634897 |
| MERTK | 86.7731638 | 0.68739864 | 0.61709592 | 1.11392512 | 0.26531131 | 0.84994492 |
| TMEM87B | 33.7498177 | 0.14092012 | 0.34464344 | 0.40888672 | 0.6826228 | 0.9673431 |
| ZC3H8 | 16.0840911 | -0.689003 | 0.44215748 | -1.5582751 | 0.11916804 | 0.70431251 |
| ZC3H6 | 24.9752548 | 0.15260272 | 0.50515522 | 0.30209074 | 0.76258289 | 0.98163473 |
| TTL | 19.5669727 | -0.0865594 | 0.60555202 | -0.142943 | 0.88633521 | 0.99019008 |
| SLC20A1 | 36.7704628 | 0.70922143 | 0.45757172 | 1.54996778 | 0.12114925 | 0.70946026 |
| PSD4 | 15.2154949 | -0.1275948 | 0.45979739 | -0.2775021 | 0.78139457 | 0.98264092 |
| PAX8 | 25.6039762 | -0.6499524 | 0.45852446 | -1.417487 | 0.15634061 | 0.74486619 |
| CBWD2 | 14.5202128 | 0.07878285 | 0.57191008 | 0.13775392 | 0.89043491 | 0.99080204 |
| SLC35F5 | 40.452372 | -0.3465647 | 0.34179973 | -1.0139409 | 0.31061095 | 0.86858542 |
| ACTR3 | 139.280416 | -0.4478701 | 0.28506876 | -1.5710951 | 0.11616056 | 0.70122655 |
| DDX18 | 19.1193571 | 0.41669239 | 0.50089918 | 0.83188875 | 0.40547174 | 0.91575086 |
| CCDC93 | 38.2139757 | 0.3295485 | 0.40791641 | 0.80788243 | 0.41915827 | 0.91985469 |
| INSIG2 | 19.3356786 | 0.92147277 | 0.44319528 | 2.07915743 | 0.03760288 | 0.47294171 |
| MARCO | 28.4351789 | 2.59085408 | 1.22477966 | 2.11536341 | 0.03439899 | 0.45650836 |
| DBI | 29.5011633 | 0.1195624 | 0.58100869 | 0.20578419 | 0.83695949 | 0.98320463 |
| PTPN4 | 67.9919926 | -0.1089421 | 0.33896183 | -0.3213992 | 0.74790791 | 0.97818861 |
| EPB41L5 | 13.5787367 | -0.210112 | 0.58451302 | -0.3594651 | 0.71924718 | 0.97348719 |
| RALB | 17.1707254 | 0.81489519 | 0.52786246 | 1.54376423 | 0.12264546 | 0.71294115 |
| CLASP1 | 146.620562 | 0.28218203 | 0.3175215 | 0.88870212 | 0.37416319 | 0.90262385 |
| NIFK-AS1 | 8.73548662 | 0.93692326 | 0.67701735 | 1.38389845 | 0.16638955 | NA |
| TSN | 9.87888925 | -0.0352782 | 0.57388459 | -0.0614726 | 0.95098285 | NA |
| GYPC | 20.7938408 | 0.35485483 | 0.6177846 | 0.57439896 | 0.56569785 | 0.95415437 |
| BIN1 | 11.0666977 | 0.10978779 | 0.54922467 | 0.19989595 | 0.84156195 | NA |
| ERCC3 | 14.4914477 | -0.5716719 | 0.50334878 | -1.1357371 | 0.2560666 | 0.8406233 |
| MAP3K2 | 91.2750189 | 0.3890523 | 0.29933199 | 1.29973512 | 0.19369177 | 0.78135737 |
| IWS1 | 45.5223428 | -0.2998843 | 0.37964844 | -0.7899001 | 0.42958612 | 0.92204028 |

| | | | | | | |
|---|---|---|---|---|---|---|
| WDR33 | 58.5835812 | 0.09794373 | 0.30302359 | 0.32322149 | 0.74652751 | 0.97805001 |
| SAP130 | 18.1936178 | 0.14684413 | 0.42823131 | 0.34290844 | 0.73166734 | 0.97605445 |
| UGGT1 | 35.206043 | 0.44249696 | 0.44218332 | 1.00070931 | 0.31696736 | 0.87263466 |
| HS6ST1 | 10.2480848 | 1.25923613 | 0.69779676 | 1.80458867 | 0.07113907 | NA |
| SMPD4 | 10.3520593 | -0.9053933 | 0.52002526 | -1.7410564 | 0.0816737 | NA |
| MZT2B | 20.3410701 | 0.91361536 | 0.57050416 | 1.60141753 | 0.10928447 | 0.68725031 |
| PTPN18 | 28.4300046 | 1.18085257 | 0.54384815 | 2.17129096 | 0.02990919 | 0.42727094 |
| FAM168B | 30.321372 | -0.2943835 | 0.44145427 | -0.6668494 | 0.50486833 | 0.94166217 |
| PLEKHB2 | 29.3584731 | 0.94133432 | 0.43286984 | 2.17463597 | 0.02965741 | 0.42629633 |
| MZT2A | 10.1466507 | 0.13806293 | 0.63362264 | 0.21789457 | 0.82751125 | NA |
| NCKAP5 | 54.8835342 | 0.40616336 | 0.6780042 | 0.5990573 | 0.54913467 | 0.95415437 |
| MGAT5 | 175.407897 | -0.3593553 | 0.29033506 | -1.237726 | 0.21581766 | 0.80634897 |
| TMEM163 | 25.6230399 | 1.54135951 | 0.90651456 | 1.70031412 | 0.08907186 | 0.64845882 |
| CCNT2 | 48.5436427 | 0.15228239 | 0.33603119 | 0.45317933 | 0.65041961 | 0.96084839 |
| RAB3GAP1 | 69.2974375 | -0.192289 | 0.29519786 | -0.6513901 | 0.51479471 | 0.94596835 |
| ZRANB3 | 22.0681286 | -0.2379355 | 0.49320108 | -0.482431 | 0.62949978 | 0.95877428 |
| R3HDM1 | 67.6200284 | 0.29699284 | 0.32966812 | 0.90088433 | 0.36764982 | 0.89868152 |
| UBXN4 | 55.4149952 | 0.15696462 | 0.28759628 | 0.54578112 | 0.5852164 | 0.95415437 |
| DARS | 47.9915486 | 0.14460121 | 0.34010482 | 0.42516661 | 0.67071522 | 0.96412305 |
| CXCR4 | 72.4402183 | -0.1873131 | 0.50030257 | -0.3743997 | 0.70810699 | 0.97184211 |
| HNMT | 27.903789 | 0.32476096 | 0.56587254 | 0.57391185 | 0.56602746 | 0.95415437 |
| SPOPL | 54.7322535 | -0.1609705 | 0.33768209 | -0.4766926 | 0.63358105 | 0.95877428 |
| KYNU | 172.199647 | 0.84225862 | 0.56376472 | 1.49398959 | 0.13517834 | 0.73044824 |
| ARHGAP15 | 592.629446 | 0.02348811 | 0.27240373 | 0.08622538 | 0.93128726 | 0.99514836 |
| GTDC1 | 93.1720335 | 0.59719776 | 0.30959949 | 1.92893652 | 0.05373874 | 0.55218699 |
| ZEB2 | 534.127513 | -0.10767 | 0.25057184 | -0.4296971 | 0.66741602 | 0.96412305 |
| ACVR2A | 29.2629688 | -0.7030452 | 0.49563108 | -1.418485 | 0.15604923 | 0.74486619 |
| ORC4 | 31.6509827 | -0.1236068 | 0.35150735 | -0.3516479 | 0.72510231 | 0.97546503 |
| MBD5 | 176.207435 | 0.21690743 | 0.32716944 | 0.66298193 | 0.50734213 | 0.94305491 |
| EPC2 | 60.1715261 | -0.1426365 | 0.33422373 | -0.4267695 | 0.66954719 | 0.96412305 |
| MMADHC | 13.2368767 | 0.18159292 | 0.49123388 | 0.36966692 | 0.71163068 | 0.97317955 |
| NMI | 14.0748506 | 1.51504756 | 0.70035375 | 2.16326044 | 0.03052115 | 0.43032651 |
| RIF1 | 48.2447818 | 0.04385189 | 0.34972245 | 0.12539054 | 0.90021437 | 0.99194856 |
| ARL5A | 17.8699748 | -0.1692299 | 0.62638692 | -0.2701683 | 0.78703074 | 0.98320463 |
| STAM2 | 32.4430112 | -0.5232499 | 0.40484203 | -1.2924793 | 0.19619122 | 0.78135737 |
| FMNL2 | 223.635221 | 0.44992403 | 0.52553414 | 0.85612712 | 0.39192747 | 0.90910959 |
| PRPF40A | 54.9925972 | -0.1662867 | 0.29330397 | -0.5669432 | 0.57075277 | 0.95415437 |
| ARL6IP6 | 23.5870512 | 0.63149681 | 0.45001983 | 1.40326441 | 0.16053801 | 0.74887162 |
| NR4A2 | 20.5016705 | 1.40120137 | 1.07005164 | 1.3094708 | 0.19037492 | 0.78046165 |
| GPD2 | 65.0624287 | 0.48833315 | 0.38016489 | 1.28452985 | 0.19895663 | 0.78142991 |
| CYTIP | 120.776662 | -0.1209277 | 0.47365671 | -0.2553066 | 0.79848631 | 0.98320463 |
| ACVR1 | 28.8260791 | -0.1434805 | 0.37151809 | -0.3862005 | 0.69934815 | 0.9694491 |
| PKP4 | 40.8319066 | 0.14690953 | 0.53642513 | 0.27386772 | 0.78418628 | 0.98264092 |
| WDSUB1 | 18.2550924 | -0.2248707 | 0.45817327 | -0.4907983 | 0.6235691 | 0.95803203 |
| BAZ2B | 156.968476 | 0.17503534 | 0.26971643 | 0.64896062 | 0.51636383 | 0.94596835 |
| 7-Mar | 89.9794622 | -0.2888434 | 0.24795255 | -1.1649138 | 0.24405388 | 0.83666131 |
| ITGB6 | 10.1887631 | -1.8897689 | 0.74146133 | -2.5487087 | 0.01081226 | NA |
| RBMS1 | 183.4935 | 0.57953021 | 0.38330933 | 1.51191263 | 0.13055609 | 0.72931666 |
| TANK | 141.393594 | 0.33533562 | 0.29298407 | 1.1445524 | 0.25239462 | 0.84008912 |
| PSMD14 | 28.4715323 | -0.5841823 | 0.38131297 | -1.5320285 | 0.12551541 | 0.71904999 |
| DPP4 | 14.4786234 | -0.4053525 | 0.57072465 | -0.7102418 | 0.47755418 | 0.9318939 |
| IFIH1 | 18.5020225 | 0.76265681 | 0.90883873 | 0.83915527 | 0.40138219 | 0.91528253 |
| GCA | 30.5610438 | 0.53834843 | 0.53322866 | 1.00960144 | 0.31268628 | 0.87058449 |
| COBLL1 | 35.1281004 | -0.910694 | 0.63206093 | -1.4408326 | 0.14963197 | 0.73934224 |

| | | | | | | |
|---|---|---|---|---|---|---|
| AC019197.1 | 18.8067927 | 0.7698333 | 0.86799446 | 0.88691039 | 0.37512715 | 0.90337408 |
| GALNT3 | 14.9260757 | -0.0541864 | 0.51761312 | -0.1046851 | 0.91662572 | 0.99498068 |
| TTC21B | 23.6999933 | -0.2998101 | 0.41882127 | -0.7158427 | 0.4740885 | 0.9314919 |
| STK39 | 82.1018752 | 0.36886427 | 0.3879566 | 0.95078744 | 0.34171229 | 0.88510795 |
| CERS6 | 103.30343 | 0.61569897 | 0.41493412 | 1.48384752 | 0.13784938 | 0.73044824 |
| LRP2 | 19.1609398 | 2.29514522 | 1.0047009 | 2.28440644 | 0.02234766 | 0.38788767 |
| FASTKD1 | 18.1962268 | -0.6898937 | 0.56953476 | -1.2113285 | 0.22576951 | 0.81875277 |
| PPIG | 55.0997053 | 0.08898465 | 0.29228758 | 0.30444211 | 0.76079109 | 0.98117406 |
| SSB | 15.3829053 | 0.55498631 | 0.50310935 | 1.1031127 | 0.26997823 | 0.84994492 |
| UBR3 | 100.433183 | 0.00609693 | 0.33847726 | 0.01801283 | 0.98562862 | 0.99811561 |
| GORASP2 | 18.5061054 | 0.46428968 | 0.46784646 | 0.99239755 | 0.32100363 | 0.87363761 |
| TLK1 | 99.1477028 | 0.05413974 | 0.26536954 | 0.20401642 | 0.83834067 | 0.98331031 |
| METTL8 | 18.767157 | 0.04636822 | 0.46656595 | 0.09938191 | 0.92083504 | 0.99514836 |
| DCAF17 | 12.0501781 | -0.2956521 | 0.51852455 | -0.5701795 | 0.56855593 | NA |
| CYBRD1 | 12.0415952 | 1.81888403 | 0.68715778 | 2.6469671 | 0.00812173 | NA |
| DYNC1I2 | 31.793974 | -0.2086441 | 0.35010227 | -0.595952 | 0.55120733 | 0.95415437 |
| SLC25A12 | 26.081528 | 1.01750418 | 0.48703794 | 2.0891682 | 0.03669259 | 0.46928949 |
| HAT1 | 20.5706646 | 0.17494661 | 0.51343246 | 0.34073928 | 0.73329986 | 0.97605445 |
| ITGA6 | 27.5762262 | 0.92736361 | 0.38991847 | 2.37835261 | 0.01739019 | 0.34827529 |
| AC078883.1 | 11.3319867 | 1.56435783 | 0.71308412 | 2.19379142 | 0.02825041 | NA |
| PDK1 | 26.7901608 | 0.01942322 | 0.50287407 | 0.03862443 | 0.96918983 | 0.99662129 |
| MAP3K20 | 28.3922248 | -0.2125249 | 0.55151399 | -0.3853482 | 0.69997944 | 0.96979939 |
| SP3 | 68.4369794 | 0.01130776 | 0.29492236 | 0.03834147 | 0.96941542 | 0.99662129 |
| OLA1 | 53.0025148 | -0.3977298 | 0.32019258 | -1.242158 | 0.21417829 | 0.80634897 |
| CIR1 | 27.8753912 | -0.0431644 | 0.42692692 | -0.1011049 | 0.91946715 | 0.99498068 |
| GPR155 | 52.5821234 | 0.62641885 | 0.60031625 | 1.04348142 | 0.29672538 | 0.8608685 |
| AC010894.3 | 61.2837676 | -0.3906856 | 0.46830435 | -0.8342558 | 0.40413684 | 0.91528253 |
| WIPF1 | 173.809365 | -0.0869554 | 0.25301141 | -0.3436818 | 0.73108562 | 0.97605445 |
| ATF2 | 38.5176432 | -0.3252281 | 0.34281153 | -0.9487081 | 0.34276909 | 0.88530691 |
| ATP5MC3 | 26.1447898 | 0.59570966 | 0.57419025 | 1.03747785 | 0.29951321 | 0.86225566 |
| LNPK | 21.510378 | 0.04709795 | 0.56504415 | 0.08335268 | 0.93357111 | 0.99514836 |
| MTX2 | 26.9875683 | -0.1139639 | 0.43870981 | -0.2597705 | 0.79504084 | 0.98320463 |
| HNRNPA3 | 73.3696643 | -0.167305 | 0.3272039 | -0.5113172 | 0.60912898 | 0.95688177 |
| NFE2L2 | 112.221037 | 0.19789246 | 0.29874984 | 0.6624019 | 0.50771369 | 0.94305491 |
| AGPS | 42.9200366 | -0.2678362 | 0.39009168 | -0.6865982 | 0.49233599 | 0.93678212 |
| RBM45 | 9.71768661 | -0.0743522 | 0.57250541 | -0.1298715 | 0.89666806 | NA |
| AC009948.1 | 12.7855636 | 1.51846417 | 0.64320172 | 2.36078998 | 0.01823606 | NA |
| PRKRA | 13.1405855 | 0.0300815 | 0.50452304 | 0.05962365 | 0.95245538 | 0.99514836 |
| PLEKHA3 | 12.5264058 | -0.5210011 | 0.51725454 | -1.0072431 | 0.31381796 | NA |
| TTN-AS1 | 35.9281853 | 0.38654022 | 0.37858145 | 1.02102261 | 0.30724373 | 0.8673137 |
| TTN | 64.0620665 | -2.1245174 | 0.64058508 | -3.3165265 | 0.00091144 | 0.06269654 |
| CCDC141 | 30.4885426 | -2.8064118 | 0.76431537 | -3.6717982 | 0.00024085 | 0.02990804 |
| SESTD1 | 59.9875951 | 0.65726194 | 0.40869494 | 1.60819694 | 0.10779205 | 0.68018384 |
| CWC22 | 18.1181073 | -0.4036715 | 0.50588164 | -0.7979563 | 0.42489583 | 0.91985469 |
| UBE2E3 | 43.1273428 | -0.5061297 | 0.47013705 | -1.0765577 | 0.28167791 | 0.85171619 |
| LINC01934 | 73.9256619 | 0.91886787 | 0.59452479 | 1.54555014 | 0.12221325 | 0.71192434 |
| ITGA4 | 165.09909 | 0.27131051 | 0.46822294 | 0.57944728 | 0.56228741 | 0.95415437 |
| SSFA2 | 24.0252111 | 0.30623726 | 0.43500582 | 0.70398428 | 0.48144256 | 0.933206 |
| PDE1A | 15.4688363 | -0.0256646 | 0.83186677 | -0.0308518 | 0.97538772 | 0.9973754 |
| DNAJC10 | 22.8143499 | -0.1346677 | 0.44027844 | -0.3058694 | 0.75970409 | 0.9808103 |
| NCKAP1 | 10.4909244 | -1.1916133 | 0.57859278 | -2.0595026 | 0.03944612 | NA |
| ZNF804A | 34.3040647 | 1.40604573 | 0.6792214 | 2.07008455 | 0.03844443 | 0.47809317 |
| ZC3H15 | 20.4446199 | -0.0902795 | 0.40098055 | -0.2251469 | 0.82186497 | 0.98320463 |
| ITGAV | 37.751946 | 0.36359818 | 0.3823854 | 0.95086837 | 0.3416712 | 0.88510795 |

| | | | | | | |
|---|---|---|---|---|---|---|
| AC007319.1 | 18.0504371 | 1.75610182 | 1.16986894 | 1.50110987 | 0.13332715 | 0.73044824 |
| TFPI | 21.3411632 | 0.359772 | 0.86000048 | 0.41833931 | 0.67569905 | 0.96623219 |
| WDR75 | 17.1150954 | -0.16221 | 0.46139499 | -0.3515643 | 0.72516501 | 0.97546503 |
| SLC40A1 | 12.7152492 | 1.1092788 | 0.94070852 | 1.17919502 | 0.23832053 | NA |
| ANKAR | 22.8665251 | -0.1424551 | 0.68166465 | -0.2089813 | 0.83446286 | 0.98320463 |
| ORMDL1 | 33.0595205 | 0.46583572 | 0.45522585 | 1.02330683 | 0.3061628 | 0.8673137 |
| PMS1 | 26.3410543 | 0.32675742 | 0.41694812 | 0.78368843 | 0.43322295 | 0.92284736 |
| HIBCH | 39.9091346 | 0.00820452 | 0.49138307 | 0.0166968 | 0.9866785 | 0.99811561 |
| MFSD6 | 23.4062566 | 0.29329021 | 0.46556059 | 0.62997215 | 0.52871281 | 0.95182065 |
| NAB1 | 17.4558524 | 0.64762577 | 0.4653812 | 1.39160277 | 0.16404272 | 0.75321592 |
| GLS | 137.213683 | -0.4543103 | 0.33733957 | -1.3467447 | 0.17806247 | 0.76504408 |
| STAT1 | 65.1010597 | 0.8942911 | 0.68373381 | 1.30795214 | 0.19088954 | 0.78122755 |
| STAT4 | 143.329864 | 1.1059085 | 0.66119405 | 1.67259293 | 0.09440746 | 0.65925012 |
| NABP1 | 59.3583852 | 0.38760372 | 0.47450345 | 0.81686176 | 0.41400743 | 0.91894074 |
| SLC39A10 | 33.0860098 | 0.78144485 | 0.52621072 | 1.48504167 | 0.13753279 | 0.73044824 |
| STK17B | 169.56801 | -0.4875279 | 0.32808751 | -1.4859691 | 0.13728731 | 0.73044824 |
| AC114760.2 | 13.4199949 | 0.13718922 | 0.60679063 | 0.22608987 | 0.82113152 | 0.98320463 |
| GTF3C3 | 13.197995 | -0.5710558 | 0.53855175 | -1.0603545 | 0.28898337 | 0.85703857 |
| ANKRD44 | 530.007821 | -0.1882918 | 0.33541642 | -0.5613673 | 0.57454718 | 0.95415437 |
| SF3B1 | 63.8511135 | 0.0974097 | 0.35312601 | 0.2758497 | 0.78266351 | 0.98264092 |
| COQ10B | 11.60419 | 0.14542472 | 0.52986013 | 0.27445871 | 0.78373213 | NA |
| HSPD1 | 30.9893752 | 0.46189083 | 0.44133827 | 1.04656873 | 0.29529853 | 0.8608685 |
| PLCL1 | 61.3266496 | -0.3776174 | 0.46950011 | -0.8042968 | 0.42122561 | 0.91985469 |
| SPATS2L | 39.9094471 | 0.51423158 | 0.67073234 | 0.76667181 | 0.44327667 | 0.92899684 |
| BZW1 | 13.779663 | -0.7894495 | 0.58495564 | -1.3495887 | 0.17714796 | 0.76409729 |
| CLK1 | 52.2972005 | 0.19263538 | 0.39494132 | 0.48775697 | 0.62572199 | 0.95876809 |
| ORC2 | 28.7418527 | 0.14404837 | 0.36824958 | 0.3911705 | 0.69567121 | 0.9685713 |
| FAM126B | 49.5081221 | 0.24263711 | 0.36188403 | 0.67048307 | 0.5025499 | 0.94115436 |
| NDUFB3 | 12.8627244 | -0.0623686 | 0.61900454 | -0.1007562 | 0.91974397 | NA |
| CFLAR | 147.41786 | 0.51043906 | 0.28059815 | 1.81911057 | 0.06889456 | 0.59946936 |
| CFLAR-AS1 | 14.2878425 | -0.132952 | 0.63013623 | -0.2109893 | 0.83289564 | 0.98320463 |
| CASP10 | 24.3515024 | 1.523954 | 0.55945392 | 2.72400271 | 0.0064496 | 0.19837198 |
| CASP8 | 47.6292298 | -0.0010134 | 0.40965196 | -0.0024739 | 0.99802616 | 0.99892869 |
| TRAK2 | 21.8540563 | -0.0207413 | 0.44240379 | -0.0468831 | 0.96260641 | 0.99653147 |
| ALS2 | 11.9384318 | 0.63342999 | 0.6143425 | 1.03106979 | 0.30250809 | NA |
| SUMO1 | 18.3633264 | -0.0826864 | 0.46404344 | -0.1781868 | 0.85857625 | 0.98751288 |
| NOP58 | 29.8049276 | -0.7807783 | 0.39165933 | -1.9935139 | 0.04620521 | 0.52613091 |
| BMPR2 | 73.8854255 | -0.2125598 | 0.37890053 | -0.5609911 | 0.57480362 | 0.95415437 |
| FAM117B | 30.5461167 | -0.5526177 | 0.37354604 | -1.479383 | 0.13903798 | 0.73044824 |
| WDR12 | 9.7995878 | 0.01903001 | 0.56491394 | 0.03368656 | 0.9731271 | NA |
| CARF | 13.7373601 | -0.7422373 | 0.49478403 | -1.5001237 | 0.13358235 | 0.73044824 |
| NBEAL1 | 95.8974222 | -0.5129683 | 0.31454592 | -1.6308218 | 0.10292792 | 0.67250824 |
| CYP20A1 | 33.5627955 | -0.1803254 | 0.39278851 | -0.4590902 | 0.64616937 | 0.96049083 |
| ABI2 | 26.0653266 | 0.07079802 | 0.48903905 | 0.14476967 | 0.88489273 | 0.99019008 |
| RAPH1 | 30.6190217 | 0.31844228 | 0.65880007 | 0.4833671 | 0.62883511 | 0.95877428 |
| CD28 | 12.4313278 | -0.6908215 | 0.75529521 | -0.9146377 | 0.36038186 | NA |
| PARD3B | 12.0586278 | -0.5823251 | 0.6192531 | -0.9403668 | 0.34702942 | NA |
| NRP2 | 25.3253385 | -0.6828143 | 0.66948249 | -1.0199136 | 0.30776943 | 0.8673137 |
| INO80D | 84.652268 | 0.01230161 | 0.71133366 | 0.01729373 | 0.98620229 | 0.99811561 |
| NDUFS1 | 32.3889581 | 0.36305813 | 0.38919117 | 0.93285294 | 0.35089591 | 0.89076052 |
| EEF1B2 | 32.761796 | -0.2439227 | 0.58845188 | -0.414516 | 0.67849626 | 0.96723295 |
| FASTKD2 | 11.0115692 | -0.3486412 | 0.62201892 | -0.5604993 | 0.57513892 | NA |
| KLF7 | 47.2503733 | 0.26785959 | 0.35417033 | 0.7563016 | 0.44946839 | 0.92914597 |
| CREB1 | 57.9889058 | 0.28198215 | 0.30762395 | 0.91664561 | 0.35932838 | 0.8957272 |

| | | | | | | |
|---|---|---|---|---|---|---|
| METTL21A | 14.4168056 | -0.1324508 | 0.56026536 | -0.2364072 | 0.81311668 | 0.98320463 |
| CCNYL1 | 14.851578 | -0.4554106 | 0.45113285 | -1.0094823 | 0.3127434 | 0.87058449 |
| PLEKHM3 | 37.1012242 | 0.41181942 | 0.35980018 | 1.14457815 | 0.25238395 | 0.84008912 |
| IDH1 | 18.8323957 | 0.88652661 | 0.52596773 | 1.6855152 | 0.09188922 | 0.65902576 |
| PIKFYVE | 71.3283739 | -0.9033433 | 0.31065478 | -2.9078685 | 0.00363901 | 0.15372746 |
| PTH2R | 15.8132278 | -0.493136 | 0.57880723 | -0.8519865 | 0.39422158 | 0.90940709 |
| RPE | 17.3557945 | -0.422194 | 0.44552506 | -0.9476324 | 0.34331665 | 0.88574094 |
| KANSL1L | 38.4482439 | 0.70955078 | 0.50929073 | 1.39321363 | 0.1635552 | 0.75321592 |
| AC007038.1 | 11.7690764 | -0.0177689 | 0.61107478 | -0.0290781 | 0.97680229 | NA |
| LANCL1 | 13.264449 | -0.1133349 | 0.47154821 | -0.2403463 | 0.81006177 | 0.98320463 |
| ERBB4 | 29.6248466 | -0.9644454 | 0.69407719 | -1.3895362 | 0.16466975 | 0.75321592 |
| IKZF2 | 46.6069589 | 0.74870179 | 0.68368181 | 1.09510268 | 0.27347165 | 0.84994492 |
| SPAG16 | 10.4294886 | -0.2024325 | 0.64706533 | -0.3128471 | 0.75439686 | NA |
| BARD1 | 18.7749906 | -0.0572403 | 0.47308754 | -0.120993 | 0.90369661 | 0.99324672 |
| ATIC | 9.20994954 | -1.1226935 | 0.58446419 | -1.9208936 | 0.05474512 | NA |
| XRCC5 | 92.3231895 | -0.1368579 | 0.27513086 | -0.4974285 | 0.6188869 | 0.95688177 |
| SMARCAL1 | 15.6558614 | -0.310709 | 0.47164925 | -0.6587713 | 0.51004267 | 0.94332619 |
| RPL37A | 58.7188098 | -0.3666506 | 0.4549734 | -0.8058727 | 0.42031626 | 0.91985469 |
| IGFBP5 | 52.3943393 | -0.5676326 | 1.39747003 | -0.4061859 | 0.684606 | 0.9673431 |
| DIRC3 | 65.0129771 | -0.5537877 | 0.76309497 | -0.7257126 | 0.46801495 | 0.9314919 |
| TNS1 | 21.9034568 | -0.9766636 | 0.61752483 | -1.5815779 | 0.11374597 | 0.69838966 |
| ARPC2 | 120.21059 | -0.6514931 | 0.30137333 | -2.1617477 | 0.03063763 | 0.43032651 |
| PNKD | 16.4563367 | 1.02009715 | 0.54146884 | 1.88394434 | 0.0595725 | 0.57281427 |
| TMBIM1 | 19.1944861 | 0.14115886 | 0.543647 | 0.25965169 | 0.79513246 | 0.98320463 |
| SLC11A1 | 113.75396 | 0.97669593 | 0.53344983 | 1.83090493 | 0.06711473 | 0.58951068 |
| CTDSP1 | 15.7418461 | 0.14549087 | 0.52241834 | 0.27849496 | 0.78063244 | 0.98264092 |
| USP37 | 31.3169949 | -0.4500872 | 0.4188396 | -1.0746051 | 0.28255155 | 0.85176121 |
| CNOT9 | 15.6238454 | -0.6883005 | 0.53032542 | -1.2978833 | 0.19432745 | 0.78135737 |
| RETREG2 | 11.4577338 | 0.13346254 | 0.52515932 | 0.25413724 | 0.79938953 | NA |
| ANKZF1 | 11.0766243 | -0.2969448 | 0.51330317 | -0.5784978 | 0.56292808 | NA |
| TUBA4A | 14.0980478 | -0.1679357 | 0.62768469 | -0.2675479 | 0.78904736 | 0.98320463 |
| FARSB | 14.0744642 | -0.6859141 | 0.54343618 | -1.2621797 | 0.20688412 | 0.79234878 |
| ACSL3 | 51.458336 | 0.83715875 | 0.53462863 | 1.56586965 | 0.11737914 | 0.70122655 |
| AP1S3 | 10.0485863 | -0.4151702 | 0.62354948 | -0.6658175 | 0.50552774 | NA |
| WDFY1 | 36.881423 | 0.30259865 | 0.38150172 | 0.79317767 | 0.42767432 | 0.92129717 |
| CUL3 | 92.5046585 | -0.154416 | 0.29608637 | -0.5215235 | 0.60200214 | 0.95688177 |
| DOCK10 | 275.049861 | -0.300807 | 0.33522614 | -0.8973256 | 0.36954522 | 0.90130596 |
| RHBDD1 | 44.5161833 | -0.4143239 | 0.50318357 | -0.8234051 | 0.41027766 | 0.91773508 |
| COL4A3 | 9.87196292 | -0.5515218 | 0.64890914 | -0.8499214 | 0.39536879 | NA |
| MFF | 24.0259575 | -0.1836254 | 0.43383659 | -0.4232593 | 0.67210612 | 0.96412305 |
| AGFG1 | 131.915383 | -0.2431563 | 0.4562598 | -0.5329339 | 0.59407937 | 0.9553365 |
| PID1 | 125.742242 | 0.32329252 | 0.87998469 | 0.36738426 | 0.71333241 | 0.97348719 |
| TRIP12 | 119.432795 | -0.1198707 | 0.23133387 | -0.518172 | 0.6043383 | 0.95688177 |
| SP110 | 63.0313322 | -0.4955839 | 0.58022467 | -0.8541241 | 0.39303623 | 0.90930707 |
| SP140 | 42.8125619 | -0.4721389 | 0.41195642 | -1.1460895 | 0.25175811 | 0.84008912 |
| SP140L | 36.585538 | -0.4416317 | 0.37830616 | -1.1673924 | 0.24305195 | 0.83507265 |
| SP100 | 165.859056 | -0.1287164 | 0.34361062 | -0.3745995 | 0.70795836 | 0.97184211 |
| CAB39 | 139.152233 | -0.0342684 | 0.25503339 | -0.1343685 | 0.89311123 | 0.99080204 |
| ITM2C | 9.79699798 | -0.9514638 | 0.78014176 | -1.2196038 | 0.22261512 | NA |
| PSMD1 | 48.2831385 | -0.3556364 | 0.3342181 | -1.0640847 | 0.28729032 | 0.85677513 |
| ARMC9 | 31.1888338 | -0.1572371 | 0.57680453 | -0.2726003 | 0.78516047 | 0.98264092 |
| NCL | 31.0859226 | -0.4856762 | 0.41902224 | -1.1590701 | 0.2464276 | 0.83817091 |
| PTMA | 165.252537 | -0.6457302 | 0.49514939 | -1.3041118 | 0.19219547 | 0.78135737 |
| PDE6D | 16.8573477 | 0.92441997 | 0.49480826 | 1.86823874 | 0.0617288 | 0.57509627 |

| | | | | | | |
|---|---|---|---|---|---|---|
| DIS3L2 | 47.2856189 | -0.2871797 | 0.37791568 | -0.7599042 | 0.44731185 | 0.92914597 |
| EIF4E2 | 13.5790253 | 0.8709397 | 0.49207867 | 1.76991964 | 0.07674053 | 0.62507515 |
| GIGYF2 | 73.2667806 | 0.13165857 | 0.30331157 | 0.43407038 | 0.66423733 | 0.96384486 |
| INPP5D | 182.034177 | 0.14356163 | 0.32639363 | 0.439842 | 0.66005155 | 0.96298563 |
| ATG16L1 | 21.153729 | -0.1785691 | 0.41093337 | -0.4345451 | 0.66389264 | 0.96379377 |
| DGKD | 92.0946676 | -1.3807649 | 0.48848187 | -2.8266452 | 0.00470384 | 0.17239111 |
| USP40 | 19.7504031 | 0.84599415 | 0.50930232 | 1.66108441 | 0.0966965 | 0.66178928 |
| ARL4C | 50.4768456 | -0.0977812 | 0.45006543 | -0.2172599 | 0.8280058 | 0.98320463 |
| AGAP1 | 31.6897367 | 0.02827112 | 0.44714444 | 0.06322591 | 0.94958661 | 0.99514836 |
| LRRFIP1 | 171.873164 | -0.4014808 | 0.3095632 | -1.2969267 | 0.19465641 | 0.78135737 |
| ILKAP | 21.5095221 | -0.3344567 | 0.42055701 | -0.7952707 | 0.42645607 | 0.92112288 |
| HDAC4 | 80.0829226 | 0.41710954 | 0.32136997 | 1.29791074 | 0.19431801 | 0.78135737 |
| NDUFA10 | 32.9570493 | -0.5634457 | 0.40206732 | -1.4013715 | 0.161103 | 0.74887162 |
| COPS9 | 10.6882181 | -0.5834077 | 0.63805043 | -0.9143598 | 0.3605278 | NA |
| ANKMY1 | 23.5934852 | -0.5615895 | 0.63080072 | -0.8902804 | 0.37331534 | 0.9025457 |
| RNPEPL1 | 16.0249206 | 0.45320584 | 0.62431276 | 0.72592755 | 0.46788318 | 0.9314919 |
| SNED1 | 19.8874866 | -0.5780659 | 0.75162515 | -0.769088 | 0.44184104 | 0.92899684 |
| MTERF4 | 19.8776318 | -0.5140334 | 0.45100871 | -1.1397416 | 0.25439398 | 0.84026145 |
| PPP1R7 | 11.3342936 | -0.231322 | 0.56283241 | -0.4109963 | 0.68107526 | NA |
| HDLBP | 50.3395873 | -0.0515251 | 0.34285923 | -0.1502807 | 0.88054319 | 0.99019008 |
| 2-Sep | 68.9878748 | -0.0039079 | 0.33334265 | -0.0117232 | 0.99064642 | 0.99827179 |
| FARP2 | 31.1891158 | -1.4256918 | 0.39669709 | -3.5939054 | 0.00032576 | 0.03466819 |
| STK25 | 10.3631655 | -0.6676646 | 0.50907183 | -1.3115332 | 0.18967769 | NA |
| ATG4B | 23.2408121 | -0.0429741 | 0.3829986 | -0.1122043 | 0.9106614 | 0.99471296 |
| ING5 | 14.9348252 | -0.2490976 | 0.50290836 | -0.4953141 | 0.62037844 | 0.95688177 |
| D2HGDH | 19.3479093 | 0.26434033 | 0.53508027 | 0.49401995 | 0.62129209 | 0.95702817 |
| TRNT1 | 16.9936868 | 0.60123081 | 0.67527854 | 0.89034491 | 0.37328071 | 0.9025457 |
| CRBN | 29.1198863 | 0.75007049 | 0.54034635 | 1.38812908 | 0.16509774 | 0.75321592 |
| SUMF1 | 127.134303 | 0.15208462 | 0.32169319 | 0.47276295 | 0.63638231 | 0.95933464 |
| ITPR1 | 94.8022677 | 0.82776435 | 0.40170316 | 2.0606369 | 0.03933769 | 0.48300729 |
| AC018816.1 | 12.1314286 | 1.00089947 | 0.73370941 | 1.36416332 | 0.17251617 | NA |
| BHLHE40 | 21.4777617 | -0.5372032 | 0.70320157 | -0.7639392 | 0.44490351 | 0.92914597 |
| ARL8B | 65.148572 | -0.2373891 | 0.34398718 | -0.6901104 | 0.49012473 | 0.93633431 |
| AC026202.2 | 13.562478 | 0.62363967 | 0.84557599 | 0.73753238 | 0.46079866 | 0.9314919 |
| EDEM1 | 34.428918 | -0.7692679 | 0.5481768 | -1.4033208 | 0.16052121 | 0.74887162 |
| RAD18 | 11.1495839 | -1.0420794 | 0.59666079 | -1.746519 | 0.08072081 | NA |
| SRGAP3 | 20.8138354 | 1.51129387 | 0.58318801 | 2.59143508 | 0.00955766 | 0.26141219 |
| THUMPD3-A: | 30.9411438 | -0.0153088 | 0.4510268 | -0.033942 | 0.97292339 | 0.9973754 |
| SETD5 | 96.3102835 | -0.0179921 | 0.24793379 | -0.072568 | 0.94214992 | 0.99514836 |
| MTMR14 | 31.9608521 | 0.14595759 | 0.40495497 | 0.36042918 | 0.71852621 | 0.97348719 |
| OGG1 | 10.0681822 | -0.2140337 | 0.6025301 | -0.3552249 | 0.72242111 | NA |
| TADA3 | 8.38250103 | -0.3371958 | 0.61404576 | -0.5491378 | 0.58291087 | NA |
| EMC3 | 14.0712942 | -0.3813943 | 0.54659135 | -0.6977687 | 0.48532188 | 0.93417265 |
| BRK1 | 19.0937709 | 0.01063793 | 0.54400904 | 0.01955469 | 0.98439861 | 0.99811561 |
| VHL | 12.647085 | -0.7227286 | 0.48407679 | -1.4930041 | 0.13543611 | NA |
| IRAK2 | 19.3508037 | 0.14537054 | 0.51487619 | 0.28234077 | 0.77768223 | 0.98264092 |
| SEC13 | 18.6648651 | 0.5715442 | 0.46565404 | 1.22740092 | 0.21967194 | 0.81042354 |
| ATG7 | 201.844858 | 0.20389016 | 0.44083601 | 0.46250795 | 0.6437171 | 0.95992767 |
| VGLL4 | 40.8530438 | 0.16230134 | 0.33546829 | 0.48380531 | 0.62852405 | 0.95877428 |
| TAMM41 | 19.2358871 | -0.5054832 | 0.41165985 | -1.2279148 | 0.21947897 | 0.81042354 |
| PPARG | 37.9913141 | -0.6092478 | 0.84413797 | -0.7217396 | 0.4704546 | 0.9314919 |
| RAF1 | 64.67508 | -0.0369112 | 0.25592385 | -0.1442272 | 0.88532103 | 0.99019008 |
| RPL32 | 111.327657 | -0.5945082 | 0.52976544 | -1.1222103 | 0.26177304 | 0.84837688 |
| IQSEC1 | 75.0131465 | -0.8722162 | 0.36829801 | -2.368235 | 0.01787318 | 0.34827529 |

| | | | | | | |
|---|---|---|---|---|---|---|
| NUP210 | 49.0220879 | -0.6269778 | 0.31334733 | -2.0009036 | 0.04540278 | 0.52128421 |
| XPC | 20.4180871 | 0.06653968 | 0.47958513 | 0.13874426 | 0.88965224 | 0.99080204 |
| SLC6A6 | 42.6584919 | 0.16946926 | 0.36347976 | 0.46624128 | 0.64104281 | 0.95992767 |
| CCDC174 | 12.0250514 | 0.51797055 | 0.55253292 | 0.93744741 | 0.34852846 | NA |
| FGD5-AS1 | 17.696495 | -0.0750163 | 0.5090334 | -0.1473702 | 0.88283985 | 0.99019008 |
| NR2C2 | 52.7159852 | 0.32183303 | 0.30638572 | 1.05041788 | 0.29352603 | 0.86027209 |
| MRPS25 | 12.2399955 | 0.82746621 | 0.55471868 | 1.49168622 | 0.13578142 | NA |
| RBSN | 13.278352 | 0.49882458 | 0.51009537 | 0.97790453 | 0.32812154 | 0.87848313 |
| CAPN7 | 47.448539 | -0.2563082 | 0.42776228 | -0.5991838 | 0.54905029 | 0.95415437 |
| SH3BP5 | 53.0261614 | -0.5470218 | 0.38208808 | -1.4316642 | 0.15223995 | 0.74315008 |
| METTL6 | 9.48080495 | 0.27021763 | 0.63489059 | 0.42561291 | 0.67038994 | NA |
| HACL1 | 13.5229217 | 0.59685445 | 0.55166391 | 1.08191681 | 0.2792895 | 0.84994492 |
| ANKRD28 | 88.4331585 | 0.21829074 | 0.79905255 | 0.27318696 | 0.7847095 | 0.98264092 |
| OXNAD1 | 57.2138719 | -0.4681613 | 0.38556812 | -1.2142117 | 0.2246669 | 0.81875277 |
| RFTN1 | 103.276603 | -1.4846514 | 0.46360183 | -3.2024278 | 0.00136274 | 0.07695335 |
| PLCL2 | 152.536609 | 0.10384844 | 0.25442008 | 0.40817707 | 0.68314368 | 0.9673431 |
| TBC1D5 | 284.420781 | 0.04327876 | 0.2668455 | 0.16218658 | 0.87115893 | 0.98751288 |
| RAB5A | 33.3087871 | 0.01436121 | 0.38304313 | 0.03749241 | 0.9700924 | 0.9967641 |
| KAT2B | 98.8159104 | 0.28485215 | 0.31427135 | 0.90638918 | 0.3647299 | 0.89868152 |
| UBE2E2 | 183.950551 | -0.5842396 | 0.38583637 | -1.5142159 | 0.12997109 | 0.72931666 |
| UBE2E1 | 81.9581087 | 0.06175381 | 0.24705293 | 0.24996187 | 0.80261684 | 0.98320463 |
| RPL15 | 128.826296 | -0.1916601 | 0.55550534 | -0.3450193 | 0.73007984 | 0.97605445 |
| NR1D2 | 19.0050278 | -0.113348 | 0.42982392 | -0.2637081 | 0.79200488 | 0.98320463 |
| THRB | 35.0194911 | 0.62139892 | 0.82001036 | 0.75779398 | 0.44857433 | 0.92914597 |
| TOP2B | 65.5479878 | 0.37948443 | 0.38408702 | 0.9880168 | 0.32314442 | 0.87489296 |
| NGLY1 | 49.4520107 | -0.0798592 | 0.36809985 | -0.2169497 | 0.82824753 | 0.98320463 |
| NEK10 | 20.5671139 | 1.40263845 | 1.23676598 | 1.13411791 | 0.25674509 | 0.84072794 |
| SLC4A7 | 95.4553391 | -0.1055955 | 0.47793402 | -0.2209416 | 0.82513794 | 0.98320463 |
| CMC1 | 43.2900166 | 0.51681882 | 0.52018953 | 0.99352023 | 0.3204565 | 0.87363761 |
| AZI2 | 25.2042879 | -0.0579045 | 0.46484585 | -0.1245672 | 0.90086621 | 0.99194856 |
| TGFBR2 | 126.184444 | -0.3268542 | 0.33380845 | -0.9791668 | 0.32749755 | 0.87723689 |
| STT3B | 83.7456304 | 0.20375947 | 0.31550073 | 0.64582885 | 0.5183902 | 0.94668603 |
| OSBPL10 | 38.6423816 | -2.64801 | 0.57643452 | -4.5937741 | 4.35E-06 | 0.00200746 |
| GPD1L | 10.5251846 | -0.1680192 | 0.53101423 | -0.3164118 | 0.75168994 | NA |
| CMTM7 | 51.3683686 | 0.54655643 | 0.41334634 | 1.32227234 | 0.18607748 | 0.77798634 |
| CMTM6 | 57.3554057 | 0.38563328 | 0.50848455 | 0.75839724 | 0.44821321 | 0.92914597 |
| DYNC1LI1 | 24.2305076 | 0.06115775 | 0.45166045 | 0.13540648 | 0.8922905 | 0.99080204 |
| CNOT10 | 29.1058922 | -0.300331 | 0.40334563 | -0.7445997 | 0.45651377 | 0.9314919 |
| GLB1 | 36.2244787 | 0.63418857 | 0.38525937 | 1.64613406 | 0.09973616 | 0.6683775 |
| CRTAP | 12.8261597 | 0.11258824 | 0.4678664 | 0.24064185 | 0.80983271 | NA |
| UBP1 | 29.8101788 | 0.06213517 | 0.38937086 | 0.15957837 | 0.87321322 | 0.98813417 |
| CLASP2 | 100.037614 | 0.07783475 | 0.31254151 | 0.24903814 | 0.80333128 | 0.98320463 |
| PDCD6IP | 73.0195429 | 0.16379826 | 0.29952344 | 0.54686291 | 0.58447292 | 0.95415437 |
| TRANK1 | 40.0435908 | 0.37493295 | 0.48046223 | 0.78035884 | 0.43517969 | 0.92377614 |
| EPM2AIP1 | 15.869975 | 0.34073562 | 0.46479867 | 0.73308219 | 0.4635083 | 0.9314919 |
| MLH1 | 12.7072156 | -0.504641 | 0.48381574 | -1.0430437 | 0.29692806 | NA |
| LRRFIP2 | 61.815233 | 0.2749652 | 0.31772853 | 0.86540922 | 0.38681422 | 0.90910959 |
| GOLGA4 | 122.46856 | 0.01178754 | 0.21835172 | 0.0539842 | 0.95694775 | 0.99514836 |
| ITGA9 | 25.0889717 | -0.1453592 | 0.57086695 | -0.2546289 | 0.79900977 | 0.98320463 |
| ACAA1 | 17.2194281 | 1.2171102 | 0.64097443 | 1.8988436 | 0.05758504 | 0.56804925 |
| MYD88 | 11.2425007 | 0.39834942 | 0.76725321 | 0.51918899 | 0.60362895 | NA |
| OXSR1 | 43.9650404 | 0.08856103 | 0.31865081 | 0.27792502 | 0.78106992 | 0.98264092 |
| EXOG | 14.9038194 | -0.7970883 | 0.43098581 | -1.8494537 | 0.06439233 | 0.58706281 |
| WDR48 | 20.4611415 | 0.53735945 | 0.51620315 | 1.04098445 | 0.29788277 | 0.8608685 |

| | | | | | | |
|---|---|---|---|---|---|---|
| CX3CR1 | 18.4400913 | 2.70856451 | 1.04565632 | 2.5903009 | 0.00958921 | 0.26141219 |
| RPSA | 62.9938338 | -0.2761249 | 0.57860025 | -0.4772291 | 0.63319902 | 0.95877428 |
| EIF1B | 14.0724724 | -0.1181598 | 0.50375207 | -0.2345595 | 0.81455065 | 0.98320463 |
| RPL14 | 115.521505 | -0.208391 | 0.67186231 | -0.3101691 | 0.75643233 | 0.97989151 |
| ZNF621 | 9.87278524 | 0.23393533 | 0.56996995 | 0.4104345 | 0.68148724 | NA |
| CTNNB1 | 99.4840368 | -0.002738 | 0.42781568 | -0.0064001 | 0.99489353 | 0.99859588 |
| ULK4 | 44.375055 | -0.1488655 | 0.33025043 | -0.4507654 | 0.65215865 | 0.96178254 |
| TRAK1 | 82.1308469 | -0.7301904 | 0.30394649 | -2.402365 | 0.01628945 | 0.33511451 |
| NKTR | 132.045676 | 0.139482 | 0.27207454 | 0.51266097 | 0.60818849 | 0.95688177 |
| AC006059.1 | 20.4248264 | 0.37570349 | 0.52706417 | 0.71282305 | 0.47595526 | 0.9314919 |
| SNRK | 54.3198116 | 0.78648845 | 0.33270622 | 2.36391263 | 0.01808308 | 0.34868204 |
| ANO10 | 35.1636187 | 0.56845734 | 0.40312396 | 1.41013039 | 0.15850118 | 0.74650686 |
| ABHD5 | 14.7133768 | 1.11007777 | 0.66215279 | 1.67646771 | 0.09364662 | 0.6590317 |
| TCAIM | 25.7657256 | -0.0141052 | 0.36803048 | -0.0383262 | 0.96942762 | 0.99662129 |
| ZNF445 | 13.3107656 | 0.42537146 | 0.52222919 | 0.81453023 | 0.41534126 | 0.91894074 |
| ZDHHC3 | 16.7195698 | -0.2488025 | 0.49041894 | -0.5073265 | 0.61192573 | 0.95688177 |
| CDCP1 | 12.3076589 | 0.49409986 | 1.04689508 | 0.47196693 | 0.63695039 | NA |
| LARS2 | 19.9286844 | -0.2925912 | 0.45626664 | -0.6412723 | 0.52134576 | 0.94895469 |
| LIMD1 | 60.8971575 | -0.5016515 | 0.38501298 | -1.3029468 | 0.19259291 | 0.78135737 |
| SACM1L | 48.4575438 | 0.82113135 | 0.45314002 | 1.81209189 | 0.069972 | 0.60598598 |
| FYCO1 | 17.4640088 | 0.4970964 | 0.50079751 | 0.99260958 | 0.32090025 | 0.87363761 |
| CCR1 | 14.5326682 | 1.15028951 | 0.84795041 | 1.35655281 | 0.17492334 | 0.76214163 |
| LTF | 22.128374 | 2.19588057 | 0.90677804 | 2.42162962 | 0.01545109 | 0.32566073 |
| CCDC12 | 31.1424089 | -0.7987182 | 0.48352823 | -1.6518543 | 0.09856425 | 0.66681487 |
| NBEAL2 | 27.4479929 | -1.4375153 | 0.67556273 | -2.1278784 | 0.03334717 | 0.45010547 |
| SETD2 | 123.598832 | -0.3793322 | 0.24694398 | -1.5361065 | 0.12451227 | 0.71741321 |
| KIF9-AS1 | 13.686665 | -0.0948481 | 0.59170897 | -0.1602952 | 0.87264855 | 0.98813417 |
| KLHL18 | 20.3723099 | -0.5820223 | 0.47179475 | -1.2336345 | 0.21733914 | 0.80640512 |
| SCAP | 15.5518834 | 0.23881256 | 0.50375934 | 0.47406081 | 0.63545654 | 0.95933464 |
| SMARCC1 | 73.5918947 | -0.0434161 | 0.32259594 | -0.1345837 | 0.89294106 | 0.99080204 |
| DHX30 | 10.5605287 | -0.670887 | 0.58881615 | -1.1393829 | 0.25454349 | NA |
| MAP4 | 92.5315952 | 0.05859557 | 0.32957431 | 0.17779167 | 0.85888659 | 0.98751288 |
| ZNF589 | 10.0854869 | -0.240524 | 0.5403533 | -0.4451236 | 0.65623046 | NA |
| TMA7 | 24.0350828 | -0.3659152 | 0.71341188 | -0.5129087 | 0.60801515 | 0.95688177 |
| SHISA5 | 20.9224049 | 0.8678601 | 0.44679871 | 1.94239613 | 0.05208916 | 0.54907299 |
| PFKFB4 | 16.6277018 | 1.03943678 | 0.81166177 | 1.28062799 | 0.20032436 | 0.78345938 |
| UQCRC1 | 8.69190593 | 0.89693958 | 0.72572604 | 1.23592034 | 0.21648818 | NA |
| IP6K2 | 16.896699 | -0.5353078 | 0.50827176 | -1.053192 | 0.292253 | 0.85950405 |
| PRKAR2A | 44.7880203 | 0.22766716 | 0.39460421 | 0.57695064 | 0.5639728 | 0.95415437 |
| SLC25A20 | 8.9437772 | 1.45277071 | 0.79522837 | 1.82685976 | 0.06772086 | NA |
| ARIH2 | 43.7660065 | 0.31554563 | 0.34127051 | 0.92462025 | 0.35516348 | 0.89303957 |
| WDR6 | 13.3311884 | -0.3551668 | 0.5071162 | -0.7003658 | 0.48369889 | 0.93399425 |
| QRICH1 | 29.125554 | -0.389145 | 0.35418901 | -1.0986932 | 0.27190192 | 0.84994492 |
| USP4 | 43.240622 | -0.568174 | 0.4240205 | -1.3399683 | 0.18025566 | 0.76851683 |
| RHOA | 92.9577533 | -0.5595401 | 0.352964 | -1.5852611 | 0.11290702 | 0.69735209 |
| IP6K1 | 32.4386422 | 0.47914719 | 0.36951718 | 1.29668447 | 0.19473978 | 0.78135737 |
| RBM6 | 158.904411 | -0.22296 | 0.30973892 | -0.7198321 | 0.47162837 | 0.9314919 |
| RBM5 | 52.5725553 | 0.14428812 | 0.29009512 | 0.49738208 | 0.61891964 | 0.95688177 |
| GNAI2 | 76.3058003 | 0.39505979 | 0.58311568 | 0.67749813 | 0.49808995 | 0.93916044 |
| RASSF1 | 31.4685717 | -0.0160193 | 0.45954721 | -0.0348589 | 0.97219225 | 0.99734828 |
| MAPKAPK3 | 18.3170622 | 0.7436001 | 0.60793089 | 1.22316552 | 0.22126719 | 0.81199775 |
| MANF | 11.3721988 | 0.18144142 | 0.72713078 | 0.24953065 | 0.80295033 | NA |
| DCAF1 | 21.7849065 | 0.21781227 | 0.43232091 | 0.5038208 | 0.61438731 | 0.95688177 |
| RAD54L2 | 19.9888259 | -0.5031591 | 0.47334053 | -1.0629959 | 0.28778379 | 0.85677513 |

| | | | | | | |
|---|---|---|---|---|---|---|
| TEX264 | 8.83889156 | 0.66237725 | 0.60069448 | 1.10268576 | 0.27016365 | NA |
| RPL29 | 140.850963 | -0.4117152 | 0.56921804 | -0.7232996 | 0.46949581 | 0.9314919 |
| WDR82 | 30.6753604 | 0.33235004 | 0.44814666 | 0.74160999 | 0.45832367 | 0.9314919 |
| DNAH1 | 10.8337232 | -0.7771749 | 0.49977656 | -1.5550447 | 0.11993543 | NA |
| NISCH | 18.0439047 | 0.31216035 | 0.45604332 | 0.68449714 | 0.4936613 | 0.93720536 |
| STAB1 | 41.2572311 | -0.2743755 | 0.56222896 | -0.4880138 | 0.6255401 | 0.95876809 |
| SMIM4 | 11.4846614 | -0.2045349 | 0.73801743 | -0.277141 | 0.78167184 | NA |
| PBRM1 | 91.9905441 | 0.03760914 | 0.26471555 | 0.14207378 | 0.88702173 | 0.99019008 |
| GNL3 | 11.8270508 | -0.3502633 | 0.50599122 | -0.6922319 | 0.48879172 | NA |
| SPCS1 | 20.0203597 | 0.2824344 | 0.53243212 | 0.53046086 | 0.59579244 | 0.9553365 |
| NEK4 | 12.9688567 | 0.43921739 | 0.67796817 | 0.64784368 | 0.51708606 | 0.94596835 |
| SFMBT1 | 15.8773464 | -1.3196565 | 0.54201747 | -2.4347122 | 0.01490364 | 0.32436474 |
| PRKCD | 25.8573601 | 0.42762477 | 0.48172391 | 0.88769681 | 0.37470386 | 0.90299244 |
| TKT | 29.3821254 | 0.74898095 | 0.49991972 | 1.49820244 | 0.13408066 | 0.73044824 |
| DCP1A | 36.2624615 | -0.2932309 | 0.42343157 | -0.6925106 | 0.48861673 | 0.93596573 |
| SELENOK | 14.2013908 | 0.53608435 | 0.59054592 | 0.90777759 | 0.36399574 | 0.89868152 |
| CACNA2D3 | 12.325075 | -1.0967037 | 0.54752916 | -2.0030051 | 0.04517674 | NA |
| CCDC66 | 31.9360507 | 0.17262735 | 0.48655522 | 0.35479498 | 0.72274319 | 0.97441549 |
| FAM208A | 48.058658 | -0.3480225 | 0.39281433 | -0.8859719 | 0.37563266 | 0.90341205 |
| ARHGEF3 | 102.935753 | 0.22637199 | 0.30915587 | 0.73222608 | 0.46403059 | 0.9314919 |
| APPL1 | 31.0671268 | 0.5648095 | 0.38282493 | 1.47537281 | 0.14011234 | 0.73044824 |
| ARF4 | 21.6172813 | 0.18499458 | 0.4459836 | 0.41480131 | 0.67828736 | 0.96718429 |
| DENND6A | 50.6857957 | -0.1095118 | 0.29141397 | -0.3757946 | 0.70706961 | 0.97184211 |
| DENND6A-AS | 11.4813672 | -0.055768 | 0.66478113 | -0.0838893 | 0.93314442 | NA |
| SLMAP | 76.3906951 | -0.2174412 | 0.26060177 | -0.834381 | 0.4040663 | 0.91528253 |
| FLNB | 18.8492612 | 0.34814087 | 0.49753483 | 0.69973166 | 0.4840949 | 0.93399425 |
| ABHD6 | 17.5361685 | -1.2148998 | 0.5927394 | -2.0496356 | 0.0404 | 0.48881259 |
| PXK | 38.8905785 | -1.1187938 | 0.33047954 | -3.3853647 | 0.00071084 | 0.05315912 |
| FHIT | 171.227666 | -0.2642315 | 0.3100133 | -0.8523233 | 0.3940347 | 0.90940709 |
| PTPRG | 31.1146133 | 0.28728121 | 0.84403023 | 0.34036839 | 0.73357912 | 0.97605445 |
| THOC7 | 10.2600015 | 0.08873592 | 0.6438149 | 0.13782832 | 0.8903761 | NA |
| ATXN7 | 10.968241 | -0.1782064 | 0.53842732 | -0.3309759 | 0.74066271 | NA |
| ADAMTS9-AS | 10.2453985 | -0.5875778 | 0.83211052 | -0.7061295 | 0.48010761 | NA |
| MAGI1 | 41.0002612 | -1.0350768 | 0.70460411 | -1.4690189 | 0.14182766 | 0.73044824 |
| SLC25A26 | 31.529475 | -0.2841644 | 0.38762007 | -0.7331002 | 0.46349729 | 0.9314919 |
| LRIG1 | 18.8613392 | 0.93950283 | 0.73020772 | 1.28662408 | 0.19822535 | 0.78142991 |
| SUCLG2 | 118.052291 | -0.0701836 | 0.25894773 | -0.2710338 | 0.78636501 | 0.98300067 |
| FAM19A1 | 20.1349692 | -0.0190927 | 0.9072865 | -0.0210438 | 0.98321074 | 0.99763261 |
| TMF1 | 35.3349361 | 0.02229163 | 0.35128191 | 0.06345796 | 0.94940183 | 0.99514836 |
| UBA3 | 35.3111772 | 0.30976689 | 0.32596524 | 0.95030651 | 0.34195653 | 0.88510795 |
| ARL6IP5 | 42.1601429 | -0.1262339 | 0.51060194 | -0.2472257 | 0.80473352 | 0.98320463 |
| FRMD4B | 134.967574 | 0.1786616 | 0.7166146 | 0.24931337 | 0.80311839 | 0.98320463 |
| MITF | 59.0344998 | -0.1859625 | 0.64599899 | -0.2878681 | 0.77344774 | 0.98264092 |
| FOXP1 | 461.609556 | 0.24424461 | 0.27285241 | 0.89515285 | 0.37070538 | 0.90226557 |
| EIF4E3 | 25.4160346 | 0.63555327 | 0.53404932 | 1.19006475 | 0.23402095 | 0.82817204 |
| RYBP | 57.4031204 | 0.78402898 | 0.32944288 | 2.37986317 | 0.01731907 | 0.34827529 |
| SHQ1 | 14.2815971 | -0.2197311 | 0.50036353 | -0.4391429 | 0.66055803 | 0.96298563 |
| GXYLT2 | 10.6495715 | 0.70837511 | 0.61536059 | 1.15115449 | 0.24966868 | NA |
| PPP4R2 | 44.9611783 | 0.15291728 | 0.34394011 | 0.4446044 | 0.65660567 | 0.96230291 |
| GBE1 | 57.9630622 | -0.4676893 | 0.4667373 | -1.0020397 | 0.3163244 | 0.87263466 |
| CHMP2B | 10.8897807 | 0.46997479 | 0.5324151 | 0.88272251 | 0.37738622 | NA |
| CGGBP1 | 39.6440111 | 0.12106525 | 0.3357992 | 0.3605287 | 0.7184518 | 0.97348719 |
| ZNF654 | 45.0131986 | 0.68490439 | 0.33741944 | 2.02983088 | 0.04237373 | 0.50429299 |
| C3orf38 | 11.987341 | -0.7449838 | 0.50941082 | -1.4624421 | 0.14362009 | NA |

| | | | | | | |
|---|---|---|---|---|---|---|
| ARL13B | 12.9751474 | 0.2882876 | 0.55000341 | 0.52415602 | 0.60017003 | 0.95688177 |
| NSUN3 | 24.8225905 | -0.1621557 | 0.40448506 | -0.4008941 | 0.68849813 | 0.9673431 |
| MTRNR2L12 | 58.282646 | -0.0803113 | 0.56992675 | -0.1409151 | 0.88793703 | 0.99019008 |
| CRYBG3 | 19.1956939 | 0.55538064 | 0.50662123 | 1.09624432 | 0.27297187 | 0.84994492 |
| CLDND1 | 20.425447 | -0.0380462 | 0.58939554 | -0.0645513 | 0.94853127 | 0.99514836 |
| ST3GAL6 | 27.2621711 | 1.49430288 | 0.70581973 | 2.11711689 | 0.03424993 | 0.45650836 |
| CMSS1 | 52.3281336 | -0.3613075 | 0.42716812 | -0.8458204 | 0.39765293 | 0.91429865 |
| FILIP1L | 24.7094307 | 0.06973266 | 0.56970415 | 0.12240154 | 0.90258102 | 0.99262388 |
| TBC1D23 | 27.3123104 | -0.4357756 | 0.37224972 | -1.1706539 | 0.24173793 | 0.83493679 |
| TOMM70 | 12.0764984 | -0.3048835 | 0.49500103 | -0.615925 | 0.53794404 | NA |
| TFG | 31.5919516 | 0.25443522 | 0.33294523 | 0.76419544 | 0.44475078 | 0.92914597 |
| SENP7 | 78.6931068 | 0.16342888 | 0.32625131 | 0.50092941 | 0.6164208 | 0.95688177 |
| PCNP | 31.8649755 | -0.1301301 | 0.33993692 | -0.3828067 | 0.70186306 | 0.97078484 |
| ZBTB11 | 21.9599017 | 0.11419125 | 0.46896886 | 0.24349432 | 0.80762248 | 0.98320463 |
| RPL24 | 43.4108708 | -0.6349853 | 0.54368238 | -1.1679342 | 0.2428333 | 0.83507265 |
| CEP97 | 9.066315 | -0.3899922 | 0.58288225 | -0.6690755 | 0.50344734 | NA |
| NXPE3 | 15.6859653 | 0.75768326 | 0.70032681 | 1.08189954 | 0.27929717 | 0.84994492 |
| NFKBIZ | 38.5460791 | 0.24907866 | 0.53109195 | 0.46899347 | 0.6390743 | 0.95933464 |
| ALCAM | 94.0772415 | -0.075598 | 0.53559094 | -0.1411488 | 0.88775242 | 0.99019008 |
| CBLB | 284.90565 | -0.076826 | 0.39066681 | -0.1966535 | 0.8440987 | 0.9842451 |
| BBX | 133.973626 | -0.1522333 | 0.26341033 | -0.5779322 | 0.5633099 | 0.95415437 |
| CD47 | 87.8663955 | -0.041672 | 0.31038124 | -0.1342606 | 0.8931965 | 0.99080204 |
| IFT57 | 12.2268583 | -1.4049452 | 0.60161742 | -2.3352801 | 0.01952879 | NA |
| DZIP3 | 13.0009878 | -0.254346 | 0.52165273 | -0.4875773 | 0.6258493 | 0.95876809 |
| TRAT1 | 12.9788603 | -0.3121489 | 0.72529707 | -0.4303739 | 0.66692366 | 0.96412305 |
| CD96 | 92.6210055 | 0.8342339 | 0.50841931 | 1.64083835 | 0.10083097 | 0.6683775 |
| PHLDB2 | 12.2359345 | 0.33776245 | 0.587066 | 0.57533983 | 0.56506149 | NA |
| ATG3 | 26.0491952 | -0.1166984 | 0.46159495 | -0.2528157 | 0.80041066 | 0.98320463 |
| SLC35A5 | 7.55846031 | -0.3552996 | 0.71527689 | -0.4967301 | 0.61937937 | NA |
| CD200R1 | 16.4180441 | -0.7610063 | 0.6346007 | -1.1991892 | 0.2304544 | 0.82393202 |
| NEPRO | 12.4919944 | -0.2621166 | 0.55872266 | -0.4691355 | 0.6389728 | NA |
| CFAP44 | 9.24965728 | 0.3029822 | 0.58615316 | 0.51689937 | 0.60522641 | NA |
| SPICE1 | 13.6619456 | -0.4046954 | 0.5893033 | -0.6867353 | 0.49224955 | 0.93678212 |
| SIDT1 | 36.3740091 | 0.91982581 | 0.66222346 | 1.38899611 | 0.16483393 | 0.75321592 |
| USF3 | 26.0320512 | -0.562455 | 0.51758889 | -1.0866829 | 0.27717699 | 0.84994492 |
| NAA50 | 51.6970926 | -0.4931657 | 0.45847113 | -1.0756745 | 0.28207284 | 0.85171619 |
| ATP6V1A | 44.4341952 | 0.44244845 | 0.36762576 | 1.20352952 | 0.22877147 | 0.82329316 |
| QTRT2 | 9.7777861 | -0.5105631 | 0.55728116 | -0.9161678 | 0.35957887 | NA |
| TIGIT | 8.55308601 | -0.4028449 | 0.83648409 | -0.4815931 | 0.63009506 | NA |
| ZBTB20 | 401.231086 | 0.23835808 | 0.26361612 | 0.90418629 | 0.36589663 | 0.89868152 |
| ARHGAP31 | 45.1210657 | -0.0996865 | 0.5480811 | -0.1818827 | 0.85567481 | 0.98751288 |
| TMEM39A | 19.5428349 | 0.22268218 | 0.57403446 | 0.38792475 | 0.69807172 | 0.96861929 |
| POGLUT1 | 14.0045747 | -0.2685598 | 0.69211957 | -0.3880251 | 0.69799742 | 0.96861929 |
| TIMMDC1 | 14.273679 | 0.21323605 | 0.50804431 | 0.41971939 | 0.67469046 | 0.96574154 |
| COX17 | 10.375074 | 0.52431096 | 0.59538347 | 0.88062735 | 0.37851955 | NA |
| GSK3B | 133.38086 | -0.0534405 | 0.26046363 | -0.2051744 | 0.83743587 | 0.98320463 |
| LRRC58 | 19.2818748 | 0.87860918 | 0.466857 | 1.8819664 | 0.05984058 | 0.57281427 |
| NDUFB4 | 10.3351967 | 0.07845062 | 0.71826495 | 0.10922239 | 0.9130261 | NA |
| HCLS1 | 78.3071558 | 0.08887289 | 0.33731183 | 0.26347397 | 0.79218528 | 0.98320463 |
| GOLGB1 | 56.0304407 | -0.2211965 | 0.33270669 | -0.6648394 | 0.50615322 | 0.94166217 |
| IQCB1 | 44.7081811 | -0.693329 | 0.38433856 | -1.8039537 | 0.07123856 | 0.60903299 |
| EAF2 | 35.9534925 | 0.31335377 | 0.52911068 | 0.59222726 | 0.55369842 | 0.95415437 |
| SLC15A2 | 17.1318876 | 0.29961186 | 0.48153495 | 0.62220168 | 0.53380926 | 0.95293563 |
| CD86 | 61.4468971 | 0.84536057 | 0.75658058 | 1.11734374 | 0.26384738 | 0.84928231 |

| | | | | | | |
|---|---|---|---|---|---|---|
| FAM162A | 15.8629091 | 0.38796194 | 0.48456981 | 0.80063169 | 0.4233449 | 0.91985469 |
| KPNA1 | 46.605819 | -0.1094962 | 0.30720708 | -0.3564246 | 0.7215226 | 0.97417403 |
| PARP9 | 51.7778262 | 0.1961796 | 0.59193099 | 0.3314231 | 0.74032492 | 0.97605445 |
| DTX3L | 19.1364358 | -0.5868547 | 0.68283524 | -0.8594383 | 0.39009877 | 0.90910959 |
| PARP15 | 37.436694 | -1.7781494 | 0.41833443 | -4.250545 | 2.13E-05 | 0.00621121 |
| PARP14 | 148.967803 | -1.1115751 | 0.67053438 | -1.6577451 | 0.09736893 | 0.66301053 |
| HSPBAP1 | 27.5820312 | 0.89075137 | 0.51243921 | 1.73825763 | 0.08216543 | 0.63740875 |
| DIRC2 | 41.98656 | 0.5601905 | 0.59514685 | 0.94126434 | 0.34656941 | 0.88790825 |
| SEC22A | 32.9150243 | 0.08180329 | 0.37809601 | 0.21635587 | 0.82871036 | 0.98320463 |
| HACD2 | 18.9431372 | -0.0419184 | 0.45054042 | -0.0930403 | 0.92587156 | 0.99514836 |
| CCDC14 | 28.802435 | 0.39145616 | 0.55187782 | 0.70931671 | 0.47812796 | 0.9318939 |
| ITGB5 | 16.9989377 | -0.1656353 | 0.64929624 | -0.2550997 | 0.79864607 | 0.98320463 |
| HEG1 | 26.4280983 | 0.40072112 | 0.47034556 | 0.85197173 | 0.39422978 | 0.90940709 |
| ZNF148 | 116.604701 | -0.024188 | 0.2770184 | -0.0873154 | 0.9304208 | 0.99514836 |
| SNX4 | 27.4040702 | -0.240152 | 0.36466978 | -0.6585464 | 0.51018708 | 0.94332619 |
| OSBPL11 | 31.498072 | 0.13586156 | 0.4500904 | 0.30185395 | 0.7627634 | 0.98163473 |
| ZXDC | 38.641072 | -0.0787209 | 0.47114684 | -0.1670835 | 0.86730431 | 0.98751288 |
| MGLL | 17.2595328 | 0.47644442 | 0.59387598 | 0.80226249 | 0.42240114 | 0.91985469 |
| SEC61A1 | 26.6505277 | -0.2502692 | 0.58893886 | -0.4249494 | 0.67087354 | 0.96412305 |
| RUVBL1 | 12.6953202 | -0.5272914 | 0.47464195 | -1.1109245 | 0.26660083 | NA |
| EEFSEC | 37.7212483 | -0.5828768 | 0.43302701 | -1.3460519 | 0.17828579 | 0.76504408 |
| RPN1 | 23.7423561 | 0.45525313 | 0.51908214 | 0.87703487 | 0.3804677 | 0.90676497 |
| RAB7A | 95.904748 | -0.2127324 | 0.28784601 | -0.7390492 | 0.4598771 | 0.9314919 |
| ACAD9 | 8.20355605 | 0.599224 | 0.6124923 | 0.97833719 | 0.32790758 | NA |
| CNBP | 46.3770429 | 0.09157141 | 0.47664404 | 0.19211697 | 0.84765058 | 0.98545782 |
| COPG1 | 13.933036 | -0.0342188 | 0.53980524 | -0.063391 | 0.94945519 | 0.99514836 |
| HMCES | 10.2462202 | -0.1591293 | 0.60923386 | -0.2611957 | 0.7939416 | NA |
| H1FX | 22.6278213 | 0.25368993 | 0.59935538 | 0.4232713 | 0.67209733 | 0.96412305 |
| MBD4 | 20.0665367 | 0.24299288 | 0.48378087 | 0.5022788 | 0.61547142 | 0.95688177 |
| PLXND1 | 28.3679432 | 0.67049388 | 0.48414864 | 1.38489263 | 0.1660853 | 0.75461088 |
| TMCC1 | 103.390116 | -0.2499883 | 0.2420966 | -1.0325974 | 0.30179234 | 0.86225566 |
| PIK3R4 | 18.6470748 | -0.3884289 | 0.65141854 | -0.5962816 | 0.55098714 | 0.95415437 |
| ATP2C1 | 54.0688382 | 0.49930962 | 0.33069959 | 1.50985858 | 0.13107951 | 0.72977266 |
| MRPL3 | 15.8639302 | -0.2059491 | 0.44696604 | -0.4607712 | 0.64496277 | 0.96049083 |
| ACPP | 11.6149494 | 0.65757756 | 0.84669295 | 0.7766423 | 0.43736984 | NA |
| DNAJC13 | 56.8052603 | 0.30405907 | 0.38931906 | 0.78100225 | 0.43480117 | 0.92343564 |
| UBA5 | 12.7738874 | -0.4056735 | 0.66836718 | -0.606962 | 0.54387612 | NA |
| CDV3 | 37.2668669 | -0.4976037 | 0.42731613 | -1.1644862 | 0.24422703 | 0.83666131 |
| TOPBP1 | 24.4507432 | -0.4227361 | 0.37413883 | -1.1298909 | 0.25852219 | 0.84404826 |
| RYK | 36.0132088 | 0.07702164 | 0.372829 | 0.20658705 | 0.83633238 | 0.98320463 |
| CEP63 | 22.7677676 | 0.0081195 | 0.47699311 | 0.01702227 | 0.98641885 | 0.99811561 |
| PPP2R3A | 15.5077986 | 0.6414547 | 0.58178553 | 1.10256213 | 0.27021736 | 0.84994492 |
| MSL2 | 35.9559164 | 0.13590485 | 0.3718366 | 0.36549615 | 0.71474108 | 0.97348719 |
| PCCB | 11.5249208 | -0.3960622 | 0.48722647 | -0.8128915 | 0.41628029 | NA |
| STAG1 | 226.71539 | -0.3758978 | 0.22788772 | -1.6494868 | 0.09904795 | 0.66763865 |
| NCK1 | 39.6513744 | 0.28344568 | 0.41603824 | 0.68129719 | 0.49568346 | 0.93805563 |
| ARMC8 | 53.6762787 | -0.2032433 | 0.30267441 | -0.6714916 | 0.50190743 | 0.94042029 |
| PIK3CB | 70.8438456 | 0.7707816 | 0.41967327 | 1.83662305 | 0.06626556 | 0.58951068 |
| COPB2 | 25.4010486 | 0.53293456 | 0.41029106 | 1.29891828 | 0.19397198 | 0.78135737 |
| AC097103.2 | 13.2516977 | -0.2369032 | 0.69534048 | -0.3407009 | 0.73332873 | 0.97605445 |
| SLC25A36 | 32.3765445 | 0.59435482 | 0.45898963 | 1.29491994 | 0.19534785 | 0.78135737 |
| ZBTB38 | 37.1054613 | -0.1688251 | 0.33571884 | -0.5028765 | 0.6150511 | 0.95688177 |
| RASA2 | 186.305347 | 0.20708469 | 0.38504687 | 0.53781684 | 0.59070351 | 0.95415437 |
| RNF7 | 13.7769426 | -0.1088866 | 0.56978126 | -0.1911025 | 0.84844532 | 0.98545782 |

| | | | | | | |
|---|---|---|---|---|---|---|
| ATP1B3 | 94.2403625 | -0.2055387 | 0.61155121 | -0.336094 | 0.73679997 | 0.97605445 |
| TFDP2 | 62.0364493 | 0.22422967 | 0.48863894 | 0.45888621 | 0.64631588 | 0.96049083 |
| GK5 | 28.743946 | 1.20316314 | 0.5486728 | 2.19286091 | 0.0283174 | 0.42013003 |
| XRN1 | 115.887243 | -0.1827282 | 0.44616386 | -0.409554 | 0.68213315 | 0.9673431 |
| ATR | 61.9355471 | -0.3162858 | 0.26419999 | -1.1971456 | 0.23124981 | 0.82457247 |
| U2SURP | 43.9251732 | -0.1422141 | 0.35763722 | -0.3976491 | 0.69088887 | 0.96818917 |
| SLC9A9 | 213.703036 | 0.2590454 | 0.48670064 | 0.5322479 | 0.59455431 | 0.9553365 |
| C3orf58 | 11.2382454 | -0.4464503 | 0.57322885 | -0.7788343 | 0.43607735 | NA |
| PLSCR1 | 81.2810067 | 0.95405298 | 0.66346518 | 1.437985 | 0.15043831 | 0.74043418 |
| GYG1 | 13.837395 | 0.66065969 | 0.51096737 | 1.29295867 | 0.19602535 | 0.78135737 |
| HLTF | 11.6135486 | 0.00456529 | 0.58868063 | 0.00775513 | 0.99381237 | NA |
| HPS3 | 26.0094444 | 0.49608527 | 0.49493475 | 1.0023246 | 0.31618684 | 0.87263466 |
| WWTR1 | 12.7473098 | -0.9842043 | 0.61882541 | -1.5904393 | 0.11173582 | NA |
| ANKUB1 | 48.7822151 | 0.8701175 | 0.79754163 | 1.09099946 | 0.27527312 | 0.84994492 |
| RNF13 | 131.417733 | 0.65338473 | 0.47406266 | 1.37826661 | 0.16812099 | 0.7567539 |
| TSC22D2 | 52.0947393 | 0.18367902 | 0.31675457 | 0.57987805 | 0.56199686 | 0.95415437 |
| SERP1 | 46.4106345 | 0.302556 | 0.56393359 | 0.53650998 | 0.59160614 | 0.95415437 |
| EIF2A | 19.2088773 | -0.2188267 | 0.49604366 | -0.441144 | 0.65910872 | 0.96280444 |
| SELENOT | 30.3482919 | -0.2202239 | 0.39633914 | -0.5556452 | 0.57845342 | 0.95415437 |
| SIAH2 | 12.9592969 | 0.50338704 | 0.69814515 | 0.72103494 | 0.47088802 | 0.9314919 |
| GPR171 | 15.9142055 | 0.88595954 | 0.59977944 | 1.47714224 | 0.13963751 | 0.73044824 |
| MBNL1 | 850.449995 | 0.17943023 | 0.2469213 | 0.7266697 | 0.46742832 | 0.9314919 |
| RAP2B | 18.7925465 | -0.0556789 | 0.4675228 | -0.1190935 | 0.90520125 | 0.99324672 |
| DHX36 | 56.8032809 | -0.1531749 | 0.36300539 | -0.4219632 | 0.67305188 | 0.96444048 |
| GMPS | 19.7377121 | -0.2571096 | 0.41068005 | -0.6260581 | 0.53127682 | 0.95265314 |
| SSR3 | 21.4376458 | 0.3724076 | 0.5314775 | 0.70070248 | 0.48348871 | 0.93399425 |
| TIPARP | 14.777378 | 0.7751339 | 0.59057712 | 1.31250241 | 0.18935067 | 0.77850418 |
| LINC00886 | 10.7280851 | -0.2228702 | 0.68567188 | -0.3250392 | 0.74515143 | NA |
| CCNL1 | 64.3694993 | 0.20563054 | 0.42584806 | 0.48287302 | 0.6291859 | 0.95877428 |
| RSRC1 | 99.2296454 | 0.05004603 | 0.26960835 | 0.18562492 | 0.85273891 | 0.98751288 |
| GFM1 | 17.0231693 | -0.8512722 | 0.47725701 | -1.7836766 | 0.07447622 | 0.61929783 |
| MFSD1 | 38.0573351 | 0.4062008 | 0.54017559 | 0.75197919 | 0.45206358 | 0.92931643 |
| SMC4 | 26.6272824 | 0.60984908 | 0.60410874 | 1.00950217 | 0.31273386 | 0.87058449 |
| KPNA4 | 54.6128673 | -0.0222125 | 0.37332727 | -0.0594988 | 0.95255481 | 0.99514836 |
| PPM1L | 85.2453512 | 0.81164921 | 0.54908803 | 1.47817684 | 0.13936045 | 0.73044824 |
| NMD3 | 17.7489075 | 0.51890439 | 0.63123224 | 0.82204989 | 0.4110485 | 0.91823911 |
| PDCD10 | 34.6992732 | -0.2986022 | 0.38925357 | -0.767115 | 0.44301315 | 0.92899684 |
| GOLIM4 | 22.2875412 | 0.49679965 | 0.51840174 | 0.95832943 | 0.33789667 | 0.88287071 |
| MECOM | 67.261176 | -1.0523417 | 0.51370874 | -2.0485182 | 0.04050925 | 0.48881259 |
| SEC62 | 42.144197 | -0.5401766 | 0.32922454 | -1.6407543 | 0.10084843 | 0.6683775 |
| GPR160 | 8.75251039 | -0.6924899 | 0.66886742 | -1.0353171 | 0.30052083 | NA |
| PHC3 | 124.036744 | 0.1343658 | 0.29620046 | 0.45363132 | 0.65009421 | 0.96084839 |
| PRKCI | 19.3184909 | -0.2504027 | 0.44352408 | -0.5645752 | 0.57236274 | 0.95415437 |
| SKIL | 42.8902529 | -0.0924873 | 0.43324972 | -0.2134735 | 0.83095763 | 0.98320463 |
| RPL22L1 | 9.48846823 | -0.5193309 | 0.65416793 | -0.7938801 | 0.42726526 | NA |
| TNIK | 107.726812 | 1.44680684 | 0.39793137 | 3.63581999 | 0.0002771 | 0.03066918 |
| PLD1 | 38.0234859 | 1.65810955 | 0.72040624 | 2.3016313 | 0.02135597 | 0.37758452 |
| FNDC3B | 344.399906 | 1.17899522 | 0.44489529 | 2.650051 | 0.00804796 | 0.23440749 |
| TNFSF10 | 36.9319645 | 0.57965467 | 0.50683188 | 1.14368234 | 0.2527554 | 0.84008912 |
| NCEH1 | 22.7087086 | 0.41031715 | 0.49167012 | 0.8345375 | 0.40397816 | 0.91528253 |
| NLGN1 | 10.3215 | 0.40016925 | 0.67018334 | 0.59710415 | 0.55043785 | NA |
| NAALADL2 | 30.4839278 | 1.39025454 | 0.64285993 | 2.16260881 | 0.03057128 | 0.43032651 |
| TBL1XR1 | 189.6259 | -0.1945913 | 0.33070217 | -0.5884185 | 0.55625143 | 0.95415437 |
| ZMAT3 | 19.0110934 | -0.2896332 | 0.45839109 | -0.6318473 | 0.52748669 | 0.95136878 |

| | | | | | | |
|---|---|---|---|---|---|---|
| PIK3CA | 54.8263167 | 0.35983272 | 0.35741502 | 1.00676439 | 0.31404801 | 0.87152625 |
| MFN1 | 26.2294247 | -0.401141 | 0.43152236 | -0.9295949 | 0.35258089 | 0.89109867 |
| GNB4 | 35.7951351 | 0.95352226 | 0.52320327 | 1.82247001 | 0.0683837 | 0.59777078 |
| NDUFB5 | 10.4222541 | 0.18415702 | 0.58037356 | 0.31730774 | 0.75101011 | NA |
| USP13 | 8.22233243 | -0.1653015 | 0.58575766 | -0.2822012 | 0.77778924 | NA |
| TTC14 | 48.1552556 | 0.11586371 | 0.31305782 | 0.37010323 | 0.71130557 | 0.97317955 |
| FXR1 | 42.6240551 | -0.1130751 | 0.38106672 | -0.296733 | 0.76667036 | 0.98264092 |
| ATP11B | 99.1118398 | 0.02031901 | 0.27125222 | 0.07490818 | 0.94028777 | 0.99514836 |
| DCUN1D1 | 27.6073024 | 0.15260833 | 0.37748846 | 0.40427284 | 0.6860121 | 0.9673431 |
| MCCC1 | 21.844993 | -0.6365165 | 0.41351295 | -1.5392905 | 0.1237334 | 0.71625587 |
| MCF2L2 | 15.647443 | 0.98604639 | 0.50068183 | 1.96940719 | 0.04890635 | 0.53486612 |
| B3GNT5 | 14.0282817 | 0.35794869 | 0.63838229 | 0.56071213 | 0.5749938 | 0.95415437 |
| KLHL6 | 34.0141433 | -0.0773226 | 0.49982858 | -0.1546982 | 0.87705921 | 0.98939075 |
| KLHL24 | 48.2497176 | 0.6500923 | 0.39020629 | 1.66602211 | 0.09570901 | 0.66124053 |
| YEATS2 | 48.476692 | 0.18713559 | 0.45872952 | 0.4079432 | 0.68331538 | 0.9673431 |
| ABCC5 | 28.2161702 | -0.1305077 | 0.47540155 | -0.2745209 | 0.78368432 | 0.98264092 |
| DVL3 | 9.5272426 | 0.21502431 | 0.62799815 | 0.34239641 | 0.73205259 | NA |
| AP2M1 | 17.2583235 | 0.40281049 | 0.5091353 | 0.7911659 | 0.42884719 | 0.92200479 |
| PSMD2 | 11.4514248 | 0.11803758 | 0.52158276 | 0.22630653 | 0.82096302 | NA |
| EIF4G1 | 24.8320435 | -0.6400026 | 0.53870712 | -1.1880343 | 0.23481988 | 0.82903329 |
| POLR2H | 9.92160449 | -0.935774 | 0.58407008 | -1.6021604 | 0.10912015 | NA |
| VPS8 | 91.7564653 | 0.36349366 | 0.30960771 | 1.17404591 | 0.24037664 | 0.83493679 |
| MAP3K13 | 16.4636563 | 0.21961503 | 0.68603515 | 0.32012213 | 0.74887575 | 0.97818861 |
| SENP2 | 19.4745396 | -0.1141165 | 0.46215716 | -0.2469214 | 0.80496905 | 0.98320463 |
| IGF2BP2 | 16.4939513 | 3.38051168 | 1.05432974 | 3.20631349 | 0.00134447 | 0.07670434 |
| TRA2B | 47.436118 | 0.33280681 | 0.3271538 | 1.01727937 | 0.30902055 | 0.8673137 |
| ETV5 | 16.101558 | -0.4910798 | 0.75212702 | -0.6529214 | 0.51380694 | 0.94596835 |
| DGKG | 24.1633722 | -1.9242772 | 0.87898822 | -2.1891957 | 0.02858262 | 0.42168851 |
| EIF4A2 | 41.7127123 | 0.15957104 | 0.36766089 | 0.4340169 | 0.66427617 | 0.96384486 |
| ST6GAL1 | 181.055742 | -0.6670124 | 0.38436871 | -1.7353453 | 0.08267966 | 0.63740875 |
| BCL6 | 56.8834395 | -0.1065816 | 0.68516669 | -0.1555557 | 0.87638322 | 0.98939075 |
| LPP | 319.171814 | 0.15924218 | 0.30726763 | 0.51825238 | 0.60428219 | 0.95688177 |
| TPRG1 | 40.572276 | 0.17855743 | 0.61532317 | 0.29018479 | 0.77167487 | 0.98264092 |
| P3H2 | 15.0309802 | 0.07986135 | 0.62442086 | 0.12789667 | 0.89823075 | 0.99141995 |
| IL1RAP | 33.1251506 | 0.65026326 | 0.75604282 | 0.86008788 | 0.3897406 | 0.90910959 |
| CCDC50 | 30.5844203 | -0.8696839 | 0.40341144 | -2.1558236 | 0.03109744 | 0.43457887 |
| MB21D2 | 35.9940128 | 0.41466113 | 0.92127783 | 0.45009346 | 0.65264305 | 0.96178254 |
| OPA1 | 47.7677187 | -0.2986262 | 0.32485882 | -0.9192491 | 0.3579653 | 0.8957272 |
| ATP13A3 | 71.6779194 | -0.0173153 | 0.41326292 | -0.041899 | 0.96657924 | 0.99653147 |
| LSG1 | 9.65200368 | -0.439774 | 0.53215915 | -0.8263956 | 0.40857969 | NA |
| XXYLT1 | 11.7860814 | 0.19430258 | 0.54804397 | 0.3545383 | 0.72293551 | NA |
| ACAP2 | 191.568577 | -0.0796278 | 0.22705525 | -0.3506979 | 0.72581498 | 0.9754739 |
| PPP1R2 | 38.714643 | 0.25753014 | 0.46554934 | 0.55317474 | 0.58014376 | 0.95415437 |
| MUC20-OT1 | 72.0774889 | 0.41196238 | 0.41624969 | 0.98970016 | 0.3223207 | 0.87433122 |
| TNK2 | 25.4675387 | 0.68777104 | 0.41887705 | 1.64194015 | 0.10060241 | 0.6683775 |
| TFRC | 69.8291986 | 0.30688979 | 1.09529781 | 0.28018845 | 0.77933293 | 0.98264092 |
| PCYT1A | 17.1012208 | -0.0623447 | 0.42703585 | -0.1459941 | 0.88392607 | 0.99019008 |
| UBXN7 | 33.8997788 | -0.2287724 | 0.40579833 | -0.5637588 | 0.57291831 | 0.95415437 |
| RNF168 | 32.8298369 | -0.0223316 | 0.35616577 | -0.0627001 | 0.9500053 | 0.99514836 |
| PIGX | 19.8090109 | 0.04419538 | 0.4642311 | 0.09520124 | 0.92415498 | 0.99514836 |
| PAK2 | 93.443467 | -0.0299235 | 0.29717806 | -0.1006922 | 0.91979478 | 0.99511227 |
| SENP5 | 51.8413835 | -0.673164 | 0.44707444 | -1.505709 | 0.1321419 | 0.73044824 |
| NCBP2 | 12.9536164 | -0.8532935 | 0.48244224 | -1.7686957 | 0.07694467 | 0.62507515 |
| DLG1 | 105.180261 | -0.0675669 | 0.26930971 | -0.2508894 | 0.80189966 | 0.98320463 |

| | | | | | | |
|---|---|---|---|---|---|---|
| RUBCN | 36.7132228 | -0.4747709 | 0.37409947 | -1.2691035 | 0.20440416 | 0.78882329 |
| FYTTD1 | 16.9739784 | 0.08316897 | 0.48005135 | 0.17325015 | 0.8624548 | 0.98751288 |
| LRCH3 | 76.6515511 | 0.03668335 | 0.25647874 | 0.14302685 | 0.88626898 | 0.99019008 |
| IQCG | 12.268366 | -0.2075539 | 0.55872707 | -0.3714763 | 0.71028282 | NA |
| RPL35A | 67.455313 | -0.3312726 | 0.50261211 | -0.6591018 | 0.50983037 | 0.94332619 |
| ZNF595 | 10.459181 | 0.61239362 | 0.85885314 | 0.71303649 | 0.47582318 | NA |
| ZNF718 | 28.5549827 | -0.6109701 | 0.39745587 | -1.5372023 | 0.12424377 | 0.71741321 |
| ZNF141 | 25.0698415 | -0.1630753 | 0.37164551 | -0.4387925 | 0.66081189 | 0.96298563 |
| ZNF721 | 42.0432541 | -0.5084481 | 0.32438538 | -1.5674198 | 0.11701659 | 0.70122655 |
| PIGG | 16.2177586 | -0.734754 | 0.45098909 | -1.6292057 | 0.10326948 | 0.67250824 |
| ATP5ME | 8.53008741 | -0.163209 | 0.63378025 | -0.2575167 | 0.79677992 | NA |
| PCGF3 | 21.6266329 | 0.36134241 | 0.43069917 | 0.83896703 | 0.40148781 | 0.91528253 |
| AC139887.2 | 32.6130762 | 0.1905031 | 0.43563509 | 0.4372997 | 0.66189401 | 0.96298563 |
| GAK | 42.3592141 | -0.2799007 | 0.35041576 | -0.7987674 | 0.42442527 | 0.91985469 |
| SPON2 | 19.208608 | 1.85166242 | 0.56880731 | 3.25534216 | 0.00113256 | 0.07001544 |
| CTBP1 | 19.254417 | -0.7878477 | 0.41463944 | -1.9000789 | 0.05742276 | 0.56745994 |
| MAEA | 16.7771179 | -0.171333 | 0.44386668 | -0.3860011 | 0.69949589 | 0.9694491 |
| UVSSA | 16.6630102 | -0.0247455 | 0.47069581 | -0.0525721 | 0.95807283 | 0.99514836 |
| SLBP | 18.9937241 | -0.3366458 | 0.43222593 | -0.7788653 | 0.43605907 | 0.92457889 |
| TACC3 | 14.8597132 | 0.46194972 | 0.47615986 | 0.97015678 | 0.33196835 | 0.88047896 |
| NSD2 | 40.8874604 | -0.097208 | 0.36481283 | -0.2664599 | 0.78988504 | 0.98320463 |
| NELFA | 11.4857928 | -0.348936 | 0.53222333 | -0.6556195 | 0.51206896 | NA |
| MXD4 | 16.8168771 | 0.17941041 | 0.54279158 | 0.33053278 | 0.74099742 | 0.97605445 |
| ZFYVE28 | 38.7225302 | 0.7945741 | 0.44969374 | 1.76692277 | 0.07724116 | 0.62520507 |
| RNF4 | 28.8827539 | 0.03086921 | 0.37802666 | 0.08165883 | 0.93491802 | 0.99514836 |
| FAM193A | 52.5370183 | -0.6637009 | 0.27708514 | -2.395296 | 0.01660695 | 0.33912498 |
| TNIP2 | 11.8963935 | -0.1328902 | 0.61028669 | -0.2177505 | 0.8276235 | NA |
| SH3BP2 | 39.5274082 | -0.3613727 | 0.49543775 | -0.7294008 | 0.46575651 | 0.9314919 |
| ADD1 | 84.4105816 | -0.3000339 | 0.34167745 | -0.8781202 | 0.37987848 | 0.9065319 |
| MFSD10 | 10.5801702 | 0.95934051 | 0.65367771 | 1.46760474 | 0.14221162 | NA |
| NOP14 | 7.03661123 | -0.4304363 | 0.69502458 | -0.6193109 | 0.53571157 | NA |
| HTT | 68.6977751 | -0.2762014 | 0.27441185 | -1.0065216 | 0.31416473 | 0.87152625 |
| RGS12 | 12.9257258 | 0.98472172 | 0.65655954 | 1.49982091 | 0.1336608 | NA |
| LRPAP1 | 17.238197 | 0.56177893 | 0.51934988 | 1.08169648 | 0.27938742 | 0.84994492 |
| LYAR | 8.59580848 | -0.9465437 | 0.61654126 | -1.535248 | 0.12472292 | NA |
| STX18 | 40.0064186 | -0.6435025 | 0.33534727 | -1.9189138 | 0.05499525 | 0.5553717 |
| STX18-AS1 | 12.9023232 | 0.65854018 | 0.64689061 | 1.01800856 | 0.30867389 | NA |
| JAKMIP1 | 20.1730955 | 0.93452061 | 0.80432948 | 1.16186294 | 0.24529114 | 0.83810223 |
| MAN2B2 | 10.7865282 | 1.12095382 | 0.60955246 | 1.83897841 | 0.06591836 | NA |
| MRFAP1 | 14.120458 | 0.22542063 | 0.53634123 | 0.4202933 | 0.6742712 | 0.96568758 |
| KIAA0232 | 54.9383096 | 0.24043282 | 0.30711178 | 0.78288373 | 0.43369539 | 0.92311128 |
| TBC1D14 | 80.5713212 | 0.56944861 | 0.48239497 | 1.18046132 | 0.23781679 | 0.83190778 |
| AFAP1 | 9.25189665 | 0.00149371 | 0.68192099 | 0.00219044 | 0.99825228 | NA |
| SH3TC1 | 30.041759 | -0.2986275 | 0.49470932 | -0.6036423 | 0.54608149 | 0.95415437 |
| ACOX3 | 9.10490148 | 0.04408497 | 0.68681266 | 0.06418776 | 0.94882072 | NA |
| SLC2A9 | 23.0781418 | 1.01054903 | 0.60125615 | 1.68072964 | 0.09281544 | 0.6590317 |
| WDR1 | 34.7674175 | -0.0456527 | 0.39075844 | -0.116831 | 0.90699394 | 0.99368779 |
| RAB28 | 34.4984019 | -0.4623166 | 0.32683748 | -1.4145153 | 0.15721065 | 0.74486619 |
| BOD1L1 | 48.0251827 | -0.4629015 | 0.37455865 | -1.2358586 | 0.21651114 | 0.80634897 |
| AC092546.1 | 19.8668411 | -0.838903 | 0.7768341 | -1.0798998 | 0.28018681 | 0.84994492 |
| CPEB2 | 31.4606114 | 0.03497537 | 0.40641803 | 0.08605763 | 0.93142061 | 0.99514836 |
| AC098829.1 | 13.428729 | 1.47946254 | 0.80225483 | 1.84413043 | 0.06516413 | 0.58951068 |
| FBXL5 | 71.7750342 | 0.48975972 | 0.39505363 | 1.23972971 | 0.21507539 | 0.80634897 |
| FAM200B | 10.0231831 | 0.05583077 | 0.60749284 | 0.09190358 | 0.92677465 | NA |

| | | | | | | |
|---|---|---|---|---|---|---|
| CD38 | 43.0218922 | 1.12196656 | 0.81656009 | 1.37401593 | 0.16943674 | 0.75745073 |
| TAPT1 | 32.7851252 | 0.63002677 | 0.39398092 | 1.59913016 | 0.10979168 | 0.68862129 |
| TAPT1-AS1 | 11.8612681 | -0.0064878 | 0.61270371 | -0.0105888 | 0.99155155 | NA |
| LDB2 | 11.9366744 | 0.96352517 | 0.80563898 | 1.19597635 | 0.23170579 | NA |
| LAP3 | 26.6073154 | 0.68424867 | 0.669338 | 1.02227674 | 0.30664994 | 0.8673137 |
| MED28 | 17.6862541 | 0.18272384 | 0.42857676 | 0.42635032 | 0.6698526 | 0.96412305 |
| DCAF16 | 11.5503844 | 0.24199185 | 0.66723359 | 0.36267936 | 0.71684442 | NA |
| LCORL | 75.9735716 | 0.63302415 | 0.30209913 | 2.09541865 | 0.03613379 | 0.46830073 |
| AC024230.1 | 11.4734672 | 1.06059384 | 0.90007879 | 1.17833445 | 0.2386633 | NA |
| PACRGL | 11.2306169 | -0.2877551 | 0.56223182 | -0.5118086 | 0.60878496 | NA |
| KCNIP4 | 41.2594958 | -0.9806427 | 0.9866438 | -0.9939177 | 0.32026295 | 0.87363761 |
| DHX15 | 45.6775763 | -0.079219 | 0.31255993 | -0.253452 | 0.79991891 | 0.98320463 |
| CCDC149 | 10.4319216 | 0.2921524 | 0.59931784 | 0.4874749 | 0.62592183 | NA |
| SEPSECS | 11.0636482 | 1.20627431 | 0.68318867 | 1.76565328 | 0.07745404 | NA |
| ANAPC4 | 20.4221138 | 0.0849962 | 0.47120383 | 0.18038097 | 0.8568535 | 0.98751288 |
| SEL1L3 | 87.5809626 | -1.0072002 | 0.29809526 | -3.3787863 | 0.00072807 | 0.05372155 |
| RBPJ | 315.102998 | 0.50148763 | 0.67152673 | 0.74678729 | 0.45519197 | 0.9314919 |
| TBC1D19 | 18.3610359 | -0.2855927 | 0.52794566 | -0.5409509 | 0.58854145 | 0.95415437 |
| STIM2 | 81.851012 | -0.2098497 | 0.36185746 | -0.5799238 | 0.56196603 | 0.95415437 |
| ARAP2 | 163.553677 | -0.4126758 | 0.37685749 | -1.0950447 | 0.27349707 | 0.84994492 |
| DTHD1 | 38.3316177 | -0.897047 | 0.47596584 | -1.8846878 | 0.059472 | 0.57281427 |
| C4orf19 | 12.3877189 | 1.0324672 | 0.77536125 | 1.33159505 | 0.18299329 | NA |
| RELL1 | 29.6181134 | 1.0619502 | 0.56779663 | 1.87030028 | 0.06144213 | 0.57339082 |
| TBC1D1 | 102.310407 | -0.4396004 | 0.25578687 | -1.71862 | 0.08568359 | 0.64676224 |
| KLF3 | 38.1192221 | -0.2532347 | 0.42482938 | -0.5960857 | 0.551118 | 0.95415437 |
| TLR1 | 25.8368493 | 0.50001489 | 0.50317353 | 0.99372255 | 0.32035796 | 0.87363761 |
| TMEM156 | 31.0767771 | -0.861835 | 0.43847637 | -1.9655221 | 0.04935384 | 0.53564799 |
| KLHL5 | 43.7464225 | -0.4676907 | 0.44647048 | -1.0475289 | 0.2948557 | 0.8608685 |
| RFC1 | 28.3636192 | -0.2860987 | 0.39306759 | -0.7278612 | 0.46669854 | 0.9314919 |
| SMIM14 | 33.4151836 | -0.0716967 | 0.38544264 | -0.1860114 | 0.85243584 | 0.98751288 |
| UBE2K | 113.662036 | -0.0743102 | 0.22504369 | -0.3302035 | 0.74124624 | 0.97605445 |
| PDS5A | 117.550254 | -0.4212122 | 0.29160271 | -1.4444728 | 0.14860603 | 0.7383468 |
| N4BP2 | 29.9357259 | -0.4315696 | 0.46480761 | -0.9284907 | 0.35315308 | 0.89109867 |
| RHOH | 96.7607428 | -1.2408718 | 0.42608956 | -2.9122324 | 0.00358856 | 0.15276206 |
| RBM47 | 249.898191 | 0.26294386 | 0.48821581 | 0.5385812 | 0.59017586 | 0.95415437 |
| APBB2 | 14.0431271 | -0.3590677 | 0.8548969 | -0.4200128 | 0.67447607 | 0.96573107 |
| TMEM33 | 22.1361897 | 0.25007426 | 0.4518647 | 0.5534273 | 0.57997085 | 0.95415437 |
| SLC30A9 | 36.8870908 | -0.068491 | 0.39002213 | -0.1756079 | 0.860602 | 0.98751288 |
| ATP8A1 | 102.853545 | -0.0767595 | 0.53265896 | -0.1441062 | 0.88541659 | 0.99019008 |
| GUF1 | 10.4044022 | -0.2164241 | 0.57021047 | -0.3795513 | 0.7042785 | NA |
| ATP10D | 28.644789 | 0.37615667 | 0.41775063 | 0.90043351 | 0.36788959 | 0.89868152 |
| NFXL1 | 18.3839188 | 0.35546548 | 0.73477318 | 0.4837758 | 0.62854499 | 0.95877428 |
| TXK | 58.9253032 | 1.10639739 | 0.35611354 | 3.10686696 | 0.00189081 | 0.09861218 |
| TEC | 25.5201655 | 1.10621214 | 0.49513076 | 2.23418181 | 0.02547111 | 0.40044642 |
| SLAIN2 | 39.8068108 | 0.28328463 | 0.33008908 | 0.85820662 | 0.39077838 | 0.90910959 |
| FRYL | 153.496859 | 0.23964752 | 0.28755085 | 0.83340919 | 0.404614 | 0.91551819 |
| OCIAD1 | 38.1202566 | 0.26247286 | 0.36783061 | 0.71356993 | 0.47549316 | 0.9314919 |
| DCUN1D4 | 20.6340968 | -0.1024769 | 0.45706765 | -0.2242052 | 0.82259763 | 0.98320463 |
| SCFD2 | 70.6560323 | -0.5010956 | 0.50460825 | -0.9930389 | 0.32069097 | 0.87363761 |
| AC058822.1 | 19.6433902 | 0.67861929 | 0.74177358 | 0.91486043 | 0.36026491 | 0.89725742 |
| CHIC2 | 57.0901527 | 0.28179337 | 0.35163534 | 0.80137954 | 0.42291196 | 0.91985469 |
| TMEM165 | 75.8665467 | 0.04381282 | 0.25801172 | 0.16980944 | 0.86516001 | 0.98751288 |
| CLOCK | 37.7558153 | -0.1249709 | 0.30761306 | -0.4062602 | 0.68455145 | 0.9673431 |
| EXOC1 | 43.2618609 | -0.185499 | 0.36444245 | -0.508994 | 0.61075646 | 0.95688177 |

| | | | | | | |
|---|---|---|---|---|---|---|
| CEP135 | 26.1757685 | -0.351402 | 0.49876906 | -0.7045384 | 0.48109752 | 0.93286394 |
| AASDH | 11.4670977 | 1.10929622 | 0.58070694 | 1.91025136 | 0.05610086 | NA |
| SRP72 | 20.6853156 | 0.08679403 | 0.47322225 | 0.18341072 | 0.85447577 | 0.98751288 |
| REST | 21.3686522 | 0.16456714 | 0.49590989 | 0.33184887 | 0.74000338 | 0.97605445 |
| POLR2B | 37.2577837 | -0.5386006 | 0.33042376 | -1.63003 | 0.10309516 | 0.67250824 |
| IGFBP7 | 49.7887295 | 0.8091819 | 0.48535081 | 1.66721038 | 0.09547258 | 0.66124053 |
| CENPC | 34.3826422 | -0.649826 | 0.35098023 | -1.8514603 | 0.06410336 | 0.5853927 |
| UBA6 | 52.9250572 | 0.05602617 | 0.32497477 | 0.17240161 | 0.86312181 | 0.98751288 |
| UBA6-AS1 | 24.7587418 | -1.0906918 | 0.82892605 | -1.3157891 | 0.18824485 | 0.77798634 |
| YTHDC1 | 54.1246645 | -0.2882035 | 0.32307565 | -0.8920619 | 0.37235975 | 0.9025457 |
| UGT2B7 | 20.0060297 | 0.32736824 | 0.73943469 | 0.44272773 | 0.65796268 | 0.9627619 |
| RUFY3 | 61.049524 | -0.0511889 | 0.3138281 | -0.1631114 | 0.87043075 | 0.98751288 |
| GRSF1 | 14.4300258 | -0.1706742 | 0.46618054 | -0.3661117 | 0.71428173 | 0.97348719 |
| MOB1B | 50.1205778 | 0.16869006 | 0.35432461 | 0.47608902 | 0.63401096 | 0.95916256 |
| DCK | 9.17417533 | -0.0994913 | 0.5936746 | -0.1675856 | 0.8669093 | NA |
| SLC4A4 | 17.1488656 | 0.80189177 | 0.53273974 | 1.50522237 | 0.13226692 | 0.73044824 |
| ANKRD17 | 173.231241 | 0.09103698 | 0.24136115 | 0.37718159 | 0.70603866 | 0.97184211 |
| MTHFD2L | 12.7507849 | 1.49686109 | 0.76887276 | 1.9468255 | 0.05155566 | NA |
| RCHY1 | 21.3067824 | 0.53561963 | 0.45431395 | 1.17896365 | 0.23841265 | 0.83267279 |
| G3BP2 | 33.0344915 | -0.4601304 | 0.31409653 | -1.4649332 | 0.14293915 | 0.73044824 |
| USO1 | 47.1345772 | -0.0386827 | 0.32842779 | -0.1177816 | 0.90624073 | 0.99368779 |
| NAAA | 38.5082427 | 0.86710403 | 0.70600617 | 1.22818195 | 0.21937867 | 0.81042354 |
| SDAD1 | 13.1648314 | -0.4125798 | 0.53980405 | -0.7643141 | 0.44468011 | 0.92914597 |
| NUP54 | 20.8856908 | -0.2901966 | 0.4745131 | -0.611567 | 0.54082425 | 0.95415437 |
| SCARB2 | 41.3983868 | 0.57733873 | 0.36707324 | 1.57281616 | 0.11576139 | 0.70122655 |
| 11-Sep | 37.5462053 | 0.32748487 | 0.58971464 | 0.55532769 | 0.57867054 | 0.95415437 |
| CCNI | 77.0774023 | -0.1895506 | 0.33638379 | -0.5634951 | 0.57309783 | 0.95415437 |
| CCNG2 | 17.2861637 | -0.3201057 | 0.51656253 | -0.6196843 | 0.53546565 | 0.95378558 |
| CNOT6L | 167.952736 | 0.08677994 | 0.52173945 | 0.16632811 | 0.86789875 | 0.98751288 |
| MRPL1 | 22.5922411 | -0.6073722 | 0.51318427 | -1.1835362 | 0.2365967 | 0.83171006 |
| BMP2K | 146.940194 | -0.335176 | 0.41117027 | -0.8151757 | 0.41497173 | 0.91894074 |
| PAQR3 | 8.66341666 | -0.1423753 | 0.55237978 | -0.2577489 | 0.79660071 | NA |
| ANTXR2 | 74.7377082 | 0.84552773 | 0.47545043 | 1.77837197 | 0.07534278 | 0.6223089 |
| RASGEF1B | 183.87793 | -0.3172343 | 0.62123548 | -0.5106507 | 0.60959568 | 0.95688177 |
| HNRNPD | 62.1477082 | -0.1972726 | 0.26932017 | -0.7324834 | 0.46387356 | 0.9314919 |
| HNRNPDL | 55.1966881 | -0.4589733 | 0.39293051 | -1.1680776 | 0.24277545 | 0.83507265 |
| ENOPH1 | 11.4527542 | -0.1899884 | 0.62244734 | -0.305228 | 0.7601925 | NA |
| SEC31A | 59.0858865 | 0.39863669 | 0.35615503 | 1.11927858 | 0.26302131 | 0.84921816 |
| LIN54 | 19.8509803 | 1.22163082 | 0.46938532 | 2.6026183 | 0.00925149 | 0.25727508 |
| PLAC8 | 37.5797729 | 0.3356685 | 0.6155516 | 0.54531335 | 0.58553803 | 0.95415437 |
| HELQ | 14.8549741 | -0.0004924 | 0.48037224 | -0.001025 | 0.99918216 | 0.99936274 |
| GPAT3 | 24.691434 | -0.6795284 | 0.71278165 | -0.9533473 | 0.34041414 | 0.8844375 |
| WDFY3 | 132.775556 | 1.32393463 | 0.54175281 | 2.44379835 | 0.01453355 | 0.32121732 |
| ARHGAP24 | 203.177144 | -1.0796919 | 0.33894531 | -3.1854458 | 0.00144531 | 0.08079146 |
| MAPK10 | 34.0832308 | -3.613571 | 0.8356084 | -4.324479 | 1.53E-05 | 0.0047006 |
| PTPN13 | 20.5939586 | 1.81935159 | 0.90378095 | 2.01304485 | 0.04410991 | 0.51607664 |
| AFF1 | 223.682189 | -0.1348979 | 0.44987268 | -0.299858 | 0.76428549 | 0.98222198 |
| KLHL8 | 20.1462279 | 0.75296293 | 0.48001945 | 1.56860922 | 0.11673902 | 0.70122655 |
| HSD17B11 | 25.501707 | 0.19874475 | 0.38550667 | 0.51554166 | 0.60617457 | 0.95688177 |
| SPP1 | 477.097064 | 0.77832467 | 0.74639614 | 1.04277691 | 0.29705163 | 0.8608685 |
| PKD2 | 39.8622555 | 0.68072896 | 0.65705887 | 1.0360243 | 0.3001908 | 0.86225566 |
| PPM1K | 30.7813289 | -0.6613895 | 0.42100765 | -1.570968 | 0.11619008 | 0.70122655 |
| HERC6 | 13.6822171 | -0.7123123 | 0.96784558 | -0.7359772 | 0.46174455 | 0.9314919 |
| HERC5 | 25.3808106 | 0.42431242 | 0.92632682 | 0.45805909 | 0.64690999 | 0.96049083 |

| | | | | | | |
|---|---|---|---|---|---|---|
| HERC3 | 29.3575131 | -0.1434405 | 0.36477781 | -0.393227 | 0.6941518 | 0.9685713 |
| FAM13A-AS1 | 11.0580712 | 0.13709399 | 0.58105241 | 0.23594084 | 0.81347858 | NA |
| FAM13A | 62.8574049 | 0.81023679 | 0.75404199 | 1.07452475 | 0.28258756 | 0.85176121 |
| GPRIN3 | 131.510595 | 0.28314882 | 0.41925414 | 0.67536322 | 0.49944503 | 0.93956733 |
| SNCA | 21.6511035 | 1.25239362 | 0.79510603 | 1.57512781 | 0.11522695 | 0.70122655 |
| CCSER1 | 96.1129834 | -0.3967363 | 0.92702547 | -0.427967 | 0.66867518 | 0.96412305 |
| SMARCAD1 | 26.4760773 | -0.2450654 | 0.40185857 | -0.60983 | 0.54197441 | 0.95415437 |
| PDLIM5 | 69.7599629 | -0.2060817 | 0.45950189 | -0.4484893 | 0.65380012 | 0.96178254 |
| RAP1GDS1 | 97.2568893 | 0.1590773 | 0.29068407 | 0.54725151 | 0.58420595 | 0.95415437 |
| TSPAN5 | 52.4446675 | 0.16537157 | 0.38627995 | 0.42811326 | 0.66856867 | 0.96412305 |
| EIF4E | 35.9976908 | -0.8695701 | 0.41478128 | -2.0964546 | 0.03604188 | 0.46820605 |
| METAP1 | 18.3493436 | -0.0060274 | 0.44764041 | -0.0134649 | 0.9892569 | 0.99811561 |
| ADH5 | 12.9109749 | 0.35078916 | 0.50490107 | 0.69476811 | 0.48720064 | NA |
| DAPP1 | 32.6402549 | -2.289845 | 0.71091158 | -3.2209983 | 0.00127745 | 0.0752064 |
| DNAJB14 | 62.9224178 | -0.0346435 | 0.27320885 | -0.1268024 | 0.89909681 | 0.99179975 |
| H2AFZ | 25.2467328 | -0.3198114 | 0.66556483 | -0.4805113 | 0.6308639 | 0.95877428 |
| PPP3CA | 212.486858 | -0.4344659 | 0.38663737 | -1.1237039 | 0.26113866 | 0.8475436 |
| BANK1 | 134.434464 | -3.2472542 | 0.86729244 | -3.7441283 | 0.00018102 | 0.02594752 |
| SLC39A8 | 41.7811245 | 2.53848592 | 0.69720638 | 3.64093902 | 0.00027165 | 0.03066918 |
| NFKB1 | 82.5116718 | -0.1158776 | 0.41659176 | -0.2781562 | 0.78089244 | 0.98264092 |
| MANBA | 97.3927467 | 0.53057879 | 0.39484339 | 1.3437702 | 0.17902271 | 0.76545642 |
| UBE2D3 | 118.401818 | 0.21510686 | 0.23045499 | 0.93340074 | 0.3506131 | 0.89048867 |
| CISD2 | 9.7290814 | 0.66785468 | 0.60675424 | 1.10070048 | 0.27102704 | NA |
| TET2 | 127.148996 | 0.58503344 | 0.27054287 | 2.16244262 | 0.03058408 | 0.43032651 |
| PPA2 | 48.1152901 | 0.63682638 | 0.39419133 | 1.61552611 | 0.10619681 | 0.67559944 |
| GSTCD | 13.7454745 | 0.16588881 | 0.53332811 | 0.31104457 | 0.75576674 | 0.97989151 |
| TBCK | 70.2847289 | 0.37165201 | 0.39034121 | 0.95212087 | 0.34103569 | 0.88510795 |
| AIMP1 | 13.0561719 | -0.2558208 | 0.50628631 | -0.5052888 | 0.613356 | 0.95688177 |
| PAPSS1 | 32.7954957 | 0.66373085 | 0.4682405 | 1.41749988 | 0.15633683 | 0.74486619 |
| SGMS2 | 14.0971071 | 1.48456469 | 0.73693396 | 2.01451523 | 0.04395547 | 0.51607664 |
| LEF1 | 27.1759892 | -0.6392929 | 0.70090838 | -0.9120919 | 0.36172034 | 0.89853007 |
| RPL34 | 65.5988504 | -0.3384601 | 0.54761385 | -0.6180635 | 0.53653347 | 0.95415437 |
| OSTC | 14.7696451 | -0.1095666 | 0.67181327 | -0.1630908 | 0.87044692 | 0.98751288 |
| SEC24B | 68.6702353 | -0.1489971 | 0.27696336 | -0.5379668 | 0.59059998 | 0.95415437 |
| MCUB | 55.4292641 | 0.18903122 | 0.38800701 | 0.48718506 | 0.62612719 | 0.95876809 |
| ELOVL6 | 13.0234684 | -0.122804 | 0.57395072 | -0.2139627 | 0.83057617 | 0.98320463 |
| FAM241A | 12.9624565 | 0.99034417 | 0.65326085 | 1.5160011 | 0.12951907 | 0.72931666 |
| AP1AR | 13.0413359 | -0.3058309 | 0.53363829 | -0.5731052 | 0.56657348 | 0.95415437 |
| ALPK1 | 43.757152 | 0.19611068 | 0.51294225 | 0.38232506 | 0.70222027 | 0.97086717 |
| LARP7 | 19.8348121 | -0.8001711 | 0.47986609 | -1.6674883 | 0.09541735 | 0.66124053 |
| ANK2 | 28.7083149 | -2.6123228 | 0.74165151 | -3.522305 | 0.00042781 | 0.04012728 |
| CAMK2D | 167.370713 | -1.4977089 | 0.41103551 | -3.6437458 | 0.0002687 | 0.03066918 |
| SNHG8 | 10.2175161 | -0.4559291 | 0.52189359 | -0.8736054 | 0.38233318 | NA |
| METTL14 | 11.530071 | 0.62944257 | 0.59449344 | 1.05878808 | 0.2896963 | NA |
| SEC24D | 47.4895332 | 0.42418632 | 0.43548126 | 0.97406332 | 0.3300251 | 0.88037558 |
| USP53 | 54.1833366 | -0.6499069 | 0.44580803 | -1.4578178 | 0.14489077 | 0.73251288 |
| C4orf3 | 23.5047795 | 0.21317919 | 0.48365794 | 0.44076436 | 0.6593836 | 0.96280444 |
| PDE5A | 9.85430783 | 0.93118992 | 0.73271085 | 1.27088322 | 0.2037702 | NA |
| ANXA5 | 44.6993236 | 0.45252672 | 0.57466956 | 0.78745553 | 0.43101525 | 0.92284736 |
| KIAA1109 | 115.312655 | 0.13014283 | 0.32592387 | 0.39930438 | 0.68966894 | 0.9673431 |
| SPATA5 | 63.6598926 | -0.6493387 | 0.31205711 | -2.0808327 | 0.03744922 | 0.47208194 |
| MFSD8 | 17.670169 | -0.1090412 | 0.49512161 | -0.2202311 | 0.82569116 | 0.98320463 |
| ABHD18 | 44.4908428 | 0.30408975 | 0.32548561 | 0.93426482 | 0.35016731 | 0.89048867 |
| LARP1B | 22.0558274 | -0.0032609 | 0.54722646 | -0.0059589 | 0.99524553 | 0.99859588 |

| | | | | | | |
|---|---|---|---|---|---|---|
| PGRMC2 | 10.8358037 | 0.38579003 | 0.54705757 | 0.70520919 | 0.48068006 | NA |
| JADE1 | 20.9785965 | 0.24565664 | 0.45444925 | 0.54055902 | 0.58881157 | 0.95415437 |
| SCLT1 | 112.952029 | 0.1239791 | 0.42911773 | 0.28891627 | 0.77264545 | 0.98264092 |
| AC096711.2 | 13.3808823 | -0.3859658 | 0.53372713 | -0.7231519 | 0.46958657 | 0.9314919 |
| ELF2 | 152.728751 | 0.55536834 | 0.42866016 | 1.2955912 | 0.19511636 | 0.78135737 |
| NDUFC1 | 9.70631113 | -0.5120787 | 0.71877311 | -0.7124344 | 0.4761958 | NA |
| NAA15 | 42.8533684 | -0.1250037 | 0.30913379 | -0.4043675 | 0.68594251 | 0.9673431 |
| AC097376.2 | 26.9944125 | 0.1709776 | 0.54218073 | 0.31535168 | 0.75249465 | 0.97875726 |
| SETD7 | 10.0955996 | 0.63762812 | 0.68731689 | 0.92770617 | 0.35356001 | NA |
| MGST2 | 13.2847873 | 1.81182603 | 0.80506725 | 2.25052755 | 0.02441548 | 0.39492781 |
| MAML3 | 162.005381 | 0.5836907 | 0.37189527 | 1.56950288 | 0.11653081 | 0.70122655 |
| ELMOD2 | 9.69981101 | 0.09786277 | 0.57371705 | 0.17057672 | 0.86455661 | NA |
| TBC1D9 | 46.4950815 | -1.3140888 | 0.31346171 | -4.1921828 | 2.76E-05 | 0.00694978 |
| RNF150 | 22.492939 | 0.43518854 | 0.72631261 | 0.59917525 | 0.54905603 | 0.95415437 |
| ZNF330 | 11.7629747 | 0.68754654 | 0.65514819 | 1.04945194 | 0.29397016 | NA |
| IL15 | 32.5895463 | 0.78317498 | 0.4832995 | 1.62047545 | 0.10513018 | 0.67519097 |
| INPP4B | 121.126013 | 0.39957478 | 0.53703229 | 0.74404238 | 0.45685084 | 0.9314919 |
| USP38 | 12.4053395 | 0.21342346 | 0.51994683 | 0.4104717 | 0.68145996 | NA |
| GAB1 | 53.223786 | -0.1574081 | 0.40343665 | -0.3901681 | 0.69641222 | 0.9685713 |
| SMARCA5 | 32.9264672 | 0.14675828 | 0.42390607 | 0.34620471 | 0.72918887 | 0.97605445 |
| ANAPC10 | 23.4366766 | 0.22290419 | 0.40075806 | 0.55620638 | 0.57806978 | 0.95415437 |
| ABCE1 | 12.0819835 | -0.4119609 | 0.52688082 | -0.7818862 | 0.43428143 | NA |
| OTUD4 | 28.5068839 | 0.36674289 | 0.37779514 | 0.97074539 | 0.33167508 | 0.88047896 |
| ZNF827 | 20.2646494 | 0.66484786 | 0.61366544 | 1.08340443 | 0.27862895 | 0.84994492 |
| LSM6 | 10.9131678 | -0.1793709 | 0.5594925 | -0.3205956 | 0.74851684 | NA |
| SLC10A7 | 43.4573991 | 0.01002524 | 0.41362581 | 0.02423747 | 0.98066319 | 0.99758514 |
| TMEM184C | 12.8463507 | 0.40397834 | 0.52362229 | 0.77150715 | 0.44040638 | NA |
| PRMT9 | 13.7466508 | 0.59594731 | 0.57472019 | 1.0369347 | 0.29976629 | 0.86225566 |
| ARHGAP10 | 36.9240497 | 0.47623505 | 0.59337916 | 0.80258136 | 0.42221674 | 0.91985469 |
| NR3C2 | 53.2741013 | 0.04383072 | 0.69291197 | 0.06325583 | 0.94956279 | 0.99514836 |
| LRBA | 238.243002 | -0.3989547 | 0.26692729 | -1.4946192 | 0.13501384 | 0.73044824 |
| RPS3A | 48.0673243 | 0.04897029 | 0.62265837 | 0.07864713 | 0.9373133 | 0.99514836 |
| FBXW7 | 218.90369 | 0.08003237 | 0.41340818 | 0.19359165 | 0.84649564 | 0.98517495 |
| TMEM154 | 26.3642636 | -1.3505213 | 0.55568266 | -2.4303823 | 0.0150829 | 0.32478127 |
| ARFIP1 | 34.6007946 | 0.2692139 | 0.33908856 | 0.79393387 | 0.42723393 | 0.92129717 |
| TMEM131L | 139.055851 | -0.0765689 | 0.34713268 | -0.2205754 | 0.82542305 | 0.98320463 |
| TLR2 | 115.624626 | 0.90280247 | 0.53009422 | 1.70309812 | 0.08854971 | 0.64733696 |
| FGB | 8.9282793 | 3.82312625 | 2.08827171 | 1.83076092 | 0.06713624 | NA |
| PDGFC | 59.8680669 | 1.31855036 | 0.88325273 | 1.49283474 | 0.13548045 | 0.73044824 |
| FAM198B | 18.9780732 | 1.51039353 | 0.93086713 | 1.62256619 | 0.10468217 | 0.67519097 |
| FAM198B-AS | 14.0091379 | -0.008958 | 0.7991352 | -0.0112096 | 0.99105623 | 0.99827179 |
| TMEM144 | 12.9444671 | 0.32191461 | 0.83819884 | 0.38405518 | 0.70093755 | NA |
| ETFDH | 16.7036923 | -0.2083986 | 0.67149312 | -0.3103511 | 0.75629396 | 0.97989151 |
| PPID | 10.4102955 | -0.1729754 | 0.52969721 | -0.3265552 | 0.74400434 | NA |
| FNIP2 | 67.6180363 | 0.33891822 | 0.43502282 | 0.77908146 | 0.43593173 | 0.92457889 |
| RAPGEF2 | 104.058848 | 0.97276648 | 0.38601145 | 2.52004566 | 0.01173396 | 0.29101961 |
| NAF1 | 10.3104156 | 0.48656835 | 0.59031084 | 0.82425786 | 0.40979306 | NA |
| TMA16 | 9.81588589 | -0.6591187 | 0.56358809 | -1.1695043 | 0.24220052 | NA |
| 1-Mar | 172.594947 | -1.1042044 | 0.46494329 | -2.3749228 | 0.01755262 | 0.34827529 |
| TMEM192 | 12.1618823 | 0.41286018 | 0.56577314 | 0.72972742 | 0.46555681 | NA |
| KLHL2 | 63.7960179 | -0.3310769 | 0.53418354 | -0.6197812 | 0.53540188 | 0.95378558 |
| DDX60 | 29.8106526 | -0.4746088 | 0.97868821 | -0.4849438 | 0.6277162 | 0.95877428 |
| DDX60L | 82.1221148 | -0.4813508 | 0.73975789 | -0.6506869 | 0.51524861 | 0.94596835 |
| PALLD | 18.5831666 | -0.1643429 | 0.43722481 | -0.3758773 | 0.70700814 | 0.97184211 |

| | | | | | | |
|---|---|---|---|---|---|---|
| CBR4 | 37.0907508 | 0.02379211 | 0.40230495 | 0.0591395 | 0.952841 | 0.99514836 |
| SH3RF1 | 26.2511742 | -1.6638992 | 0.50846912 | -3.2723702 | 0.0010665 | 0.07001544 |
| NEK1 | 33.4816607 | -0.1512858 | 0.38507111 | -0.3928775 | 0.69440996 | 0.9685713 |
| CLCN3 | 31.9422962 | 0.18546761 | 0.37716062 | 0.49174703 | 0.62289819 | 0.95783378 |
| GALNT7 | 40.5280088 | 0.12846146 | 0.35782014 | 0.35901125 | 0.71958667 | 0.97348719 |
| HMGB2 | 32.4871498 | 0.06411176 | 0.58577625 | 0.10944753 | 0.91284754 | 0.99498068 |
| SAP30 | 14.1007903 | 0.86291128 | 0.67689797 | 1.27480257 | 0.20237913 | 0.78704575 |
| CEP44 | 13.0491896 | -0.8099326 | 0.57986522 | -1.3967602 | 0.1624857 | 0.751836 |
| SPCS3 | 35.8943388 | 0.11768798 | 0.44872402 | 0.26227251 | 0.79311135 | 0.98320463 |
| DCTD | 10.2380605 | 0.15882297 | 0.55704803 | 0.28511541 | 0.77555573 | NA |
| WWC2 | 25.8682216 | 0.47380081 | 0.41735277 | 1.1352526 | 0.2562695 | 0.8406233 |
| TRAPPC11 | 28.8404705 | 0.39584234 | 0.39923941 | 0.99149116 | 0.3214458 | 0.87363761 |
| IRF2 | 53.9479984 | -0.1835838 | 0.30837914 | -0.5953185 | 0.55163059 | 0.95415437 |
| CASP3 | 21.3253835 | 0.42319838 | 0.66937221 | 0.63223178 | 0.52723543 | 0.95136478 |
| PRIMPOL | 14.6556169 | 0.78520718 | 0.72574157 | 1.08193772 | 0.2792802 | 0.84994492 |
| ACSL1 | 173.812103 | 0.40020387 | 0.54235751 | 0.7378968 | 0.46057716 | 0.9314919 |
| CFAP97 | 12.9757136 | 0.35129649 | 0.49154493 | 0.71467828 | 0.47480786 | 0.9314919 |
| SNX25 | 36.0667301 | -0.8963077 | 0.39868981 | -2.2481329 | 0.02456772 | 0.39492781 |
| UFSP2 | 10.6800476 | -0.4003186 | 0.54808651 | -0.730393 | 0.46514997 | NA |
| SDHA | 25.1731987 | 0.41675329 | 0.39640831 | 1.05132328 | 0.29311014 | 0.86027209 |
| PDCD6 | 14.6488442 | 0.25361621 | 0.50421502 | 0.50299218 | 0.61496977 | 0.95688177 |
| EXOC3 | 15.3992827 | 0.16744257 | 0.44615264 | 0.37530333 | 0.70743489 | 0.97184211 |
| BRD9 | 11.6557168 | -0.569532 | 0.51943296 | -1.0964494 | 0.27288214 | NA |
| SLC12A7 | 14.1326976 | 1.03182889 | 0.69767449 | 1.4789546 | 0.13915245 | 0.73044824 |
| CLPTM1L | 18.0102898 | -0.6840742 | 0.69225998 | -0.9881753 | 0.32306679 | 0.87489296 |
| LPCAT1 | 35.8596979 | 0.09137747 | 0.45298003 | 0.20172516 | 0.84013159 | 0.98356002 |
| NDUFS6 | 17.9457835 | -0.2949271 | 0.49572597 | -0.5949399 | 0.55188366 | 0.95415437 |
| ICE1 | 28.3383503 | -0.0374895 | 0.34826337 | -0.1076469 | 0.91427583 | 0.99498068 |
| MED10 | 11.2298152 | -0.4698945 | 0.50516152 | -0.9301867 | 0.35227443 | NA |
| NSUN2 | 19.6691479 | -0.0144009 | 0.48968802 | -0.0294083 | 0.97653891 | 0.99758514 |
| SRD5A1 | 13.369455 | 0.80150881 | 0.70327836 | 1.13967507 | 0.2544217 | 0.84026145 |
| TENT4A | 41.4329575 | 0.36306277 | 0.5077727 | 0.71501041 | 0.47460261 | 0.9314919 |
| MTRR | 12.8239586 | 0.12043878 | 0.50395065 | 0.23898924 | 0.81111393 | NA |
| CCT5 | 15.8524488 | 0.12519878 | 0.53323553 | 0.23479077 | 0.81437113 | 0.98320463 |
| 6-Mar | 77.0870039 | -0.3099356 | 0.33201658 | -0.9334943 | 0.35056483 | 0.89048867 |
| DAP | 26.2509788 | -0.3228891 | 0.51060846 | -0.6323614 | 0.52715073 | 0.95136478 |
| TRIO | 140.51239 | -0.2367607 | 0.33055752 | -0.7162466 | 0.47383911 | 0.9314919 |
| OTULINL | 25.4148354 | 1.67666899 | 0.50531672 | 3.31805565 | 0.00090646 | 0.06269654 |
| OTULIN | 26.1556729 | -0.2133595 | 0.39612337 | -0.5386187 | 0.59014998 | 0.95415437 |
| ANKH | 74.5167015 | -1.1219404 | 0.57725393 | -1.943582 | 0.05194588 | 0.54907299 |
| RETREG1 | 27.3496617 | -0.079077 | 0.56444073 | -0.140098 | 0.88858254 | 0.99061558 |
| MYO10 | 11.9989306 | -0.4721153 | 0.5240176 | -0.9009532 | 0.36761321 | NA |
| BASP1 | 49.6877688 | 0.67364745 | 0.67167345 | 1.00293892 | 0.31589033 | 0.87263466 |
| LINC02100 | 13.695485 | 0.52836022 | 0.54830731 | 0.9636206 | 0.33523618 | 0.88047896 |
| LINC02241 | 8.96699672 | 0.1783213 | 0.6400682 | 0.27859733 | 0.78055386 | NA |
| CDH6 | 8.09958154 | 2.9590463 | 1.03449541 | 2.86037646 | 0.00423138 | NA |
| DROSHA | 23.3967393 | 0.40140169 | 0.44375387 | 0.90455929 | 0.36569891 | 0.89868152 |
| PDZD2 | 12.8938307 | 1.52564076 | 0.93602638 | 1.62991214 | 0.10312007 | NA |
| GOLPH3 | 59.5245613 | -0.0276598 | 0.3170414 | -0.0872435 | 0.93047795 | 0.99514836 |
| MTMR12 | 20.4782549 | -0.2707174 | 0.45622343 | -0.5933878 | 0.55292162 | 0.95415437 |
| ZFR | 81.4520893 | -0.38717 | 0.26400392 | -1.4665311 | 0.14250366 | 0.73044824 |
| SUB1 | 64.044472 | -0.0830787 | 0.53521279 | -0.1552255 | 0.87664351 | 0.98939075 |
| TARS | 12.1050565 | -0.0127064 | 0.50812815 | -0.0250062 | 0.98004998 | NA |
| C1QTNF3-AN | 23.7521962 | 0.32411647 | 0.56640365 | 0.57223584 | 0.56716221 | 0.95415437 |

| | | | | | | |
|---|---|---|---|---|---|---|
| DNAJC21 | 13.5268716 | -0.8057619 | 0.53495491 | -1.5062239 | 0.1320097 | 0.73044824 |
| IL7R | 97.7139425 | 0.73379383 | 0.63723499 | 1.15152783 | 0.24951515 | 0.83865855 |
| LMBRD2 | 17.2874566 | 0.79483828 | 0.53332962 | 1.49033215 | 0.13613692 | 0.73044824 |
| NADK2 | 9.68688424 | -0.0705453 | 0.72839813 | -0.0968499 | 0.92284559 | NA |
| SLC1A3 | 79.3473452 | 0.70961427 | 0.93896869 | 0.75573794 | 0.44980633 | 0.92914597 |
| NIPBL | 214.413354 | 0.0800296 | 0.20569344 | 0.38907221 | 0.69722273 | 0.96861929 |
| CPLANE1 | 23.5616542 | 0.28869731 | 0.3830387 | 0.75370271 | 0.45102775 | 0.92914597 |
| NUP155 | 11.717961 | -0.7752312 | 0.51312457 | -1.510805 | 0.13083815 | NA |
| WDR70 | 117.571363 | -0.0049136 | 0.3386969 | -0.0145073 | 0.98842522 | 0.99811561 |
| RICTOR | 124.677839 | -0.0126822 | 0.29159203 | -0.0434928 | 0.96530868 | 0.99653147 |
| FYB1 | 222.440889 | 1.30728685 | 0.40064251 | 3.26297587 | 0.00110249 | 0.07001544 |
| TTC33 | 18.0986104 | 0.05282845 | 0.63806847 | 0.08279433 | 0.93401508 | 0.99514836 |
| PTGER4 | 17.2308119 | 0.36919022 | 0.60414023 | 0.61110021 | 0.54113324 | 0.95415437 |
| PRKAA1 | 25.8356403 | 0.05021605 | 0.43049513 | 0.11664721 | 0.90713963 | 0.99368779 |
| RPL37 | 42.8223759 | -0.386228 | 0.43172916 | -0.8946072 | 0.37099708 | 0.90226557 |
| C7 | 11.3256973 | 0.72749558 | 0.73006979 | 0.99647402 | 0.31901988 | NA |
| OXCT1 | 25.056435 | 0.10213568 | 0.44838127 | 0.22778756 | 0.8198114 | 0.98320463 |
| SELENOP | 54.3893194 | 0.31828155 | 0.79128296 | 0.40223481 | 0.68751123 | 0.9673431 |
| ZNF131 | 46.5613076 | -0.1883927 | 0.35566517 | -0.5296912 | 0.59632603 | 0.9553365 |
| HMGCS1 | 13.067852 | -0.3286319 | 0.47725825 | -0.6885829 | 0.49108577 | 0.93633431 |
| TMEM267 | 13.9526723 | 0.04074472 | 0.45496115 | 0.08955649 | 0.92863966 | 0.99514836 |
| PAIP1 | 13.0436424 | -0.6821214 | 0.51349651 | -1.3283856 | 0.18405076 | 0.77694741 |
| NNT-AS1 | 9.98432891 | -0.2938225 | 0.58905942 | -0.4987994 | 0.61792071 | NA |
| NNT | 44.4237909 | -0.1711871 | 0.31595749 | -0.5418041 | 0.58795348 | 0.95415437 |
| EMB | 107.497467 | 1.00941077 | 0.58905253 | 1.71361758 | 0.08659898 | 0.64676224 |
| PARP8 | 353.715122 | 0.75037753 | 0.52879088 | 1.41904401 | 0.15588619 | 0.74486619 |
| ITGA1 | 16.5998786 | 1.36530722 | 0.5678982 | 2.40414075 | 0.01621053 | 0.33473536 |
| NDUFS4 | 37.6624003 | 0.0018364 | 0.32441874 | 0.00566059 | 0.99548353 | 0.99859588 |
| ARL15 | 227.375134 | 0.36635304 | 0.28132873 | 1.30222406 | 0.1928398 | 0.78135737 |
| SNX18 | 26.9075817 | -0.0593154 | 0.56265072 | -0.1054213 | 0.91604153 | 0.99498068 |
| GZMK | 10.0742143 | -1.5665841 | 0.87894514 | -1.7823457 | 0.07469287 | NA |
| GZMA | 23.9616988 | 0.03402732 | 0.63141027 | 0.05389099 | 0.95702202 | 0.99514836 |
| DHX29 | 20.7692663 | 0.12730641 | 0.46201111 | 0.27554836 | 0.78289498 | 0.98264092 |
| MTREX | 47.8521118 | -0.3144073 | 0.30975799 | -1.0150093 | 0.31010138 | 0.86847218 |
| PLPP1 | 20.9353535 | 0.37112748 | 0.53902933 | 0.68851073 | 0.49113122 | 0.93633431 |
| SLC38A9 | 38.510185 | 0.01282068 | 0.3497649 | 0.03665515 | 0.97075997 | 0.99676559 |
| IL6ST | 47.9263826 | 0.67879417 | 0.42531037 | 1.59599724 | 0.11048941 | 0.69012233 |
| MAP3K1 | 143.835456 | 0.35029641 | 0.27163281 | 1.28959536 | 0.19719119 | 0.78135737 |
| GPBP1 | 104.890498 | -0.164172 | 0.31849779 | -0.5154573 | 0.60623351 | 0.95688177 |
| PDE4D | 212.72001 | -0.6638019 | 0.52196018 | -1.2717481 | 0.20346263 | 0.78882329 |
| ERCC8 | 19.7590386 | -0.3610803 | 0.62384156 | -0.5788012 | 0.56272329 | 0.95415437 |
| NDUFAF2 | 34.9641032 | -0.1561895 | 0.690078 | -0.2263359 | 0.82094015 | 0.98320463 |
| ZSWIM6 | 281.590894 | 0.59685604 | 0.33455884 | 1.78400914 | 0.07442218 | 0.61929783 |
| KIF2A | 81.6393812 | -0.1517795 | 0.27844736 | -0.5450922 | 0.58569011 | 0.95415437 |
| IPO11 | 14.802917 | -0.2467888 | 0.46251922 | -0.5335753 | 0.5936354 | 0.9549937 |
| SREK1IP1 | 15.2856904 | 0.276725 | 0.44539466 | 0.62130291 | 0.53440034 | 0.95368316 |
| CWC27 | 37.635473 | 0.17113597 | 0.342493 | 0.49967728 | 0.61730233 | 0.95688177 |
| CENPK | 11.3005887 | -0.1536859 | 0.79780361 | -0.1926362 | 0.84724386 | NA |
| PPWD1 | 21.7333138 | 0.13077163 | 0.44019648 | 0.2970756 | 0.76640879 | 0.98264092 |
| TRIM23 | 13.2263431 | 0.08463761 | 0.58724833 | 0.14412576 | 0.88540116 | 0.99019008 |
| TRAPPC13 | 13.5574184 | -0.0974301 | 0.47977593 | -0.2030741 | 0.83907712 | 0.98342625 |
| SGTB | 17.8070269 | -0.2718088 | 0.46010015 | -0.59076 | 0.55468125 | 0.95415437 |
| NLN | 14.9919563 | -0.0357538 | 0.65768303 | -0.0543633 | 0.9566457 | 0.99514836 |
| ERBIN | 173.097944 | 0.35041735 | 0.2669205 | 1.31281545 | 0.18924514 | 0.77850418 |

| | | | | | | |
|---|---|---|---|---|---|---|
| SREK1 | 33.4776885 | -0.251051 | 0.42283362 | -0.5937347 | 0.55268956 | 0.95415437 |
| MAST4 | 41.8356715 | -0.5645325 | 0.48260075 | -1.1697714 | 0.24209298 | 0.83493679 |
| PIK3R1 | 128.495747 | 0.40618024 | 0.50105638 | 0.81064776 | 0.41756798 | 0.91954684 |
| SLC30A5 | 17.2507745 | 1.4841678 | 0.53079633 | 2.79611542 | 0.00517209 | 0.17966878 |
| CDK7 | 11.5555595 | 0.06036747 | 0.59142147 | 0.10207183 | 0.91869966 | NA |
| CCDC125 | 10.3911672 | 0.22836952 | 0.56652637 | 0.40310484 | 0.6868711 | NA |
| RAD17 | 11.2151303 | 0.08155432 | 0.51422126 | 0.15859772 | 0.87398583 | NA |
| GTF2H2C | 9.1771133 | 0.5240651 | 0.6488187 | 0.80772193 | 0.41925067 | NA |
| NAIP | 134.106822 | 0.52974231 | 0.59688741 | 0.88750793 | 0.3748055 | 0.90299244 |
| GTF2H2 | 8.93745032 | 0.28326367 | 0.75376838 | 0.3757967 | 0.70706804 | NA |
| BDP1 | 57.1691033 | -0.4880471 | 0.44212285 | -1.1038721 | 0.26964863 | 0.84994492 |
| MCCC2 | 18.4972398 | -0.6357267 | 0.43686863 | -1.4551896 | 0.14561677 | 0.73251288 |
| MRPS27 | 26.305937 | -0.9348729 | 0.47482947 | -1.9688602 | 0.04896914 | 0.53486612 |
| TNPO1 | 56.9999982 | -0.1438434 | 0.38962798 | -0.3691815 | 0.71199245 | 0.97317955 |
| FCHO2 | 82.0972025 | 0.32512111 | 0.52641604 | 0.61761247 | 0.53683082 | 0.95415437 |
| BTF3 | 30.5759329 | -0.2565602 | 0.57861971 | -0.4434004 | 0.65747613 | 0.96270373 |
| ANKRA2 | 9.64025999 | 0.43965537 | 0.64170408 | 0.68513725 | 0.49325732 | NA |
| HEXB | 24.2745889 | 0.27830625 | 0.44021868 | 0.6322 | 0.5272562 | 0.95136478 |
| GFM2 | 12.4057947 | 0.58920228 | 0.53108102 | 1.10943955 | 0.26724061 | NA |
| NSA2 | 17.710729 | -0.3564453 | 0.54625717 | -0.6525229 | 0.51406393 | 0.94596835 |
| HMGCR | 9.53434243 | -0.7275437 | 0.55238932 | -1.317085 | 0.18781013 | NA |
| COL4A3BP | 95.1700735 | 0.36191981 | 0.2595012 | 1.39467489 | 0.1631139 | 0.75321592 |
| POLK | 52.0343887 | -0.3966436 | 0.30677987 | -1.2929258 | 0.19603672 | 0.78135737 |
| POC5 | 16.5635358 | 0.04101472 | 0.45313056 | 0.09051414 | 0.92787866 | 0.99514836 |
| IQGAP2 | 265.158475 | 0.22599108 | 0.27435086 | 0.82373017 | 0.4100929 | 0.91773508 |
| F2R | 10.7900196 | 0.54558908 | 0.66482582 | 0.82064965 | 0.41184586 | NA |
| S100Z | 30.4771951 | 2.28387778 | 0.89030416 | 2.56527811 | 0.01030931 | 0.2796654 |
| WDR41 | 26.1929761 | 0.35854557 | 0.4792124 | 0.74819759 | 0.45434098 | 0.9314919 |
| TBCA | 57.9821805 | -0.5043624 | 0.2954747 | -1.7069563 | 0.08783018 | 0.64676224 |
| AP3B1 | 128.097341 | 0.47338479 | 0.24119671 | 1.9626503 | 0.04968682 | 0.53809564 |
| SCAMP1 | 25.0299861 | 0.39951903 | 0.40603271 | 0.98395775 | 0.32513629 | 0.87685391 |
| LHFPL2 | 31.665166 | 0.72085786 | 0.65945056 | 1.09311887 | 0.27434161 | 0.84994492 |
| ARSB | 37.0787937 | -0.5743141 | 0.38933385 | -1.47512 | 0.1401803 | 0.73044824 |
| JMY | 20.7253206 | 0.69671842 | 0.49238005 | 1.41500132 | 0.15706811 | 0.74486619 |
| TENT2 | 139.330233 | -0.3414609 | 0.22012525 | -1.5512118 | 0.12085092 | 0.70921423 |
| SERINC5 | 85.4416139 | 1.05968904 | 0.51466652 | 2.05898189 | 0.03949597 | 0.48300729 |
| ZFYVE16 | 45.0395058 | 0.37549648 | 0.36341417 | 1.03324666 | 0.30148848 | 0.86225566 |
| DHFR | 86.0616458 | -0.0441543 | 0.35395168 | -0.1247468 | 0.90072402 | 0.99194856 |
| MSH3 | 48.2744334 | -0.2678874 | 0.32116586 | -0.8341092 | 0.40421946 | 0.91528253 |
| RASGRF2 | 21.2111207 | 0.63406171 | 0.64672444 | 0.9804202 | 0.32687874 | 0.87715548 |
| CKMT2-AS1 | 12.4441233 | 0.29780808 | 0.49823208 | 0.59772963 | 0.55002035 | NA |
| SSBP2 | 100.074527 | 0.35213917 | 0.49617678 | 0.70970505 | 0.47788706 | 0.9318939 |
| ATG10 | 28.3062114 | 0.34462594 | 0.44540758 | 0.77373164 | 0.4390895 | 0.92780501 |
| RPS23 | 107.379124 | -0.4019841 | 0.60650854 | -0.662784 | 0.50746892 | 0.94305491 |
| TMEM167A | 20.1597463 | -0.0330536 | 0.5570482 | -0.0593371 | 0.95268363 | 0.99514836 |
| XRCC4 | 42.5848378 | -0.0534409 | 0.39102367 | -0.1366692 | 0.89129225 | 0.99080204 |
| VCAN | 311.556393 | 1.80454424 | 1.76530524 | 1.02222788 | 0.30667306 | 0.8673137 |
| COX7C | 28.512443 | 0.37182828 | 0.59968282 | 0.62004157 | 0.53523042 | 0.95378558 |
| RASA1 | 124.498258 | -0.3137611 | 0.30807129 | -1.0184691 | 0.3084551 | 0.8673137 |
| CCNH | 109.191676 | 0.59424047 | 0.36649039 | 1.62143535 | 0.10492431 | 0.67519097 |
| TMEM161B | 21.0880214 | -0.7425779 | 0.49018832 | -1.5148828 | 0.12980209 | 0.72931666 |
| TMEM161B-. | 24.9341171 | -0.2568125 | 0.40656087 | -0.6316705 | 0.52760224 | 0.95136878 |
| MEF2C | 220.52322 | -1.6283156 | 0.37324777 | -4.3625594 | 1.29E-05 | 0.00444621 |
| MEF2C-AS1 | 49.4440015 | -2.9190436 | 0.76350165 | -3.8232315 | 0.00013171 | 0.02143839 |

| | | | | | | |
|---|---|---|---|---|---|---|
| LYSMD3 | 9.57864186 | -0.1265662 | 0.62690813 | -0.2018896 | 0.84000307 | NA |
| ARRDC3 | 38.6836455 | 0.33303612 | 0.77026572 | 0.43236523 | 0.66547599 | 0.96406914 |
| FAM172A | 165.288884 | -0.0918314 | 0.2368282 | -0.3877554 | 0.69819703 | 0.96861929 |
| KIAA0825 | 33.3311052 | 0.34834958 | 0.48559432 | 0.7173675 | 0.47314737 | 0.9314919 |
| SLF1 | 26.2894358 | -0.0618066 | 0.43178914 | -0.1431406 | 0.88617915 | 0.99019008 |
| MCTP1 | 145.883678 | 0.23718029 | 0.34919012 | 0.67922966 | 0.49699235 | 0.93900843 |
| TTC37 | 42.3743475 | -0.1816759 | 0.36651511 | -0.4956846 | 0.62011692 | 0.95688177 |
| RHOBTB3 | 26.8318111 | 0.29607895 | 0.5653059 | 0.52374997 | 0.60045245 | 0.95688177 |
| GLRX | 29.7840415 | 1.37334213 | 0.57272585 | 2.39790493 | 0.01648914 | 0.33796638 |
| ELL2 | 119.741924 | 0.63477149 | 0.74167114 | 0.85586651 | 0.39207162 | 0.90910959 |
| AC104123.1 | 17.4941051 | 0.35128088 | 0.66582539 | 0.52758709 | 0.59778596 | 0.95538391 |
| CAST | 93.6453365 | -0.0855018 | 0.34114037 | -0.2506353 | 0.8020961 | 0.98320463 |
| ERAP1 | 29.0656343 | 0.24819746 | 0.45623242 | 0.5440154 | 0.58643087 | 0.95415437 |
| ERAP2 | 36.4585201 | -0.1318647 | 0.47475427 | -0.2777535 | 0.78120159 | 0.98264092 |
| LNPEP | 106.660972 | 0.2586071 | 0.24978862 | 1.03530375 | 0.30052708 | 0.86225566 |
| CHD1 | 101.621613 | -0.2014648 | 0.39852649 | -0.5055242 | 0.61319071 | 0.95688177 |
| ST8SIA4 | 122.488355 | 0.55141462 | 0.41182408 | 1.33895673 | 0.18058476 | 0.76886756 |
| PAM | 82.0928225 | 0.68884243 | 0.44264228 | 1.55620568 | 0.11965919 | 0.70431251 |
| PPIP5K2 | 52.7311376 | -0.5018947 | 0.41682511 | -1.2040894 | 0.22855503 | 0.82329316 |
| EFNA5 | 18.3872477 | -1.0113514 | 0.76820009 | -1.3165209 | 0.18799928 | 0.77798634 |
| FBXL17 | 161.24509 | -0.0488966 | 0.20414556 | -0.2395184 | 0.81070361 | 0.98320463 |
| FER | 85.3065421 | -0.5045202 | 0.24476143 | -2.0612731 | 0.03927699 | 0.48300729 |
| PJA2 | 62.6212185 | -0.0227757 | 0.28045416 | -0.0812099 | 0.93527499 | 0.99514836 |
| MAN2A1 | 101.341147 | -0.2899387 | 0.32846591 | -0.8827056 | 0.37739534 | 0.90489854 |
| SLC25A46 | 21.8643522 | -0.9091011 | 0.38321391 | -2.3723071 | 0.01767739 | 0.34827529 |
| WDR36 | 17.2487947 | -0.0289333 | 0.44710914 | -0.064712 | 0.94840332 | 0.99514836 |
| CAMK4 | 83.2805783 | 0.48577444 | 0.60145988 | 0.80765893 | 0.41928695 | 0.91985469 |
| STARD4 | 10.5151052 | -0.2142342 | 0.55443674 | -0.3863998 | 0.69920061 | NA |
| EPB41L4A | 15.0812336 | -0.2532793 | 0.63666948 | -0.3978191 | 0.69076352 | 0.96818917 |
| APC | 46.9967978 | 0.10152414 | 0.36013231 | 0.28190789 | 0.77801414 | 0.98264092 |
| SRP19 | 13.329531 | -0.0382554 | 0.53667135 | -0.0712826 | 0.9431728 | 0.99514836 |
| REEP5 | 37.6322162 | -0.1750217 | 0.42012843 | -0.416591 | 0.67697761 | 0.96718429 |
| DCP2 | 47.9220062 | 0.30141924 | 0.36500037 | 0.82580531 | 0.40891454 | 0.91716068 |
| MCC | 16.5492769 | -0.2001988 | 0.71188977 | -0.2812216 | 0.77854046 | 0.98264092 |
| YTHDC2 | 49.7137475 | -0.2598233 | 0.34489008 | -0.7533511 | 0.45123898 | 0.92914597 |
| PGGT1B | 26.4455019 | -0.8047009 | 0.42938973 | -1.8740571 | 0.06092256 | 0.57281427 |
| CCDC112 | 12.402253 | -0.2744701 | 0.71409747 | -0.3843594 | 0.7007121 | NA |
| FEM1C | 10.6473719 | -0.1565487 | 0.58607131 | -0.2671154 | 0.78938028 | NA |
| ATG12 | 20.3252735 | 0.12389672 | 0.41467205 | 0.29878242 | 0.76510607 | 0.98238909 |
| AP3S1 | 27.0709494 | 0.63236705 | 0.51268578 | 1.23343982 | 0.21741172 | 0.80640512 |
| COMMD10 | 107.946729 | -0.6002309 | 0.30449074 | -1.9712617 | 0.04869395 | 0.53486612 |
| SEMA6A-AS1 | 12.2810626 | -1.2343481 | 0.5072486 | -2.4334185 | 0.014957 | NA |
| DTWD2 | 14.123663 | 1.31589695 | 0.63352295 | 2.07711016 | 0.03779139 | 0.47316192 |
| DMXL1 | 88.53972 | -0.4405339 | 0.26142203 | -1.6851445 | 0.0919607 | 0.65902576 |
| TNFAIP8 | 165.510855 | 0.40260022 | 0.30534625 | 1.31850389 | 0.18733502 | 0.77798634 |
| HSD17B4 | 38.8608087 | 1.47744349 | 0.72664877 | 2.0332292 | 0.04202937 | 0.50341641 |
| SRFBP1 | 16.236356 | -0.4987894 | 0.51395128 | -0.9704993 | 0.33179766 | 0.88047896 |
| SNX2 | 62.4266931 | -1.0582796 | 0.40541539 | -2.6103588 | 0.00904473 | 0.25407889 |
| SNX24 | 34.0291628 | 0.17946004 | 0.46224368 | 0.38823687 | 0.69784075 | 0.96861929 |
| CEP120 | 23.5189634 | -0.0223606 | 0.38877074 | -0.0575161 | 0.95413407 | 0.99514836 |
| CSNK1G3 | 44.9006932 | -0.3265832 | 0.4010936 | -0.8142318 | 0.41551217 | 0.91894074 |
| GRAMD2B | 17.3935011 | -0.1770911 | 0.58708313 | -0.3016458 | 0.76292211 | 0.98163473 |
| LMNB1 | 13.4857542 | 0.18681632 | 0.68059394 | 0.27449014 | 0.78370798 | 0.98264092 |
| 3-Mar | 36.3654243 | -1.1518423 | 0.48454038 | -2.3771854 | 0.01744532 | 0.34827529 |

| | | | | | | |
|---|---|---|---|---|---|---|
| PRRC1 | 11.44125 | 0.43504041 | 0.59270109 | 0.7339963 | 0.46295099 | NA |
| LINC01184 | 30.3412614 | -0.8786793 | 0.44919085 | -1.956138 | 0.0504489 | 0.54421873 |
| SLC12A2 | 16.1767163 | 0.02186746 | 0.46396174 | 0.04713202 | 0.96240801 | 0.99653147 |
| HINT1 | 50.1620642 | 0.24620845 | 0.45747149 | 0.53819408 | 0.59044307 | 0.95415437 |
| CDC42SE2 | 263.003013 | -0.0028379 | 0.2505542 | -0.0113265 | 0.99096299 | 0.99827179 |
| AC116366.3 | 65.6290148 | -0.1650512 | 0.4856093 | -0.3398849 | 0.73394324 | 0.97605445 |
| IRF1 | 35.249491 | 0.40831061 | 0.47519328 | 0.85925164 | 0.3902017 | 0.90910959 |
| KIF3A | 10.9462042 | -0.3055649 | 0.57741172 | -0.5291976 | 0.59666837 | NA |
| UQCRQ | 13.0547446 | 0.56837227 | 0.75032316 | 0.7575033 | 0.44874839 | 0.92914597 |
| AFF4 | 72.805894 | 0.04884923 | 0.30912939 | 0.15802195 | 0.8744395 | 0.98899412 |
| ZCCHC10 | 17.9913614 | -0.8187368 | 0.4800816 | -1.7054118 | 0.08811765 | 0.64676224 |
| HSPA4 | 26.2238947 | -0.1903917 | 0.35972107 | -0.529276 | 0.59661398 | 0.95534194 |
| C5orf15 | 13.7219398 | -0.1868398 | 0.587637 | -0.3179511 | 0.75052202 | 0.97818861 |
| VDAC1 | 18.9007447 | 0.26166911 | 0.46494966 | 0.56279019 | 0.57357776 | 0.95415437 |
| TCF7 | 18.7841717 | 0.17487917 | 0.46713524 | 0.37436518 | 0.70813264 | 0.97184211 |
| SKP1 | 27.165429 | -0.239739 | 0.43192092 | -0.555053 | 0.57885844 | 0.95415437 |
| UBE2B | 48.8146655 | 0.15943145 | 0.28708178 | 0.55535205 | 0.57865388 | 0.95415437 |
| JADE2 | 18.6165731 | 0.01439384 | 0.44507474 | 0.03234028 | 0.97420069 | 0.9973754 |
| SAR1B | 17.976626 | 0.16934478 | 0.52150588 | 0.32472266 | 0.74539098 | 0.97805001 |
| SEC24A | 24.7724466 | -0.3391288 | 0.65705003 | -0.5161384 | 0.60575772 | 0.95688177 |
| CAMLG | 9.99677074 | 0.10318707 | 0.56570183 | 0.1824054 | 0.85526459 | NA |
| DDX46 | 40.6591139 | -0.453943 | 0.3273556 | -1.3866969 | 0.1655342 | 0.75337545 |
| C5orf24 | 10.2334736 | -0.0664019 | 0.55241303 | -0.1202033 | 0.90432212 | NA |
| TXNDC15 | 14.8711369 | -0.8983123 | 0.47615504 | -1.8865961 | 0.05921467 | 0.57281427 |
| PCBD2 | 21.847962 | -0.2660911 | 0.55489798 | -0.4795315 | 0.63156057 | 0.95877428 |
| H2AFY | 68.921302 | 0.63192943 | 0.33939672 | 1.86191967 | 0.06261441 | 0.58041569 |
| CXCL14 | 12.9787264 | 2.59714377 | 1.07167566 | 2.42344198 | 0.01537421 | 0.32566073 |
| TGFBI | 63.60344 | -0.1514372 | 0.66004082 | -0.2294362 | 0.81852992 | 0.98320463 |
| SMAD5 | 21.9558648 | 0.17513162 | 0.42894983 | 0.40827996 | 0.68306815 | 0.9673431 |
| HNRNPA0 | 14.0339002 | -1.0146241 | 0.47640612 | -2.1297461 | 0.03319258 | 0.44911423 |
| FAM13B | 95.7543884 | -0.2076592 | 0.26421126 | -0.7859587 | 0.43189166 | 0.92284736 |
| BRD8 | 10.21043 | -0.3672171 | 0.72068662 | -0.5095378 | 0.6103753 | NA |
| FAM53C | 11.6571776 | -0.1429599 | 0.59692987 | -0.2394919 | 0.81072415 | NA |
| KDM3B | 49.0944672 | -0.0605796 | 0.30638061 | -0.1977267 | 0.84325893 | 0.9842451 |
| ETF1 | 24.0871665 | -0.1595514 | 0.45688507 | -0.3492156 | 0.72692746 | 0.97570133 |
| HSPA9 | 33.7528502 | 0.05912839 | 0.35477996 | 0.16666214 | 0.8676359 | 0.98751288 |
| CTNNA1 | 67.7011554 | 0.1780826 | 0.32789503 | 0.54310856 | 0.58705506 | 0.95415437 |
| SIL1 | 48.3383728 | 0.61410863 | 0.3683642 | 1.66712352 | 0.09548985 | 0.66124053 |
| PAIP2 | 30.8417923 | 0.12813027 | 0.51478257 | 0.24890173 | 0.80343679 | 0.98320463 |
| TMEM173 | 10.2034109 | 1.01357674 | 0.65765612 | 1.54119562 | 0.12326918 | NA |
| UBE2D2 | 86.1359194 | -0.3444365 | 0.38667251 | -0.8907707 | 0.37305221 | 0.9025457 |
| CXXC5 | 8.34050135 | 0.76298689 | 0.64446632 | 1.183905 | 0.23645067 | NA |
| PURA | 22.944631 | -0.2967604 | 0.41813179 | -0.7097294 | 0.47787196 | 0.9318939 |
| CYSTM1 | 41.1144688 | 0.11286781 | 0.40148824 | 0.28112357 | 0.77861562 | 0.98264092 |
| PFDN1 | 15.9238163 | 0.84396611 | 0.57561885 | 1.46618915 | 0.14259676 | 0.73044824 |
| SRA1 | 10.716302 | 0.90815473 | 0.59791099 | 1.51887947 | 0.12879284 | NA |
| CD14 | 70.4148548 | 1.03694976 | 0.83855638 | 1.2365892 | 0.21623963 | 0.80634897 |
| NDUFA2 | 12.2469557 | 0.44257726 | 0.57733906 | 0.76658119 | 0.44333056 | NA |
| IK | 7.30561481 | -0.5193086 | 0.69336688 | -0.7489666 | 0.45387736 | NA |
| HARS | 14.2440285 | 0.04689231 | 0.47296621 | 0.09914517 | 0.92102301 | 0.99514836 |
| TAF7 | 15.2841977 | 0.10621254 | 0.53081883 | 0.20009188 | 0.84140872 | 0.98380297 |
| DIAPH1 | 111.302607 | -0.0741559 | 0.29432763 | -0.2519503 | 0.80107946 | 0.98320463 |
| HDAC3 | 12.6605335 | -0.4027166 | 0.50549339 | -0.7966802 | 0.4256368 | NA |
| DELE1 | 13.8762717 | -0.5527116 | 0.5682286 | -0.9726924 | 0.33070621 | 0.88037558 |

| | | | | | | |
|---|---|---|---|---|---|---|
| RNF14 | 8.43376327 | -0.168396 | 0.57584911 | -0.2924308 | 0.76995727 | NA |
| GNPDA1 | 10.9912594 | -0.0157555 | 0.57303866 | -0.0274946 | 0.97806525 | NA |
| NDFIP1 | 52.9475344 | 0.02916211 | 0.32228162 | 0.09048642 | 0.92790068 | 0.99514836 |
| ARHGAP26 | 342.549961 | 0.60613863 | 0.44868006 | 1.3509373 | 0.17671552 | 0.76409729 |
| NR3C1 | 173.975557 | 0.36173706 | 0.27552858 | 1.31288396 | 0.18922205 | 0.77850418 |
| LARS | 26.0701869 | 0.5480586 | 0.43594576 | 1.25717154 | 0.20869153 | 0.79538492 |
| TCERG1 | 41.9229327 | -0.4629542 | 0.33952937 | -1.3635173 | 0.17271954 | 0.76081596 |
| PPP2R2B | 91.1469028 | -1.0828646 | 0.6623561 | -1.6348677 | 0.10207679 | 0.67089422 |
| JAKMIP2 | 21.5604767 | 0.91495587 | 0.61864659 | 1.47896373 | 0.13915001 | 0.73044824 |
| FBXO38 | 32.1807841 | 0.24753137 | 0.44559561 | 0.55550676 | 0.57854809 | 0.95415437 |
| CSNK1A1 | 135.79668 | 0.05997061 | 0.25515222 | 0.23503856 | 0.8141788 | 0.98320463 |
| PPARGC1B | 13.3565696 | 0.2472358 | 0.6330698 | 0.39053482 | 0.69614112 | 0.9685713 |
| SLC26A2 | 9.70930105 | 1.17608761 | 0.57711843 | 2.0378618 | 0.04156376 | NA |
| HMGXB3 | 17.6737581 | -0.5856675 | 0.44499734 | -1.3161146 | 0.18813558 | 0.77798634 |
| CSF1R | 55.0565548 | 1.06500142 | 0.61059084 | 1.74421454 | 0.08112169 | 0.63497512 |
| TCOF1 | 8.85705298 | -0.627796 | 0.59788284 | -1.0500319 | 0.29370346 | NA |
| CD74 | 276.786657 | -1.3293962 | 0.65585761 | -2.0269586 | 0.04266665 | 0.504524 |
| RPS14 | 107.639091 | -0.2246403 | 0.46514269 | -0.4829492 | 0.6291318 | 0.95877428 |
| DCTN4 | 44.6458627 | -0.2008127 | 0.39407583 | -0.5095788 | 0.61034658 | 0.95688177 |
| GPX3 | 150.810931 | 1.11580384 | 0.76035911 | 1.46746954 | 0.14224837 | 0.73044824 |
| TNIP1 | 34.4871374 | 0.28041652 | 0.42185128 | 0.66472838 | 0.50622424 | 0.94166217 |
| ANXA6 | 33.4528983 | 0.43593988 | 0.46538999 | 0.93671951 | 0.34890286 | 0.89048867 |
| CCDC69 | 29.4699639 | -0.0600538 | 0.55028136 | -0.109133 | 0.91309704 | 0.99498068 |
| GM2A | 11.1742595 | 0.82626677 | 0.71935413 | 1.14862309 | 0.25071143 | NA |
| SLC36A1 | 22.9923802 | 0.29389496 | 0.60543423 | 0.48542838 | 0.62737251 | 0.95877428 |
| SPARC | 10.2928456 | 0.00347289 | 0.92505467 | 0.00375425 | 0.99700455 | NA |
| ATOX1 | 17.5831061 | 0.70478827 | 0.49042751 | 1.43708961 | 0.15069253 | 0.74061499 |
| G3BP1 | 39.9868831 | 0.36420965 | 0.31950178 | 1.13992996 | 0.25431548 | 0.84026145 |
| MFAP3 | 9.94038387 | 0.62917031 | 0.57490819 | 1.09438399 | 0.2737866 | NA |
| GALNT10 | 55.723987 | 0.42540155 | 0.39820995 | 1.06828457 | 0.28539217 | 0.85509489 |
| SAP30L-AS1 | 12.8625334 | 0.39983682 | 0.50781501 | 0.78736707 | 0.43106702 | NA |
| LARP1 | 46.4264126 | 0.23719326 | 0.33836023 | 0.7010081 | 0.48329796 | 0.93399425 |
| CNOT8 | 17.3969723 | -0.3831279 | 0.47764452 | -0.8021193 | 0.42248397 | 0.91985469 |
| HAVCR2 | 38.6184479 | 0.87792101 | 0.58378809 | 1.50383508 | 0.13262384 | 0.73044824 |
| ITK | 67.4755133 | 0.1160796 | 0.37461552 | 0.30986331 | 0.75666491 | 0.9799167 |
| AC010609.1 | 9.8014504 | 0.11409441 | 0.81905854 | 0.13929945 | 0.88921352 | NA |
| CYFIP2 | 69.9246345 | -0.5841465 | 0.31452662 | -1.8572242 | 0.06327926 | 0.58364573 |
| ADAM19 | 54.8652208 | -1.0788008 | 0.58465958 | -1.8451777 | 0.06501169 | 0.58951068 |
| CLINT1 | 55.4567833 | 0.10201448 | 0.33025835 | 0.30889297 | 0.75740295 | 0.98023104 |
| EBF1 | 39.3294236 | -2.6856185 | 0.60839884 | -4.41424 | 1.01E-05 | 0.00373971 |
| RNF145 | 67.2014085 | 0.07517376 | 0.31856569 | 0.23597569 | 0.81345153 | 0.98320463 |
| UBLCP1 | 14.3949132 | -0.0511438 | 0.47354009 | -0.1080032 | 0.91399317 | 0.99498068 |
| TTC1 | 20.3760227 | -0.1858221 | 0.42970962 | -0.4324366 | 0.66542416 | 0.96406914 |
| PWWP2A | 25.7774942 | 0.20557309 | 0.40425757 | 0.50852009 | 0.61108865 | 0.95688177 |
| SLU7 | 9.48763136 | 0.13589416 | 0.60114251 | 0.22605981 | 0.8211549 | NA |
| CCNG1 | 10.8744684 | 0.192825 | 0.63101919 | 0.30557708 | 0.75992667 | NA |
| MAT2B | 17.0518928 | -0.1584755 | 0.4495794 | -0.3524972 | 0.72446544 | 0.97499799 |
| RARS | 21.1238506 | -0.1156099 | 0.42883227 | -0.2695924 | 0.78747385 | 0.98320463 |
| PANK3 | 18.5232168 | 0.36630389 | 0.45535584 | 0.80443438 | 0.42114616 | 0.91985469 |
| DOCK2 | 321.693287 | -0.1028498 | 0.29411377 | -0.3496939 | 0.72656841 | 0.9754739 |
| FAM196B | 15.5415854 | -0.403539 | 0.74007877 | -0.545265 | 0.5855713 | 0.95415437 |
| LCP2 | 105.775632 | 0.72119814 | 0.35237658 | 2.04666875 | 0.04069062 | 0.48952585 |
| NPM1 | 22.5091521 | -0.2997134 | 0.41332569 | -0.7251264 | 0.46837451 | 0.9314919 |
| FBXW11 | 79.0378033 | -0.4555152 | 0.31342089 | -1.4533657 | 0.14612223 | 0.73379351 |

| | | | | | | |
|---|---|---|---|---|---|---|
| STK10 | 157.168087 | -0.0022488 | 0.35425118 | -0.0063479 | 0.99493514 | 0.99859588 |
| UBTD2 | 14.4560322 | -0.0515113 | 0.50269825 | -0.1024697 | 0.91838385 | 0.99498068 |
| SH3PXD2B | 38.8660009 | 0.88623981 | 0.7128768 | 1.2431879 | 0.2137986 | 0.80634897 |
| DUSP1 | 52.1321504 | 0.90448271 | 0.68044099 | 1.32925958 | 0.18376234 | 0.77688373 |
| ERGIC1 | 44.3329882 | 0.65973332 | 0.30551374 | 2.15942275 | 0.03081738 | 0.43175545 |
| ATP6V0E1 | 65.1629107 | -0.3035219 | 0.27329715 | -1.1105932 | 0.2667435 | 0.84994492 |
| CREBRF | 51.6403788 | 0.168048 | 0.37948789 | 0.44282837 | 0.65788988 | 0.9627619 |
| CPEB4 | 99.8632804 | -0.9792455 | 0.46848822 | -2.0902243 | 0.03659766 | 0.46928949 |
| SFXN1 | 23.8602297 | 0.09998743 | 0.47717144 | 0.20954193 | 0.8340252 | 0.98320463 |
| HRH2 | 24.7677766 | 1.91096586 | 0.66416002 | 2.87726723 | 0.00401136 | 0.15732988 |
| THOC3 | 24.2712553 | -0.5129622 | 0.46767534 | -1.0968339 | 0.27271402 | 0.84994492 |
| SIMC1 | 27.262123 | 0.16021173 | 0.35651166 | 0.44938706 | 0.65315246 | 0.96178254 |
| KIAA1191 | 9.64592881 | 0.58007047 | 0.59417933 | 0.97625487 | 0.32893818 | NA |
| CLTB | 9.71509268 | -0.1812792 | 0.60095362 | -0.3016526 | 0.76291687 | NA |
| FAF2 | 24.2266895 | -0.1978629 | 0.37507583 | -0.5275277 | 0.59782715 | 0.95538391 |
| RNF44 | 10.3333801 | 0.25039625 | 0.83682637 | 0.29922127 | 0.76477122 | NA |
| HK3 | 11.4040488 | 1.22798262 | 1.01341868 | 1.2117229 | 0.22561847 | NA |
| UIMC1 | 47.450386 | 0.52407219 | 0.3521234 | 1.48831968 | 0.13666661 | 0.73044824 |
| ZNF346 | 14.5462922 | -0.0425302 | 0.50255347 | -0.0846283 | 0.93255693 | 0.99514836 |
| NSD1 | 102.236038 | -0.0675937 | 0.24339404 | -0.2777131 | 0.78123257 | 0.98264092 |
| PRELID1 | 21.5103297 | 1.20998215 | 0.70779931 | 1.70949891 | 0.08735857 | 0.64676224 |
| LMAN2 | 18.0773882 | -0.0263309 | 0.63305291 | -0.0415934 | 0.9668228 | 0.99653147 |
| RGS14 | 9.90270921 | -0.1721073 | 0.66654928 | -0.2582064 | 0.79624762 | NA |
| GRK6 | 12.9681026 | -0.015144 | 0.55881113 | -0.0271005 | 0.9783796 | 0.99758514 |
| DOK3 | 17.7762454 | 0.35271261 | 0.47573187 | 0.7414105 | 0.45844458 | 0.9314919 |
| FAM193B | 25.8575736 | 0.07963047 | 0.35958401 | 0.22145163 | 0.82474079 | 0.98320463 |
| TMED9 | 17.7547943 | 0.50564894 | 0.66327618 | 0.76235052 | 0.44585083 | 0.92914597 |
| RMND5B | 9.74304522 | -0.6675988 | 0.5587407 | -1.1948277 | 0.23215437 | NA |
| HNRNPAB | 16.2299193 | -0.140275 | 0.5519757 | -0.2541326 | 0.79939316 | 0.98320463 |
| PHYKPL | 52.9999603 | 0.30618114 | 0.52685979 | 0.5811435 | 0.56114374 | 0.95415437 |
| CLK4 | 16.1604974 | -0.0791564 | 0.48044023 | -0.1647581 | 0.86913437 | 0.98751288 |
| ADAMTS2 | 20.4375237 | -0.0469391 | 1.14100159 | -0.0411385 | 0.96718548 | 0.99653147 |
| RUFY1 | 32.3852824 | 0.0756537 | 0.5035822 | 0.15023109 | 0.8805823 | 0.99019008 |
| HNRNPH1 | 102.007701 | -0.0944005 | 0.31479432 | -0.2998801 | 0.76426864 | 0.98222198 |
| CANX | 50.6069074 | 0.10233423 | 0.43525641 | 0.23511253 | 0.8141214 | 0.98320463 |
| MAML1 | 16.2763517 | -0.0319505 | 0.43404632 | -0.0736107 | 0.94132014 | 0.99514836 |
| SQSTM1 | 53.6204665 | 0.23147817 | 0.41611905 | 0.55627872 | 0.57802033 | 0.95415437 |
| MRNIP | 9.57602712 | 0.77823802 | 0.69933343 | 1.11282828 | 0.26578219 | NA |
| TBC1D9B | 19.9262457 | 0.16553395 | 0.44842363 | 0.36914635 | 0.71201864 | 0.97317955 |
| RNF130 | 116.376545 | 0.41286958 | 0.43730276 | 0.94412754 | 0.34510448 | 0.88711459 |
| MAPK9 | 17.6024099 | -0.6189917 | 0.41266184 | -1.4999975 | 0.13361506 | 0.73044824 |
| CNOT6 | 26.8519239 | -0.3690102 | 0.38959687 | -0.947159 | 0.34355779 | 0.88595005 |
| MGAT1 | 59.6037243 | 0.32252314 | 0.40346404 | 0.79938511 | 0.42406714 | 0.91985469 |
| RACK1 | 57.2372627 | -0.0673917 | 0.5411452 | -0.1245353 | 0.90089148 | 0.99194856 |
| TRIM52 | 10.0683811 | -0.5320533 | 0.52973082 | -1.0043843 | 0.31519342 | NA |
| DUSP22 | 20.3519505 | -0.2108829 | 0.45514744 | -0.4633288 | 0.64312867 | 0.95992767 |
| EXOC2 | 58.8653258 | -0.1162823 | 0.2937467 | -0.3958592 | 0.69220891 | 0.9685713 |
| GMDS | 113.678727 | 0.72605985 | 0.38728141 | 1.87476037 | 0.0608257 | 0.57281427 |
| GMDS-DT | 84.9383141 | 0.32808125 | 0.55770429 | 0.58827099 | 0.55635042 | 0.95415437 |
| WRNIP1 | 14.3705477 | 0.19240177 | 0.50196496 | 0.38329722 | 0.7014994 | 0.97078484 |
| SERPINB1 | 33.6796482 | 0.86837059 | 0.41711438 | 2.08185246 | 0.03735595 | 0.47208194 |
| SERPINB9 | 56.0630352 | 0.35644535 | 0.65315192 | 0.54573115 | 0.58525076 | 0.95415437 |
| SERPINB6 | 31.8127046 | 0.19489814 | 0.48364928 | 0.40297412 | 0.68696726 | 0.9673431 |
| NQO2 | 14.1271867 | -0.4203248 | 0.56235098 | -0.7474421 | 0.45479676 | 0.9314919 |

| | | | | | | |
|---|---|---|---|---|---|---|
| RIPK1 | 25.0942529 | 0.08380004 | 0.39982868 | 0.20958987 | 0.83398778 | 0.98320463 |
| PSMG4 | 9.65375637 | -0.5007876 | 0.54231336 | -0.9234285 | 0.35578393 | NA |
| SLC22A23 | 16.8641315 | 0.45440259 | 0.72389211 | 0.62772143 | 0.53018645 | 0.95245713 |
| PRPF4B | 60.0628411 | -0.368307 | 0.31345189 | -1.1750034 | 0.23999335 | 0.83493679 |
| CDYL | 69.0327727 | -0.1847815 | 0.28407218 | -0.6504739 | 0.51538617 | 0.94596835 |
| LYRM4 | 24.4709348 | -1.0006232 | 0.45298607 | -2.2089491 | 0.02717818 | 0.40712515 |
| FARS2 | 89.6367305 | -0.1149424 | 0.33278733 | -0.3453929 | 0.729799 | 0.97605445 |
| F13A1 | 193.342038 | 1.57778417 | 0.94127863 | 1.67621373 | 0.09369634 | 0.6590317 |
| LY86 | 26.8860999 | 0.52528042 | 0.88596199 | 0.59289273 | 0.55325295 | 0.95415437 |
| RREB1 | 115.461927 | 0.37722002 | 0.32558532 | 1.15859037 | 0.2466232 | 0.83817091 |
| SSR1 | 27.6068213 | 0.41922452 | 0.54927961 | 0.76322607 | 0.44532858 | 0.92914597 |
| SNRNP48 | 10.0491848 | -0.143618 | 0.57066636 | -0.2516673 | 0.80129825 | NA |
| SLC35B3 | 11.3044193 | -0.1920498 | 0.55001709 | -0.3491706 | 0.72696126 | NA |
| GCNT2 | 22.2221592 | -0.5896589 | 0.41237282 | -1.429917 | 0.15274084 | 0.74341937 |
| MAK | 9.04459796 | 0.27817291 | 0.66206156 | 0.42016169 | 0.67436734 | NA |
| AL139807.1 | 11.7691419 | -0.3855148 | 0.55106616 | -0.6995799 | 0.48418972 | NA |
| NEDD9 | 150.100015 | -0.811786 | 0.43172347 | -1.8803379 | 0.06006204 | 0.57281427 |
| TMEM170B | 29.1821359 | 1.0605564 | 0.76138636 | 1.39292803 | 0.16364155 | 0.75321592 |
| HIVEP1 | 56.031392 | -0.6582562 | 0.40282821 | -1.6340866 | 0.10224067 | 0.67109695 |
| PHACTR1 | 44.9512261 | 0.28014498 | 0.57819991 | 0.48451232 | 0.62802232 | 0.95877428 |
| GFOD1 | 36.5097265 | 0.53579197 | 0.48429295 | 1.10633856 | 0.26858001 | 0.84994492 |
| SIRT5 | 12.8028912 | 0.27837834 | 0.64094713 | 0.43432341 | 0.66405361 | NA |
| NOL7 | 15.5218572 | -0.1049412 | 0.54776486 | -0.1915808 | 0.84807056 | 0.98545782 |
| RANBP9 | 68.7024527 | -0.1034735 | 0.32885849 | -0.3146444 | 0.75303167 | 0.97883059 |
| CD83 | 52.7932256 | -0.1493621 | 0.69660014 | -0.2144158 | 0.8302228 | 0.98320463 |
| JARID2 | 184.817755 | 0.16119183 | 0.26248431 | 0.61410082 | 0.53914871 | 0.95415437 |
| DTNBP1 | 56.4304262 | 0.04099471 | 0.40481388 | 0.10126805 | 0.91933768 | 0.99498068 |
| MYLIP | 13.5608595 | 0.96479634 | 0.57671328 | 1.67292201 | 0.09434265 | 0.65925012 |
| ATXN1 | 343.279912 | 0.04873001 | 0.30216405 | 0.16127003 | 0.87188072 | 0.98751288 |
| NUP153 | 50.8943357 | -0.1585531 | 0.39133274 | -0.4051617 | 0.68535863 | 0.9673431 |
| KIF13A | 53.9942778 | 0.83598057 | 0.42118254 | 1.9848415 | 0.04716211 | 0.52845055 |
| TPMT | 9.92525782 | 0.33572283 | 0.66385981 | 0.50571345 | 0.61305783 | NA |
| KDM1B | 26.2010485 | 0.86258976 | 0.52514442 | 1.64257628 | 0.10047063 | 0.6683775 |
| DEK | 46.1853345 | 0.18016393 | 0.30988258 | 0.58139419 | 0.56097481 | 0.95415437 |
| RNF144B | 123.274427 | 1.67849799 | 0.76442107 | 2.19577672 | 0.02810793 | 0.41814322 |
| MBOAT1 | 18.8568605 | 0.727307 | 0.5169984 | 1.40678771 | 0.15949035 | 0.7479827 |
| E2F3 | 54.1602167 | 0.01877425 | 0.31168054 | 0.06023555 | 0.95196803 | 0.99514836 |
| CDKAL1 | 162.665284 | -0.4519296 | 0.28462412 | -1.5878122 | 0.11232882 | 0.695333 |
| SOX4 | 17.5898257 | -0.8891849 | 1.08421013 | -0.8201223 | 0.41214638 | 0.91894074 |
| DCDC2 | 9.38478051 | -0.4567325 | 0.70953626 | -0.6437057 | 0.5197663 | NA |
| TDP2 | 16.6894155 | 0.84849145 | 0.48898923 | 1.73519453 | 0.08270635 | 0.63740875 |
| C6orf62 | 53.8844743 | -0.3015857 | 0.34243923 | -0.8806985 | 0.37848102 | 0.9051267 |
| RIPOR2 | 325.675058 | -0.0426795 | 0.35851673 | -0.1190446 | 0.90524003 | 0.99324672 |
| CARMIL1 | 65.3431689 | -1.3109751 | 0.41939388 | -3.1258803 | 0.00177274 | 0.09433005 |
| TRIM38 | 56.8299347 | -0.2810434 | 0.39347712 | -0.714256 | 0.47506892 | 0.9314919 |
| HIST1H1C | 10.1446942 | -0.0073393 | 0.86956722 | -0.0084402 | 0.99326576 | NA |
| HIST1H4C | 10.0342285 | 0.25155262 | 0.73860948 | 0.34057594 | 0.73342285 | NA |
| HIST1H2AC | 83.9940107 | 0.70225405 | 0.69795895 | 1.00615379 | 0.31434159 | 0.87152625 |
| HIST1H1E | 10.6633259 | -0.712103 | 0.93928718 | -0.7581312 | 0.44837242 | NA |
| HIST1H1D | 10.5060449 | -0.4533571 | 0.73010252 | -0.6209499 | 0.5346326 | NA |
| BTN3A2 | 30.7711559 | 1.04576945 | 0.41145165 | 2.54165819 | 0.0110328 | 0.2861945 |
| BTN3A1 | 30.9167704 | 0.70673253 | 0.61207854 | 1.15464353 | 0.24823643 | 0.83817091 |
| BTN3A3 | 12.5050576 | -0.2910629 | 0.50832634 | -0.5725906 | 0.5669219 | NA |
| BTN2A1 | 16.690995 | -0.2897633 | 0.44870187 | -0.6457814 | 0.51842096 | 0.94668603 |

| | | | | | | |
|---|---|---|---|---|---|---|
| HMGN4 | 8.50976436 | 0.28998734 | 0.68321678 | 0.42444411 | 0.67124195 | NA |
| TRIM27 | 24.8658022 | -0.0648757 | 0.39308178 | -0.1650439 | 0.86890943 | 0.98751288 |
| GABBR1 | 13.5769344 | -0.304557 | 0.8584964 | -0.3547563 | 0.72277217 | 0.97441549 |
| HLA-F | 30.5772487 | 0.0946923 | 0.49634141 | 0.19078058 | 0.8486975 | 0.98545782 |
| HLA-A | 231.457775 | -0.0160926 | 0.49865903 | -0.0322717 | 0.97425536 | 0.9973754 |
| TRIM26 | 17.9694165 | -0.3325034 | 0.44939598 | -0.7398895 | 0.45936704 | 0.9314919 |
| HCG17 | 14.4619163 | -0.1583578 | 0.6234697 | -0.2539944 | 0.79949992 | 0.98320463 |
| HLA-E | 155.794746 | -0.1248378 | 0.40693524 | -0.3067757 | 0.75901413 | 0.98071076 |
| GNL1 | 9.9458403 | -0.8762411 | 0.49194857 | -1.7811641 | 0.07488564 | NA |
| ABCF1 | 12.7305663 | -0.4357212 | 0.53439771 | -0.8153501 | 0.41487192 | NA |
| PPP1R10 | 22.7625118 | 0.02894283 | 0.44645455 | 0.06482816 | 0.94831082 | 0.99514836 |
| PPP1R18 | 21.7249989 | 0.030505 | 0.60219736 | 0.05065615 | 0.95959952 | 0.99595344 |
| TUBB | 25.948939 | -0.400277 | 0.61392722 | -0.6519942 | 0.51440494 | 0.94596835 |
| FLOT1 | 31.099321 | 0.16386853 | 0.49580751 | 0.33050837 | 0.74101587 | 0.97605445 |
| IER3 | 23.3354584 | 0.41508078 | 0.66068918 | 0.62825424 | 0.5298374 | 0.95245713 |
| HLA-C | 139.906178 | 0.15604028 | 0.59686727 | 0.26143212 | 0.79375929 | 0.98320463 |
| HLA-B | 274.103497 | 0.01021861 | 0.5420607 | 0.01885142 | 0.98495963 | 0.99811561 |
| LTB | 31.6892296 | -0.1124222 | 0.75951274 | -0.1480188 | 0.88232794 | 0.99019008 |
| LST1 | 27.9245944 | 2.02747646 | 0.83692551 | 2.42252917 | 0.01541289 | 0.32566073 |
| AIF1 | 31.0703139 | 1.3518949 | 1.02320462 | 1.32123612 | 0.18642265 | 0.77798634 |
| PRRC2A | 18.3931077 | 0.08181423 | 0.49135179 | 0.16650846 | 0.86775683 | 0.98751288 |
| DDAH2 | 11.403356 | -0.4210429 | 0.60855043 | -0.6918784 | 0.48901368 | NA |
| CLIC1 | 71.5999122 | -0.0642559 | 0.63307126 | -0.1014987 | 0.91915463 | 0.99498068 |
| HSPA1A | 66.2806653 | 1.02542107 | 0.74425484 | 1.3777822 | 0.16827054 | 0.7567539 |
| HSPA1B | 31.4375959 | 0.02740807 | 0.82019427 | 0.03341656 | 0.97334241 | 0.9973754 |
| C6orf48 | 9.04378791 | -0.5931421 | 0.63102473 | -0.9399665 | 0.34723477 | NA |
| ATF6B | 15.3722359 | -0.0452335 | 0.49985584 | -0.0904932 | 0.92789532 | 0.99514836 |
| PBX2 | 10.0918715 | -0.0487782 | 0.56972686 | -0.0856168 | 0.93177104 | NA |
| GPSM3 | 23.5426477 | 0.39100426 | 0.759523 | 0.5148024 | 0.60669111 | 0.95688177 |
| HLA-DRA | 173.660105 | -1.0604443 | 0.67954886 | -1.5605123 | 0.11863887 | 0.70431251 |
| HLA-DRB5 | 46.5360586 | -1.097964 | 0.74270484 | -1.4783316 | 0.13931905 | 0.73044824 |
| HLA-DRB1 | 125.576694 | -1.8372413 | 0.68128877 | -2.6967146 | 0.00700273 | 0.20947616 |
| HLA-DQA1 | 41.5763087 | -1.8487154 | 0.6513537 | -2.8382665 | 0.00453593 | 0.16960699 |
| HLA-DQB1 | 55.694168 | -1.2173045 | 0.8602792 | -1.4150109 | 0.1570653 | 0.74486619 |
| PSMB8 | 20.2129663 | 0.34070685 | 0.5057274 | 0.67369665 | 0.50050419 | 0.94001041 |
| PSMB9 | 26.2450417 | 0.45687232 | 0.5294647 | 0.86289477 | 0.38819533 | 0.90910959 |
| TAP1 | 21.6100413 | -0.120111 | 0.58489303 | -0.2053555 | 0.8372944 | 0.98320463 |
| HLA-DMB | 16.1601636 | -1.2666892 | 0.58373779 | -2.1699626 | 0.03000968 | 0.42727094 |
| BRD2 | 30.9205622 | -0.2633363 | 0.42841423 | -0.6146769 | 0.53876815 | 0.95415437 |
| HLA-DOA | 10.6173011 | -2.1237153 | 0.67250115 | -3.1579355 | 0.00158891 | NA |
| HLA-DPA1 | 70.8631977 | -0.8074247 | 0.70415863 | -1.1466517 | 0.25152561 | 0.84008912 |
| HLA-DPB1 | 82.2602683 | -1.1473104 | 0.65578066 | -1.7495337 | 0.08019881 | 0.63385609 |
| RPS18 | 63.357152 | -0.224946 | 0.69506442 | -0.3236333 | 0.74621571 | 0.97805001 |
| RGL2 | 9.15591392 | 0.03871285 | 0.5781119 | 0.06696429 | 0.94661013 | NA |
| TAPBP | 55.9863611 | 0.33082846 | 0.39657469 | 0.83421476 | 0.40415998 | 0.91528253 |
| CUTA | 9.14972782 | 0.70341381 | 0.650702 | 1.08100761 | 0.27969373 | NA |
| ITPR3 | 10.8451592 | -1.116377 | 0.58565026 | -1.9062178 | 0.05662195 | NA |
| LEMD2 | 12.434982 | -0.3724738 | 0.50257591 | -0.7411294 | 0.45861501 | NA |
| NUDT3 | 13.8388643 | -0.169056 | 0.696573 | -0.2426967 | 0.80824033 | 0.98320463 |
| C6orf106 | 48.798264 | -0.2598958 | 0.40512277 | -0.6415235 | 0.52118264 | 0.94895469 |
| SNRPC | 8.40009153 | 0.27106446 | 0.62596691 | 0.4330332 | 0.66499065 | NA |
| UHRF1BP1 | 26.7314659 | -0.0510954 | 0.4915767 | -0.103942 | 0.9172154 | 0.99498068 |
| TAF11 | 13.2976316 | -0.4329257 | 0.50430015 | -0.8584683 | 0.39063392 | 0.90910959 |
| ANKS1A | 93.0658466 | 0.55225878 | 0.30566993 | 1.80671609 | 0.07080656 | 0.60656887 |

| | | | | | | |
|---|---|---|---|---|---|---|
| ZNF76 | 13.7706737 | -0.0343467 | 0.48403273 | -0.0709594 | 0.94343005 | 0.99514836 |
| DEF6 | 26.3369467 | -0.1367765 | 0.40526786 | -0.3374966 | 0.73574255 | 0.97605445 |
| PPARD | 54.6021352 | 0.09711509 | 0.59520374 | 0.16316278 | 0.87039027 | 0.98751288 |
| RPL10A | 57.6201127 | 0.01863653 | 0.68167049 | 0.02733949 | 0.97818896 | 0.99758514 |
| FKBP5 | 354.201408 | 1.01956942 | 0.65126812 | 1.56551409 | 0.11746242 | 0.70122655 |
| SRPK1 | 31.8925238 | 0.3002433 | 0.37518658 | 0.80025064 | 0.4235656 | 0.91985469 |
| MAPK14 | 88.4743746 | 0.70944889 | 0.32085605 | 2.21111271 | 0.02702803 | 0.40712515 |
| KCTD20 | 33.565393 | 0.55551946 | 0.43877578 | 1.26606684 | 0.20548915 | 0.78970624 |
| STK38 | 109.813994 | 0.98570644 | 0.35631713 | 2.76637401 | 0.00566835 | 0.19011297 |
| SRSF3 | 35.9649468 | 0.0771152 | 0.37723045 | 0.20442465 | 0.83802167 | 0.98320463 |
| CDKN1A | 13.2501926 | -0.2198552 | 0.82083577 | -0.267843 | 0.78882014 | 0.98320463 |
| C6orf89 | 21.6367049 | 0.1135514 | 0.39989742 | 0.28395133 | 0.77644768 | 0.98264092 |
| MTCH1 | 27.7645884 | -0.1108322 | 0.4445919 | -0.2492897 | 0.8031367 | 0.98320463 |
| FGD2 | 52.1403389 | 0.11633157 | 0.5051241 | 0.23030296 | 0.81785636 | 0.98320463 |
| PIM1 | 20.2000084 | 0.42645441 | 0.59518136 | 0.71651169 | 0.47367545 | 0.9314919 |
| TBC1D22B | 15.7099945 | 0.20637001 | 0.5253789 | 0.39280224 | 0.69446555 | 0.9685713 |
| CMTR1 | 11.73743 | -0.4009826 | 0.56192664 | -0.7135854 | 0.47548361 | NA |
| ZFAND3 | 381.809562 | 0.1775661 | 0.18987997 | 0.93514919 | 0.34971142 | 0.89048867 |
| BTBD9 | 163.854821 | -0.1725192 | 0.29835136 | -0.5782418 | 0.56310088 | 0.95415437 |
| GLO1 | 15.4986924 | 0.09254584 | 0.77790887 | 0.11896746 | 0.90530113 | 0.99324672 |
| OARD1 | 11.9513725 | 0.86781519 | 0.59586673 | 1.45639142 | 0.14528445 | NA |
| NFYA | 10.2448875 | 0.22262135 | 0.56818967 | 0.39180816 | 0.69519996 | NA |
| TFEB | 30.2566198 | -2.0059593 | 0.62522347 | -3.2083877 | 0.00133481 | 0.07670434 |
| CCND3 | 255.810263 | -0.7728023 | 0.55843861 | -1.3838626 | 0.16640053 | 0.75542292 |
| TAF8 | 11.2816815 | -0.1299291 | 0.59278927 | -0.2191826 | 0.82650782 | NA |
| TRERF1 | 89.5238297 | 0.07198469 | 0.32604 | 0.22078484 | 0.82525997 | 0.98320463 |
| UBR2 | 111.898197 | -0.1695868 | 0.27487959 | -0.6169495 | 0.53726804 | 0.95415437 |
| BICRAL | 51.989455 | 0.20677376 | 0.41616918 | 0.49685024 | 0.61929466 | 0.95688177 |
| RPL7L1 | 14.4684242 | 0.01507201 | 0.48653811 | 0.03097806 | 0.97528703 | 0.9973754 |
| CNPY3 | 22.2493107 | -0.6206304 | 0.57906385 | -1.0717824 | 0.28381779 | 0.85176121 |
| PEX6 | 8.45340706 | 0.16203912 | 0.57650433 | 0.28107181 | 0.77865532 | NA |
| KLHDC3 | 8.13581382 | 0.0780904 | 0.63067652 | 0.12382006 | 0.90145775 | NA |
| RRP36 | 11.3507988 | -0.0216521 | 0.51985043 | -0.0416506 | 0.96677722 | NA |
| CUL9 | 8.71602597 | -1.0994431 | 0.5544037 | -1.9831093 | 0.04735522 | NA |
| ZNF318 | 12.9515105 | -2.8185828 | 0.68313393 | -4.1259593 | 3.69E-05 | 0.00863983 |
| ABCC10 | 8.96824014 | 0.09164355 | 0.58622846 | 0.15632737 | 0.87577499 | NA |
| POLR1C | 10.1209358 | -0.4452536 | 0.65370567 | -0.6811225 | 0.49579398 | NA |
| XPO5 | 51.5639993 | -0.5992403 | 0.50285792 | -1.1916693 | 0.23339096 | 0.82817204 |
| POLH | 10.4997037 | -1.6013127 | 0.60583792 | -2.6431371 | 0.00821418 | NA |
| GTPBP2 | 10.2222222 | -0.2571837 | 0.59619131 | -0.4313778 | 0.66619372 | NA |
| VEGFA | 19.517127 | 0.62586682 | 0.93643151 | 0.66835301 | 0.50390828 | 0.94166217 |
| MRPL14 | 8.32777617 | 0.17613912 | 0.8393912 | 0.20984151 | 0.83379137 | NA |
| HSP90AB1 | 61.2725793 | -0.895172 | 0.39572339 | -2.2621154 | 0.02369027 | 0.39492781 |
| CDC5L | 16.5053064 | 0.5762203 | 0.58793516 | 0.98007457 | 0.32704931 | 0.87715548 |
| SUPT3H | 78.0337595 | -0.0232717 | 0.46939268 | -0.0495784 | 0.96045834 | 0.99614757 |
| RUNX2 | 63.694542 | 1.10074288 | 0.38067903 | 2.89152484 | 0.00383377 | 0.15732988 |
| ENPP4 | 11.1393299 | 0.35280265 | 0.66918802 | 0.52721005 | 0.59804773 | NA |
| CD2AP | 55.0171207 | -0.5217348 | 0.37125494 | -1.4053275 | 0.1599239 | 0.74874693 |
| PKHD1 | 20.5590595 | -0.2051659 | 0.82078188 | -0.2499639 | 0.80261525 | 0.98320463 |
| TRAM2 | 15.8191475 | 1.65831712 | 0.76096231 | 2.17923686 | 0.02931408 | 0.42316846 |
| FBXO9 | 21.1353681 | 0.45697914 | 0.41181178 | 1.10967961 | 0.26713711 | 0.84994492 |
| ELOVL5 | 71.6379408 | -0.1614132 | 0.4438802 | -0.3636414 | 0.71612584 | 0.97348719 |
| GCLC | 18.6936941 | -0.5397373 | 0.44523584 | -1.2122504 | 0.22541653 | 0.81875277 |
| DST | 53.8809453 | 0.3985165 | 0.52990325 | 0.75205521 | 0.45201786 | 0.92931643 |

| | | | | | | |
|---|---|---|---|---|---|---|
| ZNF451 | 52.9658139 | 0.18071137 | 0.34994816 | 0.51639469 | 0.60557878 | 0.95688177 |
| PRIM2 | 70.7219652 | -0.1375397 | 0.30225487 | -0.4550455 | 0.64907652 | 0.96084839 |
| KHDRBS2 | 36.6730213 | -1.1076202 | 0.63793264 | -1.7362651 | 0.08251698 | 0.63740875 |
| PTP4A1 | 18.5993254 | 0.45493394 | 0.51659434 | 0.88064058 | 0.37851239 | 0.9051267 |
| PHF3 | 73.0144529 | 0.06143977 | 0.30507419 | 0.20139288 | 0.84039138 | 0.98365608 |
| LMBRD1 | 67.8578206 | -0.1888019 | 0.26706654 | -0.7069469 | 0.47959948 | 0.93257966 |
| COL19A1 | 14.2177862 | 0.66552061 | 0.6656931 | 0.99974089 | 0.31743592 | 0.87265846 |
| FAM135A | 45.2120537 | -0.5044326 | 0.43926251 | -1.1483625 | 0.25081896 | 0.83984169 |
| SMAP1 | 64.2436271 | 0.11096079 | 0.3710653 | 0.29903306 | 0.76491482 | 0.98237146 |
| OGFRL1 | 41.896639 | 0.77390246 | 0.44269073 | 1.7481786 | 0.08043311 | 0.63385609 |
| KCNQ5 | 79.6902776 | -0.8681232 | 0.57918209 | -1.4988779 | 0.13390531 | 0.73044824 |
| OOEP | 81.5071698 | -0.5775744 | 0.64121585 | -0.9007487 | 0.36772193 | 0.89868152 |
| CGAS | 17.130529 | 0.63968217 | 0.4801153 | 1.33235113 | 0.18274483 | 0.77495012 |
| MTO1 | 12.5083637 | -0.625934 | 0.50731057 | -1.233828 | 0.21726699 | NA |
| EEF1A1 | 261.597146 | -0.028401 | 0.53338039 | -0.0532473 | 0.95753491 | 0.99514836 |
| SLC17A5 | 24.9540198 | 0.22772955 | 0.47294766 | 0.4815111 | 0.63015329 | 0.95877428 |
| COX7A2 | 15.3125793 | -0.1479621 | 0.51102629 | -0.2895391 | 0.77216883 | 0.98264092 |
| TMEM30A | 29.8226187 | 0.46558279 | 0.53367614 | 0.87240699 | 0.38298639 | 0.90868001 |
| SENP6 | 118.178278 | -0.2217158 | 0.26474557 | -0.8374674 | 0.4023299 | 0.91528253 |
| MYO6 | 21.6115182 | -0.3257597 | 0.46078755 | -0.7069628 | 0.47958957 | 0.93257966 |
| PHIP | 144.233459 | 0.30944814 | 0.28274518 | 1.09444179 | 0.27376126 | 0.84994492 |
| HMGN3 | 29.4037872 | 0.39409724 | 0.43717327 | 0.90146692 | 0.36734012 | 0.89868152 |
| BCKDHB | 27.2579302 | 0.45710958 | 0.46967212 | 0.97325251 | 0.33042782 | 0.88037558 |
| TENT5A | 28.5660385 | -0.7532595 | 0.78400803 | -0.9607803 | 0.33666263 | 0.88143379 |
| IBTK | 47.0249074 | 0.02209605 | 0.32836024 | 0.06729211 | 0.94634916 | 0.99514836 |
| UBE3D | 19.9787546 | -0.4452976 | 0.44795486 | -0.9940681 | 0.32018973 | 0.87363761 |
| DOPEY1 | 46.5752019 | -0.3773748 | 0.34015702 | -1.1094135 | 0.26725182 | 0.84994492 |
| PGM3 | 10.0773219 | 0.01781697 | 0.75425334 | 0.023622 | 0.98115412 | NA |
| ME1 | 37.2145287 | 0.23943658 | 0.76000583 | 0.31504572 | 0.75272694 | 0.97875726 |
| CYB5R4 | 54.3982789 | 0.21625415 | 0.38975744 | 0.5548429 | 0.57900212 | 0.95415437 |
| CEP162 | 17.3351438 | 0.58416045 | 0.46216944 | 1.263953 | 0.2062469 | 0.79151895 |
| SNX14 | 47.4250846 | 0.05041327 | 0.33524906 | 0.15037557 | 0.88046831 | 0.99019008 |
| SYNCRIP | 17.878435 | -0.0579206 | 0.54261234 | -0.1067439 | 0.91499215 | 0.99498068 |
| ZNF292 | 133.471666 | 0.40657013 | 0.3955294 | 1.02791381 | 0.30399037 | 0.86537178 |
| SMIM8 | 10.747061 | -0.370492 | 0.6129435 | -0.6044472 | 0.54554639 | NA |
| RARS2 | 38.3062242 | 0.29525767 | 0.43045842 | 0.68591449 | 0.49276703 | 0.93678212 |
| ORC3 | 20.4827317 | 0.38857276 | 0.44607216 | 0.87109844 | 0.38370041 | 0.90910959 |
| AKIRIN2 | 26.7832739 | -0.1246796 | 0.3914017 | -0.3185463 | 0.75007055 | 0.97818861 |
| RNGTT | 75.7801108 | -0.5119059 | 0.31496896 | -1.6252583 | 0.10410752 | 0.6746265 |
| PNRC1 | 39.9688555 | 0.40855681 | 0.33293761 | 1.22712724 | 0.21977477 | 0.81042354 |
| UBE2J1 | 25.7751662 | 0.33902877 | 0.52145124 | 0.65016389 | 0.51558637 | 0.94596835 |
| ANKRD6 | 15.5554637 | 0.62658061 | 0.49829969 | 1.25743728 | 0.20859534 | 0.79538492 |
| LYRM2 | 10.4519041 | 0.03058816 | 0.61277874 | 0.04991714 | 0.96018842 | NA |
| MDN1 | 28.34198 | -0.4233571 | 0.34434807 | -1.2294453 | 0.2189049 | 0.80977253 |
| CASP8AP2 | 21.3963986 | -0.5319676 | 0.38753102 | -1.3727097 | 0.16984261 | 0.75745073 |
| BACH2 | 173.540538 | -2.6237081 | 0.74858266 | -3.504901 | 0.00045678 | 0.04077104 |
| MAP3K7 | 34.128355 | -0.0162663 | 0.38973627 | -0.0417367 | 0.96670857 | 0.99653147 |
| UFL1 | 24.8566682 | 0.03217023 | 0.52643944 | 0.06110908 | 0.95127233 | 0.99514836 |
| MMS22L | 23.9914699 | -0.431718 | 0.51697295 | -0.8350882 | 0.40366805 | 0.91528253 |
| FBXL4 | 31.5092488 | 0.32738974 | 0.44472095 | 0.73616891 | 0.46162791 | 0.9314919 |
| PNISR | 102.091478 | -0.2670002 | 0.38405886 | -0.6952063 | 0.486926 | 0.93497601 |
| USP45 | 15.2787601 | -0.6196721 | 0.49751952 | -1.2455232 | 0.21293951 | 0.80545265 |
| CCNC | 21.5512718 | 0.04935677 | 0.58640995 | 0.0841677 | 0.9329231 | 0.99514836 |
| ASCC3 | 97.4502034 | 0.12199956 | 0.29139285 | 0.41867728 | 0.675452 | 0.96612856 |

| | | | | | | |
|---|---|---|---|---|---|---|
| HACE1 | 21.264732 | 1.05539039 | 0.58375165 | 1.80794416 | 0.07061519 | 0.60640882 |
| PREP | 31.0862316 | 0.4774021 | 0.38676487 | 1.2343471 | 0.21707359 | 0.80640512 |
| ATG5 | 61.5641523 | 0.00322056 | 0.34189189 | 0.00941982 | 0.99248418 | 0.99859588 |
| PRDM1 | 31.7360859 | 1.15552614 | 0.67725872 | 1.70618128 | 0.08797433 | 0.64676224 |
| CRYBG1 | 61.5443679 | 0.7652774 | 0.51827651 | 1.47658129 | 0.13978791 | 0.73044824 |
| QRSL1 | 10.0695277 | -0.7136366 | 0.5648135 | -1.2634907 | 0.2064129 | NA |
| CD24 | 31.2074438 | -0.1893 | 0.39624461 | -0.4777351 | 0.63283876 | 0.95877428 |
| PDSS2 | 76.739964 | -0.1227533 | 0.27284437 | -0.4499021 | 0.652781 | 0.96178254 |
| SCML4 | 37.0388675 | 0.8836309 | 0.60385079 | 1.46332656 | 0.14337804 | 0.73044824 |
| SEC63 | 52.1770991 | -0.2752867 | 0.29340043 | -0.9382628 | 0.34810935 | 0.88939849 |
| OSTM1 | 20.18335 | -0.1163807 | 0.47601553 | -0.2444894 | 0.80685183 | 0.98320463 |
| SNX3 | 43.2505473 | -0.5494931 | 0.414164 | -1.3267524 | 0.18459061 | 0.77694741 |
| AFG1L | 15.3702207 | 0.22198436 | 0.50770505 | 0.43723095 | 0.66194386 | 0.96298563 |
| FOXO3 | 127.333784 | 0.19212975 | 0.5311204 | 0.36174425 | 0.71754315 | 0.97348719 |
| ARMC2 | 17.8471475 | -0.4412615 | 0.63798194 | -0.6916521 | 0.48915585 | 0.93633431 |
| SESN1 | 62.0757455 | 0.32875733 | 0.42249819 | 0.77812719 | 0.43649403 | 0.92489992 |
| CD164 | 49.4389027 | 0.29160075 | 0.31217675 | 0.93408861 | 0.35025819 | 0.89048867 |
| MICAL1 | 14.4294749 | -1.0957065 | 0.55017157 | -1.9915724 | 0.046418 | 0.52641679 |
| AK9 | 13.8374609 | -0.958223 | 0.6681582 | -1.434126 | 0.15153631 | 0.74212563 |
| FIG4 | 32.0265029 | -0.1824655 | 0.35525499 | -0.5136184 | 0.60751878 | 0.95688177 |
| CDC40 | 21.3423606 | 0.18527446 | 0.46780727 | 0.39604869 | 0.69206911 | 0.9685713 |
| CDK19 | 82.0119567 | 0.76679624 | 0.39560072 | 1.9383085 | 0.0525856 | 0.54907299 |
| AMD1 | 27.7393822 | 0.64584364 | 0.48255668 | 1.33837881 | 0.18077298 | 0.76894519 |
| GTF3C6 | 11.5606112 | -0.505907 | 0.60449425 | -0.8369095 | 0.40264344 | NA |
| SLC16A10 | 152.796587 | -0.0611891 | 1.04386898 | -0.0586176 | 0.95325668 | 0.99514836 |
| REV3L | 91.0439167 | 0.21206977 | 0.39513 | 0.53670885 | 0.59146874 | 0.95415437 |
| TRAF3IP2-AS | 41.9148729 | -0.546579 | 0.36500316 | -1.4974637 | 0.13427265 | 0.73044824 |
| FYN | 317.404468 | 0.73456325 | 0.5317636 | 1.38137183 | 0.16716466 | 0.7567539 |
| MARCKS | 38.9891842 | 1.18293738 | 0.61481564 | 1.92405221 | 0.05434804 | 0.55278922 |
| HDAC2 | 17.983914 | 0.23278448 | 0.43510655 | 0.53500568 | 0.59264593 | 0.95423409 |
| NT5DC1 | 48.7824889 | -0.1954276 | 0.38629219 | -0.5059061 | 0.61292256 | 0.95688177 |
| DSE | 19.6693959 | -0.0736344 | 0.62306333 | -0.1181812 | 0.90592409 | 0.99368779 |
| TSPYL1 | 10.2562087 | 0.63178384 | 0.58608167 | 1.07797918 | 0.28104305 | NA |
| RWDD1 | 8.82463584 | 0.62566277 | 0.65827219 | 0.9504621 | 0.3418775 | NA |
| ZUP1 | 11.1787499 | -0.5194628 | 0.52782234 | -0.9841621 | 0.32503581 | NA |
| KPNA5 | 30.8452414 | -0.1186917 | 0.45370222 | -0.2616069 | 0.79362449 | 0.98320463 |
| GOPC | 19.7148506 | 0.07734823 | 0.40484842 | 0.1910548 | 0.84848266 | 0.98545782 |
| NUS1 | 20.2254045 | -0.1610902 | 0.41018136 | -0.3927292 | 0.69451949 | 0.9685713 |
| CEP85L | 117.84348 | 0.29050724 | 0.26748087 | 1.08608606 | 0.27744092 | 0.84994492 |
| MCM9 | 30.8973975 | 0.3708892 | 0.33730338 | 1.09957154 | 0.27151885 | 0.84994492 |
| MAN1A1 | 156.483051 | -0.1689916 | 0.464866 | -0.3635276 | 0.71621079 | 0.97348719 |
| TBC1D32 | 22.6283023 | -0.33533 | 0.52741202 | -0.6358027 | 0.52490505 | 0.95007724 |
| SERINC1 | 38.997569 | -0.2956737 | 0.35129987 | -0.8416562 | 0.39998041 | 0.91508218 |
| RNF217 | 18.9510752 | 1.06867525 | 0.51737185 | 2.06558445 | 0.03886773 | 0.48119472 |
| NCOA7 | 51.4944326 | 0.28284941 | 0.41322773 | 0.68448797 | 0.49366708 | 0.93720536 |
| TRMT11 | 13.1045175 | -1.1839579 | 0.63888257 | -1.8531699 | 0.06385801 | 0.58463007 |
| RNF146 | 14.8929803 | 0.13512285 | 0.49619871 | 0.272316 | 0.78537906 | 0.98264092 |
| ECHDC1 | 35.1780956 | 0.06782393 | 0.36347221 | 0.18660004 | 0.85197424 | 0.98751288 |
| THEMIS | 161.679702 | 0.28318292 | 0.51942645 | 0.54518386 | 0.58562707 | 0.95415437 |
| PTPRK | 13.9397183 | -0.768529 | 0.594678 | -1.2923448 | 0.19623775 | 0.78135737 |
| ARHGAP18 | 105.544359 | 0.02385975 | 0.39660398 | 0.06016013 | 0.9520281 | 0.99514836 |
| L3MBTL3 | 31.9551724 | -0.803257 | 0.45002149 | -1.7849304 | 0.07427261 | 0.61929783 |
| SAMD3 | 74.8149188 | 0.66681251 | 0.50340737 | 1.32459821 | 0.18530445 | 0.77694741 |
| EPB41L2 | 56.2732656 | -0.6691471 | 0.52923702 | -1.2643619 | 0.20610017 | 0.79150473 |

| | | | | | | |
|---|---|---|---|---|---|---|
| AKAP7 | 26.0155559 | 0.05447039 | 0.59605129 | 0.0913854 | 0.92718636 | 0.99514836 |
| MED23 | 26.6105653 | -0.4373954 | 0.45787377 | -0.955275 | 0.33943864 | 0.8844375 |
| STX7 | 101.403181 | -0.7072836 | 0.3755933 | -1.8831102 | 0.05968544 | 0.57281427 |
| SLC18B1 | 12.5098618 | 0.33310049 | 0.6218499 | 0.5356606 | 0.59219314 | NA |
| RPS12 | 66.6271368 | -0.0565715 | 0.5711429 | -0.0990496 | 0.92109891 | 0.99514836 |
| TBPL1 | 12.7329345 | -0.1103231 | 0.55831309 | -0.1976007 | 0.84335746 | NA |
| SGK1 | 42.8804506 | -2.8989055 | 0.66781936 | -4.3408528 | 1.42E-05 | 0.00462026 |
| HBS1L | 28.1856069 | -0.0198088 | 0.36341697 | -0.0545071 | 0.95653111 | 0.99514836 |
| AHI1 | 40.3934999 | -0.7439516 | 0.35246825 | -2.1106912 | 0.03479886 | 0.45961076 |
| PDE7B | 26.5804866 | 0.08701471 | 0.77179479 | 0.11274332 | 0.91023405 | 0.99471296 |
| BCLAF1 | 66.887017 | -0.4098536 | 0.2771197 | -1.4789768 | 0.13914652 | 0.73044824 |
| MAP7 | 12.7429578 | 1.12555168 | 0.87530761 | 1.28589271 | 0.19848051 | NA |
| MAP3K5 | 158.757317 | 0.43491588 | 0.40545184 | 1.07266965 | 0.28341936 | 0.85176121 |
| IFNGR1 | 67.2867174 | 0.40965363 | 0.42424992 | 0.96559506 | 0.33424685 | 0.88047896 |
| AL357060.1 | 18.5458353 | 0.83480152 | 0.72577325 | 1.1502236 | 0.25005179 | 0.83865855 |
| TNFAIP3 | 43.2769913 | 0.95054507 | 0.64769335 | 1.46758504 | 0.14221697 | 0.73044824 |
| HEBP2 | 11.7031681 | -0.3638423 | 0.50767631 | -0.7166816 | 0.47357058 | NA |
| NHSL1 | 54.3326888 | 1.33136704 | 0.73585621 | 1.8092761 | 0.07040812 | 0.60640882 |
| REPS1 | 50.2966575 | 0.44137261 | 0.43277029 | 1.01987734 | 0.30778664 | 0.8673137 |
| ABRACL | 15.476891 | 0.49346348 | 0.57410676 | 0.8595326 | 0.39004674 | 0.90910959 |
| HECA | 37.9505633 | -0.1179132 | 0.36868047 | -0.319825 | 0.74910103 | 0.97818861 |
| AL031772.1 | 12.6735398 | 0.90556484 | 1.03454456 | 0.87532705 | 0.38139598 | NA |
| VTA1 | 24.6159534 | 0.6473773 | 0.3974431 | 1.62885531 | 0.10334365 | 0.67250824 |
| HIVEP2 | 103.645534 | -0.118968 | 0.40308035 | -0.2951472 | 0.76788143 | 0.98264092 |
| AIG1 | 25.4808491 | -0.0236096 | 0.53021775 | -0.0445282 | 0.96448337 | 0.99653147 |
| PEX3 | 12.4768302 | 0.35401766 | 0.50505734 | 0.70094548 | 0.48333704 | NA |
| PHACTR2 | 116.695797 | -0.0025076 | 0.38053816 | -0.0065896 | 0.99474232 | 0.99859588 |
| PLAGL1 | 30.2484518 | 0.49331593 | 0.62104961 | 0.79432613 | 0.4270056 | 0.92126666 |
| SF3B5 | 15.5457554 | 0.70170991 | 0.63296977 | 1.1085994 | 0.26760304 | 0.84994492 |
| STX11 | 22.521771 | 0.76209129 | 0.70811407 | 1.07622673 | 0.28182587 | 0.85171619 |
| UTRN | 550.428259 | -0.5293587 | 0.39957618 | -1.3248005 | 0.18523733 | 0.77694741 |
| SHPRH | 50.2722312 | -0.4466745 | 0.35276432 | -1.2662123 | 0.20543709 | 0.78970624 |
| STXBP5 | 75.7486006 | 0.39853544 | 0.44084466 | 0.90402691 | 0.36598113 | 0.89868152 |
| SASH1 | 67.4809935 | 0.24886809 | 0.64967621 | 0.3830648 | 0.70167172 | 0.97078484 |
| UST | 61.4658283 | -1.287623 | 0.41731383 | -3.0855027 | 0.00203208 | 0.10223227 |
| TAB2 | 99.5315679 | -0.1646808 | 0.28476511 | -0.5783041 | 0.5630588 | 0.95415437 |
| ZC3H12D | 13.0233029 | -2.538718 | 0.78644522 | -3.2280926 | 0.00124619 | 0.07415478 |
| PPIL4 | 20.7721774 | -0.3051274 | 0.46045991 | -0.6626579 | 0.50754965 | 0.94305491 |
| GINM1 | 13.6687906 | -0.3886087 | 0.45297967 | -0.8578943 | 0.39095084 | 0.90910959 |
| KATNA1 | 12.3440098 | -0.8611953 | 0.4635336 | -1.8578919 | 0.06318437 | NA |
| LATS1 | 31.3391957 | -0.3111771 | 0.38705333 | -0.8039644 | 0.42141754 | 0.91985469 |
| NUP43 | 12.9567655 | -0.4572776 | 0.46225068 | -0.9892416 | 0.32254494 | 0.87433122 |
| PCMT1 | 36.5378254 | -0.2165317 | 0.3206528 | -0.675284 | 0.49949532 | 0.93956733 |
| PLEKHG1 | 35.9424111 | -1.796335 | 0.48357672 | -3.7146846 | 0.00020346 | 0.02814832 |
| MTHFD1L | 38.9858573 | -0.0425692 | 0.65037216 | -0.0654536 | 0.94781282 | 0.99514836 |
| AKAP12 | 11.6703669 | 0.10536917 | 0.70168931 | 0.15016499 | 0.88063445 | NA |
| RMND1 | 13.317476 | -0.6824633 | 0.56124582 | -1.2159792 | 0.22399284 | 0.81875277 |
| ESR1 | 31.1089517 | -1.3545422 | 0.39641096 | -3.4170149 | 0.00063312 | 0.05005251 |
| SYNE1 | 130.216463 | 0.0090566 | 0.36314934 | 0.02493906 | 0.98010357 | 0.99758514 |
| OPRM1 | 10.9892542 | 0.24842379 | 0.69911034 | 0.35534276 | 0.72233283 | NA |
| IPCEF1 | 49.1181695 | 0.09692727 | 0.39872174 | 0.24309502 | 0.80793179 | 0.98320463 |
| SCAF8 | 91.5407352 | -0.0764012 | 0.28768556 | -0.2655717 | 0.79056905 | 0.98320463 |
| TIAM2 | 41.53556 | 0.12881786 | 0.40385774 | 0.31896842 | 0.74975046 | 0.97818861 |
| TFB1M | 22.0613219 | -0.0449827 | 0.40063232 | -0.1122793 | 0.91060197 | 0.99471296 |

| | | | | | | |
|---|---|---|---|---|---|---|
| ARID1B | 300.417401 | -0.0598316 | 0.2517771 | -0.237637 | 0.81216264 | 0.98320463 |
| ZDHHC14 | 46.063387 | 0.11494109 | 0.41265309 | 0.27854169 | 0.78059657 | 0.98264092 |
| SNX9 | 88.2524926 | -1.1415175 | 0.52743132 | -2.164296 | 0.03044164 | 0.43032651 |
| SYNJ2 | 16.968342 | -1.430368 | 0.71178148 | -2.0095605 | 0.04447772 | 0.5181889 |
| TULP4 | 67.7319532 | 0.20351953 | 0.30015002 | 0.67805934 | 0.49773407 | 0.93916044 |
| TMEM181 | 59.9854386 | 0.30221564 | 0.32321404 | 0.93503252 | 0.34977154 | 0.89048867 |
| DYNLT1 | 30.4310859 | 0.86732647 | 0.49614539 | 1.74812965 | 0.08044158 | 0.63385609 |
| SYTL3 | 148.875997 | 0.86585619 | 0.66958242 | 1.29312862 | 0.19596658 | 0.78135737 |
| EZR | 82.7816352 | -1.4748769 | 0.43418706 | -3.3968697 | 0.00068161 | 0.05238962 |
| AL035530.2 | 11.8248447 | -0.5487237 | 0.70906753 | -0.7738667 | 0.43900965 | NA |
| TAGAP | 17.1577001 | -0.1324698 | 0.53959419 | -0.245499 | 0.8060701 | 0.98320463 |
| WTAP | 48.754519 | -0.4851437 | 0.28580434 | -1.6974678 | 0.08960825 | 0.64907336 |
| TCP1 | 12.9482562 | -0.2550488 | 0.49106008 | -0.5193841 | 0.60349293 | NA |
| IGF2R | 104.694134 | 0.1942149 | 0.23795041 | 0.81619908 | 0.41438628 | 0.91894074 |
| MAP3K4 | 33.604401 | -0.3967649 | 0.37008121 | -1.0721023 | 0.28367409 | 0.85176121 |
| AGPAT4 | 40.7809614 | -1.0231754 | 0.6502072 | -1.5736144 | 0.11557663 | 0.70122655 |
| PRKN | 72.7912435 | -0.7767883 | 0.45256759 | -1.7164028 | 0.08608834 | 0.64676224 |
| QKI | 282.656087 | 0.5433273 | 0.44821654 | 1.21219824 | 0.22543651 | 0.81875277 |
| SFT2D1 | 22.3447302 | -0.4290006 | 0.38709252 | -1.1082638 | 0.26774789 | 0.84994492 |
| MPC1 | 21.1717573 | 0.32060449 | 0.42990253 | 0.74576087 | 0.45581188 | 0.9314919 |
| RPS6KA2 | 53.6176053 | -0.3024528 | 0.53027891 | -0.5703656 | 0.56842976 | 0.95415437 |
| AL159163.1 | 10.4242872 | 0.16717738 | 0.68783981 | 0.24304697 | 0.80796901 | NA |
| Z94721.1 | 17.9126541 | 0.22151373 | 0.53313761 | 0.41549071 | 0.67778271 | 0.96718429 |
| FGFR1OP | 18.6954311 | -0.4130835 | 0.49743308 | -0.8304303 | 0.40629554 | 0.9169561 |
| AFDN | 21.1280077 | 0.09188473 | 0.41717351 | 0.22025542 | 0.82567224 | 0.98320463 |
| WDR27 | 15.4230481 | -0.5835445 | 0.49142385 | -1.1874566 | 0.23504757 | 0.82903329 |
| PHF10 | 15.6852318 | -0.7064371 | 0.47058066 | -1.5012031 | 0.13330304 | 0.73044824 |
| FAM120B | 63.4791095 | -0.2614386 | 0.46722509 | -0.559556 | 0.57578235 | 0.95415437 |
| PSMB1 | 25.4624506 | -0.2331701 | 0.52095979 | -0.447578 | 0.6544578 | 0.96178254 |
| PDCD2 | 8.73151104 | -0.3910869 | 0.58827551 | -0.6648023 | 0.50617693 | NA |
| FAM20C | 16.7797279 | -0.8681492 | 0.65585021 | -1.3237005 | 0.18560254 | 0.77694741 |
| SUN1 | 20.4721164 | -1.770058 | 0.59827533 | -2.9586009 | 0.00309039 | 0.13681775 |
| COX19 | 14.3108656 | 0.0122175 | 0.50117357 | 0.02437778 | 0.98055128 | 0.99758514 |
| C7orf50 | 13.730529 | -0.1266469 | 0.47298659 | -0.26776 | 0.78888409 | 0.98320463 |
| INTS1 | 10.5198896 | -0.0807844 | 0.61695657 | -0.1309402 | 0.8958226 | NA |
| MAD1L1 | 49.3865923 | -0.4137894 | 0.38434655 | -1.0766048 | 0.28165684 | 0.85171619 |
| SNX8 | 36.6109995 | -2.1153434 | 0.64821534 | -3.2633344 | 0.0011011 | 0.07001544 |
| EIF3B | 21.9599128 | 0.49776409 | 0.49771662 | 1.00009538 | 0.31726435 | 0.87263466 |
| CHST12 | 16.6516004 | 0.72384801 | 0.50982902 | 1.41978582 | 0.15567004 | 0.74486619 |
| LFNG | 16.6003358 | -0.4248957 | 0.63651338 | -0.6675361 | 0.50442973 | 0.94166217 |
| TTYH3 | 34.3947665 | -0.0065159 | 0.42049664 | -0.0154956 | 0.98763679 | 0.99811561 |
| GNA12 | 45.1369463 | 0.50291538 | 0.65444098 | 0.7684656 | 0.44221062 | 0.92899684 |
| CARD11 | 100.534427 | -0.8200968 | 0.30210351 | -2.7146218 | 0.00663515 | 0.20175229 |
| SDK1 | 37.4795697 | -0.0259318 | 0.62552945 | -0.0414558 | 0.96693251 | 0.99653147 |
| FOXK1 | 38.6541482 | -0.1894571 | 0.39556777 | -0.4789499 | 0.63197427 | 0.95877428 |
| AP5Z1 | 9.55238086 | -0.2101777 | 0.54403024 | -0.3863345 | 0.69924892 | NA |
| RBAK-RBAKI | 18.7250909 | -0.6230545 | 0.45103263 | -1.3813956 | 0.16715734 | 0.7567539 |
| WIPI2 | 26.3929304 | 0.38674236 | 0.40531887 | 0.95416816 | 0.33999853 | 0.8844375 |
| TNRC18 | 66.6666362 | 0.53146196 | 0.41038239 | 1.29504085 | 0.19530614 | 0.78135737 |
| FBXL18 | 7.04032171 | 0.3088937 | 0.69265097 | 0.44595866 | 0.65562711 | NA |
| ACTB | 468.761614 | -0.0159242 | 0.73168502 | -0.0217637 | 0.98263646 | 0.99759864 |
| RNF216 | 64.2009039 | 0.15283059 | 0.28452793 | 0.5371374 | 0.5911727 | 0.95415437 |
| EIF2AK1 | 20.9445843 | 0.19789078 | 0.41784486 | 0.4735987 | 0.6357861 | 0.95933464 |
| USP42 | 19.6105695 | -0.1437182 | 0.41881676 | -0.3431529 | 0.73148342 | 0.97605445 |

| | | | | | | |
|---|---|---|---|---|---|---|
| CYTH3 | 21.2420618 | 1.22275152 | 0.46467421 | 2.63141681 | 0.00850297 | 0.24255374 |
| RAC1 | 62.9226408 | -0.2510991 | 0.37497783 | -0.6696372 | 0.50308912 | 0.94152694 |
| DAGLB | 16.1258793 | -0.8758801 | 0.43689362 | -2.0047902 | 0.04498548 | 0.52128421 |
| ZNF12 | 10.266526 | -0.2194663 | 0.53543476 | -0.4098844 | 0.68189078 | NA |
| CCZ1B | 12.5864569 | 0.10561143 | 0.57461662 | 0.18379459 | 0.8541746 | NA |
| C1GALT1 | 46.926596 | -0.504524 | 0.38108029 | -1.323931 | 0.18552596 | 0.77694741 |
| MIOS | 24.684081 | -0.8878246 | 0.39349975 | -2.2562266 | 0.02405644 | 0.39492781 |
| RPA3 | 9.9487264 | 0.62868822 | 0.65708568 | 0.95678271 | 0.33867694 | NA |
| UMAD1 | 75.4648583 | 0.086018 | 0.29296087 | 0.29361601 | 0.76905134 | 0.98264092 |
| GLCCI1 | 103.674995 | 0.10986909 | 0.51572003 | 0.21304017 | 0.83129563 | 0.98320463 |
| ICA1 | 13.5141585 | 0.78408992 | 0.55357893 | 1.4164013 | 0.15665804 | 0.74486619 |
| NDUFA4 | 27.5014176 | 0.03040771 | 0.47839419 | 0.06356203 | 0.94931897 | 0.99514836 |
| PHF14 | 71.5841917 | -0.3116762 | 0.36020996 | -0.8652626 | 0.38689468 | 0.90910959 |
| THSD7A | 14.6640495 | -1.2843858 | 0.70057941 | -1.8333193 | 0.06675509 | 0.58951068 |
| TMEM106B | 17.3224727 | -0.470616 | 0.4494161 | -1.0471721 | 0.29502021 | 0.8608685 |
| ANKMY2 | 10.8747675 | -0.1257379 | 0.57204067 | -0.2198058 | 0.82602238 | NA |
| BZW2 | 11.0276007 | -0.7089722 | 0.66249594 | -1.0701533 | 0.2845503 | NA |
| SNX13 | 83.2189445 | -0.2658699 | 0.34995225 | -0.759732 | 0.44741483 | 0.92914597 |
| HDAC9 | 206.97598 | -2.5554857 | 0.5501879 | -4.6447508 | 3.40E-06 | 0.00171296 |
| MACC1 | 13.9458433 | 0.00252728 | 0.54344298 | 0.0046505 | 0.99628945 | 0.99859588 |
| ITGB8 | 13.1237508 | 1.72954188 | 0.85491128 | 2.02306592 | 0.04306635 | 0.50816459 |
| SP4 | 27.6572779 | 0.10458519 | 0.45768685 | 0.22850819 | 0.81925119 | 0.98320463 |
| STEAP1B | 49.7288494 | -0.5181484 | 0.3952955 | -1.3107874 | 0.18992959 | 0.77993359 |
| TOMM7 | 24.2335045 | -0.2808728 | 0.54480821 | -0.5155444 | 0.60617263 | 0.95688177 |
| FAM126A | 54.3936081 | -1.8248207 | 0.69416553 | -2.6287977 | 0.00856873 | 0.24317625 |
| KLHL7 | 11.3576884 | 0.65495388 | 0.5845936 | 1.12035759 | 0.26256141 | NA |
| NUPL2 | 17.4256935 | -0.0188674 | 0.44301619 | -0.0425885 | 0.96602957 | 0.99653147 |
| TRA2A | 74.6660383 | 0.26196853 | 0.30887578 | 0.84813554 | 0.39636249 | 0.91333232 |
| CCDC126 | 13.89594 | -0.2737638 | 0.59010936 | -0.4639205 | 0.64270472 | 0.95992767 |
| MPP6 | 24.3956436 | -1.9820265 | 0.63932055 | -3.1002077 | 0.00193385 | 0.09883988 |
| OSBPL3 | 83.4405151 | 0.64987521 | 0.44176691 | 1.47108168 | 0.14126903 | 0.73044824 |
| CYCS | 15.1147806 | -0.6202954 | 0.54890045 | -1.1300691 | 0.2584471 | 0.84404826 |
| HNRNPA2B1 | 155.534989 | -0.1478741 | 0.22473851 | -0.657983 | 0.51054903 | 0.94368014 |
| CBX3 | 27.2433447 | 0.05990075 | 0.46100949 | 0.12993387 | 0.89661874 | 0.99118739 |
| SNX10 | 43.8924036 | 0.7296936 | 0.43415864 | 1.6807073 | 0.09281978 | 0.6590317 |
| SKAP2 | 111.8013 | 0.00992589 | 0.36543246 | 0.02716202 | 0.9783305 | 0.99758514 |
| HOTAIRM1 | 8.37172766 | -0.4482421 | 0.72792068 | -0.6157843 | 0.5380369 | NA |
| HIBADH | 31.7140122 | 0.70373857 | 0.55566658 | 1.26647632 | 0.2053426 | 0.78970624 |
| TAX1BP1 | 92.2074507 | 0.39628722 | 0.25781396 | 1.53710535 | 0.12426751 | 0.71741321 |
| JAZF1 | 187.068294 | -0.7883637 | 0.45187481 | -1.7446506 | 0.0810457 | 0.63497512 |
| CREB5 | 41.2028354 | -0.2614178 | 0.59944132 | -0.4361023 | 0.66276249 | 0.96360881 |
| CPVL | 82.5661244 | 0.69671723 | 0.68280813 | 1.02037042 | 0.30755282 | 0.8673137 |
| SCRN1 | 10.5859471 | -0.4441908 | 0.58548424 | -0.7586726 | 0.44804844 | NA |
| PLEKHA8 | 12.5887614 | -0.3849814 | 0.49269706 | -0.7813754 | 0.43458175 | NA |
| MTURN | 10.1545712 | 0.4526248 | 0.65678515 | 0.68915199 | 0.49072763 | NA |
| ZNRF2 | 45.6502595 | 0.43788064 | 0.34293864 | 1.27684838 | 0.20165578 | 0.78572563 |
| AC006978.2 | 38.412694 | -0.0675277 | 0.46097056 | -0.1464902 | 0.88353444 | 0.99019008 |
| NOD1 | 13.8562107 | 1.38712445 | 0.5654405 | 2.4531749 | 0.01416015 | 0.3205889 |
| AC005154.6 | 17.2199544 | -0.0133309 | 0.52498801 | -0.0253928 | 0.97974165 | 0.99758514 |
| GARS | 13.3412568 | -0.1702717 | 0.4930032 | -0.3453764 | 0.72981142 | 0.97605445 |
| AQP1 | 10.2007199 | 1.82137965 | 0.93768233 | 1.94242719 | 0.05208541 | NA |
| AVL9 | 72.7698172 | -0.0267314 | 0.27543538 | -0.0970514 | 0.92268559 | 0.99514836 |
| KBTBD2 | 23.4049688 | 0.76298219 | 0.4717496 | 1.61734572 | 0.10580368 | 0.67519097 |
| NT5C3A | 43.419011 | 0.35168641 | 0.44779454 | 0.78537448 | 0.43223404 | 0.92284736 |

| | | | | | | |
|---|---|---|---|---|---|---|
| BBS9 | 44.8895496 | 0.70379179 | 0.48732829 | 1.44418417 | 0.14868718 | 0.7383468 |
| DPY19L1 | 18.4102106 | 0.33346905 | 0.57961402 | 0.5753295 | 0.56506847 | 0.95415437 |
| HERPUD2 | 38.6563104 | 0.22427668 | 0.33688338 | 0.66573981 | 0.50557742 | 0.94166217 |
| 7-Sep | 103.645611 | -0.2703361 | 0.2567075 | -1.0530901 | 0.29229971 | 0.85950405 |
| EEPD1 | 28.9668866 | -0.0035457 | 0.68750739 | -0.0051574 | 0.99588503 | 0.99859588 |
| AOAH | 390.017759 | 1.064928 | 0.67538546 | 1.5767707 | 0.1148483 | 0.70122655 |
| ELMO1 | 325.231941 | 0.10376001 | 0.26392791 | 0.39313768 | 0.69421779 | 0.9685713 |
| ELMO1-AS1 | 11.1314287 | -0.0173512 | 0.59869164 | -0.0289818 | 0.9768791 | NA |
| STARD3NL | 15.587312 | 0.9936 | 0.50553718 | 1.96543408 | 0.04936402 | 0.53564799 |
| TRGC2 | 22.2830989 | -0.169218 | 0.55450587 | -0.305169 | 0.76023745 | 0.9808103 |
| TRGC1 | 8.74110884 | 2.08888266 | 1.0396861 | 2.00914745 | 0.0445215 | NA |
| TRG-AS1 | 50.7712876 | 1.00895354 | 0.60725043 | 1.66151144 | 0.09661077 | 0.66178928 |
| VPS41 | 62.2885213 | -0.4026081 | 0.28806148 | -1.3976465 | 0.16221923 | 0.75177046 |
| RALA | 40.4925217 | 0.58679408 | 0.36993208 | 1.58622112 | 0.11268916 | 0.69678413 |
| CDK13 | 125.093416 | -0.1692355 | 0.24654796 | -0.6864204 | 0.49244807 | 0.93678212 |
| STK17A | 66.5703035 | 0.23476817 | 0.4107675 | 0.57153541 | 0.56763676 | 0.95415437 |
| COA1 | 61.2244715 | -0.1508328 | 0.3140178 | -0.4803319 | 0.63099141 | 0.95877428 |
| BLVRA | 15.2893775 | -0.2898067 | 0.52738399 | -0.5495174 | 0.58265044 | 0.95415437 |
| DBNL | 23.2279221 | -0.0829328 | 0.49570991 | -0.1673011 | 0.86713315 | 0.98751288 |
| NUDCD3 | 37.0053743 | -0.3133433 | 0.31324756 | -1.0003057 | 0.31716259 | 0.87263466 |
| TMED4 | 11.4541659 | -0.157843 | 0.50893967 | -0.3101408 | 0.75645389 | NA |
| OGDH | 70.5562408 | -0.7130549 | 0.43584005 | -1.6360472 | 0.10182971 | 0.67049522 |
| ZMIZ2 | 18.5995743 | 0.41950156 | 0.52973084 | 0.79191454 | 0.42841051 | 0.92178217 |
| PPIA | 64.4570445 | -0.4015233 | 0.50332607 | -0.79774 | 0.4250214 | 0.91985469 |
| H2AFV | 31.6282265 | -0.2466826 | 0.40585739 | -0.6078062 | 0.54331605 | 0.95415437 |
| PURB | 13.563221 | 0.90206011 | 0.48003001 | 1.87917442 | 0.06022068 | 0.57281427 |
| MYO1G | 31.8751891 | -0.0347159 | 0.61283928 | -0.0566476 | 0.9548259 | 0.99514836 |
| CCM2 | 87.4695615 | -0.3327176 | 0.34555252 | -0.9628568 | 0.3356194 | 0.88047896 |
| TNS3 | 99.8683414 | -0.3213002 | 0.47173912 | -0.6810971 | 0.49581003 | 0.93805563 |
| HUS1 | 16.64676 | -0.1504686 | 0.50778564 | -0.296323 | 0.76698344 | 0.98264092 |
| UPP1 | 25.4974702 | -0.8873393 | 0.64603926 | -1.3735068 | 0.16959486 | 0.75745073 |
| IKZF1 | 254.883712 | 0.52728606 | 0.3551612 | 1.48463868 | 0.13763956 | 0.73044824 |
| GRB10 | 19.2896115 | 2.04374673 | 0.87790034 | 2.32799401 | 0.01991242 | 0.36765113 |
| VOPP1 | 128.697309 | -1.0257911 | 0.37412776 | -2.7418203 | 0.00610998 | 0.1932149 |
| NIPSNAP2 | 21.5732838 | 0.07072538 | 0.45464043 | 0.15556334 | 0.87637723 | 0.98939075 |
| CCT6A | 14.0676028 | -0.210581 | 0.47087278 | -0.4472141 | 0.65472046 | 0.96178254 |
| SUMF2 | 16.725033 | -0.8165404 | 0.42358433 | -1.9276926 | 0.05389337 | 0.55230721 |
| CHCHD2 | 37.2500718 | -0.4250826 | 0.53261651 | -0.7981026 | 0.42481094 | 0.91985469 |
| ZNF736 | 18.3155209 | -0.3248754 | 0.5173884 | -0.6279141 | 0.53006025 | 0.95245713 |
| ZNF680 | 16.2373929 | 0.30662126 | 0.56000591 | 0.54753219 | 0.58401316 | 0.95415437 |
| ZNF107 | 23.9369803 | -0.8864303 | 0.60637874 | -1.4618427 | 0.14378433 | 0.73044824 |
| ZNF138 | 11.2574203 | 1.02513495 | 0.57947804 | 1.76906608 | 0.07688285 | NA |
| ZNF117 | 9.08003272 | -0.1034546 | 0.71077355 | -0.1455521 | 0.88427502 | NA |
| VKORC1L1 | 27.5791756 | -0.2952916 | 0.38073746 | -0.775578 | 0.43799821 | 0.92656044 |
| GUSB | 13.5228519 | 0.07378291 | 0.52141069 | 0.14150632 | 0.88746996 | 0.99019008 |
| TPST1 | 38.5390534 | 0.58273847 | 0.67812885 | 0.85933295 | 0.39015685 | 0.90910959 |
| AC027644.4 | 17.2291918 | 0.17595893 | 0.63322839 | 0.27787594 | 0.7811076 | 0.98264092 |
| TMEM248 | 27.1238973 | -0.3232013 | 0.36181505 | -0.8932776 | 0.37170853 | 0.90230329 |
| SBDS | 12.2194539 | -0.4932877 | 0.54572809 | -0.9039075 | 0.36604444 | NA |
| TYW1 | 44.7899173 | -0.239043 | 0.36483048 | -0.6552168 | 0.51232821 | 0.94507477 |
| AUTS2 | 164.023202 | -0.3285125 | 0.41506676 | -0.791469 | 0.42867038 | 0.92198284 |
| TYW1B | 26.2861653 | 0.26343899 | 0.43955581 | 0.59933001 | 0.54895284 | 0.95415437 |
| POM121 | 32.1112361 | 0.12876174 | 0.39057465 | 0.32967255 | 0.7416474 | 0.97605445 |
| BAZ1B | 41.8143033 | 0.18406239 | 0.34402476 | 0.53502656 | 0.59263149 | 0.95423409 |

| | | | | | | |
|---|---|---|---|---|---|---|
| BUD23 | 10.5962812 | 0.46475463 | 0.59570192 | 0.78017984 | 0.43528503 | NA |
| LIMK1 | 12.0561862 | -0.716486 | 0.69393738 | -1.0324937 | 0.30184089 | NA |
| EIF4H | 22.4226656 | 0.35372429 | 0.43956002 | 0.80472352 | 0.42097924 | 0.91985469 |
| LAT2 | 27.8926134 | -0.3936498 | 0.56155919 | -0.7009943 | 0.48330658 | 0.93399425 |
| CLIP2 | 18.0573283 | -0.1454037 | 0.71571458 | -0.2031587 | 0.83901095 | 0.98342625 |
| GTF2I | 96.5389669 | -0.2980719 | 0.29242737 | -1.0193025 | 0.30805938 | 0.8673137 |
| NCF1 | 22.3838485 | -1.0798111 | 0.61119526 | -1.7667203 | 0.07727508 | 0.62520507 |
| RCC1L | 9.47760866 | 0.08156654 | 0.60501111 | 0.13481825 | 0.89275558 | NA |
| GTF2IRD2B | 8.17786232 | -0.4196474 | 0.70786443 | -0.5928359 | 0.553291 | NA |
| POM121C | 29.0723974 | 0.21900543 | 0.38764142 | 0.56496913 | 0.57209476 | 0.95415437 |
| HIP1 | 67.8373982 | 0.95859085 | 0.47268047 | 2.02798909 | 0.04256136 | 0.50435669 |
| POR | 19.3792649 | 0.14112084 | 0.49963863 | 0.28244581 | 0.77760169 | 0.98264092 |
| STYXL1 | 14.387177 | 0.8853343 | 0.60576176 | 1.46152227 | 0.14387217 | 0.73044824 |
| MDH2 | 11.0536869 | -0.0097234 | 0.51858855 | -0.0187497 | 0.98504081 | NA |
| HSPB1 | 21.4510932 | -0.4180379 | 0.51839725 | -0.8064046 | 0.42000958 | 0.91985469 |
| YWHAG | 26.8258194 | -0.3621154 | 0.39495591 | -0.9168502 | 0.35922114 | 0.8957272 |
| DTX2 | 23.2044814 | -0.2126272 | 0.46361542 | -0.4586284 | 0.64650104 | 0.96049083 |
| CCDC146 | 29.7940289 | -0.5442543 | 0.49413415 | -1.1014303 | 0.27070943 | 0.84994492 |
| FGL2 | 46.5031029 | 0.81489507 | 0.89918072 | 0.90626394 | 0.36479617 | 0.89868152 |
| GSAP | 79.7942678 | 0.13862632 | 0.41156223 | 0.33682954 | 0.7362454 | 0.97605445 |
| PTPN12 | 101.622347 | 0.36283754 | 0.3595869 | 1.00903993 | 0.31295548 | 0.8706069 |
| RSBN1L | 56.0480463 | 0.30943171 | 0.29919095 | 1.03422816 | 0.30102951 | 0.86225566 |
| PHTF2 | 91.8920903 | -0.5530634 | 0.3887897 | -1.4225259 | 0.15487365 | 0.74486619 |
| MAGI2 | 15.2878277 | -0.8988485 | 0.56483979 | -1.5913336 | 0.11153452 | 0.69385269 |
| CD36 | 47.7170259 | 1.3189992 | 1.10595201 | 1.19263693 | 0.2330116 | 0.82817204 |
| DMTF1 | 54.1479049 | -0.4199594 | 0.31936934 | -1.3149646 | 0.18852181 | 0.77798634 |
| TMEM243 | 21.3820686 | -0.8112913 | 0.44146915 | -1.8377079 | 0.06610546 | 0.58951068 |
| ABCB1 | 14.8994895 | 0.84335132 | 0.63649493 | 1.32499299 | 0.18517348 | 0.77694741 |
| SLC25A40 | 18.4441669 | 0.75835892 | 0.45632988 | 1.66186557 | 0.09653973 | 0.66178928 |
| DBF4 | 12.8335899 | -0.7226382 | 0.65149589 | -1.1091984 | 0.26734461 | NA |
| SRI | 11.9303938 | -0.4705055 | 0.52330818 | -0.8990984 | 0.36860025 | NA |
| GTPBP10 | 12.494159 | -0.1628971 | 0.57869737 | -0.2814893 | 0.77833515 | NA |
| CDK14 | 175.087786 | -2.6543765 | 0.70661287 | -3.7564791 | 0.00017232 | 0.02594752 |
| MTERF1 | 18.6970339 | -0.4338417 | 0.55477476 | -0.7820142 | 0.43420622 | 0.9231261 |
| AKAP9 | 107.173479 | 0.01456816 | 0.37225082 | 0.03913533 | 0.9687825 | 0.99653147 |
| KRIT1 | 36.778843 | -0.3014321 | 0.37452918 | -0.8048293 | 0.42091818 | 0.91985469 |
| ANKIB1 | 75.9026118 | 0.18792732 | 0.31189508 | 0.60253375 | 0.5468189 | 0.95415437 |
| GATAD1 | 11.1682456 | 0.49947633 | 0.63112648 | 0.79140449 | 0.42870799 | NA |
| PEX1 | 9.56433456 | -0.0248564 | 0.57124437 | -0.0435127 | 0.96529287 | NA |
| FAM133B | 25.4138185 | -0.4521679 | 0.40147377 | -1.1262702 | 0.26005117 | 0.8455672 |
| CDK6 | 84.5828291 | 1.32511036 | 0.45952861 | 2.88362975 | 0.00393121 | 0.15732988 |
| SAMD9 | 24.001693 | 0.65938641 | 0.79679142 | 0.82755209 | 0.40792421 | 0.9169561 |
| SAMD9L | 44.2568486 | 0.55152167 | 0.67699357 | 0.81466308 | 0.41526519 | 0.91894074 |
| VPS50 | 43.8577956 | -0.3563265 | 0.3272831 | -1.0887408 | 0.2762682 | 0.84994492 |
| BET1 | 8.89820149 | 0.33399114 | 0.71208117 | 0.46903521 | 0.63904447 | NA |
| CASD1 | 30.1108147 | 0.20159367 | 0.46370147 | 0.43474882 | 0.66374476 | 0.96379377 |
| PPP1R9A | 10.5517141 | -1.6152079 | 0.5927797 | -2.724803 | 0.00643399 | NA |
| PON2 | 10.0681934 | 0.3471011 | 0.57599774 | 0.60260845 | 0.5467692 | NA |
| PDK4 | 65.618899 | 1.46297684 | 1.14693271 | 1.27555595 | 0.20211254 | 0.78656172 |
| SLC25A13 | 63.0902397 | -0.4323569 | 0.36970615 | -1.1694608 | 0.24221801 | 0.83493679 |
| SEM1 | 24.8976679 | -0.1859378 | 0.39963746 | -0.4652662 | 0.64174082 | 0.95992767 |
| LMTK2 | 23.7181219 | -0.610529 | 0.38836999 | -1.5720292 | 0.11594377 | 0.70122655 |
| TECPR1 | 17.3391644 | 0.44489647 | 0.47256678 | 0.94144676 | 0.34647596 | 0.88790825 |
| BRI3 | 55.0629227 | 0.48806337 | 0.43087123 | 1.13273604 | 0.25732512 | 0.84162955 |

| | | | | | | |
|---|---|---|---|---|---|---|
| BAIAP2L1 | 90.0513472 | -0.0686901 | 0.69811962 | -0.0983931 | 0.92162016 | 0.99514836 |
| TRRAP | 48.3214633 | -0.0234031 | 0.33919197 | -0.0689968 | 0.9449922 | 0.99514836 |
| SMURF1 | 55.951556 | 0.10853291 | 0.29581686 | 0.36689225 | 0.71369939 | 0.97348719 |
| ARPC1B | 25.817225 | 0.26180341 | 0.68070991 | 0.38460349 | 0.70053121 | 0.96991236 |
| PDAP1 | 10.0116901 | -0.6232687 | 0.58502063 | -1.065379 | 0.28670447 | NA |
| ZNF789 | 13.8548601 | 0.23620716 | 0.57897966 | 0.40797143 | 0.68329465 | 0.9673431 |
| ZNF394 | 14.7450576 | -0.1508406 | 0.48903129 | -0.3084478 | 0.75774165 | 0.98030982 |
| TRIM4 | 16.4630761 | 0.11276835 | 0.49380421 | 0.22836653 | 0.81936131 | 0.98320463 |
| ZKSCAN1 | 33.401985 | 0.15885935 | 0.38487814 | 0.41275233 | 0.67978808 | 0.9673431 |
| COPS6 | 8.88668987 | -0.182702 | 0.78209565 | -0.2336057 | 0.81529108 | NA |
| LAMTOR4 | 30.4052967 | -0.0822052 | 0.46868168 | -0.1753966 | 0.86076798 | 0.98751288 |
| CASTOR3 | 19.9713684 | -0.8428896 | 0.43126608 | -1.9544537 | 0.05064759 | 0.5452991 |
| PILRB | 21.2511596 | -0.1572996 | 0.41834949 | -0.3760004 | 0.7069166 | 0.97184211 |
| PILRA | 39.3841503 | 1.33904063 | 0.63274567 | 2.11623832 | 0.03432454 | 0.45650836 |
| TSC22D4 | 19.1204581 | 0.19866327 | 0.44800821 | 0.44343667 | 0.65744992 | 0.96270373 |
| LRCH4 | 9.94320415 | -0.0470176 | 0.55363853 | -0.0849246 | 0.9323213 | NA |
| GNB2 | 24.1858141 | -0.1606375 | 0.62510419 | -0.2569771 | 0.79719646 | 0.98320463 |
| GIGYF1 | 12.8790112 | -0.5706914 | 0.46768289 | -1.2202529 | 0.22236902 | NA |
| SLC12A9 | 16.7461313 | -0.7964937 | 0.49383889 | -1.6128613 | 0.10677463 | 0.67742699 |
| SRRT | 9.53967272 | -1.4292795 | 0.59953592 | -2.3839765 | 0.0171267 | NA |
| TRIM56 | 42.5172885 | 0.13229048 | 0.38227754 | 0.34605873 | 0.72929858 | 0.97605445 |
| PLOD3 | 7.79025075 | 0.12095957 | 0.64234055 | 0.18831066 | 0.85063313 | NA |
| ZNHIT1 | 10.7334799 | 0.02482602 | 0.59770682 | 0.04153545 | 0.96686903 | NA |
| FIS1 | 17.0840023 | -0.5008566 | 0.43338234 | -1.1556921 | 0.24780711 | 0.83817091 |
| CUX1 | 199.712738 | -0.0139669 | 0.39302235 | -0.0355372 | 0.97165137 | 0.99724011 |
| SH2B2 | 11.3955427 | -1.3933075 | 0.73947946 | -1.8841734 | 0.05954153 | NA |
| PRKRIP1 | 41.035335 | -0.103753 | 0.35425703 | -0.2928749 | 0.76961774 | 0.98264092 |
| ORAI2 | 18.0800181 | -1.0156359 | 0.79239193 | -1.2817342 | 0.19993589 | 0.78249307 |
| AC105052.3 | 9.479229 | -0.0122339 | 0.85060344 | -0.0143826 | 0.98852477 | NA |
| FAM185A | 15.7392134 | 0.43034641 | 0.60898166 | 0.70666564 | 0.47977429 | 0.93257966 |
| ARMC10 | 10.0378295 | -0.6094979 | 0.56674569 | -1.0754346 | 0.28218021 | NA |
| NAPEPLD | 11.3517196 | -0.0296829 | 0.52546192 | -0.0564892 | 0.95495213 | NA |
| PMPCB | 11.5440275 | -0.4482146 | 0.53533302 | -0.8372632 | 0.40244466 | NA |
| DNAJC2 | 15.3535599 | -0.4888331 | 0.4534824 | -1.0779538 | 0.28105438 | 0.85131633 |
| PSMC2 | 10.9186003 | 0.23973464 | 0.55540376 | 0.43164029 | 0.66600287 | NA |
| ORC5 | 20.5304108 | -0.5217205 | 0.42060496 | -1.2404051 | 0.2148256 | 0.80634897 |
| AC007384.1 | 18.9161813 | -0.7478066 | 0.76245902 | -0.9807827 | 0.32669992 | 0.87715548 |
| LINC01004 | 38.8324997 | -0.3568645 | 0.50339206 | -0.7089196 | 0.47837437 | 0.9318939 |
| KMT2E | 185.601379 | -0.1371416 | 0.25549844 | -0.5367609 | 0.59143277 | 0.95415437 |
| SRPK2 | 183.60364 | -0.0620497 | 0.22211019 | -0.2793646 | 0.77996505 | 0.98264092 |
| RINT1 | 10.1829985 | -1.1093032 | 0.51195878 | -2.1667822 | 0.03025147 | NA |
| ATXN7L1 | 63.2247607 | 0.60444855 | 0.45403102 | 1.33129352 | 0.18309245 | 0.77523613 |
| SYPL1 | 16.6794908 | -0.9341492 | 0.5386994 | -1.7340826 | 0.08290343 | 0.63740875 |
| NAMPT | 282.047856 | 0.53823834 | 0.75036525 | 0.71730179 | 0.4731879 | 0.9314919 |
| PIK3CG | 21.6478616 | 1.02927244 | 0.50253186 | 2.04817349 | 0.040543 | 0.48881259 |
| HBP1 | 33.6615824 | 0.20102042 | 0.38556024 | 0.52137229 | 0.60210745 | 0.95688177 |
| COG5 | 145.037774 | 0.21550071 | 0.31781672 | 0.67806601 | 0.49772984 | 0.93916044 |
| BCAP29 | 16.6364856 | 0.06964629 | 0.46624513 | 0.14937699 | 0.88125617 | 0.99019008 |
| SLC26A3 | 104.871665 | -0.1359344 | 0.77846038 | -0.1746196 | 0.86137854 | 0.98751288 |
| DLD | 19.0898078 | 0.60760214 | 0.47302221 | 1.28451081 | 0.19896328 | 0.78142991 |
| PNPLA8 | 33.6005917 | 0.07355154 | 0.41454254 | 0.17742821 | 0.85917205 | 0.98751288 |
| IMMP2L | 178.40668 | -0.6694504 | 0.372888 | -1.7953121 | 0.07260398 | 0.61222939 |
| DOCK4 | 326.417723 | 0.90791014 | 0.64889273 | 1.3991683 | 0.16176252 | 0.7510015 |
| ZNF277 | 33.0534613 | 0.07156134 | 0.39073876 | 0.1831437 | 0.85468527 | 0.98751288 |

| | | | | | | |
|---|---|---|---|---|---|---|
| IFRD1 | 17.6935099 | -0.22615 | 0.44802026 | -0.5047763 | 0.61371599 | 0.95688177 |
| TMEM168 | 27.5729326 | -0.6122361 | 0.37183138 | -1.6465423 | 0.09965215 | 0.6683775 |
| BMT2 | 35.0927966 | 0.37467871 | 0.40402371 | 0.92736813 | 0.35373543 | 0.89109867 |
| MDFIC | 96.3503119 | -0.5471818 | 0.31339807 | -1.7459642 | 0.08081717 | 0.63497512 |
| TFEC | 69.4689336 | -1.7323154 | 0.5051162 | -3.4295384 | 0.00060461 | 0.04993888 |
| TES | 42.0291838 | -0.8961379 | 0.40673302 | -2.2032583 | 0.02757655 | 0.41134396 |
| CAPZA2 | 62.6094713 | -0.10873 | 0.36839968 | -0.2951413 | 0.76788591 | 0.98264092 |
| ST7 | 32.8365477 | -0.6210682 | 0.51441135 | -1.2073377 | 0.2273021 | 0.82088564 |
| CTTNBP2 | 13.4186515 | 1.40527642 | 0.7883054 | 1.78265482 | 0.07464251 | 0.61929783 |
| LSM8 | 20.0399995 | -0.5021102 | 0.40199274 | -1.249053 | 0.2116457 | 0.80277403 |
| CPED1 | 27.802782 | 0.07980423 | 0.6252715 | 0.12763133 | 0.89844074 | 0.99141995 |
| FAM3C | 53.5794336 | -2.7717732 | 0.7018335 | -3.9493316 | 7.84E-05 | 0.01536239 |
| CADPS2 | 16.5906902 | 0.2220322 | 0.70175335 | 0.31639635 | 0.75170169 | 0.97857378 |
| NDUFA5 | 12.4700024 | 1.03729189 | 0.51711853 | 2.00590742 | 0.04486613 | NA |
| WASL | 25.1123195 | -0.1395952 | 0.39100009 | -0.357021 | 0.72107611 | 0.97417403 |
| POT1 | 16.4887795 | -0.7275937 | 0.44540227 | -1.6335653 | 0.10235017 | 0.67109695 |
| ZNF800 | 45.8361078 | -0.0358231 | 0.4084559 | -0.0877038 | 0.93011213 | 0.99514836 |
| SND1 | 147.249327 | -0.1356239 | 0.25747967 | -0.5267362 | 0.59837679 | 0.95567595 |
| RBM28 | 12.0387622 | -0.459425 | 0.535773 | -0.8574994 | 0.39116897 | NA |
| IMPDH1 | 10.1335132 | 0.64183103 | 0.64477974 | 0.9954268 | 0.31952873 | NA |
| METTL2B | 9.39369729 | 0.00237138 | 0.63366609 | 0.00374232 | 0.99701407 | NA |
| CALU | 19.3154507 | -0.3576651 | 0.47139375 | -0.7587395 | 0.44800841 | 0.92914597 |
| ATP6V1F | 18.8861899 | -0.0800198 | 0.44811574 | -0.1785694 | 0.8582758 | 0.98751288 |
| TNPO3 | 52.1533239 | -0.036264 | 0.33025248 | -0.109807 | 0.91256244 | 0.99498068 |
| TSPAN33 | 10.8047395 | -1.8161632 | 0.770045 | -2.3585157 | 0.01834818 | NA |
| AHCYL2 | 67.9091901 | 0.73079258 | 0.31592033 | 2.31321796 | 0.02071066 | 0.37333155 |
| NRF1 | 54.8054197 | 0.05957286 | 0.29706312 | 0.20053939 | 0.84105875 | 0.98380297 |
| UBE2H | 91.7704253 | -0.4286445 | 0.25806595 | -1.6609884 | 0.09671578 | 0.66178928 |
| KLHDC10 | 35.8746985 | 0.47935616 | 0.37804136 | 1.26799926 | 0.20479822 | 0.78924326 |
| TMEM209 | 9.17253471 | -0.1290291 | 0.58034942 | -0.22233 | 0.82405702 | NA |
| COPG2 | 14.859375 | -0.3030379 | 0.51233549 | -0.5914833 | 0.55419665 | 0.95415437 |
| AC016831.7 | 378.079526 | -0.0422141 | 0.4242397 | -0.0995054 | 0.92073702 | 0.99514836 |
| LINC00513 | 32.5970449 | -0.5483414 | 0.50449883 | -1.0869032 | 0.27707958 | 0.84994492 |
| MKLN1 | 163.491379 | -0.1853474 | 0.24469423 | -0.7574655 | 0.44877104 | 0.92914597 |
| CHCHD3 | 125.308294 | -0.3564491 | 0.34077979 | -1.0459809 | 0.29556987 | 0.8608685 |
| EXOC4 | 227.045469 | 0.26254591 | 0.23270608 | 1.12822969 | 0.25922293 | 0.84484505 |
| SLC35B4 | 10.5714983 | -0.0078739 | 0.63625883 | -0.0123753 | 0.99012621 | NA |
| AKR1B1 | 11.0547723 | 0.07296551 | 0.55471688 | 0.13153649 | 0.89535093 | NA |
| CALD1 | 10.9628622 | 0.56338668 | 0.62714018 | 0.89834251 | 0.36900298 | NA |
| CYREN | 16.1647156 | -1.2880188 | 0.60144067 | -2.1415559 | 0.03222924 | 0.44147672 |
| TMEM140 | 15.5714397 | -0.4936712 | 0.50156006 | -0.9842714 | 0.32498209 | 0.87685391 |
| WDR91 | 11.4322788 | 0.32544816 | 0.58232628 | 0.55887596 | 0.57624638 | NA |
| CNOT4 | 83.6433718 | -0.1245265 | 0.26471482 | -0.4704174 | 0.6380568 | 0.95933464 |
| NUP205 | 22.5480414 | -0.4774063 | 0.44564878 | -1.0712614 | 0.28405193 | 0.85200183 |
| STMP1 | 17.2714212 | 0.09096152 | 0.56077807 | 0.16220591 | 0.8711437 | 0.98751288 |
| MTPN | 52.3710769 | -0.2550775 | 0.55928237 | -0.45608 | 0.64833245 | 0.96049083 |
| AC024084.1 | 12.3447615 | 0.33767911 | 0.66722785 | 0.50609265 | 0.61279161 | NA |
| AC078845.1 | 12.9757772 | -0.4387305 | 0.93003299 | -0.4717365 | 0.63711491 | 0.95933464 |
| CREB3L2 | 45.2681295 | 0.69012845 | 0.58587743 | 1.17793999 | 0.23882053 | 0.83267279 |
| TRIM24 | 27.8626866 | 0.42327017 | 0.5512899 | 0.76778148 | 0.44261702 | 0.92899684 |
| ZC3HAV1 | 114.587447 | 0.64299695 | 0.45870958 | 1.40175173 | 0.1609894 | 0.74887162 |
| UBN2 | 27.9906231 | 0.06065599 | 0.37119898 | 0.16340561 | 0.87019909 | 0.98751288 |
| HIPK2 | 110.337407 | 0.78687976 | 0.33539546 | 2.34612522 | 0.01896973 | 0.35951542 |
| TBXAS1 | 204.6286 | 1.66047979 | 0.51035041 | 3.25360725 | 0.0011395 | 0.07001544 |

| | | | | | | |
|---|---|---|---|---|---|---|
| PARP12 | 29.8113161 | 0.10584347 | 0.77121315 | 0.13724283 | 0.89083886 | 0.99080204 |
| KDM7A | 55.6191358 | -0.1232275 | 0.38338991 | -0.3214157 | 0.74789536 | 0.97818861 |
| SLC37A3 | 12.0664959 | -0.2312536 | 0.47703579 | -0.4847721 | 0.62783805 | NA |
| MKRN1 | 21.653761 | -0.1039573 | 0.46382467 | -0.2241306 | 0.82265572 | 0.98320463 |
| NDUFB2 | 19.5771151 | 0.92434928 | 0.50295489 | 1.83783735 | 0.06608638 | 0.58951068 |
| BRAF | 125.024952 | -0.2468518 | 0.24713198 | -0.9988661 | 0.31785957 | 0.87297016 |
| AGK | 19.0784428 | -0.1502584 | 0.49085606 | -0.3061151 | 0.75951702 | 0.9808103 |
| KIAA1147 | 20.3739457 | 0.0732087 | 0.43249374 | 0.16927113 | 0.86558339 | 0.98751288 |
| SSBP1 | 19.7116955 | -0.5395704 | 0.49275047 | -1.0950175 | 0.27350896 | 0.84994492 |
| TRBC1 | 11.568159 | 0.46562105 | 0.92143971 | 0.50531906 | 0.61333476 | NA |
| TRBC2 | 16.9543578 | -0.3251131 | 0.65518465 | -0.4962159 | 0.61974206 | 0.95688177 |
| GSTK1 | 24.7627258 | 0.88265858 | 0.44654459 | 1.97664153 | 0.04808216 | 0.53331354 |
| CASP2 | 12.5468939 | -0.69781 | 0.48514286 | -1.4383598 | 0.150332 | NA |
| ZYX | 20.1409761 | -0.4273804 | 0.49487874 | -0.8636062 | 0.38780425 | 0.90910959 |
| EPHA1-AS1 | 14.7764131 | -0.0790411 | 0.6481276 | -0.1219529 | 0.90293629 | 0.9928173 |
| TCAF2 | 21.9003012 | 0.75548296 | 0.49931337 | 1.51304373 | 0.13026855 | 0.72931666 |
| TCAF1 | 12.0954597 | 0.44538427 | 0.70611825 | 0.63075027 | 0.52820383 | NA |
| TPK1 | 67.4788766 | 0.07732614 | 0.30441084 | 0.25401901 | 0.79948087 | 0.98320463 |
| CUL1 | 50.2218008 | -0.211151 | 0.37302281 | -0.5660538 | 0.57135718 | 0.95415437 |
| EZH2 | 23.3812378 | -0.0142866 | 0.70506191 | -0.0202628 | 0.9838337 | 0.99789878 |
| PDIA4 | 13.3137024 | -0.5310759 | 0.66216584 | -0.8020285 | 0.42253646 | 0.91985469 |
| ZNF398 | 12.7216623 | 0.02110353 | 0.52027928 | 0.04056194 | 0.96764513 | NA |
| ZNF862 | 11.9745205 | 1.20978171 | 0.64785239 | 1.86737245 | 0.06184959 | NA |
| GIMAP8 | 20.2314485 | 0.72706281 | 0.75706509 | 0.96037027 | 0.33686889 | 0.88143379 |
| GIMAP7 | 34.9977759 | 0.7726047 | 0.54886014 | 1.40765315 | 0.1592338 | 0.74741294 |
| GIMAP4 | 26.4020116 | 1.04170956 | 0.53727704 | 1.93886858 | 0.05251734 | 0.54907299 |
| GIMAP2 | 11.2534165 | 1.25309269 | 0.62804896 | 1.99521496 | 0.04601944 | NA |
| TMEM176B | 21.5333707 | 2.04238468 | 0.7435208 | 2.74690996 | 0.00601596 | 0.19198507 |
| AGAP3 | 15.474571 | 0.56164188 | 0.54446404 | 1.03154999 | 0.30228298 | 0.86273028 |
| NUB1 | 22.2889058 | -0.1571962 | 0.56551659 | -0.2779693 | 0.78103596 | 0.98264092 |
| RHEB | 36.4949506 | 0.15484078 | 0.34129513 | 0.45368588 | 0.65005493 | 0.96084839 |
| PRKAG2 | 128.534028 | 0.90963787 | 0.49701081 | 1.83021747 | 0.06721743 | 0.58951068 |
| GALNT11 | 43.1585204 | 0.53347566 | 0.43157378 | 1.23611693 | 0.2164151 | 0.80634897 |
| KMT2C | 256.761643 | -0.0777469 | 0.17633499 | -0.4409045 | 0.6592821 | 0.96280444 |
| INSIG1 | 18.041322 | 0.1856733 | 0.72254265 | 0.25697209 | 0.79720032 | 0.98320463 |
| RBM33 | 83.8262247 | -0.049989 | 0.23948182 | -0.208738 | 0.83465274 | 0.98320463 |
| LMBR1 | 58.0058722 | 0.35346762 | 0.39151811 | 0.90281297 | 0.36662517 | 0.89868152 |
| UBE3C | 81.7386033 | -0.0876708 | 0.27477485 | -0.3190641 | 0.74967793 | 0.97818861 |
| DNAJB6 | 62.3859903 | 0.15958076 | 0.30845203 | 0.51736007 | 0.60490483 | 0.95688177 |
| ESYT2 | 92.4011889 | -0.0414392 | 0.48510306 | -0.0854235 | 0.93192468 | 0.99514836 |
| WDR60 | 17.7606518 | -0.2649584 | 0.58550964 | -0.4525261 | 0.65089002 | 0.96105266 |
| GTPBP6 | 12.8346344 | -0.1311895 | 0.52985971 | -0.247593 | 0.80444935 | NA |
| CSF2RA | 43.4489752 | 0.72618654 | 0.70849603 | 1.0249691 | 0.30537777 | 0.8673137 |
| IL3RA | 14.0697458 | -1.6309849 | 0.78502913 | -2.0776106 | 0.03774524 | 0.47316192 |
| SLC25A6 | 39.8226773 | 0.01984687 | 0.60245934 | 0.03294308 | 0.97371998 | 0.9973754 |
| LINC00106 | 8.26337099 | -0.9286311 | 0.69564192 | -1.3349268 | 0.18190029 | NA |
| ASMTL | 10.6400726 | -0.085759 | 0.51582511 | -0.166256 | 0.86795547 | NA |
| P2RY8 | 63.4247368 | -0.0081699 | 0.35746244 | -0.0228553 | 0.98176572 | 0.99758514 |
| AKAP17A | 16.8044039 | -0.1239791 | 0.46439498 | -0.266969 | 0.78949303 | 0.98320463 |
| DHRSX | 68.342027 | 0.38519895 | 0.59435451 | 0.64809629 | 0.51692268 | 0.94596835 |
| CD99 | 58.8349555 | -0.2710078 | 0.58624707 | -0.4622758 | 0.64388354 | 0.95992767 |
| PRKX | 56.7821221 | -0.56911 | 0.35574668 | -1.599762 | 0.10965138 | 0.68862129 |
| PUDP | 10.9482619 | 0.1788003 | 0.53351578 | 0.33513592 | 0.73752256 | NA |
| TBL1X | 125.42744 | -1.8364143 | 0.43778848 | -4.1947524 | 2.73E-05 | 0.00694978 |

| | | | | | | |
|---|---|---|---|---|---|---|
| WWC3 | 22.1288505 | -0.7108174 | 0.4793242 | -1.4829574 | 0.13808574 | 0.73044824 |
| ARHGAP6 | 52.2752315 | 0.18244724 | 0.44406767 | 0.41085459 | 0.68117916 | 0.9673431 |
| MSL3 | 13.3459328 | 0.10123826 | 0.49221432 | 0.20567924 | 0.83704148 | 0.98320463 |
| TMSB4X | 636.948678 | -0.0467203 | 0.67998414 | -0.0687079 | 0.94522216 | 0.99514836 |
| RAB9A | 17.6354864 | 0.3264813 | 0.48003355 | 0.68012184 | 0.49642732 | 0.93890252 |
| OFD1 | 30.9847898 | -0.1339731 | 0.42606112 | -0.3144457 | 0.75318258 | 0.97883059 |
| MOSPD2 | 14.1730312 | 1.04432831 | 0.53748088 | 1.9430055 | 0.05201549 | 0.54907299 |
| CA5B | 30.4898532 | -0.3738783 | 0.56223858 | -0.6649815 | 0.50606231 | 0.94166217 |
| ZRSR2 | 24.326071 | -0.2423674 | 0.50809968 | -0.4770076 | 0.63335671 | 0.95877428 |
| AP1S2 | 32.3622375 | 0.21335311 | 0.49556902 | 0.4305215 | 0.66681633 | 0.96412305 |
| CTPS2 | 13.970722 | 0.50631 | 0.5553579 | 0.91168236 | 0.36193595 | 0.89853007 |
| SYAP1 | 24.8696257 | 0.283567 | 0.53550712 | 0.52952984 | 0.59643795 | 0.9553365 |
| TXLNG | 11.7622394 | -0.4198876 | 0.57997284 | -0.723978 | 0.46907926 | NA |
| RBBP7 | 17.7568519 | -0.829325 | 0.58346258 | -1.4213851 | 0.15520485 | 0.74486619 |
| REPS2 | 16.5926263 | -0.1039779 | 0.64144435 | -0.1620997 | 0.87122735 | 0.98751288 |
| NHS | 18.2418404 | -0.0590673 | 0.45370122 | -0.1301898 | 0.89641622 | 0.99118739 |
| SCML1 | 11.514231 | -0.0147744 | 0.76991451 | -0.0191896 | 0.98468984 | NA |
| CDKL5 | 17.8032905 | -0.8975019 | 0.47882852 | -1.8743703 | 0.06087941 | 0.57281427 |
| PHKA2 | 16.5591977 | -0.4596643 | 0.52670789 | -0.8727121 | 0.38282005 | 0.90868001 |
| PDHA1 | 9.82809085 | 0.34776403 | 0.58048176 | 0.59909553 | 0.54910918 | NA |
| SH3KBP1 | 172.162502 | 0.6276979 | 0.35733979 | 1.75658551 | 0.07898848 | 0.62793067 |
| BCLAF3 | 18.5674803 | -0.3016218 | 0.54571474 | -0.5527097 | 0.58046219 | 0.95415437 |
| EIF1AX | 26.1093557 | 0.23136918 | 0.44672076 | 0.51792798 | 0.60450852 | 0.95688177 |
| RPS6KA3 | 150.629634 | -0.0518767 | 0.27955242 | -0.1855705 | 0.85278158 | 0.98751288 |
| SMS | 26.1173819 | -0.3455417 | 0.40279083 | -0.8578688 | 0.39096493 | 0.90910959 |
| ACOT9 | 25.2717276 | 0.14031754 | 0.48308979 | 0.2904585 | 0.77146549 | 0.98264092 |
| SAT1 | 300.203218 | 0.61661265 | 0.48566481 | 1.26962596 | 0.2042179 | 0.78882329 |
| APOO | 133.694376 | -0.1411463 | 0.57749927 | -0.2444095 | 0.80691368 | 0.98320463 |
| EIF2S3 | 23.8181843 | -0.2343377 | 0.46583888 | -0.5030446 | 0.61493295 | 0.95688177 |
| ZFX | 39.1912055 | -0.3266691 | 0.4533674 | -0.7205394 | 0.47119292 | 0.9314919 |
| PDK3 | 34.5937793 | 0.51431408 | 0.49750837 | 1.03377977 | 0.30123913 | 0.86225566 |
| POLA1 | 26.1696558 | -0.3395055 | 0.51377765 | -0.6608024 | 0.50873907 | 0.94316985 |
| GK | 67.8341178 | 0.90054203 | 0.65893337 | 1.3666663 | 0.17172993 | 0.75906822 |
| TAB3 | 19.0837087 | -0.0712444 | 0.48627511 | -0.1465106 | 0.88351835 | 0.99019008 |
| DMD | 11.5189014 | -0.7077466 | 0.53207175 | -1.3301714 | 0.1834618 | NA |
| RPGR | 9.88201077 | 0.44890156 | 0.65535257 | 0.68497718 | 0.49335832 | NA |
| BCOR | 19.2085222 | -0.0397559 | 0.44732337 | -0.0888751 | 0.92918115 | 0.99514836 |
| ATP6AP2 | 27.541671 | 0.6954892 | 0.37184713 | 1.87036323 | 0.06143339 | 0.57339082 |
| CXorf38 | 19.1837701 | 0.53969513 | 0.5182545 | 1.04137085 | 0.29770346 | 0.8608685 |
| MED14 | 30.2793823 | 0.07075261 | 0.36292029 | 0.19495359 | 0.84542928 | 0.98458147 |
| USP9X | 89.9091464 | -0.1711523 | 0.39862611 | -0.4293556 | 0.66766449 | 0.96412305 |
| DDX3X | 55.6639919 | -0.1207393 | 0.41373117 | -0.2918302 | 0.77041645 | 0.98264092 |
| CASK | 100.374276 | -0.0656736 | 0.39534863 | -0.1661158 | 0.86806585 | 0.98751288 |
| KDM6A | 132.236593 | -0.0684969 | 0.41317697 | -0.165781 | 0.86832929 | 0.98751288 |
| KRBOX4 | 9.32228663 | 0.28752145 | 0.58643547 | 0.4902866 | 0.62393111 | NA |
| SLC9A7 | 19.9206833 | -1.2947121 | 0.57918239 | -2.2354134 | 0.02539022 | 0.40031189 |
| RP2 | 23.3937554 | 0.09564517 | 0.55023958 | 0.17382459 | 0.86200332 | 0.98751288 |
| NDUFB11 | 13.4486601 | -0.3524918 | 0.69103472 | -0.5100928 | 0.60998644 | 0.95688177 |
| RBM10 | 14.6188953 | -0.1045118 | 0.48610148 | -0.215 | 0.82976729 | 0.98320463 |
| UBA1 | 21.0837959 | -0.066962 | 0.47411316 | -0.1412362 | 0.88768332 | 0.99019008 |
| ZNF41 | 8.18836138 | 0.24716937 | 0.6097998 | 0.40532872 | 0.6852359 | NA |
| TIMP1 | 59.665006 | 0.07009115 | 0.80410851 | 0.08716628 | 0.93053934 | 0.99514836 |
| ZNF81 | 16.7496965 | 0.16922445 | 0.47443428 | 0.35668681 | 0.72132628 | 0.97417403 |
| RBM3 | 22.5825537 | -0.1740239 | 0.5367846 | -0.3241969 | 0.74578894 | 0.97805001 |

| | | | | | | |
|---|---|---|---|---|---|---|
| WAS | 15.1588606 | 0.29352253 | 0.51705936 | 0.56767665 | 0.57025455 | 0.95415437 |
| OTUD5 | 20.070814 | -0.3819207 | 0.42713393 | -0.8941475 | 0.37124299 | 0.90226557 |
| GRIPAP1 | 20.0253209 | 0.37983466 | 0.47511319 | 0.79946142 | 0.42402291 | 0.91985469 |
| TFE3 | 11.6313765 | 0.14162304 | 0.58703585 | 0.24125109 | 0.80936052 | NA |
| PLP2 | 11.4684337 | -0.4458912 | 0.70131279 | -0.6357951 | 0.52490999 | NA |
| CLCN5 | 25.3327926 | -0.2321942 | 0.52982562 | -0.4382464 | 0.66120769 | 0.96298563 |
| MAGED1 | 33.7334219 | 0.05742158 | 0.37736593 | 0.15216419 | 0.87905744 | 0.99019008 |
| KANTR | 21.3479435 | 0.0867175 | 0.41058178 | 0.21120639 | 0.83272622 | 0.98320463 |
| KDM5C | 16.7274587 | -0.6249852 | 0.60528563 | -1.0325459 | 0.30181647 | 0.86225566 |
| IQSEC2 | 11.0178747 | 0.79978938 | 0.63430842 | 1.26088407 | 0.20735062 | NA |
| SMC1A | 21.8387514 | -0.6266468 | 0.39025643 | -1.6057309 | 0.10833304 | 0.68204217 |
| HUWE1 | 72.5978359 | -0.0926185 | 0.31352773 | -0.2954077 | 0.76768242 | 0.98264092 |
| PHF8 | 19.3225459 | -0.134458 | 0.41264674 | -0.3258429 | 0.74454322 | 0.97805001 |
| FAM120C | 10.7342769 | -0.6902247 | 0.53323638 | -1.2944067 | 0.19552499 | NA |
| RRAGB | 12.1141931 | -0.1248535 | 0.49296508 | -0.2532704 | 0.80005924 | NA |
| AL445472.1 | 13.6068548 | -1.4076459 | 0.55832651 | -2.5211877 | 0.01169595 | 0.29101961 |
| NBDY | 39.3817352 | -0.2382083 | 0.36092504 | -0.6599939 | 0.50925775 | 0.94332619 |
| FAAH2 | 14.444326 | -0.6278878 | 0.67736295 | -0.9269592 | 0.35394769 | 0.89109867 |
| ARHGEF9 | 17.001384 | -0.4420682 | 0.5445432 | -0.8118148 | 0.4168979 | 0.91953487 |
| MTMR8 | 11.5687981 | -0.6125532 | 0.54685481 | -1.1201387 | 0.26265466 | NA |
| MSN | 130.908948 | -0.1350009 | 0.3376639 | -0.3998086 | 0.68929749 | 0.9673431 |
| AL034397.3 | 35.3661141 | 1.8851024 | 0.6548342 | 2.87874764 | 0.00399258 | 0.15732988 |
| VSIG4 | 28.0281792 | 1.08540674 | 0.83611051 | 1.29816182 | 0.19423174 | 0.78135737 |
| OPHN1 | 17.4929606 | 0.5204141 | 0.56097821 | 0.92769039 | 0.35356819 | 0.89109867 |
| YIPF6 | 15.6252925 | -0.3470977 | 0.49860577 | -0.6961365 | 0.48634335 | 0.93459582 |
| EDA | 41.4069162 | -0.5927641 | 0.70186103 | -0.8445605 | 0.39835625 | 0.91435233 |
| IGBP1 | 15.9909424 | -0.4318976 | 0.44822285 | -0.9635779 | 0.3352576 | 0.88047896 |
| NONO | 21.5776155 | -0.5171749 | 0.41292962 | -1.2524528 | 0.21040492 | 0.79971212 |
| TAF1 | 45.874878 | -0.2136718 | 0.40264779 | -0.5306669 | 0.59564964 | 0.9553365 |
| OGT | 96.842484 | -0.2944374 | 0.350237 | -0.8406804 | 0.40052699 | 0.91508218 |
| NHSL2 | 16.6160376 | -0.232977 | 0.52972984 | -0.4398034 | 0.66007953 | 0.96298563 |
| PIN4 | 14.2208781 | -0.4888032 | 0.62125763 | -0.7867962 | 0.43140117 | 0.92284736 |
| RPS4X | 97.6230462 | -0.3644423 | 0.51602449 | -0.7062501 | 0.48003262 | 0.93257966 |
| HDAC8 | 70.5810469 | 0.25085499 | 0.32053898 | 0.78260369 | 0.43385987 | 0.92311128 |
| CHIC1 | 21.3828549 | -0.8998083 | 0.60892 | -1.4777119 | 0.13948491 | 0.73044824 |
| XIST | 90.1917128 | 0.73168215 | 4.35691172 | 0.16793596 | 0.86663366 | 0.98751288 |
| JPX | 63.0652257 | -0.0663577 | 0.5563892 | -0.1192649 | 0.90506548 | 0.99324672 |
| FTX | 332.137486 | -0.1364683 | 0.29293373 | -0.4658676 | 0.64131025 | 0.95992767 |
| RLIM | 14.8426259 | -0.0275506 | 0.51887582 | -0.0530967 | 0.95765489 | 0.99514836 |
| ABCB7 | 29.1888477 | -0.2250665 | 0.37689261 | -0.5971634 | 0.55039831 | 0.95415437 |
| ATRX | 139.832794 | -0.0792969 | 0.24403792 | -0.3249367 | 0.745229 | 0.97805001 |
| MAGT1 | 24.1672687 | -0.2395163 | 0.44724865 | -0.5355328 | 0.59228146 | 0.9542025 |
| COX7B | 18.9568894 | 0.69152447 | 0.64358557 | 1.07448722 | 0.28260437 | 0.85176121 |
| PGK1 | 57.7090015 | 0.35862405 | 0.41098148 | 0.87260392 | 0.38287901 | 0.90868001 |
| CYSLTR1 | 11.221438 | -0.3733849 | 0.67310354 | -0.5547214 | 0.57908524 | NA |
| GPR174 | 32.0595454 | -0.845892 | 0.39414539 | -2.1461421 | 0.03186165 | 0.43969912 |
| BRWD3 | 52.2383951 | 0.10728738 | 0.3370898 | 0.31827536 | 0.75027607 | 0.97818861 |
| SH3BGRL | 92.6087937 | -0.0177176 | 0.26617191 | -0.0665644 | 0.94692846 | 0.99514836 |
| APOOL | 12.4156653 | 0.14042329 | 0.55113547 | 0.25478906 | 0.79888603 | NA |
| CHM | 32.5177019 | 0.08982941 | 0.40459027 | 0.22202564 | 0.82429393 | 0.98320463 |
| DIAPH2 | 186.466993 | 0.36863078 | 0.29399565 | 1.25386475 | 0.20989117 | 0.79885676 |
| TRMT2B | 13.4223625 | 0.06130133 | 0.52890517 | 0.11590231 | 0.90772996 | 0.99389264 |
| BTK | 14.9749131 | -0.3889927 | 0.64366554 | -0.6043398 | 0.54561774 | 0.95415437 |
| ARMCX4 | 12.8918635 | 0.24096051 | 0.64615084 | 0.37291681 | 0.70921036 | NA |

| | | | | | | |
|---|---|---|---|---|---|---|
| ARMCX3 | 12.1275305 | 0.15820898 | 0.50567367 | 0.31286774 | 0.75438115 | NA |
| ZMAT1 | 13.2956285 | 0.18173799 | 0.64323195 | 0.2825388 | 0.7775304 | 0.98264092 |
| MORF4L2 | 23.397522 | -0.4815377 | 0.51204725 | -0.9404164 | 0.34700399 | 0.88790825 |
| SLC25A53 | 14.5823135 | -0.4144309 | 0.4823668 | -0.8591614 | 0.39025149 | 0.90910959 |
| FAM199X | 13.0632451 | 0.30452695 | 0.49878715 | 0.61053488 | 0.54150755 | 0.95415437 |
| RBM41 | 16.1347864 | -0.4113365 | 0.46168671 | -0.8909429 | 0.37295982 | 0.9025457 |
| TSC22D3 | 54.4923482 | -0.2712673 | 0.61919756 | -0.4380949 | 0.66131751 | 0.96298563 |
| ACSL4 | 83.6072734 | -0.032459 | 0.25379543 | -0.1278942 | 0.89823273 | 0.99141995 |
| TMEM164 | 42.1400235 | 1.36235396 | 0.65081068 | 2.09331838 | 0.03632074 | 0.46855712 |
| AMMECR1 | 21.7919822 | -0.1545687 | 0.49586774 | -0.3117135 | 0.75525826 | 0.97989151 |
| ALG13 | 52.7355093 | -0.1749566 | 0.40733726 | -0.4295129 | 0.66754999 | 0.96412305 |
| WDR44 | 40.0474504 | -0.2504572 | 0.41202933 | -0.6078626 | 0.5432786 | 0.95415437 |
| DOCK11 | 148.726415 | -0.0830517 | 0.22390089 | -0.3709305 | 0.71068932 | 0.97302195 |
| IL13RA1 | 29.0180797 | 0.2752455 | 0.447387 | 0.6152291 | 0.53840344 | 0.95415437 |
| SLC25A43 | 21.2470257 | 0.45469227 | 0.63549316 | 0.71549515 | 0.47430314 | 0.9314919 |
| SLC25A5 | 17.9198026 | 0.47279551 | 0.63769262 | 0.741416 | 0.45844124 | 0.9314919 |
| UBE2A | 12.193701 | 0.37764312 | 0.63692675 | 0.59291453 | 0.55323836 | NA |
| 6-Sep | 90.3782773 | -0.0984371 | 0.35877157 | -0.2743725 | 0.78379837 | 0.98264092 |
| UPF3B | 14.4533631 | -0.4021714 | 0.52498452 | -0.7660634 | 0.44363861 | 0.92899684 |
| NDUFA1 | 16.101788 | 0.32472456 | 0.69889483 | 0.46462578 | 0.64219947 | 0.95992767 |
| LAMP2 | 33.5677603 | 0.69504839 | 0.37755054 | 1.84094131 | 0.06563016 | 0.58951068 |
| CUL4B | 13.6502582 | -0.0076359 | 0.54520983 | -0.0140055 | 0.98882558 | 0.99811561 |
| MCTS1 | 7.84133564 | 0.26146668 | 0.61288374 | 0.42661709 | 0.66965825 | NA |
| THOC2 | 58.1678716 | 0.42876987 | 0.30300746 | 1.41504724 | 0.15705465 | 0.74486619 |
| XIAP | 40.5661781 | -0.2590772 | 0.34837851 | -0.7436658 | 0.45707867 | 0.9314919 |
| STAG2 | 100.398618 | 0.25234189 | 0.29237966 | 0.86306241 | 0.38810315 | 0.90910959 |
| SASH3 | 30.6266036 | -0.5912461 | 0.47710604 | -1.2392342 | 0.21525877 | 0.80634897 |
| ELF4 | 14.5642754 | -0.4246369 | 0.55931147 | -0.7592137 | 0.44772476 | 0.92914597 |
| AIFM1 | 8.05418777 | 1.32123934 | 0.70635922 | 1.87049209 | 0.06141551 | NA |
| ENOX2 | 33.8529256 | -0.1062834 | 0.43889687 | -0.2421603 | 0.80865598 | 0.98320463 |
| STK26 | 17.2385933 | 0.89041163 | 0.56625826 | 1.57244793 | 0.11584671 | 0.70122655 |
| RAP2C | 13.5807828 | -1.2783167 | 0.6138425 | -2.0824832 | 0.03729835 | 0.47208194 |
| RAP2C-AS1 | 20.9677352 | 0.22993148 | 0.57362478 | 0.40083952 | 0.68853828 | 0.9673431 |
| MBNL3 | 23.2505687 | -0.127115 | 0.58540601 | -0.21714 | 0.82809926 | 0.98320463 |
| PHF6 | 14.615915 | -0.8050774 | 0.59555261 | -1.3518158 | 0.17643426 | 0.76409729 |
| HPRT1 | 12.5931275 | -0.018053 | 0.57108137 | -0.0316119 | 0.97478157 | NA |
| FAM122B | 12.3065991 | 0.1491592 | 0.58089467 | 0.25677495 | 0.79735251 | NA |
| ZNF75D | 23.6959113 | -0.4327046 | 0.48835444 | -0.8860462 | 0.37559261 | 0.90341205 |
| INTS6L | 29.6554297 | -0.7567709 | 0.4179014 | -1.8108838 | 0.07015884 | 0.60640882 |
| SLC9A6 | 10.5521851 | 0.05639182 | 0.53288193 | 0.10582422 | 0.91572182 | NA |
| HTATSF1 | 9.16673061 | 0.82080627 | 0.7004122 | 1.17189031 | 0.2412411 | NA |
| ARHGEF6 | 50.0888732 | 0.3807122 | 0.35155595 | 1.08293488 | 0.27883733 | 0.84994492 |
| RBMX | 36.0945077 | 0.0544059 | 0.35184966 | 0.15462826 | 0.87711439 | 0.98939075 |
| ATP11C | 43.8296911 | 0.17024997 | 0.4558522 | 0.37347626 | 0.70879401 | 0.97186969 |
| FMR1 | 20.6788513 | 0.4065432 | 0.48387582 | 0.84018087 | 0.40080698 | 0.91508218 |
| IDS | 18.7092802 | -0.5237797 | 0.46890086 | -1.1170371 | 0.26397846 | 0.84928231 |
| LINC00894 | 21.6839349 | 0.32470159 | 0.48303701 | 0.67220851 | 0.50145096 | 0.94005069 |
| MTM1 | 43.7508865 | 0.29240886 | 0.40489209 | 0.72218961 | 0.47017791 | 0.9314919 |
| MTMR1 | 30.4436179 | -0.4838714 | 0.45913473 | -1.0538767 | 0.29193936 | 0.85935767 |
| CD99L2 | 15.5623704 | 0.76714413 | 0.68696391 | 1.11671678 | 0.26411544 | 0.84928231 |
| BCAP31 | 16.7016286 | 0.77648473 | 0.66381444 | 1.1697316 | 0.242109 | 0.83493679 |
| SSR4 | 14.8403147 | 0.50348856 | 0.70979843 | 0.7093402 | 0.47811339 | 0.9318939 |
| ARHGAP4 | 20.7441962 | -0.3148534 | 0.48536928 | -0.6486883 | 0.51653989 | 0.94596835 |
| IRAK1 | 10.4252233 | -0.2220715 | 0.65447952 | -0.3393102 | 0.73437609 | NA |

| | | | | | | |
|---|---|---|---|---|---|---|
| MECP2 | 71.5250922 | -0.3081896 | 0.27952704 | -1.1025393 | 0.27022727 | 0.84994492 |
| FLNA | 71.8600148 | -0.2330994 | 0.55174346 | -0.4224779 | 0.67267623 | 0.96433967 |
| EMD | 8.6548973 | -0.4430641 | 0.61280176 | -0.7230138 | 0.46967143 | NA |
| RPL10 | 176.600465 | -0.30241 | 0.60660922 | -0.4985253 | 0.61811384 | 0.95688177 |
| TAZ | 15.6146043 | -0.350587 | 0.43068481 | -0.8140222 | 0.41563221 | 0.91894074 |
| ATP6AP1 | 16.8877165 | 0.89045824 | 0.48896667 | 1.82110212 | 0.06859133 | 0.59777078 |
| GDI1 | 11.5922076 | -0.448687 | 0.52595558 | -0.8530892 | 0.39360986 | NA |
| GAB3 | 40.7785459 | 0.63111341 | 0.53052702 | 1.18959712 | 0.23420478 | 0.82817204 |
| MPP1 | 16.4992546 | 0.8216793 | 0.58331975 | 1.40862587 | 0.15894582 | 0.74686546 |
| BRCC3 | 12.8922369 | -0.0329784 | 0.55425316 | -0.0595007 | 0.95255334 | NA |
| TMLHE-AS1 | 12.3473684 | 0.74124868 | 0.66201507 | 1.1196855 | 0.26284781 | NA |
| TMLHE | 33.6111424 | 0.18295736 | 0.35863086 | 0.5101551 | 0.6099428 | 0.95688177 |
| VAMP7 | 12.1397713 | 0.56485021 | 0.52124368 | 1.08365863 | 0.27851619 | NA |
| FBXO25 | 10.6824595 | 0.62458105 | 0.59802132 | 1.04441268 | 0.2962945 | NA |
| ERICH1 | 58.1415942 | 0.09513721 | 0.38313931 | 0.24830971 | 0.80389478 | 0.98320463 |
| CSMD1 | 17.7926918 | 0.29452258 | 1.22622582 | 0.24018625 | 0.81018587 | 0.98320463 |
| MCPH1 | 61.3804221 | -0.1850074 | 0.30483151 | -0.6069169 | 0.54390606 | 0.95415437 |
| ANGPT2 | 10.4859843 | 0.40035277 | 0.76364601 | 0.52426487 | 0.60009433 | NA |
| AGPAT5 | 18.2990311 | -0.0523597 | 0.46525303 | -0.1125404 | 0.91039496 | 0.99471296 |
| DEFB1 | 24.0952247 | 0.53208955 | 0.65786639 | 0.80881096 | 0.41862389 | 0.91985469 |
| FAM85B | 18.6142956 | -1.6999918 | 0.69027114 | -2.4627885 | 0.01378612 | 0.31861834 |
| PRAG1 | 28.5074417 | -1.9410736 | 0.47084248 | -4.1225542 | 3.75E-05 | 0.00863983 |
| MFHAS1 | 36.3995638 | -0.2843387 | 0.43418776 | -0.654875 | 0.51254823 | 0.94516558 |
| ERI1 | 18.1079715 | 0.97417336 | 0.54277296 | 1.79480821 | 0.07268426 | 0.61222939 |
| TNKS | 80.7404117 | 0.05333306 | 0.27313118 | 0.19526538 | 0.8451852 | 0.98458147 |
| MSRA | 55.4201732 | -0.1598497 | 0.48783948 | -0.3276686 | 0.74316221 | 0.97711087 |
| XKR6 | 86.785894 | -0.5076542 | 0.30217894 | -1.6799788 | 0.09296144 | 0.6590317 |
| MTMR9 | 23.0339944 | 0.20262335 | 0.43584048 | 0.46490255 | 0.64200124 | 0.95992767 |
| BLK | 32.700154 | -3.0406885 | 0.85515723 | -3.555707 | 0.00037696 | 0.03630623 |
| FDFT1 | 22.9284843 | -0.2959467 | 0.40525433 | -0.7302741 | 0.46522265 | 0.9314919 |
| CTSB | 195.121873 | 0.70498489 | 0.47579819 | 1.48168889 | 0.1384231 | 0.73044824 |
| LONRF1 | 14.0068463 | -1.2108261 | 0.62667429 | -1.9321458 | 0.05334151 | 0.551221 |
| MSR1 | 124.434204 | 0.48340704 | 0.60102605 | 0.80430297 | 0.42122202 | 0.91985469 |
| ZDHHC2 | 26.5197784 | -0.5610596 | 0.59795629 | -0.9382953 | 0.34809267 | 0.88939849 |
| CNOT7 | 11.8408788 | -0.1206298 | 0.51037544 | -0.236355 | 0.8131572 | NA |
| VPS37A | 22.7216987 | 0.09028235 | 0.45917974 | 0.19661658 | 0.84412759 | 0.9842451 |
| MTUS1 | 11.0188376 | 0.28424834 | 0.68634396 | 0.41414853 | 0.67876533 | NA |
| PCM1 | 121.166311 | -0.3234544 | 0.25041703 | -1.2916628 | 0.19647396 | 0.78135737 |
| ASAH1 | 65.8874034 | 1.11057255 | 0.49457266 | 2.24551952 | 0.0247348 | 0.39492781 |
| PSD3 | 150.788093 | -3.0527465 | 0.85125031 | -3.5861914 | 0.00033554 | 0.03499569 |
| AC100849.1 | 39.6201773 | -1.6681359 | 1.1713688 | -1.4240911 | 0.15442009 | 0.74486619 |
| CSGALNACT1 | 82.6051347 | -0.2794201 | 0.47089035 | -0.5933869 | 0.55292228 | 0.95415437 |
| INTS10 | 32.2940995 | 0.23290442 | 0.348744 | 0.66783778 | 0.50423715 | 0.94166217 |
| ATP6V1B2 | 36.5291077 | 0.71076887 | 0.46322483 | 1.53439286 | 0.12493303 | 0.71776265 |
| DOK2 | 14.5413341 | 1.60071044 | 0.87092353 | 1.83794602 | 0.06607036 | 0.58951068 |
| XPO7 | 32.3486276 | -0.1759052 | 0.3511629 | -0.5009221 | 0.61642596 | 0.95688177 |
| SLC39A14 | 10.9095588 | -0.3915121 | 0.62499541 | -0.6264239 | 0.53103692 | NA |
| PPP3CC | 86.8634175 | -1.2185676 | 0.39716366 | -3.0681749 | 0.0021537 | 0.10651853 |
| BIN3 | 12.8481535 | 0.03436762 | 0.78865338 | 0.04357759 | 0.96524111 | NA |
| TNFRSF10B | 25.7335385 | 0.0776001 | 0.60436502 | 0.12839939 | 0.89783291 | 0.99141995 |
| TNFRSF10D | 8.99597939 | -0.869503 | 0.59947013 | -1.4504525 | 0.14693237 | NA |
| TNFRSF10A | 11.5225679 | 0.70257421 | 0.71264503 | 0.9858684 | 0.3241977 | NA |
| CHMP7 | 10.9484456 | -0.7088659 | 0.51688988 | -1.3714061 | 0.17024839 | NA |
| ENTPD4 | 27.9972093 | -0.1183684 | 0.40235709 | -0.2941874 | 0.76861468 | 0.98264092 |

| | | | | | | |
|---|---|---|---|---|---|---|
| SLC25A37 | 34.9021642 | 0.62684349 | 0.5553993 | 1.12863572 | 0.25905154 | 0.84484505 |
| ADAM28 | 69.0305779 | 0.0866188 | 0.66655893 | 0.1299492 | 0.89660662 | 0.99118739 |
| AC120193.1 | 24.4325809 | -1.3834799 | 0.48570297 | -2.8484073 | 0.00439386 | 0.16654554 |
| DOCK5 | 101.475295 | 1.13779763 | 0.44899379 | 2.53410552 | 0.01127348 | 0.28883068 |
| KCTD9 | 15.1572976 | 0.26465228 | 0.46569478 | 0.56829557 | 0.56983429 | 0.95415437 |
| PPP2R2A | 83.9297025 | 0.19131553 | 0.34583885 | 0.5531927 | 0.58013147 | 0.95415437 |
| BNIP3L | 45.6834957 | 0.26742601 | 0.46959422 | 0.56948317 | 0.56902829 | 0.95415437 |
| DPYSL2 | 25.3970533 | -0.1478198 | 0.51857606 | -0.2850495 | 0.77560624 | 0.98264092 |
| PTK2B | 113.113142 | -0.0334696 | 0.35523394 | -0.0942185 | 0.92493558 | 0.99514836 |
| CLU | 21.6022357 | 0.72419289 | 0.43326303 | 1.67148554 | 0.09462581 | 0.65925012 |
| ELP3 | 30.525126 | -0.0523481 | 0.36520814 | -0.1433377 | 0.88602348 | 0.99019008 |
| INTS9 | 23.3436304 | -0.1749745 | 0.38629179 | -0.4529593 | 0.65057805 | 0.96084839 |
| HMBOX1 | 82.9616876 | -0.4928889 | 0.27437768 | -1.7963885 | 0.07243275 | 0.61222939 |
| KIF13B | 76.9429764 | -0.5007398 | 0.27180904 | -1.8422484 | 0.06543883 | 0.58951068 |
| SARAF | 93.1606815 | 0.19944235 | 0.52613792 | 0.37906858 | 0.70463694 | 0.97155642 |
| LEPROTL1 | 48.2195919 | 0.30837368 | 0.54019383 | 0.57085746 | 0.56809627 | 0.95415437 |
| RBPMS | 21.0154879 | -0.649035 | 0.5195711 | -1.2491745 | 0.21160126 | 0.80277403 |
| GTF2E2 | 24.0086654 | -0.7746764 | 0.37395819 | -2.071559 | 0.03830659 | 0.47745196 |
| GSR | 19.4258717 | 0.29135008 | 0.44580805 | 0.65353257 | 0.513413 | 0.94596835 |
| PPP2CB | 22.1803089 | 0.42730284 | 0.49778199 | 0.85841362 | 0.39066411 | 0.90910959 |
| WRN | 30.5783326 | -0.1156007 | 0.35025561 | -0.3300467 | 0.7413647 | 0.97605445 |
| NRG1 | 28.2011549 | -0.9980488 | 0.64567506 | -1.5457448 | 0.12216622 | 0.71192434 |
| FUT10 | 9.54177108 | -0.0631553 | 0.62431268 | -0.1011597 | 0.91942367 | NA |
| RAB11FIP1 | 39.0895138 | -1.0572079 | 0.42138211 | -2.5089054 | 0.01211059 | 0.29654868 |
| ASH2L | 15.5053551 | 0.29702996 | 0.49150384 | 0.60432887 | 0.54562502 | 0.95415437 |
| BAG4 | 11.1895829 | -0.098737 | 0.54953271 | -0.1796744 | 0.85740819 | NA |
| DDHD2 | 11.9405345 | 0.57878865 | 0.53941101 | 1.07300117 | 0.28327059 | NA |
| NSD3 | 82.7055927 | 0.13841882 | 0.26322632 | 0.52585479 | 0.5989891 | 0.95582632 |
| TACC1 | 86.4883776 | -0.4234173 | 0.39741184 | -1.065437 | 0.28667823 | 0.85677513 |
| PLEKHA2 | 108.725474 | -0.5394043 | 0.37945193 | -1.4215352 | 0.15516122 | 0.74486619 |
| ADAM9 | 47.3245467 | -0.3470895 | 0.59724227 | -0.5811537 | 0.56113689 | 0.95415437 |
| GOLGA7 | 10.443836 | 0.38651897 | 0.73136572 | 0.52848932 | 0.59715976 | NA |
| GPAT4 | 21.9885347 | 0.31028048 | 0.51934797 | 0.59744237 | 0.55021208 | 0.95415437 |
| KAT6A | 143.967165 | -0.1441117 | 0.26768931 | -0.5383544 | 0.5903324 | 0.95415437 |
| AP3M2 | 9.5855388 | 0.61178315 | 0.60458149 | 1.01191182 | 0.31158022 | NA |
| IKBKB | 52.7754957 | -0.0640314 | 0.31239178 | -0.2049714 | 0.83759448 | 0.98320463 |
| POLB | 13.0569286 | -0.0042518 | 0.50571959 | -0.0084075 | 0.99329189 | 0.99859588 |
| VDAC3 | 20.8735319 | 0.0380483 | 0.41312789 | 0.09209812 | 0.92662008 | 0.99514836 |
| SLC20A2 | 36.7707356 | -0.6308707 | 0.45358561 | -1.3908526 | 0.16427012 | 0.75321592 |
| RNF170 | 14.2722395 | -0.153534 | 0.50764053 | -0.3024463 | 0.76231186 | 0.98163473 |
| HOOK3 | 49.8538386 | 0.15023513 | 0.33095043 | 0.45395055 | 0.64986442 | 0.96084839 |
| FNTA | 10.7359945 | 0.74785724 | 0.54912137 | 1.36191611 | 0.17322436 | NA |
| HGSNAT | 32.8137123 | 1.00214153 | 0.43089085 | 2.32574338 | 0.02003224 | 0.36765113 |
| SPIDR | 235.703833 | -0.0214432 | 0.35001333 | -0.0612639 | 0.95114905 | 0.99514836 |
| CEBPD | 55.0980663 | 0.40562439 | 0.58110964 | 0.69801696 | 0.48516659 | 0.93417265 |
| PRKDC | 63.8063299 | -0.0638912 | 0.30797801 | -0.2074537 | 0.83565558 | 0.98320463 |
| UBE2V2 | 19.4597049 | -0.3488408 | 0.41860685 | -0.8333375 | 0.40465441 | 0.91551819 |
| PCMTD1 | 68.0650913 | 0.28911245 | 0.30831699 | 0.93771171 | 0.34839258 | 0.88971138 |
| ST18 | 22.8418209 | 1.63818404 | 0.89491022 | 1.83055685 | 0.06716671 | 0.58951068 |
| RB1CC1 | 66.0150994 | 0.09812777 | 0.31600687 | 0.31052417 | 0.75616238 | 0.97989151 |
| ATP6V1H | 50.1146827 | 0.14791097 | 0.37943021 | 0.38982393 | 0.69666675 | 0.96861929 |
| TCEA1 | 21.8932967 | -0.5780582 | 0.37471051 | -1.5426795 | 0.12290857 | 0.71297276 |
| LYPLA1 | 38.9100304 | -0.3255423 | 0.3427892 | -0.9496865 | 0.3422716 | 0.88510795 |
| TMEM68 | 8.81738787 | -0.1176019 | 0.59999324 | -0.1960054 | 0.84460593 | NA |

| | | | | | | |
|---|---|---|---|---|---|---|
| TGS1 | 16.2994681 | 0.43446751 | 0.47771756 | 0.90946524 | 0.3631046 | 0.89868152 |
| LYN | 278.687201 | 0.2735368 | 0.44325952 | 0.61710305 | 0.53716676 | 0.95415437 |
| RPS20 | 45.5694883 | 0.18529295 | 0.47160489 | 0.3928987 | 0.6943943 | 0.9685713 |
| IMPAD1 | 14.8906578 | 0.13274023 | 0.48429138 | 0.27409167 | 0.78401417 | 0.98264092 |
| FAM110B | 22.0900044 | 1.52602814 | 0.96381629 | 1.58331848 | 0.11334889 | 0.69801324 |
| UBXN2B | 28.3942903 | 0.40838129 | 0.50064684 | 0.81570732 | 0.41466755 | 0.91894074 |
| SDCBP | 74.0355038 | 0.56879572 | 0.33352694 | 1.70539662 | 0.08812048 | 0.64676224 |
| NSMAF | 40.7190644 | -0.3532004 | 0.60733875 | -0.5815542 | 0.56086703 | 0.95415437 |
| TOX | 183.944743 | -1.1633375 | 0.46621729 | -2.4952689 | 0.01258617 | 0.30389677 |
| RAB2A | 105.975689 | -0.1196962 | 0.25063062 | -0.47758 | 0.63294916 | 0.95877428 |
| CHD7 | 41.8551585 | 0.02389849 | 0.34811321 | 0.06865148 | 0.94526704 | 0.99514836 |
| ASPH | 63.6107251 | 0.40273063 | 0.41602241 | 0.96805032 | 0.33301924 | 0.88047896 |
| YTHDF3 | 44.3899909 | 0.34899419 | 0.36770467 | 0.94911546 | 0.34256189 | 0.88530691 |
| ARMC1 | 12.9647793 | -0.4207314 | 0.53797957 | -0.7820583 | 0.43418032 | 0.9231261 |
| MTFR1 | 27.3718378 | 0.10758989 | 0.50110101 | 0.21470699 | 0.82999577 | 0.98320463 |
| PDE7A | 179.228621 | -0.3193571 | 0.35148564 | -0.9085922 | 0.36356544 | 0.89868152 |
| MYBL1 | 45.1912715 | 0.64392302 | 0.71483404 | 0.90080072 | 0.36769428 | 0.89868152 |
| VCPIP1 | 20.4263162 | 0.5364748 | 0.42259724 | 1.26947067 | 0.20427325 | 0.78882329 |
| SGK3 | 56.024821 | 1.077539 | 0.45251687 | 2.38121286 | 0.01725574 | 0.34827529 |
| CSPP1 | 31.9246956 | -0.0127131 | 0.47113275 | -0.0269842 | 0.97847234 | 0.99758514 |
| ARFGEF1 | 118.025396 | 0.2058477 | 0.26697749 | 0.77103017 | 0.44068905 | 0.92880096 |
| NCOA2 | 220.308738 | 0.28538356 | 0.24299755 | 1.17442982 | 0.24022291 | 0.83493679 |
| TRAM1 | 35.8659646 | -0.2746638 | 0.46608749 | -0.5892966 | 0.55566231 | 0.95415437 |
| LACTB2 | 8.82996205 | 1.60243794 | 0.66660973 | 2.40386223 | 0.01622289 | NA |
| MSC-AS1 | 14.9013682 | 0.85094641 | 0.62044174 | 1.37151702 | 0.17021384 | 0.75745073 |
| TERF1 | 18.381824 | -0.3791038 | 0.45871839 | -0.8264413 | 0.4085538 | 0.9169561 |
| RPL7 | 65.9913937 | 0.01095151 | 0.65959429 | 0.01660341 | 0.98675301 | 0.99811561 |
| STAU2 | 54.8158962 | -0.5766032 | 0.54024682 | -1.0672958 | 0.28583827 | 0.85570689 |
| ELOC | 15.4381727 | -0.0039929 | 0.47038212 | -0.0084885 | 0.99322721 | 0.99859588 |
| LY96 | 21.5913039 | -0.7665032 | 0.52689208 | -1.4547631 | 0.14573485 | 0.73251288 |
| GDAP1 | 12.9739059 | -0.212518 | 0.6660432 | -0.3190754 | 0.7496693 | 0.97818861 |
| PEX2 | 11.8663259 | -0.4760317 | 0.51695597 | -0.9208361 | 0.357136 | NA |
| ZC2HC1A | 9.57874015 | 0.64128736 | 0.63729093 | 1.00627096 | 0.31428525 | NA |
| IL7 | 14.2546305 | 0.23967549 | 0.60652295 | 0.3951631 | 0.69272253 | 0.9685713 |
| TPD52 | 44.1120645 | -0.5798336 | 0.41306905 | -1.4037208 | 0.160402 | 0.74887162 |
| ZNF704 | 9.12170005 | 0.47952666 | 0.74905334 | 0.64017692 | 0.52205759 | NA |
| PAG1 | 183.745015 | 1.45089936 | 0.50813089 | 2.85536537 | 0.00429873 | 0.16520275 |
| IMPA1 | 9.82358786 | -0.0345456 | 0.57952698 | -0.0596101 | 0.95246619 | NA |
| ZFAND1 | 9.67475706 | 0.37457297 | 0.65470262 | 0.57212689 | 0.56723601 | NA |
| C8orf59 | 10.9391744 | -0.168266 | 0.59849508 | -0.2811485 | 0.7785965 | NA |
| WWP1 | 89.7844187 | 0.18073213 | 0.38358804 | 0.47116206 | 0.63752501 | 0.95933464 |
| RMDN1 | 20.3882044 | -0.1936102 | 0.44231088 | -0.4377242 | 0.66158619 | 0.96298563 |
| CPNE3 | 17.1441488 | -0.1345082 | 0.45226028 | -0.2974132 | 0.76615103 | 0.98264092 |
| AF117829.1 | 18.9997648 | -1.438858 | 0.48934993 | -2.9403457 | 0.00327846 | 0.14285834 |
| RIPK2 | 22.3011829 | 0.21347688 | 0.72534637 | 0.29431026 | 0.76852083 | 0.98264092 |
| NBN | 20.8316204 | 0.40472941 | 0.39846065 | 1.01573244 | 0.30975681 | 0.86794644 |
| DECR1 | 46.73362 | 0.1055857 | 0.3558841 | 0.29668563 | 0.76670653 | 0.98264092 |
| PIP4P2 | 14.2273292 | -0.5893059 | 0.56648605 | -1.0402832 | 0.29820833 | 0.8608685 |
| LRRC69 | 13.1376627 | -0.9760589 | 0.53170532 | -1.8357139 | 0.06639998 | 0.58951068 |
| TRIQK | 12.7435403 | 0.46544562 | 0.67365118 | 0.69092972 | 0.48960971 | NA |
| RBM12B | 15.0913361 | 0.47512074 | 0.58266089 | 0.8154327 | 0.41482467 | 0.91894074 |
| VIRMA | 22.5519287 | -0.4542591 | 0.459981 | -0.9875605 | 0.32336795 | 0.87507005 |
| DPY19L4 | 15.0654697 | -0.3158193 | 0.57508874 | -0.5491662 | 0.58289141 | 0.95415437 |
| INTS8 | 23.3539841 | -0.0022401 | 0.45356104 | -0.0049389 | 0.99605937 | 0.99859588 |

| | | | | | | |
|---|---|---|---|---|---|---|
| NDUFAF6 | 29.2863626 | 0.12830706 | 0.59657701 | 0.21507208 | 0.82971112 | 0.98320463 |
| TP53INP1 | 28.7326467 | -0.098119 | 0.71106318 | -0.1379892 | 0.89024896 | 0.99080204 |
| PLEKHF2 | 11.2350102 | -0.6667869 | 0.51136441 | -1.3039369 | 0.19225511 | NA |
| UQCRB | 33.3647757 | 0.19229216 | 0.57925502 | 0.3319646 | 0.73991599 | 0.97605445 |
| PTDSS1 | 17.9358866 | -0.3957574 | 0.44797296 | -0.8834405 | 0.37699833 | 0.90433844 |
| SDC2 | 25.597924 | 0.45542538 | 0.73497844 | 0.61964455 | 0.53549183 | 0.95378558 |
| CPQ | 103.222608 | -0.4538301 | 0.40515631 | -1.1201358 | 0.2626559 | 0.84921816 |
| MTDH | 102.383769 | -0.0110645 | 0.24971946 | -0.0443077 | 0.96465912 | 0.99653147 |
| RPL30 | 99.4904182 | -0.4295785 | 0.6382599 | -0.6730464 | 0.50091775 | 0.94001041 |
| NIPAL2 | 14.5660541 | -0.9517113 | 0.71240479 | -1.3359137 | 0.18157749 | 0.77059034 |
| STK3 | 82.3878421 | 0.07540793 | 0.34901611 | 0.21605861 | 0.82894206 | 0.98320463 |
| AC104986.2 | 9.75967033 | 0.13893401 | 0.62339448 | 0.22286693 | 0.82363908 | NA |
| VPS13B | 224.877123 | 0.06586026 | 0.21152554 | 0.31135841 | 0.75552817 | 0.97989151 |
| COX6C | 27.0051264 | 0.32169133 | 0.47680417 | 0.67468229 | 0.49987764 | 0.93996699 |
| RNF19A | 89.5454717 | -0.3338335 | 0.33350114 | -1.0009965 | 0.3168285 | 0.87263466 |
| ANKRD46 | 9.62428186 | 1.12464653 | 0.66999437 | 1.678591 | 0.09323178 | NA |
| PABPC1 | 144.219305 | -0.3698026 | 0.42980746 | -0.8603913 | 0.38957339 | 0.90910959 |
| YWHAZ | 110.68789 | -0.2131605 | 0.3368538 | -0.6327982 | 0.52686544 | 0.95136478 |
| ZNF706 | 36.2105234 | 0.63954933 | 0.46845017 | 1.36524517 | 0.17217601 | 0.75982617 |
| NCALD | 75.0804825 | 0.61704204 | 0.43808361 | 1.40850292 | 0.1589822 | 0.74686546 |
| RRM2B | 11.6399964 | 0.20188843 | 0.62257049 | 0.32428204 | 0.74572451 | NA |
| UBR5 | 134.538895 | 0.07050633 | 0.29758005 | 0.23693232 | 0.8127093 | 0.98320463 |
| KLF10 | 11.9175254 | 1.35637813 | 0.71597942 | 1.89443732 | 0.05816699 | NA |
| AZIN1 | 28.4071238 | 0.72417912 | 0.40954748 | 1.76824216 | 0.07702043 | 0.62507515 |
| ATP6V1C1 | 33.7965507 | 0.40009398 | 0.47773173 | 0.83748672 | 0.40231904 | 0.91528253 |
| SLC25A32 | 12.7094006 | -0.1501748 | 0.51325103 | -0.2925951 | 0.76983162 | NA |
| OXR1 | 69.2427127 | -0.3625218 | 0.2903792 | -1.2484426 | 0.211869 | 0.8030706 |
| EIF3E | 38.3968556 | 0.0333786 | 0.46086415 | 0.07242612 | 0.9422628 | 0.99514836 |
| EMC2 | 14.7441528 | -0.5523646 | 0.49231371 | -1.1219768 | 0.26187231 | 0.84837688 |
| NUDCD1 | 12.6962655 | -0.2902343 | 0.48943615 | -0.5929973 | 0.55318298 | NA |
| ENY2 | 19.6694131 | 0.02973152 | 0.46360715 | 0.06413085 | 0.94886604 | 0.99514836 |
| TRPS1 | 297.281327 | -0.1818138 | 0.50990798 | -0.3565619 | 0.72141979 | 0.97417403 |
| EIF3H | 58.5318123 | -0.084197 | 0.39299176 | -0.2142463 | 0.83035496 | 0.98320463 |
| RAD21 | 39.7884391 | 0.09192332 | 0.41280889 | 0.22267766 | 0.82378639 | 0.98320463 |
| EXT1 | 174.996923 | 0.82359485 | 0.76688486 | 1.07394851 | 0.28284576 | 0.85176121 |
| SAMD12 | 29.9650903 | -0.2694161 | 0.69227442 | -0.3891752 | 0.69714652 | 0.96861929 |
| ENPP2 | 12.0586306 | 0.49481189 | 0.77338932 | 0.63979665 | 0.52230481 | NA |
| TAF2 | 39.1104146 | -0.6407343 | 0.34581206 | -1.8528396 | 0.06390535 | 0.58463007 |
| DEPTOR | 11.8694732 | -0.2961681 | 0.70031233 | -0.4229086 | 0.67236194 | NA |
| SNTB1 | 87.6502987 | 0.21414408 | 0.61989989 | 0.34544946 | 0.7297565 | 0.97605445 |
| ZHX2 | 116.546017 | -1.001973 | 0.40174922 | -2.4940259 | 0.01263033 | 0.30389677 |
| DERL1 | 19.5542923 | 0.22065266 | 0.52662017 | 0.41899773 | 0.67521779 | 0.96604324 |
| TBC1D31 | 16.7975161 | 0.3634209 | 0.59653253 | 0.60922226 | 0.54237713 | 0.95415437 |
| ATAD2 | 20.7791225 | -0.0283445 | 0.61558359 | -0.0460449 | 0.96327448 | 0.99653147 |
| FBXO32 | 10.090127 | 0.83034947 | 0.80448295 | 1.03215297 | 0.30200046 | NA |
| FAM91A1 | 42.1095697 | 0.05312472 | 0.34323325 | 0.15477732 | 0.87699687 | 0.98939075 |
| TMEM65 | 55.3802207 | -0.2252109 | 0.38316851 | -0.5877595 | 0.55669374 | 0.95415437 |
| TATDN1 | 14.8156929 | -0.3754692 | 0.53847063 | -0.6972882 | 0.48562243 | 0.93417265 |
| NDUFB9 | 18.791396 | -0.68469 | 0.63338676 | -1.0809983 | 0.27969787 | 0.84994492 |
| MTSS1 | 313.609248 | -1.9239904 | 0.70363652 | -2.7343526 | 0.00625031 | 0.19652953 |
| WASHC5 | 19.3003722 | 0.79377461 | 0.44482508 | 1.78446458 | 0.07434821 | 0.61929783 |
| NSMCE2 | 131.869931 | -0.0506951 | 0.23516902 | -0.2155688 | 0.82932387 | 0.98320463 |
| LINC00861 | 29.3083804 | -0.1665394 | 0.73142272 | -0.2276924 | 0.81988538 | 0.98320463 |
| PCAT1 | 44.2638582 | -0.1950874 | 0.59177504 | -0.3296648 | 0.74165323 | 0.97605445 |

| | | | | | | |
|---|---|---|---|---|---|---|
| PVT1 | 65.1951124 | 0.61831455 | 0.53550699 | 1.15463396 | 0.24824035 | 0.83817091 |
| CCDC26 | 30.9560183 | 1.00106013 | 0.76217202 | 1.3134307 | 0.18903786 | 0.77850418 |
| FAM49B | 350.328357 | 0.03737633 | 0.33139037 | 0.1127864 | 0.9101999 | 0.99471296 |
| ASAP1 | 205.764546 | 0.20205162 | 0.35827369 | 0.56395885 | 0.57278214 | 0.95415437 |
| EFR3A | 77.058748 | -0.0920112 | 0.36164874 | -0.2544216 | 0.79916987 | 0.98320463 |
| KCNQ3 | 28.9637163 | 3.180673 | 1.02660917 | 3.09823162 | 0.00194679 | 0.09883988 |
| TMEM71 | 35.7531695 | -0.2745125 | 0.49124035 | -0.5588151 | 0.57628789 | 0.95415437 |
| PHF20L1 | 59.2389612 | -0.4677528 | 0.44696644 | -1.0465054 | 0.29532773 | 0.8608685 |
| TG | 31.1826151 | 0.63407996 | 0.59159476 | 1.07181471 | 0.28380327 | 0.85176121 |
| SLA | 104.510473 | 0.74516141 | 0.32739735 | 2.27601542 | 0.02284509 | 0.39019972 |
| NDRG1 | 28.1627715 | 1.44524101 | 0.51289494 | 2.81781103 | 0.00483523 | 0.17334548 |
| ST3GAL1 | 106.225114 | 0.00554486 | 0.39308778 | 0.0141059 | 0.98874549 | 0.99811561 |
| ZFAT | 22.8646037 | -0.4751627 | 0.45647344 | -1.0409427 | 0.29790217 | 0.8608685 |
| TRAPPC9 | 140.20633 | 0.19998508 | 0.23801382 | 0.84022465 | 0.40078244 | 0.91508218 |
| AGO2 | 54.2005202 | -0.6238031 | 0.30157179 | -2.0685061 | 0.03859246 | 0.47885804 |
| PTK2 | 62.6868592 | -1.8019158 | 0.55600293 | -3.2408387 | 0.00119179 | 0.07168852 |
| DENND3 | 51.825641 | 0.4122051 | 0.56692635 | 0.72708756 | 0.46717232 | 0.9314919 |
| SLC45A4 | 18.0709422 | 0.358753 | 0.51986272 | 0.6900918 | 0.49013646 | 0.93633431 |
| LY6E | 33.9432352 | 0.13054104 | 0.71771491 | 0.18188426 | 0.85567356 | 0.98751288 |
| ZC3H3 | 16.7940292 | 0.44606863 | 0.51960365 | 0.85847864 | 0.39062822 | 0.90910959 |
| GSDMD | 13.6933994 | 0.87324877 | 0.51790617 | 1.68611385 | 0.09177387 | 0.65902576 |
| EEF1D | 64.8522624 | -0.2133559 | 0.40106284 | -0.5319764 | 0.59474237 | 0.9553365 |
| PUF60 | 9.07045072 | -0.018209 | 0.62450429 | -0.0291575 | 0.97673895 | NA |
| PLEC | 36.7578561 | -0.923035 | 0.53153407 | -1.736549 | 0.08246681 | 0.63740875 |
| PARP10 | 12.4905332 | 0.49183779 | 0.62757278 | 0.78371434 | 0.43320774 | NA |
| GRINA | 20.5492088 | 1.71803756 | 0.7137428 | 2.40708217 | 0.01608055 | 0.33329503 |
| CYC1 | 8.32429709 | 0.4443172 | 0.61204863 | 0.72595081 | 0.46786893 | NA |
| MAF1 | 10.3041611 | -0.7069237 | 0.66824002 | -1.057889 | 0.29010606 | NA |
| MROH1 | 23.9118136 | 0.45166311 | 0.41815008 | 1.08014594 | 0.2800772 | 0.84994492 |
| HSF1 | 16.8570925 | 0.01325123 | 0.47969244 | 0.02762444 | 0.97796169 | 0.99758514 |
| VPS28 | 18.4966376 | -0.0071801 | 0.44485613 | -0.0161403 | 0.98712246 | 0.99811561 |
| CYHR1 | 8.76099707 | -0.1386196 | 0.56747075 | -0.2442763 | 0.80701687 | NA |
| ZNF251 | 13.154837 | -0.6057783 | 0.51854216 | -1.1682335 | 0.2427126 | 0.83507265 |
| RPL8 | 96.1107168 | -0.3443212 | 0.58389765 | -0.5896945 | 0.55539547 | 0.95415437 |
| WASHC1 | 29.7031283 | 0.07569908 | 0.44753328 | 0.16914737 | 0.86568073 | 0.98751288 |
| DOCK8 | 345.114105 | 0.15274849 | 0.27680788 | 0.5518213 | 0.58107079 | 0.95415437 |
| KANK1 | 21.8373522 | 0.36168959 | 0.64860677 | 0.55764078 | 0.5770897 | 0.95415437 |
| SMARCA2 | 104.529069 | 0.52705097 | 0.29890111 | 1.76329547 | 0.07785067 | 0.62717836 |
| PUM3 | 28.5409314 | -0.5370174 | 0.37604446 | -1.428069 | 0.15327201 | 0.74419469 |
| RFX3 | 109.949954 | -0.5635844 | 0.3346143 | -1.6842808 | 0.09212742 | 0.65902576 |
| RFX3-AS1 | 32.8041129 | -0.5451747 | 0.56125437 | -0.9713505 | 0.33137375 | 0.88037558 |
| GLIS3 | 28.1660618 | -0.1226298 | 0.53112587 | -0.2308865 | 0.817403 | 0.98320463 |
| CDC37L1 | 11.5406941 | -0.0078894 | 0.53301825 | -0.0148013 | 0.98819068 | NA |
| AK3 | 22.5284536 | -0.2197307 | 0.48170732 | -0.4561498 | 0.64828228 | 0.96049083 |
| RCL1 | 13.3034253 | -0.6053937 | 0.49901945 | -1.2131664 | 0.22506619 | 0.81875277 |
| JAK2 | 98.8163932 | 0.03502251 | 0.29381741 | 0.11919822 | 0.90511832 | 0.99324672 |
| PLGRKT | 10.2984163 | -0.6274102 | 0.49972685 | -1.2555063 | 0.20929501 | NA |
| RIC1 | 88.715504 | 0.08246671 | 0.37259985 | 0.22132782 | 0.82483719 | 0.98320463 |
| ERMP1 | 14.7596705 | 0.02495693 | 0.46986045 | 0.05311562 | 0.95763979 | 0.99514836 |
| KIAA2026 | 75.3970507 | 0.33206095 | 0.29761798 | 1.11572881 | 0.26453823 | 0.84994492 |
| UHRF2 | 53.2939448 | -0.9262496 | 0.30209709 | -3.066066 | 0.00216895 | 0.10651853 |
| KDM4C | 179.036192 | -0.0679198 | 0.26481431 | -0.2564807 | 0.79757966 | 0.98320463 |
| NFIB | 13.4988378 | 0.45272362 | 0.7248928 | 0.62453872 | 0.53227385 | 0.95265314 |
| ZDHHC21 | 45.7921993 | -1.3316284 | 0.44496268 | -2.9926743 | 0.00276545 | 0.12544207 |

| | | | | | | |
|---|---|---|---|---|---|---|
| TTC39B | 30.1108222 | 0.54341296 | 0.4888237 | 1.11167475 | 0.26627802 | 0.84994492 |
| SNAPC3 | 27.9474088 | 0.45389735 | 0.41387998 | 1.09668834 | 0.27277766 | 0.84994492 |
| PSIP1 | 25.7540592 | 0.57678066 | 0.49193863 | 1.17246466 | 0.24101055 | 0.83493679 |
| CCDC171 | 11.1099275 | 0.82494933 | 0.71738263 | 1.14994327 | 0.25016724 | NA |
| BNC2 | 66.1861389 | 0.28281223 | 0.49594382 | 0.57025054 | 0.56850778 | 0.95415437 |
| CNTLN | 35.7339863 | 0.64708994 | 0.4255506 | 1.5205946 | 0.1283616 | 0.72782081 |
| HAUS6 | 14.8955723 | -0.9380813 | 0.53779774 | -1.7443014 | 0.08110655 | 0.63497512 |
| PLIN2 | 50.3064808 | -0.1029586 | 0.5664298 | -0.1817677 | 0.85576503 | 0.98751288 |
| DENND4C | 83.5310567 | -0.3787169 | 0.38940145 | -0.9725617 | 0.33077121 | 0.88037558 |
| RPS6 | 124.133819 | -0.3163764 | 0.61034401 | -0.5183576 | 0.6042088 | 0.95688177 |
| MLLT3 | 82.9695523 | 0.12022533 | 0.36258605 | 0.33157738 | 0.7402084 | 0.97605445 |
| FOCAD | 39.6551188 | 0.49037505 | 0.47962612 | 1.02241107 | 0.30658638 | 0.8673137 |
| HACD4 | 14.0632405 | 1.02269461 | 0.56492407 | 1.81032223 | 0.07024583 | 0.60640882 |
| CAAP1 | 16.614219 | -3.87E-01 | 0.42171766 | -0.9183078 | 0.35845776 | 0.8957272 |
| PLAA | 16.901148 | -0.0439919 | 0.45191304 | -0.097346 | 0.92245162 | 0.99514836 |
| IFT74 | 8.47800397 | 0.78366317 | 0.66543029 | 1.17767883 | 0.23892467 | NA |
| MOB3B | 53.0742456 | 0.68017382 | 0.43273747 | 1.57179322 | 0.11599852 | 0.70122655 |
| C9orf72 | 57.7784394 | 0.40855253 | 0.5683958 | 0.71878176 | 0.4722754 | 0.9314919 |
| ACO1 | 11.8159481 | 1.2679996 | 0.64650348 | 1.96131907 | 0.04984181 | NA |
| DDX58 | 42.7970903 | -0.3670707 | 0.76735154 | -0.4783606 | 0.63239358 | 0.95877428 |
| APTX | 15.5787989 | -0.3160277 | 0.52976895 | -0.5965387 | 0.55081544 | 0.95415437 |
| DNAJA1 | 30.9978333 | 0.29025645 | 0.50186383 | 0.57835697 | 0.56302314 | 0.95415437 |
| SMU1 | 15.8915042 | -0.0988613 | 0.44283043 | -0.2232486 | 0.82334202 | 0.98320463 |
| B4GALT1 | 70.8984798 | -0.4213087 | 0.39646731 | -1.0626568 | 0.28793762 | 0.85677513 |
| BAG1 | 9.88429627 | -0.2477938 | 0.54711787 | -0.4529075 | 0.65061535 | NA |
| CHMP5 | 13.0019671 | 0.4227463 | 0.66508091 | 0.63563138 | 0.52501671 | 0.95007724 |
| NFX1 | 49.1363824 | -0.7852827 | 0.31178703 | -2.5186509 | 0.01178054 | 0.29101961 |
| UBE2R2 | 106.636696 | -0.3107232 | 0.31158455 | -0.9972355 | 0.31865019 | 0.87298176 |
| UBAP2 | 38.9959484 | -0.2052599 | 0.3640555 | -0.5638148 | 0.57288019 | 0.95415437 |
| DCAF12 | 13.8642054 | 0.86273627 | 0.54994053 | 1.56878104 | 0.11669897 | 0.70122655 |
| UBAP1 | 53.274089 | 0.07901848 | 0.35861041 | 0.2203463 | 0.82560146 | 0.98320463 |
| FAM219A | 12.775082 | -0.0205575 | 0.7088911 | -0.0289996 | 0.97686495 | NA |
| VCP | 26.8667963 | -0.4326972 | 0.40685781 | -1.0635096 | 0.28755091 | 0.85677513 |
| UNC13B | 14.9669879 | 1.37955931 | 0.71056648 | 1.94149225 | 0.0521986 | 0.54907299 |
| RUSC2 | 12.0638979 | 1.15519787 | 0.71034261 | 1.6262545 | 0.10389553 | NA |
| CD72 | 14.2909528 | -2.3392139 | 0.85192502 | -2.7457979 | 0.00603639 | 0.19198507 |
| CCDC107 | 11.8324002 | 0.00588977 | 0.56074186 | 0.01050353 | 0.99161955 | NA |
| TLN1 | 43.2364748 | -0.2849081 | 0.48796336 | -0.5838718 | 0.55930655 | 0.95415437 |
| GBA2 | 9.03587989 | 1.63678664 | 0.68895165 | 2.37576415 | 0.01751265 | NA |
| RECK | 14.1193224 | -0.5510166 | 0.64492267 | -0.8543918 | 0.39288795 | 0.90930707 |
| GLIPR2 | 24.3873637 | 0.09112326 | 0.49961595 | 0.18238661 | 0.85527933 | 0.98751288 |
| CLTA | 41.1227898 | -0.0232387 | 0.36480337 | -0.063702 | 0.9492075 | 0.99514836 |
| GNE | 11.7020002 | -0.2409478 | 0.5543123 | -0.4346788 | 0.66379561 | NA |
| RNF38 | 100.886205 | -0.1333466 | 0.2827895 | -0.47154 | 0.63725518 | 0.95933464 |
| EBLN3P | 15.9706499 | -0.7662115 | 0.4489001 | -1.7068643 | 0.08784729 | 0.64676224 |
| ZCCHC7 | 209.530987 | -0.9948735 | 0.3918459 | -2.5389407 | 0.01111887 | 0.2861945 |
| GRHPR | 7.98812698 | 0.29872769 | 0.58523946 | 0.51043668 | 0.60974556 | NA |
| AL138752.2 | 17.0904284 | -0.5954157 | 0.58179897 | -1.0234045 | 0.30611665 | 0.8673137 |
| TRMT10B | 16.4973801 | 0.61925971 | 0.45157272 | 1.37133994 | 0.17026901 | 0.75745073 |
| DCAF10 | 38.7035702 | 0.07064978 | 0.42044035 | 0.16803758 | 0.86655372 | 0.98751288 |
| CBWD5 | 16.2605418 | -0.7629718 | 0.52355515 | -1.4572902 | 0.14503629 | 0.73251288 |
| PIP5K1B | 24.5332057 | -2.957399 | 0.77047677 | -3.8384013 | 0.00012384 | 0.02107273 |
| APBA1 | 13.9591774 | -0.6026446 | 0.61987137 | -0.9722091 | 0.33094655 | 0.88037558 |
| PTAR1 | 35.7580547 | 0.0594628 | 0.3686699 | 0.16129008 | 0.87186494 | 0.98751288 |

| | | | | | | |
|---|---|---|---|---|---|---|
| MAMDC2 | 83.7225794 | 0.75358227 | 0.99850985 | 0.75470689 | 0.45042487 | 0.92914597 |
| SMC5 | 64.9897155 | -0.0583554 | 0.31505124 | -0.1852252 | 0.85305242 | 0.98751288 |
| KLF9 | 21.2070681 | -0.7615588 | 0.63473218 | -1.1998112 | 0.23021266 | 0.82393202 |
| CEMIP2 | 149.173308 | -1.1183785 | 0.33965843 | -3.2926564 | 0.00099246 | 0.06639798 |
| ABHD17B | 21.3874248 | -0.2727494 | 0.48123728 | -0.5667669 | 0.57087253 | 0.95415437 |
| C9orf85 | 19.9334157 | 0.30893803 | 0.41965808 | 0.73616605 | 0.46162965 | 0.9314919 |
| ZFAND5 | 46.8198784 | -0.0442545 | 0.36869421 | -0.1200304 | 0.9044591 | 0.99324672 |
| ALDH1A1 | 11.206652 | 2.26869032 | 0.87845226 | 2.58259944 | 0.00980591 | NA |
| ANXA1 | 55.4394285 | -0.1434983 | 0.59119123 | -0.2427274 | 0.80821654 | 0.98320463 |
| CARNMT1 | 19.9493509 | -0.2395939 | 0.40556752 | -0.5907621 | 0.55467985 | 0.95415437 |
| NMRK1 | 16.501314 | 0.14519397 | 0.62778004 | 0.23128159 | 0.81709605 | 0.98320463 |
| OSTF1 | 50.7445504 | 0.29621808 | 0.38671289 | 0.76598968 | 0.44368244 | 0.92899684 |
| PCSK5 | 25.6778945 | 0.22865213 | 0.49145726 | 0.46525333 | 0.64175005 | 0.95992767 |
| GCNT1 | 20.0379608 | -0.3673265 | 0.58445216 | -0.6284972 | 0.5296783 | 0.95245713 |
| PRUNE2 | 12.4371873 | -0.2519599 | 0.6415311 | -0.3927478 | 0.69450577 | NA |
| VPS13A | 41.6138207 | -0.0229033 | 0.31956044 | -0.0716712 | 0.94286357 | 0.99514836 |
| GNAQ | 292.661169 | 1.13242362 | 0.24352225 | 4.65018538 | 3.32E-06 | 0.00171296 |
| CEP78 | 14.9869989 | 0.83305774 | 0.61673545 | 1.35075378 | 0.17677432 | 0.76409729 |
| TLE4 | 84.6514601 | 0.58338981 | 0.35195338 | 1.65757696 | 0.09740288 | 0.66301053 |
| TLE1 | 11.8607831 | 0.80150285 | 0.72922141 | 1.0991214 | 0.27171512 | NA |
| FRMD3 | 19.9153087 | 0.87550887 | 0.59494733 | 1.47157375 | 0.14113602 | 0.73044824 |
| UBQLN1 | 38.6764436 | -0.2215783 | 0.31908966 | -0.6944077 | 0.48742654 | 0.93530459 |
| GKAP1 | 12.0333835 | -0.3443482 | 0.54474063 | -0.6321324 | 0.52730038 | NA |
| HNRNPK | 59.050504 | 0.21465346 | 0.36078285 | 0.59496581 | 0.55186632 | 0.95415437 |
| AGTPBP1 | 164.360939 | -0.1606519 | 0.33945313 | -0.4732669 | 0.63602274 | 0.95933464 |
| NAA35 | 25.743836 | 0.15511398 | 0.43215869 | 0.35892829 | 0.71964873 | 0.97348719 |
| TUT7 | 84.69431 | -0.1344325 | 0.35356149 | -0.3802238 | 0.70377928 | 0.9714177 |
| DAPK1 | 114.529955 | -0.0994715 | 0.31721407 | -0.3135784 | 0.75384123 | 0.97905594 |
| CTSL | 58.2092674 | -0.2118614 | 0.77970272 | -0.2717208 | 0.78583671 | 0.98278427 |
| SPIN1 | 41.9342735 | 0.01364738 | 0.37206327 | 0.03668026 | 0.97073995 | 0.99676559 |
| SECISBP2 | 44.9900475 | -0.2566476 | 0.38968808 | -0.6585975 | 0.51015429 | 0.94332619 |
| SEMA4D | 93.9806884 | 0.93548032 | 0.35994471 | 2.59895558 | 0.00935079 | 0.25873629 |
| SYK | 113.704075 | -0.3933966 | 0.33558923 | -1.1722562 | 0.2410942 | 0.83493679 |
| AL158071.4 | 15.8863765 | 0.73026092 | 0.78604607 | 0.92903069 | 0.35287318 | 0.89109867 |
| AUH | 53.2371807 | 0.74519789 | 0.44460698 | 1.6760823 | 0.09372207 | 0.6590317 |
| NFIL3 | 15.7596441 | 0.40056254 | 0.62106486 | 0.64496089 | 0.51895253 | 0.94668603 |
| SPTLC1 | 29.805997 | 0.01643042 | 0.37407598 | 0.04392268 | 0.96496604 | 0.99653147 |
| IARS | 26.7021705 | -0.6527542 | 0.5176713 | -1.2609433 | 0.20732927 | 0.79343028 |
| NOL8 | 15.9371589 | -0.4813995 | 0.47834885 | -1.0063775 | 0.314234 | 0.87152625 |
| CENPP | 51.5313699 | -0.7607395 | 0.51798589 | -1.468649 | 0.14192802 | 0.73044824 |
| IPPK | 8.8450192 | 0.01829159 | 0.74879275 | 0.02442811 | 0.98051113 | NA |
| BICD2 | 17.974613 | 0.2470304 | 0.47272944 | 0.52256191 | 0.60127916 | 0.95688177 |
| FGD3 | 62.6870577 | 0.38992531 | 0.36695569 | 1.06259508 | 0.28796562 | 0.85677513 |
| CARD19 | 9.70862007 | -0.6164979 | 0.56528861 | -1.0905897 | 0.27545346 | NA |
| NINJ1 | 16.4195973 | 1.29089932 | 0.57609034 | 2.2407932 | 0.02503947 | 0.39704426 |
| FAM120A | 73.2111854 | -0.1951836 | 0.27149905 | -0.7189106 | 0.47219599 | 0.9314919 |
| PHF2 | 16.0858402 | -0.7907868 | 0.48558813 | -1.6285134 | 0.10341607 | 0.67250824 |
| AL158152.2 | 12.699406 | -0.5289465 | 0.52566319 | -1.006246 | 0.31429726 | NA |
| ZNF169 | 11.7493212 | -0.110032 | 0.59743468 | -0.184174 | 0.85387692 | NA |
| MFSD14B | 30.7734053 | 0.07516161 | 0.43161603 | 0.17414 | 0.86175544 | 0.98751288 |
| FBP1 | 11.9550649 | 1.83525303 | 0.78083782 | 2.3503639 | 0.01875506 | NA |
| C9orf3 | 58.1021198 | -0.04427 | 0.43182751 | -0.1025178 | 0.9183457 | 0.99498068 |
| FANCC | 17.409787 | 0.30660089 | 0.56409834 | 0.54352383 | 0.58676919 | 0.95415437 |
| LINC00476 | 14.5193225 | 0.06453647 | 0.6874661 | 0.09387586 | 0.92520777 | 0.99514836 |

| | | | | | | |
|---|---|---|---|---|---|---|
| ERCC6L2 | 35.4404619 | -0.1208364 | 0.38330493 | -0.3152488 | 0.75257276 | 0.97875726 |
| HABP4 | 11.9244025 | -0.6575147 | 0.4934866 | -1.332386 | 0.18273336 | NA |
| ZNF782 | 17.1793889 | 0.40970328 | 0.54670837 | 0.74940004 | 0.45361612 | 0.9314919 |
| MFSD14C | 59.1178521 | -0.4527419 | 0.35185386 | -1.2867328 | 0.19818744 | 0.78142991 |
| TDRD7 | 18.0924705 | -0.509208 | 0.50726121 | -1.0038379 | 0.31545675 | 0.87263466 |
| TSTD2 | 13.2317378 | -0.0648679 | 0.77202704 | -0.0840228 | 0.93303828 | 0.99514836 |
| NCBP1 | 12.6139409 | 0.34951846 | 0.55148944 | 0.63377181 | 0.52622975 | NA |
| XPA | 13.2604704 | -0.4523651 | 0.62656547 | -0.7219758 | 0.47030938 | 0.9314919 |
| ANP32B | 27.3145645 | -0.2466554 | 0.39100331 | -0.630827 | 0.52815365 | 0.95174286 |
| NANS | 9.49193179 | -0.0315171 | 0.60677242 | -0.0519422 | 0.95857473 | NA |
| TRIM14 | 47.5018104 | 0.81968511 | 0.41212014 | 1.98894697 | 0.04670706 | 0.52744812 |
| CORO2A | 8.31421849 | 0.30688069 | 0.57792389 | 0.53100537 | 0.59541506 | NA |
| TBC1D2 | 21.1986322 | 0.92205655 | 0.49202759 | 1.87399357 | 0.06093131 | 0.57281427 |
| TGFBR1 | 58.9247157 | -0.0319873 | 0.45561799 | -0.0702065 | 0.94402933 | 0.99514836 |
| SEC61B | 31.6611543 | 0.01282755 | 0.52313204 | 0.02452067 | 0.98043729 | 0.99758514 |
| STX17 | 17.1189335 | -0.4365872 | 0.45957103 | -0.9499886 | 0.34211804 | 0.88510795 |
| ERP44 | 46.4417477 | -0.0397315 | 0.32846125 | -0.1209624 | 0.90372079 | 0.99324672 |
| INVS | 60.2937049 | 0.693265 | 0.29573291 | 2.34422676 | 0.01906658 | 0.3601175 |
| TEX10 | 10.9455476 | -1.1312789 | 0.56869044 | -1.9892701 | 0.0466714 | NA |
| ALDOB | 29.7691237 | 1.74492857 | 1.07716475 | 1.61992729 | 0.1052479 | 0.67519097 |
| RNF20 | 10.4867362 | -0.2180655 | 0.54560939 | -0.3996733 | 0.68939719 | NA |
| ABCA1 | 93.6942928 | 0.31660774 | 0.67354793 | 0.4700597 | 0.63831236 | 0.95933464 |
| SLC44A1 | 48.6281755 | 0.1461323 | 0.46761157 | 0.31250787 | 0.75465458 | 0.97942271 |
| FSD1L | 15.6568246 | -0.4671595 | 0.53517319 | -0.8729127 | 0.38271068 | 0.90868001 |
| FKTN | 11.6853483 | -0.5639024 | 0.51951697 | -1.085436 | 0.2777286 | NA |
| RAD23B | 44.5714801 | -0.1304468 | 0.32534443 | -0.40095 | 0.68845696 | 0.9673431 |
| ELP1 | 13.4303343 | 0.64763319 | 0.51898347 | 1.24788788 | 0.21207212 | 0.80329028 |
| TMEM245 | 41.8941835 | -0.0282238 | 0.34253126 | -0.0823976 | 0.93433053 | 0.99514836 |
| TXN | 21.4615143 | 0.17986515 | 0.53804117 | 0.33429626 | 0.73815601 | 0.97605445 |
| LPAR1 | 14.5072781 | 1.40084022 | 0.97889229 | 1.43104632 | 0.15241694 | 0.74315008 |
| ECPAS | 50.5610947 | -0.0163849 | 0.3809397 | -0.0430118 | 0.96569211 | 0.99653147 |
| PTGR1 | 12.2163683 | 0.24258837 | 0.72858018 | 0.33296043 | 0.73916415 | NA |
| C9orf84 | 16.3478758 | 0.78022376 | 1.03041183 | 0.75719604 | 0.44893242 | 0.92914597 |
| UGCG | 62.4906285 | 0.03115723 | 0.64042205 | 0.04865109 | 0.96119735 | 0.99614757 |
| SUSD1 | 43.223059 | -0.2856384 | 0.45673849 | -0.6253871 | 0.53171702 | 0.95265314 |
| PTBP3 | 112.268931 | -0.0495739 | 0.25236031 | -0.1964409 | 0.84426512 | 0.9842451 |
| HSDL2 | 26.7684743 | 0.67242786 | 0.44136164 | 1.52353035 | 0.12762607 | 0.72782081 |
| KIAA1958 | 29.55564 | 0.95437037 | 0.50498619 | 1.88989399 | 0.05877214 | 0.57281427 |
| INIP | 13.6525755 | 0.29150437 | 0.55219946 | 0.52789688 | 0.59757091 | 0.95538391 |
| SNX30 | 33.5300538 | 0.45925765 | 0.37456086 | 1.22612289 | 0.22015243 | 0.81059452 |
| FKBP15 | 44.2829069 | 0.51417046 | 0.42378208 | 1.21328976 | 0.22501906 | 0.81875277 |
| SLC31A1 | 16.7792883 | 0.77566514 | 0.61905869 | 1.25297512 | 0.21021476 | 0.79953848 |
| CDC26 | 8.59231158 | 0.73841082 | 0.8507153 | 0.86798817 | 0.38540081 | NA |
| RGS3 | 10.7098347 | 0.22563937 | 0.68635225 | 0.32875156 | 0.74234348 | NA |
| ZNF618 | 15.0614546 | 1.0679274 | 0.72605237 | 1.47086828 | 0.14132674 | 0.73044824 |
| AKNA | 93.7709809 | -0.7206732 | 0.41780853 | -1.7248886 | 0.08454755 | 0.64676224 |
| ATP6V1G1 | 28.6925473 | -0.1767757 | 0.52368032 | -0.3375641 | 0.73569172 | 0.97605445 |
| TMEM268 | 12.9196881 | -1.3193123 | 0.74343359 | -1.7746202 | 0.07596061 | NA |
| TNFSF8 | 19.0949341 | 0.89568216 | 0.54515544 | 1.6429849 | 0.10038606 | 0.6683775 |
| ASTN2 | 11.3075622 | -0.433447 | 0.65338972 | -0.663382 | 0.50708596 | NA |
| CDK5RAP2 | 43.5804668 | -0.2391036 | 0.29329501 | -0.8152325 | 0.41493927 | 0.91894074 |
| MEGF9 | 78.4103772 | 0.46489679 | 0.42575956 | 1.0919233 | 0.27486681 | 0.84994492 |
| FBXW2 | 23.7196581 | -0.7287542 | 0.44229771 | -1.6476554 | 0.09942341 | 0.6683775 |
| PSMD5 | 17.7465756 | -0.0475246 | 0.40643974 | -0.1169291 | 0.90691621 | 0.99368779 |

| | | | | | | |
|---|---|---|---|---|---|---|
| TRAF1 | 17.513269 | -0.0940345 | 0.47941847 | -0.1961429 | 0.84449833 | 0.98429944 |
| CNTRL | 74.3840827 | -0.2697389 | 0.28543226 | -0.945019 | 0.34464916 | 0.8871109 |
| RAB14 | 36.1930928 | 0.49607085 | 0.34156261 | 1.45235704 | 0.14640236 | 0.73411592 |
| GSN | 39.3651204 | -0.2626218 | 0.54984131 | -0.4776321 | 0.6329121 | 0.95877428 |
| STOM | 38.9386042 | 0.36493773 | 0.33642787 | 1.08474288 | 0.27803555 | 0.84994492 |
| MRRF | 12.1865809 | -0.0965328 | 0.50645468 | -0.1906049 | 0.84883513 | NA |
| RC3H2 | 46.6295014 | -0.0229427 | 0.29651302 | -0.0773749 | 0.93832532 | 0.99514836 |
| RABGAP1 | 63.0633255 | -0.223664 | 0.30039984 | -0.7445543 | 0.4565412 | 0.9314919 |
| STRBP | 47.0375394 | -1.2287644 | 0.66540103 | -1.8466524 | 0.06479754 | 0.58881703 |
| DENND1A | 186.976339 | 0.50982278 | 0.25471669 | 2.00152874 | 0.04533544 | 0.52128421 |
| NEK6 | 26.5577246 | 0.25802147 | 0.50357093 | 0.51238357 | 0.60838258 | 0.95688177 |
| PSMB7 | 25.7808864 | 0.23512318 | 0.42714271 | 0.5504558 | 0.58200679 | 0.95415437 |
| NR6A1 | 28.6066666 | 0.41633817 | 0.55144447 | 0.75499564 | 0.4502516 | 0.92914597 |
| RPL35 | 45.3148299 | -0.5609379 | 0.57647601 | -0.9730463 | 0.33053029 | 0.88037558 |
| ARPC5L | 16.5098644 | -1.0476031 | 0.50335638 | -2.0812355 | 0.03741236 | 0.47208194 |
| GOLGA1 | 15.2386347 | 0.9398743 | 0.47570949 | 1.97573166 | 0.04818518 | 0.53331354 |
| SCAI | 27.0115478 | 0.89857921 | 0.43036741 | 2.08793505 | 0.03680369 | 0.46928949 |
| PPP6C | 25.361445 | -0.3173705 | 0.41539301 | -0.7640248 | 0.44485248 | 0.92914597 |
| HSPA5 | 39.3932389 | 0.22366368 | 0.65178801 | 0.34315403 | 0.73148259 | 0.97605445 |
| GAPVD1 | 48.4553481 | -0.2870303 | 0.28388636 | -1.0110748 | 0.31198062 | 0.87021207 |
| MAPKAP1 | 83.7746551 | 0.01632227 | 0.24223577 | 0.06738174 | 0.94627781 | 0.99514836 |
| PBX3 | 108.072153 | 1.61151442 | 0.53893508 | 2.99018281 | 0.00278811 | 0.12544207 |
| MVB12B | 61.8080545 | 0.86503278 | 0.49888473 | 1.73393317 | 0.08292994 | 0.63740875 |
| ZBTB43 | 19.2424824 | -0.1812773 | 0.50596656 | -0.3582793 | 0.72013433 | 0.97390601 |
| RALGPS1 | 16.8632922 | -0.4749633 | 0.49792185 | -0.9538913 | 0.34013866 | 0.8844375 |
| RPL12 | 20.9261003 | -0.6550803 | 0.46231112 | -1.4169685 | 0.15649213 | 0.74486619 |
| LRSAM1 | 19.7313358 | -0.2316829 | 0.40928585 | -0.5660661 | 0.57134884 | 0.95415437 |
| FAM129B | 14.8913625 | 0.15254329 | 0.64587547 | 0.23618065 | 0.8132925 | 0.98320463 |
| SH2D3C | 35.6090262 | 0.22000769 | 0.51992099 | 0.42315602 | 0.67218143 | 0.96412305 |
| CDK9 | 9.90111094 | -0.4816766 | 0.6166079 | -0.7811716 | 0.43470155 | NA |
| FPGS | 11.5052376 | -0.6905724 | 0.67629263 | -1.0211148 | 0.30720007 | NA |
| ENG | 24.250327 | -0.009244 | 0.43749475 | -0.0211295 | 0.98314239 | 0.99763261 |
| FAM102A | 15.2461841 | 0.306831 | 0.60658398 | 0.50583434 | 0.61297295 | 0.95688177 |
| SLC25A25 | 17.8859977 | -0.2135477 | 0.51891212 | -0.4115296 | 0.68068421 | 0.9673431 |
| C9orf16 | 13.3062774 | -0.2198328 | 0.55214909 | -0.3981403 | 0.69052679 | 0.9681721 |
| GOLGA2 | 12.2140481 | 0.32267746 | 0.48353346 | 0.66733222 | 0.50455995 | NA |
| SLC27A4 | 12.4903926 | -0.5323744 | 0.54778661 | -0.9718645 | 0.33111796 | NA |
| URM1 | 9.10114036 | -0.7349626 | 0.57209232 | -1.2846923 | 0.19889984 | NA |
| ODF2 | 8.09880437 | -0.5956772 | 0.5806388 | -1.0258998 | 0.30493881 | NA |
| GLE1 | 13.4551417 | -0.050277 | 0.4797034 | -0.1048085 | 0.91652779 | 0.99498068 |
| SPTAN1 | 34.9932286 | -0.2142231 | 0.39371482 | -0.5441073 | 0.58636765 | 0.95415437 |
| SET | 34.9839309 | -0.1187311 | 0.48139922 | -0.2466375 | 0.80518877 | 0.98320463 |
| ZER1 | 19.268749 | -0.0919991 | 0.51131361 | -0.179927 | 0.85720991 | 0.98751288 |
| TBC1D13 | 8.18380401 | 0.04577194 | 0.58527437 | 0.07820595 | 0.93766423 | NA |
| NUP188 | 11.7343635 | -0.1959803 | 0.51386338 | -0.3813859 | 0.7029169 | NA |
| PTPA | 11.0414015 | 0.43631177 | 0.58300065 | 0.74838985 | 0.45422504 | NA |
| C9orf78 | 16.1696661 | 0.06568921 | 0.45614814 | 0.14400851 | 0.88549375 | 0.99019008 |
| USP20 | 14.7009951 | -0.2923062 | 0.47202018 | -0.6192664 | 0.53574087 | 0.95392212 |
| FNBP1 | 250.413522 | 0.01903992 | 0.27487931 | 0.06926647 | 0.94477751 | 0.99514836 |
| GPR107 | 37.2600933 | 0.28504734 | 0.31971813 | 0.89155824 | 0.37262976 | 0.9025457 |
| FUBP3 | 26.1319697 | -0.1824378 | 0.35199247 | -0.5183003 | 0.60424874 | 0.95688177 |
| ABL1 | 62.9014684 | 0.03262944 | 0.28194116 | 0.11573139 | 0.90786543 | 0.99389264 |
| NUP214 | 42.8747024 | -0.5819145 | 0.372954 | -1.5602849 | 0.11869257 | 0.70431251 |
| AL157938.2 | 9.90859799 | 0.6822689 | 0.62275353 | 1.0955681 | 0.27326783 | NA |

| | | | | | | |
|---|---|---|---|---|---|---|
| FAM78A | 14.7366654 | 0.23959882 | 0.51198717 | 0.46797818 | 0.6398002 | 0.95952691 |
| PRRC2B | 72.5999112 | 0.0389642 | 0.30143206 | 0.12926364 | 0.89714905 | 0.99137525 |
| RAPGEF1 | 232.111138 | -0.3701965 | 0.31938406 | -1.1590951 | 0.24641744 | 0.83817091 |
| MED27 | 42.2260339 | -0.0676799 | 0.37616127 | -0.1799226 | 0.85721333 | 0.98751288 |
| SETX | 125.007781 | -0.0033903 | 0.2656926 | -0.0127603 | 0.98981902 | 0.99811561 |
| TTF1 | 16.3843526 | 0.17614105 | 0.44729339 | 0.39379309 | 0.69373381 | 0.9685713 |
| DDX31 | 8.72822157 | 0.13893845 | 0.64284847 | 0.2161294 | 0.82888688 | NA |
| TSC1 | 15.8203003 | -0.4973974 | 0.47664473 | -1.043539 | 0.29669873 | 0.8608685 |
| RALGDS | 10.6027876 | -0.2180048 | 0.57788098 | -0.3772486 | 0.7059889 | NA |
| RPL7A | 58.5298024 | -0.2775294 | 0.57968287 | -0.4787608 | 0.63210879 | 0.95877428 |
| SURF4 | 21.9050917 | -0.1413551 | 0.44445308 | -0.3180429 | 0.75045243 | 0.97818861 |
| VAV2 | 21.1845055 | -0.3407958 | 0.55643125 | -0.6124671 | 0.54022879 | 0.95415437 |
| BRD3 | 10.6524949 | 0.43398631 | 0.59299908 | 0.73184989 | 0.4642602 | NA |
| RXRA | 27.1017968 | 0.75585329 | 0.58314399 | 1.29616922 | 0.19491719 | 0.78135737 |
| FCN1 | 71.8768484 | 2.42109823 | 1.46755485 | 1.64974974 | 0.09899413 | 0.66763865 |
| CAMSAP1 | 15.8315534 | -0.3651492 | 0.56656295 | -0.644499 | 0.51925193 | 0.94680071 |
| UBAC1 | 11.3981183 | 1.45053064 | 0.66736302 | 2.17352564 | 0.02974078 | NA |
| NACC2 | 21.3621911 | 0.32025638 | 0.51238378 | 0.62503223 | 0.53194991 | 0.95265314 |
| SEC16A | 20.2835349 | 0.20513548 | 0.44766113 | 0.4582383 | 0.64678125 | 0.96049083 |
| NOTCH1 | 12.9861156 | 0.32957818 | 0.83541533 | 0.39450818 | 0.69320589 | 0.9685713 |
| AGPAT2 | 10.4572115 | 0.89091667 | 0.62754693 | 1.41968135 | 0.15570047 | NA |
| SNHG7 | 11.0589069 | -0.1787226 | 0.56258016 | -0.3176838 | 0.75072481 | NA |
| RABL6 | 22.4855706 | 0.26344826 | 0.44001423 | 0.59872668 | 0.54935516 | 0.95415437 |
| PHPT1 | 13.6646892 | 0.21219832 | 0.58642761 | 0.36184914 | 0.71746477 | 0.97348719 |
| EDF1 | 22.4098363 | 0.73990263 | 0.61069158 | 1.21158152 | 0.22567261 | 0.81875277 |
| FBXW5 | 17.2089119 | 0.28392437 | 0.52963558 | 0.53607495 | 0.59190675 | 0.95415437 |
| PAXX | 9.65036867 | -0.8025074 | 0.64653 | -1.2412531 | 0.21451227 | NA |
| MAN1B1 | 11.380595 | -0.3984726 | 0.47516068 | -0.8386061 | 0.40169041 | NA |
| DPP7 | 13.5100586 | -0.3316168 | 0.57893244 | -0.5728075 | 0.56677506 | 0.95415437 |
| SSNA1 | 8.58888747 | -0.1600659 | 0.78143797 | -0.2048351 | 0.83770098 | NA |
| TUBB4B | 11.5639386 | 1.01017373 | 0.64937536 | 1.55560834 | 0.11980126 | NA |
| NELFB | 14.3543758 | -0.8523135 | 0.49943778 | -1.7065459 | 0.0879065 | 0.64676224 |
| EXD3 | 10.2281196 | -0.5767475 | 0.57519399 | -1.0027008 | 0.31600522 | NA |
| MRPL41 | 18.2319104 | 0.20942677 | 0.48422928 | 0.43249506 | 0.66538164 | 0.96406914 |
| ARRDC1 | 12.0095773 | 0.25893324 | 0.53039493 | 0.48818951 | 0.62541561 | NA |
| EHMT1 | 116.185995 | -0.6226267 | 0.33881492 | -1.8376603 | 0.06611248 | 0.58951068 |
| PSMD13 | 16.2151344 | 0.16961603 | 0.49781816 | 0.34071886 | 0.73331524 | 0.97605445 |
| PGGHG | 14.1591887 | -0.106856 | 0.55888879 | -0.1911936 | 0.8483739 | 0.98545782 |
| IFITM2 | 88.1629998 | 0.43821861 | 0.60735717 | 0.72151713 | 0.4705914 | 0.9314919 |
| IFITM3 | 71.4424707 | 0.62733311 | 0.55057932 | 1.13940552 | 0.25453406 | 0.84026145 |
| SIGIRR | 13.9954731 | 0.33597832 | 0.50306526 | 0.66786229 | 0.5042215 | 0.94166217 |
| RNH1 | 16.132513 | 1.08249257 | 0.51843009 | 2.08802033 | 0.036796 | 0.46928949 |
| PHRF1 | 15.6596236 | -0.964483 | 0.58338655 | -1.6532487 | 0.09828027 | 0.66570745 |
| DEAF1 | 16.5308674 | -1.1356051 | 0.4196833 | -2.7058621 | 0.00681273 | 0.20602004 |
| TALDO1 | 27.4542924 | 0.05062153 | 0.59922105 | 0.08447888 | 0.93267569 | 0.99514836 |
| RPLP2 | 64.9045331 | -0.2049196 | 0.54469526 | -0.3762096 | 0.70676108 | 0.97184211 |
| PNPLA2 | 9.4579923 | 0.80399732 | 0.62953957 | 1.2771196 | 0.20156003 | NA |
| CD151 | 13.695014 | 0.32224178 | 0.62139313 | 0.51857957 | 0.60405396 | 0.95688177 |
| POLR2L | 18.9480775 | -0.1633345 | 0.60027208 | -0.2721008 | 0.78554453 | 0.98264092 |
| AP2A2 | 63.5249469 | -0.2904837 | 0.43688053 | -0.6649042 | 0.50611176 | 0.94166217 |
| TOLLIP | 12.3994076 | -0.7386767 | 0.62831458 | -1.1756479 | 0.23973561 | NA |
| MOB2 | 9.33779823 | 0.0347451 | 0.52437338 | 0.06626022 | 0.94717065 | NA |
| LSP1 | 27.1955662 | -0.5573338 | 0.56164255 | -0.9923283 | 0.3210374 | 0.87363761 |
| TSPAN32 | 10.9265434 | 0.57488961 | 0.74519047 | 0.77146667 | 0.44043037 | NA |

| | | | | | | |
|---|---|---|---|---|---|---|
| CD81 | 48.7072052 | 0.09266408 | 0.42999414 | 0.21550081 | 0.82937688 | 0.98320463 |
| KCNQ1 | 53.1912948 | 0.0726961 | 0.41521401 | 0.17508103 | 0.86101596 | 0.98751288 |
| KCNQ1OT1 | 25.6234715 | 0.32581994 | 0.82001213 | 0.39733552 | 0.69112006 | 0.96826795 |
| CDKN1C | 13.9929982 | 1.49682588 | 0.94571787 | 1.58274041 | 0.11348064 | 0.69801324 |
| NAP1L4 | 73.8464478 | 0.23870173 | 0.28832366 | 0.82789506 | 0.40772993 | 0.9169561 |
| CARS | 18.8733728 | -0.0761147 | 0.45674045 | -0.1666476 | 0.86764737 | 0.98751288 |
| OSBPL5 | 22.4306874 | 3.3614895 | 0.97088874 | 3.46228084 | 0.00053562 | 0.04491075 |
| ZNF195 | 12.6989053 | -1.8466364 | 0.48896296 | -3.7766387 | 0.00015896 | NA |
| NUP98 | 60.1304534 | -0.1528091 | 0.26156385 | -0.5842133 | 0.5590768 | 0.95415437 |
| RHOG | 22.8120314 | 0.15861896 | 0.70363533 | 0.22542779 | 0.8216465 | 0.98320463 |
| STIM1 | 119.54472 | -0.2649047 | 0.25696978 | -1.0308787 | 0.30259773 | 0.86318342 |
| HBB | 29.4108007 | -1.3129032 | 1.861801 | -0.7051791 | 0.48069878 | 0.93274442 |
| TRIM5 | 13.0413301 | 0.75959224 | 0.67616863 | 1.12337693 | 0.26127744 | 0.8475436 |
| TRIM22 | 58.5262252 | 0.00082272 | 0.5098952 | 0.00161351 | 0.99871261 | 0.9992543 |
| AC084337.2 | 9.80916429 | 0.49915333 | 0.56458957 | 0.88409945 | 0.37664253 | NA |
| DNHD1 | 16.3338852 | -0.0069917 | 0.54498112 | -0.0128293 | 0.98976397 | 0.99811561 |
| ILK | 13.3973283 | 0.56690633 | 0.57220061 | 0.99074751 | 0.32180889 | 0.87363761 |
| TAF10 | 9.50702856 | -0.2251909 | 0.58011795 | -0.3881812 | 0.69788197 | NA |
| TPP1 | 25.7382299 | 0.11978907 | 0.51045417 | 0.23467154 | 0.81446368 | 0.98320463 |
| PPFIBP2 | 21.1213621 | -1.0121814 | 0.57915931 | -1.7476736 | 0.08052057 | 0.63385609 |
| EIF3F | 17.6571297 | 0.90998619 | 0.53510735 | 1.70056754 | 0.08902422 | 0.64845882 |
| TRIM66 | 17.6521152 | 0.21826842 | 0.50456612 | 0.43258636 | 0.6653153 | 0.96406914 |
| RPL27A | 72.5501931 | -0.0174841 | 0.61748992 | -0.0283148 | 0.97741111 | 0.99758514 |
| TMEM9B | 11.6664384 | 0.59824034 | 0.56299442 | 1.06260439 | 0.2879614 | NA |
| DENND5A | 74.0800182 | 0.35784452 | 0.4350597 | 0.82251818 | 0.41078204 | 0.91823911 |
| TMEM41B | 22.9090824 | -0.3150625 | 0.3963295 | -0.7949508 | 0.42664212 | 0.92112288 |
| IPO7 | 36.3158839 | -0.6798044 | 0.39328654 | -1.728522 | 0.08389468 | 0.64303763 |
| ZNF143 | 18.9980404 | -0.4248266 | 0.46614935 | -0.911353 | 0.36210938 | 0.89853007 |
| SWAP70 | 54.4727897 | -0.923778 | 0.48970184 | -1.886409 | 0.05923986 | 0.57281427 |
| SBF2 | 205.755622 | 0.69688746 | 0.40131524 | 1.73650884 | 0.0824739 | 0.63740875 |
| AMPD3 | 22.9668917 | -0.0210103 | 0.53201007 | -0.0394924 | 0.96849783 | 0.99653147 |
| MTRNR2L8 | 19.7981106 | 0.85377537 | 0.81829543 | 1.04335835 | 0.29678236 | 0.8608685 |
| RNF141 | 10.7796801 | -1.4888387 | 0.65842309 | -2.2612189 | 0.02374571 | NA |
| LYVE1 | 28.9710757 | 0.40471832 | 1.13583115 | 0.35631909 | 0.72160161 | 0.97417403 |
| CTR9 | 18.688953 | 0.30144689 | 0.52816331 | 0.5707456 | 0.5681721 | 0.95415437 |
| EIF4G2 | 48.3921259 | -0.081696 | 0.41200367 | -0.1982896 | 0.84281849 | 0.9842451 |
| ZBED5 | 16.3540819 | -0.9573596 | 0.43885092 | -2.1815145 | 0.02914539 | 0.42316846 |
| USP47 | 91.9293535 | -0.2099084 | 0.25734254 | -0.8156772 | 0.41468479 | 0.91894074 |
| MICAL2 | 23.9390476 | 0.66417548 | 0.68941048 | 0.96339627 | 0.3353487 | 0.88047896 |
| ARNTL | 34.9611898 | -0.0335651 | 0.42194669 | -0.0795482 | 0.93659659 | 0.99514836 |
| BTBD10 | 30.5654658 | 0.19790293 | 0.40444025 | 0.48932551 | 0.62461126 | 0.95876809 |
| FAR1 | 77.9521152 | 0.12310652 | 0.30445987 | 0.40434399 | 0.68595979 | 0.9673431 |
| RRAS2 | 30.3406086 | -0.2416597 | 0.50264988 | -0.4807714 | 0.63067901 | 0.95877428 |
| COPB1 | 35.6025756 | -0.113425 | 0.35826678 | -0.3165938 | 0.75155186 | 0.97857378 |
| PSMA1 | 41.4630038 | 0.35145307 | 0.57156448 | 0.61489662 | 0.538623 | 0.95415437 |
| PDE3B | 240.194385 | 0.43370033 | 0.5491618 | 0.78974964 | 0.429674 | 0.92204028 |
| SOX6 | 10.5094022 | -1.0019802 | 0.55925924 | -1.7916203 | 0.0731938 | NA |
| C11orf58 | 35.7659616 | 0.05123161 | 0.31665538 | 0.1617898 | 0.87147139 | 0.98751288 |
| PLEKHA7 | 91.3959531 | -1.1438181 | 0.54071677 | -2.1153738 | 0.0343981 | 0.45650836 |
| RPS13 | 58.6653076 | -0.6831491 | 0.62224913 | -1.0978707 | 0.27226097 | 0.84994492 |
| PIK3C2A | 51.147669 | -0.0117304 | 0.32077653 | -0.0365688 | 0.97082879 | 0.99676559 |
| NUCB2 | 15.5016216 | 0.45338432 | 0.71669196 | 0.63260695 | 0.52699035 | 0.95136478 |
| SERGEF | 40.1437086 | 0.91428579 | 0.3889232 | 2.3508132 | 0.01873244 | 0.3571212 |
| HPS5 | 22.1433742 | 0.03433555 | 0.59164998 | 0.05803355 | 0.95372191 | 0.99514836 |

| | | | | | | |
|---|---|---|---|---|---|---|
| GTF2H1 | 20.0634029 | 0.10623916 | 0.5288078 | 0.20090316 | 0.84077429 | 0.98380297 |
| LDHA | 41.6651762 | 0.31383284 | 0.39992045 | 0.78473817 | 0.4326071 | 0.92284736 |
| TSG101 | 26.4178048 | -0.4999079 | 0.36525288 | -1.3686625 | 0.1711048 | 0.75751515 |
| UEVLD | 18.7189932 | 0.23123995 | 0.4686329 | 0.49343516 | 0.62170514 | 0.95702817 |
| SPTY2D1 | 11.9098668 | 0.43720043 | 0.54087685 | 0.80831788 | 0.41890761 | NA |
| ZDHHC13 | 16.3909532 | -0.0272919 | 0.45307118 | -0.0602376 | 0.95196638 | 0.99514836 |
| NAV2 | 28.2803388 | 0.7406299 | 0.71190389 | 1.04035097 | 0.29817687 | 0.8608685 |
| PRMT3 | 25.2612897 | 0.00189492 | 0.45426373 | 0.00417141 | 0.99667171 | 0.99865675 |
| LGR4 | 11.0171174 | -0.139898 | 0.73242066 | -0.1910077 | 0.84851956 | NA |
| BDNF-AS | 30.8365353 | -0.6586123 | 0.58031002 | -1.1349318 | 0.25640389 | 0.8406233 |
| METTL15 | 52.3152205 | 0.05583279 | 0.39184798 | 0.14248586 | 0.88669625 | 0.99019008 |
| ARL14EP | 12.0440059 | -0.2468084 | 0.56801022 | -0.434514 | 0.66391521 | NA |
| DNAJC24 | 16.3429971 | 0.03098132 | 0.47553461 | 0.0651505 | 0.94805417 | 0.99514836 |
| IMMP1L | 19.6238725 | 0.08086253 | 0.45468963 | 0.17784115 | 0.85884773 | 0.98751288 |
| ELP4 | 55.125005 | 0.32525315 | 0.29443082 | 1.10468443 | 0.26929635 | 0.84994492 |
| EIF3M | 20.8307825 | -0.4200043 | 0.52221319 | -0.8042775 | 0.42123671 | 0.91985469 |
| QSER1 | 30.4534441 | -0.450076 | 0.39403779 | -1.1422154 | 0.2533645 | 0.84026145 |
| TCP11L1 | 16.2814903 | -0.0185629 | 0.46772372 | -0.0396878 | 0.96834205 | 0.99653147 |
| CSTF3 | 25.5795898 | 0.1191942 | 0.42923878 | 0.27768741 | 0.78125233 | 0.98264092 |
| HIPK3 | 86.635501 | 0.73055481 | 0.32974181 | 2.2155359 | 0.0267233 | 0.40712515 |
| FBXO3 | 16.3898811 | 0.29707027 | 0.45024252 | 0.65980057 | 0.50938182 | 0.94332619 |
| LMO2 | 13.5551984 | 1.66474737 | 0.77640979 | 2.14416072 | 0.03202001 | 0.43969912 |
| CAPRIN1 | 42.7429485 | 0.23094822 | 0.36485119 | 0.63299292 | 0.52673826 | 0.95136478 |
| NAT10 | 10.1367977 | -0.7843355 | 0.57428845 | -1.3657518 | 0.17201687 | NA |
| ABTB2 | 21.2544662 | -1.0767066 | 0.93513987 | -1.1513856 | 0.24957363 | 0.83865855 |
| CAT | 34.6969021 | 1.40839108 | 0.66572721 | 2.11556784 | 0.03438158 | 0.45650836 |
| APIP | 12.325192 | -0.7306411 | 0.55893877 | -1.3071935 | 0.191147 | NA |
| PDHX | 18.6434621 | -0.5322423 | 0.41516081 | -1.2820149 | 0.19983743 | 0.78249307 |
| CD44 | 207.2445 | 0.07314087 | 0.36594519 | 0.19986837 | 0.84158353 | 0.98380297 |
| TRIM44 | 60.4650922 | -0.5419932 | 0.40116453 | -1.3510496 | 0.17667956 | 0.76409729 |
| LDLRAD3 | 35.0017938 | 2.11650059 | 0.59565328 | 3.55324255 | 0.00038051 | 0.03630623 |
| PRR5L | 31.7421878 | 0.42999653 | 0.58208991 | 0.73871155 | 0.46008217 | 0.9314919 |
| TRAF6 | 10.0176128 | 0.04696277 | 0.54525887 | 0.08612931 | 0.93136362 | NA |
| API5 | 22.1935161 | -0.2376067 | 0.44750286 | -0.5309614 | 0.59544554 | 0.9553365 |
| TTC17 | 136.553433 | -0.0274904 | 0.26896578 | -0.1022077 | 0.9185918 | 0.99498068 |
| HSD17B12 | 54.8854018 | -0.8380728 | 0.34340429 | -2.4404843 | 0.01466758 | 0.32121732 |
| EXT2 | 22.5376479 | -0.1182158 | 0.3870892 | -0.3053969 | 0.76006392 | 0.9808103 |
| CD82 | 25.124635 | -1.0429477 | 0.70414689 | -1.4811507 | 0.13856642 | 0.73044824 |
| PHF21A | 181.669308 | 0.41103446 | 0.25266235 | 1.62681329 | 0.10377676 | 0.67327148 |
| DGKZ | 27.4847585 | 0.56785682 | 0.46017485 | 1.23400229 | 0.21720205 | 0.80640512 |
| AMBRA1 | 153.296796 | 0.01946526 | 0.36718264 | 0.05301247 | 0.95772197 | 0.99514836 |
| ATG13 | 18.6123364 | -0.5364294 | 0.51325814 | -1.0451454 | 0.29595576 | 0.8608685 |
| ARHGAP1 | 15.0052745 | -0.450994 | 0.54139613 | -0.8330204 | 0.40483323 | 0.91554846 |
| CKAP5 | 25.6167062 | -0.101457 | 0.394537 | -0.2571546 | 0.79705943 | 0.98320463 |
| LRP4 | 17.2704205 | 1.02949193 | 0.63500783 | 1.62122716 | 0.10496893 | 0.67519097 |
| C11orf49 | 37.3469915 | -0.0781371 | 0.46250821 | -0.1689421 | 0.8658422 | 0.98751288 |
| ARFGAP2 | 11.4744179 | -0.4713781 | 0.61209777 | -0.7701027 | 0.44123897 | NA |
| DDB2 | 24.7312936 | 0.24253282 | 0.58995627 | 0.41110304 | 0.68099698 | 0.9673431 |
| MADD | 24.1366359 | -0.1943212 | 0.40531853 | -0.4794283 | 0.63163398 | 0.95877428 |
| SPI1 | 50.9852978 | 0.3369201 | 0.59349964 | 0.56768376 | 0.57024972 | 0.95415437 |
| CELF1 | 122.173295 | -0.1168184 | 0.24499771 | -0.4768142 | 0.6334944 | 0.95877428 |
| MTCH2 | 19.714644 | -0.2365713 | 0.48584156 | -0.4869311 | 0.62630719 | 0.95877428 |
| FNBP4 | 84.6356647 | -0.280298 | 0.27255241 | -1.0284189 | 0.30375283 | 0.86514059 |
| NUP160 | 39.7182801 | -0.779268 | 0.39426084 | -1.9765291 | 0.04809488 | 0.53331354 |

| | | | | | | |
|---|---|---|---|---|---|---|
| PTPRJ | 294.113539 | 0.92323863 | 0.54723112 | 1.68710914 | 0.09158237 | 0.65902576 |
| SERPING1 | 14.6354721 | 0.44056862 | 0.69452692 | 0.63434347 | 0.52585669 | 0.95101011 |
| ZDHHC5 | 18.3641163 | 0.3186157 | 0.51123945 | 0.62322206 | 0.53313861 | 0.95293563 |
| CTNND1 | 23.4366228 | 1.05651162 | 0.52796627 | 2.00109679 | 0.04538196 | 0.52128421 |
| LPXN | 40.6123798 | -0.3188495 | 0.37407538 | -0.8523671 | 0.39401037 | 0.90940709 |
| AP001636.3 | 13.1103054 | -0.0103929 | 1.18447935 | -0.0087743 | 0.99299925 | 0.99859588 |
| FAM111A | 21.6758835 | 0.24921607 | 0.40508675 | 0.61521655 | 0.53841172 | 0.95415437 |
| MPEG1 | 38.5443434 | 0.3927365 | 0.97701145 | 0.40197738 | 0.68770067 | 0.9673431 |
| OSBP | 23.172587 | -0.3548309 | 0.4934237 | -0.7191201 | 0.47206692 | 0.9314919 |
| PATL1 | 22.2276768 | 0.26440495 | 0.42022056 | 0.62920516 | 0.52921475 | 0.95241444 |
| STX3 | 15.6422278 | 0.4632249 | 0.53942144 | 0.85874395 | 0.3904818 | 0.90910959 |
| MS4A6A | 66.8576735 | 1.3869674 | 0.73499579 | 1.88704128 | 0.05915477 | 0.57281427 |
| MS4A4E | 52.7640839 | 3.20819238 | 1.08920781 | 2.94543645 | 0.00322499 | 0.14164378 |
| MS4A4A | 37.458857 | 0.66701563 | 0.7503203 | 0.88897453 | 0.37401677 | 0.90262385 |
| MS4A14 | 10.9046405 | 0.91049244 | 0.81219595 | 1.12102558 | 0.26227697 | NA |
| MS4A1 | 33.978681 | -4.3527956 | 0.83692565 | -5.2009346 | 1.98E-07 | 0.00027433 |
| PRPF19 | 7.87166271 | -0.5829094 | 0.62289531 | -0.9358063 | 0.34937293 | NA |
| SLC15A3 | 11.853577 | -0.6028187 | 0.78525109 | -0.7676763 | 0.44267951 | NA |
| CD6 | 26.4890846 | -0.0302291 | 0.51105327 | -0.0591507 | 0.95283211 | 0.99514836 |
| CD5 | 6.86562219 | 1.28085845 | 0.80925732 | 1.58275793 | 0.11347665 | NA |
| VPS37C | 9.76064455 | 0.04974765 | 0.76633116 | 0.06491665 | 0.94824037 | NA |
| DDB1 | 17.6880753 | -0.0838489 | 0.42600885 | -0.1968243 | 0.84396504 | 0.9842451 |
| CPSF7 | 19.0315012 | 0.09537744 | 0.46096444 | 0.20690846 | 0.83608135 | 0.98320463 |
| AP003108.2 | 11.5819701 | 0.10533853 | 0.64834973 | 0.16247177 | 0.87093436 | NA |
| TMEM258 | 9.79425577 | -0.0108948 | 0.87735693 | -0.0124178 | 0.9900923 | NA |
| FADS1 | 10.2732568 | 0.13220513 | 0.58376292 | 0.22647058 | 0.82083544 | NA |
| FTH1 | 499.540758 | 1.07804095 | 0.48359996 | 2.22919984 | 0.02580061 | 0.40178254 |
| AHNAK | 98.7596863 | -0.2807656 | 0.39333703 | -0.7138042 | 0.47534825 | 0.9314919 |
| B3GAT3 | 10.2673002 | -0.2601638 | 0.58951222 | -0.4413205 | 0.65898101 | NA |
| GANAB | 19.4530807 | 0.03144697 | 0.42466482 | 0.07405128 | 0.94096958 | 0.99514836 |
| UBXN1 | 16.5387861 | 0.08090956 | 0.54989594 | 0.14713613 | 0.88302457 | 0.99019008 |
| TTC9C | 10.6205146 | -0.0063835 | 0.50936977 | -0.0125322 | 0.99000102 | NA |
| TMEM179B | 11.6243087 | 0.17094194 | 0.52532903 | 0.32539977 | 0.74487853 | NA |
| NXF1 | 10.4085945 | -0.8576427 | 0.58706767 | -1.4608923 | 0.14404499 | NA |
| STX5 | 20.4735842 | -0.0313232 | 0.45009486 | -0.0695925 | 0.94451797 | 0.99514836 |
| SLC3A2 | 20.1730804 | -0.1606655 | 0.49536471 | -0.3243377 | 0.74568236 | 0.97805001 |
| RARRES3 | 22.3924387 | 1.63007929 | 0.85255957 | 1.91198286 | 0.05587839 | 0.55970424 |
| PLA2G16 | 10.0576013 | 1.8647702 | 0.60391248 | 3.08781529 | 0.00201634 | NA |
| ATL3 | 22.4975343 | -0.0031274 | 0.54077945 | -0.0057832 | 0.99538571 | 0.99859588 |
| RTN3 | 63.6541245 | 0.63086958 | 0.36238499 | 1.74088219 | 0.08170423 | 0.63740875 |
| MARK2 | 56.3584417 | 0.07483139 | 0.30254972 | 0.24733585 | 0.80464831 | 0.98320463 |
| OTUB1 | 11.2100285 | -0.2598846 | 0.55580283 | -0.4675841 | 0.64008204 | NA |
| STIP1 | 16.6005842 | 0.03444512 | 0.47275995 | 0.07285963 | 0.94191782 | 0.99514836 |
| FERMT3 | 26.9116261 | -0.0429108 | 0.56130355 | -0.0764485 | 0.93906227 | 0.99514836 |
| VEGFB | 10.0400676 | -0.882625 | 0.55301797 | -1.5960151 | 0.11048541 | NA |
| FKBP2 | 9.4955837 | -0.0747351 | 0.68774052 | -0.1086675 | 0.91346619 | NA |
| ESRRA | 16.3049069 | -0.377949 | 0.60490757 | -0.6248045 | 0.53209938 | 0.95265314 |
| TRMT112 | 14.5915862 | -1.1125368 | 0.59885473 | -1.8577741 | 0.0632011 | 0.58364573 |
| PRDX5 | 11.6050058 | -0.2898751 | 0.5595774 | -0.518025 | 0.6044408 | NA |
| CCDC88B | 23.9033333 | -1.1676851 | 0.57873426 | -2.0176533 | 0.04362738 | 0.5136892 |
| RASGRP2 | 19.6070719 | -1.5706245 | 0.48316894 | -3.2506735 | 0.00115132 | 0.07001544 |
| SF1 | 54.9089694 | -0.7141792 | 0.29219791 | -2.4441627 | 0.01451888 | 0.32121732 |
| MAP4K2 | 10.4322647 | 0.77092063 | 0.54941837 | 1.40315774 | 0.16056981 | NA |
| EHD1 | 16.6121386 | 0.22797034 | 0.50698463 | 0.44965926 | 0.65295615 | 0.96178254 |

| | | | | | | |
|---|---|---|---|---|---|---|
| ATG2A | 9.72775461 | 0.65117435 | 0.67315312 | 0.96734954 | 0.33336933 | NA |
| VPS51 | 19.0242325 | -0.0861047 | 0.44221843 | -0.1947108 | 0.84561937 | 0.98458147 |
| FAU | 67.4821751 | -0.2626173 | 0.65590944 | -0.4003866 | 0.68887181 | 0.9673431 |
| CAPN1 | 15.1309031 | 0.69279422 | 0.57967833 | 1.19513562 | 0.23203405 | 0.82637384 |
| POLA2 | 11.528863 | -0.1077023 | 0.53774911 | -0.2002836 | 0.84125878 | NA |
| DPF2 | 11.9578444 | -0.3649528 | 0.48484775 | -0.7527163 | 0.45162039 | NA |
| FRMD8 | 10.4129658 | -0.0987313 | 0.56779857 | -0.1738844 | 0.86195634 | NA |
| NEAT1 | 1691.76238 | -0.1795622 | 0.50702656 | -0.3541475 | 0.72322834 | 0.97451805 |
| MALAT1 | 12108.6361 | -0.3761678 | 0.28514391 | -1.319221 | 0.18709526 | 0.77798634 |
| SCYL1 | 10.9542533 | -0.3558571 | 0.53594515 | -0.6639805 | 0.5067028 | NA |
| LTBP3 | 9.30468755 | -1.0044461 | 0.57461437 | -1.7480351 | 0.08045796 | NA |
| EHBP1L1 | 42.7486213 | -0.6064618 | 0.60804146 | -0.9974021 | 0.3185694 | 0.87298176 |
| MAP3K11 | 14.0575521 | -0.3778964 | 0.68080949 | -0.5550692 | 0.57884735 | 0.95415437 |
| SIPA1 | 17.924987 | -0.7083493 | 0.57090075 | -1.2407574 | 0.21469539 | 0.80634897 |
| RELA | 7.97516388 | -0.364537 | 0.57040614 | -0.6390832 | 0.5227688 | NA |
| CFL1 | 83.4393768 | 0.15913353 | 0.66820916 | 0.23814927 | 0.81176532 | 0.98320463 |
| CTSW | 9.29236555 | 0.44860026 | 0.76158392 | 0.58903589 | 0.5558372 | NA |
| CCDC85B | 16.7700557 | -0.4814801 | 0.49164365 | -0.9793273 | 0.32741827 | 0.87723689 |
| DRAP1 | 12.7723837 | -1.2374043 | 0.58794721 | -2.104618 | 0.03532457 | NA |
| SART1 | 7.80204666 | -0.2856367 | 0.58816924 | -0.485637 | 0.62722456 | NA |
| BANF1 | 9.37711287 | 0.87453434 | 0.60924489 | 1.43543976 | 0.15116182 | NA |
| SF3B2 | 21.6298707 | -0.3848756 | 0.45810297 | -0.8401508 | 0.40082385 | 0.91508218 |
| PACS1 | 160.583737 | 0.07581703 | 0.32794591 | 0.2311876 | 0.81716906 | 0.98320463 |
| RAB1B | 10.7567609 | 0.2421259 | 0.74150907 | 0.32653128 | 0.74402243 | NA |
| ZDHHC24 | 7.92327442 | -0.7175239 | 0.57258497 | -1.2531309 | 0.21015809 | NA |
| CCS | 11.2854579 | 0.47890966 | 0.54372444 | 0.8807948 | 0.37842889 | NA |
| RBM14-RBM | 16.1253492 | 0.16379708 | 0.51849614 | 0.315908 | 0.75207234 | 0.97859589 |
| RBM4 | 10.7382345 | -0.0236852 | 0.64750149 | -0.0365794 | 0.97082038 | NA |
| RBM4B | 7.70718361 | 0.78434091 | 0.73102854 | 1.0729279 | 0.28330347 | NA |
| C11orf80 | 21.9683792 | 0.0181609 | 0.58462757 | 0.03106405 | 0.97521846 | 0.9973754 |
| KDM2A | 174.270427 | 0.0033566 | 0.21858099 | 0.01535633 | 0.98774791 | 0.99811561 |
| GRK2 | 36.5723665 | 0.13116592 | 0.38559813 | 0.34016223 | 0.73373436 | 0.97605445 |
| ANKRD13D | 43.570106 | 0.24271178 | 0.4226818 | 0.57421867 | 0.56581983 | 0.95415437 |
| RAD9A | 14.8322939 | 0.14438855 | 0.50246143 | 0.28736246 | 0.7738348 | 0.98264092 |
| PPP1CA | 19.5439482 | 1.03848708 | 0.62144718 | 1.67107861 | 0.09470615 | 0.65925012 |
| TBC1D10C | 16.6759739 | -0.7969334 | 0.51192889 | -1.5567268 | 0.11953536 | 0.70431251 |
| CORO1B | 9.48313667 | 0.35058856 | 0.59809959 | 0.58617088 | 0.55776069 | NA |
| AIP | 8.61274135 | -0.6491059 | 0.64256126 | -1.0101852 | 0.31240659 | NA |
| CDK2AP2 | 10.4012968 | 0.09747436 | 0.76563886 | 0.12731114 | 0.89869414 | NA |
| GSTP1 | 42.8756514 | 0.01054386 | 0.66123452 | 0.01594571 | 0.9872777 | 0.99811561 |
| NDUFV1 | 12.4156296 | 0.53378665 | 0.51031034 | 1.04600399 | 0.29555919 | NA |
| UNC93B1 | 21.5889842 | -0.3576424 | 0.53707704 | -0.6659051 | 0.50547175 | 0.94166217 |
| NDUFS8 | 12.4057138 | 0.33506614 | 0.67030866 | 0.49986843 | 0.61716772 | NA |
| TCIRG1 | 18.3910208 | 0.18830624 | 0.64962838 | 0.28986763 | 0.77191751 | 0.98264092 |
| CHKA | 31.9200147 | -0.3997669 | 0.47174141 | -0.847428 | 0.39675658 | 0.91333232 |
| KMT5B | 38.9823755 | -0.150921 | 0.35023345 | -0.4309154 | 0.66652988 | 0.96412305 |
| PPP6R3 | 146.542505 | -0.1817621 | 0.24890774 | -0.730239 | 0.46524413 | 0.9314919 |
| CPT1A | 32.6326335 | 0.96985513 | 0.638161 | 1.51976559 | 0.1285699 | 0.72800883 |
| TPCN2 | 12.6671235 | -0.5128421 | 0.69443595 | -0.7385017 | 0.46020965 | NA |
| ORAOV1 | 8.55316742 | -0.3775576 | 0.60622595 | -0.6228001 | 0.5334159 | NA |
| PPFIA1 | 53.3927642 | -0.2638794 | 0.2898015 | -0.9105521 | 0.36253139 | 0.89853007 |
| AP002336.2 | 20.1580518 | -0.1644662 | 0.54912551 | -0.2995056 | 0.76455426 | 0.98222198 |
| NADSYN1 | 15.4286949 | -0.4722074 | 0.44416996 | -1.0631232 | 0.2877261 | 0.85677513 |
| AP002495.2 | 11.9930823 | -0.6234926 | 0.52650913 | -1.1842009 | 0.23633354 | NA |

| | | | | | | |
|---|---|---|---|---|---|---|
| RNF121 | 12.7655376 | 0.43118518 | 0.5821103 | 0.74072764 | 0.4588586 | NA |
| NUMA1 | 29.9715424 | -0.1979419 | 0.36921059 | -0.5361221 | 0.5918742 | 0.95415437 |
| LAMTOR1 | 19.5877714 | 0.79335162 | 0.42018431 | 1.88810384 | 0.05901201 | 0.57281427 |
| INPPL1 | 14.4128962 | -0.8429271 | 0.51984577 | -1.6214945 | 0.10491162 | 0.67519097 |
| CLPB | 12.2996516 | -0.1958323 | 0.61945215 | -0.3161379 | 0.75189781 | NA |
| ARAP1 | 40.7165692 | -0.1721447 | 0.54784774 | -0.31422 | 0.75335393 | 0.97883059 |
| ATG16L2 | 34.2024782 | 0.28027341 | 0.47085381 | 0.59524507 | 0.55167967 | 0.95415437 |
| FCHSD2 | 324.913026 | -0.5896262 | 0.34392312 | -1.7144129 | 0.08645293 | 0.64676224 |
| RELT | 14.1472544 | -0.0132075 | 0.56788913 | -0.0232571 | 0.98144516 | 0.99758514 |
| FAM168A | 69.9449134 | 0.31759863 | 0.3879647 | 0.81862765 | 0.41299889 | 0.91894074 |
| AP000763.3 | 9.4041461 | 0.93420761 | 0.89180137 | 1.04755121 | 0.29484542 | NA |
| RAB6A | 56.4418228 | 0.2779272 | 0.26824324 | 1.03610139 | 0.30015484 | 0.86225566 |
| MRPL48 | 8.68202183 | -1.0113786 | 0.65579917 | -1.5422079 | 0.12302309 | NA |
| COA4 | 8.82071172 | -0.887616 | 0.69726682 | -1.2729934 | 0.20302039 | NA |
| PAAF1 | 10.5950507 | -0.0650562 | 0.59688324 | -0.1089931 | 0.91320795 | NA |
| UCP2 | 22.6185426 | -0.5208424 | 0.73160384 | -0.7119187 | 0.47651513 | 0.9314919 |
| C2CD3 | 23.9751028 | -0.1132967 | 0.38881867 | -0.291387 | 0.77075532 | 0.98264092 |
| PPME1 | 14.6544365 | 0.00267108 | 0.51371485 | 0.00519953 | 0.99585139 | 0.99859588 |
| PGM2L1 | 10.4933189 | 0.26396551 | 0.63000458 | 0.41898983 | 0.67522357 | NA |
| POLD3 | 13.6440804 | 0.12078637 | 0.52722403 | 0.22909875 | 0.81879216 | 0.98320463 |
| RNF169 | 37.7808124 | -0.1206837 | 0.32135632 | -0.3755449 | 0.70725523 | 0.97184211 |
| XRRA1 | 29.846073 | -0.6750166 | 0.72661087 | -0.9289933 | 0.35289258 | 0.89109867 |
| SPCS2 | 33.153687 | -0.2430913 | 0.53019929 | -0.4584904 | 0.64660017 | 0.96049083 |
| NEU3 | 9.78717557 | -0.0242383 | 0.64495335 | -0.0375815 | 0.97002138 | NA |
| SLCO2B1 | 71.7884739 | -0.0307767 | 0.62955204 | -0.0488867 | 0.96100959 | 0.99614757 |
| ARRB1 | 59.167821 | 1.54688409 | 0.55896523 | 2.76740665 | 0.00565042 | 0.19011297 |
| RPS3 | 139.428798 | -0.1299168 | 0.65401885 | -0.1986438 | 0.84254142 | 0.9842451 |
| UVRAG | 133.345136 | -0.4870558 | 0.29026455 | -1.6779719 | 0.09335259 | 0.6590317 |
| THAP12 | 10.8008962 | -0.1900934 | 0.60557547 | -0.3139053 | 0.75359293 | NA |
| EMSY | 40.6276194 | -0.579949 | 0.35077004 | -1.6533596 | 0.09825769 | 0.66570745 |
| ACER3 | 81.6699412 | 0.36048542 | 0.44553877 | 0.80910002 | 0.41845762 | 0.91985469 |
| PAK1 | 39.1553995 | 0.08249802 | 0.6206078 | 0.13293101 | 0.89424794 | 0.99115168 |
| CLNS1A | 22.780045 | 0.22017156 | 0.4712399 | 0.46721757 | 0.64034422 | 0.95963232 |
| RSF1 | 90.1830556 | 0.07147601 | 0.29356655 | 0.24347464 | 0.80763773 | 0.98320463 |
| AAMDC | 9.89970576 | 0.32779908 | 0.60220878 | 0.54432797 | 0.5862158 | NA |
| INTS4 | 27.2630868 | -0.1269374 | 0.36272698 | -0.349953 | 0.72637398 | 0.9754739 |
| GAB2 | 155.891101 | 0.7808774 | 0.49750161 | 1.56959775 | 0.11650872 | 0.70122655 |
| AP003086.2 | 43.3188119 | 0.95930131 | 0.70470415 | 1.36128233 | 0.17342448 | 0.76214163 |
| NARS2 | 21.9911029 | -0.1815682 | 0.44706167 | -0.4061367 | 0.68464213 | 0.9673431 |
| TENM4 | 17.5842809 | 0.65152687 | 1.14752613 | 0.56776648 | 0.57019354 | 0.95415437 |
| PRCP | 20.3217855 | 0.37491724 | 0.58994036 | 0.63551719 | 0.52509116 | 0.95007724 |
| RAB30 | 29.9612445 | -1.6459971 | 0.61175288 | -2.6906243 | 0.00713185 | 0.21105689 |
| RAB30-AS1 | 19.5403165 | 0.2146107 | 0.53467028 | 0.40138887 | 0.68813384 | 0.9673431 |
| PCF11 | 43.4969468 | -0.1141156 | 0.5028219 | -0.2269502 | 0.82046243 | 0.98320463 |
| DLG2 | 35.7681781 | -1.4658332 | 0.86798175 | -1.6887834 | 0.09126095 | 0.65760166 |
| CREBZF | 23.1887814 | -0.0799296 | 0.55247405 | -0.1446757 | 0.8849669 | 0.99019008 |
| SYTL2 | 45.5755952 | 0.84786936 | 0.70352688 | 1.20516981 | 0.22813775 | 0.82248491 |
| PICALM | 350.92711 | 0.24293299 | 0.2687546 | 0.90392123 | 0.36603717 | 0.89868152 |
| EED | 15.4128711 | -0.6058383 | 0.48600308 | -1.246573 | 0.2125541 | 0.80456523 |
| HIKESHI | 13.3880376 | 0.63132622 | 0.53964269 | 1.16989674 | 0.24204252 | 0.83493679 |
| TMEM135 | 49.1992175 | 0.30728416 | 0.40392145 | 0.76075228 | 0.44680504 | 0.92914597 |
| CTSC | 59.9893883 | 0.40378439 | 0.40502881 | 0.99692758 | 0.31879966 | 0.87298176 |
| CHORDC1 | 20.075711 | 0.32300632 | 0.64501037 | 0.50077694 | 0.61652811 | 0.95688177 |
| SLC36A4 | 38.3464049 | 0.68820554 | 0.45722404 | 1.50518231 | 0.13227721 | 0.73044824 |

| | | | | | | |
|---|---|---|---|---|---|---|
| SMCO4 | 15.9645985 | 1.25034764 | 0.77570736 | 1.61188059 | 0.10698793 | 0.67742699 |
| CEP295 | 19.2971279 | -0.443607 | 0.54857332 | -0.8086557 | 0.4187132 | 0.91985469 |
| TAF1D | 51.2289843 | -0.2519903 | 0.37464289 | -0.6726147 | 0.50119245 | 0.94005069 |
| MRE11 | 23.3556316 | -0.552916 | 0.42597266 | -1.2980081 | 0.19428454 | 0.78135737 |
| CWC15 | 9.47985271 | 0.74629177 | 0.571834 | 1.30508463 | 0.19186403 | NA |
| AP000787.1 | 11.0892177 | -1.747011 | 0.70904922 | -2.4638783 | 0.01374428 | NA |
| SESN3 | 50.2623685 | -0.3369273 | 0.45244082 | -0.7446882 | 0.45646023 | 0.9314919 |
| FAM76B | 12.1179881 | 0.32982619 | 0.55408213 | 0.59526588 | 0.55166576 | NA |
| CEP57 | 27.8350108 | 0.44179997 | 0.45291787 | 0.97545271 | 0.32933575 | 0.88004256 |
| MTMR2 | 14.1701674 | -0.4610864 | 0.5817581 | -0.7925742 | 0.42802597 | 0.92167149 |
| MAML2 | 350.659079 | 0.30007439 | 0.4187617 | 0.71657554 | 0.47363604 | 0.9314919 |
| CCDC82 | 21.2164481 | -0.3771085 | 0.41353484 | -0.9119146 | 0.36181367 | 0.89853007 |
| BIRC3 | 44.6760079 | -1.189527 | 0.50946034 | -2.3348766 | 0.01954987 | 0.36550326 |
| BIRC2 | 19.7178892 | 0.47510979 | 0.4364829 | 1.08849576 | 0.27637631 | 0.84994492 |
| TMEM123 | 50.8638144 | 0.77355202 | 0.34386789 | 2.24956167 | 0.02447678 | 0.39492781 |
| DCUN1D5 | 10.1929041 | -0.2701483 | 0.6041255 | -0.4471725 | 0.65475052 | NA |
| DYNC2H1 | 12.4412912 | 0.34811343 | 0.71504944 | 0.48683827 | 0.62637295 | NA |
| PDGFD | 33.5319131 | -0.2297746 | 0.66077446 | -0.3477353 | 0.72803899 | 0.97605445 |
| CASP4 | 31.0199322 | 0.42591867 | 0.37494633 | 1.1359457 | 0.25597929 | 0.8406233 |
| CASP1 | 17.3579447 | -0.0638717 | 0.57041617 | -0.1119738 | 0.91084417 | 0.99471296 |
| AASDHPPT | 11.985012 | 0.09413413 | 0.58810611 | 0.16006317 | 0.87283131 | NA |
| CWF19L2 | 36.5465527 | -0.3939609 | 0.37770395 | -1.0430416 | 0.29692901 | 0.8608685 |
| CUL5 | 58.0773256 | -0.1329973 | 0.29453882 | -0.4515441 | 0.65159745 | 0.96158408 |
| ACAT1 | 17.2346972 | 0.50145995 | 0.52591759 | 0.9534953 | 0.34033917 | 0.8844375 |
| NPAT | 29.2063056 | 0.13811947 | 0.38396273 | 0.35972103 | 0.71905576 | 0.97348719 |
| ATM | 186.706374 | -0.3633528 | 0.3351851 | -1.0840362 | 0.27834876 | 0.84994492 |
| C11orf65 | 13.1129499 | 1.17980688 | 0.98509302 | 1.19766038 | 0.23104926 | 0.82438852 |
| DDX10 | 44.7014399 | -0.6486834 | 0.33708452 | -1.9243939 | 0.05430523 | 0.55278922 |
| RDX | 34.8593137 | 0.12413154 | 0.45612802 | 0.27214188 | 0.78551293 | 0.98264092 |
| FDX1 | 15.643731 | -0.9001604 | 0.52588069 | -1.7117198 | 0.08694832 | 0.64676224 |
| SIK2 | 53.7658701 | 0.54903667 | 0.38523681 | 1.42519266 | 0.15410152 | 0.74480157 |
| PPP2R1B | 11.1300689 | -0.0526482 | 0.54785752 | -0.0960983 | 0.9234425 | NA |
| CRYAB | 93.9784124 | 2.8761503 | 0.99473464 | 2.89137441 | 0.00383561 | 0.15732988 |
| IL18 | 28.1502284 | -0.0443052 | 0.52313036 | -0.0846924 | 0.93250593 | 0.99514836 |
| PTS | 15.1179416 | -0.670207 | 0.61605512 | -1.0879011 | 0.27663878 | 0.84994492 |
| NCAM1 | 25.5859209 | 2.03977683 | 0.72358564 | 2.81898468 | 0.00481758 | 0.17334548 |
| USP28 | 22.7241015 | -0.0134467 | 0.5182098 | -0.0259484 | 0.9792985 | 0.99758514 |
| ZBTB16 | 60.0290831 | 0.46717098 | 0.61824378 | 0.75564202 | 0.44986385 | 0.92914597 |
| NNMT | 16.1230424 | 1.73549528 | 0.88160014 | 1.96857418 | 4.90E-02 | 0.53486612 |
| CADM1 | 44.4330697 | -0.0146983 | 0.49919246 | -0.0294441 | 0.97651042 | 0.99758514 |
| SIK3 | 308.97692 | 0.10694882 | 0.30088861 | 0.35544323 | 0.72225757 | 0.97441549 |
| PAFAH1B2 | 27.2003625 | -0.2428926 | 0.43643911 | -0.5565326 | 0.57784683 | 0.95415437 |
| SIDT2 | 21.5709404 | -0.3000758 | 0.70651808 | -0.4247249 | 0.67103721 | 0.96412305 |
| TAGLN | 14.2946133 | -0.1817962 | 0.72666299 | -0.2501795 | 0.80244854 | 0.98320463 |
| PCSK7 | 56.3707804 | 0.00635878 | 0.26520161 | 0.02397715 | 0.98087083 | 0.99758514 |
| RNF214 | 13.2025623 | -0.2865632 | 0.48784892 | -0.5874015 | 0.55693407 | 0.95415437 |
| CEP164 | 9.23208195 | 0.10242726 | 0.59256341 | 0.17285451 | 0.86276579 | NA |
| IL10RA | 61.3747827 | -0.36564 | 0.36333521 | -1.0063434 | 0.31425042 | 0.87152625 |
| JAML | 43.8286055 | 0.51544557 | 0.79435567 | 0.64888512 | 0.51641264 | 0.94596835 |
| MPZL3 | 10.9773518 | -0.4982501 | 0.5841062 | -0.8530129 | 0.39365217 | NA |
| CD3E | 17.6212111 | 0.74932251 | 0.57493219 | 1.30332329 | 0.19246441 | 0.78135737 |
| CD3D | 14.7856099 | 0.48078586 | 0.8340226 | 0.57646623 | 0.56430009 | 0.95415437 |
| CD3G | 16.3484636 | 0.55725651 | 0.64510464 | 0.86382345 | 0.38768489 | 0.90910959 |
| UBE4A | 35.7510062 | -0.0649095 | 0.35737256 | -0.1816297 | 0.85587331 | 0.98751288 |

| | | | | | | |
|---|---|---|---|---|---|---|
| ATP5MG | 34.0542136 | -0.3355663 | 0.46824109 | -0.7166528 | 0.47358835 | 0.9314919 |
| KMT2A | 84.3572191 | 0.28755454 | 0.25336828 | 1.13492717 | 0.25640584 | 0.8406233 |
| ARCN1 | 16.548551 | 0.36613018 | 0.5444947 | 0.67242193 | 0.50131512 | 0.94005069 |
| AP002954.1 | 8.80365072 | 0.33827206 | 0.86020793 | 0.39324453 | 0.69413888 | NA |
| DDX6 | 65.6513929 | -0.055506 | 0.27415756 | -0.2024603 | 0.83955692 | 0.98342625 |
| BCL9L | 11.5104924 | -0.7783106 | 0.52686307 | -1.477254 | 0.13960756 | NA |
| CCDC84 | 25.9808698 | -0.5691905 | 0.38662658 | -1.472197 | 0.14096769 | 0.73044824 |
| RPS25 | 76.8717382 | -0.4492375 | 0.55229689 | -0.8133986 | 0.41598954 | 0.91923465 |
| CBL | 89.1790442 | 0.22350222 | 0.27588235 | 0.81013598 | 0.41786203 | 0.91967721 |
| ARHGEF12 | 40.05255 | 0.09570191 | 0.35686768 | 0.26817198 | 0.78856694 | 0.98320463 |
| SORL1 | 118.954679 | -0.2173534 | 0.34881385 | -0.6231216 | 0.53320463 | 0.95293563 |
| UBASH3B | 116.756925 | -1.4396735 | 0.71992079 | -1.9997666 | 0.04552548 | 0.52161072 |
| HSPA8 | 41.130633 | 0.13108889 | 0.4297361 | 0.3050451 | 0.7603318 | 0.9808103 |
| GRAMD1B | 42.3179731 | 1.13495384 | 0.63200922 | 1.79578683 | 0.07252843 | 0.61222939 |
| TBRG1 | 24.7498811 | 0.49469999 | 0.4410309 | 1.12169008 | 0.26199423 | 0.84837688 |
| MSANTD2 | 12.7598694 | 0.63107656 | 0.53525547 | 1.17901935 | 0.23839047 | NA |
| EI24 | 9.21788688 | 0.69903988 | 0.69284088 | 1.00894722 | 0.31299995 | NA |
| STT3A | 11.7583988 | 0.18210982 | 0.59910335 | 0.30397062 | 0.76115027 | NA |
| FAM118B | 14.3298403 | 0.42211394 | 0.54053755 | 0.78091512 | 0.43485242 | 0.92343564 |
| SRPRA | 11.3959305 | -0.1136587 | 0.59645344 | -0.1905576 | 0.84887224 | NA |
| ETS1 | 132.915534 | 0.35967855 | 0.37691487 | 0.95427001 | 0.33994698 | 0.8844375 |
| FLI1 | 176.741366 | 0.75252783 | 0.26883178 | 2.79925175 | 0.00512212 | 0.17940383 |
| ARHGAP32 | 24.8586122 | 0.48625005 | 0.55910984 | 0.86968608 | 0.38447198 | 0.90910959 |
| NFRKB | 10.0371195 | -0.1857558 | 0.58453564 | -0.3177836 | 0.7506491 | NA |
| PRDM10 | 15.1580069 | -1.1898099 | 0.47268898 | -2.5171096 | 0.0118322 | 0.29101961 |
| APLP2 | 75.3007451 | 0.37735854 | 0.33058979 | 1.14147064 | 0.25367412 | 0.84026145 |
| ST14 | 12.4959964 | -1.4720651 | 0.54391902 | -2.7064049 | 0.00680161 | NA |
| ZBTB44 | 40.6513137 | -0.0267621 | 0.39870281 | -0.0671229 | 0.9464839 | 0.99514836 |
| SNX19 | 24.0268598 | -1.3147189 | 0.44140962 | -2.9784555 | 0.00289705 | 0.1292926 |
| NCAPD3 | 13.6382067 | 0.02646216 | 0.53704598 | 0.04927354 | 0.96070131 | 0.99614757 |
| VPS26B | 9.07995806 | 0.50617677 | 0.56826488 | 0.89074088 | 0.3730682 | NA |
| ZMYND11 | 45.5367965 | -0.0302044 | 0.3368973 | -0.0896546 | 0.92856167 | 0.99514836 |
| DIP2C | 14.7731663 | -0.1818742 | 0.44975524 | -0.4043848 | 0.68592975 | 0.9673431 |
| LARP4B | 102.61368 | -0.1554516 | 0.26823083 | -0.579544 | 0.56222215 | 0.95415437 |
| GTPBP4 | 11.2061372 | -0.7109773 | 0.54542449 | -1.3035302 | 0.1923938 | NA |
| IDI1 | 13.7241967 | 0.53217326 | 0.69555571 | 0.76510515 | 0.44420893 | 0.92914597 |
| WDR37 | 55.7102626 | 0.37991961 | 0.33026537 | 1.1503465 | 0.25000119 | 0.83865855 |
| PFKP | 13.2334267 | 0.38987675 | 0.49750935 | 0.78365714 | 0.43324132 | 0.92284736 |
| PITRM1 | 10.2206173 | 0.601484 | 0.59888748 | 1.00433557 | 0.31521689 | NA |
| KLF6 | 74.2693303 | -0.1002825 | 0.4859908 | -0.2063464 | 0.83652032 | 0.98320463 |
| FAM208B | 77.4620944 | -0.4334598 | 0.25437307 | -1.7040319 | 0.08837512 | 0.64691526 |
| GDI2 | 71.9797383 | -1.0654467 | 0.45822494 | -2.3251608 | 0.02006336 | 0.36765113 |
| FBH1 | 23.2572161 | 0.50881456 | 0.4846184 | 1.04992828 | 0.29375109 | 0.86027209 |
| IL15RA | 11.0545694 | -0.1853746 | 0.65599874 | -0.2825837 | 0.77749597 | NA |
| IL2RA | 28.9366188 | -2.6524643 | 0.70891404 | -3.741588 | 0.00018286 | 0.02594752 |
| RBM17 | 25.1593702 | -0.6909496 | 0.38401848 | -1.7992612 | 0.07197737 | 0.61092445 |
| PFKFB3 | 89.7685199 | 0.42066609 | 0.70935247 | 0.5930283 | 0.55316221 | 0.95415437 |
| AL137145.2 | 9.39325928 | -0.0573068 | 0.6969287 | -0.0822276 | 0.93446574 | NA |
| PRKCQ | 70.0020341 | 0.22948856 | 0.39129521 | 0.58648446 | 0.55755 | 0.95415437 |
| SFMBT2 | 188.661269 | 0.65230712 | 0.54805149 | 1.19022962 | 0.23395615 | 0.82817204 |
| KIN | 13.4377174 | -0.1749836 | 0.51091547 | -0.3424903 | 0.73198191 | 0.97605445 |
| ATP5F1C | 22.7048773 | 0.31412131 | 0.42090365 | 0.74630218 | 0.4554849 | 0.9314919 |
| TAF3 | 58.0617016 | 0.03697747 | 0.36120717 | 0.10237192 | 0.91846147 | 0.99498068 |
| GATA3 | 10.5882197 | 0.70141578 | 0.6793885 | 1.03242221 | 0.30187437 | NA |

| | | | | | | |
|---|---|---|---|---|---|---|
| CELF2 | 435.950668 | 0.04712819 | 0.26707673 | 0.17645937 | 0.85993306 | 0.98751288 |
| USP6NL | 43.3649396 | -1.9011428 | 0.54124796 | -3.5125173 | 0.00044388 | 0.04075978 |
| UPF2 | 49.1434301 | -0.0102596 | 0.33723962 | -0.0304224 | 0.97573021 | 0.99738871 |
| DHTKD1 | 17.2483888 | -1.1017535 | 0.45297527 | -2.4322596 | 0.01500495 | 0.32436474 |
| NUDT5 | 13.8840535 | 0.72787966 | 0.47905232 | 1.5194158 | 0.12865787 | 0.72800883 |
| CDC123 | 28.4608776 | -0.287502 | 0.45834442 | -0.6272619 | 0.5304876 | 0.95245713 |
| CAMK1D | 209.443803 | -0.8356641 | 0.5314107 | -1.5725392 | 0.11582557 | 0.70122655 |
| OPTN | 21.3218233 | 0.43880436 | 0.40611452 | 1.08049415 | 0.27992219 | 0.84994492 |
| AL157392.5 | 10.6404755 | -0.176974 | 0.53098526 | -0.3332936 | 0.73891265 | NA |
| FRMD4A | 118.553706 | 0.8200187 | 0.63589468 | 1.28955114 | 0.19720655 | 0.78135737 |
| FAM107B | 100.158049 | -1.0069135 | 0.45961593 | -2.1907715 | 0.02846833 | 0.42123996 |
| DCLRE1C | 21.8903987 | -0.2282211 | 0.56018643 | -0.4074021 | 0.68371268 | 0.9673431 |
| RPP38 | 12.1725558 | -0.0499469 | 0.54005594 | -0.0924847 | 0.92631292 | NA |
| NMT2 | 12.5843786 | 0.82755468 | 0.59945166 | 1.38051947 | 0.16742676 | NA |
| MINDY3 | 38.329643 | 0.25679021 | 0.35276121 | 0.72794346 | 0.4666482 | 0.9314919 |
| PTER | 15.2681099 | 0.25213562 | 0.47490004 | 0.53092355 | 0.59547175 | 0.9553365 |
| RSU1 | 57.5058743 | 0.15854922 | 0.2847458 | 0.5568097 | 0.57765746 | 0.95415437 |
| CUBN | 23.6484157 | 0.5396026 | 0.68135529 | 0.79195481 | 0.42838703 | 0.92178217 |
| VIM-AS1 | 10.326345 | 0.02851016 | 0.67577844 | 0.04218861 | 0.96634834 | NA |
| VIM | 141.514884 | 0.03250298 | 0.6148191 | 0.05286592 | 0.95783874 | 0.99514836 |
| ST8SIA6 | 33.1495335 | 3.10834262 | 0.89409815 | 3.47651164 | 0.00050798 | 0.04392458 |
| STAM | 31.2304531 | 0.90206612 | 0.4206432 | 2.14449235 | 0.03199346 | 0.43969912 |
| MRC1 | 131.571996 | 2.41492727 | 1.05426983 | 2.29061593 | 0.02198564 | 0.38502694 |
| NSUN6 | 22.1045114 | 0.2918498 | 0.45046796 | 0.64788136 | 0.51706169 | 0.94596835 |
| ARL5B | 8.27088033 | -0.3406627 | 0.61928563 | -0.5500898 | 0.58225781 | NA |
| PLXDC2 | 420.925667 | 1.33813527 | 0.55316443 | 2.41905515 | 0.01556088 | 0.32607581 |
| NEBL | 10.7422982 | -0.598968 | 1.01091441 | -0.5925012 | 0.55351499 | NA |
| MLLT10 | 104.231134 | -0.0946991 | 0.25714143 | -0.3682764 | 0.71266718 | 0.97332186 |
| DNAJC1 | 101.42457 | 0.35763771 | 0.43658349 | 0.81917368 | 0.41268733 | 0.91894074 |
| PIP4K2A | 308.018225 | -0.0931508 | 0.27040725 | -0.3444834 | 0.73048275 | 0.97605445 |
| ARHGAP21 | 52.7660543 | 0.30447861 | 0.40308941 | 0.75536247 | 0.45003153 | 0.92914597 |
| APBB1IP | 207.400496 | -0.0360402 | 0.32322631 | -0.1115016 | 0.91121861 | 0.99471296 |
| ABI1 | 109.60893 | 0.11482493 | 0.23006971 | 0.49908754 | 0.61771771 | 0.95688177 |
| ANKRD26 | 17.9424312 | -0.8169063 | 0.48705852 | -1.6772242 | 0.09349866 | 0.6590317 |
| YME1L1 | 45.4112294 | 0.08682643 | 0.32984605 | 0.2632332 | 0.79237084 | 0.98320463 |
| ACBD5 | 18.6874283 | 0.20896287 | 0.51592171 | 0.40502825 | 0.68545675 | 0.9673431 |
| RAB18 | 35.3442509 | 0.53292525 | 0.40877196 | 1.3037226 | 0.19232818 | 0.78135737 |
| MPP7 | 44.2739824 | 0.9032163 | 0.51474365 | 1.75469149 | 0.07931209 | 0.62881535 |
| WAC-AS1 | 12.722039 | 0.09654126 | 0.64457834 | 0.14977429 | 0.88094269 | NA |
| WAC | 179.596476 | -0.0883345 | 0.21629489 | -0.4083986 | 0.68298108 | 0.9673431 |
| SVIL | 64.164446 | 0.80629441 | 0.56819402 | 1.41904769 | 0.15588511 | 0.74486619 |
| MAP3K8 | 73.0512653 | -0.2849059 | 0.38599905 | -0.7381 | 0.46045368 | 0.9314919 |
| ZNF438 | 68.6054661 | 0.81369284 | 0.44667972 | 1.82164716 | 0.06850854 | 0.59777078 |
| ZEB1 | 92.0983966 | -0.596837 | 0.38311394 | -1.5578577 | 0.119267 | 0.70431251 |
| ARHGAP12 | 50.4914727 | 0.0424075 | 0.43076842 | 0.09844616 | 0.92157802 | 0.99514836 |
| KIF5B | 55.4798384 | -0.243111 | 0.31885814 | -0.7624426 | 0.44579591 | 0.92914597 |
| EPC1 | 102.840349 | -0.0534074 | 0.24121756 | -0.2214075 | 0.82477519 | 0.98320463 |
| CCDC7 | 31.6799451 | -0.0667256 | 0.59680023 | -0.1118057 | 0.9109775 | 0.99471296 |
| ITGB1 | 74.5054367 | 0.02979354 | 0.32117102 | 0.09276535 | 0.92608998 | 0.99514836 |
| NRP1 | 79.6093975 | -0.673079 | 0.61135906 | -1.1009552 | 0.27091614 | 0.84994492 |
| PARD3 | 28.6976012 | 0.91124707 | 0.72333796 | 1.25978052 | 0.20774855 | 0.79343028 |
| CUL2 | 25.1339977 | 0.19380173 | 0.38757627 | 0.50003509 | 0.61705037 | 0.95688177 |
| CREM | 82.3466407 | -0.1129765 | 0.82315592 | -0.137248 | 0.89083477 | 0.99080204 |
| CCNY | 145.887266 | 0.1847053 | 0.2906925 | 0.63539752 | 0.52516919 | 0.95007724 |

| | | | | | | |
|---|---|---|---|---|---|---|
| ZNF248 | 13.6881126 | -0.6523989 | 0.50085026 | -1.3025828 | 0.19271723 | 0.78135737 |
| ZNF33A | 58.7853583 | -0.429718 | 0.30897717 | -1.3907759 | 0.16429339 | 0.75321592 |
| ZNF33B | 21.6676286 | 0.23952788 | 0.54305435 | 0.44107534 | 0.65915846 | 0.96280444 |
| BMS1 | 13.3197247 | -0.567412 | 0.58713907 | -0.9664014 | 0.33384335 | 0.88047896 |
| CSGALNACT2 | 68.6381157 | 0.17887832 | 0.34690066 | 0.51564711 | 0.6061009 | 0.95688177 |
| HNRNPF | 35.0198857 | -0.0026769 | 0.36274282 | -0.0073797 | 0.99411188 | 0.99859588 |
| RASSF4 | 33.6142596 | 2.48595539 | 0.664254 | 3.7424771 | 0.00018222 | 0.02594752 |
| ALOX5 | 60.3776068 | 1.423014 | 0.58777346 | 2.42102462 | 0.01547683 | 0.32566073 |
| 8-Mar | 32.0063234 | -0.2013237 | 0.42178036 | -0.4773188 | 0.63313513 | 0.95877428 |
| ZFAND4 | 10.7714275 | 0.5569922 | 0.54982112 | 1.01304257 | 0.31103983 | NA |
| WASHC2C | 19.1293139 | -0.6010688 | 0.49962085 | -1.2030499 | 0.22895703 | 0.82329316 |
| AGAP4 | 10.0323273 | 0.94739239 | 0.61689874 | 1.53573403 | 0.12460363 | NA |
| TIMM23 | 12.0832711 | -0.2916885 | 0.60480252 | -0.4822871 | 0.629602 | NA |
| NCOA4 | 28.4138755 | 0.3089426 | 0.46999468 | 0.65733211 | 0.51096739 | 0.94382295 |
| MAPK8 | 58.8193422 | 0.37231518 | 0.38689873 | 0.96230656 | 0.33589563 | 0.88055255 |
| ARHGAP22 | 27.7392377 | 0.08823087 | 0.58505908 | 0.15080677 | 0.88012815 | 0.99019008 |
| WDFY4 | 81.0409948 | -2.193457 | 0.48326944 | -4.5387868 | 5.66E-06 | 0.00240852 |
| TMEM273 | 9.85939077 | 0.62539457 | 0.72883816 | 0.85807056 | 0.3908535 | NA |
| ERCC6 | 15.4937147 | -0.9570741 | 0.48056767 | -1.991549 | 0.04642056 | 0.52641679 |
| PARG | 40.3420588 | -0.1196126 | 0.38876564 | -0.3076728 | 0.75833129 | 0.98051527 |
| TIMM23B | 40.753842 | -0.7114978 | 0.48406091 | -1.4698518 | 0.1416019 | 0.73044824 |
| AGAP6 | 12.3334174 | -0.2927079 | 0.51666322 | -0.5665352 | 0.57103002 | NA |
| WASHC2A | 9.56558904 | 0.37845371 | 0.66887909 | 0.56580287 | 0.5715278 | NA |
| SGMS1 | 121.523842 | -0.1087482 | 0.30280301 | -0.3591383 | 0.71949163 | 0.97348719 |
| PRKG1 | 11.934853 | -0.0956037 | 0.58446574 | -0.1635746 | 0.87006606 | NA |
| AC074327.1 | 44.6230582 | 2.02176646 | 1.00683871 | 2.00803411 | 0.04463967 | 0.51898302 |
| IPMK | 34.1374706 | 0.21616088 | 0.38505087 | 0.56138267 | 0.5745367 | 0.95415437 |
| CISD1 | 10.0235869 | 0.33327583 | 0.69932309 | 0.47656918 | 0.63366894 | NA |
| UBE2D1 | 18.4198254 | 0.06452288 | 0.5699041 | 0.11321709 | 0.90985845 | 0.99471296 |
| BICC1 | 20.6266598 | -0.2502932 | 0.60314834 | -0.4149778 | 0.67815813 | 0.96718429 |
| CCDC6 | 41.2822883 | -0.4391901 | 0.41295197 | -1.0635379 | 0.28753809 | 0.85677513 |
| ANK3 | 179.607043 | -0.3679181 | 0.62182141 | -0.5916781 | 0.55406614 | 0.95415437 |
| ARID5B | 198.788429 | 0.19057575 | 0.31806536 | 0.59917166 | 0.54905842 | 0.95415437 |
| NRBF2 | 10.0983257 | 0.1956832 | 0.5943986 | 0.32921208 | 0.74199539 | NA |
| JMJD1C | 269.693332 | -0.5249426 | 0.24092003 | -2.1789082 | 0.02933849 | 0.42316846 |
| REEP3 | 28.1862459 | -0.0836557 | 0.39076862 | -0.2140798 | 0.83048479 | 0.98320463 |
| SIRT1 | 19.4440563 | -0.2152353 | 0.46427867 | -0.4635907 | 0.64294104 | 0.95992767 |
| HERC4 | 158.567893 | -0.1060634 | 0.28090865 | -0.3775725 | 0.70574819 | 0.97184211 |
| HNRNPH3 | 41.6189852 | -0.214633 | 0.40802955 | -0.5260233 | 0.59887205 | 0.95582632 |
| RUFY2 | 19.0604874 | 0.02469951 | 0.42335865 | 0.0583418 | 0.95347637 | 0.99514836 |
| SLC25A16 | 19.911504 | -0.5075576 | 0.46035009 | -1.102547 | 0.27022394 | 0.84994492 |
| CCAR1 | 52.7462732 | -0.6295038 | 0.36567352 | -1.7214914 | 0.08516169 | 0.64676224 |
| DDX50 | 19.3240424 | -0.2785619 | 0.43022963 | -0.6474726 | 0.51732615 | 0.94609482 |
| DDX21 | 29.5069101 | -0.4041384 | 0.38281238 | -1.0557088 | 0.29110129 | 0.85872202 |
| SRGN | 340.429049 | 0.93891706 | 0.62531897 | 1.501501 | 0.13322603 | 0.73044824 |
| VPS26A | 42.0185327 | -0.0373582 | 0.35964818 | -0.1038743 | 0.91726912 | 0.99498068 |
| SUPV3L1 | 9.93943824 | -0.0453015 | 0.74066869 | -0.0611629 | 0.95122946 | NA |
| HK1 | 48.7012756 | 0.61651577 | 0.44937014 | 1.37195536 | 0.17007733 | 0.75745073 |
| SAR1A | 20.5338593 | 0.03597229 | 0.47347285 | 0.0759754 | 0.93943867 | 0.99514836 |
| PPA1 | 19.9177488 | 0.60208406 | 0.61068534 | 0.98591537 | 0.32417465 | 0.87663738 |
| EIF4EBP2 | 20.5168478 | -0.0953978 | 0.44922237 | -0.212362 | 0.83182464 | 0.98320463 |
| PALD1 | 13.3765952 | 2.01869969 | 1.05617794 | 1.91132538 | 0.05596278 | 0.55970424 |
| PRF1 | 34.5329804 | 0.64188906 | 0.70974712 | 0.90439122 | 0.36578799 | 0.89868152 |
| SGPL1 | 31.3606757 | -0.3967785 | 0.36542273 | -1.0858069 | 0.27756443 | 0.84994492 |

| | | | | | | |
|---|---|---|---|---|---|---|
| CDH23 | 16.0329648 | 0.41028792 | 0.60596353 | 0.67708352 | 0.49835297 | 0.93916044 |
| VSIR | 56.6648039 | 0.22868047 | 0.38077231 | 0.60057011 | 0.54812635 | 0.95415437 |
| PSAP | 205.929678 | 0.88386977 | 0.63031305 | 1.40227109 | 0.16083431 | 0.74887162 |
| SPOCK2 | 30.1855225 | 0.08171932 | 0.45619497 | 0.17913245 | 0.8578337 | 0.98751288 |
| ASCC1 | 29.1075946 | -0.4731944 | 0.40682913 | -1.163128 | 0.24477757 | 0.83763924 |
| ANAPC16 | 53.4543308 | 0.00903185 | 0.29726148 | 0.03038351 | 0.97576119 | 0.99738871 |
| DDIT4 | 35.4602481 | 0.11803307 | 0.49159543 | 0.24010205 | 8.10E-01 | 0.98320463 |
| DNAJB12 | 9.41682669 | -0.6627388 | 0.57457909 | -1.1534336 | 0.24873244 | NA |
| MICU1 | 78.0625073 | -0.6627869 | 0.44322643 | -1.4953687 | 0.13481824 | 0.73044824 |
| MCU | 79.8227244 | -2.164942 | 0.56454948 | -3.8348135 | 0.00012566 | 0.02107273 |
| P4HA1 | 64.6946525 | -0.0007498 | 0.59298249 | -0.0012645 | 0.99899109 | 0.99935226 |
| ECD | 13.9272792 | -0.4998074 | 0.47081042 | -1.0615895 | 0.2884221 | 0.85703857 |
| FAM149B1 | 14.2310509 | 0.4825218 | 0.64570144 | 0.74728314 | 0.45489267 | 0.9314919 |
| DNAJC9 | 14.0736058 | 0.22961282 | 0.61072355 | 0.37596851 | 0.70694031 | 0.97184211 |
| MRPS16 | 8.92092467 | -0.2763281 | 0.58374667 | -0.4733699 | 0.63594929 | NA |
| ANXA7 | 25.9810419 | -0.2631551 | 0.45326216 | -0.5805803 | 0.56152337 | 0.95415437 |
| PPP3CB | 52.3579637 | -0.7900641 | 0.33734418 | -2.342012 | 0.0191801 | 0.36102949 |
| USP54 | 11.1824747 | 0.62691212 | 0.88399314 | 0.70918211 | 0.47821148 | NA |
| SEC24C | 13.5917962 | 0.22687324 | 0.46736606 | 0.48542943 | 0.62737176 | 0.95877428 |
| FUT11 | 10.2115006 | -0.2396742 | 0.55453053 | -0.432211 | 0.6655881 | NA |
| ZSWIM8 | 15.3506911 | 0.32798442 | 0.44595705 | 0.7354619 | 0.46205824 | 0.9314919 |
| CAMK2G | 27.3486453 | 0.28711952 | 0.41492781 | 0.69197462 | 0.48895327 | 0.93628629 |
| VCL | 50.8659324 | 0.86417089 | 0.43540651 | 1.98474496 | 0.04717285 | 0.52845055 |
| AP3M1 | 14.4051193 | -0.4323433 | 0.63349556 | -0.6824725 | 0.49494021 | 0.93805563 |
| ADK | 172.818377 | -0.4071782 | 0.43734136 | -0.9310307 | 0.35183769 | 0.89109867 |
| KAT6B | 78.0277712 | 0.06399126 | 0.33130657 | 0.19314818 | 0.84684292 | 0.98537189 |
| SAMD8 | 31.0347788 | -0.0677943 | 0.37969539 | -0.1785492 | 0.85829172 | 0.98751288 |
| VDAC2 | 12.2076801 | 0.26349869 | 0.6107344 | 0.43144564 | 0.66614437 | NA |
| LRMDA | 544.179214 | 0.97477032 | 0.55510585 | 1.75600802 | 0.07908704 | 0.62793067 |
| KCNMA1 | 154.987474 | 3.01109065 | 0.86514557 | 3.48044395 | 0.00050058 | 0.04392458 |
| POLR3A | 11.7615671 | 0.11331366 | 0.52738481 | 0.21485955 | 0.82987682 | NA |
| RPS24 | 119.154228 | -0.0382128 | 0.59966881 | -0.0637231 | 0.94919071 | 0.99514836 |
| ZMIZ1 | 52.7452795 | 0.828427 | 0.42463506 | 1.95091522 | 0.05106713 | 0.54780456 |
| NUTM2B-AS | 72.541594 | -0.0181994 | 0.33145457 | -0.0549076 | 0.95621205 | 0.99514836 |
| ANXA11 | 42.4581029 | 0.22879383 | 0.31059697 | 0.73662607 | 0.46134977 | 0.9314919 |
| TSPAN14 | 63.6716935 | 0.77872793 | 0.45037726 | 1.72905695 | 0.0837989 | 0.64303763 |
| GHITM | 17.3339635 | 0.05492257 | 0.49069682 | 0.11192771 | 0.91088072 | 0.99471296 |
| CCSER2 | 100.931127 | 0.35098518 | 0.26934353 | 1.30311345 | 0.19253603 | 0.78135737 |
| WAPL | 71.2147149 | -0.0503635 | 0.27879181 | -0.1806492 | 0.85664292 | 0.98751288 |
| BMPR1A | 25.4064371 | 0.15856322 | 0.49422589 | 0.32083147 | 0.74833811 | 0.97818861 |
| GLUD1 | 41.741087 | -0.3854998 | 0.354106 | -1.0886565 | 0.27630541 | 0.84994492 |
| SHLD2 | 48.4411563 | -0.6603618 | 0.38511086 | -1.7147318 | 0.08639441 | 0.64676224 |
| NUTM2A-AS | 62.1684793 | -0.1723679 | 0.33648713 | -0.512257 | 0.60847119 | 0.95688177 |
| PAPSS2 | 39.7654871 | 1.14739625 | 0.99612147 | 1.15186378 | 0.24937705 | 0.83865855 |
| ATAD1 | 29.807244 | -0.0844977 | 0.34196449 | -0.2470951 | 0.80483464 | 0.98320463 |
| PTEN | 151.617228 | 0.31361517 | 0.32362635 | 0.96906563 | 0.33251245 | 0.88047896 |
| RNLS | 12.8891173 | 0.21409867 | 0.66401182 | 0.32243201 | 0.74712544 | NA |
| STAMBPL1 | 24.016776 | -1.4823757 | 0.47715627 | -3.106688 | 0.00189196 | 0.09861218 |
| FAS | 9.72522271 | 0.52112748 | 0.70249485 | 0.74182391 | 0.45819403 | NA |
| LIPA | 27.009692 | 1.20430824 | 0.72995153 | 1.64984685 | 0.09897426 | 0.66763865 |
| KIF20B | 17.9312569 | -1.220938 | 0.54364596 | -2.2458329 | 0.02471471 | 0.39492781 |
| PCGF5 | 78.2174428 | 0.39334325 | 0.37248519 | 1.05599701 | 0.29096958 | 0.85872202 |
| HECTD2 | 11.7286127 | -0.3172275 | 0.65016623 | -0.4879175 | 0.62560828 | NA |
| TNKS2 | 57.3894761 | -0.2990136 | 0.29850263 | -1.0017119 | 0.31648276 | 0.87263466 |

| | | | | | | |
|---|---|---|---|---|---|---|
| BTAF1 | 65.2075622 | -0.3116784 | 0.29674453 | -1.0503258 | 0.29356835 | 0.86027209 |
| 5-Mar | 26.4434515 | -0.7890636 | 0.39853161 | -1.9799273 | 0.0477117 | 0.53233176 |
| CPEB3 | 64.1784379 | 0.08690694 | 0.3411389 | 0.25475528 | 0.79891212 | 0.98320463 |
| IDE | 24.3446459 | 0.02893378 | 0.40686936 | 0.0711132 | 0.94330766 | 0.99514836 |
| EXOC6 | 62.2062658 | -0.238209 | 0.30629839 | -0.7777023 | 0.43674455 | 0.92489992 |
| MYOF | 44.0005655 | -1.7619934 | 0.96046391 | -1.8345233 | 0.06657636 | 0.58951068 |
| FRA10AC1 | 16.4510529 | -0.1853727 | 0.46076773 | -0.4023126 | 0.687454 | 0.9673431 |
| TBC1D12 | 26.4340134 | -0.5311835 | 0.4977606 | -1.0671465 | 0.28590568 | 0.85570689 |
| TCTN3 | 12.5395978 | -0.5354081 | 0.53326469 | -1.0040194 | 0.31536926 | NA |
| ENTPD1 | 74.1185713 | -0.1442915 | 0.61805777 | -0.2334596 | 0.81540454 | 0.98320463 |
| ENTPD1-AS1 | 53.0689925 | 0.21369435 | 0.37136647 | 0.57542716 | 0.56500244 | 0.95415437 |
| AL365273.2 | 15.6997981 | -0.9832084 | 0.50724224 | -1.9383409 | 0.05258165 | 0.54907299 |
| ZNF518A | 43.9965345 | 0.47879296 | 0.42947194 | 1.11484107 | 0.26491853 | 0.84994492 |
| BLNK | 24.5504564 | -0.8310318 | 0.4290152 | -1.9370684 | 0.05273699 | 0.54961673 |
| TM9SF3 | 60.9174604 | 0.00144041 | 0.3171354 | 0.00454193 | 0.99637608 | 0.99859588 |
| PIK3AP1 | 113.125889 | -0.3629994 | 0.58922473 | -0.6160627 | 0.53785314 | 0.95415437 |
| LCOR | 115.897157 | -0.2756753 | 0.25523006 | -1.0801051 | 0.2800954 | 0.84994492 |
| PGAM1 | 29.2838431 | 0.80161207 | 0.45768187 | 1.75146128 | 0.07986649 | 0.63230491 |
| EXOSC1 | 12.791866 | 0.32459484 | 0.5358218 | 0.60578879 | 0.54465502 | NA |
| MMS19 | 21.6933193 | -0.0337333 | 0.40861242 | -0.0825557 | 0.93420485 | 0.99514836 |
| ZFYVE27 | 11.6350306 | -0.4561536 | 0.49312378 | -0.9250286 | 0.35495105 | NA |
| R3HCC1L | 54.1889076 | 0.17379152 | 0.2919115 | 0.5953569 | 0.55160493 | 0.95415437 |
| HPS1 | 16.7683968 | -0.1078809 | 0.42633104 | -0.2530449 | 0.80023352 | 0.98320463 |
| SLC25A28 | 11.1502681 | 1.02121689 | 0.6867499 | 1.48702882 | 0.1370072 | NA |
| CUTC | 15.2514171 | -0.2631412 | 0.47333134 | -0.5559344 | 0.57825572 | 0.95415437 |
| COX15 | 11.8780571 | 0.17417191 | 0.52530319 | 0.33156455 | 0.74021809 | NA |
| DNMBP | 21.8902101 | -1.2434685 | 0.57979671 | -2.1446629 | 0.03197981 | 0.43969912 |
| ERLIN1 | 12.1658026 | 1.44307326 | 0.87664842 | 1.64612543 | 0.09973794 | NA |
| CHUK | 11.6827267 | 0.06218214 | 0.55198918 | 0.11265102 | 0.91030724 | NA |
| CWF19L1 | 19.2795474 | -0.098979 | 0.39908073 | -0.2480174 | 0.80412096 | 0.98320463 |
| BLOC1S2 | 10.2149129 | 0.24172568 | 0.6278616 | 0.38499835 | 0.70023864 | NA |
| HIF1AN | 8.99497025 | 0.12057968 | 0.56972519 | 0.21164533 | 0.83238374 | NA |
| SLF2 | 28.1104398 | -0.1754937 | 0.36696098 | -0.4782354 | 0.63248268 | 0.95877428 |
| BTRC | 45.1363302 | 0.45910519 | 0.33744059 | 1.36055117 | 0.17365557 | 0.76214163 |
| FBXW4 | 47.3578121 | -0.414885 | 0.32871135 | -1.262156 | 0.20689266 | 0.79234878 |
| OGA | 105.929508 | 0.15765927 | 0.30970604 | 0.509061 | 0.61070947 | 0.95688177 |
| ARMH3 | 79.6243858 | 0.5207633 | 0.2938298 | 1.77232977 | 0.07633983 | 0.62402456 |
| GBF1 | 73.6989879 | -0.1506748 | 0.30355102 | -0.496374 | 0.61963057 | 0.95688177 |
| NFKB2 | 9.23789251 | 0.01821312 | 0.82524311 | 0.02207001 | 0.98239211 | NA |
| ACTR1A | 12.2709578 | -0.2458912 | 0.51335392 | -0.4789897 | 0.63194598 | NA |
| SUFU | 30.5088134 | 0.36328661 | 0.38194655 | 0.95114517 | 0.34153069 | 0.88510795 |
| TRIM8 | 15.9736668 | 0.14700733 | 0.57462028 | 0.25583388 | 0.79807911 | 0.98320463 |
| ARL3 | 10.5878255 | 0.46572236 | 0.53953217 | 0.86319666 | 0.38802935 | NA |
| WBP1L | 44.0593175 | -0.1109885 | 0.34816794 | -0.3187786 | 0.74989439 | 0.97818861 |
| CNNM2 | 23.6310546 | 1.25574745 | 0.57507428 | 2.18362652 | 0.0289897 | 0.42316846 |
| NT5C2 | 115.172169 | 0.15394392 | 0.26304639 | 0.58523487 | 0.5583898 | 0.95415437 |
| ATP5MD | 8.55805682 | 0.41479832 | 0.70273325 | 0.59026425 | 0.5550135 | NA |
| PDCD11 | 9.37455642 | -0.6922853 | 0.57047025 | -1.2135345 | 0.22492553 | NA |
| SH3PXD2A | 16.3603993 | -0.3115879 | 0.53290549 | -0.5846964 | 0.55875188 | 0.95415437 |
| STN1 | 10.2471343 | -0.0854889 | 0.53191385 | -0.1607194 | 0.87231438 | NA |
| SLK | 31.2530914 | 0.45874704 | 0.3970587 | 1.15536327 | 0.2479417 | 0.83817091 |
| GSTO1 | 26.0167695 | 0.53489565 | 0.51508752 | 1.03845586 | 0.29905787 | 0.86225566 |
| ITPRIP | 11.5231135 | -0.26612 | 0.52056154 | -0.5112171 | 0.60919905 | NA |
| XPNPEP1 | 18.324709 | 0.56087861 | 0.45710438 | 1.22702523 | 0.21981311 | 0.81042354 |

| | | | | | | |
|---|---|---|---|---|---|---|
| ADD3 | 102.56975 | 0.02346036 | 0.31494229 | 0.07449099 | 0.94061971 | 0.99514836 |
| MXI1 | 24.3939158 | -0.1630375 | 0.41757229 | -0.3904413 | 0.69621024 | 0.9685713 |
| SMNDC1 | 11.7024522 | -0.6342294 | 0.51792356 | -1.2245618 | 0.22074036 | NA |
| SMC3 | 23.1714069 | 0.33408458 | 0.39825759 | 0.83886556 | 0.40154476 | 0.91528253 |
| PDCD4 | 73.5946304 | 0.80573749 | 0.47437113 | 1.69853819 | 0.08940623 | 0.64845882 |
| BBIP1 | 12.8367259 | -1.2873664 | 0.47582481 | -2.7055469 | 0.0068192 | NA |
| SHOC2 | 60.5441151 | -0.1501723 | 0.32477946 | -0.4623823 | 0.64380716 | 0.95992767 |
| ACSL5 | 16.4119306 | 0.09037169 | 0.49489594 | 0.18260745 | 0.85510604 | 0.98751288 |
| ZDHHC6 | 11.2449571 | 0.20322898 | 0.53790784 | 0.37781374 | 0.70556897 | NA |
| VTI1A | 187.273962 | -0.2033214 | 0.20728764 | -0.980866 | 0.3266588 | 0.87715548 |
| TCF7L2 | 161.670607 | 0.05759432 | 0.68282201 | 0.08434749 | 0.93278016 | 0.99514836 |
| CASP7 | 11.6846821 | -0.3587224 | 0.54929777 | -0.6530564 | 0.51371993 | NA |
| NHLRC2 | 30.0087873 | -0.5095609 | 0.41048681 | -1.2413576 | 0.21447366 | 0.80634897 |
| CCDC186 | 30.365426 | -0.3272471 | 0.57145446 | -0.5726566 | 0.56687726 | 0.95415437 |
| ABLIM1 | 30.6558322 | -0.4241072 | 0.5019559 | -0.8449092 | 0.39816153 | 0.91429865 |
| FAM160B1 | 20.757955 | 1.110587 | 0.40328509 | 2.7538509 | 0.00588986 | 0.19198507 |
| SHTN1 | 37.5783194 | 0.31404383 | 0.58819878 | 0.53390765 | 0.59340543 | 0.95490132 |
| PDZD8 | 39.7538333 | 0.79568324 | 0.60877087 | 1.30703238 | 0.19120171 | 0.78122755 |
| RAB11FIP2 | 29.4848336 | -0.0149131 | 0.39197831 | -0.0380456 | 0.96965129 | 0.99666609 |
| FAM204A | 20.4838029 | -0.0231175 | 0.41420853 | -0.0558112 | 0.95549219 | 0.99514836 |
| CACUL1 | 56.7661257 | 0.16489234 | 0.29261116 | 0.56352033 | 0.57308062 | 0.95415437 |
| EIF3A | 67.4845181 | -0.5520744 | 0.31059608 | -1.7774675 | 0.07549135 | 0.62260674 |
| FAM45A | 14.9011521 | 0.91526181 | 0.61673638 | 1.48404056 | 0.13779816 | 0.73044824 |
| PRDX3 | 13.5780794 | 0.21018079 | 0.51267971 | 0.40996509 | 0.68183155 | 0.9673431 |
| GRK5 | 75.1479382 | -1.317863 | 0.35915752 | -3.6693176 | 0.0002432 | 0.02990804 |
| RGS10 | 24.8576583 | 0.58003458 | 0.48052306 | 1.20709 | 0.22739749 | 0.82088564 |
| TIAL1 | 36.3318679 | -0.6901686 | 0.34673084 | -1.9905024 | 0.04653562 | 0.52664239 |
| INPP5F | 30.095531 | -1.2764108 | 0.51888632 | -2.4599046 | 0.0138974 | 0.31861834 |
| MCMBP | 24.4767268 | -0.1240323 | 0.43798243 | -0.2831901 | 0.77703114 | 0.98264092 |
| SEC23IP | 21.3215555 | -0.0272311 | 0.42551359 | -0.0639958 | 0.94897358 | 0.99514836 |
| WDR11 | 40.2503515 | -0.3864629 | 0.3356813 | -1.1512794 | 0.24961733 | 0.83865855 |
| ATE1 | 50.3596546 | 0.35428056 | 0.30936376 | 1.14519088 | 0.25213009 | 0.84008912 |
| PLEKHA1 | 39.823377 | -0.086399 | 0.37858235 | -0.2282172 | 0.81947742 | 0.98320463 |
| IKZF5 | 9.71050465 | 0.82209403 | 0.56497913 | 1.45508743 | 0.14564506 | NA |
| ACADSB | 9.7337183 | 1.25086707 | 0.64002004 | 1.9544186 | 0.05065173 | NA |
| BUB3 | 17.8401171 | 0.10322578 | 0.45336163 | 0.22768973 | 0.81988746 | 0.98320463 |
| CHST15 | 92.1387092 | -0.4356249 | 0.61629756 | -0.7068419 | 0.47966475 | 0.93257966 |
| OAT | 8.70175066 | 0.7481213 | 0.71022965 | 1.05335128 | 0.29218003 | NA |
| LHPP | 8.60384247 | 0.83535198 | 0.69878549 | 1.19543407 | 0.23191748 | NA |
| ABRAXAS2 | 9.6916314 | -0.2460155 | 0.63099911 | -0.3898825 | 0.69662342 | NA |
| ZRANB1 | 22.2572125 | -0.4864117 | 0.38323858 | -1.2692139 | 0.20436479 | 0.78882329 |
| CTBP2 | 89.9641338 | -0.36452 | 0.42151542 | -0.8647844 | 0.38715712 | 0.90910959 |
| EDRF1 | 19.313306 | 0.27318838 | 0.42346811 | 0.64512149 | 0.51884846 | 0.94668603 |
| BCCIP | 10.7346434 | -0.5627428 | 0.59262656 | -0.949574 | 0.34232873 | NA |
| DHX32 | 16.5666061 | 0.19149632 | 0.58224823 | 0.3288912 | 0.74223793 | 0.9765917 |
| DOCK1 | 24.2571656 | 0.20612851 | 0.63361406 | 0.32532186 | 0.74493748 | 0.97805001 |
| PTPRE | 106.204828 | 1.07488549 | 0.46278327 | 2.32265417 | 0.02019774 | 0.36889203 |
| MGMT | 33.7072586 | 0.00664563 | 0.44292222 | 0.01500405 | 0.98802895 | 0.99811561 |
| GLRX3 | 17.1599221 | -0.8897751 | 0.4252833 | -2.0921939 | 0.03642117 | 0.46873201 |
| PPP2R2D | 40.4816617 | -0.0308694 | 0.3035105 | -0.1017078 | 0.91898858 | 0.99498068 |
| BNIP3 | 12.002386 | 1.32832978 | 0.89467657 | 1.48470388 | 0.13762228 | NA |
| INPP5A | 28.9152577 | -0.3260776 | 0.50869673 | -0.6410058 | 0.5215189 | 0.94895469 |
| ADAM8 | 11.9502168 | 0.49861456 | 0.69845096 | 0.71388628 | 0.4752975 | NA |
| TUBGCP2 | 22.8009677 | 0.3029953 | 0.38044603 | 0.79642125 | 0.42578723 | 0.92046401 |

| | | | | | | |
|---|---|---|---|---|---|---|
| ECHS1 | 10.784611 | 0.25459047 | 0.60435756 | 0.42125801 | 0.67356669 | NA |
| KDM5A | 78.0086247 | -0.0804362 | 0.28766312 | -0.2796196 | 0.77976939 | 0.98264092 |
| NINJ2 | 8.0807273 | -0.5460848 | 0.6619309 | -0.8249877 | 0.4093786 | NA |
| WNK1 | 171.780233 | 0.08392893 | 0.24903721 | 0.33701362 | 0.73610663 | 0.97605445 |
| RAD52 | 9.44349386 | -0.609596 | 0.58936659 | -1.0343239 | 0.30098475 | NA |
| ERC1 | 132.290433 | -0.1487337 | 0.26002622 | -0.5719952 | 0.56732524 | 0.95415437 |
| ADIPOR2 | 38.6948558 | 0.43013326 | 0.36112544 | 1.191091 | 0.23361786 | 0.82817204 |
| DCP1B | 11.0563397 | 1.41646464 | 0.64059684 | 2.21116396 | 0.02702449 | NA |
| ITFG2 | 23.4412275 | -0.2592842 | 0.51140458 | -0.507004 | 0.61215203 | 0.95688177 |
| TULP3 | 8.69112521 | -0.6082348 | 0.55004713 | -1.1057866 | 0.26881889 | NA |
| CRACR2A | 23.5227557 | -0.4154609 | 0.43152139 | -0.9627818 | 0.33565705 | 0.88047896 |
| PARP11 | 25.1203648 | 0.26236023 | 0.51039209 | 0.51403662 | 0.60722639 | 0.95688177 |
| CCND2 | 31.1182271 | 0.41109782 | 0.44250661 | 0.92902075 | 0.35287833 | 0.89109867 |
| TIGAR | 15.0248864 | 0.10584281 | 0.59161158 | 0.17890591 | 0.85801158 | 0.98751288 |
| C12orf4 | 14.5820111 | -0.1976146 | 0.48888654 | -0.4042135 | 0.68605571 | 0.9673431 |
| CD9 | 19.0523639 | -0.1926125 | 0.54479034 | -0.3535535 | 0.72367351 | 0.97475123 |
| TNFRSF1A | 41.0070689 | 0.29819003 | 0.41642923 | 0.71606412 | 0.47395176 | 0.9314919 |
| LTBR | 13.0526459 | 0.80066886 | 0.57729458 | 1.38693292 | 0.16546222 | 0.75337545 |
| MRPL51 | 10.8510646 | -0.3314486 | 0.63588845 | -0.521237 | 0.60220171 | NA |
| NCAPD2 | 12.630865 | -0.8403473 | 0.68310319 | -1.2301909 | 0.21862562 | NA |
| GAPDH | 119.11859 | 0.2245037 | 0.55742502 | 0.40275138 | 0.68713113 | 0.9673431 |
| IFFO1 | 8.98895195 | 1.43498057 | 0.61522451 | 2.33245026 | 0.01967702 | NA |
| CHD4 | 9.95126687 | 0.35830948 | 0.57285995 | 0.62547482 | 0.53165947 | NA |
| ING4 | 9.96143569 | 0.81523296 | 0.66385524 | 1.22802822 | 0.21943638 | NA |
| MLF2 | 11.0818419 | -0.7371108 | 0.57031865 | -1.2924544 | 0.19619983 | NA |
| PTMS | 15.4440215 | -0.7756934 | 0.44066437 | -1.7602817 | 0.07836006 | 0.62756088 |
| CD4 | 34.8343123 | 1.0563433 | 0.60090702 | 1.75791471 | 0.07876202 | 0.62793067 |
| TPI1 | 29.6395869 | 0.84451113 | 0.57681719 | 1.464088 | 0.14316991 | 0.73044824 |
| PTPN6 | 21.5272419 | -0.7213082 | 0.74801868 | -0.9642917 | 0.33489973 | 0.88047896 |
| PHB2 | 13.3537097 | 0.19124764 | 0.6786105 | 0.2818224 | 0.77807969 | 0.98264092 |
| LPCAT3 | 22.293551 | -0.4770227 | 0.54858732 | -0.8695474 | 0.38454782 | 0.90910959 |
| C1S | 12.7929686 | 0.40423886 | 0.66211654 | 0.61052525 | 0.54151392 | NA |
| C1RL | 15.6589839 | 0.97154212 | 0.57702937 | 1.68369613 | 0.09224041 | 0.65902576 |
| CLSTN3 | 9.80404072 | 0.07575334 | 0.58989865 | 0.12841756 | 0.89781854 | NA |
| CD163L1 | 30.5737189 | 0.55557062 | 0.69356681 | 0.80103403 | 0.42311194 | 0.91985469 |
| CD163 | 128.074147 | 0.3910937 | 0.77688039 | 0.50341559 | 0.61467212 | 0.95688177 |
| SLC2A3 | 165.384405 | -0.2576181 | 0.76695635 | -0.3358967 | 0.73694875 | 0.97605445 |
| C3AR1 | 18.5962644 | 0.95923202 | 0.56196828 | 1.70691487 | 0.08783788 | 0.64676224 |
| NECAP1 | 12.0953275 | -0.2197027 | 0.51325647 | -0.4280563 | 0.66861014 | NA |
| LINC00937 | 14.1882076 | 1.36680593 | 0.77830222 | 1.75613778 | 0.07906489 | 0.62793067 |
| RIMKLB | 12.148865 | -1.0447161 | 0.48512943 | -2.1534791 | 0.03128104 | NA |
| M6PR | 34.9043121 | 0.14691592 | 0.40711546 | 0.36087041 | 0.71819632 | 0.97348719 |
| KLRG1 | 17.2825829 | -0.5263137 | 0.87414769 | -0.6020879 | 0.54711565 | 0.95415437 |
| A2M | 30.6051363 | -0.5107227 | 0.64672123 | -0.7897107 | 0.42969676 | 0.92204028 |
| PZP | 15.6039415 | -1.1018613 | 1.59832109 | -0.6893867 | 0.49057995 | 0.93633431 |
| KLRB1 | 11.5096093 | 1.27202519 | 1.07740021 | 1.18064316 | 0.23774451 | NA |
| CLEC2D | 123.060291 | -0.634563 | 0.2807733 | -2.2600545 | 0.02381787 | 0.39492781 |
| CD69 | 45.2674441 | 1.10427664 | 0.64600144 | 1.70940275 | 0.08737637 | 0.64676224 |
| KLRF1 | 19.4238809 | 1.98095125 | 1.03582391 | 1.91244017 | 0.05581976 | 0.55970424 |
| CLEC2B | 39.9734058 | 0.49402312 | 0.67004484 | 0.73729859 | 0.46094079 | 0.9314919 |
| CLEC12A | 29.5892859 | 0.75116528 | 0.89935989 | 0.83522213 | 0.40359265 | 0.91528253 |
| CLEC7A | 48.6739755 | 0.60688059 | 0.68922151 | 0.88053055 | 0.37857196 | 0.9051267 |
| GABARAPL1 | 18.5506554 | -0.0717583 | 0.50341093 | -0.1425442 | 0.88665016 | 0.99019008 |
| KLRD1 | 45.1752489 | 1.00656415 | 0.86047044 | 1.16978353 | 0.24208809 | 0.83493679 |

| | | | | | | |
|---|---|---|---|---|---|---|
| AC022075.1 | 20.3731863 | 0.50225167 | 0.60172844 | 0.83468162 | 0.40389699 | 0.91528253 |
| KLRC4-KLRK1 | 13.1028794 | 1.10566615 | 0.81893395 | 1.3501286 | 0.17697474 | 0.76409729 |
| YBX3 | 34.7195947 | -1.2658398 | 0.52403431 | -2.4155667 | 0.01571075 | 0.3268544 |
| AC244131.2 | 16.3966231 | -0.0538139 | 0.94563831 | -0.0569075 | 0.95461891 | 0.99514836 |
| ETV6 | 354.033378 | 0.73594083 | 0.44837743 | 1.64134227 | 0.10072639 | 0.6683775 |
| BORCS5 | 34.7271446 | -0.0954125 | 0.42647619 | -0.2237228 | 0.82297297 | 0.98320463 |
| DUSP16 | 46.9014728 | 0.71705101 | 0.48584294 | 1.47589056 | 0.13997327 | 0.73044824 |
| CREBL2 | 21.2725265 | 0.58116334 | 0.47992802 | 1.21093856 | 0.22591895 | 0.81875277 |
| CDKN1B | 24.3605193 | 0.68159287 | 0.61433761 | 1.10947606 | 0.26722487 | 0.84994492 |
| APOLD1 | 20.4150583 | 0.54284611 | 0.79586031 | 0.68208718 | 0.49518383 | 0.93805563 |
| ATF7IP | 169.03557 | 0.24346377 | 0.39720878 | 0.61293652 | 0.53991832 | 0.95415437 |
| PLBD1 | 16.9030988 | 0.66334142 | 0.91086974 | 0.72825058 | 0.46646021 | 0.9314919 |
| H2AFJ | 18.7469486 | -0.4898258 | 0.65868065 | -0.7436469 | 0.45709012 | 0.9314919 |
| WBP11 | 12.4400828 | -0.0633042 | 0.58306849 | -0.1085708 | 0.91354291 | NA |
| MGP | 22.3096795 | -0.4726782 | 0.83997487 | -0.5627289 | 0.57361948 | 0.95415437 |
| ARHGDIB | 61.3711271 | 0.25399563 | 0.55568467 | 0.4570859 | 0.64760931 | 0.96049083 |
| PTPRO | 14.1183658 | -1.2704971 | 0.70465441 | -1.8030073 | 0.07138706 | 0.60903299 |
| EPS8 | 50.310638 | -0.419466 | 0.3950844 | -1.0617123 | 0.28836631 | 0.85703857 |
| STRAP | 16.8718042 | 0.0782347 | 0.45490946 | 0.17197862 | 0.86345434 | 0.98751288 |
| DERA | 31.8737855 | 0.08889904 | 0.36791549 | 0.24162897 | 0.80906767 | 0.98320463 |
| MGST1 | 23.4379326 | 1.25920334 | 0.84181373 | 1.49582181 | 0.1347001 | 0.73044824 |
| PLEKHA5 | 47.195009 | 0.53082008 | 0.45890083 | 1.15672069 | 0.24738649 | 0.83817091 |
| AEBP2 | 62.4231149 | 0.24168935 | 0.39635681 | 0.60977721 | 0.5420094 | 0.95415437 |
| PYROXD1 | 14.6786561 | -0.4076623 | 0.45304506 | -0.8998272 | 0.3682122 | 0.89868152 |
| RECQL | 20.7452105 | -0.3586239 | 0.42040133 | -0.8530513 | 0.39363088 | 0.90940709 |
| GOLT1B | 11.7110793 | 0.10797067 | 0.56495502 | 0.19111374 | 0.84843648 | NA |
| LDHB | 32.4488356 | 1.08489581 | 0.69395416 | 1.56335371 | 0.11796942 | 0.70279044 |
| ST8SIA1 | 25.5662597 | 1.46926151 | 0.69536653 | 2.112931 | 0.03460667 | 0.45816587 |
| C2CD5 | 63.8446585 | -0.1682793 | 0.46652996 | -0.3607042 | 0.71832058 | 0.97348719 |
| ETNK1 | 85.4822305 | 0.04207514 | 0.30684013 | 0.13712397 | 0.89093281 | 0.99080204 |
| SOX5 | 230.82018 | -5.7257762 | 0.86896652 | -6.5891794 | 4.42E-11 | 2.45E-07 |
| BCAT1 | 35.7466402 | 0.65034436 | 0.67451662 | 0.96416358 | 0.33496392 | 0.88047896 |
| LRMP | 63.309734 | -2.0376428 | 0.5181409 | -3.9326036 | 8.40E-05 | 0.01550087 |
| KRAS | 39.3781 | -0.5766541 | 0.3626916 | -1.5899295 | 0.1118507 | 0.69385269 |
| RASSF8 | 15.8589951 | -0.1359323 | 0.59582879 | -0.2281399 | 0.81953748 | 0.98320463 |
| ITPR2 | 266.154236 | 0.38189303 | 0.33055367 | 1.15531323 | 0.24796218 | 0.83817091 |
| INTS13 | 7.49706823 | -0.478873 | 0.629244 | -0.7610291 | 0.44663966 | NA |
| FGFR1OP2 | 43.6616823 | 0.36077344 | 0.38040234 | 0.94839964 | 0.34292604 | 0.88530691 |
| TM7SF3 | 26.2116374 | 0.78461176 | 0.56869929 | 1.37966017 | 0.1676913 | 0.7567539 |
| MED21 | 9.73666193 | 0.55263321 | 0.63835927 | 0.86570877 | 0.38664989 | NA |
| STK38L | 32.0463215 | -0.2366891 | 0.42738103 | -0.5538127 | 0.57970702 | 0.95415437 |
| PPFIBP1 | 11.4247558 | 0.40780453 | 0.74088615 | 0.55042806 | 0.58202581 | NA |
| MRPS35 | 11.8870963 | 0.5750098 | 0.57031602 | 1.00823015 | 0.31334399 | NA |
| KLHL42 | 10.829643 | -0.5748914 | 0.61936277 | -0.9281981 | 0.35330481 | NA |
| CCDC91 | 176.971996 | -0.2233158 | 0.26944844 | -0.8287888 | 0.40722394 | 0.9169561 |
| FAR2 | 53.0613825 | 0.53211105 | 0.53241668 | 0.99942596 | 0.31758839 | 0.87265846 |
| ERGIC2 | 16.2990271 | -0.7083033 | 0.43036989 | -1.6458011 | 0.09980471 | 0.6683775 |
| TMTC1 | 24.138884 | 0.94536241 | 1.09873484 | 0.86040997 | 0.38956309 | 0.90910959 |
| IPO8 | 24.5208461 | -0.0109233 | 0.42763564 | -0.0255436 | 0.97962138 | 0.99758514 |
| CAPRIN2 | 11.5200856 | 1.18313952 | 0.57729191 | 2.04946493 | 0.04041667 | NA |
| SINHCAF | 25.4332151 | 0.25299758 | 0.4639194 | 0.54534814 | 0.5855141 | 0.95415437 |
| DENND5B | 36.7242187 | -0.5932943 | 0.86161871 | -0.688581 | 0.49108698 | 0.93633431 |
| AMN1 | 13.0789916 | -1.0897415 | 0.5639082 | -1.9324803 | 0.05330025 | 0.551221 |
| KIAA1551 | 108.533985 | 0.06548036 | 0.32943171 | 0.19876762 | 0.84244453 | 0.9842451 |

| | | | | | | |
|---|---|---|---|---|---|---|
| BICD1 | 43.8839926 | -0.8134077 | 0.61308225 | -1.3267513 | 0.18459096 | 0.77694741 |
| FGD4 | 190.389197 | -0.3152753 | 0.55639599 | -0.5666383 | 0.57095993 | 0.95415437 |
| DNM1L | 41.7608632 | -0.7483963 | 0.33134873 | -2.2586363 | 0.02390602 | 0.39492781 |
| YARS2 | 10.8388538 | -0.3666713 | 0.60711266 | -0.6039593 | 0.5458707 | NA |
| CPNE8 | 59.136108 | 0.49761699 | 0.43231347 | 1.15105594 | 0.24970922 | 0.83865855 |
| KIF21A | 17.656156 | 0.32381987 | 0.5585799 | 0.57971988 | 0.56210353 | 0.95415437 |
| SLC2A13 | 38.6420489 | 0.80601081 | 0.42039376 | 1.91727588 | 0.05520289 | 0.55645321 |
| LRRK2 | 94.7023692 | 0.44709832 | 0.63365492 | 0.70558644 | 0.48044536 | 0.93257966 |
| GXYLT1 | 17.1664065 | 0.00668162 | 0.53909291 | 0.01239419 | 0.99011112 | 0.99822826 |
| YAF2 | 50.1776112 | -0.5400904 | 0.2826524 | -1.9107936 | 0.05603111 | 0.55970424 |
| PPHLN1 | 70.6623148 | 0.05406119 | 0.30143958 | 0.17934337 | 0.8576681 | 0.98751288 |
| PUS7L | 13.4752229 | -1.0711494 | 0.47003895 | -2.2788525 | 0.02267584 | 0.38988793 |
| IRAK4 | 24.2891001 | -0.2188624 | 0.44029324 | -0.4970833 | 0.61913032 | 0.95688177 |
| TMEM117 | 58.3702531 | -0.4061453 | 0.4194478 | -0.9682857 | 0.3329017 | 0.88047896 |
| NELL2 | 26.2497151 | 0.28886822 | 1.08133326 | 0.26714079 | 0.78936076 | 0.98320463 |
| ANO6 | 67.4805513 | 0.29288986 | 0.33327155 | 0.87883248 | 0.37949211 | 0.90600059 |
| ARID2 | 114.093641 | 0.18861767 | 0.25010036 | 0.75416794 | 0.45074839 | 0.92914597 |
| SCAF11 | 139.563999 | -0.1627828 | 0.1981722 | -0.8214207 | 0.41140665 | 0.91823911 |
| SLC38A1 | 115.235202 | -1.1877138 | 0.34692967 | -3.4235001 | 0.0006182 | 0.05005251 |
| SLC38A2 | 59.9660047 | -0.0859807 | 0.59614979 | -0.1442267 | 0.88532148 | 0.99019008 |
| AC008014.1 | 66.2987795 | 0.05883406 | 0.60670215 | 0.09697355 | 0.9227474 | 0.99514836 |
| PCED1B | 94.0198239 | 0.6059459 | 0.41204039 | 1.47059831 | 0.14139978 | 0.73044824 |
| RPAP3 | 13.9849733 | -0.5174109 | 0.50125178 | -1.0322375 | 0.30196087 | 0.86225566 |
| HDAC7 | 19.2033218 | -0.2274356 | 0.53236875 | -0.4272144 | 0.66922317 | 0.96412305 |
| VDR | 10.7379188 | 0.8726715 | 0.79811658 | 1.09341357 | 0.27421226 | NA |
| SENP1 | 16.6435781 | -0.2778683 | 0.51780287 | -0.5366295 | 0.59152354 | 0.95415437 |
| ZNF641 | 10.4138175 | -0.5437478 | 0.57177615 | -0.9509802 | 0.34161442 | NA |
| CCNT1 | 18.6903719 | -0.2267136 | 0.50856043 | -0.4457949 | 0.65574541 | 0.96211819 |
| DDX23 | 9.97504432 | -0.0724963 | 0.68548204 | -0.1057595 | 0.91577314 | NA |
| CCDC65 | 10.1616795 | 0.37550808 | 0.7155893 | 0.52475364 | 0.59975447 | NA |
| PRKAG1 | 13.6083492 | -0.303291 | 0.58209267 | -0.5210356 | 0.60234197 | 0.95688177 |
| KMT2D | 23.3204157 | -0.5759675 | 0.47985366 | -1.2002982 | 0.23002356 | 0.82393202 |
| LMBR1L | 13.5034391 | 1.05094181 | 0.56536694 | 1.85886675 | 0.06304603 | 0.58343933 |
| TUBA1B | 40.2051413 | 0.42330558 | 0.58588732 | 0.7225034 | 0.46998504 | 0.9314919 |
| TUBA1A | 20.497914 | 1.22559692 | 0.88751798 | 1.38092631 | 0.16730162 | 0.7567539 |
| TUBA1C | 13.8662418 | 0.89215381 | 0.6068966 | 1.47002606 | 0.1415547 | 0.73044824 |
| SPATS2 | 38.6462933 | 0.44455115 | 0.62925786 | 0.70646896 | 0.47989655 | 0.93257966 |
| PRPF40B | 11.041084 | 0.33907682 | 0.62938134 | 0.53874623 | 0.59006197 | NA |
| FMNL3 | 36.1843291 | -0.2447825 | 0.43516296 | -0.5625077 | 0.57377014 | 0.95415437 |
| TMBIM6 | 71.3793207 | -0.0264957 | 0.41587744 | -0.0637103 | 0.94920094 | 0.99514836 |
| NCKAP5L | 11.5343204 | -0.081422 | 0.52626919 | -0.1547156 | 0.87704555 | NA |
| AQP2 | 14.7191696 | -2.3852012 | 0.83837538 | -2.8450277 | 0.00444076 | 0.16717788 |
| AC074032.1 | 25.2920407 | -0.1231061 | 0.47978099 | -0.2565881 | 0.79749676 | 0.98320463 |
| CERS5 | 21.2833597 | -0.0019853 | 0.44377038 | -0.0044737 | 0.99643051 | 0.99859588 |
| LIMA1 | 13.5835396 | -0.0501832 | 0.57694137 | -0.0869815 | 0.93068623 | 0.99514836 |
| LARP4 | 40.0398471 | -0.1424545 | 0.35512291 | -0.4011414 | 0.68831602 | 0.9673431 |
| DIP2B | 133.149432 | -0.6748829 | 0.41109764 | -1.6416608 | 0.10066031 | 0.6683775 |
| ATF1 | 19.5856082 | -0.3022154 | 0.49485497 | -0.6107151 | 0.54138818 | 0.95415437 |
| METTL7A | 17.9604216 | -0.1892881 | 0.49188245 | -0.3848238 | 0.70036798 | 0.96991236 |
| SLC11A2 | 33.3305218 | 0.31612087 | 0.4342088 | 0.72803885 | 0.46658981 | 0.9314919 |
| LETMD1 | 15.9816572 | -0.0461663 | 0.47439411 | -0.0973164 | 0.92247515 | 0.99514836 |
| TFCP2 | 64.6974 | -0.0763946 | 0.31031895 | -0.2461809 | 0.8055422 | 0.98320463 |
| DAZAP2 | 36.5560483 | 0.15917181 | 0.47238811 | 0.33695136 | 0.73615357 | 0.97605445 |
| BIN2 | 60.6014369 | 0.98940293 | 0.39943487 | 2.47700687 | 0.01324894 | 0.31333166 |

| | | | | | | |
|---|---|---|---|---|---|---|
| GALNT6 | 9.79269939 | 0.18822631 | 0.57131001 | 0.32946441 | 0.74180469 | NA |
| SLC4A8 | 18.2062629 | -0.2670015 | 0.4666475 | -0.5721696 | 0.56720708 | 0.95415437 |
| ACVR1B | 18.7128525 | 0.38540698 | 0.55950997 | 0.68882952 | 0.49093055 | 0.93633431 |
| NR4A1 | 24.8151388 | -0.985046 | 0.69267589 | -1.4220879 | 0.15500073 | 0.74486619 |
| EIF4B | 46.7401926 | -0.2078548 | 0.32287108 | -0.6437702 | 0.51972445 | 0.94735016 |
| AC068888.1 | 14.2103063 | -0.3993366 | 0.56076971 | -0.7121222 | 0.47638908 | 0.9314919 |
| CSAD | 11.082133 | -0.1692224 | 0.59698381 | -0.2834623 | 0.77682247 | NA |
| PFDN5 | 46.0794665 | 0.17444538 | 0.63516863 | 0.2746442 | 0.7835896 | 0.98264092 |
| SP1 | 23.9173869 | -0.4088274 | 0.38906466 | -1.0507955 | 0.29335251 | 0.86027209 |
| PRR13 | 23.8162963 | -0.03213 | 0.49351987 | -0.0651037 | 0.94809145 | 0.99514836 |
| PCBP2 | 57.7409567 | -0.474605 | 0.26926293 | -1.7626081 | 0.07796661 | 0.62717836 |
| ATP5MC2 | 37.2144865 | -0.560302 | 0.40924044 | -1.3691266 | 0.17095972 | 0.75751515 |
| CALCOCO1 | 11.0526425 | 0.04697124 | 0.52915301 | 0.08876683 | 0.92926722 | NA |
| CBX5 | 17.7614896 | 0.34612803 | 0.45955774 | 0.75317637 | 0.45134394 | 0.92914597 |
| HNRNPA1 | 24.562582 | -0.407013 | 0.53942909 | -0.7545254 | 0.45053377 | 0.92914597 |
| COPZ1 | 12.7122105 | -0.1734392 | 0.46560614 | -0.3725019 | 0.70951917 | NA |
| ITGA5 | 14.9989772 | 0.81804113 | 0.60573593 | 1.35049135 | 0.17685843 | 0.76409729 |
| NCKAP1L | 83.3902078 | 0.35570359 | 0.34111924 | 1.04275439 | 0.29706206 | 0.8608685 |
| TESPA1 | 15.5354211 | 0.65982021 | 0.7378896 | 0.8941991 | 0.37121538 | 0.90226557 |
| CD63 | 40.9538052 | 0.00941387 | 0.57513541 | 0.0163681 | 0.98694073 | 0.99811561 |
| DGKA | 19.9790476 | 0.02335841 | 0.52925029 | 0.0441349 | 0.96479687 | 0.99653147 |
| RAB5B | 12.3594018 | -0.2798844 | 0.50874654 | -0.550145 | 0.58221993 | NA |
| RPS26 | 10.9377068 | -0.6974541 | 0.52420285 | -1.3305042 | 0.18335219 | NA |
| PA2G4 | 8.73068306 | -0.8678766 | 0.69023552 | -1.257363 | 0.20862223 | NA |
| RPL41 | 172.679934 | -0.3222549 | 0.5384398 | -0.5984976 | 0.54950795 | 0.95415437 |
| ESYT1 | 15.0297878 | 0.01113227 | 0.46323565 | 0.02403155 | 0.98082744 | 0.99758514 |
| MYL6 | 68.528614 | 0.71341754 | 0.59433988 | 1.2003528 | 0.23000235 | 0.82393202 |
| SMARCC2 | 30.8558552 | -0.7131441 | 0.4219706 | -1.6900326 | 0.09102172 | 0.65685853 |
| RNF41 | 8.6263469 | 0.83184436 | 0.64119871 | 1.29732694 | 0.19451872 | NA |
| CS | 12.726024 | -0.6395485 | 0.55007913 | -1.1626481 | 0.24497229 | NA |
| STAT2 | 37.1341733 | 1.00490058 | 0.50946476 | 1.97246335 | 0.04855674 | 0.53486612 |
| RBMS2 | 13.7488589 | 0.29635214 | 0.58425994 | 0.50722653 | 0.61199588 | 0.95688177 |
| BAZ2A | 31.9398086 | -0.15778 | 0.36827518 | -0.4284296 | 0.66833835 | 0.96412305 |
| ATP5F1B | 31.0468928 | 1.07761476 | 0.45583365 | 2.36405269 | 0.01807624 | 0.34868204 |
| PTGES3 | 66.5083289 | 0.17293892 | 0.34691051 | 0.49851164 | 0.61812347 | 0.95688177 |
| NACA | 49.1332802 | -0.280134 | 0.56617825 | -0.4947806 | 0.62075501 | 0.95688177 |
| NEMP1 | 10.0785448 | 0.04812559 | 0.70684544 | 0.06808503 | 0.94571795 | NA |
| STAT6 | 46.740252 | -0.1781631 | 0.34215035 | -0.5207158 | 0.60256475 | 0.95688177 |
| LRP1 | 46.8415416 | 1.07897843 | 0.60259132 | 1.79056418 | 0.07336326 | 0.61513982 |
| R3HDM2 | 97.6232469 | -0.271373 | 0.35310098 | -0.7685422 | 0.44216512 | 0.92899684 |
| ARHGAP9 | 22.3387597 | 0.71663007 | 0.67654092 | 1.05925605 | 0.28948318 | 0.85760168 |
| MARS | 22.9824912 | -0.5518314 | 0.40993895 | -1.3461306 | 0.1782604 | 0.76504408 |
| DCTN2 | 10.9298123 | -0.4810626 | 0.57150022 | -0.8417541 | 0.39992563 | NA |
| OS9 | 30.7319174 | 0.51220643 | 0.47079864 | 1.08795222 | 0.2766162 | 0.84994492 |
| CTDSP2 | 16.1068804 | -0.0479518 | 0.55170944 | -0.086915 | 0.93073904 | 0.99514836 |
| SLC16A7 | 63.0593778 | 1.07426656 | 0.49935182 | 2.151322 | 0.03145079 | 0.43730826 |
| FAM19A2 | 40.1042713 | 2.22118812 | 0.91030422 | 2.44005033 | 0.01468522 | 0.32121732 |
| USP15 | 251.578195 | 0.44074688 | 0.27258984 | 1.6168867 | 0.10590274 | 0.67519097 |
| MON2 | 107.366945 | -0.1743977 | 0.27325286 | -0.6382284 | 0.52332501 | 0.94953463 |
| PPM1H | 12.6274328 | 1.48378298 | 0.89002062 | 1.66713326 | 0.09548791 | NA |
| SRGAP1 | 59.4127409 | -0.3777881 | 0.46381425 | -0.8145245 | 0.41534452 | 0.91894074 |
| XPOT | 10.1818868 | 0.01496264 | 0.57948751 | 0.02582047 | 0.97940054 | NA |
| TBK1 | 39.7704566 | -0.0689698 | 0.32203818 | -0.2141666 | 0.83041716 | 0.98320463 |
| RASSF3 | 107.447937 | 0.32831092 | 0.43072716 | 0.76222479 | 0.44592585 | 0.92914597 |

| | | | | | | |
|---|---|---|---|---|---|---|
| GNS | 46.5758734 | 0.73960352 | 0.48907106 | 1.5122619 | 0.13046725 | 0.72931666 |
| LEMD3 | 44.4984096 | 0.25166646 | 0.390143 | 0.64506211 | 0.51888694 | 0.94668603 |
| IRAK3 | 146.113412 | 1.40976654 | 0.51654846 | 2.72920482 | 0.00634873 | 0.19738118 |
| HELB | 33.1219986 | 0.98911907 | 0.40753121 | 2.42710016 | 0.01522005 | 0.32566073 |
| GRIP1 | 14.114319 | -0.5358997 | 0.68957883 | -0.7771406 | 0.43707585 | 0.92496281 |
| CAND1 | 29.7069131 | -0.1085609 | 0.41260125 | -0.2631133 | 0.79246323 | 0.98320463 |
| DYRK2 | 13.8223144 | 0.51860843 | 0.70109388 | 0.73971325 | 0.45947401 | 0.9314919 |
| IFNG-AS1 | 47.6580617 | -0.0818464 | 0.87638723 | -0.0933907 | 0.92559317 | 0.99514836 |
| LINC02384 | 25.0566593 | 0.2833739 | 0.94422289 | 0.30011335 | 0.76409069 | 0.98222198 |
| RAP1B | 198.694431 | 0.35533659 | 0.23399539 | 1.51856236 | 0.12887269 | 0.72847954 |
| NUP107 | 28.7101316 | -0.1700966 | 0.3846215 | -0.4422441 | 0.65831261 | 0.96280444 |
| SLC35E3 | 23.7251523 | -0.1868376 | 0.510383 | -0.3660732 | 0.71431043 | 0.97348719 |
| MDM2 | 55.5448123 | -0.0733192 | 0.39928433 | -0.1836265 | 0.85430644 | 0.98751288 |
| CPM | 34.4421139 | 1.18593366 | 0.60295834 | 1.96685838 | 0.04919954 | 0.53564799 |
| CPSF6 | 44.8970853 | -0.4352909 | 0.43148975 | -1.0088093 | 0.31306609 | 0.8706069 |
| LYZ | 161.380421 | 1.78744495 | 1.56591677 | 1.14146868 | 0.25367494 | 0.84026145 |
| AC020656.1 | 49.2695519 | 0.51190037 | 2.21874003 | 0.2307167 | 0.8175349 | 0.98320463 |
| FRS2 | 45.8248079 | -0.0208119 | 0.45354846 | -0.0458867 | 0.96340052 | 0.99653147 |
| CCT2 | 13.8350526 | -0.3320928 | 0.48122544 | -0.6900981 | 0.4901325 | 0.93633431 |
| CNOT2 | 85.7268003 | 0.11836157 | 0.38577891 | 0.30681192 | 0.75898653 | 0.98071076 |
| ZFC3H1 | 82.1714179 | 0.08908407 | 0.26872689 | 0.3315041 | 0.74026374 | 0.97605445 |
| RAB21 | 37.2975143 | 0.23562791 | 0.38002343 | 0.62003521 | 0.53523461 | 0.95378558 |
| TBC1D15 | 41.0580627 | 0.51263876 | 0.37756327 | 1.35775591 | 0.17454115 | 0.76214163 |
| ATXN7L3B | 10.1396828 | 0.51934178 | 0.54385242 | 0.95493145 | 0.33961237 | NA |
| CAPS2 | 12.8491298 | -0.0999464 | 0.69090154 | -0.1446608 | 0.8849787 | NA |
| GLIPR1 | 28.8803274 | -0.580273 | 0.49427575 | -1.1739864 | 0.24040046 | 0.83493679 |
| KRR1 | 11.7543477 | -0.4302878 | 0.55222808 | -0.7791849 | 0.43587078 | NA |
| NAP1L1 | 51.0191716 | -0.351659 | 0.38589762 | -0.9112755 | 0.36215024 | 0.89853007 |
| OSBPL8 | 151.891095 | 0.05236788 | 0.34225856 | 0.15300678 | 0.87839293 | 0.99019008 |
| ZDHHC17 | 40.9221191 | 0.20086338 | 0.32159559 | 0.62458374 | 0.53224429 | 0.95265314 |
| SYT1 | 105.625889 | -4.162282 | 1.12835544 | -3.6888039 | 2.25E-04 | 0.02990804 |
| PAWR | 7.78018253 | -1.1758669 | 0.6454229 | -1.821855 | 0.068477 | NA |
| PPP1R12A | 155.61179 | 0.1912308 | 0.25701763 | 0.74403769 | 0.45685367 | 0.9314919 |
| METTL25 | 20.8240562 | -0.4698771 | 0.42018736 | -1.1182562 | 0.26345758 | 0.84928231 |
| TMTC2 | 66.8789465 | 1.18960547 | 0.33898642 | 3.5093013 | 0.00044929 | 0.04075978 |
| CEP290 | 20.8028595 | -0.7019604 | 0.42291524 | -1.6598134 | 0.09695199 | 0.66178928 |
| TMTC3 | 12.6335707 | -0.4326497 | 0.68284003 | -0.6336032 | 0.52633978 | NA |
| DUSP6 | 36.5431244 | 0.92086743 | 0.72467514 | 1.27073136 | 0.20382424 | 0.78882329 |
| POC1B | 28.2889945 | 0.13507837 | 0.34764589 | 0.38855161 | 0.69760787 | 0.96861929 |
| ATP2B1 | 165.947216 | -1.1671245 | 0.29539668 | -3.9510414 | 7.78E-05 | 0.01536239 |
| BTG1 | 123.763998 | 0.28061335 | 0.30830321 | 0.91018627 | 0.36272428 | 0.89853007 |
| LINC02391 | 16.8389316 | -1.2501838 | 0.57178252 | -2.1864674 | 0.02878143 | 0.42210257 |
| EEA1 | 59.3902327 | -0.9360084 | 0.37522898 | -2.4944993 | 0.0126135 | 0.30389677 |
| NUDT4 | 19.1509355 | -1.0546383 | 0.43210235 | -2.440714 | 0.01465826 | 0.32121732 |
| UBE2N | 22.1042169 | -0.3506721 | 0.40552617 | -0.8647336 | 0.38718503 | 0.90910959 |
| MRPL42 | 14.0375001 | -0.4836484 | 0.51338696 | -0.9420737 | 0.34615489 | 0.88774757 |
| CRADD | 59.3471254 | -0.2162539 | 0.30782413 | -0.7025243 | 0.48235227 | 0.93382901 |
| PLXNC1 | 93.1132088 | -0.7553219 | 0.55702291 | -1.355998 | 0.17509981 | 0.76214163 |
| CEP83 | 18.6291791 | -1.0337767 | 0.4275681 | -2.417806 | 0.0156144 | 0.32607581 |
| TMCC3 | 27.161074 | 0.8440356 | 0.68994438 | 1.22333861 | 0.22120183 | 0.81199775 |
| NDUFA12 | 14.2678449 | -0.008997 | 0.64278286 | -0.0139969 | 0.98883246 | 0.99811561 |
| NR2C1 | 18.8657801 | -0.629783 | 0.44219882 | -1.4242078 | 0.15438633 | 0.74486619 |
| FGD6 | 14.6924513 | 0.57287788 | 0.68861466 | 0.83192809 | 0.40544953 | 0.91575086 |
| VEZT | 32.1430977 | 0.24138079 | 0.34733525 | 0.69495046 | 0.48708635 | 0.93497601 |

| | | | | | | |
|---|---|---|---|---|---|---|
| METAP2 | 23.3162523 | 0.14189525 | 0.49913982 | 0.28427955 | 0.77619616 | 0.98264092 |
| LTA4H | 36.4372739 | 1.52008176 | 0.79967085 | 1.9008843 | 0.05731717 | 0.56742972 |
| ELK3 | 27.3529889 | -0.9425777 | 0.47809158 | -1.9715421 | 0.0486619 | 0.53486612 |
| CDK17 | 93.921937 | 0.06071265 | 0.3209201 | 0.18918308 | 0.84994933 | 0.98653147 |
| NEDD1 | 17.9480256 | -0.1762352 | 0.48596239 | -0.3626519 | 0.71686492 | 0.97348719 |
| TMPO | 46.8168803 | -0.1354147 | 0.49176334 | -0.2753656 | 0.78303541 | 0.98264092 |
| SLC25A3 | 36.2059479 | 0.05612777 | 0.42455026 | 0.13220525 | 0.89482195 | 0.99115168 |
| IKBIP | 17.5584825 | -0.4183992 | 0.60949251 | -0.6864714 | 0.49241587 | 0.93678212 |
| APAF1 | 42.0345645 | -0.291527 | 0.42554787 | -0.6850627 | 0.49330434 | 0.93715971 |
| UHRF1BP1L | 36.657604 | -0.5113095 | 0.34140145 | -1.4976783 | 0.13421685 | 0.73044824 |
| ACTR6 | 9.03443644 | -0.1687821 | 0.61569761 | -0.2741315 | 0.7839836 | NA |
| SCYL2 | 36.4475371 | 0.02556561 | 0.32190067 | 0.07942081 | 0.93669792 | 0.99514836 |
| UTP20 | 11.7665867 | -1.7340039 | 0.51934645 | -3.3388191 | 0.00084135 | NA |
| ARL1 | 10.6456525 | -0.0162408 | 0.54265988 | -0.0299282 | 0.97612433 | NA |
| CHPT1 | 29.8909316 | -0.884465 | 0.36225292 | -2.4415677 | 0.01462365 | 0.32121732 |
| GNPTAB | 55.1390097 | 0.47837055 | 0.29554942 | 1.61858057 | 0.10553754 | 0.67519097 |
| DRAM1 | 38.3124547 | 0.21004604 | 0.33940168 | 0.61887153 | 0.53600099 | 0.95407832 |
| WASHC3 | 22.8391619 | 0.01346188 | 0.394154 | 0.03415385 | 0.97275447 | 0.9973754 |
| C12orf42 | 17.2080108 | -0.8781623 | 0.71053854 | -1.2359108 | 0.21649173 | 0.80634897 |
| HSP90B1 | 59.0772011 | -0.1141138 | 0.63774953 | -0.1789321 | 0.85799104 | 0.98751288 |
| TDG | 16.7334449 | 0.25175331 | 0.45531842 | 0.55291705 | 0.58032021 | 0.95415437 |
| HCFC2 | 12.9862866 | -0.4235952 | 0.56959832 | -0.7436735 | 0.45707402 | 0.9314919 |
| TXNRD1 | 150.975559 | -0.1913952 | 0.52426173 | -0.3650756 | 0.71505495 | 0.97348719 |
| CHST11 | 421.83079 | 0.45120731 | 0.27023476 | 1.66968642 | 0.09498142 | 0.66033566 |
| SLC41A2 | 19.4599582 | 0.0349472 | 0.64555401 | 0.05413521 | 0.95682744 | 0.99514836 |
| WASHC4 | 90.3028352 | -0.2451456 | 0.34890753 | -0.7026091 | 0.4822994 | 0.93382901 |
| APPL2 | 33.5308723 | -0.0932454 | 0.44905715 | -0.207647 | 0.83550462 | 0.98320463 |
| C12orf75 | 21.9636411 | -0.035232 | 0.60826549 | -0.0579221 | 0.95381064 | 0.99514836 |
| TCP11L2 | 11.0605051 | 0.84412235 | 0.66666849 | 1.26618006 | 0.20544862 | NA |
| RIC8B | 20.8340453 | 0.82713177 | 0.44642792 | 1.85277788 | 0.0639142 | 0.58463007 |
| TMEM263 | 9.66259747 | -0.4279724 | 0.70722707 | -0.6051414 | 0.54508505 | NA |
| CRY1 | 17.3334664 | -0.3105999 | 0.55420397 | -0.5604434 | 0.57517706 | 0.95415437 |
| BTBD11 | 31.1538837 | 1.51737224 | 0.54425292 | 2.787991 | 0.0053036 | 0.18175221 |
| PWP1 | 10.3887011 | -0.0780554 | 0.57417922 | -0.1359425 | 0.89186673 | NA |
| PRDM4 | 11.9248309 | -0.6099586 | 0.50283466 | -1.2130401 | 0.22511449 | NA |
| CMKLR1 | 27.9122208 | -0.034571 | 0.59386454 | -0.0582137 | 0.95357842 | 0.99514836 |
| SART3 | 15.8837869 | -0.2903907 | 0.46586407 | -0.6233378 | 0.53306259 | 0.95293563 |
| ISCU | 16.4222034 | 0.19395198 | 0.48465267 | 0.40018759 | 0.68901836 | 0.9673431 |
| SELPLG | 18.4540902 | 0.15771008 | 0.7402291 | 0.21305576 | 0.83128347 | 0.98320463 |
| CORO1C | 45.3028191 | -0.4302272 | 0.38434708 | -1.1193717 | 0.2629816 | 0.84921816 |
| SSH1 | 37.6472248 | 0.22569934 | 0.52992631 | 0.42590703 | 0.67017559 | 0.96412305 |
| ACACB | 10.0507644 | -0.0707046 | 0.58645818 | -0.120562 | 0.90403796 | NA |
| UBE3B | 13.4096307 | 0.1683529 | 0.51949752 | 0.32406873 | 0.745886 | 0.97805001 |
| GLTP | 11.3132455 | 0.14905849 | 0.58125293 | 0.25644342 | 0.79760847 | NA |
| TCHP | 21.0771218 | -0.1981694 | 0.59282986 | -0.334277 | 0.73817054 | 0.97605445 |
| GIT2 | 64.2014545 | 0.11121315 | 0.3351075 | 0.33187306 | 0.73998511 | 0.97605445 |
| ANKRD13A | 32.205054 | -0.3086281 | 0.392066 | -0.7871842 | 0.43117407 | 0.92284736 |
| ATP2A2 | 58.248362 | -0.0796212 | 0.33813302 | -0.235473 | 0.8138416 | 0.98320463 |
| ANAPC7 | 16.750205 | -0.2244634 | 0.47348769 | -0.4740639 | 0.63545432 | 0.95933464 |
| ARPC3 | 83.0484967 | -0.1085277 | 0.44136257 | -0.2458924 | 0.80576553 | 0.98320463 |
| VPS29 | 30.1887328 | 0.18431373 | 0.38321882 | 0.48096211 | 0.63054343 | 0.95877428 |
| PPTC7 | 29.5139794 | -0.3374362 | 0.40956841 | -0.8238823 | 0.41000645 | 0.91773508 |
| HVCN1 | 16.3465852 | -2.1072753 | 0.59117902 | -3.5645299 | 0.00036451 | 0.0361345 |
| PPP1CC | 21.4024572 | -0.2089707 | 0.41270658 | -0.5063421 | 0.6126165 | 0.95688177 |

| | | | | | | |
|---|---|---|---|---|---|---|
| SH2B3 | 64.9418839 | 0.04559212 | 0.4887768 | 0.09327801 | 0.9256827 | 0.99514836 |
| ATXN2 | 82.2388423 | -0.182239 | 0.32936431 | -0.5533054 | 0.58005433 | 0.95415437 |
| BRAP | 13.8159243 | -0.4556165 | 0.50466008 | -0.9028187 | 0.36662213 | 0.89868152 |
| ACAD10 | 9.13476425 | -0.0340977 | 0.62192044 | -0.0548265 | 0.95627669 | NA |
| ALDH2 | 19.3697127 | 1.75353828 | 0.67271406 | 2.60666216 | 0.00914295 | 0.25554089 |
| MAPKAPK5 | 16.6471153 | -1.0184682 | 0.5738142 | -1.7749094 | 0.07591286 | 0.62329636 |
| TMEM116 | 14.0788587 | -0.7432589 | 0.6192438 | -1.2002686 | 0.23003502 | 0.82393202 |
| ERP29 | 27.8700248 | -0.5687938 | 0.48279521 | -1.1781264 | 0.23874621 | 0.83267279 |
| NAA25 | 30.1153423 | -1.3224314 | 0.36922817 | -3.5816102 | 0.00034148 | 0.03499569 |
| TRAFD1 | 10.5643066 | 0.30156492 | 0.62560203 | 0.48203955 | 0.62977785 | NA |
| HECTD4 | 60.0780963 | -0.3635568 | 0.37405571 | -0.9719322 | 0.3310843 | 0.88037558 |
| RPL6 | 79.243972 | -0.1746163 | 0.63318922 | -0.2757727 | 0.78272268 | 0.98264092 |
| PTPN11 | 31.2913537 | -0.2623406 | 0.36856681 | -0.7117858 | 0.47659743 | 0.9314919 |
| OAS1 | 19.2220461 | -0.5172601 | 0.90559131 | -0.5711849 | 0.56787432 | 0.95415437 |
| OAS2 | 14.4901927 | 0.20921164 | 0.75994977 | 0.27529667 | 0.78308832 | 0.98264092 |
| DDX54 | 8.80499286 | -0.1305624 | 0.64683382 | -0.2018485 | 0.84003515 | NA |
| TPCN1 | 33.1591716 | 0.11733306 | 0.46383365 | 0.25296366 | 0.80029629 | 0.98320463 |
| SLC8B1 | 17.9928691 | 0.46221834 | 0.55339927 | 0.83523483 | 0.4035855 | 0.91528253 |
| PLBD2 | 9.83374614 | 0.18510165 | 0.58370183 | 0.31711678 | 0.75115499 | NA |
| RBM19 | 13.5757455 | -0.2356927 | 0.49647363 | -0.4747336 | 0.63497686 | 0.95933464 |
| MED13L | 401.428493 | 0.0512579 | 0.38201003 | 0.13417946 | 0.89326066 | 0.99080204 |
| FBXW8 | 20.4015238 | -0.7211445 | 0.44493419 | -1.6207891 | 0.10506288 | 0.67519097 |
| TESC | 10.4042534 | 0.47474893 | 0.60535075 | 0.78425431 | 0.4328909 | NA |
| FBXO21 | 15.2232287 | -0.0396126 | 0.48703519 | -0.0813342 | 0.93517619 | 0.99514836 |
| WSB2 | 10.9820898 | 0.7005497 | 0.65124651 | 1.07570588 | 0.28205882 | NA |
| PEBP1 | 36.9702718 | 0.39000634 | 0.50905678 | 0.76613524 | 0.44359584 | 0.92899684 |
| TAOK3 | 237.451345 | 0.22969384 | 0.23048191 | 0.99658078 | 0.31896804 | 0.87298176 |
| SUDS3 | 10.211453 | -0.43233 | 0.5279801 | -0.8188377 | 0.41287899 | NA |
| BICDL1 | 25.8142284 | -1.4857105 | 0.43815561 | -3.3908285 | 0.00069682 | 0.05282445 |
| RAB35 | 11.5942589 | -0.6653066 | 0.5674056 | -1.1725415 | 0.2409797 | NA |
| GCN1 | 13.866392 | -0.7064193 | 0.51418306 | -1.3738673 | 0.16948289 | 0.75745073 |
| RPLP0 | 88.5927128 | 0.1233977 | 0.60313118 | 0.20459512 | 0.83788847 | 0.98320463 |
| PXN | 32.3667406 | 0.71410991 | 0.46055133 | 1.55055445 | 0.12100849 | 0.70938666 |
| SRSF9 | 23.9649836 | 0.5064464 | 0.60043182 | 0.84347029 | 0.39896547 | 0.91499168 |
| DYNLL1 | 27.343396 | 0.42312253 | 0.55883242 | 0.75715458 | 0.44895726 | 0.92914597 |
| RNF10 | 34.6383557 | 0.26132437 | 0.37475669 | 0.69731742 | 0.48560417 | 0.93417265 |
| MLEC | 20.1998949 | 1.53274465 | 0.52509453 | 2.91898802 | 0.0035117 | 0.15064909 |
| SPPL3 | 123.282934 | -0.1877022 | 0.27774638 | -0.6758045 | 0.49916479 | 0.93956733 |
| P2RX7 | 21.6692115 | 0.88974476 | 0.60136507 | 1.4795418 | 0.13899557 | 0.73044824 |
| P2RX4 | 18.190941 | 0.09547252 | 0.42964333 | 0.22221343 | 0.82414774 | 0.98320463 |
| CAMKK2 | 19.0704689 | 0.17057747 | 0.76073191 | 0.2242281 | 0.82257982 | 0.98320463 |
| ANAPC5 | 39.2520157 | 0.15651437 | 0.41183625 | 0.38004031 | 0.7039155 | 0.9714177 |
| RNF34 | 19.9779037 | -0.4857102 | 0.52246751 | -0.9296466 | 0.35255408 | 0.89109867 |
| KDM2B | 45.5300572 | 0.47695639 | 0.42301352 | 1.12752042 | 0.25952251 | 0.84531935 |
| ORAI1 | 10.1199005 | -0.1259997 | 0.73395209 | -0.1716729 | 0.86369466 | NA |
| TMEM120B | 14.5175261 | -0.5375559 | 0.49363299 | -1.0889789 | 0.27616319 | 0.84994492 |
| RHOF | 10.1666506 | -1.0894946 | 0.52729137 | -2.0662097 | 0.03880868 | NA |
| SETD1B | 11.4278595 | -0.5462164 | 0.47840309 | -1.1417494 | 0.25355821 | NA |
| MLXIP | 41.1463769 | 0.41115411 | 0.34269779 | 1.19975711 | 0.23023369 | 0.82393202 |
| CLIP1 | 62.0695158 | -0.8205537 | 0.3527439 | -2.3262024 | 0.02000776 | 0.36765113 |
| ZCCHC8 | 21.2930214 | -0.0370949 | 0.36910811 | -0.1004987 | 0.91994839 | 0.99511227 |
| RSRC2 | 36.2668022 | -0.194831 | 0.32351168 | -0.6022378 | 0.54701584 | 0.95415437 |
| KNTC1 | 11.7312453 | -0.8782893 | 0.69924309 | -1.2560572 | 0.20909523 | NA |
| DENR | 24.1745602 | 0.35201423 | 0.43798482 | 0.80371332 | 0.42156256 | 0.91985469 |

| | | | | | | |
|---|---|---|---|---|---|---|
| VPS37B | 50.9183219 | 0.34819835 | 0.64518781 | 0.53968525 | 0.58941411 | 0.95415437 |
| ARL6IP4 | 25.1331628 | -0.0079084 | 0.52867718 | -0.0149588 | 0.98806503 | 0.99811561 |
| MPHOSPH9 | 30.3195404 | 0.21529275 | 0.49069444 | 0.43875115 | 0.66084186 | 0.96298563 |
| C12orf65 | 11.0547689 | -0.6755127 | 0.63115553 | -1.0702794 | 0.28449358 | NA |
| SBNO1 | 35.0000338 | -0.2230504 | 0.34303186 | -0.6502324 | 0.51554211 | 0.94596835 |
| RILPL2 | 33.1091721 | 0.03909597 | 0.47471228 | 0.08235718 | 0.93436268 | 0.99514836 |
| SNRNP35 | 8.07450508 | -0.7825714 | 0.55289124 | -1.4154165 | 0.15694642 | NA |
| TMED2 | 13.4229809 | 0.63615109 | 0.6313502 | 1.00760417 | 0.31364453 | 0.87152625 |
| EIF2B1 | 8.22167902 | 0.69902186 | 0.66516863 | 1.05089421 | 0.29330718 | NA |
| ATP6V0A2 | 18.8061946 | 0.29016692 | 0.59513631 | 0.48756379 | 0.62585885 | 0.95876809 |
| CCDC92 | 15.2376084 | 0.33047908 | 0.51842097 | 0.63747243 | 0.52381716 | 0.94980478 |
| NCOR2 | 67.8455062 | -0.3285308 | 0.35239885 | -0.9322698 | 0.3511971 | 0.89109867 |
| SCARB1 | 20.3339132 | -0.8478103 | 0.87263072 | -0.9715568 | 0.33127109 | 0.88037558 |
| UBC | 105.349317 | -0.2018598 | 0.30739205 | -0.6566852 | 0.51138337 | 0.94427613 |
| BRI3BP | 12.2596478 | -0.2558291 | 0.64274734 | -0.3980243 | 0.69061225 | NA |
| LINC00944 | 11.8197797 | 0.00292654 | 0.97371509 | 0.00300554 | 0.99760193 | NA |
| SLC15A4 | 22.4782684 | -1.0556869 | 0.61851209 | -1.7068169 | 0.08785608 | 0.64676224 |
| GLT1D1 | 27.3208562 | -1.6732689 | 0.79934726 | -2.093294 | 0.03632291 | 0.46855712 |
| STX2 | 15.6528689 | 0.78222478 | 0.71361851 | 1.09613857 | 0.27301814 | 0.84994492 |
| RAN | 20.4417099 | -0.5059007 | 0.5233317 | -0.9666923 | 0.33369789 | 0.88047896 |
| SFSWAP | 41.9033491 | -0.1471091 | 0.34738696 | -0.4234732 | 0.67195004 | 0.96412305 |
| ULK1 | 14.5046413 | 0.59361586 | 0.64448118 | 0.92107556 | 0.35701098 | 0.8957272 |
| EP400 | 38.2449425 | -0.3837674 | 0.31579336 | -1.2152485 | 0.22427133 | 0.81875277 |
| FBRSL1 | 19.6174429 | -0.1292961 | 0.41856034 | -0.3089066 | 0.75739257 | 0.98023104 |
| ANKLE2 | 28.9349162 | -0.4254299 | 0.45132398 | -0.9426264 | 0.34587201 | 0.88774757 |
| GOLGA3 | 25.224231 | -1.0482938 | 0.45711253 | -2.2932949 | 0.02183103 | 0.38353313 |
| CHFR | 38.1977985 | -0.3306076 | 0.36293894 | -0.9109178 | 0.36233866 | 0.89853007 |
| ZNF26 | 12.4691642 | -0.6575079 | 0.50905936 | -1.2916134 | 0.19649105 | NA |
| ZNF84 | 11.5485057 | -0.2822452 | 0.62850523 | -0.4490737 | 0.6533785 | NA |
| ZNF140 | 8.01332601 | 0.34902311 | 0.64352708 | 0.54235963 | 0.58757078 | NA |
| MPHOSPH8 | 47.0161383 | 0.37000597 | 0.32600761 | 1.13496114 | 0.2563916 | 0.8406233 |
| PSPC1 | 86.4685917 | -0.1964486 | 0.27573939 | -0.7124431 | 0.47619046 | 0.9314919 |
| ZMYM5 | 25.8390205 | -0.5762325 | 0.39982948 | -1.4411956 | 0.14952944 | 0.73934224 |
| ZMYM2 | 149.378066 | -0.0956559 | 0.29532634 | -0.323899 | 0.74601451 | 0.97805001 |
| CRYL1 | 18.6403404 | 0.5933734 | 0.43671597 | 1.35871697 | 0.17423629 | 0.76214163 |
| IFT88 | 16.7700127 | -0.8480062 | 0.49515659 | -1.7126021 | 0.08678577 | 0.64676224 |
| XPO4 | 45.8640772 | -0.3479264 | 0.35428576 | -0.9820503 | 0.32607508 | 0.87715548 |
| LATS2 | 23.1717043 | 0.86104097 | 0.57201088 | 1.50528776 | 0.13225011 | 0.73044824 |
| SAP18 | 15.1173819 | 0.1531954 | 0.53014796 | 0.28896724 | 0.77260644 | 0.98264092 |
| MRPL57 | 11.5859034 | 0.71119535 | 0.54374058 | 1.30796814 | 0.19088412 | NA |
| ZDHHC20 | 89.9536062 | -0.0361644 | 0.33341736 | -0.1084657 | 0.91362627 | 0.99498068 |
| MICU2 | 75.5428431 | -0.3280097 | 0.26489932 | -1.2382429 | 0.21562601 | 0.80634897 |
| SACS | 12.3049795 | -1.35844 | 0.49924196 | -2.7210053 | 0.00650837 | NA |
| MIPEP | 14.4393455 | -1.608659 | 0.65786485 | -2.4452727 | 0.01447426 | 0.32121732 |
| PARP4 | 85.7834609 | -0.0018651 | 0.3907355 | -0.0047733 | 0.99619143 | 0.99859588 |
| MTMR6 | 23.4245323 | -0.481938 | 0.43948032 | -1.0966089 | 0.27281241 | 0.84994492 |
| NUP58 | 32.3387428 | -0.2074662 | 0.42982144 | -0.48268 | 0.62932296 | 0.95877428 |
| CDK8 | 53.4654405 | 0.04346216 | 0.32870065 | 0.13222414 | 0.89480701 | 0.99115168 |
| USP12 | 36.4282524 | -0.3408286 | 0.32760189 | -1.0403744 | 0.29816601 | 0.8608685 |
| RPL21 | 40.3152037 | -0.3034364 | 0.64524416 | -0.4702661 | 0.63816492 | 0.95933464 |
| GTF3A | 20.5541654 | -0.9136615 | 0.44442631 | -2.0558222 | 0.03979966 | 0.48300729 |
| MTIF3 | 18.8144899 | -0.2226302 | 0.4453888 | -0.4998558 | 0.61717659 | 0.95688177 |
| LNX2 | 11.7695641 | 0.67129414 | 0.61340653 | 1.09437071 | 0.27379243 | NA |
| POLR1D | 19.7881336 | -0.4465018 | 0.42865792 | -1.0416273 | 0.29758448 | 0.8608685 |

| | | | | | | |
|---|---|---|---|---|---|---|
| FLT3 | 40.6103678 | 2.10358009 | 1.56723146 | 1.34222681 | 0.17952247 | 0.76657201 |
| PAN3 | 233.49035 | 0.01851072 | 0.30220105 | 0.06125299 | 0.95115773 | 0.99514836 |
| FLT1 | 18.970013 | -0.6055156 | 0.66057647 | -0.9166473 | 0.3593275 | 0.8957272 |
| POMP | 19.06428 | -0.1577043 | 0.49661354 | -0.3175594 | 0.75081921 | 0.97829887 |
| SLC46A3 | 11.7554405 | 0.17651763 | 0.61704005 | 0.28607159 | 0.77482329 | NA |
| SLC7A1 | 10.2046408 | -0.2067367 | 0.58394943 | -0.3540319 | 0.72331497 | NA |
| UBL3 | 57.9056762 | 0.44461449 | 0.33084973 | 1.34385628 | 0.17899486 | 0.76545642 |
| KATNAL1 | 22.0706726 | -0.255316 | 0.60360828 | -0.4229829 | 0.67230771 | 0.96412305 |
| LINC00426 | 16.4469957 | 0.75695868 | 0.78609162 | 0.96293951 | 0.33557788 | 0.88047896 |
| HMGB1 | 62.0447311 | -0.3314414 | 0.29204004 | -1.1349176 | 0.25640985 | 0.8406233 |
| USPL1 | 25.1185292 | -0.3372996 | 0.38275244 | -0.8812475 | 0.37818389 | 0.9051267 |
| ALOX5AP | 27.7496675 | 0.1020635 | 0.50499829 | 0.20210663 | 0.83983337 | 0.98342625 |
| HSPH1 | 37.7892182 | -0.9542288 | 0.6786011 | -1.4061704 | 0.15967352 | 0.74820769 |
| FRY | 60.4307538 | 2.84283949 | 0.58066202 | 4.89585918 | 9.79E-07 | 0.00077379 |
| N4BP2L1 | 31.512158 | 0.70684144 | 0.47391335 | 1.49149933 | 0.13583045 | 0.73044824 |
| N4BP2L2 | 208.014628 | -0.0730555 | 0.25790127 | -0.2832691 | 0.77697057 | 0.98264092 |
| PDS5B | 94.8940734 | -0.1911064 | 0.31022988 | -0.6160156 | 0.53788425 | 0.95415437 |
| STARD13 | 111.128321 | 0.47880212 | 0.70265111 | 0.68142228 | 0.49560433 | 0.93805563 |
| NBEA | 26.3462643 | 0.20723627 | 0.45451649 | 0.45594884 | 0.64842677 | 0.96049083 |
| SPART | 24.3464042 | 0.89679824 | 0.48987191 | 1.83067905 | 0.06714846 | 0.58951068 |
| ALG5 | 12.5488156 | -0.0549866 | 0.47615866 | -0.1154795 | 0.90806504 | NA |
| EXOSC8 | 8.13011476 | -0.7266703 | 0.76143836 | -0.954339 | 0.33991208 | NA |
| SUPT20H | 33.2519742 | -0.0638794 | 0.45772352 | -0.139559 | 0.88900843 | 0.99080204 |
| UFM1 | 23.1075837 | -0.1605448 | 0.52164669 | -0.3077654 | 0.75826088 | 0.98051527 |
| PROSER1 | 12.3368957 | -1.080919 | 0.50976554 | -2.1204239 | 0.03397031 | NA |
| NHLRC3 | 25.8635659 | 1.3769531 | 0.47720602 | 2.88544785 | 0.00390857 | 0.15732988 |
| COG6 | 24.4610269 | 0.03607777 | 0.47790865 | 0.07549092 | 0.93982412 | 0.99514836 |
| FOXO1 | 130.34143 | -0.7339041 | 0.33017691 | -2.2227601 | 0.02623198 | 0.40477881 |
| MRPS31 | 22.2243791 | 0.68642056 | 0.43700543 | 1.57073692 | 0.11624377 | 0.70122655 |
| ELF1 | 190.24259 | 0.0402539 | 0.28398619 | 0.14174598 | 0.88728066 | 0.99019008 |
| WBP4 | 12.1832247 | -0.506041 | 0.55280741 | -0.9154021 | 0.3599806 | NA |
| NAA16 | 26.6785373 | 0.12438934 | 0.51579329 | 0.24116121 | 0.80943017 | 0.98320463 |
| VWA8 | 66.4315842 | -0.4720659 | 0.34709017 | -1.3600669 | 0.17380875 | 0.76214163 |
| DGKH | 43.1283501 | -0.0170658 | 0.40506757 | -0.0421308 | 0.96639439 | 0.99653147 |
| AKAP11 | 33.5656677 | -0.3200303 | 0.36649169 | -0.8732267 | 0.38253953 | 0.90868001 |
| EPSTI1 | 88.3244323 | 0.27929806 | 0.85890529 | 0.32517912 | 0.74504551 | 0.97805001 |
| DNAJC15 | 23.3272199 | -0.2735814 | 0.48061801 | -0.5692284 | 0.56920113 | 0.95415437 |
| TSC22D1 | 53.2199633 | 0.06101763 | 0.40480397 | 0.15073376 | 0.88018574 | 0.99019008 |
| GPALPP1 | 14.4275735 | 0.45610728 | 0.4647063 | 0.98149578 | 0.32634831 | 0.87715548 |
| GTF2F2 | 42.6918209 | -0.2375811 | 0.32582475 | -0.7291684 | 0.46589866 | 0.9314919 |
| TPT1 | 222.846728 | 0.2164142 | 0.49581203 | 0.43648437 | 0.66248534 | 0.96352007 |
| COG3 | 24.6055871 | 0.08229045 | 0.39249039 | 0.20966234 | 0.83393122 | 0.98320463 |
| ZC3H13 | 41.5474412 | 0.00847215 | 0.30475719 | 0.02779967 | 0.97782193 | 0.99758514 |
| LCP1 | 183.479778 | 0.45003511 | 0.43084851 | 1.04453212 | 0.29623926 | 0.8608685 |
| RUBCNL | 27.7041667 | 0.91173788 | 0.69007034 | 1.32122456 | 0.1864265 | 0.77798634 |
| LRCH1 | 75.1239981 | 0.9881648 | 0.4363724 | 2.26449885 | 0.02354345 | 0.3948379 |
| ESD | 9.64375382 | 0.47636419 | 0.55422233 | 0.85951822 | 0.39005467 | NA |
| SUCLA2 | 22.3803641 | -0.0020196 | 0.57422159 | -0.0035171 | 0.99719375 | 0.99881813 |
| MED4 | 20.8862785 | 0.38716994 | 0.47384995 | 0.81707287 | 0.41388679 | 0.91894074 |
| ITM2B | 132.155795 | 0.50011019 | 0.32065611 | 1.55964654 | 0.11884343 | 0.70431251 |
| RB1 | 161.825124 | -0.7413429 | 0.33535975 | -2.21059 | 0.02706424 | 0.40712515 |
| LPAR6 | 40.4158545 | 0.15664391 | 0.58300255 | 0.26868478 | 0.78817226 | 0.98320463 |
| RCBTB2 | 34.3666128 | -0.4791397 | 0.44696313 | -1.0719894 | 0.28372479 | 0.85176121 |
| FNDC3A | 190.648453 | -0.3452202 | 0.43020435 | -0.8024563 | 0.42228905 | 0.91985469 |

| | | | | | | |
|---|---|---|---|---|---|---|
| CAB39L | 11.29334 | -0.7459651 | 0.58298239 | -1.2795671 | 0.20069744 | NA |
| SETDB2 | 28.9559963 | 0.23615018 | 0.52131845 | 0.45298643 | 0.65055851 | 0.96084839 |
| PHF11 | 23.6108701 | -0.4309308 | 0.46893325 | -0.9189598 | 0.35811662 | 0.8957272 |
| RCBTB1 | 11.6559128 | -0.3912722 | 0.52398084 | -0.7467301 | 0.45522651 | NA |
| KPNA3 | 42.6736236 | 0.09054335 | 0.32010412 | 0.28285593 | 0.77728728 | 0.98264092 |
| DLEU2 | 288.420682 | 1.14430738 | 0.41403175 | 2.76381555 | 0.00571298 | 0.19045569 |
| TRIM13 | 24.9752233 | 0.6064569 | 0.41002291 | 1.47908053 | 0.13911879 | 0.73044824 |
| RNASEH2B | 37.9479653 | 0.06533558 | 0.39132374 | 0.16696042 | 0.86740119 | 0.98751288 |
| INTS6 | 73.4584026 | 0.36863748 | 0.3801557 | 0.96970133 | 0.33219539 | 0.88047896 |
| WDFY2 | 92.1292358 | 0.71541467 | 0.29522631 | 2.42327546 | 0.01538126 | 0.32566073 |
| VPS36 | 35.4837296 | 0.37133954 | 0.33543463 | 1.10703995 | 0.26827666 | 0.84994492 |
| SUGT1 | 18.9338757 | 0.34728979 | 0.4560092 | 0.76158507 | 0.44630769 | 0.92914597 |
| TDRD3 | 41.0593419 | -0.0436165 | 0.35437665 | -0.1230796 | 0.90204406 | 0.99262388 |
| PCDH9 | 31.1460419 | -4.5578587 | 0.75204523 | -6.0606178 | 1.36E-09 | 3.75E-06 |
| DACH1 | 34.9328219 | 0.75998153 | 0.87599107 | 0.86756768 | 0.38563105 | 0.90910959 |
| DIS3 | 17.1142416 | -0.4838563 | 0.47282488 | -1.0233308 | 0.30615148 | 0.8673137 |
| PIBF1 | 69.7675727 | 0.14567332 | 0.32860175 | 0.44331267 | 0.65753959 | 0.96270373 |
| KLF12 | 136.594313 | 0.13222746 | 0.30936746 | 0.42741231 | 0.66907905 | 0.96412305 |
| TBC1D4 | 34.9742267 | -0.5741537 | 0.60883468 | -0.9430371 | 0.34566192 | 0.88765341 |
| COMMD6 | 18.7697408 | 0.34361044 | 0.57803494 | 0.5944458 | 0.55221398 | 0.95415437 |
| UCHL3 | 15.7375721 | 0.62469009 | 0.67564476 | 0.92458364 | 0.35518253 | 0.89303957 |
| LMO7 | 17.3008032 | -1.4973625 | 0.49774524 | -3.008291 | 0.00262721 | 0.12115836 |
| KCTD12 | 36.926383 | -0.0244293 | 0.61127357 | -0.0399646 | 0.96812136 | 0.99653147 |
| AC001226.2 | 14.5179254 | 0.01990503 | 0.87168177 | 0.02283521 | 0.98178173 | 0.99758514 |
| FBXL3 | 23.6243118 | 0.09797935 | 0.38559545 | 0.25409882 | 0.79941921 | 0.98320463 |
| MYCBP2 | 222.511534 | -0.3676081 | 0.18356401 | -2.0026154 | 0.04521859 | 0.52128421 |
| RNF219 | 12.9996876 | -1.019266 | 0.48441607 | -2.1041127 | 0.03536862 | 0.46381501 |
| RBM26 | 92.4309579 | -0.0933034 | 0.33013118 | -0.2826253 | 0.77746405 | 0.98264092 |
| NDFIP2 | 9.90916523 | -0.6279977 | 0.57663084 | -1.089081 | 0.27611819 | NA |
| GPC5 | 19.4979937 | 1.61623913 | 1.10362057 | 1.46448804 | 0.14306065 | 0.73044824 |
| GPC6 | 23.4454653 | 1.85994786 | 1.78359427 | 1.04280883 | 0.29703684 | 0.8608685 |
| ABCC4 | 32.5777368 | 0.84523101 | 0.61708483 | 1.36971607 | 0.17077555 | 0.75751515 |
| DNAJC3 | 51.0082544 | 0.37418224 | 0.40731541 | 0.91865476 | 0.35827618 | 0.8957272 |
| UGGT2 | 36.3840625 | -0.1208513 | 0.38616557 | -0.3129521 | 0.75431704 | 0.97921428 |
| MBNL2 | 76.1069497 | 0.36182135 | 0.40188168 | 0.90031811 | 0.36795099 | 0.89868152 |
| RAP2A | 16.3696301 | 0.30202469 | 0.49744839 | 0.60714779 | 0.54375284 | 0.95415437 |
| IPO5 | 14.3216807 | -0.815765 | 0.57893022 | -1.4090904 | 0.15880844 | 0.74686546 |
| FARP1 | 31.2108899 | -0.6873435 | 0.49909418 | -1.3771819 | 0.16845602 | 0.7567539 |
| STK24 | 65.6866809 | -0.1432664 | 0.30957222 | -0.4627883 | 0.64351615 | 0.95992767 |
| DOCK9 | 44.7804756 | 0.01351476 | 0.36064128 | 0.03747425 | 0.97010687 | 0.9967641 |
| UBAC2 | 132.239646 | -0.1232754 | 0.22855727 | -0.5393634 | 0.58963614 | 0.95415437 |
| GPR183 | 28.0944864 | -1.2881198 | 0.74971866 | -1.7181376 | 0.08577152 | 0.64676224 |
| TM9SF2 | 59.3442317 | 0.09414412 | 0.30066555 | 0.31311909 | 0.75419018 | 0.97921428 |
| CLYBL | 32.1167663 | -0.2724925 | 0.55713773 | -0.4890936 | 0.6247754 | 0.95876809 |
| PCCA | 94.71634 | 0.16711352 | 0.2868561 | 0.58256915 | 0.56018338 | 0.95415437 |
| GGACT | 18.7639383 | -1.1578951 | 0.42013017 | -2.756039 | 0.0058506 | 0.19198507 |
| TMTC4 | 13.0900598 | 0.20922619 | 0.53345596 | 0.39220892 | 0.69490385 | 0.9685713 |
| TPP2 | 54.5822482 | 0.30757861 | 0.35748912 | 0.86038595 | 0.38957633 | 0.90910959 |
| ARGLU1 | 73.3251309 | -0.1944821 | 0.34532955 | -0.5631783 | 0.57331347 | 0.95415437 |
| ABHD13 | 14.0177039 | 0.55516817 | 0.51026009 | 1.08801018 | 0.27659061 | 0.84994492 |
| TNFSF13B | 46.2158207 | 2.2783263 | 0.89019925 | 2.55934421 | 0.01048698 | 0.2803622 |
| IRS2 | 27.8045117 | 1.51736367 | 0.95466156 | 1.58942576 | 0.1119643 | 0.69385269 |
| RAB20 | 36.03018 | 0.56075928 | 0.83316999 | 0.67304306 | 0.50091989 | 0.94001041 |
| CARS2 | 24.9508503 | 0.40308963 | 0.4387672 | 0.91868679 | 0.35825942 | 0.8957272 |

| | | | | | | |
|---|---|---|---|---|---|---|
| ANKRD10 | 73.6786192 | -0.0706824 | 0.25291563 | -0.2794703 | 0.77988389 | 0.98264092 |
| ARHGEF7 | 60.2667182 | -0.1386663 | 0.31109159 | -0.4457412 | 0.65578421 | 0.96211819 |
| TUBGCP3 | 39.4326388 | -0.6177319 | 0.37620483 | -1.6420094 | 0.10058806 | 0.6683775 |
| ATP11A | 84.4043414 | 0.23894847 | 0.32718986 | 0.73030523 | 0.46520363 | 0.9314919 |
| PCID2 | 17.2968289 | -0.0351916 | 0.46277532 | -0.0760446 | 0.93938359 | 0.99514836 |
| CUL4A | 52.9090955 | -0.1148692 | 0.40926995 | -0.2806686 | 0.77896459 | 0.98264092 |
| LAMP1 | 34.0401053 | 0.32709045 | 0.44608342 | 0.7332495 | 0.46340627 | 0.9314919 |
| DCUN1D2 | 13.2208241 | 0.67848572 | 0.52860016 | 1.28355185 | 0.19929881 | 0.78165811 |
| TMCO3 | 21.3304673 | 0.75552613 | 0.49302885 | 1.53241768 | 0.1254194 | 0.71904999 |
| TFDP1 | 12.2603426 | -0.5991872 | 0.59842552 | -1.0012727 | 0.31669497 | NA |
| GAS6 | 13.2284852 | 0.54710978 | 0.63209861 | 0.86554498 | 0.38673974 | 0.90910959 |
| RASA3 | 73.4089287 | 0.19375493 | 0.38749061 | 0.50002485 | 0.61705758 | 0.95688177 |
| CDC16 | 23.9411511 | -0.1456018 | 0.38637449 | -0.3768412 | 0.70629164 | 0.97184211 |
| UPF3A | 22.47426 | -0.7319433 | 0.43501777 | -1.6825595 | 0.09246039 | 0.65902576 |
| TEP1 | 18.7448289 | -0.2144075 | 0.45778126 | -0.4683623 | 0.63952554 | 0.95952691 |
| OSGEP | 10.0411817 | -0.3273505 | 0.54689935 | -0.5985571 | 0.54946825 | NA |
| APEX1 | 9.61985683 | -0.4882486 | 0.55675137 | -0.8769599 | 0.38050843 | NA |
| PNP | 9.52460911 | 1.4113005 | 0.79285258 | 1.78002889 | 0.07507123 | NA |
| RNASE6 | 10.3129454 | 0.74884002 | 0.85675437 | 0.87404284 | 0.38209492 | NA |
| RNASE1 | 50.5296098 | -0.0839875 | 0.90449499 | -0.0928557 | 0.92601822 | 0.99514836 |
| HNRNPC | 150.417454 | -0.2048229 | 0.2305703 | -0.8883318 | 0.37436229 | 0.90271063 |
| SUPT16H | 19.2831236 | 0.13734971 | 0.50821665 | 0.27025818 | 0.78696164 | 0.98320463 |
| CHD8 | 36.1147594 | 0.07039181 | 0.34253717 | 0.20550124 | 0.83718053 | 0.98320463 |
| RAB2B | 11.1338112 | 0.21235051 | 0.64962295 | 0.32688272 | 0.74375659 | NA |
| TOX4 | 16.8361081 | 0.35008397 | 0.45287178 | 0.7730311 | 0.43950398 | 0.92818763 |
| TRAC | 18.4401283 | 0.94907291 | 0.6584949 | 1.44127602 | 0.14950672 | 0.73934224 |
| DAD1 | 20.952942 | -0.1864335 | 0.44611363 | -0.4179059 | 0.67601593 | 0.96643558 |
| OXA1L | 11.1696093 | -0.3958968 | 0.60057083 | -0.6592009 | 0.50976677 | NA |
| SLC7A7 | 31.3290644 | 0.15199073 | 0.71764564 | 0.21179078 | 0.83227026 | 0.98320463 |
| LRP10 | 15.0343268 | 0.07684198 | 0.52530023 | 0.14628203 | 0.88369875 | 0.99019008 |
| RBM23 | 17.0149716 | 0.17350293 | 0.52008132 | 0.33360731 | 0.7386759 | 0.97605445 |
| PSMB5 | 9.52249699 | 0.03128326 | 0.56165828 | 0.05569803 | 0.95558237 | NA |
| ACIN1 | 42.3364885 | 0.47319777 | 0.31085134 | 1.52226388 | 0.12794297 | 0.72782081 |
| SLC7A8 | 15.6312412 | 0.95776311 | 0.82945807 | 1.15468538 | 0.24821929 | 0.83817091 |
| PABPN1 | 12.9072079 | -0.1733025 | 0.60124642 | -0.2882387 | 0.77316401 | NA |
| AP1G2 | 7.46734879 | -0.7457271 | 0.69569467 | -1.0719173 | 0.28375719 | NA |
| DHRS4L2 | 8.52689017 | 0.44796405 | 0.62949052 | 0.71162953 | 0.47669422 | NA |
| PSME1 | 21.0760716 | 0.36817938 | 0.61472914 | 0.59892944 | 0.54921994 | 0.95415437 |
| LTB4R | 10.8384815 | 0.49997329 | 0.68534421 | 0.72952143 | 0.46568276 | NA |
| GZMH | 12.4095114 | 0.82526085 | 1.03270406 | 0.79912618 | 0.42421725 | NA |
| GZMB | 19.7700534 | 0.69593336 | 0.91623258 | 0.75955971 | 0.44751781 | 0.92914597 |
| PRKD1 | 15.4769534 | -0.8189268 | 0.6950266 | -1.1782669 | 0.23869022 | 0.83267279 |
| G2E3 | 24.5902633 | -0.5848158 | 0.43501158 | -1.3443682 | 0.17882934 | 0.76545642 |
| SCFD1 | 84.1587903 | -0.5594389 | 0.28894604 | -1.9361364 | 0.052851 | 0.54976963 |
| STRN3 | 97.8243322 | -0.0842423 | 0.29987882 | -0.280921 | 0.77877096 | 0.98264092 |
| AP4S1 | 9.68305108 | 0.19023494 | 0.58044018 | 0.32774255 | 0.74310633 | NA |
| HECTD1 | 79.2935222 | -0.2148076 | 0.33603273 | -0.6392461 | 0.52266283 | 0.94895469 |
| HEATR5A | 13.8123522 | 0.72978564 | 0.50689735 | 1.43971091 | 0.14994921 | 0.7402488 |
| NUBPL | 39.4600788 | -0.2810284 | 0.44596961 | -0.6301514 | 0.52859554 | 0.95182065 |
| ARHGAP5 | 61.4382281 | -1.0938602 | 0.40504513 | -2.7005884 | 0.00692169 | 0.20817747 |
| NPAS3 | 20.5078118 | -0.4233522 | 0.94729783 | -0.4469051 | 0.65494361 | 0.96178254 |
| EAPP | 14.7847366 | 0.0203923 | 0.46035249 | 0.04429714 | 0.96466755 | 0.99653147 |
| SNX6 | 81.7317246 | -0.2907548 | 0.35302412 | -0.8236118 | 0.41016018 | 0.91773508 |
| BAZ1A | 114.551639 | -0.123598 | 0.29155797 | -0.4239227 | 0.67162217 | 0.96412305 |

| | | | | | | |
|---|---|---|---|---|---|---|
| SRP54 | 35.8899019 | -0.3193707 | 0.45649442 | -0.6996159 | 0.48416722 | 0.93399425 |
| FAM177A1 | 23.1949706 | 0.515538 | 0.41924724 | 1.22967535 | 0.2188187 | 0.80977253 |
| PPP2R3C | 39.5994942 | -0.0459584 | 0.44520316 | -0.1032302 | 0.91778027 | 0.99498068 |
| NFKBIA | 38.5624533 | 0.56517645 | 0.64420322 | 0.87732633 | 0.38030941 | 0.90676497 |
| RALGAPA1 | 86.6152786 | -0.3233478 | 0.33580848 | -0.9628936 | 0.3356009 | 0.88047896 |
| SEC23A | 34.8341194 | -0.1991914 | 0.40011821 | -0.4978314 | 0.61860291 | 0.95688177 |
| TRAPPC6B | 15.8455529 | -0.0300579 | 0.47193851 | -0.0636903 | 0.9492168 | 0.99514836 |
| PNN | 32.2990768 | -0.5650939 | 0.33438666 | -1.6899416 | 0.09103912 | 0.65685853 |
| MIA2 | 39.4508687 | -0.0035404 | 0.33250079 | -0.0106477 | 0.99150454 | 0.99854161 |
| FBXO33 | 35.4412063 | -0.0674935 | 0.34771621 | -0.1941051 | 0.8460936 | 0.98491417 |
| KLHL28 | 23.7737703 | -0.0729411 | 0.43218116 | -0.1687743 | 0.86597415 | 0.98751288 |
| TOGARAM1 | 22.9084634 | -0.516138 | 0.48104049 | -1.0729616 | 0.28328836 | 0.85176121 |
| PRPF39 | 13.6319524 | -0.0633192 | 0.51458541 | -0.1230489 | 0.90206837 | 0.99262388 |
| FKBP3 | 11.0605186 | -0.4980542 | 0.498884 | -0.9983367 | 0.31811611 | NA |
| FANCM | 11.9923684 | -0.7105757 | 0.64935703 | -1.0942759 | 0.273834 | NA |
| MIS18BP1 | 59.485716 | 0.11278956 | 0.39343446 | 0.28667942 | 0.77435779 | 0.98264092 |
| RPS29 | 48.3643652 | -0.2603875 | 0.50065597 | -0.5200928 | 0.60299893 | 0.95688177 |
| RPL36AL | 60.5038687 | -0.2111776 | 0.48959762 | -0.4313289 | 0.66622921 | 0.96412305 |
| KLHDC1 | 10.4752662 | 0.30613007 | 0.88064888 | 0.34761876 | 0.72812652 | NA |
| KLHDC2 | 12.914517 | 0.41779683 | 0.60569878 | 0.68977658 | 0.4903347 | NA |
| NEMF | 67.856932 | -0.3987595 | 0.27044817 | -1.4744395 | 0.1403633 | 0.73044824 |
| ARF6 | 22.5742078 | 0.12986225 | 0.6576274 | 0.19747086 | 0.84345908 | 0.9842451 |
| LINC01588 | 18.2116067 | -1.3746109 | 0.43543725 | -3.1568518 | 0.00159482 | 0.0873837 |
| VCPKMT | 10.6721711 | 0.24443224 | 0.58204187 | 0.41995645 | 0.67451727 | NA |
| SOS2 | 79.5266727 | 0.49200187 | 0.26077378 | 1.88669992 | 0.0592007 | 0.57281427 |
| ATP5S | 10.7763305 | -0.683785 | 0.5223672 | -1.3090122 | 0.19053022 | NA |
| MAP4K5 | 49.9601532 | -0.043869 | 0.37466264 | -0.1170892 | 0.90678935 | 0.99368779 |
| NIN | 89.1876268 | -0.2352164 | 0.31778169 | -0.7401823 | 0.45918941 | 0.9314919 |
| PYGL | 23.0942417 | 0.45158296 | 0.66146283 | 0.68270346 | 0.49479424 | 0.93805563 |
| TMX1 | 15.1155251 | -0.574004 | 0.45187754 | -1.2702646 | 0.2039904 | 0.78882329 |
| GNG2 | 148.473155 | 0.10150245 | 0.54908024 | 0.18485905 | 0.85333959 | 0.98751288 |
| RTRAF | 13.90957 | -0.7081047 | 0.5017423 | -1.4112916 | 0.15815866 | 0.74650686 |
| PTGDR | 19.6545684 | 0.34318378 | 0.67329574 | 0.50970733 | 0.61025652 | 0.95688177 |
| TXNDC16 | 15.8494935 | 0.0404629 | 0.66321337 | 0.06101038 | 0.95135095 | 0.99514836 |
| ERO1A | 39.0163244 | 0.81328885 | 0.49342095 | 1.64826574 | 0.09929815 | 0.6683775 |
| PSMC6 | 25.5236625 | -0.0163529 | 0.41808888 | -0.0391133 | 0.96880002 | 0.99653147 |
| STYX | 13.6913694 | 0.15669928 | 0.61942553 | 0.25297517 | 0.8002874 | 0.98320463 |
| DDHD1 | 50.9646325 | 0.35622803 | 0.36015872 | 0.98908623 | 0.32262095 | 0.87433122 |
| CNIH1 | 10.8822654 | -0.2877955 | 0.64504512 | -0.4461634 | 0.65547921 | NA |
| GMFB | 11.6130296 | -0.20849 | 0.5360428 | -0.3889429 | 0.69731839 | NA |
| SAMD4A | 47.4443866 | -1.1047751 | 0.58028843 | -1.903838 | 0.05693129 | 0.56563334 |
| GCH1 | 38.6982752 | 0.71713289 | 0.67599761 | 1.06085122 | 0.28875752 | 0.85703857 |
| SOCS4 | 10.9266164 | 0.15681296 | 0.57379875 | 0.27328913 | 0.78463097 | NA |
| MAPK1IP1L | 34.1764221 | -0.5566328 | 0.41047588 | -1.3560669 | 0.17507788 | 0.76214163 |
| LGALS3 | 19.3358167 | 0.60030313 | 0.60299573 | 0.99553463 | 0.31947631 | 0.87307748 |
| FBXO34 | 62.3379346 | 0.88817543 | 0.53077969 | 1.67334101 | 0.09426018 | 0.65925012 |
| ATG14 | 15.0855251 | 0.12402106 | 0.64975829 | 0.19087262 | 0.84862539 | 0.98545782 |
| KTN1 | 74.8822068 | -0.1835453 | 0.27105631 | -0.6771481 | 0.49831201 | 0.93916044 |
| PELI2 | 58.3275315 | 0.98761032 | 0.44607906 | 2.21398046 | 0.02683012 | 0.40712515 |
| TMEM260 | 28.2298548 | 0.62409334 | 0.37281097 | 1.67402089 | 0.09412649 | 0.65925012 |
| EXOC5 | 33.268343 | 0.0056622 | 0.36869133 | 0.01535755 | 0.98774693 | 0.99811561 |
| AP5M1 | 15.0965797 | 0.19362667 | 0.44557989 | 0.43454983 | 0.66388922 | 0.96379377 |
| ARMH4 | 15.971099 | 0.13749367 | 0.50242634 | 0.27365936 | 0.78434641 | 0.98264092 |
| ACTR10 | 23.6176292 | 0.0667406 | 0.42860966 | 0.15571417 | 0.87625833 | 0.98939075 |

| | | | | | | |
|---|---|---|---|---|---|---|
| PSMA3 | 21.7870964 | 0.5972326 | 0.49454496 | 1.20764066 | 0.22718551 | 0.82088564 |
| ARID4A | 54.2159095 | 0.62309719 | 0.3746802 | 1.66301069 | 0.0963103 | 0.66178928 |
| KIAA0586 | 20.5555435 | 0.56359481 | 0.40806644 | 1.38113494 | 0.16723748 | 0.7567539 |
| DAAM1 | 28.7458096 | -0.8085133 | 0.42041268 | -1.923142 | 0.05446222 | 0.55278922 |
| JKAMP | 14.6324744 | 0.28881518 | 0.49411763 | 0.58450694 | 0.5588793 | 0.95415437 |
| RTN1 | 61.5004982 | 0.69493003 | 0.87157869 | 0.79732334 | 0.42526327 | 0.92001834 |
| PCNX4 | 43.2302406 | -0.1572828 | 0.3933406 | -0.399864 | 0.68925665 | 0.9673431 |
| DHRS7 | 11.2526346 | 0.17064858 | 0.51114142 | 0.33385785 | 0.73848683 | NA |
| PPM1A | 27.4547975 | 0.26400141 | 0.46764217 | 0.56453722 | 0.57238858 | 0.95415437 |
| MNAT1 | 55.0306799 | 0.28883871 | 0.37430321 | 0.77167041 | 0.44030966 | 0.92861039 |
| SLC38A6 | 23.6984303 | 0.7188099 | 0.52507163 | 1.36897495 | 0.17100711 | 0.75751515 |
| PRKCH | 225.217575 | -0.0415455 | 0.25714163 | -0.1615667 | 0.87164711 | 0.98751288 |
| HIF1A | 255.329779 | -0.4687574 | 0.59411473 | -0.7890014 | 0.43011118 | 0.9224707 |
| HIF1A-AS2 | 41.8765512 | 0.33303869 | 0.85333995 | 0.39027669 | 0.69633196 | 0.9685713 |
| SNAPC1 | 11.0412677 | -0.2611689 | 0.58112672 | -0.4494181 | 0.65313006 | NA |
| PPP2R5E | 89.2662683 | -0.2036527 | 0.23890919 | -0.8524271 | 0.39397709 | 0.90940709 |
| SGPP1 | 39.9498788 | -0.8384192 | 0.46349628 | -1.8089018 | 0.07046627 | 0.60640882 |
| SYNE2 | 165.996714 | -0.2589423 | 0.26270986 | -0.985659 | 0.32430049 | 0.87663738 |
| ESR2 | 14.7319714 | 0.18941729 | 0.78447899 | 0.24145617 | 0.80920158 | 0.98320463 |
| ZBTB25 | 21.4236781 | 0.60631991 | 0.52373102 | 1.15769333 | 0.24698921 | 0.83817091 |
| ZBTB1 | 43.1798354 | 0.16867506 | 0.28294163 | 0.59614792 | 0.55107641 | 0.95415437 |
| MAX | 55.2105169 | 0.34256183 | 0.35696335 | 0.95965548 | 0.33722864 | 0.88195807 |
| FUT8 | 92.7566688 | 0.37711122 | 0.39874472 | 0.945746 | 0.34427816 | 0.8871109 |
| GPHN | 83.4434306 | 0.46604293 | 0.42377017 | 1.09975397 | 0.27143933 | 0.84994492 |
| MPP5 | 18.1415174 | 0.14110442 | 0.45564902 | 0.30967788 | 0.75680593 | 0.9799167 |
| ATP6V1D | 14.3471839 | -0.4766632 | 0.54496782 | -0.8746631 | 0.38175725 | 0.90866436 |
| EIF2S1 | 15.8844658 | 0.44323166 | 0.45846653 | 0.96676994 | 0.33365906 | 0.88047896 |
| VTI1B | 9.48432348 | -0.4839775 | 0.6307042 | -0.7673604 | 0.44286724 | NA |
| ZFYVE26 | 16.6264764 | -0.4079582 | 0.57270071 | -0.7123411 | 0.4762536 | 0.9314919 |
| RAD51B | 172.070097 | 0.91988193 | 0.47756042 | 1.92621056 | 0.05407809 | 0.55278922 |
| ZFP36L1 | 107.397662 | -0.2869461 | 0.48157099 | -0.5958542 | 0.55127265 | 0.95415437 |
| ACTN1 | 22.8622353 | 0.56221013 | 0.49918483 | 1.12625644 | 0.26005699 | 0.8455672 |
| DCAF5 | 56.2781272 | 0.00671881 | 0.30283306 | 0.02218651 | 0.98229918 | 0.99759864 |
| ERH | 16.8319506 | -0.153723 | 0.51991115 | -0.2956718 | 0.76748077 | 0.98264092 |
| SLC39A9 | 20.1785659 | 0.62456894 | 0.43125705 | 1.44825215 | 0.14754655 | 0.73693374 |
| SUSD6 | 91.5158373 | 0.64778705 | 0.43840113 | 1.47761264 | 0.13951149 | 0.73044824 |
| SRSF5 | 56.1188605 | -0.082443 | 0.40626145 | -0.2029309 | 0.83918902 | 0.98342625 |
| MED6 | 10.0806711 | -0.2188637 | 0.52044819 | -0.4205293 | 0.67409884 | NA |
| PCNX1 | 227.196337 | 0.19545192 | 0.32164892 | 0.60765607 | 0.54341561 | 0.95415437 |
| SIPA1L1 | 453.184659 | -0.7024849 | 0.55343601 | -1.2693156 | 0.20432853 | 0.78882329 |
| ZFYVE1 | 14.8788841 | 0.27456934 | 0.53102493 | 0.51705547 | 0.60511744 | 0.95688177 |
| RBM25 | 58.0263428 | -0.5285713 | 0.29646112 | -1.7829365 | 0.07459665 | 0.61929783 |
| PSEN1 | 61.905939 | 0.01002978 | 0.3090675 | 0.03245175 | 0.97411179 | 0.9973754 |
| NUMB | 154.914125 | 0.604818 | 0.37108134 | 1.62987989 | 0.10312689 | 0.67250824 |
| ELMSAN1 | 24.5764115 | 0.12700567 | 0.5151566 | 0.24653798 | 0.80526581 | 0.98320463 |
| LIN52 | 25.7992363 | 0.39716788 | 0.5075019 | 0.78259387 | 0.43386564 | 0.92311128 |
| ABCD4 | 14.8310434 | -0.3314582 | 0.44704934 | -0.7414354 | 0.4584295 | 0.9314919 |
| NPC2 | 41.4937057 | 0.57390078 | 0.51181519 | 1.12130469 | 0.26215819 | 0.84841135 |
| AREL1 | 19.6107725 | -0.5695179 | 0.40362706 | -1.4110002 | 0.15824456 | 0.74650686 |
| YLPM1 | 39.8977185 | -0.5562831 | 0.31023549 | -1.7930996 | 0.07295701 | 0.61359282 |
| DLST | 13.9993463 | -0.4606825 | 0.46040383 | -1.0006053 | 0.31701766 | 0.87263466 |
| MLH3 | 17.0643062 | -0.3123954 | 0.45977907 | -0.6794469 | 0.49685476 | 0.93900843 |
| NEK9 | 24.2119581 | -0.4881833 | 0.38910241 | -1.2546396 | 0.20960961 | 0.79833418 |
| TMED10 | 34.7968931 | -0.0097505 | 0.38059708 | -0.025619 | 0.9795612 | 0.99758514 |

| | | | | | | |
|---|---|---|---|---|---|---|
| FOS | 43.1826593 | 0.16687279 | 0.68628556 | 0.24315358 | 8.08E-01 | 0.98320463 |
| JDP2 | 24.2180172 | 0.72070695 | 0.76305208 | 0.94450558 | 0.34491135 | 0.88711459 |
| BATF | 9.7225494 | 0.30693176 | 0.60044691 | 0.51117218 | 0.6092305 | NA |
| FLVCR2 | 18.0315753 | 0.99806776 | 0.76701038 | 1.30124414 | 0.1931749 | 0.78135737 |
| TTLL5 | 32.3736336 | -0.3696983 | 0.40874047 | -0.9044817 | 0.36574004 | 0.89868152 |
| IFT43 | 12.4449525 | 0.40755469 | 0.63058933 | 0.64630761 | 0.51808016 | NA |
| GPATCH2L | 54.3276046 | -0.2067219 | 0.31261681 | -0.6612628 | 0.50844379 | 0.94305491 |
| VASH1 | 13.0390917 | 1.50868917 | 0.92233066 | 1.63573568 | 0.10189492 | 0.67049522 |
| ANGEL1 | 9.25883693 | 0.70651717 | 0.61096738 | 1.15639099 | 0.24752126 | NA |
| TMED8 | 19.9816857 | -0.3310823 | 0.41606989 | -0.7957372 | 0.42618481 | 0.92093196 |
| AHSA1 | 10.4062046 | -0.1407075 | 0.64446681 | -0.2183316 | 0.82717077 | NA |
| SPTLC2 | 79.1766939 | 0.10218474 | 0.43119125 | 0.2369824 | 0.81267044 | 0.98320463 |
| ALKBH1 | 10.2761882 | -0.0387821 | 0.58060622 | -0.0667958 | 0.94674427 | NA |
| SLIRP | 14.9439939 | 0.01301902 | 0.53941461 | 0.02413546 | 0.98074456 | 0.99758514 |
| SNW1 | 22.7226655 | -0.1763059 | 0.45771554 | -0.3851867 | 0.70009913 | 0.96979939 |
| CEP128 | 69.6699633 | 0.45534449 | 0.80334804 | 0.5668085 | 0.5708443 | 0.95415437 |
| GTF2A1 | 19.8530482 | 0.09122267 | 0.5596269 | 0.16300623 | 0.87051353 | 0.98751288 |
| STON2 | 20.7836658 | 1.33477281 | 0.7042915 | 1.89519936 | 0.05806599 | 0.57075882 |
| SEL1L | 44.58525 | 0.28856888 | 0.34046496 | 0.84757293 | 0.39667586 | 0.91333232 |
| LINC02328 | 46.5788726 | 0.4077155 | 0.44905643 | 0.90793822 | 0.36391086 | 0.89868152 |
| GALC | 23.0190084 | 0.25819305 | 0.66020825 | 0.39107819 | 0.69573944 | 0.9685713 |
| GPR65 | 19.0148735 | 1.0669453 | 0.96547299 | 1.10510114 | 0.26911577 | 0.84994492 |
| LINC01146 | 12.5398219 | 0.53780087 | 0.63329816 | 0.84920643 | 0.39576644 | NA |
| ZC3H14 | 21.2853235 | 0.71747199 | 0.5358822 | 1.3388614 | 0.1806158 | 0.76886756 |
| FOXN3 | 376.765009 | 0.16143289 | 0.27950632 | 0.57756436 | 0.56355828 | 0.95415437 |
| TDP1 | 14.9844603 | -0.1228879 | 0.53286432 | -0.2306176 | 0.81761186 | 0.98320463 |
| KCNK13 | 11.2656386 | 0.61112028 | 0.82292087 | 0.74262338 | 0.45770973 | NA |
| PSMC1 | 13.528524 | -0.0733871 | 0.58697584 | -0.1250258 | 0.90050316 | 0.99194856 |
| NRDE2 | 11.5804283 | -0.0526278 | 0.52082211 | -0.1010476 | 0.91951268 | NA |
| CALM1 | 70.4737095 | -0.3418884 | 0.3770579 | -0.9067266 | 0.3645514 | 0.89868152 |
| TTC7B | 20.4300697 | 0.77765223 | 0.5449875 | 1.42691755 | 0.15360366 | 0.74419469 |
| RPS6KA5 | 62.5313806 | 0.14335827 | 0.33677218 | 0.42568324 | 0.67033868 | 0.96412305 |
| DGLUCY | 56.1804509 | -0.0807951 | 0.47820621 | -0.1689545 | 0.86583241 | 0.98751288 |
| CCDC88C | 84.8653157 | 0.0793266 | 0.27432329 | 0.28917195 | 0.7724498 | 0.98264092 |
| PPP4R3A | 61.4054302 | 0.24136098 | 0.29726355 | 0.81194274 | 0.41682449 | 0.91953487 |
| CATSPERB | 16.5852666 | 0.03342803 | 0.81535398 | 0.04099818 | 0.96729734 | 0.99653147 |
| TC2N | 65.2094166 | -0.1700676 | 0.57821872 | -0.2941232 | 0.76866374 | 0.98264092 |
| TRIP11 | 39.6653237 | 0.41104755 | 0.35938091 | 1.14376568 | 0.25272083 | 0.84008912 |
| ATXN3 | 32.4556963 | 0.01830991 | 0.38278562 | 0.04783334 | 0.96184907 | 0.99641945 |
| NDUFB1 | 11.3532345 | -0.027094 | 0.52017806 | -0.052086 | 0.95846015 | NA |
| CPSF2 | 21.5714041 | -0.2176695 | 0.43317914 | -0.502493 | 0.61532075 | 0.95688177 |
| SLC24A4 | 16.0689121 | 1.71656231 | 0.98686765 | 1.73940479 | 0.08196359 | 0.63740875 |
| RIN3 | 132.057212 | -0.0601524 | 0.39987524 | -0.1504279 | 0.88042703 | 0.99019008 |
| LGMN | 76.560986 | -1.0113153 | 0.92059461 | -1.0985458 | 0.27196625 | 0.84994492 |
| GOLGA5 | 11.6882495 | 0.70164277 | 0.66406654 | 1.05658504 | 0.29070101 | NA |
| ITPK1 | 35.8529205 | 0.46568893 | 0.44963328 | 1.03570831 | 0.30033824 | 0.86225566 |
| BTBD7 | 66.0675661 | -0.0505587 | 0.34974775 | -0.1445577 | 0.88506008 | 0.99019008 |
| DDX24 | 36.402098 | -0.4810639 | 0.31059244 | -1.548859 | 0.12141562 | 0.71026852 |
| SERPINA1 | 37.2192432 | 0.97829802 | 0.79309051 | 1.23352633 | 0.21737946 | 0.80640512 |
| DICER1 | 75.1707012 | 0.38578564 | 0.32423218 | 1.18984376 | 0.2341078 | 0.82817204 |
| CLMN | 19.8957132 | 0.86729925 | 0.44970653 | 1.92858941 | 0.05378186 | 0.55218699 |
| SYNE3 | 23.1292421 | -1.2965545 | 0.52441372 | -2.4723887 | 0.01342135 | 0.31605855 |
| ATG2B | 28.4626047 | 0.12603304 | 0.47268462 | 0.2666324 | 0.7897522 | 0.98320463 |
| PAPOLA | 71.6202855 | 0.0724693 | 0.37014444 | 0.19578655 | 0.84477724 | 0.98441719 |

| | | | | | | |
|---|---|---|---|---|---|---|
| VRK1 | 21.1533751 | -0.6371646 | 0.44097329 | -1.4449052 | 0.1484845 | 0.7383468 |
| BCL11B | 75.6486739 | 0.24262336 | 0.33081478 | 0.7334115 | 0.46330749 | 0.9314919 |
| SETD3 | 40.9342499 | -0.0249257 | 0.33173642 | -0.0751371 | 0.94010564 | 0.99514836 |
| CCNK | 17.9719827 | 0.18708736 | 0.45233596 | 0.41360268 | 0.67916511 | 0.9673431 |
| EVL | 91.4901308 | 0.17584894 | 0.46953062 | 0.37452071 | 0.70801695 | 0.97184211 |
| YY1 | 44.4220415 | 0.04022529 | 0.33489173 | 0.1201143 | 0.9043926 | 0.99324672 |
| SLC25A29 | 11.4037468 | -0.6542524 | 0.59258812 | -1.1040592 | 0.26956746 | NA |
| WARS | 48.2030743 | -0.3120382 | 0.74378917 | -0.4195251 | 0.6748324 | 0.96574154 |
| WDR25 | 13.6231918 | 0.3593453 | 0.73725907 | 0.4874071 | 0.62596986 | 0.95876809 |
| PPP2R5C | 191.167542 | 0.10702775 | 0.40841663 | 0.26205532 | 0.79327879 | 0.98320463 |
| DYNC1H1 | 71.2417624 | 0.1561854 | 0.30234359 | 0.51658246 | 0.60544766 | 0.95688177 |
| HSP90AA1 | 150.967638 | 0.08290827 | 0.51928408 | 0.1596588 | 0.87314986 | 0.98813417 |
| WDR20 | 48.9081795 | -0.1245642 | 0.33352102 | -0.3734824 | 0.70878947 | 0.97186969 |
| TECPR2 | 16.7750077 | 0.5291752 | 0.58120954 | 0.91047231 | 0.36257348 | 0.89853007 |
| RCOR1 | 75.5515074 | 0.45631269 | 0.30789771 | 1.48202691 | 0.13833313 | 0.73044824 |
| TRAF3 | 80.6793876 | -0.7649801 | 0.34421901 | -2.2223644 | 0.02625869 | 0.40477881 |
| CDC42BPB | 20.8159432 | -0.2647345 | 0.43516966 | -0.6083478 | 0.54295685 | 0.95415437 |
| TNFAIP2 | 28.3748207 | 1.1376357 | 0.83767084 | 1.35809394 | 0.17443388 | 0.76214163 |
| EIF5 | 29.3693085 | 0.45756509 | 0.34565217 | 1.32377323 | 0.18557837 | 0.77694741 |
| MARK3 | 113.808426 | -0.1009311 | 0.21521964 | -0.4689677 | 0.63909272 | 0.95933464 |
| CKB | 9.40986197 | 1.84702592 | 0.66085854 | 2.79488848 | 0.00519176 | NA |
| BAG5 | 9.14575633 | 0.13855121 | 0.60245178 | 0.22997892 | 0.81810815 | NA |
| KLC1 | 32.6547019 | -0.5027205 | 0.40678824 | -1.2358285 | 0.21652233 | 0.80634897 |
| PPP1R13B | 15.7031532 | -0.3679728 | 0.50922483 | -0.7226136 | 0.46991734 | 0.9314919 |
| ATP5MPL | 15.4828723 | 0.57232174 | 0.62457361 | 0.91633992 | 0.35948863 | 0.8957272 |
| SIVA1 | 12.7145754 | 0.4081761 | 0.57602752 | 0.7086052 | 0.47856951 | NA |
| AKT1 | 12.5184122 | 0.50877334 | 0.51914511 | 0.98002144 | 0.32707554 | NA |
| GPR132 | 15.3892404 | 0.72010691 | 0.5352221 | 1.34543568 | 0.17848458 | 0.76509192 |
| BRF1 | 14.0575067 | 0.39267553 | 0.59386285 | 0.6612226 | 0.50846957 | 0.94305491 |
| PACS2 | 18.1122629 | 0.10247634 | 0.49594887 | 0.20662682 | 0.83630131 | 0.98320463 |
| IGHG3 | 46.3137805 | -1.7877734 | 1.88528593 | -0.948277 | 0.34298844 | 0.88530691 |
| IGHM | 188.599027 | -5.3971356 | 1.86498002 | -2.8939375 | 0.00380444 | 0.15732988 |
| NIPA1 | 8.5465588 | -0.2143987 | 0.6903567 | -0.3105623 | 0.75613342 | NA |
| NIPA2 | 19.5648117 | -1.205986 | 0.39989995 | -3.0157193 | 0.0025637 | 0.11922303 |
| CYFIP1 | 35.5344103 | 0.50601098 | 0.51500025 | 0.98254512 | 0.32583137 | 0.87715548 |
| AC124312.1 | 24.944095 | 0.12519759 | 0.49727451 | 0.25176757 | 0.80122073 | 0.98320463 |
| UBE3A | 64.945464 | 0.08682537 | 0.34805366 | 0.24945971 | 0.8030052 | 0.98320463 |
| ATP10A | 48.9587531 | -1.2366204 | 0.81873538 | -1.5104031 | 0.1309406 | 0.72973343 |
| HERC2 | 59.7825195 | -0.4836652 | 0.37201437 | -1.3001251 | 0.19355811 | 0.78135737 |
| APBA2 | 31.8323235 | 0.43820054 | 0.74476104 | 0.58837736 | 0.55627903 | 0.95415437 |
| AC091057.6 | 9.5864029 | 0.07561403 | 0.6123666 | 0.12347837 | 0.9017283 | NA |
| FAN1 | 8.32843233 | 0.20437496 | 0.59761573 | 0.34198391 | 0.732363 | NA |
| MTMR10 | 36.4481944 | -0.5598021 | 0.38189196 | -1.4658651 | 0.14268504 | 0.73044824 |
| KLF13 | 46.0309561 | 0.07349906 | 0.37750413 | 0.19469736 | 0.84562988 | 0.98458147 |
| LINC02256 | 30.7787544 | 0.7210486 | 0.54428768 | 1.32475643 | 0.18525195 | 0.77694741 |
| FMN1 | 189.97215 | 1.31359562 | 0.67505323 | 1.94591414 | 0.05166505 | 0.54907299 |
| AVEN | 18.8038945 | 1.34500765 | 0.55290116 | 2.43263671 | 0.01498933 | 0.32436474 |
| KATNBL1 | 44.0820067 | -0.3654244 | 0.34654835 | -1.0544687 | 0.2916684 | 0.85935767 |
| SLC12A6 | 69.8481032 | -1.344246 | 0.4179207 | -3.2165096 | 0.0012976 | 0.07558873 |
| NOP10 | 21.6583805 | 0.55784126 | 0.650086 | 0.85810379 | 0.39083515 | 0.90910959 |
| GOLGA8A | 18.0116573 | -0.3106491 | 0.46969292 | -0.6613877 | 0.50836374 | 0.94305491 |
| GOLGA8B | 19.0126407 | 0.60079681 | 0.4597496 | 1.30679137 | 0.19128358 | 0.78122755 |
| AQR | 34.4163192 | -0.3008442 | 0.3640208 | -0.826448 | 0.40855003 | 0.9169561 |
| ZNF770 | 11.1203899 | 0.41344097 | 0.54622858 | 0.75690102 | 0.44910917 | NA |

| | | | | | | |
|---|---|---|---|---|---|---|
| DPH6 | 33.4966241 | -0.3686076 | 0.49954294 | -0.7378897 | 0.46058146 | 0.9314919 |
| C15orf41 | 15.3154627 | -0.0243473 | 0.47893919 | -0.050836 | 0.95945621 | 0.99595344 |
| SPRED1 | 103.221487 | -0.5831028 | 0.68054063 | -0.8568229 | 0.39154274 | 0.90910959 |
| FAM98B | 8.91194349 | -0.3542954 | 0.53396274 | -0.6635208 | 0.50699709 | NA |
| RASGRP1 | 59.0871592 | 0.33894335 | 0.43964291 | 0.77095148 | 0.44073569 | 0.92880096 |
| C15orf53 | 28.0171567 | 2.0489832 | 0.95743239 | 2.14008134 | 0.03234819 | 0.44201211 |
| THBS1 | 79.8729285 | 1.24438908 | 1.39133716 | 0.89438356 | 0.37111671 | 0.90226557 |
| EIF2AK4 | 28.6521987 | -0.2396308 | 0.3951273 | -0.6064649 | 0.54420611 | 0.95415437 |
| SRP14 | 46.2426128 | 0.21587373 | 0.5047684 | 0.42766886 | 0.66889223 | 0.96412305 |
| PLCB2 | 20.3506008 | 0.61368173 | 0.56569484 | 1.08482823 | 0.27799774 | 0.84994492 |
| IVD | 15.2548253 | 0.38887178 | 0.46753905 | 0.8317418 | 0.4055547 | 0.91575086 |
| CCDC32 | 17.4418769 | -0.4058072 | 0.50749538 | -0.7996273 | 0.42392674 | 0.91985469 |
| DNAJC17 | 10.3364226 | -0.4421172 | 0.62824027 | -0.7037391 | 0.48159527 | NA |
| INO80 | 44.3057317 | -0.197956 | 0.33971329 | -0.5827148 | 0.56008529 | 0.95415437 |
| CHP1 | 15.5097667 | 0.55994428 | 0.51498514 | 1.08730182 | 0.27690345 | 0.84994492 |
| RTF1 | 24.0059889 | 0.1006398 | 0.41735132 | 0.24113929 | 0.80944717 | 0.98320463 |
| MGA | 47.832938 | -0.1368079 | 0.43106254 | -0.3173736 | 0.75096017 | 0.97829887 |
| MAPKBP1 | 14.0960009 | 0.03864704 | 0.49330069 | 0.07834378 | 0.93755459 | 0.99514836 |
| EHD4 | 19.082159 | -0.4595421 | 0.44744335 | -1.0270398 | 0.30440174 | 0.86609729 |
| VPS39 | 28.1113669 | 0.02480308 | 0.35954503 | 0.06898462 | 0.94500187 | 0.99514836 |
| TMEM87A | 47.2822159 | -0.0761167 | 0.35838466 | -0.2123882 | 0.83180421 | 0.98320463 |
| GANC | 23.0502122 | -0.3966992 | 0.50510533 | -0.7853791 | 0.43223131 | 0.92284736 |
| ZNF106 | 52.5478895 | 0.14209303 | 0.38434805 | 0.36969884 | 0.7116069 | 0.97317955 |
| SNAP23 | 38.6284053 | 0.12386383 | 0.35422441 | 0.34967615 | 0.72658175 | 0.9754739 |
| STARD9 | 12.6982825 | 0.55147497 | 0.75911764 | 0.72646839 | 0.46755168 | NA |
| TTBK2 | 52.0960884 | 0.41165388 | 0.4712276 | 0.87357761 | 0.38234833 | 0.90868001 |
| UBR1 | 71.733329 | -0.1857003 | 0.26307896 | -0.7058728 | 0.48026726 | 0.93257966 |
| AC068724.3 | 14.3117372 | -0.1181045 | 0.61773387 | -0.1911899 | 0.84837681 | 0.98545782 |
| TMEM62 | 20.564664 | -0.5038118 | 0.43590786 | -1.155776 | 0.2477728 | 0.83817091 |
| CCNDBP1 | 11.8483758 | -0.5232495 | 0.58049003 | -0.9013928 | 0.36737953 | NA |
| TUBGCP4 | 9.84430329 | 0.56487478 | 0.58589737 | 0.964119 | 0.33498627 | NA |
| TP53BP1 | 17.0768367 | 0.62952533 | 0.49004668 | 1.2846232 | 0.19892399 | 0.78142991 |
| CATSPER2 | 9.19748651 | -0.9002335 | 0.69120926 | -1.3024037 | 0.19277842 | NA |
| PDIA3 | 42.9732309 | -0.0723858 | 0.38834203 | -0.1863971 | 0.85213336 | 0.98751288 |
| SERF2 | 121.833928 | 0.28952838 | 0.50423101 | 0.57419788 | 0.5658339 | 0.95415437 |
| MFAP1 | 8.4184274 | -0.1350152 | 0.85268843 | -0.1583406 | 0.87418845 | NA |
| CASC4 | 58.9003625 | 0.03154712 | 0.33016828 | 0.09554862 | 0.92387907 | 0.99514836 |
| CTDSPL2 | 53.2626612 | -0.2247114 | 0.33186329 | -0.6771205 | 0.4983295 | 0.93916044 |
| EIF3J | 20.7849102 | 0.32915299 | 0.43287589 | 0.76038652 | 0.44702358 | 0.92914597 |
| SPG11 | 100.462931 | -0.2741208 | 0.39380906 | -0.6960754 | 0.48638163 | 0.93459582 |
| PATL2 | 16.3850216 | -1.0219598 | 0.74480487 | -1.3721175 | 0.17002686 | 0.75745073 |
| B2M | 817.639008 | 0.27225763 | 0.50967642 | 0.53417741 | 0.5932188 | 0.95487866 |
| GATM | 9.99052841 | 1.71251115 | 0.7685179 | 2.22832955 | 0.02585855 | NA |
| SQOR | 14.1004792 | 1.5456985 | 0.64974253 | 2.37894001 | 0.0173625 | 0.34827529 |
| MYEF2 | 12.9913415 | -1.4870912 | 0.59064789 | -2.5177288 | 0.01181142 | 0.29101961 |
| SLC12A1 | 24.8538436 | -1.0188273 | 0.82161608 | -1.2400285 | 0.21496487 | 0.80634897 |
| DUT | 11.9153429 | 0.17538919 | 0.61153935 | 0.28679951 | 0.77426584 | NA |
| CEP152 | 14.8094168 | 0.65691147 | 0.64601627 | 1.01686522 | 0.30921755 | 0.8673137 |
| EID1 | 18.4530578 | 0.42433973 | 0.4577743 | 0.92696277 | 0.35394587 | 0.89109867 |
| SECISBP2L | 34.4405879 | 0.14166085 | 0.37105139 | 0.38178228 | 0.70262286 | 0.97110762 |
| COPS2 | 15.0486556 | -0.7965226 | 0.5167349 | -1.5414532 | 0.12320654 | 0.71395284 |
| GALK2 | 39.9995371 | 0.36585297 | 0.48545646 | 0.75362674 | 0.45107338 | 0.92914597 |
| DTWD1 | 17.6619504 | 0.08199488 | 0.58995597 | 0.13898475 | 0.8894622 | 0.99080204 |
| ATP8B4 | 198.669451 | 2.87565129 | 0.80692703 | 3.56370674 | 0.00036565 | 0.0361345 |

| | | | | | | |
|---|---|---|---|---|---|---|
| GABPB1 | 37.9964883 | 0.04404818 | 0.37963441 | 0.11602789 | 0.90763044 | 0.99389264 |
| GABPB1-AS1 | 33.1091711 | 0.67962774 | 0.62596617 | 1.085726 | 0.27760023 | 0.84994492 |
| USP8 | 52.0636926 | -0.3405739 | 0.3021906 | -1.1270168 | 0.25973537 | 0.84551502 |
| TRPM7 | 93.3395335 | -0.3140792 | 0.31672678 | -0.9916407 | 0.32137281 | 0.87363761 |
| SPPL2A | 56.2095194 | -0.3622152 | 0.36321507 | -0.9972473 | 0.31864449 | 0.87298176 |
| AP4E1 | 18.3951718 | -0.3544865 | 0.48073624 | -0.7373826 | 0.46088972 | 0.9314919 |
| DMXL2 | 166.214139 | 1.75222415 | 0.52863492 | 3.31462049 | 0.00091768 | 0.06269654 |
| TMOD2 | 13.2637687 | 1.04894487 | 0.49887586 | 2.10261703 | 0.03549926 | 0.46442767 |
| TMOD3 | 69.6311404 | 0.18225288 | 0.34278385 | 0.53168458 | 0.59494448 | 0.9553365 |
| MAPK6 | 26.1893482 | 0.34152001 | 0.40767366 | 0.83772892 | 0.40218297 | 0.91528253 |
| GNB5 | 15.9162466 | -0.4761802 | 0.47586037 | -1.0006722 | 0.31698532 | 0.87263466 |
| MYO5A | 146.371838 | -0.371696 | 0.31189384 | -1.1917389 | 0.23336365 | 0.82817204 |
| ARPP19 | 27.6782365 | -0.0621205 | 0.37514998 | -0.1655883 | 0.86848094 | 0.98751288 |
| FAM214A | 73.9162291 | 0.23841532 | 0.50330758 | 0.47369706 | 0.63571595 | 0.95933464 |
| RSL24D1 | 12.4107168 | 0.02424786 | 0.59480617 | 0.04076599 | 0.96748246 | NA |
| RAB27A | 71.1809395 | 0.74752243 | 0.35003817 | 2.1355455 | 0.03271648 | 0.44484764 |
| PIGB | 17.6694991 | 0.17130207 | 0.50901996 | 0.33653311 | 0.73646889 | 0.97605445 |
| CCPG1 | 38.4508583 | 0.19041201 | 0.3744594 | 0.5084984 | 0.61110386 | 0.95688177 |
| NEDD4 | 10.7576258 | 1.53560438 | 0.6446717 | 2.3819944 | 0.01721916 | NA |
| RFX7 | 80.2548982 | 0.59976227 | 0.30022279 | 1.99772397 | 0.04574659 | 0.52198279 |
| ZNF280D | 18.0017505 | 0.19807701 | 0.48525957 | 0.40818776 | 0.68313584 | 0.9673431 |
| AC090517.4 | 14.2219364 | 0.97618014 | 0.70104649 | 1.39246135 | 0.16378274 | 0.75321592 |
| TCF12 | 268.250966 | 0.13033223 | 0.25374949 | 0.51362558 | 0.60751379 | 0.95688177 |
| CGNL1 | 11.5393344 | 0.53298216 | 0.61703212 | 0.86378349 | 0.38770684 | NA |
| ALDH1A2 | 18.5461291 | 0.40577238 | 0.75243843 | 0.53927653 | 0.58969606 | 0.95415437 |
| AQP9 | 27.9679035 | 0.94212001 | 0.93002505 | 1.01300498 | 0.31105778 | 0.86895193 |
| ADAM10 | 143.593326 | -0.0576488 | 0.32621354 | -0.1767209 | 0.8597276 | 0.98751288 |
| MINDY2 | 29.4161156 | -0.1003145 | 0.41991471 | -0.2388926 | 0.81118885 | 0.98320463 |
| RNF111 | 78.3373147 | 0.3865587 | 0.30742732 | 1.25739867 | 0.20860931 | 0.79538492 |
| SLTM | 87.6301405 | 0.26376103 | 0.27951144 | 0.94365022 | 0.34534842 | 0.88726006 |
| MYO1E | 38.5086741 | -0.0177481 | 0.74916592 | -0.0236905 | 0.98109951 | 0.99758514 |
| GTF2A2 | 11.5252935 | 0.25072256 | 0.5469543 | 0.45839765 | 0.64666678 | NA |
| BNIP2 | 33.6124833 | -0.3784188 | 0.3617353 | -1.0461208 | 0.29550526 | 0.8608685 |
| ANXA2 | 66.7390824 | 0.07674924 | 0.5131154 | 0.14957502 | 0.88109992 | 0.99019008 |
| ICE2 | 23.8754654 | 0.12922187 | 0.44923923 | 0.287646 | 0.77361773 | 0.98264092 |
| RORA-AS1 | 17.799255 | 0.20251136 | 0.51727697 | 0.39149502 | 0.69543137 | 0.9685713 |
| RORA | 109.339664 | 0.27595283 | 0.41847483 | 0.65942517 | 0.50962279 | 0.94332619 |
| AC009554.1 | 21.9072855 | 0.31463041 | 0.66501095 | 0.47312065 | 0.63612711 | 0.95933464 |
| VPS13C | 172.557502 | 0.12449253 | 0.26500808 | 0.46976882 | 0.63852019 | 0.95933464 |
| TLN2 | 16.0602523 | 2.00471217 | 0.81604303 | 2.45662556 | 0.01402488 | 0.31939784 |
| TPM1 | 13.4736105 | 0.12780056 | 0.48201909 | 0.26513589 | 0.79090478 | 0.98320463 |
| LACTB | 16.8774936 | 0.54254319 | 0.58911996 | 0.92093838 | 0.3570826 | 0.8957272 |
| RPS27L | 23.3727607 | -0.0210183 | 0.42260666 | -0.0497348 | 0.96033371 | 0.99614757 |
| RAB8B | 113.036043 | 0.22481691 | 0.36389843 | 0.61780127 | 0.53670634 | 0.95415437 |
| APH1B | 17.464737 | -0.8138578 | 0.47889351 | -1.6994547 | 0.08923354 | 0.64845882 |
| CA12 | 16.0984677 | 0.84670492 | 0.73373427 | 1.1539667 | 0.24851382 | 0.83858261 |
| USP3 | 134.063533 | -0.0685402 | 0.2640958 | -0.2595279 | 0.79522794 | 0.98320463 |
| USP3-AS1 | 26.2393022 | -0.3052118 | 0.52186359 | -0.5848497 | 0.55864877 | 0.95415437 |
| HERC1 | 152.452888 | -0.0095154 | 0.26606655 | -0.0357634 | 0.97147104 | 0.99723998 |
| DAPK2 | 10.3222929 | 0.40974801 | 0.66033732 | 0.62051318 | 0.53491998 | NA |
| FAM96A | 10.0760521 | -0.3782027 | 0.5402261 | -0.7000822 | 0.48387595 | NA |
| SNX1 | 50.5941352 | -0.1327228 | 0.35656556 | -0.3722256 | 0.70972489 | 0.97211298 |
| PPIB | 38.3060728 | -0.1033082 | 0.66242532 | -0.1559545 | 0.8760689 | 0.98939075 |
| TRIP4 | 25.3157772 | 0.18037939 | 0.39300038 | 0.4589802 | 0.64624839 | 0.96049083 |

| | | | | | | |
|---|---|---|---|---|---|---|
| ZNF609 | 153.790484 | 0.14066047 | 0.34123674 | 0.41220789 | 0.68018706 | 0.9673431 |
| OAZ2 | 15.2625457 | 0.29838281 | 0.57242014 | 0.52126539 | 0.60218191 | 0.95688177 |
| AC069368.1 | 14.0242111 | -0.8338595 | 0.68647897 | -1.2146905 | 0.22448415 | 0.81875277 |
| ANKDD1A | 11.3366411 | -1.6883168 | 0.79049767 | -2.1357644 | 0.03269862 | NA |
| SPG21 | 20.9572362 | 0.38267125 | 0.45154256 | 0.8474755 | 0.39673014 | 0.91333232 |
| MTFMT | 9.62917741 | -0.6775465 | 0.53353299 | -1.2699242 | 0.20411162 | NA |
| PDCD7 | 9.12709917 | 0.09241319 | 0.57708168 | 0.16013884 | 0.87277171 | NA |
| CLPX | 16.2970147 | -0.271293 | 0.46510941 | -0.5832886 | 0.55969901 | 0.95415437 |
| DPP8 | 44.3579202 | 0.01667817 | 0.30782047 | 0.05418147 | 0.95679058 | 0.99514836 |
| HACD3 | 11.9997388 | 0.61683743 | 0.5795741 | 1.06429432 | 0.28719539 | NA |
| DENND4A | 126.486788 | -0.3727535 | 0.28508757 | -1.3075051 | 0.19104122 | 0.78122755 |
| RAB11A | 58.9560797 | 0.34629305 | 0.53018439 | 0.65315588 | 0.51365579 | 0.94596835 |
| DIS3L | 12.1249002 | -0.381305 | 0.5145163 | -0.7410941 | 0.45863639 | NA |
| MAP2K1 | 88.9123511 | -0.1425433 | 0.44083405 | -0.323349 | 0.74643093 | 0.97805001 |
| RPL4 | 61.5090147 | 0.00305302 | 0.49060087 | 0.00622302 | 0.99503478 | 0.99859588 |
| SMAD3 | 59.2976226 | 0.62602194 | 0.48746823 | 1.28423124 | 0.19906106 | 0.78142991 |
| AAGAB | 27.4844323 | 0.63408291 | 0.42446686 | 1.49383372 | 0.13521909 | 0.73044824 |
| IQCH-AS1 | 9.3862015 | 0.28334275 | 0.60480495 | 0.46848616 | 0.63943697 | NA |
| MAP2K5 | 69.6716021 | 0.39901555 | 0.32574878 | 1.224918 | 0.22060612 | 0.81064692 |
| PIAS1 | 188.954785 | -0.1760061 | 0.28148611 | -0.6252746 | 0.53179086 | 0.95265314 |
| FEM1B | 13.3654598 | 0.51034041 | 0.56717628 | 0.89979153 | 0.3682312 | 0.89868152 |
| ANP32A | 41.8369819 | 0.25622047 | 0.37586646 | 0.68167952 | 0.49544162 | 0.93805563 |
| GLCE | 11.6225574 | 0.58458146 | 0.606252 | 0.9642549 | 0.33491815 | NA |
| RPLP1 | 156.10966 | -0.5841803 | 0.6295564 | -0.9279237 | 0.35344717 | 0.89109867 |
| TLE3 | 27.2672176 | -0.4214359 | 0.61270231 | -0.6878315 | 0.4915589 | 0.93673794 |
| UACA | 14.9372535 | 0.82216855 | 0.59216583 | 1.38840931 | 0.16501244 | 0.75321592 |
| THSD4-AS1 | 22.5297667 | 0.22569765 | 0.52182018 | 0.43251996 | 0.66536355 | 0.96406914 |
| MYO9A | 76.710956 | -0.5124899 | 0.35120166 | -1.4592467 | 0.14449721 | 0.73227798 |
| PKM | 79.8617556 | -0.206794 | 0.4378979 | -0.4722426 | 0.63675365 | 0.95933464 |
| PARP6 | 14.0391401 | 0.10658811 | 0.48311786 | 0.22062548 | 0.82538406 | 0.98320463 |
| AC009690.1 | 24.6034308 | 0.76310399 | 0.61605104 | 1.23870256 | 0.21545567 | 0.80634897 |
| ARIH1 | 128.677959 | -0.0244286 | 0.22721983 | -0.1075108 | 0.91438379 | 0.99498068 |
| BBS4 | 7.95066931 | 0.60756355 | 0.6610711 | 0.9190593 | 0.35806456 | NA |
| ADPGK | 42.0577224 | 0.03706176 | 0.43466526 | 0.08526507 | 0.93205066 | 0.99514836 |
| NEO1 | 12.1709379 | 0.65304726 | 0.68645341 | 0.95133516 | 0.34143426 | NA |
| NPTN | 46.8703779 | 0.25992913 | 0.35594181 | 0.73025738 | 0.46523287 | 0.9314919 |
| PML | 14.9601984 | 0.77714442 | 0.64881495 | 1.19779056 | 0.23099856 | 0.82438852 |
| ARID3B | 18.4732695 | 0.54410564 | 0.49482002 | 1.09960313 | 0.27150508 | 0.84994492 |
| CLK3 | 9.87909739 | -0.6842378 | 0.56406336 | -1.2130513 | 0.2251102 | NA |
| EDC3 | 8.1736213 | -0.2327989 | 0.59249891 | -0.3929102 | 0.69438577 | NA |
| CSK | 36.8278765 | -1.2068442 | 0.60728822 | -1.9872676 | 0.04689275 | 0.52744812 |
| SCAMP2 | 19.6894728 | -0.3587359 | 0.48927977 | -0.7331918 | 0.46344143 | 0.9314919 |
| COX5A | 13.2564689 | 0.64450932 | 0.54415236 | 1.18442807 | 0.23624366 | 0.83171006 |
| PPCDC | 22.8179382 | -0.4133073 | 0.4084334 | -1.011933 | 0.31157007 | 0.86950518 |
| MAN2C1 | 15.0012874 | -0.1248271 | 0.51920502 | -0.2404197 | 0.81000487 | 0.98320463 |
| SIN3A | 31.0434761 | -0.2158572 | 0.38552478 | -0.559905 | 0.57554426 | 0.95415437 |
| PTPN9 | 39.1309208 | -0.14878 | 0.40979935 | -0.3630557 | 0.71656325 | 0.97348719 |
| UBE2Q2 | 28.8759151 | -0.5049495 | 0.5492427 | -0.9193558 | 0.3579095 | 0.8957272 |
| FBXO22 | 19.1226017 | 0.06024035 | 0.52375247 | 0.11501683 | 0.90843179 | 0.99423609 |
| ETFA | 42.2333873 | 0.35282728 | 0.32647323 | 1.08072347 | 0.27982014 | 0.84994492 |
| SCAPER | 84.9991885 | 0.38970653 | 0.34680299 | 1.12371158 | 0.2611354 | 0.8475436 |
| RCN2 | 14.06159 | -0.2543945 | 0.55284109 | -0.4601584 | 0.6454025 | 0.96049083 |
| PSTPIP1 | 22.7414224 | -0.0115998 | 0.41035475 | -0.0282679 | 0.97744852 | 0.99758514 |
| TSPAN3 | 16.9604988 | -1.4282277 | 0.48878763 | -2.9219799 | 0.00347814 | 0.15037519 |

| | | | | | | |
|---|---|---|---|---|---|---|
| PEAK1 | 107.237853 | 0.3949771 | 0.44325549 | 0.89108226 | 0.37288504 | 0.9025457 |
| HMG20A | 21.475871 | -0.1400861 | 0.41981811 | -0.3336829 | 0.73861884 | 0.97605445 |
| LINGO1 | 197.23615 | -0.2429812 | 0.58475268 | -0.4155281 | 0.67775534 | 0.96718429 |
| TBC1D2B | 53.5609551 | 0.68126215 | 0.40746117 | 1.67196829 | 0.09453058 | 0.65925012 |
| IDH3A | 11.300316 | -0.1082143 | 0.67817435 | -0.159567 | 0.87322217 | NA |
| IREB2 | 39.3326725 | 0.11605431 | 0.37637536 | 0.30834726 | 0.75781811 | 0.98030982 |
| PSMA4 | 22.3264392 | 0.02336783 | 0.4982111 | 0.04690347 | 0.96259016 | 0.99653147 |
| MORF4L1 | 53.6616353 | -0.0433672 | 0.2791024 | -0.1553808 | 0.8765211 | 0.98939075 |
| CTSH | 28.0978557 | 0.83017509 | 0.50736919 | 1.63623474 | 0.10179047 | 0.67049522 |
| TMED3 | 12.8298809 | -0.8188412 | 0.71485149 | -1.1454704 | 0.25201435 | NA |
| BCL2A1 | 14.5357791 | 0.73802115 | 0.95580502 | 0.77214613 | 0.44002788 | 0.92861039 |
| ZFAND6 | 97.8902713 | 0.01786372 | 0.31743538 | 0.05627514 | 0.95512263 | 0.99514836 |
| MESD | 16.8338446 | 0.2473975 | 0.42102956 | 0.58760127 | 0.55679996 | 0.95415437 |
| IL16 | 16.8500512 | 0.28241335 | 0.51485684 | 0.54852792 | 0.58332947 | 0.95415437 |
| EFL1 | 21.6789818 | 0.2885582 | 0.41249465 | 0.69954411 | 0.48421206 | 0.93399425 |
| WHAMM | 17.465854 | -0.6613187 | 0.42712413 | -1.5483057 | 0.12154871 | 0.71029626 |
| BTBD1 | 30.631134 | -0.550894 | 0.32738757 | -1.6826968 | 0.0924338 | 0.65902576 |
| TM6SF1 | 12.0251187 | 0.07176632 | 0.58808366 | 0.12203421 | 0.90287193 | NA |
| WDR73 | 12.265804 | 0.42160444 | 0.57743558 | 0.73013242 | 0.46530925 | NA |
| SEC11A | 47.8624714 | 0.16841219 | 0.30295875 | 0.5558915 | 0.57828503 | 0.95415437 |
| ZNF592 | 27.8906528 | -0.5775492 | 0.33756101 | -1.7109477 | 0.08709076 | 0.64676224 |
| PDE8A | 84.7218239 | -0.4264453 | 0.38749903 | -1.1005067 | 0.2711114 | 0.84994492 |
| AKAP13 | 398.73628 | -0.6979545 | 0.27228391 | -2.5633334 | 0.01036724 | 0.27986499 |
| AC087286.2 | 21.4572746 | -0.4098389 | 0.58620837 | -0.6991352 | 0.48446757 | 0.93416151 |
| ISG20 | 42.8057982 | -1.0348217 | 0.45565254 | -2.2710763 | 0.02314236 | 0.39285221 |
| ABHD2 | 53.339091 | 0.51449431 | 0.32329757 | 1.59139557 | 0.11152058 | 0.69385269 |
| POLG | 12.6374572 | 0.21662677 | 0.54266653 | 0.39918949 | 0.68975359 | NA |
| ANPEP | 24.5856241 | 0.62047596 | 0.65001476 | 0.95455672 | 0.33980191 | 0.8844375 |
| ZNF710 | 87.5855456 | 0.48749621 | 0.44048092 | 1.10673628 | 0.26840796 | 0.84994492 |
| IDH2 | 21.4114116 | 0.69116606 | 0.43673539 | 1.58257398 | 0.1135186 | 0.69801324 |
| SEMA4B | 18.2297936 | -1.1525648 | 0.59742945 | -1.9292065 | 0.05370523 | 0.55218699 |
| CIB1 | 25.0638411 | -0.5924567 | 0.57313233 | -1.0337172 | 0.3012684 | 0.86225566 |
| IQGAP1 | 164.937948 | 0.13081874 | 0.30351867 | 0.43100722 | 0.66646312 | 0.96412305 |
| CRTC3 | 59.6539227 | -0.1011629 | 0.2986774 | -0.3387029 | 0.73483355 | 0.97605445 |
| FES | 15.129853 | 0.73751898 | 0.53196487 | 1.38640543 | 0.16562314 | 0.75337545 |
| MAN2A2 | 12.0564353 | -0.2817869 | 0.51844356 | -0.5435248 | 0.58676854 | NA |
| UNC45A | 8.77325256 | 0.79153508 | 0.59876167 | 1.32195349 | 0.18618364 | NA |
| SLCO3A1 | 128.139175 | 0.07861332 | 0.36111941 | 0.21769343 | 0.82766799 | 0.98320463 |
| AC013394.1 | 54.7760221 | -0.0442616 | 0.34123958 | -0.1297084 | 0.89679712 | 0.99118739 |
| CHD2 | 87.9259931 | -0.2591443 | 0.25096523 | -1.0325903 | 0.30179565 | 0.86225566 |
| MCTP2 | 133.784781 | 0.05515918 | 0.41685159 | 0.13232331 | 0.89472857 | 0.99115168 |
| IGF1R | 60.5689685 | -0.0970092 | 0.47463562 | -0.2043866 | 0.83805137 | 0.98320463 |
| LRRC28 | 22.1929634 | 0.65175755 | 0.39842026 | 1.63585444 | 0.10187006 | 0.67049522 |
| MEF2A | 253.652034 | -0.8797881 | 0.39154159 | -2.2469852 | 0.02464097 | 0.39492781 |
| AC090825.1 | 17.3838983 | -0.181079 | 0.52530638 | -0.3447113 | 0.73031146 | 0.97605445 |
| ADAMTS17 | 15.6252069 | 0.4441367 | 0.51035988 | 0.8702422 | 0.38416806 | 0.90910959 |
| LINS1 | 10.9040293 | 0.99329699 | 0.63039614 | 1.57567112 | 0.11510162 | NA |
| ASB7 | 13.6104934 | -0.2448162 | 0.50458537 | -0.4851829 | 0.62754663 | 0.95877428 |
| LRRK1 | 46.2787564 | -0.7880022 | 0.50268297 | -1.5675928 | 0.1169762 | 0.70122655 |
| CHSY1 | 50.6305476 | 0.35457943 | 0.44976489 | 0.78836619 | 0.43048254 | 0.92265313 |
| SELENOS | 19.4970619 | 0.5435983 | 0.53177725 | 1.02222931 | 0.30667238 | 0.8673137 |
| SNRPA1 | 12.8055624 | -0.6532258 | 0.51144013 | -1.2772283 | 0.20152166 | NA |
| TM2D3 | 13.8982002 | 0.1268389 | 0.52898591 | 0.23977747 | 0.81050277 | 0.98320463 |
| TARSL2 | 24.3667458 | 0.6695917 | 0.63635415 | 1.0522312 | 0.29269349 | 0.85998836 |

| | | | | | | |
|---|---|---|---|---|---|---|
| MPG | 11.3135766 | -0.5090958 | 0.51403543 | -0.9903904 | 0.32198331 | NA |
| NPRL3 | 15.2897071 | 0.09633915 | 0.50432141 | 0.19102729 | 0.84850421 | 0.98545782 |
| LUC7L | 38.3450321 | -0.4928652 | 0.34860571 | -1.4138186 | 0.15741517 | 0.74519724 |
| AXIN1 | 18.8270044 | -0.3441074 | 0.47821322 | -0.719569 | 0.47179038 | 0.9314919 |
| TMEM8A | 9.45245564 | -0.1526976 | 0.57476861 | -0.2656679 | 0.79049498 | NA |
| RAB11FIP3 | 24.1174185 | -0.1482178 | 0.42629314 | -0.3476897 | 0.7280732 | 0.97605445 |
| CAPN15 | 10.8984287 | -0.3357569 | 0.61974204 | -0.5417689 | 0.58797774 | NA |
| RAB40C | 10.6753573 | 0.6105083 | 0.71466559 | 0.85425731 | 0.39296244 | NA |
| RHOT2 | 11.1746443 | 0.71750138 | 0.60137161 | 1.19310816 | 0.23282701 | NA |
| STUB1 | 11.2295653 | 1.09904199 | 0.59013426 | 1.86235925 | 0.06255247 | NA |
| LMF1 | 17.4553797 | 0.11272097 | 0.63536961 | 0.17741007 | 0.8591863 | 0.98751288 |
| UBE2I | 31.9753248 | -0.9772553 | 0.39352148 | -2.4833596 | 0.01301496 | 0.30911937 |
| GNPTG | 9.09043461 | 0.59380285 | 0.63626067 | 0.93326977 | 0.3506807 | NA |
| UNKL | 11.8002188 | 0.44397211 | 0.51837966 | 0.85646127 | 0.39174268 | NA |
| CLCN7 | 19.4310289 | 0.61566654 | 0.47619213 | 1.29289525 | 0.19604729 | 0.78135737 |
| MAPK8IP3 | 30.5880976 | -1.0493376 | 0.39714705 | -2.6421891 | 0.0082372 | 0.23742025 |
| NME3 | 15.0153298 | 0.1514676 | 0.47835616 | 0.31664189 | 0.75151535 | 0.97857378 |
| MRPS34 | 11.5883989 | -0.2129786 | 0.80281911 | -0.2652884 | 0.79078732 | NA |
| SPSB3 | 13.9834168 | -0.5867102 | 0.47893251 | -1.2250374 | 0.22056112 | 0.81064692 |
| HAGH | 16.1458215 | -0.115908 | 0.51590114 | -0.224671 | 0.82223524 | 0.98320463 |
| NDUFB10 | 12.530701 | -0.1775395 | 0.55972801 | -0.3171889 | 0.75110027 | NA |
| RPS2 | 89.8855591 | -0.1007942 | 0.54475753 | -0.1850258 | 0.8532088 | 0.98751288 |
| TSC2 | 15.1449597 | -0.0380291 | 0.45786742 | -0.0830571 | 0.93380614 | 0.99514836 |
| PKD1 | 10.7684132 | -0.1691564 | 0.59421548 | -0.2846719 | 0.77589554 | NA |
| PGP | 13.217688 | -0.3498188 | 0.48950493 | -0.7146379 | 0.4748328 | 0.9314919 |
| ECI1 | 11.926597 | -0.3638497 | 0.57874076 | -0.6286921 | 0.52955065 | NA |
| RNPS1 | 19.9473542 | 0.2676982 | 0.45980433 | 0.58220025 | 0.5604318 | 0.95415437 |
| PDPK1 | 50.1431392 | -0.322383 | 0.3207672 | -1.0050371 | 0.31487896 | 0.87257896 |
| SRRM2 | 109.41223 | -0.3832145 | 0.25404238 | -1.5084667 | 0.13143511 | 0.73044824 |
| ELOB | 18.4460224 | 0.02499668 | 0.67881374 | 0.03682406 | 0.97062529 | 0.99676559 |
| MMP25-AS1 | 7.38207502 | 0.02443684 | 0.93874167 | 0.02603149 | 0.97923223 | NA |
| IL32 | 34.7149218 | 0.91173846 | 0.59361363 | 1.53591228 | 0.1245599 | 0.71741321 |
| ZNF263 | 9.08176495 | -0.4764535 | 0.63929572 | -0.7452787 | 0.45610326 | NA |
| ZNF75A | 12.0173644 | -0.953147 | 0.74127693 | -1.2858177 | 0.19850668 | NA |
| NLRC3 | 17.1231418 | 0.39054854 | 0.50532983 | 0.77285866 | 0.43960603 | 0.92818763 |
| DNASE1 | 19.459753 | 0.40401349 | 0.50037458 | 0.80742208 | 0.41942335 | 0.91985469 |
| TRAP1 | 9.03743192 | 0.21791806 | 0.55249867 | 0.39442278 | 0.69326893 | NA |
| CREBBP | 91.1425699 | -0.1471851 | 0.29434148 | -0.5000489 | 0.61704065 | 0.95688177 |
| ADCY9 | 26.1566517 | 0.29178564 | 0.47950734 | 0.60851131 | 0.54284841 | 0.95415437 |
| DNAJA3 | 10.6969128 | -0.6056603 | 0.53878126 | -1.1241301 | 0.26095782 | NA |
| NMRAL1 | 9.27658866 | 0.00789722 | 0.56536618 | 0.01396832 | 0.98885526 | NA |
| HMOX2 | 15.3183426 | 0.74444427 | 0.50736162 | 1.46728533 | 0.14229845 | 0.73044824 |
| MGRN1 | 25.7992515 | 0.52322147 | 0.3978336 | 1.31517668 | 0.18845053 | 0.77798634 |
| ANKS3 | 9.24886415 | -0.6596679 | 0.53465368 | -1.2338228 | 0.21726893 | NA |
| GLYR1 | 34.7416904 | -0.347136 | 0.44634668 | -0.7777274 | 0.43672974 | 0.92489992 |
| UBN1 | 18.4986177 | 0.10445084 | 0.44146016 | 0.23660309 | 0.81296472 | 0.98320463 |
| METTL22 | 8.52760462 | 0.06525319 | 0.64439958 | 0.10126199 | 0.91934249 | NA |
| ABAT | 16.3189161 | 1.15484534 | 0.55343076 | 2.0867025 | 0.03691504 | 0.46962714 |
| PMM2 | 24.135294 | -0.2652251 | 0.35592394 | -0.7451736 | 0.45616676 | 0.9314919 |
| CARHSP1 | 18.498971 | -0.8119088 | 0.45448132 | -1.7864514 | 0.0740262 | 0.61929783 |
| USP7 | 33.85513 | -0.333712 | 0.361772 | -0.9224373 | 0.35630053 | 0.89503727 |
| C16orf72 | 29.1819604 | 0.12321314 | 0.36535767 | 0.33723978 | 0.73593615 | 0.97605445 |
| ATF7IP2 | 51.4023391 | 0.22453045 | 0.38307591 | 0.58612522 | 0.55779137 | 0.95415437 |
| NUBP1 | 8.60013813 | -0.4918344 | 0.66797144 | -0.7363105 | 0.46154178 | NA |

| | | | | | | |
|---|---|---|---|---|---|---|
| CIITA | 71.0047228 | -0.7908845 | 0.58536818 | -1.351089 | 0.17666693 | 0.76409729 |
| DEXI | 7.83709498 | 0.19484854 | 0.56873439 | 0.34260024 | 0.73189922 | NA |
| CLEC16A | 59.1790151 | -0.3302426 | 0.39063067 | -0.8454088 | 0.39788262 | 0.91429865 |
| LITAF | 85.8936517 | 0.35856265 | 0.45456906 | 0.78879688 | 0.43023073 | 0.9224707 |
| TXNDC11 | 43.7980565 | -0.2977795 | 0.50174679 | -0.5934856 | 0.55285625 | 0.95415437 |
| ZC3H7A | 41.7418562 | -0.1087348 | 0.31563072 | -0.3445002 | 0.73047017 | 0.97605445 |
| RSL1D1 | 11.8604417 | -0.1850611 | 0.5633779 | -0.3284849 | 0.74254505 | NA |
| AC007216.4 | 12.6996026 | -0.3151969 | 0.50356862 | -0.6259264 | 0.53136323 | NA |
| GSPT1 | 32.5340724 | -0.0741952 | 0.33517377 | -0.2213633 | 0.82480955 | 0.98320463 |
| SNX29 | 134.473497 | 0.38527864 | 0.31816242 | 1.21094953 | 0.22591475 | 0.81875277 |
| CPPED1 | 54.449002 | 0.73687691 | 0.49592226 | 1.48587182 | 0.13731303 | 0.73044824 |
| MKL2 | 27.5264883 | -0.2067929 | 0.416419 | -0.4965982 | 0.61947242 | 0.95688177 |
| PARN | 44.9986262 | -0.1775406 | 0.2952631 | -0.6012963 | 0.54764262 | 0.95415437 |
| BFAR | 13.6201292 | -0.0177864 | 0.56460296 | -0.0315024 | 0.97486887 | 0.9973754 |
| PDXDC1 | 66.698101 | 0.1350015 | 0.35385079 | 0.38152097 | 0.70281672 | 0.97113301 |
| RRN3 | 14.8504352 | -0.3526992 | 0.45510064 | -0.7749916 | 0.43834461 | 0.92687256 |
| C16orf45 | 13.6920464 | -2.3718618 | 0.49415998 | -4.7997853 | 1.59E-06 | 0.00109875 |
| MARF1 | 45.1028889 | -1.717159 | 0.56231869 | -3.0537114 | 0.00226029 | 0.10783163 |
| NDE1 | 25.867259 | -0.9875505 | 0.43658629 | -2.2619825 | 0.02369848 | 0.39492781 |
| ABCC1 | 60.1860691 | 1.21133068 | 0.4304012 | 2.81442217 | 0.0048865 | 0.17334548 |
| XYLT1 | 133.299798 | -0.3438405 | 0.53320188 | -0.6448599 | 0.51901796 | 0.94668603 |
| RPS15A | 23.8567166 | 0.10708358 | 0.89882874 | 0.1191368 | 0.90516698 | 0.99324672 |
| SMG1 | 99.7943194 | -0.3640005 | 0.23020516 | -1.5812004 | 0.11383221 | 0.69838966 |
| VPS35L | 36.9345478 | -0.248822 | 0.38177667 | -0.6517475 | 0.51456406 | 0.94596835 |
| UMOD | 48.0203775 | -0.4505227 | 0.83793559 | -0.5376579 | 0.59081328 | 0.95415437 |
| ACSM3 | 39.7773711 | 5.28590017 | 1.31919031 | 4.00692768 | 6.15E-05 | 0.01361667 |
| THUMPD1 | 13.8639461 | 0.51340224 | 0.53187146 | 0.96527503 | 0.33440707 | 0.88047896 |
| METTL9 | 37.0334553 | 0.22101052 | 0.39904406 | 0.55384992 | 0.57968156 | 0.95415437 |
| IGSF6 | 15.1923673 | 1.59708384 | 0.67348408 | 2.37137577 | 0.017722 | 0.34827529 |
| NPIPB4 | 14.6369978 | -0.5272152 | 0.58435721 | -0.9022139 | 0.36694326 | 0.89868152 |
| UQCRC2 | 28.9795412 | 0.18631742 | 0.37738798 | 0.49370257 | 0.62151625 | 0.95702817 |
| MOSMO | 35.5515454 | -0.026411 | 0.46391308 | -0.0569309 | 0.95460026 | 0.99514836 |
| EEF2K | 17.6467605 | 0.91558196 | 0.49877492 | 1.83566156 | 0.06640773 | 0.58951068 |
| POLR3E | 9.65598965 | -0.5325547 | 0.51112562 | -1.0419253 | 0.29744634 | NA |
| CDR2 | 53.0237233 | -0.1116117 | 0.5247588 | -0.2126913 | 0.83156772 | 0.98320463 |
| NPIPB5 | 42.149498 | -0.2121421 | 0.33127368 | -0.6403833 | 0.52192341 | 0.94895469 |
| USP31 | 8.74324519 | 0.39598753 | 0.65433736 | 0.60517335 | 0.54506382 | NA |
| COG7 | 15.2466061 | 0.05092139 | 0.482777 | 0.10547601 | 0.9159981 | 0.99498068 |
| GGA2 | 44.7883963 | -1.1354063 | 0.34493042 | -3.2916966 | 0.00099585 | 0.06639798 |
| NDUFAB1 | 8.47469825 | 0.98351013 | 0.62102307 | 1.58369338 | 0.11326351 | NA |
| DCTN5 | 14.8222671 | 0.74890548 | 0.51412092 | 1.45667186 | 0.14520698 | 0.73251288 |
| PRKCB | 314.215269 | 0.38669904 | 0.33317809 | 1.16063769 | 0.24578927 | 0.83817091 |
| RBBP6 | 30.645798 | 0.24488363 | 0.42638637 | 0.5743233 | 0.56574905 | 0.95415437 |
| TNRC6A | 98.1227314 | 0.05599987 | 0.24070058 | 0.23265365 | 0.81603037 | 0.98320463 |
| ARHGAP17 | 46.4446814 | -0.2248349 | 0.39200339 | -0.5735534 | 0.56627006 | 0.95415437 |
| LCMT1 | 15.4488473 | -0.0123027 | 0.5113493 | -0.0240593 | 0.98080533 | 0.99758514 |
| NSMCE1 | 7.97318889 | 0.66672828 | 0.73086381 | 0.91224695 | 0.36163873 | NA |
| IL4R | 40.5766923 | 0.04705158 | 0.48581671 | 0.09685048 | 0.92284513 | 0.99514836 |
| IL21R | 30.2149698 | -2.0125876 | 0.70600368 | -2.8506758 | 0.00436264 | 0.16650253 |
| GTF3C1 | 17.5255838 | -0.366209 | 0.46651305 | -0.784992 | 0.43245825 | 0.92284736 |
| KIAA0556 | 17.3358809 | 0.56853375 | 0.47635069 | 1.19351931 | 0.23266606 | 0.82802183 |
| XPO6 | 78.7443323 | -0.0226783 | 0.39640216 | -0.0572103 | 0.95437764 | 0.99514836 |
| NUPR1 | 11.8397982 | 1.31415388 | 1.11148139 | 1.18234448 | 0.23706904 | NA |
| ATXN2L | 23.3634746 | -0.3438377 | 0.42450865 | -0.8099663 | 0.41795956 | 0.91967721 |

| | | | | | | |
|---|---|---|---|---|---|---|
| TUFM | 17.1873377 | 0.39590794 | 0.55633067 | 0.71164141 | 0.47668686 | 0.9314919 |
| RABEP2 | 19.172835 | -2.2010475 | 0.7193014 | -3.0599794 | 0.00221352 | 0.10651853 |
| NFATC2IP | 45.8787783 | 0.00282321 | 0.348482 | 0.00810144 | 0.99353606 | 0.99859588 |
| SPNS1 | 9.38522982 | -0.437097 | 0.59270733 | -0.7374584 | 0.46084365 | NA |
| LAT | 8.10566508 | 0.39302207 | 0.75719059 | 0.51905303 | 0.60372376 | NA |
| AC009093.2 | 66.7651017 | 0.22887557 | 0.47341916 | 0.48345226 | 0.62877465 | 0.95877428 |
| MAZ | 10.3841631 | -0.1394564 | 0.58864509 | -0.2369109 | 0.8127259 | NA |
| MVP | 30.052073 | 0.38076099 | 0.44073605 | 0.86392068 | 0.38763147 | 0.90910959 |
| KCTD13 | 15.2005551 | 0.63085823 | 0.50997181 | 1.2370453 | 0.21607026 | 0.80634897 |
| TMEM219 | 12.7766188 | 0.86200355 | 0.67090004 | 1.28484647 | 0.19884594 | NA |
| PPP4C | 14.0580613 | 0.08250632 | 0.49225575 | 0.16760865 | 0.86689117 | 0.98751288 |
| YPEL3 | 24.5128438 | -0.5187216 | 0.47764341 | -1.0860019 | 0.27747817 | 0.84994492 |
| CORO1A | 42.1636793 | 0.06936876 | 0.61983898 | 0.11191416 | 0.91089147 | 0.99471296 |
| NPIPB13 | 12.8880345 | -0.5867842 | 0.54518348 | -1.0763059 | 0.28179046 | NA |
| CD2BP2 | 9.05268062 | 0.50792531 | 0.71822657 | 0.70719371 | 0.47944611 | NA |
| 1-Sep | 10.0468282 | -0.0389336 | 0.56171227 | -0.0693123 | 0.94474103 | NA |
| SEPHS2 | 11.9216895 | 1.38400555 | 0.70389238 | 1.96621754 | 0.04927349 | NA |
| ITGAL | 92.5147781 | 0.92333003 | 0.52007595 | 1.77537537 | 0.07583593 | 0.62329636 |
| FBRS | 11.1257218 | 0.66271389 | 0.59921051 | 1.10597841 | 0.26873586 | NA |
| RNF40 | 7.56012179 | 0.12702248 | 0.59467395 | 0.21360021 | 0.83085884 | NA |
| BCL7C | 9.37471381 | -0.5493487 | 0.55997529 | -0.9810231 | 0.32658134 | NA |
| STX4 | 10.4764571 | 0.09366293 | 0.66731709 | 0.14035745 | 0.88837757 | NA |
| KAT8 | 13.1418389 | -0.7396418 | 0.52908414 | -1.3979663 | 0.16212319 | 0.75177046 |
| FUS | 74.6533159 | -0.2370448 | 0.33591014 | -0.7056793 | 0.48038762 | 0.93257966 |
| PYCARD | 18.7611667 | -0.2195443 | 0.75256229 | -0.2917291 | 0.77049379 | 0.98264092 |
| ITGAM | 40.9343572 | 0.90640203 | 0.69990026 | 1.29504458 | 0.19530485 | 0.78135737 |
| ITGAX | 62.3770137 | -0.662526 | 0.62756696 | -1.0557056 | 0.29110273 | 0.85872202 |
| ZNF720 | 14.2722911 | -0.7319925 | 0.45510918 | -1.6083888 | 0.10775005 | 0.68018384 |
| ZNF267 | 26.0192135 | 0.54236186 | 0.55711232 | 0.97352336 | 0.33029325 | 0.88037558 |
| VPS35 | 30.2954939 | 0.72910001 | 0.37887458 | 1.92438356 | 0.05430652 | 0.55278922 |
| C16orf87 | 9.6431534 | 1.03773761 | 0.56575321 | 1.83425846 | 0.06661564 | NA |
| DNAJA2 | 9.65144725 | -0.7273174 | 0.59932136 | -1.2135683 | 0.22491261 | NA |
| NETO2 | 12.6777809 | 0.63975547 | 0.65668554 | 0.9742189 | 0.32994786 | NA |
| ITFG1 | 126.747485 | -0.1313002 | 0.28368747 | -0.4628339 | 0.64348341 | 0.95992767 |
| PHKB | 105.127012 | 0.01286769 | 0.23864401 | 0.05392001 | 0.9569989 | 0.99514836 |
| LONP2 | 55.0803608 | 0.07749618 | 0.28448032 | 0.27241316 | 0.78530435 | 0.98264092 |
| SIAH1 | 13.9786884 | -0.7607989 | 0.51076042 | -1.4895416 | 0.13634482 | 0.73044824 |
| N4BP1 | 61.6077138 | 0.2699144 | 0.49549505 | 0.54473682 | 0.58593454 | 0.95415437 |
| CNEP1R1 | 9.84259165 | -0.2647039 | 0.56836634 | -0.4657276 | 0.64141049 | NA |
| HEATR3 | 26.6465182 | -0.1496612 | 0.55746833 | -0.2684659 | 0.78834071 | 0.98320463 |
| TENT4B | 45.1262192 | -0.0896318 | 0.32635823 | -0.2746424 | 0.78359096 | 0.98264092 |
| ADCY7 | 52.3675878 | 0.61643217 | 0.43643081 | 1.41243965 | 0.15782055 | 0.74647771 |
| BRD7 | 20.1184531 | 0.02171293 | 0.39080731 | 0.05555916 | 0.955693 | 0.99514836 |
| SNX20 | 9.34094176 | 0.46024315 | 0.59651867 | 0.7715486 | 0.44038183 | NA |
| NOD2 | 12.7239879 | 0.43545666 | 0.72909423 | 0.59725703 | 0.55033579 | NA |
| CYLD | 57.6046719 | -0.0642153 | 0.31157289 | -0.2061004 | 0.83671249 | 0.98320463 |
| CHD9 | 133.331932 | -0.1582764 | 0.26490396 | -0.5974859 | 0.55018302 | 0.95415437 |
| RBL2 | 73.9415399 | 0.5335757 | 0.27351333 | 1.95082161 | 0.05107827 | 0.54780456 |
| FTO | 98.9790458 | 0.61921175 | 0.38406984 | 1.61223738 | 0.10691029 | 0.67742699 |
| LPCAT2 | 36.4623398 | 0.57715507 | 0.40212466 | 1.43526405 | 0.15121187 | 0.74119263 |
| GNAO1 | 10.5525533 | 0.62167469 | 0.66561648 | 0.93398333 | 0.35031249 | NA |
| AMFR | 21.6390722 | -0.3633275 | 0.4878262 | -0.7447888 | 0.45639941 | 0.9314919 |
| NUDT21 | 12.0567891 | -0.1714815 | 0.50604606 | -0.3388653 | 0.73471118 | NA |
| BBS2 | 11.8131841 | -0.2061312 | 0.67821406 | -0.3039324 | 0.76117937 | NA |

| | | | | | | |
|---|---|---|---|---|---|---|
| MT2A | 141.052476 | 0.01302615 | 0.72764178 | 0.01790187 | 0.98571714 | 0.99811561 |
| MT1E | 30.662057 | -0.1534709 | 0.93177201 | -0.1647087 | 0.86917329 | 0.98751288 |
| MT1G | 100.674739 | -0.4323642 | 1.06669831 | -0.4053294 | 0.68523542 | 0.9673431 |
| MT1H | 33.4269541 | -0.2514931 | 1.03258342 | -0.2435572 | 0.8075738 | 0.98320463 |
| MT1X | 36.5590114 | 0.73236976 | 0.76785978 | 0.9537806 | 0.34019471 | 0.8844375 |
| NUP93 | 21.496669 | 0.00599542 | 0.45490039 | 0.01317963 | 0.98948448 | 0.99811561 |
| HERPUD1 | 80.3640584 | -0.331644 | 0.44062007 | -0.7526756 | 0.45164486 | 0.92914597 |
| NLRC5 | 72.4327906 | -0.8731387 | 0.40033237 | -2.1810345 | 0.02918086 | 0.42316846 |
| FAM192A | 25.2268584 | 0.20213987 | 0.4279909 | 0.47229947 | 0.63671305 | 0.95933464 |
| RSPRY1 | 17.8161599 | 0.30496039 | 0.5122651 | 0.59531752 | 0.55163124 | 0.95415437 |
| ARL2BP | 10.1771515 | -0.2697537 | 0.55654165 | -0.4846963 | 0.6278918 | NA |
| ADGRG5 | 14.6781854 | -0.0635683 | 0.62562147 | -0.1016083 | 0.91906759 | 0.99498068 |
| ADGRG1 | 13.0626531 | 1.19867555 | 0.65164075 | 1.83947298 | 0.06584565 | 0.58951068 |
| KIFC3 | 14.919397 | 1.08796615 | 0.84805883 | 1.28288995 | 0.19953063 | 0.78201313 |
| USB1 | 8.84937916 | 0.63323021 | 0.62960146 | 1.00576357 | 0.31452931 | NA |
| CSNK2A2 | 14.8686559 | -0.4048525 | 0.48322176 | -0.8378193 | 0.40213219 | 0.91528253 |
| CNOT1 | 66.1607863 | -0.0146029 | 0.30038138 | -0.0486144 | 0.9612266 | 0.99614757 |
| GOT2 | 9.29716035 | 0.58681743 | 0.6104977 | 0.96121154 | 0.33644582 | NA |
| CMTM3 | 11.2729574 | -0.1713199 | 0.6170512 | -0.277643 | 0.78128645 | NA |
| CMTM4 | 10.2972922 | 0.15106248 | 0.7136534 | 0.21167485 | 0.83236071 | NA |
| DYNC1LI2 | 30.0605427 | 0.18695574 | 0.40885058 | 0.45727156 | 0.64747587 | 0.96049083 |
| NAE1 | 10.609639 | -0.2682454 | 0.51548258 | -0.5203772 | 0.60280068 | NA |
| CDH16 | 12.5805883 | 0.76689016 | 0.61811141 | 1.24069893 | 0.21471699 | NA |
| FAM96B | 10.6139717 | 0.04564473 | 0.56735129 | 0.08045232 | 0.93587752 | NA |
| CBFB | 67.2132337 | -0.3028475 | 0.25263516 | -1.1987542 | 0.23062354 | 0.82393202 |
| C16orf70 | 13.3601929 | 0.45908695 | 0.56104045 | 0.81827782 | 0.41319857 | 0.91894074 |
| TRADD | 8.41138078 | 0.61728852 | 0.66306982 | 0.93095554 | 0.35187657 | NA |
| E2F4 | 8.1009764 | -0.2483484 | 0.61281185 | -0.4052605 | 0.68528607 | NA |
| FHOD1 | 8.43928363 | 0.04180028 | 0.67877066 | 0.06158233 | 0.95089545 | NA |
| ATP6V0D1 | 28.5557498 | -0.3491599 | 0.49466615 | -0.7058495 | 0.48028174 | 0.93257966 |
| AC009061.2 | 10.0719641 | 0.10161078 | 0.76573894 | 0.13269637 | 0.89443351 | NA |
| RIPOR1 | 14.1167326 | -1.2849389 | 0.57549642 | -2.2327486 | 0.02556553 | 0.40079212 |
| CTCF | 36.6063931 | -0.1298682 | 0.35345638 | -0.3674237 | 0.713303 | 0.97348719 |
| RANBP10 | 20.8715584 | -0.1564815 | 0.45493362 | -0.3439657 | 0.73087211 | 0.97605445 |
| DPEP2 | 16.7298256 | 1.33522372 | 0.54182349 | 2.46431495 | 0.01372754 | 0.31861834 |
| DUS2 | 10.3598793 | -1.128322 | 0.57677104 | -1.9562738 | 0.05043291 | NA |
| NFATC3 | 117.94565 | -0.3368919 | 0.26135132 | -1.2890384 | 0.19738474 | 0.78135737 |
| SLC7A6 | 27.1063751 | 0.15388768 | 0.40605822 | 0.37897936 | 0.70470319 | 0.97155642 |
| SLC7A6OS | 14.9577631 | -0.1470503 | 0.52223848 | -0.2815768 | 0.778268 | 0.98264092 |
| PRMT7 | 11.7304684 | 0.99558797 | 0.67035272 | 1.48517033 | 0.13749871 | NA |
| ZFP90 | 11.9301533 | -0.5453455 | 0.47566128 | -1.1464998 | 0.25158842 | NA |
| TANGO6 | 39.8595563 | -0.3981497 | 0.31855817 | -1.2498493 | 0.21135459 | 0.80277027 |
| CHTF8 | 8.62046196 | 1.11951473 | 0.6685905 | 1.6744401 | 0.09404414 | NA |
| SNTB2 | 20.2016467 | 0.16639627 | 0.55040587 | 0.30231558 | 0.7624115 | 0.98163473 |
| TERF2 | 16.0975895 | -0.9273004 | 0.4640317 | -1.9983557 | 0.04567812 | 0.52198279 |
| CYB5B | 24.1470399 | 0.68711991 | 0.39952395 | 1.71984659 | 0.08546033 | 0.64676224 |
| NFAT5 | 159.248643 | -0.3408203 | 0.23891071 | -1.4265592 | 0.153707 | 0.74419469 |
| WWP2 | 63.0167097 | -0.6150211 | 0.38030942 | -1.6171598 | 0.1058438 | 0.67519097 |
| PDPR | 20.4278875 | 0.00379295 | 0.47314638 | 0.00801644 | 0.99360387 | 0.99859588 |
| AARS | 8.6280916 | -0.5457091 | 0.59373998 | -0.9191045 | 0.35804092 | NA |
| ST3GAL2 | 25.7845599 | 0.56415699 | 0.52929382 | 1.06586734 | 0.28648361 | 0.85677513 |
| COG4 | 17.7301073 | -0.0297358 | 0.43601929 | -0.0681983 | 0.9456278 | 0.99514836 |
| SF3B3 | 27.862 | 0.5654165 | 0.38637379 | 1.46339251 | 0.14336001 | 0.73044824 |
| VAC14 | 27.9056782 | 0.43472483 | 0.52514331 | 0.82782132 | 0.4077717 | 0.9169561 |

| | | | | | | |
|---|---|---|---|---|---|---|
| CMTR2 | 8.41195138 | -0.0379917 | 0.59084571 | -0.0643006 | 0.94873088 | NA |
| PHLPP2 | 11.3671269 | 0.41084289 | 0.53993765 | 0.76090802 | 0.44671201 | NA |
| AP1G1 | 46.4806273 | -0.3070938 | 0.31297343 | -0.9812136 | 0.32648742 | 0.87715548 |
| IST1 | 38.6398887 | 0.20474761 | 0.32714014 | 0.62587126 | 0.53139938 | 0.95265314 |
| DHX38 | 9.51124068 | 0.02307144 | 0.55179023 | 0.04181198 | 0.96664858 | NA |
| LINC01572 | 16.8056021 | 0.40433346 | 0.74977753 | 0.53927124 | 0.58969971 | 0.95415437 |
| ZFHX3 | 84.0416346 | 0.17049362 | 0.56941628 | 0.29941824 | 0.76462094 | 0.98222198 |
| PSMD7 | 14.8331381 | -0.0347428 | 0.57802857 | -0.0601057 | 0.95207145 | 0.99514836 |
| GLG1 | 130.169714 | -0.2811425 | 0.31203382 | -0.9010001 | 0.36758824 | 0.89868152 |
| RFWD3 | 15.6512927 | 0.15195167 | 0.45922251 | 0.33088898 | 0.74072835 | 0.97605445 |
| MLKL | 19.5025852 | -0.1414216 | 0.56930724 | -0.2484099 | 0.80381724 | 0.98320463 |
| WDR59 | 26.131096 | 0.79477865 | 0.46466773 | 1.71042359 | 0.08718757 | 0.64676224 |
| ZNRF1 | 19.6402787 | 1.73608916 | 0.77114981 | 2.25129946 | 0.02436658 | 0.39492781 |
| CFDP1 | 42.190445 | -0.1435787 | 0.35247623 | -0.4073429 | 0.68375614 | 0.9673431 |
| GABARAPL2 | 16.7706306 | -0.2799768 | 0.49470148 | -0.5659509 | 0.57142713 | 0.95415437 |
| ADAT1 | 10.0392374 | -0.2067628 | 0.64835592 | -0.3189032 | 0.7497999 | NA |
| KARS | 11.8214013 | -0.3439112 | 0.60559951 | -0.5678855 | 0.57011274 | NA |
| TERF2IP | 36.2136877 | -0.1664254 | 0.40550581 | -0.4104144 | 0.68150197 | 0.9673431 |
| WWOX | 310.529732 | -1.5366211 | 0.36395692 | -4.2219863 | 2.42E-05 | 0.00670053 |
| MAF | 37.4454637 | 1.00854775 | 0.67962854 | 1.48396909 | 0.13781712 | 0.73044824 |
| CDYL2 | 33.7640508 | -0.1149766 | 0.39112319 | -0.2939653 | 0.76878441 | 0.98264092 |
| CMC2 | 19.7602169 | -0.2951964 | 0.48226367 | -0.6121057 | 0.54046781 | 0.95415437 |
| GAN | 12.8692279 | 1.00032353 | 0.51520057 | 1.9416196 | 0.05218317 | NA |
| CMIP | 236.53457 | 0.27990477 | 0.50642803 | 0.55270394 | 0.58046616 | 0.95415437 |
| PLCG2 | 120.500549 | -0.72472 | 0.31226716 | -2.3208332 | 0.02029585 | 0.36946457 |
| MPHOSPH6 | 13.8719544 | 0.03736332 | 0.61050072 | 0.0612011 | 0.95119905 | 0.99514836 |
| HSBP1 | 15.4079177 | 0.38773938 | 0.57299078 | 0.67669392 | 0.49860018 | 0.93916044 |
| MBTPS1 | 30.1626865 | 0.31453376 | 0.36761293 | 0.85561125 | 0.39221285 | 0.90910959 |
| TLDC1 | 9.81749577 | 1.01434638 | 0.61692098 | 1.64420795 | 0.10013326 | NA |
| COTL1 | 104.587118 | 0.58033401 | 0.76140533 | 0.76218801 | 0.4459478 | 0.92914597 |
| KLHL36 | 11.7915861 | -0.2378773 | 0.52248329 | -0.4552821 | 0.64890634 | NA |
| USP10 | 43.0234963 | -0.7905159 | 0.35228527 | -2.2439653 | 0.02483464 | 0.39492781 |
| ZDHHC7 | 25.0597105 | 0.62211007 | 0.46943051 | 1.32524421 | 0.18509017 | 0.77694741 |
| KIAA0513 | 20.8820089 | 1.40319264 | 0.6677856 | 2.1012622 | 0.03561796 | 0.46488152 |
| GSE1 | 45.8526666 | 0.19422362 | 0.47654721 | 0.40756428 | 0.68359359 | 0.9673431 |
| COX4I1 | 63.3693065 | -0.1163846 | 0.55270674 | -0.2105721 | 0.83322118 | 0.98320463 |
| IRF8 | 29.5786236 | -0.1127045 | 0.66498253 | -0.1694849 | 0.86541527 | 0.98751288 |
| MAP1LC3B | 21.2767893 | 0.81311647 | 0.44425228 | 1.83030343 | 0.06720458 | 0.58951068 |
| ZCCHC14 | 18.6976223 | -0.1288359 | 0.42698833 | -0.3017316 | 0.7628567 | 0.98163473 |
| KLHDC4 | 29.7824555 | -0.2316182 | 0.43645995 | -0.5306746 | 0.5956443 | 0.9553365 |
| SLC7A5 | 14.9947806 | 0.73873519 | 0.74114864 | 0.99674364 | 0.31888896 | 0.87298176 |
| BANP | 44.757167 | 0.39714356 | 0.30149847 | 1.31723242 | 0.18776073 | 0.77798634 |
| ZFPM1 | 19.4191622 | -1.9409202 | 0.83757725 | -2.3173028 | 0.02048724 | 0.37172586 |
| ZC3H18 | 21.3326919 | -0.2114467 | 0.45834006 | -0.4613315 | 0.6445608 | 0.96049083 |
| CYBA | 134.045435 | -0.1401321 | 0.65429049 | -0.2141741 | 0.83041126 | 0.98320463 |
| MVD | 8.47598226 | 0.00513972 | 0.93909624 | 0.00547305 | 0.99563316 | NA |
| RNF166 | 24.8197349 | 0.19714078 | 0.45850064 | 0.42996839 | 0.66721864 | 0.96412305 |
| PIEZO1 | 23.4721368 | 0.57756193 | 0.48870243 | 1.18182741 | 0.23727419 | 0.83171006 |
| APRT | 17.4532038 | 0.3892446 | 0.64368601 | 0.60471192 | 0.54537043 | 0.95415437 |
| GALNS | 9.81070825 | 0.05175881 | 0.58211412 | 0.08891523 | 0.92914928 | NA |
| CBFA2T3 | 17.0090526 | 2.06843243 | 0.95284466 | 2.17079711 | 0.02994651 | 0.42727094 |
| ACSF3 | 16.8234615 | 0.05582963 | 0.46343183 | 0.12047 | 0.90411084 | 0.99324672 |
| ANKRD11 | 221.079001 | -0.1827878 | 0.26155534 | -0.6988493 | 0.48464623 | 0.93417265 |
| SPG7 | 40.7084301 | -0.4310802 | 0.41665564 | -1.0346198 | 0.30084649 | 0.86225566 |

| | | | | | | |
|---|---|---|---|---|---|---|
| RPL13 | 202.973126 | -0.139707 | 0.54944711 | -0.2542683 | 0.79928831 | 0.98320463 |
| CHMP1A | 8.64387847 | 0.50184972 | 0.71467428 | 0.70220761 | 0.4825497 | NA |
| CDK10 | 8.39051703 | -0.188234 | 0.6533024 | -0.288127 | 0.77324954 | NA |
| VPS9D1 | 14.3235417 | 0.80384862 | 0.52775234 | 1.52315501 | 0.12771993 | 0.72782081 |
| ZNF276 | 16.2727427 | -0.4882679 | 0.46809564 | -1.0430944 | 0.29690459 | 0.8608685 |
| TCF25 | 80.848036 | -0.4346651 | 0.2684291 | -1.6192922 | 0.10538442 | 0.67519097 |
| DEF8 | 8.94421157 | 0.50464826 | 0.60477555 | 0.83443892 | 0.40403369 | NA |
| FAM157C | 26.7426551 | 0.96729505 | 0.73200159 | 1.32143847 | 0.18635521 | 0.77798634 |
| RPH3AL | 13.2430837 | 0.21375757 | 0.76131897 | 0.28077268 | 0.77888476 | 0.98264092 |
| RFLNB | 10.3843102 | 1.65958236 | 0.79776781 | 2.08028244 | 0.03749963 | NA |
| VPS53 | 68.2348959 | 0.0876516 | 0.35600699 | 0.24620752 | 0.80552159 | 0.98320463 |
| GLOD4 | 8.93339215 | -0.2538098 | 0.57188525 | -0.4438125 | 0.65717817 | NA |
| ABR | 117.443902 | 0.08934328 | 0.39441309 | 0.22652211 | 0.82079537 | 0.98320463 |
| YWHAE | 56.3189311 | 0.21971611 | 0.32624045 | 0.67347906 | 0.50064257 | 0.94001041 |
| CRK | 35.0742738 | 0.25082317 | 0.46594049 | 0.5383159 | 0.59035898 | 0.95415437 |
| MYO1C | 11.5329839 | -1.262241 | 0.74008582 | -1.7055333 | 0.088095 | NA |
| PITPNA | 39.2248296 | 0.19905996 | 0.46398968 | 0.42901808 | 0.66791007 | 0.96412305 |
| SLC43A2 | 42.8864452 | 0.23315337 | 0.52433894 | 0.44466156 | 0.65656435 | 0.96230291 |
| PRPF8 | 32.9372943 | 0.37194575 | 0.37926722 | 0.98069575 | 0.3267428 | 0.87715548 |
| WDR81 | 11.5520919 | 0.5691787 | 0.6510799 | 0.87420715 | 0.38200545 | NA |
| SMYD4 | 11.1146031 | -0.1534798 | 0.52661416 | -0.2914465 | 0.77070987 | NA |
| RPA1 | 23.2346754 | 0.15955322 | 0.43286687 | 0.36859652 | 0.71242849 | 0.97332186 |
| SMG6 | 71.7245823 | -0.4474174 | 0.26339205 | -1.6986747 | 0.08938048 | 0.64845882 |
| SGSM2 | 10.3781743 | 0.52116798 | 0.54683992 | 0.95305403 | 0.3405627 | NA |
| METTL16 | 26.5149216 | -0.0072749 | 0.39449579 | -0.018441 | 0.98528704 | 0.99811561 |
| PAFAH1B1 | 120.019358 | -0.1198814 | 0.26539658 | -0.4517068 | 0.65148024 | 0.96158408 |
| RAP1GAP2 | 101.371149 | 0.70964821 | 0.48971904 | 1.44909255 | 0.14731174 | 0.73693374 |
| ITGAE | 28.5663568 | -1.8626153 | 0.50760659 | -3.6694073 | 0.00024311 | 0.02990804 |
| NCBP3 | 21.2440338 | -0.4227408 | 0.51070112 | -0.8277656 | 0.40780328 | 0.9169561 |
| P2RX1 | 23.7079952 | 0.89462828 | 0.65987844 | 1.35574711 | 0.17517964 | 0.76214163 |
| ATP2A3 | 31.1170282 | -0.2503286 | 0.50934645 | -0.4914702 | 0.6230939 | 0.95783378 |
| ZZEF1 | 55.4452077 | 0.29496525 | 0.2789447 | 1.05743267 | 0.29031417 | 0.85822878 |
| ANKFY1 | 51.0829451 | 0.70107003 | 0.41126314 | 1.70467508 | 0.08825504 | 0.64689191 |
| UBE2G1 | 57.9897554 | -0.0066665 | 0.45665478 | -0.0145986 | 0.98835241 | 0.99811561 |
| ARRB2 | 40.5870375 | 0.39278935 | 0.44614195 | 0.8804134 | 0.3786354 | 0.9051267 |
| CXCL16 | 14.9521865 | 0.33564838 | 0.73207288 | 0.4584904 | 0.64660016 | 0.96049083 |
| MINK1 | 25.3721995 | 0.4231667 | 0.40777506 | 1.03774543 | 0.29938859 | 0.86225566 |
| RNF167 | 9.04709035 | -0.3314837 | 0.53199472 | -0.6230959 | 0.53322149 | NA |
| PFN1 | 93.3244932 | -0.188775 | 0.65128288 | -0.289851 | 0.77193021 | 0.98264092 |
| SPAG7 | 11.479003 | -0.6931439 | 0.58698928 | -1.1808459 | 0.23766394 | NA |
| CAMTA2 | 11.1363772 | -0.4197751 | 0.51734177 | -0.8114078 | 0.41713153 | NA |
| AC087500.1 | 18.625725 | -2.0187402 | 0.59241111 | -3.4076677 | 0.00065521 | 0.05106918 |
| SCIMP | 29.7809919 | -2.0765672 | 0.6782202 | -3.0617891 | 0.00220018 | 0.10651853 |
| RABEP1 | 77.5039097 | -1.0038009 | 0.37256754 | -2.694279 | 0.00705411 | 0.2098787 |
| NUP88 | 24.1551317 | -0.5960008 | 0.43186822 | -1.3800524 | 0.16757051 | 0.7567539 |
| RPAIN | 13.9619495 | -0.0506743 | 0.52055456 | -0.0973467 | 0.92245107 | 0.99514836 |
| DERL2 | 9.68073694 | 0.30740428 | 0.69494243 | 0.44234495 | 0.65823961 | NA |
| NLRP1 | 59.2747437 | 0.48645126 | 0.40039051 | 1.21494203 | 0.2243882 | 0.81875277 |
| KIAA0753 | 10.7386979 | -0.259067 | 0.59198492 | -0.4376243 | 0.66165864 | NA |
| XAF1 | 53.7915549 | 1.18011705 | 0.98069918 | 1.20334255 | 0.22884379 | 0.82329316 |
| ACADVL | 29.1654817 | 0.68788886 | 0.54575873 | 1.26042666 | 0.20751549 | 0.79343028 |
| EIF5A | 13.2903405 | -0.3570792 | 0.60439322 | -0.590806 | 0.55465039 | 0.95415437 |
| GPS2 | 9.35655503 | 0.3091907 | 0.79105803 | 0.39085717 | 0.69590281 | NA |
| ACAP1 | 42.7354216 | -0.3984642 | 0.32670651 | -1.2196396 | 0.22260152 | 0.81581246 |

| | | | | | | |
|---|---|---|---|---|---|---|
| ZBTB4 | 17.9664987 | -0.3819105 | 0.42825841 | -0.8917758 | 0.37251311 | 0.9025457 |
| POLR2A | 27.6708989 | 0.12055258 | 0.49462162 | 0.24372687 | 0.80744236 | 0.98320463 |
| EIF4A1 | 13.7739963 | 0.4619151 | 0.66017756 | 0.69968313 | 0.48412521 | 0.93399425 |
| CD68 | 29.062268 | 2.0330183 | 0.74637263 | 2.72386502 | 0.00645229 | 0.19837198 |
| SAT2 | 15.16974 | 0.29272022 | 0.59988671 | 0.48795916 | 0.62557877 | 0.95876809 |
| TP53 | 10.580805 | -0.488575 | 0.59761961 | -0.8175351 | 0.41362271 | NA |
| KDM6B | 11.6815827 | -0.0413812 | 0.570245 | -0.0725674 | 0.9421504 | NA |
| NAA38 | 19.5974305 | 0.01387881 | 0.44073936 | 0.03148984 | 0.97487889 | 0.9973754 |
| CHD3 | 24.0168637 | 0.0688316 | 0.4042076 | 0.17028774 | 0.86478385 | 0.98751288 |
| PER1 | 17.0938602 | -0.6205006 | 0.53934655 | -1.1504674 | 0.24995142 | 0.83865855 |
| CTC1 | 11.4784116 | -0.6579732 | 0.50152161 | -1.3119538 | 0.18953573 | NA |
| NDEL1 | 39.0665509 | -0.2454536 | 0.47503219 | -0.5167094 | 0.60535904 | 0.95688177 |
| PIK3R5 | 135.10719 | 0.13019324 | 0.26779285 | 0.48617146 | 0.6268456 | 0.95877428 |
| STX8 | 90.6625245 | -0.0114139 | 0.2650206 | -0.0430679 | 0.96564741 | 0.99653147 |
| GAS7 | 134.723574 | -1.5784144 | 0.43181284 | -3.6553205 | 0.00025686 | 0.03066918 |
| SCO1 | 9.44714246 | -0.9514756 | 0.56535674 | -1.6829649 | 0.09238189 | NA |
| MAP2K4 | 56.1022272 | 0.08306444 | 0.31005125 | 0.26790551 | 0.78877205 | 0.98320463 |
| COX10 | 18.7153159 | 0.24836874 | 0.47303831 | 0.52504996 | 0.59954846 | 0.95624942 |
| HS3ST3B1 | 11.8761195 | 0.78857227 | 0.76240848 | 1.03431729 | 0.30098786 | NA |
| PMP22 | 16.5039603 | 0.97017613 | 0.75452414 | 1.28581192 | 0.19850871 | 0.78142991 |
| ZSWIM7 | 15.8205856 | -0.2145616 | 0.46275989 | -0.4636563 | 0.64289398 | 0.95992767 |
| TTC19 | 32.6453337 | 0.35181132 | 0.43335309 | 0.81183528 | 0.41688616 | 0.91953487 |
| NCOR1 | 123.862902 | -0.1625504 | 0.21551148 | -0.7542539 | 0.45069678 | 0.92914597 |
| PIGL | 26.4382309 | -0.1618126 | 0.36173647 | -0.4473217 | 0.65464277 | 0.96178254 |
| UBB | 47.5151179 | 0.37161407 | 0.39693176 | 0.93621652 | 0.34916172 | 0.89048867 |
| TRPV2 | 11.4869229 | 0.09431553 | 0.56897375 | 0.16576429 | 0.86834245 | NA |
| AC098850.3 | 18.5274718 | 0.7760775 | 0.82902132 | 0.93613696 | 0.34920268 | 0.89048867 |
| MPRIP | 38.5369294 | -0.7725833 | 0.43416629 | -1.7794641 | 0.07516371 | 0.6217578 |
| COPS3 | 10.8415012 | -0.6406494 | 0.54222654 | -1.1815161 | 0.23739776 | NA |
| RAI1 | 21.0202742 | 0.08514175 | 0.50961809 | 0.16706971 | 0.8673152 | 0.98751288 |
| SREBF1 | 16.2301142 | 0.06786711 | 0.65006406 | 0.10440066 | 0.9168514 | 0.99498068 |
| TOM1L2 | 31.7017459 | 0.54276189 | 0.37256741 | 1.45681526 | 0.14516738 | 0.73251288 |
| ALKBH5 | 15.8925619 | 0.48239221 | 0.44849001 | 1.07559187 | 0.28210983 | 0.85171619 |
| FLII | 15.6573851 | -0.177006 | 0.5136298 | -0.3446178 | 0.73038173 | 0.97605445 |
| TOP3A | 16.2516971 | 0.19118105 | 0.46020672 | 0.41542428 | 0.67783133 | 0.96718429 |
| SHMT1 | 9.70790288 | 0.20082533 | 0.60730115 | 0.33068491 | 0.7408825 | NA |
| PRPSAP2 | 23.0450446 | -0.7569697 | 0.50090185 | -1.5112136 | 0.13073403 | 0.72931666 |
| GRAPL | 12.6725812 | -2.1292746 | 0.7128812 | -2.9868575 | 0.00281861 | NA |
| EPN2 | 15.5656873 | -0.6388483 | 0.56097732 | -1.1388131 | 0.25478112 | 0.84026145 |
| AC004448.2 | 8.61045723 | -0.6744268 | 0.65201217 | -1.0343776 | 0.30095968 | NA |
| SLC47A1 | 9.73380042 | 0.54898271 | 0.7564631 | 0.72572306 | 0.46800857 | NA |
| ALDH3A2 | 9.289399 | 1.2230538 | 0.6859792 | 1.78293132 | 0.07459748 | NA |
| ULK2 | 19.472319 | 1.40655063 | 0.59355777 | 2.36969456 | 0.01780279 | 0.34827529 |
| AKAP10 | 78.9867038 | -0.0288747 | 0.3455581 | -0.0835596 | 0.93340659 | 0.99514836 |
| SPECC1 | 21.8881213 | -0.3280747 | 0.87806919 | -0.3736319 | 0.70867819 | 0.97186969 |
| USP22 | 29.7125492 | 0.24564719 | 0.3498504 | 0.70214923 | 0.48258611 | 0.93382901 |
| DHRS7B | 18.6554137 | 0.42954151 | 0.67440867 | 0.63691577 | 0.52417971 | 0.95007724 |
| MAP2K3 | 22.3484426 | 0.50630996 | 0.48005993 | 1.05468075 | 0.29157136 | 0.85935767 |
| WSB1 | 169.536846 | 0.27251799 | 0.4708308 | 0.57880238 | 0.56272252 | 0.95415437 |
| KSR1 | 39.854937 | 1.26444702 | 0.4601887 | 2.7476707 | 0.00600203 | 0.19198507 |
| LGALS9 | 15.7642172 | 0.23673709 | 0.83433227 | 0.28374438 | 0.77660629 | 0.98264092 |
| NLK | 83.9333685 | 0.78692308 | 0.49297301 | 1.59628025 | 0.11042624 | 0.69012233 |
| SARM1 | 10.4201711 | -0.1794841 | 0.68092687 | -0.2635879 | 0.79209746 | NA |
| KIAA0100 | 22.7202977 | 0.4817992 | 0.44075692 | 1.09311773 | 0.27434211 | 0.84994492 |

| | | | | | | |
|---|---|---|---|---|---|---|
| SDF2 | 9.85679013 | -0.2193766 | 0.56661106 | -0.3871732 | 0.69862796 | NA |
| SUPT6H | 27.6959631 | -0.0421633 | 0.39023498 | -0.1080459 | 0.91395932 | 0.99498068 |
| RPL23A | 78.6029374 | -0.7098836 | 0.57943754 | -1.2251253 | 0.220528 | 0.81064692 |
| FAM222B | 43.1406826 | -0.0866126 | 0.34219755 | -0.2531071 | 0.80018548 | 0.98320463 |
| FLOT2 | 29.5190452 | 0.22709409 | 0.43779057 | 0.51872769 | 0.60395065 | 0.95688177 |
| PHF12 | 33.54183 | -0.1733071 | 0.4091005 | -0.4236297 | 0.67183586 | 0.96412305 |
| MYO18A | 13.9622739 | 0.36602513 | 0.6168313 | 0.59339585 | 0.55291626 | 0.95415437 |
| NUFIP2 | 71.700363 | 0.30124564 | 0.28839282 | 1.04456705 | 0.29622312 | 0.8608685 |
| TAOK1 | 126.400324 | 0.1922923 | 0.25421111 | 0.75642759 | 0.44939287 | 0.92914597 |
| ABHD15-AS1 | 44.6346318 | 0.52783511 | 0.64930693 | 0.8129208 | 0.41626347 | 0.91923465 |
| SSH2 | 430.968303 | -0.0454023 | 0.32028394 | -0.1417564 | 0.88727243 | 0.99019008 |
| NSRP1 | 43.7780911 | -0.1342427 | 0.2961002 | -0.4533693 | 0.65028284 | 0.96084839 |
| BLMH | 10.8822485 | -0.2158197 | 0.70766964 | -0.3049724 | 0.76038718 | NA |
| CPD | 42.4107604 | 0.28414919 | 0.60507636 | 0.46960881 | 0.63863453 | 0.95933464 |
| GOSR1 | 26.6102597 | 0.13124912 | 0.35893453 | 0.36566311 | 0.71461647 | 0.97348719 |
| CRLF3 | 57.130684 | 0.05293114 | 0.36747968 | 0.14403829 | 0.88547024 | 0.99019008 |
| TEFM | 8.94572285 | 0.23326082 | 0.58758915 | 0.39697946 | 0.69138261 | NA |
| ADAP2 | 41.6053871 | -0.1598673 | 0.52736219 | -0.3031452 | 0.76177918 | 0.98163473 |
| RNF135 | 15.016846 | 0.10225519 | 0.49899507 | 0.20492225 | 0.83763287 | 0.98320463 |
| NF1 | 127.799594 | -0.2619686 | 0.32299067 | -0.8110718 | 0.41732443 | 0.91954684 |
| EVI2B | 19.8616569 | -0.3520099 | 0.58250193 | -0.6043068 | 0.54563969 | 0.95415437 |
| RAB11FIP4 | 9.39044598 | -1.2393157 | 0.60507839 | -2.0481903 | 0.04054136 | NA |
| UTP6 | 18.4133314 | -0.0262938 | 0.44793452 | -0.0587001 | 0.95319099 | 0.99514836 |
| SUZ12 | 51.8620864 | 0.14169581 | 0.35083408 | 0.40388268 | 0.686299 | 0.9673431 |
| LRRC37B | 21.3739384 | 0.21341404 | 0.43062945 | 0.49558626 | 0.62018635 | 0.95688177 |
| RHOT1 | 51.6855644 | 0.6215472 | 0.3906673 | 1.59098855 | 0.11161215 | 0.69385269 |
| ZNF207 | 93.0610763 | 0.11529015 | 0.28365395 | 0.40644648 | 0.68441457 | 0.9673431 |
| PSMD11 | 20.7495278 | -0.8792697 | 0.5291059 | -1.6618028 | 0.09655231 | 0.66178928 |
| MYO1D | 54.3469529 | 0.40888777 | 0.66346264 | 0.61629358 | 0.53770078 | 0.95415437 |
| SLFN5 | 37.1419911 | 1.26793489 | 0.44098174 | 2.87525482 | 0.00403702 | 0.15732988 |
| SLFN11 | 19.4296036 | 0.73331698 | 0.45519582 | 1.61099235 | 0.10718139 | 0.67787635 |
| SLFN12L | 110.040358 | 0.40645198 | 0.47397599 | 0.85753707 | 0.39114814 | 0.90910959 |
| AP2B1 | 49.1536021 | 0.01759935 | 0.33646492 | 0.05230665 | 0.95828435 | 0.99514836 |
| TAF15 | 86.5603894 | 0.10785893 | 0.24975832 | 0.43185322 | 0.6658481 | 0.96412305 |
| CCL5 | 68.0787518 | 0.26318076 | 0.58655674 | 0.44868764 | 0.653657 | 0.96178254 |
| CCL4 | 26.261983 | 0.87017886 | 0.93431029 | 0.9313596 | 0.35166758 | 0.89109867 |
| AC243829.2 | 16.3598429 | -0.7717123 | 0.54713018 | -1.410473 | 0.15840008 | 0.74650686 |
| MYO19 | 12.870713 | -0.622255 | 0.50430086 | -1.2338963 | 0.21724155 | NA |
| GGNBP2 | 47.977263 | -0.1961767 | 0.35669528 | -0.5499839 | 0.5823304 | 0.95415437 |
| AATF | 38.7046668 | -0.8225482 | 0.32913095 | -2.4991516 | 0.0124491 | 0.30349487 |
| ACACA | 35.2264541 | -0.7832072 | 0.41172893 | -1.9022399 | 0.0571398 | 0.56668757 |
| TADA2A | 17.011827 | -0.4480871 | 0.44260525 | -1.0123855 | 0.31135376 | 0.8693399 |
| SYNRG | 80.5709332 | 0.16136643 | 0.24108611 | 0.6693311 | 0.50328429 | 0.94157379 |
| DDX52 | 14.2251639 | 0.12345545 | 0.50741643 | 0.24330204 | 0.80777142 | 0.98320463 |
| SOCS7 | 9.11435934 | 1.22688829 | 0.80804882 | 1.51833437 | 0.12893013 | NA |
| MLLT6 | 37.4919506 | 0.05390365 | 0.35386084 | 0.15233009 | 0.87892659 | 0.99019008 |
| PSMB3 | 14.3190727 | -0.4777218 | 0.53100871 | -0.8996496 | 0.36830677 | 0.89868152 |
| PIP4K2B | 10.5525956 | 0.96531815 | 0.62952309 | 1.53341181 | 0.12517442 | NA |
| CWC25 | 8.97316463 | -0.2497831 | 0.54356012 | -0.4595317 | 0.64585237 | NA |
| RPL23 | 40.3967547 | -0.0810423 | 0.58613236 | -0.1382663 | 0.89002997 | 0.99080204 |
| LASP1 | 31.7836884 | 0.13783473 | 0.47468414 | 0.29037146 | 0.77153208 | 0.98264092 |
| RPL19 | 117.977894 | -0.4354359 | 0.59436767 | -0.7326037 | 0.4638002 | 0.9314919 |
| FBXL20 | 76.5773502 | 0.57904139 | 0.37039314 | 1.56331563 | 0.11797837 | 0.70279044 |
| MED1 | 14.6504099 | 0.80284074 | 0.58796328 | 1.36546068 | 0.1721083 | 0.75982617 |

| | | | | | | |
|---|---|---|---|---|---|---|
| CDK12 | 42.7407522 | 0.42802521 | 0.35815449 | 1.19508542 | 0.23205366 | 0.82637384 |
| STARD3 | 14.5606936 | 0.93598254 | 0.49972651 | 1.87298956 | 0.06106983 | 0.57281427 |
| IKZF3 | 94.9123518 | -1.2264578 | 0.40048098 | -3.0624621 | 0.00219524 | 0.10651853 |
| PSMD3 | 8.75341539 | -0.6019357 | 0.58248917 | -1.0333852 | 0.30142367 | NA |
| MED24 | 9.18382699 | -0.8835822 | 0.65128855 | -1.3566679 | 0.17488677 | NA |
| MSL1 | 18.4367203 | -0.0796878 | 0.432574 | -0.1842178 | 0.8538426 | 0.98751288 |
| CASC3 | 22.7135702 | -0.1438706 | 0.399108 | -0.3604803 | 0.71848802 | 0.97348719 |
| WIPF2 | 36.0251982 | -0.3736782 | 0.32086839 | -1.164584 | 0.24418743 | 0.83666131 |
| RARA | 16.1120058 | 1.34341805 | 0.69877796 | 1.92252494 | 0.05453974 | 0.55278922 |
| IGFBP4 | 14.042449 | 0.73735768 | 0.56031606 | 1.31596742 | 0.18818498 | 0.77798634 |
| KRT10 | 10.9211212 | -0.4846632 | 0.57307001 | -0.8457312 | 0.39770271 | NA |
| EIF1 | 85.3215234 | -0.0728982 | 0.44076195 | -0.1653913 | 0.86863603 | 0.98751288 |
| ACLY | 16.1267511 | -0.1003785 | 0.50080095 | -0.2004358 | 0.84113973 | 0.98380297 |
| DNAJC7 | 40.4150288 | -0.4500128 | 0.41109478 | -1.0946693 | 0.27366154 | 0.84994492 |
| STAT5B | 74.7100866 | 0.40563753 | 0.29495173 | 1.37526751 | 0.16904853 | 0.75745073 |
| STAT5A | 19.5295363 | 0.68519737 | 0.48583808 | 1.410341 | 0.15843902 | 0.74650686 |
| STAT3 | 102.411156 | 0.05024159 | 0.3067959 | 0.16376227 | 0.86991829 | 0.98751288 |
| ATP6V0A1 | 44.2361217 | -0.9178414 | 0.4898387 | -1.8737626 | 0.06096316 | 0.57281427 |
| MLX | 10.5328011 | 0.37344137 | 0.57843727 | 0.64560392 | 0.5185359 | NA |
| RETREG3 | 26.8852425 | -0.0137901 | 0.39709921 | -0.034727 | 0.97229743 | 0.99734828 |
| EZH1 | 20.803381 | -0.7912948 | 0.4377747 | -1.8075389 | 0.0706783 | 0.60640882 |
| BECN1 | 11.1974075 | 0.37258108 | 0.58600989 | 0.63579316 | 0.52491125 | NA |
| RPL27 | 32.8687191 | 0.0742586 | 0.65614921 | 0.11317334 | 0.90989313 | 0.99471296 |
| IFI35 | 11.9952059 | 0.43984857 | 0.83019117 | 0.529816 | 0.59623951 | NA |
| BRCA1 | 10.1352067 | -0.1384174 | 0.66999395 | -0.2065949 | 0.83632621 | NA |
| NBR1 | 24.2662901 | 0.3917549 | 0.38231014 | 1.02470446 | 0.30550266 | 0.8673137 |
| TMEM106A | 18.9514239 | 0.37883485 | 0.49208281 | 0.76985996 | 0.44138297 | 0.92899684 |
| LINC00854 | 24.7102579 | -0.2064166 | 0.55333188 | -0.3730431 | 0.70911639 | 0.97207087 |
| DHX8 | 14.7249545 | -0.0355533 | 0.50947376 | -0.0697844 | 0.94436527 | 0.99514836 |
| DUSP3 | 8.82332471 | 0.14036189 | 0.56097052 | 0.25021259 | 0.80242295 | NA |
| LSM12 | 12.8615983 | 0.75256384 | 0.50339737 | 1.49496976 | 0.13492234 | NA |
| HDAC5 | 10.5397633 | 0.20631998 | 0.73995483 | 0.27882781 | 0.78037697 | NA |
| TMUB2 | 9.02961031 | -0.5348838 | 0.59896806 | -0.8930089 | 0.37185243 | NA |
| UBTF | 13.0877866 | -0.8626878 | 0.59965902 | -1.4386305 | 0.15025523 | 0.74043418 |
| SLC25A39 | 9.70331983 | -0.043285 | 0.56430542 | -0.0767049 | 0.93885834 | NA |
| GRN | 54.942761 | 0.13261942 | 0.6169424 | 0.2149624 | 0.82979663 | 0.98320463 |
| GPATCH8 | 118.736898 | -0.1994669 | 0.26492025 | -0.7529321 | 0.45149075 | 0.92914597 |
| EFTUD2 | 20.1061666 | 0.17886832 | 0.43636039 | 0.40990961 | 0.68187225 | 0.9673431 |
| DCAKD | 8.28702176 | 1.76497153 | 0.70325357 | 2.50972283 | 0.0120826 | NA |
| NMT1 | 22.8259314 | -0.4638397 | 0.37251956 | -1.2451418 | 0.21307963 | 0.80545265 |
| AC138150.1 | 12.1126889 | -0.5061681 | 0.65016266 | -0.7785254 | 0.43625935 | NA |
| AC008105.3 | 31.0341549 | 0.08451309 | 0.44125988 | 0.19152678 | 0.8481129 | 0.98545782 |
| FMNL1 | 66.3584677 | -0.0290603 | 0.42408406 | -0.0685248 | 0.94536786 | 0.99514836 |
| MAP3K14 | 40.0666072 | -0.5550464 | 0.42349343 | -1.3106377 | 0.18998021 | 0.77993359 |
| ARHGAP27 | 22.3562205 | 0.16071027 | 0.55113935 | 0.29159644 | 0.7705952 | 0.98264092 |
| PLEKHM1 | 29.9575969 | 0.0577419 | 0.39620933 | 0.14573584 | 0.88412993 | 0.99019008 |
| KANSL1 | 256.990144 | 0.07890261 | 0.22555289 | 0.34981868 | 0.72647478 | 0.9754739 |
| ARL17B | 41.3336638 | 1.04257804 | 0.98585746 | 1.05753427 | 0.29026782 | 0.85822878 |
| LRRC37A2 | 12.3115391 | 0.49525791 | 0.64215199 | 0.77124718 | 0.44056043 | NA |
| ARL17A | 15.3971789 | -0.5099742 | 0.55894998 | -0.912379 | 0.36156924 | 0.89853007 |
| NSF | 105.974276 | -0.100505 | 0.3816578 | -0.263338 | 0.79229007 | 0.98320463 |
| AC005670.2 | 8.10838976 | 0.04881398 | 0.59268666 | 0.08236052 | 0.93436003 | NA |
| CDC27 | 67.7492625 | -0.1500824 | 0.25631309 | -0.5855433 | 0.55818248 | 0.95415437 |
| EFCAB13 | 14.5742035 | 0.16421999 | 0.57144949 | 0.28737446 | 0.77382561 | 0.98264092 |

| | | | | | | |
|---|---|---|---|---|---|---|
| NPEPPS | 77.999869 | 0.10224052 | 0.25447308 | 0.40177342 | 0.68785079 | 0.9673431 |
| AC025682.1 | 9.67364198 | -0.0872271 | 0.55986106 | -0.1558013 | 0.87618964 | NA |
| KPNB1 | 66.1667193 | -0.0610936 | 0.33256307 | -0.1837054 | 0.85424458 | 0.98751288 |
| TBX21 | 9.92000195 | -0.2258441 | 0.61730271 | -0.3658563 | 0.71447228 | NA |
| SP2 | 9.78688848 | -1.1332753 | 0.60290809 | -1.8796817 | 0.06015148 | NA |
| CDK5RAP3 | 12.1379747 | -0.9555247 | 0.5645169 | -1.6926415 | 0.09052373 | NA |
| NFE2L1 | 12.8632936 | 0.84898511 | 0.60906857 | 1.39390727 | 0.16334561 | NA |
| SKAP1 | 241.149171 | 0.35626652 | 0.44526517 | 0.80012213 | 0.42364004 | 0.91985469 |
| CALCOCO2 | 32.7836203 | 0.63636304 | 0.4554367 | 1.39725904 | 0.16233569 | 0.75177046 |
| ATP5MC1 | 9.31761092 | 0.04299024 | 0.59212488 | 0.07260334 | 0.94212177 | NA |
| UBE2Z | 26.8188727 | -0.7073933 | 0.43747053 | -1.6170078 | 0.1058766 | 0.67519097 |
| SNF8 | 11.7443972 | -1.0558029 | 0.5402362 | -1.9543356 | 0.05066154 | NA |
| ABI3 | 14.763985 | 1.35949556 | 0.86188532 | 1.57735087 | 0.11471481 | 0.70122655 |
| ZNF652 | 65.4879309 | -0.4456499 | 0.34861028 | -1.2783612 | 0.2011221 | 0.78491514 |
| PHB | 13.2544397 | -0.3473439 | 0.55528406 | -0.6255246 | 0.53162678 | 0.95265314 |
| SPOP | 40.7257733 | 0.44308901 | 0.30802051 | 1.43850491 | 0.15029085 | 0.74043418 |
| FAM117A | 22.8185843 | 1.10070225 | 0.74143157 | 1.48456351 | 0.13765949 | 0.73044824 |
| KAT7 | 14.8924878 | -0.0857089 | 0.50201184 | -0.1707308 | 0.86443547 | 0.98751288 |
| PPP1R9B | 10.4381975 | 0.44273863 | 0.60927965 | 0.72665915 | 0.46743479 | NA |
| ACSF2 | 10.4602883 | 0.78760738 | 0.60428218 | 1.3033768 | 0.19244615 | NA |
| ABCC3 | 15.6747071 | 1.80416057 | 0.79932299 | 2.25711081 | 0.02400115 | 0.39492781 |
| LUC7L3 | 84.5512863 | -0.3655953 | 0.28327679 | -1.2905938 | 0.19684455 | 0.78135737 |
| TOB1 | 9.66005601 | 1.04002486 | 0.63924926 | 1.62694732 | 0.10374829 | NA |
| SPAG9 | 90.9564046 | -0.1931443 | 0.32227015 | -0.5993243 | 0.54895664 | 0.95415437 |
| MBTD1 | 47.1918582 | 0.09041769 | 0.39216843 | 0.23055829 | 0.81765797 | 0.98320463 |
| UTP18 | 14.0631192 | -0.0114417 | 0.46384959 | -0.0246668 | 0.9803207 | 0.99758514 |
| STXBP4 | 17.9662892 | 0.470468 | 0.56770168 | 0.82872398 | 0.40726062 | 0.9169561 |
| DGKE | 15.5093701 | -0.7940427 | 0.45155768 | -1.7584524 | 0.07867057 | 0.62793067 |
| TRIM25 | 20.0257453 | 0.74902054 | 0.51825622 | 1.44527072 | 0.14838186 | 0.7383468 |
| SCPEP1 | 22.1312055 | 0.40389443 | 0.60273248 | 0.67010563 | 0.50279046 | 0.94128633 |
| MSI2 | 136.071206 | 0.78446098 | 0.64871447 | 1.20925463 | 0.22656503 | 0.82055687 |
| CUEDC1 | 17.3458236 | -0.1479937 | 0.55078469 | -0.2686962 | 0.7881635 | 0.98320463 |
| VEZF1 | 14.5289683 | 0.57502743 | 0.50828263 | 1.13131433 | 0.25792281 | 0.84308615 |
| DYNLL2 | 9.43389555 | -0.455306 | 0.66837272 | -0.6812158 | 0.49573498 | NA |
| TSPOAP1-AS | 24.7669941 | -0.4926383 | 0.60331105 | -0.8165577 | 0.41418121 | 0.91894074 |
| AC004687.1 | 16.46824 | 0.41644348 | 0.77117967 | 0.54000838 | 0.58919125 | 0.95415437 |
| MTMR4 | 9.10589748 | -0.0847601 | 0.6158598 | -0.1376289 | 0.89053371 | NA |
| TRIM37 | 20.0670801 | -0.2426411 | 0.42011642 | -0.5775569 | 0.56356332 | 0.95415437 |
| SKA2 | 13.1487533 | -0.7065925 | 0.45479664 | -1.553645 | 0.12026913 | 0.70654921 |
| YPEL2 | 36.4195486 | 0.25035762 | 0.42308803 | 0.59173884 | 0.55402548 | 0.95415437 |
| DHX40 | 11.7803877 | 0.03314248 | 0.50666073 | 0.06541356 | 0.94784473 | NA |
| CLTC | 95.4112282 | 0.06063413 | 0.29310939 | 0.2068652 | 0.83611513 | 0.98320463 |
| PTRH2 | 14.9536426 | -0.4606933 | 0.53157328 | -0.8666601 | 0.38612829 | 0.90910959 |
| VMP1 | 264.050575 | 0.49803029 | 0.33567655 | 1.48366128 | 0.1378988 | 0.73044824 |
| RPS6KB1 | 18.2006375 | -0.1571506 | 0.43646014 | -0.3600572 | 0.71880435 | 0.97348719 |
| RNFT1 | 9.32449183 | -0.9409817 | 0.6727945 | -1.3986168 | 0.16192792 | NA |
| USP32 | 113.284512 | 0.14669551 | 0.22995614 | 0.63792822 | 0.52352041 | 0.94957782 |
| APPBP2 | 38.5947512 | 0.3010114 | 0.34068991 | 0.88353485 | 0.37694736 | 0.90433844 |
| PPM1D | 26.4010914 | 0.9785153 | 0.38701695 | 2.52835257 | 0.01145992 | 0.29101961 |
| BCAS3 | 153.006727 | 0.13807786 | 0.2595009 | 0.5320901 | 0.5946636 | 0.9553365 |
| INTS2 | 11.9258788 | -0.3569145 | 0.51820826 | -0.6887473 | 0.49098231 | NA |
| MED13 | 116.917942 | -0.1290884 | 0.2218369 | -0.5819069 | 0.56062939 | 0.95415437 |
| TLK2 | 73.3501069 | -0.1597789 | 0.29684269 | -0.5382613 | 0.59039666 | 0.95415437 |
| TANC2 | 81.0006235 | 1.04371722 | 0.44228326 | 2.35983888 | 0.01828287 | 0.3513105 |

| | | | | | | |
|---|---|---|---|---|---|---|
| DCAF7 | 48.0249517 | -0.1530947 | 0.39430838 | -0.3882612 | 0.69782273 | 0.96861929 |
| MAP3K3 | 63.1989415 | -0.0520942 | 0.36996709 | -0.1408076 | 0.88802193 | 0.99019008 |
| CCDC47 | 9.54999014 | -0.0451078 | 0.63439642 | -0.0711036 | 0.94331533 | NA |
| DDX42 | 28.1176043 | -0.2802194 | 0.38106918 | -0.7353505 | 0.46212608 | 0.9314919 |
| SMARCD2 | 11.3234284 | -0.0753914 | 0.582049 | -0.1295277 | 0.89694014 | NA |
| ICAM2 | 12.9198834 | -0.1032129 | 0.61930488 | -0.1666593 | 0.86763817 | NA |
| ERN1 | 61.3416927 | 0.83146188 | 0.83482379 | 0.99597291 | 0.31926331 | 0.87300562 |
| TEX2 | 16.6995996 | 0.54157922 | 0.70577781 | 0.76735088 | 0.44287292 | 0.92899684 |
| PECAM1 | 80.2495792 | 0.66051959 | 0.57083854 | 1.15710405 | 0.24722985 | 0.83817091 |
| MILR1 | 8.70799623 | -0.4196144 | 0.70111111 | -0.5984991 | 0.54950695 | NA |
| POLG2 | 10.5392754 | -1.019241 | 0.51998108 | -1.9601502 | 0.04997823 | NA |
| DDX5 | 147.553338 | -0.2489217 | 0.29737475 | -0.8370641 | 0.40255656 | 0.91528253 |
| CEP95 | 28.3787227 | 0.09612668 | 0.42820419 | 0.22448795 | 0.82237764 | 0.98320463 |
| SMURF2 | 83.0735131 | -0.0511693 | 0.28155014 | -0.1817414 | 0.85578566 | 0.98751288 |
| LRRC37A3 | 40.6557492 | -0.1022823 | 0.42648913 | -0.2398238 | 0.81046684 | 0.98320463 |
| GNA13 | 98.9300452 | -0.2324213 | 0.55909627 | -0.4157089 | 0.677623 | 0.96718429 |
| PRKCA | 145.343853 | 0.6547292 | 0.53671169 | 1.21988994 | 0.2225066 | 0.81581246 |
| HELZ | 139.281526 | 0.2591314 | 0.30697966 | 0.84413213 | 0.39859557 | 0.91452235 |
| PSMD12 | 19.8228417 | 0.11202935 | 0.40394294 | 0.27733954 | 0.7815194 | 0.98264092 |
| PITPNC1 | 227.599765 | 0.48426764 | 0.4682651 | 1.03417409 | 0.30105478 | 0.86225566 |
| NOL11 | 11.1187957 | 0.34888789 | 0.58159849 | 0.59987758 | 0.54858783 | NA |
| BPTF | 141.816943 | -0.59037 | 0.2906147 | -2.0314525 | 0.04220911 | 0.50341641 |
| ARSG | 24.816907 | 0.29189927 | 0.63358659 | 0.46070935 | 0.64500714 | 0.96049083 |
| SLC16A6 | 12.3543472 | 1.46534026 | 0.98252631 | 1.49140053 | 0.13585637 | NA |
| WIPI1 | 26.0325121 | 0.7925335 | 0.54837202 | 1.44524788 | 0.14838827 | 0.7383468 |
| PRKAR1A | 52.123791 | 0.60629527 | 0.35735232 | 1.69663171 | 0.08976631 | 0.64936831 |
| FAM20A | 33.1995737 | 0.97097086 | 0.89260426 | 1.08779547 | 0.27668541 | 0.84994492 |
| ABCA5 | 17.7054639 | -0.1264595 | 0.60525508 | -0.2089359 | 0.83449828 | 0.98320463 |
| MAP2K6 | 24.5776312 | 0.52903935 | 0.54164555 | 0.97672611 | 0.32870477 | 0.8791939 |
| KCNJ16 | 10.4370259 | -0.0150571 | 0.77563607 | -0.0194126 | 0.98451199 | NA |
| SLC39A11 | 114.146686 | 0.42581106 | 0.35468938 | 1.2005182 | 0.22993815 | 0.82393202 |
| FAM104A | 11.4240257 | -0.3354947 | 0.53222215 | -0.6303659 | 0.5284552 | NA |
| RPL38 | 29.3925877 | 0.44167636 | 0.60998072 | 0.7240825 | 0.46901509 | 0.9314919 |
| TTYH2 | 7.87144195 | 0.59713254 | 0.85369314 | 0.69946976 | 0.48425851 | NA |
| CD300A | 23.6296652 | 0.70856099 | 0.55505262 | 1.27656543 | 0.20175571 | 0.78572563 |
| CD300E | 28.7010027 | 2.32279531 | 0.8822252 | 2.63288253 | 0.00846636 | 0.24255374 |
| RAB37 | 8.94928931 | 1.09017122 | 0.77640049 | 1.40413515 | 0.16027861 | NA |
| CD300LF | 11.951535 | 1.20921028 | 0.8347398 | 1.44860743 | 0.14744724 | NA |
| SLC9A3R1 | 21.8835236 | 0.12164804 | 0.42022827 | 0.28948085 | 0.77221343 | 0.98264092 |
| TMEM104 | 11.8189888 | 0.25396358 | 0.58906242 | 0.43113187 | 0.66637249 | NA |
| NT5C | 10.0071002 | 0.06957081 | 0.63736714 | 0.10915344 | 0.91308079 | NA |
| JPT1 | 16.9752051 | 0.10672745 | 0.51743058 | 0.2062643 | 0.83658447 | 0.98320463 |
| SUMO2 | 10.0822319 | -0.1955166 | 0.52913577 | -0.3695017 | 0.71175378 | NA |
| NUP85 | 10.2919256 | -0.6230233 | 0.56411865 | -1.1044189 | 0.26941145 | NA |
| GGA3 | 19.3104418 | -0.4342475 | 0.48060794 | -0.9035379 | 0.36624049 | 0.89868152 |
| GRB2 | 163.114934 | 0.00712011 | 0.34267818 | 0.02077784 | 0.98342287 | 0.99766493 |
| AC011933.4 | 30.4236874 | 0.41210534 | 0.57712009 | 0.71407207 | 0.47518261 | 0.9314919 |
| TMEM94 | 9.41577971 | -0.230921 | 0.54808745 | -0.4213215 | 0.67352037 | NA |
| TSEN54 | 13.1785077 | 0.23610201 | 0.69684128 | 0.33881749 | 0.73474723 | 0.97605445 |
| LLGL2 | 17.2239916 | -0.3513049 | 0.48983346 | -0.7171926 | 0.47325525 | 0.9314919 |
| MYO15B | 18.9599375 | -0.0684636 | 0.49744172 | -0.1376315 | 0.89053166 | 0.99080204 |
| SAP30BP | 17.25499 | -0.102125 | 0.44249052 | -0.2307959 | 0.81747339 | 0.98320463 |
| H3F3B | 110.712598 | 0.26584386 | 0.40560027 | 0.65543313 | 0.51218893 | 0.94507477 |
| UNK | 27.4643333 | 0.09148916 | 0.43461028 | 0.2105085 | 0.83327082 | 0.98320463 |

| | | | | | | |
|---|---|---|---|---|---|---|
| UNC13D | 23.4196935 | -0.1310392 | 0.48867012 | -0.2681547 | 0.78858022 | 0.98320463 |
| WBP2 | 14.8129724 | -0.445883 | 0.52147933 | -0.8550349 | 0.39253182 | 0.90910959 |
| ACOX1 | 15.9079667 | 0.24334037 | 0.45310543 | 0.53705021 | 0.59123293 | 0.95415437 |
| SRP68 | 9.09170198 | -0.2005058 | 0.55992794 | -0.3580921 | 0.72027438 | NA |
| EXOC7 | 15.9091517 | 0.86339568 | 0.46086103 | 1.8734404 | 0.0610076 | 0.57281427 |
| RNF157 | 14.4498812 | -0.5099738 | 0.62727445 | -0.8129994 | 0.41621838 | 0.91923465 |
| UBALD2 | 20.5293629 | -0.4716335 | 0.51815761 | -0.9102124 | 0.3627105 | 0.89853007 |
| PRPSAP1 | 15.7110223 | 0.25337542 | 0.51414609 | 0.49280821 | 0.62214811 | 0.95744373 |
| UBE2O | 21.0426409 | 0.66966169 | 0.52860368 | 1.26685022 | 0.20520885 | 0.78970624 |
| RHBDF2 | 46.6733923 | -0.2296912 | 0.46527065 | -0.4936723 | 0.62153765 | 0.95702817 |
| SRSF2 | 31.5396745 | -0.6774409 | 0.38068289 | -1.7795412 | 0.07515107 | 0.6217578 |
| MFSD11 | 25.6421342 | -0.18289 | 0.41078121 | -0.4452247 | 0.65615736 | 0.96219972 |
| SEC14L1 | 75.1444351 | 0.44308304 | 0.36127715 | 1.22643528 | 0.22003492 | 0.81059452 |
| 9-Sep | 85.6867454 | 0.00361589 | 0.41429483 | 0.00872782 | 0.99303629 | 0.99859588 |
| TNRC6C | 44.9385639 | 0.45334292 | 0.42743406 | 1.06061488 | 0.28886496 | 0.85703857 |
| TMC6 | 35.7881154 | 0.37481947 | 0.49716908 | 0.75390745 | 0.4509048 | 0.92914597 |
| TMC8 | 60.2164347 | -0.2975587 | 0.3560398 | -0.8357455 | 0.40329807 | 0.91528253 |
| SYNGR2 | 21.7940896 | -0.254226 | 0.68329683 | -0.3720579 | 0.70984976 | 0.97211298 |
| AFMID | 9.2301642 | 0.32671352 | 0.57637725 | 0.56683972 | 0.57082308 | NA |
| SOCS3 | 11.2297876 | 0.3024079 | 0.67572237 | 0.44753277 | 0.65449043 | NA |
| PGS1 | 28.6643174 | 0.21264871 | 0.3762741 | 0.5651431 | 0.57197644 | 0.95415437 |
| CYTH1 | 212.520371 | 0.20309788 | 0.43384432 | 0.46813539 | 0.63968777 | 0.95952691 |
| USP36 | 40.4099263 | -0.1408462 | 0.67209125 | -0.2095641 | 0.83400786 | 0.98320463 |
| TIMP2 | 45.5843076 | 0.6280893 | 0.44274914 | 1.41861212 | 0.15601213 | 0.74486619 |
| TBC1D16 | 17.4949717 | 0.346619 | 0.79697919 | 0.434916 | 0.6636234 | 0.96379377 |
| GAA | 15.1713065 | 1.05449135 | 0.63534815 | 1.65970633 | 0.09697355 | 0.66178928 |
| RNF213 | 232.04533 | 0.29866354 | 0.51556941 | 0.57928871 | 0.56239438 | 0.95415437 |
| AC124319.2 | 18.1452752 | 0.27079672 | 0.61369425 | 0.44125674 | 0.65902714 | 0.96280444 |
| RPTOR | 51.5271549 | 0.05437378 | 0.33491549 | 0.16235074 | 0.87102966 | 0.98751288 |
| SLC38A10 | 28.6420049 | 0.25543352 | 0.41601076 | 0.61400701 | 0.53921069 | 0.95415437 |
| ACTG1 | 143.473672 | -0.2637497 | 0.54040629 | -0.4880581 | 0.62550866 | 0.95876809 |
| NPLOC4 | 34.514488 | -0.4133446 | 0.39905428 | -1.0358104 | 0.30029059 | 0.86225566 |
| CCDC137 | 10.0262683 | -0.7278974 | 0.52377486 | -1.3897142 | 0.16461568 | NA |
| HGS | 15.6705882 | 0.36502506 | 0.50479834 | 0.72311066 | 0.4696119 | 0.9314919 |
| MCRIP1 | 10.8065849 | -0.2394829 | 0.72850878 | -0.3287303 | 0.74235952 | NA |
| P4HB | 43.1336096 | 0.42924374 | 0.47333324 | 0.90685316 | 0.36448445 | 0.89868152 |
| ARHGDIA | 31.3156882 | -0.048824 | 0.61256097 | -0.0797047 | 0.93647211 | 0.99514836 |
| ANAPC11 | 13.0113627 | -0.4550835 | 0.55629246 | -0.8180653 | 0.41331991 | 0.91894074 |
| SIRT7 | 11.8571929 | 0.67394817 | 0.63191973 | 1.06650914 | 0.28619355 | NA |
| CCDC57 | 19.2345709 | -0.0130586 | 0.41897259 | -0.0311681 | 0.97513548 | 0.9973754 |
| SLC16A3 | 27.6219942 | 0.70057091 | 0.59321061 | 1.18098175 | 0.23760997 | 0.83171006 |
| CSNK1D | 34.9477445 | -0.5382807 | 0.39936956 | -1.347826 | 0.17771436 | 0.76504408 |
| CD7 | 19.5652926 | 1.24316087 | 0.81709898 | 1.52143241 | 0.12815136 | 0.72782081 |
| HEXDC | 14.372365 | -0.8270458 | 0.60513213 | -1.3667194 | 0.17171328 | 0.75906822 |
| CYBC1 | 22.384381 | 0.28691061 | 0.55501227 | 0.51694463 | 0.60519481 | 0.95688177 |
| NARF | 22.3925971 | 0.01719392 | 0.39713505 | 0.04329491 | 0.96546645 | 0.99653147 |
| FOXK2 | 47.2237687 | 0.14767102 | 0.31715157 | 0.46561656 | 0.64148999 | 0.95992767 |
| WDR45B | 23.6354726 | -0.2066218 | 0.48658067 | -0.4246404 | 0.67109886 | 0.96412305 |
| TBCD | 33.2926908 | -0.2612582 | 0.35109657 | -0.7441207 | 0.45680347 | 0.9314919 |
| B3GNTL1 | 14.7773075 | 0.19609229 | 0.47835487 | 0.40993058 | 0.68185687 | 0.9673431 |
| METRNL | 32.2839524 | -0.3367321 | 0.77821506 | -0.4326979 | 0.66523422 | 0.96406914 |
| USP14 | 28.2546803 | -0.3461123 | 0.42308112 | -0.8180754 | 0.41331414 | 0.91894074 |
| THOC1 | 32.1970336 | -0.2945662 | 0.43129586 | -0.6829794 | 0.49461987 | 0.93805563 |
| COLEC12 | 77.1246666 | -0.6767934 | 0.84663278 | -0.7993943 | 0.42406181 | 0.91985469 |

| | | | | | | |
|---|---|---|---|---|---|---|
| ENOSF1 | 28.4663279 | 0.39872287 | 0.51463965 | 0.77476129 | 0.43848073 | 0.92687256 |
| YES1 | 19.6195485 | 0.01837347 | 0.43020292 | 0.04270885 | 0.96593363 | 0.99653147 |
| METTL4 | 10.5863678 | 0.0074688 | 0.63082568 | 0.01183973 | 0.99055349 | NA |
| SMCHD1 | 301.416501 | -0.2589759 | 0.37392412 | -0.6925893 | 0.4885673 | 0.93596573 |
| AP001011.1 | 55.1143235 | -0.0443637 | 0.6683745 | -0.0663755 | 0.94707884 | 0.99514836 |
| EMILIN2 | 47.146922 | 1.57601564 | 0.39548522 | 3.98501779 | 6.75E-05 | 0.01436181 |
| LPIN2 | 59.9184395 | 0.25965092 | 0.34107109 | 0.76128096 | 0.44648927 | 0.92914597 |
| MYL12A | 61.8359591 | 0.36007068 | 0.51993136 | 0.69253503 | 0.4886014 | 0.93596573 |
| MYL12B | 54.2811961 | -0.0835543 | 0.44339152 | -0.1884436 | 0.85052893 | 0.98696312 |
| TGIF1 | 11.3621045 | 0.32063414 | 0.71956121 | 0.44559675 | 0.65588856 | NA |
| EPB41L3 | 142.471887 | 0.85143419 | 0.77369775 | 1.10047391 | 0.27112569 | 0.84994492 |
| L3MBTL4 | 52.389077 | -0.6982271 | 0.54373911 | -1.2841216 | 0.19909942 | 0.78142991 |
| PTPRM | 99.8994735 | -1.8817335 | 0.51595706 | -3.6470739 | 0.00026524 | 0.03066918 |
| RAB12 | 36.5558679 | -0.8072949 | 0.48294881 | -1.6715952 | 0.09460418 | 0.65925012 |
| NDUFV2 | 33.483899 | -1.449273 | 0.43646108 | -3.3205092 | 0.00089853 | 0.06269654 |
| ANKRD12 | 233.521574 | -0.4923552 | 0.2563807 | -1.9204068 | 0.05480654 | 0.55447783 |
| TWSG1 | 15.1945026 | -1.044885 | 0.8770482 | -1.1913656 | 0.2335101 | 0.82817204 |
| RALBP1 | 23.4566893 | -0.0497693 | 0.4789417 | -0.1039152 | 0.91723662 | 0.99498068 |
| PPP4R1 | 35.8249339 | -0.0210078 | 0.47249728 | -0.0444612 | 0.96453679 | 0.99653147 |
| RAB31 | 154.918695 | -0.1775283 | 0.61502239 | -0.2886533 | 0.77284668 | 0.98264092 |
| VAPA | 63.1142712 | -0.0551805 | 0.30040323 | -0.1836881 | 0.85425812 | 0.98751288 |
| NAPG | 22.6124621 | 0.09109008 | 0.48461834 | 0.18796251 | 0.85090604 | 0.98719372 |
| MPPE1 | 17.4819467 | -0.0400755 | 0.4627744 | -0.0865984 | 0.93099078 | 0.99514836 |
| IMPA2 | 13.8378304 | 1.05869843 | 0.73086002 | 1.44856526 | 0.14745903 | 0.73693374 |
| AFG3L2 | 16.324792 | -0.2853699 | 0.55820107 | -0.5112314 | 0.60918902 | 0.95688177 |
| SPIRE1 | 25.5252303 | -0.2784583 | 0.50414716 | -0.5523354 | 0.58071855 | 0.95415437 |
| PSMG2 | 13.0714956 | -0.2142798 | 0.52285735 | -0.4098247 | 0.68193457 | 0.9673431 |
| PTPN2 | 86.5160241 | 0.12908853 | 0.37063375 | 0.34829135 | 0.72762139 | 0.97605445 |
| SEH1L | 11.586263 | 0.05579816 | 0.53190045 | 0.10490339 | 0.91645247 | NA |
| CEP192 | 66.5192621 | -0.3771778 | 0.28611039 | -1.3182945 | 0.18740507 | 0.77798634 |
| LDLRAD4 | 235.613634 | 0.18909288 | 0.40714584 | 0.46443526 | 0.64233593 | 0.95992767 |
| FAM210A | 14.3135886 | 0.64162403 | 0.74227826 | 0.86439825 | 0.38736916 | 0.90910959 |
| RNMT | 34.6917985 | -0.2171896 | 0.39017597 | -0.5566452 | 0.57776986 | 0.95415437 |
| ROCK1 | 149.357769 | 0.41632777 | 0.19494622 | 2.13560319 | 0.03271177 | 0.44484764 |
| ESCO1 | 28.5236212 | -0.2474316 | 0.41807975 | -0.5918287 | 0.55396531 | 0.95415437 |
| SNRPD1 | 8.60266976 | -0.3290088 | 0.75963187 | -0.4331162 | 0.66493039 | NA |
| ABHD3 | 68.3424245 | 0.43997859 | 0.32749647 | 1.34346056 | 0.17912289 | 0.76545642 |
| MIB1 | 60.0933812 | 0.02360398 | 0.3744149 | 0.06304232 | 0.9497328 | 0.99514836 |
| RBBP8 | 13.1576042 | 0.42736396 | 0.64761059 | 0.65990884 | 0.50931233 | 0.94332619 |
| CABLES1 | 17.8569881 | -0.056102 | 0.69477904 | -0.0807479 | 0.9356424 | 0.99514836 |
| TMEM241 | 24.2469685 | -1.889771 | 0.47928519 | -3.9428947 | 8.05E-05 | 0.01536239 |
| RIOK3 | 40.362621 | 0.04149134 | 0.3800828 | 0.10916394 | 0.91307246 | 0.99498068 |
| RMC1 | 21.4652657 | -1.1074617 | 0.52090522 | -2.1260331 | 0.03350051 | 0.45107493 |
| NPC1 | 46.3364174 | -0.4984849 | 0.40506149 | -1.2306402 | 0.21845748 | 0.80919927 |
| TTC39C | 84.7035212 | 0.24119548 | 0.41954252 | 0.57490115 | 0.56535815 | 0.95415437 |
| OSBPL1A | 24.4707335 | 0.56024164 | 0.53807299 | 1.04120008 | 0.2977827 | 0.8608685 |
| SS18 | 44.5071086 | -0.1889841 | 0.35240542 | -0.5362689 | 0.59177274 | 0.95415437 |
| TAF4B | 11.2448919 | -0.0888802 | 0.5822638 | -0.1526459 | 0.87867749 | NA |
| TRAPPC8 | 60.34847 | -0.0465907 | 0.31292225 | -0.1488891 | 0.88164115 | 0.99019008 |
| RNF125 | 37.7885245 | 1.12138807 | 0.56586304 | 1.98173056 | 0.0475094 | 0.53114555 |
| RNF138 | 32.8427512 | 0.0297383 | 0.34380174 | 0.08649841 | 0.93107022 | 0.99514836 |
| GAREM1 | 15.3583282 | 0.24366002 | 0.48596296 | 0.50139628 | 0.61609226 | 0.95688177 |
| DTNA | 41.9284971 | 0.12436814 | 0.54289305 | 0.22908406 | 0.81880358 | 0.98320463 |
| MAPRE2 | 50.6096792 | 0.05167829 | 0.30277631 | 0.17068143 | 0.86447427 | 0.98751288 |

| | | | | | | |
|---|---|---|---|---|---|---|
| ZNF397 | 10.1390659 | 0.49948284 | 0.65152945 | 0.76663125 | 0.44330079 | NA |
| ZNF24 | 15.650759 | 0.13162765 | 0.54981035 | 0.23940554 | 0.81079113 | 0.98320463 |
| GALNT1 | 49.5527161 | -0.0656497 | 0.50302314 | -0.1305104 | 0.89616265 | 0.99118739 |
| RPRD1A | 33.9678751 | -0.26506 | 0.35415245 | -0.7484348 | 0.45419795 | 0.9314919 |
| ELP2 | 24.6921994 | -0.115441 | 0.45670225 | -0.2527708 | 0.80044531 | 0.98320463 |
| TPGS2 | 11.9264977 | 0.92402693 | 0.51222023 | 1.80396416 | 0.07123693 | NA |
| KIAA1328 | 45.3678985 | -0.7908667 | 0.4626861 | -1.7092943 | 0.08739646 | 0.64676224 |
| AC016205.1 | 26.8240103 | -0.3904321 | 0.53453556 | -0.7304138 | 0.46513731 | 0.9314919 |
| PIK3C3 | 49.8175129 | 0.15608347 | 0.34783148 | 0.44873304 | 0.65362425 | 0.96178254 |
| SETBP1 | 141.727904 | -2.8030366 | 0.60185418 | -4.657335 | 3.20E-06 | 0.00171296 |
| SLC14A1 | 10.8082204 | 1.10555154 | 0.83807243 | 1.31915991 | 0.18711566 | NA |
| EPG5 | 38.3162087 | 0.26475471 | 0.35442458 | 0.74699873 | 0.45506433 | 0.9314919 |
| PSTPIP2 | 40.7832503 | 0.25035031 | 0.50273122 | 0.49798044 | 0.61849783 | 0.95688177 |
| ATP5F1A | 26.6201054 | 0.24370004 | 0.38863481 | 0.62706696 | 0.53061535 | 0.95245713 |
| C18orf25 | 34.3590972 | -0.1533969 | 0.39199096 | -0.3913276 | 0.6955551 | 0.9685713 |
| PIAS2 | 41.9725272 | 0.26089987 | 0.37895943 | 0.68846388 | 0.49116071 | 0.93633431 |
| SMAD2 | 84.7928714 | 0.15313118 | 0.29415613 | 0.52057789 | 0.60266085 | 0.95688177 |
| CTIF | 18.9947246 | 0.41636568 | 0.51855349 | 0.80293681 | 0.42201126 | 0.91985469 |
| SMAD7 | 10.1051382 | 0.99722742 | 0.62481308 | 1.59604119 | 0.1104796 | NA |
| DYM | 120.233592 | 0.02499327 | 0.22438604 | 0.11138516 | 0.91131093 | 0.99471296 |
| ACAA2 | 10.3534201 | 1.43123407 | 0.65766689 | 2.17622946 | 0.02953811 | NA |
| MBD1 | 11.5736082 | -0.2237373 | 0.60491961 | -0.3698629 | 0.71148464 | NA |
| ME2 | 47.9377105 | -0.2640751 | 0.36253229 | -0.728418 | 0.46635774 | 0.9314919 |
| SMAD4 | 19.6379292 | -0.2630429 | 0.41112458 | -0.6398132 | 0.52229405 | 0.94895469 |
| MBD2 | 83.3368273 | 0.10107016 | 0.30827349 | 0.32785875 | 0.74301846 | 0.97711087 |
| POLI | 9.72601908 | 0.3216797 | 0.57006432 | 0.56428669 | 0.57255904 | NA |
| TCF4 | 148.528742 | 0.77076529 | 0.78632252 | 0.98021521 | 0.3269799 | 0.87715548 |
| TXNL1 | 15.6021738 | 0.05738695 | 0.43766875 | 0.1311196 | 0.8956807 | 0.99115168 |
| WDR7 | 75.4478286 | -0.1331673 | 0.25500393 | -0.5222167 | 0.60151944 | 0.95688177 |
| NARS | 15.4459631 | 0.18554096 | 0.57074761 | 0.32508407 | 0.74511745 | 0.97805001 |
| AC027097.2 | 33.9654167 | -1.1795653 | 0.42944371 | -2.7467287 | 0.00601929 | 0.19198507 |
| ATP8B1 | 26.1847946 | -0.2409384 | 0.41488604 | -0.580734 | 0.56141974 | 0.95415437 |
| NEDD4L | 41.3751974 | -0.2655056 | 0.49457218 | -0.5368389 | 0.59137889 | 0.95415437 |
| MALT1 | 86.2763312 | -0.1090715 | 0.49227156 | -0.2215677 | 0.82465041 | 0.98320463 |
| AC104365.1 | 13.91426 | -0.2214574 | 0.73844256 | -0.2998979 | 0.76425505 | 0.98222198 |
| ZNF532 | 20.463452 | -0.7184011 | 0.62885851 | -1.1423892 | 0.2532923 | 0.84026145 |
| LMAN1 | 21.4955379 | 0.38977139 | 0.60335302 | 0.64600885 | 0.51827362 | 0.94668603 |
| PIGN | 34.3021405 | -0.4486134 | 0.3323892 | -1.349663 | 0.1771241 | 0.76409729 |
| KIAA1468 | 76.5649677 | -0.4559395 | 0.29248785 | -1.5588322 | 0.11903609 | 0.70431251 |
| ZCCHC2 | 44.6423967 | 0.58883735 | 0.4339904 | 1.35679809 | 0.17484538 | 0.76214163 |
| PHLPP1 | 53.086232 | -1.3337275 | 0.56243842 | -2.3713306 | 0.01772417 | 0.34827529 |
| BCL2 | 244.09271 | -0.3362279 | 0.48766426 | -0.689466 | 0.49053009 | 0.93633431 |
| KDSR | 11.6516698 | 0.01663182 | 0.50654673 | 0.03283374 | 0.97380717 | NA |
| VPS4B | 32.9238139 | 0.35977966 | 0.41750211 | 0.86174333 | 0.38882878 | 0.90910959 |
| TMX3 | 28.071275 | -0.2923199 | 0.46156887 | -0.6333181 | 0.52652595 | 0.95136478 |
| CD226 | 42.1028225 | 1.05460317 | 0.58681087 | 1.79717729 | 0.07230748 | 0.61222939 |
| RTTN | 27.4667756 | -0.6073523 | 0.43734935 | -1.388712 | 0.16492033 | 0.75321592 |
| CYB5A | 18.8406028 | 1.66112127 | 0.7131106 | 2.32940202 | 0.01983778 | 0.36765113 |
| CNDP2 | 30.675116 | 0.43202413 | 0.46995452 | 0.91928923 | 0.35794431 | 0.8957272 |
| ZNF407 | 116.287557 | -0.3868353 | 0.28704867 | -1.3476295 | 0.17777756 | 0.76504408 |
| ZADH2 | 10.7446725 | 1.03659785 | 0.57761117 | 1.79462916 | 0.0727128 | NA |
| TSHZ1 | 19.7963652 | -0.2390819 | 0.46727435 | -0.511652 | 0.60889456 | 0.95688177 |
| ZNF516 | 46.7941635 | 0.33402077 | 0.4342377 | 0.76921182 | 0.44176757 | 0.92899684 |
| ZNF236 | 29.4046367 | -0.3773219 | 0.34158259 | -1.1046286 | 0.26932055 | 0.84994492 |

| | | | | | | |
|---|---|---|---|---|---|---|
| MBP | 144.57577 | 0.18476432 | 0.29453695 | 0.6273044 | 0.53045972 | 0.95245713 |
| ATP9B | 97.0188574 | 0.07318544 | 0.2955428 | 0.24763061 | 0.80442023 | 0.98320463 |
| NFATC1 | 27.8585817 | -0.7936852 | 0.54985706 | -1.4434391 | 0.14889683 | 0.7383468 |
| CTDP1 | 15.2368143 | 0.45229017 | 0.47851964 | 0.9451862 | 0.34456382 | 0.8871109 |
| PQLC1 | 18.4640846 | 0.42617667 | 0.48632682 | 0.87631744 | 0.38085749 | 0.9073032 |
| TXNL4A | 23.2422703 | -0.5453029 | 0.39594734 | -1.3772106 | 0.16844716 | 0.7567539 |
| ADNP2 | 14.7446603 | -0.5586209 | 0.48288649 | -1.1568369 | 0.24733899 | 0.83817091 |
| RBCK1 | 18.6697739 | -0.0615603 | 0.60546026 | -0.1016752 | 0.9190145 | 0.99498068 |
| TBC1D20 | 12.2095052 | -0.0771794 | 0.49705242 | -0.1552742 | 0.87660519 | NA |
| CSNK2A1 | 43.7106596 | 0.06257369 | 0.33082282 | 0.18914562 | 0.84997868 | 0.98653147 |
| PSMF1 | 27.3855459 | 0.36290435 | 0.43130831 | 0.84140357 | 0.40012188 | 0.91508218 |
| AL136531.2 | 9.45306385 | -0.7696158 | 0.5280896 | -1.4573583 | 0.1450175 | NA |
| SDCBP2-AS1 | 10.7842159 | -0.0544252 | 0.55809618 | -0.0975194 | 0.92231391 | NA |
| NSFL1C | 18.8471078 | 0.4507718 | 0.45189293 | 0.99751903 | 0.31851264 | 0.87298176 |
| SIRPB2 | 23.7578799 | 1.4239806 | 0.57907298 | 2.45906935 | 0.01392977 | 0.31861834 |
| SIRPA | 25.4242313 | 0.8527619 | 0.68831205 | 1.23891757 | 0.21537603 | 0.80634897 |
| STK35 | 9.21204959 | -0.1785202 | 0.65403106 | -0.2729536 | 0.78488886 | NA |
| SNRPB | 17.8853419 | -0.3756892 | 0.47366244 | -0.7931582 | 0.42768566 | 0.92129717 |
| NOP56 | 10.2616088 | -0.5863113 | 0.58377067 | -1.0043522 | 0.31520889 | NA |
| PTPRA | 83.782321 | 0.02403032 | 0.28425493 | 0.08453792 | 0.93262875 | 0.99514836 |
| ITPA | 11.1728629 | 0.23603119 | 0.53090372 | 0.44458379 | 0.65662056 | NA |
| C20orf194 | 89.291205 | 0.15849179 | 0.33087387 | 0.47900969 | 0.63193174 | 0.95877428 |
| ATRN | 54.8985084 | 0.29489957 | 0.35068278 | 0.84092971 | 0.40038731 | 0.91508218 |
| SIGLEC1 | 26.8619719 | 0.9335141 | 0.97187681 | 0.96052719 | 0.33678995 | 0.88143379 |
| CDC25B | 9.40760301 | -1.1169034 | 0.59665992 | -1.8719262 | 0.06121681 | NA |
| MAVS | 18.3981674 | -0.2758392 | 0.48515229 | -0.5685621 | 0.56965334 | 0.95415437 |
| PANK2 | 35.2143129 | -0.4155634 | 0.3248322 | -1.2793173 | 0.20078536 | 0.78415396 |
| RNF24 | 50.0833115 | 1.47509688 | 0.6382904 | 2.31101216 | 0.02083218 | 0.37430293 |
| PRNP | 21.50391 | 0.68676848 | 0.46754865 | 1.46887065 | 0.14186788 | 0.73044824 |
| RASSF2 | 52.614881 | -0.0752944 | 0.50107226 | -0.1502665 | 0.88055436 | 0.99019008 |
| SLC23A2 | 41.6922916 | 0.24704762 | 0.38290321 | 0.64519599 | 0.51880018 | 0.94668603 |
| TMEM230 | 17.2502582 | -0.603688 | 0.41571173 | -1.4521793 | 0.14645175 | 0.73411592 |
| CDS2 | 34.1726705 | -0.3435624 | 0.31667279 | -1.084913 | 0.27796021 | 0.84994492 |
| GPCPD1 | 110.170103 | -0.2032313 | 0.34324959 | -0.5920803 | 0.55379682 | 0.95415437 |
| SHLD1 | 29.1193315 | -0.9438193 | 0.4590603 | -2.0559811 | 0.03978434 | 0.48300729 |
| CRLS1 | 16.5351202 | 0.24437228 | 0.55130226 | 0.4432637 | 0.65757501 | 0.96270373 |
| TMX4 | 20.1129811 | 0.00390421 | 0.52746424 | 0.00740185 | 0.99409423 | 0.99859588 |
| PLCB1 | 126.221646 | -0.0489777 | 0.58341206 | -0.0839505 | 0.93309582 | 0.99514836 |
| SLX4IP | 35.4397586 | 0.36492813 | 0.52922808 | 0.68954795 | 0.49047851 | 0.93633431 |
| SPTLC3 | 13.5204761 | 0.13300066 | 0.71608387 | 0.18573336 | 0.85265387 | 0.98751288 |
| TASP1 | 34.6102662 | -0.2750215 | 0.40964517 | -0.6713651 | 0.50198799 | 0.94042029 |
| ESF1 | 17.4033012 | -0.0574642 | 0.53820274 | -0.1067705 | 0.91497106 | 0.99498068 |
| NDUFAF5 | 9.87405246 | -0.4522262 | 0.60550143 | -0.7468623 | 0.45514669 | NA |
| MACROD2 | 28.4335008 | -2.2103818 | 0.49099868 | -4.501808 | 6.74E-06 | 0.00266335 |
| KIF16B | 44.0435601 | 0.33338171 | 0.46557967 | 0.71605727 | 0.47395599 | 0.9314919 |
| SNRPB2 | 9.54169468 | -0.1255012 | 0.55500693 | -0.2261255 | 0.82110385 | NA |
| DSTN | 29.100281 | 0.90961575 | 0.4920614 | 1.84858181 | 0.06451822 | 0.58724315 |
| RRBP1 | 37.0042485 | 0.00759934 | 0.55471104 | 0.01369963 | 0.98906962 | 0.99811561 |
| SNX5 | 26.3098794 | -0.4639956 | 0.41890084 | -1.1076501 | 0.26801296 | 0.84994492 |
| MGME1 | 13.435447 | -0.2178186 | 0.61290343 | -0.355388 | 0.72229891 | 0.97441549 |
| SEC23B | 24.8820604 | 0.13527937 | 0.38274348 | 0.35344658 | 0.72375367 | 0.97475123 |
| SLC24A3 | 67.3683537 | 6.00110467 | 2.13022576 | 2.81712144 | 0.00484562 | 0.17334548 |
| RIN2 | 79.9469718 | -1.7643656 | 0.63307936 | -2.786958 | 0.00532054 | 0.18175221 |
| NAA20 | 10.8352797 | 0.26393462 | 0.52846985 | 0.49943174 | 0.61747526 | NA |

| | | | | | | |
|---|---|---|---|---|---|---|
| RALGAPA2 | 91.8744552 | 0.89468169 | 0.35189291 | 2.54248283 | 0.0110068 | 0.2861945 |
| KIZ | 21.3604643 | -0.0441882 | 0.46712436 | -0.0945962 | 0.92463559 | 0.99514836 |
| XRN2 | 57.7256216 | 0.02407215 | 0.34702114 | 0.06936796 | 0.94469673 | 0.99514836 |
| CD93 | 20.7446307 | 0.44731325 | 0.76954494 | 0.58126982 | 0.56105862 | 0.95415437 |
| NAPB | 14.2618271 | -0.6214397 | 0.47976339 | -1.2953046 | 0.19521518 | 0.78135737 |
| CST3 | 124.974434 | 1.1621908 | 0.67605275 | 1.71908301 | 0.08559926 | 0.64676224 |
| CST7 | 23.0549858 | 0.34453323 | 0.83906298 | 0.41061665 | 0.68135365 | 0.9673431 |
| APMAP | 25.2495291 | 0.76762268 | 0.674739 | 1.13765867 | 0.25526304 | 0.8406233 |
| ACSS1 | 18.5738991 | 0.43555032 | 0.47435479 | 0.91819525 | 0.35851666 | 0.8957272 |
| ENTPD6 | 9.44103437 | -0.2420492 | 0.55425848 | -0.4367081 | 0.66232305 | NA |
| PYGB | 12.3589656 | -0.4771829 | 0.50431673 | -0.9461969 | 0.34404819 | NA |
| ABHD12 | 30.7659662 | 0.29984756 | 0.54800612 | 0.54716097 | 0.58426815 | 0.95415437 |
| ZNF337 | 10.0701084 | 0.25320374 | 0.55454653 | 0.45659603 | 0.64796144 | NA |
| HM13 | 49.546703 | 1.03472293 | 0.4588607 | 2.25498265 | 0.02413442 | 0.39492781 |
| BCL2L1 | 37.2276256 | 0.62911868 | 0.47897002 | 1.31348239 | 0.18902045 | 0.77850418 |
| AL117381.1 | 13.2783983 | -0.598383 | 0.62998262 | -0.9498405 | 0.34219328 | 0.88510795 |
| HCK | 82.491455 | 0.28969878 | 0.68369456 | 0.42372544 | 0.67176606 | 0.96412305 |
| TM9SF4 | 34.5172373 | -0.2263137 | 0.33892297 | -0.6677437 | 0.50429718 | 0.94166217 |
| KIF3B | 15.3859208 | -0.4377267 | 0.58475159 | -0.7485686 | 0.45411727 | 0.9314919 |
| ASXL1 | 69.1629833 | -0.2586912 | 0.2790073 | -0.9271843 | 0.35383088 | 0.89109867 |
| NOL4L | 33.0353435 | 1.57806521 | 0.67046577 | 2.35368497 | 0.01858836 | 0.35594452 |
| MAPRE1 | 16.6839533 | -0.2897395 | 0.50681045 | -0.5716921 | 0.56753058 | 0.95415437 |
| CDK5RAP1 | 13.9844285 | -0.3028834 | 0.51372502 | -0.5895827 | 0.55547044 | 0.95415437 |
| CBFA2T2 | 65.3350114 | 0.40979152 | 0.30025632 | 1.36480562 | 0.17231415 | 0.75982989 |
| CHMP4B | 19.8975981 | -0.0653966 | 0.44203766 | -0.1479436 | 0.88238728 | 0.99019008 |
| RALY | 57.4088705 | 0.05644763 | 0.42089916 | 0.134112 | 0.89331401 | 0.99080204 |
| EIF2S2 | 24.6840554 | -0.1943771 | 0.41608915 | -0.4671526 | 0.6403907 | 0.95963232 |
| ITCH | 120.044523 | -0.2525528 | 0.3171196 | -0.796396 | 0.42580193 | 0.92046401 |
| DYNLRB1 | 23.1039558 | 0.07432823 | 0.45772197 | 0.16238729 | 0.87100088 | 0.98751288 |
| PIGU | 16.6913828 | -0.7558188 | 0.44055565 | -1.7156035 | 0.08623465 | 0.64676224 |
| NCOA6 | 46.9089913 | -0.4339352 | 0.28052315 | -1.5468782 | 0.12189261 | 0.71155453 |
| ACSS2 | 17.3681635 | 0.48607572 | 0.56530798 | 0.85984232 | 0.38987597 | 0.90910959 |
| TRPC4AP | 69.9745888 | 0.11545916 | 0.27458484 | 0.42048629 | 0.67413024 | 0.96568758 |
| EDEM2 | 13.4122017 | -0.2430003 | 0.47758681 | -0.5088087 | 0.61088634 | 0.95688177 |
| MMP24OS | 17.4018249 | -0.0204839 | 0.50841728 | -0.0402895 | 0.96786231 | 0.99653147 |
| MMP24 | 9.55949494 | 0.12034618 | 0.71859888 | 0.16747338 | 0.86699759 | NA |
| UQCC1 | 24.4937974 | -0.1488658 | 0.43818163 | -0.3397353 | 0.73405588 | 0.97605445 |
| CEP250 | 12.6952621 | -0.3547063 | 0.50019946 | -0.7091297 | 0.47824401 | NA |
| FO393401.1 | 16.5712132 | -0.6694959 | 0.43591962 | -1.5358242 | 0.12458151 | 0.71741321 |
| ERGIC3 | 11.5137117 | -0.0365622 | 0.56644347 | -0.064547 | 0.94853469 | NA |
| CPNE1 | 17.9893385 | 0.11317526 | 0.5461526 | 0.20722278 | 0.83583588 | 0.98320463 |
| RBM39 | 127.607542 | 0.07700252 | 0.22949109 | 0.33553601 | 0.73722079 | 0.97605445 |
| PHF20 | 91.43482 | 0.03969892 | 0.28793016 | 0.13787691 | 0.8903377 | 0.99080204 |
| SCAND1 | 16.7136842 | -0.3376272 | 0.5548652 | -0.6084851 | 0.54286576 | 0.95415437 |
| NORAD | 28.8052198 | -0.3306073 | 0.59471187 | -0.5559117 | 0.5782712 | 0.95415437 |
| DLGAP4 | 22.9293525 | -0.5570581 | 0.44850995 | -1.2420195 | 0.21422938 | 0.80634897 |
| RAB5IF | 11.2960786 | 0.59983856 | 0.7136402 | 0.84053359 | 0.40060928 | NA |
| SLA2 | 17.3309687 | 0.29566211 | 0.64426359 | 0.45891482 | 0.64629534 | 0.96049083 |
| NDRG3 | 12.4839842 | -0.3064524 | 0.52134068 | -0.587816 | 0.5566558 | NA |
| SOGA1 | 15.6765102 | -0.6615102 | 0.49150072 | -1.3458987 | 0.17833518 | 0.76504408 |
| SAMHD1 | 118.848282 | 0.2901176 | 0.45983876 | 0.63091158 | 0.52809834 | 0.95174286 |
| RBL1 | 24.9882086 | -0.3462993 | 0.39822174 | -0.8696142 | 0.38451125 | 0.90910959 |
| RPN2 | 32.832715 | 0.38198085 | 0.50143567 | 0.7617744 | 0.44619466 | 0.92914597 |
| BLCAP | 21.9799552 | -0.0374526 | 0.40600373 | -0.0922468 | 0.92650194 | 0.99514836 |

| | | | | | | |
|---|---|---|---|---|---|---|
| CTNNBL1 | 31.5730076 | -0.0567848 | 0.42466573 | -0.1337165 | 0.89362679 | 0.99084966 |
| TTI1 | 9.88174628 | 0.65800715 | 0.52586416 | 1.25128729 | 0.21082968 | NA |
| RPRD1B | 22.1238362 | 0.20346464 | 0.48180308 | 0.42229833 | 0.67280727 | 0.96433967 |
| TGM2 | 10.425232 | -0.3153784 | 0.63376022 | -0.4976305 | 0.6187445 | NA |
| RALGAPB | 48.8483162 | -0.5168178 | 0.3685673 | -1.4022346 | 0.16084521 | 0.74887162 |
| PPP1R16B | 86.609716 | -0.649055 | 0.36619819 | -1.7724146 | 0.07632577 | 0.62402456 |
| DHX35 | 10.9159186 | -0.3558344 | 0.51076046 | -0.6966756 | 0.48600582 | NA |
| MAFB | 34.8431881 | 1.35340939 | 0.4787017 | 2.82725002 | 0.00469497 | 0.17239111 |
| TOP1 | 83.5956303 | -0.0885821 | 0.29968881 | -0.2955803 | 0.7675506 | 0.98264092 |
| PLCG1 | 12.2049804 | -0.8094041 | 0.48884948 | -1.6557327 | 0.09777596 | NA |
| ZHX3 | 15.3039794 | -0.3637671 | 0.49526534 | -0.7344893 | 0.46265056 | 0.9314919 |
| CHD6 | 66.0151918 | 0.00797788 | 0.34525878 | 0.02310695 | 0.98156496 | 0.99758514 |
| SRSF6 | 14.3282836 | -0.3127658 | 0.5425113 | -0.5765147 | 0.56426731 | 0.95415437 |
| SERINC3 | 25.8218953 | 0.3815948 | 0.42848423 | 0.89056906 | 0.3731604 | 0.9025457 |
| YWHAB | 59.6012623 | 0.09375809 | 0.47293351 | 0.19824793 | 0.84285109 | 0.9842451 |
| STK4 | 163.504695 | 0.05204798 | 0.30556357 | 0.17033438 | 0.86474718 | 0.98751288 |
| SLPI | 29.7278489 | 0.19172591 | 0.94812276 | 0.20221633 | 8.40E-01 | 0.98342625 |
| SDC4 | 10.5968926 | 0.16063652 | 0.64912132 | 0.24746764 | 0.80454633 | NA |
| PIGT | 9.55173773 | 0.41735551 | 0.60768922 | 0.68679104 | 0.49221442 | NA |
| WFDC2 | 44.742615 | 0.93189326 | 0.60725004 | 1.53461209 | 1.25E-01 | 0.71776265 |
| DNTTIP1 | 9.59155637 | 1.3266941 | 0.58291917 | 2.27594866 | 0.02284908 | NA |
| CTSA | 21.1500823 | 1.35997356 | 0.50877777 | 2.67302078 | 0.00751716 | 0.22127638 |
| PLTP | 31.8274813 | 0.940794 | 0.98532967 | 0.95480125 | 0.33967822 | 0.8844375 |
| ZNF335 | 13.9314566 | 0.02517885 | 0.45138783 | 0.05578097 | 0.95551629 | 0.99514836 |
| NCOA5 | 15.3789223 | -0.6559713 | 0.50509432 | -1.2987106 | 0.19404327 | 0.78135737 |
| SLC35C2 | 14.4398156 | -0.3117847 | 0.55988117 | -0.5568765 | 0.57761181 | 0.95415437 |
| ELMO2 | 16.7171825 | 0.15263502 | 0.47974791 | 0.31815671 | 0.75036607 | 0.97818861 |
| SLC13A3 | 10.4683764 | 1.28369367 | 0.69096759 | 1.85782038 | 0.06319453 | NA |
| ZMYND8 | 78.1151147 | -0.5170116 | 0.29239985 | -1.7681663 | 0.07703311 | 0.62507515 |
| NCOA3 | 185.877345 | -0.985106 | 0.28400583 | -3.4686119 | 0.00052315 | 0.04454058 |
| SULF2 | 47.9018318 | 2.04399118 | 1.00597639 | 2.03184806 | 0.04216904 | 0.50341641 |
| PREX1 | 144.567199 | 0.31990643 | 0.35467117 | 0.90198037 | 0.36706729 | 0.89868152 |
| ARFGEF2 | 55.9539009 | -0.1992429 | 0.39778735 | -0.5008779 | 0.61645704 | 0.95688177 |
| CSE1L | 22.0401226 | -0.615645 | 0.40475337 | -1.5210374 | 0.12825045 | 0.72782081 |
| STAU1 | 50.8572022 | 0.20218778 | 0.37472291 | 0.5395661 | 0.5894963 | 0.95415437 |
| DDX27 | 13.834231 | -0.6208148 | 0.44587664 | -1.3923466 | 0.16381745 | 0.75321592 |
| ZNFX1 | 27.548261 | -0.6015947 | 0.55419596 | -1.0855271 | 0.27768827 | 0.84994492 |
| ZFAS1 | 29.0048419 | -0.3117923 | 0.35746158 | -0.8722401 | 0.38307743 | 0.90868001 |
| B4GALT5 | 73.438484 | 0.9891141 | 0.35970996 | 2.74975456 | 0.00596399 | 0.19198507 |
| SLC9A8 | 27.3603968 | -0.6063602 | 0.51489232 | -1.1776447 | 0.23893827 | 0.83267279 |
| RNF114 | 25.8553931 | 0.70451172 | 0.51950046 | 1.356133 | 0.17505685 | 0.76214163 |
| CEBPB | 81.0561923 | 0.93235174 | 0.72982375 | 1.2775026 | 0.20142486 | 0.78554277 |
| PTPN1 | 94.152081 | -0.3327506 | 0.3169621 | -1.0498121 | 0.29380453 | 0.86027209 |
| ADNP | 69.1576848 | -0.1142487 | 0.35757094 | -0.3195134 | 0.74933723 | 0.97818861 |
| DPM1 | 19.2194007 | -0.8340063 | 0.43141633 | -1.9331821 | 0.05321377 | 0.551221 |
| NFATC2 | 85.0128848 | -0.581117 | 0.44042487 | -1.3194464 | 0.18701993 | 0.77798634 |
| ZFP64 | 9.5641102 | -0.190163 | 0.56181191 | -0.3384817 | 0.73500023 | NA |
| TSHZ2 | 26.0432399 | -0.7794924 | 0.89033923 | -0.8755004 | 0.38130169 | 0.90797054 |
| ZNF217 | 38.1761567 | 0.12468752 | 0.44657953 | 0.27920563 | 0.78008702 | 0.98264092 |
| CASS4 | 13.8310665 | 0.69224432 | 0.80621647 | 0.85863331 | 0.39054285 | 0.90910959 |
| RTF2 | 18.6251502 | 0.48826411 | 0.43690983 | 1.11753977 | 0.2637636 | 0.84928231 |
| RBM38 | 12.7567846 | -1.163417 | 0.68126297 | -1.7077355 | 0.08768543 | NA |
| PCK1 | 21.4991055 | 2.51403768 | 1.29439631 | 1.94224725 | 0.05210718 | 0.54907299 |
| ZBP1 | 12.0877386 | -0.9548937 | 0.90832387 | -1.05127 | 0.29313459 | NA |

| | | | | | | |
|---|---|---|---|---|---|---|
| RAB22A | 27.8329707 | -0.3126972 | 0.42964413 | -0.727805 | 0.46673293 | 0.9314919 |
| VAPB | 28.0076965 | 0.06134162 | 0.39360149 | 0.15584704 | 0.87615359 | 0.98939075 |
| GNAS | 93.2498604 | -0.462841 | 0.30172294 | -1.5339933 | 0.1250313 | 0.71776265 |
| NELFCD | 11.6713924 | 0.33793135 | 0.53300014 | 0.63401738 | 0.52606948 | NA |
| CTSZ | 45.5670933 | 0.25527472 | 0.49218566 | 0.51865534 | 0.60400111 | 0.95688177 |
| ATP5F1E | 63.4763222 | -0.2801343 | 0.53040853 | -0.5281483 | 0.59739644 | 0.95538391 |
| ZNF831 | 38.1870236 | -0.443612 | 0.42873308 | -1.0347044 | 0.300807 | 0.86225566 |
| MIR646HG | 37.2678105 | 1.24354436 | 0.79358523 | 1.56699535 | 0.11711578 | 0.70122655 |
| TAF4 | 16.5584751 | 0.03328841 | 0.45904377 | 0.07251685 | 0.94219059 | 0.99514836 |
| PSMA7 | 30.1506063 | 0.06569964 | 0.47873323 | 0.13723644 | 0.89084391 | 0.99080204 |
| SS18L1 | 12.2524211 | -0.518655 | 0.52517732 | -0.9875808 | 0.32335799 | NA |
| MTG2 | 10.6257674 | -0.3993608 | 0.56934903 | -0.7014341 | 0.48303214 | NA |
| OSBPL2 | 15.4842528 | 0.05745995 | 0.45335597 | 0.12674357 | 0.89914336 | 0.99179975 |
| OGFR | 21.873141 | -0.1344285 | 0.57936589 | -0.2320269 | 0.81651715 | 0.98320463 |
| DIDO1 | 41.1651271 | 0.21958155 | 0.31229897 | 0.70311329 | 0.48198515 | 0.93382901 |
| GID8 | 11.7969772 | -0.4642452 | 0.51387068 | -0.903428 | 0.36629876 | NA |
| YTHDF1 | 9.64516422 | -0.7850384 | 0.56425966 | -1.3912715 | 0.16414311 | NA |
| ARFGAP1 | 11.9070313 | -0.1509635 | 0.75280258 | -0.2005352 | 0.84106199 | NA |
| PPDPF | 23.6273511 | -0.1499365 | 0.57328023 | -0.2615414 | 0.79367506 | 0.98320463 |
| ZBTB46 | 15.926921 | 0.80121844 | 0.85205458 | 0.94033699 | 0.34704473 | 0.88790825 |
| TPD52L2 | 18.221343 | 0.0411068 | 0.52331071 | 0.07855143 | 0.93738943 | 0.99514836 |
| DNAJC5 | 66.0712173 | -0.4829675 | 0.35862428 | -1.3467229 | 0.17806951 | 0.76504408 |
| PRPF6 | 17.461214 | 0.51385613 | 0.46093043 | 1.11482363 | 0.26492601 | 0.84994492 |
| RGS19 | 17.2570534 | -0.0015801 | 0.57940598 | -0.0027272 | 0.99782405 | 0.99892869 |
| PCMTD2 | 17.271212 | -0.0270298 | 0.47680786 | -0.056689 | 0.95479294 | 0.99514836 |
| BSG | 21.6307872 | 0.04330613 | 0.5329901 | 0.08125129 | 0.93524211 | 0.99514836 |
| RNF126 | 13.7750413 | -0.7247455 | 0.57049041 | -1.2703904 | 0.20394562 | 0.78882329 |
| PTBP1 | 24.2256211 | 0.1037421 | 0.48139637 | 0.21550245 | 0.8293756 | 0.98320463 |
| CFD | 19.8540108 | 2.79384883 | 0.81739378 | 3.41799618 | 0.00063084 | 0.05005251 |
| R3HDM4 | 9.23204061 | -0.1487656 | 0.74126024 | -0.2006929 | 0.84093874 | NA |
| ARID3A | 31.1685431 | 0.83656397 | 0.44594578 | 1.87593201 | 0.06066462 | 0.57281427 |
| TMEM259 | 28.5694906 | 0.36706156 | 0.39272192 | 0.93466023 | 0.34996343 | 0.89048867 |
| CNN2 | 18.0913224 | -0.2232374 | 0.49197132 | -0.453761 | 0.65000087 | 0.96084839 |
| ABCA7 | 15.6911648 | 0.50657818 | 0.60167822 | 0.84194203 | 0.3998204 | 0.91508218 |
| ARHGAP45 | 36.9819793 | -0.4309646 | 0.39303235 | -1.0965118 | 0.27285486 | 0.84994492 |
| POLR2E | 12.9969254 | 0.26397787 | 0.55892198 | 0.47229825 | 0.63671392 | 0.95933464 |
| GPX4 | 29.4224227 | 0.24397519 | 0.39857068 | 0.61212529 | 0.54045487 | 0.95415437 |
| SBNO2 | 21.3752469 | 1.19497622 | 0.52747668 | 2.2654579 | 0.02348459 | 0.3948379 |
| STK11 | 12.9356564 | -0.1585893 | 0.57998762 | -0.2734357 | 0.78451834 | NA |
| ATP5F1D | 20.5729057 | 0.29996855 | 0.48910661 | 0.6132989 | 0.53967872 | 0.95415437 |
| MIDN | 34.5318569 | -0.5095498 | 0.50835224 | -1.0023559 | 0.31617175 | 0.87263466 |
| CIRBP | 34.2750371 | -0.7864587 | 0.50921189 | -1.5444626 | 0.12247631 | 0.7127065 |
| MUM1 | 10.3758653 | 0.25657273 | 0.57858859 | 0.44344588 | 0.65744326 | NA |
| DAZAP1 | 23.2560928 | 0.11408575 | 0.41873506 | 0.2724533 | 0.78527349 | 0.98264092 |
| RPS15 | 137.962826 | -0.2973025 | 0.50782148 | -0.5854468 | 0.5582473 | 0.95415437 |
| C19orf25 | 14.441836 | 0.24502546 | 0.48182776 | 0.5085333 | 0.61107939 | 0.95688177 |
| TCF3 | 16.5579962 | -1.8541102 | 0.59725281 | -3.1043977 | 0.00190667 | 0.09861218 |
| REXO1 | 9.64883216 | 0.0719652 | 0.6543401 | 0.10998134 | 0.91242418 | NA |
| ABHD17A | 22.460893 | 0.61753701 | 0.6663915 | 0.92668802 | 0.35408854 | 0.89109867 |
| CSNK1G2 | 23.6326588 | 0.06050818 | 0.46066668 | 0.13134915 | 0.89549911 | 0.99115168 |
| BTBD2 | 10.1284332 | -0.1987893 | 0.55750082 | -0.3565722 | 0.72141213 | NA |
| MKNK2 | 12.3715167 | -0.54154 | 0.52724931 | -1.0271043 | 0.30437136 | NA |
| MOB3A | 39.2035349 | -1.1395636 | 0.50779106 | -2.2441585 | 0.0248222 | 0.39492781 |
| AP3D1 | 24.8181787 | -0.4523004 | 0.48077209 | -0.9407792 | 0.34681803 | 0.88790825 |

| | | | | | | |
|---|---|---|---|---|---|---|
| DOT1L | 18.2826363 | -0.8361947 | 0.58552702 | -1.4281061 | 0.15326131 | 0.74419469 |
| PLEKHJ1 | 10.6204347 | -0.1516251 | 0.60452112 | -0.2508185 | 0.80195447 | NA |
| AC005258.1 | 11.0547456 | -0.2963782 | 0.67677337 | -0.4379283 | 0.66143826 | NA |
| LSM7 | 17.7384027 | -0.6560621 | 0.63222378 | -1.0377055 | 0.29940716 | 0.86225566 |
| SPPL2B | 9.66666268 | -1.0418445 | 0.53120389 | -1.9612892 | 0.04984529 | NA |
| GADD45B | 18.9755892 | -0.6172694 | 0.64392593 | -0.9586031 | 0.33775872 | 0.88287071 |
| GNG7 | 31.8455337 | -0.1325821 | 0.61072155 | -0.217091 | 0.82813744 | 0.98320463 |
| SGTA | 8.24784764 | -0.5972904 | 0.58464999 | -1.0216205 | 0.30696057 | NA |
| AES | 45.7026308 | -0.7506309 | 0.45807603 | -1.6386601 | 0.10128407 | 0.67034929 |
| GNA15 | 13.1181186 | 2.31852955 | 0.82443664 | 2.81225923 | 0.00491948 | 0.17340397 |
| S1PR4 | 12.2824111 | -0.4620964 | 0.6431853 | -0.7184498 | 0.47247999 | NA |
| NCLN | 9.96929175 | 0.22343108 | 0.54780741 | 0.40786429 | 0.68337331 | NA |
| NFIC | 33.0343862 | 0.27343846 | 0.36165186 | 0.75608201 | 0.44960003 | 0.92914597 |
| MFSD12 | 19.0583935 | -0.0592814 | 0.50593622 | -0.1171718 | 0.90672395 | 0.99368779 |
| PIP5K1C | 13.2744286 | 0.04573576 | 0.61861493 | 0.07393253 | 0.94106407 | 0.99514836 |
| EEF2 | 69.6887249 | -0.1469999 | 0.57604878 | -0.2551866 | 0.79857896 | 0.98320463 |
| ZBTB7A | 28.6751624 | -0.540564 | 0.53018588 | -1.0195744 | 0.30793034 | 0.8673137 |
| MAP2K2 | 23.2743261 | -0.5921688 | 0.40099519 | -1.4767478 | 0.13974324 | 0.73044824 |
| SH3GL1 | 20.8999202 | -0.4143943 | 0.44304762 | -0.9353269 | 0.34961987 | 0.89048867 |
| MYDGF | 24.2115712 | 0.54640347 | 0.57796227 | 0.94539644 | 0.34445652 | 0.8871109 |
| DPP9 | 14.8627589 | -0.2362433 | 0.51598298 | -0.457851 | 0.64705946 | 0.96049083 |
| PLIN3 | 10.3494226 | 0.5265263 | 0.67622947 | 0.77862075 | 0.43620316 | NA |
| ARRDC5 | 10.8501382 | -0.4544831 | 0.74126904 | -0.6131149 | 0.53980038 | NA |
| KDM4B | 48.7852457 | 0.14671596 | 0.40403081 | 0.36313061 | 0.71650732 | 0.97348719 |
| SAFB2 | 25.7629781 | -0.3434537 | 0.39548372 | -0.8684395 | 0.3851538 | 0.90910959 |
| SAFB | 36.6176501 | 0.08596222 | 0.34472215 | 0.24936669 | 0.80307715 | 0.98320463 |
| RPL36 | 60.028814 | -0.1269712 | 0.56628091 | -0.2242194 | 0.82258658 | 0.98320463 |
| LONP1 | 14.3466487 | -0.2593418 | 0.47150055 | -0.5500349 | 0.58229544 | 0.95415437 |
| RANBP3 | 12.6333909 | 0.18800873 | 0.5046994 | 0.37251626 | 0.70950851 | NA |
| RFX2 | 15.7535206 | 1.361618 | 0.87520126 | 1.55577701 | 0.11976113 | 0.70431251 |
| MLLT1 | 9.19995768 | 0.54725101 | 0.55813898 | 0.98049236 | 0.32684314 | NA |
| KHSRP | 11.6077957 | -0.1326465 | 0.61386631 | -0.2160838 | 0.82892245 | NA |
| DENND1C | 14.3680275 | -0.1120806 | 0.52744975 | -0.2124953 | 0.83172061 | 0.98320463 |
| C3 | 15.8298738 | 3.86441682 | 1.18433643 | 3.2629384 | 0.00110263 | 0.07001544 |
| VAV1 | 70.9774705 | 0.06824051 | 0.3975195 | 0.17166582 | 0.86370026 | 0.98751288 |
| INSR | 37.8841666 | -0.0804841 | 0.47701788 | -0.1687235 | 0.86601411 | 0.98751288 |
| AC119396.1 | 22.8367392 | -0.0302973 | 0.56076269 | -0.0540288 | 0.95691221 | 0.99514836 |
| AC008878.3 | 11.9292478 | 0.28382358 | 0.64676168 | 0.43883797 | 0.66077895 | NA |
| MCOLN1 | 12.8522411 | 0.99293247 | 0.73348631 | 1.35371643 | 0.17582686 | NA |
| PNPLA6 | 18.0165357 | 0.3975484 | 0.49241601 | 0.80734255 | 0.41946916 | 0.91985469 |
| STXBP2 | 13.300885 | 0.62528182 | 0.87363753 | 0.71572226 | 0.47416286 | 0.9314919 |
| MAP2K7 | 10.6780099 | -0.1854727 | 0.63565726 | -0.291781 | 0.77045405 | NA |
| ELAVL1 | 19.1306291 | 0.0635781 | 0.47134354 | 0.13488697 | 0.89270124 | 0.99080204 |
| CERS4 | 17.7105441 | -0.2741189 | 0.47984643 | -0.5712639 | 0.56782078 | 0.95415437 |
| RPS28 | 24.5294714 | 0.12767389 | 0.80076338 | 0.15944023 | 0.87332205 | 0.98813417 |
| RAB11B | 10.6496307 | -0.5492272 | 0.65823295 | -0.8343964 | 0.40405762 | NA |
| 2-Mar | 12.3925211 | 0.95204436 | 0.65664369 | 1.44986446 | 0.14709632 | NA |
| HNRNPM | 59.712767 | -0.0092808 | 0.28518358 | -0.0325432 | 0.97403885 | 0.9973754 |
| PRAM1 | 22.9383041 | 2.00545918 | 0.71759095 | 2.79471081 | 0.00519462 | 0.17966878 |
| MYO1F | 140.991163 | 0.73768323 | 0.37429021 | 1.97088571 | 0.04873695 | 0.53486612 |
| ZNF317 | 8.66407895 | 0.3128242 | 0.63419719 | 0.49326015 | 0.62182878 | NA |
| ZNF266 | 14.3552891 | -0.2161484 | 0.4663393 | -0.4635002 | 0.64300586 | 0.95992767 |
| ZNF121 | 11.6680355 | -1.2305994 | 0.49416237 | -2.4902733 | 0.01276449 | NA |
| ZNF562 | 18.3839186 | -0.9635171 | 0.49500968 | -1.9464611 | 0.05159938 | 0.54907299 |

| | | | | | | |
|---|---|---|---|---|---|---|
| ZNF846 | 15.1706536 | 0.29829339 | 0.49836093 | 0.5985489 | 0.54947374 | 0.95415437 |
| UBL5 | 11.1394567 | 0.64648659 | 0.75322055 | 0.85829654 | 0.39072874 | NA |
| PIN1 | 9.26741069 | 0.0507584 | 0.59409253 | 0.08543855 | 0.93191275 | NA |
| C19orf66 | 17.5088716 | -1.0712467 | 0.58502114 | -1.8311248 | 0.06708192 | 0.58951068 |
| EIF3G | 9.75093681 | 0.5926549 | 0.62662354 | 0.94579099 | 0.3442552 | NA |
| DNMT1 | 25.611321 | -0.130669 | 0.38024657 | -0.3436427 | 0.73111498 | 0.97605445 |
| ICAM1 | 10.1917751 | 1.28539928 | 0.84598641 | 1.51940889 | 0.12865961 | NA |
| ICAM3 | 10.2870462 | 0.06331252 | 0.9110913 | 0.06949087 | 0.9445989 | NA |
| TYK2 | 17.6937253 | 0.1468148 | 0.4391434 | 0.33432087 | 0.73813745 | 0.97605445 |
| CDC37 | 22.2775737 | 0.19153417 | 0.40846421 | 0.46891297 | 0.63913184 | 0.95933464 |
| PDE4A | 23.6392764 | 1.26375429 | 0.78697606 | 1.60583574 | 0.10831 | 0.68204217 |
| KRI1 | 9.23077303 | -0.9851748 | 0.60314939 | -1.6333844 | 0.10238818 | NA |
| CDKN2D | 11.7461181 | -1.2357832 | 0.78697797 | -1.5702894 | 0.11634779 | NA |
| SLC44A2 | 27.7287732 | -0.9312702 | 0.5165774 | -1.8027699 | 0.07142436 | 0.60903299 |
| ILF3 | 35.3446452 | -0.5561141 | 0.42123537 | -1.320198 | 0.18676891 | 0.77798634 |
| DNM2 | 82.2706672 | 0.106569 | 0.31673143 | 0.33646486 | 0.73652034 | 0.97605445 |
| CARM1 | 9.81195688 | -0.5974703 | 0.69555 | -0.8589898 | 0.39034616 | NA |
| SMARCA4 | 33.5774903 | -0.4464988 | 0.42362065 | -1.0540062 | 0.2918801 | 0.85935767 |
| LDLR | 9.2352389 | -1.0189631 | 0.69108866 | -1.4744318 | 0.14036537 | NA |
| PRKCSH | 10.9500322 | 0.05732147 | 0.58854921 | 0.09739452 | 0.9224131 | NA |
| ZNF439 | 10.9077386 | -0.3497615 | 0.62248951 | -0.5618753 | 0.574201 | NA |
| ZNF44 | 29.8491221 | -0.1478188 | 0.53852633 | -0.2744876 | 0.78370994 | 0.98264092 |
| ZNF490 | 12.0547146 | 0.28254454 | 0.53555777 | 0.52757061 | 0.5977974 | NA |
| ZNF791 | 17.6530283 | 0.09218355 | 0.51165539 | 0.18016727 | 0.85702126 | 0.98751288 |
| MAN2B1 | 17.7794464 | -1.3832161 | 0.55652876 | -2.4854352 | 0.01293932 | 0.3086473 |
| TNPO2 | 11.4745299 | -0.284826 | 0.62846163 | -0.4532114 | 0.6503965 | NA |
| TRIR | 32.2942173 | -0.6862964 | 0.52169513 | -1.3155123 | 0.18833778 | 0.77798634 |
| HOOK2 | 9.56368295 | -0.293813 | 0.58800592 | -0.4996769 | 0.61730262 | NA |
| JUNB | 37.1840094 | 0.53603337 | 0.63693642 | 0.84158066 | 0.40002272 | 0.91508218 |
| CALR | 40.3513339 | 0.35477291 | 0.57683069 | 0.61503821 | 0.53852949 | 0.95415437 |
| RAD23A | 14.5056953 | -0.5872206 | 0.5126297 | -1.1455065 | 0.25199942 | 0.84008912 |
| GADD45GIP1 | 15.5148395 | 0.17296436 | 0.59061065 | 0.29285682 | 0.76963159 | 0.98264092 |
| LYL1 | 11.6223968 | -0.1390219 | 0.71252965 | -0.1951103 | 0.84530658 | NA |
| IER2 | 25.0830881 | -0.1531546 | 0.40927297 | -0.3742114 | 0.70824706 | 0.97184211 |
| CCDC130 | 12.6873443 | -0.6331395 | 0.53039625 | -1.1937103 | 0.23259129 | NA |
| C19orf53 | 17.2510834 | 0.27174446 | 0.66770882 | 0.40698048 | 0.68402232 | 0.9673431 |
| CC2D1A | 8.58761829 | -0.1180621 | 0.60024704 | -0.1966892 | 0.8440708 | NA |
| RFX1 | 12.4671913 | 0.42839317 | 0.57828032 | 0.74080537 | 0.45881146 | NA |
| IL27RA | 12.9564015 | 0.70680604 | 0.70443319 | 1.00336844 | 0.31568312 | 0.87263466 |
| PRKACA | 13.4561184 | 0.29894402 | 0.68092073 | 0.4390291 | 0.66064045 | 0.96298563 |
| ADGRE5 | 98.9967138 | -0.6622125 | 0.57306637 | -1.1555598 | 0.24786126 | 0.83817091 |
| DDX39A | 9.00485703 | 0.07762231 | 0.59530089 | 0.13039173 | 0.89625651 | NA |
| PKN1 | 32.4431989 | 0.05924027 | 0.53926537 | 0.10985364 | 0.91252545 | 0.99498068 |
| DNAJB1 | 15.0160949 | -0.1852198 | 0.84918959 | -0.2181136 | 0.8273406 | 0.98320463 |
| TECR | 12.490858 | -0.177937 | 0.59530057 | -0.2989028 | 0.76501424 | NA |
| NDUFB7 | 13.8418882 | -0.2138489 | 0.5594428 | -0.3822534 | 0.70227345 | 0.97086717 |
| ZNF333 | 10.4614181 | -0.3981123 | 0.58221714 | -0.6837866 | 0.49410996 | NA |
| ADGRE2 | 53.7870068 | 1.12421917 | 0.49527839 | 2.26987324 | 0.02321528 | 0.39288481 |
| BRD4 | 63.9911404 | -0.5989141 | 0.26349732 | -2.2729417 | 0.02302969 | 0.3921425 |
| AKAP8 | 14.5410308 | -0.6403999 | 0.45593639 | -1.4045818 | 0.16014569 | 0.74887162 |
| AKAP8L | 28.2730766 | -0.5010118 | 0.4391468 | -1.1408754 | 0.25392176 | 0.84026145 |
| RASAL3 | 27.6943892 | -0.0484235 | 0.3949869 | -0.1225952 | 0.90242766 | 0.99262388 |
| TPM4 | 58.9889862 | -0.3234624 | 0.49895068 | -0.6482853 | 0.51680045 | 0.94596835 |
| RAB8A | 11.9284289 | 0.63448693 | 0.55950174 | 1.13402136 | 0.25678559 | NA |

| | | | | | | |
|---|---|---|---|---|---|---|
| HSH2D | 29.4065699 | -0.0900379 | 0.5324144 | -0.1691124 | 0.8657082 | 0.98751288 |
| AP1M1 | 10.1604646 | -0.9292585 | 0.5724426 | -1.6233217 | 0.10452066 | NA |
| KLF2 | 45.9772739 | -1.1421368 | 0.64799352 | -1.7625744 | 0.0779723 | 0.62717836 |
| EPS15L1 | 26.5192983 | -0.2495681 | 0.45929756 | -0.543369 | 0.58687574 | 0.95415437 |
| SIN3B | 14.1496063 | 0.01274817 | 0.48004354 | 0.02655629 | 0.97881364 | 0.99758514 |
| MYO9B | 138.791321 | -0.8088812 | 0.45971428 | -1.7595303 | 0.07848748 | 0.62767299 |
| BST2 | 26.3102375 | 0.28164498 | 0.59871088 | 0.47041901 | 0.63805569 | 0.95933464 |
| SLC27A1 | 9.40381292 | -0.1025431 | 0.56445126 | -0.1816686 | 0.85584281 | NA |
| PGLS | 14.0160471 | 0.07709832 | 0.62405209 | 0.12354468 | 0.9016758 | 0.99261465 |
| COLGALT1 | 21.4801696 | 0.65787969 | 0.60647088 | 1.08476715 | 0.2780248 | 0.84994492 |
| FCHO1 | 12.1965013 | -0.5919178 | 0.63530924 | -0.9317004 | 0.3514914 | NA |
| JAK3 | 38.9378133 | 0.52810437 | 0.5157136 | 1.02402646 | 0.30582278 | 0.8673137 |
| RPL18A | 10.4889781 | 0.03441273 | 0.65926079 | 0.05219897 | 0.95837015 | NA |
| ARRDC2 | 15.2614583 | -0.267423 | 0.44014658 | -0.607577 | 0.54346804 | 0.95415437 |
| IL12RB1 | 10.2677035 | -0.4520509 | 0.59195524 | -0.7636573 | 0.4450715 | NA |
| MAST3 | 19.8963036 | -0.7919634 | 0.62864735 | -1.2597895 | 0.20774529 | 0.79343028 |
| JUND | 55.9291037 | -0.9992738 | 0.51728634 | -1.9317615 | 0.05338895 | 0.551221 |
| LSM4 | 14.78379 | -0.1204992 | 0.50246276 | -0.2398172 | 0.810472 | 0.98320463 |
| PGPEP1 | 15.3570428 | -0.3375832 | 0.57981718 | -0.5822236 | 0.56041607 | 0.95415437 |
| LRRC25 | 18.0873861 | 0.89463604 | 0.822697 | 1.08744294 | 0.2768411 | 0.84994492 |
| SSBP4 | 22.5953045 | -0.502895 | 0.47573502 | -1.0570905 | 0.29047027 | 0.85822878 |
| ELL | 30.2447053 | 0.144005 | 0.38322576 | 0.37577067 | 0.7070874 | 0.97184211 |
| FKBP8 | 14.5356069 | -0.4618006 | 0.59077076 | -0.7816918 | 0.43439573 | 0.92317434 |
| KXD1 | 8.10223498 | -0.3694916 | 0.60734257 | -0.6083743 | 0.54293928 | NA |
| UBA52 | 47.0233384 | 0.08035437 | 0.62488745 | 0.12859016 | 0.89768196 | 0.99141995 |
| REX1BD | 10.4923287 | 0.40585928 | 0.57523219 | 0.70555733 | 0.48046347 | NA |
| CRTC1 | 21.213854 | 0.32957058 | 0.51833563 | 0.63582467 | 0.5248907 | 0.95007724 |
| UPF1 | 14.2129474 | -0.1498136 | 0.48177848 | -0.3109596 | 0.75583134 | 0.97989151 |
| COPE | 14.5997853 | -0.1345402 | 0.45929551 | -0.2929273 | 0.76957771 | 0.98264092 |
| SUGP2 | 36.5059422 | 0.28000995 | 0.33877614 | 0.82653385 | 0.40850132 | 0.9169561 |
| SLC25A42 | 12.6464453 | -0.1215616 | 0.51989882 | -0.2338178 | 0.81512639 | NA |
| SUGP1 | 7.98064724 | 0.20664865 | 0.58219521 | 0.35494736 | 0.72262902 | NA |
| MAU2 | 23.4849867 | 0.13342072 | 0.37374635 | 0.35698201 | 0.72110528 | 0.97417403 |
| GATAD2A | 71.1396703 | 0.08594958 | 0.37814301 | 0.22729384 | 0.82019527 | 0.98320463 |
| PBX4 | 18.7709706 | -0.4125875 | 0.53614279 | -0.7695478 | 0.44156819 | 0.92899684 |
| GMIP | 16.0978447 | -0.1847354 | 0.5004701 | -0.3691238 | 0.71203542 | 0.97317955 |
| ZNF253 | 10.2921354 | -0.1479286 | 0.63359686 | -0.2334743 | 0.81539315 | NA |
| ZNF90 | 47.5923892 | -0.2804285 | 0.43010997 | -0.6519927 | 0.5144059 | 0.94596835 |
| ZNF430 | 26.7094122 | -0.2127313 | 0.43281721 | -0.4915037 | 0.62307024 | 0.95783378 |
| ZNF431 | 32.5218194 | -0.3584184 | 0.4643639 | -0.771848 | 0.44020445 | 0.92861039 |
| ZNF708 | 23.0441422 | 0.15082936 | 0.45289195 | 0.33303609 | 0.73910704 | 0.97605445 |
| ZNF429 | 9.93503241 | 0.24506473 | 0.85517455 | 0.28656691 | 0.77444395 | NA |
| ZNF100 | 20.6295289 | -0.1660542 | 0.54425516 | -0.3051036 | 0.76028727 | 0.9808103 |
| ZNF43 | 17.1334377 | -0.3187116 | 0.49045764 | -0.6498249 | 0.51580536 | 0.94596835 |
| ZNF91 | 37.1863256 | -0.2456584 | 0.41544159 | -0.5913189 | 0.5543068 | 0.95415437 |
| ZNF675 | 10.7008405 | -0.724281 | 0.57851011 | -1.2519763 | 0.21057849 | NA |
| ZNF254 | 34.1965525 | 0.2316398 | 0.45920192 | 0.50443996 | 0.61395225 | 0.95688177 |
| LINC00662 | 10.5357707 | 1.1302698 | 0.61490964 | 1.83810715 | 0.06604662 | NA |
| UQCRFS1 | 12.0415641 | 0.71465783 | 0.67217782 | 1.06319758 | 0.28769236 | NA |
| C19orf12 | 10.8549905 | -0.0170129 | 0.69551485 | -0.0244609 | 0.980485 | NA |
| URI1 | 64.5757537 | -0.086795 | 0.38127522 | -0.2276441 | 0.81992297 | 0.98320463 |
| TSHZ3 | 21.6061046 | 0.27616196 | 0.57826649 | 0.47756867 | 0.63295724 | 0.95877428 |
| ZNF507 | 11.7583981 | -0.8213043 | 0.64021898 | -1.282849 | 0.19954497 | NA |
| DPY19L3 | 12.2482257 | 0.39554598 | 0.51629679 | 0.76612133 | 0.44360411 | NA |

| | | | | | | |
|---|---|---|---|---|---|---|
| ANKRD27 | 21.0404994 | 0.33334008 | 0.56746123 | 0.58742353 | 0.55691929 | 0.95415437 |
| CEP89 | 10.2905248 | 0.81485514 | 0.63376206 | 1.28574301 | 0.19853277 | NA |
| PEPD | 16.3513741 | 1.08409402 | 0.65491264 | 1.65532617 | 0.09785835 | 0.66447623 |
| LSM14A | 89.8406269 | -0.452578 | 0.28337098 | -1.5971219 | 0.11023855 | 0.69011326 |
| KIAA0355 | 25.0349774 | -0.7210582 | 0.39700397 | -1.8162494 | 0.06933212 | 0.6023296 |
| GPI | 22.5418618 | 0.11132251 | 0.50334406 | 0.22116584 | 0.82496331 | 0.98320463 |
| UBA2 | 34.1096823 | -0.0435737 | 0.34571254 | -0.1260404 | 0.89969994 | 0.99182061 |
| ZNF302 | 15.5306897 | -0.1380676 | 0.5761491 | -0.2396386 | 0.81061045 | 0.98320463 |
| GRAMD1A | 21.1010735 | 0.28193346 | 0.59837895 | 0.47116206 | 0.63752501 | 0.95933464 |
| FXYD5 | 27.8369786 | -0.1492899 | 0.68093284 | -0.2192432 | 0.82646061 | 0.98320463 |
| USF2 | 19.4658883 | 0.21850444 | 0.55099167 | 0.39656579 | 0.69168769 | 0.9685713 |
| COX6B1 | 25.427546 | -0.0699425 | 0.54201844 | -0.1290408 | 0.89732538 | 0.99137525 |
| KMT2B | 14.1350097 | -0.3227389 | 0.47197195 | -0.6838096 | 0.49409543 | 0.93769688 |
| NFKBID | 25.6829705 | 0.44906484 | 0.66656121 | 0.67370382 | 0.50049963 | 0.94001041 |
| HCST | 24.6350977 | -0.1080708 | 0.6763242 | -0.1597915 | 0.87304535 | 0.98813417 |
| TYROBP | 54.8152855 | 1.32212272 | 0.77322655 | 1.70987754 | 0.08728852 | 0.64676224 |
| TBCB | 8.6479501 | -0.5576251 | 0.59341453 | -0.939689 | 0.34737711 | NA |
| CAPNS1 | 26.9450867 | -0.0963583 | 0.48498323 | -0.1986839 | 0.84251006 | 0.9842451 |
| ZNF565 | 15.1592101 | -0.0434582 | 0.58362681 | -0.0744624 | 0.94064247 | 0.99514836 |
| ZNF146 | 14.9588354 | 0.17849825 | 0.53813974 | 0.33169499 | 0.74011958 | 0.97605445 |
| ZFP14 | 17.1775262 | -0.1271354 | 0.50871148 | -0.2499165 | 0.80265189 | 0.98320463 |
| ZNF566 | 8.65375013 | -0.133173 | 0.7037471 | -0.1892342 | 0.84990927 | NA |
| ZNF529 | 12.5646466 | 1.34326703 | 0.63715391 | 2.10823006 | 0.03501109 | NA |
| ZNF420 | 9.72113664 | 0.19004015 | 0.59235613 | 0.32082077 | 0.74834622 | NA |
| HKR1 | 13.1832461 | -0.0895444 | 0.51208098 | -0.1748637 | 0.86118673 | 0.98751288 |
| SIPA1L3 | 98.4161992 | -0.8457998 | 0.46666549 | -1.8124328 | 0.06991935 | 0.60598598 |
| KCNK6 | 12.8123271 | 1.27266188 | 0.70873093 | 1.79569118 | 0.07254365 | NA |
| PSMD8 | 15.5043584 | -0.5869135 | 0.50766741 | -1.1560985 | 0.24764086 | 0.83817091 |
| RYR1 | 11.7015147 | 0.2435243 | 0.72245411 | 0.33707927 | 0.73605715 | NA |
| MAP4K1 | 10.8372612 | -1.1631216 | 0.56166822 | -2.0708339 | 0.03837432 | NA |
| EIF3K | 18.0702218 | -0.0505752 | 0.54993757 | -0.0919654 | 0.92672554 | 0.99514836 |
| ACTN4 | 65.3956298 | 0.32153078 | 0.29724819 | 1.0816913 | 0.27938972 | 0.84994492 |
| ECH1 | 11.0399771 | 0.69433621 | 0.56440708 | 1.23020463 | 0.21862049 | NA |
| HNRNPL | 23.1278837 | -0.211383 | 0.40399497 | -0.5232318 | 0.60081295 | 0.95688177 |
| SIRT2 | 12.4531332 | 0.40242977 | 0.52952186 | 0.75998708 | 0.44726231 | NA |
| GMFG | 23.5044624 | -0.2601966 | 0.39786598 | -0.6539805 | 0.51312439 | 0.94591286 |
| SAMD4B | 26.0778295 | 0.28699912 | 0.41886425 | 0.68518409 | 0.49322776 | 0.93715971 |
| MED29 | 9.50175131 | 0.83573706 | 0.7637093 | 1.09431306 | 0.2738177 | NA |
| ZFP36 | 45.509434 | -0.0108034 | 0.65592589 | -0.0164705 | 0.98685906 | 0.99811561 |
| RPS16 | 60.6286036 | -0.7604067 | 0.55234425 | -1.3766898 | 0.16860816 | 0.7567539 |
| SUPT5H | 21.1235737 | -0.6842073 | 0.39808749 | -1.7187359 | 0.08566247 | 0.64676224 |
| ZNF780B | 23.3245608 | 0.2083953 | 0.39866924 | 0.52272731 | 0.60116403 | 0.95688177 |
| AKT2 | 17.5847926 | -0.4531935 | 0.50556551 | -0.8964091 | 0.37003432 | 0.90170407 |
| PLD3 | 31.7635516 | 0.80167332 | 0.41304629 | 1.94088011 | 0.05227282 | 0.54907299 |
| BLVRB | 22.2862529 | 0.5743728 | 0.44463313 | 1.29179037 | 0.19642975 | 0.78135737 |
| SHKBP1 | 12.1251082 | -0.1403052 | 0.62596969 | -0.2241405 | 0.82264797 | NA |
| SNRPA | 11.484031 | -0.4834272 | 0.49104171 | -0.9844932 | 0.32487305 | NA |
| AXL | 12.0979598 | 0.6322153 | 0.78759767 | 0.80271352 | 0.42214033 | NA |
| HNRNPUL1 | 26.0748069 | 0.40524699 | 0.40903413 | 0.99074126 | 0.32181194 | 0.87363761 |
| TGFB1 | 58.5256443 | 0.84011979 | 0.3281505 | 2.56016613 | 0.01046221 | 0.2803622 |
| RPS19 | 254.054996 | 0.11070457 | 0.65555825 | 0.16887068 | 0.86589836 | 0.98751288 |
| ARHGEF1 | 43.2434992 | -0.2057934 | 0.32972356 | -0.6241392 | 0.5325362 | 0.95281453 |
| RABAC1 | 22.2550244 | 0.53027082 | 0.71394349 | 0.74273501 | 0.45764213 | 0.9314919 |
| POU2F2 | 126.936333 | -2.2741778 | 0.78889013 | -2.882756 | 0.00394213 | 0.15732988 |

| | | | | | | |
|---|---|---|---|---|---|---|
| LIPE-AS1 | 14.2559972 | -0.7907612 | 0.53647926 | -1.4739828 | 0.14048622 | 0.73044824 |
| PLAUR | 65.4725239 | 1.26766328 | 0.88503111 | 1.43233753 | 0.15204725 | 0.74315008 |
| SMG9 | 10.8574882 | -1.211905 | 0.65708805 | -1.8443571 | 0.06513111 | NA |
| ZNF224 | 11.7675121 | -0.1102225 | 0.50305581 | -0.219106 | 0.82656749 | NA |
| ZNF235 | 17.6768954 | 0.78687475 | 0.51975521 | 1.51393335 | 0.13004274 | 0.72931666 |
| BCL3 | 8.91640087 | -0.2056649 | 0.63457649 | -0.3240978 | 0.74586397 | NA |
| NECTIN2 | 12.4010287 | 2.0939843 | 0.66756541 | 3.13674774 | 0.00170833 | NA |
| APOE | 97.5159736 | 1.62939095 | 1.02473741 | 1.59005706 | 0.11182194 | 0.69385269 |
| CLPTM1 | 12.4210027 | -0.2388237 | 0.50019412 | -0.477462 | 0.63303314 | NA |
| RELB | 19.9762701 | -1.0207537 | 0.74011868 | -1.3791757 | 0.1678406 | 0.7567539 |
| CLASRP | 25.2740196 | -0.4949287 | 0.43411664 | -1.1400824 | 0.25425199 | 0.84026145 |
| MARK4 | 20.7439693 | 0.38198761 | 0.46352682 | 0.82408956 | 0.40988867 | 0.91773508 |
| ERCC1 | 7.75538134 | 0.29862633 | 0.68055893 | 0.4387957 | 0.66080958 | NA |
| VASP | 35.6180083 | 0.20562887 | 0.56521124 | 0.36380889 | 0.71600073 | 0.97348719 |
| OPA3 | 9.10109221 | -0.5138917 | 0.65612869 | -0.7832178 | 0.43349921 | NA |
| EML2 | 11.799935 | -0.4547687 | 0.50404677 | -0.9022351 | 0.36693197 | NA |
| SNRPD2 | 17.1561323 | 0.32590056 | 0.7093567 | 0.45943115 | 0.64592458 | 0.96049083 |
| SYMPK | 30.3610104 | -0.608231 | 0.40875144 | -1.4880218 | 0.13674515 | 0.73044824 |
| PPP5C | 12.4825686 | 0.28123517 | 0.53328933 | 0.52735946 | 0.59794399 | NA |
| CALM3 | 16.5843949 | 0.8009786 | 0.71553878 | 1.11940628 | 0.26296685 | 0.84921816 |
| PRKD2 | 28.0369487 | -0.8625734 | 0.50043038 | -1.7236631 | 0.08476868 | 0.64676224 |
| AP2S1 | 21.3821152 | 0.89355269 | 0.59974588 | 1.4898855 | 0.13625434 | 0.73044824 |
| ARHGAP35 | 39.7063777 | 0.16815196 | 0.36828069 | 0.45658642 | 0.64796834 | 0.96049083 |
| TMEM160 | 12.8383144 | 0.2345202 | 0.62094768 | 0.37768109 | 0.70566752 | NA |
| ZC3H4 | 13.1686784 | -0.3792394 | 0.51773751 | -0.7324936 | 0.46386734 | 0.9314919 |
| SAE1 | 17.6449511 | -0.8978485 | 0.55164797 | -1.6275751 | 0.10361502 | 0.67301115 |
| C5AR1 | 44.8228985 | 0.68549801 | 0.72012137 | 0.9519201 | 0.3411375 | 0.88510795 |
| DHX34 | 13.7128996 | 0.09820201 | 0.56309789 | 0.17439598 | 0.86155428 | 0.98751288 |
| NAPA | 18.4803918 | 0.28118259 | 0.61786627 | 0.45508648 | 0.64904703 | 0.96084839 |
| NOP53 | 11.82462 | -0.2000715 | 0.66962691 | -0.2987806 | 0.76510748 | NA |
| SELENOW | 19.2646005 | -0.1906715 | 0.44600714 | -0.4275076 | 0.66900962 | 0.96412305 |
| CARD8 | 74.511901 | 0.30434588 | 0.29763804 | 1.02253689 | 0.30652686 | 0.8673137 |
| EMP3 | 59.6781395 | -1.4005557 | 0.77747592 | -1.8014137 | 0.0716377 | 0.60991239 |
| KDELR1 | 9.08827787 | 0.56526109 | 0.61098833 | 0.92515856 | 0.35488343 | NA |
| RPL18 | 76.5064335 | -0.3519381 | 0.56102301 | -0.6273148 | 0.53045288 | 0.95245713 |
| NUCB1 | 20.2408472 | 0.43759828 | 0.49789212 | 0.87890179 | 0.37945453 | 0.90600059 |
| BAX | 17.8098587 | -0.2772304 | 0.55674936 | -0.4979447 | 0.61852305 | 0.95688177 |
| FTL | 806.861412 | 1.18401868 | 0.63522208 | 1.86394447 | 0.0623295 | 0.57971676 |
| GYS1 | 14.4831094 | -0.1262198 | 0.58279143 | -0.216578 | 0.82853726 | 0.98320463 |
| SNRNP70 | 46.6439867 | -0.0114183 | 0.34853446 | -0.0327608 | 0.97386536 | 0.9973754 |
| CD37 | 40.3453711 | -1.2841259 | 0.55484133 | -2.3144021 | 0.02064568 | 0.37333155 |
| PIH1D1 | 8.9167849 | 0.29335534 | 0.66739633 | 0.43955192 | 0.66026167 | NA |
| RPL13A | 106.402612 | -0.2284745 | 0.51081754 | -0.4472721 | 0.65467859 | 0.96178254 |
| RPS11 | 78.9815612 | -0.5122874 | 0.58634162 | -0.8737012 | 0.38228097 | 0.90868001 |
| FCGRT | 39.3779201 | 1.18015676 | 0.46452421 | 2.54057103 | 0.01106716 | 0.2861945 |
| NOSIP | 11.777514 | 1.14224138 | 0.56826994 | 2.01003308 | 0.04442769 | NA |
| PRR12 | 13.1745559 | -0.1282053 | 0.57625789 | -0.222479 | 0.82394105 | 0.98320463 |
| IRF3 | 21.5331142 | -0.0015263 | 0.41990437 | -0.0036348 | 0.99709986 | 0.99881813 |
| PRMT1 | 9.60679497 | -0.4439078 | 0.54109939 | -0.8203813 | 0.4119988 | NA |
| AP2A1 | 18.8874921 | 0.45248728 | 0.5899286 | 0.76702041 | 0.44306938 | 0.92899684 |
| MED25 | 11.222086 | -0.5093658 | 0.58689265 | -0.8679028 | 0.38544754 | NA |
| PNKP | 11.3555728 | -0.7399672 | 0.63351588 | -1.1680325 | 0.24279364 | NA |
| VRK3 | 11.7785315 | 0.03379723 | 0.58077685 | 0.05819314 | 0.95359479 | NA |
| NR1H2 | 9.88071188 | -0.3978958 | 0.56883062 | -0.6994979 | 0.48424093 | NA |

| | | | | | | |
|---|---|---|---|---|---|---|
| EMC10 | 18.9033772 | -0.5444879 | 0.45884802 | -1.186641 | 0.23536926 | 0.82963916 |
| SIGLEC9 | 12.3413673 | 1.24020941 | 0.77458961 | 1.60111806 | 0.10935077 | NA |
| ETFB | 14.5043032 | 1.03112229 | 0.52917411 | 1.94855016 | 0.05134917 | 0.5485836 |
| NKG7 | 50.7548383 | 0.77926313 | 0.95255913 | 0.81807324 | 0.41331537 | 0.91894074 |
| SIGLEC10 | 18.8942396 | -0.1585114 | 0.71860925 | -0.2205807 | 0.82541889 | 0.98320463 |
| FPR1 | 21.4613297 | 0.79560944 | 0.7148357 | 1.11299623 | 0.26571005 | 0.84994492 |
| FPR3 | 30.5545493 | 1.30945079 | 0.8795364 | 1.48879658 | 0.13654094 | 0.73044824 |
| ZNF577 | 8.70389996 | 1.09759958 | 0.874963 | 1.25445257 | 0.20967756 | NA |
| PPP2R1A | 22.6770142 | -0.2742979 | 0.4420965 | -0.620448 | 0.53496288 | 0.95378558 |
| ZNF766 | 9.18845973 | -0.1536137 | 0.60954643 | -0.2520131 | 0.80103091 | NA |
| ZNF83 | 32.1274541 | 0.05294472 | 0.40360513 | 0.1311795 | 0.89563331 | 0.99115168 |
| ZNF611 | 22.1328148 | -0.1300635 | 0.38179378 | -0.3406644 | 0.73335626 | 0.97605445 |
| ZNF160 | 17.832572 | 0.03433483 | 0.62627113 | 0.05482423 | 0.9562785 | 0.99514836 |
| ZNF761 | 11.3721051 | -0.0034208 | 0.50963696 | -0.0067123 | 0.99464442 | NA |
| ZNF331 | 42.748328 | 0.38112278 | 0.9543696 | 0.39934506 | 0.68963897 | 0.9673431 |
| MYADM | 14.9793476 | -1.1306965 | 0.79868533 | -1.4156971 | 0.15686421 | 0.74486619 |
| OSCAR | 11.049784 | 1.61645574 | 0.8170615 | 1.97837708 | 0.04788618 | NA |
| PRPF31 | 9.7875479 | -0.3002246 | 0.64063798 | -0.4686337 | 0.63933147 | NA |
| MBOAT7 | 19.0538917 | 0.11564969 | 0.47613383 | 0.24289323 | 0.80808811 | 0.98320463 |
| RPS9 | 163.188673 | -0.2259951 | 0.51612862 | -0.4378659 | 0.6614835 | 0.96298563 |
| LILRB3 | 16.7607133 | 0.77874968 | 0.66095392 | 1.17822083 | 0.23870858 | 0.83267279 |
| LILRA6 | 12.6979771 | 1.59467819 | 1.03943504 | 1.53417783 | 0.12498591 | NA |
| LILRB2 | 30.8553083 | 1.34547409 | 0.93585809 | 1.4376903 | 0.15052195 | 0.74043418 |
| LAIR1 | 22.5925992 | 1.1578472 | 0.54352998 | 2.13023612 | 0.03315212 | 0.44911423 |
| LENG8 | 34.2868218 | -0.5961925 | 0.43321357 | -1.3762091 | 0.16875689 | 0.75680766 |
| LILRB1 | 42.0346912 | -0.5010894 | 0.74311855 | -0.6743061 | 0.50011671 | 0.94001041 |
| LILRB4 | 10.8934043 | 1.41855128 | 0.79160169 | 1.79200133 | 0.07313275 | NA |
| PPP1R12C | 15.2015816 | -0.0261804 | 0.4863582 | -0.0538295 | 0.95707097 | 0.99514836 |
| PPP6R1 | 30.3109691 | -0.2027379 | 0.38524402 | -0.5262584 | 0.59870867 | 0.95582632 |
| RPL28 | 121.905804 | 0.18570294 | 0.48851881 | 0.38013467 | 0.70384545 | 0.9714177 |
| U2AF2 | 9.81678034 | -0.933599 | 0.55541818 | -1.6808938 | 0.09278355 | NA |
| EPN1 | 13.6342253 | 0.21070201 | 0.53978757 | 0.39034246 | 0.69628333 | 0.9685713 |
| ZNF787 | 10.3792675 | -0.6648148 | 0.52219827 | -1.2731079 | 0.20297974 | NA |
| ZSCAN5A | 16.9417725 | -0.7481198 | 0.68388197 | -1.0939311 | 0.27398518 | 0.84994492 |
| ZNF264 | 9.9597382 | 0.4070934 | 0.55732404 | 0.73044292 | 0.46511949 | NA |
| AC005261.1 | 11.3465823 | 0.34915559 | 0.54197462 | 0.64422867 | 0.51942716 | NA |
| ZNF586 | 24.2947387 | -0.1497181 | 0.54998954 | -0.2722199 | 0.78545295 | 0.98264092 |
| ZNF814 | 16.786129 | -1.1201904 | 0.44965294 | -2.4912334 | 0.01273005 | 0.30497006 |
| ZNF274 | 12.7632363 | -0.7484468 | 0.49255839 | -1.5195088 | 0.12863448 | NA |
| RPS5 | 40.477814 | -0.0617539 | 0.72940737 | -0.0846631 | 0.93252926 | 0.99514836 |
| TRIM28 | 11.7224761 | -0.5456877 | 0.64010527 | -0.8524968 | 0.3939384 | NA |
| CHMP2A | 7.59080368 | 0.02347911 | 0.7142799 | 0.03287103 | 0.97377744 | NA |
| MZF1 | 9.66460134 | -0.1045975 | 0.61595533 | -0.1698134 | 0.86515686 | NA |
| RPS4Y1 | 25.2262138 | -0.1484288 | 2.07656119 | -0.0714781 | 0.94301721 | 0.99514836 |
| ZFY | 12.2147676 | 0.07501498 | 1.71580603 | 0.04371997 | 0.96512762 | NA |
| LINC00278 | 90.8510803 | 0.1657847 | 2.32968505 | 0.07116185 | 0.94326894 | 0.99514836 |
| TTTY15 | 9.34704079 | 0.57506655 | 1.74520888 | 0.32951159 | 0.74176904 | NA |
| USP9Y | 46.2959158 | 0.59925854 | 2.11557796 | 0.28325997 | 0.77697756 | 0.98264092 |
| DDX3Y | 17.0475245 | 1.19601405 | 1.86545545 | 0.64113782 | 0.52143314 | 0.94895469 |
| UTY | 101.332309 | 0.09310145 | 2.32248064 | 0.04008707 | 0.96802371 | 0.99653147 |
| TTTY14 | 29.9801024 | 0.10556043 | 1.02631566 | 0.10285376 | 0.91807904 | 0.99498068 |
| KDM5D | 14.4716788 | -0.1009011 | 1.96758775 | -0.0512816 | 0.95910109 | 0.99580965 |
| IL17RA | 55.6728486 | 1.6167671 | 0.48689386 | 3.32057405 | 0.00089833 | 0.06269654 |
| HDHD5 | 10.120865 | 0.1103357 | 0.67971863 | 0.16232554 | 0.8710495 | NA |

| | | | | | | |
|---|---|---|---|---|---|---|
| ADA2 | 33.2988863 | 1.35962443 | 0.53249848 | 2.55329262 | 0.01067098 | 0.28288008 |
| ATP6V1E1 | 19.3102737 | 0.10073512 | 0.45995659 | 0.21901006 | 0.82664221 | 0.98320463 |
| BCL2L13 | 24.5159885 | 0.25171623 | 0.48146472 | 0.52281344 | 0.60110408 | 0.95688177 |
| BID | 30.2622214 | 0.97471997 | 0.53059246 | 1.83704076 | 0.06620388 | 0.58951068 |
| MICAL3 | 29.479556 | -0.8255665 | 0.48568829 | -1.6997867 | 0.08917105 | 0.64845882 |
| PEX26 | 11.5415801 | -0.280098 | 0.64310027 | -0.4355433 | 0.6631681 | NA |
| DGCR2 | 28.5332041 | 0.41838043 | 0.36482476 | 1.14679836 | 0.25146497 | 0.84008912 |
| HIRA | 12.0540559 | -0.3274625 | 0.68066886 | -0.4810893 | 0.63045305 | NA |
| UFD1 | 11.6241958 | -0.7918711 | 0.49998584 | -1.583787 | 0.1132422 | NA |
| TXNRD2 | 10.0669921 | -0.3068626 | 0.51438411 | -0.5965631 | 0.55079911 | NA |
| COMT | 25.3338259 | 0.51900591 | 0.4807132 | 1.07965812 | 0.28029445 | 0.84994492 |
| TANGO2 | 20.8763168 | 0.30670374 | 0.45145389 | 0.67936891 | 0.49690415 | 0.93900843 |
| DGCR8 | 8.85229157 | -0.1014551 | 0.58298625 | -0.1740265 | 0.86184461 | NA |
| KLHL22 | 19.9508234 | 0.14897332 | 0.41192301 | 0.36165331 | 0.71761112 | 0.97348719 |
| MED15 | 50.1318495 | -0.110519 | 0.33467507 | -0.3302278 | 0.74122788 | 0.97605445 |
| PI4KA | 61.2246906 | 0.3670676 | 0.28083305 | 1.30706696 | 0.19118997 | 0.78122755 |
| SNAP29 | 11.7947179 | 0.23704791 | 0.48038891 | 0.49345002 | 0.62169465 | NA |
| CRKL | 11.1781156 | 0.18985519 | 0.49952433 | 0.38007197 | 0.703892 | NA |
| UBE2L3 | 37.8627683 | 0.40486816 | 0.35259171 | 1.14826343 | 0.25085983 | 0.83984169 |
| SDF2L1 | 12.2439717 | 0.54160097 | 0.90302309 | 0.59976426 | 0.54866336 | NA |
| PPIL2 | 22.2583461 | -0.3956514 | 0.40575605 | -0.9750968 | 0.32951226 | 0.88007764 |
| YPEL1 | 15.179688 | 0.31065308 | 0.52655265 | 0.58997534 | 0.55520718 | 0.95415437 |
| MAPK1 | 103.777859 | 0.10111151 | 0.34019499 | 0.29721634 | 0.76630135 | 0.98264092 |
| PPM1F | 18.840886 | 0.60871678 | 0.76727097 | 0.79335308 | 0.42757214 | 0.92129717 |
| IGLC2 | 164.887789 | -5.9243437 | 2.56745804 | -2.3074744 | 0.02102839 | 0.37539066 |
| BCR | 37.1289888 | -0.3965441 | 0.55561315 | -0.7137055 | 0.47540933 | 0.9314919 |
| CHCHD10 | 16.7366423 | 0.57477149 | 0.51638031 | 1.11307787 | 0.26567499 | 0.84994492 |
| SMARCB1 | 13.3169225 | -0.1914422 | 0.62289908 | -0.3073407 | 0.75858408 | 0.98061301 |
| DDT | 14.4418572 | 0.81778254 | 0.69231914 | 1.18122192 | 0.23751458 | 0.83171006 |
| CABIN1 | 31.5187987 | 0.22851065 | 0.34542991 | 0.66152537 | 0.50827545 | 0.94305491 |
| SNRPD3 | 15.9696478 | 0.35140598 | 0.47833931 | 0.73463747 | 0.46256031 | 0.9314919 |
| KIAA1671 | 10.7483885 | -0.0754227 | 0.68127565 | -0.1107081 | 0.91184786 | NA |
| LRP5L | 9.64159825 | 0.30483522 | 0.55943911 | 0.54489437 | 0.58582617 | NA |
| GRK3 | 154.723383 | -0.3596546 | 0.40109812 | -0.8966749 | 0.36989244 | 0.90170407 |
| HPS4 | 11.1075952 | 0.26555546 | 0.54475981 | 0.48747256 | 0.62592349 | NA |
| TPST2 | 36.2747429 | 0.61752191 | 0.43176644 | 1.43022211 | 0.15265328 | 0.74341937 |
| MIAT | 12.5331059 | 0.26420431 | 0.65527896 | 0.40319364 | 0.68680578 | NA |
| MIATNB | 27.452459 | 0.07319612 | 0.43953012 | 0.16653265 | 0.86773779 | 0.98751288 |
| PITPNB | 42.7770066 | 0.00366048 | 0.31598489 | 0.01158437 | 0.99075722 | 0.99827179 |
| TTC28 | 40.1001122 | -1.7945959 | 0.63758961 | -2.8146567 | 0.00488294 | 0.17334548 |
| HSCB | 9.92626822 | -0.4057862 | 0.62703166 | -0.6471542 | 0.51753212 | NA |
| XBP1 | 40.400191 | 0.35018554 | 0.99728337 | 0.35113946 | 0.72548373 | 0.9754739 |
| Z93930.2 | 18.4205371 | 0.61851855 | 0.43070272 | 1.43606837 | 0.15098289 | 0.74119263 |
| EWSR1 | 46.0951872 | -0.5765477 | 0.3067784 | -1.8793622 | 0.06019505 | 0.57281427 |
| AP1B1 | 37.5778288 | -0.017741 | 0.36840138 | -0.0481567 | 0.96159137 | 0.99633901 |
| NF2 | 15.7354208 | 0.3465819 | 0.46810362 | 0.74039568 | 0.45905994 | 0.9314919 |
| UQCR10 | 26.5655589 | 0.30744285 | 0.60164121 | 0.51100697 | 0.60934618 | 0.95688177 |
| ASCC2 | 17.9298842 | 0.23075037 | 0.5226177 | 0.44152805 | 0.65883076 | 0.96280444 |
| MTMR3 | 99.3031316 | 0.09121662 | 0.27270991 | 0.33448223 | 0.7380157 | 0.97605445 |
| TBC1D10A | 9.02879587 | 0.34337924 | 0.65009526 | 0.52819836 | 0.59736167 | NA |
| SF3A1 | 11.0952523 | -0.4394908 | 0.5258102 | -0.8358355 | 0.40324746 | NA |
| DUSP18 | 11.6405569 | -0.4275943 | 0.65987021 | -0.6479976 | 0.51698653 | NA |
| RNF185 | 14.1448689 | 0.06988441 | 0.51142041 | 0.13664768 | 0.89130929 | 0.99080204 |
| LIMK2 | 38.4347045 | 0.92403445 | 0.51566585 | 1.79192485 | 0.073145 | 0.61424045 |

| | | | | | | |
|---|---|---|---|---|---|---|
| PIK3IP1 | 22.2850756 | 0.18299178 | 0.57522959 | 0.31811955 | 0.75039425 | 0.97818861 |
| PIK3IP1-AS1 | 11.4869346 | -0.3776375 | 0.59881115 | -0.6306454 | 0.52827238 | NA |
| DRG1 | 14.7985445 | -0.019883 | 0.48678348 | -0.0408457 | 0.96741894 | 0.99653147 |
| EIF4ENIF1 | 15.8800399 | -0.3172074 | 0.48616897 | -0.6524632 | 0.5141024 | 0.94596835 |
| SFI1 | 28.3203343 | -0.3121968 | 0.37780079 | -0.826353 | 0.40860386 | 0.9169561 |
| PISD | 20.4998855 | 0.00310304 | 0.48452372 | 0.00640432 | 0.99489013 | 0.99859588 |
| PRR14L | 47.5897245 | -0.357948 | 0.31429503 | -1.1388916 | 0.25474836 | 0.84026145 |
| YWHAH | 11.3270308 | 0.00610275 | 1.10543906 | 0.00552065 | 0.99559518 | NA |
| RTCB | 21.3426364 | -0.1087784 | 0.48502428 | -0.2242741 | 0.82254406 | 0.98320463 |
| FBXO7 | 25.1468136 | -0.0346386 | 0.39861917 | -0.0868964 | 0.93075383 | 0.99514836 |
| TIMP3 | 15.1075271 | -0.2289708 | 0.63092465 | -0.3629131 | 0.71666982 | 0.97348719 |
| LARGE1 | 42.1304307 | -3.6610117 | 0.72244163 | -5.0675537 | 4.03E-07 | 0.000446 |
| HMGXB4 | 13.4962199 | -0.3198867 | 0.46097182 | -0.6939399 | 0.4877199 | 0.93554313 |
| TOM1 | 22.6379274 | 0.65364332 | 0.47948433 | 1.36322144 | 0.17281273 | 0.76081596 |
| HMOX1 | 52.1977821 | -0.5333181 | 0.67107112 | -0.7947267 | 0.42677251 | 0.92112288 |
| APOL6 | 32.2931373 | 1.02712377 | 0.48952749 | 2.09819429 | 0.03588799 | 0.4673038 |
| RBFOX2 | 17.182884 | -0.0050163 | 0.52132006 | -0.0096222 | 0.99232269 | 0.99859588 |
| APOL3 | 14.1487484 | 0.74071516 | 0.57451699 | 1.28928332 | 0.19729961 | 0.78135737 |
| APOL2 | 13.0479213 | 1.00450982 | 0.64547586 | 1.55623144 | 0.11965307 | 0.70431251 |
| MYH9 | 116.192728 | -0.0080175 | 0.36487226 | -0.0219734 | 0.98246917 | 0.99759864 |
| TXN2 | 9.53007393 | 0.03607474 | 0.60622865 | 0.05950683 | 0.95254843 | NA |
| EIF3D | 20.5712259 | 0.15203292 | 0.43109941 | 0.35266324 | 0.72434092 | 0.97499799 |
| NCF4 | 16.293769 | -0.6878686 | 0.5402962 | -1.2731324 | 0.20297106 | 0.7882399 |
| IL2RB | 13.7901517 | 0.95273855 | 0.74079538 | 1.28610217 | 0.19840741 | 0.78142991 |
| RAC2 | 36.0404739 | -0.4575381 | 0.62897825 | -0.7274308 | 0.4669621 | 0.9314919 |
| CYTH4 | 49.9911606 | -0.4803709 | 0.50518638 | -0.9508786 | 0.34166601 | 0.88510795 |
| MFNG | 8.79980223 | -0.7414997 | 0.5813269 | -1.2755296 | 0.20212185 | NA |
| GGA1 | 24.7206425 | 0.0266708 | 0.42587712 | 0.06262557 | 0.95006466 | 0.99514836 |
| LGALS1 | 49.8045515 | -0.1407331 | 0.83557892 | -0.1684259 | 0.86624823 | 0.98751288 |
| TRIOBP | 15.6881113 | 0.43016641 | 0.47492148 | 0.90576323 | 0.36506119 | 0.89868152 |
| EIF3L | 28.9634477 | -0.810815 | 0.37053044 | -2.1882548 | 0.02865104 | 0.42168851 |
| POLR2F | 8.43170989 | -0.724995 | 0.58506992 | -1.2391595 | 0.21528643 | NA |
| TMEM184B | 13.4740375 | 0.18455528 | 0.62214177 | 0.29664506 | 0.76673751 | 0.98264092 |
| TPTEP2-CSNI | 36.3585052 | 0.37155538 | 0.50306293 | 0.73858629 | 0.46015825 | 0.9314919 |
| DDX17 | 181.412545 | 0.20523349 | 0.26054067 | 0.78772151 | 0.43085962 | 0.92284736 |
| JOSD1 | 19.894369 | 0.4081841 | 0.52366876 | 0.77947002 | 0.43570289 | 0.92453213 |
| GTPBP1 | 12.9785444 | -0.0935426 | 0.56563087 | -0.1653775 | 0.8686469 | 0.98751288 |
| SUN2 | 45.5913453 | -0.5671293 | 0.40776662 | -1.3908184 | 0.16428051 | 0.75321592 |
| AL021707.1 | 11.1156748 | 0.83927782 | 0.54829852 | 1.53069503 | 0.12584478 | NA |
| CBX6 | 12.9811777 | 0.00763535 | 0.57787763 | 0.01321274 | 0.98945807 | 0.99811561 |
| APOBEC3G | 12.6110093 | 0.09455348 | 0.52896651 | 0.17875135 | 0.85813295 | NA |
| CBX7 | 12.093087 | -0.2723612 | 0.55926196 | -0.4870012 | 0.62625751 | NA |
| RPL3 | 63.9369394 | -0.0636918 | 0.62873074 | -0.1013022 | 0.91931059 | 0.99498068 |
| TAB1 | 7.3832188 | 0.14670527 | 0.61056875 | 0.24027642 | 0.81011597 | NA |
| ATF4 | 19.4794558 | -0.3100467 | 0.510089 | -0.6078287 | 0.5433011 | 0.95415437 |
| RPS19BP1 | 13.2054004 | -0.1252473 | 0.59616393 | -0.2100887 | 0.83359846 | 0.98320463 |
| GRAP2 | 77.0581675 | -0.7416213 | 0.54851414 | -1.352055 | 0.17635772 | 0.76409729 |
| TNRC6B | 282.503119 | 0.04528708 | 0.21389932 | 0.21172147 | 0.83232433 | 0.98320463 |
| MKL1 | 119.767749 | 0.04265733 | 0.24782349 | 0.17212787 | 0.863337 | 0.98751288 |
| SLC25A17 | 14.6597018 | -0.0260626 | 0.56359022 | -0.0462439 | 0.96311584 | 0.99653147 |
| ST13 | 23.0536786 | 0.37081165 | 0.4728026 | 0.78428429 | 0.43287332 | 0.92284736 |
| XPNPEP3 | 12.8723861 | 1.27882513 | 0.56318604 | 2.27069749 | 0.0231653 | NA |
| RBX1 | 23.0437637 | -0.1751257 | 0.4017379 | -0.4359202 | 0.66289461 | 0.96360881 |
| EP300 | 79.8768768 | -0.2871186 | 0.2510009 | -1.1438945 | 0.25266738 | 0.84008912 |

| | | | | | | |
|---|---|---|---|---|---|---|
| L3MBTL2 | 8.08996115 | 0.21224708 | 0.64151039 | 0.33085525 | 0.74075382 | NA |
| RANGAP1 | 10.7365817 | -0.562192 | 0.5609732 | -1.0021727 | 0.31626019 | NA |
| ZC3H7B | 14.3424155 | -0.5868231 | 0.45427192 | -1.2917882 | 0.19643049 | 0.78135737 |
| TOB2 | 12.186261 | -0.810867 | 0.52770827 | -1.536582 | 0.12439571 | NA |
| ACO2 | 23.7151266 | 0.04747229 | 0.45772324 | 0.10371395 | 0.91739636 | 0.99498068 |
| DESI1 | 11.2567833 | -0.960368 | 0.56154299 | -1.7102306 | 0.08722324 | NA |
| XRCC6 | 35.0483531 | -0.3600408 | 0.37741784 | -0.953958 | 0.34010491 | 0.8844375 |
| SNU13 | 27.6025867 | -0.333333 | 0.39080073 | -0.8529487 | 0.39368773 | 0.90940709 |
| SREBF2 | 80.927733 | -1.2325472 | 0.44832519 | -2.7492258 | 0.00597362 | 0.19198507 |
| SMDT1 | 11.765362 | -0.399846 | 0.60919045 | -0.6563564 | 0.51159484 | NA |
| TCF20 | 39.4170135 | -0.0205547 | 0.34568911 | -0.05946 | 0.95258572 | 0.99514836 |
| NFAM1 | 22.1887027 | 1.94272355 | 0.79219169 | 2.45234023 | 0.01419304 | 0.3205889 |
| RRP7A | 7.72146859 | 0.10188652 | 0.6417334 | 0.15876767 | 0.87385192 | NA |
| POLDIP3 | 13.8278402 | 0.28992157 | 0.5473643 | 0.52966839 | 0.59634187 | 0.9553365 |
| CYB5R3 | 15.9621079 | 0.47942572 | 0.51664518 | 0.92795934 | 0.35342866 | 0.89109867 |
| ARFGAP3 | 32.63561 | -0.332178 | 0.32662852 | -1.0169902 | 0.30915808 | 0.8673137 |
| PACSIN2 | 67.2413057 | -0.1016193 | 0.36230245 | -0.2804818 | 0.77910786 | 0.98264092 |
| TSPO | 48.3662884 | 1.29931783 | 0.79315659 | 1.63816053 | 0.10138821 | 0.67034929 |
| PARVB | 31.1615208 | 0.39409965 | 0.35658065 | 1.10521882 | 0.26906478 | 0.84994492 |
| PARVG | 30.5391932 | -0.1929881 | 0.3804389 | -0.5072775 | 0.61196014 | 0.95688177 |
| PRR5 | 10.3603096 | -0.4951059 | 0.64029419 | -0.7732475 | 0.43937593 | NA |
| NUP50-DT | 26.7345013 | 0.06372508 | 0.41663546 | 0.15295165 | 0.8784364 | 0.99019008 |
| NUP50 | 19.5847585 | -0.1514145 | 0.41093656 | -0.3684619 | 0.71252881 | 0.97332186 |
| KIAA0930 | 23.1844766 | 0.67516714 | 0.50347979 | 1.34100145 | 0.17991998 | 0.76767708 |
| FAM118A | 21.4834869 | 1.2925971 | 0.75241032 | 1.71794176 | 0.08580724 | 0.64676224 |
| ATXN10 | 62.3586745 | -0.0061331 | 0.27692006 | -0.0221477 | 0.98233013 | 0.99759864 |
| MIRLET7BHG | 20.9623476 | 0.28065286 | 0.6894977 | 0.40703958 | 0.68397892 | 0.9673431 |
| PPARA | 23.7588524 | -0.1697541 | 0.45933626 | -0.369564 | 0.7117074 | 0.97317955 |
| TTC38 | 13.0470139 | 0.69473876 | 0.59258253 | 1.17239157 | 0.24103988 | 0.83493679 |
| TRMU | 9.13147688 | -0.98013 | 0.51990125 | -1.8852233 | 0.05939969 | NA |
| GRAMD4 | 27.0226648 | -0.4031194 | 0.51385436 | -0.7845012 | 0.43274607 | 0.92284736 |
| CERK | 26.1880796 | 0.68410331 | 0.42190029 | 1.62148102 | 0.10491452 | 0.67519097 |
| TBC1D22A | 238.383719 | -0.6819014 | 0.39831631 | -1.7119596 | 0.08690412 | 0.64676224 |
| C22orf34 | 51.3900508 | -0.0372845 | 0.52632276 | -0.0708396 | 0.94352538 | 0.99514836 |
| BRD1 | 26.263947 | 0.41716592 | 0.39201694 | 1.06415277 | 0.2872595 | 0.85677513 |
| ZBED4 | 19.4697638 | 0.11788055 | 0.45946956 | 0.2565579 | 0.79752008 | 0.98320463 |
| PIM3 | 15.8954749 | -0.0923927 | 0.65296276 | -0.1414976 | 0.88747686 | 0.99019008 |
| TRABD | 19.5503332 | -0.2814529 | 0.58657438 | -0.4798247 | 0.63135202 | 0.95877428 |
| SELENOO | 19.7505816 | 0.51678446 | 0.48310123 | 1.06972292 | 0.28474405 | 0.85361514 |
| PLXNB2 | 23.7142512 | 0.43880708 | 0.60608637 | 0.7240009 | 0.46906518 | 0.9314919 |
| PPP6R2 | 72.6568083 | -0.1225001 | 0.3453939 | -0.3546679 | 0.72283836 | 0.97441549 |
| LMF2 | 7.15909203 | 1.65129878 | 0.82839167 | 1.99337926 | 0.04621993 | NA |
| TYMP | 33.0426412 | 1.73340357 | 0.87196252 | 1.98793357 | 0.04681904 | 0.52744812 |
| FP671120.1 | 220.216089 | -0.873567 | 0.43600093 | -2.0035898 | 0.04511402 | 0.52128421 |
| FP236383.1 | 117.551931 | -1.1210689 | 0.50614321 | -2.2149242 | 0.02676527 | 0.40712515 |
| SAMSN1 | 198.139512 | 1.20365786 | 0.71671836 | 1.67940146 | 0.09307383 | 0.6590317 |
| NRIP1 | 28.7707934 | 0.63895408 | 0.43123318 | 1.48169046 | 0.13842268 | 0.73044824 |
| USP25 | 114.128004 | 0.35389135 | 0.37996997 | 0.93136663 | 0.35166394 | 0.89109867 |
| MIR99AHG | 46.7845402 | 0.84590761 | 0.59103015 | 1.43124274 | 0.15236066 | 0.74315008 |
| LINC01684 | 21.7836713 | -0.5202941 | 0.58534549 | -0.8888666 | 0.37407477 | 0.90262385 |
| ATP5PF | 15.9177267 | 0.15706019 | 0.62223445 | 0.2524132 | 0.8007217 | 0.98320463 |
| GABPA | 12.5035188 | -0.4989172 | 0.49701249 | -1.0038323 | 0.31545944 | NA |
| APP | 73.6584919 | -0.6810916 | 0.40591455 | -1.6779188 | 0.09336296 | 0.6590317 |
| ADAMTS1 | 12.1504476 | -0.7818595 | 0.6322792 | -1.2365732 | 0.21624557 | NA |

| | | | | | | |
|---|---|---|---|---|---|---|
| AF165147.1 | 30.559713 | 0.00162265 | 0.57671516 | 0.00281361 | 0.99775506 | 0.99892869 |
| LTN1 | 25.3412526 | 0.14455124 | 0.38081119 | 0.37958768 | 0.70425151 | 0.9714177 |
| USP16 | 23.78734 | -0.196891 | 0.39788716 | -0.4948413 | 0.62071212 | 0.95688177 |
| CCT8 | 16.6704596 | 0.42092034 | 0.61799391 | 0.68110758 | 0.49580342 | 0.93805563 |
| BACH1 | 215.012951 | 0.2811087 | 0.34955895 | 0.8041811 | 0.42129239 | 0.91985469 |
| TIAM1 | 43.9474999 | 0.25674499 | 0.51828385 | 0.49537525 | 0.62033526 | 0.95688177 |
| SOD1 | 19.2857676 | 0.80631888 | 0.5699814 | 1.4146407 | 0.15717387 | 0.74486619 |
| SCAF4 | 30.7669169 | -0.4014226 | 0.36179494 | -1.1095307 | 0.26720133 | 0.84994492 |
| URB1 | 9.35275023 | 0.58180372 | 0.58309333 | 0.99778833 | 0.31838201 | NA |
| EVA1C | 18.1147671 | 0.62156283 | 0.55567747 | 1.11856764 | 0.26332464 | 0.84928231 |
| SYNJ1 | 33.4600073 | 0.90659746 | 0.40035603 | 2.26447811 | 0.02354472 | 0.3948379 |
| PAXBP1 | 38.5522751 | -0.1023036 | 0.36479809 | -0.2804389 | 0.77914078 | 0.98264092 |
| IFNAR1 | 31.6315568 | -0.2392183 | 0.43968843 | -0.5440632 | 0.586398 | 0.95415437 |
| IFNGR2 | 69.8748609 | -0.0317343 | 0.44875745 | -0.0707159 | 0.94362383 | 0.99514836 |
| TMEM50B | 19.543704 | -0.1639443 | 0.51299392 | -0.3195833 | 0.7492842 | 0.97818861 |
| GART | 13.7100618 | 0.69709858 | 0.53673228 | 1.29878267 | 0.19401852 | 0.78135737 |
| SON | 92.4970806 | 0.20209219 | 0.25571841 | 0.79029192 | 0.42935731 | 0.92204028 |
| AP000311.1 | 20.7286102 | 0.05262129 | 0.89008255 | 0.05911956 | 0.95285688 | 0.99514836 |
| ITSN1 | 92.2000646 | 0.39297769 | 0.65976401 | 0.59563372 | 0.55141995 | 0.95415437 |
| LINC00649 | 15.9126566 | 0.13374809 | 0.65621782 | 0.20381661 | 0.83849681 | 0.98331031 |
| MRPS6 | 45.1401267 | -1.5673565 | 0.51848504 | -3.0229541 | 0.0025032 | 0.11739593 |
| KCNE1 | 36.431599 | 1.93381311 | 0.87128937 | 2.21948434 | 0.02645379 | 0.40665356 |
| RCAN1 | 15.1025831 | 0.39775572 | 0.69455244 | 0.57267918 | 0.56686193 | 0.95415437 |
| RUNX1 | 387.742991 | 0.95188991 | 0.47273764 | 2.01356909 | 0.04405479 | 0.51607664 |
| CBR1 | 12.1061152 | 0.6934456 | 0.67245737 | 1.03121125 | 0.30244177 | NA |
| DOPEY2 | 33.6016136 | -0.5794596 | 0.70491041 | -0.822033 | 0.4110581 | 0.91823911 |
| MORC3 | 55.031495 | 0.22967985 | 0.28762851 | 0.79852949 | 0.42456328 | 0.91985469 |
| HLCS | 38.3189814 | 0.20748987 | 0.45333695 | 0.45769458 | 0.64717188 | 0.96049083 |
| TTC3 | 85.074778 | -0.0426477 | 0.33768794 | -0.1262933 | 0.89949977 | 0.99182061 |
| VPS26C | 23.3885633 | 0.13703377 | 0.43044971 | 0.31835025 | 0.75021927 | 0.97818861 |
| DYRK1A | 163.023119 | 0.04451773 | 0.28346074 | 0.15705079 | 0.87520482 | 0.98939075 |
| KCNJ15 | 14.5082082 | 1.07903244 | 0.58722044 | 1.83752535 | 0.06613238 | 0.58951068 |
| ETS2 | 41.7777239 | 1.58770908 | 0.55541282 | 2.85861081 | 0.004255 | 0.16466571 |
| BRWD1 | 97.1391925 | 0.10404579 | 0.28550481 | 0.36442745 | 0.71553884 | 0.97348719 |
| HMGN1 | 30.5006215 | 0.06985442 | 0.5148171 | 0.13568784 | 0.89206806 | 0.99080204 |
| MX2 | 52.6007528 | 0.12000597 | 1.02129535 | 0.11750369 | 0.90646092 | 0.99368779 |
| MX1 | 52.0697485 | 0.73939214 | 1.15147627 | 0.64212539 | 0.52079177 | 0.94895469 |
| C2CD2 | 9.05950074 | 0.9351713 | 0.6597401 | 1.41748441 | 0.15634135 | NA |
| ABCG1 | 22.8497428 | 0.76697198 | 0.64573684 | 1.18774699 | 0.2349331 | 0.82903329 |
| UBASH3A | 7.83150704 | 0.37427841 | 0.78677895 | 0.47570974 | 0.63428118 | NA |
| SLC37A1 | 15.2921976 | 0.06246028 | 0.54368967 | 0.11488223 | 0.90853847 | 0.99423609 |
| NDUFV3 | 15.0356787 | 0.58298287 | 0.49754663 | 1.17171505 | 0.24131148 | 0.83493679 |
| PKNOX1 | 16.7968748 | 0.61091487 | 0.55067037 | 1.10940212 | 0.26725675 | 0.84994492 |
| RRP1B | 11.4605346 | 0.28395641 | 0.62257254 | 0.45610173 | 0.64831683 | NA |
| PDXK | 26.3338453 | 0.5880417 | 0.44564414 | 1.31953199 | 0.18699132 | 0.77798634 |
| CSTB | 39.0548795 | 0.75540001 | 0.62560788 | 1.20746563 | 0.22725287 | 0.82088564 |
| AGPAT3 | 56.4626119 | 0.52117387 | 0.38054376 | 1.36955043 | 0.17082728 | 0.75751515 |
| TRAPPC10 | 61.2842463 | -0.3240879 | 0.30805283 | -1.052053 | 0.29277522 | 0.85998836 |
| PFKL | 24.2652945 | -0.4151205 | 0.50047704 | -0.8294497 | 0.40685001 | 0.9169561 |
| TRPM2 | 29.1691869 | 0.69167718 | 0.67916163 | 1.01842794 | 0.30847463 | 0.8673137 |
| UBE2G2 | 31.3763919 | -0.2690295 | 0.38165704 | -0.7048986 | 0.48087335 | 0.93275609 |
| SUMO3 | 14.5162766 | -0.553614 | 0.54384448 | -1.0179638 | 0.30869514 | 0.8673137 |
| PTTG1IP | 34.9795905 | 0.34422993 | 0.40613183 | 0.84758176 | 0.39667093 | 0.91333232 |
| ITGB2 | 72.202681 | 0.60665929 | 0.61564105 | 0.98541073 | 0.32442236 | 0.87663738 |

| | | | | | | |
|---|---|---|---|---|---|---|
| ITGB2-AS1 | 11.344053 | -0.995932 | 0.61259061 | -1.6257709 | 0.1039984 | NA |
| ADARB1 | 33.2350593 | -1.1207299 | 0.4390048 | -2.5528875 | 0.0106834 | 0.28288008 |
| COL18A1 | 9.43687807 | -0.4254725 | 0.51847045 | -0.8206302 | 0.41185694 | NA |
| LSS | 9.55779445 | 0.74136008 | 0.63210883 | 1.17283614 | 0.24086152 | NA |
| MCM3AP | 30.6808037 | 0.30312986 | 0.39572838 | 0.76600486 | 0.44367341 | 0.92899684 |
| PCNT | 19.3937687 | 0.18978786 | 0.431012 | 0.44033079 | 0.65969754 | 0.96298563 |
| DIP2A | 67.2506565 | 0.49600139 | 0.4269483 | 1.16173644 | 0.24534254 | 0.83810223 |
| PRMT2 | 64.4217366 | 0.28851128 | 0.29312493 | 0.98426047 | 0.32498746 | 0.87685391 |
| AL592183.1 | 14.5049358 | -0.1437388 | 0.63573596 | -0.2260982 | 0.82112502 | 0.98320463 |

**Bulk RNA sequencing**

| | baseMean | log2 fold cha | lfcSE | stat | pvalue | padj |
|---|---|---|---|---|---|---|
| AL669831.5 | 95.8470735 | 0.08380638 | 0.34352386 | 0.24396088 | 0.80726112 | 0.97334138 |
| LINC00115 | 207.323489 | -0.3259006 | 0.28545395 | -1.1416926 | 0.25358183 | 0.8188642 |
| AL645608.3 | 48.6450863 | 0.41986904 | 0.55658974 | 0.75436001 | 0.45063308 | 0.89557948 |
| AL645608.1 | 809.247011 | 0.76362212 | 0.45023054 | 1.69606911 | 0.08987279 | 0.65375543 |
| SAMD11 | 1835.76659 | 0.15318622 | 0.29583189 | 0.51781512 | 0.60458727 | 0.93662382 |
| NOC2L | 4793.92796 | -0.028588 | 0.1391872 | -0.2053923 | 0.83726561 | 0.97577999 |
| KLHL17 | 1320.78855 | -0.1886725 | 0.30811351 | -0.6123474 | 0.54030792 | 0.92243385 |
| PLEKHN1 | 379.557691 | -0.5754394 | 0.31081757 | -1.8513733 | 0.06411587 | 0.61837523 |
| PERM1 | 198.376548 | -0.86406 | 0.42607172 | -2.0279684 | 0.04256347 | 0.56530166 |
| AL645608.8 | 311.826336 | -1.5998053 | 0.85814988 | -1.8642493 | 0.0622867 | 0.61308561 |
| HES4 | 543.92029 | -1.0350009 | 0.32626742 | -3.1722471 | 0.00151264 | 0.17279992 |
| ISG15 | 7996.22468 | -4.6946556 | 1.25404914 | -3.7435978 | NA | NA |
| AGRN | 25692.3903 | -0.0478438 | 0.34364145 | -0.139226 | 0.8892716 | 0.98439687 |
| AL645608.9 | 105.038932 | -1.0015877 | 0.5019358 | -1.9954499 | 0.04599384 | 0.57718395 |
| RNF223 | 65.2439681 | -1.0817633 | 0.86168228 | -1.2554085 | 0.20933051 | 0.79086734 |
| C1orf159 | 1278.42743 | -0.3015273 | 0.30991929 | -0.9729219 | 0.33059211 | 0.85384238 |
| LINC01342 | 59.4818739 | -0.100224 | 0.71394424 | -0.1403807 | 0.88835921 | 0.98423722 |
| AL390719.2 | 842.494106 | -0.0747969 | 0.3442238 | -0.2172914 | 0.82798123 | 0.97548201 |
| TTLL10 | 49.3249307 | 0.06469565 | 0.37640273 | 0.1718788 | 0.86353281 | 0.98176973 |
| TNFRSF18 | 53.1478349 | 0.70392868 | 0.80070336 | 0.87913792 | 0.3793265 | 0.87069278 |
| TNFRSF4 | 161.051812 | -0.4832066 | 0.63072892 | -0.7661082 | 0.44361194 | 0.89366134 |
| SDF4 | 10798.0249 | 0.04897945 | 0.11897537 | 0.41167722 | 0.68057602 | 0.95070301 |
| B3GALT6 | 900.41407 | -0.2913281 | 0.21886488 | -1.3310866 | 0.18316051 | 0.76723162 |
| C1QTNF12 | 278.591975 | 0.4010131 | 0.26499738 | 1.51327192 | 1.30E-01 | 7.08E-01 |
| AL162741.1 | 42.9237858 | -0.1283931 | 0.43545938 | -0.2948451 | 0.76811222 | 0.96765147 |
| UBE2J2 | 2813.28059 | -0.2760391 | 0.16365535 | -1.6867098 | 0.09165916 | 0.65518742 |
| LINC01786 | 73.8921813 | -0.1780483 | 0.48452067 | -0.3674732 | 0.71326611 | 0.95564093 |
| SCNN1D | 697.554511 | -0.2852829 | 0.39656059 | -0.7193931 | 0.47189877 | 0.90160919 |
| ACAP3 | 5191.33338 | -0.1090913 | 0.17284084 | -0.6311661 | 0.52793192 | 0.91857508 |
| PUSL1 | 843.972419 | -0.163607 | 0.20205524 | -0.8097144 | 0.41810436 | 0.88445833 |
| INTS11 | 6952.42049 | -0.2676542 | 0.16596521 | -1.6127126 | 1.07E-01 | 6.77E-01 |
| CPTP | 2513.36826 | -0.1999301 | 0.20054597 | -0.9969288 | 0.31879906 | 0.84914926 |
| TAS1R3 | 81.3649694 | -0.3578649 | 0.31841887 | -1.1238808 | 0.2610636 | 0.82424998 |
| DVL1 | 4688.87291 | -0.3290222 | 0.14474597 | -2.273101 | 0.02302009 | 0.51967919 |
| MXRA8 | 2779.68752 | 0.22315295 | 0.26905927 | 0.82938214 | 0.4068882 | 0.88132015 |
| AURKAIP1 | 4480.89603 | -0.113022 | 0.13571928 | -0.8327628 | 0.40497851 | 0.88132015 |
| CCNL2 | 13344.6843 | -0.3437185 | 0.1859941 | -1.8480074 | 0.06460127 | 0.62023173 |
| MRPL20 | 3283.16054 | -0.0144726 | 0.12746227 | -0.1135438 | 0.90959941 | 0.98662757 |
| AL391244.3 | 250.762256 | 0.34517264 | 0.26715259 | 1.29204302 | 0.19634225 | 0.77798606 |
| ANKRD65 | 218.874766 | 0.45335196 | 0.50168669 | 0.90365556 | 0.36617807 | 0.86683872 |
| LINC01770 | 75.8264124 | 0.24933725 | 0.53617203 | 0.46503219 | 0.64190841 | 0.94088547 |
| VWA1 | 4373.89732 | 0.25691696 | 0.44188562 | 0.58141053 | 0.5609638 | 0.92448946 |
| ATAD3C | 498.081607 | 0.22155058 | 0.37935962 | 0.58401203 | 0.55921222 | 0.92421175 |
| ATAD3B | 3111.29602 | -0.2484269 | 0.21127894 | -1.175824 | 2.40E-01 | 0.81177764 |
| ATAD3A | 1587.67852 | -0.1362676 | 0.18363687 | -0.7420491 | 0.45805758 | 0.8971553 |
| TMEM240 | 77.1747459 | -0.0813394 | 0.51275553 | -0.1586319 | 0.87395892 | 0.98275231 |
| SSU72 | 5451.75725 | -0.3513214 | 0.20943808 | -1.6774474 | 0.09345503 | 0.65756964 |
| FNDC10 | 244.226306 | 0.58137488 | 0.54503327 | 1.06667778 | 2.86E-01 | 0.83107779 |
| AL691432.2 | 577.671903 | -0.3228533 | 0.26273996 | -1.2287939 | 0.21914909 | 0.79847731 |
| MIB2 | 4453.14024 | -0.1645789 | 0.19282626 | -0.8535087 | 0.39337728 | 0.8783848 |
| MMP23B | 170.154493 | -0.3645568 | 0.4405325 | -0.8275367 | 0.40793295 | 0.88168279 |
| CDK11B | 1811.97341 | -0.3467901 | 0.21373805 | -1.6225004 | 0.10469625 | 0.67439401 |

*This means that positive fold changes imply higher mean value in non-COVID AKI than in COVID AKI and vice versa.

| | | | | | | |
|---|---|---|---|---|---|---|
| FO704657.1 | 652.56172 | 0.11691626 | 0.54952469 | 0.21275888 | 0.83151504 | 0.97550416 |
| SLC35E2B | 3020.91255 | -0.0572587 | 0.32325972 | -0.1771291 | 0.859407 | 0.9802651 |
| CDK11A | 1728.30647 | -0.3414727 | 0.35291719 | -0.9675718 | 0.33325827 | 0.855011 |
| SLC35E2A | 1623.53038 | -0.1482196 | 0.49344934 | -0.3003745 | 0.7638915 | 0.96678847 |
| NADK | 3848.58569 | -0.3717095 | 0.14524731 | -2.559149 | 0.01049287 | 0.40196898 |
| GNB1 | 16521.7774 | -0.1234614 | 0.19739078 | -0.625467 | 0.53166461 | 0.91962754 |
| CALML6 | 41.2203193 | 0.2421342 | 0.36990164 | 0.65459076 | 0.51273127 | 0.91568064 |
| TMEM52 | 83.8595811 | 0.76553908 | 0.60397575 | 1.2674997 | 0.20497667 | 0.78502176 |
| PRKCZ | 2496.23575 | -0.3282064 | 0.30232531 | -1.0856067 | 0.27765303 | 0.82805357 |
| AL590822.2 | 262.459335 | -0.2811772 | 0.41296589 | -0.6808727 | 0.49595206 | 0.90874439 |
| PRKCZ-AS1 | 351.384914 | -0.2788263 | 0.33423801 | -0.8342149 | 0.40415992 | 0.88132015 |
| FAAP20 | 3888.53439 | -0.2329246 | 0.12022452 | -1.9374138 | 0.05269479 | 0.58742503 |
| SKI | 4234.48665 | -0.0592229 | 0.27922737 | -0.2120955 | 0.83203254 | 0.97550416 |
| MORN1 | 667.169625 | -0.3535648 | 0.23309372 | -1.5168353 | 0.12930827 | 0.7075522 |
| RER1 | 9081.50509 | 0.00044243 | 0.11376699 | 0.00388888 | 0.99689713 | 0.99976751 |
| PEX10 | 4034.51534 | 0.03333084 | 0.11338229 | 0.29396866 | 0.76878185 | 0.96796999 |
| PLCH2 | 3303.24335 | 0.45660612 | 0.65061393 | 0.70180809 | 0.48279885 | 0.90541935 |
| PANK4 | 1707.48382 | -0.1076943 | 0.13154273 | -0.8187017 | 0.41295663 | 0.88291217 |
| AL139246.5 | 240.735767 | -0.7631861 | 0.33556348 | -2.2743418 | 0.02294544 | 0.51967919 |
| TNFRSF14-A | 4003.15233 | -0.3980326 | 0.27867642 | -1.4282967 | 0.15320647 | 0.74258845 |
| TNFRSF14 | 16005.2528 | -0.4829768 | 0.26010721 | -1.8568374 | 0.06333429 | 0.6165319 |
| FAM213B | 2802.47409 | 0.38445066 | 0.2213671 | 1.73671093 | 0.08243821 | 0.64661469 |
| MMEL1 | 826.859605 | 0.32397826 | 0.29631343 | 1.09336343 | 0.27423426 | 0.82719407 |
| TTC34 | 215.876825 | -0.2274324 | 0.52971513 | -0.4293484 | 0.66766969 | 0.94844217 |
| LINC00982 | 6453.12025 | 0.00185687 | 0.52270673 | 0.00355241 | 0.99716559 | 0.99976751 |
| PRDM16 | 1211.25173 | -0.4188726 | 0.35719085 | -1.1726858 | 0.24092182 | 0.8124487 |
| ARHGEF16 | 4364.28863 | -0.0379087 | 0.16679635 | -0.2272751 | 0.82020981 | 0.97454841 |
| MEGF6 | 1256.17272 | 0.3155043 | 0.3434779 | 0.91855778 | 0.35832692 | 0.86480228 |
| AL513320.1 | 143.156225 | 0.38723178 | 0.47279294 | 0.81903036 | 0.41276909 | 0.88286984 |
| TPRG1L | 5648.60697 | 0.2827713 | 0.31587848 | 0.89519015 | 0.37068545 | 0.86762749 |
| WRAP73 | 1482.82993 | -0.2954283 | 0.09795014 | -3.0161088 | 0.00256041 | 0.21645768 |
| TP73 | 74.9894829 | -0.0262062 | 0.29280064 | -0.0895019 | 0.92868308 | 0.98963127 |
| CCDC27 | 70.9557126 | -0.3568732 | 0.33008923 | -1.0811416 | 0.27963413 | 0.82825734 |
| SMIM1 | 190.721874 | 0.12838787 | 0.42561202 | 0.30165471 | 0.7629153 | 0.96639375 |
| LRRC47 | 3082.40814 | 0.00581701 | 0.15551293 | 0.03740534 | 0.97016182 | 0.99400331 |
| AL365330.1 | 466.147536 | -0.0206704 | 0.19941323 | -0.103656 | 0.91744236 | 0.9868044 |
| CEP104 | 1859.76302 | -0.3760939 | 0.22090663 | -1.7025019 | 0.08866133 | 0.65363785 |
| DFFB | 468.930432 | 0.0857224 | 0.19454387 | 0.44063278 | 0.65947887 | 0.94605737 |
| C1orf174 | 1546.84001 | -0.3076539 | 0.24216065 | -1.2704539 | 0.20392298 | 0.78410353 |
| LINC01134 | 88.1964439 | -0.8827291 | 0.61581185 | -1.4334396 | 0.15173226 | 0.74024851 |
| AJAP1 | 657.914835 | 0.46197752 | 0.22838336 | 2.02281606 | 0.04309212 | 0.56792499 |
| NPHP4 | 1053.69038 | 0.06979152 | 0.16337512 | 0.42718572 | 0.66924406 | 0.94873698 |
| KCNAB2 | 3739.08096 | 0.83799464 | 0.43162483 | 1.94148849 | 0.05219905 | 0.58645127 |
| CHD5 | 146.45308 | 0.17916673 | 0.63793332 | 0.28085496 | 7.79E-01 | 9.68E-01 |
| RPL22 | 13174.4003 | 0.36510674 | 0.22078862 | 1.65364833 | 0.09819899 | 0.66549286 |
| AL031847.1 | 69.2533219 | 0.4602678 | 0.51721074 | 0.8899038 | 0.37351754 | 0.86791605 |
| RNF207 | 1924.10116 | -0.069859 | 0.26942721 | -0.2592871 | 0.79541372 | 0.97101097 |
| ICMT | 5357.18417 | 0.16381135 | 0.23390882 | 0.70032138 | 0.48372663 | 0.90541935 |
| GPR153 | 903.137737 | 0.47725336 | 0.39550914 | 1.206681 | 0.22755502 | 0.80300331 |
| ACOT7 | 3133.93922 | 0.25287971 | 0.54000996 | 0.46828712 | 0.63957928 | 0.94088547 |
| HES2 | 222.58047 | -0.4877873 | 0.4746984 | -1.027573 | 0.30415072 | 0.84211658 |
| ESPN | 3682.02787 | 0.81909587 | 0.41830241 | 1.95814286 | 0.05021325 | 0.5861488 |
| AL031848.1 | 162.391442 | 0.64796793 | 0.53351132 | 1.21453455 | 0.22454367 | 0.80078694 |
| TNFRSF25 | 1168.24241 | 0.2221919 | 0.3564407 | 0.62336289 | 0.53304608 | 0.92026174 |

| | | | | | | |
|---|---|---|---|---|---|---|
| PLEKHG5 | 1422.23432 | 0.10591643 | 0.36526176 | 0.28997404 | 0.7718361 | 0.96836854 |
| NOL9 | 1102.10995 | -0.2061902 | 0.14980685 | -1.3763737 | 0.16870596 | 0.75766078 |
| ZBTB48 | 2082.02337 | 0.05260723 | 0.15778338 | 0.33341427 | 0.73882159 | 0.9602325 |
| KLHL21 | 5455.77174 | -1.0502868 | 0.50665314 | -2.0729898 | 0.03817323 | 0.55538555 |
| PHF13 | 1196.72871 | -0.2104066 | 0.18255807 | -1.1525462 | 0.24909671 | 0.81646572 |
| THAP3 | 4430.1757 | 0.03100913 | 0.16595113 | 0.18685699 | 0.85177276 | 0.97872457 |
| DNAJC11 | 6126.73621 | 0.145326 | 0.17873381 | 0.81308621 | 0.41616863 | 0.88399703 |
| CAMTA1 | 1876.03394 | 0.01448722 | 0.1672015 | 0.08664527 | 9.31E-01 | 0.98975562 |
| VAMP3 | 5175.61192 | 0.02583787 | 0.19689468 | 0.13122686 | 0.89559584 | 0.98474725 |
| PER3 | 1127.92276 | 0.07892608 | 0.47321288 | 0.16678769 | 0.86753711 | 0.98224588 |
| Z98884.1 | 56.6312159 | 0.15790939 | 0.59434202 | 0.26568774 | 0.79047971 | 0.97090793 |
| UTS2 | 287.531089 | -0.0162633 | 0.56072334 | -0.0290041 | 0.9768613 | 0.99455099 |
| PARK7 | 7609.6277 | 0.16544733 | 0.15677516 | 1.05531596 | 0.29128085 | 0.83454586 |
| AL034417.4 | 187.751086 | -0.1803093 | 0.69705007 | -0.2586748 | 0.79588614 | 0.97101097 |
| ERRFI1 | 9855.15291 | -0.682932 | 0.63693383 | -1.0722181 | 0.28362208 | 0.83107779 |
| SLC45A1 | 378.135336 | -0.0085392 | 0.38257279 | -0.0223204 | 0.98219234 | 0.99595643 |
| RERE | 11891.8609 | -0.2891493 | 0.18366537 | -1.5743267 | 0.11541193 | 0.6873836 |
| ENO1 | 93919.6014 | -0.0946623 | 0.25150139 | -0.3763889 | 0.70662782 | 0.95501548 |
| SLC2A5 | 2431.41701 | 1.25442253 | 0.64670634 | 1.93970965 | 0.05241499 | 0.58645127 |
| GPR157 | 659.393556 | -0.1757012 | 0.2360385 | -0.7443753 | 0.45664946 | 0.89688594 |
| MIR34AHG | 514.450374 | 0.55550434 | 0.50475517 | 1.10054214 | 0.27109598 | 0.82716979 |
| LINC01759 | 45.7315021 | 0.48852834 | 0.77333054 | 0.63171996 | 0.52756988 | 0.91837147 |
| H6PD | 4397.39565 | -0.1071117 | 0.21372736 | -0.5011604 | 0.61625825 | 0.93910158 |
| SPSB1 | 3960.86485 | -0.901502 | 0.48879521 | -1.8443348 | 0.06513436 | 0.6216672 |
| BX323043.1 | 72.7446701 | 1.31386331 | 0.78468341 | 1.67438651 | 0.09405466 | 0.65783452 |
| AL928921.2 | 106.250839 | 0.53644316 | 0.58315308 | 0.9199011 | 3.58E-01 | 8.64E-01 |
| SLC25A33 | 1022.08815 | 0.19652836 | 0.444969 | 0.44166753 | 0.65872981 | 0.9458503 |
| TMEM201 | 1155.29952 | -0.4068651 | 0.16823218 | -2.418474 | 0.01558576 | 0.45426553 |
| PIK3CD | 2254.1477 | -0.5852134 | 0.28666923 | -2.0414239 | 0.04120871 | 0.55882236 |
| PIK3CD-AS2 | 122.257931 | 0.31688808 | 0.34190972 | 0.92681799 | 0.35402104 | 0.86364633 |
| CLSTN1 | 10861.8659 | 0.11590021 | 0.13901838 | 0.83370419 | 0.4044477 | 0.88132015 |
| CTNNBIP1 | 1494.72088 | 0.07420009 | 0.22305747 | 0.33265008 | 0.73939843 | 0.96029564 |
| AL357140.1 | 523.596798 | 0.11208626 | 0.26184091 | 0.42807007 | 0.6686001 | 0.9486589 |
| LZIC | 670.555403 | 0.10286531 | 0.23469564 | 0.43829238 | 0.66117435 | 0.94637197 |
| NMNAT1 | 951.792527 | 0.41631781 | 0.2299569 | 1.81041667 | 0.0702312 | 0.62928077 |
| RBP7 | 526.095993 | -0.305223 | 0.62450792 | -0.4887416 | 0.62502464 | 0.94088547 |
| UBE4B | 2284.97811 | 0.11133133 | 0.16897591 | 0.65885922 | 0.50998618 | 0.91439937 |
| KIF1B | 2918.04691 | 0.04681216 | 0.19740867 | 0.23713323 | 0.81255344 | 0.97371854 |
| AL139424.2 | 145.946147 | 0.16535925 | 0.36843778 | 0.44881186 | 0.65356738 | 0.94495971 |
| PGD | 5185.10819 | -0.3009256 | 0.23085592 | -1.3035212 | 0.19239687 | 0.77442218 |
| CENPS-CORT | 879.697646 | 0.28849602 | 0.23515622 | 1.22682707 | 0.21988759 | 0.79858446 |
| CENPS | 877.02355 | 0.33925988 | 0.24983813 | 1.35791878 | 0.17448946 | 0.75974924 |
| CORT | 141.198365 | 0.02072631 | 0.18193535 | 0.1139213 | 0.90930018 | 0.98662757 |
| DFFA | 2494.754 | -0.034318 | 0.16361443 | -0.2097492 | 0.83386342 | 0.97550416 |
| AL354956.1 | 185.183087 | 0.04316413 | 0.28087561 | 0.15367704 | 0.87786439 | 0.98275231 |
| PEX14 | 1842.13721 | -0.1935509 | 0.14490625 | -1.3356975 | 0.18164817 | 0.76528574 |
| CASZ1 | 2102.91394 | -0.1542407 | 0.2331601 | -0.6615225 | 5.08E-01 | 0.91439937 |
| AL139423.1 | 920.869232 | -0.3482739 | 0.22935943 | -1.5184634 | 0.12889762 | 0.70633634 |
| TARDBP | 8287.5425 | -0.2262693 | 0.14826409 | -1.5261232 | 0.12697918 | 0.70463514 |
| AL109811.4 | 4301.04787 | -0.1686675 | 0.15647211 | -1.0779397 | 0.28106068 | 0.82937461 |
| MASP2 | 307.321069 | 0.2616836 | 0.27489299 | 0.95194713 | 0.3411238 | 0.85889221 |
| AL109811.3 | 438.609091 | 0.21159069 | 0.26852766 | 0.78796609 | 0.43071654 | 0.88966606 |
| SRM | 1926.27889 | -0.3705001 | 0.30303956 | -1.2226131 | 0.22147588 | 0.80020811 |
| EXOSC10 | 3158.851 | -0.3641429 | 0.15308668 | -2.3786711 | 0.01737517 | 0.46859699 |

| | | | | | | |
|---|---|---|---|---|---|---|
| AL109811.1 | 1327.0958 | -0.3461176 | 0.17472839 | -1.9808893 | 0.04760369 | 0.58241466 |
| AL109811.2 | 103.619506 | -0.3101079 | 0.38915776 | -0.7968693 | 0.42552696 | 0.88622485 |
| MTOR | 3048.8362 | -0.0771007 | 0.12227757 | -0.6305381 | 0.52834259 | 0.91888066 |
| ANGPTL7 | 71.5439801 | 1.98056188 | 1.19567749 | 1.65643486 | 0.09763379 | 0.66549286 |
| UBIAD1 | 896.641865 | -0.3679692 | 0.3159681 | -1.1645771 | 0.24419023 | 0.81468457 |
| FBXO2 | 1493.79174 | -0.6112691 | 0.31827659 | -1.9205595 | 0.05478727 | 0.59329767 |
| FBXO44 | 3833.44677 | -0.0537623 | 0.30008421 | -0.1791573 | 0.85781417 | 0.97964261 |
| FBXO6 | 776.521201 | -0.5224389 | 0.39173806 | -1.3336435 | 0.18232073 | 0.76611496 |
| MAD2L2 | 1232.51632 | 0.06773347 | 0.1794928 | 0.37736035 | 0.70590583 | 0.95501548 |
| AGTRAP | 3586.15426 | -0.2993406 | 0.25549504 | -1.1716101 | 0.24135365 | 0.8124487 |
| C1orf167 | 975.002872 | 0.16639331 | 0.33533792 | 0.49619591 | 0.61975619 | 0.93945324 |
| MTHFR | 3164.09107 | 0.13017684 | 0.27772866 | 0.46871949 | 0.63927015 | 0.94088547 |
| CLCN6 | 2349.45055 | -0.5544012 | 0.29714385 | -1.8657672 | 0.06207394 | 0.61236592 |
| NPPA-AS1 | 504.609394 | -1.5271759 | 0.55749328 | -2.739362 | 0.00615586 | 0.33470127 |
| NPPA | 220.440028 | -1.6202242 | 0.63071127 | -2.5688843 | 1.02E-02 | 4.00E-01 |
| KIAA2013 | 4168.08364 | 0.12978215 | 0.17197106 | 0.75467436 | 0.45044439 | 0.89557948 |
| PLOD1 | 7677.73213 | 0.30347432 | 0.29505911 | 1.02852042 | 0.30370509 | 0.84211658 |
| MFN2 | 6284.83288 | 0.22493552 | 0.2186486 | 1.02875356 | 0.30359549 | 0.84211658 |
| MIIP | 1290.35428 | 0.03356732 | 0.16821563 | 0.19954933 | 0.84183306 | 0.9762188 |
| TNFRSF8 | 40.7635893 | 0.54934663 | 0.58750157 | 0.9350556 | 0.34975965 | 0.86364633 |
| TNFRSF1B | 2258.18728 | 0.06874364 | 0.54992242 | 0.12500607 | 0.90051874 | 0.98526603 |
| VPS13D | 3318.4711 | 0.0069313 | 0.1196203 | 0.05794421 | 0.95379307 | 0.99170503 |
| DHRS3 | 5607.91446 | 0.06864845 | 0.31103412 | 0.22071035 | 0.82531797 | 0.97548201 |
| AC254633.1 | 53.5240863 | -0.7691174 | 0.45567009 | -1.6878822 | 0.09143385 | 0.65518742 |
| PDPN | 251.338438 | -0.0099189 | 0.27962226 | -0.0354726 | 0.97170288 | 0.99404213 |
| PRDM2 | 1704.81335 | -0.1858906 | 0.14515734 | -1.2806143 | 0.20032916 | 0.78134034 |
| KAZN | 3521.69651 | 0.02180426 | 0.19415664 | 0.11230243 | 0.91058361 | 0.98662757 |
| TMEM51-AS | 843.373791 | 0.08399596 | 0.25816886 | 0.32535282 | 0.74491406 | 0.96144901 |
| TMEM51 | 3325.35443 | -0.1031585 | 0.30250352 | -0.3410158 | 0.73309166 | 0.95861504 |
| FHAD1 | 97.2183708 | 0.3590034 | 0.4343229 | 0.82658177 | 0.40847415 | 0.88203324 |
| EFHD2 | 2294.67716 | -0.1804903 | 0.25821879 | -0.698982 | 0.48456326 | 0.90541935 |
| CASP9 | 1586.05272 | 0.2325213 | 0.31121454 | 0.74714153 | 0.45497814 | 0.89670027 |
| DNAJC16 | 4163.59856 | 0.61093596 | 0.37500908 | 1.62912309 | 0.10328696 | 0.67439401 |
| AL121992.3 | 135.149399 | 0.98655793 | 0.55689617 | 1.77152939 | 0.07647271 | 0.63773415 |
| AGMAT | 7171.16421 | 1.70665409 | 0.6322722 | 2.69923947 | 0.00694981 | 0.3503878 |
| DDI2 | 2335.57555 | -0.1839262 | 0.25578723 | -0.7190594 | 0.4721043 | 0.90160919 |
| RSC1A1 | 344.090087 | -0.3543768 | 0.25499024 | -1.3897661 | 0.16459991 | 0.75286112 |
| PLEKHM2 | 6031.78043 | -0.7843221 | 0.25836852 | -3.0356721 | 0.0024 | 0.21083339 |
| AL450998.3 | 888.453703 | -0.697383 | 0.26722262 | -2.6097452 | 0.00906097 | 0.38593543 |
| SLC25A34 | 847.325154 | 0.06686225 | 0.55904794 | 0.1196002 | 0.90479986 | 0.98624484 |
| SLC25A34-AS | 92.0710048 | -0.0149446 | 0.57327044 | -0.0260691 | 0.97920224 | 0.99507804 |
| TMEM82 | 893.491795 | 0.12294412 | 0.34527279 | 0.35607821 | 0.721782 | 0.95739595 |
| FBLIM1 | 3503.89 | -0.432862 | 0.38944602 | -1.1114814 | 0.26636118 | 0.8265123 |
| UQCRHL | 313.688324 | 0.37902865 | 0.37086338 | 1.02201692 | 0.30677289 | 0.84286067 |
| AL450998.2 | 284.980431 | -0.1270439 | 0.30941344 | -0.410596 | 0.68136882 | 0.95070301 |
| SPEN | 3638.61983 | -0.6799785 | 0.2336577 | -2.9101483 | 0.00361257 | 0.26039706 |
| ZBTB17 | 2021.3464 | -0.5156629 | 0.14691027 | -3.5100533 | 0.00044802 | 0.10365784 |
| SRARP | 99.7338515 | 1.08846393 | 0.41010115 | 2.65413527 | 0.00795119 | 0.37071188 |
| AL355994.3 | 69.1903056 | 1.26192597 | 0.42545709 | 2.9660476 | 0.00301654 | 0.24038155 |
| HSPB7 | 2277.23793 | -1.3613546 | 0.79374159 | -1.7151106 | 0.08632494 | 0.65077063 |
| CLCNKA | 10216.7566 | 0.80466129 | 0.6619853 | 1.21552744 | 2.24E-01 | 8.01E-01 |
| CLCNKB | 14822.9315 | 0.88065011 | 0.60804011 | 1.44834214 | 0.14752139 | 0.73412064 |
| FAM131C | 731.699305 | -0.146737 | 0.61197383 | -0.2397766 | 0.81050342 | 0.97352497 |
| EPHA2 | 2767.04201 | -0.5780491 | 0.28768665 | -2.009301 | 0.04450522 | 0.57267043 |

| | | | | | | |
|---|---|---|---|---|---|---|
| ARHGEF19-A | 97.4553775 | -0.1076004 | 0.46057952 | -0.2336195 | 0.81528038 | 0.9738447 |
| ARHGEF19 | 566.08228 | -0.0152757 | 0.39607958 | -0.0385674 | 0.96923533 | 0.99400331 |
| CPLANE2 | 271.45564 | 0.35609333 | 0.33034414 | 1.07794656 | 0.2810576 | 0.82937461 |
| FBXO42 | 1768.69898 | -0.369875 | 0.14708345 | -2.5147292 | 0.01191238 | 0.4059091 |
| SZRD1 | 7548.29439 | -0.2306553 | 0.177674 | -1.2981939 | 0.19422073 | 0.7762856 |
| SPATA21 | 2485.47826 | -0.1710806 | 0.20349877 | -0.8406962 | 0.40051814 | 0.88086294 |
| NECAP2 | 2994.52806 | -0.1467654 | 0.13624677 | -1.0772026 | 0.28138977 | 0.82957339 |
| LINC01772 | 118.029236 | 0.12613566 | 0.31012334 | 0.40672739 | 0.68420822 | 0.95070301 |
| NBPF1 | 487.44912 | 0.16174743 | 0.23097052 | 0.70029467 | 0.4837433 | 0.90541935 |
| CROCC | 2476.26101 | -0.1937671 | 0.24445462 | -0.7926506 | 0.42798142 | 0.88798629 |
| BX284668.5 | 297.144584 | 0.48258047 | 0.56977326 | 0.84696933 | 0.39701222 | 0.87988087 |
| MFAP2 | 214.944665 | 0.3120764 | 0.48177878 | 0.64775871 | 0.51714102 | 0.91636259 |
| AL049569.1 | 200.17418 | 0.00894514 | 0.23345055 | 0.03831707 | 0.96943488 | 0.99400331 |
| ATP13A2 | 2961.4693 | -0.0904915 | 0.16580537 | -0.5457696 | 0.58522435 | 0.93289552 |
| SDHB | 6823.58104 | 0.27895118 | 0.21844151 | 1.27700628 | 0.20160003 | 0.78134034 |
| PADI2 | 1189.38704 | 0.93165428 | 0.57957655 | 1.60747409 | 0.10795041 | 0.67730117 |
| AC004824.1 | 1266.99119 | -0.1180738 | 0.244588 | -0.4827458 | 0.62927621 | 0.94088547 |
| RCC2 | 3728.99278 | -0.1585628 | 0.18846012 | -0.8413599 | 4.00E-01 | 8.81E-01 |
| ARHGEF10L | 10036.69 | -0.0440949 | 0.22865604 | -0.1928437 | 8.47E-01 | 0.97684681 |
| IGSF21 | 116.534854 | -0.4806469 | 0.69693267 | -0.6896604 | 0.49040775 | 0.90600166 |
| KLHDC7A | 3612.33094 | 0.46655455 | 0.50659549 | 0.92096073 | 0.35707093 | 0.86390849 |
| AL080251.1 | 11113.4338 | 0.83371281 | 0.40210539 | 2.07336887 | 0.03813796 | 0.55538555 |
| ALDH4A1 | 25450.0278 | 0.77438705 | 0.3654017 | 2.11927596 | 0.03406715 | 0.55369025 |
| IFFO2 | 1477.6539 | 0.08781686 | 0.24382354 | 0.36016565 | 0.71872326 | 0.95648644 |
| UBR4 | 10102.0329 | -0.0526792 | 0.18576843 | -0.2835746 | 0.77673638 | 0.96836854 |
| AL035413.1 | 102.63927 | 0.0821612 | 0.27505801 | 0.29870497 | 0.76516516 | 0.96678847 |
| EMC1 | 2578.37861 | 0.03901829 | 0.16701373 | 0.23362324 | 0.81527748 | 0.9738447 |
| MRTO4 | 1444.57417 | -0.2490003 | 0.24506307 | -1.0160664 | 0.30959779 | 0.84432074 |
| AKR7A3 | 3807.15565 | 0.45379592 | 0.32117919 | 1.41290574 | 0.15768344 | 0.74522306 |
| AKR7A2 | 4294.86577 | 0.56241952 | 0.32010857 | 1.75696489 | 0.07892379 | 0.64079926 |
| PQLC2 | 1489.8663 | -0.5751217 | 0.35772407 | -1.6077243 | 0.10789557 | 0.67720384 |
| CAPZB | 9501.20464 | -0.0731204 | 0.15371892 | -0.4756761 | 0.63430515 | 0.94088547 |
| MINOS1 | 3119.07328 | 0.00400931 | 0.14758052 | 0.02716696 | 0.97832656 | 0.99476647 |
| NBL1 | 5710.7332 | -0.0833385 | 0.16682991 | -0.4995415 | 0.61739797 | 0.93910158 |
| MINOS1-NBI | 3649.78649 | -0.007084 | 0.15213255 | -0.0465647 | 0.96286018 | 0.99320711 |
| TMCO4 | 2261.08363 | -0.1005852 | 0.24493381 | -0.4106627 | 6.81E-01 | 9.51E-01 |
| RNF186 | 1310.60019 | 1.08377784 | 0.80033839 | 1.3541495 | 0.17568869 | 0.7610139 |
| AL391883.1 | 574.89038 | 1.02897164 | 0.80836966 | 1.27289741 | 0.20305445 | 0.78252625 |
| OTUD3 | 979.962004 | -0.209414 | 0.27689224 | -0.7563013 | 0.44946856 | 0.89557948 |
| PLA2G2A | 343.295016 | 0.4060009 | 1.90498149 | 0.21312591 | 0.83122876 | 0.97550416 |
| UBXN10 | 456.715187 | -0.0441553 | 0.32788215 | -0.1346681 | 0.89287431 | 0.98474725 |
| VWA5B1 | 71.6156441 | 0.66149634 | 0.67301706 | 0.98288198 | 0.32566553 | 0.85271781 |
| CAMK2N1 | 5543.52155 | 0.47188539 | 0.2684918 | 1.75754117 | 0.07882561 | 0.64079926 |
| MUL1 | 2940.14405 | -0.4902366 | 0.26363867 | -1.8595017 | 0.06295606 | 0.6147045 |
| FAM43B | 111.160875 | -0.2859918 | 0.75531403 | -0.3786396 | 0.70495551 | 0.95501548 |
| CDA | 365.73231 | -0.4735766 | 0.33856304 | -1.3987842 | 1.62E-01 | 7.50E-01 |
| PINK1 | 10604.6954 | 0.14253286 | 0.23955108 | 0.59499985 | 0.55184357 | 0.92373235 |
| PINK1-AS | 9799.59405 | 0.14866311 | 0.20148433 | 0.73783956 | 0.46061195 | 0.89755243 |
| DDOST | 9825.6671 | 0.26380145 | 0.12852529 | 2.05252559 | 0.04011861 | 0.55657615 |
| KIF17 | 277.801461 | -0.3986772 | 0.26089637 | -1.5281058 | 0.12648627 | 0.70431949 |
| SH2D5 | 503.388923 | -1.3825068 | 0.82918217 | -1.6673137 | 0.09545205 | 0.65888357 |
| HP1BP3 | 7477.62021 | -0.1682754 | 0.08099023 | -2.0777245 | 0.03773474 | 0.55538555 |
| EIF4G3 | 2993.56428 | -0.2757615 | 0.22962894 | -1.2009006 | 0.22978976 | 0.80551898 |
| ECE1 | 7032.76259 | -0.168206 | 0.20965784 | -0.8022881 | 0.42238631 | 0.88451372 |

| | | | | | | |
|---|---|---|---|---|---|---|
| NBPF3 | 311.373136 | 0.51169153 | 0.37929868 | 1.34904641 | 0.17732206 | 0.76115956 |
| ALPL | 5376.79481 | -0.1690237 | 0.46604397 | -0.3626775 | 0.71684581 | 0.95604949 |
| RAP1GAP | 15714.6781 | 0.50754822 | 0.45092573 | 1.12556943 | 0.26034781 | 0.82424998 |
| USP48 | 2307.09334 | 0.02588987 | 0.10890324 | 0.23773274 | 0.81208839 | 0.97355727 |
| LDLRAD2 | 6901.88485 | 0.11104578 | 0.55232676 | 0.20105088 | 0.84065879 | 0.97618283 |
| HSPG2 | 18825.8817 | 0.07809415 | 0.50018648 | 0.15613007 | 0.8759305 | 0.98275231 |
| CELA3A | 77.9887576 | -25.001864 | 3.67845117 | -6.7968455 | NA | NA |
| CDC42 | 12658.5122 | -0.1073555 | 0.21019933 | -0.5107317 | 0.60953891 | 0.93760173 |
| WNT4 | 85.8905606 | 0.60733424 | 0.43049862 | 1.41076932 | 0.15831264 | 0.74658713 |
| ZBTB40 | 1916.93801 | -0.372043 | 0.26914211 | -1.3823291 | 0.16687066 | 0.75558932 |
| C1QA | 4792.39079 | -0.5977027 | 0.64193902 | -0.9310895 | 0.35180726 | 0.86364633 |
| C1QC | 4406.4407 | -0.7048871 | 0.60547094 | -1.1641965 | 0.24434439 | 0.81468457 |
| C1QB | 5089.01038 | -0.625487 | 0.62543047 | -1.0000904 | 0.31726676 | 0.84819508 |
| EPHB2 | 592.257846 | -0.8992835 | 0.38952882 | -2.3086444 | 0.02096332 | 0.50728403 |
| KDM1A | 3292.88489 | -0.0095119 | 0.11508246 | -0.0826526 | 0.93412781 | 0.99029255 |
| AL031428.1 | 57.6015502 | -0.4446481 | 0.43936457 | -1.0120254 | 0.3115259 | 0.84529065 |
| LUZP1 | 1609.94641 | -0.1578496 | 0.17770361 | -0.8882743 | 0.37439321 | 0.86791605 |
| LINC01355 | 680.496414 | -0.0551327 | 0.28589231 | -0.1928444 | 0.84708082 | 0.97684681 |
| HNRNPR | 5556.06255 | -0.3528858 | 0.25465441 | -1.3857438 | 0.16582514 | 0.75515387 |
| ZNF436 | 766.986041 | 0.02956479 | 0.21368534 | 0.13835666 | 0.88995855 | 0.98459146 |
| ZNF436-AS1 | 327.801153 | 0.30330541 | 0.33977686 | 0.89266058 | 0.37203897 | 0.86772685 |
| TCEA3 | 2718.03076 | 0.61762986 | 0.36377797 | 1.69782094 | 0.08954156 | 0.65375543 |
| ASAP3 | 2763.75021 | 0.42090696 | 0.33161877 | 1.26924948 | 0.2043521 | 0.78441247 |
| ID3 | 3707.77063 | -0.4188528 | 0.45534452 | -0.919859 | 0.35764644 | 0.86403383 |
| RPL11 | 26915.5475 | 0.19098104 | 0.17789861 | 1.07353864 | 0.28302951 | 0.83078187 |
| ELOA | 2539.0328 | -0.2422803 | 0.20305146 | -1.1931965 | 0.23279242 | 0.80667958 |
| ELOA-AS1 | 762.082837 | 0.09544433 | 0.13172926 | 0.7245492 | 0.46872863 | 0.90003342 |
| PITHD1 | 2208.49071 | -0.1519797 | 0.24769356 | -0.6135796 | 0.53949315 | 0.92198621 |
| LYPLA2 | 6259.23187 | -0.0919052 | 0.11548358 | -0.7958295 | 0.42613117 | 0.88665752 |
| GALE | 3635.74425 | 0.23876004 | 0.27256508 | 0.87597443 | 0.38104394 | 0.87171941 |
| HMGCL | 5655.88022 | 0.61345058 | 0.28147705 | 2.17939823 | 0.0293021 | 0.53606221 |
| FUCA1 | 5264.90484 | 0.34087315 | 0.18097411 | 1.88354649 | 0.05962635 | 0.6066891 |
| PNRC2 | 6283.05161 | -0.3383842 | 0.15030149 | -2.2513698 | 0.02436213 | 0.51967919 |
| SRSF10 | 4541.16331 | -0.2564623 | 0.14501425 | -1.7685314 | 0.07697211 | 0.63806922 |
| AL591178.1 | 331.972386 | 0.31693667 | 0.59117059 | 0.53611711 | 0.59187762 | 0.93289552 |
| MYOM3 | 3947.45404 | 0.28409388 | 0.48013892 | 0.59169102 | 0.55405751 | 0.92373235 |
| IL22RA1 | 526.935315 | 0.55075497 | 0.70163709 | 0.78495703 | 0.43247876 | 0.88980461 |
| IFNLR1 | 544.383425 | 0.05900403 | 0.23098446 | 0.25544587 | 0.79837874 | 0.9717324 |
| GRHL3 | 42.2846249 | 0.79512639 | 0.59984028 | 1.32556352 | 0.18498432 | 0.76820626 |
| STPG1 | 734.344807 | 0.02844629 | 0.16545838 | 0.17192414 | 0.86349716 | 0.98176973 |
| NIPAL3 | 1625.30554 | 0.30512184 | 0.1951588 | 1.56345413 | 0.11794582 | 0.69069542 |
| RCAN3 | 818.17603 | 0.30293967 | 0.28855467 | 1.04985191 | 0.29378621 | 0.83567608 |
| AL445686.2 | 184.553685 | 0.27525829 | 0.32754219 | 0.84037507 | 0.40069812 | 0.88098118 |
| NCMAP | 387.861362 | 0.69912008 | 0.45045317 | 1.55203722 | 0.12065332 | 0.69467399 |
| SRRM1 | 3316.99061 | -0.0730205 | 0.14590982 | -0.5004499 | 0.61675834 | 0.93910158 |
| CLIC4 | 17250.4866 | -0.0636783 | 0.27309312 | -0.2331742 | 0.81562611 | 0.9738447 |
| RUNX3 | 265.052337 | -0.0389782 | 0.6790988 | -0.0573969 | 0.95422902 | 0.99170503 |
| SYF2 | 2843.66181 | -0.0444677 | 0.16282767 | -0.2730965 | 0.78477907 | 0.97021865 |
| RSRP1 | 8270.17342 | 0.09548849 | 0.2274292 | 0.41986031 | 0.67458751 | 0.94935413 |
| RHD | 79.7939028 | -0.6774118 | 1.32451285 | -0.5114422 | 0.60904143 | 0.93748023 |
| TMEM50A | 6295.851 | -0.1063215 | 0.12058146 | -0.88174 | 0.37791742 | 0.8699853 |
| RHCE | 256.68546 | -0.0211777 | 0.20957274 | -0.1010517 | 0.91950942 | 0.9872458 |
| MACO1 | 1837.76294 | -0.466644 | 0.26597258 | -1.7544816 | 0.07934802 | 0.64079926 |
| LDLRAP1 | 1746.5509 | 0.13136377 | 0.13666602 | 0.96120291 | 0.33645015 | 0.85679125 |

| | | | | | | |
|---|---|---|---|---|---|---|
| MAN1C1 | 3635.6175 | 0.97702351 | 0.34569258 | 2.82627853 | 0.00470923 | 0.29927103 |
| SELENON | 7749.55697 | -0.0134895 | 0.23237306 | -0.0580509 | 0.95370808 | 0.99170503 |
| AL020996.2 | 2219.13389 | 0.05643621 | 0.12827184 | 0.43997351 | 0.65995629 | 0.94626603 |
| AL020996.1 | 2009.99679 | 0.09515945 | 0.23520916 | 0.40457372 | 0.68579088 | 0.95097288 |
| MTFR1L | 5782.52181 | 0.11722539 | 0.18220003 | 0.64338844 | 0.51997208 | 0.91657872 |
| AL020996.3 | 1451.91033 | 0.20207635 | 0.17906991 | 1.12847745 | 0.25911834 | 0.82364948 |
| AUNIP | 1543.64071 | 0.19417219 | 0.17750979 | 1.09386751 | 0.27401309 | 0.82719197 |
| AL033528.2 | 1175.26737 | 0.64159087 | 0.34909772 | 1.83785466 | 0.06608383 | 0.62276463 |
| PAQR7 | 4448.76344 | 0.61762231 | 0.32932245 | 1.87543333 | 0.06073313 | 0.60964095 |
| STMN1 | 2279.83833 | 0.06506406 | 0.33737857 | 0.1928518 | 8.47E-01 | 9.77E-01 |
| PAFAH2 | 1928.46873 | 0.37898758 | 0.2449459 | 1.54722973 | 0.12180786 | 0.69725141 |
| EXTL1 | 48.7545094 | 0.16336019 | 0.47736938 | 0.3422092 | 7.32E-01 | 0.95848028 |
| SLC30A2 | 3448.11494 | -0.1353655 | 0.56699221 | -0.2387431 | 0.81130476 | 0.97352497 |
| TRIM63 | 68.6806471 | 1.66687216 | 0.56678461 | 2.94092699 | 0.00327232 | 0.25133805 |
| PDIK1L | 464.515256 | 0.3405176 | 0.24142523 | 1.41044743 | 0.15840761 | 0.7466592 |
| FAM110D | 276.399904 | -0.2879228 | 0.51916118 | -0.5545924 | 0.57917352 | 0.93139743 |
| AL391650.1 | 814.092821 | -0.153728 | 0.17708935 | -0.8680814 | 0.38534976 | 0.87368012 |
| ZNF593 | 808.819647 | -0.1536847 | 0.17652899 | -0.8705916 | 0.38397721 | 0.87278555 |
| CNKSR1 | 1182.86231 | -0.181063 | 0.27233035 | -0.6648653 | 0.50613664 | 0.91391336 |
| CEP85 | 805.061494 | -0.4777684 | 0.36276839 | -1.3170066 | 0.1878364 | 0.76970626 |
| SH3BGRL3 | 5613.072 | -0.3253659 | 0.27424856 | -1.1863906 | 2.35E-01 | 8.07E-01 |
| UBXN11 | 5646.32369 | -0.0961831 | 0.17766367 | -0.5413771 | 0.58824766 | 0.93289552 |
| CD52 | 290.090903 | 0.50919097 | 0.8268031 | 0.61585518 | 0.5379901 | 0.92127611 |
| CRYBG2 | 137.16785 | 0.06822221 | 0.67974964 | 0.10036373 | 0.92005556 | 0.98733173 |
| DHDDS | 2314.93813 | -0.0172095 | 0.24513501 | -0.0702043 | 0.94403108 | 0.99136103 |
| HMGN2 | 6186.11141 | -0.0175323 | 0.30126213 | -0.0581963 | 0.95359228 | 0.99170503 |
| RPS6KA1 | 4721.11877 | 0.07279862 | 0.26860099 | 0.27102884 | 0.78636885 | 0.97083917 |
| AL512408.1 | 41.7020517 | -0.0360567 | 0.68555032 | -0.0525953 | 0.95805438 | 0.99227301 |
| ARID1A | 3402.5306 | -0.1912015 | 0.0897945 | -2.1293234 | 0.03322751 | 0.55156489 |
| PIGV | 963.869172 | 0.13257044 | 0.25894198 | 0.51196965 | 0.60867225 | 0.93748023 |
| ZDHHC18 | 2242.09294 | -0.6139163 | 0.3691994 | -1.6628314 | 0.09634619 | 0.66233356 |
| SFN | 2596.81901 | -2.7220544 | 1.11236718 | -2.4470826 | NA | NA |
| AL034380.1 | 169.195024 | -0.2472348 | 0.22067137 | -1.1203755 | 0.26255378 | 0.82424998 |
| GPN2 | 2092.18481 | -0.1546635 | 0.15580795 | -0.992655 | 0.32087813 | 0.85106295 |
| GPATCH3 | 918.310153 | -0.3638832 | 0.28616856 | -1.2715694 | 0.20352617 | 0.78338136 |
| NUDC | 5213.45263 | -0.1084086 | 0.14036784 | -0.7723183 | 0.43992595 | 0.89208528 |
| NR0B2 | 583.712435 | -0.5811216 | 0.48803297 | -1.1907426 | 0.23375465 | 0.80725541 |
| KDF1 | 831.130468 | -0.1242175 | 0.17857159 | -0.6956175 | 0.48666837 | 0.90573694 |
| TRNP1 | 1839.00979 | 0.02877057 | 0.37635028 | 0.07644624 | 0.93906408 | 0.99113899 |
| TENT5B | 400.649859 | -1.0511254 | 0.5818889 | -1.8064022 | 0.07085554 | 0.63023427 |
| SLC9A1 | 2223.28596 | -0.2656081 | 0.17545352 | -1.5138373 | 0.13006712 | 0.70814767 |
| WDTC1 | 5411.94213 | 0.01265296 | 0.15813393 | 0.08001421 | 9.36E-01 | 0.99068685 |
| TMEM222 | 2854.38411 | -0.3034373 | 0.20233618 | -1.4996691 | 0.13370013 | 0.71505705 |
| SYTL1 | 431.908077 | 0.3424032 | 0.56749441 | 0.6033596 | 0.54626949 | 0.92345821 |
| MAP3K6 | 3358.45283 | -0.0633339 | 0.27792061 | -0.227885 | 0.81973563 | 0.97454841 |
| FCN3 | 1387.71436 | -0.5225577 | 0.73599191 | -0.7100047 | 4.78E-01 | 0.90387017 |
| GPR3 | 93.3835864 | -0.2676314 | 0.63167533 | -0.423685 | 0.67179556 | 0.94922827 |
| WASF2 | 7116.23134 | -0.018365 | 0.15882111 | -0.1156335 | 0.90794304 | 0.98662757 |
| AHDC1 | 3453.96882 | -0.3438053 | 0.33608606 | -1.022968 | 0.306323 | 0.84278367 |
| FGR | 796.303772 | -0.8215323 | 0.40741231 | -2.0164642 | 0.04375146 | 0.57059577 |
| IFI6 | 16346.2949 | -3.8897182 | 1.28173222 | -3.0347355 | 0.00240747 | 0.21083339 |
| AL445490.1 | 58.5911541 | -4.2874164 | 1.58097028 | -2.7118893 | NA | NA |
| FAM76A | 812.376625 | -0.1366418 | 0.1585098 | -0.8620401 | 0.38866545 | 0.87568585 |
| STX12 | 3551.12222 | 0.03248674 | 0.15218655 | 0.21346659 | 0.83096304 | 0.97550416 |

| | | | | | | |
|---|---|---|---|---|---|---|
| PPP1R8 | 1774.8334 | -0.2383276 | 0.18903944 | -1.2607297 | 0.20740626 | 0.78813404 |
| THEMIS2 | 1320.18574 | -0.2375352 | 0.51041952 | -0.4653725 | 0.64166471 | 0.94088547 |
| RPA2 | 1733.69454 | 0.2387317 | 0.21096187 | 1.13163435 | 0.25778819 | 0.82187662 |
| SMPDL3B | 645.801987 | 0.18963539 | 0.3342543 | 0.56733868 | 0.5704841 | 0.92759349 |
| AL512288.1 | 289.103122 | 0.15202045 | 0.35430166 | 0.42907066 | 0.66787181 | 0.94847921 |
| XKR8 | 810.550717 | -0.1943491 | 0.18903137 | -1.0281317 | 0.30388789 | 0.84211658 |
| EYA3 | 2051.08556 | 0.09901509 | 0.22648175 | 0.43718794 | 0.66197506 | 0.94642611 |
| PTAFR | 400.468983 | 0.0959659 | 0.42892142 | 0.22373773 | 8.23E-01 | 0.9753443 |
| DNAJC8 | 4216.97183 | -0.2021625 | 0.18007338 | -1.1226674 | 0.2615788 | 0.82424998 |
| ATP5IF1 | 7091.62587 | 0.31866195 | 0.15378795 | 2.07208666 | 0.03825736 | 0.55538555 |
| AL353622.1 | 421.028758 | 0.3556689 | 0.38577116 | 0.92196861 | 0.35654495 | 0.86390849 |
| SESN2 | 2768.47061 | -0.1316645 | 0.28776114 | -0.457548 | 0.6472772 | 0.94277004 |
| MED18 | 569.576468 | -0.0243883 | 0.22020667 | -0.1107521 | 0.91181294 | 0.98662757 |
| PHACTR4 | 2622.49466 | -0.3790418 | 0.16791332 | -2.2573661 | 2.40E-02 | 5.20E-01 |
| RCC1 | 1757.7312 | -0.4068688 | 0.28708822 | -1.4172257 | 0.15641696 | 0.74522306 |
| TRNAU1AP | 1068.95375 | -0.1170908 | 0.15765284 | -0.7427131 | 0.45765542 | 0.89688594 |
| SNHG12 | 1866.59497 | -0.3504112 | 0.25964569 | -1.3495747 | 0.17715246 | 0.76115956 |
| TAF12 | 757.876705 | 0.1025546 | 0.11604099 | 0.88377903 | 0.37681551 | 0.86987124 |
| RAB42 | 157.612144 | 0.75331308 | 0.79279541 | 0.95019859 | 0.34201135 | 0.85889221 |
| GMEB1 | 650.543495 | -0.422582 | 0.24806186 | -1.7035349 | 0.08846802 | 0.65363785 |
| YTHDF2 | 3545.9155 | -0.2061052 | 0.22852602 | -0.9018895 | 0.36711556 | 0.86683872 |
| EPB41 | 2744.3513 | 0.09645124 | 0.17990476 | 0.53612392 | 0.59187291 | 0.93289552 |
| TMEM200B | 1516.72293 | 0.14382683 | 0.1932917 | 0.7440921 | 0.45682076 | 0.89688594 |
| SRSF4 | 6035.49502 | -0.1668381 | 0.1447636 | -1.1524869 | 0.24912105 | 0.81646572 |
| MECR | 1189.99469 | 0.35193176 | 0.16505436 | 2.13221726 | 0.03298899 | 0.55137197 |
| PTPRU | 1321.41514 | 0.51812286 | 0.26521676 | 1.95358265 | 0.05075059 | 0.5861488 |
| MATN1 | 41.9992839 | 0.00031977 | 0.44579817 | 0.0007173 | 9.99E-01 | 1.00E+00 |
| MATN1-AS1 | 99.1415999 | 0.16813757 | 0.37824561 | 0.44451955 | 6.57E-01 | 0.94545756 |
| LAPTM5 | 4195.99163 | -0.6280731 | 0.4527617 | -1.3872047 | 0.16537936 | 0.75428926 |
| SDC3 | 3015.42901 | 0.38073675 | 0.23587349 | 1.61415657 | 0.10649347 | 0.67674128 |
| PUM1 | 4019.42357 | -0.0923785 | 0.1390665 | -0.6642759 | 0.50651375 | 0.9141539 |
| SNRNP40 | 1386.85664 | -0.0312053 | 0.15462119 | -0.2018178 | 0.84005916 | 0.97618283 |
| ZCCHC17 | 1488.89039 | -0.092496 | 0.14047971 | -0.6584293 | 0.5102623 | 0.91439937 |
| AL451070.1 | 500.194592 | 0.03919314 | 0.19282079 | 0.20326201 | 8.39E-01 | 9.76E-01 |
| FABP3 | 4530.4565 | 0.09143635 | 0.4648108 | 0.19671735 | 0.84404873 | 0.97664894 |
| SERINC2 | 13963.9978 | -0.4082604 | 0.3053501 | -1.3370238 | 0.18121486 | 0.76522169 |
| TINAGL1 | 17317.3222 | -0.2552203 | 0.27415112 | -0.9309474 | 0.3518808 | 0.86364633 |
| HCRTR1 | 558.94997 | -0.0944451 | 0.17676257 | -0.5343048 | 0.59313069 | 0.93289552 |
| PEF1 | 4558.93061 | -0.1172658 | 0.12540428 | -0.9351018 | 0.34973583 | 0.86364633 |
| AC114488.2 | 549.566334 | 0.10041631 | 0.29418465 | 0.34133769 | 7.33E-01 | 9.58E-01 |
| AC114488.3 | 158.437445 | 0.13426019 | 0.28823315 | 0.46580411 | 0.64135572 | 0.94088547 |
| COL16A1 | 3542.36902 | 0.03102892 | 0.36701087 | 0.08454496 | 0.93262316 | 0.9899349 |
| ADGRB2 | 221.841249 | 0.93485579 | 0.45138228 | 2.07109545 | 0.03834988 | 0.55538555 |
| SPOCD1 | 72.9910533 | -0.3294894 | 0.69099614 | -0.4768325 | 0.63348138 | 0.94088547 |
| PTP4A2 | 9178.81635 | -0.2490676 | 0.26055866 | -0.9558983 | 0.33912361 | 0.85733087 |
| KHDRBS1 | 5607.75529 | -0.1393027 | 0.18668276 | -0.7462002 | 0.4555465 | 0.89688594 |
| TMEM39B | 710.582748 | -0.3391533 | 0.17475518 | -1.9407336 | 0.05229059 | 0.58645127 |
| KPNA6 | 3746.12188 | -0.1902464 | 0.16556812 | -1.149052 | 0.25053453 | 0.81738655 |
| TXLNA | 4438.60149 | -0.3032073 | 0.19330885 | -1.5685123 | 0.11676161 | 0.68913995 |
| CCDC28B | 431.370212 | -0.0099276 | 0.27310873 | -0.0363505 | 0.97100289 | 0.99400331 |
| AL049795.1 | 296.212895 | 0.09852194 | 0.24012872 | 0.41028804 | 0.68159466 | 0.95070301 |
| IQCC | 332.080133 | 0.14339104 | 0.22989495 | 0.62372419 | 0.53280873 | 0.92014084 |
| DCDC2B | 455.718715 | -0.1587645 | 0.34945516 | -0.4543202 | 0.64959837 | 0.94312988 |
| TMEM234 | 1405.83077 | -0.1806373 | 0.25094129 | -0.719839 | 0.4716241 | 0.90160919 |

| | | | | | | |
|---|---|---|---|---|---|---|
| EIF3I | 7217.39682 | 0.04534733 | 0.17371878 | 0.26103872 | 0.79406264 | 0.97090793 |
| FAM167B | 828.410704 | -0.479995 | 0.52883869 | -0.9076398 | 0.36406857 | 0.8663515 |
| LCK | 299.552592 | 0.1856682 | 0.79889967 | 0.2324049 | 0.81622354 | 0.97388289 |
| HDAC1 | 4353.46695 | 0.01104184 | 0.13711686 | 0.08052868 | 0.93581678 | 0.99068685 |
| MARCKSL1 | 2226.00957 | -0.4790227 | 0.37131055 | -1.2900865 | 0.19702064 | 0.77816059 |
| TSSK3 | 578.7584 | -0.1396258 | 0.30215536 | -0.4620994 | 0.64401003 | 0.94208859 |
| FAM229A | 956.119388 | -0.1382212 | 0.26603524 | -0.5195596 | 0.60337059 | 0.93602816 |
| BSDC1 | 4827.52683 | -0.1929006 | 0.11758441 | -1.6405287 | 0.10089528 | 0.67247782 |
| AL033529.1 | 94.1399156 | -0.1704174 | 0.3350731 | -0.5085977 | 0.61103421 | 0.93833106 |
| ZBTB8A | 361.248948 | -0.2020218 | 0.26648335 | -0.7581029 | 0.44838941 | 0.8948759 |
| ZBTB8OS | 1121.96175 | -0.0469694 | 0.16068202 | -0.292313 | 0.77004731 | 0.96836854 |
| RBBP4 | 6116.6281 | -0.0332872 | 0.12702391 | -0.2620549 | 0.79327909 | 0.97090793 |
| SYNC | 944.217434 | -0.1724832 | 0.12003305 | -1.4369639 | 0.15072826 | 0.73796431 |
| KIAA1522 | 11632.2214 | -0.1636671 | 0.20850797 | -0.7849443 | 0.43248624 | 0.88980461 |
| YARS | 3489.72028 | -0.1979347 | 0.20643277 | -0.9588337 | 0.33764251 | 0.8571036 |
| S100PBP | 1445.69866 | -0.1165435 | 0.17561151 | -0.6636439 | 0.50691823 | 0.91420888 |
| FNDC5 | 71.8367202 | 0.3912925 | 0.44023888 | 0.88881858 | 0.37410059 | 0.86791605 |
| HPCA | 1007.22879 | -0.3123318 | 0.43020521 | -0.7260065 | 4.68E-01 | 9.00E-01 |
| TMEM54 | 2545.60571 | -0.282911 | 0.43706611 | -0.6472957 | 0.5174406 | 0.91636259 |
| RNF19B | 2798.0044 | -0.2253045 | 0.34633991 | -0.6505299 | 0.51534998 | 0.91636259 |
| AL020995.1 | 96.7884264 | 0.00837012 | 0.26095761 | 0.03207465 | 0.97441252 | 0.99455099 |
| AK2 | 10184.725 | 0.20588913 | 0.25108286 | 0.82000472 | 0.41221342 | 0.88268208 |
| AZIN2 | 654.91138 | -0.1310709 | 0.24533599 | -0.5342507 | 0.59316813 | 0.93289552 |
| TRIM62 | 536.617867 | -0.2203467 | 0.2427789 | -0.9076024 | 0.36408832 | 0.8663515 |
| ZNF362 | 1755.20344 | 0.13978724 | 0.23778342 | 0.58787631 | 0.55661532 | 0.92373235 |
| AL513327.1 | 558.510418 | -0.5681092 | 0.31053126 | -1.8294749 | 0.06732849 | 0.6249306 |
| PHC2 | 16756.7907 | -0.6757027 | 0.22344873 | -3.0239721 | 0.00249479 | 0.21493583 |
| ZSCAN20 | 134.322888 | -0.5329312 | 0.29315789 | -1.8178983 | 0.06907968 | 0.62779744 |
| SMIM12 | 2570.6334 | -0.3159238 | 0.26184032 | -1.2065516 | 0.22760489 | 0.80300331 |
| GJB3 | 45.5796813 | 1.8623466 | 0.72994376 | 2.55135628 | 0.01073046 | 0.40196898 |
| GJA4 | 1613.70689 | -0.7899919 | 0.45882795 | -1.7217606 | 0.08511289 | 0.64988556 |
| AL122010.1 | 157.175522 | -0.7036135 | 0.42746813 | -1.6460022 | 0.0997633 | 0.67115544 |
| DLGAP3 | 93.0119224 | -0.0351756 | 0.37437051 | -0.0939593 | 0.92514151 | 0.98869145 |
| AC114490.3 | 699.355028 | 0.35454151 | 0.22378837 | 1.5842714 | 0.11313197 | 0.68130021 |
| AC114490.2 | 1745.51031 | 0.14195468 | 0.11551985 | 1.2288336 | 0.2191342 | 0.79847731 |
| TMEM35B | 1363.43981 | 0.24034135 | 0.22084389 | 1.08828616 | 0.2764688 | 0.82730125 |
| ZMYM6 | 1329.91085 | -0.0438349 | 0.15934753 | -0.27509 | 0.78324709 | 0.96971192 |
| ZMYM1 | 474.205894 | 0.07871859 | 0.20148006 | 0.39070167 | 0.69601777 | 0.9525827 |
| SFPQ | 12770.7382 | -0.193598 | 0.19005874 | -1.0186218 | 0.30838255 | 0.84388103 |
| ZMYM4 | 2230.33434 | -0.1563726 | 0.14490591 | -1.0791317 | 0.28052904 | 0.82871922 |
| KIAA0319L | 5040.08842 | 0.11743292 | 0.19575507 | 0.5998972 | 0.54857475 | 0.92373235 |
| NCDN | 2079.13583 | -0.2908351 | 0.14708208 | -1.9773659 | 0.04800029 | 0.58413853 |
| TFAP2E | 77.0398034 | 0.29274318 | 0.35273474 | 0.82992443 | 0.40658151 | 0.88132015 |
| PSMB2 | 4143.37777 | -0.0836141 | 0.19331356 | -0.4325311 | 0.66535549 | 0.94716499 |
| C1orf216 | 589.827697 | -0.3705696 | 0.25090822 | -1.4769129 | 0.139699 | 0.72475166 |
| AGO4 | 1442.43139 | -0.1612285 | 0.1287581 | -1.2521811 | 0.21050389 | 0.79134644 |
| AGO1 | 1709.51891 | 0.14751848 | 0.14972932 | 0.98523442 | 0.32450894 | 0.85169085 |
| AGO3 | 1410.99076 | -0.1498084 | 0.11228364 | -1.3341963 | 0.18213954 | 0.76569697 |
| TEKT2 | 276.810043 | -0.1533314 | 0.60995478 | -0.2513816 | 0.8015191 | 0.97210205 |
| ADPRHL2 | 2923.88395 | -0.2340097 | 0.27448873 | -0.8525292 | 0.39392043 | 0.87884163 |
| COL8A2 | 210.386437 | 0.31326386 | 0.50097859 | 0.62530389 | 0.53177163 | 0.91962754 |
| TRAPPC3 | 3461.1684 | -0.2534045 | 0.2317921 | -1.0932404 | 0.27428825 | 0.82719407 |
| MAP7D1 | 4688.23396 | -0.5002811 | 0.31025737 | -1.6124714 | 0.1068594 | 0.67674128 |
| THRAP3 | 5469.33803 | -0.3167191 | 0.12403013 | -2.5535657 | 0.01066262 | 0.40196898 |

| | | | | | | |
|---|---|---|---|---|---|---|
| SH3D21 | 2552.63405 | 0.30860035 | 0.19268928 | 1.601544 | 0.10925648 | 0.67859123 |
| EVA1B | 1424.24687 | -0.0627694 | 0.15438151 | -0.4065864 | 0.68431178 | 0.95070301 |
| AL591845.1 | 844.382429 | 0.53738258 | 0.29726456 | 1.80775868 | 0.07064407 | 0.62981127 |
| STK40 | 5909.93652 | -0.3472824 | 0.35244282 | -0.9853581 | 3.24E-01 | 0.85169085 |
| LSM10 | 1129.60391 | 0.05281318 | 0.18861864 | 0.27999978 | 0.77947768 | 0.96836854 |
| OSCP1 | 713.422153 | 0.03147928 | 0.20321555 | 0.15490587 | 0.87689552 | 0.98275231 |
| MRPS15 | 3199.15873 | 0.01431933 | 0.17043944 | 0.08401418 | 0.93304516 | 0.99004639 |
| CSF3R | 898.343128 | -0.3908159 | 0.4687857 | -0.833677 | 0.40446301 | 0.88132015 |
| LINC01137 | 330.658231 | 0.39092207 | 0.24605877 | 1.58873454 | 0.11212034 | 0.68001961 |
| ZC3H12A | 2306.23448 | 0.17652477 | 0.53840133 | 0.32786838 | 0.74301118 | 0.96109001 |
| MEAF6 | 2542.12284 | -0.1155033 | 0.09583622 | -1.2052152 | 0.22812025 | 0.80300331 |
| SNIP1 | 793.534315 | -0.3329513 | 0.20912424 | -1.5921222 | 0.11135725 | 0.68001961 |
| DNALI1 | 1480.58462 | 0.06139745 | 0.30351522 | 0.20228787 | 0.83969168 | 0.97618283 |
| GNL2 | 1842.70465 | -0.3646676 | 0.25935125 | -1.4060763 | 1.60E-01 | 7.48E-01 |
| C1orf109 | 784.816778 | 0.02600778 | 0.24120456 | 0.10782458 | 0.91413485 | 0.98662757 |
| CDCA8 | 69.8863883 | 0.14605894 | 0.47613217 | 0.30676134 | 0.75902504 | 0.96502715 |
| EPHA10 | 54.0135085 | 1.17493176 | 0.74322757 | 1.58085061 | 0.1139122 | 0.68214043 |
| MANEAL | 274.570015 | 0.75117093 | 0.4924352 | 1.52542088 | 0.12715416 | 0.70481988 |
| AL929472.3 | 175.417097 | 0.64384407 | 0.5384311 | 1.19577801 | 0.2317832 | 0.80649152 |
| YRDC | 1006.04278 | -0.3369957 | 0.29895728 | -1.1272368 | 0.25964237 | 0.82424998 |
| C1orf122 | 2233.58959 | -0.0323964 | 0.18085929 | -0.1791247 | 0.85783981 | 0.97964261 |
| MTF1 | 985.283375 | -0.293853 | 0.22852846 | -1.2858485 | 1.98E-01 | 7.80E-01 |
| AL929472.2 | 544.836719 | 0.16483103 | 0.23773704 | 0.6933334 | 4.88E-01 | 0.90585781 |
| INPP5B | 1976.71777 | 0.17392939 | 0.20676105 | 0.84120966 | 0.40023049 | 0.88077967 |
| SF3A3 | 2909.56271 | -0.0610259 | 0.18612922 | -0.3278682 | 0.74301129 | 0.96109001 |
| FHL3 | 680.1686 | -0.4679575 | 0.4110561 | -1.1384273 | 0.25494208 | 0.81992039 |
| UTP11 | 1013.23899 | -0.0980353 | 0.2672724 | -0.3667992 | 0.71376881 | 0.95564093 |
| RRAGC | 2878.32406 | -0.7807358 | 0.46215379 | -1.689342 | 0.09115391 | 0.65495306 |
| AL139260.1 | 96.7790724 | -0.4199612 | 0.54552003 | -0.7698365 | 0.44139687 | 0.89290667 |
| MYCBP | 1384.50042 | 0.08642134 | 0.19484522 | 0.4435384 | 0.65737635 | 0.94558535 |
| AL139260.3 | 569.167615 | -0.0012076 | 0.16591997 | -0.0072785 | 0.99419265 | 0.99936518 |
| GJA9 | 57.3179343 | -1.0531335 | 0.50430178 | -2.0883002 | 0.03677076 | 0.55521179 |
| RHBDL2 | 111.328098 | -0.1438431 | 0.35955665 | -0.4000569 | 0.68911463 | 0.95158502 |
| AKIRIN1 | 4192.75855 | -0.2865936 | 0.14210225 | -2.0168125 | 4.37E-02 | 5.71E-01 |
| NDUFS5 | 5160.93439 | 0.10294096 | 0.2360635 | 0.43607317 | 0.66278364 | 0.94642611 |
| MACF1 | 9747.73738 | -0.1346706 | 0.22897394 | -0.5881483 | 0.55643277 | 0.92373235 |
| BMP8A | 239.56504 | 0.04389015 | 0.28767588 | 0.15256807 | 8.79E-01 | 0.98275231 |
| PABPC4 | 5104.61137 | -0.0386294 | 0.18717464 | -0.2063816 | 0.83649285 | 0.97577999 |
| AL365277.1 | 855.496359 | 0.11660658 | 0.24891843 | 0.46845296 | 0.6394607 | 0.94088547 |
| HEYL | 1077.4952 | -0.5947097 | 0.56706284 | -1.0487545 | 0.29429112 | 0.83615978 |
| HPCAL4 | 617.529982 | -0.3748011 | 0.72398804 | -0.5176896 | 0.60467486 | 0.93662382 |
| PPIE | 2005.93015 | 0.20099407 | 0.21824303 | 0.92096441 | 0.35706901 | 0.86390849 |
| BMP8B | 309.980864 | -0.1037667 | 0.30511036 | -0.3400957 | 0.73378445 | 0.95895327 |
| AL033527.5 | 43.4690755 | -0.9258396 | 0.54163135 | -1.7093539 | 0.08738542 | 0.65271545 |
| TRIT1 | 1259.83569 | -0.2445056 | 0.11659738 | -2.097008 | 0.03599287 | 0.55369025 |
| MYCL | 125.607139 | 0.33306964 | 0.38853274 | 0.85724987 | 0.39130681 | 0.87716276 |
| MFSD2A | 1784.70588 | 0.17589396 | 0.60974392 | 0.28847185 | 0.77298557 | 0.96836854 |
| CAP1 | 12028.5679 | -0.1376844 | 0.20876088 | -0.6595316 | 5.10E-01 | 9.14E-01 |
| PPT1 | 11898.0644 | -0.0600873 | 0.17038441 | -0.3526575 | 7.24E-01 | 0.95827709 |
| RLF | 1108.04361 | -0.24342 | 0.2170425 | -1.1215315 | 0.26206169 | 0.82424998 |
| AL050341.2 | 474.225035 | 0.00450517 | 0.36489009 | 0.01234666 | 0.99014904 | 0.99809685 |
| ZMPSTE24 | 3634.87271 | -0.1225386 | 0.22421148 | -0.5465314 | 0.58470068 | 0.93282331 |
| COL9A2 | 1553.3711 | 0.85023999 | 0.43123835 | 1.97162427 | 0.04865251 | 0.58448511 |
| SMAP2 | 2089.1472 | -0.152494 | 0.30124785 | -0.5062077 | 0.61271085 | 0.93879706 |

| | | | | | | |
|---|---|---|---|---|---|---|
| ZFP69 | 185.736721 | 0.10432145 | 0.26136953 | 0.39913392 | 0.68979453 | 0.95158502 |
| EXO5 | 319.59259 | -0.2509353 | 0.28839905 | -0.8700976 | 3.84E-01 | 8.73E-01 |
| AL603839.1 | 124.7336 | -0.1883218 | 0.35822337 | -0.5257105 | 0.59908938 | 0.93484879 |
| AL603839.2 | 70.9874295 | -0.3263885 | 0.35128196 | -0.9291353 | 3.53E-01 | 8.64E-01 |
| ZNF684 | 517.852811 | 0.59467743 | 0.2586991 | 2.29872245 | 2.15E-02 | 5.14E-01 |
| RIMS3 | 353.538819 | 0.51732751 | 0.41210047 | 1.25534316 | 2.09E-01 | 7.91E-01 |
| NFYC-AS1 | 362.144976 | -0.2401418 | 0.21477158 | -1.1181264 | 2.64E-01 | 0.82424998 |
| NFYC | 2947.0207 | -0.0668734 | 0.15648386 | -0.42735 | 0.66912441 | 0.9486589 |
| KCNQ4 | 110.823237 | -0.5131571 | 0.43451183 | -1.1809968 | 0.23760399 | 0.80962478 |
| CITED4 | 865.47705 | -0.1545238 | 0.27388359 | -0.5641952 | 0.57262131 | 0.92928554 |
| CTPS1 | 1726.77147 | -0.0428156 | 0.33954187 | -0.1260981 | 0.89965427 | 0.98506781 |
| SLFNL1-AS1 | 695.972323 | -0.0752723 | 0.19052722 | -0.3950738 | 0.69278841 | 0.9519683 |
| SLFNL1 | 132.74577 | 0.03368291 | 0.33421682 | 0.10078162 | 0.91972382 | 0.98731493 |
| SCMH1 | 3179.55019 | -0.0529438 | 0.16474114 | -0.3213756 | 0.74792574 | 0.96241535 |
| FOXO6 | 74.8059695 | -0.3094039 | 0.65290784 | -0.473886 | 0.63558122 | 0.94088547 |
| EDN2 | 361.923645 | 0.91894625 | 1.11557058 | 0.8237455 | 0.41008418 | 0.88249519 |
| HIVEP3 | 396.061328 | -0.0866544 | 0.27140654 | -0.3192791 | 7.50E-01 | 9.62E-01 |
| FOXJ3 | 2534.44992 | -0.3164332 | 0.17295511 | -1.8295686 | 6.73E-02 | 6.25E-01 |
| RIMKLA | 371.981024 | 0.44394371 | 0.47560593 | 0.93342762 | 0.35059922 | 0.86364633 |
| ZMYND12 | 332.25044 | 0.56843571 | 0.4431382 | 1.2827504 | 0.19957954 | 0.78134034 |
| PPCS | 3269.64323 | 0.48216511 | 0.24836694 | 1.94134179 | 0.05221683 | 0.58645127 |
| CCDC30 | 562.90854 | -0.1082447 | 0.30031192 | -0.3604409 | 7.19E-01 | 0.95648644 |
| PPIH | 633.411794 | -0.021079 | 0.16512391 | -0.1276555 | 0.89842161 | 0.98474725 |
| YBX1 | 16185.4997 | -0.1908604 | 0.2747612 | -0.6946411 | 0.48728026 | 0.90573694 |
| CLDN19 | 1466.61511 | 0.76031157 | 0.74208579 | 1.02456022 | 0.30557074 | 0.84270033 |
| P3H1 | 2023.38808 | -0.2559039 | 0.29159604 | -0.8775973 | 0.38016229 | 0.87090707 |
| C1orf50 | 570.665602 | 0.21486249 | 0.20180212 | 1.06471865 | 0.28700327 | 0.83155545 |
| AC098484.3 | 457.225111 | 0.23826009 | 0.2426952 | 0.98172561 | 0.32623505 | 0.85282528 |
| TMEM269 | 277.842595 | 0.13699776 | 0.22127455 | 0.61913022 | 0.53583058 | 0.92101152 |
| SVBP | 573.347361 | 0.09493767 | 0.18252599 | 0.52013236 | 0.60297132 | 0.93588323 |
| ERMAP | 1136.23002 | 0.1879119 | 0.20258 | 0.92759355 | 0.35361844 | 0.86364633 |
| AL512353.1 | 434.66456 | 0.13441884 | 0.25969304 | 0.51760664 | 0.60473275 | 0.93663607 |
| ZNF691 | 762.232576 | 0.08912421 | 0.15564025 | 0.57262955 | 0.56689555 | 0.9256133 |
| SLC2A1 | 7115.5216 | 0.00591494 | 0.4613396 | 0.01282123 | 0.98977042 | 0.99807483 |
| SLC2A1-AS1 | 55.1142386 | 0.97680357 | 0.66865081 | 1.46085753 | 0.14405454 | 0.72926154 |
| FAM183A | 54.1405586 | 0.5815341 | 0.51850506 | 1.12155916 | 2.62E-01 | 8.24E-01 |
| EBNA1BP2 | 3398.48552 | -0.0390029 | 0.22338535 | -0.1745992 | 0.86139457 | 0.98167219 |
| CFAP57 | 829.317881 | 0.03747517 | 0.2943783 | 0.12730275 | 0.89870078 | 0.98477645 |
| TMEM125 | 962.151272 | 0.00999846 | 0.33312478 | 0.03001415 | 0.97605577 | 0.99455099 |
| C1orf210 | 1451.08025 | 0.08999326 | 0.42571257 | 0.21139442 | 0.83257951 | 0.97550416 |
| TIE1 | 2518.59056 | -0.3904413 | 0.38644275 | -1.0103471 | 0.31232904 | 0.84541735 |
| CDC20 | 157.474702 | 0.61457035 | 0.43490916 | 1.4131005 | 0.15762618 | 0.74522306 |
| ELOVL1 | 4998.67283 | -0.2145344 | 0.15688962 | -1.3674224 | 1.71E-01 | 7.60E-01 |
| MED8 | 1956.6552 | -0.2120448 | 0.17133928 | -1.2375726 | 0.21587457 | 0.7954827 |
| AL139289.1 | 485.922995 | -0.1744164 | 0.19889983 | -0.8769059 | 0.38053775 | 0.87120181 |
| SZT2 | 5995.3447 | -0.051767 | 0.10326922 | -0.5012819 | 0.61617272 | 0.93910158 |
| SZT2-AS1 | 508.706431 | -0.3621732 | 0.13513194 | -2.6801447 | 0.00735904 | 0.3610358 |
| HYI | 3392.67312 | 0.1214209 | 0.15415228 | 0.78766856 | 0.4308906 | 0.88966606 |
| PTPRF | 7500.33812 | 0.05158626 | 0.15040393 | 0.34298478 | 0.73160991 | 0.95848028 |
| AL451062.3 | 1598.55802 | 0.1952918 | 0.13293262 | 1.46910361 | 0.14180469 | 0.72784886 |
| KDM4A | 1988.9221 | 0.01185751 | 0.14800282 | 0.08011679 | 0.93614437 | 0.99068685 |
| KDM4A-AS1 | 786.443996 | -0.0348636 | 0.17134585 | -0.2034693 | 0.83876822 | 0.97612168 |
| ST3GAL3 | 815.008559 | 0.3130458 | 0.19692362 | 1.58968132 | 0.11190666 | 0.68001961 |
| ARTN | 336.138241 | -0.0274804 | 0.30277192 | -0.0907628 | 0.92768106 | 0.98917108 |

| | | | | | | |
|---|---|---|---|---|---|---|
| IPO13 | 4165.37568 | -0.13497 | 0.27351201 | -0.4934702 | 0.6216804 | 0.94016713 |
| DPH2 | 1749.95037 | -0.2126802 | 0.26297503 | -0.8087467 | 0.41866088 | 0.88451372 |
| ATP6V0B | 10388.094 | -0.0486903 | 0.19082613 | -0.2551552 | 0.7986032 | 0.9717324 |
| B4GALT2 | 2138.54204 | -0.336113 | 0.18875983 | -1.7806383 | 0.07497157 | 0.63773415 |
| CCDC24 | 1845.664 | 0.04915674 | 0.35369756 | 0.13897959 | 0.88946627 | 0.98446024 |
| SLC6A9 | 817.592188 | -0.599358 | 0.20645632 | -2.9030741 | 0.00369519 | 0.26317334 |
| DMAP1 | 2495.29205 | -0.0674034 | 0.14119083 | -0.4773923 | 0.63308279 | 0.94088547 |
| ERI3 | 3695.51112 | 0.13162261 | 0.21816105 | 0.60332774 | 0.54629068 | 0.92345821 |
| RNF220 | 3367.71484 | -0.1075213 | 0.12215027 | -0.8802377 | 0.37873057 | 0.87069278 |
| TMEM53 | 1885.35272 | 0.12163232 | 0.1502931 | 0.80930074 | 0.41834218 | 0.88445833 |
| ARMH1 | 177.394918 | 0.40848726 | 0.52566453 | 0.77708736 | 0.43710723 | 0.89130865 |
| KIF2C | 80.3948761 | -0.0225573 | 0.5828494 | -0.0387018 | 0.96912812 | 0.99400331 |
| AL592166.1 | 3140.2464 | 0.15430352 | 0.24141825 | 0.63915432 | 0.52272254 | 0.91723878 |
| RPS8 | 26217.7439 | 0.17400567 | 0.20867684 | 0.83385236 | 0.40436419 | 0.88132015 |
| BEST4 | 51.5650977 | 0.29863496 | 0.4891383 | 0.61053277 | 0.54150894 | 0.92292765 |
| PLK3 | 1097.17491 | -0.0806912 | 0.50506127 | -0.1597651 | 0.87306608 | 0.98275231 |
| TCTEX1D4 | 55.477138 | 0.25449992 | 0.42812336 | 0.59445465 | 0.55220807 | 0.92373235 |
| BTBD19 | 675.755489 | -0.0619014 | 0.29816077 | -0.2076107 | 0.83553292 | 0.97550416 |
| PTCH2 | 209.15125 | -0.5620997 | 0.40174467 | -1.3991466 | 0.16176903 | 0.7498147 |
| EIF2B3 | 866.960741 | 0.26555653 | 0.16324919 | 1.62669432 | 0.10380204 | 0.67439401 |
| HECTD3 | 4047.52926 | -0.1480114 | 0.18970234 | -0.7802295 | 0.43525582 | 0.89055338 |
| UROD | 4428.88258 | 0.10245296 | 0.17247744 | 0.59400792 | 0.55250682 | 0.92373235 |
| ZSWIM5 | 497.70607 | 0.06203354 | 0.24019064 | 0.25826794 | 0.79620012 | 0.97101097 |
| LINC01144 | 100.935164 | 0.56290537 | 0.40132183 | 1.40262835 | 0.16072769 | 0.74855808 |
| MUTYH | 1348.5035 | 0.04220175 | 0.21380008 | 0.19738885 | 8.44E-01 | 9.76E-01 |
| TOE1 | 736.147015 | -0.2248813 | 0.22143649 | -1.0155567 | 3.10E-01 | 8.45E-01 |
| TESK2 | 389.71347 | -0.3078959 | 0.24626112 | -1.2502821 | 2.11E-01 | 7.91E-01 |
| CCDC163 | 184.307052 | -0.4933665 | 0.79395738 | -0.6214017 | 0.53433533 | 0.92084006 |
| MMACHC | 13134.3537 | -0.0070269 | 0.32670204 | -0.0215086 | 0.98283997 | 0.9961464 |
| PRDX1 | 28942.1181 | 0.00139524 | 0.34073681 | 0.00409477 | 0.99673285 | 0.99976751 |
| AKR1A1 | 13363.5038 | 0.68237722 | 0.29047882 | 2.34914617 | 0.01881652 | 0.48174914 |
| NASP | 3985.24575 | -0.2449549 | 0.21760341 | -1.1256941 | 0.26029502 | 0.82424998 |
| CCDC17 | 497.289243 | -0.2675827 | 0.4581036 | -0.5841095 | 0.55914662 | 0.92421175 |
| GPBP1L1 | 5890.1069 | -0.1646433 | 0.12004235 | -1.3715435 | 0.1702056 | 0.75956362 |
| TMEM69 | 1236.38822 | -0.1534221 | 0.13990542 | -1.0966132 | 0.27281053 | 0.82716979 |
| IPP | 540.164947 | 0.26409975 | 0.21744289 | 1.21457067 | 0.22452988 | 0.80078694 |
| MAST2 | 2722.61425 | -0.2222326 | 0.26803886 | -0.8291061 | 0.40704439 | 0.88132015 |
| PIK3R3 | 1769.34103 | 0.10330657 | 0.36553773 | 0.28261533 | 0.77747173 | 0.96836854 |
| AL358075.4 | 119.491479 | -0.2128474 | 0.51611503 | -0.412403 | 0.68004408 | 0.95070301 |
| TSPAN1 | 24544.1162 | 0.23428817 | 0.25218613 | 0.92902876 | 0.35287418 | 0.86364633 |
| POMGNT1 | 5741.70851 | 0.17390382 | 0.14626896 | 1.18893182 | 0.2344665 | 0.80730051 |
| LURAP1 | 314.191233 | -0.0918829 | 0.4198058 | -0.2188701 | 8.27E-01 | 0.97548201 |
| RAD54L | 373.447723 | -0.1734007 | 0.26303316 | -0.6592351 | 0.50974483 | 0.91439937 |
| LRRC41 | 5724.86038 | -0.3327412 | 0.1840551 | -1.8078346 | 0.07063225 | 0.62981127 |
| UQCRH | 6336.99083 | 0.33153103 | 0.16169932 | 2.05029335 | 0.04033581 | 0.55657615 |
| NSUN4 | 1408.02978 | -0.1807073 | 0.1508582 | -1.1978621 | 0.2309707 | 0.80610188 |
| FAAH | 3489.27283 | -0.0768595 | 0.37555817 | -0.204654 | 0.83784243 | 0.9758253 |
| MKNK1-AS1 | 447.050027 | -0.1823676 | 0.17527582 | -1.0404607 | 0.29812589 | 0.83998573 |
| MKNK1 | 2096.36304 | -0.1986929 | 0.13712024 | -1.4490411 | 0.14732612 | 0.73386234 |
| MOB3C | 1477.97669 | -0.2371535 | 0.35083692 | -0.6759651 | 0.49906281 | 0.91147492 |
| ATPAF1 | 3781.62914 | 0.38148838 | 0.22135917 | 1.72339093 | 0.08481785 | 0.64988556 |
| EFCAB14-AS | 89.4680091 | -0.3166893 | 0.4545466 | -0.6967147 | 0.48598137 | 0.90545573 |
| EFCAB14 | 6271.49795 | 0.08837427 | 0.12963141 | 0.68173496 | 0.49540656 | 0.90847703 |
| CYP4B1 | 136.314979 | 0.15446211 | 0.85902042 | 0.17981192 | 0.85730023 | 0.97953896 |

| | | | | | | |
|---|---|---|---|---|---|---|
| CYP4A11 | 27627.7766 | 1.26981516 | 0.73271198 | 1.73303453 | 0.08308953 | 0.6471083 |
| CYP4X1 | 325.959259 | 0.19742428 | 0.43193202 | 0.45707258 | 0.64761888 | 0.94282139 |
| CYP4A22-AS1 | 101.182891 | -0.5884044 | 0.97613158 | -0.6027921 | 0.54664702 | 0.92345821 |
| CYP4A22 | 4419.25676 | 1.20506932 | 1.03687254 | 1.16221548 | 0.24514795 | 0.81492642 |
| LINC00853 | 112.37541 | 0.671527 | 0.76335801 | 0.87970126 | 0.37902117 | 0.87069278 |
| PDZK1IP1 | 49030.0912 | 0.38248194 | 0.34539236 | 1.10738391 | 2.68E-01 | 8.27E-01 |
| TAL1 | 438.511214 | -0.3863903 | 0.40265155 | -0.9596145 | 0.33724927 | 0.85700184 |
| AL135960.1 | 80.0791938 | -0.2006984 | 0.45634883 | -0.4397917 | 0.66008799 | 0.94628868 |
| STIL | 140.820751 | -0.1846113 | 0.31323393 | -0.5893719 | 0.55561185 | 0.92373235 |
| CMPK1 | 7010.30245 | 0.22160873 | 0.22040542 | 1.00545951 | 0.31467564 | 0.84696964 |
| FOXD2-AS1 | 135.12106 | 0.00953267 | 0.33257688 | 0.02866305 | 0.97713332 | 0.99455099 |
| FOXD2 | 139.206818 | -0.0116848 | 0.37202178 | -0.0314089 | 0.97494341 | 0.99455099 |
| TRABD2B | 3967.90509 | 0.4739918 | 0.26881772 | 1.76324615 | 0.07785898 | 0.64079926 |
| AL691459.1 | 642.964389 | 0.74493698 | 0.29471674 | 2.52763713 | 0.0114833 | 0.40353404 |
| SLC5A9 | 1238.30459 | 0.56421903 | 0.60255578 | 0.93637643 | 0.34907941 | 0.86356499 |
| AL109659.2 | 201.57373 | 0.58735413 | 0.38457385 | 1.52728566 | 0.12668999 | 0.70443503 |
| SPATA6 | 770.521786 | 0.20269016 | 0.21986423 | 0.92188784 | 0.35658708 | 0.86390849 |
| AGBL4 | 109.266444 | 1.16590468 | 0.44820873 | 2.60125383 | 0.00928837 | 0.38664841 |
| BEND5 | 531.219248 | 0.15951368 | 0.31914242 | 0.49981973 | 6.17E-01 | 9.39E-01 |
| FAF1 | 2052.86515 | 0.18987583 | 0.11284613 | 1.68260827 | 0.09245095 | 0.65583276 |
| CDKN2C | 491.800083 | 0.34522699 | 0.33303884 | 1.03659677 | 0.29992382 | 0.84169995 |
| RNF11 | 6771.88743 | -0.0352586 | 0.21999577 | -0.1602694 | 8.73E-01 | 0.98275231 |
| TTC39A | 5396.82501 | -0.4654512 | 0.49043314 | -0.9490615 | 0.34258935 | 0.85889221 |
| TTC39A-AS1 | 98.6834459 | -0.369356 | 0.57194979 | -0.645784 | 0.51841922 | 0.91636259 |
| EPS15 | 3059.32417 | -0.0875793 | 0.09498903 | -0.9219937 | 0.35653184 | 0.86390849 |
| OSBPL9 | 7114.40151 | -0.0586145 | 0.16502452 | -0.3551866 | 0.72244981 | 0.95741464 |
| NRDC | 5789.58924 | -0.1450659 | 0.16335329 | -0.8880499 | 0.37451391 | 0.86791605 |
| AL050343.1 | 977.092675 | -0.3070369 | 0.18502859 | -1.6594025 | 0.09703472 | 0.66486533 |
| RAB3B | 125.396849 | 0.07284277 | 0.68377254 | 0.10653071 | 0.91516129 | 0.9866558 |
| TXNDC12 | 2948.76259 | 0.07484576 | 0.12481147 | 0.59967051 | 0.54872584 | 0.92373235 |
| KTI12 | 456.919077 | -0.3346594 | 0.3389717 | -0.9872784 | 0.3235062 | 0.85140144 |
| AL445685.1 | 86.563429 | -0.3015758 | 0.37516122 | -0.8038566 | 0.42147977 | 0.88451372 |
| BTF3L4 | 2039.13404 | -0.038699 | 0.1224031 | -0.3161601 | 0.75188101 | 0.96289476 |
| ZFYVE9 | 1299.09656 | -0.025377 | 0.14969005 | -0.1695304 | 0.86537947 | 0.98197545 |
| CC2D1B | 4723.78639 | -0.4993727 | 0.21663835 | -2.3050983 | 0.02116107 | 0.50998124 |
| AL513218.1 | 625.531612 | -0.4769372 | 0.23276864 | -2.0489752 | 0.04046454 | 0.55657615 |
| ORC1 | 69.5145301 | -0.0130809 | 0.33061114 | -0.0395657 | 0.96843937 | 0.99400331 |
| PRPF38A | 1887.33953 | -0.0552995 | 0.1378559 | -0.4011397 | 0.68831729 | 0.95158502 |
| TUT4 | 1777.95952 | -0.2757669 | 0.19059936 | -1.4468408 | 0.14794151 | 0.73460889 |
| GPX7 | 463.732382 | -0.2236331 | 0.25357601 | -0.8819174 | 0.37782146 | 0.86987124 |
| COA7 | 933.517524 | -0.26872 | 0.37606031 | -0.7145662 | 0.47487711 | 0.90226631 |
| ZYG11B | 2117.30047 | -0.0213608 | 0.20160077 | -0.105956 | 0.9156173 | 0.9868044 |
| ZYG11A | 222.486062 | -0.0059775 | 0.58420837 | -0.0102318 | 0.99183633 | 0.99897913 |
| ECHDC2 | 10797.3388 | 0.12743272 | 0.26804452 | 0.47541627 | 0.6344903 | 0.94088547 |
| SCP2 | 9144.75436 | 0.53275224 | 0.22696787 | 2.34725838 | 0.01891213 | 0.48353659 |
| PODN | 575.014549 | 0.32625241 | 0.62305942 | 0.52362968 | 0.60053614 | 0.93488657 |
| AL445183.1 | 163.138415 | 0.32413187 | 0.67129837 | 0.48284322 | 0.62920706 | 0.94088547 |
| SLC1A7 | 87.846003 | 2.02519483 | 0.8487913 | 2.385975 | 0.01703391 | 0.46671416 |
| CPT2 | 3242.91023 | 0.59882035 | 0.31760399 | 1.88543082 | 0.0593717 | 0.60533169 |
| AL606760.2 | 2016.85779 | 0.64404782 | 0.31720414 | 2.0303891 | 0.042317 | 0.56485966 |
| C1orf123 | 2013.41551 | 0.05946817 | 0.14620827 | 0.40673601 | 0.68420189 | 0.95070301 |
| MAGOH | 1004.15693 | -0.2515594 | 0.20236681 | -1.2430865 | 2.14E-01 | 0.7926732 |
| AL606760.1 | 104.693101 | 0.01092126 | 0.45656755 | 0.02392035 | 0.98091614 | 0.99576712 |
| LRP8 | 461.873952 | -0.3905301 | 0.48643702 | -0.8028379 | 4.22E-01 | 8.85E-01 |

| | | | | | | |
|---|---|---|---|---|---|---|
| GLIS1 | 467.028724 | 0.64185316 | 0.49358494 | 1.30039048 | 1.93E-01 | 0.77548688 |
| NDC1 | 907.349279 | -0.1265188 | 0.21445868 | -0.589945 | 5.55E-01 | 0.92373235 |
| YIPF1 | 1973.24324 | -0.0370848 | 0.16230716 | -0.2284854 | 0.81926892 | 0.97454841 |
| DIO1 | 3792.93953 | 2.22369623 | 0.54135272 | 4.10766616 | 4.00E-05 | 0.02340735 |
| HSPB11 | 808.235532 | 0.10691902 | 0.12485057 | 0.85637587 | 0.3917899 | 0.87744531 |
| LRRC42 | 1431.45666 | -0.3088401 | 0.17171585 | -1.798553 | 0.07208942 | 0.6321115 |
| TMEM59 | 18422.6724 | 0.19735135 | 0.11991335 | 1.64578296 | 0.09980845 | 0.67115544 |
| TCEANC2 | 453.082569 | 0.2034013 | 0.21525933 | 0.94491278 | 0.3447034 | 0.85999891 |
| AL357673.1 | 77.3295745 | 0.67057662 | 0.37819605 | 1.77309258 | 0.07621337 | 0.63773415 |
| CYB5RL | 307.765128 | 0.12500963 | 0.283466 | 0.44100397 | 0.65921012 | 0.94601668 |
| MRPL37 | 3329.85927 | 0.15293048 | 0.20116119 | 0.76023849 | 0.44711204 | 0.89480207 |
| SSBP3 | 3649.83904 | -0.3633194 | 0.21074626 | -1.723966 | 0.08471399 | 0.64988556 |
| SSBP3-AS1 | 48.6035412 | 0.02680855 | 0.43204084 | 0.06205096 | 9.51E-01 | 9.92E-01 |
| ACOT11 | 6125.86626 | -0.4583507 | 0.53559816 | -0.8557735 | 0.3921231 | 0.87765297 |
| FAM151A | 9629.33622 | -1.0253108 | 0.96511722 | -1.0623692 | 0.28806811 | 0.83295411 |
| MROH7 | 1025.96335 | -0.2778271 | 0.35690712 | -0.7784298 | 0.43631569 | 0.89104919 |
| MROH7-TTC4 | 2515.26728 | -0.2982154 | 0.20301252 | -1.4689507 | 0.14184616 | 0.72784886 |
| TTC4 | 1660.84891 | -0.3156352 | 0.23356433 | -1.3513844 | 0.17657232 | 0.76115956 |
| PARS2 | 433.286236 | 0.3990623 | 0.23322822 | 1.71103779 | 0.08707414 | 0.65231349 |
| TTC22 | 2379.0696 | 0.39340895 | 0.48814046 | 0.80593393 | 0.42028095 | 0.88451372 |
| DHCR24 | 7681.43511 | -0.2450311 | 0.30260426 | -0.809741 | 0.41808904 | 0.88445833 |
| AC096536.3 | 116.530301 | 0.12150322 | 0.45816369 | 0.26519609 | 0.79085841 | 0.97090793 |
| TMEM61 | 727.531143 | -0.2541172 | 0.50403082 | -0.5041699 | 0.61414198 | 0.93895494 |
| AL590440.1 | 517.739569 | -0.2157045 | 0.49767 | -0.4334287 | 0.66470334 | 0.94681198 |
| BSND | 1237.72607 | 0.43972243 | 0.49154566 | 0.89457087 | 0.37101653 | 0.86762749 |
| USP24 | 3808.65385 | -0.2648049 | 0.21671678 | -1.221894 | 0.22174771 | 0.80054995 |
| AC119674.1 | 62.6846364 | -0.1556968 | 0.52947301 | -0.29406 | 0.76871205 | 0.96796999 |
| AC119674.2 | 2753.47035 | -0.6969888 | 0.50138447 | -1.3901284 | 0.1644899 | 0.75286112 |
| PLPP3 | 13100.2233 | -0.5739093 | 0.44755798 | -1.2823127 | 0.19973298 | 0.78134034 |
| PRKAA2 | 2459.90718 | -0.3238563 | 0.15188668 | -2.1322235 | 0.03298848 | 0.55137197 |
| FYB2 | 677.765379 | 1.05270317 | 0.28316255 | 3.71766388 | 0.00020107 | 0.06978369 |
| DAB1 | 500.573251 | 0.47998761 | 0.36488653 | 1.31544349 | 0.1883609 | 0.76970626 |
| OMA1 | 1188.00458 | 0.6322809 | 0.37775523 | 1.67378465 | 0.09417293 | 0.65783452 |
| TACSTD2 | 10799.2414 | 0.16372814 | 0.55002715 | 0.29767283 | 0.76595288 | 0.96704052 |
| MYSM1 | 1765.59765 | -0.2322857 | 0.14278824 | -1.6267845 | 0.10378286 | 0.67439401 |
| AL136985.3 | 77.3232875 | 0.17162708 | 0.47388886 | 0.36216736 | 0.71722697 | 0.95611001 |
| JUN | 12293.1487 | -0.2596224 | 0.38596492 | -0.6726581 | 0.5011648 | 0.91240816 |
| AC105277.1 | 96.7499736 | 0.68041685 | 0.39790176 | 1.71001216 | 0.08726362 | 0.65256204 |
| FGGY | 1079.64185 | 0.65338525 | 0.36777004 | 1.7766136 | 0.07563184 | 0.63773415 |
| HOOK1 | 2516.03127 | 0.37841165 | 0.17556776 | 2.15535954 | 0.03113371 | 0.54466273 |
| CYP2J2 | 427.318085 | 0.7912522 | 0.89374262 | 0.88532446 | NA | NA |
| NFIA | 1506.7478 | 0.22499962 | 0.27030631 | 0.83238762 | 4.05E-01 | 0.88132015 |
| TM2D1 | 1515.13626 | -0.0009854 | 0.10386833 | -0.0094867 | 0.99243081 | 0.99906045 |
| PATJ | 5797.21 | 0.15877318 | 0.23119384 | 0.6867535 | 0.49223808 | 0.90636999 |
| KANK4 | 65.8161396 | 0.45174176 | 0.83256548 | 0.54259007 | 5.87E-01 | 0.93289552 |
| USP1 | 1314.85714 | -0.1098681 | 0.22751307 | -0.4829089 | 0.62916039 | 0.94088547 |
| DOCK7 | 1370.64885 | 0.22986144 | 0.1548845 | 1.48408287 | 0.13778694 | 0.72209298 |
| ANGPTL3 | 1614.73029 | 0.04044084 | 0.66953622 | 0.06040127 | 0.95183605 | 0.99170503 |
| ATG4C | 626.401716 | 0.08246354 | 0.18958337 | 0.43497241 | 0.66358246 | 0.94642611 |
| ALG6 | 613.726302 | 0.25986974 | 0.17176821 | 1.51290941 | 0.13030267 | 0.7084428 |
| ITGB3BP | 1062.00158 | 0.49771696 | 0.22806031 | 2.18239189 | 2.91E-02 | 5.36E-01 |
| EFCAB7 | 435.45662 | 0.48029974 | 0.23484494 | 2.04517817 | 0.0408373 | 0.55782198 |
| PGM1 | 5051.35803 | 0.08630384 | 0.21718842 | 0.3973685 | 0.69109574 | 0.95158502 |
| ROR1 | 165.203706 | 0.01515582 | 0.32591707 | 0.04650209 | 0.96291007 | 0.99320711 |

| | | | | | | |
|---|---|---|---|---|---|---|
| AL445205.1 | 56.5538101 | -0.5906128 | 0.86849822 | -0.6800392 | 0.49647967 | 0.90890243 |
| CACHD1 | 704.403505 | 0.24015513 | 0.26789688 | 0.89644617 | 0.37001452 | 0.86762749 |
| RAVER2 | 1000.8948 | 0.06851998 | 0.19548888 | 0.35050578 | 7.26E-01 | 0.95848028 |
| JAK1 | 8175.71633 | -0.0538203 | 0.13769108 | -0.3908769 | 0.69588825 | 0.9525827 |
| AK4 | 15797.0113 | 0.5612242 | 0.53534971 | 1.04833193 | 0.2944857 | 0.83630273 |
| DNAJC6 | 833.610199 | 0.19482451 | 0.4147801 | 0.46970554 | 0.63856541 | 0.94088547 |
| LEPROT | 11761.4775 | -0.3091779 | 0.17239699 | -1.7934065 | 0.07290795 | 0.63551996 |
| LEPR | 1458.24835 | 0.19747661 | 0.25936259 | 0.76139202 | 0.44642295 | 0.89480207 |
| PDE4B | 742.987128 | 0.32334801 | 0.57832131 | 0.55911481 | 0.57608337 | 0.93083012 |
| SGIP1 | 1414.78091 | -0.3974877 | 0.43385949 | -0.9161668 | 0.3595794 | 0.86480228 |
| TCTEX1D1 | 87.3786808 | 1.01108541 | 0.74997393 | 1.34816074 | 0.17760669 | 0.76115956 |
| WDR78 | 497.175292 | 0.21971088 | 0.37246539 | 0.58988266 | 0.55526932 | 0.92373235 |
| AL592161.1 | 53.2310553 | 0.32331687 | 0.49069172 | 0.65890019 | 0.50995986 | 0.91439937 |
| MIER1 | 2147.36026 | -0.2057674 | 0.13107095 | -1.5698935 | 0.1164399 | 0.68882582 |
| SLC35D1 | 1298.00674 | 0.24570221 | 0.25371457 | 0.96841979 | 0.33283476 | 0.855011 |
| IL12RB2 | 57.9278443 | 0.20239416 | 0.6164342 | 0.32833052 | 0.74266177 | 0.96109001 |
| SERBP1 | 9389.65894 | -0.0469831 | 0.14038336 | -0.3346768 | 0.7378689 | 0.9602325 |
| LINC01702 | 127.771246 | 0.56241129 | 0.70535014 | 0.7973505 | 4.25E-01 | 8.86E-01 |
| GADD45A | 8080.32013 | -0.4622874 | 0.27666096 | -1.6709528 | 0.09473101 | 0.65888357 |
| GNG12 | 7475.74138 | -0.1137069 | 0.24225472 | -0.4693692 | 0.63880575 | 0.94088547 |
| GNG12-AS1 | 312.804006 | 0.12939926 | 0.28060218 | 0.46114845 | 0.6446921 | 0.94208859 |
| DIRAS3 | 453.019131 | 0.23728942 | 0.32214703 | 0.73658734 | 0.46137333 | 0.89755243 |
| WLS | 6521.10356 | 0.02196505 | 0.20115674 | 0.10919372 | 0.91304884 | 0.98662757 |
| LRRC7 | 100.286192 | -0.1447476 | 0.34230987 | -0.4228555 | 0.67240068 | 0.94922827 |
| LRRC40 | 866.444987 | 0.19488598 | 0.15968011 | 1.22047749 | 0.22228392 | 0.80078694 |
| SRSF11 | 8389.40666 | -0.1785967 | 0.14218157 | -1.2561168 | 0.20907362 | 0.79086734 |
| ANKRD13C | 1203.99221 | -0.2992096 | 0.21128927 | -1.4161134 | 0.15674231 | 0.74522306 |
| HHLA3 | 453.005122 | 0.11596883 | 0.37817558 | 0.3066534 | 0.7591072 | 0.96502715 |
| CTH | 672.495077 | 0.33540298 | 0.45486439 | 0.73736918 | 0.46089787 | 0.89755243 |
| LINC01788 | 176.276396 | 0.80520938 | 0.6252439 | 1.28783245 | 0.19780429 | 0.77879208 |
| PTGER3 | 7061.11115 | -0.5078357 | 0.31842007 | -1.594861 | 0.11074333 | 0.68001961 |
| AL031429.1 | 64.4516384 | -2.1800658 | 0.96702162 | -2.2544127 | 0.02417022 | 0.51967919 |
| ZRANB2-AS1 | 383.676159 | -0.3995299 | 0.19765973 | -2.0213014 | 0.04324857 | 0.56886632 |
| ZRANB2 | 4352.08007 | -0.141333 | 0.18858954 | -0.7494211 | 0.45360347 | 0.89670027 |
| NEGR1 | 652.423558 | 0.2997118 | 0.4291952 | 0.69831116 | 0.48498262 | 0.90541935 |
| LRRIQ3 | 82.520245 | 0.74872063 | 0.52885265 | 1.41574526 | 0.1568501 | 0.74522306 |
| FPGT | 766.175333 | 0.11724104 | 0.19852389 | 0.5905639 | 0.55481266 | 0.92373235 |
| FPGT-TNNI3 | 414.111914 | 0.67327568 | 0.34524833 | 1.95012002 | 0.05116182 | 0.5861488 |
| TNNI3K | 265.797425 | 0.75095139 | 0.42190994 | 1.7798855 | 0.0750947 | 0.63773415 |
| AC135803.1 | 81.5063403 | -0.0153764 | 0.50684731 | -0.0303373 | 0.97579808 | 0.99455099 |
| CRYZ | 12064.7663 | 0.61588349 | 0.34468137 | 1.78681975 | 0.07396663 | 0.6374448 |
| TYW3 | 1340.30399 | -0.0271289 | 0.19441505 | -0.1395414 | 0.88902234 | 0.98423722 |
| SLC44A5 | 133.054871 | 0.96402613 | 0.98567657 | 0.97803494 | 0.32805704 | 0.85317073 |
| ACADM | 10070.426 | 0.58885832 | 0.3603067 | 1.63432521 | 0.10219058 | 0.67279543 |
| RABGGTB | 4597.44993 | -0.4583602 | 0.25770149 | -1.7786479 | 0.07529751 | 0.63773415 |
| AC092813.2 | 56.1837414 | 0.40437658 | 1.11434752 | 0.36288194 | 0.71669308 | 0.95603249 |
| ST6GALNAC3 | 788.07963 | 0.12328218 | 0.35712174 | 0.34521051 | 7.30E-01 | 0.95848028 |
| PIGK | 1956.89275 | 0.41933972 | 0.19152394 | 2.18948984 | 0.02856126 | 0.53606221 |
| AK5 | 179.397303 | 0.93163181 | 0.64082167 | 1.45380821 | 0.14599949 | 0.73247558 |
| AC118549.1 | 1657.16259 | -0.1939598 | 0.16148315 | -1.2011145 | 0.2297068 | 0.80551898 |
| USP33 | 3150.16332 | -0.0639368 | 0.13703156 | -0.4665843 | 0.64079734 | 0.94088547 |
| MIGA1 | 1122.94456 | 0.31707321 | 0.1777832 | 1.78348247 | 0.0745078 | 0.63773415 |
| NEXN-AS1 | 40.6823894 | -0.6480257 | 0.54139543 | -1.1969544 | 0.23132431 | 0.80649152 |
| NEXN | 501.898545 | -1.0679687 | 0.56640502 | -1.8855212 | 0.0593595 | 0.60533169 |

| | | | | | | |
|---|---|---|---|---|---|---|
| FUBP1 | 4038.04889 | -0.1891574 | 0.1590819 | -1.1890566 | 0.23441741 | 0.80730051 |
| DNAJB4 | 1577.05775 | -0.29478 | 0.22645591 | -1.3017105 | 0.19301536 | 0.7751938 |
| GIPC2 | 2250.86664 | 0.53184664 | 0.48093357 | 1.10586299 | 0.26878582 | 0.82683094 |
| PTGFR | 323.635618 | 0.48337675 | 0.29574129 | 1.63445812 | 0.10216269 | 0.67279543 |
| IFI44L | 3705.78327 | -3.4234123 | 1.26639954 | -2.703264 | NA | NA |
| IFI44 | 3684.98838 | -2.7282517 | 1.06429454 | -2.5634368 | NA | NA |
| ADGRL4 | 1758.30049 | -0.1464821 | 0.36172533 | -0.404954 | 0.68551135 | 0.95097288 |
| LINC01781 | 136.810484 | 0.80305037 | 0.68822661 | 1.16684004 | 0.24327497 | 0.81356465 |
| ADGRL2 | 2605.05136 | -0.0727616 | 0.2131656 | -0.3413382 | 7.33E-01 | 0.95848028 |
| TTLL7 | 469.222587 | -0.0446947 | 0.30872091 | -0.1447738 | 8.85E-01 | 0.98349622 |
| AL359504.2 | 106.060613 | 0.17973836 | 0.37755067 | 0.47606421 | 0.63402863 | 0.94088547 |
| PRKACB | 1749.2865 | 0.32752582 | 0.17606401 | 1.86026556 | 0.06284796 | 0.6147045 |
| SAMD13 | 67.4711436 | 0.96921268 | 0.47779203 | 2.02852419 | 0.04250678 | 0.56530166 |
| RPF1 | 1305.3877 | -0.148996 | 0.1476383 | -1.0091964 | 3.13E-01 | 8.46E-01 |
| GNG5 | 4772.88392 | -0.2340839 | 0.16768367 | -1.3959849 | 0.16271905 | 0.75032457 |
| SPATA1 | 1297.80961 | 0.13577424 | 0.1644778 | 0.82548674 | 0.40909531 | 0.8821603 |
| CTBS | 2053.98088 | 0.29946457 | 0.19036728 | 1.57308846 | 0.11569834 | 0.6873836 |
| SSX2IP | 1273.98035 | 0.70275346 | 0.23816291 | 2.95072588 | 0.00317028 | 0.24755935 |
| MCOLN2 | 191.95003 | -0.0284936 | 0.4746811 | -0.0600268 | 0.95213433 | 0.99170503 |
| WDR63 | 53.7855157 | 0.24287958 | 0.72801441 | 0.33361919 | 0.73866693 | 0.9602325 |
| MCOLN3 | 630.308032 | -0.0308071 | 0.28120775 | -0.1095527 | 0.91276409 | 0.98662757 |
| SYDE2 | 171.730994 | -0.2868357 | 0.25579583 | -1.1213461 | 0.26214056 | 0.82424998 |
| C1orf52 | 895.69553 | -0.1256755 | 0.13569013 | -0.9261951 | 0.3543446 | 0.86364633 |
| BCL10 | 951.276796 | -0.0390122 | 0.20344868 | -0.1917547 | 0.84793433 | 0.97711043 |
| DDAH1 | 9064.47418 | 0.37821509 | 0.21598889 | 1.75108586 | 0.07993112 | 0.64079926 |
| CYR61 | 15366.541 | -1.5022774 | 0.61768187 | -2.4321215 | 0.01501067 | 0.44724159 |
| ZNHIT6 | 912.848659 | -0.1254989 | 0.23504234 | -0.5339415 | 0.59338202 | 0.93289552 |
| COL24A1 | 105.160589 | -0.2452785 | 0.32215625 | -0.7613651 | 0.44643901 | 0.89480207 |
| AL136382.1 | 67.0357246 | -0.3931821 | 0.55391106 | -0.7098289 | 0.47781025 | 0.90387017 |
| ODF2L | 1635.04692 | -0.2550389 | 0.25279022 | -1.0088953 | 0.31302484 | 0.84566794 |
| SH3GLB1 | 4788.64316 | -0.1889441 | 0.18709924 | -1.0098603 | 0.31256224 | 0.84541735 |
| SELENOF | 7631.00083 | 0.22517664 | 0.13439273 | 1.67551206 | 0.09383381 | 0.65783452 |
| HS2ST1 | 1645.03278 | 0.08881829 | 0.16910093 | 0.52523831 | 0.59941753 | 0.93487648 |
| AC093155.3 | 362.274267 | -0.0601529 | 0.25920113 | -0.2320702 | 0.81648346 | 0.97388289 |
| LINC01140 | 150.376789 | -0.2562124 | 0.51913526 | -0.4935369 | 0.62163324 | 0.94016713 |
| LMO4 | 1874.59395 | -0.2752827 | 0.21305909 | -1.2920486 | 1.96E-01 | 7.78E-01 |
| PKN2 | 2314.15099 | -0.0458779 | 0.15001033 | -0.3058319 | 0.75973265 | 0.96510454 |
| GTF2B | 1353.61499 | -0.2201758 | 0.12813391 | -1.7183258 | 0.0857372 | 0.65049132 |
| KYAT3 | 1855.49705 | 0.35700949 | 0.22853204 | 1.56218568 | 0.11824425 | 0.69095557 |
| RBMXL1 | 950.352285 | -0.2020334 | 0.20990098 | -0.9625179 | 0.33578952 | 0.85641269 |
| GBP3 | 2504.79282 | 0.52502938 | 0.42171824 | 1.24497669 | 0.21314032 | 0.7926732 |
| GBP1 | 2270.13971 | -1.1248976 | 0.66394035 | -1.6942751 | 0.09021302 | 0.65403568 |
| GBP2 | 5262.91847 | -0.323296 | 0.37323406 | -0.8662017 | 0.38637954 | 0.87432552 |
| GBP7 | 53.4897253 | 0.03145619 | 0.74905008 | 0.04199477 | 0.96650287 | 0.99400331 |
| GBP4 | 1702.84272 | -1.0439279 | 0.55432562 | -1.8832395 | 0.05966792 | 0.6066891 |
| GBP5 | 360.728838 | -0.614522 | 0.86196987 | -0.7129274 | 0.47589067 | 0.90272018 |
| AC099063.1 | 48.2976258 | -0.6880702 | 0.88101102 | -0.7810007 | 0.4348021 | 0.89055338 |
| LRRC8B | 647.218608 | 0.07005182 | 0.14534538 | 0.48196795 | 0.62982871 | 0.94088547 |
| LRRC8C-DT | 95.8479134 | 0.22471854 | 0.29650331 | 0.75789556 | 0.44851351 | 0.8948759 |
| LRRC8C | 483.754447 | -0.1981636 | 0.28677116 | -0.6910166 | 0.48955511 | 0.90600166 |
| AC093423.3 | 129.078503 | 0.28397948 | 0.48966747 | 0.57994353 | 0.5619527 | 0.92452095 |
| AC099568.2 | 66.1396054 | 0.58403192 | 0.56042156 | 1.04212964 | 0.29735158 | 0.83898658 |
| LRRC8D | 2040.63555 | 0.44411782 | 0.3090825 | 1.43689087 | 0.150749 | 0.73796431 |
| ZNF326 | 1885.04663 | -0.1332365 | 0.16107435 | -0.827174 | 0.40813846 | 0.88181733 |

| | | | | | | |
|---|---|---|---|---|---|---|
| ZNF644 | 1155.84669 | -0.2151381 | 0.14397133 | -1.4943122 | 0.13509405 | 0.71649038 |
| HFM1 | 100.570741 | 0.19456289 | 0.4364716 | 0.445763 | 0.65576845 | 0.94532497 |
| CDC7 | 403.829081 | -0.0969591 | 0.52073615 | -0.1861961 | 0.85229096 | 0.97872457 |
| TGFBR3 | 3143.97374 | 0.04686045 | 0.21115197 | 0.22192762 | 0.82437023 | 0.97548201 |
| EPHX4 | 61.3746215 | 0.42525988 | 0.49789828 | 0.85410997 | 0.39304406 | 0.87805922 |
| BTBD8 | 188.861742 | 0.29876109 | 0.46716614 | 0.63951787 | 0.52248609 | 0.91723878 |
| GLMN | 422.22632 | 0.00538725 | 0.16082675 | 0.03349725 | 0.97327806 | 0.99445887 |
| RPAP2 | 937.8197 | 0.01743376 | 0.14340994 | 0.12156593 | 0.90324279 | 0.98617053 |
| GFI1 | 64.1492842 | 0.23387728 | 0.74344205 | 0.31458711 | 0.75307517 | 0.96301881 |
| EVI5 | 932.264355 | -0.0900698 | 0.19589647 | -0.4597824 | 0.6456724 | 0.94208859 |
| RPL5 | 26207.5449 | 0.32961963 | 0.25076749 | 1.3144432 | 0.1886971 | 0.76970626 |
| FAM69A | 611.792192 | -0.0200199 | 0.22633134 | -0.0884541 | 0.92951573 | 0.98964715 |
| MTF2 | 1332.14975 | -0.1007238 | 0.21449702 | -0.4695813 | 0.63865418 | 0.94088547 |
| TMED5 | 6714.33052 | 0.03299587 | 0.23425623 | 0.14085377 | 0.88798546 | 0.98423722 |
| CCDC18 | 912.446794 | 0.05121208 | 0.25556641 | 0.20038657 | 8.41E-01 | 9.76E-01 |
| DR1 | 2546.52513 | -0.0276714 | 0.13496109 | -0.2050321 | 0.83754704 | 0.97577999 |
| FNBP1L | 4242.04744 | 0.00056462 | 0.14498524 | 0.00389434 | 0.99689277 | 0.99976751 |
| BCAR3 | 2057.56406 | -0.3677067 | 0.3795554 | -0.9687827 | 0.33265363 | 0.855011 |
| AL109613.1 | 274.400863 | -0.1035481 | 0.34918605 | -0.2965414 | 0.76681667 | 0.96741943 |
| DNTTIP2 | 2286.94505 | -0.6727022 | 0.39342781 | -1.709849 | 8.73E-02 | 0.65256204 |
| GCLM | 3608.00428 | -0.3776732 | 0.5886263 | -0.641618 | 0.52112127 | 0.91723878 |
| ABCA4 | 357.406072 | 1.2923253 | 0.82651221 | 1.56358888 | 0.11791415 | 0.69069542 |
| ARHGAP29 | 5252.27325 | -0.0651582 | 0.29368699 | -0.2218627 | 0.82442075 | 0.97548201 |
| ABCD3 | 4723.54091 | 0.51958844 | 0.2570728 | 2.02117239 | 0.04326192 | 0.56886632 |
| F3 | 1429.82267 | -1.1441898 | 0.39924865 | -2.8658577 | 0.00415881 | 0.27935586 |
| SLC44A3 | 2615.58186 | 0.68594062 | 0.35713482 | 1.92067698 | 0.05477244 | 0.59329767 |
| CNN3 | 19958.487 | -0.2349968 | 0.17981435 | -1.3068858 | 0.19125149 | 0.77292503 |
| AC105942.1 | 754.208535 | -0.0684402 | 0.28005003 | -0.2443855 | 0.8069323 | 0.97334138 |
| ALG14 | 586.551029 | 0.17421107 | 0.28700074 | 0.60700565 | 0.54384716 | 0.92325413 |
| TMEM56 | 707.602174 | -0.0257574 | 0.28235705 | -0.0912228 | 0.92731553 | 0.98915637 |
| TMEM56-RW | 780.240269 | 0.03304076 | 0.17138991 | 0.19278124 | 0.8471303 | 0.97684681 |
| RWDD3 | 907.680939 | 0.02370391 | 0.18097801 | 0.13097672 | 0.89579372 | 0.98474725 |
| PTBP2 | 1533.49407 | -0.2446874 | 0.17277633 | -1.4162093 | 0.15671424 | 0.74522306 |
| DPYD | 1163.48809 | -0.0699056 | 0.29350271 | -0.2381771 | 8.12E-01 | 0.97355727 |
| DPYD-AS1 | 67.9261255 | -0.0212419 | 0.36555187 | -0.0581092 | 0.95366165 | 0.99170503 |
| SNX7 | 1484.93059 | 0.32077339 | 0.15647832 | 2.04995416 | 0.0403689 | 0.55657615 |
| PALMD | 640.310708 | -0.1522667 | 0.50162182 | -0.3035489 | 0.76147162 | 0.96606476 |
| FRRS1 | 123.973968 | 0.34372459 | 0.34889451 | 0.98518201 | 0.32453468 | 0.85169085 |
| AGL | 1536.11121 | 0.05471807 | 0.16415917 | 0.33332325 | 7.39E-01 | 9.60E-01 |
| AC118553.1 | 62.1102901 | 0.82582719 | 0.52499638 | 1.57301503 | 0.11571534 | 0.6873836 |
| SLC35A3 | 1790.55822 | 0.20827634 | 0.16567883 | 1.25710895 | 0.20871419 | 0.79042964 |
| AC118553.2 | 3540.84886 | -0.1257813 | 0.17223938 | -0.7302701 | 0.4652251 | 0.89921277 |
| MFSD14A | 3175.32589 | -0.1767529 | 0.18219958 | -0.9701061 | 0.33199359 | 0.85452795 |
| AC093019.2 | 54.9025456 | -0.6080602 | 0.39123315 | -1.5542144 | 0.12013329 | 0.69467399 |
| SASS6 | 214.255503 | -0.5443722 | 0.24203435 | -2.2491526 | 0.02450279 | 0.51967919 |
| TRMT13 | 943.836388 | -0.2051369 | 0.15324801 | -1.338594 | 0.18070289 | 0.76466689 |
| LRRC39 | 399.731769 | -0.2400382 | 0.14395361 | -1.6674692 | 0.09542115 | 0.65888357 |
| DBT | 1898.59713 | 0.30060671 | 0.22868073 | 1.31452574 | 0.18866934 | 0.76970626 |
| RTCA-AS1 | 75.9872741 | 0.40549001 | 0.41503792 | 0.97699508 | 0.32857159 | 0.85317073 |
| RTCA | 2507.64332 | 0.34617822 | 0.27850124 | 1.24300425 | 0.21386627 | 0.7926732 |
| CDC14A | 582.567794 | 0.14318127 | 0.31738605 | 0.45112656 | 0.65189834 | 0.9440713 |
| VCAM1 | 11582.7264 | 0.96764248 | 0.49771291 | 1.94417798 | 0.05187399 | 0.58645127 |
| EXTL2 | 936.642348 | 0.03303739 | 0.14792212 | 0.2233431 | 0.82326849 | 0.97542663 |
| SLC30A7 | 1771.44327 | -0.1073391 | 0.23113549 | -0.464399 | 0.64236192 | 0.94090101 |

| | | | | | | |
|---|---|---|---|---|---|---|
| DPH5 | 889.531751 | 0.38889816 | 0.17008551 | 2.28648609 | 0.02222584 | 0.51807763 |
| AC093157.1 | 258.80632 | 0.00738443 | 0.25375897 | 0.02910019 | 0.97678468 | 0.99455099 |
| S1PR1 | 2058.58021 | -0.4959452 | 0.35649639 | -1.3911648 | 0.16417545 | 0.752829 |
| RNPC3 | 1824.70665 | -0.0412409 | 0.21535217 | -0.1915047 | 8.48E-01 | 0.97711043 |
| AMY2B | 2956.98074 | -0.0424761 | 0.2932622 | -0.1448401 | 0.88483709 | 0.98349622 |
| PRMT6 | 1176.86543 | -0.3371863 | 0.25179196 | -1.3391466 | 0.18052295 | 0.76438786 |
| NTNG1 | 503.799935 | -0.0337949 | 0.75019536 | -0.0450481 | 0.96406897 | 0.99342622 |
| VAV3 | 6618.7996 | 0.34713163 | 0.22587439 | 1.53683484 | 0.12433376 | 0.70102567 |
| SLC25A24 | 1549.41924 | -0.0785352 | 0.22773237 | -0.3448576 | 0.7302014 | 0.95848028 |
| AL359258.2 | 106.161968 | -0.0520414 | 0.36721177 | -0.1417204 | 0.88730085 | 0.98423722 |
| FAM102B | 459.701022 | -0.0309457 | 0.35427849 | -0.0873485 | 0.93039453 | 0.98975557 |
| HENMT1 | 137.669351 | 0.11910455 | 0.3426483 | 0.34760001 | 0.7281406 | 0.95848028 |
| PRPF38B | 2616.16873 | -0.0711241 | 0.15784026 | -0.4506081 | 0.652272 | 0.9443159 |
| STXBP3 | 2977.65445 | 0.03736833 | 0.1219391 | 0.30645077 | 0.75926146 | 0.96502715 |
| GPSM2 | 1182.75911 | -0.0898563 | 0.14350824 | -0.6261401 | 0.53122306 | 0.91962754 |
| CLCC1 | 1504.42594 | 0.07199796 | 0.12213833 | 0.58947881 | 0.55554012 | 0.92373235 |
| WDR47 | 737.272435 | -0.1880849 | 0.17898171 | -1.0508609 | 0.29332246 | 0.83540174 |
| TAF13 | 1441.49752 | -0.241471 | 0.37201797 | -0.6490844 | 0.51628384 | 0.91636259 |
| AL356488.3 | 218.881295 | 0.00512075 | 0.28719253 | 0.01783038 | 0.98577417 | 0.99705651 |
| TMEM167B | 2857.3149 | 0.03675862 | 0.14775582 | 0.2487795 | 0.80353135 | 0.97274243 |
| KIAA1324 | 39.8841997 | 0.81716613 | 0.37891071 | 2.15661925 | 0.03103534 | 0.54466273 |
| SARS | 6239.51777 | -0.0617511 | 0.1514923 | -0.4076189 | 0.68355347 | 0.95070301 |
| CELSR2 | 3094.20082 | 0.06626059 | 0.3567486 | 0.18573469 | 0.85265282 | 0.97883015 |
| PSRC1 | 267.385888 | 0.32761915 | 0.36626421 | 0.89448859 | 0.37106053 | 0.86762749 |
| SORT1 | 7180.1074 | 0.12533952 | 0.20810006 | 0.60230409 | 0.54697174 | 0.92345821 |
| PSMA5 | 4265.25289 | 0.09589299 | 0.16559537 | 0.57908014 | 0.5625351 | 0.92452095 |
| SYPL2 | 1547.143 | -0.4084853 | 0.29996713 | -1.3617669 | 0.17327146 | 0.75974924 |
| ATXN7L2 | 1523.99684 | -0.2234375 | 0.26085897 | -0.8565451 | 0.39169636 | 0.87744531 |
| CYB561D1 | 1118.42557 | 0.00097037 | 0.17905436 | 0.0054194 | 0.99567597 | 0.99940889 |
| AMIGO1 | 772.826702 | -0.0025518 | 0.27355107 | -0.0093284 | 0.99255713 | 0.99906045 |
| GNAI3 | 4086.53782 | -0.2154743 | 0.19535297 | -1.1029997 | 0.27002728 | 0.82716979 |
| GNAT2 | 66.6862784 | -0.5056922 | 0.38351229 | -1.3185815 | 0.18730905 | 0.76958556 |
| AMPD2 | 4372.08612 | -0.4352735 | 0.21969013 | -1.9813066 | 0.0475569 | 0.58241466 |
| AL355310.2 | 950.271911 | -0.4531575 | 0.21968673 | -2.062744 | 0.03913696 | 0.55657615 |
| GSTM4 | 4058.91854 | 0.48776868 | 0.2196086 | 2.22108183 | 0.02634542 | 0.52678638 |
| GSTM2 | 3011.18905 | 0.42673837 | 0.38582361 | 1.10604525 | 2.69E-01 | 0.82676682 |
| GSTM1 | 413.398844 | -0.0913912 | 1.54739482 | -0.0590613 | 0.95290326 | 0.99170503 |
| AC000032.1 | 147.124916 | 0.17207446 | 1.34250299 | 0.12817436 | 0.89801099 | 0.98474725 |
| GSTM5 | 186.491215 | -0.0886182 | 0.5799917 | -0.1527923 | 0.8785621 | 0.98275231 |
| AL158847.1 | 1107.94869 | 0.15920168 | 0.40588318 | 0.39223523 | 6.95E-01 | 0.95234774 |
| GSTM3 | 5670.66055 | 0.10337262 | 0.42283411 | 0.24447559 | 0.8068625 | 0.97334138 |
| CSF1 | 2838.09187 | -0.433108 | 0.30539479 | -1.4181906 | 0.15613512 | 0.74522306 |
| AHCYL1 | 11991.7909 | 0.19523537 | 0.24278744 | 0.80414115 | 0.42131546 | 0.88451372 |
| STRIP1 | 1661.41275 | -0.233592 | 0.17292265 | -1.3508466 | 0.17674456 | 0.76115956 |
| AL160006.1 | 952.37298 | -0.1311111 | 0.19218479 | -0.6822135 | 0.49510395 | 0.90816709 |
| ALX3 | 380.813658 | -0.1520924 | 0.20681293 | -0.7354103 | 0.46208968 | 0.89762182 |
| SLC6A17 | 413.876407 | -0.6649085 | 0.47363873 | -1.4038304 | 0.16036935 | 0.74781837 |
| LINC02586 | 41.3855915 | 0.26554179 | 0.5700716 | 0.46580427 | 0.64135561 | 0.94088547 |
| KCNC4 | 1864.30307 | 0.2985756 | 0.31313569 | 0.95350231 | 0.34033563 | 0.85814041 |
| RBM15 | 499.846304 | -0.3378473 | 0.29739851 | -1.1360087 | 0.25595294 | 0.82052927 |
| SLC16A4 | 6805.67707 | 0.5800266 | 0.41769489 | 1.38863705 | 0.16494315 | 0.75299828 |
| AL355488.1 | 712.650946 | 0.06605343 | 0.26458527 | 0.24964895 | 0.80285884 | 0.97273128 |
| LAMTOR5 | 7624.57857 | 0.06171873 | 0.27077483 | 0.22793379 | 8.20E-01 | 9.75E-01 |
| AL365361.1 | 71.6633455 | 0.18579154 | 0.76403603 | 0.24317117 | 0.8078728 | 0.97334138 |

| | | | | | | |
|---|---|---|---|---|---|---|
| CD53 | 1303.73198 | -0.6157349 | 0.52791902 | -1.1663435 | 0.24347559 | 0.81376111 |
| AL360270.1 | 70.5684579 | -0.0702977 | 0.45289218 | -0.1552195 | 0.87664827 | 0.98275231 |
| LRIF1 | 571.988471 | 0.03076592 | 0.30649197 | 0.10038083 | 0.92004199 | 0.98733173 |
| AL360270.3 | 85.3349456 | -0.0699661 | 0.43972051 | -0.1591149 | 0.87357836 | 0.98275231 |
| DRAM2 | 2822.37926 | 0.26588847 | 0.23852736 | 1.11470848 | 0.26497536 | 0.82455959 |
| CEPT1 | 2684.12843 | 0.06060733 | 0.1642287 | 0.36904226 | 0.71209622 | 0.95560193 |
| AL355816.1 | 40.4377901 | -0.0572623 | 0.36408631 | -0.1572768 | 0.87502672 | 0.98275231 |
| AL355816.2 | 835.592327 | 0.06113793 | 0.2271421 | 0.26916159 | 0.78780534 | 0.97090793 |
| DENND2D | 2685.96292 | -0.0683766 | 0.26211975 | -0.2608604 | 0.7942002 | 0.97091363 |
| CHI3L2 | 889.570986 | 0.6563151 | 0.84194998 | 0.77951792 | 0.43567469 | 0.89085849 |
| PIFO | 478.172024 | 0.15951076 | 0.41493074 | 0.38442744 | 0.70066167 | 0.95360275 |
| OVGP1 | 670.801292 | 0.51448162 | 0.379458 | 1.35583286 | 0.17515235 | 0.76073329 |
| WDR77 | 2273.78239 | -0.2223784 | 0.16369393 | -1.3585012 | 0.17430469 | 0.75974924 |
| ATP5PB | 9474.17878 | 0.29135289 | 0.17169381 | 1.69693303 | 0.08970932 | 0.65375543 |
| C1orf162 | 1040.31797 | -0.6968404 | 0.53327894 | -1.3067091 | 0.19131154 | 0.77292503 |
| TMIGD3 | 137.310286 | -0.5844229 | 0.62256893 | -0.9387281 | 0.34787039 | 0.86225576 |
| ADORA3 | 203.412394 | -1.0433136 | 0.61864293 | -1.6864553 | 0.09170814 | 0.65518742 |
| RAP1A | 3967.6235 | -0.2163153 | 0.15870408 | -1.3630105 | 0.1728792 | 0.75974924 |
| LINC01160 | 56.3948076 | -0.4972805 | 0.53513232 | -0.9292664 | 0.35275103 | 0.86364633 |
| INKA2 | 642.198176 | -0.0738853 | 0.42092549 | -0.1755306 | 0.86066276 | 0.98106934 |
| AL049557.1 | 632.342306 | -0.0781668 | 0.42086344 | -0.1857296 | 0.85265679 | 0.97883015 |
| DDX20 | 779.631744 | -0.2808396 | 0.25314512 | -1.1094014 | 0.26725706 | 0.8265123 |
| KCND3 | 363.37113 | 0.2080232 | 0.34744885 | 0.59871603 | 5.49E-01 | 0.92373235 |
| CTTNBP2NL | 1474.07317 | -0.0567425 | 0.19041835 | -0.2979888 | 0.76571168 | 0.96704052 |
| WNT2B | 585.639831 | 0.10073402 | 0.16724308 | 0.60232102 | 0.54696047 | 0.92345821 |
| ST7L | 877.115306 | -0.0242967 | 0.16276175 | -0.1492774 | 0.88133474 | 0.98275231 |
| CAPZA1 | 4829.65148 | -0.1546893 | 0.11116627 | -1.391513 | 0.16406991 | 0.7527134 |
| MOV10 | 5799.86633 | -0.5345683 | 0.42387873 | -1.2611349 | 0.20726025 | 0.78790396 |
| AL603832.1 | 327.128886 | -0.4642948 | 0.41363025 | -1.1224875 | 0.26165522 | 0.82424998 |
| RHOC | 16315.6092 | -0.1175394 | 0.22261867 | -0.5279855 | 0.59750939 | 0.93416444 |
| AL603832.3 | 8475.34491 | -0.057079 | 0.23025647 | -0.2478931 | 0.80421709 | 0.97287842 |
| PPM1J | 279.922575 | 0.02959569 | 0.56743349 | 0.0521571 | 0.95840351 | 0.99237995 |
| SLC16A1 | 1403.28624 | 0.10939536 | 0.39133644 | 0.279543 | 0.77982814 | 0.96836854 |
| SLC16A1-AS1 | 96.5624061 | -0.5237085 | 0.29611159 | -1.7686186 | 0.07695754 | 0.63806922 |
| LRIG2 | 1279.08765 | -0.0862802 | 0.16885431 | -0.5109744 | 0.60936899 | 0.93748023 |
| MAGI3 | 794.740366 | 0.16123694 | 0.1958604 | 0.82322378 | 0.41038075 | 0.88249519 |
| PHTF1 | 1058.42122 | -0.2558209 | 0.23399952 | -1.093254 | 0.27428231 | 0.82719407 |
| RSBN1 | 993.781652 | -0.1864718 | 0.13669681 | -1.3641268 | 0.17252766 | 0.75974924 |
| PTPN22 | 121.673634 | -0.0123792 | 0.74109215 | -0.016704 | 0.98667279 | 0.99715899 |
| AP4B1-AS1 | 328.839496 | 0.25419266 | 0.2156951 | 1.17848138 | 2.39E-01 | 8.11E-01 |
| BCL2L15 | 137.350605 | 1.64607928 | 0.42208734 | 3.89985465 | 9.63E-05 | 4.40E-02 |
| AP4B1 | 1032.53742 | 0.23866531 | 0.15830444 | 1.50763501 | 0.13164797 | 0.7112178 |
| DCLRE1B | 254.094451 | 0.0333186 | 0.33481341 | 0.09951395 | 0.92073021 | 0.98767854 |
| HIPK1 | 2796.77362 | 0.03831753 | 0.12830517 | 0.29864369 | 0.76521193 | 0.96678847 |
| OLFML3 | 1355.95603 | 0.27665357 | 0.48246899 | 0.57341213 | 0.56636568 | 0.9256133 |
| SYT6 | 230.584633 | 0.44972363 | 0.81563764 | 0.55137675 | 0.58137544 | 0.9317705 |
| TRIM33 | 2174.27975 | -0.4233645 | 0.25181605 | -1.681245 | 0.09271535 | 0.65662464 |
| BCAS2 | 1535.26607 | -0.1146468 | 0.2198377 | -0.5215065 | 0.60201395 | 0.9355118 |
| DENND2C | 106.738136 | -0.1640835 | 0.37393101 | -0.4388069 | 0.66080145 | 0.94628868 |
| NRAS | 2744.03446 | -0.4072172 | 0.35314865 | -1.1531042 | 2.49E-01 | 8.16E-01 |
| CSDE1 | 18163.4323 | 0.03310295 | 0.20362178 | 0.1625708 | 0.87085638 | 0.98275231 |
| SIKE1 | 1624.43658 | -0.1137942 | 0.15906592 | -0.7153902 | 0.47436794 | 0.90199039 |
| TSPAN2 | 348.516646 | 0.13918907 | 0.34949919 | 0.39825292 | 0.69044376 | 0.95158502 |
| NGF | 200.21398 | 0.10677152 | 0.52951844 | 0.20163891 | 0.84019903 | 0.97618283 |

| | | | | | | |
|---|---|---|---|---|---|---|
| VANGL1 | 956.935409 | -0.297447 | 0.19691407 | -1.5105421 | 0.13090517 | 0.71007055 |
| CASQ2 | 177.716824 | -0.9194624 | 0.76477505 | -1.2022652 | 0.22926079 | 0.80535641 |
| SLC22A15 | 525.614327 | 0.13449016 | 0.29290902 | 0.45915337 | 0.64612403 | 0.94208859 |
| MAB21L3 | 78.8546632 | 0.3080091 | 0.75758907 | 0.40656487 | 0.68432761 | 0.95070301 |
| ATP1A1 | 152812.579 | -0.1436306 | 0.24057269 | -0.5970363 | 0.55048313 | 0.92373235 |
| LINC01762 | 533.92426 | -0.216481 | 0.53168292 | -0.4071619 | 0.68388909 | 0.95070301 |
| AL390066.1 | 230.188357 | 0.55320677 | 0.42709631 | 1.29527406 | 0.1952257 | 0.77723652 |
| CD58 | 1030.83211 | -0.1010461 | 0.21408922 | -0.4719814 | 0.63694005 | 0.94088547 |
| IGSF3 | 3915.81109 | -0.1121726 | 0.23027317 | -0.4871285 | 0.62616724 | 0.94088547 |
| CD2 | 204.157736 | 0.05609963 | 0.90535114 | 0.0619645 | 0.9505911 | 0.99170503 |
| PTGFRN | 1299.22068 | 0.31202381 | 0.2888864 | 1.08009173 | 0.28010133 | 0.82825734 |
| CD101 | 54.3072386 | 0.20891216 | 0.28775554 | 0.72600568 | 4.68E-01 | 0.89984974 |
| TTF2 | 551.767868 | 0.09614948 | 0.17128544 | 0.56134065 | 5.75E-01 | 0.93024203 |
| TRIM45 | 409.98731 | 0.21448137 | 0.42720656 | 0.50205541 | 0.61562854 | 0.93910158 |
| VTCN1 | 1054.81937 | 2.01786494 | 0.55717971 | 3.62156931 | 0.00029282 | 0.08492861 |
| MAN1A2 | 2114.97888 | -0.0136037 | 0.18987105 | -0.0716471 | 0.94288276 | 0.99136103 |
| TENT5C | 894.123266 | 0.20200936 | 0.46803743 | 0.43160941 | 0.66602532 | 0.9475861 |
| GDAP2 | 832.560334 | -0.0003712 | 0.10315868 | -0.003598 | 0.99712924 | 0.99976751 |
| WDR3 | 1138.03128 | -0.0702555 | 0.20279995 | -0.3464274 | 0.72902156 | 0.95848028 |
| SPAG17 | 430.133185 | -0.2175567 | 0.3772991 | -0.576616 | 0.56419886 | 0.92470993 |
| TBX15 | 56.3495369 | -1.8044875 | 1.02134659 | -1.7667729 | 0.07726627 | 0.63862831 |
| WARS2 | 878.400938 | -0.0358137 | 0.23987238 | -0.1493032 | 0.88131442 | 0.98275231 |
| AL359915.2 | 458.979537 | -0.3398352 | 0.28394649 | -1.1968282 | 0.23137352 | 0.80649152 |
| HAO2 | 5775.995 | 2.31447618 | 0.80440308 | 2.87725921 | 0.00401146 | 0.27238681 |
| HSD3B2 | 60.3738633 | 1.50642789 | 0.95731472 | 1.57359733 | 0.11558057 | 0.6873836 |
| ZNF697 | 1332.35042 | -0.4513525 | 0.46383285 | -0.9730931 | 0.33050703 | 0.85384238 |
| PHGDH | 2994.77511 | 0.14095168 | 0.34663796 | 0.40662506 | 0.68428339 | 0.95070301 |
| HMGCS2 | 3797.38911 | 1.15177833 | 0.57565907 | 2.00079941 | 0.04541401 | 0.57529633 |
| NOTCH2 | 3001.45742 | -0.1339236 | 0.11929091 | -1.1226639 | 0.26158028 | 0.82424998 |
| SEC22B | 3682.48543 | -0.1337785 | 0.1527729 | -0.8756687 | 0.38121015 | 0.87171941 |
| NBPF26 | 182.172195 | -0.2249975 | 0.41673692 | -0.539903 | 0.5892639 | 0.93289552 |
| LINC00623 | 285.524595 | 0.42447299 | 0.36226492 | 1.17171982 | 0.24130956 | 0.8124487 |
| FCGR1B | 41.9901889 | -0.0837923 | 0.81332664 | -0.1030241 | 0.91794382 | 0.9868044 |
| SRGAP2C | 538.702563 | 0.18828133 | 0.29829257 | 0.63119686 | 0.52791182 | 0.91857508 |
| AC245595.1 | 342.792293 | -1.0769473 | 0.54936734 | -1.9603409 | 0.04995596 | 0.5861488 |
| NBPF15 | 844.238526 | 0.2542858 | 0.25102609 | 1.01298555 | 3.11E-01 | 8.45E-01 |
| SRGAP2B | 236.253656 | 0.12175077 | 0.35755706 | 0.34050725 | 0.73347457 | 0.95884813 |
| NBPF20 | 227.495804 | -0.2988514 | 0.37188811 | -0.8036056 | 0.42162481 | 0.88451372 |
| GPR89A | 502.808603 | 0.0513358 | 0.22351039 | 0.22967973 | 8.18E-01 | 0.97454841 |
| PDZK1 | 18495.0832 | 0.642069 | 0.48700476 | 1.31840397 | 0.18736845 | 0.76958556 |
| CD160 | 65.4238594 | -0.0891742 | 0.25271322 | -0.3528672 | 0.72418798 | 0.95813674 |
| RNF115 | 2647.49653 | -0.2935996 | 0.14272817 | -2.0570543 | 0.039681 | 0.55657615 |
| POLR3C | 1073.07414 | -0.2639358 | 0.20556412 | -1.2839585 | 0.19915648 | 0.78134034 |
| NUDT17 | 719.472116 | -0.115988 | 0.22439133 | -0.5169004 | 0.60522569 | 0.93687349 |
| PIAS3 | 2252.70436 | -0.1545648 | 0.18278447 | -0.8456122 | 0.39776911 | 0.88054917 |
| ANKRD35 | 57.9069165 | -0.0747108 | 0.52490167 | -0.1423329 | 0.88681707 | 0.98423722 |
| ITGA10 | 352.408728 | -0.77775 | 0.42336406 | -1.8370713 | 0.06619937 | 0.6234384 |
| PEX11B | 1720.21731 | 0.10044311 | 0.18421549 | 0.54524789 | 0.58558304 | 0.93289552 |
| RBM8A | 4020.46056 | -0.0782339 | 0.15968986 | -0.4899112 | 0.62419673 | 0.94073751 |
| LIX1L-AS1 | 1473.54144 | -0.182744 | 0.16367973 | -1.1164729 | 0.26421978 | 0.82424998 |
| AC243547.3 | 1029.58937 | 0.07505215 | 0.16493484 | 0.45504121 | 0.64907959 | 0.94312988 |
| LIX1L | 1300.46801 | 0.05069128 | 0.19029285 | 0.26638564 | 0.78994222 | 0.97090793 |
| ANKRD34A | 77.1509784 | -0.477682 | 0.42044552 | -1.1361329 | 2.56E-01 | 8.21E-01 |
| POLR3GL | 1007.91875 | 0.30537283 | 0.20996485 | 1.45439982 | 1.46E-01 | 0.73247558 |

| | | | | | | |
|---|---|---|---|---|---|---|
| TXNIP | 54062.6377 | -0.1303578 | 0.48594094 | -0.2682585 | 0.78850032 | 0.97090793 |
| NBPF10 | 192.548283 | -0.0276589 | 0.19572555 | -0.1413147 | 0.88762135 | 0.98423722 |
| AC239799.1 | 102.949066 | 0.21889318 | 0.40229135 | 0.54411604 | 0.58636162 | 0.93289552 |
| NOTCH2NL | 108.717592 | 0.18298519 | 0.40996597 | 0.4463424 | 0.65534994 | 0.94532497 |
| NBPF12 | 786.269768 | -0.0986704 | 0.18168206 | -0.5430937 | 0.58706527 | 0.93289552 |
| PRKAB2 | 1358.97206 | 0.15267667 | 0.17172839 | 0.88905898 | 0.37397138 | 0.86791605 |
| AC242426.2 | 199.370893 | 0.10261308 | 0.26571071 | 0.38618348 | 0.69936079 | 0.95307625 |
| FMO5 | 1338.49994 | 0.94442245 | 0.4703167 | 2.0080564 | 0.0446373 | 0.57267043 |
| CHD1L | 1767.72135 | 0.13164673 | 0.17017332 | 0.77360381 | 0.43916512 | 0.89175645 |
| LINC00624 | 70.8031923 | 0.12028103 | 0.32621884 | 0.36871269 | 0.71234189 | 0.95560193 |
| BCL9 | 1791.53568 | -0.2532313 | 0.22880605 | -1.1067509 | 0.26840164 | 0.82676682 |
| ACP6 | 1933.26259 | 0.18634561 | 0.23169112 | 0.80428465 | 0.4212326 | 0.88451372 |
| GJA5 | 1667.44889 | -0.2417538 | 0.59357206 | -0.4072864 | 0.68379762 | 0.95070301 |
| AC241644.3 | 686.190071 | -0.2481862 | 0.593613 | -0.4180943 | 0.6758782 | 0.94959168 |
| GPR89B | 639.693969 | 0.21905115 | 0.24164033 | 0.90651732 | 0.3646621 | 0.8663515 |
| NBPF11 | 630.3787 | -0.0837609 | 0.15628981 | -0.5359334 | 0.5920046 | 0.93289552 |
| NBPF14 | 601.27837 | -0.0294991 | 0.22051639 | -0.1337729 | 0.89358215 | 0.98474725 |
| PDE4DIP | 3809.93097 | 0.0296252 | 0.26109774 | 0.11346403 | 0.90966268 | 0.98662757 |
| NBPF9 | 767.881872 | -0.3926116 | 0.26468702 | -1.483305 | 0.1379934 | 0.72209298 |
| AC245297.3 | 251.964383 | 0.10386182 | 0.41770374 | 0.24864948 | 0.80363193 | 0.97274243 |
| NBPF19 | 669.670702 | -0.2258641 | 0.59993377 | -0.3764817 | 0.70655879 | 0.95501548 |
| FCGR1A | 211.557109 | 0.12157331 | 0.71629395 | 0.16972545 | 0.86522606 | 0.98197545 |
| HIST2H2BF | 173.82246 | -0.3407297 | 0.68178865 | -0.4997585 | 0.61724512 | 0.93910158 |
| AC239868.3 | 169.591401 | -0.5138742 | 0.40463054 | -1.2699837 | 0.20409044 | 0.78419929 |
| HIST2H2BE | 1363.87247 | -0.3654755 | 0.43261783 | -0.8448 | 0.39822253 | 0.88074856 |
| BOLA1 | 707.538711 | 0.17705908 | 0.18090143 | 0.97876002 | 0.32769857 | 0.85317073 |
| SV2A | 283.920263 | -0.1917752 | 0.51271534 | -0.3740384 | 0.70837577 | 0.95501548 |
| SF3B4 | 3286.62796 | -0.3284883 | 0.25021432 | -1.3128278 | 1.89E-01 | 7.70E-01 |
| MTMR11 | 2395.95872 | 0.21356855 | 0.37058584 | 0.57629982 | 0.56441254 | 0.92474393 |
| OTUD7B | 1619.09059 | -0.5955921 | 0.26341402 | -2.2610493 | 0.0237562 | 0.51967919 |
| VPS45 | 2016.32407 | 0.12218566 | 0.2065198 | 0.59164139 | 0.55409075 | 0.92373235 |
| PLEKHO1 | 1751.51941 | -0.7682199 | 0.35755904 | -2.1485119 | 0.03167311 | 0.54507414 |
| ANP32E | 2634.46032 | 0.18289628 | 0.22531504 | 0.81173579 | 0.41694325 | 0.88445833 |
| CA14 | 49.4426918 | 0.09704199 | 0.56738867 | 0.17103266 | 0.86419809 | 0.98176973 |
| APH1A | 9043.97208 | -0.030951 | 0.10997196 | -0.2814446 | 0.77836939 | 0.96836854 |
| C1orf54 | 1576.91289 | -0.0377877 | 0.12318983 | -0.3067437 | 0.75903845 | 0.96502715 |
| CIART | 500.724351 | -0.9139047 | 0.42852057 | -2.1326975 | 0.03294955 | 0.55137197 |
| MRPS21 | 3258.0344 | -0.0042855 | 0.11209945 | -0.0382299 | 0.9695044 | 0.99400331 |
| PRPF3 | 2720.29659 | -0.2839866 | 0.19516367 | -1.4551201 | 0.14563602 | 0.73192937 |
| RPRD2 | 2641.8312 | 0.04005392 | 0.15657017 | 0.25582089 | 0.79808914 | 0.97170189 |
| TARS2 | 1707.61435 | 0.3145316 | 0.21206903 | 1.48315668 | 0.13803279 | 0.72209298 |
| ECM1 | 1182.04318 | -0.0499063 | 0.35696084 | -0.1398088 | 0.88881109 | 0.98423722 |
| AL356356.1 | 240.275991 | -0.6407001 | 0.48436513 | -1.3227627 | 0.1859143 | 0.76908034 |
| ADAMTSL4 | 1984.30906 | -0.7915756 | 0.46628525 | -1.6976208 | 0.08957935 | 0.65375543 |
| MCL1 | 17019.6565 | -0.3981107 | 0.26076891 | -1.5266802 | 0.12684056 | 0.70443503 |
| ENSA | 7165.8488 | -0.1541371 | 0.12725277 | -1.2112668 | 0.22579316 | 0.80246466 |
| GOLPH3L | 1598.47032 | 0.06149411 | 0.1738939 | 0.35363003 | 0.72361617 | 0.95810643 |
| CTSS | 1804.52241 | -0.1259443 | 0.47445054 | -0.2654529 | 0.79066056 | 0.97090793 |
| CTSK | 812.542944 | 0.63459998 | 0.28715764 | 2.20993592 | 0.02710961 | 0.52773677 |
| ARNT | 2461.33898 | -0.1349933 | 0.13958675 | -0.9670922 | 0.33349794 | 0.855011 |
| SETDB1 | 2186.54993 | -0.1315115 | 0.12491495 | -1.052808 | 0.29242902 | 0.83524572 |
| CERS2 | 21411.2009 | -0.0119757 | 0.2351919 | -0.0509189 | 0.95939014 | 0.9924381 |
| AL590133.2 | 12557.1436 | -0.0253159 | 0.24291268 | -0.104218 | 0.91699632 | 0.9868044 |
| ANXA9 | 1629.94272 | 0.33325989 | 0.19976227 | 1.66828243 | 0.09525967 | 0.65888357 |

| | | | | | | |
|---|---|---|---|---|---|---|
| MINDY1 | 2739.90581 | 0.06247078 | 0.29989078 | 0.20831176 | 0.83498555 | 0.97550416 |
| PRUNE1 | 1825.07001 | -0.0258653 | 0.23612035 | -0.1095429 | 0.91277189 | 0.98662757 |
| BNIPL | 257.168646 | -0.2787584 | 0.61142941 | -0.4559127 | 0.64845275 | 0.9431121 |
| C1orf56 | 2406.25905 | -0.1725262 | 0.16039165 | -1.075656 | 0.28208113 | 0.83003334 |
| CDC42SE1 | 6778.46496 | -0.3072068 | 0.21073395 | -1.4577944 | 0.14489722 | 0.73039059 |
| MLLT11 | 621.203348 | -0.4524149 | 0.26479187 | -1.7085678 | 0.08753104 | 0.65271545 |
| GABPB2 | 1191.93451 | 0.22975834 | 0.31832228 | 0.72177901 | 0.47043035 | 0.9008571 |
| SEMA6C | 1711.56755 | 0.17970232 | 0.33202881 | 0.54122508 | 0.58835246 | 0.93289552 |
| TNFAIP8L2 | 105.871697 | -0.0291689 | 0.55353714 | -0.0526955 | 9.58E-01 | 9.92E-01 |
| SCNM1 | 1078.20611 | -0.3036214 | 0.1538242 | -1.9738207 | 0.04840214 | 0.58440502 |
| LYSMD1 | 596.482128 | 0.05133179 | 0.22669902 | 0.22643145 | 0.82086587 | 0.97472105 |
| TMOD4 | 60.0856173 | 0.67380557 | 0.31926975 | 2.11045851 | 3.48E-02 | 5.54E-01 |
| VPS72 | 1628.11997 | -0.0324435 | 0.11462022 | -0.2830525 | 0.77713662 | 0.96836854 |
| PIP5K1A | 3407.50046 | -0.2067297 | 0.11907097 | -1.7361891 | 0.08253041 | 0.64661469 |
| PSMD4 | 5268.6025 | 0.09672268 | 0.1279242 | 0.75609371 | 0.44959301 | 0.89557948 |
| AL391069.2 | 44.6735179 | -0.0262589 | 0.30382702 | -0.086427 | 0.93112699 | 0.98975562 |
| ZNF687 | 2308.11969 | -0.3258195 | 0.14850041 | -2.1940648 | 0.02823075 | 0.53300785 |
| PI4KB | 5156.57215 | -0.1541716 | 0.13406589 | -1.1499689 | 0.2501567 | 0.81721208 |
| RFX5 | 2274.01666 | -0.0718117 | 0.22374711 | -0.3209505 | 0.74824791 | 0.96241535 |
| AL391069.1 | 416.210636 | 0.01329443 | 0.20445096 | 0.06502505 | 0.94815405 | 0.99170503 |
| SELENBP1 | 6511.68592 | 0.17668983 | 0.41638617 | 0.42434126 | 0.67131695 | 0.94922827 |
| PSMB4 | 10118.1412 | -0.1409675 | 0.18845512 | -0.7480165 | 0.45445022 | 0.89670027 |
| POGZ | 4507.82456 | -0.1089868 | 0.2330526 | -0.4676489 | 0.64003566 | 0.94088547 |
| CGN | 4226.61815 | -0.1105908 | 0.33196966 | -0.3331353 | 0.73903213 | 0.9602325 |
| TUFT1 | 1632.54514 | -0.0364244 | 0.26086149 | -0.1396312 | 0.88895138 | 0.98423722 |
| SNX27 | 3520.36528 | 0.02448521 | 0.21145799 | 0.11579232 | 0.90781714 | 0.98662757 |
| AL589765.7 | 575.789472 | -0.1681757 | 0.18461762 | -0.9109408 | 0.36232659 | 0.86561928 |
| MRPL9 | 2095.15162 | -0.1292019 | 0.14069463 | -0.9183147 | 0.35845413 | 0.86480228 |
| OAZ3 | 680.673839 | -0.0060814 | 0.18508655 | -0.0328568 | 0.97378875 | 0.99445887 |
| AL589765.4 | 274.471804 | -0.0205374 | 0.21170772 | -0.0970084 | 0.92271971 | 0.98826666 |
| TDRKH | 477.700216 | 0.25002267 | 0.19441872 | 1.28600099 | 1.98E-01 | 0.78046101 |
| TDRKH-AS1 | 60.7430643 | 0.13999477 | 0.28014343 | 0.49972533 | 0.6172685 | 0.93910158 |
| LINGO4 | 75.5396317 | -1.877814 | 0.60188136 | -3.1199072 | 0.00180908 | 0.18722055 |
| RORC | 3645.18422 | -0.3417236 | 0.45450713 | -0.7518553 | 0.45213808 | 0.89670027 |
| AL450992.2 | 93.2404384 | -0.3668192 | 0.3278841 | -1.1187465 | 0.2632483 | 0.82424998 |
| THEM5 | 130.390836 | -0.502202 | 0.23970192 | -2.0951106 | 0.03616116 | 0.55384187 |
| THEM4 | 1918.99814 | 0.02578948 | 0.29090364 | 0.08865301 | 0.92935768 | 0.98963127 |
| S100A10 | 11161.4752 | -0.4401387 | 0.37523001 | -1.1729839 | 0.24080228 | 0.8124487 |
| S100A11 | 15086.8222 | -0.4332771 | 0.34625044 | -1.2513402 | 0.2108104 | 0.79134644 |
| FLG-AS1 | 327.901303 | -0.0905455 | 0.55260596 | -0.1638519 | 0.86984774 | 0.98275231 |
| FLG | 48.8353031 | -0.1699569 | 0.61783314 | -0.2750854 | 0.78325066 | 0.96971192 |
| S100A9 | 3073.31118 | -0.5520247 | 0.66311616 | -0.8324705 | 0.40514341 | 0.88132015 |
| S100A12 | 280.451011 | -1.2417809 | 0.94207366 | -1.3181356 | 0.18745825 | 0.76958556 |
| S100A8 | 1423.06608 | -0.9762567 | 0.65296902 | -1.4951042 | 0.13488725 | 0.7163281 |
| BX470102.1 | 346.155477 | -0.10825 | 0.46585738 | -0.2323672 | 0.8162528 | 0.97388289 |
| S100A6 | 28699.992 | -0.1881285 | 0.45796835 | -0.4107894 | 0.68122697 | 0.95070301 |
| S100A4 | 2167.51802 | 0.16950916 | 0.59788314 | 0.28351554 | 0.77678167 | 0.96836854 |
| S100A3 | 224.38082 | -0.5290144 | 0.65937251 | -0.8022997 | 0.42237961 | 0.88451372 |
| S100A2 | 3910.45088 | -0.1947822 | 0.78208005 | -0.2490566 | 0.80331704 | 0.97274243 |
| S100A16 | 6492.33829 | -0.4894645 | 0.38068374 | -1.2857509 | 0.19853002 | 0.78046101 |
| S100A14 | 1301.25387 | -0.3816046 | 0.38936686 | -0.9800644 | 0.32705431 | 0.85307234 |
| S100A13 | 8110.68943 | 0.55965059 | 0.26509542 | 2.11112884 | 0.03476124 | 0.55369025 |
| S100A1 | 5060.01216 | 0.67410084 | 0.3752354 | 1.79647455 | 0.07241908 | 0.6324352 |
| AL162258.2 | 4269.14321 | 0.54906445 | 0.34426617 | 1.59488355 | 0.11073828 | 0.68001961 |

| | | | | | | |
|---|---|---|---|---|---|---|
| CHTOP | 4656.47979 | -0.2411537 | 0.15650727 | -1.5408467 | 0.12335409 | 0.69884494 |
| SNAPIN | 1504.49664 | 0.04651656 | 0.17878406 | 0.26018293 | 0.79472267 | 0.97101097 |
| ILF2 | 8117.34332 | 0.25778632 | 0.22434749 | 1.14904927 | 0.25053566 | 0.81738655 |
| NPR1 | 5727.87922 | 0.10284883 | 0.34946388 | 0.29430462 | 0.76852514 | 0.96793882 |
| INTS3 | 7569.53544 | -0.1871059 | 0.16621101 | -1.125713 | 0.260287 | 0.82424998 |
| AL513523.4 | 165.485469 | -0.1089234 | 0.38561564 | -0.2824662 | 0.77758608 | 0.96836854 |
| AL513523.3 | 115.722991 | -0.0172128 | 0.44564542 | -0.0386243 | 0.9691899 | 0.99400331 |
| SLC27A3 | 2320.37299 | -0.1787633 | 0.18935515 | -0.9440638 | 0.34513703 | 0.8605056 |
| GATAD2B | 2285.08131 | -0.2143971 | 0.12384964 | -1.7311081 | 0.08343249 | 0.64772507 |
| AL358472.5 | 174.511468 | -0.2716841 | 0.24008659 | -1.131609 | 0.25779887 | 0.82187662 |
| DENND4B | 3517.5127 | -0.299959 | 0.1137214 | -2.6376651 | 0.0083479 | 0.37698298 |
| CRTC2 | 4402.79244 | -0.3691462 | 0.17449553 | -2.1155055 | 0.03438689 | 0.55369025 |
| SLC39A1 | 11173.3058 | -0.3633592 | 0.29710787 | -1.2229874 | 0.22133446 | 0.80015993 |
| CREB3L4 | 729.099855 | 0.25004457 | 0.26222477 | 0.95355055 | 0.3403112 | 0.85814041 |
| JTB | 5396.05354 | -0.0838393 | 0.16992508 | -0.4933895 | 0.62173738 | 0.94016713 |
| AL358472.2 | 120.257638 | -0.2778822 | 0.32392267 | -0.8578658 | 0.39096655 | 0.87710956 |
| RAB13 | 6324.2278 | 0.07680475 | 0.19376001 | 0.39639112 | 6.92E-01 | 9.52E-01 |
| RPS27 | 25568.8373 | 0.25235862 | 0.1487514 | 1.69651259 | 8.98E-02 | 6.54E-01 |
| TPM3 | 11146.3911 | -0.1887346 | 0.24000321 | -0.7863835 | 0.43164283 | 0.88976671 |
| C1orf43 | 10336.876 | 0.04729983 | 0.21247129 | 0.22261751 | 0.8238332 | 0.97548201 |
| UBAP2L | 6275.5205 | -0.332582 | 0.15876928 | -2.0947504 | 0.03619319 | 0.55384187 |
| HAX1 | 3145.60675 | 0.12446481 | 0.1803014 | 0.69031529 | 0.48999594 | 0.90600166 |
| ATP8B2 | 1224.93095 | -0.156774 | 0.37761649 | -0.4151673 | 0.6780194 | 0.95018034 |
| IL6R | 2954.48868 | -0.859555 | 0.53664721 | -1.6017133 | 0.10921903 | 0.67859123 |
| SHE | 902.307661 | 0.09088164 | 0.39262316 | 0.23147295 | 0.8169474 | 0.97397142 |
| AL162591.2 | 223.145578 | 0.06849575 | 0.44181459 | 0.15503279 | 0.87679547 | 0.98275231 |
| TDRD10 | 196.907364 | -0.2455303 | 0.35119654 | -0.699125 | 0.48447389 | 0.90541935 |
| UBE2Q1 | 4402.56056 | -0.2622297 | 0.21029238 | -1.2469765 | 0.21240613 | 0.79203904 |
| UBE2Q1-AS1 | 579.500584 | -0.1523895 | 0.25581894 | -0.5956929 | 0.55138039 | 0.92373235 |
| ADAR | 10370.1999 | -0.4450377 | 0.32139873 | -1.3846905 | 0.16614714 | 0.75515387 |
| KCNN3 | 327.389811 | -0.08529 | 0.37565045 | -0.2270462 | 0.8203878 | 0.9746981 |
| PMVK | 2736.62139 | 0.67722686 | 0.27438542 | 2.46815905 | 0.013581 | 0.42608223 |
| PBXIP1 | 9175.37254 | 0.33018575 | 0.25809664 | 1.27931056 | 0.20078772 | 0.78134034 |
| PYGO2 | 4193.5722 | -0.2219496 | 0.12518783 | -1.7729324 | 0.0762399 | 0.63773415 |
| AL451085.2 | 1671.31497 | -0.2702607 | 0.2676496 | -1.0097558 | 0.3126123 | 0.84541735 |
| SHC1 | 7606.88444 | -0.3373481 | 0.22359294 | -1.50876 | 0.13136014 | 0.71081672 |
| CKS1B | 511.532119 | 0.09440534 | 0.19461404 | 0.48509008 | 0.62761245 | 0.94088547 |
| FLAD1 | 1776.87118 | -0.1590445 | 0.13050347 | -1.2186992 | 0.22295837 | 0.80078694 |
| ZBTB7B | 4251.24688 | -0.1223554 | 0.20094353 | -0.6089043 | 0.54258786 | 0.92292765 |
| DCST2 | 100.582997 | -0.150266 | 0.41696812 | -0.3603776 | 0.71856475 | 0.95648644 |
| DCST1 | 51.8883912 | 0.10447767 | 0.3323443 | 0.31436578 | 0.75324324 | 0.96301881 |
| ADAM15 | 4650.32728 | -0.3408832 | 0.32096718 | -1.0620499 | 2.88E-01 | 8.33E-01 |
| EFNA4 | 391.619379 | -0.0792475 | 0.22690601 | -0.3492524 | 7.27E-01 | 9.58E-01 |
| AL691442.1 | 191.485109 | -0.1431601 | 0.37697242 | -0.3797629 | 7.04E-01 | 9.55E-01 |
| EFNA3 | 221.218432 | -0.2060703 | 0.48456083 | -0.4252722 | 0.67063823 | 0.94922827 |
| EFNA1 | 11288.8109 | -0.593947 | 0.38840164 | -1.5292082 | 0.12621284 | 0.70418422 |
| SLC50A1 | 2729.84225 | -0.1098635 | 0.14274524 | -0.7696475 | 0.441509 | 0.89297648 |
| DPM3 | 972.132656 | 0.14538677 | 0.1713456 | 0.84850012 | 0.39615951 | 0.87967259 |
| KRTCAP2 | 3062.6022 | -0.0877219 | 0.14300555 | -0.6134163 | 0.53960112 | 0.92198621 |
| AL713999.1 | 8010.0395 | 0.09476208 | 0.35030736 | 0.27051126 | 0.78676696 | 0.97090793 |
| TRIM46 | 133.508973 | 0.2454334 | 0.44596053 | 0.55034781 | 0.58208084 | 0.9317705 |
| MUC1 | 25253.2433 | 0.16947263 | 0.48884481 | 0.34667983 | 0.72883187 | 0.95848028 |
| AC234582.1 | 1193.1853 | -0.2603593 | 0.31017484 | -0.8393954 | 0.40124749 | 0.88132015 |
| THBS3 | 2601.94841 | -0.2524387 | 0.3197104 | -0.7895855 | 0.42976986 | 0.88870396 |

| | | | | | | |
|---|---|---|---|---|---|---|
| MTX1 | 1565.16838 | -0.0176766 | 0.11175404 | -0.1581742 | 0.87431956 | 0.98275231 |
| AC234582.2 | 184.567775 | -0.0428762 | 0.17726654 | -0.2418741 | 0.8088777 | 0.97334138 |
| GBA | 3610.9977 | 0.19411443 | 0.21054826 | 0.92194743 | 0.356556 | 0.86390849 |
| FAM189B | 1887.95883 | -0.3262433 | 0.21838921 | -1.493862 | 0.13521169 | 0.71649038 |
| SCAMP3 | 3749.16966 | -0.132183 | 0.1698027 | -0.7784504 | 4.36E-01 | 8.91E-01 |
| CLK2 | 4310.10056 | -0.2327697 | 0.16300254 | -1.428013 | 0.15328812 | 0.74258845 |
| HCN3 | 1543.17011 | 0.37265569 | 0.34055629 | 1.09425577 | 0.27384282 | 0.82719197 |
| PKLR | 3204.47567 | 1.9389992 | 0.61841406 | 3.13543843 | NA | NA |
| FDPS | 5955.08371 | 0.22635969 | 0.22165521 | 1.02122431 | 0.30714818 | 0.84298407 |
| RUSC1-AS1 | 4749.16584 | 0.19417029 | 0.18456907 | 1.05201965 | 0.29279053 | 0.83524862 |
| RUSC1 | 3036.16221 | -0.2700002 | 0.13077623 | -2.064597 | 0.03896115 | 0.55613287 |
| ASH1L | 2126.65609 | -0.1357433 | 0.11870509 | -1.1435341 | 0.25281689 | 0.81835388 |
| ASH1L-AS1 | 180.075977 | -0.2647622 | 0.38249762 | -0.6921932 | 0.48881602 | 0.905872 |
| AL353807.2 | 72.3619872 | -0.4307849 | 0.4387867 | -0.9817639 | 0.32621617 | 0.85282528 |
| MSTO1 | 1435.04022 | -0.1496924 | 0.23752294 | -0.6302228 | 0.52854882 | 0.91888066 |
| YY1AP1 | 4174.68862 | -0.2687955 | 0.10661207 | -2.521248 | 0.01169394 | 0.40423902 |
| DAP3 | 4346.51091 | 0.09574772 | 0.14290834 | 0.66999388 | 0.50286169 | 0.91353729 |
| GON4L | 1794.54097 | -0.1421314 | 0.15219753 | -0.9338617 | 0.35037525 | 0.86364633 |
| SYT11 | 968.804183 | -0.2696701 | 0.30265762 | -0.8910071 | 0.37292537 | 0.86791605 |
| RIT1 | 2924.86017 | -0.3093603 | 0.24749459 | -1.2499679 | 0.21131128 | 0.79134644 |
| KHDC4 | 4080.94985 | -0.1256845 | 0.16732003 | -0.7511621 | 0.45255511 | 0.89670027 |
| ARHGEF2 | 3091.8603 | -0.2340656 | 0.26613652 | -0.8794945 | 0.3791332 | 0.87069278 |
| AL355388.2 | 70.9650986 | 0.17890145 | 0.63541063 | 0.2815525 | 0.77828666 | 0.96836854 |
| SSR2 | 8114.78912 | 0.09997133 | 0.17854768 | 0.55991392 | 0.57553815 | 0.93080432 |
| UBQLN4 | 3044.15484 | -0.0040889 | 0.13773792 | -0.0296862 | 0.97631736 | 0.99455099 |
| LAMTOR2 | 2183.45796 | 0.00100649 | 0.1382334 | 0.0072811 | 0.99419057 | 0.99936518 |
| RAB25 | 1862.9486 | 0.02107955 | 0.31919123 | 0.06604052 | 0.94734557 | 0.99166614 |
| MEX3A | 222.859887 | -0.0901271 | 0.42030156 | -0.2144343 | 0.83020843 | 0.97550416 |
| LMNA | 16749.5564 | -0.1672842 | 0.34346805 | -0.4870445 | 0.62622677 | 0.94088547 |
| SEMA4A | 2407.87213 | -0.0369683 | 0.50684316 | -0.0729383 | 0.9418552 | 0.99136103 |
| SLC25A44 | 3685.04361 | -0.0491721 | 0.2721879 | -0.1806551 | 0.85663829 | 0.97953896 |
| PMF1-BGLAF | 2167.33306 | 0.03391239 | 0.17146137 | 0.19778443 | 0.84321373 | 0.97649449 |
| PMF1 | 2950.71818 | -0.0065515 | 0.16858387 | -0.0388618 | 0.96900056 | 0.99400331 |
| BGLAP | 318.840051 | -0.0194177 | 0.25272961 | -0.076832 | 0.93875716 | 0.99113899 |
| PAQR6 | 599.27535 | -0.0522104 | 0.43441631 | -0.1201851 | 0.90433653 | 0.98624484 |
| SMG5 | 9339.27038 | -0.2552601 | 0.24453694 | -1.0438511 | 0.29655431 | 0.83831021 |
| TMEM79 | 2490.41257 | 0.00372051 | 0.29783327 | 0.01249194 | 0.99003314 | 0.99807483 |
| GLMP | 5521.54524 | 0.01364668 | 0.33943553 | 0.04020405 | 0.96793045 | 0.99400331 |
| CCT3 | 10003.1503 | 0.02071928 | 0.17352059 | 0.11940534 | 0.90495423 | 0.98624484 |
| RHBG | 1744.57591 | 1.74128692 | 0.37666315 | 4.62292883 | 3.78E-06 | 0.00506488 |
| MEF2D | 4070.40788 | -0.2481982 | 0.20246879 | -1.2258592 | 0.22025168 | 0.79901989 |
| IQGAP3 | 129.7228 | 0.36335742 | 0.53173725 | 0.68334017 | 0.49439191 | 0.90796767 |
| NAXE | 4223.02895 | -0.0197627 | 0.20671136 | -0.0956052 | 0.92383411 | 0.98869145 |
| GPATCH4 | 1407.79916 | -0.3840677 | 0.24784093 | -1.5496542 | 0.12122453 | 0.69625158 |
| HAPLN2 | 40.3932737 | -1.010271 | 0.6774972 | -1.491181 | 0.13591398 | 0.71771311 |
| BCAN | 59.2871539 | 1.0698686 | 0.5378327 | 1.98922191 | 0.04667671 | 0.58047001 |
| AL365181.3 | 68.3411895 | 1.18714402 | 0.56373938 | 2.10583838 | 0.03521839 | 0.55369025 |
| NES | 5623.93775 | -0.1428112 | 0.40937677 | -0.3488502 | 0.72720175 | 0.95848028 |
| CRABP2 | 339.682912 | 0.89576335 | 0.50345195 | 1.77924297 | 0.07519994 | 0.63773415 |
| ISG20L2 | 1492.63612 | -0.2180917 | 0.20073362 | -1.0864733 | 0.27726966 | 0.82782817 |
| RRNAD1 | 1967.87169 | -0.0868769 | 0.16519832 | -0.5258945 | 0.59896153 | 0.93484879 |
| MRPL24 | 2654.43036 | 0.28247613 | 0.148449 | 1.90284959 | 0.05706017 | 0.60049913 |
| HDGF | 17600.177 | -0.0992482 | 0.21406512 | -0.4636354 | 0.64290901 | 0.94123567 |
| PRCC | 2828.96071 | -0.4012015 | 0.20387098 | -1.9679189 | 0.04907737 | 0.58508842 |

| | | | | | | |
|---|---|---|---|---|---|---|
| SH2D2A | 182.578185 | 0.67445486 | 0.79268906 | 0.85084416 | 0.39485593 | 0.87884163 |
| NTRK1 | 238.311793 | -0.0613711 | 0.69086989 | -0.0888317 | 0.92921568 | 0.98963127 |
| INSRR | 465.450904 | 0.02151856 | 0.65134906 | 0.03303691 | 0.97364515 | 0.99445887 |
| PEAR1 | 1782.73353 | -0.1758946 | 0.35318815 | -0.4980196 | 6.18E-01 | 9.39E-01 |
| ARHGEF11 | 2949.31723 | -0.2562637 | 0.15547845 | -1.6482264 | 0.09930623 | 0.6694597 |
| ETV3 | 978.436431 | 0.01658555 | 0.22188895 | 0.07474706 | 0.94041596 | 0.99136103 |
| FCRL5 | 154.158003 | -1.0428308 | 1.03175305 | -1.0107368 | 0.31214242 | 0.84541735 |
| FCRL3 | 58.8812504 | -1.3478512 | 1.00068897 | -1.3469232 | 0.17800496 | 0.76115956 |
| FCRL2 | 43.8520225 | -0.5962432 | 1.07593097 | -0.554165 | NA | NA |
| FCRL1 | 44.3980067 | -1.094962 | 1.26694592 | -0.8642532 | NA | NA |
| KIRREL1 | 2470.03148 | 0.02353868 | 0.26579808 | 0.08855849 | 0.9294328 | 0.98963127 |
| CD1D | 183.969191 | 0.39603028 | 0.49532945 | 0.79952904 | 0.42398372 | 0.88565404 |
| CD1C | 75.4661348 | 2.19192513 | 1.21923135 | 1.79779263 | 0.07220988 | 0.6321115 |
| MNDA | 297.964201 | -0.0171375 | 0.48947545 | -0.0350119 | 0.97207025 | 0.99408319 |
| PYHIN1 | 67.8457653 | 0.31306408 | 0.8802321 | 0.35566084 | 7.22E-01 | 9.57E-01 |
| IFI16 | 2782.94805 | -0.3571158 | 0.40455978 | -0.8827269 | 3.77E-01 | 0.86987124 |
| AIM2 | 43.45411 | -1.2237583 | 0.89286725 | -1.3705938 | 0.17050162 | 0.75956362 |
| CADM3 | 391.931485 | 0.3387847 | 0.85472583 | 0.39636652 | 0.69183466 | 0.95159158 |
| CADM3-AS1 | 302.781341 | -0.0058618 | 0.90576907 | -0.0064716 | 0.99483646 | 0.99940889 |
| ACKR1 | 2774.00564 | -0.1167643 | 1.19179379 | -0.0979736 | 0.92195327 | 0.98817994 |
| FCER1A | 117.976962 | 3.27115968 | 1.21742806 | 2.68694289 | 0.00721093 | 0.35846154 |
| APCS | 256.332144 | 1.92398549 | 1.96239545 | 0.980427 | NA | NA |
| DUSP23 | 7388.47029 | -0.5036917 | 0.46557984 | -1.0818588 | 0.27931527 | 0.82825734 |
| FCRL6 | 91.261188 | 0.08527167 | 0.66491467 | 0.12824453 | 0.89795546 | 0.98474725 |
| SLAMF8 | 656.433456 | -0.2501252 | 0.33790299 | -0.7402279 | 0.45916174 | 0.89755243 |
| AL590560.1 | 876.223153 | 0.0099421 | 0.28666751 | 0.03468163 | 9.72E-01 | 0.99416401 |
| SNHG28 | 426.855817 | 0.10867448 | 0.27938037 | 0.38898395 | 0.69728802 | 0.95269909 |
| VSIG8 | 151.057751 | 0.83986068 | 0.62834722 | 1.33661876 | 0.18134711 | 0.76522169 |
| AL590560.2 | 382.735378 | -0.4818817 | 0.53894437 | -0.8941213 | 0.37125699 | 0.86762749 |
| CFAP45 | 124.223035 | -0.0577311 | 0.44789766 | -0.1288935 | 0.89744191 | 0.98474725 |
| TAGLN2 | 17399.1269 | -0.438703 | 0.36860657 | -1.1901661 | 0.23398113 | 0.80725541 |
| IGSF9 | 116.541494 | -0.0611929 | 0.66697357 | -0.0917471 | 0.92689897 | 0.98915637 |
| KCNJ10 | 2096.10839 | 0.75581695 | 0.47982423 | 1.57519548 | 0.11521133 | 0.68719782 |
| PIGM | 835.113372 | 0.02725869 | 0.13486898 | 0.20211235 | 0.83982889 | 0.97618283 |
| IGSF8 | 4327.336 | 0.37269562 | 0.15473795 | 2.40855984 | 0.0160156 | 0.45684684 |
| ATP1A2 | 294.45353 | 0.1660662 | 0.69965735 | 0.23735361 | 0.81238247 | 0.97364432 |
| AL121987.2 | 531.322774 | -0.6383744 | 0.42365633 | -1.5068212 | 0.13185649 | 0.71172882 |
| PEA15 | 9378.621 | -0.5759936 | 0.39913041 | -1.4431213 | 0.1489863 | 0.73655347 |
| DCAF8 | 8171.6012 | -0.0625719 | 0.12989435 | -0.4817142 | 0.630009 | 0.94088547 |
| AL139011.2 | 4984.62049 | 0.05840012 | 0.16118811 | 0.36231038 | 0.71712011 | 0.95611001 |
| AL139011.1 | 302.310216 | -0.0480416 | 0.3019889 | -0.1590841 | 0.87360263 | 0.98275231 |
| PEX19 | 3953.67744 | 0.2722891 | 0.14924726 | 1.82441608 | 0.06808919 | 0.62613325 |
| COPA | 7406.3711 | -0.1252351 | 0.1721672 | -0.7274038 | 0.46697861 | 0.89983399 |
| NCSTN | 6800.53735 | 0.26157114 | 0.18212718 | 1.43620047 | 0.15094531 | 0.73796431 |
| VANGL2 | 723.968901 | 0.35642837 | 0.4644256 | 0.76746064 | 0.44280769 | 0.89343541 |
| SLAMF6 | 131.350844 | -0.1277731 | 0.91555376 | -0.1395583 | 0.88900902 | 0.98423722 |
| CD84 | 221.706254 | -0.2308518 | 0.57266477 | -0.4031185 | 0.68686107 | 0.9510919 |
| SLAMF1 | 165.754785 | 0.83654882 | 0.83291538 | 1.00436232 | 0.31520401 | 0.84696964 |
| AL121985.1 | 217.285086 | 0.27472859 | 0.73605179 | 0.37324628 | 0.70896515 | 0.95501548 |
| CD48 | 301.743437 | 0.2435344 | 0.74779963 | 0.32566798 | 0.74467557 | 0.9613927 |
| SLAMF7 | 181.903385 | -0.4836098 | 0.82769998 | -0.5842816 | 5.59E-01 | 9.24E-01 |
| LY9 | 59.2326672 | -0.0315889 | 1.06142205 | -0.0297609 | 0.97625771 | 0.99455099 |
| ITLN1 | 128.506896 | 0.18553576 | 0.89009351 | 0.20844524 | 0.83488133 | 0.97550416 |
| F11R | 9983.89859 | 0.0419773 | 0.18401582 | 0.22811786 | 0.81955462 | 0.97454841 |

| | | | | | | |
|---|---|---|---|---|---|---|
| AL591806.3 | 5046.53218 | 0.10854959 | 0.1959003 | 0.5541063 | 0.5795061 | 0.93139743 |
| TSTD1 | 2280.33586 | 0.25000447 | 0.23713154 | 1.05428604 | 0.29175198 | 0.83460625 |
| USF1 | 3583.68973 | -0.3205326 | 0.16416374 | -1.9525178 | 0.05087676 | 0.5861488 |
| ARHGAP30 | 777.748114 | -0.0913585 | 0.54016717 | -0.1691301 | 0.86569432 | 0.98197545 |
| NECTIN4 | 602.230528 | 0.02930062 | 0.46160993 | 0.06347484 | 0.94938839 | 0.99170503 |
| KLHDC9 | 663.476083 | 0.56495944 | 0.41315834 | 1.36741629 | 0.17149486 | 0.75974924 |
| PFDN2 | 1659.09807 | 0.03390986 | 0.17032541 | 0.19908866 | 0.84219339 | 0.9763625 |
| NIT1 | 4705.36358 | 0.16172993 | 0.17529022 | 0.92264096 | 0.35619434 | 0.86390849 |
| DEDD | 2565.02942 | -0.2565716 | 0.10874902 | -2.3593005 | 0.01830942 | 0.47570049 |
| UFC1 | 4181.53153 | 0.10098583 | 0.15503678 | 0.65136695 | 0.51480964 | 0.91636259 |
| AL590714.1 | 944.094318 | 0.15147579 | 0.2110655 | 0.71767195 | 0.47295958 | 0.90175304 |
| USP21 | 2099.53047 | -0.0434172 | 0.18757749 | -0.2314627 | 0.81695533 | 0.97397142 |
| PPOX | 1904.37788 | 0.09606464 | 0.23684485 | 0.40560159 | 0.68503536 | 0.95097288 |
| B4GALT3 | 2687.44011 | -0.1565927 | 0.12468171 | -1.2559394 | 0.20913792 | 0.79086734 |
| ADAMTS4 | 4815.7132 | 0.68403537 | 1.17180345 | 0.58374582 | NA | NA |
| NDUFS2 | 10874.5432 | 0.11281079 | 0.13493624 | 0.83603034 | 0.40313784 | 0.88132015 |
| FCER1G | 1201.92487 | -0.4358239 | 0.57370301 | -0.7596682 | 0.44745293 | 0.89480207 |
| TOMM40L | 1517.53524 | 0.05294215 | 0.14946067 | 0.35422126 | 0.72317307 | 0.95801135 |
| NR1I3 | 1101.95192 | 0.2583138 | 0.35293843 | 0.73189481 | 0.46423278 | 0.89830907 |
| MPZ | 301.209156 | -0.1357952 | 0.28651642 | -0.4739525 | 0.63553375 | 0.94088547 |
| SDHC | 8475.99724 | 0.44784716 | 0.2500471 | 1.79105117 | 7.33E-02 | 6.37E-01 |
| CFAP126 | 344.227401 | 0.15913445 | 0.45673939 | 0.34841412 | 0.7275292 | 0.95848028 |
| AL831711.1 | 38.3262917 | 0.38794348 | 0.36657833 | 1.05828263 | 0.28992661 | 0.83401253 |
| FCGR2A | 1359.06001 | 0.71603853 | 0.48795891 | 1.46741563 | 0.14226303 | 0.72784886 |
| HSPA6 | 958.424464 | -2.0192198 | 0.77508375 | -2.6051633 | 0.00918305 | 0.38664841 |
| FCGR3A | 1479.71655 | 0.24778904 | 0.64042109 | 0.38691581 | 0.69881854 | 0.95297835 |
| FCGR3B | 133.94423 | -0.1530876 | 0.83371585 | -0.1836208 | 0.85431096 | 0.97906925 |
| FCGR2B | 293.842349 | 0.09303366 | 0.71421733 | 0.1302596 | 0.89636104 | 0.98474725 |
| AL451067.1 | 55.6918907 | 0.03177831 | 0.84415644 | 0.03764505 | 0.96997069 | 0.99400331 |
| FCRLB | 228.136698 | 1.11896763 | 0.46785949 | 2.39167456 | 0.01677171 | 0.46515254 |
| DUSP12 | 749.069514 | -0.2775938 | 0.19489815 | -1.4243018 | 0.15435914 | 0.74270721 |
| ATF6 | 3365.45455 | 0.02457746 | 0.14708916 | 0.16709226 | 0.86729746 | 0.98223481 |
| OLFML2B | 503.906684 | -0.894515 | 0.38679871 | -2.3126111 | 0.02074403 | 0.50488816 |
| NOS1AP | 107.368489 | -0.9156205 | 0.44785361 | -2.0444639 | 0.04090774 | 0.55811692 |
| AL512785.2 | 119.92003 | -0.8167804 | 0.44406593 | -1.8393224 | 0.06586779 | 0.62237346 |
| C1orf226 | 1230.84953 | -0.5162268 | 0.2329823 | -2.2157339 | 0.02670973 | 0.52678638 |
| UHMK1 | 4112.05088 | 0.05162826 | 0.15842843 | 0.32587751 | 0.74451703 | 0.9613927 |
| UAP1 | 1824.25176 | -0.2030714 | 0.25825376 | -0.786325 | 0.4316771 | 0.88976671 |
| DDR2 | 1109.68204 | -0.1128786 | 0.37836537 | -0.2983323 | 0.76544953 | 0.96698953 |
| HSD17B7 | 795.109258 | 0.05994129 | 0.21410612 | 0.27996064 | 0.7795077 | 0.96836854 |
| RGS4 | 120.896356 | 0.71674068 | 0.5122402 | 1.39922772 | 0.16174471 | 0.7498147 |
| RGS5 | 9195.00224 | -0.6218931 | 0.65481478 | -0.9497237 | 0.34225267 | 0.85889221 |
| PBX1 | 2721.84289 | 0.18536455 | 0.21966256 | 0.84386046 | 0.39874738 | 0.88074856 |
| MGST3 | 10113.9663 | -0.0646742 | 0.18276576 | -0.3538637 | 0.72344106 | 0.95810643 |
| ALDH9A1 | 7731.67903 | 0.57334856 | 0.25592807 | 2.24027226 | 0.02507325 | 0.52085426 |
| TMCO1 | 4773.94598 | 0.07017561 | 0.13264596 | 0.52904446 | 0.59677461 | 0.9337017 |
| UCK2 | 960.494881 | -0.1943893 | 0.29904428 | -0.6500353 | 0.51566942 | 0.91636259 |
| AL358115.1 | 122.833685 | -0.1686848 | 0.4457686 | -0.3784133 | 0.70512359 | 0.95501548 |
| FAM78B | 90.9666182 | 0.46016973 | 0.3736127 | 1.23167582 | 0.21807021 | 0.79708421 |
| POGK | 2594.38834 | -0.2925598 | 0.20686044 | -1.4142858 | 1.57E-01 | 0.74522306 |
| TADA1 | 866.482944 | 0.06186413 | 0.17751605 | 0.34849879 | 0.72746562 | 0.95848028 |
| ILDR2 | 570.868762 | 0.19629179 | 0.46508472 | 0.42205599 | 0.67298415 | 0.94922827 |
| MAEL | 85.5014234 | 0.8724808 | 0.72451405 | 1.20422896 | 0.22850109 | 0.80389319 |
| POU2F1 | 1126.85 | -0.2635537 | 0.17579679 | -1.4991952 | 0.13382301 | 0.71505705 |

| | | | | | | |
|---|---|---|---|---|---|---|
| CD247 | 210.134019 | 0.00796152 | 0.76683364 | 0.01038233 | 0.99171625 | 0.99891489 |
| CREG1 | 7591.89217 | -0.0159143 | 0.31687359 | -0.0502229 | 0.95994478 | 0.99255196 |
| RCSD1 | 545.462555 | -0.3412666 | 0.33323635 | -1.0240977 | 0.30578915 | 0.84270033 |
| MPZL1 | 4743.46397 | -0.2098543 | 0.1874898 | -1.1192836 | 0.26301918 | 0.82424998 |
| ADCY10 | 36.3489629 | -0.1772241 | 0.50893028 | -0.3482287 | 0.72766844 | 0.95848028 |
| MPC2 | 6444.92161 | 0.26258557 | 0.20290355 | 1.29413988 | 0.19561711 | 0.77752069 |
| DCAF6 | 6011.70299 | -0.092777 | 0.16771634 | -0.5531782 | 0.58014141 | 0.93139743 |
| GPR161 | 241.246942 | -0.1947097 | 0.30988821 | -0.6283225 | 0.52979268 | 0.91950775 |
| TIPRL | 1910.24244 | -0.1120169 | 0.20739949 | -0.5401022 | 0.58912656 | 0.93289552 |
| SFT2D2 | 5004.04643 | -0.2515738 | 0.34648058 | -0.7260833 | 0.46778769 | 0.89984974 |
| TBX19 | 349.862692 | 0.03161661 | 0.3031414 | 0.10429658 | 0.91693399 | 0.9868044 |
| DPT | 499.240732 | -0.4124867 | 0.86022239 | -0.4795117 | 0.63157462 | 0.94088547 |
| AL031726.1 | 51.6345695 | 1.97277281 | 1.01294275 | 1.94756594 | 0.05146692 | 0.58634747 |
| ATP1B1 | 53793.7217 | 0.35688441 | 0.25307817 | 1.41017462 | 0.15848813 | 0.74675717 |
| NME7 | 6476.53728 | 0.35382058 | 0.27949676 | 1.26592019 | 0.20554166 | 0.78571765 |
| BLZF1 | 1110.11928 | -0.4174479 | 0.22452434 | -1.8592546 | 0.06299105 | 0.6147045 |
| CCDC181 | 265.256002 | 0.08623209 | 0.33734492 | 0.25561994 | 0.79824432 | 0.97170189 |
| SLC19A2 | 1584.14474 | -0.2786672 | 0.20497797 | -1.3594982 | 0.17398876 | 0.75974924 |
| F5 | 280.467473 | 0.39720527 | 0.46615583 | 0.85208691 | 0.39416586 | 0.87884163 |
| SELP | 839.965883 | 0.25842158 | 0.71901688 | 0.35940961 | 0.71928869 | 0.9566409 |
| C1orf112 | 580.252457 | 0.02216167 | 0.16718107 | 0.13256087 | 0.89454068 | 0.98474725 |
| SELL | 387.358967 | -0.4605034 | 0.74954965 | -0.6143734 | 0.53896859 | 0.9215968 |
| SELE | 1530.32804 | 3.32228793 | 1.32691642 | 2.50376579 | 1.23E-02 | 0.41344377 |
| METTL18 | 273.572341 | 0.16648073 | 0.17237891 | 0.96578367 | 0.33415244 | 0.85533716 |
| SCYL3 | 1063.11936 | -0.0035776 | 0.14140538 | -0.0253007 | 0.97981515 | 0.99537078 |
| KIFAP3 | 1909.14317 | 0.20345493 | 0.17714977 | 1.14849108 | 0.25076589 | 0.81760675 |
| AL356475.1 | 102.865021 | 0.45941042 | 0.26440433 | 1.73752983 | 0.08229369 | 0.64611064 |
| GORAB | 516.81038 | -0.1614385 | 0.26934839 | -0.5993668 | 0.5489283 | 0.92373235 |
| PRRX1 | 373.853818 | 0.30641092 | 0.75315718 | 0.40683529 | 0.68412897 | 0.95070301 |
| FMO3 | 704.050561 | 0.82447486 | 0.64319451 | 1.28184374 | 0.19989747 | 0.78134034 |
| FMO2 | 1028.86934 | 0.40302789 | 0.6363206 | 0.63337238 | 0.5264905 | 0.91769856 |
| AL021026.1 | 76.1354994 | 0.58816557 | 0.77434092 | 0.75956927 | 0.44751209 | 0.89480207 |
| FMO1 | 4868.59236 | 1.39567886 | 0.47072165 | 2.96497701 | 0.00302706 | 0.24038155 |
| FMO4 | 1515.40314 | 0.46032713 | 0.58933436 | 0.78109671 | 0.43474562 | 0.89055338 |
| PRRC2C | 5166.8907 | -0.2391895 | 0.15592628 | -1.5339907 | 0.12503193 | 0.70228923 |
| MYOCOS | 89.805446 | -1.2793349 | 0.98356878 | -1.3007071 | 1.93E-01 | 7.75E-01 |
| MYOC | 61.9751619 | -0.667607 | 1.20015558 | -0.5562671 | 5.78E-01 | 9.31E-01 |
| VAMP4 | 1206.81527 | -0.0382695 | 0.1758748 | -0.217595 | 0.82774465 | 0.97548201 |
| METTL13 | 1573.81856 | -0.133864 | 0.15176666 | -0.882038 | 0.37775627 | 0.86987124 |
| DNM3 | 396.681668 | 0.1016108 | 0.41802316 | 0.24307458 | 0.80794762 | 0.97334138 |
| DNM3OS | 303.257805 | -0.5593065 | 0.32673291 | -1.7118157 | 0.08693064 | 0.65231349 |
| PIGC | 1252.79864 | -0.2189358 | 0.13157057 | -1.6640181 | 0.09610882 | 0.66169558 |
| SUCO | 2282.16038 | -0.2380072 | 0.29786149 | -0.7990533 | 0.4242595 | 0.88565404 |
| TNFSF4 | 81.6984108 | 0.46949226 | 0.36997634 | 1.26897917 | 0.20444849 | 0.78449023 |
| AL645568.1 | 75.4071159 | -0.3902393 | 0.31238854 | -1.2492115 | 2.12E-01 | 0.79149017 |
| PRDX6 | 15180.1544 | 0.15727177 | 0.21451 | 0.73316754 | 0.46345625 | 0.89830907 |
| ANKRD45 | 58.4462049 | 0.08934897 | 0.6073213 | 0.14711976 | 0.88303749 | 0.98293314 |
| KLHL20 | 790.799708 | 0.0218573 | 0.17787486 | 0.1228802 | 0.90220196 | 0.98584146 |
| CENPL | 162.778213 | -0.2659491 | 0.35780746 | -0.7432743 | 0.45731565 | 0.89688594 |
| DARS2 | 1431.54619 | 0.14045208 | 0.2360621 | 0.59497936 | 0.55185727 | 0.92373235 |
| GAS5-AS1 | 340.412329 | 0.22423607 | 0.12847655 | 1.74534624 | 0.08092462 | 0.64313548 |
| ZBTB37 | 977.188364 | 0.0299286 | 0.26129699 | 0.11453863 | 0.90881083 | 0.98662757 |
| SERPINC1 | 146.193391 | 1.41837276 | 0.71467215 | 1.98464814 | 0.04718363 | 0.58213855 |
| RC3H1 | 1694.17789 | -0.189076 | 0.15485016 | -1.2210258 | 0.22207623 | 0.80067924 |

| | | | | | | |
|---|---|---|---|---|---|---|
| RABGAP1L | 1613.99474 | -0.194699 | 0.2430947 | -0.8009183 | 0.42317892 | 0.8846631 |
| CACYBP | 2704.24575 | 0.12260154 | 0.21658925 | 0.56605552 | 0.57135604 | 0.9282864 |
| MRPS14 | 1298.11522 | 0.11371931 | 0.11650007 | 0.97613085 | 0.32899962 | 0.85317073 |
| KIAA0040 | 2547.37329 | -0.4262242 | 0.21715037 | -1.9628067 | 0.04966863 | 0.5861488 |
| COP1 | 1616.69319 | 0.00451887 | 0.14194342 | 0.03183568 | 0.97460309 | 0.99455099 |
| PAPPA2 | 1815.668 | 2.50056356 | 0.59102529 | 4.23089095 | 2.33E-05 | 0.01615665 |
| SEC16B | 1417.05719 | 0.27256159 | 0.55076863 | 0.49487494 | 0.6206884 | 0.94002099 |
| RASAL2-AS1 | 55.3016262 | -0.8703104 | 0.46631501 | -1.8663573 | 0.06199139 | 0.61210783 |
| RASAL2 | 1487.82309 | -0.4605127 | 0.21788188 | -2.1135886 | 0.03455042 | 0.55369025 |
| C1orf220 | 41.8573596 | 0.59785413 | 0.5463418 | 1.0942859 | 0.27382961 | 0.82719197 |
| RALGPS2 | 1071.94543 | 0.07586463 | 0.12265381 | 0.61852647 | 0.53622835 | 0.92102241 |
| ANGPTL1 | 609.632903 | -0.9638332 | 0.53081016 | -1.8157776 | 0.06940449 | 0.62779744 |
| FAM20B | 2337.62664 | 0.06257761 | 0.17423522 | 0.35915594 | 0.71947844 | 0.9566409 |
| TOR3A | 2003.56671 | -0.0892718 | 0.24098063 | -0.3704524 | 0.71104545 | 0.95560193 |
| ABL2 | 1293.4156 | -0.4035011 | 0.2064717 | -1.9542683 | 0.05066949 | 0.5861488 |
| SOAT1 | 2886.92249 | 0.07576417 | 0.20283583 | 0.37352458 | 0.70875806 | 0.95501548 |
| AXDND1 | 2855.54188 | 0.26801307 | 1.09526689 | 0.24470116 | 0.80668783 | 0.97334138 |
| NPHS2 | 5164.49552 | 0.26314196 | 1.32647529 | 0.19837683 | 8.43E-01 | 0.97644404 |
| TDRD5 | 106.186688 | 0.10151159 | 0.49189864 | 0.20636689 | 0.83650433 | 0.97577999 |
| TOR1AIP2 | 3512.2482 | -0.3667451 | 0.14912045 | -2.4593882 | 0.0139174 | 0.42969693 |
| AL353708.3 | 167.843928 | 0.08428906 | 0.32341887 | 0.26061887 | 0.79438644 | 0.97101097 |
| TOR1AIP1 | 2925.64231 | 0.0272757 | 0.18015666 | 0.1513999 | 0.87966027 | 0.98275231 |
| CEP350 | 1375.05149 | -0.1354961 | 0.08177523 | -1.6569333 | 0.09753296 | 0.66549286 |
| QSOX1 | 6002.68533 | 0.42839393 | 0.23272703 | 1.84075708 | 0.06565717 | 0.62237346 |
| LHX4 | 233.225366 | 0.30476946 | 0.43319754 | 0.70353459 | 0.48172266 | 0.90541935 |
| ACBD6 | 1840.2659 | 0.02641856 | 0.11749385 | 0.22485058 | 0.82209552 | 0.97493464 |
| XPR1 | 2680.70066 | -0.1696389 | 0.22275953 | -0.7615339 | 0.44633824 | 0.89480207 |
| KIAA1614 | 833.594759 | 0.14781637 | 0.31221994 | 0.47343667 | 0.63590168 | 0.94088547 |
| AL162431.2 | 412.505853 | -0.0808912 | 0.13166633 | -0.6143653 | 0.53897396 | 0.9215968 |
| KIAA1614-AS | 263.525944 | 0.15159804 | 0.3703087 | 0.40938288 | 0.6822587 | 0.95070301 |
| STX6 | 1778.90832 | -0.0368014 | 0.14472678 | -0.2542818 | 0.79927787 | 0.97192231 |
| MR1 | 1302.23985 | 0.44958667 | 0.20690659 | 2.17289683 | 0.02978808 | 0.53938016 |
| IER5 | 748.525874 | 0.03774496 | 0.26981536 | 0.13989182 | 0.88874546 | 0.98423722 |
| CACNA1E | 421.820991 | -1.0358467 | 0.66187633 | -1.5650155 | 0.11757928 | 0.69036323 |
| GLUL | 16677.0982 | -0.2169462 | 0.19350917 | -1.121116 | 0.26223849 | 0.82424998 |
| RNASEL | 786.962329 | 0.59333483 | 0.33207147 | 1.78676848 | 0.07397492 | 0.6374448 |
| RGS16 | 805.587817 | 0.70521287 | 0.63079432 | 1.11797593 | 0.26357727 | 0.82424998 |
| NPL | 2887.69103 | 1.27388759 | 0.47978735 | 2.65510878 | 0.00792828 | 0.37071188 |
| DHX9 | 7046.87073 | -0.1812476 | 0.17235257 | -1.0516095 | 0.29297873 | 0.83524862 |
| LAMC1 | 6362.48039 | 0.09321782 | 0.19349497 | 0.48175835 | 0.62997762 | 0.94088547 |
| LAMC2 | 2175.76137 | 0.10420568 | 0.47571107 | 0.21905246 | 0.82660918 | 0.97548201 |
| NMNAT2 | 43.5617401 | 0.08502965 | 0.49296096 | 0.1724876 | 0.86305421 | 0.98176973 |
| SMG7 | 3503.09886 | -0.2696725 | 0.14482319 | -1.862081 | 0.06259167 | 0.61431334 |
| NCF2 | 388.007697 | -0.0886779 | 0.43220337 | -0.2051764 | 0.83743432 | 0.97577999 |
| ARPC5 | 5124.8129 | -0.263253 | 0.20027104 | -1.3144835 | 0.18868356 | 0.76970626 |
| RGL1 | 2356.05907 | -0.1126002 | 0.29212344 | -0.3854542 | 0.6999009 | 0.95328923 |
| COLGALT2 | 95.9014019 | -0.3683875 | 0.37673281 | -0.9778481 | 0.32814946 | 0.85317073 |
| TSEN15 | 1108.24525 | 0.09811705 | 0.18387624 | 0.53360373 | 0.59361573 | 0.93304795 |
| C1orf21 | 913.71989 | -0.1441835 | 0.30969911 | -0.4655599 | 0.64153053 | 0.94088547 |
| EDEM3 | 1811.9231 | -0.1909487 | 0.17767398 | -1.0747141 | 0.28250276 | 0.83055556 |
| FAM129A | 736.027754 | -0.4143624 | 0.5069344 | -0.8173887 | 0.41370633 | 0.88316314 |
| RNF2 | 768.581672 | -0.1336163 | 0.15166463 | -0.8809982 | 0.37831882 | 0.87045602 |
| TRMT1L | 1243.00029 | 0.20744582 | 0.21701356 | 0.95591181 | 0.33911679 | 0.85733087 |
| SWT1 | 451.103346 | -0.2241406 | 0.20824058 | -1.0763539 | 0.281769 | 0.83001101 |

| | | | | | | |
|---|---|---|---|---|---|---|
| IVNS1ABP | 24312.2338 | -0.8961636 | 0.32968212 | -2.7182655 | 0.00656251 | 0.33974611 |
| HMCN1 | 236.726669 | -0.2248948 | 0.57053052 | -0.3941854 | 0.69344413 | 0.95200619 |
| PRG4 | 575.6778 | 0.08758913 | 0.24486387 | 0.35770539 | 0.7205638 | 0.95719352 |
| TPR | 3718.15215 | -0.063168 | 0.12780142 | -0.494267 | 0.62111761 | 0.94016713 |
| ODR4 | 1716.82793 | -0.1673168 | 0.13853322 | -1.2077741 | 0.22713418 | 0.80300331 |
| PTGS2 | 734.146134 | 0.81638399 | 0.79566658 | 1.02603781 | NA | NA |
| PLA2G4A | 561.943761 | 0.75335868 | 0.33817604 | 2.22771159 | 0.02589975 | 0.52502381 |
| BRINP3 | 74.3842178 | 0.01019569 | 0.56920949 | 0.01791201 | 0.98570905 | 0.99705651 |
| RGS18 | 72.9148753 | -0.3231937 | 0.72351478 | -0.4466995 | 0.65509205 | 0.94532497 |
| RGS1 | 1583.82096 | -0.775341 | 0.87071157 | -0.8904682 | 0.37321451 | 0.86791605 |
| AL136454.1 | 50.5588898 | 0.10404653 | 0.39266859 | 0.26497288 | 0.79103036 | 0.97090793 |
| RGS2 | 2162.44153 | -0.4909551 | 0.49538417 | -0.9910592 | 0.32165666 | 0.85119105 |
| UCHL5 | 1505.71197 | 0.11599866 | 0.17645784 | 0.65737322 | 0.51094097 | 0.91474443 |
| TROVE2 | 2069.36124 | -0.0598157 | 0.10846379 | -0.5514805 | 0.58130436 | 0.9317705 |
| GLRX2 | 1613.13945 | -0.0967024 | 0.16115517 | -0.6000575 | 0.54846789 | 0.92373235 |
| CDC73 | 1510.74304 | -0.1838087 | 0.14660622 | -1.2537576 | 0.20993014 | 0.79134644 |
| B3GALT2 | 59.3154819 | -0.1572727 | 0.7884266 | -0.1994766 | 0.84188992 | 0.9762188 |
| KCNT2 | 175.375175 | 0.01666589 | 0.46117155 | 0.03613815 | 0.97117221 | 0.99400331 |
| CFH | 3022.15316 | 0.13662839 | 0.40981424 | 0.33339103 | 0.73883913 | 0.9602325 |
| CFHR1 | 96.6156812 | 0.51214197 | 0.84986909 | 0.60261278 | 0.54676632 | 0.92345821 |
| ZBTB41 | 853.854276 | 0.16448225 | 0.11441496 | 1.43759393 | 0.1505493 | 0.73796431 |
| DENND1B | 763.580397 | 0.07794422 | 0.18273793 | 0.42653553 | 0.66971767 | 0.94898648 |
| C1orf53 | 92.2076714 | 0.54886404 | 0.39094951 | 1.40392561 | 0.160341 | 0.74781837 |
| NEK7 | 2270.90616 | -0.0181678 | 0.11796328 | -0.1540125 | 0.87759988 | 0.98275231 |
| ATP6V1G3 | 264.11849 | 2.19130082 | 0.50730994 | 4.31945176 | 1.56E-05 | 0.01316658 |
| PTPRC | 1126.167 | 0.01349496 | 0.64489788 | 0.02092572 | 0.98330491 | 0.9961464 |
| NR5A2 | 92.2389185 | -0.7432782 | 0.5286442 | -1.4060084 | 0.15972162 | 0.74768867 |
| ZNF281 | 685.361862 | -0.4127955 | 0.30770499 | -1.3415301 | 0.1797484 | 0.76283169 |
| DDX59 | 892.352706 | -0.1010305 | 0.2061948 | -0.489976 | 0.62415088 | 0.94073751 |
| CAMSAP2 | 1523.96514 | -0.216441 | 0.18016211 | -1.201368 | 0.22960849 | 0.80551898 |
| INAVA | 2064.16612 | -0.170866 | 0.43021601 | -0.3971632 | 0.69124712 | 0.95158502 |
| KIF21B | 253.864608 | -0.5970917 | 0.39579992 | -1.5085695 | 0.13140882 | 0.71081672 |
| TMEM9 | 5124.40774 | 0.34047357 | 0.1990034 | 1.71089318 | 0.08710083 | 0.65231349 |
| IGFN1 | 227.150884 | -0.6197278 | 0.71445475 | -0.8674137 | 0.38571538 | 0.87420019 |
| PKP1 | 57.8128873 | -0.0537379 | 0.69206476 | -0.0776487 | 0.93810753 | 0.99113899 |
| TNNT2 | 886.187638 | 0.74448699 | 0.6360039 | 1.17056985 | 0.24177172 | 0.81269742 |
| LAD1 | 4371.77334 | -0.0013632 | 0.22762508 | -0.0059888 | 0.99522162 | 0.99940889 |
| AC119427.1 | 96.1953956 | 0.14517882 | 0.44818316 | 0.32392744 | 0.74599297 | 0.96179514 |
| TNNI1 | 549.578593 | 0.47027669 | 0.51810059 | 0.9076938 | 0.36404002 | 0.8663515 |
| PHLDA3 | 989.5258 | 0.77096286 | 0.33641795 | 2.29168171 | 0.02192402 | 0.51617839 |
| CSRP1 | 11250.1388 | -0.2821103 | 0.40059873 | -0.7042218 | 0.48129467 | 0.90541935 |
| AC096677.1 | 202.172302 | 0.05061783 | 0.41639218 | 0.12156288 | 0.90324521 | 0.98617053 |
| NAV1 | 1105.46152 | -0.1533239 | 0.24322482 | -0.6303794 | 0.52844636 | 0.91888066 |
| IPO9-AS1 | 219.585239 | -0.2661653 | 0.23268938 | -1.1438652 | 0.25267954 | 0.81835388 |
| IPO9 | 2706.09016 | -0.140815 | 0.12019618 | -1.1715429 | 0.24138063 | 0.8124487 |
| SHISA4 | 677.359229 | -0.0190496 | 0.30584473 | -0.0622851 | 0.95033577 | 0.99170503 |
| AL513217.1 | 434.890617 | -0.7075503 | 0.53637633 | -1.3191304 | 0.18712551 | 0.76958556 |
| LMOD1 | 1588.2964 | -0.7029986 | 0.55077396 | -1.2763831 | 0.20182014 | 0.78134034 |
| TIMM17A | 3017.80168 | -0.0995125 | 0.23156129 | -0.4297457 | 0.66738066 | 0.94830184 |
| RNPEP | 4181.87208 | 0.13441615 | 0.14906308 | 0.90174002 | 0.36719499 | 0.86683872 |
| AC099676.1 | 657.31449 | 0.12720701 | 0.21161634 | 0.60112094 | 0.54775944 | 0.92366021 |
| ELF3-AS1 | 133.21138 | -0.8429286 | 0.40039954 | -2.1052188 | 3.53E-02 | 5.54E-01 |
| ELF3 | 16162.796 | 0.1000003 | 0.3989795 | 0.2506402 | 0.8020923 | 0.97233627 |
| GPR37L1 | 213.632585 | -0.29426 | 0.19060684 | -1.5438059 | 0.12263536 | 0.69788192 |

| | | | | | | |
|---|---|---|---|---|---|---|
| ARL8A | 3298.47845 | -0.2877141 | 0.1475604 | -1.9498058 | 0.05119927 | 0.5861488 |
| PTPN7 | 340.664508 | 0.09397488 | 0.53383325 | 0.17603789 | 0.86026418 | 0.98078908 |
| LGR6 | 123.344321 | 0.00520347 | 0.55535806 | 0.00936958 | 0.99252427 | 0.99906045 |
| UBE2T | 129.25361 | 0.31066318 | 0.27389369 | 1.13424731 | 0.25669082 | 0.82101614 |
| PPP1R12B | 2492.55727 | -0.4905364 | 0.42358711 | -1.1580532 | 0.24684235 | 0.81542272 |
| SYT2 | 102.597517 | -0.3386841 | 0.36045894 | -0.9395914 | 3.47E-01 | 0.86197176 |
| KDM5B | 2172.48417 | -0.2450483 | 0.19948094 | -1.2284295 | 2.19E-01 | 0.79847731 |
| PCAT6 | 163.286468 | -0.9454985 | 0.4503347 | -2.0995462 | 0.03576878 | 0.55369025 |
| RABIF | 705.429402 | -0.5861074 | 0.31359773 | -1.8689784 | 0.06162582 | 0.61093984 |
| KLHL12 | 2155.29683 | 0.18677953 | 0.2268765 | 0.82326522 | 0.41035719 | 0.88249519 |
| ADIPOR1 | 5519.81134 | 0.03811225 | 0.14436753 | 0.26399459 | 0.79178408 | 0.97090793 |
| CYB5R1 | 3890.02938 | 0.17025151 | 0.17995147 | 0.94609679 | 0.34409922 | 0.85961157 |
| TMEM183A | 1980.61774 | -0.0728829 | 0.17629423 | -0.4134161 | 0.67930175 | 0.95070301 |
| PPFIA4 | 671.731616 | -0.4216105 | 0.45117515 | -0.9344718 | 0.35006055 | 0.86364633 |
| ADORA1 | 508.701457 | 0.73057146 | 0.36387503 | 2.00775376 | 0.04466946 | 0.57267043 |
| MYBPH | 64.513981 | 1.20503816 | 0.8006468 | 1.50508085 | 0.13230329 | 0.71284199 |
| CHI3L1 | 3826.1809 | 0.7387148 | 0.70486224 | 1.0480272 | 0.29462608 | 0.83630273 |
| BTG2 | 8163.91235 | 0.26838155 | 0.4926264 | 0.54479734 | 0.58589291 | 0.93289552 |
| FMOD | 1615.10022 | -0.2225721 | 0.48986741 | -0.4543518 | 0.64957566 | 0.94312988 |
| PRELP | 2496.34384 | -0.3614258 | 0.5988867 | -0.6034962 | 0.54617867 | 0.92345821 |
| ATP2B4 | 2598.3436 | 0.15444963 | 0.27896416 | 0.55365401 | 5.80E-01 | 9.31E-01 |
| ZC3H11A | 7519.52865 | -0.3361098 | 0.17916771 | -1.8759505 | 0.06066207 | 0.60958066 |
| ZBED6 | 7516.25141 | -0.3353113 | 0.17919426 | -1.8712169 | 0.06131503 | 0.61013576 |
| SNRPE | 1735.38668 | -0.0409714 | 0.15655636 | -0.2617041 | 0.79354956 | 0.97090793 |
| SOX13 | 4095.47396 | 0.14104712 | 0.30601612 | 0.46091403 | 0.64486029 | 0.94208859 |
| AL592146.1 | 2581.29604 | 0.98095694 | 0.5763397 | 1.70204645 | 0.08874666 | 0.65363785 |
| ETNK2 | 6263.55682 | 1.03594418 | 0.58146455 | 1.78161194 | 0.07481254 | 0.63773415 |
| ERLNC1 | 591.151604 | 1.17545889 | 0.61894673 | 1.89912771 | 0.05754769 | 0.60129769 |
| REN | 4412.21687 | 1.54683279 | 1.0177106 | 1.5199142 | 0.12853254 | 0.70610612 |
| GOLT1A | 601.214149 | -0.0974536 | 0.35098951 | -0.2776538 | 0.7812781 | 0.96902269 |
| PLEKHA6 | 5917.295 | -0.2192707 | 0.23063935 | -0.9507082 | 0.34175252 | 0.85889221 |
| PPP1R15B | 3103.04955 | -0.2433283 | 0.22371212 | -1.0876851 | 0.27673414 | 0.82755832 |
| PIK3C2B | 1460.35931 | -0.2473016 | 0.28039848 | -0.8819647 | 0.37779587 | 0.86987124 |
| MDM4 | 4070.88232 | -0.2248347 | 0.1706627 | -1.3174215 | 0.18769737 | 0.76970626 |
| LRRN2 | 1755.32853 | 0.14215572 | 0.58330806 | 0.24370608 | 0.80745846 | 0.97334138 |
| AL512306.2 | 167.05895 | -1.2087129 | 0.51585276 | -2.3431355 | 0.01912244 | 0.48680513 |
| NFASC | 5045.4044 | 0.3890962 | 0.37709046 | 1.03183784 | 0.30214809 | 0.84211658 |
| TMEM81 | 285.291974 | 0.02242303 | 0.39165393 | 0.05725216 | 0.95434433 | 0.99170503 |
| RBBP5 | 1026.66955 | -0.022132 | 0.19602837 | -0.112902 | 0.91010822 | 0.98662757 |
| DSTYK | 1201.35199 | 0.11000586 | 0.13717141 | 0.80195911 | 0.42257661 | 0.88451372 |
| TMCC2 | 251.40581 | -0.730488 | 0.35828644 | -2.038838 | 0.0414662 | 0.55945849 |
| NUAK2 | 3128.00064 | -0.8456952 | 0.30006634 | -2.8183608 | 0.00482695 | 0.30153976 |
| LEMD1 | 99.7210901 | 1.16855159 | 0.4273526 | 2.73439682 | 0.00624947 | 0.33470127 |
| BLACAT1 | 205.948476 | 0.35921121 | 0.42242466 | 0.85035568 | 0.39512737 | 0.87915019 |
| CDK18 | 6222.88369 | -0.2173391 | 0.3750844 | -0.5794406 | 0.5622919 | 0.92452095 |
| MFSD4A | 4676.4466 | 0.96545818 | 0.6055279 | 1.59440742 | 0.11084481 | 0.68001961 |
| ELK4 | 3828.76898 | -0.3188966 | 0.21856411 | -1.4590527 | 0.1445506 | 0.73032824 |
| SLC45A3 | 916.182764 | -0.0011159 | 0.31875976 | -0.0035007 | 9.97E-01 | 1.00E+00 |
| NUCKS1 | 8625.13042 | 0.4068691 | 0.16036318 | 2.53717279 | 0.01117518 | 0.40196898 |
| RAB29 | 5092.16338 | 0.18209483 | 0.41085246 | 0.44321223 | 0.65761223 | 0.94558535 |
| SLC41A1 | 3306.75414 | 0.16347656 | 0.24323388 | 0.67209616 | 0.50152248 | 0.91259827 |
| PM20D1 | 339.520472 | 0.84573577 | 1.02948606 | 0.8215126 | 0.41135435 | 0.88268208 |
| SLC26A9 | 828.56857 | 0.45351674 | 0.42788039 | 1.05991476 | 0.28918338 | 0.83312616 |
| RAB7B | 82.5009333 | 0.33272903 | 0.72077169 | 0.46162888 | 0.64434748 | 0.94208859 |

| | | | | | | |
|---|---|---|---|---|---|---|
| RHEX | 1348.74254 | 0.10537557 | 0.46467639 | 0.22677196 | 0.82060107 | 0.97472105 |
| SRGAP2 | 1706.52386 | -0.0644695 | 0.21682577 | -0.297333 | 0.7662123 | 0.96704052 |
| IKBKE | 456.574118 | -0.1181817 | 0.45708667 | -0.2585543 | 0.79597915 | 0.97101097 |
| C1orf147 | 58.454168 | 0.13273315 | 0.50711666 | 0.26174086 | 7.94E-01 | 0.97090793 |
| RASSF5 | 608.261279 | 0.06214356 | 0.49906785 | 0.12451926 | 0.90090415 | 0.98545758 |
| EIF2D | 2547.33568 | 0.38048098 | 0.16478775 | 2.30891544 | 0.02094827 | 0.50728403 |
| DYRK3 | 302.442327 | -0.5121401 | 0.26977175 | -1.8984201 | 0.05764076 | 0.60129769 |
| MAPKAPK2 | 10657.2952 | -0.5702654 | 0.25146503 | -2.2677723 | 0.02334309 | 0.51967919 |
| IL24 | 84.9090288 | 0.45551237 | 0.77521646 | 0.58759378 | 0.55680499 | 0.92373235 |
| FCMR | 277.739875 | -0.4584355 | 0.74960857 | -0.6115665 | 0.5408246 | 0.92275982 |
| PIGR | 23787.8768 | 1.17174098 | 0.41492048 | 2.8240134 | 0.00474264 | 0.29927103 |
| FCAMR | 1873.49134 | 1.610292 | 0.3710859 | 4.33940503 | 1.43E-05 | 0.01316658 |
| C1orf116 | 924.508072 | 0.08572992 | 0.3321074 | 0.25813914 | 0.79629952 | 0.97101097 |
| PFKFB2 | 4937.164 | 0.57721989 | 0.55437882 | 1.04120119 | 0.29778218 | 0.83946087 |
| YOD1 | 1243.16985 | -0.5125986 | 0.47202141 | -1.0859648 | 0.27749458 | 0.82797737 |
| C4BPA | 111.571021 | -1.3065008 | 0.99704121 | -1.3103779 | 0.19006802 | 0.77065452 |
| CD55 | 2138.91892 | -0.0395822 | 0.24841524 | -0.159339 | 0.87340178 | 0.98275231 |
| CR2 | 283.476171 | 1.97786549 | 1.02737792 | 1.92515865 | 0.05420951 | 0.59329767 |
| CR1 | 356.73433 | 0.51877943 | 0.50143681 | 1.03458586 | 0.30086236 | 0.84206414 |
| CR1L | 55.5877016 | -1.0887492 | 0.79127692 | -1.3759395 | 0.16884037 | 0.75786012 |
| CD46 | 14632.0047 | 0.28578262 | 0.12010821 | 2.37937629 | 0.01734196 | 0.46859699 |
| MIR29B2CH( | 991.682088 | 0.09806313 | 0.37550986 | 0.26114663 | 0.79397943 | 0.97090793 |
| CD34 | 4317.96985 | -0.0823561 | 0.43152434 | -0.1908493 | 0.84864363 | 0.97741457 |
| AL356275.1 | 100.161301 | -0.6329584 | 0.45877543 | -1.3796694 | 0.16768847 | 0.7558879 |
| PLXNA2 | 1239.70714 | -0.4493051 | 0.24438489 | -1.8385143 | 0.06598666 | 0.62237346 |
| CAMK1G | 289.15371 | 1.1264627 | 0.5832218 | 1.93144821 | 0.05342765 | 0.59038182 |
| LAMB3 | 3246.64529 | -1.0872217 | 0.72127542 | -1.50736 | 0.1317184 | 0.71139325 |
| AL031316.1 | 63.6812904 | 1.19766391 | 0.71981393 | 1.66385209 | 0.09614199 | 0.66169558 |
| G0S2 | 6402.91722 | 0.62531143 | 0.52911044 | 1.18181646 | 0.23727853 | 0.80880993 |
| HSD11B1 | 242.166845 | -0.0762534 | 0.70632721 | -0.1079577 | 0.91402927 | 0.98662757 |
| TRAF3IP3 | 475.875022 | 0.44160956 | 0.54320953 | 0.81296357 | 0.41623894 | 0.88399703 |
| C1orf74 | 407.284767 | 0.47533956 | 0.38197467 | 1.24442691 | 0.21334248 | 0.7926732 |
| IRF6 | 2326.95573 | 0.12688549 | 0.17756596 | 0.71458229 | 0.47486719 | 0.90226631 |
| UTP25 | 1011.71475 | -0.4721684 | 0.31825022 | -1.4836389 | 0.13790474 | 0.72209298 |
| SERTAD4-AS | 132.420991 | 1.01013364 | 0.52716903 | 1.91614754 | 0.05534632 | 0.59617546 |
| SERTAD4 | 178.199717 | 0.65244226 | 0.63893435 | 1.02114131 | 0.30718749 | 0.84298407 |
| HHAT | 478.625209 | 0.73668164 | 0.37589697 | 1.95979672 | 0.05001956 | 0.5861488 |
| KCNH1 | 37.4027454 | 0.68665256 | 0.58468618 | 1.17439506 | 0.24023682 | 0.8124487 |
| RCOR3 | 2649.58122 | -0.0472191 | 0.1625542 | -0.2904822 | 0.77144739 | 0.96836854 |
| TRAF5 | 1084.71007 | 0.03601982 | 0.29644865 | 0.12150443 | 0.90329151 | 0.98617053 |
| LINC00467 | 456.780206 | 0.56553233 | 0.3353497 | 1.68639582 | 0.09171959 | 0.65518742 |
| SLC30A1 | 3500.89516 | -0.5060233 | 0.44416556 | -1.1392674 | 0.25459165 | 0.81992039 |
| LPGAT1 | 4432.32391 | 0.01774687 | 0.38785351 | 0.04575663 | 0.96350423 | 0.99320711 |
| INTS7 | 593.523761 | -0.3485326 | 0.2138567 | -1.6297483 | 0.1031547 | 0.67430143 |
| PPP2R5A | 3905.44756 | -0.0075382 | 0.13360644 | -0.0564212 | 9.55E-01 | 0.99185043 |
| TMEM206 | 902.420215 | 0.4288771 | 0.3924991 | 1.092683 | 2.75E-01 | 0.82719407 |
| NENF | 2383.91437 | 0.0946778 | 0.08594918 | 1.10155564 | 2.71E-01 | 8.27E-01 |
| AC092803.2 | 129.655302 | -0.1078758 | 0.39163848 | -0.2754475 | 0.78297249 | 0.96964828 |
| ATF3 | 3778.0659 | -0.717225 | 0.60016007 | -1.1950561 | 0.23206509 | 0.80649152 |
| BATF3 | 207.748967 | -0.1232611 | 0.55237149 | -0.2231489 | 0.82341961 | 0.97545556 |
| NSL1 | 1022.61463 | 0.11201352 | 0.13960682 | 0.8023499 | 4.22E-01 | 0.88451372 |
| TATDN3 | 1006.72145 | 0.24426308 | 0.20246595 | 1.2064403 | 0.22764777 | 0.80300331 |
| FLVCR1-DT | 182.274405 | 0.69503816 | 0.61132492 | 1.13693739 | 0.25556446 | 0.82052927 |
| FLVCR1 | 663.517045 | 0.12540789 | 0.17241436 | 0.7273634 | 0.46700337 | 0.89983399 |

| | | | | | | |
|---|---|---|---|---|---|---|
| VASH2 | 60.5415347 | 0.32846548 | 0.32493577 | 1.01086279 | 0.31208211 | 0.84541735 |
| ANGEL2 | 1290.56895 | -0.0532556 | 0.1551096 | -0.3433416 | 0.73134151 | 0.95848028 |
| RPS6KC1 | 891.946056 | -0.3478585 | 0.22187226 | -1.5678321 | 0.11692031 | 0.68956727 |
| PROX1 | 495.326104 | -0.2127167 | 0.70006701 | -0.3038519 | 0.76124074 | 0.96603553 |
| SMYD2 | 1764.74397 | 0.11622135 | 0.10091526 | 1.15167263 | 0.24945562 | 0.81646572 |
| PTPN14 | 2844.19193 | 0.01039335 | 0.22673479 | 0.04583923 | 0.96343839 | 0.99320711 |
| CENPF | 74.5386051 | 0.7836326 | 0.81022401 | 0.96718017 | 0.33345397 | 0.855011 |
| KCNK2 | 50.6425331 | 3.71211093 | 1.18966577 | 3.12029734 | 0.00180669 | 0.18722055 |
| KCTD3 | 3124.22839 | 0.07484478 | 0.15730325 | 0.47579932 | 0.63421735 | 0.94088547 |
| ESRRG | 1530.59686 | 0.15544178 | 0.2227565 | 0.69781031 | 0.48529583 | 0.90543654 |
| GPATCH2 | 760.102268 | -0.1458421 | 0.19714672 | -0.7397644 | 0.45944298 | 0.89755243 |
| SPATA17 | 246.097941 | 0.23814364 | 0.45018466 | 0.52899102 | 0.59681168 | 0.9337017 |
| SPATA17-AS | 56.4639475 | 0.09292175 | 0.54326279 | 0.17104382 | 0.86418931 | 0.98176973 |
| RRP15 | 715.471091 | -0.2774694 | 0.25583973 | -1.084544 | 0.27812366 | 0.82805357 |
| TGFB2 | 243.92081 | 0.1707287 | 0.482567 | 0.35379275 | 0.72349421 | 0.95810643 |
| LYPLAL1-DT | 193.869988 | 0.57808628 | 0.3510856 | 1.64656789 | 0.09964689 | 0.67098264 |
| LYPLAL1 | 2407.71534 | 0.16147341 | 0.27662209 | 0.58373287 | 0.55940005 | 0.92421175 |
| AC096642.1 | 81.7637605 | -0.3043994 | 0.36708153 | -0.8292419 | 0.40696755 | 0.88132015 |
| AL513283.1 | 64.8932586 | 1.83700287 | 0.66909753 | 2.74549344 | 0.006042 | 0.33171839 |
| EPRS | 3476.86227 | 0.03912418 | 0.15972195 | 0.24495178 | 0.80649377 | 0.97334138 |
| BPNT1 | 2150.24943 | 0.13828705 | 0.25732544 | 0.53740139 | 0.59099039 | 0.93289552 |
| IARS2 | 5121.24002 | 0.279284 | 0.20170349 | 1.38462654 | 0.16616669 | 0.75515387 |
| RAB3GAP2 | 1833.79049 | -0.155152 | 0.11799419 | -1.3149124 | 0.18853935 | 0.76970626 |
| MARK1 | 397.632262 | -0.4728025 | 0.28590233 | -1.6537202 | 0.09818438 | 0.66549286 |
| C1orf115 | 4593.79639 | 0.15773642 | 0.26092466 | 0.60452863 | 0.54549224 | 0.92345821 |
| 2-Mar | 4881.6187 | 0.21575243 | 0.31340799 | 0.68840756 | 0.49119616 | 0.90600166 |
| 1-Mar | 615.513459 | 0.19497187 | 0.28999643 | 0.67232508 | 0.50137677 | 0.91259827 |
| HLX | 434.425262 | -0.0750272 | 0.34642797 | -0.2165737 | 0.82854056 | 0.97550416 |
| DUSP10 | 132.0803 | -0.1061771 | 0.37848218 | -0.280534 | 0.77906781 | 0.96836854 |
| TAF1A | 138.081414 | 0.08912993 | 0.25469106 | 0.34995313 | 0.72637387 | 0.95848028 |
| TAF1A-AS1 | 85.6827152 | 0.99109536 | 0.50637173 | 1.95724862 | 0.05031825 | 0.5861488 |
| MIA3 | 2865.11251 | 0.06762703 | 0.16407554 | 0.4121701 | 0.68021475 | 0.95070301 |
| AL592148.3 | 588.291524 | 0.04087724 | 0.18570303 | 0.22012155 | 0.82577649 | 0.97548201 |
| AIDA | 1626.02064 | 0.00700696 | 0.16741447 | 0.04185397 | 0.96661511 | 0.99400331 |
| BROX | 2460.82043 | -0.0189195 | 0.14936576 | -0.1266656 | 0.89920503 | 0.98497876 |
| FAM177B | 102.113299 | -0.359519 | 0.56586255 | -0.6353469 | 0.52520222 | 0.91759401 |
| AL392172.1 | 533.368434 | -0.978744 | 0.67383204 | -1.4525044 | 0.14636142 | 0.73247558 |
| AC093152.1 | 178.867818 | -0.5162094 | 0.53191179 | -0.9704794 | 0.33180759 | 0.85452131 |
| DISP1 | 1146.59094 | -0.1424594 | 0.30998139 | -0.4595742 | 0.64582188 | 0.94208859 |
| TLR5 | 583.176958 | -0.3649454 | 0.37698054 | -0.9680749 | 0.33300696 | 0.855011 |
| SUSD4 | 1111.15702 | 0.14843071 | 0.42649214 | 0.34802683 | 0.72782003 | 0.95848028 |
| CAPN8 | 226.649777 | 1.69992596 | 0.79093584 | 2.14925898 | 0.03161388 | 0.54507414 |
| CAPN2 | 15465.3467 | 0.01808019 | 0.26247176 | 0.06888431 | 0.94508171 | 0.99164528 |
| TP53BP2 | 1980.21169 | -0.4733161 | 0.29172434 | -1.6224772 | 0.10470122 | 0.67439401 |
| FBXO28 | 1888.88679 | -0.1112964 | 0.24611904 | -0.4522056 | 0.65112086 | 0.94389357 |
| DEGS1 | 4309.36751 | -0.5152686 | 0.22964019 | -2.2438083 | 0.02484474 | 0.52085426 |
| AC092809.4 | 175.362156 | -0.1086852 | 0.24631178 | -0.4412503 | 0.65903177 | 0.94599528 |
| NVL | 805.590472 | -0.0018106 | 0.13043698 | -0.0138807 | 0.98892516 | 0.99792431 |
| CNIH4 | 2144.1574 | -0.3418946 | 0.19487456 | -1.7544344 | 0.0793561 | 0.64079926 |
| WDR26 | 6063.72072 | -0.2130472 | 0.17711534 | -1.2028728 | 0.22902556 | 0.80513377 |
| CNIH3 | 193.793468 | 0.84210537 | 0.32192099 | 2.61587596 | 0.00889989 | 0.38245454 |
| DNAH14 | 354.224664 | -0.1065864 | 0.22252891 | -0.4789776 | 6.32E-01 | 0.94088547 |
| LBR | 3128.17405 | -0.4236379 | 0.24129561 | -1.75568 | 0.07914307 | 0.64079926 |
| ENAH | 1756.76838 | -0.4604731 | 0.32071398 | -1.4357749 | 0.1510664 | 0.73796431 |

| | | | | | | |
|---|---|---|---|---|---|---|
| SRP9 | 7995.1628 | 0.16391498 | 0.14246904 | 1.15053048 | 0.24992545 | 0.81699857 |
| EPHX1 | 13832.4634 | 0.0537209 | 0.41488367 | 0.12948425 | 0.89697449 | 0.98474725 |
| AL591895.1 | 1241.7672 | 0.12764754 | 0.37742322 | 0.33820795 | 0.73520648 | 0.95895327 |
| TMEM63A | 7267.22147 | -0.0068234 | 0.38756078 | -0.0176059 | 0.98595326 | 0.99705792 |
| LEFTY1 | 82.4896363 | 1.04241692 | 0.59550224 | 1.75048364 | 8.00E-02 | 0.64079926 |
| AL117348.2 | 2461.91619 | 0.14777041 | 0.15690784 | 0.94176567 | 0.34631263 | 0.86135217 |
| PYCR2 | 4451.26982 | 0.12765267 | 0.14994895 | 0.85130754 | 0.39459854 | 0.87884163 |
| SDE2 | 923.68402 | -0.1795395 | 0.23771407 | -0.7552751 | 0.45008395 | 0.89557948 |
| H3F3A | 4224.87431 | 0.29892817 | 0.21436023 | 1.39451322 | 0.16316268 | 0.75131002 |
| ACBD3 | 2194.50172 | -0.1555522 | 0.21910597 | -0.7099405 | 0.47774101 | 0.90387017 |
| ACBD3-AS1 | 174.012858 | -0.2338516 | 0.40224696 | -0.5813632 | 0.56099572 | 0.92448946 |
| LIN9 | 131.96344 | 0.0214823 | 0.22602323 | 0.09504467 | 0.92427935 | 0.98869145 |
| PARP1 | 4046.40837 | -0.05066 | 0.13989457 | -0.36213 | 0.71725489 | 0.95611001 |
| STUM | 1097.93189 | -1.0072072 | 0.44434408 | -2.2667281 | 0.02340684 | 0.51967919 |
| ITPKB | 2565.59275 | -0.2477027 | 0.14745051 | -1.6799038 | 0.09297603 | 0.65753347 |
| PSEN2 | 1234.25638 | -0.1801472 | 0.24039721 | -0.749373 | 0.45363243 | 0.89670027 |
| COQ8A | 4244.40872 | -0.0847774 | 0.20679686 | -0.4099547 | 6.82E-01 | 0.95070301 |
| CDC42BPA | 3535.78629 | 0.08919446 | 0.18518542 | 0.48164947 | 0.63005497 | 0.94088547 |
| AL353689.2 | 401.347454 | 0.08741471 | 0.26607685 | 0.3285318 | 0.74250961 | 0.96109001 |
| ZNF678 | 259.052062 | -0.0067745 | 0.22596899 | -0.0299798 | 0.97608316 | 0.99455099 |
| SNAP47 | 2364.53162 | -0.1127951 | 0.12217566 | -0.9232211 | 0.35589198 | 0.86364633 |
| JMJD4 | 1168.77411 | -0.3124103 | 0.17631067 | -1.7719306 | 0.07640608 | 0.63773415 |
| WNT9A | 92.4518793 | 0.74008857 | 0.49936266 | 1.48206631 | 0.13832265 | 0.72249298 |
| ARF1 | 21702.909 | -0.1109497 | 0.16063561 | -0.690692 | 0.48975914 | 0.90600166 |
| C1orf35 | 986.007341 | 0.04186667 | 0.15154546 | 0.27626474 | 0.78234473 | 0.96957563 |
| MRPL55 | 1754.42259 | 0.13929374 | 0.13498749 | 1.03190107 | 0.30211847 | 0.84211658 |
| GUK1 | 10626.2626 | 0.07290342 | 0.14565574 | 0.50051867 | 6.17E-01 | 9.39E-01 |
| GJC2 | 70.4405706 | 0.34607347 | 0.4403947 | 0.78582569 | 0.43196961 | 0.88976671 |
| IBA57 | 695.736377 | -0.0521282 | 0.30404504 | -0.1714488 | 0.86387088 | 0.98176973 |
| OBSCN-AS1 | 223.512308 | -0.0312321 | 0.31402302 | -0.099458 | 0.92077465 | 0.98767854 |
| OBSCN | 3286.87618 | 0.08047622 | 0.21234782 | 0.378983 | 0.70470049 | 0.95501548 |
| AL353593.1 | 40.9157505 | 0.12860759 | 0.33255683 | 0.38672365 | 0.69896081 | 0.95299571 |
| TRIM11 | 1510.52353 | -0.1942719 | 0.15618873 | -1.243828 | 0.21356286 | 0.7926732 |
| AL670729.2 | 451.408323 | -0.1758482 | 0.16754664 | -1.049548 | 0.29392599 | 0.83580671 |
| AL670729.1 | 72.0692409 | -0.2488968 | 0.36936752 | -0.673846 | 0.50040923 | 0.91180773 |
| TRIM17 | 251.321492 | -0.059486 | 0.36785004 | -0.1617127 | 0.87153214 | 0.98275231 |
| HIST3H2A | 69.7937395 | 0.10895162 | 0.4364278 | 0.24964409 | 0.8028626 | 0.97273128 |
| RNF187 | 6266.94637 | -0.11377 | 0.12724256 | -0.894119 | 0.37125824 | 0.86762749 |
| RHOU | 1120.68299 | 0.59230568 | 0.30864205 | 1.91906993 | 0.05497548 | 0.59417275 |
| RAB4A | 1717.55442 | 0.27037245 | 0.17026024 | 1.58799522 | 0.11228742 | 0.68001961 |
| SPHAR | 148.600147 | 0.51122758 | 0.23504702 | 2.17500132 | 2.96E-02 | 0.53755678 |
| CCSAP | 231.423548 | 0.00846012 | 0.33151116 | 0.02551987 | 0.9796403 | 0.99530732 |
| ACTA1 | 185.323565 | 3.83476984 | 1.58081557 | 2.42581735 | NA | NA |
| NUP133 | 1743.85303 | 0.18058376 | 0.11637754 | 1.55170626 | 0.12073252 | 0.69491653 |
| ABCB10 | 1058.74426 | 0.08024577 | 0.15921091 | 0.50402179 | 0.61424606 | 0.93895494 |
| TAF5L | 929.091976 | -0.4830763 | 0.26468651 | -1.8250884 | 0.06798768 | 0.62603855 |
| URB2 | 301.77919 | -0.4880724 | 0.29362897 | -1.662208 | 0.09647108 | 0.66233356 |
| GALNT2 | 6476.60971 | -0.2926558 | 0.31753524 | -0.9216483 | 0.35671203 | 0.86390849 |
| AL136988.1 | 2562.78825 | -0.2189708 | 0.31601262 | -0.6929179 | 0.48836105 | 0.90585781 |
| PGBD5 | 326.422307 | 0.1784202 | 0.28542584 | 0.62510178 | 0.53190426 | 0.91962754 |
| COG2 | 1454.70756 | 0.0699844 | 0.14094095 | 0.49655122 | 0.61950556 | 0.93945324 |
| AGT | 11479.4159 | -0.2057533 | 0.56428657 | -0.3646256 | 0.71539093 | 0.95590747 |
| AL512328.1 | 1523.3704 | -0.025514 | 0.60694019 | -0.042037 | 0.96646918 | 0.99400331 |
| C1orf198 | 2453.94534 | -0.3509578 | 0.1972514 | -1.7792409 | 0.07520028 | 0.63773415 |

| | | | | | | |
|---|---|---|---|---|---|---|
| TTC13 | 676.776692 | -0.0549498 | 0.20006597 | -0.2746584 | 0.78357873 | 0.96978847 |
| ARV1 | 1046.2311 | 0.044167 | 0.13358953 | 0.33061724 | 0.74093362 | 0.96089108 |
| FAM89A | 690.122726 | 0.06207114 | 0.24493318 | 0.25342071 | 0.79994312 | 0.97208422 |
| C1orf131 | 522.303926 | -0.0837604 | 0.15362785 | -0.5452166 | 0.58560458 | 0.93289552 |
| GNPAT | 2440.45823 | 0.04369572 | 0.14603565 | 0.29921269 | 0.76477777 | 0.96678847 |
| EXOC8 | 694.194579 | -0.3256217 | 0.2669342 | -1.2198574 | 0.22251894 | 0.80078694 |
| SPRTN | 457.402952 | -0.4541758 | 0.34142731 | -1.3302268 | 0.18344354 | 0.76756316 |
| EGLN1 | 4837.98997 | -0.0756374 | 0.22717554 | -0.3329471 | 0.73917424 | 0.9602325 |
| AL445524.1 | 186.917559 | -0.4854774 | 0.45476054 | -1.0675451 | 0.28572575 | 0.83107779 |
| TSNAX | 2059.73007 | 0.06495831 | 0.1202179 | 0.5403381 | 0.58896389 | 0.93289552 |
| TSNAX-DISC1 | 538.325324 | 0.05026172 | 0.24851287 | 0.20224998 | 0.8397213 | 0.97618283 |
| DISC1 | 235.227523 | 0.08072848 | 0.33250721 | 0.24278716 | 0.80817028 | 0.97334138 |
| SIPA1L2 | 929.674192 | 0.09740647 | 0.37312008 | 0.2610593 | 0.79404678 | 0.97090793 |
| MAP10 | 85.8882953 | 0.36597731 | 0.39631335 | 0.92345442 | 0.35577044 | 0.86364633 |
| NTPCR | 1876.00147 | 0.38217499 | 0.15810668 | 2.41719702 | 0.01564055 | 0.45444887 |
| PCNX2 | 809.935537 | 0.01973552 | 0.23708254 | 0.08324322 | 0.93365814 | 0.99019225 |
| MAP3K21 | 1800.98801 | 0.41084366 | 0.23907371 | 1.71848114 | 0.08570889 | 0.65049132 |
| KCNK1 | 1073.36856 | 0.01304059 | 0.21113289 | 0.06176484 | 0.9507501 | 0.99170503 |
| SLC35F3 | 53.2778906 | -0.2469224 | 0.63724868 | -0.3874821 | 0.69839935 | 0.9529028 |
| AL355472.1 | 478.036829 | -0.2265376 | 0.22005343 | -1.0294664 | 0.30326055 | 0.84211658 |
| COA6 | 1055.55959 | -0.2722608 | 0.22568895 | -1.2063541 | 0.22768099 | 0.80300331 |
| TARBP1 | 1646.76008 | 0.12235918 | 0.31217654 | 0.39195507 | 0.69509141 | 0.95234774 |
| IRF2BP2 | 7537.96387 | -0.1038414 | 0.2378501 | -0.4365835 | 6.62E-01 | 0.94642611 |
| AL160408.2 | 2860.99191 | -0.2499904 | 0.21850915 | -1.1440731 | 0.25259332 | 0.81833349 |
| LINC01132 | 50.383309 | -0.1777252 | 0.53706479 | -0.3309195 | 0.74070533 | 0.96089108 |
| LINC01348 | 66.288897 | -0.5155773 | 0.44492684 | -1.1587912 | 0.24654132 | 0.81542272 |
| TOMM20 | 8162.27104 | 0.14557457 | 0.25156294 | 0.57868051 | 0.56280477 | 0.92452095 |
| RBM34 | 1640.3525 | -0.0965751 | 0.18529039 | -0.5212095 | 0.60222082 | 0.9355118 |
| ARID4B | 3306.20135 | -0.1383404 | 0.1045099 | -1.3237066 | 0.18560049 | 0.76908034 |
| GGPS1 | 1527.40658 | 0.10991292 | 0.12283683 | 0.89478795 | 0.37090045 | 0.86762749 |
| TBCE | 2717.12416 | -0.0230099 | 0.17825988 | -0.1290805 | 0.89729395 | 0.98474725 |
| B3GALNT2 | 213.075709 | -0.1036124 | 0.23584531 | -0.4393235 | 0.66042718 | 0.94628868 |
| GNG4 | 127.827187 | 0.2460281 | 0.56053713 | 0.4389149 | 0.6607232 | 0.94628868 |
| LYST | 967.004207 | -0.1266153 | 0.14168423 | -0.8936441 | 0.37151235 | 0.86762749 |
| NID1 | 5333.3669 | -0.1313505 | 0.30701806 | -0.4278266 | 0.66877738 | 0.9486589 |
| GPR137B | 3499.50971 | -0.2002645 | 0.54751466 | -0.3657701 | 0.71453662 | 0.95579447 |
| ERO1B | 1278.00442 | 0.22076735 | 0.18624866 | 1.18533657 | 0.2358844 | 0.80776058 |
| EDARADD | 96.3477036 | 0.3178669 | 0.26615007 | 1.19431452 | 0.23235496 | 0.80649152 |
| LGALS8 | 4870.453 | -0.338589 | 0.19465002 | -1.7394759 | 0.08195108 | 0.64555743 |
| LGALS8-AS1 | 105.486529 | -0.3913719 | 0.37147669 | -1.0535571 | 0.29208573 | 0.83495709 |
| AL359921.2 | 82.585225 | -0.807951 | 0.35486245 | -2.2768006 | 0.02279814 | 0.51967919 |
| AL359921.1 | 273.261568 | -0.3182483 | 0.19922586 | -1.5974247 | 0.11017108 | 0.67996643 |
| HEATR1 | 2728.0858 | -0.2721697 | 0.16859559 | -1.6143348 | 1.06E-01 | 0.67674128 |
| ACTN2 | 106.334275 | 0.02265137 | 0.56452679 | 0.04012453 | 0.96799385 | 0.99400331 |
| MTR | 4256.54951 | 0.12682216 | 0.26245143 | 0.48322146 | 0.6289385 | 0.94088547 |
| LINC01139 | 658.256243 | -0.0729863 | 0.57823896 | -0.1262217 | 0.89955645 | 0.98506781 |
| CHRM3 | 165.281646 | 0.21352051 | 0.44806385 | 0.47654035 | 6.34E-01 | 0.94088547 |
| CHRM3-AS2 | 130.064959 | 1.6288375 | 0.5820662 | 2.79837157 | 0.0051361 | 0.30843846 |
| FMN2 | 193.488254 | 0.45062342 | 0.73613738 | 0.61214582 | 0.54044129 | 0.92247611 |
| GREM2 | 60.2894473 | 1.70552398 | 0.81085962 | 2.10335295 | 0.03543493 | 0.55369025 |
| RGS7 | 46.3886599 | 0.85456413 | 0.85558785 | 0.99880349 | 3.18E-01 | 8.49E-01 |
| FH | 4563.33121 | 0.57052002 | 0.18886596 | 3.02076679 | 0.00252135 | 0.21493583 |
| KMO | 2293.30999 | 0.13456762 | 0.37494403 | 0.35890056 | 0.71966948 | 0.9566409 |
| OPN3 | 1776.77418 | -0.0484055 | 0.26881442 | -0.1800704 | 0.8570973 | 0.97953896 |

| | | | | | | |
|---|---|---|---|---|---|---|
| CHML | 622.480584 | -0.4511497 | 0.29625842 | -1.5228248 | 0.12780255 | 0.70579329 |
| CEP170 | 802.768237 | -0.3756618 | 0.18925358 | -1.9849654 | 0.04714833 | 0.58208614 |
| AL606534.1 | 44.4126166 | -0.7907243 | 0.39038103 | -2.0255192 | 0.04281408 | 0.56625172 |
| SDCCAG8 | 1387.83749 | 0.06282721 | 0.14672498 | 0.42819708 | 0.66850764 | 0.9486589 |
| AKT3 | 2360.1555 | -0.070708 | 0.24789632 | -0.2852323 | 0.77546615 | 0.96836854 |
| ZBTB18 | 1652.00383 | 0.14451186 | 0.16563115 | 0.87249202 | 0.38294002 | 0.8718674 |
| AL645465.1 | 723.820583 | -0.0121535 | 0.18776705 | -0.0647266 | 0.94839169 | 0.99170503 |
| ADSS | 2888.04357 | -0.0223057 | 0.17639244 | -0.1264547 | 0.89937198 | 0.98504648 |
| CATSPERE | 152.186255 | -0.1447655 | 0.32387977 | -0.446973 | 0.65489458 | 0.94532497 |
| DESI2 | 1609.07369 | -0.1513742 | 0.19957771 | -0.7584723 | 0.44816828 | 0.8948759 |
| COX20 | 6335.98024 | -0.0150618 | 0.13457173 | -0.1119243 | 0.91088342 | 0.98662757 |
| HNRNPU | 22595.7808 | -0.1922724 | 0.11862417 | -1.6208532 | 1.05E-01 | 0.67473222 |
| AL356512.1 | 139.714823 | 0.30581128 | 0.29194669 | 1.04749016 | 0.29487357 | 0.83630273 |
| EFCAB2 | 733.533588 | -0.0555147 | 0.27090802 | -0.2049208 | 0.83763398 | 0.97577999 |
| KIF26B | 351.031784 | 0.75844088 | 0.4219784 | 1.79734526 | 0.07228083 | 0.6321115 |
| SMYD3 | 526.828517 | 0.42181248 | 0.22986984 | 1.83500574 | 0.06650484 | 0.62378208 |
| TFB2M | 980.41828 | -0.0147016 | 0.2964992 | -0.0495838 | 0.96045404 | 0.99281834 |
| CNST | 1129.7653 | -0.3479353 | 0.12479681 | -2.7880146 | 0.00530322 | 0.31308771 |
| SCCPDH | 3758.58115 | 0.56103809 | 0.25962712 | 2.16093794 | 0.03070013 | 0.54443002 |
| AHCTF1 | 1880.82827 | -0.2534169 | 0.21931022 | -1.1555179 | 2.48E-01 | 0.81602355 |
| ZNF670 | 148.540836 | -0.180193 | 0.32175432 | -0.560033 | 0.57545695 | 0.93076883 |
| ZNF669 | 176.232561 | -0.2209459 | 0.37333655 | -0.5918142 | 5.54E-01 | 0.92373235 |
| C1orf229 | 45.1774471 | 0.20069288 | 0.37271275 | 0.5384653 | 0.59025586 | 0.93289552 |
| ZNF124 | 290.982978 | -0.0890719 | 0.19933672 | -0.4468414 | 0.6549896 | 0.94532497 |
| AL390728.6 | 702.320093 | -0.2153013 | 0.22641209 | -0.9509265 | 0.34164168 | 0.85889221 |
| ZNF496 | 1711.66447 | -0.0994649 | 0.17631227 | -0.5641406 | 0.57265843 | 0.92928554 |
| NLRP3 | 99.5633243 | -0.0416089 | 0.53485345 | -0.077795 | 0.93799113 | 0.99113899 |
| AC138089.1 | 50.7104 | 1.30757778 | 0.89086951 | 1.46775455 | 1.42E-01 | 7.28E-01 |
| AC098483.1 | 45.2793923 | 1.86402183 | 1.11029078 | 1.67885914 | 0.0931795 | 0.65756964 |
| OR2T10 | 119.393808 | 0.1273891 | 0.66490922 | 0.19158871 | 0.84806439 | 0.97711043 |
| LYPD8 | 118.798137 | -0.0366204 | 0.5817902 | -0.0629443 | 0.94981089 | 0.99170503 |
| SH3BP5L | 2631.91245 | -0.4055918 | 0.18181102 | -2.2308429 | 0.02569154 | 0.52502381 |
| ZNF672 | 2226.78744 | -0.1498629 | 0.1251708 | -1.1972672 | 0.23120242 | 0.80649152 |
| ZNF692 | 2816.57361 | -0.0311192 | 0.28618833 | -0.1087366 | 0.91341139 | 0.98662757 |
| PGBD2 | 423.122981 | -0.2490687 | 0.22306364 | -1.1165812 | 0.26417344 | 0.82424998 |
| FAM110C | 3735.22576 | 0.13579927 | 0.48587048 | 0.27949685 | 7.80E-01 | 0.96836854 |
| SH3YL1 | 5362.62508 | 0.62737769 | 0.27702629 | 2.26468648 | 0.02353193 | 0.51967919 |
| ACP1 | 4399.63544 | -0.0176242 | 0.16367277 | -0.1076797 | 9.14E-01 | 9.87E-01 |
| ALKAL2 | 490.582793 | -0.0539437 | 0.46711994 | -0.1154814 | 0.90806357 | 0.98662757 |
| LINC01874 | 174.761909 | -0.8829718 | 0.72062325 | -1.2252891 | 0.22046632 | 0.79912826 |
| TMEM18 | 1310.72525 | 0.22902291 | 0.17444414 | 1.31287247 | 0.18922593 | 0.76970626 |
| AC092159.2 | 248.386273 | 0.47400094 | 0.24027351 | 1.97275572 | 0.0485234 | 0.58440502 |
| PXDN | 2030.12952 | 0.50848276 | 0.37224843 | 1.36597691 | 0.17194621 | 0.75974924 |
| EIPR1 | 1303.81963 | 0.21171949 | 0.14643837 | 1.44579249 | 0.14823541 | 0.73474739 |
| TRAPPC12 | 2210.809 | -0.0801312 | 0.12590763 | -0.6364285 | 0.5244972 | 0.91723878 |
| TRAPPC12-AS | 70.112855 | -0.5354727 | 0.45667123 | -1.1725563 | 2.41E-01 | 0.8124487 |
| AC114810.1 | 45.2550767 | 0.29849294 | 0.47975307 | 0.62218036 | 0.53382328 | 0.92062041 |
| ADI1 | 12712.0291 | 0.41398704 | 0.43948825 | 0.94197521 | 0.34620533 | 0.86135217 |
| AC231981.1 | 1422.73892 | 0.75243325 | 0.4992931 | 1.5069971 | 0.1318114 | 0.71169042 |
| AC108488.1 | 266.39902 | 0.27418978 | 0.21682921 | 1.26454263 | 0.20603533 | 0.78673759 |
| AC108488.2 | 247.183558 | -0.1000111 | 0.26326076 | -0.3798936 | 0.7040244 | 0.95485335 |
| RNASEH1 | 521.985891 | -0.323912 | 0.28593924 | -1.1328 | 0.25729825 | 0.82115501 |
| RNASEH1-AS | 248.127662 | 0.28324522 | 0.21800213 | 1.29927734 | 0.19384877 | 0.77620852 |
| RPS7 | 22177.2013 | 0.06573307 | 0.24195628 | 0.27167335 | 0.78587319 | 0.97065795 |

| | | | | | | |
|---|---|---|---|---|---|---|
| COLEC11 | 1077.99674 | 0.50025803 | 0.43708015 | 1.1445453 | 0.25239756 | 0.81833349 |
| ALLC | 123.019814 | -1.4290291 | 0.69926727 | -2.0436093 | 0.04099216 | 0.55811692 |
| LINC00487 | 106.144239 | -4.5282242 | 1.21493255 | -3.7271404 | NA | NA |
| NRIR | 144.409012 | -4.5028231 | 1.1630432 | -3.8715871 | NA | NA |
| CMPK2 | 969.894147 | -3.7239994 | 1.16651316 | -3.1924195 | NA | NA |
| RSAD2 | 1532.45596 | -4.7843962 | 1.1730193 | -4.078702 | 4.53E-05 | 0.02496292 |
| RNF144A | 963.953344 | 0.32181511 | 0.26925969 | 1.19518488 | 0.23201481 | 0.80649152 |
| LINC01814 | 44.3645842 | 0.25231215 | 0.65688089 | 0.3841064 | 0.70089959 | 0.9538566 |
| ID2-AS1 | 42.5922827 | 0.69814502 | 0.53199522 | 1.31231446 | 0.18941406 | 0.76970626 |
| ID2 | 5417.05362 | 0.03423016 | 0.18796852 | 0.18210581 | 0.85549969 | 0.97920671 |
| KIDINS220 | 3672.15603 | -0.1005587 | 0.10583107 | -0.950181 | 0.34202028 | 0.85889221 |
| MBOAT2 | 1640.91365 | 0.56837189 | 0.13193371 | 4.30801112 | 1.65E-05 | 0.01316658 |
| AC093904.4 | 58.6727298 | 1.31158292 | 0.60162424 | 2.18006994 | 0.02925228 | 0.53606221 |
| ASAP2 | 3574.03472 | 0.10010309 | 0.16707172 | 0.59916238 | 0.5490646 | 0.92373235 |
| ITGB1BP1 | 6446.90025 | -0.0803239 | 0.24320207 | -0.3302764 | 0.74119116 | 0.96102556 |
| CPSF3 | 1260.95517 | 0.19285238 | 0.13212436 | 1.45962773 | 0.14439241 | 0.73032824 |
| IAH1 | 2629.92201 | 0.0865402 | 0.12419412 | 0.69681405 | 0.48591917 | 0.90545573 |
| ADAM17 | 1590.34634 | -0.3197923 | 0.18449313 | -1.7333561 | 0.0830324 | 0.6471083 |
| AC073195.1 | 43.2278239 | 0.00649456 | 0.43173245 | 0.01504301 | 0.98799787 | 0.99754499 |
| YWHAQ | 11385.766 | -0.2211335 | 0.22014171 | -1.0045051 | 0.3151352 | 0.84696964 |
| TAF1B | 650.63719 | -0.0949419 | 0.23248996 | -0.4083701 | 0.683002 | 0.95070301 |
| AC010969.2 | 330.880613 | 0.07360346 | 0.26384518 | 0.27896456 | 0.78027202 | 0.96864684 |
| GRHL1 | 571.830661 | 0.0740026 | 0.30527465 | 0.24241318 | 0.80846002 | 0.97334138 |
| AC104794.2 | 381.496487 | -0.3775314 | 0.35825067 | -1.0538192 | 0.29196571 | 0.83474133 |
| KLF11 | 2437.45358 | -0.1975856 | 0.1477606 | -1.3372007 | 0.18115715 | 0.76522169 |
| AC104794.4 | 817.54165 | -0.0320515 | 0.16688882 | -0.1920532 | 0.84770053 | 0.97711043 |
| CYS1 | 15332.299 | 0.22128813 | 0.3434501 | 0.64430941 | 0.51937481 | 0.91636259 |
| RRM2 | 147.972779 | -0.2705909 | 0.7119508 | -0.3800696 | 0.70389375 | 0.95481129 |
| AC007240.1 | 70.7112939 | -0.2831724 | 0.76168052 | -0.3717732 | 0.71006171 | 0.95524448 |
| HPCAL1 | 3740.90368 | 0.05548247 | 0.20177174 | 0.27497645 | 0.78333433 | 0.96971192 |
| ODC1 | 6565.95423 | 0.03512988 | 0.22979328 | 0.15287603 | 0.87849604 | 0.98275231 |
| AC007249.2 | 51.6064076 | 0.10911954 | 0.54455153 | 0.20038424 | 0.84118009 | 0.97618283 |
| NOL10 | 1219.1318 | -0.1090118 | 0.21375054 | -0.5099956 | 0.61005452 | 0.93791514 |
| AC007314.1 | 285.154569 | -0.1542084 | 0.23570838 | -0.6542338 | 0.51296118 | 0.91568064 |
| RN7SL832P | 39.7436922 | 0.27391557 | 0.56520488 | 0.48463058 | 0.62793842 | 0.94088547 |
| ATP6V1C2 | 8415.87818 | 0.43769452 | 0.18292354 | 2.39277304 | 0.01672158 | 0.46515254 |
| AC092687.3 | 158.79779 | 0.14914587 | 0.47721385 | 0.31253466 | 7.55E-01 | 9.64E-01 |
| PDIA6 | 14383.181 | 0.43033865 | 0.18830203 | 2.28536391 | 2.23E-02 | 0.51834617 |
| PQLC3 | 1301.06716 | 0.41264883 | 0.16676557 | 2.47442458 | 1.33E-02 | 4.24E-01 |
| ROCK2 | 2206.01824 | -0.2503137 | 0.16279475 | -1.5376028 | 0.12414576 | 0.70078785 |
| E2F6 | 738.765066 | -0.4785986 | 0.2824719 | -1.6943228 | 0.09020396 | 0.65403568 |
| GREB1 | 253.119468 | 0.71231949 | 0.38192216 | 1.86509073 | 0.06216869 | 0.61245813 |
| LPIN1 | 2241.71268 | -0.6628574 | 0.32332858 | -2.0501044 | 0.04035424 | 0.55657615 |
| TRIB2 | 5240.39159 | -0.380987 | 0.24644099 | -1.5459563 | 0.12211513 | 0.69785915 |
| FAM84A | 1230.84898 | 0.27193662 | 0.37720115 | 0.72093264 | 0.47095096 | 0.90089741 |
| NBAS | 1657.63476 | 0.24142813 | 0.14185308 | 1.70195902 | 0.08876305 | 0.65363785 |
| DDX1 | 5194.09353 | 0.05813953 | 0.15563806 | 0.37355601 | 0.70873466 | 0.95501548 |
| MYCNOS | 131.151726 | 1.29745249 | 0.81448882 | 1.59296537 | 0.11116796 | 0.68001961 |
| MYCN | 375.894249 | 0.68992822 | 0.50548842 | 1.36487444 | 1.72E-01 | 7.60E-01 |
| FAM49A | 288.33127 | -0.1463921 | 0.42646991 | -0.3432648 | 7.31E-01 | 9.58E-01 |
| VSNL1 | 63.0117307 | 0.80578715 | 0.57325002 | 1.40564695 | 1.60E-01 | 7.48E-01 |
| SMC6 | 988.187547 | -0.0083602 | 0.13567263 | -0.0616203 | 0.95086517 | 0.99170503 |
| GEN1 | 415.636838 | -0.6284612 | 0.50952374 | -1.2334286 | 0.21741588 | 0.79597404 |
| KCNS3 | 521.986707 | 0.43636561 | 0.31408105 | 1.38934077 | 0.16472915 | 0.75297293 |

| | | | | | | |
|---|---|---|---|---|---|---|
| RDH14 | 1273.39215 | -0.2312064 | 0.14367123 | -1.609274 | 0.10755645 | 0.67707349 |
| NT5C1B-RDH | 452.89015 | -0.293297 | 0.14847339 | -1.9754177 | 0.04822076 | 0.58440502 |
| NT5C1B | 43.5759176 | -0.7869966 | 0.41607326 | -1.8914855 | 0.05855956 | 0.60332684 |
| OSR1 | 552.095634 | -2.2747986 | 0.86814671 | -2.6202928 | 0.00878543 | 0.38201332 |
| LINC00954 | 51.8507383 | -0.3888889 | 0.51996602 | -0.7479121 | 0.45451321 | 0.89670027 |
| TTC32 | 559.2695 | -0.3977704 | 0.23403446 | -1.6996233 | 0.08920181 | 0.65375543 |
| WDR35 | 1138.12569 | 0.10491163 | 0.16732011 | 0.62701151 | 5.31E-01 | 9.20E-01 |
| MATN3 | 234.430127 | 0.73975405 | 0.38077064 | 1.94278125 | 0.05204259 | 0.58645127 |
| LAPTM4A | 24495.6525 | 0.17897443 | 0.18332426 | 0.97627247 | 0.32892946 | 0.85317073 |
| SDC1 | 21472.0934 | 0.31657813 | 0.3512058 | 0.90140349 | 0.36737383 | 0.86683872 |
| PUM2 | 4956.04787 | -0.2227263 | 0.15359638 | -1.4500754 | 1.47E-01 | 0.73337969 |
| RHOB | 22096.1387 | -0.666549 | 0.31752441 | -2.0992056 | 3.58E-02 | 0.55369025 |
| HS1BP3 | 2680.42646 | 0.04467712 | 0.20338552 | 0.21966716 | 0.82613038 | 0.97548201 |
| AC012065.3 | 42.2389333 | 0.04548151 | 0.45866747 | 0.09916009 | 0.92101116 | 0.98767854 |
| GDF7 | 1175.87911 | 0.11536526 | 0.45531727 | 0.25337336 | 0.79997971 | 0.97208422 |
| AC012065.4 | 505.212777 | 0.09015502 | 0.4998198 | 0.18037504 | 0.85685815 | 0.97953896 |
| LDAH | 1035.76065 | 0.47542778 | 0.19727168 | 2.41001537 | 0.01595185 | 0.4564177 |
| APOB | 110.34317 | -0.2056586 | 0.83359484 | -0.2467129 | 8.05E-01 | 9.73E-01 |
| AC018467.1 | 74.6251445 | -0.0313428 | 0.64592687 | -0.0485238 | 0.96129881 | 0.99320711 |
| KLHL29 | 1092.24987 | -0.7738439 | 0.52208019 | -1.4822319 | 0.13827859 | 0.72249298 |
| AC009242.1 | 77.2030247 | 0.83136209 | 0.5666248 | 1.46721797 | 0.14231677 | 0.72784886 |
| ATAD2B | 630.030436 | -0.0584363 | 0.1566788 | -0.372969 | 0.70917151 | 0.95501548 |
| UBXN2A | 902.016705 | -0.1453891 | 0.18788416 | -0.7738232 | 0.43903536 | 0.89175645 |
| MFSD2B | 308.09993 | -0.1546959 | 0.2598985 | -0.5952165 | 0.55169874 | 0.92373235 |
| WDCP | 364.013698 | -0.1914796 | 0.17363886 | -1.1027462 | 0.27013739 | 0.82716979 |
| FKBP1B | 388.069901 | -0.1817463 | 0.2834267 | -0.6412463 | 0.52136265 | 0.91723878 |
| SF3B6 | 2619.41838 | -0.1071916 | 0.11489382 | -0.9329619 | 0.35083966 | 0.86364633 |
| FAM228B | 1040.15659 | 0.29096469 | 0.24092125 | 1.20771704 | 0.22715612 | 0.80300331 |
| TP53I3 | 2653.22149 | 0.16283606 | 0.29912364 | 0.5443771 | 0.586182 | 0.93289552 |
| PFN4 | 49.827821 | 0.45888484 | 0.43617126 | 1.05207491 | 0.29276518 | 0.83524862 |
| AC008073.3 | 279.628416 | 0.55921322 | 0.34618071 | 1.61537947 | 0.10622854 | 0.67623273 |
| ITSN2 | 2561.13736 | -0.0333993 | 0.15433917 | -0.216402 | 0.82867442 | 0.97550416 |
| AC009228.1 | 127.74921 | -0.0942356 | 0.28576916 | -0.3297613 | 0.74158032 | 0.96109001 |
| NCOA1 | 2586.4443 | -0.2453286 | 0.10830356 | -2.2651939 | 0.02350078 | 0.51967919 |
| PTRHD1 | 641.208561 | 0.13032591 | 0.22258289 | 0.5855163 | 0.55820061 | 0.92421175 |
| CENPO | 1983.94993 | -0.1758295 | 0.27332169 | -0.6433059 | 0.52002561 | 0.91657872 |
| ADCY3 | 2077.44477 | -0.2071551 | 0.34562021 | -0.5993722 | 0.54892471 | 0.92373235 |
| AC012073.1 | 151.838295 | -0.0563955 | 0.39409104 | -0.1431027 | 8.86E-01 | 0.98391254 |
| DNAJC27 | 526.059782 | -0.0155276 | 0.11224352 | -0.1383387 | 0.88997275 | 0.98459146 |
| DNAJC27-AS | 98.1315808 | 0.48706673 | 0.51295522 | 0.94953071 | 0.34235076 | 0.85889221 |
| EFR3B | 120.864361 | 0.42881437 | 0.52843084 | 0.81148626 | 0.41708648 | 0.88445833 |
| POMC | 64.3175282 | -0.436234 | 0.59526575 | -0.7328391 | 0.46365659 | 0.89830907 |
| DNMT3A | 1768.2869 | -0.2036686 | 0.17141853 | -1.1881366 | 2.35E-01 | 8.07E-01 |
| AC012074.1 | 39.7292132 | 0.00640398 | 0.41971561 | 0.01525791 | 9.88E-01 | 9.98E-01 |
| DTNB | 774.654339 | 0.21566253 | 0.18213077 | 1.18410814 | 0.23637026 | 0.80806879 |
| AC104699.1 | 124.482034 | 0.18921905 | 0.27366553 | 0.69142452 | 0.4892988 | 0.90600166 |
| ASXL2 | 1565.13851 | -0.1856355 | 0.21109183 | -0.8794065 | 0.37918089 | 0.87069278 |
| KIF3C | 390.164226 | -0.047632 | 0.20067917 | -0.2373542 | 0.81238204 | 0.97364432 |
| RAB10 | 6805.73631 | -0.2665904 | 0.28511415 | -0.9350305 | 0.34977256 | 0.86364633 |
| GAREM2 | 288.849626 | 0.37720179 | 0.31121593 | 1.21202598 | 0.22550244 | 0.80214224 |
| HADHA | 16083.3727 | 0.34554438 | 0.23390724 | 1.47727102 | 0.139603 | 0.72475166 |
| HADHB | 14294.282 | 0.34928309 | 0.25865246 | 1.35039537 | 0.17688919 | 0.76115956 |
| ADGRF3 | 223.874043 | 0.13842938 | 0.45705985 | 0.30286927 | 7.62E-01 | 9.66E-01 |
| SELENOI | 1699.53012 | -0.2410548 | 0.25744141 | -0.9363483 | 3.49E-01 | 8.64E-01 |

| | | | | | | |
|---|---|---|---|---|---|---|
| C2orf70 | 82.1134995 | -0.0292138 | 0.68257016 | -0.0427997 | 9.66E-01 | 9.94E-01 |
| KCNK3 | 991.910918 | 0.09285153 | 0.51146809 | 0.18153925 | 8.56E-01 | 9.79E-01 |
| SLC35F6 | 5234.4652 | 0.21039868 | 0.38252854 | 0.55002086 | 5.82E-01 | 9.32E-01 |
| CENPA | 223.29637 | 0.31432021 | 0.35274033 | 0.89108101 | 0.37288571 | 0.86791605 |
| MAPRE3 | 2191.32066 | 0.09791448 | 0.22093658 | 0.44317914 | 6.58E-01 | 0.94558535 |
| TMEM214 | 5887.61978 | -0.1587471 | 0.15428603 | -1.0289142 | 3.04E-01 | 0.84211658 |
| AGBL5 | 3695.73289 | 0.28341969 | 0.11519548 | 2.46033694 | 0.01388066 | 0.4295825 |
| AC013472.3 | 258.807307 | 0.41838406 | 0.23810397 | 1.75714865 | 0.07889248 | 0.64079926 |
| OST4 | 4576.03669 | 0.18563055 | 0.17716098 | 1.04780725 | 0.29472742 | 0.83630273 |
| EMILIN1 | 4444.41394 | -0.1999571 | 0.52259424 | -0.382624 | 0.7019986 | 0.95431462 |
| KHK | 11223.3889 | 0.79843195 | 0.48539876 | 1.64489902 | 0.09999064 | 0.67165899 |
| CGREF1 | 5901.44422 | 0.69697301 | 0.43400986 | 1.60589212 | 0.10829762 | 0.67789099 |
| ABHD1 | 207.348497 | 0.10872669 | 0.17654352 | 0.61586337 | 0.5379847 | 0.92127611 |
| PREB | 3730.71333 | -0.0273536 | 0.13560443 | -0.2017163 | 0.84013855 | 0.97618283 |
| SLC5A6 | 2819.04029 | -0.0196094 | 0.12888087 | -0.1521511 | 0.87906776 | 0.98275231 |
| ATRAID | 4739.93433 | 0.22993244 | 0.13077308 | 1.75825516 | 0.0787041 | 0.64079926 |
| CAD | 1749.21379 | -0.1190237 | 0.14843503 | -0.8018574 | 0.42263544 | 0.88451372 |
| TRIM54 | 73.3480589 | 0.68396551 | 0.66493806 | 1.02861538 | 3.04E-01 | 8.42E-01 |
| UCN | 92.1654394 | 0.10705129 | 0.4813482 | 0.22239885 | 0.82400341 | 0.97548201 |
| MPV17 | 2605.26666 | 0.24784504 | 0.1701025 | 1.4570335 | 0.14510713 | 0.73064288 |
| GTF3C2 | 3247.45555 | -0.0154422 | 0.18575367 | -0.0831326 | 0.93374607 | 0.99022947 |
| GTF3C2-AS1 | 700.490856 | 0.00638545 | 0.28400566 | 0.02248354 | 0.98206224 | 0.99595643 |
| AC074117.1 | 1213.18558 | -0.3409122 | 0.18219624 | -1.8711263 | 0.06132758 | 0.61013576 |
| EIF2B4 | 2429.09429 | -0.2306537 | 0.15716748 | -1.4675665 | 0.14222203 | 0.72784886 |
| SNX17 | 5830.0811 | -0.0029132 | 0.08728305 | -0.0333761 | 0.97337464 | 0.99445887 |
| ZNF513 | 2390.15186 | -0.1731413 | 0.17223497 | -1.0052619 | 0.31477076 | 0.84696964 |
| PPM1G | 4527.17889 | -0.1673395 | 0.20419167 | -0.8195219 | 0.41248872 | 0.88268208 |
| NRBP1 | 6634.7404 | -0.5346175 | 0.36459295 | -1.4663406 | 0.14255551 | 0.72784886 |
| KRTCAP3 | 802.447284 | 0.08027142 | 0.26043047 | 0.30822591 | 0.75791044 | 0.96502715 |
| IFT172 | 1853.78258 | 0.18581893 | 0.29202643 | 0.63630861 | 0.52457529 | 0.91723878 |
| FNDC4 | 920.264697 | -0.4336835 | 0.44934678 | -0.9651422 | 0.33447359 | 0.85557672 |
| GCKR | 88.8825266 | -0.2514567 | 0.3608365 | -0.6968717 | 0.48588311 | 0.90545573 |
| C2orf16 | 54.6237714 | 0.07897032 | 0.3967039 | 0.19906616 | 0.84221099 | 0.9763625 |
| ZNF512 | 1774.51237 | 0.02471458 | 0.23013305 | 0.10739257 | 0.91447755 | 0.9866558 |
| CCDC121 | 207.802932 | -0.1743027 | 0.30250089 | -0.5762057 | 0.56447615 | 0.92474393 |
| GPN1 | 2020.44615 | -0.2195964 | 0.2426726 | -0.9049081 | 0.3655141 | 0.8663515 |
| SUPT7L | 3550.09004 | -0.198131 | 0.11759195 | -1.6849026 | 0.09200736 | 0.65583276 |
| SLC4A1AP | 1673.72041 | -0.0652314 | 0.15575328 | -0.4188124 | 0.67535322 | 0.94958156 |
| MRPL33 | 3610.5352 | 0.04471758 | 0.19778912 | 0.22608715 | 0.82113364 | 0.97472105 |
| RBKS | 1941.89303 | 0.15215314 | 0.32546891 | 0.46748903 | 0.64015004 | 0.94088547 |
| BABAM2 | 4214.08069 | 0.18437947 | 0.12634918 | 1.45928505 | 0.14448667 | 0.73032824 |
| AC104695.2 | 57.2129026 | -0.2810998 | 0.47865105 | -0.587275 | 0.55701902 | 0.92373235 |
| FOSL2 | 12871.3247 | -0.2878451 | 0.33878369 | -0.8496429 | 0.39552367 | 0.87927742 |
| AC104695.3 | 64.1219351 | 0.20165005 | 0.51089791 | 0.39469736 | 0.69306625 | 0.9519683 |
| PLB1 | 136.050179 | -0.0557709 | 0.58973377 | -0.0945696 | 0.92465669 | 0.98869145 |
| AC092164.1 | 50.4743356 | 0.33297023 | 0.40841163 | 0.81528097 | 0.4149115 | 0.88372047 |
| PPP1CB | 11610.3553 | -0.2563153 | 0.19251972 | -1.3313716 | 0.18306675 | 0.76723162 |
| SPDYA | 1568.94635 | -0.1451486 | 0.26221288 | -0.5535524 | 0.57988519 | 0.93139743 |
| TRMT61B | 814.359277 | -0.1318653 | 0.22313039 | -0.5909787 | 0.5545347 | 0.92373235 |
| WDR43 | 1467.63828 | -0.7337355 | 0.41250617 | -1.7787261 | 0.07528468 | 0.63773415 |
| TOGARAM2 | 104.85714 | -0.0487586 | 0.46397691 | -0.1050884 | 0.91630568 | 0.9868044 |
| C2orf71 | 49.4430273 | 0.74424148 | 0.95108015 | 0.78252236 | 0.43390765 | 0.89055338 |
| CLIP4 | 1673.10501 | -0.1557906 | 0.21606534 | -0.7210346 | 0.47088824 | 0.90089741 |
| YPEL5 | 6972.90433 | -0.2021001 | 0.12456086 | -1.6225006 | 0.1046962 | 0.67439401 |

| | | | | | | |
|---|---|---|---|---|---|---|
| LBH | 3259.76718 | -0.3845045 | 0.30337199 | -1.2674357 | 0.20499955 | 0.78502176 |
| LINC01936 | 324.789917 | -0.943759 | 0.66767177 | -1.4135074 | 0.15750659 | 0.74522306 |
| LCLAT1 | 602.823906 | 0.07325776 | 0.20504027 | 0.35728476 | 0.72087864 | 0.95728372 |
| CAPN13 | 532.738887 | -0.9264791 | 0.49923333 | -1.8558038 | 0.06348153 | 0.6173884 |
| GALNT14 | 4919.50853 | 0.26078067 | 0.34330571 | 0.75961648 | 0.44748387 | 0.89480207 |
| CAPN14 | 50.5001905 | -0.6088897 | 0.77208112 | -0.7886343 | 0.43032576 | 0.88936206 |
| EHD3 | 1758.89645 | 0.54300831 | 0.49550201 | 1.09587509 | 0.27313344 | 0.82716979 |
| XDH | 93.8552863 | 2.46684717 | 0.86412544 | 2.85473274 | 0.00430731 | 0.28584892 |
| SRD5A2 | 48.8802945 | 0.52858768 | 1.08055973 | 0.48917951 | 0.62471461 | 0.94088547 |
| MEMO1 | 952.162115 | -0.2063028 | 0.22196401 | -0.9294425 | 0.3526598 | 0.86364633 |
| DPY30 | 2379.98001 | -0.0666534 | 0.1546717 | -0.4309346 | 0.66651595 | 0.94809681 |
| SPAST | 680.288422 | -0.0632383 | 0.13929347 | -0.4539931 | 0.64983382 | 0.94312988 |
| SLC30A6 | 1411.91187 | -0.0220799 | 0.12661543 | -0.1743854 | 0.86156257 | 0.98167219 |
| NLRC4 | 111.728517 | -0.253018 | 0.3676313 | -0.6882384 | 0.49130264 | 0.90600166 |
| YIPF4 | 2788.37921 | 0.16042055 | 0.14006364 | 1.14534041 | 0.25206817 | 0.81833349 |
| BIRC6 | 3092.37617 | -0.0979721 | 0.08563644 | -1.1440464 | 0.25260441 | 0.81833349 |
| TTC27 | 707.336904 | -0.0068486 | 0.23576267 | -0.0290488 | 0.97682564 | 0.99455099 |
| LTBP1 | 2045.2908 | 0.02286099 | 0.64851405 | 0.03525135 | 0.97187932 | 0.99404213 |
| RASGRP3 | 936.206496 | -0.6735095 | 0.29771308 | -2.2622772 | 0.02368028 | 0.51967919 |
| AC020594.1 | 98.2114802 | -0.5970163 | 0.32211647 | -1.8534175 | 0.06382254 | 0.61812911 |
| FAM98A | 2542.03289 | -0.2116418 | 0.21083524 | -1.0038254 | 0.31546278 | 0.84712537 |
| LINC01320 | 2738.91059 | -0.4656543 | 0.58733201 | -0.7928298 | 0.42787701 | 0.88792416 |
| AC073218.1 | 325.340908 | -0.7345386 | 0.65852585 | -1.1154286 | 0.26466678 | 0.82424998 |
| AC009414.2 | 128.089354 | -0.2778051 | 0.29020031 | -0.9572874 | 0.33842223 | 0.85719299 |
| CRIM1 | 8698.58464 | 0.10081213 | 0.26388009 | 0.38203766 | 0.70243343 | 0.95431462 |
| AC007378.1 | 156.300538 | 0.36007408 | 0.44776144 | 0.804165 | 0.42130169 | 0.88451372 |
| FEZ2 | 2426.96251 | 0.09924359 | 0.19165233 | 0.51783136 | 0.60457593 | 0.93662382 |
| STRN | 1592.13259 | -0.3568504 | 0.17409213 | -2.0497789 | 0.04038601 | 0.55657615 |
| HEATR5B | 1348.85556 | -0.3129349 | 0.16725595 | -1.8709942 | 0.06134588 | 0.61013576 |
| GPATCH11 | 503.928473 | -0.0735367 | 0.28462189 | -0.2583662 | 0.79612426 | 0.97101097 |
| EIF2AK2 | 2845.29323 | -0.9522685 | 0.47250099 | -2.0153787 | NA | NA |
| SULT6B1 | 143.717126 | 0.36593226 | 0.22974639 | 1.59276609 | 0.11121267 | 0.68001961 |
| CEBPZOS | 3358.0044 | 0.04709171 | 0.12417372 | 0.37924052 | 0.70450926 | 0.95501548 |
| CEBPZ | 2930.37244 | -0.058936 | 0.16451609 | -0.3582387 | 0.72016467 | 0.95692068 |
| NDUFAF7 | 1291.77672 | -0.0303393 | 0.14441388 | -0.2100858 | 0.83360071 | 0.97550416 |
| PRKD3 | 1657.87061 | -0.0402347 | 0.15919419 | -0.2527398 | 0.80046933 | 0.97210205 |
| QPCT | 844.187509 | 0.16337188 | 0.47657505 | 0.3428041 | 0.73174584 | 0.95848028 |
| AC006369.1 | 416.72635 | -0.1293734 | 0.31802023 | -0.4068087 | 0.6841485 | 0.95070301 |
| CDC42EP3 | 2855.10533 | -0.1397226 | 0.23185666 | -0.6026249 | 0.54675825 | 0.92345821 |
| RMDN2 | 364.572378 | 0.14611625 | 0.23329626 | 0.62631202 | 0.5311103 | 0.91962754 |
| CYP1B1 | 2666.4958 | 0.08035397 | 0.4719299 | 0.17026675 | 0.86480036 | 0.98197545 |
| CYP1B1-AS1 | 169.383012 | 0.17469822 | 0.45830838 | 0.38118051 | 0.70306931 | 0.95431462 |
| ATL2 | 2272.33908 | -0.1168284 | 0.18255839 | -0.6399508 | 0.52220461 | 0.91723878 |
| HNRNPLL | 1408.72428 | 0.07327465 | 0.18322625 | 0.39991347 | 0.68922025 | 0.95158502 |
| GALM | 5154.08522 | 1.00749048 | 0.48281295 | 2.08670972 | 0.03691438 | 0.55521179 |
| SRSF7 | 5858.9299 | -0.0110058 | 0.13857441 | -0.0794218 | 0.93669714 | 0.99068685 |
| GEMIN6 | 540.204827 | -0.0160284 | 0.14648373 | -0.1094209 | 0.91286869 | 0.98662757 |
| DHX57 | 985.311084 | 0.01059088 | 0.1848512 | 0.05729406 | 0.95431095 | 0.99170503 |
| MORN2 | 1146.93796 | 0.25821638 | 0.13319166 | 1.93868279 | 0.05253998 | 0.58645127 |
| ARHGEF33 | 94.6384082 | -0.0098167 | 0.39096805 | -0.0251086 | 0.97996832 | 0.99537078 |
| SOS1 | 2161.7059 | -0.0720843 | 0.12587013 | -0.5726879 | 0.56685602 | 0.9256133 |
| MAP4K3 | 2214.36448 | -0.0789107 | 0.16824969 | -0.4690094 | 0.63906292 | 0.94088547 |
| AC007388.1 | 625.970691 | 0.44059662 | 0.25530554 | 1.72576206 | 0.08439023 | 0.64988556 |
| TMEM178A | 1092.51719 | 1.01069705 | 0.27624932 | 3.65864085 | 0.00025356 | 0.08192931 |

| | | | | | | |
|---|---|---|---|---|---|---|
| THUMPD2 | 720.412708 | -0.1589336 | 0.13395966 | -1.186429 | 0.23545293 | 0.80747753 |
| SLC8A1-AS1 | 252.763824 | 0.7830962 | 0.41641682 | 1.88055851 | 0.060032 | 0.6075531 |
| SLC8A1 | 1327.66604 | 1.01308478 | 0.37527745 | 2.69956209 | 0.00694308 | 0.3503878 |
| PKDCC | 2239.86459 | 0.39587238 | 0.35235572 | 1.12350208 | 0.26122432 | 0.82424998 |
| EML4 | 3172.84708 | 0.11320088 | 0.12327203 | 0.91830144 | 0.35846108 | 0.86480228 |
| COX7A2L | 5635.55863 | 0.14873808 | 0.2598497 | 0.57240044 | 0.56705072 | 0.92561704 |
| MTA3 | 1823.93544 | 0.1078543 | 0.19692595 | 0.54768961 | 0.58390505 | 0.93233498 |
| OXER1 | 694.159477 | 0.3271546 | 0.45572568 | 0.71787616 | 0.47283365 | 0.90164585 |
| HAAO | 1925.17235 | 0.34020557 | 0.36377627 | 0.93520549 | 0.34968241 | 0.86364633 |
| LINC01819 | 41.7158749 | -0.1014058 | 0.79316542 | -0.1278495 | 0.89826807 | 0.98474725 |
| ZFP36L2 | 14527.707 | -0.1904173 | 0.26521061 | -0.7179854 | 0.47276632 | 0.90160919 |
| AC010883.1 | 336.388497 | 0.02557736 | 0.2053637 | 0.12454665 | 0.90088246 | 0.98545758 |
| THADA | 1644.0569 | -0.1525525 | 0.15532567 | -0.9821463 | 0.32602777 | 0.85282528 |
| PLEKHH2 | 1651.88045 | -0.0238422 | 0.36765147 | -0.0648499 | 0.94829349 | 0.99170503 |
| C1GALT1C1L | 57.4973004 | 0.3526561 | 0.32079051 | 1.09933458 | 0.27162215 | 0.82716979 |
| DYNC2LI1 | 1771.75061 | 0.42813047 | 0.29183995 | 1.46700435 | 0.14237487 | 0.72784886 |
| LRPPRC | 6599.68823 | 0.26074202 | 0.1995465 | 1.30667299 | 0.19132379 | 0.77292503 |
| PPM1B | 2598.30772 | -0.0308643 | 0.10947512 | -0.2819297 | 0.77799743 | 0.96836854 |
| SLC3A1 | 30468.3213 | 0.66195689 | 0.36937728 | 1.79208883 | 0.07311874 | 0.63587855 |
| PREPL | 16813.7595 | 0.44247629 | 0.30935274 | 1.43032929 | 0.15262253 | 0.7422778 |
| CAMKMT | 390.728188 | 0.30413589 | 0.24537752 | 1.23946111 | 0.21517478 | 0.7949124 |
| LINC01833 | 87.9799802 | 0.91973691 | 0.79762233 | 1.15309825 | 0.24887006 | 0.81646572 |
| SRBD1 | 605.020853 | 0.294508 | 0.163644 | 1.79968715 | 0.07191005 | 0.63189503 |
| PRKCE | 1187.45265 | -0.0047065 | 0.17074961 | -0.0275638 | 0.97801005 | 0.99476647 |
| EPAS1 | 21006.4688 | -0.0738021 | 0.40284892 | -0.1832004 | 0.85464076 | 0.97910552 |
| ATP6V1E2 | 202.204756 | -0.2907611 | 0.35269067 | -0.8244082 | 0.40970765 | 0.88249519 |
| RHOQ | 6241.50371 | -0.2480206 | 0.27135255 | -0.914016 | 0.36070846 | 0.86504879 |
| AC018682.1 | 77.2002086 | -0.6901097 | 0.4848307 | -1.4234034 | 1.55E-01 | 7.43E-01 |
| PIGF | 2330.31405 | -0.1658607 | 0.2439658 | -0.6798523 | 0.49659797 | 0.90890243 |
| CRIPT | 748.277885 | 0.23735336 | 0.18615977 | 1.27499811 | 0.20230991 | 0.78191175 |
| SOCS5 | 1250.40938 | -0.1232432 | 0.21226663 | -0.5806058 | 0.56150613 | 0.92452095 |
| AC016722.2 | 306.660958 | -0.3336579 | 0.43166704 | -0.7729521 | 0.43955073 | 0.8919464 |
| MCFD2 | 5711.24448 | -0.1634701 | 0.28560362 | -0.5723669 | 0.56707341 | 0.92561704 |
| TTC7A | 2389.35635 | 0.26071529 | 0.24732391 | 1.05414512 | 0.29181648 | 0.83460625 |
| AC093732.2 | 60.962952 | 0.43855505 | 0.52030433 | 0.84288179 | 0.39929456 | 0.88077967 |
| AC073283.3 | 2721.15168 | 0.16508684 | 0.21212673 | 0.77824626 | 0.43642385 | 0.89104919 |
| CALM2 | 17651.0134 | 0.11370192 | 0.18854464 | 0.60305038 | 0.54647517 | 0.92345821 |
| AC106869.1 | 117.922765 | 0.33396039 | 0.57716947 | 0.57861756 | 0.56284726 | 0.92452095 |
| EPCAM | 14923.7665 | -0.0585381 | 0.23392158 | -0.2502465 | 0.80239669 | 0.97245417 |
| MSH2 | 687.245165 | 0.16241763 | 0.15569587 | 1.04317239 | 2.97E-01 | 8.39E-01 |
| MSH6 | 3585.49021 | -0.0670838 | 0.10962002 | -0.6119666 | 0.54055988 | 0.92247611 |
| FBXO11 | 3033.2444 | -0.12807 | 0.11306016 | -1.1327599 | 0.25731509 | 0.82115501 |
| AC079807.1 | 83.5902381 | 0.26614559 | 0.34994731 | 0.76053047 | 0.44693757 | 0.89480207 |
| FOXN2 | 586.596843 | -0.3293016 | 0.20120707 | -1.6366304 | 0.10170773 | 0.67279543 |
| PPP1R21 | 2693.45099 | 0.27408321 | 0.2330095 | 1.17627482 | 0.23948507 | 0.81163641 |
| STON1 | 1292.31047 | -0.1588749 | 0.25064329 | -0.6338684 | 0.52616668 | 0.91769856 |
| STON1-GTF2 | 191.568934 | 0.12732024 | 0.44522676 | 0.28596718 | 0.77490326 | 0.96836854 |
| GTF2A1L | 54.5431702 | 0.21091428 | 0.94642118 | 0.22285456 | 0.8236487 | 0.9754803 |
| ASB3 | 825.861786 | -0.0410278 | 0.17453552 | -0.2350685 | 0.81415558 | 0.9738447 |
| GPR75-ASB3 | 777.491259 | -0.0370238 | 0.17377009 | -0.2130617 | 0.83127884 | 0.97550416 |
| CHAC2 | 127.93881 | -0.5558733 | 0.30697559 | -1.8108063 | 7.02E-02 | 0.62928077 |
| ERLEC1 | 2709.01293 | 0.18074528 | 0.13698091 | 1.31949247 | 1.87E-01 | 0.76958556 |
| PSME4 | 2657.97166 | -0.2828164 | 0.19653656 | -1.4390013 | 0.15015014 | 0.73796431 |
| ACYP2 | 605.750868 | -0.1993166 | 0.1684271 | -1.1833999 | 0.2366507 | 0.80849753 |

| | | | | | | |
|---|---|---|---|---|---|---|
| SPTBN1 | 20182.2027 | -0.0321435 | 0.27447339 | -0.1171097 | 0.90677312 | 0.98662757 |
| AC093110.1 | 2391.5761 | -0.0768781 | 0.29283954 | -0.2625265 | 0.79291557 | 0.97090793 |
| EML6 | 290.961973 | 0.17301698 | 0.39450626 | 0.43856587 | 0.66097613 | 0.94637197 |
| RTN4 | 23747.9129 | -0.1374619 | 0.26023399 | -0.5282243 | 0.5973437 | 0.93416444 |
| CLHC1 | 365.135559 | -0.2550894 | 0.28726587 | -0.8879908 | 0.3745457 | 0.86791605 |
| RPS27A | 27631.6562 | 0.18363736 | 0.22818734 | 0.80476578 | 0.42095485 | 0.88451372 |
| MTIF2 | 1856.00365 | 0.2663746 | 0.1953706 | 1.36343235 | 0.17274629 | 0.75974924 |
| AC012358.3 | 116.447625 | 0.28313488 | 0.28499779 | 0.99346345 | 0.32048416 | 0.85028262 |
| CCDC88A | 907.283009 | -0.0684986 | 0.14883647 | -0.4602271 | 0.64535319 | 0.94208859 |
| CFAP36 | 2822.80021 | 0.14805201 | 0.13648852 | 1.0847213 | 0.27804511 | 0.82805357 |
| PPP4R3B | 4243.10241 | -0.1079817 | 0.13147348 | -0.8213196 | 0.41146424 | 0.88268208 |
| AC015982.1 | 64.6717109 | -0.1404178 | 0.38615721 | -0.3636285 | 0.71613547 | 0.95603249 |
| PNPT1 | 1890.53924 | -0.9916838 | 0.39221825 | -2.5283978 | 1.15E-02 | 0.40353404 |
| EFEMP1 | 6353.11777 | 0.20309105 | 0.38208315 | 0.53153626 | 0.59504722 | 0.9334601 |
| AC007743.1 | 61.1310322 | -0.6411361 | 0.58758275 | -1.0911418 | 0.27521049 | 0.82719407 |
| CCDC85A | 105.607613 | 0.28231053 | 0.56235807 | 0.50201206 | 0.61565904 | 0.93910158 |
| VRK2 | 993.796079 | 0.01030195 | 0.23021591 | 0.04474908 | 0.96430732 | 0.99342622 |
| FANCL | 1104.89514 | 0.07235907 | 0.24698253 | 0.29297242 | 0.76954323 | 0.96836854 |
| BCL11A | 88.7314236 | 0.18719389 | 0.53374304 | 0.35071911 | 0.72579908 | 0.95848028 |
| PAPOLG | 771.55436 | -0.1498542 | 0.17910407 | -0.8366881 | 0.4027679 | 0.88132015 |
| REL | 777.395408 | 0.31808705 | 0.28368226 | 1.12127929 | 0.262169 | 0.82424998 |
| PUS10 | 493.84725 | 0.32530408 | 0.29368968 | 1.10764559 | 2.68E-01 | 8.27E-01 |
| PEX13 | 1663.90901 | -0.1851198 | 0.21860744 | -0.8468136 | 0.397099 | 0.87988087 |
| KIAA1841 | 316.657608 | 0.11235106 | 0.15634463 | 0.71861157 | 0.47238028 | 0.90160919 |
| C2orf74 | 866.918358 | 0.13031538 | 0.38153309 | 0.34155722 | 7.33E-01 | 9.58E-01 |
| USP34 | 5917.62872 | -0.271822 | 0.10909879 | -2.4915215 | 0.01271973 | 0.41598747 |
| AC016727.1 | 875.290981 | -0.2674803 | 0.23672377 | -1.1299258 | 0.25850748 | 0.82294696 |
| XPO1 | 5569.98476 | -0.2096958 | 0.25508622 | -0.8220585 | 0.41104363 | 0.88268208 |
| FAM161A | 288.133329 | 0.32673995 | 0.25356121 | 1.28860385 | 0.19753584 | 0.77822561 |
| CCT4 | 6303.51542 | 0.245815 | 0.17220797 | 1.42743104 | 0.15345569 | 0.74258845 |
| AC107081.2 | 1068.62328 | 0.20171417 | 0.166094 | 1.21445787 | 0.22457293 | 0.80078694 |
| COMMD1 | 1184.52461 | 0.07886036 | 0.12706645 | 0.62062295 | 0.53484773 | 0.92084006 |
| AC018462.1 | 57.0039748 | -0.1627106 | 0.27865336 | -0.5839176 | 0.55927573 | 0.92421175 |
| B3GNT2 | 2235.10356 | -0.2418662 | 0.21759422 | -1.1115469 | 0.266333 | 0.8265123 |
| TMEM17 | 196.229541 | 0.11323993 | 0.33912004 | 0.33392284 | 0.73843778 | 0.9602325 |
| EHBP1 | 1485.89346 | 0.10223098 | 0.11699132 | 0.87383393 | 0.3822087 | 0.87185666 |
| AC009501.1 | 523.847595 | 0.13941637 | 0.11620139 | 1.19978233 | 0.23022389 | 0.80556869 |
| WDPCP | 261.743476 | 0.10254464 | 0.17896901 | 0.57297429 | 0.5666621 | 0.9256133 |
| MDH1 | 12637.6492 | 0.54108274 | 0.25206601 | 2.14659141 | 0.03182582 | 0.54649555 |
| UGP2 | 8189.1298 | 0.13863595 | 0.19766616 | 0.7013641 | 0.48307582 | 0.90541935 |
| VPS54 | 2125.00438 | 0.01016658 | 0.07553132 | 0.13460089 | 0.89292744 | 0.98474725 |
| PELI1 | 1621.45879 | -0.470765 | 0.25195831 | -1.8684241 | 0.06170298 | 0.61119212 |
| AC012368.1 | 387.338166 | -0.006505 | 0.72746622 | -0.008942 | 0.99286541 | 0.99906069 |
| LGALSL | 1010.4939 | -0.2643256 | 0.31541634 | -0.8380213 | 0.40201874 | 0.88132015 |
| AC008074.3 | 84.5241029 | 0.48988733 | 0.45582534 | 1.07472598 | 0.28249743 | 0.83055556 |
| AFTPH | 3673.28406 | -0.1909336 | 0.21398372 | -0.8922808 | 0.37224247 | 0.86775997 |
| SERTAD2 | 1405.22832 | -0.1170596 | 0.26242322 | -0.4460719 | 6.56E-01 | 0.94532497 |
| SLC1A4 | 623.091245 | -1.1179877 | 0.51532441 | -2.1694832 | 0.03004602 | 0.54012717 |
| CEP68 | 1404.95272 | 0.00020265 | 0.20602821 | 0.00098358 | 0.99921522 | 0.99994357 |
| RAB1A | 10988.7438 | -0.1601808 | 0.15634912 | -1.0245069 | 0.30559591 | 0.84270033 |
| ACTR2 | 7856.70109 | -0.0634753 | 0.20553029 | -0.3088367 | 0.75744572 | 0.96502715 |
| SPRED2 | 1906.8031 | -0.3248803 | 0.33324542 | -0.9748981 | 0.32961082 | 0.85356118 |
| MEIS1 | 1261.839 | 0.18499119 | 0.25280046 | 0.73176764 | 0.4643104 | 0.89830907 |
| ETAA1 | 823.373171 | -0.0707318 | 0.170854 | -0.4139898 | 0.67888155 | 0.9504646 |

| | | | | | | |
|---|---|---|---|---|---|---|
| C1D | 1296.44147 | 0.01267493 | 0.15746869 | 0.08049174 | 0.93584617 | 0.99068685 |
| WDR92 | 496.996557 | 0.05189566 | 0.13925723 | 0.37266041 | 0.7094012 | 0.95501548 |
| AC017083.3 | 852.080974 | -0.069788 | 0.22209148 | -0.3142307 | 0.75334585 | 0.96301881 |
| PNO1 | 1033.65551 | -0.2828544 | 0.33514469 | -0.843977 | 0.39868227 | 0.88074856 |
| PPP3R1 | 2666.02395 | -0.2226936 | 0.16830429 | -1.3231606 | 0.18578199 | 0.76908034 |
| AC017083.2 | 979.531029 | -0.2089541 | 0.18595131 | -1.1237035 | 0.26113885 | 0.82424998 |
| CNRIP1 | 525.922085 | -0.039527 | 0.34475105 | -0.1146537 | 0.90871961 | 0.98662757 |
| PLEK | 531.338811 | -0.1503203 | 0.60826256 | -0.2471305 | 0.8048072 | 0.97300739 |
| FBXO48 | 188.383542 | 0.25786429 | 0.22443353 | 1.14895617 | 0.25057405 | 0.81738655 |
| APLF | 168.15889 | 0.15269732 | 0.26886784 | 0.56792706 | 0.5700845 | 0.92748817 |
| ARHGAP25 | 613.55407 | -0.2158658 | 0.30162699 | -0.7156713 | 0.47419433 | 0.90199039 |
| ANTXR1 | 1394.6789 | 0.36723038 | 0.41956055 | 0.87527384 | 0.38142493 | 0.87184834 |
| GFPT1 | 4043.76872 | 0.04669001 | 0.24853521 | 0.18786075 | 0.85098581 | 0.97844748 |
| NFU1 | 1519.95319 | 0.2285867 | 0.15271625 | 1.49680658 | 0.13444359 | 0.71630293 |
| AAK1 | 3338.74097 | -0.1282096 | 0.15925177 | -0.8050749 | 0.42077644 | 0.88451372 |
| ANXA4 | 9995.93261 | 0.45107256 | 0.294718 | 1.53052261 | 0.12588742 | 0.70347499 |
| GMCL1 | 1221.83308 | -0.055082 | 0.19042797 | -0.2892535 | 0.77238736 | 0.96836854 |
| SNRNP27 | 1268.24231 | 0.07976804 | 0.11226811 | 0.71051376 | 0.4773856 | 0.90379671 |
| MXD1 | 1859.49238 | -0.4570846 | 0.21079863 | -2.1683473 | 0.03013228 | 0.54012717 |
| ASPRV1 | 158.841591 | 0.15931741 | 0.22929093 | 0.69482649 | 0.48716404 | 0.90573694 |
| PCBP1 | 16558.9593 | -0.1605501 | 0.18135652 | -0.8852734 | 0.37600921 | 0.86900834 |
| C2orf42 | 648.694634 | -0.0407072 | 0.17365674 | -0.2344121 | 0.81466509 | 0.9738447 |
| TIA1 | 4693.09014 | -0.1190791 | 0.26776773 | -0.4447106 | 6.57E-01 | 9.45E-01 |
| PCYOX1 | 5965.07204 | 0.68600887 | 0.3287976 | 2.08641692 | 0.03694087 | 0.55521179 |
| SNRPG | 1669.74908 | -0.1071301 | 0.1839286 | -0.5824551 | 0.5602602 | 0.92434754 |
| FAM136A | 2519.61351 | -0.0612822 | 0.19499201 | -0.3142805 | 0.75330804 | 0.96301881 |
| AC022201.2 | 221.497125 | 0.06848461 | 0.19857917 | 0.34487311 | 0.73018977 | 0.95848028 |
| TGFA | 1487.62882 | 0.30446588 | 0.21324611 | 1.42776756 | 0.15335877 | 0.74258845 |
| VAX2 | 4222.12075 | 2.40077334 | 0.51835658 | 4.63150933 | 3.63E-06 | 0.00506488 |
| ATP6V1B1 | 4373.62207 | 2.4160527 | 0.51987507 | 4.64737175 | 3.36E-06 | 0.00506488 |
| AC007040.2 | 483.545923 | 0.11890168 | 0.2591304 | 0.45884883 | 0.64634273 | 0.94208859 |
| ANKRD53 | 79.2569029 | -0.9222716 | 0.46362594 | -1.9892579 | 0.04667274 | 0.58047001 |
| TEX261 | 4363.0455 | 0.04943302 | 0.13456191 | 0.36736267 | 0.71334851 | 0.95564093 |
| AC007040.1 | 77.9580498 | -0.1019013 | 0.29660663 | -0.3435572 | 0.73117931 | 0.95848028 |
| NAGK | 3496.49025 | -0.1881656 | 0.15267209 | -1.2324818 | 0.21776916 | 0.79648942 |
| MCEE | 707.632164 | 0.32465108 | 0.13129833 | 2.47262149 | 0.01341261 | 0.42532276 |
| MPHOSPH10 | 1199.1413 | -0.1704924 | 0.23500608 | -0.7254807 | 0.46815719 | 0.89984974 |
| PAIP2B | 1341.43958 | -0.1649822 | 0.47777394 | -0.3453144 | 0.72985802 | 0.95848028 |
| ZNF638 | 4609.52109 | -0.0694417 | 0.13208333 | -0.5257416 | 0.59906778 | 0.93484879 |
| DYSF | 1831.06144 | -0.2666114 | 0.27069238 | -0.984924 | 3.25E-01 | 0.85169085 |
| CYP26B1 | 1782.13829 | -1.9745102 | 0.60025006 | -3.2894794 | 0.00100373 | 0.15012259 |
| EXOC6B | 1141.82525 | -0.1649774 | 0.11685559 | -1.4118061 | 0.15800705 | 0.74574881 |
| SPR | 2518.86637 | 0.38861569 | 0.21976366 | 1.76833461 | 0.07700498 | 0.63806922 |
| EMX1 | 3775.89943 | 0.80298304 | 0.3080303 | 2.60683133 | 0.00913844 | 0.38664841 |
| AC012366.1 | 1051.36476 | 0.77930566 | 0.31257102 | 2.49321149 | 0.01265934 | 0.41598747 |
| SFXN5 | 4270.803 | 0.28163853 | 0.30196538 | 0.93268481 | 0.35098273 | 0.86364633 |
| RAB11FIP5 | 8995.79131 | 0.1470373 | 0.30388596 | 0.48385685 | 0.62848747 | 0.94088547 |
| SMYD5 | 1831.10096 | -0.0263207 | 0.19360627 | -0.1359496 | 0.89186115 | 0.98474725 |
| PRADC1 | 997.214293 | 0.5204557 | 0.2943632 | 1.76807323 | 0.07704866 | 0.63806922 |
| CCT7 | 8353.1369 | 0.17044109 | 0.16767632 | 1.01648869 | 0.30939673 | 0.84414094 |
| FBXO41 | 236.236146 | 0.4496557 | 0.45943932 | 0.9787053 | 0.32772561 | 0.85317073 |
| ALMS1 | 575.690465 | 0.15824092 | 0.18921696 | 0.83629357 | 0.40298977 | 0.88132015 |
| NAT8 | 18648.0735 | 1.29818071 | 0.59140104 | 2.19509372 | 0.02815688 | 0.53300785 |
| TPRKB | 928.064588 | -0.0043564 | 0.1516655 | -0.0287237 | 0.97708494 | 0.99455099 |

| | | | | | | |
|---|---|---|---|---|---|---|
| DUSP11 | 1060.14206 | -0.1069401 | 0.12694497 | -0.842413 | 0.3995568 | 0.88077967 |
| STAMBP | 1934.45066 | 0.27643963 | 0.15493872 | 1.78418692 | 0.0743933 | 0.63773415 |
| ACTG2 | 842.354476 | -3.2622307 | 1.65957538 | -1.965702 | NA | NA |
| DGUOK | 1710.46782 | -0.1285001 | 0.11217029 | -1.1455802 | 0.2519689 | 0.81833349 |
| DGUOK-AS1 | 709.71124 | -0.1216456 | 0.13387013 | -0.9086837 | 3.64E-01 | 8.66E-01 |
| TET3 | 602.968538 | -0.4445818 | 0.20685661 | -2.149227 | 0.03161641 | 0.54507414 |
| BOLA3 | 631.212578 | 0.37569582 | 0.20519237 | 1.83094441 | 0.06710884 | 0.6249306 |
| BOLA3-AS1 | 96.1220321 | -0.2289424 | 0.31314639 | -0.7311033 | 0.46471604 | 0.89869398 |
| MOB1A | 6010.33968 | -0.0744309 | 0.13964837 | -0.532988 | 0.59404186 | 0.93317501 |
| MTHFD2 | 1627.66999 | -0.3631217 | 0.38517518 | -0.9427444 | 0.34581167 | 0.86111349 |
| AC006030.1 | 3432.00915 | -0.0627084 | 0.1448333 | -0.4329696 | 0.66503688 | 0.94692723 |
| SLC4A5 | 323.538472 | -0.2499521 | 0.32139808 | -0.7777026 | 0.43674434 | 0.89122667 |
| DCTN1 | 6947.11349 | 0.03493867 | 0.08448533 | 0.41354714 | 0.6792058 | 0.95070301 |
| DCTN1-AS1 | 101.165199 | 0.3491935 | 0.36139885 | 0.96622748 | 0.33393037 | 0.85533716 |
| AC005041.1 | 432.047086 | 0.32731524 | 0.4124205 | 0.79364444 | 0.42740245 | 0.88770346 |
| C2orf81 | 448.606815 | 0.27303877 | 0.39523813 | 0.69082091 | 0.48967809 | 0.90600166 |
| WDR54 | 1080.59723 | 0.11334235 | 0.2393571 | 0.47352826 | 0.63583634 | 0.94088547 |
| RTKN | 6537.21553 | -0.0288551 | 0.16178357 | -0.1783563 | 0.85844313 | 0.97982623 |
| INO80B | 1538.50955 | -0.1341536 | 0.1414301 | -0.9485502 | 0.34284941 | 0.85923253 |
| INO80B-WBI | 5266.97195 | 0.15621736 | 0.15341544 | 1.01826364 | 0.30855268 | 0.84392671 |
| WBP1 | 4592.92737 | 0.25398659 | 0.18581709 | 1.36686346 | 1.72E-01 | 0.75974924 |
| MOGS | 5085.30071 | -0.1598631 | 0.14262126 | -1.1208924 | 2.62E-01 | 0.82424998 |
| MRPL53 | 2045.84646 | 0.09421159 | 0.14733363 | 0.63944389 | 5.23E-01 | 0.91723878 |
| CCDC142 | 2297.47308 | 0.04479126 | 0.14563639 | 0.30755545 | 0.75842063 | 0.96502715 |
| TTC31 | 1711.29014 | -0.05521 | 0.18431838 | -0.2995363 | 0.76453089 | 0.96678847 |
| LBX2 | 254.1339 | -0.1696188 | 0.38601426 | -0.4394108 | 0.66036388 | 0.94628868 |
| LBX2-AS1 | 649.10388 | -0.3710374 | 0.4131707 | -0.8980244 | 0.36917256 | 0.86762749 |
| PCGF1 | 975.219187 | -0.2163263 | 0.22573903 | -0.9583026 | 0.33791021 | 0.8571036 |
| AUP1 | 8646.44862 | -0.2227718 | 0.16673218 | -1.3361055 | 0.18151479 | 0.76522169 |
| HTRA2 | 3192.06495 | -0.2542235 | 0.16512694 | -1.5395639 | 0.12366668 | 0.69957382 |
| LOXL3 | 1142.44749 | -0.1809056 | 0.19462955 | -0.9294868 | 0.35263685 | 0.86364633 |
| DOK1 | 562.617331 | 0.14329042 | 0.2406693 | 0.59538303 | 0.55158747 | 0.92373235 |
| M1AP | 83.2767728 | 0.42468381 | 0.51818616 | 0.81955839 | 0.4124679 | 0.88268208 |
| SEMA4F | 445.774215 | 0.14863059 | 0.31229171 | 0.47593512 | 0.6341206 | 0.94088547 |
| HK2 | 692.65776 | 0.38364679 | 0.55578498 | 0.69027916 | 0.49001865 | 0.90600166 |
| LINC01291 | 112.471198 | -0.0334968 | 1.01036083 | -0.0331533 | 0.97355233 | 0.99445887 |
| POLE4 | 1068.51577 | -0.0058944 | 0.20311087 | -0.0290206 | 0.97684814 | 0.99455099 |
| TACR1 | 154.035805 | 0.84284487 | 0.67566198 | 1.24743569 | 0.21223779 | 0.79203904 |
| AC007681.1 | 54.2085179 | -0.1575548 | 0.67621795 | -0.232994 | 0.81576604 | 0.9738447 |
| EVA1A | 2129.89161 | 0.01407874 | 0.42453471 | 0.03316275 | 0.9735448 | 0.99445887 |
| MRPL19 | 2357.35629 | -0.0100624 | 0.19948622 | -0.0504414 | 0.95977064 | 0.99255196 |
| GCFC2 | 1431.30708 | 0.13277331 | 0.09293284 | 1.42870176 | 0.15308997 | 0.74258845 |
| AC005034.3 | 371.193182 | 0.20004896 | 0.27329275 | 0.73199512 | 0.46417155 | 0.89830907 |
| REG3G | 92.9636968 | 1.44700475 | 1.35780497 | 1.06569411 | 0.28656194 | 0.83146886 |
| REG1B | 76.5527986 | 3.91397252 | 1.73860094 | 2.25121961 | NA | NA |
| REG1A | 2811.90675 | 1.77854747 | 1.61359421 | 1.10222723 | 0.2703629 | 0.82716979 |
| LRRTM1 | 70.0832136 | -3.5870265 | 1.26697873 | -2.8311655 | NA | NA |
| SUCLG1 | 13628.9616 | 0.43830999 | 0.26965818 | 1.62542816 | 0.10407136 | 0.67439401 |
| DNAH6 | 159.030781 | 0.05448626 | 0.69196276 | 0.07874161 | 0.93723815 | 0.99077727 |
| TRABD2A | 458.032737 | -0.0134041 | 0.35575246 | -0.0376782 | 0.96994422 | 0.99400331 |
| TMSB10 | 21204.4736 | -0.0673711 | 0.28821947 | -0.2337492 | 0.81517968 | 0.9738447 |
| KCMF1 | 2731.65975 | -0.2793784 | 0.28890581 | -0.9670225 | 0.33353281 | 0.855011 |
| TCF7L1 | 1351.06696 | -0.1036299 | 0.29102915 | -0.3560808 | 0.72178004 | 0.95739595 |
| TGOLN2 | 15056.8023 | 0.29546111 | 0.16998803 | 1.73812894 | 8.22E-02 | 6.46E-01 |

| | | | | | | |
|---|---|---|---|---|---|---|
| RETSAT | 7371.65312 | 0.16274488 | 0.25698193 | 0.6332931 | 5.27E-01 | 9.18E-01 |
| ELMOD3 | 1818.69806 | 0.23334408 | 0.1347052 | 1.73225734 | 0.08322775 | 0.6471966 |
| CAPG | 6216.12254 | 0.1251271 | 0.27572087 | 0.45381801 | 0.64995982 | 0.94312988 |
| SH2D6 | 84.4346442 | 0.29651493 | 0.53424101 | 0.5550209 | 0.57888037 | 0.93139743 |
| MAT2A | 16545.5414 | -0.2995224 | 0.26025781 | -1.150868 | 0.24978656 | 0.816687 |
| GGCX | 7239.76108 | -0.1394797 | 0.13953156 | -0.9996282 | 0.31749045 | 0.84819508 |
| VAMP8 | 7111.45943 | 0.0291272 | 0.17327025 | 0.1681027 | 0.86650249 | 0.98210602 |
| VAMP5 | 1393.83868 | -0.2615514 | 0.19230679 | -1.3600735 | 0.17380666 | 0.75974924 |
| RNF181 | 3413.17271 | 0.19096313 | 0.16859424 | 1.13267888 | 0.25734913 | 0.82115501 |
| TMEM150A | 3377.17912 | 0.07884475 | 0.20509694 | 0.38442675 | 0.70066218 | 0.95360275 |
| USP39 | 2194.96377 | -0.018289 | 0.15387426 | -0.1188569 | 0.90538877 | 0.98625102 |
| C2orf68 | 1733.87732 | -0.2795579 | 0.17626666 | -1.5859942 | 0.11274064 | 0.68098515 |
| SFTPB | 101.073614 | 0.57629498 | 0.80731561 | 0.713841 | 0.47532551 | 0.90226631 |
| GNLY | 377.122138 | 0.13796052 | 0.8993365 | 0.15340255 | 0.87808083 | 0.98275231 |
| ATOH8 | 814.022335 | -0.3925715 | 0.35517712 | -1.1052839 | 0.26903661 | 0.82683094 |
| ST3GAL5 | 446.589158 | 0.01873808 | 0.37440287 | 0.05004791 | 0.96008421 | 0.99255196 |
| POLR1A | 1837.84612 | -0.1478082 | 0.12991375 | -1.1377412 | 0.25522857 | 0.82017125 |
| PTCD3 | 3514.08745 | 0.08852483 | 0.12645752 | 0.70003612 | 0.48390474 | 0.90541935 |
| IMMT | 6234.9415 | 0.24997682 | 0.12125797 | 2.06152905 | 0.03925259 | 0.55657615 |
| MRPL35 | 2136.43558 | 0.17229118 | 0.1496328 | 1.15142659 | 0.24955678 | 0.81646572 |
| REEP1 | 299.889684 | -0.7670562 | 0.53495821 | -1.4338619 | 0.15161167 | 0.73993604 |
| KDM3A | 3166.17014 | -0.084107 | 0.2486299 | -0.3382821 | 0.73515063 | 0.95895327 |
| CHMP3 | 4728.42396 | 0.24549908 | 0.16379254 | 1.49884166 | 0.13391471 | 0.71521677 |
| RNF103-CHM | 2628.5012 | 0.2072443 | 0.17473082 | 1.18607751 | 0.23559169 | 0.80747753 |
| AC015971.1 | 1452.35218 | 0.01180846 | 0.2099869 | 0.05623429 | 0.95515517 | 0.99185043 |
| RNF103 | 2291.77401 | -0.199039 | 0.22737927 | -0.8753613 | 0.38137735 | 0.87184592 |
| RMND5A | 2862.45551 | -0.3339299 | 0.24457462 | -1.3653497 | 1.72E-01 | 7.60E-01 |
| CD8A | 262.40851 | 0.16041286 | 0.86262852 | 0.18595821 | 0.85247753 | 0.97883015 |
| CD8B | 63.1420975 | 0.32669672 | 0.9303195 | 0.35116615 | 0.7254637 | 0.95848028 |
| PLGLB1 | 75.816303 | 0.51259601 | 0.28526552 | 1.79690844 | 0.07235016 | 0.63220999 |
| LINC01943 | 208.55957 | -0.032444 | 0.44883679 | -0.0722846 | 0.94237542 | 0.99136103 |
| CYTOR | 386.786062 | 0.01561641 | 0.43130755 | 0.03620713 | 0.9711172 | 0.99400331 |
| RGPD2 | 60.0425732 | 0.15695867 | 0.68326479 | 0.22971866 | 0.8183104 | 0.97454841 |
| KRCC1 | 1621.77032 | 0.46223733 | 0.15842798 | 2.91764968 | 0.0035268 | 0.26021977 |
| FABP1 | 5012.4692 | 2.98627641 | 0.85191135 | 3.50538402 | 0.00045595 | 0.10420661 |
| THNSL2 | 2868.88076 | 0.77474987 | 0.64761202 | 1.19631792 | 0.23157252 | 0.80649152 |
| AC104134.1 | 352.662482 | -0.3144832 | 0.40704857 | -0.7725937 | 0.43976287 | 0.89203402 |
| EIF2AK3 | 1401.16378 | -0.2972972 | 0.31538654 | -0.942644 | 3.46E-01 | 8.61E-01 |
| AC062029.1 | 88.0712592 | 0.91701631 | 0.42503641 | 2.15750058 | 0.03096668 | 0.54466273 |
| RPIA | 767.497294 | -0.2357745 | 0.1962646 | -1.2013094 | 0.22963119 | 0.80551898 |
| IGKC | 19683.8626 | 0.9575181 | 1.24598534 | 0.76848263 | 0.4422005 | 0.89331461 |
| IGKJ5 | 382.576587 | 1.30932581 | 1.28724898 | 1.0171504 | 0.30908189 | 0.84414094 |
| IGKJ4 | 664.425121 | 0.48036741 | 0.99056002 | 0.48494528 | 0.62771516 | 0.94088547 |
| IGKJ3 | 632.725689 | 0.58950107 | 1.4499222 | 0.40657428 | 0.68432069 | 0.95070301 |
| IGKJ2 | 641.29505 | 0.82778291 | 1.1780807 | 0.70265383 | 0.48227151 | 0.90541935 |
| IGKJ1 | 1100.53355 | 1.25586731 | 1.20530479 | 1.04194999 | 0.29743487 | 0.83898658 |
| IGKV4-1 | 441.127706 | 0.32664133 | 0.98863003 | 0.33039795 | 0.74109929 | 0.96097294 |
| IGKV1-5 | 749.273471 | -0.0573645 | 1.14505407 | -0.0500976 | 0.96004459 | 0.99255196 |
| IGKV1-6 | 209.298367 | 2.76611639 | 1.23129049 | 2.24651811 | 0.02467084 | 0.52067143 |
| IGKV1-9 | 218.451876 | 1.03272295 | 1.11405181 | 0.92699724 | 0.35392797 | 0.86364633 |
| IGKV3-11 | 743.818652 | 1.69294831 | 1.24507818 | 1.35971245 | 0.17392094 | 0.75974924 |
| IGKV1-12 | 177.646056 | 0.91656579 | 1.16825069 | 0.78456259 | 0.43271007 | 0.88993419 |
| IGKV3-15 | 630.009559 | 0.20935845 | 1.38822882 | 0.15080976 | 0.88012579 | 0.98275231 |
| IGKV1-16 | 228.404862 | 0.93159699 | 1.57224463 | 0.59252675 | 0.55349792 | 0.92373235 |

| | | | | | | |
|---|---|---|---|---|---|---|
| IGKV1-17 | 357.995437 | 3.33638284 | 1.31264633 | 2.54172259 | 0.01103077 | 0.40196898 |
| IGKV3-20 | 2584.75541 | 0.285337 | 1.46051395 | 0.19536753 | 0.84510524 | 0.97675487 |
| IGKV2-24 | 528.776264 | 4.43312941 | 1.7546347 | 2.52652555 | 0.0115197 | 0.40353404 |
| IGKV1-27 | 154.790907 | 1.49264182 | 1.03140624 | 1.447191 | 1.48E-01 | 0.73455299 |
| IGKV2-28 | 190.838227 | 1.04203546 | 0.99475496 | 1.04752979 | 0.2948553 | 0.83630273 |
| IGKV2-30 | 164.174851 | 0.83270275 | 1.15908107 | 0.71841631 | 0.47250063 | 0.90160919 |
| IGKV1-33 | 151.018827 | 0.2973516 | 1.19514978 | 0.24879861 | 0.80351657 | 0.97274243 |
| IGKV1-39 | 351.34627 | 0.97879776 | 1.13240999 | 0.86434928 | 0.38739605 | 0.87460898 |
| IGKV2D-29 | 54.2756591 | 1.7350329 | 1.60913075 | 1.07824233 | 0.28092563 | 0.82937461 |
| IGKV3D-20 | 103.02034 | 0.6189722 | 1.50515427 | 0.41123506 | NA | NA |
| IGKV3D-15 | 101.82992 | 1.64507151 | 1.91070055 | 0.8609782 | NA | NA |
| AL845331.2 | 74.0420098 | -0.6462262 | 0.70557671 | -0.9158837 | 3.60E-01 | 0.86480228 |
| AC103563.7 | 82.0113004 | 0.93945746 | 0.58693155 | 1.60062524 | 1.09E-01 | 6.79E-01 |
| MAL | 9348.06807 | 0.6906114 | 0.47146114 | 1.46483208 | 0.14296674 | 0.72784886 |
| AC103563.2 | 200.774736 | 0.51078869 | 0.50503879 | 1.01138505 | 0.31183217 | 0.84537057 |
| MRPS5 | 4043.10262 | -0.082837 | 0.14767636 | -0.560936 | 0.57484114 | 0.93024927 |
| ZNF514 | 1109.67115 | 0.10757972 | 0.27282963 | 0.39431098 | 0.69335146 | 0.9519683 |
| ZNF2 | 315.117799 | 0.03658458 | 0.15832402 | 0.23107409 | 0.81725725 | 0.97419018 |
| PROM2 | 5735.12912 | 0.37777121 | 0.42583555 | 0.88712935 | 0.37500926 | 0.86819624 |
| KCNIP3 | 1337.87783 | -0.1082776 | 0.3976727 | -0.2722782 | 0.78540811 | 0.97054816 |
| FAHD2A | 1767.43335 | 0.24198086 | 0.30937009 | 0.78217277 | 0.43411304 | 0.89055338 |
| AC009237.14 | 303.076283 | 0.0624375 | 0.36692322 | 0.17016502 | 0.86488036 | 0.98197545 |
| LINC00342 | 827.450132 | 0.21623103 | 0.34182556 | 0.63257713 | 5.27E-01 | 9.18E-01 |
| ANKRD36C | 61.8411754 | -0.3557896 | 0.4377053 | -0.812852 | 0.41630295 | 0.88399703 |
| GPAT2 | 440.700133 | 0.39316563 | 0.42255494 | 0.93044855 | 0.35213889 | 0.86364633 |
| ADRA2B | 445.482797 | -0.2274468 | 0.52124377 | -0.4363539 | 0.66257998 | 0.94642611 |
| DUSP2 | 554.168854 | 0.19063838 | 0.91768068 | 0.20773934 | 0.8354325 | 0.97550416 |
| STARD7 | 7580.90768 | 0.1074719 | 0.20392512 | 0.52701649 | 0.59818214 | 0.9346978 |
| TMEM127 | 6210.21981 | -0.1943171 | 0.14770601 | -1.3155669 | 0.18831946 | 0.76970626 |
| CIAO1 | 4327.57969 | -0.1154612 | 0.19527016 | -0.5912895 | 0.55432648 | 0.92373235 |
| SNRNP200 | 7120.17695 | -0.0978734 | 0.12733408 | -0.7686344 | 0.44211039 | 0.89331461 |
| ITPRIPL1 | 52.0114015 | -0.1128039 | 0.61845582 | -0.1823961 | 0.85527192 | 0.97917572 |
| NEURL3 | 760.480611 | -0.7575327 | 0.54413665 | -1.3921737 | 0.1638698 | 0.75265041 |
| ARID5A | 2104.73318 | -0.8872599 | 0.3946882 | -2.2480022 | 0.02457605 | 0.52017724 |
| KANSL3 | 4145.88823 | -0.2803031 | 0.14866568 | -1.885459 | 0.05936789 | 0.60533169 |
| FER1L5 | 102.017784 | -0.0256734 | 0.45458486 | -0.0564766 | 0.9549621 | 0.99185043 |
| LMAN2L | 1906.26573 | 0.21615276 | 0.25103815 | 0.86103549 | 0.3892185 | 0.87568585 |
| CNNM4 | 1196.29003 | -0.2171317 | 0.25802806 | -0.8415043 | 0.40006546 | 0.88077967 |
| CNNM3-DT | 45.7755976 | 0.27928912 | 0.31956436 | 0.8739683 | 0.38213552 | 0.87185666 |
| CNNM3 | 1716.35688 | -0.0779036 | 0.12063984 | -0.6457537 | 0.51843887 | 0.91636259 |
| ANKRD23 | 1012.88887 | -0.0589099 | 0.12154765 | -0.484665 | 0.62791401 | 0.94088547 |
| ANKRD39 | 628.717895 | -0.0165809 | 0.14375466 | -0.1153414 | 0.90817453 | 0.98662757 |
| SEMA4C | 3220.66368 | 0.1065279 | 0.27944629 | 0.38121063 | 0.70304696 | 0.95431462 |
| FAHD2B | 478.377795 | 0.68402609 | 0.38779172 | 1.76390068 | 0.0777487 | 0.64069017 |
| ANKRD36 | 135.308511 | -0.229764 | 0.37287513 | -0.6161956 | 0.53776541 | 0.92123049 |
| ANKRD36B | 40.5968811 | -0.2408622 | 0.56864004 | -0.4235758 | 0.67187521 | 0.94922827 |
| COX5B | 9082.48481 | 0.17465526 | 0.1627906 | 1.07288294 | 0.28332364 | 0.83094967 |
| ACTR1B | 5117.47733 | -0.043928 | 0.14469933 | -0.3035813 | 0.76144689 | 0.96606476 |
| C2orf92 | 160.883417 | -0.3452037 | 0.29900714 | -1.1545 | 0.24829524 | 0.8162591 |
| ZAP70 | 467.720654 | 0.11699439 | 0.7168094 | 0.16321548 | 0.87034877 | 0.98275231 |
| TMEM131 | 2746.35826 | -0.0210729 | 0.13775792 | -0.1529703 | 8.78E-01 | 9.83E-01 |
| INPP4A | 1125.98234 | -0.0986897 | 0.14718486 | -0.6705153 | 0.50252937 | 0.91319792 |
| COA5 | 1755.95201 | 0.16502031 | 0.14138496 | 1.1671702 | 0.24314164 | 0.81340905 |
| UNC50 | 2208.99547 | 0.0278224 | 0.10691219 | 0.26023602 | 0.79468172 | 0.97101097 |

| | | | | | | |
|---|---|---|---|---|---|---|
| MGAT4A | 3803.6782 | 0.05746544 | 0.29562082 | 0.19438903 | 0.84587128 | 0.9767918 |
| KIAA1211L | 427.879391 | 0.20288322 | 0.33246987 | 0.61023039 | 0.5417092 | 0.92292765 |
| TSGA10 | 435.956033 | 0.33053348 | 0.3183181 | 1.03837474 | 0.29909562 | 0.84082654 |
| C2orf15 | 209.828405 | 0.11102459 | 0.25390756 | 0.4372638 | 0.66192005 | 0.94642611 |
| AC079447.1 | 1103.87049 | 0.1109101 | 0.14026166 | 0.79073707 | 0.42909744 | 0.88851358 |
| LIPT1 | 398.24565 | 0.07346851 | 0.21594208 | 0.34022322 | 0.73368843 | 0.95895327 |
| AC092587.1 | 429.014704 | 0.17831807 | 0.16090167 | 1.1082425 | 0.2677571 | 0.82676682 |
| MITD1 | 1418.13061 | -0.1097622 | 0.15857517 | -0.6921777 | 0.48882576 | 0.905872 |
| MRPL30 | 1454.12881 | 0.1737427 | 0.09483735 | 1.83200725 | 6.70E-02 | 0.6249306 |
| LYG1 | 1717.04782 | 0.10709295 | 0.44409375 | 0.24114941 | 0.80943932 | 0.97334138 |
| TXNDC9 | 1674.15078 | -0.03272 | 0.21183036 | -0.1544631 | 8.77E-01 | 0.98275231 |
| EIF5B | 5290.11908 | -0.2366918 | 0.16144955 | -1.4660418 | 0.1426369 | 0.72784886 |
| REV1 | 2215.57732 | -0.2086147 | 0.11788922 | -1.7695823 | 0.07679675 | 0.63773415 |
| AC018690.1 | 45.454923 | -0.4283624 | 0.55370576 | -0.7736283 | 0.43915063 | 0.89175645 |
| AFF3 | 241.237634 | -0.0693062 | 0.43058707 | -0.1609575 | 0.87212685 | 0.98275231 |
| LONRF2 | 564.931369 | 0.26825642 | 0.37791404 | 0.7098345 | 4.78E-01 | 0.90387017 |
| CHST10 | 397.159599 | 0.02300699 | 0.27174007 | 0.08466542 | 0.93252739 | 0.9899349 |
| PDCL3 | 1046.50596 | -0.1002259 | 0.30082328 | -0.3331722 | 0.73900434 | 0.9602325 |
| NPAS2 | 3240.04019 | -0.1899966 | 0.3677142 | -0.5166963 | 0.60536817 | 0.93692336 |
| AC016738.2 | 153.066882 | -0.1955241 | 0.35272076 | -0.5543312 | 0.57935222 | 0.93139743 |
| AC016738.1 | 1123.66234 | -0.1183941 | 0.40279093 | -0.2939343 | 0.76880813 | 0.96796999 |
| RPL31 | 19604.6591 | 0.14277753 | 0.16598927 | 0.86016119 | 0.38970019 | 0.87580901 |
| TBC1D8 | 2258.98674 | -0.1869072 | 0.23180893 | -0.8062987 | 0.42007063 | 0.88451372 |
| TBC1D8-AS1 | 49.7542965 | 0.49964838 | 0.46318416 | 1.07872509 | 0.2807103 | 0.82912399 |
| CNOT11 | 3237.82471 | -0.363716 | 0.20200262 | -1.8005507 | 0.07177372 | 0.63189503 |
| RNF149 | 3889.75322 | -0.3595944 | 0.17222088 | -2.0879838 | 0.0367993 | 0.55521179 |
| RFX8 | 47.8970831 | -0.1727818 | 0.79980487 | -0.2160299 | 0.82896444 | 0.97550416 |
| MAP4K4 | 3361.22685 | -0.28463 | 0.35599638 | -0.7995307 | 0.42398275 | 0.88565404 |
| LINC01127 | 121.413575 | -1.5685228 | 0.54235171 | -2.8920768 | 0.00382704 | 0.26845232 |
| IL1R2 | 1066.70169 | -1.4943203 | 0.47132376 | -3.1704753 | 0.0015219 | 0.17279992 |
| IL1R1 | 5004.87026 | -0.4049907 | 0.24713558 | -1.6387389 | 0.10126764 | 0.6725344 |
| IL1RL2 | 262.28833 | -0.6101399 | 0.28234961 | -2.1609376 | 0.03070016 | 0.54443002 |
| IL1RL1 | 13373.3902 | 0.88924935 | 0.84512403 | 1.05221164 | 0.29270246 | 0.83524862 |
| IL18R1 | 587.145101 | 0.34085395 | 0.35168202 | 0.96921061 | 0.33244012 | 0.85498288 |
| IL18RAP | 130.450848 | -0.3382333 | 0.41168544 | -0.8215818 | 0.41131496 | 0.88268208 |
| SLC9A4 | 598.579403 | 0.63150803 | 0.46741269 | 1.35107163 | 0.17667249 | 0.76115956 |
| SLC9A2 | 370.666276 | 0.95116409 | 0.33990199 | 2.79834812 | 0.00513647 | 0.30843846 |
| MFSD9 | 726.151631 | -0.1497774 | 0.31050961 | -0.48236 | 0.62955024 | 0.94088547 |
| TMEM182 | 167.369897 | 0.67827819 | 0.28225978 | 2.4030281 | 0.01625994 | 0.45896904 |
| POU3F3 | 5784.93032 | -0.2785748 | 0.28782972 | -0.9678459 | 0.33312132 | 0.855011 |
| AC018730.1 | 59.549903 | 0.34773361 | 0.58349302 | 0.59595162 | 0.55120755 | 0.92373235 |
| MRPS9 | 1833.76308 | 0.17631095 | 0.16186269 | 1.08926245 | 0.27603817 | 0.82719407 |
| AC104655.1 | 754.791667 | 0.17363691 | 0.16875044 | 1.0289568 | 0.30349998 | 0.84211658 |
| TGFBRAP1 | 1605.31912 | -0.2331936 | 0.20181249 | -1.1554966 | 0.24788713 | 0.81602355 |
| AC012360.3 | 242.991131 | -0.6469097 | 0.30365041 | -2.1304424 | 0.03313511 | 0.5514965 |
| C2orf49 | 986.683309 | -0.0868056 | 0.15278031 | -0.5681728 | 0.56991766 | 0.9273161 |
| FHL2 | 2208.70049 | 0.22523038 | 0.36414021 | 0.61852654 | 0.53622831 | 0.92102241 |
| NCK2 | 2578.12464 | -0.3321989 | 0.12483514 | -2.6611012 | 0.00778855 | 0.36878977 |
| AC010978.1 | 417.895485 | -0.2535379 | 0.17148359 | -1.4784964 | 0.13927497 | 0.72409321 |
| C2orf40 | 543.255279 | 0.49797329 | 0.4710796 | 1.05708947 | 0.29047076 | 0.83442894 |
| UXS1 | 3671.96055 | -0.0744403 | 0.24879626 | -0.2992019 | 0.76478599 | 0.96678847 |
| ST6GAL2 | 54.8633502 | 0.15893163 | 0.5363464 | 0.29632272 | 0.76698364 | 0.96742767 |
| SLC5A7 | 44.6442368 | 0.88069207 | 0.72368955 | 1.21694733 | 0.22362426 | 0.80078694 |
| LINC01594 | 60.7304908 | 0.62231618 | 0.55349526 | 1.12433877 | 0.26086934 | 0.82424999 |

| | | | | | | |
|---|---|---|---|---|---|---|
| SULT1C2 | 8092.95499 | 0.08856738 | 0.35312126 | 0.25081295 | 0.80195873 | 0.97233627 |
| SULT1C4 | 829.772754 | 0.89795519 | 0.40730644 | 2.20461821 | 0.0274809 | 0.52904542 |
| GCC2 | 2310.3752 | 0.0692789 | 0.14084898 | 0.49186655 | 0.62281369 | 0.94016713 |
| GCC2-AS1 | 174.499876 | 0.10731224 | 0.2194943 | 0.48890674 | 0.62490772 | 0.94088547 |
| LIMS1 | 2347.24894 | -0.2251622 | 0.26855271 | -0.8384282 | 0.40179028 | 0.88132015 |
| RANBP2 | 4212.83242 | -0.0646056 | 0.20135881 | -0.3208483 | 0.74832534 | 0.96241535 |
| CCDC138 | 134.050141 | -0.2971163 | 0.37624025 | -0.7896983 | 0.42970398 | 0.88870396 |
| EDAR | 111.347365 | 0.94440578 | 0.44363558 | 2.12878729 | 0.03327186 | 0.55181237 |
| SH3RF3-AS1 | 46.6785737 | -0.0814586 | 0.62190783 | -0.1309818 | 0.89578973 | 0.98474725 |
| SH3RF3 | 678.939763 | -0.1129423 | 0.31197995 | -0.3620178 | 0.71733874 | 0.95611001 |
| 10-Sep | 3468.20258 | -0.3610631 | 0.32065105 | -1.1260313 | 0.26015227 | 0.82424998 |
| SOWAHC | 2003.50922 | -0.979621 | 0.51648488 | -1.896708 | 0.05786648 | 0.60129769 |
| AC013271.1 | 91.24785 | -0.7587701 | 0.63204532 | -1.2004996 | 0.22994537 | 0.80551898 |
| LINC01123 | 190.347647 | 0.01682307 | 0.31543549 | 0.05333285 | 0.95746671 | 0.99208733 |
| MALL | 109.60633 | 0.44619315 | 0.40291712 | 1.10740678 | 0.2681181 | 0.82676682 |
| NPHP1 | 785.423936 | 0.24447308 | 0.36681148 | 0.66648155 | 0.50510335 | 0.91365534 |
| SMIM37 | 859.567592 | -0.0522532 | 0.17996405 | -0.2903534 | 0.77154586 | 0.96836854 |
| LINC01106 | 239.256297 | 0.28855028 | 0.28528186 | 1.0114568 | 0.31179785 | 0.84537057 |
| AC112229.3 | 97.7700541 | -0.1402087 | 0.5957478 | -0.2353491 | 0.81393778 | 0.9738447 |
| BUB1 | 57.4288335 | -0.1143382 | 0.78266324 | -0.1460886 | 0.88385144 | 0.98317437 |
| MIR4435-2HG | 1091.83087 | -0.4289548 | 0.3671404 | -1.1683671 | 0.24265873 | 0.81335097 |
| BCL2L11 | 1705.89176 | -0.332261 | 0.250619 | -1.3257615 | 0.1849187 | 0.76820626 |
| ANAPC1 | 1494.30445 | -0.1270628 | 0.11786055 | -1.0780778 | 0.28099902 | 0.82937461 |
| MERTK | 1423.57868 | -0.3031891 | 0.26432665 | -1.1470243 | 0.25137157 | 0.81815815 |
| TMEM87B | 1715.86234 | -0.133876 | 0.22537938 | -0.5940028 | 0.55251024 | 0.92373235 |
| FBLN7 | 111.602398 | -0.9043407 | 0.67296004 | -1.3438252 | 0.17900491 | 0.76120583 |
| ZC3H8 | 656.376699 | -0.0026164 | 0.13044987 | -0.0200564 | 0.98399835 | 0.9961464 |
| ZC3H6 | 655.345235 | 0.45319809 | 0.3146248 | 1.44043981 | 0.14974301 | 0.73734464 |
| RGPD8 | 64.4298487 | -0.1252194 | 0.47758133 | -0.2621949 | 0.79317122 | 0.97090793 |
| TTL | 1835.26108 | -0.0833714 | 0.29663966 | -0.2810529 | 0.77866985 | 0.96836854 |
| POLR1B | 960.244217 | -0.270429 | 0.21136479 | -1.2794418 | 0.20074152 | 0.78134034 |
| CHCHD5 | 943.124722 | 0.35977883 | 0.20847584 | 1.72575796 | 0.08439097 | 0.64988556 |
| AC079922.2 | 91.6964741 | -0.027716 | 0.38425111 | -0.0721299 | 0.94249856 | 0.99136103 |
| SLC20A1 | 3401.26681 | -0.4748936 | 0.3799822 | -1.2497787 | 0.21138041 | 0.79134644 |
| IL1B | 164.826606 | 0.94719519 | 0.79612657 | 1.18975452 | 0.23414289 | 0.80725541 |
| IL1RN | 169.261088 | -1.5260803 | 0.71824918 | -2.1247227 | 0.03360976 | 0.55369025 |
| PSD4 | 2761.95184 | -0.5549001 | 0.25377368 | -2.1865943 | 0.02877215 | 0.53606221 |
| PAX8 | 42315.1657 | -0.2969137 | 0.31990088 | -0.9281427 | 0.35333355 | 0.86364633 |
| CBWD2 | 1040.33631 | -0.0846602 | 0.10435241 | -0.8112916 | 0.41719822 | 0.88445833 |
| RABL2A | 1001.62157 | 0.01304996 | 0.19485187 | 0.06697374 | 0.94660261 | 0.99166614 |
| AC017074.1 | 68.784229 | -0.5720569 | 0.46921476 | -1.2191793 | 2.23E-01 | 0.80078694 |
| SLC35F5 | 4053.78214 | 0.05720779 | 0.19982872 | 0.2862841 | 0.77466054 | 0.96836854 |
| ACTR3 | 8260.4371 | -0.0254807 | 0.20037607 | -0.1271646 | 0.89881015 | 0.98477645 |
| DDX18 | 2506.68068 | -0.1674093 | 0.21339274 | -0.7845125 | 0.43273946 | 0.88993419 |
| AC009404.1 | 173.29967 | -0.2945382 | 0.42532726 | -0.6924978 | 0.48862476 | 0.90585781 |
| CCDC93 | 2652.11626 | -0.3868111 | 0.25524907 | -1.5154262 | 0.12966451 | 0.70814767 |
| INSIG2 | 2182.97305 | -0.0265053 | 0.20444395 | -0.1296459 | 0.8968466 | 0.98474725 |
| MARCO | 393.526555 | -0.0245617 | 0.9704745 | -0.025309 | 0.97980852 | 0.99537078 |
| STEAP3 | 1089.94674 | -0.4615201 | 0.25598504 | -1.8029181 | 0.07140109 | 0.6314902 |
| STEAP3-AS1 | 96.5944965 | -0.4636529 | 0.34847473 | -1.3305208 | 0.18334674 | 0.76732946 |
| C2orf76 | 292.198995 | 0.50019661 | 0.20712008 | 2.41500787 | 0.01573487 | 0.45577618 |
| DBI | 12557.2577 | 0.11462004 | 0.34452495 | 0.3326901 | 0.73936822 | 0.96029564 |
| TMEM37 | 9972.69893 | -0.8607782 | 0.45546632 | -1.8898833 | 0.05877356 | 0.60332684 |
| SCTR | 1196.28448 | -0.1934317 | 0.76933887 | -0.2514259 | 0.80148482 | 0.97210205 |

| | | | | | | |
|---|---|---|---|---|---|---|
| AC013275.1 | 493.576661 | -0.0644264 | 0.69940033 | -0.0921166 | 0.92660536 | 0.98914964 |
| CFAP221 | 1215.23854 | 0.17445066 | 0.48052702 | 0.36304027 | 0.7165748 | 0.95603249 |
| TMEM177 | 921.554748 | -0.1002563 | 0.23731786 | -0.4224559 | 0.67269232 | 0.94922827 |
| PTPN4 | 1150.21444 | -0.1378893 | 0.16531039 | -0.8341237 | 0.4042113 | 0.88132015 |
| EPB41L5 | 3794.02304 | 0.10592973 | 0.18933072 | 0.55949571 | 0.57582346 | 0.93083012 |
| AC012363.1 | 52.3774587 | -0.4787462 | 0.50211046 | -0.9534678 | 0.34035311 | 0.85814041 |
| TMEM185B | 736.687373 | -0.3667614 | 0.27671469 | -1.3254134 | 0.18503407 | 0.76820626 |
| RALB | 5447.67284 | 0.0903221 | 0.26423107 | 0.34182999 | 0.73247884 | 0.95848028 |
| INHBB | 580.985531 | -0.0489823 | 0.5669206 | -0.0864007 | 0.9311479 | 0.98975562 |
| GLI2 | 167.334725 | -0.198537 | 0.40664542 | -0.4882312 | 0.62538608 | 0.94088547 |
| TFCP2L1 | 10962.2178 | 0.3712419 | 0.28508745 | 1.30220359 | 0.1928468 | 0.77506795 |
| CLASP1 | 2514.97158 | -0.08809 | 0.16815771 | -0.5238535 | 0.60038042 | 0.93487648 |
| NIFK-AS1 | 673.301747 | -0.0167873 | 0.17929274 | -0.0936306 | 0.92540263 | 0.98876686 |
| NIFK | 1217.44839 | -0.1967187 | 0.26671203 | -0.7375698 | 0.46077592 | 0.89755243 |
| TSN | 3123.03774 | 0.04180358 | 0.18486173 | 0.22613429 | 0.82109697 | 0.97472105 |
| GYPC | 2113.84897 | -0.1016059 | 0.22227481 | -0.4571185 | 0.6475859 | 0.94282139 |
| BIN1 | 8585.16841 | 0.2499289 | 0.425482 | 0.58740181 | 0.55693388 | 0.92373235 |
| ERCC3 | 2394.55079 | -0.2837704 | 0.13202774 | -2.149324 | 0.03160873 | 0.54507414 |
| MAP3K2 | 2409.86261 | -0.095914 | 0.23481168 | -0.408472 | 0.68292721 | 0.95070301 |
| AC068282.1 | 84.6528632 | 0.37427634 | 0.50059557 | 0.74766211 | 0.45466399 | 0.89670027 |
| PROC | 1883.6603 | -0.0698348 | 0.33916364 | -0.2059031 | 0.83686659 | 0.97577999 |
| IWS1 | 2269.42014 | -0.0827335 | 0.13239318 | -0.6249076 | 0.53203168 | 0.91964636 |
| AC010976.1 | 254.350764 | -0.1426391 | 0.22262044 | -0.6407279 | 0.52169946 | 0.91723878 |
| MYO7B | 3997.73798 | 0.40223594 | 0.48686035 | 0.8261834 | 0.40870006 | 0.88203324 |
| AC010976.2 | 252.678561 | -0.4957348 | 0.44769131 | -1.1073139 | 0.26815825 | 0.82676682 |
| LIMS2 | 3526.92683 | -0.0272768 | 0.39228769 | -0.0695327 | 0.94456557 | 0.99143676 |
| GPR17 | 365.686991 | 0.63253841 | 0.59108088 | 1.0701385 | 0.28455697 | 0.83107779 |
| WDR33 | 4536.0589 | -0.1977438 | 0.12635966 | -1.5649279 | 0.11759981 | 0.69036323 |
| SFT2D3 | 704.498178 | -0.1392492 | 0.21837409 | -0.6376637 | 0.52369261 | 0.91723878 |
| POLR2D | 1277.40218 | -0.2687888 | 0.21025311 | -1.2784059 | 0.20110633 | 0.78134034 |
| AMMECR1L | 1501.28999 | -0.2950645 | 0.16246376 | -1.8161864 | 0.06934178 | 0.62779744 |
| AC012306.2 | 90.0394138 | -0.1869748 | 0.26992083 | -0.6927023 | 0.48849642 | 0.90585781 |
| SAP130 | 1086.26786 | -0.1467272 | 0.19581716 | -0.7493071 | 0.45367213 | 0.89670027 |
| UGGT1 | 2452.76555 | 0.12132634 | 0.16600199 | 0.73087282 | 0.46485684 | 0.89887351 |
| HS6ST1 | 6563.54589 | 0.70746061 | 0.28838486 | 2.45318219 | 0.01415986 | 0.43503269 |
| CCDC74B | 273.444151 | -0.0275321 | 0.56297175 | -0.0489049 | 0.9609951 | 0.99316252 |
| SMPD4 | 5336.70017 | -0.0746938 | 0.12597846 | -0.5929095 | 0.55324172 | 0.92373235 |
| MZT2B | 2807.41653 | 0.12187378 | 0.16835137 | 0.72392509 | 0.46911172 | 0.9003198 |
| CCDC115 | 2549.08162 | 0.47104997 | 0.16317765 | 2.88673087 | 0.00389267 | 0.26919756 |
| IMP4 | 3055.83302 | -0.1095674 | 0.20613486 | -0.5315328 | 0.59504964 | 0.9334601 |
| PTPN18 | 4185.20861 | 0.03038203 | 0.11624214 | 0.2613685 | 7.94E-01 | 0.97090793 |
| ARHGEF4 | 116.793243 | 0.27543572 | 0.33907199 | 0.81232224 | 4.17E-01 | 8.84E-01 |
| FAM168B | 6434.95571 | -0.1786071 | 0.33337004 | -0.5357624 | 5.92E-01 | 0.93289552 |
| PLEKHB2 | 14013.9719 | 0.2569485 | 0.38479234 | 0.66775887 | 0.50428753 | 0.91356552 |
| MZT2A | 1040.78641 | 0.11855645 | 0.21349998 | 0.55529957 | 0.57868977 | 0.93139743 |
| CCDC74A | 457.615493 | 0.19348931 | 0.48243529 | 0.4010679 | 6.88E-01 | 0.95158502 |
| GPR39 | 899.573928 | -0.0647258 | 0.44650136 | -0.1449621 | 0.88474083 | 0.98344768 |
| LYPD1 | 916.498885 | -0.1648205 | 0.48145945 | -0.3423352 | 0.73209862 | 0.95848028 |
| NCKAP5 | 333.871653 | 0.86609113 | 0.54426108 | 1.59131557 | 0.11153858 | 0.68001961 |
| MGAT5 | 2489.20542 | -0.0568296 | 0.2027633 | -0.2802758 | 0.77926593 | 0.96836854 |
| TMEM163 | 309.935324 | -0.0044455 | 0.47997511 | -0.009262 | 9.93E-01 | 0.99906045 |
| CCNT2-AS1 | 973.859587 | 0.87658543 | 0.59770323 | 1.46658976 | 0.14248769 | 0.72784886 |
| ACMSD | 4773.03822 | 0.81672263 | 0.62747988 | 1.30159173 | 0.19305599 | 0.7751938 |
| CCNT2 | 2253.53345 | -0.2224793 | 0.16559451 | -1.3435187 | 0.17910409 | 0.76120583 |

| | | | | | | |
|---|---|---|---|---|---|---|
| RAB3GAP1 | 2859.22804 | -0.1011935 | 0.12396808 | -0.8162866 | 0.41433624 | 0.88364943 |
| ZRANB3 | 388.054347 | 0.05027448 | 0.19031955 | 0.26415827 | 0.79165797 | 0.97090793 |
| R3HDM1 | 967.067067 | 0.15408127 | 0.07853593 | 1.96192082 | 0.0497717 | 0.5861488 |
| UBXN4 | 7082.84379 | -0.1856118 | 0.19526833 | -0.9505472 | 0.3418343 | 0.85889221 |
| MCM6 | 822.24828 | 0.32467413 | 0.24690435 | 1.31497938 | 0.18851683 | 0.76970626 |
| DARS | 4539.48057 | 0.14955222 | 0.17440432 | 0.85750296 | 0.39116698 | 0.87710956 |
| DARS-AS1 | 421.265347 | -0.0218262 | 0.29300929 | -0.0744897 | 0.94062074 | 0.99136103 |
| CXCR4 | 1789.34887 | -0.1880999 | 0.72335366 | -0.2600386 | 0.794834 | 0.97101097 |
| THSD7B | 86.2825353 | 0.26393962 | 0.73560319 | 0.35880706 | 0.71973943 | 0.9566409 |
| HNMT | 3276.57249 | 0.53152814 | 0.3509445 | 1.51456467 | 0.12988268 | 0.70814767 |
| SPOPL | 1241.56945 | -0.1705921 | 0.19630025 | -0.8690365 | 0.38482718 | 0.87312908 |
| NXPH2 | 65.9799546 | -0.2996728 | 0.98647877 | -0.3037803 | 0.7612953 | 0.96603936 |
| LRP1B | 87.6027152 | -0.0229434 | 0.42527982 | -0.053949 | 0.95697584 | 0.99200096 |
| KYNU | 756.797263 | 0.40834827 | 0.41286268 | 0.98906559 | 0.32263105 | 0.85119105 |
| ARHGAP15 | 269.87816 | -0.0544857 | 0.61127382 | -0.0891347 | 0.92897487 | 0.98963127 |
| GTDC1 | 900.921185 | 0.15736198 | 0.15736502 | 0.99998072 | 0.31731984 | 0.84819508 |
| ZEB2 | 2119.29319 | -0.1863129 | 0.24076913 | -0.7738238 | 0.439035 | 0.89175645 |
| ZEB2-AS1 | 90.7735997 | 0.23186316 | 0.42450842 | 0.54619214 | 0.58493387 | 0.93289552 |
| TEX41 | 200.417659 | 0.3331889 | 0.40308672 | 0.8265936 | 0.40846745 | 0.88203324 |
| ACVR2A | 897.774739 | -0.0779048 | 0.10326243 | -0.7544349 | 0.45058814 | 0.89557948 |
| ORC4 | 2150.60293 | 0.0150693 | 0.13605808 | 0.11075638 | 0.91180953 | 0.98662757 |
| MBD5 | 804.125424 | -0.0069651 | 0.16363974 | -0.0425636 | 0.96604938 | 0.99393925 |
| EPC2 | 1081.9766 | -0.2866231 | 0.20913683 | -1.3705052 | 1.71E-01 | 0.75956362 |
| KIF5C | 379.84399 | -0.907211 | 0.53875131 | -1.6839143 | 0.09219824 | 0.65583276 |
| LYPD6B | 390.617625 | 0.40424064 | 0.53694386 | 0.75285457 | 0.45153731 | 0.89639459 |
| LYPD6 | 106.286968 | 0.03950035 | 0.49604865 | 0.07962999 | 0.93653154 | 0.99068685 |
| MMADHC | 4219.63777 | -0.0211144 | 0.18128026 | -0.116474 | 0.9072769 | 0.98662757 |
| AC007364.1 | 479.220229 | 0.04203876 | 0.24853763 | 0.16914443 | 0.86568304 | 0.98197545 |
| RND3 | 1784.25595 | -1.2902581 | 0.53560836 | -2.4089582 | 0.01599813 | 0.45684684 |
| RBM43 | 576.777898 | 0.03369265 | 0.37667835 | 0.08944674 | 0.92872688 | 0.98963127 |
| NMI | 1066.80254 | -0.7784783 | 0.51419874 | -1.5139639 | NA | NA |
| TNFAIP6 | 98.2009637 | 0.9326258 | 0.82967477 | 1.12408602 | 0.26097654 | 0.82424998 |
| RIF1 | 1774.34349 | -0.2498295 | 0.17563384 | -1.4224453 | 1.55E-01 | 7.44E-01 |
| NEB | 253.642288 | 0.70627465 | 0.43194936 | 1.63508669 | 0.10203088 | 0.67279543 |
| ARL5A | 2040.12316 | -0.0763416 | 0.18108458 | -0.4215798 | 0.67333176 | 0.94935413 |
| AC068547.1 | 897.319751 | -0.1576376 | 0.25179998 | -0.6260429 | 0.53128677 | 0.91962754 |
| CACNB4 | 210.061841 | -0.1013966 | 0.46095743 | -0.2199695 | 0.82589494 | 0.97548201 |
| STAM2 | 1750.24554 | -0.1444485 | 0.13338776 | -1.0829216 | 0.27884322 | 0.82825734 |
| FMNL2 | 3582.68977 | -0.3302027 | 0.33974097 | -0.971925 | 0.33108789 | 0.85384238 |
| PRPF40A | 2995.26381 | -0.0707933 | 0.12489203 | -0.5668363 | 0.57082541 | 0.92790694 |
| ARL6IP6 | 615.30558 | -0.1435935 | 0.1532689 | -0.9368729 | 0.34882392 | 0.86356499 |
| RPRM | 72.362804 | -0.1120983 | 0.74012372 | -0.1514589 | 0.87961374 | 0.98275231 |
| KCNJ3 | 743.767486 | 0.30842435 | 0.35636837 | 0.86546501 | 0.38678362 | 0.87439225 |
| NR4A2 | 939.646683 | 0.49373401 | 0.82829613 | 0.59608393 | 0.55111916 | 0.92373235 |
| GPD2 | 1216.56026 | 0.07230382 | 0.20365725 | 0.35502699 | 0.72256937 | 0.95741464 |
| CYTIP | 517.045065 | -0.43633 | 0.47239264 | -0.9236596 | 0.35566356 | 0.86364633 |
| ACVR1C | 68.4343046 | 0.84661707 | 0.43541065 | 1.94441057 | 0.05184596 | 0.58645127 |
| ACVR1 | 2100.66907 | -0.2928362 | 0.2470991 | -1.1850963 | 0.23597936 | 0.80776058 |
| UPP2 | 1843.08969 | 3.21198829 | 0.69065198 | 4.65066105 | 3.31E-06 | 0.00506488 |
| CCDC148 | 291.135851 | -0.0983735 | 0.45625144 | -0.2156124 | 0.82928985 | 0.97550416 |
| PKP4 | 9300.46284 | 0.03081701 | 0.26158361 | 0.1178094 | 0.90621868 | 0.98662757 |
| PKP4-AS1 | 555.739161 | 0.24702549 | 0.25905662 | 0.95355792 | 0.34030747 | 0.85814041 |
| TANC1 | 2170.14367 | -0.1083181 | 0.21092696 | -0.5135339 | 6.08E-01 | 9.37E-01 |
| WDSUB1 | 1149.44276 | -0.0163804 | 0.2230837 | -0.0734272 | 9.41E-01 | 9.91E-01 |

| | | | | | | |
|---|---|---|---|---|---|---|
| BAZ2B | 1459.87126 | -0.0521235 | 0.2040923 | -0.255392 | 0.79842032 | 0.9717324 |
| AC009506.1 | 164.211011 | 0.27919998 | 0.30921399 | 0.90293452 | 0.36656065 | 0.86683872 |
| 7-Mar | 4527.09388 | -0.1404321 | 0.14656952 | -0.9581261 | 3.38E-01 | 8.57E-01 |
| CD302 | 1719.80982 | 0.11560834 | 0.32740743 | 0.35310237 | 0.7240117 | 0.95810643 |
| LY75-CD302 | 504.284851 | 0.01720471 | 0.31669505 | 0.0543258 | 0.9566756 | 0.99198394 |
| LY75 | 701.012405 | -0.090837 | 0.40237561 | -0.2257517 | 0.82139451 | 0.97472105 |
| PLA2R1 | 2657.08189 | 0.4613741 | 0.5287203 | 0.87262415 | 0.38286798 | 0.8718674 |
| ITGB6 | 11229.1933 | -0.0585044 | 0.54101717 | -0.1081377 | 0.91388644 | 0.98662757 |
| LINC02478 | 61.4824189 | 0.46278084 | 0.72308637 | 0.64000769 | 0.5221676 | 0.91723878 |
| RBMS1 | 4127.854 | -0.2145629 | 0.22646549 | -0.9474418 | 3.43E-01 | 8.59E-01 |
| TANK | 1982.88916 | -0.0196172 | 0.09792467 | -0.2003299 | 0.84122259 | 0.97618283 |
| LINC01806 | 66.4321876 | -0.4061399 | 0.85912987 | -0.472734 | 0.63640298 | 0.94088547 |
| PSMD14 | 3001.66875 | -0.075206 | 0.29718565 | -0.2530607 | 0.80022135 | 0.97210205 |
| DPP4 | 7737.08829 | -0.2672656 | 0.6433692 | -0.4154156 | 0.67783769 | 0.95018034 |
| AC008063.1 | 198.746764 | -0.4828787 | 0.89995312 | -0.5365599 | NA | NA |
| AC007750.1 | 47.654767 | 0.64478796 | 0.6960853 | 0.92630597 | 0.35428699 | 0.86364633 |
| FAP | 102.738332 | 0.81548794 | 0.68502981 | 1.19044154 | 0.23387289 | 0.80725541 |
| IFIH1 | 1251.94896 | -1.1880389 | 0.65949321 | -1.8014422 | NA | NA |
| GCA | 1581.54035 | 0.05766929 | 0.22097769 | 0.26097334 | 0.79411307 | 0.97090793 |
| FIGN | 430.215869 | -0.247358 | 0.32416883 | -0.7630531 | 0.44543172 | 0.89453029 |
| GRB14 | 2062.58181 | 0.26022102 | 0.25692662 | 1.01282235 | 0.31114502 | 0.84476049 |
| COBLL1 | 6344.2846 | 0.06388867 | 0.27099403 | 0.23575675 | 0.81362143 | 0.9738447 |
| AC019197.1 | 478.663066 | 0.69114687 | 0.34290587 | 2.0155586 | 0.04384615 | 0.57103595 |
| SLC38A11 | 541.424786 | 0.56977007 | 0.4689932 | 1.21487916 | 0.22441218 | 0.80078694 |
| SCN3A | 357.880119 | -0.2899616 | 0.36798617 | -0.7879686 | 0.43071506 | 0.88966606 |
| SCN2A | 851.928463 | 1.2551137 | 0.33861529 | 3.70660663 | 0.00021005 | 0.07157522 |
| CSRNP3 | 230.317329 | 0.43018808 | 0.46174965 | 0.93164788 | 0.35151853 | 0.86364633 |
| GALNT3 | 1301.8126 | 0.05177494 | 0.24288553 | 0.21316602 | 0.83119747 | 0.97550416 |
| TTC21B | 847.095395 | 0.25143168 | 0.18204484 | 1.38115249 | 0.16723208 | 0.75558932 |
| SCN9A | 303.913484 | 0.50871211 | 0.43913948 | 1.15842945 | 0.24668883 | 0.81542272 |
| SCN7A | 234.412173 | -0.2301377 | 0.64812747 | -0.355081 | NA | NA |
| STK39 | 1614.83097 | 0.17905274 | 0.22751516 | 0.78699257 | 0.43128622 | 0.88976671 |
| CERS6 | 1569.48108 | 0.14855061 | 0.1999294 | 0.74301533 | 0.4574724 | 0.89688594 |
| NOSTRIN | 2113.61588 | 0.04495676 | 0.46074958 | 0.09757308 | 9.22E-01 | 9.88E-01 |
| SPC25 | 69.0930803 | 0.69115041 | 0.81331537 | 0.84979386 | 0.3954397 | 0.87927742 |
| LRP2 | 17582.5875 | 1.4322793 | 0.37704049 | 3.79874136 | 0.00014543 | 0.05828957 |
| BBS5 | 1017.78872 | -0.0050343 | 0.24984497 | -0.0201495 | 0.98392409 | 0.9961464 |
| AC093899.2 | 299.847163 | 0.28967742 | 0.27381469 | 1.05793236 | 0.29008628 | 0.83407594 |
| KLHL41 | 77.6195163 | 1.38959239 | 0.72281681 | 1.92246828 | 5.45E-02 | 0.59329767 |
| FASTKD1 | 1039.47655 | -0.0347539 | 0.19130276 | -0.1816695 | 0.85584211 | 0.97929708 |
| PPIG | 2128.30474 | -0.0062199 | 0.1219635 | -0.0509982 | 0.95932699 | 0.9924381 |
| PHOSPHO2 | 149.470942 | -0.4112739 | 0.25873513 | -1.5895556 | 0.11193501 | 0.68001961 |
| KLHL23 | 514.392875 | -0.4341658 | 0.24705911 | -1.7573355 | 7.89E-02 | 0.64079926 |
| SSB | 4217.86921 | -0.1690717 | 0.12906456 | -1.3099776 | 0.19020342 | 0.77089149 |
| METTL5 | 2768.33595 | -0.1138244 | 0.12584585 | -0.9044751 | 0.36574355 | 0.86665822 |
| UBR3 | 1892.48741 | -0.0257655 | 0.12448236 | -0.2069815 | 0.8360243 | 0.97576953 |
| MYO3B | 427.725539 | 1.22324723 | 0.52159717 | 2.34519531 | 0.01901711 | 0.48489761 |
| AC007405.1 | 39.9882101 | 0.17202582 | 0.58941421 | 0.29185896 | 0.77039447 | 0.96836854 |
| AC007405.3 | 197.006238 | 0.18458839 | 0.28323972 | 0.65170378 | 0.51459229 | 0.91636259 |
| ERICH2 | 55.0179073 | -0.0957239 | 0.33833353 | -0.2829277 | 0.77723226 | 0.96836854 |
| GAD1 | 579.921347 | -0.9507927 | 0.27456375 | -3.4629215 | 0.00053434 | 0.11251851 |
| GORASP2 | 4030.43182 | -0.0869721 | 0.20775089 | -0.4186363 | 0.67548197 | 0.94958156 |
| TLK1 | 2548.11896 | -0.1728155 | 0.17314276 | -0.9981097 | 0.31822615 | 0.8490262 |
| METTL8 | 582.98036 | 0.06149614 | 0.21483657 | 0.28624615 | 0.7746896 | 0.96836854 |

| | | | | | | |
|---|---|---|---|---|---|---|
| DCAF17 | 810.960669 | -0.0433329 | 0.12008689 | -0.360846 | 0.7182146 | 0.95648644 |
| CYBRD1 | 3016.40904 | 0.17605492 | 0.40589233 | 0.43374784 | 0.66447156 | 0.94676967 |
| DYNC1I2 | 5193.18883 | 0.1520657 | 0.10965392 | 1.38677849 | 0.16550932 | 0.75469833 |
| SLC25A12 | 2025.28209 | 0.20290541 | 0.24705417 | 0.82129925 | 0.41147583 | 0.88268208 |
| HAT1 | 1779.18353 | -0.0745335 | 0.19542649 | -0.381389 | 0.70291462 | 0.95431462 |
| METAP1D | 430.537861 | 0.34281945 | 0.30804246 | 1.11289673 | 0.26575278 | 0.82611274 |
| ITGA6 | 7099.45985 | -0.2751889 | 0.52533014 | -0.5238399 | 0.60038988 | 0.93487648 |
| AC078883.3 | 65.5823046 | 0.04369357 | 0.44769293 | 0.09759718 | 0.92225216 | 0.98826666 |
| AC078883.1 | 415.46503 | -0.2573047 | 0.46280286 | -0.5559705 | 0.57823105 | 0.93130183 |
| PDK1 | 1261.34442 | -0.011923 | 0.3762277 | -0.031691 | 0.9747185 | 0.99455099 |
| RAPGEF4 | 477.234141 | -0.038323 | 0.33774437 | -0.1134674 | 0.90966 | 0.98662757 |
| MAP3K20 | 953.367398 | -0.4338836 | 0.41766467 | -1.0388324 | 0.29888267 | 0.84056272 |
| MAP3K20-AS | 53.020472 | -0.7079151 | 0.45031387 | -1.5720483 | 0.11593936 | 0.6873836 |
| CDCA7 | 44.853757 | 0.05504947 | 0.49663386 | 0.11084518 | 0.91173912 | 0.98662757 |
| SP3 | 2774.0944 | -0.2326601 | 0.16053972 | -1.4492369 | 0.14727145 | 0.73386234 |
| AC016737.1 | 39.7683784 | 0.22752303 | 0.50476129 | 0.45075372 | 0.65216706 | 0.9443159 |
| OLA1 | 3058.56923 | 0.21031572 | 0.19696291 | 1.06779354 | 0.28561365 | 0.83107779 |
| CIR1 | 1369.32279 | -0.1209157 | 0.19801706 | -0.6106325 | 0.54144289 | 0.92292765 |
| SCRN3 | 716.995507 | 0.28799869 | 0.24117716 | 1.19413752 | 0.23242418 | 0.80649152 |
| GPR155 | 2114.74616 | 0.32592952 | 0.44843944 | 0.72680832 | 0.46734339 | 0.89983399 |
| WIPF1 | 2216.14779 | -0.1198496 | 0.2814677 | -0.4258023 | 0.67025195 | 0.949185 |
| AC010894.4 | 85.5453459 | -0.2865103 | 0.41052888 | -0.6979054 | 0.48523639 | 0.90543654 |
| CHN1 | 330.934536 | -0.1421585 | 0.35720414 | -0.3979755 | 0.69064821 | 0.95158502 |
| ATF2 | 2245.05615 | -0.1751174 | 0.16064149 | -1.0901129 | 0.2756634 | 0.82719407 |
| AC096649.1 | 45.2556396 | -0.09555 | 0.457354 | -0.2089191 | 0.83451136 | 0.97550416 |
| ATP5MC3 | 14368.3192 | 0.31523633 | 0.19998028 | 1.57633705 | 0.11494815 | 0.68606472 |
| LNPK | 1169.79723 | -0.0301944 | 0.25497237 | -0.1184222 | 0.90573315 | 0.9863644 |
| HOXD11 | 426.980178 | 0.05788151 | 0.44846445 | 0.12906598 | 0.89730544 | 0.98474725 |
| HOXD10 | 1305.82926 | 0.18739902 | 0.38979096 | 0.480768 | 0.6306814 | 0.94088547 |
| HOXD-AS2 | 417.863934 | 0.2263634 | 0.35101911 | 0.64487485 | 0.51900829 | 0.91636259 |
| HOXD9 | 1867.95623 | 0.2094608 | 0.40767093 | 0.51379871 | 0.60739273 | 0.93739614 |
| HOXD8 | 2904.55166 | 0.28162118 | 0.27148017 | 1.0373545 | 0.29957068 | 0.84123067 |
| HOXD3 | 1199.89734 | 0.01822571 | 0.33629683 | 0.05419531 | 0.95677956 | 0.99198394 |
| AC009336.2 | 1786.68765 | -0.0130156 | 0.31942887 | -0.0407465 | 0.96749801 | 0.99400331 |
| HOXD4 | 904.918205 | -0.0380267 | 0.2916147 | -0.1304003 | 0.8962497 | 0.98474725 |
| HAGLR | 1861.37011 | 0.01486687 | 0.2403049 | 0.0618667 | 9.51E-01 | 0.99170503 |
| AC009336.1 | 52.989554 | 0.13405865 | 0.32327605 | 0.41468785 | 0.67837043 | 0.95031695 |
| HAGLROS | 110.831073 | -0.0374506 | 0.45208635 | -0.0828394 | 0.93397926 | 0.99025262 |
| HOXD1 | 448.719026 | 0.45446057 | 0.36906238 | 1.23139228 | 0.21817618 | 0.79708421 |
| MTX2 | 2616.23464 | 0.06422505 | 0.10537827 | 0.60947149 | 0.54221196 | 0.92292765 |
| AC017048.3 | 52.6080029 | 0.33349264 | 0.48848569 | 0.68270708 | 0.49479195 | 0.90816709 |
| LINC01116 | 563.361472 | 0.18009247 | 0.33545736 | 0.53685653 | 5.91E-01 | 9.33E-01 |
| LINC01117 | 56.2966796 | 0.04788495 | 0.414417 | 0.11554776 | 0.90801097 | 0.98662757 |
| HNRNPA3 | 14485.3619 | -0.1184163 | 0.17905531 | -0.6613391 | 0.50839489 | 0.91439937 |
| NFE2L2 | 8736.26792 | -0.068851 | 0.24223533 | -0.284232 | 0.77623257 | 0.96836854 |
| AC079305.1 | 276.335293 | -0.118437 | 0.32387214 | -0.3656905 | 0.71459602 | 0.95579447 |
| AGPS | 2150.67888 | 0.25351528 | 0.15889822 | 1.59545702 | 0.11061007 | 0.68001961 |
| TTC30B | 354.581314 | 0.09824992 | 0.23793482 | 0.41292786 | 0.67965947 | 0.95070301 |
| AC073834.1 | 53.7626948 | -0.1092216 | 0.35592898 | -0.3068635 | 0.75894727 | 0.96502715 |
| TTC30A | 529.056898 | 0.06214989 | 0.23112187 | 0.26890526 | 0.78800259 | 0.97090793 |
| PDE11A | 156.216169 | 0.6564155 | 0.38615427 | 1.69987891 | 0.0891537 | 0.65375543 |
| RBM45 | 585.6517 | 0.06587759 | 0.17750657 | 0.37112761 | 0.71054249 | 0.95546221 |
| OSBPL6 | 230.625104 | 0.47107302 | 0.49607211 | 0.94960594 | 0.34231252 | 0.85889221 |
| AC009948.1 | 865.722521 | -0.4056979 | 0.32639978 | -1.2429478 | 0.21388705 | 0.7926732 |

| | | | | | | |
|---|---|---|---|---|---|---|
| PRKRA | 2243.66596 | 0.04917973 | 0.1432916 | 0.34321429 | 0.73143725 | 0.95848028 |
| PJVK | 272.353462 | 0.1958634 | 0.33818607 | 0.57915867 | 0.56248211 | 0.92452095 |
| FKBP7 | 330.87797 | 0.12033473 | 0.34491353 | 0.34888377 | 0.72717657 | 0.95848028 |
| PLEKHA3 | 1342.85855 | -0.0099177 | 0.16040292 | -0.0618299 | 0.95069826 | 0.99170503 |
| TTN-AS1 | 568.984077 | -0.1055484 | 0.23229033 | -0.4543813 | 0.64955441 | 0.94312988 |
| TTN | 989.519924 | -0.1095868 | 0.23391023 | -0.4684995 | 0.63942742 | 0.94088547 |
| AC010680.2 | 80.0787821 | -0.6351572 | 0.4737253 | -1.3407711 | 0.17999479 | 0.76318604 |
| SESTD1 | 2697.83861 | 0.0775632 | 0.16669588 | 0.46529768 | 0.6417183 | 0.94088547 |
| ZNF385B | 763.485785 | 0.1611691 | 0.42057953 | 0.38320718 | 0.70156615 | 0.95424422 |
| CWC22 | 1017.89049 | -0.0967556 | 0.17297362 | -0.5593663 | 0.57591173 | 0.93083012 |
| UBE2E3 | 2929.52767 | -0.0384861 | 0.16145543 | -0.2383699 | 0.81159423 | 0.97352497 |
| AC068196.1 | 44.5030088 | -1.0726575 | 0.57493463 | -1.8657034 | 0.06208288 | 0.61236592 |
| ITGA4 | 252.93501 | 0.01673152 | 0.56270659 | 0.02973401 | 0.97627919 | 0.99455099 |
| CERKL | 617.85496 | 0.24257299 | 0.28961206 | 0.83757903 | 0.40226717 | 0.88132015 |
| SSFA2 | 5370.59141 | 0.03404121 | 0.1589948 | 0.21410268 | 0.83046699 | 0.97550416 |
| PPP1R1C | 59.2643018 | -0.372792 | 0.45072467 | -0.8270947 | 0.40818341 | 0.88181733 |
| PDE1A | 3169.68657 | -0.5818568 | 0.42059805 | -1.3834035 | 0.16654118 | 0.75558932 |
| DNAJC10 | 4155.30904 | 0.13163667 | 0.13500694 | 0.97503634 | 0.32954223 | 0.85350346 |
| FRZB | 1018.39022 | -0.0384561 | 0.54983076 | -0.0699416 | 0.94424011 | 0.99136103 |
| NCKAP1 | 8312.26336 | -0.1228271 | 0.19667945 | -0.6245038 | 0.53229675 | 0.91985 |
| DUSP19 | 180.879253 | 0.69384915 | 0.25234563 | 2.74959846 | 0.00596683 | 0.32986555 |
| NUP35 | 654.748583 | -0.2884833 | 0.16910992 | -1.7058921 | 0.08802817 | 0.65363785 |
| FSIP2 | 98.5759032 | -0.305353 | 0.33462463 | -0.912524 | 0.36149293 | 0.86528137 |
| LINC01473 | 45.4269219 | 0.10572476 | 0.50926222 | 0.20760378 | 0.83553835 | 0.97550416 |
| ZC3H15 | 3127.99786 | -0.1210273 | 0.1908985 | -0.6339878 | 0.52608881 | 0.91769856 |
| ITGAV | 7749.38495 | -0.1359454 | 0.3261669 | -0.416797 | 6.77E-01 | 9.50E-01 |
| FAM171B | 220.589669 | 0.33408558 | 0.29433184 | 1.13506435 | 0.25634836 | 0.82067614 |
| AC007319.1 | 730.084258 | -0.4491458 | 0.45416592 | -0.9889465 | 0.32268932 | 0.85119105 |
| CALCRL | 1139.42835 | -0.1158364 | 0.31603277 | -0.366533 | 0.71396737 | 0.95564093 |
| TFPI | 11730.8722 | -0.3297009 | 0.36676997 | -0.8989309 | 0.36868946 | 0.86762749 |
| GULP1 | 1813.34807 | -0.8078703 | 0.49651747 | -1.6270733 | 0.10372153 | 0.67439401 |
| COL3A1 | 10240.6102 | -0.3144065 | 0.51292564 | -0.612967 | 0.53989815 | 0.92210255 |
| COL5A2 | 1076.78846 | -0.1141419 | 0.42062278 | -0.271364 | 0.78611109 | 0.97075604 |
| WDR75 | 1754.50817 | -0.0323537 | 0.12763688 | -0.253482 | 0.79989573 | 0.97208422 |
| SLC40A1 | 5100.55612 | 0.34171413 | 0.24177089 | 1.41337995 | 0.15754404 | 0.74522306 |
| ASNSD1 | 1926.074 | -0.1256922 | 0.27043087 | -0.4647851 | 0.64208535 | 0.94088547 |
| ANKAR | 203.823318 | 0.15443216 | 0.27918068 | 0.55316206 | 0.58015245 | 0.93139743 |
| OSGEPL1 | 435.949926 | 0.37502972 | 0.2854263 | 1.31392839 | 0.1888703 | 0.76970626 |
| OSGEPL1-AS | 54.4062192 | 0.43739258 | 0.31373458 | 1.39414846 | 0.16327278 | 0.75149188 |
| ORMDL1 | 3044.78014 | -0.1346426 | 0.14934816 | -0.9015353 | 0.36730376 | 0.86683872 |
| PMS1 | 815.68645 | -0.0109786 | 0.16568438 | -0.0662624 | 0.94716893 | 0.99166614 |
| C2orf88 | 525.097068 | -0.178164 | 0.26933302 | -0.6615008 | 0.50829118 | 0.91439937 |
| MSTN | 54.4329652 | 0.42950172 | 0.48665846 | 0.88255265 | 0.37747802 | 0.86987124 |
| HIBCH | 4623.59928 | 0.71228477 | 0.3405067 | 2.09183771 | 0.03645304 | 0.55384187 |
| INPP1 | 1624.09963 | -0.2629421 | 0.1463351 | -1.7968492 | 0.07235956 | 0.63220999 |
| MFSD6 | 1328.17872 | 0.01647487 | 0.21835117 | 0.07545127 | 0.93985566 | 0.99135146 |
| NEMP2 | 320.667571 | -0.1053086 | 0.15482543 | -0.6801766 | 0.49639266 | 0.90890243 |
| NAB1 | 1640.30033 | 0.094285 | 0.17819681 | 0.529106 | 0.59673192 | 0.9337017 |
| AC006460.1 | 206.975564 | -0.1531469 | 0.30465481 | -0.5026899 | 0.61518232 | 0.93895494 |
| AC005540.1 | 199.3842 | -0.2201637 | 0.55999046 | -0.3931561 | 0.69420421 | 0.95222846 |
| GLS | 20700.6617 | 0.05465802 | 0.30568589 | 0.17880453 | 0.85809119 | 0.97974211 |
| STAT1 | 16188.601 | -0.3299497 | 0.58740447 | -0.5617079 | NA | NA |
| STAT4 | 254.60747 | -0.0984055 | 0.63388012 | -0.1552432 | 0.87662963 | 0.98275231 |
| MYO1B | 3162.23876 | 0.15408145 | 0.2175315 | 0.70831791 | 0.47874786 | 0.90444322 |

| | | | | | | |
|---|---|---|---|---|---|---|
| NABP1 | 2770.6678 | -0.1363094 | 0.36866819 | -0.3697346 | 7.12E-01 | 9.56E-01 |
| CAVIN2 | 1953.29447 | -0.3989589 | 0.3176186 | -1.2560942 | 0.20908183 | 0.79086734 |
| AC098617.1 | 263.103598 | -0.1855627 | 0.39467817 | -0.470162 | 0.63823929 | 0.94088547 |
| SLC39A10 | 1099.43036 | 0.4796053 | 0.21663195 | 2.21391763 | 0.02683445 | 0.52685774 |
| DNAH7 | 248.169786 | 0.04188347 | 0.4944122 | 0.08471368 | 0.93248902 | 0.9899349 |
| STK17B | 875.217216 | -0.1446496 | 0.33719602 | -0.4289779 | 0.66793934 | 0.94847921 |
| HECW2 | 301.549748 | 0.27194664 | 0.5164848 | 0.52653367 | 0.59851746 | 0.9346978 |
| CCDC150 | 219.374833 | 0.37921708 | 0.50167987 | 0.75589456 | 0.44971242 | 0.89557948 |
| GTF3C3 | 1391.87441 | 0.09714341 | 0.16872493 | 0.5757502 | 0.56478404 | 0.92477055 |
| PGAP1 | 550.014793 | -0.064834 | 0.25070962 | -0.258602 | 0.79594233 | 0.97101097 |
| ANKRD44 | 706.411986 | 0.14829985 | 0.28264405 | 0.52468765 | 0.59980034 | 0.93487648 |
| SF3B1 | 13207.7248 | -0.0189477 | 0.09201592 | -0.2059173 | 0.8368555 | 0.97577999 |
| COQ10B | 2216.46046 | -0.2200491 | 0.25742458 | -0.8548101 | 0.39265628 | 0.8778303 |
| HSPD1 | 22133.8568 | 0.35891966 | 0.32025497 | 1.12073097 | 0.2624024 | 0.82424998 |
| HSPE1 | 6248.13389 | 0.32393818 | 0.213165 | 1.51965929 | 0.12859663 | 0.70610612 |
| HSPE1-MOB4 | 5804.7654 | 0.32635985 | 0.20938324 | 1.55867225 | 0.11907397 | 0.69233212 |
| MOB4 | 1531.55774 | -0.0823736 | 0.13511318 | -0.6096639 | 0.54208448 | 0.92292765 |
| RFTN2 | 210.365628 | -0.4720619 | 0.38896384 | -1.2136395 | 0.2248854 | 0.80125043 |
| MARS2 | 445.786323 | -0.2375753 | 0.36071541 | -0.6586226 | 0.51013815 | 0.91439937 |
| PLCL1 | 2019.94362 | -0.0431322 | 0.36071441 | -0.1195742 | 0.90482043 | 0.98624484 |
| SATB2 | 678.009021 | -0.398059 | 0.279433 | -1.424524 | 0.15429483 | 0.74258845 |
| SATB2-AS1 | 44.100784 | -0.1759571 | 0.52307319 | -0.3363911 | 0.73657599 | 0.95988947 |
| FTCDNL1 | 133.433316 | 1.12317798 | 0.37184499 | 3.02055428 | 0.00252312 | 0.21493583 |
| C2orf69 | 1036.80344 | -0.0058623 | 0.2165568 | -0.0270703 | 0.97840366 | 0.99476647 |
| TYW5 | 677.253123 | -0.0706098 | 0.15804095 | -0.4467818 | 6.55E-01 | 9.45E-01 |
| MAIP1 | 1460.14431 | 0.18715109 | 0.28719524 | 0.65165108 | 0.51462629 | 0.91636259 |
| SPATS2L | 5930.59833 | 0.12979369 | 0.32821797 | 0.39544969 | 0.69251104 | 0.9519683 |
| KCTD18 | 857.554515 | 0.11999579 | 0.21512508 | 0.55779545 | 0.57698407 | 0.93129843 |
| SGO2 | 93.315023 | 0.05993523 | 0.39121124 | 0.15320427 | 0.87823719 | 0.98275231 |
| AOX1 | 13196.0924 | 1.15671459 | 0.458195 | 2.52450288 | 0.01158621 | 0.40423902 |
| AC007163.1 | 248.425144 | -0.3592839 | 0.53731504 | -0.6686653 | 0.503709 | 0.91356552 |
| BZW1 | 5754.7603 | -0.3173883 | 0.40162068 | -0.7902689 | 0.42937076 | 0.88851358 |
| CLK1 | 9422.8375 | -0.0712424 | 0.17369706 | -0.4101529 | 0.68169378 | 0.95070301 |
| PPIL3 | 834.3614 | 0.11340615 | 0.16681091 | 0.6798485 | 0.49660039 | 0.90890243 |
| NIF3L1 | 1011.31539 | 0.15651796 | 0.1950076 | 0.80262493 | 0.42219155 | 0.88451372 |
| ORC2 | 1326.41035 | -0.1237338 | 0.13620369 | -0.9084465 | 0.36364236 | 0.8663515 |
| AC005037.1 | 83.4575008 | -0.102088 | 0.31926281 | -0.3197617 | 0.74914896 | 0.96249332 |
| FAM126B | 1294.05948 | -0.2781883 | 0.13014263 | -2.137565 | 0.03255206 | 0.55104386 |
| NDUFB3 | 2339.82932 | 0.32613631 | 0.21135627 | 1.54306425 | 0.12281519 | 0.69788192 |
| CFLAR | 11597.2395 | -0.6057686 | 0.2351915 | -2.5756398 | 0.01000548 | 0.39989496 |
| CFLAR-AS1 | 555.267568 | -0.6222237 | 0.26900577 | -2.3130497 | 0.0207199 | 0.50488816 |
| CASP10 | 1744.80218 | -0.0500818 | 0.32682945 | -0.1532353 | 8.78E-01 | 0.98275231 |
| CASP8 | 1348.71004 | -0.1625125 | 0.16619687 | -0.9778316 | 0.32815764 | 0.85317073 |
| ALS2CR12 | 48.0991442 | 0.03589673 | 0.42441705 | 0.08457891 | 0.93259617 | 0.9899349 |
| TRAK2 | 3461.0869 | -0.2669173 | 0.25686989 | -1.0391149 | 0.29875128 | 0.84056272 |
| STRADB | 3998.32062 | 0.43791876 | 0.41283259 | 1.06076597 | 0.28879627 | 0.83295411 |
| TMEM237 | 625.440836 | 0.09052584 | 0.30523707 | 0.29657549 | 0.76679062 | 0.96741943 |
| ALS2 | 1752.21798 | -0.2029237 | 0.15104506 | -1.3434646 | 0.17912159 | 0.76120583 |
| FZD7 | 775.671966 | 0.55324791 | 0.34516336 | 1.60285816 | 0.10896597 | 0.67844893 |
| SUMO1 | 4857.68611 | -0.1087972 | 0.0882263 | -1.2331606 | 0.21751587 | 0.7960866 |
| NOP58 | 1877.17967 | -0.2920908 | 0.20846077 | -1.4011785 | 0.16116069 | 0.74908546 |
| AC064836.3 | 42.2157092 | -0.280998 | 0.51875593 | -0.5416767 | 0.58804127 | 0.93289552 |
| BMPR2 | 3571.06678 | 0.07551029 | 0.23007187 | 0.32820306 | 0.74275813 | 0.96109001 |
| FAM117B | 864.927414 | -0.1340194 | 0.17462723 | -0.7674599 | 0.44280811 | 0.89343541 |

| | | | | | | |
|---|---|---|---|---|---|---|
| ICA1L | 399.786694 | -0.0844717 | 0.35081263 | -0.2407887 | 0.80971886 | 0.97334138 |
| WDR12 | 1506.52107 | 0.35165041 | 0.25470451 | 1.38062104 | 0.16739551 | 0.75558932 |
| CARF | 551.547205 | 0.17178341 | 0.30916908 | 0.55562932 | 0.57846429 | 0.93139743 |
| NBEAL1 | 470.896507 | -0.0196965 | 0.18554273 | -0.1061563 | 0.91545836 | 0.9868044 |
| CYP20A1 | 717.878463 | -0.1103488 | 0.17516772 | -0.629961 | 0.52872009 | 0.91900791 |
| ABI2 | 4558.06174 | 0.02564388 | 0.18294894 | 0.14016958 | 0.888526 | 0.98423722 |
| AC080075.1 | 49.0324248 | -0.2900309 | 0.42377266 | -0.6844022 | 0.49372125 | 0.90749607 |
| RAPH1 | 2052.2365 | -0.069253 | 0.21122133 | -0.3278692 | 0.74301058 | 0.96109001 |
| CD28 | 77.4239952 | -0.0928275 | 0.82496081 | -0.1125236 | NA | NA |
| CTLA4 | 55.4160306 | 0.30991861 | 1.08351119 | 0.28603176 | 0.7748538 | 0.96836854 |
| PARD3B | 898.867039 | 0.37286552 | 0.18166065 | 2.05253871 | 0.04011734 | 0.55657615 |
| NRP2 | 985.343827 | -0.5218699 | 0.33759711 | -1.5458363 | 0.12214411 | 0.69785915 |
| INO80D | 1802.62515 | -0.406058 | 0.20342292 | -1.996127 | 0.0459201 | 0.57687945 |
| AC007383.2 | 256.551704 | 0.10307127 | 0.19229439 | 0.53600769 | 0.59195324 | 0.93289552 |
| NDUFS1 | 8117.64709 | 0.36519018 | 0.21461051 | 1.70164167 | 0.08882256 | 0.65375543 |
| AC007383.3 | 1092.12699 | 0.40782624 | 0.20706234 | 1.96958192 | 0.04888631 | 0.58504358 |
| EEF1B2 | 11227.5259 | 0.28624645 | 0.24042196 | 1.1906003 | 0.23381053 | 0.80725541 |
| ZDBF2 | 527.317632 | 0.03915008 | 0.27847023 | 0.14058981 | 0.888194 | 0.98423722 |
| ADAM23 | 88.4663263 | -0.596673 | 0.54933278 | -1.0861777 | 0.2774004 | 0.82782817 |
| MDH1B | 379.396151 | 0.30463284 | 0.57197151 | 0.53260142 | 0.59430952 | 0.93345246 |
| FASTKD2 | 1364.00626 | -0.0548768 | 0.13765665 | -0.39865 | 0.69015112 | 0.95158502 |
| CPO | 93.7646281 | -0.1373279 | 0.32063298 | -0.4283024 | 0.66843097 | 0.9486589 |
| KLF7 | 1399.00869 | -0.4126648 | 0.25481949 | -1.6194397 | 0.1053527 | 0.67473222 |
| CREB1 | 1551.15713 | -0.0649931 | 0.10378804 | -0.6262097 | 0.53117738 | 0.91962754 |
| METTL21A | 546.567369 | -0.1256313 | 0.17937825 | -0.7003706 | 0.48369586 | 0.90541935 |
| CCNYL1 | 1382.86727 | -0.3130031 | 0.31916468 | -0.9806947 | 0.32674334 | 0.85282528 |
| AC096772.1 | 315.548218 | -0.2796338 | 0.35831577 | -0.7804118 | 0.43514854 | 0.89055338 |
| FZD5 | 1682.43286 | 0.08012593 | 0.32842572 | 0.24396972 | 0.80725427 | 0.97334138 |
| PLEKHM3 | 592.789524 | 0.19823779 | 0.24633715 | 0.80474176 | 0.42096872 | 0.88451372 |
| IDH1 | 5860.2325 | 0.69237477 | 0.43285674 | 1.59954715 | 0.10969908 | 0.67859564 |
| PIKFYVE | 1925.11905 | -0.0349995 | 0.11625862 | -0.3010487 | 0.76337739 | 0.96671773 |
| PTH2R | 946.100539 | -0.2290893 | 0.32805065 | -0.6983353 | 0.48496753 | 0.90541935 |
| MAP2 | 478.695131 | -0.5065932 | 0.37388748 | -1.3549349 | 1.75E-01 | 0.76073329 |
| RPE | 2981.77625 | 0.18108161 | 0.23016055 | 0.78676218 | 0.4314211 | 0.88976671 |
| KANSL1L | 2889.90347 | 0.14331938 | 0.29738562 | 0.48193112 | 0.62985487 | 0.94088547 |
| AC007038.2 | 238.088313 | 0.03506469 | 0.27507291 | 0.12747416 | 0.89856513 | 0.98474725 |
| AC007038.1 | 121.888644 | -0.223218 | 0.30289319 | -0.7369528 | 0.46115107 | 0.89755243 |
| ACADL | 1556.87279 | 0.72083081 | 0.28305627 | 2.54659895 | 0.01087784 | 0.40196898 |
| LANCL1 | 2882.18758 | 0.21268007 | 0.20554264 | 1.03472482 | 0.30079744 | 0.84200819 |
| CPS1 | 181.387771 | 0.49370902 | 0.42171345 | 1.17072155 | 0.24171072 | 0.81269742 |
| ERBB4 | 1319.66138 | -0.0224459 | 0.29481196 | -0.0761364 | 0.93931058 | 0.99113899 |
| IKZF2 | 254.639112 | 0.45294076 | 0.49473691 | 0.91551844 | 0.35991953 | 0.86480228 |
| SPAG16 | 978.256393 | 0.33176132 | 0.21143485 | 1.56909475 | 0.11662586 | 0.6888681 |
| BARD1 | 149.4027 | 0.32995533 | 0.29467317 | 1.11973319 | 0.26282748 | 0.82424998 |
| ATIC | 3282.53883 | 0.11789851 | 0.20844316 | 0.56561467 | 5.72E-01 | 9.28E-01 |
| FN1 | 8853.14891 | -0.1091452 | 0.52412192 | -0.2082439 | 8.35E-01 | 9.76E-01 |
| MREG | 475.894448 | 0.0111826 | 0.30544765 | 0.03661053 | 9.71E-01 | 0.99400331 |
| PECR | 1700.29893 | 0.97810742 | 0.35225 | 2.77674213 | 5.49E-03 | 3.15E-01 |
| TMEM169 | 157.037725 | 1.4427223 | 0.40923971 | 3.52537223 | 0.00042289 | 0.09906683 |
| XRCC5 | 11205.6384 | -0.0315314 | 0.16905668 | -0.1865135 | 0.8520421 | 0.97872457 |
| LINC01963 | 483.910559 | 0.15800708 | 0.48647279 | 0.32480148 | 0.74533133 | 0.96176407 |
| SMARCAL1 | 812.439646 | -0.0834101 | 0.1759905 | -0.4739464 | 0.63553812 | 0.94088547 |
| RPL37A | 21809.0823 | 0.11856815 | 0.1607147 | 0.73775549 | 0.46066305 | 0.89755243 |
| IGFBP2 | 5083.51029 | 0.42707943 | 0.39659821 | 1.07685667 | 0.2815443 | 0.82975653 |

| | | | | | | |
|---|---|---|---|---|---|---|
| IGFBP5 | 47814.9359 | -0.1176408 | 0.45000815 | -0.2614192 | 0.79376927 | 0.97090793 |
| AC007563.2 | 1035.21331 | -0.3090203 | 0.47347397 | -0.6526658 | 0.51397173 | 0.91636259 |
| DIRC3 | 49.3139833 | 0.31313077 | 0.3545605 | 0.88315187 | 0.37715422 | 0.86987124 |
| TNS1 | 23799.9229 | -0.051257 | 0.25817612 | -0.1985351 | 0.84262639 | 0.97644404 |
| AC010136.1 | 79.6857899 | -0.293215 | 0.61626839 | -0.475791 | 0.63422326 | 0.94088547 |
| RUFY4 | 175.623124 | -5.3146201 | 1.36411906 | -3.896009 | NA | NA |
| CXCR2 | 53.9244839 | -0.5005084 | 0.93581603 | -0.5348363 | 0.59276303 | 0.93289552 |
| CXCR1 | 62.4390398 | -0.5861462 | 0.80317591 | -0.7297856 | 0.46552124 | 0.89933527 |
| ARPC2 | 9947.38489 | -0.1554353 | 0.18810369 | -0.8263276 | 0.40861827 | 0.88203324 |
| GPBAR1 | 131.314832 | -0.8647316 | 0.75360963 | -1.147453 | 0.25119445 | 0.81815815 |
| AAMP | 7250.90429 | 0.03704894 | 0.16374706 | 0.22625714 | 0.82100143 | 0.97472105 |
| PNKD | 7660.7581 | 0.08576686 | 0.11514011 | 0.74489122 | 0.45633748 | 0.89688594 |
| TMBIM1 | 18350.6322 | -0.1170547 | 0.22468302 | -0.5209771 | 0.60238271 | 0.9355118 |
| CATIP | 317.649895 | 0.02941672 | 0.41393186 | 0.07106657 | 0.94334478 | 0.99136103 |
| SLC11A1 | 1348.69023 | -0.895685 | 0.51851938 | -1.7273897 | 0.0840977 | 0.64942281 |
| CTDSP1 | 10209.6385 | -0.1976159 | 0.10938529 | -1.8066039 | 0.07082406 | 0.63023427 |
| AC021016.3 | 119.634125 | -0.0822181 | 0.25122348 | -0.3272709 | 0.74346304 | 0.96124462 |
| VIL1 | 2781.46593 | 0.1891089 | 0.26321207 | 0.7184659 | 0.47247006 | 0.90160919 |
| USP37 | 897.267032 | -0.1007556 | 0.16720489 | -0.6025878 | 0.54678293 | 0.92345821 |
| CNOT9 | 2139.35536 | -0.229314 | 0.14940942 | -1.5348029 | 0.12483224 | 0.70228923 |
| PLCD4 | 1519.57386 | -0.4238401 | 0.4808582 | -0.8814243 | 0.37808823 | 0.87016475 |
| AC012510.1 | 503.414501 | -1.1830311 | 0.57801635 | -2.0467087 | 0.04068669 | 0.55657615 |
| ZNF142 | 1237.03741 | -0.5106528 | 0.28791306 | -1.7736355 | 0.07612347 | 0.63773415 |
| BCS1L | 2134.84283 | 0.04970078 | 0.15846417 | 0.31364051 | 0.75379409 | 0.96330412 |
| RNF25 | 1382.33327 | -0.3331138 | 0.20784707 | -1.6026868 | 0.10900381 | 0.67845911 |
| STK36 | 2485.58029 | -0.3158012 | 0.2387774 | -1.3225759 | 0.18597646 | 0.76908034 |
| TTLL4 | 1051.20466 | -0.3274756 | 0.2130827 | -1.5368473 | 0.12433071 | 0.70102567 |
| CYP27A1 | 4279.13737 | 0.78903628 | 0.34459703 | 2.28973614 | 0.02203662 | 0.51639767 |
| WNT6 | 44.939666 | -0.3458989 | 0.62934789 | -0.5496148 | 0.58258364 | 0.93182555 |
| WNT10A | 113.548865 | 1.35523857 | 0.8426459 | 1.60831325 | 0.10776659 | 0.67708979 |
| CFAP65 | 94.3324383 | 0.82721991 | 0.57390239 | 1.44139478 | 1.49E-01 | 0.73670625 |
| IHH | 432.709358 | 0.98244605 | 0.62317025 | 1.57652914 | 0.11490391 | 0.68601916 |
| AC097468.3 | 171.229611 | -0.0009932 | 0.49694059 | -0.0019987 | 0.99840527 | 0.99994357 |
| NHEJ1 | 1453.11544 | 0.08866322 | 0.4029068 | 0.22005888 | 0.8258253 | 0.97548201 |
| AC068946.1 | 4677.52487 | -0.1801728 | 0.48577236 | -0.3708997 | 0.71071223 | 0.95560193 |
| SLC23A3 | 4331.98566 | -0.6269094 | 0.57778413 | -1.0850236 | 0.27791121 | 0.82805357 |
| CNPPD1 | 9960.03838 | -0.0607452 | 0.17387256 | -0.3493662 | 0.7268144 | 0.95848028 |
| RETREG2 | 10265.4392 | -0.1884976 | 0.20867569 | -0.903304 | 0.36636459 | 0.86683872 |
| ZFAND2B | 2716.53294 | -0.0525589 | 0.10432588 | -0.5037954 | 0.61440513 | 0.93895494 |
| ABCB6 | 2808.57153 | -0.0105956 | 0.37296042 | -0.0284093 | 0.97733568 | 0.99458341 |
| AC068946.2 | 6179.59196 | -0.1460465 | 0.22843183 | -0.6393437 | 0.52259934 | 0.91723878 |
| ATG9A | 4722.10463 | -0.2552955 | 0.13497368 | -1.8914466 | 0.05856476 | 0.60332684 |
| ANKZF1 | 4940.46881 | -0.2068296 | 0.34088901 | -0.6067358 | 0.54402624 | 0.92343047 |
| GLB1L | 2034.20753 | -0.0931251 | 0.29596489 | -0.3146491 | 0.75302809 | 0.96301881 |
| STK16 | 2291.47696 | -0.042588 | 0.15459684 | -0.275478 | 0.78294902 | 0.96964828 |
| TUBA4A | 5616.38517 | -0.1895075 | 0.32473675 | -0.5835728 | 0.55950775 | 0.92426044 |
| TUBA4B | 129.989081 | 0.22073191 | 0.41564362 | 0.53106051 | 0.59537685 | 0.9334601 |
| DNAJB2 | 8194.0728 | -0.2989245 | 0.18252925 | -1.6376802 | 0.10148842 | 0.67279543 |
| DNPEP | 3425.63893 | -0.0029539 | 0.24512521 | -0.0120504 | 0.99038538 | 0.99814875 |
| DES | 4209.10534 | -3.4695624 | 1.50997807 | -2.2977568 | NA | NA |
| AC053503.4 | 1144.06914 | -3.3763884 | 1.54868456 | -2.1801653 | NA | NA |
| SPEG | 1083.98869 | -0.4147675 | 0.58527076 | -0.7086763 | 0.4785254 | 0.90440142 |
| GMPPA | 2714.98855 | 0.05464982 | 0.18573839 | 0.29423011 | 7.69E-01 | 9.68E-01 |
| CHPF | 3961.27426 | 0.36740504 | 0.21797582 | 1.68553115 | 0.09188614 | 0.6555151 |

| | | | | | | |
|---|---|---|---|---|---|---|
| TMEM198 | 437.193462 | 0.52658543 | 0.24535329 | 2.14623337 | 0.03185436 | 0.54649555 |
| AC009955.3 | 49.5744292 | 0.58400898 | 0.4102207 | 1.42364581 | 0.15454903 | 0.74300499 |
| OBSL1 | 10220.1183 | 0.06867743 | 0.28058385 | 0.24476617 | 0.80663749 | 0.97334138 |
| AC009955.4 | 398.813313 | -0.1068613 | 0.3297709 | -0.3240472 | 0.74590228 | 0.96176407 |
| INHA | 165.336808 | 0.49160389 | 0.33838202 | 1.45280734 | 0.14627725 | 0.73247558 |
| STK11IP | 1813.44712 | -0.0787443 | 0.1726837 | -0.4560033 | 0.64838757 | 0.9431121 |
| SLC4A3 | 1084.84293 | -0.5160707 | 0.42244729 | -1.2216214 | 0.22185083 | 0.80054995 |
| EPHA4 | 508.550288 | 0.15938477 | 0.40976908 | 0.38896241 | 0.69730395 | 0.95269909 |
| SGPP2 | 1395.14046 | 0.07162951 | 0.27711701 | 0.25848112 | 0.79603562 | 0.97101097 |
| AC097461.1 | 41.3416392 | 0.05347335 | 0.53824574 | 0.09934746 | 0.92086239 | 0.98767854 |
| FARSB | 1586.75918 | -0.1021944 | 0.16231058 | -0.6296227 | 0.52894145 | 0.91907723 |
| MOGAT1 | 47.3815913 | -1.1623787 | 0.47459278 | -2.4492128 | 0.01431688 | 0.43699139 |
| ACSL3 | 3562.02842 | -0.2417584 | 0.17656896 | -1.3692009 | 0.1709365 | 0.75956362 |
| KCNE4 | 590.286215 | -0.5758441 | 0.62272974 | -0.9247095 | 0.35511704 | 0.86364633 |
| SCG2 | 59.2445418 | 0.66678053 | 0.47596841 | 1.4008924 | 0.16124625 | 0.74924826 |
| AP1S3 | 392.760009 | 0.14817996 | 0.269873 | 0.54907294 | 0.5829554 | 0.93191649 |
| WDFY1 | 2131.64748 | -0.3705685 | 0.19323898 | -1.9176696 | 0.05515292 | 0.59574695 |
| MRPL44 | 2245.31731 | -0.0118008 | 0.23746859 | -0.0496941 | 0.96036619 | 0.99278629 |
| AC073641.1 | 291.082592 | -0.1090043 | 0.26133223 | -0.4171101 | 0.67659788 | 0.94989294 |
| SERPINE2 | 11469.1567 | -1.079772 | 0.79755597 | -1.353851 | 0.17578391 | 0.7610139 |
| FAM124B | 53.1814721 | -0.3681976 | 0.76773254 | -0.479591 | 0.63151825 | 0.94088547 |
| AC073052.2 | 1227.32533 | -0.1111328 | 0.16686851 | -0.6659905 | 0.50541717 | 0.91368371 |
| CUL3 | 3760.72403 | -0.0903748 | 0.19388566 | -0.4661245 | 0.64112641 | 0.94088547 |
| DOCK10 | 421.26382 | -0.2549925 | 0.41466827 | -0.6149312 | 0.53860014 | 0.9215968 |
| IRS1 | 1722.32978 | -0.4826241 | 0.26284091 | -1.8361832 | 0.06633056 | 0.62373361 |
| RHBDD1 | 1217.80037 | 0.15870213 | 0.20640843 | 0.76887424 | 0.44196797 | 0.89331461 |
| COL4A4 | 4041.36863 | 0.24332752 | 0.22936469 | 1.06087611 | 0.28874621 | 0.83295411 |
| COL4A3 | 5344.40035 | 0.38044093 | 0.21452707 | 1.77339357 | 0.07616351 | 0.63773415 |
| AC097662.1 | 141.190868 | 0.12842111 | 0.19795273 | 0.64874635 | 0.51650234 | 0.91636259 |
| MFF | 4113.17112 | -0.0365717 | 0.11137138 | -0.328376 | 0.74262741 | 0.96109001 |
| AGFG1 | 3742.67351 | -0.2440424 | 0.21397026 | -1.1405435 | 0.25405992 | 0.8196483 |
| SLC19A3 | 768.150657 | -0.0227029 | 0.37334148 | -0.0608101 | 0.95151044 | 0.99170503 |
| CCL20 | 229.83353 | 1.79422798 | 0.73429837 | 2.44345903 | 0.01454722 | 0.43901686 |
| PID1 | 387.888202 | -0.2689428 | 0.46516089 | -0.5781716 | 0.56314828 | 0.92452095 |
| DNER | 1439.88661 | 0.25194412 | 0.28364948 | 0.88822348 | 0.37442055 | 0.86791605 |
| TRIP12 | 6720.50119 | -0.1477451 | 0.15404846 | -0.9590822 | 0.33751733 | 0.8571036 |
| FBXO36 | 364.538205 | 0.46950629 | 0.20436009 | 2.29744608 | 0.02159334 | 0.51394743 |
| SLC16A14 | 212.421559 | 0.41700435 | 0.42669988 | 0.97727787 | 0.32843161 | 0.85317073 |
| AC009950.1 | 141.993632 | -1.2864621 | 0.57532651 | -2.2360556 | NA | NA |
| SP110 | 1128.19757 | -1.1069533 | 0.54591717 | -2.0276946 | 0.04259143 | 0.56530166 |
| SP140 | 137.986598 | -0.7903479 | 0.78798396 | -1.003 | 0.31586088 | 0.84750116 |
| SP140L | 792.3203 | -0.3598021 | 0.37327211 | -0.9639136 | 0.33508925 | 0.85581042 |
| SP100 | 3266.58242 | -0.7208393 | 0.37848268 | -1.9045503 | 0.05683855 | 0.60049913 |
| CAB39 | 4616.23078 | -0.0191945 | 0.17317615 | -0.110838 | 0.91174479 | 0.98662757 |
| ITM2C | 13731.9393 | 0.38782855 | 0.3941605 | 0.9839356 | 0.32514718 | 0.85224941 |
| C2orf72 | 337.804867 | 0.12930473 | 0.65488452 | 0.19744661 | 0.84347806 | 0.97649449 |
| PSMD1 | 3501.81291 | -0.0544063 | 0.22925548 | -0.2373175 | 0.8124105 | 0.97364432 |
| HTR2B | 175.284291 | 0.30509583 | 0.3444981 | 0.88562412 | 0.37582011 | 0.86889275 |
| ARMC9 | 506.827725 | -0.161521 | 0.29840652 | -0.5412783 | 0.58831579 | 0.93289552 |
| B3GNT7 | 294.813912 | 0.04557576 | 0.3183301 | 0.14317138 | 0.88615483 | 0.98391254 |
| NCL | 16938.7777 | -0.1873799 | 0.23336981 | -0.8029313 | 0.42201447 | 0.88451372 |
| NMUR1 | 83.5010665 | 0.42618558 | 0.53332319 | 0.79911316 | 0.4242248 | 0.88565404 |
| PTMA | 30735.8025 | -0.1407415 | 0.28594597 | -0.492196 | 0.62258076 | 0.94016713 |
| PDE6D | 858.882499 | -0.0199113 | 0.17732364 | -0.1122882 | 0.91059492 | 0.98662757 |

| | | | | | | |
|---|---|---|---|---|---|---|
| COPS7B | 2195.65592 | -0.1714605 | 0.19781189 | -0.8667857 | 0.38605946 | 0.87432552 |
| DIS3L2 | 1515.11181 | -0.1382776 | 0.19859896 | -0.6962657 | 0.48626246 | 0.90573694 |
| AC019130.1 | 66.1072564 | 0.12417383 | 0.30041365 | 0.41334283 | 0.67935546 | 0.95070301 |
| ALPI | 523.446869 | -0.5272908 | 0.94686699 | -0.5568795 | 0.57760975 | 0.93130183 |
| ECEL1 | 279.29343 | -1.6385161 | 0.30304094 | -5.4069133 | 6.41E-08 | 3.00E-04 |
| TIGD1 | 470.0784 | -0.0889759 | 0.27053834 | -0.3288846 | 0.74224293 | 0.96109001 |
| EIF4E2 | 4537.08015 | 0.0396802 | 0.20827608 | 0.1905173 | 0.84890379 | 0.97741457 |
| AC073254.1 | 930.694498 | 0.20300765 | 0.32922731 | 0.61661848 | 0.5374864 | 0.92123049 |
| EFHD1 | 8522.63902 | 0.08756538 | 0.31330314 | 0.27949092 | 0.7798681 | 0.96836854 |
| GIGYF2 | 2972.25761 | -0.1169743 | 0.12938151 | -0.9041039 | 0.36594032 | 0.86672412 |
| KCNJ13 | 110.13187 | 1.54146789 | 0.46063416 | 3.34640377 | 0.00081867 | 0.13341488 |
| SNORC | 620.663686 | -0.0227071 | 0.1061743 | -0.2138665 | 0.83065118 | 0.97550416 |
| NGEF | 1024.34004 | 0.60808726 | 0.4863221 | 1.25037966 | 2.11E-01 | 0.79134644 |
| INPP5D | 1125.13878 | -0.1751639 | 0.46221972 | -0.3789624 | 7.05E-01 | 9.55E-01 |
| ATG16L1 | 1759.25325 | -0.3543576 | 0.27158795 | -1.3047619 | 0.19197395 | 0.77371694 |
| DGKD | 1520.46396 | -0.723448 | 0.25208431 | -2.8698652 | 0.00410647 | 0.2778315 |
| USP40 | 2692.55778 | 0.06136802 | 0.26968356 | 0.22755565 | 0.8199917 | 0.97454841 |
| UGT1A8 | 33858.6952 | 0.98996526 | 0.51840388 | 1.90964089 | 0.05617947 | 0.59895387 |
| UGT1A10 | 34061.7343 | 0.98218081 | 0.51827916 | 1.89508064 | 0.05808172 | 0.60193532 |
| UGT1A9 | 41221.7428 | 1.08599088 | 0.53611393 | 2.02567181 | 0.04279843 | 0.56625172 |
| UGT1A7 | 33923.3525 | 0.98639653 | 0.51820426 | 1.9034898 | 0.05697666 | 0.60049913 |
| UGT1A6 | 36578.6949 | 0.89105596 | 0.51976991 | 1.71432771 | 0.08646856 | 0.65132927 |
| AC114812.2 | 2074.22506 | 0.03241847 | 0.58697627 | 0.05522961 | 0.95595554 | 0.99185043 |
| UGT1A5 | 33846.466 | 0.9896134 | 0.51834002 | 1.90919737 | 5.62E-02 | 0.59895387 |
| UGT1A4 | 33955.608 | 0.98543943 | 0.51834458 | 1.90112806 | 0.05728524 | 0.60077875 |
| UGT1A3 | 33856.7217 | 0.98892063 | 0.51846864 | 1.90738753 | 0.05647042 | 0.5996103 |
| UGT1A1 | 34398.9484 | 0.97515998 | 0.5177896 | 1.88331317 | 0.05965794 | 0.6066891 |
| MROH2A | 74.0212771 | 1.65096585 | 0.48084747 | 3.4334502 | 0.00059595 | 0.11756558 |
| ARL4C | 1496.77369 | -0.1759017 | 0.54041801 | -0.325492 | 0.74480874 | 0.9613927 |
| SH3BP4 | 3241.82236 | -0.6431644 | 0.32783522 | -1.9618526 | 0.04977965 | 0.5861488 |
| AGAP1 | 2674.02748 | -0.0262198 | 0.35189215 | -0.0745108 | 0.94060397 | 0.99136103 |
| IQCA1 | 654.004147 | -0.2482822 | 0.29075079 | -0.8539347 | 0.39314118 | 0.87817149 |
| AC093915.1 | 84.6896815 | -0.037437 | 0.32356241 | -0.1157027 | 0.90788815 | 0.98662757 |
| ACKR3 | 853.347612 | -0.2786834 | 0.45410592 | -0.6136968 | 0.53941568 | 0.92195069 |
| COPS8 | 2539.14875 | 0.28370084 | 0.14328307 | 1.98000253 | 0.04770324 | 0.58241466 |
| COL6A3 | 2672.98232 | -0.1614955 | 0.51281681 | -0.3149185 | 0.7528235 | 0.96301881 |
| MLPH | 390.99786 | 1.15338018 | 0.63427308 | 1.81842841 | 0.06899868 | 0.62779744 |
| RAB17 | 8954.36489 | -0.0804706 | 0.25223601 | -0.3190289 | 0.74970458 | 0.96249332 |
| AC104667.2 | 134.713696 | -0.2106378 | 0.35030005 | -0.6013067 | 0.54763571 | 0.92361227 |
| LRRFIP1 | 6198.5838 | -0.3916036 | 0.20713165 | -1.8906026 | 0.05867742 | 0.60332684 |
| RBM44 | 55.5481072 | -0.4637365 | 0.52989884 | -0.8751416 | 0.38149686 | 0.87185666 |
| RAMP1 | 548.535319 | -0.7044847 | 0.62616394 | -1.1250803 | 0.26055503 | 0.82424998 |
| UBE2F | 1616.80207 | -0.1202738 | 0.1451462 | -0.8286392 | 0.40730861 | 0.88132015 |
| UBE2F-SCLY | 2482.25071 | -0.0239457 | 0.18382723 | -0.1302619 | 0.89635925 | 0.98474725 |
| SCLY | 1599.34213 | 0.01794459 | 0.21029115 | 0.0853321 | 0.93199737 | 0.98987582 |
| ESPNL | 80.5357576 | -1.398436 | 0.76532043 | -1.8272555 | 0.06766136 | 0.62557548 |
| KLHL30 | 173.885051 | -0.4294184 | 0.37890702 | -1.1333081 | 0.2570849 | 0.82107009 |
| ERFE | 875.165594 | -0.2204319 | 0.5727254 | -0.3848824 | 0.70032458 | 0.95355954 |
| ILKAP | 1748.48394 | -0.2405083 | 0.1604347 | -1.499104 | 0.13384665 | 0.71505705 |
| HES6 | 331.047014 | -0.2724107 | 0.28964661 | -0.9404933 | 0.34696459 | 0.86150045 |
| PER2 | 751.468123 | -0.2713886 | 0.39649861 | -0.6844629 | 0.49368292 | 0.90749607 |
| TRAF3IP1 | 937.61436 | -0.0971003 | 0.15167695 | -0.6401785 | 0.52205653 | 0.91723878 |
| ASB1 | 1563.08721 | 0.04513712 | 0.20390055 | 0.2213683 | 0.82480567 | 0.97548201 |
| TWIST2 | 49.5555314 | 0.31492651 | 0.62979864 | 0.50004316 | 0.61704468 | 0.93910158 |

| | | | | | | |
|---|---|---|---|---|---|---|
| HDAC4 | 858.429158 | -0.4256233 | 0.17086915 | -2.4909312 | 0.01274088 | 0.41598747 |
| AC062017.1 | 49.4544078 | 0.67782769 | 0.44914317 | 1.50915729 | 0.13125859 | 0.71081672 |
| NDUFA10 | 6338.89601 | 0.40942236 | 0.16723339 | 2.44820934 | 0.01435682 | 0.43749787 |
| COPS9 | 1359.32985 | -0.0656905 | 0.23074407 | -0.2846898 | 0.77588177 | 0.96836854 |
| GPC1 | 5657.33051 | -0.4004365 | 0.34529365 | -1.1596983 | 0.24617166 | 0.81542272 |
| ANKMY1 | 986.23843 | 0.08937696 | 0.1541896 | 0.57965626 | 0.56214645 | 0.92452095 |
| DUSP28 | 304.267589 | -0.0584522 | 0.20888319 | -0.279832 | 0.77960643 | 0.96836854 |
| RNPEPL1 | 5742.0542 | -0.0721429 | 0.17874273 | -0.4036131 | 0.68649724 | 0.95097288 |
| CAPN10-DT | 276.189292 | -0.0818244 | 0.3250254 | -0.2517477 | 0.80123605 | 0.97210205 |
| CAPN10 | 2620.74572 | -0.248774 | 0.19237715 | -1.2931576 | 0.19595656 | 0.77791948 |
| GPR35 | 498.113971 | -0.3449037 | 0.28168397 | -1.224435 | 0.22078815 | 0.7995731 |
| KIF1A | 234.229465 | -2.1567609 | 1.01481594 | -2.125273 | NA | NA |
| AGXT | 1736.95971 | -2.1988098 | 1.15688713 | -1.900626 | 0.05735102 | 0.60112719 |
| C2orf54 | 1043.16037 | 0.92615712 | 0.9108168 | 1.01684238 | 0.30922842 | 0.84414094 |
| CROCC2 | 123.09667 | -0.9183702 | 0.91139429 | -1.0076541 | 0.31362056 | 0.8464135 |
| SNED1 | 2345.16836 | -0.1377671 | 0.20550903 | -0.6703701 | 0.50262187 | 0.91319792 |
| MTERF4 | 2604.16202 | -0.059935 | 0.15067346 | -0.3977809 | 6.91E-01 | 0.95158502 |
| PASK | 799.225428 | 0.29292871 | 0.24764762 | 1.18284483 | 0.23687065 | 0.80849753 |
| PPP1R7 | 2579.44931 | 0.2294318 | 0.13453539 | 1.70536388 | 0.08812658 | 0.65363785 |
| ANO7 | 144.653445 | -0.4586497 | 0.34464817 | -1.3307766 | 0.18326253 | 0.76723162 |
| HDLBP | 17355.9958 | -0.0677933 | 0.18734391 | -0.3618657 | 0.71745238 | 0.95611001 |
| 2-Sep | 17686.598 | 0.04441364 | 0.18906322 | 0.23491421 | 0.81427532 | 0.9738447 |
| AC005104.1 | 1916.12096 | -0.0151884 | 0.19747927 | -0.0769113 | 0.93869408 | 0.99113899 |
| FARP2 | 2017.99924 | -0.1712204 | 0.14572107 | -1.1749869 | 0.23999994 | 0.81209487 |
| STK25 | 7790.4707 | -0.1276988 | 0.11531312 | -1.1074091 | 0.26811707 | 0.82676682 |
| BOK-AS1 | 57.9441978 | -0.2434423 | 0.51942658 | -0.4686751 | 0.63930188 | 0.94088547 |
| BOK | 3253.72614 | -0.1657846 | 0.25394175 | -0.652845 | 0.51385618 | 0.91636259 |
| THAP4 | 4268.36098 | -0.1641575 | 0.10958159 | -1.498039 | 0.13412311 | 0.71560922 |
| ATG4B | 7906.74336 | -0.4810839 | 0.23328933 | -2.062177 | 0.03919089 | 0.55657615 |
| DTYMK | 843.501581 | 0.05358049 | 0.19702876 | 0.2719425 | 0.78566624 | 0.97065795 |
| ING5 | 2625.98777 | -0.192208 | 0.15666176 | -1.2268979 | 0.21986097 | 0.79858446 |
| AC114730.3 | 185.348977 | -0.0993873 | 0.24325634 | -0.4085704 | 0.68285497 | 0.95070301 |
| D2HGDH | 5713.34301 | 0.81363744 | 0.37026126 | 2.19746846 | 0.02798701 | 0.53262541 |
| AC114730.1 | 383.237824 | 0.85130461 | 0.41759556 | 2.03858633 | 0.04149133 | 0.55945849 |
| AC114730.2 | 1044.28741 | 0.872696 | 0.45084833 | 1.93567534 | 0.05290748 | 0.58742503 |
| AC131097.3 | 114.361249 | -0.7293802 | 0.74917298 | -0.9735805 | 0.33026489 | 0.85384238 |
| NEU4 | 392.753646 | -1.4414404 | 0.59484526 | -2.4232191 | 0.01538364 | 0.45243543 |
| PDCD1 | 92.5914798 | 0.04917952 | 0.82606409 | 0.05953475 | 0.95252619 | 0.99170503 |
| LINC01238 | 124.601653 | -0.269536 | 0.54415919 | -0.4953256 | 0.6203703 | 0.9398661 |
| CHL1 | 1340.40671 | 0.44633641 | 0.47901682 | 0.93177608 | 0.35145226 | 0.86364633 |
| LINC01266 | 41.6200657 | 0.66846796 | 0.71861299 | 0.9302197 | 0.35225735 | 0.86364633 |
| CNTN4 | 324.221225 | 0.16167234 | 0.28145638 | 0.5744135 | 0.56568802 | 0.92513982 |
| TRNT1 | 1557.32292 | -0.0486262 | 0.16667775 | -0.2917376 | 0.77048723 | 0.96836854 |
| CRBN | 2351.48507 | 0.05691449 | 0.17480812 | 0.32558266 | 0.74474013 | 0.9613927 |
| AC024060.1 | 405.004315 | 0.07740768 | 0.17050733 | 0.45398447 | 0.64984001 | 0.94312988 |
| SUMF1 | 3249.50614 | 0.66207002 | 0.22728955 | 2.91289245 | 0.00358098 | 0.26033637 |
| LRRN1 | 82.7902596 | 0.04678421 | 0.53746113 | 0.08704669 | 0.9306344 | 0.98975562 |
| SETMAR | 518.558357 | 0.19947663 | 0.18974024 | 1.05131434 | 0.29311424 | 0.8352732 |
| ITPR1-DT | 67.1122153 | -0.1792977 | 0.56366411 | -0.3180932 | 0.75041426 | 0.96255347 |
| ITPR1 | 2472.114 | -0.1150306 | 0.16617999 | -0.6922048 | 0.48880875 | 0.905872 |
| EGOT | 237.717085 | 0.70622414 | 0.64679325 | 1.09188546 | 0.27488345 | 0.82719407 |
| AC018816.1 | 361.910926 | -0.1310585 | 0.19623046 | -0.6678807 | 0.50420975 | 0.91356552 |
| BHLHE40-AS | 805.446908 | 0.03896233 | 0.5765444 | 0.06757907 | 0.94612072 | 0.99166614 |
| BHLHE40 | 21124.6217 | 0.09457778 | 0.56631568 | 0.1670054 | 0.8673658 | 0.98224199 |

| | | | | | | |
|---|---|---|---|---|---|---|
| ARL8B | 5421.37487 | -0.4731107 | 0.30545149 | -1.5488897 | 0.12140823 | 0.69687955 |
| EDEM1 | 2638.58185 | -0.0423407 | 0.26967834 | -0.1570043 | 0.87524144 | 0.98275231 |
| LMCD1 | 3249.62415 | -1.8563291 | 0.61721249 | -3.0076013 | 0.00263318 | 0.21835619 |
| SSUH2 | 69.6673424 | 0.30258762 | 0.65458099 | 0.46226156 | 6.44E-01 | 9.42E-01 |
| OXTR | 60.4660545 | 0.31959208 | 0.54080689 | 0.59095416 | 0.55455113 | 0.92373235 |
| RAD18 | 343.785323 | -0.2014988 | 0.27105069 | -0.7433988 | 0.45724027 | 0.89688594 |
| SRGAP3 | 711.913789 | 1.01811085 | 0.38353195 | 2.6545659 | 0.00794105 | 0.37071188 |
| AC026191.1 | 135.882658 | 0.0561251 | 0.2995593 | 0.18735889 | 8.51E-01 | 9.79E-01 |
| THUMPD3-A: | 1533.03755 | -0.0983512 | 0.18919496 | -0.5198404 | 6.03E-01 | 9.36E-01 |
| THUMPD3 | 1816.8082 | -0.1219479 | 0.16779339 | -0.726774 | 4.67E-01 | 0.89983399 |
| SETD5 | 4162.17557 | -0.3283283 | 0.18747969 | -1.7512737 | 0.07989877 | 0.64079926 |
| MTMR14 | 2584.06248 | 0.03055625 | 0.17644317 | 0.173179 | 8.63E-01 | 9.82E-01 |
| BRPF1 | 1136.24329 | -0.3305341 | 0.22881764 | -1.4445305 | 1.49E-01 | 0.73553131 |
| OGG1 | 4379.20854 | 0.23801284 | 0.29012604 | 0.82037737 | 4.12E-01 | 8.83E-01 |
| CAMK1 | 2348.1462 | 0.04331858 | 0.27487497 | 0.15759375 | 0.87477693 | 0.98275231 |
| TADA3 | 4443.24439 | -0.0472339 | 0.08917986 | -0.5296472 | 0.59635654 | 0.9334601 |
| ARPC4 | 8034.91013 | 0.11852573 | 0.1331773 | 0.88998445 | 0.37347424 | 0.86791605 |
| ARPC4-TTLL3 | 6193.36936 | 0.10635734 | 0.13260927 | 0.80203551 | 0.42253242 | 0.88451372 |
| TTLL3 | 3857.92742 | 0.05122419 | 0.32773898 | 0.15629569 | 0.87579995 | 0.98275231 |
| AC022382.1 | 125.357007 | 0.13863392 | 0.3781386 | 0.36662197 | 0.71390102 | 0.95564093 |
| RPUSD3 | 1556.46615 | 0.09246002 | 0.17344581 | 0.53307728 | 0.59398009 | 0.93317501 |
| CIDEC | 114.729841 | 4.2054721 | 1.24572717 | 3.37591747 | 0.0007357 | 0.12885748 |
| JAGN1 | 1829.26372 | -0.1947706 | 0.15288786 | -1.2739439 | 2.03E-01 | 7.82E-01 |
| IL17RE | 541.938717 | 0.14138579 | 0.49404884 | 0.28617777 | 0.77474197 | 0.96836854 |
| IL17RC | 2456.94523 | 0.10569589 | 0.28874519 | 0.36605248 | 0.71432592 | 0.95579447 |
| CRELD1 | 3324.64522 | 0.05176015 | 0.37652776 | 0.13746702 | 0.89066166 | 0.98474725 |
| PRRT3 | 233.474886 | -0.0312607 | 0.29733757 | -0.1051354 | 0.9162684 | 0.9868044 |
| AC018809.2 | 140.007996 | 0.00143501 | 0.26343897 | 0.00544722 | 0.99565377 | 0.99940889 |
| EMC3 | 3309.76067 | 0.02503448 | 0.18209402 | 0.13748109 | 0.89065054 | 0.98474725 |
| AC022007.1 | 190.805226 | 0.41050235 | 0.33264538 | 1.23405397 | 0.2171828 | 0.79574248 |
| FANCD2 | 185.777217 | 0.29936088 | 0.39123455 | 0.76516984 | 0.44417042 | 0.8938278 |
| FANCD2OS | 49.6467915 | 0.25552011 | 0.50815207 | 0.50284181 | 0.6150755 | 0.93895494 |
| BRK1 | 6502.02535 | 0.30337315 | 0.18822873 | 1.61172612 | 1.07E-01 | 6.77E-01 |
| VHL | 2261.64144 | -0.2003666 | 0.1482866 | -1.351212 | 0.17662752 | 0.76115956 |
| IRAK2 | 817.61219 | -0.0564536 | 0.28205758 | -0.2001492 | 0.84136392 | 0.97618641 |
| TATDN2 | 2604.05157 | -0.3146413 | 0.14871475 | -2.1157372 | 0.03436716 | 0.55369025 |
| AC022384.1 | 1360.61637 | -0.237296 | 0.21711768 | -1.0929373 | 0.27442132 | 0.82719407 |
| LINC00852 | 73.286578 | -0.2786382 | 0.60430889 | -0.4610856 | 0.64473716 | 0.94208859 |
| GHRL | 81.9485327 | -0.1066109 | 0.37533466 | -0.2840422 | 0.77637801 | 0.96836854 |
| GHRLOS | 79.7488025 | -0.1004847 | 0.36247529 | -0.277218 | 0.78161272 | 0.96913822 |
| SEC13 | 5559.96047 | -0.0343537 | 0.25763818 | -0.1333408 | 0.89392384 | 0.98474725 |
| SLC6A1 | 159.789132 | -0.4586793 | 0.66610943 | -0.6885946 | 0.49107845 | 0.90600166 |
| HRH1 | 472.048486 | 0.18432216 | 0.39957527 | 0.46129522 | 0.64458682 | 0.94208859 |
| ATG7 | 3976.95838 | 0.0770426 | 0.1133423 | 0.67973384 | 0.49667301 | 0.90890243 |
| VGLL4 | 4822.18579 | 0.09437821 | 0.11896934 | 0.79329865 | 0.42760384 | 0.88790684 |
| TAMM41 | 579.376504 | 0.35265811 | 0.16665677 | 2.11607433 | 0.03433849 | 0.55369025 |
| SYN2 | 274.813438 | 0.58239931 | 0.38730218 | 1.50373364 | 0.13264996 | 0.71328342 |
| TIMP4 | 121.258672 | 0.25071672 | 0.63577115 | 0.39435058 | 0.69332222 | 0.9519683 |
| PPARG | 1224.63608 | -0.1711158 | 0.24158522 | -0.7083041 | 0.4787564 | 0.90444322 |
| TSEN2 | 610.181136 | -0.0114804 | 0.17356432 | -0.066145 | 0.94726236 | 0.99166614 |
| MKRN2OS | 83.5661578 | 0.33220199 | 0.35395866 | 0.9385333 | 0.34797041 | 0.86226544 |
| MKRN2 | 1854.24996 | -0.0703618 | 0.12536106 | -0.5612735 | 0.5746111 | 0.93024203 |
| RAF1 | 6206.64914 | -0.2308826 | 0.16340377 | -1.4129576 | 0.15766818 | 0.74522306 |
| CAND2 | 327.939907 | -0.1662923 | 0.23709933 | -0.7013613 | 0.48307758 | 0.90541935 |

| | | | | | | |
|---|---|---|---|---|---|---|
| RPL32 | 19169.3015 | 0.15840316 | 0.16569217 | 0.95600871 | 0.33906783 | 0.85733087 |
| IQSEC1 | 2556.62509 | 0.07079272 | 0.21265159 | 0.33290473 | 0.7392062 | 0.9602325 |
| NUP210 | 2088.52429 | 0.19255359 | 0.26121113 | 0.7371569 | 0.46102694 | 0.89755243 |
| HDAC11 | 2528.86639 | 0.41905451 | 0.33427144 | 1.2536354 | 0.20997456 | 0.79134644 |
| FBLN2 | 1474.06971 | 0.44878449 | 0.80464622 | 0.55774137 | 0.577021 | 0.93129843 |
| CHCHD4 | 957.314379 | 0.22901371 | 0.27550238 | 0.83125854 | 0.40582759 | 0.88132015 |
| TMEM43 | 4607.94238 | -0.1244777 | 0.13929955 | -0.8935975 | 0.37153729 | 0.86762749 |
| AC090004.1 | 2319.37694 | -0.0362601 | 0.12763119 | -0.2841007 | 0.77633318 | 0.96836854 |
| AC093495.1 | 2882.31938 | -0.0159702 | 0.20378852 | -0.0783665 | 0.93753653 | 0.99082908 |
| XPC | 3944.54067 | 0.01821926 | 0.23004009 | 0.07920039 | 0.93687324 | 0.99068685 |
| LSM3 | 2009.77142 | 0.29561919 | 0.19978514 | 1.47968558 | 0.13895717 | 0.72399122 |
| AC093496.1 | 80.7878292 | -1.8216903 | 0.97268817 | -1.8728411 | 0.06109034 | 0.60989294 |
| SLC6A6 | 7746.84915 | -0.4884771 | 0.47374676 | -1.0310932 | 0.3024971 | 0.84211658 |
| GRIP2 | 1199.89149 | -0.6690743 | 0.39565702 | -1.6910461 | 0.09082801 | 0.65495306 |
| CCDC174 | 1255.31496 | -0.0022379 | 0.15691314 | -0.0142623 | 0.98862074 | 0.99788078 |
| FGD5 | 2485.8767 | -0.011688 | 0.31131189 | -0.0375444 | 0.97005097 | 0.99400331 |
| FGD5-AS1 | 6920.43228 | -0.1246605 | 0.17856376 | -0.6981291 | 0.48509648 | 0.90543654 |
| NR2C2 | 4015.42829 | -0.4286627 | 0.18399244 | -2.3297841 | 0.01981756 | 0.49477123 |
| MRPS25 | 4871.38399 | -0.2227041 | 0.16025869 | -1.3896536 | 0.16463408 | 0.75286112 |
| RBSN | 1333.39091 | -0.0830419 | 0.13365021 | -0.6213379 | 0.5343773 | 0.92084006 |
| CAPN7 | 1692.01054 | -0.0092683 | 0.13688046 | -0.0677105 | 0.94601605 | 0.99166614 |
| SH3BP5-AS1 | 6282.39799 | -0.4658635 | 0.3294347 | -1.4141299 | 0.15732376 | 0.74522306 |
| SH3BP5 | 6403.36671 | -0.450402 | 0.34688816 | -1.298407 | 0.19414752 | 0.7762856 |
| METTL6 | 657.132493 | -0.1429889 | 0.19279506 | -0.7416629 | 0.45829163 | 0.8971553 |
| EAF1 | 1604.55533 | 0.04158181 | 0.16156621 | 0.25736702 | 0.79689546 | 0.97135726 |
| EAF1-AS1 | 550.461853 | 0.0307969 | 0.23264936 | 0.13237475 | 8.95E-01 | 9.85E-01 |
| COLQ | 359.785707 | 0.08613375 | 0.34657491 | 0.24852852 | 0.80372551 | 0.97274243 |
| HACL1 | 1406.77978 | 0.53927851 | 0.25024206 | 2.15502743 | 0.03115969 | 0.54466273 |
| BTD | 1658.26751 | 0.29773325 | 0.21843505 | 1.36302873 | 1.73E-01 | 0.75974924 |
| ANKRD28 | 2303.14409 | -0.1331235 | 0.16534486 | -0.8051264 | 0.42074676 | 0.88451372 |
| GALNT15 | 1205.04746 | -0.0004735 | 0.4347178 | -0.0010891 | 0.99913099 | 0.99994357 |
| DPH3 | 1320.0913 | -0.2230258 | 0.29389435 | -0.7588638 | 0.44793402 | 0.89480207 |
| OXNAD1 | 880.980278 | 0.12851312 | 0.14920138 | 0.86134001 | 0.3890508 | 0.87568585 |
| RFTN1 | 1503.19804 | 0.06728808 | 0.32955098 | 0.2041811 | 0.83821198 | 0.97599451 |
| PLCL2 | 1168.00743 | -0.0841376 | 0.20704277 | -0.4063777 | 0.68446511 | 0.95075309 |
| PLCL2-AS1 | 117.511173 | 0.23912741 | 0.26748981 | 0.89396828 | 0.37133886 | 0.86762749 |
| TBC1D5 | 2179.18699 | 0.33036167 | 0.22682371 | 1.45646883 | 0.14526306 | 0.73103518 |
| SATB1 | 1381.78773 | 0.23235056 | 0.28856043 | 0.80520589 | 0.42070088 | 0.88451372 |
| EFHB | 153.075882 | -0.0757223 | 0.51228135 | -0.1478139 | 0.88248965 | 0.98275231 |
| RAB5A | 3533.8504 | -0.186995 | 0.17272534 | -1.0826149 | 0.27897938 | 0.82825734 |
| PP2D1 | 608.495988 | -0.2248833 | 0.22566062 | -0.9965554 | 0.31898035 | 0.84914926 |
| KAT2B | 1572.34716 | -0.1495625 | 0.14600175 | -1.0243884 | 0.30565185 | 0.84270033 |
| ZNF385D | 321.33289 | -0.333309 | 0.62129419 | -0.5364754 | 0.59163003 | 0.93289552 |
| UBE2E2 | 1515.01404 | -0.0117247 | 0.2163664 | -0.0541889 | 0.95678468 | 0.99198394 |
| UBE2E1 | 3347.9164 | -0.0750884 | 0.13851589 | -0.5420922 | 0.58775497 | 0.93289552 |
| NKIRAS1 | 836.043722 | 0.01641881 | 0.21523116 | 0.07628453 | 0.93919274 | 0.99113899 |
| RPL15 | 30532.4467 | 0.21503082 | 0.22609912 | 0.95104666 | 0.34158069 | 0.85889221 |
| NR1D2 | 1647.72415 | -0.1954458 | 0.52022537 | -0.3756945 | 0.70714402 | 0.95501548 |
| THRB | 866.821759 | -0.0279727 | 0.27137701 | -0.1030769 | 0.91790197 | 0.9868044 |
| RARB | 824.35283 | -0.6263214 | 0.27069235 | -2.3137757 | 0.02068003 | 0.50488816 |
| TOP2B | 4805.09143 | 0.06266992 | 0.16236504 | 0.38598163 | 0.69951027 | 0.95307625 |
| NGLY1 | 1849.50924 | -0.1233402 | 0.1815132 | -0.6795108 | 0.49681428 | 0.90890243 |
| OXSM | 986.230591 | 0.11719708 | 0.23374261 | 0.50139372 | 0.61609406 | 0.93910158 |
| NEK10 | 122.057234 | 0.3768783 | 0.53577743 | 0.70342324 | 0.48179203 | 0.90541935 |

| | | | | | | |
|---|---|---|---|---|---|---|
| SLC4A7 | 1418.88854 | -0.0813745 | 0.38362122 | -0.212122 | 0.83201182 | 0.97550416 |
| EOMES | 42.6531677 | -0.0459085 | 0.89443671 | -0.0513267 | 0.95906519 | 0.99242674 |
| CMC1 | 3035.12347 | -0.1393097 | 0.12895926 | -1.0802613 | 0.28002585 | 0.82825734 |
| AZI2 | 2666.20228 | -0.1337432 | 0.18773758 | -0.7123945 | 0.47622049 | 0.90305051 |
| ZCWPW2 | 76.559488 | 0.40168428 | 0.33400818 | 1.20261809 | 0.22912414 | 0.80521698 |
| AC098650.1 | 159.214583 | -0.2801675 | 0.46745344 | -0.5993485 | 0.54894052 | 0.92373235 |
| RBMS3 | 979.953281 | -0.200431 | 0.38476913 | -0.5209123 | 0.60242785 | 0.9355118 |
| TGFBR2 | 10859.8183 | -0.0205342 | 0.30992485 | -0.0662555 | 9.47E-01 | 9.92E-01 |
| STT3B | 5919.80173 | 0.13126884 | 0.11251047 | 1.1667255 | 0.24332124 | 0.8135742 |
| OSBPL10 | 2348.83618 | -0.0239614 | 0.22400706 | -0.106967 | 0.91481514 | 0.9866558 |
| ZNF860 | 92.1730486 | 0.00067896 | 0.28000694 | 0.00242481 | 0.99806529 | 0.99990633 |
| GPD1L | 2373.29363 | 0.66490884 | 0.28045647 | 2.3708094 | 0.01774918 | 0.4711578 |
| CMTM8 | 397.94718 | 0.12216248 | 0.30694027 | 0.39800081 | 0.69062959 | 0.95158502 |
| CMTM7 | 1690.6272 | 0.22291314 | 0.20210869 | 1.10293696 | 0.27005454 | 0.82716979 |
| CMTM6 | 4489.40228 | 0.08862276 | 0.16961742 | 0.5224862 | 0.60133186 | 0.93531084 |
| DYNC1LI1 | 1944.112 | 0.00553126 | 0.2111263 | 0.02619883 | 9.79E-01 | 0.99502683 |
| CNOT10 | 1145.34105 | -0.0454161 | 0.14684984 | -0.3092693 | 0.75711671 | 0.96499587 |
| CNOT10-AS1 | 90.668114 | 0.1116919 | 0.21600642 | 0.51707678 | 0.60510256 | 0.93687349 |
| TRIM71 | 226.69915 | -0.3131032 | 0.76207549 | -0.4108559 | 0.68117823 | 0.95070301 |
| CCR4 | 44.8419706 | 0.53911718 | 0.92055357 | 0.58564455 | 0.5581144 | 0.92421175 |
| GLB1 | 5820.17157 | 0.37024114 | 0.19094726 | 1.93897066 | 0.05250491 | 0.58645127 |
| TMPPE | 82.8142859 | -0.2670553 | 0.4180295 | -0.6388431 | 0.52292501 | 0.91723878 |
| CRTAP | 8166.29274 | 0.42279564 | 0.2105057 | 2.00847596 | 0.04459274 | 0.57267043 |
| AC112211.1 | 538.179327 | 0.5027237 | 0.3436967 | 1.46269573 | 0.14355066 | 0.72905668 |
| SUSD5 | 125.23669 | -1.1073123 | 0.50224988 | -2.2047039 | 0.02747488 | 0.52904542 |
| FBXL2 | 750.906703 | 0.02456707 | 0.30857837 | 0.07961371 | 0.93654449 | 0.99068685 |
| UBP1 | 3116.08256 | -0.2491117 | 0.11052891 | -2.2538144 | 0.02420784 | 0.51967919 |
| CLASP2 | 1738.52686 | 0.15910409 | 0.12962023 | 1.22746349 | 0.21964844 | 0.79858446 |
| AC112220.3 | 54.4036825 | -0.871695 | 0.49011998 | -1.778534 | 0.0753162 | 0.63773415 |
| AC112220.4 | 80.6672495 | -1.0130924 | 0.5221274 | -1.9403165 | 0.05234124 | 0.58645127 |
| PDCD6IP | 7700.46271 | -0.1796481 | 0.18601366 | -0.965779 | 0.33415478 | 0.85533716 |
| ARPP21 | 48.1099323 | -0.1913995 | 0.59104661 | -0.3238316 | 0.74606556 | 0.96182257 |
| STAC | 239.741568 | -0.0675169 | 0.34590955 | -0.1951864 | 0.84524699 | 0.97675487 |
| TRANK1 | 985.586305 | -1.1148274 | 0.71454246 | -1.5601975 | 0.11871321 | 0.69200756 |
| EPM2AIP1 | 1910.35264 | -0.0702049 | 0.1772826 | -0.3960058 | 0.69210073 | 0.95169563 |
| MLH1 | 1438.76808 | 0.14142447 | 0.1250547 | 1.13090091 | 0.2580968 | 0.822338 |
| LRRFIP2 | 3163.5848 | -0.1150759 | 0.21756816 | -0.5289187 | 0.59686182 | 0.9337017 |
| AC126118.1 | 40.9653068 | 0.43084213 | 0.41782105 | 1.03116426 | 0.3024638 | 0.84211658 |
| AC097359.2 | 214.09378 | -0.0377078 | 0.48658311 | -0.0774951 | 0.93822967 | 0.99113899 |
| GOLGA4 | 4487.15573 | -0.2900439 | 0.24120499 | -1.2024787 | 0.22917813 | 0.80521698 |
| ITGA9 | 857.597039 | -0.2638535 | 0.39651137 | -0.6654374 | 0.50577078 | 0.91386909 |
| CTDSPL | 5016.68956 | 0.03361379 | 0.1603329 | 0.20964996 | 0.83394088 | 0.97550416 |
| VILL | 520.292886 | 0.19058746 | 0.34271118 | 0.55611685 | 0.57813097 | 0.93130183 |
| PLCD1 | 1655.15218 | 0.36160882 | 0.31140925 | 1.16120129 | 0.24556005 | 0.81542272 |
| DLEC1 | 4897.21512 | 0.83809885 | 0.36755409 | 2.28020551 | 0.0225955 | 0.51958567 |
| ACAA1 | 14212.4182 | 0.94837622 | 0.39471065 | 2.40271249 | 0.01627398 | 0.45896904 |
| MYD88 | 2406.70033 | -0.4049945 | 0.26832274 | -1.5093558 | 0.13120788 | 0.71081672 |
| OXSR1 | 3922.32423 | -0.7239432 | 0.3338972 | -2.168162 | 0.03014636 | 0.54012717 |
| SLC22A13 | 804.893728 | 0.205655 | 0.78077032 | 0.26340013 | 7.92E-01 | 0.97090793 |
| SLC22A14 | 69.4007257 | 0.04141201 | 0.62526472 | 0.06623116 | 0.94719379 | 0.99166614 |
| XYLB | 461.693559 | 0.5696304 | 0.50979867 | 1.11736345 | 0.26383896 | 0.82424998 |
| ACVR2B-AS1 | 137.159786 | -0.0309015 | 0.35509307 | -0.0870238 | 0.9306526 | 0.98975562 |
| ACVR2B | 790.169413 | 0.05076349 | 0.14849273 | 0.34185839 | 0.73245747 | 0.95848028 |
| EXOG | 510.135798 | -0.1862483 | 0.21269395 | -0.8756632 | 0.38121314 | 0.87171941 |

| | | | | | | |
|---|---|---|---|---|---|---|
| SCN11A | 55.540316 | 0.41132746 | 0.65133012 | 0.63151918 | 0.52770111 | 0.91842929 |
| WDR48 | 2497.08194 | -0.030845 | 0.17424349 | -0.1770225 | 0.85949075 | 0.9802651 |
| GORASP1 | 4046.25522 | -0.1684594 | 0.08258214 | -2.0399005 | 0.04136024 | 0.55943411 |
| TTC21A | 582.308252 | -0.2262137 | 0.37919431 | -0.596564 | 0.55079848 | 0.92373235 |
| CSRNP1 | 3527.58533 | -0.6615472 | 0.39165629 | -1.6891014 | 9.12E-02 | 6.55E-01 |
| AC092053.2 | 82.5986943 | -0.745455 | 0.48321961 | -1.5426836 | 0.12290755 | 0.69788192 |
| CX3CR1 | 152.484562 | 1.12668063 | 0.57409353 | 1.96253846 | 0.04969983 | 0.5861488 |
| SLC25A38 | 2592.09745 | -0.206473 | 0.24226891 | -0.8522473 | 0.39407687 | 0.87884163 |
| RPSA | 26185.7675 | 0.24867125 | 0.20388905 | 1.21964002 | 0.22260137 | 0.80078694 |
| MYRIP | 225.238742 | -0.5686886 | 0.42813369 | -1.3282969 | 0.18408005 | 0.76816707 |
| EIF1B | 3325.91143 | -0.4098846 | 0.27595201 | -1.4853473 | 0.13745186 | 0.72209298 |
| ENTPD3-AS1 | 334.91086 | 0.40844171 | 0.26651914 | 1.53250424 | 0.12539805 | 0.70277658 |
| ENTPD3 | 112.074979 | 0.86151361 | 0.48345744 | 1.78198439 | 0.07475178 | 0.63773415 |
| RPL14 | 19194.0318 | 0.11308061 | 0.22009113 | 0.51378992 | 0.60739887 | 0.93739614 |
| ZNF619 | 207.477758 | -0.0266355 | 0.24007642 | -0.1109461 | 0.91165909 | 0.98662757 |
| ZNF620 | 170.133844 | -0.2823086 | 0.28973993 | -0.9743517 | 0.32988193 | 0.85371079 |
| ZNF621 | 1272.77862 | -0.1934868 | 0.13859475 | -1.3960614 | 0.162696 | 0.75032457 |
| CTNNB1 | 12846.4003 | -0.1661086 | 0.14074216 | -1.1802336 | 0.2379073 | 0.80962787 |
| ULK4 | 311.034841 | 0.22425363 | 0.37596 | 0.59648268 | 5.51E-01 | 9.24E-01 |
| TRAK1 | 3707.19266 | -0.1470486 | 0.17077127 | -0.8610849 | 0.38919126 | 0.87568585 |
| VIPR1 | 162.716701 | 1.01142914 | 0.60416397 | 1.6740971 | 0.09411152 | 0.65783452 |
| SEC22C | 2264.23579 | -0.0478217 | 0.18838242 | -0.2538546 | 0.79960789 | 0.97207132 |
| SS18L2 | 763.78161 | -0.0128911 | 0.16359817 | -0.0787975 | 0.93719372 | 0.99077727 |
| NKTR | 4463.93793 | -0.0709335 | 0.17053146 | -0.4159553 | 0.67744274 | 0.95018034 |
| ZBTB47 | 1455.2812 | -0.2553266 | 0.31723313 | -0.8048548 | 0.42090349 | 0.88451372 |
| CCDC13 | 293.95944 | 0.04492864 | 0.32965695 | 0.13628908 | 0.89159276 | 0.98474725 |
| CCDC13-AS1 | 135.820908 | 0.02325051 | 0.46258001 | 0.05026267 | 9.60E-01 | 9.93E-01 |
| AC099329.3 | 3531.42205 | 0.2495345 | 0.1978241 | 1.26139589 | 2.07E-01 | 7.88E-01 |
| HIGD1A | 8847.4171 | 0.21677241 | 0.17116739 | 1.2664352 | 2.05E-01 | 7.86E-01 |
| AC099329.1 | 4961.16026 | 0.25373016 | 0.19210766 | 1.32077063 | 0.18657786 | 0.76915147 |
| ACKR2 | 118.556509 | 0.1946797 | 0.43472917 | 0.44781836 | 0.65428429 | 0.94532497 |
| AC092042.3 | 134.318984 | 0.08848447 | 0.48080194 | 0.18403517 | 0.85398586 | 0.97906925 |
| KRBOX1 | 114.000489 | 0.29565629 | 0.45211127 | 0.65394586 | 0.51314668 | 0.91580631 |
| CYP8B1 | 1371.51929 | 1.04262195 | 0.55145275 | 1.89068228 | 0.05866677 | 0.60332684 |
| ZNF662 | 477.925537 | 0.180371 | 0.33451956 | 0.5391942 | 0.58975287 | 0.93289552 |
| FAM198A | 161.703213 | -0.2368776 | 0.43729072 | -0.5416936 | 0.58802961 | 0.93289552 |
| POMGNT2 | 1364.99154 | -0.0540344 | 0.11219746 | -0.4816013 | 0.63008918 | 0.94088547 |
| SNRK | 3586.60172 | -0.1907789 | 0.21815259 | -0.8745203 | 0.38183496 | 0.87185666 |
| AC104184.1 | 218.314372 | -0.1628365 | 0.34718391 | -0.4690209 | 0.63905469 | 0.94088547 |
| SNRK-AS1 | 678.52989 | -0.1365709 | 0.22774403 | -0.5996685 | 0.54872718 | 0.92373235 |
| ANO10 | 2736.71996 | -0.0102323 | 0.22246143 | -0.0459957 | 0.96331371 | 0.99320711 |
| ABHD5 | 2103.75809 | -0.5418173 | 0.45288403 | -1.1963709 | 0.23155185 | 0.80649152 |
| AC006058.1 | 412.655657 | -0.7298805 | 0.6273061 | -1.1635157 | 0.24462034 | 0.81473945 |
| AC104187.1 | 49.181628 | -0.1985048 | 0.43269088 | -0.4587682 | 6.46E-01 | 0.94208859 |
| TCAIM | 2357.10643 | 0.17394972 | 0.21850317 | 0.79609699 | 0.42597566 | 0.88643237 |
| ZNF445 | 1291.82517 | -0.0949995 | 0.17917142 | -0.5302156 | 0.59596245 | 0.9334601 |
| ZNF852 | 56.6720239 | -0.1661584 | 0.3571154 | -0.4652794 | 0.64173142 | 0.94088547 |
| ZKSCAN7 | 255.703166 | -0.1527237 | 0.29636506 | -0.5153228 | 0.60632749 | 0.93708311 |
| ZNF660 | 118.645006 | 0.31588578 | 0.30371627 | 1.04006869 | 0.29830799 | 0.84005861 |
| ZNF197 | 657.732461 | 0.0215651 | 0.18170579 | 0.1186814 | 0.90552777 | 0.98625536 |
| ZNF35 | 266.459995 | -0.4159467 | 0.32399297 | -1.2838139 | 0.19920709 | 0.78134034 |
| AC124045.1 | 138.031848 | -0.7881039 | 0.41219734 | -1.9119578 | 0.05588161 | 0.59889467 |
| ZNF502 | 314.386627 | -0.2501472 | 0.25723929 | -0.9724302 | 0.3308366 | 0.85384238 |
| ZNF501 | 117.907066 | 0.03517466 | 0.32284957 | 0.10895062 | 0.91324165 | 0.98662757 |

| | | | | | | |
|---|---|---|---|---|---|---|
| KIAA1143 | 1003.64077 | 0.036565 | 0.18678271 | 0.19576222 | 0.84479628 | 0.97675487 |
| TMEM42 | 935.346433 | 0.19553647 | 0.31648009 | 0.61784763 | 0.53667578 | 0.92105614 |
| ZDHHC3 | 4133.56783 | 0.12682913 | 0.09269709 | 1.36821041 | 0.17124622 | 0.75956362 |
| EXOSC7 | 3357.84168 | 0.40841999 | 0.22646018 | 1.8034958 | 0.07131038 | 0.63128386 |
| CLEC3B | 2471.11007 | 0.67615096 | 0.36849097 | 1.83491868 | 0.06651774 | 0.62378208 |
| CDCP1 | 998.561549 | 0.23381285 | 0.31948864 | 0.73183463 | 0.46426951 | 0.89830907 |
| TMEM158 | 122.392917 | -0.706035 | 0.60620723 | -1.1646761 | 0.24415014 | 0.81468457 |
| LARS2 | 1670.57414 | 0.0688369 | 0.17329443 | 0.39722512 | 0.69120146 | 0.95158502 |
| LIMD1 | 1429.67259 | -0.1545744 | 0.1226082 | -1.2607184 | 0.20741033 | 0.78813404 |
| SACM1L | 3135.95538 | -0.0475284 | 0.10406713 | -0.4567086 | 0.64788051 | 0.94285781 |
| SLC6A20 | 1651.93136 | -0.4743391 | 0.40102768 | -1.1828089 | 0.2368849 | 0.80849753 |
| LZTFL1 | 1411.43689 | -0.1521274 | 0.19235005 | -0.7908884 | 0.42900911 | 0.88851358 |
| CCR9 | 52.4906114 | -0.6867603 | 0.76615963 | -0.8963671 | 0.37005673 | 0.86762749 |
| FYCO1 | 1944.69878 | 0.01637027 | 0.15922542 | 0.10281193 | 0.91811224 | 0.9868044 |
| CXCR6 | 61.0701536 | 0.07060326 | 0.74707209 | 0.09450663 | 0.92470672 | 0.98869145 |
| CCR1 | 501.007878 | -0.5507619 | 0.5181956 | -1.0628456 | 0.28785199 | 0.83266626 |
| CCR2 | 122.505676 | -0.5559007 | 0.79486075 | -0.6993687 | 0.48432164 | 0.90541935 |
| CCR5 | 144.50902 | -0.3664807 | 0.6355779 | -0.5766101 | 0.56420285 | 0.92470993 |
| CCRL2 | 140.88592 | -0.7547147 | 0.28249619 | -2.6715926 | 0.00754922 | 0.36652851 |
| LINC02009 | 541.862821 | -0.1544298 | 0.67826467 | -0.2276836 | 0.81989219 | 0.97454841 |
| LTF | 67621.8477 | 0.35350282 | 1.0442939 | 0.33850894 | 0.73497969 | 0.95895327 |
| LRRC2 | 919.662678 | -0.7308978 | 0.39478083 | -1.8514014 | 0.06411183 | 0.61837523 |
| TDGF1 | 1877.21882 | -1.8706407 | 0.76039968 | -2.4600756 | 0.01389077 | 0.4295825 |
| AC104304.1 | 261.647387 | -1.6584116 | 0.69097462 | -2.400105 | 0.01639037 | 0.45915089 |
| ALS2CL | 8850.79745 | -0.1571479 | 0.30692148 | -0.5120132 | 0.60864176 | 0.93748023 |
| TMIE | 70.9200908 | -0.1705116 | 0.32166439 | -0.5300915 | 0.59604849 | 0.9334601 |
| PRSS50 | 258.536945 | -0.3014635 | 0.4687917 | -0.6430649 | 0.52018202 | 0.91657872 |
| PRSS45 | 60.2678208 | -0.5373828 | 0.54352098 | -0.9887066 | 0.3228067 | 0.85119105 |
| AC109583.3 | 128.761466 | -0.5822124 | 0.48125231 | -1.2097861 | 0.22636096 | 0.80246466 |
| PRSS42 | 42.6756375 | 0.60234651 | 0.69760658 | 0.86344729 | 0.38789159 | 0.87510248 |
| MYL3 | 1103.26418 | 0.76796414 | 0.28116191 | 2.73139469 | 0.00630669 | 0.33470127 |
| PTH1R | 17592.2569 | 0.91028179 | 0.46395494 | 1.96200472 | 0.04976194 | 0.5861488 |
| CCDC12 | 1897.22141 | -0.0492947 | 0.2028187 | -0.2430483 | 0.80796797 | 0.97334138 |
| NBEAL2 | 4899.7175 | -0.3588315 | 0.21316114 | -1.6833815 | 0.09230127 | 0.65583276 |
| SETD2 | 3222.50348 | -0.3704399 | 0.14485444 | -2.557325 | 0.01054806 | 0.40196898 |
| KIF9-AS1 | 295.207958 | 0.11278671 | 0.30919494 | 0.36477539 | 0.71527908 | 0.95590747 |
| KIF9 | 552.745897 | 0.22816077 | 0.15869072 | 1.4377701 | 0.1504993 | 0.73796431 |
| KLHL18 | 934.33447 | -0.3701805 | 0.18045814 | -2.051337 | 0.04023414 | 0.55657615 |
| AC099778.1 | 107.549194 | -0.2129135 | 0.34708192 | -0.6134388 | 0.53958621 | 0.92198621 |
| PTPN23 | 6209.69855 | -0.6247873 | 0.20938589 | -2.9839037 | 0.00284596 | 0.23089261 |
| SCAP | 5828.29613 | -0.0494258 | 0.09202578 | -0.5370864 | 0.59120793 | 0.93289552 |
| ELP6 | 1249.39421 | -0.0865293 | 0.16336613 | -0.5296651 | 0.59634414 | 0.9334601 |
| SMARCC1 | 3715.8969 | 0.08799395 | 0.15335515 | 0.57379195 | 0.5661086 | 0.92553792 |
| DHX30 | 4507.20264 | -0.2226463 | 0.10517175 | -2.1169777 | 0.03426174 | 0.55369025 |
| MAP4 | 12538.3668 | -0.0131346 | 0.12431472 | -0.105656 | 0.91585528 | 0.9868044 |
| CDC25A | 95.7767479 | 0.16042768 | 0.32833653 | 0.48860747 | 0.62511963 | 0.94088547 |
| ZNF589 | 934.432126 | -0.1371503 | 0.23647998 | -0.5799658 | 5.62E-01 | 9.25E-01 |
| NME6 | 1203.41622 | -0.2055951 | 0.17045605 | -1.2061473 | 0.22776068 | 0.80300331 |
| PLXNB1 | 19765.2301 | -0.2028284 | 0.17376135 | -1.1672815 | 2.43E-01 | 0.81340905 |
| CCDC51 | 3080.78901 | 0.21264403 | 0.19631148 | 1.08319713 | 2.79E-01 | 0.82820819 |
| TMA7 | 3438.35177 | 0.27840924 | 0.21879086 | 1.27249025 | 0.20319899 | 0.78268834 |
| AC134772.1 | 2924.68631 | 0.26435962 | 0.21754195 | 1.21521216 | 0.22428518 | 0.80078694 |
| ATRIP | 1613.92729 | -0.6527806 | 0.46865451 | -1.3928824 | 0.16365537 | 0.75228482 |
| TREX1 | 1227.39912 | -0.9025395 | 0.54526619 | -1.6552273 | 0.09787841 | 0.66549286 |

| | | | | | | |
|---|---|---|---|---|---|---|
| SHISA5 | 11155.7163 | -0.217164 | 0.29234679 | -0.7428301 | 0.45758454 | 0.89688594 |
| PFKFB4 | 1223.02417 | -0.612656 | 0.48674465 | -1.2586806 | 0.20814572 | 0.78934595 |
| UCN2 | 93.1595115 | -2.1035564 | 0.91667308 | -2.2947728 | 0.02174615 | 0.51522706 |
| COL7A1 | 6169.71938 | -1.0810028 | 0.53927374 | -2.004553 | 0.04501086 | 0.5734524 |
| UQCRC1 | 15174.1572 | 0.1717176 | 0.16026462 | 1.07146296 | 0.28396132 | 0.83107779 |
| SLC26A6 | 4060.09951 | -0.0356897 | 0.39883654 | -0.0894845 | 0.92869686 | 0.98963127 |
| CELSR3 | 154.198095 | -0.0855521 | 0.40940258 | -0.2089681 | 0.83447314 | 0.97550416 |
| NCKIPSD | 2260.0263 | -0.1294611 | 0.27440404 | -0.4717899 | 0.63707673 | 0.94088547 |
| IP6K2 | 6090.86867 | -0.241689 | 0.16788128 | -1.4396424 | 0.1499686 | 0.73768021 |
| PRKAR2A | 2217.27303 | -0.0214184 | 0.23536489 | -0.091001 | 0.92749182 | 0.98915637 |
| PRKAR2A-AS | 81.7469012 | -0.2337293 | 0.34071571 | -0.685995 | 0.49271629 | 0.90696683 |
| SLC25A20 | 2017.16328 | 0.21629451 | 0.51184046 | 0.42258188 | 0.67260035 | 0.94922827 |
| ARIH2OS | 188.592234 | -0.3012817 | 0.22947466 | -1.3129193 | 0.18921013 | 0.76970626 |
| ARIH2 | 4750.63413 | -0.2108719 | 0.10626878 | -1.9843256 | 0.04721955 | 0.58218854 |
| AC137630.2 | 500.625447 | -0.1516377 | 0.14555897 | -1.0417612 | 0.29752241 | 0.83910722 |
| AC137630.1 | 955.648695 | -0.0569448 | 0.13907344 | -0.4094582 | 0.68220347 | 0.95070301 |
| AC137630.4 | 66.8379388 | -0.1459808 | 0.3750556 | -0.3892243 | 0.69711021 | 0.95269909 |
| P4HTM | 3840.761 | 0.01682271 | 0.1553321 | 0.10830155 | 0.91375649 | 0.98662757 |
| WDR6 | 14608.9727 | -0.3984425 | 0.16099136 | -2.4749312 | 0.01332619 | 0.42389917 |
| DALRD3 | 6248.91715 | -0.1568941 | 0.12097583 | -1.2969042 | 0.19466416 | 0.7762856 |
| NDUFAF3 | 3154.81751 | 0.25048158 | 0.19814469 | 1.26413471 | 2.06E-01 | 0.78697574 |
| IMPDH2 | 5798.4803 | 0.00569336 | 0.21252637 | 0.02678897 | 0.97862805 | 0.99483482 |
| AC137630.3 | 555.85414 | 0.06546111 | 0.2320154 | 0.28214125 | 0.77783521 | 0.96836854 |
| QRICH1 | 3706.61914 | -0.1605514 | 0.12465554 | -1.2879608 | 0.19775961 | 0.77877977 |
| QARS | 8470.55906 | 0.25078554 | 0.10638623 | 2.35731212 | 0.01840777 | 0.47570049 |
| USP19 | 4157.74788 | -0.1737287 | 0.14826629 | -1.171734 | 2.41E-01 | 0.8124487 |
| LAMB2 | 19172.5221 | -0.0204391 | 0.23565881 | -0.0867316 | 0.93088482 | 0.98975562 |
| CCDC71 | 1006.54743 | -0.2104005 | 0.17163372 | -1.2258695 | 0.22024777 | 0.79901989 |
| KLHDC8B | 1636.90027 | -0.1076925 | 0.27932407 | -0.3855467 | 0.69983239 | 0.95328923 |
| C3orf62 | 825.054301 | -0.3808329 | 0.20948809 | -1.8179215 | 0.06907614 | 0.62779744 |
| USP4 | 2952.38558 | -0.1056246 | 0.1026167 | -1.0293122 | 0.30333301 | 0.84211658 |
| RHOA | 23935.4669 | -0.069964 | 0.23695439 | -0.2952634 | 7.68E-01 | 0.96762687 |
| TCTA | 3652.20545 | 0.32811076 | 0.16671281 | 1.96811966 | 4.91E-02 | 0.58508842 |
| AMT | 2445.78233 | 0.59282262 | 0.33874709 | 1.75004489 | 0.08011057 | 0.64079926 |
| AC104452.1 | 3757.99907 | 0.45051657 | 0.33297064 | 1.35302192 | 0.17604863 | 0.76115956 |
| NICN1 | 1339.83824 | 0.22879343 | 0.37342539 | 0.61268848 | 0.54008234 | 0.92224904 |
| DAG1 | 6364.8384 | -0.0737841 | 0.21265629 | -0.3469639 | 0.72861846 | 0.95848028 |
| BSN | 163.48926 | 0.35201968 | 0.36794884 | 0.95670822 | 0.33871455 | 0.85733087 |
| APEH | 8468.80477 | 0.39133573 | 0.19087253 | 2.05024649 | 0.04034038 | 0.55657615 |
| MST1 | 9426.29026 | 0.17052507 | 0.34473808 | 0.4946511 | 0.62084643 | 0.94014891 |
| RNF123 | 3786.24263 | 0.06630398 | 0.17391405 | 0.38124567 | 0.70302097 | 0.95431462 |
| AMIGO3 | 337.245958 | -0.2639265 | 0.183385 | -1.4391933 | 1.50E-01 | 0.73796431 |
| GMPPB | 3624.13885 | -0.0401992 | 0.09018113 | -0.4457612 | 0.65576975 | 0.94532497 |
| IP6K1 | 3709.28061 | -0.4862639 | 0.19178186 | -2.5355052 | 0.01122853 | 0.40196898 |
| INKA1 | 106.780341 | -0.2251442 | 0.39939316 | -0.5637158 | 0.57294757 | 0.92928554 |
| UBA7 | 4264.07755 | -0.8384236 | 0.68157907 | -1.2301194 | 2.19E-01 | 0.7978514 |
| TRAIP | 90.5481861 | 0.03667445 | 0.46603887 | 0.07869398 | 9.37E-01 | 9.91E-01 |
| MST1R | 287.805245 | 0.07560262 | 0.34758386 | 0.21750901 | 0.82781169 | 0.97548201 |
| MON1A | 820.513127 | 0.03971466 | 0.25404286 | 0.15633053 | 0.87577249 | 0.98275231 |
| RBM6 | 8281.36957 | -0.1674702 | 0.23843874 | -0.7023615 | 0.48245377 | 0.90541935 |
| RBM5 | 7818.31236 | -0.1278102 | 0.16180064 | -0.7899239 | 0.42957221 | 0.88858861 |
| RBM5-AS1 | 147.186319 | -0.0275071 | 0.35849841 | -0.0767288 | 0.93883932 | 0.99113899 |
| SEMA3F-AS1 | 2847.95159 | -0.1182838 | 0.18388689 | -0.6432422 | 0.52006698 | 0.91657872 |
| SEMA3F | 4329.12378 | -0.314202 | 0.2850072 | -1.1024355 | 0.2702724 | 0.82716979 |

| | | | | | | |
|---|---|---|---|---|---|---|
| SLC38A3 | 1799.61664 | 4.37260773 | 1.18289115 | 3.69654276 | NA | NA |
| GNAI2 | 11521.0355 | 0.01727147 | 0.20120379 | 0.0858407 | 0.93159305 | 0.98979451 |
| U73166.1 | 72.5996099 | 0.35366557 | 0.53104562 | 0.66597964 | 0.50542412 | 0.91368371 |
| SEMA3B-AS1 | 94.7571955 | 0.58998023 | 0.44031766 | 1.3398968 | 0.1802789 | 0.76387222 |
| SEMA3B | 9849.14102 | 0.30735922 | 0.26526802 | 1.15867423 | 0.246589 | 0.81542272 |
| LSMEM2 | 668.610541 | -0.1254997 | 0.27398972 | -0.4580452 | 0.64691994 | 0.94246942 |
| IFRD2 | 3379.62136 | -0.1528886 | 0.24288562 | -0.6294676 | 0.52904298 | 0.91907723 |
| HYAL3 | 227.575555 | 0.30955906 | 0.24507789 | 1.26310482 | 0.20655151 | 0.7877456 |
| NAA80 | 689.252672 | 0.29593485 | 0.26172004 | 1.13073057 | 0.25816851 | 0.82239817 |
| HYAL1 | 5082.76377 | 0.264838 | 0.39934212 | 0.66318574 | 0.5072116 | 0.91430909 |
| HYAL2 | 6053.31769 | -0.3451034 | 0.22080134 | -1.5629589 | 0.11806226 | 0.69079142 |
| TUSC2 | 3958.62327 | -0.2217966 | 0.13461035 | -1.6476938 | 0.09941551 | 0.66980808 |
| RASSF1 | 2221.67371 | -0.3438691 | 0.25407667 | -1.3534067 | 0.17592573 | 0.76108589 |
| RASSF1-AS1 | 52.5577141 | -0.0976842 | 0.41599168 | -0.2348226 | 0.81434646 | 0.9738447 |
| ZMYND10-AS | 43.968678 | 0.40756725 | 0.69608152 | 0.58551655 | 0.55820044 | 0.92421175 |
| ZMYND10 | 224.039578 | 0.49944629 | 0.57208477 | 0.87302846 | 3.83E-01 | 8.72E-01 |
| NPRL2 | 1605.02971 | 0.08954101 | 0.10106303 | 0.88599182 | 3.76E-01 | 8.69E-01 |
| CYB561D2 | 2513.26121 | 0.12231236 | 0.15692848 | 0.77941467 | 0.43573548 | 0.89085849 |
| Z84492.1 | 556.160979 | 0.33048647 | 0.23106942 | 1.43024749 | 0.152646 | 0.7422778 |
| TMEM115 | 3325.29664 | -0.2208594 | 0.18150151 | -1.2168463 | 0.22366271 | 0.80078694 |
| CACNA2D2 | 1062.1387 | 0.98867812 | 0.35869841 | 2.75629356 | 0.00584605 | 0.32704725 |
| C3orf18 | 896.661894 | 0.30958455 | 0.29179124 | 1.06097962 | 0.28869916 | 0.83295411 |
| HEMK1 | 2315.03562 | -0.0410219 | 0.31314537 | -0.1309994 | 0.89577575 | 0.98474725 |
| CISH | 5950.72014 | 0.00077855 | 0.29244031 | 0.00266226 | 0.99787582 | 0.99989813 |
| MAPKAPK3 | 4192.13085 | -0.1009892 | 0.16586193 | -0.6088751 | 0.54260724 | 0.92292765 |
| LINC02019 | 143.685889 | -0.59255 | 0.42361078 | -1.3988076 | 0.16187068 | 0.7498147 |
| DOCK3 | 221.777118 | -0.2634552 | 0.45526692 | -0.5786829 | 0.56280314 | 0.92452095 |
| MANF | 2256.54188 | 0.54045253 | 0.31497081 | 1.71588133 | 0.08618377 | 0.65049132 |
| RBM15B | 2858.44856 | -0.4044732 | 0.23824563 | -1.6977149 | 0.08956158 | 0.65375543 |
| DCAF1 | 2007.18364 | -0.2499565 | 0.19224377 | -1.3002061 | 0.19353034 | 0.77548688 |
| RAD54L2 | 1388.67908 | -0.0636187 | 0.10958802 | -0.5805259 | 0.56156 | 0.92452095 |
| TEX264 | 5168.66374 | 0.10390113 | 0.16941975 | 0.61327636 | 0.53969362 | 0.9220053 |
| RRP9 | 1098.25865 | -0.2293163 | 0.26368875 | -0.8696476 | 0.38449303 | 0.87309542 |
| PARP3 | 2624.05663 | -0.0574213 | 0.42167854 | -0.1361732 | 0.89168437 | 0.98474725 |
| PCBP4 | 2491.90576 | 0.06550902 | 0.26853528 | 0.24394941 | 0.80727 | 0.97334138 |
| ABHD14B | 6836.037 | 0.39159053 | 0.20372345 | 1.9221672 | 0.05458472 | 0.59329767 |
| ABHD14A | 2562.66038 | 0.31882412 | 0.27219261 | 1.17131807 | 0.24147095 | 0.81260677 |
| ABHD14A-AC | 27107.1461 | 0.80358135 | 0.41404767 | 1.94079427 | 0.05228324 | 0.58645127 |
| ACY1 | 26039.7579 | 0.83311124 | 0.4245232 | 1.96246341 | 0.04970856 | 0.5861488 |
| RPL29 | 19170.263 | 0.21756413 | 0.13621767 | 1.59717994 | 0.1102256 | 0.67996643 |
| DUSP7 | 1115.35762 | -0.0175176 | 0.36002843 | -0.0486561 | 0.96119338 | 0.99320711 |
| POC1A | 182.542421 | 0.24009512 | 0.34083165 | 0.70443905 | 0.48115939 | 0.90541935 |
| ALAS1 | 4362.97048 | 0.43566459 | 0.20760665 | 2.0985098 | 0.03586013 | 0.55369025 |
| TLR9 | 67.8229921 | -0.0270959 | 0.49804784 | -0.0544042 | 0.95661317 | 0.99198394 |
| AC097637.1 | 1443.33938 | 0.19256157 | 0.1907664 | 1.00941033 | 0.31277788 | 0.84548829 |
| TWF2 | 2635.12472 | 0.15402662 | 0.20137294 | 0.7648824 | 0.44434157 | 0.8938278 |
| PPM1M | 1246.82451 | 0.06711509 | 0.30906287 | 0.21715675 | 0.82808619 | 0.97548201 |
| WDR82 | 3880.76052 | -0.1451131 | 0.15683764 | -0.9252443 | 0.35483885 | 0.86364633 |
| GLYCTK | 2694.63169 | 0.10215452 | 0.42061634 | 0.24286865 | 0.80810715 | 0.97334138 |
| DNAH1 | 946.682858 | 0.05908377 | 0.21946999 | 0.26921115 | 0.7877672 | 0.97090793 |
| BAP1 | 5477.69377 | -0.1279653 | 0.1492188 | -0.8575683 | 0.39113086 | 0.87710956 |
| PHF7 | 322.702338 | -0.174858 | 0.27973976 | -0.6250737 | 0.53192266 | 0.91962754 |
| SEMA3G | 3121.65825 | 0.19097171 | 0.45755053 | 0.41737841 | 0.67640164 | 0.94968858 |
| TNNC1 | 212.6143 | 0.82564284 | 0.55776876 | 1.4802601 | 0.13880385 | 0.72399122 |

| | | | | | | |
|---|---|---|---|---|---|---|
| NISCH | 7171.85733 | -0.0498537 | 0.22658517 | -0.2200218 | 0.82585421 | 0.97548201 |
| STAB1 | 10175.7517 | -0.4877073 | 0.40916445 | -1.1919592 | 0.23327726 | 0.80725541 |
| NT5DC2 | 2585.98491 | -0.1805312 | 0.36457773 | -0.4951789 | 0.62047381 | 0.93988843 |
| SMIM4 | 819.543019 | -0.3217783 | 0.3003925 | -1.0711927 | 0.28408278 | 0.83107779 |
| PBRM1 | 1559.22542 | -0.3263502 | 0.15728564 | -2.0748885 | 0.03799687 | 0.55538555 |
| GNL3 | 2693.23609 | -0.3919831 | 0.27021389 | -1.4506399 | 0.14688015 | 0.73337969 |
| GLT8D1 | 2582.09367 | 0.27359906 | 0.0878245 | 3.11529325 | 0.00183762 | 0.18819049 |
| SPCS1 | 5347.20429 | 0.1861258 | 0.09520334 | 1.95503435 | 0.05057902 | 0.5861488 |
| NEK4 | 1135.05491 | 0.07162014 | 0.13416072 | 0.53383841 | 0.59345334 | 0.93289552 |
| ITIH3 | 157.744039 | -0.571525 | 0.4559779 | -1.2534051 | 0.21005832 | 0.79134644 |
| ITIH4 | 1209.4454 | -1.1021136 | 0.3483262 | -3.1640273 | 0.00155602 | 0.17357983 |
| AC006254.1 | 1855.3847 | -0.8425818 | 0.34334866 | -2.4540122 | 0.01412722 | 0.43474248 |
| ITIH4-AS1 | 113.815731 | -1.4258441 | 0.41878556 | -3.4047116 | 0.00066234 | 0.12385487 |
| MUSTN1 | 545.069613 | -0.5221289 | 0.48832933 | -1.0692147 | 0.28497295 | 0.83107779 |
| TMEM110-M | 1914.0253 | -0.1536525 | 0.22893383 | -0.6711658 | 0.50211494 | 0.91280785 |
| STIMATE | 1314.93538 | 0.05187326 | 0.22755942 | 0.2279548 | 0.81968138 | 0.97454841 |
| SFMBT1 | 605.097523 | -0.5642693 | 0.1990571 | -2.8347109 | 0.00458672 | 0.29641268 |
| AC096887.1 | 292.036342 | 0.23256388 | 0.23771352 | 0.97833677 | 0.32790779 | 0.85317073 |
| RFT1 | 880.619122 | 0.21679893 | 0.18110374 | 1.19709799 | 0.23126836 | 0.80649152 |
| PRKCD | 2883.0651 | -0.0153704 | 0.20448345 | -0.0751669 | 0.94008194 | 0.99136103 |
| TKT | 19598.0965 | -0.0722002 | 0.34055229 | -0.2120091 | 0.83209994 | 0.97550416 |
| DCP1A | 1284.25344 | -0.2406538 | 0.19076735 | -1.2615041 | 0.2071273 | 0.78790396 |
| CACNA1D | 2005.98865 | 0.26727544 | 0.3237669 | 0.82551809 | 0.40907751 | 0.8821603 |
| AC012467.1 | 48.9668112 | -0.3994413 | 0.94608081 | -0.4222063 | 0.67287444 | 0.94922827 |
| CHDH | 7529.75234 | 0.26453207 | 0.4332391 | 0.61059141 | 0.54147011 | 0.92292765 |
| IL17RB | 12555.0294 | 0.67349524 | 0.60261403 | 1.1176229 | 0.26372809 | 0.82424998 |
| AC012467.2 | 2318.74182 | 0.54442337 | 0.52541426 | 1.03617929 | 0.3001185 | 0.84186812 |
| ACTR8 | 1306.1749 | 0.27319775 | 0.19221351 | 1.42132439 | 0.15522248 | 0.74407586 |
| SELENOK | 2681.38534 | 0.03384198 | 0.1976265 | 0.17124213 | 0.86403339 | 0.98176973 |
| CACNA2D3 | 536.582506 | -0.4251425 | 0.34101068 | -1.2467133 | 0.21250265 | 0.79203904 |
| WNT5A | 353.268317 | 1.84808326 | 0.62131538 | 2.97446887 | 0.00293496 | 0.23606923 |
| ERC2 | 75.5455814 | 0.01149272 | 0.391106 | 0.02938517 | 0.9765574 | 0.99455099 |
| CCDC66 | 1220.06068 | -0.0445368 | 0.20422715 | -0.2180749 | 0.82737075 | 0.97548201 |
| FAM208A | 2967.33748 | -0.0109746 | 0.11705968 | -0.0937526 | 0.92530571 | 0.98871967 |
| ARHGEF3 | 2573.31752 | -0.0923624 | 0.19363417 | -0.4769944 | 0.63336611 | 0.94088547 |
| IL17RD | 1010.61377 | 0.14520089 | 0.24539346 | 0.59170643 | 0.55404719 | 0.92373235 |
| HESX1 | 85.7286298 | -0.4989195 | 0.68571407 | -0.7275911 | NA | NA |
| APPL1 | 2091.02766 | -0.0225834 | 0.09545769 | -0.2365802 | 0.81298248 | 0.9738447 |
| ASB14 | 166.041172 | -0.3178741 | 0.21271576 | -1.4943609 | 0.13508132 | 0.71649038 |
| PDE12 | 1660.13932 | -0.1761776 | 0.27832001 | -0.6330038 | 0.52673116 | 0.91769856 |
| ARF4 | 8857.26908 | -0.557717 | 0.35230461 | -1.5830534 | 0.11340929 | 0.68193708 |
| ARF4-AS1 | 46.3680776 | 0.1895287 | 0.5445059 | 0.34807466 | 0.72778412 | 0.95848028 |
| DENND6A | 1504.72366 | -0.1048704 | 0.1760571 | -0.5956615 | 0.55140139 | 0.92373235 |
| DENND6A-AS | 69.1550588 | 0.31165497 | 0.37310303 | 0.83530539 | 0.40354578 | 0.88132015 |
| SLMAP | 2398.80338 | -0.2375338 | 0.13978382 | -1.6992936 | 0.08926387 | 0.65375543 |
| FLNB | 10822.014 | -0.0443958 | 0.13323933 | -0.3332036 | 0.73898061 | 0.9602325 |
| FLNB-AS1 | 1951.28865 | -0.1046953 | 0.12136146 | -0.862673 | 0.38831726 | 0.87553582 |
| DNASE1L3 | 1311.33904 | 0.08786256 | 0.46776299 | 0.18783565 | 0.85100549 | 0.97844748 |
| ABHD6 | 1953.73276 | 0.38094447 | 0.48626589 | 0.78340775 | 0.43338771 | 0.89055338 |
| RPP14 | 1778.29793 | 0.04613777 | 0.21444392 | 0.21515077 | 0.82964977 | 0.97550416 |
| HTD2 | 820.584661 | 0.04145706 | 0.26500339 | 0.15643973 | 0.87568642 | 0.98275231 |
| PXK | 1552.62873 | -0.1065065 | 0.18534252 | -0.5746467 | 0.56553025 | 0.92513982 |
| PDHB | 5204.42184 | 0.12631976 | 0.14616932 | 0.86420162 | 0.38747715 | 0.87460898 |
| AC135507.1 | 1724.12029 | 0.14366375 | 0.15136003 | 0.94915248 | 0.34254307 | 0.85889221 |

| | | | | | | |
|---|---|---|---|---|---|---|
| KCTD6 | 1225.2726 | 0.01386818 | 0.19895643 | 0.06970459 | 9.44E-01 | 0.99136103 |
| ACOX2 | 3374.80327 | 1.03955199 | 0.46134611 | 2.25330174 | 0.02424013 | 0.51967919 |
| FAM107A | 3716.39219 | 0.64746163 | 0.36902799 | 1.75450549 | 0.07934393 | 0.64079926 |
| FAM3D-AS1 | 100.580468 | 2.72332266 | 0.76137238 | 3.57686033 | 3.48E-04 | 0.08914272 |
| FAM3D | 110.987949 | 3.63026063 | 0.83208129 | 4.36286774 | 1.28E-05 | 0.01316658 |
| C3orf67 | 66.3837322 | 0.25048237 | 0.31265312 | 0.80115105 | 0.4230442 | 0.88455556 |
| FHIT | 906.431551 | 0.37226498 | 0.3716783 | 1.00157845 | 0.31654723 | 0.84783414 |
| PTPRG | 3325.24786 | 0.14666059 | 0.19746449 | 0.74271881 | 0.45765194 | 0.89688594 |
| C3orf14 | 316.240232 | 0.33073569 | 0.39606664 | 0.83505061 | 0.40368921 | 0.88132015 |
| THOC7 | 1573.93129 | 0.17305599 | 0.1476349 | 1.17218887 | 0.24112123 | 0.8124487 |
| ATXN7 | 1341.55149 | -0.3854254 | 0.23835737 | -1.6170064 | 0.1058769 | 0.67537064 |
| PSMD6-AS2 | 170.38852 | -0.3979985 | 0.24030079 | -1.6562515 | 0.09767089 | 0.66549286 |
| PSMD6 | 3865.36604 | -0.1561505 | 0.19580537 | -0.7974782 | 0.42517337 | 0.88584481 |
| PSMD6-AS1 | 81.268743 | -0.3142904 | 0.28711803 | -1.0946382 | 0.27367517 | 0.82719197 |
| AC092040.1 | 163.261809 | -0.0816672 | 0.3151177 | -0.2591643 | 0.7955085 | 0.97101097 |
| PRICKLE2 | 1020.51073 | 0.01272415 | 0.16938696 | 0.0751188 | 0.94012019 | 0.99136103 |
| ADAMTS9 | 4223.65504 | 0.09060548 | 0.36209628 | 0.25022482 | 0.80241349 | 0.97245417 |
| ADAMTS9-A! | 7511.57972 | -0.392734 | 0.46409303 | -0.8462397 | 0.39741903 | 0.88006972 |
| ADAMTS9-A! | 247.392297 | 0.32743273 | 0.27531958 | 1.18928241 | 0.23432856 | 0.80730051 |
| MAGI1 | 2106.78404 | 0.03185409 | 0.1418713 | 0.22452811 | 0.8223464 | 0.97504706 |
| SLC25A26 | 1588.28391 | -0.1857497 | 0.20901125 | -0.8887066 | 0.37416078 | 0.86791605 |
| LRIG1 | 1371.57294 | -0.0540783 | 0.15420773 | -0.3506849 | 0.72582477 | 0.95848028 |
| KBTBD8 | 64.1088977 | -0.4663791 | 0.50982345 | -0.9147854 | 0.36030429 | 0.86480228 |
| SUCLG2 | 7658.87386 | 0.44319863 | 0.25259932 | 1.75455197 | 0.07933597 | 0.64079926 |
| SUCLG2-AS1 | 73.1810689 | -0.010455 | 0.38532293 | -0.0271331 | 0.97835357 | 0.99476647 |
| EOGT | 824.340183 | 0.00363947 | 0.2642616 | 0.01377223 | 0.9890117 | 0.99795769 |
| AC109587.1 | 896.21372 | -0.1175096 | 0.13914354 | -0.8445205 | 0.39837863 | 0.88074856 |
| TMF1 | 1973.83479 | -0.1769876 | 0.16389891 | -1.0798586 | 2.80E-01 | 8.28E-01 |
| UBA3 | 2773.57058 | -0.0604396 | 0.10194441 | -0.5928685 | 5.53E-01 | 9.24E-01 |
| ARL6IP5 | 6480.77461 | 0.04961176 | 0.24395803 | 0.20336189 | 0.83885219 | 0.97612168 |
| FRMD4B | 2014.05635 | 0.04661668 | 0.16408061 | 0.28410841 | 0.7763273 | 0.96836854 |
| MITF | 1821.49958 | -0.1341643 | 0.21387586 | -0.6273001 | 0.53046257 | 0.91962754 |
| FOXP1 | 1435.38035 | -0.3153285 | 0.18365902 | -1.7169236 | 0.08599314 | 0.65049132 |
| EIF4E3 | 1598.01326 | 0.35765843 | 0.14341623 | 2.49384902 | 0.01263663 | 0.41598747 |
| GPR27 | 275.326006 | 0.55756621 | 0.47342167 | 1.17773698 | 0.23890148 | 0.81124801 |
| PROK2 | 42.588757 | 0.16611933 | 0.99501854 | 0.16695099 | 0.86740862 | 0.98224199 |
| RYBP | 2162.93735 | -0.095724 | 0.1974837 | -0.4847184 | 0.62787611 | 0.94088547 |
| SHQ1 | 748.96662 | -0.0519049 | 0.23732756 | -0.2187058 | 0.8268792 | 0.97548201 |
| GXYLT2 | 107.025695 | 0.04845537 | 0.33586355 | 0.14427099 | 0.88528648 | 0.98356741 |
| PPP4R2 | 2525.40246 | -0.2857756 | 0.17616096 | -1.6222416 | 0.10475163 | 0.67439401 |
| PDZRN3 | 868.207522 | 0.46179939 | 0.31843462 | 1.45021729 | 1.47E-01 | 7.33E-01 |
| CNTN3 | 424.898787 | 0.38705827 | 0.29284348 | 1.32172407 | 1.86E-01 | 7.69E-01 |
| LINC00960 | 146.330748 | 0.48237031 | 0.65247316 | 0.73929526 | 0.45972773 | 0.89755243 |
| ZNF717 | 418.687531 | 0.29136863 | 0.34707785 | 0.8394907 | 0.401194 | 0.88132015 |
| ROBO2 | 101.933218 | 0.95376012 | 0.6204337 | 1.53724745 | 0.12423272 | 0.70102567 |
| ROBO1 | 450.292273 | 0.43487264 | 0.37377555 | 1.16345931 | 0.2446432 | 0.81473945 |
| LINC02027 | 109.897332 | 0.43693085 | 0.70951246 | 0.61581844 | 5.38E-01 | 9.21E-01 |
| GBE1 | 4259.65261 | 0.02699746 | 0.41104245 | 0.06568047 | 0.94763222 | 0.99170503 |
| VGLL3 | 287.865699 | -0.8693777 | 0.62871239 | -1.3827908 | 0.16672902 | 0.75558932 |
| CHMP2B | 2987.23557 | -0.2956818 | 0.29578562 | -0.999649 | 0.31748039 | 0.84819508 |
| CGGBP1 | 4231.69089 | -0.1467787 | 0.10904721 | -1.3460107 | 0.17829906 | 0.76115956 |
| ZNF654 | 775.701006 | -0.0650541 | 0.16647942 | -0.3907634 | 0.6959721 | 0.9525827 |
| C3orf38 | 635.240528 | -0.2273489 | 0.1741622 | -1.3053862 | 0.19176137 | 0.77347283 |
| EPHA3 | 106.549232 | -0.3084038 | 0.57518514 | -0.5361818 | 0.5918329 | 0.93289552 |

| | | | | | | |
|---|---|---|---|---|---|---|
| PROS1 | 4482.83948 | 0.51881753 | 0.25785234 | 2.01207224 | 4.42E-02 | 0.57267043 |
| ARL13B | 584.997572 | 0.10129592 | 0.21608748 | 0.46877276 | 0.63923207 | 0.94088547 |
| DHFR2 | 464.528577 | -0.043254 | 0.18472401 | -0.2341545 | 0.81486505 | 0.9738447 |
| NSUN3 | 386.688411 | -0.039293 | 0.17751736 | -0.2213473 | 0.82482201 | 0.97548201 |
| MTRNR2L12 | 385.063728 | -0.256605 | 0.3806989 | -0.6740366 | 0.50028804 | 0.91180773 |
| ARL6 | 609.463847 | -0.0682172 | 0.15747289 | -0.4331997 | 0.6648697 | 0.94692723 |
| AC110491.3 | 83.5842824 | -0.124497 | 0.33026385 | -0.3769621 | 0.70620176 | 0.95501548 |
| CRYBG3 | 1866.51101 | -0.2309359 | 0.1697443 | -1.360493 | 0.17367397 | 0.75974924 |
| RIOX2 | 2891.3339 | -0.1870265 | 0.13872214 | -1.3482094 | 0.17759105 | 0.76115956 |
| CLDND1 | 3075.23492 | -0.1363012 | 0.15101634 | -0.9025596 | 0.36675968 | 0.86683872 |
| CPOX | 1335.69016 | 0.07576979 | 0.11418955 | 0.66354396 | 0.50698223 | 0.91420888 |
| ST3GAL6-AS | 131.329481 | -0.7409147 | 0.57039066 | -1.2989601 | 0.19395763 | 0.77620852 |
| ST3GAL6 | 1397.40974 | -0.3041106 | 0.35203286 | -0.8638699 | 0.38765939 | 0.87468392 |
| DCBLD2 | 1712.86899 | 0.16982939 | 0.25669844 | 0.66159103 | 0.50823336 | 0.91439937 |
| COL8A1 | 1093.3575 | 0.36108571 | 0.5971438 | 0.60468804 | 0.5453863 | 0.92345821 |
| CMSS1 | 583.871972 | -0.0510622 | 0.21761684 | -0.2346428 | 8.14E-01 | 9.74E-01 |
| FILIP1L | 1345.79319 | -1.424597 | 0.69418289 | -2.0521926 | 0.04015095 | 0.55657615 |
| TBC1D23 | 1834.68398 | -0.1674386 | 0.16646077 | -1.0058741 | 0.31447615 | 0.84696964 |
| NIT2 | 5460.09438 | 0.40961366 | 0.30024181 | 1.36427923 | 0.1724797 | 0.75974924 |
| TOMM70 | 3911.96223 | 0.23286027 | 0.19695855 | 1.18228054 | 0.23709441 | 0.80857441 |
| LNP1 | 210.877381 | 0.35720696 | 0.28901142 | 1.23596138 | 0.21647292 | 0.7954827 |
| TMEM45A | 248.723069 | 0.67467015 | 0.38067245 | 1.77231145 | 0.07634287 | 0.63773415 |
| TFG | 5408.61604 | -0.2222467 | 0.17317413 | -1.2833715 | 0.19936195 | 0.78134034 |
| ABI3BP | 1192.22428 | 1.06630973 | 0.4349473 | 2.45158376 | 0.01422291 | 0.43625454 |
| IMPG2 | 44.7273237 | -1.127172 | 0.56327558 | -2.0011022 | 0.04538138 | 0.57529633 |
| SENP7 | 1112.30482 | 0.09434328 | 0.2447554 | 0.38545943 | 0.69989706 | 0.95328923 |
| TRMT10C | 877.216424 | -0.1829383 | 0.22992513 | -0.7956427 | 0.42623972 | 0.88678491 |
| PCNP | 6262.22279 | -0.0906123 | 0.12157161 | -0.7453407 | 0.4560658 | 0.89688594 |
| ZBTB11 | 1104.59423 | -0.1257561 | 0.16540439 | -0.7602946 | 0.44707849 | 0.89480207 |
| ZBTB11-AS1 | 249.712725 | -0.1548455 | 0.28923124 | -0.5353693 | 0.59239451 | 0.93289552 |
| RPL24 | 15468.4855 | 0.18395367 | 0.17820224 | 1.03227472 | 0.30194344 | 0.84211658 |
| CEP97 | 378.403893 | -0.1229883 | 0.20149866 | -0.6103679 | 0.54161815 | 0.92292765 |
| NXPE3 | 495.811917 | -0.0703149 | 0.19736815 | -0.3562625 | 0.72164403 | 0.95739595 |
| NFKBIZ | 4923.14639 | -0.0324999 | 0.41827761 | -0.0776993 | 0.93806726 | 0.99113899 |
| ZPLD1 | 180.989933 | -0.6061252 | 0.82392932 | -0.735652 | 0.46194254 | 0.89762182 |
| ALCAM | 2023.55132 | 0.32427481 | 0.21499131 | 1.5083159 | 0.1314737 | 0.7108257 |
| CBLB | 1317.79604 | -0.1936545 | 0.17328557 | -1.1175457 | 0.26376107 | 0.82424998 |
| LINC00882 | 54.3662673 | 0.52947267 | 0.47312856 | 1.11908838 | 0.26310244 | 0.82424998 |
| DUBR | 197.559049 | -0.0839649 | 0.3622195 | -0.2318067 | 0.81668819 | 0.97388889 |
| BBX | 2576.5269 | 0.04346841 | 0.18465958 | 0.23539754 | 0.8139002 | 0.9738447 |
| CD47 | 3593.70912 | 0.0026978 | 0.19176848 | 0.01406799 | 0.98877574 | 0.99788078 |
| IFT57 | 2587.76435 | 0.17720403 | 0.15413689 | 1.14965359 | 0.25028658 | 0.81721208 |
| HHLA2 | 574.910482 | 0.74912234 | 0.56918928 | 1.31612166 | 0.18813321 | 0.76970626 |
| CIP2A | 103.383987 | -0.207953 | 0.45534412 | -0.4566942 | 0.64789087 | 0.94285781 |
| DZIP3 | 1133.9756 | 0.53515487 | 0.39112525 | 1.36824424 | 0.17123564 | 0.75956362 |
| C3orf85 | 230.457835 | 1.774141 | 0.8866732 | 2.00089616 | NA | NA |
| LINC00488 | 39.7236773 | 0.55055953 | 0.65131409 | 0.84530572 | 0.39794017 | 0.88054917 |
| NECTIN3 | 683.898047 | -0.2475841 | 0.2343395 | -1.056519 | 0.29073118 | 0.83442894 |
| CD96 | 147.578254 | 0.18957235 | 0.90975787 | 0.20837671 | 0.83493484 | 0.97550416 |
| PLCXD2 | 554.190524 | -0.0063281 | 0.36042547 | -0.0175574 | 0.98599191 | 0.99705792 |
| PHLDB2 | 4697.39549 | 0.04486433 | 0.17768017 | 0.25250048 | 0.80065424 | 0.97210205 |
| ABHD10 | 2498.12366 | 0.36641762 | 0.2618138 | 1.39953518 | 0.16165256 | 0.74969823 |
| TAGLN3 | 157.417422 | 0.71870758 | 0.59096641 | 1.21615641 | 0.22392534 | 0.80078694 |
| C3orf52 | 481.65903 | 0.19573677 | 0.40314428 | 0.48552535 | 0.62730373 | 0.94088547 |

| | | | | | | |
|---|---|---|---|---|---|---|
| GCSAM | 138.415702 | -0.3243111 | 0.52666465 | -0.6157829 | 0.53803783 | 0.92127611 |
| AC128688.2 | 160.493961 | 0.19323188 | 0.3525754 | 0.54805832 | 5.84E-01 | 9.32E-01 |
| SLC9C1 | 47.3077183 | 0.18923292 | 0.42881752 | 0.44129009 | 0.659003 | 0.94599528 |
| CD200 | 1009.23652 | 0.28848342 | 0.2897556 | 0.99560948 | 0.31943993 | 0.84928695 |
| AC092894.1 | 87.9460072 | 0.05333637 | 0.40909635 | 0.13037606 | 0.89626891 | 0.98474725 |
| ATG3 | 2562.47634 | 0.03297326 | 0.1693092 | 0.19475172 | 0.84558732 | 0.97675487 |
| SLC35A5 | 1701.94934 | -0.0186609 | 0.19581274 | -0.0952998 | 9.24E-01 | 9.89E-01 |
| CCDC80 | 1688.30456 | -0.1667571 | 0.73324764 | -0.2274227 | 0.82009509 | 0.97454841 |
| CD200R1 | 49.8986215 | 0.59977232 | 0.72923151 | 0.82247176 | 0.41080846 | 0.88268208 |
| AC074044.1 | 89.8162946 | 0.12131728 | 0.28203447 | 0.43015055 | 0.66708613 | 0.94830184 |
| GTPBP8 | 461.054053 | 0.04871741 | 0.19782825 | 0.24626116 | 0.80548008 | 0.97321269 |
| NEPRO | 2259.51619 | -0.0099165 | 0.17800216 | -0.05571 | 0.9555728 | 0.99185043 |
| AC078785.1 | 85.8155617 | -0.0470377 | 0.33652491 | -0.1397749 | 0.88883787 | 0.98423722 |
| BOC | 1113.47024 | 0.13869189 | 0.29901893 | 0.46382309 | 0.6427745 | 0.94111226 |
| CFAP44 | 861.005321 | -0.0256009 | 0.29928462 | -0.0855403 | 0.93183188 | 0.9898238 |
| SPICE1 | 589.212539 | -0.0443489 | 0.20920988 | -0.2119829 | 0.83212034 | 0.97550416 |
| SIDT1 | 152.242982 | -0.3782692 | 0.57423095 | -0.6587406 | 0.51006237 | 0.91439937 |
| USF3 | 1165.49284 | 0.12934953 | 0.17543084 | 0.73732495 | 0.46092477 | 0.89755243 |
| NAA50 | 5178.37916 | -0.3318502 | 0.25313488 | -1.3109619 | 0.18987065 | 0.7704467 |
| AC108693.2 | 49.5313561 | 0.14018369 | 0.64856028 | 0.21614597 | 0.82887397 | 0.97550416 |
| ATP6V1A | 8798.21092 | 0.25141805 | 0.20306643 | 1.23810737 | 0.21567625 | 0.7954827 |
| GRAMD1C | 2203.02223 | -0.1174103 | 0.38156507 | -0.3077072 | 0.75830512 | 0.96502715 |
| AC128687.2 | 166.793492 | -0.1749438 | 0.38945671 | -0.4491996 | 0.65328766 | 0.94493429 |
| ZDHHC23 | 1191.23433 | -0.0653769 | 0.2187076 | -0.2989238 | 0.76499817 | 0.96678847 |
| CCDC191 | 638.595529 | 0.21978963 | 0.44086319 | 0.49854384 | 0.61810078 | 0.93928244 |
| QTRT2 | 1002.56144 | -0.2442807 | 0.21388826 | -1.1420948 | 0.2534146 | 0.81869384 |
| TIGIT | 74.867753 | 0.42214693 | 1.00913053 | 0.41832738 | 0.67570777 | 0.94959168 |
| ZBTB20 | 4179.58732 | -0.1521532 | 0.24325049 | -0.6255002 | 0.53164283 | 0.91962754 |
| LSAMP | 165.862442 | 0.44134173 | 0.54795543 | 0.80543362 | 0.4205695 | 0.88451372 |
| IGSF11 | 362.985961 | 0.86615491 | 0.49933836 | 1.73460517 | 0.08281076 | 0.64680474 |
| AC083800.1 | 61.961994 | 0.93454046 | 0.64138201 | 1.45707308 | 0.14509621 | 0.73064288 |
| UPK1B | 356.073294 | 0.70788626 | 0.58221526 | 1.21584973 | 0.22404217 | 0.80078694 |
| B4GALT4 | 1917.26321 | 0.3749059 | 0.25153899 | 1.49044847 | 0.13610635 | 0.71852653 |
| B4GALT4-AS | 174.6705 | 0.48116262 | 0.34331147 | 1.40153378 | 0.16105451 | 0.74877764 |
| ARHGAP31 | 1108.82598 | 0.07473819 | 0.23222673 | 0.32183286 | 0.74757932 | 0.96231266 |
| TMEM39A | 1179.05339 | -0.1812851 | 0.16419472 | -1.1040862 | 0.26955573 | 0.82716979 |
| POGLUT1 | 826.230957 | 0.02591245 | 0.17043235 | 0.15203952 | 0.87915577 | 0.98275231 |
| TIMMDC1 | 3460.74541 | 0.13215013 | 0.15766624 | 0.83816376 | 0.40193874 | 0.88132015 |
| ADPRH | 572.435016 | -0.2107603 | 0.15643039 | -1.3473105 | 0.17788027 | 0.76115956 |
| PLA1A | 1370.29371 | -0.9503281 | 0.50082679 | -1.8975185 | 0.05775955 | 0.60129769 |
| POPDC2 | 201.571714 | 0.01245919 | 0.31301286 | 0.03980407 | 0.96824933 | 0.99400331 |
| COX17 | 1947.72623 | -0.1467255 | 0.25594619 | -0.573267 | 0.56646394 | 0.9256133 |
| MAATS1 | 625.869746 | 0.42275642 | 0.41777728 | 1.01191819 | 0.31157717 | 0.84529065 |
| GSK3B | 2194.05233 | -0.143517 | 0.23351125 | -0.6146041 | 0.5388162 | 0.9215968 |
| AC092910.3 | 63.7915393 | 0.03603197 | 0.35785895 | 0.10068763 | 0.91979843 | 0.98733173 |
| LRRC58 | 1965.16708 | -0.1238222 | 0.34689363 | -0.3569458 | 0.7211324 | 0.95731494 |
| FSTL1 | 6583.05758 | 0.08286128 | 0.39669662 | 0.20887822 | 0.83454331 | 0.97550416 |
| NDUFB4 | 5344.48379 | 0.17850857 | 0.2474221 | 0.72147381 | 0.47061805 | 0.9008571 |
| HGD | 7558.0687 | 0.56354915 | 0.43342537 | 1.30022188 | 0.19352493 | 0.77548688 |
| RABL3 | 1288.05367 | 0.15714408 | 0.15660059 | 1.00347057 | 0.31563387 | 0.84734197 |
| GTF2E1 | 561.67754 | 0.09328742 | 0.24076878 | 0.38745648 | 0.6984183 | 0.9529028 |
| HCLS1 | 2132.35519 | -0.2678071 | 0.42499538 | -0.6301413 | 0.52860213 | 0.9188881 |
| GOLGB1 | 6581.01608 | -0.1695607 | 0.15422244 | -1.0994557 | 0.27156933 | 0.82716979 |
| IQCB1 | 1239.212 | -0.007833 | 0.17118799 | -0.0457569 | 0.96350403 | 0.99320711 |

| | | | | | | |
|---|---|---|---|---|---|---|
| EAF2 | 374.659447 | 0.35010193 | 0.39787917 | 0.87992023 | 0.37890253 | 0.87069278 |
| SLC15A2 | 1029.20805 | 0.14289051 | 0.51763859 | 0.27604301 | 0.78251504 | 0.96964828 |
| ILDR1 | 967.155072 | 0.31257305 | 0.29902392 | 1.0453112 | 0.29587916 | 0.83750889 |
| CD86 | 274.723961 | -0.6125868 | 0.73565643 | -0.8327078 | 0.40500956 | 0.88132015 |
| CASR | 2551.31413 | -0.068464 | 0.4510177 | -0.1517989 | 0.87934557 | 0.98275231 |
| CSTA | 58.2073317 | 0.47931591 | 0.74087023 | 0.64696338 | 0.51765565 | 0.91636259 |
| CCDC58 | 923.606424 | 0.05353733 | 0.2950343 | 0.18146138 | 0.85600545 | 0.97929708 |
| FAM162A | 3681.72309 | 0.17033682 | 0.2398483 | 0.71018562 | 0.47758904 | 0.90387017 |
| WDR5B | 460.664896 | 0.09117891 | 0.25642397 | 0.35557873 | 0.72215607 | 0.95739595 |
| AC083798.2 | 1588.69754 | -0.2941833 | 0.22012438 | -1.3364413 | 0.18140509 | 0.76522169 |
| KPNA1 | 3112.43125 | -0.2207124 | 0.20334523 | -1.0854072 | 0.27774135 | 0.82805357 |
| PARP9 | 3522.20977 | -1.061083 | 0.5114318 | -2.0747303 | NA | NA |
| DTX3L | 2475.75347 | -0.6560342 | 0.42371203 | -1.548302 | 0.1215496 | 0.69725141 |
| PARP15 | 156.824461 | -0.7065026 | 0.71705664 | -0.9852815 | 0.32448584 | 0.85169085 |
| PARP14 | 7062.86583 | -1.4786392 | 0.62900406 | -2.3507626 | NA | NA |
| HSPBAP1 | 1084.07838 | -0.3071236 | 0.1810081 | -1.6967396 | 0.08974591 | 0.65375543 |
| DIRC2 | 1469.96122 | -0.1330105 | 0.22388303 | -0.5941072 | 0.55244039 | 0.92373235 |
| LINC02035 | 703.45513 | -0.0961151 | 0.1965425 | -0.4890296 | 0.62482076 | 0.94088547 |
| SEMA5B | 500.493915 | 0.57360555 | 0.37981054 | 1.51024125 | 0.13098188 | 0.7100756 |
| PDIA5 | 1820.23027 | 0.07117717 | 0.18542731 | 0.38385483 | 0.70108605 | 0.95400919 |
| SEC22A | 804.944165 | 0.0318425 | 0.125528 | 0.25366848 | 0.79975169 | 0.97208422 |
| ADCY5 | 2075.0631 | 0.0985867 | 0.35581684 | 0.2770715 | 0.78172521 | 0.96919239 |
| AC112503.1 | 190.497637 | 0.23739858 | 0.3961958 | 0.5991951 | 0.54904279 | 0.92373235 |
| HACD2 | 2147.15111 | 0.12139451 | 0.17370204 | 0.69886634 | 0.48463557 | 0.90541935 |
| MYLK-AS1 | 326.202133 | -0.1317198 | 0.33212853 | -0.3965929 | 0.69166769 | 0.95159158 |
| MYLK | 5686.6118 | -0.5306349 | 0.46338532 | -1.1451266 | 0.25215671 | 0.81833349 |
| CCDC14 | 2199.31949 | 0.0244095 | 0.20080902 | 0.12155581 | 0.90325081 | 0.98617053 |
| KALRN | 1010.83228 | -0.3178194 | 0.31735896 | -1.0014509 | 0.31660887 | 0.84783414 |
| UMPS | 1104.1083 | 0.08179013 | 0.20327249 | 0.40236692 | 0.68741401 | 0.95153453 |
| ITGB5 | 7824.99944 | 0.16940449 | 0.23268112 | 0.72805432 | 0.46658034 | 0.89960721 |
| MUC13 | 1658.97201 | 1.67374271 | 1.14205877 | 1.46554867 | 1.43E-01 | 0.72784886 |
| HEG1 | 2412.01255 | -0.0711065 | 0.3506096 | -0.2028082 | 8.39E-01 | 0.97618283 |
| SLC12A8 | 515.286275 | -0.1733286 | 0.35594183 | -0.4869577 | 0.6262883 | 0.94088547 |
| ZNF148 | 1866.77445 | -0.1029664 | 0.12591893 | -0.8177198 | 0.41351722 | 0.88316314 |
| SNX4 | 3403.37147 | 0.23871533 | 0.27261996 | 0.8756341 | 0.38122898 | 0.87171941 |
| OSBPL11 | 895.564521 | -0.0996229 | 0.20867779 | -0.4774004 | 0.63307705 | 0.94088547 |
| ALG1L | 69.1636055 | 0.34377961 | 0.51127147 | 0.67240131 | 0.50132825 | 0.91259827 |
| SLC41A3 | 2783.06779 | -0.0529951 | 0.157966 | -0.335484 | 0.73726002 | 0.96023056 |
| AC079848.1 | 47.906401 | 0.50977413 | 0.4867204 | 1.04736544 | 0.29493106 | 0.83630273 |
| ALDH1L1 | 14507.176 | 0.54625583 | 0.4948788 | 1.1038174 | 0.26967236 | 0.82716979 |
| ALDH1L1-AS1 | 1836.55438 | 0.37405667 | 0.44978731 | 0.83163012 | 0.40561776 | 0.88132015 |
| ALDH1L1-AS2 | 811.684267 | 0.45395311 | 0.53232219 | 0.85277886 | 0.39378195 | 0.87884163 |
| AC016924.1 | 167.397554 | 1.11869192 | 0.8109521 | 1.37947966 | NA | NA |
| AC063919.1 | 71.6468342 | 0.5224925 | 0.71555393 | 0.73019304 | 0.4652722 | 0.89921277 |
| KLF15 | 2384.06472 | -0.0312951 | 0.49146553 | -0.0636772 | 0.94922726 | 0.99170503 |
| CFAP100 | 46.8845728 | -0.0216571 | 0.66942433 | -0.0323518 | 0.97419153 | 0.99455099 |
| ZXDC | 2553.72455 | -0.2633419 | 0.23577654 | -1.1169132 | 0.26403143 | 0.82424998 |
| CHST13 | 284.404048 | 0.11234307 | 0.53388148 | 0.21042698 | 8.33E-01 | 9.76E-01 |
| TXNRD3NB | 185.353181 | 0.2322849 | 0.24628121 | 0.94316939 | 0.34559427 | 0.86111349 |
| TXNRD3 | 454.458856 | 0.31467766 | 0.15692348 | 2.00529365 | 0.04493166 | 0.57322281 |
| CHCHD6 | 492.735646 | 0.39441309 | 0.22395719 | 1.76110927 | 0.07821991 | 0.64079926 |
| PLXNA1 | 2334.13756 | -0.3249629 | 0.1709116 | -1.9013507 | 0.05725609 | 0.60077875 |
| TPRA1 | 2357.15172 | -0.2116361 | 0.1981043 | -1.0683067 | 0.28538218 | 0.83107779 |
| MCM2 | 439.806023 | 0.26955732 | 0.38488717 | 0.70035414 | 0.48370617 | 0.90541935 |

| | | | | | | |
|---|---|---|---|---|---|---|
| PODXL2 | 523.000752 | 0.53746723 | 0.37830008 | 1.42074311 | 0.15539146 | 0.74442517 |
| ABTB1 | 3830.44963 | -0.1102866 | 0.19809933 | -0.5567239 | 0.5777161 | 0.93130183 |
| MGLL | 15505.7041 | -0.2828934 | 0.19547472 | -1.4472121 | 0.14783752 | 0.73455299 |
| KBTBD12 | 171.16297 | -0.2415437 | 0.32678785 | -0.7391452 | 0.45981881 | 0.89755243 |
| SEC61A1 | 14457.6398 | -0.0482631 | 0.17314031 | -0.2787515 | 0.78043552 | 0.96864684 |
| RUVBL1 | 2646.98642 | 0.16567924 | 0.14824511 | 1.11760339 | 0.26373642 | 0.82424998 |
| EEFSEC | 1351.4624 | 0.3562396 | 0.23538276 | 1.51344812 | 0.13016587 | 0.70814767 |
| GATA2 | 2037.84408 | 0.42066401 | 0.37401279 | 1.12473159 | 0.26070279 | 0.82424998 |
| GATA2-AS1 | 182.79346 | -0.4428539 | 0.40838644 | -1.0843991 | 0.2781879 | 0.82805357 |
| RPN1 | 14284.8188 | -0.0826517 | 0.18280768 | -0.4521238 | 0.65117982 | 0.94390603 |
| RAB7A | 13623.6096 | -0.2074282 | 0.20872362 | -0.9937936 | 0.32032337 | 0.85016134 |
| AC112484.3 | 44.6627514 | 0.32595352 | 0.37031926 | 0.88019597 | 0.37875316 | 0.87069278 |
| AC112484.1 | 42.2031392 | -0.2515436 | 0.35740184 | -0.7038118 | 0.48155 | 0.90541935 |
| ACAD9 | 2778.92545 | 0.11232842 | 0.10757801 | 1.04415784 | 0.29641237 | 0.83817467 |
| KIAA1257 | 497.973197 | 0.09680485 | 0.12839963 | 0.75393403 | 0.45088883 | 0.89590699 |
| EFCC1 | 210.736596 | 0.04743246 | 0.30994909 | 0.15303307 | 0.87837219 | 0.98275231 |
| RAB43 | 1901.25017 | -0.075823 | 0.22261893 | -0.3405955 | 0.73340813 | 0.95884813 |
| ISY1-RAB43 | 2612.36689 | -0.0479336 | 0.15075685 | -0.3179528 | 0.75052072 | 0.96255347 |
| ISY1 | 1376.65537 | -0.1261398 | 0.14937476 | -0.8444516 | 0.39841711 | 0.88074856 |
| AC108673.3 | 61.5372593 | 0.19576744 | 0.30682138 | 0.63805017 | 0.52344102 | 0.91723878 |
| CNBP | 14212.619 | -0.130109 | 0.15426627 | -0.8434051 | 0.39900191 | 0.88077967 |
| COPG1 | 7974.75435 | -0.0277891 | 0.15001974 | -0.185236 | 0.85304392 | 0.97886533 |
| HMCES | 2116.68322 | 0.20141614 | 0.145856 | 1.38092456 | 0.16730216 | 0.75558932 |
| H1FX | 2909.26176 | -0.358458 | 0.26270731 | -1.3644766 | 0.17241763 | 0.75974924 |
| H1FX-AS1 | 1702.51057 | -0.2500344 | 0.26457709 | -0.9450341 | 0.34464148 | 0.85999891 |
| EFCAB12 | 206.42537 | 0.06889458 | 0.56408463 | 0.12213519 | 0.90279195 | 0.98608873 |
| MBD4 | 2411.67378 | -0.1091644 | 0.18597543 | -0.5869832 | 0.55721501 | 0.92373235 |
| IFT122 | 2592.73258 | -0.0050449 | 0.18270416 | -0.0276123 | 0.97797137 | 0.99476647 |
| PLXND1 | 4945.02952 | -0.2194483 | 0.31633002 | -0.693732 | 0.48785027 | 0.90584365 |
| TMCC1 | 1714.45815 | 0.14371248 | 0.39834836 | 0.36077086 | 0.71827075 | 0.95648644 |
| TMCC1-AS1 | 90.5533671 | 0.23853185 | 0.44166329 | 0.54007623 | 0.58914446 | 0.93289552 |
| LINC02021 | 81.4451282 | 0.18355143 | 0.68522164 | 0.26787163 | 0.78879813 | 0.97090793 |
| PIK3R4 | 1262.1158 | -0.0337249 | 0.12910877 | -0.2612132 | 0.79392813 | 0.97090793 |
| ATP2C1 | 4619.17121 | 0.20597188 | 0.17013577 | 1.21063247 | 0.2260363 | 0.80246466 |
| ASTE1 | 599.816778 | -0.0426913 | 0.129663 | -0.3292482 | 0.74196812 | 0.96109001 |
| NEK11 | 681.998036 | -0.2128184 | 0.28368787 | -0.7501852 | 0.45314317 | 0.89670027 |
| NUDT16 | 3863.85531 | 0.43236449 | 0.23567558 | 1.83457485 | 0.06656871 | 0.62378208 |
| MRPL3 | 3692.00391 | 0.15090602 | 0.18058042 | 0.83567213 | 0.40333938 | 0.88132015 |
| CPNE4 | 84.2054637 | 1.88400417 | 0.84300688 | 2.23486214 | 0.0254264 | 0.52250144 |
| ACPP | 1148.28622 | 0.61661979 | 0.50794685 | 1.21394551 | 0.22476853 | 0.80113867 |
| DNAJC13 | 2731.04538 | 0.01571275 | 0.18735629 | 0.0838656 | 0.93316329 | 0.99011568 |
| ACAD11 | 5988.72762 | 0.61300359 | 0.53659911 | 1.14238652 | 0.25329339 | 0.81848561 |
| NPHP3-ACAD | 6422.12706 | 0.52764464 | 0.49235867 | 1.07166721 | 0.28386954 | 0.83107779 |
| ACKR4 | 717.03885 | 0.93655464 | 0.55063794 | 1.70085382 | 0.08897044 | 0.65375543 |
| UBA5 | 3437.14874 | 0.18770317 | 0.20578198 | 0.9121458 | 0.36169197 | 0.86528137 |
| NPHP3 | 1505.73788 | -0.0144463 | 0.22333618 | -0.064684 | 0.94842564 | 0.99170503 |
| CDV3 | 8189.6614 | -0.6025823 | 0.38411762 | -1.5687442 | 0.11670754 | 0.689104 |
| TOPBP1 | 930.213146 | -0.1822414 | 0.18291651 | -0.9963091 | 0.31909996 | 0.84922641 |
| TF | 96.1790112 | 0.4459428 | 0.66388181 | 0.67172016 | 0.50176186 | 0.91259827 |
| SRPRB | 3249.27967 | -0.0252059 | 0.21109451 | -0.119406 | 0.90495372 | 0.98624484 |
| RAB6B | 865.219097 | -0.2716109 | 0.30777425 | -0.8825005 | 0.3775062 | 0.86987124 |
| C3orf36 | 68.0045116 | 0.8313631 | 0.6331662 | 1.31302507 | 0.1891745 | 0.76970626 |
| SLCO2A1 | 6009.31856 | 0.33977191 | 0.34076292 | 0.99709181 | 0.31871995 | 0.84914926 |
| RYK | 2581.70789 | -0.0909082 | 0.11999057 | -0.7576282 | 4.49E-01 | 8.95E-01 |

| | | | | | | |
|---|---|---|---|---|---|---|
| AMOTL2 | 4962.20379 | -0.377647 | 0.28355278 | -1.3318402 | 1.83E-01 | 0.76723162 |
| ANAPC13 | 3082.71721 | 0.18884803 | 0.18584782 | 1.01614336 | 3.10E-01 | 0.84432074 |
| CEP63 | 1367.32617 | -0.0010453 | 0.16788283 | -0.0062263 | 0.99503219 | 0.99940889 |
| EPHB1 | 115.486277 | -0.6754567 | 0.52652322 | -1.2828621 | 0.19954041 | 0.78134034 |
| PPP2R3A | 4137.76772 | -0.1309116 | 0.2268897 | -0.5769836 | 5.64E-01 | 9.25E-01 |
| AC072039.2 | 123.52617 | -0.249777 | 0.33868519 | -0.7374903 | 0.46082424 | 0.89755243 |
| MSL2 | 1309.31956 | -0.2905377 | 0.21453511 | -1.3542662 | 0.17565146 | 0.7610139 |
| PCCB | 4782.33829 | 0.42355 | 0.17961709 | 2.35807183 | 0.01837014 | 0.47570049 |
| STAG1 | 1757.66376 | -0.2290973 | 0.23617727 | -0.9700226 | 0.33203525 | 0.85452795 |
| SLC35G2 | 400.535168 | 0.08689806 | 0.24347954 | 0.35690087 | 0.72116602 | 0.95731494 |
| AC096992.2 | 74.5229419 | 0.31513607 | 0.40736832 | 0.77359002 | 0.43917328 | 0.89175645 |
| NCK1-DT | 96.6472756 | 0.31621187 | 0.37492848 | 0.84339249 | 0.39900897 | 0.88077967 |
| NCK1 | 1147.55922 | -0.0748005 | 0.18265846 | -0.4095104 | 0.68216514 | 0.95070301 |
| IL20RB | 295.248013 | -0.2026548 | 0.21168799 | -0.9573278 | 0.33840182 | 0.85719299 |
| DZIP1L | 465.733139 | -0.2306659 | 0.31407878 | -0.7344206 | 0.46269244 | 0.89796156 |
| A4GNT | 37.9102136 | 1.92535249 | 0.7534813 | 2.55527575 | 0.01061037 | 0.40196898 |
| DBR1 | 732.823296 | -0.1916111 | 0.1726739 | -1.10967 | 0.26714124 | 0.8265123 |
| ARMC8 | 2306.20094 | -0.2647076 | 0.12975004 | -2.0401347 | 0.04133691 | 0.55943411 |
| NME9 | 66.287369 | 0.33097062 | 0.4733258 | 0.69924483 | 0.48439904 | 0.90541935 |
| MRAS | 3464.86335 | -0.457457 | 0.35866298 | -1.2754509 | 0.20214968 | 0.78163854 |
| ESYT3 | 250.255466 | -0.0304699 | 0.30156575 | -0.1010391 | 0.91951943 | 0.9872458 |
| CEP70 | 1335.75198 | -0.131087 | 0.3084701 | -0.4249584 | 6.71E-01 | 0.94922827 |
| FAIM | 685.862717 | -0.0218303 | 0.16363758 | -0.1334062 | 0.89387215 | 0.98474725 |
| PIK3CB | 2163.07242 | 0.1634265 | 0.22328921 | 0.73190505 | 0.46422653 | 0.89830907 |
| MRPS22 | 1831.05952 | -0.0840941 | 0.18294907 | -0.4596587 | 6.46E-01 | 0.94208859 |
| COPB2 | 6313.00645 | -0.0803175 | 0.18974507 | -0.4232914 | 6.72E-01 | 9.49E-01 |
| AC097103.2 | 82.9265465 | 0.15577211 | 0.32383388 | 0.48102477 | 0.6304989 | 0.94088547 |
| RBP1 | 564.736847 | 0.08345677 | 0.4201466 | 0.19863727 | 0.8425465 | 0.97644404 |
| NMNAT3 | 498.986735 | 0.40959796 | 0.28098552 | 1.45771911 | 0.14491798 | 0.73039059 |
| CLSTN2 | 906.268184 | 1.39506964 | 0.54197667 | 2.57404002 | 1.01E-02 | 0.39989496 |
| SLC25A36 | 4346.84061 | -0.1506108 | 0.24522814 | -0.614166 | 0.53910564 | 0.92167294 |
| PXYLP1 | 724.440548 | 0.41190483 | 0.22523666 | 1.82876461 | 0.06743488 | 0.62502327 |
| ZBTB38 | 4040.30442 | -0.4268425 | 0.35617999 | -1.1983899 | 0.23076526 | 0.80610188 |
| RASA2 | 1122.08104 | -0.0870829 | 0.17237574 | -0.5051924 | 0.6134237 | 0.9388166 |
| RNF7 | 3171.70185 | -0.1985311 | 0.21498976 | -0.9234442 | 0.35577574 | 0.86364633 |
| ATP1B3 | 6555.11779 | -0.5921678 | 0.3604468 | -1.6428715 | 0.10040952 | 0.67206241 |
| TFDP2 | 3061.73059 | 0.16006856 | 0.14716497 | 1.08768107 | 0.27673593 | 0.82755832 |
| GK5 | 1325.99268 | 0.23530692 | 0.20319179 | 1.15805332 | 0.24684228 | 0.81542272 |
| XRN1 | 1629.23681 | -0.4438094 | 0.30288369 | -1.46528 | 0.14284453 | 0.72784886 |
| ATR | 1413.78482 | -0.141802 | 0.1446618 | -0.9802312 | 0.32697202 | 0.85297643 |
| PLS1 | 1995.01491 | 0.6354233 | 0.28211824 | 2.25232975 | 0.02430144 | 0.51967919 |
| TRPC1 | 476.488673 | 0.22112041 | 0.20094216 | 1.1004182 | 0.27114995 | 0.82716979 |
| PCOLCE2 | 890.486044 | 0.58548782 | 0.5520295 | 1.06060966 | 0.28886733 | 0.83295411 |
| PAQR9-AS1 | 94.6063062 | 0.103953 | 0.29844003 | 0.34832124 | 0.72759895 | 0.95848028 |
| U2SURP | 3367.55398 | -0.0864999 | 0.13826357 | -0.625616 | 0.53156687 | 0.91962754 |
| AC026304.1 | 92.5887608 | 0.0936204 | 0.31093159 | 0.30109646 | 0.76334094 | 0.96671773 |
| CHST2 | 1110.70561 | -0.7508872 | 0.53890336 | -1.3933616 | NA | NA |
| SLC9A9 | 313.691827 | -0.1452005 | 0.32708753 | -0.4439194 | 0.65710087 | 0.94545756 |
| C3orf58 | 799.046548 | -0.2261094 | 0.20437336 | -1.1063544 | 0.26857317 | 0.82676682 |
| PLOD2 | 4706.13388 | -0.0678188 | 0.32325533 | -0.2097996 | 8.34E-01 | 0.97550416 |
| PLSCR4 | 1773.2919 | -0.0433536 | 0.21917886 | -0.1978001 | 0.84320145 | 0.97649449 |
| PLSCR2 | 52.6602341 | 0.37151769 | 0.45922993 | 0.80900148 | 0.4185143 | 0.88445833 |
| PLSCR1 | 6472.55647 | -0.429007 | 0.45493759 | -0.943002 | 0.34567989 | 0.86111349 |
| AGTR1 | 543.428133 | -0.1894943 | 0.34980806 | -0.5417094 | 0.58801872 | 0.93289552 |

| | | | | | | |
|---|---|---|---|---|---|---|
| CPA3 | 216.512218 | 1.16194906 | 1.06948303 | 1.08645862 | 0.27727614 | 0.82782817 |
| GYG1 | 2013.76866 | -0.1974736 | 0.21654638 | -0.9119229 | 0.36180931 | 0.86528137 |
| HLTF | 1405.83761 | 0.41616156 | 0.20371179 | 2.04289389 | 4.11E-02 | 5.58E-01 |
| HPS3 | 2558.04953 | 0.40366355 | 0.25844391 | 1.56190007 | 1.18E-01 | 6.91E-01 |
| CP | 2445.93856 | 0.58453514 | 0.43507518 | 1.34352675 | 0.17910147 | 0.76120583 |
| AC093001.1 | 77.0288847 | 0.6711693 | 0.37795115 | 1.77580967 | 0.07576429 | 0.63773415 |
| TM4SF18 | 1433.66701 | 0.43957684 | 0.30396488 | 1.44614351 | 0.14813695 | 0.73474739 |
| TM4SF1 | 6338.31544 | 0.00453984 | 0.40288569 | 0.01126831 | 0.99100938 | 0.99846819 |
| WWTR1 | 5613.3355 | -0.14512 | 0.24826638 | -0.5845334 | 0.55886149 | 0.92421175 |
| WWTR1-AS1 | 311.965905 | -0.0819848 | 0.27854864 | -0.2943286 | 0.76850679 | 0.96793882 |
| COMMD2 | 1395.38595 | -0.1615982 | 0.14548776 | -1.1107338 | 0.26668295 | 0.8265123 |
| ANKUB1 | 1108.42997 | 0.27577973 | 0.23957131 | 1.15113838 | 2.50E-01 | 8.16E-01 |
| RNF13 | 3905.61172 | -0.090932 | 0.16973622 | -0.5357254 | 0.59214836 | 0.93289552 |
| PFN2 | 6290.781 | 0.33664595 | 0.18462875 | 1.82336693 | 0.06824784 | 0.62648383 |
| TSC22D2 | 2217.06044 | -0.2142997 | 0.20387964 | -1.0511087 | 0.29320867 | 0.8352732 |
| SERP1 | 6173.3019 | -0.0775216 | 0.18556927 | -0.4177499 | 6.76E-01 | 0.94959168 |
| EIF2A | 3755.75996 | 0.10325791 | 0.23911734 | 0.43182947 | 0.66586536 | 0.94753096 |
| SELENOT | 5129.13984 | 0.00667073 | 0.14314746 | 0.04660041 | 0.96283171 | 0.99320711 |
| ERICH6 | 50.877635 | 0.38823096 | 0.37807568 | 1.02686045 | 0.30448618 | 0.84211658 |
| ERICH6-AS1 | 113.058491 | 0.21870586 | 0.22861913 | 0.95663849 | 3.39E-01 | 8.57E-01 |
| SIAH2 | 1538.83726 | -0.3441311 | 0.2103661 | -1.6358675 | 0.10186732 | 0.67279543 |
| SIAH2-AS1 | 175.102807 | -0.2561874 | 0.26009419 | -0.9849793 | 0.32463425 | 0.85169085 |
| MED12L | 75.0692395 | 0.45072666 | 0.33863799 | 1.33099849 | 0.18318951 | 0.76723162 |
| GPR171 | 73.0798063 | -0.5216603 | 0.81906751 | -0.6368953 | 0.52419304 | 0.91723878 |
| P2RY14 | 67.1733583 | -0.1701519 | 0.60246557 | -0.282426 | 0.77761688 | 0.96836854 |
| P2RY13 | 68.3947634 | 0.88505226 | 0.61387959 | 1.44173592 | 0.14937689 | 0.73670625 |
| IGSF10 | 96.0028578 | 0.6380512 | 0.39536554 | 1.61382604 | 0.10656516 | 0.67674128 |
| SUCNR1 | 2105.94662 | 0.54767241 | 0.414335 | 1.32181063 | 0.18623122 | 0.76908034 |
| MBNL1 | 4015.65738 | -0.1760429 | 0.16062567 | -1.0959825 | 0.27308642 | 0.82716979 |
| MBNL1-AS1 | 175.480178 | -0.7975044 | 0.55644137 | -1.4332227 | 0.1517942 | 0.74024851 |
| P2RY1 | 90.8270611 | -0.6651723 | 0.3981782 | -1.6705393 | 0.09481272 | 0.65888357 |
| RAP2B | 1597.81337 | -0.0835459 | 0.33078685 | -0.2525671 | 0.80060274 | 0.97210205 |
| ARHGEF26 | 344.559398 | -0.0350973 | 0.43957451 | -0.0798438 | 0.93636148 | 0.99068685 |
| DHX36 | 2186.30089 | -0.1908834 | 0.18525504 | -1.0303818 | 3.03E-01 | 0.84211658 |
| MME | 12667.018 | 0.81822741 | 0.72198236 | 1.13330666 | 0.2570855 | 0.82107009 |
| MME-AS1 | 2136.70714 | 1.04977505 | 0.71081695 | 1.47685708 | 0.13971395 | 0.72475166 |
| DWORF | 75.7895117 | 2.13321652 | 1.1167611 | 1.91018161 | 0.05610983 | 0.59895387 |
| PLCH1 | 222.50371 | 0.71146377 | 0.48666213 | 1.4619255 | 0.14376163 | 0.72920837 |
| C3orf33 | 453.014869 | 0.22809064 | 0.24960458 | 0.91380789 | 0.3608178 | 0.86504879 |
| AC104472.3 | 1305.2657 | 0.32533451 | 0.22216154 | 1.46440521 | 0.14308327 | 0.72784886 |
| SLC33A1 | 2247.49297 | 0.32328914 | 0.18829389 | 1.7169391 | 8.60E-02 | 6.50E-01 |
| GMPS | 1848.50635 | -0.0331063 | 0.15565395 | -0.2126914 | 0.83156764 | 0.97550416 |
| KCNAB1 | 258.89059 | -0.0360493 | 0.42533881 | -0.0847543 | 0.93245676 | 0.9899349 |
| AC084036.1 | 172.648144 | 0.02127144 | 0.51828974 | 0.0410416 | 0.96726273 | 0.99400331 |
| SSR3 | 5854.45094 | 0.07734878 | 0.23125436 | 0.33447491 | 0.73802122 | 0.9602325 |
| TIPARP-AS1 | 60.1950397 | 0.4111565 | 0.45076644 | 0.91212759 | 0.36170156 | 0.86528137 |
| TIPARP | 1368.79155 | -0.1605052 | 0.30949058 | -0.518611 | 0.60403202 | 0.93655508 |
| LINC00886 | 224.367406 | 0.61087381 | 0.52874084 | 1.1553369 | 0.24795249 | 0.81602355 |
| LEKR1 | 73.3360385 | 0.1233504 | 0.56769596 | 0.2172825 | 0.8279882 | 0.97548201 |
| CCNL1 | 7608.79189 | -0.2954088 | 0.19873864 | -1.4864184 | 0.13716849 | 0.72209298 |
| VEPH1 | 1068.67162 | 0.7922748 | 0.51840331 | 1.52829811 | 0.12643854 | 0.70431949 |
| PTX3 | 839.255618 | -0.5964719 | 1.10605828 | -0.5392771 | 0.58969569 | 0.93289552 |
| PQLC2L | 97.3794315 | 1.11560755 | 0.44270766 | 2.5199644 | 0.01173667 | 0.40423902 |
| RSRC1 | 895.820273 | -0.0151253 | 0.10378385 | -0.1457386 | 0.88412772 | 0.98317437 |

| | | | | | | |
|---|---|---|---|---|---|---|
| AC106707.1 | 190.596051 | 0.0459459 | 0.22563087 | 0.20363303 | 0.83864028 | 0.97612168 |
| MLF1 | 768.529338 | 0.32118341 | 0.20955302 | 1.53270716 | 0.12534802 | 0.70270635 |
| GFM1 | 2986.23219 | 0.1525934 | 0.22929224 | 0.66549746 | 0.50573237 | 0.91386909 |
| LXN | 908.106293 | 0.14839753 | 0.25086426 | 0.59154512 | 0.55415523 | 0.92373235 |
| AC080013.4 | 64.4695448 | 0.49188391 | 0.45454371 | 1.08214875 | 0.27918644 | 0.82825734 |
| RARRES1 | 2284.69528 | 0.43381796 | 0.46679869 | 0.92934698 | 0.35270929 | 0.86364633 |
| MFSD1 | 3850.41567 | 0.02521216 | 0.17982829 | 0.14020131 | 0.88850094 | 0.98423722 |
| AC080013.1 | 163.674781 | 0.16402925 | 0.47322373 | 0.34662094 | 0.72887611 | 0.95848028 |
| IQCJ-SCHIP1 | 1171.36374 | -0.6068567 | 0.22547358 | -2.6914759 | 0.00711366 | 0.35456697 |
| SCHIP1 | 1090.9973 | -0.5747578 | 0.21955983 | -2.6177729 | 0.00885057 | 0.38245454 |
| AC079594.2 | 781.010286 | 0.18652021 | 0.27923642 | 0.66796517 | 0.50415582 | 0.91356552 |
| IFT80 | 1635.3288 | 0.21490534 | 0.29018797 | 0.74057289 | 0.45895245 | 0.89755243 |
| SMC4 | 835.873886 | -0.0191927 | 0.1712361 | -0.1120835 | 0.91075717 | 0.98662757 |
| TRIM59 | 702.479615 | 0.02383457 | 0.15889958 | 0.14999767 | 0.88076645 | 0.98275231 |
| KPNA4 | 3854.45163 | -0.3675125 | 0.26856877 | -1.3684111 | 0.17118342 | 0.75956362 |
| AC069224.1 | 85.5310988 | 0.03267587 | 0.45951045 | 0.07111017 | 0.94331007 | 0.99136103 |
| PPM1L | 900.543564 | -0.0620694 | 0.25522 | -0.2431996 | 0.80785077 | 0.97334138 |
| B3GALNT1 | 1276.4627 | 0.21300463 | 0.22972972 | 0.92719665 | 0.35382444 | 0.86364633 |
| NMD3 | 2344.02224 | -0.1246255 | 0.15398653 | -0.8093274 | 0.41832684 | 0.88445833 |
| LINC02067 | 136.419608 | 0.20884967 | 0.26417821 | 0.7905636 | 0.42919869 | 0.88851358 |
| AC112491.1 | 143.646685 | 0.27397778 | 0.28612979 | 0.95752973 | 0.33829995 | 0.85711496 |
| BCHE | 164.845946 | 1.97831403 | 0.49484634 | 3.99783503 | 6.39E-05 | 0.030786 |
| ZBBX | 47.3845512 | 0.35676643 | 1.01028334 | 0.35313502 | 0.72398722 | 0.95810643 |
| PDCD10 | 1774.9107 | -0.0786386 | 0.13494372 | -0.5827508 | 5.60E-01 | 9.24E-01 |
| SERPINI1 | 1193.47099 | 1.70584276 | 0.47198767 | 3.61416807 | 0.00030131 | 0.08492861 |
| GOLIM4 | 2398.97408 | 0.21564629 | 0.26671507 | 0.80852682 | 0.41878737 | 0.88451372 |
| MECOM | 4425.88856 | -0.2944608 | 0.29488673 | -0.9985558 | 3.18E-01 | 8.49E-01 |
| ACTRT3 | 160.396165 | 0.06864666 | 0.24081971 | 0.28505418 | 0.77560264 | 0.96836854 |
| MYNN | 901.794678 | -0.0697559 | 0.15792632 | -0.441699 | 0.658707 | 0.9458503 |
| AC078795.1 | 293.508693 | 0.03304129 | 0.23881462 | 0.13835538 | 0.88995956 | 0.98459146 |
| LRRC34 | 221.874952 | 0.11708685 | 0.30613343 | 0.38246997 | 0.7021128 | 0.95431462 |
| AC078795.2 | 81.7758791 | 0.0355108 | 0.38737693 | 0.09166989 | 0.92696032 | 0.98915637 |
| SEC62 | 7334.56758 | -0.0133171 | 0.14121226 | -0.0943054 | 0.92486657 | 0.98869145 |
| GPR160 | 2171.27633 | 0.5355175 | 0.27614049 | 1.93929362 | 0.0524656 | 0.58645127 |
| PHC3 | 2501.51456 | 0.01365589 | 0.14004079 | 0.09751366 | 0.92231849 | 0.98826666 |
| AC008040.5 | 1619.0353 | 0.01908434 | 0.15660707 | 0.12186132 | 9.03E-01 | 9.86E-01 |
| PRKCI | 2435.12432 | -0.2268095 | 0.17490871 | -1.2967309 | 0.19472379 | 0.7762856 |
| SKIL | 1885.77988 | -0.1807306 | 0.24600631 | -0.7346582 | 0.4625477 | 0.89792898 |
| CLDN11 | 401.342401 | 1.25494622 | 0.31394747 | 3.99731278 | 6.41E-05 | 0.030786 |
| AC026316.5 | 102.20127 | 1.27240085 | 0.45889964 | 2.77272138 | 0.00555897 | 0.31762392 |
| RPL22L1 | 850.214778 | -0.2526522 | 0.19123762 | -1.3211429 | 0.18645371 | 0.76908034 |
| EIF5A2 | 412.383709 | 0.11236667 | 0.39775242 | 0.28250406 | 0.77755703 | 0.96836854 |
| SLC2A2 | 1404.83219 | 0.78498558 | 0.46621455 | 1.68374319 | 0.09223131 | 0.65583276 |
| TNIK | 903.362457 | 0.45094518 | 0.36380749 | 1.23951591 | 0.2151545 | 0.7949124 |
| PLD1 | 1240.03305 | 1.07471251 | 0.26720437 | 4.02206187 | 5.77E-05 | 0.02922122 |
| FNDC3B | 3347.04638 | 0.16972012 | 0.2050558 | 0.82767773 | 0.40785303 | 0.88161173 |
| TNFSF10 | 17953.0297 | -0.3046497 | 0.40995487 | -0.7431299 | 0.45740302 | 0.89688594 |
| LINC02068 | 36.3357369 | -0.2798438 | 0.68929909 | -0.4059832 | 0.68475497 | 0.95097288 |
| NCEH1 | 2254.90224 | -0.0223309 | 0.34653389 | -0.0644408 | 0.94861926 | 0.99170503 |
| ECT2 | 621.005739 | -0.1821565 | 0.31934697 | -0.5704031 | 0.56840431 | 0.92638187 |
| NLGN1 | 342.565528 | 0.59612676 | 0.47639049 | 1.2513406 | 0.21081024 | 0.79134644 |
| NAALADL2 | 411.47943 | 0.49066198 | 0.45311527 | 1.08286348 | 0.27886902 | 0.82825734 |
| TBL1XR1 | 5947.89495 | -0.4192165 | 0.20028277 | -2.0931229 | 0.03633819 | 0.55384187 |
| LINC00501 | 67.2767487 | -0.4445064 | 0.72181829 | -0.6158148 | 0.53801677 | 0.92127611 |

| | | | | | | |
|---|---|---|---|---|---|---|
| KCNMB2 | 202.723663 | 0.38781601 | 0.60529791 | 0.6407027 | 0.52171586 | 0.91723878 |
| AC117457.1 | 143.129584 | 0.3831415 | 0.55647885 | 0.68851044 | 0.4911314 | 0.90600166 |
| ZMAT3 | 1440.82462 | 0.4714558 | 0.21284614 | 2.21500757 | 0.02675955 | 0.52678638 |
| PIK3CA | 1496.21466 | -0.2006955 | 0.13442555 | -1.4929863 | 0.13544079 | 0.71649038 |
| KCNMB3 | 692.971466 | -0.2861167 | 0.12603738 | -2.2700936 | 0.0232019 | 0.51967919 |
| ZNF639 | 1359.98491 | -0.3426113 | 0.18048643 | -1.8982661 | 0.05766104 | 0.60129769 |
| MFN1 | 2451.84934 | -0.17768 | 0.15534552 | -1.1437731 | 0.25271774 | 0.81835388 |
| GNB4 | 1091.46197 | -0.1924608 | 0.31581622 | -0.6094076 | 0.54225431 | 0.92292765 |
| AC007620.2 | 514.063048 | -0.1973664 | 0.30798113 | -0.6408392 | 0.52162713 | 0.91723878 |
| ACTL6A | 1484.93068 | -0.0817648 | 0.22760438 | -0.3592411 | 0.71941475 | 0.9566409 |
| AC090425.2 | 381.626261 | -0.0339069 | 0.26398163 | -0.1284443 | 0.8977974 | 0.98474725 |
| MRPL47 | 1651.99617 | -0.1016721 | 0.1919493 | -0.5296821 | 0.59633238 | 0.9334601 |
| NDUFB5 | 4551.56095 | 0.47102949 | 0.20118867 | 2.34123275 | 0.01922018 | 0.48808321 |
| USP13 | 589.694025 | -0.0489387 | 0.17840798 | -0.2743077 | 0.7838482 | 0.96978847 |
| TTC14 | 2624.98578 | -0.0526299 | 0.23956037 | -0.2196939 | 0.82610957 | 0.97548201 |
| CCDC39 | 849.842864 | -0.1495801 | 0.21722342 | -0.6886002 | 0.49107488 | 0.90600166 |
| FXR1 | 4311.10523 | 0.04405615 | 0.10591012 | 0.41597673 | 0.67742703 | 0.95018034 |
| DNAJC19 | 2245.74198 | 0.00965661 | 0.18569101 | 0.05200363 | 0.9585258 | 0.99237995 |
| ATP11B | 2279.16804 | -0.0419724 | 0.1553722 | -0.2701407 | 0.787052 | 0.97090793 |
| AC092953.2 | 286.752237 | 0.18767764 | 0.21189315 | 0.88571829 | 0.37576935 | 0.86889275 |
| DCUN1D1 | 1709.51755 | -0.2251855 | 0.12844937 | -1.7531068 | 0.07958367 | 0.64079926 |
| MCCC1 | 3690.98508 | 0.48650071 | 0.22228842 | 2.18860127 | 0.02862583 | 0.53606221 |
| MCCC1-AS1 | 863.46048 | 0.44405588 | 0.24356152 | 1.82317749 | 0.06827651 | 0.62648383 |
| LAMP3 | 452.091539 | -3.0494918 | 1.38783412 | -2.1973028 | NA | NA |
| MCF2L2 | 459.728371 | -0.4097004 | 0.33120136 | -1.2370132 | 0.21608219 | 0.7954827 |
| B3GNT5 | 690.7449 | -0.4874846 | 0.34673312 | -1.4059361 | 0.1597431 | 0.74768867 |
| KLHL6 | 214.1357 | -0.0219951 | 0.53460824 | -0.0411424 | 0.96718233 | 0.99400331 |
| KLHL24 | 3101.63255 | -0.2778074 | 0.19086913 | -1.4554862 | 0.14553471 | 0.73161643 |
| YEATS2 | 1677.61386 | -0.4036007 | 0.34138652 | -1.1822397 | 0.23711059 | 0.80857441 |
| YEATS2-AS1 | 323.809724 | -0.42294 | 0.41183912 | -1.0269543 | 0.30444198 | 0.84211658 |
| AC131160.1 | 1770.65727 | 0.07684122 | 0.15893721 | 0.48346907 | 0.62876272 | 0.94088547 |
| MAP6D1 | 112.666152 | -0.2619204 | 0.28826666 | -0.9086046 | 0.36355887 | 0.8663515 |
| PARL | 1949.7516 | 0.06618258 | 0.14075404 | 0.47020019 | 0.638212 | 0.94088547 |
| ABCC5 | 4015.8001 | -0.1904691 | 0.33415838 | -0.5699966 | 0.56867999 | 0.92642835 |
| EIF2B5 | 3646.3639 | -0.1465485 | 0.09677211 | -1.5143674 | 0.12993268 | 0.70814767 |
| DVL3 | 6712.27148 | -0.3397439 | 0.17796333 | -1.9090671 | 0.05625343 | 0.59895387 |
| AP2M1 | 17263.8847 | -0.0213248 | 0.13467698 | -0.15834 | 0.87418887 | 0.98275231 |
| ABCF3 | 3794.18244 | -0.2081495 | 0.12905721 | -1.6128469 | 0.10677778 | 0.67674128 |
| VWA5B2 | 45.0695342 | -0.1245597 | 0.52749731 | -0.2361333 | 0.81332921 | 0.9738447 |
| ALG3 | 1915.69161 | -0.2104477 | 0.20946646 | -1.0046845 | 0.31504879 | 0.84696964 |
| EEF1AKMT4 | 352.759964 | -0.5977549 | 0.37718659 | -1.5847724 | 0.11301807 | 0.68130021 |
| EEF1AKMT4- | 235.849709 | -0.5670501 | 0.37888757 | -1.4966184 | 0.13449258 | 0.71630293 |
| PSMD2 | 9138.78728 | -0.3040258 | 0.3407845 | -0.8921352 | 0.37232047 | 0.86775997 |
| EIF4G1 | 17043.85 | -0.1470954 | 0.21925437 | -0.670889 | 0.50229122 | 0.91295119 |
| FAM131A | 1648.70592 | -0.0233682 | 0.22675586 | -0.1030543 | 0.91791984 | 0.9868044 |
| CLCN2 | 918.876665 | 0.19331213 | 0.18420992 | 1.04941216 | 0.29398847 | 0.83580671 |
| POLR2H | 4178.6889 | -0.1386311 | 0.14354934 | -0.9657381 | 0.33417527 | 0.85533716 |
| THPO | 501.931899 | -0.3118424 | 0.33416483 | -0.9331992 | 0.35071712 | 0.86364633 |
| CHRD | 3367.03539 | -0.1551423 | 0.41280724 | -0.3758226 | 0.70704879 | 0.95501548 |
| EPHB3 | 422.410201 | -0.1003867 | 0.66685692 | -0.150537 | 0.88034094 | 0.98275231 |
| MAGEF1 | 2368.96073 | -0.2215867 | 0.18535267 | -1.195487 | 0.23189681 | 0.80649152 |
| VPS8 | 1562.39774 | 0.00580152 | 0.12893132 | 0.044997 | 0.9641097 | 0.99342622 |
| C3orf70 | 97.751958 | -0.1429468 | 0.6012672 | -0.2377426 | 0.81208073 | 0.97355727 |
| EHHADH-AS1 | 1292.66231 | 1.58359213 | 0.61244556 | 2.58568635 | 0.00971853 | 0.39368232 |

| | | | | | | |
|---|---|---|---|---|---|---|
| EHHADH | 7773.94965 | 1.49473846 | 0.61291258 | 2.43874656 | 0.0147383 | 0.44193678 |
| MAP3K13 | 2082.19902 | 0.18072679 | 0.21482223 | 0.84128534 | 0.4001881 | 0.88077967 |
| TMEM41A | 2016.84674 | 0.07796902 | 0.31605672 | 0.24669313 | 0.80514572 | 0.97303568 |
| LIPH | 811.803999 | 0.45920749 | 0.45401858 | 1.01142886 | 0.31181121 | 0.84537057 |
| SENP2 | 1900.15418 | -0.1975911 | 0.18409558 | -1.0733069 | 0.28313344 | 0.83078187 |
| IGF2BP2 | 1275.511 | -0.0877284 | 0.30855728 | -0.284318 | 0.77616668 | 0.96836854 |
| TRA2B | 4683.64213 | -0.1387185 | 0.19683237 | -0.7047546 | 0.48096297 | 0.90541935 |
| ETV5 | 1343.66663 | 0.54202536 | 0.30193496 | 1.79517254 | 0.07262621 | 0.63365357 |
| DGKG | 331.658304 | -0.3276562 | 0.44654941 | -0.7337513 | 0.46310033 | 0.89807153 |
| CRYGS | 415.137089 | 0.16752491 | 0.17162294 | 0.97612193 | 0.32900405 | 0.85317073 |
| TBCCD1 | 491.161554 | 0.0316884 | 0.15746024 | 0.20124696 | 0.84050547 | 0.97618283 |
| DNAJB11 | 2987.70082 | 0.05822957 | 0.25319658 | 0.22997773 | 0.81810908 | 0.97452375 |
| AC068631.1 | 7465.65251 | 0.30469948 | 0.62592879 | 0.48679575 | 0.62640308 | 0.94088547 |
| AC068631.2 | 1547.99928 | 0.07266464 | 0.27181652 | 0.26732972 | 0.78921531 | 0.97090793 |
| FETUB | 54.6931807 | 0.29908967 | 0.58219861 | 0.51372447 | 0.60744464 | 0.93739614 |
| HRG | 1121.57653 | 1.13105137 | 0.58824132 | 1.92276764 | 0.05450924 | 0.59329767 |
| KNG1 | 15932.6161 | 0.29567832 | 0.64892905 | 0.45564044 | 0.64864856 | 0.94312988 |
| AC112907.3 | 685.403193 | 0.31769007 | 0.25208837 | 1.26023297 | 0.20758533 | 0.78834607 |
| EIF4A2 | 19498.772 | 0.12533104 | 0.14612613 | 0.85769085 | 0.39106319 | 0.87710956 |
| RFC4 | 499.900584 | -0.0147753 | 0.22525603 | -0.0655933 | 0.94770166 | 0.99170503 |
| ADIPOQ | 223.569291 | 6.2747336 | 3.12219361 | 2.00971957 | NA | NA |
| ST6GAL1 | 8537.04627 | -0.3809357 | 0.27133919 | -1.4039097 | 0.16034574 | 0.74781837 |
| RPL39L | 102.828807 | 0.2682918 | 0.25939353 | 1.0343041 | 0.30099402 | 0.84211658 |
| MASP1 | 600.488803 | 0.11368131 | 0.52534481 | 0.2163937 | 0.82868088 | 0.97550416 |
| RTP4 | 523.054704 | -1.7383075 | 0.8372417 | -2.0762314 | NA | NA |
| SST | 49.0759135 | -1.0604168 | 1.11619611 | -0.9500273 | 0.34209837 | 0.85889221 |
| AC072022.1 | 258.761916 | -0.4369718 | 0.43577822 | -1.002739 | 0.3159868 | 0.84750116 |
| BCL6 | 8008.60969 | -0.5980915 | 0.39158298 | -1.5273685 | 0.1266694 | 0.70443503 |
| LPP-AS2 | 123.124207 | -0.1648059 | 0.32326518 | -0.5098164 | 0.6101801 | 0.93792127 |
| LPP | 6195.54295 | -0.0740439 | 0.32716103 | -0.2263226 | 0.82095055 | 0.97472105 |
| TPRG1 | 71.2193103 | -0.0503644 | 0.45654034 | -0.1103175 | 0.91215755 | 0.98662757 |
| P3H2 | 5830.84771 | -0.1707683 | 0.25265564 | -0.6758937 | 0.49910815 | 0.91147492 |
| P3H2-AS1 | 49.7758732 | 0.62861163 | 0.64065585 | 0.98120017 | 0.32649404 | 0.85282528 |
| CLDN1 | 2799.23722 | 0.29016674 | 0.34332023 | 0.84517808 | 0.39801142 | 0.88054917 |
| CLDN16 | 2042.73452 | 0.50765407 | 0.4276569 | 1.18705923 | 0.23520426 | 0.80747753 |
| TMEM207 | 227.476585 | 1.6553286 | 0.75825597 | 2.18307362 | 0.02903039 | 0.53606221 |
| IL1RAP | 349.341425 | 0.14449114 | 0.30597001 | 0.47223955 | 0.63675582 | 0.94088547 |
| GMNC | 151.519721 | 2.10262435 | 0.64089642 | 3.28075533 | 0.0010353 | 0.15012259 |
| UTS2B | 131.236936 | 0.0619317 | 0.41093949 | 0.15070759 | 0.88020639 | 0.98275231 |
| CCDC50 | 5634.57736 | -0.1098308 | 0.24738778 | -0.4439621 | 0.65706996 | 0.94545756 |
| FGF12 | 96.252231 | -0.0345216 | 0.54848301 | -0.0629401 | 0.94981423 | 0.99170503 |
| MB21D2 | 341.722109 | 0.35014252 | 0.28172171 | 1.24286666 | 0.21391697 | 0.7926732 |
| ATP13A4 | 44.198484 | 0.44493777 | 0.68738831 | 0.64728737 | 0.51744597 | 0.91636259 |
| OPA1 | 2775.72291 | 0.17647659 | 0.15939303 | 1.10717884 | 0.26821661 | 0.82676682 |
| HES1 | 3635.39237 | 0.10942438 | 0.51467812 | 0.2126074 | 0.8316332 | 0.97550416 |
| LINC00887 | 55.8511523 | 0.80306977 | 0.61912175 | 1.29711122 | 0.19459292 | 0.7762856 |
| CPN2 | 2972.50483 | 0.27091846 | 0.64571205 | 0.41956545 | 0.67480294 | 0.94935413 |
| ATP13A3 | 5621.18755 | -0.2767438 | 0.334954 | -0.8262143 | 0.40868253 | 0.88203324 |
| LINC00884 | 80.1622548 | 0.17290006 | 0.39763222 | 0.43482407 | 0.66369014 | 0.94642611 |
| TMEM44-AS | 332.937194 | -0.1722725 | 0.30901732 | -0.5574848 | 0.57719624 | 0.93130183 |
| TMEM44 | 732.298275 | -0.264774 | 0.21581911 | -1.2268329 | 0.21988542 | 0.79858446 |
| AC046143.1 | 230.541267 | -0.1780523 | 0.29401557 | -0.605588 | 0.54478841 | 0.92345821 |
| LSG1 | 2055.28776 | -0.1436752 | 0.25226455 | -0.5695417 | 0.56898857 | 0.92660886 |
| FAM43A | 1538.02951 | -1.1030421 | 0.68608046 | -1.6077446 | 0.10789113 | 0.67720384 |

| | | | | | | |
|---|---|---|---|---|---|---|
| XXYLT1 | 553.87531 | -0.1800281 | 0.16484451 | -1.0921088 | 0.2747853 | 0.82719407 |
| ACAP2 | 1821.09263 | -0.0923402 | 0.15929426 | -0.5796831 | 0.56212833 | 0.92452095 |
| PPP1R2 | 2631.49032 | 0.03677239 | 0.14355294 | 0.25615905 | 0.79782802 | 0.97154613 |
| APOD | 1870.78321 | -0.4110397 | 0.73228231 | -0.5613132 | 0.57458408 | 0.93024203 |
| MUC20-OT1 | 6759.12435 | -0.1019087 | 0.39725179 | -0.2565343 | 0.79753827 | 0.97148232 |
| MUC20 | 4590.9845 | 0.16991634 | 0.42198965 | 0.40265523 | 0.68720187 | 0.95138142 |
| MUC4 | 456.248932 | 0.63128017 | 0.41679136 | 1.51461914 | 0.12986888 | 0.70814767 |
| LINC01983 | 36.1895428 | 1.55700205 | 0.81943953 | 1.9000817 | 0.0574224 | 0.60129769 |
| TNK2 | 2958.64812 | -0.1157838 | 0.29073956 | -0.3982389 | 0.6904541 | 0.95158502 |
| TNK2-AS1 | 95.8142822 | -0.0957572 | 0.4243673 | -0.2256471 | 0.82147593 | 0.974755 |
| TFRC | 3965.87623 | -0.2477714 | 0.48989203 | -0.5057674 | 0.61301994 | 0.93879706 |
| SLC51A | 751.182533 | 1.13778906 | 0.46099447 | 2.46811869 | 0.01358253 | 0.42608223 |
| PCYT1A | 3065.69318 | 0.08917867 | 0.17299519 | 0.51549796 | 0.60620509 | 0.93708311 |
| AC069257.3 | 458.897644 | 0.28481506 | 0.17563958 | 1.62158807 | 0.10489158 | 0.67459611 |
| TCTEX1D2 | 389.612847 | 0.41563926 | 0.37348184 | 1.11287675 | 0.26576136 | 0.82611274 |
| TM4SF19-TC | 220.869328 | 0.42513797 | 0.33143968 | 1.28270088 | 0.19959689 | 0.78134034 |
| UBXN7 | 1575.39272 | -0.090167 | 0.14812788 | -0.6087106 | 0.54271631 | 0.92292765 |
| RNF168 | 851.533555 | -0.2038744 | 0.22415912 | -0.9095076 | 0.36308223 | 0.8663515 |
| WDR53 | 323.226487 | -0.249155 | 0.24545452 | -1.0150762 | 0.31006951 | 0.84456632 |
| FBXO45 | 449.714281 | -0.3438629 | 0.27812888 | -1.2363438 | 0.21633079 | 0.7954827 |
| NRROS | 331.780408 | -0.4237924 | 0.34517768 | -1.2277514 | 0.21954032 | 0.79858446 |
| PIGX | 1069.89933 | 0.3093989 | 0.14040046 | 2.20368864 | 0.02754625 | 0.52904542 |
| CEP19 | 202.748731 | 0.0549267 | 0.34399034 | 0.15967512 | 0.873137 | 0.98275231 |
| PAK2 | 3830.64184 | -0.2546903 | 0.24595285 | -1.0355249 | 0.30042386 | 0.84200819 |
| SENP5 | 1638.07255 | -0.2491312 | 0.13961129 | -1.7844635 | 0.07434838 | 0.63773415 |
| NCBP2 | 3811.03135 | -0.1507272 | 0.15697573 | -0.9601944 | 0.33695737 | 0.85679125 |
| NCBP2-AS1 | 255.408325 | -0.078216 | 0.24329329 | -0.3214887 | 0.74784007 | 0.96238898 |
| NCBP2-AS2 | 734.849245 | -0.3318858 | 0.19792919 | -1.6767907 | 0.09358341 | 0.65756964 |
| PIGZ | 260.499936 | 0.30040743 | 0.46845579 | 0.64127168 | 0.52134619 | 0.91723878 |
| MELTF | 910.473848 | 1.09703776 | 0.74388946 | 1.47473223 | 0.14028455 | 0.725307 |
| MELTF-AS1 | 264.507638 | 0.18572328 | 0.59945603 | 0.30981969 | 0.75669808 | 0.9648441 |
| DLG1 | 2751.61906 | 0.17672339 | 0.12916634 | 1.36818455 | 0.17125432 | 0.75956362 |
| LINC02012 | 69.1898777 | 1.1625791 | 0.45327199 | 2.56485977 | 0.01032175 | 0.40019066 |
| BDH1 | 2158.88724 | 0.71920893 | 0.28215477 | 2.54898731 | 0.01080362 | 0.40196898 |
| AC024560.1 | 70.3870108 | -0.0750243 | 1.63602478 | -0.0458577 | 0.96342367 | 0.99320711 |
| RUBCN | 1891.91679 | -0.4192094 | 0.17021866 | -2.4627701 | 0.01378683 | 0.42920097 |
| FYTTD1 | 2944.19799 | -0.1594176 | 0.16288678 | -0.9787017 | 0.32772738 | 0.85317073 |
| LRCH3 | 1783.41618 | -0.2598289 | 0.15089538 | -1.7219143 | 0.08508505 | 0.64988556 |
| IQCG | 491.351452 | 0.15545751 | 0.23580319 | 0.65926806 | 0.50972365 | 0.91439937 |
| RPL35A | 11314.3417 | 0.12953788 | 0.18519043 | 0.69948475 | 0.48424914 | 0.90541935 |
| LMLN | 621.845155 | 0.10380991 | 0.24071989 | 0.43124776 | 0.66628823 | 0.94784483 |
| ZNF595 | 416.803213 | -0.2298196 | 0.22809351 | -1.0075675 | 0.31366212 | 0.8464135 |
| ZNF718 | 297.27169 | -0.1227129 | 0.22952612 | -0.534636 | 0.5929016 | 0.93289552 |
| ZNF141 | 497.740178 | 0.1145855 | 0.20721969 | 0.55296627 | 0.58028651 | 0.93141054 |
| ZNF721 | 1248.74989 | -0.0812279 | 0.1918471 | -0.423399 | 0.67200417 | 0.94922827 |
| PIGG | 2906.41412 | -0.2521219 | 0.14038733 | -1.7959023 | 0.07251005 | 0.63293474 |
| PDE6B | 508.728651 | 0.28740273 | 0.31760935 | 0.90489379 | 0.36552165 | 0.8663515 |
| AC107464.3 | 84.3506332 | 0.47236933 | 0.57725316 | 0.81830532 | 0.41318287 | 0.88316314 |
| ATP5ME | 4115.5682 | 0.16146446 | 0.15840136 | 1.0193376 | 0.30804272 | 0.8437043 |
| MYL5 | 5021.50922 | 0.16593333 | 0.11413058 | 1.45389021 | 0.14597675 | 0.73247558 |
| SLC49A3 | 1064.09805 | -0.1004712 | 0.29748594 | -0.3377344 | 0.73556338 | 0.95895327 |
| PCGF3 | 3496.3924 | -0.3966394 | 0.25087868 | -1.5810008 | 0.11387785 | 0.68214043 |
| AC139887.2 | 489.626031 | -0.271242 | 0.26679409 | -1.0166718 | 0.30930959 | 0.84414094 |
| AC139887.1 | 168.534497 | -0.5534218 | 0.30062953 | -1.8408764 | 0.06563967 | 0.62237346 |

| | | | | | | |
|---|---|---|---|---|---|---|
| CPLX1 | 416.714885 | -0.2864363 | 0.47865633 | -0.5984174 | 0.54956149 | 0.92373235 |
| GAK | 9665.76172 | -0.30463 | 0.18063048 | -1.6864818 | 0.09170303 | 0.65518742 |
| TMEM175 | 1762.47513 | 0.07563493 | 0.29725041 | 0.25444854 | 0.79914906 | 0.97189179 |
| DGKQ | 2173.47981 | -0.2544467 | 0.1784678 | -1.4257288 | 0.15394663 | 0.74258845 |
| SLC26A1 | 1842.42302 | 0.17928098 | 0.48267573 | 0.37143153 | 0.71031615 | 0.95530618 |
| IDUA | 2153.60268 | -0.1441407 | 0.1561066 | -0.9233477 | 0.35582603 | 0.86364633 |
| FGFRL1 | 3518.20712 | 0.19935418 | 0.18031025 | 1.10561753 | 0.2688921 | 0.82683094 |
| RNF212 | 374.154874 | -0.2670319 | 0.35475521 | -0.7527217 | 0.45161719 | 0.89639459 |
| SPON2 | 4094.24813 | -0.6074055 | 0.5206963 | -1.1665254 | 0.24340207 | 0.81369928 |
| AC092535.4 | 364.542812 | -0.9086331 | 0.59022693 | -1.5394639 | 0.12369108 | 0.69957382 |
| CTBP1-AS | 3430.80348 | -0.2196141 | 0.07784713 | -2.8210941 | 0.00478602 | 0.29998235 |
| CTBP1 | 7695.30866 | -0.233207 | 0.09064031 | -2.5728835 | 0.01008551 | 0.39989496 |
| CTBP1-DT | 715.974852 | -0.104096 | 0.18890303 | -0.5510554 | 0.5815957 | 0.9317705 |
| MAEA | 5715.66439 | -0.2168478 | 0.10310654 | -2.1031429 | 0.03545328 | 0.55369025 |
| UVSSA | 2026.69563 | -0.0518641 | 0.19237468 | -0.2695997 | 0.78746825 | 0.97090793 |
| FAM53A | 99.3080904 | -0.1427417 | 0.39725746 | -0.3593179 | 0.7193573 | 0.9566409 |
| SLBP | 2016.95505 | 0.01064417 | 0.17678307 | 0.06021033 | 0.95198812 | 0.99170503 |
| AC016773.1 | 74.8222764 | 0.07728587 | 0.49448824 | 0.15629467 | 0.87580076 | 0.98275231 |
| TMEM129 | 5442.66242 | -0.0079548 | 0.1967659 | -0.0404275 | 0.9677523 | 0.99400331 |
| TACC3 | 544.432169 | -0.1800142 | 0.36096828 | -0.498698 | 0.61799218 | 0.93928244 |
| FGFR3 | 6021.69705 | 0.26247725 | 0.6142249 | 0.42733086 | 0.66913836 | 0.9486589 |
| LETM1 | 3355.80642 | -0.0765728 | 0.24120981 | -0.317453 | 0.75089991 | 0.96272571 |
| NSD2 | 1966.7165 | -0.1974231 | 0.12937305 | -1.5259983 | 0.12701028 | 0.70463514 |
| NELFA | 2447.03102 | -0.1228584 | 0.18217249 | -0.6744068 | 0.50005273 | 0.91180773 |
| C4orf48 | 140.468381 | -0.5448374 | 0.57430614 | -0.9486881 | 0.34277926 | 0.85917162 |
| NAT8L | 1551.818 | 0.46396313 | 0.60164738 | 0.77115458 | NA | NA |
| POLN | 299.405072 | -0.2798251 | 0.1639934 | -1.7063195 | 0.08794861 | 0.65363785 |
| HAUS3 | 664.421304 | -0.0262188 | 0.19830199 | -0.1322164 | 0.89481309 | 0.98474725 |
| MXD4 | 6477.20782 | -0.1850259 | 0.22577503 | -0.8195145 | 0.41249291 | 0.88268208 |
| ZFYVE28 | 1525.24416 | -0.1370054 | 0.29003424 | -0.4723766 | 0.636658 | 0.94088547 |
| CFAP99 | 104.170789 | -0.0403261 | 0.50274516 | -0.0802118 | 0.9360688 | 0.99068685 |
| RNF4 | 3882.33184 | -0.2337404 | 0.20126479 | -1.1613577 | 0.24549645 | 0.81542272 |
| FAM193A | 1539.34543 | -0.3461177 | 0.11971822 | -2.8911031 | 0.00383892 | 0.26845232 |
| TNIP2 | 3500.81385 | -0.1039603 | 0.1878842 | -0.5533211 | 0.58004357 | 0.93139743 |
| SH3BP2 | 9786.45669 | 0.15109958 | 0.23232274 | 0.65038655 | 0.51544257 | 0.91636259 |
| ADD1 | 12344.8502 | -0.1902185 | 0.10965509 | -1.7346985 | 0.08279422 | 0.64680474 |
| MFSD10 | 4982.55236 | -0.2178745 | 0.17268609 | -1.2616798 | 0.20706405 | 0.78790396 |
| NOP14-AS1 | 1488.1681 | -0.1303016 | 0.18582932 | -0.7011894 | 0.4831848 | 0.90541935 |
| NOP14 | 1947.29809 | -0.1492421 | 0.21723385 | -0.6870113 | 0.4920756 | 0.90624891 |
| GRK4 | 239.102876 | -0.2561581 | 0.27302365 | -0.9382269 | 0.34812782 | 0.86254144 |
| HTT | 3468.86301 | 0.10100715 | 0.11092994 | 0.91054901 | 0.36253305 | 0.86572781 |
| MSANTD1 | 61.2545679 | -0.0822759 | 0.25653074 | -0.3207253 | 0.74841855 | 0.96241535 |
| RGS12 | 2795.2926 | -0.1552306 | 0.19143418 | -0.8108824 | 0.41743323 | 0.88445833 |
| HGFAC | 104.539524 | 0.57015316 | 0.5105852 | 1.11666605 | 0.26413714 | 0.82424998 |
| DOK7 | 231.311492 | 0.24247216 | 0.42304515 | 0.57315906 | 0.566537 | 0.9256133 |
| LRPAP1 | 10510.2011 | 0.33362386 | 0.19059194 | 1.75046151 | 0.08003871 | 0.64079926 |
| AL121796.1 | 37.1904029 | 2.0864372 | 0.98373496 | 2.12093427 | 0.03392733 | 0.55369025 |
| AC141928.1 | 352.479777 | 0.33594016 | 0.40852283 | 0.82232899 | 0.41088968 | 0.88268208 |
| ADRA2C | 1167.43446 | 0.36710024 | 0.34714192 | 1.05749326 | 0.29028653 | 0.83439568 |
| TMEM128 | 1226.08076 | 0.40743893 | 0.1838177 | 2.21653804 | 0.02665468 | 0.52678638 |
| LYAR | 334.485408 | -0.4689062 | 0.25017802 | -1.8742901 | 0.06089045 | 0.60989294 |
| ZBTB49 | 280.393823 | -0.1954926 | 0.21081579 | -0.927315 | 0.35376299 | 0.86364633 |
| NSG1 | 179.495408 | 1.05836133 | 0.62659935 | 1.68905591 | 0.09120872 | 0.65495306 |
| STX18 | 2845.8456 | 0.04692724 | 0.1338936 | 0.35048156 | 0.72597733 | 0.95848028 |

| | | | | | | |
|---|---|---|---|---|---|---|
| STX18-AS1 | 100.66197 | 0.18692794 | 0.40659501 | 0.45973987 | 0.64570295 | 0.94208859 |
| MSX1 | 128.011876 | 0.28106618 | 0.21372686 | 1.31507186 | 0.18848575 | 0.76970626 |
| STK32B | 1257.50856 | 0.36632848 | 0.44180913 | 0.82915553 | 0.40701641 | 0.88132015 |
| EVC2 | 573.382763 | -0.7400572 | 0.25857817 | -2.8620249 | 0.00420944 | 0.28174673 |
| EVC | 3386.97465 | -0.2833744 | 0.2517349 | -1.125686 | 0.26029847 | 0.82424998 |
| CRMP1 | 2895.24983 | -0.3125983 | 0.21199322 | -1.4745675 | 0.14032886 | 0.725307 |
| WFS1 | 2189.92414 | 0.10908187 | 0.1507961 | 0.72337328 | 4.69E-01 | 0.90050905 |
| MAN2B2 | 3547.07056 | 0.63100548 | 0.28026793 | 2.25143663 | 2.44E-02 | 5.20E-01 |
| MRFAP1 | 16291.6364 | -0.1625335 | 0.18524522 | -0.8773965 | 3.80E-01 | 0.87101747 |
| LINC02482 | 111.945113 | 0.22610403 | 0.37783687 | 0.59841706 | 0.54956169 | 0.92373235 |
| AC093323.1 | 757.103339 | -0.0093943 | 0.25685913 | -0.0365737 | 0.9708249 | 0.99400331 |
| MRFAP1L1 | 3428.31073 | -0.1865684 | 0.13426498 | -1.3895535 | 0.16466452 | 0.75286112 |
| BLOC1S4 | 688.071098 | -0.4167433 | 0.25665734 | -1.6237342 | 0.10443254 | 0.67439401 |
| KIAA0232 | 2378.87792 | -0.0209264 | 0.18825223 | -0.1111616 | 0.91148819 | 0.98662757 |
| TBC1D14 | 5541.24041 | 0.24494763 | 0.20164209 | 1.21476438 | 2.24E-01 | 0.80078694 |
| AC097382.2 | 4694.95228 | 0.25206773 | 0.18536291 | 1.3598607 | 0.17387401 | 0.75974924 |
| CCDC96 | 168.561525 | -0.152355 | 0.3562587 | -0.4276527 | 0.66890397 | 0.9486589 |
| TADA2B | 2255.21746 | -0.2287122 | 0.20087477 | -1.1385809 | 2.55E-01 | 0.81992039 |
| GRPEL1 | 3705.49045 | -0.4058745 | 0.31938483 | -1.2708009 | 2.04E-01 | 0.7837895 |
| LINC02447 | 70.6455636 | 0.85650246 | 0.47486102 | 1.80369081 | 7.13E-02 | 6.31E-01 |
| SORCS2 | 1266.94799 | 0.16050639 | 0.50915412 | 0.31524126 | 7.53E-01 | 9.63E-01 |
| AFAP1 | 1053.16024 | -0.1249615 | 0.18676999 | -0.6690664 | 0.50345313 | 0.91356552 |
| ABLIM2 | 1012.31587 | 0.04017326 | 0.42116985 | 0.09538495 | 0.92400907 | 0.98869145 |
| SH3TC1 | 5652.32446 | 0.01185545 | 0.21236554 | 0.05582566 | 0.95548069 | 0.99185043 |
| HTRA3 | 364.471075 | -0.5160552 | 0.5231849 | -0.9863725 | 0.32395036 | 0.85161364 |
| AC104825.1 | 522.755347 | 0.05823986 | 0.1816867 | 0.32055101 | 0.74855067 | 0.96247232 |
| ACOX3 | 907.135826 | 0.11318499 | 0.23497003 | 0.48169968 | 0.63001931 | 0.94088547 |
| TRMT44 | 459.266259 | -0.2419832 | 0.19456171 | -1.2437348 | 0.21359717 | 0.7926732 |
| AC105345.1 | 149.050311 | -0.0292078 | 0.3834447 | -0.0761722 | 0.9392821 | 0.99113899 |
| GPR78 | 230.63258 | 0.61914173 | 0.53545191 | 1.15629754 | 0.24755947 | 0.81602355 |
| CPZ | 231.1531 | 0.6077641 | 0.54047959 | 1.12449037 | 0.26080505 | 0.82424998 |
| SLC2A9 | 3185.87929 | 0.72836391 | 0.39909588 | 1.8250349 | 0.06799576 | 0.62603855 |
| WDR1 | 18574.9166 | 0.07384193 | 0.17774103 | 0.41544675 | 0.67781489 | 0.95018034 |
| ZNF518B | 622.87196 | -0.3995111 | 0.2342067 | -1.7058055 | 0.0880443 | 0.65363785 |
| CLNK | 421.974506 | 0.58206667 | 0.64420039 | 0.90354907 | 0.36623456 | 0.86683872 |
| HS3ST1 | 2010.62193 | 0.57282341 | 0.53537973 | 1.06993855 | 0.28464697 | 0.83107779 |
| AC005699.1 | 94.9674425 | 0.34647875 | 0.70912929 | 0.48859743 | 0.62512673 | 0.94088547 |
| RAB28 | 1002.39936 | -0.0523523 | 0.14429537 | -0.3628132 | 0.71674442 | 0.95603249 |
| BOD1L1 | 1847.68829 | -0.4691677 | 0.25059266 | -1.8722323 | 0.06117448 | 0.60989294 |
| CPEB2 | 2237.35833 | -0.4000067 | 0.25906967 | -1.5440119 | 0.12258544 | 0.69788192 |
| AC098829.1 | 87.2093306 | -0.5900244 | 0.33623222 | -1.754812 | 0.07929146 | 0.64079926 |
| C1QTNF7 | 171.052763 | 1.11393345 | 0.5711743 | 1.95025135 | 0.05114617 | 0.5861488 |
| CC2D2A | 1215.64968 | 0.12087591 | 0.31175356 | 0.38772904 | 0.69821656 | 0.9529028 |
| FBXL5 | 7807.47776 | 0.16345893 | 0.18606959 | 0.87848279 | 0.37968177 | 0.8708011 |
| FAM200B | 1540.01964 | 0.22573758 | 0.15437657 | 1.4622528 | 0.14367195 | 0.72909721 |
| BST1 | 421.455592 | 0.56226546 | 0.28385316 | 1.98083216 | 0.0476101 | 0.58241466 |
| CD38 | 330.424912 | -1.5102538 | 0.62283748 | -2.424796 | 0.01531699 | 0.45205627 |
| FGFBP2 | 706.221388 | 0.42104187 | 0.46478685 | 0.90588163 | 0.36499851 | 0.8663515 |
| PROM1 | 5288.77559 | 0.41933398 | 0.4581813 | 0.91521409 | 0.36007925 | 0.86480228 |
| TAPT1 | 2050.94452 | -0.0468461 | 0.14893011 | -0.314551 | 0.75310262 | 0.96301881 |
| TAPT1-AS1 | 132.138456 | -0.0374626 | 0.51005454 | -0.0734482 | 0.94144944 | 0.99136103 |
| AC097515.1 | 276.9279 | 0.02627074 | 0.44189825 | 0.05944975 | 0.95259389 | 0.99170503 |
| LDB2 | 2030.29879 | -0.0719278 | 0.44857349 | -0.1603478 | 0.87260709 | 0.98275231 |
| QDPR | 4088.5539 | 0.60461361 | 0.34157154 | 1.77009365 | 0.07671154 | 0.63773415 |

| | | | | | | |
|---|---|---|---|---|---|---|
| LAP3 | 9080.19406 | -0.298163 | 0.54335826 | -0.5487411 | 0.58318311 | 0.93214795 |
| MED28 | 2984.04896 | -0.2716281 | 0.12928597 | -2.1009864 | 0.03564216 | 0.55369025 |
| FAM184B | 902.556696 | -0.0972841 | 0.14523293 | -0.6698486 | 0.50295429 | 0.91353729 |
| DCAF16 | 1349.36046 | -0.2075204 | 0.22218733 | -0.9339883 | 0.35030995 | 0.86364633 |
| NCAPG | 201.947828 | -0.0837498 | 0.21601255 | -0.3877081 | 6.98E-01 | 9.53E-01 |
| LCORL | 602.499604 | 0.0355783 | 0.15538343 | 0.228971 | 0.81889146 | 0.97454841 |
| SLIT2 | 1205.43467 | 0.24097009 | 0.51456288 | 0.46830058 | 0.63956965 | 0.94088547 |
| PACRGL | 574.016102 | -0.0259791 | 0.17435536 | -0.1490007 | 0.88155307 | 0.98275231 |
| KCNIP4 | 851.040466 | 0.16525565 | 0.44213784 | 0.37376501 | 0.70857916 | 0.95501548 |
| ADGRA3 | 1565.20377 | -0.1479028 | 0.17363764 | -0.8517898 | 0.39433075 | 0.87884163 |
| ERVH-1 | 81.702128 | -1.4977387 | 0.93094756 | -1.6088325 | 1.08E-01 | 0.67707349 |
| PPARGC1A | 3335.79232 | -0.7361226 | 0.22407374 | -3.2851803 | 0.00101917 | 0.15012259 |
| DHX15 | 5789.80079 | -0.343239 | 0.19328184 | -1.7758469 | 7.58E-02 | 6.38E-01 |
| SOD3 | 6554.6589 | 0.95582158 | 0.57905133 | 1.65066815 | 0.09880636 | 0.6678501 |
| CCDC149 | 831.25336 | -0.467247 | 0.20751185 | -2.251664 | 0.02434351 | 0.51967919 |
| LGI2 | 465.561169 | 0.26875229 | 0.61620415 | 0.43614165 | 6.63E-01 | 9.46E-01 |
| SEPSECS | 1511.82952 | 0.45970323 | 0.30635209 | 1.50057155 | 0.13346642 | 0.71505705 |
| SEPSECS-AS1 | 92.6756156 | -0.0138791 | 0.39327521 | -0.0352911 | 0.97184766 | 0.99404213 |
| AC104662.2 | 340.704293 | -0.058954 | 0.41445044 | -0.1422462 | 0.88688556 | 0.98423722 |
| PI4K2B | 1245.38311 | -0.2511347 | 0.31215481 | -0.8045197 | 0.42109689 | 0.88451372 |
| ZCCHC4 | 411.242362 | -0.0081767 | 0.21332861 | -0.0383293 | 0.96942509 | 0.99400331 |
| ANAPC4 | 2363.08619 | -0.644402 | 0.20126477 | -3.2017624 | 0.0013659 | 0.16515003 |
| SLC34A2 | 18718.1502 | -0.4471966 | 0.54540198 | -0.8199394 | 0.41225063 | 0.88268208 |
| SEL1L3 | 3410.58937 | -0.1193923 | 0.26806004 | -0.4453938 | 0.65603519 | 0.94537209 |
| SMIM20 | 1407.58097 | 0.25032332 | 0.2159925 | 1.15894451 | 0.24647881 | 0.81542272 |
| RBPJ | 2956.91718 | -0.1319848 | 0.09807818 | -1.3457097 | 0.17839616 | 0.76115956 |
| TBC1D19 | 593.099481 | 0.1701483 | 0.32589352 | 0.52209785 | 0.60160221 | 0.93549842 |
| STIM2 | 1766.32263 | 0.27865086 | 0.29619767 | 0.94075977 | 0.34682798 | 0.86137069 |
| PCDH7 | 205.530682 | 0.21867338 | 0.45208213 | 0.48370276 | 0.62859683 | 0.94088547 |
| ARAP2 | 812.490676 | -0.1318429 | 0.25618328 | -0.5146429 | 0.60680258 | 0.93720843 |
| C4orf19 | 1380.75878 | -0.6127109 | 0.32966862 | -1.8585662 | 0.06308865 | 0.61484368 |
| AC027607.1 | 105.048049 | -0.5721765 | 0.36237542 | -1.5789606 | 0.11434509 | 0.68355388 |
| RELL1 | 659.128141 | -0.5496267 | 0.29876883 | -1.8396386 | 0.06582131 | 0.62237346 |
| PGM2 | 1441.05839 | -0.0755272 | 0.18630255 | -0.4054006 | 0.68518304 | 0.95097288 |
| TBC1D1 | 8909.87665 | -0.066917 | 0.22335583 | -0.2995984 | 0.76448352 | 0.96678847 |
| KLF3 | 2899.63908 | -0.1611812 | 0.20173069 | -0.7989918 | 0.42429517 | 0.88565404 |
| TLR1 | 292.0695 | -0.4003493 | 0.41391807 | -0.9672188 | 0.33343468 | 0.855011 |
| TLR6 | 71.5989966 | -0.4658691 | 0.4756153 | -0.9795083 | 0.32732888 | 0.85317073 |
| FAM114A1 | 1444.90115 | -0.240575 | 0.2358784 | -1.0199111 | 0.30777062 | 0.8434443 |
| TMEM156 | 43.9610959 | -0.1588379 | 0.6657285 | -0.2385926 | 0.8114215 | 0.97352497 |
| KLHL5 | 1444.88456 | 0.1104078 | 0.17695386 | 0.62393556 | 0.53266991 | 0.92011707 |
| AC079921.1 | 288.262754 | 0.10665546 | 0.21802976 | 0.48917846 | 0.62471536 | 0.94088547 |
| WDR19 | 2022.0375 | -0.1661421 | 0.20732175 | -0.8013734 | 0.42291554 | 0.88453836 |
| RFC1 | 1966.51313 | 0.07813727 | 0.14165657 | 0.55159648 | 0.58122485 | 0.9317705 |
| KLB | 39.2172604 | 0.94005721 | 0.73805998 | 1.27368674 | 0.20277446 | 0.78193335 |
| RPL9 | 22781.9744 | 0.32165181 | 0.17400875 | 1.84848071 | 0.06453283 | 0.61989227 |
| LIAS | 4233.13441 | 0.22967163 | 0.17424256 | 1.31811443 | 0.18746535 | 0.76958556 |
| UGDH | 3816.47329 | -0.3152511 | 0.46645437 | -0.6758456 | 0.49913871 | 0.91147492 |
| UGDH-AS1 | 188.468956 | 0.60707266 | 0.30841028 | 1.96839306 | 0.04902283 | 0.58508842 |
| SMIM14 | 3073.63176 | 0.08855524 | 0.19738922 | 0.44863259 | 0.65369672 | 0.94505036 |
| AC108471.2 | 55.7475319 | 0.60762457 | 0.53249811 | 1.14108305 | 0.25383536 | 0.81906483 |
| UBE2K | 4000.21537 | -0.2968998 | 0.19156309 | -1.5498802 | 0.12117028 | 0.69615335 |
| PDS5A | 3742.51452 | -0.1759919 | 0.13298293 | -1.323417 | 0.18569676 | 0.76908034 |
| N4BP2 | 321.4094 | -0.1128404 | 0.298818 | -0.3776227 | 0.70571093 | 0.95501548 |

| | | | | | | |
|---|---|---|---|---|---|---|
| RHOH | 122.98936 | -0.4690819 | 0.84829501 | -0.5529703 | 0.58028375 | 0.93141054 |
| RBM47 | 8468.5238 | 0.014896 | 0.22904043 | 0.06503656 | 0.94814489 | 0.99170503 |
| AC098869.2 | 232.030896 | -0.0766858 | 0.4200205 | -0.1825764 | 0.85513041 | 0.97910552 |
| NSUN7 | 672.907433 | 0.13516955 | 0.28014522 | 0.48249815 | 0.62945211 | 0.94088547 |
| APBB2 | 2270.50581 | -0.1057477 | 0.15319671 | -0.6902738 | 0.490022 | 0.90600166 |
| AC131953.1 | 218.398778 | -0.1563532 | 0.18832945 | -0.8302111 | 0.40641945 | 0.88132015 |
| UCHL1 | 5030.01055 | -0.2437284 | 0.37529464 | -0.649432 | 0.51605916 | 0.91636259 |
| LIMCH1 | 2794.26347 | 0.47023346 | 0.18588176 | 2.52974506 | 0.01141454 | 0.40353404 |
| TMEM33 | 2991.6545 | -0.0304593 | 0.19098705 | -0.1594838 | 0.87328772 | 0.98275231 |
| DCAF4L1 | 118.825263 | -1.0121385 | 0.69156862 | -1.4635402 | 0.14331963 | 0.72849284 |
| SLC30A9 | 5262.43052 | 0.38750447 | 0.16925244 | 2.28950597 | 0.02204997 | 0.51639767 |
| SHISA3 | 1523.39314 | 0.04648018 | 0.38258008 | 0.12149139 | 0.90330184 | 0.98617053 |
| ATP8A1 | 675.623565 | 0.19340233 | 0.27188004 | 0.7113517 | 4.77E-01 | 0.90335153 |
| KCTD8 | 75.6672733 | 0.23620716 | 0.26189518 | 0.90191491 | 0.36710207 | 0.86683872 |
| GUF1 | 1323.61886 | 0.18664899 | 0.16171901 | 1.15415616 | 0.24843615 | 0.8162591 |
| GNPDA2 | 1095.16915 | -0.1155332 | 0.18276045 | -0.6321564 | 0.52728467 | 0.91813082 |
| AC096586.1 | 56.1156467 | -0.2516283 | 0.40090678 | -0.6276479 | 0.5302346 | 0.91962754 |
| GABRA2 | 308.900822 | 0.15806836 | 0.44780702 | 0.35298322 | 0.72410102 | 0.95813674 |
| COMMD8 | 1475.88139 | 0.33568152 | 0.19688122 | 1.70499515 | 0.08819533 | 0.65363785 |
| ATP10D | 1798.44241 | -0.0306405 | 0.26228545 | -0.1168212 | 0.90700177 | 0.98662757 |
| AC107398.2 | 43.9199412 | -0.3235705 | 0.38584331 | -0.8386061 | 0.40169041 | 0.88132015 |
| CORIN | 101.584783 | 0.10093218 | 0.47236041 | 0.2136762 | 0.83079957 | 0.97550416 |
| AC107068.2 | 65.9280528 | -0.0173888 | 0.30502975 | -0.0570067 | 0.95453983 | 0.99170503 |
| AC107068.1 | 166.44116 | 0.06160237 | 0.28584705 | 0.21550815 | 0.82937116 | 0.97550416 |
| NFXL1 | 502.004696 | 0.07286654 | 0.17063539 | 0.42703063 | 0.66935702 | 0.94876933 |
| NIPAL1 | 268.890255 | 0.03475716 | 0.39013401 | 0.0890903 | 0.92901015 | 0.98963127 |
| CNGA1 | 233.16707 | 1.19803305 | 0.54132288 | 2.21315797 | 0.02688676 | 0.52685774 |
| TXK | 76.4618779 | -0.0437036 | 0.29803715 | -0.1466382 | 0.88341764 | 0.98293342 |
| TEC | 259.777537 | -0.1655749 | 0.25529586 | -0.6485608 | 0.51662231 | 0.91636259 |
| SLAIN2 | 2168.1071 | -0.0858262 | 0.17441453 | -0.4920816 | 0.62266166 | 0.94016713 |
| FRYL | 1664.46317 | 0.09007703 | 0.12212476 | 0.73758202 | 0.46076848 | 0.89755243 |
| OCIAD1 | 8008.44787 | -0.0528377 | 0.09293176 | -0.5685644 | 0.56965176 | 0.92719921 |
| OCIAD1-AS1 | 66.6800039 | 0.0995854 | 0.28818733 | 0.34555786 | 0.72967502 | 0.95848028 |
| OCIAD2 | 5540.40363 | -0.1205553 | 0.43005537 | -0.280325 | 0.77922816 | 0.96836854 |
| CWH43 | 401.398604 | 0.69109932 | 0.39042052 | 1.77014086 | 0.07670368 | 0.63773415 |
| DCUN1D4 | 2645.00067 | -0.0914181 | 0.18143988 | -0.5038481 | 0.61436815 | 0.93895494 |
| LRRC66 | 167.831094 | -0.1028697 | 0.49971055 | -0.2058586 | 8.37E-01 | 0.97577999 |
| SGCB | 3580.58577 | 0.20641074 | 0.16857386 | 1.22445281 | 0.22078146 | 0.7995731 |
| SPATA18 | 694.132546 | 0.57392922 | 0.38654065 | 1.4847836 | 0.13760116 | 0.72209298 |
| USP46 | 1110.4745 | 0.08193105 | 0.28153233 | 0.29101825 | 0.77103736 | 0.96836854 |
| USP46-AS1 | 96.8058428 | -0.2251388 | 0.34156053 | -0.6591476 | 0.50980097 | 0.91439937 |
| ERVMER34-1 | 232.141443 | -0.5219201 | 0.45491464 | -1.1472924 | 0.25126078 | 0.81815815 |
| RASL11B | 867.661917 | -0.1130124 | 0.46514127 | -0.2429637 | 0.80803351 | 0.97334138 |
| SCFD2 | 733.95635 | 0.15923495 | 0.21993334 | 0.72401459 | 0.46905678 | 0.9003198 |
| AC023154.1 | 358.763908 | 0.07744499 | 0.87336986 | 0.08867377 | 0.92934118 | 0.98963127 |
| FIP1L1 | 1883.69237 | -0.1959482 | 0.12587134 | -1.5567341 | 0.11953363 | 0.69355411 |
| AC058822.1 | 1004.97927 | -0.3259768 | 0.29245924 | -1.1146058 | 0.26501939 | 0.82455959 |
| LNX1 | 1602.77217 | -0.2269593 | 0.25115526 | -0.9036611 | 0.36617511 | 0.86683872 |
| CHIC2 | 657.410267 | -0.2386703 | 0.22309516 | -1.0698138 | 0.28470311 | 0.83107779 |
| PDGFRA | 3437.39412 | -0.4818636 | 0.4066465 | -1.1849694 | 0.23602956 | 0.80778488 |
| KIT | 1053.88212 | 0.08035831 | 0.53746204 | 0.14951439 | 0.88114775 | 0.98275231 |
| KDR | 2574.42517 | 0.25496311 | 0.40157265 | 0.63491154 | 0.52548611 | 0.91759401 |
| SRD5A3 | 598.102505 | 0.24816621 | 0.26514349 | 0.93596946 | 0.34928891 | 0.86364633 |
| TMEM165 | 5026.24555 | -0.227841 | 0.19770959 | -1.1524025 | 2.49E-01 | 8.16E-01 |

| | | | | | | |
|---|---|---|---|---|---|---|
| CLOCK | 1388.37728 | -0.0148513 | 0.14546243 | -0.1020974 | 0.91867934 | 0.98709788 |
| EXOC1 | 2153.88999 | -0.0357468 | 0.13204123 | -0.2707244 | 0.78660305 | 0.97090793 |
| CEP135 | 541.058441 | -0.2480594 | 0.22706717 | -1.0924493 | 0.27463567 | 0.82719407 |
| KIAA1211 | 63.1806125 | 1.25574631 | 0.63503681 | 1.97743863 | 0.04799207 | 0.58413853 |
| AASDH | 735.296769 | 0.19326756 | 0.19473164 | 0.99248157 | 0.32096266 | 0.85116192 |
| PPAT | 590.56752 | -0.0762254 | 0.1857659 | -0.4103306 | 0.68156346 | 0.95070301 |
| AC068620.2 | 78.8910771 | 0.38933911 | 0.2626661 | 1.48225872 | 0.13827147 | 0.72249298 |
| PAICS | 3915.3738 | 0.11525382 | 0.15858194 | 0.7267777 | 0.46736214 | 0.89983399 |
| SRP72 | 3035.59726 | -0.0023449 | 0.18211826 | -0.0128755 | 0.98972712 | 0.99807483 |
| HOPX | 532.911875 | -0.2762543 | 0.51616514 | -0.5352053 | 0.59250792 | 0.93289552 |
| REST | 1958.26443 | -0.0297495 | 0.13674813 | -0.2175493 | 0.82778031 | 0.97548201 |
| NOA1 | 1191.49403 | 0.12429949 | 0.17843221 | 0.69662025 | 0.48604048 | 0.90545573 |
| POLR2B | 13547.4099 | 0.05385405 | 0.14288337 | 0.37690917 | 0.70624111 | 0.95501548 |
| IGFBP7 | 66871.9668 | 0.09911321 | 0.24791187 | 0.39979213 | 0.68930963 | 0.95158502 |
| IGFBP7-AS1 | 19263.0406 | 0.15579272 | 0.27250417 | 0.57170765 | 0.56752005 | 0.92614913 |
| ADGRL3 | 156.924903 | -0.5702067 | 0.56956523 | -1.0011261 | 0.31676582 | 0.84807262 |
| CENPC | 869.561552 | 0.00253528 | 0.20208541 | 0.01254556 | 0.98999035 | 0.99807483 |
| STAP1 | 348.203374 | 0.06463771 | 0.65800188 | 0.09823333 | 0.92174702 | 0.98801538 |
| UBA6 | 1843.06296 | -0.2594935 | 0.1906397 | -1.3611726 | 0.17345916 | 0.75974924 |
| UBA6-AS1 | 673.074395 | -0.0789486 | 0.22149777 | -0.3564308 | 0.72151798 | 0.95739595 |
| YTHDC1 | 3833.79579 | -0.1518128 | 0.1232206 | -1.2320404 | 0.217934 | 0.79678134 |
| UGT2A3 | 5149.23913 | 0.60477194 | 0.47993209 | 1.26011982 | 0.20762614 | 0.78834607 |
| UGT2B7 | 36966.4419 | 1.36456009 | 0.49357922 | 2.76462223 | 0.00569888 | 0.3236443 |
| AC111000.4 | 241.936282 | -0.895581 | 0.65392118 | -1.369555 | 0.17082587 | 0.75956362 |
| UGT2B11 | 64.0830568 | -1.1471648 | 0.61033956 | -1.8795517 | 0.06016919 | 0.60821514 |
| UGT2B28 | 148.723923 | 1.6419124 | 1.24099387 | 1.32306246 | 0.18581461 | 0.76908034 |
| UGT2A1 | 186.438431 | 0.81870506 | 1.26092676 | 0.64928835 | 0.51615201 | 0.91636259 |
| UGT2A2 | 110.252755 | 0.84719779 | 1.25451806 | 0.67531733 | 0.49947418 | 0.91161831 |
| SULT1B1 | 94.9942945 | -0.5185237 | 0.78965578 | -0.6566453 | 5.11E-01 | 0.91507116 |
| ENAM | 993.786609 | -0.2104316 | 0.52985651 | -0.3971483 | 0.69125812 | 0.95158502 |
| JCHAIN | 1884.92179 | -0.7371502 | 1.2570873 | -0.5863954 | 0.55760986 | 0.92413922 |
| UTP3 | 1977.17835 | -0.1431392 | 0.22472247 | -0.6369599 | 0.52415098 | 0.91723878 |
| AC009570.1 | 122.351414 | -0.6698617 | 0.698162 | -0.9594646 | 0.33732476 | 0.85707745 |
| RUFY3 | 3116.94079 | -0.1465289 | 0.26229949 | -0.5586321 | 0.57641282 | 0.9311568 |
| GRSF1 | 4979.68307 | 0.03084765 | 0.15363135 | 0.20079006 | 8.41E-01 | 9.76E-01 |
| MOB1B | 3044.80768 | 0.16525468 | 0.15827225 | 1.0441166 | 0.29643145 | 0.83817467 |
| DCK | 738.995878 | 0.11176295 | 0.16154294 | 0.69184668 | 0.48903362 | 0.90600166 |
| SLC4A4 | 10641.7068 | 1.33879901 | 0.50329316 | 2.66007788 | 0.00781226 | 0.36878977 |
| GC | 2289.57405 | -1.5285518 | 1.16485697 | -1.3122227 | 0.18944501 | 0.76970626 |
| ADAMTS3 | 186.655071 | 0.27368887 | 0.4533343 | 0.60372417 | 0.54602704 | 0.92345821 |
| COX18 | 1061.54208 | 0.33862769 | 0.15719892 | 2.15413498 | 0.03122959 | 0.54466273 |
| ANKRD17 | 3974.90897 | -0.2131585 | 0.12587575 | -1.6934037 | 0.09037864 | 0.65447686 |
| ALB | 8513.69931 | 0.06175606 | 1.45374084 | 0.04248079 | 0.96611542 | 0.99393925 |
| AFP | 86.0774083 | 0.15124124 | 0.85404113 | 0.17708894 | 0.85943853 | 0.9802651 |
| AFM | 1588.32831 | 0.32446378 | 1.37135396 | 0.23660105 | 0.81296631 | 0.9738447 |
| LINC02499 | 203.648136 | 1.22475099 | 1.16390741 | 1.05227527 | 0.29267327 | 0.83524862 |
| RASSF6 | 1395.35201 | 0.45554357 | 0.37991654 | 1.19906222 | 0.23050375 | 0.80584899 |
| CXCL8 | 1451.26977 | 1.75357153 | 1.14949312 | 1.52551721 | 0.12713014 | 0.70481988 |
| CXCL6 | 316.747757 | 2.77938135 | 0.81684105 | 3.40259752 | 0.00066749 | 0.12385487 |
| CXCL1 | 1550.03934 | 2.01952452 | 0.78030935 | 2.58810755 | 0.00965048 | 0.39368232 |
| CXCL3 | 261.841742 | 1.42667509 | 0.97875758 | 1.45763887 | 0.14494011 | 0.73039059 |
| CXCL2 | 1798.11316 | 1.19350902 | 0.8043813 | 1.48376028 | 0.13787253 | 0.72209298 |
| MTHFD2L | 771.267202 | -0.0431429 | 0.1987329 | -0.2170898 | 0.8281384 | 0.97548201 |
| AC093677.2 | 137.926511 | -0.0884619 | 0.27871309 | -0.3173943 | 0.75094443 | 0.96272571 |

| | | | | | | |
|---|---|---|---|---|---|---|
| AREG | 99.4683732 | 0.19349888 | 0.83403655 | 0.23200287 | 0.81653578 | 0.97388289 |
| BTC | 431.483829 | -0.3952869 | 0.53186431 | -0.7432101 | 0.4573545 | 0.89688594 |
| PARM1 | 5059.71186 | 0.0344507 | 0.31460371 | 0.10950505 | 0.91280192 | 0.98662757 |
| RCHY1 | 1363.74421 | -0.0711124 | 0.1529425 | -0.4649614 | 6.42E-01 | 0.94088547 |
| THAP6 | 725.557584 | 0.03230902 | 0.1732737 | 0.18646233 | 8.52E-01 | 9.79E-01 |
| CDKL2 | 420.542465 | 0.03304918 | 0.33568016 | 0.09845436 | 0.92157151 | 0.98794026 |
| G3BP2 | 5772.90483 | 0.07417174 | 0.26619628 | 0.27863553 | 0.78052455 | 0.96864684 |
| USO1 | 4051.175 | 0.20406664 | 0.15014658 | 1.35911617 | 0.17410978 | 0.75974924 |
| NAAA | 1388.86516 | 0.80034206 | 0.24695753 | 3.24080849 | 0.00119191 | 0.15779959 |
| SDAD1 | 1656.75451 | 0.0018557 | 0.15540696 | 0.01194088 | 0.99047278 | 0.99814875 |
| CXCL9 | 401.087088 | 0.46701243 | 1.2631692 | 0.36971487 | 0.71159495 | 0.95560193 |
| CXCL10 | 2034.87952 | -4.562951 | 1.47856542 | -3.0860663 | NA | NA |
| CXCL11 | 966.161953 | -6.09991 | 1.84733798 | -3.302 | NA | NA |
| NUP54 | 1440.90971 | -0.129662 | 0.12948023 | -1.001404 | 0.31663151 | 0.84783414 |
| SCARB2 | 9721.2128 | -0.1180815 | 0.20425695 | -0.5781026 | 0.56319488 | 0.92452095 |
| FAM47E | 551.198712 | 0.36670202 | 0.30859895 | 1.18828021 | 0.23472303 | 0.80747753 |
| FAM47E-STB | 1708.30217 | 0.04116217 | 0.30726817 | 0.13396172 | 0.89343285 | 0.98474725 |
| STBD1 | 1577.45333 | -0.0030248 | 0.35240148 | -0.0085833 | 0.99315162 | 0.99906069 |
| CCDC158 | 53.0726231 | 1.05372698 | 0.52928049 | 1.99086686 | 0.04649553 | 0.57983846 |
| SHROOM3 | 2536.3417 | -0.1720017 | 0.23285176 | -0.7386748 | 0.46010446 | 0.89755243 |
| SOWAHB | 547.539003 | 0.58425381 | 0.39384974 | 1.48344344 | 0.13795663 | 0.72209298 |
| 11-Sep | 5776.6294 | 0.1085018 | 0.24546877 | 0.44201874 | 0.65847565 | 0.9458503 |
| CCNI | 15298.7087 | 0.02494934 | 0.21448783 | 0.11632056 | 0.90739849 | 0.98662757 |
| CCNG2 | 4786.53551 | 0.22583998 | 0.29930575 | 0.75454608 | 0.45052139 | 0.89557948 |
| CNOT6L | 1974.40796 | 0.11875296 | 0.10758338 | 1.10382259 | 0.2696701 | 0.82716979 |
| MRPL1 | 1046.3139 | 0.18038417 | 0.17700424 | 1.01909523 | 0.30815776 | 0.84370848 |
| FRAS1 | 1926.56324 | 0.99203025 | 0.39619452 | 2.50389694 | 0.01228338 | 0.41344377 |
| ANXA3 | 1463.22043 | 0.26778355 | 0.35836604 | 0.74723472 | 0.45492189 | 0.89670027 |
| LINC01094 | 54.7995952 | 0.13290124 | 0.61824272 | 0.21496612 | 0.82979373 | 0.97550416 |
| BMP2K | 1725.08141 | -0.0782387 | 0.31114568 | -0.2514537 | 0.80146334 | 0.97210205 |
| PAQR3 | 1893.89386 | 0.05850288 | 0.24217061 | 0.24157711 | 0.80910786 | 0.97334138 |
| LINC00989 | 41.0190204 | -0.6605655 | 0.63920653 | -1.0334148 | 0.30140982 | 0.84211658 |
| ANTXR2 | 1382.64084 | 0.0805457 | 0.26594301 | 0.30286829 | 0.76199024 | 0.96607083 |
| PRDM8 | 102.367177 | -0.0199712 | 0.45182066 | -0.0442016 | 0.96474372 | 0.99357855 |
| FGF5 | 100.483228 | -0.7266154 | 0.9545671 | -0.7611989 | 0.4465383 | 0.89480207 |
| PRKG2 | 102.159847 | 0.97209934 | 0.52656416 | 1.84611754 | 0.06487514 | 0.62075686 |
| RASGEF1B | 879.036602 | -0.5893151 | 0.26665963 | -2.2099899 | 0.02710586 | 0.52773677 |
| HNRNPD | 9319.48777 | -0.130883 | 0.11730883 | -1.1157128 | 0.2645451 | 0.82424998 |
| AC124016.1 | 59.5213723 | 0.03626055 | 0.38897845 | 0.09321995 | 0.92572883 | 0.98876911 |
| HNRNPDL | 16776.988 | -0.2533125 | 0.16552824 | -1.5303279 | 0.12593557 | 0.70347499 |
| ENOPH1 | 2144.08189 | 0.13865343 | 0.18502652 | 0.74937057 | 0.45363388 | 0.89670027 |
| TMEM150C | 1328.3656 | 0.10618961 | 0.37845699 | 0.28058568 | 0.7790282 | 0.96836854 |
| SCD5 | 2940.40372 | 0.1357038 | 0.39123473 | 0.34686031 | 0.72869627 | 0.95848028 |
| SEC31A | 7909.9376 | -0.1157014 | 0.21253437 | -0.5443891 | 0.58617372 | 0.93289552 |
| THAP9-AS1 | 2612.74836 | 0.16820426 | 0.27791739 | 0.60523113 | 0.54502543 | 0.92345821 |
| THAP9 | 304.177864 | 0.08052023 | 0.37038437 | 0.21739642 | 0.82789942 | 0.97548201 |
| LIN54 | 658.010035 | -0.0535646 | 0.31315865 | -0.1710463 | 0.86418737 | 0.98176973 |
| COPS4 | 1742.96672 | 0.18982244 | 0.11005475 | 1.72480003 | 0.08456352 | 0.64988556 |
| PLAC8 | 121.049226 | -0.7830722 | 0.64209796 | -1.2195525 | 0.22263458 | 0.80078694 |
| COQ2 | 421.311189 | 0.06163015 | 0.23033355 | 0.26756912 | 0.789031 | 0.97090793 |
| HPSE | 373.841913 | 0.02286223 | 0.36430337 | 0.06275602 | 0.94996079 | 0.99170503 |
| HELQ | 732.355181 | -0.0151658 | 0.11373754 | -0.1333402 | 0.89392433 | 0.98474725 |
| MRPS18C | 1408.87374 | -0.0456125 | 0.17217133 | -0.2649252 | 0.79106712 | 0.97090793 |
| ABRAXAS1 | 1429.26995 | -0.0192558 | 0.17190179 | -0.1120166 | 0.91081028 | 0.98662757 |

| | | | | | | |
|---|---|---|---|---|---|---|
| GPAT3 | 5455.13499 | -0.9610358 | 0.69852265 | -1.375812 | 0.16887985 | 0.75786012 |
| CDS1 | 1907.14686 | -0.0398561 | 0.1464586 | -0.2721324 | 7.86E-01 | 9.71E-01 |
| WDFY3 | 3710.37415 | -0.0274492 | 0.22275578 | -0.1232256 | 0.90192847 | 0.98572048 |
| ARHGAP24 | 5134.84467 | 0.14092554 | 0.31415878 | 0.44858063 | 0.65373421 | 0.94505036 |
| MAPK10 | 3428.49759 | 0.18627128 | 0.33370949 | 0.55818396 | 0.57671877 | 0.93127944 |
| PTPN13 | 2684.17025 | 0.33884703 | 0.21328672 | 1.58869257 | 0.11212982 | 0.68001961 |
| C4orf36 | 96.2335571 | 0.21960717 | 0.29056843 | 0.75578471 | 0.44977829 | 0.89557948 |
| AC093827.5 | 193.955791 | -0.080044 | 0.29842898 | -0.2682179 | 0.78853162 | 0.97090793 |
| AC093827.4 | 286.910189 | -0.2132274 | 0.36260317 | -0.5880461 | 0.55650135 | 0.92373235 |
| AFF1 | 7351.70209 | -0.2391739 | 0.18958812 | -1.2615447 | 0.20711268 | 0.78790396 |
| KLHL8 | 996.925843 | 0.11992275 | 0.28308252 | 0.42363178 | 0.67183437 | 0.94922827 |
| HSD17B11 | 3326.1456 | 0.26438787 | 0.26727172 | 0.98921007 | 0.32256037 | 0.85119105 |
| NUDT9 | 2899.31546 | -0.0511589 | 0.21008083 | -0.2435201 | 0.80760251 | 0.97334138 |
| SPARCL1 | 9966.66302 | -0.4096738 | 0.50534699 | -0.8106783 | 0.41755044 | 0.88445833 |
| SPP1 | 623220.032 | -0.2701339 | 0.40467595 | -0.6675313 | 0.50443285 | 0.91356552 |
| PKD2 | 4081.84401 | 0.05720103 | 0.1709374 | 0.33463144 | 0.73790313 | 0.9602325 |
| ABCG2 | 292.512024 | 0.18336159 | 0.44087898 | 0.41590005 | 0.67748314 | 0.95018034 |
| PPM1K | 2414.08284 | -0.2943852 | 0.41478011 | -0.7097379 | 0.47786669 | 0.90388572 |
| HERC6 | 1428.5055 | -1.9487522 | 0.81266683 | -2.3979719 | NA | NA |
| HERC5 | 1086.79535 | -2.0059128 | 0.6817909 | -2.9421231 | NA | NA |
| PYURF | 3273.90583 | 0.1079466 | 0.12839508 | 0.84073777 | 0.40049486 | 0.88086294 |
| PIGY | 1251.56596 | 0.1542588 | 0.14758116 | 1.04524724 | 0.29590872 | 0.83750889 |
| AC098582.1 | 51.3336621 | 0.47286205 | 0.42880186 | 1.10275185 | 0.27013494 | 0.82716979 |
| HERC3 | 1801.09391 | -0.0239334 | 0.16769336 | -0.1427211 | 0.88651044 | 0.98418886 |
| NAP1L5 | 393.01404 | 0.12548716 | 0.30857734 | 0.40666356 | 0.68425511 | 0.95070301 |
| FAM13A-AS1 | 1708.51008 | -0.3845688 | 0.24476074 | -1.5712031 | 0.11613547 | 0.68754779 |
| FAM13A | 3482.77917 | -0.2536688 | 0.29098153 | -0.8717693 | 0.38333425 | 0.87195864 |
| TIGD2 | 380.517919 | 0.12442749 | 0.25726825 | 0.48364884 | 0.62863511 | 0.94088547 |
| GPRIN3 | 1189.51499 | -0.2365882 | 0.37911595 | -0.6240523 | 0.53259322 | 0.92011707 |
| AC097478.1 | 105.31022 | 0.3497076 | 0.39257503 | 0.8908045 | 0.37303406 | 0.86791605 |
| SNCA | 286.444344 | 0.40651345 | 0.40845595 | 0.9952443 | 0.31961746 | 0.84951792 |
| MMRN1 | 260.933335 | -1.384709 | 0.61535907 | -2.2502455 | 0.02443337 | 0.51967919 |
| CCSER1 | 241.819084 | -0.24541 | 0.29374185 | -0.8354616 | 0.40345785 | 0.88132015 |
| SMARCAD1 | 1387.91948 | -0.1793965 | 0.14513622 | -1.2360558 | 0.21643781 | 0.7954827 |
| HPGDS | 56.2093586 | 0.71934174 | 0.73851971 | 0.97403188 | 0.33004071 | 0.85377783 |
| PDLIM5 | 4515.55868 | -0.0277566 | 0.17525038 | -0.1583825 | 0.87415539 | 0.98275231 |
| BMPR1B | 572.180531 | 0.14040877 | 0.29220468 | 0.48051512 | 0.63086116 | 0.94088547 |
| UNC5C | 168.68641 | 0.13969573 | 0.27383242 | 0.51015046 | 0.60994606 | 0.93791514 |
| STPG2 | 38.5463004 | -0.5270404 | 0.58831676 | -0.8958447 | 0.37033573 | 0.86762749 |
| AC019077.1 | 41.7714996 | 0.27098793 | 0.39318114 | 0.68921904 | 0.49068544 | 0.90600166 |
| RAP1GDS1 | 1619.90158 | 0.06631382 | 0.16509582 | 0.4016687 | 0.68792786 | 0.95158502 |
| TSPAN5 | 565.674777 | -0.467815 | 0.28871486 | -1.6203357 | 0.10516019 | 0.67473222 |
| EIF4E | 2210.87662 | -0.604913 | 0.37074569 | -1.6316118 | 0.1027613 | 0.67335541 |
| METAP1 | 2295.66005 | 0.11775013 | 0.17626268 | 0.66803776 | 0.50410949 | 0.91356552 |
| AC019131.2 | 64.8997529 | 0.50273107 | 0.50005031 | 1.00536098 | 0.31472306 | 0.84696964 |
| ADH5 | 8428.8208 | 0.36606775 | 0.21945023 | 1.66811285 | 0.09529333 | 0.65888357 |
| AP002026.1 | 95.7591204 | -0.2719543 | 0.61849516 | -0.4397032 | 0.66015205 | 0.94628868 |
| ADH4 | 51.3080093 | -0.5952402 | 0.64841376 | -0.9179944 | 0.35862181 | 0.86480228 |
| ADH6 | 1966.77753 | 2.183536 | 0.52907368 | 4.12709249 | 3.67E-05 | 0.02220982 |
| ADH1B | 5686.3141 | 0.27650011 | 0.40390294 | 0.68457069 | 0.49361487 | 0.90749607 |
| ADH1C | 725.17418 | 2.15352154 | 0.86136347 | 2.50013104 | 0.01241474 | 0.4154725 |
| TRMT10A | 336.103796 | 0.30519366 | 0.2859694 | 1.06722486 | 0.28587031 | 0.83107779 |
| MTTP | 666.988575 | 1.86828162 | 0.72371759 | 2.58150643 | 0.00983702 | 0.39561269 |
| C4orf54 | 139.32211 | 0.28645507 | 0.70158593 | 0.40829648 | 0.68305602 | 0.95070301 |

| | | | | | | |
|---|---|---|---|---|---|---|
| DAPP1 | 152.441796 | 0.4471779 | 0.73711728 | 0.60665774 | 0.54407808 | 0.92343481 |
| LAMTOR3 | 2788.52595 | -0.1348008 | 0.18497462 | -0.7287527 | 0.46615295 | 0.89960479 |
| DNAJB14 | 1848.98895 | -0.1069141 | 0.1052961 | -1.015366 | 0.30993139 | 0.84455826 |
| H2AFZ | 5389.83673 | -0.6135661 | 0.34850079 | -1.7605875 | 0.07830825 | 0.64079926 |
| AC097460.1 | 1527.75165 | -0.562415 | 0.36984453 | -1.5206795 | 0.12834028 | 0.70596572 |
| DDIT4L | 1915.87512 | -1.4532172 | 0.67362641 | -2.1573044 | 0.03098195 | 0.54466273 |
| EMCN | 3433.0157 | 0.85442842 | 0.54274616 | 1.57426894 | 0.11542529 | 0.6873836 |
| PPP3CA | 1360.71018 | -0.0313509 | 0.22412124 | -0.1398836 | 0.88875193 | 0.98423722 |
| AP001816.1 | 190.883735 | -0.0963223 | 0.18024142 | -0.5344073 | 0.59305979 | 0.93289552 |
| BANK1 | 252.10329 | 0.0074821 | 0.40603319 | 0.01842731 | 0.98529796 | 0.99692004 |
| SLC39A8 | 1803.74443 | -0.2695596 | 0.35577633 | -0.7576659 | 0.448651 | 0.8948759 |
| NFKB1 | 2490.38803 | 0.08708093 | 0.25270279 | 0.34459824 | 0.73039643 | 0.95848028 |
| MANBA | 1599.63506 | 0.34912199 | 0.16044052 | 2.17602137 | 0.02955366 | 0.53721163 |
| UBE2D3 | 16108.6342 | -0.3791115 | 0.23906926 | -1.5857812 | 0.11278896 | 0.68098515 |
| AC018797.2 | 73.2514853 | -0.2013073 | 0.43078169 | -0.467307 | 0.64028026 | 0.94088547 |
| CISD2 | 2594.16736 | -0.0303188 | 0.29440484 | -0.1029833 | 0.91797624 | 0.9868044 |
| SLC9B1 | 860.480902 | 0.02301048 | 0.31273254 | 0.0735788 | 0.94134554 | 0.99136103 |
| SLC9B2 | 5865.77045 | 0.5025222 | 0.22198265 | 2.26379043 | 0.02358701 | 0.51967919 |
| BDH2 | 9909.83527 | 0.57666496 | 0.24817835 | 2.3235909 | 0.02014743 | 0.49813061 |
| CXXC4 | 67.7192733 | 0.46229776 | 0.58679322 | 0.78783759 | 0.43079171 | 0.88966606 |
| AC096577.1 | 108.43803 | 1.33174252 | 0.91154665 | 1.46097023 | 1.44E-01 | 7.29E-01 |
| TET2 | 999.284381 | -0.0815581 | 0.20441371 | -0.3989855 | 0.68990386 | 0.95158502 |
| PPA2 | 4143.94873 | 0.3749952 | 0.19157366 | 1.95744649 | 5.03E-02 | 5.86E-01 |
| ARHGEF38 | 1496.52791 | -0.1936117 | 0.24451751 | -0.7918114 | 0.42847068 | 0.88820133 |
| INTS12 | 1055.38639 | -0.2307016 | 0.14101646 | -1.6359906 | 0.10184154 | 0.67279543 |
| GSTCD | 392.348507 | 0.21760166 | 0.16879641 | 1.28913675 | 0.19735055 | 0.77816059 |
| NPNT | 5902.49432 | 0.38510855 | 0.41733396 | 0.92278267 | 0.35612048 | 0.86384336 |
| AC109361.2 | 127.131868 | -0.1882912 | 0.58517274 | -0.3217703 | 0.74762672 | 0.96231266 |
| TBCK | 1855.11108 | 0.06884982 | 0.24149172 | 0.28510221 | 0.77556584 | 0.96836854 |
| AIMP1 | 2537.38163 | 0.04231762 | 0.15175483 | 0.2788552 | 0.78035595 | 0.96864684 |
| PAPSS1 | 2171.24883 | 0.24637169 | 0.14154565 | 1.74058118 | 0.08175702 | 0.64541209 |
| SGMS2 | 1686.129 | -0.400739 | 0.32088436 | -1.248858 | 0.21171701 | 0.79177972 |
| AC096564.1 | 108.949174 | -0.7735571 | 0.43567599 | -1.775533 | 0.07580991 | 0.63773415 |
| CYP2U1 | 491.355016 | 0.25287241 | 0.21629726 | 1.16909668 | 0.24236468 | 0.81319028 |
| HADH | 6095.13683 | 0.89424509 | 0.37515567 | 2.38366403 | 0.01714124 | 0.4678237 |
| LEF1 | 294.56096 | 0.12567804 | 0.39424821 | 0.31877898 | 0.74989412 | 0.96249332 |
| RPL34 | 12594.2953 | 0.22170427 | 0.18156159 | 1.2210968 | 0.22204937 | 0.80067924 |
| OSTC | 2811.12139 | 0.07721025 | 0.19560143 | 0.39473255 | 0.69304027 | 0.9519683 |
| ETNPPL | 181.750877 | 0.71209931 | 0.81457703 | 0.87419517 | 0.38201197 | 0.87185666 |
| COL25A1 | 189.676199 | -0.4605141 | 0.614897 | -0.7489288 | 4.54E-01 | 8.97E-01 |
| SEC24B-AS1 | 75.1033998 | 0.16703278 | 0.46169732 | 0.36177984 | 0.71751656 | 0.95612755 |
| SEC24B | 2039.03513 | -0.2064816 | 0.13413884 | -1.5393128 | 0.12372796 | 0.69957382 |
| MCUB | 429.168507 | -0.6868166 | 0.42305256 | -1.6234782 | 0.10448721 | 0.67439401 |
| CASP6 | 1062.12704 | 0.05140707 | 0.28874791 | 0.17803442 | 0.85869594 | 0.97983564 |
| PLA2G12A | 2520.83817 | 0.11781365 | 0.23787719 | 0.49527089 | 0.62040892 | 0.9398661 |
| AC126283.2 | 16635.3961 | 0.35229494 | 0.23065934 | 1.52733874 | 0.1266768 | 0.70443503 |
| CFI | 18420.7853 | 0.31156593 | 0.23318458 | 1.33613437 | 0.18150536 | 0.76522169 |
| GAR1 | 738.713748 | -0.4088274 | 0.30067211 | -1.3597119 | 0.17392111 | 0.75974924 |
| LRIT3 | 59.1069416 | -0.4251354 | 0.4439279 | -0.9576676 | 0.33823039 | 0.8571036 |
| EGF | 5375.88535 | -0.2132097 | 0.44227659 | -0.4820733 | 0.62975387 | 0.94088547 |
| ELOVL6 | 946.213289 | 0.32348608 | 0.22177203 | 1.45864236 | 0.14466357 | 0.73032824 |
| ENPEP | 11521.7836 | 0.61114674 | 0.42104343 | 1.45150523 | 0.14663923 | 0.73323526 |
| AC098798.1 | 540.867462 | 0.84486885 | 0.55551169 | 1.52088402 | 0.12828894 | 0.70596572 |
| PITX2 | 107.795538 | 0.0008035 | 0.66936163 | 0.00120039 | 0.99904223 | 0.99994357 |

| | | | | | | |
|---|---|---|---|---|---|---|
| FAM241A | 773.109701 | -0.1253406 | 0.32117668 | -0.3902542 | 0.69634858 | 0.9525827 |
| AC109347.1 | 69.9680234 | 0.41610789 | 0.59035482 | 0.70484373 | 4.81E-01 | 0.90541935 |
| AP1AR | 1698.67004 | -0.2145523 | 0.4494041 | -0.4774152 | 0.63306649 | 0.94088547 |
| TIFA | 569.611849 | -0.0061986 | 0.30040991 | -0.0206339 | 0.98353774 | 0.9961464 |
| ALPK1 | 1664.65788 | -0.10934 | 0.3284961 | -0.3328502 | 0.73924735 | 0.9602325 |
| ZGRF1 | 142.334473 | -0.5517417 | 0.40657847 | -1.3570361 | 0.17476974 | 0.76029706 |
| LARP7 | 1839.21319 | -0.2090823 | 0.15489087 | -1.3498688 | 1.77E-01 | 0.76115956 |
| AC106864.1 | 460.575105 | -0.1941225 | 0.20268002 | -0.9577784 | 0.33817452 | 0.8571036 |
| ANK2 | 6413.3897 | 0.3733397 | 0.47649804 | 0.78350732 | 0.43332925 | 0.89055338 |
| CAMK2D | 1689.09268 | -0.0157345 | 0.13223935 | -0.1189847 | 0.90528749 | 0.98625102 |
| ARSJ | 505.830918 | 0.2838585 | 0.21048176 | 1.34861325 | 0.17746122 | 0.76115956 |
| UGT8 | 2509.12848 | -0.1798409 | 0.3068206 | -0.5861434 | 0.55777918 | 0.92421175 |
| TRAM1L1 | 334.797642 | -0.0757286 | 0.54809343 | -0.1381674 | 0.89010812 | 0.98462839 |
| SNHG8 | 1958.4826 | 0.11156757 | 0.23831599 | 0.46814975 | 0.6396775 | 0.94088547 |
| PRSS12 | 107.567073 | -0.0519527 | 0.7227971 | -0.0718773 | 0.94269959 | 0.99136103 |
| METTL14 | 1151.94027 | -0.1156271 | 0.16011081 | -0.7221692 | 0.47019047 | 0.9008571 |
| SEC24D | 1527.43537 | 0.05648992 | 0.28113332 | 0.2009364 | 0.84074831 | 0.97618283 |
| SYNPO2 | 2264.2906 | -0.9913228 | 0.61293549 | -1.6173362 | 0.10580572 | 0.67526487 |
| MYOZ2 | 50.6282851 | 0.10421782 | 0.8062124 | 0.12926844 | 0.89714524 | 0.98474725 |
| USP53 | 2688.20158 | -0.3122496 | 0.28659895 | -1.0895001 | 2.76E-01 | 0.82719407 |
| C4orf3 | 8474.8265 | 0.06051552 | 0.18176308 | 0.33293625 | 0.7391824 | 0.9602325 |
| PDE5A | 991.781412 | 0.08519409 | 0.25895272 | 0.32899479 | 0.74215963 | 0.96109001 |
| MAD2L1 | 145.945051 | 0.76654494 | 0.42388331 | 1.80838669 | 0.07054634 | 0.62981127 |
| PRDM5 | 273.332302 | 0.15495839 | 0.33388364 | 0.46410895 | 0.64256969 | 0.94090101 |
| NDNF | 755.174037 | 0.59789703 | 0.70031993 | 0.85374841 | 0.3932444 | 0.87825756 |
| QRFPR | 105.917838 | 1.18481328 | 0.61289446 | 1.93314404 | 0.05321846 | 0.58946046 |
| ANXA5 | 18006.317 | -0.3324611 | 0.33541937 | -0.9911804 | 0.3215975 | 0.85119105 |
| EXOSC9 | 1001.69739 | -0.0472954 | 0.17678328 | -0.267533 | 0.7890588 | 0.97090793 |
| CCNA2 | 315.596302 | -0.0417719 | 0.3247841 | -0.1286144 | 0.89766273 | 0.98474725 |
| BBS7 | 628.42667 | -0.0774746 | 0.16550862 | -0.4681003 | 0.63971284 | 0.94088547 |
| KIAA1109 | 3251.82492 | 0.15638538 | 0.16697399 | 0.93658527 | 0.34897193 | 0.86356499 |
| BBS12 | 759.658168 | 0.13018096 | 0.23477418 | 0.55449437 | 0.57924056 | 0.93139743 |
| FGF2 | 1033.16674 | -0.4800347 | 0.33547486 | -1.4309111 | 1.52E-01 | 0.7422778 |
| AC021205.3 | 40.8134832 | 0.34405981 | 0.47537597 | 0.72376358 | 0.46921089 | 0.90041792 |
| NUDT6 | 654.251289 | 0.05632711 | 0.24074498 | 0.23397004 | 0.81500824 | 0.9738447 |
| SPATA5 | 360.412511 | 0.07118123 | 0.21809556 | 0.32637636 | 0.74413962 | 0.9613927 |
| SPRY1 | 2616.75677 | -0.1969242 | 0.36619743 | -0.5377541 | 0.59074684 | 0.93289552 |
| LINC01091 | 39.8195645 | 1.27112895 | 0.5773014 | 2.2018463 | 0.02767617 | 0.53038691 |
| ANKRD50 | 702.094134 | -0.2872724 | 0.31568914 | -0.909985 | 3.63E-01 | 0.86610681 |
| FAT4 | 513.174986 | -0.0826735 | 0.34466951 | -0.2398631 | 0.81043641 | 0.97352497 |
| INTU | 805.489429 | -0.0884384 | 0.41511013 | -0.213048 | 0.83128954 | 0.97550416 |
| HSPA4L | 1802.90914 | 0.40173059 | 0.29017112 | 1.38446097 | 0.16621735 | 0.75517075 |
| AC093591.2 | 165.136348 | 0.60527262 | 0.39019547 | 1.55120359 | 0.1208529 | 0.69496908 |
| PLK4 | 67.257293 | 0.31736804 | 0.46268128 | 0.68593232 | 0.49275579 | 0.90696683 |
| MFSD8 | 1302.03343 | -0.1696402 | 0.16447187 | -1.0314235 | 0.30234226 | 0.84211658 |
| ABHD18 | 1143.6906 | -0.0473039 | 0.15841447 | -0.2986085 | 0.76523879 | 0.96678847 |
| LARP1B | 1571.98782 | -0.1161456 | 0.28984123 | -0.4007215 | 0.68862521 | 0.95158502 |
| PGRMC2 | 5601.2243 | -0.2911629 | 0.18665567 | -1.5598929 | 0.11878519 | 0.69221182 |
| AC108062.1 | 242.821755 | 0.04706732 | 0.28154392 | 0.16717577 | 0.86723175 | 0.98223481 |
| JADE1 | 3234.75329 | -0.2670102 | 0.17446313 | -1.5304677 | 0.125901 | 0.70347499 |
| SCLT1 | 733.469104 | -0.2259589 | 0.22897 | -0.9868492 | 0.32371659 | 0.85154042 |
| C4orf33 | 770.078743 | 0.05537703 | 0.4536761 | 0.12206292 | 0.90284919 | 0.98608873 |
| PCDH10 | 260.919892 | 0.26178772 | 0.43068121 | 0.60784569 | 5.43E-01 | 9.23E-01 |
| PABPC4L | 86.6089616 | -0.2690903 | 0.3779621 | -0.7119504 | 0.47649552 | 0.90309179 |

| | | | | | | |
|---|---|---|---|---|---|---|
| PCDH18 | 490.46561 | -0.0935339 | 0.415846 | -0.2249243 | 0.8220382 | 0.97493464 |
| LINC02172 | 27.2156184 | -0.3084946 | 1.82738875 | -0.1688172 | 0.86594045 | 0.98197545 |
| SLC7A11 | 396.696414 | -3.0011754 | 1.08127712 | -2.775584 | NA | NA |
| NOCT | 386.376398 | -0.0320994 | 0.53865148 | -0.0595922 | 0.95248044 | 0.99170503 |
| ELF2 | 1796.73291 | -0.0136917 | 0.1181425 | -0.1158916 | 0.90773844 | 0.98662757 |
| MGARP | 47.717793 | 0.65546335 | 0.49702909 | 1.31876255 | 0.18724851 | 0.76958556 |
| NDUFC1 | 3411.08967 | 0.18547978 | 0.16212034 | 1.14408704 | 0.25258754 | 0.81833349 |
| NAA15 | 1343.94574 | -0.1237428 | 0.20426688 | -0.6057901 | 0.54465416 | 0.92345821 |
| AC097376.2 | 241.783366 | -0.1622693 | 0.29405017 | -0.5518421 | 0.58105652 | 0.9317705 |
| RAB33B | 583.004959 | -0.1888047 | 0.23439525 | -0.8054972 | 0.42053284 | 0.88451372 |
| SETD7 | 1885.85624 | -0.061574 | 0.29002977 | -0.2123024 | 0.83187114 | 0.97550416 |
| MGST2 | 1850.06588 | 0.36943825 | 0.18300385 | 2.01874581 | 0.04351365 | 0.56947573 |
| MAML3 | 444.589813 | 0.14079563 | 0.17188605 | 0.81912194 | 0.41271685 | 0.88285886 |
| SCOC | 5179.28025 | 0.25038971 | 0.15515713 | 1.6137815 | 0.10657483 | 0.67674128 |
| SCOC-AS1 | 40.6364146 | 0.49713476 | 0.45850677 | 1.08424737 | 0.27825513 | 0.82805357 |
| ELMOD2 | 1379.14488 | 0.35211195 | 0.135315 | 2.60216484 | 0.00926373 | 0.38664841 |
| TBC1D9 | 2282.64413 | -0.2450123 | 0.31995846 | -0.7657629 | 0.44381738 | 0.8938278 |
| RNF150 | 1234.73459 | 0.25370083 | 0.39262252 | 0.64616984 | 0.51816937 | 0.91636259 |
| ZNF330 | 2160.0597 | -0.0579952 | 0.2121656 | -0.2733489 | 0.78458501 | 0.97021865 |
| LINC02432 | 289.379505 | -0.5107595 | 0.52399273 | -0.9747454 | 0.32968657 | 0.85356118 |
| IL15 | 451.160491 | -0.400144 | 0.25161127 | -1.5903264 | 0.11176125 | 0.68001961 |
| INPP4B | 632.684243 | -0.0302369 | 0.27893487 | -0.1084011 | 0.9136775 | 0.98662757 |
| USP38 | 1487.30399 | -0.241585 | 0.27494718 | -0.8786598 | 3.80E-01 | 0.8708011 |
| GAB1 | 1491.02383 | -0.3294838 | 0.22024319 | -1.4959997 | 0.13465374 | 0.71630293 |
| SMARCA5 | 3512.64195 | -0.0924606 | 0.12637233 | -0.7316522 | 0.4643809 | 0.89832395 |
| SMARCA5-A! | 77.6640144 | 0.19995031 | 0.5087472 | 0.39302489 | 0.69430109 | 0.95222846 |
| GYPA | 75.374413 | 0.35914007 | 0.84174705 | 0.42666032 | 0.66962676 | 0.94898648 |
| HHIP-AS1 | 81.0508442 | 0.46167533 | 0.30324095 | 1.52247024 | 0.1278913 | 0.70579329 |
| HHIP | 133.173637 | 0.96937703 | 0.45384207 | 2.13593472 | 0.03268474 | 0.55104386 |
| ANAPC10 | 586.553633 | 0.12865263 | 0.12806393 | 1.00459695 | 0.31509097 | 0.84696964 |
| ABCE1 | 2505.27361 | -0.1250606 | 0.25009306 | -0.5000564 | 0.61703535 | 0.93910158 |
| OTUD4 | 2956.57776 | -0.1848604 | 0.16954381 | -1.0903402 | 0.2755633 | 0.82719407 |
| SMAD1 | 740.797799 | -0.0130533 | 0.23813145 | -0.0548155 | 0.95628544 | 0.99185043 |
| SMAD1-AS1 | 158.638767 | 0.02336271 | 0.33523835 | 0.06968985 | 0.94444052 | 0.99136103 |
| MMAA | 952.72076 | 0.21695939 | 0.23497324 | 0.92333659 | 0.35583182 | 0.86364633 |
| AC093864.1 | 101.415634 | 0.30619598 | 0.29517395 | 1.03734082 | 0.29957705 | 0.84123067 |
| ZNF827 | 1051.76353 | 0.33491179 | 0.28182875 | 1.18835211 | 0.23469472 | 0.80747753 |
| LSM6 | 509.288129 | -0.0008932 | 0.10474709 | -0.0085274 | 0.99319617 | 0.99906069 |
| REELD1 | 75.8471641 | 0.46619512 | 0.57881212 | 0.80543427 | 0.42056912 | 0.88451372 |
| SLC10A7 | 381.119613 | 0.07856135 | 0.15627167 | 0.50272288 | 0.61515912 | 0.93895494 |
| EDNRA | 497.334133 | -0.267988 | 0.33388709 | -0.8026307 | 0.4221882 | 0.88451372 |
| TMEM184C | 1933.90919 | -0.2090055 | 0.20812573 | -1.004227 | 0.31526921 | 0.84696964 |
| PRMT9 | 754.592183 | 0.20060748 | 0.20786674 | 0.96507732 | 3.35E-01 | 8.56E-01 |
| ARHGAP10 | 963.6987 | -0.347828 | 0.25250757 | -1.3774954 | 0.16835912 | 0.75735257 |
| NR3C2 | 1113.02135 | 0.24491566 | 0.23503602 | 1.04203456 | 0.29739566 | 0.83898658 |
| DCLK2 | 543.359216 | -0.3330207 | 0.39390158 | -0.8454414 | 0.39786445 | 0.88054917 |
| LRBA | 3246.7116 | 0.16758392 | 0.12275897 | 1.36514608 | 0.17220714 | 0.75974924 |
| RPS3A | 28397.6735 | 0.40683856 | 0.26092532 | 1.55921456 | 0.11894561 | 0.69233212 |
| SH3D19 | 20581.5371 | 0.39874808 | 0.24535499 | 1.6251884 | 0.10412242 | 0.67439401 |
| AC095055.1 | 63.3364154 | -2.1510725 | 0.98554269 | -2.1826274 | NA | NA |
| FAM160A1 | 156.608877 | -0.7743354 | 0.30815928 | -2.5127766 | 0.01197852 | 0.40742171 |
| AC092611.2 | 82.3961536 | -0.2956278 | 0.30398092 | -0.972521 | 0.33079142 | 0.85384238 |
| GATB | 1584.28759 | 0.45541265 | 0.20354198 | 2.23743842 | 0.0252577 | 0.52140487 |
| AC092611.1 | 50.5514033 | -0.941256 | 0.59536562 | -1.5809714 | 0.11388458 | 0.68214043 |

| | | | | | | |
|---|---|---|---|---|---|---|
| FBXW7 | 863.747308 | -0.2045014 | 0.14317334 | -1.4283481 | 0.1531917 | 0.74258845 |
| TMEM154 | 157.403879 | 0.3641523 | 0.4883621 | 0.74566045 | 0.45587256 | 0.89688594 |
| TIGD4 | 44.608036 | 0.65318836 | 0.36438927 | 1.79255651 | 0.07304387 | 0.63580072 |
| ARFIP1 | 2051.66743 | 0.14974585 | 0.11249461 | 1.33113805 | 0.18314359 | 0.76723162 |
| FHDC1 | 444.630687 | -0.3260752 | 0.29606083 | -1.1013791 | 0.2707317 | 0.82716979 |
| TRIM2 | 3696.39539 | -0.1512172 | 0.24081933 | -0.6279281 | 5.30E-01 | 0.91962754 |
| TMEM131L | 509.848298 | -0.1029873 | 0.27949723 | -0.3684733 | 7.13E-01 | 9.56E-01 |
| TLR2 | 1893.788 | -0.8815314 | 0.51615915 | -1.7078674 | 0.08766095 | 0.65322219 |
| RNF175 | 105.224093 | -0.7502447 | 0.54752104 | -1.3702573 | 0.17060659 | 0.75956362 |
| SFRP2 | 236.718155 | 2.36660185 | 1.12805802 | 2.09794339 | 0.03591015 | 0.55369025 |
| PLRG1 | 2597.66203 | -0.1078406 | 0.17018936 | -0.6336509 | 0.52630869 | 0.91769856 |
| FGB | 18974.6434 | 0.73846522 | 1.19395104 | 0.61850544 | 0.53624221 | 0.92102241 |
| FGA | 9052.51907 | 0.93143133 | 1.27906403 | 0.72821322 | 0.46648308 | 0.89960479 |
| FGG | 5015.37965 | 1.95777634 | 1.51046641 | 1.29614028 | 0.19492716 | 0.77660075 |
| LRAT | 59.4643281 | 0.12964758 | 0.59377281 | 0.21834543 | 0.82715998 | 0.97548201 |
| AC097467.3 | 67.8439742 | 0.17555404 | 0.28837366 | 0.60877281 | 0.54267504 | 0.92292765 |
| MAP9 | 694.342759 | 0.03124418 | 0.19047331 | 0.16403441 | 8.70E-01 | 0.98275231 |
| GUCY1A1 | 1025.95474 | -0.3363424 | 0.2484502 | -1.3537619 | 0.17581237 | 0.7610139 |
| GUCY1B1 | 1331.02278 | 0.1125359 | 0.21415865 | 0.52547916 | 0.59925014 | 0.93487648 |
| TDO2 | 52.2845659 | 0.50994157 | 1.20628571 | 0.42273698 | 0.67248717 | 0.94922827 |
| CTSO | 2703.37404 | -0.1500332 | 0.30469943 | -0.4923973 | 0.62243852 | 0.94016713 |
| PDGFC | 1736.95432 | -0.0908242 | 0.32972265 | -0.2754564 | 0.78296561 | 0.96964828 |
| GLRB | 295.49348 | -0.4397856 | 0.34499765 | -1.2747495 | 0.20239793 | 0.78193335 |
| FAM198B | 1366.27739 | -0.2076881 | 0.53636622 | -0.3872132 | 0.69859839 | 0.9529028 |
| FAM198B-AS | 136.143836 | -0.1273336 | 0.52897126 | -0.2407193 | 0.80977266 | 0.97334138 |
| TMEM144 | 469.259231 | 0.51250714 | 0.32482171 | 1.5778106 | 0.11460913 | 0.68447727 |
| RXFP1 | 185.379044 | -0.4589774 | 0.58860715 | -0.7797686 | 0.43552707 | 0.89077954 |
| C4orf46 | 226.944264 | -0.0812896 | 0.23979818 | -0.3389917 | 0.73461596 | 0.95895327 |
| ETFDH | 3459.96346 | 0.18817829 | 0.27834002 | 0.67607342 | 0.49899403 | 0.91147492 |
| PPID | 2574.15908 | -0.1597269 | 0.15944415 | -1.0017736 | 0.31645296 | 0.84783414 |
| FNIP2 | 6016.64976 | -1.0380807 | 0.44510835 | -2.3321978 | 0.01969029 | 0.49477123 |
| RAPGEF2 | 2080.75767 | -0.3137624 | 0.20063425 | -1.5638527 | 0.11785217 | 0.69069542 |
| NAF1 | 391.071103 | -0.1666013 | 0.22301229 | -0.7470499 | 4.55E-01 | 0.89670027 |
| NPY1R | 2633.04559 | -0.2975693 | 0.37754367 | -0.7881718 | 0.43059624 | 0.88966606 |
| NPY5R | 203.628438 | -0.4996397 | 0.31912562 | -1.5656522 | 0.11743006 | 0.6903252 |
| TMA16 | 539.295488 | 0.18160046 | 0.2011372 | 0.90286858 | 3.67E-01 | 8.67E-01 |
| 1-Mar | 434.705005 | -0.2415801 | 0.38139826 | -0.6334065 | 0.5264682 | 0.91769856 |
| AC093788.1 | 79.7951713 | -0.2191927 | 0.39362274 | -0.5568599 | 0.57762317 | 0.93130183 |
| APELA | 64.8220866 | -0.1334935 | 0.79184871 | -0.1685846 | 0.8661234 | 0.98197545 |
| FAM218A | 73.1804006 | 0.17435498 | 0.51291816 | 0.33992749 | 0.73391113 | 0.95895327 |
| TMEM192 | 3036.64771 | 0.5978522 | 0.36754557 | 1.62660701 | 0.10382059 | 0.67439401 |
| KLHL2 | 2114.41688 | 0.1693972 | 0.270578 | 0.62605681 | 0.53127768 | 0.91962754 |
| MSMO1 | 3838.03409 | 0.43908831 | 0.38364094 | 1.14452932 | 0.25240419 | 0.81833349 |
| CPE | 4063.52976 | -0.0687834 | 0.51729705 | -0.132967 | 0.89421949 | 0.98474725 |
| TLL1 | 247.670584 | -0.5974037 | 0.56018562 | -1.0664389 | 0.2862253 | 0.83107779 |
| DDX60 | 1217.59993 | -1.1443075 | 0.75971107 | -1.5062404 | NA | NA |
| DDX60L | 1320.4563 | -2.1974562 | 0.74372681 | -2.9546551 | NA | NA |
| PALLD | 6018.25856 | -0.344846 | 0.44572032 | -0.7736825 | 0.43911859 | 0.89175645 |
| CBR4 | 2286.75046 | 0.24015975 | 0.17385511 | 1.38137869 | 0.16716255 | 0.75558932 |
| SH3RF1 | 1591.95099 | 0.15523649 | 0.22138267 | 0.70121339 | 0.48316985 | 0.90541935 |
| NEK1 | 690.041147 | 0.05756594 | 0.20116569 | 0.28616184 | 0.77475417 | 0.96836854 |
| CLCN3 | 3584.78638 | 0.20058608 | 0.15251941 | 1.31515117 | 0.1884591 | 0.76970626 |
| HPF1 | 667.519176 | 0.23731859 | 0.17897502 | 1.32598724 | 0.18484392 | 0.76820626 |
| LINC02275 | 202.893944 | -0.0403484 | 0.69272199 | -0.0582462 | 0.9535525 | 0.99170503 |

| | | | | | | |
|---|---|---|---|---|---|---|
| MFAP3L | 3182.44255 | -0.0981347 | 0.24038022 | -0.4082479 | 0.68309172 | 0.95070301 |
| AADAT | 320.747525 | 1.06243471 | 0.44448322 | 2.39026954 | 0.01683601 | 0.46606159 |
| AC105285.1 | 84.6370932 | 0.20325809 | 0.34784104 | 0.58434188 | 0.55899032 | 0.92421175 |
| GALNT7 | 1051.79254 | 0.4861634 | 0.13960274 | 3.48247744 | 0.0004968 | 0.10826133 |
| AC097534.1 | 240.870483 | 0.38837202 | 0.19551997 | 1.98635472 | 0.04699395 | 0.58171004 |
| HMGB2 | 2671.13346 | -0.0835203 | 0.35966376 | -0.2322178 | 0.81636885 | 0.97388289 |
| AC097534.2 | 113.104721 | -0.3136838 | 0.44445219 | -0.7057762 | 0.48032732 | 0.90541935 |
| SAP30 | 425.973438 | -0.3624723 | 0.27138926 | -1.3356179 | 0.1816742 | 0.76528574 |
| SCRG1 | 78.4659801 | -1.926404 | 1.16362351 | -1.6555217 | NA | NA |
| FBXO8 | 1356.82937 | 0.0799655 | 0.16860264 | 0.47428381 | 0.63529754 | 0.94088547 |
| CEP44 | 717.541275 | -0.0090839 | 0.13743854 | -0.066094 | 0.94730295 | 0.99166614 |
| HPGD | 1559.00658 | 0.65498298 | 0.48237991 | 1.35781562 | 0.1745222 | 0.75974924 |
| GPM6A | 599.099413 | -0.4494495 | 0.50613516 | -0.8880029 | 0.37453918 | 0.86791605 |
| WDR17 | 218.5492 | 0.32964784 | 0.41705808 | 0.79041231 | 0.42928702 | 0.88851358 |
| SPATA4 | 56.9801532 | 0.80505105 | 0.53790535 | 1.49664071 | 0.13448677 | 0.71630293 |
| AC019163.1 | 266.827288 | -0.0134375 | 0.31917588 | -0.0421005 | 0.96641859 | 0.99400331 |
| SPCS3 | 4508.60687 | 0.04730718 | 0.20314941 | 0.23286889 | 0.81586322 | 0.9738447 |
| VEGFC | 488.892424 | -0.1340227 | 0.52105905 | -0.2572122 | 0.79701498 | 0.9713715 |
| AGA | 989.035905 | 0.61259203 | 0.1606383 | 3.81348675 | 0.00013702 | 0.05706421 |
| TENM3 | 212.499556 | 0.60190301 | 0.28973946 | 2.07739398 | 0.03776521 | 0.55538555 |
| DCTD | 3979.75642 | -0.0704568 | 0.1135767 | -0.6203452 | 0.53503056 | 0.92084006 |
| WWC2-AS2 | 66.4969324 | 0.42095092 | 0.39304967 | 1.07098658 | 0.28417546 | 0.83107779 |
| WWC2 | 2211.88828 | -0.067896 | 0.16193134 | -0.4192887 | 0.67500518 | 0.94943501 |
| CLDN22 | 303.459021 | 0.01802784 | 0.20957829 | 0.08601961 | 0.93145083 | 0.98975562 |
| CDKN2AIP | 1404.74237 | -0.1135521 | 0.22996798 | -0.4937737 | 0.62146602 | 0.94016713 |
| ING2 | 468.094781 | 0.03606831 | 0.26771821 | 0.13472489 | 0.8928294 | 0.98474725 |
| RWDD4 | 964.542442 | 0.18131954 | 0.21565894 | 0.84076989 | 0.40047686 | 0.88086294 |
| TRAPPC11 | 1595.25915 | 0.17783425 | 0.15042551 | 1.18220807 | 0.23712315 | 0.80857441 |
| STOX2 | 757.911944 | 0.10883648 | 0.3172393 | 0.34307376 | 0.73154297 | 0.95848028 |
| ENPP6 | 1741.57821 | 0.98067127 | 0.5629294 | 1.74208574 | 0.08149344 | 0.64441714 |
| IRF2 | 1673.26182 | -0.0789665 | 0.19828387 | -0.3982497 | 0.69044611 | 0.95158502 |
| CASP3 | 1062.94295 | -0.0962257 | 0.25474932 | -0.377727 | 0.70563341 | 0.95501548 |
| PRIMPOL | 500.419618 | -0.002609 | 0.18126383 | -0.0143933 | 0.98851621 | 0.99788078 |
| CENPU | 228.02317 | 0.09411071 | 0.31180808 | 0.30182254 | 0.76278734 | 0.96636231 |
| ACSL1 | 19343.3054 | -0.0136052 | 0.25742772 | -0.0528505 | 0.95785105 | 0.99223683 |
| LINC02437 | 173.759486 | -0.692481 | 0.83511646 | -0.8292029 | 0.40698958 | 0.88132015 |
| HELT | 32.1818703 | -0.3844778 | 0.80336764 | -0.4785826 | 0.6322356 | 0.94088547 |
| SLC25A4 | 5855.58177 | 0.21725364 | 0.19203074 | 1.1313482 | 0.25790856 | 0.82187798 |
| CFAP97 | 1541.23895 | 0.1804949 | 0.17491405 | 1.03190622 | 0.30211605 | 0.84211658 |
| SNX25 | 1167.20604 | -0.1388656 | 0.17146885 | -0.8098589 | 0.41802125 | 0.88445833 |
| LRP2BP | 388.566651 | -0.0948359 | 0.29543637 | -0.3210027 | 0.74820838 | 0.96241535 |
| AC112722.1 | 42.0987377 | -0.4042544 | 0.41672054 | -0.970085 | 0.33200411 | 0.85452795 |
| ANKRD37 | 2049.08635 | 0.10631832 | 0.55945436 | 0.19003931 | 0.84927832 | 0.97754115 |
| UFSP2 | 3022.39243 | 0.30109326 | 0.34249648 | 0.87911345 | 0.37933976 | 0.87069278 |
| C4orf47 | 198.179824 | -0.2286904 | 0.4507177 | -0.5073916 | 6.12E-01 | 9.39E-01 |
| CCDC110 | 95.5170219 | 0.12850739 | 0.37071366 | 0.34664864 | 0.7288553 | 0.95848028 |
| PDLIM3 | 1065.79024 | -1.3850926 | 0.72992622 | -1.897579 | NA | NA |
| SORBS2 | 1870.8782 | 0.54532577 | 0.32106166 | 1.6985079 | 0.08941194 | 0.65375543 |
| AC093797.1 | 95.7219366 | 0.55438698 | 0.31964807 | 1.73436675 | 0.08285303 | 0.64680474 |
| TLR3 | 795.232516 | -0.5294234 | 0.48713991 | -1.0867994 | 0.27712548 | 0.82779863 |
| FAM149A | 2014.51096 | 0.47661266 | 0.23604238 | 2.01918254 | 0.04346825 | 0.56927916 |
| CYP4V2 | 4568.8847 | 0.22315242 | 0.29948472 | 0.74512121 | 0.45619845 | 0.89688594 |
| KLKB1 | 1037.61488 | -0.1561438 | 0.36790897 | -0.4244087 | 0.67126778 | 0.94922827 |
| F11 | 885.348056 | 0.32650747 | 0.55123322 | 0.59232183 | 0.5536351 | 0.92373235 |

| | | | | | | |
|---|---|---|---|---|---|---|
| F11-AS1 | 405.231663 | 0.25657465 | 0.56507944 | 0.45405058 | 0.64979242 | 0.94312988 |
| FAT1 | 6760.04196 | 0.35135902 | 0.16957022 | 2.07205619 | 0.0382602 | 0.55538555 |
| LINC01060 | 90.1879567 | 0.95418043 | 0.99052298 | 0.96330974 | 0.33539211 | 0.85611325 |
| FRG1 | 972.097729 | 0.08630762 | 0.19106733 | 0.45171314 | 0.65147565 | 0.94393131 |
| PLEKHG4B | 180.297886 | 1.02989526 | 0.80508178 | 1.27924304 | 0.20081148 | 0.78134034 |
| CCDC127 | 1003.47044 | -0.2271223 | 0.20680122 | -1.098264 | 0.27208921 | 0.82716979 |
| SDHA | 14437.6595 | 0.47137847 | 0.16602375 | 2.83922305 | 0.00452235 | 0.29501922 |
| PDCD6 | 3966.61815 | 0.1251712 | 0.16817845 | 0.74427607 | 0.45670948 | 0.89688594 |
| AHRR | 764.121339 | 0.31047347 | 0.2161216 | 1.43656847 | 0.15084065 | 0.73796431 |
| EXOC3-AS1 | 268.551851 | 0.21084956 | 0.32227929 | 0.65424484 | 0.51295408 | 0.91568064 |
| EXOC3 | 11185.9321 | 0.45233798 | 0.24378485 | 1.85548024 | 0.06352768 | 0.61751671 |
| SLC9A3 | 17600.0883 | 0.81088 | 0.2981744 | 2.71948231 | 6.54E-03 | 3.39E-01 |
| AC106772.2 | 40.1394341 | 0.86657583 | 0.83144467 | 1.04225316 | 0.29729432 | 0.83898658 |
| CEP72 | 223.498039 | 0.02219836 | 0.27261344 | 0.08142796 | 9.35E-01 | 0.99068685 |
| TPPP | 882.125613 | 0.40965519 | 0.35740589 | 1.14619035 | 0.2517164 | 0.81833349 |
| ZDHHC11B | 580.25863 | 0.87137585 | 0.4924081 | 1.76962128 | 0.07679025 | 0.63773415 |
| ZDHHC11 | 557.35719 | -0.7012155 | 0.37717407 | -1.8591296 | 0.06300877 | 0.6147045 |
| BRD9 | 1486.52449 | 0.00161215 | 0.19041113 | 0.0084667 | 0.99324463 | 0.99906069 |
| TRIP13 | 140.937091 | -0.0549329 | 0.41233658 | -0.1332233 | 0.89401676 | 0.98474725 |
| AC116351.1 | 55.0733666 | 1.06064851 | 0.57520523 | 1.84394794 | 0.06519073 | 0.6216672 |
| NKD2 | 622.22405 | 0.62456218 | 0.31824133 | 1.96254268 | 0.04969934 | 0.5861488 |
| SLC12A7 | 11127.6543 | -0.0799806 | 0.25522311 | -0.3133752 | 0.75399562 | 0.96339067 |
| AC114291.1 | 1154.62009 | 0.75531229 | 0.7426335 | 1.01707274 | 0.30911883 | 0.84414094 |
| SLC6A19 | 11527.4147 | 0.7340728 | 0.67946115 | 1.08037494 | 0.27997525 | 0.82825734 |
| SLC6A18 | 1189.99193 | -2.5904927 | 0.8724435 | -2.9692384 | 0.00298539 | 0.23909904 |
| CLPTM1L | 7461.22114 | 0.41369959 | 0.19975617 | 2.0710229 | 0.03835666 | 0.55538555 |
| LPCAT1 | 4027.34275 | -0.6097983 | 0.33625634 | -1.8134924 | 0.0697559 | 0.62779744 |
| AC091849.2 | 44.2866314 | 0.2079913 | 0.62026011 | 0.33532915 | 0.73737681 | 0.96023056 |
| MRPL36 | 1010.67099 | -0.3820908 | 0.21787011 | -1.7537552 | 7.95E-02 | 6.41E-01 |
| NDUFS6 | 2505.70058 | 0.01172518 | 0.17609499 | 0.0665844 | 0.94691257 | 0.99166614 |
| IRX2 | 1648.6185 | -0.6328295 | 0.40960654 | -1.5449693 | 0.1223537 | 0.69788192 |
| C5orf38 | 419.623698 | -0.5978893 | 0.44904307 | -1.3314743 | 0.18303301 | 0.76723162 |
| IRX1 | 1551.40477 | -0.6944985 | 0.4828477 | -1.4383387 | 0.15033797 | 0.73796431 |
| AC016595.1 | 490.331317 | -0.7621642 | 0.50252768 | -1.5166612 | 0.12935224 | 0.70755752 |
| LINC01020 | 202.364588 | 2.35818075 | 1.08085864 | 2.18176612 | 0.0291268 | 0.53606221 |
| LINC02121 | 63.9549896 | -0.6163064 | 1.31538728 | -0.4685361 | 0.63940129 | 0.94088547 |
| AC022424.1 | 162.033161 | 1.29814608 | 0.47521602 | 2.73169682 | 0.00630091 | 0.33470127 |
| ADAMTS16 | 455.782227 | 1.11205204 | 0.42199458 | 2.63522824 | 0.00840807 | 0.37787934 |
| ICE1 | 1938.03972 | -0.2812203 | 0.1840052 | -1.5283277 | 0.12643119 | 0.70431949 |
| LINC02145 | 50.7872302 | 0.17748141 | 0.6206601 | 0.28595588 | 0.77491191 | 0.96836854 |
| MED10 | 1190.14309 | -0.3730549 | 0.27536526 | -1.3547639 | 0.1754928 | 0.76079356 |
| UBE2QL1 | 222.63363 | 0.77740869 | 0.50085512 | 1.5521628 | 0.12062327 | 0.69467399 |
| LINC01018 | 52.1685168 | 0.09796851 | 0.58734847 | 0.16679794 | 0.86752904 | 0.98224588 |
| NSUN2 | 2803.44532 | -0.2095747 | 0.2483082 | -0.8440105 | 0.39866351 | 0.88074856 |
| SRD5A1 | 966.475449 | 0.53218749 | 0.29038456 | 1.83269897 | 0.06684735 | 0.6249306 |
| TENT4A | 1838.66896 | -0.4987711 | 0.1547148 | -3.2238097 | 0.00126497 | 0.16018166 |
| ADCY2 | 60.6651975 | 0.6203738 | 0.55158712 | 1.12470683 | 0.26071329 | 0.82424998 |
| C5orf49 | 262.462486 | 0.09767105 | 0.4237047 | 0.23051679 | 0.81769022 | 0.97438078 |
| MTRR | 2619.27912 | 0.14457573 | 0.21345024 | 0.67732757 | 0.49819814 | 0.91054529 |
| FASTKD3 | 760.437446 | 0.06428935 | 0.24050589 | 0.26730884 | 0.78923138 | 0.97090793 |
| MIR4458HG | 610.436275 | -0.4649447 | 0.31747922 | -1.4644886 | 0.1430605 | 0.72784886 |
| AC021088.1 | 48.170766 | 0.22074266 | 0.58771407 | 0.37559532 | 7.07E-01 | 0.95501548 |
| SEMA5A | 1887.21942 | 0.46083037 | 0.49722705 | 0.92680069 | 0.35403003 | 0.86364633 |
| SNHG18 | 786.427805 | -0.2886259 | 0.15291799 | -1.8874554 | 5.91E-02 | 6.05E-01 |

| | | | | | | |
|---|---|---|---|---|---|---|
| FAM173B | 556.369251 | 0.27951368 | 0.24132112 | 1.15826451 | 0.24675611 | 0.81542272 |
| CCT5 | 7711.30097 | -0.0639957 | 0.21500163 | -0.2976523 | 0.76596857 | 0.96704052 |
| AC012640.4 | 2168.04065 | -0.0334494 | 0.21136903 | -0.1582514 | 0.8742587 | 0.98275231 |
| CMBL | 16889.2327 | 1.54624646 | 0.71785802 | 2.15397253 | 0.03124233 | 0.54466273 |
| AC012640.2 | 74.2344062 | 0.40686809 | 0.3958863 | 1.02773977 | 0.30407225 | 0.84211658 |
| 6-Mar | 7450.11051 | 0.05292011 | 0.14319342 | 0.36957083 | 0.71170229 | 0.95560193 |
| ROPN1L | 44.8576974 | 0.30618357 | 0.57317436 | 0.53418922 | 0.59321063 | 0.93289552 |
| AC092336.1 | 227.702156 | -0.7073227 | 0.78376863 | -0.9024636 | 0.36681062 | 0.86683872 |
| LINC02212 | 440.201211 | -0.7412735 | 0.79150708 | -0.9365342 | 0.34899819 | 0.86356499 |
| ANKRD33B | 1580.39193 | 0.15750732 | 0.40388574 | 0.38997988 | 0.69655142 | 0.9525827 |
| DAP | 9744.1809 | 0.08186583 | 0.22312854 | 0.36689985 | 0.71369372 | 0.95564093 |
| AC012629.2 | 372.772056 | 0.19827692 | 0.34728748 | 0.57093024 | 0.56804693 | 0.92618483 |
| DNAH5 | 896.893746 | -0.1449548 | 0.73619109 | -0.1968983 | 0.84390712 | 0.97664894 |
| TRIO | 3451.34712 | -0.6994581 | 0.26399039 | -2.649559 | 0.00805969 | 0.37112203 |
| OTULINL | 426.266657 | 0.3142635 | 0.40368335 | 0.77849013 | 0.43628012 | 0.89104919 |
| AC010491.1 | 52.5244868 | 0.16888204 | 0.56515724 | 0.2988231 | 0.76507502 | 0.96678847 |
| OTULIN | 1118.6682 | -0.2839475 | 0.21072724 | -1.3474646 | 0.17783066 | 0.76115956 |
| ANKH | 2010.85709 | -0.1714713 | 0.30373767 | -0.5645376 | 5.72E-01 | 9.29E-01 |
| FBXL7 | 598.824977 | -0.4618161 | 0.29854919 | -1.5468676 | 1.22E-01 | 6.98E-01 |
| ZNF622 | 1477.81644 | -0.2860221 | 0.31706024 | -0.9021064 | 0.36700033 | 0.86683872 |
| RETREG1 | 6561.95511 | 0.19680048 | 0.41237335 | 0.47723861 | 0.63319222 | 0.94088547 |
| MYO10 | 4216.02825 | -0.3726421 | 0.21280528 | -1.7510941 | 0.07992971 | 0.64079926 |
| BASP1 | 583.080666 | -0.3091838 | 0.48741008 | -0.6343401 | 0.52585887 | 0.91769856 |
| AC026785.2 | 98.2674075 | -0.049333 | 0.53154953 | -0.0928097 | 0.92605472 | 0.98876911 |
| CDH9 | 305.457311 | 1.05742241 | 0.58464054 | 1.80867102 | 0.07050213 | 0.62981127 |
| AC113386.1 | 148.771744 | 0.18863119 | 0.56833231 | 0.33190299 | 0.73996251 | 0.96043876 |
| CDH6 | 7287.68876 | 0.41795141 | 0.55014389 | 0.7597129 | 0.44742621 | 0.89480207 |
| DROSHA | 1585.81653 | -0.0537229 | 0.14962072 | -0.3590607 | 0.71954966 | 0.9566409 |
| C5orf22 | 929.387709 | -0.0942029 | 0.16704508 | -0.563937 | 0.57279698 | 0.92928554 |
| PDZD2 | 1342.37919 | 0.28249754 | 0.47568091 | 0.59388033 | 0.55259215 | 0.92373235 |
| AC025178.1 | 40.7950353 | -0.6450556 | 0.47275683 | -1.3644554 | 0.17242429 | 0.75974924 |
| GOLPH3 | 5596.86671 | -0.3248068 | 0.14442317 | -2.2489935 | 0.02451291 | 0.51967919 |
| AC025181.2 | 158.85896 | -0.1206867 | 0.25501288 | -0.4732571 | 0.63602976 | 0.94088547 |
| MTMR12 | 2265.19998 | -0.0129404 | 0.11685312 | -0.1107406 | 0.91182202 | 0.98662757 |
| ZFR | 3836.58613 | -0.0700733 | 0.13501536 | -0.5190025 | 0.60375901 | 0.93628858 |
| SUB1 | 5482.29505 | -0.0268224 | 0.09886728 | -0.2712975 | 0.78616222 | 0.97075604 |
| LINC02061 | 130.39022 | -2.5439465 | 2.15537632 | -1.1802795 | 0.23788907 | 0.80962787 |
| NPR3 | 5808.97701 | 0.46003884 | 0.40519272 | 1.13535811 | 0.25622531 | 0.82067614 |
| TARS | 3368.70653 | -0.2016115 | 0.2486679 | -0.8107659 | 0.41750011 | 0.88445833 |
| ADAMTS12 | 128.101668 | 0.11766167 | 0.38819176 | 0.30310193 | 0.76181218 | 0.96607083 |
| SLC45A2 | 55.3011226 | 0.58085149 | 0.45043962 | 1.2895213 | 0.19721692 | 0.77816059 |
| AMACR | 11552.235 | 0.65888899 | 0.47535935 | 1.38608608 | 0.16572062 | 0.75515387 |
| C1QTNF3-AN | 11551.9853 | 0.63981776 | 0.46070644 | 1.38877536 | 0.16490107 | 0.75299828 |
| C1QTNF3 | 1011.47256 | 0.2791355 | 0.55216955 | 0.50552498 | 0.61319016 | 0.93879706 |
| RAI14 | 2193.38951 | -0.1312648 | 0.13505405 | -0.9719427 | 0.33107908 | 0.85384238 |
| AC026801.2 | 126.40127 | 0.59922667 | 0.38794207 | 1.54462924 | 0.12243597 | 0.69788192 |
| TTC23L | 76.9954277 | 0.25457875 | 0.46960469 | 0.54211288 | 0.58774075 | 0.93289552 |
| RAD1 | 1192.44234 | -0.2213631 | 0.21018843 | -1.0531648 | 0.29226546 | 0.83524572 |
| BRIX1 | 1104.29744 | -0.3003615 | 0.20236854 | -1.4842301 | 0.13774789 | 0.72209298 |
| DNAJC21 | 1952.5478 | -0.0300248 | 0.15208659 | -0.197419 | 0.8434997 | 0.97649449 |
| AGXT2 | 7801.44367 | 1.82826821 | 0.58055166 | 3.14919128 | 0.00163723 | 0.18012143 |
| PRLR | 1843.37165 | 0.93211055 | 0.5061231 | 1.84166768 | 0.06552377 | 0.62237346 |
| SPEF2 | 958.385016 | 0.30124787 | 0.48029334 | 0.62721642 | 0.53051738 | 0.91962754 |
| IL7R | 549.94663 | 0.09586981 | 0.89124587 | 0.10756831 | 0.91433814 | 0.9866558 |

| | | | | | | |
|---|---|---|---|---|---|---|
| UGT3A1 | 4252.40144 | 0.31213229 | 0.4634003 | 0.67356947 | 0.50058507 | 0.91197285 |
| UGT3A2 | 146.278136 | 0.22412694 | 0.66187478 | 0.33862438 | 7.35E-01 | 9.59E-01 |
| LMBRD2 | 900.193705 | 0.33192259 | 0.17242152 | 1.92506477 | 0.05422126 | 0.59329767 |
| SKP2 | 370.415184 | 0.13739479 | 0.15149366 | 0.90693427 | 0.36444156 | 0.8663515 |
| NADK2 | 1965.68773 | -0.2589762 | 0.28731407 | -0.9013696 | 0.36739186 | 0.86683872 |
| RANBP3L | 823.507829 | -0.2486598 | 0.63810828 | -0.3896827 | 0.69677115 | 0.9525827 |
| SLC1A3 | 365.334125 | -0.546657 | 0.58426352 | -0.9356342 | 0.34946154 | 0.86364633 |
| NIPBL | 2483.05555 | -0.361067 | 0.16775176 | -2.1523889 | 0.03136674 | 0.54507414 |
| CPLANE1 | 734.769599 | 0.07069664 | 0.27007075 | 0.26177083 | 0.79349813 | 0.97090793 |
| NUP155 | 748.799566 | 0.10775398 | 0.14928649 | 0.72179327 | 0.47042159 | 0.9008571 |
| WDR70 | 1021.47367 | 0.05518985 | 0.15513147 | 0.35576182 | 0.72201895 | 0.95739595 |
| GDNF-AS1 | 114.848667 | 0.56319236 | 0.57940527 | 0.97201802 | 0.33104159 | 0.85384238 |
| GDNF | 70.5016122 | 0.41276839 | 0.52638887 | 0.78415106 | 0.43295148 | 0.89017484 |
| EGFLAM | 347.328185 | 0.48536186 | 0.53011823 | 0.91557286 | 0.35989097 | 0.86480228 |
| LIFR | 3338.44043 | -0.404723 | 0.23686962 | -1.708632 | 0.08751914 | 0.65271545 |
| LIFR-AS1 | 232.733339 | 0.20928725 | 0.4215826 | 0.49643237 | 0.61958939 | 0.93945324 |
| OSMR-AS1 | 47.4965337 | 0.22389393 | 0.44078974 | 0.50793816 | 0.61149671 | 0.93834929 |
| OSMR | 6385.14601 | -0.2338512 | 0.34671757 | -0.6744718 | 0.5000114 | 0.91180773 |
| RICTOR | 1934.50316 | -0.4711079 | 0.18922068 | -2.4897274 | 1.28E-02 | 4.16E-01 |
| FYB1 | 746.393636 | 0.21944188 | 0.41030999 | 0.53481973 | 0.59277452 | 0.93289552 |
| C9 | 1721.08483 | -0.0391452 | 0.41612169 | -0.0940715 | 0.92505233 | 0.98869145 |
| DAB2 | 22170.9235 | -0.1896297 | 0.28605877 | -0.6629047 | 0.50739162 | 0.91430909 |
| TTC33 | 666.77367 | -0.0074315 | 0.18429243 | -0.0403246 | 0.96783431 | 0.99400331 |
| PTGER4 | 1001.33063 | -0.1856883 | 0.30105679 | -0.6167882 | 0.53737442 | 0.92123049 |
| PRKAA1 | 3555.61052 | -0.0345642 | 0.12339319 | -0.2801143 | 0.77938984 | 0.96836854 |
| RPL37 | 19594.3085 | 0.15611599 | 0.14771105 | 1.05690128 | 0.29055664 | 0.83442894 |
| CARD6 | 546.479239 | -0.2352986 | 0.34590812 | -0.6802344 | 0.49635605 | 0.90890243 |
| C7 | 42369.1967 | 0.32933476 | 0.3556948 | 0.92589143 | 0.35450241 | 0.86364633 |
| C6 | 590.133341 | 0.40032202 | 0.63280234 | 0.63261778 | 0.52698327 | 0.9178228 |
| PLCXD3 | 668.614907 | -0.3353194 | 0.51738469 | -0.6481046 | 0.51691728 | 0.91636259 |
| OXCT1 | 2879.35715 | -0.2052505 | 0.44142154 | -0.4649762 | 0.64194851 | 0.94088547 |
| C5orf51 | 1457.79062 | -0.0759005 | 0.23177094 | -0.3274805 | 0.74330447 | 0.96117222 |
| FBXO4 | 402.740108 | 0.31574119 | 0.20155614 | 1.5665174 | 0.11722754 | 0.68956727 |
| GHR | 736.513281 | 0.21664592 | 0.25279466 | 0.85700353 | 0.39144293 | 0.8772797 |
| CCDC152 | 19025.9142 | 0.26160998 | 0.36457929 | 0.71756676 | 0.47302446 | 0.90175304 |
| SELENOP | 24247.7912 | 0.25676074 | 0.36194207 | 0.70939733 | 0.47807794 | 0.90404077 |
| AC008945.1 | 3466.78396 | 0.30772206 | 0.38634112 | 0.79650352 | 0.42573943 | 0.88640673 |
| ANXA2R | 101.471537 | 0.20130399 | 0.50552244 | 0.3982098 | 0.69047554 | 0.95158502 |
| AC025171.2 | 40.4279925 | 0.32919537 | 0.40590148 | 0.81102283 | 0.41735256 | 0.88445833 |
| ZNF131 | 1208.19822 | -0.2271782 | 0.17164167 | -1.3235606 | 0.18564902 | 0.76908034 |
| NIM1K | 56.7813374 | -0.5108761 | 0.47318012 | -1.0796651 | 0.28029135 | 0.82835218 |
| AC114947.2 | 1555.06911 | 0.07312854 | 0.4056126 | 0.1802916 | 0.85692365 | 0.97953896 |
| HMGCS1 | 3791.72484 | -0.0271344 | 0.40906431 | -0.066333 | 0.94711274 | 0.99166614 |
| CCL28 | 481.957201 | 0.42100228 | 0.47068767 | 0.89444084 | 0.37108607 | 0.86762749 |
| TMEM267 | 694.972241 | 0.15861927 | 0.19023728 | 0.83379696 | 0.40439541 | 0.88132015 |
| C5orf34 | 86.5787641 | 0.45609652 | 0.35674025 | 1.27851153 | 0.20106912 | 0.78134034 |
| PAIP1 | 3873.39897 | -0.0373884 | 0.15756696 | -0.2372855 | 0.81243529 | 0.97364432 |
| NNT-AS1 | 1013.15346 | 0.07234197 | 0.16793674 | 0.43076918 | 0.66663621 | 0.94819591 |
| NNT | 5692.94813 | 0.62713206 | 0.17513829 | 3.58078208 | 0.00034257 | 0.08914272 |
| MRPS30-DT | 64.1201711 | -0.3216783 | 0.48337153 | -0.6654886 | 0.50573802 | 0.91386909 |
| MRPS30 | 1851.1559 | 0.0438389 | 0.24279415 | 0.18055996 | 0.85671298 | 0.97953896 |
| AC093297.2 | 641.002919 | 0.19392424 | 0.12433142 | 1.55973635 | 0.1188222 | 0.69221226 |
| EMB | 466.976686 | 0.11368941 | 0.6066221 | 0.1874139 | 0.85133613 | 0.97855304 |
| PARP8 | 1225.73988 | -0.1113075 | 0.30288539 | -0.3674904 | 0.71325324 | 0.95564093 |

| | | | | | | |
|---|---|---|---|---|---|---|
| ITGA1 | 4390.80785 | -0.040396 | 0.2663913 | -0.1516416 | 0.87946965 | 0.98275231 |
| PELO | 2372.97467 | -0.4983306 | 0.24184652 | -2.0605242 | 0.03934846 | 0.55657615 |
| ITGA2 | 2329.86787 | -0.6946684 | 0.50738788 | -1.3691071 | 0.17096579 | 0.75956362 |
| MOCS2 | 3371.02659 | 0.36842687 | 0.161006 | 2.2882804 | 0.0221212 | 0.51647233 |
| AC008966.1 | 206.560236 | 0.15107648 | 0.37771772 | 0.39997193 | 0.6891772 | 0.95158502 |
| FST | 203.512722 | 0.86481101 | 0.47785512 | 1.8097766 | 0.07033044 | 0.62936167 |
| NDUFS4 | 2545.54094 | 0.28429599 | 0.15759749 | 1.80393736 | 0.07124113 | 0.63126712 |
| AC025175.1 | 69.5783076 | -0.1074288 | 0.53773201 | -0.1997813 | 0.84165166 | 0.9762188 |
| ARL15 | 1564.74402 | -0.1647774 | 0.27955944 | -0.5894182 | 0.55558079 | 0.92373235 |
| SNX18 | 1777.13894 | 0.07121152 | 0.2038867 | 0.34927004 | 0.72688659 | 0.95848028 |
| ESM1 | 1193.0158 | 0.46764412 | 0.62871451 | 0.74380997 | 0.45699145 | 0.89688594 |
| GZMK | 138.470024 | -0.0175737 | 0.96157644 | -0.018276 | 0.98541871 | 0.99696242 |
| GZMA | 138.493241 | 0.20385984 | 0.83813083 | 0.24323153 | 0.80782604 | 0.97334138 |
| CDC20B | 48.3264127 | 1.65564381 | 1.26755681 | 1.30616932 | 0.19149498 | 0.77328076 |
| GPX8 | 630.362555 | 0.06779186 | 0.34934999 | 0.19405142 | 0.84613562 | 0.97680348 |
| CCNO | 216.896 | 0.48194179 | 0.6000944 | 0.80310995 | 0.42191119 | 0.88451372 |
| DHX29 | 1942.20993 | -0.0279909 | 0.15542784 | -0.1800896 | 0.85708224 | 0.97953896 |
| MTREX | 4929.99819 | -0.1653686 | 0.20552896 | -0.8045999 | 0.42105063 | 0.88451372 |
| PLPP1 | 6173.43723 | -0.2686418 | 0.35108966 | -0.7651657 | 0.44417285 | 0.8938278 |
| SLC38A9 | 650.183807 | -0.0104116 | 0.17719493 | -0.0587577 | 0.95314508 | 0.99170503 |
| IL6ST | 11285.6616 | -0.186455 | 0.21785162 | -0.8558809 | 0.39206368 | 0.87765297 |
| AC008914.1 | 197.297952 | -0.4380023 | 0.39193122 | -1.1175489 | 0.26375971 | 0.82424998 |
| C5orf67 | 212.188135 | -0.5977985 | 0.82712067 | -0.7227464 | 0.46983573 | 0.9008571 |
| MAP3K1 | 1844.32893 | -0.0173044 | 0.26476102 | -0.0653585 | 0.94788854 | 0.99170503 |
| SETD9 | 345.067392 | 0.44600168 | 0.35069296 | 1.27177253 | 0.20345395 | 0.78338136 |
| MIER3 | 1005.38251 | -0.1151793 | 0.15805893 | -0.7287112 | 0.46617835 | 0.89960479 |
| GPBP1 | 4120.18028 | -0.2757714 | 0.15734548 | -1.7526487 | 0.07966233 | 0.64079926 |
| PLK2 | 1278.69743 | -0.4095569 | 0.28469031 | -1.438605 | 0.15026248 | 0.73796431 |
| GAPT | 53.2469688 | 0.32201736 | 0.81147728 | 0.39682856 | 0.69149389 | 0.95159158 |
| RAB3C | 40.2125928 | -0.2926385 | 0.94544338 | -0.3095251 | 0.75692211 | 0.96493281 |
| PDE4D | 1894.06989 | 0.13318677 | 0.25814296 | 0.51594192 | 0.60589498 | 0.93698713 |
| PART1 | 79.532609 | 0.69542691 | 0.61744291 | 1.12630156 | 0.2600379 | 0.82424998 |
| DEPDC1B | 168.748572 | -1.1474282 | 0.72986556 | -1.572109 | 0.11592528 | 0.6873836 |
| ELOVL7 | 1713.28654 | -0.2451571 | 0.20192266 | -1.2141139 | 0.22470424 | 0.80106184 |
| ERCC8 | 430.024859 | 0.22698796 | 0.14222178 | 1.596014 | 0.11048567 | 0.68001961 |
| AC022445.1 | 45.0351237 | 0.44733287 | 0.27733918 | 1.61294512 | 0.10675643 | 0.67674128 |
| NDUFAF2 | 599.043956 | -0.0516317 | 0.20278391 | -0.2546143 | 0.79902105 | 0.97186224 |
| SMIM15 | 2417.72626 | -0.0203359 | 0.17573106 | -0.1157215 | 9.08E-01 | 0.98662757 |
| SMIM15-AS1 | 315.775371 | 0.03990423 | 0.28139086 | 0.14181069 | 0.88722954 | 0.98423722 |
| ZSWIM6 | 1043.59457 | -0.246816 | 0.19306262 | -1.2784244 | 0.20109981 | 0.78134034 |
| KIF2A | 719.29564 | -0.1516207 | 0.21366398 | -0.7096221 | 0.47793852 | 0.90393034 |
| DIMT1 | 1148.62384 | -0.0737594 | 0.1552449 | -0.4751167 | 0.63470381 | 0.94088547 |
| IPO11 | 1104.71582 | -0.0490836 | 0.12571983 | -0.3904209 | 0.69622535 | 0.9525827 |
| LRRC70 | 210.877783 | -0.6177511 | 0.32289766 | -1.9131484 | 0.05572906 | 0.59817776 |
| RNF180 | 681.099826 | 0.21476558 | 0.23665417 | 0.90750813 | 0.36413815 | 0.8663515 |
| RGS7BP | 46.4154754 | -0.6775019 | 0.58380586 | -1.1604918 | 0.24584862 | 0.81542272 |
| SREK1IP1 | 859.499548 | -0.1272725 | 0.14530069 | -0.8759249 | 0.38107085 | 0.87171941 |
| CWC27 | 709.000837 | 0.02273622 | 0.13886949 | 0.16372366 | 0.86994869 | 0.98275231 |
| ADAMTS6 | 84.5447798 | -0.7528571 | 0.46800374 | -1.6086562 | 0.10769154 | 0.67707349 |
| CENPK | 56.590955 | 0.14728186 | 0.55271378 | 0.2664704 | 0.78987695 | 0.97090793 |
| PPWD1 | 1360.32358 | -0.0665886 | 0.23343198 | -0.2852589 | 0.77544579 | 0.96836854 |
| TRIM23 | 687.008385 | -0.0732236 | 0.21436283 | -0.341587 | 0.7326617 | 0.95848028 |
| TRAPPC13 | 1355.44309 | 0.07518626 | 0.11342589 | 0.66286679 | 0.50741588 | 0.91430909 |
| SHLD3 | 215.764921 | -0.4142277 | 0.2539008 | -1.6314548 | 0.1027944 | 0.67335541 |

| | | | | | | |
|---|---|---|---|---|---|---|
| SGTB | 552.637939 | -0.0232945 | 0.26086176 | -0.0892982 | 0.92884491 | 0.98963127 |
| NLN | 1395.36629 | 0.04440292 | 0.3285036 | 0.13516722 | 0.89247966 | 0.98474725 |
| ERBIN | 3663.0417 | -0.0459238 | 0.15029359 | -0.3055605 | 0.75993932 | 0.96514582 |
| SREK1 | 3532.11083 | -0.3435817 | 0.13288802 | -2.5854977 | 0.00972385 | 0.39368232 |
| MAST4 | 2210.50065 | -0.6399674 | 0.23904777 | -2.6771527 | 0.00742508 | 0.36301578 |
| CD180 | 97.1497777 | -0.8208797 | 0.57165613 | -1.4359676 | 0.15101155 | 0.73796431 |
| PIK3R1 | 3429.0712 | 0.00655584 | 0.35541054 | 0.01844583 | 0.98528319 | 0.99692004 |
| SLC30A5 | 2491.04777 | 0.05042974 | 0.1565172 | 0.32219935 | 0.74730168 | 0.96226645 |
| AC010273.1 | 682.004552 | 0.12940386 | 0.17496262 | 0.73960862 | 0.45953751 | 0.89755243 |
| CCNB1 | 309.664753 | 0.32228205 | 0.30612416 | 1.05278215 | 0.29244085 | 0.83524572 |
| CENPH | 90.7452788 | 0.07361943 | 0.33418413 | 0.22029602 | 0.82564062 | 0.97548201 |
| MRPS36 | 1704.13156 | 0.23111264 | 0.23161465 | 0.99783253 | 0.31836057 | 0.8490262 |
| CDK7 | 1418.6769 | -0.1702785 | 0.18910314 | -0.9004533 | 0.36787908 | 0.8674411 |
| CCDC125 | 817.491457 | 0.19877618 | 0.32672165 | 0.60839611 | 0.54292479 | 0.92297267 |
| AK6 | 1827.95668 | -0.1639433 | 0.22952048 | -0.714286 | 0.47505036 | 0.90226631 |
| TAF9 | 2270.12829 | -0.3945368 | 0.21605342 | -1.8261077 | 0.06783405 | 0.62599695 |
| RAD17 | 2087.44604 | -0.0550487 | 0.16006866 | -0.3439066 | 0.73091656 | 0.95848028 |
| MARVELD2 | 1075.94001 | 0.11477317 | 0.17868109 | 0.64233527 | 0.52065552 | 0.91680964 |
| OCLN | 1105.52296 | -0.0820643 | 0.24712182 | -0.3320802 | 0.73982869 | 0.96043876 |
| GTF2H2C | 635.142077 | -0.1616436 | 0.25266494 | -0.6397549 | 0.52233199 | 0.91723878 |
| SMN2 | 202.949669 | -0.1371104 | 0.86751209 | -0.1580501 | 0.87441733 | 0.98275231 |
| SMN1 | 342.106881 | -0.0772213 | 0.13692939 | -0.5639495 | 0.5727885 | 0.92928554 |
| NAIP | 371.117277 | -0.3422898 | 0.33669592 | -1.0166142 | 0.30933701 | 0.84414094 |
| GTF2H2 | 548.908363 | -0.020886 | 0.49996947 | -0.0417745 | 0.96667843 | 0.99400331 |
| BDP1 | 1260.72628 | -0.2296448 | 0.13811611 | -1.662694 | 0.0963737 | 0.66233356 |
| MCCC2 | 5651.23598 | 0.38405072 | 0.19685105 | 1.95097112 | 0.05106048 | 0.5861488 |
| MAP1B | 1926.85338 | -1.3351148 | 0.52985504 | -2.5197737 | 0.01174303 | 0.40423902 |
| MRPS27 | 2754.37989 | 0.27248023 | 0.12269045 | 2.22087569 | 0.02635938 | 0.52678638 |
| PTCD2 | 561.699104 | 0.01993936 | 0.21777533 | 0.09155929 | 0.92704819 | 0.98915637 |
| ZNF366 | 46.5174336 | -0.3094114 | 0.39200701 | -0.7893006 | 0.42993635 | 0.88894938 |
| TNPO1 | 5092.87258 | -0.3444413 | 0.28547714 | -1.2065459 | 0.22760709 | 0.80300331 |
| FCHO2 | 2132.77346 | 0.07205382 | 0.14492424 | 0.49718265 | 0.61906026 | 0.93945324 |
| TMEM171 | 994.649866 | -0.3216974 | 0.39410956 | -0.816264 | 0.41434916 | 0.88364943 |
| AC116345.3 | 114.195071 | 3.20832906 | 1.0062407 | 3.18843101 | NA | NA |
| TMEM174 | 3860.12776 | 2.13684731 | 0.9860498 | 2.16707849 | 0.03022887 | 0.5405936 |
| AC116345.1 | 51.4055319 | 0.99844556 | 0.63811933 | 1.56466903 | 0.11766053 | 0.69038073 |
| FOXD1 | 209.54498 | 0.08820362 | 0.32154203 | 0.27431443 | 0.78384299 | 0.96978847 |
| FOXD1-AS1 | 132.903328 | 0.17916833 | 0.32284549 | 0.55496618 | 0.57891779 | 0.93139743 |
| BTF3 | 12386.4358 | 0.1691958 | 0.1812641 | 0.93342146 | 0.3506024 | 0.86364633 |
| ANKRA2 | 755.459625 | 0.22169413 | 0.21650784 | 1.02395429 | 0.30585687 | 0.84270033 |
| UTP15 | 550.982807 | -0.2128067 | 0.25402652 | -0.8377342 | 0.40218 | 0.88132015 |
| ARHGEF28 | 4077.44166 | -0.0285516 | 0.22445204 | -0.1272059 | 0.89877743 | 0.98477645 |
| ENC1 | 641.973825 | 0.70569213 | 0.4229239 | 1.6686031 | 0.09519606 | 0.65888357 |
| HEXB | 8236.7835 | 0.25849938 | 0.30017089 | 0.86117406 | 0.38914219 | 0.87568585 |
| GFM2 | 2356.32376 | 0.20954044 | 0.15210286 | 1.3776233 | 0.16831962 | 0.75735257 |
| NSA2 | 3118.66633 | 0.15401352 | 0.1601917 | 0.96143258 | 0.33633471 | 0.85665246 |
| FAM169A | 450.607998 | 0.32253427 | 0.31510554 | 1.02357538 | 0.30603588 | 0.84270033 |
| AC010501.1 | 40.6951075 | 0.73259541 | 0.79195343 | 0.92504859 | 0.35494063 | 0.86364633 |
| GCNT4 | 400.085083 | -0.0838933 | 0.33579111 | -0.2498377 | 0.8027129 | 0.97269122 |
| HMGCR | 2376.23553 | -0.071754 | 0.40356157 | -0.177802 | 0.85887849 | 0.97992012 |
| COL4A3BP | 2563.07623 | -0.1968267 | 0.26658125 | -0.7383368 | 0.46030982 | 0.89755243 |
| POLK | 1049.64786 | -0.0674849 | 0.20674251 | -0.3264199 | 0.74410672 | 0.9613927 |
| AC010245.1 | 323.096906 | -0.0869728 | 0.18772942 | -0.4632878 | 0.64315808 | 0.9413797 |
| AC010245.2 | 115.468812 | 0.23773813 | 0.30060195 | 0.79087355 | 0.42901778 | 0.88851358 |

| | | | | | | |
|---|---|---|---|---|---|---|
| ANKDD1B | 258.16848 | 0.6039002 | 0.58049611 | 1.04031739 | 0.29819247 | 0.83998573 |
| POC5 | 405.749372 | 0.00864327 | 0.18371321 | 0.04704761 | 0.96247528 | 0.99320711 |
| IQGAP2 | 2580.00193 | 0.91814173 | 0.33053942 | 2.77770718 | 0.00547439 | 0.314681 |
| F2RL2 | 53.1133752 | -2.3940181 | 0.86676473 | -2.762016 | 0.00574457 | 0.32369964 |
| F2R | 1726.92224 | -0.0906706 | 0.34505679 | -0.26277 | 0.79272785 | 0.97090793 |
| F2RL1 | 2905.11164 | 0.26432234 | 0.18206352 | 1.45181387 | 0.14655337 | 0.7330015 |
| CRHBP | 1262.32463 | 1.62931472 | 0.76262344 | 2.13646032 | 0.03264191 | 0.55104386 |
| AGGF1 | 1606.53617 | 0.12368148 | 0.16980454 | 0.72837557 | 0.46638371 | 0.89960479 |
| AC008581.2 | 783.905186 | 0.17213845 | 0.23161917 | 0.74319606 | 0.45736299 | 0.89688594 |
| ZBED3 | 1755.14249 | 0.1542883 | 0.26710218 | 0.57763775 | 0.56350871 | 0.92452095 |
| AC008581.1 | 772.106372 | 0.11139267 | 0.28078836 | 0.39671401 | 0.69157837 | 0.95159158 |
| ZBED3-AS1 | 220.810767 | 0.28982996 | 0.4199766 | 0.69010978 | 0.49012515 | 0.90600166 |
| AC022414.1 | 355.254975 | 0.21913051 | 0.28448752 | 0.77026405 | 0.44114328 | 0.89290667 |
| PDE8B | 667.149541 | 0.16814468 | 0.336576 | 0.49957417 | 0.61737495 | 0.93910158 |
| WDR41 | 1427.38343 | 0.14526073 | 0.18858111 | 0.77028248 | 0.44113235 | 0.89290667 |
| TBCA | 3381.63437 | -0.20044 | 0.14592893 | -1.3735455 | 0.16958281 | 0.7586159 |
| AP3B1 | 2842.32903 | 0.25792622 | 0.13963205 | 1.84718496 | 0.06472034 | 0.62073893 |
| SCAMP1-AS1 | 580.403023 | -0.597416 | 0.43363027 | -1.3777082 | 0.1682934 | 0.75735257 |
| SCAMP1 | 2199.32281 | -0.0512903 | 0.15540608 | -0.3300403 | 0.74136953 | 0.96109001 |
| LHFPL2 | 1116.41121 | -0.6476174 | 0.39101957 | -1.6562277 | 0.09767572 | 0.66549286 |
| ARSB | 1520.14522 | 0.38654698 | 0.23613624 | 1.63696593 | 0.10163759 | 0.67279543 |
| DMGDH | 9779.83592 | 1.07713519 | 0.48614487 | 2.2156671 | 0.02671431 | 0.52678638 |
| BHMT2 | 18282.474 | 0.76101487 | 0.39044193 | 1.94911157 | 0.0512821 | 0.58634747 |
| BHMT | 19547.8415 | 1.04764932 | 0.48961583 | 2.13973742 | 0.03237599 | 0.55014009 |
| JMY | 1456.86834 | -0.0652023 | 0.16223149 | -0.4019088 | 0.68775111 | 0.95158502 |
| HOMER1 | 517.984579 | -0.6407964 | 0.35900529 | -1.7849218 | 0.074274 | 0.63773415 |
| TENT2 | 2556.82289 | 0.17021551 | 0.16201387 | 1.05062308 | 0.29343174 | 0.83540174 |
| CMYA5 | 916.467947 | 0.08333747 | 0.31459249 | 0.26490609 | 0.79108181 | 0.97090793 |
| MTX3 | 1229.53047 | 0.06774323 | 0.15081495 | 0.44918116 | 0.65330098 | 0.94493429 |
| THBS4 | 215.606155 | 1.29247101 | 0.79342847 | 1.62896979 | 0.10331942 | 0.67439401 |
| AC012636.1 | 54.7177108 | 0.90862332 | 0.75249548 | 1.20748011 | 0.2272473 | 0.80300331 |
| SERINC5 | 1138.82033 | 0.03442701 | 0.12673065 | 0.27165495 | 0.78588734 | 0.97065795 |
| ZFYVE16 | 1659.13765 | -0.0532776 | 0.16019648 | -0.3325768 | 0.73945373 | 0.9603009 |
| AC008771.1 | 113.977922 | 0.18605187 | 0.29445711 | 0.6318471 | 0.52748679 | 0.91831212 |
| FAM151B | 104.738759 | -0.5916825 | 0.26705357 | -2.2155948 | 0.02671927 | 0.52678638 |
| ANKRD34B | 88.9884118 | 0.58697775 | 0.54002308 | 1.08694939 | 0.27705918 | 0.82779863 |
| DHFR | 746.77874 | 0.48013856 | 0.40356878 | 1.18973165 | 0.23415188 | 0.80725541 |
| MSH3 | 809.62097 | 0.35266033 | 0.24661348 | 1.43001239 | 0.15271346 | 0.7423959 |
| RASGRF2 | 987.104417 | -0.0482164 | 0.28773462 | -0.1675725 | 0.8669196 | 0.98222237 |
| CKMT2-AS1 | 367.758881 | -0.0251663 | 0.26733294 | -0.0941384 | 0.92499919 | 0.98869145 |
| CKMT2 | 754.375091 | 0.70821031 | 0.35110166 | 2.017109 | 0.04368414 | 0.57051182 |
| ZCCHC9 | 1015.63322 | 0.06807501 | 0.12567952 | 0.54165556 | 0.58805581 | 0.93289552 |
| ACOT12 | 120.295576 | 1.39940613 | 0.44622824 | 3.13607704 | 1.71E-03 | 1.83E-01 |
| SSBP2 | 1113.00063 | 0.08072998 | 0.27281145 | 0.2959186 | 0.76729224 | 0.96749893 |
| ATG10 | 18926.873 | 0.17904913 | 0.24304364 | 0.73669538 | 0.46130762 | 0.89755243 |
| RPS23 | 22953.3241 | 0.15511784 | 0.24889313 | 0.6232307 | 5.33E-01 | 9.20E-01 |
| ATP6AP1L | 312.919267 | 0.48710852 | 0.38610343 | 1.26160113 | 0.20709235 | 0.78790396 |
| AC104118.1 | 46.2732469 | -0.5516446 | 0.54821811 | -1.0062503 | 0.3142952 | 0.84696964 |
| TMEM167A | 3830.91352 | -0.097489 | 0.22675325 | -0.4299342 | 6.67E-01 | 9.48E-01 |
| XRCC4 | 386.9988 | 0.08119321 | 0.15835155 | 0.51274023 | 0.60813305 | 0.93748023 |
| VCAN | 2049.66282 | 0.53031921 | 0.5812168 | 0.91242927 | 3.62E-01 | 0.86528137 |
| VCAN-AS1 | 400.817631 | 0.55076873 | 0.58210102 | 0.94617379 | 0.34405995 | 0.85961157 |
| HAPLN1 | 44.4563411 | -0.108339 | 1.15488563 | -0.0938093 | 0.92526063 | 0.98871967 |
| EDIL3 | 784.840669 | 0.25490704 | 0.43949984 | 0.57999347 | 0.56191902 | 0.92452095 |

| | | | | | | |
|---|---|---|---|---|---|---|
| COX7C | 13753.1713 | 0.14788759 | 0.15442255 | 0.95768134 | 0.33822347 | 0.8571036 |
| RASA1 | 1754.64883 | -0.0595186 | 0.1221991 | -0.4870627 | 0.62621393 | 0.94088547 |
| CCNH | 2195.2485 | 0.08172137 | 0.11549452 | 0.70757794 | 0.4792074 | 0.90480914 |
| AC018754.1 | 328.303871 | 0.1485364 | 0.23423083 | 0.63414538 | 0.52598595 | 0.91769856 |
| TMEM161B | 1144.62544 | -0.0462824 | 0.15834045 | -0.2922966 | 0.77005985 | 0.96836854 |
| TMEM161B-. | 720.716626 | 0.0814592 | 0.31677344 | 0.25715288 | 0.79706076 | 0.9713715 |
| MEF2C-AS2 | 77.3215495 | -0.2741806 | 0.37632238 | -0.7285789 | 0.46625926 | 0.89960479 |
| MEF2C | 1177.31359 | -0.2407723 | 0.25361103 | -0.9493764 | 0.3424292 | 0.85889221 |
| MEF2C-AS1 | 43.1659098 | -0.5648902 | 0.48938538 | -1.154285 | 0.24838333 | 0.8162591 |
| CETN3 | 607.058609 | 0.33198828 | 0.21209743 | 1.56526308 | 0.11752124 | 0.69036323 |
| MBLAC2 | 317.523049 | 0.1898904 | 0.26516871 | 0.71611166 | 0.47392241 | 0.90199039 |
| POLR3G | 122.568004 | -0.0114161 | 0.28017565 | -0.0407463 | 0.96749812 | 0.99400331 |
| LYSMD3 | 1221.77372 | -0.0965311 | 0.23917926 | -0.403593 | 0.68651203 | 0.95097288 |
| ADGRV1 | 746.873916 | 0.18389102 | 0.46010708 | 0.39967005 | 0.68939955 | 0.95158502 |
| LUCAT1 | 79.1701583 | -1.1161237 | 0.91485967 | -1.2199944 | 0.222467 | 0.80078694 |
| ARRDC3 | 9211.70743 | 0.22893113 | 0.43576387 | 0.52535592 | 0.59933579 | 0.93487648 |
| NR2F1-AS1 | 764.712898 | -0.0908145 | 0.28316155 | -0.3207162 | 0.74842545 | 0.96241535 |
| NR2F1 | 3137.43613 | -0.2525297 | 0.36078026 | -0.6999543 | 0.48395583 | 0.90541935 |
| FAM172A | 995.377478 | 0.25010344 | 0.1997596 | 1.25202212 | 0.2105618 | 0.79134644 |
| KIAA0825 | 64.3268899 | 0.06981498 | 0.36825521 | 0.18958314 | 0.8496358 | 0.97783251 |
| SLF1 | 259.971712 | -0.1199971 | 0.23691499 | -0.5064987 | 6.13E-01 | 9.39E-01 |
| MCTP1 | 303.348081 | -0.4667532 | 0.6775494 | -0.6888844 | 4.91E-01 | 9.06E-01 |
| FAM81B | 57.1705626 | 0.14448688 | 0.88558898 | 0.16315343 | 0.87039763 | 0.98275231 |
| TTC37 | 2993.40895 | 4.40E-05 | 0.10983459 | 0.00040037 | 0.99968055 | 0.99994357 |
| ARSK | 468.95948 | 0.03573362 | 0.20634604 | 0.1731733 | 0.86251521 | 0.98176973 |
| RFESD | 79.4057536 | 0.07481719 | 0.37109286 | 0.20161313 | 0.84021918 | 0.97618283 |
| SPATA9 | 64.1825969 | 0.05491952 | 0.37012593 | 0.14838063 | 8.82E-01 | 0.98275231 |
| AC008840.1 | 61.2030161 | -0.3166375 | 0.56905199 | -0.5564298 | 0.57791707 | 0.93130183 |
| RHOBTB3 | 3030.1329 | -0.0792751 | 0.32121756 | -0.2467958 | 0.80506628 | 0.97303568 |
| GLRX | 9070.03045 | 0.23777713 | 0.38802826 | 0.61278303 | 0.54001982 | 0.92222629 |
| AC008592.4 | 79.3016372 | 1.14308004 | 0.71461753 | 1.59956898 | 0.10969423 | 0.67859564 |
| LINC01554 | 343.758687 | 1.24213032 | 0.68347344 | 1.81737907 | 0.06915909 | 0.62779744 |
| ELL2 | 2983.55295 | -0.5707636 | 0.27844777 | -2.0498051 | 0.04038346 | 0.55657615 |
| AC099509.1 | 41.7930159 | 0.1954784 | 0.87166113 | 0.22425963 | 0.82255529 | 0.97519297 |
| CAST | 8965.86717 | -0.056458 | 0.13430198 | -0.4203808 | 0.67420726 | 0.94935413 |
| AC020900.1 | 539.070995 | -0.0475722 | 0.25751009 | -0.184739 | 0.85343375 | 0.97895715 |
| ERAP1 | 3753.98999 | 0.01915472 | 0.16273316 | 0.1177063 | 0.90630038 | 0.98662757 |
| AC008906.1 | 175.931249 | -0.4355635 | 0.34391768 | -1.2664761 | 0.20534269 | 0.78571765 |
| AC009126.1 | 126.322941 | 0.13195244 | 0.34673899 | 0.38055265 | 0.70353522 | 0.95467045 |
| ERAP2 | 1332.79064 | 1.54464056 | 0.71444809 | 2.1620053 | 0.03061777 | 0.54443002 |
| LNPEP | 3291.11753 | 0.09426351 | 0.18195029 | 0.51807287 | 0.60440743 | 0.93659663 |
| LIX1-AS1 | 49.8513329 | 0.01751929 | 0.31813648 | 0.05506847 | 0.95608392 | 0.99185043 |
| LIX1 | 171.824395 | 2.77108231 | 0.83126937 | 3.33355519 | 0.00085744 | 0.13734375 |
| RIOK2 | 789.015434 | -0.0472754 | 0.21587428 | -0.2189952 | 0.82665378 | 0.97548201 |
| RGMB | 688.126526 | -0.1222833 | 0.21596955 | -0.5662062 | 0.57125358 | 0.92828088 |
| AC008522.1 | 461.797995 | -0.1363994 | 0.22953993 | -0.5942295 | 0.5523586 | 0.92373235 |
| CHD1 | 1807.7876 | -0.1665741 | 0.15151543 | -1.0993873 | 0.27159919 | 0.82716979 |
| LINC02062 | 82.4922095 | 0.26381689 | 0.41903227 | 0.62958609 | 0.52896543 | 0.91907723 |
| FAM174A | 1213.07732 | 0.19909187 | 0.15848615 | 1.25620985 | 0.2090399 | 0.79086734 |
| ST8SIA4 | 240.597529 | -0.2746738 | 0.50472194 | -0.5442081 | 0.58629828 | 0.93289552 |
| SLCO4C1 | 3065.84356 | 0.24022277 | 0.35144392 | 0.68353087 | 0.49427145 | 0.90796767 |
| PAM | 10393.246 | 0.26024985 | 0.19409852 | 1.34081318 | 0.17998112 | 0.76318604 |
| GIN1 | 387.703971 | 0.10669698 | 0.14227599 | 0.74992963 | 0.45329709 | 0.89670027 |
| PPIP5K2 | 1665.97751 | 0.07201099 | 0.17979799 | 0.40051056 | 0.68878051 | 0.95158502 |

| | | | | | | |
|---|---|---|---|---|---|---|
| C5orf30 | 586.198183 | 0.34736376 | 0.27905759 | 1.24477445 | 0.21321467 | 0.7926732 |
| NUDT12 | 1375.52114 | 0.3277091 | 0.22476856 | 1.45798464 | 0.14484478 | 0.73039059 |
| EFNA5 | 1299.98492 | -0.3867268 | 0.3048385 | -1.2686286 | 0.20457358 | 0.78460928 |
| FBXL17 | 1236.26612 | 0.3337154 | 0.16487463 | 2.02405553 | 0.04296444 | 0.56664077 |
| LINC01023 | 135.979265 | -0.1312 | 0.339721 | -0.3861993 | 0.69934911 | 0.95307625 |
| FER | 809.734932 | -0.0196104 | 0.18023329 | -0.1088055 | 0.91335674 | 0.98662757 |
| AC008467.1 | 80.5576506 | -0.3697991 | 0.43394803 | -0.8521738 | 0.39411765 | 0.87884163 |
| PJA2 | 5395.87053 | 0.02580223 | 0.16415439 | 0.15718269 | 0.87510087 | 0.98275231 |
| MAN2A1 | 3280.35348 | 0.01347299 | 0.23363824 | 0.05766604 | 0.95401464 | 0.99170503 |
| TMEM232 | 176.139133 | 0.28041599 | 0.50595388 | 0.55423232 | 0.57941986 | 0.93139743 |
| SLC25A46 | 2857.63106 | -0.0873872 | 0.21880605 | -0.3993819 | 0.68961185 | 0.95158502 |
| TSLP | 49.1309384 | 0.13141163 | 0.50560134 | 0.25991155 | 0.794932 | 0.97101097 |
| WDR36 | 1249.30666 | -0.1777674 | 0.15550335 | -1.1431743 | 0.25296625 | 0.81835388 |
| CAMK4 | 119.435457 | 0.18795111 | 0.56786916 | 0.33097609 | 0.74066255 | 0.96089108 |
| STARD4 | 1094.36569 | -0.2766696 | 0.29230159 | -0.9465209 | 0.34388294 | 0.85961157 |
| STARD4-AS1 | 929.367341 | -0.2312319 | 0.26918181 | -0.8590175 | 0.39033085 | 0.87650906 |
| NREP | 2104.47889 | -0.2497704 | 0.29076933 | -0.8589983 | 0.39034147 | 0.87650906 |
| EPB41L4A | 1935.92326 | -0.2030903 | 0.14432664 | -1.4071574 | 0.15938073 | 0.74762924 |
| EPB41L4A-A! | 2389.68859 | -0.2670899 | 0.21844175 | -1.2227053 | 0.22144101 | 0.80020811 |
| AC010261.2 | 105.995407 | -0.0641651 | 0.29972713 | -0.2140785 | 0.83048585 | 0.97550416 |
| AC104126.1 | 59.1864573 | 2.44298727 | 1.08539147 | 2.25078908 | 0.0243989 | 0.51967919 |
| EPB41L4A-D' | 116.459294 | -0.6843272 | 0.31935315 | -2.142854 | 0.03212482 | 0.54881614 |
| APC | 1153.59306 | -0.4318181 | 0.27926841 | -1.5462475 | 0.1220448 | 0.69785915 |
| AC008575.1 | 1110.93889 | -0.2932509 | 0.24722133 | -1.1861878 | 2.36E-01 | 0.80747753 |
| SRP19 | 1615.58792 | -0.3140776 | 0.22486258 | -1.3967533 | 0.16248778 | 0.75026273 |
| REEP5 | 11805.1515 | 0.0271983 | 0.1531658 | 0.17757427 | 0.85905734 | 0.98000934 |
| DCP2 | 1735.76058 | -0.0582319 | 0.13786967 | -0.4223691 | 0.67275566 | 0.94922827 |
| MCC | 890.629343 | 0.29850784 | 0.20753887 | 1.43832257 | 0.15034255 | 0.73796431 |
| YTHDC2 | 1827.70374 | 0.05276481 | 0.09949112 | 0.53034696 | 0.5958714 | 0.9334601 |
| KCNN2 | 88.7411798 | -0.1631629 | 0.58366866 | -0.2795472 | 0.77982495 | 0.96836854 |
| TRIM36 | 51.2759095 | -0.0209438 | 0.65658912 | -0.0318979 | 0.97455347 | 0.99455099 |
| PGGT1B | 1256.79674 | 0.0009643 | 0.15697721 | 0.00614293 | 0.99509868 | 0.99940889 |
| AC008494.3 | 64.7110525 | 0.25146755 | 0.4036802 | 0.62293753 | 0.53332558 | 0.92049136 |
| CCDC112 | 186.094385 | 0.1684397 | 0.25969892 | 0.64859606 | 0.5165995 | 0.91636259 |
| FEM1C | 1462.14735 | -0.4910364 | 0.29058785 | -1.6898036 | 0.09106553 | 0.65495306 |
| TMED7-TICA | 968.447817 | 0.03429011 | 0.22935734 | 0.14950516 | 0.88115504 | 0.98275231 |
| TICAM2 | 361.393505 | -0.027048 | 0.30791494 | -0.0878426 | 9.30E-01 | 9.90E-01 |
| AC010226.1 | 414.385428 | 0.05365476 | 0.28996173 | 0.18504083 | 0.85319701 | 0.97886533 |
| TMED7 | 4602.02958 | -0.0776188 | 0.20745092 | -0.3741548 | 0.70828912 | 0.95501548 |
| CDO1 | 180.411862 | 0.48524808 | 0.57181529 | 0.84860983 | 0.39609843 | 0.87967259 |
| ATG12 | 1793.70316 | -0.1141706 | 0.12407558 | -0.9201696 | 0.35748414 | 0.86390849 |
| AP3S1 | 2200.3584 | 0.07251659 | 0.18493644 | 0.39211631 | 0.69497228 | 0.95234774 |
| AC034236.2 | 264.116381 | 0.2179181 | 0.24460083 | 0.89091317 | 0.37297576 | 0.86791605 |
| COMMD10 | 1245.67509 | 0.34195722 | 0.18607065 | 1.83778158 | 0.0660946 | 0.62276463 |
| SEMA6A | 2342.22113 | -0.7010041 | 0.42387797 | -1.6537875 | 0.0981707 | 0.66549286 |
| SEMA6A-AS1 | 199.160304 | -0.5813607 | 0.31296645 | -1.8575815 | 0.06322847 | 0.61588607 |
| DTWD2 | 401.529961 | 0.33571724 | 0.18441233 | 1.82047068 | 0.06868736 | 0.62750387 |
| DMXL1 | 2234.40779 | 0.17233746 | 0.22988718 | 0.74966104 | 0.45345888 | 0.89670027 |
| TNFAIP8 | 1236.17502 | 0.1204572 | 0.2542291 | 0.47381356 | 0.63563286 | 0.94088547 |
| HSD17B4 | 7744.23983 | 0.30060963 | 0.28046429 | 1.07182853 | 0.28379705 | 0.83107779 |
| PRR16 | 119.941717 | -0.8726418 | 0.59882605 | -1.4572542 | 0.14504624 | 0.73064288 |
| SRFBP1 | 431.764773 | -0.073976 | 0.25875854 | -0.2858882 | 0.77496374 | 0.96836854 |
| LOX | 732.738953 | 0.03687363 | 0.37288389 | 0.0988877 | 0.92122743 | 0.98779251 |
| SNCAIP | 166.010137 | -0.1166303 | 0.43187248 | -0.2700573 | 0.78711619 | 0.97090793 |

| | | | | | | |
|---|---|---|---|---|---|---|
| SNX2 | 4565.99851 | 0.0102674 | 0.14428428 | 0.07116093 | 0.94326968 | 0.99136103 |
| SNX24 | 1620.51207 | 0.04636965 | 0.1766868 | 0.26243984 | 0.79298236 | 0.97090793 |
| PPIC | 2207.27805 | -0.057429 | 0.24954122 | -0.2301383 | 0.81798434 | 0.97446554 |
| AC106786.2 | 510.522118 | 0.00212665 | 0.24687653 | 0.00861422 | 0.99312693 | 0.99906069 |
| PRDM6 | 131.551546 | -0.1358247 | 0.29956755 | -0.4534027 | 0.6502588 | 0.94322083 |
| CEP120 | 851.191739 | 0.06641701 | 0.18352619 | 0.36189392 | 0.7174313 | 0.95611001 |
| CSNK1G3 | 1108.61509 | -0.0712112 | 0.12135463 | -0.5868029 | 0.55733609 | 0.92376719 |
| ZNF608 | 836.569687 | -0.0723135 | 0.25132275 | -0.2877315 | 0.77355228 | 0.96836854 |
| GRAMD2B | 3010.89536 | -0.0826396 | 0.22093649 | -0.3740421 | 0.70837298 | 0.95501548 |
| AC093535.1 | 537.831685 | -0.071104 | 0.2767744 | -0.2569022 | 0.79725425 | 0.9713715 |
| ALDH7A1 | 6617.07554 | 0.57257358 | 0.27511412 | 2.08122206 | 0.03741359 | 0.55532957 |
| PHAX | 1255.57 | -0.2015242 | 0.23786637 | -0.8472161 | 0.39687467 | 0.87988087 |
| LMNB1 | 303.376893 | 0.52806619 | 0.53858563 | 0.9804684 | 0.32685496 | 0.8529085 |
| 3-Mar | 220.217576 | 0.40622522 | 0.48802779 | 0.83238134 | 0.40519373 | 0.88132015 |
| C5orf63 | 244.508523 | 0.06437194 | 0.28822802 | 0.22333687 | 0.82327333 | 0.97542663 |
| PRRC1 | 2494.90308 | -0.1648429 | 0.29551931 | -0.5578077 | 0.57697572 | 0.93129843 |
| CTXN3 | 1074.90125 | 1.25664483 | 0.7991842 | 1.5724095 | 0.11585561 | 0.6873836 |
| CCDC192 | 67.3118003 | 0.47174535 | 0.36386131 | 1.29649768 | 0.19480408 | 0.77644051 |
| LINC01184 | 804.326577 | 0.2050026 | 0.19912836 | 1.02949973 | 0.30324491 | 0.84211658 |
| SLC12A2 | 874.049485 | 0.0298325 | 0.2562582 | 0.1164158 | 0.90732302 | 0.98662757 |
| FBN2 | 50.6452668 | 0.88022889 | 0.57876258 | 1.52088078 | 0.12828976 | 0.70596572 |
| ISOC1 | 1690.28192 | 0.09124139 | 0.19724376 | 0.4625819 | 0.64366409 | 0.94197319 |
| ADAMTS19 | 47.1919137 | 0.21888271 | 1.00757877 | 0.21723633 | 0.82802418 | 0.97548201 |
| CHSY3 | 48.8193894 | 0.28198499 | 0.43841532 | 0.64319146 | 0.52009987 | 0.91657872 |
| HINT1 | 12077.6778 | 0.24479394 | 0.16653681 | 1.46990893 | 0.14158642 | 0.72784886 |
| LYRM7 | 817.669461 | 0.20801488 | 0.19163428 | 1.08547844 | 0.27770981 | 0.82805357 |
| CDC42SE2 | 5935.11805 | -0.079988 | 0.19863389 | -0.4026905 | 0.68717589 | 0.95138142 |
| RAPGEF6 | 624.074989 | -0.1982963 | 0.16397049 | -1.2093416 | 0.22653165 | 0.80269044 |
| AC008695.1 | 159.725654 | -0.1979293 | 0.28201803 | -0.7018322 | 0.48278384 | 0.90541935 |
| FNIP1 | 1357.67869 | -0.6608476 | 0.3328338 | -1.9855184 | 0.04708682 | 0.58171004 |
| AC026398.1 | 103.419802 | 0.77069369 | 0.45827379 | 1.68173199 | 0.09262082 | 0.65662464 |
| ACSL6 | 236.127577 | 0.69565532 | 0.49005394 | 1.41954847 | 0.15573918 | 0.7451386 |
| P4HA2-AS1 | 535.12813 | -0.1416678 | 0.31757344 | -0.4460946 | 0.65552893 | 0.94532497 |
| P4HA2 | 3554.23169 | -0.0871032 | 0.35868989 | -0.242837 | 8.08E-01 | 0.97334138 |
| PDLIM4 | 963.080747 | -0.495227 | 0.49995548 | -0.9905423 | 0.32190914 | 0.85119105 |
| SLC22A4 | 827.646989 | -0.2163813 | 0.363487 | -0.595293 | 0.55164762 | 0.92373235 |
| SLC22A5 | 6598.53576 | 0.37160347 | 0.53146205 | 0.6992098 | 0.48442093 | 0.90541935 |
| AC116366.3 | 3053.32414 | -0.0899003 | 0.17343944 | -0.5183385 | 0.60422212 | 0.93655508 |
| AC116366.1 | 141.582407 | -0.9451544 | 0.51101235 | -1.8495724 | 0.0643752 | 0.61964848 |
| IRF1 | 3292.40526 | -0.1758785 | 0.46977257 | -0.3743908 | 0.70811358 | 0.95501548 |
| RAD50 | 2352.64303 | -0.0256973 | 0.15362875 | -0.1672688 | 0.86715855 | 0.98223481 |
| TH2LCRR | 452.808435 | 0.07782039 | 0.18576016 | 0.41892942 | 0.67526772 | 0.94958156 |
| KIF3A | 760.757626 | -0.0326892 | 0.17490145 | -0.1869009 | 0.85173831 | 0.97872457 |
| CCNI2 | 314.872295 | 0.48118808 | 0.32022133 | 1.50267341 | 0.13292328 | 0.71419589 |
| 8-Sep | 3788.81024 | 0.19896884 | 0.29788224 | 0.66794463 | 0.50416894 | 0.91356552 |
| SOWAHA | 556.179371 | 0.70023486 | 0.34199279 | 2.04751348 | 0.04060769 | 0.55657615 |
| SHROOM1 | 1427.38482 | 0.12460444 | 0.53007373 | 0.23507001 | 0.81415439 | 0.9738447 |
| GDF9 | 2937.6614 | 0.35261192 | 0.20111138 | 1.75331658 | 0.07954768 | 0.64079926 |
| UQCRQ | 6275.48126 | 0.278417 | 0.17992089 | 1.54744119 | 0.1217569 | 0.69725141 |
| LEAP2 | 392.285719 | 0.27759779 | 0.52365444 | 0.53011637 | 0.59603125 | 0.9334601 |
| AFF4 | 4870.88856 | -0.3116355 | 0.16374078 | -1.9032248 | 5.70E-02 | 6.00E-01 |
| ZCCHC10 | 729.180312 | -0.0952893 | 0.12422315 | -0.767082 | 0.44303275 | 0.89352768 |
| HSPA4 | 5303.57889 | -0.0556376 | 0.27030855 | -0.2058299 | 0.83692379 | 0.97577999 |
| FSTL4 | 214.715191 | 0.51003992 | 0.6381345 | 0.79926711 | 0.42413554 | 0.88565404 |

| | | | | | | |
|---|---|---|---|---|---|---|
| C5orf15 | 4942.46191 | 0.1622978 | 0.17561862 | 0.92414916 | 0.35540866 | 0.86364633 |
| VDAC1 | 13657.0212 | 0.18603188 | 0.22840827 | 0.81447087 | 0.41537526 | 0.88381712 |
| AC008608.2 | 59.4571997 | -0.1719268 | 0.36993072 | -0.4647541 | 0.64210754 | 0.94088547 |
| TCF7 | 410.931758 | -0.1993894 | 0.57166105 | -0.3487896 | 0.7272473 | 0.95848028 |
| SKP1 | 14839.4798 | 0.14217607 | 0.14370387 | 0.98936839 | 0.32248294 | 0.85119105 |
| AC104109.3 | 6860.53287 | 0.15126048 | 0.17684716 | 0.85531758 | 3.92E-01 | 8.78E-01 |
| PPP2CA | 7069.90468 | -0.2372605 | 0.1853157 | -1.2803044 | 0.20043809 | 0.78134034 |
| CDKL3 | 182.984911 | 0.24362462 | 0.20124779 | 1.21057041 | 0.22606009 | 0.80246466 |
| UBE2B | 3502.82166 | -0.2560473 | 0.12837776 | -1.994483 | 0.04609929 | 0.57789088 |
| CDKN2AIPNL | 732.107057 | 0.03064704 | 0.1477265 | 0.20745794 | 0.83565224 | 0.97550416 |
| JADE2 | 3069.14438 | 0.00200867 | 0.2948036 | 0.00681359 | 0.99456359 | 0.99940889 |
| SAR1B | 3915.15425 | 0.10616697 | 0.21752055 | 0.48807788 | 0.62549468 | 0.94088547 |
| SEC24A | 1445.53561 | -0.2761679 | 0.27500354 | -1.0042341 | 0.3152658 | 0.84696964 |
| CAMLG | 1841.58718 | 0.06548743 | 0.15072011 | 0.43449695 | 0.66392761 | 0.94642611 |
| DDX46 | 2563.94481 | -0.1320167 | 0.14770113 | -0.8938095 | 0.37142381 | 0.86762749 |
| C5orf24 | 2856.09843 | -0.048026 | 0.12731567 | -0.3772198 | 0.70601028 | 0.95501548 |
| TXNDC15 | 2758.38214 | 0.15228739 | 0.17892842 | 0.85110793 | 0.3947094 | 0.87884163 |
| PCBD2 | 763.501719 | 0.26295935 | 0.25915389 | 1.01468418 | 0.3102564 | 0.84456632 |
| CATSPER3 | 129.19453 | 0.34396135 | 0.29541593 | 1.16432905 | 0.24429069 | 0.81468457 |
| C5orf66 | 1620.31271 | -0.1087966 | 0.15857997 | -0.6860675 | 0.49267054 | 0.90696683 |
| H2AFY | 5897.88435 | -0.1741799 | 0.15877633 | -1.0970142 | 0.27263521 | 0.82716979 |
| AC026691.1 | 43.1572791 | 0.69074803 | 0.62744938 | 1.10088248 | 0.27094781 | 0.82716979 |
| CXCL14 | 22939.7904 | 0.85151256 | 0.46368946 | 1.83638541 | 0.06630068 | 0.62373361 |
| SLC25A48 | 561.637682 | 0.95644306 | 0.69886067 | 1.36857475 | 0.17113224 | 0.75956362 |
| LECT2 | 90.6147723 | -0.4536397 | 0.60835857 | -0.7456782 | 0.4558618 | 0.89688594 |
| TGFBI | 5098.89981 | 0.08147728 | 0.51135699 | 0.15933542 | 0.87340462 | 0.98275231 |
| SMAD5 | 2154.06102 | 0.01265134 | 0.13908274 | 0.09096271 | 0.92752222 | 0.98915637 |
| SMIM32 | 2369.09338 | 0.50762087 | 0.48983074 | 1.03631895 | 3.00E-01 | 8.42E-01 |
| TRPC7 | 127.156366 | -0.2552065 | 0.82344336 | -0.309926 | 0.75661727 | 0.9648441 |
| TRPC7-AS1 | 62.5021463 | -0.1203961 | 0.72260694 | -0.1666136 | 0.86767411 | 0.98224588 |
| SPOCK1 | 2284.09836 | 0.47581296 | 0.50215144 | 0.94754872 | 0.34335924 | 0.85938376 |
| KLHL3 | 882.044936 | 0.37099445 | 0.352404 | 1.05275323 | 0.29245411 | 0.83524572 |
| HNRNPA0 | 6681.84598 | -0.2229279 | 0.19971567 | -1.1162262 | 0.26432534 | 0.82424998 |
| AC106791.1 | 41.2121933 | -0.2187903 | 0.54539405 | -0.40116 | 0.6883023 | 0.95158502 |
| MYOT | 87.4354139 | 0.19856819 | 0.61238173 | 0.32425558 | 0.74574455 | 0.96176407 |
| PKD2L2 | 348.69281 | -0.5198557 | 0.26464676 | -1.9643383 | 0.04949088 | 0.5861488 |
| FAM13B | 1612.81459 | -0.4972518 | 0.20537883 | -2.4211441 | 0.01547174 | 0.45305611 |
| NME5 | 671.029151 | 0.24234244 | 0.47952523 | 0.50537995 | 0.613292 | 0.9388166 |
| BRD8 | 3314.13911 | -0.1595318 | 0.1853726 | -0.8606009 | 0.38945789 | 0.87568585 |
| KIF20A | 72.5418622 | 0.27077232 | 0.56661814 | 0.47787442 | 0.63273959 | 0.94088547 |
| CDC23 | 1566.52451 | 0.05636296 | 0.14818879 | 0.38034562 | 0.70368888 | 0.9547407 |
| CDC25C | 48.5460742 | -0.6895746 | 0.55117716 | -1.2510944 | 0.21090006 | 0.79134644 |
| FAM53C | 2076.80657 | -0.5872214 | 0.24684827 | -2.3788759 | 0.01736552 | 0.46859699 |
| AC104116.1 | 606.443276 | -0.6184933 | 0.22398934 | -2.7612621 | 0.00575784 | 0.32369964 |
| KDM3B | 3577.21124 | -0.0093912 | 0.10078782 | -0.0931776 | 0.92576246 | 0.98876911 |
| REEP2 | 113.399446 | 0.30617474 | 0.27883428 | 1.09805273 | 0.27218147 | 0.82716979 |
| EGR1 | 7728.44628 | -0.1368218 | 0.75896716 | -0.1802736 | 0.85693777 | 0.97953896 |
| ETF1 | 5792.86583 | -0.319296 | 0.25064462 | -1.2738993 | 0.20269912 | 0.78193335 |
| HSPA9 | 20623.5022 | -0.1483031 | 0.33445207 | -0.4434211 | 0.65746116 | 0.94558535 |
| CTNNA1 | 19868.4829 | -0.1021915 | 0.18262479 | -0.5595706 | 0.57577236 | 0.93083012 |
| SIL1 | 2751.50335 | 0.05329825 | 0.18039528 | 0.29545256 | 0.76764819 | 0.96754958 |
| AC011405.1 | 889.058575 | 0.1427777 | 0.18175689 | 0.78554213 | 0.43213578 | 0.88976671 |
| MATR3 | 11010.0552 | -0.0416877 | 0.1128689 | -0.3693466 | 0.71186939 | 0.95560193 |
| SNHG4 | 153.057666 | -0.4602304 | 0.40208422 | -1.144612 | 0.25236991 | 0.81833349 |

| | | | | | | |
|---|---|---|---|---|---|---|
| PAIP2 | 5439.99812 | -0.179349 | 0.15313222 | -1.1712038 | 0.24151688 | 0.81261543 |
| AC135457.1 | 3482.93867 | -0.1725223 | 0.16142754 | -1.0687292 | 0.2851917 | 0.83107779 |
| SLC23A1 | 2758.05424 | 0.08258757 | 0.48983403 | 0.16860316 | 0.86610879 | 0.98197545 |
| MZB1 | 325.482548 | -0.8447578 | 0.95188979 | -0.8874534 | 0.37483484 | 0.86791605 |
| PROB1 | 648.335599 | -0.6549877 | 0.32043302 | -2.0440707 | 0.04094657 | 0.55811692 |
| SPATA24 | 214.652023 | 0.12562046 | 0.25649432 | 0.48975921 | 0.6243043 | 0.94075157 |
| DNAJC18 | 455.590791 | -0.1143459 | 0.24800913 | -0.4610553 | 0.64475896 | 0.94208859 |
| AC142391.1 | 970.03652 | -0.2270859 | 0.39860981 | -0.5696947 | 0.56888477 | 0.92660886 |
| ECSCR | 1026.50445 | -0.2386228 | 0.3994788 | -0.5973353 | 0.55028353 | 0.92373235 |
| TMEM173 | 3066.78027 | -0.0428584 | 0.27433196 | -0.1562284 | 0.87585299 | 0.98275231 |
| UBE2D2 | 4420.0462 | -0.1528465 | 0.18430152 | -0.8293287 | 0.40691843 | 0.88132015 |
| CXXC5 | 5804.34336 | 0.15694936 | 0.17587568 | 0.89238809 | 3.72E-01 | 0.86775997 |
| CXXC5-AS1 | 90.1117258 | 0.58689483 | 0.36577977 | 1.60450325 | 0.10860317 | 0.67800851 |
| NRG2 | 102.12635 | 0.04963346 | 0.51745903 | 0.09591767 | 0.92358596 | 0.98858365 |
| MALINC1 | 190.152427 | 0.16333592 | 0.26545161 | 0.61531337 | 0.53834779 | 0.92155425 |
| PURA | 1713.86117 | -0.2306866 | 0.16347086 | -1.411179 | 1.58E-01 | 0.74620522 |
| IGIP | 1501.17513 | 0.36461207 | 0.24874759 | 1.46579133 | 1.43E-01 | 7.28E-01 |
| CYSTM1 | 12197.5055 | -0.0980767 | 0.15405946 | -0.6366161 | 0.52437493 | 0.91723878 |
| PFDN1 | 2444.17808 | -0.1127012 | 0.09809651 | -1.1488813 | 2.51E-01 | 8.17E-01 |
| HBEGF | 947.126434 | -0.2964972 | 0.49617727 | -0.5975631 | 0.55013149 | 0.92373235 |
| SLC4A9 | 1685.4212 | 1.75452714 | 0.55195086 | 3.17877417 | 0.00147899 | 0.17236139 |
| AC008438.1 | 227.347861 | 1.28856548 | 0.49742026 | 2.59049658 | 0.00958376 | 0.39368232 |
| ANKHD1 | 4623.28699 | -0.1344283 | 0.13079728 | -1.0277608 | 0.30406236 | 0.84211658 |
| ANKHD1-EIF4 | 5988.09257 | -0.0401434 | 0.14013564 | -0.2864614 | 0.77452477 | 0.96836854 |
| SRA1 | 2900.42 | -0.0218674 | 0.16713869 | -0.1308338 | 0.89590677 | 0.98474725 |
| EIF4EBP3 | 1518.65368 | 0.21140414 | 0.22054196 | 0.95856652 | 0.33777716 | 0.8571036 |
| APBB3 | 2346.28847 | 0.01663756 | 0.29797246 | 0.0558359 | 0.95547253 | 0.99185043 |
| SLC35A4 | 5826.32788 | -0.0308745 | 0.12517466 | -0.2466517 | 0.80517781 | 0.97303568 |
| CD14 | 3120.59165 | 0.04477112 | 0.5875533 | 0.07619925 | 0.93926058 | 0.99113899 |
| NDUFA2 | 3073.34762 | 0.21923105 | 0.12559514 | 1.74553765 | 0.08089132 | 0.64313548 |
| TMCO6 | 733.208991 | 0.08605142 | 0.21348608 | 0.40307746 | 0.68689124 | 0.9510919 |
| IK | 6886.99304 | 0.0020091 | 0.09552877 | 0.02103134 | 0.98322065 | 0.9961464 |
| WDR55 | 2999.83149 | -0.3210802 | 0.16872985 | -1.9029245 | 0.0570504 | 0.60049913 |
| DND1 | 902.392171 | -0.3833905 | 0.1766298 | -2.1705878 | 0.02996234 | 0.54012717 |
| HARS | 3398.59647 | -0.1359267 | 0.12338154 | -1.1016778 | 0.27060177 | 0.82716979 |
| HARS2 | 2520.28326 | 0.2093572 | 0.16354031 | 1.28015657 | 0.20049008 | 0.78134034 |
| ZMAT2 | 3853.00762 | 0.11555359 | 0.12479255 | 0.9259654 | 0.35446396 | 0.86364633 |
| PCDHA1 | 683.740513 | 0.28270388 | 0.37420738 | 0.75547382 | 0.44996474 | 0.89557948 |
| PCDHA2 | 693.651406 | 0.26001236 | 0.36834417 | 0.70589516 | 0.48025334 | 0.90541935 |
| PCDHA3 | 714.940808 | 0.2651486 | 0.37428545 | 0.70841278 | 0.47868896 | 0.90444322 |
| PCDHA4 | 706.968583 | 0.26263212 | 0.36855137 | 0.71260657 | 0.47608925 | 0.90298437 |
| PCDHA5 | 684.860925 | 0.27439311 | 0.37296766 | 0.73570214 | 0.46191199 | 0.89762182 |
| PCDHA6 | 693.175695 | 0.26682535 | 0.37258647 | 0.71614341 | 4.74E-01 | 9.02E-01 |
| PCDHA7 | 690.650146 | 0.26127366 | 0.36865516 | 0.70872103 | 0.47849761 | 0.90440142 |
| PCDHA8 | 675.729703 | 0.26620291 | 0.37193445 | 0.71572534 | 0.47416096 | 0.90199039 |
| PCDHA9 | 675.104932 | 0.27031053 | 0.37072434 | 0.72914156 | 4.66E-01 | 0.89960479 |
| PCDHA10 | 815.954053 | 0.24919289 | 0.35878555 | 0.69454551 | 0.48734017 | 0.90573694 |
| PCDHA11 | 701.439369 | 0.2664721 | 0.36834133 | 0.72343796 | 0.46941086 | 0.90050905 |
| PCDHA12 | 690.810636 | 0.26522176 | 0.37159366 | 0.71374134 | 0.47538714 | 0.90229191 |
| PCDHA13 | 675.366051 | 0.27209407 | 0.37130184 | 0.73281098 | 0.46367372 | 0.89830907 |
| PCDHAC1 | 684.550641 | 0.27077566 | 0.37258412 | 0.7267504 | 0.46737887 | 0.89983399 |
| PCDHAC2 | 682.166912 | 0.26410944 | 0.36722352 | 0.71920622 | 0.47201387 | 0.90160919 |
| PCDHB2 | 105.913947 | 0.72295789 | 0.51699388 | 1.39838771 | 0.16199667 | 0.75010861 |
| AC244517.2 | 65.4523075 | -0.5567697 | 0.59876011 | -0.9298711 | 0.35243785 | 0.86364633 |

| | | | | | | |
|---|---|---|---|---|---|---|
| PCDHB3 | 120.534025 | -0.3288466 | 0.45777692 | -0.7183555 | 0.47253811 | 0.90160919 |
| PCDHB4 | 185.531942 | -0.3433402 | 0.42355584 | -0.8106137 | 0.41758752 | 0.88445833 |
| PCDHB5 | 254.5091 | 0.25604017 | 0.3565505 | 0.71810352 | 0.47269346 | 0.90160919 |
| AC244517.11 | 61.4150509 | 0.19023662 | 0.40652537 | 0.46795755 | 0.63981495 | 0.94088547 |
| PCDHB6 | 68.0515312 | 0.06930489 | 0.50854746 | 0.13628008 | 0.89159987 | 0.98474725 |
| PCDHB7 | 108.623455 | -0.5619974 | 0.67791066 | -0.8290139 | 0.40709654 | 0.88132015 |
| PCDHB16 | 232.755774 | -0.3321713 | 0.48140623 | -0.6900022 | 0.49019283 | 0.90600166 |
| AC244517.4 | 58.0999394 | -0.4491435 | 0.52096507 | -0.8621374 | 0.38861189 | 0.87568585 |
| PCDHB9 | 54.1856768 | 0.03452384 | 0.37267084 | 0.09263896 | 0.92619039 | 0.98876911 |
| PCDHB10 | 65.0495507 | -0.1844527 | 0.56124772 | -0.3286476 | 0.74242203 | 0.96109001 |
| PCDHB11 | 87.9015924 | -0.0292534 | 0.52561822 | -0.0556553 | 0.95561645 | 0.99185043 |
| PCDHB12 | 61.1881741 | -0.0644911 | 0.58872023 | -0.1095446 | 0.91277057 | 0.98662757 |
| PCDHB13 | 103.205407 | 0.40692138 | 0.37648177 | 1.08085281 | 0.27976259 | 0.82825734 |
| PCDHB14 | 245.878197 | 0.07769613 | 0.36086761 | 0.2153037 | 0.82953055 | 0.97550416 |
| AC244517.10 | 44.4332751 | -0.1880679 | 0.4455645 | -0.4220891 | 0.67295995 | 0.94922827 |
| PCDHB15 | 251.638103 | -0.2285387 | 0.42721611 | -0.5349486 | 0.59268539 | 0.93289552 |
| TAF7 | 4428.00898 | -0.1215565 | 0.11165201 | -1.0887084 | 0.27628247 | 0.82719407 |
| PCDHGA1 | 4167.87317 | 0.14811936 | 0.27455343 | 0.53949193 | 0.58954747 | 0.93289552 |
| PCDHGA2 | 4241.65523 | 0.14209693 | 0.27405052 | 0.51850632 | 0.60410505 | 0.93655508 |
| PCDHGA3 | 4176.38575 | 0.14724173 | 0.2731193 | 0.53911143 | 0.58980997 | 0.93289552 |
| PCDHGB1 | 4176.52849 | 0.14656782 | 0.27449514 | 0.53395416 | 0.59337325 | 0.93289552 |
| PCDHGA4 | 4174.96621 | 0.14576896 | 0.27487351 | 0.53031287 | 0.59589503 | 0.9334601 |
| PCDHGB2 | 4177.85294 | 0.1486054 | 0.27417268 | 0.54201389 | 0.58780894 | 0.93289552 |
| PCDHGA5 | 4177.30167 | 0.14672213 | 0.27484655 | 0.5338329 | 0.59345715 | 0.93289552 |
| PCDHGB3 | 4157.71215 | 0.14584572 | 0.27433652 | 0.53163071 | 0.59498179 | 0.9334601 |
| PCDHGA6 | 4271.27325 | 0.13921842 | 0.27396142 | 0.50816796 | 0.61133556 | 0.93834929 |
| PCDHGA7 | 4212.03055 | 0.14613017 | 0.27364213 | 0.53401927 | 0.59332821 | 0.93289552 |
| PCDHGB4 | 4312.98561 | 0.13736224 | 0.26896671 | 0.5107035 | 0.60955869 | 0.93760173 |
| PCDHGA8 | 4261.11368 | 0.14002439 | 0.27186476 | 0.51505165 | 0.60651693 | 0.93708311 |
| PCDHGB5 | 4290.12444 | 0.14465972 | 0.2690645 | 0.53763958 | 0.5908259 | 0.93289552 |
| PCDHGA9 | 4196.97323 | 0.14064629 | 0.27441854 | 0.51252475 | 0.6082838 | 0.93748023 |
| PCDHGB6 | 4255.11436 | 0.14056573 | 0.2751017 | 0.51095916 | 0.60937965 | 0.93748023 |
| PCDHGA10 | 4205.60081 | 0.14014781 | 0.27308798 | 0.51319656 | 0.60781383 | 0.93748023 |
| PCDHGB7 | 4311.83081 | 0.14810465 | 0.27248345 | 0.54353633 | 0.58676058 | 0.93289552 |
| PCDHGA11 | 4186.15618 | 0.14561901 | 0.2722682 | 0.53483665 | 0.59276282 | 0.93289552 |
| PCDHGA12 | 4211.34927 | 0.13997828 | 0.27355908 | 0.51169306 | 0.60886584 | 0.93748023 |
| PCDHGC3 | 4477.48521 | 0.1334557 | 0.27501621 | 0.48526484 | 0.62748849 | 0.94088547 |
| PCDHGC4 | 4146.58513 | 0.14618064 | 0.27474537 | 0.53205862 | 0.5946854 | 0.9334601 |
| PCDHGC5 | 4151.12445 | 0.1455467 | 0.27469455 | 0.52984925 | 0.59621646 | 0.9334601 |
| DIAPH1 | 9304.61286 | -0.1882099 | 0.14312241 | -1.3150276 | 0.18850061 | 0.76970626 |
| AC008781.1 | 511.325914 | 0.23972755 | 0.13104968 | 1.82928757 | 0.06735654 | 0.6249306 |
| HDAC3 | 2787.00115 | 0.15586622 | 0.1115641 | 1.39710014 | 0.16238346 | 0.75026273 |
| RELL2 | 1366.64568 | -0.2264098 | 0.30171644 | -0.7504059 | 0.45301031 | 0.89670027 |
| FCHSD1 | 2126.07718 | -0.2596298 | 0.21861076 | -1.1876351 | 0.2349772 | 0.80747753 |
| ARAP3 | 2097.51356 | -0.3481005 | 0.2896739 | -1.2016978 | 0.22948064 | 0.80551898 |
| PCDH1 | 4377.08894 | 0.1014379 | 0.23290312 | 0.43553688 | 0.66317278 | 0.94642611 |
| DELE1 | 4503.33902 | -0.01383 | 0.11184166 | -0.1236567 | 0.90158712 | 0.98572048 |
| PCDH12 | 1255.01027 | -0.224306 | 0.27159446 | -0.8258859 | 0.40886883 | 0.88207556 |
| RNF14 | 3431.41217 | 0.19566814 | 0.2388738 | 0.81912765 | 4.13E-01 | 0.88285886 |
| AC005740.3 | 176.273001 | 0.17726676 | 0.35780634 | 0.49542654 | 0.62029906 | 0.93985162 |
| GNPDA1 | 5793.18472 | -0.0469383 | 0.55059702 | -0.0852498 | 0.93206277 | 0.98987582 |
| NDFIP1 | 9966.67583 | 0.14042411 | 0.12180851 | 1.15282679 | 0.24898149 | 0.81646572 |
| SPRY4 | 2069.38108 | 0.16680133 | 0.2802631 | 0.59515981 | 0.55173665 | 0.92373235 |
| FGF1 | 3498.0706 | 0.75697524 | 0.66790966 | 1.13334973 | 0.25706741 | 0.82107009 |

| | | | | | | |
|---|---|---|---|---|---|---|
| ARHGAP26 | 1628.1206 | 0.02178002 | 0.20114734 | 0.10827894 | 0.91377443 | 0.98662757 |
| NR3C1 | 2058.10617 | -0.2051414 | 0.16724276 | -1.2266086 | 0.21996972 | 0.79858446 |
| YIPF5 | 2169.22004 | -0.2812329 | 0.27547129 | -1.0209154 | 0.30729454 | 0.8430694 |
| KCTD16 | 172.341376 | -0.6048695 | 0.61978933 | -0.9759275 | 0.32910038 | 0.85327551 |
| PRELID2 | 84.7018513 | -0.2518082 | 0.36254102 | -0.6945647 | 0.48732816 | 0.90573694 |
| SH3RF2 | 217.167698 | -0.6824374 | 0.73468285 | -0.928887 | 0.35294763 | 0.86364633 |
| AC091887.1 | 653.327584 | 0.09771445 | 0.19388449 | 0.50398281 | 6.14E-01 | 0.93895494 |
| LARS | 4385.99598 | 0.10341183 | 0.11122787 | 0.92972951 | 0.35251115 | 0.86364633 |
| AC091959.3 | 187.887365 | -0.0900628 | 0.26262303 | -0.3429355 | 0.73164701 | 0.95848028 |
| RBM27 | 1117.61064 | -0.2028043 | 0.16979124 | -1.1944333 | 0.23230853 | 0.80649152 |
| TCERG1 | 2261.41644 | -0.2662572 | 0.18809637 | -1.4155359 | 0.15691143 | 0.74522306 |
| PPP2R2B | 259.682529 | 0.76083883 | 0.34660464 | 2.19512015 | 0.02815498 | 0.53300785 |
| STK32A | 395.353563 | 1.40952617 | 0.52797654 | 2.66967575 | 0.00759245 | 0.3667272 |
| DPYSL3 | 1181.87383 | -0.9279129 | 0.6815249 | -1.3615245 | NA | NA |
| SPINK1 | 1176.23025 | 0.09894903 | 1.08744046 | 0.0909926 | 0.92749847 | 0.98915637 |
| SPINK5 | 36.1640669 | 0.56648127 | 0.68569016 | 0.82614758 | 0.40872038 | 0.88203324 |
| AC091948.1 | 43.0050081 | 0.35366403 | 0.5272795 | 0.67073352 | 0.5023903 | 0.91304273 |
| FBXO38 | 1646.64016 | -0.0663266 | 0.09747812 | -0.6804259 | 4.96E-01 | 0.90890243 |
| ADRB2 | 89.2636051 | -0.6209767 | 0.40797078 | -1.5221108 | 0.12798132 | 0.70580262 |
| ABLIM3 | 4193.70965 | -0.3113812 | 0.54496658 | -0.5713768 | 0.56774427 | 0.92618483 |
| AC012613.2 | 143.578464 | -0.1272902 | 0.70922121 | -0.1794788 | 0.85756174 | 0.97959852 |
| AFAP1L1 | 949.243771 | -0.5255639 | 0.26954007 | -1.9498546 | 0.05119346 | 0.5861488 |
| GRPEL2 | 960.386606 | -0.5920108 | 0.29860036 | -1.9826193 | 0.04740997 | 0.58224788 |
| GRPEL2-AS1 | 45.714089 | -0.4725062 | 0.45423748 | -1.0402184 | 0.29823847 | 0.83998904 |
| PCYOX1L | 436.846351 | 0.39516587 | 0.3629729 | 1.0886925 | 0.27628951 | 0.82719407 |
| CARMN | 2668.99636 | -0.1291267 | 0.59221487 | -0.2180402 | 0.82739777 | 0.97548201 |
| AC131025.2 | 40.0817737 | -0.4769194 | 0.76221118 | -0.6257051 | 0.53150839 | 0.91962754 |
| CSNK1A1 | 8457.58634 | -0.1382032 | 0.13875159 | -0.9960475 | 3.19E-01 | 0.84928695 |
| ARHGEF37 | 1140.54363 | 0.68786675 | 0.22200431 | 3.09843871 | 0.00194543 | 0.18989241 |
| PPARGC1B | 282.936475 | 0.28009848 | 0.42124508 | 0.66492997 | 0.50609529 | 0.91391336 |
| SLC26A2 | 1020.3736 | -0.4880191 | 0.47836117 | -1.0201896 | 0.30763854 | 0.8434443 |
| TIGD6 | 387.972547 | 0.2101333 | 0.23622643 | 0.88954187 | 0.37371193 | 0.86791605 |
| HMGXB3 | 2901.74931 | -0.1915234 | 0.14128689 | -1.3555636 | 0.17523806 | 0.76073329 |
| CSF1R | 1868.29823 | -0.0801051 | 0.5081794 | -0.1576315 | 0.87474717 | 0.98275231 |
| PDGFRB | 7126.62054 | -0.1344376 | 0.32275017 | -0.4165376 | 0.67701665 | 0.95012499 |
| CAMK2A | 148.91853 | 1.39005684 | 0.55219385 | 2.51733487 | 0.01182464 | 0.40438968 |
| ARSI | 121.864097 | 0.31358516 | 0.47609625 | 0.65865917 | 0.51011466 | 0.91439937 |
| TCOF1 | 1679.3997 | 0.06478605 | 0.19413781 | 0.33371167 | 0.73859715 | 0.9602325 |
| CD74 | 86086.9911 | -0.2061284 | 0.38277784 | -0.5385066 | 0.59022732 | 0.93289552 |
| RPS14 | 19846.8601 | 0.33306849 | 0.20270134 | 1.64314892 | 0.10035213 | 0.67191826 |
| NDST1 | 4633.48361 | -0.3216372 | 0.14514193 | -2.2160182 | 0.02669025 | 0.52678638 |
| SYNPO | 8665.93634 | -0.157199 | 0.3260557 | -0.482123 | 0.62971858 | 0.94088547 |
| MYOZ3 | 43.3841363 | 0.06206127 | 0.48973349 | 0.12672459 | 0.89915839 | 0.98497876 |
| RBM22 | 2434.79586 | -0.220295 | 0.17934984 | -1.2282978 | 0.21933521 | 0.79847731 |
| DCTN4 | 2492.98625 | -0.0584315 | 0.14852152 | -0.3934212 | 0.69400845 | 0.95222846 |
| SMIM3 | 2293.4492 | -0.2492148 | 0.30861428 | -0.8075283 | 0.4193622 | 0.88451372 |
| ZNF300 | 518.003367 | -0.2280541 | 0.27314959 | -0.8349053 | 0.40377102 | 0.88132015 |
| GPX3 | 616637.432 | 0.37727124 | 0.39663845 | 0.95117163 | 0.34151726 | 0.85889221 |
| TNIP1 | 12116.718 | -0.0614915 | 0.12144728 | -0.5063228 | 0.61263004 | 0.93879706 |
| ANXA6 | 7285.17114 | 0.2169893 | 0.3301751 | 0.65719461 | 0.51105579 | 0.9148626 |
| CCDC69 | 1233.95048 | 0.07577251 | 0.37509696 | 0.20200779 | 0.83991063 | 0.97618283 |
| GM2A | 1653.01386 | 0.63557473 | 0.19264939 | 3.29912659 | 0.00096986 | 0.14829062 |
| SLC36A2 | 6410.91038 | 0.21385282 | 0.74932448 | 0.28539414 | 0.7753422 | 0.96836854 |
| SLC36A1 | 990.792947 | -0.3761575 | 0.41133382 | -0.9144822 | 0.36046353 | 0.86497402 |

| | | | | | | |
|---|---|---|---|---|---|---|
| FAT2 | 96.548209 | -0.0373493 | 0.47126458 | -0.0792533 | 0.93683111 | 0.99068685 |
| AC011337.1 | 68.6637031 | -0.3625375 | 0.43601039 | -0.8314882 | 0.40569787 | 0.88132015 |
| SPARC | 25130.15 | -0.070387 | 0.36402784 | -0.193356 | 0.84668019 | 0.97684681 |
| CLMAT3 | 1209.60123 | -0.0613736 | 0.41078109 | -0.149407 | 0.88123249 | 0.98275231 |
| ATOX1 | 2566.37075 | 0.16036868 | 0.17591906 | 0.91160492 | 0.36197673 | 0.86528137 |
| AC091982.3 | 78.0531958 | 0.58771689 | 0.37405236 | 1.5712156 | 0.11613258 | 0.68754779 |
| G3BP1 | 5977.10243 | -0.1693901 | 0.15556396 | -1.0888774 | 0.27620793 | 0.82719407 |
| LINC01933 | 42.9962016 | 0.36776672 | 0.58087936 | 0.63312066 | 0.52665485 | 0.91769856 |
| FAM114A2 | 1138.54485 | 0.02991997 | 0.15757785 | 0.18987419 | 0.84940772 | 0.97763005 |
| MFAP3 | 358.504241 | -0.278003 | 0.26797456 | -1.0374233 | 0.29953865 | 0.84123067 |
| GALNT10 | 1817.32555 | 0.09454019 | 0.13905008 | 0.67990029 | 0.4965676 | 0.90890243 |
| SAP30L | 1980.22089 | -0.1109623 | 0.17967913 | -0.6175579 | 0.53686678 | 0.92105614 |
| LARP1 | 8717.49826 | -0.2800291 | 0.20584049 | -1.360418 | 1.74E-01 | 7.60E-01 |
| FAXDC2 | 1989.68656 | 0.24910069 | 0.30360157 | 0.82048553 | 0.41193938 | 0.88268208 |
| CNOT8 | 2804.49785 | -0.1545292 | 0.1579319 | -0.9784544 | 0.32784963 | 0.85317073 |
| GEMIN5 | 687.119979 | -0.0661763 | 0.1437604 | -0.4603239 | 0.64528378 | 0.94208859 |
| MRPL22 | 1214.61516 | -0.1134246 | 0.1451897 | -0.7812165 | 0.43467515 | 0.89055338 |
| SGCD | 110.800061 | 0.44969436 | 0.47891478 | 0.93898618 | 0.34773784 | 0.86225576 |
| TIMD4 | 159.644043 | 0.10389609 | 0.52285364 | 0.1987097 | 0.84248983 | 0.97644404 |
| HAVCR1 | 360.625335 | -0.1142356 | 0.48717415 | -0.2344862 | 8.15E-01 | 9.74E-01 |
| HAVCR2 | 2756.90645 | 0.8202125 | 0.48319261 | 1.69748561 | 0.08960489 | 0.65375543 |
| MED7 | 698.091836 | -0.1079606 | 0.14110817 | -0.7650911 | 0.44421729 | 0.8938278 |
| ITK | 170.75472 | 0.1248365 | 0.84432784 | 0.14785312 | 0.88245869 | 0.98275231 |
| AC009185.1 | 43.0646643 | 0.42818524 | 0.58562651 | 0.73115753 | 0.46468294 | 0.89869398 |
| CYFIP2 | 13756.0498 | 0.25502223 | 0.32830626 | 0.77678151 | 0.43728769 | 0.89130865 |
| ADAM19 | 553.521779 | -0.6771135 | 0.42214151 | -1.6039964 | 1.09E-01 | 0.67800851 |
| THG1L | 441.331968 | -0.1567683 | 0.21055416 | -0.7445507 | 0.45654336 | 0.89688594 |
| LSM11 | 325.243653 | -0.2518902 | 0.24929846 | -1.0103962 | 0.31230553 | 0.84541735 |
| CLINT1 | 6996.09907 | -0.0030759 | 0.14217661 | -0.0216346 | 0.98273945 | 0.99613392 |
| EBF1 | 534.898082 | -0.1715452 | 0.35332896 | -0.4855112 | 0.62731376 | 0.94088547 |
| LINC02202 | 89.7869216 | 0.0797408 | 0.55715604 | 0.14312112 | 0.88619452 | 0.98391254 |
| RNF145 | 3399.1741 | -0.0421214 | 0.31459078 | -0.1338927 | 0.89348742 | 0.98474725 |
| UBLCP1 | 1237.22143 | -0.1083836 | 0.21252355 | -0.5099839 | 0.61006272 | 0.93791514 |
| ADRA1B | 234.171966 | -0.4909525 | 0.59132825 | -0.8302537 | 0.40639534 | 0.88132015 |
| TTC1 | 2178.6497 | -0.0544902 | 0.21944147 | -0.2483132 | 0.80389208 | 0.97274243 |
| PWWP2A | 1016.86719 | 0.02106657 | 0.12258984 | 0.17184597 | 0.86355862 | 0.98176973 |
| CCNJL | 371.320397 | -0.3930705 | 0.39134917 | -1.0043984 | 0.31518663 | 0.84696964 |
| C1QTNF2 | 56.2209081 | -0.6887132 | 0.66601801 | -1.0340759 | 0.30110068 | 0.84211658 |
| ZBED8 | 335.288243 | 0.11390987 | 0.31203903 | 0.36505005 | 0.71507404 | 0.95590747 |
| SLU7 | 1862.56526 | -0.1001985 | 0.13568321 | -0.738474 | 0.46022648 | 0.89755243 |
| PTTG1 | 98.5542473 | -0.2618101 | 0.48935896 | -0.5350062 | 0.59264554 | 0.93289552 |
| MIR3142HG | 102.901902 | 0.88121433 | 0.56905003 | 1.54857092 | 0.1214849 | 0.6971061 |
| ATP10B | 47.889547 | 1.78548409 | 0.96381799 | 1.85251168 | NA | NA |
| GABRB2 | 92.0079666 | 1.61175268 | 0.60107652 | 2.68144342 | 0.00733053 | 0.3610358 |
| CCNG1 | 6587.6247 | 0.53018609 | 0.19028649 | 2.78625189 | 0.00533214 | 0.31308771 |
| NUDCD2 | 1211.35611 | 0.00320764 | 0.12756466 | 0.02514522 | 0.97993913 | 0.99537078 |
| HMMR | 61.5078415 | 0.66228386 | 0.60413808 | 1.09624586 | 0.2729712 | 0.82716979 |
| MAT2B | 5340.99764 | 0.14330899 | 0.13658992 | 1.04919151 | 0.29408998 | 0.83596851 |
| TENM2 | 130.504357 | 0.32654674 | 0.49658361 | 0.65758664 | 0.51080378 | 0.91474443 |
| AC008637.1 | 137.338317 | 0.28291517 | 0.40315544 | 0.70175208 | 0.48283379 | 0.90541935 |
| WWC1 | 10958.7244 | -0.2242978 | 0.2502961 | -0.8961296 | 0.37018353 | 0.86762749 |
| RARS | 2815.12001 | -0.0943128 | 0.22766898 | -0.414254 | 0.6786881 | 0.95040677 |
| PANK3 | 2990.81492 | -0.1444391 | 0.24510967 | -0.5892836 | 0.55567104 | 0.92373235 |
| AC011365.2 | 642.203753 | -0.1735235 | 0.45967423 | -0.3774924 | 0.70580771 | 0.95501548 |

| | | | | | | |
|---|---|---|---|---|---|---|
| SLIT3 | 3552.24265 | 0.01738426 | 0.45002794 | 0.03862928 | 0.96918595 | 0.99400331 |
| AC011365.1 | 240.096131 | -0.1187503 | 0.44838093 | -0.2648424 | 0.79113084 | 0.97090793 |
| SPDL1 | 330.645524 | 0.07366167 | 0.22891595 | 0.32178477 | 0.74761575 | 0.96231266 |
| DOCK2 | 432.153018 | -0.049281 | 0.58512994 | -0.0842224 | 0.93287964 | 0.98999825 |
| FOXI1 | 1895.76998 | -0.1279814 | 0.54451984 | -0.2350354 | 0.81418129 | 0.9738447 |
| LINC01187 | 1014.69203 | 0.40744394 | 0.56210242 | 0.72485713 | 0.46853968 | 0.89984974 |
| C5orf58 | 106.274059 | -0.4974724 | 0.51563431 | -0.9647775 | 0.33465629 | 0.85557672 |
| LCP2 | 693.65022 | -0.3412005 | 0.4944172 | -0.6901065 | 0.49012724 | 0.90600166 |
| KCNIP1 | 107.526658 | 1.25065319 | 0.24406571 | 5.12424775 | 2.99E-07 | 0.00093308 |
| KCNMB1 | 247.730196 | -0.941329 | 0.69291133 | -1.3585129 | 0.174301 | 0.75974924 |
| AC008514.1 | 74.9153168 | 0.95979845 | 0.70931569 | 1.35313298 | 0.17601315 | 0.76115956 |
| GABRP | 346.163895 | 1.7056829 | 0.64408473 | 2.64822751 | 0.00809151 | 0.37167377 |
| RANBP17 | 168.615062 | 0.39881311 | 0.32734732 | 1.21831795 | 0.22310317 | 0.80078694 |
| NPM1 | 17690.3337 | -0.0403387 | 0.23099845 | -0.1746275 | 0.86137236 | 0.98167219 |
| FBXW11 | 2877.13339 | -0.1684221 | 0.17794281 | -0.9464956 | 0.34389587 | 0.85961157 |
| STK10 | 1498.26697 | -0.4486006 | 0.31221048 | -1.436853 | 0.15075977 | 0.73796431 |
| UBTD2 | 1544.11763 | -0.2203747 | 0.21100963 | -1.044382 | 0.29630869 | 0.83817467 |
| SH3PXD2B | 639.849927 | -0.0950805 | 0.31027512 | -0.3064392 | 0.75927024 | 0.96502715 |
| NEURL1B | 1861.28574 | -0.3858852 | 0.50574075 | -0.7630099 | 0.4454575 | 0.89453029 |
| AC027309.1 | 553.152107 | -0.3625596 | 0.52327804 | -0.6928623 | 0.488396 | 0.90585781 |
| DUSP1 | 18873.4704 | -0.75647 | 0.46312549 | -1.6334017 | 0.10238454 | 0.67279543 |
| ERGIC1 | 10448.284 | 0.08953911 | 0.2396757 | 0.37358443 | 0.70871352 | 0.95501548 |
| AC008429.1 | 77.3291476 | -0.856243 | 0.56617882 | -1.512319 | 0.13045273 | 0.7084713 |
| RPL26L1 | 791.297663 | 0.0796618 | 0.17923022 | 0.44446636 | 0.65670544 | 0.94545756 |
| ATP6V0E1 | 12127.7378 | -0.1535314 | 0.19316126 | -0.7948353 | 0.42670932 | 0.88709241 |
| CREBRF | 2291.96335 | -0.3266509 | 0.18211097 | -1.7936917 | 7.29E-02 | 0.63541838 |
| AC008378.1 | 44.945043 | 0.37180416 | 0.33883294 | 1.09730818 | 0.2725067 | 0.82716979 |
| BNIP1 | 559.793805 | 0.00395224 | 0.13200155 | 0.02994083 | 9.76E-01 | 0.99455099 |
| STC2 | 1132.02569 | 0.1633121 | 0.53861547 | 0.30320721 | 0.76173195 | 0.96607083 |
| BOD1 | 1859.70658 | -0.1196982 | 0.14968763 | -0.799653 | 0.42391189 | 0.88565404 |
| LINC01485 | 110.479177 | 0.79184345 | 0.71329139 | 1.11012619 | 0.26694465 | 0.8265123 |
| CPEB4 | 3846.7052 | -0.4142614 | 0.24355232 | -1.7009132 | 0.08895929 | 0.65375543 |
| SFXN1 | 3907.03458 | -0.0232718 | 0.36110764 | -0.0644456 | 0.94861541 | 0.99170503 |
| HRH2 | 376.94904 | -0.1886929 | 0.33533514 | -0.5626996 | 0.57363948 | 0.92978516 |
| THOC3 | 1045.16496 | -0.0670231 | 0.22145986 | -0.302642 | 0.76216267 | 0.96622409 |
| AC139491.7 | 162.076833 | -0.5435476 | 0.55900195 | -0.9723536 | 0.33087466 | 0.85384238 |
| SIMC1 | 357.731229 | 0.17986546 | 0.17035002 | 1.05585812 | 0.29103304 | 0.83454586 |
| KIAA1191 | 15470.7573 | 0.46073989 | 0.39827719 | 1.15683222 | 0.24734091 | 0.81566357 |
| AC138956.1 | 1717.04582 | 0.50095618 | 0.4410912 | 1.1357202 | 2.56E-01 | 8.21E-01 |
| ARL10 | 978.546786 | -0.3606346 | 0.1547295 | -2.3307423 | 0.01976695 | 0.49477123 |
| NOP16 | 3074.96587 | -0.2122439 | 0.21266614 | -0.9980147 | 0.31827221 | 0.8490262 |
| HIGD2A | 3323.74374 | -0.104188 | 0.15912475 | -0.6547567 | 0.51262439 | 0.91568064 |
| CLTB | 3388.77091 | -0.1829868 | 0.17993172 | -1.0169789 | 0.30916346 | 0.84414094 |
| FAF2 | 2337.25685 | -0.3542258 | 0.24219472 | -1.462566 | 0.14358616 | 0.72905668 |
| RNF44 | 5232.09291 | -0.2049167 | 0.160218 | -1.2789865 | 0.2009018 | 0.78134034 |
| CDHR2 | 3582.35338 | 0.5168162 | 0.65372351 | 0.79057306 | 0.42919317 | 0.88851358 |
| GPRIN1 | 290.21962 | -0.1794201 | 0.5510117 | -0.3256194 | 0.74471232 | 0.9613927 |
| TSPAN17 | 3152.17113 | -0.0587709 | 0.16302903 | -0.3604934 | 0.71847819 | 0.95648644 |
| HK3 | 469.299761 | -0.6275233 | 0.64234064 | -0.9769322 | 0.32860271 | 0.85317073 |
| UIMC1 | 1623.62945 | -0.3855224 | 0.18329686 | -2.1032679 | 0.03544236 | 0.55369025 |
| ZNF346 | 771.00558 | -0.2089996 | 0.19189703 | -1.0891238 | 0.27609929 | 0.82719407 |
| FGFR4 | 5336.55326 | 0.55967 | 0.28375547 | 1.97236725 | 0.0485677 | 0.58440502 |
| NSD1 | 2279.01368 | -0.1540617 | 0.10421254 | -1.4783416 | 0.13931637 | 0.72409321 |
| RAB24 | 3328.33017 | -0.1471326 | 0.2935823 | -0.501163 | 0.61625642 | 0.93910158 |

| | | | | | | |
|---|---|---|---|---|---|---|
| MXD3 | 8720.24729 | -0.1369598 | 0.25253039 | -0.5423498 | 0.58757758 | 0.93289552 |
| PRELID1 | 6910.85439 | -0.1540177 | 0.31317538 | -0.4917938 | 0.62286511 | 0.94016713 |
| LMAN2 | 8455.00255 | 0.06717136 | 0.14190645 | 0.47334961 | 0.63596377 | 0.94088547 |
| RGS14 | 3332.2948 | 0.13222324 | 0.31252272 | 0.4230836 | 0.67223427 | 0.94922827 |
| SLC34A1 | 11431.0944 | 0.19722717 | 0.82376524 | 0.23942158 | 0.8107787 | 0.97352497 |
| F12 | 689.73207 | -0.1118729 | 0.49646033 | -0.2253411 | 0.8217139 | 0.97487098 |
| GRK6 | 1415.56912 | -0.1184726 | 0.15236013 | -0.7775827 | 0.43681506 | 0.89127323 |
| PRR7 | 204.686076 | -0.4527834 | 0.35498346 | -1.2755057 | 0.2021303 | 0.78163854 |
| DBN1 | 5586.03689 | -0.3699421 | 0.26869888 | -1.3767907 | 0.16857695 | 0.7576272 |
| PDLIM7 | 4558.32736 | -0.6572214 | 0.43216066 | -1.5207804 | 0.12831496 | 0.70596572 |
| AC145098.1 | 52.6833504 | -0.7990839 | 0.36115362 | -2.2125872 | 0.02692612 | 0.52685774 |
| DOK3 | 492.627032 | -0.7540166 | 0.28131844 | -2.6802956 | 0.00735572 | 0.3610358 |
| DDX41 | 4223.19313 | 0.01680734 | 0.17239598 | 0.09749265 | 0.92233517 | 0.98826666 |
| FAM193B | 5560.47383 | -0.3249636 | 0.22673434 | -1.4332349 | 0.15179073 | 0.74024851 |
| TMED9 | 8645.70778 | -0.1396037 | 0.21966212 | -0.6355381 | 0.52507751 | 0.91759401 |
| B4GALT7 | 1797.40345 | 0.18341698 | 0.17799792 | 1.03044451 | 0.30280139 | 0.84211658 |
| AC139795.2 | 116.744118 | -0.320393 | 0.3358447 | -0.9539916 | 0.34008789 | 0.85814041 |
| FAM153A | 181.528761 | -0.5250179 | 0.74860251 | -0.7013307 | NA | NA |
| AC106795.2 | 41.711645 | -0.4700005 | 0.38753051 | -1.212809 | 0.22520287 | 0.80176092 |
| FAM153C | 45.3578856 | -0.3964942 | 0.84911546 | -0.4669497 | 0.64053587 | 0.94088547 |
| N4BP3 | 692.017404 | -0.3073799 | 0.25407331 | -1.2098079 | 0.2263526 | 0.80246466 |
| RMND5B | 3584.18706 | -0.0211419 | 0.13253242 | -0.1595222 | 0.8732575 | 0.98275231 |
| NHP2 | 2471.31826 | 0.0107387 | 0.20491937 | 0.05240449 | 0.95820639 | 0.99235997 |
| HNRNPAB | 6233.8256 | -0.2379648 | 0.19735146 | -1.2057918 | 0.22789778 | 0.80300331 |
| PHYKPL | 5031.12046 | -0.2746745 | 0.16198232 | -1.6957064 | 0.08994149 | 0.65375543 |
| COL23A1 | 424.674535 | 0.25990332 | 0.5083468 | 0.51127167 | 0.60916084 | 0.93748023 |
| CLK4 | 2198.38374 | -0.0078479 | 0.19470547 | -0.0403063 | 0.96784891 | 0.99400331 |
| ZNF354A | 967.366029 | -0.413148 | 0.20546058 | -2.0108385 | 0.04434252 | 0.57267043 |
| ZNF354B | 681.138353 | -0.1574183 | 0.18885041 | -0.8335609 | 0.40452849 | 0.88132015 |
| ZFP2 | 224.133677 | 0.2118324 | 0.31827234 | 0.66556963 | 0.50568622 | 0.91386909 |
| ZNF454 | 55.7865306 | 0.00748062 | 0.406642 | 0.01839608 | 0.98532288 | 0.99692004 |
| GRM6 | 39.3829945 | -0.1587759 | 0.48346498 | -0.3284124 | 0.7425999 | 0.96109001 |
| ZNF879 | 183.493767 | -0.2435734 | 0.2194953 | -1.1096976 | 0.26712938 | 0.8265123 |
| ZNF354C | 243.390684 | -0.3074517 | 0.18994673 | -1.6186206 | 0.10552893 | 0.6752194 |
| ADAMTS2 | 1084.80507 | -0.5381692 | 0.391646 | -1.3741216 | 0.16940394 | 0.75852708 |
| RUFY1 | 3066.77332 | 0.06579583 | 0.17460462 | 0.37682757 | 0.70630175 | 0.95501548 |
| AC136604.2 | 321.849083 | -0.0981351 | 0.21954283 | -0.4469974 | 0.65487693 | 0.94532497 |
| HNRNPH1 | 23499.1636 | -0.2927182 | 0.15233969 | -1.9214833 | 0.0546708 | 0.59329767 |
| CANX | 28596.7489 | 0.20997078 | 0.16228858 | 1.29381117 | 0.19573065 | 0.77765279 |
| MAML1 | 2295.5714 | -0.2808528 | 0.1842147 | -1.5245951 | 0.12736013 | 0.70557946 |
| LTC4S | 315.154433 | 0.01767585 | 0.53855133 | 0.0328211 | 0.97381725 | 0.99445887 |
| MGAT4B | 15675.4058 | -0.2240651 | 0.39801913 | -0.5629505 | 0.57346862 | 0.92970376 |
| SQSTM1 | 59325.5067 | -0.9217676 | 0.59677279 | -1.5445872 | 0.12244615 | 0.69788192 |
| MRNIP | 36852.6155 | -0.8992785 | 0.55605406 | -1.6172502 | 0.10582428 | 0.67526487 |
| AC008393.1 | 110.50474 | 0.32755122 | 0.39410586 | 0.83112496 | 0.40590304 | 0.88132015 |
| TBC1D9B | 9061.69694 | -0.127598 | 0.18279271 | -0.6980476 | 4.85E-01 | 9.05E-01 |
| RNF130 | 4218.91456 | -0.0517299 | 0.19415506 | -0.266436 | 0.78990344 | 0.97090793 |
| MAPK9 | 2066.74889 | 0.11274408 | 0.10972499 | 1.02751498 | 0.30417803 | 0.84211658 |
| AC008610.1 | 105.700569 | 0.63341069 | 0.42582222 | 1.48750034 | 0.13688271 | 0.72140578 |
| GFPT2 | 918.647988 | -0.3504087 | 0.44456362 | -0.7882083 | 0.4305749 | 0.88966606 |
| CNOT6 | 1721.14794 | -0.2641088 | 0.14792975 | -1.7853665 | 0.07420189 | 0.63773415 |
| FLT4 | 3462.55752 | -0.4796294 | 0.34483728 | -1.3908863 | 0.16425992 | 0.75286112 |
| MGAT1 | 8530.15095 | -0.4756673 | 0.14965453 | -3.1784357 | 0.00148072 | 0.17236139 |
| LINC00847 | 1515.42068 | 0.01559182 | 0.22129461 | 0.07045729 | 0.9438297 | 0.99136103 |

| | | | | | | |
|---|---|---|---|---|---|---|
| ZFP62 | 887.908803 | -0.0868331 | 0.19128705 | -0.4539415 | 0.64987092 | 0.94312988 |
| BTNL9 | 1782.42603 | 0.41303899 | 0.53319488 | 0.77464921 | 0.43854697 | 0.89175645 |
| AC008443.5 | 84.4035272 | 0.53090721 | 0.42989927 | 1.23495723 | 0.21684642 | 0.7954827 |
| TRIM7 | 238.62924 | 0.36501264 | 0.41953397 | 0.87004312 | 0.38427684 | 0.87298595 |
| TRIM41 | 4058.01365 | 0.03457943 | 0.14433506 | 0.2395775 | 0.81065781 | 0.97352497 |
| AC008443.2 | 517.980387 | 0.12112291 | 0.23125363 | 0.52376654 | 6.00E-01 | 9.35E-01 |
| RACK1 | 38247.4872 | 0.15870977 | 0.21396577 | 0.74175307 | 0.45823696 | 0.8971553 |
| TRIM52 | 1964.68389 | -0.172992 | 0.18797425 | -0.9202961 | 0.35741804 | 0.86390849 |
| AC138035.1 | 66.7314113 | -0.6884924 | 0.48274764 | -1.4261953 | 0.15381197 | 0.74258845 |
| DUSP22 | 2297.46434 | -0.0493064 | 0.16322121 | -0.3020832 | 0.76258863 | 0.96630779 |
| IRF4 | 106.037039 | -0.5553772 | 0.74793587 | -0.7425465 | 0.45775626 | 0.89698977 |
| EXOC2 | 1322.88581 | -0.0371946 | 0.14054277 | -0.2646499 | 0.79127919 | 0.97090793 |
| FOXQ1 | 1058.33098 | -0.1590088 | 0.42578133 | -0.3734519 | 0.70881216 | 0.95501548 |
| FOXCUT | 78.673892 | 0.80713467 | 0.61060646 | 1.32185739 | 0.18621564 | 0.76908034 |
| FOXC1 | 2105.74381 | -0.3036008 | 0.29500996 | -1.0291206 | 0.303423 | 0.84211658 |
| GMDS | 1579.91133 | 0.2664452 | 0.24981099 | 1.06658722 | 0.28615828 | 0.83107779 |
| GMDS-DT | 348.957073 | 0.15248584 | 0.29822819 | 0.51130592 | 0.60913686 | 0.93748023 |
| MYLK4 | 424.16198 | 0.91578859 | 0.74189625 | 1.23438903 | 0.21705798 | 0.79574248 |
| WRNIP1 | 3290.66982 | -0.0761105 | 0.10495977 | -0.7251399 | 0.46836624 | 0.89984974 |
| SERPINB1 | 3649.04626 | 0.06085338 | 0.19713261 | 0.3086926 | 0.75755537 | 0.96502715 |
| SERPINB9P1 | 48.0126912 | -2.1189444 | 1.14844784 | -1.8450506 | NA | NA |
| SERPINB9 | 909.977879 | -0.0958923 | 0.37733449 | -0.2541307 | 0.79939455 | 0.97200113 |
| SERPINB6 | 6898.21121 | 0.05059153 | 0.20972857 | 0.24122385 | 0.80938163 | 0.97334138 |
| LINC01011 | 321.357922 | 0.72811401 | 0.45324704 | 1.60643962 | 0.10817735 | 0.67789099 |
| NQO2 | 7639.83791 | 0.41521202 | 0.55025847 | 0.75457633 | 0.45050323 | 0.89557948 |
| AL133351.1 | 140.492728 | 1.40923859 | 0.74051407 | 1.90305444 | 0.05703344 | 0.60049913 |
| RIPK1 | 2460.64949 | -0.3228422 | 0.25830777 | -1.2498355 | 0.21135964 | 0.79134644 |
| AL031963.3 | 41.1065447 | -0.4225116 | 0.47889448 | -0.8822645 | 0.3776338 | 0.86987124 |
| BPHL | 3933.09868 | 0.62950238 | 0.37486363 | 1.67928369 | 0.09309677 | 0.65756964 |
| AL031963.1 | 1746.04701 | 0.663432 | 0.36848685 | 1.80042247 | 0.07179396 | 0.63189503 |
| TUBB2A | 1232.12738 | -0.273653 | 0.38977781 | -0.7020743 | 0.4826328 | 0.90541935 |
| TUBB2B | 689.990989 | -0.1755909 | 0.47533351 | -0.3694057 | 0.71182536 | 0.95560193 |
| PSMG4 | 1687.34238 | 0.08027498 | 0.23437559 | 0.34250572 | 0.73197034 | 0.95848028 |
| SLC22A23 | 1780.05808 | 0.11635515 | 0.36174268 | 0.32165171 | 0.74771657 | 0.96233424 |
| PXDC1 | 2940.48735 | -0.5031818 | 0.29318161 | -1.7162802 | 0.08611077 | 0.65049132 |
| AL391422.3 | 649.046967 | -0.1685863 | 0.28958826 | -0.5821585 | 5.60E-01 | 9.24E-01 |
| FAM50B | 856.17782 | 0.11310976 | 0.21768997 | 0.51959106 | 6.03E-01 | 9.36E-01 |
| PRPF4B | 3664.69583 | -0.3315014 | 0.16132198 | -2.0549054 | 3.99E-02 | 5.57E-01 |
| C6orf201 | 4139.15635 | 0.31691937 | 0.35482465 | 0.89317179 | 0.37176518 | 0.86762749 |
| ECI2 | 7490.43345 | 0.25254863 | 0.32265941 | 0.78270965 | 0.43379764 | 0.89055338 |
| AL136309.4 | 624.55885 | 0.35713451 | 0.41188589 | 0.86707149 | 0.38590284 | 0.87420019 |
| CDYL | 1027.17594 | -0.5851704 | 0.36816902 | -1.5894069 | 0.11196856 | 0.68001961 |
| RPP40 | 191.330711 | -0.2788411 | 0.30790195 | -0.9056164 | 0.36513892 | 0.8663515 |
| AL359643.3 | 46.4816965 | -0.5229357 | 0.38717725 | -1.3506364 | 0.17681195 | 0.76115956 |
| PPP1R3G | 223.463215 | 0.37195176 | 0.5635106 | 0.6600617 | 0.50921424 | 0.91439937 |
| LYRM4 | 842.858144 | -0.1640315 | 0.205345 | -0.7988095 | 0.42440085 | 0.88565404 |
| FARS2 | 801.362449 | 0.2313243 | 0.18645287 | 1.24065829 | 0.21473201 | 0.79468653 |
| NRN1 | 407.650702 | -0.154546 | 0.55474168 | -0.2785909 | 0.78055878 | 0.96864684 |
| F13A1 | 3918.16783 | 0.11205617 | 0.73953623 | 0.15152222 | 0.87956378 | 0.98275231 |
| LY86 | 134.412022 | -0.1224316 | 0.71757703 | -0.170618 | 0.86452413 | 0.98188271 |
| AL031123.2 | 113.647306 | 0.1825302 | 0.52243967 | 0.34938043 | 0.72680372 | 0.95848028 |
| AL031123.1 | 127.628596 | 0.1225606 | 0.61567667 | 0.19906651 | 0.84221072 | 0.9763625 |
| RREB1 | 2371.01274 | -0.1178133 | 0.16392493 | -0.7187027 | 4.72E-01 | 9.02E-01 |
| SSR1 | 6637.19972 | 0.136815 | 0.17691997 | 0.77331575 | 0.43933554 | 0.89175645 |

| | | | | | | |
|---|---|---|---|---|---|---|
| AL139095.4 | 421.116958 | 0.05895435 | 0.23496101 | 0.25091119 | 0.80188277 | 0.97233627 |
| RIOK1 | 646.875265 | -0.2101809 | 0.24323672 | -0.8641002 | 0.38753284 | 0.87460898 |
| AL031058.1 | 75.3566947 | -0.1602816 | 0.30657041 | -0.5228214 | 0.60109857 | 0.93518083 |
| DSP | 5021.21832 | -0.2423266 | 0.20429427 | -1.1861647 | 0.23555727 | 0.80747753 |
| SNRNP48 | 770.657854 | 0.01182817 | 0.15896392 | 0.07440789 | 0.94068583 | 0.99136103 |
| BMP6 | 882.154838 | -0.6761613 | 0.55748003 | -1.2128889 | 0.22517231 | 0.80176092 |
| TXNDC5 | 9830.36184 | 0.10267669 | 0.27337922 | 0.37558339 | 7.07E-01 | 0.95501548 |
| BLOC1S5-TXI | 10031.7643 | 0.10764643 | 0.2680869 | 0.40153559 | 0.68802584 | 0.95158502 |
| BLOC1S5 | 1496.03023 | 0.1974867 | 0.19260526 | 1.02534426 | 0.30520078 | 0.84268008 |
| EEF1E1-BLO( | 1419.83729 | -0.0976359 | 0.2190802 | -0.4456626 | 0.65584095 | 0.94532497 |
| EEF1E1 | 1676.74207 | -0.1610305 | 0.24076493 | -0.6688287 | 0.50360472 | 0.91356552 |
| SLC35B3 | 1374.63643 | 0.14228972 | 0.15139655 | 0.93984787 | 0.3472956 | 0.86197176 |
| TFAP2A | 941.396836 | 0.50084054 | 0.31858622 | 1.57207221 | 0.1159338 | 0.6873836 |
| AL138885.3 | 261.250244 | 0.46968881 | 0.3513781 | 1.33670485 | 0.181319 | 0.76522169 |
| TFAP2A-AS1 | 87.7309722 | 0.12471934 | 0.53771979 | 0.23194114 | 0.81658373 | 0.97388289 |
| GCNT2 | 1396.23535 | -0.4453676 | 0.15437889 | -2.8848998 | 0.00391538 | 0.26977276 |
| C6orf52 | 51.2569714 | 0.62616219 | 0.39975964 | 1.56634669 | 0.11726748 | 0.68958574 |
| PAK1IP1 | 658.791581 | -0.2348677 | 0.30296049 | -0.7752419 | 0.43819673 | 0.89152136 |
| TMEM14C | 4635.01855 | 0.29177363 | 0.14777834 | 1.97440058 | 0.04833621 | 0.58440502 |
| TMEM14B | 2649.45345 | 0.21819779 | 0.17444908 | 1.25078215 | 0.21101397 | 0.79134644 |
| AL024498.2 | 968.425918 | 0.30785692 | 0.21649349 | 1.42201469 | 0.15502199 | 0.74379596 |
| MAK | 146.439823 | -0.1547041 | 0.34421743 | -0.4494373 | 0.65311623 | 0.94481291 |
| SYCP2L | 68.9489091 | 0.12318857 | 0.39049546 | 0.31546736 | 0.75240683 | 0.96301881 |
| SMIM13 | 581.854575 | -0.1172945 | 0.20871175 | -0.561993 | 0.5741208 | 0.93011739 |
| AL139807.1 | 417.680904 | -1.0181354 | 0.33539628 | -3.0356192 | 0.00240042 | 0.21083339 |
| NEDD9 | 5105.51676 | -0.9280662 | 0.28673328 | -3.2366881 | 0.00120926 | 0.15785077 |
| TMEM170B | 604.779136 | 0.12119073 | 0.21668797 | 0.55928683 | 0.57596598 | 0.93083012 |
| ADTRP | 258.706897 | 0.45712946 | 0.64163899 | 0.71244028 | 0.47619218 | 0.90305051 |
| AL157373.2 | 183.101249 | 0.06377258 | 0.3744419 | 0.17031368 | 0.86476346 | 0.98197545 |
| HIVEP1 | 846.211929 | -0.3378662 | 0.19203214 | -1.7594253 | 0.07850529 | 0.64079926 |
| EDN1 | 677.00184 | 0.02651421 | 0.3740455 | 0.070885 | 0.94348928 | 0.99136103 |
| PHACTR1 | 950.552852 | -0.6009648 | 0.17762701 | -3.3832964 | 0.00071621 | 0.12783376 |
| AL008729.1 | 261.987428 | -0.6701195 | 0.57519743 | -1.1650253 | NA | NA |
| TBC1D7 | 1110.21496 | -0.7492919 | 0.52038828 | -1.4398708 | NA | NA |
| AL008729.2 | 259.844293 | -0.7719568 | 0.30972492 | -2.4923948 | 0.01268849 | 0.41598747 |
| GFOD1 | 503.697319 | -0.3962836 | 0.43247307 | -0.9163196 | 0.35949929 | 0.86480228 |
| SIRT5 | 1337.20584 | 0.11868205 | 0.33817396 | 0.35094971 | 0.72562608 | 0.95848028 |
| AL441883.1 | 103.950211 | -0.0862562 | 0.28026417 | -0.3077676 | 0.75825921 | 0.96502715 |
| NOL7 | 2938.33082 | -0.1397395 | 0.18148506 | -0.7699781 | 0.44131291 | 0.89290667 |
| RANBP9 | 2361.01486 | -0.0294103 | 0.10685377 | -0.2752392 | 0.78313251 | 0.96971192 |
| MCUR1 | 2399.10186 | 0.37967468 | 0.28244342 | 1.34425041 | 0.17886743 | 0.76120583 |
| RNF182 | 111.739443 | 0.07635765 | 0.34675055 | 0.22020917 | 0.82570825 | 0.97548201 |
| CD83 | 981.954737 | -0.0545455 | 0.30470879 | -0.1790085 | 0.85793103 | 0.97967862 |
| JARID2 | 611.974436 | -0.2925071 | 0.19532151 | -1.4975674 | 0.13424567 | 0.71576049 |
| DTNBP1 | 1535.90532 | -0.2893776 | 0.21343045 | -1.3558401 | 0.17515005 | 0.76073329 |
| MYLIP | 1834.60262 | -0.0817214 | 0.20559516 | -0.3974869 | 0.69100848 | 0.95158502 |
| GMPR | 2343.09343 | -0.895086 | 0.54771224 | -1.6342267 | 0.10221125 | 0.67279543 |
| ATXN1 | 1549.90082 | 0.08505654 | 0.21155369 | 0.40205653 | 6.88E-01 | 0.95158502 |
| AL137003.1 | 39.9475048 | 0.17322537 | 0.46089991 | 0.37584161 | 0.70703465 | 0.95501548 |
| AL137003.2 | 201.014613 | 0.47917698 | 0.40299868 | 1.18902866 | 0.23442839 | 0.80730051 |
| RBM24 | 102.466982 | -0.7365899 | 0.51586551 | -1.427872 | 0.15332871 | 0.74258845 |
| CAP2 | 680.32689 | 0.68584669 | 0.23296871 | 2.94394333 | 0.00324059 | 0.24992576 |
| FAM8A1 | 2603.13543 | 0.06458675 | 0.20361046 | 0.31720745 | 0.7510862 | 0.96272571 |
| NUP153 | 1944.76874 | -0.3682577 | 0.21186086 | -1.7382053 | 0.08217465 | 0.64573009 |

| | | | | | | |
|---|---|---|---|---|---|---|
| AL138724.1 | 76.0536857 | -0.8376761 | 0.52531576 | -1.5946144 | 0.11079849 | 0.68001961 |
| KIF13A | 1432.62029 | -0.1191637 | 0.14869524 | -0.8013956 | 0.42290269 | 0.88453836 |
| NHLRC1 | 64.6975277 | -0.0893627 | 0.56875987 | -0.1571185 | 0.87515148 | 0.98275231 |
| TPMT | 4794.16485 | 0.60875205 | 0.25625163 | 2.37560264 | 0.01752032 | 0.46859699 |
| KDM1B | 980.777518 | 0.33361081 | 0.21140123 | 1.57809308 | 0.11454423 | 0.68430774 |
| DEK | 3869.72279 | -0.3060142 | 0.20731636 | -1.4760736 | 0.13992413 | 0.72475166 |
| RNF144B | 2876.66105 | -0.0724893 | 0.41941549 | -0.1728342 | 0.86278176 | 0.98176973 |
| AL022068.1 | 84.3606534 | -0.4215887 | 0.48458259 | -0.8700039 | 0.38429828 | 0.87298595 |
| ID4 | 1887.40004 | -0.28573 | 0.28287424 | -1.0100954 | 0.3124496 | 0.84541735 |
| MBOAT1 | 223.099723 | 0.5506157 | 0.32064777 | 1.71719798 | 0.085943 | 0.65049132 |
| E2F3 | 1123.05234 | -0.2681836 | 0.20783639 | -1.2903592 | 0.19692599 | 0.77816059 |
| CDKAL1 | 624.197577 | -0.0488927 | 0.17469513 | -0.2798745 | 0.77957376 | 0.96836854 |
| SOX4 | 2320.54198 | -0.342524 | 0.38596856 | -0.8874402 | 0.37484193 | 0.86791605 |
| CASC15 | 300.272597 | -0.7914613 | 0.51986062 | -1.5224489 | 0.12789663 | 0.70579329 |
| DCDC2 | 4429.62286 | 0.31861382 | 0.23802583 | 1.33856829 | 0.18071126 | 0.76466689 |
| KAAG1 | 585.056338 | -0.0360329 | 0.29418299 | -0.1224846 | 0.90251527 | 0.98601135 |
| MRS2 | 2104.62298 | 0.03026286 | 0.15537637 | 0.19477133 | 0.84557197 | 0.97675487 |
| GPLD1 | 390.858434 | 0.19840182 | 0.15459828 | 1.28333782 | 0.19937375 | 0.78134034 |
| ALDH5A1 | 1897.08713 | 1.05564044 | 0.39659207 | 2.66177899 | 0.00777289 | 0.36878977 |
| TDP2 | 5766.13192 | 0.40825576 | 0.61985221 | 0.65863403 | 0.51013081 | 0.91439937 |
| ACOT13 | 2874.57493 | 0.2752348 | 0.30084743 | 0.91486506 | 0.36026248 | 0.86480228 |
| AL031775.1 | 814.276965 | 0.50388792 | 0.39919857 | 1.2622488 | 0.20685927 | 0.78790396 |
| C6orf62 | 7352.02116 | -0.1662553 | 0.19529304 | -0.8513118 | 0.39459616 | 0.87884163 |
| AL031775.2 | 53.7076868 | -0.2723907 | 0.30572113 | -0.8909775 | 0.37294124 | 0.86791605 |
| GMNN | 663.166703 | -0.1505044 | 0.18309892 | -0.821984 | 4.11E-01 | 8.83E-01 |
| RIPOR2 | 526.759895 | -0.5049981 | 0.47338205 | -1.0667876 | 0.28606774 | 0.83107779 |
| CARMIL1 | 1696.45594 | 0.03657342 | 0.17271431 | 0.21175677 | 8.32E-01 | 0.97550416 |
| SCGN | 73.3737889 | 2.04160292 | 0.96712839 | 2.1109947 | 0.03477277 | 0.55369025 |
| SLC17A4 | 418.117974 | 1.60814403 | 0.59583185 | 2.69898972 | 0.00695503 | 0.3503878 |
| SLC17A1 | 6835.60523 | 0.52596882 | 0.63278932 | 0.83119106 | 4.06E-01 | 0.88132015 |
| SLC17A3 | 9989.71082 | 2.30114215 | 0.67889483 | 3.38954145 | 7.00E-04 | 1.27E-01 |
| SLC17A2 | 81.0855783 | 0.93046106 | 0.53308842 | 1.745416 | 0.08091248 | 0.64313548 |
| TRIM38 | 4912.46581 | 0.32467195 | 0.30588146 | 1.06143061 | 0.28849425 | 0.83295411 |
| U91328.2 | 465.238246 | 0.06681996 | 0.30177025 | 0.22142661 | 0.82476027 | 0.97548201 |
| U91328.1 | 355.4609 | -0.0542589 | 0.22715498 | -0.2388628 | 0.811212 | 0.97352497 |
| HIST1H1C | 2210.66576 | -0.6802617 | 0.34737447 | -1.9582951 | 0.0501954 | 0.5861488 |
| HFE | 671.437254 | 0.21200882 | 0.28017671 | 0.75669679 | 0.44923154 | 0.89546252 |
| HIST1H2BC | 296.337088 | -2.0058642 | 0.80570308 | -2.4895824 | 0.01278933 | 0.41611943 |
| HIST1H2AC | 2404.43101 | -0.6381193 | 0.36648537 | -1.7411862 | 0.08165095 | 0.64505626 |
| HIST1H2BD | 1111.72202 | -0.6671735 | 0.36956344 | -1.8053016 | 0.07102749 | 0.63056668 |
| AL353759.1 | 55.1807997 | -0.2495349 | 0.45494975 | -0.5484889 | 0.58335628 | 0.93220705 |
| HIST1H2BE | 86.6263357 | -0.3657124 | 0.47577947 | -0.7686595 | 0.44209549 | 0.89331461 |
| AL031777.3 | 63.7270757 | -1.4687347 | 0.64377921 | -2.2814262 | 2.25E-02 | 5.20E-01 |
| HIST1H4E | 34.8436461 | -0.2291299 | 0.71801808 | -0.3191144 | 7.50E-01 | 9.62E-01 |
| HIST1H2BG | 97.596844 | -1.0299153 | 0.40748996 | -2.5274618 | 0.01148903 | 0.40353404 |
| HIST1H2AE | 48.148992 | -1.1895249 | 0.46251242 | -2.5718766 | 0.01011489 | 0.39989496 |
| HIST1H3E | 100.605324 | 0.03778621 | 0.51451437 | 0.07344054 | 0.94145556 | 0.99136103 |
| HIST1H2BH | 47.8390538 | -1.5232129 | 0.77323534 | -1.9699214 | 0.04884738 | 0.58504358 |
| HIST1H4H | 164.449464 | -0.5593238 | 0.38809335 | -1.4412094 | 0.14952553 | 0.73670625 |
| BTN3A2 | 1843.00731 | 0.16838499 | 0.54883559 | 0.30680407 | 7.59E-01 | 0.96502715 |
| BTN2A2 | 937.900264 | 0.09446895 | 0.30844691 | 0.30627296 | 0.75939683 | 0.96502715 |
| BTN3A1 | 1469.55764 | -0.335061 | 0.61831793 | -0.5418911 | 0.58789351 | 0.93289552 |
| BTN3A3 | 1105.73973 | -0.5762119 | 0.58399 | -0.9866812 | 0.32379896 | 0.85156215 |
| BTN2A1 | 2218.0366 | -0.2215832 | 0.18169659 | -1.2195232 | 0.22264566 | 0.80078694 |

| | | | | | | |
|---|---|---|---|---|---|---|
| HCG11 | 465.953308 | 0.12708388 | 0.19477579 | 0.65246241 | 0.51410292 | 0.91636259 |
| HMGN4 | 2599.08126 | -0.0317666 | 0.13524081 | -0.2348892 | 0.81429476 | 0.9738447 |
| ABT1 | 1183.60022 | -0.2279199 | 0.24046412 | -0.9478334 | 0.34321426 | 0.85936336 |
| ZNF322 | 294.223841 | 0.21668508 | 0.27580912 | 0.78563424 | 0.4320818 | 0.88976671 |
| LINC00240 | 165.294506 | -0.6769596 | 0.39550414 | -1.7116371 | 0.08696356 | 0.65231349 |
| HIST1H4I | 197.952264 | -0.0370376 | 0.33779466 | -0.1096454 | 0.91269061 | 0.98662757 |
| HIST1H2BK | 367.682014 | -0.1010627 | 0.32808945 | -0.3080339 | 0.75805651 | 0.96502715 |
| PRSS16 | 327.457517 | 0.64649037 | 0.41672467 | 1.55136092 | 0.12081521 | 0.69496908 |
| ZNF391 | 243.999802 | -0.0336481 | 0.27102323 | -0.1241522 | 0.90119479 | 0.98553843 |
| AL031118.1 | 42.4785547 | 0.82757339 | 0.75857901 | 1.09095213 | 0.27529395 | 0.82719407 |
| ZNF184 | 397.988363 | -0.1134803 | 0.2181226 | -0.5202595 | 0.60288275 | 0.93585899 |
| HIST1H2BN | 92.6974632 | -0.1953779 | 0.35677072 | -0.5476287 | 0.58394686 | 0.93233498 |
| AL121944.1 | 106.459993 | -0.6664121 | 0.55914126 | -1.1918493 | 0.23332033 | 0.80725541 |
| ZNF165 | 464.444855 | -1.4297415 | 0.68123368 | -2.0987535 | 0.03583864 | 0.55369025 |
| ZSCAN16-AS | 519.72998 | 0.26147228 | 0.17048909 | 1.53365992 | 0.12511334 | 0.70228923 |
| ZSCAN16 | 310.651372 | -0.3012461 | 0.22389286 | -1.3454923 | 0.17846631 | 0.76115956 |
| ZKSCAN8 | 1462.24747 | -0.2759875 | 0.1576565 | -1.750562 | 0.08002139 | 0.64079926 |
| ZSCAN9 | 701.903216 | -0.2090835 | 0.21285168 | -0.9822969 | 0.32595362 | 0.85282528 |
| ZKSCAN4 | 265.367205 | 0.06553069 | 0.17722884 | 0.36975185 | 0.7115674 | 0.95560193 |
| NKAPL | 45.5028729 | -0.1369465 | 0.4887429 | -0.2802015 | 0.77932294 | 0.96836854 |
| ZSCAN26 | 792.180285 | -0.0221359 | 0.23314043 | -0.0949465 | 0.92435734 | 0.98869145 |
| AL021997.3 | 111.645507 | -0.0533543 | 0.28245116 | -0.1888973 | 0.85017331 | 0.97816532 |
| PGBD1 | 344.891869 | -0.1132719 | 0.18967025 | -0.5972042 | 0.55037103 | 0.92373235 |
| ZSCAN31 | 366.338581 | 0.86221327 | 0.44131836 | 1.95372172 | 5.07E-02 | 0.5861488 |
| ZKSCAN3 | 364.732376 | 0.26277563 | 0.24148271 | 1.08817575 | 0.27651753 | 0.82730125 |
| ZSCAN12 | 271.701903 | -0.0975384 | 0.16338435 | -0.5969872 | 0.5505159 | 0.92373235 |
| ZBED9 | 112.620861 | 0.05475159 | 0.33457901 | 0.16364324 | 0.870012 | 0.98275231 |
| TRIM27 | 3323.68644 | -0.1254791 | 0.16922315 | -0.7415007 | 0.45838994 | 0.89719956 |
| ZNF311 | 83.9377531 | 0.04225339 | 0.42767137 | 0.09879874 | 0.92129807 | 0.98779251 |
| OR2I1P | 1622.37387 | 1.98E-01 | 0.47622795 | 0.41590444 | 0.67747993 | 0.95018034 |
| UBD | 1172.48195 | 0.04966855 | 0.4685389 | 0.1060073 | 9.16E-01 | 9.87E-01 |
| GABBR1 | 4329.88751 | -0.0225854 | 0.33151933 | -0.0681269 | 9.46E-01 | 9.92E-01 |
| HLA-F | 7432.61787 | -0.6597523 | 0.59156704 | -1.1152621 | 0.26473813 | 0.82424998 |
| HLA-G | 2163.62326 | 0.05538748 | 0.64731033 | 0.08556558 | 0.93181176 | 0.9898238 |
| HLA-A | 78072.8067 | -0.4138864 | 0.37413819 | -1.1062395 | 0.26862288 | 0.82676682 |
| ZNRD1 | 818.807399 | 0.18690875 | 0.13611979 | 1.37311961 | 0.16971517 | 0.7586159 |
| PPP1R11 | 3706.99426 | -0.017299 | 0.12911117 | -0.1339849 | 0.89341449 | 0.98474725 |
| RNF39 | 357.678227 | -0.180505 | 0.14225539 | -1.2688796 | 0.20448401 | 0.78449023 |
| TRIM10 | 277.03069 | 1.14334589 | 0.50370715 | 2.26986232 | 0.02321594 | 0.51967919 |
| TRIM15 | 577.816228 | 1.14261645 | 0.44868429 | 2.54659339 | 0.01087801 | 0.40196898 |
| TRIM26 | 3481.47753 | -0.0790816 | 0.25185353 | -0.3139986 | 0.75352213 | 0.96308792 |
| HCG18 | 1036.3591 | -0.2895858 | 0.29051316 | -0.9968079 | 0.31885775 | 0.84914926 |
| TRIM39 | 1188.44437 | -0.2203677 | 0.22850676 | -0.9643814 | 0.33485475 | 0.85579065 |
| TRIM39-RPP: | 1161.70234 | -0.1949932 | 0.19241665 | -1.0133905 | 0.31087368 | 0.84476049 |
| RPP21 | 939.806062 | -0.205492 | 0.18245918 | -1.1262356 | 0.26006579 | 0.82424998 |
| HLA-E | 47004.8997 | -0.4661033 | 0.26569123 | -1.7543046 | 0.07937833 | 0.64079926 |
| GNL1 | 4530.20878 | -0.1036032 | 0.14247734 | -0.7271555 | 0.4671307 | 0.89983399 |
| PRR3 | 784.406867 | -0.00717 | 0.19141393 | -0.0374579 | 0.97011987 | 0.99400331 |
| ABCF1 | 4451.24829 | -0.2608128 | 0.21533829 | -1.2111771 | 0.22582753 | 0.80246466 |
| PPP1R10 | 3647.25457 | -0.214679 | 0.22292832 | -0.9629955 | 0.33554978 | 0.85616588 |
| MRPS18B | 4378.94503 | 0.34594888 | 0.18632934 | 1.85665278 | 0.06336057 | 0.6165319 |
| ATAT1 | 1027.05491 | -0.1658895 | 0.18239281 | -0.9095178 | 0.36307687 | 0.8663515 |
| C6orf136 | 2163.43531 | 0.07152052 | 0.15042876 | 0.47544445 | 0.63447022 | 0.94088547 |
| DHX16 | 3077.72394 | -0.1887326 | 0.15385144 | -1.2267197 | 0.21992797 | 0.79858446 |

| | | | | | | |
|---|---|---|---|---|---|---|
| PPP1R18 | 2843.09257 | -0.3636938 | 0.37006221 | -0.9827911 | 0.32571026 | 0.85271781 |
| NRM | 655.726321 | -0.1752324 | 0.36814555 | -0.4759866 | 0.63408391 | 0.94088547 |
| MDC1 | 2044.9804 | 0.04641032 | 0.1505699 | 0.30823103 | 0.75790654 | 0.96502715 |
| MDC1-AS1 | 277.464479 | -0.0029137 | 0.21261128 | -0.0137044 | 0.98906585 | 0.9979586 |
| TUBB | 16932.3043 | -0.300446 | 0.38377017 | -0.7828799 | 4.34E-01 | 0.89055338 |
| AL662797.1 | 6763.38436 | -0.3733615 | 0.4081764 | -0.9147063 | 0.36034582 | 0.86480228 |
| FLOT1 | 14100.3803 | -0.1069152 | 0.22582949 | -0.4734333 | 0.63590406 | 0.94088547 |
| IER3-AS1 | 2887.06724 | 0.29764561 | 0.5034202 | 0.59124684 | 0.55435504 | 0.92373235 |
| IER3 | 9408.15072 | 0.25478623 | 0.5218158 | 0.48826852 | 0.62535966 | 0.94088547 |
| DDR1 | 19555.684 | -0.0472539 | 0.17743091 | -0.2663231 | 0.7899904 | 0.97090793 |
| GTF2H4 | 1716.99264 | -0.1449003 | 0.16923269 | -0.8562193 | 0.3918765 | 0.87744531 |
| VARS2 | 4812.8779 | 0.01707242 | 0.16574649 | 0.10300321 | 0.91796042 | 0.9868044 |
| SFTA2 | 78.7085947 | 0.57806772 | 0.51461794 | 1.12329492 | 0.26131226 | 0.82424998 |
| PSORS1C1 | 207.294737 | -0.3696661 | 0.78639278 | -0.4700782 | 0.63829918 | 0.94088547 |
| PSORS1C2 | 50.6279184 | -0.6364406 | 0.94756603 | -0.6716583 | 0.50180125 | 0.91259827 |
| CCHCR1 | 2097.03232 | -0.0583237 | 0.34643662 | -0.1683532 | 8.66E-01 | 0.98200144 |
| TCF19 | 855.730522 | -0.0618283 | 0.37312127 | -0.1657057 | 8.68E-01 | 0.98246119 |
| POU5F1 | 2736.15424 | -0.3239724 | 0.56738041 | -0.5709968 | 0.56800181 | 0.92618483 |
| AL662844.4 | 231.071025 | -0.206445 | 0.43206107 | -0.4778144 | 0.63278233 | 0.94088547 |
| HCG27 | 201.614603 | 0.23996551 | 0.46935191 | 0.51126992 | 0.60916206 | 0.93748023 |
| HLA-C | 68538.1282 | -0.47321 | 0.46028018 | -1.0280912 | 0.30390694 | 0.84211658 |
| HLA-B | 100294.191 | -0.608982 | 0.49453531 | -1.2314226 | 0.21816483 | 0.79708421 |
| AL645933.2 | 182.071052 | 0.33017511 | 0.43045092 | 0.76704473 | 0.44305492 | 0.89352768 |
| MICA | 2138.65437 | 0.17485877 | 0.19470213 | 0.89808352 | 0.36914102 | 0.86762749 |
| MICB | 270.281703 | 0.20527669 | 0.43896574 | 0.46763715 | 0.64004409 | 0.94088547 |
| MCCD1 | 1419.64064 | 1.50391144 | 0.68086342 | 2.20882984 | 0.02718648 | 0.52852884 |
| DDX39B | 18695.3646 | -0.2511052 | 0.18864057 | -1.3311303 | 0.18314615 | 0.76723162 |
| ATP6V1G2-D | 11748.7322 | -0.2308267 | 0.17905629 | -1.289129 | 0.19735325 | 0.77816059 |
| ATP6V1G2 | 495.946365 | -0.1545321 | 0.15880631 | -0.9730852 | 0.33051098 | 0.85384238 |
| NFKBIL1 | 1333.86863 | -0.2100624 | 0.11764906 | -1.7855003 | 0.07418021 | 0.63773415 |
| TNF | 89.2018017 | 0.45327856 | 0.52667388 | 0.86064371 | 0.3894343 | 0.87568585 |
| LTB | 310.846478 | 0.70470901 | 0.69904572 | 1.00810145 | 0.31340576 | 0.84638821 |
| LST1 | 291.519498 | 0.29442655 | 0.55960836 | 0.52612965 | 0.59879813 | 0.93484879 |
| AIF1 | 867.443496 | 0.33905465 | 0.44876285 | 0.75553191 | 4.50E-01 | 0.89557948 |
| PRRC2A | 15078.179 | -0.2235964 | 0.13943516 | -1.6035872 | 0.10880507 | 0.67800851 |
| BAG6 | 14068.9749 | -0.1082754 | 0.16336189 | -0.662795 | 0.50746183 | 0.91430909 |
| APOM | 5352.05935 | 0.70455409 | 0.69106506 | 1.0195192 | 0.30795655 | 0.84365058 |
| C6orf47 | 2194.40889 | -0.2555503 | 0.17388875 | -1.4696193 | 0.14166489 | 0.72784886 |
| C6orf47-AS1 | 685.897192 | -0.2521978 | 0.17552139 | -1.4368491 | 0.15076088 | 0.73796431 |
| GPANK1 | 2295.15502 | 0.03949599 | 0.14028311 | 0.28154484 | 7.78E-01 | 9.68E-01 |
| CSNK2B | 6052.33274 | 0.01390926 | 0.10135133 | 0.1372381 | 0.8908426 | 0.98474725 |
| AL662899.2 | 6747.11077 | -0.0094534 | 0.10000657 | -0.0945274 | 9.25E-01 | 0.98869145 |
| LY6G5B | 1392.80425 | -0.2034 | 0.23399198 | -0.8692604 | 0.38470473 | 0.87312908 |
| LY6G5C | 101.361183 | -0.2141501 | 0.34502889 | -0.6206728 | 0.5348149 | 0.92084006 |
| ABHD16A | 3311.31358 | 0.04575036 | 0.20741064 | 0.22057867 | 0.82542051 | 0.97548201 |
| AL662899.1 | 3234.75905 | 0.04915874 | 0.20886591 | 0.23536028 | 0.81392911 | 0.9738447 |
| DDAH2 | 4093.13229 | 0.05019456 | 0.20150886 | 0.24909357 | 0.8032884 | 0.97274243 |
| CLIC1 | 18358.3879 | -0.2050293 | 0.22033545 | -0.9305327 | 0.35209534 | 0.86364633 |
| MSH5 | 1473.95031 | 0.14449954 | 0.30080987 | 0.48036833 | 0.63096551 | 0.94088547 |
| MSH5-SAPCI | 2256.44074 | 0.15262938 | 0.26766566 | 0.570224 | 0.56852578 | 0.92642835 |
| SAPCD1 | 1079.67828 | 0.14535923 | 0.2820132 | 0.51543413 | 0.60624969 | 0.93708311 |
| SAPCD1-AS1 | 737.327902 | 0.07651006 | 0.28275826 | 0.27058469 | 0.78671047 | 0.97090793 |
| VWA7 | 1010.98575 | -0.2347413 | 0.29542935 | -0.7945766 | 0.42685982 | 0.88709241 |
| VARS | 3891.58982 | -0.0488731 | 0.18307121 | -0.2669625 | 0.78949806 | 0.97090793 |

| | | | | | | |
|---|---|---|---|---|---|---|
| LSM2 | 966.612016 | 0.14705809 | 0.20477618 | 0.71814058 | 0.47267061 | 0.90160919 |
| HSPA1L | 112.423664 | 0.3253784 | 0.32090038 | 1.01395454 | 0.31060445 | 0.84476049 |
| HSPA1A | 7483.74624 | -0.9396497 | 0.40351731 | -2.3286477 | 0.01987774 | 0.49477123 |
| HSPA1B | 8171.29099 | -0.559587 | 0.37953224 | -1.4744123 | 0.14037062 | 0.725307 |
| C6orf48 | 4783.47945 | 0.16222384 | 0.13791311 | 1.17627568 | 0.23948473 | 0.81163641 |
| NEU1 | 7717.08614 | 0.10213708 | 0.29714105 | 0.34373264 | 0.73104736 | 0.95848028 |
| SLC44A4 | 7748.73251 | 0.17107133 | 0.32632111 | 0.52424229 | 0.60011003 | 0.93487648 |
| EHMT2-AS1 | 1173.03748 | 0.07400975 | 0.16698963 | 0.44319971 | 0.65762129 | 0.94558535 |
| EHMT2 | 2830.82701 | 0.05670212 | 0.14353842 | 0.39503098 | 0.69282002 | 0.9519683 |
| C2 | 2729.2912 | 0.2070057 | 0.57650456 | 0.35907036 | 0.71954246 | 0.9566409 |
| ZBTB12 | 192.248145 | 0.06125508 | 0.2857305 | 0.21438062 | 0.83025025 | 0.97550416 |
| AL645922.1 | 14378.9427 | -0.2498804 | 0.51852899 | -0.4819025 | 0.62987518 | 0.94088547 |
| C2-AS1 | 76.8752487 | 0.50960115 | 0.78768299 | 0.64696223 | 0.51765639 | 0.91636259 |
| CFB | 12580.4937 | -0.3341328 | 0.56852033 | -0.5877236 | 0.55671786 | 0.92373235 |
| NELFE | 3441.0069 | -0.0509195 | 0.1455105 | -0.349937 | 0.72638601 | 0.95848028 |
| SKIV2L | 3877.19336 | 0.08628079 | 0.1580958 | 0.54575005 | 0.58523776 | 0.93289552 |
| DXO | 1979.74784 | -0.2489822 | 0.14866434 | -1.6747945 | 0.09397456 | 0.65783452 |
| STK19 | 2095.97876 | -0.0587405 | 0.16276632 | -0.3608886 | 0.71818275 | 0.95648644 |
| C4A | 5823.25213 | 0.46121911 | 0.40395114 | 1.14176956 | 0.25354982 | 0.8188642 |
| C4A-AS1 | 48.3994924 | -0.3532805 | 0.77118888 | -0.4580986 | 0.64688161 | 0.94246942 |
| C4B | 6659.0037 | 0.15384443 | 0.59597776 | 0.25813788 | 0.7963005 | 0.97101097 |
| C4B-AS1 | 47.1468004 | -0.388342 | 0.79193459 | -0.4903713 | 0.62387116 | 0.94054939 |
| CYP21A2 | 869.38743 | 0.50530379 | 0.46986446 | 1.07542458 | 0.28218468 | 0.8300774 |
| TNXB | 3633.10354 | 0.09614762 | 0.48186898 | 0.19953062 | 0.84184769 | 0.9762188 |
| ATF6B | 5663.23429 | 0.05681352 | 0.18525706 | 0.30667396 | 0.75909155 | 0.96502715 |
| FKBPL | 334.880535 | -0.3869484 | 0.18895481 | -2.0478354 | 0.04057613 | 0.55657615 |
| PRRT1 | 373.898646 | 0.2181927 | 0.45748056 | 0.4769442 | 0.63340186 | 0.94088547 |
| AL662884.4 | 369.947609 | 0.23059887 | 0.29316125 | 0.78659396 | 0.4315196 | 0.88976671 |
| PPT2 | 1943.64581 | -0.0031882 | 0.20852131 | -0.0152895 | 0.9878012 | 0.9975136 |
| PPT2-EGFL8 | 6263.8963 | 0.09429388 | 0.2079985 | 0.45333924 | 0.65030448 | 0.94322083 |
| EGFL8 | 2190.12767 | -0.0048301 | 0.28315579 | -0.0170583 | 0.98639014 | 0.99715899 |
| AGPAT1 | 5395.68927 | 0.19229647 | 0.21232568 | 0.90566753 | 0.36511185 | 0.8663515 |
| RNF5 | 4484.24278 | 0.17520062 | 0.12445362 | 1.4077583 | 0.15920265 | 0.7475863 |
| AGER | 892.14847 | 0.09457314 | 0.36886305 | 0.25639094 | 0.79764898 | 0.97148232 |
| PBX2 | 6050.76832 | -0.2213194 | 0.2149875 | -1.0294524 | 3.03E-01 | 8.42E-01 |
| GPSM3 | 1232.51422 | -0.268685 | 0.39566326 | -0.679075 | 4.97E-01 | 9.09E-01 |
| NOTCH4 | 3770.91448 | -0.1398937 | 0.37926611 | -0.3688538 | 7.12E-01 | 9.56E-01 |
| AL662796.1 | 54.386231 | 0.32739767 | 0.42732064 | 0.76616396 | 0.44357875 | 0.89366134 |
| HLA-DRA | 20397.1684 | 0.17184789 | 0.40804053 | 0.42115396 | 0.67364266 | 0.94935413 |
| HLA-DRB5 | 6428.44171 | 0.87717039 | 0.47609368 | 1.84243235 | 0.06541193 | 0.62237346 |
| HLA-DRB1 | 17116.7617 | 0.11665115 | 0.41215781 | 0.28302546 | 0.77715732 | 0.96836854 |
| HLA-DQA1 | 2489.5304 | 0.54526908 | 0.57221335 | 0.95291221 | 0.34063456 | 0.85849463 |
| HLA-DQB1 | 4356.99765 | 0.72606596 | 0.65216387 | 1.11331828 | 2.66E-01 | 8.26E-01 |
| HLA-DQB1-A | 1086.12477 | 0.57611198 | 0.69235509 | 0.83210478 | 4.05E-01 | 8.81E-01 |
| HLA-DQA2 | 606.994837 | 2.18599117 | 1.1839593 | 1.84633979 | 0.06484288 | 0.62075686 |
| HLA-DQB2 | 423.760193 | 1.50999236 | 0.72402561 | 2.08555102 | 0.03701931 | 0.55522536 |
| HLA-DOB | 137.363826 | 0.53276302 | 0.6820698 | 0.7810975 | 0.43474515 | 0.89055338 |
| AL669918.1 | 3239.05211 | -1.0823678 | 0.52256081 | -2.0712763 | 3.83E-02 | 5.55E-01 |
| TAP2 | 4701.38048 | -1.0323893 | 0.49574265 | -2.0825106 | 0.03729585 | 0.55532957 |
| PSMB8 | 3956.76305 | -0.2696455 | 0.40738418 | -0.6618949 | 0.50803861 | 0.91439937 |
| PSMB8-AS1 | 3020.14098 | -0.6959257 | 0.51335787 | -1.3556347 | 0.17521544 | 0.76073329 |
| PSMB9 | 1534.21378 | -0.6666464 | 0.60684747 | -1.0985402 | 0.27196866 | 0.82716979 |
| TAP1 | 6025.33547 | -0.7680182 | 0.53953379 | -1.4234849 | 0.15459563 | 0.74300499 |
| HLA-DMB | 2405.60017 | 0.31676268 | 0.42124709 | 0.75196408 | 0.45207266 | 0.89670027 |

| | | | | | | |
|---|---|---|---|---|---|---|
| AL645941.2 | 1689.13218 | 0.31252945 | 0.38017378 | 0.82206999 | 0.41103706 | 0.88268208 |
| HLA-DMA | 4231.56297 | 0.02192933 | 0.39853128 | 0.05502537 | 0.95611825 | 0.99185043 |
| BRD2 | 13707.0866 | -0.3438489 | 0.27107858 | -1.2684474 | 0.20463823 | 0.78460928 |
| AL645941.3 | 1213.67198 | -0.4377366 | 0.4039482 | -1.0836453 | 0.27852211 | 0.82805357 |
| HLA-DOA | 1223.3237 | 0.27712564 | 0.43780673 | 0.63298625 | 5.27E-01 | 0.91769856 |
| HLA-DPA1 | 10587.4409 | 0.32141617 | 0.50633055 | 0.63479513 | 0.52556204 | 0.91759401 |
| HLA-DPB1 | 5884.17229 | 0.45796818 | 0.50408862 | 0.90850727 | 0.36361027 | 0.8663515 |
| COL11A2 | 230.107754 | 0.07437731 | 0.31121607 | 0.2389893 | 0.81111388 | 0.97352497 |
| RXRB | 4299.39179 | -0.0259747 | 0.13321023 | -0.1949902 | 0.84540065 | 0.97675487 |
| SLC39A7 | 9541.20417 | -0.2368798 | 0.27720813 | -0.8545197 | 0.39281712 | 0.8778303 |
| HSD17B8 | 1230.0833 | 0.48180163 | 0.29272256 | 1.64593271 | 0.09977761 | 0.67115544 |
| RING1 | 3872.7692 | -0.3447981 | 0.13662393 | -2.5237023 | 0.01161262 | 0.40423902 |
| HCG25 | 1505.74686 | 0.05119464 | 0.19131879 | 0.26758817 | 0.78901634 | 0.97090793 |
| VPS52 | 3449.61468 | 0.09304567 | 0.16866969 | 0.5516443 | 0.58119208 | 0.9317705 |
| RPS18 | 31784.0663 | 0.25107173 | 0.21565747 | 1.16421532 | 0.24433676 | 0.81468457 |
| B3GALT4 | 488.28101 | -0.2128784 | 0.26509964 | -0.8030128 | 0.42196733 | 0.88451372 |
| WDR46 | 2301.24264 | -0.1612839 | 0.21244007 | -0.7591972 | 0.44773458 | 0.89480207 |
| PFDN6 | 2004.82036 | -0.1550795 | 0.1135781 | -1.3653994 | 0.17212756 | 0.75974924 |
| RGL2 | 6420.73575 | -0.1789414 | 0.2412735 | -0.7416538 | 0.4582971 | 0.8971553 |
| TAPBP | 10332.7877 | -0.1660364 | 0.25723926 | -0.6454552 | 0.51863221 | 0.91636259 |
| ZBTB22 | 1397.50057 | -0.0983972 | 0.15127212 | -0.6504647 | 5.15E-01 | 0.91636259 |
| DAXX | 2695.40793 | -0.2738717 | 0.19912337 | -1.3753868 | 0.16901156 | 0.75786012 |
| AL662820.1 | 361.010555 | -0.2764786 | 0.28065978 | -0.9851024 | 0.32457378 | 0.85169085 |
| KIFC1 | 74.3525637 | -0.3137944 | 0.60334199 | -0.5200937 | 0.60299824 | 0.93588323 |
| PHF1 | 7184.00629 | -0.2373702 | 0.16396402 | -1.447697 | 0.14770181 | 0.73443345 |
| CUTA | 4756.08656 | 0.1708901 | 0.18492796 | 0.92409012 | 0.3554394 | 0.86364633 |
| SYNGAP1 | 3022.32863 | -0.4825222 | 0.25958586 | -1.8588154 | 0.06305331 | 0.61481896 |
| SYNGAP1-AS | 253.997391 | -0.3818818 | 0.336888 | -1.1335572 | 0.25698034 | 0.82107009 |
| ZBTB9 | 547.289475 | -0.1600395 | 0.26949329 | -0.5938532 | 0.5526103 | 0.92373235 |
| BAK1 | 1730.75193 | -0.3790026 | 0.19409138 | -1.952702 | 0.05085492 | 0.5861488 |
| ITPR3 | 4768.27068 | 0.01206107 | 0.2586304 | 0.04663439 | 0.96280462 | 0.99320711 |
| UQCC2 | 3872.68413 | 0.06147165 | 0.20767655 | 0.29599707 | 0.76723232 | 0.96749893 |
| IP6K3 | 354.003911 | 0.02490008 | 0.68364264 | 0.03642266 | 0.97094535 | 0.99400331 |
| LEMD2 | 3622.05662 | -0.1990344 | 0.14589109 | -1.3642669 | 0.17248359 | 0.75974924 |
| HMGA1 | 5486.92399 | -1.1857537 | 0.63157737 | -1.877448 | 0.06045671 | 0.60849584 |
| SMIM29 | 1058.26411 | -0.1694456 | 0.28311338 | -0.5985078 | 0.54950115 | 0.92373235 |
| NUDT3 | 5411.84732 | 0.05085021 | 0.21801311 | 0.23324383 | 0.81557208 | 0.9738447 |
| RPS10-NUDT | 19085.0191 | 0.13363718 | 0.18148492 | 0.73635416 | 0.46151519 | 0.89755243 |
| RPS10 | 18058.2567 | 0.14849701 | 0.19063123 | 0.77897526 | 0.43599429 | 0.89095725 |
| PACSIN1 | 206.394977 | 0.258155 | 0.63961995 | 0.40360685 | 0.68650185 | 0.95097288 |
| C6orf106 | 8645.61316 | -0.1026117 | 0.30422094 | -0.3372934 | 0.7358957 | 0.95920305 |
| AL451165.2 | 323.504809 | 0.13106661 | 0.42027126 | 0.31186193 | 0.75514545 | 0.96360109 |
| SNRPC | 2403.62974 | -0.0896789 | 0.17889634 | -0.5012898 | 0.61616717 | 0.93910158 |
| UHRF1BP1 | 1388.66343 | -0.1248177 | 0.21575461 | -0.5785171 | 0.56291502 | 0.92452095 |
| TAF11 | 1399.56873 | -0.0585984 | 0.14482437 | -0.4046167 | 0.68575927 | 0.95097288 |
| ANKS1A | 2740.92966 | -0.1653993 | 0.18521359 | -0.8930191 | 0.37184697 | 0.86762749 |
| SCUBE3 | 367.245929 | 0.78361386 | 0.57052685 | 1.37349164 | NA | NA |
| Z97832.2 | 185.427327 | 0.09651823 | 0.40078211 | 0.24082469 | 0.809691 | 0.97334138 |
| ZNF76 | 3616.70811 | -0.0385513 | 0.15771889 | -0.2444303 | 0.80689758 | 0.97334138 |
| DEF6 | 482.087369 | -0.1988979 | 0.58838948 | -0.3380378 | 0.73533473 | 0.95895327 |
| PPARD | 3070.29535 | -0.0151854 | 0.19610004 | -0.0774372 | 0.93827578 | 0.99113899 |
| FANCE | 169.954253 | -0.213817 | 0.28402666 | -0.752806 | 0.45156649 | 0.89639459 |
| RPL10A | 19567.0651 | 0.28843011 | 0.22566416 | 1.27813874 | 0.20120051 | 0.78134034 |
| TEAD3 | 2865.46555 | -0.3727817 | 0.2714526 | -1.3732845 | 0.16966392 | 0.7586159 |

| | | | | | | |
|---|---|---|---|---|---|---|
| FKBP5 | 8912.78926 | -0.5121467 | 0.68438812 | -0.7483279 | 0.45426243 | 0.89670027 |
| ARMC12 | 79.3805502 | -0.9413089 | 0.53923175 | -1.7456482 | 0.0808721 | 0.64313548 |
| LHFPL5 | 667.741875 | 0.18874988 | 0.22548209 | 0.83709478 | 0.40253929 | 0.88132015 |
| SRPK1 | 2677.09228 | 0.19144578 | 0.2404516 | 0.79619255 | 0.42592013 | 0.88641523 |
| MAPK14 | 4015.74721 | -0.2126787 | 0.23584887 | -0.9017586 | 0.36718513 | 0.86683872 |
| MAPK13 | 2722.20317 | 0.08508771 | 0.15991629 | 0.53207657 | 0.59467297 | 0.9334601 |
| Z84485.1 | 58.0554467 | -0.1119966 | 0.68232868 | -0.1641388 | 0.86962186 | 0.98275231 |
| BRPF3 | 3187.44476 | -0.3602512 | 0.29762871 | -1.2104048 | 0.22612361 | 0.80246466 |
| PNPLA1 | 242.229079 | 1.23051398 | 0.68888872 | 1.7862304 | 0.07406196 | 0.63749647 |
| ETV7 | 290.077759 | -1.2412247 | 0.95410764 | -1.3009273 | NA | NA |
| KCTD20 | 2699.65713 | 0.04418656 | 0.19664148 | 0.22470619 | 0.82220785 | 0.97500616 |
| STK38 | 3190.89884 | 0.19372093 | 0.16430091 | 1.17906177 | 0.23837358 | 0.81092018 |
| SRSF3 | 9478.1463 | -0.1051888 | 0.18939113 | -0.5554052 | 0.57861752 | 0.93139743 |
| CDKN1A | 12250.2459 | 0.02968337 | 0.45283041 | 0.06555074 | 0.94773551 | 0.99170503 |
| CPNE5 | 305.077804 | -0.2726379 | 0.2580051 | -1.0567152 | 0.29064158 | 0.83442894 |
| PPIL1 | 1122.40314 | 0.03014365 | 0.19496255 | 0.1546125 | 0.87712682 | 0.98275231 |
| C6orf89 | 5670.21102 | 0.00073024 | 0.16095245 | 0.00453702 | 0.99637999 | 0.99976751 |
| PI16 | 162.381178 | -0.3323582 | 1.01318641 | -0.3280326 | 0.74288698 | 0.96109001 |
| MTCH1 | 15008.5653 | -0.3148806 | 0.15731071 | -2.0016478 | 0.04532262 | 0.57529633 |
| FGD2 | 627.833833 | -0.8969997 | 0.60836052 | -1.4744541 | 1.40E-01 | 7.25E-01 |
| PIM1 | 2818.6801 | -0.3655086 | 0.34828928 | -1.0494396 | 0.29397584 | 0.83580671 |
| TMEM217 | 177.543952 | -1.315651 | 0.5511727 | -2.3870033 | 1.70E-02 | 0.46671416 |
| TBC1D22B | 1461.71231 | -0.3926731 | 0.19384469 | -2.0257097 | 0.04279454 | 0.56625172 |
| RNF8 | 1204.88982 | -0.1738144 | 0.17929682 | -0.9694224 | 0.33233448 | 0.85496237 |
| CMTR1 | 2920.57006 | -0.5157024 | 0.3688821 | -1.3980143 | 0.16210877 | 0.75026273 |
| CCDC167 | 332.592685 | 0.56372737 | 0.23474423 | 2.40145363 | 0.01633008 | 0.45915089 |
| MDGA1 | 302.545043 | -0.1605576 | 0.48951261 | -0.3279948 | 0.74291557 | 0.96109001 |
| ZFAND3 | 5053.19998 | -0.2143653 | 0.13348361 | -1.6059294 | 0.10828943 | 0.67789099 |
| BTBD9 | 1276.46574 | -0.1451834 | 0.1675972 | -0.866264 | 0.38634539 | 0.87432552 |
| GLO1 | 4839.95308 | 0.16555296 | 0.15447427 | 1.07171865 | 0.28384642 | 0.83107779 |
| SAYSD1 | 1001.34611 | -0.1096567 | 0.1579903 | -0.6940724 | 0.48763679 | 0.90573694 |
| KCNK5 | 6286.99577 | 1.25625195 | 0.48207332 | 2.60593544 | 0.00916237 | 0.38664841 |
| KIF6 | 118.02768 | 0.32051208 | 0.75271801 | 0.42580632 | 0.67024898 | 0.949185 |
| DAAM2 | 1928.07018 | -0.2102043 | 0.33711339 | -0.6235418 | 0.53292855 | 0.92017817 |
| AL590999.1 | 465.543557 | -0.1627866 | 0.42114578 | -0.3865328 | 0.69910213 | 0.95299633 |
| MOCS1 | 4159.03151 | 0.20513962 | 0.2808852 | 0.73033263 | 0.46518689 | 0.89921277 |
| UNC5CL | 9800.63475 | 0.38473438 | 0.48922424 | 0.78641724 | 0.43162309 | 0.88976671 |
| OARD1 | 3203.53722 | 0.27757367 | 0.23031383 | 1.20519759 | 0.22812703 | 0.80300331 |
| NFYA | 929.11169 | -0.1064148 | 0.18131859 | -0.5868941 | 0.55727484 | 0.92374738 |
| TREM2 | 81.5638629 | -0.2142866 | 0.69012671 | -0.3105033 | 0.75617827 | 0.96450325 |
| TREM1 | 50.8830274 | 0.1219934 | 0.76646323 | 0.15916406 | 0.87353962 | 0.98275231 |
| FOXP4 | 3417.9179 | -0.1121645 | 0.20373712 | -0.5505353 | 0.58195227 | 0.9317705 |
| MDFI | 327.590504 | -0.079727 | 0.46291563 | -0.1722279 | 0.86325837 | 0.98176973 |
| TFEB | 1809.72356 | 0.12399365 | 0.27285847 | 0.45442478 | 0.64952312 | 0.94312988 |
| FRS3 | 539.546658 | 0.17094499 | 0.31212808 | 0.54767578 | 0.58391454 | 0.93233498 |
| AL365205.1 | 5335.68629 | 0.04248332 | 0.12567722 | 0.33803514 | 0.73533671 | 0.95895327 |
| PRICKLE4 | 1125.54047 | -0.0786249 | 0.3699314 | -0.2125393 | 0.83168635 | 0.97550416 |
| TOMM6 | 4054.15133 | 0.07064378 | 0.1495668 | 0.47232262 | 0.63669653 | 0.94088547 |
| USP49 | 420.655985 | -0.0125922 | 0.21636102 | -0.0581999 | 0.9535894 | 0.99170503 |
| MED20 | 1063.51851 | -0.0712791 | 0.15714026 | -0.4536015 | 0.6501157 | 0.943166 |
| BYSL | 668.600395 | -0.3292663 | 0.31683193 | -1.0392459 | 0.29869038 | 0.84056272 |
| CCND3 | 3570.47292 | -0.3787298 | 0.30407251 | -1.2455246 | 0.21293896 | 0.7926732 |
| TAF8 | 1565.80857 | -0.1911409 | 0.13171163 | -1.4512073 | 0.14672214 | 0.73337969 |
| AL512274.1 | 62.0219671 | 0.58230808 | 0.4849779 | 1.20068991 | 0.22987151 | 0.80551898 |

| | | | | | | |
|---|---|---|---|---|---|---|
| C6orf132 | 610.977395 | 0.30348419 | 0.33271108 | 0.91215532 | 0.36168696 | 0.86528137 |
| GUCA1B | 91.7987396 | 0.21584163 | 0.3739762 | 0.57715339 | 5.64E-01 | 9.25E-01 |
| MRPS10 | 2010.10829 | 0.14779451 | 0.17823781 | 0.82919844 | 0.40699213 | 0.88132015 |
| TRERF1 | 603.642339 | 0.21702918 | 0.3621178 | 0.5993331 | 0.54895078 | 0.92373235 |
| UBR2 | 2569.50606 | -0.1000455 | 0.13801919 | -0.7248667 | 0.46853379 | 0.89984974 |
| PRPH2 | 96.5838409 | 0.55811645 | 0.47494908 | 1.17510797 | 0.23995152 | 0.81209487 |
| TBCC | 839.966063 | -0.1020984 | 0.15562261 | -0.656064 | 0.51178297 | 0.91512475 |
| BICRAL | 866.110963 | -0.007878 | 0.10414851 | -0.0756416 | 0.93970425 | 0.99134327 |
| RPL7L1 | 5052.14421 | -1.43E-01 | 0.17101739 | -0.8372758 | 0.40243756 | 0.88132015 |
| C6orf226 | 276.094282 | -0.5482802 | 0.28989932 | -1.8912779 | 0.05858726 | 0.60332684 |
| CNPY3 | 2267.14853 | -0.1349833 | 0.18702564 | -0.721737 | 0.47045621 | 0.9008571 |
| AL035587.1 | 89.9645319 | -0.155739 | 0.27008201 | -0.5766361 | 5.64E-01 | 0.92470993 |
| GNMT | 48.6177175 | -0.7090982 | 0.44513496 | -1.5929959 | 0.11116111 | 0.68001961 |
| PEX6 | 2858.23442 | 0.29024423 | 0.32897921 | 0.88225706 | 0.37763781 | 0.86987124 |
| PPP2R5D | 2647.4781 | 0.00260984 | 0.15332794 | 0.01702128 | 0.98641964 | 0.99715899 |
| MEA1 | 3902.58487 | -0.0304628 | 0.12760517 | -0.2387268 | 0.81131746 | 0.97352497 |
| KLHDC3 | 4865.21856 | -0.1755289 | 0.12443992 | -1.4105516 | 0.15837687 | 0.7466592 |
| RRP36 | 1747.84584 | -0.1426691 | 0.19888961 | -0.7173279 | 0.4731718 | 0.90179314 |
| AL136304.1 | 94.5166543 | -0.1274389 | 0.25083984 | -0.5080488 | 0.61141912 | 0.93834929 |
| CUL7 | 2722.72331 | -0.328767 | 0.14472959 | -2.2715948 | 0.023111 | 0.51967919 |
| KLC4 | 3866.18976 | 0.28689553 | 0.19661925 | 1.45914263 | 0.14452586 | 0.73032824 |
| MRPL2 | 2382.17093 | 0.1751062 | 0.19346392 | 0.90511034 | 0.36540693 | 0.8663515 |
| AL355385.1 | 544.733164 | 0.35992843 | 0.21205179 | 1.69736091 | 0.08962845 | 0.65375543 |
| PTK7 | 2705.82688 | 0.12294638 | 0.33434493 | 0.36772319 | 0.71307965 | 0.95564093 |
| SRF | 2671.45982 | -0.2983603 | 0.21856795 | -1.3650687 | 0.17223147 | 0.75974924 |
| CUL9 | 2434.87555 | -0.2069102 | 0.13532402 | -1.5289984 | 0.12626485 | 0.70426472 |
| AL133375.1 | 670.72375 | -0.1149692 | 0.17279282 | -0.6653587 | 0.50582107 | 0.91387185 |
| DNPH1 | 3118.71481 | 0.36301944 | 0.1855425 | 1.95652987 | 0.05040277 | 0.5861488 |
| TTBK1 | 192.035774 | 0.46802263 | 0.6301067 | 0.74276726 | 0.4576226 | 0.89688594 |
| SLC22A7 | 8126.99411 | 1.79058576 | 0.71097436 | 2.51849555 | 0.01178574 | 0.40423902 |
| CRIP3 | 1255.59481 | 0.28109197 | 0.5977517 | 0.47024871 | 0.63817734 | 0.94088547 |
| ZNF318 | 1353.01635 | -0.3065998 | 0.19095972 | -1.6055731 | 0.10836775 | 0.67800851 |
| ABCC10 | 1666.00003 | -0.2772797 | 0.18113613 | -1.5307806 | 0.12582363 | 0.70347499 |
| DLK2 | 105.751014 | -0.6811089 | 0.31201069 | -2.1829665 | 0.02903828 | 0.53606221 |
| TJAP1 | 2689.02553 | -0.2690219 | 0.14899869 | -1.8055322 | 0.07099144 | 0.63056668 |
| LRRC73 | 219.056129 | -0.1176368 | 0.28305483 | -0.4155971 | 0.67770484 | 0.95018034 |
| POLR1C | 2550.29787 | -0.2693113 | 0.1863499 | -1.4451916 | 0.14840409 | 0.73519456 |
| YIPF3 | 6436.22927 | -0.1702645 | 0.12137918 | -1.4027484 | 0.16069187 | 0.74855808 |
| AL355802.2 | 391.77722 | -0.4434557 | 0.28759143 | -1.5419643 | 1.23E-01 | 6.98E-01 |
| XPO5 | 1889.9299 | -0.2346144 | 0.15023117 | -1.5616893 | 0.1183612 | 0.69103031 |
| POLH | 428.50942 | 0.09830089 | 0.18936155 | 0.51911746 | 0.60367884 | 0.93626289 |
| GTPBP2 | 4603.70733 | -0.655565 | 0.23561235 | -2.7823881 | 0.00539605 | 0.31308771 |
| MAD2L1BP | 1155.66331 | -0.1603049 | 0.25621082 | -0.6256757 | 0.5315277 | 0.91962754 |
| RSPH9 | 1425.3648 | -0.0695002 | 0.2020242 | -0.3440194 | 0.7308317 | 0.95848028 |
| MRPS18A | 1982.62929 | -0.0388184 | 0.19883093 | -0.1952331 | 0.84521043 | 0.97675487 |
| VEGFA | 21084.7265 | -0.1442812 | 0.35069564 | -0.4114143 | 0.68076876 | 0.95070301 |
| AL136131.3 | 914.467052 | -0.0625801 | 0.49970713 | -0.1252336 | 0.90033858 | 0.98526603 |
| AL109615.3 | 134.500859 | -0.2332864 | 0.3604365 | -0.6472329 | 0.5174812 | 0.91636259 |
| C6orf223 | 449.69156 | -0.4510174 | 0.44051297 | -1.0238459 | 3.06E-01 | 0.84270033 |
| MRPL14 | 1580.78065 | -0.1018329 | 0.24593656 | -0.4140616 | 6.79E-01 | 0.95046197 |
| TMEM63B | 3213.01056 | -0.0150898 | 0.18666167 | -0.0808404 | 0.93556892 | 0.99068685 |
| SLC29A1 | 4067.708 | -0.0557179 | 0.2893315 | -0.1925747 | 0.84729204 | 0.97693491 |
| HSP90AB1 | 41927.6273 | -0.0266934 | 0.17714094 | -0.1506902 | 0.88022014 | 0.98275231 |
| SLC35B2 | 3878.00031 | -0.0606975 | 0.16662277 | -0.3642807 | 0.71564845 | 0.95590747 |

| | | | | | | |
|---|---|---|---|---|---|---|
| NFKBIE | 1385.44304 | 0.16902712 | 0.26621335 | 0.63493105 | 5.25E-01 | 0.91759401 |
| TMEM151B | 105.259742 | -0.0390701 | 0.29847882 | -0.1308975 | 0.89585641 | 0.98474725 |
| AL353588.1 | 103.354967 | 0.00419816 | 0.32180465 | 0.01304569 | 0.98959134 | 0.99800524 |
| AARS2 | 1168.00893 | 0.09705037 | 0.21124621 | 0.45941828 | 0.64593382 | 0.94208859 |
| CDC5L | 1652.76526 | -0.1684311 | 0.15573026 | -1.0815568 | 0.27944952 | 0.82825734 |
| SUPT3H | 404.40702 | 0.10362367 | 0.25308128 | 0.40944818 | 0.68221079 | 0.95070301 |
| RUNX2 | 231.693507 | -0.020044 | 0.33624664 | -0.059611 | 0.95246547 | 0.99170503 |
| CLIC5 | 3863.60246 | 0.53622018 | 0.74204511 | 0.72262477 | 0.46991044 | 0.9008571 |
| ENPP4 | 2051.50782 | 0.63704764 | 0.14757996 | 4.3166271 | 1.58E-05 | 0.01316658 |
| ENPP5 | 1784.78535 | 0.5905277 | 0.24594696 | 2.40103681 | 0.01634869 | 0.45915089 |
| RCAN2 | 1772.82295 | 0.34287658 | 0.39341804 | 0.87153244 | 0.3834635 | 0.87214678 |
| CYP39A1 | 335.99013 | 0.61798716 | 0.43722097 | 1.41344355 | 0.15752535 | 0.74522306 |
| SLC25A27 | 1724.14248 | -0.0459174 | 0.43731137 | -0.1049994 | 0.9163763 | 0.9868044 |
| TDRD6 | 280.600071 | 0.0977613 | 0.45664893 | 0.21408415 | 0.83048144 | 0.97550416 |
| PLA2G7 | 54.655639 | 0.65843305 | 0.78553991 | 0.83819172 | 0.40192304 | 0.88132015 |
| ADGRF5 | 6374.31 | 0.04215463 | 0.45939644 | 0.0917609 | 0.92688801 | 0.98915637 |
| ADGRF1 | 3641.22091 | 0.16586744 | 0.37178869 | 0.44613363 | 0.65550072 | 0.94532497 |
| TNFRSF21 | 3999.15277 | 0.21857653 | 0.35219663 | 0.62060938 | 0.53485667 | 0.92084006 |
| AL355353.1 | 101.208785 | 0.63947268 | 0.48827636 | 1.30965315 | 0.1903132 | 0.77100296 |
| CD2AP | 4136.34424 | 0.16095211 | 0.1963879 | 0.81956225 | 0.4124657 | 0.88268208 |
| PTCHD4 | 74.4998999 | 1.36768602 | 0.575346 | 2.37715396 | 0.0174468 | 0.46859699 |
| MUT | 3473.42417 | 0.57571641 | 0.2421265 | 2.37775051 | 0.01741861 | 0.46859699 |
| CENPQ | 133.532716 | 0.19876655 | 0.23485635 | 0.84633248 | 0.39736729 | 0.88006972 |
| C6orf141 | 36.544876 | 1.09094242 | 0.74288347 | 1.46852429 | 0.14196186 | 0.72784886 |
| AL138826.1 | 470.892791 | 1.10803638 | 1.23922079 | 0.8941396 | 0.3712472 | 0.86762749 |
| TFAP2B | 1302.07502 | -0.2770658 | 0.32421268 | -0.8545805 | 0.39278343 | 0.8778303 |
| PKHD1 | 3984.46403 | 0.34414637 | 0.45241218 | 0.7606921 | 0.44684099 | 0.89480207 |
| MCM3 | 2204.47028 | -0.0572893 | 0.23516705 | -0.2436112 | 0.80753197 | 0.97334138 |
| PAQR8 | 634.182664 | 0.02722361 | 0.4479146 | 0.06077857 | 0.95153556 | 0.99170503 |
| EFHC1 | 1488.02015 | 0.0295424 | 0.26033871 | 0.11347678 | 0.90965257 | 0.98662757 |
| TRAM2 | 1378.35363 | -0.321743 | 0.18449062 | -1.7439532 | 0.08116724 | 0.6439041 |
| TRAM2-AS1 | 438.271834 | 0.28797318 | 0.28214285 | 1.02066447 | 0.30741343 | 0.84327213 |
| TMEM14A | 1074.29611 | 0.58730264 | 0.2290722 | 2.56383205 | 0.01035236 | 0.40019066 |
| GSTA2 | 7207.83336 | 1.42231009 | 1.11412322 | 1.2766183 | 0.20173704 | 0.78134034 |
| GSTA1 | 8686.75108 | 1.56590411 | 0.74254268 | 2.10884054 | 0.03495835 | 0.55369025 |
| GSTA4 | 1031.53733 | 0.62644617 | 0.2749312 | 2.27855612 | 0.02269346 | 0.51967919 |
| ICK | 595.69919 | 0.09217304 | 0.20412159 | 0.45155947 | 0.65158638 | 0.94393131 |
| FBXO9 | 3806.88554 | 0.01510146 | 0.14813059 | 0.10194695 | 0.91879879 | 0.98709788 |
| GCM1 | 119.740659 | 0.22087903 | 0.5198689 | 0.42487449 | 6.71E-01 | 9.49E-01 |
| ELOVL5 | 3872.97918 | 0.27386146 | 0.2863402 | 0.95641988 | 0.33886015 | 0.85733087 |
| GCLC | 4170.57718 | -0.6360682 | 0.58643579 | -1.0846339 | 0.27808382 | 0.82805357 |
| AL033397.2 | 233.33843 | -0.6073957 | 0.652702 | -0.9305866 | 0.35206747 | 0.86364633 |
| AL033397.1 | 119.452405 | 1.17699415 | 0.895156 | 1.31484808 | 0.18856096 | 0.76970626 |
| LINC01564 | 121.051867 | -0.8509083 | 0.88209972 | -0.9646395 | 0.33472539 | 0.85557672 |
| KLHL31 | 74.0650521 | 0.0622171 | 0.42589326 | 0.14608614 | 0.88385339 | 0.98317437 |
| LRRC1 | 856.446636 | 0.24685697 | 0.17866661 | 1.38166264 | 0.16707531 | 0.75558932 |
| AL033384.2 | 169.525008 | 0.28158822 | 0.21674273 | 1.29918184 | 0.19388153 | 0.77620852 |
| TINAG | 3155.50855 | 0.87570062 | 0.3973717 | 2.20373173 | 0.02754322 | 0.52904542 |
| FAM83B | 98.8067824 | -0.3207419 | 0.40271762 | -0.7964438 | 4.26E-01 | 0.88640673 |
| BMP5 | 112.283396 | 0.57400013 | 0.84968121 | 0.67554764 | 0.4993279 | 0.91161831 |
| COL21A1 | 117.538768 | 1.12224945 | 0.71217498 | 1.57580578 | 0.11507057 | 0.68657674 |
| DST | 5227.60719 | -0.0765787 | 0.27980869 | -0.2736825 | 0.78432864 | 0.97017379 |
| BEND6 | 51.3848395 | 0.4007766 | 0.36313558 | 1.10365555 | 0.26974258 | 0.82716979 |
| KIAA1586 | 270.506577 | -0.1813856 | 0.21320065 | -0.850774 | 0.3948949 | 0.87884163 |

| | | | | | | |
|---|---|---|---|---|---|---|
| ZNF451 | 1605.6561 | -0.121658 | 0.11756216 | -1.0348398 | 0.30074374 | 0.84200819 |
| ZNF451-AS1 | 200.208098 | -0.1663628 | 0.23644943 | -0.7035874 | 0.48168975 | 0.90541935 |
| BAG2 | 1008.53458 | -0.4603902 | 0.29589948 | -1.5559008 | 0.11973169 | 0.69405865 |
| RAB23 | 667.174373 | -0.2388891 | 0.27572401 | -0.8664066 | 0.38626721 | 0.87432552 |
| PRIM2 | 267.905226 | 0.03674914 | 0.16253465 | 0.22610034 | 0.82112338 | 0.97472105 |
| AL021368.2 | 199.783889 | 0.01144271 | 0.19495891 | 0.05869294 | 0.95319668 | 0.99170503 |
| KHDRBS2 | 42.4507966 | 0.74948113 | 0.45184028 | 1.65873021 | 0.09717017 | 0.66486533 |
| LGSN | 152.236042 | -0.7821599 | 0.7144697 | -1.0947418 | 0.27362975 | 0.82719197 |
| PTP4A1 | 9301.05334 | -0.1762979 | 0.29269383 | -0.6023288 | 0.54695529 | 0.92345821 |
| AL135905.1 | 62.7169019 | 0.02583244 | 0.33919622 | 0.07615781 | 0.93929355 | 0.99113899 |
| PHF3 | 2632.39613 | -0.1260212 | 0.12437658 | -1.0132231 | 0.31095361 | 0.84476049 |
| EYS | 53.3249002 | -0.0617118 | 0.28507646 | -0.2164744 | 0.82861794 | 0.97550416 |
| ADGRB3 | 57.5181701 | -0.1465977 | 0.59251475 | -0.247416 | 0.80458626 | 0.97287947 |
| LMBRD1 | 5912.81336 | 0.33522077 | 0.11983235 | 2.79741458 | 0.00515134 | 0.30843846 |
| COL19A1 | 70.5557103 | 0.30051004 | 0.50213083 | 0.5984696 | 0.54952664 | 0.92373235 |
| COL9A1 | 45.8235043 | 0.56474961 | 0.87508578 | 0.64536486 | 0.51869077 | 0.91636259 |
| FAM135A | 1417.24623 | -0.0763878 | 0.23093081 | -0.3307821 | 0.74080909 | 0.96089108 |
| SDHAF4 | 643.207232 | 0.66725096 | 0.20674501 | 3.22741014 | 0.00124916 | 0.1592555 |
| SMAP1 | 1784.35954 | -0.0439782 | 0.200629 | -0.2192017 | 0.82649296 | 0.97548201 |
| B3GAT2 | 1615.91602 | -0.0612036 | 0.19314167 | -0.3168844 | 0.75133134 | 0.96284879 |
| OGFRL1 | 2269.67675 | -1.0025498 | 0.51268881 | -1.9554744 | 0.05052711 | 0.5861488 |
| AL136164.2 | 246.01828 | -0.6491835 | 0.35668203 | -1.8200624 | 0.06874951 | 0.62750387 |
| LINC00472 | 558.257735 | -0.4419016 | 0.43472352 | -1.0165119 | 0.30938567 | 0.84414094 |
| KHDC1 | 63.1278793 | 0.39821121 | 0.40721605 | 0.97788683 | 0.3281303 | 0.85317073 |
| AC019205.1 | 74.224918 | 0.08482585 | 0.39877112 | 0.21271814 | 0.83154682 | 0.97550416 |
| DDX43 | 84.8354145 | 0.06978595 | 0.80941345 | 0.08621793 | 0.93129318 | 0.98975562 |
| CGAS | 175.640284 | -0.042301 | 0.4001923 | -0.1057016 | 0.91581912 | 0.9868044 |
| MTO1 | 1495.27454 | 0.09385715 | 0.15762909 | 0.59543043 | 5.52E-01 | 9.24E-01 |
| AL603910.1 | 208.050624 | 0.22744724 | 0.19296849 | 1.17867556 | 2.39E-01 | 0.81095554 |
| EEF1A1 | 332840.473 | 0.36123476 | 0.27343828 | 1.32108338 | 0.18647356 | 0.76908034 |
| SLC17A5 | 3600.22293 | 0.02867184 | 0.46346274 | 0.06186439 | 0.95067083 | 0.99170503 |
| CD109 | 1018.31726 | 0.11927587 | 0.34464187 | 0.3460864 | 0.72927778 | 0.95848028 |
| COL12A1 | 2496.68726 | 0.52867106 | 0.47768424 | 1.10673749 | 0.26840744 | 0.82676682 |
| COX7A2 | 5525.51709 | 0.09614546 | 0.1789723 | 0.53720861 | 0.59112352 | 0.93289552 |
| TMEM30A | 7992.85936 | 0.10846149 | 0.15768408 | 0.68784046 | 0.49155326 | 0.90600166 |
| AL080250.1 | 55.6676702 | -0.1033026 | 0.28073299 | -0.3679746 | 0.71289219 | 0.95564093 |
| FILIP1 | 223.042393 | -0.6397834 | 0.44334015 | -1.4430982 | 0.14899281 | 0.73655347 |
| SENP6 | 3978.17642 | -0.2367466 | 0.17984563 | -1.3163877 | 0.18804396 | 0.76970626 |
| MYO6 | 8988.45781 | 0.06358272 | 0.23880114 | 0.26625801 | 0.79004051 | 0.97090793 |
| IMPG1 | 77.538995 | 0.13026459 | 0.34004323 | 0.38308243 | 0.70165865 | 0.95424422 |
| MEI4 | 262.121149 | 1.14588167 | 0.39031193 | 2.93581002 | 0.00332678 | 0.25241775 |
| IRAK1BP1 | 345.434137 | 0.39947645 | 0.32266061 | 1.23807008 | 0.21569008 | 0.7954827 |
| PHIP | 1936.96249 | 7.37E-05 | 0.13747312 | 0.00053633 | 1.00E+00 | 1.00E+00 |
| HMGN3 | 5229.70706 | 0.2820674 | 0.1640942 | 1.71893584 | 0.08562606 | 0.65049132 |
| HMGN3-AS1 | 1297.76885 | 0.2588964 | 0.16075121 | 1.61054093 | 0.10727982 | 0.67707349 |
| LCA5 | 290.969827 | 0.17061872 | 0.20369876 | 0.83760312 | 4.02E-01 | 8.81E-01 |
| SH3BGRL2 | 1631.38606 | 0.27067606 | 0.2013223 | 1.34449123 | 0.17878959 | 0.76120583 |
| ELOVL4 | 105.625227 | -0.1827802 | 0.62020706 | -0.2947084 | 0.76821666 | 0.96768037 |
| BCKDHB | 1449.04889 | 0.68087661 | 0.2655845 | 2.56369102 | 0.01035657 | 0.40019066 |
| AL359715.3 | 61.4050257 | 1.12392774 | 0.34195859 | 3.28673637 | 0.00101356 | 0.15012259 |
| AL359715.1 | 83.7780338 | 0.59314915 | 0.57680805 | 1.02833023 | 0.30379452 | 0.84211658 |
| AL359715.2 | 90.2118162 | 0.30140596 | 0.4410891 | 0.68332217 | 0.49440329 | 0.90796767 |
| TENT5A | 798.581887 | -0.7420883 | 0.35165269 | -2.1102877 | 0.03483359 | 0.55369025 |
| IBTK | 2770.02191 | -0.1910557 | 0.21316257 | -0.8962911 | 0.37009733 | 0.86762749 |

| | | | | | | |
|---|---|---|---|---|---|---|
| TPBG | 1297.05783 | 0.01192048 | 0.34304569 | 0.03474894 | 0.97227993 | 0.99416401 |
| UBE3D | 237.583271 | 0.13343416 | 0.18911054 | 0.70558814 | 0.4804443 | 0.90541935 |
| DOPEY1 | 2004.77452 | -0.1609008 | 0.19828638 | -0.8114564 | 0.41710363 | 0.88445833 |
| PGM3 | 2631.72394 | -0.2329686 | 0.24890694 | -0.9359666 | 0.3492904 | 0.86364633 |
| RWDD2A | 652.694477 | 0.21312087 | 0.17043139 | 1.25047896 | 2.11E-01 | 0.79134644 |
| ME1 | 666.756934 | 0.55670087 | 0.33191227 | 1.67725307 | 0.09349301 | 0.65756964 |
| PRSS35 | 126.714351 | 0.6910313 | 0.75393148 | 0.91657043 | 0.35936779 | 0.86480228 |
| CYB5R4 | 971.85968 | -0.120196 | 0.26873755 | -0.4472617 | 0.65468614 | 0.94532497 |
| MRAP2 | 237.655741 | 0.08944029 | 0.47693085 | 0.18753303 | 0.85124273 | 0.97854014 |
| CEP162 | 549.740416 | -0.0455429 | 0.26244314 | -0.1735344 | 0.86223141 | 0.98176973 |
| TBX18 | 68.027544 | -0.6014806 | 0.81547408 | -0.737584 | 0.46076728 | 0.89755243 |
| LINC02535 | 389.746033 | 1.53037647 | 0.94230421 | 1.62407898 | 0.10435895 | 0.67439401 |
| NT5E | 1685.2505 | 0.56874903 | 0.27063349 | 2.10154709 | 0.03559297 | 0.55369025 |
| SNX14 | 3746.30062 | 0.16184559 | 0.14862285 | 1.08896843 | 0.27616781 | 0.82719407 |
| AL589666.1 | 1307.15234 | -0.218617 | 0.26906622 | -0.8125027 | 0.41650322 | 0.88419652 |
| SYNCRIP | 6280.90587 | -0.3062019 | 0.2391622 | -1.2803106 | 0.20043591 | 0.78134034 |
| ZNF292 | 1892.61403 | -0.7037065 | 0.34325692 | -2.0500868 | 4.04E-02 | 5.57E-01 |
| SMIM8 | 351.794692 | 0.12486975 | 0.26701189 | 0.46765615 | 0.6400305 | 0.94088547 |
| C6orf163 | 59.1470448 | -0.1403445 | 0.37983108 | -0.369492 | 0.71176107 | 0.95560193 |
| CFAP206 | 114.901392 | 0.11101266 | 0.35112462 | 0.31616314 | 0.75187868 | 0.96289476 |
| AL049697.1 | 1127.43121 | 0.06698015 | 0.1353567 | 0.4948418 | 0.6207118 | 0.94002099 |
| SLC35A1 | 1581.61621 | 0.00396578 | 0.16979327 | 0.02335654 | 0.98136587 | 0.99595643 |
| RARS2 | 1631.71136 | -0.0297308 | 0.14873285 | -0.1998937 | 0.84156375 | 0.97621603 |
| ORC3 | 1063.92435 | 0.10084415 | 0.22933949 | 0.43971562 | 0.66014309 | 0.94628868 |
| AKIRIN2 | 1841.90452 | -0.2156536 | 0.19829501 | -1.0875394 | 0.27679849 | 0.82760008 |
| CNR1 | 138.638768 | 0.76967063 | 0.98434402 | 0.78191223 | 0.43426616 | 0.89055338 |
| RNGTT | 675.525995 | 0.17485711 | 0.18085026 | 0.96686121 | 0.33361342 | 0.855011 |
| PNRC1 | 9170.95787 | -0.1624668 | 0.25979482 | -0.6253658 | 0.53173099 | 0.91962754 |
| SRSF12 | 202.417621 | -0.0683661 | 0.31820029 | -0.2148524 | 0.82988236 | 0.97550416 |
| PM20D2 | 1416.33594 | -0.1666164 | 0.20103448 | -0.8287953 | 0.40722027 | 0.88132015 |
| UBE2J1 | 3342.86288 | -0.1989739 | 0.20805502 | -0.9563524 | 0.33889422 | 0.85733087 |
| RRAGD | 3506.34943 | -0.0617421 | 0.32249966 | -0.1914485 | 0.84817419 | 0.97711043 |
| ANKRD6 | 1552.78489 | -0.2027009 | 0.12259153 | -1.6534658 | 0.0982361 | 0.66549286 |
| LYRM2 | 2296.6845 | -0.1687282 | 0.14770205 | -1.142355 | 2.53E-01 | 8.18E-01 |
| MDN1 | 1735.51559 | -0.1592846 | 0.11333662 | -1.4054115 | 1.60E-01 | 7.48E-01 |
| CASP8AP2 | 447.978691 | -0.1136827 | 0.12921417 | -0.8798008 | 0.37896721 | 0.87069278 |
| BACH2 | 175.952189 | -0.0922749 | 0.29024989 | -0.3179153 | 0.75054922 | 0.96255347 |
| MAP3K7 | 2328.98793 | -0.0653292 | 0.15337836 | -0.4259347 | 0.67015545 | 0.949185 |
| EPHA7 | 741.051611 | 0.49012555 | 0.48380743 | 1.01305917 | 0.3110319 | 0.84476049 |
| MANEA-DT | 42.0663502 | -0.0535345 | 0.41063085 | -0.1303712 | 0.89627272 | 0.98474725 |
| MANEA | 839.340406 | 0.0569963 | 0.15260114 | 0.3734985 | 0.70877746 | 0.95501548 |
| UFL1 | 1696.77311 | 0.17928912 | 0.19990184 | 0.89688576 | 0.36977988 | 0.86762749 |
| FHL5 | 252.259141 | -0.9384123 | 0.66877689 | -1.403177 | 0.16056407 | 0.74816787 |
| GPR63 | 51.7696612 | -0.1429279 | 0.36894298 | -0.3873983 | 0.69846136 | 0.9529028 |
| NDUFAF4 | 1089.98114 | -0.1635016 | 0.20874191 | -0.7832714 | 0.43346776 | 0.89055338 |
| KLHL32 | 142.486038 | 0.60638942 | 0.37994008 | 1.59601329 | 0.11048583 | 0.68001961 |
| MMS22L | 279.755602 | 0.18158652 | 0.34177429 | 0.53130539 | 0.59520717 | 0.9334601 |
| FBXL4 | 1562.44214 | -0.0301594 | 0.14692804 | -0.2052663 | 0.83736406 | 0.97577999 |
| FAXC | 132.884941 | -0.2740132 | 0.3413602 | -0.8027099 | 0.42214245 | 0.88451372 |
| COQ3 | 444.087829 | 0.39126565 | 0.2281638 | 1.71484542 | 0.08637357 | 0.65087539 |
| PNISR | 6610.73587 | -0.0564323 | 0.18319944 | -0.3080374 | 0.75805387 | 0.96502715 |
| AL513550.1 | 234.804591 | -0.1208 | 0.27579957 | -0.4379991 | 0.66138691 | 0.94642611 |
| USP45 | 820.94074 | 0.08284105 | 0.15689809 | 0.52799272 | 0.59750439 | 0.93416444 |
| TSTD3 | 139.782013 | 0.25336765 | 0.23800458 | 1.06454945 | 0.28707987 | 0.83155545 |

| | | | | | | |
|---|---|---|---|---|---|---|
| CCNC | 3847.70146 | -0.0505684 | 0.16215009 | -0.3118619 | 0.75514545 | 0.96360109 |
| AL137784.2 | 332.455182 | -0.1975472 | 0.31885682 | -0.6195484 | 0.53555512 | 0.92101152 |
| SIM1 | 1311.3517 | -0.2860702 | 0.36008479 | -0.7944522 | 0.4269322 | 0.88714229 |
| ASCC3 | 1110.61537 | -0.0702524 | 0.19675228 | -0.3570602 | 0.72104675 | 0.95729223 |
| AL133338.1 | 67.2006393 | 0.04590487 | 0.34575218 | 0.13276813 | 0.89437676 | 0.98474725 |
| GRIK2 | 98.938871 | -0.4035264 | 0.64150054 | -0.6290352 | 0.52932602 | 0.91920857 |
| HACE1 | 702.555479 | 0.06646749 | 0.19863244 | 0.33462557 | 0.73790755 | 0.9602325 |
| BVES | 150.792518 | -0.680919 | 0.56808087 | -1.1986304 | 0.23067168 | 0.80610188 |
| PREP | 1321.77628 | 0.18604551 | 0.13495158 | 1.37860931 | 0.16801524 | 0.75673485 |
| AL133406.2 | 137.468904 | 0.05191125 | 0.18654994 | 0.27826999 | 0.78080511 | 0.96875661 |
| ATG5 | 1889.00326 | -0.1975568 | 0.15116369 | -1.3069064 | 1.91E-01 | 7.73E-01 |
| PRDM1 | 294.199094 | 0.05991734 | 0.50227704 | 0.11929141 | 0.90504449 | 0.98624484 |
| CRYBG1 | 2164.48538 | 0.48482562 | 0.35223861 | 1.3764125 | 0.16869396 | 0.75766078 |
| AL109920.1 | 80.4343482 | 1.59948263 | 0.89515153 | 1.78682891 | 0.07396514 | 0.6374448 |
| RTN4IP1 | 500.546913 | 0.49658703 | 0.23990171 | 2.06996033 | 0.03845606 | 0.55565329 |
| QRSL1 | 1230.48566 | 0.03588468 | 0.19999185 | 0.17943069 | 0.85759954 | 0.97959852 |
| LINC02532 | 5098.98221 | -0.7644383 | 0.48986048 | -1.5605225 | 0.11863645 | 0.69194841 |
| CD24 | 65552.2531 | -0.1102015 | 0.22060575 | -0.4995403 | 0.61739878 | 0.93910158 |
| C6orf203 | 908.741429 | 0.21311605 | 0.20104211 | 1.06005679 | 0.28911876 | 0.83308345 |
| BEND3 | 182.761793 | -0.1233074 | 0.28740889 | -0.4290314 | 0.66790037 | 0.94847921 |
| PDSS2 | 1644.69585 | 0.38958316 | 0.25114123 | 1.55125134 | 0.12084146 | 0.69496908 |
| SOBP | 538.676323 | 0.04973931 | 0.26847557 | 0.18526567 | 0.85302066 | 0.97886533 |
| SCML4 | 70.9381239 | 0.38580269 | 0.75675067 | 0.50981479 | 0.61018122 | 0.93792127 |
| SEC63 | 3887.99113 | 0.07380957 | 0.15658921 | 0.47135798 | 0.63738512 | 0.94088547 |
| AL024507.2 | 99.7540288 | 0.19607468 | 0.42111625 | 0.46560703 | 0.64149681 | 0.94088547 |
| OSTM1 | 2064.25192 | 0.03731181 | 0.23653603 | 0.1577426 | 0.87465963 | 0.98275231 |
| SNX3 | 7137.03225 | -0.0009935 | 0.12231714 | -0.0081225 | 0.99351927 | 0.99920627 |
| AFG1L | 359.783355 | 0.38396137 | 0.25280915 | 1.51877953 | 0.128818 | 0.70633634 |
| FOXO3 | 2723.11002 | -0.1867541 | 0.12705146 | -1.4699088 | 0.14158644 | 0.72784886 |
| ARMC2 | 213.362704 | 0.32831132 | 0.3937421 | 0.83382326 | 0.40438059 | 0.88132015 |
| SESN1 | 1569.01131 | 0.24467378 | 0.25643674 | 0.95412918 | 0.34001826 | 0.85814041 |
| AL390208.1 | 212.722622 | 0.41518057 | 0.30819107 | 1.34715317 | 0.17793091 | 0.76115956 |
| CEP57L1 | 368.001922 | 0.13105205 | 0.23771992 | 0.5512876 | 0.58143654 | 0.9317705 |
| CD164 | 19986.6758 | -0.1377865 | 0.15452543 | -0.8916756 | 0.37256686 | 0.86791605 |
| PPIL6 | 391.277117 | 0.1601472 | 0.21483739 | 0.74543449 | 0.4560091 | 0.89688594 |
| SMPD2 | 909.310041 | -0.2037769 | 0.15337678 | -1.3286032 | 0.18397891 | 0.76791734 |
| MICAL1 | 5379.40049 | -0.0991922 | 0.22683601 | -0.4372859 | 0.661904 | 0.94642611 |
| ZBTB24 | 715.428914 | -0.2583472 | 0.19909344 | -1.2976178 | 0.1944187 | 0.7762856 |
| AK9 | 385.396268 | -0.0227039 | 0.18435988 | -0.1231497 | 9.02E-01 | 0.98572048 |
| FIG4 | 1021.52718 | 0.29774136 | 0.12339227 | 2.41296616 | 0.01582329 | 0.4564177 |
| WASF1 | 1034.0353 | -0.2883122 | 0.24805445 | -1.1622941 | 0.24511604 | 0.81492642 |
| CDC40 | 1050.1713 | -0.0009477 | 0.13864322 | -0.0068353 | 0.99454629 | 0.99940889 |
| METTL24 | 78.4009043 | 0.50110578 | 0.35031845 | 1.43042929 | 0.15259385 | 0.7422778 |
| DDO | 536.797165 | 0.60696272 | 0.56923663 | 1.06627488 | 0.2862994 | 0.83107779 |
| CDK19 | 884.055925 | 0.18963224 | 0.13850528 | 1.36913365 | 0.1709575 | 0.75956362 |
| AMD1 | 5329.79528 | -0.5151927 | 0.34738051 | -1.483079 | 0.13805342 | 0.72209298 |
| GTF3C6 | 2256.30016 | 0.00530464 | 0.17731929 | 0.02991577 | 0.97613423 | 0.99455099 |
| RPF2 | 1184.81566 | -0.1086286 | 0.17997347 | -0.603581 | 0.54612222 | 0.92345821 |
| SLC16A10 | 848.362878 | 0.05468761 | 0.31481592 | 0.17371298 | 0.86209104 | 0.98176973 |
| MFSD4B | 347.984757 | -0.189812 | 0.28353958 | -0.6694373 | 0.50321659 | 0.91356552 |
| AL080317.1 | 207.604181 | -0.3255997 | 0.3363589 | -0.9680129 | 0.33303794 | 0.855011 |
| AL080317.2 | 92.5425661 | -0.2559076 | 0.59516886 | -0.4299747 | 0.66721404 | 0.94830184 |
| REV3L | 1455.78876 | -0.0828991 | 0.18671338 | -0.4439914 | 0.65704883 | 0.94545756 |
| AL080317.3 | 798.418447 | -0.0296427 | 0.21495765 | -0.1379002 | 0.89031927 | 0.98466501 |

| | | | | | | |
|---|---|---|---|---|---|---|
| TRAF3IP2-AS | 244.37129 | 0.10568776 | 0.226183 | 0.46726658 | 0.64030917 | 0.94088547 |
| TRAF3IP2 | 1365.19305 | 0.11854378 | 0.18717247 | 0.63333983 | 0.52651175 | 0.91769856 |
| Z97989.1 | 145.646865 | 0.28944124 | 0.36324881 | 0.79681262 | 0.42555986 | 0.88622485 |
| FYN | 1859.01134 | 0.15669292 | 0.28195517 | 0.55573701 | 0.57839065 | 0.93139743 |
| WISP3 | 214.969108 | -0.2196825 | 0.41539648 | -0.5288501 | 0.59690945 | 0.9337017 |
| TUBE1 | 1183.02832 | 0.02223221 | 0.23963355 | 0.09277586 | 0.92608163 | 0.98876911 |
| FAM229B | 585.076274 | 0.29571657 | 0.3362866 | 0.87935876 | 0.37920678 | 0.87069278 |
| LAMA4 | 1082.15527 | -0.1850578 | 0.31859743 | -0.5808515 | 0.56134056 | 0.92452095 |
| RFPL4B | 44.9399123 | 1.68614522 | 1.14048767 | 1.47844231 | 0.13928943 | 0.72409321 |
| MARCKS | 3528.16626 | -0.1084583 | 0.29596379 | -0.366458 | 0.71402335 | 0.95564093 |
| LINC01268 | 142.050651 | 0.2360696 | 0.61061779 | 0.38660779 | 0.69904659 | 0.95299571 |
| HDAC2 | 3083.6823 | -0.0428781 | 0.14881555 | -0.2881291 | 0.77324791 | 0.96836854 |
| FRK | 2359.57477 | 0.60169027 | 0.21993487 | 2.73576563 | 0.00622353 | 0.33470127 |
| NT5DC1 | 2785.13328 | 0.75167018 | 0.22741226 | 3.3053195 | 0.00094868 | 0.14816039 |
| AL050331.2 | 307.31944 | 0.73349452 | 0.30245237 | 2.42515716 | 0.01530176 | 0.45205627 |
| TSPYL4 | 2084.78702 | -0.157346 | 0.25490191 | -0.6172804 | 0.53704978 | 0.92105614 |
| DSE | 3025.97592 | -0.0442918 | 0.26155702 | -0.1693391 | 0.86552991 | 0.98197545 |
| TSPYL1 | 7373.70784 | 0.18900839 | 0.17902447 | 1.05576849 | 0.291074 | 0.83454586 |
| Z84488.2 | 229.02273 | -0.5548504 | 0.45752821 | -1.2127129 | 0.22523959 | 0.80176092 |
| Z84488.1 | 45.0962322 | -0.9520221 | 0.87027599 | -1.0939313 | 0.27398513 | 0.82719197 |
| CALHM6 | 128.897547 | -0.9532026 | 0.82906776 | -1.1497282 | 0.25025585 | 0.81721208 |
| CALHM5 | 374.378898 | -0.3799009 | 0.32864372 | -1.1559657 | 0.24769518 | 0.81602355 |
| RWDD1 | 1745.17116 | 0.03653122 | 0.15958491 | 0.22891401 | 0.81893575 | 0.97454841 |
| RSPH4A | 86.4007752 | -0.1667775 | 0.49510736 | -0.3368512 | 0.73622908 | 0.95955556 |
| ZUP1 | 482.920555 | -0.0976342 | 0.20136305 | -0.4848667 | 6.28E-01 | 9.41E-01 |
| KPNA5 | 732.261004 | -0.1853936 | 0.16772801 | -1.1053227 | 0.2690198 | 0.82683094 |
| FAM162B | 86.6455652 | -0.092769 | 0.4979055 | -0.1863185 | 0.85219498 | 0.97872457 |
| AL132671.2 | 118.705528 | -0.0215057 | 0.37202115 | -0.0578077 | 0.95390183 | 0.99170503 |
| DCBLD1 | 687.53505 | -0.4352927 | 0.19437152 | -2.239488 | 2.51E-02 | 5.21E-01 |
| GOPC | 1731.17504 | 0.04887083 | 0.10822901 | 0.45155021 | 0.65159305 | 0.94393131 |
| NUS1 | 2437.0693 | 0.00838039 | 0.25160228 | 0.0333081 | 0.97342889 | 0.99445887 |
| SLC35F1 | 55.7209663 | -0.4815975 | 0.65204327 | -0.7385974 | 0.46015149 | 0.89755243 |
| CEP85L | 308.498327 | -0.0314697 | 0.22296233 | -0.1411437 | 0.88775641 | 0.98423722 |
| PLN | 829.588389 | -0.7703741 | 0.65104633 | -1.1832862 | 0.23669575 | 0.80849753 |
| MCM9 | 919.960808 | -0.1879782 | 0.17895198 | -1.0504396 | 0.29351604 | 0.83540174 |
| ASF1A | 711.904273 | 0.05842195 | 0.21182237 | 0.27580635 | 0.78269681 | 0.96964828 |
| AL365275.1 | 43.5376413 | 0.01507767 | 0.33497445 | 0.04501141 | 0.96409821 | 0.99342622 |
| FAM184A | 474.294328 | 0.1610636 | 0.32023862 | 0.50294872 | 0.61500032 | 0.93895494 |
| MAN1A1 | 6520.57438 | 0.18503188 | 0.19901641 | 0.92973177 | 0.35250998 | 0.86364633 |
| TBC1D32 | 208.108241 | -0.1157042 | 0.31151829 | -0.3714203 | 7.10E-01 | 0.95530618 |
| GJA1 | 2481.66861 | 0.32215505 | 0.31241944 | 1.03116196 | 3.02E-01 | 8.42E-01 |
| HSF2 | 691.901569 | -0.242973 | 0.14663823 | -1.6569551 | 0.09752856 | 0.66549286 |
| SERINC1 | 14561.5375 | 0.04797996 | 0.15991954 | 0.30002563 | 0.7641576 | 0.96678847 |
| PKIB | 186.337352 | 0.00179805 | 0.64814465 | 0.00277416 | 0.99778655 | 0.99989813 |
| SMPDL3A | 2244.69579 | 0.51778198 | 0.53819312 | 0.96207469 | 0.33601209 | 0.85641269 |
| RNF217 | 928.182606 | -0.3267564 | 0.16853963 | -1.9387509 | 0.05253167 | 0.58645127 |
| TPD52L1 | 2780.85566 | 0.14001527 | 0.34232824 | 0.40900884 | 0.68253318 | 0.95070301 |
| HDDC2 | 2851.78818 | 0.25650175 | 0.14640082 | 1.75205137 | 0.07976498 | 0.64079926 |
| HEY2 | 230.393059 | 0.16357935 | 0.29879766 | 0.5474586 | 0.58406371 | 0.93236269 |
| NCOA7 | 7298.57981 | 0.15388859 | 0.36532231 | 0.42124061 | 6.74E-01 | 9.49E-01 |
| HINT3 | 2009.0445 | 0.33092291 | 0.13037569 | 2.53822552 | 0.01114162 | 0.40196898 |
| TRMT11 | 975.535164 | 0.07050936 | 0.14497838 | 0.48634397 | 0.62672331 | 0.94088547 |
| CENPW | 110.075049 | -0.3721051 | 0.42130988 | -0.88321 | 0.37712284 | 0.86987124 |
| RSPO3 | 184.988231 | 0.04360796 | 0.63088908 | 0.06912143 | 0.94489296 | 0.99164528 |

| | | | | | | |
|---|---|---|---|---|---|---|
| RNF146 | 2527.97104 | 0.13550308 | 0.13001055 | 1.04224683 | 0.29729726 | 0.83898658 |
| ECHDC1 | 4061.79103 | 0.21872582 | 0.19133197 | 1.14317443 | 0.25296618 | 0.81835388 |
| AL096711.2 | 208.945074 | 1.11081631 | 0.49676037 | 2.23612104 | 0.02534384 | 0.52250144 |
| KIAA0408 | 178.659352 | 1.11598591 | 0.50262949 | 2.22029531 | 0.02639873 | 0.52678638 |
| PTPRK | 2219.3753 | -0.0439277 | 0.12008442 | -0.3658065 | 0.71450946 | 0.95579447 |
| LAMA2 | 1179.54568 | 0.20992185 | 0.38377235 | 0.54699576 | 0.58438164 | 0.9326319 |
| AL356124.2 | 79.1513301 | 0.32241105 | 0.47400366 | 0.68018684 | 0.49638616 | 0.90890243 |
| ARHGAP18 | 2293.91481 | 0.22122196 | 0.19736719 | 1.12086492 | 0.26234536 | 0.82424998 |
| L3MBTL3 | 638.3605 | 0.21893715 | 0.20974615 | 1.04381964 | 0.29656884 | 0.83831021 |
| AL355581.1 | 155.824251 | 0.07809243 | 0.29411562 | 0.2655161 | 0.79061191 | 0.97090793 |
| SAMD3 | 97.1543083 | -0.0202407 | 0.56558029 | -0.0357875 | 0.97145179 | 0.99404213 |
| TMEM200A | 1117.64673 | 0.21385526 | 0.312039 | 0.68534786 | 0.49312444 | 0.90711083 |
| SMLR1 | 176.244608 | 0.50662966 | 0.75432097 | 0.67163672 | 0.50181499 | 0.91259827 |
| EPB41L2 | 2773.35134 | 0.05737437 | 0.25402326 | 0.22586266 | 0.82130824 | 0.97472105 |
| AKAP7 | 498.079862 | 0.22369532 | 0.34000199 | 0.65792355 | 0.51058726 | 0.91472286 |
| MED23 | 1550.10177 | -0.0990638 | 0.1250004 | -0.7925081 | 0.42806445 | 0.88800332 |
| ENPP3 | 609.703024 | 0.36988299 | 0.41221499 | 0.89730603 | 0.36955564 | 0.86762749 |
| ENPP1 | 1280.46787 | 0.22881745 | 0.50885883 | 0.44966784 | 0.65294997 | 0.9447282 |
| CTGF | 13760.8213 | -1.2179241 | 0.47640227 | -2.5565035 | 0.010573 | 0.40196898 |
| AL133346.1 | 195.707095 | -1.020456 | 0.4983754 | -2.047565 | 0.04060265 | 0.55657615 |
| MOXD1 | 2198.33862 | 0.70266458 | 0.4527293 | 1.5520634 | 0.12064705 | 0.69467399 |
| STX7 | 4703.2512 | 0.14530873 | 0.15782447 | 0.92069836 | 0.35720793 | 0.86390849 |
| VNN1 | 3639.06938 | 1.05001028 | 1.02778608 | 1.02162338 | 0.30695919 | 0.84286067 |
| VNN2 | 256.433079 | 0.07817993 | 0.51089973 | 0.15302401 | 0.87837934 | 0.98275231 |
| SLC18B1 | 1541.43691 | -0.1386972 | 0.26886096 | -0.5158696 | 0.60594547 | 0.93698713 |
| RPS12 | 24695.596 | 0.23123934 | 0.1924069 | 1.20182456 | 0.22943151 | 0.80551898 |
| EYA4 | 394.546054 | 0.22566733 | 0.44893963 | 0.50266744 | 0.61519811 | 0.93895494 |
| TARID | 1318.86655 | -0.0416019 | 0.41079415 | -0.1012719 | 0.91933465 | 0.9872458 |
| TCF21 | 1848.63768 | 0.04043132 | 0.47613848 | 0.08491503 | 0.93232894 | 0.9899349 |
| TBPL1 | 875.638078 | -0.1870419 | 0.13620889 | -1.3731986 | 0.16969061 | 0.7586159 |
| SLC2A12 | 440.153602 | 0.36927519 | 0.34451982 | 1.07185468 | 0.28378531 | 0.83107779 |
| SGK1 | 9537.02264 | -0.4442658 | 0.46538836 | -0.9546131 | 0.3397734 | 0.85802452 |
| AL078590.3 | 42.297622 | 0.1688819 | 0.55745541 | 0.30295141 | 0.76192689 | 0.96607083 |
| ALDH8A1 | 4556.22947 | 0.64969709 | 0.48496811 | 1.33966972 | 0.18035275 | 0.7640124 |
| AL445190.1 | 294.924031 | 0.94718515 | 0.59046706 | 1.60412869 | 0.10868569 | 0.67800851 |
| HBS1L | 1686.73179 | 0.16492689 | 0.22144174 | 0.74478685 | 0.45640059 | 0.89688594 |
| AHI1 | 1195.3896 | -0.1000046 | 0.19879246 | -0.5030602 | 0.61492194 | 0.93895494 |
| AL049552.1 | 76.8506174 | -0.3494026 | 0.41778222 | -0.8363271 | 0.40297089 | 0.88132015 |
| LINC00271 | 139.872326 | 0.68228351 | 0.51641376 | 1.32119545 | 0.1864362 | 0.76908034 |
| PDE7B | 929.87329 | -0.1229806 | 0.23782806 | -0.5170989 | 0.60508715 | 0.93687349 |
| AL138828.1 | 592.712958 | -1.8195449 | 0.66587773 | -2.732551 | 0.00628459 | 0.33470127 |
| MTFR2 | 52.9211574 | -0.0098657 | 0.4812047 | -0.0205021 | 0.98364283 | 0.9961464 |
| BCLAF1 | 4624.55488 | -0.0120103 | 0.13896853 | -0.0864248 | 0.93112874 | 0.98975562 |
| AL023284.4 | 283.396055 | -0.0478959 | 0.35299127 | -0.1356859 | 0.89206959 | 0.98474725 |
| MAP7 | 2916.94027 | 0.1238288 | 0.24332611 | 0.50890058 | 6.11E-01 | 0.93808191 |
| AL024508.2 | 47.5837056 | 0.02301248 | 0.51304443 | 0.04485476 | 0.96422308 | 0.99342622 |
| MAP3K5 | 1642.37318 | 0.34106559 | 0.25657933 | 1.3292793 | 0.18375584 | 0.76781833 |
| PEX7 | 652.64648 | 0.56319507 | 0.16829684 | 3.3464387 | 0.00081857 | 0.13341488 |
| IFNGR1 | 3515.73211 | -0.1042959 | 0.15300834 | -0.6816353 | 0.49546957 | 0.90848207 |
| AL357060.1 | 57.4804755 | 0.04859705 | 0.4552291 | 0.10675295 | 0.91498497 | 0.9866558 |
| TNFAIP3 | 2756.72545 | 0.49846728 | 0.72447818 | 0.6880363 | 0.49142992 | 0.90600166 |
| PERP | 3979.62589 | 0.163825 | 0.18827152 | 0.87015288 | 0.38421686 | 0.87298595 |
| ARFGEF3 | 647.522016 | -0.1266766 | 0.27209372 | -0.4655626 | 0.64152865 | 0.94088547 |
| HEBP2 | 3188.98449 | 0.01664215 | 0.1379379 | 0.12064961 | 0.90396857 | 0.98624484 |

| | | | | | | |
|---|---|---|---|---|---|---|
| NHSL1 | 1315.47594 | 0.09854737 | 0.2245105 | 0.43894328 | 0.66070264 | 0.94628868 |
| AL590617.2 | 178.779987 | -0.1937482 | 0.3883203 | -0.4989391 | 0.61782229 | 0.93924932 |
| CCDC28A | 1869.52236 | 0.36890903 | 0.1737204 | 2.12357926 | 0.03370534 | 0.55369025 |
| REPS1 | 1624.6739 | -0.0290127 | 0.1185751 | -0.2446782 | 0.80670562 | 0.97334138 |
| ABRACL | 1697.02781 | -0.4681119 | 0.43950638 | -1.0650855 | 0.28683723 | 0.83155545 |
| HECA | 1558.69135 | -0.3278079 | 0.2260058 | -1.45044 | 0.14693585 | 0.73337969 |
| CITED2 | 6255.60553 | -0.7120133 | 0.244529 | -2.9117746 | 0.00359382 | 0.26033637 |
| FILNC1 | 218.918741 | -1.4018096 | 0.92926277 | -1.508518 | 0.13142201 | 0.71081672 |
| AL050338.1 | 43.6018084 | -1.5186608 | 0.97658256 | -1.5550767 | 0.11992781 | 0.69433886 |
| VTA1 | 1847.80556 | -0.2346613 | 0.2416405 | -0.9711174 | 0.33148981 | 0.85408791 |
| ADGRG6 | 928.132146 | 0.61063003 | 0.37199468 | 1.64150206 | 0.10069324 | 0.67247782 |
| AL023584.1 | 217.943236 | -0.44231 | 0.40278493 | -1.0981294 | 0.272148 | 0.82716979 |
| HIVEP2 | 1634.42125 | -0.2913959 | 0.29512963 | -0.9873488 | 0.32347165 | 0.85140144 |
| LINC01277 | 41.9926026 | -0.2585772 | 0.55618756 | -0.46491 | 0.64199591 | 0.94088547 |
| AIG1 | 4331.93504 | 0.54950075 | 0.21393827 | 2.56850138 | 0.01021393 | 0.40019066 |
| ADAT2 | 609.891034 | -0.1277121 | 0.27193347 | -0.4696447 | 0.63860889 | 0.94088547 |
| PEX3 | 1369.74375 | -0.0108476 | 0.21374658 | -0.0507499 | 0.95952478 | 0.99246392 |
| FUCA2 | 4769.03966 | 0.16073658 | 0.24363437 | 0.65974511 | 0.50941741 | 0.91439937 |
| PHACTR2 | 969.621729 | 0.00533631 | 0.16934723 | 0.03151105 | 0.97486198 | 0.99455099 |
| AL049844.1 | 153.75256 | -0.2245282 | 0.27819731 | -0.8070827 | 0.41961884 | 0.88451372 |
| LTV1 | 1086.73673 | -0.1665453 | 0.27020963 | -0.6163558 | 0.53765974 | 0.92123049 |
| AL049844.3 | 424.092247 | -0.1172238 | 0.3179524 | -0.3686836 | 0.71236356 | 0.95560193 |
| PLAGL1 | 1781.32439 | -0.3443602 | 0.20788463 | -1.6564967 | 0.09762127 | 0.66549286 |
| SF3B5 | 3751.96036 | 0.05404701 | 0.14454884 | 0.37390141 | 0.70847767 | 0.95501548 |
| STX11 | 250.918323 | -0.6175158 | 0.37110817 | -1.6639779 | 0.09611684 | 0.66169558 |
| UTRN | 2954.94426 | -0.0084778 | 0.32190885 | -0.0263359 | 0.97898941 | 0.99499459 |
| EPM2A | 807.841441 | -0.2082128 | 0.31626157 | -0.6583565 | 0.51030911 | 0.91439937 |
| FBXO30 | 952.50282 | -0.6467077 | 0.36771739 | -1.7587086 | 0.07862702 | 0.64079926 |
| AL356599.1 | 933.762 | -0.0753771 | 0.14227445 | -0.5298009 | 0.59624999 | 0.9334601 |
| SHPRH | 954.212473 | -0.0106851 | 0.17755944 | -0.0601777 | 0.95201413 | 0.99170503 |
| GRM1 | 86.5202286 | -0.742553 | 0.77841467 | -0.9539298 | 0.34011916 | 0.85814041 |
| RAB32 | 1181.88864 | -0.225365 | 0.38829594 | -0.5803951 | 0.56164823 | 0.92452095 |
| STXBP5-AS1 | 69.9627195 | -0.3530429 | 0.22289752 | -1.5838798 | 0.11322108 | 0.68151616 |
| STXBP5 | 736.79671 | 0.01986874 | 0.17958328 | 0.11063805 | 0.91190337 | 0.98662757 |
| SAMD5 | 2229.56288 | -0.8980827 | 0.52977581 | -1.6952128 | 0.09003506 | 0.65393309 |
| SASH1 | 3856.33266 | -0.4041058 | 0.37631405 | -1.0738526 | 0.28288875 | 0.83078187 |
| UST | 328.056484 | -0.2277466 | 0.27877611 | -0.8169515 | 0.41395616 | 0.88349303 |
| TAB2 | 3311.41876 | -0.2794218 | 0.17430775 | -1.6030373 | 0.10892642 | 0.67842804 |
| ZC3H12D | 195.751134 | -0.1193796 | 0.44145289 | -0.2704243 | 0.78683384 | 0.97090793 |
| PPIL4 | 1360.09275 | 0.04842196 | 0.12582067 | 0.38484901 | 0.70034928 | 0.95355954 |
| GINM1 | 5048.76465 | 0.19780396 | 0.16265879 | 1.21606687 | 0.22395945 | 0.80078694 |
| AL078581.3 | 1073.42858 | 0.17151573 | 0.16425886 | 1.04417945 | 0.29640237 | 0.83817467 |
| KATNA1 | 980.082528 | 0.04520797 | 0.15666533 | 0.28856397 | 0.77291507 | 0.96836854 |
| LATS1 | 1424.14316 | -0.1659161 | 0.15342077 | -1.0814451 | 0.27949916 | 0.82825734 |
| NUP43 | 1586.20213 | 0.10951104 | 0.12228564 | 0.8955347 | 0.37050133 | 0.86762749 |
| PCMT1 | 3116.75521 | -0.1227401 | 0.23630428 | -0.5194153 | 0.60347118 | 0.93607461 |
| AL355312.2 | 897.241049 | -0.0563034 | 0.28412964 | -0.1981609 | 0.8429192 | 0.97649449 |
| LRP11 | 2003.45714 | 0.37719987 | 0.14832048 | 2.5431409 | 0.01098609 | 0.40196898 |
| AL355312.4 | 91.0383132 | 0.41607033 | 0.42748004 | 0.97330937 | 0.33039957 | 0.85384238 |
| AL355312.3 | 53.2871617 | 0.83964992 | 0.43350294 | 1.93689557 | 0.05275811 | 0.58742503 |
| RAET1E | 67.039323 | 0.25450862 | 0.69029203 | 0.36869703 | 0.71235356 | 0.95560193 |
| RAET1G | 41.0716607 | 0.12964714 | 0.41406013 | 0.31311187 | 0.75419567 | 0.96348234 |
| ULBP2 | 59.2065977 | 0.19549739 | 0.40622731 | 0.48125122 | 0.63033796 | 0.94088547 |
| PPP1R14C | 740.869988 | 0.27853722 | 0.28840674 | 0.96577914 | 0.33415471 | 0.85533716 |

| | | | | | | |
|---|---|---|---|---|---|---|
| IYD | 1745.14567 | 1.62053783 | 0.67896559 | 2.38677459 | 0.01699691 | 0.46671416 |
| PLEKHG1 | 920.385991 | -0.0499003 | 0.29708907 | -0.167964 | 0.86661157 | 0.98217027 |
| MTHFD1L | 1025.39494 | 0.04256687 | 0.3151531 | 0.13506729 | 0.89255867 | 0.98474725 |
| AKAP12 | 6323.03163 | -0.0499265 | 0.26188166 | -0.1906454 | 0.84880344 | 0.97741457 |
| ZBTB2 | 841.319358 | -0.4420951 | 0.30148157 | -1.4664083 | 0.14253708 | 0.72784886 |
| RMND1 | 1644.92175 | 0.22262094 | 0.34657598 | 0.64234384 | 0.52064995 | 0.91680964 |
| ARMT1 | 2102.40581 | 0.19030063 | 0.23845169 | 0.79806787 | 0.42483111 | 0.88573606 |
| CCDC170 | 79.970106 | 1.17546321 | 0.61524511 | 1.91056084 | 0.05606104 | 0.59895387 |
| ESR1 | 271.317522 | 0.68676168 | 0.36859909 | 1.86316703 | 0.06243877 | 0.61393757 |
| SYNE1 | 6638.38092 | 0.1139321 | 0.24763122 | 0.46008777 | 0.64545322 | 0.94208859 |
| MYCT1 | 585.782125 | -0.5801209 | 0.27805382 | -2.086362 | 0.03694584 | 0.55521179 |
| VIP | 68.8202939 | 0.2931998 | 0.61664179 | 0.47547831 | 0.63444609 | 0.94088547 |
| FBXO5 | 237.330082 | -0.5065357 | 0.26630202 | -1.9021101 | 0.05715677 | 0.60069729 |
| AL080276.2 | 401.199611 | -0.1853569 | 0.33034743 | -0.5610969 | 0.57473147 | 0.93024927 |
| MTRF1L | 870.727055 | -0.195187 | 0.26199167 | -0.7450122 | 0.45626434 | 0.89688594 |
| RGS17 | 91.6683969 | 0.31652327 | 0.39834181 | 0.79460217 | 4.27E-01 | 0.88709241 |
| IPCEF1 | 357.849627 | 0.54543471 | 0.43982011 | 1.24013136 | 0.21492681 | 0.79487183 |
| CNKSR3 | 3028.93959 | -0.0555507 | 0.17608194 | -0.3154819 | 0.75239579 | 0.96301881 |
| SCAF8 | 1897.31448 | -0.4585088 | 0.23960673 | -1.9135889 | 0.05567271 | 0.59817776 |
| TIAM2 | 1203.03827 | -0.3400007 | 0.16766738 | -2.0278283 | 4.26E-02 | 5.65E-01 |
| AL596202.1 | 142.405389 | 0.30415854 | 0.40231706 | 0.75601699 | 0.44963901 | 0.89557948 |
| TFB1M | 516.238863 | -0.0214786 | 0.20109811 | -0.1068068 | 0.91494225 | 0.9866558 |
| AL355297.4 | 49.3439315 | 0.06255403 | 0.36246642 | 0.17257883 | 0.8629825 | 0.98176973 |
| ARID1B | 2510.30206 | -0.1749332 | 0.10986887 | -1.5922001 | 0.11133974 | 0.68001961 |
| TMEM242 | 1245.83204 | 0.19883262 | 0.19583326 | 1.01531589 | 0.30995527 | 0.84455826 |
| ZDHHC14 | 1483.88909 | -0.2153024 | 0.21789923 | -0.9880822 | 0.32311238 | 0.85119105 |
| SNX9 | 5117.78885 | -0.2095448 | 0.28428653 | -0.73709 | 4.61E-01 | 8.98E-01 |
| AL391863.1 | 217.606478 | -0.020917 | 0.49145987 | -0.0425609 | 0.96605158 | 0.99393925 |
| SYNJ2 | 1049.66898 | -0.5696845 | 0.35471964 | -1.6060135 | 0.10827094 | 0.67789099 |
| SERAC1 | 773.380229 | -0.4258862 | 0.24336301 | -1.7500038 | 0.08011765 | 0.64079926 |
| GTF2H5 | 1152.93649 | 0.23623592 | 0.17341291 | 1.36227413 | 1.73E-01 | 0.75974924 |
| TULP4 | 1268.08309 | -0.0731145 | 0.20112422 | -0.363529 | 0.71620977 | 0.95603249 |
| TMEM181 | 1512.8348 | 0.02251395 | 0.16700958 | 0.13480632 | 0.89276501 | 0.98474725 |
| DYNLT1 | 4972.04734 | -0.2442699 | 0.24707774 | -0.9886359 | 0.32284133 | 0.85119105 |
| SYTL3 | 733.406859 | 0.00922071 | 0.34038869 | 0.02708878 | 0.97838892 | 0.99476647 |
| EZR | 21031.1422 | 0.11013396 | 0.20934542 | 0.52608728 | 0.59882756 | 0.93484879 |
| EZR-AS1 | 826.949189 | 0.13817881 | 0.36304089 | 0.380615 | 0.70348895 | 0.95467045 |
| RSPH3 | 1093.41365 | 0.11633004 | 0.17987406 | 0.64673051 | 0.51780637 | 0.91636259 |
| AL035530.2 | 120.559186 | 0.09506485 | 0.3309431 | 0.28725437 | 7.74E-01 | 0.96836854 |
| TAGAP | 150.059259 | -0.2705344 | 0.60091846 | -0.4502015 | 0.65256513 | 0.94437526 |
| FNDC1 | 100.886893 | 0.06462116 | 0.55754216 | 0.11590364 | 0.90772891 | 0.98662757 |
| AL078604.2 | 73.9255001 | -0.2014694 | 0.66657875 | -0.3022439 | 0.76246615 | 0.96630779 |
| SOD2 | 166660.659 | -0.157631 | 0.45580617 | -0.345829 | 7.29E-01 | 0.95848028 |
| WTAP | 6750.4528 | -0.2941165 | 0.18929991 | -1.5537066 | 0.12025441 | 0.69467399 |
| SOD2-OT1 | 218.735041 | 0.2098734 | 0.40749411 | 0.51503419 | 0.60652913 | 0.93708311 |
| ACAT2 | 2680.94221 | 0.04008964 | 0.33435042 | 0.11990305 | 0.90455995 | 0.98624484 |
| TCP1 | 9822.95181 | -0.0060163 | 0.26258767 | -0.0229115 | 0.98172088 | 0.99595643 |
| MRPL18 | 2228.34754 | -0.0310906 | 0.19589748 | -0.1587087 | 0.87389839 | 0.98275231 |
| IGF2R | 7360.2056 | -0.205425 | 0.27562421 | -0.7453084 | 0.45608531 | 0.89688594 |
| SLC22A1 | 103.698517 | 0.04230037 | 0.55165228 | 0.0766794 | 0.93887859 | 0.99113899 |
| SLC22A2 | 7218.04321 | 0.98374302 | 0.48472758 | 2.02947607 | 0.04240982 | 0.56525418 |
| AL162582.1 | 41.0667357 | 0.84087378 | 0.67747032 | 1.24119647 | 0.21453318 | 0.79426438 |
| SLC22A3 | 773.990057 | -0.4321063 | 0.42393507 | -1.0192746 | 0.3080726 | 0.8437043 |
| LPA | 117.058974 | -0.6721772 | 1.08684495 | -0.6184665 | 0.53626789 | 0.92102241 |

| | | | | | | |
|---|---|---|---|---|---|---|
| PLG | 15002.1088 | 0.25532268 | 0.80795141 | 0.31601242 | 0.75199308 | 0.96297248 |
| MAP3K4 | 1187.03505 | -0.0557936 | 0.10127969 | -0.5508863 | 0.58171161 | 0.9317705 |
| AGPAT4 | 883.758104 | -0.362821 | 0.36812652 | -0.9855878 | 0.32433544 | 0.85169085 |
| PRKN | 646.935587 | 0.5691364 | 0.26525219 | 2.14564257 | 0.03190151 | 0.54649555 |
| PACRG | 314.0752 | 0.36070944 | 0.37269242 | 0.96784755 | 0.33312051 | 0.855011 |
| CAHM | 68.3311823 | 0.0903748 | 0.39378916 | 0.22950048 | 0.81847995 | 0.97454841 |
| QKI | 4365.21497 | -0.2062021 | 0.14834765 | -1.3899925 | 0.16453114 | 0.75286112 |
| PDE10A | 844.057678 | -0.2552575 | 0.23611691 | -1.081064 | 0.27966866 | 0.82825734 |
| LINC00473 | 581.994216 | -1.6899462 | 1.21100483 | -1.3954908 | 1.63E-01 | 0.75050573 |
| LINC00602 | 106.176078 | -1.2323823 | 0.98144896 | -1.2556764 | 0.20923333 | 0.79086734 |
| SFT2D1 | 2333.64851 | -0.0034999 | 0.10657777 | -0.0328387 | 0.97380323 | 0.99445887 |
| MPC1 | 6127.0022 | 0.36615171 | 0.22566993 | 1.62250998 | 0.1046942 | 0.67439401 |
| RPS6KA2 | 2474.15996 | 0.34625604 | 0.23877102 | 1.45015943 | 0.14701407 | 0.73337969 |
| AL159163.1 | 1375.23851 | -0.3577284 | 0.3842627 | -0.9309474 | 0.35188075 | 0.86364633 |
| RNASET2 | 3115.78668 | -0.5233203 | 0.46745821 | -1.1195018 | 0.26292613 | 0.82424998 |
| FGFR1OP | 1567.02793 | 0.15067809 | 0.22839811 | 0.65971688 | 0.50943553 | 0.91439937 |
| CCR6 | 83.9185381 | -0.1395572 | 0.44518973 | -0.313478 | 0.75391757 | 0.96339067 |
| UNC93A | 64.0357121 | 2.23690199 | 0.98430968 | 2.27255918 | 0.02305276 | 0.51967919 |
| LINC01558 | 177.218257 | -0.2230368 | 0.52821591 | -0.4222455 | 0.67284583 | 0.94922827 |
| AL009178.2 | 107.997598 | 0.12233612 | 0.55715601 | 0.21957247 | 0.82620413 | 0.97548201 |
| AL009178.3 | 75.7233526 | 0.6197378 | 0.58644597 | 1.0567688 | 0.29061712 | 0.83442894 |
| AFDN-DT | 132.013458 | 0.59163884 | 0.60015555 | 0.98580915 | 0.32422677 | 0.85169085 |
| AFDN | 5397.2395 | -0.0770405 | 0.12372655 | -0.6226678 | 0.53350286 | 0.92049136 |
| KIF25-AS1 | 36.9714445 | -1.0401595 | 1.56381357 | -0.6651429 | 0.50595908 | 0.91390261 |
| KIF25 | 33.4805054 | 0.36972937 | 0.81902361 | 0.451427 | 6.52E-01 | 0.94393131 |
| FRMD1 | 1544.07117 | 0.79959248 | 0.59879731 | 1.33533078 | 0.18176811 | 0.76543911 |
| DACT2 | 425.683047 | 1.22559498 | 0.56732091 | 2.16032048 | 0.03074787 | 0.54443002 |
| SMOC2 | 946.71546 | -0.3835543 | 0.60564363 | -0.6333003 | 0.52653755 | 0.91769856 |
| BX322234.1 | 70.49883 | -0.1737322 | 0.39181645 | -0.443402 | 0.657475 | 0.94558535 |
| THBS2 | 2855.7114 | -0.00276 | 0.42535954 | -0.0064885 | 9.95E-01 | 9.99E-01 |
| WDR27 | 1321.42359 | -0.0260908 | 0.24463777 | -0.1066509 | 0.91506592 | 0.9866558 |
| C6orf120 | 1751.49928 | -0.1288761 | 0.15460047 | -0.8336077 | 0.40450209 | 0.88132015 |
| PHF10 | 3987.26832 | 0.03105922 | 0.09783681 | 0.31745947 | 0.75089499 | 0.96272571 |
| AL354892.2 | 68.5140518 | 0.22698429 | 0.27720218 | 0.8188402 | 0.41287759 | 0.88290038 |
| TCTE3 | 330.443488 | -0.0657357 | 0.27205535 | -0.2416261 | 0.80906986 | 0.97334138 |
| ERMARD | 1195.99352 | 0.05350875 | 0.25435055 | 0.21037402 | 0.83337577 | 0.97550416 |
| AL596442.2 | 65.3995573 | -0.6140535 | 0.56812792 | -1.0808366 | 0.27976979 | 0.82825734 |
| DLL1 | 959.749996 | -0.3392789 | 0.37225223 | -0.9114221 | 0.36207302 | 0.86540116 |
| FAM120B | 1745.7911 | -0.0759235 | 0.11420867 | -0.6647786 | 0.50619209 | 0.91392543 |
| PSMB1 | 7041.71991 | -0.0737623 | 0.20475021 | -0.3602549 | 0.71865653 | 0.95648644 |
| TBP | 886.219568 | -0.1144177 | 0.12309327 | -0.9295207 | 0.35261931 | 0.86364633 |
| PDCD2 | 2287.25861 | -0.1471579 | 0.16662141 | -0.8831872 | 0.37713513 | 0.86987124 |
| AC093627.5 | 77.118301 | -0.234988 | 0.46927862 | -0.5007429 | 0.61655205 | 0.93910158 |
| AC093627.4 | 381.880163 | 0.33346533 | 0.3951004 | 0.8440015 | 0.39866857 | 0.88074856 |
| AC093627.7 | 494.480784 | 0.58909798 | 0.70787555 | 0.83220557 | 0.40529292 | 0.88132015 |
| AC093627.2 | 80.0302074 | -1.3653437 | 1.45979819 | -0.9352962 | 0.34963567 | 0.86364633 |
| FAM20C | 8017.6828 | 0.27240258 | 0.22936686 | 1.1876283 | 0.23497988 | 0.80747753 |
| PDGFA | 1948.3512 | -0.1125717 | 0.25769187 | -0.4368462 | 0.66222287 | 0.94642611 |
| AC147651.1 | 148.383275 | -0.0875063 | 0.33289535 | -0.2628642 | 0.79265527 | 0.97090793 |
| PRKAR1B | 1609.1551 | -0.1406686 | 0.24311243 | -0.5786154 | 0.56284869 | 0.92452095 |
| AC147651.2 | 49.6774315 | -0.0407052 | 0.75994207 | -0.0535636 | 0.95728288 | 0.99206141 |
| AC147651.4 | 49.848248 | -0.5587416 | 0.37608238 | -1.4856894 | 0.13736129 | 0.72209298 |
| DNAAF5 | 1630.01532 | -0.3369856 | 0.1923959 | -1.7515215 | 7.99E-02 | 6.41E-01 |
| SUN1 | 11155.091 | -0.1976319 | 0.14306748 | -1.3813894 | 0.16715926 | 0.75558932 |

| | | | | | | |
|---|---|---|---|---|---|---|
| GET4 | 6043.76859 | -0.3323988 | 0.22384074 | -1.484979 | 0.1375494 | 0.72209298 |
| AC073957.3 | 2243.87475 | -0.1841967 | 0.2760728 | -0.6672033 | 0.50464229 | 0.91365534 |
| ADAP1 | 1491.01611 | 0.21557975 | 0.31656223 | 0.68100275 | 0.49586975 | 0.90868241 |
| COX19 | 1783.48247 | 0.05475774 | 0.14633847 | 0.37418553 | 0.70826629 | 0.95501548 |
| C7orf50 | 3264.06363 | 0.00732073 | 0.20458086 | 0.03578405 | 0.97145455 | 0.99404213 |
| GPR146 | 677.190977 | -0.3934044 | 0.2967778 | -1.3255857 | 0.18497695 | 0.76820626 |
| AC073957.2 | 589.895964 | -0.3668688 | 0.30722451 | -1.1941391 | 0.23242355 | 0.80649152 |
| GPER1 | 847.749413 | -0.0684526 | 0.51820309 | -0.1320961 | 0.89490827 | 0.98474725 |
| ZFAND2A | 854.390844 | -0.4066392 | 0.47030774 | -0.8646237 | 0.38724539 | 0.87460898 |
| AC091729.3 | 266.10153 | 0.5120294 | 0.46077536 | 1.11123435 | 0.26646749 | 0.8265123 |
| UNCX | 51.5702788 | -0.7492006 | 0.85784671 | -0.8733502 | 0.38247221 | 0.8718674 |
| MICALL2 | 4380.6589 | -0.2765522 | 0.32255832 | -0.857371 | 0.39123986 | 0.87716276 |
| AC102953.2 | 248.358169 | -0.0015796 | 0.25785287 | -0.0061259 | 0.99511226 | 0.99940889 |
| INTS1 | 6650.7828 | -0.1129402 | 0.14711345 | -0.767708 | 0.44266068 | 0.89343541 |
| MAFK | 5666.59346 | -0.2831942 | 0.1983255 | -1.4279264 | 0.15331305 | 0.74258845 |
| TMEM184A | 3555.92569 | -0.1428297 | 0.11774209 | -1.2130724 | 0.22510213 | 0.80176092 |
| PSMG3 | 1113.83788 | 0.08967641 | 0.16568094 | 0.54125964 | 0.58832863 | 0.93289552 |
| PSMG3-AS1 | 507.020688 | 0.09774675 | 0.24774931 | 0.39453893 | 0.69318319 | 0.9519683 |
| ELFN1 | 410.945102 | -0.4706479 | 0.54874334 | -0.8576832 | 0.39106744 | 0.87710956 |
| MAD1L1 | 1065.08914 | -0.0246114 | 0.20954824 | -0.1174496 | 0.90650378 | 0.98662757 |
| MRM2 | 1142.01255 | -0.0789995 | 0.18088387 | -0.4367416 | 6.62E-01 | 0.94642611 |
| NUDT1 | 217.438755 | 0.07192893 | 0.26481087 | 0.27162379 | 7.86E-01 | 0.97065795 |
| SNX8 | 2199.84874 | -0.2650336 | 0.35377996 | -0.7491482 | 0.45376787 | 0.89670027 |
| EIF3B | 7334.89647 | -0.2690766 | 0.21068538 | -1.277149 | 0.20154964 | 0.78134034 |
| CHST12 | 1258.03267 | -0.1241183 | 0.21072907 | -0.5889948 | 5.56E-01 | 0.92373235 |
| LFNG | 535.841797 | 0.22812229 | 0.41976474 | 0.54345272 | 0.58681814 | 0.93289552 |
| BRAT1 | 3018.05515 | 0.01516923 | 0.13375263 | 0.11341253 | 0.9097035 | 0.98662757 |
| IQCE | 1990.07824 | -0.1868605 | 0.18274029 | -1.0225467 | 0.30652221 | 0.84278367 |
| TTYH3 | 12289.812 | 0.05434318 | 0.36571724 | 0.14859343 | 0.88187446 | 0.98275231 |
| GNA12 | 2437.40195 | -0.056265 | 0.24244686 | -0.2320717 | 0.81648236 | 0.97388289 |
| CARD11 | 279.500283 | 0.43415285 | 0.57798267 | 0.75115202 | 0.45256117 | 0.89670027 |
| SDK1 | 842.907 | 1.147891 | 0.19221873 | 5.97179581 | 2.35E-09 | 2.20E-05 |
| AC017000.1 | 79.3104546 | 0.55263709 | 0.60561954 | 0.91251529 | 0.36149752 | 0.86528137 |
| FOXK1 | 2571.85368 | -0.0822853 | 0.17934901 | -0.4587997 | 0.64637798 | 0.94208859 |
| AP5Z1 | 3879.82512 | -0.7658296 | 0.29008186 | -2.6400463 | 0.00828947 | 0.37565563 |
| RADIL | 89.9772326 | -0.0320983 | 0.55933827 | -0.0573861 | 0.95423762 | 0.99170503 |
| RBAK-RBAKI | 130.697412 | -0.6561184 | 0.29171593 | -2.249169 | 0.02450174 | 0.51967919 |
| RBAK | 898.027128 | -0.275489 | 0.23104447 | -1.1923635 | 0.23311876 | 0.80695949 |
| WIPI2 | 5208.3944 | -0.3131514 | 0.15074323 | -2.0773827 | 0.03776625 | 0.55538555 |
| SLC29A4 | 700.504234 | -0.0579015 | 0.44852922 | -0.129092 | 0.89728485 | 0.98474725 |
| TNRC18 | 7992.41665 | 0.01722047 | 0.24475452 | 0.07035812 | 0.94390863 | 0.99136103 |
| AC092171.4 | 57.3086913 | 0.52904222 | 0.54753457 | 0.96622615 | 3.34E-01 | 0.85533716 |
| AC092171.3 | 109.503197 | 0.39580141 | 0.35075266 | 1.12843451 | 0.25913646 | 0.82364948 |
| AC092171.5 | 54.7262066 | 0.51743854 | 0.47863626 | 1.0810684 | 0.27966669 | 0.82825734 |
| FBXL18 | 184.943197 | -0.3445968 | 0.32963759 | -1.0453808 | 0.29584701 | 0.83750889 |
| AC092171.2 | 598.187666 | -0.0370867 | 0.33918383 | -0.109341 | 0.91293201 | 0.98662757 |
| ACTB | 128601.298 | -0.0748631 | 0.30882436 | -0.2424131 | 0.80846006 | 0.97334138 |
| FSCN1 | 1562.3073 | 0.03824562 | 0.26659416 | 0.1434601 | 0.88592682 | 0.98385457 |
| RNF216 | 2330.09091 | -0.4904994 | 0.17431658 | -2.8138427 | 0.00489532 | 0.30278265 |
| CCZ1 | 661.108099 | -0.0792803 | 0.33360524 | -0.237647 | 8.12E-01 | 9.74E-01 |
| RSPH10B | 260.795914 | -0.0926962 | 0.4067602 | -0.2278891 | 0.81973245 | 0.97454841 |
| PMS2 | 865.81621 | 0.30544244 | 0.25453516 | 1.20000098 | 0.23013896 | 0.80556869 |
| AIMP2 | 1655.91966 | -0.0190025 | 0.21684271 | -0.0876324 | 0.93016884 | 0.98973492 |
| EIF2AK1 | 6721.26089 | 0.05218047 | 0.10470345 | 0.49836441 | 0.61822722 | 0.93929439 |

| | | | | | | |
|---|---|---|---|---|---|---|
| USP42 | 751.660842 | -0.3162659 | 0.17530605 | -1.8040786 | 0.07121898 | 0.63126712 |
| CYTH3 | 1584.47673 | -0.18664 | 0.24587976 | -0.7590701 | 0.4478106 | 0.89480207 |
| FAM220A | 629.432277 | -0.0239329 | 0.11502052 | -0.2080749 | 0.83517049 | 0.97550416 |
| RAC1 | 13091.9498 | -0.1922077 | 0.12951951 | -1.4840055 | 0.13780747 | 0.72209298 |
| DAGLB | 1762.47035 | -0.2359395 | 0.24700526 | -0.9552005 | 0.33947633 | 0.85759056 |
| KDELR2 | 8579.47201 | 0.0319608 | 0.1738561 | 0.18383482 | 0.85414304 | 0.97906925 |
| GRID2IP | 76.3094889 | 0.0996211 | 0.40948635 | 0.24328308 | 0.80778611 | 0.97334138 |
| ZDHHC4 | 1988.43979 | 0.0570145 | 0.11569144 | 0.49281523 | 0.62214315 | 0.94016713 |
| AC079742.1 | 836.176984 | 0.03228768 | 0.11352609 | 0.28440754 | 0.77609808 | 0.96836854 |
| C7orf26 | 1420.11209 | -0.3174427 | 0.1536027 | -2.066648 | 0.03876733 | 0.55608529 |
| ZNF853 | 471.154641 | -0.0670105 | 0.40415617 | -0.1658034 | 0.86831168 | 0.98243355 |
| ZNF316 | 3759.17455 | -0.4672946 | 0.16084783 | -2.9051968 | 0.00367022 | 0.26253301 |
| ZNF12 | 1279.61293 | -0.0507121 | 0.13721671 | -0.3695765 | 0.71169806 | 0.95560193 |
| RSPH10B2 | 189.395569 | -0.0978567 | 0.47410511 | -0.2064029 | 8.36E-01 | 0.97577999 |
| CCZ1B | 822.056085 | -0.1754238 | 0.33462116 | -0.5242461 | 0.60010738 | 0.93487648 |
| C1GALT1 | 2241.70975 | -0.2855428 | 0.27429664 | -1.0410001 | 0.29787552 | 0.83957554 |
| COL28A1 | 688.834294 | -0.3326909 | 0.44551248 | -0.74676 | 0.45520847 | 0.89670027 |
| AC004982.1 | 214.104029 | -0.1189498 | 0.22477296 | -0.5291996 | 0.59666699 | 0.9337017 |
| AC004982.2 | 468.888084 | -0.0194619 | 0.27227361 | -0.0714791 | 0.94301645 | 0.99136103 |
| MIOS | 1448.57222 | -0.1637682 | 0.15890196 | -1.0306244 | 0.30271697 | 0.84211658 |
| RPA3 | 1040.05005 | 0.31339449 | 0.17314245 | 1.81003842 | 0.07028983 | 0.62936167 |
| UMAD1 | 1135.75228 | 0.0400906 | 0.14155704 | 0.28321163 | 7.77E-01 | 9.68E-01 |
| GLCCI1 | 688.125796 | 0.00681712 | 0.22364083 | 0.03048245 | 0.97568229 | 0.99455099 |
| ICA1 | 1732.14288 | 0.12973634 | 0.2164247 | 0.59945259 | 0.54887112 | 0.92373235 |
| AC006042.1 | 442.974644 | 0.01895087 | 0.30401793 | 0.06233472 | 0.95029628 | 0.99170503 |
| AC007009.1 | 49.0250172 | 0.29155337 | 0.404315 | 0.72110451 | 0.47084522 | 0.90089741 |
| AC007128.1 | 49.0968275 | 0.29851349 | 0.40228678 | 0.74204152 | 0.45806218 | 0.8971553 |
| NDUFA4 | 10708.1945 | 0.25313958 | 0.17633538 | 1.43555748 | 0.1511283 | 0.73796431 |
| PHF14 | 1094.48209 | 0.01960228 | 0.15312768 | 0.12801266 | 0.89813895 | 0.98474725 |
| THSD7A | 1503.46969 | -0.5893505 | 0.30683159 | -1.920762 | 0.05476171 | 0.59329767 |
| TMEM106B | 3449.29595 | -0.1683042 | 0.21471486 | -0.7838499 | 0.43312819 | 0.89044047 |
| AC013470.2 | 48.3206268 | 1.14991935 | 0.68078011 | 1.68912006 | 0.09119642 | 0.65495306 |
| AC005281.1 | 258.134508 | 0.19262241 | 0.32810299 | 0.58707911 | 0.55715058 | 0.92373235 |
| SCIN | 5920.72654 | 0.40821466 | 0.2243824 | 1.81928112 | 0.06886855 | 0.62775561 |
| ARL4A | 525.224538 | -0.8025133 | 0.39354031 | -2.0392151 | 0.04142857 | 0.55945849 |
| ETV1 | 316.452464 | 0.96285114 | 0.29682852 | 3.2437959 | 0.00117948 | 0.15779959 |
| DGKB | 58.036466 | 0.46625346 | 0.70661235 | 0.65984335 | 0.50935436 | 0.91439937 |
| AGMO | 295.687106 | 0.76450198 | 0.40429761 | 1.89093868 | 5.86E-02 | 6.03E-01 |
| MEOX2 | 34.4839165 | 0.18809367 | 0.88297815 | 0.21302189 | 0.83130989 | 0.97550416 |
| ISPD | 171.603497 | 0.16029477 | 0.38520889 | 0.41612428 | 0.67731907 | 0.95018034 |
| SOSTDC1 | 662.500146 | 0.75373535 | 0.55379767 | 1.3610302 | 0.17350415 | 0.75974924 |
| ANKMY2 | 1115.16265 | 0.20529841 | 0.23640894 | 0.86840374 | 0.38517335 | 0.8735972 |
| BZW2 | 1911.24139 | 0.05798943 | 0.31351968 | 0.18496265 | 0.85325833 | 0.97887576 |
| AC073333.1 | 388.684805 | 0.11768273 | 0.3481382 | 0.33803454 | 0.73533716 | 0.95895327 |
| TSPAN13 | 809.370704 | 0.24075342 | 0.37698151 | 0.63863456 | 0.52306069 | 0.91723878 |
| AGR2 | 700.771464 | 1.81118212 | 0.57601357 | 3.14433932 | 0.00166462 | 0.18012143 |
| AGR3 | 94.6670429 | 1.81419428 | 0.56111794 | 3.23317819 | 0.00122421 | 0.15785077 |
| AHR | 1689.80989 | -0.1159389 | 0.22312948 | -0.5196036 | 0.60333989 | 0.93602816 |
| AC019117.3 | 173.125331 | 0.03345122 | 0.38106632 | 0.08778321 | 0.93004899 | 0.98973492 |
| SNX13 | 2054.93455 | -0.0139902 | 0.09860085 | -0.1418877 | 0.8871687 | 0.98423722 |
| HDAC9 | 270.36217 | -0.1660766 | 0.3468551 | -0.478807 | 0.63207593 | 0.94088547 |
| TWIST1 | 48.6172753 | 0.54558915 | 0.52029829 | 1.04860839 | 0.29435839 | 0.83622414 |
| TWISTNB | 664.948137 | -0.0015216 | 0.20637008 | -0.0073732 | 9.94E-01 | 0.99936518 |
| MACC1 | 1676.72493 | 0.12242617 | 0.23059939 | 0.53090412 | 0.59548522 | 0.9334601 |

| | | | | | | |
|---|---|---|---|---|---|---|
| AC004130.1 | 92.6514576 | 0.87797453 | 0.42744454 | 2.05400807 | 0.03997491 | 0.55657615 |
| ITGB8 | 5285.35629 | 0.43512708 | 0.26524124 | 1.64049555 | 0.10090217 | 0.67247782 |
| SP4 | 276.751944 | -0.0306615 | 0.22108665 | -0.1386853 | 0.88969884 | 0.9845209 |
| DNAH11 | 200.416746 | -0.6048881 | 0.49791792 | -1.2148349 | 2.24E-01 | 0.80078694 |
| CDCA7L | 277.863068 | -0.25216 | 0.38473139 | -0.6554182 | 0.51219851 | 0.91551505 |
| RAPGEF5 | 2231.7529 | -0.2790282 | 0.28442362 | -0.9810305 | 0.32657771 | 0.85282528 |
| AC002480.2 | 59.9539671 | -0.3649342 | 0.73913173 | -0.4937336 | 6.21E-01 | 9.40E-01 |
| AC073072.1 | 354.764565 | 3.27290647 | 1.52241035 | 2.14981885 | NA | NA |
| IL6 | 1339.70068 | 3.16674047 | 1.54427597 | 2.05063118 | NA | NA |
| TOMM7 | 5496.43977 | 0.31708257 | 0.19161535 | 1.6547869 | 0.09796773 | 0.66549286 |
| FAM126A | 626.946673 | -0.0721275 | 0.2284715 | -0.3156958 | 0.75223342 | 0.96301881 |
| KLHL7 | 1205.35009 | 0.04546218 | 0.13944899 | 0.32601297 | 0.74441454 | 0.9613927 |
| NUPL2 | 1440.174 | -0.1212817 | 0.13550922 | -0.8950072 | 0.37078325 | 0.86762749 |
| AC005082.1 | 502.053753 | -0.1791313 | 0.47477497 | -0.3772973 | 0.70595271 | 0.95501548 |
| GPNMB | 3228.34893 | 0.30422154 | 0.51658677 | 0.58890695 | 0.5559237 | 0.92373235 |
| MALSU1 | 1073.05433 | 0.11270339 | 0.14515315 | 0.77644466 | 0.43748649 | 0.89130865 |
| TRA2A | 4279.12283 | -0.0954933 | 0.22491875 | -0.4245678 | 0.67115179 | 0.94922827 |
| CCDC126 | 411.943495 | 0.07633464 | 0.23168344 | 0.32947818 | 0.74179429 | 0.96109001 |
| FAM221A | 470.470381 | -0.0674721 | 0.21885003 | -0.3083031 | 7.58E-01 | 9.65E-01 |
| AC004485.1 | 112.403137 | -0.3185413 | 0.84605609 | -0.3765014 | 0.70654419 | 0.95501548 |
| MPP6 | 1043.58853 | -0.1634023 | 0.27376532 | -0.5968701 | 5.51E-01 | 0.92373235 |
| GSDME | 1050.0749 | 0.32853331 | 0.24184238 | 1.35846043 | 0.17431763 | 0.75974924 |
| OSBPL3 | 2911.40956 | -0.3239701 | 0.33111065 | -0.9784346 | 0.32785941 | 0.85317073 |
| CYCS | 7970.0312 | 0.0664928 | 0.1904647 | 0.34910824 | 0.72700805 | 0.95848028 |
| C7orf31 | 204.050694 | 0.21941824 | 0.22371716 | 0.98078414 | 0.3266992 | 0.85282528 |
| AC010719.1 | 61.6803543 | 0.89543598 | 0.43534363 | 2.05684871 | 0.03970078 | 0.55657615 |
| NFE2L3 | 1121.29 | -0.3331226 | 0.22670965 | -1.4693801 | 0.14172973 | 0.72784886 |
| AC004520.1 | 202.611281 | -0.3937876 | 0.34599537 | -1.1381297 | 0.25506631 | 0.819931 |
| HNRNPA2B1 | 40817.3473 | -0.1526619 | 0.137712 | -1.1085593 | 0.26762033 | 0.82676682 |
| CBX3 | 7701.76285 | -0.3698325 | 0.21922492 | -1.6870002 | 0.09160332 | 0.65518742 |
| SNX10 | 4544.86296 | 0.07589551 | 0.30188547 | 0.25140497 | 0.80150102 | 0.97210205 |
| AC004540.2 | 560.945759 | 0.16790947 | 0.23237177 | 0.72258982 | 0.46993193 | 0.9008571 |
| AC004540.1 | 181.386257 | 0.21973827 | 0.3160951 | 0.69516506 | 0.48695187 | 0.90573694 |
| SKAP2 | 2610.99418 | 0.25693589 | 0.20703186 | 1.24104517 | 2.15E-01 | 0.79431436 |
| HOXA1 | 89.7756068 | -0.0062267 | 0.24146314 | -0.0257875 | 0.97942684 | 0.99524212 |
| HOTAIRM1 | 1569.49083 | -0.0482727 | 0.29960676 | -0.1611203 | 0.87199862 | 0.98275231 |
| HOXA2 | 186.301573 | 0.0665384 | 0.43818868 | 0.15184875 | 0.87930623 | 0.98275231 |
| HOXA3 | 1225.78472 | 0.13199001 | 0.30350514 | 0.4348856 | 0.66364547 | 0.94642611 |
| HOXA-AS2 | 1411.61842 | 0.0296989 | 0.34388453 | 0.08636302 | 9.31E-01 | 0.98975562 |
| HOXA4 | 573.329499 | -0.0107627 | 0.29431498 | -0.0365685 | 0.97082906 | 0.99400331 |
| HOXA-AS3 | 1159.98362 | 0.27542124 | 0.30326299 | 0.90819274 | 0.3637764 | 0.8663515 |
| HOXA5 | 1303.68093 | 0.14662657 | 0.25758276 | 0.56924063 | 0.56919285 | 0.92686099 |
| HOXA6 | 173.227572 | 0.21324265 | 0.39942475 | 0.53387439 | 0.59342844 | 0.93289552 |
| HOXA7 | 1107.53269 | 0.33332495 | 0.36523092 | 0.91264165 | 0.36143104 | 0.86528137 |
| HOXA9 | 2454.18786 | 0.22010634 | 0.19550996 | 1.12580628 | 0.26024753 | 0.82424998 |
| AC004080.3 | 903.882171 | 0.20345559 | 0.19773584 | 1.0289262 | 0.30351436 | 0.84211658 |
| HOXA10-AS | 794.032015 | -0.0452147 | 0.2280134 | -0.1982985 | 0.84281153 | 0.97645468 |
| HOXA10 | 1193.13451 | 0.00010158 | 0.22660048 | 0.00044827 | 0.99964233 | 0.99994357 |
| HOXA11 | 250.048135 | -0.1222146 | 0.36023361 | -0.3392649 | 0.73441021 | 0.95895327 |
| HOXA11-AS | 211.514528 | -0.0641713 | 0.40813305 | -0.1572313 | 0.87506259 | 0.98275231 |
| HIBADH | 6577.41991 | 0.65080158 | 0.3384994 | 1.92260777 | 0.05452933 | 0.59329767 |
| AC005091.1 | 230.903536 | 0.34800691 | 0.34997647 | 0.99437232 | 0.32004164 | 0.84966452 |
| TAX1BP1 | 7218.55096 | 0.09043386 | 0.16494049 | 0.54828176 | 0.58349845 | 0.93225443 |
| JAZF1 | 617.831556 | 0.09108431 | 0.29941331 | 0.30420929 | 0.76096845 | 0.9658862 |

| | | | | | | |
|---|---|---|---|---|---|---|
| CREB5 | 753.05126 | 0.03299144 | 0.40474116 | 0.08151245 | 0.93503443 | 0.99068685 |
| TRIL | 681.602487 | 0.07605761 | 0.47736143 | 0.15932919 | 0.87340953 | 0.98275231 |
| AC005013.1 | 67.4031481 | 0.19410737 | 0.37087733 | 0.52337352 | 0.60071435 | 0.93489351 |
| AC005162.3 | 180.613297 | -0.021495 | 0.4805792 | -0.0447272 | 0.96432473 | 0.99342622 |
| CPVL | 5430.65977 | 0.47909393 | 0.31805989 | 1.50630099 | 0.13198992 | 0.7121526 |
| AC004593.3 | 1196.29713 | 0.21545765 | 0.24194259 | 0.89053212 | 0.37318023 | 0.86791605 |
| CHN2 | 749.511233 | -0.0269742 | 0.26259586 | -0.1027212 | 0.91818422 | 0.9868044 |
| AC007255.1 | 91.1340021 | -0.1248554 | 0.34950558 | -0.3572344 | 7.21E-01 | 0.95728372 |
| PRR15 | 1118.49203 | -0.1846617 | 0.19774514 | -0.9338366 | 0.35038817 | 0.86364633 |
| WIPF3 | 520.837237 | 0.18607577 | 0.36682413 | 0.50726152 | 0.61197133 | 0.93856019 |
| SCRN1 | 3250.85106 | 0.24948155 | 0.18808247 | 1.32644761 | 0.18469148 | 0.76820626 |
| FKBP14 | 568.602309 | 0.09148369 | 0.2430583 | 0.37638579 | 0.70663011 | 0.95501548 |
| PLEKHA8 | 935.905386 | 0.15122643 | 0.26484335 | 0.57100329 | 0.56799741 | 0.92618483 |
| MTURN | 4936.52837 | -0.0026619 | 0.37880586 | -0.0070271 | 0.99439326 | 0.99940602 |
| ZNRF2 | 1685.27488 | 0.16209 | 0.13879589 | 1.16782989 | 0.24287539 | 0.8133895 |
| AC006978.2 | 668.635263 | 0.1858425 | 0.14017113 | 1.32582576 | 0.18489742 | 0.76820626 |
| LINC01176 | 256.948609 | -0.4179953 | 0.32402058 | -1.2900271 | 0.19704124 | 0.77816059 |
| NOD1 | 875.344608 | 0.00011895 | 0.27030855 | 0.00044007 | 0.99964888 | 0.99994357 |
| AC006027.1 | 454.987108 | 0.04504734 | 0.34228563 | 0.13160744 | 0.8952948 | 0.98474725 |
| GGCT | 1699.01451 | 0.56671648 | 0.28239171 | 2.0068453 | 0.04476614 | 0.57267043 |
| AC005154.6 | 1191.74147 | 0.56164214 | 0.27774137 | 2.02217677 | 0.04315809 | 0.56839482 |
| AC005154.4 | 48.7871679 | 0.07608194 | 0.42275332 | 0.17996769 | 0.85717793 | 0.97953896 |
| GARS | 5517.9516 | -0.391118 | 0.346117 | -1.1300168 | 0.25846916 | 0.82294696 |
| INMT | 1097.76025 | -0.1077733 | 0.55371251 | -0.1946377 | 0.84567657 | 0.97675487 |
| INMT-MINDY | 629.739627 | 0.13010411 | 0.26923788 | 0.48323109 | 0.62893166 | 0.94088547 |
| MINDY4 | 564.169094 | 0.09214699 | 0.29786157 | 0.30936181 | 0.75704632 | 0.96499587 |
| AC004691.2 | 19846.4491 | 1.30242542 | 0.46137364 | 2.82292984 | 0.0047587 | 0.29927103 |
| AQP1 | 63002.4957 | 1.26976006 | 0.43604622 | 2.91198503 | 0.0035914 | 0.26033637 |
| GHRHR | 55.7159506 | 0.3812628 | 0.96042188 | 0.3969743 | NA | NA |
| PDE1C | 425.353383 | 0.13681763 | 0.50746973 | 0.26960748 | 0.78746225 | 0.97090793 |
| LSM5 | 1071.55773 | 0.22088996 | 0.16429013 | 1.34451141 | 0.17878307 | 0.76120583 |
| AVL9 | 2144.16246 | -0.0536825 | 0.17473205 | -0.3072275 | 0.75867022 | 0.96502715 |
| AC018645.2 | 83.5186981 | -0.1347659 | 0.24178561 | -0.5573776 | 0.57726946 | 0.93130183 |
| LINC00997 | 261.803951 | -0.1802662 | 0.24961476 | -0.7221777 | 0.47018526 | 0.9008571 |
| KBTBD2 | 2511.00382 | -0.2285893 | 0.17269415 | -1.3236656 | 0.18561413 | 0.76908034 |
| FKBP9 | 3803.65426 | 0.40746108 | 0.12474035 | 3.26647366 | 0.00108896 | 0.15578769 |
| NT5C3A | 1992.82465 | -1.1266165 | 0.4741146 | -2.3762535 | 0.01748944 | 0.46859699 |
| RP9 | 404.953398 | -0.0614544 | 0.13479308 | -0.4559165 | 0.64845001 | 0.9431121 |
| BBS9 | 691.447626 | 0.048696 | 0.24228259 | 0.20098846 | 0.8407076 | 0.97618283 |
| BMPER | 40.4239849 | 0.12982657 | 0.46747806 | 0.27771692 | 0.78122967 | 0.96902269 |
| DPY19L1 | 1182.73797 | -0.2541918 | 0.24469397 | -1.0388153 | 0.29889067 | 0.84056272 |
| HERPUD2 | 1594.6473 | -0.1849409 | 0.10755392 | -1.719518 | 0.08552009 | 0.6504594 |
| AC018647.2 | 464.560015 | -0.4349873 | 0.20698081 | -2.1015826 | 0.03558985 | 0.55369025 |
| SEPT7-AS1 | 130.86405 | 0.29968458 | 0.34273816 | 0.87438346 | 3.82E-01 | 8.72E-01 |
| 7-Sep | 5937.22008 | -0.1502556 | 0.15772411 | -0.952648 | 0.34076845 | 0.85849463 |
| EEPD1 | 505.863094 | -0.3176012 | 0.30746577 | -1.0329644 | 0.30162054 | 0.84211658 |
| KIAA0895 | 885.717746 | -0.1333083 | 0.15978959 | -0.8342738 | 0.40412668 | 0.88132015 |
| ANLN | 62.3378805 | 0.33718575 | 0.64293467 | 0.52444792 | 0.59996704 | 0.93487648 |
| AOAH | 308.078594 | -0.1511729 | 0.59123314 | -0.2556908 | 0.79818962 | 0.97170189 |
| ELMO1 | 529.812158 | -0.0537314 | 0.36050593 | -0.1490445 | 0.88151853 | 0.98275231 |
| EPDR1 | 1603.84742 | 0.14314144 | 0.2763488 | 0.51797382 | 0.60447653 | 0.93662382 |
| SFRP4 | 245.244237 | 0.58683307 | 0.63525517 | 0.92377536 | 0.35560329 | 0.86364633 |
| STARD3NL | 2143.85854 | -0.0563332 | 0.11475856 | -0.4908845 | 0.62350817 | 0.94045607 |
| TRGC2 | 68.0098763 | 0.0591852 | 0.63418615 | 0.09332466 | 0.92564564 | 0.98876911 |

| | | | | | | |
|---|---|---|---|---|---|---|
| TRGC1 | 61.6848107 | -0.7136235 | 0.67638164 | -1.0550605 | 0.29139766 | 0.83454586 |
| TRG-AS1 | 51.5885802 | 0.38362372 | 0.71453429 | 0.53688638 | 0.5913461 | 0.93289552 |
| AMPH | 145.995551 | -0.6635063 | 0.47604261 | -1.3937961 | 0.16337919 | 0.75149188 |
| VPS41 | 2432.84432 | 0.09460498 | 0.17715275 | 0.53403057 | 0.59332039 | 0.93289552 |
| YAE1D1 | 411.65617 | -0.2676134 | 0.15646199 | -1.7104054 | 0.08719093 | 0.65231349 |
| RALA | 2179.20913 | -0.3530499 | 0.34408695 | -1.0260485 | 0.30486872 | 0.842333 |
| LINC00265 | 254.59839 | 0.01590295 | 0.34718075 | 0.04580598 | 0.9634649 | 0.99320711 |
| AC072061.1 | 72.1984467 | -0.1191422 | 0.32277602 | -0.3691172 | 0.71204036 | 0.95560193 |
| CDK13 | 2524.49405 | -0.1934756 | 0.11125198 | -1.7390751 | 0.08202155 | 0.64559676 |
| MPLKIP | 1357.02724 | -0.1104556 | 0.18609712 | -0.5935376 | 0.55282143 | 0.92373235 |
| SUGCT | 1043.65657 | 0.73580242 | 0.35379968 | 2.07971477 | 0.0375517 | 0.55538555 |
| INHBA | 388.350768 | -1.1697682 | 0.84451672 | -1.3851333 | 0.16601171 | 0.75515387 |
| GLI3 | 299.601084 | 0.5914756 | 0.38049756 | 1.55447936 | 0.12007012 | 0.69467399 |
| C7orf25 | 728.120292 | 0.0309778 | 0.16212688 | 0.19107131 | 0.84846972 | 0.97733073 |
| AC010132.3 | 5286.46362 | -0.0135804 | 0.12767243 | -0.1063694 | 0.91528923 | 0.98673697 |
| PSMA2 | 4933.06269 | -0.0214943 | 0.12348553 | -0.1740634 | 0.86181567 | 0.98176973 |
| MRPL32 | 2659.09565 | -0.1699438 | 0.15793019 | -1.076069 | 0.28189638 | 0.83001101 |
| HECW1 | 736.195584 | 2.87720332 | 0.78740638 | 3.65402593 | 0.00025816 | 0.08200305 |
| LUARIS | 93.0364661 | 2.33141524 | 0.84452184 | 2.76063343 | 0.00576894 | 0.32369964 |
| STK17A | 1253.75508 | -0.6207816 | 0.32194452 | -1.9282253 | 0.0538271 | 0.59266174 |
| COA1 | 1892.01231 | 0.1706427 | 0.13159218 | 1.29675412 | 0.1947158 | 0.7762856 |
| BLVRA | 2339.11867 | 0.0273913 | 0.13573967 | 0.20179286 | 0.84007866 | 0.97618283 |
| MRPS24 | 2761.3758 | -0.1603344 | 0.17415425 | -0.920646 | 3.57E-01 | 0.86390849 |
| URGCP-MRP | 2889.0527 | -0.1365866 | 0.17289554 | -0.789995 | 0.4295307 | 0.88858861 |
| URGCP | 4273.96762 | -0.109115 | 0.16368696 | -0.666608 | 0.50502256 | 0.91365534 |
| UBE2D4 | 1296.57023 | 0.02865059 | 0.1872775 | 0.1529847 | 0.87841034 | 0.98275231 |
| LINC00957 | 340.624448 | 0.32039945 | 0.36050146 | 0.88876048 | 0.37413182 | 0.86791605 |
| DBNL | 5282.63268 | 0.02118073 | 0.14233664 | 0.14880731 | 0.88170569 | 0.98275231 |
| PGAM2 | 366.096391 | 0.15753995 | 0.33208655 | 0.47439426 | 0.63521878 | 0.94088547 |
| POLM | 2665.84056 | 0.06010402 | 0.15875584 | 0.37859405 | 0.70498934 | 0.95501548 |
| AEBP1 | 15469.6314 | -0.0754357 | 0.46229608 | -0.1631763 | 0.87037966 | 0.98275231 |
| POLD2 | 3628.56127 | 0.25010502 | 0.13183145 | 1.89715745 | 0.05780716 | 0.60129769 |
| YKT6 | 5179.98548 | -0.391416 | 0.30928538 | -1.2655495 | 0.2056744 | 0.78583974 |
| CAMK2B | 195.671758 | 0.41244028 | 0.35734176 | 1.15418998 | 0.24842229 | 0.8162591 |
| NUDCD3 | 4181.50634 | -0.1312183 | 0.13641652 | -0.9618944 | 0.33610265 | 0.85641741 |
| NPC1L1 | 174.31541 | 0.71959494 | 0.43641671 | 1.64887118 | 0.09917403 | 0.66904984 |
| DDX56 | 5515.08107 | 0.05087146 | 0.16995765 | 0.29931846 | 0.76469707 | 0.96678847 |
| TMED4 | 7156.69376 | 0.47559979 | 0.15785171 | 3.01295302 | 0.00258719 | 0.21645768 |
| OGDH | 14079.3277 | 0.21795327 | 0.19770111 | 1.1024383 | 0.27027117 | 0.82716979 |
| ZMIZ2 | 10510.3871 | -0.0486095 | 0.20543289 | -0.2366198 | 0.81295174 | 0.9738447 |
| PPIA | 35136.6251 | -0.0542027 | 0.20686205 | -0.2620232 | 0.79330357 | 0.97090793 |
| H2AFV | 5350.8031 | 0.03286784 | 0.25183771 | 0.13051197 | 0.89616138 | 0.98474725 |
| PURB | 2348.33699 | -0.1337054 | 0.12280109 | -1.0887969 | 0.27624346 | 0.82719407 |
| AC004854.2 | 46.6174457 | 0.101574 | 0.34605415 | 0.29352055 | 0.7691243 | 0.96823796 |
| MYO1G | 442.504354 | -0.2551545 | 0.65477641 | -0.3896819 | 0.69677176 | 0.9525827 |
| SNHG15 | 1212.56639 | -0.2103853 | 0.28477368 | -0.7387805 | 0.46004027 | 0.89755243 |
| CCM2 | 2207.86874 | 0.05836538 | 0.16972546 | 0.34388115 | 0.73093567 | 0.95848028 |
| NACAD | 362.839067 | -0.2407745 | 0.38497094 | -0.6254356 | 0.53168522 | 0.91962754 |
| TBRG4 | 4019.79498 | 0.0581462 | 0.2052703 | 0.28326649 | 0.77697256 | 0.96836854 |
| RAMP3 | 3104.08596 | -0.3734661 | 0.31274988 | -1.1941368 | 0.23242447 | 0.80649152 |
| ADCY1 | 340.765158 | 0.38223992 | 0.51905525 | 0.7364147 | 0.46147836 | 0.89755243 |
| IGFBP1 | 67.8756897 | 0.56433557 | 1.07316808 | 0.52585945 | 0.59898587 | 0.93484879 |
| IGFBP3 | 7096.93695 | -0.6295724 | 0.39218591 | -1.6052906 | 0.10842987 | 0.67800851 |
| TNS3 | 11113.8612 | -0.2409832 | 0.273458 | -0.8812438 | 0.37818586 | 0.87028257 |

| | | | | | | |
|---|---|---|---|---|---|---|
| HUS1 | 717.167804 | -0.2140008 | 0.15490805 | -1.3814696 | 0.16713462 | 0.75558932 |
| UPP1 | 4908.4391 | -0.3799502 | 0.48398498 | -0.7850455 | 4.32E-01 | 0.88980461 |
| ABCA13 | 97.2558857 | -0.0031728 | 0.77766539 | -0.0040799 | 0.99674474 | 0.99976751 |
| IKZF1 | 317.251987 | -0.1241104 | 0.64273645 | -0.1930968 | 0.84688312 | 0.97684681 |
| FIGNL1 | 261.990737 | 0.02094918 | 0.27210585 | 0.07698907 | 0.93863224 | 0.99113899 |
| DDC | 11388.7492 | 1.09337762 | 0.5521413 | 1.98024966 | 0.04767548 | 0.58241466 |
| DDC-AS1 | 1548.74773 | 1.20808048 | 0.61197245 | 1.97407657 | 0.04837304 | 0.58440502 |
| GRB10 | 3861.05937 | -0.276174 | 0.31036748 | -0.8898289 | 0.37355774 | 0.86791605 |
| AC004830.1 | 272.526937 | -0.2144288 | 0.24131051 | -0.888601 | 0.37421757 | 0.86791605 |
| COBL | 2114.90413 | -0.1864184 | 0.24196705 | -0.7704289 | 0.44104553 | 0.89290667 |
| SEC61G | 1988.4423 | -0.0347659 | 0.23660231 | -0.1469381 | 0.88318088 | 0.98293314 |
| EGFR | 2971.56619 | -0.0213002 | 0.14264041 | -0.1493283 | 0.88129461 | 0.98275231 |
| LANCL2 | 1156.67025 | 0.21021033 | 0.34775213 | 0.60448322 | 0.54552242 | 0.92345821 |
| VOPP1 | 2701.54759 | -0.0921862 | 0.2197814 | -0.419445 | 0.67489094 | 0.94935413 |
| AC092647.5 | 567.678267 | -0.1127551 | 0.19383202 | -0.5817155 | 0.56075831 | 0.9244521 |
| ZNF713 | 85.2484708 | 0.13497764 | 0.3344059 | 0.40363414 | 0.68648178 | 0.95097288 |
| NIPSNAP2 | 4085.95994 | 0.17515947 | 0.15830743 | 1.10645138 | 0.2685312 | 0.82676682 |
| MRPS17 | 844.716754 | -0.1265558 | 0.20765439 | -0.6094541 | 0.54222351 | 0.92292765 |
| PSPH | 1010.34355 | 0.24758997 | 0.4230129 | 0.58530123 | 0.55834519 | 0.92421175 |
| CCT6A | 8481.96353 | -0.164307 | 0.32648108 | -0.5032665 | 6.15E-01 | 0.93895494 |
| SUMF2 | 8947.20756 | 0.2625926 | 0.1502416 | 1.74780218 | 0.08049829 | 0.64196529 |
| PHKG1 | 1618.2682 | 0.15905868 | 0.18576024 | 0.85625796 | 0.39185511 | 0.87744531 |
| CHCHD2 | 11715.1527 | 0.03232146 | 0.22005773 | 0.14687718 | 0.88322897 | 0.98293314 |
| NUPR2 | 186.156089 | 0.33986214 | 0.30128331 | 1.12804835 | 0.2592995 | 0.82364948 |
| ZNF736 | 894.405269 | 0.14984537 | 0.27006013 | 0.5548593 | 0.5789909 | 0.93139743 |
| ZNF680 | 718.63511 | -0.0038433 | 0.23519461 | -0.0163407 | 0.98696257 | 0.99726698 |
| ZNF107 | 309.594456 | -0.533397 | 0.42913711 | -1.2429524 | 0.21388537 | 0.7926732 |
| ZNF138 | 404.071385 | 0.07597781 | 0.26415466 | 0.28762625 | 0.77363285 | 0.96836854 |
| ZNF273 | 203.066025 | 0.02396375 | 0.40762309 | 0.05878898 | 0.95312019 | 0.99170503 |
| ZNF117 | 1076.66195 | -0.0978607 | 0.24123922 | -0.4056585 | 0.68499355 | 0.95097288 |
| ERV3-1 | 361.099422 | 0.03610443 | 0.39735229 | 0.09086253 | 0.92760183 | 0.98915637 |
| ZNF92 | 326.471479 | -0.0890529 | 0.23640697 | -0.376693 | 0.70640174 | 0.95501548 |
| VKORC1L1 | 1304.71894 | -0.0036536 | 0.16164774 | -0.0226022 | 0.98196761 | 0.99595643 |
| GUSB | 3659.65552 | 0.18594311 | 0.16584569 | 1.12118145 | 0.26221063 | 0.82424998 |
| ASL | 6696.00834 | 0.30399733 | 0.18382626 | 1.65372088 | 0.09818424 | 0.66549286 |
| AC068533.4 | 4006.98736 | 0.2732512 | 0.16330772 | 1.6732289 | 0.09428224 | 0.65783452 |
| CRCP | 1902.50761 | -0.0509716 | 0.15026625 | -0.3392089 | 0.7344524 | 0.95895327 |
| TPST1 | 1779.13103 | -0.0359536 | 0.31817813 | -0.1129983 | 0.91003188 | 0.98662757 |
| AC008267.5 | 99.2801593 | 0.43842962 | 0.47563564 | 0.92177621 | 0.35664532 | 0.86390849 |
| KCTD7 | 850.087322 | -0.0234769 | 0.2084494 | -0.1126262 | 0.91032689 | 0.98662757 |
| AC027644.4 | 1965.55368 | -0.2179203 | 0.20246811 | -1.0763192 | 0.28178451 | 0.83001101 |
| RABGEF1 | 2695.43434 | -0.1501649 | 0.15560583 | -0.9650338 | 0.33452787 | 0.85557672 |
| AC027644.3 | 519.440388 | 0.11722637 | 0.30231218 | 0.38776597 | 0.69818923 | 0.9529028 |
| AC073335.2 | 900.013515 | 0.09713096 | 0.38832121 | 0.25013045 | 0.80248647 | 0.97247973 |
| TMEM248 | 5291.7508 | 0.02313853 | 0.10326456 | 0.22407042 | 0.82270251 | 0.97519922 |
| SBDS | 6035.26317 | 0.00946639 | 0.20665997 | 0.04580658 | 0.96346442 | 0.99320711 |
| TYW1 | 2149.54494 | 0.06672486 | 0.13508106 | 0.49396158 | 0.62133331 | 0.94016713 |
| AC006480.2 | 80.0862136 | -0.0665114 | 0.37182332 | -0.178879 | 0.85803274 | 0.97973506 |
| LINC01372 | 107.13147 | -0.0486769 | 0.36540012 | -0.1332153 | 0.89402308 | 0.98474725 |
| AUTS2 | 1855.24762 | -0.073811 | 0.2322085 | -0.3178651 | 0.75058729 | 0.96255347 |
| GALNT17 | 300.705003 | 1.08219294 | 0.49842205 | 2.17123809 | 0.02991318 | 0.54012717 |
| TYW1B | 226.074347 | 0.60182647 | 0.49779874 | 1.20897548 | 2.27E-01 | 0.80288508 |
| POM121 | 4790.46617 | -0.1060407 | 0.10328306 | -1.0266998 | 0.30456183 | 0.84211658 |
| TRIM74 | 58.4440099 | 0.05848919 | 0.4976586 | 0.11752874 | 0.90644107 | 0.98662757 |

| | | | | | | |
|---|---|---|---|---|---|---|
| NSUN5 | 1672.97025 | -0.1329654 | 0.10933664 | -1.21611 | 0.22394302 | 0.80078694 |
| TRIM50 | 380.02561 | -1.2922665 | 1.01797407 | -1.2694493 | 0.20428085 | 0.7843531 |
| BAZ1B | 2644.49466 | -0.1987325 | 0.11586218 | -1.7152495 | 0.0862995 | 0.65077063 |
| BCL7B | 3560.49468 | -0.3889674 | 0.22891686 | -1.6991643 | 0.08928823 | 0.65375543 |
| TBL2 | 1613.47139 | -0.2440177 | 0.15161071 | -1.609502 | 0.10750662 | 0.67707349 |
| MLXIPL | 8200.95942 | -0.0087497 | 0.61480761 | -0.0142316 | 0.9886452 | 0.99788078 |
| VPS37D | 796.059202 | 0.20421413 | 0.26065138 | 0.78347612 | 0.43334757 | 0.89055338 |
| DNAJC30 | 899.976872 | -0.0028747 | 0.22032876 | -0.0130472 | 0.98959013 | 0.99800524 |
| BUD23 | 3977.36625 | -0.0514893 | 0.13058689 | -0.3942916 | 0.69336574 | 0.9519683 |
| STX1A | 389.618019 | -0.3280108 | 0.28194127 | -1.1634012 | 0.24466677 | 0.81473945 |
| ABHD11 | 2985.87944 | 0.01444369 | 0.20517813 | 0.07039586 | 0.94387859 | 0.99136103 |
| CLDN3 | 2867.9914 | -0.0125239 | 0.32475394 | -0.0385641 | 0.96923789 | 0.99400331 |
| CLDN4 | 13892.2949 | -0.3550527 | 0.41883105 | -0.847723 | 0.39659226 | 0.8796941 |
| METTL27 | 336.398598 | 0.35242393 | 0.53385099 | 0.66015413 | 0.50915493 | 0.91439937 |
| ELN | 1886.76124 | -0.4842638 | 0.71242117 | -0.6797437 | 0.49666677 | 0.90890243 |
| ELN-AS1 | 184.66431 | -0.3337922 | 0.68752031 | -0.4855015 | 0.62732062 | 0.94088547 |
| LIMK1 | 1593.14847 | -0.158273 | 0.22637665 | -0.6991578 | 0.48445343 | 0.90541935 |
| EIF4H | 12635.7485 | -0.0206283 | 0.16491902 | -0.1250811 | 0.90045935 | 0.98526603 |
| LAT2 | 339.968517 | -0.2174525 | 0.43829303 | -0.496135 | 0.61979914 | 0.93945324 |
| RFC2 | 860.150839 | 0.0480484 | 0.19446148 | 0.24708442 | 0.80484289 | 0.97300739 |
| CLIP2 | 1957.07018 | -0.413074 | 0.3095537 | -1.3344178 | 0.18206696 | 0.76569697 |
| GTF2IRD1 | 2933.41974 | -0.640614 | 0.27793316 | -2.3049213 | 0.02117098 | 0.50998124 |
| GTF2I | 7756.78707 | 0.02745181 | 0.26138419 | 0.10502476 | 0.91635616 | 0.9868044 |
| NCF1 | 252.471572 | -0.9934752 | 0.61115084 | -1.6255809 | 0.10403883 | 0.67439401 |
| GTF2IRD2 | 405.911907 | 0.16799797 | 0.33685065 | 0.49873131 | 0.61796868 | 0.93928244 |
| CASTOR2 | 86.166562 | -0.7283246 | 0.55120063 | -1.3213421 | 0.18638733 | 0.76908034 |
| RCC1L | 3769.56732 | 0.13588221 | 0.15882706 | 0.85553564 | 0.39225468 | 0.87765581 |
| GTF2IRD2B | 783.138232 | 0.05738823 | 0.27637536 | 0.20764596 | 0.83550542 | 0.97550416 |
| TRIM73 | 148.789271 | 0.48476444 | 0.28156237 | 1.72169472 | 0.08512483 | 0.64988556 |
| POM121C | 2669.85182 | -0.2333316 | 0.16549915 | -1.4098657 | 1.59E-01 | 0.74675717 |
| SPDYE5 | 48.7231657 | -0.2099316 | 0.38791843 | -0.5411746 | 0.58838727 | 0.93289552 |
| HIP1 | 2376.42368 | -0.262726 | 0.33365016 | -0.7874295 | 0.4310305 | 0.88966606 |
| RHBDD2 | 6863.03361 | -0.0576042 | 0.15386445 | -0.3743831 | 0.70811933 | 0.95501548 |
| POR | 13324.1947 | -0.4636301 | 0.41175615 | -1.1259822 | 2.60E-01 | 8.24E-01 |
| TMEM120A | 3408.73951 | 0.0769735 | 0.20090287 | 0.38313786 | 7.02E-01 | 0.95424422 |
| STYXL1 | 1037.4834 | 0.48659666 | 0.32402077 | 1.50174528 | 0.1331629 | 0.71486852 |
| MDH2 | 11209.528 | 0.34253619 | 0.21035886 | 1.62834213 | 0.10345236 | 0.67439401 |
| SRRM3 | 148.560404 | -0.1344539 | 0.54798628 | -0.2453599 | 0.80617778 | 0.97334138 |
| HSPB1 | 22074.6755 | -0.0432642 | 0.27812385 | -0.1555574 | 0.87638191 | 0.98275231 |
| YWHAG | 7015.96474 | -0.0976772 | 0.20868759 | -0.4680548 | 0.63974542 | 0.94088547 |
| SSC4D | 385.734006 | 0.69376982 | 0.35950072 | 1.92981482 | 0.05362979 | 0.59191745 |
| ZP3 | 183.351332 | 0.16628492 | 0.4568506 | 0.36398096 | 0.71587223 | 0.95590747 |
| DTX2 | 954.740163 | -0.2106802 | 0.15142935 | -1.3912773 | 0.16414135 | 0.752829 |
| UPK3B | 42.1229145 | -0.1936929 | 0.55360035 | -0.3498785 | 0.72642989 | 0.95848028 |
| POMZP3 | 548.404371 | -0.6099579 | 0.39496385 | -1.5443385 | 0.12250636 | 0.69788192 |
| CCDC146 | 815.393985 | 0.28737081 | 0.44438114 | 0.64667643 | 0.51784138 | 0.91636259 |
| FGL2 | 3060.25326 | 0.75289916 | 0.41469574 | 1.81554593 | 0.06944005 | 0.62779744 |
| GSAP | 1433.05778 | 0.117607 | 0.36284056 | 0.3241286 | 0.74584068 | 0.96176407 |
| PTPN12 | 3661.58568 | -0.2942806 | 0.18523245 | -1.5887095 | 0.11212599 | 0.68001961 |
| APTR | 614.153699 | 0.40464673 | 0.20705474 | 1.95429833 | 0.05066595 | 0.5861488 |
| RSBN1L | 1117.09002 | -0.1243237 | 0.15441595 | -0.805122 | 0.4207493 | 0.88451372 |
| TMEM60 | 746.094958 | -0.3623647 | 0.20431032 | -1.7735994 | 0.07612944 | 0.63773415 |
| PHTF2 | 809.619436 | -0.0041276 | 0.19447871 | -0.0212239 | 0.98306702 | 0.9961464 |
| MAGI2 | 680.591108 | 0.32119916 | 0.27975967 | 1.14812532 | 2.51E-01 | 8.18E-01 |

| | | | | | | |
|---|---|---|---|---|---|---|
| GNAI1 | 3643.28898 | -0.3720893 | 0.19353055 | -1.9226384 | 0.05452548 | 0.59329767 |
| CD36 | 726.830613 | 1.21904764 | 0.82694905 | 1.47415085 | 0.14044098 | 0.72546979 |
| SEMA3C | 558.736044 | 0.05744255 | 0.48343032 | 0.1188228 | 0.90541574 | 0.98625102 |
| HGF | 1058.96434 | -0.4649917 | 0.40570578 | -1.1461304 | 0.25174122 | 0.81833349 |
| CACNA2D1 | 876.142232 | 0.21711051 | 0.13119885 | 1.65482019 | 0.09796098 | 0.66549286 |
| PCLO | 743.840667 | -0.0633518 | 0.48753019 | -0.1299443 | 0.89661046 | 0.98474725 |
| SEMA3A | 105.189805 | -0.8712476 | 0.43775673 | -1.9902552 | 0.04656283 | 0.57983846 |
| AC003984.1 | 45.8426777 | 0.37224746 | 0.67401393 | 0.55228452 | 0.58075344 | 0.9317705 |
| SEMA3D | 235.93739 | 0.17658058 | 0.45116726 | 0.39138606 | 0.6955119 | 0.9525827 |
| KIAA1324L | 296.164095 | -0.3791454 | 0.29587542 | -1.281436 | 0.20004055 | 0.78134034 |
| AC005076.1 | 39.7548369 | 1.00676841 | 0.51892984 | 1.94008579 | 0.05236926 | 0.58645127 |
| DMTF1 | 3812.52915 | -0.082439 | 0.20072721 | -0.4107015 | 0.68129146 | 0.95070301 |
| TMEM243 | 1731.90836 | 0.31902386 | 0.14653381 | 2.17713481 | 0.02947051 | 0.53682638 |
| TP53TG1 | 621.677199 | 0.54100719 | 0.24845179 | 2.17751377 | 0.02944226 | 0.53682638 |
| CROT | 1200.25156 | 0.67027152 | 0.17982601 | 3.72733365 | 0.00019352 | 0.06842806 |
| ABCB4 | 69.2797307 | 0.24116389 | 0.37695734 | 0.63976441 | 5.22E-01 | 9.17E-01 |
| ABCB1 | 3301.37771 | 0.22648602 | 0.3623288 | 0.62508422 | 5.32E-01 | 9.20E-01 |
| RUNDC3B | 185.317802 | 0.16127303 | 0.41412094 | 0.38943462 | 6.97E-01 | 9.53E-01 |
| SLC25A40 | 897.60984 | 0.29087205 | 0.13874608 | 2.09643434 | 0.03604368 | 0.55369025 |
| DBF4 | 342.136003 | -0.0199007 | 0.28872262 | -0.0689268 | 0.94504785 | 0.99164528 |
| ADAM22 | 587.511101 | 0.63094566 | 0.35397383 | 1.78246416 | 0.07467357 | 0.63773415 |
| SRI | 3387.85943 | 0.13498604 | 0.20758805 | 0.65025922 | 0.51552479 | 0.91636259 |
| AC003991.2 | 718.826291 | 0.07942392 | 0.2480293 | 0.32021991 | 0.74880163 | 0.96249332 |
| AC003991.1 | 788.982322 | 0.09949349 | 0.64790246 | 0.15356245 | 0.87795474 | 0.98275231 |
| STEAP4 | 2307.66901 | 0.12842267 | 0.62931242 | 0.20406823 | 0.83830018 | 0.97599451 |
| STEAP1 | 341.222634 | -0.0428414 | 0.33141689 | -0.1292673 | 0.89714617 | 0.98474725 |
| STEAP2 | 936.27894 | 0.2094489 | 0.33928424 | 0.61732573 | 0.5370199 | 0.92105614 |
| CFAP69 | 788.444201 | 0.02398219 | 0.3247598 | 0.07384592 | 0.94113298 | 0.99136103 |
| FAM237B | 43.7361868 | 0.411337 | 0.37680561 | 1.09164246 | 0.27499028 | 0.82719407 |
| GTPBP10 | 1064.67734 | -0.0124355 | 0.15536946 | -0.0800381 | 0.93620692 | 0.99068685 |
| CLDN12 | 3189.33885 | -0.1131723 | 0.2607489 | -0.4340278 | 0.66426822 | 0.94655191 |
| CDK14 | 1441.87502 | 0.15794146 | 0.33502344 | 0.47143406 | 0.6373308 | 0.94088547 |
| FZD1 | 2391.91747 | 0.10031343 | 0.23318784 | 0.43018295 | 0.66706257 | 0.94830184 |
| MTERF1 | 492.747381 | -0.0461525 | 0.19250506 | -0.239747 | 0.81052638 | 0.97352497 |
| AKAP9 | 2740.21787 | -0.0153137 | 0.14093929 | -0.1086546 | 0.91347644 | 0.98662757 |
| CYP51A1 | 3855.81154 | -0.042962 | 0.31934376 | -0.1345321 | 0.8929818 | 0.98474725 |
| LRRD1 | 3759.10532 | -0.0494549 | 0.31890917 | -0.1550753 | 0.87676194 | 0.98275231 |
| KRIT1 | 1926.26328 | 0.13602058 | 0.13536985 | 1.00480702 | 0.31498978 | 0.84696964 |
| ANKIB1 | 2631.61116 | -0.2503695 | 0.22365459 | -1.1194472 | 0.26294939 | 0.82424998 |
| GATAD1 | 2743.23259 | -0.0135438 | 0.12559968 | -0.1078333 | 0.91412791 | 0.98662757 |
| AC007566.1 | 609.435046 | 0.00098102 | 0.14710504 | 0.00666885 | 0.99467907 | 0.99940889 |
| PEX1 | 1733.95313 | 0.1203307 | 0.1611398 | 0.7467472 | 0.45521617 | 0.89670027 |
| RBM48 | 776.886928 | -0.1484322 | 0.13657021 | -1.0868563 | 0.27710034 | 0.82779863 |
| FAM133B | 528.491128 | -0.140571 | 0.15519853 | -0.9057497 | 0.36506835 | 0.8663515 |
| CDK6 | 1200.75114 | 0.36408389 | 0.16490558 | 2.20783248 | 0.02725595 | 0.52878242 |
| SAMD9 | 718.116397 | -1.6345137 | 0.6463696 | -2.5287601 | NA | NA |
| SAMD9L | 1586.55463 | -1.8931324 | 0.80418477 | -2.3541013 | NA | NA |
| HEPACAM2 | 428.019801 | 2.03907893 | 0.40741 | 5.0049801 | 5.59E-07 | 0.00130877 |
| VPS50 | 1238.29225 | 0.09946206 | 0.17850638 | 0.55719053 | 0.57739726 | 0.93130183 |
| CALCR | 189.409814 | -0.3373446 | 0.47288785 | -0.7133713 | 0.47561602 | 0.90257104 |
| GNGT1 | 150.74329 | 0.5197959 | 0.68880662 | 0.75463256 | 0.45046948 | 0.89557948 |
| TFPI2 | 1103.02467 | 0.56374232 | 0.66940481 | 0.84215457 | 4.00E-01 | 0.88077967 |
| GNG11 | 2276.94288 | 0.11818104 | 0.29685198 | 0.39811437 | 0.69054588 | 0.95158502 |
| BET1 | 1366.30435 | 0.22276048 | 0.14259753 | 1.56216222 | 0.11824978 | 0.69095557 |

| | | | | | | |
|---|---|---|---|---|---|---|
| COL1A2 | 10460.5931 | -0.2920384 | 0.45237012 | -0.645574 | 0.51855525 | 0.91636259 |
| CASD1 | 1165.20458 | 0.03577707 | 0.17381578 | 0.20583329 | 0.83692114 | 0.97577999 |
| SGCE | 1637.46124 | 0.08187971 | 0.26212989 | 0.31236312 | 0.75476458 | 0.96351771 |
| PEG10 | 710.566151 | 0.93065854 | 0.27888103 | 3.33711668 | 0.00084652 | 0.13676471 |
| PPP1R9A | 882.024036 | 0.25649261 | 0.21107841 | 1.2151532 | 2.24E-01 | 0.80078694 |
| PON3 | 125.991281 | -0.0041096 | 0.52587542 | -0.0078147 | 0.99376484 | 0.99925673 |
| PON2 | 5344.65953 | -0.2794573 | 0.18331709 | -1.5244477 | 0.12739692 | 0.70557946 |
| AC005021.1 | 390.475855 | -0.00244 | 0.26064042 | -0.0093615 | 0.99253075 | 0.99906045 |
| PDK4 | 16160.2653 | -0.462359 | 0.76341005 | -0.6056496 | 0.54474746 | 0.92345821 |
| DYNC1I1 | 187.811643 | 0.78838625 | 0.45088638 | 1.74852531 | 0.08037311 | 0.64123984 |
| SLC25A13 | 2691.3339 | 0.01614008 | 0.26713552 | 0.06041907 | 0.95182187 | 0.99170503 |
| SEM1 | 2295.1474 | -0.1549748 | 0.20252988 | -0.7651948 | 0.44415558 | 0.8938278 |
| SDHAF3 | 696.281134 | 0.21386538 | 0.17880358 | 1.19609117 | 0.23166098 | 0.80649152 |
| TAC1 | 323.948413 | -1.5237459 | 0.96174129 | -1.5843616 | 0.11311146 | 0.68130021 |
| ASNS | 1547.80228 | 0.04553026 | 0.21761299 | 0.20922582 | 0.83427196 | 0.97550416 |
| LMTK2 | 2143.55556 | -0.3533291 | 0.30631062 | -1.1534993 | 0.24870552 | 0.81646572 |
| TECPR1 | 2172.11337 | -0.1541064 | 0.13674707 | -1.1269448 | 0.25976583 | 0.82424998 |
| BRI3 | 5392.24203 | -0.4192719 | 0.31074397 | -1.3492521 | 0.17725601 | 0.76115956 |
| BAIAP2L1 | 1589.99247 | -0.1399607 | 0.23211626 | -0.6029769 | 0.54652408 | 0.92345821 |
| NPTX2 | 98.0639272 | -0.0902624 | 0.82235967 | -0.1097602 | 0.91259952 | 0.98662757 |
| TMEM130 | 296.686122 | 1.00146116 | 0.48314533 | 2.07279487 | 0.03819137 | 0.55538555 |
| TRRAP | 2617.57451 | -0.2007495 | 0.12011671 | -1.6712875 | 0.09466491 | 0.65888357 |
| SMURF1 | 2427.37172 | -0.4405365 | 0.22911333 | -1.9227885 | 0.05450662 | 0.59329767 |
| AC004922.1 | 9410.47655 | 0.00615706 | 0.23105062 | 0.02664812 | 0.97874039 | 0.99483482 |
| ARPC1A | 4068.01325 | -0.0114227 | 0.20284974 | -0.0563111 | 0.955094 | 0.99185043 |
| ARPC1B | 6056.63486 | -0.0052504 | 0.31442058 | -0.0166985 | 0.98667714 | 0.99715899 |
| PDAP1 | 6310.3036 | -0.1453039 | 0.16907016 | -0.8594297 | 0.39010351 | 0.87645794 |
| BUD31 | 2786.33079 | -0.0727833 | 0.14470187 | -0.5029882 | 0.61497254 | 0.93895494 |
| PTCD1 | 4420.19463 | -0.0268377 | 0.12170688 | -0.2205113 | 0.825473 | 0.97548201 |
| ATP5MF-PTC | 4276.93508 | 0.00472576 | 0.13087984 | 0.03610761 | 0.97119655 | 0.99400331 |
| CPSF4 | 1598.27917 | -0.3403835 | 0.1350553 | -2.5203271 | 0.01172458 | 0.40423902 |
| ATP5MF | 5004.98512 | -0.0108515 | 0.12563815 | -0.0863711 | 0.93117142 | 0.98975562 |
| ZNF789 | 858.727512 | 0.19832694 | 0.30250719 | 0.65561066 | 0.51207466 | 0.91551505 |
| ZNF394 | 1183.55638 | -0.1134627 | 0.15243415 | -0.7443391 | 0.45667134 | 0.89688594 |
| ZKSCAN5 | 742.075155 | -0.1671333 | 0.14512717 | -1.1516333 | 0.24947179 | 0.81646572 |
| FAM200A | 544.570571 | 0.19852354 | 0.17187393 | 1.15505322 | 0.24806863 | 0.81605305 |
| ZNF655 | 4030.07182 | -0.2916536 | 0.15847315 | -1.8403975 | 0.06570991 | 0.62237346 |
| AC005020.2 | 384.960834 | -0.054303 | 0.2312844 | -0.234789 | 0.81437251 | 0.9738447 |
| TMEM225B | 134.626708 | 0.6191587 | 0.47425315 | 1.30554472 | 0.19170743 | 0.77347283 |
| ZSCAN25 | 740.63444 | -0.2974831 | 0.17522148 | -1.6977549 | 0.08955404 | 0.65375543 |
| CYP3A5 | 1308.05136 | 0.89487073 | 0.56237735 | 1.59122825 | 0.11155822 | 0.68001961 |
| CYP3A7 | 284.441633 | 1.44681792 | 0.6405017 | 2.25888223 | 0.02389071 | 0.51967919 |
| CYP3A7-CYP3 | 238.399039 | 1.35940987 | 0.6446448 | 2.10877349 | 0.03496414 | 0.55369025 |
| CYP3A4 | 132.830607 | 0.13429701 | 0.94195892 | 0.14257204 | 0.88662818 | 0.98420297 |
| TRIM4 | 1793.26818 | -0.1602484 | 0.2036609 | -0.7868391 | 0.43137609 | 0.88976671 |
| GJC3 | 51.2845944 | 1.41095988 | 0.95134883 | 1.48311517 | 0.13804381 | 0.72209298 |
| AZGP1 | 10714.8068 | 0.42642889 | 0.57805859 | 0.73769146 | 0.46070196 | 0.89755243 |
| ZKSCAN1 | 3393.6535 | 0.07042611 | 0.20607363 | 0.34175217 | 0.73253741 | 0.95848028 |
| ZSCAN21 | 966.082456 | 0.00457617 | 0.20414895 | 0.02241582 | 0.98211626 | 0.99595643 |
| ZNF3 | 1912.68973 | -0.0685457 | 0.17795531 | -0.3851851 | 0.70010026 | 0.95346115 |
| COPS6 | 4020.74426 | 0.08001624 | 0.12931408 | 0.61877443 | 0.53606497 | 0.92101152 |
| MCM7 | 2888.51196 | -0.2008233 | 0.22622229 | -0.8877257 | 0.37468833 | 0.86791605 |
| AP4M1 | 3372.10666 | -0.10625 | 0.1358296 | -0.7822302 | 0.43407931 | 0.89055338 |
| TAF6 | 3999.50839 | -0.0967606 | 0.14548212 | -0.6651033 | 0.50598439 | 0.91390261 |

| | | | | | | |
|---|---|---|---|---|---|---|
| AC073842.2 | 52.4695868 | 0.10678884 | 0.34460601 | 0.30988676 | 0.75664707 | 0.9648441 |
| CNPY4 | 702.011001 | 0.05780617 | 0.25893352 | 0.22324717 | 0.82334314 | 0.97542663 |
| MBLAC1 | 130.216581 | 0.02398395 | 0.34801615 | 0.06891619 | 0.94505633 | 0.99164528 |
| LAMTOR4 | 3891.68813 | 0.1145736 | 0.13286385 | 0.86233846 | 0.38850129 | 0.87568585 |
| C7orf43 | 2103.06388 | -0.1151214 | 0.19350961 | -0.594913 | 0.55190163 | 0.92373235 |
| GAL3ST4 | 127.806842 | -0.1202167 | 0.51623972 | -0.23287 | 0.81586239 | 0.9738447 |
| GPC2 | 65.0337789 | -0.0074267 | 0.33753332 | -0.0220027 | 0.98244578 | 0.99605174 |
| STAG3 | 1151.05474 | -0.3463944 | 0.32439711 | -1.0678098 | 0.28560632 | 0.83107779 |
| CASTOR3 | 1445.7057 | -0.3613259 | 0.29211499 | -1.2369302 | 0.21611299 | 0.7954827 |
| PVRIG | 124.129706 | -0.283343 | 0.51476243 | -0.5504345 | 0.5820214 | 0.9317705 |
| SPDYE3 | 418.570034 | 0.40312918 | 0.23442238 | 1.71967019 | 0.08549241 | 0.6504594 |
| PILRB | 11449.8825 | -0.5108815 | 0.34520032 | -1.4799565 | 0.13888485 | 0.72399122 |
| PILRA | 2026.00445 | -0.4458363 | 0.38678712 | -1.1526659 | 0.24904754 | 0.81646572 |
| ZCWPW1 | 402.381809 | 0.09752976 | 0.30982058 | 0.31479433 | 0.75291781 | 0.96301881 |
| MEPCE | 3758.82832 | -0.4019497 | 0.35111387 | -1.1447845 | 2.52E-01 | 0.81833349 |
| AC092849.2 | 602.510574 | -0.2899739 | 0.19421915 | -1.4930245 | 0.13543078 | 0.71649038 |
| PPP1R35 | 1167.96636 | -0.3334068 | 0.18544713 | -1.7978536 | 0.07220021 | 0.6321115 |
| AC092849.1 | 228.816078 | -0.3783234 | 0.24710425 | -1.5310276 | 0.12576257 | 0.70347499 |
| C7orf61 | 49.0577781 | 0.03492162 | 0.51347723 | 0.06801007 | 0.94577762 | 0.99166614 |
| TSC22D4 | 5325.32399 | -0.3298633 | 0.2496645 | -1.3212265 | 0.18642587 | 0.76908034 |
| NYAP1 | 152.043541 | 0.00167022 | 0.78203648 | 0.00213573 | 0.99829594 | 0.99990633 |
| AGFG2 | 2427.28888 | -0.0193044 | 0.3243528 | -0.0595167 | 0.95254055 | 0.99170503 |
| SAP25 | 790.35176 | -0.0288139 | 0.32720767 | -0.0880599 | 0.92982904 | 0.98973492 |
| LRCH4 | 5508.69452 | -0.1702056 | 0.18561448 | -0.9169843 | 0.35915088 | 0.86480228 |
| FBXO24 | 152.796059 | 1.06133891 | 0.48783245 | 2.17562179 | 0.02958355 | 0.53723387 |
| PCOLCE-AS1 | 774.189154 | 0.28192559 | 0.60370183 | 0.46699475 | 0.64050363 | 0.94088547 |
| PCOLCE | 2622.02926 | 0.11201216 | 0.48672342 | 0.23013514 | 0.81798676 | 0.97446554 |
| MOSPD3 | 1005.13094 | 0.07084836 | 0.13112137 | 0.54032655 | 0.58897185 | 0.93289552 |
| GNB2 | 10934.7305 | -0.2071511 | 0.16243283 | -1.2753031 | 0.20220197 | 0.78165577 |
| GIGYF1 | 8561.12388 | -0.2838006 | 0.22549455 | -1.2585699 | 0.20818573 | 0.78934595 |
| POP7 | 839.52907 | -0.2402813 | 0.23637244 | -1.0165368 | 0.30937384 | 0.84414094 |
| EPO | 121.026989 | 0.51437132 | 1.28093387 | 0.40155963 | 0.68800815 | 0.95158502 |
| EPHB4 | 4654.68184 | -0.1022132 | 0.28168692 | -0.3628611 | 0.71670863 | 0.95603249 |
| SLC12A9 | 3572.39653 | 0.07860042 | 0.15937785 | 0.49317031 | 0.62189225 | 0.94016713 |
| TRIP6 | 6499.11185 | -0.344791 | 0.15732333 | -2.1916073 | 0.02840788 | 0.53506736 |
| SRRT | 7414.83706 | -0.4532097 | 0.1945294 | -2.3297749 | 0.01981805 | 0.49477123 |
| UFSP1 | 66.0758578 | -0.16874 | 0.42510453 | -0.3969378 | 0.69141336 | 0.95158829 |
| ACHE | 442.625985 | 0.01541277 | 0.4464606 | 0.03452214 | 0.97246079 | 0.99416401 |
| MUC3A | 302.95823 | 0.62704814 | 0.49045281 | 1.27850861 | 0.20107015 | 0.78134034 |
| AC254629.1 | 137.177824 | 0.70171964 | 0.49589606 | 1.41505389 | 0.1570527 | 0.74522306 |
| TRIM56 | 4446.12659 | -0.2678272 | 0.23107565 | -1.1590458 | 0.24643753 | 0.81542272 |
| SERPINE1 | 9493.88585 | -0.0611758 | 0.96627938 | -0.0633107 | 0.94951912 | 0.99170503 |
| AP1S1 | 3040.87737 | 0.43130404 | 0.12515145 | 3.44625689 | 0.00056841 | 0.11578877 |
| VGF | 712.155369 | -1.6449595 | 1.64165522 | -1.0020128 | NA | NA |
| MOGAT3 | 169.606897 | 1.31842097 | 0.72616952 | 1.81558292 | 0.06943438 | 0.62779744 |
| PLOD3 | 4446.44434 | -0.1663986 | 0.18858857 | -0.8823367 | 0.37759474 | 0.86987124 |
| ZNHIT1 | 4189.38848 | 0.17766608 | 0.11402838 | 1.55808647 | 0.11921275 | 0.69233536 |
| CLDN15 | 1300.5343 | -0.4907772 | 0.31884601 | -1.5392295 | 0.12374828 | 0.69957382 |
| FIS1 | 4921.83321 | 0.19438998 | 0.10154793 | 1.9142683 | 0.05558588 | 0.59801093 |
| AC006329.1 | 68.3898415 | -0.1920029 | 0.40427293 | -0.4749338 | 0.63483417 | 0.94088547 |
| IFT22 | 1751.67185 | 0.10622271 | 0.11577745 | 0.91747327 | 0.35889469 | 0.86480228 |
| COL26A1 | 342.466861 | 0.90172742 | 0.91158251 | 0.98918904 | 0.32257066 | 0.85119105 |
| CUX1 | 6542.61257 | 0.18102166 | 0.16899881 | 1.07114168 | 0.28410573 | 0.83107779 |
| SH2B2 | 612.601944 | -0.0554473 | 0.36942678 | -0.1500902 | 0.88069344 | 0.98275231 |

| | | | | | | |
|---|---|---|---|---|---|---|
| SPDYE6 | 40.4152605 | -0.5080421 | 0.64111183 | -0.7924391 | 0.42810469 | 0.88800332 |
| PRKRIP1 | 3230.91323 | -0.0919112 | 0.17226385 | -0.5335489 | 0.59365369 | 0.93304795 |
| ORAI2 | 2298.47374 | -0.4436517 | 0.29075589 | -1.5258564 | 1.27E-01 | 0.70463514 |
| ALKBH4 | 727.475833 | -0.1244093 | 0.12017979 | -1.0351931 | 3.01E-01 | 0.84200819 |
| LRWD1 | 1494.54698 | -0.4280708 | 0.20282693 | -2.1105224 | 0.03481339 | 0.55369025 |
| POLR2J | 1498.93334 | -0.0509832 | 0.19299054 | -0.2641744 | 0.79164553 | 0.97090793 |
| RASA4B | 217.652403 | 0.0344158 | 0.56980144 | 0.06039964 | 0.95183735 | 0.99170503 |
| AC093668.2 | 631.047584 | -0.2595026 | 0.45561553 | -0.569565 | 0.56897279 | 0.92660886 |
| POLR2J3 | 1107.71141 | -0.3608725 | 0.40290896 | -0.8956675 | 0.37043036 | 0.86762749 |
| AC093668.3 | 494.800312 | -0.3381523 | 0.4370365 | -0.7737392 | 0.43908502 | 0.89175645 |
| AC093668.1 | 558.569255 | -0.1842904 | 0.39325649 | -0.4686266 | 0.63933658 | 0.94088547 |
| RASA4 | 562.092608 | 0.52704093 | 0.55649193 | 0.94707739 | 0.34359934 | 0.85950284 |
| AC105052.1 | 732.20007 | 0.13354374 | 0.44560207 | 0.29969283 | 0.76441147 | 0.96678847 |
| UPK3BL1 | 1362.50849 | -0.193425 | 0.46390459 | -0.41695 | 6.77E-01 | 0.9499862 |
| AC105052.3 | 1380.67335 | -0.2125517 | 0.46665237 | -0.455482 | 0.64876254 | 0.94312988 |
| POLR2J2 | 43.8199552 | -1.1151754 | 0.69913476 | -1.5950793 | 0.11069451 | 0.68001961 |
| FAM185A | 375.40358 | 0.1491142 | 0.18071953 | 0.82511393 | 0.40930691 | 0.88221056 |
| FBXL13 | 56.1311324 | -0.3199305 | 0.29071188 | -1.100507 | 0.27111128 | 0.82716979 |
| LRRC17 | 132.7589 | -0.1907607 | 0.39736464 | -0.4800645 | 0.63118152 | 0.94088547 |
| ARMC10 | 1791.84615 | 0.08081971 | 0.13398868 | 0.60318311 | 0.54638688 | 0.92345821 |
| NAPEPLD | 2060.84742 | 0.15418261 | 0.24758545 | 0.62274505 | 0.53345208 | 0.92049136 |
| PMPCB | 4974.62407 | 0.05227509 | 0.11707093 | 0.44652494 | 0.6552181 | 0.94532497 |
| DNAJC2 | 1403.56037 | -0.3085206 | 0.21606258 | -1.4279224 | 0.15331419 | 0.74258845 |
| PSMC2 | 4018.6283 | -0.1213878 | 0.1889411 | -0.6424639 | 0.52057201 | 0.91680964 |
| RELN | 166.737763 | 0.42167481 | 0.49706873 | 0.84832294 | 0.39625814 | 0.87967259 |
| ORC5 | 699.521056 | 0.11816322 | 0.19915782 | 0.59331449 | 0.5529707 | 0.92373235 |
| LHFPL3 | 98.5761042 | 0.2091003 | 0.36503794 | 0.57281797 | 0.56676795 | 0.9256133 |
| LINC01004 | 355.526866 | 0.08437077 | 0.22559555 | 0.37399131 | 0.70841078 | 0.95501548 |
| KMT2E-AS1 | 421.445269 | -0.5034829 | 0.27473609 | -1.8326058 | 0.06686122 | 0.6249306 |
| KMT2E | 2687.77948 | -0.195243 | 0.18161327 | -1.0750479 | 0.2823533 | 0.83044305 |
| SRPK2 | 1529.25511 | -0.1647834 | 0.12132317 | -1.3582185 | 0.17439437 | 0.75974924 |
| PUS7 | 585.208401 | -0.6585516 | 0.33131626 | -1.9876826 | 0.0468468 | 0.58171004 |
| RINT1 | 1093.73154 | -0.1176264 | 0.16304494 | -0.7214357 | 4.71E-01 | 9.01E-01 |
| EFCAB10 | 515.968406 | -0.1344651 | 0.16048887 | -0.8378469 | 4.02E-01 | 0.88132015 |
| AC073073.2 | 87.1481505 | -0.2097364 | 0.27636479 | -0.7589115 | 0.44790548 | 0.89480207 |
| ATXN7L1 | 698.416489 | -0.1569405 | 0.39587532 | -0.3964393 | 6.92E-01 | 9.52E-01 |
| CDHR3 | 463.983878 | 0.45998912 | 0.44273475 | 1.03897225 | 0.29881764 | 0.84056272 |
| SYPL1 | 8762.85714 | 0.01036706 | 0.11028388 | 0.09400341 | 0.92510645 | 0.98869145 |
| NAMPT | 17895.304 | 0.6533582 | 0.48720555 | 1.34103193 | 0.17991009 | 0.76317223 |
| AC007032.1 | 50.3677266 | 0.17845381 | 0.41740918 | 0.42752727 | 0.66899533 | 0.9486589 |
| CCDC71L | 1464.66078 | -0.0818191 | 0.38060589 | -0.2149707 | 0.82979013 | 0.97550416 |
| PIK3CG | 50.4049662 | -0.2845454 | 0.60399218 | -0.4711078 | 0.63756374 | 0.94088547 |
| PRKAR2B | 734.258176 | 0.49816746 | 0.38196453 | 1.3042244 | 0.19215708 | 0.7739557 |
| HBP1 | 4562.35429 | -0.1036943 | 0.13534666 | -0.766139 | 0.44359363 | 0.89366134 |
| COG5 | 3523.75029 | -0.1140866 | 0.10935624 | -1.0432561 | 0.29682969 | 0.83877759 |
| DUS4L | 493.981927 | 0.43789002 | 0.25832924 | 1.695085 | 0.0900593 | 0.65393309 |
| BCAP29 | 1973.14566 | -0.1570071 | 0.14154596 | -1.1092307 | 0.26733068 | 0.82660356 |
| SLC26A4 | 285.689767 | 0.63208195 | 0.49484778 | 1.27732601 | 2.01E-01 | 7.81E-01 |
| AC002467.1 | 97.4651912 | 0.10645848 | 0.33974483 | 0.31334834 | 0.75401603 | 0.96339067 |
| CBLL1 | 1032.28527 | -0.2256403 | 0.2140759 | -1.0540201 | 0.29187373 | 0.83460625 |
| DLD | 9448.26561 | 0.16557402 | 0.12170183 | 1.36048913 | 0.17367519 | 0.75974924 |
| LAMB1 | 17851.653 | -0.2078731 | 0.19879112 | -1.0456862 | 0.29570595 | 0.83750889 |
| AC005046.1 | 122.208543 | -1.172792 | 0.54559452 | -2.149567 | 0.03158948 | 0.54507414 |
| LAMB4 | 97.6164884 | -0.1651789 | 0.43530074 | -0.3794592 | 0.70434688 | 0.95488362 |

| | | | | | | |
|---|---|---|---|---|---|---|
| NRCAM | 1396.51819 | -0.2094249 | 0.34044474 | -0.615151 | 0.53845499 | 0.9215968 |
| PNPLA8 | 2718.42792 | -0.3228632 | 0.30693587 | -1.0518913 | 0.29284944 | 0.83524862 |
| THAP5 | 1076.80426 | 0.20693933 | 0.14175941 | 1.4597925 | 0.14434711 | 0.73032824 |
| DNAJB9 | 4179.24317 | 0.25257822 | 0.26828728 | 0.94144686 | 0.34647591 | 0.86135217 |
| IMMP2L | 1276.9343 | 0.19485776 | 0.35527523 | 0.54846987 | 0.58336931 | 0.93220705 |
| DOCK4 | 992.457086 | -0.3652272 | 0.21896427 | -1.6679762 | 0.09532045 | 0.65888357 |
| ZNF277 | 1013.50092 | 0.14790651 | 0.14758384 | 1.00218636 | 0.3162536 | 0.84756299 |
| IFRD1 | 3479.05973 | -0.5005057 | 0.25129076 | -1.9917396 | 0.04639964 | 0.57943708 |
| LSMEM1 | 503.689157 | -0.4065669 | 0.4079609 | -0.9965831 | 0.31896691 | 0.84914926 |
| TMEM168 | 1447.1458 | 0.15000946 | 0.13225613 | 1.1342345 | 0.2566962 | 0.82101614 |
| BMT2 | 570.32942 | 0.09882334 | 0.22250089 | 0.44414806 | 0.65693554 | 0.94545756 |
| GPR85 | 43.6812814 | 0.42474517 | 0.6974207 | 0.60902289 | 0.54250926 | 0.92292765 |
| SMIM30 | 2053.78185 | 0.08820156 | 0.10661268 | 0.82730837 | 0.4080623 | 0.88181557 |
| FOXP2 | 280.807085 | 0.39010377 | 0.41049416 | 0.95032721 | 0.34194602 | 0.85889221 |
| MDFIC | 1183.62504 | 0.48144979 | 0.21552972 | 2.23379768 | 0.02549639 | 0.52304994 |
| TFEC | 1458.56695 | 0.26652183 | 0.56171478 | 0.47447894 | 0.63515841 | 0.94088547 |
| TES | 2486.67464 | -0.3351277 | 0.25443455 | -1.3171468 | 0.18778942 | 0.76970626 |
| CAV2 | 1461.32696 | 0.15159931 | 0.32039599 | 0.47316233 | 0.63609737 | 0.94088547 |
| CAV1 | 1235.80077 | -0.2684871 | 0.54014165 | -0.4970679 | 0.61914116 | 0.93945324 |
| AC006159.2 | 231.687095 | 1.88900571 | 0.73052167 | 2.58583117 | 0.00971445 | 0.39368232 |
| LINC01510 | 426.795692 | 2.33034936 | 0.73574069 | 3.16735147 | 0.00153834 | 0.17279992 |
| MET | 5281.72662 | -0.2970724 | 0.35751154 | -0.830945 | 0.40600471 | 0.88132015 |
| CAPZA2 | 8269.89468 | -0.0322052 | 0.17854352 | -0.1803774 | 0.85685632 | 0.97953896 |
| ST7-AS1 | 200.739802 | -0.1583594 | 0.24692697 | -0.6413207 | 5.21E-01 | 9.17E-01 |
| ST7 | 1802.5282 | -0.0867225 | 0.16756912 | -0.5175329 | 0.60478422 | 0.93663838 |
| ST7-AS2 | 222.756128 | 0.14360725 | 0.30828145 | 0.46583164 | 0.64133601 | 0.94088547 |
| CFTR | 301.465869 | -0.0611222 | 0.6703059 | -0.0911855 | 0.92734517 | 0.98915637 |
| CTTNBP2 | 631.910688 | 0.19879197 | 0.21335607 | 0.93173807 | 0.3514719 | 0.86364633 |
| LSM8 | 1323.35647 | -0.3395581 | 0.21120608 | -1.6077098 | 0.10789876 | 0.67720384 |
| TSPAN12 | 5567.25485 | 0.66598095 | 0.3051934 | 2.18216038 | 0.0290977 | 0.53606221 |
| ING3 | 668.818809 | -0.2591251 | 0.1864643 | -1.3896766 | 0.16462711 | 0.75286112 |
| CPED1 | 535.441348 | -0.2517599 | 0.45216752 | -0.5567846 | 0.57767461 | 0.93130183 |
| FAM3C | 3981.09593 | -0.1034737 | 0.2128088 | -0.4862285 | 6.27E-01 | 0.94088547 |
| AASS | 786.733495 | 0.23710594 | 0.34642393 | 0.68443869 | 0.49369819 | 0.90749607 |
| CADPS2 | 2026.03582 | 0.50604419 | 0.22617792 | 2.23737218 | 0.02526203 | 0.52140487 |
| SLC13A1 | 3882.81355 | 1.20273953 | 0.5919871 | 2.03169888 | 0.04218415 | 0.56485966 |
| IQUB | 182.414189 | 0.28284644 | 0.36239131 | 0.78050007 | 0.43509659 | 0.89055338 |
| AC073323.1 | 47.5621453 | 0.21186783 | 0.39398745 | 0.53775275 | 0.59074776 | 0.93289552 |
| NDUFA5 | 5918.20303 | 0.31106382 | 0.13035262 | 2.38632579 | 1.70E-02 | 0.46671416 |
| ASB15 | 200.617868 | 3.18779106 | 1.16969443 | 2.72531953 | 0.00642393 | 0.33628727 |
| WASL | 4599.96828 | 0.10057426 | 0.19575893 | 0.5137659 | 0.60741567 | 0.93739614 |
| AC006333.2 | 362.325076 | -0.3560941 | 0.21927734 | -1.6239438 | 0.1043878 | 0.67439401 |
| GPR37 | 61.9364294 | 0.74435986 | 0.51335279 | 1.44999671 | 0.14705944 | 0.73337969 |
| POT1 | 992.604752 | 0.16985318 | 0.19720062 | 0.86132174 | 0.38906087 | 0.87568585 |
| POT1-AS1 | 41.1292079 | 0.69747281 | 0.3858912 | 1.80743382 | 0.07069466 | 0.62981127 |
| AC073486.1 | 322.725217 | 0.11828472 | 0.85270991 | 0.13871625 | 0.88967438 | 0.9845209 |
| GRM8 | 48.2964101 | 0.32768028 | 0.66002215 | 0.49646861 | 0.61956383 | 0.93945324 |
| ZNF800 | 808.458106 | -0.3732953 | 0.25150263 | -1.4842599 | 0.13774 | 0.72209298 |
| AC000123.1 | 93.3252856 | -0.3546215 | 0.26473352 | -1.3395415 | 0.18039445 | 0.76401635 |
| GCC1 | 2737.16199 | -0.0682496 | 0.11230552 | -0.6077135 | 0.54337753 | 0.92297267 |
| ARF5 | 5764.44686 | 0.00372108 | 0.1333733 | 0.02789973 | 0.97774212 | 0.99476647 |
| FSCN3 | 1977.92039 | -0.0178935 | 0.14134618 | -0.1265933 | 0.89926232 | 0.98498393 |
| SND1 | 6768.83654 | 0.10234678 | 0.17602876 | 0.58142077 | 0.5609569 | 0.92448946 |
| LRRC4 | 377.105346 | -0.2548904 | 0.42678684 | -0.5972312 | 0.55035303 | 0.92373235 |

| | | | | | | |
|---|---|---|---|---|---|---|
| RBM28 | 1763.27102 | -0.3437277 | 0.19457564 | -1.7665504 | 0.07730356 | 0.63862831 |
| IMPDH1 | 1710.27391 | 0.02889093 | 0.27977513 | 0.10326481 | 0.9177528 | 0.9868044 |
| HILPDA | 4608.14159 | -0.8497924 | 0.86744357 | -0.9796515 | 0.32725817 | 0.85312914 |
| METTL2B | 1123.11239 | 0.15133011 | 0.17664903 | 0.85667105 | 0.39162671 | 0.87744531 |
| AC090114.2 | 355.339996 | -0.3394394 | 0.37581788 | -0.9032019 | 0.36641877 | 0.86683872 |
| AC018638.7 | 158.157024 | 0.17385413 | 0.49861275 | 0.34867566 | 0.72733282 | 0.95848028 |
| FAM71F2 | 314.355092 | -0.1289604 | 0.3069842 | -0.4200882 | 0.67442102 | 0.94935413 |
| AC018638.6 | 90.1127766 | -0.145891 | 0.40573954 | -0.3595681 | 0.71917013 | 0.9566409 |
| CALU | 4804.41678 | -0.132873 | 0.30394131 | -0.4371668 | 0.66199039 | 0.94642611 |
| OPN1SW | 133.380257 | -0.5391602 | 0.68538572 | -0.7866522 | 0.43148548 | 0.88976671 |
| CCDC136 | 58.1976244 | -0.477885 | 0.38567676 | -1.2390815 | 0.21531531 | 0.79511806 |
| FLNC | 2114.22395 | -2.086129 | 0.69233161 | -3.0131934 | 0.00258514 | 0.21645768 |
| FLNC-AS1 | 105.499203 | -1.8651143 | 0.72184314 | -2.5838222 | 0.00977122 | 0.39381151 |
| KCP | 6503.79787 | 0.12861816 | 0.27396014 | 0.46947764 | 0.63872827 | 0.94088547 |
| ATP6V1F | 6593.01651 | 0.06306804 | 0.26106216 | 0.24158246 | 0.80910372 | 0.97334138 |
| ATP6V1FNB | 1568.37161 | 0.43871405 | 0.50080966 | 0.87600956 | 0.38102484 | 0.87171941 |
| IRF5 | 1866.49028 | 0.36125041 | 0.2361495 | 1.52975304 | 0.12607787 | 0.70364066 |
| TNPO3 | 2600.81955 | -0.1512754 | 0.18928298 | -0.7992021 | 0.42417323 | 0.88565404 |
| TSPAN33 | 7994.36598 | 0.14088631 | 0.29960423 | 0.47024139 | 0.63818256 | 0.94088547 |
| SMO | 2362.84423 | -0.0257309 | 0.31204798 | -0.0824582 | 0.93428236 | 0.99034987 |
| AC011005.4 | 238.644639 | -0.1522228 | 0.39165847 | -0.388662 | 0.69752621 | 0.95272493 |
| AHCYL2 | 2196.61913 | 0.40506107 | 0.17804956 | 2.27499063 | 0.02290649 | 0.51967919 |
| STRIP2 | 755.721043 | -0.0659979 | 0.85806792 | -0.0769145 | 0.93869155 | 0.99113899 |
| AC078846.1 | 197.340489 | -0.3915507 | 0.22729441 | -1.7226586 | 0.08495029 | 0.64988556 |
| NRF1 | 1109.78244 | -0.0217033 | 0.11820109 | -0.1836137 | 0.85431652 | 0.97906925 |
| UBE2H | 7637.92547 | -0.4197028 | 0.13655766 | -3.0734476 | 0.00211601 | 0.19927695 |
| AC073320.1 | 162.237197 | 0.19855959 | 0.34477951 | 0.5759031 | 0.56468068 | 0.92477055 |
| ZC3HC1 | 1234.26902 | -0.0513791 | 0.26456782 | -0.1942 | 0.84601925 | 0.9767918 |
| KLHDC10 | 2553.34446 | 0.1035829 | 0.16303872 | 0.63532699 | 0.52521517 | 0.91759401 |
| TMEM209 | 753.204051 | 0.08201089 | 0.2109983 | 0.38868033 | 0.69751263 | 0.95272493 |
| AC087071.2 | 94.0375811 | 0.12382484 | 0.3255206 | 0.38039018 | 7.04E-01 | 9.55E-01 |
| CPA4 | 63.2320258 | -0.872459 | 0.75789304 | -1.1511638 | 0.24966485 | 0.81646572 |
| CEP41 | 398.223207 | 0.04954798 | 0.21121641 | 0.23458393 | 0.81453169 | 0.9738447 |
| MEST | 1526.05015 | 0.25115023 | 0.50160265 | 0.50069558 | 0.61658538 | 0.93910158 |
| AC007938.2 | 265.42535 | 0.49754341 | 0.46870414 | 1.06152981 | 0.2884492 | 0.83295411 |
| COPG2 | 1165.34509 | 0.01249516 | 0.21952985 | 0.0569178 | 0.95461068 | 0.99170503 |
| AC007938.3 | 314.321134 | 0.03407429 | 0.25709862 | 0.13253394 | 0.89456198 | 0.98474725 |
| AC016831.7 | 792.836913 | -0.6508026 | 0.16900636 | -3.850758 | 0.00011775 | 0.05015467 |
| LINC00513 | 91.9815007 | -0.1954813 | 0.82019197 | -0.238336 | 0.81162051 | 0.97352497 |
| AC016831.1 | 341.474768 | -1.2119211 | 0.37300292 | -3.2490928 | 0.00115774 | 0.15697311 |
| MKLN1 | 4469.16807 | -0.4666686 | 0.15653361 | -2.9812675 | 0.00287058 | 0.23188586 |
| AC008264.2 | 2846.56449 | -0.3025721 | 0.1299366 | -2.3286134 | 0.01987955 | 0.49477123 |
| PODXL | 16604.3485 | 0.29731397 | 0.5010863 | 0.59333886 | 0.5529544 | 0.92373235 |
| PLXNA4 | 206.378038 | 0.33089632 | 0.49750907 | 0.66510611 | 0.50598263 | 0.91390261 |
| CHCHD3 | 3204.53085 | 0.1633515 | 0.19393713 | 0.84229104 | 0.39962507 | 0.88077967 |
| EXOC4 | 2131.36807 | 0.22302614 | 0.20048417 | 1.11243762 | 0.26595004 | 0.82642508 |
| LRGUK | 80.7061997 | -0.2743976 | 0.30123266 | -0.9109158 | 0.36233973 | 0.86561928 |
| SLC35B4 | 1679.50663 | 0.03421793 | 0.18956937 | 0.18050349 | 0.85675731 | 0.97953896 |
| AKR1B1 | 12675.2079 | -0.2375174 | 0.55378867 | -0.4288953 | 0.66799942 | 0.94847921 |
| AKR1B10 | 13615.7054 | 3.61898431 | 1.56371447 | 2.3143511 | 0.02064847 | 0.50488816 |
| AKR1B15 | 293.704922 | 6.6772527 | 1.72808954 | 3.86395064 | NA | NA |
| BPGM | 2088.63271 | 0.11591122 | 0.26536065 | 0.43680638 | 0.66225177 | 0.94642611 |
| CALD1 | 12417.7628 | -0.0759792 | 0.33241777 | -0.2285655 | 0.81920662 | 0.97454841 |
| AGBL3 | 461.235093 | 0.00782309 | 0.3364511 | 0.02325178 | 9.81E-01 | 9.96E-01 |

| | | | | | | |
|---|---|---|---|---|---|---|
| CYREN | 4899.09459 | -0.6359468 | 0.24091822 | -2.639679 | 0.00829846 | 0.37565563 |
| TMEM140 | 4862.27158 | -0.6547637 | 0.37285773 | -1.7560684 | 0.07907673 | 0.64079926 |
| AC083862.2 | 1446.74346 | -0.756718 | 0.26802861 | -2.8232732 | 0.0047536 | 0.29927103 |
| WDR91 | 6266.68881 | 0.03466201 | 0.33858216 | 0.10237399 | 9.18E-01 | 9.87E-01 |
| AC009542.1 | 276.245447 | 0.09204937 | 0.38300827 | 0.24033259 | 0.81007243 | 0.97334138 |
| STRA8 | 80.7365964 | 0.46424675 | 0.67560407 | 0.68715801 | 4.92E-01 | 9.06E-01 |
| CNOT4 | 1137.96489 | -0.1697331 | 0.14800345 | -1.1468188 | 0.25145653 | 0.81829255 |
| NUP205 | 1464.97878 | -0.0648114 | 0.10380813 | -0.6243387 | 0.5324052 | 0.91995259 |
| STMP1 | 2244.22933 | -0.255236 | 0.20689638 | -1.2336419 | 0.21733639 | 0.79592123 |
| SLC13A4 | 199.803682 | 0.93078926 | 0.32939569 | 2.8257482 | 0.00471703 | 0.29927103 |
| FAM180A | 177.828217 | 0.52273901 | 0.66529009 | 0.78573094 | 0.43202513 | 0.88976671 |
| MTPN | 5331.95165 | -0.0614052 | 0.14091433 | -0.4357626 | 0.66300895 | 0.94642611 |
| LUZP6 | 1316.78578 | -0.0703804 | 0.13081469 | -0.5380159 | 0.59056604 | 0.93289552 |
| PTN | 985.834518 | -0.0279813 | 0.470197 | -0.0595098 | 0.95254604 | 0.99170503 |
| DGKI | 158.257077 | -1.2737215 | 0.59775758 | -2.1308328 | 0.03310292 | 0.55145048 |
| CREB3L2 | 3606.50146 | 0.09590002 | 0.2147252 | 0.44661746 | 0.65515129 | 0.94532497 |
| TRIM24 | 1081.80352 | -0.325908 | 0.2797354 | -1.1650581 | 0.24399546 | 0.81468457 |
| SVOPL | 158.751549 | -0.1292173 | 0.61023821 | -0.2117489 | 8.32E-01 | 0.97550416 |
| ATP6V0A4 | 5025.94955 | 0.37004325 | 0.39019443 | 0.94835605 | 0.34294822 | 0.85923855 |
| TMEM213 | 5485.08266 | 0.59751448 | 0.38928195 | 1.53491442 | 0.12480485 | 0.70228923 |
| KIAA1549 | 1355.88695 | -0.0196448 | 0.34486464 | -0.0569639 | 0.95457393 | 0.99170503 |
| ZC3HAV1L | 46.4757511 | -0.1810625 | 0.36181471 | -0.5004287 | 0.61677327 | 0.93910158 |
| ZC3HAV1 | 2592.16341 | -0.1759709 | 0.26650945 | -0.6602801 | 0.50907413 | 0.91439937 |
| TTC26 | 507.159112 | 0.15611021 | 0.17010636 | 0.9177212 | 0.35876485 | 0.86480228 |
| UBN2 | 1587.73281 | -0.0717153 | 0.19636719 | -0.3652103 | 0.71495447 | 0.95590747 |
| FMC1 | 1104.72119 | 0.33188032 | 0.28724697 | 1.15538316 | 0.24793355 | 0.81602355 |
| LUC7L2 | 5262.48682 | -0.0595839 | 0.09622473 | -0.6192157 | 0.53577427 | 0.92101152 |
| C7orf55-LUC | 4718.2461 | 0.03557245 | 0.10182469 | 0.34934991 | 0.72682663 | 0.95848028 |
| KLRG2 | 146.296376 | 0.61659939 | 0.69475217 | 0.88750984 | 0.37480447 | 0.86791605 |
| HIPK2 | 9365.59439 | 0.05507685 | 0.23975455 | 0.2297218 | 0.81830795 | 0.97454841 |
| TBXAS1 | 847.269009 | -0.335955 | 0.28676145 | -1.1715488 | 0.24137827 | 0.8124487 |
| PARP12 | 2637.52575 | -1.3123222 | 0.59938542 | -2.1894463 | NA | NA |
| KDM7A | 1328.27952 | -0.2047741 | 0.27098677 | -0.7556609 | 0.44985251 | 0.89557948 |
| SLC37A3 | 2011.17114 | -0.3048004 | 0.19496718 | -1.5633421 | 0.11797215 | 0.69069542 |
| RAB19 | 41.3853509 | -0.6376172 | 0.44869617 | -1.4210444 | 0.15530385 | 0.74419572 |
| MKRN1 | 5447.54979 | -0.1217368 | 0.1270625 | -0.958086 | 0.33801938 | 0.8571036 |
| DENND2A | 1276.11069 | 0.13152484 | 0.33487347 | 0.3927598 | 0.69449689 | 0.95222846 |
| ADCK2 | 1018.62897 | 0.14983987 | 0.14884767 | 1.00666588 | 0.31409536 | 0.84685098 |
| NDUFB2 | 5440.16847 | 0.1773249 | 0.13381495 | 1.32515021 | 0.18512134 | 0.76820626 |
| NDUFB2-AS1 | 2478.78162 | 0.23048247 | 0.15940172 | 1.44592214 | 0.14819904 | 0.73474739 |
| BRAF | 2056.8744 | -0.0860817 | 0.11052484 | -0.7788446 | 0.43607124 | 0.89101735 |
| MRPS33 | 1473.6144 | 0.4966283 | 0.1705791 | 2.91142523 | 0.00359784 | 0.26033637 |
| TMEM178B | 443.751672 | 0.79620082 | 0.49600836 | 1.60521652 | 0.10844616 | 0.67800851 |
| AGK | 2028.08841 | 0.15378902 | 0.17261018 | 0.8909615 | 0.37294983 | 0.86791605 |
| AC004918.1 | 562.178879 | 0.05667498 | 0.1592986 | 0.3557783 | 0.7220066 | 0.95739595 |
| KIAA1147 | 2432.18499 | 0.10272572 | 0.22557771 | 0.45538949 | 0.64882906 | 0.94312988 |
| WEE2-AS1 | 130.380578 | -0.1019179 | 0.35706072 | -0.2854357 | 0.77531035 | 0.96836854 |
| WEE2 | 45.312624 | 0.46839408 | 0.52827967 | 0.88664037 | 0.37527255 | 0.86848393 |
| SSBP1 | 2392.08926 | -0.1407503 | 0.20151013 | -0.6984777 | 0.48487851 | 0.90541935 |
| TAS2R4 | 41.5075038 | -0.5402447 | 0.40692699 | -1.3276207 | 0.18430346 | 0.76820626 |
| TAS2R5 | 43.7419512 | -0.8313049 | 0.45470849 | -1.828215 | 0.0675173 | 0.62547785 |
| MGAM | 2563.45492 | 0.25569664 | 0.57619009 | 0.44377133 | 0.65720792 | 0.94547736 |
| CLEC5A | 76.7179563 | -0.1744415 | 0.79757213 | -0.2187157 | 0.82687154 | 0.97548201 |
| TRBC1 | 276.05139 | 0.95560754 | 1.01477474 | 0.94169425 | 0.3463492 | 0.86135217 |

| | | | | | | |
|---|---|---|---|---|---|---|
| TRBC2 | 460.88638 | 0.48263313 | 0.92659569 | 0.52086701 | 0.60245942 | 0.9355118 |
| EPHB6 | 729.305891 | 0.17685686 | 0.36684387 | 0.4821039 | 0.62973214 | 0.94088547 |
| TRPV6 | 971.314332 | 0.56347985 | 0.33897134 | 1.662323 | 0.09644803 | 0.66233356 |
| AC245427.1 | 67.0665556 | 0.64819364 | 0.45645543 | 1.42005899 | 0.15559051 | 0.74477742 |
| TRPV5 | 139.648066 | 1.77311074 | 0.51123769 | 3.46827078 | 0.00052382 | 0.11155563 |
| KEL | 48.1863616 | 0.00251196 | 0.44527487 | 0.00564136 | 0.99549887 | 0.99940889 |
| GSTK1 | 11126.6599 | 0.32301515 | 0.17864159 | 1.8081744 | 7.06E-02 | 6.30E-01 |
| AC073342.1 | 1876.13931 | 0.25577912 | 0.45492821 | 0.56224063 | 0.57395208 | 0.92999713 |
| TMEM139 | 3068.51879 | 0.14140018 | 0.44651364 | 0.31667605 | 0.75148943 | 0.96285385 |
| CASP2 | 1176.89623 | -0.1824945 | 0.15894337 | -1.1481728 | 0.25089724 | 0.81781433 |
| FAM131B | 58.8682823 | 0.26963342 | 0.56490432 | 0.47730812 | 0.63314273 | 0.94088547 |
| AC093673.1 | 322.8407 | -0.6593135 | 0.34096453 | -1.9336717 | 0.0531535 | 0.58908917 |
| ZYX | 9048.2771 | -0.4175983 | 0.27506755 | -1.5181662 | 0.1289725 | 0.70654009 |
| EPHA1 | 2804.94207 | -0.2403041 | 0.23137623 | -1.0385858 | 0.29899741 | 0.84073674 |
| TCAF2 | 373.60961 | -0.0316134 | 0.42525993 | -0.074339 | 0.94074065 | 0.99136103 |
| TCAF1 | 894.148808 | 0.162324 | 0.24966885 | 0.65015719 | 0.51559069 | 0.91636259 |
| CTAGE4 | 66.7879673 | -1.2194256 | 0.81649901 | -1.4934808 | 0.13531136 | 0.71649038 |
| ARHGEF35 | 561.707187 | -0.2941609 | 0.8914443 | -0.3299824 | 0.74141329 | 0.96109001 |
| AC004889.1 | 155.44464 | -0.0625057 | 0.4755714 | -0.1314328 | 0.89543294 | 0.98474725 |
| OR2A42 | 36.8445298 | 0.652633 | 0.42859222 | 1.52273648 | 0.12782465 | 0.70579329 |
| AC074386.1 | 296.515076 | 0.04372597 | 0.39214413 | 0.11150484 | 0.91121603 | 0.98662757 |
| OR2A7 | 72.2992284 | -0.1859619 | 0.45725127 | -0.4066951 | 0.68423194 | 0.95070301 |
| CTAGE8 | 45.5122577 | -0.2389234 | 0.5536177 | -0.4315675 | 0.6660558 | 0.9475861 |
| OR2A1-AS1 | 163.13509 | -0.0418331 | 0.43619214 | -0.0959051 | 0.92359592 | 0.98858365 |
| ARHGEF5 | 2312.02254 | -0.3875032 | 0.22675391 | -1.708915 | 0.0874667 | 0.65271545 |
| TPK1 | 406.364055 | 0.15755781 | 0.19143465 | 0.82303703 | 0.41048694 | 0.88252101 |
| AC005229.4 | 134.098354 | -0.0129461 | 0.29909807 | -0.043284 | 0.96547518 | 0.99377925 |
| CUL1 | 2592.44471 | -0.2924866 | 0.1842924 | -1.5870793 | 0.1124947 | 0.68030433 |
| EZH2 | 317.579961 | -0.3426839 | 0.2573206 | -1.3317392 | 0.18294589 | 0.76723162 |
| PDIA4 | 7645.97395 | 0.4752008 | 0.24465693 | 1.94231492 | 0.05209899 | 0.58645127 |
| ZNF786 | 358.651107 | -0.179601 | 0.21379955 | -0.8400438 | 0.40088383 | 0.88119978 |
| ZNF425 | 188.448389 | 0.1647755 | 0.26707291 | 0.61696824 | 0.53725567 | 0.92119932 |
| ZNF398 | 601.712256 | -0.2037715 | 0.24062212 | -0.8468529 | 0.39707713 | 0.87988087 |
| ZNF282 | 1701.22357 | -0.2850969 | 0.16593892 | -1.7180832 | 0.08578145 | 0.65049132 |
| ZNF212 | 894.946941 | -0.2615873 | 0.20429022 | -1.280469 | 0.20038023 | 0.78134034 |
| ZNF783 | 1582.25864 | -0.1035699 | 0.19874598 | -0.5211172 | 0.60228515 | 0.9355118 |
| ZNF777 | 1006.88788 | -0.2663616 | 0.24793717 | -1.0743108 | 0.2826834 | 0.83076205 |
| ZNF746 | 1985.9639 | -0.5051542 | 0.34473295 | -1.4653493 | 0.14282564 | 0.72784886 |
| KRBA1 | 1096.98072 | -0.1687348 | 0.23477709 | -0.7187023 | 0.47232437 | 0.90160919 |
| ZNF467 | 1296.63596 | -0.3894074 | 0.40824648 | -0.9538538 | 0.34015767 | 0.85814041 |
| SSPO | 464.900882 | -0.8748289 | 0.38794683 | -2.2550226 | 0.02413191 | 0.51967919 |
| ZNF862 | 1448.82739 | 0.11111644 | 0.2441069 | 0.45519583 | 0.64896836 | 0.94312988 |
| AC004877.1 | 141.074829 | -0.445376 | 0.40109567 | -1.1103983 | 0.26682742 | 0.8265123 |
| ATP6V0E2-A | 1191.93566 | 0.79534005 | 0.29780037 | 2.67071548 | 0.00756898 | 0.366538 |
| ATP6V0E2 | 3574.4913 | 1.06068322 | 0.34406186 | 3.08282709 | 0.00205044 | 0.1948632 |
| ACTR3C | 713.918652 | -0.415201 | 0.27892274 | -1.4885878 | 0.13659594 | 0.72070508 |
| LRRC61 | 1529.30964 | 0.12776016 | 0.3701612 | 0.34514736 | 0.72998359 | 0.95848028 |
| ZBED6CL | 820.053261 | -1.4810195 | 0.58166205 | -2.5461855 | 0.01089073 | 0.40196898 |
| RARRES2 | 5761.1821 | -0.2461252 | 0.40597316 | -0.6062598 | 0.54434222 | 0.92345821 |
| AC005586.1 | 458.578871 | -0.0367696 | 0.20006459 | -0.1837887 | 0.85417921 | 0.97906925 |
| REPIN1 | 6958.78071 | -0.0640684 | 0.12216689 | -0.5244337 | 6.00E-01 | 0.93487648 |
| AC073111.3 | 228.253484 | -0.0466329 | 0.1952454 | -0.2388424 | 0.8112278 | 0.97352497 |
| ZNF775 | 254.968953 | -0.1148017 | 0.20875856 | -0.5499257 | 0.58237035 | 0.93177403 |
| AC073111.5 | 753.852524 | -0.193526 | 0.44293953 | -0.4369129 | 0.66217453 | 0.94642611 |

| | | | | | | |
|---|---|---|---|---|---|---|
| GIMAP8 | 1077.00887 | -0.0516096 | 0.38699886 | -0.1333586 | 0.89390978 | 0.98474725 |
| GIMAP7 | 846.55754 | -0.2659254 | 0.30346669 | -0.8762919 | 0.38087139 | 0.87165941 |
| GIMAP4 | 1630.64161 | -0.436162 | 0.31375135 | -1.3901519 | 0.16448276 | 0.75286112 |
| GIMAP6 | 1447.27207 | -0.4152494 | 0.25786333 | -1.6103471 | 0.10732211 | 0.67707349 |
| GIMAP2 | 421.009129 | -0.3102265 | 0.37023675 | -0.837914 | 0.40207902 | 0.88132015 |
| GIMAP1 | 620.226183 | -0.1571865 | 0.41046643 | -0.382946 | 0.7017598 | 0.95426502 |
| GIMAP1-GIM | 1819.59641 | -0.567116 | 0.21452641 | -2.6435719 | 0.00820363 | 0.37407374 |
| GIMAP5 | 1740.18588 | -0.5987545 | 0.20636193 | -2.9014773 | 0.00371408 | 0.26317334 |
| TMEM176B | 20799.511 | 0.23989633 | 0.21983891 | 1.09123691 | 0.27516865 | 0.82719407 |
| TMEM176A | 17860.7916 | 0.15723708 | 0.20410678 | 0.77036678 | 0.44108235 | 0.89290667 |
| AOC1 | 13687.563 | 0.28803034 | 0.54331981 | 0.53013038 | 0.59602154 | 0.9334601 |
| KCNH2 | 136.636897 | -0.381725 | 0.51644489 | -0.7391398 | 0.45982212 | 0.89755243 |
| NOS3 | 1938.54722 | -0.4243178 | 0.33946041 | -1.2499773 | 0.21130786 | 0.79134644 |
| ATG9B | 794.325027 | -0.3560787 | 0.35777163 | -0.9952682 | 0.31960582 | 0.84951792 |
| ABCB8 | 3244.92874 | -0.05177 | 0.08556604 | -0.6050294 | 0.54515948 | 0.92345821 |
| ASIC3 | 332.589578 | -0.948912 | 0.45502586 | -2.0854023 | 0.0370328 | 0.55522536 |
| CDK5 | 609.450644 | 0.42199778 | 0.19618434 | 2.15102688 | 0.03147408 | 0.54507414 |
| SLC4A2 | 10765.5874 | -0.6024538 | 0.24400724 | -2.4689999 | 0.01354913 | 0.42608223 |
| AC010973.2 | 4009.17822 | -0.5765599 | 0.26133377 | -2.2062203 | 0.02736858 | 0.52904542 |
| FASTK | 5508.16586 | -0.1292524 | 0.14413737 | -0.8967306 | 0.36986267 | 0.86762749 |
| TMUB1 | 2639.03593 | -0.3054886 | 0.15960867 | -1.913985 | 5.56E-02 | 0.59805701 |
| AGAP3 | 3768.61937 | -0.1116036 | 0.18327295 | -0.6089474 | 0.54255933 | 0.92292765 |
| ABCF2 | 2809.28621 | -0.2484742 | 0.20431484 | -1.2161339 | 0.22393392 | 0.80078694 |
| CHPF2 | 3418.70789 | -0.5193144 | 0.3100607 | -1.6748798 | 0.09395782 | 0.65783452 |
| SMARCD3 | 3636.41564 | 0.07851348 | 0.27357354 | 0.28699225 | 0.77411826 | 0.96836854 |
| AC021097.1 | 1031.48021 | 0.08713021 | 0.31949327 | 0.27271376 | 0.78507326 | 0.97032632 |
| NUB1 | 3586.03788 | -0.3785197 | 0.34329635 | -1.1026034 | 0.27019944 | 0.82716979 |
| WDR86 | 853.698229 | -0.1969299 | 0.3630618 | -0.5424143 | 0.58753314 | 0.93289552 |
| RHEB | 4560.50444 | -0.1756565 | 0.22143888 | -0.7932507 | 0.42763178 | 0.88790684 |
| PRKAG2 | 2686.42049 | -0.5438231 | 0.27076497 | -2.0084693 | 0.04459345 | 0.57267043 |
| PRKAG2-AS1 | 195.532219 | 0.30203761 | 0.37231259 | 0.81124736 | 0.41722363 | 0.88445833 |
| GALNT11 | 9548.7076 | 0.70994624 | 0.31182632 | 2.27673611 | 2.28E-02 | 0.51967919 |
| AC006017.1 | 2321.05656 | 0.7337497 | 0.30648784 | 2.39405816 | 0.01666311 | 0.46470725 |
| KMT2C | 2446.48572 | -0.2095269 | 0.11051344 | -1.8959405 | 0.05796791 | 0.60187073 |
| LINC01003 | 248.414788 | -0.2391717 | 0.37312819 | -0.6409908 | 0.52152867 | 0.91723878 |
| ACTR3B | 536.635972 | 0.1667256 | 0.22687723 | 0.73487143 | 0.4624178 | 0.89776981 |
| DPP6 | 359.281282 | 1.54947203 | 0.77449056 | 2.00063384 | 0.04543186 | 0.57529633 |
| PAXIP1-AS2 | 1062.2093 | -0.1743432 | 0.1874256 | -0.9301996 | 0.35226776 | 0.86364633 |
| PAXIP1 | 910.468084 | -0.2661275 | 0.21614172 | -1.2312639 | 0.21822418 | 0.79708421 |
| AC093726.1 | 207.59101 | -0.0616394 | 0.37124897 | -0.1660324 | 0.86813143 | 0.98237277 |
| AC093726.2 | 66.7513036 | -0.199414 | 0.41470768 | -0.4808545 | 0.63061992 | 0.94088547 |
| PAXIP1-AS1 | 726.63637 | 0.0641026 | 0.25368461 | 0.25268621 | 0.80051071 | 0.97210205 |
| AC099552.3 | 36.023853 | -0.9706037 | 1.04649635 | -0.9274793 | 0.35367775 | 0.86364633 |
| AC144652.1 | 46.6317002 | 0.42063959 | 0.67932095 | 0.61920597 | 0.53578068 | 0.92101152 |
| INSIG1 | 3492.21609 | -0.0575921 | 0.51129866 | -0.1126389 | 0.91031683 | 0.98662757 |
| AC009403.1 | 485.88443 | -0.0267288 | 0.32312038 | -0.0827207 | 0.93407359 | 0.99029255 |
| RBM33 | 3747.71654 | -0.1694769 | 0.13435109 | -1.261448 | 0.20714747 | 0.78790396 |
| SHH | 627.378321 | -0.0038601 | 0.40298578 | -0.0095788 | 9.92E-01 | 0.99906045 |
| AC021218.1 | 216.637086 | -0.2980601 | 0.72781879 | -0.4095252 | 0.6821543 | 0.95070301 |
| LINC01006 | 522.854429 | -0.1364271 | 0.17774849 | -0.7675288 | 0.4427672 | 0.89343541 |
| RNF32 | 257.671937 | -0.3982785 | 0.24229899 | -1.643748 | 0.10022827 | 0.67191826 |
| AC005534.1 | 50.2494934 | -0.4766183 | 0.42619075 | -1.1183215 | 0.26342969 | 0.82424998 |
| LMBR1 | 1980.30779 | 0.12908206 | 0.13783833 | 0.93647437 | 0.349029 | 0.86356499 |
| NOM1 | 1083.5662 | -0.2103064 | 0.21766983 | -0.9661718 | 0.33395823 | 0.85533716 |

| | | | | | | |
|---|---|---|---|---|---|---|
| UBE3C | 3594.63648 | -0.1618184 | 0.16896808 | -0.9576864 | 0.3382209 | 0.8571036 |
| DNAJB6 | 4764.39591 | -0.2825012 | 0.19598442 | -1.4414472 | 0.14945839 | 0.73670625 |
| PTPRN2 | 656.957567 | 0.6361445 | 0.31413984 | 2.02503605 | 0.04286366 | 0.56634074 |
| NCAPG2 | 266.270971 | 0.21726307 | 0.21598044 | 1.00593865 | 0.31444508 | 0.84696964 |
| ESYT2 | 4595.52335 | 0.14497201 | 0.19640645 | 0.73812244 | 0.46044005 | 0.89755243 |
| WDR60 | 1580.32156 | 0.04977038 | 0.22403893 | 0.22215058 | 0.82419666 | 0.97548201 |
| VIPR2 | 150.612639 | 0.59907557 | 0.31988687 | 1.87277324 | 0.06109971 | 0.60989294 |
| PLCXD1 | 697.597695 | 0.05833186 | 0.34495451 | 0.16910014 | 0.86571788 | 0.98197545 |
| GTPBP6 | 4230.94701 | -0.1522242 | 0.180796 | -0.8419668 | 0.39980654 | 0.88077967 |
| LINC00685 | 81.8124334 | 0.52076743 | 0.32786735 | 1.58834794 | 0.11220768 | 0.68001961 |
| PPP2R3B | 1014.04014 | -0.1084003 | 0.2352576 | -0.4607728 | 0.6449616 | 0.94208859 |
| CSF2RA | 281.300838 | -0.5603408 | 0.61521874 | -0.9107992 | 0.36240116 | 0.86561928 |
| IL3RA | 890.956085 | 0.0028567 | 0.32655242 | 0.00874805 | 0.99302016 | 0.99906069 |
| SLC25A6 | 28541.5459 | 0.17827189 | 0.23427727 | 0.76094401 | 0.44669051 | 0.89480207 |
| LINC00106 | 390.608109 | -0.1251977 | 0.26611463 | -0.4704653 | 0.63802261 | 0.94088547 |
| ASMTL-AS1 | 1649.19136 | -0.0568987 | 0.1929735 | -0.2948526 | 0.76810647 | 0.96765147 |
| ASMTL | 2709.09088 | -0.0095112 | 0.19206007 | -0.0495222 | 0.96050314 | 0.99281834 |
| P2RY8 | 397.275582 | -0.1946447 | 0.48544402 | -0.4009622 | 6.88E-01 | 0.95158502 |
| AKAP17A | 5069.09503 | -0.1765826 | 0.14384265 | -1.2276098 | 0.21959347 | 0.79858446 |
| DHRSX | 1426.387 | 0.06096439 | 0.2089565 | 0.29175638 | 0.7704729 | 0.96836854 |
| ZBED1 | 2795.60498 | -0.1611524 | 0.1752027 | -0.9198057 | 0.35767432 | 0.86403383 |
| CD99 | 4302.29619 | 0.132336 | 0.26367464 | 0.50189128 | 0.615744 | 0.93910158 |
| GYG2 | 182.7731 | 1.11372001 | 0.38408273 | 2.89968777 | 0.00373535 | 0.26317334 |
| ARSD | 5222.55811 | 0.45252491 | 0.35014788 | 1.29238225 | 0.1962248 | 0.77798606 |
| ARSD-AS1 | 1416.58339 | 0.45196452 | 0.34289566 | 1.31808177 | 0.18747628 | 0.76958556 |
| ARSE | 2367.55891 | 0.57230743 | 0.48946033 | 1.16926213 | 0.24229803 | 0.81319028 |
| ARSF | 370.572554 | 0.12616259 | 0.6460587 | 0.19528038 | 0.84517346 | 0.97675487 |
| MXRA5 | 1018.47099 | 0.1848866 | 0.58680508 | 0.31507327 | 0.75270602 | 0.96301881 |
| PRKX | 2793.63966 | -0.6075658 | 0.46436594 | -1.3083771 | 0.19074544 | 0.77208646 |
| BX890604.1 | 545.625719 | -0.577765 | 0.35552872 | -1.6250866 | 1.04E-01 | 0.67439401 |
| FAM239A | 104.751313 | -1.9953977 | 1.93060613 | -1.0335602 | NA | NA |
| FAM239B | 53.2735413 | -7.8218522 | 1.66656955 | -4.6933848 | 2.69E-06 | 0.00503611 |
| NLGN4X | 56.9062614 | 0.77098702 | 0.52755782 | 1.46142657 | 0.14389841 | 0.72920837 |
| PUDP | 1127.41878 | -0.1410236 | 0.33566058 | -0.4201376 | 0.67438494 | 0.94935413 |
| STS | 707.244696 | 0.23793922 | 0.30576648 | 0.77817299 | 0.43646703 | 0.89104919 |
| PNPLA4 | 1391.80333 | 0.55950004 | 0.35582528 | 1.57240104 | 0.11585757 | 0.6873836 |
| ANOS1 | 85.793479 | -0.2979977 | 0.59768855 | -0.4985836 | 0.61807273 | 0.93928244 |
| TBL1X | 2685.62781 | -0.0851268 | 0.18041433 | -0.4718405 | 6.37E-01 | 9.41E-01 |
| GPR143 | 419.908809 | 0.52768071 | 0.40297004 | 1.30947875 | 0.19037223 | 0.77107542 |
| SHROOM2 | 705.097815 | -0.0227634 | 0.32046364 | -0.0710328 | 0.94337165 | 0.99136103 |
| WWC3 | 1510.55407 | -0.0517465 | 0.14416052 | -0.3589506 | 0.71963204 | 0.9566409 |
| CLCN4 | 482.385217 | 0.01198924 | 0.33595229 | 0.03568733 | 0.97153167 | 0.99404213 |
| MID1 | 1258.35776 | 0.48026633 | 0.32696679 | 1.46885354 | 0.14187252 | 0.72784886 |
| HCCS | 1233.67217 | -0.0016313 | 0.184852 | -0.0088249 | 0.99295885 | 0.99906069 |
| ARHGAP6 | 1063.5753 | 0.02954208 | 0.23015154 | 0.12835923 | 0.89786469 | 0.98474725 |
| AC004554.2 | 80.1235063 | 0.11575196 | 0.49202121 | 0.23525807 | 0.81400844 | 0.9738447 |
| MSL3 | 1314.99279 | -0.2560712 | 0.14313342 | -1.7890386 | 0.0736086 | 0.6374448 |
| PRPS2 | 1758.17134 | -0.0123794 | 0.22565504 | -0.05486 | 0.95625003 | 0.99185043 |
| TLR7 | 68.7831188 | -0.4289145 | 0.69243787 | -0.6194267 | 0.53563531 | 0.92101152 |
| TLR8 | 73.6345596 | -0.3016205 | 0.75004764 | -0.4021351 | 0.68758459 | 0.95158502 |
| TMSB4X | 39706.3502 | -0.0195487 | 0.34151006 | -0.0572419 | 0.95435253 | 0.99170503 |
| TCEANC | 171.4676 | 0.23401167 | 0.31239506 | 0.7490889 | 0.45380363 | 0.89670027 |
| RAB9A | 2124.38847 | -0.5340932 | 0.32770423 | -1.6298027 | 0.1031432 | 0.67430143 |
| TRAPPC2 | 833.75528 | -0.121529 | 0.13745318 | -0.8841486 | 0.376616 | 0.86970773 |

| | | | | | | |
|---|---|---|---|---|---|---|
| OFD1 | 1809.90886 | -0.2253578 | 0.20837357 | -1.0815084 | 0.27947101 | 0.82825734 |
| GPM6B | 581.582404 | -0.0005178 | 0.38046258 | -0.0013611 | 0.99891401 | 0.99994357 |
| GEMIN8 | 997.804025 | 0.03949635 | 0.19663978 | 0.20085635 | 0.8408109 | 0.97618283 |
| MOSPD2 | 721.705033 | 0.07357853 | 0.14014 | 0.5250359 | 6.00E-01 | 0.93487648 |
| ASB9 | 491.599989 | 0.7302667 | 0.38781196 | 1.88304327 | 0.0596945 | 0.6066891 |
| PIGA | 771.69255 | -0.1226604 | 0.26965287 | -0.4548827 | 0.64919362 | 0.94312988 |
| VEGFD | 132.829457 | 1.26356487 | 0.52556028 | 2.40422446 | 0.01620682 | 0.45896904 |
| PIR | 896.03018 | 0.70112739 | 0.55226645 | 1.26954552 | 0.20424657 | 0.7843531 |
| BMX | 77.0894738 | -0.2243629 | 0.64281388 | -0.3490325 | 0.72706492 | 0.95848028 |
| ACE2 | 3582.74812 | 1.43496701 | 0.65405981 | 2.19393851 | 0.02823983 | 0.53300785 |
| CLTRN | 14602.0953 | -0.2157569 | 0.53264765 | -0.4050649 | 0.68542981 | 0.95097288 |
| CA5B | 780.219609 | -0.216223 | 0.36282589 | -0.5959414 | 0.55121436 | 0.92373235 |
| INE2 | 254.144484 | -0.0145235 | 0.39869515 | -0.0364276 | 0.97094138 | 0.99400331 |
| ZRSR2 | 832.562464 | -0.0736558 | 0.26238904 | -0.2807123 | 0.77893105 | 0.96836854 |
| AP1S2 | 1214.59268 | -0.2477981 | 0.2372083 | -1.0446433 | 0.29618787 | 0.83799168 |
| CTPS2 | 822.59416 | 0.22921628 | 0.18795152 | 1.21955004 | 0.22263549 | 0.80078694 |
| SYAP1 | 3224.64781 | 0.12721323 | 0.17961578 | 0.70825194 | 0.47878882 | 0.90444322 |
| TXLNG | 1072.08113 | -0.1880518 | 0.1639181 | -1.1472301 | 0.25128652 | 0.81815815 |
| RBBP7 | 4641.07621 | 0.17017638 | 0.1093642 | 1.55605188 | 0.11969576 | 0.69405865 |
| REPS2 | 1784.66882 | 0.53670469 | 0.47674615 | 1.12576616 | 2.60E-01 | 8.24E-01 |
| NHS | 1274.72914 | 0.13976604 | 0.23214095 | 0.60207407 | 0.54712483 | 0.92345821 |
| SCML1 | 1219.04042 | 0.07296659 | 0.26377591 | 0.27662341 | 0.78206929 | 0.96942659 |
| RAI2 | 1384.66962 | 0.27574201 | 0.25768782 | 1.07006222 | 0.2845913 | 0.83107779 |
| SCML2 | 84.2746173 | -0.1607737 | 0.29212507 | -0.5503591 | 0.58207307 | 0.9317705 |
| CDKL5 | 680.394762 | -0.165802 | 0.19900805 | -0.8331421 | 0.40476462 | 0.88132015 |
| PHKA2-AS1 | 846.693385 | 0.32554108 | 0.23895404 | 1.36235855 | 0.17308476 | 0.75974924 |
| PHKA2 | 1953.16527 | 0.31293031 | 0.24107209 | 1.29807773 | 1.94E-01 | 0.7762856 |
| ADGRG2 | 46.3120596 | 0.22841063 | 0.48465069 | 0.4712892 | 0.63743423 | 0.94088547 |
| PDHA1 | 9411.82113 | 0.28334374 | 0.24270475 | 1.16744212 | 0.24303187 | 0.81340905 |
| MAP3K15 | 1496.68413 | 0.00059694 | 0.28111293 | 0.00212348 | 0.99830571 | 0.99990633 |
| SH3KBP1 | 1333.64027 | -0.212382 | 0.25175345 | -0.8436111 | 0.39888676 | 0.88074856 |
| BCLAF3 | 926.926336 | -0.1920247 | 0.20404448 | -0.9410923 | 0.34665758 | 0.86135217 |
| MAP7D2 | 424.774596 | -0.2282352 | 0.56223869 | -0.4059401 | 0.68478661 | 0.95097288 |
| EIF1AX | 3632.97297 | -0.1284895 | 0.25137891 | -0.5111388 | 0.60925386 | 0.93748023 |
| RPS6KA3 | 2964.31875 | -0.2826888 | 0.3090444 | -0.9147191 | 0.36033911 | 0.86480228 |
| MBTPS2 | 878.329408 | 0.21242448 | 0.18301678 | 1.1606831 | 0.2457708 | 0.81542272 |
| YY2 | 73.0933563 | -0.3835777 | 0.3906659 | -0.981856 | 0.32617077 | 0.85282528 |
| SMS | 7226.69167 | -0.2981899 | 0.2788585 | -1.0693233 | 0.28492403 | 0.83107779 |
| PRDX4 | 2229.79725 | 0.00400458 | 0.26610928 | 0.01504865 | 0.98799337 | 0.99754499 |
| ACOT9 | 1914.09052 | -0.5019571 | 0.29907779 | -1.6783497 | 0.09327885 | 0.65756964 |
| SAT1 | 40069.7911 | -0.2713319 | 0.27304631 | -0.9937214 | 0.32035854 | 0.85016134 |
| APOO | 1123.87452 | 0.33725584 | 0.15900846 | 2.12099307 | 0.03392239 | 0.55369025 |
| CXorf58 | 127.41656 | 0.39710053 | 0.22569692 | 1.75944152 | 0.07850254 | 0.64079926 |
| KLHL15 | 460.663828 | -0.1762487 | 0.17498618 | -1.007215 | 0.31383146 | 0.84646403 |
| EIF2S3 | 6305.55293 | 0.17619524 | 0.17295184 | 1.01875317 | 0.30832016 | 0.84388103 |
| ZFX | 1240.79154 | -0.3564582 | 0.2287118 | -1.5585473 | 0.11910355 | 0.69233212 |
| PDK3 | 362.508292 | 0.35121396 | 0.31807382 | 1.10419009 | 0.26951069 | 0.82716979 |
| POLA1 | 386.524204 | 0.08185735 | 0.2044312 | 0.40041515 | 6.89E-01 | 0.95158502 |
| CXorf21 | 52.0572766 | -0.6053761 | 0.73866574 | -0.8195535 | 0.4124707 | 0.88268208 |
| GK | 3255.02637 | 0.01349226 | 0.56459472 | 0.02389725 | 0.98093457 | 0.99576712 |
| GK-AS1 | 398.678031 | -0.1384754 | 0.56651554 | -0.2444336 | 0.80689502 | 0.97334138 |
| TAB3 | 2157.50054 | 0.04153628 | 0.17887268 | 0.23221142 | 0.81637381 | 0.97388289 |
| DMD | 645.14915 | -0.460953 | 0.22727613 | -2.0281627 | 0.04254364 | 0.56530166 |
| TMEM47 | 2440.49438 | 0.02341394 | 0.2632643 | 0.08893699 | 0.92913198 | 0.98963127 |

| | | | | | | |
|---|---|---|---|---|---|---|
| PRRG1 | 1239.66286 | -0.0585021 | 0.32277279 | -0.1812486 | 0.85617248 | 0.97934134 |
| AF241726.2 | 1095.57266 | 0.66690748 | 0.42407604 | 1.57261299 | 0.11580846 | 0.6873836 |
| XK | 79.047141 | 0.02804625 | 0.28429974 | 0.0986503 | 0.92141593 | 0.98782999 |
| CYBB | 971.689667 | 0.15838259 | 0.59099581 | 0.26799275 | 0.7887049 | 0.97090793 |
| DYNLT3 | 2948.1683 | -0.1651692 | 0.273623 | -0.6036379 | 0.54608443 | 0.92345821 |
| SYTL5 | 130.772106 | -0.1675553 | 0.50915408 | -0.3290857 | 0.74209089 | 0.96109001 |
| SRPX | 471.096151 | -0.0133857 | 0.59404698 | -0.0225331 | 0.98202274 | 0.99595643 |
| RPGR | 626.592044 | -0.1169376 | 0.23303898 | -0.5017941 | 0.61581238 | 0.93910158 |
| TSPAN7 | 2323.9334 | 0.75405377 | 0.42699502 | 1.76595447 | 0.07740349 | 0.63862831 |
| MID1IP1 | 3071.4266 | -0.3683257 | 0.24798247 | -1.4852893 | 0.1374672 | 0.72209298 |
| MID1IP1-AS1 | 93.7019627 | 0.12485617 | 0.28748871 | 0.43429937 | 0.66407106 | 0.94648686 |
| BCOR | 1403.19857 | 0.20868343 | 0.23163706 | 0.9009069 | 0.36763782 | 0.86719956 |
| ATP6AP2 | 9964.66406 | 0.29717487 | 0.16638434 | 1.78607478 | 0.07408715 | 0.63749647 |
| CXorf38 | 1401.1832 | -0.0267247 | 0.21151448 | -0.1263492 | 0.89945553 | 0.98506781 |
| MED14 | 2172.87483 | -0.0418101 | 0.16520264 | -0.253084 | 0.80020332 | 0.97210205 |
| MED14OS | 42.0506591 | -0.1030636 | 0.39494288 | -0.2609584 | 0.79412463 | 0.97090793 |
| USP9X | 5016.89165 | 0.00713784 | 0.13855344 | 0.05151684 | 0.95891368 | 0.99242674 |
| DDX3X | 13257.3846 | -0.3668358 | 0.27847939 | -1.3172817 | 0.18774421 | 0.76970626 |
| NYX | 50.2508437 | 0.64730785 | 0.76766615 | 0.84321532 | 0.39910803 | 0.88077967 |
| CASK | 1428.43183 | 0.4342726 | 0.15809665 | 2.74688044 | 0.00601651 | 0.33163337 |
| CASK-AS1 | 240.736811 | 0.55302237 | 0.21263431 | 2.6008144 | 0.00930028 | 0.38664841 |
| GPR34 | 236.657694 | -0.4942357 | 0.47599355 | -1.0383244 | 0.29911907 | 0.84082654 |
| MAOA | 12284.4262 | -0.2319582 | 0.48539913 | -0.477871 | 0.632742 | 0.94088547 |
| MAOB | 6126.17517 | 0.6306436 | 0.44985284 | 1.40188867 | 0.16094849 | 0.74874703 |
| EFHC2 | 191.360693 | 0.28224778 | 0.50069725 | 0.56370948 | 0.57295187 | 0.92928554 |
| FUNDC1 | 715.706552 | 0.08748214 | 0.15316374 | 0.57116746 | 0.56788613 | 0.92618483 |
| KDM6A | 1182.40987 | -0.2926696 | 0.30839876 | -0.9489973 | 0.34262196 | 0.85889221 |
| CXorf36 | 896.128099 | -0.8175736 | 0.42126136 | -1.9407752 | 0.05228555 | 0.58645127 |
| MIR222HG | 397.474356 | -0.5926091 | 0.478096 | -1.2395191 | 0.21515334 | 0.7949124 |
| KRBOX4 | 800.955532 | -0.1114257 | 0.16725095 | -0.6662187 | 0.50527129 | 0.91368371 |
| ZNF674 | 179.025585 | -0.2555 | 0.23475248 | -1.0883803 | 0.27642727 | 0.82730125 |
| ZNF674-AS1 | 252.59718 | -0.1743215 | 0.30813372 | -0.5657334 | 0.57157502 | 0.92841021 |
| CHST7 | 260.998122 | -0.3780594 | 0.38546367 | -0.9807913 | 0.32669566 | 0.85282528 |
| SLC9A7 | 845.937564 | 0.03951291 | 0.1401239 | 0.28198552 | 0.77795461 | 0.96836854 |
| RP2 | 584.555108 | -0.4431127 | 0.32092369 | -1.3807415 | 0.16735845 | 0.75558932 |
| JADE3 | 603.2353 | -0.1762484 | 0.17356314 | -1.0154716 | 0.30988106 | 0.84455826 |
| RGN | 2701.09871 | 0.61130708 | 0.49443877 | 1.23636561 | 0.21632269 | 0.7954827 |
| NDUFB11 | 3334.86019 | 0.18357827 | 0.13641667 | 1.34571729 | 0.17839371 | 0.76115956 |
| RBM10 | 3566.94797 | -0.1294661 | 0.13549336 | -0.9555159 | 0.33931685 | 0.85759056 |
| UBA1 | 19093.0817 | -0.1266847 | 0.08968037 | -1.4126244 | 0.15776619 | 0.74532297 |
| CDK16 | 6796.20072 | -0.121127 | 0.14263229 | -0.849226 | 0.39575557 | 0.87939946 |
| USP11 | 4321.55439 | -0.0179606 | 0.22709669 | -0.0790879 | 0.93696274 | 0.99068685 |
| ZNF41 | 507.013708 | 0.11666579 | 0.1341776 | 0.86948782 | 0.38458037 | 0.87309759 |
| LINC01560 | 104.404706 | 0.16312933 | 0.33920585 | 0.48091543 | 0.63057661 | 0.94088547 |
| ARAF | 5395.29494 | -0.065783 | 0.10495818 | -0.6267544 | 0.53082021 | 0.91962754 |
| SYN1 | 119.297479 | -0.2784677 | 0.2958951 | -0.9411029 | 0.34665213 | 0.86135217 |
| TIMP1 | 15591.8267 | -0.2717325 | 0.54733402 | -0.4964655 | 0.61956599 | 0.93945324 |
| CFP | 160.368211 | 0.09304518 | 0.47797432 | 0.19466565 | 0.8456547 | 0.97675487 |
| ELK1 | 1589.54398 | -0.3031313 | 0.12751548 | -2.3772118 | 0.01744407 | 0.46859699 |
| UXT | 1993.63252 | -0.0136229 | 0.19527707 | -0.0697617 | 0.94438333 | 0.99136103 |
| UXT-AS1 | 883.910614 | -0.0363347 | 0.27533897 | -0.1319634 | 0.89501321 | 0.98474725 |
| ZNF81 | 151.814995 | -0.1561444 | 0.17116226 | -0.9122597 | 0.36163202 | 0.86528137 |
| ZNF182 | 665.529601 | 0.0112907 | 0.15344295 | 0.07358242 | 0.94134266 | 0.99136103 |
| ZNF630 | 224.155924 | 0.15723284 | 0.21160489 | 0.74304916 | 0.45745192 | 0.89688594 |

| | | | | | | |
|---|---|---|---|---|---|---|
| SLC38A5 | 295.800854 | -1.1003641 | 0.48170306 | -2.2843204 | 0.02235271 | 0.51866171 |
| FTSJ1 | 1549.68527 | -0.1628056 | 0.2322222 | -0.7010769 | 0.48325504 | 0.90541935 |
| PORCN | 511.226517 | 0.01812554 | 0.25978323 | 0.06977177 | 0.94437531 | 0.99136103 |
| EBP | 2688.72028 | 0.40229181 | 0.26276922 | 1.5309701 | 0.12577678 | 0.70347499 |
| TBC1D25 | 1181.26261 | -0.366723 | 0.15264581 | -2.4024442 | 0.01628592 | 0.45896904 |
| AC115618.1 | 1109.87758 | 0.1810514 | 0.52404312 | 0.34548951 | 0.7297264 | 0.95848028 |
| RBM3 | 8956.92834 | 0.12241881 | 0.42468078 | 0.28826077 | 0.77314713 | 0.96836854 |
| AC115618.2 | 337.011692 | 0.39873942 | 0.20446701 | 1.95014063 | 0.05115936 | 0.5861488 |
| WDR13 | 8700.96767 | -0.1280894 | 0.16447961 | -0.7787557 | 0.43612362 | 0.89102723 |
| WAS | 520.160887 | -0.1945333 | 0.51993215 | -0.3741513 | 0.70829177 | 0.95501548 |
| SUV39H1 | 304.248788 | 0.14920753 | 0.37976881 | 0.39289043 | 0.69440041 | 0.95222846 |
| AC231533.1 | 66.3689268 | 0.0246993 | 0.45300495 | 0.05452326 | 0.95651828 | 0.99198394 |
| GLOD5 | 179.762742 | 0.68126276 | 0.49069725 | 1.38835657 | 0.16502849 | 0.75303284 |
| HDAC6 | 8016.73638 | 0.34127231 | 0.30262141 | 1.12772033 | 0.25943805 | 0.82394992 |
| PCSK1N | 2872.96163 | 0.05029483 | 0.35446051 | 0.14189121 | 0.88716594 | 0.98423722 |
| TIMM17B | 2306.26268 | 0.07748218 | 0.11089907 | 0.69867293 | 0.48475646 | 0.90541935 |
| PQBP1 | 2911.66787 | -0.1652602 | 0.18332256 | -0.9014723 | 0.36733727 | 0.86683872 |
| SLC35A2 | 2086.27723 | -0.2327974 | 0.13160923 | -1.7688534 | 0.07691834 | 0.63806922 |
| PIM2 | 1033.78613 | 0.09447158 | 0.45769135 | 0.20640892 | 0.8364715 | 0.97577999 |
| OTUD5 | 4714.5254 | -0.1073005 | 0.07679561 | -1.3972213 | 0.16234704 | 0.75026273 |
| KCND1 | 251.039757 | -0.0852027 | 0.45243097 | -0.188322 | 0.85062422 | 0.97839793 |
| GRIPAP1 | 3753.73779 | -0.153117 | 0.18298049 | -0.8367942 | 0.40270827 | 0.88132015 |
| TFE3 | 6616.97727 | -0.4886796 | 0.23519343 | -2.0777774 | 0.03772986 | 0.55538555 |
| CCDC120 | 2251.78261 | 0.15731451 | 0.26327952 | 0.59751897 | 0.55016095 | 0.92373235 |
| PRAF2 | 1397.48167 | -0.4059904 | 0.35031832 | -1.1589186 | 0.24648938 | 0.81542272 |
| WDR45 | 6382.93067 | -0.1742517 | 0.17849407 | -0.9762327 | 0.32894916 | 0.85317073 |
| GPKOW | 1351.92052 | 0.02800811 | 0.13870043 | 0.20193243 | 8.40E-01 | 9.76E-01 |
| MAGIX | 535.265022 | 0.1534171 | 0.31092196 | 0.49342641 | 0.62171133 | 0.94016713 |
| PLP2 | 2947.88655 | -0.239347 | 0.35634318 | -0.6716757 | 5.02E-01 | 9.13E-01 |
| PRICKLE3 | 607.864609 | -0.1709483 | 0.20469123 | -0.8351522 | 0.40363199 | 0.88132015 |
| SYP | 323.95761 | -0.01278 | 0.49903252 | -0.0256096 | 0.97956872 | 0.99528855 |
| CCDC22 | 1442.13516 | 0.01562965 | 0.12961998 | 0.12058054 | 0.90402328 | 0.98624484 |
| FOXP3 | 133.370882 | 0.24859315 | 0.35260226 | 0.70502426 | 0.48079514 | 0.90541935 |
| PPP1R3F | 450.181983 | -0.1904177 | 0.22888233 | -0.8319456 | 0.40543963 | 0.88132015 |
| AC232271.1 | 233.626683 | -0.0060042 | 0.3348578 | -0.0179305 | 9.86E-01 | 0.99705651 |
| USP27X-AS1 | 115.885404 | 0.41799157 | 0.36460308 | 1.14642906 | 0.25161766 | 0.81833349 |
| USP27X | 276.66296 | 0.13547105 | 0.20619936 | 0.65699063 | 0.51118694 | 0.91492259 |
| CLCN5 | 5507.77442 | 0.52714811 | 0.29682846 | 1.77593521 | 0.0757436 | 0.63773415 |
| SHROOM4 | 1057.77017 | 0.01589959 | 0.15128883 | 0.10509425 | 0.91630102 | 0.9868044 |
| NUDT10 | 234.626061 | 0.76295016 | 0.6806695 | 1.12088194 | 0.26233812 | 0.82424998 |
| NUDT11 | 41.8818926 | -0.1933059 | 0.540976 | -0.3573281 | 0.72084618 | 0.95728372 |
| GSPT2 | 521.178203 | 0.08150065 | 0.13516797 | 0.60295831 | 0.54653643 | 0.92345821 |
| MAGED1 | 6701.28171 | -0.1149495 | 0.23346667 | -0.4923592 | 0.62246543 | 0.94016713 |
| GPR173 | 376.698803 | 0.33678676 | 0.36407551 | 0.92504645 | 0.35494174 | 0.86364633 |
| TSPYL2 | 3263.05143 | -0.383461 | 0.28937591 | -1.325131 | 0.1851277 | 0.76820626 |
| KANTR | 935.13237 | 0.00623804 | 0.25713313 | 0.02425997 | 0.98064524 | 0.99573612 |
| KDM5C | 6477.67874 | -0.507974 | 0.21962942 | -2.3128688 | 0.02072986 | 0.50488816 |
| IQSEC2 | 1779.27055 | 0.23916928 | 0.23338831 | 1.02476977 | 0.30547184 | 0.84270033 |
| SMC1A | 2743.41677 | -0.2081785 | 0.17269122 | -1.2054954 | 2.28E-01 | 0.80300331 |
| RIBC1 | 158.502333 | -0.0512537 | 0.46472182 | -0.110289 | 0.91218019 | 0.98662757 |
| HSD17B10 | 4116.81164 | 0.31197175 | 0.16419979 | 1.89995217 | 0.0574394 | 0.60129769 |
| HUWE1 | 8592.23553 | -0.0301733 | 0.09599595 | -0.3143187 | 0.75327896 | 0.96301881 |
| PHF8 | 2360.04899 | -0.4539718 | 0.21934984 | -2.0696244 | 0.03848753 | 0.55565329 |
| FAM120C | 669.769327 | 0.01953248 | 0.14384475 | 0.13578863 | 0.89198838 | 0.98474725 |

| | | | | | | |
|---|---|---|---|---|---|---|
| WNK3 | 243.907955 | 0.16879607 | 0.27615489 | 0.61123694 | 0.54104273 | 0.92292765 |
| TSR2 | 2665.90502 | 0.12869122 | 0.14451282 | 0.89051764 | 0.373188 | 0.86791605 |
| FGD1 | 887.64435 | -0.0771751 | 0.17518177 | -0.4405429 | 0.65954397 | 0.94607834 |
| GNL3L | 198.628392 | -0.2680176 | 0.32774106 | -0.8177724 | 0.41348716 | 0.88316314 |
| MAGED2 | 12888.6281 | 0.02591493 | 0.13396442 | 0.19344636 | 0.84660941 | 0.97684681 |
| TRO | 1380.4764 | 0.06424352 | 0.32714281 | 0.19637761 | 0.84431462 | 0.97675487 |
| APEX2 | 1665.84312 | 0.38511378 | 0.23597039 | 1.63204277 | 0.10267048 | 0.67335541 |
| ALAS2 | 66.8159607 | -2.8263683 | 0.53108752 | -5.3218505 | 1.03E-07 | 0.000385 |
| FAM104B | 629.811341 | 0.34327319 | 0.18692972 | 1.8363757 | 0.06630211 | 0.62373361 |
| MTRNR2L10 | 43.8614957 | -0.7419357 | 0.6509869 | -1.1397091 | 0.2544075 | 0.81992039 |
| MAGEH1 | 1277.5123 | -0.1022288 | 0.31601789 | -0.3234904 | 0.74632387 | 0.96195706 |
| USP51 | 255.887548 | 0.36459211 | 0.39984992 | 0.91182239 | 0.36186222 | 0.86528137 |
| RRAGB | 1081.16925 | 0.17064175 | 0.156998 | 1.08690397 | 0.27707925 | 0.82779863 |
| AL445472.1 | 72.1178595 | 0.03330192 | 0.32713405 | 0.101799 | 0.91891622 | 0.9871265 |
| KLF8 | 919.028083 | -0.0837059 | 0.42517263 | -0.196875 | 0.84392536 | 0.97664894 |
| UBQLN2 | 2546.73618 | -0.1998192 | 0.22297981 | -0.8961315 | 0.37018253 | 0.86762749 |
| NBDY | 1620.79275 | 0.31021989 | 0.12395521 | 2.50267723 | 0.01232579 | 0.41397424 |
| SPIN3 | 747.934018 | 0.06196918 | 0.19813981 | 0.3127548 | 0.75446696 | 0.96350701 |
| SPIN2B | 197.595789 | 0.18560142 | 0.22308922 | 0.8319605 | 0.40543124 | 0.88132015 |
| FAAH2 | 905.077355 | 0.0988808 | 0.24206696 | 0.40848532 | 0.6829174 | 0.95070301 |
| ZXDB | 536.606414 | -0.0517869 | 0.18682439 | -0.2771957 | 0.78162981 | 0.96913822 |
| ZXDA | 162.24544 | -0.0707086 | 0.25908261 | -0.2729191 | 7.85E-01 | 0.97025923 |
| SPIN4 | 176.542696 | 0.03095418 | 0.16259211 | 0.19037934 | 8.49E-01 | 0.97741457 |
| ARHGEF9 | 1299.87817 | -0.0250898 | 0.19725663 | -0.1271938 | 0.89878701 | 0.98477645 |
| AMER1 | 344.017435 | -0.1822382 | 0.21179973 | -0.8604269 | 0.38955377 | 0.87577651 |
| MTMR8 | 46.1175821 | -0.1744569 | 0.26460882 | -0.659301 | 0.50970248 | 0.91439937 |
| ZC4H2 | 581.53082 | -0.127736 | 0.20597214 | -0.6201615 | 0.53515145 | 0.92084006 |
| ZC3H12B | 161.0484 | 0.4080634 | 0.25720523 | 1.58652839 | 0.1126195 | 0.6806198 |
| LAS1L | 1902.38037 | 0.02962235 | 0.22975128 | 0.12893225 | 0.89741126 | 0.98474725 |
| MSN | 25924.0125 | -0.2015552 | 0.2336254 | -0.8627281 | 0.38828698 | 0.87553582 |
| AL034397.3 | 194.706866 | -0.0467111 | 0.66957429 | -0.0697623 | 0.94438283 | 0.99136103 |
| VSIG4 | 1649.77554 | -0.4856287 | 0.46991669 | -1.0334357 | 0.30140005 | 0.84211658 |
| HEPH | 672.392143 | -0.1797933 | 0.47038235 | -0.3822279 | 0.70229231 | 0.95431462 |
| EDA2R | 170.344989 | 0.91764155 | 0.50335264 | 1.82305897 | 0.06829446 | 0.62648383 |
| AR | 693.680481 | 0.26534632 | 0.37696445 | 0.70390276 | 0.48149333 | 0.90541935 |
| AL157700.1 | 39.9382416 | 0.49989549 | 0.40212872 | 1.24312308 | 0.21382248 | 0.7926732 |
| OPHN1 | 533.562719 | 0.20237006 | 0.2611979 | 0.77477677 | 0.43847158 | 0.89175645 |
| YIPF6 | 2671.4426 | -0.009461 | 0.11600888 | -0.0815538 | 0.93500154 | 0.99068685 |
| STARD8 | 2650.21914 | -0.1814856 | 0.23037669 | -0.7877777 | 0.43082675 | 0.88966606 |
| EFNB1 | 3975.21705 | -0.0053471 | 0.19627356 | -0.027243 | 0.97826592 | 0.99476647 |
| PJA1 | 1278.31636 | -0.0261477 | 0.18817488 | -0.1389541 | 0.88948644 | 0.98446024 |
| FAM155B | 220.616933 | 0.45105965 | 0.43332767 | 1.04092048 | 0.29791245 | 0.83957554 |
| EDA | 391.477705 | 0.18215085 | 0.22825675 | 0.79800858 | 0.42486551 | 0.88573606 |
| IGBP1 | 2729.30801 | 0.30967893 | 0.14882528 | 2.08082207 | 0.0374502 | 0.55532957 |
| IGBP1-AS2 | 721.998536 | 0.36935166 | 0.16754778 | 2.20445572 | 0.02749231 | 0.52904542 |
| PDZD11 | 1206.44689 | 0.15421204 | 0.12675658 | 1.21659984 | 0.2237565 | 0.80078694 |
| GDPD2 | 53.3387803 | 2.71939209 | 0.63900843 | 4.25564354 | 2.08E-05 | 0.01556334 |
| DLG3 | 2298.62728 | 0.02330735 | 0.20936255 | 0.11132528 | 0.91135841 | 0.98662757 |
| SNX12 | 2834.71083 | -0.1475936 | 0.19703852 | -0.7490595 | 0.45382133 | 0.89670027 |
| FOXO4 | 1519.74285 | 0.00296946 | 0.22759576 | 0.01304706 | 0.98959025 | 0.99800524 |
| AL590764.2 | 474.909854 | -0.2234754 | 0.74523793 | -0.2998712 | 0.76427542 | 0.96678847 |
| IL2RG | 584.48062 | -0.2800222 | 0.75528177 | -0.370752 | 0.71082225 | 0.95560193 |
| MED12 | 1841.38332 | -0.1251077 | 0.2076525 | -0.6024857 | 0.54685087 | 0.92345821 |
| NLGN3 | 75.6569664 | 0.16777913 | 0.37978987 | 0.44176832 | 0.65865687 | 0.9458503 |

| | | | | | | |
|---|---|---|---|---|---|---|
| GJB1 | 2558.23053 | 1.61624542 | 0.53209688 | 3.03750218 | 0.00238548 | 0.21083339 |
| ZMYM3 | 2656.05718 | 0.18755758 | 0.34448443 | 0.54445881 | 0.58612578 | 0.93289552 |
| NONO | 11615.5446 | -0.0215125 | 0.1709732 | -0.1258236 | 0.89987158 | 0.98514342 |
| ITGB1BP2 | 59.2669623 | -0.4285716 | 0.45792245 | -0.9359043 | 0.34932245 | 0.86364633 |
| TAF1 | 1429.40581 | 0.09079427 | 0.14257247 | 0.63682887 | 5.24E-01 | 0.91723878 |
| OGT | 9272.08563 | -0.1974773 | 0.1802449 | -1.0956052 | 0.27325159 | 0.82716979 |
| GCNA | 226.815149 | 0.13096652 | 0.33584134 | 0.38996546 | 0.69656209 | 0.9525827 |
| CXCR3 | 49.8801231 | -0.4853308 | 0.83148743 | -0.5836899 | 0.55942896 | 0.92421175 |
| NHSL2 | 361.983026 | -0.0994111 | 0.41534311 | -0.2393469 | 0.81083661 | 0.97352497 |
| RTL5 | 684.55089 | -0.054759 | 0.26352989 | -0.2077903 | 0.83539269 | 0.97550416 |
| PIN4 | 928.454318 | 0.35256014 | 0.18113845 | 1.94635727 | 0.05161184 | 0.58645127 |
| RPS4X | 35261.0494 | 0.29269896 | 0.26694302 | 1.09648477 | 0.27286669 | 0.82716979 |
| CITED1 | 109.668455 | 2.3091132 | 0.64351674 | 3.58827217 | 0.00033288 | 0.08914272 |
| HDAC8 | 1023.85001 | 0.18420093 | 0.15651214 | 1.17691148 | 0.23923084 | 0.81163641 |
| PHKA1 | 546.224179 | -0.1485372 | 0.35470966 | -0.418757 | 0.67539374 | 0.94958156 |
| NAP1L2 | 458.010997 | 0.51449821 | 0.34349722 | 1.49782349 | 0.13417912 | 0.71560922 |
| CHIC1 | 936.178157 | -0.1640144 | 0.11460366 | -1.4311448 | 0.15238873 | 0.7422778 |
| TSIX | 7144.07983 | -1.7759338 | 2.74137467 | -0.647826 | 0.51709751 | 0.91636259 |
| XIST | 7296.25256 | -1.77652 | 3.11193557 | -0.570873 | 0.56808573 | 0.92618483 |
| JPX | 1371.03151 | -0.1383096 | 0.28035982 | -0.4933287 | 6.22E-01 | 0.94016713 |
| FTX | 4277.65542 | -0.1152172 | 0.20984159 | -0.5490674 | 0.58295918 | 0.93191649 |
| SLC16A2 | 1816.52019 | 0.62589886 | 0.26988916 | 2.31909597 | 0.02038983 | 0.50082026 |
| RLIM | 2210.3218 | -0.1005663 | 0.14511821 | -0.6929958 | 0.48831218 | 0.90585781 |
| NEXMIF | 205.547014 | 0.57678381 | 0.31663616 | 1.82159801 | 0.068516 | 0.62698165 |
| ABCB7 | 1282.4717 | 0.08948363 | 0.13003404 | 0.6881554 | 0.49135493 | 0.90600166 |
| UPRT | 691.090323 | 0.14002777 | 0.13066157 | 1.07168289 | 0.28386249 | 0.83107779 |
| ZDHHC15 | 196.561977 | 0.28578306 | 0.25542756 | 1.11884189 | 0.2632076 | 0.82424998 |
| PBDC1 | 1097.46274 | -0.5673002 | 0.31491211 | -1.8014557 | 0.07163108 | 0.63189503 |
| MAGEE1 | 350.785841 | -0.0209655 | 0.17169834 | -0.1221064 | 0.90281472 | 0.98608873 |
| ATRX | 2157.28645 | -0.1106953 | 0.1068484 | -1.0360034 | 0.30020054 | 0.84197221 |
| MAGT1 | 4430.6215 | 0.34689227 | 0.10644082 | 3.25901551 | 1.12E-03 | 0.15680011 |
| COX7B | 7241.59326 | 0.06179882 | 0.16033975 | 0.3854242 | 0.69992315 | 0.95328923 |
| ATP7A | 624.40679 | 0.12155194 | 0.13236893 | 0.91828153 | 0.3584715 | 0.86480228 |
| PGK1 | 29168.5233 | 0.20110353 | 0.21374012 | 0.94087872 | 3.47E-01 | 0.86135217 |
| TAF9B | 1415.10233 | 0.03654489 | 0.19882473 | 0.18380457 | 0.85416677 | 0.97906925 |
| CYSLTR1 | 86.750502 | 0.01436004 | 0.54590206 | 0.02630516 | 0.97901394 | 0.99499459 |
| GPR174 | 81.8952794 | 0.19771961 | 0.9312618 | 0.21231367 | 0.83186233 | 0.97550416 |
| ITM2A | 840.082085 | 0.28472108 | 0.5554009 | 0.51264065 | 0.60820271 | 0.93748023 |
| BRWD3 | 776.791681 | -0.0732332 | 0.155497 | -0.4709623 | 0.63766763 | 0.94088547 |
| HMGN5 | 431.207335 | 0.73201869 | 0.25436153 | 2.87786716 | 0.00400374 | 0.27238681 |
| SH3BGRL | 5546.07756 | 0.26252187 | 0.23700408 | 1.10766813 | 0.26800517 | 0.82676682 |
| POU3F4 | 99.6905291 | 0.66352242 | 0.67746373 | 0.97942133 | 0.32737184 | 0.85317073 |
| RPS6KA6 | 837.880293 | -0.0378272 | 0.2487776 | -0.1520523 | 0.87914571 | 0.98275231 |
| HDX | 229.779344 | -0.3753217 | 0.30080474 | -1.2477255 | 0.21213159 | 0.79203904 |
| APOOL | 1732.05304 | 0.37506561 | 0.25666199 | 1.46132121 | 0.14392731 | 0.72920837 |
| SATL1 | 589.701011 | 0.51736114 | 0.23637654 | 2.18871605 | 0.02861748 | 0.53606221 |
| ZNF711 | 543.203196 | -0.4524211 | 0.32004919 | -1.4135986 | 0.15747978 | 0.74522306 |
| POF1B | 154.342148 | 0.42894532 | 0.47195151 | 0.90887583 | 0.36341567 | 0.8663515 |
| CHM | 1007.68395 | -0.126148 | 0.16437955 | -0.7674191 | 0.44283238 | 0.89343541 |
| DACH2 | 62.9336817 | 1.3006993 | 0.66369266 | 1.95979161 | 0.05002015 | 0.5861488 |
| KLHL4 | 108.637791 | 0.73235832 | 0.38381754 | 1.90808975 | 0.05637962 | 0.59932524 |
| PABPC5 | 45.5884377 | -0.1980102 | 0.32706156 | -0.6054219 | 5.45E-01 | 9.23E-01 |
| NAP1L3 | 141.087169 | 0.26740505 | 0.40518177 | 0.65996316 | 0.50927747 | 0.91439937 |
| DIAPH2 | 597.948245 | 0.09182621 | 0.19711668 | 0.46584696 | 0.64132505 | 0.94088547 |

| | | | | | | |
|---|---|---|---|---|---|---|
| PCDH19 | 36.0487631 | 0.53882499 | 0.48887313 | 1.10217756 | 0.27038449 | 0.82716979 |
| TSPAN6 | 2899.23626 | 0.48277781 | 0.18136351 | 2.66193471 | 0.00776929 | 0.36878977 |
| SRPX2 | 1492.20856 | -0.2915285 | 0.49549074 | -0.5883631 | 0.5562886 | 0.92373235 |
| SYTL4 | 1214.5286 | -0.1916305 | 0.40559325 | -0.4724696 | 0.63659161 | 0.94088547 |
| CSTF2 | 529.608858 | -0.1430645 | 0.16215495 | -0.8822702 | 0.3776307 | 0.86987124 |
| TRMT2B | 1078.61653 | -0.064334 | 0.12771689 | -0.5037235 | 0.61445568 | 0.93895494 |
| TIMM8A | 307.432656 | 0.00896199 | 0.24693691 | 0.03629263 | 9.71E-01 | 0.99400331 |
| BTK | 178.821439 | -0.3012433 | 0.55682912 | -0.5409977 | 0.58850915 | 0.93289552 |
| RPL36A | 8508.13337 | 0.15370651 | 0.17810958 | 0.86298845 | 0.38814382 | 0.87553582 |
| RPL36A-HNR | 9538.94132 | 0.14137726 | 0.17069563 | 0.82824184 | 0.40753356 | 0.88132776 |
| GLA | 2297.11307 | -1.1775597 | 0.47199787 | -2.4948412 | 0.01260135 | 0.41598747 |
| HNRNPH2 | 6582.97436 | -0.1454976 | 0.16946358 | -0.8585776 | 0.39057362 | 0.87677931 |
| ARMCX4 | 985.02405 | -0.0776142 | 0.2704552 | -0.2869761 | 0.77413065 | 0.96836854 |
| ARMCX1 | 1372.94055 | -0.6700485 | 0.30777502 | -2.1770722 | 0.02947518 | 0.53682638 |
| ARMCX6 | 769.260801 | -0.1744733 | 0.12735234 | -1.3700047 | 0.17068544 | 0.75956362 |
| ARMCX3 | 4087.46265 | -0.1137453 | 0.12866444 | -0.8840459 | 0.37667144 | 0.86970773 |
| ARMCX3-AS1 | 350.149793 | -0.0577242 | 0.28501011 | -0.2025339 | 0.83949936 | 0.97618283 |
| ARMCX2 | 2884.50827 | -0.0895388 | 0.21432725 | -0.4177669 | 0.67611757 | 0.94959168 |
| ZMAT1 | 1273.45481 | 0.34717214 | 0.47657765 | 0.72846919 | 0.46632642 | 0.89960479 |
| TCEAL2 | 392.245886 | -0.2001031 | 0.38976361 | -0.5133959 | 0.60767439 | 0.93739614 |
| BEX5 | 290.804054 | 0.25762041 | 0.34468261 | 0.74741343 | 0.45481404 | 0.89670027 |
| ARMCX5 | 741.241876 | 0.09792175 | 0.22363218 | 0.43786968 | 0.66148075 | 0.94642611 |
| AL035427.1 | 87.0859845 | 0.1516346 | 0.27389696 | 0.55361915 | 5.80E-01 | 0.93139743 |
| GPRASP1 | 1034.03271 | 0.27501911 | 0.45232487 | 0.60801236 | 0.54317928 | 0.92297267 |
| GPRASP2 | 969.894323 | 0.37587818 | 0.26444907 | 1.42136322 | 0.15521119 | 0.74407586 |
| BHLHB9 | 383.29262 | -0.1789044 | 0.23207376 | -0.7708945 | 0.44076946 | 0.89290667 |
| LINC00630 | 61.339365 | -0.3336058 | 0.29757986 | -1.1210632 | 0.26226098 | 0.82424998 |
| BEX1 | 236.624492 | -0.7501903 | 0.62144527 | -1.2071704 | 0.22736654 | 0.80300331 |
| NXF3 | 66.7646225 | 0.19954657 | 0.61477876 | 0.32458273 | 0.7454969 | 0.96176407 |
| BEX4 | 3491.64142 | -0.0891104 | 0.20850071 | -0.4273865 | 0.66909784 | 0.9486589 |
| TCEAL8 | 2132.61647 | -0.0233781 | 0.09382694 | -0.2491615 | 0.80323583 | 0.97274243 |
| BEX2 | 1555.12255 | 0.19870882 | 0.34443477 | 0.57691279 | 0.56399837 | 0.92470993 |
| TCEAL7 | 122.238659 | -0.4207893 | 0.27651345 | -1.521768 | 0.12806722 | 0.70580262 |
| TCEAL9 | 3071.98569 | -0.2422317 | 0.19349365 | -1.2518847 | 0.21061189 | 0.79134644 |
| BEX3 | 5312.06652 | 0.11858223 | 0.23662535 | 0.50113917 | 0.61627318 | 0.93910158 |
| Z69733.1 | 46.7298493 | -0.2094211 | 0.49667243 | -0.4216483 | 0.67328174 | 0.94935413 |
| RAB40A | 98.2738571 | -0.2076179 | 0.38518949 | -0.5390021 | 0.58988542 | 0.93289552 |
| TCEAL4 | 4523.0607 | -0.0555337 | 0.18656503 | -0.2976643 | 0.76595942 | 0.96704052 |
| TCEAL3 | 1869.06652 | -0.0920608 | 0.25404429 | -0.3623808 | 0.71706748 | 0.95611001 |
| TCEAL1 | 1173.76238 | -0.2101597 | 0.19664009 | -1.068753 | 0.28518097 | 0.83107779 |
| MORF4L2 | 9671.07005 | -0.2674279 | 0.16920793 | -1.5804692 | 0.11399946 | 0.68214043 |
| MORF4L2-AS | 220.338387 | 0.05490647 | 0.22776278 | 0.24106868 | 0.80950188 | 0.97334138 |
| PLP1 | 79.6699684 | -1.5560923 | 0.96512146 | -1.612328 | 0.10689059 | 0.67674128 |
| RAB9B | 44.624632 | -0.5950146 | 0.59645556 | -0.9975841 | 0.31848109 | 0.8490262 |
| SLC25A53 | 127.079235 | 0.0732085 | 0.26068338 | 0.280833 | 0.77883849 | 0.96836854 |
| ZCCHC18 | 42.1340054 | 0.19514907 | 0.58854756 | 0.33157741 | 0.74020838 | 0.96054877 |
| FAM199X | 3464.67216 | -0.0258583 | 0.14279353 | -0.1810891 | 0.85629766 | 0.97942475 |
| NRK | 809.279541 | -0.0478001 | 0.35860264 | -0.1332954 | 0.8939598 | 0.98474725 |
| MUM1L1 | 115.368886 | 0.28815619 | 0.58497527 | 0.49259551 | 0.62229842 | 0.94016713 |
| CXorf57 | 270.009764 | 0.12134984 | 0.25516232 | 0.47557898 | 0.63437435 | 0.94088547 |
| RNF128 | 2341.79281 | 0.52208078 | 0.21870195 | 2.3871793 | 0.01697821 | 0.46671416 |
| TBC1D8B | 745.275968 | 0.2285002 | 0.35901074 | 0.63647178 | 0.52446897 | 0.91723878 |
| MORC4 | 923.801482 | 0.10316785 | 0.13868255 | 0.74391372 | 0.45692867 | 0.89688594 |
| RIPPLY1 | 66.6238305 | 1.15771856 | 0.56852971 | 2.03633782 | 0.04171644 | 0.56071666 |

| | | | | | | |
|---|---|---|---|---|---|---|
| CLDN2 | 14966.8355 | 0.74125999 | 0.46143332 | 1.60642927 | 0.10817963 | 0.67789099 |
| RBM41 | 368.619354 | -0.0995013 | 0.27308453 | -0.3643609 | 7.16E-01 | 0.95590747 |
| NUP62CL | 208.039494 | 0.22965041 | 0.40542666 | 0.56644132 | 0.57109381 | 0.92818222 |
| PRPS1 | 1618.75774 | 0.03018149 | 0.23927044 | 0.12613967 | 0.89962137 | 0.98506781 |
| TSC22D3 | 7765.76995 | -0.4770791 | 0.50426271 | -0.9460923 | 3.44E-01 | 0.85961157 |
| NCBP2L | 78.8419604 | 0.13896677 | 0.32863979 | 0.42285436 | 0.67240152 | 0.94922827 |
| MID2 | 340.738027 | -0.6127983 | 0.2993994 | -2.0467584 | 0.04068181 | 0.55657615 |
| PSMD10 | 1434.00858 | 0.08636957 | 0.10890265 | 0.79308972 | 0.42772555 | 0.88791007 |
| ATG4A | 1110.24881 | 0.06545769 | 0.15549492 | 0.42096354 | 0.67378171 | 0.94935413 |
| COL4A6 | 488.891744 | 0.43845275 | 0.34086966 | 1.28627686 | 0.19834646 | 0.78046101 |
| COL4A5 | 1634.331 | 0.21747144 | 0.29412829 | 0.73937614 | 0.45967863 | 0.89755243 |
| NXT2 | 388.47447 | 0.06731818 | 0.22433596 | 0.30007754 | 0.76411801 | 0.96678847 |
| KCNE5 | 45.2373967 | -0.2611026 | 0.56642886 | -0.4609627 | 0.64482537 | 0.94208859 |
| ACSL4 | 9487.60766 | 0.32338859 | 0.30310822 | 1.06690803 | 0.28601337 | 0.83107779 |
| TMEM164 | 2323.50094 | 0.03493784 | 0.14800432 | 0.23605961 | 0.81338642 | 0.9738447 |
| AMMECR1 | 574.41953 | -0.3164959 | 0.19452061 | -1.6270556 | 0.10372529 | 0.67439401 |
| CHRDL1 | 1234.80017 | 0.73335574 | 0.53316775 | 1.37546904 | 0.16898608 | 0.75786012 |
| PAK3 | 58.8967017 | -0.2273678 | 0.49806639 | -0.456501 | 0.64802972 | 0.94298664 |
| CAPN6 | 570.901328 | 0.40102266 | 0.67435397 | 0.59467679 | 0.55205954 | 0.92373235 |
| ALG13 | 1906.5967 | -0.0477624 | 0.16919694 | -0.2822886 | 0.77772226 | 0.96836854 |
| AMOT | 1414.42389 | 0.7117359 | 0.242311 | 2.93728262 | 0.00331102 | 0.25241775 |
| IL13RA2 | 63.1366566 | 1.846756 | 0.84000599 | 2.19850338 | 0.02791325 | 0.53262541 |
| LRCH2 | 165.413841 | -0.4623901 | 0.39518294 | -1.1700659 | 0.24197443 | 0.81313301 |
| PLS3-AS1 | 43.0424669 | -0.0846615 | 0.48504218 | -0.1745447 | 0.86143742 | 0.98167219 |
| PLS3 | 7559.09409 | 0.19933203 | 0.2043716 | 0.9753411 | 0.32939109 | 0.85350346 |
| AGTR2 | 26.9957978 | 2.04262061 | 1.62743997 | 1.25511272 | 0.20943784 | 0.79086734 |
| KLHL13 | 1464.54376 | 0.56112223 | 0.16937265 | 3.31294478 | 0.00092319 | 0.14539111 |
| WDR44 | 743.341148 | -0.1339718 | 0.20810996 | -0.6437548 | 0.51973443 | 0.91657872 |
| DOCK11 | 852.941245 | -0.2771638 | 0.35033851 | -0.7911313 | 0.42886739 | 0.88851358 |
| IL13RA1 | 4633.0534 | -0.029261 | 0.22913362 | -0.1277029 | 0.89838408 | 0.98474725 |
| LONRF3 | 127.183422 | -1.1802288 | 0.39237639 | -3.0078996 | 0.0026306 | 0.21835619 |
| PGRMC1 | 10832.6622 | 0.23438393 | 0.23095634 | 1.01484084 | 0.3101817 | 0.84456632 |
| SLC25A43 | 1033.14547 | -0.0189556 | 0.23753176 | -0.0798022 | 0.93639457 | 0.99068685 |
| SLC25A5 | 22547.9863 | 0.33575428 | 0.28312845 | 1.18587263 | 0.2356726 | 0.80747753 |
| CXorf56 | 960.813345 | 0.02605906 | 0.16006799 | 0.16279998 | 0.87067593 | 0.98275231 |
| UBE2A | 3708.64358 | -0.234452 | 0.13362939 | -1.7544944 | 0.07934582 | 0.64079926 |
| NKRF | 698.209988 | -0.1125787 | 0.12505009 | -0.9002691 | 0.36797707 | 0.86745387 |
| 6-Sep | 2235.34724 | -0.0976428 | 0.29317531 | -0.3330525 | 0.73909464 | 0.9602325 |
| SOWAHD | 43.3852089 | 0.24878687 | 0.40771236 | 0.61020193 | 0.54172805 | 0.92292765 |
| RPL39 | 5840.24876 | 0.13396581 | 0.17257991 | 0.77625381 | 0.43759914 | 0.89130865 |
| UPF3B | 790.824522 | -0.0781986 | 0.13261578 | -0.5896625 | 0.55541691 | 0.92373235 |
| RNF113A | 640.352462 | 0.08501019 | 0.12354743 | 0.68807736 | 0.49140407 | 0.90600166 |
| NDUFA1 | 5297.25308 | 0.19178943 | 0.14954962 | 1.28244685 | 0.19968594 | 0.78134034 |
| NKAP | 863.629249 | 0.09587883 | 0.12636057 | 0.75877172 | 0.44798913 | 0.89480207 |
| RHOXF1 | 44.0502089 | -0.1637737 | 0.54060203 | -0.3029468 | 0.76193038 | 0.96607083 |
| ZBTB33 | 777.028572 | -0.2246869 | 0.19401232 | -1.1581062 | 0.24682072 | 0.81542272 |
| TMEM255A | 211.194833 | -0.7973272 | 0.99835914 | -0.7986377 | 0.42450053 | 0.88565404 |
| LAMP2 | 12294.6634 | 0.02081804 | 0.20045492 | 0.10385396 | 0.91728525 | 0.9868044 |
| CUL4B | 2341.00592 | -0.0887187 | 0.13185084 | -0.6728714 | 0.50102914 | 0.91233842 |
| MCTS1 | 1183.20627 | 0.16379645 | 0.14768114 | 1.10912231 | 0.26737742 | 0.82661173 |
| C1GALT1C1 | 1602.03937 | -0.0910527 | 0.14657763 | -0.6211912 | 0.53447383 | 0.92084006 |
| GLUD2 | 157.545051 | 0.30514696 | 0.31194978 | 0.97819257 | 0.32797909 | 0.85317073 |
| GRIA3 | 141.225229 | -0.5987437 | 0.58875257 | -1.0169701 | 0.30916766 | 0.84414094 |
| THOC2 | 2246.84192 | -0.293099 | 0.15461812 | -1.8956317 | 0.05800876 | 0.60193532 |

| | | | | | | |
|---|---|---|---|---|---|---|
| XIAP | 2428.0057 | 0.02345621 | 0.17943432 | 0.13072308 | 8.96E-01 | 0.98474725 |
| STAG2 | 3034.45407 | -0.03967 | 0.10570147 | -0.3753022 | 0.70743573 | 0.95501548 |
| SH2D1A | 54.3813493 | 0.55591458 | 1.00890327 | 0.5510088 | 0.58162764 | 0.9317705 |
| TENM1 | 338.887305 | 0.94431128 | 0.44092849 | 2.14164271 | 0.03222224 | 0.54925606 |
| DCAF12L1 | 255.328846 | -0.4497819 | 0.38361996 | -1.1724675 | 0.2410094 | 0.8124487 |
| SMARCA1 | 2672.12174 | 0.06768773 | 0.20742584 | 0.32632254 | 0.74418034 | 0.9613927 |
| OCRL | 1570.13826 | 0.10934858 | 0.14297479 | 0.76481018 | 0.44438458 | 0.8938278 |
| APLN | 787.294086 | -0.3609852 | 0.80890107 | -0.4462662 | 0.655405 | 0.94532497 |
| XPNPEP2 | 13693.8922 | 0.56256713 | 0.77604978 | 0.72491114 | NA | NA |
| SASH3 | 537.471628 | -0.461875 | 0.60697897 | -0.7609407 | 0.44669247 | 0.89480207 |
| ZDHHC9 | 4486.63753 | -0.0115237 | 0.24908867 | -0.0462634 | 0.96310029 | 0.99320711 |
| UTP14A | 721.938484 | -0.1881807 | 0.20932351 | -0.8989946 | 0.36865554 | 0.86762749 |
| BCORL1 | 999.447853 | -0.0211009 | 0.29844044 | -0.0707038 | 0.94363346 | 0.99136103 |
| ELF4 | 1058.98996 | -0.0834221 | 0.28097035 | -0.2969071 | 0.76653741 | 0.96729533 |
| AIFM1 | 9917.9447 | 0.86444404 | 0.33700955 | 2.56504313 | 0.0103163 | 0.40019066 |
| RAB33A | 43.9532069 | -0.6873949 | 0.56178479 | -1.2235912 | 0.2211065 | 0.79979866 |
| ZNF280C | 184.221539 | 0.11754207 | 0.21612381 | 0.54386452 | 0.58653471 | 0.93289552 |
| SLC25A14 | 544.104466 | -0.0446876 | 0.17680886 | -0.2527452 | 0.80046512 | 0.97210205 |
| RBMX2 | 726.581969 | -0.0414035 | 0.13857663 | -0.2987771 | 7.65E-01 | 0.96678847 |
| ENOX2 | 438.828214 | 0.04352692 | 0.23840389 | 0.18257639 | 0.85513041 | 0.97910552 |
| IGSF1 | 69.8939098 | -0.1349217 | 0.60518833 | -0.2229417 | 0.82358088 | 0.9754753 |
| STK26 | 371.080896 | -0.3175961 | 0.22010801 | -1.44291 | 0.14904582 | 0.73662124 |
| FRMD7 | 78.3025565 | -0.1417543 | 0.92468438 | -0.1533002 | 0.87816158 | 0.98275231 |
| RAP2C | 2955.7922 | -0.2792925 | 0.17697123 | -1.5781801 | 1.15E-01 | 6.84E-01 |
| RAP2C-AS1 | 253.315556 | -0.2715854 | 0.24001501 | -1.131535 | 0.25782999 | 0.82187662 |
| MBNL3 | 923.009224 | 0.35738809 | 0.22458148 | 1.59135158 | 0.11153048 | 0.68001961 |
| HS6ST2 | 1052.79891 | -0.7673365 | 0.3655232 | -2.0992826 | 0.035792 | 0.55369025 |
| GPC4 | 5407.80463 | -0.3858848 | 0.19226896 | -2.0070052 | 0.0447491 | 0.57267043 |
| GPC3 | 4167.74169 | 0.06752837 | 0.61211604 | 0.11031955 | 9.12E-01 | 9.87E-01 |
| CCDC160 | 266.467498 | -0.2903851 | 0.45065858 | -0.6443572 | 0.51934384 | 0.91636259 |
| PHF6 | 989.835802 | -0.0021193 | 0.20420413 | -0.0103786 | 0.99171925 | 0.99891489 |
| HPRT1 | 1849.32893 | 0.28202786 | 0.20043152 | 1.4071033 | 0.15939676 | 0.74762924 |
| MIR503HG | 826.325523 | 0.18388851 | 0.4008903 | 0.45870033 | 0.64644938 | 0.94208859 |
| FAM122B | 1708.9121 | 0.00210456 | 0.15162077 | 0.01388044 | 0.98892537 | 0.99792431 |
| FAM122C | 278.477004 | -0.2133843 | 0.40033261 | -0.5330175 | 0.59402146 | 0.93317501 |
| MOSPD1 | 643.940197 | -0.1476044 | 0.25957477 | -0.5686392 | 0.56960103 | 0.92719921 |
| SMIM10 | 249.409335 | 0.03561524 | 0.28305453 | 0.12582464 | 0.89987073 | 0.98514342 |
| RTL8B | 886.878179 | -0.1174876 | 0.18863496 | -0.6228303 | 0.53339604 | 0.92049136 |
| RTL8C | 5859.80865 | 0.01067421 | 0.22318795 | 0.04782613 | 9.62E-01 | 9.93E-01 |
| RTL8A | 2288.44562 | -0.1811261 | 0.14514212 | -1.2479225 | 2.12E-01 | 0.79203904 |
| SMIM10L2B | 382.549193 | 0.36061225 | 0.37480177 | 0.96214125 | 0.33597865 | 0.85641269 |
| ZNF75D | 1116.51562 | 0.04778112 | 0.18737584 | 0.25500152 | 0.79872192 | 0.97175068 |
| ZNF449 | 484.794764 | 0.04926888 | 0.17059883 | 0.28879962 | 0.77273472 | 0.96836854 |
| SMIM10L2A | 1858.30304 | 0.21290279 | 0.56893885 | 0.37421032 | 0.70824784 | 0.95501548 |
| INTS6L | 676.893759 | -0.0086 | 0.18633148 | -0.0461544 | 0.96318721 | 0.99320711 |
| MMGT1 | 1534.05225 | 0.05040982 | 0.2250821 | 0.22396191 | 0.82278694 | 0.97519922 |
| SLC9A6 | 995.434121 | -0.0041058 | 0.11035528 | -0.0372055 | 0.97032112 | 0.99400331 |
| FHL1 | 3496.02484 | 0.05710357 | 0.43358908 | 0.13169975 | 0.89522178 | 0.98474725 |
| MAP7D3 | 570.720099 | -0.0497399 | 0.17482352 | -0.2845148 | 0.77601591 | 0.96836854 |
| HTATSF1 | 2993.2823 | 0.17849611 | 0.1780547 | 1.00247906 | 0.31611227 | 0.84756299 |
| VGLL1 | 125.702716 | 1.46335787 | 1.0243464 | 1.42857717 | 0.1531258 | 0.74258845 |
| ARHGEF6 | 785.767893 | 0.12723309 | 0.31162203 | 0.40829299 | 0.68305859 | 0.95070301 |
| AL683813.1 | 189.718609 | -0.1309064 | 0.29384039 | -0.4455017 | 0.65595721 | 0.94534713 |
| RBMX | 7361.07795 | -0.1034684 | 0.169727 | -0.6096169 | 5.42E-01 | 0.92292765 |

| | | | | | | |
|---|---|---|---|---|---|---|
| FGF13 | 449.831277 | -0.2059806 | 0.35004156 | -0.5884462 | 0.55623281 | 0.92373235 |
| MCF2 | 63.8397339 | 0.38043372 | 0.50765611 | 0.74939258 | 0.45362062 | 0.89670027 |
| ATP11C | 1269.28695 | 0.14779418 | 0.19212 | 0.76928055 | 0.44172678 | 0.89314342 |
| LDOC1 | 1842.40114 | 0.12230267 | 0.22661513 | 0.53969332 | 0.58940855 | 0.93289552 |
| SLITRK4 | 453.297109 | -0.0186616 | 0.44960288 | -0.0415069 | 0.96689176 | 0.99400331 |
| SLITRK2 | 156.682948 | 0.14193547 | 0.50129978 | 0.28313491 | 0.77707342 | 0.96836854 |
| AL589669.1 | 262.757729 | 0.59171309 | 0.65800257 | 0.8992565 | 0.36851605 | 0.86762749 |
| FMR1 | 3761.60343 | 0.03768021 | 0.28367121 | 0.13283058 | 8.94E-01 | 9.85E-01 |
| AFF2 | 69.4183796 | 0.77693773 | 0.64251633 | 1.20921087 | 0.22658184 | 0.8027165 |
| IDS | 5787.27064 | -0.0124843 | 0.2049016 | -0.0609285 | 0.95141615 | 0.99170503 |
| AC244197.3 | 1460.13092 | -0.2562817 | 0.25077368 | -1.0219642 | 0.30679783 | 0.84286067 |
| AC244197.2 | 52.7878956 | -0.5636342 | 0.69448941 | -0.8115807 | 0.41703225 | 0.88445833 |
| LINC00893 | 566.604784 | 0.25100873 | 0.35155913 | 0.71398723 | 0.47523508 | 0.90226631 |
| CXorf40A | 1028.71889 | 0.09537745 | 0.12893592 | 0.73972755 | 0.45946533 | 0.89755243 |
| TMEM185A | 646.816422 | -0.181749 | 0.13504451 | -1.3458449 | 0.17835255 | 0.76115956 |
| CXorf40B | 1175.68814 | 0.08881783 | 0.20840528 | 0.42617842 | 0.66997784 | 0.949185 |
| HSFX4 | 58.4119977 | -0.0729781 | 0.31337556 | -0.2328774 | 0.81585659 | 0.9738447 |
| LINC00894 | 342.314363 | -0.0541025 | 0.40020575 | -0.1351866 | 0.89246433 | 0.98474725 |
| MAMLD1 | 497.024035 | -0.289959 | 0.45531592 | -0.6368304 | 0.52423531 | 0.91723878 |
| MTM1 | 991.505765 | 0.11181433 | 0.19841143 | 0.56354783 | 0.5730619 | 0.92928554 |
| MTMR1 | 1394.43127 | -0.2027364 | 0.1156387 | -1.7531882 | 0.0795697 | 0.64079926 |
| CD99L2 | 3562.59455 | 0.25266615 | 0.17420427 | 1.45040155 | 1.47E-01 | 7.33E-01 |
| HMGB3 | 1171.56162 | 0.03871881 | 0.43830072 | 0.08833845 | 0.92960768 | 0.9896488 |
| VMA21 | 1856.03286 | 0.09461563 | 0.17503576 | 0.54055026 | 0.58881761 | 0.93289552 |
| GABRE | 1400.12819 | 0.0735699 | 0.47142621 | 0.15605815 | 0.87598718 | 0.98275231 |
| CETN2 | 2713.88621 | 0.236511 | 0.17497692 | 1.3516697 | 0.176481 | 0.76115956 |
| NSDHL | 1152.9823 | -0.0364141 | 0.19375023 | -0.1879437 | 0.85092076 | 0.97844748 |
| ZNF185 | 1047.72819 | 0.56956936 | 0.26509499 | 2.14854815 | 0.03167024 | 0.54507414 |
| PNMA3 | 479.608433 | 0.07443659 | 0.69245736 | 0.10749629 | 9.14E-01 | 0.9866558 |
| PNMA6A | 405.085595 | 0.2659236 | 0.34378663 | 0.77351351 | 0.43921854 | 0.89175645 |
| ZNF275 | 971.209754 | -0.0854819 | 0.17639709 | -0.4845993 | 0.62796061 | 0.94088547 |
| ZFP92 | 195.612264 | -0.3598332 | 0.44442816 | -0.8096544 | 0.41813882 | 0.88445833 |
| TREX2 | 1207.02946 | 0.25825859 | 0.28708496 | 0.89958941 | 0.3683388 | 0.86762749 |
| HAUS7 | 1209.99485 | 0.25215739 | 0.28361379 | 0.88908719 | 0.37395622 | 0.86791605 |
| BGN | 16305.3104 | -0.2170648 | 0.48199749 | -0.4503442 | 0.65246225 | 0.94437526 |
| ATP2B3 | 41.3268469 | 0.18131637 | 1.09707318 | 0.16527281 | 0.86872926 | 0.98272083 |
| CCNQ | 948.870748 | -0.1890768 | 0.25245263 | -0.7489597 | 0.4538815 | 0.89670027 |
| DUSP9 | 3686.31513 | 0.31920362 | 0.59706352 | 0.53462255 | 0.59291089 | 0.93289552 |
| PNCK | 422.917551 | -0.2010263 | 0.33990976 | -0.5914109 | 0.55424515 | 0.92373235 |
| SLC6A8 | 16138.1311 | -0.311877 | 0.34464098 | -0.904933 | 0.36550086 | 0.8663515 |
| BCAP31 | 12054.9807 | 0.05365138 | 0.13545354 | 0.39608694 | 0.69204089 | 0.95168318 |
| ABCD1 | 1131.55106 | -0.4099335 | 0.1702804 | -2.4074027 | 0.01606644 | 0.45716266 |
| PLXNB3 | 311.765485 | -0.179019 | 0.48814483 | -0.3667334 | 0.71381792 | 0.95564093 |
| SRPK3 | 286.362347 | -0.1401925 | 0.45267475 | -0.3096981 | 0.75679057 | 0.9648851 |
| IDH3G | 4731.71846 | 0.24778408 | 0.15164543 | 1.63396999 | 0.10226515 | 0.67279543 |
| SSR4 | 6415.73546 | 0.0920145 | 0.170779 | 0.53879285 | 0.5900298 | 0.93289552 |
| PDZD4 | 399.06827 | 0.13142061 | 0.57623387 | 0.22806818 | 0.81959324 | 0.97454841 |
| L1CAM | 3575.42308 | 0.44232601 | 0.33739315 | 1.31101064 | 0.18985417 | 0.7704467 |
| U52112.2 | 582.150492 | 0.15337624 | 0.37555062 | 0.40840365 | 0.68297735 | 0.95070301 |
| AVPR2 | 457.793223 | 0.81284499 | 0.42715675 | 1.90291967 | 0.05705103 | 0.60049913 |
| ARHGAP4 | 5402.45768 | -0.038242 | 0.2997466 | -0.1275811 | 0.89848049 | 0.98474725 |
| NAA10 | 3569.5044 | -0.1901189 | 0.18158135 | -1.0470176 | 0.29509147 | 0.83653142 |
| RENBP | 3677.59149 | 0.15871758 | 0.4189346 | 0.37886004 | 0.7047918 | 0.95501548 |
| HCFC1 | 3840.64554 | -0.4496935 | 0.17004261 | -2.644593 | 0.00817892 | 0.37396013 |

| | | | | | | |
|---|---|---|---|---|---|---|
| TMEM187 | 513.734209 | 0.36307007 | 0.22460892 | 1.61645439 | 0.10599611 | 0.6756711 |
| IRAK1 | 6821.43394 | -0.2811555 | 0.29428566 | -0.9553831 | 0.33938399 | 0.85759056 |
| MECP2 | 3538.94691 | -0.2918232 | 0.10822383 | -2.6964782 | 0.0070077 | 0.35209453 |
| FLNA | 40051.0975 | -0.6668246 | 0.5619551 | -1.1866154 | 0.23537934 | 0.80747753 |
| EMD | 3465.20828 | -0.0467037 | 0.15446877 | -0.3023501 | 0.76238516 | 0.96630779 |
| RPL10 | 48801.931 | 0.32887069 | 0.20825896 | 1.57914304 | 0.11430325 | 0.68352177 |
| AC245140.2 | 13510.7827 | 0.39066984 | 0.23467764 | 1.66470837 | 0.09597095 | 0.66168746 |
| DNASE1L1 | 2133.00235 | 0.03456712 | 0.11943309 | 0.28942668 | 0.77225488 | 0.96836854 |
| TAZ | 2743.3625 | -0.0161794 | 0.19072403 | -0.0848313 | 0.93239553 | 0.9899349 |
| AC244090.1 | 167.313402 | 0.73853187 | 0.26683586 | 2.76773848 | 0.00564467 | 0.32154048 |
| ATP6AP1 | 11232.9807 | 0.19366445 | 0.11550741 | 1.67664097 | 0.09361271 | 0.65756964 |
| GDI1 | 9827.12311 | -0.1246029 | 0.15128152 | -0.8236494 | 4.10E-01 | 0.88249519 |
| FAM50A | 4638.01351 | -0.3127349 | 0.21525157 | -1.452881 | 0.14625679 | 0.73247558 |
| PLXNA3 | 2728.63699 | -0.3199684 | 0.24765567 | -1.291989 | 0.19636096 | 0.77798606 |
| LAGE3 | 388.544232 | -0.197054 | 0.26256618 | -0.7504926 | 0.45295805 | 0.89670027 |
| UBL4A | 2415.03935 | -0.1381086 | 0.23171116 | -0.5960376 | 0.55115014 | 0.92373235 |
| SLC10A3 | 1792.39428 | -0.2781134 | 0.22225089 | -1.2513487 | 0.2108073 | 0.79134644 |
| FAM3A | 3192.85407 | 0.02272388 | 0.1601257 | 0.14191278 | 0.8871489 | 0.98423722 |
| G6PD | 3174.69587 | -0.5823797 | 0.29616225 | -1.9664211 | 0.04924999 | 0.5861488 |
| IKBKG | 1486.94819 | -0.38804 | 0.19935511 | -1.9464762 | 0.05159757 | 0.58645127 |
| GAB3 | 304.296452 | -0.2677189 | 0.31735113 | -0.8436048 | 0.39889029 | 0.88074856 |
| DKC1 | 2006.41228 | -0.2681859 | 0.24307983 | -1.1032833 | 0.26990414 | 0.82716979 |
| MPP1 | 3279.84779 | 0.33450362 | 0.32967457 | 1.01464793 | 0.31027368 | 0.84456632 |
| F8 | 1686.40595 | 0.43403941 | 0.35353911 | 1.22769844 | 0.21956019 | 0.79858446 |
| F8A1 | 738.63041 | -0.5440431 | 0.25948467 | -2.096629 | 0.03602643 | 0.55369025 |
| FUNDC2 | 4464.07344 | -0.3249999 | 0.2322124 | -1.3995803 | 0.16163905 | 0.74969823 |
| CMC4 | 749.155357 | 0.01040485 | 0.16310285 | 0.0637932 | 0.94913489 | 0.99170503 |
| MTCP1 | 836.112154 | -0.0025617 | 0.16374241 | -0.0156446 | 0.98751789 | 0.997408 |
| BRCC3 | 1145.23156 | 0.00526217 | 0.1266313 | 0.04155508 | 0.96685338 | 0.99400331 |
| VBP1 | 1737.76462 | 0.09364078 | 0.16571669 | 0.56506547 | 0.57202923 | 0.92874623 |
| RAB39B | 64.7548421 | 0.05909185 | 0.36723004 | 0.16091236 | 0.87216243 | 0.98275231 |
| CLIC2 | 514.142862 | 0.02776537 | 0.31736997 | 0.08748581 | 0.93028537 | 0.9897375 |
| TMLHE | 614.655645 | 0.3558731 | 0.22375275 | 1.59047475 | 0.11172783 | 0.68001961 |
| SPRY3 | 147.928263 | -0.089829 | 0.28356545 | -0.3167842 | 0.75140738 | 0.96285385 |
| VAMP7 | 2764.32022 | 0.11026538 | 0.15274795 | 0.72187792 | 0.47036954 | 0.9008571 |
| ZNF596 | 408.450241 | 0.04114645 | 0.2071736 | 0.19860858 | 0.84256894 | 0.97644404 |
| AC004908.1 | 81.1928799 | 0.02270313 | 0.4085014 | 0.05557662 | 0.95567909 | 0.99185043 |
| FBXO25 | 2007.61752 | -0.0961845 | 0.14955941 | -0.6431192 | 0.52014678 | 0.91657872 |
| AC083964.1 | 335.323935 | -0.0926644 | 0.20528203 | -0.4514005 | 0.65170094 | 0.94393131 |
| TDRP | 2216.6482 | -0.2103308 | 0.19896572 | -1.0571208 | 0.29045646 | 0.83442894 |
| ERICH1 | 1317.52625 | -0.4253165 | 0.20935863 | -2.0315212 | 0.04220216 | 0.56485966 |
| DLGAP2 | 180.199761 | -0.4587126 | 0.93952867 | -0.4882369 | 0.62538204 | 0.94088547 |
| CLN8 | 1881.91732 | 0.05366446 | 0.26731059 | 0.20075696 | 0.84088862 | 0.97618283 |
| AC100810.1 | 778.972183 | 0.37510149 | 0.31627276 | 1.18600631 | 0.23561981 | 0.80747753 |
| AC019257.8 | 1464.40649 | -0.0705562 | 0.16202571 | -0.4354629 | 0.66322646 | 0.94642611 |
| ARHGEF10 | 1300.06926 | -0.0533439 | 0.20948345 | -0.254645 | 0.79899732 | 0.97186224 |
| KBTBD11-OT | 67.4016998 | -0.6624051 | 0.51642116 | -1.2826839 | 0.19960283 | 0.78134034 |
| AC019257.1 | 119.976106 | -0.0572775 | 0.45128878 | -0.1269198 | 0.89900389 | 0.98487356 |
| KBTBD11 | 3152.36248 | -0.1886002 | 0.32291903 | -0.5840479 | 0.55918808 | 0.92421175 |
| MYOM2 | 682.721522 | -0.034988 | 0.43662733 | -0.0801324 | 0.93613193 | 0.99068685 |
| CSMD1 | 337.916449 | -0.8363729 | 0.6403836 | -1.3060498 | 0.19153563 | 0.77328076 |
| AC016065.1 | 199.559734 | -0.0266271 | 0.30453148 | -0.0874363 | 0.93032474 | 0.9897375 |
| MCPH1 | 817.694279 | -0.2509715 | 0.14481305 | -1.7330721 | 0.08308285 | 0.6471083 |
| ANGPT2 | 391.08314 | -0.1088531 | 0.5423371 | -0.2007111 | 0.84092452 | 0.97618283 |

| | | | | | | |
|---|---|---|---|---|---|---|
| AF287957.1 | 43.6780634 | 0.49321115 | 0.46172137 | 1.06820083 | 0.28542993 | 0.83107779 |
| MCPH1-AS1 | 143.073254 | -0.1474241 | 0.25396423 | -0.5804916 | 0.56158313 | 0.92452095 |
| AGPAT5 | 1788.81252 | -0.0074878 | 0.30896218 | -0.0242354 | 0.98066481 | 0.99573612 |
| DEFB1 | 17030.6175 | 0.2976767 | 0.41695244 | 0.71393442 | 0.47526773 | 0.90226631 |
| DEFA3 | 201.175213 | -1.793758 | 1.10492537 | -1.62342 | 0.10449965 | 0.67439401 |
| PRAG1 | 873.085191 | -0.1379883 | 0.20256186 | -0.6812154 | 0.4957352 | 0.90861354 |
| CLDN23 | 602.418013 | -0.5555015 | 0.28403441 | -1.9557543 | 0.0504941 | 0.5861488 |
| MFHAS1 | 3403.88829 | -0.0760706 | 0.14914031 | -0.5100605 | 0.61000906 | 0.93791514 |
| ERI1 | 1040.88451 | -0.2556733 | 0.26612819 | -0.9607148 | 0.33669557 | 0.85679125 |
| PPP1R3B | 2089.41868 | 0.21209524 | 0.40435044 | 0.52453322 | 0.59990772 | 0.93487648 |
| AC022784.3 | 46.3719643 | 0.06112687 | 0.37422725 | 0.16334157 | 0.87024951 | 0.98275231 |
| AC021242.3 | 40.707931 | -0.3031522 | 0.35898098 | -0.8444798 | 0.39840137 | 0.88074856 |
| TNKS | 1808.51909 | -0.1265486 | 0.13826979 | -0.9152295 | 0.36007119 | 0.86480228 |
| MSRA | 4373.88288 | 0.67019126 | 0.38117243 | 1.75823642 | 0.07870729 | 0.64079926 |
| AC104964.4 | 220.643085 | -0.2742432 | 0.3688598 | -0.7434891 | 0.45718564 | 0.89688594 |
| SOX7 | 710.483653 | 0.63039165 | 0.42046455 | 1.49927418 | 0.13380252 | 0.71505705 |
| PINX1 | 765.535932 | 0.10740543 | 0.2950509 | 0.3640234 | 0.71584054 | 0.95590747 |
| XKR6 | 72.1146675 | -0.0027739 | 0.37072144 | -0.0074823 | 0.99403003 | 0.99936252 |
| MTMR9 | 845.887739 | -0.0813248 | 0.15755332 | -0.5161733 | 0.6057334 | 0.93698713 |
| AF131216.1 | 90.9167712 | -0.1072968 | 0.24340753 | -0.4408114 | 0.65934955 | 0.94601668 |
| FAM167A | 856.046705 | -0.0001284 | 0.46462527 | -0.0002763 | 0.99977952 | 0.99994357 |
| BLK | 98.6254664 | -0.5750103 | 1.05229224 | -0.5464359 | 0.58476633 | 0.93284864 |
| NEIL2 | 1064.79288 | 0.47166104 | 0.22370549 | 2.10840175 | 0.03499625 | 0.55369025 |
| FDFT1 | 5565.33215 | 0.15438815 | 0.25384839 | 0.60819037 | 0.54306122 | 0.92297267 |
| AC069185.1 | 53.0475851 | 0.61896906 | 0.32303494 | 1.91610564 | 0.05535165 | 0.59617546 |
| CTSB | 98804.4629 | 0.08801357 | 0.53877147 | 0.16335975 | 0.87023519 | 0.98275231 |
| FAM86B1 | 277.818924 | 0.35502839 | 0.45796005 | 0.77523879 | 4.38E-01 | 8.92E-01 |
| AC145124.1 | 79.5919054 | 0.84376002 | 0.46484661 | 1.81513643 | 0.06950295 | 0.62779744 |
| LONRF1 | 1412.23347 | -0.2213473 | 0.2561014 | -0.8642955 | 0.38742556 | 0.87460898 |
| TRMT9B | 1803.60555 | 0.06020103 | 0.38294177 | 0.15720674 | 0.87508192 | 0.98275231 |
| DLC1 | 2010.50163 | -0.1882378 | 0.32531084 | -0.5786399 | 0.56283216 | 0.92452095 |
| C8orf48 | 58.4487567 | -0.5127234 | 0.55889902 | -0.9173811 | 0.35894297 | 0.86480228 |
| TUSC3 | 2641.54538 | 0.22584158 | 0.23892915 | 0.94522407 | 0.34454449 | 0.85999891 |
| MSR1 | 849.999702 | -0.5692638 | 0.4054642 | -1.4039805 | 0.16032465 | 0.74781837 |
| MICU3 | 381.757047 | 0.06774479 | 0.25176249 | 0.26908213 | 0.78786648 | 0.97090793 |
| ZDHHC2 | 2618.08013 | 0.12619724 | 0.16849557 | 0.74896471 | 0.45387848 | 0.89670027 |
| CNOT7 | 3327.85496 | -0.0722195 | 0.174062 | -0.4149065 | 0.67821037 | 0.95023479 |
| VPS37A | 2591.38508 | -0.1796052 | 0.21642358 | -0.8298784 | 0.40660755 | 0.88132015 |
| MTMR7 | 180.281973 | -0.3384278 | 0.30301933 | -1.116852 | 0.26405759 | 0.82424998 |
| SLC7A2 | 2068.99552 | 0.0040016 | 0.33240834 | 0.01203822 | 0.99039512 | 0.99814875 |
| PDGFRL | 939.704066 | -0.2826286 | 0.238334 | -1.1858508 | 0.23568124 | 0.80747753 |
| MTUS1 | 3570.6639 | -0.3851059 | 0.21395321 | -1.799954 | 0.0718679 | 0.63189503 |
| FGL1 | 183.541104 | 0.07993447 | 1.13155378 | 0.07064134 | 0.94368321 | 0.99136103 |
| AC087273.1 | 50.5224771 | 0.5686237 | 1.30106648 | 0.43704431 | 0.66207921 | 0.94642611 |
| PCM1 | 4009.93954 | -0.0843539 | 0.13370802 | -0.6308815 | 0.52811803 | 0.91881312 |
| ASAH1 | 20173.5853 | 0.29827351 | 0.30770812 | 0.96933908 | 0.33237604 | 0.85496237 |
| AC124242.1 | 65.318738 | 0.01964461 | 0.39241291 | 0.05006108 | 0.96007371 | 0.99255196 |
| NAT1 | 366.67363 | 0.19854692 | 0.34461002 | 0.57614958 | 0.56451408 | 0.92474393 |
| NAT2 | 45.9316868 | 0.49183224 | 0.43905334 | 1.12021067 | 0.262624 | 0.82424998 |
| PSD3 | 475.549685 | -0.0594845 | 0.34419385 | -0.1728227 | 0.86279081 | 0.98176973 |
| SH2D4A | 971.527786 | 0.15306351 | 0.21853555 | 0.70040556 | 0.48367407 | 0.90541935 |
| CSGALNACT1 | 1284.98048 | -0.4649904 | 0.24764331 | -1.8776618 | 0.06042744 | 0.60849584 |
| INTS10 | 3338.97885 | -0.010298 | 0.1609506 | -0.0639824 | 0.94898427 | 0.99170503 |
| LPL | 1412.61414 | 1.18152404 | 0.77614076 | 1.52230638 | 0.12793233 | 0.70579329 |

| | | | | | | |
|---|---|---|---|---|---|---|
| ATP6V1B2 | 9857.83182 | -0.3050895 | 0.47340009 | -0.6444643 | 0.51927437 | 0.91636259 |
| LZTS1 | 410.889337 | -0.2686293 | 0.39532649 | -0.6795125 | 0.49681322 | 0.90890243 |
| AC018541.1 | 40.2704752 | 3.659092 | 1.20287971 | 3.0419434 | 0.00235056 | 0.21083339 |
| GFRA2 | 97.4834305 | 1.44625869 | 0.6086242 | 2.37627535 | 0.01748841 | 0.46859699 |
| DOK2 | 280.109063 | -0.3595718 | 0.61223749 | -0.5873078 | 0.55699704 | 0.92373235 |
| XPO7 | 3211.518 | 0.08297053 | 0.19653962 | 0.42215677 | 6.73E-01 | 9.49E-01 |
| NPM2 | 393.41159 | -0.1640806 | 0.33184332 | -0.494452 | 0.62098703 | 0.94016713 |
| FGF17 | 192.959394 | -0.5224182 | 0.38245435 | -1.3659621 | 0.17195085 | 0.75974924 |
| DMTN | 6461.68074 | -0.0703754 | 0.24524243 | -0.2869627 | 0.77414085 | 0.96836854 |
| FAM160B2 | 5749.26702 | -0.1895004 | 0.19874015 | -0.9535083 | 0.3403326 | 0.85814041 |
| NUDT18 | 485.803963 | 0.04545696 | 0.24474054 | 0.18573532 | 0.85265233 | 0.97883015 |
| HR | 174.087239 | 0.5052224 | 0.59860709 | 0.84399668 | 0.39867126 | 0.88074856 |
| REEP4 | 887.864919 | 0.05228566 | 0.27284564 | 0.19163092 | 0.84803132 | 0.97711043 |
| LGI3 | 68.0483692 | 1.14985329 | 0.93991731 | 1.2233558 | 0.22119534 | 0.79996563 |
| BMP1 | 3026.33597 | -0.1032666 | 0.2258796 | -0.4571752 | 0.64754511 | 0.94282139 |
| PHYHIP | 681.698924 | -0.0536845 | 0.36005574 | -0.1491006 | 0.88147424 | 0.98275231 |
| POLR3D | 929.701389 | -0.312709 | 0.17918458 | -1.7451781 | 0.08095388 | 0.64313548 |
| PIWIL2 | 55.9577887 | 0.94466559 | 0.43343988 | 2.17946165 | 0.02929739 | 0.53606221 |
| SLC39A14 | 10979.4559 | -0.6864879 | 0.41505949 | -1.6539505 | 0.09813756 | 0.66549286 |
| PPP3CC | 1192.40335 | -0.2284487 | 0.16512781 | -1.383466 | 0.16652203 | 0.75558932 |
| AC037459.2 | 230.240134 | -0.1774625 | 0.21217725 | -0.836388 | 0.40293664 | 0.88132015 |
| SORBS3 | 6185.9131 | -0.1985664 | 0.20685625 | -0.9599247 | 0.33709312 | 0.85688344 |
| AC037459.4 | 1174.63593 | -0.0785842 | 0.22324586 | -0.3520074 | 0.72483272 | 0.95847868 |
| PDLIM2 | 3052.2111 | 0.13579945 | 0.38681855 | 0.35106756 | 0.72553766 | 0.95848028 |
| AC037459.1 | 1621.02892 | -0.0163573 | 0.3135018 | -0.052176 | 0.95838845 | 0.99237995 |
| C8orf58 | 1005.50828 | -0.3247463 | 0.17619968 | -1.8430583 | 0.0653205 | 0.62203839 |
| CCAR2 | 6460.66 | -0.2897628 | 0.14749061 | -1.9646188 | 0.04945838 | 0.5861488 |
| AC037459.3 | 1745.24089 | -0.2156478 | 0.22290083 | -0.9674608 | 0.33331374 | 0.855011 |
| BIN3 | 1935.14378 | -0.3221153 | 0.14194683 | -2.2692676 | 0.02325206 | 0.51967919 |
| EGR3 | 257.882509 | -0.3183637 | 0.92332289 | -0.3448021 | 0.73024317 | 0.95848028 |
| PEBP4 | 82.4974595 | 0.53395764 | 0.50316692 | 1.06119384 | 0.28860182 | 0.83295411 |
| AC107959.1 | 738.443846 | 0.10401999 | 0.39257891 | 0.26496581 | 0.7910358 | 0.97090793 |
| RHOBTB2 | 2579.07444 | 0.14962939 | 0.35450598 | 0.42207859 | 0.67296765 | 0.94922827 |
| TNFRSF10B | 4443.00276 | 0.04231296 | 0.19300585 | 0.21923147 | 0.82646974 | 0.97548201 |
| AC107959.4 | 95.0661835 | 0.18582195 | 0.36737395 | 0.50581145 | 0.61298902 | 0.93879706 |
| AC107959.5 | 78.7658765 | 0.57605936 | 0.39418662 | 1.46138741 | 0.14390915 | 0.72920837 |
| TNFRSF10C | 173.091879 | 0.49737073 | 0.3641887 | 1.36569512 | 0.17203468 | 0.75974924 |
| TNFRSF10D | 1785.52957 | -0.1121033 | 0.28323908 | -0.3957904 | 0.69225963 | 0.95177446 |
| TNFRSF10A- | 126.042837 | -0.3299638 | 0.35321611 | -0.9341697 | 0.35021638 | 0.86364633 |
| TNFRSF10A | 593.700704 | -0.6068305 | 0.40779443 | -1.4880793 | 0.13672998 | 0.72102859 |
| AC100861.2 | 58.0032313 | 0.17819537 | 0.3675732 | 0.48478879 | 0.62782618 | 0.94088547 |
| CHMP7 | 2318.62871 | -0.246709 | 0.14430575 | -1.7096268 | 0.0873349 | 0.65260901 |
| R3HCC1 | 1576.01237 | 1.98E-05 | 0.18004553 | 0.00010988 | 0.99991233 | 0.99996568 |
| LOXL2 | 635.799638 | 0.39478108 | 0.41032427 | 0.96211975 | 0.33598946 | 0.85641269 |
| ENTPD4 | 2355.5759 | -0.0554568 | 0.1664583 | -0.3331575 | 0.73901543 | 0.9602325 |
| SLC25A37 | 4149.33232 | 0.03014827 | 0.35407544 | 0.08514647 | 0.93214495 | 0.98987582 |
| NKX3-1 | 134.909225 | -0.2923104 | 0.47763785 | -0.6119918 | 0.5405432 | 0.92247611 |
| STC1 | 2275.76788 | 0.56914153 | 0.5216052 | 1.09113468 | 0.27521362 | 0.82719407 |
| ADAM28 | 521.473566 | 0.17053008 | 0.38616496 | 0.44159907 | 0.65877936 | 0.9458503 |
| AF106564.1 | 56.5150338 | 1.46191905 | 1.1054991 | 1.32240637 | 0.18603287 | 0.76908034 |
| NEFM | 202.416336 | 1.5711478 | 1.08155158 | 1.45267949 | NA | NA |
| NEFL | 350.700117 | 3.28494492 | 0.9039253 | 3.63408893 | 0.00027896 | 0.08298534 |
| DOCK5 | 1291.49485 | -0.0856453 | 0.21009358 | -0.407653 | 0.68352842 | 0.95070301 |
| GNRH1 | 277.982392 | 0.17176342 | 0.30478809 | 0.5635503 | 0.57306022 | 0.92928554 |

| | | | | | | |
|---|---|---|---|---|---|---|
| KCTD9 | 1533.72639 | -0.0818419 | 0.17163827 | -0.476828 | 0.63348463 | 0.94088547 |
| EBF2 | 64.2817401 | 0.10439359 | 0.83393039 | 0.12518262 | 0.90037898 | 0.98526603 |
| PPP2R2A | 2444.1762 | -0.2749046 | 0.14613013 | -1.8812312 | 0.05994048 | 0.6075531 |
| BNIP3L | 6982.95321 | 0.02805579 | 0.2535732 | 0.11064178 | 0.91190041 | 0.98662757 |
| PNMA2 | 838.368758 | 0.0642575 | 0.41851074 | 0.15353848 | 0.87797365 | 0.98275231 |
| DPYSL2 | 3704.83032 | 0.2714855 | 0.26442752 | 1.02669157 | 0.30456572 | 0.84211658 |
| AC090150.1 | 180.326501 | -0.0555119 | 0.82373235 | -0.0673907 | 0.94627067 | 0.99166614 |
| TRIM35 | 1213.80075 | -0.2304709 | 0.32396789 | -0.7114005 | 0.47683607 | 0.90335153 |
| PTK2B | 2610.15654 | -0.0125341 | 0.28592017 | -0.0438378 | 0.9650337 | 0.99357855 |
| EPHX2 | 7280.53133 | 1.18151372 | 0.4149856 | 2.84711983 | 0.00441168 | 0.29060638 |
| CLU | 80559.0511 | 0.02643398 | 0.4272389 | 0.06187165 | 0.95066504 | 0.99170503 |
| SCARA3 | 1225.89844 | 1.23444682 | 0.55387264 | 2.22875571 | 0.02583016 | 0.52502381 |
| CCDC25 | 2112.28921 | 0.13849242 | 0.11625981 | 1.19123218 | 0.23356245 | 0.80725541 |
| AC013643.2 | 190.895108 | 0.18940454 | 0.21153717 | 0.89537237 | 0.37058806 | 0.86762749 |
| SCARA5 | 177.330226 | -0.3343821 | 1.17859112 | -0.2837134 | 7.77E-01 | 0.96836854 |
| NUGGC | 360.575276 | 0.79823172 | 0.46162901 | 1.72916281 | 0.08377996 | 0.64854202 |
| ELP3 | 1702.48618 | 0.14204253 | 0.17363764 | 0.81803999 | 0.41333435 | 0.88316314 |
| ZNF395 | 5061.32357 | -0.432039 | 0.28368653 | -1.5229451 | 0.12777245 | 0.70579329 |
| FBXO16 | 890.155143 | -0.40056 | 0.44940826 | -0.8913053 | 0.3727654 | 0.86791605 |
| FZD3 | 678.150829 | -0.2155871 | 0.12851671 | -1.6775029 | 0.09344419 | 0.65756964 |
| EXTL3 | 2116.992 | -0.0834748 | 0.19660713 | -0.4245766 | 0.67114538 | 0.94922827 |
| EXTL3-AS1 | 69.4315439 | 0.05803084 | 0.40433634 | 0.1435212 | 0.88587857 | 0.98385457 |
| INTS9 | 1052.8378 | -0.1555517 | 0.11174695 | -1.3919997 | 0.1639225 | 0.75265041 |
| HMBOX1 | 1493.91513 | -0.1784686 | 0.16255803 | -1.0978763 | 0.2722585 | 0.82716979 |
| KIF13B | 6239.80352 | 0.13677358 | 0.12452957 | 1.0983221 | 0.27206387 | 0.82716979 |
| AC108449.3 | 2086.72648 | 0.19144081 | 0.14216691 | 1.34659195 | 0.17811169 | 0.76115956 |
| DUSP4 | 173.504159 | 0.32786136 | 0.56983698 | 0.57535993 | 0.5650479 | 0.92493342 |
| SARAF | 16875.3299 | 0.35384584 | 0.21249186 | 1.66522072 | 0.09586873 | 0.66151541 |
| AC044849.1 | 1351.6427 | 0.4824457 | 0.279558 | 1.72574457 | 0.08439338 | 0.64988556 |
| LEPROTL1 | 3410.57425 | 0.20820342 | 0.19096798 | 1.09025303 | 0.2756017 | 0.82719407 |
| AC026979.2 | 40.451686 | -0.2641148 | 0.44918983 | -0.5879803 | 0.5565455 | 0.92373235 |
| DCTN6 | 1708.23454 | 0.08986738 | 0.15506622 | 0.57954195 | 0.56222355 | 0.92452095 |
| RBPMS-AS1 | 1772.57734 | -0.5267122 | 0.27042959 | -1.9476869 | 0.05145244 | 0.58634747 |
| RBPMS | 14955.259 | -0.2748397 | 0.23912598 | -1.1493513 | 2.50E-01 | 8.17E-01 |
| GTF2E2 | 1268.87654 | -0.1594019 | 0.20057388 | -0.794729 | 0.42677113 | 0.88709241 |
| GSR | 6385.17173 | 0.30961819 | 0.37210194 | 0.83207893 | 0.4053644 | 0.88132015 |
| UBXN8 | 705.067818 | 0.43072092 | 0.13239771 | 3.25323551 | 0.00114099 | 0.15697311 |
| PPP2CB | 7380.52278 | 0.01607681 | 0.20380198 | 0.07888444 | 0.93712454 | 0.99077727 |
| WRN | 419.012531 | -0.2260619 | 0.28419846 | -0.7954369 | 0.42635942 | 0.8869355 |
| AC009563.1 | 95.3786 | -0.2702185 | 0.3641941 | -0.741963 | 0.45810978 | 0.8971553 |
| NRG1 | 872.202876 | -0.1565722 | 0.48559491 | -0.3224338 | 0.74712411 | 0.96224616 |
| AC090204.1 | 59.8183553 | 0.48832584 | 0.46054794 | 1.06031491 | 0.28900136 | 0.83308345 |
| AC067838.1 | 74.4452866 | 0.59571554 | 0.38912969 | 1.53089203 | 1.26E-01 | 0.70347499 |
| FUT10 | 290.331229 | 0.42142042 | 0.23796668 | 1.770922 | 0.07657367 | 0.63773415 |
| TTI2 | 787.18528 | -0.2054073 | 0.16589115 | -1.2382055 | 0.21563989 | 0.7954827 |
| MAK16 | 940.888195 | -0.3027174 | 0.21023328 | -1.439912 | 0.14989229 | 0.73749842 |
| RNF122 | 703.749208 | -0.0812573 | 0.33606162 | -0.2417928 | 0.80894071 | 0.97334138 |
| DUSP26 | 124.43501 | -0.661336 | 0.52393656 | -1.2622444 | 2.07E-01 | 7.88E-01 |
| UNC5D | 237.389107 | -0.0323382 | 0.56144434 | -0.0575982 | 9.54E-01 | 9.92E-01 |
| AC124067.4 | 134.524611 | 1.18997984 | 0.76072796 | 1.56426463 | 0.11775543 | 0.69069313 |
| ZNF703 | 561.423796 | -0.2973129 | 0.38288897 | -0.7764989 | 0.43745448 | 0.89130865 |
| AC138356.1 | 88.4918579 | 0.53782137 | 0.40539255 | 1.32666813 | 0.18461849 | 0.76820626 |
| ERLIN2 | 3611.84421 | 0.2046893 | 0.14826868 | 1.38052962 | 0.16742364 | 0.75558932 |
| PLPBP | 3354.62856 | 0.31975465 | 0.21931986 | 1.4579375 | 0.14485777 | 0.73039059 |

| | | | | | | |
|---|---|---|---|---|---|---|
| ADGRA2 | 2098.03511 | -0.1675271 | 0.42037786 | -0.3985156 | 0.69025016 | 0.95158502 |
| BRF2 | 1210.22847 | -0.0852145 | 0.27114533 | -0.314276 | 0.75331145 | 0.96301881 |
| RAB11FIP1 | 2576.03682 | 0.16841404 | 0.17152599 | 0.98185725 | 3.26E-01 | 8.53E-01 |
| EIF4EBP1 | 1143.0618 | 0.01513018 | 0.29046618 | 0.0520893 | 9.58E-01 | 9.92E-01 |
| ASH2L | 1651.51811 | 0.07651829 | 0.19207652 | 0.39837401 | 6.90E-01 | 9.52E-01 |
| AC084024.3 | 862.661614 | -0.2177734 | 0.14227564 | -1.530644 | 0.12585741 | 0.70347499 |
| LSM1 | 1516.20073 | -0.1862789 | 0.13223806 | -1.4086633 | 0.15893475 | 0.74726448 |
| BAG4 | 1251.3098 | -0.0029894 | 0.16250895 | -0.0183952 | 0.98532362 | 0.99692004 |
| DDHD2 | 2529.9783 | -0.1331398 | 0.10412199 | -1.2786901 | 0.20100622 | 0.78134034 |
| PLPP5 | 2548.81412 | -0.0560148 | 0.23817641 | -0.235182 | 0.81406745 | 0.9738447 |
| NSD3 | 3136.30605 | -0.2941943 | 0.16000124 | -1.8387001 | 0.06595931 | 0.62237346 |
| AC087362.1 | 79.0081174 | -0.2548337 | 0.40199235 | -0.6339267 | 0.52612866 | 0.91769856 |
| LETM2 | 390.192478 | -0.3312612 | 0.34578249 | -0.9580047 | 0.3380604 | 0.8571036 |
| AC087623.4 | 106.956556 | -0.4439256 | 0.39803165 | -1.1153023 | 0.26472091 | 0.82424998 |
| FGFR1 | 5863.96318 | -0.7337446 | 0.35627085 | -2.0595135 | 0.03944507 | 0.55657615 |
| AC087623.2 | 212.28042 | -0.7611553 | 0.44990704 | -1.6918057 | 0.09068303 | 0.65495306 |
| TACC1 | 5427.09818 | -0.3265716 | 0.26956795 | -1.2114629 | 0.22571804 | 0.80238655 |
| PLEKHA2 | 2815.84409 | -0.3368344 | 0.23192648 | -1.4523327 | 0.14640913 | 0.73247558 |
| AC108863.1 | 731.852941 | -0.2492294 | 0.21865859 | -1.1398106 | 0.25436523 | 0.81992039 |
| TM2D2 | 2591.89715 | 0.12806013 | 0.21330095 | 0.60037299 | 0.54825769 | 0.92373235 |
| ADAM9 | 6669.51522 | 0.17423957 | 0.29453324 | 0.59157864 | 0.55413278 | 0.92373235 |
| ADAM32 | 45.3679007 | 0.16865803 | 0.33400107 | 0.50496254 | 0.61358515 | 0.93886343 |
| IDO1 | 194.500424 | -1.9865344 | 1.09451229 | -1.814995 | 0.06952467 | 0.62779744 |
| IDO2 | 81.6447813 | 0.57102702 | 0.77892296 | 0.7330982 | 0.46349854 | 0.89830907 |
| TCIM | 12721.6411 | -0.3291157 | 0.42904363 | -0.7670914 | 0.44302716 | 0.89352768 |
| SFRP1 | 9125.91855 | 0.08674385 | 0.32338119 | 0.26824026 | 0.78851439 | 0.97090793 |
| GOLGA7 | 2716.12388 | -0.0406131 | 0.20738485 | -0.1958344 | 0.8447398 | 0.97675487 |
| GINS4 | 110.324861 | 0.16174762 | 0.37157948 | 0.4352975 | 0.6633465 | 0.94642611 |
| AC009630.2 | 43.9066412 | 0.25429187 | 0.40870634 | 0.62218723 | 0.53381876 | 0.92062041 |
| GPAT4 | 7364.958 | -0.2652396 | 0.32020961 | -0.8283311 | 0.40748301 | 0.88132015 |
| AC009630.3 | 3165.22367 | -0.2662981 | 0.30711362 | -0.8670996 | 0.38588742 | 0.87420019 |
| ANK1 | 116.554646 | -0.7493674 | 0.55557428 | -1.348816 | 0.17739609 | 0.76115956 |
| KAT6A | 1597.55388 | -0.2219645 | 0.13126984 | -1.6909027 | 0.09085539 | 0.65495306 |
| AP3M2 | 1151.16817 | 0.00649325 | 0.12227847 | 0.05310219 | 0.95765048 | 0.9922229 |
| PLAT | 6971.59122 | -0.394968 | 0.41902998 | -0.9425769 | 0.34589736 | 0.86111349 |
| IKBKB | 3067.32352 | -0.2354997 | 0.11046731 | -2.13185 | 0.03301918 | 0.55137197 |
| POLB | 661.136592 | 0.20644284 | 0.15024365 | 1.37405366 | 0.16942502 | 0.75852708 |
| VDAC3 | 4965.22447 | 0.04456022 | 0.16573723 | 0.26886063 | 0.78803694 | 0.97090793 |
| SLC20A2 | 3276.89697 | -0.5377666 | 0.1894595 | -2.8384254 | 0.00453367 | 0.29501922 |
| SMIM19 | 1507.91725 | 0.12604886 | 0.26886232 | 0.46882309 | 0.63919609 | 0.94088547 |
| THAP1 | 441.397671 | -0.3217405 | 0.25301464 | -1.2716282 | 0.20350527 | 0.78338136 |
| RNF170 | 1362.98379 | 0.35773693 | 0.20629508 | 1.7341031 | 0.08289979 | 0.64680474 |
| HOOK3 | 1812.83657 | -0.0011226 | 0.14283299 | -0.0078593 | 0.99372929 | 0.99925673 |
| AC110275.1 | 1038.81461 | -0.1821448 | 0.16500146 | -1.1038981 | 0.26963736 | 0.82716979 |
| FNTA | 3304.21635 | -0.277115 | 0.14195831 | -1.9520871 | 0.05092787 | 0.5861488 |
| HGSNAT | 4800.41355 | 0.09167178 | 0.15274486 | 0.60016277 | 0.54839776 | 0.92373235 |
| AC120036.4 | 42.9433549 | -0.1397873 | 0.41103419 | -0.3400867 | 0.73379124 | 0.95895327 |
| SPIDR | 2817.48953 | -0.4642335 | 0.21435003 | -2.1657731 | 0.03032853 | 0.54183693 |
| CEBPD | 10284.5055 | -0.5253099 | 0.40993878 | -1.2814349 | 0.20004095 | 0.78134034 |
| PRKDC | 3226.91752 | 0.03616558 | 0.17655627 | 0.20483882 | 0.83769805 | 0.97577999 |
| MCM4 | 1136.75666 | -0.3900447 | 0.35801188 | -1.0894741 | 0.27594486 | 0.82719407 |
| UBE2V2 | 1895.35335 | -0.0369688 | 0.17239971 | -0.2144368 | 0.83020644 | 0.97550416 |
| SNAI2 | 467.734053 | -0.3017852 | 0.46682158 | -0.6464679 | 0.51797635 | 0.91636259 |
| PXDNL | 93.0478285 | 1.61621074 | 0.75641385 | 2.13667525 | 0.03262441 | 0.55104386 |

| | | | | | | |
|---|---|---|---|---|---|---|
| PCMTD1 | 3836.43187 | 0.0557425 | 0.23698781 | 0.23521254 | 0.81404377 | 0.9738447 |
| AC064807.1 | 114.321928 | 0.25905197 | 0.36865711 | 0.70269084 | 0.48224843 | 0.90541935 |
| AC064807.2 | 97.3472401 | 1.25311151 | 0.63383159 | 1.97704174 | 0.04803691 | 0.58420491 |
| ST18 | 47.5583176 | 0.74534552 | 0.86056945 | 0.86610735 | 3.86E-01 | 8.74E-01 |
| RB1CC1 | 2223.78937 | -0.2665475 | 0.2978869 | -0.8947942 | 0.37089709 | 0.86762749 |
| ATP6V1H | 4528.39134 | 0.17519506 | 0.16376593 | 1.0697894 | 0.28471411 | 0.83107779 |
| TCEA1 | 3118.30248 | -0.003217 | 0.1692415 | -0.0190084 | 0.98483443 | 0.99674796 |
| LYPLA1 | 3754.89536 | 0.15000296 | 0.16433974 | 0.91276135 | 0.36136807 | 0.86528137 |
| MRPL15 | 2478.07168 | -0.1660147 | 0.17765463 | -0.93448 | 0.35005635 | 0.86364633 |
| SOX17 | 321.088595 | -0.7920502 | 0.65657919 | -1.2063285 | 0.22769085 | 0.80300331 |
| RP1 | 105.968977 | -0.2002192 | 0.71085249 | -0.2816607 | 0.77820371 | 0.96836854 |
| XKR4 | 123.850397 | 1.07979112 | 0.44500736 | 2.42645678 | 0.01524707 | 0.45205627 |
| TMEM68 | 1068.79958 | 0.11630499 | 0.15799632 | 0.73612466 | 0.46165484 | 0.89759034 |
| TGS1 | 908.314536 | -0.0007136 | 0.19238016 | -0.0037094 | 0.99704035 | 0.99976751 |
| LYN | 1409.75702 | -0.0346162 | 0.26661523 | -0.1298357 | 0.89669638 | 0.98474725 |
| RPS20 | 22046.5536 | 0.02289685 | 0.15459379 | 0.14810975 | 0.88225615 | 0.98275231 |
| PLAG1 | 300.638495 | 0.15119823 | 0.31928355 | 0.47355472 | 0.63581747 | 0.94088547 |
| CHCHD7 | 1553.25882 | 0.10560926 | 0.24189225 | 0.43659631 | 0.66240414 | 0.94642611 |
| AC107952.2 | 59.7205847 | 0.72005117 | 0.34188508 | 2.10612049 | 0.03519389 | 0.55369025 |
| IMPAD1 | 4320.04791 | -0.0669845 | 0.21507219 | -0.3114512 | 0.75545765 | 0.96391829 |
| LINC01606 | 761.440871 | -0.4408764 | 0.50506372 | -0.8729124 | 0.38271083 | 0.8718674 |
| FAM110B | 770.189425 | 0.43569425 | 0.20180513 | 2.15898508 | 0.03085132 | 0.54443002 |
| UBXN2B | 1873.95947 | -0.1322755 | 0.14301056 | -0.9249356 | 0.35499943 | 0.86364633 |
| SDCBP | 11600.7275 | -0.3300648 | 0.28378775 | -1.1630691 | 0.24480147 | 0.81484785 |
| NSMAF | 1291.49422 | -0.2502957 | 0.21137954 | -1.1841057 | 0.23637122 | 0.80806879 |
| TOX | 451.820784 | 0.40732434 | 0.32047128 | 1.27101667 | 0.20372272 | 0.78366254 |
| AC090152.1 | 123.504902 | 0.44085378 | 0.2931935 | 1.50362738 | 1.33E-01 | 7.13E-01 |
| CA8 | 807.763338 | -0.2989845 | 0.55185818 | -0.5417777 | 0.58797166 | 0.93289552 |
| RAB2A | 6461.75977 | -0.0610815 | 0.16319818 | -0.3742782 | 0.70819732 | 0.95501548 |
| CHD7 | 936.556619 | -0.1936957 | 0.12038907 | -1.6089142 | 0.1076351 | 0.67707349 |
| CLVS1 | 828.985096 | 0.3352031 | 0.22801178 | 1.47011308 | 0.14153113 | 0.72784886 |
| ASPH | 5474.12344 | 0.19887934 | 0.19243242 | 1.03350227 | 0.30136891 | 0.84211658 |
| GGH | 2480.98979 | 0.57917253 | 0.47783579 | 1.21207441 | 0.2254839 | 0.80214224 |
| TTPA | 53.273337 | 1.7510996 | 0.79565724 | 2.20082156 | 0.02774866 | 0.53065062 |
| YTHDF3-AS1 | 48.9380382 | 0.47257922 | 0.44610855 | 1.05933685 | 0.2894464 | 0.83338685 |
| YTHDF3 | 2876.50448 | -0.2684781 | 0.23872109 | -1.1246519 | 2.61E-01 | 0.82424998 |
| AC011978.2 | 58.2713681 | -0.0405139 | 0.2887904 | -0.1402882 | 0.88843226 | 0.98423722 |
| CYP7B1 | 146.421602 | 0.28265256 | 0.32356473 | 0.87355802 | 0.38235899 | 0.87185666 |
| AC100814.1 | 59.9290749 | 0.59603815 | 0.37317923 | 1.59719005 | 0.11022335 | 0.67996643 |
| ARMC1 | 1479.6663 | -0.2254796 | 0.22804273 | -0.9887604 | 0.3227804 | 0.85119105 |
| MTFR1 | 2263.9437 | 0.099885 | 0.30888131 | 0.32337663 | 0.74641003 | 0.96200195 |
| AC055822.1 | 44.0739822 | -0.592006 | 0.4527965 | -1.3074439 | 0.19106199 | 0.77292503 |
| PDE7A | 1693.88008 | -0.17019 | 0.38293319 | -0.4444379 | 0.65672604 | 0.94545756 |
| TRIM55 | 420.301516 | 0.23625879 | 0.45057024 | 0.52435506 | 0.60003161 | 0.93487648 |
| RRS1 | 860.039323 | -0.5159268 | 0.37068404 | -1.3918235 | 0.16397586 | 0.75265041 |
| ADHFE1 | 1628.43077 | 0.64132879 | 0.30446682 | 2.1063996 | 0.03516966 | 0.55369025 |
| VXN | 680.989077 | 0.38958084 | 0.35339089 | 1.10240768 | 0.27028447 | 0.82716979 |
| MYBL1 | 137.838862 | 0.34508406 | 0.33345401 | 1.03487751 | 0.30072612 | 0.84200819 |
| VCPIP1 | 1230.20298 | -0.2522274 | 0.18773954 | -1.3434965 | 0.17911127 | 0.76120583 |
| SGK3 | 1082.25921 | 0.36516723 | 0.26077641 | 1.40030776 | 0.16142118 | 0.74936693 |
| C8orf44 | 442.154015 | 0.09723032 | 0.3418604 | 0.28441527 | 0.77609216 | 0.96836854 |
| MCMDC2 | 680.947103 | -0.0135027 | 0.16256481 | -0.0830606 | 9.34E-01 | 9.90E-01 |
| PPP1R42 | 950.524283 | 0.14132003 | 0.12658548 | 1.11640002 | 0.26425094 | 0.82424998 |
| COPS5 | 3276.5924 | -0.0087413 | 0.10381086 | -0.0842041 | 0.93289415 | 0.98999825 |

| | | | | | | |
|---|---|---|---|---|---|---|
| CSPP1 | 991.575718 | -0.2610362 | 0.20850432 | -1.2519461 | 0.21058952 | 0.79134644 |
| ARFGEF1 | 3926.31427 | -0.001748 | 0.12822045 | -0.0136325 | 0.98912314 | 0.99796268 |
| AC021321.1 | 39.0881602 | 0.14547668 | 0.51564913 | 0.28212339 | 0.7778489 | 0.96836854 |
| PREX2 | 399.531453 | -0.0677852 | 0.46928475 | -0.1444437 | 0.8851501 | 0.98356741 |
| SULF1 | 1370.5602 | -0.0935832 | 0.39172021 | -0.2389033 | 0.81118059 | 0.97352497 |
| NCOA2 | 2325.45711 | -0.1238185 | 0.16108596 | -0.7686486 | 0.44210195 | 0.89331461 |
| TRAM1 | 13002.0443 | -0.1695659 | 0.2040563 | -0.8309759 | 0.40598726 | 0.88132015 |
| LACTB2-AS1 | 1352.33427 | 0.63614828 | 0.29200204 | 2.17857474 | 0.02936327 | 0.53635197 |
| LACTB2 | 3237.79189 | 0.58744695 | 0.28895914 | 2.03297587 | 0.04205496 | 0.56438286 |
| XKR9 | 49.4468268 | 0.38931949 | 0.85188593 | 0.45700895 | 0.64766461 | 0.94282139 |
| MSC-AS1 | 448.553649 | 1.08490346 | 0.3704757 | 2.92840649 | 0.00340704 | 0.25438802 |
| MSC | 585.996742 | 1.12332542 | 0.5458933 | 2.05777469 | 3.96E-02 | 5.57E-01 |
| TRPA1 | 119.620001 | 0.85723165 | 0.47110681 | 1.81961209 | 0.0688181 | 0.62760097 |
| TERF1 | 1536.4272 | 0.14796887 | 0.17235139 | 0.85853017 | 0.39059978 | 0.87677931 |
| AC022893.1 | 114.794878 | 0.2362217 | 0.42540537 | 0.55528613 | 0.57869896 | 0.93139743 |
| SBSPON | 1187.88197 | 0.17883118 | 0.32241281 | 0.55466524 | 0.57912366 | 0.93139743 |
| RPL7 | 34207.3044 | 0.32420981 | 0.27379333 | 1.18414064 | 0.23635739 | 0.80806879 |
| RDH10 | 5780.42273 | -0.8381419 | 0.42260374 | -1.9832808 | 0.04733607 | 0.58218854 |
| RDH10-AS1 | 144.653128 | -1.3585335 | 0.57203306 | -2.3749212 | 0.0175527 | 0.46859699 |
| STAU2 | 1863.92862 | -0.0108096 | 0.17506913 | -0.0617449 | 0.950766 | 0.99170503 |
| AC022826.2 | 430.969215 | 0.12663065 | 0.23918364 | 0.52942855 | 0.5965082 | 0.93361952 |
| UBE2W | 1997.47391 | 0.11993417 | 0.12666278 | 0.94687775 | 0.34370108 | 0.85952787 |
| ELOC | 2191.58904 | -0.3162573 | 0.25214265 | -1.2542793 | 0.20974052 | 0.79134644 |
| TMEM70 | 1397.22086 | -0.0853087 | 0.24154109 | -0.3531852 | 0.72394957 | 0.95810643 |
| LY96 | 147.251465 | -0.4582795 | 0.50267285 | -0.9116854 | 0.36193437 | 0.86528137 |
| JPH1 | 157.82823 | 0.33078641 | 0.53352719 | 0.61999916 | 0.53525834 | 0.92084006 |
| GDAP1 | 473.576193 | 0.0170905 | 0.26888089 | 0.06356159 | 0.94931932 | 0.99170503 |
| PI15 | 657.147028 | 0.59071386 | 0.76146933 | 0.77575529 | 0.43789349 | 0.89143622 |
| CRISPLD1 | 107.187975 | 0.56759443 | 0.52466323 | 1.08182622 | 0.27932976 | 0.82825734 |
| HNF4G | 820.630601 | 0.96530819 | 0.51622921 | 1.86992169 | 0.06149469 | 0.61037866 |
| ZFHX4 | 51.9446813 | 0.99962154 | 0.54991834 | 1.81776359 | 0.06910027 | 0.62779744 |
| PEX2 | 2176.12183 | 0.15356424 | 0.09056323 | 1.69565774 | 0.08995072 | 0.65375543 |
| PKIA | 337.320179 | -0.1904343 | 0.38784166 | -0.4910105 | 0.623419 | 0.94045607 |
| ZC2HC1A | 526.010177 | 0.08688418 | 0.21072508 | 0.41231056 | 0.68011181 | 0.95070301 |
| IL7 | 308.62123 | 0.18799029 | 0.25908575 | 0.72559103 | 0.46808952 | 0.89984974 |
| AC083837.1 | 44.8396088 | -0.5577623 | 0.85085656 | -0.6555303 | 0.51212638 | 0.91551505 |
| HEY1 | 219.695892 | -0.2568197 | 0.61009967 | -0.4209472 | NA | NA |
| MRPS28 | 1475.15742 | 0.2602128 | 0.14073351 | 1.84897538 | 0.06446137 | 0.61989227 |
| AC036214.3 | 2214.473 | 0.23160745 | 0.14112197 | 1.64118635 | 0.10075874 | 0.67247782 |
| TPD52 | 3743.83174 | 0.22114396 | 0.15902951 | 1.39058445 | 0.16435147 | 0.75286112 |
| AC009812.1 | 198.253765 | 0.19375405 | 0.35164659 | 0.55099084 | 0.58163995 | 0.9317705 |
| ZBTB10 | 1692.17849 | -0.0150008 | 0.1481922 | -0.1012256 | 0.91937139 | 0.9872458 |
| AC009812.4 | 158.375745 | 0.15769283 | 0.2937109 | 0.5368981 | 0.591338 | 0.93289552 |
| ZNF704 | 1816.87704 | 0.34640605 | 0.18043031 | 1.91988837 | 0.054872 | 0.59342538 |
| PAG1 | 548.708581 | -0.2560034 | 0.24185953 | -1.0584798 | 0.28983676 | 0.83399826 |
| AC009902.2 | 99.8874952 | -0.1854708 | 0.64994322 | -0.2853646 | 0.77536481 | 0.96836854 |
| FABP5 | 216.375855 | -0.2442535 | 0.59328108 | -0.4116994 | 0.68055974 | 0.95070301 |
| AC018616.1 | 71.1492906 | -0.3230677 | 0.46953681 | -0.6880562 | 0.49141737 | 0.90600166 |
| FABP4 | 474.950731 | 3.21591098 | 0.89029439 | 3.6121883 | 0.00030362 | 0.08492861 |
| AC023644.1 | 71.4120571 | -0.3295746 | 0.47259275 | -0.6973754 | 0.48556786 | 0.90543654 |
| IMPA1 | 1465.50684 | -0.1776353 | 0.17538423 | -1.0128353 | 0.31113882 | 0.84476049 |
| SLC10A5 | 42.8040873 | 0.30140057 | 0.3253594 | 0.92636197 | 3.54E-01 | 0.86364633 |
| ZFAND1 | 1983.66328 | 0.15178551 | 0.12297658 | 1.23426361 | 0.21710469 | 0.79574248 |
| CHMP4C | 1058.76365 | -0.05668 | 0.21064074 | -0.2690836 | 0.78786535 | 0.97090793 |

| | | | | | | |
|---|---|---|---|---|---|---|
| AC132219.1 | 55.2100067 | -0.5263664 | 0.29605506 | -1.7779341 | 0.07541469 | 0.63773415 |
| SNX16 | 612.277835 | -0.6119651 | 0.19718032 | -3.103581 | 0.00191194 | 0.18915498 |
| RALYL | 390.60879 | 0.80114984 | 0.56331807 | 1.42219799 | 0.15496878 | 0.74373109 |
| LRRCC1 | 390.105416 | 0.34466243 | 0.25382636 | 1.35786694 | 0.17450591 | 0.75974924 |
| E2F5 | 1860.88247 | -0.085332 | 0.13056811 | -0.6535442 | 0.51340551 | 0.91609375 |
| C8orf59 | 1696.33641 | -0.108108 | 0.153409 | -0.7047045 | 0.48099415 | 0.90541935 |
| CA13 | 652.356782 | -0.270212 | 0.27249753 | -0.9916125 | 0.3213866 | 0.85119105 |
| CA1 | 69.5744016 | -3.815913 | 0.52944026 | -7.2074477 | 5.70E-13 | 1.07E-08 |
| CA3 | 250.01537 | 0.75334214 | 0.42162308 | 1.78676684 | 0.07397518 | 0.6374448 |
| CA3-AS1 | 118.675305 | 0.49716086 | 0.32243752 | 1.54188281 | 1.23E-01 | 6.98E-01 |
| CA2 | 9979.91937 | 0.25417221 | 0.26221755 | 0.96931806 | 0.33238652 | 0.85496237 |
| ATP6V0D2 | 2240.13767 | 0.62541095 | 0.58735958 | 1.06478377 | 0.28697379 | 0.83155545 |
| AC023194.3 | 892.913858 | 0.65947621 | 0.59051388 | 1.11678359 | 0.26408687 | 0.82424998 |
| SLC7A13 | 947.322858 | 2.33223323 | 1.96890284 | 1.18453444 | 0.23620157 | 0.80802614 |
| WWP1 | 3561.42357 | 0.09441514 | 0.17152909 | 0.55043221 | 0.58202296 | 0.9317705 |
| RMDN1 | 2625.64124 | 0.29165701 | 0.18529903 | 1.57398024 | 1.15E-01 | 0.6873836 |
| CPNE3 | 4186.55934 | 0.09414039 | 0.14766512 | 0.63752624 | 0.52378212 | 0.91723878 |
| AF117829.1 | 304.033866 | -0.2505192 | 0.29921447 | -0.8372562 | 0.4024486 | 0.88132015 |
| RIPK2 | 754.97264 | -0.4221562 | 0.31857247 | -1.3251495 | 0.18512156 | 0.76820626 |
| OSGIN2 | 1651.07871 | -0.4771751 | 0.40188619 | -1.187339 | 0.23509394 | 0.80747753 |
| NBN | 2141.15024 | -0.137941 | 0.19463555 | -0.7087143 | 0.4785018 | 0.90440142 |
| DECR1 | 6529.00737 | 0.26657045 | 0.22915734 | 1.16326386 | 0.24472246 | 0.81477059 |
| CALB1 | 7108.89616 | 2.55028175 | 0.56395888 | 4.52210584 | 6.12E-06 | 0.00764975 |
| TMEM64 | 1577.38975 | -0.1850978 | 0.2632171 | -0.7032136 | 0.48192263 | 0.90541935 |
| NECAB1 | 463.355002 | 0.00746832 | 0.25019942 | 0.02984947 | 0.97618711 | 0.99455099 |
| C8orf88 | 218.466872 | -0.043515 | 0.29417636 | -0.1479215 | 0.8824047 | 0.98275231 |
| PIP4P2 | 894.620948 | -0.1582002 | 0.18176321 | -0.8703644 | 0.38410131 | 0.87296175 |
| AC087439.1 | 195.256861 | -0.1961906 | 0.2256838 | -0.8693162 | 0.38467422 | 0.87312908 |
| OTUD6B-AS1 | 1438.1879 | 0.35920467 | 0.18685071 | 1.92241528 | 0.05455352 | 0.59329767 |
| OTUD6B | 486.220652 | 0.00766401 | 0.2083472 | 0.03678482 | 0.97065658 | 0.99400331 |
| LRRC69 | 75.6978418 | -0.4632926 | 0.31011486 | -1.4939386 | 0.13519167 | 0.71649038 |
| SLC26A7 | 1505.11494 | 2.47792748 | 0.59322892 | 4.17701733 | 2.95E-05 | 0.01976885 |
| RUNX1T1 | 216.039785 | 0.10739421 | 0.45945781 | 0.23374118 | 0.81518592 | 0.9738447 |
| TRIQK | 1196.60317 | 0.25270871 | 0.13849288 | 1.82470545 | 0.06804549 | 0.62603855 |
| FAM92A | 1760.35911 | -0.2417808 | 0.16645243 | -1.4525519 | 0.1463482 | 0.73247558 |
| AC120053.1 | 213.653667 | 0.30061296 | 0.35982179 | 0.83544958 | 0.40346462 | 0.88132015 |
| RBM12B | 856.781251 | -0.1609548 | 0.17106034 | -0.940924 | 0.3467438 | 0.86135217 |
| AC010834.1 | 246.948784 | -0.0812243 | 0.18413989 | -0.441101 | 6.59E-01 | 9.46E-01 |
| TMEM67 | 644.698814 | -0.1407265 | 0.22104529 | -0.6366411 | 0.52435863 | 0.91723878 |
| PDP1 | 1402.05324 | -0.3956126 | 0.40259863 | -0.9826477 | 0.32578088 | 0.85271781 |
| GEM | 2654.09215 | -0.4995453 | 0.48882574 | -1.0219291 | 0.30681444 | 0.84286067 |
| RAD54B | 287.288804 | -0.432501 | 0.59978097 | -0.7210983 | 0.47084903 | 0.90089741 |
| FSBP | 265.120134 | -0.4394928 | 0.61559865 | -0.7139275 | 4.75E-01 | 0.90226631 |
| VIRMA | 1620.13913 | -0.054514 | 0.13367045 | -0.4078239 | 6.83E-01 | 9.51E-01 |
| AC108860.2 | 84.7789851 | 0.18062129 | 0.54416557 | 0.3319234 | 7.40E-01 | 9.60E-01 |
| ESRP1 | 1201.52215 | 0.03872076 | 0.28490596 | 0.13590715 | 8.92E-01 | 9.85E-01 |
| DPY19L4 | 1424.0847 | 0.2563061 | 0.15701084 | 1.63241024 | 0.10259309 | 0.67335541 |
| AP003692.1 | 521.141121 | 0.3392873 | 0.17356857 | 1.95477387 | 0.05060977 | 0.5861488 |
| INTS8 | 1216.07897 | 0.02270379 | 0.20155942 | 0.11264066 | 0.91031545 | 0.98662757 |
| CCNE2 | 313.716684 | -0.043858 | 0.24583431 | -0.1784048 | 0.85840504 | 0.97982623 |
| NDUFAF6 | 593.736932 | 0.24127406 | 0.17457567 | 1.38206006 | 0.16695326 | 0.75558932 |
| TP53INP1 | 2068.79587 | -0.0358962 | 0.23412616 | -0.15332 | 0.87814592 | 0.98275231 |
| PLEKHF2 | 1362.56121 | 0.15309332 | 0.31788832 | 0.48159467 | 0.63009391 | 0.94088547 |
| C8orf37 | 186.721535 | 0.11618878 | 0.24930646 | 0.46604802 | 0.64118113 | 0.94088547 |

| | | | | | | |
|---|---|---|---|---|---|---|
| UQCRB | 10525.2984 | 0.20624072 | 0.17657438 | 1.16801043 | 0.24280256 | 0.8133895 |
| AP003465.1 | 2992.22294 | 0.27140411 | 0.17587741 | 1.54314368 | 0.12279592 | 0.69788192 |
| MTERF3 | 845.137081 | -0.0176687 | 0.11621263 | -0.1520377 | 0.87915721 | 0.98275231 |
| PTDSS1 | 3097.54694 | 0.27405476 | 0.21988116 | 1.24637671 | 0.21262613 | 0.79203904 |
| SDC2 | 5462.05184 | -0.1131185 | 0.19303121 | -0.5860114 | 0.55786784 | 0.92421175 |
| CPQ | 3803.10131 | 0.50176321 | 0.21202238 | 2.36655777 | 0.01795437 | 0.47458799 |
| TSPYL5 | 946.75502 | -0.2613624 | 0.22894144 | -1.1416124 | 0.25361517 | 0.8188642 |
| MTDH | 3852.24963 | -0.1897587 | 0.2632923 | -0.7207152 | 0.47108479 | 0.90106144 |
| LAPTM4B | 8199.43205 | 0.02562124 | 0.27637699 | 0.09270396 | 0.92613875 | 0.98876911 |
| MATN2 | 2569.30094 | 0.6472753 | 0.46163456 | 1.40213787 | 0.16087408 | 0.74874703 |
| RPL30 | 16841.666 | 0.13051748 | 0.18921243 | 0.68979335 | 0.49032415 | 0.90600166 |
| AP003352.1 | 11586.8168 | 0.13399432 | 0.18855044 | 0.71065505 | 0.47729802 | 0.90372319 |
| ERICH5 | 346.273883 | 0.07044974 | 0.35984033 | 0.19578055 | 0.84478193 | 0.97675487 |
| RIDA | 6800.19934 | 0.45508523 | 0.39319728 | 1.15739667 | 0.24711033 | 0.8156466 |
| POP1 | 354.392749 | 0.05443867 | 0.23873438 | 0.22803027 | 0.81962271 | 0.97454841 |
| NIPAL2 | 856.136006 | 0.19472775 | 0.25339026 | 0.76848949 | 0.44219643 | 0.89331461 |
| STK3 | 1616.37682 | -0.229953 | 0.226614 | -1.0147343 | 0.31023248 | 0.84456632 |
| OSR2 | 105.010317 | 0.05568091 | 0.8275047 | 0.06728773 | 0.94635265 | 0.99166614 |
| AC104986.2 | 83.749375 | 0.62277772 | 0.31919453 | 1.95109141 | 0.05104617 | 0.5861488 |
| VPS13B | 1218.84686 | -0.0504915 | 0.1161908 | -0.434557 | 0.66388401 | 0.94642611 |
| COX6C | 6094.21299 | 0.0898041 | 0.19467479 | 0.46130316 | 0.64458112 | 0.94208859 |
| POLR2K | 1795.49917 | -0.0067844 | 0.0814106 | -0.0833353 | 0.93358495 | 0.99019225 |
| SPAG1 | 1026.71892 | -0.4524128 | 0.26557259 | -1.7035373 | 0.08846757 | 0.65363785 |
| RNF19A | 4567.94602 | -0.2840141 | 0.21045354 | -1.3495335 | 0.17716566 | 0.76115956 |
| AP000424.1 | 91.7046404 | -0.1511675 | 0.71082523 | -0.2126648 | 0.83158841 | 0.97550416 |
| AP000424.2 | 72.3737763 | -0.3322565 | 0.9835505 | -0.3378134 | 0.73550382 | 0.95895327 |
| ANKRD46 | 1425.80748 | 0.39608412 | 0.19185639 | 2.0644823 | 0.03897201 | 0.55613287 |
| PABPC1 | 19703.079 | -0.0017448 | 0.30891171 | -0.0056482 | 0.99549339 | 0.99940889 |
| YWHAZ | 21356.5852 | -0.1896316 | 0.22263151 | -0.8517734 | 0.39433987 | 0.87884163 |
| AP003469.4 | 402.743899 | 0.14357585 | 0.29375669 | 0.48875772 | 0.62501323 | 0.94088547 |
| ZNF706 | 2351.11295 | -0.1139052 | 0.20365699 | -0.5592994 | 0.57595744 | 0.93083012 |
| GRHL2 | 678.303181 | 0.14738124 | 0.41814855 | 0.35246144 | 0.72449223 | 0.95828829 |
| NCALD | 898.676677 | 0.23462202 | 0.3595524 | 0.65253916 | 0.51405342 | 0.91636259 |
| AP000426.1 | 129.176967 | 0.32788143 | 0.38516028 | 0.85128568 | 0.39461068 | 0.87884163 |
| RRM2B | 2019.65332 | 0.42507965 | 0.16643642 | 2.55400623 | 0.01064914 | 0.40196898 |
| UBR5-AS1 | 420.316931 | 0.23811747 | 0.24842759 | 0.95849849 | 3.38E-01 | 0.8571036 |
| UBR5 | 3186.38849 | -0.1100579 | 0.11622441 | -0.9469427 | 0.34366796 | 0.85952787 |
| AP002907.1 | 490.281069 | -0.106937 | 0.15114023 | -0.7075349 | 0.47923414 | 0.90480914 |
| KLF10 | 4331.11016 | 0.16166222 | 0.37163556 | 0.43500203 | 0.66356095 | 0.94642611 |
| AZIN1 | 3893.40547 | -0.0073791 | 0.19079026 | -0.0386767 | 0.96914816 | 0.99400331 |
| AZIN1-AS1 | 243.837574 | -0.1206596 | 0.37606499 | -0.3208478 | 0.74832575 | 0.96241535 |
| AP003354.2 | 57.7337653 | -0.3768745 | 0.41683861 | -0.9041257 | 0.36592874 | 0.86672412 |
| AP003550.1 | 269.197455 | -0.4743761 | 0.35236157 | -1.3462766 | 0.17821332 | 0.76115956 |
| ATP6V1C1 | 5884.65672 | -0.4483237 | 0.31850945 | -1.407568 | 0.15925904 | 0.74762924 |
| BAALC | 199.165426 | -0.3582175 | 0.33938667 | -1.0554849 | 0.29120362 | 0.83454586 |
| BAALC-AS1 | 88.2288995 | 0.18348686 | 0.28907592 | 0.63473586 | 0.5256007 | 0.91759401 |
| AC025370.1 | 66.4909433 | -0.2929763 | 0.43778548 | -0.6692235 | 0.50335291 | 0.91356552 |
| FZD6 | 1601.71489 | -0.1375448 | 0.14767059 | -0.9314298 | 0.3516313 | 0.86364633 |
| CTHRC1 | 151.660925 | 0.26840156 | 0.40075336 | 0.66974252 | 0.50302194 | 0.91356552 |
| SLC25A32 | 1156.56541 | -0.2869369 | 0.29309799 | -0.9789793 | 0.32759021 | 0.85317073 |
| DCAF13 | 2019.98982 | -0.0521813 | 0.28539643 | -0.182838 | 0.85492517 | 0.97910552 |
| RIMS2 | 54.7691186 | 0.19353465 | 0.64511701 | 0.29999929 | 7.64E-01 | 9.67E-01 |
| DPYS | 8950.87612 | 1.26595214 | 0.63555061 | 1.99189824 | 4.64E-02 | 5.79E-01 |
| LRP12 | 507.090171 | -0.1904988 | 0.22022919 | -0.8650023 | 3.87E-01 | 8.75E-01 |

| | | | | | | |
|---|---|---|---|---|---|---|
| ZFPM2 | 151.112454 | -0.060086 | 0.35688567 | -0.168362 | 0.86629852 | 0.98200144 |
| AC027031.2 | 542.634687 | -1.1093249 | 0.33041467 | -3.3573718 | 0.00078687 | 0.13166762 |
| OXR1 | 2071.93116 | -0.2510928 | 0.19390354 | -1.2949366 | 0.19534211 | 0.77730222 |
| ANGPT1 | 434.430578 | 0.25198894 | 0.50271077 | 0.50126026 | 0.61618797 | 0.93910158 |
| EIF3E | 8833.08263 | 0.31918585 | 0.20195601 | 1.58047214 | 1.14E-01 | 0.68214043 |
| EMC2 | 2144.88814 | 0.31492463 | 0.22236695 | 1.41623845 | 0.1567057 | 0.74522306 |
| NUDCD1 | 1217.47752 | -0.0786102 | 0.15370329 | -0.5114412 | 0.60904217 | 0.93748023 |
| ENY2 | 2701.05551 | -0.1779006 | 0.13108015 | -1.3571896 | 0.17472099 | 0.76026144 |
| PKHD1L1 | 103.656703 | 0.85546447 | 0.83544629 | 1.02396106 | 0.30585367 | 0.84270033 |
| EBAG9 | 1239.9437 | 0.13282289 | 0.17429887 | 0.76204102 | 0.44603552 | 0.89480207 |
| SYBU | 328.225195 | -0.1398447 | 0.47760575 | -0.2928037 | 0.7696722 | 0.96836854 |
| KCNV1 | 46.0390387 | -0.1390273 | 0.65352505 | -0.2127345 | 0.83153409 | 0.97550416 |
| TRPS1 | 915.559519 | -0.0723737 | 0.22883078 | -0.3162759 | 0.75179313 | 0.96289476 |
| EIF3H | 7027.10165 | 0.24397982 | 0.16090904 | 1.51625929 | 0.1294538 | 0.70764335 |
| UTP23 | 888.700005 | -0.010705 | 0.16696734 | -0.0641145 | 0.94887908 | 0.99170503 |
| RAD21 | 5772.28121 | -0.0575104 | 0.12922508 | -0.4450402 | 0.65629072 | 0.94545756 |
| RAD21-AS1 | 66.73702 | 0.32867022 | 0.44930007 | 0.73151607 | 0.46446399 | 0.89839196 |
| AARD | 55.9432058 | -0.4188392 | 0.46565241 | -0.8994674 | 0.36840374 | 0.86762749 |
| SLC30A8 | 150.032388 | 0.89731394 | 0.88221834 | 1.01711096 | 0.30910064 | 0.84414094 |
| MED30 | 303.699417 | 0.09721566 | 0.2411511 | 0.40313174 | 0.68685131 | 0.9510919 |
| EXT1 | 1851.08213 | 0.04831885 | 0.13458299 | 0.35902645 | 0.7195753 | 0.9566409 |
| SAMD12 | 930.928476 | 0.09214912 | 0.23832246 | 0.38665732 | 0.69900992 | 0.95299571 |
| TNFRSF11B | 3553.64425 | 0.64523725 | 0.38141699 | 1.69168459 | 0.09070613 | 0.65495306 |
| COLEC10 | 40.7603825 | 0.85686038 | 0.46846498 | 1.82908094 | 0.06738748 | 0.6249306 |
| MAL2 | 6491.71403 | 0.07537549 | 0.2566862 | 0.29364838 | 0.7690266 | 0.96818001 |
| MAL2-AS1 | 2000.90803 | 0.09217205 | 0.252827 | 0.36456569 | 0.71543563 | 0.95590747 |
| NOV | 1175.94671 | -1.1078262 | 0.63788084 | -1.736729 | 0.08243501 | 0.64661469 |
| ENPP2 | 7132.44004 | -0.0800597 | 0.54923505 | -0.1457658 | 0.88410627 | 0.98317437 |
| TAF2 | 1120.15925 | -0.0287183 | 0.15067374 | -0.1905992 | 0.84883964 | 0.97741457 |
| DSCC1 | 71.0638769 | 0.38119747 | 0.36983588 | 1.03072064 | 0.30267184 | 0.84211658 |
| AP005717.1 | 673.198796 | -0.4794744 | 0.35874003 | -1.336551 | 0.18136923 | 0.76522169 |
| DEPTOR | 6432.96845 | -0.4192119 | 0.36493621 | -1.1487264 | 0.25066883 | 0.81743249 |
| AC091563.1 | 641.762528 | -0.2645939 | 0.29516942 | -0.8964135 | 0.37003194 | 0.86762749 |
| COL14A1 | 2200.70523 | 0.4358926 | 0.82974937 | 0.52533044 | 0.5993535 | 0.93487648 |
| MRPL13 | 1771.26626 | 0.31828322 | 0.17781603 | 1.78995797 | 0.07346067 | 0.6374448 |
| MTBP | 312.347192 | 0.05333196 | 0.29180553 | 0.18276542 | 0.85498208 | 0.97910552 |
| SNTB1 | 2141.33912 | 0.12322029 | 0.22489805 | 0.54789397 | 0.58376471 | 0.93233498 |
| AC104958.2 | 164.13774 | -0.0802318 | 0.5370239 | -0.1494008 | 0.88123741 | 0.98275231 |
| HAS2 | 94.5519356 | 0.75452708 | 0.92768399 | 0.81334494 | 0.41602032 | 0.88399703 |
| ZHX2 | 1787.76361 | 0.14506038 | 0.16633264 | 0.87211014 | 0.3831483 | 0.8718674 |
| AC104316.1 | 385.431433 | -0.1096294 | 0.2754929 | -0.3979389 | 0.6906752 | 0.95158502 |
| DERL1 | 3862.55671 | -0.1039124 | 0.16179838 | -0.6422338 | 0.52072136 | 0.91683945 |
| TBC1D31 | 600.351163 | -0.0664279 | 0.19221995 | -0.3455825 | 0.72965649 | 0.95848028 |
| C8orf76 | 684.067603 | -0.1929399 | 0.17563307 | -1.0985397 | 2.72E-01 | 0.82716979 |
| ZHX1-C8orf7( | 634.583809 | -0.1526126 | 0.17832898 | -0.8557927 | 0.39211247 | 0.87765297 |
| ZHX1 | 1471.31357 | -0.0080861 | 0.12533679 | -0.0645151 | 9.49E-01 | 0.99170503 |
| ATAD2 | 249.427465 | 0.03215584 | 0.31328415 | 0.10264113 | 0.9182478 | 0.9868044 |
| WDYHV1 | 474.573916 | 0.41149045 | 0.25398497 | 1.62013702 | 0.10520285 | 0.67473222 |
| FBXO32 | 900.624179 | -0.5889682 | 0.23243209 | -2.5339367 | 0.01127891 | 0.40196898 |
| ANXA13 | 145.030487 | -0.0183037 | 0.76962843 | -0.0237825 | 0.98102611 | 0.99576712 |
| FAM91A1 | 2497.71117 | -0.1350097 | 0.22913102 | -0.5892251 | 0.55571029 | 0.92373235 |
| FER1L6 | 87.477777 | 0.97975827 | 0.69505982 | 1.4096028 | 0.158657 | 0.74685158 |
| TMEM65 | 1423.32389 | 0.2037602 | 0.14408948 | 1.41412267 | 0.15732589 | 0.74522306 |
| TRMT12 | 531.329438 | -0.2766882 | 0.159535 | -1.734342 | 0.08285742 | 0.64680474 |

| | | | | | | |
|---|---|---|---|---|---|---|
| RNF139-AS1 | 313.746413 | -0.0149404 | 0.32555941 | -0.0458915 | 0.96339669 | 0.99320711 |
| RNF139 | 1383.85762 | -0.0627542 | 0.22165902 | -0.2831116 | 7.77E-01 | 9.68E-01 |
| TATDN1 | 3203.74518 | 0.20335322 | 0.21727856 | 0.93591021 | 0.34931942 | 0.86364633 |
| AC090198.1 | 424.707161 | 0.00441008 | 0.12315357 | 0.03580962 | 0.97143416 | 0.99404213 |
| NDUFB9 | 7324.98245 | 0.08335455 | 0.22102391 | 0.3771291 | 0.70607767 | 0.95501548 |
| MTSS1 | 3451.96021 | -0.0777292 | 0.34406143 | -0.2259166 | 0.82126628 | 0.97472105 |
| ZNF572 | 113.217639 | 0.32501859 | 0.32924622 | 0.98715968 | 0.32356436 | 0.85143496 |
| SQLE | 1768.79147 | -0.0515474 | 0.48211906 | -0.1069184 | 0.91485373 | 0.9866558 |
| WASHC5 | 2087.74101 | 0.07317865 | 0.23466284 | 0.31184591 | 0.75515763 | 0.96360109 |
| WASHC5-AS1 | 317.5629 | 0.10288667 | 0.28260071 | 0.36407082 | 0.71580512 | 0.95590747 |
| NSMCE2 | 748.455809 | -0.0233763 | 0.16787847 | -0.1392456 | 0.88925605 | 0.98439687 |
| TRIB1 | 2897.72305 | -0.2915106 | 0.31725901 | -0.9188411 | 0.35817867 | 0.86480228 |
| LINC00861 | 94.5549329 | 0.62357131 | 0.88782299 | 0.70235995 | 0.48245472 | 0.90541935 |
| FAM84B | 5336.44423 | -0.7391127 | 0.44170652 | -1.6733118 | 0.09426593 | 0.65783452 |
| PCAT1 | 550.853067 | -0.7457989 | 0.42077323 | -1.7724485 | 0.07632014 | 0.63773415 |
| POU5F1B | 84.1468358 | 0.13817347 | 0.52165452 | 0.26487544 | 0.79110542 | 0.97090793 |
| MYC | 2287.11732 | -0.8935546 | 0.48794331 | -1.8312673 | 0.06706066 | 0.6249306 |
| PVT1 | 169.15083 | -0.0251264 | 0.54823848 | -0.0458312 | 0.96344482 | 0.99320711 |
| AC103718.1 | 104.945497 | 1.52420998 | 0.78521743 | 1.94113111 | 0.05224238 | 0.58645127 |
| AC022973.4 | 84.040903 | 0.07906384 | 0.30258726 | 0.26129269 | 0.7938668 | 0.97090793 |
| FAM49B | 2093.31995 | -0.0467061 | 0.14948848 | -0.3124395 | 0.75470652 | 0.96351771 |
| ASAP1 | 1403.92605 | -0.3408793 | 0.18197292 | -1.873242 | 0.06103498 | 0.60989294 |
| EFR3A | 2947.63308 | 0.10420598 | 0.16668763 | 0.62515722 | 0.53186787 | 0.91962754 |
| KCNQ3 | 175.680497 | 0.5848721 | 0.41151456 | 1.42126706 | 0.15523913 | 0.74407586 |
| LRRC6 | 615.674236 | 0.21120085 | 0.30363613 | 0.69557219 | 0.48669679 | 0.90573694 |
| TMEM71 | 104.181641 | 0.26123363 | 0.61441702 | 0.42517316 | 0.67071045 | 0.94922827 |
| PHF20L1 | 1613.83438 | -0.0562621 | 0.1417016 | -0.3970462 | 0.69133342 | 0.95158502 |
| AF230666.1 | 140.088868 | -0.2106204 | 0.25805263 | -0.8161916 | 0.41439058 | 0.88364943 |
| TG | 314.447775 | 0.88494731 | 0.49528529 | 1.78674256 | 0.07397911 | 0.6374448 |
| SLA | 955.393103 | -0.6210877 | 0.41932091 | -1.4811752 | 0.13855988 | 0.72333593 |
| WISP1 | 1557.87856 | 0.0639277 | 0.48893176 | 0.13074973 | 0.89597329 | 0.98474725 |
| NDRG1 | 52382.0116 | 0.4535205 | 0.53740802 | 0.84390349 | 0.39872334 | 0.88074856 |
| ST3GAL1 | 5564.88326 | 0.07300984 | 0.48612093 | 0.15018864 | 0.88061579 | 0.98275231 |
| AC103706.1 | 50.1886943 | -0.092531 | 0.369643 | -0.2503252 | 0.80233587 | 0.97245417 |
| ZFAT | 1016.64662 | 0.30460825 | 0.44619379 | 0.6826815 | 0.49480812 | 0.90816709 |
| ZFAT-AS1 | 93.6155917 | 0.69115258 | 0.55720947 | 1.24038198 | 0.21483414 | 0.79487183 |
| AC083843.3 | 1005.63508 | -0.6258879 | 0.37727954 | -1.65895 | 0.09712586 | 0.66486533 |
| KHDRBS3 | 1349.0892 | -0.3428497 | 0.34645236 | -0.9896012 | 0.32236908 | 0.85119105 |
| AC046195.1 | 54.2198272 | -0.1614118 | 1.11101626 | -0.145283 | 0.88448745 | 0.98339935 |
| FAM135B | 93.7966508 | 2.00747036 | 0.81072327 | 2.47614744 | 0.01328087 | 0.42371784 |
| COL22A1 | 196.669323 | 0.21076098 | 0.6830673 | 0.30855082 | 0.75766323 | 0.96502715 |
| TRAPPC9 | 1853.62212 | 0.16600853 | 0.25295092 | 0.65628749 | 0.51163916 | 0.91510941 |
| AC107375.1 | 229.978112 | -0.4873536 | 0.27076496 | -1.7999137 | 0.07187426 | 0.63189503 |
| CHRAC1 | 1349.39672 | -0.3990717 | 0.22552229 | -1.7695445 | 0.07680305 | 0.63773415 |
| AGO2 | 2054.06949 | -0.3474792 | 0.20840668 | -1.6673131 | 0.09545216 | 0.65888357 |
| ERICD | 65.9950379 | -0.3906784 | 0.29958988 | -1.3040441 | 0.19221853 | 0.7740178 |
| PTK2 | 4653.25699 | -0.1224065 | 0.09490425 | -1.2897893 | 0.19712382 | 0.77816059 |
| DENND3 | 3087.19603 | -0.294423 | 0.36368845 | -0.8095473 | 0.41820041 | 0.88445833 |
| AC040970.1 | 50.9579975 | -0.4359315 | 0.60062476 | -0.7257968 | 0.46796337 | 0.89984974 |
| SLC45A4 | 3421.47321 | -0.4886033 | 0.3333971 | -1.4655296 | 0.14277648 | 0.72784886 |
| PTP4A3 | 4729.90625 | -0.4522626 | 0.37911159 | -1.1929537 | 0.23288749 | 0.80667958 |
| AC100803.3 | 585.635841 | -0.3654697 | 0.55104607 | -0.6632289 | 0.50718399 | 0.91430909 |
| TSNARE1 | 1227.28037 | -0.248215 | 0.1722308 | -1.4411768 | 0.14953474 | 0.73670625 |
| ADGRB1 | 131.417096 | 0.07924516 | 0.61094625 | 0.12970889 | 0.89679675 | 0.98474725 |

| | | | | | | |
|---|---|---|---|---|---|---|
| ARC | 150.391216 | -0.0417641 | 0.58644197 | -0.071216 | 0.94322583 | 0.99136103 |
| JRK | 1113.12827 | -0.0895514 | 0.17791512 | -0.5033376 | 0.61472697 | 0.93895494 |
| PSCA | 89.3506188 | 1.14862099 | 1.0080296 | 1.13947149 | 0.25450655 | 0.81992039 |
| LY6K | 106.674324 | -0.3764051 | 0.46648117 | -0.8069031 | 0.4197223 | 0.88451372 |
| AC108002.1 | 81.3663605 | -0.2274851 | 0.42877149 | -0.5305508 | 0.59573009 | 0.9334601 |
| THEM6 | 2300.91456 | 0.19887988 | 0.14506257 | 1.37099377 | 0.1703769 | 0.75956362 |
| AC108002.2 | 101.664435 | 0.39743396 | 0.3880844 | 1.02409155 | 0.30579204 | 0.84270033 |
| LYNX1-SLUR | 237.354002 | 0.46264293 | 0.52542619 | 0.88050984 | 3.79E-01 | 0.87066232 |
| LYNX1 | 2131.90855 | 0.71180599 | 0.50208691 | 1.41769477 | 0.1562799 | 0.74522306 |
| LY6E-DT | 122.842249 | -0.1617439 | 0.74125913 | -0.2182015 | 0.82727211 | 0.97548201 |
| LY6E | 15167.8899 | -1.2434215 | 0.60290741 | -2.0623755 | 0.039172 | 0.55657615 |
| GPIHBP1 | 537.781909 | -0.2224128 | 0.81762917 | -0.2720216 | NA | NA |
| ZFP41 | 805.240574 | -0.3164499 | 0.21373601 | -1.480564 | 0.13872279 | 0.7237761 |
| AC138696.1 | 792.448087 | -0.1832496 | 0.18420024 | -0.9948389 | 0.31981462 | 0.84960209 |
| GLI4 | 1470.4372 | -0.1845329 | 0.15784769 | -1.1690568 | 0.24238076 | 0.81319028 |
| MINCR | 275.728878 | 0.38403447 | 0.2762765 | 1.39003668 | 0.16451774 | 0.75286112 |
| ZNF696 | 398.034618 | -0.0254651 | 0.23203183 | -0.1097481 | 0.91260912 | 0.98662757 |
| TOP1MT | 1205.66853 | 0.10658605 | 0.20043984 | 0.53176078 | 0.59489169 | 0.9334601 |
| RHPN1-AS1 | 58.7305867 | 0.26035543 | 0.44091342 | 0.59049106 | 0.55486148 | 0.92373235 |
| RHPN1 | 5257.58374 | -0.1488255 | 0.34620676 | -0.4298746 | 0.66728689 | 0.94830184 |
| ZC3H3 | 1940.42526 | 0.09675951 | 0.25788808 | 0.37519964 | 0.707512 | 0.95501548 |
| AC067930.4 | 72.6253165 | 1.84512174 | 0.89304614 | 2.06609901 | 0.03881913 | 0.55608529 |
| AC067930.1 | 55.9261829 | 1.87183113 | 0.93560164 | 2.00067107 | 0.04542785 | 0.57529633 |
| GSDMD | 6877.0179 | -0.2076545 | 0.35650991 | -0.5824649 | 0.56025355 | 0.92434754 |
| MROH6 | 708.643931 | 0.3437606 | 0.41317909 | 0.83198935 | 0.40541496 | 0.88132015 |
| AC067930.5 | 2894.83243 | 0.16322511 | 0.19587158 | 0.8333272 | 0.40466022 | 0.88132015 |
| NAPRT | 4638.08249 | 0.16792538 | 0.19865597 | 0.84530749 | 0.39793918 | 0.88054917 |
| AC067930.2 | 8592.57478 | -0.0119734 | 0.18560217 | -0.0645109 | 0.94856346 | 0.99170503 |
| EEF1D | 17700.6386 | -0.0886521 | 0.17340131 | -0.5112538 | 0.60917332 | 0.93748023 |
| TIGD5 | 664.906959 | -0.3856769 | 0.24328035 | -1.5853188 | 0.11289392 | 0.68124529 |
| PYCR3 | 855.171152 | -0.0492861 | 0.21225827 | -0.2321988 | 0.81638358 | 0.97388289 |
| TSTA3 | 2592.68228 | -0.1423701 | 0.22086105 | -0.6446138 | 0.51917746 | 0.91636259 |
| ZNF623 | 902.62208 | -0.1474237 | 0.15643077 | -0.9424214 | 0.34597692 | 0.86119716 |
| ZNF707 | 746.104476 | -0.2435183 | 0.18009704 | -1.3521504 | 0.1763272 | 0.76115956 |
| MAPK15 | 1478.36806 | 0.0167037 | 0.58007177 | 0.02879593 | 0.97702735 | 0.99455099 |
| AC105219.2 | 206.603893 | 0.09150948 | 0.58458016 | 0.15653881 | 0.87560833 | 0.98275231 |
| FAM83H | 2991.43396 | -0.0583232 | 0.18417212 | -0.3166777 | 0.75148814 | 0.96285385 |
| IQANK1 | 288.245616 | 0.06749857 | 0.44115951 | 0.15300265 | 0.87839619 | 0.98275231 |
| SCRIB | 6228.47301 | -0.0450108 | 0.14772111 | -0.3047011 | 0.76059381 | 0.96560687 |
| PUF60 | 6733.36524 | -0.2728042 | 0.17329591 | -1.5742101 | 0.1154389 | 0.6873836 |
| NRBP2 | 10817.9033 | -0.1240123 | 0.42550679 | -0.291446 | 0.77071023 | 0.96836854 |
| EPPK1 | 354.471683 | 0.29037006 | 0.53669405 | 0.54103462 | 0.58848372 | 0.93289552 |
| PLEC | 17137.7801 | -0.2520992 | 0.25439319 | -0.9909827 | 0.32169404 | 0.85119105 |
| PARP10 | 9094.89016 | -0.9322389 | 0.57907687 | -1.6098707 | 0.10742609 | 0.67707349 |
| GRINA | 14528.8155 | -0.2027728 | 0.17786143 | -1.1400605 | 0.25426111 | 0.81992039 |
| SPATC1 | 61.2854558 | -0.2496366 | 0.6448694 | -0.3871119 | 0.69867336 | 0.9529028 |
| OPLAH | 3598.78619 | 0.15391005 | 0.28622827 | 0.53771787 | 0.59077184 | 0.93289552 |
| AC109322.1 | 392.496996 | -0.3884786 | 0.26122219 | -1.4871577 | 0.13697317 | 0.72167955 |
| EXOSC4 | 956.73409 | -0.4828665 | 0.23905613 | -2.0198875 | 0.04339506 | 0.56911599 |
| GPAA1 | 7121.22313 | 0.16078122 | 0.14434062 | 1.11390145 | 0.26532147 | 0.82529288 |
| CYC1 | 8102.53675 | 0.13176864 | 0.18904814 | 0.69701103 | 0.48579589 | 0.90545573 |
| SHARPIN | 3997.98239 | -0.0553366 | 0.11713121 | -0.4724323 | 0.63661824 | 0.94088547 |
| MAF1 | 6277.6801 | -0.317131 | 0.15698222 | -2.0201714 | 0.04336562 | 0.56911599 |
| WDR97 | 337.366824 | -0.3797765 | 0.25183546 | -1.5080343 | 0.13154575 | 0.71087052 |

| | | | | | | |
|---|---|---|---|---|---|---|
| HGH1 | 1243.57935 | 0.04425757 | 0.12712909 | 0.34813095 | 0.72774184 | 0.95848028 |
| MROH1 | 4830.04108 | -0.5105205 | 0.23915185 | -2.1347128 | 0.03278448 | 0.55104386 |
| BOP1 | 2709.3879 | -0.4271339 | 0.23038319 | -1.8540152 | 0.06373699 | 0.61812911 |
| SCX | 195.933258 | -0.4713453 | 0.37187699 | -1.2674765 | 0.20498496 | 0.78502176 |
| HSF1 | 5481.33239 | -0.2837991 | 0.12359537 | -2.2961953 | 0.02166472 | 0.51394743 |
| DGAT1 | 3086.93053 | -0.018992 | 0.1795279 | -0.1057888 | 0.91574991 | 0.9868044 |
| AC233992.1 | 621.892214 | -0.1516649 | 0.12600714 | -1.2036211 | 0.22873607 | 0.80434391 |
| AC233992.2 | 231.244177 | -0.1015972 | 0.20463346 | -0.4964838 | 6.20E-01 | 0.93945324 |
| SLC52A2 | 2765.49935 | -0.2437686 | 0.24944926 | -0.977227 | 0.32845677 | 0.85317073 |
| FBXL6 | 1918.32504 | -0.1429089 | 0.18388906 | -0.7771474 | 0.43707178 | 0.89130865 |
| ADCK5 | 709.294658 | 0.06795455 | 0.16998186 | 0.39977532 | 0.68932201 | 0.95158502 |
| CPSF1 | 6597.04706 | -0.2181787 | 0.30782789 | -0.7087686 | 0.4784681 | 0.90440142 |
| SLC39A4 | 3584.10583 | 0.51199598 | 0.27843382 | 1.83884266 | 0.06593833 | 0.62237346 |
| VPS28 | 7449.00716 | -0.0049594 | 0.15885913 | -0.0312191 | 0.9750948 | 0.99455099 |
| TONSL | 546.968975 | -0.0411582 | 0.21242658 | -0.1937526 | 0.84636957 | 0.97681836 |
| CYHR1 | 4490.20387 | -0.0074442 | 0.11500339 | -0.0647304 | 0.94838867 | 0.99170503 |
| AC084125.1 | 714.7042 | 0.43933089 | 0.26972133 | 1.62883259 | 0.10334846 | 0.67439401 |
| KIFC2 | 4402.91515 | -0.0768964 | 0.24067649 | -0.319501 | 0.74934661 | 0.96249332 |
| FOXH1 | 1626.87067 | 0.01457318 | 0.25479429 | 0.05719587 | 0.95438917 | 0.99170503 |
| PPP1R16A | 6929.13111 | -0.0935888 | 0.21426075 | -0.4367987 | 0.66225732 | 0.94642611 |
| GPT | 2974.85391 | 0.01666857 | 0.55090137 | 0.03025691 | 0.97586216 | 0.99455099 |
| MFSD3 | 1732.62669 | 0.18692289 | 0.20626635 | 0.90622098 | 0.3648189 | 0.8663515 |
| RECQL4 | 555.45399 | -0.3092929 | 0.27165978 | -1.1385304 | 0.25489907 | 0.81992039 |
| AC084125.4 | 141.718758 | -0.1194013 | 0.32130047 | -0.3716188 | 0.71017669 | 0.95524448 |
| LRRC14 | 2002.03065 | -0.0331305 | 0.14618729 | -0.2266307 | 0.82071089 | 0.97472105 |
| LRRC24 | 973.021184 | 0.11744123 | 0.18257077 | 0.64326418 | 0.5200527 | 0.91657872 |
| C8orf82 | 1595.47128 | 0.13123183 | 0.25026283 | 0.52437601 | 0.60001704 | 0.93487648 |
| ARHGAP39 | 841.856514 | -0.3464108 | 0.25695712 | -1.3481269 | 0.17761757 | 0.76115956 |
| ZNF251 | 1517.77824 | 0.00708253 | 0.16160819 | 0.04382529 | 0.96504367 | 0.99357855 |
| ZNF34 | 398.845109 | 0.2914556 | 0.24747158 | 1.17773362 | 0.23890282 | 0.81124801 |
| RPL8 | 41053.7755 | 0.01163285 | 0.20102742 | 0.05786699 | 0.95385458 | 0.99170503 |
| ZNF517 | 874.239748 | -0.0002753 | 0.16637247 | -0.0016545 | 0.99867989 | 0.99994357 |
| ZNF7 | 1271.18282 | -0.1050534 | 0.16125681 | -0.6514664 | 0.51474543 | 0.91636259 |
| COMMD5 | 1400.2169 | -0.2712182 | 0.26940461 | -1.0067318 | 0.31406366 | 0.84685098 |
| ZNF250 | 975.373359 | 0.21097242 | 0.26077533 | 0.80901987 | 0.41850372 | 0.88445833 |
| ZNF16 | 463.81786 | 0.1440425 | 0.22179982 | 0.6494257 | 0.51606326 | 0.91636259 |
| ZNF252P-AS | 95.4869376 | 0.22209281 | 0.37040866 | 0.59958859 | 0.54878045 | 0.92373235 |
| C8orf33 | 2283.04001 | -0.0619472 | 0.26044549 | -0.2378509 | 0.81199676 | 0.97355727 |
| WASHC1 | 4556.99697 | -0.1005708 | 0.21468592 | -0.4684556 | 0.63945883 | 0.94088547 |
| FOXD4 | 41.2554146 | -0.0057567 | 0.5293811 | -0.0108744 | 0.99132368 | 0.99873026 |
| CBWD1 | 1373.94233 | 0.32284861 | 0.24371032 | 1.32472276 | 0.18526312 | 0.7684852 |
| C9orf66 | 965.213029 | 1.40661985 | 0.81974083 | 1.7159324 | 0.08617442 | 0.65049132 |
| DOCK8 | 1672.45933 | 0.28279407 | 0.29854563 | 0.94723905 | 0.34351698 | 0.85941153 |
| KANK1 | 3179.24969 | 0.15050033 | 0.18667939 | 0.8061968 | 0.42012938 | 0.88451372 |
| LINC01230 | 432.872547 | 0.40581463 | 0.59242524 | 0.68500563 | 0.49334037 | 0.90715187 |
| DMRT2 | 1594.28597 | 0.25901841 | 0.6679785 | 0.38776459 | 0.69819025 | 0.9529028 |
| SMARCA2 | 4221.64488 | 0.28233445 | 0.18239972 | 1.5478886 | 0.12164912 | 0.69725141 |
| VLDLR-AS1 | 275.259296 | -0.0848716 | 0.34976511 | -0.2426531 | 0.80827413 | 0.97334138 |
| VLDLR | 3060.79504 | -0.2212889 | 0.27687937 | -0.7992251 | 0.4241599 | 0.88565404 |
| PUM3 | 2201.44928 | -0.2250116 | 0.28837937 | -0.7802624 | 0.43523643 | 0.89055338 |
| RFX3 | 584.901398 | -0.0877831 | 0.24998027 | -0.3511603 | 0.7254681 | 0.95848028 |
| RFX3-AS1 | 47.1640212 | 0.20246722 | 0.49649775 | 0.40779081 | 0.68342726 | 0.95070301 |
| GLIS3 | 1765.01525 | -0.4351531 | 0.20583257 | -2.1141121 | 3.45E-02 | 0.55369025 |
| SLC1A1 | 4495.56096 | 0.71387613 | 0.33203664 | 2.14999204 | 3.16E-02 | 5.45E-01 |

| | | | | | | |
|---|---|---|---|---|---|---|
| SPATA6L | 386.939769 | -0.3252955 | 0.35426471 | -0.9182271 | 0.35849996 | 0.86480228 |
| PLPP6 | 571.829431 | -0.1002087 | 0.18454842 | -0.5429941 | 0.58713388 | 0.93289552 |
| CDC37L1-DT | 65.7732663 | 0.6254599 | 0.3904841 | 1.60175509 | 0.10920978 | 0.67859123 |
| CDC37L1 | 1576.30621 | -0.1351099 | 0.15281081 | -0.8841645 | 0.37660742 | 0.86970773 |
| AK3 | 7708.31046 | 0.07738361 | 0.19988658 | 0.3871376 | 0.69865434 | 0.9529028 |
| RCL1 | 937.664643 | -0.2448483 | 0.24252986 | -1.0095592 | 0.31270653 | 0.84541735 |
| JAK2 | 825.565205 | -0.275682 | 0.25445679 | -1.083414 | 0.27862472 | 0.82805357 |
| PLGRKT | 1086.73304 | 0.54790927 | 0.17300172 | 3.16707417 | 0.00153981 | 0.17279992 |
| CD274 | 480.964449 | -1.1133012 | 0.50184019 | -2.2184378 | 0.026525 | 0.52678638 |
| PDCD1LG2 | 114.251367 | -0.707717 | 0.51403414 | -1.3767899 | 0.16857721 | 0.7576272 |
| RIC1 | 1177.64104 | -0.2725869 | 0.15546283 | -1.7533898 | 0.07953512 | 0.64079926 |
| ERMP1 | 4943.12645 | 0.21295353 | 0.44003137 | 0.48395079 | 0.62842079 | 0.94088547 |
| KIAA2026 | 1320.86386 | -0.2208665 | 0.12269786 | -1.8000842 | 0.07184734 | 0.63189503 |
| RANBP6 | 1342.72905 | 0.02573296 | 0.13615545 | 0.18899689 | 0.85009526 | 0.97816532 |
| IL33 | 853.806609 | -0.1879295 | 0.79512729 | -0.2363515 | 0.81315991 | 0.9738447 |
| UHRF2 | 1923.13005 | -0.1536438 | 0.13875419 | -1.107309 | 0.26816037 | 0.82676682 |
| GLDC | 4723.17101 | 0.93316132 | 0.34149347 | 2.73258905 | 0.00628387 | 0.33470127 |
| AL162411.1 | 67.4689421 | 0.20054866 | 0.33116106 | 0.6055925 | 0.54478538 | 0.92345821 |
| AL354707.1 | 64.6149539 | 0.75116001 | 0.64046598 | 1.17283359 | 0.24086255 | 0.8124487 |
| KDM4C | 1320.78693 | -0.0129837 | 0.15526969 | -0.0836201 | 0.93335851 | 0.99017182 |
| DMAC1 | 1800.13742 | 0.22528265 | 0.19847596 | 1.13506262 | 0.25634908 | 0.82067614 |
| PTPRD | 1731.05584 | 0.19367343 | 0.40559758 | 0.47750144 | 0.6330051 | 0.94088547 |
| PTPRD-AS1 | 82.7481793 | 0.39409425 | 0.60829564 | 0.64786631 | 0.51707142 | 0.91636259 |
| TYRP1 | 553.694523 | -1.9251866 | 0.53726418 | -3.5833146 | 0.00033926 | 0.08914272 |
| LURAP1L | 733.70965 | -1.8520572 | 0.70310694 | -2.6341046 | 0.00843595 | 0.3779007 |
| MPDZ | 1829.48304 | -0.097303 | 0.24868321 | -0.3912729 | 0.69559549 | 0.9525827 |
| LINC01235 | 42.0863962 | -1.2549962 | 0.73128695 | -1.7161474 | 0.08613507 | 0.65049132 |
| NFIB | 2652.01004 | 0.10975789 | 0.22961219 | 0.47801422 | 0.63264009 | 0.94088547 |
| AL159169.2 | 88.0137641 | -0.4638085 | 0.36414441 | -1.2736939 | 0.20277192 | 0.78193335 |
| ZDHHC21 | 1444.97751 | -0.2363247 | 0.1831303 | -1.2904731 | 0.19688643 | 0.77816059 |
| FREM1 | 1204.67121 | 1.08777687 | 0.52911553 | 2.05583999 | 0.03979794 | 0.55657615 |
| TTC39B | 1926.31584 | 0.10143999 | 0.22422847 | 0.45239568 | 0.65098396 | 0.9438312 |
| SNAPC3 | 2159.10483 | -0.128986 | 0.17775345 | -0.7256456 | 0.46805607 | 0.89984974 |
| PSIP1 | 2707.43308 | 0.10218887 | 0.14984305 | 0.68197268 | 0.49525622 | 0.9083575 |
| CCDC171 | 129.529986 | 0.57447352 | 0.36671059 | 1.56655832 | 0.11721797 | 0.68956727 |
| BNC2 | 935.983057 | 0.49838984 | 0.27400377 | 1.81891599 | 0.06892425 | 0.62779744 |
| BNC2-AS1 | 48.5648239 | 0.27224905 | 0.38814584 | 0.70140915 | 0.48304771 | 0.90541935 |
| CNTLN | 374.550583 | -0.0946834 | 0.1412309 | -0.6704155 | 0.50259294 | 0.91319792 |
| SH3GL2 | 770.647311 | -0.1211888 | 0.32971427 | -0.367557 | 0.71320355 | 0.95564093 |
| ADAMTSL1 | 407.748037 | 0.12995876 | 0.40808773 | 0.31845791 | 0.75013762 | 0.96250371 |
| RRAGA | 4232.94743 | -0.2006804 | 0.1624981 | -1.2349704 | 0.21684151 | 0.7954827 |
| HAUS6 | 735.377257 | -0.0703642 | 0.21700597 | -0.3242502 | 0.74574862 | 0.96176407 |
| PLIN2 | 35679.241 | -1.0801002 | 0.64159736 | -1.6834548 | 0.09228708 | 0.65583276 |
| DENND4C | 1562.47146 | 0.09777282 | 0.16607345 | 0.58873239 | 0.5560408 | 0.92373235 |
| AL391834.1 | 137.362438 | 0.11326774 | 0.27172226 | 0.41685115 | 0.67678729 | 0.95000096 |
| AL391834.2 | 103.602085 | 0.30598679 | 0.2672839 | 1.14480068 | 2.52E-01 | 0.81833349 |
| RPS6 | 45124.0921 | 0.27922893 | 0.24725303 | 1.12932464 | 0.25876091 | 0.82361383 |
| ACER2 | 57.8715135 | 0.58197063 | 0.42508116 | 1.36908122 | 0.17097389 | 0.75956362 |
| MLLT3 | 789.642445 | 0.23829019 | 0.2130173 | 1.11864244 | 0.26329271 | 0.82424998 |
| FOCAD | 1179.33414 | 0.16398537 | 0.23861368 | 0.68724212 | 0.49193017 | 0.90615916 |
| HACD4 | 318.223731 | 0.45579691 | 0.38433993 | 1.18592128 | 0.23565339 | 0.80747753 |
| KLHL9 | 1720.48886 | -0.163052 | 0.11772977 | -1.3849683 | 0.16606215 | 0.75515387 |
| MTAP | 1421.17301 | -0.1201105 | 0.14330233 | -0.8381616 | 4.02E-01 | 0.88132015 |
| AL359922.1 | 162.644537 | 0.17276812 | 0.22764536 | 0.75893537 | 0.44789122 | 0.89480207 |

| | | | | | | |
|---|---|---|---|---|---|---|
| AL359922.2 | 138.026389 | -0.08363 | 0.29032205 | -0.2880594 | 0.77330126 | 0.96836854 |
| CDKN2B | 154.091769 | -1.4391658 | 0.57399313 | -2.5072876 | 0.01216617 | 0.41257332 |
| DMRTA1 | 501.833268 | 0.29554107 | 0.29309066 | 1.00836059 | 0.31328138 | 0.84623903 |
| LINC01239 | 50.0416952 | 1.78097168 | 0.70460679 | 2.52761073 | 0.01148416 | 0.40353404 |
| TUSC1 | 932.430198 | -0.2749694 | 0.20116963 | -1.3668536 | 0.1716712 | 0.75974924 |
| CAAP1 | 1678.10585 | 0.01037789 | 0.14493327 | 0.07160464 | 0.94291655 | 0.99136103 |
| PLAA | 1678.83046 | -0.1231416 | 0.20326762 | -0.6058104 | 0.54464067 | 0.92345821 |
| IFT74 | 1125.14844 | 0.22816408 | 0.2047134 | 1.11455369 | 0.26504173 | 0.82455959 |
| LRRC19 | 2849.32903 | 0.67533618 | 0.52396498 | 1.28889564 | 1.97E-01 | 0.77819208 |
| TEK | 1971.16062 | -0.041886 | 0.3489752 | -0.1200257 | 0.90446279 | 0.98624484 |
| MOB3B | 1053.03018 | 0.00884611 | 0.15928344 | 0.05553693 | 9.56E-01 | 9.92E-01 |
| C9orf72 | 896.353743 | -0.2823469 | 0.27155191 | -1.0397529 | 0.29845473 | 0.84021932 |
| ACO1 | 8469.58141 | 0.59361401 | 0.45642137 | 1.30058331 | 0.19340112 | 0.77548688 |
| DDX58 | 1772.77697 | -1.4498696 | 0.680516 | -2.1305445 | NA | NA |
| TOPORS | 1258.25286 | -0.074491 | 0.21812363 | -0.3415081 | 0.73272111 | 0.95848028 |
| SMIM27 | 340.241938 | -0.08019 | 0.19137486 | -0.4190207 | 0.67520098 | 0.94956788 |
| NDUFB6 | 2291.83117 | 0.22098624 | 0.19195461 | 1.15124214 | 0.24963263 | 0.81646572 |
| APTX | 1427.87814 | 0.0860772 | 0.07561815 | 1.13831399 | 0.2549894 | 0.81992039 |
| DNAJA1 | 8648.08602 | -0.2456449 | 0.17575224 | -1.3976775 | 0.16220995 | 0.75026273 |
| SMU1 | 2776.40552 | 0.01037139 | 0.10268038 | 0.10100656 | 0.91954525 | 0.9872458 |
| B4GALT1 | 9519.91136 | -0.0356716 | 0.19850994 | -0.1796968 | 0.85739059 | 0.97953896 |
| B4GALT1-AS | 147.603281 | 0.15876261 | 0.4033482 | 0.3936118 | 0.69386766 | 0.95222846 |
| BAG1 | 5211.56479 | 0.13965091 | 0.18797498 | 0.74292286 | 0.45752838 | 0.89688594 |
| CHMP5 | 4347.96061 | -0.0819104 | 0.20860382 | -0.3926601 | 0.69457053 | 0.95222846 |
| NFX1 | 2322.98192 | -0.2792448 | 0.22333683 | -1.2503302 | 0.21117894 | 0.79134644 |
| AQP7 | 3426.08095 | 0.23244923 | 0.41507389 | 0.56001891 | 0.57546654 | 0.93076883 |
| AQP3 | 17776.2631 | -0.1133413 | 0.40820272 | -0.2776593 | 0.78127393 | 0.96902269 |
| NOL6 | 2147.61158 | -0.3512844 | 0.21612356 | -1.6253867 | 0.10408019 | 0.67439401 |
| PRSS3 | 76.7481152 | 0.35841742 | 0.70010088 | 0.5119511 | 6.09E-01 | 0.93748023 |
| UBE2R2 | 3693.50912 | -0.1610476 | 0.20146572 | -0.7993798 | 0.4240702 | 0.88565404 |
| UBAP2 | 1780.58214 | -0.3763941 | 0.16181442 | -2.3260849 | 0.02001402 | 0.49637604 |
| DCAF12 | 1961.01396 | -0.0401596 | 0.12782225 | -0.3141835 | 0.75338167 | 0.96301881 |
| UBAP1 | 5349.64682 | -0.5349208 | 0.20358076 | -2.6275606 | 8.60E-03 | 3.80E-01 |
| KIF24 | 362.560027 | -0.2741921 | 0.19074017 | -1.4375162 | 1.51E-01 | 7.38E-01 |
| NUDT2 | 638.214542 | 0.50996602 | 0.2562329 | 1.99024411 | 4.66E-02 | 5.80E-01 |
| MYORG | 2103.37452 | 0.87637008 | 0.39105557 | 2.24103723 | 2.50E-02 | 5.21E-01 |
| C9orf24 | 102.111309 | -0.0020384 | 0.37036504 | -0.0055038 | 0.99560864 | 0.99940889 |
| FAM219A | 2803.51878 | -0.3210379 | 0.21275787 | -1.5089353 | 0.13131532 | 0.71081672 |
| DNAI1 | 175.46093 | 0.04511351 | 0.53535919 | 0.08426774 | 0.93284356 | 0.98999825 |
| CNTFR | 203.980284 | -0.441208 | 0.64392884 | -0.6851814 | 0.49322949 | 0.90715187 |
| RPP25L | 945.976607 | 0.14799348 | 0.18648807 | 0.79358152 | 0.42743909 | 0.88770346 |
| DCTN3 | 2295.9555 | 0.29563593 | 0.1300437 | 2.27335843 | 0.02300459 | 0.51967919 |
| SIGMAR1 | 2647.01604 | 0.17527279 | 0.22296833 | 0.78608827 | 0.43181578 | 0.88976671 |
| GALT | 2868.38621 | 0.19231309 | 0.38001357 | 0.506069 | 0.61280821 | 0.93879706 |
| AL162231.3 | 1600.96891 | 0.28219464 | 0.32252401 | 0.87495701 | 0.3815973 | 0.87185666 |
| IL11RA | 1179.68653 | 0.07654832 | 0.29689061 | 0.25783343 | 0.79653547 | 0.97117111 |
| AL162231.1 | 198.2716 | 0.23697715 | 0.20352814 | 1.16434588 | 0.24428387 | 0.81468457 |
| AL162231.2 | 227.123359 | 0.61646792 | 0.2029229 | 3.03794167 | 0.002382 | 0.21083339 |
| CCL19 | 823.236371 | 0.7710791 | 1.35267841 | 0.57003874 | 0.56865142 | 0.92642835 |
| CCL21 | 1955.30584 | 0.13264422 | 0.82129448 | 0.16150629 | 0.87169466 | 0.98275231 |
| PHF24 | 70.987128 | -1.5646105 | 0.89632678 | -1.7455804 | 0.08088389 | 0.64313548 |
| DNAJB5 | 966.102429 | -0.5118937 | 0.39556513 | -1.2940819 | 0.19563713 | 0.77752069 |
| VCP | 15735.9713 | -0.0606906 | 0.267951 | -0.226499 | 0.82081331 | 0.97472105 |
| FANCG | 580.622047 | 0.23246519 | 0.25788881 | 0.90141637 | 0.36736698 | 0.86683872 |

| | | | | | | |
|---|---|---|---|---|---|---|
| PIGO | 2629.67252 | 0.12312492 | 0.15061446 | 0.81748406 | 0.41365186 | 0.88316314 |
| STOML2 | 5076.72311 | 0.01972397 | 0.16488805 | 0.11962037 | 0.90478388 | 0.98624484 |
| FAM214B | 2377.73173 | -0.1932542 | 0.28367808 | -0.6812449 | 0.49571655 | 0.90861354 |
| UNC13B | 3417.37854 | 0.32027943 | 0.17680532 | 1.81148067 | 0.07006648 | 0.62928077 |
| RUSC2 | 4475.74013 | -0.677897 | 0.26911354 | -2.5189998 | 0.01176887 | 0.40423902 |
| FAM166B | 140.459608 | 0.23339277 | 0.37031154 | 0.63026059 | 0.52852411 | 0.91888066 |
| TESK1 | 3509.81947 | -0.2999885 | 0.1886876 | -1.5898689 | 0.11186436 | 0.68001961 |
| CD72 | 1418.83752 | -0.45731 | 0.17965976 | -2.5454226 | 0.01091456 | 0.40196898 |
| SIT1 | 53.906594 | 0.2225349 | 0.86800671 | 0.25637463 | 0.79766158 | 0.97148232 |
| CCDC107 | 1603.08839 | -0.0600877 | 0.17803451 | -0.337506 | 0.73573551 | 0.95906094 |
| ARHGEF39 | 1681.60064 | -0.0277575 | 0.17913515 | -0.1549528 | 0.87685852 | 0.98275231 |
| CA9 | 83.4422529 | 0.42822865 | 0.94564533 | 0.45284277 | 0.65066196 | 0.94346674 |
| TPM2 | 7850.36125 | -0.659185 | 0.52269248 | -1.2611335 | 0.20726074 | 0.78790396 |
| TLN1 | 16086.8422 | -0.1913178 | 0.17933052 | -1.0668447 | 0.28604198 | 0.83107779 |
| CREB3 | 3244.45832 | -0.0791157 | 0.15804428 | -0.5005919 | 0.61665836 | 0.93910158 |
| GBA2 | 8015.66946 | -0.034056 | 0.13899968 | -0.2450075 | 0.80645062 | 0.97334138 |
| RGP1 | 2933.6944 | -0.0697114 | 0.22440135 | -0.310655 | 0.75606288 | 0.96449353 |
| MSMP | 645.259263 | -0.145407 | 0.20847908 | -0.6974655 | 0.48551153 | 0.90543654 |
| AL133410.2 | 527.129007 | -0.0617032 | 0.23032408 | -0.2678975 | 0.78877825 | 0.97090793 |
| AL133410.1 | 75.13153 | 0.36870683 | 0.45185166 | 0.81599087 | 0.41450536 | 0.88364943 |
| NPR2 | 1310.59026 | -0.4352923 | 0.26758726 | -1.6267302 | 0.10379442 | 0.67439401 |
| SPAG8 | 445.93141 | -0.1090911 | 0.21332604 | -0.5113818 | 0.60908375 | 0.93748023 |
| HINT2 | 1828.3215 | 0.31430496 | 0.21463343 | 1.46438024 | 0.14309008 | 0.72784886 |
| TMEM8B | 2486.2461 | 0.31059797 | 0.268426 | 1.15710837 | 0.24722809 | 0.81566357 |
| HRCT1 | 256.508628 | -0.3246381 | 0.50218862 | -0.6464465 | 0.51799025 | 0.91636259 |
| SPAAR | 172.495747 | 0.19566541 | 0.3731247 | 0.52439682 | 0.60000257 | 0.93487648 |
| RECK | 643.813121 | 0.14966298 | 0.31310453 | 0.47799686 | 0.63265244 | 0.94088547 |
| GLIPR2 | 576.439577 | -0.3780894 | 0.52439813 | -0.7209968 | 0.4709115 | 0.90089741 |
| CLTA | 7109.3924 | -0.0125326 | 0.1972511 | -0.0635363 | 0.94933945 | 0.99170503 |
| GNE | 2153.67038 | -0.0789493 | 0.29696456 | -0.2658543 | 0.79035139 | 0.97090793 |
| RNF38 | 2553.01264 | -0.3084788 | 0.24062459 | -1.2819922 | 0.19984539 | 0.78134034 |
| MELK | 50.6597836 | 0.27817065 | 0.69659188 | 0.39933088 | 6.90E-01 | 9.52E-01 |
| EBLN3P | 2956.81216 | -0.0800722 | 0.13167762 | -0.608093 | 0.54312578 | 0.92297267 |
| ZCCHC7 | 634.027084 | -0.0963987 | 0.11552811 | -0.8344178 | 0.40404559 | 0.88132015 |
| GRHPR | 8699.326 | 0.26308416 | 0.32663475 | 0.80543836 | 0.42056676 | 0.88451372 |
| ZBTB5 | 1357.44115 | -0.3199195 | 0.18386514 | -1.7399681 | 0.08186462 | 0.64544584 |
| POLR1E | 464.714689 | 0.44110415 | 0.24155375 | 1.82611179 | 0.06783343 | 0.62599695 |
| AL513165.1 | 81.6257426 | -0.2360573 | 0.45095186 | -0.5234646 | 0.60065098 | 0.93488657 |
| FBXO10 | 789.798436 | 0.19610068 | 0.34833079 | 0.56297256 | 0.57345357 | 0.92970376 |
| AL513165.2 | 2118.07228 | 0.00133479 | 0.30866246 | 0.00432444 | 0.9965496 | 0.99976751 |
| TOMM5 | 2356.78093 | -0.0723846 | 0.27664385 | -0.2616525 | 0.79358938 | 0.97090793 |
| AL138752.2 | 3859.2852 | -0.0688475 | 0.21444533 | -0.3210493 | 0.74817304 | 0.96241535 |
| FRMPD1 | 63.7717608 | 0.6437952 | 0.45959932 | 1.40077493 | 0.16128139 | 0.74924826 |
| TRMT10B | 631.419791 | 0.00518132 | 0.22856696 | 0.02266872 | 0.98191453 | 0.99595643 |
| EXOSC3 | 764.979275 | -0.1993209 | 0.14530775 | -1.3717158 | 0.17015192 | 0.75956362 |
| DCAF10 | 1690.87038 | -0.2148415 | 0.20599599 | -1.0429401 | 0.29697605 | 0.8389551 |
| SLC25A51 | 532.571015 | 0.02852697 | 0.30006721 | 0.09506861 | 0.92426033 | 0.98869145 |
| SHB | 2225.47876 | -0.0915545 | 0.25231213 | -0.3628621 | 0.71670794 | 0.95603249 |
| ALDH1B1 | 6660.89955 | 0.47404127 | 0.67252219 | 0.70487083 | 0.48089063 | 0.90541935 |
| FAM95C | 862.851774 | 0.04525646 | 0.23628806 | 0.19153086 | 0.84810971 | 0.97711043 |
| ANKRD18A | 112.531981 | -0.3403126 | 0.34832106 | -0.9770084 | 0.32856498 | 0.85317073 |
| FAM201A | 205.208705 | 0.61113664 | 0.24112822 | 2.53448822 | 0.01126117 | 0.40196898 |
| CNTNAP3 | 319.456432 | 0.14396093 | 0.38424965 | 0.37465469 | 0.7079173 | 0.95501548 |
| GLIDR | 248.680269 | -0.3464547 | 0.28154219 | -1.2305604 | 0.21848732 | 0.79772711 |

| | | | | | | |
|---|---|---|---|---|---|---|
| FAM95B1 | 64.1375665 | -0.4315724 | 0.73018918 | -0.5910419 | 0.55449236 | 0.92373235 |
| CBWD6 | 364.386861 | 0.03329794 | 0.25920572 | 0.12846142 | 0.89778383 | 0.98474725 |
| CNTNAP3B | 413.942548 | 0.80236606 | 0.49100886 | 1.63411726 | 0.10223423 | 0.67279543 |
| BX088651.4 | 143.103922 | -0.2428322 | 0.59463317 | -0.4083731 | 0.68299976 | 0.95070301 |
| FAM27C | 60.8620183 | -0.8254206 | 0.46457489 | -1.7767224 | 0.07561393 | 0.63773415 |
| AL590399.1 | 57.6642974 | 0.24582243 | 0.43302132 | 0.56769128 | 0.57024461 | 0.92752597 |
| BX005266.2 | 42.8544534 | -0.4885049 | 0.8074451 | -0.6050008 | 0.54517845 | 0.92345821 |
| LINC01410 | 85.8793973 | -0.1859497 | 0.54127928 | -0.3435374 | 0.73119423 | 0.95848028 |
| AL512625.3 | 148.513936 | 0.84168771 | 0.78992588 | 1.06552746 | 0.28663731 | 0.83155545 |
| AL512625.1 | 157.541764 | 0.29946056 | 0.516918 | 0.57931927 | 0.56237377 | 0.92452095 |
| CBWD5 | 723.963844 | -0.131577 | 0.31315622 | -0.4201639 | 0.67436569 | 0.94935413 |
| ZNF658 | 138.154185 | 0.22198778 | 0.22384045 | 0.99172323 | 0.32133256 | 0.85119105 |
| CBWD3 | 177.877696 | 0.04499216 | 0.24522633 | 0.18347199 | 0.85442769 | 0.97907853 |
| PGM5 | 1311.89834 | 0.18886627 | 0.46167069 | 0.40909306 | 0.68247137 | 0.95070301 |
| TMEM252 | 3422.36105 | 1.44865141 | 0.61153789 | 2.36886614 | 0.01784271 | 0.47230257 |
| AL353616.2 | 277.215443 | 0.71527623 | 0.67349498 | 1.06203647 | 0.28821913 | 0.83295411 |
| LINC01506 | 85.0226347 | 1.11579265 | 0.63640879 | 1.75326404 | 0.07955669 | 0.64079926 |
| PIP5K1B | 569.578169 | 0.0647909 | 0.36172751 | 0.17911522 | 0.85784723 | 0.97964261 |
| FAM122A | 1327.97343 | -0.0571693 | 0.12540114 | -0.4558912 | 0.6484682 | 0.9431121 |
| FXN | 516.419339 | -0.026354 | 0.17114031 | -0.1539908 | 0.87761702 | 0.98275231 |
| AL358113.1 | 8354.93586 | 0.02392864 | 0.21324698 | 0.11221091 | 0.91065618 | 0.98662757 |
| TJP2 | 7425.24844 | 0.05402071 | 0.22712343 | 0.23784739 | 0.81199946 | 0.97355727 |
| FAM189A2 | 387.720464 | 0.04801195 | 0.26062938 | 0.18421542 | 0.85384446 | 0.97906925 |
| APBA1 | 284.431386 | 0.49424238 | 0.22294213 | 2.21690886 | 0.02662932 | 0.52678638 |
| PTAR1 | 2407.50254 | -0.2285617 | 0.17709776 | -1.2905964 | 0.19684366 | 0.77816059 |
| C9orf135 | 49.4240476 | 1.89873625 | 0.74328464 | 2.55452103 | 0.0106334 | 0.40196898 |
| MAMDC2-AS | 143.493736 | 0.24136046 | 0.39976158 | 0.60376103 | 0.54600253 | 0.92345821 |
| MAMDC2 | 198.840944 | 0.7581631 | 0.5786303 | 1.31027202 | 0.19010383 | 0.77065452 |
| SMC5-AS1 | 89.090149 | 0.27129114 | 0.37877255 | 0.71623759 | 0.47384465 | 0.90199039 |
| SMC5 | 1664.75686 | -0.1312091 | 0.17200886 | -0.7628042 | 0.44558021 | 0.89464524 |
| KLF9 | 5233.7842 | -0.022733 | 0.35034362 | -0.0648877 | 0.94826343 | 0.99170503 |
| TRPM3 | 1470.97638 | 0.36512004 | 0.3632525 | 1.00514116 | 0.31482888 | 0.84696964 |
| CEMIP2 | 3976.1906 | 0.03562625 | 0.308232 | 0.11558258 | 0.90798337 | 0.98662757 |
| ABHD17B | 852.684527 | -0.1167584 | 0.22552605 | -0.5177156 | 0.60465668 | 0.93662382 |
| C9orf85 | 438.617486 | 0.12644373 | 0.17151265 | 0.73722682 | 0.46098443 | 0.89755243 |
| GDA | 5111.48029 | 1.33118569 | 0.50155706 | 2.6541062 | 0.00795188 | 0.37071188 |
| AL135924.2 | 119.67068 | 0.3335352 | 0.88513675 | 0.37681771 | 0.70630908 | 0.95501548 |
| LINC01504 | 89.9822651 | 0.38366288 | 0.91744574 | 0.41818591 | 6.76E-01 | 9.50E-01 |
| ZFAND5 | 13269.7717 | -0.2178263 | 0.23862282 | -0.9128478 | 0.36132261 | 0.86528137 |
| TMC1 | 65.2799217 | 1.87912141 | 0.73878755 | 2.54352069 | 0.01097416 | 0.40196898 |
| ALDH1A1 | 21956.6075 | 1.19850978 | 0.43663594 | 2.74487206 | 0.00605345 | 0.33171839 |
| ANXA1 | 4036.59836 | 0.51393245 | 0.4258781 | 1.20675952 | 0.22752477 | 0.80300331 |
| RORB | 51.1462157 | 0.97837599 | 0.43939002 | 2.22666869 | 2.60E-02 | 0.52533244 |
| TRPM6 | 271.17246 | -0.0352508 | 0.59014573 | -0.0597323 | 0.95236886 | 0.99170503 |
| C9orf40 | 266.318493 | 0.1503047 | 0.1684763 | 0.89214152 | 0.37231709 | 0.86775997 |
| CARNMT1 | 921.517934 | 0.16715232 | 0.15234198 | 1.09721778 | 0.2725462 | 0.82716979 |
| NMRK1 | 1970.55695 | 0.23575209 | 0.31108109 | 0.75784769 | 0.44854217 | 0.8948759 |
| OSTF1 | 1799.6134 | 0.16547002 | 0.16273086 | 1.01683248 | 0.30923313 | 0.84414094 |
| PCSK5 | 884.338284 | -0.3783476 | 0.41754661 | -0.9061206 | 0.364872 | 0.8663515 |
| RFK | 2390.05436 | 0.19775416 | 0.19152055 | 1.03254801 | 0.30181547 | 0.84211658 |
| GCNT1 | 246.204097 | -0.1198054 | 0.31028744 | -0.386111 | 0.69941447 | 0.95307625 |
| PRUNE2 | 3406.06647 | 0.21849933 | 0.36178252 | 0.60395213 | 0.54587547 | 0.92345821 |
| PCA3 | 75.336797 | -0.9831827 | 0.6332001 | -1.5527204 | 0.12048996 | 0.69467399 |
| VPS13A | 2541.18073 | -0.0402598 | 0.23993065 | -0.1677978 | 0.86674237 | 0.9821997 |

| | | | | | | |
|---|---|---|---|---|---|---|
| GNA14 | 662.531166 | 0.35643844 | 0.4629263 | 0.769968 | 0.44131887 | 0.89290667 |
| GNAQ | 3520.01063 | -0.0794701 | 0.1850453 | -0.4294627 | 0.66758653 | 0.94839594 |
| CEP78 | 635.475754 | 0.11574336 | 0.20120659 | 0.57524634 | 0.56512471 | 0.92497835 |
| PSAT1 | 5782.16801 | 0.81175104 | 0.59940508 | 1.35426121 | 0.17565306 | 0.7610139 |
| TLE4 | 978.807242 | -0.2868291 | 0.23618456 | -1.2144278 | 0.2245844 | 0.80078694 |
| LINC01507 | 72.7286989 | -0.0708278 | 0.46408599 | -0.1526179 | 0.87869963 | 0.98275231 |
| TLE1 | 1809.88604 | -0.2746665 | 0.16983195 | -1.6172838 | 0.10581704 | 0.67526487 |
| RASEF | 690.945551 | -0.1159266 | 0.27112004 | -0.4275839 | 0.66895408 | 0.9486589 |
| FRMD3 | 1201.07475 | -0.1501596 | 0.29719879 | -0.5052496 | 0.61338355 | 0.9388166 |
| AL137847.2 | 36.6344525 | -0.1696499 | 0.48385375 | -0.3506222 | 0.72587178 | 0.95848028 |
| IDNK | 898.249783 | 0.44553298 | 0.42421871 | 1.05024359 | 0.29360613 | 0.83540174 |
| UBQLN1 | 5814.74302 | -0.0798875 | 0.19655388 | -0.4064407 | 0.68441885 | 0.95075309 |
| AL354920.1 | 258.968405 | 0.35330203 | 0.40765846 | 0.86666183 | 0.38612733 | 0.87432552 |
| GKAP1 | 365.517963 | -0.0260234 | 0.16771142 | -0.1551674 | 0.87668932 | 0.98275231 |
| AL354733.1 | 97.9269638 | -0.1635284 | 0.3035899 | -0.538649 | 0.59012907 | 0.93289552 |
| AL354733.2 | 90.4622091 | -0.1108187 | 0.31071253 | -0.3566599 | 0.72134643 | 0.95739595 |
| KIF27 | 162.7315 | -0.1235081 | 0.25059476 | -0.4928599 | 0.6221116 | 0.94016713 |
| C9orf64 | 911.306141 | 0.33214811 | 0.19487502 | 1.70441602 | 0.08830339 | 0.65363785 |
| HNRNPK | 24839.581 | -0.1353365 | 0.1424646 | -0.9499656 | 0.34212972 | 0.85889221 |
| AL354733.3 | 2241.10877 | -0.0753339 | 0.21135972 | -0.356425 | 0.72152228 | 0.95739595 |
| RMI1 | 331.31325 | 0.07511136 | 0.15374546 | 0.48854364 | 0.62516482 | 0.94088547 |
| NTRK2 | 2494.72498 | -0.3375039 | 0.44938754 | -0.7510308 | 0.45263409 | 0.89670027 |
| AGTPBP1 | 962.775092 | -0.0628117 | 0.10311431 | -0.6091466 | 0.54242725 | 0.92292765 |
| NAA35 | 955.512678 | -0.1582086 | 0.16545987 | -0.9561751 | 0.33898377 | 0.85733087 |
| GOLM1 | 7981.49296 | -0.2143124 | 0.23985222 | -0.8935187 | 0.37157946 | 0.86762749 |
| ISCA1 | 3539.70114 | -0.0129944 | 0.29193912 | -0.0445106 | 0.96449743 | 0.99349491 |
| TUT7 | 949.099775 | -0.2187656 | 0.16281854 | -1.343616 | 0.17907259 | 0.76120583 |
| GAS1 | 670.014246 | 0.1713062 | 0.54767128 | 0.31279018 | 0.75444008 | 0.96350701 |
| AL513318.2 | 169.698379 | 0.2269084 | 0.44193232 | 0.51344603 | 0.60763935 | 0.93739614 |
| DAPK1 | 4875.42854 | -0.545553 | 0.2203551 | -2.4757901 | 0.01329417 | 0.42371784 |
| CTSL | 25256.5391 | -1.0959009 | 0.62541671 | -1.7522731 | 0.07972685 | 0.64079926 |
| AL772337.2 | 51.4212111 | 1.97441799 | 0.7410092 | 2.6644986 | 0.00771032 | 0.36878977 |
| AL772337.1 | 167.397512 | 0.58149889 | 0.78615915 | 0.7396707 | 0.45949983 | 0.89755243 |
| CDK20 | 1306.48764 | -0.0397007 | 0.19348727 | -0.2051851 | 0.83742753 | 0.97577999 |
| SPIN1 | 4496.18537 | -0.0205115 | 0.15648262 | -0.1310784 | 0.89571332 | 0.98474725 |
| NXNL2 | 261.126946 | 0.84862127 | 0.43707097 | 1.94160977 | 0.05218436 | 0.58645127 |
| C9orf47 | 246.301584 | -1.4747687 | 0.78494723 | -1.8788126 | 0.06027009 | 0.60825814 |
| S1PR3 | 2252.11244 | -0.2175641 | 0.49200308 | -0.4422006 | 0.65834402 | 0.9458503 |
| SHC3 | 262.17477 | -0.3634395 | 0.43159857 | -0.8420777 | 0.39974448 | 0.88077967 |
| CKS2 | 672.60041 | -0.327042 | 0.47934367 | -0.6822704 | 0.49506797 | 0.90816709 |
| SECISBP2 | 3491.65375 | 0.13367273 | 0.10625512 | 1.25803562 | 0.20837887 | 0.78989249 |
| SEMA4D | 5487.49588 | -0.5106385 | 0.27622842 | -1.8486096 | 0.0645142 | 0.61989227 |
| GADD45G | 665.485384 | -0.0912579 | 0.41541088 | -0.2196811 | 0.82611949 | 0.97548201 |
| LINC01508 | 60.9360656 | 0.97244505 | 0.51454432 | 1.88991504 | 0.05876932 | 0.60332684 |
| DIRAS2 | 87.2754708 | 0.87952055 | 0.45873796 | 1.91726136 | 0.05520474 | 0.59596312 |
| SYK | 1559.57802 | 0.187052 | 0.22996457 | 0.81339488 | 0.41599169 | 0.88399703 |
| AL158071.3 | 87.8021417 | -0.7736987 | 0.54670993 | -1.4151904 | 0.15701267 | 0.74522306 |
| AL158071.5 | 73.081693 | -0.0120474 | 0.37529719 | -0.0321008 | 0.97439163 | 0.99455099 |
| AUH | 2637.9041 | 0.08990017 | 0.18493715 | 0.48611203 | 0.62688774 | 0.94088547 |
| NFIL3 | 4050.08594 | -0.8196087 | 0.39579237 | -2.0708047 | 0.03837705 | 0.55538555 |
| ROR2 | 259.807269 | 0.38834131 | 0.31173399 | 1.24574579 | 0.21285774 | 0.79260219 |
| SPTLC1 | 4040.4054 | 0.00568257 | 0.166194 | 0.03419241 | 0.97272372 | 0.99426317 |
| IARS | 4713.85023 | 0.189865 | 0.26765207 | 0.70937245 | 0.47809338 | 0.90404077 |
| NOL8 | 1153.01913 | -0.0481659 | 0.12713242 | -0.3788638 | 0.70478899 | 0.95501548 |

| | | | | | | |
|---|---|---|---|---|---|---|
| CENPP | 937.699039 | -0.2851247 | 0.13360104 | -2.1341506 | 0.03283046 | 0.55132224 |
| OGN | 301.395 | -0.7729209 | 0.81955004 | -0.9431039 | 0.34562775 | 0.86111349 |
| OMD | 77.4300178 | 0.21238356 | 0.66776348 | 0.31805207 | 0.75044544 | 0.96255347 |
| ASPN | 355.044929 | -0.4361603 | 0.48130185 | -0.9062095 | 0.36482495 | 0.8663515 |
| ECM2 | 784.699262 | 0.27588704 | 0.43590412 | 0.63290762 | 0.52679397 | 0.91770272 |
| IPPK | 1000.17084 | -0.2438972 | 0.16036898 | -1.5208502 | 0.12829742 | 0.70596572 |
| BICD2 | 1903.16826 | 0.01944963 | 0.22497037 | 0.0864542 | 0.93110536 | 0.98975562 |
| ZNF484 | 186.532238 | -0.143777 | 0.29693176 | -0.4842089 | 6.28E-01 | 9.41E-01 |
| FGD3 | 901.199753 | 0.36759904 | 0.40186849 | 0.9147247 | 0.36033618 | 0.86480228 |
| SUSD3 | 1082.80531 | 0.31975769 | 0.56751423 | 0.56343554 | 0.57313834 | 0.92932909 |
| CARD19 | 2582.61873 | 0.10952571 | 0.21937265 | 0.49926783 | 0.61759072 | 0.93924106 |
| NINJ1 | 6025.69709 | 0.29066752 | 0.15237831 | 1.90753863 | 0.05645087 | 0.5996103 |
| WNK2 | 200.748602 | -0.1418807 | 0.38564439 | -0.3679054 | 0.71294375 | 0.95564093 |
| FAM120AOS | 1874.58974 | 0.08870316 | 0.14711042 | 0.60296998 | 0.54652866 | 0.92345821 |
| FAM120A | 7418.91241 | 0.18470358 | 0.14366428 | 1.28566116 | 0.19856134 | 0.78046101 |
| PHF2 | 2151.89012 | -0.0238297 | 0.15933458 | -0.1495577 | 8.81E-01 | 0.98275231 |
| PTPDC1 | 587.683555 | -0.3041159 | 0.23353577 | -1.3022242 | 0.19283977 | 0.77506795 |
| AL158152.2 | 63.2012226 | 0.20532248 | 0.33957172 | 0.60465129 | 0.54541072 | 0.92345821 |
| AL158152.1 | 43.0987494 | -0.133867 | 0.70773252 | -0.1891492 | 0.84997589 | 0.97816384 |
| MIRLET7DHG | 64.609206 | 0.04243687 | 0.43140972 | 0.0983679 | 0.92164017 | 0.98795735 |
| ZNF169 | 263.780468 | -0.113694 | 0.27697043 | -0.4104913 | 0.68144556 | 0.95070301 |
| MFSD14B | 2366.3738 | -0.1764406 | 0.22434778 | -0.7864604 | 0.43159781 | 0.88976671 |
| PCAT7 | 5389.62083 | 0.60304092 | 0.50972667 | 1.18306723 | 0.23678251 | 0.80849753 |
| FBP1 | 11812.1816 | 0.68663287 | 0.53559103 | 1.28200965 | 0.19983926 | 0.78134034 |
| C9orf3 | 2911.86568 | -0.4063532 | 0.23671062 | -1.7166664 | 0.08604014 | 0.65049132 |
| FANCC | 1076.35606 | 0.08826939 | 0.19780827 | 0.44623713 | 0.65542596 | 0.94532497 |
| PTCH1 | 908.920904 | -0.2455565 | 0.18393644 | -1.3350072 | 0.18187399 | 0.76543911 |
| LINC00476 | 223.511814 | -0.1903359 | 0.26863454 | -0.7085311 | 0.47861548 | 0.90444322 |
| ERCC6L2 | 455.571079 | -0.1270715 | 0.19051603 | -0.6669857 | 0.50478129 | 0.91365534 |
| LINC00092 | 40.1659159 | 0.74050311 | 0.67697294 | 1.09384447 | 2.74E-01 | 0.82719197 |
| HSD17B3 | 248.429091 | -0.2528609 | 0.30823252 | -0.8203576 | 0.41201231 | 0.88268208 |
| AL160269.1 | 2746.34561 | 0.09970146 | 0.21956698 | 0.45408224 | 0.64976964 | 0.94312988 |
| HSD17B3-AS | 64.6890374 | -0.213472 | 0.39188936 | -0.5447252 | 0.58594253 | 0.93289552 |
| SLC35D2 | 2620.6878 | 0.13289827 | 0.22487038 | 0.59099943 | 0.5545208 | 0.92373235 |
| ZNF367 | 212.562829 | -0.3965497 | 0.5399681 | -0.7343947 | 0.46270822 | 0.89796156 |
| HABP4 | 2405.51707 | -0.0070921 | 0.23430183 | -0.030269 | 0.9758525 | 0.99455099 |
| CDC14B | 2500.474 | 0.20519978 | 0.24770618 | 0.82839991 | 0.40744406 | 0.88132015 |
| AAED1 | 768.059428 | -0.4683115 | 0.26318445 | -1.7794043 | 0.0751735 | 0.63773415 |
| ZNF510 | 435.173396 | -0.0668493 | 0.13272783 | -0.5036572 | 0.61450226 | 0.93895494 |
| ZNF782 | 264.408407 | -0.3830293 | 0.24895506 | -1.5385481 | 0.12391466 | 0.6996941 |
| MFSD14C | 1399.59478 | -0.2087206 | 0.18242721 | -1.144131 | 0.25256933 | 0.81833349 |
| CTSV | 642.014753 | -0.4694622 | 0.54991725 | -0.853696 | 0.39327343 | 0.87825756 |
| CCDC180 | 562.696142 | 0.21889283 | 0.3821697 | 0.57276344 | 0.56680488 | 0.9256133 |
| TDRD7 | 1029.00585 | -0.5237505 | 0.35623807 | -1.470226 | 0.14150056 | 0.72784886 |
| TMOD1 | 1587.9253 | -0.4734055 | 0.22890002 | -2.0681757 | 0.03862351 | 0.5559736 |
| TSTD2 | 1612.31926 | -0.3086253 | 0.12421165 | -2.4846729 | 0.01296705 | 0.41899224 |
| NCBP1 | 1476.60426 | -0.0622595 | 0.11249541 | -0.5534405 | 0.57996184 | 0.93139743 |
| XPA | 790.236745 | -0.0752485 | 0.18582403 | -0.404945 | 0.68551795 | 0.95097288 |
| TRMO | 601.089052 | -0.2333073 | 0.32097191 | -0.7268776 | 0.46730096 | 0.89983399 |
| ANP32B | 4394.71917 | 0.0295311 | 0.16171242 | 0.18261492 | 0.85510018 | 0.97910552 |
| NANS | 1926.9598 | 0.00499487 | 0.19409994 | 0.02573349 | 0.97946991 | 0.99524212 |
| TRIM14 | 2723.84003 | -0.2924827 | 0.42494825 | -0.6882785 | 0.49127745 | 0.90600166 |
| CORO2A | 247.226024 | 1.15549726 | 0.4789639 | 2.41249342 | 0.01584382 | 0.4564177 |
| TBC1D2 | 754.952765 | 0.01458415 | 0.16653858 | 0.08757222 | 0.93021668 | 0.98973492 |

| | | | | | | |
|---|---|---|---|---|---|---|
| ANKS6 | 1950.45062 | -0.0058105 | 0.14378226 | -0.0404119 | 0.96776473 | 0.99400331 |
| GALNT12 | 475.009195 | 0.27182029 | 0.35708809 | 0.76121355 | 0.44652953 | 0.89480207 |
| AL136084.2 | 236.339929 | 0.36349573 | 0.371628 | 0.97811716 | 0.32801638 | 0.85317073 |
| COL15A1 | 2413.1613 | -0.4603779 | 0.70774149 | -0.6504888 | 0.51537654 | 0.91636259 |
| TGFBR1 | 2756.10124 | -0.5052577 | 0.42804745 | -1.1803778 | 0.23784997 | 0.80962787 |
| ALG2 | 2574.06172 | -0.112016 | 0.22001927 | -0.5091192 | 0.61066868 | 0.93807505 |
| SEC61B | 3543.558 | -0.1526005 | 0.22193903 | -0.6875784 | 0.49171829 | 0.90612511 |
| NR4A3 | 830.436163 | 0.16712797 | 1.11868348 | 0.14939701 | 0.88124037 | 0.98275231 |
| STX17 | 480.174671 | -0.2726588 | 0.26884755 | -1.0141762 | 0.31049868 | 0.84476049 |
| ERP44 | 3993.7523 | -0.0609802 | 0.11504591 | -0.530051 | 0.59607655 | 0.9334601 |
| INVS | 667.462571 | -0.1295492 | 0.26791603 | -0.4835439 | 0.62870959 | 0.94088547 |
| TEX10 | 1146.34442 | -0.2633961 | 0.15937841 | -1.6526463 | 0.09840287 | 0.66624572 |
| MSANTD3 | 1038.07281 | -0.6674815 | 0.4168331 | -1.6013161 | 0.10930693 | 0.67859123 |
| MSANTD3-TI | 298.666558 | -0.3071588 | 0.38107238 | -0.8060378 | 0.42022104 | 0.88451372 |
| TMEFF1 | 88.0350994 | 0.24869105 | 0.40828072 | 0.60911777 | 0.54244638 | 0.92292765 |
| PLPPR1 | 730.653189 | 0.55217018 | 0.45816317 | 1.20518236 | 0.22813291 | 0.80300331 |
| MRPL50 | 1458.53706 | -0.1168902 | 0.17116746 | -0.6828999 | 0.4946701 | 0.90816709 |
| ZNF189 | 1389.10456 | -0.1821456 | 0.23997371 | -0.7590233 | 0.44783862 | 0.89480207 |
| ALDOB | 200931.914 | 0.43967203 | 0.67754302 | 0.64892121 | 5.16E-01 | 0.91636259 |
| TMEM246-A: | 203.87339 | 0.54351885 | 0.48570434 | 1.1190323 | 0.26312636 | 0.82424998 |
| TMEM246 | 974.36602 | 0.08037203 | 0.25756167 | 0.31204964 | 0.7550028 | 0.96360109 |
| RNF20 | 1493.80981 | 0.33417186 | 0.19440359 | 1.71895934 | 0.08562178 | 0.65049132 |
| SMC2 | 514.600723 | 0.14089811 | 0.18576865 | 0.75846009 | 0.4481756 | 0.8948759 |
| NIPSNAP3A | 1415.37406 | 0.18613226 | 0.26098144 | 0.71320114 | 0.4757213 | 0.90257104 |
| NIPSNAP3B | 219.448992 | -0.0439663 | 0.39917392 | -0.1101432 | 0.91229579 | 0.98662757 |
| ABCA1 | 1823.47687 | -0.1661855 | 0.42208284 | -0.3937272 | 0.69378246 | 0.95222846 |
| SLC44A1 | 3456.89869 | -0.1953395 | 0.16505197 | -1.1835028 | 0.23660996 | 0.80849753 |
| FSD1L | 482.525551 | 0.01885944 | 0.23651743 | 0.07973806 | 0.93644559 | 0.99068685 |
| FKTN | 772.198175 | 0.07632082 | 0.16252182 | 0.46960356 | 0.63863828 | 0.94088547 |
| TMEM38B | 1859.328 | -0.5968971 | 0.33019727 | -1.8076985 | 0.07065344 | 0.62981127 |
| ZNF462 | 714.368539 | -0.1791811 | 0.15227684 | -1.1766797 | 0.23932338 | 0.81163641 |
| RAD23B | 7146.81475 | -0.1235446 | 0.19839533 | -0.6227192 | 0.53346908 | 0.92049136 |
| KLF4 | 1245.10274 | -0.4095907 | 0.39053076 | -1.0488051 | 0.29426782 | 0.83615978 |
| ELP1 | 2020.46166 | 0.11294551 | 0.17235196 | 0.65531897 | 0.51226241 | 0.91551505 |
| FAM206A | 1058.08744 | -0.043223 | 0.15011063 | -0.2879413 | 0.77339168 | 0.96836854 |
| CTNNAL1 | 3216.9453 | -0.4040926 | 0.31302209 | -1.2909396 | 0.19672461 | 0.77816059 |
| TMEM245 | 5582.65531 | 0.04866998 | 0.27908729 | 0.1743898 | 0.86155913 | 0.98167219 |
| EPB41L4B | 731.880446 | 0.372031 | 0.18998766 | 1.95818504 | 0.0502083 | 0.5861488 |
| PTPN3 | 5762.82128 | -0.0557123 | 0.2318538 | -0.2402905 | 0.81010505 | 0.97334138 |
| PALM2-AKAF | 5278.66022 | -0.3117874 | 0.26872993 | -1.1602257 | 0.24595694 | 0.81542272 |
| AKAP2 | 5275.2048 | -0.3124209 | 0.2688222 | -1.1621843 | 2.45E-01 | 8.15E-01 |
| TXN | 11661.6018 | 0.13171584 | 0.54641848 | 0.24105304 | 0.80951401 | 0.97334138 |
| SVEP1 | 1581.45443 | 0.92349222 | 0.44627989 | 2.06931174 | 3.85E-02 | 5.56E-01 |
| LPAR1 | 1079.54436 | -0.0449678 | 0.4099585 | -0.1096886 | 9.13E-01 | 0.98662757 |
| ECPAS | 4834.60949 | -0.1039377 | 0.16930662 | -0.6139024 | 5.39E-01 | 9.22E-01 |
| ZNF483 | 67.5035706 | -0.7462132 | 0.3799621 | -1.9639149 | 0.04953997 | 0.5861488 |
| PTGR1 | 14043.8941 | 0.13938217 | 0.57794126 | 0.24117013 | 0.80942326 | 0.97334138 |
| DNAJC25 | 1258.70879 | -0.5986119 | 0.36290893 | -1.6494824 | 0.09904884 | 0.66868673 |
| DNAJC25-GN | 1722.28604 | -0.2765993 | 0.2908544 | -0.9509889 | 0.34161002 | 0.85889221 |
| GNG10 | 1642.60538 | -0.2870115 | 0.29620634 | -0.9689581 | 0.33256609 | 0.855011 |
| C9orf84 | 144.349468 | 0.05699134 | 0.71184616 | 0.08006132 | 0.93618849 | 0.99068685 |
| UGCG | 8698.23254 | -0.4158571 | 0.37175585 | -1.1186296 | 0.2632982 | 0.82424998 |
| SUSD1 | 1102.29411 | 0.03161229 | 0.25126677 | 0.12581164 | 0.89988102 | 0.98514342 |
| PTBP3 | 5074.35842 | -0.2535445 | 0.32447338 | -0.781403 | 0.43456552 | 0.89055338 |

| | | | | | | |
|---|---|---|---|---|---|---|
| HSDL2 | 5157.86541 | 0.45841672 | 0.38189613 | 1.20037016 | 0.22999561 | 0.80551898 |
| C9orf147 | 47.7332299 | -0.1682771 | 0.357548 | -0.470642 | 0.63789639 | 0.94088547 |
| KIAA1958 | 943.272473 | 0.16392126 | 0.25733479 | 0.63699614 | 0.52412736 | 0.91723878 |
| INIP | 1269.20885 | 0.00230142 | 0.10779229 | 0.02135049 | 0.98296607 | 0.9961464 |
| SNX30 | 2641.93145 | 0.02720136 | 0.27308032 | 0.09960938 | 0.92065445 | 0.98767854 |
| ZNF883 | 114.230063 | 0.13137839 | 0.46542729 | 0.28227479 | 0.77773282 | 0.96836854 |
| ZFP37 | 158.123929 | 0.05429739 | 0.31117751 | 0.17449009 | 0.86148032 | 0.98167219 |
| SLC31A2 | 1614.90607 | -0.2754336 | 0.33024835 | -0.8340195 | 0.40427001 | 0.88132015 |
| FKBP15 | 3641.96286 | -0.2280188 | 0.1877065 | -1.2147622 | 0.22445679 | 0.80078694 |
| SLC31A1 | 2797.77893 | 0.15581611 | 0.33476576 | 0.46544817 | 0.64161054 | 0.94088547 |
| CDC26 | 716.698171 | -0.1380804 | 0.15543605 | -0.888342 | 3.74E-01 | 8.68E-01 |
| PRPF4 | 1919.82647 | -0.2792645 | 0.26305827 | -1.0616068 | 0.28841421 | 0.83295411 |
| RNF183 | 810.120407 | -0.2669462 | 0.41047247 | -0.6503389 | 0.51547335 | 0.91636259 |
| WDR31 | 266.472159 | 0.15005575 | 0.38011251 | 0.39476668 | 0.69301509 | 0.9519683 |
| BSPRY | 1765.03667 | -0.3420956 | 0.23468561 | -1.4576761 | 0.14492984 | 0.73039059 |
| HDHD3 | 4756.62887 | -0.03167 | 0.31305591 | -0.1011642 | 0.91942015 | 0.9872458 |
| ALAD | 6034.63495 | 0.34050928 | 0.30183162 | 1.1281432 | 0.25925945 | 0.82364948 |
| POLE3 | 1569.6813 | -0.4011646 | 0.28887334 | -1.3887213 | 1.65E-01 | 7.53E-01 |
| C9orf43 | 226.159236 | -0.2931341 | 0.32280182 | -0.908093 | 0.36382909 | 0.8663515 |
| RGS3 | 5172.69149 | 0.3167576 | 0.25338143 | 1.25012158 | 0.21125514 | 0.79134644 |
| ZNF618 | 1636.28312 | 0.13251576 | 0.29713827 | 0.44597339 | 0.65561647 | 0.94532497 |
| AMBP | 987.25125 | 0.37548512 | 0.6257658 | 0.60004097 | 0.54847893 | 0.92373235 |
| KIF12 | 24625.2549 | -0.1989893 | 0.38074616 | -0.5226297 | 0.60123195 | 0.93529286 |
| COL27A1 | 8797.96549 | -0.5178625 | 0.35288603 | -1.4675064 | 0.14223834 | 0.72784886 |
| AKNA | 1632.86886 | -0.0941069 | 0.38004454 | -0.2476207 | 0.80442791 | 0.97287947 |
| WHRN | 1343.22875 | -0.4740781 | 0.2398482 | -1.9765757 | 0.04808962 | 0.58440502 |
| ATP6V1G1 | 6326.23331 | 0.0715718 | 0.20569919 | 0.347944 | 0.72788224 | 0.95848028 |
| TMEM268 | 834.113142 | -0.3805924 | 0.36040432 | -1.0560152 | 0.29096128 | 0.83454586 |
| TNFSF15 | 1485.33138 | -0.3763991 | 0.50117081 | -0.7510395 | 0.4526289 | 0.89670027 |
| TNFSF8 | 50.5629608 | -0.1833088 | 0.64878233 | -0.2825429 | 0.77752728 | 0.96836854 |
| TNC | 8130.50845 | 0.27691318 | 0.65178563 | 0.42485315 | 0.67094373 | 0.94922827 |
| PAPPA | 2185.59029 | -0.1736807 | 0.43489383 | -0.3993634 | 0.68962547 | 0.95158502 |
| PAPPA-AS1 | 730.324552 | -0.2275028 | 0.48251884 | -0.47149 | 0.63729083 | 0.94088547 |
| ASTN2 | 581.204061 | -0.1310458 | 0.30084338 | -0.4355949 | 0.6631307 | 0.94642611 |
| TRIM32 | 591.44299 | -0.2639317 | 0.23455689 | -1.1252354 | 0.26048928 | 0.82424998 |
| TLR4 | 1342.84376 | -0.0399034 | 0.34558459 | -0.1154663 | 0.90807554 | 0.98662757 |
| AL160272.1 | 41.115189 | 0.01862028 | 0.42424608 | 0.04389028 | 0.96499187 | 0.99357855 |
| BRINP1 | 149.044917 | 0.34899331 | 0.53803031 | 0.6486499 | 0.51656469 | 0.91636259 |
| CDK5RAP2 | 1348.57865 | -0.0081758 | 0.14882615 | -0.0549351 | 0.95619015 | 0.99185043 |
| MEGF9 | 1805.43967 | 0.27772182 | 0.23064683 | 1.20409984 | 0.22855099 | 0.80391781 |
| FBXW2 | 3469.89299 | -0.0532394 | 0.12828782 | -0.415 | 0.67814192 | 0.95020993 |
| B3GNT10 | 144.00544 | 0.16602173 | 0.2351514 | 0.70602056 | 0.48017535 | 0.90541935 |
| PSMD5 | 1502.08379 | 0.2014047 | 0.20068306 | 1.00359593 | 0.31557342 | 0.84730106 |
| PHF19 | 648.395042 | 0.08274961 | 0.34416233 | 0.24043773 | 0.80999093 | 0.97334138 |
| TRAF1 | 1040.47408 | 0.03156998 | 0.39927602 | 0.07906806 | 0.93697848 | 0.99068685 |
| C5 | 421.449503 | 0.90548187 | 0.39491947 | 2.29282664 | 0.02185799 | 0.51592008 |
| CNTRL | 551.638001 | 0.13166829 | 0.23116706 | 0.56958066 | 0.56896215 | 0.92660886 |
| RAB14 | 6128.99856 | -0.0585379 | 0.120167 | -0.4871378 | 0.62616068 | 0.94088547 |
| GSN | 19744.9188 | -0.2547198 | 0.37364483 | -0.6817163 | 0.49541835 | 0.90847703 |
| GSN-AS1 | 628.07177 | -0.1390145 | 0.34403705 | -0.4040683 | 0.68616252 | 0.95097288 |
| STOM | 16465.2151 | -0.3761078 | 0.23038088 | -1.6325479 | 0.10256412 | 0.67335541 |
| DAB2IP | 8311.31955 | -0.3054705 | 0.13479212 | -2.2662337 | 0.02343708 | 0.51967919 |
| TTLL11 | 180.184386 | -0.5829394 | 0.23839429 | -2.4452742 | 0.0144742 | 0.43901686 |
| NDUFA8 | 2226.08109 | 0.18713499 | 0.17130011 | 1.09243939 | 0.27464002 | 0.82719407 |

| | | | | | | |
|---|---|---|---|---|---|---|
| LHX6 | 130.171742 | 0.302224 | 0.6009297 | 0.50292738 | 0.61501533 | 0.93895494 |
| RBM18 | 1439.59721 | -0.021333 | 0.14776189 | -0.1443741 | 0.88520506 | 0.98356741 |
| MRRF | 2670.52295 | -0.1847046 | 0.11163182 | -1.6545873 | 0.09800824 | 0.66549286 |
| PTGS1 | 598.054785 | 0.30286939 | 0.67973283 | 0.44557123 | 0.655907 | 0.94534713 |
| PDCL | 688.749406 | -0.211294 | 0.19756906 | -1.0694692 | 0.28485829 | 0.83107779 |
| RC3H2 | 2069.54817 | -0.1126096 | 0.12603953 | -0.8934468 | 0.37161797 | 0.86762749 |
| ZBTB6 | 396.425777 | -0.0798339 | 0.20240945 | -0.3944179 | 0.69327252 | 0.9519683 |
| ZBTB26 | 498.449987 | -0.1836833 | 0.22873492 | -0.8030399 | 0.42195165 | 0.88451372 |
| RABGAP1 | 2410.96932 | -0.1518874 | 0.0984 | -1.5435712 | 0.12269225 | 0.69788192 |
| STRBP | 1357.32612 | -0.0587839 | 0.11340884 | -0.5183358 | 0.60422398 | 0.93655508 |
| CRB2 | 1202.00863 | 0.09841284 | 0.36728546 | 0.26794645 | 0.78874054 | 0.97090793 |
| DENND1A | 2723.36963 | 0.05056405 | 0.1950257 | 0.25926867 | 0.79542795 | 0.97101097 |
| NEK6 | 2127.04675 | 0.05134434 | 0.47499669 | 0.1080941 | 0.91392105 | 0.98662757 |
| PSMB7 | 5182.39587 | -0.1261313 | 0.29024035 | -0.4345753 | 0.66387072 | 0.94642611 |
| NR6A1 | 502.631333 | -0.4115028 | 0.25398244 | -1.620202 | 0.1051889 | 0.67473222 |
| OLFML2A | 517.498766 | 0.61534468 | 0.49487626 | 1.24343139 | 0.21370891 | 0.7926732 |
| WDR38 | 530.644556 | 0.03100501 | 0.17241775 | 0.17982493 | 0.85729001 | 0.97953896 |
| RPL35 | 11849.1915 | 0.01844025 | 0.1535115 | 0.1201229 | 0.90438579 | 0.98624484 |
| ARPC5L | 2521.799 | -0.3405985 | 0.23860999 | -1.4274278 | 0.15345661 | 0.74258845 |
| GOLGA1 | 1267.50591 | -0.0665451 | 0.21566257 | -0.308561 | 0.7576555 | 0.96502715 |
| SCAI | 431.863064 | 0.32971171 | 0.25416352 | 1.29724246 | 0.19454777 | 0.7762856 |
| PPP6C | 3421.44944 | -0.0952355 | 0.14392173 | -0.6617171 | 5.08E-01 | 9.14E-01 |
| RABEPK | 814.186358 | 0.33059318 | 0.20282078 | 1.62997686 | 0.10310639 | 0.67430143 |
| HSPA5 | 27123.9882 | 0.44929866 | 0.32080655 | 1.40052835 | 0.16135516 | 0.74924826 |
| GAPVD1 | 2018.43511 | -0.1386834 | 0.1473917 | -0.9409175 | 3.47E-01 | 8.61E-01 |
| MAPKAP1 | 4975.76396 | 0.09874249 | 0.35303462 | 0.27969633 | 0.7797105 | 0.96836854 |
| PBX3 | 619.225673 | -8.42E-05 | 0.19130548 | -0.0004403 | 0.99964867 | 0.99994357 |
| MVB12B | 641.715438 | -0.2023845 | 0.21363202 | -0.9473509 | 0.34346003 | 0.85938376 |
| LMX1B | 342.135891 | 0.44015405 | 0.85900509 | 0.51239981 | 6.08E-01 | 9.37E-01 |
| ZBTB43 | 1129.95286 | -0.6580505 | 0.34452361 | -1.9100303 | 5.61E-02 | 0.59895387 |
| ZBTB34 | 814.848508 | -0.2884832 | 0.2517081 | -1.146102 | 0.25175296 | 0.81833349 |
| RALGPS1 | 1376.29995 | 0.14374072 | 0.1707634 | 0.84175369 | 0.39992584 | 0.88077967 |
| ANGPTL2 | 1365.00286 | -1.1564338 | 0.59640185 | -1.9390178 | 0.05249917 | 0.58645127 |
| AL450263.1 | 132.401914 | 0.11285787 | 0.23881428 | 0.47257588 | 0.6365158 | 0.94088547 |
| GARNL3 | 350.210571 | 0.22091818 | 0.24999732 | 0.88368217 | 0.37686781 | 0.86987124 |
| SLC2A8 | 1266.28029 | 0.22613457 | 0.18283684 | 1.23681074 | 0.21615735 | 0.7954827 |
| ZNF79 | 205.881692 | -0.0093384 | 0.31793856 | -0.0293716 | 0.97656821 | 0.99455099 |
| RPL12 | 23695.1852 | 0.08558325 | 0.16575213 | 0.51633271 | 0.60562205 | 0.93698713 |
| LRSAM1 | 2045.0302 | -0.2719702 | 0.17358537 | -1.5667807 | 0.11716596 | 0.68956727 |
| FAM129B | 12526.3961 | -0.4923439 | 0.27969163 | -1.7603098 | 0.07835529 | 0.64079926 |
| STXBP1 | 2218.1665 | -0.1020008 | 0.26420259 | -0.3860703 | 0.69944462 | 0.95307625 |
| AL162426.1 | 191.437543 | -0.2351597 | 0.29593868 | -0.794623 | 0.42683281 | 0.88709241 |
| PTRH1 | 857.242567 | 0.04698472 | 0.22978272 | 0.20447456 | 0.83798267 | 0.97592799 |
| CFAP157 | 841.641796 | 0.03773365 | 0.27448572 | 0.13747034 | 0.89065903 | 0.98474725 |
| TTC16 | 93.7447237 | 0.13405726 | 0.37459271 | 0.35787473 | 0.72043706 | 0.95709301 |
| TOR2A | 852.444841 | 0.06626479 | 0.17694498 | 0.37449376 | 0.708037 | 0.95501548 |
| SH2D3C | 1381.83135 | -0.2939933 | 0.36628846 | -0.8026277 | 0.42218995 | 0.88451372 |
| CDK9 | 4174.57244 | -0.1108982 | 0.13445477 | -0.8247996 | 0.40948537 | 0.88239224 |
| FPGS | 3423.81962 | -0.0808195 | 0.19267233 | -0.4194663 | 0.6748754 | 0.94935413 |
| ENG | 19658.6556 | 0.2403824 | 0.52634262 | 0.45670328 | 0.64788434 | 0.94285781 |
| AL162586.1 | 4928.19016 | 0.22144608 | 0.48903331 | 0.45282412 | 0.6506754 | 0.94346674 |
| AK1 | 2268.33025 | 0.05122711 | 0.26734034 | 0.19161757 | 0.84804178 | 0.97711043 |
| AL157935.2 | 7058.12648 | 0.05188511 | 0.21075462 | 0.2461873 | 0.80553725 | 0.97321903 |
| ST6GALNAC( | 5182.24621 | 0.03242017 | 0.20769169 | 0.15609758 | 0.87595611 | 0.98275231 |

| | | | | | | |
|---|---|---|---|---|---|---|
| ST6GALNAC4 | 1959.36245 | 0.221086 | 0.48737612 | 0.45362501 | 0.65009875 | 0.943166 |
| PIP5KL1 | 123.857458 | -0.4569369 | 0.36647611 | -1.2468394 | 0.21245639 | 0.79203904 |
| AL157935.1 | 885.859081 | -1.3234 | 0.79922218 | -1.65586 | NA | NA |
| DPM2 | 3206.74165 | -0.7209975 | 0.48349212 | -1.4912291 | NA | NA |
| FAM102A | 9451.68315 | -0.2335743 | 0.52035143 | -0.4488779 | 0.65351972 | 0.94495971 |
| NAIF1 | 470.59059 | -0.0681067 | 0.21586744 | -0.3155025 | 0.75238018 | 0.96301881 |
| SLC25A25 | 4498.90223 | -0.1565996 | 0.42349112 | -0.3697825 | 0.71154458 | 0.95560193 |
| SLC25A25-AS | 2544.64466 | -0.1292628 | 0.23726916 | -0.5447941 | 0.58589516 | 0.93289552 |
| PTGES2 | 4090.84222 | 0.065182 | 0.14748936 | 0.44194374 | 0.65852991 | 0.9458503 |
| AL590708.1 | 63.7623327 | 0.47540886 | 0.49358598 | 0.96317335 | 3.35E-01 | 0.85616588 |
| LCN2 | 7435.77427 | 0.45480764 | 0.70884983 | 0.64161353 | 0.52112415 | 0.91723878 |
| C9orf16 | 1999.26229 | 0.2944984 | 0.21488774 | 1.37047554 | 0.17053851 | 0.75956362 |
| CIZ1 | 4228.85877 | -0.0927688 | 0.1206238 | -0.7690751 | 0.44184871 | 0.89318162 |
| DNM1 | 2325.25545 | -0.2008605 | 0.3391118 | -0.5923135 | 0.55364064 | 0.92373235 |
| GOLGA2 | 4765.80876 | -0.3119347 | 0.24052415 | -1.2968956 | 0.1946671 | 0.7762856 |
| SWI5 | 807.958147 | 0.20221846 | 0.19936607 | 1.01430732 | 0.31043613 | 0.84476049 |
| TRUB2 | 2765.92146 | 0.00963436 | 0.16887497 | 0.05705027 | 0.95450515 | 0.99170503 |
| AL359091.4 | 39.6668091 | 0.28327132 | 0.39369779 | 0.71951462 | 0.4718239 | 0.90160919 |
| COQ4 | 4902.9931 | 0.07415575 | 0.18495908 | 0.40093059 | 0.68847123 | 0.95158502 |
| SLC27A4 | 3537.79797 | 0.18501744 | 0.27446367 | 0.67410537 | 0.50024432 | 0.91180773 |
| URM1 | 2826.52281 | -0.0592786 | 0.13943053 | -0.4251481 | 0.67072872 | 0.94922827 |
| AL359091.3 | 99.0122816 | 0.03894934 | 0.22740632 | 0.17127641 | 0.86400643 | 0.98176973 |
| CERCAM | 1722.39861 | 0.17959392 | 0.43789354 | 0.41013146 | 0.68170951 | 0.95070301 |
| ODF2 | 1757.05633 | -0.1640674 | 0.18147082 | -0.9040978 | 0.36594357 | 0.86672412 |
| ODF2-AS1 | 99.018286 | -0.0766231 | 0.3743628 | -0.204676 | 0.8378253 | 0.9758253 |
| GLE1 | 1718.98671 | 0.0269617 | 0.17322806 | 0.1556428 | 0.87631459 | 0.98275231 |
| AL356481.1 | 270.339007 | 0.05829415 | 0.20378629 | 0.28605533 | 0.77483574 | 0.96836854 |
| SPTAN1 | 11246.2939 | 0.00695003 | 0.10824322 | 0.06420749 | 0.94880501 | 0.99170503 |
| AL356481.3 | 1220.92028 | -0.0507844 | 0.1416271 | -0.3585785 | 0.71991044 | 0.95680034 |
| WDR34 | 3563.95493 | 0.12422093 | 0.26683249 | 0.46553901 | 0.64154551 | 0.94088547 |
| SET | 10701.6569 | -0.0963238 | 0.19901464 | -0.4840036 | 0.62838334 | 0.94088547 |
| PKN3 | 972.704531 | -0.4510498 | 0.29803032 | -1.513436 | 0.13016895 | 0.70814767 |
| ZDHHC12 | 1383.52542 | 0.00292701 | 0.18684715 | 0.01566525 | 0.98750145 | 0.997408 |
| AL441992.1 | 109.773822 | 0.26444723 | 0.36476804 | 0.72497368 | 0.46846818 | 0.89984974 |
| ZER1 | 4623.46958 | -0.0120928 | 0.17653065 | -0.0685023 | 0.94538577 | 0.99166614 |
| TBC1D13 | 4344.21685 | 0.03555025 | 0.26450468 | 0.1344031 | 0.89308383 | 0.98474725 |
| ENDOG | 1746.42013 | -0.2321042 | 0.16557101 | -1.4018407 | 0.16096281 | 0.74874703 |
| SPOUT1 | 3258.89949 | -0.2408024 | 0.13508252 | -1.7826315 | 0.07464632 | 0.63773415 |
| KYAT1 | 778.641958 | 0.30628017 | 0.31787261 | 0.96353117 | 0.33528103 | 0.85599307 |
| LRRC8A | 3696.18791 | -0.4867057 | 0.31338689 | -1.5530506 | 0.12041106 | 0.69467399 |
| PHYHD1 | 2808.34943 | 0.15892014 | 0.28393951 | 0.55969718 | 0.57568601 | 0.93083012 |
| AL672142.1 | 792.390439 | 0.13502652 | 0.28773393 | 0.46927563 | 0.63887263 | 0.94088547 |
| DOLK | 938.790835 | 0.04731991 | 0.23001461 | 0.20572567 | 0.83700521 | 0.97577999 |
| NUP188 | 2148.45895 | -0.2209719 | 0.12326349 | -1.7926796 | 0.07302417 | 0.63580072 |
| AL592211.1 | 44.3269385 | 0.17361261 | 0.39871655 | 0.43542866 | 6.63E-01 | 0.94642611 |
| SH3GLB2 | 9685.11149 | -0.0828264 | 0.27937863 | -0.2964665 | 0.76687386 | 0.96741943 |
| MIGA2 | 3042.93559 | -0.1143767 | 0.17625463 | -0.6489287 | 0.51638445 | 0.91636259 |
| DOLPP1 | 754.329758 | 0.27593133 | 0.21753587 | 1.26844064 | 0.20464064 | 0.78460928 |
| CRAT | 5926.1857 | 0.33845762 | 0.16457616 | 2.05654102 | 0.0397304 | 0.55657615 |
| AL158151.2 | 540.753854 | 0.3681861 | 0.17627145 | 2.08874493 | 0.03673069 | 0.55521179 |
| PTPA | 9096.57099 | 0.21888567 | 0.11612387 | 1.88493254 | 0.05943895 | 0.60540505 |
| IER5L | 2075.43327 | -0.2387474 | 0.42742221 | -0.5585752 | 0.57645169 | 0.9311568 |
| AL158151.3 | 275.379102 | 0.0070834 | 0.42005764 | 0.01686293 | 0.98654597 | 0.99715899 |
| AL158151.1 | 1039.57482 | -0.1790442 | 0.43781938 | -0.4089454 | 0.68257976 | 0.95070301 |

| | | | | | | |
|---|---|---|---|---|---|---|
| AL161785.1 | 91.2142432 | -0.0289213 | 0.46812431 | -0.0617812 | 0.95073709 | 0.99170503 |
| LINC01503 | 2422.67102 | -0.4196635 | 0.47208915 | -0.8889497 | 0.37403011 | 0.86791605 |
| AL353803.2 | 44.4156509 | 0.18218517 | 0.62980772 | 0.2892711 | 0.77237393 | 0.96836854 |
| AL353803.1 | 67.9063159 | 0.07184916 | 0.67005194 | 0.10722924 | 9.15E-01 | 9.87E-01 |
| NTMT1 | 2214.4976 | -0.1578327 | 0.1530224 | -1.0314354 | 0.30233669 | 0.84211658 |
| ASB6 | 3440.91291 | -0.2567787 | 0.13211742 | -1.9435643 | 0.05194802 | 0.58645127 |
| AL590369.1 | 172.95112 | -0.2508477 | 0.25377851 | -0.9884512 | 0.32293171 | 0.85119105 |
| PRRX2 | 52.6145636 | -1.2625813 | 0.6843107 | -1.845041 | 0.06503157 | 0.62154765 |
| PTGES | 677.258644 | 0.35211205 | 0.55604921 | 0.633239 | 0.52657758 | 0.91769856 |
| TOR1B | 1279.51226 | -0.4206917 | 0.17440181 | -2.4121981 | 1.59E-02 | 0.4564177 |
| TOR1A | 1929.46264 | -0.1189509 | 0.12018885 | -0.9897003 | 0.32232062 | 0.85119105 |
| C9orf78 | 3030.01902 | 0.0611448 | 0.17949044 | 0.34065768 | 7.33E-01 | 0.95884813 |
| USP20 | 2946.55918 | 0.04087961 | 0.12105613 | 0.33769136 | 0.73559579 | 0.95895327 |
| FNBP1 | 2593.54337 | -0.1952736 | 0.20126245 | -0.9702435 | 0.33192511 | 0.85452795 |
| GPR107 | 4453.55365 | 0.03933144 | 0.14606394 | 0.2692755 | 0.78771768 | 0.97090793 |
| NCS1 | 4077.25468 | -0.3757951 | 0.35732307 | -1.0516955 | 0.29293928 | 0.83524862 |
| HMCN2 | 647.885762 | -1.7569516 | 0.91277866 | -1.9248386 | NA | NA |
| ASS1 | 39558.8131 | 0.38875781 | 0.48325823 | 0.8044515 | 0.42113627 | 0.88451372 |
| FUBP3 | 2851.56318 | -0.2326719 | 0.23584525 | -0.9865446 | 0.32386594 | 0.85156215 |
| EXOSC2 | 1067.48757 | -0.1152599 | 0.18971065 | -0.6075563 | 0.5434818 | 0.92297267 |
| ABL1 | 5125.22908 | -0.234897 | 0.17304424 | -1.3574389 | 0.17464179 | 0.76009329 |
| LAMC3 | 2619.47672 | 0.098735 | 0.58956006 | 0.16747234 | 0.86699841 | 0.98223481 |
| AIF1L | 27930.2994 | 0.34570554 | 0.31897943 | 1.08378634 | 0.27845955 | 0.82805357 |
| NUP214 | 3814.46051 | 0.09977304 | 0.19150792 | 0.52098648 | 0.60237619 | 0.9355118 |
| AL157938.2 | 46.1844544 | -0.0511805 | 0.5245573 | -0.0975689 | 0.92227463 | 0.98826666 |
| AL157938.3 | 74.518502 | -0.0632028 | 0.33912896 | -0.186368 | 0.8521562 | 0.97872457 |
| FAM78A | 294.427871 | -0.2117527 | 0.45057943 | -0.4699565 | 0.63838613 | 0.94088547 |
| PLPP7 | 279.140191 | 0.09658675 | 0.28100135 | 0.34372342 | 0.7310543 | 0.95848028 |
| PRRC2B | 10379.8031 | -0.1868006 | 0.10545619 | -1.7713574 | 0.07650128 | 0.63773415 |
| AL358781.1 | 6409.25366 | -0.0942368 | 0.13361945 | -0.7052623 | 0.48064699 | 0.90541935 |
| POMT1 | 2744.75661 | 0.04085806 | 0.08164403 | 0.50044152 | 0.61676423 | 0.93910158 |
| AL358781.2 | 345.220942 | 0.00137374 | 0.13818867 | 0.00994102 | 0.99206834 | 0.99906045 |
| UCK1 | 2684.06194 | -0.1428642 | 0.17243697 | -0.8285012 | 0.40738672 | 0.88132015 |
| PRRT1B | 38.1444899 | -0.7771846 | 0.6107883 | -1.2724288 | 2.03E-01 | 7.83E-01 |
| RAPGEF1 | 5378.32554 | -0.4703334 | 0.27831976 | -1.6899031 | 9.10E-02 | 6.55E-01 |
| MED27 | 738.265343 | 0.13462898 | 0.18572552 | 0.72488144 | 0.46852477 | 0.89984974 |
| NTNG2 | 83.7886226 | -0.6854965 | 0.45186315 | -1.5170445 | 0.12925544 | 0.70746969 |
| SETX | 3347.29403 | -0.2670753 | 0.1959103 | -1.3632532 | 0.17280274 | 0.75974924 |
| TTF1 | 536.990328 | -0.0592286 | 0.14387214 | -0.4116749 | 0.68057769 | 0.95070301 |
| DDX31 | 1034.99858 | -0.4807339 | 0.20542949 | -2.3401404 | 1.93E-02 | 0.48835495 |
| GTF3C4 | 1212.76617 | -0.3503982 | 0.21247005 | -1.6491649 | 0.09911385 | 0.66888465 |
| AK8 | 246.658738 | 0.49499363 | 0.36311228 | 1.36319715 | 0.17282039 | 0.75974924 |
| SPACA9 | 713.092132 | 0.33840945 | 0.29840058 | 1.13407773 | 2.57E-01 | 8.21E-01 |
| TSC1 | 2931.57389 | -0.2218077 | 0.13677676 | -1.6216769 | 0.10487256 | 0.67459611 |
| GTF3C5 | 3384.80261 | -0.5066778 | 0.24065613 | -2.1054018 | 0.03525635 | 0.55369025 |
| CEL | 567.407781 | -0.1531098 | 0.61711815 | -0.2481044 | 0.80405359 | 0.97274342 |
| RALGDS | 3397.59954 | -0.2415467 | 0.24891616 | -0.9703937 | 0.33185028 | 0.85452131 |
| AL162417.1 | 2999.36327 | -0.2820118 | 0.24329853 | -1.1591182 | 0.24640801 | 0.81542272 |
| GBGT1 | 640.220183 | -0.2125451 | 0.37583518 | -0.5655274 | 0.57171513 | 0.92846735 |
| ABO | 2680.01552 | -0.0464305 | 0.41630172 | -0.111531 | 0.91119531 | 0.98662757 |
| SURF6 | 2217.7883 | -0.0157403 | 0.19293958 | -0.0815817 | 0.93497933 | 0.99068685 |
| MED22 | 2464.80619 | -0.3691211 | 0.18288586 | -2.0183139 | 0.04355858 | 0.56966602 |
| RPL7A | 41382.6453 | 0.24282191 | 0.21333057 | 1.13824244 | 0.25501926 | 0.81992039 |
| SURF1 | 2823.99727 | 0.0957685 | 0.16250594 | 0.58932306 | 5.56E-01 | 9.24E-01 |

| | | | | | | |
|---|---|---|---|---|---|---|
| SURF2 | 732.905952 | -0.1354768 | 0.22117213 | -0.61254 | 5.40E-01 | 9.22E-01 |
| SURF4 | 12497.2186 | -0.2295482 | 0.25432163 | -0.90259 | 0.36674356 | 0.86683872 |
| STKLD1 | 114.046577 | 0.29584224 | 0.29929582 | 0.98846099 | 0.32292693 | 0.85119105 |
| REXO4 | 1365.67498 | -0.0758166 | 0.17919213 | -0.4231024 | 0.67222056 | 0.94922827 |
| ADAMTS13 | 670.335838 | -0.0839247 | 0.43595465 | -0.1925078 | 0.84734447 | 0.97693526 |
| CACFD1 | 4032.79258 | 0.29242555 | 0.26503388 | 1.10335158 | 0.26987451 | 0.82716979 |
| SLC2A6 | 460.949803 | -0.2229709 | 0.44468578 | -0.5014123 | 0.61608097 | 0.93910158 |
| ADAMTSL2 | 3453.22529 | -0.2301813 | 0.58698359 | -0.3921427 | 0.69495277 | 0.95234774 |
| DBH | 329.909702 | 0.41235799 | 0.44558847 | 0.92542339 | 0.35474572 | 0.86364633 |
| DBH-AS1 | 788.118984 | 0.37900267 | 0.40281793 | 0.94087834 | 0.34676721 | 0.86135217 |
| SARDH | 1289.881 | 0.96759077 | 0.51244696 | 1.8881774 | 0.05900214 | 0.60423993 |
| VAV2 | 5039.21343 | -0.2941951 | 0.18456169 | -1.5940203 | 0.11093148 | 0.68001961 |
| BRD3OS | 2160.79988 | -0.0252665 | 0.17716804 | -0.1426134 | 0.8865955 | 0.98420297 |
| BRD3 | 1872.43806 | -0.0714129 | 0.19026843 | -0.3753272 | 0.70741715 | 0.95501548 |
| BX649632.1 | 41.3203812 | 0.40089339 | 0.42089496 | 0.95247849 | 0.34085437 | 0.85859566 |
| WDR5 | 2535.93806 | -0.1897606 | 0.13594353 | -1.395878 | 0.16275123 | 0.75032457 |
| RXRA | 5763.45297 | -0.0570393 | 0.30617473 | -0.1862965 | 0.85221224 | 0.97872457 |
| COL5A1 | 2007.31558 | -0.3750739 | 0.45929032 | -0.8166379 | 4.14E-01 | 0.88364943 |
| FCN2 | 163.241094 | 0.26042964 | 0.62033285 | 0.41982242 | 0.67461518 | 0.94935413 |
| FCN1 | 394.875139 | 0.96114366 | 1.07104873 | 0.89738556 | 0.36951321 | 0.86762749 |
| OLFM1 | 531.917381 | 0.94163218 | 0.54116258 | 1.74001719 | 0.081856 | 0.64544584 |
| PPP1R26-AS1 | 275.020056 | 0.10837734 | 0.40477738 | 0.26774554 | 0.78889519 | 0.97090793 |
| PPP1R26 | 2135.15394 | -0.1897621 | 0.17890717 | -1.0606735 | 0.2888383 | 0.83295411 |
| C9orf116 | 278.613334 | 0.1353259 | 0.19570122 | 0.69149235 | 0.48925619 | 0.90600166 |
| MRPS2 | 3067.17568 | 0.04854181 | 0.1451118 | 0.33451315 | 0.73799237 | 0.9602325 |
| AL161452.1 | 1531.3824 | 0.09723443 | 0.15520965 | 0.62647155 | 0.53100569 | 0.91962754 |
| CAMSAP1 | 1217.85015 | -0.2903763 | 0.29350325 | -0.9893463 | 0.32249374 | 0.85119105 |
| UBAC1 | 2232.63585 | -0.0189079 | 0.15724021 | -0.1202484 | 0.90428635 | 0.98624484 |
| NACC2 | 2703.61752 | -0.040939 | 0.26598418 | -0.1539151 | 0.87767668 | 0.98275231 |
| TMEM250 | 2240.12247 | -0.4547064 | 0.2322006 | -1.958248 | 0.05020092 | 0.5861488 |
| QSOX2 | 1205.18852 | -0.1679418 | 0.13328142 | -1.2600544 | 0.20764974 | 0.78834607 |
| CCDC187 | 75.3553621 | 0.62547344 | 0.57015643 | 1.09702077 | 0.27263231 | 0.82716979 |
| GPSM1 | 1352.14556 | -0.0790507 | 0.32439894 | -0.2436835 | 0.80747594 | 0.97334138 |
| DNLZ | 434.490799 | -0.3856475 | 0.30468912 | -1.265708 | 0.20561763 | 0.78578305 |
| CARD9 | 647.998544 | -0.2880893 | 0.32894342 | -0.8758019 | 0.38113774 | 0.87171941 |
| SNAPC4 | 2037.45715 | -0.6319424 | 0.27119756 | -2.330192 | 0.01979601 | 0.49477123 |
| ENTR1 | 2146.42748 | -0.4032772 | 0.22898902 | -1.7611202 | 0.07821807 | 0.64079926 |
| PMPCA | 3400.70994 | 0.15271682 | 0.20384847 | 0.7491683 | 0.45375577 | 0.89670027 |
| INPP5E | 2106.20523 | -0.1369661 | 0.16791158 | -0.8157039 | 4.15E-01 | 8.84E-01 |
| SEC16A | 4900.30159 | -0.178983 | 0.19148551 | -0.9347077 | 0.34993894 | 0.86364633 |
| C9orf163 | 59.7422338 | 0.34895362 | 0.41272184 | 0.84549346 | 0.39783538 | 0.88054917 |
| NOTCH1 | 4601.74093 | -0.1252922 | 0.17361739 | -0.7216573 | 0.47050523 | 0.9008571 |
| NALT1 | 152.548808 | 0.95835359 | 0.53752491 | 1.78290079 | 0.07460246 | 0.63773415 |
| AL590226.2 | 47.9334355 | 0.36570129 | 0.76711653 | 0.47672195 | 0.63356013 | 0.94088547 |
| AL590226.1 | 72.2500478 | 0.03005097 | 0.4943591 | 0.06078774 | 0.95152825 | 0.99170503 |
| EGFL7 | 6951.24279 | -0.0588689 | 0.30401911 | -0.1936356 | 0.84646124 | 0.97681836 |
| AGPAT2 | 3977.48785 | 0.19157167 | 0.24448595 | 0.78356922 | 0.43329292 | 0.89055338 |
| FAM69B | 1128.14642 | -0.0238804 | 0.37250571 | -0.0641075 | 0.94888462 | 0.99170503 |
| SNHG7 | 2642.28766 | -0.280434 | 0.21894187 | -1.2808603 | 0.20024273 | 0.78134034 |
| LCN10 | 173.046165 | -1.7442818 | 0.87943706 | -1.9834072 | 0.04732196 | 0.58218854 |
| AL355987.1 | 206.696132 | -1.86722 | 0.84201881 | -2.2175515 | 0.02658543 | 0.52678638 |
| LCN6 | 73.936436 | -2.5563344 | 0.92814624 | -2.7542366 | 0.00588292 | 0.32813047 |
| TMEM141 | 3458.02167 | 0.53241423 | 0.27343219 | 1.94715274 | 0.05151643 | 0.58645127 |
| AL355987.3 | 2519.3197 | 0.55199062 | 0.27034463 | 2.04180351 | 0.04117103 | 0.55871557 |

| | | | | | | |
|---|---|---|---|---|---|---|
| CCDC183 | 734.923755 | 0.33434872 | 0.2908269 | 1.14964853 | 0.25028866 | 0.81721208 |
| CCDC183-AS: | 667.894599 | 0.50611002 | 0.34989246 | 1.44647307 | 0.14804456 | 0.73474739 |
| RABL6 | 6420.37559 | 0.04510182 | 0.12103596 | 0.37263155 | 0.70942269 | 0.95501548 |
| AJM1 | 780.015626 | 0.01397109 | 0.43694137 | 0.03197474 | 0.97449219 | 0.99455099 |
| PHPT1 | 4620.39605 | 0.28583292 | 0.21254149 | 1.34483352 | 0.178679 | 0.76120583 |
| MAMDC4 | 2142.15732 | -0.2068666 | 0.40594946 | -0.509587 | 0.61034083 | 0.93792127 |
| EDF1 | 9567.02561 | 0.0493113 | 0.13568171 | 0.36343365 | 0.71628097 | 0.95603249 |
| TRAF2 | 1055.01431 | 0.01676573 | 0.18536436 | 0.09044745 | 0.92793165 | 0.9892705 |
| FBXW5 | 7570.65013 | -0.1363915 | 0.15656283 | -0.8711615 | 0.38366599 | 0.87239556 |
| C8G | 164.314452 | 1.47871468 | 0.55117698 | 2.68283098 | 0.00730019 | 0.3610358 |
| LCN12 | 1991.07622 | 0.44755383 | 0.59688498 | 0.74981587 | 0.45336561 | 0.89670027 |
| PTGDS | 6681.52965 | 1.19263394 | 0.43718702 | 2.72797197 | 0.0063725 | 0.33501741 |
| AL807752.7 | 6393.61366 | 1.22460057 | 0.4376677 | 2.79801448 | 0.00514178 | 0.30843846 |
| LCNL1 | 199.122117 | 1.86457098 | 0.66919553 | 2.78628726 | 0.00533156 | 0.31308771 |
| PAXX | 1724.61459 | 0.05008234 | 0.2037613 | 0.24578926 | 0.80584537 | 0.97334138 |
| ABCA2 | 4769.95548 | 0.38329417 | 0.33362819 | 1.14886628 | 0.25061113 | 0.81738655 |
| C9orf139 | 90.8333882 | 0.36793961 | 0.50914862 | 0.7226566 | 0.46989088 | 0.9008571 |
| AL807752.6 | 1172.63578 | -0.0178901 | 0.44616476 | -0.0400975 | 0.96801537 | 0.99400331 |
| NPDC1 | 4186.19284 | -0.0468015 | 0.45305181 | -0.1033027 | 0.91772272 | 0.9868044 |
| ENTPD2 | 817.954893 | 0.76535123 | 0.4529102 | 1.68985206 | 0.09105626 | 0.65495306 |
| SAPCD2 | 136.443917 | 1.34299386 | 0.55163728 | 2.43455963 | 0.01490992 | 0.44565688 |
| AL807752.3 | 43.619453 | 1.48961225 | 0.70858898 | 2.10222327 | 0.03553372 | 0.55369025 |
| UAP1L1 | 3366.74216 | -0.6482525 | 0.32173248 | -2.0148806 | 0.04391716 | 0.57116687 |
| MAN1B1-DT | 142.797314 | -0.0281372 | 0.38517297 | -0.0730509 | 0.94176558 | 0.99136103 |
| MAN1B1 | 4497.10742 | 0.25597564 | 0.17036862 | 1.50248112 | 0.1329729 | 0.7142577 |
| DPP7 | 7686.67632 | -0.0723602 | 0.10971559 | -0.6595252 | 0.50955858 | 0.91439937 |
| ANAPC2 | 3518.80283 | -0.0882982 | 0.14326838 | -0.6163132 | 0.53768786 | 0.92123049 |
| SSNA1 | 2357.0929 | -0.1567284 | 0.21855223 | -0.717121 | 0.47329945 | 0.90189171 |
| TPRN | 3601.50174 | -0.1944373 | 0.29989884 | -0.6483429 | 0.5167632 | 0.91636259 |
| TMEM203 | 1811.53469 | -0.0056503 | 0.11200112 | -0.0504488 | 0.95976477 | 0.99255196 |
| NDOR1 | 1594.18533 | -0.3354581 | 0.15968261 | -2.1007806 | 0.03566023 | 0.55369025 |
| BX255925.3 | 1453.52948 | -0.2209405 | 0.14773529 | -1.4955162 | 0.13477977 | 0.71630293 |
| RNF208 | 452.136363 | -0.1049295 | 0.30348686 | -0.3457466 | 7.30E-01 | 9.58E-01 |
| CYSRT1 | 237.294707 | 0.62714783 | 0.48478712 | 1.29365613 | 0.19578423 | 0.77770076 |
| RNF224 | 216.893028 | 0.68208357 | 0.62171079 | 1.0971075 | 0.2725944 | 0.82716979 |
| SLC34A3 | 1609.9857 | -0.3884273 | 0.69580804 | -0.5582391 | 0.57668112 | 0.93127944 |
| TUBB4B | 13607.4258 | -0.2892198 | 0.30193562 | -0.9578856 | 0.33812044 | 0.8571036 |
| FAM166A | 1340.27952 | -0.3539859 | 0.30738722 | -1.1515959 | 2.49E-01 | 0.81646572 |
| STPG3-AS1 | 779.744722 | 0.4342881 | 0.75551458 | 0.57482425 | 0.56541017 | 0.92512239 |
| STPG3 | 733.540138 | 0.31654862 | 0.76596852 | 0.41326584 | 0.67941186 | 0.95070301 |
| NELFB | 4101.85559 | -0.120061 | 0.13737813 | -0.8739458 | 0.38214776 | 0.87185666 |
| TOR4A | 1349.43373 | -0.1095526 | 0.29089225 | -0.3766088 | 0.70646437 | 0.95501548 |
| NRARP | 271.109433 | -0.0030997 | 0.46643095 | -0.0066455 | 0.99469772 | 0.99940889 |
| EXD3 | 2384.68397 | 0.02936305 | 0.31056579 | 0.09454695 | 0.92467469 | 0.98869145 |
| NOXA1 | 2038.37192 | 0.27586754 | 0.27296577 | 1.01063051 | 3.12E-01 | 8.45E-01 |
| ENTPD8 | 389.403752 | 0.77526439 | 0.28321569 | 2.7373639 | 0.00619337 | 0.33470127 |
| NSMF | 6273.33959 | 0.06942986 | 0.23484161 | 0.29564548 | 0.76750084 | 0.96749893 |
| PNPLA7 | 2024.61025 | 0.45550059 | 0.40952568 | 1.11226381 | 0.26602474 | 0.8265123 |
| MRPL41 | 2507.58275 | 0.08636213 | 0.15000559 | 0.57572604 | 0.56480037 | 0.92477055 |
| DPH7 | 1797.22691 | -0.2531662 | 0.14860318 | -1.7036391 | 0.08844853 | 0.65363785 |
| ZMYND19 | 1099.19549 | -0.3989739 | 0.28135101 | -1.4180646 | 0.15617191 | 0.74522306 |
| ARRDC1 | 3871.21322 | -0.1355383 | 0.13149654 | -1.0307368 | 0.30266426 | 0.84211658 |
| ARRDC1-AS1 | 1067.76068 | -0.1105667 | 0.19141006 | -0.5776433 | 0.56350496 | 0.92452095 |
| EHMT1 | 2554.34302 | -0.3230862 | 0.09938907 | -3.2507214 | 0.00115113 | 0.15697311 |

| | | | | | | |
|---|---|---|---|---|---|---|
| LINC01001 | 73.6634231 | -0.5671843 | 0.49826778 | -1.1383122 | 0.25499014 | 0.81992039 |
| BET1L | 4402.15089 | -0.0105848 | 0.14587115 | -0.0725627 | 0.94215412 | 0.99136103 |
| AC069287.2 | 1685.35299 | 0.00852789 | 0.15550459 | 0.0548401 | 0.95626586 | 0.99185043 |
| RIC8A | 4972.70839 | -0.3792732 | 0.21391475 | -1.7730109 | 0.0762269 | 0.63773415 |
| SIRT3 | 1842.4014 | 0.18663011 | 0.20338855 | 0.91760382 | 0.35882631 | 0.86480228 |
| PSMD13 | 5208.83274 | -0.0875959 | 0.15198473 | -0.5763468 | 5.64E-01 | 0.92474393 |
| NLRP6 | 246.132369 | 0.17687281 | 0.66772009 | 0.26489066 | 7.91E-01 | 9.71E-01 |
| AC136475.3 | 582.684179 | 0.47616854 | 0.91303847 | 0.52152079 | 0.60200403 | 0.9355118 |
| PGGHG | 11324.5892 | 0.20885234 | 0.62339032 | 0.3350266 | 0.73760502 | 0.9602325 |
| IFITM2 | 24158.6462 | -0.8385049 | 0.48680191 | -1.7224766 | 0.08498322 | 0.64988556 |
| AC136475.2 | 150.645579 | 0.44415755 | 0.58086979 | 0.7646422 | 0.44448463 | 0.8938278 |
| IFITM1 | 15621.3105 | -1.5877138 | 0.6914536 | -2.2961972 | 0.02166461 | 0.51394743 |
| AC136475.1 | 91.4641516 | 0.47699024 | 0.3862664 | 1.23487375 | 0.2168775 | 0.7954827 |
| IFITM3 | 38388.3241 | -0.7308637 | 0.41676984 | -1.7536386 | 0.07949245 | 0.64079926 |
| AC136475.5 | 59.2964383 | -1.0069523 | 0.60580257 | -1.662179 | 0.09647689 | 0.66233356 |
| AC136475.9 | 46.8956263 | -1.2556183 | 0.66795036 | -1.8798078 | 0.06013428 | 0.60819023 |
| B4GALNT4 | 85.2182234 | 0.31642603 | 0.71618066 | 0.44182431 | 0.65861634 | 0.9458503 |
| PKP3 | 303.851608 | 0.06787584 | 0.45561082 | 0.14897767 | 0.88157125 | 0.98275231 |
| SIGIRR | 5554.61153 | -0.0498453 | 0.19992289 | -0.2493226 | 0.80311122 | 0.97273362 |
| ANO9 | 2729.4574 | -0.3104331 | 0.35050542 | -0.8856726 | 0.37579397 | 0.86889275 |
| PTDSS2 | 2955.23934 | -0.0394165 | 0.11882084 | -0.3317306 | 0.7400927 | 0.96046516 |
| AC138230.1 | 48.7386824 | 0.1903767 | 0.40732866 | 0.46737861 | 0.64022902 | 0.94088547 |
| RNH1 | 12184.7888 | -0.1050044 | 0.13331686 | -0.7876306 | 0.43091283 | 0.88966606 |
| HRAS | 1918.03403 | -0.3593987 | 0.21461293 | -1.6746368 | 0.09400552 | 0.65783452 |
| LRRC56 | 470.883902 | -0.3034884 | 0.21743687 | -1.3957542 | 0.16278852 | 0.75032457 |
| LMNTD2 | 3652.90266 | -0.2732838 | 0.41135336 | -0.6643528 | 0.50646453 | 0.9141539 |
| AP006284.1 | 2831.88186 | -0.2720647 | 0.42705393 | -0.6370733 | 0.52407707 | 0.91723878 |
| RASSF7 | 6594.38766 | -0.0225701 | 0.29926722 | -0.0754179 | 0.93988223 | 0.99135146 |
| MIR210HG | 1585.32007 | -0.3159328 | 0.51395208 | -0.6147126 | 0.53874453 | 0.9215968 |
| PHRF1 | 4009.85537 | -0.2690254 | 0.09742459 | -2.7613704 | 0.00575594 | 0.32369964 |
| IRF7 | 3690.58885 | -1.9276048 | 0.64529651 | -2.9871613 | NA | NA |
| CDHR5 | 10893.2076 | 0.80341168 | 0.54165257 | 1.48326017 | 0.1380053 | 0.72209298 |
| DRD4 | 95.4457701 | 0.30395875 | 0.45220862 | 0.67216487 | 0.50147874 | 0.91259827 |
| DEAF1 | 2482.54243 | -0.1314939 | 0.14152386 | -0.9291287 | 0.3528224 | 0.86364633 |
| EPS8L2 | 21258.3664 | -0.0177835 | 0.23192364 | -0.0766781 | 0.93887965 | 0.99113899 |
| TMEM80 | 1803.10173 | -0.3031125 | 0.18541793 | -1.634753 | 0.10210084 | 0.67279543 |
| AP006621.4 | 1644.85285 | 0.18337801 | 0.21511641 | 0.85245941 | 0.39395916 | 0.87884163 |
| TALDO1 | 10631.0626 | 0.06171529 | 0.28771575 | 0.21450092 | 0.83015645 | 0.97550416 |
| GATD1 | 5169.63814 | -0.2776914 | 0.14877962 | -1.8664616 | 0.06197681 | 0.61210783 |
| AP006621.3 | 597.845812 | -0.2413105 | 0.4554797 | -0.5297942 | 0.5962546 | 0.9334601 |
| AP006621.2 | 131.641651 | -0.3022057 | 0.52604176 | -0.57449 | 0.56563626 | 0.92513982 |
| CEND1 | 48.9323942 | -0.7848861 | 0.55023633 | -1.4264526 | 0.15373773 | 0.74258845 |
| SLC25A22 | 2367.14583 | -0.2225243 | 0.200576 | -1.1094264 | 0.26724627 | 0.8265123 |
| PANO1 | 74.4443604 | -0.1536012 | 0.25406142 | -0.6045829 | 0.5454562 | 0.92345821 |
| PIDD1 | 2220.30115 | -0.0389741 | 0.22203686 | -0.1755298 | 0.86066334 | 0.98106934 |
| RPLP2 | 20920.9429 | -0.0654292 | 0.21785837 | -0.3003289 | 0.76392631 | 0.96678847 |
| PNPLA2 | 12009.8524 | -0.112074 | 0.18174366 | -0.61666 | 0.53745904 | 0.92123049 |
| AP006621.1 | 4176.07966 | -0.1034732 | 0.19677083 | -0.5258562 | 0.59898813 | 0.93484879 |
| CRACR2B | 1386.22106 | -0.286354 | 0.44944612 | -0.6371264 | 0.52404254 | 0.91723878 |
| CD151 | 17515.5694 | -0.0242909 | 0.2285505 | -0.1062823 | 0.91535834 | 0.98675471 |
| POLR2L | 8977.23211 | 0.03853032 | 0.19869763 | 0.19391432 | 0.84624297 | 0.97680707 |
| TSPAN4 | 3701.2847 | 0.20320513 | 0.20319959 | 1.00002725 | 0.31729732 | 0.84819508 |
| AP006623.1 | 233.528356 | 0.03434085 | 0.20008257 | 0.17163339 | 0.86372575 | 0.98176973 |
| CHID1 | 3944.78468 | 0.2708972 | 0.1504492 | 1.80058915 | 7.18E-02 | 6.32E-01 |

| | | | | | | |
|---|---|---|---|---|---|---|
| AP2A2 | 7839.1381 | -0.0307578 | 0.17272405 | -0.1780746 | 0.85866442 | 0.97983564 |
| MUC6 | 370.384932 | -0.8802549 | 0.89834316 | -0.9798649 | 0.32715283 | 0.85308232 |
| AC139749.1 | 98.8807766 | 0.16219257 | 0.52729903 | 0.30759126 | 0.75839337 | 0.96502715 |
| TOLLIP | 6762.50552 | -0.2936529 | 0.22025318 | -1.3332514 | 0.18244933 | 0.7664835 |
| TOLLIP-AS1 | 94.6031131 | 0.2138393 | 0.4310552 | 0.49608334 | 0.61983561 | 0.93945324 |
| BRSK2 | 213.053612 | -0.4214924 | 0.53443682 | -0.7886665 | 0.43030695 | 0.88936206 |
| MOB2 | 2092.27739 | -0.4092258 | 0.13122805 | -3.1184321 | 0.00181816 | 0.18722055 |
| DUSP8 | 1211.10729 | -0.5377265 | 0.35952876 | -1.4956427 | 0.1347468 | 0.71630293 |
| KRTAP5-1 | 76.9038586 | -0.6730341 | 0.62687823 | -1.0736282 | 0.28298937 | 0.83078187 |
| IFITM10 | 1079.81222 | 0.31251078 | 0.65870826 | 0.47442973 | 6.35E-01 | 0.94088547 |
| AC068580.4 | 84317.9941 | 0.28904227 | 0.26666497 | 1.0839154 | 0.27840231 | 0.82805357 |
| CTSD | 135622.734 | 0.2522396 | 0.25072963 | 1.0060223 | 0.31440484 | 0.84696964 |
| AC068580.3 | 17647.907 | 0.21917078 | 0.27178428 | 0.80641449 | 0.42000389 | 0.88451372 |
| LSP1 | 660.499212 | 0.45949392 | 0.76529097 | 0.60041728 | 0.54822818 | 0.92373235 |
| PRR33 | 244.790895 | 0.38907876 | 0.79922998 | 0.48681702 | 0.62638801 | 0.94088547 |
| TNNT3 | 57.4034493 | 2.28427338 | 0.84275754 | 2.71047515 | 0.00671869 | 0.34592016 |
| MRPL23 | 490.150647 | 1.0442162 | 0.52588393 | 1.98564006 | 0.0470733 | 0.58171004 |
| IGF2 | 4504.85919 | 2.09597132 | 0.4934294 | 4.24776339 | 2.16E-05 | 0.01556334 |
| INS-IGF2 | 411.50994 | 2.21052403 | 0.53943322 | 4.09786408 | 4.17E-05 | 0.02368067 |
| ASCL2 | 104.535098 | -0.5551192 | 0.65257417 | -0.8506606 | 0.3949579 | 0.87887746 |
| C11orf21 | 118.021991 | 0.49586596 | 0.74259551 | 0.66774705 | 5.04E-01 | 9.14E-01 |
| TSPAN32 | 135.416116 | 0.6535605 | 0.57042221 | 1.14574871 | 0.25189914 | 0.81833349 |
| CD81-AS1 | 1463.77565 | 0.04973598 | 0.22877277 | 0.21740343 | 0.82789396 | 0.97548201 |
| CD81 | 27184.2889 | -0.0330147 | 0.21395125 | -0.1543097 | 0.87736557 | 0.98275231 |
| TSSC4 | 1939.54369 | -0.4664763 | 0.2288736 | -2.0381394 | 0.041536 | 0.55945849 |
| TRPM5 | 169.216941 | 0.47108855 | 0.52429182 | 0.89852357 | 0.36890649 | 0.86762749 |
| AC124057.1 | 50.4210297 | 0.51314895 | 0.62301065 | 0.82365999 | 0.41013278 | 0.88249519 |
| KCNQ1 | 3774.49411 | 0.11981275 | 0.33995059 | 0.35244167 | 0.72450706 | 0.95828829 |
| KCNQ1OT1 | 1351.39956 | -0.0256376 | 0.29923655 | -0.0856769 | 0.9317233 | 0.98982066 |
| CDKN1C | 3960.67806 | 0.1931008 | 0.33719666 | 0.57266521 | 0.5668714 | 0.9256133 |
| SLC22A18AS | 1024.37564 | 0.95083085 | 0.42683953 | 2.22760729 | 0.02590671 | 0.52502381 |
| SLC22A18 | 4938.52348 | 0.59485194 | 0.28542694 | 2.0840778 | 0.03715309 | 0.55527244 |
| PHLDA2 | 255.505419 | -0.2805566 | 0.41285585 | -0.6795511 | 0.49678874 | 0.90890243 |
| NAP1L4 | 6372.36049 | 0.12297827 | 0.12270916 | 1.00219305 | 0.31625036 | 0.84756299 |
| CARS | 2445.58856 | -0.3751319 | 0.22470494 | -1.6694423 | 0.09502977 | 0.65888357 |
| CARS-AS1 | 362.407993 | -0.303175 | 0.30318184 | -0.9999776 | 0.31732134 | 0.84819508 |
| OSBPL5 | 2251.11761 | -0.3033934 | 0.31583923 | -0.9605945 | 0.33675609 | 0.85679125 |
| ZNF195 | 1198.66869 | -0.3262952 | 0.18293793 | -1.7836389 | 0.07448236 | 0.63773415 |
| ART5 | 54.8008558 | -0.8110555 | 0.87166396 | -0.930468 | 0.3521288 | 0.86364633 |
| CHRNA10 | 105.768927 | -0.3425375 | 0.27852569 | -1.2298239 | 0.21876307 | 0.79794445 |
| NUP98 | 3520.72749 | -0.2705659 | 0.18724966 | -1.4449472 | 0.14847272 | 0.73534016 |
| PGAP2 | 1105.5513 | 0.2400082 | 0.23696193 | 1.01285554 | 3.11E-01 | 0.84476049 |
| RHOG | 1896.90393 | -0.434284 | 0.2166988 | -2.0040906 | 0.04506035 | 0.573693 |
| STIM1 | 3015.78536 | 0.41069267 | 0.16436376 | 2.49868147 | 0.01246563 | 0.41598747 |
| RRM1 | 1177.77018 | 0.36438471 | 0.21728906 | 1.67695833 | 0.09355063 | 0.65756964 |
| RRM1-AS1 | 176.995887 | 0.48687928 | 0.24953738 | 1.95112766 | 0.05104186 | 0.5861488 |
| TRIM21 | 1286.96875 | -0.8236524 | 0.45326779 | -1.8171431 | 0.06919521 | 0.62779744 |
| TRIM68 | 1112.83074 | 0.17422319 | 0.24751324 | 0.70389444 | 0.48149852 | 0.90541935 |
| OR51E1 | 47.3382619 | -0.3486646 | 0.54855902 | -0.6356008 | 0.52503664 | 0.91759401 |
| OR51E2 | 47.6973069 | -1.2764943 | 0.51413263 | -2.4828113 | 0.01303501 | 0.42046322 |
| HBB | 7344.54616 | -1.1189327 | 0.74815848 | -1.4955825 | 1.35E-01 | 0.71630293 |
| HBD | 74.5344411 | -2.6438325 | 0.61443716 | -4.3028525 | 1.69E-05 | 0.01316658 |
| AC104389.4 | 329.772331 | -0.4983837 | 0.50914446 | -0.9788649 | 0.32764672 | 0.85317073 |
| UBQLNL | 61.2423727 | -1.1789926 | 0.64054888 | -1.8405974 | 0.06568058 | 0.62237346 |

| | | | | | | |
|---|---|---|---|---|---|---|
| TRIM6 | 1524.56603 | 0.94047671 | 0.68705178 | 1.36885857 | 0.17104349 | 0.75956362 |
| TRIM6-TRIM | 876.1959 | 0.20366305 | 0.55210111 | 0.36888723 | 0.71221178 | 0.95560193 |
| TRIM34 | 450.212544 | -0.2703098 | 0.48825121 | -0.5536285 | 5.80E-01 | 9.31E-01 |
| TRIM5 | 1689.2108 | -0.5948374 | 0.50567548 | -1.1763223 | 0.23946609 | 0.81163641 |
| TRIM22 | 4566.03853 | -1.4193431 | 0.79188658 | -1.7923566 | 0.07307586 | 0.63580072 |
| AC022762.2 | 162.502384 | 0.48190047 | 0.30301137 | 1.59037092 | 0.11175122 | 0.68001961 |
| FAM160A2 | 2982.94549 | -0.0321377 | 0.19607483 | -0.1639052 | 0.86980577 | 0.98275231 |
| CAVIN3 | 852.305737 | -0.0181447 | 0.2729967 | -0.0664648 | 0.94700777 | 0.99166614 |
| SMPD1 | 7608.52556 | 0.08174512 | 0.27902349 | 0.29296861 | 0.76954614 | 0.96836854 |
| APBB1 | 3728.25951 | -0.2336103 | 0.19645566 | -1.1891249 | 0.23439053 | 0.80730051 |
| HPX | 52.4518225 | 0.48082834 | 0.62174833 | 0.77334882 | 0.43931597 | 0.89175645 |
| TRIM3 | 1432.73682 | -0.0642246 | 0.17477861 | -0.3674628 | 0.71327387 | 0.95564093 |
| ARFIP2 | 4153.85193 | -0.2902945 | 0.16841625 | -1.7236726 | 0.08476697 | 0.64988556 |
| TIMM10B | 1931.87913 | -0.0307212 | 0.16924637 | -0.1815175 | 0.85596138 | 0.97929708 |
| AC084337.2 | 431.430161 | -0.1098986 | 0.21830847 | -0.5034098 | 0.61467619 | 0.93895494 |
| DNHD1 | 1309.14357 | -0.481476 | 0.38445325 | -1.2523655 | 0.21043671 | 0.79134644 |
| RRP8 | 1051.86842 | -0.1762032 | 0.13278721 | -1.3269595 | 0.18452209 | 0.76820626 |
| AC091564.2 | 1121.02736 | -0.1153352 | 0.23746995 | -0.4856835 | 0.62719155 | 0.94088547 |
| ILK | 8580.12917 | -0.1479119 | 0.15655374 | -0.9447994 | 0.34476132 | 0.85999891 |
| TAF10 | 10295.1088 | -0.1214304 | 0.14001854 | -0.8672453 | 0.38580765 | 0.87420019 |
| TPP1 | 13347.57 | -0.1359156 | 0.24363161 | -0.5578734 | 0.57693085 | 0.93129843 |
| AC091564.6 | 652.875996 | 0.05092085 | 0.35322218 | 0.14416097 | 0.88537336 | 0.98356741 |
| DCHS1 | 1537.22247 | -0.0358904 | 0.32968192 | -0.1088639 | 0.91331048 | 0.98662757 |
| AC091564.4 | 579.862575 | -0.1338287 | 0.35541359 | -0.3765436 | 0.70651282 | 0.95501548 |
| MRPL17 | 2430.25123 | -0.1708646 | 0.13128521 | -1.3014761 | 0.19309555 | 0.7751938 |
| ZNF215 | 245.122217 | 0.04728796 | 0.34643999 | 0.13649682 | 0.89142854 | 0.98474725 |
| ZNF214 | 199.43263 | 0.2430087 | 0.3676962 | 0.66089532 | 0.50867945 | 0.91439937 |
| SYT9 | 177.661265 | 0.38268362 | 0.46187189 | 0.82854927 | 4.07E-01 | 0.88132015 |
| AC107884.1 | 167.28425 | -0.1151927 | 0.32762779 | -0.3515962 | 0.72514111 | 0.95848028 |
| OLFML1 | 318.467323 | -0.1151074 | 0.49433698 | -0.2328522 | 0.81587618 | 0.9738447 |
| PPFIBP2 | 2476.12994 | -0.2059366 | 0.32003862 | -0.6434743 | 0.51991637 | 0.91657872 |
| CYB5R2 | 335.559241 | 0.39137919 | 0.322452 | 1.21375952 | 0.22483957 | 0.80123947 |
| OVCH2 | 169.502751 | 1.17149815 | 0.48684874 | 2.40628774 | 0.01611557 | 0.45760882 |
| EIF3F | 6156.17314 | 0.32661308 | 0.17380953 | 1.87914369 | 0.06022488 | 0.60825814 |
| TUB | 1906.72742 | 0.50732706 | 0.47039404 | 1.07851505 | 2.81E-01 | 8.29E-01 |
| RIC3 | 1162.86744 | -0.0881421 | 0.29766076 | -0.296116 | 0.76714153 | 0.96749893 |
| LMO1 | 54.5662103 | -0.0797816 | 0.6187943 | -0.1289307 | 0.89741251 | 0.98474725 |
| STK33 | 765.93046 | 0.23663945 | 0.2712529 | 0.87239416 | 0.38299339 | 0.8718674 |
| TRIM66 | 1123.69921 | 0.06508648 | 0.32171693 | 0.20230979 | 0.83967455 | 0.97618283 |
| RPL27A | 21963.6153 | 0.16139343 | 0.18199536 | 0.88679972 | 0.37518674 | 0.86839257 |
| ST5 | 3980.47323 | -0.1092844 | 0.31379513 | -0.3482667 | 0.72763986 | 0.95848028 |
| AC091053.1 | 1860.54598 | -0.0641414 | 0.32690406 | -0.1962087 | 0.84444684 | 0.97675487 |
| AKIP1 | 881.057194 | 0.15281522 | 0.18234166 | 0.83807079 | 0.40199095 | 0.88132015 |
| TMEM9B | 3381.34772 | 0.02101851 | 0.09620636 | 0.21847323 | 8.27E-01 | 0.97548201 |
| TMEM9B-AS | 221.77131 | -0.0061508 | 0.34711902 | -0.0177196 | 0.98586251 | 0.99705651 |
| NRIP3 | 92.0437966 | -0.7601151 | 0.6503881 | -1.1687101 | 0.24252045 | 0.81333856 |
| SCUBE2 | 104.435226 | 0.4577287 | 0.37505394 | 1.22043432 | 0.22230027 | 0.80078694 |
| DENND5A | 3305.65709 | -0.3604763 | 0.18474708 | -1.9511878 | 0.05103471 | 0.5861488 |
| TMEM41B | 2268.16655 | -0.1572697 | 0.20277476 | -0.775588 | 0.43799228 | 0.89152136 |
| IPO7 | 5453.22215 | -0.17612 | 0.23395801 | -0.7527848 | 0.45157922 | 0.89639459 |
| AC132192.2 | 164.776278 | -0.0009039 | 0.34742071 | -0.0026018 | 0.99792405 | 0.99989813 |
| ZNF143 | 925.531543 | -0.1992492 | 0.26320428 | -0.7570135 | 0.44904178 | 0.89545563 |
| WEE1 | 1194.48346 | -0.2271996 | 0.2602389 | -0.8730424 | 0.38263997 | 0.8718674 |
| SWAP70 | 2978.48137 | -0.30004 | 0.26059844 | -1.15135 | 0.24958827 | 0.81646572 |

| | | | | | | |
|---|---|---|---|---|---|---|
| SBF2-AS1 | 1076.92224 | -0.1088201 | 0.19561581 | -0.556295 | 5.78E-01 | 0.93130183 |
| SBF2 | 1942.29833 | 0.09855505 | 0.14799961 | 0.66591424 | 0.50546592 | 0.91368371 |
| ADM | 9586.19377 | -0.4626469 | 0.66228475 | -0.698562 | 0.48482582 | 0.90541935 |
| AMPD3 | 1002.0691 | 0.0473285 | 0.19350047 | 0.24459112 | 0.80677304 | 0.97334138 |
| MTRNR2L8 | 177.316142 | -0.5102571 | 0.4750949 | -1.0740109 | 0.28281778 | 0.83076616 |
| RNF141 | 1895.44414 | 0.34866126 | 0.15363936 | 2.26934858 | 0.02324714 | 0.51967919 |
| LYVE1 | 2084.33283 | -0.8529874 | 0.41871687 | -2.0371461 | 0.0416354 | 0.5601501 |
| MRVI1 | 1142.92817 | -0.6069971 | 0.5650932 | -1.0741539 | 0.28275369 | 0.83076616 |
| CTR9 | 2212.69851 | -0.0873417 | 0.19902421 | -0.4388496 | 0.66077049 | 0.94628868 |
| EIF4G2 | 28233.6218 | -0.0566617 | 0.270251 | -0.2096633 | 0.83393044 | 0.97550416 |
| ZBED5 | 2547.67783 | -0.2658579 | 0.19429991 | -1.3682861 | 0.17122254 | 0.75956362 |
| ZBED5-AS1 | 642.577925 | 0.97982419 | 0.34421255 | 2.84656733 | 0.00441934 | 0.29060638 |
| GALNT18 | 2496.87851 | 0.73241385 | 0.30867961 | 2.37273157 | 0.01765709 | 0.47014911 |
| USP47 | 3082.48071 | -0.0498051 | 0.13714459 | -0.3631576 | 0.71648717 | 0.95603249 |
| DKK3 | 1979.88218 | 0.1179307 | 0.48130673 | 0.24502193 | 0.80643945 | 0.97334138 |
| MICAL2 | 980.782648 | -0.4445534 | 0.47747753 | -0.9310457 | 0.35182991 | 0.86364633 |
| AC079329.1 | 323.37168 | -0.3888803 | 0.50660849 | -0.767615 | 0.44271592 | 0.89343541 |
| PARVA | 3182.30978 | 0.01569663 | 0.17562201 | 0.08937735 | 0.92878202 | 0.98963127 |
| TEAD1 | 2827.32342 | -0.2631719 | 0.20546766 | -1.2808436 | 0.2002486 | 0.78134034 |
| LINC00958 | 578.204266 | -0.6018238 | 0.29288835 | -2.054789 | 0.03989939 | 0.55657615 |
| RASSF10 | 248.34793 | -0.2222686 | 0.34374869 | -0.6466021 | 0.5178895 | 0.91636259 |
| ARNTL | 969.484162 | -0.3505975 | 0.33247021 | -1.054523 | 0.29164352 | 0.83460625 |
| BTBD10 | 1103.89843 | -0.1694394 | 0.18498527 | -0.9159617 | 0.35968699 | 0.86480228 |
| FAR1 | 2437.05032 | 0.03957008 | 0.12067962 | 0.32789365 | 0.74299208 | 0.96109001 |
| SPON1 | 2249.19394 | -0.3353142 | 0.41746743 | -0.8032106 | 0.42185304 | 0.88451372 |
| SPON1-AS1 | 144.323649 | -0.5252829 | 0.49545804 | -1.0601965 | 0.28905523 | 0.83308345 |
| RRAS2 | 2831.25582 | -0.8636173 | 0.49173573 | -1.756263 | 0.07904351 | 0.64079926 |
| COPB1 | 6505.38283 | -0.1787146 | 0.21087809 | -0.8474784 | 0.39672853 | 0.87978812 |
| AC018523.2 | 4759.04228 | -0.0370432 | 0.22721791 | -0.1630294 | 0.8704953 | 0.98275231 |
| PSMA1 | 4940.09087 | -0.0562377 | 0.21662515 | -0.2596084 | 0.79516586 | 0.97101097 |
| PDE3B | 355.287125 | 0.13440307 | 0.39378448 | 0.34131124 | 0.73286928 | 0.95848028 |
| CYP2R1 | 606.06907 | 0.2100562 | 0.13582144 | 1.54656146 | 0.12196903 | 0.69774774 |
| CALCA | 237.937188 | 0.3968302 | 0.84403947 | 0.47015598 | 0.63824358 | 0.94088547 |
| INSC | 112.031282 | -0.2827862 | 0.84633652 | -0.3341297 | 0.73828167 | 0.9602325 |
| AC073172.1 | 56.0063043 | -0.2564054 | 0.95168446 | -0.2694228 | 0.78760438 | 0.97090793 |
| AC087379.2 | 84.139434 | -0.3630392 | 0.95674721 | -0.3794516 | 0.70435257 | 0.95488362 |
| SOX6 | 1162.69764 | 0.27076299 | 0.32459961 | 0.83414454 | 0.40419954 | 0.88132015 |
| C11orf58 | 5957.40604 | -0.013119 | 0.09336563 | -0.1405123 | 0.88825524 | 0.98423722 |
| PLEKHA7 | 4213.40991 | 0.01501523 | 0.20245274 | 0.0741666 | 0.94087782 | 0.99136103 |
| RPS13 | 13185.3392 | 0.14734051 | 0.22271662 | 0.66156046 | 0.50825296 | 0.91439937 |
| PIK3C2A | 3328.80262 | -0.024328 | 0.13106879 | -0.1856127 | 0.85274851 | 0.97883015 |
| NUCB2 | 1842.62132 | 0.214408 | 0.20914695 | 1.0251548 | 0.30529015 | 0.84268008 |
| NCR3LG1 | 51.2404952 | 0.22606266 | 0.47809352 | 0.47284193 | 0.63632596 | 0.94088547 |
| AC124798.1 | 551.79516 | -0.0099699 | 0.39546166 | -0.0252109 | 0.97988678 | 0.99537078 |
| KCNJ11 | 426.380273 | -0.4223516 | 0.3408125 | -1.2392491 | 0.21525327 | 0.79504564 |
| ABCC8 | 83.5101084 | -0.9219692 | 0.69996778 | -1.3171594 | 0.18778519 | 0.76970626 |
| USH1C | 5576.31183 | 0.75429913 | 0.43101612 | 1.75004854 | 0.08010994 | 0.64079926 |
| SERGEF | 1160.6767 | 0.06446311 | 0.20381921 | 0.31627593 | 0.75179308 | 0.96289476 |
| TPH1 | 55.5554086 | -0.3736994 | 0.58685701 | -0.6367811 | 0.52426744 | 0.91723878 |
| SAAL1 | 368.351626 | -0.0621296 | 0.1693881 | -0.3667882 | 0.71377699 | 0.95564093 |
| SAA4 | 87.3967356 | 1.17719478 | 0.94439917 | 1.24650129 | 0.21258042 | 0.79203904 |
| SAA2-SAA4 | 2332.36916 | 1.54705975 | 1.31879196 | 1.17308855 | 0.2407603 | 0.8124487 |
| SAA2 | 3114.79705 | 1.54562758 | 1.31164474 | 1.17838888 | 0.23864161 | 0.81095554 |
| SAA1 | 6996.43746 | 1.67563681 | 1.34906659 | 1.24207124 | 0.21421028 | 0.79353921 |

| | | | | | | |
|---|---|---|---|---|---|---|
| HPS5 | 972.833385 | 0.04526184 | 0.1654898 | 0.27350229 | 0.78446714 | 0.97021865 |
| GTF2H1 | 1973.20486 | -0.2673199 | 0.31523827 | -0.8479932 | 0.39644174 | 0.87967259 |
| LDHA | 54013.359 | -0.127213 | 0.34320861 | -0.370658 | 0.71089228 | 0.95560193 |
| LDHC | 72.1925159 | 2.74895871 | 0.97093362 | 2.83125298 | 0.0046366 | 0.29749322 |
| TSG101 | 3901.53151 | -0.0729206 | 0.16199661 | -0.4501363 | 0.65261213 | 0.94437526 |
| UEVLD | 1244.9512 | 0.0788073 | 0.15616706 | 0.50463461 | 0.6138155 | 0.93895494 |
| SPTY2D1OS | 192.459197 | -0.4214427 | 0.26513789 | -1.5895226 | 0.11194246 | 0.68001961 |
| SPTY2D1 | 1217.82692 | -0.3764997 | 0.35349263 | -1.0650849 | 0.2868375 | 0.83155545 |
| TMEM86A | 1501.93852 | 0.00212942 | 0.38685732 | 0.00550441 | 0.99560814 | 0.99940889 |
| IGSF22 | 289.726881 | -0.1104269 | 0.31963646 | -0.3454766 | 0.7297361 | 0.95848028 |
| AC103974.1 | 110.401628 | 0.12169686 | 0.34167251 | 0.35617985 | 0.72170588 | 0.95739595 |
| ZDHHC13 | 1076.92029 | -0.1494609 | 0.15909823 | -0.9394256 | 0.34751229 | 0.86204208 |
| NAV2 | 2379.15625 | -0.5192619 | 0.26139872 | -1.9864745 | 0.04698066 | 0.58171004 |
| AC009549.1 | 50.0835323 | -1.5106897 | 0.66693749 | -2.2651144 | 0.02350566 | 0.51967919 |
| NAV2-AS1 | 747.661709 | -0.4645079 | 0.2842669 | -1.6340554 | 0.10224722 | 0.67279543 |
| HTATIP2 | 2738.14509 | 0.26303762 | 0.40896826 | 0.64317369 | 0.5201114 | 0.91657872 |
| PRMT3 | 753.498629 | 0.05903781 | 0.2023624 | 0.291743 | 0.77048313 | 0.96836854 |
| NELL1 | 617.243315 | -0.3255074 | 0.50841903 | -0.6402346 | 0.5220201 | 0.91723878 |
| ANO5 | 461.526703 | 0.13380871 | 0.28832208 | 0.46409454 | 0.64258001 | 0.94090101 |
| FANCF | 729.087192 | 0.11643681 | 0.17655238 | 0.65950294 | 0.50957286 | 0.91439937 |
| GAS2 | 286.857138 | 0.29860527 | 0.31384398 | 0.95144495 | 0.34137855 | 0.85889221 |
| SVIP | 1604.064 | 0.37493007 | 0.17721905 | 2.11563078 | 0.03437622 | 0.55369025 |
| ANO3 | 349.87288 | -0.038621 | 0.51067717 | -0.0756271 | 0.93971577 | 0.99134327 |
| MUC15 | 1074.11263 | -0.2541587 | 0.63003944 | -0.4034013 | 0.68665306 | 0.95097288 |
| SLC5A12 | 14999.4217 | 0.93456833 | 0.57777835 | 1.61752051 | 0.10576597 | 0.67526487 |
| FIBIN | 1237.18509 | 0.39776766 | 0.49085512 | 0.81035655 | 0.41773528 | 0.88445833 |
| BBOX1 | 11501.827 | 0.8217095 | 0.52803019 | 1.55617902 | 0.11966553 | 0.69405865 |
| BBOX1-AS1 | 45.896987 | -0.7267591 | 0.67056096 | -1.0838077 | 0.27845007 | 0.82805357 |
| CCDC34 | 166.817506 | 0.50647214 | 0.26865339 | 1.88522517 | 0.05939944 | 0.60533169 |
| LGR4 | 6095.10176 | -0.3057947 | 0.18734023 | -1.6322961 | 0.10261712 | 0.67335541 |
| LIN7C | 1447.62743 | -0.1945651 | 0.11046288 | -1.7613618 | 0.07817719 | 0.64079926 |
| BDNF-AS | 280.008085 | 0.69222338 | 0.35654703 | 1.94146441 | 0.05220197 | 0.58645127 |
| BDNF | 124.04953 | -0.1033944 | 0.48959238 | -0.2111847 | 0.83274315 | 0.97550416 |
| METTL15 | 679.369011 | 0.22074503 | 0.20721708 | 1.06528397 | 0.28674745 | 0.83155545 |
| ARL14EP | 1138.21957 | 0.2288072 | 0.11863934 | 1.92859478 | 0.05378119 | 0.59266174 |
| MPPED2 | 608.266341 | 0.58068637 | 0.32846593 | 1.76787401 | 0.07708197 | 0.63806922 |
| DCDC1 | 170.158555 | 0.15279983 | 0.44131079 | 0.34624087 | 0.7291617 | 0.95848028 |
| DNAJC24 | 440.794211 | -0.1099831 | 0.24466038 | -0.4495336 | 0.65304675 | 0.94478532 |
| IMMP1L | 567.785952 | 0.13568163 | 0.13200808 | 1.0278282 | 0.30403065 | 0.84211658 |
| ELP4 | 892.006963 | 0.28683307 | 0.20130723 | 1.42485231 | 0.1541999 | 0.74258845 |
| PAX6 | 131.471892 | 0.53965467 | 0.31357756 | 1.72096076 | 0.08525794 | 0.65020388 |
| AL035078.4 | 2690.16057 | -0.2367093 | 0.28330575 | -0.8355261 | 0.40342156 | 0.88132015 |
| RCN1 | 5550.54373 | -0.2654697 | 0.23799333 | -1.1154501 | 0.26465761 | 0.82424998 |
| AL078612.1 | 1364.00226 | -0.2462413 | 0.24726228 | -0.9958707 | 0.31931297 | 0.84928695 |
| WT1 | 778.582668 | 0.47447911 | 0.66449051 | 0.7140495 | 0.47519657 | 0.90226631 |
| EIF3M | 5601.07216 | 0.08244687 | 0.19617228 | 0.42027786 | 0.67428248 | 0.94935413 |
| CCDC73 | 1357.84926 | 0.08370291 | 0.16608277 | 0.50398307 | 0.61427327 | 0.93895494 |
| PRRG4 | 2116.64186 | -0.209003 | 0.23401408 | -0.8931216 | 0.37179205 | 0.86762749 |
| QSER1 | 1031.86609 | -0.0971108 | 0.12924022 | -0.7513976 | 0.45241338 | 0.89670027 |
| DEPDC7 | 826.40243 | 0.58532566 | 0.49830361 | 1.17463661 | 0.24014013 | 0.81235851 |
| TCP11L1 | 724.252023 | 0.02202511 | 0.17216028 | 0.12793378 | 0.89820138 | 0.98474725 |
| LINC00294 | 1315.77011 | -0.1531817 | 0.24143493 | -0.6344637 | 0.52577826 | 0.91769856 |
| CSTF3 | 1459.94816 | 0.04522133 | 0.17651722 | 0.25618652 | 0.79780681 | 0.97154613 |
| HIPK3 | 5309.56289 | -0.2461368 | 0.24060587 | -1.0229873 | 0.30631384 | 0.84278367 |

| | | | | | | |
|---|---|---|---|---|---|---|
| KIAA1549L | 216.704271 | 0.64366968 | 0.19820492 | 3.24749605 | 0.00116425 | 0.15697311 |
| AL049629.1 | 192.276242 | 0.56044185 | 0.27831126 | 2.01372324 | 0.0440386 | 0.57234908 |
| AL049629.2 | 17594.8048 | -0.0221253 | 0.2540738 | -0.087082 | 0.93060633 | 0.98975562 |
| C11orf91 | 95.990536 | -0.7220473 | 0.64581704 | -1.118037 | 0.2635512 | 0.82424998 |
| CD59 | 44979.9422 | -0.0825657 | 0.26100264 | -0.3163404 | 0.75174416 | 0.96289476 |
| FBXO3 | 3011.36609 | 0.14508431 | 0.14085111 | 1.03005442 | 0.30298446 | 0.84211658 |
| LMO2 | 2233.31391 | 0.15280628 | 0.30317199 | 0.50402505 | 0.61424377 | 0.93895494 |
| CAPRIN1 | 7549.26665 | -0.024791 | 0.13648416 | -0.18164 | 0.85586527 | 0.97929708 |
| NAT10 | 1847.12699 | -0.1904878 | 0.15916903 | -1.196764 | 0.23139853 | 0.80649152 |
| ABTB2 | 2241.10096 | -0.2842741 | 0.3008337 | -0.9449545 | 0.34468212 | 0.85999891 |
| CAT | 15711.7327 | 0.45776033 | 0.32922281 | 1.39042714 | 0.16439921 | 0.75286112 |
| ELF5 | 515.868782 | 1.08322879 | 0.59322184 | 1.82600963 | 6.78E-02 | 6.26E-01 |
| EHF | 483.55308 | 0.23647992 | 0.42065196 | 0.56217477 | 0.57399695 | 0.92999713 |
| APIP | 1542.12472 | -0.2128324 | 0.21547858 | -0.9877196 | 0.32329 | 0.85119105 |
| PDHX | 2104.29198 | 0.17496151 | 0.17006605 | 1.02878562 | 0.30358042 | 0.84211658 |
| CD44 | 5185.77451 | -0.3457496 | 0.54421089 | -0.6353228 | 0.52521791 | 0.91759401 |
| CD44-AS1 | 442.133321 | -0.5789267 | 0.57467832 | -1.0073927 | 0.31374611 | 0.84646403 |
| PAMR1 | 367.517351 | 1.25843803 | 0.48875685 | 2.57477318 | 0.01003058 | 0.39989496 |
| FJX1 | 439.406552 | -0.2639855 | 0.4529349 | -0.5828331 | 0.56000565 | 0.92434754 |
| TRIM44 | 5263.80188 | -0.0598688 | 0.11157897 | -0.5365596 | 0.59157187 | 0.93289552 |
| LDLRAD3 | 510.953711 | 0.12310434 | 0.37522927 | 0.32807765 | 0.74285295 | 0.96109001 |
| COMMD9 | 2306.49466 | -0.0381105 | 0.17609966 | -0.2164142 | 0.82866489 | 0.97550416 |
| PRR5L | 544.61175 | 0.57465895 | 0.48760944 | 1.17852302 | 0.23858816 | 0.81095554 |
| TRAF6 | 700.755397 | -0.1933339 | 0.16671676 | -1.1596551 | 0.24618927 | 0.81542272 |
| RAG1 | 162.727024 | 0.67920736 | 0.34310281 | 1.97960302 | 0.04774815 | 0.58258343 |
| RAG2 | 82.4040976 | -0.0715548 | 0.2578043 | -0.2775545 | 0.78135433 | 0.9690531 |
| C11orf74 | 493.710891 | 0.05188951 | 0.23049007 | 0.22512686 | 0.82188058 | 0.97492651 |
| API5 | 4158.70635 | 0.02247647 | 0.1457291 | 0.15423459 | 0.87742477 | 0.98275231 |
| AC087276.2 | 378.371276 | 0.17958891 | 0.3350885 | 0.5359447 | 0.59199677 | 0.93289552 |
| TTC17 | 3718.599 | -0.2006679 | 0.18413135 | -1.0898083 | 0.27579761 | 0.82719407 |
| HSD17B12 | 5435.41162 | 0.12197446 | 0.25526157 | 0.47784108 | 0.63276332 | 0.94088547 |
| AC087521.2 | 407.405736 | 0.16247144 | 0.32067509 | 0.50665438 | 0.61239734 | 0.9387918 |
| AC087521.4 | 2152.66667 | 0.13382852 | 0.28449384 | 0.47040921 | 0.63806268 | 0.94088547 |
| ALKBH3 | 781.12794 | -0.0613119 | 0.20300809 | -0.3020172 | 0.76263899 | 0.96630779 |
| C11orf96 | 5252.62821 | -0.7663732 | 0.69865052 | -1.0969336 | 0.27267042 | 0.82716979 |
| AC087521.1 | 1291.60435 | -0.7077075 | 0.64919946 | -1.0901233 | 0.27565883 | 0.82719407 |
| ACCS | 1875.80921 | 0.3731106 | 0.52422518 | 0.71173727 | 0.47662748 | 0.90313654 |
| EXT2 | 3216.95242 | -0.0365965 | 0.18612876 | -0.196619 | 0.84412569 | 0.97664894 |
| CD82 | 2470.28767 | 0.86187758 | 0.40078969 | 2.15044847 | 0.03151976 | 0.54507414 |
| AC010768.1 | 373.642539 | 0.84187125 | 0.43762057 | 1.92374697 | 0.05438631 | 0.59329767 |
| TSPAN18 | 2519.62476 | 0.29865588 | 0.35307683 | 0.84586654 | 0.3976272 | 0.88042667 |
| TP53I11 | 3436.77908 | -0.1519402 | 0.31117846 | -0.4882734 | 0.62535619 | 0.94088547 |
| PRDM11 | 736.988674 | -0.3965975 | 0.20014362 | -1.9815647 | 0.04752798 | 0.58241466 |
| SYT13 | 1300.41136 | -0.1556602 | 0.40797691 | -0.3815417 | 0.70280135 | 0.95431462 |
| CHST1 | 288.948804 | 0.10374104 | 0.47557981 | 0.21813593 | 0.82732321 | 0.97548201 |
| SLC35C1 | 1691.57124 | -0.1727292 | 0.29403183 | -0.5874506 | 0.55690111 | 0.92373235 |
| CRY2 | 3459.23018 | 0.20013985 | 0.24948597 | 0.80220884 | 0.42243216 | 0.88451372 |
| MAPK8IP1 | 1630.35901 | 0.29061639 | 0.23558756 | 1.23358123 | 0.21735899 | 0.79592123 |
| AC068385.1 | 395.682438 | 0.22973951 | 0.23925165 | 0.96024209 | 0.33693339 | 0.85679125 |
| PEX16 | 1645.8914 | 0.08381277 | 0.21010925 | 0.39890092 | 0.68996621 | 0.95158502 |
| LARGE2 | 2008.00928 | 0.40332188 | 0.40851074 | 0.98729811 | 0.32349651 | 0.85140144 |
| PHF21A | 2040.52188 | -0.2505795 | 0.23743683 | -1.0553522 | 0.29126427 | 0.83454586 |
| CREB3L1 | 337.827462 | 0.23626395 | 0.43979206 | 0.53721741 | 0.59111744 | 0.93289552 |
| DGKZ | 5635.76478 | -0.0697827 | 0.19236447 | -0.3627628 | 0.71678206 | 0.95603249 |

| | | | | | | |
|---|---|---|---|---|---|---|
| MDK | 2136.80777 | -1.5466631 | 0.75960942 | -2.0361295 | 0.04173735 | 0.56071666 |
| AMBRA1 | 1799.39161 | -0.1528565 | 0.19949407 | -0.7662206 | 0.44354506 | 0.89366134 |
| HARBI1 | 295.761777 | -0.177299 | 0.32834397 | -0.5399796 | 0.58921113 | 0.93289552 |
| ATG13 | 4045.44747 | -0.0532493 | 0.14413388 | -0.3694433 | 0.71179733 | 0.95560193 |
| ARHGAP1 | 6052.19402 | -0.0324975 | 0.14807999 | -0.2194594 | 0.8262922 | 0.97548201 |
| ZNF408 | 654.356113 | -0.272908 | 0.20799293 | -1.3121021 | 0.18948569 | 0.76970626 |
| CKAP5 | 1979.53397 | -0.0524382 | 0.12802636 | -0.4095889 | 0.68210755 | 0.95070301 |
| LRP4 | 221.655946 | -0.0202324 | 0.32164526 | -0.0629027 | 0.94984397 | 0.99170503 |
| C11orf49 | 2255.38672 | 0.08786738 | 0.23382222 | 0.37578711 | 0.70707517 | 0.95501548 |
| ARFGAP2 | 4961.91805 | 0.00904031 | 0.22028747 | 0.04103868 | 0.96726506 | 0.99400331 |
| PACSIN3 | 1461.05034 | -0.1466708 | 0.20972673 | -0.6993423 | 0.48433815 | 0.90541935 |
| DDB2 | 1031.10661 | 0.31730027 | 0.25192055 | 1.25952514 | 0.20784072 | 0.7886501 |
| ACP2 | 3507.77981 | -0.0971814 | 0.20832895 | -0.4664805 | 0.64087163 | 0.94088547 |
| NR1H3 | 2072.61041 | 0.27809653 | 0.3054926 | 0.91032163 | 0.36265291 | 0.86590307 |
| MADD | 3073.48816 | -0.1447696 | 0.16392599 | -0.8831401 | 0.37716058 | 0.86987124 |
| SPI1 | 748.465901 | -0.3667519 | 0.49435852 | -0.7418743 | 0.45816353 | 0.8971553 |
| AC090559.1 | 84.7391789 | 0.08345399 | 0.34976894 | 0.2385975 | 0.81141771 | 0.97352497 |
| SLC39A13 | 8273.94133 | -0.4892626 | 0.1938174 | -2.524348 | 0.01159131 | 0.40423902 |
| PSMC3 | 4993.75062 | -0.2117392 | 0.19858327 | -1.0662488 | 0.28631119 | 0.83107779 |
| RAPSN | 64.6105069 | -0.1212659 | 0.69990852 | -0.1732596 | 0.86244736 | 0.98176973 |
| CELF1 | 5408.33188 | -0.3307652 | 0.13844373 | -2.3891668 | 0.01688663 | 0.46671416 |
| NDUFS3 | 5221.7463 | 0.12306475 | 0.12213325 | 1.00762691 | 0.31363361 | 0.8464135 |
| PTPMT1 | 2378.72534 | -0.0506611 | 0.12105777 | -0.4184869 | 0.67559114 | 0.94958156 |
| KBTBD4 | 1534.53274 | -0.1791902 | 0.20940596 | -0.8557072 | 0.39215976 | 0.87765297 |
| KF459542.1 | 156.994703 | -0.205552 | 0.31155063 | -0.6597709 | 0.50940084 | 0.91439937 |
| MTCH2 | 7150.13352 | 0.40554069 | 0.25980061 | 1.56096893 | 0.11853108 | 0.69159122 |
| AGBL2 | 199.995394 | -0.1961839 | 0.64125897 | -0.3059356 | 0.75965371 | 0.96510454 |
| FNBP4 | 4445.82355 | -0.3053833 | 0.21684753 | -1.4082858 | 0.15904645 | 0.7475863 |
| NUP160 | 1943.0099 | -0.2983163 | 0.15996057 | -1.8649368 | 0.06219027 | 0.61245813 |
| PTPRJ | 3456.21439 | 0.00034465 | 0.15809004 | 0.00218007 | 0.99826056 | 0.99990633 |
| FOLH1 | 840.25001 | -0.0601414 | 0.49052179 | -0.122607 | 0.90241831 | 0.98599252 |
| APLNR | 507.752595 | 0.49798009 | 0.72124457 | 0.69044554 | 4.90E-01 | 9.06E-01 |
| TNKS1BP1 | 10503.0143 | -0.0799694 | 0.26124202 | -0.3061121 | 0.75951926 | 0.96502715 |
| AP000781.1 | 770.304537 | -0.0097174 | 0.18723735 | -0.051899 | 0.95860915 | 0.99237995 |
| SSRP1 | 4865.24244 | -0.0142873 | 0.16920378 | -0.0844387 | 0.93270763 | 0.9899685 |
| PRG2 | 58.1654399 | -0.5641939 | 0.82884278 | -0.6807007 | NA | NA |
| AP000781.2 | 1100.59633 | -0.3510598 | 0.39907971 | -0.8796734 | 0.37903628 | 0.87069278 |
| SLC43A3 | 1659.39365 | -0.3442375 | 0.39463434 | -0.8722948 | 0.3830476 | 0.8718674 |
| RTN4RL2 | 227.303958 | -0.9739604 | 0.42407735 | -2.2966575 | 0.02163832 | 0.51394743 |
| AP002893.1 | 118.33617 | -0.5986326 | 0.43723739 | -1.3691248 | 0.17096026 | 0.75956362 |
| SLC43A1 | 731.109102 | -0.3183141 | 0.32964651 | -0.9656226 | 0.33423305 | 0.85536824 |
| TIMM10 | 1190.16698 | 0.02169323 | 0.23850339 | 0.09095564 | 0.92752784 | 0.98915637 |
| SMTNL1 | 48.7406823 | -2.8854391 | 1.03316886 | -2.792805 | NA | NA |
| UBE2L6 | 3576.29617 | -1.2541939 | 0.61320687 | -2.045303 | 0.04082499 | 0.55782198 |
| SERPING1 | 38524.7315 | -0.358697 | 0.32716736 | -1.0963716 | 2.73E-01 | 8.27E-01 |
| YPEL4 | 89.6251682 | -0.4534985 | 0.42796256 | -1.0596686 | 0.28929539 | 0.83312616 |
| CLP1 | 569.740091 | -0.3074146 | 0.33226623 | -0.9252057 | 0.35485891 | 0.86364633 |
| ZDHHC5 | 6227.48989 | -0.2852905 | 0.21414996 | -1.3321994 | 0.18279466 | 0.76723162 |
| MED19 | 554.094481 | -0.3368936 | 0.21967249 | -1.5336177 | 0.12512373 | 0.70228923 |
| TMX2 | 5489.0343 | 0.0467568 | 0.14298934 | 0.32699502 | 0.74367165 | 0.96137808 |
| TMX2-CTNNI | 1105.18308 | 0.08330636 | 0.1786733 | 0.46624962 | 0.64103684 | 0.94088547 |
| SELENOH | 3171.63268 | 0.12894178 | 0.14218652 | 0.90684954 | 0.36448637 | 0.8663515 |
| AP001931.1 | 1623.0479 | 0.16586165 | 0.1558237 | 1.06441866 | 0.28713908 | 0.83159845 |
| CTNND1 | 15197.6533 | 0.20256058 | 0.12708896 | 1.59384868 | 0.11096993 | 0.68001961 |

| | | | | | | |
|---|---|---|---|---|---|---|
| LPXN | 772.26751 | 0.18548486 | 0.20874845 | 0.88855686 | 3.74E-01 | 0.86791605 |
| ZFP91 | 3410.711 | -0.3033123 | 0.21590079 | -1.404869 | 0.16006023 | 0.74781837 |
| ZFP91-CNTF | 269.772413 | -0.6749312 | 0.50719103 | -1.3307239 | 0.18327988 | 0.76723162 |
| AP001350.1 | 59.2899444 | -0.7272396 | 0.6122837 | -1.1877494 | 0.23493217 | 0.80747753 |
| CNTF | 42.4460824 | -0.9708879 | 0.7189024 | -1.3505141 | 0.17685112 | 0.76115956 |
| GLYAT | 7586.17482 | 0.94455594 | 0.74180934 | 1.27331363 | 0.20290677 | 0.78228262 |
| GLYATL1 | 9614.14105 | 1.72745853 | 0.62763985 | 2.75230857 | 0.00591767 | 0.32908936 |
| FAM111A-DT | 221.536385 | 0.13397968 | 0.33047102 | 0.40542036 | 0.68516854 | 0.95097288 |
| FAM111A | 2022.85866 | -0.0745919 | 0.34052099 | -0.2190521 | 0.82660943 | 0.97548201 |
| DTX4 | 2069.42128 | 0.26343737 | 0.3898755 | 0.67569613 | 0.49923359 | 0.91154879 |
| MPEG1 | 957.31628 | 0.15207065 | 0.55556217 | 0.27372391 | 0.7842968 | 0.97017379 |
| OSBP | 5254.64495 | -0.0674164 | 0.17293277 | -0.3898417 | 0.69665359 | 0.9525827 |
| AP000442.2 | 295.644433 | -0.7076514 | 0.33436725 | -2.1163896 | 0.03431168 | 0.55369025 |
| PATL1 | 3481.86732 | -0.7239103 | 0.25491841 | -2.8397728 | 0.00451457 | 0.29501922 |
| STX3 | 7751.62043 | -1.1098829 | 0.55995291 | -1.9821004 | 0.04746801 | 0.58241466 |
| MRPL16 | 2235.31462 | 0.0556635 | 0.16280281 | 0.34190752 | 0.73242049 | 0.95848028 |
| TCN1 | 51.6531096 | 0.43472668 | 0.64509719 | 0.6738933 | 0.50037915 | 0.91180773 |
| MS4A3 | 72.242174 | 2.22020342 | 1.29431518 | 1.7153499 | 0.08628109 | 0.65077063 |
| MS4A6A | 1306.1359 | -0.0436781 | 0.61656789 | -0.0708407 | 0.94352453 | 0.99136103 |
| MS4A4E | 54.502486 | 0.33910094 | 0.78020757 | 0.43462913 | 0.66383165 | 0.94642611 |
| MS4A4A | 750.782352 | -0.291806 | 0.55412016 | -0.5266114 | 0.59846347 | 0.9346978 |
| MS4A7 | 909.289867 | -0.0145483 | 0.50794383 | -0.0286415 | 0.97715051 | 0.99455099 |
| MS4A14 | 190.769963 | 0.00055912 | 0.65680798 | 0.00085127 | 0.99932079 | 0.99994357 |
| MS4A1 | 251.969791 | -1.0627616 | 1.09752828 | -0.9683228 | NA | NA |
| MS4A10 | 86.1181651 | 5.84783757 | 1.75621169 | 3.32980222 | NA | NA |
| AP000777.2 | 345.379534 | -0.3326524 | 0.28135692 | -1.1823147 | 0.23708084 | 0.80857441 |
| AP000777.3 | 334.270611 | -0.3211635 | 0.25092245 | -1.2799315 | 0.20056924 | 0.78134034 |
| CCDC86 | 1327.76365 | -0.3667159 | 0.25175647 | -1.4566293 | 1.45E-01 | 7.31E-01 |
| PTGDR2 | 166.317114 | -0.2538935 | 0.18666129 | -1.3601828 | 0.17377207 | 0.75974924 |
| PRPF19 | 4774.39803 | -0.2698332 | 0.22470343 | -1.2008414 | 0.22981272 | 0.80551898 |
| AP003721.3 | 38.5123699 | 0.94302471 | 0.47550925 | 1.983189 | 0.04734632 | 0.58218854 |
| TMEM109 | 5605.52071 | 0.05841383 | 0.1607533 | 0.36337563 | 0.7163243 | 0.95603249 |
| TMEM132A | 1526.15359 | -0.1691067 | 0.4271984 | -0.3958505 | 0.69221533 | 0.95177446 |
| SLC15A3 | 1286.93054 | -1.3382512 | 0.59003942 | -2.2680708 | 0.02332489 | 0.51967919 |
| CD6 | 214.258419 | 0.12139588 | 0.82777903 | 0.14665252 | 0.8834063 | 0.98293342 |
| CD5 | 150.720966 | 0.38268232 | 0.87951563 | 0.43510576 | 0.66348566 | 0.94642611 |
| VPS37C | 2126.52293 | -0.1580778 | 0.16386498 | -0.9646832 | 0.3347035 | 0.85557672 |
| VWCE | 134.243904 | 0.77996614 | 0.31471204 | 2.47834861 | 0.01319921 | 0.42221215 |
| DDB1 | 11124.0605 | 0.01470552 | 0.187228 | 0.07854339 | 0.93739582 | 0.99079211 |
| TKFC | 6546.01678 | 0.33449552 | 0.31227775 | 1.07114748 | 0.28410312 | 0.83107779 |
| CYB561A3 | 2394.80996 | -0.111809 | 0.18791922 | -0.5949845 | 0.55185385 | 0.92373235 |
| TMEM138 | 1734.38036 | -0.1501845 | 0.15187668 | -0.9888583 | 0.32273249 | 0.85119105 |
| TMEM216 | 554.785247 | -0.0376195 | 0.15872824 | -0.2370055 | 0.81265252 | 0.97371854 |
| CPSF7 | 5787.90253 | -0.2943307 | 0.16446887 | -1.7895831 | 7.35E-02 | 6.37E-01 |
| AP003108.2 | 1608.79119 | -0.2998079 | 0.18329654 | -1.6356442 | 1.02E-01 | 6.73E-01 |
| SDHAF2 | 1678.9796 | -0.3648769 | 0.19515089 | -1.8697169 | 0.06152315 | 0.61037866 |
| PPP1R32 | 495.0503 | 0.54923556 | 0.32574115 | 1.68611048 | 0.09177452 | 0.65518742 |
| LRRC10B | 130.091979 | -1.1471584 | 0.61884656 | -1.8537041 | 0.0637815 | 0.61812911 |
| SYT7 | 1255.66428 | -0.2588667 | 0.40507358 | -0.639061 | 0.52278324 | 0.91723878 |
| DAGLA | 600.118394 | -0.1169201 | 0.34616164 | -0.3377615 | 0.73554292 | 0.95895327 |
| DKFZP434K0 | 107.440463 | -0.2184777 | 0.37449625 | -0.5833908 | 0.55963022 | 0.92429981 |
| MYRF | 3235.41963 | 0.61806083 | 0.36077694 | 1.7131384 | 0.08668708 | 0.65228863 |
| TMEM258 | 2008.60506 | 0.1541068 | 0.14106523 | 1.09245065 | 0.27463507 | 0.82719407 |
| FEN1 | 614.678316 | -0.2290065 | 0.39875923 | -0.5742977 | 0.56576637 | 0.92513982 |

| | | | | | | |
|---|---|---|---|---|---|---|
| FADS2 | 2294.88602 | 0.09594166 | 0.49707607 | 0.19301203 | 0.84694955 | 0.97684681 |
| FADS1 | 2206.85767 | -0.4460615 | 0.59794922 | -0.7459856 | 0.45567611 | 0.89688594 |
| FADS3 | 5319.85894 | -0.5788312 | 0.34632018 | -1.6713758 | 0.09464748 | 0.65888357 |
| RAB3IL1 | 1903.24711 | -0.7252816 | 0.34563718 | -2.09839 | 0.03587071 | 0.55369025 |
| BEST1 | 171911.879 | -0.010117 | 0.42487227 | -0.0238119 | 0.98100267 | 0.99576712 |
| FTH1 | 193912.136 | 0.01160136 | 0.42247728 | 0.02746031 | 0.97809259 | 0.99476647 |
| INCENP | 428.591632 | -0.4129347 | 0.2527727 | -1.6336206 | 0.10233854 | 0.67279543 |
| SCGB1D2 | 68.3811542 | 0.60177701 | 0.62801652 | 0.95821844 | 0.33795262 | 0.8571036 |
| AP003306.2 | 550.305241 | 0.02040826 | 0.46514055 | 0.04387547 | 0.96500367 | 0.99357855 |
| ASRGL1 | 4274.62694 | -0.115547 | 0.45194672 | -0.2556651 | 0.79820947 | 0.97170189 |
| AP003064.1 | 1984.82507 | -0.0873289 | 0.47886869 | -0.1823651 | 0.85529622 | 0.97917572 |
| AHNAK | 21539.8504 | 0.13884864 | 0.21655143 | 0.64118088 | 0.52140517 | 0.91723878 |
| EEF1G | 52076.847 | 0.32046033 | 0.25107379 | 1.27635914 | 0.20182859 | 0.78134034 |
| AP002990.1 | 53458.5804 | 0.3131406 | 0.24189287 | 1.29454248 | 0.19547811 | 0.77743338 |
| TUT1 | 2248.12776 | 0.05517431 | 0.25307547 | 0.21801526 | 0.82741722 | 0.97548201 |
| MTA2 | 4510.88679 | -0.3421211 | 0.19758598 | -1.7315048 | 0.08336178 | 0.647557 |
| EML3 | 4780.54629 | -0.2626294 | 0.22979739 | -1.1428737 | 0.25309104 | 0.81835388 |
| AP001458.1 | 383.656735 | -0.0760119 | 0.26069951 | -0.2915689 | 0.77061624 | 0.96836854 |
| ROM1 | 460.770834 | -0.1442959 | 0.19319462 | -0.7468941 | 0.45512751 | 0.89670027 |
| B3GAT3 | 3158.0415 | -0.0233547 | 0.14713199 | -0.1587329 | 0.87387934 | 0.98275231 |
| GANAB | 18184.0166 | 0.22025986 | 0.11847159 | 1.85917878 | 0.0630018 | 0.6147045 |
| INTS5 | 1470.63101 | -0.1321317 | 0.15025104 | -0.8794061 | 0.37918111 | 0.87069278 |
| AP001458.2 | 1267.36175 | -0.0892021 | 0.16208819 | -0.5503308 | 0.58209251 | 0.9317705 |
| C11orf98 | 1529.97568 | -0.1050619 | 0.16071203 | -0.6537274 | 0.51328742 | 0.91597025 |
| LBHD1 | 2470.89032 | -0.1233202 | 0.1358552 | -0.9077323 | 0.36401965 | 0.8663515 |
| CSKMT | 328.046858 | -0.3899188 | 0.20071872 | -1.942613 | 0.05206294 | 0.58645127 |
| UQCC3 | 991.814896 | -0.1139927 | 0.18467157 | -0.6172725 | 0.53705499 | 0.92105614 |
| UBXN1 | 4359.13971 | 0.04366664 | 0.13853805 | 0.31519599 | 0.75261285 | 0.96301881 |
| LRRN4CL | 63.6902686 | 0.66695679 | 0.81979292 | 0.81356739 | 0.41589283 | 0.88399703 |
| BSCL2 | 4774.2064 | 0.14729037 | 0.14295141 | 1.03035271 | 0.30284446 | 0.84211658 |
| HNRNPUL2-E | 9976.8049 | 0.08185795 | 0.0757509 | 1.08062028 | 0.27986606 | 0.82825734 |
| GNG3 | 123.323544 | 0.03895048 | 0.21105554 | 0.18455086 | 0.85358133 | 0.97906651 |
| HNRNPUL2 | 7715.85035 | 0.04271747 | 0.11570811 | 0.36918305 | 7.12E-01 | 0.95560193 |
| TTC9C | 930.09881 | 0.03531634 | 0.15427309 | 0.22892091 | 0.81893039 | 0.97454841 |
| ZBTB3 | 227.765671 | 0.11842185 | 0.27550944 | 0.42982866 | 0.66732029 | 0.94830184 |
| POLR2G | 2006.00474 | 0.35776212 | 0.15904111 | 2.24949467 | 0.02448104 | 0.51967919 |
| AP001160.2 | 128.99068 | -0.2707846 | 0.3162583 | -0.8562135 | 0.39187969 | 0.87744531 |
| TAF6L | 1650.49511 | -0.1561178 | 0.19410579 | -0.8042923 | 0.42122816 | 0.88451372 |
| TMEM223 | 2477.73276 | -0.0260988 | 0.11923427 | -0.2188864 | 0.82673851 | 0.97548201 |
| AP001160.3 | 589.924625 | -0.1139749 | 0.21860541 | -0.5213726 | 0.60210726 | 0.9355118 |
| TMEM179B | 2639.68574 | 0.0470887 | 0.15334154 | 0.30708378 | 0.7587796 | 0.96502715 |
| NXF1 | 7962.20368 | -0.6084686 | 0.18478505 | -3.2928452 | 0.00099179 | 0.14989637 |
| STX5 | 2413.27937 | -0.0585088 | 0.10793917 | -0.5420534 | 0.58778174 | 0.93289552 |
| AP001160.1 | 827.076617 | -0.3821309 | 0.19574985 | -1.9521391 | 0.05092169 | 0.5861488 |
| WDR74 | 1943.87609 | -0.3480318 | 0.18051458 | -1.9279983 | 0.05385534 | 0.59266174 |
| SLC3A2 | 20973.8779 | -0.7709487 | 0.52537422 | -1.4674277 | 0.14225973 | 0.72784886 |
| SLC22A6 | 25155.2995 | 1.82610236 | 0.84716959 | 2.15553342 | 0.03112012 | 0.54466273 |
| SLC22A8 | 20321.1527 | 1.32900308 | 1.11966046 | 1.18696973 | 0.23523956 | 0.80747753 |
| SLC22A24 | 105.904289 | 0.88321724 | 0.6399106 | 1.38021972 | 0.167519 | 0.75558932 |
| HRASLS5 | 113.514392 | 1.87871002 | 0.65209421 | 2.88104079 | 0.00396364 | 0.27110453 |
| LGALS12 | 55.1815749 | 2.53451499 | 1.17947009 | 2.14885906 | NA | NA |
| RARRES3 | 5921.30613 | -0.1907873 | 0.48882055 | -0.3903014 | 0.69631371 | 0.9525827 |
| HRASLS2 | 512.216264 | 0.17537817 | 0.86691078 | 0.20230244 | NA | NA |
| PLA2G16 | 4494.73355 | 0.02616968 | 0.35173314 | 0.07440208 | 0.94069045 | 0.99136103 |

| | | | | | | |
|---|---|---|---|---|---|---|
| ATL3 | 3881.69923 | -0.0344812 | 0.2121025 | -0.1625684 | 0.87085829 | 0.98275231 |
| AP000753.2 | 76.9879658 | -0.5208138 | 0.47466448 | -1.0972252 | 0.27254294 | 0.82716979 |
| RTN3 | 7777.28776 | 0.0356789 | 0.18013677 | 0.1980656 | 0.84299374 | 0.97649449 |
| C11orf95 | 1821.44795 | 0.24104915 | 0.32379115 | 0.74445872 | 0.45659901 | 0.89688594 |
| SPINDOC | 1019.93824 | -0.0323153 | 0.18789307 | -0.1719877 | 0.86344724 | 0.98176973 |
| MARK2 | 3152.28293 | -0.279031 | 0.11606653 | -2.4040612 | 0.01621406 | 0.45896904 |
| RCOR2 | 51.2056981 | 0.5814554 | 0.6276027 | 0.92647051 | 0.35420152 | 0.86364633 |
| NAA40 | 2481.20197 | -0.2735346 | 0.22476921 | -1.2169575 | 0.22362037 | 0.80078694 |
| COX8A | 9209.85905 | 0.20120805 | 0.13245381 | 1.51908095 | 0.12874212 | 0.70633634 |
| AP000721.1 | 7117.32861 | 0.19379816 | 0.13687496 | 1.41587742 | 0.1568114 | 0.74522306 |
| OTUB1 | 5145.61116 | -0.2444193 | 0.12303076 | -1.986652 | 0.04696098 | 0.58171004 |
| MACROD1 | 2026.67409 | 0.17246331 | 0.23564917 | 0.73186472 | 0.46425114 | 0.89830907 |
| FLRT1 | 242.105327 | -0.1566079 | 0.48918254 | -0.320142 | 0.74886072 | 0.96249332 |
| AP006333.1 | 56.6342737 | -0.0279074 | 0.70056674 | -0.0398355 | 0.96822425 | 0.99400331 |
| STIP1 | 5090.99749 | -0.2026141 | 0.15469807 | -1.3097388 | 0.1902842 | 0.77100296 |
| FERMT3 | 984.119317 | -0.228005 | 0.38667815 | -0.5896506 | 0.55542495 | 0.92373235 |
| TRPT1 | 2783.37915 | 0.0191163 | 0.13674617 | 0.13979407 | 0.8888227 | 0.98423722 |
| NUDT22 | 4189.44449 | 0.03493812 | 0.1172571 | 0.29796167 | 0.76573242 | 0.96704052 |
| AP001453.1 | 1553.17906 | 0.029821 | 0.14633601 | 0.20378441 | 0.83852198 | 0.97612168 |
| DNAJC4 | 2442.35168 | 0.0119022 | 0.1736044 | 0.06855934 | 0.94534038 | 0.99166614 |
| VEGFB | 4111.89405 | -0.0036982 | 0.26214708 | -0.0141075 | 0.98874423 | 0.99788078 |
| FKBP2 | 5323.6035 | 0.25026929 | 0.15604829 | 1.60379386 | 0.1087595 | 0.67800851 |
| PPP1R14B | 2722.39296 | -0.1856972 | 0.25556731 | -0.7266079 | 0.46746618 | 0.89983399 |
| AP001453.2 | 215.684786 | -0.1432832 | 0.25835516 | -0.5545978 | 0.57916977 | 0.93139743 |
| AP001453.4 | 154.222586 | -0.1637845 | 0.28465041 | -0.5753884 | 0.56502864 | 0.92493342 |
| PLCB3 | 2177.41147 | -0.016498 | 0.2870852 | -0.0574673 | 0.95417292 | 0.99170503 |
| BAD | 2476.58452 | -0.0130518 | 0.13424079 | -0.097227 | 0.92254611 | 0.98826666 |
| GPR137 | 4216.36001 | -0.1276946 | 0.11979464 | -1.0659458 | 0.28644812 | 0.8312673 |
| ESRRA | 7590.83678 | 0.13604483 | 0.21641896 | 0.6286179 | 0.52959924 | 0.91931447 |
| TRMT112 | 5589.58884 | 0.03003753 | 0.14292079 | 0.2101691 | 0.83353569 | 0.97550416 |
| PRDX5 | 12787.6509 | 0.06355111 | 0.17649029 | 0.36008276 | 0.71878525 | 0.95648644 |
| CCDC88B | 1123.35538 | -0.2801275 | 0.46763442 | -0.599031 | 0.54915218 | 0.92373235 |
| RPS6KA4 | 2075.47693 | -0.0809573 | 0.20161444 | -0.4015453 | 0.68801866 | 0.95158502 |
| SLC22A11 | 8453.1817 | 0.30025742 | 0.48147288 | 0.62362271 | 0.53287539 | 0.92017117 |
| SLC22A12 | 27082.965 | 0.74751382 | 0.53705036 | 1.39188774 | 0.1639564 | 0.75265041 |
| NRXN2 | 833.306005 | 0.76786705 | 0.30955596 | 2.48054358 | 0.01311822 | 0.42169573 |
| RASGRP2 | 1292.62748 | -0.4327008 | 0.4740525 | -0.9127698 | 0.36136362 | 0.86528137 |
| PYGM | 262.982454 | -0.393095 | 0.48975384 | -0.8026379 | 0.42218406 | 0.88451372 |
| SF1 | 13788.4324 | -0.3382471 | 0.15495106 | -2.1829288 | 0.02904105 | 0.53606221 |
| MAP4K2 | 1316.28267 | 0.18394755 | 0.27551531 | 0.6676491 | 0.50435761 | 0.91356552 |
| MEN1 | 2878.49606 | -0.1967848 | 0.17313368 | -1.1366059 | 0.25570308 | 0.82052927 |
| CDC42BPG | 1672.26903 | 0.42517922 | 0.1781528 | 2.38659861 | 0.01700505 | 0.46671416 |
| EHD1 | 5564.01518 | -0.1909718 | 0.19847814 | -0.9621806 | 0.33595891 | 0.85641269 |
| MIR194-2HG | 200.89248 | 1.36357714 | 0.51446292 | 2.65048672 | 0.00803759 | 0.37112203 |
| ATG2A | 4465.91489 | -0.4489537 | 0.26463849 | -1.6964792 | 0.08979517 | 0.65375543 |
| PPP2R5B | 2368.47078 | -0.1314224 | 0.22870738 | -0.5746312 | 0.56554072 | 0.92513982 |
| MAJIN | 64.895714 | 0.20851823 | 0.73667033 | 0.28305502 | 0.77713466 | 0.96836854 |
| BATF2 | 763.231051 | -1.058362 | 0.58163812 | -1.8196228 | NA | NA |
| ARL2 | 3878.57436 | 0.13214953 | 0.17436346 | 0.75789695 | 0.44851268 | 0.8948759 |
| ARL2-SNX15 | 3812.79783 | 0.03887997 | 0.11926357 | 0.3260004 | 0.74442405 | 0.9613927 |
| SNX15 | 1788.60463 | -0.0980021 | 0.17227993 | -0.5688537 | 0.56945546 | 0.9271275 |
| SAC3D1 | 300.671598 | -0.0438733 | 0.20811703 | -0.2108106 | 0.83303507 | 0.97550416 |
| NAALADL1 | 220.991877 | -0.0688103 | 0.26536544 | -0.2593038 | 7.95E-01 | 0.97101097 |
| CDCA5 | 96.9290931 | -0.117427 | 0.57221353 | -0.2052154 | 0.83740384 | 0.97577999 |

| | | | | | | |
|---|---|---|---|---|---|---|
| ZFPL1 | 1803.31077 | -0.1125856 | 0.18524223 | -0.6077749 | 0.54333677 | 0.92297267 |
| TMEM262 | 1664.58642 | -0.1200613 | 0.17923679 | -0.6698477 | 0.5029549 | 0.91353729 |
| VPS51 | 5196.73847 | 0.13746931 | 0.16004943 | 0.85891784 | 0.39038585 | 0.87650906 |
| AP003068.1 | 1640.9731 | 0.14167745 | 0.15436945 | 0.91778169 | 0.35873317 | 0.86480228 |
| TM7SF2 | 1814.61118 | 0.39012821 | 0.28088497 | 1.38892521 | 0.16485549 | 0.75299828 |
| ZNHIT2 | 587.878023 | -0.2387328 | 0.16975311 | -1.4063528 | 0.15961938 | 0.74768867 |
| AP003068.3 | 199.45972 | -0.1551468 | 0.20580683 | -0.7538467 | 0.45094129 | 0.89590699 |
| FAU | 12974.5639 | 0.05864554 | 0.18176863 | 0.32263841 | 0.7469691 | 0.96224616 |
| SYVN1 | 11996.0139 | 0.00429664 | 0.13668818 | 0.03143389 | 0.97492352 | 0.99455099 |
| MRPL49 | 5022.55407 | 0.01722647 | 0.11672067 | 0.14758717 | 0.88266858 | 0.98276701 |
| AP003068.2 | 343.365824 | 0.32145782 | 0.34653779 | 0.92762701 | 0.35360108 | 0.86364633 |
| CAPN1 | 11039.5527 | 0.04074175 | 0.14801028 | 0.27526294 | 0.78311423 | 0.96971192 |
| POLA2 | 557.682024 | -0.1545598 | 0.16765063 | -0.9219162 | 0.3565723 | 0.86390849 |
| CDC42EP2 | 1079.17032 | -0.0354436 | 0.23203605 | -0.1527504 | 0.87859515 | 0.98275231 |
| DPF2 | 2323.16424 | -0.1588719 | 0.11975936 | -1.3265926 | 0.18464347 | 0.76820626 |
| AP000944.1 | 94.2453904 | 0.29917958 | 0.39454075 | 0.75829828 | 0.44827243 | 0.8948759 |
| SLC25A45 | 1263.68945 | 0.39256247 | 0.38864038 | 1.01009181 | 0.31245131 | 0.84541735 |
| FRMD8 | 2626.11669 | -0.2971271 | 0.17081628 | -1.7394543 | 0.08195488 | 0.64555743 |
| NEAT1 | 157860.337 | -0.3587491 | 0.36936455 | -0.9712602 | 0.33141875 | 0.85408791 |
| MALAT1 | 46455.467 | -0.0626827 | 0.15015579 | -0.4174512 | 0.6763484 | 0.94968498 |
| SCYL1 | 6206.33621 | -0.1727147 | 0.16971842 | -1.0176543 | 0.30884226 | 0.84414094 |
| LTBP3 | 8605.10786 | -0.0444205 | 0.27580891 | -0.1610553 | 0.87204981 | 0.98275231 |
| SSSCA1 | 2447.54953 | -0.2101119 | 0.23325853 | -0.9007683 | 0.3677115 | 0.86726419 |
| FAM89B | 1545.62311 | -0.2084547 | 0.26976873 | -0.7727164 | 0.43969023 | 0.89203402 |
| EHBP1L1 | 4485.03135 | -0.2617414 | 0.34751453 | -0.7531813 | 0.45134099 | 0.89639459 |
| MAP3K11 | 5888.10634 | -0.138287 | 0.10963464 | -1.2613437 | 2.07E-01 | 0.78790396 |
| PCNX3 | 4350.21528 | -0.1680051 | 0.13677148 | -1.2283633 | 0.21931062 | 0.79847731 |
| SIPA1 | 3172.01409 | -0.2063863 | 0.31393632 | -0.6574144 | 0.51091446 | 0.91474443 |
| RELA | 6162.30505 | -0.4111491 | 0.18857907 | -2.1802478 | 0.0292391 | 0.53606221 |
| KAT5 | 4063.14256 | -0.1896738 | 0.20299272 | -0.9343872 | 0.35010418 | 0.86364633 |
| RNASEH2C | 5064.75671 | -0.2015448 | 0.11133046 | -1.8103297 | 0.07024467 | 0.62928077 |
| AP5B1 | 1336.98202 | -0.5709334 | 0.25441941 | -2.244064 | 0.02482829 | 0.52085426 |
| OVOL1 | 819.674846 | -0.2133463 | 0.32396546 | -0.6585465 | 0.51018706 | 0.91439937 |
| OVOL1-AS1 | 62.8341586 | 1.45404297 | 0.92315528 | 1.57507952 | NA | NA |
| AP001266.1 | 385.708919 | -0.2394309 | 0.30782978 | -0.7778031 | 0.43668512 | 0.89122667 |
| CFL1 | 27902.0373 | -0.1702894 | 0.22346122 | -0.7620535 | 0.44602805 | 0.89480207 |
| SNX32 | 8054.03216 | -0.0980272 | 0.23806072 | -0.411774 | 0.68050506 | 0.95070301 |
| MUS81 | 3140.51465 | -0.1528011 | 0.14943554 | -1.022522 | 0.3065339 | 0.84278367 |
| EFEMP2 | 3660.4087 | 0.13371287 | 0.27643787 | 0.48369954 | 0.62859912 | 0.94088547 |
| CTSW | 178.384669 | -0.3618465 | 0.77781349 | -0.4652098 | 0.64178119 | 0.94088547 |
| FIBP | 2461.9994 | 0.06216556 | 0.12808281 | 0.48535443 | 0.62742495 | 0.94088547 |
| CCDC85B | 811.110902 | -0.430165 | 0.4150814 | -1.0363388 | 0.3000441 | 0.84181141 |
| FOSL1 | 888.991673 | -2.3498888 | 1.12214637 | -2.0941018 | 0.03625091 | 0.55384187 |
| C11orf68 | 1862.96644 | -0.0523278 | 0.16309669 | -0.3208392 | 0.74833224 | 0.96241535 |
| DRAP1 | 2504.53366 | 0.02226165 | 0.15385434 | 0.14469302 | 0.88495325 | 0.9835088 |
| SART1 | 3208.68017 | -0.1910708 | 0.15272693 | -1.2510618 | 0.21091191 | 0.79134644 |
| EIF1AD | 1146.38842 | -0.153727 | 0.21680026 | -0.709072 | 0.47827979 | 0.904302 |
| BANF1 | 3271.65424 | -0.0164011 | 0.18295697 | -0.0896444 | 0.92856982 | 0.98963127 |
| CST6 | 69.2398542 | 0.6673494 | 0.88705063 | 0.75232391 | 0.45185629 | 0.89658431 |
| GAL3ST3 | 173.528767 | -0.5070123 | 0.74948967 | -0.6764767 | NA | NA |
| SF3B2 | 10623.7837 | -0.0923553 | 0.10867187 | -0.8498545 | 0.39540599 | 0.87927742 |
| AP006287.2 | 63.9825164 | -0.2505644 | 0.31305975 | -0.8003726 | 0.42349497 | 0.88520178 |
| PACS1 | 4926.58553 | 0.10315835 | 0.13333413 | 0.77368299 | 0.43911828 | 0.89175645 |
| AP000759.1 | 70.8869024 | 0.67968986 | 0.30479517 | 2.22998893 | 0.02574818 | 0.52502381 |

| | | | | | | |
|---|---|---|---|---|---|---|
| KLC2 | 3269.6485 | -0.3198088 | 0.18632803 | -1.716375 | 0.08609343 | 0.65049132 |
| AP001107.4 | 384.486297 | -0.2605455 | 0.19786229 | -1.3168022 | 0.18790492 | 0.76970626 |
| AP001107.8 | 420.301539 | -0.4062535 | 0.17789282 | -2.2836983 | 0.02238927 | 0.51866171 |
| AP001107.7 | 102.984136 | -0.172786 | 0.29109116 | -0.5935804 | 0.5527928 | 0.92373235 |
| RAB1B | 11866.9477 | -0.0428009 | 0.15030333 | -0.2847635 | 0.77582536 | 0.96836854 |
| AP001107.1 | 2508.04407 | 0.06957598 | 0.19682891 | 0.35348458 | 0.72372519 | 0.95810643 |
| AP001107.2 | 5796.36771 | -0.0912642 | 0.13182827 | -0.6922961 | 0.48875141 | 0.905872 |
| CNIH2 | 56.1493934 | -0.0890337 | 0.36353423 | -0.2449114 | 8.07E-01 | 9.73E-01 |
| YIF1A | 3204.15883 | -0.0893009 | 0.19897389 | -0.4488073 | 0.65357066 | 0.94495971 |
| CD248 | 1077.16112 | 0.25343162 | 0.60381488 | 0.41971742 | 0.6746919 | 0.94935413 |
| RIN1 | 405.297836 | -0.1290373 | 0.5524611 | -0.2335682 | 8.15E-01 | 9.74E-01 |
| AP001107.6 | 56.1359847 | -0.0957191 | 0.52777195 | -0.1813644 | 0.85608153 | 0.97929708 |
| BRMS1 | 2396.76328 | -0.1753465 | 0.2144641 | -0.8176029 | 0.41358395 | 0.88316314 |
| B4GAT1 | 2102.45236 | 0.44571604 | 0.22392634 | 1.99045833 | 0.04654047 | 0.57983846 |
| AP001107.9 | 471.753433 | -0.2591197 | 0.31056314 | -0.8343543 | 0.40408135 | 0.88132015 |
| SLC29A2 | 1678.73396 | -0.2756772 | 0.30103563 | -0.9157627 | 0.35979135 | 0.86480228 |
| MRPL11 | 1391.87267 | 0.21869146 | 0.14424719 | 1.51608823 | 0.12949704 | 0.70764335 |
| PELI3 | 717.859064 | 0.02914467 | 0.21687793 | 0.13438285 | 0.89309984 | 0.98474725 |
| AP002748.3 | 267.382954 | 0.14165567 | 0.25333014 | 0.55917417 | 0.57604286 | 0.93083012 |
| DPP3 | 3062.09343 | 0.04865147 | 0.22208792 | 0.21906402 | 0.82660018 | 0.97548201 |
| AP002748.4 | 4619.13009 | 0.23298042 | 0.22451702 | 1.03769601 | 0.2994116 | 0.84123067 |
| BBS1 | 4212.43272 | 0.26818107 | 0.27983444 | 0.95835618 | 0.33788318 | 0.8571036 |
| ZDHHC24 | 1818.39412 | 0.25145056 | 0.13251077 | 1.89758585 | 0.05775066 | 0.60129769 |
| CTSF | 4310.82317 | 0.61031744 | 0.25086133 | 2.43288769 | 0.01497895 | 0.44700706 |
| CCDC87 | 40.3829879 | 0.00925637 | 0.38885107 | 0.02380441 | 0.98100862 | 0.99576712 |
| CCS | 2903.66629 | 0.16395578 | 0.23517093 | 0.69717707 | 4.86E-01 | 9.05E-01 |
| RBM14 | 3227.21981 | -0.1431628 | 0.10925953 | -1.3103001 | 1.90E-01 | 7.71E-01 |
| RBM14-RBM | 4561.74391 | -0.1777856 | 0.15361686 | -1.1573316 | 0.24713691 | 0.8156466 |
| RBM4 | 4903.49986 | -0.1811361 | 0.14498707 | -1.2493258 | 0.21154595 | 0.79149017 |
| RBM4B | 1215.00656 | -0.0200122 | 0.16742292 | -0.119531 | 0.90485467 | 0.98624484 |
| AP001157.1 | 804.027881 | -0.0005611 | 0.21210434 | -0.0026455 | 9.98E-01 | 1.00E+00 |
| SPTBN2 | 5782.48123 | 0.58381091 | 0.32872806 | 1.77596922 | 0.07573799 | 0.63773415 |
| C11orf80 | 1383.1043 | 0.83953191 | 0.22200284 | 3.78162689 | 0.00015581 | 0.05959132 |
| RCE1 | 1948.95346 | -0.0508908 | 0.20621557 | -0.2467843 | 0.80507513 | 0.97303568 |
| PC | 8841.0661 | 0.50064735 | 0.59235454 | 0.84518192 | 0.39800927 | 0.88054917 |
| LRFN4 | 713.896782 | -0.5288754 | 0.27029502 | -1.9566596 | 0.0503875 | 0.5861488 |
| C11orf86 | 271.446695 | -2.4973818 | 1.62739684 | -1.5345869 | 0.12488534 | 0.70228923 |
| SYT12 | 79.9274594 | 0.37309218 | 0.77245965 | 0.48299245 | 0.6291011 | 0.94088547 |
| RHOD | 982.391617 | -0.1320459 | 0.296507 | -0.4453381 | 6.56E-01 | 0.94537209 |
| KDM2A | 5433.70428 | -0.3175764 | 0.13428773 | -2.364895 | 0.01803518 | 0.47478681 |
| GRK2 | 6913.29461 | -0.1526214 | 0.23471773 | -0.6502339 | 0.51554114 | 0.91636259 |
| ANKRD13D | 2875.0317 | -0.1760471 | 0.29069522 | -0.6056071 | 0.54477567 | 0.92345821 |
| SSH3 | 3370.34871 | 0.01388542 | 0.22314557 | 0.06222586 | 0.95038297 | 0.99170503 |
| RAD9A | 3083.36058 | -0.2531913 | 0.16419027 | -1.5420601 | 0.12305899 | 0.69788192 |
| POLD4 | 5167.00951 | 0.13031967 | 0.22624083 | 0.57602189 | 0.56460039 | 0.92476629 |
| AP003419.1 | 4473.91288 | 0.14286371 | 0.23761604 | 0.60123763 | 0.54768172 | 0.92361227 |
| CLCF1 | 1243.15188 | -0.6194008 | 0.2764684 | -2.2404035 | 0.02506474 | 0.52085426 |
| AP003419.3 | 59.446069 | 0.1947178 | 0.49751703 | 0.39137916 | 0.695517 | 0.9525827 |
| PPP1CA | 8036.70197 | -0.0074064 | 0.15453298 | -0.0479276 | 0.96177395 | 0.99320711 |
| TBC1D10C | 507.411484 | 0.08962463 | 0.5848196 | 0.15325175 | 0.87819975 | 0.98275231 |
| CARNS1 | 135.548139 | 0.26867219 | 0.31914806 | 0.84184183 | 0.39987649 | 0.88077967 |
| RPS6KB2 | 2460.60819 | 0.01233736 | 0.13799412 | 0.08940499 | 0.92876006 | 0.98963127 |
| RPS6KB2-AS | 751.729297 | 0.06268545 | 0.14696713 | 0.426527 | 0.66972388 | 0.94898648 |
| CORO1B | 6752.17172 | 0.01957898 | 0.11907546 | 0.16442501 | 0.86939658 | 0.98275231 |

| | | | | | | |
|---|---|---|---|---|---|---|
| PTPRCAP | 760.89844 | 0.04126332 | 0.36435927 | 0.113249 | 9.10E-01 | 0.98662757 |
| CABP4 | 73.4678201 | -0.1311732 | 0.33032057 | -0.3971087 | 0.69128734 | 0.95158502 |
| TMEM134 | 2155.02677 | 0.0504639 | 0.18784322 | 0.26864903 | 0.78819978 | 0.97090793 |
| AIP | 2906.82381 | 0.06407287 | 0.11647685 | 0.55009106 | 0.58225692 | 0.93177403 |
| PITPNM1 | 7598.75552 | -0.390702 | 0.27752052 | -1.407831 | 0.15918113 | 0.7475863 |
| CDK2AP2 | 5052.26177 | -0.0785757 | 0.27866521 | -0.2819717 | 0.77796518 | 0.96836854 |
| GSTP1 | 22032.6586 | 0.05027981 | 0.32453955 | 0.15492661 | 0.87687918 | 0.98275231 |
| NDUFV1 | 15138.3378 | 0.32198103 | 0.17487093 | 1.84124966 | 0.06558498 | 0.62237346 |
| NUDT8 | 666.273907 | 0.33453701 | 0.32527148 | 1.02848553 | 0.30372149 | 0.84211658 |
| ACY3 | 2971.50931 | 1.29157957 | 0.59436991 | 2.17302316 | 0.02977857 | 0.53938016 |
| ALDH3B2 | 94.9999644 | 1.69766595 | 0.80808732 | 2.10084469 | 0.0356546 | 0.55369025 |
| AP003716.1 | 272.413648 | 0.81265415 | 0.75544803 | 1.07572475 | 0.28205037 | 0.83003334 |
| UNC93B1 | 2578.67283 | -0.8609271 | 0.48299713 | -1.7824683 | 0.07467289 | 0.63773415 |
| ALDH3B1 | 2133.91993 | 0.40588551 | 0.50149586 | 0.80934967 | 0.41831404 | 0.88445833 |
| AC004923.4 | 41.2069452 | -0.223027 | 0.50782154 | -0.4391839 | 6.61E-01 | 0.94628868 |
| NDUFS8 | 6031.27854 | 0.03903539 | 0.16019124 | 0.24367993 | 0.80747872 | 0.97334138 |
| TCIRG1 | 5044.90416 | -0.5284011 | 0.42741711 | -1.2362657 | 0.2163598 | 0.7954827 |
| AP002807.1 | 2117.8626 | -0.5674369 | 0.20797325 | -2.7284132 | 0.00636398 | 0.33501741 |
| CHKA | 2990.69426 | -0.6159128 | 0.26537609 | -2.3209054 | 0.02029195 | 0.5002876 |
| KMT5B | 2285.37979 | -0.1162764 | 0.1519631 | -0.7651621 | 0.44417503 | 0.8938278 |
| C11orf24 | 1695.61496 | -0.0315977 | 0.24005753 | -0.1316256 | 0.89528047 | 0.98474725 |
| LRP5 | 4890.4188 | 0.38603283 | 0.27224777 | 1.41794671 | 0.15620632 | 0.74522306 |
| PPP6R3 | 4352.81622 | -0.1437517 | 0.09721227 | -1.4787407 | 0.13920963 | 0.72409321 |
| AP003096.1 | 374.922232 | -0.0698484 | 0.19033405 | -0.3669782 | 0.7136353 | 0.95564093 |
| TESMIN | 419.147812 | 0.19581103 | 0.19597569 | 0.99915976 | 3.18E-01 | 0.84855921 |
| CPT1A | 7141.27721 | -0.1360597 | 0.32966519 | -0.4127209 | 0.67981112 | 0.95070301 |
| AP000808.1 | 49.9233475 | -0.2844666 | 0.80375522 | -0.3539219 | 0.72339742 | 0.95810643 |
| MRPL21 | 1473.67874 | 0.0204021 | 0.14414641 | 0.14153733 | 0.88744547 | 0.98423722 |
| IGHMBP2 | 1549.37483 | -0.3163295 | 0.14726013 | -2.1480999 | 0.03170582 | 0.5451365 |
| AP003071.3 | 367.696875 | 0.18855253 | 0.32250761 | 0.58464521 | 0.5587863 | 0.92421175 |
| MRGPRF | 894.083549 | 0.11072193 | 0.32421657 | 0.34150607 | 0.73272263 | 0.95848028 |
| MRGPRF-AS | 105.620955 | 0.2415772 | 0.34851197 | 0.69316757 | 0.48820439 | 0.90585781 |
| TPCN2 | 2423.95168 | 0.25360508 | 0.2899943 | 0.87451746 | 0.38183651 | 0.87185666 |
| AP003071.5 | 110.363642 | 0.48462525 | 0.75940579 | 0.63816375 | 0.52336709 | 0.91723878 |
| CCND1 | 3382.82146 | 0.04695872 | 0.36833306 | 0.12748984 | 0.89855271 | 0.98474725 |
| ORAOV1 | 2229.4574 | -0.1506071 | 0.22224508 | -0.6776623 | 0.49798584 | 0.91033482 |
| ANO1 | 727.875194 | -0.6364367 | 0.41273104 | -1.5420131 | 0.12307042 | 0.69788192 |
| ANO1-AS1 | 239.37294 | -0.646289 | 0.4444295 | -1.454199 | 0.14589113 | 0.73247558 |
| FADD | 1457.78406 | -0.0633783 | 0.16260359 | -0.3897721 | 6.97E-01 | 0.9525827 |
| AP000879.1 | 1247.87857 | -0.099875 | 0.15359781 | -0.6502369 | 5.16E-01 | 9.16E-01 |
| PPFIA1 | 2818.01811 | -0.427538 | 0.27670666 | -1.5450948 | 0.12232334 | 0.69788192 |
| AP000487.1 | 521.666104 | -0.3165571 | 0.35589125 | -0.889477 | 0.37374675 | 0.86791605 |
| CTTN | 17913.6961 | -0.3052684 | 0.21403254 | -1.4262711 | 0.15379011 | 0.74258845 |
| SHANK2 | 1432.55275 | -0.0648329 | 0.21688126 | -0.2989329 | 0.76499124 | 0.96678847 |
| DHCR7 | 2174.91099 | 0.10784317 | 0.36184906 | 0.29803357 | 0.76567754 | 0.96704052 |
| AP002387.2 | 116.503143 | -0.174089 | 0.33678933 | -0.5169078 | 0.60522052 | 0.93687349 |
| NADSYN1 | 4403.3241 | 0.05452539 | 0.16693753 | 0.32662151 | 0.74395417 | 0.9613927 |
| KRTAP5-8 | 117.020568 | 0.55431831 | 0.90603356 | 0.6118077 | 0.540665 | 0.92257149 |
| FAM86C1 | 339.787246 | 0.09642748 | 0.22478053 | 0.42898504 | 0.66793411 | 0.94847921 |
| AP002495.2 | 100.256567 | 0.14757549 | 0.4189331 | 0.35226506 | 0.72463949 | 0.95836477 |
| RNF121 | 1748.62576 | -0.0356341 | 0.159386 | -0.2235714 | 0.82309085 | 0.97537437 |
| IL18BP | 5858.63403 | -0.4096559 | 0.32650335 | -1.2546759 | 0.20959646 | 0.79128274 |
| NUMA1 | 13981.7535 | -0.0637899 | 0.229437 | -0.2780278 | 0.780991 | 0.96885896 |
| AP002490.1 | 1030.13422 | -0.1573711 | 0.37652284 | -0.4179589 | 0.6759772 | 0.94959168 |

| | | | | | | |
|---|---|---|---|---|---|---|
| LRTOMT | 1478.79603 | 0.3488305 | 0.20218847 | 1.72527398 | 0.08447811 | 0.64988556 |
| AP000812.5 | 1165.31009 | 0.43227046 | 0.2342923 | 1.84500498 | 0.06503681 | 0.62154765 |
| LAMTOR1 | 5936.11506 | -0.052319 | 0.20408329 | -0.2563612 | 0.79767195 | 0.97148232 |
| ANAPC15 | 1009.17688 | 0.4086788 | 0.29298611 | 1.39487433 | 0.16305374 | 0.75117751 |
| FOLR3 | 256.245806 | 0.18365756 | 0.33301268 | 0.55150321 | 0.58128877 | 0.9317705 |
| FOLR1 | 5278.81116 | -0.0254499 | 0.24002509 | -0.1060301 | 0.91555847 | 0.9868044 |
| FOLR2 | 643.576178 | 0.27749045 | 0.65701469 | 0.42235044 | 0.67276924 | 0.94922827 |
| INPPL1 | 10405.8419 | -0.1161958 | 0.18227139 | -0.6374881 | 0.52380697 | 0.91723878 |
| CLPB | 1420.51858 | -0.158507 | 0.17845681 | -0.8882095 | 0.37442809 | 0.86791605 |
| AP000593.3 | 77.5460459 | -0.2213414 | 0.27737404 | -0.7979889 | 0.42487694 | 0.88573606 |
| LINC01537 | 68.3417963 | 0.21715106 | 0.49261394 | 0.44081388 | 0.65934775 | 0.94601668 |
| PDE2A | 2225.35295 | -0.0078881 | 0.39342588 | -0.0200497 | 0.98400371 | 0.9961464 |
| AP005019.1 | 123.70269 | 0.10564161 | 0.40690124 | 0.25962469 | 0.79515329 | 0.97101097 |
| ARAP1 | 7538.82889 | -0.1894175 | 0.22484879 | -0.8424219 | 0.39955186 | 0.88077967 |
| ARAP1-AS1 | 1688.28469 | -0.1931277 | 0.23363842 | -0.8266093 | 4.08E-01 | 0.88203324 |
| ARAP1-AS2 | 324.743937 | -0.1663714 | 0.30235104 | -0.5502592 | 0.58214164 | 0.9317705 |
| STARD10 | 2409.56188 | -0.2716286 | 0.20151399 | -1.3479393 | 0.17767791 | 0.76115956 |
| ATG16L2 | 3583.27914 | 0.06396307 | 0.30646688 | 0.2087112 | 0.83467369 | 0.97550416 |
| FCHSD2 | 1520.78498 | 0.02338143 | 0.15910327 | 0.14695757 | 0.88316551 | 0.98293314 |
| P2RY2 | 133.038254 | 0.02390985 | 0.35236744 | 0.06785488 | 0.94590116 | 0.99166614 |
| P2RY6 | 573.987869 | 0.47223838 | 0.50523598 | 0.93468873 | 0.34994874 | 0.86364633 |
| AP002761.3 | 157.672734 | -0.4816057 | 0.35541286 | -1.3550599 | 0.17539846 | 0.76073329 |
| ARHGEF17 | 6743.87141 | -0.2552537 | 0.309331 | -0.8251799 | 0.40926946 | 0.88221056 |
| RELT | 839.872708 | -0.5619168 | 0.50236732 | -1.1185377 | 0.26333742 | 0.82424998 |
| AP000763.4 | 372.484506 | -0.4884499 | 0.51859192 | -0.9418772 | 0.3462555 | 0.86135217 |
| FAM168A | 2423.15107 | 0.12469954 | 0.14429585 | 0.86419355 | 0.38748158 | 0.87460898 |
| AP000763.3 | 55.5181667 | -0.081867 | 0.44835674 | -0.1825935 | 0.85511698 | 0.97910552 |
| PLEKHB1 | 849.611081 | 0.32481117 | 0.33122864 | 0.98062527 | 0.32677757 | 0.85282528 |
| RAB6A | 6520.31563 | -0.0320388 | 0.13255278 | -0.2417062 | 0.8090078 | 0.97334138 |
| AP002770.1 | 562.249032 | 0.06799902 | 0.33871286 | 0.20075714 | 0.84088848 | 0.97618283 |
| MRPL48 | 959.562584 | 0.14015457 | 0.18122504 | 0.7733731 | 0.43930161 | 0.89175645 |
| COA4 | 2545.23025 | 0.08229655 | 0.20589464 | 0.39970225 | 0.68937583 | 0.95158502 |
| PAAF1 | 1338.16428 | 0.59658049 | 0.31818769 | 1.87493269 | 0.06080198 | 0.60989294 |
| DNAJB13 | 102.075501 | -0.4997763 | 0.67844271 | -0.7366522 | 0.4613339 | 0.89755243 |
| AP003717.1 | 79.4911046 | -0.5187118 | 0.73456012 | -0.7061529 | 0.48009305 | 0.90537463 |
| UCP2 | 1898.54382 | 0.2975198 | 0.44148508 | 0.67390682 | 0.50037055 | 0.91180773 |
| UCP3 | 133.755329 | -0.4751221 | 0.30704269 | -1.5474137 | 0.12176352 | 0.69725141 |
| C2CD3 | 1043.53423 | -0.4564033 | 0.20499268 | -2.2264372 | 0.02598491 | 0.52533244 |
| PPME1 | 2913.71827 | 0.052522 | 0.16023321 | 0.32778471 | 0.74307445 | 0.96109001 |
| P4HA3 | 280.650355 | 0.46798863 | 0.31543619 | 1.48362377 | 0.13790876 | 0.72209298 |
| PGM2L1 | 481.055871 | -0.2324858 | 0.31701359 | -0.7333622 | 0.46333757 | 0.89830907 |
| KCNE3 | 1031.68699 | 0.76596373 | 0.38360922 | 1.99672921 | 4.59E-02 | 5.77E-01 |
| AP001372.3 | 420.71106 | 0.77595746 | 0.37874044 | 2.04878426 | 0.04048322 | 0.55657615 |
| LIPT2 | 111.236224 | 0.4939736 | 0.40502048 | 1.21962623 | 0.2226066 | 0.80078694 |
| POLD3 | 1248.4109 | 0.14000749 | 0.33759659 | 0.41471831 | 0.67834813 | 0.95031695 |
| CHRDL2 | 72.3374033 | 0.28863379 | 0.44789821 | 0.64441826 | 0.51930425 | 0.91636259 |
| RNF169 | 1342.90364 | -0.4417083 | 0.20741091 | -2.1296289 | 0.03320226 | 0.55156489 |
| XRRA1 | 1619.26416 | -0.4685507 | 0.25309635 | -1.8512739 | 0.06413016 | 0.61837523 |
| SPCS2 | 2968.04561 | 0.04017508 | 0.16178511 | 0.24832371 | 0.80388396 | 0.97274243 |
| NEU3 | 730.062531 | -0.0755354 | 0.16273242 | -0.4641691 | 0.64252662 | 0.94090101 |
| SLCO2B1 | 3602.02406 | 0.05793047 | 0.45599474 | 0.12704197 | 0.89890718 | 0.98482517 |
| AP001972.1 | 257.358561 | 0.15509459 | 0.49553646 | 0.31298321 | 0.75429342 | 0.96348234 |
| ARRB1 | 811.114447 | -0.1863044 | 0.28724515 | -0.6485904 | 5.17E-01 | 9.16E-01 |
| AP001972.2 | 41.2440659 | -0.1483762 | 0.4252817 | -0.3488892 | 7.27E-01 | 9.58E-01 |

| | | | | | | |
|---|---|---|---|---|---|---|
| RPS3 | 27214.5974 | 0.20774515 | 0.21575727 | 0.96286511 | 0.33561522 | 0.85621629 |
| GDPD5 | 688.526246 | -0.3252636 | 0.39767735 | -0.8179083 | 0.41340955 | 0.88316314 |
| SERPINH1 | 6287.97483 | -0.0088264 | 0.28953233 | -0.0304851 | 0.9756802 | 0.99455099 |
| MAP6 | 648.369437 | 0.5405573 | 0.31389939 | 1.72207182 | 0.08505651 | 0.64988556 |
| MOGAT2 | 141.206909 | 3.72503811 | 1.07744697 | 3.4572821 | 0.00054565 | 0.11362321 |
| DGAT2 | 448.439001 | 1.47111083 | 0.6647138 | 2.21314924 | 0.02688736 | 0.52685774 |
| AP003031.1 | 163.038445 | 1.41607271 | 0.72698514 | 1.94787023 | NA | NA |
| UVRAG | 1198.78612 | -0.2161235 | 0.17020322 | -1.2697966 | 0.20415711 | 0.78419929 |
| WNT11 | 286.9369 | -1.8200849 | 0.87642365 | -2.0767182 | 0.03782758 | 0.55538555 |
| AP000785.1 | 44.1031508 | -1.9143775 | 0.97645993 | -1.9605285 | 0.04993405 | 0.5861488 |
| THAP12 | 1811.7893 | -0.0295132 | 0.18383854 | -0.1605386 | 0.87245685 | 0.98275231 |
| GVQW3 | 77.4128233 | 0.84451808 | 0.54193133 | 1.55834887 | 0.11915057 | 0.69233212 |
| AP002360.3 | 43.5454937 | 0.22534749 | 0.59837605 | 0.37659845 | 0.70647204 | 0.95501548 |
| AP002360.1 | 274.00492 | 1.04748834 | 0.37483072 | 2.79456378 | 0.00519698 | 0.3091955 |
| EMSY | 920.469557 | -0.1911119 | 0.17115399 | -1.1166081 | 0.26416194 | 0.82424998 |
| AP001189.3 | 68.8528226 | 0.03134904 | 0.52733008 | 0.05944861 | 0.9525948 | 0.99170503 |
| AP001189.1 | 472.804193 | -0.092829 | 0.43939122 | -0.2112673 | 0.83267869 | 0.97550416 |
| LRRC32 | 6722.43508 | -0.1012136 | 0.40551383 | -0.2495936 | 0.80290167 | 0.97273128 |
| TSKU | 8163.81685 | 0.54129866 | 0.73603701 | 0.73542316 | 0.46208182 | 0.89762182 |
| ACER3 | 1344.44625 | 0.31044614 | 0.16428768 | 1.88964958 | 0.05880484 | 0.60332684 |
| CAPN5 | 2990.9867 | 0.78845712 | 0.37376601 | 2.10949388 | 0.03490197 | 0.55369025 |
| MYO7A | 1172.92796 | 0.0754918 | 0.35398287 | 0.213264 | 0.83112105 | 0.97550416 |
| PAK1 | 2446.48111 | 0.0764853 | 0.13248158 | 0.57732778 | 0.56371805 | 0.92453312 |
| AP003680.1 | 45.9087874 | 0.06699849 | 0.43334897 | 0.15460632 | 0.87713169 | 0.98275231 |
| CLNS1A | 2421.94601 | 0.07548761 | 0.13109215 | 0.57583624 | 0.56472588 | 0.92477055 |
| AQP11 | 362.993863 | 0.92599217 | 0.50810121 | 1.82245612 | 0.06838581 | 0.62657129 |
| RSF1 | 1859.94377 | -0.1014618 | 0.1235858 | -0.8209823 | 0.41165636 | 0.88268208 |
| AAMDC | 796.249579 | 0.38907578 | 0.30175197 | 1.28938936 | 0.19726276 | 0.77816059 |
| AP002812.2 | 324.870194 | 0.35052457 | 0.33892158 | 1.03423503 | 0.3010263 | 0.84211658 |
| AP002812.5 | 486.840473 | 0.40588182 | 0.31761845 | 1.27789121 | 0.20128779 | 0.78134034 |
| INTS4 | 1123.74625 | -0.0694563 | 0.13590295 | -0.5110729 | 0.6093 | 0.93748023 |
| KCTD14 | 793.026873 | 0.34055801 | 0.25312773 | 1.34539984 | 0.17849615 | 0.76115956 |
| AP003032.2 | 487.073531 | 0.30990442 | 0.27608739 | 1.12248668 | 0.26165557 | 0.82424998 |
| NDUFC2-KCT | 3201.04889 | 0.30943247 | 0.14900385 | 2.0766743 | 0.03783163 | 0.55538555 |
| THRSP | 199.915162 | 4.95477685 | 1.07395724 | 4.61356996 | NA | NA |
| NDUFC2 | 3710.12028 | 0.30438465 | 0.1510036 | 2.01574433 | 0.04382672 | 0.57103595 |
| ALG8 | 1295.53022 | -0.1107861 | 0.14470776 | -0.7655852 | 0.44392317 | 0.8938278 |
| KCTD21-AS1 | 254.948934 | -0.1906 | 0.37237722 | -0.5118466 | 0.60875839 | 0.93748023 |
| KCTD21 | 726.169119 | 0.08861441 | 0.22796834 | 0.38871364 | 0.69748799 | 0.95272493 |
| USP35 | 1071.48585 | -0.0623031 | 0.34872144 | -0.1786616 | 0.85820344 | 0.97975089 |
| GAB2 | 1683.22311 | -0.5044778 | 0.34208753 | -1.4747039 | 0.14029218 | 0.725307 |
| AP003086.2 | 207.462089 | 0.26453976 | 0.24831909 | 1.06532187 | 0.2867303 | 0.83155545 |
| NARS2 | 1092.25796 | 0.263877 | 0.2416034 | 1.09219076 | 0.27474926 | 0.82719407 |
| TENM4 | 524.00139 | -0.4394082 | 0.38718526 | -1.1348785 | 0.25642623 | 0.82078293 |
| FAM181B | 70.005388 | 0.50401522 | 0.44242629 | 1.13920721 | 0.25461674 | 0.81992039 |
| PRCP | 9955.39586 | 0.51555113 | 0.21852578 | 2.35922342 | 0.01831323 | 0.47570049 |
| RAB30 | 395.110504 | -0.0968031 | 0.21039251 | -0.460107 | 0.64543943 | 0.94208859 |
| RAB30-AS1 | 378.769689 | -0.3225331 | 0.26253109 | -1.2285521 | 0.21923979 | 0.79847731 |
| AP001767.3 | 211.050145 | -0.1768928 | 0.29745437 | -0.5946889 | 0.55205145 | 0.92373235 |
| PCF11 | 3935.07094 | -0.1575306 | 0.19272463 | -0.8173869 | 0.41370734 | 0.88316314 |
| PCF11-AS1 | 318.154347 | -0.1295312 | 0.20438116 | -0.6337729 | 0.52622905 | 0.91769856 |
| ANKRD42 | 1299.31757 | 0.09983908 | 0.16940779 | 0.58934173 | 0.55563206 | 0.92373235 |
| AP000873.3 | 63.298848 | -0.1566307 | 0.29644148 | -0.5283698 | 0.59724269 | 0.93414499 |
| CCDC90B | 1523.09556 | 0.07789835 | 0.13627301 | 0.57163446 | 0.56756965 | 0.92614913 |

| | | | | | | |
|---|---|---|---|---|---|---|
| AP000446.1 | 54.3436245 | 0.0856142 | 0.297845 | 0.2874455 | 0.77377123 | 0.96836854 |
| DLG2 | 177.620565 | 0.56501364 | 0.35111657 | 1.60919105 | 0.10757457 | 0.67707349 |
| TMEM126B | 2005.29214 | -0.0368666 | 0.1386921 | -0.2658161 | 0.79038087 | 0.97090793 |
| TMEM126A | 714.547546 | -0.0430162 | 0.15828762 | -0.27176 | 0.78580658 | 0.97065795 |
| CREBZF | 4850.84702 | -0.1998836 | 0.12186438 | -1.6402137 | 0.10096074 | 0.6725344 |
| CCDC89 | 67.7108877 | 0.048124 | 0.35820104 | 0.13434914 | 0.8931265 | 0.98474725 |
| SYTL2 | 2220.67779 | 0.52867146 | 0.25082646 | 2.10771812 | 0.03505538 | 0.55369025 |
| PICALM | 12358.8925 | 0.0660918 | 0.12750562 | 0.51834423 | 0.60421812 | 0.93655508 |
| EED | 1040.90488 | -0.1645394 | 0.13009457 | -1.2647674 | 0.20595472 | 0.78659006 |
| HIKESHI | 1146.76048 | 0.0957268 | 0.21019527 | 0.45541842 | 0.64880825 | 0.94312988 |
| ME3 | 1797.65148 | -0.271408 | 0.20480054 | -1.3252309 | 0.18509459 | 0.76820626 |
| PRSS23 | 4328.99988 | -0.13667 | 0.34749554 | -0.3932999 | 0.69409796 | 0.95222846 |
| AP001528.2 | 103.811681 | 1.39228744 | 0.54648494 | 2.54771422 | 0.01084313 | 0.40196898 |
| FZD4 | 4804.37145 | -0.0692293 | 0.30783587 | -0.2248903 | 0.82206463 | 0.97493464 |
| TMEM135 | 1037.81475 | 0.34717081 | 0.28136772 | 1.23386868 | 0.21725184 | 0.79583987 |
| RAB38 | 279.372752 | -0.1659583 | 0.25600658 | -0.648258 | 0.5168181 | 0.91636259 |
| CTSC | 12988.1805 | 0.24003774 | 0.19836004 | 1.2101114 | 0.22623615 | 0.80246466 |
| GRM5-AS1 | 47.9848976 | 0.65341432 | 0.40501854 | 1.61329482 | 0.10668047 | 0.67674128 |
| GRM5 | 53.9848261 | 0.4918266 | 0.37763827 | 1.30237487 | 0.19278827 | 0.77506795 |
| NOX4 | 4643.64987 | 1.14866672 | 0.57652768 | 1.99238781 | 0.04632852 | 0.57943708 |
| NAALAD2 | 167.129154 | 0.20795217 | 0.41777453 | 0.49776173 | 0.61865199 | 0.93945324 |
| CHORDC1 | 1383.7909 | -0.281194 | 0.13997327 | -2.0089121 | 0.04454646 | 0.57267043 |
| FAT3 | 48.9300562 | -0.636444 | 0.71286587 | -0.8927963 | 0.37196626 | 0.86772685 |
| SLC36A4 | 1247.07747 | -0.5490697 | 0.26756446 | -2.0521025 | 0.0401597 | 0.55657615 |
| SMCO4 | 927.234691 | -0.2880275 | 0.26582931 | -1.0835056 | 0.27858408 | 0.82805357 |
| CEP295 | 943.229854 | -0.1380903 | 0.1525411 | -0.9052661 | 0.36532443 | 0.8663515 |
| TAF1D | 5093.36293 | -0.1886374 | 0.20422711 | -0.9236648 | 0.35566088 | 0.86364633 |
| C11orf54 | 15357.8327 | 0.87739525 | 0.52423798 | 1.67365833 | 0.09419777 | 0.65783452 |
| AP001273.2 | 8938.16517 | 0.70543126 | 0.45813409 | 1.5397921 | 0.12361104 | 0.69957382 |
| MED17 | 1584.97988 | -0.1288516 | 0.18096919 | -0.7120083 | 0.47645965 | 0.90309179 |
| VSTM5 | 116.503038 | 0.41116677 | 0.58873786 | 0.69838685 | 0.4849353 | 0.90541935 |
| PANX1 | 700.686099 | -0.4297842 | 0.25222149 | -1.7039951 | 0.088382 | 0.65363785 |
| MRE11 | 1016.0825 | -0.1326379 | 0.09783274 | -1.3557616 | 0.17517504 | 0.76073329 |
| ANKRD49 | 720.796836 | -0.2793006 | 0.12510406 | -2.2325465 | 0.02557887 | 0.5239055 |
| FUT4 | 663.858907 | -0.115629 | 0.32337047 | -0.3575745 | 0.72066179 | 0.95725584 |
| PIWIL4 | 399.981085 | 0.26910568 | 0.36215643 | 0.74306478 | 0.45744246 | 0.89688594 |
| AP000943.3 | 223.38771 | -0.2171789 | 0.32875315 | -0.660614 | 0.50885988 | 0.91439937 |
| AMOTL1 | 1552.50262 | -0.0518641 | 0.26804172 | -0.1934928 | 0.84657306 | 0.97684681 |
| AP002383.2 | 170.203256 | -0.1195025 | 0.35198848 | -0.3395068 | 0.734228 | 0.95895327 |
| CWC15 | 2221.95603 | 0.04526137 | 0.10829727 | 0.41793642 | 0.6759936 | 0.94959168 |
| KDM4D | 59.173837 | 0.09615431 | 0.34311301 | 0.28024093 | 0.77929267 | 0.96836854 |
| SRSF8 | 3031.58846 | -0.1670205 | 0.2275317 | -0.7340536 | 0.46291605 | 0.89807153 |
| ENDOD1 | 4520.7093 | -0.4342294 | 0.30979 | -1.4016895 | 0.16100798 | 0.74874703 |
| AP000787.1 | 110.808484 | 0.30830416 | 0.35766137 | 0.86200016 | 0.38868743 | 0.87568585 |
| SESN3 | 1547.496 | 0.31802302 | 0.26611389 | 1.1950636 | 0.23206218 | 0.80649152 |
| FAM76B | 843.231991 | -0.0091295 | 0.15761412 | -0.0579229 | 0.95381007 | 0.99170503 |
| CEP57 | 1740.61015 | 0.34311236 | 0.11064079 | 3.10113803 | 0.00192778 | 0.18915498 |
| MTMR2 | 2067.3697 | 0.05776107 | 0.18782116 | 0.30753225 | 0.75843828 | 0.96502715 |
| MAML2 | 1346.07012 | -0.0174089 | 0.27374981 | -0.0635942 | 0.94929338 | 0.99170503 |
| CCDC82 | 1872.91404 | -0.3645703 | 0.2667221 | -1.3668546 | 0.17167089 | 0.75974924 |
| JRKL | 762.411396 | -0.032125 | 0.22028355 | -0.1458348 | 0.8840518 | 0.98317437 |
| ARHGAP42 | 927.276064 | 0.35939778 | 0.31099566 | 1.155636 | 0.24783008 | 0.81602355 |
| PGR | 225.510428 | 0.20535707 | 0.28253709 | 0.72683225 | 0.46732873 | 0.89983399 |
| TRPC6 | 150.665698 | -0.1388972 | 0.55155149 | -0.25183 | 0.80117244 | 0.97210205 |

| | | | | | | |
|---|---|---|---|---|---|---|
| CEP126 | 355.625415 | 0.22263221 | 0.34965627 | 0.63671733 | 0.52430898 | 0.91723878 |
| CFAP300 | 122.007996 | 0.65559307 | 0.45260513 | 1.44848792 | 0.14748064 | 0.73412064 |
| YAP1 | 5769.65537 | -0.1709779 | 0.16106315 | -1.0615582 | 0.2884363 | 0.83295411 |
| AP000942.2 | 718.353811 | -0.2770237 | 0.20219534 | -1.3700795 | 1.71E-01 | 0.75956362 |
| BIRC3 | 2353.81733 | 0.40860065 | 0.52512526 | 0.7781013 | 0.43650929 | 0.89104919 |
| BIRC2 | 3098.99582 | -0.0409898 | 0.17686709 | -0.2317546 | 0.81672861 | 0.97388889 |
| TMEM123 | 8219.11839 | 0.03647239 | 0.14118889 | 0.2583234 | 7.96E-01 | 0.97101097 |
| MMP7 | 10790.5312 | 0.93931329 | 0.51284177 | 1.83158501 | 0.06701327 | 0.6249306 |
| AP000851.2 | 125.61429 | -0.9714655 | 2.0754287 | -0.4680794 | 6.40E-01 | 9.41E-01 |
| MMP8 | 96.7881064 | -0.7373055 | 0.97664046 | -0.7549405 | 0.45028466 | 0.89557948 |
| MMP1 | 121.223096 | -1.0325318 | 0.69039292 | -1.4955712 | 0.13476543 | 0.71630293 |
| MMP3 | 107.031333 | -4.2228995 | 2.14443088 | -1.9692402 | NA | NA |
| DCUN1D5 | 836.786278 | -0.2858538 | 0.2475519 | -1.1547226 | 0.24820404 | 0.81621194 |
| DYNC2H1 | 858.728189 | 0.60984088 | 0.23381376 | 2.60823347 | 0.00910109 | 0.38664841 |
| PDGFD | 577.220477 | 0.39021257 | 0.4014963 | 0.97189581 | 0.33110239 | 0.85384238 |
| CASP4 | 3385.27418 | -0.1058735 | 0.333064 | -0.3178774 | 0.75057794 | 0.96255347 |
| CASP1 | 902.169407 | -0.1136351 | 0.52255764 | -0.2174594 | 0.82785033 | 0.97548201 |
| CARD16 | 206.368945 | -0.3337097 | 0.57988228 | -0.5754784 | 0.5649678 | 0.92493342 |
| GRIA4 | 85.6912793 | 1.74222819 | 0.9206782 | 1.89233131 | 0.05844685 | 0.60332684 |
| MSANTD4 | 841.21922 | -0.1397344 | 0.16301192 | -0.8572038 | 0.39133226 | 0.87716276 |
| KBTBD3 | 511.546242 | 0.24694631 | 0.16447725 | 1.50140102 | 0.13325187 | 0.71493653 |
| AASDHPPT | 2376.2809 | 0.16921844 | 0.22162079 | 0.76354949 | 0.44513576 | 0.89445558 |
| AP001001.1 | 372.561815 | 0.19891641 | 0.23622676 | 0.84205705 | 0.39975602 | 0.88077967 |
| GUCY1A2 | 488.924504 | -0.3156885 | 0.58572495 | -0.5389705 | 0.58990722 | 0.93289552 |
| AP000766.1 | 138.186028 | -0.093175 | 0.32181037 | -0.289534 | 0.77217278 | 0.96836854 |
| CWF19L2 | 657.885897 | -0.081531 | 0.19055446 | -0.4278619 | 0.66875163 | 0.9486589 |
| ALKBH8 | 397.796689 | -0.2239281 | 0.23664918 | -0.9462449 | 0.34402369 | 0.85961157 |
| ELMOD1 | 188.712852 | -0.3376145 | 0.56571412 | -0.5967936 | 0.55064522 | 0.92373235 |
| SLN | 71.9940321 | -3.2651748 | 2.27602744 | -1.4345938 | NA | NA |
| SLC35F2 | 1295.95847 | 0.0741729 | 0.24009935 | 0.30892586 | 0.75737793 | 0.96502715 |
| CUL5 | 2073.01461 | 0.03962259 | 0.14850972 | 0.26680131 | 0.78962214 | 0.97090793 |
| AP002433.1 | 2316.85722 | 0.70528783 | 0.35495771 | 1.986963 | 0.0469265 | 0.58171004 |
| ACAT1 | 20221.3801 | 0.5960884 | 0.33085545 | 1.80165811 | 0.07159921 | 0.63189503 |
| NPAT | 660.486304 | -0.1076994 | 0.14214738 | -0.7576602 | 0.44865445 | 0.8948759 |
| ATM | 1994.7499 | -0.2249947 | 0.20806807 | -1.0813515 | 0.2795408 | 0.82825734 |
| C11orf65 | 67.3303746 | 0.10572372 | 0.46281657 | 0.22843547 | 0.81930772 | 0.97454841 |
| KDELC2 | 1405.99244 | 0.11083402 | 0.15494283 | 0.71532203 | 0.47441008 | 0.90199039 |
| EXPH5 | 514.599343 | -0.1730649 | 0.33445175 | -0.5174586 | 0.60483609 | 0.93664131 |
| DDX10 | 1145.00733 | 0.00636913 | 0.30161529 | 0.02111675 | 0.98315252 | 0.9961464 |
| RDX | 3664.2917 | 0.08749731 | 0.19712811 | 0.44386014 | 0.6571437 | 0.94545756 |
| ZC3H12C | 738.671914 | 0.23108373 | 0.21615839 | 1.06904818 | 0.28504796 | 0.83107779 |
| AP000924.1 | 67.8776448 | 0.46763347 | 0.90884812 | 0.51453423 | 0.60687853 | 0.9372485 |
| FDX1 | 2131.58735 | 0.03882126 | 0.20272324 | 0.19149881 | 0.84813481 | 0.97711043 |
| ARHGAP20 | 52.6927667 | -0.4985935 | 0.69349775 | -0.7189547 | 0.47216885 | 0.90160919 |
| COLCA1 | 927.965606 | 0.2256057 | 0.43319624 | 0.52079331 | 0.60251076 | 0.93551402 |
| COLCA2 | 525.445939 | -0.000697 | 0.30159918 | -0.002311 | 0.99815607 | 0.99990633 |
| POU2AF1 | 178.9755 | -0.6189 | 0.89619954 | -0.6905828 | 0.48982774 | 0.90600166 |
| LAYN | 258.501188 | 0.08116011 | 0.29920657 | 0.27125109 | 0.78619792 | 0.97075604 |
| SIK2 | 3705.29712 | 0.00378306 | 0.24023941 | 0.01574705 | 0.98743619 | 0.997408 |
| PPP2R1B | 2314.90429 | -0.3308817 | 0.25959981 | -1.2745836 | 0.20245665 | 0.78193335 |
| ALG9 | 1187.51792 | 0.02312486 | 0.11801502 | 0.19594842 | 0.84465054 | 0.97675487 |
| AP001781.2 | 942.132658 | 0.07211717 | 0.13747117 | 0.5245985 | 0.59986233 | 0.93487648 |
| FDXACB1 | 174.253811 | 0.08698286 | 0.2314104 | 0.37588138 | 0.70700509 | 0.95501548 |
| C11orf1 | 928.257501 | 0.74626393 | 0.26304089 | 2.83706439 | 0.00455304 | 0.29525459 |

| | | | | | | |
|---|---|---|---|---|---|---|
| CRYAB | 100130.73 | 0.50284763 | 0.67804225 | 0.74161695 | 0.45831945 | 0.8971553 |
| HSPB2 | 810.343259 | -0.295569 | 0.42780094 | -0.690903 | 0.4896265 | 0.90600166 |
| HSPB2-C11or | 3021.03391 | 0.24281563 | 0.41314992 | 0.58771796 | 0.55672162 | 0.92373235 |
| C11orf52 | 2254.30577 | 0.44141465 | 0.45202295 | 0.97653151 | 0.32880114 | 0.85317073 |
| DIXDC1 | 1239.34721 | -0.0259435 | 0.39302403 | -0.06601 | 0.94736984 | 0.99166614 |
| DLAT | 2916.30327 | 0.16726185 | 0.14882433 | 1.12388776 | 2.61E-01 | 0.82424998 |
| PIH1D2 | 340.383774 | 0.18270537 | 0.20530175 | 0.88993579 | 0.37350037 | 0.86791605 |
| NKAPD1 | 1545.8592 | -0.1486985 | 0.10791692 | -1.3778979 | 0.16823481 | 0.75735257 |
| TIMM8B | 1603.30693 | 0.08649608 | 0.1814411 | 0.47671714 | 0.63356355 | 0.94088547 |
| SDHD | 9874.6475 | 0.30568535 | 0.17795623 | 1.7177558 | 8.58E-02 | 0.65049132 |
| AP002884.3 | 4274.27155 | 0.36814125 | 0.21329393 | 1.72598087 | 0.08435085 | 0.64988556 |
| IL18 | 1198.68021 | -0.0390759 | 0.46443087 | -0.0841372 | 0.93294732 | 0.98999863 |
| BCO2 | 251.223754 | 0.36761264 | 0.34030621 | 1.08024078 | 0.28003497 | 0.82825734 |
| PTS | 1378.55878 | 0.16406261 | 0.16244246 | 1.00997371 | 0.31250789 | 0.84541735 |
| AP002884.1 | 208.424616 | -0.1230568 | 0.27847373 | -0.4418972 | 0.65856359 | 0.9458503 |
| NCAM1 | 238.52498 | -0.5816794 | 0.84466093 | -0.6886543 | 0.49104081 | 0.90600166 |
| TTC12 | 820.80646 | -6.38E-05 | 0.26400276 | -0.0002415 | 0.99980732 | 0.99994357 |
| ANKK1 | 68.6663301 | -0.1860292 | 0.50779605 | -0.3663464 | 0.71410664 | 0.95566071 |
| TMPRSS5 | 81.3117285 | -0.2858469 | 0.52922293 | -0.5401258 | 0.58911031 | 0.93289552 |
| ZW10 | 847.050377 | -0.0008872 | 0.19763567 | -0.0044893 | 0.99641805 | 0.99976751 |
| USP28 | 959.102477 | 0.10985802 | 0.25338008 | 0.43357006 | 0.66460068 | 0.94680968 |
| HTR3A | 82.594511 | -2.2213062 | 1.13599499 | -1.9553838 | 0.05053779 | 0.5861488 |
| ZBTB16 | 355.040708 | -0.5380464 | 0.59590497 | -0.9029063 | 0.36657563 | 0.86683872 |
| NNMT | 17691.7576 | 0.06885789 | 0.71743076 | 0.09597844 | 0.92353769 | 0.98858365 |
| C11orf71 | 906.775803 | 0.31998093 | 0.3197939 | 1.00058483 | 0.31702757 | 0.84819508 |
| RBM7 | 1870.74681 | -0.0453357 | 0.085288 | -0.5315606 | 0.59503038 | 0.9334601 |
| AP002373.1 | 1701.26999 | -0.0966105 | 0.19643731 | -0.4918136 | 6.23E-01 | 0.94016713 |
| REXO2 | 3617.65388 | -0.1768191 | 0.18407892 | -0.9605616 | 0.33677265 | 0.85679125 |
| CADM1 | 2237.6479 | -0.106854 | 0.325415 | -0.3283622 | 0.74263778 | 0.96109001 |
| BUD13 | 825.005683 | -0.1234458 | 0.13349356 | -0.9247326 | 0.355105 | 0.86364633 |
| ZPR1 | 1845.41339 | -0.2512887 | 0.24660329 | -1.0189999 | 0.308203 | 0.84370908 |
| APOC3 | 133.553316 | -0.3408553 | 1.25472301 | -0.2716578 | 0.78588514 | 0.97065795 |
| SIK3 | 2412.9578 | -0.3828596 | 0.15837762 | -2.4173847 | 1.56E-02 | 0.45444887 |
| PAFAH1B2 | 4147.13559 | -0.1685864 | 0.19472243 | -0.8657781 | 0.38661188 | 0.87432552 |
| SIDT2 | 2690.20924 | -0.1848224 | 0.2409612 | -0.7670214 | 0.44306882 | 0.89352768 |
| TAGLN | 24613.395 | -0.8193281 | 0.64239178 | -1.2754336 | 0.20215581 | 0.78163854 |
| PCSK7 | 13866.1954 | -0.6837802 | 0.55295745 | -1.2365873 | 0.21624033 | 0.7954827 |
| RNF214 | 1430.21748 | 0.03380509 | 0.11804752 | 0.28636852 | 0.77459588 | 0.96836854 |
| BACE1 | 2427.04191 | 0.23363691 | 0.09920711 | 2.35504204 | 0.01852061 | 0.47570049 |
| BACE1-AS | 490.95961 | 0.09910131 | 0.13628239 | 0.72717622 | 0.46711801 | 0.89983399 |
| CEP164 | 2125.97059 | -0.2535596 | 0.24656073 | -1.0283861 | 0.30376825 | 0.84211658 |
| DSCAML1 | 623.001327 | 0.27289833 | 0.43524413 | 0.6270006 | 0.53065884 | 0.91962754 |
| FXYD2 | 60784.4902 | 0.05859792 | 0.25129802 | 0.23318098 | 0.81562087 | 0.9738447 |
| AP000757.2 | 27196.952 | 0.12111964 | 0.23162213 | 0.5229191 | 0.60103055 | 0.93515264 |
| FXYD6-FXYD2 | 55539.6728 | 0.1189362 | 0.23482243 | 0.50649418 | 0.61250977 | 0.9387918 |
| AP000757.1 | 2689.64086 | -0.3569767 | 0.27222428 | -1.3113331 | 0.18974525 | 0.77036736 |
| FXYD6 | 3717.13212 | -0.4636901 | 0.25434319 | -1.8230881 | 0.06829004 | 0.62648383 |
| TMPRSS13 | 136.90389 | -0.2914049 | 0.76796913 | -0.3794487 | 0.70435468 | 0.95488362 |
| IL10RA | 1007.30291 | -0.2685766 | 0.53692797 | -0.5002098 | 0.61692737 | 0.93910158 |
| SMIM35 | 118.891251 | 1.33875008 | 0.98058536 | 1.36525603 | 0.17217259 | 0.75974924 |
| TMPRSS4 | 2359.39567 | 0.70872257 | 0.49673936 | 1.42674937 | 0.15365215 | 0.74258845 |
| SCN4B | 298.443126 | -0.4543283 | 0.73957765 | -0.6143078 | 0.53901193 | 0.9215968 |
| JAML | 449.659732 | 0.82211134 | 0.82222891 | 0.99985701 | 0.31737971 | 0.84819508 |
| MPZL3 | 543.982289 | -0.0854339 | 0.23840448 | -0.3583568 | 0.72007631 | 0.95688507 |

| | | | | | | |
|---|---|---|---|---|---|---|
| MPZL2 | 2855.19617 | 0.78725342 | 0.40238167 | 1.95648427 | 0.05040813 | 0.5861488 |
| CD3E | 312.545664 | 0.43706163 | 0.90127435 | 0.48493739 | 0.62772076 | 0.94088547 |
| CD3D | 107.528872 | 0.46682108 | 0.93199418 | 0.50088411 | 0.61645269 | 0.93910158 |
| CD3G | 63.531536 | 0.13746825 | 0.99496798 | 0.13816349 | 0.89011121 | 0.98462839 |
| UBE4A | 2918.59331 | -0.0488066 | 0.12354226 | -0.3950595 | 0.69279894 | 0.9519683 |
| AP001267.1 | 6767.47984 | 0.10134963 | 0.13430395 | 0.75462878 | 0.45047175 | 0.89557948 |
| ATP5MG | 8506.51171 | 0.14891963 | 0.16047149 | 0.92801299 | 0.35340083 | 0.86364633 |
| KMT2A | 2442.96387 | -0.1646801 | 0.11265751 | -1.4617761 | 0.14380259 | 0.72920837 |
| AP001267.3 | 283.587188 | 0.11644045 | 0.48788739 | 0.23866254 | 0.81136726 | 0.97352497 |
| TTC36 | 296.569392 | 0.26210601 | 0.58902833 | 0.44498031 | 0.65633399 | 0.94545756 |
| TMEM25 | 2214.6673 | 0.14402649 | 0.22775805 | 0.63236618 | 0.52714763 | 0.91797748 |
| IFT46 | 1352.51998 | 0.01249003 | 0.21143485 | 0.0590727 | 0.9528942 | 0.99170503 |
| ARCN1 | 5859.71539 | -0.2128148 | 0.22904147 | -0.9291542 | 0.35280918 | 0.86364633 |
| PHLDB1 | 6943.47674 | -0.2733671 | 0.25224854 | -1.0837212 | 0.27848842 | 0.82805357 |
| TREH | 3622.34361 | -0.1560043 | 0.22932545 | -0.6802748 | 0.49633048 | 0.90890243 |
| AP002954.1 | 128.546127 | -0.1039373 | 0.37004981 | -0.2808738 | 0.77880718 | 0.96836854 |
| DDX6 | 4350.12971 | -0.071583 | 0.13586643 | -0.5268633 | 0.59828851 | 0.9346978 |
| AP004609.3 | 136.399835 | -0.1371251 | 0.28932437 | -0.4739494 | 0.635536 | 0.94088547 |
| CXCR5 | 2857.19584 | -0.6820771 | 0.27078598 | -2.5188788 | 0.01177292 | 0.40423902 |
| BCL9L | 6371.4339 | -0.7803363 | 0.30026728 | -2.5988056 | 0.00935487 | 0.38701912 |
| UPK2 | 88.3768905 | -0.7074224 | 0.43485579 | -1.6267977 | 0.10378007 | 0.67439401 |
| AP003392.4 | 58.0502683 | 0.09806218 | 0.39301011 | 0.24951567 | 0.80296192 | 0.97273362 |
| CCDC84 | 3324.46269 | -0.1170618 | 0.13207504 | -0.8863281 | 0.37544073 | 0.86876586 |
| RPS25 | 15565.1587 | 0.12002169 | 0.18168707 | 0.66059567 | 0.50887164 | 0.91439937 |
| TRAPPC4 | 6303.60714 | 0.32300741 | 0.23126899 | 1.39667413 | 0.16251159 | 0.75026273 |
| SLC37A4 | 8535.74694 | 0.47121097 | 0.38723693 | 1.21685441 | 0.22365961 | 0.80078694 |
| AP003392.3 | 2458.02333 | 0.41127236 | 0.29291468 | 1.40406877 | 0.16029837 | 0.74781837 |
| HYOU1 | 8112.96894 | 0.46036547 | 0.24265551 | 1.89719767 | 0.05780185 | 0.60129769 |
| VPS11 | 2770.42465 | -0.0748392 | 0.13450408 | -0.5564083 | 0.57793176 | 0.93130183 |
| HMBS | 986.57573 | 0.15369668 | 0.2045107 | 0.75153368 | 0.45233154 | 0.89670027 |
| H2AFX | 1069.00531 | -0.5640609 | 0.21192402 | -2.6616187 | 0.00777659 | 0.36878977 |
| DPAGT1 | 2019.39465 | -0.0381374 | 0.17007694 | -0.2242362 | 0.82257353 | 0.97519297 |
| C2CD2L | 1775.88127 | 0.41799499 | 0.24701847 | 1.69216083 | 0.09061531 | 0.65495306 |
| HINFP | 1732.53165 | -0.281929 | 0.14045444 | -2.0072628 | 0.04472169 | 0.57267043 |
| ABCG4 | 46.8607493 | -0.1870821 | 0.69172607 | -0.270457 | 0.78680872 | 0.97090793 |
| NLRX1 | 1442.46137 | 0.21885568 | 0.20197091 | 1.08360002 | 0.27854218 | 0.82805357 |
| PDZD3 | 3958.2437 | 1.45210073 | 0.74961744 | 1.93712239 | 0.05273038 | 0.58742503 |
| CCDC153 | 878.791805 | 1.27606408 | 0.63954063 | 1.99528227 | 0.0460121 | 0.57718395 |
| CBL | 1263.51798 | -0.4783759 | 0.22814874 | -2.0967721 | 0.03601376 | 0.55369025 |
| MCAM | 5909.87558 | -0.6390496 | 0.41159797 | -1.5526063 | 0.12051722 | 0.69467399 |
| RNF26 | 1883.26268 | -0.1118302 | 0.10458817 | -1.0692431 | 0.28496014 | 0.83107779 |
| AP003396.1 | 726.647407 | -0.1404089 | 0.12005404 | -1.1695473 | 0.24218321 | 0.81319028 |
| C1QTNF5 | 1551.83905 | -0.2536509 | 0.3585877 | -0.7073608 | 0.47934232 | 0.90485036 |
| MFRP | 1553.04323 | -0.2558131 | 0.3586169 | -0.7133326 | 0.47563999 | 0.90257104 |
| USP2 | 2478.66566 | 0.04894702 | 0.50425853 | 0.09706732 | 0.92267293 | 0.98826666 |
| AP003396.3 | 82.2705528 | -0.1489333 | 0.56668812 | -0.2628135 | 0.7926943 | 0.97090793 |
| USP2-AS1 | 79.9439944 | 0.34402663 | 0.51506003 | 0.66793503 | 0.50417507 | 0.91356552 |
| THY1 | 8837.75512 | -0.3304026 | 0.58622457 | -0.563611 | 0.5730189 | 0.92928554 |
| NECTIN1 | 913.161795 | -0.0153803 | 0.26360511 | -0.058346 | 0.95347306 | 0.99170503 |
| TRIM29 | 145.093965 | 1.99819141 | 0.6357618 | 3.14298754 | 0.00167233 | 0.18012143 |
| OAF | 6318.96081 | 0.01793795 | 0.30593767 | 0.05863271 | 0.95324466 | 0.99170503 |
| POU2F3 | 40.0590305 | 0.86057229 | 0.67753222 | 1.27015701 | 0.20402871 | 0.78418828 |
| TMEM136 | 588.087185 | 0.05782381 | 0.23024471 | 0.25114069 | 0.80170533 | 0.97222465 |
| ARHGEF12 | 9972.14113 | 0.12333773 | 0.14827336 | 0.83182666 | 0.40550679 | 0.88132015 |

| | | | | | | |
|---|---|---|---|---|---|---|
| TBCEL | 1025.46946 | -0.1585475 | 0.19254116 | -0.8234471 | 0.4102538 | 0.88249519 |
| AP000646.1 | 119.08926 | -0.1958123 | 0.36920817 | -0.5303573 | 0.59586421 | 0.9334601 |
| TECTA | 124.387326 | -0.1047848 | 0.35059374 | -0.2988781 | 0.76503307 | 0.96678847 |
| SC5D | 2692.02627 | 0.03299591 | 0.31982643 | 0.10316818 | 0.9178295 | 0.9868044 |
| SORL1 | 3565.01984 | 0.72892243 | 0.34115169 | 2.136652 | 0.03262631 | 0.55104386 |
| UBASH3B | 373.95409 | -0.0938685 | 0.45381341 | -0.2068439 | 0.83613175 | 0.97577341 |
| JHY | 529.064069 | -0.3282716 | 0.27023247 | -1.2147748 | 0.22445198 | 0.80078694 |
| HSPA8 | 49797.2969 | -0.0325239 | 0.27803722 | -0.1169768 | 0.90687841 | 0.98662757 |
| CLMP | 589.691437 | -0.861537 | 0.42711376 | -2.0171138 | 0.04368364 | 0.57051182 |
| GRAMD1B | 1522.68784 | 0.593871 | 0.56165644 | 1.05735635 | 0.29034899 | 0.83442894 |
| SCN3B | 146.123429 | -0.737995 | 0.30620733 | -2.4101154 | 0.01594748 | 0.4564177 |
| ZNF202 | 707.784125 | -0.2319986 | 0.20706221 | -1.1204293 | 0.26253089 | 0.82424998 |
| VWA5A | 2669.80503 | 0.1725251 | 0.23410513 | 0.73695563 | 0.46114933 | 0.89755243 |
| TBRG1 | 3260.23164 | -0.2108141 | 0.11514465 | -1.8308629 | 0.06712101 | 0.6249306 |
| SIAE | 7221.55758 | 0.40893617 | 0.20669723 | 1.97843082 | 0.04788013 | 0.58372668 |
| SPA17 | 507.149144 | 0.26776163 | 0.2027474 | 1.3206662 | 0.18661269 | 0.76915147 |
| NRGN | 501.615155 | -0.2660032 | 0.49508301 | -0.5372901 | 0.59106725 | 0.93289552 |
| AP000866.5 | 110.786189 | -0.1186031 | 0.53927522 | -0.2199305 | 8.26E-01 | 0.97548201 |
| VSIG2 | 345.171634 | -0.2545048 | 0.33905461 | -0.7506307 | 0.45287495 | 0.89670027 |
| ESAM | 4861.74578 | -0.1840167 | 0.4159809 | -0.4423682 | 0.65822275 | 0.9458503 |
| AP000866.2 | 105.918563 | -0.1080053 | 0.32142654 | -0.3360185 | 0.73685688 | 0.96009463 |
| MSANTD2 | 1305.31817 | 0.01420144 | 0.20338694 | 0.06982476 | 0.94433314 | 0.99136103 |
| AP000866.1 | 54.2761114 | 0.02751857 | 0.3755031 | 0.07328453 | 0.9415797 | 0.99136103 |
| ROBO3 | 1093.32426 | -0.2491907 | 0.4188589 | -0.5949276 | 0.55189189 | 0.92373235 |
| AP003501.1 | 311.430353 | -0.2309937 | 0.41347291 | -0.5586671 | 0.57638891 | 0.9311568 |
| ROBO4 | 2561.31801 | -0.2685616 | 0.41584225 | -0.6458258 | 0.5183922 | 0.91636259 |
| AP003501.2 | 259.238776 | -0.3669699 | 0.41260908 | -0.8893888 | 0.37379414 | 0.86791605 |
| SLC37A2 | 190.377108 | -0.0864815 | 0.42792099 | -0.2020968 | 0.83984103 | 0.97618283 |
| TMEM218 | 715.775825 | 0.21101823 | 0.21977805 | 0.96014245 | 0.33698353 | 0.85679125 |
| PKNOX2 | 299.595612 | -0.7058409 | 0.53248276 | -1.3255658 | 0.18498356 | 0.76820626 |
| FEZ1 | 307.970707 | 0.01862253 | 0.45665693 | 0.04078013 | 0.96747118 | 0.99400331 |
| EI24 | 4595.24089 | -0.0432625 | 0.16659126 | -0.2596922 | 0.79510118 | 0.97101097 |
| STT3A | 5643.75465 | 0.14119474 | 0.16919868 | 0.83449082 | 0.40400446 | 0.88132015 |
| CHEK1 | 287.537263 | 0.11112161 | 0.27195798 | 0.40859846 | 0.68283436 | 0.95070301 |
| HYLS1 | 270.640629 | -0.1624585 | 0.29745651 | -0.5461588 | 0.58495677 | 0.93289552 |
| PUS3 | 520.58076 | 0.19648584 | 0.24173023 | 0.81283108 | 0.41631491 | 0.88399703 |
| DDX25 | 147.983376 | -0.1886291 | 0.48522614 | -0.3887448 | 0.69746495 | 0.95272493 |
| VSIG10L2 | 128.30517 | 0.04115274 | 0.5775082 | 0.07125915 | 0.94319151 | 0.99136103 |
| CDON | 203.356513 | 0.19004031 | 0.51316078 | 0.37033289 | 0.71113447 | 0.95560193 |
| RPUSD4 | 1347.90946 | -0.0712986 | 0.21484532 | -0.3318603 | 0.73999476 | 0.96043876 |
| FAM118B | 1123.99907 | -0.0123077 | 0.21001157 | -0.058605 | 0.95326671 | 0.99170503 |
| SRPRA | 8088.67708 | -0.2312943 | 0.27490008 | -0.841376 | 0.40013734 | 0.88077967 |
| FOXRED1 | 2012.2456 | 0.12921899 | 0.15011945 | 0.86077448 | 0.38936226 | 0.87568585 |
| TIRAP | 369.03742 | 0.12030534 | 0.29437924 | 0.40867467 | 0.68277843 | 0.95070301 |
| AP001318.1 | 118.947222 | 0.21963804 | 0.37003349 | 0.5935626 | 0.5528047 | 0.92373235 |
| AP001318.2 | 262.053793 | -0.074887 | 0.16944519 | -0.4419544 | 0.6585222 | 0.9458503 |
| DCPS | 1726.78981 | -1.1697549 | 0.2329089 | -5.022371 | 5.10E-07 | 0.00130877 |
| GSEC | 172.139049 | -1.0478495 | 0.34319934 | -3.0531805 | 0.0022643 | 0.20649641 |
| ST3GAL4 | 2511.97796 | -0.9300342 | 0.28760936 | -3.2336716 | 0.0012221 | 0.15785077 |
| ETS1 | 5075.21188 | -0.3933455 | 0.35836387 | -1.0976148 | 0.27237271 | 0.82716979 |
| FLI1 | 971.055391 | -0.0885616 | 0.31537963 | -0.2808094 | 0.77885657 | 0.96836854 |
| KCNJ1 | 9907.08426 | 0.36477548 | 0.44676009 | 0.81649075 | 0.41421951 | 0.88364943 |
| KCNJ5 | 573.781796 | -0.3492815 | 0.5060873 | -0.6901605 | 0.49009324 | 0.90600166 |
| C11orf45 | 138.51949 | 0.12984049 | 0.41559878 | 0.31241788 | 0.75472296 | 0.96351771 |

| | | | | | | |
|---|---|---|---|---|---|---|
| TP53AIP1 | 43.7620114 | 0.21822673 | 0.5337924 | 0.40882323 | 0.68266939 | 0.95070301 |
| ARHGAP32 | 1328.9926 | 0.03725157 | 0.2024772 | 0.18397908 | 0.85402987 | 0.97906925 |
| BARX2 | 289.679509 | 0.40479532 | 0.3540755 | 1.14324577 | 0.25293657 | 0.81835388 |
| TMEM45B | 1602.67384 | 0.18586184 | 0.40523456 | 0.45865249 | 0.64648374 | 0.94208859 |
| NFRKB | 1738.26388 | -0.1336306 | 0.10602387 | -1.260382 | 0.20753159 | 0.78834607 |
| PRDM10 | 515.442945 | -0.3505443 | 0.27064963 | -1.295196 | 0.19525263 | 0.77723652 |
| APLP2 | 41059.7204 | 0.36174575 | 0.14766768 | 2.44972871 | 0.01429639 | 0.43699139 |
| ST14 | 8626.50332 | -0.0858869 | 0.19025986 | -0.451419 | 0.65168756 | 0.94393131 |
| ZBTB44 | 3534.59714 | -0.1691996 | 0.20696703 | -0.8175195 | 0.41363164 | 0.88316314 |
| AP002986.1 | 60.0127926 | 0.19999143 | 0.68501344 | 0.29195256 | 0.7703229 | 0.96836854 |
| ADAMTS8 | 143.679542 | -0.839599 | 0.78220843 | -1.0733699 | 0.28310516 | 0.83078187 |
| ADAMTS15 | 1069.75857 | -0.4646505 | 0.46253391 | -1.0045762 | 0.31510098 | 0.84696964 |
| AP003486.1 | 569.599388 | 0.10637728 | 0.16923162 | 0.62858987 | 5.30E-01 | 0.91931447 |
| SNX19 | 3745.74315 | -0.097005 | 0.14217695 | -0.6822833 | 0.49505983 | 0.90816709 |
| NTM | 95.6812608 | 0.19218696 | 0.57293734 | 0.33544149 | 7.37E-01 | 9.60E-01 |
| OPCML | 95.8847419 | -0.402972 | 0.77564734 | -0.5195299 | 0.60339129 | 0.93602816 |
| IGSF9B | 44.1200714 | -0.5018449 | 0.76080613 | -0.6596226 | 0.50949605 | 0.91439937 |
| JAM3 | 1172.30696 | -0.2342055 | 0.32764071 | -0.7148243 | 0.47471761 | 0.90226631 |
| NCAPD3 | 936.504828 | 0.05067057 | 0.16617283 | 0.30492692 | 7.60E-01 | 9.66E-01 |
| VPS26B | 2210.61108 | 0.08069431 | 0.11012913 | 0.73272448 | 0.46372649 | 0.89830907 |
| THYN1 | 1452.69817 | 0.42348458 | 0.18738302 | 2.25999448 | 0.02382159 | 0.51967919 |
| ACAD8 | 3203.67239 | 0.32879303 | 0.15751756 | 2.087342 | 0.03685723 | 0.55521179 |
| GLB1L3 | 71.756521 | -0.5332909 | 0.64688611 | -0.8243969 | 0.40971409 | 0.88249519 |
| GLB1L2 | 2217.09079 | 0.63912145 | 0.35731838 | 1.78866102 | 0.07366942 | 0.6374448 |
| B3GAT1 | 413.97407 | -0.0335842 | 0.72733926 | -0.0461741 | 0.96317152 | 0.99320711 |
| ZMYND11 | 4028.9436 | 0.06007202 | 0.1548785 | 0.38786541 | 0.69811563 | 0.9529028 |
| DIP2C | 2587.68873 | 0.04823884 | 0.27482585 | 0.17552511 | 0.86066704 | 0.98106934 |
| PRR26 | 544.04361 | -0.3177067 | 0.54664546 | -0.5811933 | 0.56111021 | 0.92448946 |
| LARP4B | 3317.2323 | -0.1326176 | 0.1363065 | -0.9729366 | 0.33058482 | 0.85384238 |
| GTPBP4 | 2022.61727 | -0.226595 | 0.26292907 | -0.8618103 | 0.38879191 | 0.87568585 |
| IDI2 | 162.669831 | -0.1129291 | 0.41439729 | -0.2725142 | 0.78522671 | 0.97038798 |
| IDI2-AS1 | 422.172703 | -0.1590685 | 0.54213669 | -0.2934103 | 0.76920859 | 0.96827902 |
| IDI1 | 3634.27744 | -0.0011697 | 0.39489235 | -0.0029621 | 0.9976366 | 0.99989813 |
| WDR37 | 1694.18755 | -0.1650124 | 0.22149845 | -0.7449821 | 0.45628257 | 0.89688594 |
| ADARB2 | 60.0761033 | 0.92291223 | 0.503276 | 1.83380935 | 0.0666823 | 0.62391062 |
| AL731533.2 | 74.3073089 | -0.2169514 | 0.67343468 | -0.3221566 | 0.74733409 | 0.96226645 |
| PFKP | 8561.16959 | -0.3062102 | 0.40277956 | -0.7602427 | 0.44710953 | 0.89480207 |
| AL451164.3 | 1900.82283 | -0.2611368 | 0.44237923 | -0.5903007 | 0.55498904 | 0.92373235 |
| PITRM1 | 5330.42902 | -0.2686269 | 0.26130239 | -1.0280309 | 0.3039353 | 0.84211658 |
| PITRM1-AS1 | 1454.83794 | -0.2320197 | 0.28262662 | -0.8209408 | 0.41167998 | 0.88268208 |
| AL139280.1 | 82.5473595 | 1.17399341 | 0.76108991 | 1.54251606 | 0.12294824 | 0.69788192 |
| KLF6 | 11499.212 | -0.7438983 | 0.424775 | -1.751276 | 0.07989838 | 0.64079926 |
| LINC00702 | 46.0244425 | -1.8331502 | 1.02998715 | -1.7797796 | 0.07511203 | 0.63773415 |
| AKR1E2 | 73.3053583 | -0.4217077 | 0.33824148 | -1.2467651 | 2.12E-01 | 7.92E-01 |
| AKR1C1 | 25770.3816 | -0.65316 | 1.09239023 | -0.5979182 | 5.50E-01 | 0.92373235 |
| AKR1C2 | 4110.41851 | -2.3075379 | 0.85454075 | -2.7003252 | 0.00692717 | 0.3503878 |
| AL391427.1 | 88.2328676 | -0.4222159 | 0.64546078 | -0.654131 | 0.51302741 | 0.91568064 |
| AKR1C3 | 11245.7033 | -0.0066143 | 0.57276004 | -0.0115481 | 0.99078619 | 0.9983285 |
| AKR1C4 | 47.3829421 | -1.0205684 | 0.9299462 | -1.0974488 | 0.27244523 | 0.82716979 |
| TUBAL3 | 503.186602 | 0.67612353 | 0.71101962 | 0.95092105 | 3.42E-01 | 8.59E-01 |
| NET1 | 3067.99618 | 0.29700127 | 0.32714101 | 0.90786928 | 0.36394729 | 0.8663515 |
| CALML3-AS1 | 152.816985 | 1.24730707 | 0.91608049 | 1.36156929 | 0.17333385 | 0.75974924 |
| CALML3 | 741.059498 | 1.04036953 | 1.25511658 | 0.82890271 | 0.40715946 | 0.88132015 |
| AL732437.1 | 537.599048 | 1.06734678 | 1.3681556 | 0.78013552 | 0.43531111 | 0.89055338 |

| | | | | | | |
|---|---|---|---|---|---|---|
| ASB13 | 3444.2178 | 0.39506101 | 0.31489574 | 1.25457717 | 0.20963229 | 0.79128274 |
| FAM208B | 1586.4513 | -0.3509046 | 0.16094685 | -2.1802518 | 0.0292388 | 0.53606221 |
| GDI2 | 9687.25042 | 0.10834183 | 0.21891707 | 0.49489897 | 0.62067144 | 0.94002099 |
| ANKRD16 | 401.627976 | 0.17690973 | 0.16138435 | 1.09620132 | 0.27299068 | 0.82716979 |
| FBH1 | 3968.62577 | -0.037041 | 0.12977499 | -0.2854249 | 0.7753186 | 0.96836854 |
| AL137186.2 | 1534.16176 | -0.0979734 | 0.16992045 | -0.5765839 | 0.56422059 | 0.92470993 |
| IL15RA | 923.312645 | -0.1387666 | 0.48586989 | -0.2856045 | 0.77518106 | 0.96836854 |
| IL2RA | 124.553971 | 0.07234885 | 0.688607 | 0.10506551 | 0.91632382 | 0.9868044 |
| RBM17 | 5020.29779 | -0.1640633 | 0.16540851 | -0.9918674 | 0.32126223 | 0.85119105 |
| PFKFB3 | 17809.42 | 0.06372027 | 0.5650284 | 0.11277357 | 0.91021008 | 0.98662757 |
| AL137145.2 | 74.8508645 | 0.39479435 | 0.62524708 | 0.63142134 | 5.28E-01 | 0.91845529 |
| AL137145.1 | 83.401157 | 0.57122934 | 0.57368607 | 0.99571764 | 0.31938736 | 0.84928695 |
| PRKCQ | 734.25678 | -0.4915806 | 0.30979807 | -1.5867774 | 0.11256307 | 0.6804982 |
| SFMBT2 | 536.832506 | -0.1611085 | 0.23992433 | -0.6714971 | 0.50190393 | 0.91259827 |
| ITIH5 | 2196.53153 | 0.95902128 | 0.25254241 | 3.79746624 | 0.00014618 | 0.05828957 |
| ITIH2 | 61.8508766 | 0.59813814 | 0.36365971 | 1.64477427 | 0.10001637 | 0.67165899 |
| KIN | 621.299499 | -0.1244861 | 0.1362208 | -0.9138555 | 0.36079278 | 0.86504879 |
| ATP5F1C | 11907.9967 | 0.24626484 | 0.16475479 | 1.49473557 | 0.13498347 | 0.71649038 |
| TAF3 | 539.431004 | -0.0556596 | 0.17204064 | -0.3235258 | 0.74629709 | 0.96195706 |
| GATA3 | 1988.28087 | 0.22506291 | 0.26877653 | 0.8373607 | 0.40238985 | 0.88132015 |
| GATA3-AS1 | 382.577986 | 0.17499658 | 0.35075132 | 0.49891924 | 0.61783628 | 0.93924932 |
| AL390294.1 | 153.17817 | 0.28864194 | 0.38569232 | 0.74837356 | 0.45423486 | 0.89670027 |
| CELF2 | 679.083365 | -0.2781004 | 0.39459866 | -0.7047678 | 0.48095473 | 0.90541935 |
| USP6NL | 803.355947 | -0.1603962 | 0.15961936 | -1.0048671 | 0.31496084 | 0.84696964 |
| ECHDC3 | 4975.12023 | 0.81786254 | 0.37295823 | 2.19290652 | 0.02831411 | 0.53383785 |
| PROSER2 | 1150.34366 | -0.404176 | 0.36683176 | -1.1018021 | 0.2705477 | 0.82716979 |
| PROSER2-AS | 181.887122 | -0.3891356 | 0.34698117 | -1.1214892 | 0.2620797 | 0.82424998 |
| UPF2 | 1500.873 | -0.3610013 | 0.12423069 | -2.9058948 | 0.00366205 | 0.26253301 |
| DHTKD1 | 3309.87727 | 0.3991137 | 0.34066143 | 1.17158462 | 0.24136387 | 0.8124487 |
| SEC61A2 | 1123.94369 | -0.2271082 | 0.16776687 | -1.3537133 | 0.17582787 | 0.7610139 |
| NUDT5 | 2815.51407 | -0.0623658 | 0.20369302 | -0.3061753 | 0.75947121 | 0.96502715 |
| CDC123 | 2279.33805 | -0.1383172 | 0.17606263 | -0.7856138 | 0.43209377 | 0.88976671 |
| AL512770.1 | 836.9494 | -0.1410691 | 0.17291554 | -0.8158264 | 0.41459943 | 0.88364943 |
| CAMK1D | 366.160714 | -0.1233228 | 0.28799717 | -0.4282084 | 0.66849937 | 0.9486589 |
| CCDC3 | 1149.99174 | -0.2436698 | 0.76508811 | -0.318486 | 0.75011635 | 0.96250371 |
| OPTN | 6702.03729 | -0.1035893 | 0.25939426 | -0.399351 | 0.68963463 | 0.95158502 |
| PHYH | 6192.77317 | 0.81307019 | 0.32087816 | 2.53389074 | 0.01128039 | 0.40196898 |
| SEPHS1 | 3065.54108 | 0.14992079 | 0.19291892 | 0.77711813 | 0.43708908 | 0.89130865 |
| BEND7 | 2547.28984 | -0.1783142 | 0.30880844 | -0.5774266 | 0.56365128 | 0.92453312 |
| PRPF18 | 899.863457 | -0.1277435 | 0.16092725 | -0.7937963 | 0.42731402 | 0.88769902 |
| AL157392.5 | 1217.08889 | -0.1308091 | 0.1197988 | -1.0919064 | 0.27487423 | 0.82719407 |
| FRMD4A | 2624.7866 | -0.0326796 | 0.25270621 | -0.1293185 | 0.89710562 | 0.98474725 |
| FAM107B | 7603.98696 | 0.24398026 | 0.25109758 | 0.97165515 | 0.33122214 | 0.85384238 |
| CDNF | 204.362435 | 0.59928648 | 0.3872877 | 1.54739351 | 0.12176839 | 0.69725141 |
| HSPA14 | 1293.78547 | 0.07228296 | 0.20368255 | 0.35488049 | 0.72267912 | 0.95749236 |
| AC069544.1 | 68.3654643 | -0.2102422 | 0.3472935 | -0.6053733 | 0.54493098 | 0.92345821 |
| SUV39H2 | 240.173945 | -0.0441008 | 0.20320936 | -0.2170215 | 0.82819155 | 0.97548201 |
| DCLRE1C | 281.997336 | -0.1132246 | 0.20595277 | -0.5497599 | 0.58248405 | 0.93177403 |
| OLAH | 50.5428526 | -0.3485495 | 0.68582703 | -0.5082178 | 0.6113006 | 0.93834929 |
| RPP38 | 791.050569 | -0.2017014 | 0.17917098 | -1.1257479 | 0.26027224 | 0.82424998 |
| NMT2 | 1288.49502 | 0.19595198 | 0.19832588 | 0.98803032 | 0.3231378 | 0.85119105 |
| FAM171A1 | 3224.83877 | 0.2213735 | 0.31934367 | 0.693214 | 0.48817526 | 0.90585781 |
| ITGA8 | 1093.40755 | -0.3862862 | 0.49552452 | -0.7795502 | 4.36E-01 | 8.91E-01 |
| MINDY3 | 1359.66357 | 0.17883031 | 0.15244998 | 1.17304252 | 0.24077876 | 0.8124487 |

| | | | | | | |
|---|---|---|---|---|---|---|
| PTER | 2352.91415 | 0.57601834 | 0.41661226 | 1.38262457 | 0.16678001 | 0.75558932 |
| C1QL3 | 104.796061 | 0.06020474 | 0.81423162 | 0.07394056 | 0.94105768 | 0.99136103 |
| RSU1 | 3111.48327 | -0.1526483 | 0.2369885 | -0.644117 | 0.51949954 | 0.91649637 |
| CUBN | 18046.6095 | 1.49168862 | 0.60446789 | 2.46777148 | 0.01359571 | 0.42608223 |
| TRDMT1 | 249.274081 | 0.18728722 | 0.22110034 | 0.84706889 | 0.39695673 | 0.87988087 |
| VIM-AS1 | 9069.92273 | -0.344738 | 0.4033972 | -0.854587 | 0.39277985 | 0.8778303 |
| VIM | 32545.2475 | -0.349927 | 0.40390408 | -0.8663615 | 0.38629193 | 0.87432552 |
| AL133415.1 | 12870.8055 | -0.3265317 | 0.41896327 | -0.7793801 | 0.43575582 | 0.89085849 |
| HACD1 | 253.432166 | -0.4715207 | 0.40709435 | -1.1582591 | 0.24675831 | 0.81542272 |
| STAM-AS1 | 59.717113 | 0.34433124 | 0.41804551 | 0.82366926 | 4.10E-01 | 0.88249519 |
| STAM | 1266.33629 | -0.3428381 | 0.24760197 | -1.3846338 | 0.16616448 | 0.75515387 |
| AC069542.1 | 99.3037941 | -0.2621159 | 0.42019742 | -0.6237923 | 0.53276397 | 0.92014084 |
| TMEM236 | 45.6561361 | -0.2176601 | 0.41770675 | -0.5210834 | 0.60230865 | 0.9355118 |
| MRC1 | 1334.71275 | 0.27215829 | 0.54140796 | 0.50268617 | 0.61518494 | 0.93895494 |
| CACNB2 | 409.365371 | -0.2249639 | 0.29210107 | -0.7701577 | 0.44120635 | 0.89290667 |
| AL450384.2 | 208.395262 | -0.0603596 | 0.29560591 | -0.2041895 | 0.83820541 | 0.97599451 |
| NSUN6 | 688.480508 | 0.16814918 | 0.3038393 | 0.55341483 | 5.80E-01 | 0.93139743 |
| ARL5B | 1341.05956 | -0.1890002 | 0.2956335 | -0.6393059 | 0.52262397 | 0.91723878 |
| MALRD1 | 64.4144075 | -0.6952533 | 0.5787551 | -1.201291 | 2.30E-01 | 0.80551898 |
| PLXDC2 | 2455.95375 | 0.2343416 | 0.25860296 | 0.90618298 | 3.65E-01 | 0.8663515 |
| NEBL | 2348.74803 | 0.35121646 | 0.25858554 | 1.35822158 | 0.17439339 | 0.75974924 |
| NEBL-AS1 | 41.8253919 | 0.36738471 | 0.76624044 | 0.479464 | 0.63160857 | 0.94088547 |
| CASC10 | 318.786299 | -0.5187427 | 0.46818756 | -1.1079806 | 0.26787018 | 0.82676682 |
| SKIDA1 | 154.369683 | 0.17999663 | 0.36138667 | 0.49807214 | 0.6184332 | 0.93943514 |
| MLLT10 | 1827.96078 | -0.1610712 | 0.12571015 | -1.28129 | 0.20009183 | 0.78134034 |
| DNAJC1 | 2302.40142 | 0.00587029 | 0.18067862 | 0.03249026 | 0.97408109 | 0.99452354 |
| COMMD3 | 1603.41335 | 0.33388456 | 0.22096399 | 1.5110361 | 0.13077925 | 0.70979843 |
| COMMD3-BI | 1932.52384 | 0.18577277 | 0.18520778 | 1.00305062 | 0.31583644 | 0.84750116 |
| BMI1 | 2231.60506 | -0.2114905 | 0.13346294 | -1.5846382 | 0.11304856 | 0.68130021 |
| SPAG6 | 51.03581 | 1.04160714 | 0.91619139 | 1.13688815 | 0.25558505 | 0.82052927 |
| PIP4K2A | 1969.75826 | -0.1360174 | 0.2476128 | -0.5493148 | 0.58278944 | 0.93183662 |
| ARMC3 | 82.20135 | 0.33689413 | 0.77222847 | 0.43626225 | 0.66264647 | 0.94642611 |
| MSRB2 | 1487.48829 | 0.38755995 | 0.26516501 | 1.46158026 | 0.14385627 | 0.72920837 |
| C10orf67 | 168.3865 | 0.35751818 | 0.48053608 | 0.74399863 | 0.4568773 | 0.89688594 |
| OTUD1 | 1486.20117 | -0.148007 | 0.23527585 | -0.6290787 | 0.52929755 | 0.91920857 |
| KIAA1217 | 3755.5819 | -0.2889534 | 0.1567351 | -1.8435779 | 0.06524467 | 0.6216672 |
| ARHGAP21 | 2458.60542 | -0.4382204 | 0.24633166 | -1.7789851 | 0.0752422 | 0.63773415 |
| PRTFDC1 | 217.838842 | 0.11748022 | 0.30817293 | 0.38121525 | 0.70304354 | 0.95431462 |
| THNSL1 | 565.233587 | 0.64237812 | 0.32976675 | 1.94797723 | 0.05141769 | 0.58634747 |
| MYO3A | 50.9278164 | 1.35421481 | 0.53445134 | 2.53384118 | 0.01128199 | 0.40196898 |
| APBB1IP | 601.167542 | 0.26480213 | 0.47610703 | 0.55618192 | 0.5780865 | 0.93130183 |
| PDSS1 | 415.527843 | 0.40851801 | 0.30734308 | 1.32919217 | 0.18378458 | 0.76781833 |
| ABI1 | 3388.46189 | -0.0863793 | 0.19001885 | -0.4545829 | 0.64940937 | 0.94312988 |
| ANKRD26 | 310.794047 | -0.1785887 | 0.31413623 | -0.5685072 | 0.56969063 | 0.92719921 |
| YME1L1 | 5472.95345 | -0.0562953 | 0.16671016 | -0.3376835 | 0.73560174 | 0.95895327 |
| MASTL | 516.283484 | -0.4332168 | 0.25769757 | -1.6811056 | 0.09274241 | 0.65662464 |
| ACBD5 | 2789.06115 | 0.0993696 | 0.19796683 | 0.50195074 | 0.61570217 | 0.93910158 |
| RAB18 | 5264.01473 | -0.3665308 | 0.25691556 | -1.4266587 | 0.15367829 | 0.74258845 |
| MPP7 | 938.672189 | 0.34751808 | 0.16911196 | 2.05495867 | 0.039883 | 0.55657615 |
| WAC-AS1 | 1286.76513 | -0.1051787 | 0.16945411 | -0.6206913 | 0.53480277 | 0.92084006 |
| WAC | 6732.65067 | -0.3503951 | 0.15562128 | -2.2515884 | 0.02434829 | 0.51967919 |
| BAMBI | 1244.41866 | -0.00601 | 0.24843559 | -0.0241914 | 0.98069992 | 0.99573612 |
| SVIL | 2947.06059 | -0.0287878 | 0.25647716 | -0.1122432 | 0.91063055 | 0.98662757 |
| JCAD | 1710.48061 | -0.1234499 | 0.44805203 | -0.2755259 | 0.78291225 | 0.96964828 |

| | | | | | | |
|---|---|---|---|---|---|---|
| MTPAP | 1168.65655 | -0.2660834 | 0.165675 | -1.6060564 | 0.10826153 | 0.67789099 |
| MAP3K8 | 1126.08363 | -0.17465 | 0.29040953 | -0.6013921 | 0.54757887 | 0.92361227 |
| ZNF438 | 453.805063 | -0.1121141 | 0.12219713 | -0.9174854 | 0.35888832 | 0.86480228 |
| ZEB1-AS1 | 467.039245 | 0.11467047 | 0.29169392 | 0.39311916 | 0.69423148 | 0.95222846 |
| ZEB1 | 1301.84568 | -0.44708 | 0.28561899 | -1.565302 | 0.1175121 | 0.69036323 |
| ARHGAP12 | 2198.71991 | -0.1014567 | 0.15910928 | -0.6376542 | 0.52369882 | 0.91723878 |
| KIF5B | 4898.15042 | -0.0667339 | 0.15276456 | -0.4368416 | 0.66222625 | 0.94642611 |
| AL158834.2 | 446.322464 | -0.3018423 | 0.17127401 | -1.7623355 | 0.07801263 | 0.64079926 |
| EPC1 | 1429.81269 | -0.3156496 | 0.19191644 | -1.6447242 | 0.10002669 | 0.67165899 |
| AL391839.2 | 64.9955139 | -0.4811679 | 0.57146264 | -0.8419936 | 0.39979155 | 0.88077967 |
| CCDC7 | 177.15646 | 0.05100674 | 0.26251058 | 0.19430357 | 0.84593819 | 0.9767918 |
| AL365203.2 | 359.765195 | -0.2216817 | 0.58035764 | -0.3819744 | 0.70248039 | 0.95431462 |
| ITGB1 | 11036.9243 | -0.0922993 | 0.3315648 | -0.2783748 | 0.78072468 | 0.96872095 |
| NRP1 | 10922.295 | 0.14769801 | 0.25452194 | 0.5802958 | 0.56171516 | 0.92452095 |
| AL121748.2 | 690.875595 | 0.14542476 | 0.24660898 | 0.58969775 | 0.5553933 | 0.92373235 |
| PARD3 | 2726.80825 | -0.1452692 | 0.24823243 | -0.5852143 | 0.55840363 | 0.92421175 |
| CUL2 | 1457.07446 | -0.0683677 | 0.18567562 | -0.3682107 | 0.71271614 | 0.95564093 |
| CREM | 2369.2592 | -0.5609559 | 0.30688784 | -1.8278856 | 0.06756672 | 0.62557548 |
| CCNY | 3639.21865 | -0.1625816 | 0.18972341 | -0.85694 | 0.39147805 | 0.8772797 |
| AL117336.3 | 193.844853 | 0.15192456 | 0.37656518 | 0.40344825 | 0.6866185 | 0.95097288 |
| FZD8 | 1305.59447 | 0.20391258 | 0.24554157 | 0.83046053 | 0.40627846 | 0.88132015 |
| ZNF248 | 1002.26648 | 0.1574451 | 0.26592134 | 0.59207396 | 0.55380106 | 0.92373235 |
| ZNF25 | 722.118575 | 0.28534426 | 0.15651193 | 1.823147 | 0.06828113 | 0.62648383 |
| ZNF33A | 2286.83442 | -0.0517063 | 0.20270625 | -0.25508 | 0.79866127 | 0.97173998 |
| ZNF37A | 1212.34134 | -0.2863158 | 0.1766399 | -1.6209009 | 0.10503889 | 0.67473222 |
| AL117339.5 | 179.213994 | 0.14335022 | 0.39189368 | 0.36578855 | 0.71452286 | 0.95579447 |
| AL117339.4 | 50.395967 | 0.44955673 | 0.64649723 | 0.69537302 | 0.48682156 | 0.90573694 |
| LINC00839 | 655.484781 | 0.51906125 | 0.51889804 | 1.00031454 | 0.31715831 | 0.84819508 |
| ZNF33B | 1997.0436 | 0.14124874 | 0.23718761 | 0.59551484 | 5.51E-01 | 0.92373235 |
| BMS1 | 2234.18623 | -0.176636 | 0.15722956 | -1.1234273 | 0.26125605 | 0.82424998 |
| RET | 111.916659 | 0.38972928 | 0.51697418 | 0.75386604 | 0.45092967 | 0.89590699 |
| AC010864.1 | 42.1392696 | 0.56561809 | 0.44531303 | 1.27015843 | 0.2040282 | 0.78418828 |
| CSGALNACT2 | 1349.90626 | -0.417917 | 0.34816589 | -1.2003387 | 0.23000781 | 0.80551898 |
| RASGEF1A | 212.582772 | 0.46357313 | 0.33943178 | 1.36573286 | 1.72E-01 | 0.75974924 |
| FXYD4 | 6802.23558 | 1.50795717 | 0.62255409 | 2.42221068 | 0.0154264 | 0.45243543 |
| HNRNPF | 10554.7368 | -0.3650443 | 0.27618671 | -1.32173 | 0.18625808 | 0.76908034 |
| AL450326.1 | 67.7354138 | 0.52803738 | 0.46079316 | 1.14593145 | 0.25182352 | 0.81833349 |
| ZNF487 | 304.351855 | 0.05526001 | 0.18576023 | 0.2974803 | 0.76609985 | 0.96704052 |
| ZNF239 | 228.536015 | 0.08146945 | 0.30311057 | 0.26877799 | 0.78810053 | 0.97090793 |
| ZNF485 | 114.584373 | -0.0312765 | 0.25786703 | -0.1212894 | 0.90346178 | 0.98617053 |
| ZNF32 | 1538.2827 | 0.0495976 | 0.12514694 | 0.39631491 | 6.92E-01 | 9.52E-01 |
| ZNF32-AS1 | 379.701007 | 0.01991255 | 0.18417411 | 0.10811807 | 9.14E-01 | 0.98662757 |
| ZNF32-AS2 | 323.451953 | -0.0630299 | 0.16474092 | -0.3826004 | 0.70201605 | 0.95431462 |
| CXCL12 | 6401.04525 | -0.0782147 | 0.54529838 | -0.1434348 | 8.86E-01 | 0.98385457 |
| TMEM72-AS | 78.468147 | 0.76310805 | 0.41471445 | 1.84008071 | 0.06575639 | 0.62237346 |
| TMEM72 | 5189.22027 | 1.06691034 | 0.50273887 | 2.12219586 | 0.03382129 | 0.55369025 |
| RASSF4 | 13831.8698 | 0.46269592 | 0.32030353 | 1.44455455 | 0.14858305 | 0.73553131 |
| DEPP1 | 11578.1834 | -0.4438552 | 0.3177476 | -1.3968797 | 0.16244975 | 0.75026273 |
| C10orf25 | 121.624147 | -0.0421188 | 0.28452662 | -0.1480312 | 8.82E-01 | 0.98275231 |
| ZNF22 | 1725.47211 | -0.248518 | 0.14513608 | -1.71231 | 0.08683956 | 0.65228863 |
| ALOX5 | 1011.78759 | 0.83951997 | 0.51538791 | 1.62890894 | 0.1033323 | 0.67439401 |
| 8-Mar | 2629.57666 | -0.1358584 | 0.1890904 | -0.7184837 | 0.4724591 | 0.90160919 |
| ZFAND4 | 353.076519 | -0.0974681 | 0.16332787 | -0.5967637 | 0.55066518 | 0.92373235 |
| WASHC2C | 2888.41313 | -0.1808335 | 0.25608859 | -0.7061367 | 0.48010315 | 0.90537463 |

| | | | | | | |
|---|---|---|---|---|---|---|
| AGAP4 | 583.848405 | -0.0689924 | 0.36759639 | -0.1876852 | 0.85112345 | 0.97846305 |
| TIMM23 | 2627.02592 | -0.1744814 | 0.19277921 | -0.905084 | 0.3654209 | 0.8663515 |
| NCOA4 | 9516.30763 | 0.12335518 | 0.19967108 | 0.61779193 | 0.5367125 | 0.92105614 |
| GPRIN2 | 267.066382 | -0.3743918 | 0.55264766 | -0.6774511 | 0.49811978 | 0.91049087 |
| SYT15 | 52.1148648 | -0.2146871 | 0.544219 | -0.3944867 | 0.69322177 | 0.9519683 |
| AL356056.3 | 38.2086887 | -0.4628593 | 0.47676106 | -0.9708412 | 0.33162737 | 0.85429945 |
| PTPN20 | 263.796076 | 0.26898793 | 0.36195964 | 0.74314342 | 0.45739485 | 0.89688594 |
| ZNF488 | 71.9911141 | 2.65402098 | 0.74410032 | 3.56675154 | 0.00036143 | 0.08914272 |
| AL591684.1 | 174.509982 | 0.82373815 | 0.70835922 | 1.16288195 | 0.2448774 | 0.81485217 |
| AGAP9 | 1249.53994 | 0.87583588 | 0.56793517 | 1.54214058 | 0.12303944 | 0.69788192 |
| BX547991.1 | 165.15056 | 1.34508488 | 0.65309175 | 2.05956497 | 0.03944015 | 0.55657615 |
| MAPK8 | 2025.81947 | -0.2394023 | 0.2662342 | -0.8992168 | 0.36853719 | 0.86762749 |
| ARHGAP22 | 200.956489 | -0.1583711 | 0.43266562 | -0.3660359 | 0.71433831 | 0.95579447 |
| WDFY4 | 351.037631 | -0.1912559 | 0.57444272 | -0.3329417 | 0.73917828 | 0.9602325 |
| AC060234.1 | 54.7665076 | -0.3115632 | 0.52870877 | -0.5892907 | 0.5556663 | 0.92373235 |
| VSTM4 | 1208.8302 | 0.04700365 | 0.39334765 | 0.11949644 | 0.90488206 | 0.98624484 |
| TMEM273 | 402.22555 | -0.1490024 | 0.59864369 | -0.2488999 | 0.80343818 | 0.97274243 |
| ERCC6 | 665.113968 | -0.2831011 | 0.17035822 | -1.6617987 | 0.09655314 | 0.66243513 |
| OGDHL | 15119.5939 | 1.13930106 | 0.31758939 | 3.58733985 | 0.00033407 | 0.08914272 |
| PARG | 1128.81386 | 0.18572462 | 0.23958245 | 0.77520128 | 0.43822073 | 0.89152136 |
| TIMM23B | 391.956332 | -0.2020631 | 0.27378264 | -0.7380421 | 0.46048886 | 0.89755243 |
| LINC00843 | 102.949003 | -0.191307 | 0.29859637 | -0.6406878 | 0.52172554 | 0.91723878 |
| AGAP6 | 1432.52414 | -0.1266838 | 0.14118034 | -0.8973188 | 0.36954882 | 0.86762749 |
| WASHC2A | 3078.88287 | -0.5022994 | 0.23794334 | -2.1110041 | 0.03477196 | 0.55369025 |
| ASAH2 | 60.0140374 | -0.6068966 | 0.59646631 | -1.0174869 | 0.30892187 | 0.84414094 |
| SGMS1 | 2419.49724 | 0.11485316 | 0.18087941 | 0.63497093 | 0.52544737 | 0.91759401 |
| SGMS1-AS1 | 399.321957 | 0.12484858 | 0.33525379 | 0.37240021 | 0.7095949 | 0.95501548 |
| ASAH2B | 415.069063 | 0.43645212 | 0.28645881 | 1.52361214 | 0.12760563 | 0.70557946 |
| A1CF | 1106.40985 | 1.35868703 | 0.56812481 | 2.39152912 | 1.68E-02 | 4.65E-01 |
| PRKG1 | 499.508439 | -0.1972891 | 0.39374751 | -0.5010549 | 0.6163325 | 0.93910158 |
| CSTF2T | 1592.79441 | -0.0420216 | 0.16205316 | -0.2593075 | 0.79539802 | 0.97101097 |
| PRKG1-AS1 | 96.9573074 | 0.08618724 | 0.39441332 | 0.21852011 | 8.27E-01 | 9.75E-01 |
| AC073174.1 | 65.7645261 | -2.3052619 | 1.72447736 | -1.3367887 | 0.18129162 | 0.76522169 |
| PCDH15 | 70.6171112 | 2.16504967 | 0.88525507 | 2.44567893 | 0.01445797 | 0.43901686 |
| ZWINT | 331.790462 | 0.22006399 | 0.43304865 | 0.50817384 | 0.61133144 | 0.93834929 |
| IPMK | 664.554481 | -0.0534039 | 0.23451701 | -0.2277188 | 0.81986482 | 0.97454841 |
| CISD1 | 2380.95706 | 0.09482915 | 0.21319769 | 0.44479444 | 0.65646831 | 0.94545756 |
| UBE2D1 | 1115.56235 | -0.6660567 | 0.30912192 | -2.1546733 | 0.03118741 | 0.54466273 |
| TFAM | 1734.15296 | -0.0802165 | 0.19730053 | -0.4065699 | 0.6843239 | 0.95070301 |
| BICC1 | 5105.68533 | 0.05408774 | 0.26818004 | 0.20168443 | 0.84016343 | 0.97618283 |
| LINC00844 | 53.2312757 | 1.08544501 | 0.69784877 | 1.55541581 | 1.20E-01 | 6.94E-01 |
| PHYHIPL | 1575.51656 | 0.12421417 | 0.38958842 | 0.31883436 | 0.74985212 | 0.96249332 |
| FAM13C | 568.236428 | 0.34354323 | 0.41051958 | 0.83684979 | 0.402677 | 0.88132015 |
| SLC16A9 | 7395.03248 | 1.41045022 | 0.76483449 | 1.84412475 | 0.06516496 | 0.6216672 |
| MRLN | 84.4282656 | 2.08870946 | 0.79582486 | 2.62458435 | 0.00867548 | 0.37987657 |
| CCDC6 | 2818.27225 | 0.00104107 | 0.17804068 | 0.00584736 | 0.9953345 | 0.99940889 |
| ANK3 | 5665.46412 | 0.37104269 | 0.31013851 | 1.19637734 | 0.23154934 | 0.80649152 |
| AL592430.1 | 1134.92713 | 0.48722382 | 0.37274741 | 1.30711524 | 0.19117358 | 0.77292503 |
| AL592430.2 | 106.576769 | 0.27469081 | 0.52845662 | 0.51979822 | 0.60320422 | 0.93602816 |
| CDK1 | 175.902011 | 0.6640602 | 0.6024518 | 1.10226279 | 0.27034744 | 0.82716979 |
| RHOBTB1 | 2523.80029 | 0.51225967 | 0.48299085 | 1.06059911 | 0.28887213 | 0.83295411 |
| TMEM26 | 244.252284 | 0.40649474 | 0.30429005 | 1.33587917 | 0.18158877 | 0.76526988 |
| CABCOCO1 | 71.5913398 | 0.36055307 | 0.46062977 | 0.7827394 | 0.43378016 | 0.89055338 |
| ARID5B | 4968.45806 | -0.6875939 | 0.21947581 | -3.1328914 | 0.00173093 | 0.18431503 |

| | | | | | | |
|---|---|---|---|---|---|---|
| RTKN2 | 70.5244812 | 0.05530008 | 0.40662912 | 0.13599637 | 0.89182415 | 0.98474725 |
| ZNF365 | 46.5718036 | -0.2318295 | 0.88846188 | -0.2609336 | 0.79414373 | 0.97090793 |
| AL133417.1 | 120.670942 | 0.0074458 | 0.33113263 | 0.02248586 | 0.98206039 | 0.99595643 |
| ADO | 1182.01213 | -0.0138642 | 0.17020762 | -0.0814548 | 0.93508027 | 0.99068685 |
| EGR2 | 156.522848 | 0.61233738 | 0.8124616 | 0.75368163 | 4.51E-01 | 0.89600895 |
| NRBF2 | 927.329934 | -0.1306222 | 0.29294795 | -0.4458888 | 0.65567757 | 0.94532497 |
| JMJD1C | 2374.92544 | -0.28184 | 0.18041225 | -1.5621996 | 0.11824097 | 0.69095557 |
| REEP3 | 2158.97654 | -0.2871885 | 0.22266633 | -1.2897706 | 0.19713033 | 0.77816059 |
| LINC01515 | 93.2204399 | 0.15961275 | 0.55889854 | 0.28558449 | 0.77519639 | 0.96836854 |
| DNAJC12 | 630.365588 | 0.70106629 | 0.39173529 | 1.78964294 | 0.07351133 | 0.6374448 |
| SIRT1 | 1070.82976 | -0.3875061 | 0.28093383 | -1.37935 | 0.16778687 | 0.7558879 |
| HERC4 | 2290.891 | -0.1895615 | 0.11489852 | -1.6498166 | 0.09898045 | 0.66846583 |
| ATOH7 | 91.5886374 | 0.36819686 | 0.47565181 | 0.77408907 | 0.43887812 | 0.89175645 |
| AC016395.1 | 82.2520105 | 0.15111263 | 0.64065586 | 0.23587176 | 8.14E-01 | 0.9738447 |
| PBLD | 8499.18375 | 0.66440863 | 0.33130126 | 2.00545159 | 0.04491479 | 0.57322281 |
| HNRNPH3 | 8251.33645 | -0.2205435 | 0.07727371 | -2.8540565 | 0.00431649 | 0.28584892 |
| RUFY2 | 2343.89631 | -0.1878814 | 0.10737098 | -1.7498342 | 0.08014692 | 0.64079926 |
| DNA2 | 470.534408 | -0.5098101 | 0.37244565 | -1.3688175 | 0.17105632 | 0.75956362 |
| SLC25A16 | 2533.38944 | -0.129414 | 0.23499599 | -0.5507072 | 0.58183444 | 0.9317705 |
| TET1 | 56.9147426 | 0.04368405 | 0.52169093 | 0.08373549 | 0.93326674 | 0.99016938 |
| CCAR1 | 3060.96041 | -0.2932747 | 0.20577819 | -1.4251982 | 0.15409992 | 0.74258845 |
| STOX1 | 165.161972 | 0.1087594 | 0.40507981 | 0.26848883 | 0.78832307 | 0.97090793 |
| DDX50 | 1798.36818 | -0.0039132 | 0.14626381 | -0.0267543 | 0.97865569 | 0.99483482 |
| DDX21 | 3669.76423 | -0.3524313 | 0.35680805 | -0.9877336 | 0.32328313 | 0.85119105 |
| KIF1BP | 1762.50325 | -0.0760726 | 0.13698746 | -0.5553254 | 0.57867209 | 0.93139743 |
| SRGN | 5996.07334 | -0.4321886 | 0.4083902 | -1.0582736 | 0.28993072 | 0.83401253 |
| VPS26A | 3094.66138 | -0.2795898 | 0.22620735 | -1.2359892 | 0.21646259 | 0.7954827 |
| SUPV3L1 | 1460.33202 | -0.1498322 | 0.18508958 | -0.8095117 | 0.4182209 | 0.88445833 |
| HKDC1 | 4652.95193 | -0.5131118 | 0.68900456 | -0.7447147 | 0.45644422 | 0.89688594 |
| AL596223.2 | 662.540997 | -0.5046549 | 0.67891083 | -0.7433301 | 0.45728184 | 0.89688594 |
| HK1 | 6001.45067 | -0.3250178 | 0.29453095 | -1.1035099 | 0.2698058 | 0.82716979 |
| TSPAN15 | 1902.90307 | 0.51717229 | 0.28723704 | 1.80050694 | 0.07178063 | 0.63189503 |
| FAM241B | 451.652357 | 0.05523147 | 0.27567207 | 0.20035207 | 0.84120524 | 0.97618283 |
| COL13A1 | 227.548105 | 0.37976115 | 0.58786014 | 0.64600595 | 0.5182755 | 0.91636259 |
| H2AFY2 | 986.604419 | -0.0341613 | 0.30894673 | -0.1105735 | 0.91195454 | 0.98662757 |
| AIFM2 | 2001.98674 | -0.1301278 | 0.27420015 | -0.4745722 | 0.63509193 | 0.94088547 |
| TYSND1 | 1441.92328 | 0.17884956 | 0.19843595 | 0.90129614 | 0.36743088 | 0.86683872 |
| SAR1A | 5695.03106 | -0.2829769 | 0.22627738 | -1.2505753 | 0.21108948 | 0.79134644 |
| PPA1 | 7402.44181 | 0.0579747 | 0.20219511 | 0.28672651 | 0.77432174 | 0.96836854 |
| LRRC20 | 301.563331 | 0.84794894 | 0.27656274 | 3.06602734 | 0.00216923 | 0.20321981 |
| EIF4EBP2 | 9510.77732 | 0.2146567 | 0.31822194 | 0.6745503 | 0.49996152 | 0.91180773 |
| PALD1 | 678.589172 | -0.3835693 | 0.32733149 | -1.171807 | 2.41E-01 | 8.12E-01 |
| PRF1 | 411.542734 | -0.3088759 | 0.70092692 | -0.4406678 | 0.65945353 | 0.94605737 |
| ADAMTS14 | 50.9292077 | 0.6206896 | 0.64191593 | 0.96693285 | 0.3335776 | 0.855011 |
| SGPL1 | 3619.93601 | -0.1001784 | 0.13124038 | -0.7633203 | 0.4452724 | 0.89452291 |
| PCBD1 | 5659.94086 | -0.131341 | 0.42213911 | -0.3111321 | 0.75570022 | 0.96416215 |
| UNC5B | 1396.06167 | 0.18240908 | 0.38140428 | 0.47825652 | 0.63246764 | 0.94088547 |
| SLC29A3 | 371.815456 | -0.2825324 | 0.33177198 | -0.8515862 | 0.39444382 | 0.87884163 |
| CDH23 | 202.249886 | -0.2437901 | 0.49556721 | -0.4919416 | 0.62276063 | 0.94016713 |
| VSIR | 2297.8809 | -0.3487893 | 0.31888559 | -1.0937756 | 0.27405341 | 0.82719197 |
| PSAP | 85648.7123 | 0.048474 | 0.26967911 | 0.17974696 | 0.85735123 | 0.97953896 |
| CHST3 | 1752.34154 | -0.0161546 | 0.23171546 | -0.0697172 | 0.94441874 | 0.99136103 |
| SPOCK2 | 6571.2199 | 0.2053455 | 0.44503158 | 0.46141781 | 0.64449887 | 0.94208859 |
| ASCC1 | 1948.6211 | 0.0773237 | 0.15053208 | 0.51366927 | 0.60748324 | 0.93739614 |

| | | | | | | |
|---|---|---|---|---|---|---|
| ANAPC16 | 6858.55488 | 0.22378846 | 0.17348603 | 1.28995092 | 0.1970677 | 0.77816059 |
| DDIT4 | 16607.4229 | 0.07952489 | 0.65589068 | 0.12124717 | 0.90349526 | 0.98617053 |
| DDIT4-AS1 | 11849.9827 | 0.08882132 | 0.65950632 | 0.13467849 | 0.89286608 | 0.98474725 |
| DNAJB12 | 3442.45485 | -0.114068 | 0.15697561 | -0.7266606 | 0.46743389 | 0.89983399 |
| MICU1 | 4404.01834 | 0.19590773 | 0.22575597 | 0.86778539 | 0.38551183 | 0.87394184 |
| MCU | 1334.24906 | 0.089903 | 0.16346429 | 0.54998555 | 0.58232929 | 0.93177403 |
| OIT3 | 173.787242 | 1.70451277 | 0.7432817 | 2.29322578 | 0.02183501 | 0.51592008 |
| PLA2G12B | 277.72931 | 1.4489796 | 0.74334303 | 1.94927448 | 0.05126265 | 0.58634747 |
| P4HA1 | 3718.45449 | 0.02280415 | 0.37423034 | 0.06093614 | 0.95141007 | 0.99170503 |
| AL731563.2 | 164.847868 | 0.2483092 | 0.47919437 | 0.51818055 | 0.6043323 | 0.93659663 |
| AL731563.3 | 87.0395919 | -0.0771639 | 0.33210729 | -0.2323463 | 0.81626903 | 0.97388289 |
| NUDT13 | 461.288872 | 0.40740912 | 0.289401 | 1.4077668 | 0.15920013 | 0.7475863 |
| AC016394.2 | 103.461918 | -0.1393262 | 0.32006646 | -0.4353041 | 0.6633417 | 0.94642611 |
| ECD | 1574.76372 | -0.0657602 | 0.10266216 | -0.6405491 | 0.52181567 | 0.91723878 |
| FAM149B1 | 2297.75413 | -0.0945096 | 0.10761854 | -0.8781911 | 3.80E-01 | 8.71E-01 |
| DNAJC9 | 1515.4814 | -0.3259833 | 0.22256456 | -1.4646685 | 0.14301139 | 0.72784886 |
| DNAJC9-AS1 | 1102.26187 | -0.0856579 | 0.14931335 | -0.5736791 | 0.56618498 | 0.92558207 |
| MRPS16 | 3761.13248 | -0.0779121 | 0.21376663 | -0.3644728 | 0.71550497 | 0.95590747 |
| AC016394.1 | 290.826125 | 0.31882097 | 0.44365986 | 0.71861577 | 0.4723777 | 0.90160919 |
| CFAP70 | 801.579837 | 0.30160021 | 0.57284444 | 0.52649583 | 0.59854375 | 0.9346978 |
| ANXA7 | 9559.12155 | 0.09075209 | 0.173223 | 0.52390321 | 0.60034586 | 0.93487648 |
| AL512656.1 | 2015.93807 | 0.20520994 | 0.20761534 | 0.98841415 | 0.32294986 | 0.85119105 |
| MSS51 | 228.3999 | 0.05930065 | 0.35686017 | 0.16617335 | 0.86802053 | 0.98237277 |
| PPP3CB | 2458.82517 | 0.01291994 | 0.14731961 | 0.08770009 | 0.93011505 | 0.98973492 |
| USP54 | 1185.77237 | -0.3366743 | 0.15610606 | -2.1567024 | 0.03102886 | 0.54466273 |
| AC073389.3 | 39.9936447 | -0.374924 | 0.46556638 | -0.8053074 | 0.42064233 | 0.88451372 |
| MYOZ1 | 124.495668 | -0.4257278 | 0.55080704 | -0.7729163 | 0.4395719 | 0.8919464 |
| AC073389.2 | 68.7237278 | 0.22473846 | 0.30368205 | 0.74004526 | 0.45927253 | 0.89755243 |
| AGAP5 | 341.166796 | -0.0660214 | 0.25910382 | -0.2548067 | 7.99E-01 | 9.72E-01 |
| AC022400.6 | 57.4506708 | -0.1896926 | 0.32444134 | -0.5846746 | 0.55876651 | 0.92421175 |
| SEC24C | 6414.83934 | -0.0252845 | 0.1565029 | -0.1615591 | 0.87165305 | 0.98275231 |
| FUT11 | 1449.71213 | -0.1936605 | 0.331343 | -0.5844713 | 0.55890329 | 0.92421175 |
| CHCHD1 | 1467.30441 | -0.0023207 | 0.14452561 | -0.0160576 | 0.98718842 | 0.99732542 |
| ZSWIM8 | 9368.05314 | -0.4339647 | 0.19482109 | -2.2275039 | 0.02591361 | 0.52502381 |
| ZSWIM8-AS1 | 3227.40556 | -0.4344075 | 0.18211465 | -2.3853519 | 0.01706279 | 0.46682305 |
| AC022400.7 | 5840.75696 | -0.346633 | 0.12734443 | -2.7220117 | 0.00648858 | 0.33778486 |
| NDST2 | 1898.31875 | -0.1750285 | 0.13209209 | -1.3250491 | 0.18515488 | 0.76820626 |
| CAMK2G | 2547.49338 | 0.09288409 | 0.21002044 | 0.44226216 | 0.65829951 | 0.9458503 |
| AC022400.1 | 389.692832 | 0.01076078 | 0.26463065 | 0.04066341 | 0.96756423 | 0.99400331 |
| PLAU | 4385.4884 | -0.6916345 | 0.5714455 | -1.2103244 | 0.22615442 | 0.80246466 |
| C10orf55 | 1544.54244 | -0.710296 | 0.58388222 | -1.2165055 | 0.2237924 | 0.80078694 |
| VCL | 6376.2847 | -0.1729051 | 0.27807614 | -0.6217903 | 0.53407974 | 0.92083983 |
| AP3M1 | 2061.33273 | -0.1489519 | 0.13407115 | -1.1109911 | 0.26657217 | 0.8265123 |
| ADK | 1485.7684 | 0.33818961 | 0.17392731 | 1.94443076 | 0.05184353 | 0.58645127 |
| KAT6B | 1053.51967 | -0.2188541 | 0.24183969 | -0.9049552 | 0.36548912 | 0.8663515 |
| SAMD8 | 1456.17984 | -0.1603377 | 0.15069482 | -1.0639897 | 0.28733339 | 0.83203261 |
| VDAC2 | 6545.4008 | 0.09573412 | 0.19804997 | 0.48338363 | 0.62882337 | 0.94088547 |
| COMTD1 | 1171.41024 | -0.1574858 | 0.21987898 | -0.7162387 | 4.74E-01 | 0.90199039 |
| ZNF503-AS1 | 69.6157381 | 0.29778879 | 0.42168408 | 0.70618932 | 0.48007041 | 0.90537463 |
| AC010997.3 | 395.978772 | -0.1515063 | 0.32180227 | -0.4708056 | 0.63777954 | 0.94088547 |
| ZNF503 | 2827.20581 | -0.182546 | 0.25715389 | -0.7098705 | 4.78E-01 | 0.90387017 |
| ZNF503-AS2 | 477.404652 | -0.2044738 | 0.28933514 | -0.7067024 | 0.47975145 | 0.90525794 |
| AC010997.4 | 143.287381 | 0.24851947 | 0.34019122 | 0.73052878 | 0.46506703 | 0.89918717 |
| LRMDA | 499.671044 | 0.46216885 | 0.31317755 | 1.47574067 | 0.14001352 | 0.72486007 |

| | | | | | | |
|---|---|---|---|---|---|---|
| KCNMA1 | 534.045271 | -0.3172852 | 0.44909851 | -0.7064936 | 0.47988122 | 0.90532051 |
| AL450306.1 | 1513.85397 | -0.2722666 | 0.26356086 | -1.0330311 | 0.30158934 | 0.84211658 |
| DLG5 | 2148.60234 | -0.2973362 | 0.30201809 | -0.984498 | 0.32487072 | 0.85188222 |
| POLR3A | 1066.40187 | -0.0715502 | 0.14356954 | -0.4983663 | 0.61822587 | 0.93929439 |
| RPS24 | 23009.5637 | 0.10620084 | 0.18794739 | 0.56505623 | 0.57203552 | 0.92874623 |
| ZMIZ1-AS1 | 178.294171 | -0.3163761 | 0.49398833 | -0.6404526 | 0.52187837 | 0.91723878 |
| ZMIZ1 | 6113.52359 | -0.1347823 | 0.20862997 | -0.6460352 | 0.51825654 | 0.91636259 |
| PPIF | 6232.61171 | -0.6082125 | 0.33540034 | -1.8133927 | 0.06977127 | 0.62779744 |
| ZCCHC24 | 1449.70296 | -0.0265579 | 0.32892264 | -0.0807421 | 0.93564708 | 0.99068685 |
| AL133481.1 | 308.799007 | -0.0337375 | 0.42669344 | -0.0790674 | 0.93697905 | 0.99068685 |
| EIF5AL1 | 61.3530888 | -0.6125795 | 0.69791663 | -0.8777259 | 0.38009251 | 0.87090707 |
| AL132656.2 | 133.428214 | 0.00648519 | 0.30305314 | 0.02139952 | 0.98292695 | 0.9961464 |
| NUTM2B-AS | 740.855663 | -0.1523464 | 0.22632892 | -0.6731192 | 0.50087143 | 0.91222852 |
| AL135925.1 | 239.338508 | -0.2521563 | 0.14342799 | -1.7580688 | 0.07873581 | 0.64079926 |
| SFTPD | 106.913664 | 0.33691288 | 0.42173924 | 0.79886539 | 0.42436847 | 0.88565404 |
| SFTPD-AS1 | 47.2913926 | 0.64029278 | 0.36167053 | 1.77037587 | 0.07666455 | 0.63773415 |
| TMEM254-A | 195.01239 | 0.21376168 | 0.39958522 | 0.53495894 | 5.93E-01 | 9.33E-01 |
| TMEM254 | 1362.73331 | 0.57055335 | 0.28098798 | 2.0305258 | 0.04230312 | 0.56485966 |
| PLAC9 | 365.907961 | 0.06954119 | 0.53216187 | 0.13067677 | 0.89603101 | 0.98474725 |
| ANXA11 | 16693.6067 | 0.08205596 | 0.16671952 | 0.49217969 | 0.62259232 | 0.94016713 |
| LINC00857 | 93.5331309 | 0.18590896 | 0.31805571 | 0.58451696 | 0.55887256 | 0.92421175 |
| MAT1A | 100.88389 | 1.92871113 | 0.56080138 | 3.43920541 | 0.00058342 | 0.11756558 |
| FAM213A | 6902.84898 | 0.34240752 | 0.22812568 | 1.50095996 | 0.13336592 | 0.71505705 |
| TSPAN14 | 6868.4953 | -0.2178536 | 0.12676334 | -1.7185849 | 0.08568998 | 0.65049132 |
| NRG3 | 87.746249 | 0.27337088 | 0.52174275 | 0.52395722 | 0.6003083 | 0.93487648 |
| AL603756.1 | 1441.07793 | 0.60039013 | 0.3629231 | 1.65431776 | 0.09806297 | 0.66549286 |
| GHITM | 19733.4185 | 0.32722656 | 0.26320566 | 1.24323527 | 0.21378115 | 0.7926732 |
| CDHR1 | 306.614204 | -2.0468498 | 0.80258149 | -2.5503326 | 0.01076202 | 0.40196898 |
| CCSER2 | 1708.74428 | -0.1616107 | 0.16859249 | -0.9585875 | 0.33776657 | 0.8571036 |
| GRID1 | 67.5869228 | 0.42692469 | 0.61007725 | 0.69978792 | NA | NA |
| WAPL | 2430.27733 | -0.1840456 | 0.1901874 | -0.9677067 | 0.33319085 | 0.855011 |
| AL731569.1 | 111.33599 | 0.23877713 | 0.31926934 | 0.74788621 | 0.4545288 | 0.89670027 |
| LDB3 | 251.648322 | -0.6536543 | 0.70619453 | -0.9256009 | 0.35465343 | 0.86364633 |
| AC067750.1 | 116.959515 | -0.0362964 | 0.30351244 | -0.1195879 | 0.90480963 | 0.98624484 |
| BMPR1A | 1482.78249 | -0.2422993 | 0.12979449 | -1.8667922 | 0.06193061 | 0.61210783 |
| MMRN2 | 3311.58612 | 0.03294454 | 0.37118602 | 0.08875479 | 0.92927679 | 0.98963127 |
| SNCG | 543.917447 | 0.29107158 | 0.52061689 | 0.55908978 | 0.57610045 | 0.93083012 |
| ADIRF | 4990.87033 | 0.21103578 | 0.55065196 | 0.38324713 | 0.70153654 | 0.95424422 |
| AL136982.7 | 4509.52504 | 0.22353042 | 0.55394596 | 0.40352388 | 0.68656287 | 0.95097288 |
| AL136982.2 | 38.2947092 | 0.04949679 | 0.64175124 | 0.07712769 | 0.93852197 | 0.99113899 |
| GLUD1 | 14030.2326 | 0.67276407 | 0.30236911 | 2.22497621 | 0.02608284 | 0.52561121 |
| SHLD2 | 3180.02205 | 0.20590524 | 0.29259853 | 0.70371248 | 0.48161184 | 0.90541935 |
| NUTM2A-AS | 779.890579 | -0.2627885 | 0.17127908 | -1.5342708 | 0.12496304 | 0.70228923 |
| NUTM2A | 53.2911309 | 0.36090889 | 0.86438086 | 0.41753457 | 0.67628744 | 0.94967053 |
| NUTM2D | 160.603065 | -0.1653448 | 0.40321484 | -0.4100662 | 0.68175737 | 0.95070301 |
| LINC00864 | 119.291409 | 1.49206953 | 0.56482178 | 2.6416643 | 0.00824998 | 0.37527389 |
| MINPP1 | 1354.57901 | 0.07628854 | 0.27068853 | 0.28183144 | 0.77807276 | 0.96836854 |
| AL138767.1 | 42.6884664 | 1.01298589 | 0.60199451 | 1.68271615 | 0.09243005 | 0.65583276 |
| PAPSS2 | 2417.75256 | 0.82892205 | 0.24592641 | 3.37061017 | 0.00075002 | 0.12895512 |
| ATAD1 | 2757.21928 | -0.1133327 | 0.11603216 | -0.9767353 | 0.32870022 | 0.85317073 |
| KLLN | 69.6480888 | 0.74345413 | 0.43719687 | 1.70050195 | 0.08903655 | 0.65375543 |
| PTEN | 4573.29453 | -0.0638696 | 0.17960935 | -0.3556028 | 0.72213805 | 0.95739595 |
| RNLS | 752.16365 | 0.63389205 | 0.20373288 | 3.11138809 | 0.0018621 | 0.18905527 |
| ANKRD22 | 182.527536 | -0.1547804 | 0.43123259 | -0.3589256 | 7.20E-01 | 0.9566409 |

| | | | | | | |
|---|---|---|---|---|---|---|
| STAMBPL1 | 187.442213 | 0.3715863 | 0.43258348 | 0.85899327 | 0.39034423 | 0.87650906 |
| ACTA2-AS1 | 8462.95568 | -0.412289 | 0.61068774 | -0.6751225 | 0.49959794 | 0.91161831 |
| ACTA2 | 13580.1403 | -0.4535986 | 0.61145325 | -0.741837 | 0.45818611 | 0.8971553 |
| FAS | 1519.19899 | 0.63911409 | 0.24288045 | 2.63139366 | 0.00850355 | 0.3779007 |
| CH25H | 184.50267 | 0.14797829 | 0.68162814 | 0.21709533 | 0.82813405 | 0.97548201 |
| LIPA | 4394.49985 | -0.1234233 | 0.19994702 | -0.61728 | 0.53705006 | 0.92105614 |
| IFIT2 | 3932.66773 | -2.8696873 | 0.83474623 | -3.437796 | NA | NA |
| AL353751.1 | 411.564494 | -0.8138195 | 0.6943161 | -1.1721168 | 0.24115018 | 0.8124487 |
| IFIT3 | 5473.43393 | -2.9004544 | 0.80599434 | -3.598604 | 0.00031993 | 0.08817364 |
| IFIT1 | 6493.64201 | -3.7158824 | 1.10052188 | -3.376473 | NA | NA |
| IFIT5 | 1288.97419 | -0.5413652 | 0.317857 | -1.7031722 | 0.08853584 | 0.65363785 |
| SLC16A12 | 4674.79511 | -0.1865074 | 0.53577488 | -0.3481078 | 0.7277592 | 0.95848028 |
| SLC16A12-AS | 111.393767 | 0.12957341 | 0.57099818 | 0.22692439 | 0.82048254 | 0.97472105 |
| PANK1 | 2174.61372 | 0.52170938 | 0.53057862 | 0.98328383 | 0.32546776 | 0.85244476 |
| KIF20B | 154.960952 | -0.0786823 | 0.27873749 | -0.282281 | 0.77772804 | 0.96836854 |
| LINC00865 | 217.576655 | 0.3048438 | 0.56181784 | 0.54260256 | 5.87E-01 | 0.93289552 |
| RPP30 | 920.284 | -0.0733367 | 0.14345949 | -0.5112017 | 0.6092098 | 0.93748023 |
| ANKRD1 | 122.495747 | -3.4538811 | 0.83533744 | -4.1347136 | 3.55E-05 | 0.02220169 |
| PCGF5 | 3560.69966 | -0.5361053 | 0.25974686 | -2.0639529 | 0.03902218 | 0.55613287 |
| HECTD2 | 695.515277 | 0.00843193 | 0.23905037 | 0.03527262 | 0.97186236 | 0.99404213 |
| PPP1R3C | 1530.22209 | 0.41008048 | 0.6828021 | 0.60058468 | 5.48E-01 | 9.24E-01 |
| TNKS2 | 3025.2812 | -0.3802861 | 0.1891637 | -2.0103544 | 0.04439369 | 0.57267043 |
| FGFBP3 | 80.4486304 | 0.00588383 | 0.49585695 | 0.01186597 | 0.99053254 | 0.99814875 |
| BTAF1 | 1918.68065 | -0.0336539 | 0.18849033 | -0.1785444 | 0.85829543 | 0.97979623 |
| CPEB3 | 568.681311 | 0.01757709 | 0.30511734 | 0.05760763 | 0.95406118 | 0.99170503 |
| 5-Mar | 2649.38061 | -0.2464931 | 0.14600964 | -1.6881971 | 0.0913734 | 0.65518742 |
| IDE | 1565.1834 | 0.1004534 | 0.17039014 | 0.58954936 | 0.55549281 | 0.92373235 |
| KIF11 | 108.507397 | 0.13241941 | 0.53442834 | 0.24777767 | 0.80430643 | 0.97287947 |
| HHEX | 341.731176 | -0.1394751 | 0.33971063 | -0.4105704 | 0.6813876 | 0.95070301 |
| EXOC6 | 636.202963 | 0.10563901 | 0.22172701 | 0.47643727 | 6.34E-01 | 0.94088547 |
| MYOF | 3468.12136 | -0.0063194 | 0.30123286 | -0.0209786 | 0.98326274 | 0.9961464 |
| CEP55 | 41.0229286 | -0.0188917 | 0.90228407 | -0.0209376 | 9.83E-01 | 0.9961464 |
| RBP4 | 7635.20685 | -0.0019027 | 0.52130471 | -0.0036499 | 9.97E-01 | 0.99976751 |
| PDE6C | 45.219293 | -0.2431409 | 0.34537883 | -0.7039833 | 0.4814432 | 0.90541935 |
| FRA10AC1 | 925.235041 | 0.17083766 | 0.12844511 | 1.33004415 | 0.18350373 | 0.7676436 |
| SLC35G1 | 143.553314 | 0.37166674 | 0.28559296 | 1.30138621 | 0.19312629 | 0.7751938 |
| PLCE1 | 846.134528 | 0.3492396 | 0.34675941 | 1.00715247 | 0.31386151 | 0.84646403 |
| NOC3L | 685.112838 | -0.0909079 | 0.23157589 | -0.392562 | 0.69464303 | 0.95225698 |
| TBC1D12 | 758.117054 | -0.1283286 | 0.1460335 | -0.8787611 | 3.80E-01 | 0.8708011 |
| HELLS | 368.764685 | 0.46911498 | 0.31177483 | 1.50465958 | 0.13241162 | 0.71284199 |
| CYP2C9 | 163.868645 | 0.34894273 | 0.72073238 | 0.4841502 | 0.62827928 | 0.94088547 |
| CYP2C8 | 133.187603 | 0.07203935 | 0.53565373 | 0.13448865 | 0.89301618 | 0.98474725 |
| PDLIM1 | 3836.88598 | 0.22410545 | 0.37245347 | 0.60170053 | 0.5473735 | 0.92359114 |
| SORBS1 | 2951.79827 | -0.3294049 | 0.4148197 | -0.7940918 | 0.42714199 | 0.88747982 |
| ALDH18A1 | 3275.2311 | 0.08537529 | 0.21207944 | 0.40256277 | 0.6872699 | 0.95140532 |
| TCTN3 | 2844.39854 | 0.3001416 | 0.08413067 | 3.56756446 | 0.00036031 | 0.08914272 |
| ENTPD1 | 1404.71084 | -0.1371838 | 0.3444709 | -0.3982449 | 0.69044964 | 0.95158502 |
| ENTPD1-AS1 | 321.348716 | -0.2161367 | 0.26242963 | -0.8235988 | 0.41016758 | 0.88249519 |
| AL365273.2 | 80.7571481 | -0.4901709 | 0.5858536 | -0.8366782 | 0.40277348 | 0.88132015 |
| CC2D2B | 84.1376643 | -0.0739863 | 0.32524486 | -0.2274788 | 0.82005149 | 0.97454841 |
| CCNJ | 434.607034 | -0.2035254 | 0.31249236 | -0.6512972 | 0.51485463 | 0.91636259 |
| ZNF518A | 1721.04959 | 0.09262104 | 0.25420947 | 0.36434931 | 0.71559718 | 0.95590747 |
| BLNK | 2172.10414 | 0.06876202 | 0.28912913 | 0.2378246 | 0.81201713 | 0.97355727 |
| TLL2 | 88.9090404 | 0.57534299 | 0.52292686 | 1.10023608 | 0.27122928 | 0.82716979 |

| | | | | | | |
|---|---|---|---|---|---|---|
| TM9SF3 | 10354.1977 | -0.0107431 | 0.15938428 | -0.0674039 | 0.94626016 | 0.99166614 |
| PIK3AP1 | 2027.80409 | -0.0756699 | 0.38433296 | -0.1968863 | 0.84391654 | 0.97664894 |
| LCOR | 1719.8751 | -0.1398216 | 0.13130005 | -1.0649016 | 0.28692048 | 0.83155545 |
| ARHGAP19 | 586.674654 | 0.12801451 | 0.18671224 | 0.68562463 | 4.93E-01 | 9.07E-01 |
| FRAT1 | 201.417286 | -0.0139292 | 0.32459488 | -0.0429125 | 0.96577127 | 0.99387827 |
| FRAT2 | 570.420978 | -0.2136327 | 0.29896386 | -0.7145771 | 4.75E-01 | 0.90226631 |
| RRP12 | 1728.74116 | -0.3122642 | 0.33232966 | -0.9396218 | 0.3474116 | 0.86197176 |
| AL355490.2 | 47.6343911 | -0.2765201 | 0.47299531 | -0.5846149 | 0.55880671 | 0.92421175 |
| PGAM1 | 5189.1959 | 0.06319554 | 0.21419274 | 0.29504054 | 0.76796291 | 0.96762687 |
| EXOSC1 | 902.018949 | -0.0395596 | 0.12905486 | -0.3065329 | 0.75919892 | 0.96502715 |
| ZDHHC16 | 2305.63078 | -0.302866 | 0.1741961 | -1.7386495 | 0.08209643 | 0.6456652 |
| MMS19 | 3612.95947 | -0.2002819 | 0.1091648 | -1.8346749 | 0.06655388 | 0.62378208 |
| UBTD1 | 1377.51249 | -0.3842871 | 0.24809216 | -1.548969 | 0.12138916 | 0.69687955 |
| ANKRD2 | 384.156148 | 0.06136521 | 0.77577581 | 0.07910174 | 0.9369517 | 0.99068685 |
| HOGA1 | 3929.79933 | 0.58297187 | 0.51886427 | 1.12355371 | 0.2612024 | 0.82424998 |
| AL355315.1 | 1839.6806 | 0.40333828 | 0.32195616 | 1.25277394 | 0.21028799 | 0.79134644 |
| MORN4 | 883.23149 | 0.23238686 | 0.29751272 | 0.7810989 | 0.43474433 | 0.89055338 |
| PI4K2A | 2435.74562 | -0.2315058 | 0.21647057 | -1.0694563 | 0.28486413 | 0.83107779 |
| AVPI1 | 581.445413 | 0.01236995 | 0.30927701 | 0.03999633 | 0.96809605 | 0.99400331 |
| MARVELD1 | 1066.35274 | -0.0785163 | 0.27000106 | -0.2908 | 0.77120426 | 0.96836854 |
| ZFYVE27 | 3610.80846 | -0.0765742 | 0.15712792 | -0.4873364 | 0.62601993 | 0.94088547 |
| GOLGA7B | 121.068048 | 0.12764813 | 0.30126836 | 0.42370241 | 0.67178286 | 0.94922827 |
| CRTAC1 | 484.856412 | 0.95745635 | 0.52610332 | 1.81990175 | 0.06877397 | 0.62750387 |
| R3HCC1L | 1045.75246 | -0.4862555 | 0.19695512 | -2.4688647 | 0.01355425 | 0.42608223 |
| LOXL4 | 832.238259 | 0.59095562 | 0.4376613 | 1.35025788 | 0.17693328 | 0.76115956 |
| AL139241.1 | 59.8354703 | 1.04331728 | 0.55707065 | 1.87286349 | 6.11E-02 | 6.10E-01 |
| PYROXD2 | 775.556682 | 0.35431567 | 0.35934428 | 0.98600614 | 3.24E-01 | 0.85169085 |
| HPS1 | 4448.66058 | -0.0834216 | 0.13662424 | -0.6105915 | 0.54147004 | 0.92292765 |
| HPSE2 | 87.776704 | 2.17791245 | 0.84108021 | 2.58942302 | 0.00961369 | 0.39368232 |
| CNNM1 | 398.288136 | -1.0799949 | 0.87914111 | -1.2284659 | 0.21927213 | 0.79847731 |
| GOT1 | 11087.5571 | 0.1536155 | 0.26540118 | 0.5788049 | 0.56272083 | 0.92452095 |
| AL391684.1 | 227.979831 | 0.14796989 | 0.45867392 | 0.32260367 | 0.74699541 | 0.96224616 |
| SLC25A28 | 3117.32163 | -0.5517112 | 0.38704338 | -1.4254506 | 1.54E-01 | 7.43E-01 |
| ENTPD7 | 328.6172 | -0.2517977 | 0.29635383 | -0.8496524 | 0.39551835 | 0.87927742 |
| CUTC | 954.856213 | 0.10204706 | 0.22938413 | 0.4448741 | 0.65641074 | 0.94545756 |
| COX15 | 2447.60484 | 0.22134783 | 0.20981396 | 1.0549719 | 0.29143818 | 0.83454586 |
| ABCC2 | 7489.52143 | -1.0874476 | 1.09620528 | -0.992011 | 0.32119218 | 0.85119105 |
| DNMBP | 2103.0621 | -0.205784 | 0.23543625 | -0.8740539 | 0.38208891 | 0.87185666 |
| ERLIN1 | 2312.57782 | 0.31768028 | 0.21614584 | 1.46974966 | 1.42E-01 | 7.28E-01 |
| CHUK | 1800.0577 | -0.1132099 | 0.26696761 | -0.4240587 | 0.671523 | 0.94922827 |
| AL138921.1 | 225.61612 | 0.00601608 | 0.19577464 | 0.0307296 | 0.97548519 | 0.99455099 |
| CWF19L1 | 1142.72731 | -0.049963 | 0.16699462 | -0.2991894 | 0.76479552 | 0.96678847 |
| BLOC1S2 | 2029.45577 | -0.3425511 | 0.33345753 | -1.0272706 | 0.30429306 | 0.84211658 |
| SCD | 4748.472 | 0.51641577 | 0.77224867 | 0.66871695 | 5.04E-01 | 9.14E-01 |
| OLMALINC | 374.014005 | 0.60346996 | 0.482547 | 1.25059311 | 0.21108297 | 0.79134644 |
| SEC31B | 1341.64433 | 0.10164561 | 0.23187131 | 0.4383708 | 0.66111751 | 0.94637197 |
| AL133352.1 | 6112.59669 | 0.17136099 | 0.12815175 | 1.33717248 | 0.18116634 | 0.76522169 |
| NDUFB8 | 6369.0242 | 0.15791849 | 0.12665494 | 1.24684039 | 0.21245603 | 0.79203904 |
| HIF1AN | 5366.64209 | 0.12809261 | 0.13095309 | 0.97815646 | 0.32799695 | 0.85317073 |
| PAX2 | 11621.4386 | -0.0623267 | 0.42511574 | -0.1466111 | 0.88343901 | 0.98293342 |
| SLF2 | 1500.70669 | -0.0377861 | 0.14284879 | -0.2645181 | 0.79138074 | 0.97090793 |
| MRPL43 | 3405.1095 | 0.1685583 | 0.14493501 | 1.1629923 | 0.24483263 | 0.81484785 |
| SEMA4G | 1967.69109 | 0.36007468 | 0.33352327 | 1.07960886 | 0.28031639 | 0.82835218 |
| AL133215.1 | 236.285481 | 0.39670982 | 0.43100071 | 0.9204389 | 0.35734345 | 0.86390849 |

| | | | | | | |
|---|---|---|---|---|---|---|
| TWNK | 1095.18632 | -0.1209394 | 0.23150862 | -0.5223967 | 0.60139412 | 0.93533006 |
| LZTS2 | 7867.92155 | -0.3555416 | 0.15875917 | -2.2395028 | 0.02512322 | 0.52085426 |
| PDZD7 | 738.746976 | -0.1496829 | 0.22636951 | -0.6612326 | 0.50846319 | 0.91439937 |
| SFXN3 | 3257.47781 | -0.0377416 | 0.24680642 | -0.1529198 | 0.8784615 | 0.98275231 |
| KAZALD1 | 173.423142 | 0.11891925 | 0.45304851 | 0.2624868 | 0.79294616 | 0.97090793 |
| BTRC | 1339.50238 | -0.1526583 | 0.19844633 | -0.7692675 | 0.44173451 | 0.89314342 |
| DPCD | 1355.75169 | 0.08569773 | 0.18678158 | 0.45881252 | 0.6463688 | 0.94208859 |
| POLL | 2073.40281 | -0.0223891 | 0.21416755 | -0.1045399 | 0.91674087 | 0.9868044 |
| FBXW4 | 4085.94774 | -0.0222002 | 0.1311245 | -0.1693063 | 0.86555576 | 0.98197545 |
| NPM3 | 655.103612 | 0.09889098 | 0.23502879 | 0.42076115 | 0.67392951 | 0.94935413 |
| OGA | 10185.316 | -0.1593985 | 0.18795546 | -0.8480653 | 0.39640161 | 0.87967259 |
| KCNIP2-AS1 | 59.4606036 | -0.0334113 | 0.5289848 | -0.0631611 | 0.94963821 | 0.99170503 |
| KCNIP2 | 273.501616 | 0.35703525 | 0.4801188 | 0.74363939 | 0.45709467 | 0.89688594 |
| ARMH3 | 1576.33142 | -0.065112 | 0.11973865 | -0.5437845 | 0.58658975 | 0.93289552 |
| HPS6 | 1113.76194 | 0.01115663 | 0.17593034 | 0.06341504 | 0.94943601 | 0.99170503 |
| LDB1 | 4130.18754 | -0.0176551 | 0.15993048 | -0.1103922 | 0.91209834 | 0.98662757 |
| PPRC1 | 2012.01695 | -0.6102801 | 0.37186977 | -1.6411124 | 0.10077409 | 0.67247782 |
| NOLC1 | 3265.07443 | -0.1864839 | 0.25998117 | -0.7172978 | 0.47319035 | 0.90179314 |
| GBF1 | 5213.30699 | -0.190109 | 0.10432169 | -1.8223342 | 6.84E-02 | 6.27E-01 |
| NFKB2 | 5882.3294 | -0.0152278 | 0.31782034 | -0.0479132 | 0.9617854 | 0.99320711 |
| PSD | 556.454492 | 0.12896031 | 0.17712641 | 0.72806933 | 0.46657115 | 0.89960721 |
| FBXL15 | 1598.8138 | 0.18503543 | 0.18995105 | 0.97412164 | 0.32999615 | 0.85377783 |
| CUEDC2 | 2636.5988 | 0.11771501 | 0.1673587 | 0.70336953 | 0.48182549 | 0.90541935 |
| C10orf95 | 253.128006 | 0.31588059 | 0.23207324 | 1.36112458 | 0.17347432 | 0.75974924 |
| MFSD13A | 357.379037 | 0.02142071 | 0.24815553 | 0.0863197 | 0.93121228 | 0.98975562 |
| ACTR1A | 6766.74515 | -0.1942544 | 0.17748968 | -1.0944548 | 0.27375557 | 0.82719197 |
| AL121928.1 | 592.262232 | -0.0984783 | 0.20006399 | -0.4922338 | 0.62255408 | 0.94016713 |
| SUFU | 1384.53957 | 0.19657334 | 0.20698656 | 0.94969134 | 0.34226911 | 0.85889221 |
| TRIM8 | 9427.98468 | 0.04742094 | 0.12045673 | 0.39367615 | 0.69382015 | 0.95222846 |
| AL391121.1 | 278.89791 | 0.26802938 | 0.25402905 | 1.05511311 | 0.2913736 | 0.83454586 |
| ARL3 | 1870.5671 | 0.45583434 | 0.19302211 | 2.36156546 | 0.01819796 | 0.47570049 |
| SFXN2 | 3207.8057 | 0.64432472 | 0.31029052 | 2.07652079 | 0.03784581 | 0.55538555 |
| WBP1L | 3715.28241 | -0.1677426 | 0.11039122 | -1.5195284 | 0.12862955 | 0.70610612 |
| CYP17A1 | 2805.46468 | -1.0079856 | 0.69221945 | -1.4561648 | 0.14534705 | 0.73126152 |
| CYP17A1-AS1 | 74.3879008 | -0.1564015 | 0.45117924 | -0.3466505 | 0.72885387 | 0.95848028 |
| AL358790.1 | 340.274609 | 0.06019136 | 0.3008673 | 0.20005951 | 0.84143404 | 0.97618641 |
| BORCS7 | 1981.00434 | 0.08838297 | 0.30236338 | 0.29230713 | 0.77005181 | 0.96836854 |
| BORCS7-ASN | 1490.48349 | 0.12297062 | 0.31955948 | 0.38481293 | 0.70037602 | 0.95355954 |
| AS3MT | 952.185981 | 0.15606929 | 0.43841764 | 0.35598314 | 0.72185319 | 0.95739595 |
| CNNM2 | 5023.71524 | -0.2212123 | 0.17475005 | -1.2658781 | 0.20555674 | 0.78571765 |
| NT5C2 | 4880.35958 | -0.2656233 | 0.22311678 | -1.1905124 | 0.23384505 | 0.80725541 |
| PCGF6 | 451.414208 | -0.016724 | 0.21095257 | -0.0792785 | 0.93681107 | 0.99068685 |
| TAF5 | 458.518737 | -0.2093584 | 0.2142526 | -0.9771567 | 0.32849161 | 0.85317073 |
| ATP5MD | 4033.59467 | 0.00744281 | 0.19913355 | 0.037376 | 0.97018521 | 0.99400331 |
| PDCD11 | 1652.00325 | -0.2226835 | 0.1539988 | -1.4460078 | 0.148175 | 0.73474739 |
| CALHM2 | 657.484351 | -0.3569476 | 0.4017637 | -0.8884515 | 0.37429794 | 0.86791605 |
| AL139339.2 | 68.3290869 | -0.277131 | 0.4212781 | -0.6578339 | 0.51064485 | 0.9147386 |
| NEURL1 | 138.340464 | 0.83191874 | 0.38768896 | 2.14584069 | 3.19E-02 | 0.54649555 |
| SH3PXD2A | 1816.79662 | 0.19323256 | 0.33645978 | 0.57431103 | 0.56575735 | 0.92513982 |
| STN1 | 945.763026 | 0.07253153 | 0.22480098 | 0.32264775 | 0.74696203 | 0.96224616 |
| SLK | 3431.4065 | -0.0724992 | 0.18154605 | -0.3993432 | 0.68964034 | 0.95158502 |
| COL17A1 | 52.2318851 | 1.42471199 | 0.52708786 | 2.70298768 | 6.87E-03 | 3.50E-01 |
| SFR1 | 246.715564 | 0.06399048 | 0.25036327 | 0.25559054 | 0.79826703 | 0.97170189 |
| CFAP43 | 376.338549 | 0.25004547 | 0.48799254 | 0.51239608 | 0.60837383 | 0.93748023 |

| | | | | | | |
|---|---|---|---|---|---|---|
| GSTO1 | 5876.0535 | -0.1108739 | 0.3466886 | -0.3198083 | 0.74911364 | 0.96249332 |
| GSTO2 | 1155.09323 | 0.34579301 | 0.26095547 | 1.32510351 | 0.18513682 | 0.76820626 |
| ITPRIP | 2654.86169 | -0.4580701 | 0.36701365 | -1.248101 | 0.21199409 | 0.79203904 |
| AL162742.2 | 302.411193 | -0.3988209 | 0.38050642 | -1.0481318 | 0.29457788 | 0.83630273 |
| CFAP58 | 43.7645583 | -0.1401825 | 0.33421614 | -0.4194367 | 0.67489702 | 0.94935413 |
| SORCS3 | 87.5038601 | 0.06967131 | 0.46993969 | 0.14825585 | 0.88214086 | 0.98275231 |
| SORCS1 | 334.802203 | 0.40590088 | 0.58449551 | 0.69444652 | 0.48740223 | 0.90573694 |
| XPNPEP1 | 3488.08131 | 0.32493327 | 0.24072266 | 1.34982418 | 1.77E-01 | 7.61E-01 |
| ADD3 | 5803.1164 | 0.56042978 | 0.23643982 | 2.37028508 | 0.01777437 | 0.4711592 |
| MXI1 | 3133.94891 | -0.042575 | 0.21789667 | -0.1953909 | 0.84508692 | 0.97675487 |
| SMNDC1 | 1567.62678 | -0.1361073 | 0.15145823 | -0.8986457 | 0.36884143 | 0.86762749 |
| DUSP5 | 2369.46825 | -0.414007 | 0.42387194 | -0.9767265 | 0.32870458 | 0.85317073 |
| SMC3 | 1954.21301 | -0.2117979 | 0.15131353 | -1.3997285 | 0.16159462 | 0.74969823 |
| RBM20 | 176.272448 | -0.7566807 | 0.29557143 | -2.5600604 | 0.0104654 | 0.40196898 |
| PDCD4-AS1 | 58.4145578 | 0.13722996 | 0.35508583 | 0.38646983 | 6.99E-01 | 0.95299633 |
| PDCD4 | 3490.84829 | -0.0028638 | 0.1854572 | -0.0154418 | 0.98767969 | 0.99747292 |
| BBIP1 | 1804.8478 | 0.04594728 | 0.11492092 | 0.39981651 | 0.68929167 | 0.95158502 |
| AL158163.1 | 42.2442234 | 0.0231792 | 0.41875856 | 0.05535218 | 0.95585789 | 0.99185043 |
| SHOC2 | 2036.20651 | -0.1552577 | 0.16557274 | -0.937701 | 0.34839809 | 0.86288527 |
| ADRA2A | 378.916869 | 0.53957824 | 0.33210534 | 1.62472015 | 0.1042222 | 0.67439401 |
| GPAM | 714.21538 | 1.29004028 | 0.53262223 | 2.4220549 | NA | NA |
| ACSL5 | 1031.76376 | 0.01628714 | 0.30835465 | 0.05281951 | 0.95787572 | 0.99223683 |
| AL157786.1 | 210.142717 | 0.04994183 | 0.24062138 | 0.20755357 | 0.83557756 | 0.97550416 |
| ZDHHC6 | 3753.14835 | 0.05287722 | 0.24803221 | 0.21318691 | 0.83118118 | 0.97550416 |
| VTI1A | 1023.99048 | -0.2125815 | 0.23950743 | -0.8875777 | 0.37476796 | 0.86791605 |
| TCF7L2 | 1471.29805 | -0.147021 | 0.18794191 | -0.7822682 | 0.43405699 | 0.89055338 |
| AL158212.2 | 82.3509753 | -0.1392379 | 0.35717544 | -0.3898305 | 0.69666191 | 0.9525827 |
| HABP2 | 2547.50181 | -0.0315557 | 0.67875655 | -0.0464904 | 0.96291937 | 0.99320711 |
| NRAP | 675.619592 | 0.0042481 | 0.72206972 | 0.00588322 | 0.9953059 | 0.99940889 |
| CASP7 | 1956.20759 | -0.442342 | 0.24875966 | -1.7781903 | 0.0753726 | 0.63773415 |
| PLEKHS1 | 91.248695 | 1.3252746 | 0.75855056 | 1.74711438 | 0.0806175 | 0.64264251 |
| DCLRE1A | 475.532922 | 0.21522788 | 0.29722744 | 0.72411846 | 0.46899301 | 0.9003198 |
| NHLRC2 | 1708.89835 | 0.06127739 | 0.12707068 | 0.48223073 | 0.62964205 | 0.94088547 |
| ADRB1 | 71.0515577 | 0.23077737 | 0.61861035 | 0.37305773 | 0.70910547 | 0.95501548 |
| CCDC186 | 1470.21986 | -0.1090866 | 0.13001155 | -0.8390528 | 0.40143966 | 0.88132015 |
| VWA2 | 57.1357 | 0.91076646 | 0.67145602 | 1.35640524 | 0.17497027 | 0.76073329 |
| AFAP1L2 | 1890.44601 | 0.69645801 | 0.3842607 | 1.81246224 | 0.0699148 | 0.62873 |
| ABLIM1 | 8809.15161 | 0.11843586 | 0.14053659 | 0.8427404 | 0.39937365 | 0.88077967 |
| FAM160B1 | 1408.15276 | -0.2509073 | 0.14392215 | -1.7433543 | 0.08127175 | 0.6439041 |
| TRUB1 | 1013.52204 | 0.19780118 | 0.18687139 | 1.05848833 | 0.28983287 | 0.83399826 |
| ATRNL1 | 341.961397 | -0.1417989 | 0.28909359 | -0.4904948 | 0.62378381 | 0.9405038 |
| GFRA1 | 944.56161 | 0.96001132 | 0.63397935 | 1.51426275 | 0.12995921 | 0.70814767 |
| C10orf82 | 62.1590837 | -0.8122431 | 0.44792365 | -1.8133516 | 0.0697776 | 0.62779744 |
| AC016825.1 | 747.960228 | 0.08056565 | 0.12225226 | 0.65901154 | 5.10E-01 | 0.91439937 |
| HSPA12A | 3221.54799 | 0.06695291 | 0.13912616 | 0.48123886 | 0.63034674 | 0.94088547 |
| ENO4 | 137.370415 | -0.1843518 | 0.29011022 | -0.6354544 | 0.52513207 | 0.91759401 |
| SHTN1 | 2853.48989 | -0.1823092 | 0.21179235 | -0.8607921 | 0.38935254 | 0.87568585 |
| SLC18A2 | 1145.01498 | -0.0563159 | 0.43330144 | -0.1299693 | 0.89659072 | 0.98474725 |
| AL391988.1 | 125.984685 | 0.0200443 | 0.34876572 | 0.05747211 | 0.95416912 | 0.99170503 |
| PDZD8 | 2992.99342 | -0.2973445 | 0.41532784 | -0.7159271 | 4.74E-01 | 0.90199039 |
| AC005871.2 | 48.8876169 | 0.06887736 | 0.52241939 | 0.13184303 | 8.95E-01 | 9.85E-01 |
| EMX2OS | 5172.14638 | 0.2176105 | 0.26197264 | 0.8306612 | 0.40616505 | 0.88132015 |
| EMX2 | 3578.4359 | 0.18880424 | 0.23414192 | 0.80636664 | 0.42003148 | 0.88451372 |
| RAB11FIP2 | 1599.25384 | 0.22344154 | 0.13877903 | 1.61005261 | 0.10738637 | 0.67707349 |

| | | | | | | |
|---|---|---|---|---|---|---|
| CASC2 | 222.093994 | 0.44603243 | 0.43460217 | 1.02630053 | 0.30474995 | 0.84230231 |
| FAM204A | 1770.45821 | -0.1263626 | 0.15838642 | -0.7978124 | 0.42497939 | 0.88573606 |
| CACUL1 | 2414.75999 | -0.190334 | 0.1765474 | -1.0780903 | 0.28099347 | 0.82937461 |
| NANOS1 | 94.6625712 | -0.0600157 | 0.47070495 | -0.1275018 | 0.89854327 | 0.98474725 |
| EIF3A | 7124.85416 | -0.1804291 | 0.28655389 | -0.6296515 | 0.52892261 | 0.91907723 |
| FAM45A | 1506.76037 | -0.0668544 | 0.22053315 | -0.3031489 | 0.7617764 | 0.96607083 |
| SFXN4 | 1657.93837 | 0.06733821 | 0.28457428 | 0.23662789 | 0.81294548 | 0.9738447 |
| PRDX3 | 12587.1024 | 0.28343221 | 0.16623806 | 1.70497788 | 0.08819855 | 0.65363785 |
| GRK5 | 897.591343 | 0.16539756 | 0.28970286 | 0.57092141 | 0.56805292 | 0.92618483 |
| RGS10 | 380.69318 | 0.10539451 | 0.42401897 | 0.24856085 | 0.8037005 | 0.97274243 |
| TIAL1 | 4445.31247 | -0.2236452 | 0.14675797 | -1.5239047 | 0.12753253 | 0.70557946 |
| BAG3 | 5028.91024 | -0.2086546 | 0.2843189 | -0.7338754 | 0.46302471 | 0.89807153 |
| INPP5F | 1381.56137 | 0.13562559 | 0.14333511 | 0.94621333 | 0.34403979 | 0.85961157 |
| MCMBP | 2629.01984 | -0.1884306 | 0.18251417 | -1.0324163 | 0.30187713 | 0.84211658 |
| SEC23IP | 2097.43513 | -0.1932808 | 0.17440173 | -1.1082506 | 0.2677536 | 0.82676682 |
| PLPP4 | 275.361775 | 0.37753516 | 0.34388287 | 1.09785975 | 0.27226573 | 0.82716979 |
| LINC01561 | 54.8041042 | 0.32013642 | 0.64136451 | 0.49914895 | 0.61767446 | 0.93924932 |
| WDR11 | 3651.82927 | 0.16222794 | 0.12067471 | 1.34434088 | 0.17883818 | 0.76120583 |
| FGFR2 | 2787.30975 | -0.1525763 | 0.4075788 | -0.3743479 | 0.70814548 | 0.95501548 |
| ATE1 | 1818.78165 | 0.08111476 | 0.19722413 | 0.41128213 | 0.68086567 | 0.95070301 |
| NSMCE4A | 1449.70544 | 0.03356089 | 0.18617059 | 0.18026957 | 0.85694095 | 0.97953896 |
| AL731566.1 | 263.454893 | 0.04632377 | 0.22224543 | 0.20843521 | 0.83488916 | 0.97550416 |
| TACC2 | 3174.08861 | -0.1533284 | 0.30229729 | -0.5072108 | 6.12E-01 | 0.93856019 |
| BTBD16 | 52.8782419 | 0.78946283 | 0.46132374 | 1.71129897 | 0.08702594 | 0.65231349 |
| PLEKHA1 | 9202.89957 | -0.0759302 | 0.26787613 | -0.2834528 | 0.77682977 | 0.96836854 |
| HTRA1 | 5497.25353 | 0.5675288 | 0.29451752 | 1.92697805 | 0.05398237 | 0.59329767 |
| CUZD1 | 301.487721 | 0.22040581 | 0.25241291 | 0.87319545 | 0.38255656 | 0.8718674 |
| FAM24B | 272.602997 | 0.47655586 | 0.32381323 | 1.4716998 | 0.14110196 | 0.72711201 |
| C10orf88 | 481.199125 | -0.3969182 | 0.16595146 | -2.3917729 | 0.01676721 | 0.46515254 |
| PSTK | 288.281565 | -0.3816779 | 0.31183363 | -1.2239793 | 0.22096003 | 0.79965146 |
| IKZF5 | 1251.17645 | -0.2553096 | 0.1904682 | -1.3404317 | 1.80E-01 | 0.76344164 |
| ACADSB | 7473.99677 | 1.31571227 | 0.61963863 | 2.12335416 | 0.03372419 | 0.55369025 |
| HMX2 | 102.221271 | -1.4493594 | 1.03300542 | -1.4030511 | NA | NA |
| BUB3 | 3272.05245 | -0.1820977 | 0.14741618 | -1.2352627 | 0.21673274 | 0.7954827 |
| CPXM2 | 1441.48926 | -0.3020904 | 0.55569833 | -0.5436231 | 0.58670089 | 0.93289552 |
| CHST15 | 1445.0968 | -0.1414385 | 0.32351817 | -0.4371888 | 0.6619744 | 0.94642611 |
| OAT | 5536.99484 | 0.11307175 | 0.30668661 | 0.36868825 | 0.71236011 | 0.95560193 |
| LHPP | 1887.76041 | 0.56707855 | 0.32739903 | 1.73207158 | 0.08326082 | 0.6471966 |
| AC068896.1 | 832.059712 | 0.11898298 | 0.22261513 | 0.53447838 | 0.5930106 | 0.93289552 |
| FAM53B | 2089.34958 | -0.0686935 | 0.21545348 | -0.3188322 | 0.74985374 | 0.96249332 |
| AL513190.1 | 114.282397 | -0.1116887 | 0.33301316 | -0.3353882 | 0.73733225 | 0.96023056 |
| EEF1AKMT2 | 889.353106 | 0.16499486 | 0.26523916 | 0.62206072 | 5.34E-01 | 0.92067136 |
| ABRAXAS2 | 1064.06996 | -0.1870484 | 0.25029245 | -0.7473195 | 0.4548707 | 0.89670027 |
| ZRANB1 | 2656.07697 | -0.4296625 | 0.24651068 | -1.7429772 | 0.0813376 | 0.6439041 |
| CTBP2 | 6547.7546 | -0.2788679 | 0.18530505 | -1.5049127 | 0.13234652 | 0.71284199 |
| AL731571.1 | 617.362436 | -0.3693328 | 0.29058185 | -1.2710112 | 0.20372466 | 0.78366254 |
| AL158835.1 | 109.620079 | 0.79144541 | 0.50026945 | 1.58203826 | 0.11364084 | 0.68214043 |
| AL158835.4 | 152.07102 | 0.35527535 | 0.26953159 | 1.3181214 | 0.18746301 | 0.76958556 |
| EDRF1 | 715.900377 | -0.137427 | 0.11513005 | -1.1936672 | 0.23260818 | 0.80653282 |
| EDRF1-AS1 | 150.399043 | -0.0800845 | 0.19779125 | -0.4048939 | 0.68555547 | 0.95097288 |
| UROS | 2011.98747 | 0.38558355 | 0.18308432 | 2.10604358 | 0.03520057 | 0.55369025 |
| BCCIP | 3263.96834 | -0.1870285 | 0.17702777 | -1.0564923 | 0.29074334 | 0.83442894 |
| DHX32 | 2414.35039 | -0.3100608 | 0.23262449 | -1.3328811 | 0.18257083 | 0.76682205 |
| FANK1 | 509.81727 | 0.06303405 | 0.36523634 | 0.17258429 | 0.8629782 | 0.98176973 |

| | | | | | | |
|---|---|---|---|---|---|---|
| ADAM12 | 148.126893 | 0.00798731 | 0.47671051 | 0.01675505 | 0.98663203 | 0.99715899 |
| DOCK1 | 3436.81341 | -0.0074968 | 0.1473857 | -0.0508652 | 0.95943293 | 0.9924381 |
| AL359094.1 | 69.9992998 | -0.1367077 | 0.39601196 | -0.3452112 | 0.72993563 | 0.95848028 |
| FAM196A | 43.1233166 | -0.2537176 | 0.57293968 | -0.4428347 | 0.65788531 | 0.94579845 |
| FOXI2 | 140.337441 | 0.57569988 | 0.8275498 | 0.69566796 | 0.48663679 | 0.90573694 |
| CLRN3 | 980.318717 | 1.09109992 | 0.50716525 | 2.15136961 | 0.03144704 | 0.54507414 |
| PTPRE | 788.312771 | -0.0328137 | 0.35243611 | -0.0931054 | 0.92581986 | 0.98876911 |
| MKI67 | 118.041906 | 0.02479129 | 0.84572495 | 0.02931366 | 0.97661443 | 0.99455099 |
| MGMT | 1566.86636 | 0.4454291 | 0.26011683 | 1.71241939 | 0.08681941 | 0.65228863 |
| EBF3 | 91.2484371 | -0.0680195 | 0.41714265 | -0.1630606 | 0.8704707 | 0.98275231 |
| C10orf143 | 223.345345 | 0.4179683 | 0.29180105 | 1.43237424 | 0.15203675 | 0.74123847 |
| GLRX3 | 2649.04723 | -0.3332297 | 0.33506354 | -0.9945268 | 0.31996647 | 0.84960209 |
| PPP2R2D | 2260.62862 | -0.4422298 | 0.27737246 | -1.5943537 | 0.11085683 | 0.68001961 |
| BNIP3 | 8878.59325 | -0.1469043 | 0.42211974 | -0.3480156 | 0.72782844 | 0.95848028 |
| AL162274.2 | 60.0105688 | -0.4944391 | 0.46184346 | -1.0705771 | 0.2843596 | 0.83107779 |
| DPYSL4 | 71.436974 | -0.9292938 | 0.76489204 | -1.2149345 | 0.22439106 | 0.80078694 |
| STK32C | 753.96364 | -0.168416 | 0.28925536 | -0.5822399 | 0.5604051 | 0.92434754 |
| LRRC27 | 764.097772 | -0.0210482 | 0.30932469 | -0.0680457 | 0.94574924 | 0.99166614 |
| PWWP2B | 986.603062 | -0.1129166 | 0.20701807 | -0.5454434 | 0.58544863 | 0.93289552 |
| INPP5A | 1463.63569 | -0.0212334 | 0.22887178 | -0.0927743 | 0.92608284 | 0.98876911 |
| NKX6-2 | 329.14977 | 0.08955272 | 0.31246518 | 0.28660065 | 7.74E-01 | 0.96836854 |
| CFAP46 | 379.152774 | 0.24681312 | 0.43115078 | 0.57245199 | 0.56701581 | 0.92561704 |
| KNDC1 | 313.930298 | -0.6448792 | 0.46700515 | -1.3808824 | 0.16731512 | 0.75558932 |
| VENTX | 168.686884 | 0.26811345 | 0.528645 | 0.50717107 | 0.61203479 | 0.93856019 |
| ADAM8 | 781.863082 | -0.0268655 | 0.36456405 | -0.0736921 | 0.9412554 | 0.99136103 |
| TUBGCP2 | 5334.53899 | 0.09914124 | 0.09881155 | 1.00333655 | 0.31569851 | 0.84739412 |
| AL360181.2 | 115.261895 | -0.0228084 | 0.3699469 | -0.0616532 | 0.95083902 | 0.99170503 |
| ZNF511 | 1382.98051 | -0.1779429 | 0.15048261 | -1.1824814 | 0.23701475 | 0.80857441 |
| ZNF511-PRA | 6080.55712 | -0.3635582 | 0.61761833 | -0.5886454 | 0.55609918 | 0.92373235 |
| PRAP1 | 5342.27177 | -0.4102302 | 0.76194491 | -0.5383987 | 0.59030181 | 0.93289552 |
| FUOM | 1119.81304 | 0.28696311 | 0.26626727 | 1.07772579 | 0.28115615 | 0.82939514 |
| ECHS1 | 19541.1403 | 0.64261541 | 0.32407195 | 1.98294052 | 0.04737407 | 0.58218854 |
| PAOX | 547.108353 | -0.2568343 | 0.19743053 | -1.3008844 | 0.19329803 | 0.77548688 |
| AL360181.3 | 3225.65499 | -0.5429681 | 0.18338894 | -2.9607463 | 0.00306895 | 0.24166015 |
| MTG1 | 3959.85495 | -0.6255374 | 0.20589489 | -3.0381395 | 0.00238044 | 0.21083339 |
| SPRN | 929.070697 | -0.9027581 | 0.28920832 | -3.1214804 | 0.00179944 | 0.18722055 |
| SCART1 | 314.85818 | -0.0397959 | 0.35600382 | -0.111785 | 0.91099392 | 0.98662757 |
| CYP2E1 | 327.542966 | -0.2841831 | 0.41995538 | -0.6766984 | 0.49859734 | 0.91100837 |
| SYCE1 | 57.7643959 | -0.6027063 | 0.84413209 | -0.7139952 | 0.47523013 | 0.90226631 |
| IQSEC3 | 779.40959 | 1.18650839 | 0.66382219 | 1.78738886 | 0.07387466 | 0.6374448 |
| AC026369.1 | 189.602185 | 0.72990001 | 0.89501495 | 0.81551711 | 0.41477637 | 0.88364943 |
| AC026369.2 | 71.2901248 | 0.71205563 | 0.58108067 | 1.22539894 | 0.22042495 | 0.79912826 |
| SLC6A12 | 8961.1631 | -0.9623268 | 0.51438453 | -1.8708315 | 0.06136844 | 0.61013576 |
| AC007406.1 | 49.6773692 | -0.6109566 | 0.44387581 | -1.3764133 | 0.1686937 | 0.75766078 |
| AC007406.4 | 420.602696 | -0.814932 | 0.50310725 | -1.6197977 | 0.10527573 | 0.67473222 |
| SLC6A13 | 9104.25869 | 0.78454416 | 0.4021187 | 1.9510263 | 0.05105392 | 0.5861488 |
| KDM5A | 1567.72332 | -0.1146226 | 0.14183357 | -0.8081484 | 4.19E-01 | 0.88451372 |
| CCDC77 | 293.632096 | 0.05145452 | 0.26382625 | 0.19503184 | 0.84536802 | 0.97675487 |
| B4GALNT3 | 1459.9244 | -0.0989883 | 0.34478142 | -0.2871044 | 0.77403237 | 0.96836854 |
| NINJ2 | 109.007733 | 0.00635307 | 0.31003632 | 0.02049136 | 0.9836514 | 0.9961464 |
| AC021054.1 | 388.301467 | 0.10362167 | 0.35656641 | 0.29060972 | 0.77134982 | 0.96836854 |
| WNK1 | 9158.1319 | 0.11223764 | 0.24445558 | 0.45913308 | 0.64613861 | 0.94208859 |
| RAD52 | 848.956752 | -0.0542418 | 0.33479235 | -0.1620163 | 0.87129304 | 0.98275231 |
| AC004803.1 | 105.86507 | 0.11953082 | 0.27606406 | 0.43298219 | 0.66502771 | 0.94692723 |

| | | | | | | |
|---|---|---|---|---|---|---|
| ERC1 | 2087.44854 | -0.1917192 | 0.15731366 | -1.2187066 | 0.22295556 | 0.80078694 |
| WNT5B | 141.128375 | -0.2745352 | 0.44471592 | -0.6173271 | 0.53701898 | 0.92105614 |
| FBXL14 | 619.645241 | -0.1572955 | 0.17803702 | -0.8834988 | 0.37696682 | 0.86987124 |
| ADIPOR2 | 3731.60749 | -0.2853747 | 0.17234757 | -1.6558091 | 0.09776048 | 0.66549286 |
| CACNA2D4 | 167.187268 | -0.688698 | 0.29479452 | -2.3361966 | 0.019481 | 0.49203956 |
| DCP1B | 777.556331 | 0.04010303 | 0.20773526 | 0.19304873 | 0.8469208 | 0.97684681 |
| CACNA1C | 660.91818 | -0.5508537 | 0.40402486 | -1.3634155 | 0.17275161 | 0.75974924 |
| CACNA1C-AS | 126.168623 | -0.545368 | 0.48407932 | -1.1266088 | 0.25990793 | 0.82424998 |
| FKBP4 | 7975.03519 | 0.02140986 | 0.13529769 | 0.15824259 | 0.87426565 | 0.98275231 |
| ITFG2-AS1 | 193.531714 | -0.0966621 | 0.2309987 | -0.4184528 | 0.67561606 | 0.94958156 |
| AC005841.1 | 2006.33778 | 0.05062667 | 0.15370164 | 0.32938279 | 0.74186638 | 0.96109001 |
| ITFG2 | 1746.50744 | 0.14448269 | 0.21821883 | 0.66210002 | 0.50790712 | 0.91439937 |
| NRIP2 | 757.219432 | 0.3614049 | 0.3148822 | 1.14774636 | 0.25107329 | 0.81815815 |
| TEX52 | 62.3516312 | 0.03690224 | 0.42986317 | 0.08584648 | 0.93158846 | 0.98979451 |
| FOXM1 | 486.981253 | 0.02423593 | 0.24217731 | 0.10007514 | 0.92028467 | 0.9874645 |
| RHNO1 | 744.045911 | 0.41215445 | 0.21745114 | 1.89538879 | 0.05804091 | 0.60193532 |
| TULP3 | 2461.14826 | -0.1076478 | 0.20878329 | -0.5155959 | 0.60613666 | 0.93708311 |
| TEAD4 | 1406.6452 | -0.4894359 | 0.34659727 | -1.4121171 | 0.1579155 | 0.74565239 |
| AC125807.2 | 396.527396 | -0.5936221 | 0.48965453 | -1.2123283 | 2.25E-01 | 8.02E-01 |
| TSPAN9 | 9831.28325 | 0.0285006 | 0.14694418 | 0.19395528 | 0.8462109 | 0.97680707 |
| CRACR2A | 60.9778561 | -0.181636 | 0.38350033 | -0.4736266 | 0.63576621 | 0.94088547 |
| PARP11 | 386.366349 | 0.03906499 | 0.25750147 | 0.15170784 | 0.87941737 | 0.98275231 |
| CCND2 | 1520.92401 | 0.30968043 | 0.50601881 | 0.6119939 | 0.54054179 | 0.92247611 |
| TIGAR | 462.484778 | -0.3410391 | 0.31156188 | -1.0946112 | 0.27368702 | 0.82719197 |
| C12orf4 | 683.515627 | -0.0336458 | 0.17369671 | -0.1937044 | 0.84640734 | 0.97681836 |
| RAD51AP1 | 66.8990715 | 0.22886427 | 0.44915006 | 0.50954967 | 0.61036699 | 0.93792127 |
| DYRK4 | 864.690387 | -0.0328605 | 0.15232284 | -0.2157295 | 0.82919863 | 0.97550416 |
| AC005832.4 | 2834.12518 | 0.24794399 | 0.13902537 | 1.78344422 | 0.07451402 | 0.63773415 |
| AKAP3 | 134.217091 | 0.71948584 | 0.62973932 | 1.14251377 | 0.25324053 | 0.81848561 |
| NDUFA9 | 4792.73316 | 0.22153114 | 0.14682514 | 1.50880933 | 0.13134752 | 0.71081672 |
| AC005833.2 | 3509.415 | 0.25391728 | 0.15167602 | 1.67407662 | 0.09411554 | 0.65783452 |
| KCNA5 | 103.047676 | -0.9600142 | 0.4959338 | -1.9357708 | 0.05289578 | 0.58742503 |
| NTF3 | 154.067448 | -1.0876189 | 0.47827308 | -2.2740541 | 0.02296273 | 0.51967919 |
| ANO2 | 75.1160601 | -0.1688122 | 0.47045695 | -0.3588261 | 0.71972518 | 0.9566409 |
| VWF | 4250.60131 | 0.39089036 | 0.74130858 | 0.52729777 | 0.59798682 | 0.93468482 |
| CD9 | 12035.9978 | -0.0451454 | 0.30257682 | -0.149203 | 0.88139346 | 0.98275231 |
| PLEKHG6 | 261.748729 | -0.4003102 | 0.36681368 | -1.0913177 | 0.27513313 | 0.82719407 |
| TNFRSF1A | 10620.7411 | -0.2224264 | 0.25265744 | -0.8803477 | 0.378671 | 0.87069278 |
| SCNN1A | 20784.0771 | 0.03426965 | 0.25249869 | 0.13572208 | 0.89204099 | 0.98474725 |
| LTBR | 10577.737 | -0.3335839 | 0.23890874 | -1.3962817 | 0.16262968 | 0.75032457 |
| AC005840.2 | 215.161013 | -0.9523589 | 0.5143824 | -1.851461 | 0.06410326 | 0.61837523 |
| CD27 | 604.443287 | 0.22375375 | 0.40228544 | 0.55620643 | 0.57806974 | 0.93130183 |
| TAPBPL | 2475.97263 | 0.20189776 | 0.37785936 | 0.53431985 | 0.59312026 | 0.93289552 |
| VAMP1 | 1006.25142 | 0.09279784 | 0.26599741 | 0.34886745 | 0.72718882 | 0.95848028 |
| AC005840.4 | 72.084665 | -0.5483266 | 0.47889879 | -1.1449739 | 0.25221996 | 0.81833349 |
| MRPL51 | 3083.20014 | 0.02612726 | 0.17349988 | 0.15058947 | 0.88029957 | 0.98275231 |
| NCAPD2 | 1367.6007 | 0.0390442 | 0.13971497 | 0.27945612 | 0.77989481 | 0.96836854 |
| GAPDH | 134460.915 | 0.31279828 | 0.32830061 | 0.95278008 | 0.34070151 | 0.85849463 |
| AC006064.4 | 67134.4575 | 0.28139001 | 0.32976567 | 0.85330292 | 0.39349134 | 0.87853481 |
| IFFO1 | 1688.41702 | -0.1568544 | 0.18847968 | -0.8322083 | 0.40529139 | 0.88132015 |
| NOP2 | 2071.67099 | -0.5454598 | 0.33383724 | -1.6339095 | 1.02E-01 | 0.67279543 |
| AC006064.6 | 5683.01291 | -0.1714711 | 0.17604899 | -0.9739967 | 0.33005818 | 0.85377783 |
| CHD4 | 7580.00352 | 0.06102592 | 0.09725107 | 0.62750904 | 0.53032561 | 0.91962754 |
| AC006064.2 | 938.577592 | 0.22288842 | 0.18719055 | 1.19070337 | 2.34E-01 | 8.07E-01 |

| | | | | | | |
|---|---|---|---|---|---|---|
| LPAR5 | 80.2949236 | -0.0520322 | 0.54239649 | -0.0959302 | 0.92357601 | 0.98858365 |
| ACRBP | 139.369353 | -0.1927097 | 0.38544257 | -0.49997 | 0.61709623 | 0.93910158 |
| ING4 | 1799.47216 | -0.03403 | 0.27595541 | -0.1233171 | 0.90185598 | 0.98572048 |
| ZNF384 | 3464.93754 | -0.3498356 | 0.19160226 | -1.8258427 | 0.06787396 | 0.62599695 |
| PIANP | 42.0104367 | 0.33593249 | 0.57699062 | 0.58221482 | 0.56042199 | 0.92434754 |
| COPS7A | 3781.64976 | 0.10485487 | 0.14248015 | 0.73592613 | 0.46177565 | 0.89762182 |
| AC125494.2 | 52.1162126 | -0.0548251 | 0.45840958 | -0.1195984 | 0.90480127 | 0.98624484 |
| MLF2 | 10997.9512 | -0.1305197 | 0.09927431 | -1.3147376 | 0.18859812 | 0.76970626 |
| PTMS | 7460.25715 | -0.1734784 | 0.29108712 | -0.5959672 | 0.55119711 | 0.92373235 |
| LAG3 | 178.725913 | -1.4011226 | 0.6486806 | -2.1599575 | 0.03077596 | 0.54443002 |
| CD4 | 1421.1181 | 0.15321922 | 0.53827946 | 0.28464623 | 0.77591519 | 0.96836854 |
| GPR162 | 549.161145 | -0.4487282 | 0.48388276 | -0.927349 | 0.35374539 | 0.86364633 |
| P3H3 | 923.671871 | -0.0286863 | 0.36884373 | -0.0777737 | 0.93800805 | 0.99113899 |
| GNB3 | 260.892915 | 0.21665869 | 0.46490595 | 0.46602692 | 0.64119623 | 0.94088547 |
| CDCA3 | 271.585929 | 0.17159238 | 0.37975742 | 0.45184734 | 0.65137897 | 0.94393131 |
| USP5 | 4087.44024 | 0.14856488 | 0.12156077 | 1.22214489 | 0.22165284 | 0.80054995 |
| TPI1 | 37576.7162 | 0.02951256 | 0.18679698 | 0.15799272 | 0.87446254 | 0.98275231 |
| SPSB2 | 1214.10795 | -0.2719916 | 0.32609039 | -0.8340987 | 0.40422538 | 0.88132015 |
| LRRC23 | 766.617371 | 0.14948981 | 0.194056 | 0.77034364 | 0.44109607 | 0.89290667 |
| ENO2 | 3122.18443 | -0.1793788 | 0.32226335 | -0.5566218 | 0.57778586 | 0.93130183 |
| ATN1 | 19593.1098 | -0.6238102 | 0.27201864 | -2.2932627 | 0.02183288 | 0.51592008 |
| C12orf57 | 3577.5818 | 0.14807503 | 0.30637986 | 0.48330537 | 0.62887893 | 0.94088547 |
| U47924.2 | 2968.88899 | 0.1581524 | 0.31281334 | 0.50558073 | 0.61315101 | 0.93879706 |
| PTPN6 | 2070.11039 | -0.060685 | 0.26499682 | -0.2290027 | 0.81886679 | 0.97454841 |
| MIR200CHG | 77.0559143 | 0.33014085 | 0.36467804 | 0.90529401 | 0.36530965 | 0.8663515 |
| U47924.1 | 75.7996895 | 0.26499929 | 0.257756 | 1.02810135 | 0.30390216 | 0.84211658 |
| PHB2 | 7154.92404 | 0.08019621 | 0.10541429 | 0.76077174 | 4.47E-01 | 8.95E-01 |
| EMG1 | 6593.74326 | -0.0969516 | 0.2136553 | -0.4537759 | 6.50E-01 | 0.94312988 |
| LPCAT3 | 6600.15896 | -0.0739499 | 0.24192884 | -0.305668 | 0.75985741 | 0.96512963 |
| C1S | 14937.6901 | 0.05245165 | 0.44245403 | 0.11854713 | 0.90563416 | 0.98631391 |
| C1R | 22975.0552 | -0.0969474 | 0.4050219 | -0.2393633 | 0.81082387 | 0.97352497 |
| C1RL | 9920.33358 | -0.0673354 | 0.3460717 | -0.1945708 | 0.84572899 | 0.97675522 |
| C1RL-AS1 | 1778.38223 | 0.1848741 | 0.29810665 | 0.62016093 | 0.53515184 | 0.92084006 |
| AC233309.1 | 330.398495 | 0.60418955 | 0.52422629 | 1.15253577 | 0.24910098 | 0.81646572 |
| RBP5 | 10813.6922 | 1.32183613 | 0.38751073 | 3.4110956 | 0.00064702 | 0.12385487 |
| AC018653.3 | 456.489036 | 0.32148014 | 0.34523165 | 0.93120126 | 0.35174947 | 0.86364633 |
| CLSTN3 | 8556.71669 | 0.39862508 | 0.20638513 | 1.93146216 | 0.05342592 | 0.59038182 |
| PEX5 | 3740.94331 | 0.19552413 | 0.23510942 | 0.83163036 | 0.40561762 | 0.88132015 |
| CD163L1 | 828.489572 | -0.3744566 | 0.54418899 | -0.6881003 | 0.49138964 | 0.90600166 |
| CD163 | 4829.64172 | -0.5811934 | 0.56409606 | -1.0303093 | 0.30286481 | 0.84211658 |
| GDF3 | 60.7735056 | -0.7530467 | 1.07102709 | -0.703107 | 0.48198904 | 0.90541935 |
| SLC2A3 | 3704.55559 | 0.23348263 | 0.69019558 | 0.33828474 | 7.35E-01 | 9.59E-01 |
| FOXJ2 | 1279.25858 | -0.0504236 | 0.15424374 | -0.3269085 | 0.74373711 | 0.96137808 |
| C3AR1 | 480.845881 | -0.302144 | 0.39025776 | -0.7742164 | 0.43880283 | 0.89175645 |
| NECAP1 | 1813.96909 | -0.1274627 | 0.15363853 | -0.8296271 | 0.40674966 | 0.88132015 |
| CLEC4A | 112.369638 | 0.18641751 | 0.33075277 | 0.56361587 | 5.73E-01 | 9.29E-01 |
| AC092111.3 | 47.3137049 | 0.10841468 | 0.35572482 | 0.30477119 | 0.76054042 | 0.96560687 |
| FAM66C | 122.863809 | 0.14574416 | 0.35489617 | 0.41066706 | 0.68131669 | 0.95070301 |
| AC092111.1 | 66.1592017 | 0.52952087 | 0.71204396 | 0.74366317 | 0.45708027 | 0.89688594 |
| FAM90A1 | 53.3295215 | 0.03692998 | 0.53770945 | 0.06868017 | 0.9452442 | 0.99166614 |
| LINC02449 | 172.723844 | 0.12616186 | 0.36403713 | 0.34656317 | 0.72891952 | 0.95848028 |
| LINC00937 | 108.235532 | -0.1502386 | 0.71525233 | -0.2100497 | 0.83362886 | 0.97550416 |
| CLEC4E | 241.350142 | -1.1637711 | 0.71933665 | -1.6178393 | 0.10569724 | 0.67526487 |
| MFAP5 | 138.890424 | -0.3633746 | 1.19087074 | -0.3051335 | 0.76026446 | 0.96545035 |

| | | | | | | |
|---|---|---|---|---|---|---|
| RIMKLB | 2952.99686 | -0.6840116 | 0.24338487 | -2.8104113 | 0.00494782 | 0.30302984 |
| A2ML1-AS1 | 84.3240762 | -0.682646 | 0.3664547 | -1.8628388 | 0.06248495 | 0.61406941 |
| PHC1 | 2718.53142 | -0.1660279 | 0.12119426 | -1.3699319 | 0.17070815 | 0.75956362 |
| M6PR | 4789.39164 | -0.1496071 | 0.19766774 | -0.7568614 | 0.44913292 | 0.89546252 |
| KLRG1 | 79.8742543 | -0.2046749 | 0.50661219 | -0.404007 | 0.68620755 | 0.95097288 |
| A2M-AS1 | 4253.50314 | 0.20674477 | 0.45166667 | 0.4577375 | 0.64714104 | 0.94264497 |
| A2M | 47170.2164 | 0.20594159 | 0.46964445 | 0.43850533 | 0.66102001 | 0.94637197 |
| PZP | 48.7399783 | -0.6172327 | 0.50527733 | -1.221572 | 0.22186951 | 0.80054995 |
| LINC00987 | 113.161548 | -0.502848 | 0.34366649 | -1.4631859 | 0.14341652 | 0.72859015 |
| KLRB1 | 88.7326993 | 0.41137039 | 0.86567078 | 0.4752042 | 0.63464143 | 0.94088547 |
| CLEC2D | 726.728079 | -0.0200818 | 0.40127443 | -0.050045 | 0.96008654 | 0.99255196 |
| CD69 | 237.705209 | -0.1163344 | 0.78676781 | -0.1478637 | 0.88245033 | 0.98275231 |
| KLRF1 | 68.6820477 | 0.43801095 | 0.64113453 | 0.68318103 | 0.49449246 | 0.90802304 |
| CLEC2B | 517.423367 | -0.3887962 | 0.32744365 | -1.1873683 | 0.23508239 | 0.80747753 |
| AC091814.1 | 51.7858883 | -0.425917 | 0.30119643 | -1.4140838 | 0.1573373 | 0.74522306 |
| CLEC12A | 57.6265536 | -0.1430873 | 0.87102255 | -0.1642751 | 0.86951456 | 0.98275231 |
| CLEC1A | 348.065991 | -0.2798147 | 0.3790451 | -0.7382096 | 0.46038711 | 0.89755243 |
| CLEC7A | 549.985192 | -0.9942427 | 0.48533542 | -2.0485683 | 0.04050435 | 0.55657615 |
| OLR1 | 456.297734 | 0.30678965 | 0.82452205 | 0.3720818 | 0.70983194 | 0.95512518 |
| TMEM52B | 4663.45534 | 1.22591989 | 0.66794413 | 1.83536293 | 0.06645193 | 0.62378208 |
| GABARAPL1 | 26882.9971 | -0.3319316 | 0.26496495 | -1.2527377 | 0.21030117 | 0.79134644 |
| AC115676.1 | 234.904546 | -0.5478402 | 0.34953647 | -1.5673335 | 0.11703677 | 0.68956727 |
| KLRD1 | 197.394106 | -0.6851312 | 0.45686199 | -1.4996459 | 1.34E-01 | 0.71505705 |
| AC022075.1 | 60.6903779 | 0.5058734 | 0.42410386 | 1.19280546 | 0.23294557 | 0.80667958 |
| KLRK1 | 224.441295 | 0.19749224 | 0.72415366 | 0.27272146 | 0.78506734 | 0.97032632 |
| KLRC4-KLRK1 | 220.799935 | 0.25029356 | 0.71405494 | 0.35052423 | 7.26E-01 | 0.95848028 |
| MAGOHB | 443.295187 | 0.21688772 | 0.21407042 | 1.01316064 | 0.31098344 | 0.84476049 |
| STYK1 | 72.7391464 | 0.63351113 | 0.60634367 | 1.04480539 | 0.29611292 | 0.83790613 |
| YBX3 | 12675.7956 | -0.2743784 | 0.29560191 | -0.9282023 | 0.35330264 | 0.86364633 |
| PRH1 | 485.008586 | -0.0006453 | 0.16194063 | -0.003985 | 0.99682045 | 0.99976751 |
| PRR4 | 193.268045 | -0.0995623 | 0.30776963 | -0.3234963 | 0.74631938 | 0.96195706 |
| AC018630.2 | 577.44904 | -0.021286 | 0.1606761 | -0.1324775 | 0.89460663 | 0.98474725 |
| AC006518.7 | 71.7046541 | -0.3704534 | 0.4358755 | -0.8499065 | 0.39537708 | 0.87927742 |
| TAS2R14 | 473.443625 | -0.0109906 | 0.14313045 | -0.076787 | 0.93879298 | 0.99113899 |
| TAS2R20 | 69.3802205 | -0.2261429 | 0.41837668 | -0.5405247 | 0.58883525 | 0.93289552 |
| SMIM10L1 | 1385.03052 | 0.01374188 | 0.19032626 | 0.0722017 | 0.94244139 | 0.99136103 |
| LINC01252 | 76.9986462 | 0.23094433 | 0.63250581 | 0.36512603 | 0.71501733 | 0.95590747 |
| ETV6 | 2172.23373 | -0.2647107 | 0.23685588 | -1.1176026 | 0.26373677 | 0.82424998 |
| BCL2L14 | 34.9579446 | -1.0159937 | 0.80794348 | -1.257506 | 0.20857048 | 0.79029911 |
| LRP6 | 2037.30887 | -0.0503304 | 0.22105491 | -0.2276828 | 0.81989286 | 0.97454841 |
| MANSC1 | 1400.79963 | 0.16333845 | 0.21844811 | 0.74772198 | 0.45462787 | 0.89670027 |
| LOH12CR2 | 76.2564371 | 0.56562366 | 0.48091316 | 1.17614511 | 0.2395369 | 0.81163641 |
| BORCS5 | 346.646913 | 0.12803084 | 0.18442526 | 0.69421532 | 0.48754719 | 0.90573694 |
| DUSP16 | 1848.27902 | -0.165287 | 0.19576868 | -0.8442973 | 0.39850327 | 0.88074856 |
| CREBL2 | 4044.96946 | 0.21942514 | 0.17806028 | 1.23230818 | 0.21783399 | 0.79657107 |
| CDKN1B | 3010.64934 | 0.26319937 | 0.17392203 | 1.51331811 | 0.13019887 | 0.70814767 |
| APOLD1 | 4463.12238 | -0.2359413 | 0.25725774 | -0.9171396 | 0.3590695 | 0.86480228 |
| DDX47 | 2602.70818 | -0.1265934 | 0.14817002 | -0.8543795 | 0.39289475 | 0.8778303 |
| GPRC5A | 1965.81343 | 0.83623937 | 0.51183169 | 1.6338171 | 0.10229726 | 0.67279543 |
| GPRC5D-AS1 | 222.498991 | -0.1250845 | 0.32650219 | -0.3831048 | 0.70164209 | 0.95424422 |
| HEBP1 | 2720.23232 | -0.0418441 | 0.15812854 | -0.2646208 | 0.79130156 | 0.97090793 |
| FAM234B | 716.015284 | -0.0202176 | 0.2964739 | -0.0681934 | 0.94563166 | 0.99166614 |
| EMP1 | 6786.15616 | -0.6992193 | 0.51051762 | -1.3696281 | 0.17080303 | 0.75956362 |
| LINC01559 | 96.6121085 | -2.2443805 | 0.92747676 | -2.4198779 | 0.01552572 | 0.45322041 |

| | | | | | | |
|---|---|---|---|---|---|---|
| ATF7IP | 2174.66738 | -0.0500444 | 0.17616022 | -0.2840844 | 0.77634573 | 0.96836854 |
| PLBD1 | 1820.40897 | 0.46703564 | 0.10810923 | 4.32003486 | 1.56E-05 | 0.01316658 |
| PLBD1-AS1 | 88.90043 | 0.34744123 | 0.40648541 | 0.85474465 | 0.39269253 | 0.8778303 |
| GUCY2C | 89.1346749 | 0.39090557 | 0.36022952 | 1.08515697 | 0.27785214 | 0.82805357 |
| HIST4H4 | 49.9754947 | -0.406017 | 0.43486783 | -0.9336561 | 0.35048132 | 0.86364633 |
| H2AFJ | 3111.33474 | -0.1647118 | 0.17619659 | -0.9348185 | 3.50E-01 | 8.64E-01 |
| WBP11 | 3543.52709 | -0.0449893 | 0.18188136 | -0.2473553 | 0.80463325 | 0.97287947 |
| C12orf60 | 11640.7503 | -0.1267389 | 0.48699953 | -0.2602445 | 0.79467519 | 0.97101097 |
| SMCO3 | 136.249714 | -0.4506456 | 0.76005556 | -0.5929114 | 0.55324048 | 0.92373235 |
| ART4 | 201.807565 | 0.51384394 | 0.61342614 | 0.83766227 | 0.40222041 | 0.88132015 |
| MGP | 27580.2153 | -0.1958293 | 0.49145909 | -0.3984651 | 0.69028739 | 0.95158502 |
| ERP27 | 1518.00732 | 0.61042297 | 0.30983901 | 1.9701295 | 0.04882353 | 0.58504358 |
| ARHGDIB | 3344.91026 | 0.21133543 | 0.40504231 | 0.52176137 | 0.60183649 | 0.9355118 |
| RERG | 2072.79286 | -0.0589423 | 0.25732278 | -0.2290596 | 0.81882259 | 0.97454841 |
| PTPRO | 1563.49264 | 0.73102733 | 0.64243469 | 1.13790139 | 0.25516166 | 0.82009684 |
| EPS8 | 4995.28255 | 0.13215121 | 0.30112842 | 0.43885332 | 0.66076782 | 0.94628868 |
| STRAP | 6947.17438 | 0.00901249 | 0.27022871 | 0.03335133 | 0.97339442 | 0.99445887 |
| DERA | 3239.87989 | 0.37727661 | 0.29856419 | 1.26363651 | 0.20636052 | 0.78731087 |
| MGST1 | 14413.4705 | 0.16851225 | 0.48022766 | 0.35090075 | 0.72566281 | 0.95848028 |
| LMO3 | 287.812921 | 0.47654222 | 0.25524041 | 1.8670328 | 0.06189701 | 0.61210783 |
| RERGL | 364.755518 | -0.6792585 | 0.55647711 | -1.2206405 | 0.22222218 | 0.80078694 |
| PIK3C2G | 294.484288 | 0.87233335 | 0.48288378 | 1.80650788 | 0.07083904 | 0.63023427 |
| PLEKHA5 | 2482.12068 | 0.08791479 | 0.24826559 | 0.35411587 | 0.72325205 | 0.95804826 |
| AEBP2 | 1694.10182 | -0.1097134 | 0.13191091 | -0.8317233 | 0.40556514 | 0.88132015 |
| PDE3A | 727.960688 | 0.03052386 | 0.2698592 | 0.11311032 | 0.90994309 | 0.98662757 |
| PYROXD1 | 2108.28342 | -0.0839504 | 0.21254065 | -0.3949849 | 0.69285401 | 0.9519683 |
| RECQL | 2017.64177 | -0.1204942 | 0.22603762 | -0.5330712 | 0.59398427 | 0.93317501 |
| GOLT1B | 1849.74472 | -0.1304964 | 0.18822095 | -0.6933147 | 0.48811208 | 0.90585781 |
| SPX | 2071.79788 | 0.51896938 | 0.32130476 | 1.61519354 | 0.10626879 | 0.67625921 |
| GYS2 | 70.2702846 | -0.0567349 | 0.53750784 | -0.1055518 | 0.91593799 | 0.9868044 |
| LDHB | 58054.139 | 0.27136141 | 0.25570562 | 1.06122585 | 2.89E-01 | 0.83295411 |
| AC010185.1 | 186.919056 | -0.2334243 | 0.3157502 | -0.7392689 | 0.45974375 | 0.89755243 |
| KCNJ8 | 790.514813 | -0.3720608 | 0.28232761 | -1.3178335 | 0.18755939 | 0.76966292 |
| ABCC9 | 658.031991 | 0.20754924 | 0.33675743 | 0.61631673 | 0.5376855 | 0.92123049 |
| CMAS | 1995.49387 | 0.20223646 | 0.185298 | 1.09141201 | 0.27509163 | 0.82719407 |
| ST8SIA1 | 78.201505 | 0.96717696 | 0.61133457 | 1.58207471 | 0.11363252 | 0.68214043 |
| C2CD5 | 2404.4477 | 0.05765434 | 0.19251436 | 0.29948076 | 0.76457325 | 0.96678847 |
| ETNK1 | 4032.11489 | -0.0160712 | 0.1418373 | -0.1133075 | 0.9097868 | 0.98662757 |
| SOX5 | 123.15963 | 0.09730999 | 0.41776944 | 0.23292749 | 0.81581771 | 0.9738447 |
| BCAT1 | 402.19414 | -1.1444159 | 0.51097873 | -2.2396548 | 0.02511335 | 0.52085426 |
| LRMP | 784.230564 | 0.38854234 | 0.36995054 | 1.05025484 | 2.94E-01 | 8.35E-01 |
| AC023510.1 | 101.195001 | 0.57144306 | 0.40101873 | 1.42497849 | 0.15416342 | 0.74258845 |
| AC023510.2 | 120.886422 | 0.48417506 | 0.41443096 | 1.16828882 | 0.24269028 | 0.81335097 |
| CASC1 | 160.811545 | 0.59234325 | 0.57118849 | 1.03703639 | 0.2997189 | 0.84142109 |
| ETFRF1 | 1430.60062 | 0.25105634 | 0.20002551 | 1.25512158 | 0.20943463 | 0.79086734 |
| KRAS | 2338.91602 | -0.0974362 | 0.1644353 | -0.5925501 | 0.55348226 | 0.92373235 |
| AC092794.2 | 42.0951757 | -0.3414214 | 0.34499422 | -0.9896439 | 0.32234821 | 0.85119105 |
| RASSF8-AS1 | 1390.63643 | -0.2848802 | 0.30704991 | -0.9277978 | 0.35351244 | 0.86364633 |
| RASSF8 | 1811.13316 | -0.5201343 | 0.29382362 | -1.7702263 | 0.07668945 | 0.63773415 |
| BHLHE41 | 1275.9589 | 0.33883746 | 0.58325399 | 0.58094324 | 5.61E-01 | 0.92452095 |
| SSPN | 533.158552 | 0.3565759 | 0.4080042 | 0.87395154 | 0.38214464 | 0.87185666 |
| AC022509.2 | 106.612908 | 0.17523092 | 0.25188162 | 0.69568762 | 0.48662448 | 0.90573694 |
| AC022509.1 | 58.0557746 | 0.48201747 | 0.56152357 | 0.85841005 | 0.39066608 | 0.87682312 |
| ITPR2 | 1988.62087 | 0.04964592 | 0.17382934 | 0.28560149 | 0.77518336 | 0.96836854 |

| | | | | | | |
|---|---|---|---|---|---|---|
| INTS13 | 992.117898 | -0.045575 | 0.18919354 | -0.2408911 | 8.10E-01 | 9.73E-01 |
| FGFR1OP2 | 2260.78689 | -0.1001021 | 0.11214741 | -0.8925938 | 0.37207476 | 0.86772685 |
| AC024896.1 | 835.21245 | 0.70903298 | 0.33860646 | 2.09397361 | 0.03626233 | 0.55384187 |
| TM7SF3 | 6583.4969 | 0.78711981 | 0.33126023 | 2.37613739 | 0.01749495 | 0.46859699 |
| MED21 | 1493.48169 | -0.1298304 | 0.16603041 | -0.7819675 | 0.43423366 | 0.89055338 |
| AC092747.4 | 561.895938 | -0.028618 | 0.17369088 | -0.164764 | 0.86912974 | 0.98275231 |
| STK38L | 1521.21587 | -0.5774755 | 0.31708934 | -1.8211759 | 0.06858013 | 0.62726218 |
| ARNTL2 | 225.951897 | -0.199976 | 0.47056346 | -0.4249714 | 0.67085753 | 0.94922827 |
| PPFIBP1 | 3693.78442 | -0.0100423 | 0.13725098 | -0.0731676 | 0.94167272 | 0.99136103 |
| AC009509.1 | 98.5069736 | -0.1819847 | 0.61302014 | -0.2968657 | 0.76656903 | 0.96729533 |
| REP15 | 92.9198491 | -0.3242351 | 0.55705938 | -0.5820477 | 0.56053454 | 0.92434754 |
| MRPS35 | 3661.94628 | 0.23840409 | 0.22808327 | 1.04525019 | 0.29590735 | 0.83750889 |
| AC009511.2 | 84.3738695 | -1.2104003 | 0.53538634 | -2.2607979 | 2.38E-02 | 0.51967919 |
| KLHL42 | 1821.30117 | -0.262591 | 0.22628178 | -1.1604603 | 0.24586146 | 0.81542272 |
| CCDC91 | 1083.4226 | 0.1700882 | 0.09379334 | 1.81343576 | 0.06976464 | 0.62779744 |
| AC022364.1 | 197.244547 | 0.16139264 | 0.3840245 | 0.42026653 | 0.67429076 | 0.94935413 |
| FAR2 | 571.509734 | 0.13262669 | 0.22850526 | 0.58040977 | 0.56163832 | 0.92452095 |
| AC009318.1 | 79.6501953 | 0.01703893 | 0.39304355 | 0.04335125 | 0.96542154 | 0.99377925 |
| AC009318.2 | 50.974645 | -0.2294295 | 0.29653321 | -0.7737059 | 0.43910474 | 0.89175645 |
| ERGIC2 | 1620.19502 | -0.0001293 | 0.11548299 | -0.0011197 | 0.99910664 | 0.99994357 |
| OVCH1-AS1 | 43.9832784 | -0.3549147 | 0.58377563 | -0.6079643 | 0.54321115 | 0.92297267 |
| TMTC1 | 2235.68271 | -0.0227292 | 0.35454556 | -0.0641081 | 0.94888418 | 0.99170503 |
| IPO8 | 2412.30394 | 0.14648019 | 0.11812693 | 1.24002369 | 0.21496663 | 0.79487183 |
| CAPRIN2 | 1483.31945 | 0.09237594 | 0.10890146 | 0.84825252 | 0.39629735 | 0.87967259 |
| TSPAN11 | 127.130061 | 0.37206802 | 0.59507684 | 0.62524367 | 0.53181115 | 0.91962754 |
| DDX11 | 1304.11371 | -0.4054678 | 0.38184102 | -1.061876 | 0.28829197 | 0.83295411 |
| SINHCAF | 2913.81318 | -0.1768154 | 0.1663643 | -1.0628208 | 0.28786322 | 0.83266626 |
| DENND5B | 1008.52479 | -0.0367849 | 0.29622587 | -0.1241786 | 0.90117388 | 0.98553843 |
| ETFBKMT | 772.236206 | 0.11434218 | 0.18524635 | 0.61724388 | 0.53707388 | 0.92105614 |
| AMN1 | 602.350357 | 0.07880014 | 0.19366046 | 0.40689845 | 0.68408257 | 0.95070301 |
| AC023157.3 | 117.153085 | 0.04741306 | 0.36111271 | 0.13129714 | 0.89554026 | 0.98474725 |
| KIAA1551 | 1227.64122 | -0.0814428 | 0.24800099 | -0.3283973 | 0.7426113 | 0.96109001 |
| BICD1 | 734.299019 | 0.16591832 | 0.21478525 | 0.77248468 | 0.43982741 | 0.89203402 |
| FGD4 | 1915.1881 | -0.1422123 | 0.21773744 | -0.6531366 | 0.51366823 | 0.91636259 |
| DNM1L | 3459.63767 | -0.0951891 | 0.0879761 | -1.0819882 | 0.27925779 | 0.82825734 |
| AC084824.5 | 66.1424992 | 0.01875867 | 0.32479654 | 0.05775516 | 0.95394366 | 0.99170503 |
| YARS2 | 690.261673 | -0.2407801 | 0.16825856 | -1.4310126 | 0.15242659 | 0.7422778 |
| PKP2 | 615.607174 | 0.4123649 | 0.38659496 | 1.06665877 | 0.28612595 | 0.83107779 |
| ALG10 | 130.410515 | -0.3905256 | 0.26520345 | -1.4725511 | 0.14087211 | 0.72669535 |
| ALG10B | 460.960416 | 0.02193725 | 0.2184208 | 0.10043574 | 0.9199984 | 0.98733173 |
| CPNE8 | 1413.2996 | -0.1773539 | 0.17262405 | -1.0273999 | 0.30423221 | 0.84211658 |
| KIF21A | 1892.35989 | 0.04597307 | 0.29446014 | 0.15612664 | 0.8759332 | 0.98275231 |
| SLC2A13 | 1157.36977 | 0.27551955 | 0.21974355 | 1.25382311 | 0.20990631 | 0.79134644 |
| LINC02471 | 185.707963 | -0.0816949 | 0.7005741 | -0.1166114 | 0.90716802 | 0.98662757 |
| AC079630.1 | 469.252754 | 0.03202528 | 0.52455607 | 0.06105215 | 0.95131768 | 0.99170503 |
| LRRK2 | 3391.31786 | 0.16958623 | 0.44048652 | 0.38499756 | 0.70023923 | 0.95355954 |
| MUC19 | 59.7159702 | 2.16963353 | 0.64443891 | 3.36670165 | 0.00076073 | 0.12960754 |
| CNTN1 | 637.573747 | 0.75788357 | 0.48487161 | 1.56306031 | 0.11803841 | 0.69079142 |
| PDZRN4 | 51.913499 | -0.4245758 | 0.81008754 | -0.524111 | NA | NA |
| GXYLT1 | 1091.77901 | -0.0688813 | 0.27349012 | -0.2518602 | 0.80114915 | 0.97210205 |
| YAF2 | 811.894607 | -0.2952067 | 0.14402737 | -2.0496567 | 0.04039794 | 0.55657615 |
| PPHLN1 | 2180.60676 | -0.1309797 | 0.14094799 | -0.9292769 | 0.35274561 | 0.86364633 |
| ZCRB1 | 1632.24631 | 0.13067574 | 0.14595184 | 0.89533461 | 0.37060825 | 0.86762749 |
| PRICKLE1 | 476.694509 | 0.42835681 | 0.28034369 | 1.52797022 | 1.27E-01 | 7.04E-01 |

| | | | | | | |
|---|---|---|---|---|---|---|
| AC079601.1 | 275.473707 | 0.36408092 | 0.3260393 | 1.11667802 | 0.26413202 | 0.82424998 |
| PUS7L | 715.493071 | 0.10315076 | 0.19181139 | 0.53777182 | 0.59073459 | 0.93289552 |
| IRAK4 | 1385.16005 | 0.1930255 | 0.15319723 | 1.25998033 | 0.20767646 | 0.78834607 |
| TWF1 | 3410.19378 | -0.0412067 | 0.23362451 | -0.1763799 | 0.85999546 | 0.98063093 |
| TMEM117 | 232.290289 | 0.43430006 | 0.24411979 | 1.77904486 | 0.07523241 | 0.63773415 |
| NELL2 | 72.8276457 | 1.91540302 | 0.87625509 | 2.18589659 | 0.02882317 | 0.53606221 |
| ANO6 | 4655.05144 | -0.168312 | 0.33394903 | -0.5040051 | 0.61425782 | 0.93895494 |
| AC008124.1 | 1101.26393 | 0.07384169 | 0.22057552 | 0.33476827 | 0.7377999 | 0.9602325 |
| ARID2 | 1041.37062 | -0.1031689 | 0.11956968 | -0.8628352 | 0.38822808 | 0.87553582 |
| SCAF11 | 3920.05617 | -0.0209194 | 0.09701683 | -0.2156266 | 0.8292788 | 0.97550416 |
| SLC38A1 | 4223.43986 | -0.2144333 | 0.23522538 | -0.9116077 | 0.36197529 | 0.86528137 |
| SLC38A2 | 10619.1888 | -0.7090581 | 0.33854369 | -2.094436 | 0.03622116 | 0.55384187 |
| AC008014.1 | 189.926089 | 0.57407089 | 0.28738346 | 1.99757803 | 0.04576243 | 0.57687945 |
| AC008035.1 | 67.7236578 | 0.56960302 | 0.3517932 | 1.61914167 | 0.10541678 | 0.67473222 |
| SLC38A4 | 560.138845 | 0.82229242 | 0.75288403 | 1.09219001 | 0.27474959 | 0.82719407 |
| AMIGO2 | 845.342732 | -0.7623065 | 0.40940983 | -1.8619644 | 0.0626081 | 0.61431334 |
| PCED1B | 376.594124 | -0.0670625 | 0.33033604 | -0.2030129 | 0.83912497 | 0.97618283 |
| RPAP3 | 986.432477 | -0.0191994 | 0.12373877 | -0.1551609 | 0.87669448 | 0.98275231 |
| AC004241.1 | 3994.88936 | 0.10872749 | 0.19324924 | 0.5626283 | 0.57368802 | 0.92978516 |
| ENDOU | 77.2849995 | -0.3704266 | 0.72865372 | -0.5083713 | 0.61119297 | 0.93834929 |
| RAPGEF3 | 15426.8538 | 0.09854412 | 0.16694952 | 0.59026295 | 0.55501437 | 0.92373235 |
| SLC48A1 | 5494.42205 | 0.0492846 | 0.17945167 | 0.27463997 | 0.78359286 | 0.96978847 |
| HDAC7 | 10722.6239 | -0.2908576 | 0.26079089 | -1.1152906 | 0.26472589 | 0.82424998 |
| AC004466.1 | 1473.35823 | -0.263026 | 0.23691738 | -1.1102012 | 0.26691235 | 0.8265123 |
| AC004466.3 | 37.7434887 | -0.5416299 | 0.41612881 | -1.3015919 | 0.19305593 | 0.7751938 |
| VDR | 2146.79109 | 0.58128185 | 0.42454284 | 1.36919479 | 0.17093839 | 0.75956362 |
| AC121338.2 | 77.9163323 | 0.42838722 | 0.43192168 | 0.99181688 | 0.32128686 | 0.85119105 |
| TMEM106C | 3878.23218 | 0.64438567 | 0.25373574 | 2.53959359 | 0.01109813 | 0.40196898 |
| SENP1 | 561.738853 | -0.1287311 | 0.19506842 | -0.6599279 | 0.50930009 | 0.91439937 |
| PFKM | 5004.54118 | 0.10669649 | 0.21637237 | 0.49311513 | 0.62193124 | 0.94016713 |
| ASB8 | 1836.85864 | 0.10091751 | 0.21168147 | 0.47674228 | 0.63354565 | 0.94088547 |
| CCDC184 | 56.1303758 | 0.2686302 | 0.58452093 | 0.45957328 | 0.64582255 | 0.94208859 |
| AC024257.3 | 42.3785513 | 0.07870764 | 0.52415757 | 0.15016027 | 0.88063817 | 0.98275231 |
| ZNF641 | 1219.98572 | -0.0811507 | 0.12224621 | -0.6638302 | 0.50679898 | 0.91420888 |
| KANSL2 | 1557.13107 | 0.00367933 | 0.15928678 | 0.02309879 | 0.98157147 | 0.99595643 |
| CCNT1 | 1508.15486 | -0.0973982 | 0.13861705 | -0.7026423 | 0.48227871 | 0.90541935 |
| AC117498.1 | 2054.26278 | 0.07575678 | 0.29681932 | 0.25522859 | 0.79854655 | 0.9717324 |
| ADCY6 | 5050.32157 | 0.05513608 | 0.25174527 | 0.21901538 | 0.82663807 | 0.97548201 |
| CACNB3 | 1016.64523 | 0.0201772 | 0.28748184 | 0.07018601 | 0.94404561 | 0.99136103 |
| DDX23 | 4344.00677 | -0.126146 | 0.14707537 | -0.8576966 | 0.39105999 | 0.87710956 |
| RND1 | 351.39539 | 1.54855775 | 0.67858789 | 2.28202976 | 0.02248758 | 0.51958567 |
| AC073610.3 | 2590.81605 | -0.1028483 | 0.17153656 | -0.5995707 | 5.49E-01 | 0.92373235 |
| CCDC65 | 263.585098 | 0.04706701 | 0.23977737 | 0.19629463 | 0.84437956 | 0.97675487 |
| FKBP11 | 1508.85278 | -0.5106198 | 0.26077209 | -1.9581077 | 0.05021738 | 0.5861488 |
| AC073610.2 | 1765.47298 | -0.2822492 | 0.16646152 | -1.695582 | 0.08996506 | 0.65375543 |
| ARF3 | 11991.3844 | -0.0739889 | 0.12514488 | -0.5912263 | 0.55436878 | 0.92373235 |
| DDN | 1096.69918 | 2.84780038 | 0.92047532 | 3.09383676 | 0.00197586 | 0.19087435 |
| AC011603.3 | 166.385381 | 3.06060847 | 1.0511192 | 2.91176155 | 0.00359397 | 0.26033637 |
| DDN-AS1 | 478.437002 | 0.00738871 | 0.13371322 | 0.05525789 | 0.95593301 | 0.99185043 |
| PRKAG1 | 3156.94879 | 0.08342493 | 0.1366423 | 0.61053519 | 0.54150734 | 0.92292765 |
| KMT2D | 4604.20731 | -0.4290939 | 0.17557743 | -2.4439014 | 1.45E-02 | 4.39E-01 |
| RHEBL1 | 70.8286075 | -1.2438789 | 0.52771595 | -2.3570993 | 0.01841832 | 0.47570049 |
| LMBR1L | 3020.31287 | -0.0567234 | 0.21683361 | -0.2615985 | 0.79363097 | 0.97090793 |
| AC011603.2 | 16815.3347 | -0.0973541 | 0.4311712 | -0.2257898 | 0.82136488 | 0.97472105 |

| | | | | | | |
|---|---|---|---|---|---|---|
| TUBA1B | 19461.2009 | -0.1085112 | 0.43050344 | -0.2520565 | 0.80099739 | 0.97210205 |
| TUBA1A | 4947.10137 | 0.01878489 | 0.47116353 | 0.03986915 | 0.96819745 | 0.99400331 |
| TUBA1C | 10044.9347 | -0.470687 | 0.49866797 | -0.9438886 | 0.34522655 | 0.8605867 |
| AC125611.3 | 6441.32459 | -0.5011351 | 0.50332089 | -0.9956573 | 0.31941667 | 0.84928695 |
| PRPH | 41.1729578 | -0.3268376 | 0.65845762 | -0.4963684 | 0.61963449 | 0.93945324 |
| TROAP | 71.9127559 | -0.0943511 | 0.7792103 | -0.1210855 | 9.04E-01 | 9.86E-01 |
| DNAJC22 | 1247.8816 | 0.71157769 | 0.35598231 | 1.99891303 | 0.04561776 | 0.57670328 |
| SPATS2 | 1220.50457 | 0.10418329 | 0.16517839 | 0.63073194 | 0.52821581 | 0.91881312 |
| KCNH3 | 265.010908 | 0.57423943 | 0.48069161 | 1.19461089 | 0.23223909 | 0.80649152 |
| MCRS1 | 3502.22389 | -0.0503052 | 0.17295887 | -0.2908507 | 0.77116548 | 0.96836854 |
| PRPF40B | 3253.02795 | -0.0277461 | 0.26328692 | -0.1053834 | 0.9160716 | 0.9868044 |
| FAM186B | 55.6750822 | 0.198284 | 0.36473958 | 0.54363172 | 0.58669493 | 0.93289552 |
| FMNL3 | 3614.47201 | -0.048469 | 0.24163605 | -0.200587 | 0.84102155 | 0.97618283 |
| TMBIM6 | 75350.3257 | 0.22179262 | 0.30043233 | 0.73824486 | 0.46036567 | 0.89755243 |
| NCKAP5L | 3187.58434 | -0.3999984 | 0.25459141 | -1.5711385 | 0.11615049 | 0.68754779 |
| BCDIN3D-AS | 282.058779 | -0.2267284 | 0.27870576 | -0.8135045 | 0.41592887 | 0.88399703 |
| BCDIN3D | 427.617243 | -0.2732285 | 0.28358069 | -0.9634946 | 0.33529935 | 0.85599307 |
| FAIM2 | 168.121105 | -0.1737821 | 0.5521155 | -0.3147567 | 0.75294637 | 0.96301881 |
| AQP2 | 52635.1034 | -0.4055886 | 0.48956638 | -0.8284649 | 0.40740727 | 0.88132015 |
| AC025154.2 | 4189.58759 | -0.2101913 | 0.45214212 | -0.4648787 | 0.64201831 | 0.94088547 |
| AC025154.1 | 9105.38323 | -0.251724 | 0.47405342 | -0.5310034 | 0.5954164 | 0.9334601 |
| AQP5 | 118.888448 | 0.69365171 | 0.57234277 | 1.21195155 | 0.22553093 | 0.80214224 |
| AQP6 | 3313.23173 | 0.99164094 | 0.56152232 | 1.76598671 | 0.07739808 | 0.63862831 |
| RACGAP1 | 816.467324 | 0.4559294 | 0.28774112 | 1.58451247 | 0.11307715 | 0.68130021 |
| ASIC1 | 203.259378 | -0.528563 | 0.44782512 | -1.1802889 | 0.23788531 | 0.80962787 |
| SMARCD1 | 3659.13719 | -0.2313149 | 0.16751152 | -1.3808895 | 0.16731293 | 0.75558932 |
| GPD1 | 11185.7582 | 2.01966584 | 0.58215392 | 3.46929869 | 0.00052182 | 0.11155563 |
| COX14 | 2399.63369 | 0.05700597 | 0.20819281 | 0.27381334 | 0.78422807 | 0.97017379 |
| AC074032.1 | 1071.89461 | 0.16765307 | 0.21098863 | 0.79460713 | 0.42684207 | 0.88709241 |
| CERS5 | 2356.80755 | -0.0692729 | 0.17560474 | -0.3944817 | 0.69322543 | 0.9519683 |
| LIMA1 | 4880.66059 | 0.03927201 | 0.20353737 | 0.19294742 | 0.84700015 | 0.97684681 |
| FAM186A | 60.6194837 | 0.2603068 | 0.51602193 | 0.50444911 | 0.61394582 | 0.93895494 |
| LARP4 | 3652.63199 | -0.1156475 | 0.20477376 | -0.5647576 | 0.57223864 | 0.92899552 |
| DIP2B | 1539.85973 | -0.4197691 | 0.24733773 | -1.6971494 | 0.08966843 | 0.65375543 |
| ATF1 | 1038.66901 | -0.3861248 | 0.22997078 | -1.6790167 | 0.0931488 | 0.65756964 |
| METTL7A | 10695.2851 | 0.65425216 | 0.37875675 | 1.72736765 | 0.08410166 | 0.64942281 |
| SLC11A2 | 4928.33787 | -0.3717922 | 0.22999873 | -1.6164966 | 0.10598699 | 0.6756711 |
| LETMD1 | 3959.57861 | 0.15205988 | 0.15025956 | 1.01198141 | 0.31154694 | 0.84529065 |
| CSRNP2 | 1945.96977 | -0.3191728 | 0.14078787 | -2.2670479 | 0.0233873 | 0.51967919 |
| TFCP2 | 1644.28129 | 0.30579894 | 0.13651109 | 2.24010334 | 0.02508421 | 0.52085426 |
| POU6F1 | 1250.52725 | -0.3095505 | 0.19430898 | -1.5930836 | 0.11114143 | 0.68001961 |
| DAZAP2 | 21661.2842 | -0.0206923 | 0.22485165 | -0.0920264 | 0.92667706 | 0.98915637 |
| SMAGP | 1187.57155 | -0.1816336 | 0.23561418 | -0.7708943 | 0.44076961 | 0.89290667 |
| BIN2 | 382.626358 | -0.1161159 | 0.53851342 | -0.2156231 | 0.82928153 | 0.97550416 |
| GALNT6 | 454.504569 | -0.1589116 | 0.40953877 | -0.3880259 | 0.69799688 | 0.9529028 |
| SLC4A8 | 436.72203 | -0.9306234 | 0.52590144 | -1.7695776 | 0.07679753 | 0.63773415 |
| SCN8A | 393.075233 | -0.1487214 | 0.42943799 | -0.3463163 | 0.72910505 | 0.95848028 |
| AC068987.2 | 56.6139623 | -0.3252863 | 0.40576483 | -0.8016623 | 0.42274836 | 0.88453836 |
| AC068987.3 | 76.6526324 | -0.1290692 | 0.31240147 | -0.4131516 | 0.67949553 | 0.95070301 |
| FIGNL2 | 1226.23136 | 0.02403233 | 0.22973182 | 0.10461035 | 9.17E-01 | 0.9868044 |
| AC068987.4 | 924.159625 | 0.04870805 | 0.22773971 | 0.21387599 | 8.31E-01 | 9.76E-01 |
| AC068987.1 | 65.4064005 | -0.1298905 | 0.51659478 | -0.251436 | 0.80147706 | 0.97210205 |
| ANKRD33 | 110.717137 | -2.5174586 | 0.95517312 | -2.6356045 | 0.00839875 | 0.37787934 |
| ACVRL1 | 2345.33287 | -0.2826191 | 0.34482417 | -0.8196034 | 0.41244223 | 0.88268208 |

| | | | | | | |
|---|---|---|---|---|---|---|
| ACVR1B | 7490.55113 | -0.1233604 | 0.17737337 | -0.6954845 | 0.48675173 | 0.90573694 |
| GRASP | 1344.22348 | -0.079479 | 0.329118 | -0.2414909 | 0.80917468 | 0.97334138 |
| NR4A1 | 12178.4916 | -0.6515325 | 0.72019295 | -0.9046638 | 0.36564352 | 0.86653075 |
| AC025259.3 | 3530.88305 | -0.5484208 | 0.72101041 | -0.7606281 | 0.4468792 | 0.89480207 |
| ATG101 | 2222.33394 | -0.3971057 | 0.27606656 | -1.4384419 | 0.1503087 | 0.73796431 |
| AC025259.1 | 1272.82598 | -0.3854669 | 0.27283287 | -1.4128315 | 0.15770528 | 0.74522306 |
| AC078864.2 | 53.3876958 | -0.178973 | 0.47972921 | -0.3730708 | 0.70909574 | 0.95501548 |
| AC078864.1 | 147.718821 | -0.0239471 | 0.55390801 | -0.0432329 | 0.96551585 | 0.99377925 |
| KRT80 | 467.116004 | -0.2026558 | 0.47866267 | -0.4233792 | 0.67201864 | 0.94922827 |
| KRT7 | 9784.49368 | 0.33596267 | 0.47951299 | 0.70063309 | 0.48353202 | 0.90541935 |
| KRT7-AS | 3645.85666 | 0.35202185 | 0.49142911 | 0.71632275 | 4.74E-01 | 9.02E-01 |
| KRT8 | 32609.3342 | 0.28311702 | 0.3033184 | 0.93339877 | 3.51E-01 | 8.64E-01 |
| KRT18 | 20073.7243 | 0.30818376 | 0.28629157 | 1.07646815 | 2.82E-01 | 8.30E-01 |
| EIF4B | 16914.1299 | 0.1999452 | 0.28252567 | 0.70770629 | 0.47912767 | 0.90480914 |
| AC068888.1 | 4565.05147 | 0.14747346 | 0.263161 | 0.56039256 | 0.57521171 | 0.93051727 |
| TNS2 | 15111.1555 | -0.1365335 | 0.25467339 | -0.5361122 | 0.59188099 | 0.93289552 |
| SPRYD3 | 4125.11432 | -0.0697328 | 0.14490747 | -0.481223 | 0.63035801 | 0.94088547 |
| IGFBP6 | 1479.20743 | -0.0702471 | 0.49469958 | -0.1419995 | 0.88708037 | 0.98423722 |
| SOAT2 | 69.2696863 | 1.53965844 | 0.73676462 | 2.08975621 | 0.03663971 | 0.55521179 |
| CSAD | 2976.29438 | 0.1089268 | 0.23215457 | 0.46919945 | 0.63892708 | 0.94088547 |
| AC073573.1 | 838.136019 | 0.03383816 | 0.24334457 | 0.13905452 | 0.88940707 | 0.98446024 |
| ZNF740 | 2619.09594 | -0.0535848 | 0.1315985 | -0.4071839 | 0.68387294 | 0.95070301 |
| ITGB7 | 831.921385 | 0.4452605 | 0.36608707 | 1.21626939 | 0.22388232 | 0.80078694 |
| RARG | 1181.14991 | -0.1200298 | 0.3577698 | -0.3354944 | 0.73725217 | 0.96023056 |
| MFSD5 | 1842.81485 | -0.3971179 | 0.26394689 | -1.5045373 | 0.13244308 | 0.71284199 |
| ESPL1 | 573.502502 | 0.6573247 | 0.71771403 | 0.91585879 | 0.35974096 | 0.86480228 |
| PFDN5 | 10187.029 | 0.18273864 | 0.10322087 | 1.77036526 | 0.07666631 | 0.63773415 |
| AC073611.1 | 690.938057 | 0.11647661 | 0.15138257 | 0.76941886 | 4.42E-01 | 0.89314342 |
| C12orf10 | 2073.74148 | 0.07929002 | 0.15443244 | 0.51342851 | 0.60765161 | 0.93739614 |
| AAAS | 3377.44369 | 0.06246112 | 0.23128359 | 0.27006292 | 0.78711185 | 0.97090793 |
| SP1 | 5000.41601 | 0.04846311 | 0.19247742 | 0.25178596 | 0.80120651 | 0.97210205 |
| PRR13 | 11754.3221 | 0.53452827 | 0.24619724 | 2.17113838 | 0.02992071 | 0.54012717 |
| PCBP2 | 29974.6596 | -0.0572596 | 0.14431188 | -0.3967766 | 0.69153219 | 0.95159158 |
| MAP3K12 | 5564.11868 | -0.0707977 | 0.11518022 | -0.6146691 | 0.53877327 | 0.9215968 |
| AC023509.4 | 93.8904942 | -0.1530757 | 0.31923928 | -0.4795014 | 0.63158195 | 0.94088547 |
| TARBP2 | 1664.72518 | -0.1755628 | 0.13750632 | -1.2767615 | 2.02E-01 | 7.81E-01 |
| AC023509.3 | 707.019702 | -0.2323923 | 0.20206117 | -1.1501087 | 0.2500991 | 0.81721208 |
| NPFF | 325.010726 | -0.2047076 | 0.30652031 | -0.6678436 | 0.50423343 | 0.91356552 |
| ATF7 | 2036.65012 | -0.1040107 | 0.11906356 | -0.8735729 | 0.38235087 | 0.87185666 |
| AC023509.2 | 236.115126 | -0.0087228 | 0.28413667 | -0.0306992 | 0.97550941 | 0.99455099 |
| ATP5MC2 | 9643.77019 | 0.26551051 | 0.07349978 | 3.61239863 | 0.00030338 | 0.08492861 |
| CALCOCO1 | 7725.29004 | -0.1397905 | 0.25299249 | -0.552548 | 0.58057299 | 0.931631 |
| HOTAIR | 42.9531573 | -0.1517774 | 0.72641909 | -0.2089392 | 0.83449573 | 0.97550416 |
| HOXC11 | 83.4194923 | -0.0654749 | 0.51367443 | -0.1274638 | 0.89857333 | 0.98474725 |
| HOXC10 | 2147.4065 | 0.06135439 | 0.2356618 | 0.26034932 | 0.79459433 | 0.97101097 |
| AC012531.3 | 505.299402 | 0.07499116 | 0.26835257 | 0.27945012 | 0.77989941 | 0.96836854 |
| HOXC6 | 1326.726 | -0.2070973 | 0.27393645 | -0.7560048 | 0.44964633 | 0.89557948 |
| HOXC9 | 434.252312 | -0.1412331 | 0.19532406 | -0.7230707 | 0.46963647 | 0.90077342 |
| HOXC-AS1 | 114.489952 | 0.13938835 | 0.28466197 | 0.48966271 | 0.62437259 | 0.94077559 |
| HOXC8 | 294.312786 | -0.4231234 | 0.35065496 | -1.206666 | 0.22756078 | 0.80300331 |
| HOXC4 | 1330.74624 | -0.2081087 | 0.38652144 | -0.5384144 | 0.59029101 | 0.93289552 |
| HOXC5 | 157.625355 | 0.13804067 | 0.4323803 | 0.31925754 | 0.74953123 | 0.96249332 |
| AC023794.1 | 212.206542 | 0.2410486 | 0.49600818 | 0.48597707 | 0.62698342 | 0.94088547 |
| AC023794.3 | 50.9089321 | 0.00821791 | 0.54717475 | 0.0150188 | 0.98801718 | 0.99754499 |

| | | | | | | |
|---|---|---|---|---|---|---|
| AC023794.4 | 97.4690868 | 0.4447785 | 0.52029117 | 0.8548646 | 0.39262611 | 0.8778303 |
| SMUG1 | 1630.6364 | -0.1798105 | 0.16452965 | -1.0928757 | 0.27444839 | 0.82719407 |
| LINC02381 | 2210.04527 | -0.3143354 | 0.32936846 | -0.954358 | 0.33990247 | 0.85814041 |
| SMUG1-AS1 | 51.238209 | -0.2411838 | 0.58016029 | -0.4157192 | 0.67761549 | 0.95018034 |
| AC023794.7 | 47.7137059 | -0.2107822 | 0.54609822 | -0.3859786 | 0.6995125 | 0.95307625 |
| CBX5 | 4520.07204 | 0.36910341 | 0.16006119 | 2.30601449 | 0.02110982 | 0.50981853 |
| AC078778.1 | 91.3025802 | -0.1354914 | 0.42917361 | -0.3157031 | 7.52E-01 | 9.63E-01 |
| HNRNPA1 | 30024.2236 | 0.13538776 | 0.21001402 | 0.64466057 | 0.51914718 | 0.91636259 |
| NFE2 | 123.287375 | -0.231717 | 0.47144064 | -0.4915083 | 0.62306698 | 0.94030676 |
| COPZ1 | 6228.03569 | 0.08970821 | 0.09880998 | 0.90788612 | 0.36393839 | 0.8663515 |
| AC079313.1 | 63.4830956 | -0.6923972 | 0.63030803 | -1.0985061 | 0.27198355 | 0.82716979 |
| GPR84 | 80.2169317 | 0.13085483 | 0.77239298 | 0.16941484 | 0.86547035 | 0.98197545 |
| ZNF385A | 1113.84982 | -0.1041829 | 0.38672655 | -0.2693967 | 0.78762444 | 0.97090793 |
| ITGA5 | 4299.44231 | -0.9755422 | 0.53042754 | -1.839162 | 0.06589137 | 0.62237346 |
| NCKAP1L | 604.351061 | -0.2471829 | 0.55272762 | -0.4472056 | 0.65472663 | 0.94532497 |
| PDE1B | 491.165924 | -0.5572367 | 0.46679387 | -1.1937533 | 0.23257448 | 0.80653282 |
| PPP1R1A | 10840.8156 | -0.7219317 | 0.46329727 | -1.5582473 | 0.11917463 | 0.69233212 |
| AC079310.1 | 92.5034987 | 2.2073349 | 1.39652459 | 1.5805915 | 0.11397147 | 0.68214043 |
| TESPA1 | 64.3292641 | 0.40560338 | 0.90474802 | 0.44830536 | 0.65393283 | 0.94510853 |
| METTL7B | 2014.57917 | 2.00160734 | 0.59085588 | 3.38764053 | 0.00070497 | 0.12703622 |
| ITGA7 | 3184.52128 | -0.2261082 | 0.51514878 | -0.4389183 | 0.66072072 | 0.94628868 |
| AC009779.3 | 3767.20549 | 0.1755484 | 0.22902908 | 0.76648957 | 4.43E-01 | 0.89366134 |
| BLOC1S1 | 2871.43656 | 0.10140184 | 0.15619586 | 0.64919673 | 0.51621122 | 0.91636259 |
| RDH5 | 1249.34051 | 0.40838138 | 0.42459662 | 0.96181025 | 0.33614493 | 0.85641741 |
| CD63 | 33207.8275 | -0.2569169 | 0.32509529 | -0.7902818 | 0.42936321 | 0.88851358 |
| AC009779.2 | 601.929223 | 0.04972656 | 0.24926893 | 0.19948962 | 0.84187976 | 0.9762188 |
| GDF11 | 729.864983 | -0.2161906 | 0.2599801 | -0.831566 | 0.40565394 | 0.88132015 |
| SARNP | 2271.93292 | 0.05483196 | 0.14033873 | 0.39071152 | 0.69601048 | 0.9525827 |
| AC023055.1 | 3992.51633 | -0.0398151 | 0.14155054 | -0.2812784 | 0.77849688 | 0.96836854 |
| ORMDL2 | 2273.08995 | -0.182045 | 0.207565 | -0.8770508 | 0.38045907 | 0.87118022 |
| DNAJC14 | 2467.98017 | -0.1820264 | 0.16071304 | -1.1326173 | 0.25737498 | 0.82115501 |
| MMP19 | 1452.13116 | -0.2317943 | 0.46002976 | -0.5038681 | 0.61435405 | 0.93895494 |
| PYM1 | 1093.95518 | -0.0726421 | 0.1552576 | -0.4678813 | 0.63986951 | 0.94088547 |
| DGKA | 1082.70985 | 0.07698905 | 0.53655645 | 0.14348732 | 8.86E-01 | 9.84E-01 |
| PMEL | 572.010113 | 0.02082972 | 0.18459077 | 0.11284268 | 0.91015528 | 0.98662757 |
| CDK2 | 1716.08724 | -0.1899613 | 0.24844001 | -0.7646166 | 0.44449989 | 0.8938278 |
| AC025162.1 | 263.436421 | -0.3052805 | 0.28222502 | -1.0816917 | 0.27938955 | 0.82825734 |
| RAB5B | 8670.9794 | -0.0609803 | 0.10306964 | -0.5916416 | 5.54E-01 | 9.24E-01 |
| SUOX | 4014.98076 | 0.8211581 | 0.38346054 | 2.14144094 | 0.0322385 | 0.54925606 |
| IKZF4 | 637.273402 | -0.0620471 | 0.20672395 | -0.3001449 | 0.76406664 | 0.96678847 |
| RPS26 | 7493.3985 | -0.0909252 | 0.26140765 | -0.3478291 | 0.72796851 | 0.95848028 |
| ERBB3 | 5840.60824 | 0.26491536 | 0.3588342 | 0.73826676 | 0.46035236 | 0.89755243 |
| AC034102.2 | 2095.67684 | 0.02302483 | 0.21322907 | 0.10798169 | 0.91401022 | 0.98662757 |
| PA2G4 | 5278.52196 | -0.079416 | 0.20327647 | -0.3906798 | 0.69603391 | 0.9525827 |
| AC034102.4 | 1208.19812 | -0.0877017 | 0.24694155 | -0.3551517 | 0.72247598 | 0.95741464 |
| RPL41 | 16699.7328 | 0.17915162 | 0.1552868 | 1.15368221 | 0.24863048 | 0.81646572 |
| ZC3H10 | 573.889928 | -0.2561102 | 0.17769722 | -1.4412732 | 0.14950752 | 0.73670625 |
| ESYT1 | 6000.26633 | -0.1024454 | 0.18491649 | -0.5540091 | 0.57957264 | 0.93139743 |
| AC034102.6 | 394.616052 | -0.115727 | 0.23671442 | -0.4888888 | 6.25E-01 | 0.94088547 |
| AC034102.5 | 379.367425 | 0.03937579 | 0.19964713 | 0.19722693 | 0.84364995 | 0.97652052 |
| MYL6B | 21541.9775 | -0.0524179 | 0.24346603 | -0.2152985 | 0.8295346 | 0.97550416 |
| MYL6 | 45596.1512 | -0.086141 | 0.23650831 | -0.3642196 | 0.71569402 | 0.95590747 |
| SMARCC2 | 5168.79965 | -0.1209407 | 0.15094065 | -0.8012468 | 0.42298878 | 0.88453836 |
| RNF41 | 2074.89359 | -0.1435359 | 0.15667209 | -0.9161549 | 0.35958568 | 0.86480228 |

| | | | | | | |
|---|---|---|---|---|---|---|
| NABP2 | 1227.11139 | -0.0109196 | 0.19626706 | -0.0556366 | 0.95563134 | 0.99185043 |
| SLC39A5 | 10145.9168 | 0.56659421 | 0.53287872 | 1.06327047 | 0.28765931 | 0.83259046 |
| ANKRD52 | 5332.7048 | -0.1259737 | 0.16853771 | -0.7474509 | 0.45479145 | 0.89670027 |
| COQ10A | 988.511243 | 0.08241844 | 0.20698292 | 0.39818955 | 0.69049047 | 0.95158502 |
| AC073896.5 | 157.620425 | 0.23240157 | 0.25977396 | 0.89462995 | 0.37098494 | 0.86762749 |
| CS | 10063.862 | 0.02754819 | 0.20505369 | 0.13434622 | 0.89312881 | 0.98474725 |
| AC073896.1 | 4598.15594 | -0.0359798 | 0.1477329 | -0.2435463 | 8.08E-01 | 9.73E-01 |
| AC073896.2 | 761.042915 | 0.07874944 | 0.2554428 | 0.30828599 | 7.58E-01 | 0.96502715 |
| CNPY2 | 4203.81816 | -0.0482279 | 0.12950738 | -0.372395 | 0.70959877 | 0.95501548 |
| PAN2 | 3477.40388 | 0.04766913 | 0.21516591 | 0.22154591 | 0.82466739 | 0.97548201 |
| IL23A | 70.8220849 | -0.6484143 | 0.52583297 | -1.2331184 | 0.21753159 | 0.7960866 |
| STAT2 | 6486.41714 | -0.9265144 | 0.47644895 | -1.9446248 | NA | NA |
| TIMELESS | 911.662648 | -0.4599926 | 0.20081284 | -2.2906533 | 0.02198347 | 0.51639767 |
| MIP | 303.012397 | 0.005325 | 0.19732563 | 0.02698587 | 0.97847101 | 0.99476647 |
| SPRYD4 | 1845.10625 | -0.0616487 | 0.158174 | -0.3897527 | 0.69671945 | 0.9525827 |
| GLS2 | 385.56503 | 0.32366197 | 0.21007863 | 1.54067064 | 0.12339697 | 0.69887656 |
| RBMS2 | 3476.49905 | -0.1176085 | 0.2586224 | -0.4547497 | 0.64928931 | 0.94312988 |
| BAZ2A | 5869.8226 | -0.2946993 | 0.19006053 | -1.5505549 | 0.12100838 | 0.69543637 |
| ATP5F1B | 52477.6977 | 0.47095487 | 0.16803854 | 2.80265978 | 0.00506831 | 0.30823792 |
| PTGES3 | 12614.5568 | 0.05240686 | 0.10935744 | 0.4792254 | 0.63177829 | 0.94088547 |
| NACA | 21227.9991 | 0.26169847 | 0.17278427 | 1.51459658 | 0.1298746 | 0.70814767 |
| PRIM1 | 127.32941 | 0.36748269 | 0.27524684 | 1.3351023 | 0.18184287 | 0.76543911 |
| HSD17B6 | 92.7139264 | -0.3425218 | 0.47250402 | -0.7249077 | 0.46850864 | 0.89984974 |
| RDH16 | 55.4677913 | 0.33548049 | 0.40052319 | 0.83760566 | 0.40225221 | 0.88132015 |
| GPR182 | 157.300484 | -0.4914367 | 0.56771431 | -0.8656408 | 0.38668715 | 0.87432552 |
| ZBTB39 | 294.655868 | -0.0125248 | 0.16888829 | -0.0741605 | 0.94088266 | 0.99136103 |
| NEMP1 | 652.393852 | -0.0126763 | 0.18913863 | -0.0670211 | 0.94656491 | 0.99166614 |
| NAB2 | 2657.70465 | -0.1885687 | 0.22933378 | -0.8222455 | 0.41093718 | 0.88268208 |
| STAT6 | 16930.2194 | -0.2172146 | 0.14032179 | -1.5479745 | 1.22E-01 | 6.97E-01 |
| LRP1 | 7107.54056 | 0.03635479 | 0.37649694 | 0.09656064 | 9.23E-01 | 9.89E-01 |
| NXPH4 | 39.6521782 | 0.48671258 | 0.55435927 | 0.87797319 | 0.37995827 | 0.87090707 |
| SHMT2 | 4216.62809 | 0.15932955 | 0.18437852 | 0.86414378 | 0.38750891 | 0.87460898 |
| AC137834.2 | 344.186462 | 0.33536743 | 0.24113285 | 1.39079942 | 0.16428626 | 0.75286112 |
| NDUFA4L2 | 936.029038 | -0.135156 | 0.58583393 | -0.2307071 | 0.81754235 | 0.97433259 |
| STAC3 | 239.090277 | -0.0972129 | 0.4329121 | -0.2245557 | 0.82232495 | 0.97504706 |
| AC137834.1 | 1676.19115 | 0.17221049 | 0.22881703 | 0.7526122 | 0.45168299 | 0.89643025 |
| R3HDM2 | 3261.21507 | 0.16220612 | 0.21333341 | 0.76034092 | 0.44705083 | 0.89480207 |
| INHBE | 54.2538265 | 0.79082025 | 0.83997167 | 0.94148442 | 0.34645667 | 0.86135217 |
| GLI1 | 157.073713 | 0.23661653 | 0.6203652 | 0.38141491 | 0.7028954 | 0.95431462 |
| ARHGAP9 | 757.358674 | -0.4411693 | 0.58437486 | -0.7549423 | 0.45028359 | 0.89557948 |
| MARS | 6669.17125 | -0.3534991 | 0.21644242 | -1.6332248 | 0.10242172 | 0.67279543 |
| DDIT3 | 3749.79547 | -0.5566518 | 0.48110574 | -1.157026 | 0.24726175 | 0.81566357 |
| AC022506.1 | 725.028446 | -0.5210013 | 0.54858188 | -0.9497239 | 0.34225257 | 0.85889221 |
| MBD6 | 8138.29918 | -0.3955749 | 0.24365849 | -1.6234809 | 0.10448664 | 0.67439401 |
| DCTN2 | 7579.37136 | -0.0562322 | 0.16565945 | -0.3394447 | 0.73427472 | 0.95895327 |
| KIF5A | 81.5510841 | 0.04929326 | 0.39722579 | 0.12409381 | 0.901241 | 0.98553843 |
| PIP4K2C | 3974.12157 | 0.0111852 | 0.21098764 | 0.05301354 | 0.95772112 | 0.99223378 |
| DTX3 | 3190.67102 | -0.0392186 | 0.2810062 | -0.1395648 | 0.88900387 | 0.98423722 |
| ARHGEF25 | 1112.47873 | -0.3424907 | 0.47808782 | -0.7163762 | 0.47375911 | 0.90199039 |
| AC025165.1 | 658.584097 | -0.3220363 | 0.47819023 | -0.673448 | 0.50066229 | 0.91202488 |
| SLC26A10 | 289.703389 | -0.4503092 | 0.43926764 | -1.0251363 | 0.30529887 | 0.84268008 |
| B4GALNT1 | 289.789139 | -0.5516703 | 0.45160151 | -1.2215865 | 0.22186405 | 0.80054995 |
| OS9 | 16480.5562 | 0.27692889 | 0.10196195 | 2.71600237 | 0.00660754 | 0.34113496 |
| AC025165.2 | 6235.33365 | 0.24055918 | 0.13367138 | 1.79963114 | 0.0719189 | 0.63189503 |

| | | | | | | |
|---|---|---|---|---|---|---|
| AGAP2 | 350.648411 | -0.1821406 | 0.39245637 | -0.464104 | 0.64257327 | 0.94090101 |
| AGAP2-AS1 | 233.466903 | -0.2312072 | 0.3403074 | -0.679407 | 0.49688001 | 0.90893395 |
| TSPAN31 | 4816.51467 | -0.1385397 | 0.13511003 | -1.0253844 | 0.30518185 | 0.84268008 |
| CDK4 | 5023.3688 | -0.3372048 | 0.24030471 | -1.4032386 | 0.16054571 | 0.74816787 |
| 9-Mar | 2282.50746 | -0.0426612 | 0.16938387 | -0.2518612 | 8.01E-01 | 0.97210205 |
| CYP27B1 | 1906.09896 | 0.30188448 | 0.62255156 | 0.48491482 | 0.62773677 | 0.94088547 |
| METTL1 | 1206.48243 | 0.13249698 | 0.27657287 | 0.47906717 | 0.63189084 | 0.94088547 |
| EEF1AKMT3 | 1101.4545 | 0.21973301 | 0.26513135 | 0.82877038 | 0.40723435 | 0.88132015 |
| AC025165.3 | 927.000113 | 0.18707574 | 0.17591063 | 1.06347034 | 0.28756871 | 0.83245678 |
| TSFM | 2387.75317 | 0.13864315 | 0.14867374 | 0.93253287 | 0.35106121 | 0.86364633 |
| AVIL | 723.282468 | -0.1740742 | 0.26826423 | -0.6488909 | 0.5164089 | 0.91636259 |
| AC025165.5 | 47.6409322 | 0.30345236 | 0.39091069 | 0.7762703 | 0.4375894 | 0.89130865 |
| CTDSP2 | 10619.8585 | -0.2117141 | 0.17804414 | -1.1891103 | 2.34E-01 | 8.07E-01 |
| AC084033.3 | 210.279014 | 0.42830559 | 0.19921226 | 2.14999613 | 0.03155552 | 0.54507414 |
| ATP23 | 217.976565 | 0.34422023 | 0.24430582 | 1.40897266 | 0.15884325 | 0.74706632 |
| LRIG3 | 758.092672 | 0.04633642 | 0.32634133 | 0.14198759 | 0.88708981 | 0.98423722 |
| SLC16A7 | 1879.00339 | -0.2034997 | 0.32289784 | -0.6302294 | 0.52854449 | 0.91888066 |
| FAM19A2 | 63.5891034 | 0.42014179 | 0.58990189 | 0.71222317 | 0.47632659 | 0.90309179 |
| USP15 | 2272.74221 | -0.3596346 | 0.17512328 | -2.053608 | 0.04001365 | 0.55657615 |
| MON2 | 2457.63616 | -0.1496003 | 0.09851949 | -1.5184843 | 0.12889235 | 0.70633634 |
| LINC01465 | 59.9745221 | -1.2572666 | 0.55296171 | -2.2736956 | 0.02298429 | 0.51967919 |
| AC048341.1 | 880.036783 | -0.8111837 | 0.34574995 | -2.3461571 | 0.01896811 | 0.48430706 |
| AC048341.2 | 382.025022 | -0.4481613 | 0.36804548 | -1.217679 | 0.22334598 | 0.80078694 |
| PPM1H | 1810.94562 | 0.14765912 | 0.29110097 | 0.50724365 | 0.61198387 | 0.93856019 |
| AVPR1A | 458.923008 | -0.5262085 | 0.62541944 | -0.841369 | 0.40014123 | 0.88077967 |
| DPY19L2 | 1804.8795 | 0.43027208 | 0.29750235 | 1.44628129 | 0.14809832 | 0.73474739 |
| RXYLT1 | 1047.1922 | 0.08904033 | 0.21561495 | 0.4129599 | 0.679636 | 0.95070301 |
| RXYLT1-AS1 | 224.997486 | 0.05497003 | 0.23529253 | 0.2336242 | 0.81527674 | 0.9738447 |
| SRGAP1 | 1027.65846 | -0.1493523 | 0.15492835 | -0.964009 | 0.33504143 | 0.85581042 |
| AC025576.1 | 288.75528 | -0.0132319 | 0.19295952 | -0.0685734 | 0.94532918 | 0.99166614 |
| C12orf66 | 422.83825 | 0.23261474 | 0.23814165 | 0.97679149 | 0.32867239 | 0.85317073 |
| C12orf56 | 70.4073917 | 0.52387527 | 0.42407228 | 1.23534428 | 0.2167024 | 0.7954827 |
| XPOT | 2699.43203 | -0.2178397 | 0.2557885 | -0.85164 | 0.39441394 | 0.87884163 |
| TBK1 | 1533.54249 | -0.2706364 | 0.20622839 | -1.3123142 | 0.18941415 | 0.76970626 |
| RASSF3 | 971.279009 | -0.2547149 | 0.27961951 | -0.9109338 | 0.36233027 | 0.86561928 |
| GNS | 15751.406 | 0.25494526 | 0.24187905 | 1.05401963 | 0.29187393 | 0.83460625 |
| TBC1D30 | 489.59004 | -0.063151 | 0.21097729 | -0.299326 | 0.76469133 | 0.96678847 |
| LEMD3 | 1189.65264 | -0.0231985 | 0.13915888 | -0.1667049 | 0.86760228 | 0.98224588 |
| MSRB3 | 1062.10645 | -0.1158028 | 0.47003646 | -0.2463699 | 0.8053959 | 0.97317374 |
| HMGA2 | 50.0160518 | -1.0926326 | 0.87815701 | -1.2442338 | NA | NA |
| AC090673.1 | 75.8729111 | 0.12112286 | 0.58796108 | 0.2060049 | 8.37E-01 | 0.97577999 |
| LLPH | 1094.56121 | -0.3049659 | 0.2358471 | -1.293066 | 0.19598823 | 0.77791948 |
| AC078927.1 | 3800.23798 | 0.09102797 | 0.14672025 | 0.62041857 | 0.53498225 | 0.92084006 |
| TMBIM4 | 3992.66481 | 0.18667942 | 0.1997835 | 0.93440861 | 0.35009316 | 0.86364633 |
| IRAK3 | 892.102464 | -0.1577625 | 0.42652668 | -0.3698772 | 0.71147397 | 0.95560193 |
| AC078889.1 | 119.763078 | -0.0800173 | 0.46436972 | -0.1723139 | 0.8631908 | 0.98176973 |
| HELB | 476.837324 | -0.1234035 | 0.36375851 | -0.3392457 | 0.73442467 | 0.95895327 |
| GRIP1 | 295.243948 | -0.0661279 | 0.21117023 | -0.3131498 | 0.75416686 | 0.96348234 |
| CAND1 | 3996.29142 | -0.1077701 | 0.10924434 | -0.9865051 | 0.32388533 | 0.85156215 |
| DYRK2 | 1198.49915 | 0.15462128 | 0.13021294 | 1.18744935 | 0.23505042 | 0.80747753 |
| MDM1 | 557.156619 | 0.06206742 | 0.19322264 | 0.3212223 | 0.74804194 | 0.96241535 |
| LINC02384 | 91.8420204 | 0.39024133 | 0.54796535 | 0.71216424 | 0.47636308 | 0.90309179 |
| RAP1B | 4367.74263 | -0.1139024 | 0.26647223 | -0.4274457 | 0.66905472 | 0.9486589 |
| AC090061.1 | 39.7598174 | 0.28534533 | 0.46796123 | 0.60976276 | 0.54201897 | 0.92292765 |

| | | | | | | |
|---|---|---|---|---|---|---|
| NUP107 | 1265.68531 | 0.01928995 | 0.23634255 | 0.08161863 | 0.93494999 | 0.99068685 |
| SLC35E3 | 964.287729 | -0.3746552 | 0.30446124 | -1.2305515 | 0.21849065 | 0.79772711 |
| MDM2 | 4312.66057 | 0.21323562 | 0.16864795 | 1.26438309 | 0.20609256 | 0.78679582 |
| AC025423.4 | 1108.91491 | 0.24788219 | 0.200686 | 1.23517431 | 0.21676564 | 0.7954827 |
| CPM | 5247.74545 | -0.3442544 | 0.35820672 | -0.9610495 | 0.33652726 | 0.85679125 |
| CPSF6 | 3755.57835 | -0.2766645 | 0.17510655 | -1.579978 | 0.1141119 | 0.68259531 |
| AC020656.2 | 38.3652191 | 0.74949298 | 0.81630333 | 0.91815499 | 0.35853773 | 0.86480228 |
| LYZ | 1847.09382 | 1.33746998 | 0.87113474 | 1.53531931 | 0.12470541 | 0.70225485 |
| AC020656.1 | 865.829631 | 1.32984156 | 0.86895494 | 1.53039185 | 0.12591976 | 0.70347499 |
| YEATS4 | 559.588534 | -0.0411191 | 0.30747999 | -0.1337294 | 0.89361654 | 0.98474725 |
| FRS2 | 908.368604 | -0.3468915 | 0.1915974 | -1.8105232 | 0.07021469 | 0.62928077 |
| CCT2 | 6459.71521 | -0.1995365 | 0.2435286 | -0.8193554 | 0.41258369 | 0.88277555 |
| BEST3 | 52.0579343 | 1.0053835 | 0.84684166 | 1.18721545 | 0.23514265 | 0.80747753 |
| RAB3IP | 3187.76119 | 0.10853175 | 0.15410466 | 0.70427302 | 0.48126277 | 0.90541935 |
| AC025263.2 | 518.044116 | 0.02370047 | 0.21976786 | 0.10784319 | 0.91412008 | 0.98662757 |
| MYRFL | 111.893411 | 0.24734363 | 0.53997863 | 0.45806188 | 0.64690798 | 0.94246942 |
| AC025159.1 | 277.192647 | 0.02244626 | 0.33501802 | 0.06700015 | 0.94658159 | 0.99166614 |
| CNOT2 | 2523.00566 | -0.0853918 | 0.09763489 | -0.8746038 | 0.38178954 | 0.87185666 |
| KCNMB4 | 500.367993 | 0.34806369 | 0.257759 | 1.35034547 | 0.17690519 | 0.76115956 |
| AC025569.1 | 875.778604 | 0.24878742 | 0.42901347 | 0.57990585 | 0.56197811 | 0.92452095 |
| PTPRB | 2914.70602 | 0.12487359 | 0.41337064 | 0.30208626 | 0.76258631 | 0.96630779 |
| TSPAN8 | 2231.90739 | -0.2812172 | 0.46203443 | -0.6086498 | 0.5427566 | 0.92292765 |
| LGR5 | 43.460278 | 0.19117296 | 1.23235079 | 0.15512869 | 0.87671986 | 0.98275231 |
| ZFC3H1 | 2201.02642 | -0.2658914 | 0.13333942 | -1.9940942 | 0.04614176 | 0.57803664 |
| THAP2 | 398.724392 | 0.10346625 | 0.22445542 | 0.46096569 | 0.64482322 | 0.94208859 |
| AC073612.1 | 518.512125 | 1.46E-01 | 0.34977908 | 4.18E-01 | 0.67605197 | 0.94959168 |
| TMEM19 | 1894.9883 | 0.24582416 | 0.26361876 | 0.93249875 | 0.35107883 | 0.86364633 |
| RAB21 | 4788.19609 | -0.0968383 | 0.14756972 | -0.656221 | 0.51168195 | 0.91510941 |
| AC089984.1 | 175.509535 | 0.16539779 | 0.38214548 | 0.43281368 | 0.66515013 | 0.94694459 |
| TBC1D15 | 2903.62574 | -0.2588135 | 0.22766328 | -1.136826 | 0.25561103 | 0.82052927 |
| TRHDE | 921.916664 | -0.0265921 | 0.28116443 | -0.0945786 | 0.92464956 | 0.98869145 |
| ATXN7L3B | 4899.54739 | -0.2241476 | 0.13543924 | -1.6549679 | 0.09793101 | 0.66549286 |
| AC025257.1 | 444.849566 | -0.0305927 | 0.39748669 | -0.0769652 | 0.93865121 | 0.99113899 |
| CAPS2 | 303.818234 | 0.2567534 | 0.3905367 | 0.65743732 | 0.51089976 | 0.91474443 |
| GLIPR1L1 | 38.9126493 | 0.18840918 | 0.39950801 | 0.47160301 | 0.63721018 | 0.94088547 |
| GLIPR1L2 | 70.5017439 | 0.94501752 | 0.71817371 | 1.31586204 | 0.18822035 | 0.76970626 |
| GLIPR1 | 1326.2173 | -0.3622242 | 0.25521953 | -1.4192652 | 0.15582172 | 0.74522306 |
| AC121761.1 | 118.292345 | -1.0501374 | 0.77300734 | -1.358509 | 0.17430224 | 0.75974924 |
| KRR1 | 1989.20941 | -0.1831377 | 0.14665461 | -1.2487688 | 0.21174964 | 0.79177972 |
| PHLDA1 | 2768.05034 | -0.2620962 | 0.4128499 | -0.6348463 | 0.52552865 | 0.91759401 |
| AC011611.3 | 767.001146 | -0.5050878 | 0.49271366 | -1.0251142 | 0.30530929 | 0.84268008 |
| NAP1L1 | 9433.3143 | 0.08599594 | 0.2311646 | 0.3720117 | 0.70988414 | 0.95512518 |
| BBS10 | 704.892744 | 0.03676776 | 0.13936601 | 0.26382158 | 0.7919174 | 0.97090793 |
| OSBPL8 | 2688.06153 | 0.22294726 | 0.24912934 | 0.89490566 | 0.37083752 | 0.86762749 |
| ZDHHC17 | 1799.32771 | -0.1847816 | 0.1471123 | -1.2560584 | 0.20909479 | 0.79086734 |
| CSRP2 | 2287.1386 | 0.06336531 | 0.36628592 | 0.17299411 | 0.86265606 | 0.98176973 |
| NAV3 | 103.250053 | 0.06907927 | 0.66003908 | 0.10465936 | 0.91664611 | 0.9868044 |
| SYT1 | 163.037694 | 0.78719746 | 0.56546541 | 1.39212309 | 0.16388513 | 0.75265041 |
| AC027288.3 | 236.363174 | 1.31339369 | 0.83515063 | 1.57264288 | 0.11580153 | 0.6873836 |
| PAWR | 3296.9086 | -0.4406674 | 0.17365698 | -2.5375736 | 0.01116239 | 0.40196898 |
| AC073569.2 | 112.389619 | -0.7268245 | 0.38476561 | -1.889006 | 0.05889102 | 0.60343175 |
| PPP1R12A | 2171.12556 | -0.1353952 | 0.13985687 | -0.9680983 | 0.33299529 | 0.855011 |
| OTOGL | 379.788764 | 1.98622967 | 0.90371189 | 2.19785719 | 0.02795928 | 0.53262541 |
| PTPRQ | 439.051405 | -0.3995441 | 0.89340099 | -0.4472169 | 0.65471843 | 0.94532497 |

| | | | | | | |
|---|---|---|---|---|---|---|
| LIN7A | 1015.00032 | 0.47094917 | 0.24868519 | 1.89375642 | 0.05825735 | 0.60320496 |
| ACSS3 | 2960.0638 | 0.45307427 | 0.29614655 | 1.52989888 | 0.12604176 | 0.70364066 |
| PPFIA2 | 168.514164 | 0.11463881 | 0.54422163 | 0.21064729 | 0.83316251 | 0.97550416 |
| CCDC59 | 699.928893 | -0.1192029 | 0.15191132 | -0.7846876 | 0.43263674 | 0.88991825 |
| METTL25 | 382.726804 | 0.00618876 | 0.20841562 | 0.02969433 | 0.97631084 | 0.99455099 |
| TMTC2 | 1091.76246 | 0.78271656 | 0.38549356 | 2.0304271 | 0.04231314 | 0.56485966 |
| LRRIQ1 | 106.057803 | 0.40418262 | 0.56493441 | 0.71545052 | 0.4743307 | 0.90199039 |
| ALX1 | 78.0981516 | -0.470875 | 0.55575088 | -0.847277 | 0.39684072 | 0.87988087 |
| RASSF9 | 210.645067 | 0.10689261 | 0.36938603 | 0.28937914 | 0.77229126 | 0.96836854 |
| C12orf29 | 1058.66132 | -0.0426249 | 0.20322478 | -0.2097428 | 0.83386842 | 0.97550416 |
| CEP290 | 910.552897 | -0.0957484 | 0.17234572 | -0.5555603 | 0.5785115 | 0.93139743 |
| TMTC3 | 1140.5093 | 0.03281372 | 0.1509398 | 0.2173961 | 0.82789967 | 0.97548201 |
| KITLG | 2683.57329 | 0.73147272 | 0.34315591 | 2.13160461 | 0.03303936 | 0.55137197 |
| DUSP6 | 5106.15409 | -0.1373616 | 0.3356386 | -0.4092545 | 0.68235293 | 0.95070301 |
| AC024909.2 | 992.647423 | -0.1622116 | 0.35813503 | -0.4529343 | 0.65059606 | 0.94346674 |
| AC010201.2 | 130.18141 | 0.36763313 | 0.53413075 | 0.68828303 | 0.49127456 | 0.90600166 |
| AC010201.1 | 45.2636566 | 0.42122852 | 0.60283344 | 0.69874777 | 4.85E-01 | 9.05E-01 |
| POC1B | 1193.94212 | 0.18742405 | 0.14284225 | 1.31210514 | 0.18948466 | 0.76970626 |
| GALNT4 | 653.985354 | -0.0879932 | 0.19603817 | -0.4488573 | 0.65353463 | 0.94495971 |
| POC1B-GALN | 735.235004 | -0.0732652 | 0.18276495 | -0.4008714 | 0.68851483 | 0.95158502 |
| POC1B-AS1 | 160.125979 | -0.3202189 | 0.23868237 | -1.3416109 | 0.17972218 | 0.76283169 |
| AC025034.1 | 70.1023868 | -0.5605111 | 0.4062859 | -1.3795976 | 0.16771057 | 0.7558879 |
| ATP2B1 | 1714.60072 | -0.1294834 | 0.22402542 | -0.5779854 | 0.56327398 | 0.92452095 |
| ATP2B1-AS1 | 104.941002 | 0.470767 | 0.46992454 | 1.00179277 | 0.31644369 | 0.84783414 |
| LUM | 8602.40037 | 0.84946691 | 0.52799164 | 1.60886433 | 0.10764601 | 0.67707349 |
| DCN | 16411.4063 | 0.62466076 | 0.41554248 | 1.50324164 | 0.13277674 | 0.71361312 |
| BTG1 | 9604.59676 | -0.1094425 | 0.19797355 | -0.5528138 | 0.5803909 | 0.93142344 |
| AC025164.1 | 552.023543 | -0.1915569 | 0.23141995 | -0.827746 | 4.08E-01 | 8.82E-01 |
| PLEKHG7 | 102.613002 | -0.8800745 | 0.78238999 | -1.124854 | 0.26065091 | 0.82424998 |
| EEA1 | 1192.11853 | -0.3265692 | 0.18118719 | -1.8023856 | 0.07148477 | 0.63163413 |
| AC138123.1 | 70.7848469 | -0.7358137 | 0.37769964 | -1.9481451 | 0.0513976 | 0.58634747 |
| NUDT4 | 6160.9166 | -0.4919783 | 0.37897564 | -1.2981791 | 0.1942258 | 0.7762856 |
| UBE2N | 3885.78565 | -0.2074589 | 0.1687779 | -1.2291827 | 0.21900333 | 0.79847731 |
| MRPL42 | 2844.58997 | -0.1300643 | 0.10488039 | -1.2401203 | 0.2149309 | 0.79487183 |
| SOCS2 | 1579.40168 | -0.4260631 | 0.2603845 | -1.6362846 | 0.10178004 | 0.67279543 |
| CRADD | 844.844487 | 0.28854722 | 0.20537965 | 1.40494554 | 0.16003747 | 0.74781837 |
| PLXNC1 | 436.271017 | 0.07364976 | 0.31505062 | 0.23377121 | 8.15E-01 | 9.74E-01 |
| AC073655.2 | 47.341578 | 0.02709029 | 0.39666185 | 0.06829568 | 0.94555027 | 0.99166614 |
| CEP83 | 527.278686 | 0.14444729 | 0.22371121 | 0.6456864 | 0.51848247 | 0.91636259 |
| TMCC3 | 935.8553 | 0.05090587 | 0.32966721 | 0.15441592 | 0.8772818 | 0.98275231 |
| NDUFA12 | 1865.08253 | 0.11342697 | 0.17482647 | 0.64879747 | 0.51646929 | 0.91636259 |
| NR2C1 | 1555.41132 | 0.04719256 | 0.16121259 | 0.29273494 | 0.76972476 | 0.96836854 |
| FGD6 | 712.537826 | 0.2712085 | 0.20671414 | 1.31199784 | 0.18952086 | 0.76970626 |
| VEZT | 2464.33974 | 0.05902258 | 0.16141133 | 0.36566563 | 0.7146146 | 0.95579447 |
| METAP2 | 3682.04349 | 0.43188668 | 0.23791514 | 1.81529718 | 0.06947825 | 0.62779744 |
| USP44 | 141.067694 | 0.3370506 | 0.41884607 | 0.80471234 | 0.4209857 | 0.88451372 |
| NTN4 | 7161.45299 | 0.42153126 | 0.36046217 | 1.16941885 | 0.24223492 | 0.81319028 |
| SNRPF | 1124.63868 | 0.0871204 | 0.15898832 | 0.54796729 | 0.58371437 | 0.93233498 |
| AMDHD1 | 515.887315 | 0.59168364 | 0.46670364 | 1.26779309 | 0.20487185 | 0.78501397 |
| HAL | 71.2443003 | 0.05998569 | 0.57892643 | 0.1036154 | 0.91747457 | 0.9868044 |
| AC007298.2 | 42.5057714 | -0.3302292 | 0.48977179 | -0.6742512 | 0.50015165 | 0.91180773 |
| LTA4H | 4050.46115 | 0.17962121 | 0.20749907 | 0.86564826 | 0.38668308 | 0.87432552 |
| AC007298.1 | 402.116377 | 0.28023348 | 0.21444742 | 1.30677016 | 0.19129078 | 0.77292503 |
| ELK3 | 1866.66321 | -0.4338887 | 0.37681415 | -1.151466 | 0.24954057 | 0.81646572 |

| | | | | | | |
|---|---|---|---|---|---|---|
| AC008149.1 | 83.7712846 | -0.5163814 | 0.47730669 | -1.0818649 | 0.27931255 | 0.82825734 |
| CDK17 | 1482.09114 | -0.4263294 | 0.22557938 | -1.8899306 | 0.05876724 | 0.60332684 |
| CFAP54 | 63.751996 | -0.1499476 | 0.68412461 | -0.2191817 | 0.82650852 | 0.97548201 |
| NEDD1 | 686.944366 | 0.03131105 | 0.13025971 | 0.240374 | 0.81004033 | 0.97334138 |
| TMPO-AS1 | 175.139694 | 0.20302696 | 0.27599516 | 0.73561784 | 0.4619633 | 0.89762182 |
| TMPO | 2358.37278 | 0.08983933 | 0.21270339 | 0.42236907 | 0.67275565 | 0.94922827 |
| SLC25A3 | 26365.6419 | 0.22841173 | 0.2599306 | 0.87874121 | 0.37954161 | 0.8708011 |
| IKBIP | 894.368461 | -0.5591572 | 0.37576169 | -1.4880633 | 0.1367342 | 0.72102859 |
| APAF1 | 828.949515 | 0.34903079 | 0.22833594 | 1.52858457 | 0.12636746 | 0.70431949 |
| ANKS1B | 501.57278 | 0.55404439 | 0.29585249 | 1.87270487 | 0.06110915 | 0.60989294 |
| UHRF1BP1L | 973.057768 | -0.0009819 | 0.25763229 | -0.0038112 | 0.99695914 | 0.99976751 |
| ACTR6 | 1277.16859 | 0.43418741 | 0.19216782 | 2.25941794 | 0.0238574 | 0.51967919 |
| DEPDC4 | 57.5406473 | 0.5900623 | 0.40536793 | 1.45562154 | 0.14549727 | 0.73161643 |
| SCYL2 | 2557.28486 | -0.1161462 | 0.21274459 | -0.5459418 | 0.58510594 | 0.93289552 |
| NR1H4 | 3058.4717 | -0.1832082 | 0.4018506 | -0.4559113 | 0.64845376 | 0.9431121 |
| GAS2L3 | 362.251267 | 0.09324624 | 0.35593489 | 0.26197557 | 7.93E-01 | 9.71E-01 |
| SLC5A8 | 1657.39523 | -0.9689175 | 1.12902586 | -0.8581889 | 0.39078816 | 0.87699209 |
| UTP20 | 755.691307 | -0.3401529 | 0.29961569 | -1.1352973 | 0.25625078 | 0.82067614 |
| ARL1 | 4530.81485 | 0.18175433 | 0.14354345 | 1.26619732 | 0.20544245 | 0.78571765 |
| CHPT1 | 4173.39806 | 0.33006911 | 0.32130179 | 1.02728687 | 0.3042854 | 0.84211658 |
| SYCP3 | 788.255106 | 0.20233594 | 0.33909873 | 0.59668741 | 0.5507161 | 0.92373235 |
| GNPTAB | 1567.84576 | 0.10941132 | 0.10888286 | 1.00485343 | 0.31496743 | 0.84696964 |
| DRAM1 | 3638.79334 | -0.3440076 | 0.35239848 | -0.9761891 | 0.32897075 | 0.85317073 |
| AC084398.2 | 592.247044 | -0.3461887 | 0.33858243 | -1.0224651 | 0.30656082 | 0.84278367 |
| WASHC3 | 1321.04799 | 0.20482465 | 0.19980381 | 1.02512885 | 0.30530239 | 0.84268008 |
| AC079907.1 | 150.135102 | 0.17506942 | 0.22991105 | 0.76146588 | 0.44637885 | 0.89480207 |
| NUP37 | 578.437026 | 0.32597377 | 0.14720508 | 2.21441938 | 0.02679994 | 0.52685774 |
| PARPBP | 81.207456 | 0.31110425 | 0.399739 | 0.77826843 | 0.43641078 | 0.89104919 |
| PMCH | 39.7365415 | 0.14920789 | 0.52075745 | 0.28652089 | 0.77447919 | 0.96836854 |
| IGF1 | 315.858626 | 0.00300061 | 0.67740699 | 0.00442956 | 0.99646574 | 0.99976751 |
| PAH | 13986.0367 | 1.94883403 | 0.6880253 | 2.83250344 | 0.00461851 | 0.29744137 |
| STAB2 | 46.748063 | 0.04044239 | 0.37279541 | 0.10848415 | 0.91361165 | 0.98662757 |
| NT5DC3 | 469.932006 | 0.09572314 | 0.25951606 | 0.36885247 | 0.7122377 | 0.95560193 |
| HSP90B1 | 37335.7031 | 0.71659072 | 0.28903047 | 2.47929132 | 0.01316437 | 0.42221215 |
| C12orf73 | 521.543993 | 0.14943583 | 0.21725767 | 0.68782765 | 0.49156132 | 0.90600166 |
| TDG | 1081.2708 | -0.2760087 | 0.18526286 | -1.4898221 | 0.13627101 | 0.71919319 |
| GLT8D2 | 236.408253 | 0.70725384 | 0.50935445 | 1.38852981 | 0.16497577 | 0.75299828 |
| HCFC2 | 687.552932 | -0.0582185 | 0.10546256 | -0.5520305 | 0.58092748 | 0.9317705 |
| NFYB | 1677.33607 | 0.11052039 | 0.12648889 | 0.87375573 | 0.3822513 | 0.87185666 |
| TXNRD1 | 14068.4301 | -0.9219862 | 0.69267535 | -1.3310509 | 0.18317225 | 0.76723162 |
| EID3 | 309.553459 | -2.9610243 | 0.87581058 | -3.3808958 | NA | NA |
| CHST11 | 887.188861 | -0.2562658 | 0.30867451 | -0.8302136 | 0.40641804 | 0.88132015 |
| SLC41A2 | 839.026745 | -0.4102896 | 0.33490282 | -1.2251004 | 0.2205374 | 0.79912826 |
| C12orf45 | 387.249019 | -0.021533 | 0.22763869 | -0.094593 | 0.92463812 | 0.98869145 |
| ALDH1L2 | 121.061372 | -0.6745181 | 0.45699424 | -1.4759882 | 0.13994705 | 0.72475166 |
| WASHC4 | 2910.28236 | -0.2914161 | 0.1896517 | -1.5365857 | 0.1243948 | 0.70104934 |
| APPL2 | 2151.51883 | 0.17609784 | 0.11093144 | 1.58744749 | 0.11241133 | 0.68001961 |
| C12orf75 | 3046.62545 | 0.48381925 | 0.19134484 | 2.52851994 | 0.01145446 | 0.40353404 |
| NUAK1 | 746.225256 | -0.2178097 | 0.28197572 | -0.7724414 | 0.43985307 | 0.89203402 |
| CKAP4 | 3030.46694 | 0.13180665 | 0.22189156 | 0.59401383 | 0.55250286 | 0.92373235 |
| AC079174.2 | 108.592956 | 0.7691403 | 0.54282937 | 1.41690989 | 0.15650928 | 0.74522306 |
| TCP11L2 | 234.259209 | -0.3723925 | 0.23346028 | -1.5951001 | 0.11068985 | 0.68001961 |
| POLR3B | 525.324832 | 0.20848325 | 0.20365826 | 1.02369162 | 0.30598096 | 0.84270033 |
| AC079385.1 | 84.6172439 | 0.07281713 | 0.28346543 | 0.25688187 | 7.97E-01 | 0.9713715 |

| | | | | | | |
|---|---|---|---|---|---|---|
| RIC8B | 1047.55656 | -0.0397534 | 0.15262406 | -0.2604664 | 7.95E-01 | 9.71E-01 |
| AC007541.1 | 121.943042 | 0.05551919 | 0.40572898 | 0.13683811 | 0.89115876 | 0.98474725 |
| TMEM263 | 3414.8711 | -0.0993799 | 0.17050461 | -0.5828577 | 0.55998908 | 0.92434754 |
| MTERF2 | 1528.8022 | 0.15519723 | 0.28646116 | 0.54177408 | 0.58797415 | 0.93289552 |
| CRY1 | 1660.91456 | -0.7074092 | 0.26750213 | -2.6444993 | 0.00818119 | 0.37396013 |
| AC078929.1 | 60.1287833 | -0.7752854 | 0.5652298 | -1.3716286 | 0.17017909 | 0.75956362 |
| BTBD11 | 549.093841 | -0.3556892 | 0.62091324 | -0.5728485 | 0.56674725 | 0.9256133 |
| PWP1 | 2372.92761 | -0.2057926 | 0.22645398 | -0.9087613 | 0.36347612 | 0.8663515 |
| PRDM4 | 2959.23424 | -0.4976491 | 0.43075766 | -1.1552878 | 0.24797258 | 0.81602355 |
| AC007622.2 | 304.37508 | -0.6883221 | 0.54140915 | -1.2713529 | 0.20360313 | 0.78351667 |
| WSCD2 | 105.601522 | 0.23085513 | 0.63545676 | 0.36329006 | 0.71638822 | 0.95603249 |
| CMKLR1 | 556.320727 | -0.3799853 | 0.42722483 | -0.8894271 | 0.37377359 | 0.86791605 |
| FICD | 692.17452 | 0.07592243 | 0.29829203 | 0.25452382 | 0.7990909 | 0.97188413 |
| SART3 | 2261.97436 | -0.1085861 | 0.13390239 | -0.8109346 | 0.41740324 | 0.88445833 |
| ISCU | 11153.8181 | 0.28569107 | 0.11189215 | 2.55327181 | 0.01067162 | 0.40196898 |
| TMEM119 | 343.816441 | -0.721698 | 0.61148252 | -1.180243 | 0.23790356 | 0.80962787 |
| SELPLG | 552.934913 | -0.019796 | 0.54481586 | -0.0363352 | 0.97101508 | 0.99400331 |
| CORO1C | 4428.35266 | -0.4410869 | 0.29160409 | -1.5126223 | 0.13037561 | 0.7084713 |
| SSH1 | 2272.26701 | -0.2781209 | 0.17815181 | -1.5611456 | 0.11848941 | 0.6915634 |
| DAO | 2102.14288 | 1.62437258 | 0.68699455 | 2.36446212 | 0.01805627 | 0.47478681 |
| USP30 | 1092.15896 | 0.08797223 | 0.29193855 | 0.30133819 | 0.76315662 | 0.96656878 |
| USP30-AS1 | 89.5674322 | -1.1309289 | 0.83517532 | -1.3541215 | NA | NA |
| ALKBH2 | 488.151931 | 0.00211764 | 0.1764012 | 0.01200465 | 0.9904219 | 0.99814875 |
| UNG | 2001.47233 | -0.0109009 | 0.20300947 | -0.0536965 | 0.95717701 | 0.99205545 |
| ACACB | 4057.22915 | -0.3240345 | 0.40067619 | -0.8087192 | 0.41867666 | 0.88451372 |
| KCTD10 | 3971.49617 | -0.139153 | 0.15797258 | -0.8808679 | 0.37838934 | 0.87045602 |
| UBE3B | 2412.5592 | 0.11263053 | 0.1949509 | 0.57773794 | 0.56344106 | 0.92452095 |
| MMAB | 2085.26371 | 0.0921672 | 0.15287178 | 0.60290528 | 0.5465717 | 0.92345821 |
| MVK | 4780.30058 | -0.0405947 | 0.27834746 | -0.145842 | 0.88404613 | 0.98317437 |
| FAM222A | 794.084276 | 0.05586602 | 0.25690864 | 0.21745483 | 0.82785391 | 0.97548201 |
| TRPV4 | 1967.70289 | 0.54674241 | 0.33911558 | 1.61225975 | 0.10690543 | 0.67674128 |
| GLTP | 3143.38193 | 0.05897295 | 0.27760495 | 0.21243478 | 0.83176785 | 0.97550416 |
| TCHP | 3086.95157 | -0.1006441 | 0.13648426 | -0.7374042 | 0.46087661 | 0.89755243 |
| GIT2 | 2670.4339 | -0.1281794 | 0.1533255 | -0.835995 | 0.40315771 | 0.88132015 |
| AC084876.1 | 168.729037 | -0.1097336 | 0.37500811 | -0.2926165 | 0.76981527 | 0.96836854 |
| ANKRD13A | 3942.16577 | -0.2140729 | 0.19477908 | -1.099055 | 0.27174409 | 0.82716979 |
| C12orf76 | 906.477806 | -0.0090504 | 0.19395922 | -0.0466613 | 9.63E-01 | 0.99320711 |
| IFT81 | 991.857717 | -0.0667163 | 0.24207053 | -0.2756068 | 7.83E-01 | 9.70E-01 |
| ATP2A2 | 9550.23748 | -0.0972365 | 0.19608014 | -0.4959018 | 0.61996367 | 0.93949536 |
| ANAPC7 | 2731.91039 | -0.219023 | 0.12735818 | -1.7197406 | 0.08547961 | 0.6504594 |
| ARPC3 | 9155.67287 | -0.0460433 | 0.14053437 | -0.32763 | 0.74319143 | 0.96109622 |
| GPN3 | 1212.78444 | 0.15878394 | 0.1448963 | 1.09584541 | 0.27314643 | 0.82716979 |
| FAM216A | 279.093346 | -0.1643489 | 0.23871955 | -0.6884602 | 0.49116304 | 0.90600166 |
| VPS29 | 3625.26538 | 0.10597502 | 0.11660247 | 0.90885738 | 0.36342541 | 0.8663515 |
| PPTC7 | 2363.89239 | 0.0321114 | 0.19525234 | 0.16446102 | 0.86936823 | 0.98275231 |
| TCTN1 | 2491.43197 | 0.22911616 | 0.28502071 | 0.80385793 | 0.42147903 | 0.88451372 |
| HVCN1 | 358.702889 | 0.03599235 | 0.28894659 | 0.12456404 | 0.9008687 | 0.98545758 |
| PPP1CC | 4145.31032 | -0.0983014 | 0.18254053 | -0.5385182 | 5.90E-01 | 9.33E-01 |
| MYL2 | 53.425036 | 7.56878844 | 2.78422536 | 2.71845396 | NA | NA |
| PHETA1 | 1176.18271 | 0.51607978 | 0.24978709 | 2.06607866 | 0.03882105 | 0.55608529 |
| SH2B3 | 2039.53775 | -0.1897626 | 0.30781848 | -0.6164755 | 0.53758071 | 0.92123049 |
| ATXN2 | 2796.2195 | -0.2583935 | 0.11317191 | -2.2831949 | 0.02241889 | 0.51870674 |
| BRAP | 1032.85329 | -0.1409324 | 0.17820686 | -0.7908359 | 0.42903977 | 0.88851358 |
| ACAD10 | 4126.29289 | 0.19192723 | 0.14525962 | 1.32127041 | 0.18641122 | 0.76908034 |

| | | | | | | |
|---|---|---|---|---|---|---|
| AC002996.1 | 10167.9874 | 0.82888711 | 0.43086417 | 1.9237782 | 0.05438239 | 0.59329767 |
| ALDH2 | 24732.2195 | 0.75000056 | 0.41728034 | 1.79735417 | 0.07227941 | 0.6321115 |
| MAPKAPK5-A | 833.28868 | -0.0174953 | 0.16399477 | -0.1066821 | 0.9150412 | 0.9866558 |
| MAPKAPK5 | 2190.12151 | 0.02491514 | 0.13191001 | 0.18887981 | 0.85018702 | 0.97816532 |
| TMEM116 | 1525.73764 | 0.13214826 | 0.22386584 | 0.59030113 | 0.55498878 | 0.92373235 |
| ERP29 | 8464.84217 | 0.24019789 | 0.12951916 | 1.85453566 | 0.06366256 | 0.61812911 |
| AC073575.2 | 92.511154 | -0.352584 | 0.35694016 | -0.9877957 | 0.32325272 | 0.85119105 |
| NAA25 | 1139.0019 | -0.391584 | 0.23199885 | -1.6878705 | 0.09143608 | 0.65518742 |
| TRAFD1 | 2225.26489 | -0.2234489 | 0.31236742 | -0.71534 | 0.47439897 | 0.90199039 |
| HECTD4 | 2974.29151 | -0.2363137 | 0.13614601 | -1.7357372 | 0.08261032 | 0.64680474 |
| RPL6 | 24681.5029 | 0.29353503 | 0.25039584 | 1.17228394 | 0.24108308 | 0.8124487 |
| PTPN11 | 4614.27629 | -0.1132525 | 0.26046945 | -0.4348016 | 0.66370644 | 0.94642611 |
| OAS1 | 3946.84516 | -2.7210678 | 1.01197992 | -2.6888555 | NA | NA |
| AC004551.1 | 552.355868 | -2.297182 | 0.86679797 | -2.6501931 | NA | NA |
| OAS3 | 3050.71394 | -3.0512254 | 1.12452935 | -2.7133355 | NA | NA |
| OAS2 | 2071.93387 | -3.0290842 | 1.02347481 | -2.959608 | NA | NA |
| DTX1 | 1341.2457 | -0.297363 | 0.35603733 | -0.8352018 | 0.40360412 | 0.88132015 |
| RASAL1 | 3728.29899 | -0.1857175 | 0.30100026 | -0.6170012 | 0.53723391 | 0.92119932 |
| CFAP73 | 1356.1111 | -0.0952974 | 0.16326195 | -0.5837087 | 0.55941631 | 0.92421175 |
| DDX54 | 4374.81919 | -0.0968111 | 0.15428892 | -0.6274663 | 0.53035365 | 0.91962754 |
| RITA1 | 1451.11412 | 0.08013848 | 0.24046153 | 0.33326944 | 0.73893091 | 0.9602325 |
| AC089999.1 | 267.368048 | 0.03984955 | 0.30849536 | 0.12917392 | 0.89722004 | 0.98474725 |
| IQCD | 234.903931 | 0.1752912 | 0.3595686 | 0.48750418 | 0.62590108 | 0.94088547 |
| TPCN1 | 11594.4428 | 0.65044464 | 0.28575814 | 2.27620683 | 0.02283363 | 0.51967919 |
| SLC8B1 | 3404.27785 | -0.0695493 | 0.19610424 | -0.3546548 | 0.72284823 | 0.95764871 |
| PLBD2 | 3079.30549 | -0.0216067 | 0.12033502 | -0.1795549 | 0.857502 | 0.97959852 |
| SDS | 87.3715419 | -0.5124825 | 0.63640014 | -0.8052834 | 0.42065615 | 0.88451372 |
| SDSL | 1630.2279 | 0.53763177 | 0.39435137 | 1.36333181 | 0.17277796 | 0.75974924 |
| RBM19 | 1961.3198 | -0.2934291 | 0.11282901 | -2.6006531 | 0.00930465 | 0.38664841 |
| AC073863.1 | 122.738714 | -0.4261738 | 0.2511026 | -1.6972098 | 8.97E-02 | 6.54E-01 |
| TBX3 | 1214.83155 | 0.22520046 | 0.33321741 | 0.67583643 | 4.99E-01 | 0.91147492 |
| MED13L | 2526.14045 | -0.2587788 | 0.11210391 | -2.3083836 | 0.02097781 | 0.50728403 |
| AC012157.1 | 185.055345 | -0.3429192 | 0.32526265 | -1.0542841 | 0.29175288 | 0.83460625 |
| MAP1LC3B2 | 44.1210283 | -0.0636417 | 0.27424942 | -0.2320579 | 0.81649306 | 0.97388289 |
| C12orf49 | 3921.28339 | -0.8036871 | 0.39690904 | -2.0248648 | 0.04288125 | 0.56634074 |
| RNFT2 | 153.928255 | 0.27544559 | 0.37389908 | 0.73668433 | 0.46131434 | 0.89755243 |
| HRK | 89.2758618 | -2.3757307 | 0.98351982 | -2.4155392 | 0.01571193 | 0.45577618 |
| FBXW8 | 682.902079 | -0.044957 | 0.18839585 | -0.2386303 | 0.81139228 | 0.97352497 |
| AC127164.1 | 51.4840271 | 0.11104484 | 0.35349128 | 0.31413742 | 0.75341667 | 0.96301881 |
| TESC | 1050.48045 | 0.56134234 | 0.39263053 | 1.42969611 | 0.15280426 | 0.74246943 |
| FBXO21 | 3895.6614 | 0.13631294 | 0.2736775 | 0.49807871 | 0.61842857 | 0.93943514 |
| NOS1 | 273.551798 | 1.36800414 | 0.69335765 | 1.97301368 | 0.048494 | 0.58440502 |
| AC073864.1 | 56.8851296 | -0.5100842 | 0.66711197 | -0.7646156 | 0.4445005 | 0.8938278 |
| KSR2 | 307.006128 | -0.6373449 | 0.51258828 | -1.2433855 | 0.21372579 | 0.7926732 |
| RFC5 | 1132.58714 | 0.06854011 | 0.17974302 | 0.38132278 | 0.70296375 | 0.95431462 |
| AC131159.2 | 98.2024101 | 0.13013351 | 0.22348149 | 0.58230105 | 0.56036392 | 0.92434754 |
| WSB2 | 4073.08797 | 0.13127677 | 0.30849216 | 0.42554329 | 0.67044067 | 0.94922827 |
| VSIG10 | 1264.78236 | 0.0730417 | 0.20929053 | 0.34899666 | 0.72709182 | 0.95848028 |
| AC131238.1 | 77.2445344 | -0.0670517 | 0.32082323 | -0.2089989 | 0.83444907 | 0.97550416 |
| PEBP1 | 41259.1095 | 0.34153708 | 0.28484614 | 1.19902302 | 2.31E-01 | 0.80584899 |
| TAOK3 | 1576.4647 | 0.02061923 | 0.11073709 | 0.18619981 | 0.85228807 | 0.97872457 |
| SUDS3 | 2526.60558 | -0.0345116 | 0.16289583 | -0.2118627 | 0.83221413 | 0.97550416 |
| AC026367.3 | 79.1961105 | -0.2463716 | 0.40919089 | -0.6020947 | 0.54711113 | 0.92345821 |
| AC026367.2 | 51.3589072 | -0.0284479 | 0.38718005 | -0.0734746 | 0.94142843 | 0.99136103 |

| | | | | | | |
|---|---|---|---|---|---|---|
| HSPB8 | 7270.02675 | 0.10489479 | 0.39121784 | 0.26812375 | 0.78860407 | 0.97090793 |
| AC002070.1 | 386.079989 | 0.35142364 | 0.31512265 | 1.11519637 | 0.26476628 | 0.82424998 |
| TMEM233 | 85.3717393 | -0.1224848 | 0.62084633 | -0.1972868 | 0.84360311 | 0.97652052 |
| PRKAB1 | 7713.50325 | 0.17571122 | 0.22316254 | 0.78736879 | 0.43106601 | 0.88966606 |
| CIT | 3125.2697 | -0.3958885 | 0.34048278 | -1.1627269 | 0.24494031 | 0.81487261 |
| AC002563.1 | 83.2671229 | -0.2773535 | 0.3568551 | -0.777216 | 0.43703133 | 0.89130865 |
| BICDL1 | 14223.8726 | -0.7133471 | 0.45521362 | -1.5670601 | 0.11710064 | 0.68956727 |
| RAB35 | 3357.54442 | -0.4826511 | 0.19740478 | -2.4449817 | 0.01448595 | 0.43901686 |
| AC004812.2 | 295.048004 | -0.2479121 | 0.45556992 | -0.5441801 | 0.58631755 | 0.93289552 |
| GCN1 | 4099.93266 | -0.2258323 | 0.1295594 | -1.7430796 | 0.08131971 | 0.6439041 |
| RPLP0 | 47105.9914 | 0.13944346 | 0.2744781 | 0.50803129 | 0.6114314 | 0.93834929 |
| PXN-AS1 | 5465.93752 | -0.2519323 | 0.15262972 | -1.6506111 | 0.09881801 | 0.6678501 |
| PXN | 11993.0841 | -0.3132178 | 0.15881413 | -1.9722289 | 0.04858348 | 0.58440502 |
| SIRT4 | 177.205565 | -0.1338733 | 0.4759644 | -0.2812675 | 0.77850526 | 0.96836854 |
| MSI1 | 111.499572 | -0.3391917 | 0.54792298 | -0.61905 | 0.53588344 | 0.92101152 |
| COX6A1 | 9981.76589 | 0.04034597 | 0.18582105 | 0.21712272 | 0.82811271 | 0.97548201 |
| AL021546.1 | 8283.93211 | 0.06134736 | 0.19057029 | 0.32191458 | 0.74751741 | 0.96231266 |
| TRIAP1 | 785.975147 | -0.0534706 | 0.19347337 | -0.276372 | 0.78226235 | 0.96953765 |
| GATC | 5162.44811 | -0.0731608 | 0.09607558 | -0.7614917 | 4.46E-01 | 0.89480207 |
| SRSF9 | 6781.92592 | -0.0470215 | 0.10441719 | -0.4503236 | 0.65247709 | 0.94437526 |
| DYNLL1 | 6409.6787 | 0.46386508 | 0.22096798 | 2.09924113 | 0.03579565 | 0.55369025 |
| NRAV | 527.604511 | -4.69E-05 | 0.22932759 | -0.0002045 | 0.99983686 | 0.99994357 |
| COQ5 | 1846.77455 | 0.1328561 | 0.22025419 | 0.60319444 | 0.54637935 | 0.92345821 |
| RNF10 | 11476.2158 | -0.190809 | 0.16233843 | -1.175378 | 0.23984352 | 0.81209487 |
| POP5 | 1445.44993 | 0.19345044 | 0.10561964 | 1.83157631 | 0.06701457 | 0.6249306 |
| AC125616.1 | 37.0243944 | 0.14142332 | 0.52541915 | 0.26916287 | 0.78780435 | 0.97090793 |
| CABP1 | 705.685456 | -0.4909666 | 0.51630626 | -0.9509213 | 0.34164435 | 0.85889221 |
| MLEC | 10384.9963 | 0.50724068 | 0.19266939 | 2.63269985 | 0.00847092 | 0.3779007 |
| AC069234.2 | 1080.11823 | 0.55793764 | 0.35608657 | 1.56685951 | 0.11714753 | 0.68956727 |
| UNC119B | 3746.13922 | 0.20114755 | 0.19030452 | 1.05697728 | 0.29052196 | 0.83442894 |
| AC069234.3 | 792.39521 | 0.27037115 | 0.21749322 | 1.24312448 | 0.21382197 | 0.7926732 |
| ACADS | 4506.82561 | 0.36879548 | 0.22599958 | 1.63184148 | 0.10271288 | 0.67335541 |
| SPPL3 | 2901.84153 | -0.2087062 | 0.19023071 | -1.0971216 | 0.27258824 | 0.82716979 |
| HNF1A | 2505.82736 | 0.92636509 | 0.553586 | 1.67338966 | 0.09425061 | 0.65783452 |
| C12orf43 | 2162.57185 | 0.45020101 | 0.32497548 | 1.38533838 | 0.16594902 | 0.75515387 |
| OASL | 1272.73822 | -4.8601234 | 1.32808699 | -3.6594917 | NA | NA |
| P2RX7 | 116.226814 | 0.16164929 | 0.35362045 | 0.45712654 | 0.6475801 | 0.94282139 |
| P2RX4 | 2208.44895 | -0.2413108 | 0.33103552 | -0.7289575 | 0.46602763 | 0.89960479 |
| CAMKK2 | 2303.99647 | 0.16640312 | 0.2100718 | 0.79212498 | 0.42828781 | 0.88820133 |
| ANAPC5 | 5646.21556 | 0.0076551 | 0.11036819 | 0.06935961 | 0.94470338 | 0.99152587 |
| RNF34 | 1567.12578 | -0.2488955 | 0.14175415 | -1.7558253 | 0.07911825 | 0.64079926 |
| KDM2B | 1366.57805 | -0.2716541 | 0.20454246 | -1.3281063 | 0.18414298 | 0.76816707 |
| ORAI1 | 1450.08365 | -0.3604512 | 0.30636569 | -1.1765391 | 0.2393795 | 0.81163641 |
| MORN3 | 125.635871 | 0.52275207 | 0.42789883 | 1.22167213 | 0.22183164 | 0.80054995 |
| TMEM120B | 4736.28275 | -0.3797437 | 0.35616878 | -1.0661902 | 0.28633766 | 0.83107779 |
| RHOF | 3766.79602 | -0.4977361 | 0.38618842 | -1.2888426 | 0.19745282 | 0.77819208 |
| LINC01089 | 2296.44983 | 0.34437058 | 0.38632975 | 0.89139027 | 0.37271984 | 0.86791605 |
| AC084018.1 | 422.272161 | 0.25712193 | 0.39676871 | 0.64803984 | 0.51695918 | 0.91636259 |
| AC084018.2 | 446.842654 | 0.22303947 | 0.45584138 | 0.48929184 | 0.6246351 | 0.94088547 |
| SETD1B | 2608.79434 | -0.4058766 | 0.19597272 | -2.0710872 | 0.03835065 | 0.55538555 |
| HPD | 14711.2972 | 0.8089856 | 1.10854063 | 0.72977532 | 0.46552753 | 0.89933527 |
| AC079360.1 | 4232.09298 | 0.90209598 | 1.23653381 | 0.72953604 | 0.46567383 | 0.89943247 |
| PSMD9 | 3253.00437 | -0.2377239 | 0.24749906 | -0.9605041 | 0.33680156 | 0.85679125 |
| AC069503.2 | 2920.29967 | -0.2314874 | 0.24921179 | -0.9288781 | 0.35295227 | 0.86364633 |

| | | | | | | |
|---|---|---|---|---|---|---|
| WDR66 | 633.229851 | -0.3623999 | 0.21340069 | -1.6982134 | 0.0894675 | 0.65375543 |
| AC069503.1 | 57.9598016 | 0.35871685 | 0.46731041 | 0.76762008 | 0.44271293 | 0.89343541 |
| BCL7A | 1834.53457 | -0.1580974 | 0.14245927 | -1.1097729 | 2.67E-01 | 8.27E-01 |
| MLXIP | 6476.52545 | -0.1929926 | 0.21015958 | -0.9183146 | 0.35845421 | 0.86480228 |
| LRRC43 | 291.658112 | 0.15129036 | 0.43618025 | 0.34685285 | 0.72870187 | 0.95848028 |
| B3GNT4 | 2189.41081 | 0.01292634 | 0.17761041 | 0.07277919 | 0.94198183 | 0.99136103 |
| DIABLO | 2747.41679 | 0.00962355 | 0.16412312 | 0.05863616 | 0.95324191 | 0.99170503 |
| AC048338.1 | 3203.14029 | 0.01668589 | 0.16672632 | 0.10007951 | 0.9202812 | 0.9874645 |
| VPS33A | 1220.35741 | 0.01096438 | 0.2397872 | 0.04572544 | 0.96352908 | 0.99320711 |
| CLIP1 | 3197.97295 | -0.3834792 | 0.28645976 | -1.3386843 | 0.18067345 | 0.76466689 |
| ZCCHC8 | 1520.99058 | -0.2116147 | 0.14967517 | -1.4138264 | 0.15741287 | 0.74522306 |
| RSRC2 | 4085.22903 | -0.0375256 | 0.1130672 | -0.3318875 | 0.7399742 | 0.96043876 |
| KNTC1 | 446.389156 | 0.02859578 | 0.16123537 | 0.17735423 | 0.85923016 | 0.98014684 |
| HCAR2 | 89.5503576 | 1.65420953 | 0.45499745 | 3.63564572 | 0.00027729 | 0.08298534 |
| HCAR1 | 96.5253454 | 0.81475517 | 0.4820188 | 1.6902975 | 0.09097105 | 0.65495306 |
| DENR | 2884.47633 | -0.0595242 | 0.12956622 | -0.4594115 | 0.64593872 | 0.94208859 |
| HIP1R | 10864.6458 | -0.067235 | 0.16527535 | -0.4068058 | 0.68415066 | 0.95070301 |
| VPS37B | 3288.68945 | -0.1538164 | 0.23893026 | -0.6437711 | 5.20E-01 | 0.91657872 |
| AC027290.1 | 959.192156 | -0.1354656 | 0.2340159 | -0.5788733 | 0.56267467 | 0.92452095 |
| ABCB9 | 3372.40218 | 0.11435228 | 0.09607088 | 1.19029075 | 2.34E-01 | 0.80725541 |
| OGFOD2 | 1742.49005 | -0.0886609 | 0.12567817 | -0.7054595 | 0.48052435 | 0.90541935 |
| ARL6IP4 | 6653.50229 | 0.09289705 | 0.09093736 | 1.02154989 | 0.30699398 | 0.84286067 |
| PITPNM2 | 4412.81064 | -0.473993 | 0.19405522 | -2.4425678 | 1.46E-02 | 4.39E-01 |
| MPHOSPH9 | 621.121677 | 0.13823588 | 0.17166431 | 0.80526861 | 0.42066469 | 0.88451372 |
| C12orf65 | 1053.35749 | -0.2244601 | 0.22827821 | -0.9832741 | 0.32547254 | 0.85244476 |
| AC068768.1 | 1746.7368 | -0.3557516 | 0.28772877 | -1.2364129 | 0.21630511 | 0.7954827 |
| CDK2AP1 | 2961.46635 | -0.3619577 | 0.27907951 | -1.29697 | 0.19464152 | 0.7762856 |
| SBNO1 | 2292.06341 | -0.1535323 | 0.19672174 | -0.780454 | 0.43512368 | 0.89055338 |
| AC137767.1 | 256.967303 | -0.2978599 | 0.37079677 | -0.8032969 | 0.42180317 | 0.88451372 |
| KMT5A | 1763.56759 | -0.2829141 | 0.16838741 | -1.6801381 | 0.09293046 | 0.65753347 |
| RILPL2 | 2083.91467 | 0.0550213 | 0.16966361 | 0.32429644 | 0.74571361 | 0.96176407 |
| SNRNP35 | 1793.35888 | -0.1836743 | 0.16186382 | -1.1347456 | 2.56E-01 | 0.82082102 |
| RILPL1 | 1513.18662 | -0.1466559 | 0.20416721 | -0.7183126 | 0.47256456 | 0.90160919 |
| AC055713.1 | 1296.74518 | 0.02218766 | 0.18145405 | 0.12227703 | 0.90267963 | 0.98608873 |
| TMED2 | 9216.50527 | -0.0321947 | 0.20900916 | -0.1540348 | 0.87758232 | 0.98275231 |
| DDX55 | 1035.67372 | -0.1056698 | 0.12149621 | -0.869737 | 0.38444412 | 0.87309542 |
| EIF2B1 | 2504.30661 | -0.1227381 | 0.11902827 | -1.0311674 | 0.30246231 | 0.84211658 |
| GTF2H3 | 831.307932 | 0.12062579 | 0.16160181 | 0.74643833 | 0.45540267 | 0.89688594 |
| TCTN2 | 1216.55334 | -0.0998139 | 0.2080679 | -0.4797178 | 6.31E-01 | 9.41E-01 |
| AC117503.1 | 375.946363 | -0.1583304 | 0.28369534 | -0.5581 | 0.57677607 | 0.93127944 |
| ATP6V0A2 | 936.661495 | -0.0590046 | 0.24545872 | -0.240385 | 0.81003178 | 0.97334138 |
| DNAH10 | 239.774008 | 0.15678212 | 0.33984386 | 0.46133574 | 0.64455775 | 0.94208859 |
| CCDC92 | 2555.45873 | -0.2409146 | 0.14357375 | -1.6779849 | 0.09335004 | 0.65756964 |
| AC079315.1 | 158.377118 | 0.44391929 | 0.37834162 | 1.17332925 | 2.41E-01 | 8.12E-01 |
| DNAH10OS | 346.520621 | 0.40455749 | 0.29281262 | 1.38162584 | 0.16708662 | 0.75558932 |
| ZNF664 | 6794.47553 | -0.1794809 | 0.22023875 | -0.8149377 | 0.41510796 | 0.88377697 |
| RFLNA | 233.492646 | 0.14236599 | 0.43555238 | 0.32686308 | 0.74377145 | 0.96137808 |
| AC068790.8 | 199.75097 | -0.3336986 | 0.48418433 | -0.6891975 | 0.490699 | 0.90600166 |
| NCOR2 | 7651.9429 | -0.2254475 | 0.16130941 | -1.3976094 | 0.16223041 | 0.75026273 |
| SCARB1 | 1412.53906 | 0.30656017 | 0.2871234 | 1.06769483 | 0.28565819 | 0.83107779 |
| UBC | 44178.5813 | -0.3231295 | 0.27755845 | -1.1641855 | 0.24434886 | 0.81468457 |
| DHX37 | 1404.11693 | -0.5128823 | 0.17174585 | -2.9862864 | 0.00282388 | 0.23089261 |
| BRI3BP | 1017.51273 | 0.54165352 | 0.30116142 | 1.79854884 | 7.21E-02 | 6.32E-01 |
| THRIL | 60.4546071 | 0.02287048 | 0.42448585 | 0.05387807 | 0.95703231 | 0.99200096 |

| | | | | | | |
|---|---|---|---|---|---|---|
| AACS | 1795.67248 | 0.22659793 | 0.17220783 | 1.31583988 | 0.18822779 | 0.76970626 |
| LINC00939 | 79.752477 | 1.00635385 | 0.81799742 | 1.23026531 | 0.21859777 | 0.79780737 |
| TMEM132C | 86.0657145 | 0.32214189 | 0.69138709 | 0.46593564 | 0.64126157 | 0.94088547 |
| SLC15A4 | 1808.93646 | -0.4312259 | 0.22853234 | -1.886936 | 0.05916894 | 0.60462654 |
| GLT1D1 | 105.105351 | -0.72846 | 0.57684284 | -1.2628396 | 0.20664684 | 0.78778852 |
| RIMBP2 | 598.048662 | -0.3728578 | 0.36402231 | -1.0242718 | 0.30570692 | 0.84270033 |
| STX2 | 1629.4394 | -0.1569636 | 0.21812276 | -0.7196114 | 0.4717643 | 0.90160919 |
| RAN | 9198.24339 | -0.1617719 | 0.23521516 | -0.6877616 | 0.49160295 | 0.90600166 |
| ADGRD1 | 216.110403 | -0.4417061 | 0.70195986 | -0.629247 | 0.52918737 | 0.91920857 |
| SFSWAP | 2520.58976 | -0.3022281 | 0.14155777 | -2.1350157 | 0.03275973 | 0.55104386 |
| MMP17 | 80.9475422 | -0.3267023 | 0.59736656 | -0.5469042 | 0.58444452 | 0.93265282 |
| ULK1 | 5553.20825 | -0.3271298 | 0.19187094 | -1.7049473 | 0.08820426 | 0.65363785 |
| AC131009.1 | 62.7703107 | -0.313433 | 0.35420876 | -0.8848822 | 0.37622019 | 0.86938873 |
| PUS1 | 1478.01908 | -0.4035758 | 0.21141288 | -1.9089463 | 0.05626902 | 0.59895387 |
| EP400 | 2403.67291 | -0.1739217 | 0.14778524 | -1.176854 | 0.23925377 | 0.81163641 |
| AC137590.1 | 319.084243 | -0.0328682 | 0.19129867 | -0.1718159 | 0.86358223 | 0.98176973 |
| DDX51 | 1730.42872 | -0.1054395 | 0.1321988 | -0.7975831 | 0.42511245 | 0.88581636 |
| NOC4L | 1119.44253 | -0.0636912 | 0.21543479 | -0.2956404 | 0.76750475 | 0.96749893 |
| LINC02361 | 153.301713 | 0.20510808 | 0.54870615 | 0.37380314 | 7.09E-01 | 9.55E-01 |
| GALNT9 | 842.125577 | 0.10949088 | 0.43054239 | 0.25430917 | 0.79925671 | 0.97192231 |
| AC148477.3 | 96.4616153 | 0.77648194 | 0.49074625 | 1.58224731 | 0.11359312 | 0.68214043 |
| AC148477.2 | 561.09508 | 0.54688345 | 0.51180034 | 1.06854844 | 0.28527319 | 0.83107779 |
| AC148477.4 | 271.524387 | 0.28300931 | 0.39639293 | 0.71396155 | 0.47525095 | 0.90226631 |
| FBRSL1 | 4697.13332 | -0.3462921 | 0.17144023 | -2.0198997 | 0.04339379 | 0.56911599 |
| LRCOL1 | 84.9985779 | -0.5634725 | 0.74837133 | -0.7529317 | 0.45149098 | 0.89639459 |
| P2RX2 | 94.1425174 | -0.0344501 | 0.93202322 | -0.0369627 | 0.97051471 | 0.99400331 |
| POLE | 944.472131 | -0.2123025 | 0.195402 | -1.0864909 | 2.77E-01 | 8.28E-01 |
| PXMP2 | 1635.07526 | 0.54503985 | 0.34929568 | 1.56039679 | 0.11866614 | 0.69194841 |
| AC135586.2 | 1595.36696 | 0.49319872 | 0.35256813 | 1.39887493 | 0.16185049 | 0.7498147 |
| PGAM5 | 1746.76071 | -0.3415115 | 0.25323919 | -1.3485727 | 0.17747426 | 0.76115956 |
| ANKLE2 | 3883.9563 | -0.5977215 | 0.26621385 | -2.2452682 | 0.02475092 | 0.52085426 |
| GOLGA3 | 3729.08247 | -0.4096669 | 0.25872539 | -1.5834045 | 0.1133293 | 0.68183129 |
| CHFR | 1763.55257 | -0.2782479 | 0.17491126 | -1.5907949 | 0.11165574 | 0.68001961 |
| AC127070.2 | 47.8746179 | 0.19305652 | 0.38344657 | 0.50347699 | 0.61462895 | 0.93895494 |
| ZNF605 | 945.485223 | -0.301803 | 0.21335245 | -1.4145747 | 0.15719322 | 0.74522306 |
| ZNF26 | 1068.93536 | -0.2695582 | 0.21716229 | -1.2412754 | 0.21450402 | 0.79426438 |
| ZNF84 | 1775.99065 | -0.2146415 | 0.19536904 | -1.0986463 | 0.27192236 | 0.82716979 |
| ZNF140 | 1208.34115 | -0.1051632 | 0.19624679 | -0.5358725 | 0.5920467 | 0.93289552 |
| ZNF891 | 641.835625 | -0.0511363 | 0.21019965 | -0.2432748 | 0.80779256 | 0.97334138 |
| ZNF10 | 965.642316 | 0.23582186 | 0.23959188 | 0.98426484 | 3.25E-01 | 8.52E-01 |
| AC026786.1 | 171.62691 | 0.00741134 | 0.35938657 | 0.02062219 | 9.84E-01 | 9.96E-01 |
| ZNF268 | 949.464689 | 0.06719728 | 0.21188111 | 0.31714619 | 0.75113268 | 0.96272571 |
| MPHOSPH8 | 2804.10368 | 0.12340965 | 0.12067339 | 1.02267491 | 0.30646158 | 0.84278367 |
| PSPC1-AS2 | 183.23857 | -0.2874951 | 0.32241582 | -0.8916905 | 0.37255887 | 0.86791605 |
| PSPC1 | 1896.23375 | -0.3847463 | 0.25898288 | -1.4856051 | 0.13738359 | 0.72209298 |
| ZMYM5 | 847.97638 | -0.2479923 | 0.15670883 | -1.582504 | 0.11353457 | 0.68214043 |
| AL355001.2 | 149.12405 | -0.0008797 | 0.32937622 | -0.0026707 | 0.99786909 | 0.99989813 |
| ZMYM2 | 3389.42673 | -0.1450414 | 0.14759782 | -0.98268 | 0.32576495 | 0.85271781 |
| GJA3 | 82.452668 | 0.19367639 | 0.85981361 | 0.22525392 | 0.82178175 | 0.97487098 |
| GJB2 | 968.640101 | 0.72355742 | 0.47556457 | 1.5214704 | 0.12814183 | 0.70591009 |
| CRYL1 | 14539.2472 | 0.79484027 | 0.3251709 | 2.44437701 | 0.01451025 | 0.43901686 |
| IFT88 | 1664.38105 | 0.14897489 | 0.16186055 | 0.92039034 | 0.35736881 | 0.86390849 |
| AL161772.1 | 78.9881846 | -0.0561208 | 0.38923699 | -0.1441815 | 0.88535712 | 0.98356741 |
| IL17D | 108.913241 | 0.05671501 | 0.3448489 | 0.16446338 | 0.86936638 | 0.98275231 |

| | | | | | | |
|---|---|---|---|---|---|---|
| EEF1AKMT1 | 237.142432 | 0.59822875 | 0.22977192 | 2.60357638 | 0.00922567 | 0.38664841 |
| XPO4 | 1106.49824 | -0.1531392 | 0.15704686 | -0.975118 | 0.32950172 | 0.85350346 |
| LATS2 | 1485.69713 | -0.5348268 | 0.2544143 | -2.1021886 | 3.55E-02 | 5.54E-01 |
| SAP18 | 7028.3575 | 0.07879065 | 0.14113213 | 0.55827576 | 0.57665609 | 0.93127944 |
| MRPL57 | 2089.465 | 0.09720506 | 0.15847764 | 0.61336768 | 0.53963325 | 0.92198621 |
| ZDHHC20 | 1550.36394 | -0.0094877 | 0.14191746 | -0.0668534 | 0.94669844 | 0.99166614 |
| MICU2 | 3912.92302 | 0.30563259 | 0.10115568 | 3.02140798 | 0.00251602 | 0.21493583 |
| FGF9 | 1076.84059 | 0.18885153 | 0.46728986 | 0.40414215 | 0.68610819 | 0.95097288 |
| AL136962.1 | 42.0254935 | -0.1643548 | 0.48442281 | -0.3392797 | 7.34E-01 | 0.95895327 |
| LINC00621 | 1092.64733 | -0.5851787 | 0.79088382 | -0.7399047 | 0.45935781 | 0.89755243 |
| AL157931.1 | 925.686472 | -0.2076214 | 0.7691427 | -0.2699388 | 0.78720736 | 0.97090793 |
| SACS | 493.097681 | -0.5417079 | 0.39657921 | -1.3659513 | 0.17195424 | 0.75974924 |
| TNFRSF19 | 1311.3821 | 0.71406973 | 0.32308969 | 2.21012845 | 0.02709625 | 0.52773677 |
| MIPEP | 1902.61179 | 0.01591914 | 0.27019615 | 0.05891698 | 0.95301824 | 0.99170503 |
| C1QTNF9B | 69.8378916 | -1.0806342 | 0.62532886 | -1.7281054 | 0.08396933 | 0.64920344 |
| AL359736.1 | 82.0394248 | -0.7972796 | 0.29962765 | -2.6609012 | 0.00779318 | 0.36878977 |
| SPATA13 | 3550.93226 | -0.1277792 | 0.16760559 | -0.7623804 | 0.445833 | 0.89476935 |
| PARP4 | 4070.4211 | 0.3426057 | 0.21416189 | 1.59975099 | 0.10965384 | 0.67859564 |
| ATP12A | 264.548661 | 1.8708335 | 0.68426852 | 2.73406337 | 0.0062558 | 0.33470127 |
| CENPJ | 294.257139 | -0.1336046 | 0.25947954 | -0.5148947 | 0.6066266 | 0.93708311 |
| MTMR6 | 1721.98943 | -0.0242034 | 0.18441089 | -0.1312473 | 0.89557965 | 0.98474725 |
| NUP58 | 2672.65365 | -0.0319426 | 0.20653504 | -0.1546593 | 0.87708989 | 0.98275231 |
| ATP8A2 | 70.6022513 | 0.51677739 | 0.46308317 | 1.11594939 | 0.2644438 | 0.82424998 |
| SHISA2 | 395.801968 | -0.2075268 | 0.58760215 | -0.3531757 | 0.72395673 | 0.95810643 |
| RNF6 | 1730.90668 | -0.421284 | 0.37916374 | -1.1110871 | 0.26653084 | 0.8265123 |
| CDK8 | 953.867907 | 0.00122844 | 0.20442283 | 0.00600931 | 0.9952053 | 0.99940889 |
| WASF3 | 723.204942 | 0.02481072 | 0.1551047 | 0.15996107 | 0.87291174 | 0.98275231 |
| USP12 | 2687.94538 | -0.2893206 | 0.28978124 | -0.9984103 | 0.31808046 | 0.84892422 |
| RPL21 | 20076.7554 | 0.222893 | 0.25173189 | 0.88543808 | 0.3759204 | 0.86900834 |
| RASL11A | 1639.13578 | -0.0355368 | 0.33169852 | -0.1071359 | 0.91468116 | 0.9866558 |
| GTF3A | 3797.63138 | 0.02014298 | 0.19393357 | 0.10386536 | 0.9172762 | 0.9868044 |
| MTIF3 | 2309.53641 | 0.17273054 | 0.13506869 | 1.27883485 | 0.20095522 | 0.78134034 |
| LNX2 | 946.403957 | -0.1170215 | 0.25509099 | -0.4587442 | 0.64641783 | 0.94208859 |
| POLR1D | 3616.13621 | -0.2680872 | 0.20860522 | -1.2851414 | 0.19874288 | 0.78101077 |
| LINC00543 | 74.6482075 | 0.46477034 | 0.52624199 | 0.88318749 | 0.37713498 | 0.86987124 |
| FLT3 | 114.869694 | 0.95649903 | 1.00377107 | 0.95290556 | 0.34063792 | 0.85849463 |
| PAN3-AS1 | 139.689697 | -0.0522677 | 0.32437592 | -0.1611331 | 0.87198855 | 0.98275231 |
| PAN3 | 2514.25393 | -0.2000629 | 0.1308792 | -1.5286073 | 0.12636181 | 0.70431949 |
| FLT1 | 5929.30306 | 0.58554605 | 0.44613378 | 1.31248983 | 0.18935492 | 0.76970626 |
| POMP | 4169.80524 | -0.2117673 | 0.21181543 | -0.9997727 | 0.31742053 | 0.84819508 |
| SLC46A3 | 1615.25059 | 0.4188372 | 0.15223668 | 2.75122385 | 0.00593731 | 0.32920427 |
| SLC7A1 | 1390.45985 | -0.3188811 | 0.27264654 | -1.1695771 | 0.24217122 | 0.81319028 |
| UBL3 | 2606.98032 | 0.01167989 | 0.23793075 | 0.04908947 | 0.960848 | 0.99311814 |
| KATNAL1 | 692.561741 | -0.0262976 | 0.15921688 | -0.1651685 | 0.86881137 | 0.98272083 |
| LINC00426 | 47.4134817 | 0.47398444 | 0.91763308 | 0.51652937 | 0.60548473 | 0.93698713 |
| HMGB1 | 11745.8433 | 0.05348412 | 0.14671937 | 0.36453348 | 0.71545968 | 0.95590747 |
| USPL1 | 1142.28519 | 0.10983395 | 0.12572549 | 0.87360131 | 0.38233541 | 0.87185666 |
| ALOX5AP | 498.357824 | -0.1675589 | 0.45671864 | -0.3668756 | 0.71371178 | 0.95564093 |
| MEDAG | 768.518468 | -1.441221 | 0.97894919 | -1.4722123 | 0.14096355 | 0.72696697 |
| HSPH1 | 3766.58069 | -0.4372214 | 0.28521609 | -1.532948 | 0.12528866 | 0.7025837 |
| B3GLCT | 664.016966 | 0.08063833 | 0.19659144 | 0.4101823 | 0.68167223 | 0.95070301 |
| FRY | 1574.79268 | -0.130034 | 0.28785567 | -0.4517334 | 0.65146103 | 0.94393131 |
| BRCA2 | 57.7377155 | -0.8983351 | 0.54141605 | -1.6592324 | 0.09706897 | 0.66486533 |
| N4BP2L1 | 1287.21547 | 0.00200623 | 0.26380055 | 0.00760509 | 0.99393207 | 0.99931765 |

| | | | | | | |
|---|---|---|---|---|---|---|
| N4BP2L2 | 5709.38798 | -0.2003085 | 0.14227595 | -1.4078871 | 0.1591645 | 0.7475863 |
| PDS5B | 1190.05586 | -0.142214 | 0.13944115 | -1.0198855 | 0.30778277 | 0.8434443 |
| KL | 6354.89053 | 0.38822986 | 0.48630721 | 0.79832224 | 0.42468352 | 0.88573606 |
| STARD13 | 1413.97628 | -0.1402205 | 0.24069166 | -0.582573 | 0.56018076 | 0.92434754 |
| RFC3 | 261.913484 | 0.14364365 | 0.35404486 | 0.40572159 | 0.68494718 | 0.95097288 |
| NBEA | 727.688535 | -0.1573823 | 0.18489547 | -0.8511963 | 0.39466033 | 0.87884163 |
| DCLK1 | 295.844487 | -1.3408532 | 0.77224596 | -1.7363033 | 0.08251023 | 0.64661469 |
| SOHLH2 | 44.315594 | 0.19115044 | 0.78258158 | 0.24425625 | 0.80703237 | 0.97334138 |
| CCDC169-SO | 45.7876328 | 0.28978255 | 0.64885722 | 0.44660449 | 0.65516065 | 0.94532497 |
| CCDC169 | 60.3880099 | 0.42557408 | 0.55334 | 0.76910051 | 0.44183365 | 0.89318162 |
| SPART | 4835.80658 | -0.1753017 | 0.1956371 | -0.8960553 | 0.37022321 | 0.86762749 |
| SPART-AS1 | 51.300304 | 0.03944924 | 0.35480579 | 0.11118544 | 9.11E-01 | 0.98662757 |
| RFXAP | 270.194585 | -0.1838009 | 0.21804826 | -0.8429369 | 0.39926371 | 0.88077967 |
| SMAD9 | 789.629421 | -0.4068557 | 0.39069631 | -1.0413606 | 0.29770824 | 0.83937869 |
| ALG5 | 1708.92123 | 0.13083849 | 0.13497778 | 0.96933355 | 0.33237879 | 0.85496237 |
| EXOSC8 | 1495.30726 | -0.1511598 | 0.18772058 | -0.8052385 | 0.42068209 | 0.88451372 |
| SUPT20H | 2184.96165 | -0.1234861 | 0.15061188 | -0.8198964 | 0.41227515 | 0.88268208 |
| POSTN | 1878.97932 | 0.4046753 | 0.73544591 | 0.55024481 | 0.58215147 | 0.9317705 |
| TRPC4 | 163.328614 | 0.19246432 | 0.51295172 | 0.37520943 | 0.70750472 | 0.95501548 |
| UFM1 | 4564.78254 | -0.1388528 | 0.22744709 | -0.6104838 | 0.54154134 | 0.92292765 |
| FREM2 | 1970.29522 | 0.63318869 | 0.39850055 | 1.58892802 | 0.11207664 | 0.68001961 |
| PROSER1 | 1705.82355 | -0.3134229 | 0.16320181 | -1.9204622 | 0.05479955 | 0.59329767 |
| NHLRC3 | 2099.96032 | 0.28508349 | 0.27008424 | 1.05553547 | 0.2911805 | 0.83454586 |
| LHFPL6 | 2489.19651 | -0.0292802 | 0.43044514 | -0.068023 | 0.94576733 | 0.99166614 |
| COG6 | 1224.47207 | 0.04372057 | 0.18639491 | 0.23455882 | 0.81455117 | 0.9738447 |
| LINC00598 | 63.5989189 | 0.1001459 | 0.43757279 | 0.22886683 | 0.81897242 | 0.97454841 |
| FOXO1 | 1922.87487 | -0.3144256 | 0.19528803 | -1.6100609 | 0.10738456 | 0.67707349 |
| MRPS31 | 1139.88921 | 0.01741316 | 0.17978985 | 0.09685286 | 0.92284325 | 0.98834249 |
| SLC25A15 | 264.003029 | 0.36731719 | 0.29534063 | 1.24370694 | 0.21360744 | 0.7926732 |
| ELF1 | 2558.96818 | -0.3889064 | 0.12244324 | -3.1762179 | 1.49E-03 | 1.73E-01 |
| WBP4 | 932.720088 | -0.3788256 | 0.10741156 | -3.5268608 | 0.00042052 | 0.09906683 |
| KBTBD6 | 511.204048 | 0.2201722 | 0.13273147 | 1.65877918 | 0.0971603 | 0.66486533 |
| AL354696.2 | 60.3279786 | 0.3419601 | 0.42059319 | 0.81304241 | 0.41619374 | 0.88399703 |
| KBTBD7 | 435.300665 | 0.21683648 | 0.26762028 | 0.81023934 | 0.41780263 | 0.88445833 |
| MTRF1 | 732.385098 | -0.3215612 | 0.2007113 | -1.6021079 | 0.10913175 | 0.67859123 |
| NAA16 | 950.933043 | -0.1578287 | 0.15380407 | -1.0261674 | 0.30481267 | 0.84230231 |
| RGCC | 1056.62729 | -0.6292679 | 0.53758101 | -1.1705546 | 0.24177784 | 0.81269742 |
| VWA8 | 1900.98442 | 0.1586479 | 0.27124397 | 0.58489005 | 0.55862165 | 0.92421175 |
| DGKH | 1616.02528 | -0.2360935 | 0.17614515 | -1.3403347 | 0.18013654 | 0.76344164 |
| AKAP11 | 2154.03923 | 0.24642911 | 0.11857792 | 2.07820406 | 0.03769057 | 0.55538555 |
| EPSTI1 | 1871.44285 | -1.5750391 | 0.74479075 | -2.1147404 | 0.03445208 | 0.55369025 |
| DNAJC15 | 2780.41478 | 0.1101473 | 0.15743267 | 0.69964705 | 0.48414775 | 0.90541935 |
| ENOX1 | 212.43262 | 0.24580619 | 0.39933988 | 0.61553129 | 0.53820391 | 0.9214763 |
| CCDC122 | 212.180921 | 0.4837176 | 0.26302878 | 1.83902917 | 0.0659109 | 0.62237346 |
| LACC1 | 687.801692 | 0.20130229 | 0.19932686 | 1.00991054 | 0.31253815 | 0.84541735 |
| AL512506.3 | 130.73114 | 0.3484587 | 0.49078739 | 0.70999929 | 0.47770458 | 0.90387017 |
| LINC00284 | 261.441379 | 0.01605351 | 0.79320764 | 0.02023872 | 0.98385294 | 0.9961464 |
| SMIM2-AS1 | 2048.62868 | -0.1988129 | 0.47992795 | -0.4142557 | 6.79E-01 | 0.95040677 |
| SMIM2 | 102.221554 | -1.1598795 | 0.39752287 | -2.9177679 | 0.00352547 | 0.26021977 |
| AL589745.1 | 348.791019 | -0.0669704 | 0.60651062 | -0.1104192 | 9.12E-01 | 9.87E-01 |
| SERP2 | 237.084945 | -0.016329 | 0.28615939 | -0.0570625 | 9.54E-01 | 9.92E-01 |
| TSC22D1 | 21998.24 | -0.1599342 | 0.33598333 | -0.4760182 | 0.6340614 | 0.94088547 |
| TSC22D1-AS | 164.934207 | 0.02733223 | 0.49604168 | 0.05510067 | 0.95605826 | 0.99185043 |
| NUFIP1 | 384.95764 | -0.1403661 | 0.24986577 | -0.561766 | 0.57427544 | 0.93020709 |

| | | | | | | |
|---|---|---|---|---|---|---|
| GPALPP1 | 945.666118 | 0.11707752 | 0.14785682 | 0.79183037 | 0.42845959 | 0.88820133 |
| GTF2F2 | 1230.73911 | -0.171749 | 0.1671512 | -1.0275067 | 0.30418193 | 0.84211658 |
| TPT1 | 127425.027 | 0.2064902 | 0.27894805 | 0.74024607 | 4.59E-01 | 8.98E-01 |
| AL138963.3 | 71088.416 | 0.22266014 | 0.28717841 | 0.77533731 | 0.43814037 | 0.89152136 |
| SLC25A30 | 3246.42773 | 0.09444471 | 0.28336344 | 0.33329886 | 0.7389087 | 0.9602325 |
| SLC25A30-AS | 59.0345502 | 0.2335412 | 0.37393818 | 0.62454495 | 0.53226976 | 0.91985 |
| COG3 | 1923.70059 | -0.1202793 | 0.15413831 | -0.7803338 | 0.43519441 | 0.89055338 |
| ERICH6B | 186.879114 | -0.0908575 | 0.21923647 | -0.4144269 | 0.67856146 | 0.95040677 |
| LINC01055 | 551.516116 | 1.10754471 | 0.96115956 | 1.15230058 | 0.24919758 | 0.81646572 |
| SIAH3 | 64.4601233 | 0.53828603 | 0.71842229 | 0.74926131 | 0.45369972 | 0.89670027 |
| ZC3H13 | 1950.1347 | -0.1109312 | 0.12027589 | -0.9223058 | 0.35636907 | 0.86390849 |
| LCP1 | 2072.42218 | -0.1084809 | 0.59290401 | -0.1829654 | 0.85482519 | 0.97910552 |
| RUBCNL | 166.163191 | -0.8091222 | 0.57427702 | -1.4089406 | 0.15885274 | 0.74706632 |
| LRCH1 | 2095.79402 | 0.03856272 | 0.15949816 | 0.24177533 | 0.80895425 | 0.97334138 |
| ESD | 5217.91689 | 0.18703249 | 0.12627347 | 1.48117018 | 0.13856123 | 0.72333593 |
| SUCLA2 | 3626.05922 | 0.24359721 | 0.19008485 | 1.2815183 | 0.20001168 | 0.78134034 |
| NUDT15 | 1097.88196 | -0.0585178 | 0.22100369 | -0.2647822 | 0.79117727 | 0.97090793 |
| MED4 | 2029.0778 | -0.135948 | 0.14696825 | -0.9250164 | 0.35495738 | 0.86364633 |
| MED4-AS1 | 832.583665 | -0.1162253 | 0.16234677 | -0.7159077 | 0.47404833 | 0.90199039 |
| ITM2B | 80180.9071 | 0.34788699 | 0.15742767 | 2.20982116 | 0.02711758 | 0.52773677 |
| RB1 | 2281.75967 | -0.0839314 | 0.16327428 | -0.5140516 | 0.6072159 | 0.93739614 |
| LPAR6 | 1866.12664 | -0.1915081 | 0.29709986 | -0.6445918 | 0.51919178 | 0.91636259 |
| RCBTB2 | 1651.79526 | 0.20497427 | 0.21797843 | 0.94034199 | 0.34704217 | 0.86156011 |
| CYSLTR2 | 121.039539 | -0.3448785 | 0.42051591 | -0.8201319 | 0.41214093 | 0.88268208 |
| AL161421.1 | 108.229803 | -0.2983988 | 0.37897606 | -0.7873817 | 0.43105847 | 0.88966606 |
| FNDC3A | 3501.94305 | 0.03373812 | 0.17063791 | 0.19771761 | 0.84326601 | 0.97649449 |
| CDADC1 | 837.999916 | -0.040778 | 0.26140665 | -0.1559944 | 0.87603742 | 0.98275231 |
| CAB39L | 911.611427 | -0.0443499 | 0.23591906 | -0.1879879 | 0.85088616 | 0.97844748 |
| SETDB2 | 1067.43062 | -0.1422721 | 0.20519769 | -0.6933414 | 0.48809533 | 0.90585781 |
| PHF11 | 2211.51714 | -0.6421738 | 0.32629739 | -1.968063 | 0.04906079 | 0.58508842 |
| RCBTB1 | 1929.66611 | 0.09037336 | 0.17616197 | 0.51301287 | 0.60794232 | 0.93748023 |
| EBPL | 683.326103 | 0.03651327 | 0.2648861 | 0.13784517 | 0.89036279 | 0.98466501 |
| KPNA3 | 1620.28332 | -0.3536822 | 0.25678145 | -1.3773666 | 0.16839894 | 0.75735257 |
| SPRYD7 | 859.054756 | 0.27014328 | 0.19223379 | 1.40528512 | 1.60E-01 | 0.74781837 |
| DLEU2 | 286.049281 | -0.1307818 | 0.16677715 | -0.7841708 | 0.43293992 | 0.89017484 |
| TRIM13 | 1841.75333 | -0.0729663 | 0.13237815 | -0.5511961 | 0.58149924 | 0.9317705 |
| KCNRG | 108.580577 | 0.17364416 | 0.28130118 | 0.61728913 | 0.53704404 | 0.92105614 |
| RNASEH2B | 868.061562 | 0.10640032 | 0.19824333 | 0.53671577 | 0.59146396 | 0.93289552 |
| FAM124A | 724.554318 | -0.9431029 | 0.48124755 | -1.9597043 | 0.05003036 | 0.5861488 |
| SERPINE3 | 78.5765556 | -0.0560223 | 0.26429314 | -0.2119705 | 0.83213007 | 0.97550416 |
| INTS6 | 1519.21281 | -0.2300708 | 0.14282957 | -1.6108062 | 0.10722197 | 0.67707349 |
| INTS6-AS1 | 296.835184 | 0.06698289 | 0.30290533 | 0.22113474 | 0.82498752 | 0.97548201 |
| WDFY2 | 1267.73891 | -0.0511661 | 0.19386238 | -0.2639298 | 0.79183398 | 0.97090793 |
| AL162377.3 | 1316.61687 | 0.31286239 | 0.19469075 | 1.60697101 | 0.10806073 | 0.67776643 |
| DHRS12 | 1048.21439 | 0.5349943 | 0.27852608 | 1.92080505 | 0.05475629 | 0.59329767 |
| AL162377.1 | 213.547913 | 0.44620396 | 0.33979201 | 1.31316787 | 0.18912639 | 0.76970626 |
| ATP7B | 605.906738 | 0.01433037 | 0.19906798 | 0.07198731 | 0.94261201 | 0.99136103 |
| ALG11 | 253.707079 | -0.1397561 | 0.20092439 | -0.6955657 | 0.48670087 | 0.90573694 |
| UTP14C | 1329.16162 | -0.0633273 | 0.10753601 | -0.5888936 | 0.55593267 | 0.92373235 |
| NEK5 | 126.257884 | -0.2727575 | 0.62452127 | -0.4367466 | 0.66229515 | 0.94642611 |
| NEK3 | 979.954637 | 0.09448248 | 0.30581879 | 0.30894924 | 0.75736014 | 0.96502715 |
| THSD1 | 275.327548 | -0.1838408 | 0.31319718 | -0.586981 | 0.55721649 | 0.92373235 |
| VPS36 | 2997.4286 | 0.04579691 | 0.09083293 | 0.5041884 | 0.61412899 | 0.93895494 |
| CKAP2 | 432.88031 | 0.4092896 | 0.1996681 | 2.04984969 | 0.0403791 | 0.55657615 |

| | | | | | | |
|---|---|---|---|---|---|---|
| HNRNPA1L2 | 439.649649 | 0.02936696 | 0.26134149 | 0.11237007 | 0.91052999 | 0.98662757 |
| AL139089.1 | 103.564442 | 0.26428148 | 0.38737723 | 0.68223288 | 0.49509171 | 0.90816709 |
| SUGT1 | 2727.00272 | -0.042842 | 0.11173312 | -0.3834316 | 0.70139975 | 0.95424422 |
| OLFM4 | 7054.47719 | -0.5187637 | 0.73386706 | -0.7068906 | 0.47963445 | 0.9051283 |
| PCDH17 | 464.271585 | -0.6782906 | 0.45707053 | -1.4839955 | 0.1378101 | 0.72209298 |
| DIAPH3 | 146.916316 | -0.1204429 | 0.61811256 | -0.194856 | 0.8455057 | 0.97675487 |
| TDRD3 | 1067.09588 | 0.04304758 | 0.18940027 | 0.22728362 | 0.82020321 | 0.97454841 |
| PCDH20 | 107.46002 | -0.4971742 | 0.70552228 | -0.7046896 | 0.48100345 | 0.90541935 |
| AL592490.1 | 106.031129 | -0.493563 | 0.70767508 | -0.697443 | 0.48552562 | 0.90543654 |
| PCDH9 | 495.59509 | -0.5982731 | 0.41290159 | -1.4489483 | 0.14735201 | 0.73386234 |
| DACH1 | 751.371791 | 0.32243555 | 0.33450778 | 0.96391048 | 0.33509081 | 0.85581042 |
| MZT1 | 571.143604 | 0.07101059 | 0.22031569 | 0.32231292 | 0.74721564 | 0.96224616 |
| BORA | 301.757245 | -0.2174853 | 0.19876385 | -1.0941896 | 0.27387183 | 0.82719197 |
| DIS3 | 1861.16192 | -0.235497 | 0.17861615 | -1.318453 | 1.87E-01 | 0.76958556 |
| PIBF1 | 730.892726 | -0.0033797 | 0.1742179 | -0.0193991 | 9.85E-01 | 0.99654013 |
| KLF5 | 1450.17093 | -0.1003011 | 0.33222021 | -0.3019114 | 0.76271961 | 0.96634182 |
| KLF12 | 981.381296 | -0.2431937 | 0.19478915 | -1.2484969 | 0.21184914 | 0.79199375 |
| TBC1D4 | 3866.71533 | -0.3050273 | 0.24991617 | -1.2205185 | 0.22226838 | 0.80078694 |
| COMMD6 | 2401.44281 | 0.2679983 | 0.19419578 | 1.38004183 | 0.16757377 | 0.75565447 |
| UCHL3 | 1231.56412 | -0.3081484 | 0.26326312 | -1.170496 | 0.24180144 | 0.81269742 |
| LMO7-AS1 | 424.523786 | -0.2305171 | 0.24949901 | -0.9239199 | 0.35552803 | 0.86364633 |
| LMO7 | 4118.25132 | 0.09351247 | 0.32013606 | 0.29210229 | 0.77020842 | 0.96836854 |
| KCTD12 | 3742.83664 | 0.37882432 | 0.36790576 | 1.02967763 | 0.30316136 | 0.84211658 |
| CLN5 | 4727.59649 | 0.22600815 | 0.10685248 | 2.11514181 | 0.03441786 | 0.55369025 |
| AC001226.2 | 1173.08801 | 0.34266269 | 0.148851 | 2.3020517 | 0.02133226 | 0.51189223 |
| FBXL3 | 5559.0285 | 0.05956199 | 0.12818681 | 0.46464993 | 6.42E-01 | 9.41E-01 |
| MYCBP2 | 1885.04851 | -0.0879949 | 0.12911544 | -0.6815211 | 0.49554184 | 0.9085257 |
| SLAIN1 | 433.76074 | 0.30903382 | 0.19748604 | 1.56483879 | 0.11762071 | 0.69036323 |
| EDNRB | 3466.67436 | -0.3811769 | 0.40569273 | -0.9395704 | 0.34743795 | 0.86197176 |
| RNF219-AS1 | 190.648638 | -0.1454484 | 0.26773501 | -0.5432552 | 0.58695409 | 0.93289552 |
| RNF219 | 531.516267 | -0.1273718 | 0.18582749 | -0.6854303 | 0.49307245 | 0.90711083 |
| RBM26 | 2477.26517 | -0.0011133 | 0.11091156 | -0.0100378 | 0.99199114 | 0.99906045 |
| RBM26-AS1 | 111.457068 | 0.08022896 | 0.38185142 | 0.21010517 | 0.83358559 | 0.97550416 |
| NDFIP2 | 3521.67634 | 0.05903888 | 0.28253675 | 0.20895999 | 0.83447947 | 0.97550416 |
| SPRY2 | 1386.09408 | -0.4845291 | 0.29939692 | -1.6183502 | 0.10558714 | 0.67526487 |
| MIR4500HG | 51.8459601 | 2.22385958 | 0.87831153 | 2.53197129 | 0.01134233 | 0.40335212 |
| LINC00379 | 77.7794722 | 3.17764872 | 0.99022532 | 3.2090158 | 0.0013319 | 0.16208552 |
| GPC5 | 184.281944 | 2.65637922 | 0.81631903 | 3.25409445 | 0.00113754 | 0.15697311 |
| GPC6 | 897.771169 | -0.0108318 | 0.3989547 | -0.0271503 | 0.97833982 | 0.99476647 |
| DCT | 37.3634008 | 0.77456613 | 0.68008824 | 1.13892005 | 2.55E-01 | 8.20E-01 |
| TGDS | 797.443098 | 0.01743957 | 0.12563368 | 0.13881285 | 0.88959804 | 0.9845209 |
| GPR180 | 833.376205 | -0.7202248 | 0.25577156 | -2.8158908 | 0.00486422 | 0.30193205 |
| ABCC4 | 2524.03268 | 0.33897688 | 0.26219805 | 1.29282761 | 0.19607069 | 0.77791948 |
| CLDN10 | 6293.85289 | 0.1347226 | 0.39690579 | 0.33943218 | 0.73428418 | 0.95895327 |
| CLDN10-AS1 | 108.549235 | -0.72375 | 0.53124792 | -1.3623583 | 0.17308484 | 0.75974924 |
| DZIP1 | 3487.78166 | 0.18447663 | 0.19909904 | 0.92655711 | 0.35415653 | 0.86364633 |
| DNAJC3-DT | 323.329112 | 0.44404694 | 0.39991927 | 1.11034145 | 0.26685192 | 0.8265123 |
| DNAJC3 | 4164.80225 | 0.14887045 | 0.25420922 | 0.58562176 | 0.55812972 | 0.92421175 |
| UGGT2 | 1242.866 | -0.0059714 | 0.18270698 | -0.0326829 | 9.74E-01 | 9.95E-01 |
| HS6ST3 | 97.3939293 | -0.3012047 | 0.55015057 | -0.5474951 | 0.58403863 | 0.93236269 |
| OXGR1 | 97.3366212 | 1.65295353 | 0.65051256 | 2.5410017 | 0.01105354 | 0.40196898 |
| MBNL2 | 2929.83138 | -0.0512588 | 0.1877487 | -0.273018 | 0.78483941 | 0.97022925 |
| RAP2A | 1663.56358 | -0.2146795 | 0.25308115 | -0.8482636 | 0.39629116 | 0.87967259 |
| IPO5 | 4840.18113 | 0.02324561 | 0.21127924 | 0.11002317 | 0.912391 | 0.98662757 |

| | | | | | | |
|---|---|---|---|---|---|---|
| FARP1 | 17433.8189 | -0.1986643 | 0.2235208 | -0.8887958 | 0.37411285 | 0.86791605 |
| FARP1-AS1 | 762.696413 | -0.1095218 | 0.23045919 | -0.4752328 | 0.63462104 | 0.94088547 |
| STK24 | 13696.0095 | -0.2485835 | 0.20513075 | -1.2118296 | 0.22557761 | 0.80214224 |
| SLC15A1 | 1672.59205 | 0.81728572 | 0.65547647 | 1.24685745 | 0.21244977 | 0.79203904 |
| DOCK9 | 1897.90633 | -0.3425412 | 0.28268211 | -1.2117541 | 2.26E-01 | 8.02E-01 |
| DOCK9-DT | 61.5798566 | -0.0934028 | 0.30328431 | -0.3079711 | 0.75810433 | 0.96502715 |
| UBAC2-AS1 | 52.5498709 | 0.07354018 | 0.50435794 | 0.14580951 | 0.88407177 | 0.98317437 |
| UBAC2 | 3494.29049 | 0.04043577 | 0.11762916 | 0.34375637 | 0.73102952 | 0.95848028 |
| GPR183 | 372.262158 | -0.7343985 | 0.88491573 | -0.8299078 | 0.40659092 | 0.88132015 |
| TM9SF2 | 12129.3665 | 0.09274056 | 0.18220924 | 0.50897836 | 0.6107674 | 0.93807505 |
| CLYBL | 1698.14868 | 0.33811642 | 0.43398673 | 0.7790939 | 0.4359244 | 0.89095725 |
| AL355338.1 | 278.268376 | -0.3729703 | 0.17790738 | -2.0964298 | 0.03604408 | 0.55369025 |
| PCCA | 7081.01625 | 0.56816846 | 0.27384038 | 2.07481624 | 0.03800357 | 0.55538555 |
| GGACT | 3169.10591 | 0.45553701 | 0.41330934 | 1.10216964 | 2.70E-01 | 0.82716979 |
| TMTC4 | 1259.81276 | 0.318068 | 0.12368548 | 2.57158726 | 0.01012335 | 0.39989496 |
| NALCN | 43.4804116 | -0.0159439 | 0.56052071 | -0.0284448 | 0.97730743 | 0.99458341 |
| ITGBL1 | 250.258844 | -0.5855681 | 0.69624098 | -0.8410423 | 0.40032422 | 0.88086294 |
| FGF14 | 208.403687 | -0.5537941 | 0.63327123 | -0.8744975 | 0.38184736 | 0.87185666 |
| FGF14-AS2 | 226.098753 | 0.17609202 | 0.34428112 | 0.51147742 | 0.6090168 | 0.93748023 |
| TPP2 | 1965.94654 | -0.025967 | 0.12840766 | -0.2022232 | 0.83974223 | 0.97618283 |
| TEX30 | 939.562699 | -0.1256135 | 0.23395109 | -0.5369221 | 0.59132145 | 0.93289552 |
| KDELC1 | 347.851765 | 0.12851369 | 0.25135012 | 0.51129354 | 0.60914553 | 0.93748023 |
| BIVM | 2527.35087 | 0.23371934 | 0.20905486 | 1.11798089 | 0.26357516 | 0.82424998 |
| BIVM-ERCC5 | 3680.64182 | -0.0036778 | 0.15483129 | -0.0237535 | 0.98104926 | 0.99576712 |
| ERCC5 | 3741.88036 | -0.0020446 | 0.15545291 | -0.0131528 | 0.98950587 | 0.99800524 |
| SLC10A2 | 473.623452 | 0.86364036 | 0.74176304 | 1.16430761 | 0.24429937 | 0.81468457 |
| EFNB2 | 1698.89058 | 0.05264967 | 0.37637184 | 0.13988737 | 0.88874898 | 0.98423722 |
| AL138689.2 | 271.458699 | 0.04375981 | 0.38022271 | 0.11508994 | 0.90837384 | 0.98662757 |
| ARGLU1 | 6470.25266 | -0.1145681 | 0.1572272 | -0.7286785 | 0.46619836 | 0.89960479 |
| LINC00551 | 125.95492 | 0.27429699 | 0.63817527 | 0.42981451 | 0.66733057 | 0.94830184 |
| FAM155A | 69.1028901 | -0.2341246 | 0.39985134 | -0.5855292 | 0.55819196 | 0.92421175 |
| LIG4 | 1039.18572 | -0.2976156 | 0.25658598 | -1.1599061 | 0.24608705 | 0.81542272 |
| ABHD13 | 939.045493 | -0.3647528 | 0.21912408 | -1.6645946 | 0.09599367 | 0.66168746 |
| TNFSF13B | 279.213978 | -0.5388274 | 0.62587615 | -0.8609169 | 0.38928381 | 0.87568585 |
| IRS2 | 1274.23412 | -0.4112235 | 0.36198858 | -1.1360125 | 0.25595135 | 0.82052927 |
| COL4A1 | 16002.1873 | -0.4076478 | 0.37189018 | -1.0961511 | 0.27301264 | 0.82716979 |
| COL4A2 | 24097.7559 | -0.2237816 | 0.30280876 | -0.7390195 | 0.45989518 | 0.89755243 |
| COL4A2-AS2 | 424.39384 | -0.2284093 | 0.36981452 | -0.6176319 | 0.536818 | 0.92105614 |
| COL4A2-AS1 | 1532.83136 | -0.315095 | 0.35091889 | -0.897914 | 0.36923141 | 0.86762749 |
| RAB20 | 2509.80737 | -0.0570278 | 0.37029236 | -0.1540074 | 0.87760388 | 0.98275231 |
| AL139385.1 | 201.12342 | -0.0522501 | 0.25458363 | -0.2052375 | 8.37E-01 | 9.76E-01 |
| NAXD | 4515.97165 | 0.03168343 | 0.1030435 | 0.30747625 | 7.58E-01 | 0.96502715 |
| CARS2 | 2812.95025 | 0.04587033 | 0.09589988 | 0.47831479 | 0.63242617 | 0.94088547 |
| ING1 | 914.355629 | 0.13751736 | 0.12957874 | 1.06126482 | 0.28856957 | 0.83295411 |
| LINC00346 | 89.3598434 | 0.0209614 | 0.40344333 | 0.05195625 | 0.95856356 | 0.99237995 |
| ANKRD10 | 7785.03455 | -0.1412515 | 0.15236478 | -0.9270616 | 0.35389454 | 0.86364633 |
| ARHGEF7-AS | 44.0361462 | 0.11710044 | 0.55726332 | 0.21013483 | 0.83356244 | 0.97550416 |
| ARHGEF7 | 3875.72269 | -0.1748124 | 0.1402188 | -1.2467114 | 0.21250334 | 0.79203904 |
| TUBGCP3 | 1062.15137 | 0.3231445 | 0.20375658 | 1.58593406 | 0.11275427 | 0.68098515 |
| AL139384.1 | 70.7303128 | 0.3399943 | 0.55890307 | 0.60832428 | 0.54297242 | 0.92297267 |
| ATP11A | 4021.28329 | -0.1473858 | 0.23685919 | -0.6222506 | 0.53377711 | 0.92062041 |
| MCF2L | 3280.70932 | -0.2908677 | 0.31506338 | -0.9232038 | 0.355901 | 0.86364633 |
| AL356740.1 | 63.1031262 | 0.4650169 | 0.51060198 | 0.91072287 | 0.36244141 | 0.86561928 |
| MCF2L-AS1 | 84.2905507 | 0.08999227 | 0.33428454 | 0.26920859 | 0.78776917 | 0.97090793 |

| | | | | | | |
|---|---|---|---|---|---|---|
| F7 | 43.9469406 | -0.1513535 | 0.45715259 | -0.3310787 | 7.41E-01 | 9.61E-01 |
| F10 | 288.623779 | -0.319734 | 0.31301798 | -1.0214557 | 0.30703857 | 0.84286067 |
| AL137002.2 | 132.786842 | 0.04989628 | 0.30822893 | 0.16188059 | 0.87139989 | 0.98275231 |
| PROZ | 1095.58738 | 0.12007334 | 0.53937152 | 0.22261713 | 0.8238335 | 0.97548201 |
| AL137002.1 | 174.828873 | 0.21968149 | 0.61724722 | 0.35590519 | 0.72191157 | 0.95739595 |
| PCID2 | 1868.80467 | -0.1401295 | 0.1358301 | -1.0316525 | 0.30223493 | 0.84211658 |
| CUL4A | 4269.99674 | -0.1301636 | 0.20049164 | -0.6492221 | 5.16E-01 | 0.91636259 |
| LAMP1 | 22404.6666 | 0.28987133 | 0.20602839 | 1.40694852 | 0.15944265 | 0.74762924 |
| GRTP1 | 1654.39489 | 0.15535935 | 0.3379704 | 0.45968329 | 0.64574357 | 0.94208859 |
| GRTP1-AS1 | 277.136111 | 0.30245083 | 0.37302809 | 0.81079908 | 0.41748106 | 0.88445833 |
| ADPRHL1 | 142.542634 | -0.1970463 | 0.26851821 | -0.7338285 | 0.46305324 | 0.89807153 |
| DCUN1D2 | 1026.61497 | -0.3173595 | 0.19990011 | -1.5875902 | 0.11237903 | 0.68001961 |
| TMCO3 | 3549.6368 | 0.08610433 | 0.24167593 | 0.35628014 | 0.72163078 | 0.95739595 |
| TFDP1 | 2522.88521 | -0.3714756 | 0.19511665 | -1.9038642 | 0.05692788 | 0.60049913 |
| TMEM255B | 1508.82908 | -0.2723307 | 0.51681149 | -0.5269439 | 5.98E-01 | 0.9346978 |
| GAS6-AS1 | 2373.90064 | 0.35248138 | 0.37992506 | 0.92776553 | 0.35352921 | 0.86364633 |
| GAS6 | 7990.6744 | 0.20088808 | 0.35052648 | 0.57310387 | 5.67E-01 | 9.26E-01 |
| GAS6-DT | 98.9417471 | 0.2209111 | 0.47507567 | 0.46500192 | 0.64193008 | 0.94088547 |
| RASA3 | 1326.73779 | -0.1508232 | 0.36550553 | -0.4126429 | 0.67986829 | 0.95070301 |
| CDC16 | 3544.24706 | 0.03301726 | 0.12867185 | 0.25660048 | 0.7974872 | 0.97148232 |
| UPF3A | 2580.04671 | -0.209728 | 0.24523425 | -0.855215 | 0.39243213 | 0.8778303 |
| CHAMP1 | 801.33523 | -0.1248318 | 0.2683252 | -0.4652259 | 0.6417697 | 0.94088547 |
| TTC5 | 694.823919 | 0.11757876 | 0.16849749 | 0.69780715 | 0.48529781 | 0.90543654 |
| AL356019.2 | 131.285968 | 0.39637294 | 0.27374626 | 1.44795747 | 0.14762895 | 0.73443345 |
| CCNB1IP1 | 2592.95777 | -0.0361591 | 0.22600596 | -0.1599919 | 0.87288742 | 0.98275231 |
| AL355075.4 | 97.3969259 | 0.27823796 | 0.50737743 | 0.54838458 | 0.58342787 | 0.93222113 |
| PARP2 | 1127.97251 | 0.05143127 | 0.17268907 | 0.29782583 | 0.7658361 | 0.96704052 |
| AL355075.1 | 97.7871785 | 0.19287955 | 0.30995788 | 0.62227664 | 0.53375998 | 0.92062041 |
| TEP1 | 2064.91993 | -0.2345746 | 0.13777195 | -1.7026297 | 0.08863739 | 0.65363785 |
| KLHL33 | 55.2365712 | 0.4785001 | 0.59105874 | 0.80956438 | 0.41819059 | 0.88445833 |
| OSGEP | 2017.97039 | -0.1291436 | 0.20900756 | -0.6178896 | 0.53664809 | 0.92105614 |
| AL355075.2 | 79.9744257 | -0.526934 | 0.31576386 | -1.6687597 | 0.09516502 | 0.65888357 |
| APEX1 | 5335.96418 | -0.07756 | 0.23559519 | -0.3292086 | 7.42E-01 | 9.61E-01 |
| PIP4P1 | 3000.82938 | -0.4269596 | 0.22759015 | -1.8760021 | 6.07E-02 | 6.10E-01 |
| PNP | 9726.2499 | 0.10572701 | 0.41153596 | 0.25690832 | 7.97E-01 | 9.71E-01 |
| AL355075.3 | 4258.75989 | 0.13646604 | 0.42404191 | 0.32182206 | 0.7475875 | 0.96231266 |
| RNASE4 | 4586.57486 | 0.16469212 | 0.41904093 | 0.39302157 | 0.69430355 | 0.95222846 |
| ANG | 2079.82855 | -0.0925092 | 0.42975839 | -0.2152587 | 0.8295656 | 0.97550416 |
| AL163636.2 | 4675.96 | 0.14614012 | 0.42609215 | 0.34297774 | 0.73161521 | 0.95848028 |
| AL163636.1 | 1786.9031 | 0.06367458 | 0.41891664 | 0.15199822 | 0.87918834 | 0.98275231 |
| RNASE6 | 430.382584 | -0.1258913 | 0.6038353 | -0.2084861 | 0.83484943 | 0.97550416 |
| RNASE1 | 7191.16619 | -0.6079699 | 0.39100953 | -1.5548723 | 0.11997648 | 0.6944037 |
| RNASE2 | 96.9354648 | 0.19183905 | 0.90472757 | 0.21204068 | 0.8320753 | 0.97550416 |
| METTL17 | 4083.56158 | -0.1071161 | 0.10670478 | -1.0038548 | 0.3154486 | 0.84712537 |
| AL355922.2 | 1713.79687 | -0.0862195 | 0.1389363 | -0.6205686 | 0.53488348 | 0.92084006 |
| NDRG2 | 9021.28 | -0.0291683 | 0.28309593 | -0.1030331 | 0.91793668 | 0.9868044 |
| TPPP2 | 121.941522 | 0.15073059 | 0.38554793 | 0.39095163 | 0.69583299 | 0.9525827 |
| AL161668.3 | 96.8869791 | 0.28911101 | 0.44567559 | 0.64870283 | 0.51653048 | 0.91636259 |
| ARHGEF40 | 3559.15644 | -0.044546 | 0.1189109 | -0.3746166 | 0.70794564 | 0.95501548 |
| ZNF219 | 1834.80784 | -0.2359175 | 0.29664829 | -0.7952769 | 0.42645247 | 0.8870306 |
| TMEM253 | 51.3291313 | 0.26671437 | 0.4043525 | 0.65960857 | 0.50950506 | 0.91439937 |
| HNRNPC | 17957.1842 | 0.06483429 | 0.14956357 | 0.43348986 | 0.66465893 | 0.94681198 |
| SUPT16H | 3490.16836 | -0.0396207 | 0.13951204 | -0.2839948 | 0.77641436 | 0.96836854 |
| CHD8 | 2313.75084 | -0.2713261 | 0.13489329 | -2.011413 | 0.04428185 | 0.57267043 |

| | | | | | | |
|---|---|---|---|---|---|---|
| RAB2B | 1385.62053 | -0.1153434 | 0.19542461 | -0.5902196 | 0.55504342 | 0.92373235 |
| TOX4 | 4191.74821 | -0.1156175 | 0.08531801 | -1.355136 | 0.17537423 | 0.76073329 |
| METTL3 | 3233.18769 | -0.2241307 | 0.11784453 | -1.9019186 | 5.72E-02 | 0.60069729 |
| SALL2 | 642.468941 | 0.04925878 | 0.32650209 | 0.15086818 | 0.8800797 | 0.98275231 |
| AL161747.2 | 122.794727 | -0.0063969 | 0.38187492 | -0.0167512 | 0.98663507 | 0.99715899 |
| TRAV30 | 115.426683 | 0.18336322 | 0.91268839 | 0.20090452 | 0.84077323 | 0.97618283 |
| TRAC | 407.566715 | 0.8115482 | 0.78655602 | 1.03177419 | 0.30217791 | 0.84211658 |
| DAD1 | 6288.46791 | -0.11465 | 0.24080473 | -0.4761119 | 0.63399466 | 0.94088547 |
| AC243965.2 | 152.945082 | -0.262394 | 0.32426068 | -0.8092069 | 0.41839613 | 0.88445833 |
| ABHD4 | 4933.13347 | -0.2315814 | 0.28531867 | -0.8116586 | 0.41698755 | 0.88445833 |
| OXA1L | 6847.6622 | 0.12777913 | 0.20028806 | 0.63797675 | 0.52348881 | 0.91723878 |
| SLC7A7 | 9707.23178 | 0.36394354 | 0.58331221 | 0.6239258 | 5.33E-01 | 9.20E-01 |
| MRPL52 | 2336.61207 | 0.09398805 | 0.1442136 | 0.65172803 | 5.15E-01 | 9.16E-01 |
| MMP14 | 9484.58743 | 0.01740727 | 0.24027383 | 0.07244764 | 9.42E-01 | 9.91E-01 |
| LRP10 | 22421.4599 | -0.0929328 | 0.17033012 | -0.5456041 | 0.5853381 | 0.93289552 |
| REM2 | 84.0500482 | 0.16439796 | 0.32135751 | 0.51157341 | 0.6089496 | 0.93748023 |
| RBM23 | 4592.42869 | -0.0675872 | 0.09920601 | -0.6812813 | 0.49569349 | 0.90861354 |
| PRMT5-AS1 | 2653.86616 | 0.08452185 | 0.14826411 | 0.57007624 | 0.56862599 | 0.92642835 |
| PRMT5 | 3297.22301 | 0.0931219 | 0.15281217 | 0.60938795 | 0.54226732 | 0.92292765 |
| AL132780.1 | 260.841286 | 0.54188294 | 0.33403399 | 1.62223893 | 0.1047522 | 0.67439401 |
| HAUS4 | 1828.47281 | 0.20058012 | 0.28675474 | 0.69948319 | 0.48425011 | 0.90541935 |
| AL132780.3 | 1269.43422 | 0.13254038 | 0.28431472 | 0.4661749 | 0.64109031 | 0.94088547 |
| AJUBA | 2094.43333 | -0.7520965 | 0.23410431 | -3.2126555 | 0.00131514 | 0.16147422 |
| AL132780.2 | 206.689469 | 0.28526698 | 0.22797012 | 1.25133497 | 0.21081229 | 0.79134644 |
| C14orf93 | 669.862095 | 0.21190078 | 0.21829961 | 0.97068787 | 0.33170374 | 0.85437874 |
| PSMB5 | 5490.92856 | -0.0774664 | 0.18932708 | -0.4091673 | 0.68241691 | 0.95070301 |
| CDH24 | 546.461643 | -0.2418368 | 0.29565654 | -0.8179655 | 0.4133769 | 0.88316314 |
| ACIN1 | 9630.7412 | -0.2656864 | 0.12441765 | -2.1354395 | 3.27E-02 | 5.51E-01 |
| C14orf119 | 2663.72937 | -0.1775162 | 0.14568059 | -1.2185301 | 0.22302261 | 0.80078694 |
| SLC7A8 | 15062.7218 | 0.21360299 | 0.60088728 | 0.35547931 | 0.72223055 | 0.95739595 |
| RNF212B | 402.534859 | 0.70492737 | 0.62035584 | 1.13632745 | 0.25581955 | 0.82052927 |
| HOMEZ | 1362.79207 | 0.27431209 | 0.24706045 | 1.11030356 | 0.26686824 | 0.8265123 |
| PPP1R3E | 1459.91439 | 0.21284406 | 0.36679637 | 0.58027853 | 0.5617268 | 0.92452095 |
| BCL2L2 | 4914.79472 | -0.180089 | 0.13775098 | -1.3073517 | 0.19109329 | 0.77292503 |
| BCL2L2-PABI | 6857.61155 | -0.2273782 | 0.11684387 | -1.9460005 | 0.05165468 | 0.58645127 |
| PABPN1 | 13215.2604 | -0.2281095 | 0.15011183 | -1.5195967 | 0.12861236 | 0.70610612 |
| SLC22A17 | 3455.11583 | 0.23322455 | 0.30685152 | 0.7600567 | 0.44722069 | 0.89480207 |
| EFS | 643.6827 | -0.4937791 | 0.51945751 | -0.9505668 | 0.3418243 | 0.85889221 |
| AL049829.2 | 131.96112 | -0.3420348 | 0.51091632 | -0.6694536 | 5.03E-01 | 0.91356552 |
| MYH7 | 95.3096651 | 4.67801648 | 1.64348286 | 2.84640417 | NA | NA |
| NGDN | 1458.70365 | -0.1741404 | 0.157404 | -1.1063276 | 2.69E-01 | 8.27E-01 |
| ZFHX2 | 448.842929 | -0.1313293 | 0.33399508 | -0.3932074 | 0.69416628 | 0.95222846 |
| THTPA | 2790.71159 | -0.3300177 | 0.1466906 | -2.2497535 | 0.0244646 | 0.51967919 |
| AP1G2 | 8481.78079 | -0.5326494 | 0.15612948 | -3.4115874 | 0.00064586 | 0.12385487 |
| AL135999.1 | 4150.87994 | -0.524822 | 0.16198622 | -3.2399175 | 0.00119564 | 0.15779959 |
| JPH4 | 221.533142 | -0.1595719 | 0.40742866 | -0.3916559 | 0.69531245 | 0.95248074 |
| DHRS2 | 59.913708 | 3.51635577 | 0.9283467 | 3.78776136 | 0.00015201 | 0.0593507 |
| DHRS4-AS1 | 3109.89479 | 0.84005418 | 0.39327331 | 2.13605692 | 0.03267477 | 0.55104386 |
| DHRS4 | 3268.82603 | 0.96294377 | 0.4432864 | 2.1722836 | 0.02983428 | 0.53969519 |
| DHRS4L2 | 2343.21223 | 0.76740198 | 0.37240843 | 2.0606461 | 0.03933682 | 0.55657615 |
| CARMIL3 | 212.199918 | 0.7005999 | 0.33840468 | 2.070302 | 3.84E-02 | 0.55563699 |
| CPNE6 | 271.7147 | -0.8254307 | 0.99480376 | -0.8297423 | 0.40668451 | 0.88132015 |
| NRL | 253.164388 | 0.12357611 | 0.30536301 | 0.40468592 | 0.68570839 | 0.95097288 |
| PCK2 | 12717.0993 | 0.82520829 | 0.42951044 | 1.92127641 | 0.05469687 | 0.59329767 |

| | | | | | | |
|---|---|---|---|---|---|---|
| DCAF11 | 6144.56208 | 0.22726222 | 0.17522803 | 1.29695133 | 0.19464793 | 0.7762856 |
| AL136295.3 | 1330.79302 | 0.16907729 | 0.18052695 | 0.93657647 | 0.34897646 | 0.86356499 |
| FITM1 | 93.8391953 | -0.0149713 | 0.23377524 | -0.0640416 | 0.94893708 | 0.99170503 |
| PSME1 | 8860.59303 | 0.0626747 | 0.1925087 | 0.32556814 | 0.74475112 | 0.9613927 |
| EMC9 | 1113.9498 | 0.00257706 | 0.27389728 | 0.00940887 | 0.99249292 | 0.99906045 |
| AL136295.2 | 449.379984 | -0.1049268 | 0.28076326 | -0.37372 | 0.70861265 | 0.95501548 |
| PSME2 | 6302.0554 | -0.1990752 | 0.3266359 | -0.6094712 | 5.42E-01 | 0.92292765 |
| RNF31 | 3301.70113 | -0.1442825 | 0.22954173 | -0.6285675 | 0.52963227 | 0.91931447 |
| AL136295.5 | 7684.35423 | -0.4607269 | 0.31879134 | -1.4452304 | 0.14839318 | 0.73519456 |
| IRF9 | 6397.42485 | -0.6007191 | 0.36995443 | -1.623765 | 1.04E-01 | 0.67439401 |
| REC8 | 2209.57115 | -0.2524687 | 0.41581576 | -0.6071648 | 0.54374152 | 0.92319803 |
| IPO4 | 4199.24808 | -0.1474316 | 0.20240429 | -0.7284013 | 0.46636794 | 0.89960479 |
| AL136295.4 | 5655.04933 | -0.1516401 | 0.17535436 | -0.8647637 | 0.38716852 | 0.87460898 |
| TM9SF1 | 4449.73092 | -0.0347784 | 0.10930281 | -0.3181841 | 0.75034526 | 0.96255347 |
| AL136295.1 | 6082.81332 | 0.0626256 | 0.10259418 | 0.61042056 | 0.54158325 | 0.92292765 |
| AL136295.6 | 60.6583545 | 0.05363721 | 0.35763316 | 0.14997831 | 0.88078173 | 0.98275231 |
| TSSK4 | 44.6859759 | -0.2367887 | 0.42588064 | -0.5559977 | 0.57821241 | 0.93130183 |
| CHMP4A | 2195.96173 | 0.22016687 | 0.1631926 | 1.34912289 | 0.1772975 | 0.76115956 |
| MDP1 | 574.914589 | 0.21166472 | 0.20279992 | 1.04371205 | 0.29661863 | 0.83832451 |
| NEDD8-MDP | 3975.28388 | 0.09985713 | 0.1051595 | 0.94957778 | 0.34232684 | 0.85889221 |
| NEDD8 | 4570.57846 | 0.02945691 | 0.11522768 | 0.25564093 | 0.79822811 | 0.97170189 |
| GMPR2 | 4559.28194 | 0.25943899 | 0.18561825 | 1.39770198 | 0.16220258 | 0.75026273 |
| TINF2 | 2821.30665 | -0.1703569 | 0.21259565 | -0.8013187 | 0.42294715 | 0.88453836 |
| TGM1 | 175.483794 | -0.1357918 | 0.47001839 | -0.2889075 | 0.77265216 | 0.96836854 |
| RABGGTA | 2033.29741 | -0.134173 | 0.16346539 | -0.8208036 | 0.41175814 | 0.88268208 |
| DHRS1 | 2383.47998 | 0.12548563 | 0.15428666 | 0.81332778 | 0.41603015 | 0.88399703 |
| NOP9 | 5037.01244 | -0.0598369 | 0.37805042 | -0.1582775 | 0.87423815 | 0.98275231 |
| CIDEB | 4411.55336 | -0.1084065 | 0.41111943 | -0.2636861 | 0.7920218 | 0.97090793 |
| LTB4R2 | 1093.96597 | -0.2410973 | 0.4450239 | -0.5417625 | 0.5879821 | 0.93289552 |
| LTB4R | 1229.62538 | -1.0691839 | 0.43271869 | -2.4708521 | 0.01347916 | 0.42599132 |
| ADCY4 | 1986.64423 | -0.3157921 | 0.31404286 | -1.00557 | 0.31462244 | 0.84696964 |
| AL096870.1 | 269.491753 | -0.0948671 | 0.30987269 | -0.3061487 | 0.75949143 | 0.96502715 |
| RIPK3 | 588.104104 | -0.0712786 | 0.33632343 | -0.2119346 | 0.83215808 | 0.97550416 |
| NFATC4 | 3729.24364 | -0.2971989 | 0.43159584 | -0.6886046 | 0.49107211 | 0.90600166 |
| NYNRIN | 1591.05068 | -0.7330063 | 0.49316105 | -1.4863427 | 0.1371885 | 0.72209298 |
| CBLN3 | 487.707113 | 0.12472723 | 0.41692164 | 0.29916227 | 0.76481623 | 0.96678847 |
| KHNYN | 7387.76606 | -0.2168402 | 0.10165253 | -2.1331511 | 0.03291233 | 0.55137197 |
| SDR39U1 | 5917.27565 | -0.2117231 | 0.11670015 | -1.8142484 | 0.06963948 | 0.62779744 |
| CTSG | 84.7595269 | 1.53292276 | 1.09627137 | 1.39830593 | 0.16202122 | 0.75010861 |
| GZMH | 87.345943 | 0.15712217 | 0.87922413 | 0.17870548 | 0.85816897 | 0.97975089 |
| GZMB | 125.133677 | -0.8982984 | 0.81978717 | -1.0957702 | 0.27317934 | 0.82716979 |
| STXBP6 | 139.507341 | 0.95135958 | 0.45049594 | 2.111805 | 0.03470318 | 0.55369025 |
| NOVA1 | 507.433246 | 0.13759752 | 0.25904053 | 0.53118145 | 0.59529305 | 0.9334601 |
| LINC02294 | 198.700234 | 0.85870964 | 0.80659325 | 1.06461297 | 0.28705111 | 0.83155545 |
| LINC02293 | 100.450257 | 0.74190882 | 0.80306825 | 0.9238428 | 0.35556817 | 0.86364633 |
| LINC00645 | 467.742856 | -0.0948854 | 0.43007157 | -0.220627 | 0.82538286 | 0.97548201 |
| PRKD1 | 2285.65092 | -0.3384653 | 0.32199287 | -1.0511577 | 0.29318618 | 0.8352732 |
| G2E3 | 519.056268 | 0.10189631 | 0.14317338 | 0.71169868 | 0.47665138 | 0.90313654 |
| SCFD1 | 2445.45652 | -0.0704293 | 0.1203901 | -0.5850088 | 0.55854179 | 0.92421175 |
| COCH | 491.524068 | 0.16694063 | 0.34847379 | 0.47906225 | 0.63189435 | 0.94088547 |
| AL049830.3 | 220.084957 | 0.46002105 | 0.58979172 | 0.77997204 | 0.43540733 | 0.89063183 |
| STRN3 | 1957.92857 | -0.1446441 | 0.22180294 | -0.652129 | 0.51431796 | 0.91636259 |
| AP4S1 | 460.223471 | 0.10823447 | 0.25576007 | 0.4231875 | 0.67215846 | 0.94922827 |
| HECTD1 | 5576.69447 | -0.3646819 | 0.27944257 | -1.3050336 | 0.19188141 | 0.77351031 |

| | | | | | | |
|---|---|---|---|---|---|---|
| AL136418.1 | 81.5800499 | 0.05927437 | 0.3307169 | 0.17922993 | 0.85775716 | 0.97964261 |
| HEATR5A | 1145.71144 | -0.1347943 | 0.18665952 | -0.7221397 | 4.70E-01 | 0.9008571 |
| AL139353.1 | 290.575193 | -0.5264802 | 0.3073728 | -1.7128392 | 0.08674213 | 0.65228863 |
| AL139353.2 | 51.9420154 | -0.3212434 | 0.50394208 | -0.637461 | 0.52382464 | 0.91723878 |
| DTD2 | 486.19872 | 0.10492015 | 0.18764492 | 0.55914194 | 5.76E-01 | 9.31E-01 |
| NUBPL | 782.594919 | 0.39465684 | 0.22228228 | 1.77547595 | 0.07581933 | 0.63773415 |
| ARHGAP5-AS | 241.485715 | -0.2832362 | 0.36533943 | -0.7752687 | 0.43818087 | 0.89152136 |
| ARHGAP5 | 4030.74617 | -0.3738173 | 0.25709405 | -1.4540101 | 0.14594351 | 0.73247558 |
| AKAP6 | 395.29904 | -0.2362559 | 0.35669074 | -0.662355 | 0.50774371 | 0.91439937 |
| NPAS3 | 83.6787644 | -0.3285265 | 0.31003793 | -1.0596334 | 0.2893114 | 0.83312616 |
| EGLN3 | 2040.44629 | 0.46381561 | 0.45029364 | 1.03002923 | 0.30299628 | 0.84211658 |
| SPTSSA | 2436.22352 | -0.2536512 | 0.32288473 | -0.7855782 | 0.43211465 | 0.88976671 |
| EAPP | 1522.48028 | 0.06132377 | 0.17723076 | 0.3460109 | 0.72933453 | 0.95848028 |
| SNX6 | 3045.11275 | -0.1153894 | 0.11384445 | -1.0135709 | 0.31078757 | 0.84476049 |
| CFL2 | 2180.29883 | -0.1825032 | 0.2527565 | -0.7220513 | 0.47026295 | 0.9008571 |
| BAZ1A | 1222.78787 | -0.397328 | 0.27903996 | -1.4239109 | 0.15447228 | 0.74300499 |
| AL121603.2 | 123.510302 | -0.0967836 | 0.19896658 | -0.4864314 | 0.62666136 | 0.94088547 |
| SRP54 | 3256.72844 | -0.0485355 | 0.17955868 | -0.2703045 | 0.78692599 | 0.97090793 |
| FAM177A1 | 3275.37116 | 0.04115046 | 0.21137123 | 0.19468335 | 0.84564084 | 0.97675487 |
| PPP2R3C | 961.130304 | -0.141731 | 0.2014574 | -0.7035282 | 4.82E-01 | 9.05E-01 |
| KIAA0391 | 1878.40306 | 0.28495195 | 0.15808725 | 1.802498 | 7.15E-02 | 6.32E-01 |
| AL121594.1 | 5558.64244 | -0.1031891 | 0.14366009 | -0.7182863 | 0.47258076 | 0.90160919 |
| PSMA6 | 4285.34085 | -0.244396 | 0.18152968 | -1.3463144 | 0.17820115 | 0.76115956 |
| NFKBIA | 9992.69158 | 0.15564439 | 0.45616508 | 0.3412019 | 0.73295159 | 0.95849877 |
| RALGAPA1 | 1137.65528 | -0.0131135 | 0.11378159 | -0.1152515 | 0.90824575 | 0.98662757 |
| AL162311.3 | 64.7866728 | 0.28579291 | 0.31709612 | 0.90128163 | 0.3674386 | 0.86683872 |
| BRMS1L | 604.661446 | 0.11161071 | 0.16998773 | 0.65658098 | 0.51145038 | 0.91507116 |
| PTCSC3 | 56.3356206 | -0.0291042 | 0.71610242 | -0.0406426 | 0.96758086 | 0.99400331 |
| MBIP | 1142.79115 | 0.06719006 | 0.15394964 | 0.43644181 | 0.66251621 | 0.94642611 |
| SLC25A21 | 145.492415 | 0.87125505 | 0.39974259 | 2.1795402 | 0.02929156 | 0.53606221 |
| SLC25A21-AS | 187.542697 | 0.81272166 | 0.54887708 | 1.48069885 | 0.13868684 | 0.7237761 |
| MIPOL1 | 304.718111 | -0.0316485 | 0.25365998 | -0.1247673 | 0.9007078 | 0.98541534 |
| SSTR1 | 349.848191 | 0.19746168 | 0.44883381 | 0.43994387 | 0.65997776 | 0.94626603 |
| CLEC14A | 1512.08414 | -0.1379237 | 0.33172772 | -0.415774 | 0.67757541 | 0.95018034 |
| SEC23A | 2426.10501 | 0.21398953 | 0.31350774 | 0.68256538 | 0.49488152 | 0.90816709 |
| GEMIN2 | 359.131068 | 0.06228749 | 0.1223089 | 0.50926376 | 0.61056735 | 0.93807505 |
| TRAPPC6B | 1615.83536 | -0.0794998 | 0.09458924 | -0.8404738 | 0.40064279 | 0.88096286 |
| AL132639.3 | 297.600529 | -0.2929622 | 0.21225364 | -1.380246 | 0.16751093 | 0.75558932 |
| PNN | 6595.37341 | -0.2342525 | 0.15242678 | -1.5368195 | 0.12433751 | 0.70102567 |
| MIA2 | 6340.76233 | 0.15393733 | 0.27236435 | 0.56518901 | 0.57194521 | 0.92874623 |
| AL132639.2 | 114.991423 | 0.30981782 | 0.36519212 | 0.84836941 | 0.39623227 | 0.87967259 |
| FBXO33 | 849.12187 | -0.0356574 | 0.17299722 | -0.2061154 | 0.83670076 | 0.97577999 |
| C14orf28 | 357.556911 | 0.19983242 | 0.2434794 | 0.82073645 | 0.4117964 | 0.88268208 |
| KLHL28 | 567.493979 | -0.168273 | 0.13936326 | -1.2074414 | 0.22726219 | 0.80300331 |
| TOGARAM1 | 966.248833 | -0.0476017 | 0.16093935 | -0.2957738 | 0.7674028 | 0.96749893 |
| PRPF39 | 3328.11983 | -0.1728324 | 0.14466159 | -1.1947357 | 0.23219031 | 0.80649152 |
| FKBP3 | 2742.63411 | -0.0934439 | 0.21929772 | -0.4261052 | 0.67003116 | 0.949185 |
| FANCM | 210.67601 | -0.2243984 | 0.29150752 | -0.769786 | 0.44142684 | 0.89290667 |
| MIS18BP1 | 412.489109 | -0.1460973 | 0.20116911 | -0.7262414 | 0.46769082 | 0.89984974 |
| LINC02303 | 247.154738 | 1.82684961 | 0.86786416 | 2.10499487 | 0.03529175 | 0.55369025 |
| LINC00871 | 50.3843112 | 0.91743333 | 0.68321388 | 1.34282011 | 0.17933023 | 0.76174702 |
| RPS29 | 8998.60451 | 0.03614579 | 0.18012617 | 0.20066931 | 0.84095716 | 0.97618283 |
| LRR1 | 315.159837 | -0.4924428 | 0.32960497 | -1.4940393 | 0.13516536 | 0.71649038 |
| RPL36AL | 5553.29778 | -0.1647128 | 0.25175882 | -0.6542482 | 0.51295193 | 0.91568064 |

| | | | | | | |
|---|---|---|---|---|---|---|
| MGAT2 | 1498.48756 | -0.164053 | 0.22823125 | -0.7188016 | 0.47226317 | 0.90160919 |
| AL139099.1 | 1494.37153 | -0.1614962 | 0.22827167 | -0.7074736 | 0.4792722 | 0.90480914 |
| DNAAF2 | 756.807618 | -0.1254288 | 0.20333981 | -0.6168432 | 0.53733812 | 0.92123049 |
| POLE2 | 108.570365 | 0.19073294 | 0.26264048 | 0.72621302 | 0.46770819 | 0.89984974 |
| KLHDC1 | 345.486566 | 0.34130397 | 0.25104128 | 1.3595532 | 0.17397136 | 0.75974924 |
| KLHDC2 | 6165.99751 | 0.04462325 | 0.17003674 | 0.262433 | 0.79298763 | 0.97090793 |
| NEMF | 3793.31645 | -0.0076007 | 0.12816453 | -0.0593041 | 0.95270989 | 0.99170503 |
| ARF6 | 5770.53735 | -0.0816116 | 0.17791853 | -0.4587018 | 0.64644831 | 0.94208859 |
| LINC01588 | 109.046556 | 0.10539026 | 0.3876088 | 0.27189851 | 0.78570005 | 0.97065795 |
| VCPKMT | 645.953124 | -0.1463613 | 0.2355877 | -0.6212604 | 0.53442828 | 0.92084006 |
| SOS2 | 2047.18389 | -0.017345 | 0.15314722 | -0.1132567 | 0.90982701 | 0.98662757 |
| L2HGDH | 1488.61255 | 0.66807626 | 0.3132138 | 2.13297203 | 0.03292702 | 0.55137197 |
| ATP5S | 2522.57434 | 0.14044626 | 0.20051752 | 0.70041889 | 0.48366574 | 0.90541935 |
| AL359397.2 | 516.003198 | 0.13390993 | 0.24199023 | 0.55336918 | 0.58001064 | 0.93139743 |
| CDKL1 | 716.769185 | -0.0861446 | 0.25237863 | -0.3413308 | 0.7328546 | 0.95848028 |
| MAP4K5 | 1905.74651 | -0.1517266 | 0.16158107 | -0.9390119 | 0.34772462 | 0.86225576 |
| AL118556.1 | 59.7531094 | 0.20259007 | 0.3870273 | 0.52345162 | 0.60066001 | 0.93488657 |
| ATL1 | 362.906586 | 0.00212153 | 0.24525233 | 0.00865041 | 0.99309806 | 0.99906069 |
| SAV1 | 2080.29526 | -0.1363205 | 0.12671526 | -1.0758016 | 0.28201601 | 0.83003334 |
| NIN | 1296.64331 | 0.09878482 | 0.15371282 | 0.64265827 | 0.52044586 | 0.91680964 |
| PYGL | 1700.06135 | 0.07036692 | 0.31237058 | 0.22526743 | 0.82177124 | 0.97487098 |
| AL358334.3 | 280.124437 | 0.18316184 | 0.37311228 | 0.49090274 | 0.62349524 | 0.94045607 |
| AL358334.2 | 62.0081773 | 1.84691932 | 0.88296407 | 2.09172647 | 0.03646299 | 0.55384187 |
| TRIM9 | 166.728822 | 0.28571455 | 0.57911399 | 0.49336496 | 0.62175473 | 0.94016713 |
| TMX1 | 2099.39716 | -0.1651945 | 0.18483282 | -0.893751 | 0.37145512 | 0.86762749 |
| FRMD6 | 416.74364 | -0.329679 | 0.4111541 | -0.8018381 | 0.42264662 | 0.88451372 |
| GNG2 | 539.211939 | -0.0635168 | 0.35252128 | -0.1801786 | 0.85701235 | 0.97953896 |
| AL358333.3 | 86.3575836 | -0.224381 | 0.31556509 | -0.7110451 | 0.4770563 | 0.90344707 |
| RTRAF | 4742.09617 | 0.06192725 | 0.16097503 | 0.38470098 | 0.70045897 | 0.95360275 |
| NID2 | 1398.82842 | 0.05818358 | 0.34099702 | 0.17062782 | 0.86451642 | 0.98188271 |
| PTGDR | 45.5305973 | 0.03393107 | 0.63026624 | 0.05383609 | 0.95706576 | 0.99200096 |
| PTGER2 | 320.011366 | -0.3693398 | 0.27621129 | -1.337164 | 0.18116911 | 0.76522169 |
| TXNDC16 | 379.993993 | 0.49602937 | 0.14942058 | 3.3196858 | 0.00090119 | 0.14312853 |
| GPR137C | 51.7111118 | 0.3779843 | 0.37887862 | 0.99763957 | 0.31845417 | 0.8490262 |
| ERO1A | 5034.78353 | 0.28865097 | 0.38486563 | 0.75000454 | 0.45325197 | 0.89670027 |
| AL133453.1 | 981.988186 | 0.49316016 | 0.37109752 | 1.32892336 | 0.18387325 | 0.76781833 |
| PSMC6 | 3981.22123 | 0.048819 | 0.17371981 | 0.2810215 | 0.77869391 | 0.96836854 |
| STYX | 713.236223 | -0.0810174 | 0.15062607 | -0.537871 | 0.59066611 | 0.93289552 |
| GNPNAT1 | 1682.35287 | -0.3325688 | 0.3381303 | -0.9835523 | 0.32533567 | 0.85244476 |
| AL139317.3 | 1118.38172 | -0.3054651 | 0.30984181 | -0.9858744 | 0.32419473 | 0.85169085 |
| FERMT2 | 4767.20653 | -0.424301 | 0.20551336 | -2.0645908 | 0.03896174 | 0.55613287 |
| DDHD1 | 834.136522 | 0.03290491 | 0.15363474 | 0.21417622 | 0.83040964 | 0.97550416 |
| AL352979.2 | 122.191815 | 0.09958358 | 0.27105236 | 0.3673961 | 0.71332358 | 0.95564093 |
| BMP4 | 616.811272 | -0.1766089 | 0.31893852 | -0.5537397 | 0.579757 | 0.93139743 |
| CDKN3 | 67.7730389 | 0.28630841 | 0.59180787 | 0.48378609 | 0.62853769 | 0.94088547 |
| CNIH1 | 3021.20621 | 0.18198968 | 0.18865911 | 0.96464825 | 0.33472102 | 0.85557672 |
| GMFB | 3423.73179 | -0.1926157 | 0.21781303 | -0.8843169 | 0.37652518 | 0.86970773 |
| CGRRF1 | 946.805152 | -0.2204513 | 0.253917 | -0.8682023 | 0.38528357 | 0.87368012 |
| SAMD4A | 1792.73962 | -0.4002957 | 0.30425962 | -1.3156385 | 0.18829541 | 0.76970626 |
| GCH1 | 1677.91725 | -0.0653316 | 0.38973379 | -0.1676314 | 0.8668733 | 0.98222237 |
| WDHD1 | 164.946338 | -0.3198227 | 0.30411866 | -1.0516377 | 0.29296578 | 0.83524862 |
| SOCS4 | 1286.02395 | -0.1664403 | 0.1993714 | -0.8348251 | 0.40381619 | 0.88132015 |
| MAPK1IP1L | 6784.22862 | -0.0441849 | 0.19942758 | -0.2215587 | 0.82465745 | 0.97548201 |
| LGALS3 | 5515.98057 | 0.04809221 | 0.37622176 | 0.12782942 | 0.89828396 | 0.98474725 |

| | | | | | | |
|---|---|---|---|---|---|---|
| FBXO34 | 1663.26108 | -0.1868886 | 0.09681959 | -1.9302767 | 0.05357256 | 0.59163425 |
| ATG14 | 1428.1603 | -0.087632 | 0.17284223 | -0.5070056 | 0.61215089 | 0.93858463 |
| KTN1-AS1 | 110.266256 | 0.0825349 | 0.21620811 | 0.38173822 | 0.70265555 | 0.95431462 |
| KTN1 | 9727.8805 | -0.080934 | 0.19117747 | -0.4233446 | 0.67204384 | 0.94922827 |
| PELI2 | 1368.68171 | -0.0545904 | 0.18101156 | -0.3015854 | 0.76296812 | 0.96639535 |
| TMEM260 | 1082.94617 | 0.03959528 | 0.26638357 | 0.14864009 | 8.82E-01 | 9.83E-01 |
| AL161757.2 | 179.49524 | 0.05135575 | 0.29725398 | 0.17276725 | 0.86283438 | 0.98176973 |
| AL161757.5 | 255.190536 | 0.13208442 | 0.32210351 | 0.41006824 | 0.68175589 | 0.95070301 |
| EXOC5 | 2346.78871 | 0.02366743 | 0.17176551 | 0.13778921 | 0.89040702 | 0.98466501 |
| AP5M1 | 2901.94158 | 0.09306127 | 0.12317166 | 0.75554127 | 0.44992428 | 0.89557948 |
| NAA30 | 1416.24239 | -0.1356745 | 0.25348297 | -0.535241 | 0.5924832 | 0.93289552 |
| CCDC198 | 5399.11932 | 0.56165995 | 0.38053805 | 1.47596268 | 0.13995391 | 0.72475166 |
| ARMH4 | 1142.00001 | 0.16378141 | 0.22238147 | 0.73648856 | 0.46143343 | 0.89755243 |
| ACTR10 | 3356.80211 | 0.05100358 | 0.11972188 | 0.42601718 | 0.67009533 | 0.949185 |
| PSMA3 | 4425.42211 | -0.1546387 | 0.1910133 | -0.8095702 | 0.41818725 | 0.88445833 |
| PSMA3-AS1 | 3769.56645 | 0.01044079 | 0.1563751 | 0.0667676 | 0.94676671 | 0.99166614 |
| ARID4A | 1002.69261 | -0.0753576 | 0.20090186 | -0.3750968 | 0.70758846 | 0.95501548 |
| TOMM20L | 75.4955265 | 0.13069136 | 0.28612907 | 0.45675666 | 0.64784596 | 0.94285781 |
| AL139021.1 | 260.043932 | -0.0575891 | 0.18760667 | -0.3069671 | 0.75886841 | 0.96502715 |
| TIMM9 | 1078.94045 | -0.140577 | 0.12451604 | -1.1289872 | 0.25890323 | 0.82364948 |
| KIAA0586 | 517.197828 | 0.03015438 | 0.15102141 | 0.19966955 | 0.84173903 | 0.9762188 |
| DACT1 | 683.338104 | -0.4951343 | 0.47807033 | -1.0356934 | 0.30034519 | 0.84200819 |
| DAAM1 | 943.637562 | -0.3020129 | 0.21926567 | -1.3773836 | 0.16839367 | 0.75735257 |
| GPR135 | 173.466987 | -0.3672662 | 0.25774604 | -1.4249152 | 0.15418172 | 0.74258845 |
| L3HYPDH | 1503.04287 | -0.3611358 | 0.34807298 | -1.0375289 | 0.29948945 | 0.84123067 |
| JKAMP | 2620.91392 | 0.13760819 | 0.12519697 | 1.09913361 | 0.27170979 | 0.82716979 |
| CCDC175 | 200.052432 | 0.20121869 | 0.18077868 | 1.11306647 | 0.26567988 | 0.82611274 |
| RTN1 | 247.477027 | 0.28799104 | 0.53569904 | 0.53759858 | 0.59085421 | 0.93289552 |
| PCNX4 | 3477.60016 | 0.01494199 | 0.10134403 | 0.14743833 | 0.88278605 | 0.98279039 |
| DHRS7 | 4228.69436 | 0.2060696 | 0.14571193 | 1.414226 | 0.15729556 | 0.74522306 |
| AL157756.1 | 115.36795 | 0.14368765 | 0.39826869 | 0.36078067 | 0.71826341 | 0.95648644 |
| PPM1A | 4602.81845 | -0.2430515 | 0.28539908 | -0.8516199 | 0.39442511 | 0.87884163 |
| SIX4 | 194.217697 | 0.179298 | 0.24779319 | 0.7235792 | 4.69E-01 | 0.90050905 |
| MNAT1 | 815.507759 | 0.06606951 | 0.11837861 | 0.55812036 | 5.77E-01 | 0.93127944 |
| TRMT5 | 812.837661 | -0.0969218 | 0.14815396 | -0.6541962 | 5.13E-01 | 0.91568064 |
| SLC38A6 | 608.330317 | 0.21058555 | 0.21482199 | 0.98027929 | 0.32694827 | 0.85297643 |
| PRKCH | 1246.61264 | -0.2074585 | 0.29648738 | -0.6997211 | 0.48410152 | 0.90541935 |
| TMEM30B | 1912.54024 | -0.2505319 | 0.30703113 | -0.8159822 | 0.41451029 | 0.88364943 |
| AL355916.3 | 282.448913 | -0.2533853 | 0.36989902 | -0.685012 | 0.49333635 | 0.90715187 |
| LINC01303 | 114.335121 | -0.4174724 | 0.50161742 | -0.8322527 | 0.40526635 | 0.88132015 |
| AL355916.1 | 114.470678 | -0.0613218 | 0.75618444 | -0.0810937 | 0.93536743 | 0.99068685 |
| HIF1A-AS1 | 457.268506 | 0.17176826 | 0.53909984 | 0.3186205 | 0.7500143 | 0.96249332 |
| HIF1A | 23745.196 | -0.1911733 | 0.38962547 | -0.490659 | 0.62366762 | 0.9405038 |
| SNAPC1 | 964.401183 | -0.1123327 | 0.34319894 | -0.3273108 | 0.74343279 | 0.96124462 |
| SYT16 | 82.6907235 | 0.97063013 | 0.56146964 | 1.72873128 | 8.39E-02 | 0.64887192 |
| RHOJ | 922.064468 | -0.2844147 | 0.47296512 | -0.6013438 | 5.48E-01 | 9.24E-01 |
| PPP2R5E | 1950.93602 | -0.2348405 | 0.20008269 | -1.1737175 | 0.24050822 | 0.8124487 |
| WDR89 | 621.177158 | 0.12577501 | 0.20653895 | 0.60896511 | 0.54254756 | 0.92292765 |
| AL136038.2 | 144.692283 | -0.000833 | 0.2738559 | -0.0030417 | 0.99757307 | 0.99989813 |
| AL136038.5 | 53.9308809 | -0.021715 | 0.2984079 | -0.0727694 | 0.94198961 | 0.99136103 |
| SGPP1 | 2624.51113 | -0.2685003 | 0.2453981 | -1.0941419 | 0.27389277 | 0.82719197 |
| SYNE2 | 14223.3944 | -0.2494344 | 0.31696534 | -0.7869455 | 0.4313138 | 0.88976671 |
| ESR2 | 265.603416 | -0.7970005 | 0.52207372 | -1.5266053 | 0.1268592 | 0.70443503 |
| MTHFD1 | 4674.60298 | 0.3114137 | 0.33813889 | 0.92096385 | 0.3570693 | 0.86390849 |

| | | | | | | |
|---|---|---|---|---|---|---|
| AL122035.1 | 577.069369 | 0.37837761 | 0.34031011 | 1.11186122 | 0.26619782 | 0.8265123 |
| AL122035.2 | 791.385164 | 0.27345608 | 0.31725519 | 0.86194361 | 0.38871855 | 0.87568585 |
| ZBTB25 | 1991.38009 | 0.17113149 | 0.28234926 | 0.60609859 | 0.54444929 | 0.92345821 |
| AKAP5 | 158.165036 | -0.3350133 | 0.37381072 | -0.896211 | 0.37014009 | 0.86762749 |
| ZBTB1 | 2074.45438 | 0.01655531 | 0.16086894 | 0.10291178 | 0.91803299 | 0.9868044 |
| AL049869.3 | 90.0185058 | 0.14516964 | 0.35223344 | 0.41214043 | 0.68023649 | 0.95070301 |
| HSPA2 | 4067.99982 | -0.1984641 | 0.44638464 | -0.4446035 | 0.65660635 | 0.94545756 |
| PPP1R36 | 104.53276 | 0.4273686 | 0.59865183 | 0.71388506 | 0.47529826 | 0.90226631 |
| PLEKHG3 | 6222.01416 | 0.02145529 | 0.15960125 | 0.1344306 | 0.89306208 | 0.98474725 |
| SPTB | 1588.393 | 0.18161217 | 0.20220342 | 0.89816564 | 3.69E-01 | 0.86762749 |
| CHURC1 | 2283.74639 | -0.1507483 | 0.10667886 | -1.4131035 | 0.15762528 | 0.74522306 |
| CHURC1-FNT | 1478.41117 | 0.13956666 | 0.12367298 | 1.12851373 | 0.25910302 | 0.82364948 |
| GPX2 | 667.358253 | 1.80466095 | 0.82425711 | 2.18943935 | NA | NA |
| RAB15 | 1967.57746 | 0.30632076 | 0.2774581 | 1.10402528 | 0.26958217 | 0.82716979 |
| FNTB | 1336.43216 | 0.11698064 | 0.17303396 | 0.67605593 | 0.49900514 | 0.91147492 |
| MAX | 3448.57906 | -0.2863155 | 0.13772739 | -2.0788565 | 0.03763054 | 0.55538555 |
| AL139022.1 | 489.584415 | -0.2557271 | 0.1686262 | -1.5165325 | 0.12938475 | 0.70755752 |
| FUT8 | 912.94683 | 0.18052106 | 0.28463424 | 0.63422119 | 0.52593648 | 0.91769856 |
| FUT8-AS1 | 51.9634403 | 0.30303866 | 0.57546172 | 0.5266009 | 0.59847077 | 0.9346978 |
| GPHN | 1665.18269 | 0.07589166 | 0.21026909 | 0.36092637 | 0.71815449 | 0.95648644 |
| MPP5 | 2687.48241 | 0.1235087 | 0.21175592 | 0.58325972 | 0.55971847 | 0.92434754 |
| ATP6V1D | 4371.40802 | 0.10863576 | 0.20328264 | 0.53440745 | 0.59305967 | 0.93289552 |
| EIF2S1 | 2656.62604 | -0.1852558 | 0.2487192 | -0.744839 | 0.45636905 | 0.89688594 |
| PLEK2 | 363.43879 | -0.3728411 | 0.41711496 | -0.893857 | 0.37139839 | 0.86762749 |
| TMEM229B | 278.79442 | -0.4725573 | 0.56875475 | -0.830863 | 0.40605105 | 0.88132015 |
| PLEKHH1 | 2188.12716 | -0.1640181 | 0.18893184 | -0.8681339 | 0.38532103 | 0.87368012 |
| PIGH | 2060.96796 | -0.300196 | 0.13119654 | -2.2881393 | 0.02212941 | 0.51647233 |
| ARG2 | 9154.85281 | 0.35253941 | 0.49261845 | 0.71564395 | 4.74E-01 | 0.90199039 |
| VTI1B | 9966.22593 | 0.2291364 | 0.36072218 | 0.63521572 | 0.52528773 | 0.91759401 |
| AL049779.1 | 2851.00594 | 0.1632775 | 0.20782101 | 0.78566408 | 0.43206431 | 0.88976671 |
| RDH11 | 7208.73996 | 0.10183386 | 0.22646944 | 0.44965829 | 0.65295685 | 0.9447282 |
| RDH12 | 471.624091 | 0.64038915 | 0.98211885 | 0.65204853 | 0.51436987 | 0.91636259 |
| ZFYVE26 | 1781.33328 | -0.4471684 | 0.24873694 | -1.7977561 | 0.07221566 | 0.6321115 |
| RAD51B | 162.484068 | 0.13901212 | 0.37266165 | 0.37302503 | 0.70912982 | 0.95501548 |
| ZFP36L1 | 22260.3896 | -0.4415183 | 0.26414756 | -1.6714837 | 0.09462617 | 0.65888357 |
| ACTN1 | 6598.10423 | -0.5069444 | 0.44094948 | -1.1496654 | 0.25028172 | 0.81721208 |
| DCAF5 | 4026.09351 | -0.0101963 | 0.15236992 | -0.0669179 | 0.9466471 | 0.99166614 |
| EXD2 | 1643.10236 | -0.2553268 | 0.12304624 | -2.0750473 | 0.03798215 | 0.55538555 |
| AL359317.1 | 54.3173211 | -0.2491202 | 0.48257053 | -0.5162358 | 0.60568971 | 0.93698713 |
| GALNT16 | 1330.45332 | -0.1737487 | 0.5319618 | -0.3266188 | 0.7439562 | 0.9613927 |
| ERH | 3186.53308 | -0.2063651 | 0.13619662 | -1.5152 | 1.30E-01 | 0.70814767 |
| SLC39A9 | 4409.41397 | 0.01751207 | 0.1136243 | 0.15412261 | 0.87751306 | 0.98275231 |
| PLEKHD1 | 163.955854 | 0.83728561 | 0.72223773 | 1.15929364 | 2.46E-01 | 8.15E-01 |
| SUSD6 | 5119.72087 | -9.29E-05 | 0.20760253 | -0.0004474 | 0.999643 | 0.99994357 |
| SRSF5 | 19149.7298 | -0.0673736 | 0.16970248 | -0.3970098 | 0.69136021 | 0.95158502 |
| SMOC1 | 212.90884 | -0.2327878 | 0.5013843 | -0.4642901 | 0.64243992 | 0.94090101 |
| COX16 | 1887.77982 | 0.24222997 | 0.13468595 | 1.79847992 | 0.07210099 | 0.6321115 |
| SYNJ2BP-CO) | 2003.10191 | 0.22855188 | 0.10082271 | 2.26686915 | 0.02339822 | 0.51967919 |
| SYNJ2BP | 3004.68944 | 0.2475789 | 0.08833578 | 2.80270253 | 0.00506764 | 0.30823792 |
| MED6 | 709.179181 | -0.238702 | 0.19402359 | -1.2302732 | 0.21859481 | 0.79780737 |
| AL357153.3 | 181.276726 | -0.3271057 | 0.25632575 | -1.276133 | 0.20190852 | 0.78134034 |
| TTC9 | 653.641703 | -0.4969292 | 0.49116002 | -1.0117459 | 0.31165955 | 0.84537057 |
| MAP3K9 | 791.055045 | -0.274865 | 0.28375479 | -0.9686708 | 0.33270946 | 0.855011 |
| AC004816.1 | 75.6930602 | -0.019381 | 0.3078073 | -0.0629648 | 0.94979451 | 0.99170503 |

| | | | | | | |
|---|---|---|---|---|---|---|
| PCNX1 | 2815.92881 | -0.2326083 | 0.17519897 | -1.327681 | 0.18428351 | 0.76820626 |
| AC005476.2 | 79.8899813 | 0.15189033 | 0.51893783 | 0.29269465 | 0.76975556 | 0.96836854 |
| SIPA1L1 | 3016.6436 | -0.2937208 | 0.16795862 | -1.7487691 | 0.08033094 | 0.6411764 |
| RGS6 | 184.197954 | -0.4722745 | 0.58059249 | -0.8134354 | 0.41596847 | 0.88399703 |
| DPF3 | 442.616203 | 0.99290638 | 0.56157114 | 1.76808655 | 0.07704643 | 0.63806922 |
| DCAF4 | 1055.60556 | -0.2884016 | 0.25318411 | -1.1390984 | 0.25466211 | 0.81992039 |
| ZFYVE1 | 2033.20175 | -0.3659793 | 0.26437869 | -1.3842997 | 0.16626671 | 0.75521193 |
| AL442663.3 | 84.5339837 | -0.1966638 | 0.34651195 | -0.5675528 | 0.57033869 | 0.92753481 |
| RBM25 | 4612.47382 | -0.1775003 | 0.16418249 | -1.081116 | 0.27964553 | 0.82825734 |
| PSEN1 | 4202.51791 | -0.2665048 | 0.2211464 | -1.2051058 | 0.22816246 | 0.80300331 |
| PAPLN | 5413.03588 | 0.56369116 | 0.31959846 | 1.76374806 | 0.07777441 | 0.64069017 |
| AC004846.1 | 123.482639 | -0.0718252 | 0.27774996 | -0.2585965 | 0.79594659 | 0.97101097 |
| AC004846.2 | 1779.01728 | 0.61003358 | 0.32059805 | 1.90279881 | 0.0570668 | 0.60049913 |
| NUMB | 5521.01681 | -0.0818518 | 0.15138699 | -0.5406791 | 0.58872882 | 0.93289552 |
| HEATR4 | 198.365864 | -0.1473012 | 0.26136759 | -0.5635788 | 0.57304084 | 0.92928554 |
| RIOX1 | 603.524184 | -0.1724916 | 0.27531683 | -0.6265205 | 5.31E-01 | 0.91962754 |
| ACOT1 | 827.967526 | 0.90453203 | 0.35952109 | 2.51593594 | 0.01187168 | 0.40525879 |
| ACOT2 | 1775.60597 | 0.3410348 | 0.71522145 | 0.47682407 | 6.33E-01 | 0.94088547 |
| ACOT4 | 709.850678 | 0.89175631 | 0.46979292 | 1.89819019 | 0.05767103 | 0.60129769 |
| ACOT6 | 104.745743 | 0.99756476 | 0.72889315 | 1.3686022 | 0.17112366 | 0.75956362 |
| AC005225.2 | 77.6755602 | 0.54185444 | 0.67574533 | 0.8018619 | 0.42263285 | 0.88451372 |
| DNAL1 | 1227.41385 | 0.05101032 | 0.17151404 | 0.2974119 | 0.76615206 | 0.96704052 |
| AC006146.1 | 98.241642 | 0.19839564 | 0.26002726 | 0.76298016 | 0.44547523 | 0.89453029 |
| PNMA1 | 3256.22206 | -0.1386338 | 0.19871698 | -0.6976444 | 0.48539958 | 0.90543654 |
| ELMSAN1 | 2225.05642 | -0.3518263 | 0.2820488 | -1.2473951 | 0.21225265 | 0.79203904 |
| AC005520.2 | 165.245936 | 0.62737291 | 0.83967627 | 0.74716046 | 0.45496671 | 0.89670027 |
| LINC02274 | 821.874548 | -0.314756 | 0.48802208 | -0.6449625 | 0.51895146 | 0.91636259 |
| PTGR2 | 1468.80227 | 0.37506085 | 0.27016986 | 1.38824089 | 0.16506371 | 0.75303284 |
| AC005520.1 | 3309.56443 | -0.0087504 | 0.11737727 | -0.0745497 | 0.940573 | 0.99136103 |
| AC005520.5 | 45.2996813 | 0.31827957 | 0.35610157 | 0.89378874 | 0.37143494 | 0.86762749 |
| ZNF410 | 2690.89493 | -0.0896012 | 0.11976017 | -0.748172 | 0.4543564 | 0.89670027 |
| AC005480.1 | 420.366546 | -0.0170322 | 0.19392672 | -0.087828 | 0.93001337 | 0.98973492 |
| FAM161B | 1993.13397 | -0.0410079 | 0.10311006 | -0.3977096 | 0.69084427 | 0.95158502 |
| COQ6 | 3564.37453 | 0.61911786 | 0.32013261 | 1.93394189 | 0.05312027 | 0.58906925 |
| ENTPD5 | 7787.13947 | 0.90987335 | 0.46071521 | 1.97491491 | 0.0482778 | 0.58440502 |
| BBOF1 | 3256.26986 | 0.39473293 | 0.3635505 | 1.08577191 | 0.27757992 | 0.82805357 |
| ALDH6A1 | 19174.126 | 0.74704605 | 0.49028801 | 1.52368818 | 0.12758662 | 0.70557946 |
| LIN52 | 879.232507 | 0.40286759 | 0.28419992 | 1.41754999 | 0.15632219 | 0.74522306 |
| ABCD4 | 5358.57929 | 0.13795171 | 0.14490637 | 0.95200584 | 0.34109402 | 0.85889221 |
| NPC2 | 8900.45384 | 0.43536225 | 0.2191908 | 1.986225 | 0.04700835 | 0.58171004 |
| ISCA2 | 1138.10948 | 0.13098653 | 0.16113613 | 0.8128936 | 0.41627906 | 0.88399703 |
| LTBP2 | 1158.57562 | 0.2966189 | 0.4591619 | 0.64600067 | 0.51827892 | 0.91636259 |
| AREL1 | 2251.68978 | -0.0956766 | 0.11068258 | -0.8644235 | 0.3873553 | 0.87460898 |
| FCF1 | 1452.36262 | 0.09765974 | 0.15175901 | 0.64351854 | 0.51988769 | 0.91657872 |
| YLPM1 | 2578.35844 | -0.0210172 | 0.09728058 | -0.2160472 | 0.82895093 | 0.97550416 |
| PROX2 | 84.2013774 | 0.09752823 | 0.34165847 | 0.28545533 | 0.77529532 | 0.96836854 |
| DLST | 9871.78219 | 0.22788608 | 0.23440892 | 0.97217322 | 0.33096438 | 0.85384238 |
| RPS6KL1 | 461.245476 | -0.4236426 | 0.44238684 | -0.9576293 | 0.33824973 | 0.8571036 |
| PGF | 559.196358 | -0.2969246 | 0.43137647 | -0.688319 | 0.49125193 | 0.90600166 |
| EIF2B2 | 2156.84159 | -0.1251562 | 0.21656976 | -0.5779023 | 0.56333009 | 0.92452095 |
| AL049780.1 | 545.068721 | -0.0688401 | 0.25191754 | -0.2732643 | 0.78465004 | 0.97021865 |
| MLH3 | 1301.82933 | 0.09409287 | 0.20195982 | 0.46589898 | 0.64128781 | 0.94088547 |
| ACYP1 | 565.572858 | 0.12587952 | 0.29960818 | 0.42014713 | 0.67437797 | 0.94935413 |
| ZC2HC1C | 428.524965 | 0.04133129 | 0.27165334 | 0.15214717 | 0.87907086 | 0.98275231 |

| | | | | | | |
|---|---|---|---|---|---|---|
| NEK9 | 4170.52815 | -0.0320257 | 0.20771368 | -0.154182 | 0.87746626 | 0.98275231 |
| TMED10 | 10843.7878 | 0.21288155 | 0.10212939 | 2.08442986 | 0.03712108 | 0.55527244 |
| FOS | 8853.10201 | -0.3205847 | 0.66460736 | -0.4823671 | 0.62954516 | 0.94088547 |
| AF111167.2 | 89.3218464 | 0.34728408 | 0.60410022 | 0.57487826 | 0.56537364 | 0.92512239 |
| JDP2 | 1116.90737 | -0.0999413 | 0.34930522 | -0.2861145 | 7.75E-01 | 0.96836854 |
| BATF | 69.9075949 | -0.3698924 | 0.85633667 | -0.4319474 | 0.66577963 | 0.9474809 |
| AC007182.1 | 122.091527 | -0.1496849 | 0.48341263 | -0.309642 | 0.7568332 | 0.9648851 |
| FLVCR2 | 872.201092 | -0.0719996 | 0.32515539 | -0.2214313 | 0.82475658 | 0.97548201 |
| TTLL5 | 928.00103 | -0.1006499 | 0.1307356 | -0.769874 | 0.44137464 | 0.89290667 |
| ERG28 | 1851.11373 | 0.06846931 | 0.13294792 | 0.51500849 | 0.60654709 | 0.93708311 |
| IFT43 | 1937.80928 | 0.15430507 | 0.19933696 | 0.77409162 | 0.43887662 | 0.89175645 |
| TGFB3 | 1292.74634 | -0.4526145 | 0.41940783 | -1.0791751 | 0.28050968 | 0.82871922 |
| TGFB3-AS1 | 81.6696144 | -0.211584 | 0.55408975 | -0.3818588 | 0.7025661 | 0.95431462 |
| GPATCH2L | 1948.30563 | -0.2591288 | 0.20517639 | -1.2629561 | 0.20660496 | 0.78778852 |
| ESRRB | 2215.40318 | -1.3189539 | 0.52200468 | -2.5267091 | 0.01151368 | 0.40353404 |
| VASH1 | 1753.8949 | 0.04642877 | 0.27144799 | 0.17104114 | 0.86419142 | 0.98176973 |
| AF111169.3 | 206.326743 | 0.01407239 | 0.32895094 | 0.04277961 | 0.96587722 | 0.99387827 |
| AF111169.1 | 705.838621 | 0.13697212 | 0.27883418 | 0.49123145 | 0.62326276 | 0.94038864 |
| VASH1-AS1 | 1259.1568 | 0.06220312 | 0.2521457 | 0.24669517 | 0.80514415 | 0.97303568 |
| ANGEL1 | 1237.85926 | 0.00755694 | 0.20508309 | 0.0368482 | 0.97060605 | 0.99400331 |
| IRF2BPL | 4478.17783 | -0.0836452 | 0.26949795 | -0.3103743 | 0.75627636 | 0.96450325 |
| AC007686.3 | 636.805623 | -0.0364274 | 0.22702018 | -0.160459 | 0.8725195 | 0.98275231 |
| LINC02289 | 97.968999 | -0.4212437 | 0.49559855 | -0.8499696 | 0.39534201 | 0.87927742 |
| CIPC | 1772.43042 | -0.2755758 | 0.1488403 | -1.8514865 | 0.06409959 | 0.61837523 |
| TMEM63C | 769.523854 | -0.0537333 | 0.17526142 | -0.3065896 | 0.75915581 | 0.96502715 |
| POMT2 | 2143.23715 | -0.4014275 | 0.27795345 | -1.4442258 | 1.49E-01 | 7.36E-01 |
| GSTZ1 | 1115.13713 | 0.52062705 | 0.26022209 | 2.00070276 | 0.04542443 | 0.57529633 |
| TMED8 | 1739.02432 | 0.03314955 | 0.18442368 | 0.17974674 | 0.8573514 | 0.97953896 |
| SAMD15 | 94.5666171 | 0.16679853 | 0.41726861 | 0.39973899 | 0.68934877 | 0.95158502 |
| NOXRED1 | 83.0720697 | -0.1082306 | 0.41144617 | -0.2630491 | 0.79251271 | 0.97090793 |
| VIPAS39 | 1014.84833 | 0.07715213 | 0.21375737 | 0.36093321 | 0.71814938 | 0.95648644 |
| AHSA1 | 4087.10859 | -0.1782618 | 0.12912525 | -1.3805337 | 1.67E-01 | 7.56E-01 |
| ISM2 | 76.5659738 | 0.91095615 | 0.46633938 | 1.95341887 | 0.05076998 | 0.5861488 |
| SPTLC2 | 3507.73017 | -0.2691311 | 0.17535566 | -1.5347726 | 0.12483968 | 0.70228923 |
| ALKBH1 | 735.117082 | -0.178021 | 0.2126667 | -0.8370894 | 0.40254234 | 0.88132015 |
| SLIRP | 2199.82677 | -0.0460857 | 0.1641171 | -0.2808097 | 7.79E-01 | 0.96836854 |
| SNW1 | 2534.68063 | -0.283375 | 0.1506681 | -1.8807897 | 0.06000053 | 0.6075531 |
| C14orf178 | 99.256084 | -0.1484901 | 0.28734273 | -0.5167701 | 0.60531669 | 0.93692105 |
| ADCK1 | 768.654491 | -0.1698032 | 0.26048638 | -0.6518696 | 0.51448528 | 0.91636259 |
| NRXN3 | 136.451922 | 0.23754128 | 0.41552463 | 0.57166595 | 0.56754831 | 0.92614913 |
| DIO2 | 168.604905 | -0.0817023 | 0.66377697 | -0.1230869 | 0.90203826 | 0.98572048 |
| CEP128 | 110.153016 | -0.2160972 | 0.30875502 | -0.6998987 | 0.48399059 | 0.90541935 |
| GTF2A1 | 1501.27322 | 0.20075167 | 0.19882899 | 1.00967002 | 0.31265341 | 0.84541735 |
| STON2 | 1655.63127 | -0.0966005 | 0.36416645 | -0.2652648 | 0.79080551 | 0.97090793 |
| SEL1L | 4293.60084 | 0.14479232 | 0.19315482 | 0.74961794 | 0.45348484 | 0.89670027 |
| FLRT2 | 455.71663 | 0.34814479 | 0.34317583 | 1.01447934 | 0.31035408 | 0.84466249 |
| GALC | 1752.96306 | 0.29762938 | 0.15725824 | 1.89261549 | 0.05840902 | 0.60332684 |
| GPR65 | 156.216258 | -0.3164815 | 0.57136786 | -0.5539014 | 0.5796463 | 0.93139743 |
| KCNK10 | 272.606298 | 0.61493607 | 0.69020742 | 0.89094387 | 0.37295928 | 0.86791605 |
| SPATA7 | 768.44465 | -0.0209176 | 0.22197097 | -0.0942359 | 0.92492176 | 0.98869145 |
| PTPN21 | 1371.55379 | -0.136391 | 0.14654503 | -0.9307106 | 0.35200327 | 0.86364633 |
| AL162171.1 | 41.5402497 | -0.1905361 | 0.52691863 | -0.3616043 | 0.71764773 | 0.95623434 |
| ZC3H14 | 3228.50918 | -0.3427586 | 0.2141764 | -1.6003564 | 0.10951955 | 0.67859123 |
| EML5 | 1243.80196 | -0.5315781 | 0.34489187 | -1.5412891 | 0.12324645 | 0.69844624 |

| | | | | | | |
|---|---|---|---|---|---|---|
| TTC8 | 1046.15644 | 0.06586146 | 0.16548836 | 0.39798241 | 0.69064315 | 0.95158502 |
| FOXN3 | 1829.39566 | -0.186109 | 0.22057407 | -0.8437481 | 0.39881017 | 0.88074856 |
| AL138478.1 | 835.06805 | -0.1513414 | 0.23753258 | -0.6371395 | 0.52403401 | 0.91723878 |
| FOXN3-AS1 | 61.5409523 | 0.0963537 | 0.39505039 | 0.24390229 | 0.8073065 | 0.97334138 |
| EFCAB11 | 350.350003 | 0.51571667 | 0.31411135 | 1.64182755 | 0.10062575 | 0.67247782 |
| TDP1 | 761.327388 | -0.1008759 | 0.19239969 | -0.5243037 | 0.60006735 | 0.93487648 |
| KCNK13 | 266.243039 | 0.15331818 | 0.42006457 | 0.36498717 | 0.71512098 | 0.95590747 |
| PSMC1 | 3148.48058 | -0.0862703 | 0.24863815 | -0.3469715 | 0.72861274 | 0.95848028 |
| NRDE2 | 2964.37021 | -0.0770028 | 0.19715363 | -0.3905724 | 0.69611337 | 0.9525827 |
| CALM1 | 18191.5481 | 0.3527501 | 0.28076573 | 1.25638591 | 0.20897609 | 0.79086734 |
| AL512791.1 | 2273.34434 | 0.24182547 | 0.29070668 | 0.83185382 | 0.40549146 | 0.88132015 |
| TTC7B | 1701.694 | 0.03930893 | 0.2724685 | 0.14426963 | 0.88528756 | 0.98356741 |
| AL096869.2 | 88.9951684 | 0.0114071 | 0.33711391 | 0.03383753 | 0.97300671 | 0.99444395 |
| RPS6KA5 | 355.185443 | 0.01086823 | 0.27695733 | 0.03924154 | 0.96869782 | 0.99400331 |
| DGLUCY | 4177.46557 | 0.08746374 | 0.33348197 | 0.26227426 | 0.79311001 | 0.97090793 |
| GPR68 | 174.431838 | 0.22566821 | 0.44054781 | 0.51224455 | 0.60847987 | 0.93748023 |
| CCDC88C | 1433.94674 | -0.0875069 | 0.29005827 | -0.3016873 | 0.76289046 | 0.96639375 |
| PPP4R3A | 3777.25078 | -0.353714 | 0.17927623 | -1.9730114 | 0.04849426 | 0.58440502 |
| CATSPERB | 169.802289 | -0.1418077 | 0.38799554 | -0.3654878 | 0.71474728 | 0.9558355 |
| TC2N | 1239.68139 | -0.0771176 | 0.29860902 | -0.258256 | 0.79620933 | 0.97101097 |
| FBLN5 | 8354.99626 | -0.1640561 | 0.3281262 | -0.4999788 | 0.61708999 | 0.93910158 |
| TRIP11 | 1853.97314 | 0.12060787 | 0.10782212 | 1.1185819 | 0.26331855 | 0.82424998 |
| ATXN3 | 1025.91543 | -0.1051208 | 0.13873172 | -0.7577269 | 0.44861449 | 0.8948759 |
| NDUFB1 | 2801.36871 | 0.04403016 | 0.11949675 | 0.36846326 | 0.71252784 | 0.95560193 |
| CPSF2 | 1698.3441 | -0.1734209 | 0.17251552 | -1.005248 | 0.31477743 | 0.84696964 |
| RIN3 | 1188.8773 | -0.1170657 | 0.21116198 | -0.5543881 | 0.57931326 | 0.93139743 |
| LGMN | 15705.0683 | 0.0610113 | 0.36205144 | 0.16851557 | 0.8661777 | 0.98197545 |
| GOLGA5 | 2136.08724 | -0.0565429 | 0.25948794 | -0.2179016 | 0.82750574 | 0.97548201 |
| ITPK1 | 6006.35121 | 0.03190595 | 0.17331812 | 0.18408893 | 0.85394369 | 0.97906925 |
| MOAP1 | 3609.63362 | -0.2049359 | 0.19358394 | -1.0586408 | 0.28976341 | 0.83399826 |
| TMEM251 | 1218.63088 | -0.2553447 | 0.31765876 | -0.8038334 | 0.4214932 | 0.88451372 |
| AL110118.2 | 827.468654 | -0.1826504 | 0.30785503 | -0.5933 | 0.55298037 | 0.92373235 |
| GON7 | 711.610032 | -0.0159413 | 0.16013998 | -0.0995458 | 0.92070489 | 0.98767854 |
| UBR7 | 1598.26241 | 0.04611501 | 0.19982274 | 0.23077959 | 0.81748604 | 0.97433259 |
| BTBD7 | 2166.44978 | -0.0392126 | 0.15023255 | -0.2610124 | 0.79408298 | 0.97090793 |
| UNC79 | 121.809328 | 1.05729239 | 0.41132703 | 2.57044227 | 0.01015688 | 0.39989496 |
| PRIMA1 | 128.932478 | -0.6188839 | 0.94448417 | -0.6552612 | 0.51229957 | 0.91551505 |
| ASB2 | 111.370393 | -0.6777295 | 0.7802389 | -0.868618 | 0.38505612 | 0.87343703 |
| OTUB2 | 132.417019 | 0.16184741 | 0.40016471 | 0.40445197 | 0.68588039 | 0.95097288 |
| DDX24 | 7421.2459 | -0.0644382 | 0.1089937 | -0.5912102 | 0.55437962 | 0.92373235 |
| IFI27L1 | 589.648445 | 0.06373559 | 0.29606046 | 0.21527896 | 0.82954983 | 0.97550416 |
| IFI27 | 4409.07158 | -1.6779462 | 0.75542244 | -2.2212025 | 0.02633725 | 0.52678638 |
| IFI27L2 | 712.932631 | 0.06296485 | 0.42071432 | 0.14966178 | 0.88103147 | 0.98275231 |
| SERPINA6 | 1016.85431 | 0.00202431 | 0.60853054 | 0.00332655 | 0.9973458 | 0.9998533 |
| SERPINA1 | 92074.4169 | -0.4258724 | 0.63201259 | -0.6738353 | 0.50041603 | 0.91180773 |
| SERPINA4 | 258.591774 | 0.55615102 | 0.40807312 | 1.36287099 | 0.17292317 | 0.75974924 |
| SERPINA5 | 8886.44967 | 0.6485644 | 0.35180139 | 1.84355271 | 0.06524835 | 0.6216672 |
| AL049839.2 | 40569.3012 | -0.3560109 | 0.70957892 | -0.5017214 | 0.61586353 | 0.93910158 |
| SERPINA3 | 39709.8143 | -0.3635787 | 1.24064389 | -0.2930565 | 0.76947899 | 0.96836854 |
| DICER1 | 2990.64143 | -0.0912883 | 0.12586947 | -0.7252616 | 0.46829157 | 0.89984974 |
| DICER1-AS1 | 633.146762 | 0.2220367 | 0.3581053 | 0.62003186 | 0.53523681 | 0.92084006 |
| CLMN | 6209.06101 | -0.0886625 | 0.22739188 | -0.3899104 | 0.69660281 | 0.9525827 |
| SYNE3 | 345.143709 | -0.1355205 | 0.33880327 | -0.3999977 | 0.68915818 | 0.95158502 |
| SNHG10 | 345.522575 | 0.04148704 | 0.24614724 | 0.16854563 | 0.86615405 | 0.98197545 |

| | | | | | | |
|---|---|---|---|---|---|---|
| GLRX5 | 2888.14526 | -0.000721 | 0.24409629 | -0.0029539 | 0.99764313 | 0.99989813 |
| C14orf132 | 686.311459 | -0.6626789 | 0.46835511 | -1.4149071 | 0.15709575 | 0.74522306 |
| BDKRB2 | 1687.61688 | -0.2313012 | 0.44261867 | -0.5225745 | 0.60127039 | 0.93529286 |
| AL355102.2 | 305.00555 | -0.2037065 | 0.48759616 | -0.4177771 | 0.67611009 | 0.94959168 |
| BDKRB1 | 195.110783 | -0.6276194 | 0.58233665 | -1.0777604 | 0.28114071 | 0.82939514 |
| AL355102.4 | 41.101567 | 0.4204952 | 0.73985065 | 0.56835146 | 0.56979634 | 0.92719921 |
| ATG2B | 2646.52145 | -0.1317199 | 0.15833794 | -0.8318907 | 0.40547066 | 0.88132015 |
| GSKIP | 1172.75009 | -0.1104316 | 0.2325467 | -0.4748793 | 0.634873 | 0.94088547 |
| AK7 | 415.383484 | 0.11013523 | 0.25624992 | 0.42979615 | 0.66734393 | 0.94830184 |
| AL163051.1 | 48.0903696 | -0.1837553 | 0.47299846 | -0.3884903 | 0.6976532 | 0.9527595 |
| PAPOLA | 7212.15719 | -0.1910033 | 0.13678837 | -1.3963419 | 0.16261155 | 0.75032457 |
| VRK1 | 278.772 | 0.23686919 | 0.28330823 | 0.83608299 | 0.40310822 | 0.88132015 |
| LINC01550 | 118.070298 | 0.61245345 | 0.45083131 | 1.35849803 | 0.17430571 | 0.75974924 |
| BCL11B | 133.435414 | -0.2754795 | 0.63412049 | -0.4344276 | 0.66397793 | 0.94642611 |
| SETD3 | 6657.77745 | 0.03157076 | 0.24013336 | 0.13147177 | 0.89540212 | 0.98474725 |
| CCNK | 2559.41037 | -0.3108348 | 0.14855872 | -2.092336 | 0.03640847 | 0.55384187 |
| CCDC85C | 4526.19808 | -0.1723732 | 0.18239101 | -0.9450749 | 0.34462062 | 0.85999891 |
| AL110504.1 | 976.33752 | -0.0557946 | 0.32213697 | -0.1732015 | 0.86249305 | 0.98176973 |
| HHIPL1 | 84.378292 | -0.1718139 | 0.436355 | -0.3937479 | 0.69376718 | 0.95222846 |
| CYP46A1 | 182.309678 | 0.39359974 | 0.42733777 | 0.92105068 | 0.35702397 | 0.86390849 |
| EML1 | 964.711834 | 0.04715533 | 0.30906947 | 0.15257196 | 0.87873584 | 0.98275231 |
| EVL | 4159.17628 | -0.1995237 | 0.25706237 | -0.7761686 | 0.43764946 | 0.89130865 |
| DEGS2 | 1243.72711 | 0.09432724 | 0.39569117 | 0.23838602 | 0.81158171 | 0.97352497 |
| YY1 | 5486.50971 | -0.2402008 | 0.15903911 | -1.5103255 | 0.1309604 | 0.7100756 |
| SLC25A29 | 11929.5738 | -0.4971806 | 0.36602396 | -1.3583281 | 0.17435961 | 0.75974924 |
| AL157871.1 | 4784.74258 | -0.4280268 | 0.37685282 | -1.1357929 | 0.25604323 | 0.82063559 |
| SLC25A47 | 166.355014 | -1.1234278 | 0.86074406 | -1.3051821 | 0.19183084 | 0.77347283 |
| WARS | 7823.39586 | -0.7195163 | 0.33832392 | -2.1267084 | 0.03344432 | 0.55367229 |
| AL157871.4 | 1392.47419 | -0.766052 | 0.36369621 | -2.1062965 | 0.03517861 | 0.55369025 |
| WDR25 | 523.808157 | -0.0020409 | 0.17733086 | -0.0115089 | 0.99081747 | 0.9983285 |
| BEGAIN | 129.232751 | -0.0794479 | 0.43608541 | -0.1821844 | 8.55E-01 | 9.79E-01 |
| MEG3 | 4359.13275 | 0.10494463 | 0.40519884 | 0.25899539 | 7.96E-01 | 0.97101097 |
| AL117190.2 | 626.128128 | 0.10484557 | 0.4278191 | 0.24506986 | 8.06E-01 | 0.97334138 |
| AL117190.1 | 836.503604 | 0.23177691 | 0.41190388 | 0.5626966 | 0.5736415 | 0.92978516 |
| MEG9 | 58.3429062 | 0.09502106 | 0.48121657 | 0.19746007 | 0.84346753 | 0.97649449 |
| DIO3OS | 306.911159 | 0.22005387 | 0.39718396 | 0.55403513 | 0.5795548 | 0.93139743 |
| PPP2R5C | 4023.42593 | 0.1586118 | 0.13560529 | 1.16965796 | 0.24213864 | 0.81319028 |
| AL118558.3 | 187.879571 | 0.25450506 | 0.43229312 | 0.58873262 | 5.56E-01 | 0.92373235 |
| AL118558.4 | 46.1660152 | 0.17605991 | 0.50854519 | 0.34620308 | 0.7291901 | 0.95848028 |
| DYNC1H1 | 9876.31188 | 0.0144619 | 0.12911855 | 0.11200481 | 0.9108196 | 0.98662757 |
| HSP90AA1 | 39089.5467 | 0.23866617 | 0.1988434 | 1.20027203 | 0.23003371 | 0.80551898 |
| WDR20 | 1507.7977 | -0.1032608 | 0.1909727 | -0.5407095 | 0.58870783 | 0.93289552 |
| MOK | 1215.49618 | -0.3648569 | 0.23074642 | -1.5812027 | 0.11383169 | 0.68214043 |
| ZNF839 | 1250.9544 | 0.19411501 | 0.25334046 | 0.76622192 | 0.44354427 | 0.89366134 |
| CINP | 1819.04132 | 0.22897453 | 0.17197298 | 1.33145643 | 0.18303887 | 0.76723162 |
| TECPR2 | 1547.2565 | -0.1292896 | 0.13363644 | -0.9674729 | 0.33330767 | 0.855011 |
| ANKRD9 | 2641.36974 | -0.2492911 | 0.27188849 | -0.9168873 | 0.35920171 | 0.86480228 |
| RCOR1 | 1368.22549 | -0.1150795 | 0.15205335 | -0.7568364 | 0.4491479 | 0.89546252 |
| TRAF3 | 1529.08292 | 0.10625143 | 0.16427025 | 0.64680871 | 0.51775576 | 0.91636259 |
| AMN | 10232.0836 | -0.1357562 | 0.20455977 | -0.6636503 | 0.50691417 | 0.91420888 |
| CDC42BPB | 8294.18364 | -0.1150777 | 0.13461374 | -0.8548732 | 0.39262136 | 0.8778303 |
| AL117209.1 | 1364.34415 | -0.1564439 | 0.18536803 | -0.8439636 | 0.39868977 | 0.88074856 |
| EXOC3L4 | 1107.40262 | -0.4718289 | 0.6824183 | -0.6914072 | 0.48930969 | 0.90600166 |
| TNFAIP2 | 12478.8764 | 0.48176519 | 0.38949941 | 1.236883 | 0.21613052 | 0.7954827 |

| | | | | | | |
|---|---|---|---|---|---|---|
| LINC00605 | 43.554074 | 0.32419655 | 0.69521535 | 0.46632536 | 0.64098263 | 0.94088547 |
| AL161669.3 | 197.256187 | 0.40303427 | 0.38391423 | 1.04980289 | 0.29380874 | 0.83567608 |
| AL138976.2 | 47.6688009 | 0.28101321 | 0.43473878 | 0.64639556 | 0.51802322 | 0.91636259 |
| EIF5 | 11768.7129 | -0.2920772 | 0.3640804 | -0.8022327 | 0.42241839 | 0.88451372 |
| MARK3 | 4541.90416 | -0.1676398 | 0.14466671 | -1.1588 | 0.24653773 | 0.81542272 |
| CKB | 15645.8327 | 0.1751628 | 0.32992092 | 0.53092361 | 0.59547172 | 0.9334601 |
| AL133367.1 | 468.520453 | 0.07033055 | 0.40850533 | 0.17216556 | 0.86330737 | 0.98176973 |
| TRMT61A | 1637.78355 | -0.3442054 | 0.22538933 | -1.5271593 | 0.1267214 | 0.70443503 |
| BAG5 | 2814.53693 | -0.1083498 | 0.1619967 | -0.6688397 | 5.04E-01 | 9.14E-01 |
| KLC1 | 5247.72394 | -0.2158207 | 0.21476611 | -1.0049103 | 0.31494005 | 0.84696964 |
| AL139300.1 | 2450.32775 | -0.1050007 | 0.18853606 | -0.5569263 | 0.57757779 | 0.93130183 |
| APOPT1 | 1452.70761 | -0.0213048 | 0.19242478 | -0.1107176 | 0.91184026 | 0.98662757 |
| AL049840.1 | 78.1856067 | -0.4721294 | 0.35708192 | -1.322188 | 0.18610555 | 0.76908034 |
| AL049840.2 | 199.756732 | -0.4035514 | 0.28281513 | -1.4269089 | 0.15360616 | 0.74258845 |
| XRCC3 | 3008.32235 | -0.1987506 | 0.22853482 | -0.8696731 | 0.3844791 | 0.87309542 |
| ZFYVE21 | 2975.82489 | 0.18081077 | 0.12443664 | 1.4530348 | 0.14621409 | 0.73247558 |
| PPP1R13B | 2978.88222 | -0.316897 | 0.13434146 | -2.3588922 | 0.01832958 | 0.47570049 |
| ATP5MPL | 3378.87604 | 0.15034525 | 0.17402724 | 0.86391786 | 0.38763301 | 0.87468392 |
| TDRD9 | 373.820608 | 0.4711591 | 0.60369689 | 0.7804564 | 0.43512228 | 0.89055338 |
| ASPG | 2166.63933 | -0.256658 | 0.48434719 | -0.529905 | 5.96E-01 | 0.9334601 |
| KIF26A | 653.526073 | 0.18466854 | 0.53694545 | 0.34392421 | 0.73090328 | 0.95848028 |
| C14orf180 | 106.228295 | 1.66588737 | 0.77150888 | 2.15925883 | 0.03083009 | 0.54443002 |
| BX927359.1 | 57.8543835 | 1.64331136 | 0.81746086 | 2.01026306 | 0.04440335 | 0.57267043 |
| INF2 | 8205.94747 | -0.2181655 | 0.18464234 | -1.1815575 | 0.23738134 | 0.8090132 |
| ADSSL1 | 1205.75828 | 0.17498166 | 0.43916433 | 0.39844232 | 0.69030417 | 0.95158502 |
| SIVA1 | 2554.70555 | -0.0426378 | 0.14744581 | -0.2891763 | 0.77244643 | 0.96836854 |
| AKT1 | 9013.33656 | -0.2571621 | 0.14623254 | -1.7585835 | 0.07864828 | 0.64079926 |
| AL583722.2 | 3325.47541 | -0.2231032 | 0.14640942 | -1.523831 | 0.12755093 | 0.70557946 |
| ZBTB42 | 769.328105 | -0.2763615 | 0.13377489 | -2.0658694 | 0.03884081 | 0.55608529 |
| LINC00638 | 53.8188278 | -0.0876654 | 0.43013664 | -0.2038082 | 0.83850337 | 0.97612168 |
| CEP170B | 8354.8093 | -0.4312613 | 0.16663118 | -2.5881188 | 0.00965017 | 0.39368232 |
| PLD4 | 228.254205 | 0.57598699 | 0.60929645 | 0.94533128 | 0.34448977 | 0.85999891 |
| AHNAK2 | 474.756814 | 0.30333464 | 0.68267333 | 0.44433352 | 0.65680147 | 0.94545756 |
| CLBA1 | 1454.38155 | -0.0138769 | 0.31632158 | -0.0438698 | 0.96500823 | 0.99357855 |
| CDCA4 | 280.48635 | -0.2936926 | 0.23926377 | -1.2274846 | 0.2196405 | 0.79858446 |
| GPR132 | 558.958843 | 0.70189316 | 0.47035313 | 1.49226848 | 0.13562878 | 0.7170152 |
| LINC02298 | 85.0097043 | 0.25623669 | 0.36931579 | 0.6938146 | 0.48779849 | 0.90584365 |
| JAG2 | 1500.6082 | -0.1146987 | 0.37700088 | -0.3042399 | 0.76094509 | 0.9658862 |
| NUDT14 | 1553.86308 | 0.11647646 | 0.2721963 | 0.42791344 | 0.66871414 | 0.9486589 |
| BRF1 | 2629.17619 | -0.0959882 | 0.15198346 | -0.6315698 | 0.52766804 | 0.91842929 |
| BTBD6 | 2262.05626 | -0.0930852 | 0.14433274 | -0.6449348 | 0.51896945 | 0.91636259 |
| PACS2 | 5600.58361 | -0.1899444 | 0.09256312 | -2.0520522 | 0.04016459 | 0.55657615 |
| TEX22 | 502.109082 | -0.1013469 | 0.28372172 | -0.3572052 | 0.72093818 | 0.95728372 |
| AL928654.1 | 65.8247523 | 0.24807793 | 0.31393147 | 0.79022957 | 0.42939371 | 0.88851358 |
| MTA1 | 5448.79266 | -0.2277448 | 0.27111513 | -0.84003 | 0.40089159 | 0.88119978 |
| AL928654.2 | 1062.52151 | -0.1520404 | 0.28920603 | -0.5257166 | 0.59908514 | 0.93484879 |
| CRIP2 | 7923.93674 | -0.0332707 | 0.43493098 | -0.0764965 | 0.93902412 | 0.99113899 |
| CRIP1 | 1324.85863 | 0.37639956 | 0.63834697 | 0.58964729 | 0.55542714 | 0.92373235 |
| AL928654.3 | 1362.29477 | 0.36427271 | 0.61908177 | 0.58840808 | 0.55625842 | 0.92373235 |
| TEDC1 | 392.315805 | 0.29555536 | 0.51152354 | 0.57779425 | 0.56340304 | 0.92452095 |
| TMEM121 | 94.6963962 | 0.1603739 | 0.53524279 | 0.29962832 | 0.76446068 | 0.96678847 |
| IGHA2 | 1851.48699 | 2.78395162 | 1.3039665 | 2.13498708 | 0.03276207 | 0.55104386 |
| IGHG4 | 1533.8895 | 2.40848887 | 1.38985079 | 1.73291182 | 0.08311134 | 0.6471083 |
| IGHG2 | 5639.36613 | 2.61169945 | 1.19526573 | 2.18503667 | 0.02888616 | 0.53606221 |

| | | | | | | |
|---|---|---|---|---|---|---|
| IGHGP | 1146.93447 | -0.2292298 | 1.20222189 | -0.1906718 | 0.84878277 | 0.97741457 |
| IGHA1 | 12163.461 | 1.42197098 | 1.26684397 | 1.12245155 | 0.2616705 | 0.82424998 |
| IGHG1 | 16529.7878 | -0.4544596 | 1.21835574 | -0.3730106 | 0.70914056 | 0.95501548 |
| IGHG3 | 4085.44448 | 0.31036791 | 1.27564513 | 0.24330271 | 0.80777091 | 0.97334138 |
| IGHD | 239.601129 | 0.60038511 | 1.12587839 | 0.53325929 | 0.5938541 | 0.93317501 |
| IGHM | 4649.05206 | -0.8967677 | 1.23949442 | -0.7234947 | 0.46937599 | 0.90050905 |
| IGHJ6 | 381.10181 | -0.7822116 | 1.06143678 | -0.7369366 | 4.61E-01 | 8.98E-01 |
| IGHJ5 | 159.756239 | 0.9850518 | 1.09119365 | 0.90272867 | 0.36666991 | 0.86683872 |
| IGHJ4 | 762.082051 | 0.58018578 | 1.3557013 | 0.42795989 | 0.66868032 | 0.9486589 |
| IGHJ3 | 282.39609 | 0.75801026 | 1.18223722 | 0.64116596 | 0.52141486 | 0.91723878 |
| IGHJ2 | 70.1544714 | -0.6895689 | 1.22786698 | -0.561599 | 5.74E-01 | 9.30E-01 |
| FAM30A | 87.2144718 | -0.0361106 | 1.15714545 | -0.0312066 | 9.75E-01 | 0.99455099 |
| IGHV6-1 | 82.7972974 | -1.6736523 | 1.18693058 | -1.4100676 | 0.15851973 | 0.74675717 |
| IGHV1-2 | 281.914142 | 1.2682808 | 1.54661072 | 0.8200388 | 0.41219399 | 0.88268208 |
| IGHV1-3 | 69.7243314 | 0.0493342 | 1.31008229 | 0.03765733 | 0.9699609 | 0.99400331 |
| IGHV4-4 | 64.6721901 | 0.70519711 | 1.17544149 | 0.59994233 | 0.54854467 | 0.92373235 |
| IGHV7-4-1 | 42.0238957 | -1.8286342 | 2.77729011 | -0.6584239 | NA | NA |
| IGHV2-5 | 132.499307 | -0.1531226 | 1.33588446 | -0.1146226 | 9.09E-01 | 9.87E-01 |
| IGHV3-7 | 268.601304 | 1.24092292 | 1.1795276 | 1.05205078 | 0.29277625 | 0.83524862 |
| IGHV5-10-1 | 77.7828527 | -1.0977346 | 2.30118883 | -0.4770293 | 0.63334123 | 0.94088547 |
| IGHV3-11 | 111.042409 | 0.3345895 | 1.28155168 | 0.26108155 | 0.79402962 | 0.97090793 |
| IGHV3-15 | 178.185011 | 0.46082585 | 1.13965663 | 0.404355 | 0.68595169 | 0.95097288 |
| IGHV1-18 | 245.149517 | -0.1598475 | 0.99285025 | -0.1609986 | 0.87209452 | 0.98275231 |
| IGHV3-21 | 170.985825 | 0.04624511 | 1.21224591 | 0.03814829 | 0.96956945 | 0.99400331 |
| IGHV3-23 | 504.101812 | 0.94359983 | 1.41252934 | 0.6680214 | 0.50411993 | 0.91356552 |
| IGHV1-24 | 143.545134 | -0.10757 | 1.81148781 | -0.0593821 | NA | NA |
| IGHV3-30 | 548.729921 | -1.541597 | 1.47847042 | -1.0426972 | 2.97E-01 | 0.83898658 |
| IGHV4-31 | 70.3505326 | 0.63635298 | 1.41241434 | 0.4505427 | 0.65231917 | 0.9443159 |
| IGHV3-33 | 135.890892 | -0.6809167 | 1.21049164 | -0.5625125 | 0.5737669 | 0.92978516 |
| IGHV4-34 | 235.853118 | -0.6717807 | 1.53315347 | -0.4381693 | 0.66126359 | 0.94637197 |
| IGHV4-39 | 177.552466 | -0.2942845 | 1.19765992 | -0.2457162 | 0.80590191 | 0.97334138 |
| IGHV1-46 | 81.396467 | 0.99316014 | 1.23279723 | 0.80561516 | 0.42046478 | 0.88451372 |
| IGHV3-48 | 139.290319 | 0.00045145 | 1.24983479 | 0.00036121 | 0.9997118 | 0.99994357 |
| IGHV3-49 | 152.211162 | 2.61777299 | 1.28955041 | 2.02998888 | NA | NA |
| IGHV5-51 | 954.480686 | 1.10089568 | 1.55164846 | 0.7095007 | NA | NA |
| IGHV4-59 | 183.924069 | 0.21381213 | 1.10138118 | 0.19413091 | 0.84607338 | 0.9767918 |
| IGHV4-61 | 60.1852134 | 0.33599103 | 1.25332079 | 0.26808063 | 0.78863725 | 0.97090793 |
| IGHV1-69D | 121.894676 | 1.43287124 | 1.14293309 | 1.25367902 | 2.10E-01 | 7.91E-01 |
| IGHV2-70 | 42.3616624 | -2.8056465 | 1.69944152 | -1.6509227 | 9.88E-02 | 0.6678501 |
| IGHV3-73 | 46.4900061 | 0.39592293 | 1.17985379 | 0.33556949 | 0.73719553 | 0.96023056 |
| IGHV3-74 | 138.634516 | -0.4542279 | 0.97469598 | -0.4660201 | 0.64120112 | 0.94088547 |
| AC026495.1 | 78.6567262 | 1.70343975 | 1.18529854 | 1.43713983 | 0.15067827 | 0.73796431 |
| NIPA1 | 1498.70064 | -0.1124347 | 0.1867027 | -0.6022126 | 0.54703263 | 0.92345821 |
| NIPA2 | 3164.95022 | -0.2073493 | 0.25963663 | -0.7986132 | 0.42451471 | 0.88565404 |
| CYFIP1 | 6051.16714 | 0.04171323 | 0.11724072 | 0.35579135 | 0.72199682 | 0.95739595 |
| TUBGCP5 | 822.376446 | 0.09550428 | 0.19606352 | 0.48710888 | 0.62618117 | 0.94088547 |
| NDN | 757.370131 | -0.0293358 | 0.23721945 | -0.1236653 | 0.90158032 | 0.98572048 |
| PWRN1 | 292.408261 | 0.44405041 | 0.68230758 | 0.65080679 | 0.51517122 | 0.91636259 |
| NPAP1 | 92.0358372 | 0.29680205 | 0.58755375 | 0.50514876 | 0.61345436 | 0.9388166 |
| SNRPN | 10653.6765 | 0.35971433 | 0.17650725 | 2.03795789 | 0.04155414 | 0.55945849 |
| SNURF | 9034.45981 | 0.38795703 | 0.17655516 | 2.19737009 | 0.02799403 | 0.53262541 |
| AC124312.1 | 7717.99964 | 0.35533213 | 0.16968948 | 2.09401386 | 0.03625875 | 0.55384187 |
| PWAR6 | 962.777645 | 0.35387653 | 0.17244957 | 2.05205809 | 0.04016402 | 0.55657615 |
| UBE3A | 3029.05788 | -0.0519679 | 0.14165373 | -0.3668659 | 0.71371902 | 0.95564093 |

| | | | | | | |
|---|---|---|---|---|---|---|
| ATP10A | 635.139857 | -0.5731978 | 0.37093265 | -1.5452881 | 0.1222766 | 0.69788192 |
| GABRB3 | 660.984099 | 0.79681011 | 0.34482101 | 2.3107934 | 0.02084427 | 0.50666984 |
| HERC2 | 3027.07214 | -0.1214518 | 0.11880237 | -1.022301 | 0.30663846 | 0.84286067 |
| GOLGA8M | 45.6012972 | -0.4227849 | 0.76557015 | -0.5522484 | 0.58077817 | 0.9317705 |
| APBA2 | 141.261569 | -0.086969 | 0.54211102 | -0.1604265 | 0.87254512 | 0.98275231 |
| FAM189A1 | 403.737064 | 1.20817982 | 0.46473018 | 2.59974472 | 0.00932931 | 0.38681562 |
| NSMCE3 | 1008.00097 | -0.1070836 | 0.14929061 | -0.7172828 | 0.4731996 | 0.90179314 |
| AC022613.1 | 155.168104 | 0.55088346 | 0.40117586 | 1.37317198 | 0.16969889 | 0.7586159 |
| TJP1 | 3906.69864 | -0.1530494 | 0.19373021 | -0.7900133 | 0.42952 | 0.88858861 |
| GOLGA8H | 36.0927217 | 0.28304781 | 0.57594414 | 0.4914501 | 0.62310814 | 0.94030676 |
| AC091057.2 | 45.1071933 | 0.49657041 | 0.39420084 | 1.25968889 | 0.20778161 | 0.78858551 |
| ARHGAP11B | 43.4597366 | 0.32136243 | 0.42312825 | 0.7594918 | 0.44755842 | 0.89480207 |
| FAN1 | 2363.78932 | -0.1284547 | 0.13595292 | -0.9448467 | 0.34473716 | 0.85999891 |
| MTMR10 | 6306.27322 | -0.2865119 | 0.210252 | -1.3627071 | 0.17297485 | 0.75974924 |
| LINC02352 | 63.2287761 | 0.10887926 | 0.39864802 | 0.27312129 | 7.85E-01 | 9.70E-01 |
| KLF13 | 5008.91474 | -9.87E-05 | 0.25694704 | -0.0003842 | 0.99969344 | 0.99994357 |
| OTUD7A | 271.494649 | -0.0574221 | 0.22394795 | -0.2564084 | 0.79763548 | 0.97148232 |
| AC123768.3 | 173.715354 | 0.19320653 | 0.34271605 | 0.56375103 | 0.57292358 | 0.92928554 |
| GOLGA8N | 363.54773 | 0.20709539 | 0.38127194 | 0.54316976 | 0.58701293 | 0.93289552 |
| ARHGAP11A | 121.777597 | -0.0936446 | 0.32124413 | -0.2915061 | 0.77066425 | 0.96836854 |
| SCG5 | 42.7969363 | -0.0631353 | 0.62528651 | -0.1009701 | 0.91957418 | 0.9872458 |
| GREM1 | 461.249465 | -2.3117087 | 1.41371665 | -1.6351994 | NA | NA |
| FMN1 | 465.517938 | 0.39301383 | 0.28600901 | 1.37413093 | 0.16940104 | 0.75852708 |
| RYR3 | 310.239844 | -0.2498719 | 0.32583616 | -0.7668636 | 0.44316264 | 0.8935502 |
| AC010809.2 | 77.4623913 | -0.3254444 | 0.37287281 | -0.8728026 | 0.38277068 | 0.8718674 |
| AVEN | 324.815911 | -0.1609199 | 0.27688314 | -0.5811834 | 0.56111688 | 0.92448946 |
| EMC7 | 3545.67545 | 0.0493748 | 0.15165312 | 0.32557722 | 0.74474425 | 0.9613927 |
| PGBD4 | 152.124 | -0.2689341 | 0.22225061 | -1.210049 | 0.22626008 | 0.80246466 |
| KATNBL1 | 1313.07235 | -0.1854353 | 0.12536304 | -1.479186 | 0.13909061 | 0.72409321 |
| EMC4 | 3238.4391 | -0.0579973 | 0.1646584 | -0.3522282 | 0.72466715 | 0.95836477 |
| SLC12A6 | 2982.0501 | 0.12263324 | 0.24511013 | 0.50031894 | 0.61685052 | 0.93910158 |
| NOP10 | 3052.94927 | -0.3029561 | 0.25741246 | -1.1769288 | 0.23922393 | 0.81163641 |
| LPCAT4 | 1629.39646 | 0.02498012 | 0.28020061 | 0.08915084 | 0.92896204 | 0.98963127 |
| GOLGA8A | 4094.42002 | 0.0356622 | 0.24466188 | 0.14576117 | 0.88410993 | 0.98317437 |
| GOLGA8B | 4042.04774 | 0.28266414 | 0.28323058 | 0.99800005 | 0.31827933 | 0.8490262 |
| ACTC1 | 360.884739 | -2.2260902 | 0.91747821 | -2.4263139 | 0.01525307 | 0.45205627 |
| AQR | 1752.25499 | 0.02365804 | 0.09111317 | 0.25965551 | 0.79512951 | 0.97101097 |
| ZNF770 | 2640.49808 | 0.25894738 | 0.14337402 | 1.80609699 | 0.07090319 | 0.63035896 |
| DPH6 | 424.160123 | 0.13533016 | 0.2207476 | 0.61305382 | 0.53984075 | 0.92208855 |
| DPH6-DT | 55.7125386 | 0.65270793 | 0.45094012 | 1.44743813 | 0.14777425 | 0.73455299 |
| C15orf41 | 322.189486 | -0.0538689 | 0.20534607 | -0.2623321 | 0.79306545 | 0.97090793 |
| MEIS2 | 1326.15084 | -0.1602657 | 0.31873758 | -0.502814 | 0.61509502 | 0.93895494 |
| LINC01852 | 70.3629587 | 0.70337933 | 0.48957704 | 1.43670816 | 0.15080094 | 0.73796431 |
| SPRED1 | 1641.52977 | 0.07088461 | 0.20518557 | 0.34546588 | 0.72974416 | 0.95848028 |
| FAM98B | 1619.48213 | 0.1750477 | 0.10478901 | 1.67047761 | 0.09482491 | 0.65888357 |
| RASGRP1 | 636.417253 | 0.02387846 | 0.19559448 | 0.12208145 | 0.90283451 | 0.98608873 |
| THBS1 | 15612.2569 | -0.5092292 | 0.48603955 | -1.0477114 | 0.29477161 | 0.83630273 |
| AC037198.2 | 228.133512 | -0.2702332 | 0.59217611 | -0.4563392 | 0.64814608 | 0.94308274 |
| FSIP1 | 1323.69574 | -0.4357713 | 0.4850524 | -0.8984005 | 0.36897206 | 0.86762749 |
| AC023908.3 | 122.884898 | 0.25132919 | 0.32887618 | 0.76420613 | 0.44474442 | 0.89411662 |
| GPR176 | 489.131886 | 0.12940788 | 0.44881568 | 0.2883319 | 0.77309269 | 0.96836854 |
| EIF2AK4 | 1651.0352 | -0.0450316 | 0.17139961 | -0.262729 | 0.79275947 | 0.97090793 |
| SRP14 | 11044.4803 | -0.0244868 | 0.15909254 | -0.1539156 | 0.87767625 | 0.98275231 |
| SRP14-AS1 | 480.056412 | 0.09021297 | 0.32286579 | 0.27941321 | 0.77992773 | 0.96836854 |

| | | | | | | |
|---|---|---|---|---|---|---|
| BMF | 1357.1391 | 0.24626762 | 0.4132741 | 0.59589415 | 0.55124594 | 0.92373235 |
| BMF-AS1 | 339.700503 | 0.28944866 | 0.47687742 | 0.60696658 | 0.54387309 | 0.92325413 |
| BUB1B | 49.2588062 | 0.3754648 | 0.70188235 | 0.5349398 | 0.59269149 | 0.93289552 |
| BUB1B-PAK6 | 406.680119 | 0.15693577 | 0.27298687 | 0.57488393 | 0.5653698 | 0.92512239 |
| PAK6 | 665.354257 | 0.24450437 | 0.24529176 | 0.99678997 | 0.31886647 | 0.84914926 |
| PLCB2 | 986.57406 | 0.19832054 | 0.53565367 | 0.37024023 | 0.7112035 | 0.95560193 |
| INAFM2 | 806.088323 | 0.33295131 | 0.32857737 | 1.01331175 | 0.31091128 | 0.84476049 |
| CCDC9B | 1532.44735 | -0.1687362 | 0.31662068 | -0.5329285 | 0.59408308 | 0.93317501 |
| DISP2 | 51.7131622 | 0.69903708 | 0.9158013 | 0.7633065 | 0.44528063 | 0.89452291 |
| KNSTRN | 400.577448 | 0.06821189 | 0.29955407 | 0.22771145 | 0.81987057 | 0.97454841 |
| IVD | 6190.25345 | 0.45864407 | 0.2727557 | 1.68151963 | 0.09266203 | 0.65662464 |
| BAHD1 | 2874.1776 | 0.09875267 | 0.10779728 | 0.91609611 | 0.35961649 | 0.86480228 |
| AC013356.2 | 609.251905 | 0.08702643 | 0.1533726 | 0.56741837 | 0.57042997 | 0.92758596 |
| CHST14 | 758.43858 | 0.11186514 | 0.19241961 | 0.58136038 | 0.56099759 | 0.92448946 |
| AC091045.1 | 794.337677 | -0.0842556 | 0.18343577 | -0.4593192 | 0.64600498 | 0.94208859 |
| CCDC32 | 1169.26972 | -0.1889993 | 0.14069764 | -1.3433011 | 1.79E-01 | 0.76125802 |
| RPUSD2 | 440.929759 | 0.03671492 | 0.19605904 | 0.18726462 | 0.85145316 | 0.97855304 |
| KNL1 | 237.319729 | 1.18249053 | 0.69046453 | 1.71260143 | 0.08678589 | 0.65228863 |
| RAD51 | 55.1575496 | 0.0091458 | 0.53505911 | 0.01709306 | 0.98636237 | 0.99715899 |
| RMDN3 | 2845.43707 | 0.15994958 | 0.13476022 | 1.18691987 | 0.23525923 | 0.80747753 |
| GCHFR | 1859.91663 | 0.63291766 | 0.3421613 | 1.849764 | 0.06434757 | 0.61964848 |
| DNAJC17 | 2697.35145 | 0.36230144 | 0.23414943 | 1.54730868 | 0.12178883 | 0.69725141 |
| C15orf62 | 278.461359 | 0.07797147 | 0.37524486 | 0.20778825 | 0.83539431 | 0.97550416 |
| ZFYVE19 | 2361.93236 | -0.2515023 | 0.20852586 | -1.2060966 | 0.22778026 | 0.80300331 |
| PPP1R14D | 116.696962 | 0.9557107 | 0.59922575 | 1.59490925 | 0.11073253 | 0.68001961 |
| SPINT1-AS1 | 942.312481 | 0.32351353 | 0.18092358 | 1.78812249 | 7.38E-02 | 6.37E-01 |
| SPINT1 | 12370.384 | 0.14734127 | 0.19795349 | 0.74432265 | 0.4566813 | 0.89688594 |
| RHOV | 315.154047 | 0.20660491 | 0.48590815 | 0.42519335 | 0.67069573 | 0.94922827 |
| VPS18 | 2508.12632 | -0.3162189 | 0.29946895 | -1.0559321 | 0.29099925 | 0.83454586 |
| DLL4 | 1485.40419 | -0.2927871 | 0.40205785 | -0.7282213 | 0.46647815 | 0.89960479 |
| CHAC1 | 254.67222 | -0.922905 | 0.46973058 | -1.9647539 | 0.04944272 | 0.5861488 |
| INO80 | 1546.6981 | -0.4275535 | 0.17299132 | -2.4715316 | 0.01345356 | 0.42590076 |
| CHP1 | 6771.84095 | 0.27970287 | 0.23613267 | 1.1845158 | 0.23620894 | 0.80802614 |
| NUSAP1 | 380.684715 | -0.3206798 | 0.41292947 | -0.7765971 | 0.43739654 | 0.89130865 |
| NDUFAF1 | 1073.36504 | 0.40642165 | 0.23617515 | 1.72084846 | 0.08527833 | 0.65020388 |
| RTF1 | 2169.88106 | -0.1585313 | 0.16045472 | -0.9880128 | 0.3231464 | 0.85119105 |
| LTK | 276.508654 | 0.6377898 | 0.39622795 | 1.60965376 | 0.10747347 | 0.67707349 |
| RPAP1 | 1313.90798 | 0.0574468 | 0.16518389 | 0.34777486 | 0.72800927 | 0.95848028 |
| TYRO3 | 1809.97344 | 0.18141784 | 0.58662487 | 0.30925698 | 0.75712605 | 0.96499587 |
| MGA | 1271.2359 | -0.2531608 | 0.21098817 | -1.1998816 | 0.23018534 | 0.80556869 |
| MAPKBP1 | 1716.63923 | -0.665899 | 0.24897725 | -2.6745377 | 0.00748324 | 0.3642684 |
| AC020659.2 | 697.654665 | -0.7000366 | 0.26879834 | -2.604319 | 0.0092057 | 0.38664841 |
| JMJD7 | 602.715536 | 0.43486722 | 0.25546774 | 1.70223929 | 0.08871052 | 0.65363785 |
| JMJD7-PLA2( | 2897.88261 | 0.18654012 | 0.26457835 | 0.7050468 | 0.48078111 | 0.90541935 |
| PLA2G4B | 2033.38326 | 0.14767052 | 0.28155542 | 0.5244812 | 0.59994389 | 0.93487648 |
| SPTBN5 | 762.710984 | 0.28321918 | 0.5059724 | 0.55975222 | 0.57564846 | 0.93083012 |
| AC020659.1 | 1086.27039 | 0.16794165 | 0.46373639 | 0.36214896 | 0.71724072 | 0.95611001 |
| EHD4 | 3184.73596 | -0.0616689 | 0.17578417 | -0.3508218 | 0.72572208 | 0.95848028 |
| PLA2G4F | 1723.67071 | 0.98224767 | 0.51245706 | 1.91674142 | 0.05527079 | 0.59617546 |
| VPS39 | 4166.53124 | -0.0316997 | 0.15834216 | -0.2001975 | 0.8413261 | 0.97618641 |
| AC036103.1 | 621.680406 | 0.00621406 | 0.21045564 | 0.02952669 | 0.97644453 | 0.99455099 |
| TMEM87A | 2504.79476 | 0.02336146 | 0.203364 | 0.11487512 | 0.90854411 | 0.98662757 |
| GANC | 783.681324 | 0.10748921 | 0.19731686 | 0.54475432 | 0.5859225 | 0.93289552 |
| AC012651.1 | 4577.95233 | 0.35243076 | 0.58242401 | 0.6051103 | 0.54510571 | 0.92345821 |

| | | | | | | |
|---|---|---|---|---|---|---|
| CAPN3 | 4713.8396 | 0.35322695 | 0.58562245 | 0.60316498 | 0.54639894 | 0.92345821 |
| ZNF106 | 2581.23166 | 0.05608205 | 0.16363577 | 0.34272489 | 0.73180544 | 0.95848028 |
| SNAP23 | 3334.38636 | 0.17234359 | 0.15837823 | 1.08817729 | 0.27651685 | 0.82730125 |
| AC018362.1 | 1882.59161 | 0.15327985 | 0.130638 | 1.17331751 | 0.2406685 | 0.8124487 |
| LRRC57 | 1124.8708 | -0.0674641 | 0.09625333 | -0.7009013 | 0.48336458 | 0.90541935 |
| HAUS2 | 945.445374 | -0.1563109 | 0.20300496 | -0.7699855 | 0.44130852 | 0.89290667 |
| AC018362.2 | 210.556166 | -0.1543225 | 0.2530206 | -0.6099208 | 0.54191425 | 0.92292765 |
| STARD9 | 564.350415 | -0.0757451 | 0.31646419 | -0.2393481 | 0.81083565 | 0.97352497 |
| CDAN1 | 1025.7528 | -0.1943746 | 0.13535883 | -1.4359951 | 0.15100372 | 0.73796431 |
| AC090510.3 | 325.235898 | -0.1777793 | 0.19829343 | -0.8965466 | 0.3699609 | 0.86762749 |
| TTBK2 | 935.46093 | -0.1452793 | 0.16402487 | -0.885715 | 0.37577111 | 0.86889275 |
| AC090510.1 | 356.453466 | -0.1465278 | 0.19378789 | -0.7561249 | 0.44957431 | 0.89557948 |
| UBR1 | 1510.40926 | 0.07848725 | 0.1325551 | 0.5921104 | 0.55377666 | 0.92373235 |
| TMEM62 | 1268.64519 | -0.2188163 | 0.34338582 | -0.6372316 | 0.52397397 | 0.91723878 |
| AC068724.1 | 443.719813 | -0.2462042 | 0.37833456 | -0.6507578 | 0.51520283 | 0.91636259 |
| CCNDBP1 | 2948.44561 | 0.16404872 | 0.17636926 | 0.93014345 | 0.35229681 | 0.86364633 |
| LCMT2 | 617.049843 | -0.0455244 | 0.23856637 | -0.1908248 | 0.84866286 | 0.97741457 |
| ADAL | 608.134576 | 0.24245262 | 0.25231822 | 0.96090019 | 0.33660235 | 0.85679125 |
| ZSCAN29 | 900.366774 | -0.3296553 | 0.18967506 | -1.7380003 | 0.08221077 | 0.64573009 |
| TUBGCP4 | 2512.04941 | -0.0660329 | 0.14019316 | -0.4710137 | 0.63763095 | 0.94088547 |
| TP53BP1 | 3189.85159 | -0.0485665 | 0.1807415 | -0.268707 | 0.78815519 | 0.97090793 |
| MAP1A | 420.404743 | -0.2840693 | 0.35420024 | -0.8020021 | 0.42255175 | 0.88451372 |
| PPIP5K1 | 1204.70077 | 0.31853395 | 0.17893254 | 1.78019017 | 0.07504484 | 0.63773415 |
| CKMT1B | 211.354972 | 0.0866565 | 0.38808676 | 0.22329157 | 0.82330859 | 0.97542663 |
| STRC | 48.949095 | -0.0173891 | 0.52231461 | -0.0332924 | 0.97344138 | 0.99445887 |
| AC011330.3 | 56.7675752 | 0.20604671 | 0.47115397 | 0.43732353 | 0.66187674 | 0.94642611 |
| CATSPER2 | 380.192608 | -0.1495948 | 0.43763681 | -0.3418241 | 7.32E-01 | 9.58E-01 |
| CKMT1A | 132.620603 | -0.2049579 | 0.67017096 | -0.3058292 | 0.75973466 | 0.96510454 |
| PDIA3 | 19742.6529 | 0.23119064 | 0.197777 | 1.16894601 | 0.24242538 | 0.81319028 |
| ELL3 | 374.093412 | 0.00375927 | 0.23066161 | 0.01629776 | 0.98699685 | 0.99726698 |
| AC018512.1 | 1038.91812 | 0.08207936 | 0.1140819 | 0.71947753 | 0.47184674 | 0.90160919 |
| SERF2 | 21132.5309 | -0.0338426 | 0.13781985 | -0.2455567 | 0.80602538 | 0.97334138 |
| SERINC4 | 1869.4398 | 0.08604942 | 0.13346254 | 0.64474586 | 0.51909189 | 0.91636259 |
| HYPK | 2169.95264 | -0.0985222 | 0.13915451 | -0.7080058 | 0.47894164 | 0.90464072 |
| MFAP1 | 1640.44899 | -0.0264531 | 0.10692144 | -0.2474065 | 0.80459367 | 0.97287947 |
| WDR76 | 140.764915 | -0.1662244 | 0.22891284 | -0.726147 | 0.46774866 | 0.89984974 |
| FRMD5 | 52.8336512 | 1.00442509 | 0.53965327 | 1.86124155 | 0.06271007 | 0.61467023 |
| CASC4 | 4809.92436 | 0.02552762 | 0.22081259 | 0.11560763 | 0.90796351 | 0.98662757 |
| AC025043.1 | 960.191034 | 0.03942309 | 0.28491981 | 0.13836555 | 8.90E-01 | 9.85E-01 |
| CTDSPL2 | 1330.00951 | -0.2167426 | 0.1640327 | -1.3213377 | 0.18638879 | 0.76908034 |
| EIF3J-DT | 940.784 | -0.0322708 | 0.18145103 | -0.1778487 | 0.85884176 | 0.97992012 |
| EIF3J | 3271.7271 | -0.0799422 | 0.13533328 | -0.5907059 | 0.55471752 | 0.92373235 |
| SPG11 | 3091.34546 | 0.09691511 | 0.11822911 | 0.81972289 | 0.4123741 | 0.88268208 |
| PATL2 | 125.578876 | -0.4874452 | 0.73469492 | -0.6634662 | 0.50703202 | 0.91420888 |
| B2M | 105170.069 | -0.3236267 | 0.33733595 | -0.9593604 | 0.33737722 | 0.85709455 |
| TRIM69 | 2072.24414 | -0.4610721 | 0.36996077 | -1.2462729 | 0.21266423 | 0.79203904 |
| SORD | 8039.93409 | 1.72582032 | 0.77888365 | 2.2157614 | 0.02670785 | 0.52678638 |
| AC091117.1 | 1524.68549 | 1.79194733 | 0.78951 | 2.26969554 | 0.02322606 | 0.51967919 |
| DUOX2 | 82.9542403 | 0.78372102 | 0.69039342 | 1.13518031 | 0.25629978 | 0.82067614 |
| DUOX1 | 229.689539 | 0.074459 | 0.22874031 | 0.32551762 | 0.74478935 | 0.9613927 |
| SHF | 245.610336 | -0.2114429 | 0.3925621 | -0.5386229 | 0.59014711 | 0.93289552 |
| AC051619.5 | 57.8809705 | 1.21649598 | 0.6403749 | 1.89966219 | 0.05747746 | 0.60129769 |
| SLC28A2 | 210.852285 | 0.8169466 | 0.74294189 | 1.09961036 | 0.27150192 | 0.82716979 |
| GATM | 54290.158 | 1.18979363 | 0.38248053 | 3.11072995 | 0.00186626 | 0.18905527 |

| | | | | | | |
|---|---|---|---|---|---|---|
| SPATA5L1 | 617.107577 | -0.1621675 | 0.21608311 | -0.7504865 | 0.45296174 | 0.89670027 |
| C15orf48 | 128.248072 | 0.55486476 | 0.42270838 | 1.31264198 | 0.18930362 | 0.76970626 |
| AC025580.1 | 66.6280809 | 0.54168925 | 0.58866007 | 0.92020723 | 0.35746448 | 0.86390849 |
| SLC30A4 | 825.069593 | 0.14821032 | 0.30758143 | 0.48185718 | 0.62990741 | 0.94088547 |
| BLOC1S6 | 2409.65024 | -0.041252 | 0.22248885 | -0.1854117 | 0.85290613 | 0.97886533 |
| AC090527.2 | 2716.35595 | 0.0482182 | 0.35481603 | 0.13589632 | 0.89190324 | 0.98474725 |
| SQOR | 4583.02728 | 0.00460569 | 0.32452626 | 0.01419205 | 0.98867676 | 0.99788078 |
| SEMA6D | 1155.55326 | -0.5387775 | 0.41167727 | -1.3087376 | 0.19062325 | 0.77175856 |
| SLC24A5 | 905.233344 | -0.0578333 | 0.20048288 | -0.2884699 | 0.77298707 | 0.96836854 |
| MYEF2 | 2682.46792 | -0.0445714 | 0.16805732 | -0.2652157 | 0.79084329 | 0.97090793 |
| SLC12A1 | 34363.0699 | 0.86187953 | 0.71811706 | 1.20019364 | 0.23006415 | 0.80551898 |
| AC023355.1 | 497.800437 | 0.10778017 | 0.34719167 | 0.31043419 | 0.7562308 | 0.96450325 |
| DUT | 2279.49327 | 0.02553718 | 0.23396578 | 0.10914921 | 0.91308414 | 0.98662757 |
| FBN1 | 1526.87611 | -0.14808 | 0.39750618 | -0.3725226 | 0.70950383 | 0.95501548 |
| CEP152 | 288.831328 | -0.2535817 | 0.24144067 | -1.0502856 | 0.29358681 | 0.83540174 |
| EID1 | 7836.22937 | 0.18260124 | 0.2156155 | 0.84688367 | 0.39705997 | 0.87988087 |
| SECISBP2L | 3471.39141 | -0.049756 | 0.14656817 | -0.3394731 | 0.73425333 | 0.95895327 |
| COPS2 | 3218.97663 | -0.169709 | 0.17382038 | -0.9763472 | 0.32889242 | 0.85317073 |
| GALK2 | 1908.1297 | 0.4033013 | 0.27314507 | 1.47650956 | 0.13980715 | 0.72475166 |
| FAM227B | 1042.40524 | 0.41784296 | 0.27284106 | 1.5314519 | 0.12565775 | 0.70347499 |
| AC022306.2 | 83.272591 | 0.26171955 | 0.2640053 | 0.99134206 | 0.32151858 | 0.85119105 |
| FGF7 | 810.579872 | 0.3420814 | 0.60246984 | 0.56779838 | 0.57017188 | 0.92748817 |
| DTWD1 | 909.986675 | -0.076898 | 0.15847027 | -0.4852521 | 0.62749751 | 0.94088547 |
| ATP8B4 | 1317.12466 | 1.08851075 | 0.52238391 | 2.08373715 | 0.03718408 | 0.55527244 |
| SLC27A2 | 8155.27062 | 1.15668025 | 0.53060556 | 2.17992486 | 0.02926303 | 0.53606221 |
| HDC | 106.725145 | 1.63687525 | 0.96745659 | 1.69193664 | 0.09065806 | 0.65495306 |
| GABPB1 | 732.159419 | -0.399649 | 0.23744302 | -1.6831362 | 0.09234873 | 0.65583276 |
| GABPB1-AS1 | 1445.2798 | 0.13288521 | 0.39862889 | 0.33335568 | 0.73886581 | 0.9602325 |
| USP8 | 2599.11587 | -0.0386234 | 0.13207378 | -0.2924379 | 0.76995185 | 0.96836854 |
| AC012170.3 | 174.208399 | -0.0283137 | 0.26571471 | -0.1065566 | 0.91514077 | 0.9866558 |
| AC012170.2 | 503.873952 | -0.1898033 | 0.13333779 | -1.4234768 | 0.15459799 | 0.74300499 |
| USP50 | 224.5528 | 0.02504285 | 0.17261455 | 0.14507959 | 0.88464804 | 0.98340286 |
| TRPM7 | 3136.24834 | -0.1790443 | 0.20683672 | -0.8656312 | 0.38669245 | 0.87432552 |
| AC084756.1 | 384.455644 | -0.112665 | 0.22670758 | -0.4969617 | 0.61921605 | 0.93945324 |
| SPPL2A | 4733.63052 | -0.1214384 | 0.19750057 | -0.6148762 | 0.5386365 | 0.9215968 |
| AC012100.2 | 259.982983 | -0.1120166 | 0.30952025 | -0.3619039 | 0.71742385 | 0.95611001 |
| AP4E1 | 656.017159 | -0.2307106 | 0.18680874 | -1.2350096 | 0.21682693 | 0.7954827 |
| TNFAIP8L3 | 219.402273 | -1.5619184 | 0.70328645 | -2.2208851 | 0.02635875 | 0.52678638 |
| GLDN | 95.6052485 | -1.6644378 | 0.49493634 | -3.3629331 | 0.00077119 | 0.13020612 |
| AC066613.2 | 413.533004 | 0.19130016 | 0.23060459 | 0.82955922 | 0.40678804 | 0.88132015 |
| DMXL2 | 1814.95947 | 0.15427909 | 0.25533643 | 0.60421887 | 0.54569814 | 0.92345821 |
| AC066613.1 | 175.681825 | 0.07143039 | 0.39326945 | 0.18163219 | 0.85587138 | 0.97929708 |
| LYSMD2 | 515.040745 | 0.00331785 | 0.24719685 | 0.01342189 | 0.9892912 | 0.99800524 |
| TMOD2 | 524.340807 | 0.3073485 | 0.24054233 | 1.27773147 | 0.20134412 | 0.78134034 |
| TMOD3 | 2980.92623 | -0.0184603 | 0.14692928 | -0.1256409 | 0.90001622 | 0.98517634 |
| AC090971.2 | 80.747168 | -0.0513454 | 0.44105666 | -0.1164146 | 0.907324 | 0.98662757 |
| AC090971.1 | 235.588198 | -0.2240796 | 0.31255392 | -0.7169309 | 0.47341672 | 0.90194729 |
| LEO1 | 855.426981 | 0.06302446 | 0.18161264 | 0.34702682 | 0.72857117 | 0.95848028 |
| MAPK6 | 2144.55563 | -0.3718169 | 0.43999809 | -0.8450421 | 3.98E-01 | 8.81E-01 |
| BCL2L10 | 131.927135 | 0.46050972 | 0.39390964 | 1.16907449 | 0.24237362 | 0.81319028 |
| GNB5 | 3085.91742 | 0.09275202 | 0.25177813 | 0.3683879 | 0.71258402 | 0.95560193 |
| AC023906.3 | 590.899959 | 0.13695528 | 0.26874961 | 0.50960176 | 0.61033049 | 0.93792127 |
| AC023906.4 | 416.075881 | 0.1574362 | 0.30576696 | 0.51488952 | 0.60663022 | 0.93708311 |
| CERNA1 | 134.768324 | 0.17017319 | 0.27339632 | 0.62244141 | 0.53365166 | 0.92062041 |

| | | | | | | |
|---|---|---|---|---|---|---|
| MYO5C | 1121.74201 | 0.17693844 | 0.33375671 | 0.53014196 | 0.59601351 | 0.9334601 |
| MYO5A | 1228.23173 | -0.1278079 | 0.13766785 | -0.9283784 | 0.35321134 | 0.86364633 |
| ARPP19 | 6603.45546 | -0.1944437 | 0.23844421 | -0.8154681 | 4.15E-01 | 8.84E-01 |
| AC025917.1 | 315.977415 | -0.2084382 | 0.17497522 | -1.1912442 | 0.23355773 | 0.80725541 |
| FAM214A | 1651.53802 | -0.1501143 | 0.12954922 | -1.1587431 | 0.24656092 | 0.81542272 |
| WDR72 | 7092.20892 | 0.58899834 | 0.4983726 | 1.18184335 | 0.23726787 | 0.80880993 |
| UNC13C | 41.4613661 | 0.99492883 | 0.69789761 | 1.4256086 | 0.15398135 | 0.74258845 |
| RSL24D1 | 2873.78044 | -0.0432234 | 0.20292116 | -0.2130059 | 0.83132237 | 0.97550416 |
| RAB27A | 1648.69232 | 0.0440558 | 0.27761696 | 0.15869277 | 0.87391093 | 0.98275231 |
| PIGBOS1 | 464.309016 | -0.0518972 | 0.13262907 | -0.3912959 | 0.69557854 | 0.9525827 |
| PIGB | 1366.93197 | 0.03619575 | 0.10049764 | 0.36016521 | 0.71872359 | 0.95648644 |
| CCPG1 | 5244.34946 | 0.17127394 | 0.11333802 | 1.51117816 | 0.13074306 | 0.70979843 |
| C15orf65 | 139.947869 | 0.19255994 | 0.37906079 | 0.50799224 | 0.61145878 | 0.93834929 |
| DNAAF4 | 222.196907 | -0.0424902 | 0.30297974 | -0.140241 | 0.88846957 | 0.98423722 |
| PYGO1 | 160.933532 | 0.17892699 | 0.39978345 | 0.44755978 | 0.65447093 | 0.94532497 |
| PRTG | 200.823326 | 1.14811565 | 0.45134907 | 2.54374216 | 0.0109672 | 0.40196898 |
| NEDD4 | 974.763966 | 0.16678593 | 0.31070738 | 0.53679424 | 0.59140975 | 0.93289552 |
| RFX7 | 684.623792 | 0.20655222 | 0.12209336 | 1.69175635 | 0.09069244 | 0.65495306 |
| TEX9 | 558.32864 | 0.13731527 | 0.39798071 | 0.34502998 | 0.73007183 | 0.95848028 |
| AC084782.1 | 73.3443314 | 0.52932621 | 0.4423281 | 1.1966823 | 0.23143041 | 0.80649152 |
| MNS1 | 427.836821 | 0.42496156 | 0.418 | 1.01665444 | 0.30931785 | 0.84414094 |
| ZNF280D | 1539.53058 | 0.04135646 | 0.18614426 | 0.22217427 | 0.82417823 | 0.97548201 |
| AC090517.4 | 513.559632 | -0.1397927 | 0.17704578 | -0.7895849 | 0.42977023 | 0.88870396 |
| TCF12 | 2465.24828 | -0.010624 | 0.13544556 | -0.0784373 | 0.93748018 | 0.99082541 |
| LINC00926 | 281.498892 | -0.1587804 | 0.43571704 | -0.3644116 | 0.71555066 | 0.95590747 |
| CGNL1 | 10708.295 | 0.29979926 | 0.23449321 | 1.27849868 | 0.20107365 | 0.78134034 |
| AC025271.2 | 129.622733 | 0.1172882 | 0.49202267 | 0.23837966 | 8.12E-01 | 0.97352497 |
| AC025271.4 | 180.99513 | 0.03714611 | 0.45308612 | 0.08198465 | 0.93465892 | 0.99068685 |
| GCOM1 | 830.447124 | -0.1563882 | 0.27510449 | -0.5684685 | 0.56971687 | 0.92719921 |
| MYZAP | 340.718353 | 0.09086313 | 0.5820796 | 0.15610087 | 0.87595351 | 0.98275231 |
| POLR2M | 1720.54338 | -0.1359965 | 0.14134759 | -0.9621423 | 0.33597811 | 0.85641269 |
| ALDH1A2 | 2499.75198 | -0.7921269 | 0.52974597 | -1.4952958 | 0.13483727 | 0.71630293 |
| AC025431.1 | 90.2828962 | -1.1233798 | 0.72315712 | -1.5534381 | 0.1203185 | 0.69467399 |
| AQP9 | 261.702926 | 0.1680751 | 0.61951447 | 0.27130134 | 0.78615928 | 0.97075604 |
| LIPC | 846.05546 | -0.3145025 | 0.31914033 | -0.9854678 | 0.32439434 | 0.85169085 |
| AC018904.1 | 192.416891 | -0.086768 | 0.36510367 | -0.237653 | 0.81215024 | 0.97355727 |
| ADAM10 | 4557.38625 | -0.1719187 | 0.20422972 | -0.8417909 | 0.39990499 | 0.88077967 |
| MINDY2 | 1960.6152 | -0.1621408 | 0.14980816 | -1.0823229 | 0.27910906 | 0.82825734 |
| AC090515.4 | 48.5407594 | -0.6945346 | 0.33209965 | -2.091344 | 0.03649724 | 0.55384187 |
| RNF111 | 1250.86083 | -0.1703639 | 0.17102222 | -0.9961506 | 0.31917697 | 0.84928695 |
| SLTM | 4771.6766 | -0.3001651 | 0.19358242 | -1.5505805 | 0.12100225 | 0.69543637 |
| AC025918.1 | 42.7522596 | -0.6901766 | 0.48224901 | -1.4311624 | 0.15238368 | 0.7422778 |
| CCNB2 | 73.3214699 | 0.05610921 | 0.55351217 | 0.10136943 | 0.9192572 | 0.9872458 |
| MYO1E | 3055.81912 | -0.325161 | 0.26297589 | -1.2364671 | 0.21628497 | 0.7954827 |
| FAM81A | 480.771148 | 0.51047938 | 0.29336672 | 1.74007257 | 0.08184628 | 0.64544584 |
| GCNT3 | 2820.82409 | 0.61991076 | 0.52864596 | 1.17263878 | 0.24094069 | 0.8124487 |
| GTF2A2 | 1615.26518 | -0.0436769 | 0.14993173 | -0.2913122 | 0.77081255 | 0.96836854 |
| BNIP2 | 3190.96591 | -0.109542 | 0.18945836 | -0.5781851 | 0.56313917 | 0.92452095 |
| AC092755.2 | 136.520739 | -0.0973138 | 0.28812961 | -0.337743 | 0.73555686 | 0.95895327 |
| ANXA2 | 20232.1646 | 0.0161279 | 0.34057152 | 0.04735539 | 0.96222998 | 0.99320711 |
| ICE2 | 1425.12191 | 0.20425276 | 0.16813871 | 1.21478723 | 0.22444725 | 0.80078694 |
| RORA-AS1 | 211.369707 | -0.2847289 | 0.27500183 | -1.035371 | 0.30049567 | 0.84200819 |
| RORA | 2189.31948 | -0.3078487 | 0.25175571 | -1.2228074 | 0.22140245 | 0.80020811 |
| VPS13C | 3281.36857 | -0.0307039 | 0.15538554 | -0.1975985 | 0.84335925 | 0.97649449 |

| | | | | | | |
|---|---|---|---|---|---|---|
| C2CD4A | 1391.39537 | 0.21301049 | 0.53650753 | 0.39703168 | 0.6913441 | 0.95158502 |
| C2CD4B | 160.750414 | 0.70425099 | 0.74010306 | 0.951558 | 0.34132119 | 0.85889221 |
| TLN2 | 5914.67728 | 0.30993899 | 0.47478151 | 0.6528034 | 0.51388303 | 0.91636259 |
| AC103740.1 | 932.743251 | 0.38712567 | 0.44157184 | 0.8766992 | 0.38065004 | 0.87135245 |
| TPM1 | 13803.708 | 0.31844079 | 0.30377771 | 1.04826911 | 0.29451464 | 0.83630273 |
| TPM1-AS | 649.648371 | -0.1579691 | 0.38895383 | -0.4061384 | 0.68464089 | 0.95092677 |
| AC079328.2 | 3347.17545 | 0.28185784 | 0.31479691 | 0.89536408 | 0.37059249 | 0.86762749 |
| LACTB | 1554.60774 | -0.0626316 | 0.24249189 | -0.2582834 | 0.79618818 | 0.97101097 |
| RPS27L | 4192.04416 | 0.41314941 | 0.20908006 | 1.9760345 | 0.04815087 | 0.58440502 |
| RAB8B | 1699.07052 | -0.1451723 | 0.18001249 | -0.8064569 | 0.41997944 | 0.88451372 |
| APH1B | 1019.76841 | 0.08066839 | 0.17448441 | 0.46232431 | 0.64384877 | 0.94208859 |
| CA12 | 34150.3598 | 0.78204638 | 0.41924107 | 1.8653859 | 0.06212733 | 0.61245813 |
| LINC02568 | 186.350975 | 0.66248879 | 0.38041526 | 1.74148849 | 0.08159799 | 0.6449717 |
| USP3 | 2458.31919 | -0.1514248 | 0.14674511 | -1.0318898 | 0.30212374 | 0.84211658 |
| USP3-AS1 | 1291.65353 | -0.1469196 | 0.15961289 | -0.9204747 | 0.35732475 | 0.86390849 |
| FBXL22 | 76.1336274 | 0.5253987 | 0.44407953 | 1.18311851 | 0.23676218 | 0.80849753 |
| HERC1 | 2202.76974 | -0.0753326 | 0.11188103 | -0.6733278 | 0.50073877 | 0.91207554 |
| DAPK2 | 349.400396 | 0.16160606 | 0.36841088 | 0.43865714 | 0.66090999 | 0.9463718 |
| FAM96A | 2761.2681 | 0.07988368 | 0.22695717 | 0.35197692 | 0.72485557 | 0.95847868 |
| SNX1 | 7622.65823 | -0.0215211 | 0.12979562 | -0.1658077 | 0.8683083 | 0.98243355 |
| SNX22 | 13739.3595 | 0.02609079 | 0.24419146 | 0.10684562 | 0.91491145 | 0.9866558 |
| PPIB | 17635.0528 | 0.02712787 | 0.2401704 | 0.1129526 | 0.91006814 | 0.98662757 |
| CSNK1G1 | 1036.72047 | -0.1498217 | 0.19393282 | -0.7725441 | 0.43979223 | 0.89203402 |
| AC087632.1 | 104.075572 | -0.0155286 | 0.32756405 | -0.0474063 | 0.96218938 | 0.99320711 |
| PCLAF | 55.7288223 | 0.67637562 | 0.61732919 | 1.0956482 | 0.27323276 | 0.82716979 |
| TRIP4 | 1719.85229 | 0.13100869 | 0.19714978 | 0.66451352 | 0.5063617 | 0.91414359 |
| ZNF609 | 1729.52386 | -0.2344892 | 0.16035479 | -1.4623149 | 0.14365494 | 0.72909721 |
| OAZ2 | 5324.54634 | -0.0902088 | 0.15521942 | -0.5811694 | 0.56112628 | 0.92448946 |
| RBPMS2 | 964.372438 | 0.04015708 | 0.34753178 | 0.11554937 | 0.90800969 | 0.98662757 |
| PIF1 | 73.0118275 | -0.4752161 | 0.49678413 | -0.9565848 | 0.33877688 | 0.85733087 |
| AC069368.1 | 265.62467 | 0.22385029 | 0.39226373 | 0.57066272 | 0.56822829 | 0.92633668 |
| PLEKHO2 | 2362.90386 | 0.22352844 | 0.38162853 | 0.58572257 | 0.55806197 | 0.92421175 |
| ANKDD1A | 540.15467 | -0.1422372 | 0.22279098 | -0.6384334 | 0.52319163 | 0.91723878 |
| AC103691.1 | 165.867301 | -0.2637016 | 0.35765289 | -0.7373116 | 0.46093291 | 0.89755243 |
| SPG21 | 4644.87447 | -0.0820127 | 0.19458319 | -0.4214788 | 0.67340549 | 0.94935413 |
| MTFMT | 739.655917 | 0.12612148 | 0.16481862 | 0.76521375 | 0.44414427 | 0.8938278 |
| SLC51B | 591.452354 | 0.45560548 | 0.51638632 | 0.88229581 | 0.37761686 | 0.86987124 |
| RASL12 | 507.705142 | -0.8024685 | 0.66959281 | -1.1984425 | 0.2307448 | 0.80610188 |
| UBAP1L | 564.688992 | -0.1582681 | 0.23744644 | -0.6665421 | 0.50506465 | 0.91365534 |
| PDCD7 | 1135.48196 | -0.0991279 | 0.17000042 | -0.5831036 | 0.55982358 | 0.92434754 |
| CLPX | 2822.68295 | -0.0509897 | 0.21210369 | -0.2403998 | 0.81002035 | 0.97334138 |
| CILP | 219.978681 | 1.0999698 | 0.73060054 | 1.50556939 | 0.13217775 | 0.7126419 |
| PARP16 | 693.327766 | -0.0143289 | 0.1580342 | -0.0906698 | 0.92775497 | 0.9891936 |
| IGDCC4 | 162.117615 | 0.7602755 | 0.59903308 | 1.26917115 | 0.20438003 | 0.78441247 |
| DPP8 | 1954.64702 | 0.06271026 | 0.14916491 | 0.42040892 | 0.67418675 | 0.94935413 |
| HACD3 | 5545.36496 | 0.37071221 | 0.19797799 | 1.87249202 | 0.06113856 | 0.60989294 |
| INTS14 | 1975.88369 | -0.2120879 | 0.18064607 | -1.1740522 | 2.40E-01 | 8.12E-01 |
| SLC24A1 | 697.130245 | 0.27738912 | 0.26376052 | 1.05167037 | 0.29295081 | 0.83524862 |
| AC011939.3 | 77.0163763 | 0.54099047 | 0.37973844 | 1.42463975 | 0.15426136 | 0.74258845 |
| DENND4A | 1650.48802 | -0.0252089 | 0.24376088 | -0.1034164 | 0.91763251 | 0.9868044 |
| RAB11A | 7052.87384 | 0.09659549 | 0.11573115 | 0.83465415 | 0.40391247 | 0.88132015 |
| MEGF11 | 1845.8417 | 0.58292467 | 0.62320218 | 0.93537007 | 0.34959761 | 0.86364633 |
| DIS3L | 1002.89374 | 0.32360656 | 0.1866653 | 1.73361928 | 0.08298566 | 0.6471083 |
| AC055855.1 | 60.616404 | 0.14648007 | 0.36592781 | 0.40029774 | 0.68893724 | 0.95158502 |

| | | | | | | |
|---|---|---|---|---|---|---|
| TIPIN | 230.518408 | -0.0006747 | 0.24325433 | -0.0027736 | 0.99778696 | 0.99989813 |
| MAP2K1 | 3768.81975 | -0.2499939 | 0.21488328 | -1.1633941 | 0.24466963 | 0.81473945 |
| AC116913.1 | 1264.65664 | -0.2385718 | 0.1860444 | -1.2823379 | 0.19972412 | 0.78134034 |
| SNAPC5 | 1867.60364 | -0.0520143 | 0.17934037 | -0.290031 | 0.77179252 | 0.96836854 |
| RPL4 | 43697.6333 | 0.25808302 | 0.26340914 | 0.97978007 | 0.32719469 | 0.85308232 |
| ZWILCH | 414.504763 | -0.1547295 | 0.25270149 | -0.6123016 | 0.54033823 | 0.92243385 |
| LCTL | 118.151606 | -0.1490129 | 0.27999101 | -0.532206 | 0.5945833 | 0.9334601 |
| SMAD6 | 974.052259 | 0.23553832 | 0.34109331 | 0.69053926 | 0.48985513 | 0.90600166 |
| AC087482.1 | 676.53 | 1.43250154 | 0.80780299 | 1.77333034 | 0.07617398 | 0.63773415 |
| SMAD3 | 6007.39504 | -0.6677024 | 0.20787912 | -3.2119741 | 0.00131826 | 0.16147422 |
| AAGAB | 2897.47934 | 0.00513459 | 0.1421584 | 0.0361188 | 0.97118763 | 0.99400331 |
| IQCH | 181.565522 | 0.01292193 | 0.33756281 | 0.03828007 | 0.96946438 | 0.99400331 |
| IQCH-AS1 | 652.8018 | 0.11526357 | 0.2208916 | 0.52181057 | 6.02E-01 | 0.9355118 |
| C15orf61 | 366.98341 | -0.1632846 | 0.25886964 | -0.63076 | 5.28E-01 | 9.19E-01 |
| AC016355.1 | 68.2972327 | 0.02856734 | 0.44339016 | 0.06442935 | 0.94862836 | 0.99170503 |
| MAP2K5 | 974.092633 | 0.23079097 | 0.16965966 | 1.36031734 | 0.17372952 | 0.75974924 |
| SKOR1 | 70.8084389 | -0.5665458 | 0.32626357 | -1.7364665 | 0.08248139 | 0.64661469 |
| PIAS1 | 2294.65586 | -0.0217587 | 0.14704411 | -0.1479743 | 0.88236308 | 0.98275231 |
| AC107871.1 | 2983.97718 | 0.20750534 | 0.26003966 | 0.79797575 | 0.42488456 | 0.88573606 |
| CALML4 | 1779.57324 | 0.45697402 | 0.32322991 | 1.41377391 | 0.15742829 | 0.74522306 |
| CLN6 | 1984.38844 | -0.0840675 | 0.25365842 | -0.33142 | 0.74032724 | 0.96063649 |
| FEM1B | 2435.42469 | -0.1954365 | 0.14133637 | -1.3827756 | 0.16673367 | 0.75558932 |
| ITGA11 | 748.207983 | -0.2019195 | 0.40903988 | -0.4936426 | 0.62155864 | 0.94016713 |
| CORO2B | 970.795638 | -0.0638422 | 0.38588548 | -0.1654433 | 0.86859504 | 0.98263549 |
| ANP32A | 5571.56516 | 0.33057551 | 0.1580599 | 2.09145714 | 0.0364871 | 0.55384187 |
| SPESP1 | 140.088576 | 0.1952366 | 0.50413377 | 0.38727143 | 0.69855528 | 0.9529028 |
| NOX5 | 75.5927694 | 0.25646729 | 0.55348427 | 0.4633687 | 0.64310011 | 0.94136836 |
| EWSAT1 | 81.1472223 | 1.17022243 | 0.56216761 | 2.0816255 | 0.03737669 | 0.55532957 |
| GLCE | 1494.55975 | 0.27214292 | 0.19089114 | 1.42564455 | 0.15397097 | 0.74258845 |
| AC026992.2 | 462.819506 | 0.43676032 | 0.45694799 | 0.95582064 | 0.33916286 | 0.85733087 |
| AC026992.1 | 168.052556 | 0.96299679 | 0.50549858 | 1.90504352 | 0.05677441 | 0.60049913 |
| PAQR5 | 6950.71932 | -0.0388234 | 0.40364134 | -0.0961828 | 0.92337538 | 0.98858365 |
| AC027237.3 | 948.015973 | -0.1575538 | 0.43311234 | -0.3637712 | 0.71602885 | 0.95603249 |
| KIF23 | 739.913179 | -0.4538775 | 0.54818691 | -0.8279613 | 0.40769244 | 0.88156963 |
| RPLP1 | 39993.1047 | 0.06728304 | 0.19898973 | 0.33812319 | 0.73527035 | 0.95895327 |
| DRAIC | 583.841373 | 0.31510758 | 0.48712543 | 0.64687154 | 0.51771508 | 0.91636259 |
| AC100826.1 | 153.57708 | 0.74173939 | 0.45544779 | 1.62859368 | 0.10339906 | 0.67439401 |
| TLE3 | 3718.48925 | -0.1239118 | 0.18387193 | -0.6739025 | 0.5003733 | 0.91180773 |
| UACA | 3585.38353 | 0.3266477 | 0.1755707 | 1.86049094 | 0.0628161 | 0.6147045 |
| LARP6 | 1064.97272 | -0.2110611 | 0.26271063 | -0.8033978 | 0.42174487 | 0.88451372 |
| LRRC49 | 363.893045 | 0.26535382 | 0.3135 | 0.84642364 | 0.39731645 | 0.88006972 |
| THAP10 | 155.943124 | 0.65552992 | 0.25344181 | 2.58651059 | 0.00969532 | 0.39368232 |
| THSD4 | 1723.23659 | -0.1720421 | 0.42091146 | -0.408737 | 0.68273264 | 0.95070301 |
| AC104938.1 | 908.661252 | -0.1138019 | 0.30549058 | -0.3725219 | 0.70950429 | 0.95501548 |
| MYO9A | 2835.12574 | -0.1550959 | 0.26037551 | -0.5956623 | 0.55140088 | 0.92373235 |
| SENP8 | 265.732477 | 0.25122786 | 0.27903367 | 0.90034962 | 3.68E-01 | 8.67E-01 |
| AC020779.2 | 69.5890543 | 0.10044737 | 0.41444129 | 0.24236816 | 0.8084949 | 0.97334138 |
| GRAMD2A | 674.512953 | -0.1911487 | 0.51717413 | -0.3696022 | 0.7116789 | 0.95560193 |
| PKM | 70889.36 | -0.1259677 | 0.24443261 | -0.5153474 | 0.6063103 | 0.93708311 |
| PARP6 | 4052.60424 | -0.5241316 | 0.18754368 | -2.7947175 | 0.00519451 | 0.3091955 |
| AC009690.3 | 837.432979 | -0.3547923 | 0.2582022 | -1.374087 | 0.16941469 | 0.75852708 |
| AC009690.2 | 6049.66847 | 0.11364615 | 0.21527115 | 0.52792095 | 0.5975542 | 0.93416444 |
| CELF6 | 1308.89335 | -0.0623652 | 0.2401099 | -0.2597359 | 7.95E-01 | 9.71E-01 |
| AC009690.1 | 6317.47519 | 0.13568175 | 0.18996004 | 0.71426469 | 0.47506352 | 0.90226631 |

| | | | | | | |
|---|---|---|---|---|---|---|
| HEXA | 9423.8481 | 0.0808036 | 0.20770076 | 0.38903855 | 0.69724763 | 0.95269909 |
| HEXA-AS1 | 60.7972956 | 0.12167243 | 0.36483559 | 0.33349934 | 7.39E-01 | 0.9602325 |
| TMEM202-A: | 119.06165 | -0.5805886 | 0.3060178 | -1.8972378 | 0.05779655 | 0.60129769 |
| ARIH1 | 3154.50121 | -0.1851447 | 0.1285734 | -1.439992 | 0.14986967 | 0.73749842 |
| BBS4 | 1520.57406 | 0.10527924 | 0.17127238 | 0.61468894 | 0.53876017 | 0.9215968 |
| ADPGK | 3082.80779 | -0.278558 | 0.16516789 | -1.6865144 | 0.09169677 | 0.65518742 |
| ADPGK-AS1 | 80.7164204 | -0.1975668 | 0.3489014 | -0.566254 | 0.57122114 | 0.92828088 |
| NEO1 | 3010.06384 | 0.32222854 | 0.12620206 | 2.55327485 | 1.07E-02 | 4.02E-01 |
| NPTN | 6844.06635 | 0.05107404 | 0.17365785 | 0.29410731 | 7.69E-01 | 9.68E-01 |
| CD276 | 2704.22737 | -0.6449422 | 0.34928939 | -1.8464408 | 0.06482823 | 0.62075686 |
| C15orf59 | 767.342751 | 0.83662049 | 0.52569223 | 1.59146443 | 0.11150509 | 0.68001961 |
| C15orf59-AS | 130.400348 | 1.16114551 | 0.59232059 | 1.96033284 | 0.0499569 | 0.5861488 |
| LOXL1-AS1 | 433.038216 | 0.45938762 | 0.29753149 | 1.54399664 | 0.12258915 | 0.69788192 |
| LOXL1 | 934.541994 | 0.13987863 | 0.3011187 | 0.46452986 | 0.64226817 | 0.94088547 |
| STOML1 | 889.305342 | 0.13712526 | 0.18913841 | 0.72499957 | 0.46845229 | 0.89984974 |
| PML | 5096.94785 | -0.584857 | 0.40845721 | -1.4318686 | 0.15218144 | 0.74175094 |
| ISLR | 1432.62479 | 0.28882126 | 0.52775098 | 0.54726807 | 0.58419458 | 0.93249217 |
| STRA6 | 1173.59642 | -0.5282492 | 0.70962585 | -0.7444053 | NA | NA |
| CCDC33 | 63.0433572 | 0.25708806 | 0.74181553 | 0.34656602 | 0.72891738 | 0.95848028 |
| CYP11A1 | 55.6874749 | 0.02970111 | 0.84389446 | 0.03519529 | 0.97192402 | 0.99404213 |
| SEMA7A | 300.793868 | -0.6081844 | 0.4530313 | -1.3424776 | 0.17944119 | 0.76204561 |
| UBL7 | 1861.15599 | 0.18258259 | 0.1595813 | 1.14413519 | 0.25256757 | 0.81833349 |
| UBL7-AS1 | 90.2712143 | 0.22914607 | 0.28278684 | 0.81031374 | 0.41775988 | 0.88445833 |
| ARID3B | 578.215179 | -0.5418781 | 0.29452226 | -1.8398546 | 0.06578959 | 0.62237346 |
| CLK3 | 4584.40285 | -0.198802 | 0.15950962 | -1.2463325 | 0.21264235 | 0.79203904 |
| AC100835.2 | 40.1179913 | -0.5026533 | 0.57212802 | -0.8785678 | 0.37963566 | 0.8708011 |
| EDC3 | 1531.38198 | -0.3220575 | 0.1207081 | -2.6680684 | 0.00762887 | 0.36753911 |
| CSK | 2433.1333 | -0.4235449 | 0.17502389 | -2.4199263 | 0.01552365 | 0.45322041 |
| ULK3 | 5172.07759 | -0.1357008 | 0.18364866 | -0.7389153 | 0.45995843 | 0.89755243 |
| SCAMP2 | 7393.84952 | -0.1292715 | 0.19910642 | -0.6492583 | 0.5161714 | 0.91636259 |
| MPI | 5867.12473 | 0.08857455 | 0.11713975 | 0.75614424 | 0.44956272 | 0.89557948 |
| FAM219B | 4114.39425 | -0.0289693 | 0.08686602 | -0.3334943 | 0.73876121 | 0.9602325 |
| COX5A | 5250.96381 | 0.2673906 | 0.20905052 | 1.27907167 | 0.20087182 | 0.78134034 |
| RPP25 | 669.585727 | -0.5449376 | 0.37168388 | -1.4661319 | 1.43E-01 | 7.28E-01 |
| SCAMP5 | 2544.86142 | 0.48122761 | 0.18401313 | 2.61518089 | 0.00891802 | 0.38245454 |
| PPCDC | 592.555244 | -0.0953582 | 0.25469319 | -0.3744042 | 0.70810364 | 0.95501548 |
| C15orf39 | 1585.35572 | -0.2221331 | 0.19607547 | -1.1328959 | 0.25725797 | 0.82115501 |
| AC113208.3 | 172.214935 | -0.3579788 | 0.27999957 | -1.2784978 | 0.20107396 | 0.78134034 |
| COMMD4 | 3196.44652 | 0.01855733 | 0.15391127 | 0.12057163 | 0.90403034 | 0.98624484 |
| AC068338.3 | 120.688751 | -0.0504835 | 0.2750409 | -0.1835489 | 0.85436735 | 0.97906925 |
| NEIL1 | 5984.63446 | -0.1062782 | 0.16959883 | -0.626645 | 0.530892 | 0.91962754 |
| MAN2C1 | 9487.8658 | -0.1521701 | 0.16488261 | -0.9228994 | 0.35605964 | 0.86380759 |
| AC068338.2 | 518.289445 | -0.192344 | 0.25368363 | -0.7582043 | 0.44832871 | 0.8948759 |
| SIN3A | 2503.74916 | -0.1064149 | 0.13527348 | -0.7866649 | 0.43147807 | 0.88976671 |
| PTPN9 | 2356.71063 | 0.05469282 | 0.16858518 | 0.32442245 | 0.74561822 | 0.96176407 |
| SNUPN | 1063.86983 | 0.19078086 | 0.16908706 | 1.1282996 | 0.25919341 | 0.82364948 |
| AC105020.3 | 70.9091056 | 0.43097693 | 0.28859797 | 1.49334707 | 0.13534636 | 0.71649038 |
| AC105020.2 | 551.516603 | -0.1894548 | 0.25813751 | -0.7339297 | 0.46299159 | 0.89807153 |
| IMP3 | 2433.71971 | -0.1281206 | 0.22600337 | -0.5668967 | 0.57078435 | 0.92790694 |
| AC105020.6 | 941.12489 | -0.0938169 | 0.22083294 | -0.424832 | 0.67095914 | 0.94922827 |
| SNX33 | 2386.69035 | 0.04377863 | 0.16734498 | 0.26160709 | 0.79362438 | 0.97090793 |
| CSPG4 | 1919.11052 | -1.1080032 | 0.6752381 | -1.6409074 | 0.10081665 | 0.67247782 |
| AC105020.4 | 315.071224 | -1.1435266 | 0.70442231 | -1.6233537 | 0.10451381 | 0.67439401 |
| AC105020.1 | 123.549807 | 0.14246476 | 0.27056849 | 0.52653863 | 0.59851402 | 0.9346978 |

| | | | | | | |
|---|---|---|---|---|---|---|
| UBE2Q2 | 1979.65987 | 0.2767992 | 0.15588157 | 1.77570196 | 0.07578205 | 0.63773415 |
| FBXO22 | 1560.97717 | 0.29847636 | 0.14213017 | 2.10002108 | 0.03572699 | 0.55369025 |
| NRG4 | 308.023213 | 0.40669047 | 0.251102 | 1.61962263 | 0.10531337 | 0.67473222 |
| TMEM266 | 161.908582 | 0.34839586 | 0.48375139 | 0.7201961 | 0.47140427 | 0.90140554 |
| ETFA | 7949.61172 | 0.45246876 | 0.3113577 | 1.45321208 | 0.14616488 | 0.73247558 |
| SCAPER | 730.218635 | 0.60749095 | 0.21818106 | 2.78434323 | 0.00536362 | 0.31308771 |
| RCN2 | 2831.80496 | 0.33138592 | 0.14713293 | 2.25228921 | 0.024304 | 0.51967919 |
| PSTPIP1 | 362.787157 | -0.267202 | 0.70664969 | -0.3781251 | 0.7053377 | 0.95501548 |
| TSPAN3 | 13889.8241 | -0.0243772 | 0.17813847 | -0.1368444 | 0.8911538 | 0.98474725 |
| AC090181.1 | 70.3797888 | -0.3032522 | 0.54441614 | -0.5570228 | 0.57751186 | 0.93130183 |
| PEAK1 | 1279.19928 | -0.1974162 | 0.20256874 | -0.9745639 | 0.32977661 | 0.85363861 |
| HMG20A | 1758.37537 | -0.0674287 | 0.11424006 | -0.5902369 | 0.55503183 | 0.92373235 |
| LINGO1 | 236.124695 | 0.12946934 | 0.46176333 | 0.2803803 | 0.77918575 | 0.96836854 |
| TBC1D2B | 2078.50004 | -0.1893013 | 0.20509473 | -0.9229943 | 0.35601017 | 0.86379941 |
| AC104758.2 | 418.078638 | -0.1642889 | 0.31709276 | -0.51811 | 0.60438154 | 0.93659663 |
| SH2D7 | 150.973103 | -0.2320806 | 0.3832261 | -0.6055969 | 0.54478244 | 0.92345821 |
| CIB2 | 569.886045 | -0.2064044 | 0.30960958 | -0.6666601 | 0.50498927 | 0.91365534 |
| IDH3A | 4334.81668 | -0.0141201 | 0.19240003 | -0.0733895 | 0.94149617 | 0.99136103 |
| ACSBG1 | 3073.01771 | -0.0135149 | 0.19250862 | -0.0702042 | 0.94403114 | 0.99136103 |
| DNAJA4 | 2378.06628 | 0.3324335 | 0.32977835 | 1.00805133 | 0.31342982 | 0.84638821 |
| WDR61 | 2609.04518 | 0.2684794 | 0.12897767 | 2.08159591 | 0.0373794 | 0.55532957 |
| AC090607.1 | 425.048679 | 0.43721898 | 0.20049241 | 2.18072588 | 0.0292037 | 0.53606221 |
| CRABP1 | 128.768478 | 1.02462871 | 0.96000109 | 1.06732037 | 0.28582719 | 0.83107779 |
| IREB2 | 2332.42852 | -0.1725886 | 0.14689147 | -1.1749396 | 0.24001886 | 0.81209487 |
| HYKK | 541.26644 | 0.84484467 | 0.41174807 | 2.05184854 | 0.04018438 | 0.55657615 |
| PSMA4 | 4217.41158 | -0.2497319 | 0.16667034 | -1.4983587 | 0.13404009 | 0.71560922 |
| ADAMTS7 | 499.088924 | -0.2554107 | 0.33105867 | -0.7714969 | 0.44041246 | 0.89262503 |
| MORF4L1 | 15476.999 | -0.0922404 | 0.1902568 | -0.4848204 | 0.62780374 | 0.94088547 |
| CTSH | 25558.6446 | 0.57994915 | 0.29237419 | 1.98358534 | 0.04730208 | 0.58218854 |
| RASGRF1 | 244.992928 | 0.67564353 | 0.69384027 | 0.97377388 | 0.33016883 | 0.85384238 |
| AC011944.1 | 44.352739 | 0.43819177 | 0.68763878 | 0.6372412 | 0.52396774 | 0.91723878 |
| TMED3 | 2776.17713 | -0.0405801 | 0.26992011 | -0.150341 | 0.88049556 | 0.98275231 |
| MTHFS | 2287.11064 | 0.08173797 | 0.32381131 | 0.25242468 | 0.80071282 | 0.97210205 |
| ST20-MTHFS | 2483.42298 | 0.08657481 | 0.31932997 | 0.27111394 | 0.7863034 | 0.97082232 |
| ST20 | 398.812208 | 0.04152573 | 0.30091229 | 0.13799946 | 0.89024085 | 0.98466127 |
| ST20-AS1 | 395.01797 | 0.28262999 | 0.3609115 | 0.78310052 | 0.43356809 | 0.89055338 |
| BCL2A1 | 115.943208 | -0.2484646 | 0.76622198 | -0.3242723 | 0.74573185 | 0.96176407 |
| ZFAND6 | 4224.66248 | -0.164871 | 0.16623095 | -0.9918191 | 0.32128579 | 0.85119105 |
| FAH | 4142.19481 | 0.94208154 | 0.32618187 | 2.88820941 | 0.00387442 | 0.26919756 |
| CTXND1 | 389.852037 | 0.38335528 | 0.42833362 | 0.89499228 | 0.37079122 | 0.86762749 |
| AC016705.2 | 506.804919 | 0.00705824 | 0.39383386 | 0.01792188 | 0.98570117 | 0.99705651 |
| ARNT2 | 2353.31428 | -0.1263484 | 0.37514498 | -0.3367988 | 0.73626855 | 0.95955556 |
| ABHD17C | 1080.26821 | 0.01566936 | 0.21223709 | 0.07382952 | 0.94114604 | 0.99136103 |
| CEMIP | 122.903434 | -0.3454056 | 0.38453525 | -0.8982418 | 0.36905663 | 0.86762749 |
| AC027808.1 | 43.3608011 | 0.03975896 | 0.36166452 | 0.10993325 | 0.91246231 | 0.98662757 |
| MESD | 3118.04389 | 0.01262557 | 0.17138747 | 0.07366683 | 0.94127549 | 0.99136103 |
| AC068870.2 | 64.8131844 | 0.21016705 | 0.49063393 | 0.42835816 | 0.66839038 | 0.9486589 |
| TLNRD1 | 1676.34992 | -0.4767318 | 0.34681342 | -1.374606 | 0.16925363 | 0.7584845 |
| IL16 | 586.824843 | 0.07301111 | 0.48248023 | 0.15132456 | 0.8797197 | 0.98275231 |
| STARD5 | 329.695131 | 0.06432079 | 0.45777872 | 0.14050629 | 0.88825998 | 0.98423722 |
| MEX3B | 115.738926 | 0.50313801 | 0.46704043 | 1.07729006 | 0.2813507 | 0.82957339 |
| EFL1 | 1228.04722 | -0.3497372 | 0.34918561 | -1.0015797 | 0.31654662 | 0.84783414 |
| SAXO2 | 110.225614 | 0.62538801 | 0.55737684 | 1.12202009 | 0.2618539 | 0.82424998 |
| GOLGA6L10 | 64.6820139 | -0.5366547 | 0.73281874 | -0.7323157 | 0.46397588 | 0.89830907 |

| | | | | | | |
|---|---|---|---|---|---|---|
| GOLGA6L9 | 499.07381 | 0.43735575 | 0.23254776 | 1.88071366 | 0.06001088 | 0.6075531 |
| RPS17 | 19995.391 | 0.15724483 | 0.20822275 | 0.75517603 | 0.45014337 | 0.89557948 |
| AC245033.1 | 19639.6678 | 0.1637486 | 0.21099685 | 0.77607129 | 4.38E-01 | 8.91E-01 |
| CPEB1 | 120.14748 | 0.32745745 | 0.55440869 | 0.5906427 | 0.55475985 | 0.92373235 |
| AP3B2 | 264.59295 | 0.13807575 | 0.53448626 | 0.25833359 | 0.79614947 | 0.97101097 |
| SNHG21 | 241.528315 | 0.08902848 | 0.2779328 | 0.32032377 | 0.74872291 | 0.96249332 |
| FSD2 | 308.411381 | 0.15862318 | 0.24244054 | 0.65427662 | 0.51293361 | 0.91568064 |
| WHAMM | 1825.41796 | -0.4147484 | 0.24725482 | -1.6774126 | 0.09346182 | 0.65756964 |
| HOMER2 | 813.450186 | -0.623056 | 0.40944773 | -1.5216984 | 1.28E-01 | 7.06E-01 |
| RAMMET | 595.620241 | -0.2201787 | 0.31406492 | -0.701061 | 0.48326493 | 0.90541935 |
| C15orf40 | 1700.60516 | -0.0726184 | 0.20176368 | -0.3599182 | 0.71890833 | 0.95655386 |
| BTBD1 | 3543.22907 | -0.0117209 | 0.22875475 | -0.0512377 | 0.95913614 | 0.9924381 |
| AC022558.3 | 280.319805 | 0.10829242 | 0.31668541 | 0.34195583 | 0.73238413 | 0.95848028 |
| TM6SF1 | 266.786932 | -0.3129586 | 0.38450058 | -0.8139354 | 0.41568198 | 0.88399703 |
| HDGFL3 | 1597.76313 | -0.1977489 | 0.28165778 | -0.7020894 | 0.48262341 | 0.90541935 |
| AC024270.3 | 44.3120281 | -0.6764256 | 0.52487043 | -1.2887477 | 0.19748581 | 0.77819208 |
| ADAMTSL3 | 488.832595 | 0.42284242 | 0.46219617 | 0.91485487 | 0.36026782 | 0.86480228 |
| GOLGA6L4 | 138.138103 | -0.0506517 | 0.38106583 | -0.1329212 | 0.89425567 | 0.98474725 |
| AC048382.5 | 233.757214 | 0.25877347 | 0.27892279 | 0.92776022 | 0.35353196 | 0.86364633 |
| ZSCAN2 | 730.965744 | 0.08016614 | 0.20531951 | 0.39044578 | 0.69620694 | 0.9525827 |
| AC048382.6 | 353.562652 | 0.19613096 | 0.2571604 | 0.76267947 | 0.44565458 | 0.89469871 |
| WDR73 | 3471.15844 | -0.1133804 | 0.20029597 | -0.5660645 | 0.57134996 | 0.9282864 |
| NMB | 424.980786 | -0.1784149 | 0.36274014 | -0.4918532 | 0.62282315 | 0.94016713 |
| SEC11A | 5529.84318 | 0.22041203 | 0.14063384 | 1.56727591 | 0.11705021 | 0.68956727 |
| ZNF592 | 2836.29636 | -0.3796886 | 0.18858655 | -2.0133386 | 0.04407902 | 0.57247748 |
| ALPK3 | 3742.52707 | -1.0109139 | 0.31403544 | -3.2191075 | 1.29E-03 | 1.61E-01 |
| SLC28A1 | 5352.79669 | 0.42218712 | 0.51638768 | 0.81757783 | 0.41359829 | 0.88316314 |
| PDE8A | 3260.95502 | 0.0840508 | 0.20286214 | 0.4143247 | 0.67863633 | 0.95040677 |
| AKAP13 | 4079.43017 | -0.2948189 | 0.17374507 | -1.6968475 | 0.08972549 | 0.65375543 |
| AC021739.2 | 1020.35082 | -0.243048 | 0.18159476 | -1.3384087 | 0.18076325 | 0.76471425 |
| KLHL25 | 785.670199 | -0.2517255 | 0.22788271 | -1.1046273 | 0.26932113 | 0.82716979 |
| NTRK3 | 903.535264 | -0.8823936 | 0.45379228 | -1.9444878 | 5.18E-02 | 5.86E-01 |
| MRPL46 | 1591.13365 | 0.08678795 | 0.21895159 | 0.39637964 | 0.69182498 | 0.95159158 |
| MRPS11 | 1642.98095 | -0.0460833 | 0.1113495 | -0.4138621 | 0.67897507 | 0.95052453 |
| DET1 | 404.318415 | 0.31734 | 0.32630823 | 0.97251606 | 0.33079388 | 0.85384238 |
| AEN | 1387.01146 | -0.6292377 | 0.43493762 | -1.446731 | 0.14797228 | 0.73460889 |
| ISG20 | 1214.90344 | -2.1150835 | 0.69017377 | -3.0645667 | 0.00217986 | 0.20321981 |
| ACAN | 283.790877 | -1.9359143 | 0.70767338 | -2.7356043 | 0.00622659 | 0.33470127 |
| HAPLN3 | 530.623099 | -0.150937 | 0.71277891 | -0.2117585 | 0.83229544 | 0.97550416 |
| MFGE8 | 4871.31892 | -0.4764226 | 0.35998871 | -1.3234376 | 0.18568988 | 0.76908034 |
| ABHD2 | 5029.4454 | 0.28748536 | 0.346931 | 0.82865285 | 0.40730087 | 0.88132015 |
| FANCI | 1239.40612 | -0.192859 | 0.17270174 | -1.116717 | 0.26411534 | 0.82424998 |
| POLG | 3567.89218 | -0.160498 | 0.17970782 | -0.8931051 | 0.37180089 | 0.86762749 |
| MIR9-3HG | 114.542636 | 0.26867796 | 0.51241948 | 0.52433205 | 0.60004761 | 0.93487648 |
| RHCG | 3517.0738 | 1.69652047 | 0.5317825 | 3.19025253 | 0.00142149 | 0.16968188 |
| TICRR | 134.521142 | -0.0710929 | 0.42715249 | -0.1664344 | 0.86781508 | 0.98234612 |
| KIF7 | 323.686844 | -0.0960817 | 0.42192195 | -0.2277238 | 0.819861 | 0.97454841 |
| PLIN1 | 359.854595 | 1.90987763 | 0.84494573 | 2.26035537 | 0.0237992 | 0.51967919 |
| PEX11A | 928.539947 | 0.42313317 | 0.37380917 | 1.13194967 | 0.2576556 | 0.82187662 |
| WDR93 | 98.479519 | 0.39290987 | 0.53372267 | 0.73616858 | 0.46162811 | 0.89759034 |
| MESP1 | 92.4084699 | 0.4555332 | 0.40548235 | 1.12343532 | 0.26125266 | 0.82424998 |
| ANPEP | 54879.7507 | 0.50082196 | 0.5319211 | 0.9415343 | NA | NA |
| AP3S2 | 3665.08137 | 0.23867859 | 0.15094329 | 1.58124677 | 0.11382163 | 0.68214043 |
| C15orf38-AP: | 2467.98472 | 0.230815 | 0.14264961 | 1.61805557 | 0.10565062 | 0.67526487 |

| | | | | | | |
|---|---|---|---|---|---|---|
| ARPIN | 1643.06289 | 0.34737376 | 0.19654753 | 1.76737787 | 0.07716497 | 0.63819446 |
| ZNF710 | 977.874185 | -0.1718478 | 0.27770394 | -0.6188166 | 0.53603716 | 0.92101152 |
| ZNF710-AS1 | 2545.62687 | 0.41462122 | 0.27284192 | 1.51963903 | 0.12860172 | 0.70610612 |
| IDH2 | 16826.9983 | 0.05185472 | 0.24588306 | 0.21089179 | 0.83297171 | 0.97550416 |
| SEMA4B | 11273.8451 | -0.0756154 | 0.35230331 | -0.2146315 | 0.83005466 | 0.97550416 |
| CIB1 | 10457.8527 | 0.24750957 | 0.14592373 | 1.69615702 | 0.08985615 | 0.65375543 |
| GDPGP1 | 2350.49648 | 0.1873216 | 0.17225775 | 1.08744949 | 0.27683821 | 0.82760008 |
| AC091167.7 | 208.505245 | 0.13914872 | 0.24775712 | 0.56163358 | 0.57436569 | 0.93023062 |
| TTLL13P | 55.0721896 | 0.27504131 | 0.38232596 | 0.71938957 | 0.47190092 | 0.90160919 |
| AC091167.2 | 5832.64955 | -0.0153077 | 0.17155012 | -0.0892314 | 0.92889805 | 0.98963127 |
| NGRN | 7150.52529 | -0.0296437 | 0.17217652 | -0.1721705 | 0.86330348 | 0.98176973 |
| AC091167.6 | 1112.03225 | 0.02931126 | 0.21908863 | 0.13378724 | 0.89357081 | 0.98474725 |
| ZNF774 | 179.965317 | 0.12236079 | 0.28828271 | 0.4244472 | 0.6712397 | 0.94922827 |
| IQGAP1 | 7257.78976 | -0.081272 | 0.19770628 | -0.4110745 | 0.68101793 | 0.95070301 |
| CRTC3 | 2238.61282 | -0.1087617 | 0.12811004 | -0.8489706 | 0.39589766 | 0.87961091 |
| CRTC3-AS1 | 71.5404119 | 0.2407751 | 0.39811129 | 0.60479344 | 0.54531625 | 0.92345821 |
| BLM | 43.7439736 | 0.01837948 | 0.60164992 | 0.03054847 | 0.97562964 | 0.99455099 |
| FURIN | 10326.0515 | -0.2395696 | 0.21721551 | -1.1029118 | 0.27006546 | 0.82716979 |
| FES | 1702.50146 | -0.3487418 | 0.41064551 | -0.8492528 | 0.39574066 | 0.87939946 |
| MAN2A2 | 4270.27273 | -0.0450947 | 0.11316943 | -0.3984704 | 0.69028347 | 0.95158502 |
| AC068831.4 | 1742.40603 | -0.0408454 | 0.19446249 | -0.2100424 | 0.8336346 | 0.97550416 |
| HDDC3 | 881.872114 | 0.42808248 | 0.23716707 | 1.80498283 | 7.11E-02 | 6.31E-01 |
| UNC45A | 5235.12844 | 0.08190476 | 0.13941119 | 0.58750496 | 0.55686462 | 0.92373235 |
| AC068831.1 | 867.598024 | 0.07744267 | 0.16244975 | 0.4767177 | 0.63356316 | 0.94088547 |
| RCCD1 | 941.441561 | 0.12092702 | 0.32328218 | 0.37406028 | 0.70835947 | 0.95501548 |
| PRC1 | 234.173526 | 0.43387998 | 0.38396535 | 1.12999774 | 0.25847718 | 0.82294696 |
| AC068831.7 | 1127.78225 | 0.12182213 | 0.15932417 | 0.764618 | 0.44449905 | 0.8938278 |
| PRC1-AS1 | 94.6307127 | 0.24205995 | 0.37982627 | 0.63729123 | 0.52393516 | 0.91723878 |
| VPS33B | 1020.12829 | -0.0089062 | 0.14572402 | -0.0611169 | 0.95126607 | 0.99170503 |
| SV2B | 90.6558798 | -0.7026077 | 0.6378918 | -1.1014528 | 0.27069964 | 0.82716979 |
| SLCO3A1 | 1751.72951 | 0.34394636 | 0.3565051 | 0.96477262 | 0.33465871 | 0.85557672 |
| AC116903.1 | 338.782424 | 0.20191232 | 0.3833368 | 0.52672302 | 0.59838595 | 0.9346978 |
| FAM174B | 432.625882 | 0.32064652 | 0.31122064 | 1.03028679 | 0.3028754 | 0.84211658 |
| AC013394.1 | 1695.958 | -0.0707274 | 0.17492408 | -0.4043321 | 0.68596855 | 0.95097288 |
| CHD2 | 5214.40877 | -0.2863016 | 0.19308816 | -1.4827505 | 0.13814071 | 0.72234792 |
| RGMA | 328.294611 | -0.3531067 | 0.47037846 | -0.7506865 | 0.45284137 | 0.89670027 |
| MCTP2 | 602.822674 | -0.0016209 | 0.34048321 | -0.0047604 | 0.99620173 | 0.99976751 |
| LINC01197 | 137.905825 | 0.18382431 | 0.49897932 | 0.36840065 | 0.71257451 | 0.95560193 |
| LINC00924 | 523.72216 | -0.1324958 | 0.45553287 | -0.2908589 | 0.77115922 | 0.96836854 |
| AC024337.2 | 259.020855 | 0.38462353 | 0.4132071 | 0.93082506 | 0.35194406 | 0.86364633 |
| NR2F2-AS1 | 379.110461 | 0.16464982 | 0.39738778 | 0.41433035 | 0.67863219 | 0.95040677 |
| AC012409.3 | 118.828908 | 0.24320602 | 0.45814011 | 0.5308551 | 0.59551919 | 0.9334601 |
| AC012409.1 | 70.9515208 | 1.07765393 | 0.5181217 | 2.07992435 | 0.03753247 | 0.55538555 |
| NR2F2 | 9166.55877 | -0.3433894 | 0.28531036 | -1.2035645 | 0.22875796 | 0.80434391 |
| AC103746.1 | 57.9716761 | 0.18166742 | 0.44140746 | 0.411564 | 0.68065902 | 0.95070301 |
| AC087477.2 | 202.191132 | 0.04748328 | 0.35849479 | 0.13245181 | 0.89462693 | 0.98474725 |
| ARRDC4 | 2504.22032 | 0.19277365 | 0.24549292 | 0.78525136 | 0.43230621 | 0.88980461 |
| AC015722.2 | 41.7998878 | -0.1463745 | 0.94784474 | -0.1544288 | 0.87727163 | 0.98275231 |
| FAM169B | 180.029471 | -0.8496214 | 0.55642313 | -1.5269339 | 0.12677744 | 0.70443503 |
| IGF1R | 5016.85223 | -0.1814028 | 0.19618818 | -0.924637 | 0.35515477 | 0.86364633 |
| SYNM | 886.886412 | -0.5443034 | 0.44956594 | -1.210731 | 0.22599853 | 0.80246466 |
| AC036108.2 | 89.8086685 | -0.7259244 | 0.36818285 | -1.9716411 | 0.04865059 | 0.58448511 |
| TTC23 | 1175.5454 | -0.0078165 | 0.22660321 | -0.034494 | 0.97248322 | 0.99416401 |
| AC036108.3 | 126.455514 | -0.318459 | 0.30749772 | -1.0356466 | 0.30036706 | 0.84200819 |

| | | | | | | |
|---|---|---|---|---|---|---|
| LRRC28 | 1198.48336 | 0.39969688 | 0.23104303 | 1.72996729 | 0.08363612 | 0.64783289 |
| MEF2A | 3540.31432 | -0.2162898 | 0.12389502 | -1.7457508 | 0.08085426 | 0.64313548 |
| LYSMD4 | 1688.49794 | -0.103892 | 0.13982247 | -0.7430283 | 0.45746455 | 0.89688594 |
| AC084855.1 | 49.436734 | -0.1904034 | 0.47823255 | -0.3981398 | 0.69052717 | 0.95158502 |
| ADAMTS17 | 212.159244 | -0.4880048 | 0.51462357 | -0.9482752 | 0.34298936 | 0.85923855 |
| AC027020.2 | 261.213474 | -0.6145593 | 0.34362231 | -1.7884731 | 0.07369971 | 0.6374448 |
| LINS1 | 650.154132 | 0.02499373 | 0.1276888 | 0.19573942 | 0.84481413 | 0.97675487 |
| ASB7 | 808.367712 | -0.1598295 | 0.15533085 | -1.0289617 | 0.30349768 | 0.84211658 |
| AC087762.1 | 85.5018347 | 0.28214368 | 0.62944034 | 0.44824531 | 0.65397617 | 0.94510853 |
| AC015712.1 | 50.8185379 | 0.39595646 | 0.57107705 | 0.69335034 | 0.48808972 | 0.90585781 |
| ALDH1A3 | 850.006276 | 0.94487366 | 0.49307808 | 1.91627595 | 0.05532998 | 0.59617546 |
| AC015712.2 | 573.917276 | 0.92344043 | 0.48049008 | 1.92187201 | 0.05462186 | 0.59329767 |
| LRRK1 | 538.257868 | 0.02043086 | 0.37628601 | 0.05429609 | 0.95669927 | 0.99198394 |
| AC090907.2 | 55.203478 | -0.0225305 | 0.55938624 | -0.0402771 | 0.9678722 | 0.99400331 |
| CHSY1 | 2254.19062 | 0.05073961 | 0.38271398 | 0.13257841 | 0.89452681 | 0.98474725 |
| SELENOS | 2857.17286 | -0.1453116 | 0.24051336 | -0.6041726 | 0.54572891 | 0.92345821 |
| SNRPA1 | 2003.82538 | 0.05735919 | 0.17591155 | 0.32606837 | 0.74437263 | 0.9613927 |
| AC023024.1 | 374.120362 | -0.5690341 | 0.24950458 | -2.2806562 | 0.0225688 | 0.51958567 |
| PCSK6 | 650.047501 | -0.6197048 | 0.28148506 | -2.2015548 | 0.02769677 | 0.53038691 |
| TM2D3 | 2526.4284 | -0.0328752 | 0.11730349 | -0.2802578 | 0.77927975 | 0.96836854 |
| TARSL2 | 940.825901 | 0.18471293 | 0.21953112 | 0.84139746 | 0.40012531 | 0.88077967 |
| POLR3K | 439.816069 | 1.65E-01 | 0.15598072 | 1.05E+00 | 0.29147833 | 0.83454586 |
| SNRNP25 | 1101.29052 | 0.13745822 | 0.16184361 | 0.84932746 | 0.39569911 | 0.87939946 |
| RHBDF1 | 2080.463 | -0.1338718 | 0.26724074 | -0.5009407 | 0.61641286 | 0.93910158 |
| MPG | 1708.24025 | 0.00556643 | 0.15063817 | 0.03695231 | 0.97052303 | 0.99400331 |
| NPRL3 | 3638.93332 | -0.0015619 | 0.08465393 | -0.0184506 | 0.98527935 | 0.99692004 |
| HBA2 | 1109.12918 | -0.7828564 | 0.74481073 | -1.051081 | 0.29322141 | 0.8352732 |
| HBA1 | 198.895923 | -1.6780235 | 0.75244376 | -2.2300982 | 0.02574092 | 0.52502381 |
| LUC7L | 4608.70716 | 0.01833252 | 0.19665715 | 0.09322072 | 0.92572822 | 0.98876911 |
| FAM234A | 5694.86031 | 0.0060261 | 0.28135404 | 0.0214182 | 0.98291206 | 0.9961464 |
| AC004754.1 | 2861.70673 | 0.06421859 | 0.2804037 | 0.22902189 | 0.8188519 | 0.97454841 |
| RGS11 | 373.199717 | 0.90995708 | 0.51327529 | 1.77284411 | 0.07625454 | 0.63773415 |
| ARHGDIG | 112.854716 | 0.8643832 | 0.54375665 | 1.58965083 | 1.12E-01 | 6.80E-01 |
| AXIN1 | 2621.19259 | -0.4656818 | 0.19550468 | -2.381947 | 0.01722138 | 0.46842641 |
| MRPL28 | 3352.20704 | 0.05076622 | 0.11684669 | 0.43446865 | 0.66394815 | 0.94642611 |
| TMEM8A | 9138.16259 | -0.0794496 | 0.18252833 | -0.4352729 | 0.66336436 | 0.94642611 |
| NME4 | 3841.01901 | -0.092553 | 0.27552874 | -0.3359105 | 0.73693833 | 0.96009463 |
| DECR2 | 3233.95082 | 0.45815237 | 0.25948206 | 1.76564181 | 0.07745596 | 0.63862831 |
| RAB11FIP3 | 16436.675 | 0.20761609 | 0.38506542 | 0.53917096 | 0.5897689 | 0.93289552 |
| LINC00235 | 117.606299 | -0.2214647 | 0.46265055 | -0.4786868 | 0.63216148 | 0.94088547 |
| CAPN15 | 5232.98831 | -0.5056041 | 0.20974428 | -2.4105741 | 0.01592744 | 0.4564177 |
| Z97986.1 | 952.624048 | -0.516883 | 0.23599154 | -2.1902609 | 2.85E-02 | 0.53606221 |
| PRR35 | 514.614505 | 1.11026568 | 0.81912722 | 1.3554252 | 0.17528212 | 0.76073329 |
| PIGQ | 7672.17791 | -0.0092827 | 0.21872893 | -0.0424391 | 0.96614862 | 0.99393925 |
| NHLRC4 | 474.962416 | 1.28273216 | 0.57303255 | 2.23849788 | 0.02518861 | 0.52140487 |
| RAB40C | 2535.80195 | -0.1712135 | 0.13150405 | -1.3019638 | 0.19292876 | 0.7751938 |
| WFIKKN1 | 330.531463 | -0.1422747 | 0.40906665 | -0.3478033 | 0.72798789 | 0.95848028 |
| METTL26 | 1769.23329 | 0.00616521 | 0.16585645 | 0.03717195 | 0.9703479 | 0.99400331 |
| MCRIP2 | 4307.46917 | 0.07537425 | 0.29687865 | 0.25388907 | 0.79958126 | 0.97207132 |
| AL022341.1 | 40.5598811 | -0.2823814 | 0.40476523 | -0.6976425 | 0.48540077 | 0.90543654 |
| WDR90 | 3355.44766 | -0.2996804 | 0.22548689 | -1.3290367 | 0.18383586 | 0.76781833 |
| AL022341.2 | 324.230074 | -0.3539121 | 0.24434493 | -1.4484117 | 0.14750195 | 0.73412064 |
| RHOT2 | 8415.68867 | -0.1318823 | 0.14528311 | -0.9077609 | 0.36400455 | 0.8663515 |
| RHBDL1 | 282.056745 | 0.64529006 | 0.30848688 | 2.09179092 | 0.03645722 | 0.55384187 |

| | | | | | | |
|---|---|---|---|---|---|---|
| STUB1 | 8513.62912 | -0.0012733 | 0.12025434 | -0.0105886 | 0.99155165 | 0.99885344 |
| JMJD8 | 8109.31892 | 0.00059453 | 0.12400988 | 0.00479425 | 0.99617476 | 0.99976751 |
| WDR24 | 981.922657 | -0.183821 | 0.12813862 | -1.4345478 | 0.15141599 | 0.73917351 |
| FBXL16 | 569.372969 | 0.39504971 | 0.56798261 | 0.69553134 | 0.48672238 | 0.90573694 |
| METRN | 975.761706 | 0.55942598 | 0.26146185 | 2.13960843 | 0.03238642 | 0.55014009 |
| FAM173A | 803.909044 | 0.03619712 | 0.18278983 | 0.1980259 | 0.8430248 | 0.97649449 |
| CCDC78 | 350.368376 | 0.01282581 | 0.31318632 | 0.04095267 | 0.96733363 | 0.99400331 |
| HAGHL | 1104.2973 | -0.1110332 | 0.23452311 | -0.4734425 | 0.63589753 | 0.94088547 |
| NARFL | 2738.36063 | -0.1905256 | 0.11197516 | -1.7014984 | 0.08884944 | 0.65375543 |
| MSLN | 440.276097 | 0.29959596 | 0.55740179 | 0.53748654 | 0.59093158 | 0.93289552 |
| RPUSD1 | 1239.50817 | -0.5951897 | 0.28676038 | -2.0755645 | 0.03793425 | 0.55538555 |
| CHTF18 | 898.059514 | -0.1331372 | 0.38466764 | -0.3461096 | 0.72926032 | 0.95848028 |
| LMF1 | 1517.19173 | 0.05228222 | 0.21410751 | 0.24418679 | 0.80708616 | 0.97334138 |
| LMF1-AS1 | 91.9321577 | 0.00515544 | 0.30225433 | 0.01705663 | 0.98639144 | 0.99715899 |
| AC009041.2 | 262.198612 | 0.10625323 | 0.33209613 | 0.3199472 | 0.74900836 | 0.96249332 |
| SOX8 | 79.6736442 | -0.286576 | 0.34443069 | -0.832028 | 0.40539316 | 0.88132015 |
| AC009041.3 | 378.494704 | 0.18954289 | 0.67735053 | 0.27982984 | 0.77960806 | 0.96836854 |
| SSTR5 | 160.108189 | 0.22435386 | 0.63508621 | 0.35326521 | 0.72388963 | 0.95810643 |
| CACNA1H | 4607.44932 | -0.8271064 | 0.49058749 | -1.6859509 | 0.09180525 | 0.65518742 |
| AC120498.6 | 124.192563 | -0.5782939 | 0.58121182 | -0.9949796 | 0.31974617 | 0.84960209 |
| TPSG1 | 264.571227 | -0.5440817 | 0.5541618 | -0.9818101 | 0.3261934 | 0.85282528 |
| AC120498.1C | 39.9132029 | -0.869637 | 0.73801944 | -1.1783388 | 0.23866155 | 0.81095554 |
| TPSB2 | 231.466408 | 2.13150276 | 1.07546879 | 1.98192898 | 0.04748719 | 0.58241466 |
| TPSAB1 | 266.317904 | 1.80749904 | 1.14181302 | 1.58300791 | 0.11341966 | 0.68193708 |
| TPSD1 | 44.669688 | 1.04183392 | 1.43349128 | 0.72678079 | 0.46736025 | 0.89983399 |
| AC120498.4 | 81.5663522 | -0.7795362 | 1.04294347 | -0.7474386 | 0.45479883 | 0.89670027 |
| AC120498.9 | 60.0564659 | -0.5869711 | 0.52321237 | -1.1218601 | 0.26192195 | 0.82424998 |
| UBE2I | 4622.94669 | -0.0490441 | 0.18912359 | -0.2593229 | 0.79538615 | 0.97101097 |
| AL031714.1 | 2129.99354 | -0.115864 | 0.20325125 | -0.5700529 | 0.56864184 | 0.92642835 |
| BAIAP3 | 2668.79382 | -0.0756291 | 0.21284054 | -0.3553322 | 0.72234072 | 0.95741464 |
| TSR3 | 2353.81348 | 0.00497705 | 0.14840139 | 0.03353777 | 0.97324575 | 0.99445887 |
| GNPTG | 5902.82645 | 0.17792907 | 0.07900797 | 2.25203957 | 0.02431977 | 0.51967919 |
| AL031709.1 | 157.819974 | -0.3335903 | 0.30740248 | -1.0851906 | 0.27783723 | 0.82805357 |
| UNKL | 2373.9253 | -0.0513431 | 0.13252646 | -0.3874174 | 6.98E-01 | 0.9529028 |
| C16orf91 | 563.770253 | -0.1738534 | 0.19810756 | -0.8775708 | 0.38017666 | 0.87090707 |
| CCDC154 | 403.053532 | 0.44485821 | 0.44086857 | 1.00904949 | 0.3129509 | 0.84559008 |
| CLCN7 | 11818.8513 | -0.5132372 | 0.20709441 | -2.4782766 | 0.01320187 | 0.42221215 |
| AL031600.3 | 4653.15884 | -0.4774063 | 0.20192125 | -2.3643194 | 0.01806323 | 0.47478681 |
| AL031600.1 | 497.276082 | -0.7744355 | 0.26429802 | -2.9301602 | 0.00338787 | 0.2541761 |
| TELO2 | 2987.71921 | -0.1350893 | 0.23096575 | -0.5848888 | 0.55862249 | 0.92421175 |
| IFT140 | 2365.41125 | -0.0692243 | 0.25729344 | -0.2690479 | 0.78789282 | 0.97090793 |
| TMEM204 | 1705.92091 | 0.00971099 | 0.40120228 | 0.02420472 | 0.98068931 | 0.99573612 |
| CRAMP1 | 1788.86954 | -0.3820001 | 0.13586211 | -2.8116749 | 0.00492843 | 0.30302984 |
| AL031708.1 | 853.897381 | -0.4795964 | 0.26208135 | -1.8299524 | 0.06725706 | 0.6249306 |
| JPT2 | 6073.83114 | -0.4837745 | 0.22199911 | -2.1791732 | 0.02931881 | 0.53606221 |
| MAPK8IP3 | 10781.4244 | -0.4037327 | 0.18461623 | -2.1868756 | 0.0287516 | 0.53606221 |
| AL031710.1 | 68.4473124 | 1.58022983 | 0.6680055 | 2.36559405 | 1.80E-02 | 0.47478681 |
| NME3 | 2107.18221 | 0.14889019 | 0.23401513 | 0.63624172 | 0.52461888 | 0.91723878 |
| MRPS34 | 2601.64306 | -0.1845023 | 0.22845921 | -0.8075943 | 0.4193242 | 0.88451372 |
| EME2 | 6610.56527 | -0.2392175 | 0.13897387 | -1.721313 | 0.08519405 | 0.65009026 |
| SPSB3 | 4895.73103 | -0.1951355 | 0.15943789 | -1.2238965 | 0.22099128 | 0.79965146 |
| NUBP2 | 2395.28435 | -0.0547442 | 0.14729686 | -0.3716589 | 0.71014682 | 0.95524448 |
| IGFALS | 65.2397295 | 0.58714684 | 0.44279213 | 1.3260101 | 0.18483635 | 0.76820626 |
| HAGH | 5594.32536 | 0.44109978 | 0.29942045 | 1.47317851 | 0.14070291 | 0.72614342 |

| | | | | | | |
|---|---|---|---|---|---|---|
| FAHD1 | 3549.48817 | 0.3619284 | 0.36780244 | 0.98402934 | 0.32510109 | 0.85224779 |
| MEIOB | 73.0670973 | 0.36631819 | 0.52193926 | 0.70184066 | 0.48277854 | 0.90541935 |
| MSRB1 | 5686.64234 | 0.15231306 | 0.36171531 | 0.42108548 | 0.67369267 | 0.94935413 |
| NDUFB10 | 3988.00095 | 0.23015327 | 0.14362579 | 1.60245089 | 0.10905594 | 0.67855822 |
| RPS2 | 29788.3212 | 0.02054251 | 0.18799207 | 0.1092733 | 0.91298572 | 0.98662757 |
| SNHG9 | 209.359732 | -0.3436443 | 0.37531517 | -0.9156154 | 0.35986866 | 0.86480228 |
| TBL3 | 2305.3545 | 0.04153622 | 0.13180954 | 0.31512301 | 0.75266826 | 0.96301881 |
| NOXO1 | 80.6478037 | -0.2019832 | 0.54636652 | -0.3696845 | 0.71161761 | 0.95560193 |
| GFER | 1391.82695 | 0.0216195 | 0.13572973 | 0.15928345 | 0.87344556 | 0.98275231 |
| AC005606.2 | 758.798659 | 0.02311608 | 0.15107433 | 0.1530113 | 0.87838936 | 0.98275231 |
| SYNGR3 | 198.080669 | 0.44239744 | 0.43691531 | 1.01254736 | 0.31127642 | 0.84496398 |
| AC005606.1 | 790.870169 | -0.3655265 | 0.24813564 | -1.4730915 | 0.14072637 | 0.72614342 |
| ZNF598 | 2837.88842 | -0.391715 | 0.23454457 | -1.6701091 | 0.09489778 | 0.65888357 |
| SLC9A3R2 | 5669.66802 | -0.0122982 | 0.35667621 | -0.03448 | 0.9724944 | 0.99416401 |
| NTHL1 | 436.109435 | 0.41380497 | 0.32783651 | 1.2622297 | 0.20686614 | 0.78790396 |
| TSC2 | 7858.17088 | -0.2048408 | 0.20022286 | -1.0230637 | 0.30627771 | 0.84278367 |
| PKD1 | 5558.21025 | -0.2666193 | 0.24949017 | -1.0686564 | 0.2852245 | 0.83107779 |
| AC009065.6 | 111.069752 | -0.5769167 | 0.43956392 | -1.3124752 | 1.89E-01 | 7.70E-01 |
| AC009065.3 | 62.5805774 | -0.6816075 | 0.38720767 | -1.7603149 | 0.07835442 | 0.64079926 |
| RAB26 | 81.6237787 | 1.74839883 | 0.53661173 | 3.25821955 | 0.00112114 | 0.15680011 |
| SNHG19 | 471.170469 | 0.16345767 | 0.23780268 | 0.68736681 | 0.49185161 | 0.90615916 |
| TRAF7 | 5244.86613 | -0.0543408 | 0.11575644 | -0.4694407 | 0.63875465 | 0.94088547 |
| CASKIN1 | 1240.71599 | -0.1154547 | 0.13621628 | -0.8475841 | 0.39666965 | 0.87976164 |
| MLST8 | 2334.13705 | -0.0406683 | 0.17884755 | -0.2273908 | 0.82011987 | 0.97454841 |
| BRICD5 | 945.654655 | 0.076917 | 0.25348356 | 0.30343981 | 0.76155471 | 0.96606476 |
| AC009065.7 | 486.376467 | 0.07638727 | 0.20129394 | 0.37948121 | 0.70433056 | 0.95488362 |
| PGP | 1323.94058 | 0.08722091 | 0.19078989 | 0.4571569 | 0.64755827 | 0.94282139 |
| E4F1 | 2381.02061 | -0.3538574 | 0.16945679 | -2.0881869 | 0.03678098 | 0.55521179 |
| AC009065.1 | 169.302649 | -0.0750888 | 0.28402795 | -0.2643712 | 0.79149393 | 0.97090793 |
| DNASE1L2 | 223.502423 | 0.04800968 | 0.39599694 | 0.12123751 | 0.90350291 | 0.98617053 |
| ECI1 | 4176.8514 | 0.11569677 | 0.12952812 | 0.89321741 | 0.37174075 | 0.86762749 |
| AC009065.8 | 143.426126 | -0.0920376 | 0.28714884 | -0.3205222 | 0.74857247 | 0.96247232 |
| RNPS1 | 5293.6758 | -0.2718131 | 0.1569028 | -1.7323665 | 0.08320834 | 0.6471966 |
| AC009065.4 | 105.173307 | -0.266282 | 0.38771714 | -0.6867946 | 0.49221219 | 0.90636999 |
| ABCA3 | 1936.31132 | 0.33763574 | 0.17295367 | 1.9521745 | 5.09E-02 | 0.5861488 |
| CCNF | 181.928088 | 0.10237635 | 0.24392291 | 0.41970782 | 0.67469891 | 0.94935413 |
| TEDC2 | 65.6603057 | -0.0286747 | 0.37511568 | -0.0764422 | 0.93906731 | 0.99113899 |
| TBC1D24 | 2599.03811 | 0.11436513 | 0.22733782 | 0.50306249 | 0.61492034 | 0.93895494 |
| AC093525.2 | 12036.4136 | 0.03068754 | 0.19440442 | 0.15785413 | 0.87457174 | 0.98275231 |
| ATP6V0C | 12298.2169 | 0.02798998 | 0.19658606 | 0.14238029 | 0.88677963 | 0.98423722 |
| AC093525.1 | 4981.64377 | 0.05133313 | 0.22816856 | 0.22497899 | 8.22E-01 | 9.75E-01 |
| AMDHD2 | 4198.08514 | -0.5889164 | 0.37936544 | -1.5523722 | 0.12057318 | 0.69467399 |
| CEMP1 | 1890.45743 | -0.6841102 | 0.40517186 | -1.6884445 | 0.09132594 | 0.65518742 |
| PDPK1 | 2922.4495 | 0.08492132 | 0.16222277 | 0.52348586 | 0.60063619 | 0.93488657 |
| AC093525.6 | 63.3540867 | 0.90461018 | 1.22535353 | 0.73824423 | 0.46036605 | 0.89755243 |
| AC141586.3 | 184.426015 | -0.140647 | 0.43835793 | -0.3208498 | 0.74832424 | 0.96241535 |
| AC141586.2 | 35.6734871 | -0.1247775 | 0.59988951 | -0.2080009 | 0.8352283 | 0.97550416 |
| KCTD5 | 1151.72535 | -0.0910987 | 0.2001846 | -0.4550735 | 6.49E-01 | 9.43E-01 |
| PRSS27 | 98.3627054 | -0.0975939 | 0.36871827 | -0.2646842 | 7.91E-01 | 9.71E-01 |
| SRRM2-AS1 | 103.668608 | 0.06280814 | 0.52301338 | 0.12008897 | 0.90441267 | 0.98624484 |
| SRRM2 | 39102.1484 | -0.3561773 | 0.15086686 | -2.3608714 | 0.01823205 | 0.47570049 |
| ELOB | 5121.86311 | 0.0050965 | 0.14037736 | 0.0363057 | 0.9710386 | 0.99400331 |
| AC092117.1 | 119.151406 | -0.3741774 | 0.57018367 | -0.6562401 | 0.51166965 | 0.91510941 |
| PRSS22 | 2073.97443 | -0.3667294 | 0.44880643 | -0.8171214 | 0.41385903 | 0.88338635 |

| | | | | | | |
|---|---|---|---|---|---|---|
| AC003965.2 | 155.18876 | -0.3573737 | 0.4865563 | -0.7344962 | 0.46264638 | 0.89796156 |
| FLYWCH2 | 1017.05674 | 0.34805117 | 0.20731066 | 1.678887 | 0.09317407 | 0.65756964 |
| FLYWCH1 | 3688.87155 | -0.2213488 | 0.31048658 | -0.7129095 | 0.47590179 | 0.90272018 |
| PKMYT1 | 258.965119 | 0.59743609 | 0.37994675 | 1.57242057 | 0.11585305 | 0.6873836 |
| PAQR4 | 292.458841 | 0.72356688 | 0.32905995 | 2.19889075 | 0.02788569 | 0.53262541 |
| AC004233.3 | 86.410494 | 0.3264903 | 0.51267893 | 0.6368319 | 0.52423434 | 0.91723878 |
| CLDN9 | 37.5951689 | -0.4575016 | 0.84075443 | -0.544156 | 5.86E-01 | 9.33E-01 |
| TNFRSF12A | 5172.4532 | -0.6873225 | 0.55662016 | -1.2348142 | 2.17E-01 | 7.95E-01 |
| HCFC1R1 | 2991.10182 | 0.12908558 | 0.26392059 | 0.48910765 | 0.62476548 | 0.94088547 |
| THOC6 | 1684.28351 | -0.4497518 | 0.25916375 | -1.7353962 | 0.08267066 | 0.64680474 |
| BICDL2 | 994.64289 | -0.0098776 | 0.45360743 | -0.0217756 | 0.98262698 | 0.99611734 |
| AC108134.1 | 206.090495 | 0.115279 | 0.44078553 | 0.26153082 | 0.79368319 | 0.97090793 |
| MMP25 | 466.546051 | -0.5270835 | 0.36429247 | -1.4468692 | 0.14793357 | 0.73460889 |
| MMP25-AS1 | 906.266031 | -0.4562019 | 0.31595156 | -1.4438983 | 0.14876758 | 0.73602252 |
| IL32 | 8004.8789 | 0.07094133 | 0.5334678 | 0.13298146 | 0.89420804 | 0.98474725 |
| AC108134.3 | 157.630606 | -0.1166195 | 0.4187115 | -0.2785199 | 0.78061327 | 0.96864684 |
| ZNF213-AS1 | 1019.97715 | -0.1084084 | 0.15422516 | -0.7029228 | 0.48210386 | 0.90541935 |
| ZNF205 | 1096.76602 | -0.2495926 | 0.14571706 | -1.712858 | 0.08673866 | 0.65228863 |
| ZNF213 | 948.464692 | -0.1036191 | 0.22167652 | -0.4674336 | 0.6401897 | 0.94088547 |
| ZNF200 | 375.341029 | 0.06294636 | 0.16486432 | 0.38180704 | 0.70260449 | 0.95431462 |
| MEFV | 64.9444396 | -0.3702475 | 0.66502745 | -0.5567402 | 5.78E-01 | 9.31E-01 |
| LINC00921 | 78.1709927 | -0.0564538 | 0.35578172 | -0.1586753 | 0.87392468 | 0.98275231 |
| ZNF263 | 2332.57874 | -0.2722439 | 0.18664374 | -1.4586286 | 0.14466736 | 0.73032824 |
| TIGD7 | 573.81569 | -0.3755586 | 0.27268754 | -1.377249 | 0.16843528 | 0.75735257 |
| ZNF75A | 1004.27726 | -0.4218744 | 0.29248381 | -1.4423856 | 0.14919362 | 0.73670625 |
| AC025283.3 | 850.109209 | -0.2368463 | 0.20534074 | -1.1534304 | 0.24873375 | 0.81646572 |
| ZSCAN32 | 805.712753 | -0.0420492 | 0.2098357 | -0.2003911 | 0.84117468 | 0.97618283 |
| AC025283.2 | 765.718117 | -0.0279827 | 0.19903119 | -0.1405943 | 0.88819042 | 0.98423722 |
| ZNF174 | 552.441217 | -0.0536023 | 0.16508975 | -0.324686 | 0.74541873 | 0.96176407 |
| ZNF597 | 221.299507 | 0.09476023 | 0.23053141 | 0.41105128 | 0.68103493 | 0.95070301 |
| NAA60 | 5541.94857 | -0.3629313 | 0.1559124 | -2.32779 | 0.01992326 | 0.49520126 |
| CLUAP1 | 1164.75656 | 0.17279564 | 0.28179793 | 0.61318989 | 0.53975079 | 0.92201891 |
| NLRC3 | 275.258995 | 0.0332212 | 0.45715685 | 0.07266914 | 0.94206941 | 0.99136103 |
| AC006111.1 | 109.208452 | -0.417554 | 0.24978597 | -1.6716472 | 0.09459392 | 0.65888357 |
| SLX4 | 459.427412 | -0.3685987 | 0.21029798 | -1.7527448 | 0.07964582 | 0.64079926 |
| DNASE1 | 6729.77993 | 0.29815963 | 0.21883953 | 1.36245782 | 0.17305345 | 0.75974924 |
| TRAP1 | 6551.57145 | 0.34381669 | 0.2188139 | 1.57127442 | 0.11611892 | 0.68754779 |
| CREBBP | 3343.73287 | -0.4093405 | 0.15203387 | -2.6924299 | 0.00709334 | 0.35449694 |
| ADCY9 | 2977.78148 | 0.16021204 | 0.33225799 | 0.48219168 | 0.62966979 | 0.94088547 |
| SRL | 185.159658 | -0.1100383 | 0.50068647 | -0.2197749 | 0.82604649 | 0.97548201 |
| LINC01569 | 295.891196 | 0.30349228 | 0.5256663 | 0.57734779 | 0.56370454 | 0.92453312 |
| TFAP4 | 453.498645 | 0.16612006 | 0.24425395 | 0.68011209 | 0.49643349 | 0.90890243 |
| GLIS2 | 6372.13099 | 0.2341966 | 0.31307823 | 0.74804499 | 0.45443302 | 0.89670027 |
| GLIS2-AS1 | 39.970776 | 0.9861356 | 0.58203912 | 1.69427718 | 9.02E-02 | 0.65403568 |
| PAM16 | 1177.14947 | -0.1847812 | 0.16145718 | -1.1444593 | 0.25243322 | 0.81833349 |
| CORO7-PAM | 2969.77936 | -0.1350127 | 0.15314033 | -0.8816272 | 3.78E-01 | 8.70E-01 |
| CORO7 | 2870.82392 | -0.1568477 | 0.19529079 | -0.8031494 | 0.42188839 | 0.88451372 |
| VASN | 5689.76267 | 0.45139058 | 0.30005987 | 1.50433503 | 0.13249512 | 0.71291734 |
| DNAJA3 | 3930.42219 | 0.14363087 | 0.23021654 | 0.62389465 | 0.53269678 | 0.92011707 |
| NMRAL1 | 2173.84334 | 0.28132139 | 0.2029782 | 1.38596851 | 0.16575652 | 0.75515387 |
| HMOX2 | 3457.56424 | 0.0372471 | 0.2325624 | 0.16015959 | 0.87275536 | 0.98275231 |
| CDIP1 | 4754.2349 | 0.10089167 | 0.21965572 | 0.45931729 | 0.64600633 | 0.94208859 |
| UBALD1 | 2464.68463 | -0.5132146 | 0.12672872 | -4.0497106 | 5.13E-05 | 0.02745878 |
| MGRN1 | 5846.54433 | -0.2723914 | 0.22589828 | -1.2058143 | 0.22788909 | 0.80300331 |

| | | | | | | |
|---|---|---|---|---|---|---|
| NUDT16L1 | 2013.71712 | 0.55972223 | 0.22551892 | 2.48193027 | 0.01306728 | 0.42078002 |
| ANKS3 | 2049.69346 | -0.1137399 | 0.13154121 | -0.8646713 | 0.38721924 | 0.87460898 |
| C16orf71 | 121.767717 | -0.1531347 | 0.35140833 | -0.4357743 | 0.66300048 | 0.94642611 |
| ZNF500 | 1032.39126 | -0.0002132 | 0.26147402 | -0.0008155 | 0.99934934 | 0.99994357 |
| SMIM22 | 246.173608 | 0.19060103 | 0.37678469 | 0.50586192 | 0.61295359 | 0.93879706 |
| AC020663.2 | 182.686642 | 0.2167429 | 0.34864269 | 0.62167631 | 0.53415473 | 0.92084006 |
| ROGDI | 2283.23918 | 0.57051051 | 0.27964751 | 2.04010581 | 0.04133979 | 0.55943411 |
| GLYR1 | 8265.04732 | -0.0546061 | 0.08981935 | -0.6079546 | 0.54321758 | 0.92297267 |
| UBN1 | 2252.10329 | -0.2752797 | 0.16265541 | -1.6924101 | 0.09056781 | 0.65495306 |
| PPL | 1882.91418 | 0.46603054 | 0.29902412 | 1.55850483 | 0.11911362 | 0.69233212 |
| NAGPA-AS1 | 61.4644824 | 0.02458556 | 0.39437995 | 0.06233978 | 0.95029225 | 0.99170503 |
| NAGPA | 1395.18647 | 0.08407396 | 0.19952831 | 0.42136356 | 0.67348962 | 0.94935413 |
| ALG1 | 1847.03032 | -0.1826479 | 0.18793244 | -0.9718805 | 0.33111 | 0.85384238 |
| C16orf89 | 917.280675 | 1.8619612 | 0.56773643 | 3.27962257 | 0.00103946 | 0.15012259 |
| EEF2KMT | 1309.96982 | -0.1227365 | 0.18402943 | -0.6669393 | 0.50481092 | 0.91365534 |
| METTL22 | 769.733213 | -0.3731022 | 0.15834923 | -2.3561986 | 0.01846304 | 0.47570049 |
| ABAT | 8251.01106 | 0.8816583 | 0.50326817 | 1.75186579 | 0.07979689 | 0.64079926 |
| TMEM186 | 1078.64037 | -0.0600074 | 0.28099037 | -0.2135566 | 0.83089282 | 0.97550416 |
| PMM2 | 2176.46206 | -0.4742915 | 0.29578398 | -1.6035065 | 0.10882288 | 0.67800851 |
| AC012173.1 | 703.781452 | -0.5174404 | 0.33109988 | -1.5627924 | 0.11810144 | 0.69080494 |
| AC022167.2 | 3367.04917 | -0.4042336 | 0.26178559 | -1.54414 | 0.12255443 | 0.69788192 |
| CARHSP1 | 7381.62683 | -0.3273829 | 0.32903643 | -0.9949748 | 3.20E-01 | 8.50E-01 |
| AC022167.1 | 1556.93383 | -0.2510727 | 0.3595623 | -0.6982731 | 0.48500639 | 0.90541935 |
| USP7 | 4196.2725 | -0.1541267 | 0.16741754 | -0.9206124 | 0.35725284 | 0.86390849 |
| C16orf72 | 5333.09045 | -0.2908365 | 0.25165922 | -1.1556759 | 0.24781373 | 0.81602355 |
| AC087190.1 | 74.6202626 | -0.5446429 | 0.28136466 | -1.9357189 | 0.05290214 | 0.58742503 |
| GRIN2A | 82.6844866 | 0.46770358 | 0.64774371 | 0.72205036 | NA | NA |
| ATF7IP2 | 476.213114 | 0.30118092 | 0.28398656 | 1.0605464 | 0.28889609 | 0.83295411 |
| EMP2 | 1242.999 | 0.04021674 | 0.38885506 | 0.10342347 | 0.91762688 | 0.9868044 |
| AC027277.1 | 274.986532 | 0.09834277 | 0.40790739 | 0.24109092 | 0.80948465 | 0.97334138 |
| NUBP1 | 1494.33036 | -0.0583662 | 0.21239126 | -0.2748049 | 0.78346614 | 0.96978847 |
| TVP23A | 1008.69547 | -0.1006667 | 0.19829612 | -0.5076584 | 0.61169295 | 0.93849673 |
| CIITA | 4414.96294 | 0.12866259 | 0.20572791 | 0.62540173 | 0.53170743 | 0.91962754 |
| DEXI | 2439.57981 | 0.06742079 | 0.17303918 | 0.38962733 | 0.69681214 | 0.9525827 |
| CLEC16A | 2399.19845 | -0.0264452 | 0.22521057 | -0.1174242 | 0.90652392 | 0.98662757 |
| AC007220.1 | 91.5317822 | -0.1648686 | 0.20512141 | -0.803761 | 0.42153499 | 0.88451372 |
| RMI2 | 182.123477 | -0.5417905 | 0.66834619 | -0.8106435 | NA | NA |
| SOCS1 | 1088.00796 | -0.3514446 | 0.43568815 | -0.8066426 | 0.41987245 | 0.88451372 |
| AC099489.1 | 226.033376 | 0.20804829 | 0.5033345 | 0.41334 | 0.67935752 | 0.95070301 |
| LITAF | 12426.055 | 0.17221073 | 0.33782911 | 0.50975694 | 0.61022176 | 0.93792127 |
| SNN | 2460.97721 | -0.5173583 | 0.36783739 | -1.4064864 | 0.15957973 | 0.74768867 |
| TXNDC11 | 3820.44538 | -0.0952457 | 0.14355772 | -0.6634664 | 0.50703192 | 0.91420888 |
| AC007613.1 | 108.758767 | 0.24709301 | 0.3248735 | 0.76058224 | 0.44690663 | 0.89480207 |
| ZC3H7A | 2913.67031 | 0.0253266 | 0.11885361 | 0.21309071 | 8.31E-01 | 0.97550416 |
| RSL1D1 | 3904.72707 | -0.0664296 | 0.20752057 | -0.3201107 | 0.74888439 | 0.96249332 |
| GSPT1 | 6350.85094 | -0.2041398 | 0.20981169 | -0.9729668 | 0.33056981 | 0.85384238 |
| NPIPB2 | 146.908363 | 0.06791494 | 1.15765393 | 0.05866602 | 0.95321813 | 0.99170503 |
| SNX29 | 3729.57654 | 0.0886111 | 0.32400152 | 0.27348975 | 0.78447677 | 0.97021865 |
| CPPED1 | 3667.96189 | -0.3430061 | 0.37555142 | -0.9133399 | 0.36106378 | 0.86528137 |
| SHISA9 | 273.748406 | 0.15778238 | 0.362303 | 0.43549842 | 0.66320069 | 0.94642611 |
| ERCC4 | 357.572275 | 0.0114859 | 0.19257232 | 0.0596446 | 0.9524387 | 0.99170503 |
| AC010401.1 | 45.5302472 | 0.22731584 | 0.33035915 | 0.688087 | 0.491398 | 0.90600166 |
| MKL2 | 1475.3671 | -0.1105673 | 0.17301793 | -0.6390511 | 0.52278972 | 0.91723878 |
| AC130650.2 | 74.8547835 | -1.0088017 | 0.28610549 | -3.525978 | 0.00042192 | 0.09906683 |

| | | | | | | |
|---|---|---|---|---|---|---|
| MIR193BHG | 178.742031 | -0.6232005 | 0.38749316 | -1.6082876 | 0.1077722 | 0.67708979 |
| PARN | 1554.36099 | 0.00473562 | 0.15144576 | 0.03126941 | 0.97505469 | 0.99455099 |
| BFAR | 2636.16208 | -0.2929648 | 0.20362813 | -1.4387247 | 0.15022854 | 0.73796431 |
| NOMO1 | 3174.31089 | 0.0290403 | 0.20011872 | 0.14511538 | 0.88461977 | 0.98340286 |
| NPIPA1 | 1598.89748 | -0.4588832 | 0.27795763 | -1.6509106 | 0.09875683 | 0.6678501 |
| PDXDC1 | 8995.72783 | 0.10085928 | 0.18278214 | 0.5518005 | 0.58108505 | 0.9317705 |
| AC138932.5 | 501.638194 | 0.00775698 | 0.28043627 | 0.02766041 | 0.977933 | 0.99476647 |
| NTAN1 | 1365.35818 | -0.3826947 | 0.2299613 | -1.66417 | 0.09607846 | 0.66169558 |
| RRN3 | 2206.50874 | 0.00574362 | 0.20817339 | 0.02759057 | 0.9779887 | 0.99476647 |
| NPIPA5 | 249.972823 | -0.4099417 | 0.84060637 | -0.4876738 | 0.62578094 | 0.94088547 |
| MPV17L | 868.310407 | 0.43328846 | 0.40486136 | 1.07021441 | 0.28452281 | 0.83107779 |
| AC140504.1 | 271.513368 | 0.15306284 | 0.25758247 | 0.59422848 | 5.52E-01 | 0.92373235 |
| C16orf45 | 1494.7905 | 0.08265838 | 0.22774501 | 0.36294265 | 0.71664773 | 0.95603249 |
| MARF1 | 3027.4664 | -0.0892067 | 0.13334281 | -0.6690028 | 0.50349369 | 0.91356552 |
| NDE1 | 1475.54012 | -0.6706886 | 0.57372878 | -1.1689994 | 0.24240388 | 0.81319028 |
| AC026401.3 | 89.8019779 | -0.3413474 | 0.41110208 | -0.8303227 | 0.40635638 | 0.88132015 |
| MYH11 | 7248.73496 | -1.2613402 | 1.05736892 | -1.1929046 | 0.23290675 | 0.80667958 |
| AF001548.2 | 460.91796 | -0.7770231 | 0.6712688 | -1.1575439 | 0.2470502 | 0.8156466 |
| AF001548.1 | 105.573309 | -0.7590885 | 0.86626122 | -0.8762812 | 0.38087716 | 0.87165941 |
| FOPNL | 1645.34808 | 0.06725005 | 0.1757169 | 0.38271819 | 0.70192873 | 0.95431462 |
| ABCC1 | 1741.78399 | -0.2131975 | 0.20792988 | -1.0253335 | 0.30520585 | 0.84268008 |
| ABCC6 | 3511.83479 | 0.19834933 | 0.45180444 | 0.4390159 | 0.66065002 | 0.94628868 |
| NOMO3 | 127.03679 | 0.1929162 | 0.57535975 | 0.33529665 | 0.73740132 | 0.96023056 |
| AC138969.1 | 1119.89091 | -0.8040328 | 0.2280097 | -3.5263099 | 0.00042139 | 0.09906683 |
| XYLT1 | 253.39269 | -0.3072123 | 0.44747224 | -0.6865506 | 0.49236597 | 0.90651642 |
| NPIPA8 | 37.3075968 | -0.0617341 | 1.29059549 | -0.0478338 | 0.96184866 | 0.99320711 |
| AC126755.2 | 112.557896 | -0.5636646 | 0.78115832 | -0.7215754 | 0.47055558 | 0.9008571 |
| NOMO2 | 410.625153 | 0.06222345 | 0.30182678 | 0.20615616 | 0.83666893 | 0.97577999 |
| RPS15A | 16314.4221 | 0.21797323 | 0.20846862 | 1.04559252 | 0.2957492 | 0.83750889 |
| AC138811.2 | 16507.4029 | 0.18997357 | 0.21527492 | 0.88246958 | 0.37752292 | 0.86987124 |
| ARL6IP1 | 9181.25879 | -0.2436268 | 0.23691948 | -1.0283105 | 0.30380379 | 0.84211658 |
| AC138811.1 | 45.7620195 | -1.1885397 | 0.36775319 | -3.2318952 | 0.00122972 | 0.15785077 |
| SMG1 | 4973.89346 | -0.4698917 | 0.15113238 | -3.1091398 | 0.00187633 | 0.18905527 |
| TMC7 | 79.5359989 | -0.069403 | 0.40257058 | -0.1723996 | 8.63E-01 | 9.82E-01 |
| AC099518.4 | 72.3753032 | 0.43029107 | 0.27848765 | 1.5450993 | 0.12232225 | 0.69788192 |
| COQ7 | 1013.79417 | 0.14281223 | 0.21953509 | 0.65052119 | 0.51535562 | 0.91636259 |
| ITPRIPL2 | 1412.13842 | 0.03299773 | 0.23392922 | 0.1410586 | 0.88782365 | 0.98423722 |
| AC099518.3 | 164.602599 | 0.11283705 | 0.26994667 | 0.41799755 | 0.67594891 | 0.94959168 |
| AC099518.5 | 771.955438 | 0.02518228 | 0.27061866 | 0.09305448 | 0.92586028 | 0.98876911 |
| SYT17 | 427.409942 | 0.32414828 | 0.38928132 | 0.83268386 | 0.40502305 | 0.88132015 |
| TMC5 | 44.3567051 | 2.13889197 | 0.72287623 | 2.95886332 | 0.00308776 | 0.24212433 |
| AC130456.5 | 932.763934 | -0.0408985 | 0.24591475 | -0.1663118 | 0.86791156 | 0.98237277 |
| GDE1 | 3863.71847 | 0.00338988 | 0.21745185 | 0.01558913 | 0.98756218 | 0.997408 |
| CCP110 | 711.213346 | -0.016628 | 0.24477462 | -0.0679317 | 0.94584 | 0.99166614 |
| VPS35L | 2825.63231 | 0.21879254 | 0.11320398 | 1.93272829 | 0.05326968 | 0.58967929 |
| KNOP1 | 594.102965 | -0.3969338 | 0.24991182 | -1.5882953 | 0.11221958 | 0.68001961 |
| AC002550.1 | 279.954897 | -0.2905881 | 0.32496273 | -0.8942196 | 0.37120439 | 0.86762749 |
| IQCK | 2608.19117 | 0.19932317 | 0.22855577 | 0.87209862 | 0.38315458 | 0.8718674 |
| GPRC5B | 6092.74629 | 0.12491583 | 0.2786568 | 0.44827841 | 0.65395228 | 0.94510853 |
| GP2 | 958.006338 | 0.33638206 | 0.9428208 | 0.3567826 | 0.72125456 | 0.95736467 |
| UMOD | 295077.39 | -0.6365953 | 0.55373891 | -1.1496307 | 0.250296 | 0.81721208 |
| ACSM5 | 1675.58986 | 0.49661247 | 0.71504575 | 0.69451846 | 0.48735713 | 0.90573694 |
| AC137056.1 | 583.527965 | 0.40845779 | 0.75487782 | 0.54109126 | 0.58844468 | 0.93289552 |
| ACSM2A | 15306.4691 | 0.88581027 | 0.58049261 | 1.52596303 | 0.12701907 | 0.70463514 |

| | | | | | | |
|---|---|---|---|---|---|---|
| ACSM2B | 20304.2038 | 0.52806276 | 0.57435717 | 0.91939788 | 0.3578875 | 0.86432598 |
| AC141273.1 | 53.0598332 | -0.1471731 | 0.95134303 | -0.1547004 | 0.87705753 | 0.98275231 |
| ACSM3 | 4264.94764 | 0.77265931 | 0.38165169 | 2.02451435 | 0.04291726 | 0.56641712 |
| ACSM1 | 82.4248871 | 0.76658748 | 0.52314959 | 1.46533133 | 0.14283054 | 0.72784886 |
| THUMPD1 | 2606.25352 | -0.1022712 | 0.13386947 | -0.763962 | 0.44488988 | 0.89429407 |
| ERI2 | 2026.36742 | 0.54026215 | 0.30938628 | 1.74623825 | 0.08076956 | 0.64313548 |
| REXO5 | 107.105075 | 0.54452633 | 0.30804718 | 1.76767186 | 7.71E-02 | 6.38E-01 |
| DCUN1D3 | 1398.60703 | -0.3687598 | 0.2289941 | -1.6103462 | 0.10732229 | 0.67707349 |
| LYRM1 | 2065.92938 | 0.1169118 | 0.30522461 | 0.3830353 | 0.70169359 | 0.95424422 |
| TMEM159 | 1391.31026 | -0.1549339 | 0.25744521 | -0.6018132 | 0.5472985 | 0.92354775 |
| ANKS4B | 922.616004 | 1.70250976 | 0.64683178 | 2.63207501 | 0.00848651 | 0.3779007 |
| CRYM | 2517.24755 | 1.30351123 | 0.63904811 | 2.03977011 | 0.04137323 | 0.55943411 |
| NPIPB3 | 358.182046 | -0.2595238 | 0.36496407 | -0.7110942 | 0.47702589 | 0.90344707 |
| METTL9 | 8271.23144 | 0.08651153 | 0.1721516 | 0.50253108 | 0.615294 | 0.93902482 |
| IGSF6 | 171.004345 | 0.37749242 | 0.56251239 | 0.67108286 | 0.50216775 | 0.9128153 |
| AC092375.2 | 53.6211627 | 0.39361435 | 0.46247824 | 0.8510981 | 0.39471486 | 0.87884163 |
| NPIPB4 | 911.413354 | -0.3613429 | 0.27208542 | -1.3280496 | 0.18416171 | 0.76816707 |
| AC092119.2 | 156.095073 | 0.12658318 | 0.33943904 | 0.37291875 | 0.70920891 | 0.95501548 |
| UQCRC2 | 11739.891 | 0.25753217 | 0.14645608 | 1.75842602 | 0.07867505 | 0.64079926 |
| MOSMO | 926.460388 | -0.2656084 | 0.16909916 | -1.5707257 | 0.11624637 | 0.6878981 |
| AC009019.1 | 172.381897 | -0.2229928 | 0.199601 | -1.1171926 | 0.26391197 | 0.82424998 |
| VWA3A | 70.9497963 | 0.34514738 | 0.87463804 | 0.39461739 | 0.69312528 | 0.9519683 |
| EEF2K | 1283.79871 | -0.2281199 | 0.20863055 | -1.0934156 | 0.27421135 | 0.82719407 |
| AC009034.1 | 148.276165 | -0.0348123 | 0.31498233 | -0.1105215 | 9.12E-01 | 0.98662757 |
| POLR3E | 2277.67701 | -0.194593 | 0.11476125 | -1.6956337 | 0.08995527 | 0.65375543 |
| CDR2 | 2239.80479 | -0.0347855 | 0.22113499 | -0.1573042 | 0.87500509 | 0.98275231 |
| NPIPB5 | 1096.14776 | -0.3994343 | 0.29139433 | -1.3707689 | 0.170447 | 0.75956362 |
| HS3ST2 | 54.1141925 | -0.2355001 | 1.01103281 | -0.2329302 | 0.81581559 | 0.9738447 |
| USP31 | 1445.87263 | -0.2992988 | 0.28112603 | -1.0646428 | 0.28703762 | 0.83155545 |
| SCNN1G | 3967.18709 | -0.4238438 | 0.38192116 | -1.1097677 | 0.26709913 | 0.8265123 |
| AC099482.1 | 167.644347 | -0.2872571 | 0.41551966 | -0.69132 | 0.48936445 | 0.90600166 |
| SCNN1B | 1862.42066 | 0.04776779 | 0.312763 | 0.15272838 | 0.87861248 | 0.98275231 |
| COG7 | 2255.69397 | -0.0739887 | 0.13059648 | -0.566544 | 0.57102404 | 0.92814931 |
| AC008915.2 | 239.628398 | -0.447614 | 0.38858883 | -1.1518962 | 0.24936372 | 0.81646572 |
| GGA2 | 7079.83037 | -0.6102232 | 0.24802494 | -2.4603298 | 0.01388094 | 0.4295825 |
| EARS2 | 1441.31864 | 0.16812317 | 0.20508666 | 0.81976647 | 0.41234925 | 0.88268208 |
| UBFD1 | 2654.02466 | -0.1771215 | 0.14170673 | -1.2499158 | 0.21133029 | 0.79134644 |
| NDUFAB1 | 2924.19603 | 0.1492185 | 0.17912796 | 0.83302741 | 0.40482927 | 0.88132015 |
| PALB2 | 507.767287 | -0.3442921 | 0.20718934 | -1.6617268 | 0.09656757 | 0.66243513 |
| AC008870.4 | 71.1119821 | -0.5585208 | 0.32207481 | -1.7341338 | 0.08289434 | 0.64680474 |
| DCTN5 | 3197.73661 | -0.0277214 | 0.16245724 | -0.1706379 | 0.86450851 | 0.98188271 |
| PLK1 | 209.536881 | 0.11893884 | 0.1866249 | 0.63731496 | 0.52391971 | 0.91723878 |
| CHP2 | 78.0186891 | 0.59369093 | 1.26509965 | 0.46928392 | 0.6388667 | 0.94088547 |
| PRKCB | 244.806695 | 0.01363633 | 0.59699007 | 0.0228418 | 0.98177647 | 0.99595643 |
| RBBP6 | 2511.87536 | -0.3155454 | 0.19217309 | -1.6419854 | 0.10059303 | 0.67247782 |
| TNRC6A | 3247.2016 | -0.2926553 | 0.16730884 | -1.7491918 | 0.08025787 | 0.6408661 |
| SLC5A11 | 780.910441 | 2.66864682 | 0.8642242 | 3.0879103 | 0.00201569 | 0.19273522 |
| ARHGAP17 | 2772.74594 | -0.2717195 | 0.13989987 | -1.9422425 | 0.05210775 | 0.58645127 |
| LCMT1 | 1641.07883 | 0.20027114 | 0.12163966 | 1.64642958 | 0.09967535 | 0.67098264 |
| ZKSCAN2 | 234.361244 | -0.0108049 | 0.23517032 | -0.0459448 | 0.96335426 | 0.99320711 |
| AC008741.2 | 399.410187 | 0.04496905 | 0.29645297 | 0.15169033 | 0.87943118 | 0.98275231 |
| KDM8 | 315.853999 | 0.05529315 | 0.24491088 | 0.22576847 | 0.8213815 | 0.97472105 |
| AC109449.1 | 80.7272729 | 0.13696876 | 0.27937007 | 0.49027715 | 0.62393779 | 0.94057418 |
| NSMCE1 | 2066.12033 | 0.49909049 | 0.22880459 | 2.18129579 | 0.02916155 | 0.53606221 |

| | | | | | | |
|---|---|---|---|---|---|---|
| AC106739.1 | 42.2942543 | 0.35826469 | 0.53585728 | 0.66858229 | 0.50376197 | 0.91356552 |
| AC106739.2 | 51.5692599 | -0.0782023 | 0.45625541 | -0.1714002 | 0.86390911 | 0.98176973 |
| IL4R | 5397.29997 | -0.068779 | 0.3233476 | -0.2127091 | 0.83155384 | 0.97550416 |
| IL21R | 145.689694 | 0.70326878 | 0.57394694 | 1.2253202 | 0.2204546 | 0.79912826 |
| IL21R-AS1 | 85.1493582 | 0.37249568 | 0.58330033 | 0.63860016 | 0.52308308 | 0.91723878 |
| GTF3C1 | 4265.21008 | -0.0686904 | 0.1432024 | -0.4796732 | 0.63145979 | 0.94088547 |
| KIAA0556 | 1868.91171 | -0.0777614 | 0.13181531 | -0.5899271 | 0.55523953 | 0.92373235 |
| XPO6 | 3756.3148 | -0.5094027 | 0.20968658 | -2.4293529 | 0.0151258 | 0.44995661 |
| SBK1 | 393.154349 | -0.3137274 | 0.4127983 | -0.7600016 | 0.44725364 | 0.89480207 |
| AC138894.1 | 3358.65544 | -0.122512 | 0.20166141 | -0.6075133 | 0.54351029 | 0.92297267 |
| CLN3 | 3455.52353 | -0.1172982 | 0.1940859 | -0.6043624 | 0.54560273 | 0.92345821 |
| APOBR | 331.032594 | 0.14633155 | 0.57707348 | 0.25357525 | 0.79982372 | 0.97208422 |
| NUPR1 | 5182.99297 | 0.38157193 | 0.55896569 | 0.68263928 | 0.49483481 | 0.90816709 |
| AC020765.2 | 187.617795 | -0.3310519 | 0.22610582 | -1.4641459 | 0.14315409 | 0.72784886 |
| SGF29 | 766.914706 | -0.3166774 | 0.18730797 | -1.6906778 | 0.09089835 | 0.65495306 |
| SULT1A2 | 362.857932 | 0.60173492 | 0.55946826 | 1.07554791 | 0.28212949 | 0.83004534 |
| SULT1A1 | 1972.65758 | -0.2815961 | 0.46478747 | -0.6058599 | 0.54460781 | 0.92345821 |
| EIF3C | 435.978942 | 0.09336196 | 0.25787773 | 0.36203964 | 0.71732241 | 0.95611001 |
| AC145285.6 | 239.005126 | -0.2692313 | 0.25052571 | -1.0746655 | 0.2825245 | 0.83055556 |
| ATXN2L | 9915.38767 | -0.2871583 | 0.15356257 | -1.8699761 | 0.06148714 | 0.61037866 |
| AC133550.1 | 1212.72294 | -0.1812989 | 0.14058117 | -1.2896386 | 0.19717616 | 0.77816059 |
| TUFM | 13543.0725 | 0.23201057 | 0.17857998 | 1.29919697 | 0.19387634 | 0.77620852 |
| SH2B1 | 6494.48671 | -0.0957826 | 0.2054487 | -0.4662117 | 0.64106396 | 0.94088547 |
| AC133550.2 | 46.1243691 | 0.02222974 | 0.3112891 | 0.07141189 | 0.94306995 | 0.99136103 |
| ATP2A1 | 320.682497 | -0.1112987 | 0.27022362 | -0.4118761 | 0.68043022 | 0.95070301 |
| RABEP2 | 2105.6388 | -0.1914144 | 0.23501998 | -0.8144603 | 0.41538131 | 0.88381712 |
| CD19 | 108.351145 | -1.579721 | 0.93250315 | -1.6940651 | 0.09025292 | 0.65407193 |
| NFATC2IP | 2675.09924 | -0.320523 | 0.1944501 | -1.6483559 | 0.09927965 | 0.6694597 |
| AC109460.2 | 375.733857 | -0.1075103 | 0.18587769 | -0.5783925 | 0.56299914 | 0.92452095 |
| AC109460.1 | 451.077364 | -0.3702997 | 0.27210137 | -1.3608888 | 0.17354885 | 0.75974924 |
| SPNS1 | 3675.99016 | -0.4901571 | 0.25940231 | -1.8895632 | 0.0588164 | 0.60332684 |
| LAT | 1040.56772 | -0.0895794 | 0.2372559 | -0.3775645 | 0.70575417 | 0.95501548 |
| AC109460.4 | 179.223145 | -0.045975 | 0.30813403 | -0.1492044 | 0.88139232 | 0.98275231 |
| NPIPB11 | 256.757033 | -0.4217149 | 0.39384107 | -1.0707743 | 0.28427091 | 0.83107779 |
| BOLA2 | 457.616181 | 0.07707379 | 0.38225054 | 0.20163161 | 0.84020473 | 0.97618283 |
| BOLA2-SMG | 460.364951 | 0.09102231 | 0.38261651 | 0.23789436 | 0.81196303 | 0.97355727 |
| NPIPB12 | 69.1113562 | -0.875801 | 0.47991261 | -1.8249176 | 0.06801346 | 0.62603855 |
| SPN | 226.62935 | 0.62156778 | 0.64595289 | 0.96224939 | 0.33592434 | 0.85641269 |
| AC009086.2 | 3092.1242 | 0.7228383 | 0.38715337 | 1.86705933 | 0.0618933 | 0.61210783 |
| QPRT | 10264.0338 | 0.56794136 | 0.37316716 | 1.52194893 | 0.12802188 | 0.70580262 |
| C16orf54 | 97.7852612 | -0.1834439 | 0.64110338 | -0.2861378 | 0.77477261 | 0.96836854 |
| KIF22 | 1764.6146 | 0.2201056 | 0.22302763 | 0.98689832 | 0.32369249 | 0.85154042 |
| AC009133.4 | 809.007579 | 0.26642597 | 0.21516398 | 1.23824618 | 0.21562479 | 0.7954827 |
| MAZ | 6049.05864 | -0.4463149 | 0.19237798 | -2.3199894 | 0.02034145 | 0.5002876 |
| AC009133.2 | 1063.21819 | -0.41847 | 0.20111266 | -2.080774 | 0.0374546 | 0.55532957 |
| AC009133.1 | 5389.78154 | -0.4611087 | 0.1840926 | -2.5047647 | 0.01225329 | 0.41344377 |
| PRRT2 | 882.834678 | -0.5109327 | 0.19898198 | -2.5677335 | 0.01023658 | 0.40019066 |
| AC009133.6 | 1762.58538 | -0.2785977 | 0.18830099 | -1.4795339 | 0.13899768 | 0.72400098 |
| PAGR1 | 1618.223 | -0.2411297 | 0.21769029 | -1.1076733 | 0.26800292 | 0.82676682 |
| AC120114.5 | 2207.05016 | -0.0083423 | 0.21186289 | -0.039376 | 0.96859058 | 0.99400331 |
| MVP | 25805.7696 | -0.0269542 | 0.16875479 | -0.1597239 | 0.8730986 | 0.98275231 |
| CDIPT | 5895.93796 | 0.03356713 | 0.11947738 | 0.28094966 | 0.77874901 | 0.96836854 |
| AC120114.3 | 684.860465 | 0.05258533 | 0.18492329 | 0.28436292 | 0.77613227 | 0.96836854 |
| SEZ6L2 | 1028.58974 | 1.2208562 | 0.55823842 | 2.18697987 | 0.02874399 | 0.53606221 |

| | | | | | | |
|---|---|---|---|---|---|---|
| ASPHD1 | 425.856653 | 0.61819577 | 0.41950495 | 1.47363165 | 0.1405808 | 0.72579192 |
| KCTD13 | 1024.59143 | -0.0123784 | 0.14501741 | -0.0853577 | 0.93197699 | 0.98987582 |
| TMEM219 | 5280.76886 | 0.06502 | 0.2506826 | 0.25937183 | 0.79534836 | 0.97101097 |
| TAOK2 | 5373.62256 | -0.155389 | 0.15439103 | -1.0064636 | 0.3141926 | 0.84696964 |
| HIRIP3 | 1557.9988 | 0.19481166 | 0.16563252 | 1.17616796 | 0.23952777 | 0.81163641 |
| INO80E | 5550.58041 | 0.02084508 | 0.29015839 | 0.07184036 | 0.94272895 | 0.99136103 |
| DOC2A | 1013.02724 | -0.0721456 | 0.27851271 | -0.2590389 | 0.79560523 | 0.97101097 |
| AC093512.2 | 67479.4246 | 0.08919635 | 0.20032147 | 0.44526607 | 0.65612749 | 0.94537443 |
| ALDOA | 68208.2174 | 0.08428117 | 0.19963106 | 0.42218465 | 0.67289025 | 0.94922827 |
| AC093512.1 | 1564.13556 | 0.38855014 | 0.34675595 | 1.12052912 | 0.26248835 | 0.82424998 |
| PPP4C | 4620.90366 | -0.2513456 | 0.16551652 | -1.5185531 | 0.12887503 | 0.70633634 |
| TBX6 | 211.770036 | 0.05019081 | 0.41832515 | 0.11998038 | 0.90449869 | 0.98624484 |
| YPEL3 | 4531.27088 | -0.2888355 | 0.26630359 | -1.08461 | 0.27809442 | 0.82805357 |
| AC012645.1 | 67.3345269 | -0.3339568 | 0.36238031 | -0.9215644 | 0.35675585 | 0.86390849 |
| GDPD3 | 887.204183 | 0.29173354 | 0.3738822 | 0.78028195 | 0.43522493 | 0.89055338 |
| AC012645.4 | 336.385979 | 0.32706094 | 0.42478709 | 0.76994087 | 0.44133497 | 0.89290667 |
| MAPK3 | 3651.60918 | 0.10996922 | 0.18887924 | 0.58221972 | 0.56041869 | 0.92434754 |
| CORO1A | 1677.29055 | -0.1837134 | 0.59807504 | -0.3071746 | 0.7587105 | 0.96502715 |
| AC012645.3 | 220.558311 | -0.1786788 | 0.65352421 | -0.2734081 | 0.78453953 | 0.97021865 |
| SLX1A | 125.085288 | 0.04955957 | 0.29616345 | 0.16733857 | 0.86710366 | 0.98223481 |
| CD2BP2 | 5168.42233 | -0.2455832 | 0.14566133 | -1.6859878 | 0.09179815 | 0.65518742 |
| AC106782.2 | 116.880845 | -0.2285976 | 0.35155637 | -0.6502445 | 0.51553428 | 0.91636259 |
| TBC1D10B | 3336.3437 | -0.2815582 | 0.16806389 | -1.6753046 | 0.09387448 | 0.65783452 |
| AC106782.6 | 267.338372 | -0.2611016 | 0.16660353 | -1.5672032 | 0.11706719 | 0.68956727 |
| ZNF48 | 714.846631 | -0.2171782 | 0.1815625 | -1.1961622 | 2.32E-01 | 0.80649152 |
| 1-Sep | 416.10952 | 0.33036052 | 0.56549001 | 0.58420223 | 0.55908426 | 0.92421175 |
| ZNF771 | 860.353277 | 0.14802295 | 0.24853461 | 0.59558287 | 0.55145392 | 0.92373235 |
| DCTPP1 | 1532.70898 | 0.13754779 | 0.2025279 | 0.67915475 | 0.49703982 | 0.90913397 |
| SEPHS2 | 14271.7147 | 0.11680342 | 0.39589775 | 0.29503431 | 0.76796767 | 0.96762687 |
| ITGAL | 596.867838 | -0.0414741 | 0.67842634 | -0.0611328 | 0.95125347 | 0.99170503 |
| AC116348.3 | 43.0338079 | 0.39653454 | 0.80338706 | 0.49357845 | 0.62160392 | 0.94016713 |
| ZNF768 | 3343.09015 | -0.1377994 | 0.1807926 | -0.762196 | 0.44594306 | 0.89480207 |
| AC002310.5 | 929.005509 | -0.0365702 | 0.17913802 | -0.2041455 | 0.83823977 | 0.97599451 |
| ZNF747 | 983.524201 | -0.0866406 | 0.15724357 | -0.5509964 | 0.58163614 | 0.9317705 |
| AC002310.4 | 145.449994 | 0.01963386 | 0.2357108 | 0.08329639 | 0.93361587 | 0.99019225 |
| AC002310.1 | 148.263141 | 0.08006629 | 0.23643746 | 0.33863623 | 0.73488378 | 0.95895327 |
| ZNF764 | 462.272878 | -0.1322523 | 0.13280481 | -0.9958398 | 0.31932799 | 0.84928695 |
| ZNF688 | 753.719286 | -0.0305287 | 0.16244683 | -0.1879306 | 0.85093103 | 0.97844748 |
| AC002310.2 | 340.850432 | -0.1280113 | 0.22418887 | -0.5709975 | 0.56800137 | 0.92618483 |
| ZNF785 | 851.598377 | -0.1099262 | 0.23375311 | -0.4702664 | 0.63816471 | 0.94088547 |
| ZNF689 | 456.10811 | 0.02467593 | 0.23313416 | 0.10584432 | 0.91570587 | 0.9868044 |
| PRR14 | 3093.84391 | -0.4167772 | 0.18249327 | -2.2837947 | 0.0223836 | 0.51866171 |
| FBRS | 6524.51918 | -0.309147 | 0.18517983 | -1.6694418 | 0.09502985 | 0.65888357 |
| AC093249.6 | 141.344927 | -0.0090444 | 0.25607467 | -0.0353194 | 0.97182503 | 0.99404213 |
| SRCAP | 6543.96103 | -0.6078232 | 0.19121049 | -3.1788174 | 0.00147877 | 0.17236139 |
| AC106886.5 | 6974.71519 | -0.5559828 | 0.17541013 | -3.169616 | 0.00152641 | 0.17279992 |
| TMEM265 | 1095.95606 | -0.182562 | 0.18238224 | -1.0009856 | 0.31683377 | 0.84812045 |
| PHKG2 | 3279.51952 | -0.1490825 | 0.14577087 | -1.0227177 | 0.30644134 | 0.84278367 |
| CCDC189 | 672.030607 | -0.0516863 | 0.15799845 | -0.3271318 | 0.7435682 | 0.96131427 |
| RNF40 | 5074.38812 | -0.1557004 | 0.13920304 | -1.1185131 | 0.26334792 | 0.82424998 |
| ZNF629 | 1517.82855 | -0.0733151 | 0.18828911 | -0.3893751 | 0.69699869 | 0.95263271 |
| BCL7C | 1921.431 | 0.21910881 | 0.25698604 | 0.85260978 | 0.39387574 | 0.87884163 |
| MIR762HG | 1148.12204 | 0.20692767 | 0.27088395 | 0.76389785 | 4.45E-01 | 0.89429407 |
| CTF1 | 462.530645 | 0.05938959 | 0.38690083 | 0.15350081 | 0.87800335 | 0.98275231 |

| | | | | | | |
|---|---|---|---|---|---|---|
| FBXL19-AS1 | 111.110799 | -0.9382423 | 0.45789089 | -2.049052 | 0.04045703 | 0.55657615 |
| FBXL19 | 1889.24201 | -0.2654685 | 0.23022443 | -1.1530857 | 0.24887522 | 0.81646572 |
| ORAI3 | 2112.00802 | 0.17504069 | 0.27158821 | 0.64450768 | 0.51924628 | 0.91636259 |
| SETD1A | 2814.47017 | -0.7480288 | 0.28932566 | -2.5854217 | 0.009726 | 0.39368232 |
| HSD3B7 | 3965.38687 | -0.2805693 | 0.41038549 | -0.6836725 | 0.49418201 | 0.90796767 |
| STX1B | 222.923536 | -0.0726512 | 0.45070147 | -0.1611957 | 0.87193925 | 0.98275231 |
| STX4 | 3237.49522 | -0.1688907 | 0.1576926 | -1.0710121 | 2.84E-01 | 0.83107779 |
| AC135050.3 | 70.4874552 | -0.1500504 | 0.33868324 | -0.4430404 | 6.58E-01 | 0.94565705 |
| AC135050.1 | 407.162347 | -0.3233264 | 0.29929463 | -1.0802945 | 0.28001104 | 0.82825734 |
| ZNF668 | 547.310565 | -0.3209699 | 0.27729475 | -1.1575045 | 0.2470663 | 0.8156466 |
| ZNF646 | 1333.76718 | -0.2308434 | 0.16438795 | -1.4042598 | 0.16024149 | 0.74781837 |
| PRSS53 | 721.281299 | 0.009132 | 0.21922516 | 0.04165581 | 0.96677309 | 0.99400331 |
| AC135050.2 | 4315.93956 | 0.04906131 | 0.26071946 | 0.18817662 | 0.8507382 | 0.97844029 |
| VKORC1 | 4089.0467 | 0.03149059 | 0.27826369 | 0.11316816 | 0.90989723 | 0.98662757 |
| BCKDK | 4297.67709 | -0.0353743 | 0.19845582 | -0.1782476 | 0.85852853 | 0.97982623 |
| KAT8 | 2447.10356 | -0.2383793 | 0.13463067 | -1.7706165 | 0.0766245 | 0.63773415 |
| AC009088.3 | 3612.59922 | -0.0702959 | 0.1861026 | -0.3777268 | 0.70563353 | 0.95501548 |
| PRSS8 | 10831.747 | -0.0533396 | 0.15278122 | -0.349124 | 0.72699619 | 0.95848028 |
| PRSS36 | 124.361377 | -0.2052556 | 0.25895972 | -0.7926158 | 0.42800172 | 0.88798629 |
| FUS | 14735.1603 | -0.3149836 | 0.18585886 | -1.6947462 | 0.09012357 | 0.65403568 |
| AC009088.2 | 2153.77344 | -0.432764 | 0.29793965 | -1.4525225 | 0.14635637 | 0.73247558 |
| PYCARD | 300.861345 | -0.2510161 | 0.46866243 | -0.535601 | 0.59223433 | 0.93289552 |
| PYCARD-AS1 | 142.27304 | -0.2157317 | 0.41558965 | -0.5190979 | 0.60369248 | 0.93626289 |
| ITGAM | 474.539234 | -0.3613679 | 0.56073991 | -0.6444484 | 0.51928471 | 0.91636259 |
| ITGAX | 757.364229 | -0.5491482 | 0.55114675 | -0.9963738 | 0.31906853 | 0.84922641 |
| COX6A2 | 43.56866 | 0.16439344 | 0.56547709 | 0.29071636 | 0.77126826 | 0.96836854 |
| ZNF843 | 64.7788612 | 0.98714893 | 0.31779203 | 3.10627337 | 0.00189461 | 0.18915498 |
| AC026471.4 | 121.231949 | 0.06650906 | 0.32144757 | 0.20690484 | 0.83608418 | 0.97577341 |
| AC026471.1 | 287.830202 | 0.05902598 | 0.29030354 | 0.20332504 | 0.83888099 | 0.97612168 |
| ARMC5 | 1172.71081 | -0.1674822 | 0.20734964 | -0.8077285 | 0.4192469 | 0.88451372 |
| TGFB1I1 | 1760.10526 | -0.198049 | 0.38724988 | -0.5114243 | 0.60905401 | 0.93748023 |
| SLC5A2 | 9516.51376 | 0.0303839 | 0.22826413 | 0.13310853 | 0.89410755 | 0.98474725 |
| AC026471.3 | 1604.35396 | 0.08040761 | 0.60322387 | 0.13329647 | 0.89395892 | 0.98474725 |
| C16orf58 | 8423.62612 | 0.04935484 | 0.14296803 | 0.3452159 | 0.72993207 | 0.95848028 |
| AC074050.3 | 300.121226 | 0.26088819 | 0.29913426 | 0.87214416 | 0.38312974 | 0.8718674 |
| ZNF720 | 726.671281 | 0.22405284 | 0.11715467 | 1.91245344 | 0.05581806 | 0.59879007 |
| ZNF267 | 692.299578 | 0.05632922 | 0.27598532 | 0.20410224 | 0.8382736 | 0.97599451 |
| SHCBP1 | 333.542773 | -0.0660891 | 0.44785644 | -0.1475676 | 0.88268403 | 0.98276701 |
| AC092368.3 | 327.924048 | -0.4426204 | 0.24743344 | -1.7888463 | 0.07363957 | 0.6374448 |
| VPS35 | 4837.97239 | -0.0789146 | 0.18355453 | -0.4299248 | 0.66725038 | 0.94830184 |
| ORC6 | 116.738261 | 0.0675485 | 0.35530377 | 0.19011478 | 0.84921919 | 0.97753312 |
| MYLK3 | 153.658654 | 0.90218728 | 0.77364111 | 1.16615736 | 0.24355083 | 0.81376111 |
| C16orf87 | 751.862393 | 0.47712955 | 0.33089696 | 1.44192786 | 0.14932272 | 0.73670625 |
| GPT2 | 3400.89242 | 1.03203573 | 0.56812458 | 1.81656588 | 0.06928361 | 0.62779744 |
| DNAJA2 | 3869.08815 | 0.04442177 | 0.12122292 | 0.36644699 | 0.71403156 | 0.95564093 |
| NETO2 | 244.806904 | -0.1275402 | 0.4161268 | -0.3064935 | 0.75922893 | 0.96502715 |
| ITFG1-AS1 | 865.389578 | 0.48957789 | 0.14003963 | 3.49599528 | 0.0004723 | 0.1066424 |
| ITFG1 | 4560.49997 | 0.45059885 | 0.11499538 | 3.91840838 | 8.91E-05 | 0.04176226 |
| PHKB | 3836.74843 | 0.43451954 | 0.21593824 | 2.01223989 | 0.04419466 | 0.57267043 |
| ABCC11 | 61.5753682 | 0.47053211 | 0.51481903 | 0.91397576 | 0.36072958 | 0.86504879 |
| LONP2 | 4801.59444 | 0.04483094 | 0.09007024 | 0.49773307 | 0.6186722 | 0.93945324 |
| SIAH1 | 2038.22846 | -0.2199426 | 0.16332714 | -1.3466382 | 0.17809679 | 0.76115956 |
| N4BP1 | 2195.0757 | -0.4580648 | 0.27704967 | -1.6533672 | 0.09825616 | 0.66549286 |
| ZNF423 | 204.556945 | 0.3257433 | 0.52126155 | 0.62491333 | 0.53202794 | 0.91964636 |

| | | | | | | |
|---|---|---|---|---|---|---|
| CNEP1R1 | 782.384879 | -0.0959528 | 0.14481462 | -0.6625909 | 0.50759261 | 0.91439937 |
| HEATR3 | 776.110803 | -0.0003911 | 0.18192777 | -0.0021497 | 0.99828482 | 0.99990633 |
| AC007610.2 | 74.6491859 | 0.16847001 | 0.26335627 | 0.63970381 | 0.52236518 | 0.91723878 |
| TENT4B | 954.77819 | -0.0445788 | 0.15065036 | -0.2959089 | 0.76729963 | 0.96749893 |
| ADCY7 | 758.428774 | -0.3417122 | 0.38866952 | -0.8791845 | 0.37930123 | 0.87069278 |
| BRD7 | 3366.50504 | -0.0509861 | 0.09605233 | -0.5308162 | 0.59554613 | 0.9334601 |
| AC007493.2 | 43.6826811 | 0.13284389 | 0.35191336 | 0.37749034 | 0.70580924 | 0.95501548 |
| NKD1 | 516.477257 | 0.06091961 | 0.49239382 | 0.12372131 | 0.90153594 | 0.98572048 |
| AC007608.3 | 53.5998168 | -0.0684818 | 0.76373726 | -0.0896667 | 0.92855211 | 0.98963127 |
| SNX20 | 171.595602 | 0.09087761 | 0.70596385 | 0.12872842 | 0.89757255 | 0.98474725 |
| NOD2 | 144.001183 | -0.8534815 | 0.6867298 | -1.2428199 | 0.21393421 | 0.7926732 |
| AC007728.2 | 78.8157126 | -1.1594021 | 0.55636503 | -2.0838874 | 0.03717041 | 0.55527244 |
| CYLD | 2330.23869 | -0.7487571 | 0.38844676 | -1.927567 | 0.05390901 | 0.5929042 |
| LINC02128 | 41.3828097 | -0.7397316 | 0.51529185 | -1.4355585 | 0.15112801 | 0.73796431 |
| SALL1 | 4815.8069 | -0.5671944 | 0.25451373 | -2.2285414 | 0.02584443 | 0.52502381 |
| AC087564.1 | 108.643723 | -0.4348106 | 0.26808313 | -1.6219246 | 0.10481949 | 0.67459549 |
| AC007344.1 | 160.34523 | -1.6603443 | 0.56313175 | -2.9484117 | 0.00319411 | 0.24838541 |
| HNRNPA1P4 | 162.083391 | 0.193977 | 0.27936009 | 0.69436187 | 0.4874553 | 0.90573694 |
| LINC01571 | 57.1332567 | 0.37565256 | 0.65063606 | 0.57736203 | 0.56369492 | 0.92453312 |
| TOX3 | 908.306237 | 0.33581113 | 0.2476965 | 1.35573628 | 0.17518309 | 0.76073329 |
| AC026462.3 | 149.636254 | 0.8770078 | 0.53438748 | 1.64114584 | 0.10076714 | 0.67247782 |
| AC007906.2 | 1096.67416 | 0.03794848 | 0.65103556 | 0.05828942 | 0.9535181 | 0.99170503 |
| CHD9 | 2265.53124 | -0.0513021 | 0.15384533 | -0.3334653 | 7.39E-01 | 9.60E-01 |
| AC007342.4 | 51.5903185 | 1.47771495 | 0.71420856 | 2.06902441 | 0.0385438 | 0.55565329 |
| AC007342.5 | 70.6715754 | 1.64767855 | 0.60410039 | 2.72749126 | 0.00638179 | 0.33501741 |
| RBL2 | 4859.19916 | 0.23503401 | 0.23041337 | 1.0200537 | 0.307703 | 0.8434443 |
| AKTIP | 3234.39026 | 0.1277815 | 0.15318132 | 0.8341846 | 0.40417697 | 0.88132015 |
| RPGRIP1L | 328.691301 | 0.09526138 | 0.31979763 | 0.29788019 | 0.76579461 | 0.96704052 |
| FTO | 2120.20759 | 0.12531993 | 0.19397342 | 0.64606756 | 0.5182356 | 0.91636259 |
| IRX3 | 1659.30319 | -0.6700177 | 0.54620296 | -1.2266827 | 0.21994188 | 0.79858446 |
| AC018553.2 | 105.549334 | -0.7481997 | 0.64652077 | -1.1572709 | 2.47E-01 | 0.8156466 |
| AC018553.1 | 43.0623048 | -0.6220726 | 0.4504007 | -1.3811537 | 0.1672317 | 0.75558932 |
| CRNDE | 461.49746 | -0.4260888 | 0.39711935 | -1.0729489 | 0.28329406 | 0.83094967 |
| IRX5 | 571.77946 | -0.7905773 | 0.5137956 | -1.5387 | 0.12387756 | 0.6996941 |
| IRX6 | 174.684116 | 0.03084382 | 0.67709551 | 0.04555313 | 0.96366642 | 0.99323942 |
| MMP2 | 4434.94283 | 0.55937319 | 0.40689787 | 1.37472625 | 0.16921633 | 0.7584845 |
| LPCAT2 | 1146.737 | 0.11324478 | 0.25759195 | 0.43962856 | 0.66020615 | 0.94628868 |
| CES1 | 448.416384 | -0.9235356 | 0.68485466 | -1.3485132 | 0.17749337 | 0.76115956 |
| GNAO1 | 246.483202 | -0.2208819 | 0.4537744 | -0.4867658 | 0.6264243 | 0.94088547 |
| AMFR | 10479.0892 | -0.0625223 | 0.19797044 | -0.3158166 | 0.75214173 | 0.96301881 |
| AC092140.1 | 400.165741 | 0.05413679 | 0.24524724 | 0.22074373 | 0.82529198 | 0.97548201 |
| NUDT21 | 4145.92734 | -0.0302354 | 0.13499435 | -0.2239753 | 0.82277656 | 0.97519922 |
| OGFOD1 | 1786.02337 | -0.2241909 | 0.13256434 | -1.6911857 | 9.08E-02 | 0.65495306 |
| AC092140.2 | 89.5313126 | 0.02233364 | 0.21447382 | 0.10413226 | 0.91706439 | 0.9868044 |
| BBS2 | 3803.67792 | 0.24614865 | 0.20538491 | 1.19847489 | 0.23073219 | 0.80610188 |
| MT3 | 54.987317 | 0.36971091 | 0.65875755 | 0.56122455 | 0.57464447 | 0.93024203 |
| MT2A | 28108.0712 | -1.0524445 | 0.47663122 | -2.2080898 | 2.72E-02 | 5.29E-01 |
| MT1E | 9203.30083 | -1.2683429 | 0.5922634 | -2.1415183 | 0.03223226 | 0.54925606 |
| MT1M | 1238.89206 | -2.3661093 | 0.56773233 | -4.1676494 | 3.08E-05 | 0.01988852 |
| MT1A | 368.110519 | -1.1952947 | 0.7311704 | -1.634769 | 1.02E-01 | 6.73E-01 |
| MT1F | 6314.67032 | -0.9121206 | 0.53715027 | -1.6980735 | 0.08949389 | 0.65375543 |
| MT1G | 24378.4916 | -0.8773054 | 0.66831116 | -1.31272 | 0.18927731 | 0.76970626 |
| MT1H | 6919.17782 | -1.5260756 | 0.70356046 | -2.1690752 | 0.03007697 | 0.54012717 |
| MT1X | 11738.4389 | -0.9974771 | 0.56637396 | -1.7611634 | 0.07821075 | 0.64079926 |

| | | | | | | |
|---|---|---|---|---|---|---|
| NUP93 | 1458.82508 | -0.1598894 | 0.16929776 | -0.944427 | 0.3449515 | 0.86024432 |
| SLC12A3 | 10625.3051 | 2.77871191 | 0.8165925 | 3.40281339 | 0.00066696 | 0.12385487 |
| HERPUD1 | 10828.4993 | 0.03181557 | 0.24491562 | 0.1299042 | 0.89664222 | 0.98474725 |
| CETP | 217.036042 | 0.4559619 | 0.49881749 | 0.91408562 | 0.36067186 | 0.86504879 |
| NLRC5 | 1918.04968 | -1.410466 | 0.46203083 | -3.0527529 | 0.00226753 | 0.20649641 |
| CPNE2 | 1497.29166 | -0.1298483 | 0.23752437 | -0.5466737 | 0.58460294 | 0.93282331 |
| FAM192A | 3539.02404 | -0.18397 | 0.10381933 | -1.7720204 | 0.07639117 | 0.63773415 |
| RSPRY1 | 1880.52123 | -0.0284223 | 0.15962051 | -0.1780615 | 0.85867472 | 0.97983564 |
| ARL2BP | 3317.11392 | -0.1241923 | 0.27275675 | -0.4553226 | 0.64887716 | 0.94312988 |
| AC009090.3 | 280.342122 | -0.0352979 | 0.36690619 | -0.0962041 | 0.92335846 | 0.98858365 |
| PLLP | 5539.20237 | 0.14850612 | 0.20189917 | 0.73554599 | 0.46200704 | 0.89762182 |
| CX3CL1 | 5604.06584 | -0.1838307 | 0.38798359 | -0.4738104 | 0.63563512 | 0.94088547 |
| CIAPIN1 | 2815.56833 | -0.1279596 | 0.17612762 | -0.7265165 | 0.46752222 | 0.89984944 |
| COQ9 | 5966.74248 | 0.12480003 | 0.18522137 | 0.67378851 | 0.50044578 | 0.91180773 |
| POLR2C | 4592.99114 | 0.07416632 | 0.15438751 | 0.48039069 | 0.63094962 | 0.94088547 |
| DOK4 | 5954.81995 | 0.212158 | 0.25998476 | 0.81604013 | 0.41447719 | 0.88364943 |
| CCDC102A | 494.4279 | -0.142466 | 0.36350587 | -0.3919221 | 0.69511578 | 0.95234774 |
| ADGRG5 | 1086.10922 | 0.80375535 | 0.48904034 | 1.64353587 | 0.10027211 | 0.67191826 |
| ADGRG1 | 42067.5428 | -0.5599903 | 0.41458664 | -1.3507197 | 0.17678525 | 0.76115956 |
| ADGRG3 | 139.856495 | -0.7705861 | 0.44805103 | -1.7198625 | 0.08545744 | 0.6504594 |
| DRC7 | 91.090651 | 0.04202132 | 0.7855492 | 0.05349291 | 0.95733918 | 0.99206489 |
| KATNB1 | 1143.6488 | 0.0125039 | 0.17575507 | 0.07114392 | 0.94328322 | 0.99136103 |
| KIFC3 | 11900.5841 | -0.447417 | 0.20870057 | -2.1438228 | 0.03204709 | 0.54798767 |
| AC092118.2 | 1997.2236 | -0.426997 | 0.24960911 | -1.7106628 | 0.08714338 | 0.65231349 |
| AC092118.1 | 179.10833 | -0.1461671 | 0.37320557 | -0.3916532 | 0.6953145 | 0.95248074 |
| ZNF319 | 613.736157 | -0.0463489 | 0.13498193 | -0.3433715 | 0.73131902 | 0.95848028 |
| USB1 | 2290.56259 | -0.1639682 | 0.19451768 | -0.8429474 | 0.39925784 | 0.88077967 |
| MMP15 | 2820.86848 | 0.3207557 | 0.15671117 | 2.04679546 | 0.04067817 | 0.55657615 |
| CFAP20 | 1111.21407 | 0.05460796 | 0.15860912 | 0.34429271 | 0.73062617 | 0.95848028 |
| AC026771.1 | 186.665415 | 0.23324172 | 0.2751026 | 0.8478354 | 0.39652964 | 0.8796941 |
| CSNK2A2 | 3129.0849 | -0.1458101 | 0.22268098 | -0.6547938 | 0.51260051 | 0.91568064 |
| CCDC113 | 303.76837 | 0.48106516 | 0.29706949 | 1.61936914 | 1.05E-01 | 6.75E-01 |
| GINS3 | 171.92447 | -0.5137713 | 0.2705647 | -1.898885 | 5.76E-02 | 6.01E-01 |
| NDRG4 | 336.612077 | -0.5093925 | 0.50952134 | -0.9997472 | 0.31743286 | 0.84819508 |
| SETD6 | 3212.85347 | -0.198719 | 0.15959738 | -1.2451267 | 0.21308517 | 0.7926732 |
| CNOT1 | 5755.10894 | -0.0848767 | 0.1234972 | -0.6872762 | 4.92E-01 | 9.06E-01 |
| AC009118.3 | 1247.97225 | -0.1096391 | 0.11091013 | -0.9885401 | 0.32288822 | 0.85119105 |
| SLC38A7 | 2022.7072 | -0.6644452 | 0.2972247 | -2.2354978 | 0.02538469 | 0.52250144 |
| GOT2 | 8353.86643 | 0.25962158 | 0.18419417 | 1.40949947 | 0.15868753 | 0.74685158 |
| AC092378.1 | 394.376775 | 0.43524876 | 0.35696823 | 1.2192927 | 0.22273312 | 0.80078694 |
| AC138305.1 | 46.8727475 | -0.061953 | 0.55052369 | -0.1125347 | 0.91039942 | 0.98662757 |
| CDH11 | 1042.18223 | -0.085604 | 0.42529303 | -0.2012823 | 0.84047786 | 0.97618283 |
| CDH5 | 2630.22868 | -0.0191208 | 0.39366902 | -0.0485707 | 0.9612614 | 0.99320711 |
| BEAN1 | 72.5912905 | 1.47759916 | 0.46845765 | 3.15417875 | 0.0016095 | 0.17848359 |
| AC010542.3 | 348.372527 | 0.62554436 | 0.31845519 | 1.9643089 | 0.04949428 | 0.5861488 |
| TK2 | 2908.50988 | 0.50323408 | 0.24755439 | 2.0328223 | 0.04207048 | 0.56438286 |
| AC010542.2 | 634.237409 | 0.55988311 | 0.24753936 | 2.26179427 | 0.02371012 | 0.51967919 |
| AC010542.4 | 96.1417473 | 0.44995034 | 0.34956227 | 1.2871822 | 1.98E-01 | 0.77952007 |
| CKLF | 640.010381 | 0.34349104 | 0.25494158 | 1.34733233 | 0.17787322 | 0.76115956 |
| CKLF-CMTM1 | 619.259371 | 0.34415183 | 0.26452781 | 1.30100436 | 0.19325696 | 0.77548688 |
| AC010542.5 | 359.521041 | 0.48691574 | 0.24648173 | 1.97546382 | 0.04821554 | 0.58440502 |
| CMTM1 | 69.5561723 | 0.21347071 | 0.32732719 | 0.6521631 | 0.51429596 | 0.91636259 |
| CMTM3 | 1010.23429 | -0.1407483 | 0.28856295 | -0.487756 | 0.62572268 | 0.94088547 |
| CMTM4 | 4810.65237 | 0.34879072 | 0.28596666 | 1.21969019 | 0.22258234 | 0.80078694 |

| | | | | | | |
|---|---|---|---|---|---|---|
| DYNC1LI2 | 6317.08206 | -0.0586744 | 0.14214145 | -0.4127885 | 0.67976154 | 0.95070301 |
| AC018557.1 | 3175.4706 | -0.0371341 | 0.13977103 | -0.265678 | 0.79048718 | 0.97090793 |
| NAE1 | 1833.68557 | -0.0568927 | 0.17252626 | -0.3297627 | 0.74157926 | 0.96109001 |
| PDP2 | 1311.85856 | 0.25893993 | 0.28539849 | 0.90729258 | 0.3642521 | 0.8663515 |
| CDH16 | 68838.3802 | -0.1665934 | 0.27693969 | -0.6015513 | 0.54747288 | 0.92361227 |
| RRAD | 4402.27638 | -0.3032798 | 0.4818168 | -0.6294504 | 0.52905422 | 0.91907723 |
| FAM96B | 4081.62907 | 0.00582317 | 0.20170848 | 0.02886925 | 0.97696887 | 0.99455099 |
| CES2 | 12253.3587 | 0.58357624 | 0.40647875 | 1.43568696 | 0.15109144 | 0.73796431 |
| CES3 | 1620.69102 | 0.88311284 | 0.690832 | 1.27833227 | 0.2011323 | 0.78134034 |
| CES4A | 893.008042 | 0.49912798 | 0.47055631 | 1.06071893 | 0.28881765 | 0.83295411 |
| CBFB | 1981.77369 | 0.00787731 | 0.1527913 | 0.05155604 | 0.95888245 | 0.99242674 |
| C16orf70 | 2208.45173 | -0.3062327 | 0.23963133 | -1.2779328 | 0.20127312 | 0.78134034 |
| B3GNT9 | 1175.73846 | -0.1917328 | 0.17908159 | -1.0706452 | 0.28432899 | 0.83107779 |
| TRADD | 1763.83726 | 0.02733008 | 0.24363724 | 0.1121753 | 0.91068442 | 0.98662757 |
| FBXL8 | 652.108768 | -0.0033296 | 0.25264805 | -0.0131787 | 0.98948524 | 0.99800524 |
| AC074143.1 | 798.223179 | 0.08061391 | 0.33717849 | 0.23908378 | 0.81104062 | 0.97352497 |
| HSF4 | 2378.81453 | 0.21499906 | 0.39160188 | 0.54902458 | 0.58298858 | 0.93191649 |
| NOL3 | 2276.89575 | 0.15620405 | 0.30340867 | 0.51483055 | 0.60667143 | 0.93708311 |
| KIAA0895L | 3570.26672 | -0.2221367 | 0.29597975 | -0.7505132 | 0.4529457 | 0.89670027 |
| EXOC3L1 | 556.320508 | -0.1586108 | 0.33540451 | -0.4728942 | 0.63628867 | 0.94088547 |
| E2F4 | 4236.05429 | -0.3316797 | 0.16838241 | -1.9697999 | 0.0488613 | 0.58504358 |
| ELMO3 | 2772.04893 | 0.15826066 | 0.27126601 | 0.58341501 | 0.55961396 | 0.92429981 |
| LRRC29 | 433.44513 | 0.0096289 | 0.3341891 | 0.02881273 | 0.97701395 | 0.99455099 |
| TMEM208 | 2148.46257 | 0.06731584 | 0.17222126 | 0.39086837 | 0.69589454 | 0.9525827 |
| FHOD1 | 2452.53949 | -0.1677332 | 0.35164949 | -0.4769898 | 6.33E-01 | 0.94088547 |
| SLC9A5 | 474.943787 | -0.3357712 | 0.29556886 | -1.1360167 | 0.25594958 | 0.82052927 |
| PLEKHG4 | 479.148648 | 0.00166129 | 0.32516907 | 0.005109 | 0.99592362 | 0.99960393 |
| LRRC36 | 121.330871 | 0.22324362 | 0.6245598 | 0.35744154 | 0.72076128 | 0.95728372 |
| TPPP3 | 782.287726 | 1.04945056 | 0.50568357 | 2.07531077 | 0.03795774 | 0.55538555 |
| ZDHHC1 | 1439.27562 | 0.0757717 | 0.23739099 | 0.31918523 | 0.74958606 | 0.96249332 |
| HSD11B2 | 18110.5253 | 0.6832569 | 0.63793236 | 1.07104912 | 0.28414734 | 0.83107779 |
| AC009061.1 | 274.998316 | 0.63207349 | 0.61972121 | 1.019932 | 0.30776071 | 0.8434443 |
| ATP6V0D1 | 7782.46523 | -0.0777926 | 0.2043092 | -0.3807591 | 0.703382 | 0.95466435 |
| AC009061.2 | 148.856203 | 0.09552371 | 0.45291845 | 0.2109071 | 0.83295977 | 0.97550416 |
| AC027682.6 | 160.009846 | 0.61599465 | 0.49854743 | 1.23557882 | 0.21661516 | 0.7954827 |
| RIPOR1 | 9879.67691 | -0.3120796 | 0.22113163 | -1.4112843 | 0.15816079 | 0.74620522 |
| AC027682.5 | 692.58676 | -0.3097913 | 0.34266152 | -0.9040736 | 0.36595635 | 0.86672412 |
| AC027682.2 | 1528.00226 | -0.3278894 | 0.29618383 | -1.1070468 | 0.26827369 | 0.82676682 |
| AC027682.3 | 868.441117 | -0.4363366 | 0.31074468 | -1.4041644 | 0.16026989 | 0.74781837 |
| CTCF | 2151.05433 | -0.3104998 | 0.22303628 | -1.3921494 | 0.16387717 | 0.75265041 |
| CARMIL2 | 208.456704 | -0.0145738 | 0.73259806 | -0.0198933 | 0.98412852 | 0.99619492 |
| ACD | 1199.48214 | -0.0208523 | 0.17508171 | -0.1191005 | 0.90519571 | 0.98625102 |
| PARD6A | 416.593162 | -0.0319366 | 0.1692574 | -0.1886867 | 0.85033841 | 0.97816532 |
| ENKD1 | 2353.93478 | -0.1304507 | 0.29764016 | -0.4382833 | 0.66118095 | 0.94637197 |
| C16orf86 | 506.050364 | 0.25277553 | 0.31024611 | 0.81475811 | 0.41521079 | 0.88377697 |
| GFOD2 | 1977.68007 | -0.1836104 | 0.12862881 | -1.4274434 | 0.15345212 | 0.74258845 |
| RANBP10 | 2401.40534 | -0.1403052 | 0.14202432 | -0.9878957 | 0.32320372 | 0.85119105 |
| TSNAXIP1 | 1558.57838 | -0.3200793 | 0.19002518 | -1.6844045 | 0.09210352 | 0.65583276 |
| CENPT | 4600.81685 | -0.2750568 | 0.16288042 | -1.6887039 | 0.09127619 | 0.65515398 |
| THAP11 | 1322.46707 | -0.0640882 | 0.15852786 | -0.4042711 | 0.68601338 | 0.95097288 |
| NUTF2 | 3132.89537 | -0.346395 | 0.20695535 | -1.6737667 | 0.09417646 | 0.65783452 |
| EDC4 | 4104.62175 | -0.231929 | 0.10841127 | -2.1393442 | 0.0324078 | 0.55014009 |
| AC040162.3 | 2316.76507 | -0.1752751 | 0.11817291 | -1.4832087 | 0.13801896 | 0.72209298 |
| NRN1L | 79.31224 | 0.31194151 | 0.3058647 | 1.01986765 | 0.30779124 | 0.8434443 |

| | | | | | | |
|---|---|---|---|---|---|---|
| PSKH1 | 2829.53311 | 0.36248621 | 0.22712353 | 1.59598704 | 0.11049169 | 0.68001961 |
| CTRL | 752.754483 | 0.02043028 | 0.17720817 | 0.11528971 | 0.9082155 | 0.98662757 |
| AC040162.1 | 2117.02844 | -0.2603707 | 0.38737974 | -0.6721329 | 0.50149908 | 0.91259827 |
| PSMB10 | 2560.31747 | -0.2153501 | 0.40400388 | -0.5330396 | 0.59400617 | 0.93317501 |
| LCAT | 1559.15064 | -0.2546839 | 0.28555485 | -0.8918913 | 0.37245119 | 0.86791605 |
| SLC12A4 | 4116.09123 | -0.1534901 | 0.22071454 | -0.6954236 | 0.48678985 | 0.90573694 |
| DPEP2 | 142.335001 | 0.39935387 | 0.59925018 | 0.66642262 | 0.50514101 | 0.91365534 |
| DUS2 | 776.983069 | 0.21087859 | 0.2073926 | 1.01680867 | 0.30924446 | 0.84414094 |
| DDX28 | 720.526972 | 0.09757586 | 0.33098329 | 0.29480599 | 0.76814209 | 0.96765147 |
| NFATC3 | 2835.74056 | -0.0282362 | 0.14019116 | -0.2014119 | 8.40E-01 | 9.76E-01 |
| AC020978.5 | 2590.61367 | -0.0097027 | 0.13883937 | -0.0698841 | 9.44E-01 | 9.91E-01 |
| ESRP2 | 2578.41932 | -0.1004653 | 0.22999876 | -0.4368079 | 0.66225068 | 0.94642611 |
| PLA2G15 | 4625.41842 | 0.42712823 | 0.38159372 | 1.11932721 | 0.26300057 | 0.82424998 |
| AC020978.2 | 1310.70128 | 0.43881606 | 0.38716824 | 1.13339889 | 0.25704678 | 0.82107009 |
| SLC7A6 | 2386.83673 | -0.2779577 | 0.16669176 | -1.6674949 | 9.54E-02 | 6.59E-01 |
| SLC7A6OS | 2293.26415 | -0.2672789 | 0.14769873 | -1.8096219 | 7.04E-02 | 6.29E-01 |
| PRMT7 | 1355.71564 | -0.0159057 | 0.33283303 | -0.0477888 | 0.96188458 | 0.99320711 |
| SMPD3 | 267.431784 | -0.1358848 | 0.64907076 | -0.2093529 | 0.83417276 | 0.97550416 |
| ZFP90 | 1114.17906 | 0.03930045 | 0.1957603 | 0.20075803 | 0.84088778 | 0.97618283 |
| CDH3 | 1659.17215 | 0.10426909 | 0.36184166 | 0.28816221 | 0.77322258 | 0.96836854 |
| AC126773.2 | 99.2160448 | 0.08073131 | 0.42748616 | 0.18885129 | 8.50E-01 | 0.97816532 |
| CDH1 | 5429.48009 | 0.36821422 | 0.20228228 | 1.82029889 | 6.87E-02 | 0.62750387 |
| AC099314.1 | 265.931415 | 0.26896567 | 0.3082034 | 0.87268884 | 0.3828327 | 0.8718674 |
| TANGO6 | 654.698349 | 0.14243259 | 0.20248732 | 0.70341487 | 0.48179724 | 0.90541935 |
| AC009137.1 | 53.113654 | 0.29754563 | 0.3261808 | 0.91221075 | 0.36165778 | 0.86528137 |
| HAS3 | 2296.10265 | 0.14721009 | 0.22938379 | 0.64176326 | 0.52102691 | 0.91723878 |
| CHTF8 | 6785.26887 | 0.1638394 | 0.24958267 | 0.65645342 | 0.51153243 | 0.91510397 |
| UTP4 | 4049.69312 | -0.2002528 | 0.36674607 | -0.5460257 | 0.58504827 | 0.93289552 |
| SNTB2 | 2164.30605 | -0.1508488 | 0.25917735 | -0.5820293 | 0.56054692 | 0.92434754 |
| AC026464.4 | 3568.39568 | 0.10881178 | 0.10650303 | 1.02167773 | 0.30693345 | 0.84286067 |
| VPS4A | 7006.02373 | 0.09906616 | 0.08143908 | 1.21644493 | 0.22381548 | 0.80078694 |
| COG8 | 5368.5863 | 0.12583877 | 0.11607597 | 1.08410706 | 0.27831733 | 0.82805357 |
| PDF | 1172.32388 | 0.37788181 | 0.22398013 | 1.68712201 | 0.09157989 | 0.65518742 |
| AC026464.3 | 465.608568 | -0.0571547 | 0.11057517 | -0.5168857 | 0.60523597 | 0.93687349 |
| AC026464.1 | 444.968931 | -0.3046784 | 0.3745539 | -0.8134434 | 0.41596389 | 0.88399703 |
| NIP7 | 1155.01975 | -0.2741233 | 0.28323054 | -0.967845 | 0.3331218 | 0.855011 |
| TMED6 | 320.985388 | -0.521442 | 0.67576869 | -0.771628 | 0.44033478 | 0.89262457 |
| TERF2 | 1467.90985 | -0.0982598 | 0.1002693 | -0.9799592 | 0.32710625 | 0.85308232 |
| CYB5B | 4226.43009 | -0.0005915 | 0.10363635 | -0.0057072 | 0.99544632 | 0.99940889 |
| NFAT5 | 5592.87997 | -0.2450426 | 0.18714827 | -1.3093502 | 0.19041575 | 0.77108505 |
| NQO1 | 9363.88926 | 1.02069251 | 0.63969914 | 1.59558212 | 1.11E-01 | 6.80E-01 |
| AC092115.2 | 1626.12451 | 1.29671158 | 0.68207613 | 1.90112442 | 0.05728572 | 0.60077875 |
| AC092115.3 | 949.405395 | -0.09176 | 0.11495361 | -0.7982348 | 0.42473423 | 0.88573606 |
| NOB1 | 2117.48909 | -0.1343272 | 0.12841095 | -1.0460731 | 0.29552727 | 0.83738685 |
| WWP2 | 5140.42029 | -0.3580211 | 0.11922275 | -3.0029597 | 0.00267368 | 0.2207375 |
| CLEC18A | 977.627896 | 1.22223874 | 0.74976493 | 1.63016259 | 0.10306714 | 0.67430143 |
| PDPR | 3509.68479 | -0.466383 | 0.27067408 | -1.7230429 | 0.08488077 | 0.64988556 |
| AC009060.1 | 1797.66604 | -0.44788 | 0.27470429 | -1.6304078 | 0.10301534 | 0.67430143 |
| CLEC18C | 912.35151 | 1.96800018 | 0.82608679 | 2.38231648 | 0.0172041 | 0.46842641 |
| EXOSC6 | 1803.25994 | -0.4723664 | 0.29750388 | -1.5877654 | 0.1123394 | 0.68001961 |
| AARS | 7205.36357 | -0.0971309 | 0.18363821 | -0.5289253 | 5.97E-01 | 0.9337017 |
| DDX19B | 1320.075 | -0.0255579 | 0.16430615 | -0.1555504 | 0.87638743 | 0.98275231 |
| AC012184.2 | 2014.65075 | 0.01837214 | 0.15263686 | 0.12036503 | 0.90419399 | 0.98624484 |
| DDX19A | 2328.5801 | -0.0197714 | 0.13035851 | -0.1516694 | 0.87944767 | 0.98275231 |

| | | | | | | |
|---|---|---|---|---|---|---|
| ST3GAL2 | 1764.08847 | -0.0238311 | 0.16386773 | -0.1454287 | 0.88437243 | 0.98332979 |
| AC012184.1 | 212.211201 | 0.05797023 | 0.19277185 | 0.30071939 | 0.76362848 | 0.96678847 |
| FUK | 2547.753 | 0.09118144 | 0.19265413 | 0.47329086 | 0.63600568 | 0.94088547 |
| COG4 | 3386.7741 | 0.04112356 | 0.18551953 | 0.22166703 | 8.25E-01 | 0.97548201 |
| SF3B3 | 4546.49552 | 0.03968838 | 0.13802719 | 0.28754029 | 0.77369865 | 0.96836854 |
| IL34 | 275.211684 | 0.07392075 | 0.42802021 | 0.17270388 | 0.86288419 | 0.98176973 |
| MTSS1L | 5885.35231 | -0.7886687 | 0.44422406 | -1.7753848 | 7.58E-02 | 6.38E-01 |
| VAC14 | 3412.76395 | -0.1971675 | 0.24032069 | -0.8204351 | 0.4119681 | 0.88268208 |
| VAC14-AS1 | 244.968747 | -0.1856969 | 0.38509135 | -0.4822152 | 0.62965308 | 0.94088547 |
| HYDIN | 386.638974 | -0.0221063 | 0.47220845 | -0.0468147 | 0.96266094 | 0.99320711 |
| CMTR2 | 1561.17183 | -0.4144359 | 0.35786557 | -1.158077 | 0.24683261 | 0.81542272 |
| ZNF23 | 949.634245 | 0.01834499 | 0.17095833 | 0.10730677 | 9.15E-01 | 0.9866558 |
| AC010547.4 | 975.984475 | 0.04016749 | 0.17009848 | 0.23614254 | 8.13E-01 | 0.9738447 |
| AC010547.1 | 39.7593203 | 0.18122777 | 0.39639152 | 0.45719387 | 0.6475317 | 0.94282139 |
| ZNF19 | 329.31544 | 0.2250665 | 0.32937905 | 0.68330544 | 0.49441386 | 0.90796767 |
| AC009097.2 | 135.195816 | 0.25159828 | 0.5409044 | 0.4651437 | 0.64182855 | 0.94088547 |
| MARVELD3 | 1700.87006 | -0.172841 | 0.41622908 | -0.4152544 | 0.67795569 | 0.95018034 |
| PHLPP2 | 1275.03826 | -0.2614463 | 0.25140809 | -1.0399281 | 0.2983733 | 0.84011628 |
| AP1G1 | 5972.28775 | -0.0145423 | 0.16899624 | -0.0860509 | 0.93142594 | 0.98975562 |
| ATXN1L | 2740.36791 | -0.1634968 | 0.23602103 | -0.6927214 | 0.48848441 | 0.90585781 |
| IST1 | 9384.89654 | -0.2002447 | 0.08928671 | -2.2427153 | 0.02491518 | 0.52085426 |
| ZNF821 | 643.407495 | -0.3001493 | 0.23374498 | -1.2840886 | 1.99E-01 | 0.78134034 |
| PKD1L3 | 109.506231 | 0.12522941 | 0.24928008 | 0.50236428 | 0.6154113 | 0.93910158 |
| DHODH | 1005.99473 | -0.0129968 | 0.15546453 | -0.0835997 | 0.93337471 | 0.99017182 |
| TXNL4B | 764.533496 | -0.0432875 | 0.17059201 | -0.2537486 | 0.7996898 | 0.97208422 |
| HP | 5964.97158 | -1.4403587 | 1.49398466 | -0.9641054 | 0.33499307 | 0.85581042 |
| HPR | 268.184695 | -0.8731113 | 1.03784861 | -0.8412704 | 0.40019646 | 0.88077967 |
| AC009087.1 | 125.638545 | -0.0635167 | 0.19899532 | -0.3191871 | 7.50E-01 | 9.62E-01 |
| DHX38 | 3367.83337 | -0.3428329 | 0.19290481 | -1.7772128 | 0.07553323 | 0.63773415 |
| PMFBP1 | 897.535547 | -0.3343135 | 0.21037767 | -1.5891111 | 0.11203532 | 0.68001961 |
| AC004943.2 | 210.717961 | -0.4188145 | 0.22951678 | -1.8247664 | 0.06803629 | 0.62603855 |
| ZFHX3 | 1250.7167 | -0.3764712 | 0.20234366 | -1.8605533 | 0.06280729 | 0.6147045 |
| AC116667.1 | 107.51407 | -0.0262407 | 0.49909794 | -0.0525763 | 0.95806948 | 0.99227301 |
| PSMD7 | 4476.8101 | -0.1485585 | 0.20600292 | -0.7211476 | 4.71E-01 | 0.90089741 |
| NPIPB15 | 740.440825 | 0.82666944 | 1.31933279 | 0.62658144 | 5.31E-01 | 0.91962754 |
| CLEC18B | 2018.51667 | 1.86579102 | 0.73630906 | 2.53397807 | 0.01127758 | 0.40196898 |
| AC009053.3 | 55.6995013 | 0.36744734 | 0.74488718 | 0.49329261 | 0.62180585 | 0.94016713 |
| GLG1 | 9065.45529 | 0.08429203 | 0.08223458 | 1.02501935 | 0.30535406 | 0.84268008 |
| RFWD3 | 1053.2238 | -0.1629319 | 0.21675346 | -0.7516923 | 0.45223613 | 0.89670027 |
| MLKL | 929.851004 | -0.4283966 | 0.4344328 | -0.9861056 | 0.3240813 | 0.85169085 |
| FA2H | 487.987769 | 0.3711085 | 0.39576966 | 0.93768811 | 0.34840472 | 0.86288527 |
| WDR59 | 3568.68218 | 0.03028328 | 0.17708376 | 0.17101102 | 0.8642151 | 0.98176973 |
| ZNRF1 | 1504.88021 | 0.08162185 | 0.19645794 | 0.41546731 | 0.67779984 | 0.95018034 |
| AC099508.1 | 294.591926 | 0.13312602 | 0.2440869 | 0.5454042 | 0.58547555 | 0.93289552 |
| LDHD | 4807.76455 | 0.77068657 | 0.46175336 | 1.66904376 | 0.09510871 | 0.65888357 |
| ZFP1 | 452.6765 | -0.0679952 | 0.2478423 | -0.2743487 | 0.7838167 | 0.96978847 |
| BCAR1 | 12249.0236 | -0.5576713 | 0.23992787 | -2.3243289 | 0.02010787 | 0.49780928 |
| CFDP1 | 3186.77518 | 0.26911326 | 0.2000047 | 1.34553465 | 0.17845264 | 0.76115956 |
| AC009163.4 | 789.232773 | 0.32291857 | 0.22664869 | 1.42475375 | 0.1542284 | 0.74258845 |
| AC009163.7 | 73.0480363 | 0.39715019 | 0.44829502 | 0.88591255 | 0.37566465 | 0.86889275 |
| TMEM170A | 1565.46231 | -0.314532 | 0.16168431 | -1.9453466 | 0.05173328 | 0.58645127 |
| CHST6 | 46.4524956 | 0.89535769 | 0.80549687 | 1.1115595 | 0.2663276 | 0.8265123 |
| AC009163.2 | 101.366683 | 0.49413054 | 0.28591388 | 1.72824958 | 0.08394349 | 0.64920344 |
| CHST5 | 41.3740526 | 0.41389391 | 0.38729895 | 1.06866779 | 0.28521939 | 0.83107779 |

| | | | | | | |
|---|---|---|---|---|---|---|
| TMEM231 | 715.73024 | 0.4450159 | 0.2967157 | 1.49980571 | 0.13366474 | 0.71505705 |
| GABARAPL2 | 4863.62545 | 0.14297709 | 0.18837186 | 0.75901511 | 0.44784352 | 0.89480207 |
| ADAT1 | 620.395185 | -0.0256449 | 0.17175981 | -0.1493069 | 0.88131145 | 0.98275231 |
| KARS | 5536.32777 | -0.0339874 | 0.12124099 | -0.2803292 | 0.77922496 | 0.96836854 |
| TERF2IP | 3545.82172 | 0.08578721 | 0.11650386 | 0.7363465 | 4.62E-01 | 8.98E-01 |
| MON1B | 2751.75127 | -0.1002529 | 0.23451802 | -0.4274849 | 0.66902616 | 0.9486589 |
| SYCE1L | 268.940569 | -0.04292 | 0.19647 | -0.2184558 | 8.27E-01 | 0.97548201 |
| AC009139.2 | 521.797669 | 0.37817859 | 0.28087973 | 1.34640754 | 0.17817112 | 0.76115956 |
| NUDT7 | 461.255975 | 0.79731302 | 0.35777848 | 2.2285103 | 0.02584651 | 0.52502381 |
| AC092134.1 | 369.099015 | 0.83764999 | 0.37536819 | 2.23154226 | 0.02564523 | 0.52469135 |
| VAT1L | 293.309794 | -0.0678164 | 0.66975942 | -0.1012548 | 0.91934818 | 0.9872458 |
| WWOX | 772.403728 | 0.17726486 | 0.17548249 | 1.010157 | 0.31242008 | 0.84541735 |
| MAF | 8958.51287 | 0.11852636 | 0.40844315 | 0.29019059 | 0.77167043 | 0.96836854 |
| AC009159.3 | 151.022166 | -0.0171218 | 0.4816813 | -0.0355459 | 0.97164446 | 0.99404213 |
| MAFTRR | 51.8621742 | 0.97615861 | 0.68988574 | 1.41495692 | 0.15708113 | 0.74522306 |
| DYNLRB2 | 91.6052276 | 0.25684912 | 0.41412117 | 0.62022696 | 0.53510838 | 0.92084006 |
| CDYL2 | 772.767081 | 0.04833141 | 0.19530205 | 0.24747008 | 0.80454445 | 0.97287947 |
| CMC2 | 1626.44607 | 0.05694578 | 0.1395994 | 0.40792282 | 6.83E-01 | 9.51E-01 |
| CENPN | 611.388239 | -0.1036922 | 0.20779412 | -0.4990139 | 0.61776957 | 0.93924932 |
| AC092718.2 | 47.9618473 | -0.3724461 | 0.43916439 | -0.8480789 | 0.39639402 | 0.87967259 |
| AC092718.8 | 1090.69333 | 0.71960834 | 0.36886529 | 1.95087031 | 0.05107247 | 0.5861488 |
| ATMIN | 2652.2963 | -0.2965449 | 0.2806386 | -1.0566791 | 0.29065809 | 0.83442894 |
| C16orf46 | 222.405486 | 0.2598797 | 0.18493596 | 1.40524157 | 0.15994946 | 0.74781837 |
| AC092718.3 | 1281.58634 | 0.62146358 | 0.30606238 | 2.03051282 | 0.04230444 | 0.56485966 |
| GCSH | 1165.29665 | 0.71258724 | 0.35651625 | 1.99875111 | 0.04563529 | 0.57670328 |
| BCO1 | 240.377788 | 0.48281227 | 0.45706118 | 1.05634057 | 0.29081265 | 0.83450005 |
| GAN | 534.8667 | -0.1662273 | 0.22710776 | -0.7319315 | 4.64E-01 | 8.98E-01 |
| CMIP | 4313.12415 | -0.0086418 | 0.16186382 | -0.0533894 | 9.57E-01 | 9.92E-01 |
| PLCG2 | 3957.62594 | -0.7541246 | 0.54759185 | -1.3771655 | NA | NA |
| SDR42E1 | 571.733533 | 0.15494638 | 0.1776445 | 0.87222733 | 0.38308437 | 0.8718674 |
| HSD17B2 | 107.40396 | 2.17099137 | 0.43767674 | 4.96026214 | 7.04E-07 | 0.00146592 |
| AC092142.1 | 47.0783379 | 1.95715522 | 0.54872928 | 3.56670453 | 0.0003615 | 0.08914272 |
| MPHOSPH6 | 663.376552 | 0.09697426 | 0.13698583 | 0.70791458 | 0.47899831 | 0.90465658 |
| CDH13 | 528.416373 | -0.1556401 | 0.59015451 | -0.2637277 | 0.79198976 | 0.97090793 |
| HSBP1 | 5884.30268 | -0.0625258 | 0.1335478 | -0.4681901 | 0.63964869 | 0.94088547 |
| MLYCD | 1570.11304 | 0.15055062 | 0.27954308 | 0.53855964 | 5.90E-01 | 9.33E-01 |
| AC009119.2 | 435.737205 | 0.20516556 | 0.28733721 | 0.71402362 | 0.47521258 | 0.90226631 |
| OSGIN1 | 4544.00761 | -1.1602659 | 0.83543924 | -1.3888095 | 0.1648907 | 0.75299828 |
| NECAB2 | 379.795478 | 0.96538901 | 0.71412007 | 1.35185811 | 0.17642071 | 0.76115956 |
| MBTPS1 | 5584.34697 | -0.0746574 | 0.10621163 | -0.7029116 | 0.48211082 | 0.90541935 |
| AC040169.1 | 46.2198172 | -0.5693623 | 0.42099437 | -1.3524227 | 0.17624015 | 0.76115956 |
| HSDL1 | 1294.04381 | -0.4371305 | 0.29702877 | -1.4716772 | 0.14110807 | 0.72711201 |
| DNAAF1 | 1408.59377 | -0.2313059 | 0.26974418 | -0.8575009 | 0.39116812 | 0.87710956 |
| TAF1C | 6694.38266 | -0.3508687 | 0.18516253 | -1.894923 | 0.05810261 | 0.60193532 |
| WFDC1 | 644.666017 | -0.0236094 | 0.51778122 | -0.0455972 | 0.96363132 | 0.99323942 |
| ATP2C2 | 558.746446 | -0.0177683 | 0.33552934 | -0.0529561 | 0.95776688 | 0.99223378 |
| ATP2C2-AS1 | 313.315972 | -0.0075459 | 0.37238556 | -0.0202637 | 0.98383303 | 0.9961464 |
| TLDC1 | 947.599533 | 0.05254637 | 0.22414647 | 0.23442872 | 8.15E-01 | 0.9738447 |
| AC022165.1 | 66.3232382 | -0.0443167 | 0.3388827 | -0.1307731 | 0.89595482 | 0.98474725 |
| COTL1 | 8230.13298 | -0.0477501 | 0.41436677 | -0.1152362 | 0.9082579 | 0.98662757 |
| KLHL36 | 3239.54087 | 0.21392641 | 0.20208157 | 1.05861419 | 2.90E-01 | 0.83399826 |
| USP10 | 3382.6612 | -0.2614431 | 0.18211987 | -1.4355553 | 0.15112892 | 0.73796431 |
| CRISPLD2 | 3378.43483 | 0.31187156 | 0.7226077 | 0.43159181 | 0.66603811 | 0.9475861 |
| ZDHHC7 | 4637.93612 | -0.3892304 | 0.15294115 | -2.5449686 | 0.01092876 | 0.40196898 |

| | | | | | | |
|---|---|---|---|---|---|---|
| KIAA0513 | 866.75863 | -0.1774635 | 0.23320325 | -0.760982 | 0.44666784 | 0.89480207 |
| LINC02139 | 44.2450834 | 0.54196302 | 0.64245099 | 0.84358657 | 0.39890047 | 0.88074856 |
| GSE1 | 4066.47927 | -0.0053821 | 0.16572182 | -0.0324766 | 0.97409194 | 0.99452354 |
| GINS2 | 50.8539436 | 0.77311529 | 0.57953361 | 1.33403012 | 0.18219398 | 0.76575408 |
| C16orf74 | 44.6585452 | 0.27892765 | 0.51898292 | 0.53745054 | 0.59095644 | 0.93289552 |
| EMC8 | 1292.72552 | -0.281683 | 0.21345239 | -1.3196528 | 0.18695097 | 0.76958556 |
| COX4I1 | 17076.8167 | 0.18887645 | 0.15706389 | 1.20254539 | 0.22915229 | 0.80521698 |
| IRF8 | 967.545156 | 0.38457109 | 0.52404309 | 0.73385395 | 0.46303775 | 0.89807153 |
| FOXF1 | 193.25863 | 0.59697967 | 0.53928246 | 1.10698885 | 0.26829875 | 0.82676682 |
| MTHFSD | 923.47595 | -0.0879232 | 0.22707998 | -0.3871903 | 0.6986153 | 0.9529028 |
| FOXC2-AS1 | 67.6428158 | -0.6789863 | 0.42118728 | -1.612077 | 0.10694519 | 0.67674128 |
| FOXC2 | 375.762413 | -0.6849662 | 0.3552557 | -1.9280935 | 0.0538435 | 0.59266174 |
| FOXL1 | 142.238619 | -0.442747 | 0.64573568 | -0.6856473 | 0.49293552 | 0.90705676 |
| LINC02188 | 78.7371877 | 0.39454713 | 0.50480595 | 0.78158178 | 0.4344604 | 0.89055338 |
| C16orf95 | 118.662897 | 0.15576087 | 0.33155401 | 0.46979035 | 6.39E-01 | 0.94088547 |
| AC010531.1 | 277.217059 | -0.0688813 | 0.27747047 | -0.2482474 | 8.04E-01 | 0.97274243 |
| FBXO31 | 2302.51751 | -0.3922657 | 0.20601273 | -1.9040848 | 5.69E-02 | 0.60049913 |
| AC010531.7 | 143.046732 | -0.7016062 | 0.29850101 | -2.3504315 | 0.01875166 | 0.48074536 |
| MAP1LC3B | 6862.86854 | -0.4770756 | 0.29544493 | -1.6147699 | 0.10636052 | 0.67638364 |
| ZCCHC14 | 3137.32183 | -0.551712 | 0.36648997 | -1.5053945 | 0.13222268 | 0.71267911 |
| KLHDC4 | 2304.10219 | -0.0287554 | 0.13660972 | -0.2104934 | 0.83328259 | 0.97550416 |
| SLC7A5 | 2396.63543 | -0.7098729 | 0.41705324 | -1.7021159 | 0.08873365 | 0.65363785 |
| AC126696.2 | 168.417072 | -0.5798262 | 0.44510884 | -1.3026616 | 0.19269032 | 0.77495378 |
| BANP | 876.679869 | -0.417354 | 0.18841757 | -2.2150484 | 0.02675674 | 0.52678638 |
| ZNF469 | 62.566447 | -0.2635765 | 0.3967476 | -0.6643431 | 0.50647072 | 0.9141539 |
| ZFPM1 | 306.038306 | -0.2761397 | 0.23328134 | -1.1837198 | 0.236524 | 0.8084436 |
| ZC3H18 | 2652.11467 | -0.2937621 | 0.1466264 | -2.0034737 | 0.04512647 | 0.57414472 |
| CYBA | 8364.84436 | -0.2396581 | 0.30878968 | -0.776121 | 0.43767757 | 0.89130865 |
| MVD | 2940.60586 | 0.10010137 | 0.19071517 | 0.52487369 | 0.599671 | 0.93487648 |
| SNAI3-AS1 | 185.957926 | 0.29872616 | 0.25042743 | 1.19286518 | 0.23292218 | 0.80667958 |
| SNAI3 | 78.1260895 | 0.1308383 | 0.49721718 | 0.26314114 | 0.7924418 | 0.97090793 |
| RNF166 | 1551.90113 | -0.2059775 | 0.28335932 | -0.7269129 | 0.46727933 | 0.89983399 |
| AC138028.3 | 315.805316 | -0.2160099 | 0.27185594 | -0.7945748 | 0.42686086 | 0.88709241 |
| CTU2 | 772.914777 | -0.1940377 | 0.15863298 | -1.2231861 | 0.22125941 | 0.80004295 |
| AC138028.6 | 153.259289 | -0.1272265 | 0.24095869 | -0.5280012 | 0.59749849 | 0.93416444 |
| PIEZO1 | 7708.72526 | -0.2456411 | 0.19001044 | -1.2927766 | 0.19608834 | 0.77791948 |
| AC138028.4 | 1126.84827 | -0.2101064 | 0.20629886 | -1.0184566 | 0.30846101 | 0.84388103 |
| AC138028.2 | 298.838449 | -0.3583469 | 0.39866105 | -0.898876 | 0.3687187 | 0.86762749 |
| AC138028.5 | 86.1272822 | -0.3170662 | 0.46979146 | -0.6749084 | 0.49973397 | 0.91175178 |
| CDT1 | 83.0205645 | 0.00278447 | 0.38613502 | 0.00721114 | 0.9942464 | 0.9993656 |
| APRT | 4053.06847 | 0.14648918 | 0.111144 | 1.31801246 | 0.18749948 | 0.76958556 |
| GALNS | 2029.91027 | -0.1964584 | 0.14636215 | -1.3422758 | 0.17950659 | 0.76215064 |
| TRAPPC2L | 1867.91954 | 0.02434312 | 0.1875633 | 0.12978618 | 0.8967356 | 0.98474725 |
| CBFA2T3 | 167.849219 | 0.08183026 | 0.63020993 | 0.12984603 | 0.89668825 | 0.98474725 |
| AC092384.2 | 47.6007091 | 0.02142861 | 0.62793538 | 0.03412551 | 0.97277707 | 0.99426347 |
| ACSF3 | 1989.72144 | -0.1063362 | 0.20330244 | -0.5230446 | 0.60094319 | 0.93509435 |
| AC135782.3 | 249.667235 | 0.29625407 | 0.2535546 | 1.16840345 | 0.24264407 | 0.81335097 |
| LINC00304 | 44.1162309 | -1.0819562 | 0.52651248 | -2.0549488 | 0.03988395 | 0.55657615 |
| SLC22A31 | 43.8020144 | -1.2985932 | 0.85422075 | -1.5202079 | 0.12845872 | 0.70610612 |
| AC009113.1 | 230.222957 | 0.22424568 | 0.36691162 | 0.61117083 | 0.54108649 | 0.92292765 |
| ZNF778 | 608.351503 | -0.3087406 | 0.26982477 | -1.1442263 | 0.25252978 | 0.81833349 |
| ANKRD11 | 4921.81975 | -0.284566 | 0.14522046 | -1.9595446 | 0.05004904 | 0.5861488 |
| AC137932.3 | 484.229405 | -0.4290452 | 0.19091104 | -2.2473565 | 0.02461725 | 0.52017724 |
| AC137932.1 | 43.6256253 | 0.17658205 | 0.4789998 | 0.36864745 | 0.71239053 | 0.95560193 |

| | | | | | | |
|---|---|---|---|---|---|---|
| SPG7 | 7955.34377 | -0.3274406 | 0.18424001 | -1.77725 | 0.07552712 | 0.63773415 |
| RPL13 | 35138.9301 | 0.16686696 | 0.18232904 | 0.91519686 | 0.36008829 | 0.86480228 |
| CPNE7 | 830.916451 | 0.45038812 | 0.64245326 | 0.70104419 | 0.48327544 | 0.90541935 |
| DPEP1 | 18159.4079 | -0.2727584 | 0.58911584 | -0.4629962 | 6.43E-01 | 9.42E-01 |
| CHMP1A | 5581.99499 | -0.2416477 | 0.11179247 | -2.1615741 | 3.07E-02 | 5.44E-01 |
| SPATA33 | 618.510929 | -0.0499166 | 0.12357484 | -0.4039382 | 0.68625813 | 0.95097288 |
| CDK10 | 7974.76937 | -0.0746072 | 0.2149021 | -0.3471684 | 0.72846482 | 0.95848028 |
| LINC02166 | 158.970754 | 0.38791982 | 0.39105512 | 0.99198247 | 0.32120608 | 0.85119105 |
| AC010538.1 | 76.2182949 | 0.50075037 | 0.390769 | 1.28144855 | 0.20003616 | 0.78134034 |
| SPATA2L | 1000.78156 | -0.2323156 | 0.22390802 | -1.0375492 | 0.29947997 | 0.84123067 |
| VPS9D1 | 2872.13492 | -0.421037 | 0.12483889 | -3.3726433 | 0.0007445 | 0.12895512 |
| VPS9D1-AS1 | 958.195187 | -0.3780773 | 0.17802865 | -2.1236878 | 3.37E-02 | 5.54E-01 |
| ZNF276 | 2877.10098 | -0.2640516 | 0.154626 | -1.7076794 | 0.08769584 | 0.65322246 |
| FANCA | 1844.86901 | -0.2692666 | 0.16277928 | -1.6541824 | 0.09809046 | 0.66549286 |
| SPIRE2 | 475.727131 | -0.0743444 | 0.36263267 | -0.2050128 | 0.83756209 | 0.97577999 |
| TCF25 | 7859.52272 | -0.055784 | 0.12828991 | -0.4348274 | 0.66368773 | 0.94642611 |
| MC1R | 1099.42914 | -0.2560157 | 0.3595428 | -0.7120592 | 4.76E-01 | 0.90309179 |
| AC092143.1 | 1070.60233 | -0.3226384 | 0.4542994 | -0.7101889 | 0.47758698 | 0.90387017 |
| AC092143.2 | 413.38617 | -0.1131154 | 0.33087298 | -0.3418694 | 0.73244916 | 0.95848028 |
| TUBB3 | 583.55184 | -0.5080626 | 0.70657294 | -0.7190518 | 0.47210898 | 0.90160919 |
| DEF8 | 4233.5032 | -0.1817572 | 0.12554061 | -1.4477961 | 0.14767408 | 0.73443345 |
| CENPBD1 | 500.66431 | -0.0978043 | 0.25860429 | -0.3782005 | 0.70528162 | 0.95501548 |
| DBNDD1 | 1598.53077 | 0.00664099 | 0.24598387 | 0.02699768 | 0.97846158 | 0.99476647 |
| GAS8 | 3315.87906 | -0.0015565 | 0.18398117 | -0.0084601 | 0.9932499 | 0.99906069 |
| FAM157C | 56.982182 | 0.03893135 | 0.49061156 | 0.0793527 | 0.93675209 | 0.99068685 |
| AC133919.2 | 96.4452595 | -0.1433922 | 1.11502715 | -0.1285998 | 0.89767435 | 0.98474725 |
| AC240565.1 | 40.3078774 | 0.28127287 | 0.67645867 | 0.415802 | 0.6775549 | 0.95018034 |
| DOC2B | 1574.61877 | -0.1577001 | 0.35764282 | -0.4409429 | 0.65925435 | 0.94601668 |
| RPH3AL | 1427.91834 | -0.0108024 | 0.22747418 | -0.0474884 | 0.96212397 | 0.99320711 |
| AC129507.4 | 137.966167 | 0.63681919 | 0.30595961 | 2.08138319 | 0.03739885 | 0.55532957 |
| AC129507.2 | 222.262768 | 0.50341035 | 0.44037156 | 1.14314909 | 0.2529767 | 0.81835388 |
| AC129507.3 | 96.7568306 | 0.75974459 | 0.59691627 | 1.27278251 | 0.20309523 | 0.78252625 |
| AC141424.1 | 213.798044 | -0.6126538 | 0.3434411 | -1.7838685 | 0.07444503 | 0.63773415 |
| C17orf97 | 446.842781 | -0.5899777 | 0.33886614 | -1.7410345 | 0.08167753 | 0.64505626 |
| RFLNB | 981.647428 | 0.0323411 | 0.58186592 | 0.0555817 | 0.95567504 | 0.99185043 |
| VPS53 | 2040.04374 | -0.0774548 | 0.15006205 | -0.5161521 | 0.6057482 | 0.93698713 |
| FAM57A | 980.007828 | -0.5670355 | 0.27179643 | -2.0862507 | 0.03695592 | 0.55521179 |
| GEMIN4 | 821.456771 | -0.2390997 | 0.19020529 | -1.2570612 | 0.20873146 | 0.79042964 |
| GLOD4 | 3911.72772 | 0.18002831 | 0.18136109 | 0.99265127 | 0.32087993 | 0.85106295 |
| MRM3 | 1002.00627 | -0.0977317 | 0.21294929 | -0.4589433 | 0.64627485 | 0.94208859 |
| NXN | 990.728677 | 0.1015022 | 0.39163995 | 0.25917223 | 7.96E-01 | 0.97101097 |
| TIMM22 | 1372.96286 | -0.1241979 | 0.16918337 | -0.734102 | 0.46288655 | 0.89807153 |
| ABR | 3686.60607 | 0.5356296 | 0.32225045 | 1.66215311 | 0.09648208 | 0.66233356 |
| TRARG1 | 67.583027 | 4.56770896 | 2.3346505 | 1.95648512 | NA | NA |
| YWHAE | 19875.1848 | -0.0062644 | 0.14298952 | -0.0438101 | 0.96505578 | 0.99357855 |
| CRK | 3868.27369 | -0.2555051 | 0.2848509 | -0.8969783 | 0.36973049 | 0.86762749 |
| MYO1C | 10135.3588 | -0.0398681 | 0.18314435 | -0.2176867 | 8.28E-01 | 9.75E-01 |
| INPP5K | 2146.01099 | -0.1993647 | 0.16989561 | -1.1734539 | 0.24061381 | 0.8124487 |
| PITPNA-AS1 | 371.049398 | 0.09702845 | 0.20934049 | 0.46349585 | 0.64300899 | 0.9413085 |
| PITPNA | 4840.55165 | 0.09208197 | 0.15612168 | 0.58980898 | 5.55E-01 | 0.92373235 |
| SLC43A2 | 7039.27012 | 0.45743289 | 0.35408062 | 1.29188908 | 0.19639556 | 0.77798606 |
| SCARF1 | 1826.14322 | -0.183521 | 0.22290999 | -0.8232965 | 0.41033942 | 0.88249519 |
| RILP | 1051.2293 | -0.1311266 | 0.30469377 | -0.4303554 | 0.6669371 | 0.94830184 |
| PRPF8 | 9159.71872 | 0.0912991 | 0.12915242 | 0.70690977 | 0.47962255 | 0.9051283 |

| | | | | | | |
|---|---|---|---|---|---|---|
| TLCD2 | 391.129941 | -0.2433534 | 0.14568635 | -1.6703925 | 0.09484173 | 0.65888357 |
| MIR22HG | 3794.5123 | -1.1377737 | 0.65273334 | -1.7430912 | 0.08131769 | 0.6439041 |
| WDR81 | 3598.58748 | -0.5143331 | 0.33474398 | -1.536497 | 0.12441652 | 0.70104934 |
| AC130343.1 | 500.195755 | -0.4283492 | 0.31871122 | -1.3440041 | 0.17894706 | 0.76120583 |
| SERPINF2 | 12478.8107 | 0.3180037 | 0.46844363 | 0.67885158 | 0.49723191 | 0.90913397 |
| SERPINF1 | 2687.52671 | 0.31685099 | 0.4625082 | 0.68507109 | 0.49329906 | 0.90715187 |
| SMYD4 | 555.609965 | -0.1097299 | 0.21794957 | -0.5034644 | 0.61463777 | 0.93895494 |
| RPA1 | 2409.02504 | 0.10326481 | 0.13541904 | 0.76255752 | 0.44572733 | 0.89474891 |
| RTN4RL1 | 231.486742 | 0.82502369 | 0.46817863 | 1.76219852 | 0.07803576 | 0.64079926 |
| AC099684.2 | 115.869184 | 0.55059463 | 0.62368116 | 0.88281427 | 0.37733663 | 0.86987124 |
| AC099684.1 | 47.6831314 | 1.3209778 | 0.67687325 | 1.95158813 | 0.05098712 | 0.5861488 |
| DPH1 | 4895.8624 | -0.1682847 | 0.12254137 | -1.373289 | 0.16966254 | 0.7586159 |
| OVCA2 | 2919.91783 | -0.2338099 | 0.11533218 | -2.0272741 | 0.04263439 | 0.56547141 |
| AC090617.3 | 1940.41942 | -0.2559266 | 0.13074974 | -1.9573776 | 0.05030309 | 0.5861488 |
| AC090617.5 | 79.6925849 | 0.36282847 | 0.36629466 | 0.99053718 | 0.32191163 | 0.85119105 |
| HIC1 | 3408.40349 | -0.2275785 | 0.30959525 | -0.7350838 | 0.46228844 | 0.89776981 |
| SMG6 | 2948.95851 | -0.2853249 | 0.17788853 | -1.6039532 | 0.10872437 | 0.67800851 |
| SRR | 1564.57199 | -0.2480959 | 0.17530535 | -1.4152213 | 0.15700363 | 0.74522306 |
| TSR1 | 2310.74736 | -0.2457756 | 0.2058272 | -1.1940872 | 0.23244385 | 0.80649152 |
| SGSM2 | 5866.35756 | -0.2019205 | 0.16693973 | -1.2095415 | 0.22645488 | 0.80257013 |
| AC006435.2 | 1381.26938 | -0.2339308 | 0.17955408 | -1.3028433 | 0.19262826 | 0.77495378 |
| MNT | 2456.12813 | -0.4105125 | 0.16376339 | -2.5067414 | 0.01218498 | 0.41257332 |
| AC006435.1 | 809.672387 | -0.3962273 | 0.17861726 | -2.2183037 | 0.02653413 | 0.52678638 |
| METTL16 | 1233.56184 | -0.0949839 | 0.14263779 | -0.6659094 | 0.50546898 | 0.91368371 |
| PAFAH1B1 | 6938.75401 | -0.1256275 | 0.12234906 | -1.0267958 | 0.30451661 | 0.84211658 |
| AC005696.2 | 1470.26582 | -0.0004531 | 0.11967941 | -0.003786 | 0.99697925 | 0.99976751 |
| CLUH | 15591.1913 | -0.3940303 | 0.24123009 | -1.6334212 | 1.02E-01 | 0.67279543 |
| AC005696.1 | 189.982821 | -0.2996494 | 0.31915826 | -0.9388741 | 0.34779538 | 0.86225576 |
| RAP1GAP2 | 1741.61568 | 0.00190702 | 0.22175556 | 0.00859966 | 0.99313855 | 0.99906069 |
| ASPA | 2283.57061 | 0.6255499 | 0.7048356 | 0.88751179 | 0.37480342 | 0.86791605 |
| TRPV3 | 49.1210449 | 0.4359539 | 0.36919038 | 1.18083763 | 0.23766723 | 0.80962787 |
| TRPV1 | 1443.36333 | 0.1459674 | 0.27329987 | 0.53409249 | 0.59327755 | 0.93289552 |
| AC027796.3 | 1404.58844 | 0.0937083 | 0.28507574 | 0.32871371 | 0.74237209 | 0.96109001 |
| SHPK | 1560.29299 | 0.09492213 | 0.25235842 | 0.37614014 | 0.70681271 | 0.95501548 |
| CTNS | 2544.51441 | -0.4850585 | 0.29746541 | -1.6306382 | 0.10296668 | 0.67424825 |
| AC027796.4 | 332.15381 | -0.4275338 | 0.23608645 | -1.8109204 | 0.07015317 | 0.62928077 |
| TAX1BP3 | 6886.00223 | -0.0882906 | 0.26025543 | -0.339246 | 0.7344244 | 0.95895327 |
| P2RX5-TAX1I | 7852.19431 | -0.0989623 | 0.24485728 | -0.4041632 | 0.68609272 | 0.95097288 |
| EMC6 | 1012.67851 | -0.1679004 | 0.21392244 | -0.7848657 | 0.43253233 | 0.88980461 |
| P2RX5 | 129.776108 | -1.1960755 | 0.7753323 | -1.5426617 | 0.12291287 | 0.69788192 |
| ITGAE | 767.533034 | 0.05487991 | 0.24007651 | 0.2285934 | 0.81918495 | 0.97454841 |
| AC116914.2 | 40.6950649 | -0.1236267 | 0.37387915 | -0.3306595 | 0.74090166 | 0.96089108 |
| NCBP3 | 2446.37991 | -0.0984111 | 0.10825888 | -0.9090348 | 0.36333175 | 0.8663515 |
| CAMKK1 | 457.714614 | 0.00258499 | 0.30555677 | 0.00845994 | 9.93E-01 | 9.99E-01 |
| P2RX1 | 300.126997 | -0.4142635 | 0.74640728 | -0.55501 | 0.57888784 | 0.93139743 |
| ATP2A3 | 1469.52135 | 0.10767935 | 0.42725321 | 0.25202703 | 0.80102017 | 0.97210205 |
| ZZEF1 | 2824.73231 | -0.2679796 | 0.10265713 | -2.6104334 | 0.00904276 | 0.38593543 |
| CYB5D2 | 1823.23713 | 0.09077755 | 0.20477686 | 0.44329986 | 0.65754886 | 0.94558535 |
| ANKFY1 | 3337.55251 | -0.2912811 | 0.244784 | -1.1899516 | 0.2340654 | 0.80725541 |
| AC087292.1 | 1483.16473 | -0.264438 | 0.26656179 | -0.9920327 | 0.32118158 | 0.85119105 |
| UBE2G1 | 2459.65045 | -0.1708763 | 0.19571164 | -0.8731023 | 0.38260731 | 0.8718674 |
| SPNS3 | 477.996786 | 0.36869749 | 0.34969398 | 1.05434325 | 0.29172579 | 0.83460625 |
| AC127521.1 | 221.457866 | 0.41815213 | 0.379717 | 1.10122046 | 0.27080072 | 0.82716979 |
| AC118754.1 | 65.1809697 | 0.43502929 | 0.43380388 | 1.0028248 | 0.3159454 | 0.84750116 |

| | | | | | | |
|---|---|---|---|---|---|---|
| SPNS2 | 10932.5148 | 0.125724 | 0.25031665 | 0.50225983 | 0.61548476 | 0.93910158 |
| MYBBP1A | 3685.99502 | -0.1467071 | 0.17068586 | -0.8595151 | 0.39005641 | 0.87645794 |
| GGT6 | 3331.1812 | 0.64782548 | 0.26553657 | 2.43968455 | 0.01470009 | 0.44149748 |
| SMTNL2 | 3207.70564 | -0.3572454 | 0.32352675 | -1.1042222 | 0.26949678 | 0.82716979 |
| PELP1 | 3315.42483 | -0.1112247 | 0.12867544 | -0.864382 | 0.38737808 | 0.87460898 |
| AC091153.2 | 54.8878302 | -0.4193383 | 0.33517526 | -1.2511016 | 0.2108974 | 0.79134644 |
| ARRB2 | 2268.52571 | 0.02241185 | 0.29135137 | 0.07692379 | 0.93868418 | 0.99113899 |
| MED11 | 1150.4535 | 0.10333953 | 0.12696013 | 0.81395259 | 0.41567211 | 0.88399703 |
| AC091153.4 | 489.334928 | 0.18920578 | 0.17105429 | 1.10611534 | 0.2686766 | 0.82676682 |
| CXCL16 | 3241.62974 | 0.01815209 | 0.25971897 | 0.06989127 | 0.9442802 | 0.99136103 |
| ZMYND15 | 354.877882 | -0.1460994 | 0.54873478 | -0.2662477 | 7.90E-01 | 9.71E-01 |
| TM4SF5 | 612.765072 | 0.24564333 | 0.48458233 | 0.50691763 | 0.61221261 | 0.93860247 |
| VMO1 | 179.642934 | -0.6673012 | 0.54934294 | -1.214726 | 0.22447061 | 0.80078694 |
| GLTPD2 | 862.443673 | 0.89317586 | 0.59820639 | 1.49308978 | 0.13541369 | 0.71649038 |
| PSMB6 | 5067.63121 | 0.05774174 | 0.16786979 | 0.34396743 | 0.73087078 | 0.95848028 |
| PLD2 | 1666.04004 | 0.18808166 | 0.25343654 | 0.74212529 | 0.45801143 | 0.8971553 |
| MINK1 | 10924.4881 | -0.1558397 | 0.16784192 | -0.928491 | 0.35315291 | 0.86364633 |
| CHRNE | 1117.6858 | -0.1100283 | 0.1703185 | -0.6460147 | 0.51826982 | 0.91636259 |
| C17orf107 | 247.144949 | 0.18459949 | 0.28494378 | 0.64784532 | 0.517085 | 0.91636259 |
| GP1BA | 74.698994 | -0.1590104 | 0.4596228 | -0.3459585 | 0.72937391 | 0.95848028 |
| SLC25A11 | 4772.37081 | 0.43347988 | 0.2067579 | 2.09655778 | 0.03603274 | 0.55369025 |
| RNF167 | 5607.66741 | -0.0261666 | 0.10327373 | -0.2533709 | 0.79998158 | 0.97208422 |
| PFN1 | 23009.3136 | -0.2318984 | 0.22562494 | -1.0278047 | 0.30404172 | 0.84211658 |
| ENO3 | 10354.5456 | -0.1788214 | 0.21699604 | -0.8240769 | 0.40989588 | 0.88249519 |
| SPAG7 | 3110.29094 | 0.06155028 | 0.17826733 | 0.34526952 | 0.72989176 | 0.95848028 |
| CAMTA2 | 5261.73742 | -0.3349259 | 0.17524381 | -1.9111997 | 0.05597892 | 0.59889467 |
| AC004771.4 | 1338.23693 | -0.4713799 | 0.18971277 | -2.4847033 | 0.01296594 | 0.41899224 |
| AC004771.2 | 756.331164 | -0.2151757 | 0.20147246 | -1.0680156 | 0.28551348 | 0.83107779 |
| AC004771.5 | 41.191757 | -0.4106164 | 0.43395822 | -0.9462119 | 0.34404053 | 0.85961157 |
| AC004771.3 | 59.8475517 | -0.2077943 | 0.33175216 | -0.6263541 | 0.53108273 | 0.91962754 |
| INCA1 | 263.572286 | -0.1698909 | 0.20252497 | -0.8388638 | 0.40154575 | 0.88132015 |
| KIF1C | 14445.4292 | -0.0488324 | 0.21921694 | -0.2227582 | 0.82372368 | 0.97548201 |
| AC004771.1 | 1228.99341 | 0.03645512 | 0.22293904 | 0.16352058 | 0.87010857 | 0.98275231 |
| ZFP3 | 630.698757 | -0.2830798 | 0.1990176 | -1.4223855 | 0.15491436 | 0.74373109 |
| ZNF232 | 639.639859 | -0.3034839 | 0.62675224 | -0.4842167 | NA | NA |
| USP6 | 207.209673 | -0.1275035 | 0.77096765 | -0.1653811 | NA | NA |
| ZNF594 | 357.44152 | -0.215867 | 0.26227411 | -0.8230589 | 0.4104745 | 0.88252101 |
| AC087500.1 | 302.953703 | -0.1468878 | 0.32362251 | -0.4538861 | 0.64991078 | 0.94312988 |
| SCIMP | 279.611055 | -0.5205 | 0.33103686 | -1.5723325 | 0.11587345 | 0.6873836 |
| RABEP1 | 2936.56298 | -0.1638242 | 0.16562671 | -0.9891173 | 3.23E-01 | 0.85119105 |
| NUP88 | 2110.50192 | -0.3193861 | 0.18131907 | -1.7614591 | 0.07816073 | 0.64079926 |
| RPAIN | 2105.37246 | 0.01083504 | 0.13077205 | 0.08285442 | 0.93396729 | 0.99025262 |
| AC004148.2 | 1480.25866 | 0.04386839 | 0.17805816 | 0.24637113 | 0.80539495 | 0.97317374 |
| C1QBP | 3994.67208 | 0.16084195 | 0.19670169 | 0.81769481 | 0.41353148 | 0.88316314 |
| DHX33 | 836.520652 | -0.1857342 | 0.18887018 | -0.9833962 | 0.32541247 | 0.85244476 |
| DERL2 | 2328.36563 | -0.0170947 | 0.14175522 | -0.120593 | 0.90401339 | 0.98624484 |
| MIS12 | 1066.73469 | 0.06193338 | 0.19179049 | 0.32292208 | 7.47E-01 | 0.96218092 |
| NLRP1 | 1257.10961 | 0.10593551 | 0.37028374 | 0.28609279 | 0.77480705 | 0.96836854 |
| AC007846.2 | 42.7679941 | -0.5679052 | 0.48662049 | -1.1670392 | 0.24319453 | 0.81344077 |
| WSCD1 | 903.644143 | -0.4206947 | 0.37376647 | -1.125555 | 0.26035392 | 0.82424998 |
| PIMREG | 58.9098651 | 0.24045581 | 0.45289594 | 0.53092949 | 0.59546764 | 0.9334601 |
| PITPNM3 | 419.281141 | 0.19459107 | 0.25646029 | 0.75875713 | 0.44799785 | 0.89480207 |
| KIAA0753 | 1096.99898 | -0.1166 | 0.21557749 | -0.5408728 | 0.58859528 | 0.93289552 |
| TXNDC17 | 2552.80366 | 0.23095605 | 0.21061217 | 1.09659407 | 0.27281888 | 0.82716979 |

| | | | | | | |
|---|---|---|---|---|---|---|
| MED31 | 562.072729 | -0.0875233 | 0.13143549 | -0.6659029 | 0.50547317 | 0.91368371 |
| C17orf100 | 161.031389 | 0.08014313 | 0.34655097 | 0.23125928 | 0.81711338 | 0.97408065 |
| XAF1 | 4141.10837 | -2.7294027 | 1.08168221 | -2.5232944 | NA | NA |
| ALOX12-AS1 | 308.614792 | 0.25360746 | 0.26231267 | 0.96681358 | 0.33363723 | 0.855011 |
| AC040977.2 | 864.859826 | 0.14145529 | 0.19403582 | 0.72901637 | 0.46599165 | 0.89960479 |
| AC040977.1 | 164.567595 | 0.04450642 | 0.25905092 | 0.17180569 | 0.86359029 | 0.98176973 |
| ALOX12 | 291.745263 | -0.0086593 | 0.231704 | -0.0373721 | 0.9701883 | 0.99400331 |
| RNASEK | 8533.42622 | 0.04508847 | 0.21631407 | 0.20843984 | 0.83488555 | 0.97550416 |
| RNASEK-C17 | 9798.91273 | 0.07103585 | 0.18473936 | 0.3845193 | 0.7005936 | 0.95360275 |
| C17orf49 | 2437.77207 | 0.20137512 | 0.25393994 | 0.79300294 | 0.42777611 | 0.88791007 |
| MIR497HG | 1535.52085 | 0.1096887 | 0.23082806 | 0.47519656 | 0.63464688 | 0.94088547 |
| BCL6B | 1089.12319 | -0.6044272 | 0.3926788 | -1.5392408 | 0.12374551 | 0.69957382 |
| SLC16A13 | 916.06021 | 0.66406357 | 0.60654195 | 1.09483537 | 0.27358877 | 0.82719197 |
| SLC16A11 | 220.831147 | 0.04479908 | 0.52055755 | 0.08605981 | 0.93141887 | 0.98975562 |
| CLEC10A | 253.046228 | -0.1497949 | 0.87453621 | -0.171285 | 0.86399968 | 0.98176973 |
| ASGR2 | 157.962978 | -0.2200196 | 0.48713516 | -0.4516603 | 0.65151372 | 0.94393131 |
| ASGR1 | 140.297403 | 0.06919078 | 0.47013461 | 0.14717227 | 0.88299605 | 0.98293314 |
| DLG4 | 1650.54264 | -0.1371073 | 0.42533869 | -0.3223484 | 0.74718874 | 0.96224616 |
| ACADVL | 57954.5721 | -0.1485077 | 0.32381488 | -0.4586191 | 0.64650773 | 0.94208859 |
| DVL2 | 2754.69562 | -0.2796532 | 0.16231759 | -1.7228767 | 8.49E-02 | 0.64988556 |
| PHF23 | 2527.4103 | -0.3715625 | 0.2267363 | -1.6387431 | 0.10126678 | 0.6725344 |
| GABARAP | 13446.0232 | 0.16010772 | 0.12536228 | 1.27716022 | 0.20154569 | 0.78134034 |
| AC120057.2 | 14495.9552 | 0.13995143 | 0.12025469 | 1.16379183 | 0.24450838 | 0.81473945 |
| CTDNEP1 | 5960.86045 | -0.2019478 | 0.13255391 | -1.5235145 | 0.12763003 | 0.70557946 |
| AC003688.1 | 3930.19676 | -0.055601 | 0.1510807 | -0.3680221 | 0.71285673 | 0.95564093 |
| ELP5 | 2498.83774 | -0.0872925 | 0.10612285 | -0.8225611 | 0.41075764 | 0.88268208 |
| CLDN7 | 6925.43396 | 0.05135617 | 0.19446624 | 0.26408782 | 0.79171225 | 0.97090793 |
| SLC2A4 | 329.340209 | 0.49181465 | 0.46624386 | 1.05484424 | 0.29149657 | 0.83454586 |
| AC003688.2 | 45.3413716 | 0.38652103 | 0.59068073 | 0.65436538 | 0.51287643 | 0.91568064 |
| YBX2 | 307.049135 | 0.52173001 | 0.40248133 | 1.29628376 | 0.19487774 | 0.77656894 |
| EIF5A | 13978.131 | -0.2153884 | 0.20665943 | -1.0422385 | 0.29730114 | 0.83898658 |
| GPS2 | 9787.58139 | -0.2000229 | 0.18622935 | -1.0740678 | 0.28279231 | 0.83076616 |
| AC026954.2 | 5682.30659 | -0.2460081 | 0.12322794 | -1.9963659 | 0.04589411 | 0.57687945 |
| NEURL4 | 3442.53253 | -0.1747326 | 0.11140798 | -1.5684027 | 0.11678718 | 0.68913995 |
| ACAP1 | 864.693304 | -0.1567048 | 0.5423818 | -0.2889197 | 0.77264284 | 0.96836854 |
| KCTD11 | 1105.25477 | -0.2707971 | 0.15226501 | -1.7784592 | 0.07532847 | 0.63773415 |
| AC026954.3 | 276.761955 | -0.2206199 | 0.18726794 | -1.1780976 | 0.2387577 | 0.81104912 |
| TNK1 | 975.37067 | -0.0992752 | 0.14701875 | -0.6752553 | 0.49951356 | 0.91161831 |
| PLSCR3 | 1706.9913 | 0.02250417 | 0.24510008 | 0.09181624 | 0.92684404 | 0.98915637 |
| TMEM256-PI | 3245.82235 | 0.25277663 | 0.21453105 | 1.17827529 | 0.23868688 | 0.81095554 |
| TMEM256 | 1802.42511 | 0.52799017 | 0.32045505 | 1.64762628 | 0.09942938 | 0.66980808 |
| NLGN2 | 3118.2987 | -0.5135273 | 0.37226892 | -1.3794525 | 0.16775529 | 0.7558879 |
| TMEM102 | 563.439854 | -0.0467934 | 0.23001612 | -0.2034355 | 0.83879467 | 0.97612168 |
| AC113189.1 | 147.484715 | -0.0753076 | 0.299537 | -0.2514133 | 0.80149458 | 0.97210205 |
| FGF11 | 684.330163 | -0.197423 | 0.51216055 | -0.385471 | 0.69988851 | 0.95328923 |
| AC113189.3 | 85.3960668 | -0.3817615 | 0.5094658 | -0.7493369 | 0.45365418 | 0.89670027 |
| CHRNB1 | 792.738986 | -0.0570608 | 0.27511646 | -0.207406 | 0.83569283 | 0.97550416 |
| ZBTB4 | 6666.99934 | 0.07286403 | 0.18713232 | 0.38937169 | 0.69700121 | 0.95263271 |
| POLR2A | 8198.48101 | -0.0892049 | 0.20951 | -0.4257786 | 0.67026916 | 0.949185 |
| TNFSF12 | 1946.26954 | 0.28563503 | 0.2934087 | 0.97350565 | 0.33030205 | 0.85384238 |
| TNFSF12-TN | 5355.98933 | 0.03531133 | 0.3034632 | 0.11636115 | 0.90736633 | 0.98662757 |
| AC016876.3 | 171.767703 | 0.2963905 | 0.3048834 | 0.97214376 | 0.33097904 | 0.85384238 |
| TNFSF13 | 5585.91054 | 0.00932508 | 0.31113863 | 0.02997081 | 0.97609033 | 0.99455099 |
| SENP3 | 2808.57759 | -0.157544 | 0.13890617 | -1.1341758 | 0.25672081 | 0.82101614 |

| | | | | | | |
|---|---|---|---|---|---|---|
| SENP3-EIF4A | 26348.08 | -0.1887201 | 0.23913821 | -0.7891673 | 0.43001422 | 0.88901231 |
| EIF4A1 | 25547.1041 | -0.1951933 | 0.25153051 | -0.7760225 | 0.4377357 | 0.89130865 |
| CD68 | 7320.47798 | -0.4526516 | 0.39178927 | -1.1553447 | 0.24794931 | 0.81602355 |
| AC016876.1 | 1195.76994 | -0.3246663 | 0.23435327 | -1.3853713 | 0.16593896 | 0.75515387 |
| MPDU1 | 5124.86676 | -0.1111338 | 0.1782937 | -0.6233186 | 0.53307515 | 0.92026174 |
| SOX15 | 90.2190403 | -0.3799122 | 0.3627576 | -1.0472893 | 0.29496617 | 0.83630273 |
| FXR2 | 3534.60067 | -0.2740908 | 0.16993183 | -1.6129458 | 0.10675628 | 0.67674128 |
| SHBG | 180.937514 | 0.48671255 | 0.66806827 | 0.72853715 | 0.46628484 | 0.89960479 |
| SAT2 | 9396.81755 | 0.34409808 | 0.39685324 | 0.86706633 | 0.38590567 | 0.87420019 |
| ATP1B2 | 674.007015 | 0.52940891 | 0.42317447 | 1.25104172 | 0.21091926 | 0.79134644 |
| TP53 | 1601.12368 | 0.36589023 | 0.30448709 | 1.2016609 | 0.22949493 | 0.80551898 |
| WRAP53 | 811.360428 | -0.2380184 | 0.20574035 | -1.1568871 | 0.24731849 | 0.81566357 |
| EFNB3 | 146.664931 | 0.58686805 | 0.76714958 | 0.7649982 | 0.44427262 | 0.8938278 |
| KDM6B | 4902.33758 | -0.6524694 | 0.31389967 | -2.0785921 | 0.03765486 | 0.55538555 |
| TMEM88 | 519.884356 | -0.25899 | 0.40803274 | -0.6347286 | 0.52560545 | 0.91759401 |
| NAA38 | 2165.19586 | 0.04275258 | 0.17223906 | 0.24821652 | 0.80396688 | 0.97274243 |
| CYB5D1 | 1980.87663 | -0.2125614 | 0.16637397 | -1.2776121 | 0.20138624 | 0.78134034 |
| CHD3 | 7319.50409 | 0.02158581 | 0.28619301 | 0.07542396 | 0.9398774 | 0.99135146 |
| RNF227 | 281.194232 | -0.0659868 | 0.39150434 | -0.1685469 | 0.86615304 | 0.98197545 |
| KCNAB3 | 255.122215 | -0.4847747 | 0.4090373 | -1.1851601 | 0.23595415 | 0.80776058 |
| AC104581.2 | 1438.4373 | 0.00218279 | 0.10903037 | 0.02001998 | 0.98402744 | 0.9961464 |
| TRAPPC1 | 3683.58984 | 0.04800529 | 0.1032912 | 0.4647568 | 0.64210563 | 0.94088547 |
| CNTROB | 2239.74668 | -0.1878446 | 0.12329655 | -1.5235186 | 0.127629 | 0.70557946 |
| ALOX15B | 57.5221767 | -0.6969887 | 1.16307845 | -0.599262 | 0.54899817 | 0.92373235 |
| ALOX12B | 89.1901856 | 0.22107212 | 0.6657585 | 0.33206053 | 0.73984355 | 0.96043876 |
| AC129492.1 | 94.8880373 | -0.0053388 | 0.66641984 | -0.0080112 | 9.94E-01 | 0.99920627 |
| ALOXE3 | 137.331233 | -5.0996425 | 1.37885147 | -3.6984713 | NA | NA |
| PER1 | 10715.3633 | -0.364267 | 0.38313296 | -0.9507588 | 0.34172684 | 0.85889221 |
| AC129492.3 | 1693.62832 | -0.1013017 | 0.23260574 | -0.4355082 | 6.63E-01 | 0.94642611 |
| VAMP2 | 7505.48997 | 0.00036103 | 0.23537785 | 0.00153381 | 0.99877619 | 0.99994357 |
| TMEM107 | 626.899623 | 0.21630684 | 0.26541854 | 0.81496508 | 0.4150923 | 0.88377697 |
| BORCS6 | 751.372301 | 0.03896212 | 0.15860268 | 0.24565866 | 0.80594648 | 0.97334138 |
| AURKB | 47.9789111 | -0.4329585 | 0.68851161 | -0.6288325 | 0.52945872 | 0.91931447 |
| LINC00324 | 168.75877 | 0.43927507 | 0.37996665 | 1.15608847 | 0.24764497 | 0.81602355 |
| CTC1 | 2186.36121 | -0.2015212 | 0.15068123 | -1.337401 | 1.81E-01 | 0.76522169 |
| PFAS | 1284.21602 | -0.0912236 | 0.21538556 | -0.4235361 | 0.67190417 | 0.94922827 |
| SLC25A35 | 2413.69185 | 0.0412628 | 0.17094281 | 0.24138363 | 0.8092578 | 0.97334138 |
| RANGRF | 1696.00682 | -0.0071432 | 0.15180671 | -0.0470547 | 0.96246966 | 0.99320711 |
| ARHGEF15 | 1373.70784 | -0.305786 | 0.26697701 | -1.1453644 | 0.25205824 | 0.81833349 |
| AC135178.1 | 89.7459142 | -0.3685775 | 0.27558528 | -1.3374354 | 0.18108056 | 0.76522169 |
| AC135178.3 | 77.385289 | 0.39315843 | 0.27530979 | 1.42805832 | 0.15327507 | 0.74258845 |
| KRBA2 | 177.260739 | 0.47140758 | 0.24192888 | 1.94853785 | 0.05135064 | 0.58634747 |
| AC135178.2 | 19737.672 | 0.21105434 | 0.19071761 | 1.10663266 | 0.26845278 | 0.82676682 |
| RPL26 | 20220.8077 | 0.19688939 | 0.18967676 | 1.03802586 | 0.29925801 | 0.84109095 |
| NDEL1 | 2590.66751 | -0.5099638 | 0.24677306 | -2.0665295 | 0.03877851 | 0.55608529 |
| MYH10 | 4584.65724 | 0.24278957 | 0.17098005 | 1.41998768 | 0.15561127 | 0.74477742 |
| MFSD6L | 156.833162 | 0.11999197 | 0.28268341 | 0.42447476 | 0.67121961 | 0.94922827 |
| PIK3R6 | 63.9570985 | -0.0660256 | 0.82221301 | -0.0803023 | 0.9359968 | 0.99068685 |
| PIK3R5 | 357.485288 | -0.6815252 | 0.45111769 | -1.5107481 | 0.13085265 | 0.70999117 |
| NTN1 | 363.247503 | 0.27801884 | 0.57426566 | 0.48412931 | 0.6282941 | 0.94088547 |
| STX8 | 1066.12058 | 0.23656194 | 0.14640301 | 1.61582697 | 0.10613173 | 0.67584598 |
| CFAP52 | 51.5412747 | 0.14572071 | 0.57793785 | 0.25213907 | 0.80093357 | 0.97210205 |
| USP43 | 688.893241 | -0.1361173 | 0.32590415 | -0.4176605 | 0.67619538 | 0.94961241 |
| GAS7 | 402.445925 | -0.6046527 | 0.34654202 | -1.744818 | 0.08101656 | 0.64336071 |

| | | | | | | |
|---|---|---|---|---|---|---|
| MYH8 | 265.76276 | 0.0187309 | 0.6656646 | 0.02813865 | 0.97755157 | 0.99469508 |
| MYH3 | 307.828521 | -0.582541 | 0.33234716 | -1.7528088 | 0.07963482 | 0.64079926 |
| SCO1 | 2546.99406 | -0.004517 | 0.19413469 | -0.0232675 | 0.98143689 | 0.99595643 |
| AC002347.2 | 318.003706 | 0.05985034 | 0.19396113 | 0.30856872 | 0.75764962 | 0.96502715 |
| ADPRM | 338.790871 | 0.0382613 | 0.19657008 | 0.19464457 | 0.84567121 | 0.97675487 |
| TMEM220 | 934.272993 | -0.1929362 | 0.32141577 | -0.60027 | 0.5483263 | 0.92373235 |
| TMEM220-A | 309.911237 | 0.10555422 | 0.27642737 | 0.38185154 | 0.70257149 | 0.95431462 |
| AC015908.3 | 91.3781042 | 0.32061278 | 0.38099595 | 0.8415123 | 0.400061 | 0.88077967 |
| TMEM238L | 121.67383 | -0.2433288 | 1.09361778 | -0.2224989 | 0.82392552 | 0.97548201 |
| SHISA6 | 114.562227 | 1.79605634 | 0.63459121 | 2.83025719 | 0.00465106 | 0.29749322 |
| DNAH9 | 113.689181 | 0.82682224 | 0.49951553 | 1.65524831 | 0.09787414 | 0.66549286 |
| AC005410.2 | 59.4069386 | 0.02150799 | 0.29715655 | 0.07237933 | 0.94230004 | 0.99136103 |
| ZNF18 | 450.308521 | 0.28128249 | 0.22402491 | 1.2555858 | 0.20926618 | 0.79086734 |
| MAP2K4 | 1952.01746 | -0.2450331 | 0.24963345 | -0.9815716 | 0.32631094 | 0.85282528 |
| MYOCD | 193.427563 | -1.7449838 | 0.77864943 | -2.241039 | 0.02502355 | 0.52085426 |
| ARHGAP44 | 783.847176 | -0.0380096 | 0.22327749 | -0.1702346 | 0.86482565 | 0.98197545 |
| AC005277.1 | 191.789397 | 0.0365949 | 0.28017037 | 0.13061659 | 0.89607862 | 0.98474725 |
| ELAC2 | 3695.09385 | -0.1957978 | 0.17794574 | -1.1003228 | 0.27119149 | 0.82716979 |
| HS3ST3A1 | 65.056297 | -0.2995481 | 0.45553449 | -0.657575 | 0.51081124 | 0.91474443 |
| COX10 | 899.669748 | -0.0386458 | 0.22573396 | -0.1712005 | 0.86406615 | 0.98176973 |
| HS3ST3B1 | 498.83751 | -0.344383 | 0.41545793 | -0.8289239 | 0.40714747 | 0.88132015 |
| PMP22 | 2555.8915 | -0.5306907 | 0.34037738 | -1.5591244 | 0.11896693 | 0.69233212 |
| CDRT4 | 1099.77482 | 0.0394995 | 0.24733808 | 0.15969841 | 0.87311865 | 0.98275231 |
| TVP23C-CDR | 1107.4497 | 0.04614222 | 0.23558152 | 0.19586521 | 0.84471567 | 0.97675487 |
| TVP23C | 294.898153 | 0.0132239 | 0.2410583 | 0.05485767 | 0.95625186 | 0.99185043 |
| AC005324.3 | 509.697872 | 0.05361376 | 0.49611944 | 0.10806624 | 0.91394315 | 0.98662757 |
| TRIM16 | 1099.43903 | -0.0807823 | 0.48662594 | -0.1660049 | 0.86815313 | 0.98237277 |
| ZNF286A | 357.915716 | -0.1511566 | 0.22602097 | -0.6687726 | 0.50364057 | 0.91356552 |
| AC005324.4 | 90.3451804 | 0.14835626 | 0.39459546 | 0.37597053 | 0.70693881 | 0.95501548 |
| AC015922.3 | 156.934459 | 0.14943201 | 0.22967302 | 0.65062938 | 0.51528576 | 0.91636259 |
| AC015922.4 | 295.520281 | 0.15854007 | 0.26894045 | 0.58949881 | 0.55552671 | 0.92373235 |
| ADORA2B | 45.6769467 | 0.25295842 | 0.67988569 | 0.37206022 | 0.70984802 | 0.95512518 |
| ZSWIM7 | 1327.60842 | 0.10354254 | 0.14001208 | 0.73952579 | 0.45958778 | 0.89755243 |
| TTC19 | 3634.29141 | 0.07541134 | 0.12994435 | 0.58033565 | 0.56168829 | 0.92452095 |
| AC002553.2 | 45.5877154 | -0.397028 | 0.36293435 | -1.0939388 | 0.27398182 | 0.82719197 |
| NCOR1 | 4372.87367 | -0.1002135 | 0.11575007 | -0.8657748 | 0.38661366 | 0.87432552 |
| AC002553.1 | 333.52149 | -0.0268552 | 0.14262649 | -0.1882902 | 0.85064916 | 0.97839793 |
| PIGL | 1076.13785 | 0.11317166 | 0.21343011 | 0.53025162 | 0.59593749 | 0.9334601 |
| CENPV | 726.950698 | 0.2860142 | 0.24502851 | 1.16726907 | 0.24310173 | 0.81340905 |
| UBB | 34580.4467 | 0.10900724 | 0.19545192 | 0.55771896 | 0.57703631 | 0.93129843 |
| AC093484.3 | 25330.5762 | 0.08628521 | 0.19878403 | 0.4340651 | 0.66424117 | 0.94655191 |
| TRPV2 | 453.56967 | -0.3252259 | 0.39726349 | -0.8186654 | 0.41297733 | 0.88291217 |
| LRRC75A | 5805.08473 | 0.14002975 | 0.22927912 | 0.61073924 | 0.54137223 | 0.92292765 |
| ZNF287 | 248.675878 | 0.25967919 | 0.24726486 | 1.05020659 | 0.29362314 | 0.83540174 |
| ZNF624 | 136.36152 | 0.07324162 | 0.33365495 | 0.21951306 | 0.82625041 | 0.97548201 |
| AC098850.3 | 1029.56707 | -0.48393 | 0.88578475 | -0.5463291 | 0.58483971 | 0.93288628 |
| MPRIP | 9924.50261 | -0.4108233 | 0.27322136 | -1.503628 | 0.13267717 | 0.71328342 |
| AC055811.1 | 310.343946 | -0.5494891 | 0.40701575 | -1.3500439 | 1.77E-01 | 7.61E-01 |
| PLD6 | 293.39498 | -0.2720076 | 0.41084657 | -0.6620662 | 0.50792879 | 0.91439937 |
| AC055811.2 | 981.355097 | -0.7541363 | 0.42912754 | -1.7573711 | 0.07885458 | 0.64079926 |
| FLCN | 3402.75488 | -0.7789353 | 0.38747616 | -2.0102793 | 0.04440164 | 0.57267043 |
| COPS3 | 1812.52697 | 0.01494464 | 0.22146193 | 0.06748176 | 0.94619818 | 0.99166614 |
| NT5M | 211.708042 | 0.38505863 | 0.33667962 | 1.14369449 | 0.25275036 | 0.81835388 |
| MED9 | 1772.74165 | -0.3658642 | 0.18878463 | -1.937998 | 5.26E-02 | 0.58703362 |

| | | | | | | |
|---|---|---|---|---|---|---|
| RASD1 | 13539.8447 | -0.3639256 | 0.4114917 | -0.8844058 | 0.37647722 | 0.86970773 |
| PEMT | 1103.28615 | 0.07727597 | 0.20341641 | 0.37989056 | 0.70402666 | 0.95485335 |
| AC020558.2 | 96.5320194 | -0.1553192 | 0.29689776 | -0.5231404 | 0.60087657 | 0.93506832 |
| AC020558.1 | 179.58056 | 0.15920972 | 0.20378685 | 0.7812561 | 0.43465189 | 0.89055338 |
| RAI1 | 2901.84481 | -0.1868761 | 0.20226445 | -0.9239197 | 0.35552813 | 0.86364633 |
| SREBF1 | 8079.54678 | -0.267636 | 0.34541237 | -0.7748304 | 0.4384399 | 0.89175645 |
| TOM1L2 | 6003.69496 | -0.1491484 | 0.18450556 | -0.8083682 | 0.41887865 | 0.88451372 |
| DRC3 | 830.698862 | -0.0723252 | 0.33558579 | -0.2155192 | 0.82936254 | 0.97550416 |
| ATPAF2 | 1225.01215 | 0.12263754 | 0.14929641 | 0.82143662 | 0.41139761 | 0.88268208 |
| GID4 | 819.872509 | -0.31535 | 0.1865746 | -1.6902089 | 0.090988 | 0.65495306 |
| DRG2 | 2323.09618 | -0.0364892 | 0.09730945 | -0.3749807 | 0.70767485 | 0.95501548 |
| MYO15A | 123.078633 | -0.3958294 | 0.37592677 | -1.0529427 | 0.29236724 | 0.83524572 |
| AC087164.1 | 488.503544 | -0.1079894 | 0.38953413 | -0.2772272 | 0.78160567 | 0.96913822 |
| ALKBH5 | 5871.90418 | -0.1354289 | 0.27249597 | -0.496994 | 0.61919329 | 0.93945324 |
| LLGL1 | 1678.45755 | -0.1481304 | 0.11421932 | -1.2968947 | 0.19466743 | 0.7762856 |
| FLII | 10045.4555 | -0.3259406 | 0.21250862 | -1.5337757 | 0.12508484 | 0.70228923 |
| MIEF2 | 1790.54983 | -0.3088005 | 0.23400407 | -1.3196373 | 0.18695614 | 0.76958556 |
| TOP3A | 1147.41432 | -0.3405755 | 0.14058446 | -2.422569 | 0.0154112 | 0.45243543 |
| SMCR8 | 3745.86339 | 0.06943885 | 0.22876857 | 0.3035332 | 0.76148355 | 0.96606476 |
| SHMT1 | 19568.4481 | 0.66705468 | 0.31300407 | 2.13113742 | 0.03307782 | 0.55145048 |
| EVPLL | 100.450312 | 0.51408663 | 0.39414939 | 1.30429387 | 0.19213341 | 0.7739557 |
| AL353997.2 | 81.1847628 | 0.59546473 | 0.3971272 | 1.49943072 | 0.13376193 | 0.71505705 |
| ZNF286B | 87.1543249 | 0.00698926 | 0.39355872 | 0.01775912 | 0.98583102 | 0.99705651 |
| TRIM16L | 1210.75772 | -0.5723439 | 0.63242797 | -0.9049945 | 0.36546828 | 0.8663515 |
| TVP23B | 1565.2223 | 0.1028926 | 0.15050174 | 0.68366386 | 0.49418745 | 0.90796767 |
| PRPSAP2 | 1942.53984 | 0.0414791 | 0.19769741 | 0.20981104 | 0.83381516 | 0.97550416 |
| SLC5A10 | 8205.94654 | -0.4944619 | 0.31841966 | -1.5528625 | 0.12045599 | 0.69467399 |
| AC090286.1 | 606.537262 | -0.0973978 | 0.49013263 | -0.1987172 | 0.84248396 | 0.97644404 |
| FAM83G | 3718.33822 | -0.7271566 | 0.41351077 | -1.7584949 | 0.07866333 | 0.64079926 |
| GRAP | 1282.86269 | -0.3637378 | 0.33280632 | -1.0929414 | 2.74E-01 | 8.27E-01 |
| EPN2 | 2592.94484 | -0.0712621 | 0.20954404 | -0.3400819 | 0.73379484 | 0.95895327 |
| AC124066.1 | 1472.92378 | -0.0581515 | 0.2184369 | -0.2662165 | 0.79007247 | 0.97090793 |
| B9D1 | 1118.78828 | 0.02455545 | 0.20494359 | 0.11981567 | 0.90462917 | 0.98624484 |
| MAPK7 | 2014.03665 | -0.2993975 | 0.19015524 | -1.5744901 | 0.11537418 | 0.6873836 |
| MFAP4 | 3292.06985 | 0.31058315 | 0.68383252 | 0.45418014 | 0.64969918 | 0.94312988 |
| RNF112 | 237.958816 | -0.7087336 | 0.40261291 | -1.7603351 | 0.078351 | 0.64079926 |
| SLC47A1 | 8089.33678 | 1.5369673 | 0.41592229 | 3.6953232 | 0.00021961 | 0.07349392 |
| AC025627.1 | 684.615806 | 1.58532819 | 0.54102069 | 2.93025428 | 0.00338685 | 0.2541761 |
| ALDH3A2 | 9549.24737 | 0.35511867 | 0.33408347 | 1.0629639 | 0.28779832 | 0.83266626 |
| SLC47A2 | 10914.9783 | 1.04822851 | 0.23565101 | 4.44822418 | 8.66E-06 | 0.01014159 |
| AC005722.2 | 366.89644 | 0.14923455 | 0.46840135 | 0.31860402 | 0.75002681 | 0.96249332 |
| ALDH3A1 | 132.502679 | 0.33468127 | 0.39812734 | 0.84063875 | 0.40055034 | 0.88086294 |
| ULK2 | 1161.70491 | 0.26740816 | 0.22758812 | 1.17496537 | 0.24000857 | 0.81209487 |
| AC015726.1 | 104.956391 | -0.0159872 | 0.34785315 | -0.0459597 | 0.96334239 | 0.99320711 |
| AKAP10 | 1584.3068 | -0.1755841 | 0.17855332 | -0.9833705 | 3.25E-01 | 8.52E-01 |
| SPECC1 | 1003.39041 | -0.2447265 | 0.19466816 | -1.2571471 | 0.20870036 | 0.79042964 |
| CCDC144NL-/ | 966.792875 | -0.001599 | 0.26770226 | -0.005973 | 0.99523426 | 0.99940889 |
| USP22 | 9636.39917 | 0.03480845 | 0.20826333 | 0.16713672 | 0.86726247 | 0.98223481 |
| AC087393.2 | 3256.63636 | 0.07350939 | 0.23005121 | 0.3195349 | 0.74932093 | 0.96249332 |
| DHRS7B | 1741.54101 | 0.21440193 | 0.15479486 | 1.38507132 | 0.16603066 | 0.75515387 |
| TMEM11 | 1228.12985 | -0.3300769 | 0.22393981 | -1.4739536 | 0.14049409 | 0.72554416 |
| NATD1 | 2145.42616 | -0.2535864 | 0.22466042 | -1.1287542 | 0.25900154 | 0.82364948 |
| MAP2K3 | 4851.3712 | -0.4431855 | 0.40251048 | -1.1010533 | 0.27087348 | 0.82716979 |
| KCNJ12 | 299.536727 | 0.58614643 | 0.47400014 | 1.23659548 | 0.2162373 | 0.7954827 |

| | | | | | | |
|---|---|---|---|---|---|---|
| C17orf51 | 504.34609 | -0.3279608 | 0.30644065 | -1.0702263 | 0.28451747 | 0.83107779 |
| WSB1 | 15609.6901 | -0.1975735 | 0.29138791 | -0.6780429 | 0.49774451 | 0.90998243 |
| KSR1 | 3259.49704 | 0.08280927 | 0.32736409 | 0.25295772 | 8.00E-01 | 9.72E-01 |
| AC015688.4 | 554.84466 | -0.9157146 | 0.53533178 | -1.7105553 | 0.08716323 | 0.65231349 |
| AC015688.6 | 1157.66855 | 0.08657822 | 0.36727104 | 0.23573385 | 0.8136392 | 0.9738447 |
| LGALS9 | 2284.28399 | -1.4252528 | 0.64481989 | -2.2103114 | 0.02708356 | 0.52773677 |
| NOS2 | 190.966981 | -1.4161632 | 0.6798354 | -2.0830972 | 0.03724237 | 0.55532957 |
| AC005697.1 | 260.987003 | 0.78548358 | 0.43460426 | 1.80735361 | 0.07070716 | 0.62981127 |
| AC005697.2 | 617.207785 | 0.87907724 | 0.39334815 | 2.23485797 | 0.02542667 | 0.52250144 |
| LYRM9 | 1147.07845 | 0.82911032 | 0.39745162 | 2.08606602 | 0.03697264 | 0.55521179 |
| NLK | 1675.69852 | 0.06000939 | 0.14624459 | 0.41033584 | 0.68155961 | 0.95070301 |
| ERVE-1 | 163.407851 | 0.13831245 | 0.34588856 | 0.39987576 | 6.89E-01 | 0.95158502 |
| TMEM97 | 999.118137 | 0.00788287 | 0.20235037 | 0.03895656 | 0.96892503 | 0.99400331 |
| IFT20 | 1483.07212 | -0.2368452 | 0.15091126 | -1.5694337 | 0.11654692 | 0.6888681 |
| TNFAIP1 | 5191.7217 | -0.3685306 | 0.19164317 | -1.9230043 | 5.45E-02 | 0.59329767 |
| POLDIP2 | 9156.67542 | 0.08821897 | 0.17870257 | 0.49366368 | 0.62154372 | 0.94016713 |
| TMEM199 | 1560.72888 | -0.3281762 | 0.16440287 | -1.996171 | 0.04591531 | 0.57687945 |
| AC002094.2 | 584.771472 | -0.3464815 | 0.19880169 | -1.7428498 | 0.08135985 | 0.6439041 |
| AC002094.3 | 94.5713594 | -1.1002493 | 0.78439436 | -1.4026736 | NA | NA |
| SARM1 | 2206.77305 | 0.32637137 | 0.20081575 | 1.62522797 | 0.10411399 | 0.67439401 |
| VTN | 286.126801 | -1.033095 | 0.79672866 | -1.2966711 | NA | NA |
| SLC46A1 | 2212.39117 | 0.36579043 | 0.20656072 | 1.77086154 | 0.07658373 | 0.63773415 |
| AC015917.2 | 47.6944474 | 0.91978554 | 0.53475545 | 1.72001153 | 0.08543035 | 0.6504594 |
| AC005726.1 | 2627.6506 | 0.41068478 | 0.52119561 | 0.7879667 | 4.31E-01 | 0.88966606 |
| SLC13A2 | 3709.77947 | 1.9024599 | 0.66001565 | 2.88244664 | 0.003946 | 0.27088637 |
| UNC119 | 2053.4918 | 0.15285446 | 0.25807898 | 0.59227782 | 0.55366456 | 0.92373235 |
| PIGS | 4189.45787 | 0.23833361 | 0.1516541 | 1.57156065 | 0.11605248 | 0.68754779 |
| ALDOC | 882.94072 | 0.49843982 | 0.24352185 | 2.04679709 | 0.04067801 | 0.55657615 |
| AC005726.4 | 318.779233 | 0.54656053 | 0.25294447 | 2.16079256 | 0.03071137 | 0.54443002 |
| SPAG5 | 3549.42907 | 0.49813131 | 0.52828202 | 0.94292688 | 0.34571831 | 0.86111349 |
| AC005726.5 | 282.031243 | 0.08829496 | 0.31901195 | 0.27677633 | 0.78195186 | 0.96934514 |
| SGK494 | 786.500791 | 0.0031541 | 0.36556246 | 0.00862806 | 0.99311589 | 0.99906069 |
| KIAA0100 | 10026.1757 | 0.06688386 | 0.31761495 | 0.21058159 | 0.83321378 | 0.97550416 |
| AC005726.2 | 217.363594 | 0.25016898 | 0.50383595 | 0.49652864 | 0.61952149 | 0.93945324 |
| SDF2 | 2752.44054 | -0.0778362 | 0.16884957 | -0.4609796 | 0.64481323 | 0.94208859 |
| SUPT6H | 6404.06808 | -0.3103059 | 0.18001067 | -1.7238196 | 0.08474041 | 0.64988556 |
| PROCA1 | 305.955281 | 0.33086007 | 0.36973772 | 0.89485073 | 3.71E-01 | 0.86762749 |
| RAB34 | 4005.35333 | 0.02816485 | 0.24832127 | 0.11342103 | 0.90969676 | 0.98662757 |
| RPL23A | 13690.049 | 0.18500828 | 0.1935253 | 0.95599013 | 0.33907722 | 0.85733087 |
| AC010761.1 | 9590.75009 | 0.19537329 | 0.19917003 | 0.98093723 | 0.32662369 | 0.85282528 |
| TLCD1 | 603.421713 | -0.1084744 | 0.22458493 | -0.4829997 | 6.29E-01 | 0.94088547 |
| NEK8 | 1088.65862 | 0.13534251 | 0.41901137 | 0.32300439 | 0.74669191 | 0.96216675 |
| AC010761.2 | 231.413174 | 0.19609288 | 0.47594518 | 0.41200728 | 0.68033409 | 0.95070301 |
| TRAF4 | 6517.52183 | -0.1063973 | 0.15045166 | -0.7071861 | 0.47945083 | 0.90496404 |
| AC010761.4 | 763.164582 | -0.0917578 | 0.2004058 | -0.4578597 | 0.64705321 | 0.9425903 |
| AC010761.5 | 845.605039 | -0.1094487 | 0.14187086 | -0.7714668 | 0.44043027 | 0.89262503 |
| FAM222B | 2231.86955 | -0.2414568 | 0.15085283 | -1.6006119 | 0.10946291 | 0.67859123 |
| ERAL1 | 2784.79046 | 0.0520189 | 0.16045712 | 0.32419191 | 0.74579275 | 0.96176407 |
| FLOT2 | 10986.0268 | 0.07013118 | 0.24109522 | 0.29088583 | 0.77113864 | 0.96836854 |
| AC024267.6 | 1223.04192 | -0.0457591 | 0.40051667 | -0.1142503 | 0.9090394 | 0.98662757 |
| DHRS13 | 1547.06237 | -0.0087831 | 0.38982552 | -0.0225307 | 0.98202461 | 0.99595643 |
| PHF12 | 2724.80266 | -0.0870336 | 0.1318613 | -0.660039 | 0.5092288 | 0.91439937 |
| PIPOX | 4701.27897 | 1.70849412 | 0.70974397 | 2.40719781 | 0.01607546 | 0.45716266 |
| MYO18A | 4919.5544 | -0.398654 | 0.27206291 | -1.4653009 | 0.14283883 | 0.72784886 |

| | | | | | | |
|---|---|---|---|---|---|---|
| TIAF1 | 3213.13397 | -0.3930076 | 0.23631095 | -1.6630953 | 0.09629337 | 0.66233356 |
| NUFIP2 | 4197.98948 | -0.2631957 | 0.19237907 | -1.3681098 | 0.17127773 | 0.75956362 |
| TAOK1 | 2586.56233 | -0.183093 | 0.15073489 | -1.2146689 | 0.2244924 | 0.80078694 |
| ABHD15 | 1359.22119 | 0.70261477 | 0.33973005 | 2.0681561 | 0.03862535 | 0.5559736 |
| TP53I13 | 3502.13145 | 0.21945995 | 0.2161914 | 1.0151188 | 0.3100492 | 0.84456632 |
| GIT1 | 3945.49821 | -0.2807972 | 0.20178992 | -1.3915324 | 0.16406404 | 0.7527134 |
| ANKRD13B | 343.71176 | -0.4264615 | 0.40532849 | -1.0521379 | 0.29273628 | 0.83524862 |
| CORO6 | 397.682436 | -0.5471577 | 0.39230795 | -1.3947148 | 0.16310186 | 0.75121453 |
| SSH2 | 1479.57072 | -0.089058 | 0.17909434 | -0.4972686 | 0.61899966 | 0.93945324 |
| EFCAB5 | 138.525263 | 0.05663265 | 0.35768406 | 0.1583315 | 0.87419559 | 0.98275231 |
| NSRP1 | 1569.08018 | -0.1103259 | 0.14768847 | -0.7470177 | 0.4550529 | 0.89670027 |
| SLC6A4 | 67.6480031 | 0.25025001 | 0.51695614 | 0.48408364 | 0.62832651 | 0.94088547 |
| BLMH | 2391.39501 | 0.07913196 | 0.14464936 | 0.54706058 | 0.58433711 | 0.9326319 |
| TMIGD1 | 94.4407394 | 1.88001804 | 0.80508754 | 2.3351722 | 0.01953443 | 0.49272502 |
| CPD | 7910.45637 | -0.0346955 | 0.33783266 | -0.1027002 | 0.91820089 | 0.9868044 |
| GOSR1 | 2484.00672 | -0.1975702 | 0.21048776 | -0.9386305 | 0.3479205 | 0.86225576 |
| TBC1D29 | 41.8394087 | 0.28051338 | 0.69139377 | 0.40572159 | 0.68494718 | 0.95097288 |
| CRLF3 | 1022.9632 | -0.2126616 | 0.22237133 | -0.9563356 | 0.33890272 | 0.85733087 |
| AC127024.5 | 180.20776 | -0.1893814 | 0.28822261 | -0.6570666 | 0.51113809 | 0.91492254 |
| AC127024.6 | 97.8929357 | -0.1591004 | 0.25001449 | -0.6363646 | 0.5245388 | 0.91723878 |
| ATAD5 | 126.233942 | -0.0877808 | 0.25853927 | -0.3395259 | 0.73421358 | 0.95895327 |
| TEFM | 583.321824 | -0.1306965 | 0.16645951 | -0.7851552 | 0.43236258 | 0.88980461 |
| AC130324.3 | 259.054271 | -0.0745032 | 0.24337575 | -0.3061243 | 0.75951003 | 0.96502715 |
| ADAP2 | 755.077204 | -0.5308369 | 0.35434789 | -1.4980671 | 0.13411582 | 0.71560922 |
| AC138207.1 | 132.039303 | -0.5365728 | 0.39685986 | -1.3520461 | 0.17636056 | 0.76115956 |
| AC138207.5 | 47.1539935 | -0.1316867 | 0.43602795 | -0.3020143 | 0.76264119 | 0.96630779 |
| RNF135 | 1910.45018 | 0.13025601 | 0.23909046 | 0.54479804 | 0.58589243 | 0.93289552 |
| AC138207.2 | 43.5455385 | 0.26062093 | 0.42753865 | 0.60958448 | 0.54213709 | 0.92292765 |
| NF1 | 2753.30768 | -0.1584477 | 0.09273185 | -1.7086657 | 0.08751289 | 0.65271545 |
| EVI2B | 418.790695 | -0.4593575 | 0.57508965 | -0.798758 | 0.42443074 | 0.88565404 |
| AC134669.1 | 272.818779 | -0.3936012 | 0.60935203 | -0.645934 | 0.51832212 | 0.91636259 |
| EVI2A | 255.481896 | -0.405026 | 0.61361865 | -0.6600615 | 0.50921439 | 0.91439937 |
| RAB11FIP4 | 1953.74727 | 0.30656389 | 0.20757314 | 1.47689578 | 0.13970358 | 0.72475166 |
| COPRS | 1840.11378 | -0.0953487 | 0.23557732 | -0.4047448 | 0.68566514 | 0.95097288 |
| UTP6 | 2048.59579 | -0.0762574 | 0.14344157 | -0.531627 | 0.59498438 | 0.9334601 |
| AC004253.1 | 100.723939 | -0.056722 | 0.44353395 | -0.1278866 | 0.89823875 | 0.98474725 |
| SUZ12 | 1612.52841 | -0.160419 | 0.1245123 | -1.2883789 | 0.1976141 | 0.77837029 |
| LRRC37B | 382.769798 | -0.2461881 | 0.18259584 | -1.3482679 | 0.17757224 | 0.76115956 |
| AC116407.2 | 67.6841427 | -1.2965214 | 0.83447068 | -1.5537052 | NA | NA |
| AC116407.1 | 88.9800521 | 0.31412931 | 0.25187785 | 1.24714944 | 0.21234271 | 0.79203904 |
| RHOT1 | 1601.5996 | 0.11238048 | 0.122621 | 0.91648638 | 0.35941185 | 0.86480228 |
| RHBDL3 | 48.2487002 | -0.1732741 | 0.59571738 | -0.2908663 | 0.7711536 | 0.96836854 |
| C17orf75 | 1021.40205 | 0.35345885 | 0.23572485 | 1.49945521 | 0.13375558 | 0.71505705 |
| ZNF207 | 7456.66303 | -0.3349503 | 0.14853157 | -2.2550784 | 0.02412841 | 0.51967919 |
| AC005899.6 | 43.662807 | -0.1096106 | 0.46847134 | -0.233975 | 0.81500436 | 0.9738447 |
| AC005899.8 | 248.680808 | 0.03536481 | 0.32519397 | 0.10874988 | 0.91340088 | 0.98662757 |
| PSMD11 | 4550.11299 | -0.2935656 | 0.26257629 | -1.1180204 | 0.26355829 | 0.82424998 |
| CDK5R1 | 155.303354 | -0.0933781 | 0.51077569 | -0.1828162 | 8.55E-01 | 0.97910552 |
| MYO1D | 4102.43942 | 0.15655678 | 0.24255184 | 0.64545698 | 0.51863109 | 0.91636259 |
| TMEM98 | 1795.99535 | 0.64037811 | 0.24099241 | 2.6572543 | 0.007878 | 0.37071188 |
| ASIC2 | 116.323322 | 0.05474201 | 0.67783877 | 0.08075964 | 0.93563311 | 0.99068685 |
| CCL2 | 4962.23368 | 0.28285875 | 0.81311097 | 0.34787225 | 0.72793612 | 0.95848028 |
| CCL11 | 274.459814 | -0.3904576 | 1.0860409 | -0.3595238 | 0.71920325 | 0.9566409 |
| CCL8 | 371.891726 | -2.3075912 | 1.29257467 | -1.7852672 | NA | NA |

| | | | | | | |
|---|---|---|---|---|---|---|
| CCL13 | 67.9263058 | -0.1505101 | 0.65254029 | -0.2306526 | 0.81758467 | 0.97433259 |
| TMEM132E | 528.88383 | -0.3545267 | 0.40046275 | -0.8852927 | 0.37599879 | 0.86900834 |
| CCT6B | 188.138118 | 0.12106439 | 0.59060527 | 0.20498359 | 0.83758494 | 0.97577999 |
| ZNF830 | 682.566975 | -0.0834023 | 0.11932828 | -0.6989316 | 0.48459475 | 0.90541935 |
| LIG3 | 2374.83769 | -0.2875615 | 0.22053326 | -1.3039369 | 0.1922551 | 0.7740178 |
| RFFL | 3730.59969 | -0.3123331 | 0.3619564 | -0.8629024 | 0.3881911 | 0.87553582 |
| AC004223.3 | 893.066716 | -0.3865784 | 0.41083537 | -0.940957 | 0.3467269 | 0.86135217 |
| RAD51D | 731.62502 | -0.0005904 | 0.12752564 | -0.0046294 | 0.99630626 | 0.99976751 |
| FNDC8 | 53.4691859 | -0.1772927 | 0.42548035 | -0.4166884 | 0.67690633 | 0.9500413 |
| NLE1 | 648.225701 | 0.05333579 | 0.25202538 | 0.21162865 | 0.83239676 | 0.97550416 |
| AC022916.1 | 90.6034326 | 0.01223214 | 0.42796119 | 0.02858235 | 0.97719769 | 0.99455099 |
| SLFN5 | 1533.55558 | -0.0954821 | 0.41203331 | -0.231734 | 8.17E-01 | 0.97388889 |
| SLFN11 | 842.530917 | 0.39038434 | 0.42159329 | 0.92597379 | 0.3544596 | 0.86364633 |
| AC060766.4 | 37.530262 | -0.6313483 | 0.54504856 | -1.158334 | 0.24672778 | 0.81542272 |
| SLFN12 | 180.85605 | -0.4599947 | 0.39642155 | -1.1603676 | 0.24589916 | 0.81542272 |
| SLFN13 | 564.2362 | 0.12527257 | 0.51868893 | 0.24151772 | 0.80915389 | 0.97334138 |
| SLFN12L | 48.7396258 | -0.469604 | 0.94520935 | -0.4968254 | 6.19E-01 | 0.93945324 |
| LINC02001 | 362.769726 | -0.1019952 | 0.28927578 | -0.352588 | 0.7243973 | 0.95827838 |
| PEX12 | 425.246199 | 0.10019343 | 0.17867452 | 0.56075945 | 0.57496153 | 0.93035349 |
| AP2B1 | 10397.6337 | 0.26693894 | 0.14415052 | 1.85180692 | 0.06405355 | 0.61837523 |
| TAF15 | 6090.37201 | -0.0110104 | 0.13005356 | -0.0846603 | 0.93253145 | 0.9899349 |
| RASL10B | 259.715361 | -0.5760674 | 0.51015155 | -1.1292084 | 0.25880995 | 0.82363004 |
| MMP28 | 131.521227 | 0.54361556 | 0.58692607 | 0.9262079 | 0.35433795 | 0.86364633 |
| CCL5 | 619.20221 | -0.4873369 | 0.85947622 | -0.5670162 | 0.57070318 | 0.9278692 |
| CCL14 | 1562.99192 | 0.09261156 | 0.58267755 | 0.15894136 | 0.87371508 | 0.98275231 |
| AC244100.2 | 1182.4482 | 0.09748462 | 0.59028775 | 0.16514763 | 0.86882778 | 0.98272083 |
| CCL15-CCL14 | 1944.83159 | 0.36094343 | 0.49557173 | 0.72833739 | 0.46640708 | 0.89960479 |
| CCL15 | 661.911383 | 1.38177631 | 0.62099049 | 2.2251167 | 0.0260734 | 0.52561121 |
| AC244100.3 | 308.284918 | 1.52491766 | 0.63434032 | 2.40394251 | NA | NA |
| CCL23 | 145.182446 | -0.0483682 | 0.80620332 | -0.0599951 | 0.95215954 | 0.99170503 |
| CCL18 | 46.8711144 | 0.36397072 | 0.85511062 | 0.42564168 | 0.67036896 | 0.94922827 |
| AC243829.4 | 190.592999 | -1.18432 | 0.71002547 | -1.6679965 | 0.09531642 | 0.65888357 |
| CCL3 | 218.791728 | -1.25053 | 0.7020449 | -1.7812678 | 0.07486871 | 0.63773415 |
| CCL4 | 229.818515 | -0.7499181 | 0.64509254 | -1.1624969 | 0.24503369 | 0.81492642 |
| CCL3L1 | 54.7490021 | -1.3830845 | 1.0755054 | -1.2859856 | 0.19844808 | 0.78046101 |
| CCL4L2 | 73.2151693 | -1.5870833 | 1.04119205 | -1.5242945 | 0.12743516 | 0.70557946 |
| ZNHIT3 | 2134.36097 | 0.02333586 | 0.19555491 | 0.11933149 | 0.90501274 | 0.98624484 |
| MYO19 | 2868.42793 | -0.0536624 | 0.1623971 | -0.3304394 | 7.41E-01 | 0.96097294 |
| PIGW | 508.657447 | -0.6288143 | 0.48479165 | -1.2970814 | NA | NA |
| GGNBP2 | 3542.07306 | -0.1516596 | 0.19047971 | -0.7961983 | 0.42591681 | 0.88641523 |
| DHRS11 | 1078.70685 | 0.49794751 | 0.24539 | 2.02920864 | 4.24E-02 | 0.56525418 |
| MRM1 | 470.49443 | 0.2609377 | 0.21929731 | 1.18988098 | 0.23409318 | 0.80725541 |
| LHX1-DT | 609.078383 | 0.38132166 | 0.4617576 | 0.82580483 | 4.09E-01 | 8.82E-01 |
| LHX1 | 963.688296 | 1.16747391 | 0.30032503 | 3.88736804 | 0.00010134 | 0.04521804 |
| AC243773.1 | 160.651482 | 1.05035951 | 0.32631096 | 3.21889135 | 0.00128687 | 0.16078183 |
| AATF | 2341.87095 | -0.0212135 | 0.20583014 | -0.1030631 | 0.91791291 | 0.9868044 |
| ACACA | 1423.47442 | 0.15480495 | 0.19948298 | 0.77603085 | 0.43773077 | 0.89130865 |
| TADA2A | 758.290687 | 0.03178875 | 0.15729794 | 0.2020926 | 0.83984433 | 0.97618283 |
| DUSP14 | 878.32285 | -0.262945 | 0.34304355 | -0.7665063 | 0.4433751 | 0.89366134 |
| SYNRG | 2380.1765 | -0.0279318 | 0.1427593 | -0.1956563 | 0.84487922 | 0.97675487 |
| DDX52 | 2305.74217 | -0.550804 | 0.3257597 | -1.6908291 | 0.09086944 | 0.65495306 |
| AC243585.2 | 340.716773 | -0.7603271 | 0.36107142 | -2.1057527 | 0.03522584 | 0.55369025 |
| AC243571.2 | 65.2437439 | -0.4700823 | 0.52663815 | -0.8926096 | 0.37206626 | 0.86772685 |
| HNF1B | 8146.44236 | -0.2599585 | 0.23838621 | -1.0904932 | 0.27549593 | 0.82719407 |

| | | | | | | |
|---|---|---|---|---|---|---|
| TBC1D3L | 437.234891 | 0.06838067 | 0.41697068 | 0.16399395 | 0.8697359 | 0.98275231 |
| MRPL45 | 1896.04629 | 0.27623227 | 0.19947758 | 1.38477851 | 0.16612021 | 0.75515387 |
| SOCS7 | 933.450062 | -0.0528881 | 0.22903045 | -0.2309215 | 0.81737579 | 0.97426952 |
| ARHGAP23 | 2000.33861 | -0.5228983 | 0.30183545 | -1.7323953 | 0.0832032 | 0.6471966 |
| SRCIN1 | 264.562194 | -0.1728328 | 0.46065778 | -0.375187 | 0.70752137 | 0.95501548 |
| EPOP | 188.548994 | -0.2132511 | 0.30579028 | -0.6973769 | 4.86E-01 | 9.05E-01 |
| AC006449.6 | 270.278145 | -0.0031196 | 0.2886161 | -0.0108087 | 9.91E-01 | 9.99E-01 |
| AC006449.2 | 69.9322812 | -0.1540802 | 0.34889372 | -0.441625 | 0.65876059 | 0.9458503 |
| MLLT6 | 16573.5252 | -0.4035349 | 0.20320396 | -1.9858616 | 0.04704869 | 0.58171004 |
| AC006449.3 | 942.594872 | -0.6443846 | 0.25297975 | -2.5471785 | 0.01085979 | 0.40196898 |
| CISD3 | 3674.27461 | 0.13486061 | 0.15477179 | 0.87135137 | 0.38356233 | 0.87226569 |
| PCGF2 | 4855.93597 | -0.0783638 | 0.15922018 | -0.4921723 | 0.62259754 | 0.94016713 |
| AC006449.5 | 92.9879196 | 0.20886951 | 0.35459235 | 0.58904122 | 0.55583362 | 0.92373235 |
| PSMB3 | 4441.21151 | -0.0836027 | 0.1751329 | -0.4773669 | 0.63310087 | 0.94088547 |
| PIP4K2B | 3607.20049 | 0.05544301 | 0.14129022 | 0.39240518 | 0.69475886 | 0.95234609 |
| CWC25 | 2151.728 | -0.4749463 | 0.26953114 | -1.7621204 | 0.07804896 | 0.64079926 |
| RPL23 | 22230.1584 | 0.2396952 | 0.2137976 | 1.12113137 | 0.26223195 | 0.82424998 |
| LASP1 | 12952.0111 | -0.0275695 | 0.23294014 | -0.1183544 | 0.90578682 | 0.98636553 |
| AC006441.3 | 2523.05386 | -0.0405411 | 0.23362362 | -0.1735318 | 0.86223341 | 0.98176973 |
| LINC00672 | 70.2478979 | 0.7867119 | 0.34758306 | 2.26337812 | 0.02361239 | 0.51967919 |
| AC006441.4 | 89.1592248 | 1.16103697 | 0.9277846 | 1.2514079 | 0.2107857 | 0.79134644 |
| PLXDC1 | 536.484194 | -0.0169379 | 0.63591815 | -0.0266354 | 0.97875057 | 0.99483482 |
| AC004408.1 | 68.1476562 | -0.030468 | 0.49613895 | -0.0614102 | 0.95103252 | 0.99170503 |
| CACNB1 | 536.424863 | 0.00955879 | 0.34917144 | 0.02737563 | 0.97816014 | 0.99476647 |
| RPL19 | 35150.555 | 0.13145526 | 0.1567897 | 0.83841769 | 0.40179616 | 0.88132015 |
| STAC2 | 2014.63728 | -0.7861324 | 0.40604859 | -1.936055 | 0.05286097 | 0.58742503 |
| FBXL20 | 1523.11867 | 0.04897311 | 0.16542751 | 0.29603968 | 0.76719978 | 0.96749893 |
| AC005288.1 | 398.123844 | -0.2735148 | 0.20770661 | -1.3168324 | 0.1878948 | 0.76970626 |
| MED1 | 936.923967 | -0.3138286 | 0.17157976 | -1.8290539 | 6.74E-02 | 6.25E-01 |
| CDK12 | 1683.96567 | -0.3370658 | 0.18324856 | -1.8393913 | 0.06585765 | 0.62237346 |
| AC009283.1 | 324.147461 | -0.0148096 | 0.34454867 | -0.0429826 | 0.9657154 | 0.99387827 |
| PPP1R1B | 1137.31681 | 1.38007006 | 0.53785956 | 2.56585579 | 0.01029216 | 0.40019066 |
| STARD3 | 3399.53379 | -0.1117835 | 0.12122789 | -0.922094 | 0.35647954 | 0.86390849 |
| TCAP | 269.606618 | 0.31481394 | 0.39699041 | 0.79300138 | 0.42777702 | 0.88791007 |
| PNMT | 62.0663358 | 0.39286557 | 0.5886233 | 0.66743123 | 0.50449673 | 0.91359292 |
| PGAP3 | 2688.69588 | 0.44944515 | 0.27518239 | 1.63326275 | 0.10241375 | 0.67279543 |
| ERBB2 | 16672.8104 | 0.16986121 | 0.28733781 | 0.5911551 | 0.55441651 | 0.92373235 |
| MIEN1 | 2889.66275 | -0.0084357 | 0.13505449 | -0.0624612 | 0.95019557 | 0.99170503 |
| GRB7 | 7091.66239 | -0.2810719 | 0.23611324 | -1.1904112 | 0.2338848 | 0.80725541 |
| IKZF3 | 202.845752 | -0.2923879 | 0.74454863 | -0.3927049 | 0.69453743 | 0.95222846 |
| GSDMB | 811.95858 | -0.2393394 | 0.44057783 | -0.5432399 | 0.58696467 | 0.93289552 |
| ORMDL3 | 4019.69966 | 0.26973421 | 0.13825286 | 1.95102078 | 0.05105457 | 0.5861488 |
| PSMD3 | 5780.12168 | -0.2955694 | 0.23136372 | -1.2775098 | 0.20142233 | 0.78134034 |
| CSF3 | 1181.5266 | 3.90880956 | 1.79718214 | 2.17496572 | NA | NA |
| MED24 | 4321.98871 | -0.0718366 | 0.12304665 | -0.5838159 | 0.55934421 | 0.92421175 |
| THRA | 4954.85985 | -0.0911621 | 0.15866972 | -0.5745397 | 0.56560264 | 0.92513982 |
| NR1D1 | 3262.11397 | -0.1142639 | 0.43673363 | -0.2616329 | 0.79360446 | 0.97090793 |
| MSL1 | 5524.40282 | -0.1974062 | 0.20887243 | -0.9451044 | 0.3446056 | 0.85999891 |
| CASC3 | 4746.59419 | -0.1331689 | 0.10499831 | -1.2682954 | 0.20469248 | 0.78464755 |
| RAPGEFL1 | 502.486025 | -0.4102351 | 0.36927718 | -1.1109138 | 0.26660544 | 0.8265123 |
| WIPF2 | 3155.62466 | -0.2879252 | 0.23118606 | -1.2454264 | 0.21297506 | 0.7926732 |
| CDC6 | 71.989888 | -0.5215581 | 0.65299311 | -0.7987191 | 0.42445331 | 0.88565404 |
| RARA | 3104.18971 | -0.2217122 | 0.28459668 | -0.77904 | 0.43595618 | 0.89095725 |
| RARA-AS1 | 396.678254 | -0.1500336 | 0.34013675 | -0.441098 | 0.65914205 | 0.94600866 |

| | | | | | | |
|---|---|---|---|---|---|---|
| AC080112.1 | 360.661168 | -0.0928634 | 0.28547107 | -0.3252988 | 0.74495497 | 0.96144901 |
| GJD3 | 66.5991596 | 0.107916 | 0.36246864 | 0.29772506 | 0.76591302 | 0.96704052 |
| TOP2A | 112.600679 | 0.36328161 | 0.87522844 | 0.41507062 | 0.67809021 | 0.95020851 |
| IGFBP4 | 43174.7212 | 0.02677254 | 0.26126282 | 0.10247357 | 0.91838079 | 0.98689073 |
| TNS4 | 103.976479 | 1.4497977 | 0.68456693 | 2.11783194 | 0.0341893 | 0.55369025 |
| CCR7 | 74.1575408 | -0.0523893 | 0.86103611 | -0.0608445 | 0.95148302 | 0.99170503 |
| SMARCE1 | 4806.04557 | -0.059253 | 0.1546516 | -0.3831389 | 0.7016168 | 0.95424422 |
| AC073508.2 | 4473.38909 | -0.0599123 | 0.15122505 | -0.3961797 | 0.69197244 | 0.95165887 |
| AC073508.3 | 251.524637 | 0.28219709 | 0.241089 | 1.17051003 | 0.24179578 | 0.81269742 |
| KRT222 | 60.2745168 | 1.49170082 | 0.64816515 | 2.3014209 | 0.02136785 | 0.51209068 |
| KRT10 | 722.236495 | -0.304538 | 0.32130459 | -0.947817 | 0.3432226 | 0.85936336 |
| TMEM99 | 555.248347 | 0.34839274 | 0.26388761 | 1.32023153 | 0.18675773 | 0.76957489 |
| KRT15 | 55.4990237 | 2.24526837 | 0.80430499 | 2.79156341 | 0.00524541 | 0.3110892 |
| KRT19 | 10873.6519 | 1.15141799 | 0.53422899 | 2.15528921 | 0.03113921 | 0.54466273 |
| KRT14 | 22.6778377 | -0.2227627 | 1.60138255 | -0.1391065 | NA | NA |
| KRT17 | 219.647066 | 0.4516158 | 0.7890767 | 0.57233447 | 0.5670954 | 0.92561704 |
| EIF1 | 26731.3777 | -0.0832821 | 0.2411186 | -0.3453991 | 0.72979437 | 0.95848028 |
| HAP1 | 608.826805 | -0.8627665 | 0.38390929 | -2.2473187 | 0.02461967 | 0.52017724 |
| JUP | 19766.0117 | -0.2048289 | 0.17361402 | -1.1797949 | 0.2380818 | 0.81007461 |
| P3H4 | 1025.72501 | -0.1332643 | 0.27427505 | -0.4858782 | 0.62705353 | 0.94088547 |
| FKBP10 | 2717.59538 | -0.4120813 | 0.38616493 | -1.0671122 | 0.28592115 | 0.83107779 |
| NT5C3B | 2569.27167 | -0.1548 | 0.18839777 | -0.8216657 | 0.41126719 | 0.88268208 |
| KLHL10 | 49.6585818 | -0.1729051 | 0.39558309 | -0.4370893 | 0.66204659 | 0.94642611 |
| ACLY | 9220.57622 | 0.0454757 | 0.24383745 | 0.18650008 | 0.85205262 | 0.97872457 |
| AC125257.2 | 1436.18036 | 0.03202866 | 0.23682236 | 0.1352434 | 0.89241944 | 0.98474725 |
| TTC25 | 212.775654 | -0.059094 | 0.3403636 | -0.1736203 | 0.86216385 | 0.98176973 |
| CNP | 9398.88786 | -0.4331839 | 0.27278321 | -1.5880154 | 0.11228287 | 0.68001961 |
| DNAJC7 | 4900.29838 | -0.001121 | 0.08942809 | -0.0125351 | 0.98999873 | 0.99807483 |
| NKIRAS2 | 2154.21923 | -0.1809848 | 0.10633572 | -1.7020137 | 0.0887528 | 0.65363785 |
| ZNF385C | 193.217196 | 0.03937315 | 0.3681812 | 0.10693961 | 0.91483689 | 0.9866558 |
| C17orf113 | 249.555657 | 0.12370599 | 0.50509439 | 0.24491658 | 0.80652102 | 0.97334138 |
| DHX58 | 1770.12068 | -1.1729764 | 0.68098866 | -1.722461 | NA | NA |
| KAT2A | 6005.75 | -0.1855011 | 0.30362301 | -0.6109586 | 5.41E-01 | 9.23E-01 |
| AC099811.2 | 9420.23069 | -0.0027079 | 0.16702434 | -0.0162127 | 0.98706472 | 0.99728179 |
| RAB5C | 12804.5002 | -0.0227193 | 0.1555254 | -0.1460808 | 0.88385764 | 0.98317437 |
| GHDC | 4015.68107 | 0.07433335 | 0.20382825 | 0.3646862 | 0.71534566 | 0.95590747 |
| STAT5B | 4629.77686 | -0.0226073 | 0.17268773 | -0.1309145 | 0.89584298 | 0.98474725 |
| STAT5A | 2402.16345 | -0.0147818 | 0.15391933 | -0.0960359 | 0.92349205 | 0.98858365 |
| STAT3 | 14584.5943 | -0.1697977 | 0.23159217 | -0.7331756 | 0.46345135 | 0.89830907 |
| CAVIN1 | 6953.00258 | -0.018091 | 0.43740135 | -0.0413601 | 0.96700884 | 0.99400331 |
| ATP6V0A1 | 8938.12888 | 0.23673506 | 0.29899505 | 0.79176914 | 0.4284953 | 0.88820133 |
| NAGLU | 5615.44822 | -0.2430276 | 0.32975783 | -0.7369882 | 0.46112954 | 0.89755243 |
| HSD17B1 | 1125.94149 | -0.1331691 | 0.16723634 | -0.7962929 | 0.42586181 | 0.88641523 |
| AC067852.2 | 1017.58003 | -0.1542419 | 0.16550869 | -0.931926 | 0.35137475 | 0.86364633 |
| COASY | 7642.10706 | 0.08479014 | 0.18044856 | 0.46988541 | 0.63843689 | 0.94088547 |
| MLX | 3619.39956 | 0.0718075 | 0.20222601 | 0.35508537 | 0.72252564 | 0.95741464 |
| PSMC3IP | 1039.80145 | -0.1084108 | 0.15344025 | -0.7065345 | 0.47985578 | 0.90532051 |
| RETREG3 | 3758.16574 | -0.1103443 | 0.11412963 | -0.9668331 | 0.33362745 | 0.855011 |
| TUBG1 | 2362.05485 | 0.13579501 | 0.28503803 | 0.47641017 | 0.63378219 | 0.94088547 |
| TUBG2 | 2571.4063 | -0.1453151 | 0.27745238 | -0.5237478 | 6.00E-01 | 0.93487648 |
| PLEKHH3 | 2065.87043 | -0.016183 | 0.33683705 | -0.0480441 | 9.62E-01 | 0.99320711 |
| CCR10 | 108.529286 | 0.16332968 | 0.30964921 | 0.52746679 | 0.59786947 | 0.93457934 |
| AC100793.2 | 43.2538146 | 0.25823859 | 0.46516607 | 0.55515354 | 0.57878965 | 0.93139743 |
| CNTNAP1 | 875.882007 | 0.17146345 | 0.3556785 | 0.48207425 | 0.6297532 | 0.94088547 |

| | | | | | | |
|---|---|---|---|---|---|---|
| AC100793.3 | 217.602495 | 0.28513871 | 0.46043996 | 0.61927446 | 5.36E-01 | 0.92101152 |
| EZH1 | 3993.69647 | -0.1283087 | 0.14904485 | -0.8608728 | 0.38930811 | 0.87568585 |
| RAMP2-AS1 | 262.411526 | 0.08166793 | 0.39275333 | 0.20793696 | 8.35E-01 | 0.97550416 |
| RAMP2 | 902.796531 | 0.24973273 | 0.44556611 | 0.56048411 | 0.57514928 | 0.93051727 |
| VPS25 | 3533.83695 | 0.00197414 | 0.15515289 | 0.01272381 | 0.98984814 | 0.99807483 |
| WNK4 | 5657.67399 | 0.05652757 | 0.47603931 | 0.11874558 | 0.90547692 | 0.98625536 |
| COA3 | 6752.16898 | 0.26194372 | 0.23362231 | 1.12122735 | 0.2621911 | 0.82424998 |
| CNTD1 | 2203.20256 | 0.03661064 | 0.13548808 | 0.27021299 | 0.7869964 | 0.97090793 |
| BECN1 | 4530.71694 | 0.07290116 | 0.12088718 | 0.60305116 | 0.54647466 | 0.92345821 |
| PSME3 | 5119.27663 | -0.275192 | 0.18610476 | -1.4786938 | 0.13922218 | 0.72409321 |
| AOC2 | 183.293654 | -0.5727583 | 0.52012482 | -1.101194 | 0.27081224 | 0.82716979 |
| AOC3 | 2586.90784 | -0.3172094 | 0.42932489 | -0.7388564 | 0.45999419 | 0.89755243 |
| LINC00671 | 3310.78831 | 1.66996381 | 1.02243428 | 1.63332142 | 0.10240141 | 0.67279543 |
| G6PC | 4943.13016 | 1.34889747 | 1.50327417 | 0.89730636 | 0.36955547 | 0.86762749 |
| PTGES3L-AA | 1433.99353 | 0.21299649 | 0.1347604 | 1.58055698 | 0.11397937 | 0.68214043 |
| AARSD1 | 1526.23726 | 0.19819632 | 0.12901196 | 1.53626321 | 0.12447384 | 0.70116147 |
| PTGES3L | 39.4874438 | -0.0739987 | 0.43450641 | -0.1703053 | 0.86477005 | 0.98197545 |
| RUNDC1 | 1369.31429 | -0.1894169 | 0.16619154 | -1.1397505 | 0.25439027 | 0.81992039 |
| RPL27 | 14095.7187 | 0.10247711 | 0.2283012 | 0.44886803 | 0.65352686 | 0.94495971 |
| IFI35 | 2784.47968 | -1.4684009 | 0.65640394 | -2.2370385 | NA | NA |
| VAT1 | 13146.1956 | -0.1169401 | 0.33763053 | -0.3463554 | 0.72907566 | 0.95848028 |
| RND2 | 130.455024 | 0.10754652 | 0.37040285 | 0.29035013 | 7.72E-01 | 0.96836854 |
| BRCA1 | 157.143206 | 0.19671807 | 0.38302574 | 0.51358969 | 0.60753889 | 0.93739614 |
| AC060780.1 | 881.989341 | -0.111993 | 0.31887359 | -0.3512145 | 0.72542741 | 0.95848028 |
| NBR1 | 7946.24082 | 0.29391092 | 0.22778908 | 1.29027657 | 1.97E-01 | 0.77816059 |
| TMEM106A | 1437.36146 | 0.16018698 | 0.51174304 | 0.31302229 | 0.75426373 | 0.96348234 |
| LINC00854 | 608.570436 | 0.11750524 | 0.47475267 | 0.24750833 | 0.80451485 | 0.97287947 |
| LINC00910 | 137.285307 | 0.51931135 | 0.42383207 | 1.22527619 | 0.22047118 | 0.79912826 |
| ARL4D | 2105.59567 | 0.30977688 | 0.21737335 | 1.42509133 | 0.1541308 | 0.74258845 |
| DHX8 | 1715.79894 | -0.0419908 | 0.15301394 | -0.2744245 | 7.84E-01 | 0.96978847 |
| ETV4 | 435.985733 | 0.86759578 | 0.42489222 | 2.04191968 | 0.0411595 | 0.55871557 |
| SOST | 351.147859 | 3.17871167 | 1.10891667 | 2.86650185 | 0.00415036 | 0.27935586 |
| DUSP3 | 8653.21385 | -0.1691426 | 0.27327281 | -0.6189515 | 0.53594834 | 0.92101152 |
| MPP3 | 440.919302 | -0.6535664 | 0.16969678 | -3.8513777 | 0.00011746 | 0.05015467 |
| CD300LG | 118.859857 | 0.4409896 | 0.73492968 | 0.60004325 | 0.54847741 | 0.92373235 |
| MPP2 | 228.201301 | -0.0557636 | 0.46884174 | -0.1189391 | 9.05E-01 | 0.98625102 |
| AC007993.2 | 38.5190108 | 1.30498816 | 0.9339428 | 1.39728917 | 0.16232663 | 0.75026273 |
| LINC01976 | 29.7513539 | 1.21679996 | 0.94187681 | 1.29188864 | 0.19639571 | 0.77798606 |
| PYY | 73.6575536 | 1.19447552 | 0.48939109 | 2.4407382 | 0.01465728 | 0.44091816 |
| NAGS | 1274.22796 | 0.83621136 | 0.49562099 | 1.68719926 | 0.09156504 | 0.65518742 |
| TMEM101 | 4164.53609 | 0.66980002 | 0.2874304 | 2.33030331 | 0.01979013 | 0.49477123 |
| LSM12 | 674.769669 | -0.1713569 | 0.17153859 | -0.998941 | 0.31782326 | 0.84872125 |
| G6PC3 | 3772.64631 | 0.11578479 | 0.13424326 | 0.86249981 | 0.38841254 | 0.8756453 |
| HDAC5 | 4256.45692 | -0.0829737 | 0.16388093 | -0.5063051 | 0.6126425 | 0.93879706 |
| C17orf53 | 63.6531111 | 0.56818166 | 0.43617375 | 1.30264983 | 0.19269434 | 0.77495378 |
| ASB16 | 455.432362 | -0.0100382 | 0.33098573 | -0.0303283 | 0.97580519 | 0.99455099 |
| ASB16-AS1 | 687.356601 | -0.0107872 | 0.30251048 | -0.035659 | 0.97155426 | 0.99404213 |
| TMUB2 | 3232.36597 | -0.2166416 | 0.11455121 | -1.8912206 | 0.0585949 | 0.60332684 |
| ATXN7L3 | 4022.08932 | -0.4779773 | 0.19705329 | -2.4256243 | 0.01528208 | 0.45205627 |
| AC004596.1 | 196.168741 | -0.1689528 | 0.20490549 | -0.8245403 | 0.40963265 | 0.88249519 |
| UBTF | 4861.12571 | 0.01258065 | 0.19984882 | 0.06295084 | 0.94980565 | 0.99170503 |
| AC003102.1 | 413.39334 | -0.0154678 | 0.28837731 | -0.0536372 | 0.9572242 | 0.99205545 |
| SLC4A1 | 2451.84961 | 1.61975021 | 0.57615136 | 2.81132759 | 0.00493375 | 0.30302984 |
| RUNDC3A-AS | 72.5504625 | 0.6532407 | 0.47720245 | 1.36889637 | 0.17103167 | 0.75956362 |

| | | | | | | |
|---|---|---|---|---|---|---|
| RUNDC3A | 149.795126 | 0.14600533 | 0.44564626 | 0.32762607 | 0.74319441 | 0.96109622 |
| SLC25A39 | 12472.8436 | 0.18435027 | 0.22626874 | 0.81474032 | 0.41522097 | 0.88377697 |
| GRN | 21805.138 | -0.1765221 | 0.25100217 | -0.703269 | 0.48188811 | 0.90541935 |
| FAM171A2 | 102.786553 | 0.09373092 | 0.51296563 | 0.18272359 | 0.85501491 | 0.97910552 |
| ITGA2B | 132.183449 | -0.4901165 | 0.496111 | -0.9879169 | 0.32319333 | 0.85119105 |
| GPATCH8 | 2152.58995 | -0.1016831 | 0.13655326 | -0.7446402 | 0.45648923 | 0.89688594 |
| FZD2 | 164.365371 | 0.2068135 | 0.25748691 | 0.80320007 | 0.42185911 | 0.88451372 |
| MEIOC | 448.341924 | 0.12239222 | 0.28541678 | 0.42881929 | 0.66805474 | 0.9484859 |
| CCDC43 | 924.70834 | 0.07325983 | 0.2107323 | 0.34764406 | 0.72810751 | 0.95848028 |
| DBF4B | 518.367368 | -0.1942061 | 0.41795272 | -0.4646604 | 0.64217467 | 0.94088547 |
| ADAM11 | 65.9669517 | -0.0515352 | 0.38118586 | -0.135197 | 0.89245609 | 0.98474725 |
| GJC1 | 255.283111 | -1.1008242 | 0.58538659 | -1.8805081 | 0.06003886 | 0.6075531 |
| HIGD1B | 84.1678142 | -0.516952 | 0.58800503 | -0.8791626 | 0.37931313 | 0.87069278 |
| EFTUD2 | 3822.69072 | -0.138796 | 0.15423608 | -0.8998931 | 0.36817713 | 0.86759809 |
| CCDC103 | 381.904714 | 0.01611024 | 0.28876258 | 0.05579061 | 0.95550862 | 0.99185043 |
| FAM187A | 314.137683 | 0.06539333 | 0.36268354 | 0.1803041 | 0.85691383 | 0.97953896 |
| GFAP | 224.257559 | -0.0660949 | 0.35568359 | -0.1858249 | 0.85258208 | 0.97883015 |
| KIF18B | 43.4291593 | -0.5178858 | 0.82616444 | -0.6268556 | 0.53075387 | 0.91962754 |
| C1QL1 | 205.332411 | 0.37314044 | 0.64443367 | 0.57902071 | 0.5625752 | 0.92452095 |
| DCAKD | 1721.38617 | 0.44335761 | 0.18698487 | 2.37108818 | 0.0177358 | 0.4711578 |
| NMT1 | 5868.36927 | -0.0867059 | 0.13590744 | -0.6379775 | 0.52348833 | 0.91723878 |
| PLCD3 | 1579.90573 | 0.30351853 | 0.26651947 | 1.13882311 | 0.25477694 | 0.81992039 |
| ACBD4 | 2148.39719 | 0.30079001 | 0.40150431 | 0.74915762 | 0.45376221 | 0.89670027 |
| AC142472.1 | 124.607458 | 0.41519358 | 0.29331906 | 1.41550153 | 0.1569215 | 0.74522306 |
| HEXIM1 | 4313.21539 | -0.2619779 | 0.16337746 | -1.6035132 | 0.10882139 | 0.67800851 |
| AC138150.1 | 631.492731 | -0.1992712 | 0.1861334 | -1.0705829 | 2.84E-01 | 0.83107779 |
| HEXIM2 | 676.541632 | 0.2172262 | 0.31279496 | 0.69446834 | 0.48738855 | 0.90573694 |
| AC138150.2 | 524.076696 | 0.24088907 | 0.29161234 | 0.82605926 | 0.40877048 | 0.88203324 |
| AC008105.3 | 189.265154 | 0.55355157 | 0.74440468 | 0.74361646 | 0.45710854 | 0.89688594 |
| FMNL1 | 1275.05995 | -0.1483269 | 0.53473354 | -0.2773847 | 0.78148475 | 0.96913822 |
| AC008105.1 | 226.265234 | -0.0465598 | 0.50064999 | -0.0929988 | 0.92590452 | 0.98876911 |
| MAP3K14-AS | 613.577679 | -0.0766449 | 0.30733546 | -0.2493852 | 0.80306284 | 0.97273362 |
| MAP3K14 | 2180.56966 | -0.0505941 | 0.31707802 | -0.1595636 | 0.87322486 | 0.98275231 |
| ARHGAP27 | 3923.08386 | 0.04896551 | 0.23214177 | 0.21092934 | 0.83294241 | 0.97550416 |
| AC003070.1 | 357.841095 | 0.15293347 | 0.25985868 | 0.58852555 | 0.55617959 | 0.92373235 |
| PLEKHM1 | 3179.77753 | -1.1582987 | 0.45630747 | -2.5384171 | 0.01113552 | 0.40196898 |
| LINC02210-C | 105.339039 | -0.0824347 | 0.45042188 | -0.1830167 | 0.8547849 | 0.97910552 |
| MAPT-AS1 | 115.788947 | 0.38086126 | 0.6676116 | 0.57048328 | 0.56834996 | 0.92638187 |
| MAPT | 2651.93331 | 0.35631028 | 0.51366308 | 0.69366536 | 0.4878921 | 0.90584365 |
| KANSL1 | 2735.42677 | -0.2723899 | 0.14514801 | -1.8766358 | 0.06056803 | 0.60928905 |
| KANSL1-AS1 | 182.64083 | 1.08803749 | 0.56796078 | 1.91569125 | 0.05540441 | 0.59640092 |
| ARL17B | 69.6978913 | -0.0863684 | 0.57164598 | -0.1510873 | 0.87990688 | 0.98275231 |
| LRRC37A | 99.4467716 | -0.2817758 | 0.43447984 | -0.648536 | 0.51663831 | 0.91636259 |
| LRRC37A2 | 444.62791 | -0.1898448 | 0.38261818 | -0.496173 | 0.61977239 | 0.93945324 |
| ARL17A | 165.252332 | -0.2309023 | 0.36355733 | -0.6351194 | 0.52535052 | 0.91759401 |
| NSF | 1806.91879 | -0.021967 | 0.12771698 | -0.1719975 | 0.86343946 | 0.98176973 |
| WNT3 | 186.166031 | -0.241729 | 0.44476076 | -0.5435035 | 0.58678317 | 0.93289552 |
| WNT9B | 143.947177 | 0.73744979 | 0.95535489 | 0.77191188 | 0.44016662 | 0.89238021 |
| GOSR2 | 3508.89186 | 0.05345768 | 0.22179535 | 0.24102253 | 0.80953765 | 0.97334138 |
| AC005670.2 | 1145.83013 | 0.19588943 | 0.28641268 | 0.68394121 | 0.49401229 | 0.90785294 |
| CDC27 | 2263.34608 | -0.3075635 | 0.18987267 | -1.6198409 | 0.10526645 | 0.67473222 |
| AC002558.2 | 49.768635 | -0.1424774 | 0.48763327 | -0.2921815 | 0.77014788 | 0.96836854 |
| ITGB3 | 1606.5098 | -0.1898878 | 0.33146689 | -0.572871 | 0.56673201 | 0.9256133 |
| AC068234.1 | 868.620954 | 0.06899197 | 0.26263821 | 0.26268826 | 0.79279087 | 0.97090793 |

| | | | | | | |
|---|---|---|---|---|---|---|
| AC068234.2 | 189.131489 | 0.3464949 | 0.37996699 | 0.91190792 | 0.36181719 | 0.86528137 |
| EFCAB13 | 491.889022 | 0.54477938 | 0.31138158 | 1.74955556 | 0.08019503 | 0.6408661 |
| NPEPPS | 6222.5106 | -0.1362365 | 0.13587041 | -1.0026947 | 0.3160082 | 0.84750116 |
| AC025682.1 | 1415.30727 | -0.160604 | 0.15808021 | -1.0159653 | 0.30964592 | 0.84432914 |
| KPNB1 | 7770.92528 | -0.2055081 | 0.17505179 | -1.1739844 | 0.24040128 | 0.8124487 |
| AC015674.1 | 1298.46754 | -0.2187869 | 0.18764112 | -1.1659858 | 0.24362021 | 0.81384782 |
| TBKBP1 | 2139.03591 | -0.5978544 | 0.23583893 | -2.5350115 | 0.01124436 | 0.40196898 |
| TBX21 | 85.2938077 | -0.3146065 | 0.70465757 | -0.4464671 | 0.65525984 | 0.94532497 |
| OSBPL7 | 1160.26585 | -0.0316842 | 0.27031161 | -0.1172136 | 0.9066908 | 0.98662757 |
| MRPL10 | 2419.2377 | 0.03624041 | 0.13971136 | 0.25939486 | 0.79533059 | 0.97101097 |
| LRRC46 | 235.823309 | 0.69155069 | 0.39977069 | 1.72986842 | 0.08365379 | 0.64783289 |
| SCRN2 | 6550.96698 | 0.96765258 | 0.42767665 | 2.26257985 | 0.0236616 | 0.51967919 |
| SP6 | 135.810457 | -0.2294127 | 0.33237739 | -0.6902177 | 0.49005732 | 0.90600166 |
| SP2 | 1395.86735 | -0.2765894 | 0.20839226 | -1.3272539 | 0.18442472 | 0.76820626 |
| SP2-AS1 | 59.1170753 | 0.13678175 | 0.34243768 | 0.39943545 | 0.68957238 | 0.95158502 |
| PNPO | 4571.49364 | 1.100357 | 0.36135139 | 3.04511624 | 0.0023259 | 0.21057858 |
| AC018521.2 | 1500.32763 | 1.05226713 | 0.37818527 | 2.78241176 | 0.00539565 | 0.31308771 |
| PRR15L | 1837.62666 | 0.45706151 | 0.58588771 | 0.78011793 | 0.43532146 | 0.89055338 |
| CDK5RAP3 | 10577.9754 | 0.19101089 | 0.16793325 | 1.13742154 | 0.25536211 | 0.82031903 |
| AC018521.4 | 1945.97871 | 0.26696612 | 0.17731519 | 1.50560207 | 0.13216936 | 0.7126419 |
| COPZ2 | 518.944333 | 0.16316322 | 0.27869029 | 0.5854643 | 0.55823556 | 0.92421175 |
| NFE2L1 | 24242.3326 | 0.07940243 | 0.2392898 | 0.33182538 | 0.74002112 | 0.96043876 |
| AC004477.2 | 5756.44863 | 0.0333184 | 0.18855944 | 0.17669969 | 0.85974428 | 0.98043493 |
| CBX1 | 2809.86102 | -0.0937272 | 0.15724721 | -0.5960503 | 0.55114162 | 0.92373235 |
| SNX11 | 1522.10735 | -0.2936494 | 0.16883202 | -1.7392994 | 0.08198211 | 0.64555743 |
| SKAP1 | 789.535129 | 0.35859283 | 0.28238051 | 1.26989228 | 0.204123 | 0.78419929 |
| HOXB2 | 2285.61236 | -0.0669798 | 0.33176533 | -0.2018891 | 0.84000344 | 0.97618283 |
| HOXB-AS1 | 1736.75076 | -0.0192298 | 0.30231176 | -0.0636093 | 0.94928132 | 0.99170503 |
| HOXB3 | 4344.78976 | -0.0455233 | 0.28822847 | -0.1579416 | 0.87450284 | 0.98275231 |
| HOXB-AS3 | 3206.90683 | 0.0885351 | 0.25376719 | 0.34888315 | 7.27E-01 | 9.58E-01 |
| HOXB4 | 1063.32163 | -0.0769096 | 0.22438231 | -0.3427614 | 0.73177796 | 0.95848028 |
| HOXB5 | 1266.99799 | 0.24013705 | 0.34742868 | 0.69118372 | 0.48945009 | 0.90600166 |
| HOXB6 | 3304.6837 | 0.07867796 | 0.27586212 | 0.28520756 | 0.77548513 | 0.96836854 |
| HOXB7 | 1331.89064 | -0.0205119 | 0.20323081 | -0.1009291 | 0.91960674 | 0.9872458 |
| HOXB8 | 1530.2163 | 0.28912715 | 0.30125305 | 0.95974846 | 0.33718183 | 0.85694665 |
| HOXB9 | 1745.50232 | 0.25978016 | 0.32379747 | 0.80229211 | 0.42238401 | 0.88451372 |
| AC103702.2 | 100.673427 | -0.2887621 | 0.56076554 | -0.5149426 | 0.60659312 | 0.93708311 |
| LINC02086 | 231.739288 | -0.6164556 | 0.66083408 | -0.9328447 | 0.35090014 | 0.86364633 |
| TTLL6 | 198.203304 | 0.35151214 | 0.44399307 | 0.79170638 | 0.4285319 | 0.88820133 |
| CALCOCO2 | 5649.61374 | -0.1057117 | 0.17485045 | -0.6045836 | 0.54545569 | 0.92345821 |
| ATP5MC1 | 4145.89522 | 0.41027746 | 0.23715815 | 1.7299741 | 0.0836349 | 0.64783289 |
| UBE2Z | 6025.48113 | -0.3765577 | 0.17472243 | -2.1551765 | 0.03114803 | 0.54466273 |
| SNF8 | 3415.0458 | -0.0526649 | 0.14607663 | -0.3605293 | 0.71845132 | 0.95648644 |
| AC091133.1 | 1834.07396 | 0.04548902 | 0.16329283 | 0.27857331 | 0.7805723 | 0.96864684 |
| IGF2BP1 | 201.558123 | 0.61974431 | 0.67790544 | 0.91420465 | 0.36060932 | 0.86504879 |
| B4GALNT2 | 488.640757 | 0.86346145 | 0.59848632 | 1.44274215 | 0.14909312 | 0.73666073 |
| GNGT2 | 93.0185766 | -0.495448 | 0.37578282 | -1.3184423 | 1.87E-01 | 7.70E-01 |
| ABI3 | 607.862982 | -0.4353473 | 0.32710961 | -1.3308913 | 1.83E-01 | 7.67E-01 |
| PHOSPHO1 | 48.0618753 | 0.32435582 | 0.67464611 | 0.4807792 | 0.63067343 | 0.94088547 |
| ZNF652 | 1940.89414 | -0.0503123 | 0.13418884 | -0.3749364 | 0.70770777 | 0.95501548 |
| PHB | 11289.1269 | 0.24602807 | 0.16450262 | 1.49558756 | 0.13476117 | 0.71630293 |
| AC091180.3 | 6671.96555 | 0.23983445 | 0.17124589 | 1.40052676 | 0.16135564 | 0.74924826 |
| NGFR | 775.514645 | -0.7073217 | 0.5519282 | -1.2815466 | 0.20000176 | 0.78134034 |
| AC006487.1 | 44.1599911 | -0.9469659 | 0.65198508 | -1.452435 | 0.14638069 | 0.73247558 |

| | | | | | | |
|---|---|---|---|---|---|---|
| NXPH3 | 107.782572 | -0.5419618 | 0.45389389 | -1.1940276 | 0.23246717 | 0.80649152 |
| SPOP | 2410.96066 | -0.2319005 | 0.09286899 | -2.4970719 | 0.01252236 | 0.41598747 |
| SLC35B1 | 2578.50122 | 0.12869664 | 0.18262313 | 0.70471161 | 0.48098972 | 0.90541935 |
| AC015795.1 | 145.231868 | 0.22758612 | 0.25581256 | 0.88965966 | 3.74E-01 | 0.86791605 |
| FAM117A | 1084.03875 | 0.17908667 | 0.21123641 | 0.84780208 | 0.3965482 | 0.8796941 |
| KAT7 | 2311.88128 | -0.0536579 | 0.10663328 | -0.5031999 | 0.61482375 | 0.93895494 |
| FLJ45513 | 140.097844 | -0.5194167 | 0.36664177 | -1.4166871 | 0.15657442 | 0.74522306 |
| AC009720.1 | 161.618305 | -0.4822014 | 0.44806598 | -1.076184 | 2.82E-01 | 8.30E-01 |
| ITGA3 | 22748.1346 | -0.1398162 | 0.33051658 | -0.4230232 | 0.67227835 | 0.94922827 |
| AC002401.4 | 59.129622 | -0.1243653 | 0.81835774 | -0.1519694 | 0.87921109 | 0.98275231 |
| PDK2 | 6384.14311 | 0.60178698 | 0.37603832 | 1.6003342 | 0.10952446 | 0.67859123 |
| SAMD14 | 3728.78069 | 0.47934007 | 0.32473599 | 1.4760916 | 1.40E-01 | 7.25E-01 |
| PPP1R9B | 4844.96912 | -0.2570969 | 0.22329941 | -1.1513551 | 0.24958618 | 0.81646572 |
| AC002401.1 | 1037.15723 | -0.134529 | 0.28643677 | -0.4696638 | 0.63859522 | 0.94088547 |
| SGCA | 198.815794 | -1.0995909 | 0.76714494 | -1.4333548 | NA | NA |
| COL1A1 | 9708.27104 | -0.6979801 | 0.51501363 | -1.3552653 | 0.17533304 | 0.76073329 |
| TMEM92 | 41.5287122 | 0.70267181 | 0.90429524 | 0.77703805 | 0.43713632 | 0.89130865 |
| XYLT2 | 2138.34354 | 0.13888501 | 0.27205695 | 0.51049978 | 0.60970137 | 0.93766726 |
| MRPL27 | 2376.84871 | -0.0841205 | 0.17377945 | -0.4840648 | 0.62833989 | 0.94088547 |
| EME1 | 662.605188 | -0.2585054 | 0.30034255 | -0.8607018 | 0.3894023 | 0.87568585 |
| LRRC59 | 4885.34649 | -0.2416039 | 0.32110352 | -0.7524173 | 0.45180014 | 0.89656781 |
| AC004707.1 | 427.221007 | -0.234745 | 0.42127829 | -0.5572208 | 0.57737658 | 0.93130183 |
| ACSF2 | 12066.6447 | 0.99262802 | 0.48176331 | 2.06040598 | 0.03935975 | 0.55657615 |
| CHAD | 377.38277 | 0.73216106 | 0.44661552 | 1.63935429 | 0.1011395 | 0.6725344 |
| AC021491.4 | 142.559858 | 1.29953575 | 0.63205648 | 2.05604371 | 0.0397783 | 0.55657615 |
| RSAD1 | 4198.22085 | -0.0157869 | 0.20973077 | -0.0752721 | 0.93999823 | 0.99136103 |
| MYCBPAP | 73.1451867 | 0.16826207 | 0.52171772 | 0.32251554 | 0.74706217 | 0.96224616 |
| EPN3 | 846.689308 | 0.85556582 | 0.50984659 | 1.6780848 | 0.09333055 | 0.65756964 |
| SPATA20 | 6465.57445 | 0.4584611 | 0.27542398 | 1.66456494 | 0.09599958 | 0.66168746 |
| ABCC3 | 7460.76252 | 0.62768168 | 0.4247597 | 1.4777336 | 0.1394791 | 0.7246958 |
| ANKRD40 | 2591.96507 | -0.3527714 | 0.21514817 | -1.6396674 | 0.10107435 | 0.6725344 |
| AC005921.2 | 959.689495 | -0.373958 | 0.22836865 | -1.637519 | 0.10152207 | 0.67279543 |
| LUC7L3 | 8490.33249 | -0.1513458 | 0.15834008 | -0.9558276 | 0.33915934 | 0.85733087 |
| TOB1 | 4301.82079 | 0.0533066 | 0.30603991 | 0.17418185 | 0.86172255 | 0.98176973 |
| AC005920.4 | 89.6268383 | -0.6480207 | 0.75194969 | -0.8617873 | 0.38880457 | 0.87568585 |
| SPAG9 | 6676.1727 | -0.4572429 | 0.38450328 | -1.1891781 | 2.34E-01 | 0.80730051 |
| NME1 | 2055.80645 | -0.0714781 | 0.28656392 | -0.2494317 | 0.80302687 | 0.97273362 |
| NME1-NME2 | 13223.7687 | -0.1955854 | 0.25208474 | -0.7758717 | 0.43782474 | 0.89139309 |
| NME2 | 11641.3876 | -0.2136827 | 0.25249467 | -0.8462859 | 0.39739326 | 0.88006972 |
| MBTD1 | 1133.10145 | -0.2767421 | 0.16074216 | -1.7216522 | 0.08513255 | 0.64988556 |
| UTP18 | 1663.78846 | -0.15155 | 0.25464342 | -0.5951458 | 0.55174604 | 0.92373235 |
| CA10 | 446.235338 | 0.25827825 | 0.40457366 | 0.63839611 | 0.52321586 | 0.91723878 |
| TOM1L1 | 4066.94835 | 0.26652886 | 0.14522134 | 1.83532851 | 0.06645703 | 0.62378208 |
| COX11 | 4617.9441 | 0.13058685 | 0.12694286 | 1.02870577 | 0.30361796 | 0.84211658 |
| STXBP4 | 1023.82309 | -0.0979793 | 0.29256637 | -0.3348958 | 0.73770366 | 0.9602325 |
| HLF | 1492.7228 | 0.34878466 | 0.38872124 | 0.89726164 | 0.36957932 | 0.86762749 |
| AC007638.2 | 39.2881471 | 1.07824623 | 0.89076172 | 1.21047661 | NA | NA |
| MMD | 838.557592 | 0.30245074 | 0.29567151 | 1.02292823 | 0.30634178 | 0.84278367 |
| TMEM100 | 82.2033199 | 0.46221932 | 0.31230996 | 1.48000184 | 0.13887276 | 0.72399122 |
| PCTP | 1714.2886 | 0.31125837 | 0.34297898 | 0.90751441 | 0.36413483 | 0.8663515 |
| C17orf67 | 87.839082 | -0.6404325 | 0.47970104 | -1.3350658 | 0.1818548 | 0.76543911 |
| DGKE | 467.720903 | 0.03669856 | 0.24541735 | 0.14953531 | 0.88113125 | 0.98275231 |
| TRIM25 | 2681.18329 | -0.9930997 | 0.37103528 | -2.6765641 | 0.00743813 | 0.36301578 |
| AC015912.1 | 344.523831 | -1.0913555 | 0.41798923 | -2.6109655 | 0.0090287 | 0.38593543 |

| | | | | | | |
|---|---|---|---|---|---|---|
| AC015912.3 | 57.8235983 | -2.264297 | 0.92124931 | -2.4578548 | NA | NA |
| COIL | 792.054943 | -0.1806911 | 0.20519985 | -0.8805616 | 0.37855514 | 0.87066232 |
| SCPEP1 | 5777.31357 | 0.48496305 | 0.11265502 | 4.30485077 | 1.67E-05 | 0.01316658 |
| AC007114.1 | 128.382749 | 0.50959468 | 0.46293909 | 1.10078127 | 0.27099187 | 0.82716979 |
| AKAP1 | 3590.89083 | 0.05058869 | 0.20370115 | 0.24834758 | 0.80386549 | 0.97274243 |
| MSI2 | 5541.77579 | -0.0764093 | 0.17197989 | -0.4442922 | 0.65683135 | 0.94545756 |
| AC015845.2 | 162.47735 | -2.5954446 | 1.03047182 | -2.5186954 | NA | NA |
| MRPS23 | 2369.86585 | -0.0238492 | 0.25075549 | -0.0951095 | 0.92422783 | 0.98869145 |
| CUEDC1 | 4679.66607 | -0.2456285 | 0.25538463 | -0.9617981 | 0.33615103 | 0.85641741 |
| VEZF1 | 3056.70474 | -0.2835926 | 0.17685712 | -1.6035128 | 0.10882149 | 0.67800851 |
| AC015813.2 | 2879.86449 | -0.0487682 | 0.13958485 | -0.3493802 | 0.72680386 | 0.95848028 |
| AC015813.1 | 1721.49773 | 0.08342318 | 0.23598338 | 0.35351294 | 0.72370393 | 0.95810643 |
| SRSF1 | 9416.18506 | -0.1366759 | 0.12255585 | -1.115213 | 0.26475915 | 0.82424998 |
| DYNLL2 | 8531.13769 | 0.22942077 | 0.19654584 | 1.16726338 | 0.24310402 | 0.81340905 |
| EPX | 36.5790067 | 0.44548644 | 0.55745786 | 0.79913921 | 0.42420969 | 0.88565404 |
| MKS1 | 956.977707 | 0.0571112 | 0.26499184 | 0.2155206 | 0.82936145 | 0.97550416 |
| LPO | 98.4877197 | 0.41965022 | 0.45142875 | 0.92960455 | 0.35257587 | 0.86364633 |
| MPO | 435.743228 | 1.91027975 | 1.08097016 | 1.76719008 | NA | NA |
| TSPOAP1 | 2862.90317 | 0.13311046 | 0.43750596 | 0.30424835 | 0.76093869 | 0.9658862 |
| TSPOAP1-AS | 1430.36221 | -4.96E-07 | 0.15375035 | -3.22E-06 | 0.99999743 | 0.99999743 |
| AC004687.1 | 66.1775332 | 0.95456873 | 0.909954 | 1.04902965 | 0.29416447 | 0.83605343 |
| SUPT4H1 | 2715.06134 | -0.0717283 | 0.15098326 | -0.4750748 | 0.63473369 | 0.94088547 |
| RNF43 | 765.783608 | 0.04459171 | 0.38354487 | 0.11626205 | 0.90744486 | 0.98662757 |
| MTMR4 | 2415.13835 | 0.45667968 | 0.30519945 | 1.49633193 | 0.13456718 | 0.71630293 |
| SEPT4-AS1 | 327.767098 | -0.1915548 | 0.42513835 | -0.4505705 | 0.65229913 | 0.9443159 |
| 4-Sep | 1274.58417 | -0.1058344 | 0.53346735 | -0.1983896 | 8.43E-01 | 0.97644404 |
| C17orf47 | 63.7566994 | -0.2660835 | 0.55931283 | -0.4757328 | 6.34E-01 | 0.94088547 |
| TEX14 | 62.5139487 | -0.7505327 | 0.42364092 | -1.7716246 | 0.07645689 | 0.63773415 |
| RAD51C | 506.170078 | 0.06238267 | 0.16755477 | 0.37231208 | 7.10E-01 | 0.95502998 |
| PPM1E | 203.601115 | -1.3928824 | 0.45199616 | -3.0816245 | 0.00205874 | 0.1948632 |
| TRIM37 | 1124.56952 | -0.3733537 | 0.21964321 | -1.699819 | 0.08916498 | 0.65375543 |
| SKA2 | 915.541158 | 0.09912461 | 0.23562597 | 0.42068627 | 6.74E-01 | 0.94935413 |
| AC099850.2 | 153.39386 | 0.02131786 | 0.22485553 | 0.09480692 | 9.24E-01 | 0.98869145 |
| SMG8 | 592.552098 | 0.06918679 | 0.25589008 | 0.27037698 | 7.87E-01 | 0.97090793 |
| GDPD1 | 139.268329 | 0.18944629 | 0.29400983 | 0.64435361 | 0.51934616 | 0.91636259 |
| YPEL2 | 2534.13485 | 0.02727177 | 0.23569579 | 0.11570749 | 0.90788437 | 0.98662757 |
| AC091059.1 | 275.336288 | 0.02718344 | 0.20089029 | 0.13531484 | 0.89236295 | 0.98474725 |
| DHX40 | 2887.57907 | 0.05310398 | 0.12419357 | 0.4275904 | 0.66894936 | 0.9486589 |
| AC091271.1 | 100.638493 | 0.54768869 | 0.45409314 | 1.20611533 | 0.22777303 | 0.80300331 |
| CLTC | 16085.2403 | 0.11802159 | 0.18153699 | 0.65012421 | 0.51561199 | 0.91636259 |
| PTRH2 | 1272.15496 | -0.50852 | 0.39424395 | -1.2898614 | 0.1970988 | 0.77816059 |
| VMP1 | 21202.5086 | -0.2047415 | 0.38258221 | -0.5351568 | 0.59254145 | 0.93289552 |
| TUBD1 | 413.373357 | 0.07354901 | 0.25200152 | 0.2918594 | 0.77039413 | 0.96836854 |
| RPS6KB1 | 1648.63954 | -0.2858198 | 0.1372153 | -2.0830027 | 0.03725098 | 0.55532957 |
| AC005702.1 | 218.943231 | -0.198367 | 0.18297021 | -1.0841489 | 0.2782988 | 0.82805357 |
| RNFT1 | 595.723517 | 0.01528941 | 0.17632275 | 0.08671262 | 0.93089995 | 0.98975562 |
| RNFT1-DT | 46.3711697 | 0.67672777 | 0.42533693 | 1.59103929 | 1.12E-01 | 6.80E-01 |
| HEATR6 | 480.629214 | -0.1018665 | 0.17098829 | -0.5957511 | 0.55134153 | 0.92373235 |
| CA4 | 1589.06971 | 0.9300206 | 0.3046798 | 3.05245247 | 0.0022698 | 0.20649641 |
| USP32 | 1389.29917 | -0.1851383 | 0.25300997 | -0.7317433 | 0.46432528 | 0.89830907 |
| APPBP2 | 2307.05382 | -0.0604482 | 0.13398639 | -0.4511519 | 0.65188007 | 0.9440713 |
| PPM1D | 884.923887 | -0.1444596 | 0.17202298 | -0.8397691 | 0.40103786 | 0.88132015 |
| BCAS3 | 1250.11442 | 0.2662208 | 0.24867366 | 1.07056293 | 0.284366 | 0.83107779 |
| AC005884.1 | 101.090309 | 0.39016595 | 0.28446511 | 1.37157753 | 0.17019499 | 0.75956362 |

| | | | | | | |
|---|---|---|---|---|---|---|
| AC005746.2 | 35.0087105 | -0.2994563 | 0.58902195 | -0.5083958 | 0.61117577 | 0.93834929 |
| TBX2-AS1 | 487.662976 | 0.41235978 | 0.28610013 | 1.44131282 | 0.14949633 | 0.73670625 |
| TBX2 | 5760.59185 | 0.17325688 | 0.24002569 | 0.72182639 | 0.47040122 | 0.9008571 |
| INTS2 | 530.4967 | -0.0458443 | 0.1695021 | -0.2704644 | 0.78680297 | 0.97090793 |
| MED13 | 2759.4339 | -0.3539341 | 0.18499335 | -1.9132263 | 0.05571909 | 0.59817776 |
| METTL2A | 636.716596 | -0.0433058 | 0.20348146 | -0.2128243 | 0.831464 | 0.97550416 |
| TLK2 | 1566.4447 | -0.3552046 | 0.15874359 | -2.2375996 | 0.02524718 | 0.52140487 |
| AC080038.1 | 98.1208411 | 0.58183505 | 0.55861135 | 1.04157399 | 0.29760923 | 0.83922579 |
| MRC2 | 3347.4478 | 0.13453239 | 0.33693293 | 0.39928537 | 0.68968294 | 0.95158502 |
| 10-Mar | 70.1455988 | 0.79620323 | 0.42604113 | 1.86884121 | 0.06164491 | 0.61093984 |
| TANC2 | 1307.03927 | -0.2307676 | 0.20110532 | -1.1474961 | 0.25117664 | 0.81815815 |
| CYB561 | 3419.16887 | 0.21351024 | 0.12381498 | 1.72442973 | 0.08463029 | 0.64988556 |
| ACE | 2580.0158 | 0.75908081 | 0.53547846 | 1.41757488 | 0.15631492 | 0.74522306 |
| AC113554.1 | 1383.11913 | 0.76352667 | 0.53503044 | 1.42707145 | 0.1535593 | 0.74258845 |
| KCNH6 | 1322.87209 | 1.34842513 | 0.65042776 | 2.07313588 | 0.03815964 | 0.55538555 |
| DCAF7 | 4450.16072 | 0.00849089 | 0.1072945 | 0.07913631 | 0.93692421 | 0.99068685 |
| TACO1 | 1730.42595 | 0.48141349 | 0.31223074 | 1.54185171 | 0.12310963 | 0.69788192 |
| MAP3K3 | 3397.93385 | -0.2179535 | 0.16577095 | -1.314787 | 0.18858149 | 0.76970626 |
| STRADA | 2209.05838 | 0.02002836 | 0.19045943 | 0.10515813 | 0.91625033 | 0.9868044 |
| LIMD2 | 1334.6152 | -0.1222203 | 0.37484115 | -0.3260589 | 0.74437982 | 0.9613927 |
| AC046185.1 | 2953.36428 | -0.0035164 | 0.11388523 | -0.0308766 | 0.97536795 | 0.99455099 |
| CCDC47 | 5937.38582 | -0.0620561 | 0.17445462 | -0.3557148 | 0.72205413 | 0.95739595 |
| DDX42 | 5102.05617 | -0.1567408 | 0.12861253 | -1.2187052 | 2.23E-01 | 0.80078694 |
| FTSJ3 | 5538.38163 | -0.1130472 | 0.1558627 | -0.7253002 | 0.46826789 | 0.89984974 |
| PSMC5 | 8172.02981 | -0.1060287 | 0.12656411 | -0.8377473 | 0.40217262 | 0.88132015 |
| SMARCD2 | 3318.22868 | 0.08793567 | 0.19027527 | 0.46214977 | 6.44E-01 | 0.94208859 |
| CD79B | 205.061308 | -0.4901473 | 0.5791762 | -0.8462836 | 3.97E-01 | 8.80E-01 |
| SCN4A | 81.6078203 | 0.38874515 | 0.72236127 | 0.53815891 | 0.59046734 | 0.93289552 |
| PRR29 | 1359.25803 | -0.6400424 | 0.25045312 | -2.5555377 | 0.01060239 | 0.40196898 |
| ICAM2 | 1680.80336 | -0.6152855 | 0.25633757 | -2.4002939 | 0.01638191 | 0.45915089 |
| ERN1 | 1387.47083 | -0.0645602 | 0.19797844 | -0.3260971 | 0.74435089 | 0.9613927 |
| SNHG25 | 203.916479 | -0.5053648 | 0.25299399 | -1.9975368 | 0.0457669 | 0.57687945 |
| TEX2 | 2716.29627 | -0.238203 | 0.24061316 | -0.9899835 | 0.32218221 | 0.85119105 |
| PECAM1 | 9687.82894 | 0.03018399 | 0.4206396 | 0.07175737 | 0.942795 | 0.99136103 |
| MILR1 | 109.775403 | 0.0224989 | 0.61512692 | 0.03657602 | 0.97082306 | 0.99400331 |
| POLG2 | 632.651019 | -0.2698161 | 0.14302245 | -1.88653 | 0.05922357 | 0.60485498 |
| DDX5 | 40654.1358 | -0.0403221 | 0.11660913 | -0.3457883 | 0.72950183 | 0.95848028 |
| CEP95 | 2597.28683 | -0.2546742 | 0.13840327 | -1.8400879 | 0.06575533 | 0.62237346 |
| SMURF2 | 1369.82612 | -0.0685313 | 0.12508614 | -0.5478729 | 0.58377918 | 0.93233498 |
| LRRC37A3 | 1003.17024 | -0.3642319 | 0.36789601 | -0.9900403 | 0.32215443 | 0.85119105 |
| GNA13 | 3415.00405 | -0.7247896 | 0.36816433 | -1.9686579 | 0.04899239 | 0.58508842 |
| RGS9 | 437.614859 | -0.4968853 | 0.45151471 | -1.1004852 | 0.27112076 | 0.82716979 |
| AXIN2 | 336.505646 | -0.0154413 | 0.35366151 | -0.0436613 | 0.96517437 | 0.99364606 |
| AC004805.1 | 170.888124 | 0.07288407 | 0.30053046 | 0.24251808 | 0.80837875 | 0.97334138 |
| CEP112 | 568.777232 | 0.0274748 | 0.29874767 | 0.09196656 | 0.9267246 | 0.98915637 |
| APOH | 348.519687 | -0.1128715 | 1.05329136 | -0.1071608 | 0.91466145 | 0.9866558 |
| PRKCA | 2468.00552 | 0.5071128 | 0.23852762 | 2.12601295 | 0.03350218 | 0.55367229 |
| HELZ | 2080.5474 | -0.2223638 | 0.14629498 | -1.5199688 | 0.12851883 | 0.70610612 |
| AC007448.3 | 74.4208771 | 0.5022737 | 0.50515706 | 0.99429216 | 0.32008065 | 0.84966452 |
| PSMD12 | 2797.98805 | -0.3655664 | 0.30690745 | -1.1911291 | 0.2336029 | 0.80725541 |
| PITPNC1 | 1567.34567 | -0.2213186 | 0.32439856 | -0.682243 | 0.49508533 | 0.90816709 |
| NOL11 | 1885.94864 | -0.0908053 | 0.21372244 | -0.4248748 | 0.67092793 | 0.94922827 |
| BPTF | 2215.86487 | -0.2601388 | 0.15087682 | -1.7241803 | 0.08467531 | 0.64988556 |
| C17orf58 | 446.595079 | 0.01212846 | 0.30150389 | 0.04022655 | 0.96791251 | 0.99400331 |

| | | | | | | |
|---|---|---|---|---|---|---|
| KPNA2 | 1810.45574 | -0.5292857 | 0.37672684 | -1.4049587 | 0.16003355 | 0.74781837 |
| AC005332.7 | 1651.77851 | -0.0223531 | 0.15067634 | -0.1483521 | 0.88206492 | 0.98275231 |
| AC005332.4 | 121.982854 | 0.23870525 | 0.2325771 | 1.02634889 | 0.30472716 | 0.84230231 |
| AC005332.6 | 1036.39956 | -0.2457423 | 0.22311079 | -1.1014361 | 0.27070688 | 0.82716979 |
| AC005332.5 | 174.789634 | 0.15949429 | 0.36846105 | 0.43286608 | 0.66511206 | 0.94694459 |
| AMZ2 | 2394.40796 | 0.30391736 | 0.22976721 | 1.32271857 | 0.18592899 | 0.76908034 |
| ARSG | 1366.02734 | 0.3419722 | 0.09392068 | 3.64107448 | 0.0002715 | 0.08298534 |
| SLC16A6 | 102.825578 | 0.57786323 | 0.70899599 | 0.81504442 | 0.41504688 | 0.88377697 |
| WIPI1 | 2512.27818 | 0.30921684 | 0.13894746 | 2.22542278 | 0.02605287 | 0.52561121 |
| PRKAR1A | 13892.5235 | 0.17011046 | 0.13781408 | 1.23434746 | 0.21707346 | 0.79574248 |
| FAM20A | 4693.09763 | -0.057655 | 0.63220455 | -0.0911968 | 0.92733625 | 0.98915637 |
| ABCA8 | 1248.33393 | 0.39297384 | 0.45662784 | 0.86059981 | 0.38945849 | 0.87568585 |
| ABCA9 | 528.923303 | -0.3613043 | 0.30710249 | -1.1764941 | 0.23939751 | 0.81163641 |
| ABCA6 | 419.229377 | -0.1158058 | 0.35700044 | -0.3243855 | 0.74564623 | 0.96176407 |
| ABCA10 | 388.645934 | -0.1013888 | 0.33165083 | -0.3057096 | 0.75982577 | 0.96512963 |
| ABCA5 | 1972.78779 | 0.30196335 | 0.19952324 | 1.51342447 | 0.13017187 | 0.70814767 |
| MAP2K6 | 477.704824 | 0.17935851 | 0.58450231 | 0.3068568 | 0.75895238 | 0.96502715 |
| AC005208.1 | 44.6902454 | 1.79658506 | 0.87667397 | 2.04931949 | 0.04043088 | 0.55657615 |
| KCNJ16 | 14463.7238 | 0.2228342 | 0.32198262 | 0.69206905 | 0.48889397 | 0.90590882 |
| KCNJ2-AS1 | 43.272121 | -0.0205955 | 0.56180313 | -0.0366596 | 0.97075646 | 0.99400331 |
| KCNJ2 | 131.903314 | -0.0316393 | 0.31520469 | -0.1003771 | 0.92004494 | 0.98733173 |
| LINC02097 | 75.6864164 | 1.15668308 | 0.44129221 | 2.62112732 | 0.00876395 | 0.38196566 |
| SOX9 | 3134.1179 | 1.09696635 | 0.39393033 | 2.78467094 | 0.00535821 | 0.31308771 |
| LINC00511 | 153.817816 | 0.80412772 | 0.406491 | 1.97821775 | 0.04790415 | 0.58372668 |
| SLC39A11 | 1460.1766 | 0.59337522 | 0.32597384 | 1.82031548 | 0.06871098 | 0.62750387 |
| SSTR2 | 1100.01819 | -0.7426565 | 0.51232551 | -1.4495794 | 0.14717584 | 0.73376495 |
| COG1 | 3077.82356 | -0.0447235 | 0.10125604 | -0.4416873 | 0.65871552 | 0.9458503 |
| AC097641.2 | 267.217311 | 0.09751695 | 0.16863136 | 0.57828479 | 0.56307186 | 0.92452095 |
| FAM104A | 1824.9857 | -0.1903908 | 0.14496309 | -1.3133746 | 0.18905675 | 0.76970626 |
| C17orf80 | 1073.58652 | 0.08169043 | 0.15272998 | 0.53486835 | 0.5927409 | 0.93289552 |
| CPSF4L | 198.469063 | 0.1335128 | 0.21038432 | 0.63461383 | 0.52568031 | 0.91763922 |
| CDC42EP4 | 3887.39291 | -0.0393632 | 0.21620522 | -0.1820643 | 0.85553229 | 0.97920671 |
| SDK2 | 124.01769 | 0.51342692 | 0.37844467 | 1.3566763 | 0.17488409 | 0.76061793 |
| RPL38 | 8230.74087 | 0.14756326 | 0.15365268 | 0.96036889 | 0.33686959 | 0.85679125 |
| AC100786.1 | 48.8308792 | 0.30644415 | 0.36019105 | 0.85078224 | 0.39489033 | 0.87884163 |
| TTYH2 | 307.11245 | 0.09410582 | 0.40099186 | 0.23468263 | 0.81445507 | 0.9738447 |
| AC100786.2 | 57.9861727 | -0.016684 | 0.51374151 | -0.0324755 | 0.97409285 | 0.99452354 |
| KIF19 | 49.4358709 | -1.1883182 | 0.52708215 | -2.2545218 | 0.02416336 | 0.51967919 |
| GPRC5C | 13616.496 | 0.21761351 | 0.14428548 | 1.50821492 | 0.13149953 | 0.7108257 |
| CD300A | 369.143384 | -0.4136254 | 0.62653989 | -0.6601741 | 0.5091421 | 0.91439937 |
| CD300C | 78.8946715 | -0.7428596 | 0.65537137 | -1.1334941 | 0.25700681 | 0.82107009 |
| CD300E | 260.044959 | -0.1713966 | 0.98672777 | -0.173702 | 0.86209967 | 0.98176973 |
| RAB37 | 320.677544 | 0.46668293 | 0.48963615 | 0.95312188 | 0.34052832 | 0.85846668 |
| CD300LF | 68.1617145 | 0.29021855 | 0.69343256 | 0.41852455 | 0.67556364 | 0.94958156 |
| AC016888.1 | 1790.0222 | 0.59077584 | 0.48928005 | 1.20743904 | 0.22726311 | 0.80300331 |
| SLC9A3R1 | 14549.6994 | 0.38095422 | 0.42039695 | 0.90617742 | 0.36484195 | 0.8663515 |
| NAT9 | 2617.17778 | -0.3447866 | 0.20462084 | -1.6850024 | 0.0919881 | 0.65583276 |
| TMEM104 | 1739.54878 | -0.1998739 | 0.22657987 | -0.8821343 | 0.37770417 | 0.86987124 |
| GRIN2C | 98.3640596 | -0.5831318 | 0.5846616 | -0.9973835 | 0.3185784 | 0.84904404 |
| FDXR | 829.736082 | 0.51115994 | 0.31756614 | 1.60961726 | 0.10748144 | 0.67707349 |
| FADS6 | 147.357319 | 0.24216587 | 0.83527011 | 0.28992522 | 0.77187345 | 0.96836854 |
| HID1 | 1785.54964 | -0.15117 | 0.27303599 | -0.5536634 | 0.57980921 | 0.93139743 |
| CDR2L | 1907.74732 | -0.0741117 | 0.35717252 | -0.2074956 | 0.83562285 | 0.97550416 |
| MRPL58 | 1150.99187 | 0.32979704 | 0.17557951 | 1.87833447 | 0.06033542 | 0.60825814 |

| | | | | | | |
|---|---|---|---|---|---|---|
| KCTD2 | 6479.72863 | -0.0359593 | 0.14654298 | -0.245384 | 0.80615915 | 0.97334138 |
| ATP5PD | 7392.70277 | 0.23747178 | 0.1387911 | 1.71100154 | 0.08708083 | 0.65231349 |
| SLC16A5 | 5368.79352 | 0.45713311 | 0.43616906 | 1.04806406 | 0.29460909 | 0.83630273 |
| ARMC7 | 2006.79625 | 0.32561197 | 0.33066211 | 0.98472719 | 0.32475809 | 0.85181379 |
| NT5C | 1831.024 | -0.1563909 | 0.14944089 | -1.0465066 | 0.29532718 | 0.837073 |
| JPT1 | 4026.23119 | 0.17794976 | 0.15723323 | 1.13175665 | 0.25773676 | 0.82187662 |
| SUMO2 | 6609.01086 | -0.0852945 | 0.15682629 | -0.5438788 | 0.58652489 | 0.93289552 |
| NUP85 | 1965.19248 | -0.0889415 | 0.17888158 | -0.4972086 | 0.61904192 | 0.93945324 |
| GGA3 | 2827.08502 | -0.3102114 | 0.11828413 | -2.6225954 | 0.00872628 | 0.38121041 |
| MRPS7 | 3398.44549 | 0.11164952 | 0.14286879 | 0.78148296 | 0.4345185 | 0.89055338 |
| MIF4GD | 1303.9535 | 0.11559126 | 0.16098678 | 0.71801707 | 0.47274676 | 0.90160919 |
| AC022211.2 | 348.216733 | 0.00308289 | 0.19237819 | 0.01602514 | 0.98721434 | 0.99732542 |
| SLC25A19 | 588.780165 | -0.03417 | 0.18109875 | -0.1886814 | 0.85034253 | 0.97816532 |
| GRB2 | 6582.19083 | -0.0679623 | 0.10688942 | -0.6358187 | 0.52489458 | 0.91759401 |
| TMEM94 | 5625.52201 | -0.0072661 | 0.10387508 | -0.06995 | 0.94423345 | 0.99136103 |
| CASKIN2 | 5288.6859 | -0.2955132 | 0.29987835 | -0.9854437 | 0.32440618 | 0.85169085 |
| TSEN54 | 2401.82857 | 0.00680073 | 0.16809164 | 0.04045849 | 0.9677276 | 0.99400331 |
| LLGL2 | 12746.116 | -0.0291653 | 0.23458115 | -0.124329 | 0.90105475 | 0.9855364 |
| MYO15B | 15962.8299 | 0.03591979 | 0.22675603 | 0.15840721 | 0.87413593 | 0.98275231 |
| RECQL5 | 3983.16834 | 0.00091842 | 0.34328239 | 0.00267541 | 0.99786533 | 0.99989813 |
| SMIM5 | 1891.75909 | -0.0537324 | 0.41063181 | -0.1308529 | 0.89589167 | 0.98474725 |
| SMIM6 | 316.918888 | 0.63030171 | 0.33340091 | 1.89052183 | 0.0586882 | 0.60332684 |
| SAP30BP | 3507.97124 | -0.2918982 | 0.18270933 | -1.5976097 | 0.11012987 | 0.67996643 |
| ITGB4 | 2213.68383 | 0.35196444 | 0.38851488 | 0.90592268 | 0.36497678 | 0.8663515 |
| GALK1 | 1884.85431 | 0.09040219 | 0.3134381 | 0.28842119 | 0.77302435 | 0.96836854 |
| H3F3B | 20557.2214 | -0.0998326 | 0.23777629 | -0.4198595 | 0.67458811 | 0.94935413 |
| UNK | 2346.50508 | -0.2737017 | 0.20486052 | -1.3360391 | 0.18153651 | 0.76522169 |
| AC087289.1 | 295.413058 | -0.3975224 | 0.26392072 | -1.5062189 | 0.132011 | 0.7121526 |
| UNC13D | 805.897869 | 0.01712112 | 0.33606404 | 0.05094601 | 0.95936854 | 0.9924381 |
| WBP2 | 12408.2936 | -0.086507 | 0.13185923 | -0.6560559 | 0.51178819 | 0.91512475 |
| TRIM47 | 4490.88704 | -0.0792584 | 0.23223283 | -0.3412886 | 0.73288632 | 0.95848028 |
| AC087289.5 | 1215.42502 | 0.03629223 | 0.23714546 | 0.15303783 | 0.87836845 | 0.98275231 |
| TRIM65 | 1777.2569 | 0.25327779 | 0.14632733 | 1.73089867 | 0.08346984 | 0.64774668 |
| AC087289.2 | 973.350413 | -0.0690298 | 0.1074656 | -0.6423429 | 0.52065057 | 0.91680964 |
| MRPL38 | 4347.46132 | 0.0019604 | 0.09701928 | 0.02020624 | 0.98387885 | 0.9961464 |
| AC087289.3 | 4540.61875 | 0.02987163 | 0.09499941 | 0.31444015 | 0.75318677 | 0.96301881 |
| FBF1 | 871.714011 | 0.03573267 | 0.3739553 | 0.09555333 | 0.92387533 | 0.98869145 |
| ACOX1 | 5619.00555 | 0.31395001 | 0.34758951 | 0.90322063 | 0.36640881 | 0.86683872 |
| AC087289.4 | 400.241017 | 0.40348171 | 0.39680739 | 1.01682004 | 0.30923904 | 0.84414094 |
| TEN1 | 1179.19628 | -0.3809228 | 0.27275076 | -1.3965966 | 0.16253491 | 0.75026273 |
| CDK3 | 1276.83789 | -0.3642659 | 0.14802663 | -2.4608133 | 0.01386225 | 0.4295825 |
| EVPL | 3688.22942 | -0.0425515 | 0.29564207 | -0.1439293 | 0.88555634 | 0.98365406 |
| SRP68 | 4027.72217 | 0.00801357 | 0.15475529 | 0.05178218 | 0.95870226 | 0.99237995 |
| ZACN | 5532.69683 | -0.2359768 | 0.10719317 | -2.2014163 | 0.02770657 | 0.53038691 |
| EXOC7 | 8436.82313 | -0.2128001 | 0.11898412 | -1.7884746 | 0.07369948 | 0.6374448 |
| FOXJ1 | 527.629093 | -0.4207228 | 0.6001668 | -0.7010098 | 0.4832969 | 0.90541935 |
| RNF157-AS1 | 470.100142 | 0.24834638 | 0.4007427 | 0.61971529 | 0.53544525 | 0.92095993 |
| RNF157 | 1090.63109 | 0.53056931 | 0.44429121 | 1.19419269 | 0.2324026 | 0.80649152 |
| UBALD2 | 2517.18859 | -0.4559559 | 0.24324689 | -1.8744573 | 0.06086743 | 0.60989294 |
| QRICH2 | 293.770458 | -0.5653324 | 0.23718671 | -2.3834913 | 0.01714929 | 0.4678237 |
| PRPSAP1 | 2276.7805 | -0.1559361 | 0.20607696 | -0.7566887 | 0.44923636 | 0.89546252 |
| SPHK1 | 1352.38719 | -2.2145963 | 0.8185806 | -2.7054102 | 0.00682201 | 0.34932037 |
| UBE2O | 1864.61046 | -0.343752 | 0.19936871 | -1.7242023 | 0.08467134 | 0.64988556 |
| RHBDF2 | 1973.3221 | -0.2838354 | 0.30772962 | -0.9223533 | 0.35634433 | 0.86390849 |

| | | | | | | |
|---|---|---|---|---|---|---|
| CYGB | 673.457115 | 0.86552811 | 0.47147091 | 1.83580384 | 0.06638667 | 0.62378208 |
| PRCD | 274.970526 | 0.69619418 | 0.41674443 | 1.67055424 | 0.09480976 | 0.65888357 |
| AC015802.4 | 80.1323714 | 0.69756662 | 0.38277493 | 1.82239368 | 0.06839527 | 0.62657129 |
| AC015802.6 | 2761.94636 | 0.0857249 | 0.25170996 | 0.34057018 | 0.73342718 | 0.95884813 |
| ST6GALNAC2 | 893.518226 | 1.02260311 | 0.41635735 | 2.45607075 | 0.01404655 | 0.43297104 |
| AC015802.1 | 175.539537 | 1.15505191 | 0.43912369 | 2.63035662 | 0.00852953 | 0.3779007 |
| AC005837.1 | 41.1563167 | 0.85138966 | 0.46924837 | 1.81436891 | 0.06962095 | 0.62779744 |
| MXRA7 | 4513.02133 | -0.104839 | 0.38806825 | -0.2701561 | 0.78704013 | 0.97090793 |
| JMJD6 | 2057.51912 | -0.4674411 | 0.24023383 | -1.9457756 | 0.0516817 | 0.58645127 |
| METTL23 | 1399.88349 | 0.02883991 | 0.15940088 | 0.18092693 | 8.56E-01 | 9.80E-01 |
| AC005837.2 | 763.580964 | 0.01662716 | 0.15952595 | 0.10422854 | 0.91698799 | 0.9868044 |
| SRSF2 | 10791.63 | -0.3318378 | 0.17941023 | -1.8496034 | 0.06437073 | 0.61964848 |
| MFSD11 | 5098.90038 | -0.3577151 | 0.2025902 | -1.7657077 | 0.07744491 | 0.63862831 |
| MGAT5B | 46.2041089 | -0.0612066 | 0.56475024 | -0.1083782 | 0.91369573 | 0.98662757 |
| SEC14L1 | 8133.97994 | -0.5394054 | 0.23454735 | -2.2997718 | 0.02146115 | 0.51301456 |
| 9-Sep | 11015.3564 | 0.06451968 | 0.17888703 | 0.36067277 | 0.71834408 | 0.95648644 |
| TNRC6C | 790.847616 | 0.02861754 | 0.1624908 | 0.17611791 | 0.86020131 | 0.98077707 |
| TMC6 | 2497.62481 | 0.14618426 | 0.31845935 | 0.45903585 | 0.64620842 | 0.94208859 |
| TMC8 | 851.406999 | -0.132717 | 0.67498521 | -0.1966221 | 0.84412324 | 0.97664894 |
| SYNGR2 | 7379.74653 | -0.0560476 | 0.16248783 | -0.3449339 | 0.73014409 | 0.95848028 |
| TK1 | 948.676677 | 0.2750929 | 0.30567195 | 0.89996122 | 0.36814089 | 0.86759809 |
| AFMID | 6617.53082 | 0.59376373 | 0.3171566 | 1.87214682 | 0.06118629 | 0.60989294 |
| BIRC5 | 70.1608305 | 0.2992113 | 0.7145969 | 0.4187134 | 0.6754256 | 0.94958156 |
| SOCS3 | 8819.27296 | -0.0427078 | 0.53461179 | -0.0798857 | 0.93632818 | 0.99068685 |
| AC061992.1 | 132.000142 | 0.02171344 | 0.58666064 | 0.03701193 | 0.9704755 | 0.99400331 |
| PGS1 | 2116.40908 | -0.1988678 | 0.18487872 | -1.075666 | 0.28207664 | 0.83003334 |
| DNAH17 | 934.518283 | -0.1013724 | 0.21578439 | -0.4697855 | 0.63850824 | 0.94088547 |
| CYTH1 | 3806.01756 | -0.3973661 | 0.20248631 | -1.9624345 | 0.04971193 | 0.5861488 |
| USP36 | 2952.73637 | -0.6112326 | 0.29920079 | -2.0428843 | 4.11E-02 | 5.58E-01 |
| TIMP2 | 10192.6154 | 0.08352796 | 0.31212165 | 0.26761347 | 0.78899686 | 0.97090793 |
| AC022966.1 | 1582.39443 | -0.049653 | 0.31948994 | -0.1554133 | 0.8764955 | 0.98275231 |
| CEP295NL | 653.95247 | -0.0346905 | 0.3496161 | -0.0992244 | 0.92096007 | 0.98767854 |
| LGALS3BP | 19343.3551 | -0.751543 | 0.55871273 | -1.3451332 | 0.17858224 | 0.76115956 |
| CANT1 | 2217.12765 | -0.1981677 | 0.15364551 | -1.2897722 | 0.19712975 | 0.77816059 |
| C1QTNF1 | 2448.66885 | -0.3081614 | 0.56091841 | -0.5493873 | 0.58273969 | 0.93183657 |
| ENGASE | 3985.35702 | -0.0003386 | 0.291026 | -0.0011635 | 0.99907164 | 0.99994357 |
| ENPP7 | 424.851849 | -0.3859371 | 0.7006581 | -0.5508208 | 5.82E-01 | 0.9317705 |
| CBX2 | 149.907147 | 0.1371474 | 0.58996185 | 0.23246825 | 0.81617435 | 0.97388289 |
| CBX8 | 455.830831 | -0.007481 | 0.32613359 | -0.0229385 | 0.98169931 | 0.99595643 |
| CBX4 | 1144.34127 | -0.1258124 | 0.20479301 | -0.6143395 | 0.53899101 | 0.9215968 |
| TBC1D16 | 5322.98234 | -0.1644623 | 0.33427392 | -0.4919986 | 0.6227203 | 0.94016713 |
| AC100791.3 | 54.5105867 | -0.3397469 | 0.35861509 | -0.9473859 | 0.34344217 | 0.85938376 |
| CCDC40 | 948.896781 | -0.0351463 | 0.31242512 | -0.1124951 | 0.91043082 | 0.98662757 |
| GAA | 8794.97827 | 0.17204108 | 0.24045418 | 0.71548386 | 0.47431011 | 0.90199039 |
| EIF4A3 | 3398.81491 | -0.1469426 | 0.18688873 | -0.7862574 | 0.43171669 | 0.88976671 |
| CARD14 | 831.334944 | -0.2728675 | 0.35944324 | -0.7591392 | 0.44776929 | 0.89480207 |
| AC087741.1 | 1525.7022 | -0.0855143 | 0.39235686 | -0.2179502 | 0.82746788 | 0.97548201 |
| SGSH | 3521.60663 | -0.2162827 | 0.18136535 | -1.1925248 | 0.23305551 | 0.8068896 |
| SLC26A11 | 2841.84016 | -0.2867441 | 0.24382974 | -1.1760012 | 0.23959441 | 0.81168453 |
| RNF213 | 14788.5125 | -0.6315511 | 0.47319997 | -1.334639 | 0.18199454 | 0.76569697 |
| AC124319.3 | 37.6200763 | 0.43749469 | 0.498783 | 0.87712431 | 0.38041912 | 0.87118022 |
| AC124319.2 | 233.910341 | -0.040079 | 0.31756226 | -0.1262084 | 0.89956693 | 0.98506781 |
| ENDOV | 1543.62617 | 0.00188189 | 0.19969069 | 0.00942403 | 0.99248082 | 0.99906045 |
| NPTX1 | 46.6555582 | -0.7100591 | 0.76858884 | -0.9238478 | 0.35556555 | 0.86364633 |

| | | | | | | |
|---|---|---|---|---|---|---|
| RPTOR | 1311.35035 | -0.085945 | 0.13965189 | -0.6154234 | 0.53827513 | 0.92151404 |
| AC127496.3 | 44.5616006 | -0.1974133 | 0.3591068 | -0.5497342 | 0.58250171 | 0.93177403 |
| AC127496.4 | 64.3825937 | -0.0438967 | 0.28466823 | -0.1542029 | 0.87744976 | 0.98275231 |
| CHMP6 | 1390.22136 | -0.2118581 | 0.19143483 | -1.1066851 | 0.26843012 | 0.82676682 |
| AC127496.6 | 660.510696 | -0.2498974 | 0.20419592 | -1.2238121 | 0.22102314 | 0.79965146 |
| BAIAP2-DT | 1247.31688 | 0.22981551 | 0.2855053 | 0.80494306 | 0.42085254 | 0.88451372 |
| BAIAP2 | 2548.3516 | 0.20123996 | 0.34537094 | 0.58267774 | 0.56011026 | 0.92434754 |
| AATK | 465.665873 | -0.7690823 | 0.47588176 | -1.6161205 | 0.10606827 | 0.67567147 |
| CEP131 | 913.464082 | 0.10537582 | 0.28695039 | 0.3672266 | 0.71345 | 0.95564093 |
| AC027601.2 | 91.021361 | -0.1846435 | 0.31021322 | -0.5952148 | 0.55169987 | 0.92373235 |
| TEPSIN | 3143.97561 | -0.3358864 | 0.18520336 | -1.8136087 | 0.06973799 | 0.62779744 |
| AC027601.1 | 2093.61183 | -0.3085978 | 0.18979513 | -1.6259523 | 0.10395981 | 0.67439401 |
| NDUFAF8 | 841.530946 | 0.2314484 | 0.23018549 | 1.00548651 | 0.31466264 | 0.84696964 |
| SLC38A10 | 8813.3241 | 0.23057614 | 0.20657082 | 1.11620867 | 0.26433282 | 0.82424998 |
| AC027601.4 | 690.454067 | 0.15519128 | 0.21837805 | 0.71065419 | 0.47729855 | 0.90372319 |
| LINC00482 | 229.95429 | 0.01505365 | 0.51069773 | 0.02947663 | 0.97648445 | 0.99455099 |
| TMEM105 | 116.606313 | 2.76343272 | 0.88521307 | 3.12177127 | 0.00179767 | 0.18722055 |
| AC110285.2 | 1069.49862 | 0.31019391 | 0.66041667 | 0.46969425 | 0.63857348 | 0.94088547 |
| BAHCC1 | 1993.2785 | -0.0499667 | 0.27193938 | -0.1837419 | 0.85421591 | 0.97906925 |
| AC110285.4 | 130.582182 | -0.0414615 | 0.31221344 | -0.1327986 | 0.89435263 | 0.98474725 |
| ACTG1 | 75767.0451 | 0.07478628 | 0.29840623 | 0.25061902 | 0.80210868 | 0.97233627 |
| FSCN2 | 46.4674677 | 0.98576918 | 0.41856206 | 2.35513263 | 0.0185161 | 0.47570049 |
| FAAP100 | 2919.48217 | -0.1118348 | 0.16742905 | -0.6679532 | 0.50416348 | 0.91356552 |
| NPLOC4 | 8214.04096 | -0.2580789 | 0.22586952 | -1.1426019 | 0.25320393 | 0.81848561 |
| TSPAN10 | 113.471564 | 0.27998153 | 0.40831172 | 0.68570536 | 0.49289892 | 0.90705676 |
| PDE6G | 108.820853 | 0.09802934 | 0.37228249 | 0.26331976 | 0.79230413 | 0.97090793 |
| OXLD1 | 1159.90523 | 0.03934305 | 0.10558702 | 0.37261259 | 0.7094368 | 0.95501548 |
| CCDC137 | 1752.8393 | -0.308812 | 0.21135991 | -1.4610718 | 0.14399575 | 0.72926154 |
| ARL16 | 2117.6826 | 0.13061646 | 0.19369775 | 0.6743313 | 0.50010071 | 0.91180773 |
| HGS | 9225.68937 | -0.6185203 | 0.25094922 | -2.464723 | 1.37E-02 | 0.42829183 |
| AC139530.3 | 61.0899677 | 0.14379226 | 0.41723287 | 0.34463311 | 0.73037021 | 0.95848028 |
| AC139530.1 | 3088.38304 | -0.4981196 | 0.23733132 | -2.0988365 | 0.03583131 | 0.55369025 |
| MRPL12 | 3075.21529 | 0.03026517 | 0.19835551 | 0.15258046 | 0.87872914 | 0.98275231 |
| AC139530.2 | 6571.05594 | 0.4219796 | 0.35152287 | 1.20043286 | 0.22997127 | 0.80551898 |
| SLC25A10 | 4481.63509 | 0.86871529 | 0.51626342 | 1.68269775 | 9.24E-02 | 6.56E-01 |
| GCGR | 3002.92502 | -0.0762252 | 0.54158529 | -0.1407447 | 0.88807167 | 0.98423722 |
| MCRIP1 | 3599.47767 | -0.2002322 | 0.21745163 | -0.9208127 | 0.3571482 | 0.86390849 |
| P4HB | 30464.1218 | 0.00844217 | 0.27652171 | 0.03052985 | 0.97564449 | 0.99455099 |
| AC145207.3 | 12931.9559 | -0.0277901 | 0.2857172 | -0.0972645 | 0.92251633 | 0.98826666 |
| ARHGDIA | 18388.4361 | -0.4537192 | 0.26339886 | -1.7225557 | 0.08496891 | 0.64988556 |
| AC145207.2 | 9901.92241 | -0.4833449 | 0.26055117 | -1.8550864 | 0.06358389 | 0.61774269 |
| AC145207.5 | 164.096143 | 0.43746546 | 0.41030783 | 1.06618843 | 0.28633847 | 0.83107779 |
| AC145207.6 | 106.960913 | 0.41525727 | 0.41120316 | 1.00985915 | 0.31256278 | 0.84541735 |
| ALYREF | 1332.39482 | -0.2635822 | 0.27591102 | -0.955316 | 0.33941794 | 0.85759056 |
| ANAPC11 | 2271.7199 | 0.04841424 | 0.16154827 | 0.299689 | 0.76441439 | 0.96678847 |
| NPB | 220.735043 | 0.83859937 | 0.43351475 | 1.93441944 | 0.05306158 | 0.58876675 |
| PCYT2 | 3779.17767 | 0.8102976 | 0.36904258 | 2.19567512 | 0.02811521 | 0.53300785 |
| SIRT7 | 2321.93968 | -0.3866081 | 0.31226986 | -1.2380577 | 0.21569466 | 0.7954827 |
| MAFG | 3302.99521 | -0.5742817 | 0.3660005 | -1.5690734 | 0.11663084 | 0.6888681 |
| AC145207.8 | 101.770979 | -0.7400996 | 0.52257756 | -1.4162483 | 0.15670281 | 0.74522306 |
| MAFG-DT | 121.639684 | -0.0444734 | 0.3031011 | -0.1467281 | 0.88334665 | 0.98293342 |
| PYCR1 | 282.224354 | -0.0682238 | 0.32269605 | -0.211418 | 0.8325611 | 0.97550416 |
| AC145207.4 | 113.714915 | -0.0584919 | 0.37681632 | -0.1552266 | 0.87664264 | 0.98275231 |
| NOTUM | 34.9330704 | 0.23667514 | 0.65733203 | 0.36005416 | 7.19E-01 | 0.95648644 |

| | | | | | | |
|---|---|---|---|---|---|---|
| ASPSCR1 | 2949.44634 | -0.1955438 | 0.19601724 | -0.9975848 | 0.31848072 | 0.8490262 |
| CENPX | 1230.47031 | 0.08373981 | 0.25528526 | 0.32802446 | 0.74289317 | 0.96109001 |
| LRRC45 | 859.410445 | 0.147787 | 0.22853598 | 0.64666843 | 0.51784656 | 0.91636259 |
| RAC3 | 515.987636 | 0.18834624 | 0.32883088 | 0.57277541 | 0.56679677 | 0.9256133 |
| DCXR | 7551.19042 | 0.33943773 | 0.48313282 | 0.70257642 | 0.48231976 | 0.90541935 |
| RFNG | 3696.89509 | -0.099724 | 0.13543681 | -0.7363141 | 0.46153956 | 0.89755243 |
| GPS1 | 6828.03666 | 0.02831301 | 0.14660459 | 0.19312497 | 8.47E-01 | 0.97684681 |
| DUS1L | 5882.66768 | -0.0399185 | 0.11294031 | -0.3534481 | 0.72375251 | 0.95810643 |
| FASN | 6764.86904 | 0.15219356 | 0.4153363 | 0.36643452 | 0.71404087 | 0.95564093 |
| CCDC57 | 2965.47638 | -0.2295319 | 0.23634466 | -0.9711747 | 0.33146129 | 0.85408791 |
| AC129510.1 | 297.040492 | -0.1121489 | 0.31945165 | -0.351067 | 0.72553809 | 0.95848028 |
| AC132872.2 | 62.4584253 | 0.64438241 | 0.54247912 | 1.18784739 | 0.23489354 | 0.80747753 |
| SLC16A3 | 6506.81555 | -0.577863 | 0.443447 | -1.3031162 | 0.19253508 | 0.77481211 |
| CSNK1D | 14143.2258 | -0.4241731 | 0.21528477 | -1.9702887 | 4.88E-02 | 0.58504358 |
| AC132872.3 | 1589.04608 | -0.4310014 | 0.20879003 | -2.0642816 | 3.90E-02 | 5.56E-01 |
| AC132872.1 | 401.711494 | -0.0732075 | 0.29975617 | -0.2442236 | 0.80705762 | 0.97334138 |
| CD7 | 420.432841 | 0.11408162 | 0.76424189 | 0.14927424 | 0.88133724 | 0.98275231 |
| SECTM1 | 4625.88665 | -1.1958682 | 0.57027523 | -2.0970018 | 0.03599342 | 0.55369025 |
| AC132938.1 | 40.1212435 | -0.4238385 | 0.51429709 | -0.8241122 | 0.40987578 | 0.88249519 |
| OGFOD3 | 1844.90613 | -0.002546 | 0.12586231 | -0.0202285 | 9.84E-01 | 9.96E-01 |
| HEXDC | 3250.73616 | -0.445765 | 0.34152351 | -1.3052249 | 1.92E-01 | 7.73E-01 |
| CYBC1 | 4634.64232 | -0.0811213 | 0.23293731 | -0.3482539 | 7.28E-01 | 0.95848028 |
| AC132938.3 | 80.757477 | 0.21952573 | 0.45525677 | 0.48220201 | 0.62966245 | 0.94088547 |
| NARF | 3529.34346 | -0.3357729 | 0.20417613 | -1.6445259 | 0.10006763 | 0.6716932 |
| FOXK2 | 2901.48492 | -0.257499 | 0.14414155 | -1.7864318 | 0.07402938 | 0.63749647 |
| AC124283.1 | 632.112273 | -0.2231754 | 0.29273093 | -0.7623908 | 0.44582682 | 0.89476935 |
| WDR45B | 6666.13377 | -0.2473158 | 0.20169129 | -1.2262097 | 0.22011979 | 0.79885069 |
| RAB40B | 2428.80368 | -0.1837026 | 0.31247967 | -0.5878866 | 0.55660838 | 0.92373235 |
| FN3KRP | 2175.85301 | 0.00180685 | 0.22405164 | 0.00806442 | 0.99356559 | 0.99920627 |
| FN3K | 4822.24862 | 0.02565477 | 0.38870177 | 0.06600115 | 0.94737691 | 0.99166614 |
| TBCD | 4459.44916 | 0.01686913 | 0.12980054 | 0.12996195 | 0.89659653 | 0.98474725 |
| ZNF750 | 41.0581843 | -0.1029924 | 0.66128546 | -0.1557458 | 0.8762334 | 0.98275231 |
| B3GNTL1 | 886.105033 | -0.0529272 | 0.23096281 | -0.2291592 | 0.81874522 | 0.97454841 |
| METRNL | 2902.39117 | -0.3543329 | 0.21470621 | -1.6503152 | 0.09887849 | 0.66801794 |
| AC130371.2 | 91.7964554 | 0.07368506 | 0.32230002 | 0.22862257 | 0.81916228 | 0.97454841 |
| AC144831.1 | 407.901799 | 0.15429214 | 0.39845055 | 0.38723034 | 0.69858569 | 0.9529028 |
| USP14 | 3286.01874 | -0.1752064 | 0.23838113 | -0.7349842 | 0.46234913 | 0.89776981 |
| THOC1 | 1670.51452 | -0.1488268 | 0.14488654 | -1.0271954 | 0.30432848 | 0.84211658 |
| COLEC12 | 433.002024 | 0.09351249 | 0.69236101 | 0.1350632 | 0.89256191 | 0.98474725 |
| CLUL1 | 211.981216 | 1.1269384 | 0.45706358 | 2.4656053 | 0.0136782 | 0.42795172 |
| TYMS | 1777.64129 | 0.31185206 | 0.23460074 | 1.32928846 | 0.18375282 | 0.76781833 |
| ENOSF1 | 6550.471 | 0.09848472 | 0.25454844 | 0.38689972 | 0.69883045 | 0.95297835 |
| AP001178.2 | 214.469707 | 0.44338745 | 0.37080223 | 1.19575183 | 0.23179342 | 0.80649152 |
| YES1 | 3533.47387 | -0.0691839 | 0.27502999 | -0.2515503 | 0.80138868 | 0.97210205 |
| AP001020.3 | 103.064165 | -0.1695849 | 0.5429535 | -0.3123378 | 0.75478381 | 0.96351771 |
| AP000894.4 | 51.1413448 | 0.18444556 | 0.35362685 | 0.52158245 | 6.02E-01 | 0.9355118 |
| ADCYAP1 | 49.4930318 | 0.96914505 | 0.67907255 | 1.42715981 | 0.15353384 | 0.74258845 |
| METTL4 | 568.719345 | -0.2100302 | 0.15994875 | -1.3131094 | 0.1891461 | 0.76970626 |
| NDC80 | 56.9339753 | 0.08628919 | 0.73446785 | 0.11748531 | 0.90647548 | 0.98662757 |
| SMCHD1 | 2062.80366 | -0.3726806 | 0.2653193 | -1.4046493 | 0.16012559 | 0.74781837 |
| EMILIN2 | 536.484255 | 0.17026269 | 0.45100646 | 0.37751718 | 0.7057893 | 0.95501548 |
| LPIN2 | 4267.41142 | 0.18401397 | 0.1862123 | 0.98819451 | 0.32305739 | 0.85119105 |
| AP000919.4 | 54.8199561 | 0.03436595 | 0.46894557 | 0.07328345 | 0.94158056 | 0.99136103 |
| MYOM1 | 213.829646 | -0.19195 | 0.52177508 | -0.3678787 | 0.71296366 | 0.95564093 |

| | | | | | | |
|---|---|---|---|---|---|---|
| AP005329.2 | 83.1803969 | -0.1817504 | 0.29492304 | -0.616264 | 0.5377203 | 0.92123049 |
| AP005329.3 | 69.1830822 | -0.1382422 | 0.31099546 | -0.444515 | 0.65667025 | 0.94545756 |
| MYL12A | 14268.1538 | -0.0260107 | 0.16697593 | -0.1557753 | 0.87621012 | 0.98275231 |
| AP005329.1 | 6294.26976 | 0.04669625 | 0.16450607 | 0.28385731 | 0.77651973 | 0.96836854 |
| MYL12B | 21952.4232 | 0.1057278 | 0.16283756 | 0.64928384 | 0.51615493 | 0.91636259 |
| TGIF1 | 3905.01443 | -0.4310201 | 0.26305417 | -1.6385224 | 0.10131277 | 0.6725344 |
| DLGAP1 | 295.712461 | 0.9230834 | 0.70360338 | 1.31193714 | 0.18954135 | 0.76970626 |
| DLGAP1-AS1 | 453.241122 | 0.05312407 | 0.28741551 | 0.18483368 | 0.85335949 | 0.97893189 |
| DLGAP1-AS2 | 309.480635 | -0.1448181 | 0.44037265 | -0.3288534 | 0.74226648 | 0.96109001 |
| DLGAP1-AS3 | 70.110824 | 0.22220601 | 1.03260268 | 0.21519023 | 8.30E-01 | 0.97550416 |
| AP001496.2 | 342.086312 | 0.40760904 | 0.33381459 | 1.22106419 | 0.22206172 | 0.80067924 |
| LINC00526 | 698.034423 | 0.36812542 | 0.30698973 | 1.19914573 | 0.23047128 | 0.80584899 |
| LINC00667 | 1750.06079 | 0.01997189 | 0.173285 | 0.1152546 | 0.90824332 | 0.98662757 |
| ZBTB14 | 697.052533 | 0.164653 | 0.26247784 | 0.62730246 | 0.53046099 | 0.91962754 |
| EPB41L3 | 4406.56275 | -0.0215263 | 0.30012811 | -0.0717238 | 0.94282174 | 0.99136103 |
| TMEM200C | 154.738414 | 0.65023349 | 0.53944717 | 1.2053701 | 2.28E-01 | 0.80300331 |
| L3MBTL4 | 764.78866 | -0.0255684 | 0.35160595 | -0.0727189 | 0.9420298 | 0.99136103 |
| AP001021.1 | 79.8824227 | -0.0728095 | 0.39641999 | -0.1836677 | 0.85427416 | 0.97906925 |
| ARHGAP28 | 488.060471 | 0.24172352 | 0.38877134 | 0.62176271 | 0.53409791 | 0.92083983 |
| LAMA1 | 1629.79024 | -0.4184475 | 0.32336836 | -1.2940275 | 0.19565592 | 0.77752069 |
| AP002409.1 | 249.660603 | -0.3412882 | 0.36288787 | -0.9404785 | 0.34697217 | 0.86150045 |
| PTPRM | 5116.32227 | -0.2656638 | 0.19750023 | -1.3451317 | 0.17858271 | 0.76115956 |
| AP001094.1 | 212.378314 | -0.3288516 | 0.30697769 | -1.0712557 | 0.28405448 | 0.83107779 |
| RAB12 | 2245.02328 | -0.0098116 | 0.12882895 | -0.07616 | 0.93929178 | 0.99113899 |
| AP001793.2 | 555.554623 | 0.05526773 | 0.1406431 | 0.39296441 | 0.69434577 | 0.95222846 |
| MTCL1 | 472.755632 | -0.1521562 | 0.37050961 | -0.4106673 | 0.6813165 | 0.95070301 |
| NDUFV2 | 5891.63653 | 0.17941393 | 0.12342715 | 1.45360184 | 0.14605673 | 0.73247558 |
| NDUFV2-AS1 | 145.322131 | -0.1938536 | 0.20969054 | -0.9244748 | 0.35523917 | 0.86364633 |
| ANKRD12 | 1855.62749 | -0.3875162 | 0.23223392 | -1.6686461 | 0.09518754 | 0.65888357 |
| TWSG1 | 2515.17783 | 0.39741223 | 0.23313589 | 1.7046377 | 0.08826202 | 0.65363785 |
| AP005432.2 | 129.113563 | 2.52550191 | 0.6732723 | 3.75108544 | 0.00017607 | 0.06345655 |
| RALBP1 | 3917.50615 | 0.29069453 | 0.17900032 | 1.62398885 | 0.10437819 | 0.67439401 |
| PPP4R1 | 3086.83699 | -0.2627808 | 0.22792975 | -1.1529028 | 0.24895028 | 0.81646572 |
| RAB31 | 1931.79914 | -0.3384356 | 0.38861961 | -0.870866 | 0.38382734 | 0.87255073 |
| VAPA | 7349.07813 | -0.1625335 | 0.14815181 | -1.0970739 | 0.27260907 | 0.82716979 |
| APCDD1 | 520.602692 | 0.33429026 | 0.68897271 | 0.48520102 | 0.62753376 | 0.94088547 |
| NAPG | 1958.53491 | -0.135698 | 0.17987336 | -0.7544084 | 0.45060402 | 0.89557948 |
| PIEZO2 | 168.152529 | 0.41689091 | 0.29912135 | 1.39371837 | 0.16340267 | 0.75149188 |
| GNAL | 1668.43364 | 0.15091757 | 0.16441441 | 0.91790963 | 0.35866618 | 0.86480228 |
| CHMP1B | 4491.75229 | -0.3992096 | 0.25077766 | -1.5918867 | 0.11141016 | 0.68001961 |
| CHMP1B-AS1 | 1832.98439 | -0.4842244 | 0.28087879 | -1.7239621 | 0.08471468 | 0.64988556 |
| MPPE1 | 2628.60357 | 0.1616999 | 0.1870327 | 0.8645542 | 0.38728352 | 0.87460898 |
| IMPA2 | 9913.76814 | 0.598787 | 0.31695899 | 1.88916236 | 0.05887008 | 0.60343175 |
| AP001542.3 | 180.521038 | 0.96570476 | 0.37365539 | 2.58447965 | 0.00975261 | 0.39381151 |
| TUBB6 | 6097.45587 | -0.1992187 | 0.23787963 | -0.8374767 | 0.40232465 | 0.88132015 |
| AFG3L2 | 6943.90945 | -0.0516133 | 0.26903405 | -0.1918467 | 0.84786227 | 0.97711043 |
| PRELID3A | 97.1809785 | -0.1468576 | 0.29985864 | -0.4897562 | 0.62430645 | 0.94075157 |
| AP001029.2 | 1226.06643 | -0.0578665 | 0.26077078 | -0.2219055 | 0.82438743 | 0.97548201 |
| SPIRE1 | 1931.0023 | -0.1506709 | 0.3022377 | -0.4985178 | 0.61811911 | 0.93928244 |
| PSMG2 | 2590.60487 | -0.0118879 | 0.1094598 | -0.108605 | 0.91351583 | 0.98662757 |
| CEP76 | 262.717782 | 0.01307193 | 0.19641313 | 0.06655322 | 0.94693739 | 0.99166614 |
| PTPN2 | 1968.6374 | -0.1314714 | 0.19728723 | -0.6663957 | 0.50515823 | 0.91365534 |
| SEH1L | 1697.39355 | -0.2448437 | 0.24960882 | -0.9809095 | 0.32663738 | 0.85282528 |
| AP002449.1 | 296.99488 | -0.1400017 | 0.24190249 | -0.5787526 | 0.5627561 | 0.92452095 |

| | | | | | | |
|---|---|---|---|---|---|---|
| CEP192 | 993.707235 | -0.5462614 | 0.36114602 | -1.5125777 | 0.13038696 | 0.7084713 |
| LDLRAD4 | 680.93141 | -0.2979845 | 0.43021385 | -0.6926427 | 0.48853381 | 0.90585781 |
| AP001010.1 | 89.0411034 | -0.4545016 | 0.56302341 | -0.8072517 | 0.41952147 | 0.88451372 |
| FAM210A | 1210.72621 | -0.2957424 | 0.2703732 | -1.0938303 | 0.2740294 | 0.82719197 |
| RNMT | 1997.31739 | -0.2615602 | 0.21621356 | -1.2097306 | 0.2263823 | 0.80246466 |
| ZNF519 | 111.831041 | -0.0671103 | 0.40084618 | -0.1674215 | 0.86703842 | 0.98223481 |
| ROCK1 | 2424.87589 | -0.1817771 | 0.14192002 | -1.2808415 | 0.20024933 | 0.78134034 |
| GREB1L | 68.6994355 | 1.29606106 | 0.75351151 | 1.72002821 | 0.08542731 | 0.6504594 |
| ESCO1 | 863.251634 | -0.2269953 | 0.21420749 | -1.059698 | 0.289282 | 0.83312616 |
| SNRPD1 | 1269.0447 | -0.1766859 | 0.22615115 | -0.7812736 | 0.43464161 | 0.89055338 |
| ABHD3 | 1349.05046 | -0.4824449 | 0.35864794 | -1.3451769 | 0.17856812 | 0.76115956 |
| MIB1 | 1832.54192 | -0.2147812 | 0.2586846 | -0.8302823 | 0.4063792 | 0.88132015 |
| GATA6-AS1 | 104.960172 | -0.1040629 | 0.55744521 | -0.1866782 | 0.85191293 | 0.97872457 |
| GATA6 | 369.566182 | -0.0511652 | 0.49003432 | -0.1044114 | 0.91684285 | 0.9868044 |
| AC091588.3 | 115.606835 | -0.0682362 | 0.50190035 | -0.1359556 | 0.89185635 | 0.98474725 |
| RBBP8 | 3573.82733 | -0.2419819 | 0.26166694 | -0.9247706 | 0.35508525 | 0.86364633 |
| CABLES1 | 2247.88568 | -0.094969 | 0.37283425 | -0.2547217 | 0.79893808 | 0.97186224 |
| TMEM241 | 243.153488 | 0.14692584 | 0.18140937 | 0.80991322 | 0.41799005 | 0.88445833 |
| AC011731.1 | 371.142779 | -0.0913225 | 0.39209048 | -0.2329118 | 0.81582993 | 0.9738447 |
| RIOK3 | 4307.9017 | -0.1088452 | 0.24404777 | -0.4459997 | 0.65559745 | 0.94532497 |
| RMC1 | 2827.42276 | -0.3275802 | 0.314875 | -1.0403501 | 0.29817729 | 0.83998573 |
| NPC1 | 5151.93659 | -0.8153403 | 0.46165517 | -1.7661242 | 0.07737502 | 0.63862831 |
| ANKRD29 | 320.963354 | 0.70277755 | 0.36486521 | 1.92612925 | 0.05408824 | 0.59329767 |
| LAMA3 | 525.869629 | 0.54437775 | 0.36305596 | 1.4994321 | 0.13376157 | 0.71505705 |
| TTC39C | 369.077181 | 0.00623948 | 0.35266143 | 0.01769256 | 0.98588411 | 0.99705651 |
| CABYR | 111.945359 | -0.2209844 | 0.35650146 | -0.6198696 | 0.53534364 | 0.92086969 |
| AC090772.3 | 316.947797 | 0.15815692 | 0.24322957 | 0.65023721 | 0.51553901 | 0.91636259 |
| OSBPL1A | 2430.61723 | 0.49711585 | 0.22195557 | 2.23970886 | 0.02510983 | 0.52085426 |
| IMPACT | 1247.04521 | 0.39601782 | 0.21083481 | 1.87833224 | 0.06033573 | 0.60825814 |
| ZNF521 | 319.211712 | 0.06382428 | 0.3775646 | 0.16904202 | 0.86576359 | 0.98197545 |
| SS18 | 4520.90931 | -0.1241431 | 0.16820926 | -0.7380278 | 0.46049753 | 0.89755243 |
| TAF4B | 300.782344 | 0.11360117 | 0.33015633 | 0.34408296 | 0.7307839 | 0.95848028 |
| LINC01543 | 70.7570269 | -0.6145098 | 1.11161253 | -0.5528094 | 0.58039395 | 0.93142344 |
| KCTD1 | 2069.56947 | 0.01000188 | 0.29943933 | 0.03340203 | 0.97335399 | 0.99445887 |
| AQP4-AS1 | 47.2822228 | 0.50739282 | 0.70035339 | 0.72448113 | 0.46877041 | 0.90003342 |
| AQP4 | 137.999349 | 0.78208927 | 0.77450063 | 1.0097981 | 0.31259203 | 0.84541735 |
| CHST9 | 184.018875 | 0.06226287 | 0.48037694 | 0.12961253 | 0.89687299 | 0.98474725 |
| CDH2 | 1988.29254 | -0.0502896 | 0.38981236 | -0.1290098 | 0.89734993 | 0.98474725 |
| AC110015.1 | 235.487278 | 0.02378172 | 0.42488731 | 0.05597183 | 0.95536425 | 0.99185043 |
| DSC2 | 1243.71088 | -0.1736886 | 0.2128316 | -0.8160845 | 0.41445179 | 0.88364943 |
| DSG2 | 2885.42912 | 0.25507547 | 0.30782849 | 0.82862852 | 0.40731465 | 0.88132015 |
| DSG2-AS1 | 94.3889393 | 0.10180366 | 0.43278791 | 0.2352276 | 0.81403208 | 0.9738447 |
| TTR | 61.1182451 | 0.01604553 | 0.83903828 | 0.01912372 | 0.98474241 | 0.99670866 |
| B4GALT6 | 339.190911 | -0.2736464 | 0.31569489 | -0.8668066 | 0.38604798 | 0.87432552 |
| AC017100.1 | 42.6620073 | -0.4455909 | 0.60578445 | -0.7355601 | 0.46199845 | 0.89762182 |
| TRAPPC8 | 2068.82005 | 0.04687556 | 0.16922412 | 0.27700282 | 0.78177794 | 0.96919365 |
| AC009831.1 | 58.1928148 | -0.1334291 | 0.42473836 | -0.3141442 | 0.75341151 | 0.96301881 |
| RNF125 | 215.280942 | 0.56834328 | 0.4282999 | 1.32697506 | 0.18451693 | 0.76820626 |
| AC011825.4 | 53.8326281 | -0.1492526 | 0.44838609 | -0.3328663 | 0.73923524 | 0.9602325 |
| RNF138 | 1084.31349 | -0.2065006 | 0.18968383 | -1.088657 | 0.27630515 | 0.82719407 |
| GAREM1 | 2080.12836 | 0.0341932 | 0.20935249 | 0.16332839 | 0.87025989 | 0.98275231 |
| AC015563.2 | 125.523879 | -0.4022113 | 0.33098339 | -1.215201 | 0.22428946 | 0.80078694 |
| AC021224.1 | 61.9066348 | 0.03555368 | 0.35810186 | 0.09928372 | 0.92091301 | 0.98767854 |
| KLHL14 | 543.836154 | 0.41446374 | 0.40334577 | 1.02756438 | 0.30415479 | 0.84211658 |

| | | | | | | |
|---|---|---|---|---|---|---|
| AC012123.1 | 44.118161 | 0.4457878 | 0.56778147 | 0.78513974 | 0.43237164 | 0.88980461 |
| CCDC178 | 166.75948 | 0.35997035 | 0.37911587 | 0.94949954 | 0.3423666 | 0.85889221 |
| ASXL3 | 187.327069 | 0.11729518 | 0.31191039 | 0.37605411 | 0.70687667 | 0.95501548 |
| NOL4 | 55.4316598 | 0.72129531 | 0.81084433 | 0.88956077 | 0.37370178 | 0.86791605 |
| DTNA | 394.189769 | 0.4161655 | 0.34317122 | 1.21270512 | 0.22524258 | 0.80176092 |
| MAPRE2 | 1678.13241 | -0.0367401 | 0.21184424 | -0.1734297 | 0.86231369 | 0.98176973 |
| ZNF397 | 1496.85826 | -0.1009195 | 0.14532597 | -0.6944354 | 0.48740921 | 0.90573694 |
| ZSCAN30 | 1238.15845 | -0.0001167 | 0.26004898 | -0.0004489 | 0.99964183 | 0.99994357 |
| AC011815.1 | 91.1776193 | -0.8681523 | 0.33069028 | -2.6252732 | 0.00865795 | 0.37987657 |
| ZNF24 | 3347.68789 | -0.1576842 | 0.12352696 | -1.2765164 | 0.20177304 | 0.78134034 |
| ZNF396 | 316.131413 | 0.11971945 | 0.24080536 | 0.49716271 | 0.61907431 | 0.93945324 |
| AC007998.2 | 196.609717 | -0.2214283 | 0.24313154 | -0.9107346 | 0.36243522 | 0.86561928 |
| INO80C | 726.851837 | -0.0033055 | 0.25267423 | -0.013082 | 0.98956237 | 0.99800524 |
| GALNT1 | 1975.64248 | 0.10788187 | 0.22433593 | 0.48089428 | 0.63059163 | 0.94088547 |
| C18orf21 | 556.206238 | -0.0953745 | 0.25213776 | -0.3782633 | 0.70523503 | 0.95501548 |
| RPRD1A | 2803.62295 | 0.00146832 | 0.1109111 | 0.0132387 | 0.98943736 | 0.99800524 |
| SLC39A6 | 3115.09695 | 0.06201668 | 0.18659134 | 0.33236633 | 7.40E-01 | 9.60E-01 |
| ELP2 | 3078.10456 | 0.02649231 | 0.10028801 | 0.26416233 | 0.79165483 | 0.97090793 |
| AC023043.1 | 110.936952 | -0.0330702 | 0.29819226 | -0.1109023 | 0.91169381 | 0.98662757 |
| MOCOS | 329.898616 | 0.26377496 | 0.40619713 | 0.64937672 | 5.16E-01 | 9.16E-01 |
| FHOD3 | 1974.2695 | -0.0112114 | 0.2764748 | -0.0405513 | 0.96765358 | 0.99400331 |
| TPGS2 | 3558.46737 | 0.02196395 | 0.17537481 | 0.12524007 | 0.9003335 | 0.98526603 |
| KIAA1328 | 344.510136 | -0.0066744 | 0.22488497 | -0.0296791 | 0.97632297 | 0.99455099 |
| CELF4 | 123.092504 | -0.0300395 | 0.32346452 | -0.0928681 | 0.92600832 | 0.98876911 |
| AC015961.2 | 59.3337713 | -0.1513494 | 0.32874085 | -0.4603912 | 0.64523548 | 0.94208859 |
| PIK3C3 | 2006.87387 | -0.1031747 | 0.13444072 | -0.7674367 | 0.44282193 | 0.89343541 |
| AC120049.1 | 179.276907 | 0.02152505 | 0.35161556 | 0.06121757 | 9.51E-01 | 0.99170503 |
| SETBP1 | 1441.52415 | -0.2781348 | 0.20256303 | -1.3730776 | 0.16972822 | 0.7586159 |
| SLC14A2 | 125.221004 | 3.27116969 | 1.12778952 | 2.90051434 | 0.00372551 | 0.26317334 |
| SLC14A1 | 928.604316 | 1.00278342 | 1.11345311 | 0.90060678 | 0.36779743 | 0.8673577 |
| EPG5 | 1764.57568 | 0.07357717 | 0.18516199 | 0.39736646 | 0.69109724 | 0.95158502 |
| PSTPIP2 | 448.143452 | -0.0350451 | 0.46175935 | -0.0758947 | 0.93950288 | 0.99128608 |
| ATP5F1A | 41213.8944 | 0.40265417 | 0.1494763 | 2.6937659 | 0.00706498 | 0.35402334 |
| HAUS1 | 512.848685 | -0.2674378 | 0.17043542 | -1.5691447 | 0.11661423 | 0.6888681 |
| C18orf25 | 1289.48842 | -0.2361247 | 0.14502894 | -1.628121 | 1.03E-01 | 6.74E-01 |
| RNF165 | 123.624144 | -0.0867995 | 0.63704724 | -0.1362529 | 8.92E-01 | 9.85E-01 |
| LOXHD1 | 71.9281024 | 1.15154855 | 1.07747816 | 1.06874422 | NA | NA |
| ST8SIA5 | 298.50509 | -0.0640476 | 0.38530015 | -0.1662279 | 0.86797764 | 0.98237277 |
| PIAS2 | 1771.3356 | -0.2597916 | 0.15232633 | -1.7054938 | 0.08810236 | 0.65363785 |
| KATNAL2 | 528.337844 | -0.1514919 | 0.22434731 | -0.675256 | 0.49951316 | 0.91161831 |
| AC012254.1 | 101.232031 | -0.189877 | 0.27316521 | -0.6950996 | 0.48699291 | 0.90573694 |
| AC012254.3 | 824.17297 | 0.39486923 | 0.2515687 | 1.56962782 | 0.11650172 | 0.6888681 |
| HDHD2 | 2391.09743 | 0.32889254 | 0.17931041 | 1.8342077 | 0.06662317 | 0.62390091 |
| AC012254.2 | 3339.72631 | 0.13711389 | 0.2074566 | 0.66092807 | 0.50865844 | 0.91439937 |
| IER3IP1 | 3872.38647 | -0.0114737 | 0.25300063 | -0.0453506 | 0.96382783 | 0.99335116 |
| SMAD2 | 3616.59732 | -0.1171292 | 0.1226272 | -0.9551647 | 0.33949439 | 0.85759056 |
| ZBTB7C | 251.616513 | 0.05336329 | 0.48740683 | 0.10948408 | 0.91281855 | 0.98662757 |
| CTIF | 3010.75316 | -0.2820613 | 0.19257165 | -1.4647085 | 0.14300047 | 0.72784886 |
| SMAD7 | 2213.01991 | -0.0708842 | 0.3250969 | -0.2180403 | 0.8273977 | 0.97548201 |
| DYM | 1850.66995 | 0.01060507 | 0.15821728 | 0.06702852 | 0.946559 | 0.99166614 |
| C18orf32 | 3771.70953 | 0.12603402 | 0.17591312 | 0.71645606 | 0.47370979 | 0.90199039 |
| RPL17-C18or | 23724.3103 | 0.13200297 | 0.22567299 | 0.58493028 | 0.5585946 | 0.92421175 |
| AC100778.3 | 448.490754 | -0.0689407 | 0.44944474 | -0.1533909 | 0.87809003 | 0.98275231 |
| RPL17 | 20923.8841 | 0.12357694 | 0.24467036 | 0.50507522 | 0.613506 | 0.93881898 |

| | | | | | | |
|---|---|---|---|---|---|---|
| LIPG | 2344.42491 | 0.33662805 | 0.56709619 | 0.59359956 | 0.55277997 | 0.92373235 |
| ACAA2 | 17168.2135 | 0.23995526 | 0.39331985 | 0.61007666 | 0.54181103 | 0.92292765 |
| AC090227.1 | 5798.51445 | 0.20972851 | 0.35779168 | 0.58617491 | 0.55775798 | 0.92421175 |
| SNHG22 | 511.222768 | 0.02180188 | 0.31014149 | 0.07029657 | 0.94395761 | 0.99136103 |
| MYO5B | 3621.60476 | 0.01386351 | 0.30283684 | 0.04577883 | 0.96348654 | 0.99320711 |
| CFAP53 | 167.67079 | 0.20877004 | 0.33971352 | 0.61454733 | 0.53885371 | 0.9215968 |
| MBD1 | 4197.26474 | -0.2030671 | 0.11346042 | -1.7897618 | 7.35E-02 | 0.6374448 |
| CXXC1 | 4336.00226 | -0.3062724 | 0.15084258 | -2.030411 | 4.23E-02 | 0.56485966 |
| MAPK4 | 436.075434 | 0.54965662 | 0.47220743 | 1.16401518 | 0.24441786 | 0.81473945 |
| MRO | 1275.46476 | 0.84522525 | 0.71033209 | 1.18990154 | 2.34E-01 | 8.07E-01 |
| ME2 | 1645.19194 | -0.0073187 | 0.1356117 | -0.0539677 | 0.95696088 | 0.99200096 |
| ELAC1 | 452.068505 | 0.10533248 | 0.31356822 | 0.33591566 | 0.73693447 | 0.96009463 |
| AC091551.1 | 511.946468 | -0.0292646 | 0.31068351 | -0.0941941 | 0.92495498 | 0.98869145 |
| SMAD4 | 4058.37681 | -0.1915864 | 0.13626821 | -1.4059509 | 0.15973869 | 0.74768867 |
| MEX3C | 1499.08233 | -0.1338843 | 0.19720826 | -0.6788982 | 0.49720236 | 0.90913397 |
| MBD2 | 4417.89458 | -0.1643794 | 0.18495571 | -0.8887498 | 0.37413756 | 0.86791605 |
| AC093462.1 | 58.8377339 | -0.5010122 | 0.41520567 | -1.2066602 | 0.22756304 | 0.80300331 |
| POLI | 1752.19942 | -0.1567018 | 0.19550061 | -0.8015415 | 0.42281825 | 0.88453836 |
| C18orf54 | 103.438663 | 0.09652647 | 0.28968653 | 0.33321009 | 0.7389757 | 0.9602325 |
| RAB27B | 44.8293241 | 0.97748002 | 0.49743528 | 1.96503961 | 0.04940965 | 0.5861488 |
| CCDC68 | 280.011864 | 0.4278435 | 0.37715466 | 1.13439801 | 2.57E-01 | 8.21E-01 |
| TCF4 | 1322.32339 | -0.1922917 | 0.31016266 | -0.6199704 | 0.53527728 | 0.92084006 |
| AC006305.1 | 45.9081113 | 0.60325266 | 0.75176346 | 0.8024501 | 4.22E-01 | 0.88451372 |
| TXNL1 | 3268.73258 | -0.1121876 | 0.20780542 | -0.5398685 | 0.58928773 | 0.93289552 |
| WDR7 | 755.150814 | 0.01296933 | 0.10369643 | 0.12507019 | 0.90046798 | 0.98526603 |
| FECH | 2599.39371 | 0.29473493 | 0.27221417 | 1.08273174 | 0.27892751 | 0.82825734 |
| NARS | 11005.3165 | 0.14010725 | 0.22223944 | 0.63043379 | 0.52841081 | 0.91888066 |
| AC027097.1 | 165.786809 | -0.5328972 | 0.21750241 | -2.4500747 | 0.01428266 | 0.43699139 |
| AC027097.2 | 157.169823 | -0.582625 | 0.28996628 | -2.0092853 | 0.04450689 | 0.57267043 |
| ATP8B1 | 1124.04417 | 0.33612959 | 0.26406055 | 1.27292618 | 0.20304424 | 0.78252625 |
| NEDD4L | 6468.96781 | -0.5074073 | 0.42360115 | -1.1978422 | 0.23097845 | 0.80610188 |
| AC090236.1 | 529.250654 | 0.24845979 | 0.38790295 | 0.64052051 | 0.52183426 | 0.91723878 |
| ALPK2 | 1529.74686 | 0.20950409 | 0.43645893 | 0.48000873 | 6.31E-01 | 0.94088547 |
| AC105105.1 | 213.746842 | 0.01005923 | 0.58634655 | 0.01715578 | 0.98631234 | 0.99715899 |
| MALT1 | 2134.67615 | -0.1566269 | 0.23845422 | -0.6568426 | 5.11E-01 | 9.15E-01 |
| AC104365.3 | 162.708935 | -0.2072104 | 0.30712401 | -0.6746799 | 0.49987917 | 0.91180773 |
| ZNF532 | 1838.44638 | -0.324978 | 0.20312486 | -1.5998929 | 0.10962235 | 0.67859564 |
| SEC11C | 2576.15685 | 0.28229981 | 0.27346595 | 1.0323033 | 0.30193006 | 0.84211658 |
| LMAN1 | 6517.49052 | 0.18826702 | 0.17844634 | 1.05503436 | 0.29140961 | 0.83454586 |
| CCBE1 | 309.655028 | -0.0014542 | 0.39496292 | -0.0036818 | 0.99706234 | 0.99976751 |
| PMAIP1 | 226.710359 | -1.7207787 | 0.71320332 | -2.4127463 | 0.01583284 | 0.4564177 |
| AC090409.1 | 58.5427964 | -0.265049 | 0.39766221 | -0.6665181 | 0.50508002 | 0.91365534 |
| LINC01544 | 38.4489856 | 0.06969226 | 0.86354349 | 0.08070498 | 0.93567658 | 0.99068685 |
| RNF152 | 6568.25336 | 0.58150809 | 0.49760197 | 1.16862095 | 0.24255638 | 0.81333856 |
| PIGN | 978.777246 | 0.21202049 | 0.17283764 | 1.22670321 | 2.20E-01 | 0.79858446 |
| KIAA1468 | 1715.55259 | 0.00769894 | 0.16388304 | 0.04697826 | 0.96253055 | 0.99320711 |
| TNFRSF11A | 89.3844195 | -0.1514413 | 0.51327291 | -0.2950503 | 0.76795542 | 0.96762687 |
| ZCCHC2 | 1149.59057 | -0.5053571 | 0.21843442 | -2.3135413 | 0.0206929 | 0.50488816 |
| PHLPP1 | 687.005371 | -0.2698443 | 0.20224727 | -1.3342298 | 0.18212855 | 0.76569697 |
| BCL2 | 1253.16814 | -0.17486 | 0.20196808 | -0.8657802 | 0.38661073 | 0.87432552 |
| KDSR | 3945.59734 | -0.3761834 | 0.29142733 | -1.2908309 | 0.19676232 | 0.77816059 |
| AC036176.1 | 85.2604623 | 0.5021461 | 0.3891152 | 1.29048183 | 0.19688342 | 0.77816059 |
| VPS4B | 2478.36621 | -0.1987366 | 0.20894675 | -0.951135 | 0.34153585 | 0.85889221 |
| SERPINB7 | 93.5507392 | -2.7590998 | 0.96546765 | -2.8577859 | 4.27E-03 | 2.85E-01 |

| | | | | | | |
|---|---|---|---|---|---|---|
| HMSD | 175.981077 | -1.248039 | 0.65006527 | -1.9198673 | 0.05487467 | 0.59342538 |
| AC009802.1 | 54.6321125 | -0.3636957 | 0.54327283 | -0.6694531 | 0.50320649 | 0.91356552 |
| SERPINB8 | 965.765842 | -0.949177 | 0.52688805 | -1.8014775 | 0.07162765 | 0.63189503 |
| CDH19 | 74.1722266 | -1.2806285 | 0.74500187 | -1.7189601 | NA | NA |
| AC110597.1 | 44.3173052 | 1.78450165 | 0.54067513 | 3.30050628 | 9.65E-04 | 1.48E-01 |
| DSEL | 577.23648 | 0.38000635 | 0.3169118 | 1.1990918 | 0.23049225 | 0.80584899 |
| AC110597.3 | 191.105886 | 0.02444873 | 0.34513527 | 0.07083811 | 0.9435266 | 0.99136103 |
| TMX3 | 2566.2811 | -0.1252584 | 0.1548 | -0.8091625 | 0.41842166 | 0.88445833 |
| CCDC102B | 226.144116 | -0.185822 | 0.38152202 | -0.4870544 | 0.62621981 | 0.94088547 |
| DOK6 | 423.01555 | 0.13276612 | 0.31903626 | 0.41614742 | 0.67730213 | 0.95018034 |
| CD226 | 163.210839 | 0.02554929 | 0.49334569 | 0.05178779 | 0.95869778 | 0.99237995 |
| RTTN | 539.280693 | 0.14932291 | 0.29881736 | 0.49971298 | 0.61727719 | 0.93910158 |
| SOCS6 | 1169.56662 | -0.1000246 | 0.2263446 | -0.4419131 | 0.65855213 | 0.9458503 |
| CBLN2 | 66.3789652 | 1.57362879 | 1.03869253 | 1.51500924 | NA | NA |
| FBXO15 | 147.672406 | 0.67060156 | 0.51658252 | 1.29814994 | 0.19423582 | 0.7762856 |
| TIMM21 | 1315.11106 | 0.1877394 | 0.18723545 | 1.00269154 | 0.31600972 | 0.84750116 |
| CYB5A | 15318.8366 | 0.59136568 | 0.32010801 | 1.84739417 | 0.06469003 | 0.62073893 |
| CNDP2 | 37753.7081 | 0.34310832 | 0.37764336 | 0.9085512 | 0.36358708 | 0.8663515 |
| CNDP1 | 167.818046 | 0.92805294 | 0.54167386 | 1.71330575 | 0.0866563 | 0.65228863 |
| LINC00909 | 617.149391 | 0.07708487 | 0.28857694 | 0.2671207 | 0.78937623 | 0.97090793 |
| ZNF407 | 511.694718 | -0.3741112 | 0.18272756 | -2.0473716 | 0.04062161 | 0.55657615 |
| ZADH2 | 2367.44675 | 0.02517822 | 0.1586414 | 0.15871152 | 0.87389616 | 0.98275231 |
| TSHZ1 | 1727.81499 | 0.05983312 | 0.248991 | 0.24030232 | 0.81009589 | 0.97334138 |
| ZNF516 | 1496.7293 | -0.1765038 | 0.24428934 | -0.7225194 | 0.46997523 | 0.9008571 |
| AC018413.1 | 68.5333605 | 0.04411878 | 0.30557519 | 0.14437944 | 0.88520085 | 0.98356741 |
| ZNF236-DT | 57.4166331 | 0.273867 | 0.29492857 | 0.92858755 | 0.35310288 | 0.86364633 |
| ZNF236 | 510.10718 | -0.1875953 | 0.21515041 | -0.8719263 | 0.38324859 | 0.87194871 |
| AC093330.1 | 75.9956143 | -0.3946141 | 0.4394241 | -0.8980256 | 0.36917188 | 0.86762749 |
| MBP | 3067.11899 | -0.1643764 | 0.35330008 | -0.4652601 | 0.64174521 | 0.94088547 |
| AC018529.2 | 58.031122 | -1.7696095 | 0.57170394 | -3.095325 | 0.00196597 | 0.19087435 |
| LINC01896 | 204.432572 | 1.95956827 | 0.87731257 | 2.23360332 | 0.02550919 | 0.52304994 |
| SALL3 | 164.617279 | 1.19772166 | 0.84641087 | 1.41505941 | 0.15705108 | 0.74522306 |
| ATP9B | 1086.59614 | 0.17960092 | 0.18987215 | 0.94590452 | 3.44E-01 | 8.60E-01 |
| NFATC1 | 653.851009 | -0.0489694 | 0.36181213 | -0.135345 | 0.89233915 | 0.98474725 |
| AC068473.5 | 385.293409 | 0.20438443 | 0.32929592 | 0.620671 | 0.53481612 | 0.92084006 |
| CTDP1 | 1813.09481 | -0.3460113 | 0.15566875 | -2.2227408 | 0.02623328 | 0.52678638 |
| PQLC1 | 5297.34968 | -0.0108356 | 0.3287491 | -0.03296 | 0.97370647 | 0.99445887 |
| HSBP1L1 | 2230.82001 | -0.2722698 | 0.24121288 | -1.128753 | 0.25900205 | 0.82364948 |
| TXNL4A | 2281.50544 | -0.1334086 | 0.2238788 | -0.5958967 | 0.55124421 | 0.92373235 |
| AC090360.1 | 695.754688 | 0.13360201 | 0.11951009 | 1.11791403 | 0.26360371 | 0.82424998 |
| RBFA | 894.074821 | 0.02749697 | 0.11742774 | 0.23416077 | 0.81486017 | 0.9738447 |
| RBFADN | 143.714084 | 0.76695761 | 0.40753338 | 1.8819504 | 0.05984275 | 0.60732038 |
| ADNP2 | 1322.21493 | -0.3751984 | 0.20233966 | -1.8542998 | 6.37E-02 | 0.61812911 |
| PARD6G-AS1 | 168.597518 | -0.6363403 | 0.39504408 | -1.6108082 | 1.07E-01 | 6.77E-01 |
| PARD6G | 459.040543 | 0.26828611 | 0.30929828 | 0.86740254 | 0.38572149 | 0.87420019 |
| AC139100.1 | 67.6158457 | 0.50299037 | 0.32157875 | 1.56412812 | 0.11778748 | 0.69069313 |
| C20orf96 | 977.573816 | 0.04238112 | 0.25255641 | 0.16780853 | 0.86673391 | 0.9821997 |
| ZCCHC3 | 1367.84143 | -0.1707277 | 0.13290363 | -1.2845978 | 0.19893285 | 0.78134034 |
| NRSN2-AS1 | 226.287784 | -0.1769819 | 0.29082845 | -0.6085438 | 0.54282685 | 0.92292765 |
| SOX12 | 2130.21687 | -0.2876655 | 0.25299072 | -1.1370597 | 0.25551334 | 0.82052927 |
| NRSN2 | 2962.82167 | -0.3408994 | 0.20807035 | -1.6383855 | 1.01E-01 | 6.73E-01 |
| TRIB3 | 819.211197 | -0.4279276 | 0.54063278 | -0.7915309 | 0.42863424 | 0.88821697 |
| RBCK1 | 7591.90989 | -0.5332799 | 0.26602975 | -2.0045874 | 0.04500717 | 0.5734524 |
| TBC1D20 | 3135.14036 | -0.5679618 | 0.25603845 | -2.2182674 | 0.0265366 | 0.52678638 |

| | | | | | | |
|---|---|---|---|---|---|---|
| CSNK2A1 | 5540.17682 | -0.2527062 | 0.25353719 | -0.9967224 | 0.3188993 | 0.84914926 |
| TCF15 | 40.9873935 | 0.16645043 | 0.66407644 | 0.2506495 | 0.80208511 | 0.97233627 |
| SRXN1 | 6457.62751 | -1.0105729 | 0.74110999 | -1.3635937 | 0.17269549 | 0.75974924 |
| AL121758.1 | 6447.50263 | -1.0136384 | 0.74134266 | -1.3673008 | 0.17153105 | 0.75974924 |
| SLC52A3 | 1373.25538 | 0.42462558 | 0.46383677 | 0.91546339 | 0.35994841 | 0.86480228 |
| FAM110A | 486.962153 | 0.56862783 | 0.27547138 | 2.06419929 | 0.03899882 | 0.55613287 |
| ANGPT4 | 48.9005903 | -1.6217137 | 0.44062325 | -3.6804996 | 0.00023278 | 0.07653478 |
| RSPO4 | 77.0162543 | 0.34648018 | 0.66139401 | 0.52386349 | 0.60037349 | 0.93487648 |
| PSMF1 | 5556.90953 | -0.1363183 | 0.14930773 | -0.9130023 | 0.36124134 | 0.86528137 |
| TMEM74B | 180.142498 | -0.5840835 | 0.42951839 | -1.3598567 | 0.17387528 | 0.75974924 |
| SNPH | 453.873937 | 0.31225914 | 0.28557189 | 1.09345195 | 0.27419541 | 0.82719407 |
| SDCBP2 | 820.847711 | 0.56981748 | 0.36009704 | 1.5823998 | 0.11355833 | 0.68214043 |
| AL136531.2 | 1998.3651 | -0.0212805 | 0.26813794 | -0.0793641 | 0.93674298 | 0.99068685 |
| SDCBP2-AS1 | 287.633166 | 0.07106251 | 0.33926623 | 0.20945942 | 0.83408961 | 0.97550416 |
| AL136531.1 | 39.813924 | -0.078693 | 0.4650477 | -0.1692149 | 0.86562758 | 0.98197545 |
| FKBP1A | 10675.9055 | -0.1478074 | 0.23179994 | -0.6376507 | 0.52370112 | 0.91723878 |
| NSFL1C | 3930.00646 | -0.1439823 | 0.18584942 | -0.7747255 | 0.43850187 | 0.89175645 |
| SIRPB2 | 186.551735 | 0.01019349 | 0.45576075 | 0.02236588 | 9.82E-01 | 9.96E-01 |
| AL049634.2 | 55.9198358 | -0.2548291 | 0.81943024 | -0.3109833 | 0.75581335 | 0.96424084 |
| SIRPB1 | 131.934843 | -0.3023057 | 0.5323601 | -0.5678595 | 5.70E-01 | 9.27E-01 |
| SIRPG | 47.2818345 | 0.69158008 | 1.065923 | 0.64880867 | 0.51646205 | 0.91636259 |
| AL117335.1 | 39.3218549 | 0.03649431 | 0.43782209 | 0.08335419 | 0.93356991 | 0.99019225 |
| SIRPA | 4386.17166 | -0.0063878 | 0.08891932 | -0.0718379 | 9.43E-01 | 9.91E-01 |
| STK35 | 1821.85041 | -0.1472598 | 0.15681918 | -0.939042 | 0.3477092 | 0.86225576 |
| SNRPB | 5653.15016 | -0.4184933 | 0.2787261 | -1.5014498 | 0.13323925 | 0.71493653 |
| AL049650.1 | 1882.41579 | -0.3335748 | 0.29583357 | -1.1275758 | 0.25949912 | 0.82400425 |
| ZNF343 | 789.383658 | -0.029592 | 0.13928093 | -0.2124629 | 0.83174593 | 0.97550416 |
| NOP56 | 4183.99896 | -0.5092623 | 0.30660283 | -1.6609837 | 0.09671672 | 0.66321552 |
| IDH3B | 5031.13059 | 0.13906103 | 0.15563495 | 0.89350771 | 0.37158535 | 0.86762749 |
| EBF4 | 1101.3012 | -0.3162943 | 0.30261433 | -1.0452058 | 0.29592785 | 0.83750889 |
| CPXM1 | 827.53881 | 1.71795835 | 0.63429891 | 2.70843656 | 0.0067601 | 0.34709891 |
| PCED1A | 3860.41177 | 0.06027474 | 0.27365452 | 0.22025854 | 0.82566981 | 0.97548201 |
| VPS16 | 2839.43502 | -0.126563 | 0.10564302 | -1.1980255 | 0.23090707 | 0.80610188 |
| PTPRA | 4562.82331 | 0.13446874 | 0.09965377 | 1.34935933 | 0.17722158 | 0.76115956 |
| MRPS26 | 2207.9332 | 0.20557729 | 0.18988669 | 1.08263141 | 0.27897206 | 0.82825734 |
| OXT | 47.8566743 | -0.9105482 | 1.16365374 | -0.7824907 | 0.43392626 | 0.89055338 |
| UBOX5-AS1 | 178.470617 | 0.1887486 | 0.42317575 | 0.44602886 | 0.6555764 | 0.94532497 |
| UBOX5 | 817.411697 | 0.28722028 | 0.30265377 | 0.94900612 | 0.3426175 | 0.85889221 |
| FASTKD5 | 1644.42202 | -0.1704561 | 0.39358447 | -0.4330865 | 0.66495192 | 0.94692723 |
| LZTS3 | 5647.60594 | -0.4893903 | 0.23483395 | -2.0839846 | 0.03716157 | 0.55527244 |
| DDRGK1 | 4161.37738 | -0.0723197 | 0.15534912 | -0.4655304 | 0.64155166 | 0.94088547 |
| ITPA | 2536.12266 | 0.11364852 | 0.15169306 | 0.74920052 | 0.45373636 | 0.89670027 |
| SLC4A11 | 1291.78661 | 1.51815672 | 0.58041129 | 2.6156568 | 0.0089056 | 0.38245454 |
| AL109976.1 | 90.865753 | 1.13285844 | 0.7091574 | 1.59747109 | 0.11016074 | 0.67996643 |
| C20orf194 | 2178.44824 | 0.08087965 | 0.18063893 | 0.44774206 | 0.65433936 | 0.94532497 |
| ATRN | 3697.67695 | 0.2100331 | 0.14813149 | 1.41788282 | 0.15622498 | 0.74522306 |
| ADAM33 | 1215.04847 | -1.3144407 | 0.79592339 | -1.6514663 | 0.09864339 | 0.66763302 |
| SIGLEC1 | 1137.21204 | -1.1313636 | 0.70330896 | -1.6086296 | 0.10769735 | 0.67707349 |
| HSPA12B | 832.171242 | -0.3676473 | 0.30007708 | -1.2251761 | 0.22050886 | 0.79912826 |
| C20orf27 | 1076.02137 | 0.19319935 | 0.22101108 | 0.87416139 | 0.38203037 | 0.87185666 |
| SPEF1 | 177.957267 | 0.01696744 | 0.51477117 | 0.03296113 | 9.74E-01 | 9.94E-01 |
| CENPB | 4732.80915 | -0.2735793 | 0.22700598 | -1.2051636 | 0.22814016 | 0.80300331 |
| CDC25B | 4195.26476 | 0.26605916 | 0.2962287 | 0.89815457 | 0.36910315 | 0.86762749 |
| LINC01730 | 43.6289618 | -0.3572304 | 0.7637581 | -0.4677272 | 0.6399797 | 0.94088547 |

| | | | | | | |
|---|---|---|---|---|---|---|
| AP5S1 | 845.676122 | -0.3507701 | 0.26149796 | -1.3413875 | 0.17979468 | 0.76285534 |
| MAVS | 6025.49209 | 0.1028195 | 0.16652395 | 0.61744573 | 0.53694077 | 0.92105614 |
| AL353194.1 | 52.3753541 | -0.4921019 | 0.36435282 | -1.3506191 | 0.17681748 | 0.76115956 |
| PANK2 | 1833.68846 | -0.2485627 | 0.17725357 | -1.4023 | 0.16082569 | 0.74874703 |
| AL031670.1 | 323.645314 | -0.3603812 | 0.23122261 | -1.5585899 | 0.11909348 | 0.69233212 |
| RNF24 | 2365.83297 | -0.2796174 | 0.32175462 | -0.8690393 | 0.38482562 | 0.87312908 |
| SMOX | 1397.18627 | -0.0470402 | 0.33583863 | -0.1400678 | 0.88860643 | 0.98423722 |
| PRNP | 8441.55835 | -0.0430442 | 0.35117236 | -0.1225728 | 0.90244542 | 0.98599252 |
| RASSF2 | 527.659329 | -0.0748132 | 0.42091243 | -0.1777404 | 0.85892684 | 0.97992012 |
| SLC23A2 | 1465.45443 | 0.16160167 | 0.20642102 | 0.78287409 | 0.43370105 | 0.89055338 |
| TMEM230 | 5756.85748 | 0.20081396 | 0.14142385 | 1.41994413 | 0.15562395 | 0.74477742 |
| PCNA | 1917.19841 | 0.25053357 | 0.16856857 | 1.48624123 | 0.13721532 | 0.72209298 |
| CDS2 | 4439.44674 | -0.0187781 | 0.19919044 | -0.094272 | 0.92489312 | 0.98869145 |
| LINC00654 | 54.315373 | -0.2406466 | 0.42183834 | -0.5704711 | 5.68E-01 | 0.92638187 |
| GPCPD1 | 2380.72768 | -0.0829897 | 0.3940532 | -0.2106053 | 0.83319529 | 0.97550416 |
| SHLD1 | 160.570506 | 0.23311417 | 0.25721304 | 0.90630775 | 3.65E-01 | 0.8663515 |
| CHGB | 758.817461 | -1.0251932 | 0.67016162 | -1.5297702 | 0.12607363 | 0.70364066 |
| TRMT6 | 906.661873 | -0.4856624 | 0.23414877 | -2.0741618 | 0.03806429 | 0.55538555 |
| MCM8 | 478.957194 | -0.3495182 | 0.2690762 | -1.2989563 | 1.94E-01 | 0.77620852 |
| MCM8-AS1 | 37.5989986 | -0.3352227 | 0.36112335 | -0.9282775 | 0.35326364 | 0.86364633 |
| AL035461.2 | 43.768852 | -0.1968479 | 0.40965833 | -0.4805172 | 0.63085968 | 0.94088547 |
| CRLS1 | 4247.03208 | -0.0163731 | 0.26266348 | -0.0623349 | 0.9502961 | 0.99170503 |
| LRRN4 | 658.725062 | -0.3805378 | 0.55320105 | -0.6878833 | 0.49152626 | 0.90600166 |
| AL118505.1 | 46.730699 | 0.6079741 | 0.56218957 | 1.08143968 | 0.27950158 | 0.82825734 |
| FERMT1 | 318.208274 | 0.5563164 | 0.45973436 | 1.21008229 | 0.22624732 | 0.80246466 |
| BMP2 | 465.770241 | -0.5476459 | 0.36678503 | -1.4930977 | 0.1354116 | 0.71649038 |
| TMX4 | 3713.72003 | -0.0901191 | 0.21107847 | -0.4269459 | 0.66941876 | 0.94876933 |
| PLCB1 | 952.300095 | -0.3485194 | 0.45540206 | -0.7653003 | 0.44409273 | 0.8938278 |
| PLCB4 | 503.700226 | 0.23532852 | 0.31157899 | 0.75527725 | 0.45008265 | 0.89557948 |
| LAMP5 | 49.4924057 | 1.08428806 | 0.53502077 | 2.02662799 | 0.04270047 | 0.56594734 |
| ANKEF1 | 403.549871 | 0.17672419 | 0.22589313 | 0.7823354 | 0.43401749 | 0.89055338 |
| SNAP25-AS1 | 47.9494181 | 0.57636837 | 0.3422026 | 1.68428987 | 0.09212566 | 0.65583276 |
| SNAP25 | 144.906121 | 1.81200827 | 0.53420049 | 3.39200039 | 0.00069384 | 0.12703622 |
| MKKS | 2984.65459 | 0.16712661 | 0.22907073 | 0.72958516 | 0.46564379 | 0.89943247 |
| AL034430.1 | 1007.37522 | 0.14561012 | 0.32521466 | 0.44773541 | 0.65434416 | 0.94532497 |
| SLX4IP | 580.14954 | 0.16097951 | 0.28102354 | 0.57283284 | 0.56675788 | 0.9256133 |
| JAG1 | 3682.21559 | -0.1292902 | 0.21971686 | -0.5884399 | 5.56E-01 | 0.92373235 |
| BTBD3 | 1789.00027 | -0.2834871 | 0.18508772 | -1.5316363 | 0.12561221 | 0.70347499 |
| AL035448.1 | 42.0391931 | -0.4004175 | 0.44164411 | -0.906652 | 0.36459085 | 0.8663515 |
| SPTLC3 | 2114.33886 | 0.34682315 | 0.31378395 | 1.10529282 | 0.26903273 | 0.82683094 |
| ISM1 | 217.076553 | 0.25178316 | 0.57795555 | 0.43564451 | 0.66309467 | 0.94642611 |
| TASP1 | 645.235393 | 0.1370793 | 0.147202 | 0.93123254 | 0.35173329 | 0.86364633 |
| ESF1 | 624.839671 | -0.2856421 | 0.25733593 | -1.109997 | 0.26700034 | 0.8265123 |
| NDUFAF5 | 960.776421 | 0.11809789 | 0.16948543 | 0.6968026 | 4.86E-01 | 9.05E-01 |
| MACROD2 | 283.690099 | 0.32800853 | 0.25992073 | 1.26195601 | 0.20696461 | 0.78790396 |
| FLRT3 | 2962.49354 | -0.2629939 | 0.26799053 | -0.9813551 | 0.32641768 | 0.85282528 |
| KIF16B | 1515.95813 | 0.00596283 | 0.24190789 | 0.02464918 | 0.98033479 | 0.99568904 |
| SNRPB2 | 1955.9992 | 0.18341436 | 0.20965497 | 0.87483911 | 0.38166146 | 0.87185666 |
| BFSP1 | 105.739892 | 0.96246158 | 0.58565502 | 1.64339336 | 1.00E-01 | 6.72E-01 |
| DSTN | 19182.7091 | 0.08956157 | 0.2810421 | 0.31867669 | 0.74997169 | 0.96249332 |
| RRBP1 | 16670.1881 | -0.1573688 | 0.15732411 | -1.0002843 | 0.31717294 | 0.84819508 |
| SNX5 | 7503.15759 | 0.52228258 | 0.2252731 | 2.31844181 | 2.04E-02 | 0.50103529 |
| OVOL2 | 729.974578 | 0.61195126 | 0.27200959 | 2.24974147 | 0.02446536 | 0.51967919 |
| MGME1 | 1161.06185 | 0.02382729 | 0.24514611 | 0.09719627 | 0.92257053 | 0.98826666 |

| | | | | | | |
|---|---|---|---|---|---|---|
| PET117 | 806.588579 | 0.19552952 | 0.27813344 | 0.70300614 | 0.48205193 | 0.90541935 |
| KAT14 | 1503.395 | 0.25224133 | 0.20565645 | 1.22651796 | 0.22000382 | 0.79858446 |
| ZNF133 | 1453.03355 | -0.066389 | 0.27805171 | -0.238765 | 0.81128779 | 0.97352497 |
| AL049646.2 | 83.1967096 | 0.10261161 | 0.29942026 | 0.34270095 | 0.73182345 | 0.95848028 |
| DZANK1 | 470.589687 | 0.1574946 | 0.27071923 | 0.58176362 | 0.56072591 | 0.9244521 |
| POLR3F | 642.368211 | -0.1596043 | 0.20722529 | -0.7701972 | 0.4411829 | 0.89290667 |
| RBBP9 | 1500.8991 | 0.37473216 | 0.15024083 | 2.49420983 | 0.01262379 | 0.41598747 |
| SEC23B | 4242.85685 | 0.39267812 | 0.20976982 | 1.87194768 | 0.06121384 | 0.60989294 |
| SMIM26 | 1543.34579 | -0.0446629 | 0.18743531 | -0.2382846 | 0.8116604 | 0.97352497 |
| AL121900.2 | 2152.25793 | 0.27722094 | 0.20073538 | 1.38102677 | 0.16727073 | 0.75558932 |
| DTD1 | 1770.61066 | 0.46659141 | 0.2116946 | 2.20407795 | 0.02751886 | 0.52904542 |
| AL035563.1 | 152.87111 | 0.48162155 | 0.48129646 | 1.00067545 | 0.31698374 | 0.84819508 |
| SLC24A3 | 192.093356 | 0.13514231 | 0.41205012 | 0.32797541 | 0.74293026 | 0.96109001 |
| AL121761.2 | 42.8599021 | 0.89008605 | 0.47041922 | 1.89211243 | 0.058476 | 0.60332684 |
| AL121761.1 | 51.2441447 | 0.15535781 | 0.31231105 | 0.49744578 | 0.61887473 | 0.93945324 |
| RIN2 | 2555.8964 | 0.24890471 | 0.14889781 | 1.67164787 | 0.09459378 | 0.65888357 |
| NAA20 | 3034.90643 | -0.0162785 | 0.17077614 | -0.0953207 | 0.92406011 | 0.98869145 |
| CRNKL1 | 1231.91083 | -0.1170495 | 0.14783209 | -0.7917732 | 0.4284929 | 0.88820133 |
| CFAP61 | 53.4493401 | 0.67893443 | 0.73187579 | 0.92766345 | 0.35358217 | 0.86364633 |
| RALGAPA2 | 3228.12404 | -0.1904868 | 0.22002746 | -0.8657408 | 0.38663231 | 0.87432552 |
| KIZ | 1460.77385 | 0.3671387 | 0.3234209 | 1.13517309 | 0.2563028 | 0.82067614 |
| AL121759.1 | 84.5492459 | 0.79026269 | 0.36553164 | 2.16195428 | 0.0306217 | 0.54443002 |
| KIZ-AS1 | 101.67087 | 0.50999116 | 0.35329913 | 1.44351095 | 0.14887659 | 0.73636742 |
| AL117332.1 | 133.046495 | -0.0787908 | 0.35704326 | -0.2206757 | 0.82534495 | 0.97548201 |
| XRN2 | 3910.75173 | -0.1592861 | 0.18534937 | -0.859383 | 3.90E-01 | 8.76E-01 |
| THBD | 1952.1321 | -0.2926722 | 0.49178379 | -0.5951238 | 0.55176071 | 0.92373235 |
| CD93 | 4268.61598 | -0.4558957 | 0.44712759 | -1.0196098 | 0.30791354 | 0.84365058 |
| AL118508.1 | 71.9165477 | 1.88940228 | 0.70446194 | 2.68205018 | 0.00731725 | 0.3610358 |
| NXT1 | 582.831659 | -0.2017582 | 0.27532798 | -0.7327921 | 0.46368524 | 0.89830907 |
| LINC01431 | 63.2536052 | 0.60418925 | 0.4775258 | 1.26524943 | 0.20578193 | 0.78609032 |
| GZF1 | 1090.68025 | -0.1991021 | 0.23948239 | -0.8313853 | 0.40575599 | 0.88132015 |
| NAPB | 1217.62716 | -0.1832629 | 0.24279812 | -0.7547952 | 0.45037186 | 0.89557948 |
| CST3 | 17453.3822 | -0.1679162 | 0.19722265 | -0.8514042 | 0.39454485 | 0.87884163 |
| SYNDIG1 | 384.49332 | -0.29018 | 0.4044034 | -0.717551 | 0.47303421 | 0.90175304 |
| AL035661.1 | 685.652262 | 0.17827985 | 0.33541729 | 0.53151657 | 0.59506086 | 0.9334601 |
| CST7 | 237.316744 | -0.0298554 | 0.7672466 | -0.0389124 | 0.96896027 | 0.99400331 |
| APMAP | 5528.03539 | 0.18746149 | 0.20088013 | 0.93320072 | 0.35071634 | 0.86364633 |
| ACSS1 | 7084.76158 | -0.0590643 | 0.18518291 | -0.3189512 | 0.74976355 | 0.96249332 |
| ENTPD6 | 6369.43176 | -0.0901918 | 0.21877716 | -0.4122543 | 0.68015307 | 0.95070301 |
| AL035252.3 | 126.301517 | 0.06982876 | 0.2435317 | 0.28673374 | 0.7743162 | 0.96836854 |
| PYGB | 8342.07283 | 0.35925525 | 0.28147948 | 1.27631062 | 0.20184574 | 0.78134034 |
| ABHD12 | 3802.90505 | 0.04331504 | 0.19033502 | 0.22757265 | 0.81997848 | 0.97454841 |
| GINS1 | 146.359185 | -0.2315731 | 0.35518817 | -0.6519731 | 0.51441852 | 0.91636259 |
| NINL | 2993.83164 | -0.1075153 | 0.19000289 | -0.5658611 | 0.57148818 | 0.92841021 |
| NANP | 492.953152 | -0.2951191 | 0.32255465 | -0.9149429 | 0.36022162 | 0.86480228 |
| ZNF337-AS1 | 1651.15501 | -0.1205531 | 0.25180738 | -0.4787514 | 0.63211552 | 0.94088547 |
| ZNF337 | 1671.0335 | -0.1426974 | 0.24983265 | -0.5711718 | 0.56788322 | 0.92618483 |
| FAM182B | 39.8913138 | 1.37960826 | 0.86581148 | 1.593428 | 0.11106421 | 0.68001961 |
| LINC01597 | 175.314543 | 0.742354 | 0.699538 | 1.06120612 | 0.28859624 | 0.83295411 |
| REM1 | 92.1329717 | 0.90617191 | 0.76289417 | 1.18780814 | 0.23490901 | 0.80747753 |
| HM13 | 8003.06931 | -0.0478743 | 0.21791573 | -0.2196916 | 0.82611133 | 0.97548201 |
| HM13-AS1 | 67.602002 | 0.12918889 | 0.3886762 | 0.3323818 | 0.73960098 | 0.96037421 |
| ID1 | 3769.99757 | 0.30353085 | 0.45104148 | 0.6729555 | 0.5009756 | 0.91232958 |
| COX4I2 | 91.9501438 | -0.411124 | 0.82804683 | -0.4964985 | 0.61954271 | 0.93945324 |

| | | | | | | |
|---|---|---|---|---|---|---|
| BCL2L1 | 10385.0213 | -0.3616406 | 0.26794406 | -1.3496868 | 0.17711647 | 0.76115956 |
| ABALON | 3155.09167 | -0.3073961 | 0.31024923 | -0.9908037 | 0.32178145 | 0.85119105 |
| TPX2 | 88.2109965 | 0.28781043 | 0.68391834 | 0.42082573 | 0.67388234 | 0.94935413 |
| FOXS1 | 123.077257 | -1.1068534 | 0.69419067 | -1.5944516 | 0.11083493 | 0.68001961 |
| DUSP15 | 1521.56368 | 0.35354674 | 0.50406968 | 0.70138466 | 0.48306299 | 0.90541935 |
| TTLL9 | 156.291912 | -0.1121847 | 0.49647388 | -0.225963 | 0.82123018 | 0.97472105 |
| PDRG1 | 1208.48398 | -0.2080354 | 0.2298409 | -0.9051278 | 0.36539771 | 0.8663515 |
| CCM2L | 449.260083 | 0.02131863 | 0.44138936 | 0.04829893 | 0.96147801 | 0.99320711 |
| AL031658.1 | 247.365706 | -0.0079774 | 0.45809893 | -0.0174141 | 0.98610628 | 0.99711976 |
| HCK | 529.028692 | -0.0121823 | 0.54000019 | -0.0225598 | 0.9820014 | 0.99595643 |
| TM9SF4 | 5074.47465 | -0.0831538 | 0.16128146 | -0.515582 | 0.60614637 | 0.93708311 |
| PLAGL2 | 878.943448 | 0.06137877 | 0.20196232 | 0.30391199 | 0.76119494 | 0.96603553 |
| POFUT1 | 2773.84577 | 0.32193495 | 0.12940082 | 2.48788957 | 0.01285036 | 0.41738067 |
| KIF3B | 4316.16038 | 0.26027696 | 0.13980968 | 1.86165185 | 0.06265218 | 0.61442412 |
| ASXL1 | 4560.0353 | -0.4550746 | 0.15011655 | -3.0314754 | 2.43E-03 | 2.12E-01 |
| NOL4L | 2360.56476 | 0.1140501 | 0.18380645 | 0.62049021 | 0.5349351 | 0.92084006 |
| AL034550.1 | 110.216019 | -0.0790575 | 0.29451459 | -0.2684333 | 0.78836583 | 0.97090793 |
| FO393400.1 | 1054.187 | 0.0209865 | 0.18772702 | 0.11179262 | 0.91098784 | 0.98662757 |
| COMMD7 | 2710.36039 | -0.1098657 | 0.17709634 | -0.6203726 | 0.53501254 | 0.92084006 |
| DNMT3B | 334.537703 | 0.06097699 | 0.29768528 | 0.20483711 | 0.83769939 | 0.97577999 |
| MAPRE1 | 4805.95445 | -0.2924726 | 0.21524261 | -1.3588046 | 0.17420853 | 0.75974924 |
| AL035071.1 | 820.36218 | -0.3193517 | 0.35243745 | -0.9061231 | 0.36487071 | 0.8663515 |
| CDK5RAP1 | 1672.41956 | -0.0854081 | 0.12332388 | -0.6925509 | 4.89E-01 | 0.90585781 |
| SNTA1 | 1821.01637 | 0.22018806 | 0.26786512 | 0.82201094 | 0.41107067 | 0.88268208 |
| CBFA2T2 | 1705.00571 | -0.0529049 | 0.15367363 | -0.3442676 | 0.73064503 | 0.95848028 |
| NECAB3 | 1446.72138 | 0.21454507 | 0.22210717 | 0.96595292 | 0.33406774 | 0.85533716 |
| ACTL10 | 69.831503 | -0.2018648 | 0.36675302 | -0.5504108 | 0.58203767 | 0.9317705 |
| E2F1 | 86.6727789 | 0.09423416 | 0.47490411 | 0.19842776 | 0.8427104 | 0.97644404 |
| PXMP4 | 563.260966 | 0.69259818 | 0.20655065 | 3.35316391 | 0.00079893 | 0.13250281 |
| ZNF341 | 394.517054 | -0.3245546 | 0.21996618 | -1.4754749 | 0.14008491 | 0.72502935 |
| CHMP4B | 7407.89918 | -0.2877908 | 0.2308468 | -1.2466743 | 0.21251695 | 0.79203904 |
| AL034549.1 | 131.132608 | 0.29279331 | 0.34522971 | 0.84811158 | 0.39637583 | 0.87967259 |
| RALY | 6629.73968 | -0.0518744 | 0.12050145 | -0.4304875 | 0.66684109 | 0.94830184 |
| EIF2S2 | 4470.57243 | -0.2027253 | 0.25431437 | -0.7971445 | 0.42536711 | 0.88605146 |
| AHCY | 10930.142 | 0.40503659 | 0.31992871 | 1.26602142 | 0.20550541 | 0.78571765 |
| ITCH | 2560.29932 | -0.1944557 | 0.173765 | -1.1190726 | 2.63E-01 | 0.82424998 |
| DYNLRB1 | 5474.57794 | 0.10643131 | 0.1250715 | 0.85096379 | 0.39478947 | 0.87884163 |
| MAP1LC3A | 1895.45005 | 0.06269086 | 0.30332724 | 0.20667732 | 0.83626187 | 0.97577999 |
| PIGU | 1112.7685 | 0.15686209 | 0.19064379 | 0.822802 | 4.11E-01 | 8.83E-01 |
| NCOA6 | 2395.86335 | -0.2812093 | 0.16249295 | -1.730594 | 0.08352421 | 0.64783289 |
| TP53INP2 | 4382.65572 | -0.1687223 | 0.2928621 | -0.5761151 | 0.5645374 | 0.92474393 |
| GGT7 | 2936.70404 | 0.12121037 | 0.27418706 | 0.44207182 | 0.65843723 | 0.9458503 |
| ACSS2 | 6482.57627 | 0.70879961 | 0.31351216 | 2.26083614 | 0.02376941 | 0.51967919 |
| GSS | 4481.75104 | 0.67679355 | 0.24863613 | 2.7220241 | 0.00648834 | 0.33778486 |
| MYH7B | 434.490431 | -0.2619249 | 0.2550697 | -1.0268759 | 0.30447889 | 0.84211658 |
| TRPC4AP | 5810.37551 | -0.2888455 | 0.12639039 | -2.2853438 | 0.02229267 | 0.51834617 |
| EDEM2 | 2048.31496 | -0.0239603 | 0.12763953 | -0.1877187 | 0.85109719 | 0.97846305 |
| PROCR | 995.036368 | -0.2999668 | 0.36786705 | -0.8154216 | 0.414831 | 0.88364943 |
| AL356652.1 | 323.314936 | -0.3817148 | 0.41328776 | -0.9236053 | 0.35569184 | 0.86364633 |
| MMP24OS | 5284.76526 | 0.02167818 | 0.21950202 | 0.09876071 | 0.92132827 | 0.98779251 |
| MMP24 | 3706.47877 | 0.0738467 | 0.36697384 | 0.2012315 | 0.84051756 | 0.97618283 |
| AL121753.1 | 4215.38088 | -0.1813888 | 0.36958729 | -0.4907873 | 0.62357688 | 0.94048401 |
| EIF6 | 7841.29714 | -0.2330039 | 0.36514632 | -0.6381111 | 0.52340137 | 0.91723878 |
| UQCC1 | 3076.84479 | 0.03161906 | 0.13043302 | 0.24241611 | 0.80845775 | 0.97334138 |

| | | | | | | |
|---|---|---|---|---|---|---|
| GDF5 | 55.067848 | -0.7798592 | 0.66360868 | -1.1751793 | 0.23992298 | 0.81209487 |
| CEP250 | 1407.67877 | -0.5677734 | 0.32720721 | -1.7352107 | 0.08270349 | 0.64680474 |
| ERGIC3 | 9413.00512 | 0.21067278 | 0.09624946 | 2.18882039 | 0.0286099 | 0.53606221 |
| SPAG4 | 578.131813 | -0.5445343 | 0.4843084 | -1.1243545 | 0.26086267 | 0.82424998 |
| CPNE1 | 5600.31583 | 0.04274949 | 0.17694361 | 0.24159949 | 0.80909052 | 0.97334138 |
| AL109827.1 | 1901.38238 | 0.18890676 | 0.13434326 | 1.40614988 | 0.15967962 | 0.74768867 |
| RBM12 | 2648.67053 | -0.4818688 | 0.23588071 | -2.0428494 | 0.04106735 | 0.55811692 |
| NFS1 | 2997.62401 | 0.37179087 | 0.23932919 | 1.55347068 | 0.12031073 | 0.69467399 |
| ROMO1 | 2576.81228 | -0.1424583 | 0.15550674 | -0.9160911 | 0.35961912 | 0.86480228 |
| RBM39 | 13806.2922 | -0.0778175 | 0.12426384 | -0.626228 | 0.53116543 | 0.91962754 |
| PHF20 | 1290.18546 | -0.1075307 | 0.11479948 | -0.9366828 | 0.34892173 | 0.86356499 |
| SCAND1 | 2710.65715 | -0.3920583 | 0.16735193 | -2.3427174 | 0.01914388 | 0.48680513 |
| CNBD2 | 205.784094 | -0.0243579 | 0.29914859 | -0.0814241 | 0.93510465 | 0.99068685 |
| NORAD | 12824.6961 | -0.0565793 | 0.20537554 | -0.275492 | 0.78293828 | 0.96964828 |
| EPB41L1 | 6656.7739 | 0.42302299 | 0.25743139 | 1.64324556 | 0.10033214 | 0.67191826 |
| AL121895.1 | 39.2366313 | 1.12268534 | 0.70518411 | 1.59204572 | 0.11137443 | 0.68001961 |
| AAR2 | 2002.25262 | -0.2031182 | 0.17134357 | -1.185444 | 0.23584194 | 0.80776058 |
| DLGAP4 | 4873.30429 | -0.4292048 | 0.21377887 | -2.0077046 | 0.04467469 | 0.57267043 |
| DLGAP4-AS1 | 1447.16743 | -0.3781775 | 0.18988035 | -1.9916621 | 0.04640815 | 0.57943708 |
| MYL9 | 11455.3291 | -0.4201911 | 0.54547809 | -0.7703171 | 0.44111181 | 0.89290667 |
| TGIF2 | 1410.20246 | -0.0547335 | 0.20614029 | -0.2655156 | 0.7906123 | 0.97090793 |
| TGIF2-RAB5I | 5214.94766 | -0.5260026 | 0.42024979 | -1.2516428 | 0.21070005 | 0.79134644 |
| RAB5IF | 5465.65333 | -0.5108377 | 0.42061148 | -1.214512 | 0.22455226 | 0.80078694 |
| SLA2 | 1500.28133 | -0.616474 | 0.38545283 | -1.59935 | 0.10974287 | 0.67859564 |
| NDRG3 | 1869.81412 | -0.0338386 | 0.2472494 | -0.13686 | 0.89114143 | 0.98474725 |
| DSN1 | 452.453552 | 0.01435206 | 0.14201632 | 0.10105922 | 0.91950345 | 0.9872458 |
| SOGA1 | 2104.86558 | 0.00195437 | 0.25677342 | 0.00761127 | 0.99392715 | 0.99931765 |
| TLDC2 | 554.39672 | -0.8921521 | 0.38846357 | -2.2966171 | 0.02164062 | 0.51394743 |
| SAMHD1 | 2906.29215 | -0.9982704 | 0.53926222 | -1.8511781 | 0.06414393 | 0.61837523 |
| RBL1 | 198.090442 | -0.1529125 | 0.19940538 | -0.7668422 | 4.43E-01 | 8.94E-01 |
| MROH8 | 852.583548 | 0.04648618 | 0.24941949 | 0.18637748 | 0.85214875 | 0.97872457 |
| RPN2 | 13548.0785 | 0.10600596 | 0.16912316 | 0.6267974 | 0.53079205 | 0.91962754 |
| MANBAL | 2825.98701 | 0.08115131 | 0.18919716 | 0.42892455 | 0.66797814 | 0.94847921 |
| SRC | 4742.43546 | -0.0621 | 0.14956543 | -0.4152029 | 0.67799336 | 0.95018034 |
| BLCAP | 5411.35537 | -0.0322577 | 0.12424411 | -0.2596319 | 0.79514773 | 0.97101097 |
| NNAT | 44.88257 | -0.4971321 | 0.65147003 | -0.7630928 | 0.44540804 | 0.89453029 |
| CTNNBL1 | 1641.56089 | -0.2481732 | 0.15504337 | -1.6006697 | 1.09E-01 | 0.67859123 |
| VSTM2L | 127.419468 | 0.39307859 | 0.67246983 | 0.58452971 | 0.55886398 | 0.92421175 |
| TTI1 | 1141.63867 | -0.1010537 | 0.19728972 | -0.5122096 | 0.60850436 | 0.93748023 |
| RPRD1B | 1922.73069 | -0.2505534 | 0.17717964 | -1.4141206 | 0.15732649 | 0.74522306 |
| TGM2 | 28782.9917 | -0.2538122 | 0.42398331 | -0.5986372 | 0.54941485 | 0.92373235 |
| KIAA1755 | 537.841803 | -0.1365126 | 0.7658721 | -0.1782447 | 0.85853084 | 0.97982623 |
| BPI | 151.378905 | -0.7820281 | 0.64816679 | -1.206523 | 0.2276159 | 0.80300331 |
| LBP | 2079.73988 | -1.2087722 | 0.72859554 | -1.6590442 | 0.09710689 | 0.66486533 |
| AL391095.2 | 81.8362927 | -1.1438517 | 0.87952576 | -1.3005324 | 0.19341857 | 0.77548688 |
| RALGAPB | 2844.93255 | -0.064479 | 0.09708249 | -0.6641669 | 0.50658351 | 0.91416224 |
| ARHGAP40 | 591.47588 | 0.96901069 | 0.56719642 | 1.70842174 | 0.08755812 | 0.65271545 |
| ACTR5 | 583.530474 | -0.3284854 | 0.20599756 | -1.5946081 | 0.1107999 | 0.68001961 |
| PPP1R16B | 2628.22487 | 0.70477101 | 0.71263543 | 0.98896432 | 0.3226806 | 0.85119105 |
| FAM83D | 199.494319 | 0.24749982 | 0.6583668 | 0.37592998 | 0.70696895 | 0.95501548 |
| DHX35 | 829.706905 | -0.11509 | 0.17404664 | -0.6612597 | 0.50844576 | 0.91439937 |
| MAFB | 1304.65497 | -0.2621647 | 0.22966567 | -1.1415058 | 0.25365949 | 0.8188642 |
| TOP1 | 3811.45578 | -0.1269579 | 0.17849157 | -0.7112821 | 0.47690945 | 0.90335153 |
| PLCG1-AS1 | 313.795847 | 0.10416281 | 0.32826692 | 0.31731131 | 0.7510074 | 0.96272571 |

| | | | | | | |
|---|---|---|---|---|---|---|
| PLCG1 | 5116.19548 | -0.1323508 | 0.22332368 | -0.5926411 | 0.55342138 | 0.92373235 |
| ZHX3 | 4797.78555 | -0.0932845 | 0.30674318 | -0.3041128 | 0.76104195 | 0.96591407 |
| LPIN3 | 7755.8472 | -0.2885984 | 0.51450291 | -0.5609267 | 0.57484749 | 0.93024927 |
| EMILIN3 | 2327.38193 | -0.2701377 | 0.50570356 | -0.5341819 | 0.59321572 | 0.93289552 |
| CHD6 | 1657.85442 | -0.2197141 | 0.22518247 | -0.9757156 | 0.32920539 | 0.85333863 |
| SRSF6 | 10090.9047 | -0.0093007 | 0.18971306 | -0.0490253 | 0.96089916 | 0.99311814 |
| L3MBTL1 | 1120.08454 | -0.0197051 | 0.18913382 | -0.104186 | 0.91702171 | 0.9868044 |
| Z98752.3 | 785.695516 | 0.01892075 | 0.30650902 | 0.06172982 | 0.95077799 | 0.99170503 |
| Z98752.1 | 136.44579 | -0.0256649 | 0.28974561 | -0.0885774 | 0.92941778 | 0.98963127 |
| SGK2 | 4167.72784 | 0.04582554 | 0.34485344 | 0.13288409 | 0.89428505 | 0.98474725 |
| IFT52 | 1174.96668 | 0.27202255 | 0.17008658 | 1.59931815 | 0.10974993 | 0.67859564 |
| MYBL2 | 64.9335663 | -1.1572072 | 0.80929986 | -1.4298868 | 0.15274951 | 0.7423959 |
| TOX2 | 184.368196 | -0.4391356 | 0.44293362 | -0.9914253 | 0.32147797 | 0.85119105 |
| JPH2 | 275.586274 | -0.913663 | 0.57542938 | -1.5877934 | 0.11233306 | 0.68001961 |
| OSER1 | 2452.7994 | -0.4328341 | 0.32705792 | -1.3234173 | 0.18569664 | 0.76908034 |
| OSER1-DT | 366.90301 | 0.56693978 | 0.32744146 | 1.73142332 | 0.08337629 | 0.647557 |
| FITM2 | 1198.40979 | 0.04657549 | 0.34933647 | 0.13332559 | 0.89393589 | 0.98474725 |
| R3HDML | 89.9427247 | 3.56490275 | 1.47832371 | 2.41144935 | 0.01588926 | 0.4564177 |
| AL117382.1 | 94.0093391 | 3.99608481 | 1.50750723 | 2.65078981 | 0.00803038 | 0.37112203 |
| HNF4A | 5887.38551 | 1.25159909 | 0.80634849 | 1.55218136 | 0.12061884 | 0.69467399 |
| HNF4A-AS1 | 108.612353 | 3.93036756 | 1.24857011 | 3.14789497 | 0.00164451 | 0.18012143 |
| TTPAL | 731.732392 | -0.3424879 | 0.16198384 | -2.1143336 | 0.03448679 | 0.55369025 |
| SERINC3 | 6010.79048 | 0.02989488 | 0.1161069 | 0.25747721 | 0.7968104 | 0.97131676 |
| PKIG | 3417.88102 | -0.2470342 | 0.19928487 | -1.2396035 | 0.2151221 | 0.7949124 |
| ADA | 325.722719 | -0.1010476 | 0.471166 | -0.2144629 | 0.83018613 | 0.97550416 |
| WISP2 | 74.7934065 | -0.2229875 | 1.30720108 | -0.1705839 | NA | NA |
| KCNK15 | 145.088429 | 0.02329738 | 0.59799447 | 0.03895918 | 0.96892293 | 0.99400331 |
| YWHAB | 11757.6025 | 0.02302896 | 0.1899162 | 0.12125851 | 0.90348628 | 0.98617053 |
| PABPC1L | 3339.45997 | -0.036947 | 0.36770594 | -0.1004798 | 0.91996341 | 0.98733173 |
| TOMM34 | 2399.21691 | 0.13852163 | 0.19163207 | 0.72285205 | 0.46977079 | 0.9008571 |
| STK4 | 1133.1182 | -0.2752022 | 0.17397759 | -1.5818255 | 1.14E-01 | 0.68214043 |
| SEMG2 | 71.4823332 | 1.03961542 | 0.75931811 | 1.36914345 | 0.17095444 | 0.75956362 |
| SLPI | 36277.2915 | -0.4461246 | 0.81806764 | -0.5453395 | 0.58552001 | 0.93289552 |
| SDC4 | 22450.9893 | -0.1134154 | 0.15183261 | -0.7469764 | 0.45507781 | 0.89670027 |
| SYS1 | 2869.14568 | -0.3342845 | 0.1990949 | -1.679021 | 0.09314796 | 0.65756964 |
| SYS1-DBNDD | 2126.8775 | -0.2545159 | 0.23318788 | -1.0914629 | 0.27506925 | 0.82719407 |
| TP53TG5 | 131.666743 | -0.1846232 | 0.2796015 | -0.6603082 | 0.50905609 | 0.91439937 |
| DBNDD2 | 1838.55563 | -0.2499085 | 0.28760327 | -0.8689349 | 0.38488272 | 0.87314939 |
| PIGT | 5488.7506 | 0.19090441 | 0.1891883 | 1.00907089 | 0.31294064 | 0.84559008 |
| AL031663.3 | 49.9655811 | -0.5628868 | 0.53035921 | -1.0613312 | 0.2885394 | 0.83295411 |
| WFDC2 | 29036.7666 | 0.21340719 | 0.47096038 | 0.45313195 | 0.65045373 | 0.94336429 |
| DNTTIP1 | 1103.05363 | 0.02694555 | 0.11538628 | 0.23352473 | 0.81535397 | 0.9738447 |
| UBE2C | 62.564237 | 0.10127279 | 0.80821042 | 0.12530498 | 0.90028211 | 0.98526603 |
| TNNC2 | 35.3805473 | 1.25380054 | 0.83427884 | 1.5028555 | NA | NA |
| SNX21 | 2153.53682 | 0.21697106 | 0.23707256 | 0.91520947 | 0.36008167 | 0.86480228 |
| ACOT8 | 1929.76781 | 0.19497429 | 0.20623966 | 0.94537727 | 0.3444663 | 0.85999891 |
| ZSWIM3 | 189.759704 | 0.35962281 | 0.24263024 | 1.48218461 | 1.38E-01 | 7.22E-01 |
| ZSWIM1 | 820.255581 | -0.2868159 | 0.17318127 | -1.6561598 | 9.77E-02 | 6.65E-01 |
| SPATA25 | 234.344872 | -0.3803261 | 0.34397025 | -1.1056946 | 2.69E-01 | 0.82683094 |
| NEURL2 | 482.979807 | 0.18161067 | 0.36818541 | 0.49325873 | 0.62182978 | 0.94016713 |
| CTSA | 27721.1646 | 0.0960171 | 0.28133241 | 0.34129411 | 0.73288218 | 0.95848028 |
| AL008726.1 | 6098.32341 | 0.26655687 | 0.34115666 | 0.78133275 | 0.43460681 | 0.89055338 |
| PLTP | 17341.2913 | -0.195194 | 0.4745412 | -0.411332 | 0.68082912 | 0.95070301 |
| PCIF1 | 2374.30616 | -0.1092222 | 0.19982938 | -0.5465774 | 0.58466908 | 0.93282331 |

| | | | | | | |
|---|---|---|---|---|---|---|
| ZNF335 | 2012.11917 | -0.4593306 | 0.25687361 | -1.7881579 | 0.07375054 | 0.6374448 |
| MMP9 | 402.695072 | -0.0569736 | 0.94718966 | -0.0601502 | 0.95203601 | 0.99170503 |
| SLC12A5-AS1 | 149.614625 | -0.0085656 | 0.88457363 | -0.0096833 | 0.99227393 | 0.99906045 |
| NCOA5 | 2108.97492 | -0.4763689 | 0.29978794 | -1.5890196 | 0.11205597 | 0.68001961 |
| CD40 | 1362.69701 | 0.1371403 | 0.43938969 | 0.31211542 | 0.75495281 | 0.96360109 |
| SLC35C2 | 3422.27698 | -0.0519432 | 0.15317109 | -0.3391189 | 0.73452016 | 0.95895327 |
| AL133227.1 | 327.64084 | -0.0921747 | 0.17994953 | -0.5122253 | 0.60849333 | 0.93748023 |
| ELMO2 | 2517.92598 | -0.1186398 | 0.1392226 | -0.8521589 | 0.39412589 | 0.87884163 |
| ZNF334 | 541.945775 | -0.1244299 | 0.48225119 | -0.2580189 | 0.79639229 | 0.97105972 |
| SLC13A3 | 44467.1429 | 0.42984967 | 0.48849583 | 0.87994543 | 0.37888887 | 0.87069278 |
| TP53RK | 795.575154 | -0.1984092 | 0.28771543 | -0.689602 | 0.49044448 | 0.90600166 |
| SLC2A10 | 400.999488 | 0.22078626 | 0.31989782 | 0.69017746 | 0.4900826 | 0.90600166 |
| EYA2 | 469.018977 | -0.588115 | 0.52915979 | -1.1114128 | 0.2663907 | 0.8265123 |
| ZMYND8 | 2318.78445 | 0.01048212 | 0.26173688 | 0.04004832 | 0.9680546 | 0.99400331 |
| NCOA3 | 1936.00907 | -0.3640091 | 0.12927336 | -2.8158093 | 0.00486545 | 0.30193205 |
| SULF2 | 6780.23616 | 0.1393642 | 0.30150797 | 0.46222393 | 0.64392074 | 0.94208859 |
| AL354813.1 | 646.347459 | 0.18215075 | 0.35965751 | 0.50645613 | 0.61253648 | 0.9387918 |
| LINC00494 | 111.392167 | 0.45473183 | 0.74388635 | 0.61129207 | 0.54100624 | 0.92292765 |
| PREX1 | 1719.52745 | -0.1953276 | 0.3121282 | -0.625793 | 0.53145072 | 0.91962754 |
| ARFGEF2 | 2874.00611 | -0.1187195 | 0.18136387 | -0.6545929 | 0.51272991 | 0.91568064 |
| CSE1L | 2521.13271 | 0.05312131 | 0.13701723 | 0.38769803 | 0.69823951 | 0.9529028 |
| STAU1 | 7427.66223 | -0.1351053 | 0.2513295 | -0.5375624 | 0.59087917 | 0.93289552 |
| DDX27 | 2620.62374 | -0.1470027 | 0.11952387 | -1.2299025 | 2.19E-01 | 7.98E-01 |
| ZNFX1 | 3917.37242 | -0.9377029 | 0.40253889 | -2.3294716 | 0.01983409 | 0.49477123 |
| ZFAS1 | 2236.29509 | -0.1678857 | 0.27343312 | -0.6139919 | 0.53922071 | 0.92178558 |
| KCNB1 | 75.3086457 | 0.10805681 | 0.39560122 | 0.27314578 | 0.78474115 | 0.97021865 |
| PTGIS | 821.73403 | -0.3230026 | 0.53963605 | -0.5985564 | 0.54946875 | 0.92373235 |
| B4GALT5 | 6292.54628 | -0.1496547 | 0.33004631 | -0.4534356 | 0.65023513 | 0.94322083 |
| SLC9A8 | 1898.46871 | -0.2721053 | 0.20473801 | -1.3290414 | 0.18383431 | 0.76781833 |
| SPATA2 | 999.881179 | -0.201232 | 0.18313936 | -1.0987916 | 0.27185896 | 0.82716979 |
| RNF114 | 5747.4984 | -0.0022269 | 0.23538425 | -0.0094609 | 0.99245142 | 0.99906045 |
| SNAI1 | 446.12739 | -0.1578109 | 0.52744664 | -0.2991979 | 0.76478906 | 0.96678847 |
| UBE2V1 | 6528.08449 | -0.1321913 | 0.14095196 | -0.9378467 | 0.34832319 | 0.86288527 |
| TMEM189-U | 6871.10591 | -0.128455 | 0.1434149 | -0.8956879 | 0.37041945 | 0.86762749 |
| TMEM189 | 1578.62831 | -0.1390009 | 0.17587816 | -0.7903249 | 0.42933807 | 0.88851358 |
| LINC01273 | 57.7443657 | 0.91251659 | 0.53880262 | 1.69360089 | 0.09034115 | 0.65445825 |
| CEBPB-AS1 | 242.293609 | -0.1912951 | 0.33011856 | -0.579474 | 0.56226938 | 0.92452095 |
| CEBPB | 3028.12619 | -0.3321656 | 0.37574255 | -0.8840243 | 0.37668308 | 0.86970773 |
| SMIM25 | 66.502901 | -0.1252751 | 0.76715822 | -0.1632976 | 8.70E-01 | 9.83E-01 |
| PTPN1 | 9223.92505 | -0.5074826 | 0.34558843 | -1.4684595 | 0.14197945 | 0.72784886 |
| AL133230.1 | 464.301226 | -0.5376087 | 0.51123646 | -1.0515851 | 2.93E-01 | 0.83524862 |
| RIPOR3 | 3669.67846 | 0.25143991 | 0.28282426 | 0.88903232 | 0.37398571 | 0.86791605 |
| PARD6B | 2842.48639 | -0.0315386 | 0.33264477 | -0.0948117 | 0.92446443 | 0.98869145 |
| BCAS4 | 556.5178 | -0.1160404 | 0.26863274 | -0.4319668 | 0.66576557 | 0.9474809 |
| ADNP | 3548.45889 | -0.2143921 | 0.12945587 | -1.6561015 | 0.09770128 | 0.66549286 |
| ADNP-AS1 | 91.522399 | 0.37571464 | 0.40523952 | 0.92714216 | 0.35385273 | 0.86364633 |
| DPM1 | 1707.8936 | -0.0672751 | 0.14015975 | -0.4799888 | 0.63123535 | 0.94088547 |
| MOCS3 | 396.298761 | 0.08254093 | 0.24401776 | 0.33825786 | 0.73516887 | 0.95895327 |
| NFATC2 | 350.090335 | 0.08556311 | 0.41117081 | 0.20809627 | 0.8351538 | 0.97550416 |
| ATP9A | 2231.3864 | 0.22636152 | 0.22871707 | 0.98970102 | 3.22E-01 | 8.51E-01 |
| ZFP64 | 692.35546 | 0.12045601 | 0.17940066 | 0.67143572 | 0.501943 | 0.91259827 |
| TSHZ2 | 220.297427 | 0.1046027 | 0.3924487 | 0.26653854 | 0.78982448 | 0.97090793 |
| AL157838.1 | 121.147615 | -0.1315711 | 0.36009829 | -0.3653756 | 0.71483107 | 0.95587935 |
| ZNF217 | 1908.75234 | -0.0241028 | 0.18869741 | -0.1277325 | 0.89836068 | 0.98474725 |

| | | | | | | |
|---|---|---|---|---|---|---|
| BCAS1 | 185.425257 | 1.57416821 | 0.77956237 | 2.01929732 | 0.04345633 | 0.56927916 |
| CYP24A1 | 7850.75809 | 0.49127967 | 0.68526646 | 0.71691772 | 0.47342487 | 0.90194729 |
| PFDN4 | 624.156589 | -0.1390825 | 0.18539206 | -0.7502073 | 0.45312984 | 0.89670027 |
| DOK5 | 299.978194 | 0.16885335 | 0.49985985 | 0.33780139 | 0.73551286 | 0.95895327 |
| CBLN4 | 34.3362647 | 0.1666996 | 0.98858454 | 0.16862452 | 0.86609199 | 0.98197545 |
| FAM210B | 2313.55271 | 0.26155481 | 0.1360543 | 1.92242965 | 0.05455172 | 0.59329767 |
| AURKA | 204.38678 | 0.1753479 | 0.23491481 | 0.74643187 | 4.55E-01 | 0.89688594 |
| CSTF1 | 1270.67678 | -0.1696504 | 0.13746782 | -1.23411 | 0.2171619 | 0.79574248 |
| CASS4 | 78.0256344 | -0.0237694 | 0.53928228 | -0.0440759 | 0.96484387 | 0.99357855 |
| RTF2 | 4522.05748 | 0.03048483 | 0.13027652 | 0.23400098 | 0.81498422 | 0.9738447 |
| FAM209A | 1533.6716 | 0.03992548 | 0.13513129 | 0.29545693 | 0.76764485 | 0.96754958 |
| TFAP2C | 96.0537308 | 0.83274994 | 0.81982015 | 1.01577149 | 0.30973821 | 0.84445794 |
| BMP7 | 430.529566 | 0.31617281 | 0.42332348 | 0.74688229 | 0.45513462 | 0.89670027 |
| RAE1 | 1508.67829 | -0.099204 | 0.20574348 | -0.4821732 | 0.62968294 | 0.94088547 |
| AL109955.1 | 230.320854 | 0.1476766 | 0.29577054 | 0.49929451 | 0.61757193 | 0.93924106 |
| RBM38 | 1640.80259 | 0.25044051 | 0.22996784 | 1.0890241 | 0.27614326 | 0.82719407 |
| PCK1 | 74631.8676 | 1.40100396 | 1.03816281 | 1.34950313 | 0.17717542 | 0.76115956 |
| ZBP1 | 239.07696 | -2.4272071 | 1.14922583 | -2.1120367 | 0.03468331 | 0.55369025 |
| PMEPA1 | 1243.80387 | -0.4385982 | 0.40716739 | -1.0771939 | 0.28139363 | 0.82957339 |
| NKILA | 75.3434297 | -0.133124 | 0.41763963 | -0.3187532 | 0.74991364 | 0.96249332 |
| RAB22A | 2513.4446 | -0.1841679 | 0.28815993 | -0.6391168 | 0.52274696 | 0.91723878 |
| VAPB | 4265.94743 | 0.03924622 | 0.2136286 | 0.18371239 | 0.85423909 | 0.97906925 |
| APCDD1L | 249.270189 | 0.2778412 | 0.39881126 | 0.69667341 | 4.86E-01 | 9.05E-01 |
| STX16 | 5538.70172 | -0.0847411 | 0.33587242 | -0.2523016 | 0.80080796 | 0.97210205 |
| STX16-NPEPI | 5512.14896 | 0.2312166 | 0.22532017 | 1.02616915 | 0.30481186 | 0.84230231 |
| NPEPL1 | 5666.10787 | 0.25575424 | 0.2330465 | 1.09743866 | 0.27244968 | 0.82716979 |
| GNAS-AS1 | 79.8904681 | 1.18071148 | 0.72875655 | 1.62017272 | 0.10519518 | 0.67473222 |
| GNAS | 37216.6147 | -0.0595669 | 0.17867369 | -0.3333836 | 0.73884475 | 0.9602325 |
| NELFCD | 4257.36747 | -0.0595482 | 0.12698326 | -0.4689452 | 0.6391088 | 0.94088547 |
| CTSZ | 10227.5653 | -0.0131167 | 0.20812314 | -0.0630236 | 0.94974774 | 0.99170503 |
| TUBB1 | 95.3249809 | -0.2215751 | 0.42525428 | -0.5210414 | 0.6023379 | 0.9355118 |
| ATP5F1E | 10309.9155 | 0.06825586 | 0.19840143 | 0.34402909 | 0.73082441 | 0.95848028 |
| PRELID3B | 2627.14044 | -0.1671097 | 0.20565332 | -0.8125795 | 0.41645921 | 0.88419652 |
| ZNF831 | 51.1007101 | -0.1535037 | 0.86788503 | -0.176871 | 0.85960971 | 0.98034112 |
| PHACTR3 | 145.689521 | 1.59628296 | 0.43749407 | 3.6486962 | 0.00026357 | 0.08232752 |
| SYCP2 | 208.650702 | 0.04592474 | 0.28742995 | 0.15977716 | 8.73E-01 | 9.83E-01 |
| FAM217B | 466.291214 | -0.0610865 | 0.28455707 | -0.2146722 | 8.30E-01 | 9.76E-01 |
| PPP1R3D | 477.065719 | -0.0076063 | 0.19712071 | -0.0385871 | 0.96921962 | 0.99400331 |
| CDH26 | 44.3755713 | 0.40007454 | 0.4190454 | 0.95472838 | 3.40E-01 | 8.58E-01 |
| CDH4 | 118.089816 | 0.57796027 | 0.33998854 | 1.69994045 | 8.91E-02 | 0.65375543 |
| TAF4 | 1139.666 | -0.1905527 | 0.15328172 | -1.2431533 | 0.21381134 | 0.7926732 |
| AL137077.2 | 90.6544538 | 0.11368831 | 0.24019009 | 0.47332639 | 6.36E-01 | 0.94088547 |
| LSM14B | 2236.52745 | -0.0608879 | 0.13617323 | -0.4471354 | 0.65477729 | 0.94532497 |
| PSMA7 | 6778.57123 | -0.1506148 | 0.21393504 | -0.7040213 | 0.48141953 | 0.90541935 |
| SS18L1 | 1775.53473 | -0.0359311 | 0.19388966 | -0.1853172 | 0.85298025 | 0.97886533 |
| MTG2 | 1970.81616 | -0.2835346 | 0.12039397 | -2.3550567 | 0.01851988 | 0.47570049 |
| OSBPL2 | 3373.59788 | -0.1703693 | 0.11910376 | -1.4304277 | 0.15259431 | 0.7422778 |
| ADRM1 | 5646.91373 | -0.2911955 | 0.24995602 | -1.1649868 | 2.44E-01 | 8.15E-01 |
| AL354836.1 | 1681.92594 | -0.1582877 | 0.23680178 | -0.6684395 | 0.50385311 | 0.91356552 |
| LAMA5 | 28658.6586 | -0.180753 | 0.22686089 | -0.7967569 | 0.42559221 | 0.88622485 |
| LAMA5-AS1 | 242.402605 | -0.3705766 | 0.40411247 | -0.9170135 | 0.35913558 | 0.86480228 |
| AL121832.2 | 91.6702828 | -0.2621783 | 0.25497663 | -1.0282443 | 3.04E-01 | 0.84211658 |
| RPS21 | 8366.72216 | 0.11331855 | 0.21126712 | 0.5363757 | 5.92E-01 | 0.93289552 |
| CABLES2 | 394.824936 | -0.1666212 | 0.25501101 | -0.6533882 | 0.51350602 | 0.91618586 |

| | | | | | | |
|---|---|---|---|---|---|---|
| RBBP8NL | 53.0158127 | 1.76425888 | 0.8380297 | 2.10524624 | 0.03526988 | 0.55369025 |
| GATA5 | 179.593584 | 0.27798985 | 0.48101334 | 0.57792545 | 0.56331445 | 0.92452095 |
| SLCO4A1 | 9244.8727 | -0.2391602 | 0.3649137 | -0.6553884 | 0.51221769 | 0.91551505 |
| AL357033.4 | 112.624135 | -0.699021 | 0.43591394 | -1.6035757 | 0.1088076 | 0.67800851 |
| SLCO4A1-AS | 1695.1216 | -0.2493639 | 0.37327956 | -0.6680351 | 0.50411118 | 0.91356552 |
| NTSR1 | 158.312268 | -0.2800651 | 0.41484973 | -0.6751001 | 0.49961217 | 0.91161831 |
| LINC00659 | 68.578316 | -0.4588798 | 0.30744917 | -1.4925387 | 0.13555797 | 0.71684309 |
| MRGBP | 1355.00674 | -0.4781066 | 0.16283206 | -2.9361945 | 0.00332266 | 0.25241775 |
| OGFR-AS1 | 112.518802 | -0.5910809 | 0.38808045 | -1.5230887 | 0.12773653 | 0.70579329 |
| OGFR | 5910.54475 | -0.6020292 | 0.30683243 | -1.9620783 | 0.04975337 | 0.5861488 |
| COL9A3 | 277.245907 | -0.2258264 | 0.42103488 | -0.5363603 | 0.59170953 | 0.93289552 |
| TCFL5 | 575.883249 | 0.0522492 | 0.1650304 | 0.31660348 | 0.7515445 | 0.96285859 |
| DIDO1 | 3610.31871 | -0.2276606 | 0.12760997 | -1.7840349 | 0.074418 | 0.63773415 |
| AL117379.1 | 105.443978 | -0.0173071 | 0.58161246 | -0.029757 | 0.97626083 | 0.99455099 |
| GID8 | 2643.08398 | -0.1389741 | 0.12282794 | -1.1314535 | 0.25786428 | 0.82187662 |
| SLC17A9 | 1060.93418 | -0.6389852 | 0.49265107 | -1.2970339 | 0.19461951 | 0.7762856 |
| YTHDF1 | 2968.77969 | -0.3073653 | 0.20555852 | -1.4952689 | 1.35E-01 | 7.16E-01 |
| BIRC7 | 112.825567 | -0.4934589 | 0.41660946 | -1.1844641 | 0.2362294 | 0.80802614 |
| NKAIN4 | 3534.03812 | -0.005475 | 0.7730275 | -0.0070825 | 0.994349 | 0.99940602 |
| AL121827.2 | 451.570041 | -0.1742442 | 0.86060718 | -0.2024665 | 0.83955201 | 0.97618283 |
| ARFGAP1 | 8151.07925 | -0.251943 | 0.18740508 | -1.3443768 | 0.17882659 | 0.76120583 |
| CHRNA4 | 571.807121 | 1.33752471 | 0.90916204 | 1.47116207 | 0.14124729 | 0.72762933 |
| AL121827.1 | 120.098116 | 0.73275227 | 0.94210428 | 0.77778255 | 0.43669722 | 0.89122667 |
| KCNQ2 | 75.9966204 | 0.96962959 | 1.10759815 | 0.87543446 | 0.38133756 | 0.87184592 |
| EEF1A2 | 1098.47108 | 0.00464694 | 0.74151704 | 0.0062668 | 0.99499985 | 0.99940889 |
| PPDPF | 9340.79558 | -0.1166844 | 0.28398736 | -0.4108787 | 0.68116147 | 0.95070301 |
| PTK6 | 94.4399183 | 0.87837346 | 0.51501927 | 1.70551573 | 0.08809828 | 0.65363785 |
| FNDC11 | 56.4661833 | 0.74525488 | 0.5985958 | 1.2450052 | 2.13E-01 | 0.7926732 |
| HELZ2 | 3394.92681 | -2.2761854 | 0.79387091 | -2.8671984 | NA | NA |
| GMEB2 | 1355.3815 | -0.2873214 | 0.12480953 | -2.3020789 | 0.02133072 | 0.51189223 |
| MHENCR | 635.474741 | 0.19035364 | 0.27186415 | 0.70017926 | 0.48381536 | 0.90541935 |
| STMN3 | 2139.39654 | 0.19822852 | 0.29896106 | 0.66305799 | 0.50729341 | 0.91430909 |
| RTEL1 | 2808.81362 | -0.2974867 | 0.24397337 | -1.2193407 | 0.2227149 | 0.80078694 |
| RTEL1-TNFR! | 4511.80001 | -0.1401161 | 0.27268315 | -0.5138423 | 0.60736225 | 0.93739614 |
| TNFRSF6B | 1453.80572 | 0.13770543 | 0.36277296 | 0.37959122 | 0.70424888 | 0.95488362 |
| ARFRP1 | 3317.94685 | -0.0240979 | 0.19983183 | -0.1205908 | 0.90401517 | 0.98624484 |
| ZGPAT | 4497.98657 | -0.2375585 | 0.25424999 | -0.9343503 | 0.35012325 | 0.86364633 |
| AL121845.3 | 6913.68553 | 0.0501759 | 0.33994866 | 0.14759848 | 0.88265966 | 0.98276701 |
| LIME1 | 3566.70861 | 0.42177067 | 0.40255755 | 1.04772763 | 0.29476412 | 0.83630273 |
| AL121845.2 | 2547.44538 | 0.72136081 | 0.52837752 | 1.3652375 | 0.17217842 | 0.75974924 |
| SLC2A4RG | 6777.44916 | 0.51529913 | 0.35327217 | 1.45864625 | 0.1446625 | 0.73032824 |
| ZBTB46 | 1667.22542 | 0.22347268 | 0.2057396 | 1.0861919 | 0.2773941 | 0.82782817 |
| AL121845.1 | 400.700397 | 0.27448412 | 0.27528907 | 0.99707596 | 0.31872764 | 0.84914926 |
| AL118506.1 | 627.977464 | -0.3339115 | 0.26351872 | -1.2671261 | 0.2051102 | 0.78528504 |
| ABHD16B | 293.230204 | -0.3163033 | 0.26494966 | -1.1938242 | 0.23254674 | 0.80653282 |
| TPD52L2 | 6672.52785 | -0.3031353 | 0.15043169 | -2.0151027 | 0.04389389 | 0.57116687 |
| DNAJC5 | 4943.10999 | -0.182498 | 0.16216056 | -1.1254156 | 0.26041298 | 0.82424998 |
| UCKL1 | 4075.29956 | -0.2051774 | 0.16060784 | -1.2775052 | 0.20142394 | 0.78134034 |
| UCKL1-AS1 | 554.345164 | -0.0139125 | 0.16844699 | -0.0825927 | 0.93417544 | 0.99029255 |
| ZNF512B | 4026.40168 | 0.11041196 | 0.2076076 | 0.53183007 | 0.5948437 | 0.9334601 |
| SAMD10 | 865.601985 | 0.18388414 | 0.27697114 | 0.66391081 | 0.5067474 | 0.91420888 |
| PRPF6 | 4615.50242 | 0.20698902 | 0.16181586 | 1.27916398 | 0.20083932 | 0.78134034 |
| C20orf204 | 112.90165 | -0.4709243 | 0.27215788 | -1.7303348 | 0.08357047 | 0.64783289 |
| SOX18 | 751.330171 | -0.0120839 | 0.55630554 | -0.0217217 | 0.98266994 | 0.99611734 |

| | | | | | | |
|---|---|---|---|---|---|---|
| TCEA2 | 2292.17071 | -0.0328549 | 0.21814138 | -0.1506129 | 0.8802811 | 0.98275231 |
| AL355803.1 | 38.6948459 | -0.8205644 | 0.55576525 | -1.4764586 | 0.13982082 | 0.72475166 |
| RGS19 | 479.834465 | -0.3403452 | 0.35318784 | -0.9636378 | 0.33522754 | 0.85599307 |
| OPRL1 | 69.8219336 | -0.0122133 | 0.55300562 | -0.0220854 | 0.98237987 | 0.9960388 |
| PCMTD2 | 3760.54522 | 0.05558617 | 0.19553944 | 0.28427088 | 0.7762028 | 0.96836854 |
| PLPP2 | 1183.78203 | -0.121122 | 0.52333169 | -0.2314441 | 0.81696979 | 0.97397142 |
| MIER2 | 1212.6527 | -0.3666461 | 0.2424437 | -1.5122939 | 0.13045912 | 0.7084713 |
| AC016588.1 | 283.950125 | -0.2569422 | 0.28143372 | -0.912976 | 0.36125516 | 0.86528137 |
| C2CD4C | 222.653017 | 1.01390899 | 0.746319 | 1.35854639 | 0.17429037 | 0.75974924 |
| SHC2 | 2179.18501 | -0.4871777 | 0.26563084 | -1.8340406 | 0.06664797 | 0.62390091 |
| MADCAM1 | 71.8782377 | 0.01661041 | 0.38177659 | 0.0435082 | 0.96529643 | 0.99371713 |
| AC005775.1 | 380.125663 | -0.1377655 | 0.38187823 | -0.3607578 | 0.71828053 | 0.95648644 |
| TPGS1 | 604.431961 | -0.1307137 | 0.35204597 | -0.3712971 | 0.71041623 | 0.95536097 |
| CDC34 | 2423.24393 | -0.3427194 | 0.14146734 | -2.4226041 | 0.01540971 | 0.45243543 |
| GZMM | 50.1733745 | 0.35734283 | 0.75874329 | 0.47096671 | 0.63766451 | 0.94088547 |
| BSG | 37384.6309 | 0.19501125 | 0.15061256 | 1.29478745 | 1.95E-01 | 7.77E-01 |
| HCN2 | 99.7612772 | 0.87541393 | 0.42638827 | 2.05309103 | 0.04006375 | 0.55657615 |
| POLRMT | 4481.95857 | -0.0891643 | 0.08522613 | -1.0462088 | 0.29546466 | 0.83733602 |
| RNF126 | 2232.34793 | -0.4249404 | 0.1216933 | -3.4918964 | 0.0004796 | 0.1067163 |
| FSTL3 | 3637.6188 | -0.867911 | 0.41493409 | -2.0916841 | 3.65E-02 | 5.54E-01 |
| PALM | 4406.48725 | -0.3963498 | 0.40350099 | -0.9822771 | 3.26E-01 | 8.53E-01 |
| MISP | 1615.38827 | 0.22459098 | 0.25585706 | 0.87779865 | 3.80E-01 | 8.71E-01 |
| LINC01836 | 157.959356 | -0.0412981 | 0.45303189 | -0.0911593 | 0.92736604 | 0.98915637 |
| PTBP1 | 13435.2343 | -0.4331368 | 0.16616925 | -2.6066001 | 0.00914461 | 0.38664841 |
| AZU1 | 285.397999 | 2.2595558 | 1.37744891 | 1.64039174 | NA | NA |
| PRTN3 | 100.999164 | -0.1527924 | 1.08782322 | -0.1404571 | 0.88829888 | 0.98423722 |
| ELANE | 155.136318 | 1.1318027 | 1.03439767 | 1.09416594 | 0.27388221 | 0.82719197 |
| CFD | 792.538016 | -0.3504297 | 0.89248144 | -0.3926464 | NA | NA |
| MED16 | 3473.98003 | -0.1643998 | 0.22451189 | -0.7322542 | 0.46401344 | 0.89830907 |
| R3HDM4 | 3994.52989 | -0.1013757 | 0.24152922 | -0.4197243 | 0.67468684 | 0.94935413 |
| ARID3A | 1282.91282 | -0.3139397 | 0.23158695 | -1.3556018 | 0.1752259 | 0.76073329 |
| WDR18 | 2172.70159 | -0.0822819 | 0.21529292 | -0.382186 | 0.70232342 | 0.95431462 |
| GRIN3B | 1013.37786 | -0.1249226 | 0.16699026 | -0.7480832 | 0.45440997 | 0.89670027 |
| TMEM259 | 13742.9011 | -0.1661743 | 0.10664196 | -1.5582447 | 0.11917525 | 0.69233212 |
| AC004528.2 | 4637.88137 | -0.1422119 | 0.11615765 | -1.2243009 | 0.22083875 | 0.79960183 |
| CNN2 | 4812.83333 | -0.035203 | 0.32752124 | -0.1074832 | 0.91440568 | 0.9866558 |
| ABCA7 | 2082.75692 | -0.2371584 | 0.33699462 | -0.7037454 | 4.82E-01 | 0.90541935 |
| ARHGAP45 | 1922.9258 | -0.2255108 | 0.32433983 | -0.6952916 | 0.48687256 | 0.90573694 |
| POLR2E | 7411.12136 | 0.28482218 | 0.1993611 | 1.42867477 | 0.15309773 | 0.74258845 |
| GPX4 | 15865.8276 | 0.03324689 | 0.27011012 | 0.12308644 | 0.90203865 | 0.98572048 |
| SBNO2 | 11518.1185 | -0.2976822 | 0.30054236 | -0.9904834 | 0.3219379 | 0.85119105 |
| STK11 | 4424.34222 | -0.1040972 | 0.1412818 | -0.7368055 | 0.46124061 | 0.89755243 |
| CBARP | 470.336048 | -0.4391487 | 0.38840461 | -1.1306476 | 0.25820345 | 0.82239817 |
| ATP5F1D | 5276.49416 | -0.0032379 | 0.1410344 | -0.0229581 | 0.98168367 | 0.99595643 |
| MIDN | 8217.35726 | -0.2109146 | 0.29467239 | -0.7157596 | 0.47413978 | 0.90199039 |
| CIRBP | 19961.8047 | 0.20274978 | 0.29951689 | 0.6769227 | 0.498455 | 0.9109259 |
| CIRBP-AS1 | 134.747263 | -0.7720088 | 0.36471285 | -2.1167578 | 0.03428041 | 0.55369025 |
| C19orf24 | 1410.15403 | -0.2632498 | 0.25420359 | -1.0355864 | 0.30039515 | 0.84200819 |
| MUM1 | 3410.70153 | 0.05266684 | 0.16878956 | 0.31202663 | 0.75502029 | 0.96360109 |
| AC005329.2 | 786.322439 | 0.06882524 | 0.21110061 | 0.3260305 | 0.74440128 | 0.9613927 |
| NDUFS7 | 5006.48651 | 0.11027709 | 0.14443194 | 0.76352282 | 0.44515166 | 0.89445558 |
| AC005329.3 | 2174.68474 | 0.18902537 | 0.14163951 | 1.33455251 | 0.18202285 | 0.76569697 |
| AC005329.1 | 2695.82536 | 0.11540518 | 0.16631694 | 0.69388713 | 0.487753 | 0.90584365 |
| GAMT | 2166.19925 | 0.71780826 | 0.40338263 | 1.77947241 | 0.07516234 | 0.63773415 |

| | | | | | | |
|---|---|---|---|---|---|---|
| DAZAP1 | 7274.04202 | -0.2804939 | 0.13361966 | -2.0991966 | 0.03579957 | 0.55369025 |
| RPS15 | 11140.739 | -0.0453764 | 0.13721043 | -0.3307063 | 0.74086635 | 0.96089108 |
| APC2 | 367.737236 | 0.21663339 | 0.2574455 | 0.84147282 | 0.4000831 | 0.88077967 |
| AC027307.2 | 297.117521 | 0.31376229 | 0.3051864 | 1.0281005 | 0.30390255 | 0.84211658 |
| C19orf25 | 1972.93011 | -0.125796 | 0.13497275 | -0.9320103 | 0.3513312 | 0.86364633 |
| PCSK4 | 615.718821 | -0.0480816 | 0.41119729 | -0.1169307 | 0.90691496 | 0.98662757 |
| REEP6 | 1470.04427 | 0.90753603 | 0.48473857 | 1.87221749 | 0.06117652 | 0.60989294 |
| ADAMTSL5 | 132.293618 | 0.56734692 | 0.52055025 | 1.08989847 | 0.27575787 | 0.82719407 |
| MEX3D | 530.300218 | -0.416838 | 0.29308845 | -1.4222259 | 0.15496068 | 0.74373109 |
| MBD3 | 5495.34909 | -0.3286006 | 0.13165718 | -2.4958806 | 0.01256449 | 0.41598747 |
| AC005943.1 | 5988.78964 | -0.047912 | 0.16540773 | -0.2896601 | 0.77207632 | 0.96836854 |
| UQCR11 | 5332.92924 | -0.0020854 | 0.1750012 | -0.0119162 | 0.99049244 | 0.99814875 |
| TCF3 | 2690.78275 | -0.2938412 | 0.19753317 | -1.4875537 | 0.13686864 | 0.72140578 |
| ATP8B3 | 110.290146 | -0.3838097 | 0.40655992 | -0.9440422 | 0.3451481 | 0.8605056 |
| REXO1 | 3232.80568 | -0.3340644 | 0.1298837 | -2.5720271 | 0.0101105 | 0.39989496 |
| AC012615.2 | 487.229769 | -0.2868323 | 0.14134427 | -2.0293168 | 0.04242603 | 0.56525418 |
| AC012615.3 | 433.887112 | -0.3011032 | 0.25306245 | -1.1898377 | 0.23411018 | 0.80725541 |
| KLF16 | 1369.10662 | -0.2336776 | 0.23895183 | -0.9779277 | 0.32811008 | 0.85317073 |
| AC012615.1 | 326.932695 | 0.22058386 | 0.25963941 | 0.84957774 | 0.39555989 | 0.87927742 |
| AC012615.5 | 254.037264 | 0.20500895 | 0.26950876 | 0.76067639 | 0.44685038 | 0.89480207 |
| ABHD17A | 4325.19894 | -0.1251492 | 0.13799651 | -0.9069009 | 0.36445918 | 0.8663515 |
| SCAMP4 | 6441.11181 | -0.1900613 | 0.19506988 | -0.9743242 | 0.32989561 | 0.85371079 |
| ADAT3 | 667.022249 | -0.332539 | 0.17853795 | -1.8625673 | 0.06252316 | 0.61412297 |
| CSNK1G2 | 4696.37314 | -0.3223972 | 0.14307216 | -2.2533885 | 0.02423467 | 0.51967919 |
| BTBD2 | 6017.2002 | 0.07976003 | 0.15730899 | 0.50702777 | 0.61213533 | 0.93858463 |
| MKNK2 | 8802.48253 | 0.11879368 | 0.24153404 | 0.49182997 | 0.62283955 | 0.94016713 |
| MOB3A | 2096.82861 | -0.2178608 | 0.16193755 | -1.3453384 | 0.17851597 | 0.76115956 |
| IZUMO4 | 312.311166 | 0.45873956 | 0.31447463 | 1.45874902 | 0.1446342 | 0.73032824 |
| AP3D1 | 10983.377 | -0.152621 | 0.21035253 | -0.7255487 | 0.46811546 | 0.89984974 |
| DOT1L | 5436.91086 | -0.6480909 | 0.34023776 | -1.9048177 | 0.05680377 | 0.60049913 |
| AC004490.1 | 235.192257 | -1.0678243 | 0.48672131 | -2.1939132 | 0.02824165 | 0.53300785 |
| PLEKHJ1 | 5077.79089 | -0.2437007 | 0.14963714 | -1.6286107 | 0.10339545 | 0.67439401 |
| SF3A2 | 2521.76993 | -0.2794492 | 0.21853392 | -1.2787453 | 0.20098678 | 0.78134034 |
| AMH | 108.231262 | -0.8497884 | 0.60192801 | -1.4117774 | 0.1580155 | 0.74574881 |
| JSRP1 | 5957.73157 | -0.1163358 | 0.15047922 | -0.7731019 | 0.43946207 | 0.89191668 |
| OAZ1 | 22834.3371 | -0.1800955 | 0.1001746 | -1.7978155 | 0.07220625 | 0.6321115 |
| AC005258.1 | 7682.56643 | -0.1255368 | 0.09087308 | -1.3814517 | 0.16714011 | 0.75558932 |
| LSM7 | 1776.07652 | -0.0691556 | 0.23806721 | -0.2904878 | 0.7714431 | 0.96836854 |
| SPPL2B | 7511.98991 | -0.1417841 | 0.22932517 | -0.6182665 | 0.53639966 | 0.92105614 |
| TMPRSS9 | 217.525727 | 0.0418166 | 0.40621432 | 0.1029422 | 0.91800884 | 0.9868044 |
| TIMM13 | 3114.59251 | 0.05664358 | 0.174637 | 0.32435038 | 0.74567278 | 0.96176407 |
| LMNB2 | 1991.25225 | -0.1828547 | 0.23518212 | -0.7775026 | 4.37E-01 | 8.91E-01 |
| GADD45B | 12574.2442 | -0.4528525 | 0.47749296 | -0.9483962 | 3.43E-01 | 8.59E-01 |
| GNG7 | 1440.91001 | 0.13951405 | 0.33892049 | 0.4116424 | 0.68060155 | 0.95070301 |
| DIRAS1 | 430.05417 | -0.4282539 | 0.52571973 | -0.814605 | 0.41529842 | 0.88381712 |
| AC006538.2 | 486.518662 | -0.0813972 | 0.21322303 | -0.3817468 | 0.70264916 | 0.95431462 |
| SLC39A3 | 1032.09183 | -0.0952011 | 0.19431283 | -0.4899375 | 0.62417815 | 0.94073751 |
| SGTA | 5090.98653 | -0.214243 | 0.16579485 | -1.2922174 | 0.19628186 | 0.77798606 |
| THOP1 | 2027.00109 | -0.0853185 | 0.16225239 | -0.5258381 | 0.5990007 | 0.93484879 |
| ZNF554 | 462.273387 | 0.01268553 | 0.20097437 | 0.06312015 | 0.94967083 | 0.99170503 |
| ZNF555 | 313.075348 | -0.1958065 | 0.21115037 | -0.9273321 | 0.35375412 | 0.86364633 |
| ZNF556 | 41.3932658 | 0.27253185 | 0.66358176 | 0.41069822 | 0.68129383 | 0.95070301 |
| ZNF57 | 420.795197 | -0.0773852 | 0.2281497 | -0.339186 | 0.73446959 | 0.95895327 |
| AC119403.1 | 110.994927 | -0.1317202 | 0.23727241 | -0.5551432 | 0.5787967 | 0.93139743 |

| | | | | | | |
|---|---|---|---|---|---|---|
| ZNF77 | 247.362859 | 0.25796183 | 0.31956491 | 0.80722829 | 0.41953497 | 0.88451372 |
| TLE6 | 245.963567 | 0.7952764 | 0.55706749 | 1.42761229 | 0.15340348 | 0.74258845 |
| TLE2 | 3489.31154 | 0.18141895 | 0.38202128 | 0.47489226 | 0.63486377 | 0.94088547 |
| AC005944.1 | 9742.87388 | -0.0897173 | 0.19489567 | -0.4603352 | 0.64527564 | 0.94208859 |
| AES | 20576.7409 | -0.1185217 | 0.19646426 | -0.6032737 | 0.54632664 | 0.92345821 |
| GNA11 | 6840.39301 | -0.0822235 | 0.13161015 | -0.6247504 | 0.53213489 | 0.91973993 |
| AC005262.2 | 841.337608 | -0.0180138 | 0.20694225 | -0.0870473 | 0.93063388 | 0.98975562 |
| KF456478.1 | 1372.23978 | -0.0184234 | 0.14939812 | -0.1233174 | 0.90185574 | 0.98572048 |
| GNA15 | 269.970854 | 0.5045711 | 0.53040076 | 0.95130163 | 0.34145128 | 0.85889221 |
| AC005264.1 | 51.0905554 | 0.22300419 | 0.45466058 | 0.490485 | 0.62379074 | 0.9405038 |
| S1PR4 | 98.4255904 | -0.1751276 | 0.55218388 | -0.3171545 | 0.7511264 | 0.96272571 |
| NCLN | 3821.19712 | -0.0739282 | 0.1504565 | -0.4913595 | 0.62317217 | 0.94032767 |
| NFIC | 6929.07376 | -0.3305934 | 0.20363264 | -1.6234796 | 0.10448693 | 0.67439401 |
| SMIM24 | 9727.53176 | 0.7691292 | 0.41516939 | 1.85256722 | 0.06394441 | 0.61837523 |
| DOHH | 946.186151 | -0.0654717 | 0.24024859 | -0.2725166 | 0.7852248 | 0.97038798 |
| FZR1 | 3570.04104 | -0.3735947 | 0.13681853 | -2.7305854 | 0.00632219 | 0.33470127 |
| MFSD12 | 9773.82606 | 0.00706924 | 0.28993612 | 0.02438207 | 0.98054785 | 0.99573612 |
| C19orf71 | 1069.48752 | 0.24902874 | 0.27054104 | 0.92048414 | 3.57E-01 | 8.64E-01 |
| AC005786.3 | 1844.37517 | -0.2684089 | 0.33005323 | -0.8132292 | 0.41608666 | 0.88399703 |
| HMG20B | 4477.66519 | -0.0651674 | 0.14802899 | -0.4402343 | 0.65976742 | 0.94618161 |
| GIPC3 | 411.701002 | -0.2985128 | 0.45875539 | -0.6507014 | 0.51523926 | 0.91636259 |
| TBXA2R | 290.459656 | -0.4510296 | 0.4909768 | -0.9186374 | 0.35828528 | 0.86480228 |
| CACTIN-AS1 | 886.221739 | -0.2570853 | 0.18339615 | -1.4018031 | 0.16097404 | 0.74874703 |
| CACTIN | 2031.85001 | -0.269544 | 0.1736339 | -1.5523696 | 0.12057382 | 0.69467399 |
| PIP5K1C | 3906.4733 | -0.2790122 | 0.15885495 | -1.7563962 | 0.07902078 | 0.64079926 |
| TJP3 | 1594.92171 | -0.0905571 | 0.28300992 | -0.3199784 | 0.74898471 | 0.96249332 |
| APBA3 | 1519.88998 | -0.2382129 | 0.105102 | -2.2664923 | 0.02342126 | 0.51967919 |
| AC005954.2 | 170.735313 | -0.1516262 | 0.18378654 | -0.8250126 | 4.09E-01 | 0.88223304 |
| MRPL54 | 1687.14808 | -0.0384135 | 0.2378182 | -0.1615246 | 0.87168021 | 0.98275231 |
| MATK | 121.889479 | 0.3304607 | 0.63991846 | 0.51641063 | 6.06E-01 | 9.37E-01 |
| NMRK2 | 97.9995966 | -0.9210637 | 1.02873845 | -0.8953332 | 3.71E-01 | 8.68E-01 |
| DAPK3 | 2535.45875 | -0.5338717 | 0.16772701 | -3.1829796 | 1.46E-03 | 1.72E-01 |
| EEF2 | 86742.6824 | 0.193744 | 0.27788466 | 0.69721012 | 0.48567131 | 0.90543654 |
| PIAS4 | 1275.9378 | -0.3471508 | 0.13758333 | -2.5232037 | 0.0116291 | 0.40423902 |
| ZBTB7A | 2894.98076 | -0.1404515 | 0.21048418 | -0.667278 | 0.50459455 | 0.91365534 |
| MAP2K2 | 5299.84955 | -0.170183 | 0.11330756 | -1.5019565 | 0.13310834 | 0.71478037 |
| CREB3L3 | 162.36 | 2.46837532 | 1.03077254 | 2.39468479 | 0.01663466 | 0.46460528 |
| SIRT6 | 1406.69659 | -0.3018484 | 0.19325104 | -1.5619495 | 0.11829989 | 0.69095557 |
| ANKRD24 | 358.941646 | 0.527847 | 0.5799022 | 0.91023451 | 0.36269885 | 0.86590307 |
| EBI3 | 146.141519 | 0.13571419 | 0.68052947 | 0.19942442 | 0.84193076 | 0.9762188 |
| YJU2 | 983.052793 | -0.3414885 | 0.33824015 | -1.0096038 | 0.31268515 | 0.84541735 |
| STAP2 | 2180.60826 | 0.65301029 | 0.34910034 | 1.87055185 | 0.06140722 | 0.61019766 |
| MPND | 1273.09387 | 0.57149924 | 0.24843193 | 2.30042586 | 0.02142411 | 0.51278309 |
| AC007292.3 | 211.396865 | 0.56642463 | 0.32998968 | 1.71649194 | 0.08607204 | 0.65049132 |
| AC007292.2 | 599.057664 | 0.5062398 | 0.23809414 | 2.12621692 | 0.0334852 | 0.55367229 |
| SH3GL1 | 5242.78958 | -0.3479331 | 0.17029375 | -2.043135 | 0.04103907 | 0.55811692 |
| AC007292.1 | 892.458287 | -0.3019855 | 0.18423591 | -1.6391238 | 0.10118748 | 0.6725344 |
| CHAF1A | 321.861327 | 0.00030193 | 0.32268136 | 0.00093568 | 9.99E-01 | 1.00E+00 |
| UBXN6 | 9996.81642 | 0.00868644 | 0.15587603 | 0.05572659 | 0.95555962 | 0.99185043 |
| AC011498.2 | 124.514629 | -0.3573045 | 0.31661671 | -1.1285082 | 0.25910537 | 0.82364948 |
| AC011498.6 | 842.697089 | 0.22150737 | 0.20287079 | 1.09186429 | 0.27489276 | 0.82719407 |
| HDGFL2 | 2849.08533 | -0.0386133 | 0.19122307 | -0.2019279 | 0.83997307 | 0.97618283 |
| PLIN4 | 2390.86514 | 0.10728521 | 0.44619082 | 0.24044693 | 0.8099838 | 0.97334138 |
| PLIN5 | 3995.55611 | -0.1167177 | 0.51678924 | -0.2258517 | 0.82131678 | 0.97472105 |

| | | | | | | |
|---|---|---|---|---|---|---|
| AC011498.4 | 1359.89534 | -0.0323231 | 0.47610955 | -0.06789 | 0.94587322 | 0.99166614 |
| LRG1 | 3782.80621 | -0.1677106 | 0.63909666 | -0.2624182 | 0.79299906 | 0.97090793 |
| SEMA6B | 1388.08527 | -0.4785898 | 0.49431386 | -0.9681902 | 0.3329494 | 0.855011 |
| TNFAIP8L1 | 1529.46271 | -0.0406777 | 0.28868234 | -0.1409082 | 0.88794244 | 0.98423722 |
| MYDGF | 2920.61265 | -0.0346598 | 0.21961757 | -0.1578187 | 0.87459967 | 0.98275231 |
| AC005339.1 | 458.777311 | -0.0885424 | 0.28639288 | -0.309164 | 0.75719676 | 0.96502036 |
| DPP9 | 3271.07898 | -0.2651823 | 0.16370239 | -1.619905 | 0.10525269 | 0.67473222 |
| DPP9-AS1 | 676.635649 | -0.3846679 | 0.17896307 | -2.1494263 | 0.03160062 | 0.54507414 |
| FEM1A | 1708.73158 | -0.2494719 | 0.20156887 | -1.2376509 | 0.21584552 | 0.7954827 |
| AC005523.2 | 1585.64935 | -0.2186423 | 0.20737188 | -1.0543489 | 0.29172321 | 0.83460625 |
| TICAM1 | 1995.17258 | -0.0969116 | 0.2357168 | -0.4111355 | 0.68097316 | 0.95070301 |
| AC027319.1 | 777.880695 | -0.1762885 | 0.28485934 | -0.6188616 | 0.53600757 | 0.92101152 |
| PLIN3 | 3540.91023 | -0.2596353 | 0.25623932 | -1.0132531 | 0.31093927 | 0.84476049 |
| UHRF1 | 55.5188485 | -0.2289327 | 0.58551855 | -0.3909913 | 6.96E-01 | 9.53E-01 |
| KDM4B | 2568.49205 | -0.3119394 | 0.15529683 | -2.0086654 | 0.04457263 | 0.57267043 |
| PTPRS | 2746.64653 | -0.2785892 | 0.32694437 | -0.8520996 | 3.94E-01 | 0.87884163 |
| TINCR | 93.1828255 | -1.3364924 | 0.72586407 | -1.8412434 | NA | NA |
| SAFB2 | 5771.30574 | -0.2226861 | 0.13441706 | -1.6566807 | 9.76E-02 | 6.65E-01 |
| SAFB | 5333.90659 | -0.2274357 | 0.120814 | -1.8825276 | 0.05976442 | 0.60707041 |
| RPL36 | 10301.4462 | 0.00902348 | 0.10136017 | 0.08902393 | 0.92906289 | 0.98963127 |
| C19orf70 | 2576.21344 | 0.03712451 | 0.16444915 | 0.22575072 | 0.82139531 | 0.97472105 |
| HSD11B1L | 653.203754 | -0.0543704 | 0.18612723 | -0.2921143 | 0.77019924 | 0.96836854 |
| LONP1 | 7553.61104 | -0.3878998 | 0.25387798 | -1.5278985 | 0.12653774 | 0.70431949 |
| PRR22 | 324.104414 | -0.0361428 | 0.39784484 | -0.0908466 | 0.92761449 | 0.98915637 |
| AC011499.1 | 467.645597 | 0.09164734 | 0.2338349 | 0.39193182 | 0.69510859 | 0.95234774 |
| DUS3L | 1953.35331 | 0.04719301 | 0.17774856 | 0.26550433 | 0.79062097 | 0.97090793 |
| NRTN | 179.586442 | -0.1247071 | 0.422592 | -0.2951005 | 0.76791713 | 0.96762687 |
| FUT6 | 3824.46913 | 0.97199449 | 0.54773536 | 1.77456956 | 0.07596899 | 0.63773415 |
| FUT3 | 1013.25717 | 0.89635155 | 0.55114742 | 1.6263372 | 0.10387794 | 0.67439401 |
| AC024592.2 | 78.0522151 | 0.21370708 | 0.72975412 | 0.29284806 | 0.76963829 | 0.96836854 |
| AC024592.3 | 4830.79468 | 0.11118859 | 0.09145536 | 1.215769 | 0.22407293 | 0.80078694 |
| NDUFA11 | 6126.95252 | 0.04588005 | 0.11241604 | 0.40812724 | 6.83E-01 | 0.95070301 |
| AC024592.1 | 101.571491 | 0.16284637 | 0.37162442 | 0.43820148 | 0.66124024 | 0.94637197 |
| AC104532.1 | 878.197633 | 0.79477785 | 0.39811244 | 1.99636527 | 0.04589418 | 0.57687945 |
| VMAC | 315.6798 | 0.24626195 | 0.24864428 | 0.99041871 | 0.32196951 | 0.85119105 |
| AC104532.2 | 211.880708 | 0.28719222 | 0.26667633 | 1.0769318 | 0.28151074 | 0.82975653 |
| CAPS | 2458.44489 | 0.73354783 | 0.38428677 | 1.90885529 | 5.63E-02 | 5.99E-01 |
| RANBP3 | 3135.12098 | -0.2276551 | 0.1234826 | -1.8436205 | 0.06523846 | 0.6216672 |
| RFX2 | 444.974341 | -0.2621893 | 0.34725886 | -0.7550254 | 4.50E-01 | 0.89557948 |
| ACSBG2 | 59.512175 | 1.21332732 | 0.6936371 | 1.74922494 | 0.08025214 | 0.6408661 |
| MLLT1 | 5126.3816 | -0.3173355 | 0.17876264 | -1.7751778 | 0.07586853 | 0.63773415 |
| CLPP | 2262.09211 | 0.09628038 | 0.15180993 | 0.63421661 | 0.52593947 | 0.91769856 |
| ALKBH7 | 1792.47579 | 0.40824344 | 0.14915114 | 2.73711239 | 0.00619811 | 0.33470127 |
| PSPN | 274.177627 | -0.2027417 | 0.24570211 | -0.8251523 | 0.40928513 | 0.88221056 |
| GTF2F1 | 4272.58009 | -0.1466423 | 0.12495382 | -1.1735722 | 0.24056644 | 0.8124487 |
| KHSRP | 7516.60293 | -0.318853 | 0.15033191 | -2.1209932 | 0.03392238 | 0.55369025 |
| SLC25A41 | 37.4567536 | -0.8144794 | 0.77196545 | -1.0550723 | 0.29139227 | 0.83454586 |
| SLC25A23 | 5717.51845 | 0.31703929 | 0.18056456 | 1.75582231 | 7.91E-02 | 0.64079926 |
| CRB3 | 1704.77994 | -0.2180776 | 0.22415646 | -0.9728812 | 0.33061233 | 0.85384238 |
| DENND1C | 2918.89105 | -0.1058115 | 0.22203508 | -0.4765532 | 0.63368031 | 0.94088547 |
| AC010503.4 | 1249.79989 | -0.1340813 | 0.19192291 | -0.6986209 | 0.48478899 | 0.90541935 |
| TUBB4A | 186.41103 | 0.90051348 | 0.61345058 | 1.46794789 | 0.14211838 | 0.72784886 |
| AC010503.1 | 85.896 | 0.74781051 | 0.6581501 | 1.13623095 | 2.56E-01 | 8.21E-01 |
| AC010503.2 | 61.1671083 | 0.95051861 | 0.64903786 | 1.46450409 | 0.14305627 | 0.72784886 |

| | | | | | | |
|---|---|---|---|---|---|---|
| TNFSF9 | 112.915535 | 0.15991964 | 0.69439963 | 0.23029914 | 0.81785933 | 0.97446554 |
| TNFSF14 | 188.475106 | 0.69702984 | 0.6618756 | 1.05311306 | 2.92E-01 | 0.83524572 |
| C3 | 23435.639 | 0.47060633 | 0.69535816 | 0.67678263 | 4.99E-01 | 9.11E-01 |
| GPR108 | 5652.29943 | -0.0055527 | 0.12369963 | -0.0448884 | 9.64E-01 | 0.99342622 |
| TRIP10 | 8111.58654 | -0.1043554 | 0.24698608 | -0.4225152 | 0.67264903 | 0.94922827 |
| AC008760.1 | 709.509569 | -0.134447 | 0.27624754 | -0.4866901 | 0.62647793 | 0.94088547 |
| SH2D3A | 1909.08393 | -0.4363496 | 0.26340815 | -1.6565532 | 0.09760983 | 0.66549286 |
| VAV1 | 284.41724 | -0.0815942 | 0.57282554 | -0.1424417 | 0.88673114 | 0.98423722 |
| ADGRE1 | 43.1554534 | 0.60799781 | 0.75085928 | 0.80973603 | 0.41809191 | 0.88445833 |
| ZNF557 | 383.496378 | -0.0882289 | 0.23969213 | -0.3680928 | 0.71280407 | 0.95564093 |
| INSR | 7889.26435 | -0.045771 | 0.26924095 | -0.17 | 0.86501016 | 0.98197545 |
| AC008878.3 | 3746.17174 | -0.2755562 | 0.18366681 | -1.5003047 | 0.13353549 | 0.71505705 |
| ARHGEF18 | 3991.09893 | -0.2803557 | 0.18778771 | -1.4929394 | 0.13545304 | 0.71649038 |
| PEX11G | 364.666068 | 0.79941277 | 0.43126826 | 1.85363229 | 0.06379179 | 0.61812911 |
| AC008878.1 | 823.250908 | -0.2182733 | 0.17129845 | -1.2742283 | 0.20258254 | 0.78193335 |
| ZNF358 | 2949.54999 | -0.2387459 | 0.15269136 | -1.5635846 | 0.11791516 | 0.69069542 |
| AC008878.4 | 680.622414 | -0.2099546 | 0.2509164 | -0.8367513 | 0.40273236 | 0.88132015 |
| MCOLN1 | 1881.45104 | -0.5914975 | 0.29633098 | -1.9960704 | 0.04592627 | 0.57687945 |
| AC008878.2 | 923.338077 | -0.2709837 | 0.25612919 | -1.0579963 | 0.29005714 | 0.83407594 |
| PNPLA6 | 6647.11291 | -0.2224393 | 0.27019722 | -0.8232478 | 0.41036707 | 0.88249519 |
| CAMSAP3 | 4140.63549 | -0.2479537 | 0.17502995 | -1.4166357 | 0.15658947 | 0.74522306 |
| XAB2 | 4205.52087 | -0.0085863 | 0.16466687 | -0.0521437 | 0.95841423 | 0.99237995 |
| PET100 | 1647.07099 | 0.07755599 | 0.18053183 | 0.42959731 | 0.66748859 | 0.94832869 |
| AC008763.2 | 4992.17984 | -0.012906 | 0.09918794 | -0.1301163 | 0.89647445 | 0.98474725 |
| PCP2 | 496.066565 | -0.1121773 | 0.50569995 | -0.2218258 | 0.82444946 | 0.97548201 |
| AC008763.1 | 262.889053 | -0.1706122 | 0.42180205 | -0.4044839 | 0.68585689 | 0.95097288 |
| STXBP2 | 7138.51858 | -0.0982874 | 0.13464038 | -0.7299992 | 0.4653907 | 0.89925633 |
| MCEMP1 | 69.1688176 | -0.5857682 | 0.90453143 | -0.6475929 | 0.51724828 | 0.91636259 |
| AC008763.3 | 1494.16219 | -0.3814907 | 0.17530975 | -2.1760952 | 0.02954814 | 0.53721163 |
| TRAPPC5 | 1522.01112 | -0.3780572 | 0.16996786 | -2.2242863 | 0.02612919 | 0.52592428 |
| CD209 | 368.596141 | 0.7624533 | 0.63488703 | 1.20092751 | 0.22977932 | 0.80551898 |
| EVI5L | 1721.52761 | 0.02588778 | 0.1584771 | 0.16335347 | 0.87024014 | 0.98275231 |
| PRR36 | 569.003166 | -0.2235038 | 0.4502835 | -0.4963623 | 0.6196388 | 0.93945324 |
| AC010336.3 | 150.854823 | -0.2407361 | 0.50597019 | -0.4757912 | 0.63422317 | 0.94088547 |
| LRRC8E | 340.264466 | -0.0046408 | 0.28433779 | -0.0163214 | 0.98697798 | 0.99726698 |
| AC010336.1 | 327.593816 | -0.0312671 | 0.28854022 | -0.1083632 | 9.14E-01 | 0.98662757 |
| MAP2K7 | 3406.86098 | -0.4517592 | 0.18126593 | -2.4922457 | 0.01269382 | 0.41598747 |
| AC010336.4 | 553.584351 | -0.4331941 | 0.21672492 | -1.99882 | 0.04562783 | 0.57670328 |
| TGFBR3L | 144.903135 | 0.93964818 | 0.64171697 | 1.46427199 | 0.14311965 | 0.72784886 |
| AC010336.2 | 54.0213211 | 0.66191391 | 0.57018125 | 1.16088333 | 0.24568935 | 0.81542272 |
| SNAPC2 | 964.736974 | 0.00810002 | 0.14875428 | 0.05445236 | 0.95657476 | 0.99198394 |
| CTXN1 | 514.361566 | -0.4915304 | 0.38019097 | -1.2928512 | 0.19606252 | 0.77791948 |
| TIMM44 | 2604.6013 | -0.1912856 | 0.21441166 | -0.8921419 | 0.37231689 | 0.86775997 |
| ELAVL1 | 3534.66667 | -0.116277 | 0.14371538 | -0.8090786 | 0.41846993 | 0.88445833 |
| AC010336.6 | 108.53983 | -0.1621332 | 0.39447792 | -0.4110069 | 0.68106744 | 0.95070301 |
| FBN3 | 265.797436 | 0.18705878 | 0.73286079 | 0.25524463 | 0.79853416 | 0.9717324 |
| CERS4 | 3075.20131 | -0.0633925 | 0.32568235 | -0.194645 | 0.84567084 | 0.97675487 |
| CD320 | 1715.22788 | 0.30819997 | 0.19836545 | 1.5536978 | 0.12025652 | 0.69467399 |
| AC010323.1 | 2133.2585 | 0.28653003 | 0.16675824 | 1.7182361 | 0.08575356 | 0.65049132 |
| NDUFA7 | 2049.93657 | 0.27502848 | 0.16356981 | 1.68141345 | 0.09268263 | 0.65662464 |
| RPS28 | 13697.8206 | 0.00518746 | 0.12806829 | 0.04050538 | 0.96769022 | 0.99400331 |
| KANK3 | 1516.98233 | -0.0829138 | 0.27416101 | -0.3024274 | 0.76232625 | 0.96630779 |
| ANGPTL4 | 9516.08492 | -0.2917608 | 0.76291793 | -0.3824275 | 0.7021443 | 0.95431462 |
| RAB11B-AS1 | 1228.51022 | -0.0095868 | 0.18660255 | -0.0513757 | 0.95902612 | 0.99242674 |

| | | | | | | |
|---|---|---|---|---|---|---|
| RAB11B | 8039.2593 | -0.0902176 | 0.19559338 | -0.4612508 | 0.64461868 | 0.94208859 |
| 2-Mar | 1778.50058 | 0.04447169 | 0.16196178 | 0.2745814 | 0.78363786 | 0.96978847 |
| HNRNPM | 9054.65846 | -0.2213643 | 0.21406514 | -1.0340977 | 3.01E-01 | 0.84211658 |
| PRAM1 | 147.569382 | -0.0249279 | 0.51061689 | -0.0488191 | 0.96106343 | 0.99317837 |
| ZNF414 | 630.306099 | -0.2942052 | 0.21020831 | -1.3995887 | 0.16163651 | 0.74969823 |
| MYO1F | 947.584231 | -0.1526046 | 0.54004747 | -0.2825763 | 0.77750165 | 0.96836854 |
| ADAMTS10 | 1128.1124 | 0.19175254 | 0.50171814 | 0.38219176 | 0.70231913 | 0.95431462 |
| ZNF558 | 1003.44805 | 0.05951025 | 0.13517286 | 0.44025297 | 0.6597539 | 0.94618161 |
| AC008734.1 | 42.86801 | -0.2287014 | 0.40234912 | -0.5684154 | 0.56975293 | 0.92719921 |
| ZNF317 | 1371.68217 | -0.3200982 | 0.19558223 | -1.6366424 | 0.10170521 | 0.67279543 |
| AC011451.1 | 49.7703548 | 0.17608622 | 0.26597617 | 0.66203757 | 0.50794714 | 0.91439937 |
| ZNF559 | 817.967374 | -0.0866557 | 0.11543251 | -0.7507044 | 0.45283058 | 0.89670027 |
| ZNF559-ZNF | 614.884086 | 0.25055734 | 0.27600559 | 0.90779807 | 0.36398492 | 0.8663515 |
| ZNF177 | 504.383792 | 0.35678931 | 0.31407002 | 1.13601836 | 2.56E-01 | 0.82052927 |
| ZNF266 | 1912.02825 | 0.02082331 | 0.17405171 | 0.11963867 | 0.90476939 | 0.98624484 |
| ZNF426 | 581.740976 | -0.0277483 | 0.1443099 | -0.1922827 | 0.8475208 | 0.97701835 |
| ZNF426-DT | 53.7347795 | 0.61434401 | 0.46199031 | 1.32977682 | 0.18359182 | 0.76781833 |
| ZNF121 | 1449.62407 | -0.2280954 | 0.16771368 | -1.3600289 | 0.17382078 | 0.75974924 |
| ZNF561 | 1188.28716 | 0.03231877 | 0.19280291 | 0.16762594 | 0.86687756 | 0.98222237 |
| ZNF562 | 1227.04117 | -0.1992539 | 0.16364595 | -1.2175913 | 0.22337931 | 0.80078694 |
| ZNF846 | 309.466232 | 0.19427271 | 0.29780659 | 0.65234524 | 0.51417848 | 0.91636259 |
| FBXL12 | 1256.70736 | -0.0773853 | 0.12736889 | -0.6075685 | 0.54347372 | 0.92297267 |
| UBL5 | 3956.63648 | 0.04163751 | 0.13872342 | 0.30014769 | 0.7640645 | 0.96678847 |
| PIN1 | 2532.10922 | 0.16819706 | 0.16597318 | 1.01339898 | 0.31086963 | 0.84476049 |
| OLFM2 | 267.89564 | -0.678814 | 0.6048813 | -1.1222268 | 0.26176604 | 0.82424998 |
| COL5A3 | 807.477088 | -0.1896253 | 0.59724179 | -0.3175018 | 0.75086287 | 0.96272571 |
| AC008742.1 | 43.5393689 | -0.4837915 | 0.80428393 | -0.6015183 | 0.54749483 | 0.92361227 |
| RDH8 | 429.380701 | 0.5385248 | 1.27623603 | 0.42196333 | 0.67305179 | 0.94925222 |
| C19orf66 | 4812.81639 | -1.1338311 | 0.54064728 | -2.0971734 | 0.03597823 | 0.55369025 |
| AC020931.1 | 1121.76923 | -1.2025322 | 0.57209102 | -2.1019944 | 0.03555376 | 0.55369025 |
| ANGPTL6 | 1424.81431 | -1.0429885 | 0.51958225 | -2.0073598 | 0.04471137 | 0.57267043 |
| PPAN-P2RY1 | 1822.76493 | -0.548643 | 0.22073203 | -2.4855615 | 0.01293473 | 0.41899224 |
| PPAN | 1502.99648 | -0.5045627 | 0.21265791 | -2.3726496 | 1.77E-02 | 4.70E-01 |
| P2RY11 | 1968.47306 | -0.3422599 | 0.2049703 | -1.6698026 | 0.09495842 | 0.65888357 |
| EIF3G | 7079.20603 | -0.0645773 | 0.1348486 | -0.4788872 | 0.63201887 | 0.94088547 |
| DNMT1 | 1691.20023 | -0.3379909 | 0.2202691 | -1.5344455 | 0.12492009 | 0.70228923 |
| S1PR2 | 281.226898 | 0.05013608 | 0.37305144 | 0.13439455 | 0.89309059 | 0.98474725 |
| MRPL4 | 2962.4257 | 0.00662208 | 0.11322565 | 0.05848573 | 0.95336173 | 0.99170503 |
| AC011511.2 | 1323.69713 | 0.39412989 | 0.32867057 | 1.19916393 | 0.23046421 | 0.80584899 |
| AC011511.3 | 720.538457 | -0.0155923 | 0.10493917 | -0.1485844 | 0.88188161 | 0.98275231 |
| ICAM1 | 8328.66969 | 0.62753541 | 0.59326512 | 1.05776556 | 0.29016234 | 0.83416665 |
| AC011511.5 | 3157.44246 | 0.68407373 | 0.60131502 | 1.13762953 | 0.25527521 | 0.82018048 |
| ICAM5 | 187.690951 | 0.73067606 | 0.43529757 | 1.67856684 | 0.09323649 | 0.65756964 |
| ZGLP1 | 331.081635 | -0.4490219 | 0.38322929 | -1.1716795 | 0.24132575 | 0.8124487 |
| AC011511.1 | 1300.87953 | -0.1443807 | 0.18596706 | -0.7763781 | 4.38E-01 | 0.89130865 |
| FDX2 | 1065.88787 | -0.0697067 | 0.17960416 | -0.388113 | 0.69793244 | 0.9529028 |
| AC011511.4 | 2464.78215 | -0.306105 | 0.15888513 | -1.9265806 | 0.05403192 | 0.59329767 |
| RAVER1 | 3492.98442 | -0.3322757 | 0.17096711 | -1.9435069 | 0.05195495 | 0.58645127 |
| AC114271.1 | 912.236198 | 0.51185022 | 0.35774539 | 1.43076681 | 0.15249706 | 0.7422778 |
| ICAM3 | 771.683509 | 0.64922024 | 0.40792912 | 1.59150254 | 0.11149652 | 0.68001961 |
| TYK2 | 6558.52041 | -0.1343368 | 0.16133856 | -0.8326394 | 0.40504812 | 0.88132015 |
| CDC37 | 10008.0804 | -0.1826552 | 0.17834236 | -1.0241828 | 0.30574894 | 0.84270033 |
| PDE4A | 1062.42703 | -0.0919577 | 0.37855603 | -0.2429171 | 0.80806964 | 0.97334138 |
| KEAP1 | 3628.38271 | -0.2863927 | 0.25058092 | -1.1429149 | 0.2530739 | 0.81835388 |

| | | | | | | |
|---|---|---|---|---|---|---|
| S1PR5 | 45.4686725 | -0.2282233 | 0.75535612 | -0.30214 | 0.76254533 | 0.96630779 |
| ATG4D | 2904.19229 | 0.00851205 | 0.12640282 | 0.06734064 | 0.94631053 | 0.99166614 |
| KRI1 | 3018.04559 | -0.2638341 | 0.12167295 | -2.1683878 | 0.03012919 | 0.54012717 |
| CDKN2D | 554.347522 | -0.1648269 | 0.16963482 | -0.9716572 | 0.33122113 | 0.85384238 |
| AP1M2 | 3577.73193 | 0.44952924 | 0.2802232 | 1.6041828 | 0.10867377 | 0.67800851 |
| SLC44A2 | 10531.8749 | 0.07669378 | 0.27187146 | 0.28209574 | 0.7778701 | 0.96836854 |
| ILF3-DT | 1276.0662 | 0.0092847 | 0.30525671 | 0.03041602 | 0.97573527 | 0.99455099 |
| ILF3 | 11475.9517 | -0.2624192 | 0.20234324 | -1.296901 | 0.19466524 | 0.7762856 |
| QTRT1 | 3957.69932 | 0.00534952 | 0.24550842 | 0.02178954 | 0.98261584 | 0.99611734 |
| DNM2 | 16509.5545 | 0.01063994 | 0.16036931 | 0.06634651 | 0.94710196 | 0.99166614 |
| TMED1 | 2663.25513 | -0.1417891 | 0.20884783 | -0.678911 | 0.49719424 | 0.90913397 |
| C19orf38 | 96.9283709 | -0.4459171 | 0.47976174 | -0.9294554 | 0.35265314 | 0.86364633 |
| CARM1 | 4145.9524 | -0.2080718 | 0.14789174 | -1.4069199 | 0.15945115 | 0.74762924 |
| YIPF2 | 3518.28146 | -0.101292 | 0.13833616 | -0.732216 | 0.46403671 | 0.89830907 |
| TIMM29 | 843.072193 | -0.0821222 | 0.16556959 | -0.4959982 | 0.61989571 | 0.93946834 |
| SMARCA4 | 5352.14694 | -0.0474551 | 0.13255471 | -0.3580041 | 0.72034026 | 0.95703224 |
| LDLR | 2535.46537 | -0.5332019 | 0.57475988 | -0.9276951 | 0.35356573 | 0.86364633 |
| SPC24 | 222.120407 | -0.4912376 | 0.53576382 | -0.9168922 | 0.35919915 | 0.86480228 |
| KANK2 | 4461.79953 | -0.0372002 | 0.3083622 | -0.120638 | 0.90397774 | 0.98624484 |
| DOCK6 | 4032.90414 | -0.0140154 | 0.19855534 | -0.0705867 | 0.94372669 | 0.99136103 |
| AC011472.1 | 151.442033 | -0.1493415 | 0.30911597 | -0.4831245 | 0.62900736 | 0.94088547 |
| ANGPTL8 | 382.204397 | 0.78969991 | 0.58032332 | 1.36079298 | 0.17357912 | 0.75974924 |
| RAB3D | 2919.55899 | 0.12162731 | 0.2050551 | 0.59314451 | 0.55308444 | 0.92373235 |
| TMEM205 | 6170.52963 | 0.12152199 | 0.1138552 | 1.06733808 | 0.2858192 | 0.83107779 |
| CCDC159 | 5699.54799 | 0.19182813 | 0.11215176 | 1.71043357 | 0.08718573 | 0.65231349 |
| PLPPR2 | 1971.81311 | -0.2351998 | 0.24284346 | -0.9685245 | 0.33278251 | 0.855011 |
| AC024575.1 | 93.6925572 | 0.11911491 | 0.36144183 | 0.32955485 | 0.74173635 | 0.96109001 |
| SWSAP1 | 329.566764 | 0.19566688 | 0.27098822 | 0.72204939 | 0.47026411 | 0.9008571 |
| EPOR | 3475.55593 | -0.0485682 | 0.30823769 | -0.1575673 | 0.87479776 | 0.98275231 |
| RGL3 | 8245.46804 | -0.0677208 | 0.27287784 | -0.2481726 | 0.80400087 | 0.97274243 |
| CCDC151 | 819.712093 | 0.2334847 | 0.14229988 | 1.64079342 | 0.1008403 | 0.67247782 |
| PRKCSH | 13635.4875 | 0.16901007 | 0.08785124 | 1.92382103 | 0.05437702 | 0.59329767 |
| AC008481.3 | 101.85322 | -0.1809649 | 0.24553084 | -0.7370353 | 0.46110091 | 0.89755243 |
| ZNF653 | 469.976831 | -0.3010738 | 0.18198111 | -1.6544233 | 0.09804153 | 0.66549286 |
| ECSIT | 2548.55418 | -0.0300544 | 0.23442354 | -0.1282055 | 0.89798637 | 0.98474725 |
| CNN1 | 1976.27327 | -2.2170782 | 0.88045343 | -2.5181095 | 0.01179866 | 0.40423902 |
| ELOF1 | 2213.03874 | -0.3028371 | 0.1894191 | -1.5987675 | 0.10987228 | 0.6791281 |
| ZNF627 | 1463.42488 | 0.22923093 | 0.21885462 | 1.04741192 | 0.29490964 | 0.83630273 |
| ACP5 | 5206.50007 | 0.36256877 | 0.48374684 | 0.74950106 | 0.45355526 | 0.89670027 |
| AC008543.1 | 103.922656 | -0.1648982 | 0.35944887 | -0.4587528 | 0.64641169 | 0.94208859 |
| ZNF823 | 654.691726 | -0.2452056 | 0.25991208 | -0.9434176 | 0.34546733 | 0.86095788 |
| ZNF441 | 542.570288 | 0.18165457 | 0.28839657 | 0.62987769 | 0.52877461 | 0.91901743 |
| ZNF491 | 110.403661 | -0.0282667 | 0.29451224 | -0.095978 | 0.92353805 | 0.98858365 |
| ZNF440 | 833.712469 | 0.02697002 | 0.28714427 | 0.09392497 | 0.92516876 | 0.98869145 |
| ZNF439 | 612.669771 | -0.090361 | 0.16613553 | -0.5438996 | 0.58651057 | 0.93289552 |
| ZNF69 | 505.84933 | -0.0561051 | 0.21070552 | -0.2662727 | 0.79002917 | 0.97090793 |
| ZNF700 | 1715.76048 | -0.052139 | 0.20285326 | -0.2570282 | 0.79715702 | 0.9713715 |
| AC008770.2 | 779.54774 | 0.03879605 | 0.23010988 | 0.16859795 | 0.86611289 | 0.98197545 |
| ZNF763 | 408.702814 | 0.17049472 | 0.37372579 | 0.45620271 | 0.64824421 | 0.9431121 |
| ZNF433 | 840.473952 | 0.12862175 | 0.23485029 | 0.54767548 | 0.58391475 | 0.93233498 |
| AC008770.1 | 145.695983 | 0.01562621 | 0.23729208 | 0.0658522 | 0.9474955 | 0.99170503 |
| ZNF878 | 52.7884749 | -0.5682954 | 0.49910155 | -1.1386369 | 0.25485464 | 0.81992039 |
| ZNF844 | 1082.9525 | 0.08087475 | 0.25506212 | 0.31707864 | 0.75118393 | 0.96272571 |
| ZNF788P | 330.279021 | 0.14210815 | 0.20221303 | 0.70276457 | 0.48220248 | 0.90541935 |

| | | | | | | |
|---|---|---|---|---|---|---|
| ZNF20 | 389.991401 | 0.01598623 | 0.17021362 | 0.09391864 | 0.92517379 | 0.98869145 |
| ZNF625-ZNF | 516.614076 | -0.0502616 | 0.19157481 | -0.2623603 | 0.79304371 | 0.97090793 |
| ZNF625 | 247.18328 | -0.1033873 | 0.30742597 | -0.3363 | 0.73664466 | 0.95991222 |
| ZNF136 | 538.989702 | -0.09787 | 0.19616993 | -0.4989044 | 0.61784673 | 0.93924932 |
| ZNF44 | 2319.9746 | -0.2245495 | 0.22489857 | -0.998448 | 0.31806217 | 0.84892422 |
| ZNF563 | 329.245212 | -0.0594947 | 0.23100276 | -0.2575498 | 0.79675441 | 0.97131168 |
| ZNF442 | 242.647565 | 0.16719536 | 0.32837292 | 0.50916304 | 0.61063795 | 0.93807505 |
| AC008758.4 | 74.5479731 | 0.29083709 | 0.29825742 | 0.97512105 | 0.32950022 | 0.85350346 |
| ZNF799 | 421.399196 | -0.0583204 | 0.18665588 | -0.3124485 | 0.75469969 | 0.96351771 |
| AC008758.5 | 79.4813078 | -0.1314866 | 0.34592109 | -0.3801057 | 0.70386694 | 0.95481129 |
| ZNF443 | 321.208882 | 0.16126767 | 0.19484206 | 0.82768407 | 0.40784944 | 0.88161173 |
| ZNF709 | 280.15518 | 0.00360408 | 0.22089453 | 0.01631586 | 0.9869824 | 0.99726698 |
| AC008758.1 | 289.145032 | -0.0064552 | 0.17769758 | -0.0363271 | 0.97102157 | 0.99400331 |
| ZNF564 | 845.848198 | 0.00636334 | 0.20815947 | 0.03056956 | 0.97561282 | 0.99455099 |
| AC010422.6 | 1033.3469 | 0.02936952 | 0.21036631 | 0.13961135 | 0.88896707 | 0.98423722 |
| ZNF490 | 168.576704 | 0.0447426 | 0.22834014 | 0.19594714 | 0.84465154 | 0.97675487 |
| ZNF791 | 1198.76345 | -0.0746758 | 0.14937791 | -0.4999121 | 0.61713694 | 0.93910158 |
| AC010422.3 | 1490.29453 | 0.12803466 | 0.20262863 | 0.63186857 | 0.52747276 | 0.91831212 |
| MAN2B1 | 4174.00439 | 0.1622156 | 0.16914714 | 0.95902062 | 0.33754836 | 0.8571036 |
| AC010422.5 | 2611.41281 | 0.11598084 | 0.13480848 | 0.86033789 | 0.38960281 | 0.87577651 |
| WDR83 | 4166.69853 | -0.0074685 | 0.10460073 | -0.0713997 | 0.94307968 | 0.99136103 |
| WDR83OS | 4979.3739 | -0.0168994 | 0.11009643 | -0.1534967 | 0.87800661 | 0.98275231 |
| DHPS | 3364.95085 | 0.02380585 | 0.10919298 | 0.21801628 | 0.82741643 | 0.97548201 |
| FBXW9 | 746.928377 | -0.0460741 | 0.16091746 | -0.2863212 | 0.77463211 | 0.96836854 |
| AC018761.4 | 209.300958 | 0.01971182 | 0.24367365 | 0.08089436 | 0.93552597 | 0.99068685 |
| TNPO2 | 4879.56305 | -0.114232 | 0.12645387 | -0.9033496 | 0.36634039 | 0.86683872 |
| TRIR | 5216.50604 | -0.1693363 | 0.14019684 | -1.2078469 | 0.22710616 | 0.80300331 |
| ASNA1 | 4229.6389 | 0.10793538 | 0.15363472 | 0.70254544 | 0.48233907 | 0.90541935 |
| HOOK2 | 7563.76361 | -0.1437471 | 0.23178907 | -0.6201634 | 0.53515021 | 0.92084006 |
| JUNB | 9226.99895 | -0.1731738 | 0.49235444 | -0.3517259 | 0.72504386 | 0.95848028 |
| PRDX2 | 11808.5884 | 0.33337884 | 0.19174756 | 1.73863407 | 0.08209915 | 0.6456652 |
| AC018761.2 | 4753.64806 | 0.31980638 | 0.19287955 | 1.65806261 | 0.09730483 | 0.66549286 |
| THSD8 | 298.140471 | 0.29859713 | 0.34248188 | 0.87186256 | 0.38328336 | 0.87194871 |
| RNASEH2A | 299.267343 | 0.28436759 | 0.34592475 | 0.82205042 | 0.4110482 | 0.88268208 |
| DNASE2 | 1831.34724 | 0.45909374 | 0.14071523 | 3.26257313 | 1.10E-03 | 0.15675098 |
| AC020934.2 | 373.255024 | 0.52757323 | 0.18006488 | 2.9299063 | 0.00339064 | 0.2541761 |
| GCDH | 2085.59269 | 0.62254049 | 0.4058827 | 1.53379411 | 0.12508031 | 0.70228923 |
| SYCE2 | 117.794829 | 0.27711863 | 0.36579523 | 0.75757856 | 0.44870332 | 0.8948759 |
| FARSA | 2989.42081 | -0.1045635 | 0.14443105 | -0.723968 | 0.46908538 | 0.9003198 |
| FARSA-AS1 | 280.329949 | -0.2362657 | 0.19304862 | -1.2238665 | 0.2210026 | 0.79965146 |
| CALR | 32889.3552 | 0.46306039 | 0.2498715 | 1.85319412 | 0.06385454 | 0.61812911 |
| AC092069.1 | 7167.9546 | 0.34083452 | 0.30655097 | 1.1118364 | 0.2662085 | 0.8265123 |
| RAD23A | 5958.783 | -0.2053981 | 0.1604274 | -1.2803178 | 0.20043338 | 0.78134034 |
| GADD45GIP1 | 3920.89316 | -0.1477416 | 0.11652243 | -1.2679241 | 0.20482506 | 0.78499518 |
| NFIX | 1770.97357 | 0.06273894 | 0.30216091 | 0.20763422 | 0.83551459 | 0.97550416 |
| LYL1 | 308.18414 | -0.2443891 | 0.35369498 | -0.6909601 | 0.48959058 | 0.90600166 |
| TRMT1 | 2817.82499 | -0.2108485 | 0.13312872 | -1.5837942 | 0.11324056 | 0.68151616 |
| NACC1 | 3838.35243 | -0.214866 | 0.17972557 | -1.1955229 | 0.2318828 | 0.80649152 |
| AC011446.1 | 694.291962 | -0.1877601 | 0.21191095 | -0.8860332 | 0.37559965 | 0.86889275 |
| STX10 | 2163.11695 | -0.1866242 | 0.18229625 | -1.0237414 | 0.30595745 | 0.84270033 |
| IER2 | 5234.59402 | -0.1013971 | 0.29703049 | -0.3413693 | 0.73282561 | 0.95848028 |
| AC011446.2 | 2783.69849 | -0.1302305 | 0.33675709 | -0.3867194 | 0.69896398 | 0.95299571 |
| CACNA1A | 61.2165086 | -0.3800057 | 0.50279935 | -0.7557801 | 0.44978106 | 0.89557948 |
| CCDC130 | 3092.43138 | -0.2260788 | 0.09720027 | -2.3259074 | 0.02002349 | 0.49637604 |

| | | | | | | |
|---|---|---|---|---|---|---|
| MRI1 | 2862.46671 | 0.05068797 | 0.24458973 | 0.20723669 | 0.83582502 | 0.9755977 |
| AC008686.1 | 1630.30953 | 0.09353775 | 0.2529277 | 0.3698201 | 0.71151654 | 0.95560193 |
| C19orf53 | 2717.05245 | -0.0296759 | 0.12564573 | -0.236187 | 8.13E-01 | 0.9738447 |
| ZSWIM4 | 1332.97591 | -0.3822427 | 0.29951321 | -1.2762133 | 0.20188014 | 0.78134034 |
| AC020916.1 | 1426.12087 | -0.1840857 | 0.46797374 | -0.3933676 | 0.69404798 | 0.95222846 |
| C19orf57 | 215.365279 | -0.1767945 | 0.16225312 | -1.0896217 | 0.27587982 | 0.82719407 |
| CC2D1A | 4117.35031 | -0.2592308 | 0.14618668 | -1.7732857 | 0.07618137 | 0.63773415 |
| PODNL1 | 61.7881154 | 0.14951148 | 0.4007801 | 0.37305116 | 0.70911037 | 0.95501548 |
| DCAF15 | 1814.73624 | -0.2508147 | 0.20812422 | -1.2051201 | 0.22815695 | 0.80300331 |
| RFX1 | 1471.2577 | -0.2926634 | 0.12890011 | -2.2704666 | 0.02317929 | 0.51967919 |
| IL27RA | 579.293033 | 0.28668104 | 0.47498513 | 0.60355794 | 0.54613758 | 0.92345821 |
| PALM3 | 3826.64224 | 0.01519872 | 0.50579722 | 0.03004904 | 0.97602794 | 0.99455099 |
| MISP3 | 1831.09475 | -0.0448363 | 0.26650803 | -0.1682361 | 0.86639752 | 0.98204644 |
| SAMD1 | 1811.77045 | -0.2550021 | 0.31092828 | -0.8201314 | 0.41214119 | 0.88268208 |
| PRKACA | 4234.21882 | -0.0803513 | 0.19745414 | -0.4069366 | 0.68405456 | 0.95070301 |
| ASF1B | 92.3998124 | -0.0842749 | 0.58853958 | -0.1431932 | 0.88613758 | 0.98391254 |
| AC022098.1 | 392.534435 | -0.0920519 | 0.20853423 | -0.4414234 | 0.65890649 | 0.94596036 |
| ADGRL1 | 2235.1927 | 0.01315619 | 0.36370223 | 0.03617298 | 0.97114443 | 0.99400331 |
| ADGRE5 | 3142.82436 | -0.5131649 | 0.4680103 | -1.096482 | 0.2728679 | 0.82716979 |
| AC008569.1 | 592.951102 | -0.4195798 | 0.45897967 | -0.9141576 | 0.36063404 | 0.86504879 |
| DDX39A | 3594.43129 | -1.042025 | 0.34827286 | -2.9919786 | 2.77E-03 | 0.2278311 |
| PKN1 | 6014.33057 | -0.1420639 | 0.16721497 | -0.8495886 | 3.96E-01 | 8.79E-01 |
| PTGER1 | 381.674543 | 0.28398197 | 0.58033149 | 0.48934441 | 0.62459789 | 0.94088547 |
| GIPC1 | 5539.80422 | 0.01795473 | 0.20460551 | 0.08775292 | 0.93007306 | 0.98973492 |
| DNAJB1 | 8738.1576 | -0.416548 | 0.21795874 | -1.9111322 | 0.05598759 | 0.59889467 |
| TECR | 6218.77948 | 0.07094742 | 0.11975432 | 0.59244139 | 0.55355506 | 0.92373235 |
| NDUFB7 | 4658.24749 | 0.10524672 | 0.15965951 | 0.65919481 | 0.50977068 | 0.91439937 |
| ZNF333 | 831.744459 | 0.17586825 | 0.32526049 | 0.54069969 | 0.58871459 | 0.93289552 |
| ADGRE2 | 531.133558 | 1.33594161 | 0.63938555 | 2.08941477 | 0.0366704 | 0.55521179 |
| CASP14 | 48.4407944 | -0.9142625 | 1.26496053 | -0.7227597 | NA | NA |
| SYDE1 | 1762.0836 | -0.1617807 | 0.28181897 | -0.574059 | 0.56592787 | 0.92532318 |
| ILVBL | 6560.83379 | 0.25353414 | 0.16945474 | 1.49617613 | 0.13460776 | 0.71630293 |
| NOTCH3 | 6340.23573 | -0.5847555 | 0.43942918 | -1.3307161 | 0.18328243 | 0.76723162 |
| EPHX3 | 157.703888 | 0.02988092 | 0.39472942 | 0.07569974 | 0.93965798 | 0.99134327 |
| BRD4 | 5309.97486 | -0.4324583 | 0.14340585 | -3.0156255 | 0.0025645 | 0.21645768 |
| AKAP8 | 2591.37892 | -0.3009642 | 0.1319673 | -2.2805964 | 0.02257234 | 0.51958567 |
| AKAP8L | 8586.7578 | -0.3732454 | 0.28080871 | -1.3291802 | 0.18378852 | 0.76781833 |
| WIZ | 3846.96643 | -0.4071698 | 0.15476788 | -2.6308417 | 0.00851737 | 0.3779007 |
| RASAL3 | 677.844449 | -0.3341327 | 0.62157933 | -0.5375544 | 0.59088469 | 0.93289552 |
| CYP4F3 | 4454.35212 | 0.13989347 | 0.82850296 | 0.1688509 | 0.86591392 | 0.98197545 |
| CYP4F12 | 414.219853 | 1.32125311 | 0.63608347 | 2.07716939 | 0.03778593 | 0.55538555 |
| CYP4F2 | 2315.44107 | 2.05746028 | 0.87290202 | 2.35703461 | 0.01842153 | 0.47570049 |
| CYP4F11 | 973.640675 | 0.62871115 | 0.90413508 | 0.69537303 | 0.48682156 | 0.90573694 |
| TPM4 | 13170.3141 | 0.16024992 | 0.36201326 | 0.44266311 | 0.65800943 | 0.9458503 |
| RAB8A | 3235.78947 | -0.114907 | 0.16111644 | -0.7131922 | 0.47572684 | 0.90257104 |
| AC008894.2 | 2509.1628 | -0.1260237 | 0.16482149 | -0.7646074 | 0.44450537 | 0.8938278 |
| HSH2D | 536.978195 | -1.3979886 | 0.84741474 | -1.6497101 | NA | NA |
| FAM32A | 3615.49761 | -0.3156555 | 0.21352435 | -1.4783113 | 0.13932448 | 0.72409321 |
| AP1M1 | 4213.93518 | 0.00547217 | 0.22295398 | 0.02454394 | 0.98041873 | 0.99572034 |
| KLF2 | 1579.8949 | -0.1790131 | 0.4166072 | -0.4296928 | 0.66741909 | 0.94830184 |
| EPS15L1 | 1999.74069 | 0.01069531 | 0.19823712 | 0.05395213 | 0.9569733 | 0.99200096 |
| AC008764.1 | 805.88525 | -0.346055 | 0.25557427 | -1.3540293 | 0.17572704 | 0.7610139 |
| C19orf44 | 2128.97272 | -0.2522456 | 0.11663518 | -2.1626884 | 0.03056515 | 0.54443002 |
| CHERP | 3474.73493 | -0.2818879 | 0.16648105 | -1.6932132 | 0.09041489 | 0.6544865 |

| | | | | | | |
|---|---|---|---|---|---|---|
| AC008764.6 | 90.0779638 | 0.3627311 | 0.30223791 | 1.20015091 | 0.23008074 | 0.80551898 |
| SLC35E1 | 2835.4128 | -0.1350444 | 0.21559683 | -0.6263747 | 0.53106917 | 0.91962754 |
| AC008764.3 | 517.385553 | -0.1129794 | 0.23792039 | -0.4748623 | 0.63488509 | 0.94088547 |
| AC008764.8 | 293.156442 | -0.143991 | 0.21095923 | -0.6825538 | 0.49488882 | 0.90816709 |
| MED26 | 613.497527 | -0.3957015 | 0.18859836 | -2.0981172 | 0.0358948 | 0.55369025 |
| AC008764.4 | 1300.85584 | 0.06083962 | 0.1527683 | 0.39824766 | 0.69044763 | 0.95158502 |
| AC008764.2 | 71.4663229 | 0.04778863 | 0.3879309 | 0.12318852 | 0.90195782 | 0.98572048 |
| AC024075.3 | 233.538789 | -0.1648152 | 0.24732722 | -0.6663851 | 0.50516496 | 0.91365534 |
| SMIM7 | 2664.09388 | 0.06881412 | 0.13101603 | 0.52523436 | 0.59942028 | 0.93487648 |
| TMEM38A | 687.927081 | -0.2141854 | 0.40629149 | -0.5271718 | 0.59807428 | 0.9346978 |
| NWD1 | 45.1950631 | 0.03493607 | 1.01040844 | 0.03457619 | 0.97241769 | 0.99416401 |
| SIN3B | 4791.05992 | -0.3659698 | 0.1896907 | -1.9292976 | 0.05369392 | 0.59227651 |
| F2RL3 | 755.062095 | -0.7120971 | 0.43977152 | -1.6192432 | 0.10539494 | 0.67473222 |
| CPAMD8 | 2117.48474 | 0.7509761 | 0.32365528 | 2.32029617 | 0.02032486 | 0.5002876 |
| HAUS8 | 248.975351 | -0.2430469 | 0.21031122 | -1.1556534 | 0.24782297 | 0.81602355 |
| MYO9B | 3497.88252 | -0.5435501 | 0.16480471 | -3.2981464 | 0.00097325 | 0.14829062 |
| AC020913.2 | 318.121532 | -0.5562171 | 0.17935591 | -3.1011919 | 0.00192743 | 0.18915498 |
| USE1 | 992.600239 | -0.1152829 | 0.23195276 | -0.4970101 | 0.61918191 | 0.93945324 |
| OCEL1 | 1750.0909 | 0.14649628 | 0.21407891 | 0.6843097 | 0.49377962 | 0.90751436 |
| NR2F6 | 2924.70125 | 0.13770097 | 0.23426754 | 0.58779361 | 0.55667083 | 0.92373235 |
| AC010646.1 | 534.944928 | 0.00031146 | 0.26139366 | 0.00119155 | 0.99904928 | 0.99994357 |
| USHBP1 | 704.759329 | -0.3311219 | 0.34610515 | -0.956709 | 0.33871415 | 0.85733087 |
| BABAM1 | 3323.9171 | 0.03800616 | 0.09146068 | 0.41554644 | 0.67774193 | 0.95018034 |
| AC010463.1 | 2370.50682 | 0.09869926 | 0.09793332 | 1.00782103 | 0.31354039 | 0.8464135 |
| ANKLE1 | 81.8652298 | -0.6329984 | 0.41784951 | -1.5148956 | 0.12979884 | 0.70814767 |
| ABHD8 | 979.275518 | 0.02143411 | 0.27001162 | 0.07938219 | 0.93672863 | 0.99068685 |
| MRPL34 | 3046.42941 | -0.0776646 | 0.25487081 | -0.3047214 | 0.76057834 | 0.96560687 |
| DDA1 | 2097.02225 | -0.2022925 | 0.12110822 | -1.6703448 | 0.09485116 | 0.65888357 |
| ANO8 | 1808.19267 | -0.2847757 | 0.35557006 | -0.8008991 | 0.42319005 | 0.8846631 |
| GTPBP3 | 1309.73999 | 0.04556352 | 0.1903517 | 0.2393649 | 0.81082264 | 0.97352497 |
| PLVAP | 11146.4474 | -0.1605702 | 0.54042818 | -0.2971167 | 0.76637743 | 0.9671838 |
| BST2 | 4787.84078 | -0.806815 | 0.56711422 | -1.4226675 | 0.15483255 | 0.74373109 |
| BISPR | 289.998648 | -1.1494452 | 0.62896163 | -1.8275283 | 0.06762037 | 0.62557548 |
| MVB12A | 1949.99577 | 0.04976466 | 0.25022525 | 0.19887946 | 0.84235704 | 0.97644404 |
| AC010319.4 | 115.722635 | -0.5594756 | 0.4537868 | -1.232904 | 0.21761159 | 0.79622389 |
| SLC27A1 | 1802.93426 | 0.20578508 | 0.26918672 | 0.76446966 | 0.44458741 | 0.89389686 |
| AC010618.3 | 549.77426 | 0.15444276 | 0.15275173 | 1.01107043 | 0.31198272 | 0.84541735 |
| PGLS | 2351.87956 | -0.0059106 | 0.09220588 | -0.0641022 | 0.94888884 | 0.99170503 |
| FAM129C | 77.0334144 | 0.11268758 | 0.98991808 | 0.11383526 | NA | NA |
| COLGALT1 | 3848.19749 | -0.4188841 | 0.19496282 | -2.1485332 | 0.03167142 | 0.54507414 |
| UNC13A | 67.1677302 | 0.68668124 | 0.68606243 | 1.00090198 | 0.3168742 | 0.84812045 |
| MAP1S | 1880.89291 | -0.4572469 | 0.17925333 | -2.5508416 | 0.01074632 | 0.40196898 |
| AC008761.1 | 511.661387 | -0.4133647 | 0.18447861 | -2.2407189 | 0.02504429 | 0.52085426 |
| FCHO1 | 134.300477 | 0.08288986 | 0.59103536 | 0.14024519 | 0.88846627 | 0.98423722 |
| B3GNT3 | 974.07713 | -0.1718234 | 0.31253518 | -0.5497731 | 0.58247499 | 0.93177403 |
| JAK3 | 1196.6362 | 0.23577388 | 0.54313374 | 0.43409913 | 0.66421646 | 0.94655191 |
| RPL18A | 14375.3827 | 0.01778162 | 0.18790573 | 0.09463053 | 0.9246083 | 0.98869145 |
| CCDC124 | 2462.06813 | -0.1799434 | 0.17669334 | -1.0183936 | 0.30849095 | 0.84388103 |
| ARRDC2 | 2779.8086 | -0.133862 | 0.33334812 | -0.4015683 | 0.68800176 | 0.95158502 |
| IL12RB1 | 125.88355 | -0.4365577 | 0.58280231 | -0.7490666 | 0.45381707 | 0.89670027 |
| MAST3 | 1951.48143 | 0.30833028 | 0.36186132 | 0.85206753 | 0.39417661 | 0.87884163 |
| AC007192.2 | 483.777556 | 0.4377715 | 0.33177884 | 1.31946779 | 0.18701277 | 0.76958556 |
| PIK3R2 | 2769.42265 | 0.03743836 | 0.14839942 | 0.25228106 | 0.80082383 | 0.97210205 |
| AC007192.1 | 9033.60596 | -0.3266795 | 0.24039006 | -1.358956 | 1.74E-01 | 0.75974924 |

| | | | | | | |
|---|---|---|---|---|---|---|
| IFI30 | 6335.63665 | -0.4610111 | 0.34148608 | -1.3500144 | 0.17701135 | 0.76115956 |
| MPV17L2 | 824.748334 | 0.17939108 | 0.229725 | 0.78089491 | 0.43486431 | 0.89055338 |
| RAB3A | 305.237269 | 0.63779576 | 0.42019323 | 1.51786301 | 0.12904894 | 0.70675223 |
| AC005759.1 | 152.457828 | 0.78945329 | 0.43149029 | 1.8295969 | 0.06731024 | 0.6249306 |
| AC008397.2 | 405.387259 | 0.86910435 | 0.4403554 | 1.973643 | 0.04842235 | 0.58440502 |
| PDE4C | 363.352971 | 0.75897652 | 0.49509703 | 1.53298539 | 1.25E-01 | 7.03E-01 |
| IQCN | 539.122855 | 0.83671658 | 0.49988036 | 1.67383367 | 0.09416329 | 0.65783452 |
| JUND | 7291.53211 | -0.6661046 | 0.23774363 | -2.8017769 | 0.0050822 | 0.30823792 |
| LSM4 | 3980.16497 | 0.34409819 | 0.2104018 | 1.63543371 | 0.10195816 | 0.67279543 |
| PGPEP1 | 8155.0131 | 0.6580884 | 0.32809009 | 2.00581616 | 0.04487587 | 0.57322281 |
| GDF15 | 15105.5404 | -0.2011422 | 0.47609195 | -0.422486 | 0.67267035 | 0.94922827 |
| LRRC25 | 284.072322 | 0.13625013 | 0.53001742 | 0.25706726 | 0.79712685 | 0.9713715 |
| SSBP4 | 2838.56761 | -0.1188801 | 0.30561418 | -0.3889874 | 0.69728545 | 0.95269909 |
| ISYNA1 | 4500.61292 | 0.05250927 | 0.3619322 | 0.14508042 | 0.88464738 | 0.98340286 |
| AC010335.1 | 595.359186 | -0.0677842 | 0.24121351 | -0.2810131 | 0.77870033 | 0.96836854 |
| ELL | 2623.77548 | -0.1354586 | 0.1893403 | -0.7154238 | 0.47434719 | 0.90199039 |
| AC005387.1 | 91.9838898 | 0.03961433 | 0.31871112 | 0.12429541 | 0.90108138 | 0.9855364 |
| FKBP8 | 13393.4899 | 0.03574206 | 0.1669092 | 0.21414072 | 0.83043732 | 0.97550416 |
| AC005387.2 | 44.7258954 | -0.3985408 | 0.34778958 | -1.1459251 | 0.25182613 | 0.81833349 |
| KXD1 | 3461.71332 | -0.101046 | 0.10659982 | -0.9478998 | 3.43E-01 | 8.59E-01 |
| AC005253.1 | 500.834144 | -0.0672072 | 0.15346817 | -0.437923 | 6.61E-01 | 9.46E-01 |
| AC005253.2 | 1457.91727 | -0.1007625 | 0.13121164 | -0.7679387 | 4.43E-01 | 8.93E-01 |
| UBA52 | 13281.0017 | 0.06344014 | 0.21682863 | 0.29258191 | 7.70E-01 | 9.68E-01 |
| CRLF1 | 636.87121 | -0.1591677 | 0.3585262 | -0.4439501 | 0.65707866 | 0.94545756 |
| REX1BD | 1589.4042 | 0.25805062 | 0.26245214 | 0.98322925 | 0.32549462 | 0.85244476 |
| TMEM59L | 206.534 | -0.0079625 | 0.81763798 | -0.0097384 | 9.92E-01 | 9.99E-01 |
| KLHL26 | 802.254645 | 0.01700104 | 0.25750992 | 0.06602093 | 0.94736116 | 0.99166614 |
| CRTC1 | 2310.39744 | -0.1223994 | 0.24703683 | -0.4954701 | 0.6202683 | 0.93985162 |
| COMP | 344.976123 | 1.04866711 | 0.70323282 | 1.491209 | 0.13590663 | 0.71771311 |
| UPF1 | 7058.47052 | -0.2434659 | 0.14265921 | -1.7066259 | 0.08789161 | 0.65363785 |
| CERS1 | 44.8925568 | -0.096017 | 0.46464028 | -0.2066481 | 0.83628472 | 0.97577999 |
| COPE | 6857.12282 | -0.0780189 | 0.16183909 | -0.4820772 | 0.62975107 | 0.94088547 |
| AC002985.1 | 5342.78514 | -0.1060278 | 0.24094631 | -0.4400475 | 0.6599027 | 0.94626603 |
| DDX49 | 1835.17604 | -0.1711679 | 0.21244973 | -0.8056867 | 0.42042351 | 0.88451372 |
| HOMER3 | 3572.44784 | -0.2323085 | 0.39852607 | -0.5829193 | 0.55994762 | 0.92434754 |
| HOMER3-AS | 99.8929494 | -0.4504172 | 0.53071771 | -0.8486945 | 0.3960513 | 0.87967259 |
| SUGP2 | 6058.71629 | -0.1516613 | 0.13097166 | -1.15797 | 0.24687629 | 0.81542272 |
| ARMC6 | 1562.36002 | -0.3330093 | 0.22004462 | -1.5133718 | 0.13018525 | 0.70814767 |
| SLC25A42 | 4829.00103 | 0.8445555 | 0.57209548 | 1.47624921 | 0.139877 | 0.72475166 |
| TMEM161A | 1977.9197 | 0.1618565 | 0.20105457 | 0.80503768 | 4.21E-01 | 8.85E-01 |
| BORCS8-MEI | 1109.99134 | -0.1855416 | 0.12643668 | -1.4674664 | 0.14224923 | 0.72784886 |
| MEF2B | 456.425846 | -0.2955571 | 0.15705509 | -1.8818688 | 0.05985384 | 0.60732038 |
| BORCS8 | 1906.31352 | -0.0723533 | 0.23561171 | -0.3070868 | 0.75877728 | 0.96502715 |
| RFXANK | 2266.06913 | -0.0362673 | 0.12637469 | -0.2869824 | 0.77412583 | 0.96836854 |
| NR2C2AP | 1217.2023 | -0.1393636 | 0.13371931 | -1.0422096 | 0.2973145 | 0.83898658 |
| HAPLN4 | 69.5504392 | 0.91860368 | 0.85581197 | 1.07337092 | 0.28310473 | 0.83078187 |
| TM6SF2 | 513.261775 | 0.65010754 | 0.59283248 | 1.09661256 | 0.2728108 | 0.82716979 |
| SUGP1 | 1904.16566 | -0.077314 | 0.12511491 | -0.6179438 | 5.37E-01 | 0.92105614 |
| MAU2 | 3837.18285 | -0.2386897 | 0.16940168 | -1.4090161 | 0.15883039 | 0.74706632 |
| GATAD2A | 4743.35039 | -0.3951791 | 0.1987124 | -1.9886989 | 0.04673444 | 0.58080251 |
| TSSK6 | 92.7944323 | 0.20904881 | 0.37493184 | 0.55756483 | 0.57714158 | 0.93130183 |
| NDUFA13 | 7923.00899 | 0.02735406 | 0.09706864 | 0.28180124 | 0.77809592 | 0.96836854 |
| AC011448.1 | 8272.05822 | 0.01517165 | 0.09812617 | 0.15461365 | 8.77E-01 | 0.98275231 |
| YJEFN3 | 1157.0944 | -0.4116301 | 0.31527031 | -1.3056418 | 0.19167441 | 0.77347283 |

| | | | | | | |
|---|---|---|---|---|---|---|
| PBX4 | 197.358164 | -0.0760701 | 0.40303583 | -0.1887427 | 0.85029446 | 0.97816532 |
| LPAR2 | 508.047584 | -0.0595886 | 0.31323641 | -0.1902352 | 8.49E-01 | 9.77E-01 |
| GMIP | 816.593551 | -0.4473265 | 0.31630033 | -1.4142461 | 0.15728966 | 0.74522306 |
| ATP13A1 | 5294.77634 | -0.3479476 | 0.09223769 | -3.7722933 | 0.00016175 | 0.05980058 |
| ZNF101 | 612.403124 | -0.077441 | 0.20559659 | -0.3766646 | 0.70642287 | 0.95501548 |
| ZNF14 | 329.687874 | 0.14588742 | 0.13993066 | 1.04256936 | 0.29714779 | 0.83898658 |
| LINC00663 | 335.633395 | 0.30501707 | 0.31409517 | 0.9710976 | 0.33149968 | 0.85408791 |
| AC011477.4 | 97.0753066 | 0.54038874 | 0.2863556 | 1.88712477 | 0.05914355 | 0.60462654 |
| ZNF506 | 1129.17587 | 0.1467591 | 0.16712847 | 0.87812149 | 0.37987779 | 0.87090707 |
| ZNF253 | 405.545355 | 0.15679128 | 0.2259028 | 0.69406522 | 0.48764131 | 0.90573694 |
| AC011477.3 | 73.6637136 | 0.39342272 | 0.30636735 | 1.28415355 | 0.19908823 | 0.78134034 |
| ZNF93 | 164.800575 | 0.00337081 | 0.27335969 | 0.01233105 | 0.9901615 | 0.99809685 |
| ZNF682 | 320.930135 | 0.0527579 | 0.35103119 | 0.15029404 | 0.88053263 | 0.98275231 |
| ZNF90 | 225.499908 | 0.09973127 | 0.27456498 | 0.36323374 | 7.16E-01 | 9.56E-01 |
| AC011447.3 | 109.420445 | 0.07274168 | 0.25735636 | 0.28264962 | 0.77744544 | 0.96836854 |
| ZNF486 | 783.646081 | -0.0404236 | 0.38303381 | -0.1055354 | 0.91595095 | 0.9868044 |
| AC008554.1 | 38.1844871 | 0.76755922 | 0.73853369 | 1.03930156 | 0.29866451 | 0.84056272 |
| ZNF737 | 298.761024 | 0.22564494 | 0.26944141 | 0.83745456 | 0.40233711 | 0.88132015 |
| AC010636.2 | 137.397635 | 0.66295012 | 0.30047316 | 2.2063539 | 0.02735923 | 0.52904542 |
| ZNF626 | 548.551209 | 0.51663327 | 0.1767294 | 2.92330122 | 0.00346341 | 0.25757071 |
| AC010329.1 | 148.583945 | 0.26498965 | 0.37060157 | 0.71502571 | 0.47459316 | 0.90224694 |
| ZNF85 | 170.598326 | 0.25967676 | 0.35553265 | 0.73038794 | 0.46515309 | 0.89921277 |
| ZNF430 | 277.651475 | -0.4106325 | 0.20794207 | -1.9747446 | 0.04829714 | 0.58440502 |
| ZNF714 | 111.281968 | -0.3958037 | 0.45099061 | -0.8776319 | 0.38014352 | 0.87090707 |
| ZNF431 | 928.03909 | -0.0701977 | 0.20274961 | -0.3462287 | 0.72917085 | 0.95848028 |
| ZNF708 | 257.461734 | -0.1502638 | 0.21646714 | -0.6941645 | 0.48757905 | 0.90573694 |
| ZNF738 | 249.67911 | -0.2682964 | 0.37451739 | -0.7163791 | 0.47375728 | 0.90199039 |
| ZNF493 | 868.966098 | -0.1889693 | 0.17714942 | -1.0667225 | 0.28609716 | 0.83107779 |
| ZNF429 | 325.603715 | 0.34253743 | 0.27590055 | 1.24152502 | 2.14E-01 | 7.94E-01 |
| AC123912.4 | 88.628121 | 0.84966071 | 0.57511993 | 1.47736268 | 0.13957844 | 0.72475166 |
| ZNF100 | 362.466123 | -0.0766884 | 0.36577147 | -0.209662 | 0.83393149 | 0.97550416 |
| ZNF43 | 644.02197 | 0.122177 | 0.29338813 | 0.41643472 | 0.67709192 | 0.95015947 |
| ZNF208 | 102.82944 | -0.3216591 | 0.4238964 | -0.7588153 | 0.44796304 | 0.89480207 |
| ZNF257 | 67.2799577 | -0.4670628 | 0.42509592 | -1.0987234 | 0.27188874 | 0.82716979 |
| ZNF676 | 101.868495 | -0.0515784 | 0.43409098 | -0.1188193 | 0.90541854 | 0.98625102 |
| ZNF98 | 55.1364533 | -0.1272937 | 0.46408454 | -0.2742898 | 0.7838619 | 0.96978847 |
| AC092329.3 | 127.022636 | -0.0535921 | 0.30835603 | -0.1737995 | 0.86202307 | 0.98176973 |
| AC092329.1 | 156.260141 | 0.08905911 | 0.2692742 | 0.33073763 | 0.74084267 | 0.96089108 |
| ZNF91 | 1205.03097 | -0.2617382 | 0.18012033 | -1.4531299 | 0.1461877 | 0.73247558 |
| AC010300.1 | 58.6458201 | 0.73669297 | 0.47776752 | 1.54194861 | 0.12308608 | 0.69788192 |
| ZNF675 | 276.370601 | -0.1420834 | 0.25482413 | -0.5575745 | 0.57713497 | 0.93130183 |
| ZNF681 | 106.557636 | -0.0887308 | 0.32085402 | -0.2765457 | 0.78212895 | 0.96943642 |
| AC139769.2 | 37.5301771 | -2.4018752 | 0.73247774 | -3.2791102 | 0.00104135 | 0.15012259 |
| ZNF726 | 129.611105 | 0.06130873 | 0.45807049 | 0.13384125 | 0.89352811 | 0.98474725 |
| AC011503.1 | 40.9392692 | -0.0752512 | 0.53256142 | -0.1413006 | 0.88763249 | 0.98423722 |
| ZNF254 | 902.435281 | -0.0841856 | 0.1440287 | -0.5845057 | 0.55888013 | 0.92421175 |
| AC092279.1 | 217.58642 | -1.0281625 | 0.47415877 | -2.1683928 | 0.03012881 | 0.54012717 |
| LINC00662 | 401.915728 | -0.1564612 | 0.17133811 | -0.9131721 | 0.36115205 | 0.86528137 |
| AC006504.1 | 63.7962044 | -1.7106266 | 0.61529662 | -2.7801657 | 0.00543312 | 0.31359184 |
| AC006504.5 | 429.42562 | -0.2127083 | 0.23746608 | -0.8957419 | 0.37039065 | 0.86762749 |
| LINC01532 | 75.9125448 | 0.44062737 | 0.45752576 | 0.96306569 | 0.33551456 | 0.85616588 |
| UQCRFS1 | 7905.66059 | 0.22829596 | 0.21384546 | 1.06757447 | 0.28571251 | 0.83107779 |
| POP4 | 1471.64215 | 0.03836305 | 0.21889098 | 0.17526101 | 0.86087455 | 0.98124619 |
| PLEKHF1 | 451.693261 | -0.03016 | 0.19750489 | -0.1527052 | 0.87863073 | 0.98275231 |

| | | | | | | |
|---|---|---|---|---|---|---|
| C19orf12 | 2138.17351 | 0.11161044 | 0.28037396 | 0.39807704 | 0.6905734 | 0.95158502 |
| CCNE1 | 88.7809704 | -0.1695179 | 0.29309066 | -0.5783806 | 0.56300721 | 0.92452095 |
| URI1 | 2793.5653 | -0.1055569 | 0.15784412 | -0.6687415 | 0.50366039 | 0.91356552 |
| TSHZ3 | 395.515421 | 0.0665045 | 0.39441343 | 0.16861621 | 0.86609853 | 0.98197545 |
| ZNF507 | 806.714296 | -0.1645128 | 0.16332097 | -1.0072973 | 0.31379192 | 0.84646403 |
| DPY19L3 | 1406.49019 | -0.0564112 | 0.15796801 | -0.3571049 | 0.72101325 | 0.95729223 |
| PDCD5 | 2448.62994 | -0.0615297 | 0.13543958 | -0.454296 | 0.64961581 | 0.94312988 |
| ANKRD27 | 1571.07734 | -0.2052919 | 0.15050965 | -1.3639785 | 0.17257434 | 0.75974924 |
| AC008736.1 | 43.1386553 | 0.27303711 | 0.3767102 | 0.72479352 | 0.46857871 | 0.89984974 |
| NUDT19 | 858.646053 | -0.0267075 | 0.16084239 | -0.1660474 | 0.86811962 | 0.98237277 |
| SLC7A9 | 3167.7575 | 1.67515106 | 0.5398504 | 3.10299121 | 0.00191575 | 0.18915498 |
| AC008805.2 | 516.113818 | 1.36865429 | 0.51627757 | 2.65100476 | 8.03E-03 | 0.37112203 |
| CEP89 | 1319.10363 | -0.155931 | 0.14521995 | -1.0737576 | 0.28293135 | 0.83078187 |
| FAAP24 | 62.3228197 | -0.5623336 | 0.47088227 | -1.1942126 | 0.2323948 | 0.80649152 |
| RHPN2 | 3092.80239 | 0.01720765 | 0.25098532 | 0.06856038 | 0.94533956 | 0.99166614 |
| GPATCH1 | 506.287953 | -0.0282093 | 0.22446857 | -0.1256714 | 9.00E-01 | 9.85E-01 |
| WDR88 | 52.8299107 | 0.15273384 | 0.5352644 | 0.28534279 | 7.75E-01 | 9.68E-01 |
| LRP3 | 1295.84833 | -0.1425081 | 0.36118747 | -0.3945544 | 0.69317176 | 0.9519683 |
| SLC7A10 | 51.0245703 | 1.24503439 | 0.77175498 | 1.61325087 | 0.10669002 | 0.67674128 |
| CEBPA | 558.692912 | 0.99959007 | 0.51464911 | 1.94227494 | 0.05210382 | 0.58645127 |
| AC008738.3 | 319.657491 | 0.96531637 | 0.52822946 | 1.82745651 | 0.06763116 | 0.62557548 |
| AC008738.5 | 92.196448 | 1.05380406 | 0.58905581 | 1.7889715 | 0.07361941 | 0.6374448 |
| CEBPA-DT | 57.9317694 | 0.40267442 | 0.58571381 | 0.68749347 | 0.49177182 | 0.90613466 |
| CEBPG | 1752.88395 | -0.268912 | 0.34560436 | -0.7780921 | 0.43651473 | 0.89104919 |
| PEPD | 21233.86 | 0.46088961 | 0.39465463 | 1.16783023 | 0.24287525 | 0.8133895 |
| KCTD15 | 1488.088 | -0.1273168 | 0.35176866 | -0.3619335 | 0.71740176 | 0.95611001 |
| AC008556.1 | 111.395814 | 0.0378909 | 0.5062463 | 0.07484678 | 0.94033662 | 0.99136103 |
| LSM14A | 5737.21064 | -0.2633725 | 0.15492786 | -1.6999686 | 0.08913683 | 0.65375543 |
| KIAA0355 | 1762.43721 | -0.165689 | 0.17183293 | -0.9642446 | 0.33492333 | 0.85581042 |
| GPI | 23257.1491 | 0.03075556 | 0.20933972 | 0.14691699 | 0.88319754 | 0.98293314 |
| AC092073.1 | 5047.64803 | 0.08747439 | 0.21402849 | 0.40870442 | 0.68275659 | 0.95070301 |
| PDCD2L | 208.234898 | -0.1461967 | 0.19973916 | -0.7319381 | 0.46420635 | 0.89830907 |
| UBA2 | 3380.90485 | -0.1098462 | 0.14580504 | -0.7533771 | 0.45122335 | 0.89627735 |
| WTIP | 1131.51619 | 0.01631527 | 0.35570152 | 0.04586786 | 0.96341557 | 0.99320711 |
| SCGB1B2P | 64.2555113 | -0.1030096 | 0.409738 | -0.2514036 | 8.02E-01 | 0.97210205 |
| SCGB2B2 | 47.5815783 | -0.1962773 | 0.49222656 | -0.398754 | 0.69007451 | 0.95158502 |
| ZNF302 | 1929.93422 | -0.1645729 | 0.16612036 | -0.9906845 | 0.32183964 | 0.85119105 |
| AC020910.4 | 74.9915718 | -0.0221988 | 0.34018095 | -0.065256 | 0.94797017 | 0.99170503 |
| ZNF181 | 710.830803 | -0.0314507 | 0.13420842 | -0.2343426 | 0.81471905 | 0.9738447 |
| ZNF599 | 363.421127 | 0.30514986 | 0.21904007 | 1.39312345 | 0.16358246 | 0.75213419 |
| ZNF30 | 206.367189 | 0.23081297 | 0.19760253 | 1.16806688 | 0.24277979 | 0.8133895 |
| ZNF792 | 223.383026 | 0.24966787 | 0.31903954 | 0.78256089 | 0.43388501 | 0.89055338 |
| GRAMD1A | 4343.64576 | -0.3404801 | 0.23215857 | -1.4665844 | 0.14248915 | 0.72784886 |
| AC020907.4 | 124.849892 | -0.0665316 | 0.2886819 | -0.2304668 | 0.8177291 | 0.97438078 |
| SCN1B | 536.204732 | -0.0252778 | 0.23236126 | -0.1087866 | 0.91337175 | 0.98662757 |
| HPN | 21386.7744 | 0.36835352 | 0.36075037 | 1.02107594 | 0.30721846 | 0.84298407 |
| HPN-AS1 | 2967.17017 | 0.45293279 | 0.36745808 | 1.23261078 | 0.21772101 | 0.79646877 |
| FXYD3 | 234.852094 | 0.84380441 | 0.41995494 | 2.0092737 | 0.04450812 | 0.57267043 |
| LGI4 | 522.281219 | -1.4701621 | 0.77235908 | -1.9034696 | 0.05697929 | 0.60049913 |
| FXYD1 | 323.302388 | -0.1750387 | 0.60543624 | -0.2891116 | NA | NA |
| FXYD5 | 1061.36658 | 0.20073788 | 0.46452576 | 0.43213509 | 0.66564323 | 0.94743068 |
| LSR | 4516.672 | -0.3057805 | 0.23995945 | -1.2743007 | 0.20255688 | 0.78193335 |
| USF2 | 5119.8291 | -0.1496162 | 0.16598801 | -0.9013673 | 0.36739305 | 0.86683872 |
| CD22 | 174.099467 | -0.8232407 | 0.91477408 | -0.8999388 | 0.36815282 | 0.86759809 |

| | | | | | | |
|---|---|---|---|---|---|---|
| FFAR2 | 47.6211856 | -1.1149036 | 0.7032889 | -1.5852711 | 0.11290475 | 0.68124529 |
| DMKN | 2769.1405 | 0.1544503 | 0.28105608 | 0.54953552 | 0.58263799 | 0.93183296 |
| GAPDHS | 269.340237 | 0.23579143 | 0.38746058 | 0.60855592 | 0.54281883 | 0.92292765 |
| TMEM147-A: | 1820.73764 | 0.17894181 | 0.10054595 | 1.77970191 | 0.07512476 | 0.63773415 |
| TMEM147 | 3960.58271 | 0.06002991 | 0.17526337 | 0.34251258 | 0.73196518 | 0.95848028 |
| HAUS5 | 879.134322 | 0.17403913 | 0.33920088 | 0.51308573 | 0.60789135 | 0.93748023 |
| RBM42 | 3324.45109 | -0.0962087 | 0.17166049 | -0.5604595 | 0.57516608 | 0.93051727 |
| ETV2 | 56.3117452 | 0.40843997 | 0.32689889 | 1.24943824 | 0.21150483 | 0.79149017 |
| COX6B1 | 7406.37886 | 0.2581162 | 0.15851935 | 1.6282946 | 0.10346243 | 0.67439401 |
| ZBTB32 | 40.8752261 | -1.230224 | 0.6321322 | -1.9461498 | 0.05163675 | 0.58645127 |
| KMT2B | 5802.91873 | -0.4611756 | 0.17633937 | -2.6152732 | 0.00891561 | 0.38245454 |
| IGFLR1 | 1135.82005 | -0.0173452 | 0.3447448 | -0.0503132 | 0.95987283 | 0.99255196 |
| AD000671.2 | 2288.73449 | -0.0800225 | 0.28545013 | -0.280338 | 0.77921817 | 0.96836854 |
| U2AF1L4 | 1624.21023 | -0.1856532 | 0.26701328 | -0.6952958 | 0.48686992 | 0.90573694 |
| PSENEN | 3119.31223 | -0.2005242 | 0.1428893 | -1.4033536 | 0.16051143 | 0.74816787 |
| AD000671.1 | 2972.45738 | -0.1745818 | 0.14489876 | -1.2048536 | 0.22825982 | 0.80319515 |
| AD000671.3 | 607.776823 | -0.1702069 | 0.16700159 | -1.0191933 | 0.30811121 | 0.8437043 |
| LIN37 | 787.767623 | -0.2274228 | 0.17998875 | -1.2635391 | 0.20639551 | 0.78731087 |
| AC002398.2 | 1451.98356 | -0.5656059 | 0.5029699 | -1.1245323 | 0.26078729 | 0.82424998 |
| HSPB6 | 1734.60684 | -0.5979423 | 0.50982203 | -1.1728451 | 0.24085793 | 0.8124487 |
| PROSER3 | 1194.72051 | -0.4085927 | 0.17934699 | -2.2782244 | 0.02271321 | 0.51967919 |
| AC002398.1 | 265.326204 | -0.3737352 | 0.20784759 | -1.7981213 | 0.07215778 | 0.6321115 |
| ARHGAP33 | 1398.46185 | 0.51797271 | 0.47262315 | 1.09595288 | 0.2730994 | 0.82716979 |
| PRODH2 | 9089.83719 | 1.26829591 | 0.64307636 | 1.97223221 | 0.0485831 | 0.58440502 |
| NPHS1 | 2905.43866 | 0.46246228 | 1.17230333 | 0.39449028 | 0.6932191 | 0.9519683 |
| KIRREL2 | 84.6982641 | 0.42026514 | 0.7428791 | 0.56572481 | 0.57158087 | 0.92841021 |
| APLP1 | 233.616388 | 0.34185495 | 0.43741225 | 0.7815395 | 0.43448526 | 0.89055338 |
| NFKBID | 769.359675 | -0.2429263 | 0.41754915 | -0.5817909 | 0.56070756 | 0.9244521 |
| HCST | 284.343985 | -0.2987348 | 0.57351839 | -0.5208809 | 0.60244972 | 0.9355118 |
| TYROBP | 1022.67882 | -0.3588853 | 0.54506725 | -0.658424 | 0.51026573 | 0.91439937 |
| LRFN3 | 731.769436 | -0.3184164 | 0.22585333 | -1.4098371 | 0.15858778 | 0.74675717 |
| SDHAF1 | 708.102843 | -0.0745732 | 0.14449212 | -0.5161053 | 0.60578084 | 0.93698713 |
| SYNE4 | 880.729182 | -0.0891529 | 0.21268984 | -0.4191684 | 0.67509303 | 0.94948732 |
| ALKBH6 | 851.361463 | -0.0690212 | 0.16164867 | -0.4269827 | 0.6693919 | 0.94876933 |
| AC002116.2 | 170.33738 | -0.1516074 | 0.30538078 | -0.4964536 | 0.61957438 | 0.93945324 |
| CLIP3 | 964.768136 | -0.097301 | 0.28311307 | -0.3436826 | 0.731085 | 0.95848028 |
| THAP8 | 388.964843 | 0.0999097 | 0.27221735 | 0.36702182 | 0.71360275 | 0.95564093 |
| WDR62 | 138.224745 | -0.7603343 | 0.43822902 | -1.7350159 | 0.082738 | 0.64680474 |
| POLR2I | 1283.54426 | -0.0183023 | 0.13697315 | -0.1336193 | 0.89370363 | 0.98474725 |
| TBCB | 3978.16966 | 0.03337742 | 0.15383324 | 0.21697144 | 0.8282306 | 0.97548201 |
| CAPNS1 | 20537.3389 | -0.0055469 | 0.16129568 | -0.0343897 | 0.97256638 | 0.99416401 |
| COX7A1 | 904.738655 | -0.2914398 | 0.42269288 | -0.6894835 | 0.49051904 | 0.90600166 |
| ZNF565 | 181.513541 | -0.278563 | 0.22186253 | -1.255566 | 0.20927338 | 0.79086734 |
| ZNF146 | 3064.87369 | -0.2128121 | 0.17311944 | -1.2292788 | 0.21896728 | 0.79847731 |
| AC012617.1 | 161.300324 | -0.0172156 | 0.33857137 | -0.0508477 | 0.95944685 | 0.9924381 |
| LINC00665 | 1435.07206 | -0.0528932 | 0.2126322 | -0.2487545 | 0.80355069 | 0.97274243 |
| ZFP14 | 645.99463 | 0.28535817 | 0.21853621 | 1.30577065 | 0.19163056 | 0.77347283 |
| ZFP82 | 242.839015 | -0.2365057 | 0.24823995 | -0.9527304 | 0.34072667 | 0.85849463 |
| ZNF566 | 471.627065 | -0.279234 | 0.18612183 | -1.5002754 | 0.13354307 | 0.71505705 |
| AC092295.2 | 150.589842 | -0.346727 | 0.28014282 | -1.2376796 | 0.21583489 | 0.7954827 |
| ZNF260 | 823.518677 | -0.1370033 | 0.12583057 | -1.088792 | 0.27624563 | 0.82719407 |
| ZNF529 | 962.566432 | -0.1281263 | 0.20490587 | -0.6252933 | 0.53177856 | 0.91962754 |
| ZNF529-AS1 | 199.789033 | -0.0478844 | 0.26162932 | -0.1830239 | 0.85477923 | 0.97910552 |
| ZNF382 | 277.443419 | 0.06957049 | 0.25227248 | 0.27577518 | 0.78272075 | 0.96964828 |

| | | | | | | |
|---|---|---|---|---|---|---|
| ZNF461 | 304.269342 | -0.1634854 | 0.19134751 | -0.8543901 | 0.39288888 | 0.8778303 |
| LINC01534 | 52.451016 | 0.47302165 | 0.3171803 | 1.49133364 | 0.13587392 | 0.71771311 |
| ZNF567 | 330.99011 | -0.3096184 | 0.20830409 | -1.4863772 | 0.13717937 | 0.72209298 |
| ZNF850 | 86.9162229 | -0.03429 | 0.29528831 | -0.1161236 | 0.90755456 | 0.98662757 |
| AC020928.1 | 204.464301 | -0.1007901 | 0.19801249 | -0.5090088 | 0.61074607 | 0.93807505 |
| ZNF790 | 361.718794 | 0.02256674 | 0.19435446 | 0.11611124 | 0.90756438 | 0.98662757 |
| ZNF345 | 333.317492 | -0.1590776 | 0.25606931 | -0.6212289 | 5.34E-01 | 0.92084006 |
| ZNF829 | 138.394167 | -0.1894778 | 0.34233401 | -0.5534881 | 0.57992921 | 0.93139743 |
| ZNF568 | 392.803785 | -0.1370246 | 0.22176472 | -0.6178829 | 0.53665254 | 0.92105614 |
| ZNF420 | 339.245012 | -0.2233069 | 0.19125224 | -1.1676041 | 0.24296649 | 0.81340905 |
| AC012309.1 | 252.694243 | 0.11783827 | 0.17545929 | 0.67159896 | 0.50183904 | 0.91259827 |
| ZNF585A | 567.300739 | 0.09417093 | 0.11316587 | 0.83214955 | 0.40532454 | 0.88132015 |
| AC012309.2 | 48.5987498 | -0.2709592 | 0.32997153 | -0.8211594 | 0.41155548 | 0.88268208 |
| ZNF585B | 1040.39463 | -0.0081219 | 0.17610605 | -0.0461193 | 0.96321517 | 0.99320711 |
| ZNF383 | 599.159719 | -0.0785721 | 0.15537314 | -0.5056993 | 0.61306773 | 0.93879706 |
| LINC01535 | 71.3279293 | -0.4950474 | 0.53842213 | -0.9194411 | 0.35786492 | 0.86432598 |
| AC016590.1 | 125.514683 | -0.1349056 | 0.46861915 | -0.2878789 | 0.77343942 | 0.96836854 |
| HKR1 | 3224.67231 | 0.01765435 | 0.11432597 | 0.15442118 | 8.77E-01 | 0.98275231 |
| AC016590.3 | 55.0070932 | 0.48291979 | 0.44875507 | 1.07613222 | 0.28186813 | 0.83001101 |
| ZNF527 | 302.881398 | 0.04639835 | 0.15447238 | 0.30036666 | 0.76389749 | 0.96678847 |
| ZNF569 | 287.646261 | 0.24753515 | 0.21686167 | 1.14144257 | 0.2536858 | 0.8188642 |
| ZNF570 | 339.432487 | -0.0132255 | 0.18383067 | -0.0719437 | 0.94264672 | 0.99136103 |
| ZNF793-AS1 | 137.247166 | 0.18169678 | 0.22108243 | 0.82185084 | 0.41116179 | 0.88268208 |
| ZNF793 | 502.216339 | 0.04583057 | 0.23306064 | 0.19664653 | 0.84410415 | 0.97664894 |
| ZNF571-AS1 | 90.3764367 | -0.0438904 | 0.40252199 | -0.1090386 | 0.91317185 | 0.98662757 |
| ZNF540 | 464.221204 | 0.62810005 | 0.32910775 | 1.90849368 | 0.05632744 | 0.5991104 |
| ZNF571 | 313.643304 | 0.15138723 | 0.16492589 | 0.91791067 | 0.35866563 | 0.86480228 |
| ZFP30 | 459.833295 | 0.18345151 | 0.1579361 | 1.16155528 | 0.24541616 | 0.81542272 |
| ZNF781 | 115.330373 | 0.58574191 | 0.41928409 | 1.39700488 | 0.1624121 | 0.75026273 |
| ZNF607 | 475.255635 | -0.1438562 | 0.20712007 | -0.6945548 | 0.48733435 | 0.90573694 |
| AC093227.1 | 195.28361 | 0.3525081 | 0.26815531 | 1.31456693 | 0.18865549 | 0.76970626 |
| ZNF573 | 60.3925898 | 0.15051351 | 0.27809552 | 0.54122953 | 0.58834939 | 0.93289552 |
| SIPA1L3 | 2543.45959 | -0.1686497 | 0.1778748 | -0.9481371 | 0.34305964 | 0.85929974 |
| SPINT2 | 16385.0466 | 0.00780273 | 0.15118755 | 0.05160959 | 0.95883978 | 0.99242674 |
| PPP1R14A | 712.595949 | -0.641497 | 0.62426565 | -1.0276026 | 0.30413679 | 0.84211658 |
| AC011479.1 | 6923.63302 | 0.04955961 | 0.16469483 | 0.30091782 | 0.76347716 | 0.96677875 |
| C19orf33 | 497.916036 | 0.01897938 | 0.54119758 | 0.03506922 | 0.97202454 | 0.99408319 |
| YIF1B | 1235.93734 | 0.04887816 | 0.22273276 | 0.21944757 | 0.82630142 | 0.97548201 |
| KCNK6 | 607.829679 | -0.1128121 | 0.31942658 | -0.3531708 | 0.72396038 | 0.95810643 |
| CATSPERG | 207.27743 | -0.1024278 | 0.37244169 | -0.2750171 | 0.78330311 | 0.96971192 |
| PSMD8 | 6978.86351 | 0.03127703 | 0.18453669 | 0.16948951 | 0.86541162 | 0.98197545 |
| GGN | 78.7199853 | -0.3343179 | 0.35023482 | -0.9545535 | 0.33980353 | 0.85802452 |
| SPRED3 | 389.120847 | -0.4431456 | 0.46917042 | -0.9445302 | 0.3448988 | 0.86022737 |
| FAM98C | 1009.08064 | 0.26050012 | 0.21566295 | 1.2079039 | 0.22708423 | 0.80300331 |
| RASGRP4 | 150.483355 | -0.1126547 | 0.39109989 | -0.288046 | 0.77331155 | 0.96836854 |
| RYR1 | 176.559854 | 0.03331182 | 0.56123365 | 0.05935465 | 0.95266964 | 0.99170503 |
| MAP4K1 | 290.48669 | 0.1506562 | 0.64393993 | 0.23396002 | 0.81501601 | 0.9738447 |
| AC008649.1 | 94.9289977 | 0.28524667 | 0.70547514 | 0.4043327 | 0.68596808 | 0.95097288 |
| EIF3K | 8668.8574 | 0.28506777 | 0.1430349 | 1.99299452 | 0.04626205 | 0.57915632 |
| ACTN4 | 34957.7781 | -0.0176093 | 0.20546688 | -0.0857038 | 0.93170191 | 0.98982066 |
| CAPN12 | 5547.8516 | 0.24219558 | 0.30070824 | 0.80541716 | 0.42057899 | 0.88451372 |
| AC022144.1 | 261.136781 | -0.0385829 | 0.4373441 | -0.088221 | 0.92970104 | 0.98969198 |
| LGALS4 | 203.233139 | -0.9686016 | 0.73342098 | -1.3206625 | 0.18661393 | 0.76915147 |
| ECH1 | 13010.4378 | 0.33602618 | 0.30829017 | 1.08996722 | 0.27572758 | 0.82719407 |

| | | | | | | |
|---|---|---|---|---|---|---|
| AC008982.1 | 11435.076 | 0.24819378 | 0.2142439 | 1.15846369 | 0.24667486 | 0.81542272 |
| HNRNPL | 10937.6946 | 0.0375748 | 0.16329234 | 0.23010754 | 0.81800821 | 0.97446554 |
| RINL | 500.264062 | 0.46781499 | 0.24220413 | 1.9314906 | 5.34E-02 | 0.59038182 |
| SIRT2 | 3815.86246 | 0.23892924 | 0.17880968 | 1.33622095 | 0.18147707 | 0.76522169 |
| NFKBIB | 2131.26672 | -0.1174698 | 0.27690189 | -0.4242291 | 0.67139874 | 0.94922827 |
| CCER2 | 41.5120294 | 0.34889146 | 0.4074096 | 0.85636533 | 0.39179573 | 0.87744531 |
| SARS2 | 1583.2715 | 0.05299193 | 0.19820007 | 0.26736585 | 0.78918749 | 0.97090793 |
| AC011455.2 | 2827.84603 | 0.0666635 | 0.20895519 | 0.3190325 | 0.74970187 | 0.96249332 |
| MRPS12 | 1829.45384 | 0.01460176 | 0.16144145 | 0.09044618 | 0.92793266 | 0.9892705 |
| ERVK9-11 | 379.157328 | 0.05001773 | 0.49085633 | 0.10189892 | 0.91883691 | 0.98709788 |
| FBXO17 | 4340.62587 | -0.0509994 | 0.35037533 | -0.1455566 | 0.88427146 | 0.98327586 |
| FBXO27 | 417.226153 | 0.24166596 | 0.28272092 | 0.85478624 | 0.39266949 | 0.8778303 |
| AC010605.1 | 72.7543279 | 0.34737527 | 0.39881403 | 0.87102069 | 0.38374286 | 0.8724645 |
| PAK4 | 4033.30776 | -0.0568222 | 0.20576449 | -0.2761515 | 0.78243168 | 0.96961926 |
| NCCRP1 | 77.197738 | 0.85458382 | 0.58738882 | 1.45488607 | 0.1457008 | 0.73205865 |
| LRFN1 | 65.5795052 | 0.32618784 | 0.49307985 | 0.66153147 | 0.50827154 | 0.91439937 |
| GMFG | 671.259141 | 0.04012394 | 0.32542175 | 0.12329828 | 0.9018709 | 0.98572048 |
| SAMD4B | 5502.27009 | -0.2224357 | 0.15038807 | -1.4790782 | 0.13911943 | 0.72409321 |
| PAF1 | 3473.59444 | -0.0500487 | 0.17366274 | -0.2881947 | 0.77319768 | 0.96836854 |
| MED29 | 3551.60445 | 0.17916626 | 0.19042737 | 0.94086399 | 0.34677456 | 0.86135217 |
| ZFP36 | 19343.4007 | -0.0932725 | 0.55138581 | -0.1691601 | 0.86567069 | 0.98197545 |
| PLEKHG2 | 2040.53918 | -0.4706949 | 0.37838145 | -1.2439693 | 0.21351088 | 0.7926732 |
| RPS16 | 21165.1263 | -0.0834697 | 0.2428262 | -0.3437427 | 0.73103983 | 0.95848028 |
| SUPT5H | 8659.82157 | -0.2867062 | 0.14291458 | -2.0061371 | 0.04484162 | 0.57322281 |
| TIMM50 | 2546.88489 | -0.0186808 | 0.16838868 | -0.1109386 | 0.91166503 | 0.98662757 |
| EID2B | 176.558168 | -0.1648716 | 0.18883508 | -0.8730986 | 0.38260936 | 0.8718674 |
| AC011500.3 | 57.5394757 | -0.2180793 | 0.26554436 | -0.8212536 | 0.41150183 | 0.88268208 |
| EID2 | 724.351692 | -0.1027836 | 0.20252182 | -0.5075186 | 0.61179097 | 0.93856019 |
| DYRK1B | 1667.02466 | -0.2849668 | 0.19436012 | -1.4661793 | 1.43E-01 | 7.28E-01 |
| FBL | 3178.99914 | 0.09990115 | 0.19452896 | 0.51355414 | 0.60756375 | 0.93739614 |
| FCGBP | 2329.1519 | -0.6397986 | 0.68764634 | -0.930418 | 0.35215469 | 0.86364633 |
| PSMC4 | 5074.06887 | -0.2270125 | 0.22760716 | -0.9973874 | 0.3185765 | 0.84904404 |
| ZNF546 | 417.567549 | 0.26865927 | 0.27480563 | 0.97763378 | 0.32825548 | 0.85317073 |
| ZNF780B | 989.314156 | -0.0666149 | 0.34276026 | -0.1943484 | 0.84590307 | 0.9767918 |
| ZNF780A | 629.077605 | -0.0954891 | 0.18285669 | -0.522207 | 0.60152618 | 0.93545781 |
| MAP3K10 | 1672.00207 | -0.3961729 | 0.16260632 | -2.4363932 | 0.01483455 | 0.44411232 |
| CNTD2 | 144.060257 | 0.9800294 | 0.65359536 | 1.49944364 | 0.13375858 | 0.71505705 |
| AKT2 | 10968.6413 | -0.0616271 | 0.13049845 | -0.4722438 | 0.63675281 | 0.94088547 |
| C19orf47 | 588.913183 | -0.5040513 | 0.21631172 | -2.3302081 | 0.01979516 | 0.49477123 |
| PLD3 | 9597.89453 | 0.11664506 | 0.15221796 | 0.76630287 | 0.44349611 | 0.89366134 |
| PRX | 898.593249 | 0.86326134 | 0.37711131 | 2.28914201 | 0.0220711 | 0.51639767 |
| SERTAD1 | 1383.69169 | -0.4460341 | 0.35007254 | -1.274119 | 0.20262124 | 0.78193335 |
| SERTAD3 | 1485.51918 | -0.1351705 | 0.1400658 | -0.9650502 | 0.33451969 | 0.85557672 |
| AC010271.1 | 64.1724783 | 0.10668858 | 0.28811335 | 0.37030072 | 0.71115844 | 0.95560193 |
| BLVRB | 4311.32474 | 0.36623841 | 0.28558733 | 1.28240427 | 0.19970087 | 0.78134034 |
| SPTBN4 | 206.789845 | 0.13526914 | 0.54613624 | 0.24768388 | 0.804379 | 0.97287947 |
| SHKBP1 | 3170.85229 | -0.1306234 | 0.14481706 | -0.9019891 | 0.36706263 | 0.86683872 |
| LTBP4 | 6929.6682 | 0.34530738 | 0.48505555 | 0.71189244 | 0.47653138 | 0.90309179 |
| NUMBL | 1182.19129 | -0.5093296 | 0.34148723 | -1.4915041 | 0.1358292 | 0.71771311 |
| COQ8B | 2457.28119 | -0.0504248 | 0.17717611 | -0.2846026 | 0.77594864 | 0.96836854 |
| ITPKC | 2792.50739 | -0.1624375 | 0.24823665 | -0.6543654 | 0.51287643 | 0.91568064 |
| C19orf54 | 1474.84328 | -0.1437901 | 0.14343905 | -1.0024472 | 0.31612767 | 0.84756299 |
| SNRPA | 2223.34012 | -0.0705171 | 0.13175519 | -0.535213 | 0.59250259 | 0.93289552 |
| MIA-RAB4B | 1741.84894 | 0.24041532 | 0.21309285 | 1.12821864 | 0.25922759 | 0.82364948 |

| | | | | | | |
|---|---|---|---|---|---|---|
| MIA | 41.6726943 | 0.28758358 | 0.60689396 | 0.47386134 | 0.63559879 | 0.94088547 |
| RAB4B | 1878.84268 | 0.24135888 | 0.21819641 | 1.10615422 | 0.26865977 | 0.82676682 |
| RAB4B-EGLN | 5905.70863 | -0.3091995 | 0.16723542 | -1.8488879 | 0.06447401 | 0.61989227 |
| AC008537.4 | 806.229416 | 0.1815509 | 0.22832178 | 0.79515367 | 0.42652412 | 0.88708118 |
| EGLN2 | 5121.93431 | -0.4722926 | 0.20510123 | -2.3027291 | 0.02129409 | 0.51189223 |
| AC008537.1 | 513.748854 | -0.5740138 | 0.23297764 | -2.4638149 | 0.01374671 | 0.428664 |
| CYP2B6 | 858.60617 | 0.84748526 | 0.84356787 | 1.00464384 | 0.31506838 | 0.84696964 |
| CYP2S1 | 138.669356 | 1.7094722 | 0.9013255 | 1.89662025 | 0.05787808 | 0.60129769 |
| AXL | 1667.28148 | -0.237649 | 0.38321168 | -0.6201508 | 0.5351585 | 0.92084006 |
| HNRNPUL1 | 8660.26815 | -0.0666539 | 0.19211692 | -0.3469444 | 0.72863308 | 0.95848028 |
| TGFB1 | 3971.24057 | 0.00329211 | 0.22288187 | 0.01477064 | 0.98821517 | 0.99769113 |
| CCDC97 | 1704.84507 | -0.4077061 | 0.19286811 | -2.1139112 | 0.03452286 | 0.55369025 |
| AC011462.1 | 3984.45123 | 0.40593457 | 0.24956536 | 1.62656616 | 0.10382927 | 0.67439401 |
| TMEM91 | 921.094092 | 0.14640817 | 0.29632486 | 0.49407996 | 0.62124971 | 0.94016713 |
| B9D2 | 430.683571 | -0.1408039 | 0.28639405 | -0.4916441 | 0.62297098 | 0.9402512 |
| EXOSC5 | 584.668043 | 0.24887906 | 0.21948064 | 1.13394542 | 0.25681744 | 0.82105351 |
| BCKDHA | 3686.15208 | 0.429157 | 0.25505824 | 1.68258433 | 0.09245559 | 0.65583276 |
| AC011462.2 | 115.691766 | 0.86808151 | 0.39096766 | 2.22034092 | 0.02639563 | 0.52678638 |
| B3GNT8 | 276.870911 | 0.72260429 | 0.38487573 | 1.87750027 | 0.06044956 | 0.60849584 |
| DMAC2 | 2976.19762 | -0.017307 | 0.10942423 | -0.1581643 | 0.87432737 | 0.98275231 |
| ERICH4 | 249.725882 | 3.1477117 | 0.93109461 | 3.38065719 | 0.00072313 | 0.12785022 |
| PCAT19 | 544.235006 | -0.297689 | 0.36067619 | -0.8253637 | 0.40916514 | 0.88220938 |
| LINC01480 | 66.9955758 | 0.09687211 | 0.61404863 | 0.15775967 | 0.87464618 | 0.98275231 |
| AC243960.1 | 49.3718112 | -0.0805068 | 0.71117773 | -0.1132021 | 0.90987036 | 0.98662757 |
| CEACAM21 | 56.1235022 | -0.24432 | 0.6617731 | -0.36919 | 0.71198612 | 0.95560193 |
| RPS19 | 19289.0122 | 0.01069989 | 0.15492226 | 0.06906617 | 0.94493695 | 0.99164528 |
| CD79A | 166.120748 | -0.9985437 | 1.02726651 | -0.9720396 | 0.33103085 | 0.85384238 |
| ARHGEF1 | 8169.34802 | -0.3287223 | 0.21361331 | -1.5388663 | 0.12383695 | 0.69967688 |
| RABAC1 | 4020.86456 | -0.2458388 | 0.1898648 | -1.29481 | 0.19538577 | 0.77730222 |
| AC010616.2 | 3666.87145 | -0.216408 | 0.18573495 | -1.1651443 | 0.24396057 | 0.81468457 |
| GRIK5 | 1134.09027 | -0.3827773 | 0.37767652 | -1.0135057 | 0.31081868 | 0.84476049 |
| ZNF574 | 871.62192 | -0.2152852 | 0.24076791 | -0.8941609 | 3.71E-01 | 0.86762749 |
| POU2F2 | 206.574564 | -0.3471966 | 0.52785781 | -0.6577465 | 0.51070106 | 0.91474443 |
| DEDD2 | 1979.32338 | -0.3862071 | 0.23490048 | -1.6441309 | 1.00E-01 | 6.72E-01 |
| ZNF526 | 487.112578 | -0.3890384 | 0.1818005 | -2.1399193 | 0.03236129 | 0.55014009 |
| GSK3A | 4214.65 | -0.1807626 | 0.18002285 | -1.0041094 | 0.31532586 | 0.84700043 |
| AC006486.1 | 1510.45938 | -0.1453408 | 0.22318816 | -0.6512028 | 0.5149156 | 0.91636259 |
| ERF | 4697.2924 | -0.112415 | 0.22850375 | -0.4919613 | 0.62274669 | 0.94016713 |
| CIC | 8312.27519 | -0.3275102 | 0.18060899 | -1.8133659 | 0.06977541 | 0.62779744 |
| PAFAH1B3 | 735.5751 | 0.010365 | 0.26140078 | 0.03965175 | 0.96837077 | 0.99400331 |
| PRR19 | 318.944786 | -0.1996547 | 0.2716651 | -0.7349294 | 0.46238251 | 0.89776981 |
| MEGF8 | 4347.00716 | 0.25422429 | 0.15105938 | 1.68294277 | 0.09238617 | 0.65583276 |
| LIPE-AS1 | 759.085634 | -0.1188957 | 0.24562001 | -0.4840636 | 0.62834074 | 0.94088547 |
| LIPE | 490.535834 | 0.71337051 | 0.38997671 | 1.82926439 | 6.74E-02 | 0.6249306 |
| CEACAM1 | 3741.42401 | -0.1499559 | 0.32961099 | -0.454948 | 0.64914661 | 0.94312988 |
| PSG5 | 53.2448446 | -1.0500504 | 1.34747678 | -0.7792716 | 0.43581972 | 0.89089195 |
| PSG4 | 79.5664023 | -1.6078569 | 1.50483153 | -1.068463 | 0.28531169 | 0.83107779 |
| CD177 | 111.843893 | -1.4654395 | 0.6987365 | -2.0972705 | 3.60E-02 | 5.54E-01 |
| LYPD3 | 100.225312 | 0.1617029 | 0.61042588 | 0.26490178 | 7.91E-01 | 0.97090793 |
| PHLDB3 | 689.331541 | 0.26620591 | 0.2425716 | 1.09743227 | 0.27245247 | 0.82716979 |
| ETHE1 | 1878.55499 | 0.18261477 | 0.22074595 | 0.82726215 | 0.40808849 | 0.88181557 |
| ZNF575 | 1062.13282 | 0.18807322 | 0.20945476 | 0.89791811 | 0.36922921 | 0.86762749 |
| XRCC1 | 1668.25817 | 0.18213304 | 0.17597823 | 1.03497481 | 0.30068068 | 0.84200819 |
| L34079.2 | 183.000127 | 0.22450739 | 0.19435663 | 1.15513109 | 0.24803675 | 0.81605305 |

| | | | | | | |
|---|---|---|---|---|---|---|
| L34079.1 | 327.548027 | 0.20567996 | 0.17198617 | 1.1959099 | 0.23173172 | 0.80649152 |
| L34079.3 | 231.504322 | 0.20425232 | 0.17123919 | 1.19278953 | 0.23295181 | 0.80667958 |
| PINLYP | 378.909532 | -0.0344837 | 0.22295096 | -0.1546693 | 0.87708203 | 0.98275231 |
| IRGQ | 2052.47995 | -0.0631819 | 0.19602502 | -0.3223152 | 0.74721389 | 0.96224616 |
| ZNF576 | 601.452691 | 0.0249725 | 0.15854827 | 0.15750727 | 0.87484508 | 0.98275231 |
| SRRM5 | 1072.27862 | 0.09315592 | 0.1469277 | 0.63402558 | 0.52606413 | 0.91769856 |
| ZNF428 | 1625.69856 | 0.09255922 | 0.12679134 | 0.73001217 | 0.46538274 | 0.89925633 |
| CADM4 | 2788.59067 | 0.04045026 | 0.21247045 | 0.19038063 | 0.84901087 | 0.97741457 |
| PLAUR | 992.896433 | -0.0864292 | 0.57773539 | -0.1496 | 0.88108021 | 0.98275231 |
| SMG9 | 1753.17934 | -0.2683228 | 0.149683 | -1.7926068 | 0.07303582 | 0.63580072 |
| KCNN4 | 120.326432 | 0.2805363 | 0.74938804 | 0.37435386 | 0.70814107 | 0.95501548 |
| LYPD5 | 84.3182451 | -0.168589 | 0.60350271 | -0.2793508 | 0.77997562 | 0.96836854 |
| ZNF283 | 234.258716 | -0.1972675 | 0.20990538 | -0.9397923 | 0.34732411 | 0.86197176 |
| ZNF404 | 256.565372 | -0.1468501 | 0.51498469 | -0.2851544 | 0.77552587 | 0.96836854 |
| AC006213.2 | 46.9783579 | -0.0357735 | 0.46125999 | -0.0775559 | 0.93818129 | 0.99113899 |
| ZNF45 | 542.050237 | -0.1938692 | 0.19755599 | -0.9813378 | 0.32642619 | 0.85282528 |
| ZNF155 | 406.525809 | 0.20661431 | 0.16427161 | 1.25776032 | 0.20847845 | 0.79011014 |
| AC006213.1 | 324.754161 | 0.23742546 | 0.192138 | 1.23570278 | 0.21656906 | 0.7954827 |
| ZNF230 | 276.681772 | 0.11895306 | 0.13533052 | 0.87898172 | 0.37941119 | 0.87075007 |
| ZNF222 | 189.180809 | 0.0807063 | 0.22407668 | 0.36017267 | 0.71871801 | 0.95648644 |
| AC067968.1 | 350.808933 | -0.0403696 | 0.15028925 | -0.2686128 | 0.78822765 | 0.97090793 |
| ZNF223 | 265.712366 | 0.15558628 | 0.19372273 | 0.803139 | 0.4218944 | 0.88451372 |
| ZNF284 | 272.333159 | -0.2044589 | 0.22482222 | -0.9094249 | 0.36312589 | 0.8663515 |
| ZNF224 | 1452.35433 | -0.1004854 | 0.17662612 | -0.5689158 | 0.56941328 | 0.9271275 |
| AC021092.1 | 775.521886 | -0.1088325 | 0.19777177 | -0.5502936 | 0.58211802 | 0.9317705 |
| ZNF225 | 307.907305 | 0.07691023 | 0.2133002 | 0.36057272 | 0.71841888 | 0.95648644 |
| ZNF234 | 461.84847 | -0.178582 | 0.16068173 | -1.111402 | 2.66E-01 | 0.8265123 |
| ZNF226 | 1104.34503 | 0.1421241 | 0.24663479 | 0.57625326 | 5.64E-01 | 0.92474393 |
| ZNF227 | 700.054748 | 0.02809886 | 0.11855821 | 0.23700476 | 0.8126531 | 0.97371854 |
| ZNF235 | 582.976853 | -0.0638576 | 0.13536644 | -0.4717386 | 0.63711338 | 0.94088547 |
| ZNF233 | 101.821781 | 0.53871429 | 0.33891436 | 1.58952927 | 0.11194095 | 0.68001961 |
| ZNF112 | 291.586682 | -0.2055592 | 0.19334926 | -1.0631496 | 0.2877141 | 0.83262045 |
| ZNF285 | 167.315679 | -0.0804343 | 0.25604491 | -0.3141412 | 0.7534138 | 0.96301881 |
| ZNF229 | 345.016195 | -0.1909531 | 0.28873175 | -0.6613511 | 0.50838717 | 0.91439937 |
| ZNF180 | 380.175467 | 0.03329481 | 0.22368304 | 0.14884816 | 0.88167345 | 0.98275231 |
| AC243964.2 | 129.43779 | -1.4689896 | 0.69982805 | -2.0990722 | 0.03581054 | 0.55369025 |
| PVR | 3762.35346 | -1.0574378 | 0.46134718 | -2.2920652 | 0.02190188 | 0.51617839 |
| CEACAM19 | 1660.28937 | -0.9016029 | 0.3555296 | -2.5359434 | 0.01121449 | 0.40196898 |
| BCL3 | 4855.30096 | 0.04548763 | 0.51452606 | 0.08840685 | 0.92955332 | 0.98964715 |
| CBLC | 703.866777 | 0.09704136 | 0.40132652 | 0.24180151 | 0.80893397 | 0.97334138 |
| BCAM | 45914.8034 | 0.05441161 | 0.34776864 | 0.15645923 | 0.87567106 | 0.98275231 |
| NECTIN2 | 6937.64271 | -0.2245021 | 0.21652415 | -1.0368456 | 0.2998078 | 0.84150038 |
| AC011481.2 | 390.754867 | -0.1445731 | 0.26818073 | -0.5390885 | 0.5898258 | 0.93289552 |
| TOMM40 | 2913.50405 | -0.0614621 | 0.20442642 | -0.3006563 | 0.76367656 | 0.96678847 |
| APOE | 53260.1103 | -0.5780796 | 0.48545077 | -1.1908099 | 0.23372822 | 0.80725541 |
| APOC1 | 171.505429 | -0.0225981 | 0.47739496 | -0.0473362 | 0.96224529 | 0.99320711 |
| APOC4-APOC | 322.839369 | 0.15400242 | 0.83529026 | 0.18436995 | 0.85372324 | 0.97906925 |
| APOC2 | 324.366722 | 0.17352179 | 0.8365234 | 0.20743208 | 0.83567243 | 0.97550416 |
| AC011481.1 | 36.0190278 | 0.20840925 | 0.49495213 | 0.42106951 | 0.67370433 | 0.94935413 |
| CLPTM1 | 9921.24897 | 0.04914295 | 0.23957796 | 0.20512301 | 0.83747602 | 0.97577999 |
| RELB | 1677.54235 | -0.0246666 | 0.39972033 | -0.0617097 | 0.95079402 | 0.99170503 |
| CLASRP | 4101.59823 | -0.4956582 | 0.16729836 | -2.96272 | 0.00304934 | 0.24112935 |
| ZNF296 | 112.082578 | -0.4114623 | 0.43708389 | -0.9413806 | 3.47E-01 | 0.86135217 |
| GEMIN7-AS1 | 459.853868 | -0.3757891 | 0.17919056 | -2.0971479 | 0.03598049 | 0.55369025 |

| | | | | | | |
|---|---|---|---|---|---|---|
| GEMIN7 | 801.157208 | -0.3943652 | 0.18388301 | -2.1446528 | 0.03198062 | 0.54735048 |
| MARK4 | 2021.20429 | -0.1076903 | 0.1675978 | -0.6425522 | 0.52051468 | 0.91680964 |
| PPP1R37 | 3463.77059 | -0.3055465 | 0.14733722 | -2.07379 | 0.03809882 | 0.55538555 |
| TRAPPC6A | 1294.46266 | 0.39429258 | 0.20789434 | 1.89660085 | 0.05788064 | 0.60129769 |
| BLOC1S3 | 664.497347 | -0.5395049 | 0.29009164 | -1.8597742 | 6.29E-02 | 0.6147045 |
| EXOC3L2 | 1838.35819 | 0.07369139 | 0.48997199 | 0.15039918 | 0.88044969 | 0.98275231 |
| CKM | 96.9602074 | 4.85106869 | 1.405287 | 3.45201278 | 0.00055642 | 0.11459223 |
| KLC3 | 700.513817 | 0.04839911 | 0.30149235 | 0.1605318 | 0.87246217 | 0.98275231 |
| ERCC2 | 1820.91193 | 0.31736273 | 0.21759049 | 1.45853215 | 0.14469392 | 0.73032824 |
| PPP1R13L | 1902.09419 | -0.3795842 | 0.26387761 | -1.4384858 | 0.15029626 | 0.73796431 |
| CD3EAP | 1191.8123 | -0.4819229 | 0.3325529 | -1.4491615 | 0.14729249 | 0.73386234 |
| ERCC1 | 3281.63427 | -0.3302727 | 0.27570258 | -1.1979313 | 0.23094378 | 0.80610188 |
| FOSB | 1434.35671 | -2.2359102 | 1.13882983 | -1.96334 | 0.04960668 | 0.5861488 |
| RTN2 | 276.165153 | 0.04303962 | 0.31553217 | 0.13640326 | 0.8915025 | 0.98474725 |
| PPM1N | 128.270674 | -0.0551932 | 0.37277755 | -0.1480595 | 0.88229583 | 0.98275231 |
| VASP | 4335.58264 | -0.357466 | 0.28608904 | -1.2494922 | 0.21148512 | 0.79149017 |
| OPA3 | 2404.45081 | -0.0673674 | 0.17615493 | -0.3824328 | 0.70214038 | 0.95431462 |
| GPR4 | 1183.32005 | -0.2321603 | 0.43275222 | -0.536474 | 0.591631 | 0.93289552 |
| EML2 | 3045.32229 | 0.37949189 | 0.30091662 | 1.26111976 | 0.2072657 | 0.78790396 |
| GIPR | 191.693914 | 0.73828607 | 0.40739378 | 1.81221732 | 0.06995262 | 0.6287684 |
| SNRPD2 | 3489.71371 | 0.09516371 | 0.11930381 | 0.79765863 | 4.25E-01 | 0.88581636 |
| QPCTL | 648.054042 | 0.16020302 | 0.14451588 | 1.1085496 | 0.26762453 | 0.82676682 |
| FBXO46 | 964.522915 | -0.2354874 | 0.21050012 | -1.1187044 | 0.26326625 | 0.82424998 |
| SIX5 | 1382.70236 | 0.3384282 | 0.25539111 | 1.32513695 | 1.85E-01 | 0.76820626 |
| DM1-AS | 1434.50786 | 0.02348073 | 0.1682407 | 0.1395663 | 0.88900267 | 0.98423722 |
| DMPK | 5162.73097 | -0.2789432 | 0.16514971 | -1.6890323 | 0.09121325 | 0.65495306 |
| AC011530.1 | 1229.49029 | -0.40529 | 0.19413301 | -2.0876922 | 0.0368256 | 0.55521179 |
| DMWD | 2325.47734 | -0.4838907 | 0.16432387 | -2.9447378 | 0.00323228 | 0.24992576 |
| SYMPK | 6253.02618 | -0.2299451 | 0.1421266 | -1.6178893 | 0.10568645 | 0.67526487 |
| AC092301.1 | 240.921319 | -0.2045002 | 0.21785139 | -0.9387143 | 0.34787743 | 0.86225576 |
| FOXA3 | 176.420246 | 0.2450465 | 0.55179267 | 0.44409162 | 0.65697634 | 0.94545756 |
| IRF2BP1 | 1467.44537 | -0.2530024 | 0.14849419 | -1.7037868 | 0.08842092 | 0.65363785 |
| MYPOP | 743.69001 | -0.1237316 | 0.21514451 | -0.5751092 | 5.65E-01 | 0.92504936 |
| NOVA2 | 574.287774 | -0.384397 | 0.39796637 | -0.9659032 | 0.33409264 | 0.85533716 |
| CCDC61 | 721.95719 | 0.27232898 | 0.29483975 | 0.92365084 | 0.35566813 | 0.86364633 |
| HIF3A | 1025.55936 | -0.9010759 | 0.6033401 | -1.4934792 | 0.13531178 | 0.71649038 |
| AC007193.2 | 41.9424215 | -1.2216453 | 0.78558616 | -1.5550749 | 0.11992823 | 0.69433886 |
| PPP5C | 3988.51765 | -0.2286761 | 0.12252177 | -1.8664118 | 0.06198377 | 0.61210783 |
| AC007193.1 | 123.459193 | -0.3894815 | 0.3171967 | -1.2278864 | 0.2194896 | 0.79858446 |
| AC007193.3 | 365.236161 | -0.1959836 | 0.19158368 | -1.022966 | 0.30632392 | 0.84278367 |
| CCDC8 | 1222.00367 | -0.1484874 | 0.35091632 | -0.4231418 | 0.67219179 | 0.94922827 |
| PNMA8A | 896.779109 | 0.94603578 | 0.37970246 | 2.49151871 | 0.01271983 | 0.41598747 |
| PPP5D1 | 40.4221208 | 0.55684084 | 0.38377959 | 1.45093916 | 0.1467968 | 0.73337969 |
| PNMA8B | 122.681612 | 0.16055537 | 0.38855975 | 0.41320638 | 6.79E-01 | 9.51E-01 |
| AC011484.1 | 49.728147 | 0.14460596 | 0.40892168 | 0.35362753 | 0.72361804 | 0.95810643 |
| CALM3 | 12310.2511 | 0.09060866 | 0.2382711 | 0.38027549 | 0.70374093 | 0.9547422 |
| AC093503.2 | 2575.7678 | 0.13201982 | 0.24264477 | 0.5440868 | 0.58638174 | 0.93289552 |
| PTGIR | 988.708908 | -0.0665568 | 0.51686689 | -0.1287698 | 0.89753984 | 0.98474725 |
| DACT3 | 526.572106 | -0.7608864 | 0.5963394 | -1.2759285 | 0.20198081 | 0.78144556 |
| PRKD2 | 3663.52213 | -0.4216925 | 0.29265025 | -1.4409436 | 0.14960062 | 0.73683712 |
| STRN4 | 5907.50082 | -0.4464215 | 0.13090423 | -3.4102912 | 0.00064894 | 0.12385487 |
| AC008635.1 | 572.329943 | -0.3916041 | 0.16957782 | -2.3092884 | 0.02092758 | 0.50728403 |
| FKRP | 1986.88407 | -0.0962219 | 0.17831811 | -0.5396084 | 0.58946715 | 0.93289552 |
| SLC1A5 | 6446.36272 | 0.03186802 | 0.34402926 | 0.09263173 | 0.92619614 | 0.98876911 |

| | | | | | | |
|---|---|---|---|---|---|---|
| AC008622.2 | 1278.38012 | 0.09822012 | 0.42896331 | 0.22897091 | 0.81889152 | 0.97454841 |
| AP2S1 | 2742.06225 | -0.0128707 | 0.16572526 | -0.0776627 | 0.93809639 | 0.99113899 |
| ARHGAP35 | 5416.68607 | 0.00269204 | 0.21227901 | 0.01268163 | 0.98988179 | 0.99807483 |
| NPAS1 | 84.8473646 | -0.0017997 | 0.50063664 | -0.0035948 | 0.99713174 | 0.99976751 |
| TMEM160 | 468.22114 | 0.36391372 | 0.37580814 | 0.96834974 | 0.33286973 | 0.855011 |
| ZC3H4 | 1782.4089 | -0.2479931 | 0.20486632 | -1.2105118 | 0.22608256 | 0.80246466 |
| SAE1 | 2497.04565 | -0.101114 | 0.20855726 | -0.4848263 | 0.62779955 | 0.94088547 |
| BBC3 | 539.920379 | 0.43834417 | 0.28355136 | 1.54590746 | 0.12212692 | 0.69785915 |
| CCDC9 | 2356.05851 | -0.2111507 | 0.20855369 | -1.0124524 | 0.3113218 | 0.84496478 |
| INAFM1 | 721.3158 | -0.1651014 | 0.27256566 | -0.6057307 | 0.5446936 | 0.92345821 |
| C5AR1 | 1373.5181 | -0.4218453 | 0.41851256 | -1.0079633 | 3.13E-01 | 8.46E-01 |
| C5AR2 | 192.918765 | 0.12306237 | 0.52272305 | 0.23542557 | 0.81387844 | 0.9738447 |
| DHX34 | 1754.19658 | -0.5599995 | 0.16048353 | -3.4894515 | 0.00048401 | 0.1067163 |
| MEIS3 | 292.85205 | -0.2521912 | 0.52464245 | -0.4806916 | 0.6307357 | 0.94088547 |
| KPTN | 578.799694 | -0.3884314 | 0.22635467 | -1.7160298 | 0.08615659 | 0.65049132 |
| NAPA-AS1 | 2091.38033 | -0.3477174 | 0.19786772 | -1.7573224 | 0.07886286 | 0.64079926 |
| NAPA | 10464.9492 | -0.3974404 | 0.19230486 | -2.0667207 | 0.03876048 | 0.55608529 |
| ZNF541 | 93.4647385 | 1.01196988 | 0.71179046 | 1.42172443 | 0.15510627 | 0.74400986 |
| BICRA | 1244.8227 | -0.3264596 | 0.19394059 | -1.6832968 | 0.09231765 | 0.65583276 |
| EHD2 | 3001.68969 | -0.0848083 | 0.3694182 | -0.2295726 | 0.81842387 | 0.97454841 |
| NOP53 | 9791.68112 | 0.36929901 | 0.18361481 | 2.01127029 | 0.04429691 | 0.57267043 |
| NOP53-AS1 | 89.6807608 | 0.37568274 | 0.25565136 | 1.46951196 | 0.14169398 | 0.72784886 |
| SELENOW | 6348.69318 | 0.03799444 | 0.24209851 | 0.15693793 | 0.87529377 | 0.98275231 |
| PLA2G4C | 1190.05809 | -0.6072308 | 0.37000855 | -1.6411263 | 0.1007712 | 0.67247782 |
| PLA2G4C-AS | 271.624482 | -0.7456341 | 0.36162837 | -2.0618795 | 0.03921921 | 0.55657615 |
| LIG1 | 975.580301 | 0.19600336 | 0.30761535 | 0.63717029 | 0.52401393 | 0.91723878 |
| AC011466.3 | 203.721267 | 0.16660347 | 0.33928802 | 0.49103846 | 0.62339924 | 0.94045607 |
| ZSWIM9 | 678.354686 | -0.0815815 | 0.19433955 | -0.4197886 | 0.67463988 | 0.94935413 |
| CARD8 | 1794.70503 | -0.1183823 | 0.24617047 | -0.4808954 | 0.63059082 | 0.94088547 |
| AC011466.1 | 202.58931 | 0.05726498 | 0.27540692 | 0.2079286 | 0.83528472 | 0.97550416 |
| CARD8-AS1 | 193.433888 | -0.0691075 | 0.2138178 | -0.3232076 | 0.746538 | 0.96210071 |
| CCDC114 | 116.970759 | -0.1848427 | 0.56394361 | -0.3277681 | 0.74308704 | 0.96109001 |
| EMP3 | 1241.7537 | -0.4152391 | 0.33384158 | -1.2438208 | 0.21356553 | 0.7926732 |
| TMEM143 | 840.428822 | 0.50496063 | 0.3187126 | 1.58437613 | 0.11310815 | 0.68130021 |
| KDELR1 | 10224.8913 | 0.04767329 | 0.10827001 | 0.44031855 | 0.65970641 | 0.94618161 |
| GRWD1 | 1864.96527 | -0.3629724 | 0.22667579 | -1.6012845 | 0.10931393 | 0.67859123 |
| KCNJ14 | 166.360595 | -0.6210953 | 0.43208837 | -1.4374265 | 0.15059684 | 0.73796431 |
| AC008403.2 | 482.91416 | -0.2055849 | 0.20183882 | -1.0185599 | 3.08E-01 | 8.44E-01 |
| AC008403.1 | 1129.89506 | -0.099755 | 0.21696019 | -0.4597849 | 0.64567066 | 0.94208859 |
| CYTH2 | 6269.57646 | -0.1755085 | 0.15669884 | -1.1200368 | 0.26269808 | 0.82424998 |
| LMTK3 | 501.156493 | -0.7222793 | 0.45133981 | -1.6003004 | 0.10953196 | 0.67859123 |
| SULT2B1 | 279.938557 | -0.1786439 | 0.82158082 | -0.2174393 | 0.82786603 | 0.97548201 |
| FAM83E | 85.7372509 | 1.95801641 | 0.82250429 | 2.38055465 | 0.0172866 | 0.46859699 |
| RPL18 | 17146.2452 | 0.14932279 | 0.1830881 | 0.81557887 | 0.41474104 | 0.88364943 |
| AC022154.1 | 115.116148 | -0.3941322 | 0.34129151 | -1.1548257 | 0.24816183 | 0.81621194 |
| SPHK2 | 3227.78411 | 0.26019929 | 0.31965388 | 0.81400325 | 0.41564309 | 0.88399703 |
| DBP | 1090.2361 | 0.42098954 | 0.32119859 | 1.31068303 | 0.18996487 | 0.77059128 |
| CA11 | 831.646272 | -0.0702529 | 0.24945939 | -0.2816204 | 0.77823456 | 0.96836854 |
| NTN5 | 54.9534818 | 0.70298832 | 0.56142965 | 1.25213963 | 0.21051899 | 0.79134644 |
| FUT2 | 91.5539441 | 1.25601859 | 0.58171868 | 2.15915121 | 3.08E-02 | 5.44E-01 |
| MAMSTR | 98.3216598 | -0.514329 | 0.45185358 | -1.1382647 | 0.25500998 | 0.81992039 |
| RASIP1 | 1266.17354 | -0.2501241 | 0.30980425 | -0.8073616 | 0.41945819 | 0.88451372 |
| FUT1 | 249.286184 | 0.22644015 | 0.34109987 | 0.66385293 | 0.50678444 | 0.91420888 |
| BCAT2 | 3187.16507 | 0.31820471 | 0.16145916 | 1.97080618 | 0.04874605 | 0.58504358 |

| | | | | | | |
|---|---|---|---|---|---|---|
| HSD17B14 | 2423.87474 | 0.3648889 | 0.40468114 | 0.90167014 | 0.36723212 | 0.86683872 |
| PLEKHA4 | 2502.28494 | -0.6852694 | 0.52888826 | -1.2956789 | 0.19508613 | 0.77706892 |
| PPP1R15A | 5726.59092 | -0.7272683 | 0.3326229 | -2.1864649 | 0.02878161 | 0.53606221 |
| NUCB1 | 25463.2444 | 0.12338946 | 0.13243278 | 0.93171388 | 0.35148441 | 0.86364633 |
| NUCB1-AS1 | 4581.35469 | 0.20860263 | 0.16302194 | 1.27959849 | 0.20068638 | 0.78134034 |
| DHDH | 790.769284 | 0.645531 | 0.62350989 | 1.03531798 | 0.30052043 | 0.84200819 |
| BAX | 3056.09625 | -0.0427037 | 0.19236275 | -0.2219955 | 8.24E-01 | 0.97548201 |
| AC026803.2 | 69.8985796 | -2.0177315 | 0.85683579 | -2.3548637 | 1.85E-02 | 0.47570049 |
| FTL | 323941.777 | 0.18798292 | 0.53430456 | 0.35182729 | 0.72496779 | 0.95848028 |
| GYS1 | 3602.24813 | -0.1015298 | 0.24443369 | -0.4153673 | 0.67787301 | 0.95018034 |
| RUVBL2 | 3107.6284 | 0.05316615 | 0.111155 | 0.47830643 | 0.63243212 | 0.94088547 |
| SNRNP70 | 22842.2965 | -0.0919717 | 0.22546714 | -0.4079162 | 0.6833352 | 0.95070301 |
| LIN7B | 265.246132 | 0.41937415 | 0.32504306 | 1.29021107 | 0.19697738 | 0.77816059 |
| C19orf73 | 56.4511888 | 0.56055461 | 0.34304395 | 1.63406064 | 0.10224612 | 0.67279543 |
| PPFIA3 | 433.316682 | 0.22017843 | 0.24281409 | 0.90677779 | 0.36452432 | 0.8663515 |
| HRC | 305.982064 | -0.0800352 | 0.53884945 | -0.1485298 | 0.88192463 | 0.98275231 |
| TRPM4 | 5156.08004 | 0.37665816 | 0.40200884 | 0.93693999 | 0.34878943 | 0.86356499 |
| SLC6A16 | 522.442702 | -0.3625445 | 0.33342112 | -1.0873471 | 0.27688342 | 0.82760323 |
| AC011450.1 | 214.519443 | -0.3928042 | 0.37357425 | -1.0514756 | 0.29304019 | 0.83526481 |
| CD37 | 1464.76335 | -0.4149588 | 0.38821601 | -1.0688863 | 0.28512089 | 0.83107779 |
| TEAD2 | 2362.33196 | -0.4516032 | 0.36545918 | -1.2357146 | 0.21656468 | 0.7954827 |
| SLC17A7 | 49.4758717 | -0.545274 | 0.51111974 | -1.0668224 | 0.28605205 | 0.83107779 |
| PIH1D1 | 3523.423 | -0.034191 | 0.12934647 | -0.2643363 | 0.7915208 | 0.97090793 |
| ALDH16A1 | 2796.63136 | -0.0815202 | 0.2051074 | -0.3974513 | 0.69103466 | 0.95158502 |
| AC010619.1 | 911.489536 | -0.0761806 | 0.2082493 | -0.3658146 | 0.71450343 | 0.95579447 |
| FLT3LG | 407.674887 | -0.1563213 | 0.49713105 | -0.3144469 | 0.75318165 | 0.96301881 |
| RPL13A | 55612.7333 | 0.09678942 | 0.20978364 | 0.46137734 | 0.64452791 | 0.94208859 |
| RPS11 | 27970.1539 | 0.09551694 | 0.17026819 | 0.56097936 | 0.57481161 | 0.93024927 |
| FCGRT | 11039.0263 | 0.03418563 | 0.18845739 | 0.18139713 | 0.85605587 | 0.97929708 |
| RCN3 | 1681.2071 | -0.5370168 | 0.28227596 | -1.9024533 | 5.71E-02 | 6.01E-01 |
| NOSIP | 2151.34388 | -0.155825 | 0.13402542 | -1.1626525 | 0.24497051 | 0.81487261 |
| PRRG2 | 864.96733 | -0.1734431 | 0.19378379 | -0.8950343 | 0.37076875 | 0.86762749 |
| PRR12 | 2345.71034 | -0.1460099 | 0.23075301 | -0.6327542 | 0.52689419 | 0.91779198 |
| AC011495.2 | 97.04123 | -0.173698 | 0.3091285 | -0.5618959 | 0.57418698 | 0.93014419 |
| RRAS | 2228.61814 | 0.06390444 | 0.36237753 | 0.17634768 | 0.86002081 | 0.98063093 |
| SCAF1 | 4161.5292 | -0.2782051 | 0.09320703 | -2.9848086 | 0.00283756 | 0.23089261 |
| IRF3 | 6065.97069 | -0.0432548 | 0.21634243 | -0.1999369 | 8.42E-01 | 0.97621603 |
| BCL2L12 | 1238.99751 | -0.1166903 | 0.19243256 | -0.6063956 | 5.44E-01 | 9.23E-01 |
| PRMT1 | 5429.45022 | -0.1059389 | 0.18372309 | -0.5766227 | 0.56419433 | 0.92470993 |
| ADM5 | 172.616247 | 0.10991832 | 0.36795544 | 0.29872726 | 7.65E-01 | 9.67E-01 |
| AC011495.3 | 56.9099067 | 0.19495513 | 0.37909195 | 0.51426873 | 6.07E-01 | 0.93739614 |
| CPT1C | 216.40198 | -0.7997603 | 0.44758835 | -1.786821 | 0.07396642 | 0.6374448 |
| AP2A1 | 5658.29151 | 0.03416855 | 0.15327246 | 0.22292686 | 0.82359243 | 0.9754753 |
| FUZ | 2125.93702 | -0.0499042 | 0.2521743 | -0.1978956 | 0.84312678 | 0.97649449 |
| AC006942.1 | 444.531129 | -0.1046907 | 0.33370095 | -0.3137263 | 0.75372896 | 0.96328658 |
| MED25 | 4286.83756 | -0.421246 | 0.24450953 | -1.7228202 | 0.08492105 | 0.64988556 |
| PTOV1-AS1 | 1014.69523 | -0.1711838 | 0.26141265 | -0.6548412 | 0.51257001 | 0.91568064 |
| PTOV1 | 11661.8494 | -0.3360017 | 0.13066831 | -2.5714095 | 0.01012855 | 0.39989496 |
| AC018766.1 | 916.833214 | -0.2513678 | 0.17456068 | -1.4400023 | 0.14986674 | 0.73749842 |
| PTOV1-AS2 | 4606.757 | -0.3001228 | 0.13749251 | -2.18283 | 2.90E-02 | 5.36E-01 |
| PNKP | 7125.65644 | -0.3132796 | 0.1696827 | -1.8462671 | 6.49E-02 | 6.21E-01 |
| AKT1S1 | 3300.59086 | -0.2562798 | 0.22861 | -1.1210347 | 2.62E-01 | 8.24E-01 |
| TBC1D17 | 4418.8688 | -0.0483473 | 0.14105598 | -0.3427523 | 0.73178483 | 0.95848028 |
| IL4I1 | 1195.39673 | 0.17362477 | 0.42992858 | 0.40384561 | 0.68632625 | 0.95097288 |

| | | | | | | |
|---|---|---|---|---|---|---|
| NUP62 | 3118.32564 | -0.0827031 | 0.18312855 | -0.4516125 | 0.65154817 | 0.94393131 |
| AC011452.1 | 424.518975 | -0.1907741 | 0.24306675 | -0.7848628 | 0.43253402 | 0.88980461 |
| ATF5 | 2278.21199 | 0.21812571 | 0.36925663 | 0.59071575 | 0.55471089 | 0.92373235 |
| SIGLEC11 | 99.7888345 | -1.8912281 | 0.61198854 | -3.0902999 | 0.00199955 | 0.19217166 |
| VRK3 | 3761.96838 | 0.10336881 | 0.29080703 | 0.355455 | 0.72224875 | 0.95739595 |
| ZNF473 | 533.717712 | -0.2655763 | 0.2315229 | -1.1470842 | 0.2513468 | 0.81815815 |
| AC010624.3 | 63.632174 | -0.2196221 | 0.29176821 | -0.7527281 | 0.45161335 | 0.89639459 |
| MYH14 | 3842.97823 | 0.78168319 | 0.14323698 | 5.45727228 | 4.84E-08 | 0.00030042 |
| AC008655.1 | 604.928383 | 0.68810932 | 0.21402442 | 3.21509729 | 0.001304 | 0.16147422 |
| KCNC3 | 4423.00803 | -0.3598777 | 0.32255122 | -1.1157227 | 0.26454086 | 0.82424998 |
| NR1H2 | 4881.28418 | -0.115362 | 0.13041354 | -0.8845861 | 0.37637988 | 0.86965052 |
| NAPSA | 14206.7064 | -0.743656 | 0.51726086 | -1.4376808 | 0.15052465 | 0.73796431 |
| POLD1 | 1314.67884 | -0.4690319 | 0.22269569 | -2.1061564 | 0.03519077 | 0.55369025 |
| AC020909.1 | 407.642312 | -0.3332155 | 0.31491049 | -1.0581277 | 0.28999724 | 0.83407584 |
| SPIB | 59.2870535 | 0.40578352 | 1.1533387 | 0.35183379 | NA | NA |
| MYBPC2 | 131.265183 | -0.8732389 | 0.59014934 | -1.4796913 | 0.13895565 | 0.72399122 |
| FAM71E1 | 1065.79597 | 0.27807804 | 0.20647605 | 1.34678108 | 0.17805075 | 0.76115956 |
| EMC10 | 10868.4265 | 0.18458993 | 0.16807478 | 1.09826071 | 0.27209066 | 0.82716979 |
| AC020909.3 | 3309.60721 | 0.21184248 | 0.18285205 | 1.15854586 | 0.24664135 | 0.81542272 |
| AC020909.2 | 86.5577174 | -0.5376355 | 0.48658548 | -1.1049147 | 0.26919654 | 0.82716979 |
| JOSD2 | 863.556507 | -0.2797269 | 0.24234556 | -1.1542481 | 0.24839846 | 0.8162591 |
| ASPDH | 1290.04379 | 0.23633965 | 0.58547196 | 0.40367373 | 0.68645266 | 0.95097288 |
| LRRC4B | 110.223004 | -0.5225651 | 0.60720196 | -0.8606117 | 0.38945193 | 0.87568585 |
| SYT3 | 41.0039505 | 0.18646458 | 0.46052098 | 0.40489921 | 0.6855516 | 0.95097288 |
| SHANK1 | 40.8463947 | -0.123818 | 0.46130262 | -0.2684095 | 0.78838415 | 0.97090793 |
| CLEC11A | 613.353146 | 0.01312863 | 0.29321996 | 0.04477399 | 0.96428746 | 0.99342622 |
| C19orf48 | 1451.72397 | -0.5477605 | 0.3613143 | -1.5160223 | 0.12951372 | 0.70764335 |
| KLK1 | 1650.34015 | -0.4163958 | 1.30510157 | -0.3190524 | 0.74968679 | 0.96249332 |
| AC011483.2 | 246.5167 | 1.0569811 | 0.8251667 | 1.28093039 | 2.00E-01 | 7.81E-01 |
| KLK6 | 446.311413 | 1.03396958 | 0.84296459 | 1.2265872 | 2.20E-01 | 0.79858446 |
| KLK7 | 329.55104 | 0.49646576 | 0.9620248 | 0.51606337 | 0.60581015 | 0.93698713 |
| CTU1 | 248.023302 | -0.2789374 | 0.2795787 | -0.9977062 | 0.31842184 | 0.8490262 |
| SIGLEC9 | 161.067099 | -0.5388063 | 0.50823558 | -1.0601507 | 0.28907604 | 0.83308345 |
| SIGLEC7 | 104.364372 | -0.3734326 | 0.70720329 | -0.5280413 | 0.59747066 | 0.93416444 |
| AC063977.6 | 50.6352952 | -0.4395532 | 0.68331999 | -0.6432612 | 0.52005462 | 0.91657872 |
| CD33 | 173.68656 | -0.3811629 | 0.48504597 | -0.7858283 | 0.43196806 | 0.88976671 |
| VSIG10L | 313.624438 | -0.4360172 | 0.46810256 | -0.9314566 | 0.35161745 | 0.86364633 |
| ETFB | 6187.09833 | 0.60835713 | 0.29088145 | 2.09142636 | 0.03648986 | 0.55384187 |
| AC008750.4 | 3472.09517 | 0.66083274 | 0.30097673 | 2.19562733 | 0.02811863 | 0.53300785 |
| AC008750.8 | 4543.20504 | 0.68292676 | 0.30991275 | 2.20360973 | 0.0275518 | 0.52904542 |
| CLDND2 | 83.5047785 | 0.10129 | 0.32370504 | 0.31290832 | 0.75435032 | 0.96348936 |
| NKG7 | 281.883619 | -0.2277724 | 0.63422115 | -0.3591371 | 0.71949252 | 0.9566409 |
| SIGLEC10 | 309.402355 | -1.0114318 | 0.65716058 | -1.5390939 | 0.12378138 | 0.69957382 |
| AC008750.1 | 42.1789282 | -1.0704113 | 0.79392283 | -1.3482561 | 0.17757603 | 0.76115956 |
| AC008750.2 | 70.48806 | -0.8407893 | 0.64137829 | -1.3109101 | 0.18988812 | 0.7704467 |
| ZNF175 | 683.151641 | -0.1883192 | 0.19297833 | -0.9758566 | 0.32913551 | 0.85327551 |
| SIGLEC5 | 70.7021081 | -0.2042235 | 0.63499834 | -0.3216126 | 0.74774618 | 0.96233424 |
| AC018755.2 | 52.2730192 | -0.1232725 | 0.67949577 | -0.1814176 | 0.85603978 | 0.97929708 |
| SIGLEC14 | 135.57805 | -0.6364902 | 0.76506472 | -0.8319429 | 0.40544115 | 0.88132015 |
| SPACA6P-AS | 107.285461 | -0.3036955 | 0.32799143 | -0.9259252 | 0.35448487 | 0.86364633 |
| SPACA6 | 1164.56857 | -0.2643158 | 0.28575669 | -0.9249679 | 0.35498261 | 0.86364633 |
| FPR1 | 555.443048 | -0.5526615 | 0.58558489 | -0.943777 | 3.45E-01 | 8.61E-01 |
| FPR2 | 84.2762657 | -0.5736581 | 0.74737416 | -0.7675648 | 0.4427458 | 0.89343541 |
| FPR3 | 386.161081 | -0.4300364 | 0.6488471 | -0.66277 | 0.50747789 | 0.91430909 |

| | | | | | | |
|---|---|---|---|---|---|---|
| ZNF577 | 1152.5934 | 0.1208195 | 0.20335964 | 0.5941174 | 0.55243359 | 0.92373235 |
| ZNF649-AS1 | 52.5587128 | -0.2033506 | 0.37073655 | -0.5485043 | 0.58334566 | 0.93220705 |
| ZNF649 | 433.345978 | -0.159302 | 0.24157639 | -0.6594271 | 5.10E-01 | 9.14E-01 |
| ZNF613 | 386.265378 | -0.1604508 | 0.17844757 | -0.8991479 | 0.36857388 | 0.86762749 |
| ZNF350-AS1 | 124.951938 | -1.5608269 | 0.50045429 | -3.1188202 | 0.00181577 | 0.18722055 |
| ZNF350 | 601.509338 | -0.3972913 | 0.22775406 | -1.7443874 | 0.08109155 | 0.64368353 |
| ZNF615 | 519.210727 | -0.1883681 | 0.19315199 | -0.9752326 | 0.32944491 | 0.85350346 |
| ZNF614 | 414.567673 | -0.1028785 | 0.12686949 | -0.8109005 | 0.41742283 | 0.88445833 |
| ZNF432 | 798.410846 | -0.0973342 | 0.21395762 | -0.4549228 | 0.64916481 | 0.94312988 |
| AC011468.5 | 86.8735744 | 0.11255023 | 0.37973143 | 0.29639431 | 0.76692897 | 0.96742383 |
| ZNF841 | 600.265494 | -0.2462179 | 0.24344482 | -1.0113908 | 0.31182943 | 0.84537057 |
| ZNF616 | 416.947136 | 0.01915181 | 0.19734224 | 0.09704873 | 0.9226877 | 0.98826666 |
| ZNF836 | 344.295162 | 0.01632814 | 0.20770713 | 0.07861136 | 0.93734175 | 0.99079085 |
| PPP2R1A | 18348.7798 | 0.13628701 | 0.14794009 | 0.92123107 | 0.3569298 | 0.86390849 |
| AC010320.2 | 2828.84539 | 0.17443282 | 0.15147283 | 1.15157828 | 0.24949441 | 0.81646572 |
| ZNF766 | 729.280975 | 0.00818323 | 0.21947457 | 0.03728554 | 0.97025733 | 0.99400331 |
| ZNF480 | 798.30379 | -0.3122041 | 0.28576384 | -1.0925248 | 2.75E-01 | 0.82719407 |
| AC010320.4 | 43.8849989 | -0.1473763 | 0.35834886 | -0.4112649 | 0.68087831 | 0.95070301 |
| ZNF610 | 196.659952 | 0.3340964 | 0.34805744 | 0.9598887 | 0.33711123 | 0.85688344 |
| ZNF880 | 403.302645 | 0.32134859 | 0.24632155 | 1.30458987 | 0.19203254 | 0.77378668 |
| ZNF528 | 804.455096 | -0.1060282 | 0.27030452 | -0.3922545 | 0.69487019 | 0.95234774 |
| ZNF578 | 43.8010168 | -0.1325252 | 0.60385714 | -0.2194645 | 0.82628826 | 0.97548201 |
| ZNF808 | 558.088602 | 0.09075348 | 0.20985359 | 0.43246093 | 6.65E-01 | 0.94716559 |
| ZNF701 | 355.200816 | -0.1044713 | 0.21165181 | -0.4936 | 0.62158873 | 0.94016713 |
| ZNF83 | 4252.48234 | 0.0883159 | 0.24239774 | 0.36434293 | 0.71560195 | 0.95590747 |
| ZNF611 | 542.187532 | -0.8445307 | 0.31715587 | -2.6628253 | 0.00774876 | 0.36878977 |
| ZNF600 | 579.447328 | -0.5205443 | 0.28832579 | -1.8054033 | 0.07101159 | 0.63056668 |
| ZNF28 | 490.766756 | 0.16524552 | 0.12649936 | 1.30629531 | 0.19145215 | 0.77327689 |
| ZNF468 | 2112.23381 | -0.3745492 | 0.24733348 | -1.5143491 | 0.12993731 | 0.70814767 |
| ZNF320 | 1461.87835 | -0.3794741 | 0.31355764 | -1.2102213 | 0.22619399 | 0.80246466 |
| AC010487.3 | 216.564818 | -0.3655789 | 0.37484126 | -0.9752899 | 0.32941648 | 0.85350346 |
| ZNF888 | 643.43631 | 0.18217853 | 0.37924007 | 0.48037785 | 0.63095874 | 0.94088547 |
| ZNF816-ZNF | 197.344491 | -0.1191341 | 0.39264302 | -0.3034157 | 0.76157306 | 0.96606476 |
| ZNF816 | 300.345205 | 0.37917184 | 0.26046896 | 1.45572756 | 1.45E-01 | 0.73161643 |
| ZNF160 | 2284.71478 | -0.1302844 | 0.30304354 | -0.4299196 | 0.6672541 | 0.94830184 |
| ZNF415 | 407.928897 | -0.0332023 | 0.26932269 | -0.1232806 | 0.9018849 | 0.98572048 |
| ZNF347 | 509.095263 | -0.0106745 | 0.17969821 | -0.0594024 | 0.95263162 | 0.99170503 |
| ZNF665 | 139.816819 | 0.47944184 | 0.41946843 | 1.14297476 | 0.25304907 | 0.81835388 |
| ZNF677 | 468.776263 | 0.18168106 | 0.22952621 | 0.79154822 | 0.42862415 | 0.88821697 |
| ZNF845 | 315.34272 | -0.0004354 | 0.1784465 | -0.0024402 | 0.99805303 | 0.99990633 |
| ZNF525 | 390.104125 | -0.0537627 | 0.21175309 | -0.2538936 | 8.00E-01 | 0.97207132 |
| ZNF765 | 432.516157 | -0.2416809 | 0.16667654 | -1.4499993 | 0.14705872 | 0.73337969 |
| ZNF761 | 735.153534 | -0.22565 | 0.21863771 | -1.0320724 | 0.30203818 | 0.84211658 |
| ZNF813 | 176.007259 | 0.20883785 | 0.20997097 | 0.99460342 | 3.20E-01 | 8.50E-01 |
| ZNF331 | 1350.6433 | -0.0151292 | 0.52102505 | -0.0290374 | 0.97683479 | 0.99455099 |
| MYADM | 3238.36724 | -0.2412497 | 0.57318018 | -0.4208968 | 0.67383044 | 0.94935413 |
| AC008440.2 | 1093.26234 | -0.192924 | 0.56819408 | -0.3395388 | 0.73420385 | 0.95895327 |
| OSCAR | 130.047015 | -0.0275937 | 0.73855963 | -0.0373615 | 0.97019676 | 0.99400331 |
| NDUFA3 | 2768.18855 | 0.2114244 | 0.14115126 | 1.49785703 | 1.34E-01 | 7.16E-01 |
| TFPT | 840.763623 | -0.2859612 | 0.1640215 | -1.7434372 | 0.08125727 | 0.6439041 |
| PRPF31 | 3696.18386 | -0.1035733 | 0.12694384 | -0.8158987 | 0.41455807 | 0.88364943 |
| AC245052.4 | 1692.95479 | -0.1128265 | 0.14690131 | -0.7680425 | 0.44246194 | 0.89343541 |
| CNOT3 | 3324.70189 | -0.2865154 | 0.12646113 | -2.2656398 | 0.02347344 | 0.51967919 |
| LENG1 | 942.176251 | -0.2596301 | 0.13412999 | -1.9356603 | 0.05290931 | 0.58742503 |

| | | | | | | |
|---|---|---|---|---|---|---|
| TMC4 | 4944.58599 | -0.0570371 | 0.30814478 | -0.1850985 | 0.8531518 | 0.97886533 |
| MBOAT7 | 5016.64767 | -0.2492463 | 0.18036302 | -1.3819146 | 0.16699793 | 0.75558932 |
| TSEN34 | 2548.61435 | 0.18115863 | 0.13340811 | 1.35792822 | 0.17448646 | 0.75974924 |
| RPS9 | 24265.7962 | 0.0699495 | 0.16518731 | 0.42345565 | 0.67196285 | 0.94922827 |
| LILRB3 | 459.277416 | -0.5346179 | 0.5470706 | -0.9772375 | 0.32845157 | 0.85317073 |
| LILRA6 | 215.199598 | -0.2051897 | 0.59076413 | -0.3473293 | 0.72834393 | 0.95848028 |
| LILRB5 | 702.106057 | -0.2452335 | 0.4153976 | -0.5903585 | 0.55495033 | 0.92373235 |
| LILRB2 | 600.530245 | 0.11642096 | 0.59956433 | 0.19417593 | 0.84603813 | 0.9767918 |
| LILRA5 | 224.212985 | -0.4354502 | 0.65883497 | -0.6609397 | 0.508651 | 0.91439937 |
| LAIR1 | 843.471302 | -0.2684006 | 0.57100289 | -0.4700511 | 0.63831848 | 0.94088547 |
| TTYH1 | 106.949683 | -0.5486214 | 0.41795526 | -1.312632 | 0.18930698 | 0.76970626 |
| AC245884.8 | 135.464066 | 0.23092959 | 0.52625764 | 0.43881471 | 0.6607958 | 0.94628868 |
| LENG8-AS1 | 304.753301 | 0.03513908 | 0.28100728 | 0.12504687 | 0.90048644 | 0.98526603 |
| LENG8 | 29989.538 | -0.0990812 | 0.15354578 | -0.6452876 | 0.51874085 | 0.91636259 |
| LENG9 | 1266.26748 | 0.10030125 | 0.13919021 | 0.72060565 | 0.47115218 | 0.90109838 |
| CDC42EP5 | 325.75285 | 0.11039803 | 0.42715179 | 0.25845153 | 0.79605845 | 0.97101097 |
| LILRA2 | 97.5786632 | 0.69338855 | 0.617147 | 1.12353873 | 0.26120876 | 0.82424998 |
| LILRA1 | 66.6879283 | 1.11456927 | 0.92250585 | 1.20819751 | 0.2269713 | 0.80300331 |
| LILRB1 | 357.032382 | 0.13931897 | 0.58090881 | 0.23982932 | 0.81046257 | 0.97352497 |
| LILRB4 | 388.230776 | -0.029822 | 0.69568733 | -0.0428669 | 0.96580761 | 0.99387827 |
| FCAR | 54.5323217 | -0.8640273 | 0.72820226 | -1.186521 | 0.23541662 | 0.80747753 |
| NLRP2 | 319.488281 | -0.1422957 | 0.5018927 | -0.2835183 | 0.77677959 | 0.96836854 |
| AC011476.3 | 177.683074 | -0.303959 | 0.35332881 | -0.8602722 | 0.389639 | 0.87577651 |
| RDH13 | 1687.62715 | -0.0413937 | 0.21525652 | -0.1922995 | 0.84750764 | 0.97701835 |
| EPS8L1 | 2981.57228 | -0.0102065 | 0.6409992 | -0.0159228 | 0.98729597 | 0.99735412 |
| PPP1R12C | 8086.40944 | -0.3121449 | 0.17050202 | -1.8307399 | 0.06713937 | 0.6249306 |
| TNNT1 | 49.7801498 | 4.84260903 | 1.76906844 | 2.737378 | NA | NA |
| AC010327.1 | 112.596523 | 0.45787917 | 0.46090213 | 0.99344122 | 0.32049498 | 0.85028262 |
| DNAAF3 | 257.990072 | 0.41924466 | 0.40500511 | 1.03515895 | 0.30059469 | 0.84200819 |
| AC010327.3 | 155.825677 | 0.47599589 | 0.42776964 | 1.11273885 | 2.66E-01 | 0.82615987 |
| TMEM86B | 514.310023 | -0.2152466 | 0.22235586 | -0.9680274 | 0.3330307 | 0.855011 |
| PPP6R1 | 6315.74861 | -0.3419097 | 0.22119919 | -1.5457097 | 0.12217469 | 0.69785915 |
| HSPBP1 | 1937.36648 | 0.00965679 | 0.18782283 | 0.05141434 | 0.95899536 | 0.99242674 |
| BRSK1 | 256.526046 | -0.3322495 | 0.50028824 | -0.6641162 | 0.50661592 | 0.91416224 |
| AC020922.2 | 43.2867395 | 0.13245797 | 0.53846033 | 0.24599392 | 0.80568694 | 0.97333713 |
| TMEM150B | 281.713913 | 0.0815154 | 0.53160823 | 0.15333736 | 0.87813224 | 0.98275231 |
| KMT5C | 1074.92383 | 0.18798913 | 0.31558222 | 0.59568987 | 0.55138243 | 0.92373235 |
| COX6B2 | 62.9367523 | 0.27424741 | 0.57822118 | 0.47429499 | 0.63528956 | 0.94088547 |
| IL11 | 218.393059 | -1.2785199 | 0.87202718 | -1.4661468 | 1.43E-01 | 0.72784886 |
| TMEM238 | 107.642916 | -0.0768215 | 0.39665689 | -0.1936724 | 0.84643239 | 0.97681836 |
| RPL28 | 18942.0254 | -0.0633384 | 0.18102029 | -0.3498968 | 0.72641615 | 0.95848028 |
| UBE2S | 571.052874 | -0.2828804 | 0.2524639 | -1.1204785 | 0.2625099 | 0.82424998 |
| ISOC2 | 4323.69103 | 0.15279877 | 0.23903825 | 0.63922311 | 0.5226778 | 0.91723878 |
| ZNF628 | 453.079886 | -0.5481762 | 0.17895995 | -3.0631222 | 0.00219041 | 0.20321981 |
| NAT14 | 981.817139 | 0.07935376 | 0.42377982 | 0.18725232 | 0.8514628 | 0.97855304 |
| SSC5D | 501.619987 | 0.80595896 | 0.49670556 | 1.62260909 | 0.104673 | 0.67439401 |
| ZNF579 | 918.067527 | -0.1822633 | 0.31049375 | -0.5870111 | 0.55719622 | 0.92373235 |
| FIZ1 | 1226.15118 | -0.178507 | 0.16690065 | -1.0695406 | 0.28482614 | 0.83107779 |
| ZNF524 | 753.76857 | -0.0104095 | 0.13892369 | -0.0749296 | 0.9402707 | 0.99136103 |
| ZNF865 | 1161.13581 | -0.3166697 | 0.18982162 | -1.6682489 | 0.09526633 | 0.65888357 |
| AC008735.4 | 40.0086939 | -0.2710736 | 0.33400986 | -0.8115736 | 0.41703635 | 0.88445833 |
| ZNF784 | 523.654458 | -0.0571334 | 0.25616181 | -0.2230366 | 0.82350706 | 0.9754753 |
| ZNF580 | 2229.13719 | -0.1007812 | 0.17447337 | -0.577631 | 0.5635133 | 0.92452095 |
| ZNF581 | 1061.99266 | 0.02396766 | 0.12582485 | 0.19048433 | 0.84892962 | 0.97741457 |

| | | | | | | |
|---|---|---|---|---|---|---|
| CCDC106 | 2150.60682 | -0.0432913 | 0.1158169 | -0.3737912 | 0.7085597 | 0.95501548 |
| U2AF2 | 9445.0603 | -0.3300925 | 0.13761485 | -2.398669 | 0.01645478 | 0.46026719 |
| AC008735.2 | 652.792707 | -0.2494138 | 0.15917663 | -1.5668997 | 0.11713813 | 0.68956727 |
| EPN1 | 10123.2203 | -0.1212291 | 0.11080845 | -1.0940423 | 0.27393642 | 0.82719197 |
| NLRP9 | 107.607257 | 0.14493406 | 0.41226968 | 0.35155158 | 0.72517459 | 0.95848028 |
| ZNF787 | 1337.048 | -0.509856 | 0.19421612 | -2.625199 | 0.00865983 | 0.37987657 |
| ZNF444 | 2392.4053 | -0.2024864 | 0.16611559 | -1.2189491 | 0.22286351 | 0.80078694 |
| ZSCAN5A | 344.326078 | -0.4560003 | 0.27823919 | -1.6388788 | 0.10123851 | 0.6725344 |
| AC006116.8 | 49.4304206 | -0.3228192 | 0.33023804 | -0.9775348 | 0.32830448 | 0.85317073 |
| ZNF582 | 207.662211 | -0.2360399 | 0.29267721 | -0.8064855 | 0.41996299 | 0.88451372 |
| ZNF582-AS1 | 323.311501 | -0.1907783 | 0.2161148 | -0.8827636 | 0.37736402 | 0.86987124 |
| ZNF583 | 347.759635 | -0.2704945 | 0.21630105 | -1.2505466 | 2.11E-01 | 0.79134644 |
| ZNF667 | 444.516769 | -0.2816547 | 0.2752675 | -1.0232037 | 0.30621156 | 0.84278367 |
| ZNF667-AS1 | 1115.69846 | 0.11362043 | 0.1750483 | 0.64908044 | 5.16E-01 | 9.16E-01 |
| ZNF471 | 582.451458 | -0.0375162 | 0.20389981 | -0.1839934 | 0.85401861 | 0.97906925 |
| ZFP28 | 496.695595 | 0.09526557 | 0.25326487 | 0.37614996 | 0.70680541 | 0.95501548 |
| AC005498.3 | 81.3538939 | 0.12416287 | 0.47302667 | 0.26248598 | 0.79294679 | 0.97090793 |
| ZNF470 | 466.597221 | -0.0395582 | 0.25997759 | -0.1521602 | 0.87906057 | 0.98275231 |
| ZNF71 | 403.489901 | -0.55428 | 0.21110489 | -2.6256144 | 0.00864927 | 0.37987657 |
| ZNF835 | 92.1514529 | 0.18301849 | 0.35079947 | 0.52171827 | 0.6018665 | 0.9355118 |
| AC007228.2 | 51.1538383 | 0.12249553 | 0.50036463 | 0.24481253 | 0.80660159 | 0.97334138 |
| PEG3 | 426.33468 | -0.0781581 | 0.21515506 | -0.3632639 | 0.71640773 | 0.95603249 |
| ZNF264 | 1377.44891 | -0.2185882 | 0.15155367 | -1.4423156 | 0.14921336 | 0.73670625 |
| AURKC | 157.623957 | -0.1615664 | 0.33750302 | -0.4787111 | 0.63214419 | 0.94088547 |
| ZNF805 | 51.7783563 | -0.1779578 | 0.37910073 | -0.4694209 | 0.63876879 | 0.94088547 |
| AC005261.3 | 315.936892 | -0.1594795 | 0.21298072 | -0.7487978 | 0.45397908 | 0.89670027 |
| ZNF460-AS1 | 68.3159886 | 0.66864432 | 0.56048306 | 1.19297866 | 0.23287773 | 0.80667958 |
| ZNF460 | 71.7532672 | -0.2762978 | 0.38426747 | -0.7190246 | 0.47212574 | 0.90160919 |
| AC005261.1 | 2729.99257 | -0.1366319 | 0.20466431 | -0.6675902 | 0.50439519 | 0.91356552 |
| ZNF543 | 141.453999 | 0.17888068 | 0.30066824 | 0.59494372 | 0.55188109 | 0.92373235 |
| ZNF304 | 612.254833 | 0.00135 | 0.21224565 | 0.00636058 | 0.99492503 | 0.99940889 |
| ZNF547 | 395.304045 | -0.0271666 | 0.1679454 | -0.1617588 | 0.87149584 | 0.98275231 |
| TRAPPC2B | 345.218085 | -0.3453234 | 0.25108654 | -1.3753163 | 0.16903342 | 0.75786012 |
| AC003002.1 | 244.376444 | -0.2369961 | 0.21944515 | -1.0799786 | 0.28015171 | 0.82825734 |
| ZNF548 | 688.38247 | -0.0762717 | 0.18151406 | -0.4201974 | 0.67434124 | 0.94935413 |
| AC003002.3 | 107.888767 | -0.0303691 | 0.3661331 | -0.0829456 | 0.93389478 | 0.99025262 |
| AC003002.2 | 171.663504 | 0.11827481 | 0.55086197 | 0.21470862 | 0.8299945 | 0.97550416 |
| ZNF17 | 291.603054 | 0.00199488 | 0.24858414 | 0.00802495 | 0.99359708 | 0.99920627 |
| ZNF749 | 259.949522 | 0.23385473 | 0.52371819 | 0.44652779 | 0.65521604 | 0.94532497 |
| AC004076.1 | 81.7722584 | -0.0162101 | 0.34687302 | -0.0467322 | 0.9627267 | 0.99320711 |
| VN1R1 | 122.467216 | 0.11331833 | 0.30233683 | 0.37480822 | 0.7078031 | 0.95501548 |
| ZNF772 | 343.924731 | 0.20375903 | 0.29915858 | 0.6811071 | 4.96E-01 | 9.09E-01 |
| ZNF419 | 639.202921 | 0.12637235 | 0.17112307 | 0.73848809 | 0.4602179 | 0.89755243 |
| AC003005.1 | 73.3234158 | 0.30886243 | 0.3521771 | 0.87700884 | 0.38048184 | 0.87118022 |
| ZNF773 | 280.809104 | -0.0846771 | 0.22351424 | -0.3788444 | 0.70480342 | 0.95501548 |
| ZNF549 | 415.032029 | -0.1810575 | 0.21790867 | -0.8308869 | 0.40603751 | 0.88132015 |
| ZNF550 | 520.651907 | -0.0244849 | 0.16564688 | -0.1478139 | 0.88248965 | 0.98275231 |
| ZNF416 | 174.586054 | 0.12450732 | 0.23462789 | 0.53065864 | 0.59565535 | 0.9334601 |
| ZIK1 | 226.075502 | -0.0230628 | 0.28493575 | -0.0809402 | 9.35E-01 | 0.99068685 |
| ZNF530 | 119.199771 | -0.1189523 | 0.33209773 | -0.3581845 | 0.72020523 | 0.95692068 |
| ZNF134 | 774.398741 | -0.0153028 | 0.17248326 | -0.0887205 | 0.92930401 | 0.98963127 |
| ZNF211 | 879.340271 | 0.10779319 | 0.21337616 | 0.50517916 | 0.613433 | 0.9388166 |
| ZNF551 | 282.516042 | -0.381988 | 0.23865802 | -1.6005662 | 0.10947304 | 0.67859123 |
| AC003006.1 | 95.1499686 | 0.09496335 | 0.46244556 | 0.20535034 | 0.83729842 | 0.97577999 |

| | | | | | | |
|---|---|---|---|---|---|---|
| ZNF154 | 385.23174 | -0.5570718 | 0.30578879 | -1.8217535 | 0.0684924 | 0.62698165 |
| ZNF671 | 634.623726 | -0.0361406 | 0.20950866 | -0.1725016 | 0.86304317 | 0.98176973 |
| ZNF776 | 925.889703 | -0.1159613 | 0.17960307 | -0.6456532 | 0.51850394 | 0.91636259 |
| ZNF586 | 393.098266 | -0.4376794 | 0.1750441 | -2.5003949 | 0.01240549 | 0.4154725 |
| ZNF552 | 556.039834 | -0.0388411 | 0.19098149 | -0.2033762 | 0.83884103 | 0.97612168 |
| AC010522.1 | 58.5143265 | -0.1390827 | 0.38803732 | -0.358426 | 0.72002452 | 0.95688409 |
| ZNF587B | 711.425552 | -0.3619154 | 0.33047678 | -1.0951311 | 0.27345919 | 0.82719197 |
| ZNF814 | 1504.66976 | -0.2159391 | 0.13865185 | -1.5574198 | 0.11937086 | 0.69282416 |
| ZNF587 | 1076.63457 | -0.2816271 | 0.19754501 | -1.4256353 | 0.15397366 | 0.74258845 |
| AC010326.2 | 277.77236 | 0.01488319 | 0.14465251 | 0.10288924 | 0.91805088 | 0.9868044 |
| ZNF417 | 340.908798 | -0.2319643 | 0.2080526 | -1.1149313 | 0.26487988 | 0.82446667 |
| ZNF418 | 347.000203 | -0.1498172 | 0.26632505 | -0.5625352 | 0.57375142 | 0.92978516 |
| ZNF256 | 231.343422 | -0.2303038 | 0.21136731 | -1.0895906 | 0.27589353 | 0.82719407 |
| C19orf18 | 75.6411798 | 0.233363 | 0.35097208 | 0.66490475 | 0.50611141 | 0.91391336 |
| ZNF606 | 798.351403 | 0.06847449 | 0.23091777 | 0.29653191 | 0.7668239 | 0.96741943 |
| ZSCAN1 | 115.495816 | -0.0641216 | 0.40397877 | -0.1587253 | 0.87388533 | 0.98275231 |
| ZNF135 | 3734.78282 | 0.0820727 | 0.2404506 | 0.34132874 | 7.33E-01 | 0.95848028 |
| ZSCAN18 | 5852.69344 | 0.06360872 | 0.24121989 | 0.26369601 | 0.79201417 | 0.97090793 |
| ZNF329 | 958.069795 | 0.14211773 | 0.16508801 | 0.86086043 | 0.38931492 | 0.87568585 |
| ZNF274 | 1882.15641 | -0.0787002 | 0.17907259 | -0.4394874 | 0.66030838 | 0.94628868 |
| ZNF544 | 1298.20723 | -0.1948744 | 0.10741213 | -1.8142677 | 0.06963652 | 0.62779744 |
| AC020915.5 | 1965.30013 | -0.2458068 | 0.19530937 | -1.2585509 | 0.20819257 | 0.78934595 |
| AC020915.3 | 221.40751 | -0.2756586 | 0.22649557 | -1.2170597 | 0.2235815 | 0.80078694 |
| ZNF8 | 779.833654 | -0.201906 | 0.14215261 | -1.4203468 | 0.15550675 | 0.74477742 |
| AC020915.1 | 1767.24557 | -0.2650882 | 0.20504208 | -1.2928478 | 0.19606371 | 0.77791948 |
| ERVK3-1 | 1992.544 | -0.2624452 | 0.21049895 | -1.2467766 | 0.21247943 | 0.79203904 |
| AC010642.2 | 1732.20667 | -0.3094129 | 0.24001108 | -1.289161 | 0.19734211 | 0.77816059 |
| ZSCAN22 | 291.901079 | -0.422254 | 0.2753616 | -1.5334527 | 0.12516435 | 0.7023069 |
| A1BG | 163.095407 | -0.1465879 | 0.4208775 | -0.3482912 | 0.72762148 | 0.95848028 |
| A1BG-AS1 | 298.98202 | -0.1901276 | 0.35879941 | -0.5298993 | 0.59618173 | 0.9334601 |
| ZNF497 | 413.953341 | -0.0944397 | 0.2870224 | -0.3290326 | 0.74213102 | 0.96109001 |
| AC012313.2 | 93.8724597 | -0.1006669 | 0.35750902 | -0.2815785 | 0.77826669 | 0.96836854 |
| AC012313.6 | 57.5691747 | 0.29313992 | 0.32771972 | 0.89448362 | 0.37106319 | 0.86762749 |
| ZNF837 | 208.628573 | -0.1381548 | 0.21823327 | -0.63306 | 0.52669447 | 0.91769856 |
| RPS5 | 15524.302 | 0.07656423 | 0.24078992 | 0.31797109 | 0.75050687 | 0.96255347 |
| ZNF584 | 463.837254 | 0.23198639 | 0.15196111 | 1.52661689 | 0.12685631 | 0.70443503 |
| AC012313.1 | 938.494641 | -0.3373529 | 0.30977302 | -1.0890325 | 0.27613954 | 0.82719407 |
| AC012313.5 | 135.308363 | -0.0537704 | 0.28738486 | -0.1871024 | 0.85158032 | 0.97862808 |
| ZNF132 | 341.268159 | 0.17576699 | 0.13910478 | 1.26355827 | 2.06E-01 | 7.87E-01 |
| ZNF324B | 298.038297 | -0.1132149 | 0.28456396 | -0.3978541 | 0.69073772 | 0.95158502 |
| ZNF324 | 541.781491 | -0.4069923 | 0.21382959 | -1.9033489 | 0.05699504 | 0.60049913 |
| ZNF446 | 1212.58304 | -0.1264069 | 0.15492401 | -0.8159287 | 0.41454094 | 0.88364943 |
| SLC27A5 | 1112.12009 | 0.01542869 | 0.16608548 | 0.0928961 | 0.92598611 | 0.98876911 |
| ZBTB45 | 742.677376 | -0.1296562 | 0.10825995 | -1.1976374 | 0.2310582 | 0.80623007 |
| TRIM28 | 11432.6606 | -0.282113 | 0.16855738 | -1.6736913 | 0.09419129 | 0.65783452 |
| CHMP2A | 5615.563 | 0.1336907 | 0.13087172 | 1.02153999 | 0.30699867 | 0.84286067 |
| UBE2M | 3049.52376 | -0.1688169 | 0.15214299 | -1.1095939 | 2.67E-01 | 8.27E-01 |
| MZF1-AS1 | 617.400296 | -0.2786095 | 0.21195498 | -1.3144751 | 0.18868636 | 0.76970626 |
| MZF1 | 2624.09954 | -0.2836093 | 0.12382358 | -2.2904301 | 0.0219964 | 0.51639767 |
| RPS4Y1 | 6263.86953 | 0.68279094 | 2.29600883 | 0.29738167 | 0.76617514 | 0.96704052 |
| ZFY | 402.492692 | -0.0998004 | 2.17066952 | -0.0459768 | 0.96332877 | 0.99320711 |
| ZFY-AS1 | 123.387471 | 0.47193221 | 2.27148698 | 0.20776355 | 0.83541359 | 0.97550416 |
| LINC00278 | 58.2413961 | 1.35850413 | 1.58775638 | 0.85561245 | 0.39221218 | 0.87765581 |
| TBL1Y | 38.4265996 | 0.26934562 | 1.75114197 | 0.15381141 | 0.87775843 | 0.98275231 |

| | | | | | | |
|---|---|---|---|---|---|---|
| TTTY15 | 132.759087 | 0.22912885 | 1.96339884 | 0.1167001 | 0.90709771 | 0.98662757 |
| USP9Y | 1353.07081 | 0.58732056 | 2.11166378 | 0.27813166 | 0.78091129 | 0.96882421 |
| DDX3Y | 2488.09799 | 0.41506968 | 2.38839256 | 0.17378621 | 0.86203349 | 0.98176973 |
| UTY | 587.912132 | 0.65352999 | 2.19693952 | 0.29747291 | 0.76610549 | 0.96704052 |
| TMSB4Y | 88.8036086 | -0.2019413 | 2.27620145 | -0.0887185 | 0.9293056 | 0.98963127 |
| NLGN4Y | 271.3886 | 0.36662892 | 2.46392811 | 0.14879855 | 0.8817126 | 0.98275231 |
| TTTY14 | 280.967556 | 0.73614816 | 2.13689254 | 0.3444947 | 0.73047428 | 0.95848028 |
| AC010889.1 | 78.9500825 | 0.68547656 | 1.5830447 | 0.4330115 | 0.66500641 | 0.94692723 |
| KDM5D | 2026.56893 | 0.48148718 | 2.31927593 | 0.20760237 | 0.83553945 | 0.97550416 |
| EIF1AY | 834.740227 | 0.62173552 | 2.2539283 | 0.2758453 | 0.78266689 | 0.96964828 |
| AP000547.3 | 579.96722 | -0.0558352 | 0.5399101 | -0.1034157 | 0.91763304 | 0.9868044 |
| IL17RA | 1600.68583 | -0.1489296 | 0.17796235 | -0.83686 | 0.40267126 | 0.88132015 |
| TMEM121B | 43.6739276 | 0.33346141 | 0.55825631 | 0.59732671 | 0.55028927 | 0.92373235 |
| HDHD5 | 2436.80985 | 0.01948796 | 0.18062801 | 0.10789002 | 0.91408294 | 0.98662757 |
| ADA2 | 1908.45357 | -0.0350281 | 0.47881632 | -0.0731556 | 0.94168226 | 0.99136103 |
| CECR2 | 300.316095 | -0.3380681 | 0.23707156 | -1.4260173 | 0.15386336 | 0.74258845 |
| SLC25A18 | 235.488752 | -0.3099018 | 0.59514016 | -0.5207207 | 0.60256133 | 0.93551503 |
| AC007666.1 | 61.1198227 | -0.4059147 | 0.71522082 | -0.5675375 | 0.57034903 | 0.92753481 |
| ATP6V1E1 | 9899.00407 | 0.19232352 | 0.25187889 | 0.76355554 | 0.44513215 | 0.89445558 |
| BCL2L13 | 4982.86113 | -0.2576551 | 0.36884228 | -0.6985509 | 0.48483274 | 0.90541935 |
| BID | 1029.96975 | 0.01106049 | 0.18157308 | 0.06091484 | 0.95142703 | 0.99170503 |
| MICAL3 | 3977.28377 | -0.3715759 | 0.22609395 | -1.6434579 | 0.10028822 | 0.67191826 |
| AC016027.1 | 269.4126 | -0.033908 | 0.3112298 | -0.1089484 | 0.91324341 | 0.98662757 |
| PEX26 | 3861.06685 | 0.11434296 | 0.19738801 | 0.57928018 | 0.56240014 | 0.92452095 |
| TUBA8 | 165.815434 | -0.1344078 | 0.26222386 | -0.5125688 | 0.608253 | 0.93748023 |
| AC008079.2 | 202.525135 | -0.1775649 | 0.24406116 | -0.7275426 | 0.46689363 | 0.89983399 |
| USP18 | 565.5793 | -2.5654722 | 0.92380333 | -2.7770761 | NA | NA |
| GGTLC3 | 967.073061 | 0.81324822 | 0.84646348 | 0.96075996 | 0.33667288 | 0.85679125 |
| TMEM191B | 47.8484131 | 0.10206171 | 0.8569501 | 0.11909878 | 0.90519709 | 0.98625102 |
| DGCR6 | 117.538251 | 1.86049024 | 1.13490532 | 1.6393352 | 0.10114347 | 0.6725344 |
| PRODH | 122.236036 | 2.54582482 | 1.12324625 | 2.26648859 | 0.02342148 | 0.51967919 |
| DGCR9 | 97.6154203 | 0.25912 | 0.48165582 | 0.53797752 | 0.59059257 | 0.93289552 |
| DGCR2 | 6403.40638 | -0.0865645 | 0.1232619 | -0.7022811 | 0.4825039 | 0.90541935 |
| ESS2 | 940.142451 | -0.1050601 | 0.18711693 | -0.5614678 | 0.57447868 | 0.93024203 |
| TSSK2 | 58.2437108 | -0.6046122 | 0.34694998 | -1.7426495 | 0.08139487 | 0.64390932 |
| LINC01311 | 89.2735238 | 0.09350612 | 0.36983642 | 0.25283102 | 0.8003988 | 0.97210205 |
| SLC25A1 | 6418.03759 | 0.40224719 | 0.2650949 | 1.51737052 | 1.29E-01 | 0.70722589 |
| CLTCL1 | 488.210199 | -0.2485965 | 0.24991599 | -0.9947204 | 0.31987227 | 0.84960209 |
| HIRA | 2881.69236 | -0.3266054 | 0.17837473 | -1.8310072 | 6.71E-02 | 0.6249306 |
| C22orf39 | 3019.71534 | -0.0797132 | 0.13504504 | -0.5902709 | 0.55500905 | 0.92373235 |
| MRPL40 | 1408.55904 | 0.15494433 | 0.17049889 | 0.90877031 | 0.36347138 | 0.8663515 |
| AC000068.1 | 53.1932515 | 0.21126492 | 0.41747516 | 0.50605387 | 0.61281884 | 0.93879706 |
| UFD1 | 2814.90062 | -0.3992215 | 0.30721006 | -1.2995066 | 0.19377014 | 0.77620852 |
| AC000068.2 | 737.821668 | -0.2747907 | 0.33914606 | -0.8102429 | 0.4178006 | 0.88445833 |
| CLDN5 | 1709.75898 | -0.4806384 | 0.47023203 | -1.0221304 | 0.30671919 | 0.84286067 |
| 5-Sep | 688.31866 | -0.0278485 | 0.50712565 | -0.0549144 | 0.95620662 | 0.99185043 |
| AC000093.1 | 738.34462 | -0.0485228 | 0.50550119 | -0.0959894 | 0.92352898 | 0.98858365 |
| GP1BB | 228.208606 | -0.0520865 | 0.52101752 | -0.0999707 | 0.92036757 | 0.98749691 |
| TBX1 | 59.6811216 | -0.5584054 | 0.68193242 | -0.8188575 | 0.41286775 | 0.88290038 |
| GNB1L | 275.760664 | 0.13599362 | 0.21966906 | 0.6190841 | 0.53586096 | 0.92101152 |
| RTL10 | 956.027169 | -0.1263884 | 0.1575821 | -0.8020478 | 0.42252532 | 0.88451372 |
| TXNRD2 | 2890.01256 | 0.04605578 | 0.14836135 | 0.31042979 | 0.75623414 | 0.96450325 |
| COMT | 5472.38723 | 0.1798926 | 0.25529362 | 0.70464978 | 0.48102821 | 0.90541935 |
| ARVCF | 2673.46919 | -0.2329906 | 0.21000267 | -1.1094647 | 0.26722976 | 0.8265123 |

| | | | | | | |
|---|---|---|---|---|---|---|
| TANGO2 | 2010.01054 | -0.0058372 | 0.19274682 | -0.0302841 | 0.97584044 | 0.99455099 |
| AC006547.1 | 744.426468 | -0.0074251 | 0.21596755 | -0.0343806 | 0.97257366 | 0.99416401 |
| AC006547.3 | 1023.03508 | 0.03055198 | 0.19774258 | 0.15450381 | 0.8772125 | 0.98275231 |
| DGCR8 | 2102.99365 | -0.2631355 | 0.21371415 | -1.2312497 | 0.2182295 | 0.79708421 |
| AC006547.2 | 700.437935 | -0.2328921 | 0.21027833 | -1.1075419 | 0.26805971 | 0.82676682 |
| TRMT2A | 2591.52254 | -0.1931596 | 0.13199622 | -1.463372 | 0.14336562 | 0.72852904 |
| RANBP1 | 2293.70098 | 0.04309843 | 0.13057015 | 0.33007874 | 0.74134047 | 0.96109001 |
| ZDHHC8 | 4238.59939 | -0.2490703 | 0.3217479 | -0.7741162 | 4.39E-01 | 0.89175645 |
| CCDC188 | 85.1484637 | -0.101705 | 0.54776782 | -0.1856718 | 0.85270214 | 0.97883015 |
| RTN4R | 79.2611925 | 0.01265622 | 0.28384979 | 0.04458773 | 0.96443592 | 0.99348616 |
| DGCR6L | 2061.26196 | 0.35033947 | 0.23202851 | 1.50989834 | 0.13106937 | 0.71034442 |
| ZNF74 | 515.601159 | 0.11026312 | 0.17540497 | 0.6286203 | 0.52959767 | 0.91931447 |
| SCARF2 | 962.236675 | -0.1030462 | 0.38889516 | -0.2649716 | 0.79103132 | 0.97090793 |
| AC007731.5 | 577.170729 | 0.08896355 | 0.18502067 | 0.48083034 | 0.63063709 | 0.94088547 |
| KLHL22 | 1630.531 | 0.11313029 | 0.13739512 | 0.82339383 | 0.41028407 | 0.88249519 |
| MED15 | 7325.25913 | -0.42096 | 0.15574764 | -2.7028342 | 0.0068751 | 0.35012584 |
| PI4KA | 5924.65729 | -0.1737115 | 0.10910115 | -1.592206 | 0.11133842 | 0.68001961 |
| SNAP29 | 1593.72539 | -0.1681815 | 0.18962 | -0.8869399 | 0.37511125 | 0.86832509 |
| AC007308.1 | 219.608425 | -0.3835843 | 0.24872674 | -1.5421915 | 0.12302706 | 0.69788192 |
| CRKL | 3024.25629 | -0.2269044 | 0.20474319 | -1.1082393 | 0.2677585 | 0.82676682 |
| LINC01637 | 51.1387645 | 0.98111406 | 0.58739879 | 1.67026912 | 9.49E-02 | 0.65888357 |
| AIFM3 | 159.350197 | 0.10546752 | 0.4676546 | 0.22552438 | 0.82157136 | 0.97480651 |
| LZTR1 | 4988.66509 | -0.2310299 | 0.13546941 | -1.7054024 | 0.08811939 | 0.65363785 |
| THAP7 | 1361.76224 | -0.1964685 | 0.16607453 | -1.1830143 | 0.2368035 | 0.80849753 |
| THAP7-AS1 | 280.867012 | -0.032233 | 0.38633877 | -0.0834319 | 0.93350813 | 0.99019225 |
| P2RX6 | 81.9966782 | 0.18183276 | 0.45014159 | 0.40394569 | 0.68625266 | 0.95097288 |
| GGT2 | 2379.35694 | 0.03495934 | 0.74277526 | 0.04706583 | 0.96246076 | 0.99320711 |
| HIC2 | 495.329822 | -0.0417855 | 0.25794283 | -0.1619951 | 0.87130969 | 0.98275231 |
| TMEM191C | 62.7653952 | 0.20498503 | 0.41780912 | 0.49061884 | 0.62369606 | 0.9405038 |
| UBE2L3 | 4504.10008 | -0.2294289 | 0.18676903 | -1.2284097 | 0.21929321 | 0.79847731 |
| YDJC | 1152.31052 | -0.0335 | 0.2379555 | -0.1407826 | 0.88804166 | 0.98423722 |
| SDF2L1 | 1066.36271 | 0.42899774 | 0.38078776 | 1.1266059 | 0.25990914 | 0.82424998 |
| PPIL2 | 3864.44725 | -0.1540439 | 0.23802332 | -0.6471798 | 0.51751559 | 0.91636259 |
| YPEL1 | 1167.10939 | -0.0332511 | 0.25063297 | -0.1326683 | 0.89445568 | 0.98474725 |
| MAPK1 | 5728.51052 | -0.0302019 | 0.14686809 | -0.2056395 | 0.8370725 | 0.97577999 |
| PPM1F | 2794.64301 | -0.0648146 | 0.16259402 | -0.3986285 | 0.69016694 | 0.95158502 |
| AC245452.1 | 2096.8489 | -0.2821737 | 0.28231431 | -0.9995018 | 0.31755168 | 0.84823774 |
| TOP3B | 1637.31833 | -0.3356702 | 0.31414976 | -1.0685038 | 2.85E-01 | 8.31E-01 |
| IGLV4-69 | 70.4474089 | -0.2000877 | 1.38889479 | -0.1440625 | 0.88545108 | 0.98359545 |
| IGLV6-57 | 121.789421 | -0.3864952 | 0.95583475 | -0.4043535 | 0.68595276 | 0.95097288 |
| AC245060.6 | 221.760038 | -0.2636868 | 0.4333258 | -0.6085186 | 0.54284358 | 0.92292765 |
| AC245060.5 | 104.514032 | -0.0445647 | 0.51768436 | -0.0860848 | 0.93139903 | 0.98975562 |
| AC245060.2 | 40.0761463 | -0.4343519 | 0.53370548 | -0.813842 | 0.41573546 | 0.88399703 |
| IGLV5-52 | 148.729822 | -0.569412 | 0.30598891 | -1.860891 | 0.06275957 | 0.6147045 |
| IGLV1-51 | 336.68858 | 0.37787989 | 0.86967481 | 0.43450711 | 0.66392023 | 0.94642611 |
| IGLV1-47 | 158.530702 | -0.1307776 | 1.18093172 | -0.110741 | 0.91182172 | 0.98662757 |
| IGLV7-46 | 87.3458854 | -0.1665598 | 1.47954156 | -0.1125753 | 0.91036729 | 0.98662757 |
| IGLV1-44 | 275.51428 | -0.2941563 | 1.17308342 | -0.2507548 | 0.80200367 | 0.97233627 |
| IGLV7-43 | 50.2789124 | 1.40487698 | 1.34831273 | 1.04195188 | 0.29743399 | 0.83898658 |
| IGLV1-40 | 495.98023 | -0.0882779 | 0.77076725 | -0.1145324 | 0.90881573 | 0.98662757 |
| ZNF280B | 68.8675478 | -0.3248296 | 0.45997752 | -0.7061858 | 0.48007261 | 0.90537463 |
| PRAME | 163.974286 | 0.36639466 | 0.66576471 | 0.55033656 | 0.58208855 | 0.9317705 |
| IGLV2-34 | 48.420799 | -5.2909731 | 1.68824055 | -3.1340161 | NA | NA |
| GGTLC2 | 45.102282 | -1.3067364 | 0.42732063 | -3.0579763 | 2.23E-03 | 0.20572373 |

| | | | | | | |
|---|---|---|---|---|---|---|
| IGLV3-27 | 82.5004174 | -1.3453986 | 2.09861363 | -0.6410892 | NA | NA |
| IGLV3-25 | 221.562502 | -0.3648967 | 1.11724934 | -0.3266028 | 0.74396835 | 0.9613927 |
| IGLV2-23 | 543.148452 | 1.56034082 | 1.18334051 | 1.31858988 | 0.18730626 | 0.76958556 |
| IGLV3-21 | 344.102738 | 0.85767546 | 1.44193024 | 0.59481065 | 0.55197005 | 0.92373235 |
| IGLV3-19 | 426.167231 | -1.1849779 | 1.28624535 | -0.9212689 | 0.35691003 | 0.86390849 |
| IGLV2-14 | 1505.16477 | -1.5688726 | 1.37651035 | -1.1397463 | 0.254392 | 0.81992039 |
| IGLV2-11 | 244.562112 | 0.1943695 | 1.1885765 | 0.16353133 | 0.87010011 | 0.98275231 |
| IGLV3-10 | 119.558743 | -0.1717467 | 1.3431845 | -0.1278653 | 0.89825556 | 0.98474725 |
| IGLV2-8 | 345.180692 | 1.07111067 | 1.10235375 | 0.97165784 | 0.3312208 | 0.85384238 |
| IGLV3-1 | 374.49279 | -0.7308249 | 1.11311575 | -0.6565579 | 0.51146526 | 0.91507116 |
| IGLL5 | 5197.44559 | -1.1047797 | 1.39330055 | -0.7929227 | 0.42782284 | 0.88791007 |
| IGLJ1 | 1479.57735 | -1.0619285 | 1.39911924 | -0.7589979 | 0.44785384 | 0.89480207 |
| IGLC1 | 5048.15026 | -1.1344217 | 1.39740253 | -0.8118074 | 0.41690217 | 0.88445833 |
| IGLJ2 | 666.85891 | 0.25525553 | 1.00027441 | 0.2551855 | 0.79857983 | 0.9717324 |
| IGLC2 | 3065.69843 | 0.00132644 | 1.12076923 | 0.00118351 | 0.9990557 | 0.99994357 |
| IGLJ3 | 318.086674 | -0.8577719 | 0.99095206 | -0.8656038 | 0.38670745 | 0.87432552 |
| IGLC3 | 1766.53182 | -0.8510013 | 1.14103962 | -0.7458122 | 0.45578087 | 0.89688594 |
| IGLC7 | 89.2194275 | 0.9845259 | 1.68226044 | 0.58523988 | 0.55838643 | 0.92421175 |
| RSPH14 | 290.542825 | 0.69872855 | 0.36285618 | 1.92563496 | 0.05414997 | 0.59329767 |
| GNAZ | 346.528851 | 0.36981047 | 0.37940331 | 0.974716 | 0.32970116 | 0.85356118 |
| RAB36 | 1307.31415 | 0.0624786 | 0.36576996 | 0.17081393 | 0.86437008 | 0.98188271 |
| BCR | 6200.0432 | -0.1573887 | 0.19990242 | -0.7873278 | 0.43108999 | 0.88966606 |
| PCAT14 | 37.1060511 | 0.261866 | 1.01659175 | 0.25759209 | 0.79672173 | 0.97131168 |
| AP000345.2 | 269.257195 | -1.4901756 | 0.57968003 | -2.5706864 | 0.01014972 | 0.39989496 |
| DRICH1 | 146.412426 | -1.5697475 | 0.49168939 | -3.1925592 | 0.00141018 | 0.16941145 |
| RGL4 | 1471.67814 | -0.2914314 | 0.35281382 | -0.8260203 | 0.40879257 | 0.88203324 |
| ZNF70 | 218.52863 | -0.2391313 | 0.17153443 | -1.3940715 | 0.16329601 | 0.75149188 |
| C22orf15 | 577.759119 | 0.11735443 | 0.27975999 | 0.41948253 | 0.67486352 | 0.94935413 |
| CHCHD10 | 4154.56682 | 0.07348341 | 0.24226509 | 0.30331822 | 0.76164736 | 0.96607083 |
| MMP11 | 290.830425 | 0.54110934 | 0.30795763 | 1.75709022 | 0.07890243 | 0.64079926 |
| SMARCB1 | 3849.81822 | 0.03804985 | 0.15138883 | 0.25133855 | 0.80155237 | 0.97210205 |
| DERL3 | 335.586558 | 0.82576623 | 0.43226332 | 1.91033149 | 0.05609055 | 0.59895387 |
| AP000350.5 | 96.5357749 | 0.64505181 | 0.45568769 | 1.41555681 | 0.1569053 | 0.74522306 |
| SLC2A11 | 2532.69722 | 0.50464768 | 0.27403691 | 1.84153181 | 0.06554366 | 0.62237346 |
| AP000350.4 | 11583.8576 | -0.0780514 | 0.16996984 | -0.4592071 | 0.64608544 | 0.94208859 |
| AP000350.7 | 261.910798 | 0.64177244 | 0.34218681 | 1.87550314 | 0.06072354 | 0.60964095 |
| MIF | 12307.8724 | -0.1020784 | 0.16011501 | -0.6375316 | 0.52377864 | 0.91723878 |
| MIF-AS1 | 12298.9462 | -0.1043267 | 0.16047173 | -0.6501254 | 0.51561125 | 0.91636259 |
| GSTT2B | 297.220959 | -0.2360955 | 0.58841566 | -0.4012393 | 0.68824397 | 0.95158502 |
| DDTL | 2786.2805 | 0.17675063 | 0.17330179 | 1.01990079 | 0.30777551 | 0.8434443 |
| AC253536.6 | 77.6843787 | 0.4558626 | 0.50497768 | 0.90273812 | 0.3666649 | 0.86683872 |
| DDT | 2510.52105 | 0.21523874 | 0.19531849 | 1.10198856 | 0.27046665 | 0.82716979 |
| CABIN1 | 4955.31286 | -0.2257582 | 0.19929812 | -1.1327666 | 0.2573123 | 0.82115501 |
| SUSD2 | 6099.52684 | -0.65998 | 0.56369239 | -1.1708158 | 0.24167281 | 0.81269742 |
| GGT5 | 4543.24777 | -0.2842942 | 0.2454638 | -1.158192 | 0.24678567 | 0.81542272 |
| SPECC1L | 2719.30232 | -0.3173963 | 0.16477289 | -1.9262655 | 0.05407124 | 0.59329767 |
| SPECC1L-ADO | 1638.31098 | 0.15591791 | 0.34594873 | 0.45069659 | 0.65220824 | 0.9443159 |
| ADORA2A | 1248.29611 | 0.27321583 | 0.37687484 | 0.72495109 | 0.46848203 | 0.89984974 |
| ADORA2A-AS | 891.606849 | 0.37728061 | 0.39634227 | 0.95190608 | 0.34114462 | 0.85889221 |
| UPB1 | 2825.94153 | 1.3892766 | 0.58823859 | 2.36175699 | 0.01818856 | 0.47570049 |
| GUCD1 | 9324.05672 | 0.04355169 | 0.3076921 | 0.14154307 | 0.88744094 | 0.98423722 |
| SNRPD3 | 5327.13283 | 0.14654442 | 0.19287172 | 0.75980253 | 0.44737263 | 0.89480207 |
| GGT1 | 16798.4181 | 0.43691084 | 0.40552312 | 1.07740056 | 0.28130135 | 0.82957339 |
| LRRC75B | 2825.99014 | 0.554322 | 0.43437964 | 1.27612335 | 0.20191192 | 0.78134034 |

| | | | | | | |
|---|---|---|---|---|---|---|
| AP000356.2 | 623.399861 | -0.730849 | 0.41201587 | -1.773837 | 0.07609011 | 0.63773415 |
| SGSM1 | 384.468067 | -0.1043363 | 0.23952596 | -0.4355951 | 0.66313055 | 0.94642611 |
| KIAA1671 | 2702.75475 | -0.1548576 | 0.23904635 | -0.6478139 | 0.5171053 | 0.91636259 |
| CRYBB3 | 78.8304297 | 0.02424853 | 0.55086289 | 0.04401917 | 0.96488913 | 0.99357855 |
| LRP5L | 1106.18866 | 0.04582351 | 0.271587 | 0.16872496 | 0.86601298 | 0.98197545 |
| GRK3 | 535.057235 | -0.0465355 | 0.24345465 | -0.1911464 | 0.84841087 | 0.97732301 |
| ASPHD2 | 458.645539 | -0.3842466 | 0.44215078 | -0.8690398 | 0.38482535 | 0.87312908 |
| HPS4 | 2404.51355 | -0.0013737 | 0.20704772 | -0.0066347 | 0.99470634 | 0.99940889 |
| SRRD | 1815.14685 | -0.2795695 | 0.19778556 | -1.4134979 | 0.15750938 | 0.74522306 |
| TFIP11 | 2573.26405 | -0.3539763 | 0.24163674 | -1.4649108 | 0.14294524 | 0.72784886 |
| Z95115.1 | 227.068469 | 0.32439486 | 0.31225417 | 1.0388808 | 0.29886018 | 0.84056272 |
| TPST2 | 2498.56335 | -0.1097978 | 0.15934682 | -0.6890489 | 0.49079248 | 0.90600166 |
| MIAT | 298.157333 | -0.0565513 | 0.61838519 | -0.0914499 | 0.92713512 | 0.98915637 |
| MIATNB | 127.221739 | 0.22248816 | 0.28872768 | 0.77058136 | 0.44095511 | 0.89290667 |
| LINC01422 | 44.7819386 | -0.0444128 | 0.43109877 | -0.1030224 | 0.91794522 | 0.9868044 |
| MN1 | 192.954763 | 0.37758327 | 0.42431099 | 0.8898739 | 0.3735336 | 0.86791605 |
| PITPNB | 3117.11464 | -0.2719271 | 0.16599625 | -1.638152 | 0.10138999 | 0.67261938 |
| TTC28 | 1556.82751 | 0.15847645 | 0.19447774 | 0.81488224 | 0.41513972 | 0.88377697 |
| CHEK2 | 585.621839 | -0.0085001 | 0.22689583 | -0.0374626 | 0.97011616 | 0.99400331 |
| HSCB | 643.485344 | 0.00623697 | 0.22030469 | 0.02831067 | 0.97741437 | 0.99460947 |
| CCDC117 | 1289.12395 | -0.1081484 | 0.18190395 | -0.5945356 | 0.55215393 | 0.92373235 |
| XBP1 | 8401.95977 | 0.02716913 | 0.2236093 | 0.12150268 | 0.90329289 | 0.98617053 |
| Z93930.2 | 78.5954647 | 0.41357332 | 0.29861541 | 1.38496979 | 0.16606171 | 0.75515387 |
| ZNRF3 | 1699.51462 | -0.0140814 | 0.16534813 | -0.085162 | 0.93213263 | 0.98987582 |
| KREMEN1 | 2196.79265 | 0.5597182 | 0.3340342 | 1.67563142 | 0.09381041 | 0.65783452 |
| EMID1 | 821.754204 | 0.34988894 | 0.21896188 | 1.59794458 | 0.11005531 | 0.67996643 |
| RHBDD3 | 1614.86365 | -0.4657282 | 0.20614405 | -2.2592367 | 0.02386867 | 0.51967919 |
| AL031186.1 | 122.086329 | -0.6528326 | 0.40167958 | -1.6252571 | 0.10410779 | 0.67439401 |
| EWSR1 | 12990.0432 | -0.3510177 | 0.17894557 | -1.9615892 | 0.04981033 | 0.5861488 |
| GAS2L1 | 1659.91993 | -0.1272341 | 0.26665292 | -0.4771526 | 0.63325349 | 0.94088547 |
| RASL10A | 77.5268587 | -0.3987682 | 0.32198975 | -1.2384499 | 0.21554927 | 0.7954827 |
| AP1B1 | 7197.04074 | 0.07564828 | 0.16670273 | 0.45379151 | 0.6499789 | 0.94312988 |
| NEFH | 111.955939 | -0.0184178 | 0.50676266 | -0.036344 | 0.97100803 | 0.99400331 |
| THOC5 | 2986.74597 | 0.28438232 | 0.21011198 | 1.35347979 | 0.1759024 | 0.76108589 |
| NIPSNAP1 | 6234.40923 | 0.90977817 | 0.36297895 | 2.5064213 | 0.01219602 | 0.41257332 |
| NF2 | 2357.00668 | 0.02631397 | 0.28728275 | 0.09159606 | 0.92701898 | 0.98915637 |
| CABP7 | 423.937281 | 0.16459346 | 0.17115058 | 0.96168801 | 0.33620635 | 0.85644192 |
| ZMAT5 | 801.515277 | 0.17230655 | 0.15794641 | 1.09091778 | 0.27530906 | 0.82719407 |
| UQCR10 | 3820.22086 | 0.10744984 | 0.16885721 | 0.63633555 | 0.52455774 | 0.91723878 |
| ASCC2 | 2619.71845 | -0.0715401 | 0.14244281 | -0.5022374 | 0.61550052 | 0.93910158 |
| MTMR3 | 3091.48173 | -0.3221083 | 0.18131782 | -1.7764845 | 0.0756531 | 0.63773415 |
| AC003681.1 | 2602.38321 | -0.3982052 | 0.20161607 | -1.9750666 | 0.04826059 | 0.58440502 |
| HORMAD2-A | 48.6721291 | 0.35861216 | 0.39460936 | 0.90877762 | 3.63E-01 | 8.66E-01 |
| LIF | 3517.98089 | 0.86589214 | 0.99335084 | 0.87168814 | NA | NA |
| AC004264.1 | 149.321407 | 0.5338094 | 0.88564829 | 0.60273294 | NA | NA |
| OSM | 104.70962 | 0.66647613 | 0.85872866 | 0.77611959 | 0.43767838 | 0.89130865 |
| CASTOR1 | 869.84417 | 0.06790053 | 0.42466968 | 0.15989023 | 8.73E-01 | 0.98275231 |
| AC004997.1 | 2176.57711 | -0.1139912 | 0.27080003 | -0.4209423 | 0.67379725 | 0.94935413 |
| TBC1D10A | 2562.71142 | -0.1781009 | 0.20733715 | -0.8589918 | 3.90E-01 | 0.87650906 |
| SF3A1 | 4759.34297 | -0.0835802 | 0.14628134 | -0.5713663 | 0.56775133 | 0.92618483 |
| CCDC157 | 823.161336 | -0.3408197 | 0.17395387 | -1.9592535 | 0.05008311 | 0.5861488 |
| RNF215 | 1641.66658 | -0.0640727 | 0.1623798 | -0.3945856 | 0.69314877 | 0.9519683 |
| SEC14L2 | 970.349604 | 0.1760818 | 0.23867889 | 0.73773511 | 0.46067543 | 0.89755243 |
| AC004832.3 | 2296.34297 | 0.09687425 | 0.24624982 | 0.39339823 | 0.69402537 | 0.95222846 |

| | | | | | | |
|---|---|---|---|---|---|---|
| MTFP1 | 2051.20788 | 0.07716236 | 0.27664354 | 0.2789234 | 0.78030361 | 0.96864684 |
| SEC14L4 | 46.9505882 | 0.18829247 | 0.61373448 | 0.30679793 | 0.75899718 | 0.96502715 |
| SEC14L6 | 1187.50785 | 0.58350387 | 0.54026083 | 1.08004105 | 2.80E-01 | 0.82825734 |
| GAL3ST1 | 1855.06986 | 1.14040138 | 0.69335835 | 1.64475033 | 0.10002131 | 0.67165899 |
| PES1 | 3747.69304 | -0.2421877 | 0.24166201 | -1.0021754 | 3.16E-01 | 0.84756299 |
| TCN2 | 11945.4284 | -0.220752 | 0.41013676 | -0.5382401 | 0.59041131 | 0.93289552 |
| SLC35E4 | 293.151203 | -0.0189191 | 0.23578041 | -0.0802405 | 0.93604599 | 0.99068685 |
| DUSP18 | 250.896594 | -0.0379478 | 0.15772554 | -0.2405939 | 0.80986989 | 0.97334138 |
| OSBP2 | 578.111337 | -0.1652538 | 0.20006379 | -0.8260053 | 0.40880106 | 0.88203324 |
| MORC2-AS1 | 641.695332 | -0.341081 | 0.22669663 | -1.5045704 | 0.13243457 | 0.71284199 |
| MORC2 | 1960.15846 | -0.2700666 | 0.12243676 | -2.2057638 | 0.02740054 | 0.52904542 |
| AC004542.2 | 1321.56537 | -0.2152594 | 0.15297544 | -1.4071501 | 0.15938289 | 0.74762924 |
| AC004542.1 | 1339.33768 | -0.1818038 | 0.19611319 | -0.9270349 | 0.35390842 | 0.86364633 |
| SMTN | 7549.05947 | -0.1589149 | 0.34001238 | -0.4673799 | 0.64022812 | 0.94088547 |
| AC005005.3 | 485.67098 | -0.1719669 | 0.41523622 | -0.4141423 | 0.67876987 | 0.95045025 |
| SELENOM | 3198.87795 | -0.1761454 | 0.29954542 | -0.5880423 | 0.5565039 | 0.92373235 |
| INPP5J | 2903.91474 | 0.49324548 | 0.47852171 | 1.0307693 | 0.30264902 | 0.84211658 |
| PLA2G3 | 37.0106753 | 1.18296414 | 1.01656698 | 1.16368538 | 0.24455154 | 0.81473945 |
| RNF185 | 3932.58346 | -0.2268674 | 0.23706328 | -0.9569911 | 0.33857176 | 0.85733087 |
| RNF185-AS1 | 788.200501 | -0.2302815 | 0.28861201 | -0.7978931 | 0.4249325 | 0.88573606 |
| LIMK2 | 3649.45297 | 0.12559295 | 0.18945262 | 0.66292535 | 0.50737836 | 0.91430909 |
| PIK3IP1 | 2854.82726 | 0.78800088 | 0.3317955 | 2.3749595 | 0.01755088 | 0.46859699 |
| PATZ1 | 2695.42305 | 0.34264328 | 0.2729693 | 1.25524476 | 0.20938992 | 0.79086734 |
| LINC01521 | 51.2837935 | -0.0538952 | 0.47196016 | -0.1141944 | 0.90908367 | 0.98662757 |
| DRG1 | 2091.02648 | -0.092132 | 0.18046538 | -0.5105243 | 0.60968419 | 0.93766726 |
| EIF4ENIF1 | 2099.68869 | -0.3061893 | 0.14809137 | -2.0675705 | 0.03868043 | 0.55608529 |
| AL096701.3 | 387.43285 | -0.3546603 | 0.20794284 | -1.7055665 | 0.08808882 | 0.65363785 |
| SFI1 | 2225.25029 | -0.4220313 | 0.18976993 | -2.22391 | 0.0261545 | 0.52592428 |
| PISD | 4730.37923 | -0.4867125 | 0.22435376 | -2.1693975 | 0.03005252 | 0.54012717 |
| PRR14L | 1462.92203 | -0.1506328 | 0.12171275 | -1.2376092 | 0.21586102 | 0.7954827 |
| DEPDC5 | 780.323097 | -0.1718481 | 0.14431652 | -1.1907722 | 0.23374303 | 0.80725541 |
| Z82190.2 | 944.006262 | -0.3929231 | 0.30669338 | -1.2811595 | 0.20013766 | 0.78134034 |
| YWHAH | 6431.88517 | -0.3229905 | 0.29782275 | -1.0845058 | 2.78E-01 | 8.28E-01 |
| SLC5A1 | 644.049324 | 0.4185323 | 0.6652142 | 0.62916923 | 0.52923827 | 0.91920857 |
| RFPL3 | 65.1021776 | -0.4145 | 0.24319176 | -1.7044164 | 0.08830332 | 0.65363785 |
| RFPL3S | 90.7731298 | -0.4524951 | 0.22648146 | -1.9979342 | 0.04572379 | 0.57687945 |
| RTCB | 5946.01682 | 0.29409391 | 0.27998484 | 1.05039223 | 0.29353782 | 0.83540174 |
| FBXO7 | 7576.64241 | 0.01945093 | 0.19574341 | 0.09936955 | 0.92084486 | 0.98767854 |
| TIMP3 | 28811.5484 | -0.0925045 | 0.30825428 | -0.3000916 | 0.76410725 | 0.96678847 |
| LARGE1 | 929.439875 | -0.0262306 | 0.25242259 | -0.1039155 | 0.9172364 | 0.9868044 |
| HMGXB4 | 684.02728 | -0.1813804 | 0.17908829 | -1.0127986 | 0.31115638 | 0.84476049 |
| TOM1 | 5381.51958 | -0.2227452 | 0.26159915 | -0.8514754 | 0.39450531 | 0.87884163 |
| HMOX1 | 29216.0627 | -0.733442 | 0.79063064 | -0.9276671 | 0.35358028 | 0.86364633 |
| MCM5 | 1174.59911 | -0.1027445 | 0.32198099 | -0.3191012 | 0.74964977 | 0.96249332 |
| RASD2 | 145.842954 | -0.156675 | 0.43024768 | -0.3641507 | 7.16E-01 | 0.95590747 |
| MB | 52.5861911 | 7.1253481 | 1.73972116 | 4.09568399 | NA | NA |
| APOL6 | 3110.10121 | -0.3731095 | 0.46555837 | -0.8014237 | 0.42288643 | 0.88453836 |
| RBFOX2 | 3101.39705 | -0.0201491 | 0.14601525 | -0.1379929 | 0.89024602 | 0.98466127 |
| APOL3 | 1782.92786 | -0.4198564 | 0.36790211 | -1.1412177 | 0.25377932 | 0.81902502 |
| APOL4 | 1141.04442 | -1.8537373 | 0.82402822 | -2.2496041 | 2.45E-02 | 0.51967919 |
| APOL2 | 5209.10044 | -0.7346005 | 0.43726394 | -1.6799934 | 0.09295859 | 0.65753347 |
| APOL1 | 5433.35287 | -0.7975144 | 0.42184592 | -1.8905349 | 0.05868646 | 0.60332684 |
| MYH9 | 39076.8304 | -0.3405126 | 0.28584675 | -1.191242 | 0.23355861 | 0.80725541 |
| TXN2 | 4721.25736 | 0.34225983 | 0.13412044 | 2.5518841 | 0.01071422 | 0.40196898 |

| | | | | | | |
|---|---|---|---|---|---|---|
| FOXRED2 | 936.904053 | -0.1865442 | 0.41088178 | -0.4540095 | 6.50E-01 | 9.43E-01 |
| EIF3D | 7489.85718 | 0.06181558 | 0.18291048 | 0.33795537 | 0.73539682 | 0.95895327 |
| IFT27 | 1370.56584 | 0.17050174 | 0.2153112 | 0.79188513 | 4.28E-01 | 0.88820133 |
| PVALB | 591.52199 | 0.50468077 | 0.67416818 | 0.74859773 | 0.4540997 | 0.89670027 |
| NCF4-AS1 | 63.6027786 | -0.6195827 | 0.55773993 | -1.110881 | 0.26661957 | 0.8265123 |
| NCF4 | 581.965155 | -0.6157121 | 0.44468434 | -1.3846048 | 0.16617334 | 0.75515387 |
| CSF2RB | 485.75666 | 0.13902427 | 0.52491566 | 0.26485069 | 0.79112449 | 0.97090793 |
| TST | 4951.06738 | 0.2798098 | 0.30278413 | 0.92412305 | 0.35542225 | 0.86364633 |
| MPST | 5502.25018 | 0.10127719 | 0.28699354 | 0.35289014 | 0.7241708 | 0.95813674 |
| KCTD17 | 529.80458 | -0.2888761 | 0.43266296 | -0.6676701 | 0.50434421 | 0.91356552 |
| IL2RB | 229.383497 | 0.31988521 | 0.75666898 | 0.42275449 | 0.67247439 | 0.94922827 |
| Z82188.2 | 111.210993 | 0.26571155 | 0.35917949 | 0.73977371 | 0.45943731 | 0.89755243 |
| C1QTNF6 | 836.943719 | 0.18821645 | 0.30997656 | 0.60719577 | 0.543721 | 0.92319803 |
| RAC2 | 775.548684 | -0.1768838 | 0.5981764 | -0.2957051 | 7.67E-01 | 9.67E-01 |
| CYTH4 | 885.542807 | -0.0399392 | 0.45379128 | -0.0880122 | 0.929867 | 0.98973492 |
| ELFN2 | 171.771848 | 1.48837216 | 0.68240778 | 2.18105979 | 0.02917899 | 0.53606221 |
| MFNG | 703.709611 | -0.1133736 | 0.39739001 | -0.2852955 | 0.77541779 | 0.96836854 |
| CARD10 | 4065.571 | 0.43572984 | 0.22132642 | 1.9687204 | 0.04898521 | 0.58508842 |
| CDC42EP1 | 4356.95071 | -0.275046 | 0.27101212 | -1.0148845 | 0.3101609 | 0.84456632 |
| LGALS2 | 2082.77149 | 0.5598139 | 0.40166925 | 1.39371859 | 0.1634026 | 0.75149188 |
| GGA1 | 4858.22457 | -0.2593878 | 0.14534379 | -1.7846501 | 0.07431809 | 0.63773415 |
| SH3BP1 | 505.477161 | -0.0937679 | 0.46128238 | -0.2032767 | 0.83891879 | 0.97612168 |
| Z83844.3 | 1401.6585 | 0.30011456 | 0.28216554 | 1.06361166 | 0.28750466 | 0.83239994 |
| Z83844.2 | 220.468702 | -0.0253501 | 0.45073868 | -0.0562413 | 0.95514956 | 0.99185043 |
| PDXP | 1086.55462 | 0.43800929 | 0.34592957 | 1.26618053 | 0.20544846 | 0.78571765 |
| LGALS1 | 3429.21019 | -0.0085007 | 0.38304344 | -0.0221925 | 0.98229438 | 0.99600601 |
| NOL12 | 1072.49921 | -0.0130553 | 0.1916331 | -0.0681268 | 0.94568473 | 0.99166614 |
| Z83844.1 | 689.817006 | -0.0915746 | 0.23815966 | -0.3845092 | 0.70060107 | 0.95360275 |
| TRIOBP | 5028.91769 | -0.157143 | 0.12869417 | -1.2210573 | 0.22206433 | 0.80067924 |
| H1F0 | 14319.9237 | -0.8705051 | 0.36840567 | -2.3628982 | 0.01813265 | 0.47570049 |
| GCAT | 1620.51713 | 0.51678616 | 0.40824491 | 1.26587289 | 0.2055586 | 0.78571765 |
| ANKRD54 | 1827.14552 | -0.3012767 | 0.11505504 | -2.6185444 | 0.00883058 | 0.38245454 |
| EIF3L | 10812.2795 | 0.34362977 | 0.26972132 | 1.27401782 | 0.2026571 | 0.78193335 |
| MICALL1 | 1601.5942 | -0.7550349 | 0.33671781 | -2.2423375 | 0.02493957 | 0.52085426 |
| C22orf23 | 348.63516 | -0.0954391 | 0.28226815 | -0.3381151 | 0.73527648 | 0.95895327 |
| POLR2F | 1247.4783 | -0.0648912 | 0.10061119 | -0.6449697 | 0.51894684 | 0.91636259 |
| SOX10 | 58.9546557 | -1.4853533 | 0.81346055 | -1.8259685 | 0.06785501 | 0.62599695 |
| PICK1 | 2700.97357 | -0.2036318 | 0.13073615 | -1.5575787 | 0.11933315 | 0.69281988 |
| AL031587.2 | 338.719666 | -0.2288988 | 0.17438746 | -1.3125874 | 0.18932201 | 0.76970626 |
| SLC16A8 | 75.3834029 | -0.3029875 | 0.54777571 | -0.5531234 | 0.58017894 | 0.93139743 |
| BAIAP2L2 | 3299.07999 | 0.49220083 | 0.34771772 | 1.41551841 | 0.15691655 | 0.74522306 |
| AL022322.1 | 191.011007 | 0.01136027 | 0.40725929 | 0.02789445 | 0.97774634 | 0.99476647 |
| PLA2G6 | 3070.84842 | 0.01019948 | 0.33484778 | 0.03046004 | 0.97570016 | 0.99455099 |
| AL022322.2 | 463.490377 | -0.1594105 | 0.31533735 | -0.5055236 | 0.61319112 | 0.93879706 |
| MAFF | 3006.04454 | -0.0990676 | 0.51597592 | -0.1920005 | 0.84774181 | 0.97711043 |
| TMEM184B | 4426.66244 | -0.2427607 | 0.18753947 | -1.2944515 | 0.1955095 | 0.77743338 |
| AL020993.1 | 143.953855 | -0.2570099 | 0.22141135 | -1.1607801 | 0.24573134 | 0.81542272 |
| CSNK1E | 6110.85174 | -0.205756 | 0.20896495 | -0.9846436 | 0.32479918 | 0.85181379 |
| TPTEP2-CSNI | 2573.7649 | -0.2570784 | 0.19359772 | -1.3278999 | 1.84E-01 | 0.76820236 |
| KDELR3 | 612.958287 | 0.1864922 | 0.28271688 | 0.65964296 | 0.50948298 | 0.91439937 |
| DDX17 | 33194.8381 | -0.072067 | 0.18776363 | -0.3838178 | 7.01E-01 | 9.54E-01 |
| FAM227A | 327.494376 | -0.2249019 | 0.43152439 | -0.52118 | 0.60224138 | 0.9355118 |
| CBY1 | 1770.64997 | 0.33005227 | 0.22624043 | 1.45885624 | 0.14460468 | 0.73032824 |
| AL021707.2 | 1007.25646 | 0.34519834 | 0.21066394 | 1.63862093 | 0.10129223 | 0.6725344 |

| | | | | | | |
|---|---|---|---|---|---|---|
| TOMM22 | 2521.30262 | 0.04350134 | 0.16664557 | 0.2610411 | 0.79406081 | 0.97090793 |
| JOSD1 | 2968.09444 | -0.2657589 | 0.24207951 | -1.0978166 | 0.27228458 | 0.82716979 |
| GTPBP1 | 6614.55939 | -0.0219981 | 0.21066214 | -0.1044238 | 9.17E-01 | 0.9868044 |
| AL021707.5 | 2343.91028 | 0.25531446 | 0.23796863 | 1.07289122 | 0.28331993 | 0.83094967 |
| SUN2 | 11678.0531 | 0.14845591 | 0.21807581 | 0.68075368 | 0.49602736 | 0.90879351 |
| AL021707.3 | 2163.80733 | 0.20865877 | 0.24945654 | 0.8364534 | 0.40289988 | 0.88132015 |
| AL021707.1 | 577.419378 | 0.14511702 | 0.26096753 | 0.5560731 | 0.57816088 | 0.93130183 |
| AL021707.6 | 269.07847 | -1.2084136 | 0.54617462 | -2.2125042 | 0.02693185 | 0.52685774 |
| DNAL4 | 1502.02107 | 0.10477834 | 0.19772279 | 0.52992547 | 0.5961636 | 0.9334601 |
| NPTXR | 522.881935 | 0.42154914 | 0.30539984 | 1.38031878 | 0.16748851 | 0.75558932 |
| CBX6 | 6572.27718 | 0.16853584 | 0.26869082 | 0.62724825 | 0.53049652 | 0.91962754 |
| APOBEC3A | 119.725369 | -1.1551488 | 0.63570652 | -1.8171102 | 0.06920024 | 0.62779744 |
| APOBEC3B | 89.8401442 | -0.3836249 | 0.52359585 | -0.7326737 | 0.46375748 | 0.89830907 |
| APOBEC3B-A | 43.4808632 | -0.6142184 | 0.53329699 | -1.151738 | 0.24942873 | 0.81646572 |
| APOBEC3C | 582.927237 | -0.0043768 | 0.30943533 | -0.0141446 | 0.98871462 | 0.99788078 |
| AL022318.4 | 216.627416 | -0.5043086 | 0.48631376 | -1.0370026 | 0.29973466 | 0.84142109 |
| APOBEC3D | 191.679099 | -0.5601139 | 0.55408937 | -1.0108728 | 0.3120773 | 0.84541735 |
| APOBEC3F | 230.810705 | -0.5846282 | 0.53953485 | -1.0835782 | 0.27855184 | 0.82805357 |
| APOBEC3G | 473.277971 | -0.796077 | 0.68264918 | -1.1661584 | 0.24355043 | 0.81376111 |
| APOBEC3H | 45.8697964 | -0.6210293 | 0.92418094 | -0.6719781 | 0.50159766 | 0.91259827 |
| CBX7 | 5367.14084 | 0.36325771 | 0.22114176 | 1.64264634 | 0.10045613 | 0.67213434 |
| AL031846.2 | 1354.80182 | 0.27914724 | 0.2155017 | 1.29533659 | 0.19520414 | 0.77723652 |
| PDGFB | 1564.5346 | -0.2596551 | 0.35327245 | -0.7349995 | 0.46233981 | 0.89776981 |
| RPL3 | 72010.2448 | 0.39825777 | 0.25315582 | 1.57317249 | 0.11567888 | 0.6873836 |
| SYNGR1 | 2149.38414 | 0.20938426 | 0.40222439 | 0.52056578 | 0.60266929 | 0.93560513 |
| TAB1 | 2019.58797 | -0.1635956 | 0.1761358 | -0.9288036 | 0.3529909 | 0.86364633 |
| MGAT3 | 684.214619 | 0.91869686 | 0.48052351 | 1.91186662 | 5.59E-02 | 0.59889467 |
| MIEF1 | 2328.9202 | 0.12397703 | 0.25116678 | 0.49360443 | 0.62158557 | 0.94016713 |
| AL022312.1 | 70.8686704 | -0.016121 | 0.45650948 | -0.0353136 | 0.97182971 | 0.99404213 |
| ATF4 | 18756.8682 | -0.6125398 | 0.28265904 | -2.1670625 | 0.03023009 | 0.5405936 |
| RPS19BP1 | 3957.12683 | -0.1588278 | 0.23238072 | -0.6834811 | 0.49430289 | 0.90796767 |
| GRAP2 | 101.361261 | -0.03051 | 0.80121352 | -0.0380798 | 0.96962408 | 0.99400331 |
| FAM83F | 1200.96414 | 0.2273782 | 0.33052345 | 0.68793363 | 0.49149458 | 0.90600166 |
| Z83847.1 | 50.1502342 | 0.43726778 | 0.55442457 | 0.78868759 | 0.43029462 | 0.88936206 |
| TNRC6B | 1808.31545 | -0.1066159 | 0.14040371 | -0.7593522 | 4.48E-01 | 8.95E-01 |
| ADSL | 2424.71574 | 0.00930078 | 0.16920139 | 0.05496869 | 0.9561634 | 0.99185043 |
| AL022238.4 | 7659.21754 | -0.0476963 | 0.12999264 | -0.3669157 | 0.71368191 | 0.95564093 |
| SGSM3 | 5509.19055 | -0.0781775 | 0.20029744 | -0.3903068 | 0.69630968 | 0.9525827 |
| MKL1 | 2239.50014 | -0.4148416 | 0.16175099 | -2.564693 | 0.01032671 | 0.40019066 |
| SLC25A17 | 1517.47374 | -0.0599857 | 0.13458606 | -0.4457054 | 0.65581003 | 0.94532497 |
| ST13 | 16446.5441 | 0.27082774 | 0.21597038 | 1.25400412 | 0.20984051 | 0.79134644 |
| XPNPEP3 | 1402.90944 | 0.30033867 | 0.15011758 | 2.00068949 | 0.04542586 | 0.57529633 |
| DNAJB7 | 88.5447029 | -0.2846497 | 0.52556104 | -0.5416112 | 0.58808638 | 0.93289552 |
| RBX1 | 2170.8947 | -0.0828768 | 0.13490419 | -0.6143381 | 0.53899194 | 0.9215968 |
| EP300 | 3093.83567 | -0.4085855 | 0.22584032 | -1.8091787 | 0.07042324 | 0.62967653 |
| L3MBTL2 | 1786.95986 | 0.01678402 | 0.17245058 | 0.09732652 | 0.92246709 | 0.98826666 |
| AL035681.1 | 258.78426 | -0.0956773 | 0.29134296 | -0.328401 | 0.74260845 | 0.96109001 |
| CHADL | 270.984744 | 0.10402072 | 0.28215882 | 0.36866016 | 0.71238105 | 0.95560193 |
| RANGAP1 | 5797.23082 | -0.3542658 | 0.16842181 | -2.1034435 | 0.03542702 | 0.55369025 |
| ZC3H7B | 8965.73508 | 0.10695075 | 0.15838898 | 0.67524113 | 0.49952258 | 0.91161831 |
| TEF | 2080.76786 | 0.12701467 | 0.47778196 | 0.26584233 | 0.79036064 | 0.97090793 |
| TOB2 | 4092.93943 | -0.0331166 | 0.29179032 | -0.1134945 | 0.90963851 | 0.98662757 |
| PHF5A | 1040.89402 | -0.2703286 | 0.20090195 | -1.3455749 | 0.17843966 | 0.76115956 |
| ACO2 | 20129.356 | 0.44325793 | 0.23286708 | 1.90348041 | 5.70E-02 | 0.60049913 |

| | | | | | | |
|---|---|---|---|---|---|---|
| POLR3H | 11718.1602 | 0.19337095 | 0.17490921 | 1.10555041 | 0.26892116 | 0.82683094 |
| CSDC2 | 6263.03923 | -0.2000094 | 0.42141758 | -0.474611 | 0.63506424 | 0.94088547 |
| PMM1 | 3857.52328 | 0.332283 | 0.2091951 | 1.58838808 | 1.12E-01 | 6.80E-01 |
| DESI1 | 3213.98201 | -0.1813234 | 0.22569159 | -0.8034121 | 0.4217366 | 0.88451372 |
| XRCC6 | 11791.361 | -0.0034917 | 0.17238384 | -0.0202554 | 0.98383963 | 0.9961464 |
| SNU13 | 5086.54747 | -0.0418914 | 0.10781642 | -0.388544 | 6.98E-01 | 0.9527595 |
| C22orf46 | 748.501856 | -0.0522754 | 0.26435292 | -0.1977484 | 0.84324194 | 0.97649449 |
| MEI1 | 114.273472 | 0.37341959 | 0.58125277 | 0.64243924 | 0.52058803 | 0.91680964 |
| CCDC134 | 136.834473 | -0.101084 | 0.31196172 | -0.3240271 | 0.74591755 | 0.96176407 |
| AL021453.1 | 385.360423 | 0.09372933 | 0.19340518 | 0.48462677 | 0.62794112 | 0.94088547 |
| SREBF2 | 8847.38169 | -0.0952168 | 0.24249245 | -0.3926589 | 0.69457142 | 0.95222846 |
| TNFRSF13C | 117.662094 | -0.6211172 | 0.84460935 | -0.7353899 | 0.46210209 | 0.89762182 |
| CENPM | 48.0760442 | 0.37830484 | 0.5162074 | 0.73285435 | 0.46364727 | 0.89830907 |
| 3-Sep | 50.9540255 | 0.02366203 | 0.63499325 | 0.03726344 | 0.97027496 | 0.99400331 |
| WBP2NL | 132.817313 | 0.10630563 | 0.36246849 | 0.29328241 | 0.7693063 | 0.96833698 |
| NAGA | 2286.06601 | 0.1276407 | 0.18100239 | 0.70518792 | 0.4806933 | 0.90541935 |
| PHETA2 | 541.946626 | -0.3082068 | 0.21823146 | -1.4122934 | 0.1578636 | 0.74559519 |
| SMDT1 | 2350.74607 | 0.29015899 | 0.14357555 | 2.02094985 | 0.04328496 | 0.56886632 |
| NDUFA6 | 4161.08477 | -0.1921967 | 0.20284731 | -0.9474943 | 0.34338697 | 0.85938376 |
| CYP2D6 | 312.240412 | 0.25716498 | 0.30587039 | 0.84076453 | 0.40047986 | 0.88086294 |
| AC254562.2 | 46.3062863 | 0.47012809 | 0.36680139 | 1.28169662 | 0.19994909 | 0.78134034 |
| AC254562.3 | 84.8716721 | 0.56914249 | 0.35214535 | 1.61621472 | 0.1060479 | 0.67567147 |
| TCF20 | 2177.02818 | -0.7687767 | 0.25446862 | -3.021106 | 0.00251853 | 0.21493583 |
| Z83851.1 | 64.0967705 | -0.8406958 | 0.52061742 | -1.6148053 | 0.10635287 | 0.67638364 |
| LINC01315 | 124.34876 | 0.25499541 | 0.52489222 | 0.48580529 | 0.62710523 | 0.94088547 |
| NFAM1 | 388.585249 | -0.0869043 | 0.52145438 | -0.1666576 | 0.86763951 | 0.98224588 |
| RRP7A | 2818.47306 | -0.1915007 | 0.2909187 | -0.6582619 | 0.51036983 | 0.91442074 |
| SERHL2 | 468.74819 | -0.168492 | 0.27830855 | -0.6054144 | 0.54490366 | 0.92345821 |
| POLDIP3 | 4925.098 | -0.1478909 | 0.10257368 | -1.4418019 | 0.14935825 | 0.73670625 |
| CYB5R3 | 21490.669 | 0.11915218 | 0.17746211 | 0.67142321 | 0.50195096 | 0.91259827 |
| A4GALT | 5307.67616 | 0.32046277 | 0.2715097 | 1.18029952 | 2.38E-01 | 8.10E-01 |
| ARFGAP3 | 5715.12831 | -0.0525769 | 0.1967666 | -0.2672041 | 0.78931199 | 0.97090793 |
| PACSIN2 | 8719.81535 | -0.3243354 | 0.32867398 | -0.9867999 | 0.32374076 | 0.85154042 |
| AL022476.1 | 76.9280158 | 0.47139281 | 0.35239996 | 1.3376642 | 0.18100593 | 0.76522169 |
| TTLL1 | 296.067512 | 0.55533179 | 0.263393 | 2.10837714 | 0.03499838 | 0.55369025 |
| BIK | 144.055613 | 0.2033525 | 0.33597919 | 0.60525327 | 0.54501073 | 0.92345821 |
| MCAT | 1295.10508 | 0.03385668 | 0.19734098 | 0.17156436 | 0.86378003 | 0.98176973 |
| TSPO | 3564.22404 | -0.0171993 | 0.17375672 | -0.0989852 | 0.92115001 | 0.98777092 |
| TTLL12 | 1832.70707 | 0.16108344 | 0.23503565 | 0.6853575 | 0.49311835 | 0.90711083 |
| SCUBE1 | 820.230787 | 0.99349131 | 0.39787682 | 2.49698209 | 0.01252553 | 0.41598747 |
| Z82214.1 | 38.5056908 | 1.00238377 | 0.58078782 | 1.72590357 | 0.08436476 | 0.64988556 |
| EFCAB6 | 197.025016 | 0.14966447 | 0.33373837 | 0.44844848 | 6.54E-01 | 0.94510853 |
| PNPLA3 | 523.680958 | 0.42240562 | 0.35355583 | 1.19473527 | 0.23219048 | 0.80649152 |
| SAMM50 | 2526.87503 | 0.21606364 | 0.21932085 | 0.98514863 | 0.32455107 | 0.85169085 |
| PARVB | 1164.80468 | 0.41652115 | 0.27776707 | 1.49953393 | 0.13373517 | 0.71505705 |
| PARVG | 822.640898 | -0.2219373 | 0.59714079 | -0.3716666 | 0.71014108 | 0.95524448 |
| RTL6 | 1779.09796 | -0.1082632 | 0.2198974 | -0.492335 | 0.62248256 | 0.94016713 |
| PRR5 | 1870.49482 | -0.1105434 | 0.25492253 | -0.4336353 | 0.66455326 | 0.94680968 |
| PRR5-ARHGA | 2769.23461 | 0.14297964 | 0.18514994 | 0.77223703 | 0.43997406 | 0.89208631 |
| ARHGAP8 | 2165.34474 | 0.18319607 | 0.24487334 | 0.74812582 | 0.45438427 | 0.89670027 |
| PHF21B | 231.450344 | -0.3580269 | 0.30433652 | -1.1764179 | 0.23942794 | 0.81163641 |
| NUP50-DT | 626.658769 | -0.0193139 | 0.26775661 | -0.0721324 | 0.94249655 | 0.99136103 |
| NUP50 | 2169.0185 | -0.210355 | 0.22044998 | -0.9542072 | 0.33997877 | 0.85814041 |
| KIAA0930 | 3774.65429 | -0.5654274 | 0.33715687 | -1.6770456 | 9.35E-02 | 6.58E-01 |

| | | | | | | |
|---|---|---|---|---|---|---|
| FAM118A | 1417.41085 | -1.0190563 | 0.56286798 | -1.8104713 | 0.07022274 | 0.62928077 |
| FBLN1 | 6049.16646 | 0.03903145 | 0.48530049 | 0.08042739 | 0.93589734 | 0.99068685 |
| ATXN10 | 4626.36088 | 0.10743851 | 0.17851669 | 0.60184014 | 0.54728055 | 0.92354775 |
| WNT7B | 212.130954 | 0.50767898 | 1.09088686 | 0.46538188 | 0.64165801 | 0.94088547 |
| BX537318.1 | 95.1837221 | 0.48747739 | 0.55546785 | 0.87759785 | 3.80E-01 | 0.87090707 |
| PRR34 | 198.410956 | 0.00020586 | 0.33751348 | 0.00060992 | 1.00E+00 | 0.99994357 |
| PRR34-AS1 | 247.093188 | -0.037757 | 0.32316763 | -0.116834 | 0.90699163 | 0.98662757 |
| MIRLET7BHG | 394.491644 | -0.3394516 | 0.37187095 | -0.9128211 | 0.36133665 | 0.86528137 |
| PPARA | 4075.96556 | 0.3548448 | 0.38333214 | 0.92568498 | 0.35460972 | 0.86364633 |
| CDPF1 | 481.781769 | 0.00734079 | 0.16037643 | 0.04577224 | 0.96349179 | 0.99320711 |
| TTC38 | 7961.33699 | 1.18209283 | 0.4461292 | 2.64966475 | 0.00805717 | 0.37112203 |
| GTSE1 | 49.6640214 | 0.00947603 | 0.50944988 | 0.01860051 | 0.9851598 | 0.99692004 |
| TRMU | 3400.15082 | -0.1298634 | 0.10956178 | -1.1852985 | 0.23589943 | 0.80776058 |
| CELSR1 | 4621.24737 | 0.34630157 | 0.24444688 | 1.41667411 | 0.15657823 | 0.74522306 |
| AL021392.1 | 95.8982755 | -0.2031951 | 0.36984222 | -0.5494103 | 5.83E-01 | 9.32E-01 |
| GRAMD4 | 2367.54339 | -0.2171964 | 0.2472562 | -0.8784267 | 3.80E-01 | 8.71E-01 |
| CERK | 3062.6618 | 0.2121909 | 0.28381355 | 0.7476419 | 0.45467619 | 0.89670027 |
| AL118516.1 | 240.889316 | -0.4387605 | 0.32498865 | -1.3500796 | 0.17699045 | 0.76115956 |
| TBC1D22A | 1859.95349 | 0.05013097 | 0.09299592 | 0.5390663 | 0.58984111 | 0.93289552 |
| FAM19A5 | 178.914582 | -0.567993 | 0.45294817 | -1.2539912 | 0.2098452 | 0.79134644 |
| C22orf34 | 169.327165 | -0.3944935 | 0.4791208 | -0.8233697 | 0.41029781 | 0.88249519 |
| BRD1 | 2257.63912 | -0.2665777 | 0.12342049 | -2.1599144 | 3.08E-02 | 0.54443002 |
| ZBED4 | 800.595987 | -0.420551 | 0.23183147 | -1.8140375 | 0.06967195 | 0.62779744 |
| ALG12 | 2146.87661 | -0.1732006 | 0.13759907 | -1.2587341 | 0.20812638 | 0.78934595 |
| AL671710.1 | 1208.54178 | -0.2215631 | 0.16241151 | -1.364208 | 0.17250211 | 0.75974924 |
| CRELD2 | 1929.90384 | 0.5280263 | 0.25764791 | 2.04941041 | 0.040422 | 0.55657615 |
| PIM3 | 11576.3771 | -0.3889354 | 0.28595162 | -1.360144 | 0.17378436 | 0.75974924 |
| PANX2 | 754.451074 | 0.41622325 | 0.52645365 | 0.79061709 | 0.42916747 | 0.88851358 |
| TRABD | 3679.74141 | -0.0864313 | 0.16755138 | -0.5158498 | 0.60595934 | 0.93698713 |
| AL022328.3 | 82.4694001 | -0.0513434 | 0.22473479 | -0.228462 | 0.81928708 | 0.97454841 |
| AL022328.4 | 1011.77979 | 0.03091647 | 0.13681598 | 0.22597122 | 8.21E-01 | 0.97472105 |
| SELENOO | 3497.80639 | 0.05509114 | 0.19701337 | 0.27963151 | 0.77976023 | 0.96836854 |
| AL022328.1 | 54.6251907 | -0.0599118 | 0.34562636 | -0.1733426 | 0.86238217 | 0.98176973 |
| AL022328.2 | 656.037427 | 0.01226382 | 0.23968026 | 0.0511674 | 0.95919213 | 0.9924381 |
| TUBGCP6 | 5167.2874 | -0.2129337 | 0.1539245 | -1.3833648 | 0.16655303 | 0.75558932 |
| HDAC10 | 2868.26414 | -0.110089 | 0.19043495 | -0.5780922 | 0.56320185 | 0.92452095 |
| MAPK12 | 4603.60206 | -0.0626884 | 0.16670595 | -0.3760418 | 0.70688582 | 0.95501548 |
| MAPK11 | 1771.37466 | 0.35786944 | 0.27756058 | 1.28933815 | 0.19728055 | 0.77816059 |
| PLXNB2 | 27036.4712 | 0.15538704 | 0.21356581 | 0.7275839 | 0.46686834 | 0.89983399 |
| DENND6B | 1892.48904 | 0.07290403 | 0.31386883 | 0.23227546 | 0.81632408 | 0.97388289 |
| CR559946.1 | 249.395593 | 0.04350304 | 0.33077691 | 0.13151777 | 0.89536573 | 0.98474725 |
| CR559946.2 | 331.265387 | 0.02249176 | 0.38570144 | 0.05831392 | 0.95349858 | 0.99170503 |
| PPP6R2 | 7366.65893 | -0.1461753 | 0.10693939 | -1.3668982 | 1.72E-01 | 7.60E-01 |
| SBF1 | 9894.86111 | -0.2625451 | 0.21598987 | -1.2155434 | 0.2241589 | 0.80078694 |
| ADM2 | 2055.86472 | -0.2223667 | 0.43688263 | -0.5089849 | 0.61076282 | 0.93807505 |
| MIOX | 33990.9893 | -0.5141463 | 1.1184729 | -0.4596859 | 0.64574167 | 0.94208859 |
| LMF2 | 6934.32373 | -0.0810848 | 0.11609323 | -0.6984452 | 0.48489885 | 0.90541935 |
| NCAPH2 | 2053.45236 | -0.2033371 | 0.21374535 | -0.9513055 | 0.34144931 | 0.85889221 |
| SCO2 | 2981.94848 | -0.8437356 | 0.41399078 | -2.0380542 | 0.04154451 | 0.55945849 |
| U62317.3 | 1843.49093 | -0.9390684 | 0.40130882 | -2.3400144 | 0.019283 | 0.48835495 |
| TYMP | 6606.91469 | -0.8527663 | 0.64527395 | -1.321557 | 0.1863157 | 0.76908034 |
| ODF3B | 3031.96479 | -1.2769819 | 0.70403606 | -1.8138019 | 0.06970824 | 0.62779744 |
| U62317.2 | 52.0921352 | -0.3642682 | 0.78412411 | -0.4645542 | 0.64225071 | 0.94088547 |
| KLHDC7B | 420.964329 | -0.4484374 | 0.86516453 | -0.5183262 | 0.6042307 | 0.93655508 |

| | | | | | | |
|---|---|---|---|---|---|---|
| CPT1B | 3583.50347 | -0.7723074 | 0.33288274 | -2.3200583 | 0.02033773 | 0.5002876 |
| CHKB-CPT1B | 8447.87563 | -0.3815973 | 0.20042971 | -1.9038957 | 0.05692377 | 0.60049913 |
| CHKB | 5040.8493 | -0.1120068 | 0.14352174 | -0.780417 | 0.43514549 | 0.89055338 |
| MAPK8IP2 | 265.165615 | -0.5519027 | 0.31949597 | -1.7274168 | 8.41E-02 | 6.49E-01 |
| ARSA | 4415.97485 | -0.0974703 | 0.20486997 | -0.4757667 | 0.63424059 | 0.94088547 |
| SHANK3 | 7264.64428 | -0.2899541 | 0.32091248 | -0.9035302 | 3.66E-01 | 8.67E-01 |
| ACR | 75.2831488 | 1.07802018 | 0.57838141 | 1.8638569 | 0.0623418 | 0.61330589 |
| RABL2B | 2031.78453 | -0.205388 | 0.16990287 | -1.2088556 | 0.22671833 | 0.80289651 |
| FP565260.3 | 203.42767 | -0.4546436 | 0.99914257 | -0.4550337 | 0.64908498 | 0.94312988 |
| GATD3B | 949.263755 | -0.6476793 | 0.69601746 | -0.9305503 | 0.35208624 | 0.86364633 |
| FP565260.2 | 696.308242 | -0.6464219 | 0.71266106 | -0.9070537 | 0.36437839 | 0.8663515 |
| FP565260.1 | 221.187788 | -1.0138735 | 0.83922816 | -1.2081024 | 0.22700789 | 0.80300331 |
| FP565260.6 | 261.139996 | -0.2455878 | 0.35219534 | -0.6973056 | 0.48561156 | 0.90543654 |
| SIK1B | 895.089465 | -0.7399649 | 0.54334345 | -1.3618733 | 0.17323788 | 0.75974924 |
| U2AF1L5 | 127.824539 | -0.2547395 | 1.27324115 | -0.2000717 | 0.84142454 | 0.97618641 |
| CRYAA2 | 152.419136 | 8.27305019 | 2.40882879 | 3.43446999 | 0.00059371 | 0.11756558 |
| KCNE1B | 34.9204318 | 1.54659691 | 0.71747533 | 2.15560986 | 0.03111414 | 0.54466273 |
| RBM11 | 186.908725 | -0.2463861 | 0.41636078 | -0.5917611 | 0.55401056 | 0.92373235 |
| HSPA13 | 1483.89699 | -0.1208365 | 0.36197792 | -0.3338229 | 0.73851318 | 0.9602325 |
| SAMSN1 | 389.751341 | 0.04365301 | 0.53475145 | 0.08163233 | 0.93493909 | 0.99068685 |
| AF127577.4 | 174.891185 | 0.25288304 | 0.3062887 | 0.8256362 | 0.4090105 | 0.8821603 |
| LINC02246 | 69.5563224 | 0.06934182 | 0.32731809 | 0.21184843 | 0.83222529 | 0.97550416 |
| NRIP1 | 2390.5924 | -0.7054757 | 0.38215825 | -1.8460302 | 0.06488782 | 0.62075686 |
| AF127577.1 | 795.021283 | -0.5970377 | 0.31634904 | -1.8872752 | 0.05912332 | 0.60462654 |
| USP25 | 2324.09628 | -0.3511509 | 0.14876757 | -2.3603997 | 0.01825526 | 0.47570049 |
| MIR99AHG | 462.985263 | 0.0594132 | 0.36343857 | 0.16347522 | 0.87014428 | 0.98275231 |
| CXADR | 1180.05864 | 0.3186456 | 0.20983004 | 1.51858903 | 0.12886597 | 0.70633634 |
| BTG3 | 1244.97914 | 0.08958597 | 0.26883492 | 0.33323786 | 0.73895474 | 0.9602325 |
| C21orf91 | 536.70462 | -0.0291937 | 0.14970831 | -0.1950039 | 0.8453899 | 0.97675487 |
| CHODL | 196.958063 | 1.50811039 | 0.7136413 | 2.11326109 | 0.03457843 | 0.55369025 |
| MIR155HG | 103.764821 | -1.121543 | 0.68731829 | -1.6317666 | 1.03E-01 | 6.73E-01 |
| MRPL39 | 1230.07742 | 0.24804971 | 0.13746766 | 1.80442228 | 0.07116513 | 0.6311906 |
| JAM2 | 696.015987 | 0.16572041 | 0.57744797 | 0.28698761 | 0.77412181 | 0.96836854 |
| ATP5PF | 5597.96051 | 0.12520873 | 0.18768537 | 0.66712032 | 0.50469528 | 0.91365534 |
| GABPA | 3099.2614 | 0.07708735 | 0.17265201 | 0.44648973 | 0.65524353 | 0.94532497 |
| APP | 70782.8024 | 0.26057988 | 0.20786664 | 1.2535916 | 0.20999048 | 0.79134644 |
| AP000229.1 | 42.9262707 | 1.99498672 | 0.63412444 | 3.14604925 | 0.00165492 | 0.18012143 |
| CYYR1 | 1133.74852 | 0.07944555 | 0.35465164 | 0.22401011 | 0.82274944 | 0.97519922 |
| ADAMTS1 | 25784.0734 | -0.1785991 | 0.39483795 | -0.4523353 | 6.51E-01 | 9.44E-01 |
| ADAMTS5 | 579.572012 | -0.0236782 | 0.41542658 | -0.0569973 | 0.95454731 | 0.99170503 |
| N6AMT1 | 675.984828 | 0.10835048 | 0.2133618 | 0.50782511 | 0.611576 | 0.93839412 |
| LTN1 | 1288.73287 | -0.1206765 | 0.12292154 | -0.9817359 | 0.32622996 | 0.85282528 |
| RWDD2B | 866.932005 | 0.36671897 | 0.1670676 | 2.19503343 | 0.0281612 | 0.53300785 |
| AF129075.2 | 78.0302057 | -0.1646159 | 0.48638907 | -0.338445 | 0.7350279 | 0.95895327 |
| USP16 | 2103.61062 | -0.1968994 | 0.25908751 | -0.7599725 | 0.44727103 | 0.89480207 |
| CCT8 | 6560.96577 | 0.14088116 | 0.15236257 | 0.92464419 | 0.35515102 | 0.86364633 |
| MAP3K7CL | 426.894299 | -0.5494634 | 0.50583649 | -1.086247 | 0.27736973 | 0.82782817 |
| BACH1 | 1830.89191 | -0.3625182 | 0.33741 | -1.0744145 | 0.28263694 | 0.83075577 |
| CLDN8 | 2180.53672 | 0.90539956 | 0.42596058 | 2.12554776 | 0.03354093 | 0.55369025 |
| TIAM1 | 381.882641 | 0.16192266 | 0.60770274 | 0.26645044 | 0.78989232 | 0.97090793 |
| SOD1 | 15672.4534 | 0.35361358 | 0.22773075 | 1.55277047 | 0.12047798 | 0.69467399 |
| AP000254.1 | 238.096702 | 0.08401698 | 0.16605304 | 0.50596473 | 0.61288141 | 0.93879706 |
| SCAF4 | 1687.53596 | -0.2721703 | 0.16601045 | -1.6394766 | 0.10111404 | 0.6725344 |
| HUNK | 887.324888 | 0.19608392 | 0.30341485 | 0.64625685 | 0.51811303 | 0.91636259 |

| | | | | | | |
|---|---|---|---|---|---|---|
| MIS18A | 301.235782 | 0.10989186 | 0.28798971 | 0.3815826 | 0.70277099 | 0.95431462 |
| MRAP | 272.688276 | 0.26704947 | 0.1929842 | 1.38378928 | 0.16642299 | 0.75558932 |
| URB1 | 1059.25945 | 0.08259546 | 0.16618007 | 0.49702384 | 0.61917224 | 0.93945324 |
| URB1-AS1 | 297.407087 | -0.0944367 | 0.29523294 | -0.3198719 | 0.7490654 | 0.96249332 |
| EVA1C | 873.1119 | 0.30657736 | 0.39901781 | 0.76833 | 0.44229115 | 0.89340144 |
| TCP10L | 109.704412 | -0.0764769 | 0.39143076 | -0.1953779 | 0.84509713 | 0.97675487 |
| AP000275.2 | 1862.64251 | -0.0194718 | 0.10626234 | -0.1832427 | 0.85460758 | 0.97910552 |
| CFAP298 | 2385.89767 | -0.0561808 | 0.12601159 | -0.4458384 | 0.65571398 | 0.94532497 |
| SYNJ1 | 1048.89303 | 0.09527762 | 0.1322954 | 0.72018844 | 0.47140898 | 0.90140554 |
| PAXBP1-AS1 | 445.696235 | -0.168155 | 0.15155897 | -1.1095022 | 0.26721361 | 0.8265123 |
| PAXBP1 | 2356.52355 | -0.2458172 | 0.1467244 | -1.6753672 | 0.0938622 | 0.65783452 |
| C21orf62-AS | 64.0788783 | 0.3393099 | 0.56547835 | 0.60004048 | 0.54847925 | 0.92373235 |
| C21orf62 | 99.704867 | 0.75699941 | 0.63196651 | 1.19784734 | 0.23097645 | 0.80610188 |
| OLIG1 | 142.791338 | 0.44385219 | 0.47127928 | 0.9418029 | 0.34629356 | 0.86135217 |
| IFNAR2 | 1855.25351 | 0.43514902 | 0.25935351 | 1.67782201 | 0.09338185 | 0.65756964 |
| AP000295.1 | 2369.82724 | 0.34547594 | 0.19159003 | 1.80320418 | 0.07135615 | 0.63139079 |
| IL10RB-DT | 1003.38335 | 0.34533227 | 0.31743108 | 1.08789682 | 0.27664066 | 0.82753753 |
| IL10RB | 1591.71971 | 0.2236525 | 0.18970694 | 1.17893684 | 0.23842333 | 0.81094221 |
| IFNAR1 | 3533.86819 | 0.07086986 | 0.21936017 | 0.32307534 | 0.74663818 | 0.96216367 |
| IFNGR2 | 4593.81353 | 0.11279433 | 0.1920697 | 0.58725727 | 0.55703094 | 0.92373235 |
| TMEM50B | 6202.62599 | 0.10848073 | 0.12397022 | 0.87505473 | 0.38154412 | 0.87185666 |
| DNAJC28 | 150.670028 | 0.17003888 | 0.24552659 | 0.6925477 | 0.48859344 | 0.90585781 |
| GART | 2292.08158 | -0.0284707 | 0.15359711 | -0.1853595 | 8.53E-01 | 9.79E-01 |
| SON | 9730.12574 | -0.3193136 | 0.16123264 | -1.9804527 | 0.04765269 | 0.58241466 |
| DONSON | 2931.58167 | -0.2206285 | 0.17156662 | -1.2859638 | 0.19845569 | 0.78046101 |
| AP000311.1 | 7558.73049 | 0.22205223 | 0.18173767 | 1.22182829 | 0.22177257 | 0.80054995 |
| CRYZL1 | 1162.23345 | 0.13716019 | 0.12696802 | 1.08027353 | 0.28002039 | 0.82825734 |
| ITSN1 | 1951.34916 | -0.2284434 | 0.12991611 | -1.7583918 | 0.07868087 | 0.64079926 |
| ATP5PO | 8858.2043 | 0.18594249 | 0.17782237 | 1.04566422 | 0.29571609 | 0.83750889 |
| LINC00649 | 1757.65138 | 0.21564711 | 0.18929982 | 1.13918288 | 0.25462688 | 0.81992039 |
| MRPS6 | 7507.05409 | -0.39273 | 0.38648422 | -1.0161604 | 0.30955299 | 0.84432074 |
| SLC5A3 | 8599.30733 | -0.8559568 | 0.38214528 | -2.2398729 | 0.02509918 | 0.52085426 |
| LINC00310 | 43.1022453 | -0.5092057 | 0.51366945 | -0.9913101 | 0.32153416 | 0.85119105 |
| KCNE2 | 44.2214956 | 0.72016142 | 0.44674577 | 1.61201619 | 0.10695842 | 0.67674128 |
| KCNE1 | 90.8615845 | -1.4500366 | 0.67830103 | -2.1377479 | 0.03253721 | 0.55104386 |
| RCAN1 | 5753.60963 | -0.1225175 | 0.25226115 | -0.4856771 | 0.62719611 | 0.94088547 |
| CLIC6 | 551.412168 | 0.29131279 | 0.39190946 | 0.74331657 | 0.45729004 | 0.89688594 |
| LINC01426 | 68.0069684 | 1.5808742 | 0.74219482 | 2.12999896 | 0.0331717 | 0.55156489 |
| RUNX1 | 1205.83591 | 0.10665803 | 0.50945864 | 0.20935562 | 0.83417063 | 0.97550416 |
| SETD4 | 1511.67153 | -0.1657914 | 0.14459191 | -1.146616 | 0.25154038 | 0.81833349 |
| CBR1 | 9177.55243 | 0.22532735 | 0.22652514 | 0.99471235 | 0.31987618 | 0.84960209 |
| AP000688.2 | 260.731027 | 0.48624762 | 0.37446784 | 1.29850303 | 0.19411454 | 0.7762856 |
| CBR3 | 153.509906 | -0.0673016 | 0.29516117 | -0.2280165 | 0.81963338 | 0.97454841 |
| DOPEY2 | 599.707532 | 0.26369752 | 0.1984717 | 1.32864041 | 0.18396663 | 0.76791734 |
| MORC3 | 1380.62722 | -0.1210062 | 0.17539049 | -0.6899246 | 0.49024159 | 0.90600166 |
| CHAF1B | 172.2503 | -0.2579168 | 0.36776539 | -0.7013081 | 0.48311076 | 0.90541935 |
| AP000695.2 | 92.5555544 | -0.5858864 | 0.42868325 | -1.3667117 | 0.17171568 | 0.75974924 |
| CLDN14 | 505.60446 | -0.8343217 | 0.43647437 | -1.9115021 | 0.05594009 | 0.59889467 |
| AP000696.1 | 223.162176 | -0.5945338 | 0.5191497 | -1.1452069 | 0.25212345 | 0.81833349 |
| AP000697.1 | 54.448331 | -0.7600825 | 0.85406678 | -0.8899568 | 0.37348911 | 0.86791605 |
| SIM2 | 2943.28284 | -0.484024 | 0.50404277 | -0.9602836 | 0.33691249 | 0.85679125 |
| HLCS | 1160.13898 | 0.22031175 | 0.22514776 | 0.97852073 | 0.32781684 | 0.85317073 |
| PIGP | 703.034555 | 0.52321521 | 0.18120282 | 2.88745629 | 0.00388371 | 0.26919756 |
| TTC3 | 5156.56502 | -0.015729 | 0.12172961 | -0.1292127 | 0.89718938 | 0.98474725 |

| | | | | | | |
|---|---|---|---|---|---|---|
| TTC3-AS1 | 151.420611 | -0.0443451 | 0.30058083 | -0.1475314 | 0.88271261 | 0.98276701 |
| VPS26C | 3834.21312 | 0.04624573 | 0.27185829 | 0.1701097 | 0.86492386 | 0.98197545 |
| DYRK1A | 3319.16653 | -0.3746975 | 0.16022496 | -2.3385716 | 0.01935761 | 0.48958305 |
| KCNJ15 | 15534.8452 | 0.66587489 | 0.45162072 | 1.47441174 | 0.14037076 | 0.725307 |
| ERG | 1487.32893 | -0.4641507 | 0.37890264 | -1.2249866 | 0.22058026 | 0.79912907 |
| ETS2 | 9972.43603 | -0.4698698 | 0.27583799 | -1.7034267 | 0.08848825 | 0.65363785 |
| AF064858.1 | 41.5515833 | -0.0988487 | 0.51286328 | -0.1927388 | 8.47E-01 | 9.77E-01 |
| PSMG1 | 1155.17464 | -0.2253668 | 0.28245702 | -0.7978801 | 0.42494008 | 0.88573606 |
| BRWD1 | 2621.97028 | 0.06576185 | 0.17992621 | 0.36549342 | 0.71474312 | 0.9558355 |
| AF129408.1 | 140.9657 | -0.2655305 | 0.32131755 | -0.8263803 | 0.40858841 | 0.88203324 |
| BRWD1-AS2 | 39.1141877 | 0.23122863 | 0.620397 | 0.37271075 | 0.70936373 | 0.95501548 |
| HMGN1 | 6491.80376 | -0.0909436 | 0.15226131 | -0.5972862 | 0.55031631 | 0.92373235 |
| WRB | 1861.81025 | 0.18892347 | 0.18391393 | 1.02723849 | 0.30430818 | 0.84211658 |
| LCA5L | 244.491224 | -0.0683521 | 0.23808963 | -0.2870857 | 0.77404668 | 0.96836854 |
| SH3BGR | 363.949736 | 0.61219473 | 0.28971848 | 2.1130676 | 0.03459498 | 0.55369025 |
| B3GALT5 | 64.8259014 | 1.71947347 | 0.61854637 | 2.77986188 | 0.0054382 | 0.31359184 |
| IGSF5 | 49.5172534 | 1.56701875 | 0.73699705 | 2.12622121 | 0.03348485 | 0.55367229 |
| PCP4 | 744.736526 | 0.03999385 | 0.68515095 | 0.05837232 | 0.95345206 | 0.99170503 |
| LINC00323 | 88.6138371 | -0.5481417 | 0.75172119 | -0.7291822 | 0.46589023 | 0.89960479 |
| BACE2 | 4898.53399 | -0.6350801 | 0.48118566 | -1.3198233 | 0.18689402 | 0.76958556 |
| PLAC4 | 99.141805 | -3.0149656 | 1.09205522 | -2.7608179 | NA | NA |
| FAM3B | 853.147328 | 0.45178305 | 0.29051497 | 1.5551111 | 0.11991961 | 0.69433886 |
| MX2 | 1311.97226 | -2.8839802 | 1.05239343 | -2.7404012 | NA | NA |
| MX1 | 9241.78529 | -3.237508 | 1.12667841 | -2.8734978 | NA | NA |
| AP001610.1 | 1364.40668 | -3.1255667 | 1.10075776 | -2.8394682 | NA | NA |
| TMPRSS2 | 4541.85391 | 0.14886655 | 0.25084438 | 0.59346176 | 0.55287216 | 0.92373235 |
| LINC00479 | 41.2141595 | -0.2363064 | 0.76814457 | -0.3076327 | 0.75836187 | 0.96502715 |
| RIPK4 | 2798.15011 | -0.3188019 | 0.36661819 | -0.8695748 | 0.38453282 | 0.87309542 |
| AP001615.1 | 625.481325 | -0.3517606 | 0.39635147 | -0.8874966 | 0.37481159 | 0.86791605 |
| PRDM15 | 589.141548 | -0.4473519 | 0.33924637 | -1.3186637 | 0.18728158 | 0.76958556 |
| C2CD2 | 1631.90141 | -0.2747164 | 0.21016937 | -1.307119 | 0.19117229 | 0.77292503 |
| ZBTB21 | 1025.80916 | -0.1241429 | 0.27573862 | -0.4502194 | 0.65255223 | 0.94437526 |
| ABCG1 | 1473.03844 | -0.5831921 | 0.3929772 | -1.4840354 | 0.13779953 | 0.72209298 |
| TFF3 | 131.562585 | 0.34707049 | 0.51587041 | 0.6727862 | 0.50108332 | 0.91234844 |
| TMPRSS3 | 387.441045 | -0.1428848 | 0.50203482 | -0.2846114 | 0.77594191 | 0.96836854 |
| UBASH3A | 58.815045 | 0.41147679 | 0.96635546 | 0.42580272 | 0.6702516 | 0.949185 |
| RSPH1 | 370.650903 | 0.81951785 | 0.39777273 | 2.06026655 | 0.03937307 | 0.55657615 |
| SLC37A1 | 1071.19378 | -0.0914233 | 0.21347483 | -0.4282628 | 0.66845979 | 0.9486589 |
| LINC01671 | 907.318351 | -0.7023759 | 0.5106027 | -1.375582 | 0.16895108 | 0.75786012 |
| AP001626.1 | 87.3326288 | -0.6287166 | 0.50562038 | -1.2434559 | 0.2136999 | 0.7926732 |
| PDE9A | 1808.49587 | -0.4996525 | 0.3175131 | -1.5736437 | 0.11556984 | 0.6873836 |
| WDR4 | 381.258482 | -0.3351834 | 0.19435357 | -1.7246063 | 0.08459844 | 0.64988556 |
| NDUFV3 | 2290.92524 | 0.17740914 | 0.17331034 | 1.02365009 | 3.06E-01 | 0.84270033 |
| PKNOX1 | 1046.82597 | -0.0660616 | 0.18605298 | -0.355069 | 0.72253788 | 0.95741464 |
| CBS | 66.9888587 | 0.52749256 | 0.8757696 | 0.60231888 | 0.5469619 | 0.92345821 |
| U2AF1 | 129.612297 | -0.7657484 | 1.10045522 | -0.6958469 | 0.4865247 | 0.90573694 |
| CRYAA | 302.329984 | -0.3928966 | 1.75701347 | -0.2236161 | 0.82305601 | 0.97537437 |
| LINC01679 | 119.383425 | 0.45155645 | 0.70708946 | 0.63861291 | 0.52307478 | 0.91723878 |
| SIK1 | 243.745287 | 2.59620224 | 1.15141395 | 2.25479485 | NA | NA |
| RRP1B | 1020.64781 | -0.1788063 | 0.18834799 | -0.9493402 | 0.3424476 | 0.85889221 |
| PDXK | 9490.84992 | -0.0201118 | 0.18248783 | -0.1102092 | 0.91224344 | 0.98662757 |
| CSTB | 5572.93376 | -0.0568103 | 0.17952266 | -0.3164518 | 0.75165958 | 0.96289476 |
| RRP1 | 1658.21975 | -0.415864 | 0.18366474 | -2.2642558 | 0.02355839 | 0.51967919 |
| AGPAT3 | 9093.59908 | 0.26341497 | 0.29304822 | 0.89887928 | 0.36871696 | 0.86762749 |

| | | | | | | |
|---|---|---|---|---|---|---|
| TRAPPC10 | 687.846198 | -0.0345566 | 0.35494576 | -0.0973574 | 0.9224426 | 0.98826666 |
| PWP2 | 130.176389 | 1.83289128 | 1.35731882 | 1.35037638 | 0.17689528 | 0.76115956 |
| GATD3A | 473.578675 | 2.33047017 | 1.69445928 | 1.3753474 | 0.16902377 | 0.75786012 |
| ICOSLG | 269.106928 | -0.7528288 | 0.93100245 | -0.8086217 | 0.41873275 | 0.88451372 |
| DNMT3L | 55.6990085 | -0.5662635 | 0.6548325 | -0.8647456 | 0.38717842 | 0.87460898 |
| AIRE | 96.3188737 | -0.0575654 | 0.56436329 | -0.1020006 | 0.91875619 | 0.98709788 |
| PFKL | 24545.3244 | -0.1812834 | 0.15639707 | -1.1591231 | 0.24640603 | 0.81542272 |
| C21orf2 | 4145.07192 | -0.2401591 | 0.19710849 | -1.218411 | 2.23E-01 | 0.80078694 |
| AP001062.1 | 2382.46848 | -0.2177779 | 0.22188654 | -0.981483 | 0.3263546 | 0.85282528 |
| TRPM2 | 300.880345 | -0.3698155 | 0.47303806 | -0.7817881 | 0.43433912 | 0.89055338 |
| LRRC3 | 133.7067 | -0.5304927 | 0.4624021 | -1.1472542 | 0.25127658 | 0.81815815 |
| UBE2G2 | 5417.25448 | -0.0970777 | 0.122303 | -0.7937479 | 0.42734222 | 0.88769902 |
| SUMO3 | 4530.98273 | -0.2819205 | 0.22736452 | -1.2399493 | 0.21499415 | 0.79487183 |
| PTTG1IP | 19586.089 | -0.1043358 | 0.16688803 | -0.6251842 | 0.53185015 | 0.91962754 |
| ITGB2 | 2121.84889 | -0.1212426 | 0.57436729 | -0.2110889 | 0.83281786 | 0.97550416 |
| ITGB2-AS1 | 265.724356 | -0.3639199 | 0.67485856 | -0.5392537 | 0.58971184 | 0.93289552 |
| LINC01547 | 139.73175 | 0.17283005 | 0.37666595 | 0.45884172 | 0.64634783 | 0.94208859 |
| FAM207A | 802.2484 | -0.2027962 | 0.15331988 | -1.3226998 | 0.18593523 | 0.76908034 |
| AP001505.1 | 61.559263 | 0.23103575 | 0.3603847 | 0.6410809 | 0.52147012 | 0.91723878 |
| ADARB1 | 1559.61218 | -0.482261 | 0.256694 | -1.878739 | 6.03E-02 | 6.08E-01 |
| LINC00334 | 830.796936 | -0.3743971 | 0.19213166 | -1.9486485 | 0.05133741 | 0.58634747 |
| POFUT2 | 3637.09681 | -0.4554401 | 0.17608411 | -2.5864918 | 0.00969585 | 0.39368232 |
| BX322562.1 | 366.824594 | -0.7324134 | 0.33572641 | -2.1815782 | 0.02914068 | 0.53606221 |
| COL18A1 | 41753.0512 | 0.04656331 | 0.21619308 | 0.21537836 | 0.82947234 | 0.97550416 |
| COL18A1-AS | 207.44821 | -0.5800495 | 0.79223635 | -0.7321673 | 0.46406647 | 0.89830907 |
| SLC19A1 | 19011.5205 | 0.07224916 | 0.18978176 | 0.38069599 | 0.70342885 | 0.95466435 |
| PCBP3 | 201.993653 | -0.4507172 | 0.65306642 | -0.6901552 | 0.49009659 | 0.90600166 |
| COL6A1 | 14643.1064 | 0.07043386 | 0.28354791 | 0.24840198 | 0.8038234 | 0.97274243 |
| COL6A2 | 10547.7184 | -0.4899632 | 0.47349805 | -1.0347734 | 0.30077476 | 0.84200819 |
| FTCD | 10647.5385 | 0.72237955 | 0.49079092 | 1.47186821 | 0.14105647 | 0.72711201 |
| FTCD-AS1 | 1206.24582 | 0.88014656 | 0.50271339 | 1.75079196 | 0.07998175 | 0.64079926 |
| SPATC1L | 351.126419 | -0.1552134 | 0.38832044 | -0.3997045 | 0.68937414 | 0.95158502 |
| AP001468.1 | 64.5787321 | 0.31016832 | 0.39713346 | 0.78101785 | 4.35E-01 | 8.91E-01 |
| LSS | 3866.13234 | 0.04266136 | 0.23042322 | 0.1851435 | 0.85311649 | 0.97886533 |
| AP001469.1 | 114.957443 | 0.13311966 | 0.43572068 | 0.30551604 | 0.75997315 | 0.96514582 |
| MCM3AP-AS | 2962.88734 | -0.1453801 | 0.11413735 | -1.2737292 | 0.2027594 | 0.78193335 |
| MCM3AP | 3946.27782 | -0.1704367 | 0.11619824 | -1.4667753 | 0.1424372 | 0.72784886 |
| AP001469.3 | 340.123119 | -0.0077719 | 0.18996125 | -0.0409133 | 0.96736502 | 0.99400331 |
| YBEY | 408.848984 | -0.6631439 | 0.38396687 | -1.7270862 | 0.08415218 | 0.64954534 |
| C21orf58 | 162.095115 | 0.4140896 | 0.38746523 | 1.06871421 | 0.28519847 | 0.83107779 |
| PCNT | 1350.49742 | -0.1778748 | 0.15353845 | -1.1585035 | 0.24665862 | 0.81542272 |
| DIP2A | 1812.37972 | -0.2326146 | 0.12411 | -1.8742614 | 0.0608944 | 0.60989294 |
| S100B | 70.5688435 | -0.2397717 | 0.95783286 | -0.2503273 | 0.80233425 | 0.97245417 |
| PRMT2 | 4436.47499 | 0.12083432 | 0.14669983 | 0.82368415 | 0.41011905 | 0.88249519 |
| MT-ND1 | 506074.065 | 0.05487317 | 0.50216061 | 0.10927415 | 0.91298505 | 0.98662757 |
| MT-ND2 | 497597.951 | -0.1283354 | 0.47573226 | -0.2697639 | 0.78734187 | 0.97090793 |
| MT-CO1 | 1967945.24 | 1.3395035 | 0.35523212 | 3.7707837 | 0.00016274 | 0.05980058 |
| MT-CO2 | 670189.307 | 0.30053817 | 0.31372424 | 0.95796923 | 0.33807828 | 0.8571036 |
| MT-ATP8 | 335983.065 | 0.03586074 | 0.41619548 | 0.08616322 | 0.93133667 | 0.98975562 |
| MT-ATP6 | 831631.519 | 0.12846964 | 0.41765365 | 0.30759852 | 0.75838785 | 0.96502715 |
| MT-CO3 | 1276195.01 | 0.38628941 | 0.32546071 | 1.18690029 | 0.23526695 | 0.80747753 |
| MT-ND3 | 148258.448 | -0.0071366 | 0.35564508 | -0.0200666 | 0.98399024 | 0.9961464 |
| MT-ND4L | 296783.762 | 0.10188619 | 0.40491365 | 0.25162448 | 0.80133134 | 0.97210205 |
| MT-ND4 | 1385339.17 | 0.21014646 | 0.36097613 | 0.58216166 | 0.5604578 | 0.92434754 |

| | | | | | | |
|---|---|---|---|---|---|---|
| MT-ND5 | 247438.436 | 0.241465 | 0.45808624 | 0.5271169 | 0.59811241 | 0.9346978 |
| MT-ND6 | 70037.9947 | 0.80138123 | 0.50261954 | 1.59440923 | 0.11084441 | 0.68001961 |
| MT-CYB | 701070.424 | 0.11403139 | 0.39231053 | 0.29066615 | 0.77130667 | 0.96836854 |
| AC145212.1 | 682.505015 | -0.3922106 | 0.44526529 | -0.8808471 | 0.37840058 | 0.87045602 |
| MAFIP | 638.077877 | -0.4141954 | 0.43938182 | -0.9426776 | 3.46E-01 | 0.86111349 |
| AC011043.1 | 651.508405 | 0.48788783 | 0.64398822 | 0.75760366 | 0.44868829 | 0.8948759 |
| AL592183.1 | 309.427976 | 0.26416097 | 0.43877576 | 0.60204093 | 0.5471469 | 0.92345821 |
| AC007325.4 | 69.5119305 | -0.2397005 | 0.52069147 | -0.4603503 | 0.64526478 | 0.94208859 |
| AC007325.2 | 655.131259 | -0.099357 | 0.54535451 | -0.1821879 | 0.85543523 | 0.97920671 |
| AL354822.1 | 49.6382944 | -1.2756493 | 0.5119786 | -2.4916068 | 0.01271667 | 0.41598747 |
| AC004556.1 | 120.072435 | -5.8137937 | 1.43903651 | -4.0400599 | 5.34E-05 | 0.0278187 |
| AC233755.1 | 88.3958587 | -1.0165305 | 2.00958536 | -0.5058409 | NA | NA |
| AC240274.1 | 328.811538 | 0.18534843 | 0.45457952 | 0.40773599 | 0.68346751 | 0.95070301 |
